# Intergeneric chromosomal transfer in yeast results in improved phenotypes and widespread transcriptional responses

**DOI:** 10.1101/2025.01.24.634819

**Authors:** Yilin Lyu, Yi Shi, Kunfeng Song, Jungang Zhou, Hao Chen, Xueying C. Li, Yao Yu, Hong Lu

## Abstract

Interspecific genetic exchanges caused by natural hybridization or horizontal gene transfer can lead to enhanced phenotypes, which are often of interest for industrial applications and evolutionary research. However, transferring genetic materials between distantly related species, such as intergeneric yeasts, presents technical challenges. In this study, we established a method to transfer individual chromosomes from *Saccharomyces cerevisiae* (*Sc*) into *Kluyveromyces marxianus* (*Km*), an emerging model for bioproduction. The *Sc* chromosome of interest was circularized, genetically modified to carry *Km* centromeres and replication origins, and transformed into *Km* via protoplast transformation. Using this method, we generated two synthetic strains, each containing a full set of *Km* chromosomes and either *Sc* chromosome I or III. The *Sc* chromosomes exhibited normal replication, segregation, and active transcription after the transfer. The synthetic strains displayed enhanced phenotypes in flocculation and salt tolerance, which were found to be caused by transgressive expression of *FLO9* and *SPS22* on the transferred *Sc* chromosomes, respectively. Transcriptomic analysis revealed that transgressive expression was prevalent among the transferred *Sc* genes, suggesting evolution of lineage-specific *cis-* and *trans*-regulatory interactions across a long evolutionary timescale. Our strategy has potential applications in optimizing cell factories, constructing synthetic genomes, and advancing evolutionary research.

## Introduction

The transfer of genetic materials between species is one of the major driving forces for adaptation ^1,2^. Chromosomes, which carry packs of genes, can introduce abundant genetic variation and contribute to evolution of adaptive traits when transferred between species. For example, horizontal transfer of plastid chromosomes between tobacco plants (*Nicotiana*) aided chloroplast capture ^3^. Horizontal transfer of chromosomes between *Fusarium* species, a group of filamentous fungi, led to new pathogenic lineages ^4^. Furthermore, hybridization is by nature a process of interspecific chromosome transfers. Interspecific hybridization often leads to enhanced phenotypes, giving rise to industrially important strains such as the lager yeast *Saccharomyces pastorianus* ^5^ and hybrid crops like triticale ^6^. Therefore, introduction of genetic materials from a distantly related species may hold great potential for phenotypic improvement in agricultural and industrial practices. In addition, understanding the phenotypic and molecular consequences of interspecific genetic exchanges may shed light on genome evolution, such as the molecular bases for heterosis ^7^ and evolution of gene regulation ^8,9^.

In this study, we explore the phenotypic and molecular consequences of chromosomal transfers between *Kluyveromyces marxianus* (*Km*) and *Saccharomyces cerevisiae* (*Sc*). *K. marxianus* is a yeast species that belongs to the *Saccharomycetaceae* family but only distantly related to *S. cerevisiae* (**Fig. 1a**). It has been previously argued that it may serve as an emerging model for bioproduction, including heterologous proteins, bioethanol and bulk chemicals ^10,11^. It has a number of traits different from *S. cerevisiae*, including a high growth rate, thermotolerance, and the ability to assimilate a wide variety of sugars ^12–14^. On the other hand, *S. cerevisiae* is more tolerant to ethanol than *K. marxianus* ^11^. It is widely used in industrial fermentation and has arguably the best-studied genome among eukaryotes ^15^. Mixing the genetic material between the two species is expected to provide novel phenotypes for industrial development, as well as new insights for understanding their evolution.

**Fig. 1.**
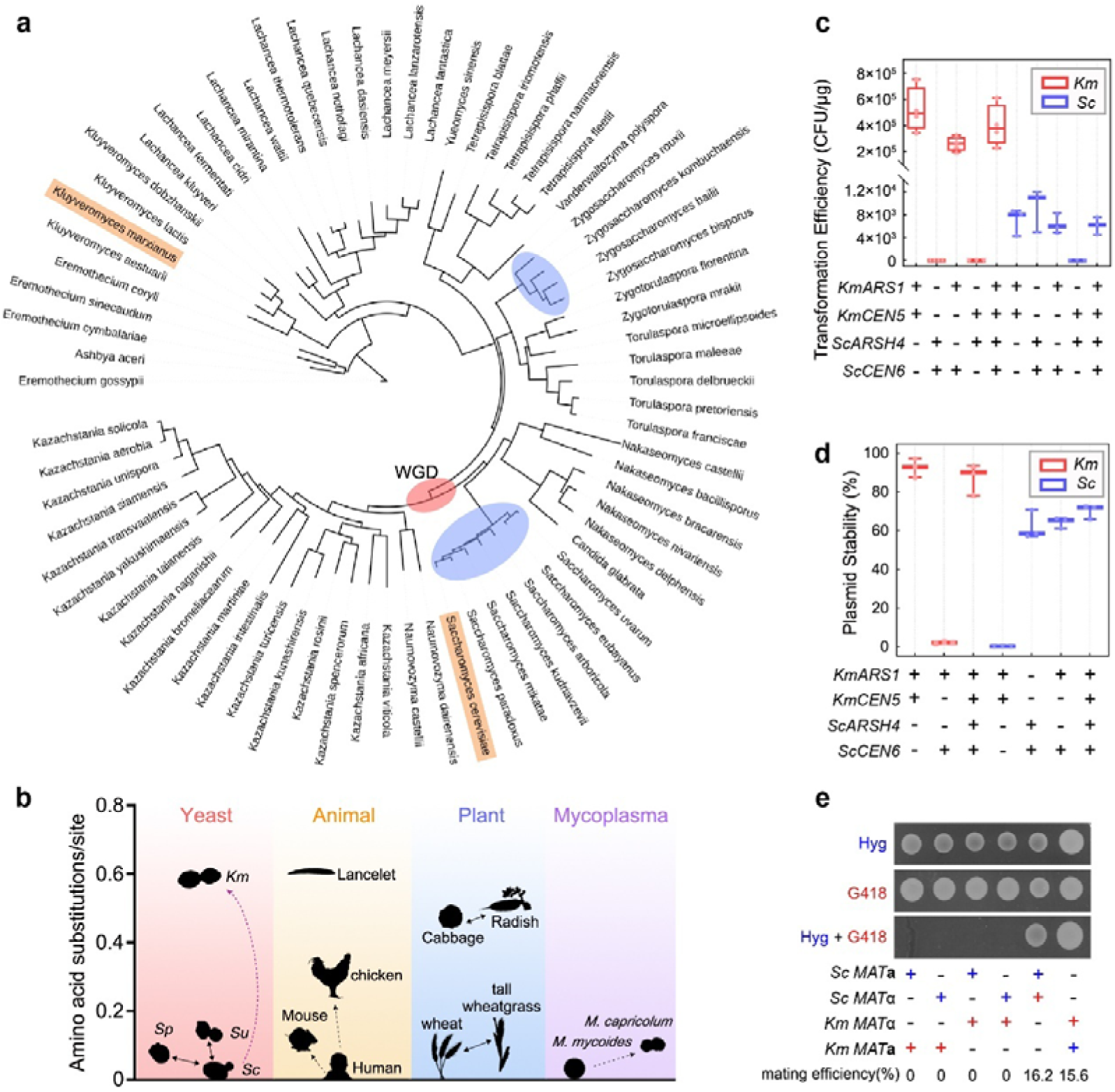
Functionality of *ARS* and *CEN* in *Sc* and *Km*. **(a)** Phylogenetic tree of the *Saccharomycetaceae* family based on 71 selected species. Blue ellipses show species capable of interspecific hybridization within the genus, while red ellipse marks the event of whole-genome duplication. *Sc* and *Km* are highlighted in orange. The tree was constructed with iqtree2 ^39^ with concatenated coding sequences of 2,408 orthologous groups ^23^. **(b)** The evolutionary distance between *Km* and *Sc*, compared to species in other taxa that are capable of interspecific hybridization (solid lines) or tolerant to chromosomal transfer (dashed lines). The magenta dashed line shows findings from this study that *Km* was tolerant to *Sc* chrI and III. Amino acid substitutions per site were generated by Orthofinder2 ^40^. *Sp*, *Saccharomyces paradoxus*. *Su*, *S. uvarum*. **(c, d)** Transformation efficiency **(c)** and stability **(d)** of plasmids containing different combinations of *ARS* and *CEN* from *Sc* (blue) and *Km* (red). Stability refers to the percentage of plasmid-containing cells after being grown in non-selective YPD medium for 24 h. Lines in the boxplots and whiskers show the median, maximum and minimum, respectively. **(e)** Mating assay between *Km* and *Sc*. Haploid *Km* or *Sc* cells containing a *kanR* (red cross) or *hygR* (blue cross) plasmid were mixed and spotted onto plates containing hygromycin (Hyg), G418 (Kan), or both. Mating efficiency was quantified by counting double-resistant colonies (see **Methods**), averaged across three replicates.

Stable allodiploid hybrids between *K. marxianus* and *S. cerevisiae* haven’t been made available, although partial integration of *S. cerevisiae* genome fragments into *K. marxianus* has been achieved by protoplast fusion ^16,17^. In order to introduce *S. cerevisiae* genetic materials in a systematic, controllable manner, we developed a genetic-engineering strategy to artificially transfer chromosomes between the two species. Artificial chromosome transfer was first established in 1977, when murine chromosomes were transferred into human cells via microcells ^18^. To date, artificial transfer of functional chromosomes has primarily been achieved between closely related species (dashed lines, **Fig. 1b**), including transfers of human chromosome 21 into mouse and chicken cells ^19–21^, and between bacteria species *Mycoplasma mycoides* and *M. capricolum* ^22^. *S. cerevisiae* (*Sc*) and *K. marxianus* (*Km*) diverged 114 million years ago ^23^, significantly exceeding the evolutionary distance between the species that have been shown capable of natural or artificial chromosome transfers (**Fig. 1b**). Previous efforts of protoplast fusion showed that the *Sc* genome was often unstable after the fusion with *Km* ^17^. One possible explanation is the incompatibility of replicating elements such as centromere (*CEN*) and autonomously replicating sequence (*ARS*), as previously shown in hybrid studies ^24,25^. In this study, we engineered the *Sc* chromosome with a series of genetic modifications to resolve potential incompatibilities, including circularization of the *Sc* chromosome and insertion of *ARS* and *CEN* from the host species ^26^. The engineered *Sc* chromosome was transformed into *Km*, creating a “monochromosomal hybrid”. Through phenotypic and gene expression analysis, we showed that the chromosome transfer led to beneficial phenotypes associated with overexpression of genes from the *Sc* chromosome. The transfer triggered widespread transcriptional responses in the transferred and host genomes, suggesting cross-species regulation. The pervasive transgressive expression found among the transferred genes provided new evidence for regulatory evolution across a long evolutionary timescale, as well as potential implications for synthetic biology.

## Results

### Compatibility of *ARS* and *CEN* between *Sc* and *Km*

It is critical to ensure proper replication and segregation of the heterologous chromosome upon chromosome transfer. Therefore, we started by investigating the compatibility of *ARS* and *CEN* between *Sc* and *Km*. In *S. cerevisiae*, there are 228 functional *ARS* sites distributed across the genome ^27^, sharing an 11-bp consensus sequence ^28^. The distribution of *ARS* in the *Km* genome is less characterized, and only 15 *KmARS*s have been experimentally validated ^29–32^. Different from canonical *ARS* in other yeasts, *KmARS*s are diverse and do not share a common consensus sequence ^32^. There is one *CEN* per chromosome in both *Sc* and *Km*, each consisting of three conserved centromere DNA elements (CDEI, II and III) ^33^. The *Km* CDEs are similar to their *Sc* counterparts, with the exception that the *Km* CDEII was approximately twice the length of that in *Sc* ^34^. We selected two *ARS-CEN* pairs from each species for our experiments, based on their chromosomal locations and previously-demonstrated functionality in plasmids ^31,32,35–38^. Pairs on the same chromosome were tested together to maximize intraspecific *ARS-CEN* compatibility: *ScARSH4* and *ScCEN6* ^35,36^ (*Sc* chrII), *ScARS1* ^37^ and *ScCEN4* (*Sc* chrIV), *KmARS1* and *KmCEN5* ^31,38^ (*Km* chrV), and *KmARS18* ^32^ and *KmCEN3* (*Km* chrIII).

We examined transformation success and stability of plasmids carrying different combinations of *KmARS1*, *KmCEN5*, *ScARSH4* and *ScCEN6*. We found that *KmARS1* and *KmCEN5* were respectively essential for successful transformation (**Fig. 1c**, left) and plasmid maintenance in *Km* (**Fig. 1d**, left). Neither could be replaced by their *Sc* counterparts, suggesting incompatibility. Plasmids carrying *KmCEN5+KmARS1* produced transformants in *Sc* (**Fig. 1c**, right), but with poor stability (**Fig. 1d**, right), suggesting that *KmARS1,* but not *KmCEN5,* can function in *Sc*. Similar results were obtained using combinations of *KmARS18*, *KmCEN3*, *ScARS1* and *ScCEN4* (**Fig. S1**). The observed incompatibility could explain the rapid loss of *Sc* chromosomes in previous protoplast fusion between *Km* and *Sc* ^17^. Finally, the plasmid containing both *ARS*s and *CEN*s from *Sc* and *Km*, known as “double *CEN/ARS* plasmid”, could replicate and segregate stably in both species (**Fig. 1c**, left). Therefore, the issue of *Sc* chromosomal instability in *Km* may be resolved by engineering *KmCEN* and *KmARS* into *Sc* chromosomes.

Using double *CEN/ARS* plasmids carrying different antibiotic markers, we examined the mating rate between *Km* and *Sc*. We transformed *Km* and *Sc* haploids with plasmids carrying a *kan^R^* or *hyg^R^* marker, mixed them in different combinations (see **Methods**) and selected for zygotes on double antibiotic medium (**Fig. 1e**). We found that the mating efficiency between *Km MAT***a** and *MAT*α cells, as well as between *Sc MAT***a** and *MAT*α cells, was around 16%. However, mixing 1.6×10^7^ *Km* cells with 1.6×10^6^ *Sc* cells, regardless of their mating types, failed to generate any zygotes, based on both double selection (**Fig. 1e**) and microscopic observation (**Fig. S2**). These results demonstrated a prezygotic reproductive barrier between the two species, which urges a synthetic method for generating *Km-Sc* hybrids.

### Engineering and transferring *Sc* chromosomes into *Km*

We selected the smallest *Sc* chromosome, chrI (252□kb, 85 genes based W303 annotation), and the third smallest chromosome, chrIII (341□kb, 156 genes), for proof-of-principle experiments of *Sc-Km* chromosomal transfer. The experimental pipeline is shown in **Fig. 2a**. First, we circularized the chromosomes of interest to avoid potential incompatibility of telomeres. *Sc* telomeres (TELs) are composed of TG_1−3_ repeat sequences, while *Km* TELs consist of a long repeat motif (25 nt) ^41^. We removed the TELs from *Sc* chrI and chrIII, respectively, and joined the ends with *KmURA3* with CRISPR/Cas9 ^42^. To avoid homologous recombination at the telomeres, we also removed the telomere-associated long repetitive sequence on chrI during the circularization ^42^. Next, to ensure stable maintenance of *Sc* chromosomes in *Km*, we inserted *KmCEN5* /*ARS1* adjacent to the native *CEN*s on chrI and chrIII. Given that chrI and chrIII contain 5 and 12 *ARS*s, respectively ^27^, we placed another copy of *KmARS1* close to a native *ScARS*, which was about ∼100 kb away from the *KmCEN5* /*ARS1* in the circular chromosomes (see **Methods** for details). The engineered circular chrI and chrIII were named R1 and R3, respectively. *Sc* cells with R1 (“*Sc*-R1”) or R3 (“*Sc*-R3”) showed no fitness defect under normal culture conditions as well as in the presence of microtubule-depolymerizing and DNA-damaging agents (**Fig. S3**), indicating that the introduction of *KmARS1* and *KmCEN5* did not affect chromosome replication in *Sc*. The circularization was confirmed by pulsed-field gel electrophoresis (PFGE), which showed an absence of linear chrI or chrIII in the gel (**Fig. 2b**). Following a protocol developed by Noskov et al. ^43^, we extracted and column-purified R1 and R3, with the undesired linear chromosomes removed by exonuclease. Finally, the purified R1 and R3 were separately transformed into *Km* using protoplast transformation ^44^, yielding four R1 transformants (*Kluyveromyces-Saccharomyces*-R1, “KS-R1” hereafter) and one R3 transformant (“KS-R3”). The low number of transformants indicates a need for further optimization of the transformation protocol, especially for transferring larger chromosomes.

**Fig. 2.**
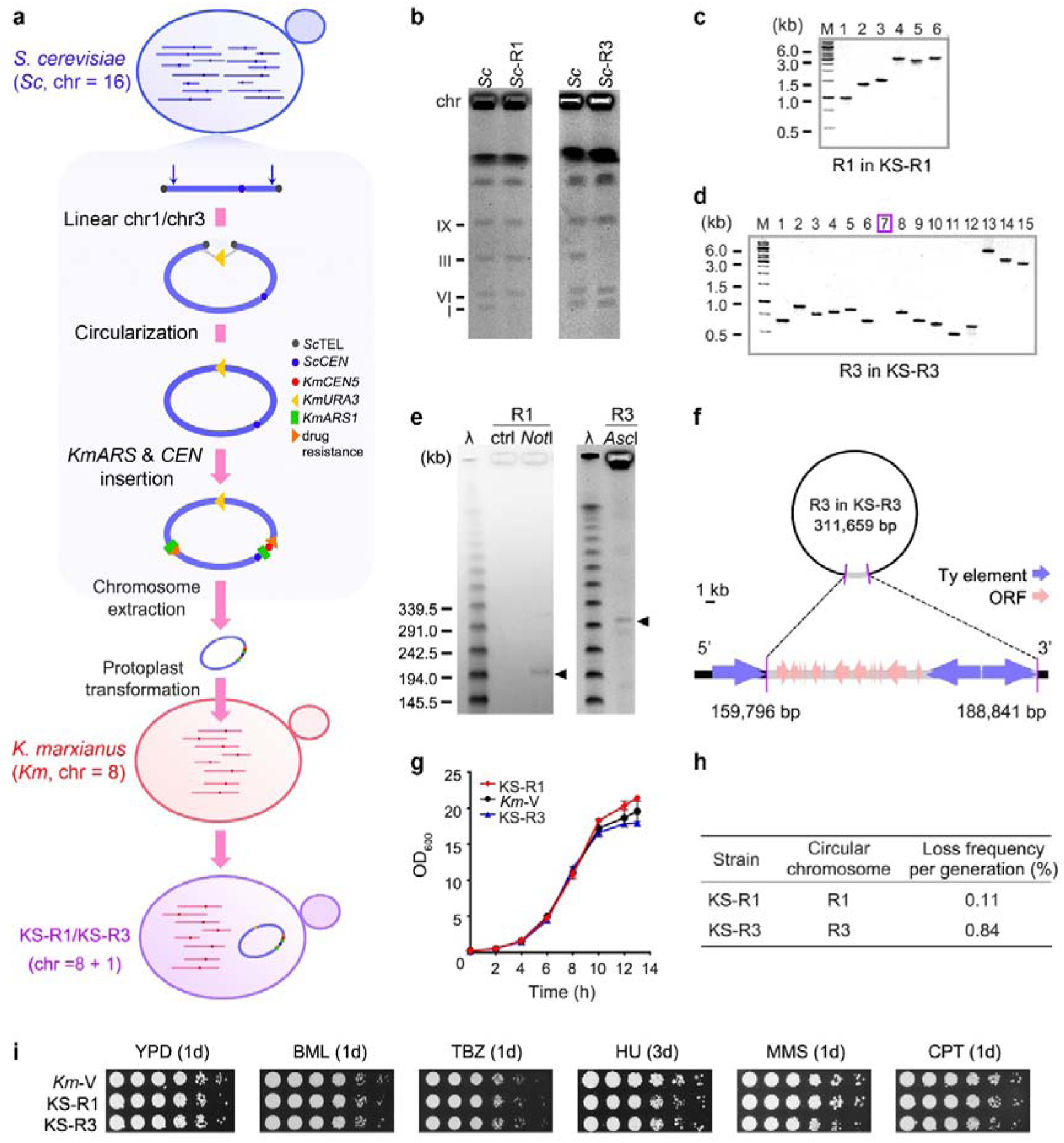
Engineering and transferring *Sc* chromosomes into *Km*. **(a)** Experimental pipeline. **(b)** PFGE of chromosomal extracts from *Sc*-R1, *Sc*-R3 and the parental *Sc* strain. **(c, d)** PCR of markers in the circular chromosomes in KS-R1 **(c,** represented by one transformant) and KS-R3 **(d)**. Purple box labels the missing marker “7” in KS-R3. See **Table S1** for marker positions. **(e)** PFGE of R1 and R3 (arrows) in KS, linearized with NotI and AscI respectively. Ctrl: non-digested KS-R1 chromosomal extracts. λ: Lambda PFG Ladder. **(f)** Illustration of the 29 kb deletion in KS-R3. Coordinates are based on a manually curated R3 sequence (**Table S1**). **(g)** Growth curves of KS-R1, KS-R3, and *Km*-V (*Km* with an empty vector) in YPD. The values represent mean ± SD (n=3). **(h)** Stability of circular chromosomes in KS. The values represent the mean (n=3). **(i)** Spot assay of KS-R1 and KS-R3 in the presence of benomyl (BML), thiabendazole (TBZ), hydroxyurea (HU), methyl methane sulfonate (MMS), camptothecin (CPT) or cycloheximide (CHX). The plates were incubated at 30 degrees for 1 day (1d) or 3 days (3d) as indicated.

We next examined the integrity of the transferred chromosomes. All four KS-R1 transformants retained six *Sc*-specific markers in R1 (**Fig. 2c**, see **Table S1** for marker positions), indicating successful transfer. The transfer was further confirmed by restriction digest (**Fig. 2e**) and whole genome sequencing, which found no mutations in the transferred R1. In the PCR analysis of KS-R3, marker 7 was absent (**Fig. 2d**). PFGE showed that the size of linearized R3 was smaller than the expected 340 kb (**Fig. 2e**). Genome sequencing revealed a 29 kb deletion between two direct repeats of putative Ty elements in R3 (**Figs. 2f & S4**). The deletion was potentially due to homologous recombination between Ty elements, the rate of which has been shown to significantly increase in circular plasmids during transformation ^45^. The deleted region contained 13 genes (*MAK32*, *PET18*, *MAK31*, *HTL1*, *HSP30*, *YCR022C*, *YCR023C*, *SLM5*, *YCR024C-B*, *PMP1*, *YCR025C*, *NPP1*, *RHB1*). We introduced the deleted region, in six individual segments via plasmids, into *Km* cells (**Fig. S4**). None of the segments affected growth (**Fig. S4**), so it was unlikely that the deletion was an adaptive response to overcome incompatibility caused by single genes in this region.

We next evaluated the *Km* genome integrity after transformation. The PFGE patterns of *Km* chromosomes in KS-R1 and KS-R3 were consistent with the parent (**Fig. S5**). Genome sequencing revealed 33 mutations in 11 ORFs, as well as 82 SNPs and indels in intergenic sequences in *Km* chromosomes in KS-R1 (**Table S2**). KS-R3 contained 4 mutations in 4 ORFs, and 47 SNPs and indels in intergenic sequences (**Table S2**). These mutations might have been induced during the protoplast transformation. Overall, the transfer of R1 or R3 into *Km* did not result in fusions, translocations, or other large-scale rearrangements in the *Km* chromosomes.

Subsequently, we examined the KS strains for any fitness defects. In rich medium (YPD), the growth curves of KS-R1 and KS-R3 were indistinguishable from *Km*-V, the parental *Km* strain with an empty vector (**Fig. 2g**). Under non-selective conditions, R1 and R3 were highly stable in the KS strains. The loss rate per generation for R1 and R3 in YPD was 0.11% and 0.84%, respectively (**Fig. 2h**), a level comparable to that of circular YACs in *Km* (0.33-0.64%) ^46^. Furthermore, KS-R1 and KS-R3 exhibited robust growth in the presence of the microtubule-depolymerizing agents benomyl (BML) and thiabendazole (TBZ), DNA replication inhibitor hydroxyurea (HU), DNA-damaging agents methyl methane sulfonate (MMS) and camptothecin (CPT) ^47^, or protein synthesis inhibitor cycloheximide (CHX) (**Fig. 2i**). Taken together, the introduction of R1 and R3 did not affect growth, chromosome segregation, DNA replication, or protein synthesis in rich medium. The *Km* cells were tolerant to the transplantation of *Sc* chromosomes, reflecting cellular plasticity.

### Enhanced phenotypes due to interactions between *Km*-encoded *trans*-factors and *Sc cis*-regulatory sequences

We next investigated if the transferred chromosomes conferred novel phenotypes. We treated the KS strains and their parents with 28 environmental conditions, chosen to represent common stressors for yeast growth (**Figs. 3a & S6-7**). For KS-R1, the phenotypes of the four independent transformants were generally consistent (**Fig. S7**), so we primarily analyzed one representative transformant (**Fig. 3a & S6**). We primarily compared the KS strains to *Sc* and *Km* strains with an empty double *CEN/ARS* plasmid (referred to as *Sc*-V and *Km*-V respectively), which carries the same antibiotic and *KmURA3* markers as the circularized chromosomes, to account for phenotypic effects associated with the vector (see **Fig. S8** for details). *Sc*-R1 and *Sc*-R3 were also examined to account for effects of chromosomal circularization and other genetic modifications, but their phenotypes were for the most times consistent with *Sc*-V (**Fig. 3a & S6**). For each strain, we semi-quantified the growth on solid media under different conditions, normalized the data to YPD growth at 30 °C, and then adjusted the data in reference to *Km*-V to reflect inter-strain differences (**Fig. 3a & Fig. S6b**). *Sc*-V and *Km*-V showed significant growth differences under 21 out of the 28 conditions (2% ethanol, 3% glycerol, 0.02% glucose, Ser, 1/4N, 1/2AA1/2N, 20, 37 and 42°C, 5-FU, 0.016% and 0.024% H_2_O_2_, and all three AcOH, TM and NaCl treatments; p < 0.05, t-test, **Table S3**), reflecting phenotypic divergence between species. KS-R1 and KS-R3 exhibited comparable growth to *Km*-V under 25 and 18 conditions, respectively (p> 0.05, t-test; **Fig. 3a & Table S3**), suggesting that the small number of genes in R1 and R3 did not trigger a widespread metabolic reprogramming in *Km*. Under a few conditions, R1 and R3 caused increased sensitivity in KS compared to *Km*, including the sensitivities to 37 □ and 42 □ for KS-R1, and to tunicamycin (TM), 1/4N, 1/2AA1/2N, and 0.024% H_2_O_2_ for KS-R3 (p < 0.05, t-test, **Fig. 3a-b & Table S3**). Under these conditions, the KS phenotypes were intermediate to those of *Km* and *Sc*, consistent with dominant effects exerted by the introduced *Sc* chromosomes. The defective phenotypes were not shared by the two strains, indicating that the defects were caused by specific genes in R1 or R3, rather than a general effect caused by chromosomal transfers. Additionally, we ruled out that the mutations in the *Km* genome caused the phenotypes, because the strains no longer exhibited the phenotypes after losing the *Sc* chromosomes (**Fig. S9**).

**Fig. 3.**
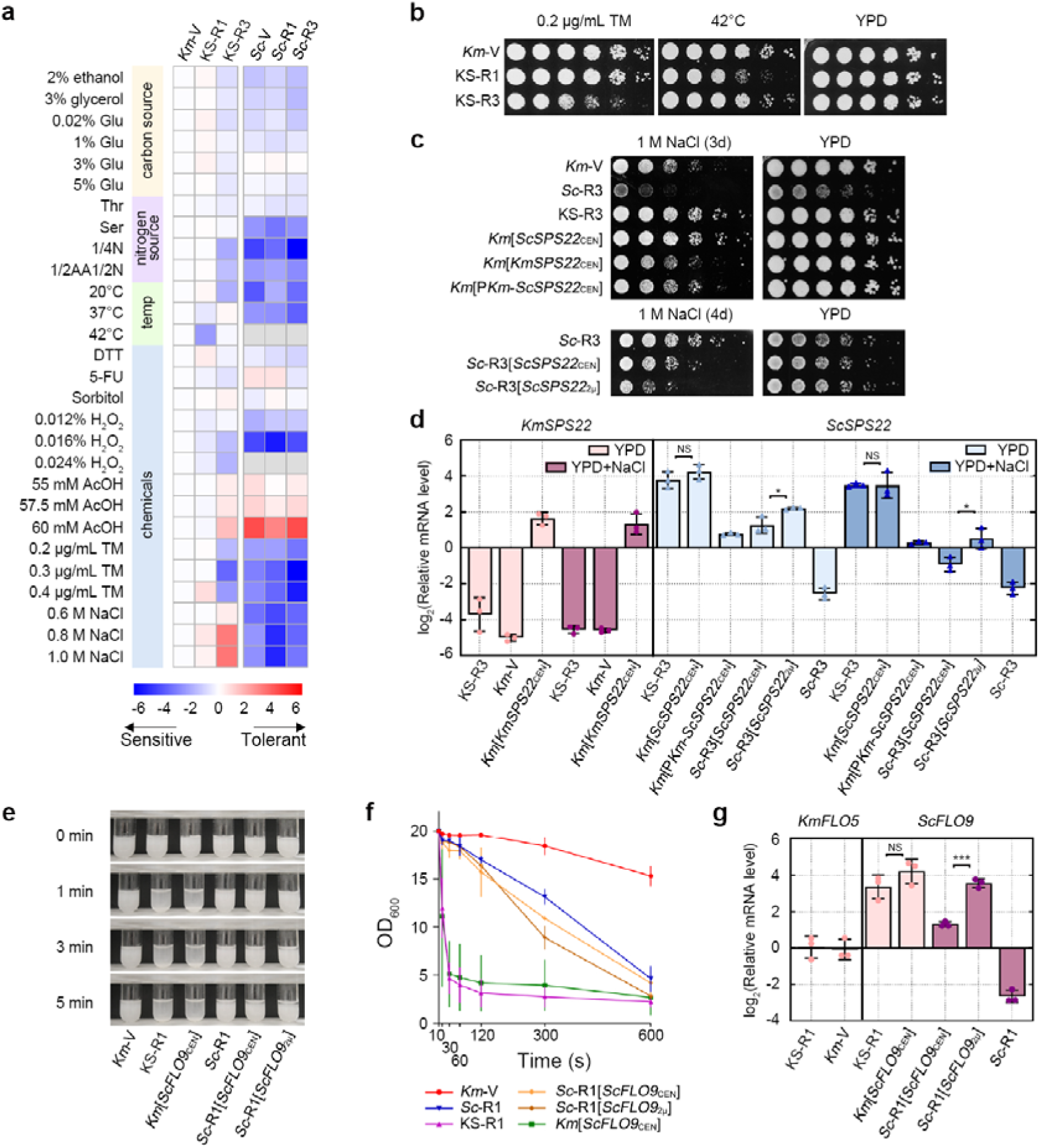
Chromosomal transfers resulted in enhanced phenotypes. **(a)** Growth phenotypes of the synthetic strains and the parental species, relative to *Km*-Vector (*Km*-V). *Km*/*Sc*-V, *Km*/*Sc* with an empty vector. KS-R1/R3, *Km* transformed with a circularized *Sc* chrI/III. *Sc*-R1/R3, *Sc* with circularized chrI or III. Relative growth (RG) was quantified by normalization to YPD growth at 30°C for each strain. Log2(RG) was averaged cross replicates, and then subtracted by the value of *Km*-V of for each condition. A value of 0 in the heatmap means equal growth to *Km*-V. n = 3 except for three H_2_O_2_ treatments and 60mM AcOH, where n=2 (see **Methods** for details). Original images and data are provided in **Fig. S6** and **Table S3**. Grey denotes no growth. Glu, glucose; 1/4 N, one-quarter ammonium sulfate; 1/2 AA, half amino acids; 1/2 N, half ammonium sulfate; TM, tunicamycin; 5-FU, 5-fluorouracil. **(b)** Growth defects under tunicamycin or 42°C treatment associated with the transferred *Sc* chromosomes. Serial dilutions were performed at 1:5 for all spot assays in this study. Growth was after 1 day unless indicated otherwise. **(c)** Improved salt (NaCl) tolerance associated with the promoter sequence of *ScSPS22*. *ScSPS22*, *KmSPS22* and P*Km-ScSPS22* alleles were expressed with a centromeric plasmid (indicated by the subscript CEN) or a 2µ plasmid (*ScSPS22*_2µ_) in either *Km* (upper panel) or *Sc*-R3 (lower panel). **(d)** Relative mRNA levels of *ScSPS22* and *KmSPS22* measured by qPCR. mRNA levels in KS and *Km* were normalized to the average of three housekeeping genes, *KmSWC4*, *KmTRK1*, and *KmMPE1*. mRNA levels of *ScSPS22* in *Sc* were normalized to the average of *ScSWC4*, *ScTRK1*, and *ScMPE1*. The values represent the mean ± SD (n=3). **(e, f)** Enhanced flocculation in KS-R1 associated with *ScFLO9*. *ScFLO9* was expressed with either a centromeric plasmid (*ScFLO9* _CEN_) or a 2µ plasmid (*ScFLO9*_2µ_) in *Km* or *Sc*-R1. Overnight cultures were vortexed vigorously and left still, before being imaged **(e)** and examined for the OD_600_ of the supernatant **(f)** at designated timepoints. The values in **(f)** represent the mean ± SD (n=3). **(g)** Relative expression levels of *ScFLO9* and *KmFLO5* measured by qPCR, normalized as described above. The values represent the mean ± SD (n=3). *, p<0.05; ***, p<0.001. NS, not significant. P-values were based on t-tests.

Importantly, the chromosomal transfer resulted in enhanced phenotypes that were superior to both parents. KS-R1 showed a mild growth increase in 0.02% glucose, while KS-R3 exhibited enhanced tolerance to 0.8M and 1M sodium chloride (NaCl) compared to both parents (p < 0.05, t-test; **Fig. 3a & S6b, Table S3**). Of these phenotypes, the increase in high-salt tolerance of KS-R3 is the most pronounced (**Fig. 3c**). High salt is a common stress in industrial processes. Therefore, increased salt tolerance is a beneficial phenotype for industrial production ^48^. We next investigated the molecular mechanisms by which R3 increased salt tolerance in KS. There are three genes in R3 that have been associated with salt tolerance based on previous large-scale studies: *SPS22* ^49^, *STE50* ^50^ and *HMLALPHA2* ^49^. We expressed the three candidate *Sc* genes in *Km* with a centromeric plasmid, including their coding sequences (CDSs) and 1000 bp-upstream sequences (promoters). We found that only *ScSPS22* enhanced salt tolerance to the same level as KS-R3 (**Fig. S10, Fig. 3c**, upper panel), indicating that it was the causal gene. *ScSPS22* is involved in β-glucan synthesis and related to cell wall function ^51^. In *S. cerevisiae*, overexpression of *ScSPS22* was previously found to cause reduced salt tolerance ^49^, which was recapitulated in our study (**Fig. 3c**, lower panel). It remains to be understood why overexpression of *ScSPS22* had an opposite effect in *Km* than *Sc*, possibility related to the difference in their cell-wall composition ^52^.

In order to understand the contribution of *cis*-regulatory and coding sequences (CDS) to the phenotypic effect of *ScSPS22*, we constructed a chimeric allele consisting of the promoter of *KmSPS22* (*FIM1_814*) and the CDS of *ScSPS22*, namely P*km-ScSPS22*. There was no substantial difference between P*km-ScSPS22* and full-length *KmSPS22* when expressed with a centromeric plasmid (**Fig. 3c**, upper panel), suggesting a lack of functional divergence in the CDS. On the contrary, P*km-ScSPS22* conferred a much lower level of salt tolerance than *ScSPS22* driven by its endogenous promoter, indicating that the *Sc* promoter was causal for the increase in salt tolerance.

The different phenotypic effects of *Sc* and *Km* promoters suggest potential divergence in gene expression. Therefore, we analyzed the transcription level of *ScSPS22* and *KmSPS22* in KS-R3 and the parental strains by qPCR (**Fig. 3d**). The relative mRNA level of *ScSPS22* in KS-R3 was respectively 53 and 80 times higher than those in Sc-R3 with and without NaCl (**Fig. 3d**, right panel), suggesting that (1) a significant difference in *trans*-acting regulators for *ScSPS22* between *Sc* and KS-R3; and (2) the up-regulation of *ScSPS22* in KS-R3 is constitutive. The relative mRNA level of *KmSPS22* did not significantly differ between KS-R3 and *Km*-V (**Fig. 3d**, left panel), indicating little difference in the *trans*-acting factors for *KmSPS22* between *Km* and KS-R3.

When transformed into *Km* on a centromeric plasmid, the *ScSPS22* allele was expressed at the same level as *ScSPS22* in KS-R3 (p > 0.05 for both YPD and NaCl conditions, Student’s t-test, **Fig. 3d**), suggesting that the promoter sequence of *ScSPS22* was causal for its up-regulation in KS-R3. The chimeric allele *Pkm-ScSPS22* drove lower expression than *ScSPS22* in *Km*, consistent with *cis*-regulatory divergence. In *Sc*-R3, expressing *ScSPS22* with centromeric or high-copy 2μ plasmids increased its relative mRNA level, but failed to match the level of *ScSPS22* in KS-R3, suggesting that the upregulation of *ScSPS22* in KS-R3 resulted from induction or de-repression by *Km*-specific *trans*-acting factors, rather than a dosage effect. Taken together, the phenotypic and expression analysis showed that the increased salt resistance of KS-R3 was caused by specific interactions between *Km*-encoded *trans*-acting regulators and the *Sc* promoter of *SPS22*.

KS-R1 showed a statistically significant growth increase in 0.02% glucose compared to *Km* and *Sc*, but the effect was very mild. However, we additionally found that it exhibited a flocculation phenotype surpassing that of *Km*-V and *Sc*-R1 (**Fig. 3e&f**). The flocculation phenotype disappeared after the loss of R1, indicating that R1 was responsible for the enhanced flocculation (**Fig. S11**). Flocculation facilitates cell sedimentation, reducing separation costs in high-density fermentation ^53^. The *FLO* family genes control flocculation ^54^, among which *ScFLO9* is present on R1 and has been shown to cause flocculation in *Km* ^55^. Introducing *ScFLO9* with its 1000 bp upstream sequence into *Km* via a centromeric plasmid enhanced flocculation to the same level as KS-R1 (**Fig. 3e&f**). This result suggests that *ScFLO9* was responsible for the increased flocculation phenotype in KS-R1. Overexpressing *ScFLO9* with a centromeric or 2μ plasmid in *Sc* slightly enhanced flocculation (**Fig. 3e&f**), consistent with its known role in *Sc* but suggesting its effect is background-dependent.

We similarly analyzed the expression level of *FLO9* to understand the molecular mechanisms of the enhanced phenotype. qPCR showed that the relative mRNA level of *ScFLO9* in KS-R1 was 64.5 times higher than in *Sc*-R1 (**Fig. 3g**), again suggesting a significant change in the *trans*-regulatory environment. The high expression in KS-R1 was recapitulated by *ScFLO9* expression from a centromeric plasmid in *Km* (light pink bars in **Fig. 3g**, p > 0.05, Student’s t-test). The expression level of *KmFLO5* (*FIM1_6*), the *ScFLO9* ortholog, was not increased in KS-R1, consistent with a lack of change in *trans* environment for *Km* genes (**Fig. 3g**, left panel). Overexpressing *ScFLO9* in *Sc*-R1 with centromeric and 2μ plasmids led to an increase in the mRNA level of *ScFLO9* (purple bars in **Fig. 3g**), with the higher level of *ScFLO9*_2μ_ correlating with faster flocculation (**Fig. 3f**). Taken together, the enhanced flocculation of KS-R1 was associated with *ScFLO9* being exposed to an alien *trans*-regulatory environment encoded by the *Km* genome. In both cases of *SPS22* and *FLO9*, the genetic combinations between *Km*-encoded *trans*-factors and *Sc*-encoded *cis*-factors could never be found in nature due to reproductive isolation, demonstrating the power of cross-species synthetic biology.

### Pervasive transcriptional responses triggered by chromosomal transfer

The experiments with *ScSPS22* and *ScFLO9* revealed significant changes in *Sc* gene expression after being transferred to *Km*. To determine if this is a general pattern, we analyzed transcriptomes of KS and the parental strains in YPD and under two stress conditions, 1 μg/mL tunicamycin (TM) and 1 M NaCl. The genes on R1 and on R3 were actively transcribed in KS under the three conditions (read counts >5) (**Supplementary Data File 2**). However, their expression in KS was poorly correlated with that in *Sc*, with an average correlation coefficient (R²) of 0.38 for R1 genes and 0.32 for R3 genes (**Fig. 4a**). This correlation was substantially lower than the correlation between the expression of human genes on Hsa21 transplanted into mouse cells and their counterparts in human cells (R²=0.90) ^21^, which aligns with the remote evolutionary distance between *Km* and *Sc* (**Fig. 1b**). Compared to *Sc*, on average, 22.7% and 18.3% of the genes in R1 and R3, respectively, were up-regulated in KS, while 27.3% and 19.3% of the genes in R1 and R3 were down-regulated in KS, respectively (FDR-adjusted p < 0.05 by Wald test and absolute log2FoldChange >1; **Fig. 4b, Table S4**). The median log2-fold change for up- and down-regulated genes was 1.81 and −1.57, respectively. Consistent with the case studies of *SPS22* and *FLO9*, the high proportion and fold change of differentially expressed *Sc* genes reflect dramatic differences in the *trans*-regulatory environment between *Sc* and the KS strains.

**Fig. 4.**
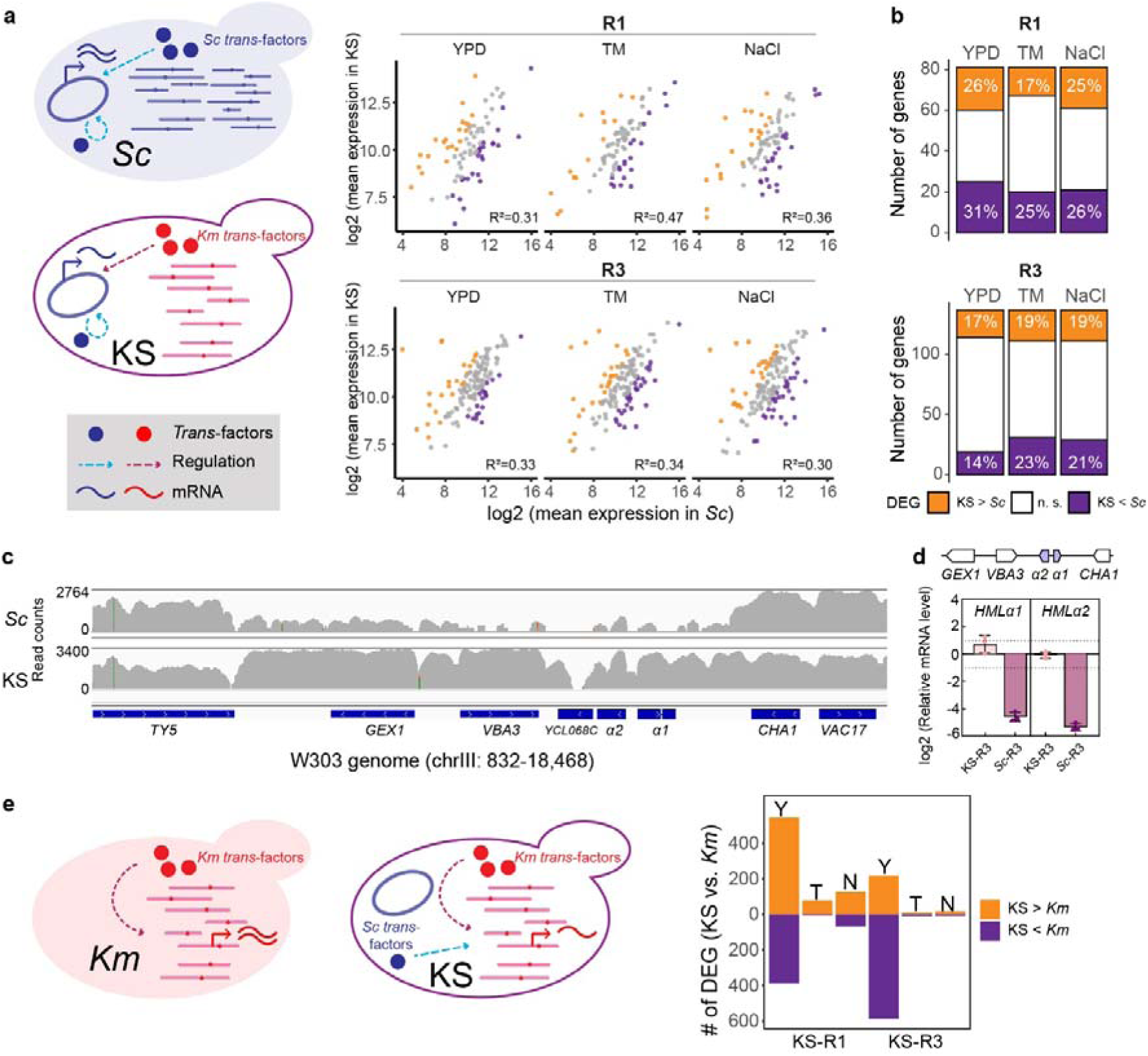
Transferred *Sc* chromosomes were actively transcribed. **(a)** Comparison of *Sc* gene expression between *Sc* and KS strains. Left, diagram showing changes in *trans*-environment. Right, correlation of *Sc* gene expression. The expression level was represented by log2 (average normalized read counts). Each point represents the average of three replicates. Adjusted R^2^ were derived from linear regression. N = 81 genes for R1, and 137 genes for R3 (see **Methods** for data filters). DEG, differentially expressed genes. n.s., not significant. **(b)** Proportion of significantly up- and down-regulated R1/R3 genes in KS. **(c)** De-repression of the *HML* locus in KS-R1, shown by raw read counts (grey bars) covering the *GEX1-CHA1* region in the W303 genome, visualized by Integrative Genomics Viewer. Data were from one representative biological replicate in YPD. Grey bars are in log scale. **(d)** The relative mRNA levels of *HML*α1 and *HML*α2 in KS-R3 and *Sc*-R3 determined by qPCR, normalized as described in **Fig. 3d**. The values represent mean ± SD (n=3). **(e)** Comparison of *Km* gene expression between *Km* and KS strains. Left, diagram showing the effects of *Sc trans*-acting factors. Right, number of DEGs between KS strains and *Km*-V. Y, YPD; T, TM; N, NaCl. In this figure, all DEGs are defined as an FDR-adjusted p-value < 0.05 by Wald test and abs(log2FoldChange) > 1.

The genes showing differential expression between *Sc* and KS were in general uniformly distributed across the transferred chromosomes (**Fig. S12**), but we found a region around *HML* that showed de-repression across multiple genes in KS-R3, providing an interesting example for chromatin-level regulatory divergence. The *HML* locus, along with its flanking genes *GEX1* and *VBA3*, is repressed in *Sc* by *SIR* complex ^56^. Consistent with this previous knowledge, we found little expression of these genes in *Sc* (**Fig. 4c**, upper panel). Interestingly, *GEX1*, *VBA3* and *HML* genes were highly expressed in KS-R3 (**Fig. 4c**, lower panel), indicating de-repression. qPCR confirmed that the relative mRNA levels of α*1* and α*2* were upregulated 42-fold and 39-fold, respectively, in KS-R3 compared with *Sc*-R3 (**Fig. 4d**). The differential expression might be explained by different composition of silencers between *Kluyveromyces* and *Saccharomyces* ^57^. The silencer of *HML* in KS-R3 might not be able to recruit essential proteins such as Rap1 and ORC to the locus ^58^, contributing to the de-repression of α*1* and α*2*, and the flanking genes *GEX1* and *VBA3*.

While the expression of *Sc* genes underwent significant changes in KS, R1 and R3 also affected the expression of *Km* genes (**Fig. 4e**). Out of 4,975 *Km* genes, 931 (18.7%) and 803 (16.1%) genes in KS-R1 and KS-R3, respectively, exhibited significant expression differences compared to *Km*-Vector in YPD (FDR-adjusted p < 0.05 by Wald test and absolute log2FoldChange >1; **Fig. 4e, Table S4**). R1 encodes a transcription factor, Oaf1, that regulates at least 97 *Sc* target genes ^59^. Among the 73 *Km* orthologs of the known *Sc* target genes (see **Methods** for ortholog assignment), 27 (37.0%) showed differential expression in KS-R1. R3 encodes four transcription factors regulating at least 305 *Sc* target genes ^59^, with 44/160 (27.2%) of their *Km* orthologs differentially expressed in KS-R3 (**Table S4**). This suggests that transcription factors encoded by R1 and R3 might directly alter gene expression in *Km*, but a large fraction of the observed transcriptional changes might be explained by indirect or uncharacterized regulation. Under TM and NaCl stresses, the number of differentially expressed genes was substantially lower than in YPD (**Fig. 4e**, **Table S4**). These results indicate that the heterologous chromosomes induced pervasive changes in expression of the endogenous genome, potentially via novel *Sc-Km* regulatory interactions. Interestingly, regulation from the *Km* genome became dominant under stress conditions, removing much of the effects associated with foreign chromosomes found in YPD.

### Co-evolution of c*is-* and *trans*-regulatory factors

Transferring genetic materials between remotely related species provides a unique opportunity to examine evolution of gene expression. Evolutionary divergence of gene expression can be caused by evolution of *cis*- or *trans*-regulatory factors, or both. *Cis*-regulatory elements such as promoters or enhancers are physically linked to a target gene (e.g., “promA” for “Gene A” in **Fig. 5a**). The regulatory activity of *cis*-elements may evolve through changes in transcription factor binding sites or other elements that interact with the transcription machinery. *Trans*-acting factors often refer to regulatory proteins such as transcription factors (TFs) encoded elsewhere in the genome (e.g., transcription factor “B” in **Fig. 5a**). They may include any direct or indirect regulators, which may evolve through coding or non-coding changes ^60^, even including turnover of transcription factors (e.g., “B” may be replaced by “C” over time, **Fig. 5a**). It has been proposed that *cis*-acting changes and *cis-trans co*-evolution plays an increasingly important role as species diverge ^61^, but this idea has been rarely tested in species as diverged as *S. cerevisiae* and *K. marxianus*. The KS strains allowed us to interrogate this question (**Fig. 5b**) ^62,63^. If the transferred genes showed the same expression level in KS (“*KSsc*”) as that in *S. cerevisiae* (“*Psc*”), it means that the expression divergence is entirely encoded by the *Sc cis*-regulatory elements (the change in *trans*-environment from *Sc* to KS had no effect; “donor-like” in **Fig. 5c**). If the *Sc* allele was expressed at the same level as its orthologous *Km* allele in KS (“*KSkm*”), the expression divergence is entirely determined by *trans*-differences (the *cis*-regulatory elements of *Sc* and *Km* had equivalent activities when exposed to the same *trans*-environment; “recipient-like” in **Fig. 5c**). An intermediate or transgressive expression level of *Sc* alleles in KS indicates contributions from both factors. In particular, transgressive expression, defined by a *KSsc* level outside the range of *Psc* and *KSkm*, may suggest novel *cis-trans* interactions specific to the synthetic background (**Fig. 5c**).

**Fig. 5.**
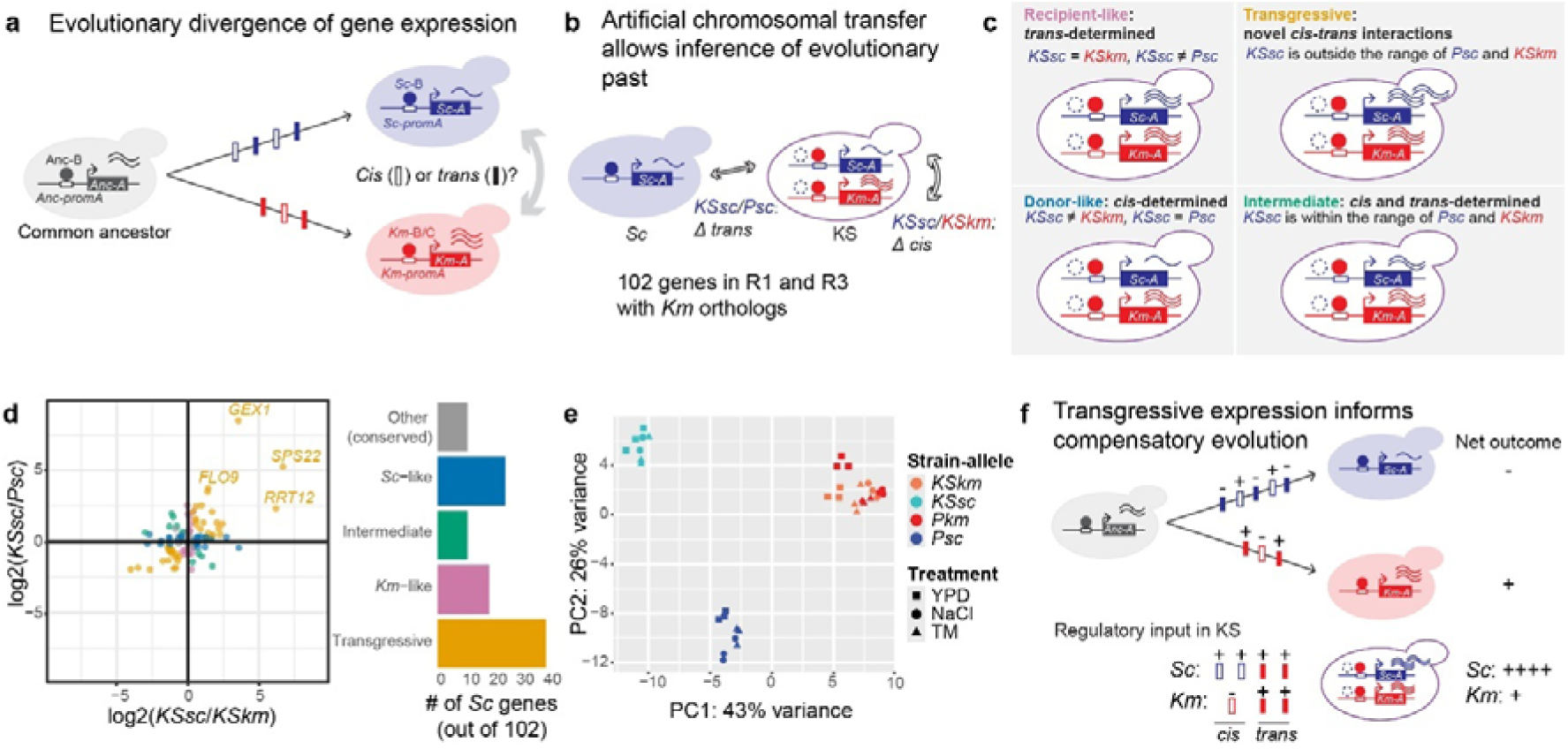
C*is* and *trans* co-evolution in *Km* and *Sc*. **(a)** Divergence of gene expression can be caused by accumulation of *cis* (unfilled vertical bars) and/or *trans* (filled vertical bars) changes during species’ divergence. In the diagram, a focal gene “*A*” is under the regulation of a transcription factor “B” which binds to A’s promoter (“*promA*”). Both A and B undergo sequence divergence in the two species, represented by different colors. In some cases, B can be replaced by other TFs such as “C” during evolution, as exemplified in *Km*. Transcripts are represented by wavy lines. **(b-c)** Comparing allelic expression of the transferred genes to donor (*Sc*, blue) and recipient (*Km* allele in KS, red) reveals contributing factors to expression divergence. In KS, *Km*-B/C is present but *Sc*-B may or may not be present, represented by dashed circles. Anc, ancestor. **(d)** 102 genes showing divergent expression were classified into five categories according to (**c**). Log2 fold changes were extracted from DESeq models, adjusted with the ashr method. Genes showing the most prominent transgressive expression were labeled. Data were from the YPD treatment. (**e)** PCA of gene expression of 162 transferred genes across treatments, strains and alleles. **(f)** Transgressive expression suggests that the *cis* and *trans* factors often evolved in opposite directions, represented by “+” and “-”. The order and number of changes was arbitrary.

In order to characterize *cis-* and *trans-* regulatory divergence between *Sc* and *Km*, we identified *Km* orthologs for 62/85 genes on R1 and 100/156 genes on R3, with an average amino-acid sequence similarity of 48.0% (BLOSUM62-based similarity, see **Methods** and **Table S5**). 106 out of the 162 genes showed significant expression divergence between *Sc* and *Km* (FDR-adjusted p < 0.05, Wald test). What factors determined their divergence? Focusing on the *Sc* allele of these genes, we classified them into *Sc*-like, intermediate, *Km*-like, transgressive and others based on the criteria in **Fig. 5c**, using the transcriptomic data in YPD (**Fig. 5d, Table S6**). Interestingly, we found that most transferred genes showed transgressive expression (40/106 genes, 37.7%, **Fig. 5d**), more than those determined by *cis*-alone (*Sc*-like) or *trans*-alone (*Km*-like). *SPS22* and *FLO9* analyzed in our phenotypic studies (**Fig. 3**) were among the most prominent examples in this class (**Fig. 5d**), suggesting that transgressive expression might be an important source of phenotypic improvement in these engineered strains. Transgressive expression was also the most abundant class under NaCl and TM stresses (54/132 and 36/122 genes respectively, **Fig. S13, Table S6**). Principal Component Analysis (PCA) of the 162 transferred genes across treatments and strains showed that *KSsc* levels were distinct from *Pkm*, *KSkm* and *Psc*, consistent with pervasive transgressive expression of *Sc* alleles (**Fig. 5e**). The second most abundant class in YPD was *Sc*-like expression (25/106, **Fig. 5d**), supporting an important role of *cis*-regulatory evolution. However, there were more *Km*-like genes than *Sc*-like genes under NaCl and TM stresses (**Fig. S13**), consistent with a predominant role of *trans*-evolution in environmental responses^64^.

In the analysis above, we defined “recipient-like” by comparing *KSsc* to *KSkm*, reasoning that this was a conservative estimation of *cis*-effects, when the two alleles were exposed to the same *trans*-environment in KS. This was different from previous analyses of introgression lines^62^ or enhancer swaps^63^, where the transferred allele was compared to the recipient parent (*Pkm* in this case). The methodological difference was largely negligible because the *trans*-environment in KS was highly similar to *Km*, as shown in PCA (*Pkm* and *KSkm* clustered together, **Fig. 5e**). However, to account for the minority of cases where the transferred *Sc* genes influenced *Km* allelic expression (**Fig. 4e**), we performed the same analysis using *KSsc/Pkm* instead of *KSsc/KSkm* (**Fig. S14**). This second method was less conservative with *cis* effects, but it provided a more authentic estimate for transgressive expression: in the cases where *KSkm* showed transgressive expression in KS, transgressive expression of *KSSc* might be missed or overestimated by the first method. The results still supported the prevalence of transgressive expression of the transferred genes (37/106 genes in YPD), although there were more genes classified into *Sc*-like, as expected (**Fig. S14**).

The prevalence of transgressive expression in KS suggests that the *cis*- and *trans*-regulatory factors often evolved in opposite directions within a lineage ^65^. For example, when an *Sc* allele showed transgressively high expression in KS (*KSsc* > *KSkm* and *KSsc* > *PSc*, **Fig. 5c & f**), it might indicate that (1) the *Sc* promoter accumulated more changes to increase gene expression compared to the *Km* promoter, via either stronger activator activity or weaker repressor activity (“+” effects for *cis*-changes in the *Sc* lineage and correspondingly “-” effects in the *Km* lineage, **Fig. 5f**), such that *KSsc > KSkm*; and (2) a *trans*-acting factor that decreased gene expression must have co-evolved with this promoter in *S. cerevisiae* (“-” effects for *trans*-changes in the *Sc* lineage and correspondingly “+” effects in the *Km* lineage, **Fig. 5f**), such that *KSsc > Psc*.

Compensatory evolution may also be identified by examination of genes showing conserved level of expression between the two species (**Fig. S15**) ^66^. For two genomes as diverged as *Sc* and *Km*, it is as important to understand the conservation of gene expression as to understand their divergence. Across 4,307 genes with identifiable orthologs between *Km* and *Sc-*R1 (see **Methods** for ortholog assignment), 29.7% showed no difference in expression levels between the parental strains in YPD (FDR-adjusted P > 0.05, Wald test; **Fig. S15b & c, Table S6**). This ratio was 21.9% and 12.8% under NaCl and TM conditions respectively. The expression level of the 4,307 orthologs was correlated genome-wide between species (Spearman’s rank correlation rho = 0.647 for YPD data, P < 2.2e-16, **Fig. S15b**), suggesting a certain level of regulatory conservation despite the deep divergence. The genes showing conserved expression in YPD were significantly enriched for rRNA processing (GO: 0006364, corrected P = 2.07e-20) and ribosome biogenesis (GO: 0042254, corrected P = 2.61e-19), consistent with a housekeeping role. There was a similar proportion of genes showing conserved expression among the transferred genes (56 out of 162, 34.6% in YPD; 18.5% in NaCl and 24.7% in TM, **Fig. S15d**), providing us with a small sample to peek into the driving force of regulatory conservation – did the *cis* and *trans* regulatory factors both maintained a conserved level of activity, or they evolved in a compensatory manner under stabilizing selection? In the latter case, we would expect to find *cis*-regulatory divergence in the KS strains. Indeed, we found that 42 out of the 56 (75%) conserved genes showed significant *cis-*divergence in the KS strains in YPD (23/30 in NaCl and 29/40 in TM; FDR-adjusted P < 0.05, **Fig. S15d-f**), consistent with compensatory *cis-trans* evolution. The top genes showing this trend in YPD were *CDC19*, *TRX3*, and *HIS4* (**Fig. S15e**). In all three cases, the *Sc* allele was down-regulated in KS strains, suggesting evolution of *Sc*-specific activators. We note that the amount of *cis*-divergence found in KS may be specific to the given *trans* environment, e.g., the *cis*-effects in KS may or may not manifest themselves in an F1 hybrid. However, such dependency itself indicates co-evolution of *cis-trans* interactions (see **Discussion**).

Taken together, our results support that lineage-specific co-evolution of *cis*- and *trans*-regulatory factors was predominantly involved in evolution of gene expression between *S. cerevisiae* and *K. marxianus*. Transgressive expression was the most prevalent pattern among the transferred genes in KS. *Cis*-only and *trans*-only evolution also occurred, with the latter predominantly contributing to environmental responses ^64^. At the same time, around one-third of yeast genome showed conserved gene expression despite the long evolutionary distance. Compensatory *cis*- and *trans*-evolution might underlie such conservation, consistent with stabilizing selection.

## Discussion

Interspecific hybridization often leads to novel phenotypes that are of interest for industrial purposes and evolutionary studies^67^. For diverged species that cannot mate naturally, artificial chromosomal transfer provides an opportunity to explore the industrial potential of mixing diverged genetic materials, as well as to understand its molecular and phenotypic consequences. In this study, we, for the first time, successfully introduced chromosomes of *S. cerevisiae* to *K. marxianus*, two species both with great industrial potential but genetically as diverged as human and lancelet (**Fig. 1b**). We show that the transferred *Sc* chromosomes were stably maintained in *Km*. The synthetic strains exhibited enhanced phenotypes, demonstrating the technology’s potential in future phenotypic screens. The improved salt resistance and flocculation phenotypes were associated with *Sc* promoters being activated by *Km*-encoded *trans*-acting factors, to a greater level than the native regulation in *Km* or *Sc*. Finally, we examined the molecular consequences of the chromosomal transfer with transcriptomic analysis, revealing broad transcriptional responses upon chromosomal transfer, as well as prevalent compensatory evolution of *cis*- and *trans*-regulatory changes during the divergence of the two species.

One of the technological advances in our study is the method to enable stable maintenance of *Sc* chromosomes in *Km*. There are several technical factors of consideration. First, we removed *Sc* TELs and circularized the chromosomes of interest. The circularization made it possible to easily remove unwanted linear chromosomes by exonuclease ^43^, which may increase transformation efficiency. In the case of *CEN* engineering, it was important to ensure that the heterologous *CEN* was not partially functional, which has been shown to disrupt chromosome segregation and lead to dicentric breakage, thereby causing genome rearrangements and instability ^68^. We found that *Sc* and *Km CEN*s were non-functional in the other species. Therefore, we may use double *CEN*s for maintaining the heterologous chromosomes. Next, we considered the density of *ARS*s. The requirement of *ARS* density for YACs varied in previous reports, from one *ARS* per 30 kb ^69^, 51 kb ^70^, to 1 Mb ^71^. In our study, *KmARS1* was positioned at a density of one *ARS* per 110 kb in R1 and 150 kb in R3, which proved to be sufficient for replication of *Sc* chromosomes in KS. Finally, we note that one potential technical pitfall was the difficulty in acquiring independent transformants, as exemplified by R3. Only one transformant of *Km*-R3 was acquired, which contained a deletion potentially mediated by Ty elements during transformation (**Fig. 2**). Future efforts are needed to optimize the transformation efficiency to ensure scalability.

There are several advantages in interspecific chromosomal transfers for synthetic biology purposes. First, it enables combination of genetic materials from remotely related species, partly circumventing the genomic incompatibility that prevented interspecific hybridization. The synthetic “monochromosomal hybrids” were viable, providing novel materials for phenotypic screens or downstream experiments such as directed evolution ^72^. In particular, directed evolution may help resolve any residual incompatibility, by selecting for mutations that improve the desired traits and/or compensate for the incompatibility.

Second, our strategy of chromosomal transfers may provide a complementary approach to whole-genome transformation (WGT) ^73,74^ or genomic libraries, in terms of (1) the ability to screen for polygenic effects and (2) a medium-level of experimental throughput. Both WGT and genomic libraries may be used for genome-wide interspecific genetic screens, where foreign genes were individually introduced to the host, avoiding incompatibility. Compared to WGT and genomic libraries, the strategy of chromosomal transfers is limited in its ability to screen the entire genome but has the strength of screening for polygenic effects. Although our experiments with chr1 and chr3 mainly found single-gene effects, future transfers with larger chromosomes might potentially find beneficial phenotypes necessitated by interactions among multiple genes in the transferred chromosome. Additionally, as demonstrated in our study, cross-species regulation may be an important source for novel phenotypes (**Figs. 3 & 5**). Our approach enables screening for such interactions by introducing full-length *cis*-regulatory elements, which are often missed from WGT or genomic libraries. Furthermore, our strategy allows for possibilities of transferring multiple chromosomes into *Km*. This can be done either by marker recycling, i.e., popping out *URA3* in the (evolved) synthetic strains and transforming another chromosome subsequently, or by mating with another (evolved) synthetic strain containing a different chromosome. Therefore, chromosomal transfer is particularly advantageous for screening for polygenic effects, including interactions within and among foreign chromosomes, as well as between host and foreign genes.

Our strategy provides a medium-throughput screening capability, between that of WGT and genomic libraries. In WGT studies, genomic DNA of a foreign species was transformed into the host species without a stable vector ^73,74^. The foreign genes were expected to randomly integrate into the host genome, and only the transformants that conferred desired phenotypes were selected. The success rate of such strategy depends on the efficiency of homologous recombination and screening conditions. Also, random integration of foreign genes can disrupt the host genome, lowering the success rate even more. From this perspective, the experimental throughput of such WGT approach can be low. Compared to WGT, chromosomal transfers allow screening for hundreds of genes at a time once a stable strain is established, so it might provide a higher throughput than the WGT approach. Genomic libraries on vectors allow generation of stable, pooled transformants for high-throughput screens with a genome-wide coverage, such as the MoBY-ORF library in *S. cerevisiae* ^75^. However, direct application of *S. cerevisiae*-based molecular libraries in non-model yeasts such as *Km* requires technical optimization. In this regard, our findings about *CEN* and *ARS* compatibility may inform future studies to convert genomic libraries on *Sc*-based vectors into *Km*-compatible versions, advancing interspecific genetic screens.

The synthetic KS strains showed interesting phenotypic and molecular characteristics, demonstrating that the transferred chromosomes were functional, even across a remote evolutionary distance such as between yeast genera. Previous interspecific chromosomal transfers in yeast have been successful in using *S. cerevisiae* as a container to maintain heterologous genetic materials from bacteria ^22,71,76,77^, and the epigenetic status of the heterologous chromosomes has been characterized only recently ^78^. In our study, we found that the *Sc* chromosomes were “alive” in the *Km* background. Genes on the transferred R1 and R3 chromosomes were actively transcribed, despite a high level of sequence divergence. Genes on R1 and R3 showed significant expression differences between KS and *Sc* backgrounds, and the same for *Km* genes between KS and *Km* background. The transcriptional responses indicate active interspecific regulation, which can give rise to transgressive phenotypes, as shown with *SPS22* and *FLO9*. It is of future interest to investigate what proportion of the regulatory interactions between the two species’ genomes are associated with beneficial phenotypes, or, on the other hand, indicative of molecular incompatibility.

The KS strains allowed for exploring patterns of regulatory evolution at an unprecedented long evolutionary timescale. It has been long debated whether evolution of gene expression primarily occurs through *cis*, *trans*, or both types of changes. A number of examples, especially from the evo-devo field, have argued for an important role of *cis*-regulatory changes ^79,80^. That is, when a gene was transferred to a different species, its expression pattern was often determined by its *cis*-regulatory sequences (donor-like), suggesting that the *trans* environment was relatively conserved ^63^. Recent transcriptomic and experimental studies showed that compensatory evolution of *cis* and *trans*-acting factors is common, possibly driven by stabilizing selection ^9,61,63,64,81^. However, most systematic studies so far were restricted to intrageneric hybrids ^61,64,81,82^ or introgression lines ^62^. The furthest phylogenetic distance examined so far by genome-wide analysis was probably between *Saccharomyces cerevisiae* and *S. uvarum* ^61,83,84^, two species that share 80% nucleotide identity in coding regions ^85^. Studies on the *Saccharomyces* species found that gene expression divergence reached a plateau as the phylogenetic distance increases within the genus, with the contribution from *cis*-regulatory differences and compensatory changes becoming increasingly important ^61^. Still, little is known for regulatory divergence beyond this phylogenetic scope. It is imaginable that both *cis* and *trans*-acting factors continuously accumulate nucleotide substitutions over a long evolutionary timespan, but it remains unknown whether they evolve in a compensatory manner under stabilizing selection, like what has been found within a genus, or they tend to change in the same direction to increase expression divergence. Evidence from enhancer-swaps showed a higher proportion of misregulation in cross-genus experiments than within *Drosophila* flies or *Caenorhabditis* worms, indicating that lineage-specific *cis-trans* co-evolution becomes abundant beyond the genus level, but the number of genes cataloged in this study was relatively small ^63^. Our study examined the expression pattern of over a hundred *Sc* genes transferred to *Km*, providing new evidence for long-term regulatory divergence at a relatively systematic scale. We found that transgressive expression, rather than *cis*-determined, was the most common pattern (37.7%) among the transferred genes. It is consistent with the model where both *cis* and *trans* changes accumulate with time, and they tend to act in opposite directions. At the same time, around one third of genes showed conserved expression between *Km* and *Sc*, with contributions from compensatory *cis-trans* co-evolution as well. The data together point to that evolution of gene expression is under stabilizing selection in *Saccharomycetaceae* yeasts, across an evolutionary timespan of 114 million years. From a practical point of view, the prevalent transgression and its association with enhanced phenotypes demonstrate the potential of cross-species regulation in generating transgressive phenotypes, providing an alternative route to previous approaches that focus on optimizing individual proteins.

We note a few limitations in our *cis-trans* analysis. First, our sample size of 162 genes provided a statistically sufficient examination of regulatory evolution, but it was a small subset of the yeast genome. Future chromosomal transfers with larger chromosomes may provide more data. Second, we used the differences between *KSsc* and *KSkm* as *cis*-effects, reasoning that the two alleles were exposed to the same *trans*-environment, similar to previous studies with F1 hybrids ^61,86^. However, we note that the estimation of *cis* effects might depend on *trans*-environment. For example, we might find more *cis*-divergence in our “monochromosomal hybrids” than in F1 hybrids due to unavailability of certain *Sc*-specific *trans-*factors. Furthermore, unlike F1 analysis, we cannot properly estimate *trans*-divergence by subtracting *cis* effects from parental differences, because *Sc* and *Km* alleles experienced different levels of *trans*-changes. Therefore, we did not quantitatively compare our conclusions to previous F1 studies. We reasoned that our analysis, similar to previous ones with introgression lines and enhancer swaps, still informs about *cis-trans* co-evolution, and the abundant transgressive expression provides novel insights for both regulatory evolution and future phenotypic studies.

In summary, our study established a strategy for engineering the structural elements and *ARS* of a functional chromosome before transferring it to a remotely related species. This strategy might be extended to other microbes as well as to plants. The resultant synthetic strains provide valuable resources for cell factories, synthetic biology, and evolutionary genomics.

## Materials and Methods

### Strains and media

Yeast strains used in this study are listed in **Table S7**. *Sc* strain W303-1A (*MATa leu2-3,112 trp1-1 can1-100 ura3-1 ade2-1 his3-11,15*) was used for chromosome engineering. The engineered *Sc* chromosome was transferred into a *ura3*Δ *Km* strain, FIM-1ΔU ^87^.

The following media were used in this study: YPD (1% yeast extract, 2% peptone, 2% dextrose)^88^, synthetic medium lacking uracil or leucine (SD-Ura or SD-Leu) □0.17% yeast nitrogen base without amino acids and ammonium sulfate, 2% glucose, 1 g/L sodium glutamate, 2 g/L DO Supplement-Ura (630416, Takara) or DO Supplement*-*Leu (630414, Takara), 2% agar for plates], SD (same recipe as SD*-*Ura, with the addition of 20 mg/L uracil), and ME (2% malt extract, 3% agar). G418 (A600958, Sangon) and hygromycin (H8080, Solarbio) were added to YPD at final concentrations of 200 mg/L and 300 mg/L, respectively, to prepare YPDG and YPDH media. Both antibiotics were supplemented into YPD and SD*-*Ura to prepare YPDGH and SD*-*Ura+GH, respectively. Unless otherwise indicated, cells were grown at 30 □.

### Plasmids

All plasmids used in this study are listed in **Table S8**. All primers are listed in **Table S1**. A previously published vector LHZ626 served as a control for circular chromosomes ^44^, which contains *CEN* and *ARS* elements from both *Sc* and *Km* (double CEN/ARS plasmid), as well as *KmURA3*, *kanMX6* and *hphMX4*.

To construct plasmids for diploid selection, a fragment containing *ScARSH4* and *ScCEN6* was amplified from LHZ626 and inserted into the HpaI and SpeI sites of a *Km*-centromeric plasmid, LHZ882 ^34^, yielding LHZ1493. KanMX6 (*kan^R^*) was amplified from pFA6a*-*KanMX6 and used to replace hphMX4 (*hyg^R^*) in LHZ1493, producing LHZ1494.

To create plasmids with different combinations of *ARS* and *CEN*, a BamHI site was first introduced between *KmARS1* and *KmCEN5* of LHZ882 to generate LHZ1495. Then, *ScCEN6*, *ScARSH4, ScARSH4*-*ScCEN6* were inserted into of LHZ1495 to replace their *Km* counterparts, generating LHZ1496-1498, respectively. Next, *KmARS18* and *KmCEN3* were inserted between the HindIII and SalI sites of LHZ1495 to replace *KmARS1* and *KmCEN5*, giving rise to LHZ1783. *ScARS1*-*ScCEN4, ScARS1* and *ScCEN4* was inserted into LHZ1783 to replace their *Km* counterparts, generating LHZ1784-1786, respectively (see **Table S8** for restriction sites). Finally, *ScARS1* and *ScCEN4* were inserted into the SalI site of LHZ1783 to obtain LHZ1787, a plasmid with *ARS* and *CEN* from both species.

In order to remove the telomeres and join the chromosomal ends, we constructed CRISPR plasmids expressing two guide RNAs (gRNAs), each targeting one end of the chromosome of interest. This is done by cloning the two gRNAs into the SapI and NotI sites of pRS425*-*Cas9*-*2xSapI, respectively. The double-gRNA plasmids, LHZ1499 (for chr1) and LHZ1500 (for chr3), were subsequently transformed into yeast (see below). In order to insert *KmARS1* and *KmCEN5* into the circularized chromosomes, we cloned the respective gDNAs into the SapI sites of pRS425-Cas9-2xSapI to construct LHZ1501_∼_1504. The homologous repair templates (“donor DNA”, see below) was assembled into pMD-18T with In-Fusion Snap Assembly (Clontech), according to the designs in **Supplementary Data File 1**.

For flocculation analysis, the *ScFLO9* cassette, including 1000 bp upstream, the ORF of *ScFLO9*, and 200 bp downstream sequence was amplified from the genome of W303*-*1A. This cassette was inserted into the NotI site of LHZ626 to produce LHZ1505, and into the SmaI and SpeI sites of pRS315 (centromeric) and pRS425 (2-micron) to produce LHZ1509 and LHZ1510, respectively.

For NaCl tolerance analysis, the *ScSPS22* cassette, including 1000 bp upstream sequence, the ORF of *ScSPS22*, and 200 bp downstream sequence was amplified from the genome of W303*-*1A. This cassette was inserted into the NotI site of LHZ626 to produce LHZ1506, and into the SmaI and SpeI sites of pRS315 and pRS425 to produce LHZ1511 and 1512, respectively. The 1000 bp upstream sequence of *ScSPS22* in LHZ1506 was replaced by the 1000 bp upstream sequence of *KmSPS22* to produce LHZ1507. The *KmSPS22* cassette, including 1000 bp upstream sequence, the ORF of *KmSPS22*, and 200 bp downstream sequence was amplified from the genome of FIM-1ΔU and inserted into the NotI site of LHZ626 to produce LHZ1508. The full sequences of LHZ626, LHZ1495, and pRS425-Cas9-2xSapI are listed in **Table S8**.

### Transformation efficiency and plasmid stability

Fim-1ΔU and W303-1A were grown in liquid YPD overnight. Cells from 1 mL-culture were pelleted and transformed with LHZ626, LHZ1495_∼_LHZ1498, respectively, using the lithium acetate (LiAc) method ^89,90^. Colony-forming units (CFUs) on YPDH plates were counted after 3 days, to evaluate transformation efficiency (CFUs/μg DNA). To measure the stability of the plasmid, transformants were grown in liquid YPD for 24 h. The cells were then diluted and spread onto YPD and YPDH plates. Stability was determined by dividing the CFU count on YPDH plates by that on YPD plates. The experiments were replicated three times.

### Mating assay

The quantitative mating assay was performed as previously described ^91^ with modifications. Strains transformed with LHZ1493 (*hyg^R^*) were used as experimental cells, and strains transformed with LHZ1494 (*kan^R^*) served as tester cells. The experimental and tester cells were cultured overnight in YPDH and YPDG liquid media, respectively. Cells were then washed and resuspended in water to an OD_600_ of 1. In each cross, 100 μL experimental cells was mixed with 500 μL tester cells. The mixture resulted in a 1:5 ratio of experimental to tester cells when both cell types were *Km* or *Sc*. It resulted in a 1:10 ratio of *Sc* experimental cells to *Km* tester cells, as CFU per OD_600_ of *Km* was twice that of *Sc*. The mixed cells were pelleted, resuspended in 20 μL ddH_2_O, and spotted onto ME plates. After incubation at 30 □ for 24 h, the cells were washed off the ME plates with ddH_2_O and then pelleted and resuspended in 1 mL ddH_2_O. To examine the mating result, 3 μL cell suspension was spotted on YPDH, YPDG, and YPDGH (**Fig.1e**). These plates were incubated at 30 □ for 24 h. To quantify mating efficiency, the cells were diluted and plated onto YPD and YPDGH plates. The mating efficiency was calculated as the number of colonies on YPDGH plates divided by one-sixth of the number on YPD plates, reflecting the 1:5 ratio of experimental to tester cells. The experiments were replicated three times.

### Engineering of *Sc* chromosome I and III

Each step of engineering *Sc* chromosome I (chr1) or III (chr3) involved transforming a CRISPR plasmid (**Table S8**) and a donor, which is a PCR product for homologous recombination. The composition and full sequences of the donors are listed in **Supplementary Data File 1**.

Chr1 and chr3 were circularized as previously described ^92^. First, LHZ1499 or LHZ1500 was transformed into W303-1A to induce double strand breaks in both telomeres in chr1 or chr3. A DNA fragment “chr1-cir” or “chr3-cir” containing *KmURA3* with homologous arms to the chromosomal ends was simultaneously transformed to connect the two chromosomal ends with *KmURA3*. The constructs above were designed to remove the left telomere (0-1111 bp) and the right telomere along with the repetitive sequence (208917-252221 bp) of chr1, and the left telomere (0-831 bp) and the right telomere (334198-341087 bp) of chr3. The donors contain direct repeat (DR) sequences ^93^, allowing marker recycling by popping out *KmURA3* in future studies. Successful circularization was selected by SD-Ura-Leu medium. The transformants with circularized chromosomes were named *Sc*-ring1 and *Sc*-ring3.

In order to insert *KmARS1* and *KmCEN5* into the circularized chromosomes, we transformed LHZ1501 and a donor “chr1-ARS1” into *Sc*-ring1 to insert *KmARS1* and *hyg^R^* at 66090 bp of chr1, resulting in a strain named *Sc*-ring1L. LHZ1502 and a donor “chr1-ARS1/CEN5” were transformed into *Sc*-ring1L to insert *KmARS1*/*KmCEN5* and *kan^R^* at 157346 bp of chr1, resulting in a strain named *Sc*-R1. Similarly, LHZ1503 and a donor “chr3-ARS1/CEN5” were transformed into *Sc*-ring3 to insert *KmARS1*/*KmCEN5* and *hyg^R^* at 108718 bp of chr3, resulting in a strain named *Sc*-ring3L. LHZ1504 and a donor “chr3-ARS1” were transformed into *Sc*-ring3L to insert *KmARS1* and *kan^R^* at 243183 bp of chr3, resulting in a strain named *Sc*-R3. All transformations described above followed the LiAc method ^89,90^.

### Extraction and protoplast transformation of R1 and R3

R1 and R3 were extracted from *Sc*-R1 and *Sc*-R3 following a previously described method ^43^, which was developed for purifying yeast artificial chromosomes up to 600 kb in size. A total of 10 μL R1 or R3 was transformed into *Km* by protoplast transformation ^45^. The transformants were selected on SD-Ura+GH plates.

### Pulsed-Field Gel Electrophoresis (PFGE)

Plugs containing the genome of *Sc*, *Km*, and KS strains were prepared by using the CHEF Yeast Genomic DNA Plug Kit (170-3593, Biorad). To separate circular R1 from linear chromosomes in KS-R1, chromosomes were separated on the CHEF MAPPER^TM^ XA System in a 1% pulsed field certified agarose gel (162-0137, Biorad) in 0.5×TBE (diluted from 10×TBE, T1051, Solarbio) at 14 □. The running time was 24 h at 6.0 V cm^-1^, with a 60_∼_120 sec switch time ramp at an included angle of 120 °. The plug containing the entrapped R1 was cut out and linearized with NotI as described in the manufacturer’s protocol. Similarly, R3 in KS-R3 was separated from linear chromosomes and then linearized with AscI. Chromosomes of W303-1A, *Sc*-R1, *Sc*-R3, linearized R1 and R3 were separated in a 1% pulsed field certified agarose gel in 0.5×TBE at 14 □. The running time was 16 h at 6.7 V cm^-1^, with a 10_∼_40 sec switch time ramp at an included angle of 120 °. Lambda PFG Ladder (N0341S, NEB) was used as a size marker for PFGE.

### Growth curves of KS-R1 and KS-R3 and the stabilities of transferred chromosomes

As a control, Fim-1ΔU was transformed with LHZ626 to produce *Km*-V (*Km*-Vector), which confers G418 and hygromycin resistance, as well as Ura+ growth. KS-R1, KS-R3, and *Km*-V were cultured in YPDGH liquid medium overnight. The overnight culture (referred to as day 0 culture) was diluted into liquid YPD to an initial OD_600_ of 0.2 and grown at 30 □. To monitor growth curves, the OD_600_ of the culture was recorded every 2 hours for the first 14 hours. To assess the stability of the transferred chromosomes, the culture was diluted into to an OD_600_ of 0.2 with fresh YPD every 24 hours, a period corresponding to approximately 7 generations. Cultures from day 0 and after 5 days of growth were diluted and plated onto YPD and YPDGH plates. Stability was determined by dividing the CFU count on YPDGH plates by that on YPD plates. The loss rate of transferred chromosomes per generation was calculated as previously described ^34^. The experiments were replicated three times.

### Genome Sequencing

*Km*-V, *Sc*-R1, *Sc*-R3, KS-R1, and KS-R3 were cultured overnight in SD-Ura+GH liquid medium. The cultures were diluted to an OD_600_ of 0.2 and cultured until the OD_600_ reached 0.6. The cells were collected and washed once with ddH_2_O. Genomic DNA was extracted using the E.Z.N.A. Fungal DNA Kit (D3390, OMEGA Bio-Tek) and sequenced on the Illumina NovaSeq 6000 platform using the 150-bp pair-end sequencing strategy (BIOZERON, Shanghai, China).

To construct the reference sequences for R1 and R3, sequences of chr1 and chr3 were extracted from the W303-1A genome (GenBank assembly: GCA002163515.1). We then manually modified the sequences to reflect our genome edits, including the insertions of *Km* elements and deletions of telomere sequences, giving rise to reference sequences R1 and R3 (provided in **Table S1**). Annotations for R1 and R3 were transferred from S288C annotations (R64) with SnapGene. The whole-genome sequencing data were processed by BIOZERON biotechnology. To identify *de novo* variants during circular chromosome transfer, the raw sequencing data of *Sc* and KS strains were aligned to reference sequences for R1 or R3 using BWA ^94^ with default settings. Picard tools were used to remove PCR duplicates. The average sequencing depth of KS-R1, *Sc*-R1, KS-R3, and *Sc*-R3 was 298, 1,183, 1,241 and 1,134 for the respective R1 or R3 chromosome. SNPs and indels were called following the best practices of GATK HaplotypeCaller ^95^. Then, all heterozygous variants were removed from the VCF files of KS-R1 and KS-R3. All variants coexisting in both KS-R1 and *Sc*-R1 or KS-R3 and *Sc*-R3 were filtered out from the KS-R1 and KS-R3 VCF files as well. The remaining variants were considered to be *de novo* mutations generated during the chromosome transfer. In the case of R1, no *de novo* mutation was found.

To identify variants in *Km*, raw sequencing data of KS-R1, KS-R3, and *Km*-V were aligned to the reference genome of FIM1 (GenBank assembly: GCA_001854445.2) ^96^ combined with either R1 or R3 sequence, using BWA with default parameters. Picard tools were used to remove PCR duplicates. The average sequencing depth of the *Km* genome for KS-R1, KS-R3 and *Km*-V was 1,036, 1,102 and 615, respectively. GATK HaplotypeCaller was used to identify the variants in these strains, following its best practices. VCFtools ^97^ was used to remove variants shared between *Km*-V and KS-R1 or KS-R3, as well as heterozygous calls. The remaining unique, homozygous variants in KS-R1 and KS-R3 were listed in **Table S2**.

### Spot assay and quantification of growth phenotypes under various conditions

To investigate whether the chromosomal transfer resulted in enhanced phenotypes, W303-1A was transformed with LHZ626 to construct a control strain *Sc*-V (*Sc*-Vector). *Km*-V, *Sc*-V, *Sc*-R1, *Sc*-R3, KS-R1, and KS-R3 were cultured overnight in liquid SD-Ura+GH medium. The cultures were diluted to an OD_600_ of 1.0 and subjected to five serial 5-fold dilutions. These dilutions were spotted on plates using a 48-pin replicator. To evaluate the growth in different carbon sources, we used YP in combination with 2% ethanol, 3% glycerol, or different concentrations of glucose (0.02%, 1%, 3%, and 5%). To evaluate the growth in various nitrogen sources, we replaced 1 g/L sodium glutamate in the SD medium with 1 g/L threonine (Thr), 1 g/L serine (Ser), or 1.25 g of ammonium sulfate (1/4 N). For the condition of 1/2 AA and 1/2 N, the amino acids supplemented in the SD medium were halved, and 2.5 g/L ammonium sulfate was added. For chemical treatments, YPD plates were supplemented with H_2_O_2_ (0.012%, 0.016% and 0.024%), acetic acid (AcOH, 55, 57.5 and 60 mM), TM (0.2, 0.3 and 0.4 μg/mL), NaCl (0.6, 0.8 and 1M), 20 mM DTT, 15 μg/mL 5-FU, or 1 M sorbitol. The plates were incubated at 30 □ or at other specified temperatures for 1-3 days before the pictures were taken.

Spot quantification was performed using the ‘gitter’ package ^98^ in R. There were three biological replicates for each condition, except for the three H_2_O_2_ treatments and the 60mM AcOH treatment, where one biological replicate was removed due to technical problems in quantification. Source data are provided in **Table S3**. The quantified growth (*G*) under a specific condition (*condition_x*) was normalized to YPD growth at 30°C of the same strain (*strain_i*), to generate a score for relative growth (*RG*):

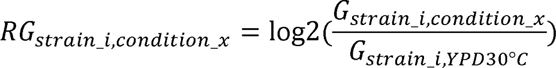

In order to visualize inter-strain differences, data across three replicates were averaged and the average *RG* for a given strain under a given condition was subtracted by that of *Km*-V under the same condition, to generate *RG’* for visualization (**Fig. 3a**):

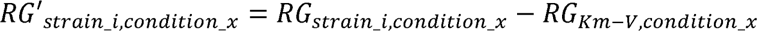

To evaluate chromosomal stability, spot dilution was performed as above on YPD supplemented with 20 μg/mL benomyl (BML), 10 μg/mL thiabendazole (TBZ), 0.05 M hydroxyurea (HU), 0.01% methyl methane sulfonate (MMS), 5 μg/mL camptothecin (CPT), or 0.05 μg/mL cycloheximide (CHX).

To perform spot assays of *SPS22*-expressing cells, *Sc*-R3 was transformed with LHZ1511 or LHZ1512, and FIM-1ΔU was transformed with LHZ1506, LHZ1507, or LHZ1508. Transformants were cultured overnight in SD-Leu medium. The culture was diluted and spotted onto plates with or without 1 M NaCl, as described above.

### Flocculation analysis

*Km*-V, KS-R1, and *Sc*-R1 were cultured overnight in SD-Ura medium. *Sc*-R1 carrying LHZ1509 (*ScFLO9_CEN_*) or LHZ1510 (*ScFLO9_2µ_*) and FIM-1ΔU carrying LHZ1505 (*ScFLO9_CEN_*) were cultured overnight in SD-Leu medium. Cells were harvested and washed with ddH_2_O and 250 mM EDTA. After two subsequent washes with ddH_2_O to ensure complete removal of EDTA, cells equivalent to an OD_600_ of 40 were pelleted and resuspended in 2 mL of ddH_2_O in a 15 mL tube. 100 μL 1 M Tris-HCl (pH 7.5) was added to the cell suspension, followed by 1 min of agitation. The tube was then left undisturbed, and pictures were taken every minute during a 5 min time window. Samples of the supernatant were taken from the surface of the cultures at various time points for OD_600_ measurement.

### RNAseq and qPCR

Three biological replicates of *Km*-V, *Sc*-R1, *Sc*-R3, KS-R1, and KS-R3 were cultured overnight in SD-Ura+GH liquid medium. The cultures were diluted into YPD to achieve an OD_600_ of 0.2. Once the OD_600_ of the cultures reached 0.6, cells were collected directly for the YPD treatment. For stress treatment, the culture was supplemented with tunicamycin (T8480, Solarbio) at a final concentration of 1 μg/mL, or with NaCl at a final concentration of 1 M, and cells were collected after 1 hour. Total RNA was extracted using the ZR Fungal/Bacterial RNA MiniPrep kit (R2014, ZymoResearch). Samples were reversed transcribed using TruSeqTM RNA sample preparation Kit (Illumina, California, USA) and sequenced by Illumina HiSeq X Ten (BIOZERON, Shanghai, China). Batch information was provided in **Supplementary Data File 2**.

The reads were mapped to corresponding reference genomes with BWA (0.7.17-r1188) ^94^ under the default settings. Different reference genomes were used for different samples: *Km*-V samples were mapped to the FIM1 genome (GCA_001854445.2), *S. cerevisiae* samples were mapped to a W303 genome ^99^ (GCA_002163515.1, ScRAPdb^100^ code W303.asm02.HP0), and KS samples were mapped to a combined genome of FIM1 and chrI and chrIII of W303. They were then processed with samtools ^101^ to remove duplicated reads. Reads with mapping quality lower than or equal to 2 were also removed with samtools, as well as those with 30 or more soft-clipped bases. The number of reads mapped to each allele (with a minimal mapping quality of 10) was counted with htseq-count ^102^, with the setting “-idattr ID -t CDS”. Counts from exons were summed up to generate counts per gene. The gene models for the RNA-seq analysis were based on the W303.asm02.HP0 annotation downloaded from ScRAPdb. There are 5,680 annotated genes in this genome, but not all genes were annotated to a known systematic name as in the S288C genome (R64), due to known intraspecific variation ^99^. We performed an all-vs-all BLAST search between W303 and S288C genes (R64 assembly) and selected the top S288C hit for each W303 gene, based on E-value and query coverage. 5,347 out of the 5,680 (94%) W303 annotations matched S288C genes unambiguously. The rest W303 gene models were kept in the analysis but flagged with “ambiguous” in the dataset. The raw read counts are provided in **Supplementary Data File 2**.

Differential expression was analyzed with DEseq2 ^103^. To examine changes in expression levels before and after chromosomal transfers (*Sc* vs. KS; *Km* vs KS), the data were fitted to a model of *count ∼ group,* where *group* represented a combination between the strain factor (*Sc*/KS/*Km*) and the treatment factor (YPD/TM/NaCl). In the *Sc* analysis (**Fig. 4a-d**), R1 and R3 data were treated separately. Data from different treatments were combined in one data frame and normalized together. In the *Km* analysis, data from KS-R1, KS-R3 and *Km*-V across three treatments were combined and normalized together. Genes with an average read count lower than 5 of any *group* were removed from the analysis, leaving 218 out of 241 *Sc* genes (chrI and chrIII) and 4,975 out of 5,081 *Km* genes in the dataset. The median of read counts across groups was 1,413 for *Sc* genes and *1,267* for Km genes, prior to normalization. Adjusted p-values for pairwise comparisons between *groups* (e.g., *Sc*-YPD vs. KS-YPD) were derived from FDR-corrected Wald significance tests, with an FDR cutoff of 0.05. Log2 fold changes were shrunken with the ashr method ^104^. Differentially expressed genes (DEG) were defined as an adjusted p-value lower than 0.05 and an absolute log2 fold change higher than 1.

For qPCR, RNA samples were reverse-transcribed using a PrimeScript RT Reagent Kit (RR037A, Takara, China). The qPCR was performed using ChamQ Universal SYBR qPCR Master Mix (Q711-02, Vazyme). The mRNA level was normalized to the average of three housekeeping genes, including *MPE1*, *TRK1*, and *SWC4*. Primers used in qPCR are listed in **Table S1.**

### *Cis* and *trans* effects

Orthologs were identified with OrthoFinder (3.0.1b1)^40^. Animo acid sequences of six species were provided to OrthoFinder, including *S. cerevisiae* (W303, GenBank #GCA_002163515.1), *K. marxianus* (FIM1, GCA_001854445.2), *Candida albicans* (SC5314, GCF_000182965.3), *Lachancea thermotolerans* (CBS6340, GCF_000142805.1), *Naumovozyma castellii* (CBS4309, GCF_000237345.1), and *Torulaspora delbrueckii* (CBS1146, GCF_000243375.1), along with a species tree according to Shen et al. (2018) ^23^. There was a total of 32,488 genes across the six genomes, 93.5% of which were placed into orthogroups. There was a total of 5,056 orthogroups, of which 3,630 were with all species present and 2,847 were single-copy orthogroups. When focusing on single-copy orthologues, the median of amino acid sequence similarity between *S. cerevisiae* and *K. marxianus* genes was 51.0% (using BLOSUM62 scoring matrix), between that of *S. cerevisiae*-*T. delbrueckii* (58.7%) and *S. cerevisiae*-*C. albicans* (37.0%), consistent with expectation (**Fig. S16a**). The high level of evolutionary divergence resulted in loss of synteny (**Fig. S16b**), so we did not further take into account of synteny in the ortholog analysis.

Between *S. cerevisiae* and *K. marxianus*, there were 4,851 orthologs, of which 3,813 were one-to-one orthologs. In the cases where multiple orthologs were identified for one gene (one-to-many or many-to-one) and many-to-many orthologous relationships, we performed pair-wise sequence alignments within the orthogroup and selected the reciprocal best match based on amino-acid sequence similarity, identifying 4,330 orthologous pairs. The median amino acid sequence similarity was 48.4%. The length of orthologs did not differ by more than two fold for 4,149/4,330 (95.8%) orthologous pairs, ruling out partial match due to local similarity. The orthologous genes were listed in **Table S4**.

Differential gene expression was identified with DESeq2^103^. R1 and R3 data were combined. YPD, TM and NaCl data were analyzed separately. Out of 241 genes on chrI and chrIII of the W303 genome, we identified *Km* orthologs for 173 genes, 162 of which had an average read count above 5 across replicates and were retained in the analysis. First, we focused on genes showing significant divergence between *Sc* and *Km* (FDR-adjusted p < 0.05 by Wald test), which included 102 genes in YPD, 132 genes in NaCl and 122 genes in TM. We designated the expression level of *Km* alleles in Km-V as *Pkm*, *Sc* alleles in *Sc*-R1 or *Sc*-R3 as *Psc*, *Km* alleles in KS as *KSkm*, and *Sc* alleles in KS as *KSsc*. We used Wald test in DESeq2 to identify significant differences between (1) *KSsc* and *KSkm* (*cis*-effects), (2) *KSsc* and *Psc* (*trans*-effects), (3) *Pkm* and *Psc* (species divergence) and (4) *KSsc* and *Pkm* (similar to *KSsc-KSkm* comparison but accounting for *trans*-differences between *K. marxianus* and KS), using an FDR-adjusted p-value cutoff of 0.05. Effect sizes were adjusted with the ashr method.

Genes were classified according to **Fig. 5c**. “*Km*-like” genes were defined as no significant difference between *KSsc* and *KSkm* but a significant difference between *KSsc* and *Psc*. “*Sc*-like” genes were defined as no significant difference between *KSsc* and *Psc* but a significant difference between *KSsc* and *KSkm*. If *KSsc* was significantly different from both *KSkm* and *Psc*, the gene was further classified into “intermediate” and “transgressive”: “intermediate” means that *KSsc* was between *Psc* and *KSkm*, and “transgressive” means that *KSsc* was outside the range of *Psc* and *KSkm*. If *KSsc* was not significantly different from either *Psc* or *KSkm*, the gene was classified as “other (conserved)”.

The above analysis of *cis*-regulatory differences primarily considered allelic differences in KS, where the two alleles share the same *trans*-environment, similar to previous studies on F1 hybrids ^64,86,105,106^. Comparing *KSsc* to *Pkm* also reveals *cis*-differences, but they might be sometimes affected by the slight differences between the *trans*-environment of KS and *Km*, i.e., reflecting *cis-* and *trans-*effects for some genes (**Fig. S14a**). In order to address if the *trans*-difference between KS and *Km* changed our conclusion, we performed the same classification analysis as above, but used *Pkm* in place of *KSkm* (**Fig. S14**).

In order to understand the major source of variance in the transcriptomic data, we performed a PCA with data from three treatments combined and normalized together. The normalized data were transformed with the VarianceStabilizingTransformation function in DESeq2 prior to the PCA.

Genome-wide expression divergence between *Km* and *Sc* orthologs was analyzed for 4,307 genes whose average read counts across replicates were higher than 5 for every condition. *Sc* data were from *Sc*-R1. YPD, NaCl and TM data were analyzed separately. DEG was identified by comparing *Psc* and *Pkm* with Wald test in DESeq2, with an FDR-adjusted p-value cutoff of 0.05. GO analysis was performed with SGD Gene Ontology Term Finder (Version 0.86, GO version 2025-07-22), with 4,307 genes as the background set. *Cis*-divergence for the transferred genes was from comparison between *KSsc* and *KSkm* described above (FDR-adjusted p-value < 0.05). The results for *cis-trans* analysis and genome-wide divergence are provided in **Table S6**.

## Author contributions

Conceptualization: HL, YY

Methodology: HL, YY, YL

Investigation: YL, KS, JZ, HC, XCL

Writing—original draft: YL, YY

Writing—review & editing: YL, YY, YW, XH, XCL

## Data and materials availability

DNA-seq and RNA-seq data are available in NCBI. (https://dataview.ncbi.nlm.nih.gov/object/PRJNA1102743?reviewer=98okerqm686mrj3ddq79r4l18u)

Source Data are provided with this paper.

## Supplementary Materials

Fig. S1. Transformation efficiency and stability of plasmids containing different combinations of *ARS* and *CEN* in *Km* and *Sc*.

Fig. S2. Zygotes were found in intraspecific but not interspecific crosses.

Fig. S3. Growth of *Sc*-R1, *Sc*-R3 and *Sc*-Vector in the presence of microtubule-depolymerizing and DNA-damaging agents.

Fig. S4. Identification of a 29 kb deletion in R3.

Fig. S5. PFGE of *Km* chromosomes in KS-R1 and KS-R3.

Fig. S6. Growth of KS and their parental strains on YPD and under 28 environmental conditions.

Fig. S7. Four independent transformants of KS-R1 showed consistent phenotypes.

Fig. S8. Effect of empty vector.

Fig. S9. Growth of KS-R1 and KS-R3 after loss of *Sc* chromosomes.

Fig. S10. Screen three candidate genes.

Fig. S11. Flocculation of *Km* and KS-R1ΔR1.

Fig. S12. Expression differences along R1 and R3 chromosomes under YPD, TM and NaCl conditions.

Fig. S13. Divergence of gene expression under NaCl and TM.

Fig. S14. Comparing expression levels of transferred genes to donor and recipient.

Fig. S15. Compensatory *cis-trans* evolution underlying conserved expression levels.

Fig. S16. Divergence of orthologous genes.

Table S1. Primers used in this study, along with manually curated R1 and R3 sequences.

Table S2. SNPs and INDELs on *Km* chromosomes of KS-R1 & KS-R3.

Table S3. Quantification of spot assays.

Table S4. Differential expression of *Sc* and *Km* genes upon chromosomal transfers.

Table S5. Orthologous genes in *Km* and *Sc* (W303).

Table S6. Classification of the transferred genes based on *cis* and *trans* effects.

Table S7. Strains used in this study.

Table S8. Plasmids used in this study, and sequences of LHZ626, LHZ1495 and pRS425-Cas9-2xSapI.

Supplementary Data File 1. Donor DNA sequences for engineering *Sc* chr1 and chr3. Supplementary Data File 2. Raw read counts of RNA-seq data and batch information.

