## Supplemental data file2 for "Intergeneric chromosomal transfer in yeast results in improved phenotypes and widespread transcriptional responses"

**Sc_counts**

| | protein_id | chr | SC-RI-1 | SC-RI-1_NaCl | SC-RI-1_TM | SC-RI-2 | SC-RI-2_NaCl | SC-RI-2_TM | SC-RI-3 | SC-RI-3_NaCl | SC-RI-3_TM | SC-RIII-1 | SC-RIII-1_NaCl | SC-RIII-1_TM | SC-RIII-2 | SC-RIII-2_NaCl | SC-RIII-2_TM | SC-RIII-3 | SC-RIII-3_NaCl | SC-RIII-3_TM |
|---|---|---|---|---|---|---|---|---|---|---|---|---|---|---|---|---|---|---|---|---|
| 1 | W303.asm02.HP0_G0000010 | chrI | 0 | 1 | 0 | 0 | 0 | 0 | 0 | 0 | 0 | 0 | 1 | 0 | 0 | 5 | 0 | 0 | 3 | 0 |
| 2 | W303.asm02.HP0_G0000020 | chrI | 7 | 2 | 1 | 0 | 2 | 0 | 3 | 2 | 1 | 9 | 5 | 1 | 4 | 2 | 1 | 4 | 1 | 1 |
| 3 | W303.asm02.HP0_G0000030 | chrI | 4 | 2 | 0 | 5 | 0 | 0 | 1 | 2 | 0 | 4 | 3 | 1 | 1 | 0 | 1 | 1 | 1 | 1 |
| 4 | W303.asm02.HP0_G0000040 | chrI | 1370 | 201 | 270 | 700 | 247 | 209 | 1001 | 225 | 289 | 452 | 124 | 142 | 561 | 112 | 106 | 449 | 121 | 122 |
| 5 | W303.asm02.HP0_G0000050 | chrI | 82 | 11 | 10 | 50 | 21 | 17 | 82 | 6 | 13 | 123 | 14 | 32 | 102 | 6 | 18 | 88 | 10 | 16 |
| 6 | W303.asm02.HP0_G0000060 | chrI | 90 | 14 | 17 | 69 | 16 | 16 | 74 | 8 | 15 | 166 | 12 | 44 | 145 | 10 | 32 | 72 | 10 | 28 |
| 7 | W303.asm02.HP0_G0000070 | chrI | 200 | 24 | 41 | 183 | 54 | 49 | 243 | 17 | 41 | 511 | 59 | 148 | 362 | 52 | 109 | 260 | 66 | 130 |
| 8 | W303.asm02.HP0_G0000080 | chrI | 43 | 14 | 9 | 45 | 13 | 6 | 40 | 10 | 11 | 45 | 18 | 15 | 57 | 13 | 9 | 49 | 14 | 20 |
| 9 | W303.asm02.HP0_G0000090 | chrI | 86 | 32 | 33 | 96 | 35 | 33 | 74 | 26 | 50 | 187 | 47 | 56 | 193 | 23 | 34 | 180 | 31 | 50 |
| 10 | W303.asm02.HP0_G0000100 | chrI | 2830 | 588 | 756 | 2642 | 716 | 823 | 2647 | 544 | 831 | 2708 | 675 | 852 | 2995 | 544 | 735 | 1897 | 600 | 770 |
| 11 | W303.asm02.HP0_G0000110 | chrI | 1348 | 264 | 227 | 1032 | 219 | 284 | 1220 | 193 | 316 | 960 | 440 | 362 | 518 | 327 | 295 | 426 | 408 | 339 |
| 12 | W303.asm02.HP0_G0000120 | chrI | 4382 | 355 | 259 | 5077 | 228 | 238 | 4259 | 265 | 268 | 1035 | 485 | 424 | 903 | 336 | 391 | 736 | 484 | 451 |
| 13 | W303.asm02.HP0_G0000130 | chrI | 15010 | 1600 | 1102 | 13371 | 1033 | 1182 | 13061 | 1621 | 1570 | 4484 | 2489 | 2013 | 3759 | 1804 | 1951 | 3319 | 2334 | 1799 |
| 14 | W303.asm02.HP0_G0000140 | chrI | 938 | 1154 | 1770 | 923 | 2082 | 1560 | 1060 | 1266 | 2183 | 6082 | 1077 | 1272 | 6679 | 659 | 752 | 6206 | 961 | 1139 |
| 15 | W303.asm02.HP0_G0000150 | chrI | 1255 | 348 | 376 | 1060 | 388 | 365 | 1283 | 343 | 355 | 2634 | 561 | 730 | 1967 | 407 | 450 | 1804 | 510 | 585 |
| 16 | W303.asm02.HP0_G0000160 | chrI | 3474 | 965 | 1115 | 3315 | 1191 | 1114 | 3271 | 991 | 1214 | 7433 | 1703 | 2198 | 5721 | 1082 | 1687 | 5526 | 1533 | 1984 |
| 17 | W303.asm02.HP0_G0000170 | chrI | 1343 | 667 | 540 | 1048 | 497 | 560 | 1054 | 678 | 663 | 1047 | 500 | 434 | 1190 | 328 | 397 | 1029 | 482 | 388 |
| 18 | W303.asm02.HP0_G0000180 | chrI | 1179 | 438 | 489 | 1310 | 436 | 464 | 1195 | 387 | 583 | 1347 | 355 | 406 | 1586 | 243 | 310 | 1155 | 343 | 369 |
| 19 | W303.asm02.HP0_G0000190 | chrI | 8394 | 1387 | 1659 | 7940 | 1425 | 1586 | 8152 | 1420 | 1886 | 11357 | 2611 | 3715 | 14690 | 2769 | 3227 | 7507 | 2439 | 3255 |
| 20 | W303.asm02.HP0_G0000200 | chrI | 4176 | 370 | 567 | 4353 | 482 | 580 | 3966 | 329 | 659 | 4591 | 722 | 1154 | 2784 | 447 | 726 | 2858 | 723 | 999 |
| 21 | W303.asm02.HP0_G0000210 | chrI | 3531 | 1458 | 1474 | 3041 | 1402 | 1551 | 3135 | 1412 | 1690 | 3716 | 1319 | 1665 | 3766 | 923 | 1338 | 3448 | 1261 | 1539 |
| 22 | W303.asm02.HP0_G0000220 | chrI | 1575 | 210 | 234 | 1525 | 220 | 243 | 1457 | 175 | 272 | 2068 | 337 | 520 | 1231 | 266 | 353 | 1136 | 347 | 498 |
| 23 | W303.asm02.HP0_G0000230 | chrI | 1026 | 465 | 357 | 1051 | 397 | 372 | 1107 | 401 | 384 | 2529 | 446 | 576 | 2264 | 288 | 381 | 2234 | 477 | 495 |
| 24 | W303.asm02.HP0_G0000240 | chrI | 743 | 501 | 689 | 526 | 646 | 731 | 622 | 483 | 788 | 1306 | 394 | 585 | 1520 | 258 | 445 | 1237 | 337 | 449 |
| 25 | W303.asm02.HP0_G0000250 | chrI | 1357 | 877 | 944 | 1179 | 874 | 1009 | 1276 | 933 | 1096 | 2040 | 746 | 927 | 2583 | 620 | 739 | 1929 | 654 | 806 |
| 26 | W303.asm02.HP0_G0000260 | chrI | 1595 | 1098 | 1462 | 986 | 1212 | 1377 | 1288 | 1255 | 1782 | 2391 | 1100 | 1298 | 2846 | 769 | 889 | 2242 | 941 | 984 |
| 27 | W303.asm02.HP0_G0000270 | chrI | 2085 | 511 | 546 | 1882 | 414 | 701 | 2182 | 335 | 769 | 4117 | 1226 | 1513 | 945 | 785 | 864 | 1057 | 1085 | 1341 |
| 28 | W303.asm02.HP0_G0000280 | chrI | 8476 | 2546 | 2579 | 7320 | 2041 | 2539 | 8372 | 2147 | 3133 | 9699 | 4095 | 4801 | 4814 | 3064 | 3043 | 4774 | 3690 | 4165 |
| 29 | W303.asm02.HP0_G0000290 | chrI | 2556 | 618 | 792 | 2568 | 732 | 909 | 2397 | 588 | 923 | 5572 | 1020 | 1529 | 3283 | 685 | 1094 | 3384 | 979 | 1448 |
| 30 | W303.asm02.HP0_G0000300 | chrI | 938 | 290 | 775 | 928 | 642 | 898 | 1028 | 195 | 1060 | 7348 | 661 | 1771 | 1973 | 475 | 1059 | 1534 | 538 | 1343 |
| 31 | W303.asm02.HP0_G0000310 | chrI | 12496 | 1469 | 1290 | 12186 | 1224 | 1320 | 11628 | 1267 | 1891 | 5254 | 2580 | 2361 | 3792 | 2407 | 2216 | 2561 | 2542 | 2103 |
| 32 | W303.asm02.HP0_G0000320 | chrI | 97911 | 20196 | 22986 | 96577 | 18274 | 26032 | 106829 | 16188 | 29574 | 100827 | 46196 | 53412 | 68142 | 37930 | 42356 | 68581 | 41269 | 50570 |
| 33 | W303.asm02.HP0_G0000330 | chrI | 188 | 149 | 105 | 121 | 119 | 94 | 188 | 138 | 122 | 372 | 137 | 111 | 271 | 62 | 80 | 303 | 132 | 87 |
| 34 | W303.asm02.HP0_G0000340 | chrI | 465 | 1074 | 1581 | 427 | 1739 | 1431 | 470 | 1177 | 1779 | 8920 | 1604 | 2232 | 7408 | 1169 | 1327 | 7471 | 1559 | 2055 |
| 35 | W303.asm02.HP0_G0000350 | chrI | 16301 | 2688 | 2744 | 15434 | 2855 | 3049 | 16808 | 2475 | 3526 | 28663 | 6004 | 7936 | 20121 | 4425 | 5746 | 18310 | 5029 | 7149 |
| 36 | W303.asm02.HP0_G0000360 | chrI | 743 | 412 | 443 | 707 | 435 | 468 | 706 | 427 | 449 | 1268 | 345 | 430 | 1347 | 239 | 343 | 1221 | 306 | 398 |
| 37 | W303.asm02.HP0_G0000370 | chrI | 3151 | 317 | 231 | 3635 | 204 | 312 | 2948 | 245 | 314 | 1284 | 591 | 421 | 817 | 335 | 371 | 578 | 581 | 377 |
| 38 | W303.asm02.HP0_G0000380 | chrI | 750 | 527 | 650 | 693 | 600 | 628 | 753 | 556 | 868 | 1877 | 486 | 533 | 2265 | 324 | 396 | 2007 | 437 | 455 |
| 39 | W303.asm02.HP0_G0000390 | chrI | 692 | 221 | 222 | 798 | 205 | 313 | 626 | 191 | 278 | 1333 | 325 | 560 | 1111 | 229 | 313 | 912 | 303 | 424 |
| 40 | W303.asm02.HP0_G0000400 | chrI | 1890 | 329 | 398 | 1884 | 380 | 475 | 1705 | 284 | 463 | 3384 | 664 | 999 | 1795 | 460 | 731 | 1743 | 615 | 859 |
| 41 | W303.asm02.HP0_G0000410 | chrI | 1093 | 473 | 487 | 980 | 425 | 514 | 925 | 481 | 517 | 1302 | 454 | 573 | 1572 | 302 | 469 | 1243 | 395 | 515 |
| 42 | W303.asm02.HP0_G0000420 | chrI | 5596 | 1179 | 1328 | 5358 | 1520 | 1371 | 5556 | 1176 | 1550 | 12486 | 1602 | 2719 | 10068 | 1253 | 2037 | 9353 | 1455 | 2411 |
| 43 | W303.asm02.HP0_G0000430 | chrI | 1096 | 72 | 85 | 1055 | 57 | 94 | 1073 | 63 | 109 | 503 | 113 | 120 | 274 | 70 | 105 | 256 | 121 | 109 |
| 44 | W303.asm02.HP0_G0000440 | chrI | 1823 | 725 | 810 | 1569 | 725 | 713 | 1556 | 707 | 851 | 2083 | 562 | 629 | 2276 | 349 | 479 | 2200 | 572 | 538 |
| 45 | W303.asm02.HP0_G0000450 | chrI | 3656 | 812 | 965 | 3390 | 806 | 1101 | 3514 | 802 | 1227 | 7531 | 1460 | 2364 | 4656 | 1088 | 1655 | 4738 | 1321 | 2134 |
| 46 | W303.asm02.HP0_G0000460 | chrI | 1768 | 2713 | 3212 | 1768 | 4186 | 3030 | 2084 | 3175 | 3323 | 14629 | 3011 | 3582 | 14739 | 1902 | 2156 | 13022 | 2382 | 3302 |
| 47 | W303.asm02.HP0_G0000470 | chrI | 781 | 175 | 313 | 793 | 341 | 351 | 808 | 183 | 359 | 2549 | 265 | 747 | 2235 | 232 | 503 | 2142 | 258 | 544 |
| 48 | W303.asm02.HP0_G0000480 | chrI | 6601 | 2563 | 4507 | 5403 | 2585 | 5441 | 6848 | 2325 | 5555 | 18191 | 4891 | 10374 | 9833 | 3902 | 6118 | 9547 | 4380 | 9236 |
| 49 | W303.asm02.HP0_G0000490 | chrI | 1600 | 815 | 790 | 1284 | 661 | 829 | 1615 | 725 | 913 | 4600 | 1363 | 1656 | 3061 | 968 | 1218 | 2859 | 1189 | 1423 |
| 50 | W303.asm02.HP0_G0000500 | chrI | 11471 | 871 | 1222 | 10632 | 1036 | 1385 | 11737 | 656 | 1706 | 4971 | 1549 | 2746 | 1072 | 1286 | 1417 | 1051 | 1210 | 2174 |
| 51 | W303.asm02.HP0_G0000510 | chrI | 1276 | 380 | 441 | 892 | 398 | 407 | 1151 | 324 | 528 | 1511 | 629 | 594 | 599 | 428 | 382 | 691 | 566 | 532 |
| 52 | W303.asm02.HP0_G0000520 | chrI | 1482 | 383 | 779 | 1395 | 720 | 873 | 1687 | 381 | 1056 | 8798 | 1050 | 1924 | 4399 | 767 | 1162 | 3975 | 872 | 1681 |
| 53 | W303.asm02.HP0_G0000530 | chrI | 428 | 32 | 69 | 376 | 54 | 42 | 399 | 40 | 74 | 143 | 42 | 55 | 239 | 26 | 38 | 130 | 35 | 52 |
| 54 | W303.asm02.HP0_G0000540 | chrI | 9023 | 414 | 593 | 8446 | 490 | 739 | 8094 | 364 | 745 | 2685 | 687 | 1358 | 1126 | 613 | 1073 | 1019 | 548 | 1060 |
| 55 | W303.asm02.HP0_G0000550 | chrI | 8829 | 3203 | 3056 | 8456 | 2825 | 2858 | 8772 | 3086 | 2970 | 12533 | 3502 | 4120 | 11430 | 2352 | 3240 | 11413 | 3264 | 3784 |
| 56 | W303.asm02.HP0_G0000560 | chrI | 3238 | 1448 | 854 | 2888 | 829 | 867 | 3121 | 1423 | 1027 | 3405 | 1699 | 1279 | 3977 | 1110 | 998 | 2893 | 1618 | 1093 |
| 57 | W303.asm02.HP0_G0000570 | chrI | 2050 | 1014 | 978 | 1824 | 925 | 924 | 2018 | 1057 | 1038 | 3282 | 1063 | 1286 | 3794 | 743 | 1111 | 3158 | 931 | 1141 |
| 58 | W303.asm02.HP0_G0000580 | chrI | 4362 | 821 | 521 | 4099 | 550 | 602 | 4099 | 608 | 578 | 4031 | 1443 | 1308 | 2846 | 911 | 1016 | 2510 | 1335 | 1205 |
| 59 | W303.asm02.HP0_G0000590 | chrI | 17761 | 2501 | 2663 | 13445 | 2133 | 3590 | 17457 | 1932 | 3613 | 16559 | 5766 | 7259 | 10863 | 4995 | 5558 | 10160 | 5156 | 6236 |
| 60 | W303.asm02.HP0_G0000600 | chrI | 416 | 227 | 241 | 428 | 259 | 276 | 425 | 201 | 303 | 1960 | 345 | 476 | 667 | 212 | 344 | 746 | 329 | 448 |
| 61 | W303.asm02.HP0_G0000610 | chrI | 1996 | 644 | 450 | 2013 | 455 | 484 | 1884 | 547 | 470 | 2963 | 661 | 797 | 2611 | 442 | 577 | 2497 | 736 | 721 |
| 62 | W303.asm02.HP0_G0000620 | chrI | 1014 | 746 | 484 | 752 | 480 | 486 | 822 | 730 | 583 | 2086 | 738 | 593 | 1915 | 472 | 494 | 1948 | 741 | 551 |
| 63 | W303.asm02.HP0_G0000630 | chrI | 3783 | 1115 | 989 | 3504 | 780 | 902 | 3246 | 1241 | 1198 | 2145 | 1049 | 938 | 2793 | 773 | 885 | 2088 | 999 | 813 |
| 64 | W303.asm02.HP0_G0000640 | chrI | 2661 | 1956 | 3312 | 2579 | 2312 | 3671 | 2710 | 2124 | 3576 | 7743 | 1907 | 3761 | 8355 | 1432 | 2505 | 7807 | 1694 | 3318 |
| 65 | W303.asm02.HP0_G0000650 | chrI | 46284 | 18303 | 12950 | 44249 | 11618 | 15671 | 45023 | 11467 | 12664 | 44896 | 21398 | 24054 | 27986 | 17734 | 20734 | 32780 | 21828 | 21832 |
| 66 | W303.asm02.HP0_G0000660 | chrI | 11335 | 14953 | 15032 | 11426 | 14277 | 15514 | 12920 | 14039 | 14779 | 34234 | 14923 | 16921 | 31194 | 11440 | 11695 | 31160 | 12529 | 15155 |
| 67 | W303.asm02.HP0_G0000670 | chrI | 2646 | 444 | 471 | 2343 | 453 | 527 | 2510 | 413 | 609 | 3797 | 862 | 1174 | 2347 | 524 | 907 | 2500 | 864 | 1045 |
| 68 | W303.asm02.HP0_G0000680 | chrI | 4345 | 742 | 613 | 4431 | 617 | 606 | 4216 | 736 | 725 | 5176 | 1295 | 1251 | 3480 | 758 | 1041 | 4124 | 1310 | 1231 |
| 69 | W303.asm02.HP0_G0000690 | chrI | 1 | 0 | 0 | 0 | 0 | 1 | 0 | 1 | 0 | 0 | 2 | 0 | 1 | 2 | 0 | 0 | 0 | 1 |
| 70 | W303.asm02.HP0_G0000700 | chrI | 3536 | 1495 | 1078 | 3670 | 1167 | 1142 | 3931 | 1237 | 940 | 6232 | 1303 | 1915 | 5225 | 1057 | 1131 | 5116 | 1171 | 1620 |
| 71 | W303.asm02.HP0_G0000710 | chrI | 4868 | 3281 | 4553 | 4342 | 3413 | 4770 | 4735 | 3483 | 5589 | 8942 | 2696 | 4367 | 10667 | 2018 | 3120 | 9140 | 2487 | 3672 |
| 72 | W303.asm02.HP0_G0000720 | chrI | 3137 | 2176 | 2420 | 2525 | 1949 | 2309 | 3041 | 2208 | 2583 | 4346 | 1125 | 1510 | 4627 | 807 | 1088 | 4110 | 1066 | 1298 |
| 73 | W303.asm02.HP0_G0000730 | chrI | 4308 | 1820 | 1690 | 4416 | 1538 | 1629 | 4406 | 1812 | 1749 | 8663 | 2342 | 2786 | 7598 | 1488 | 2082 | 7879 | 2144 | 2461 |
| 74 | W303.asm02.HP0_G0000740 | chrI | 422 | 783 | 956 | 451 | 1025 | 936 | 475 | 829 | 1142 | 3510 | 874 | 859 | 3830 | 514 | 632 | 3391 | 684 | 728 |
| 75 | W303.asm02.HP0_G0000750 | chrI | 2441 | 1094 | 984 | 2247 | 1033 | 974 | 2683 | 1084 | 1097 | 9651 | 2118 | 2326 | 6364 | 1345 | 1515 | 6242 | 1707 | 2077 |
| 76 | W303.asm02.HP0_G0000760 | chrI | 4326 | 1158 | 1009 | 2913 | 1097 | 934 | 3915 | 1225 | 969 | 4572 | 1525 | 1599 | 4191 | 947 | 992 | 3925 | 1184 | 1269 |
| 77 | W303.asm02.HP0_G0000770 | chrI | 2266 | 622 | 482 | 2121 | 533 | 515 | 2390 | 586 | 525 | 3592 | 980 | 1141 | 3425 | 886 | 1011 | 2868 | 865 | 940 |
| 78 | W303.asm02.HP0_G0000780 | chrI | 1842 | 161 | 208 | 1998 | 178 | 258 | 1618 | 112 | 279 | 1163 | 238 | 455 | 715 | 203 | 275 | 608 | 218 | 378 |
| 79 | W303.asm02.HP0_G0000790 | chrI | 167 | 21 | 31 | 105 | 30 | 30 | 115 | 25 | 29 | 122 | 26 | 35 | 124 | 19 | 41 | 63 | 41 | 32 |
| 80 | W303.asm02.HP0_G0000800 | chrI | 385 | 151 | 129 | 342 | 124 | 132 | 339 | 115 | 174 | 395 | 127 | 145 | 467 | 85 | 113 | 361 | 141 | 104 |
| 81 | W303.asm02.HP0_G0000810 | chrI | 4140 | 1338 | 1472 | 3559 | 1186 | 1351 | 3501 | 1279 | 1678 | 3169 | 1072 | 1038 | 3640 | 687 | 1031 | 3356 | 1038 | 866 |
| 82 | W303.asm02.HP0_G0000820 | chrI | 1488 | 419 | 440 | 1308 | 403 | 391 | 1346 | 403 | 460 | 1595 | 418 | 385 | 1565 | 272 | 312 | 1483 | 384 | 392 |
| 83 | W303.asm02.HP0_G0000830 | chrI | 2368 | 575 | 581 | 2212 | 583 | 634 | 2112 | 572 | 600 | 2247 | 549 | 604 | 2056 | 327 | 461 | 2045 | 486 | 564 |
| 84 | W303.asm02.HP0_G0000840 | chrI | 630 | 107 | 197 | 561 | 187 | 200 | 589 | 117 | 236 | 511 | 223 | 349 | 366 | 137 | 157 | 337 | 173 | 240 |
| 85 | W303.asm02.HP0_G0000850 | chrI | 5302 | 339 | 593 | 5320 | 619 | 790 | 5293 | 302 | 832 | 2865 | 526 | 1266 | 1277 | 466 | 673 | 941 | 350 | 922 |
| 86 | W303.asm02.HP0_G0000870 | chrII | 2053 | 368 | 379 | 1679 | 377 | 496 | 2227 | 241 | 536 | 1360 | 392 | 666 | 549 | 244 | 342 | 567 | 293 | 595 |
| 87 | W303.asm02.HP0_G0000880 | chrII | 66 | 31 | 20 | 54 | 25 | 37 | 62 | 19 | 36 | 70 | 21 | 30 | 22 | 9 | 10 | 42 | 16 | 29 |
| 88 | W303.asm02.HP0_G0000890 | chrII | 3 | 2 | 1 | 0 | 0 | 1 | 0 | 5 | 1 | 2 | 3 | 0 | 0 | 1 | 1 | 1 | 3 | 1 |
| 89 | W303.asm02.HP0_G0000900 | chrII | 21 | 5 | 2 | 41 | 0 | 1 | 14 | 1 | 2 | 5 | 1 | 2 | 5 | 1 | 1 | 4 | 2 | 0 |
| 90 | W303.asm02.HP0_G0000910 | chrII | 4334 | 1477 | 2193 | 4575 | 1618 | 2251 | 4118 | 1580 | 2677 | 3669 | 1135 | 1757 | 4886 | 863 | 1412 | 3948 | 1011 | 1590 |
| 91 | W303.asm02.HP0_G0000920 | chrII | 1384 | 186 | 233 | 1164 | 185 | 255 | 1396 | 174 | 264 | 2218 | 367 | 595 | 878 | 271 | 400 | 1050 | 319 | 563 |

| | | | | | | | | | | | | | | | | | | | | |
|---|---|---|---|---|---|---|---|---|---|---|---|---|---|---|---|---|---|---|---|---|
| 92 | W303.asm02.HP0_G0000930 | chrII | 3210 | 306 | 483 | 3289 | 469 | 507 | 3071 | 245 | 552 | 5631 | 720 | 1266 | 2974 | 559 | 834 | 3006 | 668 | 1078 |
| 93 | W303.asm02.HP0_G0000940 | chrII | 1117 | 275 | 410 | 1240 | 405 | 476 | 1076 | 283 | 490 | 4037 | 608 | 1117 | 2360 | 395 | 676 | 2599 | 565 | 969 |
| 94 | W303.asm02.HP0_G0000950 | chrII | 1942 | 632 | 779 | 2177 | 839 | 732 | 1869 | 630 | 806 | 4619 | 809 | 1118 | 4228 | 512 | 854 | 4092 | 750 | 1066 |
| 95 | W303.asm02.HP0_G0000960 | chrII | 2125 | 1175 | 1819 | 1812 | 1508 | 1733 | 1835 | 1261 | 2108 | 4270 | 1090 | 1417 | 5355 | 784 | 1358 | 4484 | 992 | 1328 |
| 96 | W303.asm02.HP0_G0000970 | chrII | 2714 | 151 | 255 | 2818 | 188 | 322 | 2513 | 111 | 303 | 2108 | 264 | 584 | 1295 | 249 | 581 | 986 | 246 | 576 |
| 97 | W303.asm02.HP0_G0000980 | chrII | 11789 | 3959 | 3897 | 10206 | 3776 | 4331 | 11166 | 3968 | 4962 | 27957 | 8241 | 10462 | 24468 | 7411 | 8066 | 22503 | 7045 | 8197 |
| 98 | W303.asm02.HP0_G0000990 | chrII | 1443 | 1515 | 1617 | 1116 | 1262 | 1320 | 1225 | 1494 | 1537 | 3807 | 970 | 1123 | 3453 | 606 | 781 | 3870 | 881 | 1016 |
| 99 | W303.asm02.HP0_G0001000 | chrII | 371 | 84 | 147 | 378 | 189 | 138 | 375 | 70 | 133 | 1794 | 161 | 357 | 1202 | 102 | 274 | 1117 | 142 | 295 |
| 100 | W303.asm02.HP0_G0001010 | chrII | 367 | 530 | 441 | 371 | 411 | 496 | 349 | 416 | 553 | 1398 | 444 | 544 | 1159 | 275 | 432 | 1089 | 446 | 475 |
| 101 | W303.asm02.HP0_G0001020 | chrII | 2339 | 1578 | 1817 | 2374 | 1543 | 1872 | 2198 | 1739 | 1800 | 5165 | 1452 | 2146 | 6207 | 1037 | 1459 | 5054 | 1443 | 1817 |
| 102 | W303.asm02.HP0_G0001030 | chrII | 12179 | 9202 | 11574 | 11174 | 11382 | 11764 | 11896 | 9438 | 13004 | 24596 | 8819 | 10240 | 25868 | 6733 | 8629 | 24499 | 7817 | 9168 |
| 103 | W303.asm02.HP0_G0001040 | chrII | 1273 | 548 | 472 | 1132 | 411 | 531 | 1159 | 598 | 546 | 991 | 405 | 496 | 1183 | 330 | 413 | 921 | 406 | 431 |
| 104 | W303.asm02.HP0_G0001050 | chrII | 12345 | 1209 | 760 | 11294 | 752 | 1044 | 12985 | 930 | 990 | 6119 | 1707 | 2075 | 4049 | 1337 | 1441 | 3654 | 1315 | 1744 |
| 105 | W303.asm02.HP0_G0001060 | chrII | 4356 | 1407 | 1960 | 4048 | 1780 | 2050 | 3978 | 1496 | 2863 | 4073 | 1138 | 1722 | 5447 | 862 | 1515 | 4199 | 1077 | 1506 |
| 106 | W303.asm02.HP0_G0001070 | chrII | 681 | 867 | 816 | 620 | 812 | 837 | 640 | 918 | 848 | 2774 | 729 | 843 | 2573 | 462 | 630 | 2634 | 739 | 747 |
| 107 | W303.asm02.HP0_G0001080 | chrII | 3208 | 392 | 666 | 2820 | 634 | 668 | 2935 | 422 | 743 | 5301 | 510 | 1357 | 3914 | 360 | 1073 | 4224 | 446 | 1146 |
| 108 | W303.asm02.HP0_G0001090 | chrII | 5194 | 5172 | 6814 | 4821 | 6261 | 6644 | 5175 | 5189 | 8069 | 14180 | 4931 | 5693 | 15479 | 3498 | 4676 | 14437 | 4223 | 5084 |
| 109 | W303.asm02.HP0_G0001100 | chrII | 5455 | 649 | 291 | 5365 | 368 | 344 | 5215 | 433 | 254 | 2034 | 569 | 573 | 1741 | 319 | 404 | 1612 | 588 | 493 |
| 110 | W303.asm02.HP0_G0001110 | chrII | 2631 | 547 | 824 | 2997 | 750 | 925 | 2521 | 548 | 892 | 8263 | 1352 | 1898 | 4962 | 934 | 1410 | 4681 | 1183 | 1626 |
| 111 | W303.asm02.HP0_G0001120 | chrII | 850 | 479 | 571 | 875 | 650 | 565 | 803 | 470 | 629 | 3333 | 650 | 920 | 2764 | 354 | 680 | 2830 | 605 | 820 |
| 112 | W303.asm02.HP0_G0001130 | chrII | 1370 | 446 | 581 | 1062 | 407 | 645 | 1530 | 474 | 697 | 3737 | 1177 | 1406 | 2983 | 851 | 921 | 2586 | 983 | 1318 |
| 113 | W303.asm02.HP0_G0001140 | chrII | 728 | 289 | 766 | 680 | 660 | 798 | 916 | 250 | 901 | 5963 | 722 | 1766 | 1441 | 504 | 885 | 1448 | 468 | 1400 |
| 114 | W303.asm02.HP0_G0001150 | chrII | 1663 | 388 | 344 | 1552 | 381 | 387 | 1501 | 399 | 376 | 2004 | 581 | 796 | 1494 | 431 | 540 | 1416 | 569 | 676 |
| 115 | W303.asm02.HP0_G0001160 | chrII | 3336 | 996 | 1646 | 3222 | 1613 | 1680 | 3413 | 1043 | 2018 | 13061 | 1631 | 3088 | 7801 | 1160 | 2160 | 7894 | 1428 | 2804 |
| 116 | W303.asm02.HP0_G0001170 | chrII | 3159 | 595 | 749 | 3053 | 531 | 743 | 2866 | 554 | 964 | 1197 | 452 | 496 | 2075 | 400 | 665 | 1275 | 414 | 408 |
| 117 | W303.asm02.HP0_G0001180 | chrII | 19724 | 6029 | 5382 | 17568 | 5629 | 5485 | 19562 | 5736 | 6496 | 39134 | 9289 | 11215 | 28686 | 6817 | 7947 | 30473 | 8510 | 10446 |
| 118 | W303.asm02.HP0_G0001190 | chrII | 2317 | 472 | 355 | 2218 | 1699 | 2059 | 2328 | 502 | 444 | 2408 | 938 | 1032 | 1300 | 478 | 546 | 1539 | 985 | 911 |
| 119 | W303.asm02.HP0_G0001200 | chrII | 431 | 107 | 86 | 360 | 92 | 115 | 330 | 101 | 118 | 906 | 239 | 283 | 554 | 138 | 163 | 608 | 248 | 223 |
| 120 | W303.asm02.HP0_G0001210 | chrII | 9906 | 8731 | 10695 | 10079 | 9982 | 10311 | 10632 | 8897 | 13125 | 21825 | 8980 | 9533 | 23449 | 6516 | 7430 | 22020 | 7677 | 8683 |
| 121 | W303.asm02.HP0_G0001220 | chrII | 465 | 1356 | 1104 | 396 | 1392 | 1206 | 443 | 1486 | 1294 | 2776 | 1237 | 894 | 3350 | 803 | 588 | 2800 | 1094 | 813 |
| 122 | W303.asm02.HP0_G0001230 | chrII | 1515 | 501 | 418 | 1179 | 423 | 483 | 1388 | 401 | 531 | 2706 | 669 | 711 | 1181 | 453 | 457 | 1284 | 642 | 658 |
| 123 | W303.asm02.HP0_G0001240 | chrII | 1750 | 1565 | 1552 | 1677 | 1856 | 1387 | 1868 | 1685 | 1577 | 6222 | 1527 | 1609 | 6561 | 1044 | 1093 | 6179 | 1327 | 1385 |
| 124 | W303.asm02.HP0_G0001250 | chrII | 2056 | 913 | 1185 | 2277 | 1180 | 1009 | 2094 | 1073 | 1277 | 8327 | 1248 | 1522 | 8149 | 851 | 1252 | 7317 | 1075 | 1384 |
| 125 | W303.asm02.HP0_G0001260 | chrII | 1742 | 186 | 296 | 1795 | 301 | 396 | 1652 | 171 | 375 | 2409 | 437 | 775 | 683 | 274 | 430 | 666 | 318 | 583 |
| 126 | W303.asm02.HP0_G0001270 | chrII | 5515 | 2794 | 2335 | 4379 | 1968 | 2501 | 4650 | 2429 | 2584 | 2946 | 2134 | 1897 | 5081 | 1784 | 2011 | 2735 | 2092 | 1538 |
| 127 | W303.asm02.HP0_G0001280 | chrII | 898 | 303 | 457 | 958 | 412 | 433 | 926 | 318 | 432 | 2468 | 334 | 644 | 1935 | 285 | 539 | 2191 | 264 | 633 |
| 128 | W303.asm02.HP0_G0001290 | chrII | 2375 | 968 | 1131 | 2226 | 1187 | 1222 | 2279 | 943 | 1298 | 8471 | 1599 | 2419 | 7101 | 1356 | 1940 | 5471 | 1427 | 2125 |
| 129 | W303.asm02.HP0_G0001300 | chrII | 1088 | 458 | 585 | 1066 | 592 | 536 | 1152 | 491 | 612 | 3326 | 556 | 831 | 3118 | 423 | 639 | 2873 | 632 | 741 |
| 130 | W303.asm02.HP0_G0001310 | chrII | 1994 | 1042 | 1234 | 1944 | 1161 | 1329 | 1792 | 1051 | 1779 | 3024 | 829 | 1037 | 4270 | 607 | 971 | 3288 | 734 | 981 |
| 131 | W303.asm02.HP0_G0001320 | chrII | 821 | 402 | 450 | 722 | 412 | 437 | 770 | 453 | 551 | 957 | 312 | 345 | 1244 | 225 | 303 | 946 | 287 | 301 |
| 132 | W303.asm02.HP0_G0001330 | chrII | 9366 | 3419 | 2266 | 8206 | 2177 | 2359 | 8933 | 3025 | 2496 | 11291 | 3923 | 3921 | 9213 | 2856 | 3226 | 9142 | 3849 | 3546 |
| 133 | W303.asm02.HP0_G0001340 | chrII | 1548 | 1478 | 1207 | 1486 | 1210 | 1181 | 1497 | 1507 | 1266 | 4272 | 1492 | 1374 | 3904 | 888 | 988 | 3748 | 1206 | 1232 |
| 134 | W303.asm02.HP0_G0001350 | chrII | 9940 | 4751 | 4684 | 9451 | 4234 | 4452 | 9586 | 4883 | 4712 | 13870 | 4254 | 4393 | 14460 | 2883 | 3677 | 13215 | 3934 | 3869 |
| 135 | W303.asm02.HP0_G0001360 | chrII | 1717 | 1211 | 1600 | 1580 | 1649 | 1435 | 1727 | 1330 | 1676 | 5887 | 1215 | 1460 | 6403 | 770 | 907 | 5102 | 1059 | 1243 |
| 136 | W303.asm02.HP0_G0001370 | chrII | 1026 | 383 | 628 | 1359 | 972 | 735 | 1501 | 464 | 799 | 7369 | 624 | 1103 | 7467 | 580 | 543 | 5544 | 504 | 920 |
| 137 | W303.asm02.HP0_G0001380 | chrII | 843 | 620 | 708 | 812 | 814 | 702 | 844 | 699 | 787 | 4963 | 815 | 1022 | 4395 | 549 | 866 | 4314 | 718 | 1058 |
| 138 | W303.asm02.HP0_G0001390 | chrII | 6836 | 615 | 691 | 7134 | 697 | 755 | 7236 | 577 | 904 | 7610 | 1413 | 1705 | 6162 | 960 | 1252 | 5435 | 1219 | 1564 |
| 139 | W303.asm02.HP0_G0001400 | chrII | 5183 | 2735 | 2934 | 5275 | 2374 | 3217 | 4856 | 2692 | 3563 | 6285 | 2295 | 2648 | 6769 | 1438 | 2049 | 6447 | 2091 | 2382 |
| 140 | W303.asm02.HP0_G0001410 | chrII | 2071 | 242 | 206 | 1965 | 286 | 272 | 1883 | 234 | 225 | 718 | 217 | 253 | 880 | 175 | 212 | 625 | 237 | 233 |
| 141 | W303.asm02.HP0_G0001420 | chrII | 20989 | 1129 | 1754 | 20011 | 1429 | 1982 | 21543 | 1021 | 2135 | 10712 | 2276 | 3918 | 4236 | 2201 | 2939 | 3193 | 1777 | 3336 |
| 142 | W303.asm02.HP0_G0001430 | chrII | 2509 | 851 | 618 | 2458 | 665 | 652 | 2428 | 734 | 579 | 3464 | 905 | 985 | 3335 | 516 | 689 | 2999 | 770 | 849 |
| 143 | W303.asm02.HP0_G0001440 | chrII | 18246 | 6643 | 5309 | 18084 | 5132 | 5047 | 17260 | 7118 | 6154 | 25858 | 9595 | 9721 | 23891 | 8037 | 7934 | 21141 | 8301 | 7555 |
| 144 | W303.asm02.HP0_G0001450 | chrII | 8 | 47 | 8 | 5 | 6 | 9 | 9 | 34 | 9 | 13 | 32 | 4 | 14 | 15 | 5 | 28 | 26 | 4 |
| 145 | W303.asm02.HP0_G0001460 | chrII | 113 | 258 | 393 | 103 | 419 | 345 | 96 | 274 | 372 | 1274 | 417 | 441 | 1921 | 226 | 343 | 1019 | 477 | 342 |
| 146 | W303.asm02.HP0_G0001470 | chrII | 802 | 1196 | 1664 | 819 | 2152 | 1606 | 804 | 1264 | 1906 | 13492 | 2021 | 2984 | 8967 | 1528 | 1847 | 8684 | 1736 | 2414 |
| 147 | W303.asm02.HP0_G0001480 | chrII | 8660 | 5565 | 5056 | 8692 | 4772 | 5059 | 8367 | 5436 | 5439 | 10642 | 4758 | 4593 | 11173 | 3218 | 3739 | 10181 | 4266 | 4100 |
| 148 | W303.asm02.HP0_G0001490 | chrII | 3041 | 2424 | 3176 | 2994 | 2224 | 3259 | 2903 | 2566 | 4146 | 6145 | 1887 | 2709 | 7537 | 1452 | 2105 | 6843 | 1793 | 2371 |
| 149 | W303.asm02.HP0_G0001500 | chrII | 268 | 346 | 397 | 293 | 286 | 405 | 209 | 348 | 540 | 701 | 252 | 269 | 753 | 168 | 212 | 762 | 202 | 223 |
| 150 | W303.asm02.HP0_G0001510 | chrII | 900 | 2814 | 2637 | 876 | 3381 | 2377 | 1187 | 2876 | 2979 | 20480 | 4560 | 4792 | 14950 | 2617 | 2370 | 17111 | 3815 | 4317 |
| 151 | W303.asm02.HP0_G0001520 | chrII | 3304 | 1746 | 1287 | 2860 | 1399 | 1392 | 2819 | 1593 | 1448 | 5033 | 1713 | 1819 | 4658 | 1278 | 1612 | 4335 | 1692 | 1621 |
| 152 | W303.asm02.HP0_G0001530 | chrII | 2160 | 316 | 517 | 2045 | 477 | 593 | 2014 | 290 | 664 | 3700 | 541 | 1022 | 2033 | 453 | 772 | 1921 | 487 | 967 |
| 153 | W303.asm02.HP0_G0001540 | chrII | 3468 | 2482 | 2410 | 3078 | 2321 | 2299 | 3272 | 2516 | 3031 | 7050 | 2079 | 2185 | 7544 | 1311 | 1647 | 7672 | 1980 | 1992 |
| 154 | W303.asm02.HP0_G0001550 | chrII | 845 | 382 | 370 | 741 | 352 | 430 | 860 | 303 | 451 | 3468 | 594 | 1041 | 2144 | 388 | 642 | 2341 | 516 | 898 |
| 155 | W303.asm02.HP0_G0001560 | chrII | 1426 | 338 | 492 | 1489 | 490 | 534 | 1514 | 371 | 617 | 3741 | 489 | 869 | 2873 | 369 | 649 | 2397 | 408 | 760 |
| 156 | W303.asm02.HP0_G0001570 | chrII | 3716 | 977 | 1156 | 3702 | 902 | 1415 | 3451 | 743 | 1448 | 3446 | 1304 | 2304 | 2725 | 992 | 1544 | 2370 | 1291 | 1855 |
| 157 | W303.asm02.HP0_G0001580 | chrII | 2256 | 663 | 826 | 1619 | 742 | 875 | 2063 | 559 | 1038 | 6899 | 1297 | 1845 | 3819 | 920 | 1513 | 4254 | 1199 | 1706 |
| 158 | W303.asm02.HP0_G0001590 | chrII | 515 | 544 | 545 | 526 | 617 | 489 | 515 | 550 | 462 | 1877 | 458 | 477 | 2108 | 287 | 368 | 1677 | 414 | 438 |
| 159 | W303.asm02.HP0_G0001600 | chrII | 11814 | 4950 | 5448 | 10433 | 5134 | 6052 | 11243 | 4730 | 7108 | 28588 | 9039 | 10613 | 25166 | 8132 | 9094 | 21780 | 7389 | 8595 |
| 160 | W303.asm02.HP0_G0001610 | chrII | 1734 | 993 | 1054 | 1529 | 710 | 1069 | 1345 | 948 | 1219 | 1555 | 719 | 801 | 1725 | 487 | 608 | 1667 | 772 | 638 |
| 161 | W303.asm02.HP0_G0001620 | chrII | 1309 | 433 | 314 | 1029 | 348 | 309 | 1022 | 433 | 361 | 1178 | 392 | 270 | 1615 | 301 | 301 | 1137 | 387 | 223 |
| 162 | W303.asm02.HP0_G0001630 | chrII | 798 | 1525 | 2088 | 766 | 2125 | 2094 | 858 | 1748 | 2959 | 5825 | 1536 | 1604 | 6667 | 882 | 1009 | 6667 | 1282 | 1391 |
| 163 | W303.asm02.HP0_G0001640 | chrII | 1003 | 549 | 483 | 926 | 409 | 460 | 893 | 571 | 430 | 1246 | 527 | 513 | 1452 | 371 | 452 | 1147 | 499 | 411 |
| 164 | W303.asm02.HP0_G0001650 | chrII | 1079 | 901 | 1210 | 1142 | 978 | 1165 | 1067 | 864 | 1703 | 2773 | 682 | 909 | 3581 | 490 | 738 | 3259 | 642 | 806 |
| 165 | W303.asm02.HP0_G0001660 | chrII | 594 | 721 | 498 | 561 | 454 | 452 | 574 | 714 | 543 | 1714 | 628 | 487 | 1934 | 389 | 384 | 1652 | 625 | 447 |
| 166 | W303.asm02.HP0_G0001670 | chrII | 3952 | 1340 | 1479 | 3790 | 1431 | 1455 | 4322 | 1309 | 1604 | 15072 | 2426 | 3457 | 11448 | 1721 | 2093 | 11351 | 2004 | 2996 |
| 167 | W303.asm02.HP0_G0001680 | chrII | 2118 | 662 | 722 | 1989 | 726 | 777 | 1989 | 649 | 937 | 6117 | 1262 | 1774 | 4013 | 983 | 1258 | 3502 | 1183 | 1615 |
| 168 | W303.asm02.HP0_G0001690 | chrII | 18093 | 1628 | 2022 | 18292 | 1857 | 2537 | 17883 | 1496 | 2505 | 16049 | 3383 | 5806 | 9313 | 2878 | 4427 | 8358 | 3091 | 5158 |
| 169 | W303.asm02.HP0_G0001700 | chrII | 928 | 853 | 746 | 800 | 757 | 755 | 872 | 916 | 931 | 1902 | 752 | 624 | 2037 | 432 | 431 | 1860 | 655 | 546 |
| 170 | W303.asm02.HP0_G0001710 | chrII | 848 | 1239 | 1667 | 867 | 1345 | 1478 | 976 | 1340 | 1645 | 6875 | 1528 | 1856 | 6577 | 959 | 1153 | 6220 | 1317 | 1587 |
| 171 | W303.asm02.HP0_G0001720 | chrII | 1987 | 546 | 564 | 1858 | 523 | 535 | 1808 | 580 | 546 | 2306 | 558 | 676 | 2261 | 309 | 518 | 2043 | 583 | 574 |
| 172 | W303.asm02.HP0_G0001730 | chrII | 134 | 348 | 477 | 110 | 438 | 445 | 128 | 312 | 685 | 1156 | 322 | 374 | 1264 | 221 | 242 | 1328 | 293 | 300 |
| 173 | W303.asm02.HP0_G0001740 | chrII | 5798 | 900 | 1300 | 5112 | 1141 | 1403 | 5307 | 918 | 1493 | 7826 | 1597 | 2736 | 3877 | 1329 | 1905 | 4059 | 1389 | 2420 |
| 174 | W303.asm02.HP0_G0001750 | chrII | 4590 | 986 | 991 | 4189 | 937 | 1082 | 4638 | 827 | 1269 | 5414 | 1719 | 2120 | 4454 | 1067 | 1379 | 3950 | 1476 | 1841 |
| 175 | W303.asm02.HP0_G0001760 | chrII | 4566 | 415 | 383 | 3907 | 421 | 468 | 4415 | 302 | 494 | 2647 | 934 | 894 | 829 | 843 | 742 | 713 | 752 | 704 |
| 176 | W303.asm02.HP0_G0001770 | chrII | 786 | 343 | 521 | 823 | 656 | 498 | 756 | 327 | 604 | 5174 | 652 | 1015 | 3791 | 410 | 611 | 3773 | 678 | 880 |
| 177 | W303.asm02.HP0_G0001780 | chrII | 237 | 33 | 45 | 236 | 36 | 62 | 199 | 25 | 53 | 288 | 74 | 105 | 118 | 50 | 72 | 111 | 51 | 95 |
| 178 | W303.asm02.HP0_G0001790 | chrII | 7632 | 838 | 1041 | 7173 | 964 | 1211 | 7728 | 767 | 1370 | 7378 | 1988 | 2262 | 2897 | 1467 | 1657 | 3073 | 1671 | 2128 |
| 179 | W303.asm02.HP0_G0001800 | chrII | 652 | 279 | 258 | 670 | 249 | 275 | 752 | 263 | 252 | 1597 | 350 | 483 | 1426 | 241 | 360 | 1448 | 295 | 433 |
| 180 | W303.asm02.HP0_G0001810 | chrII | 1176 | 838 | 810 | 1160 | 909 | 756 | 1412 | 853 | 756 | 6109 | 1167 | 1396 | 5106 | 751 | 1016 | 5304 | 1082 | 1306 |
| 181 | W303.asm02.HP0_G0001820 | chrII | 14 | 0 | 3 | 9 | 5 | 3 | 9 | 3 | 2 | 21 | 3 | 1 | 22 | 4 | 3 | 24 | 5 | 2 |
| 182 | W303.asm02.HP0_G0001830 | chrII | 1480 | 310 | 451 | 1456 | 367 | 518 | 1557 | 342 | 519 | 4007 | 701 | 1444 | 2162 | 548 | 840 | 2301 | 608 | 1325 |
| 183 | W303.asm02.HP0_G0001840 | chrII | 11651 | 1291 | 1509 | 10088 | 1238 | 1701 | 10979 | 1046 | 1798 | 5327 | 1889 | 2843 | 3150 | 1606 | 2185 | 2525 | 1507 | 2451 |
| 184 | W303.asm02.HP0_G0001850 | chrII | 2696 | 1411 | 1775 | 2472 | 1526 | 1660 | 2571 | 1408 | 1943 | 4492 | 1439 | 1797 | 5439 | 919 | 1390 | 4655 | 1270 | 1739 |

| | | | | | | | | | | | | | | | | | | | | |
|---|---|---|---|---|---|---|---|---|---|---|---|---|---|---|---|---|---|---|---|---|
| 185 | W303.asm02.HP0_G0001860 | chrII | 2685 | 988 | 1025 | 2553 | 897 | 1077 | 2763 | 902 | 1204 | 4920 | 1212 | 1639 | 3443 | 680 | 1121 | 3935 | 1147 | 1593 |
| 186 | W303.asm02.HP0_G0001870 | chrII | 2349 | 1059 | 1547 | 2199 | 2119 | 1506 | 2441 | 1210 | 2079 | 18941 | 1921 | 3701 | 13963 | 1252 | 2117 | 15167 | 1556 | 3268 |
| 187 | W303.asm02.HP0_G0001880 | chrII | 4206 | 3701 | 7302 | 3487 | 5715 | 6967 | 3930 | 3751 | 9302 | 12005 | 3322 | 5525 | 12518 | 2229 | 4210 | 12563 | 2842 | 4596 |
| 188 | W303.asm02.HP0_G0001890 | chrII | 5631 | 4086 | 4572 | 5137 | 4052 | 4805 | 5276 | 4411 | 4438 | 11098 | 3988 | 5262 | 9843 | 2678 | 3739 | 10359 | 3457 | 4520 |
| 189 | W303.asm02.HP0_G0001900 | chrII | 2640 | 1369 | 1831 | 2029 | 1570 | 1799 | 2087 | 1418 | 2539 | 4058 | 1302 | 1581 | 5533 | 936 | 1510 | 4612 | 1273 | 1392 |
| 190 | W303.asm02.HP0_G0001910 | chrII | 3547 | 876 | 690 | 3593 | 676 | 814 | 3146 | 812 | 867 | 4332 | 1487 | 1528 | 3038 | 1111 | 1414 | 3132 | 1373 | 1405 |
| 191 | W303.asm02.HP0_G0001920 | chrII | 1373 | 980 | 1154 | 1147 | 899 | 1265 | 1361 | 910 | 1265 | 4841 | 1166 | 2032 | 4130 | 786 | 1316 | 4093 | 1224 | 1899 |
| 192 | W303.asm02.HP0_G0001930 | chrII | 5509 | 1005 | 1020 | 5021 | 878 | 1219 | 4987 | 900 | 1277 | 5619 | 1535 | 2553 | 5252 | 1255 | 1834 | 4213 | 1536 | 2193 |
| 193 | W303.asm02.HP0_G0001940 | chrII | 2027 | 1196 | 1236 | 2247 | 1182 | 1157 | 1877 | 1217 | 1551 | 4760 | 1066 | 1047 | 6072 | 708 | 784 | 5039 | 924 | 894 |
| 194 | W303.asm02.HP0_G0001950 | chrII | 2629 | 891 | 687 | 3031 | 648 | 685 | 2276 | 900 | 712 | 1898 | 890 | 826 | 2729 | 691 | 685 | 1611 | 844 | 696 |
| 195 | W303.asm02.HP0_G0001960 | chrII | 3076 | 635 | 572 | 2813 | 466 | 588 | 2647 | 549 | 705 | 2093 | 786 | 803 | 2715 | 567 | 723 | 1845 | 830 | 650 |
| 196 | W303.asm02.HP0_G0001970 | chrII | 2292 | 337 | 412 | 2306 | 331 | 401 | 2397 | 309 | 446 | 3391 | 609 | 887 | 3452 | 443 | 647 | 2407 | 559 | 743 |
| 197 | W303.asm02.HP0_G0001980 | chrII | 925 | 985 | 1199 | 918 | 984 | 1133 | 917 | 860 | 1214 | 3211 | 2910 | 2098 | 2008 | 2279 | 1660 | 2096 | 2234 | 1424 |
| 198 | W303.asm02.HP0_G0001990 | chrII | 5228 | 6696 | 9649 | 4728 | 8641 | 9463 | 4940 | 7051 | 11350 | 15412 | 6356 | 7662 | 17065 | 4163 | 5977 | 16301 | 5576 | 6588 |
| 199 | W303.asm02.HP0_G0002000 | chrII | 2070 | 3584 | 5406 | 1641 | 4767 | 5109 | 1787 | 3431 | 5957 | 9153 | 2728 | 3803 | 9672 | 1881 | 2803 | 9405 | 2604 | 3379 |
| 200 | W303.asm02.HP0_G0002010 | chrII | 21621 | 8385 | 6556 | 20631 | 6297 | 6190 | 22327 | 8279 | 5947 | 21969 | 8665 | 10048 | 21757 | 7300 | 7297 | 19543 | 7565 | 8622 |
| 201 | W303.asm02.HP0_G0002020 | chrII | 2135 | 2028 | 2092 | 1949 | 1643 | 2155 | 1967 | 2144 | 2868 | 4726 | 1636 | 1675 | 5965 | 1159 | 1265 | 5148 | 1537 | 1536 |
| 202 | W303.asm02.HP0_G0002030 | chrII | 6418 | 1223 | 1492 | 5421 | 1231 | 1664 | 6496 | 1010 | 1831 | 11041 | 2652 | 3823 | 6480 | 2236 | 2631 | 5945 | 2359 | 3316 |
| 203 | W303.asm02.HP0_G0002040 | chrII | 5086 | 870 | 857 | 4713 | 680 | 908 | 4532 | 788 | 1064 | 3564 | 1373 | 1569 | 2273 | 846 | 1173 | 2428 | 1288 | 1471 |
| 204 | W303.asm02.HP0_G0002050 | chrII | 5708 | 1765 | 1684 | 5454 | 1804 | 1652 | 5824 | 1795 | 1920 | 14657 | 2966 | 3503 | 10568 | 1947 | 2469 | 10936 | 2678 | 3187 |
| 205 | W303.asm02.HP0_G0002060 | chrII | 215 | 79 | 58 | 210 | 58 | 59 | 181 | 58 | 66 | 208 | 67 | 64 | 185 | 22 | 49 | 213 | 83 | 70 |
| 206 | W303.asm02.HP0_G0002070 | chrII | 378 | 112 | 138 | 402 | 140 | 156 | 371 | 95 | 173 | 932 | 142 | 279 | 631 | 88 | 188 | 622 | 126 | 240 |
| 207 | W303.asm02.HP0_G0002080 | chrII | 299 | 117 | 92 | 231 | 84 | 78 | 333 | 62 | 108 | 536 | 145 | 168 | 324 | 77 | 114 | 450 | 128 | 179 |
| 208 | W303.asm02.HP0_G0002090 | chrII | 710 | 1018 | 654 | 611 | 821 | 614 | 695 | 858 | 667 | 4320 | 1295 | 1109 | 3664 | 862 | 880 | 3217 | 1217 | 973 |
| 209 | W303.asm02.HP0_G0002100 | chrII | 1243 | 583 | 935 | 1162 | 804 | 985 | 1145 | 610 | 1222 | 2344 | 473 | 646 | 2539 | 349 | 588 | 2497 | 451 | 646 |
| 210 | W303.asm02.HP0_G0002110 | chrII | 2156 | 1107 | 1361 | 2106 | 1270 | 1312 | 2281 | 1068 | 1591 | 12843 | 2287 | 3031 | 12868 | 2361 | 2667 | 7603 | 2027 | 2776 |
| 211 | W303.asm02.HP0_G0002120 | chrII | 4387 | 525 | 403 | 4049 | 427 | 473 | 3896 | 481 | 464 | 2150 | 567 | 580 | 2275 | 430 | 569 | 1823 | 548 | 567 |
| 212 | W303.asm02.HP0_G0002130 | chrII | 19593 | 13592 | 9822 | 18991 | 10068 | 9709 | 19662 | 12107 | 8292 | 31965 | 12193 | 13226 | 27717 | 8958 | 9577 | 28405 | 10461 | 11663 |
| 213 | W303.asm02.HP0_G0002140 | chrII | 9655 | 2023 | 1509 | 9150 | 1453 | 1541 | 9189 | 1817 | 1647 | 6586 | 1838 | 2102 | 7471 | 1562 | 2081 | 5991 | 1925 | 1899 |
| 214 | W303.asm02.HP0_G0002150 | chrII | 1799 | 1363 | 1253 | 1488 | 1242 | 1191 | 1705 | 1392 | 1110 | 5532 | 1471 | 1779 | 4568 | 854 | 1235 | 4986 | 1336 | 1643 |
| 215 | W303.asm02.HP0_G0002160 | chrII | 3975 | 2453 | 2677 | 4402 | 2756 | 2452 | 4389 | 2690 | 2900 | 11712 | 2998 | 3240 | 11145 | 2052 | 2667 | 11182 | 2634 | 2952 |
| 216 | W303.asm02.HP0_G0002170 | chrII | 845 | 533 | 439 | 837 | 435 | 420 | 1039 | 491 | 439 | 3820 | 732 | 894 | 2760 | 487 | 565 | 2731 | 689 | 789 |
| 217 | W303.asm02.HP0_G0002180 | chrII | 70 | 27 | 27 | 59 | 23 | 21 | 52 | 21 | 23 | 198 | 66 | 25 | 157 | 21 | 18 | 165 | 55 | 23 |
| 218 | W303.asm02.HP0_G0002190 | chrII | 884 | 400 | 448 | 820 | 382 | 485 | 779 | 408 | 471 | 1531 | 415 | 521 | 1361 | 286 | 405 | 1228 | 415 | 446 |
| 219 | W303.asm02.HP0_G0002200 | chrII | 1640 | 2917 | 5912 | 1633 | 6473 | 5406 | 1796 | 3271 | 6033 | 17013 | 2798 | 4517 | 19405 | 1940 | 3076 | 15984 | 2520 | 3732 |
| 220 | W303.asm02.HP0_G0002210 | chrII | 4407 | 2393 | 2067 | 4057 | 1969 | 2003 | 3967 | 2265 | 2277 | 5239 | 2068 | 2079 | 5548 | 1460 | 1728 | 5074 | 1866 | 1918 |
| 221 | W303.asm02.HP0_G0002220 | chrII | 9023 | 1924 | 2101 | 8653 | 1864 | 1910 | 8967 | 1847 | 2082 | 9143 | 2396 | 3189 | 8480 | 1786 | 2728 | 7487 | 2257 | 3027 |
| 222 | W303.asm02.HP0_G0002230 | chrII | 3116 | 888 | 659 | 2700 | 605 | 642 | 2713 | 835 | 811 | 2955 | 1183 | 1182 | 2659 | 946 | 1159 | 2281 | 1124 | 1183 |
| 223 | W303.asm02.HP0_G0002240 | chrII | 4304 | 877 | 964 | 3625 | 900 | 1072 | 4252 | 739 | 1329 | 4938 | 1370 | 1954 | 1718 | 1097 | 1423 | 1572 | 1083 | 1711 |
| 224 | W303.asm02.HP0_G0002250 | chrII | 5891 | 5204 | 5532 | 6247 | 5587 | 5809 | 6157 | 5526 | 5872 | 15673 | 5215 | 6503 | 15918 | 4192 | 5098 | 14178 | 4282 | 5262 |
| 225 | W303.asm02.HP0_G0002260 | chrII | 23 | 27 | 20 | 10 | 17 | 20 | 14 | 18 | 25 | 62 | 23 | 13 | 47 | 8 | 7 | 59 | 18 | 14 |
| 226 | W303.asm02.HP0_G0002270 | chrII | 3821 | 1152 | 1210 | 3544 | 1149 | 1176 | 3642 | 1219 | 1222 | 7213 | 1421 | 1835 | 7320 | 1121 | 1520 | 6294 | 1247 | 1734 |
| 227 | W303.asm02.HP0_G0002280 | chrII | 1513 | 1316 | 1929 | 1324 | 1584 | 1759 | 1480 | 1399 | 2087 | 5373 | 1168 | 1322 | 5968 | 766 | 1032 | 5879 | 973 | 1213 |
| 228 | W303.asm02.HP0_G0002290 | chrII | 1453 | 559 | 747 | 1471 | 753 | 698 | 1429 | 515 | 777 | 3988 | 1153 | 1304 | 2250 | 712 | 835 | 2208 | 952 | 1151 |
| 229 | W303.asm02.HP0_G0002300 | chrII | 4260 | 977 | 778 | 3989 | 771 | 849 | 4349 | 941 | 822 | 4974 | 1279 | 1436 | 4048 | 886 | 1207 | 3668 | 1196 | 1261 |
| 230 | W303.asm02.HP0_G0002310 | chrII | 0 | 0 | 0 | 0 | 0 | 0 | 0 | 0 | 0 | 0 | 0 | 0 | 0 | 0 | 0 | 0 | 0 | 0 |
| 231 | W303.asm02.HP0_G0002320 | chrII | 0 | 0 | 0 | 1 | 0 | 0 | 0 | 1 | 0 | 0 | 0 | 0 | 1 | 0 | 0 | 0 | 1 | 0 |
| 232 | W303.asm02.HP0_G0002330 | chrII | 0 | 1 | 0 | 0 | 0 | 0 | 0 | 0 | 0 | 0 | 0 | 0 | 0 | 0 | 0 | 1 | 0 | 0 |
| 233 | W303.asm02.HP0_G0002340 | chrII | 0 | 0 | 0 | 0 | 0 | 0 | 0 | 0 | 0 | 0 | 0 | 0 | 0 | 0 | 0 | 0 | 0 | 0 |
| 234 | W303.asm02.HP0_G0002350 | chrII | 0 | 0 | 0 | 0 | 0 | 0 | 0 | 0 | 0 | 0 | 0 | 0 | 0 | 0 | 0 | 0 | 0 | 0 |
| 235 | W303.asm02.HP0_G0002360 | chrII | 0 | 0 | 0 | 0 | 0 | 0 | 0 | 0 | 0 | 0 | 0 | 0 | 0 | 0 | 0 | 0 | 0 | 0 |
| 236 | W303.asm02.HP0_G0002370 | chrII | 0 | 0 | 0 | 0 | 0 | 0 | 0 | 0 | 0 | 0 | 0 | 0 | 0 | 0 | 0 | 0 | 0 | 0 |
| 237 | W303.asm02.HP0_G0002380 | chrII | 0 | 0 | 0 | 0 | 0 | 0 | 0 | 0 | 0 | 0 | 0 | 0 | 0 | 0 | 0 | 0 | 0 | 0 |
| 238 | W303.asm02.HP0_G0002390 | chrII | 0 | 0 | 0 | 0 | 0 | 0 | 0 | 0 | 0 | 0 | 0 | 0 | 0 | 0 | 0 | 0 | 0 | 0 |
| 239 | W303.asm02.HP0_G0002400 | chrII | 88 | 32 | 24 | 110 | 24 | 33 | 80 | 24 | 42 | 112 | 33 | 51 | 98 | 24 | 36 | 82 | 23 | 38 |
| 240 | W303.asm02.HP0_G0002410 | chrII | 0 | 0 | 0 | 0 | 0 | 0 | 0 | 0 | 0 | 0 | 0 | 0 | 0 | 0 | 0 | 0 | 0 | 0 |
| 241 | W303.asm02.HP0_G0002420 | chrII | 14 | 17 | 26 | 54 | 24 | 17 | 42 | 17 | 29 | 114 | 19 | 26 | 126 | 21 | 9 | 108 | 19 | 31 |
| 242 | W303.asm02.HP0_G0002430 | chrII | 69 | 85 | 78 | 100 | 90 | 74 | 74 | 82 | 110 | 447 | 79 | 137 | 304 | 52 | 118 | 406 | 72 | 122 |
| 243 | W303.asm02.HP0_G0002440 | chrII | 811 | 221 | 99 | 895 | 118 | 114 | 637 | 189 | 95 | 270 | 199 | 114 | 249 | 118 | 69 | 238 | 225 | 105 |
| 244 | W303.asm02.HP0_G0002450 | chrII | 5077 | 1810 | 1523 | 4158 | 1252 | 1552 | 4113 | 1587 | 1692 | 3520 | 1436 | 1753 | 3745 | 1132 | 1820 | 3187 | 1482 | 1519 |
| 245 | W303.asm02.HP0_G0002460 | chrII | 5852 | 1587 | 1146 | 5398 | 1023 | 1144 | 5198 | 1465 | 1246 | 3009 | 1264 | 1248 | 4131 | 1108 | 1292 | 2994 | 1442 | 1142 |
| 246 | W303.asm02.HP0_G0002470 | chrII | 9733 | 4196 | 4441 | 7080 | 3764 | 4273 | 7557 | 4369 | 5433 | 7620 | 3856 | 3933 | 7096 | 2513 | 3531 | 7252 | 3170 | 3315 |
| 247 | W303.asm02.HP0_G0002480 | chrII | 2149 | 688 | 775 | 1907 | 807 | 751 | 2192 | 749 | 845 | 5492 | 1030 | 1277 | 4369 | 641 | 891 | 4422 | 946 | 1103 |
| 248 | W303.asm02.HP0_G0002490 | chrII | 7403 | 1317 | 863 | 6305 | 693 | 820 | 6197 | 1188 | 897 | 3556 | 1442 | 1166 | 3981 | 1118 | 1408 | 2848 | 1505 | 971 |
| 249 | W303.asm02.HP0_G0002500 | chrII | 970 | 206 | 155 | 1344 | 123 | 157 | 893 | 235 | 212 | 306 | 157 | 113 | 431 | 88 | 75 | 348 | 185 | 99 |
| 250 | W303.asm02.HP0_G0002510 | chrII | 2246 | 1131 | 1194 | 2112 | 1211 | 1203 | 2259 | 1083 | 1201 | 4752 | 952 | 1321 | 5181 | 614 | 979 | 4691 | 909 | 1127 |
| 251 | W303.asm02.HP0_G0002520 | chrII | 3624 | 789 | 743 | 3646 | 727 | 732 | 3955 | 805 | 788 | 7398 | 1309 | 1542 | 5393 | 892 | 1110 | 5410 | 1221 | 1415 |
| 252 | W303.asm02.HP0_G0002530 | chrII | 2220 | 1221 | 1610 | 1595 | 1354 | 1633 | 1773 | 1317 | 2189 | 3514 | 1181 | 1299 | 4017 | 750 | 1234 | 4243 | 1052 | 1192 |
| 253 | W303.asm02.HP0_G0002540 | chrII | 5532 | 567 | 716 | 5259 | 616 | 880 | 5400 | 550 | 830 | 6200 | 1200 | 1777 | 3833 | 1040 | 1569 | 3075 | 1051 | 1630 |
| 254 | W303.asm02.HP0_G0002550 | chrII | 2325 | 306 | 377 | 2532 | 368 | 466 | 2377 | 257 | 446 | 3722 | 622 | 931 | 2232 | 412 | 588 | 2470 | 592 | 775 |
| 255 | W303.asm02.HP0_G0002560 | chrII | 108 | 70 | 109 | 108 | 93 | 98 | 83 | 50 | 91 | 315 | 52 | 88 | 315 | 37 | 45 | 244 | 50 | 64 |
| 256 | W303.asm02.HP0_G0002570 | chrII | 55 | 21 | 30 | 34 | 32 | 36 | 43 | 26 | 27 | 91 | 15 | 20 | 94 | 10 | 17 | 75 | 17 | 13 |
| 257 | W303.asm02.HP0_G0002580 | chrII | 1338 | 1119 | 1203 | 1285 | 1313 | 1156 | 1432 | 1133 | 1210 | 5613 | 1094 | 1434 | 4842 | 649 | 820 | 5310 | 931 | 1223 |
| 258 | W303.asm02.HP0_G0002590 | chrII | 3062 | 925 | 1063 | 2941 | 919 | 1042 | 2746 | 887 | 1298 | 2141 | 643 | 761 | 2472 | 446 | 668 | 2194 | 593 | 649 |
| 259 | W303.asm02.HP0_G0002600 | chrII | 987 | 334 | 382 | 942 | 392 | 428 | 1001 | 339 | 530 | 2401 | 402 | 623 | 1976 | 313 | 436 | 2201 | 392 | 570 |
| 260 | W303.asm02.HP0_G0002610 | chrII | 1000 | 255 | 350 | 1004 | 373 | 290 | 907 | 283 | 305 | 2598 | 438 | 607 | 2241 | 251 | 414 | 2209 | 429 | 548 |
| 261 | W303.asm02.HP0_G0002620 | chrII | 1116 | 640 | 892 | 1295 | 615 | 883 | 887 | 556 | 1063 | 2204 | 567 | 918 | 2325 | 331 | 733 | 2215 | 525 | 814 |
| 262 | W303.asm02.HP0_G0002630 | chrII | 19412 | 2003 | 1741 | 19086 | 1952 | 2033 | 20213 | 2180 | 1877 | 8716 | 2496 | 2698 | 10640 | 2117 | 3146 | 8097 | 2101 | 2577 |
| 263 | W303.asm02.HP0_G0002640 | chrII | 5651 | 3404 | 3600 | 6252 | 3432 | 3575 | 5500 | 3354 | 4281 | 19599 | 6463 | 6886 | 12624 | 4067 | 4824 | 13471 | 5628 | 6020 |
| 264 | W303.asm02.HP0_G0002650 | chrII | 1033 | 3364 | 3940 | 968 | 4405 | 3523 | 1008 | 3682 | 3721 | 20516 | 4742 | 5467 | 15354 | 2662 | 3168 | 17271 | 4274 | 4562 |
| 265 | W303.asm02.HP0_G0002660 | chrII | 992 | 808 | 973 | 798 | 763 | 975 | 929 | 845 | 1196 | 2450 | 662 | 837 | 2664 | 458 | 607 | 2546 | 617 | 759 |
| 266 | W303.asm02.HP0_G0002670 | chrII | 3974 | 1748 | 1594 | 4321 | 1555 | 1581 | 4130 | 1999 | 1731 | 5242 | 2100 | 2336 | 8992 | 1796 | 2281 | 5297 | 1700 | 2131 |
| 267 | W303.asm02.HP0_G0002680 | chrII | 8871 | 1114 | 1025 | 7862 | 2332 | 2872 | 7909 | 690 | 650 | 737 | 506 | 530 | 757 | 249 | 331 | 682 | 520 | 434 |
| 268 | W303.asm02.HP0_G0002690 | chrII | 8 | 4 | 4 | 3 | 5 | 6 | 7 | 2 | 3 | 14 | 7 | 4 | 13 | 1 | 1 | 21 | 6 | 1 |
| 269 | W303.asm02.HP0_G0002700 | chrII | 5773 | 1145 | 1517 | 5635 | 1538 | 1505 | 6154 | 1145 | 1680 | 9982 | 1695 | 2682 | 8362 | 1353 | 2111 | 7559 | 1484 | 2423 |
| 270 | W303.asm02.HP0_G0002710 | chrII | 2423 | 791 | 866 | 2141 | 862 | 812 | 2341 | 870 | 1061 | 7702 | 1304 | 1628 | 5271 | 906 | 1236 | 5773 | 1209 | 1544 |
| 271 | W303.asm02.HP0_G0002720 | chrII | 1120 | 112 | 121 | 1035 | 98 | 106 | 1017 | 107 | 100 | 472 | 133 | 73 | 557 | 76 | 86 | 461 | 117 | 75 |
| 272 | W303.asm02.HP0_G0002730 | chrII | 1539 | 1066 | 889 | 1394 | 834 | 870 | 1443 | 969 | 936 | 2319 | 839 | 841 | 2574 | 617 | 799 | 2322 | 758 | 722 |
| 273 | W303.asm02.HP0_G0002740 | chrII | 21057 | 15381 | 15555 | 20951 | 15393 | 14682 | 21467 | 16500 | 13325 | 49306 | 17845 | 19667 | 49351 | 13690 | 15884 | 45282 | 14718 | 18145 |
| 274 | W303.asm02.HP0_G0002750 | chrII | 27896 | 3220 | 3480 | 27032 | 3214 | 3796 | 30705 | 2855 | 4386 | 36038 | 8400 | 9760 | 22220 | 5826 | 6172 | 21795 | 7153 | 8651 |
| 275 | W303.asm02.HP0_G0002760 | chrII | 7113 | 1390 | 1214 | 6463 | 1110 | 1367 | 6822 | 1254 | 1294 | 8875 | 2108 | 3025 | 6793 | 1521 | 2178 | 6490 | 1788 | 2645 |
| 276 | W303.asm02.HP0_G0002770 | chrII | 4175 | 776 | 836 | 4185 | 849 | 862 | 4061 | 813 | 889 | 7473 | 1113 | 1774 | 5987 | 699 | 1193 | 6045 | 965 | 1678 |
| 277 | W303.asm02.HP0_G0002780 | chrII | 11021 | 10042 | 10715 | 10035 | 9378 | 10596 | 9625 | 9841 | 10606 | 18807 | 8215 | 9804 | 22696 | 6169 | 8178 | 19549 | 7169 | 8990 |

| | | | | | | | | | | | | | | | | | | | | |
|---|---|---|---|---|---|---|---|---|---|---|---|---|---|---|---|---|---|---|---|---|
| 278 | W303.asm02.HP0_G0002790 | chrII | 1448 | 865 | 671 | 1499 | 707 | 868 | 1513 | 712 | 862 | 5673 | 1438 | 1653 | 4659 | 981 | 1218 | 4032 | 1255 | 1379 |
| 279 | W303.asm02.HP0_G0002800 | chrII | 6715 | 1927 | 1822 | 5982 | 2026 | 2030 | 7225 | 1838 | 2276 | 21472 | 4043 | 4709 | 12864 | 2920 | 2541 | 12984 | 3440 | 3934 |
| 280 | W303.asm02.HP0_G0002810 | chrII | 16805 | 11855 | 14314 | 16375 | 13997 | 14178 | 17156 | 11965 | 16306 | 32548 | 11221 | 12532 | 35617 | 8396 | 10469 | 32764 | 9940 | 11132 |
| 281 | W303.asm02.HP0_G0002820 | chrII | 524 | 170 | 141 | 412 | 174 | 131 | 466 | 93 | 90 | 593 | 99 | 78 | 598 | 94 | 93 | 492 | 77 | 85 |
| 282 | W303.asm02.HP0_G0002830 | chrII | 5003 | 3852 | 3612 | 4713 | 3174 | 3713 | 4422 | 3667 | 4526 | 5264 | 3334 | 3334 | 8237 | 3229 | 3576 | 5737 | 3253 | 3007 |
| 283 | W303.asm02.HP0_G0002840 | chrII | 12921 | 2523 | 3452 | 12586 | 3241 | 3666 | 12875 | 2730 | 4110 | 21734 | 4894 | 5779 | 21262 | 3851 | 5451 | 18195 | 4434 | 5410 |
| 284 | W303.asm02.HP0_G0002850 | chrII | 2179 | 1697 | 2006 | 2067 | 1875 | 1932 | 2147 | 1666 | 2347 | 5440 | 1273 | 1449 | 6317 | 842 | 1177 | 5937 | 1132 | 1354 |
| 285 | W303.asm02.HP0_G0002860 | chrII | 1238 | 1879 | 2482 | 1206 | 2458 | 2628 | 1418 | 2117 | 2557 | 6731 | 1779 | 2379 | 7298 | 1106 | 1573 | 6639 | 1379 | 2059 |
| 286 | W303.asm02.HP0_G0002870 | chrII | 4766 | 2284 | 3231 | 4736 | 2744 | 3262 | 4549 | 2346 | 3806 | 5281 | 1831 | 2165 | 7167 | 1419 | 2266 | 5617 | 1594 | 1860 |
| 287 | W303.asm02.HP0_G0002880 | chrII | 859 | 763 | 959 | 740 | 662 | 990 | 747 | 741 | 1275 | 1837 | 644 | 966 | 2153 | 459 | 649 | 1847 | 607 | 801 |
| 288 | W303.asm02.HP0_G0002890 | chrII | 5395 | 2828 | 2601 | 4738 | 2903 | 2320 | 5955 | 3535 | 3148 | 17013 | 5593 | 5382 | 18616 | 4319 | 4325 | 14499 | 4820 | 4967 |
| 289 | W303.asm02.HP0_G0002900 | chrII | 6113 | 6013 | 4670 | 5215 | 5052 | 4627 | 6495 | 6118 | 4136 | 7806 | 5648 | 3456 | 10638 | 5983 | 4436 | 7025 | 3660 | 2805 |
| 290 | W303.asm02.HP0_G0002910 | chrII | 3155 | 737 | 655 | 3012 | 801 | 648 | 3183 | 803 | 820 | 5940 | 1124 | 1175 | 5031 | 836 | 985 | 4565 | 1014 | 1136 |
| 291 | W303.asm02.HP0_G0002920 | chrII | 1361 | 881 | 1242 | 1561 | 856 | 1300 | 1386 | 775 | 1201 | 3328 | 667 | 1351 | 3595 | 426 | 882 | 3085 | 684 | 1163 |
| 292 | W303.asm02.HP0_G0002930 | chrII | 1568 | 1774 | 1685 | 1385 | 1391 | 1715 | 1519 | 1852 | 2005 | 5085 | 1473 | 1595 | 6198 | 1121 | 1350 | 5532 | 1311 | 1410 |
| 293 | W303.asm02.HP0_G0002940 | chrII | 2050 | 341 | 676 | 1931 | 590 | 690 | 1895 | 366 | 852 | 3451 | 317 | 913 | 3097 | 245 | 817 | 3298 | 324 | 743 |
| 294 | W303.asm02.HP0_G0002950 | chrII | 1275 | 497 | 490 | 1255 | 529 | 541 | 1301 | 494 | 438 | 2510 | 609 | 703 | 2213 | 354 | 578 | 2234 | 551 | 640 |
| 295 | W303.asm02.HP0_G0002960 | chrII | 10134 | 6166 | 3515 | 11999 | 4183 | 4162 | 9244 | 4806 | 2885 | 8588 | 4168 | 3201 | 8011 | 2695 | 2778 | 9161 | 4357 | 2952 |
| 296 | W303.asm02.HP0_G0002970 | chrII | 2223 | 363 | 485 | 1988 | 457 | 548 | 2192 | 356 | 521 | 4355 | 729 | 1115 | 2547 | 578 | 761 | 2553 | 633 | 972 |
| 297 | W303.asm02.HP0_G0002980 | chrII | 2063 | 680 | 436 | 1966 | 497 | 494 | 2154 | 585 | 425 | 3068 | 858 | 897 | 2578 | 633 | 718 | 2425 | 761 | 834 |
| 298 | W303.asm02.HP0_G0002990 | chrII | 388 | 1207 | 1715 | 310 | 1739 | 1605 | 431 | 1258 | 1994 | 5665 | 1063 | 1354 | 5907 | 743 | 904 | 5700 | 974 | 1258 |
| 299 | W303.asm02.HP0_G0003000 | chrII | 1374 | 554 | 801 | 1385 | 939 | 880 | 1224 | 435 | 1024 | 3884 | 908 | 1237 | 3037 | 487 | 694 | 2548 | 1013 | 878 |
| 300 | W303.asm02.HP0_G0003010 | chrII | 7441 | 6893 | 8963 | 7039 | 8066 | 9098 | 7823 | 6989 | 9406 | 16830 | 5730 | 7948 | 17556 | 4352 | 6074 | 16516 | 4808 | 7010 |
| 301 | W303.asm02.HP0_G0003020 | chrII | 1077 | 598 | 636 | 959 | 463 | 671 | 988 | 529 | 591 | 1808 | 494 | 967 | 1760 | 375 | 569 | 1592 | 519 | 835 |
| 302 | W303.asm02.HP0_G0003030 | chrII | 4883 | 192 | 288 | 4160 | 236 | 337 | 4474 | 184 | 369 | 335 | 178 | 334 | 316 | 127 | 205 | 290 | 122 | 239 |
| 303 | W303.asm02.HP0_G0003040 | chrII | 6523 | 4876 | 5037 | 6554 | 4489 | 5186 | 6760 | 4908 | 6329 | 9821 | 4355 | 4408 | 12605 | 3265 | 3811 | 10031 | 3725 | 4093 |
| 304 | W303.asm02.HP0_G0003050 | chrII | 2336 | 1218 | 1224 | 2465 | 1061 | 1156 | 2412 | 1296 | 1234 | 5853 | 1476 | 1680 | 6107 | 1028 | 1403 | 5390 | 1358 | 1561 |
| 305 | W303.asm02.HP0_G0003060 | chrII | 4814 | 2754 | 3026 | 4316 | 2556 | 3094 | 4384 | 2680 | 3928 | 5764 | 2342 | 2475 | 7264 | 1572 | 2123 | 6031 | 2035 | 2133 |
| 306 | W303.asm02.HP0_G0003070 | chrII | 830 | 383 | 528 | 722 | 410 | 469 | 765 | 354 | 729 | 1297 | 280 | 336 | 1363 | 156 | 315 | 1428 | 251 | 295 |
| 307 | W303.asm02.HP0_G0003080 | chrII | 5910 | 510 | 815 | 5598 | 802 | 1032 | 5656 | 386 | 1076 | 6968 | 926 | 2082 | 2581 | 751 | 1292 | 2064 | 663 | 1625 |
| 308 | W303.asm02.HP0_G0003090 | chrII | 3013 | 191 | 476 | 2750 | 480 | 512 | 2687 | 188 | 627 | 3909 | 453 | 807 | 1958 | 361 | 564 | 1664 | 437 | 575 |
| 309 | W303.asm02.HP0_G0003100 | chrII | 3297 | 1194 | 1422 | 3015 | 1545 | 1606 | 3216 | 1034 | 1803 | 9175 | 1880 | 3421 | 2958 | 1524 | 1760 | 2975 | 1529 | 2804 |
| 310 | W303.asm02.HP0_G0003110 | chrII | 1024 | 223 | 123 | 858 | 173 | 191 | 1008 | 206 | 157 | 689 | 373 | 234 | 426 | 197 | 137 | 363 | 384 | 220 |
| 311 | W303.asm02.HP0_G0003120 | chrII | 24333 | 10001 | 7613 | 25176 | 7672 | 8515 | 27189 | 8697 | 9134 | 31464 | 15464 | 16189 | 25573 | 14005 | 12428 | 24949 | 13285 | 15434 |
| 312 | W303.asm02.HP0_G0003130 | chrII | 2947 | 719 | 755 | 2734 | 600 | 739 | 2689 | 713 | 761 | 2036 | 627 | 793 | 2196 | 410 | 595 | 2083 | 622 | 728 |
| 313 | W303.asm02.HP0_G0003140 | chrII | 2758 | 917 | 1235 | 2567 | 1057 | 1257 | 2306 | 986 | 1745 | 2353 | 773 | 1037 | 3355 | 536 | 912 | 2584 | 737 | 850 |
| 314 | W303.asm02.HP0_G0003150 | chrII | 22497 | 8919 | 8728 | 21900 | 9094 | 7793 | 22915 | 9658 | 8607 | 42032 | 11649 | 12859 | 39842 | 7960 | 10064 | 40176 | 10242 | 11677 |
| 315 | W303.asm02.HP0_G0003160 | chrII | 4365 | 2047 | 2382 | 3979 | 2178 | 2367 | 4110 | 2114 | 2833 | 5499 | 1763 | 2264 | 6360 | 1361 | 1853 | 5511 | 1681 | 1901 |
| 316 | W303.asm02.HP0_G0003170 | chrII | 2247 | 466 | 475 | 2224 | 491 | 505 | 2157 | 400 | 604 | 3935 | 735 | 1086 | 2567 | 531 | 751 | 2746 | 761 | 975 |
| 317 | W303.asm02.HP0_G0003180 | chrII | 2900 | 811 | 719 | 2731 | 779 | 734 | 2954 | 709 | 821 | 5137 | 1152 | 1374 | 4293 | 822 | 1064 | 4063 | 1029 | 1242 |
| 318 | W303.asm02.HP0_G0003190 | chrII | 28079 | 1984 | 1075 | 24431 | 900 | 1515 | 24981 | 1197 | 1462 | 5957 | 3063 | 2892 | 3071 | 2276 | 2613 | 2709 | 2786 | 2461 |
| 319 | W303.asm02.HP0_G0003200 | chrII | 3319 | 1514 | 1739 | 2423 | 1375 | 1837 | 2601 | 1478 | 2032 | 2518 | 1209 | 1290 | 3288 | 912 | 1290 | 2654 | 1155 | 1126 |
| 320 | W303.asm02.HP0_G0003210 | chrII | 32952 | 7876 | 4404 | 30666 | 4892 | 4639 | 33908 | 7303 | 3772 | 28033 | 9394 | 9701 | 25212 | 7573 | 7184 | 23586 | 8207 | 8647 |
| 321 | W303.asm02.HP0_G0003220 | chrII | 1104 | 309 | 247 | 1009 | 191 | 243 | 946 | 307 | 267 | 640 | 227 | 180 | 772 | 120 | 217 | 726 | 217 | 170 |
| 322 | W303.asm02.HP0_G0003230 | chrII | 949 | 942 | 1316 | 806 | 1025 | 1220 | 778 | 879 | 1531 | 3070 | 646 | 1061 | 3211 | 365 | 729 | 3264 | 622 | 940 |
| 323 | W303.asm02.HP0_G0003240 | chrII | 2656 | 1132 | 1170 | 2508 | 1148 | 1140 | 2686 | 1018 | 1044 | 5734 | 1015 | 1568 | 4937 | 712 | 1072 | 5397 | 939 | 1339 |
| 324 | W303.asm02.HP0_G0003250 | chrII | 391 | 253 | 295 | 372 | 247 | 275 | 372 | 227 | 295 | 2165 | 344 | 506 | 1719 | 182 | 311 | 1742 | 367 | 508 |
| 325 | W303.asm02.HP0_G0003260 | chrII | 1281 | 218 | 239 | 1601 | 200 | 279 | 1079 | 178 | 321 | 1038 | 389 | 431 | 632 | 314 | 432 | 468 | 371 | 386 |
| 326 | W303.asm02.HP0_G0003270 | chrII | 2373 | 822 | 759 | 2127 | 742 | 891 | 2638 | 719 | 868 | 7077 | 1585 | 1977 | 4524 | 1066 | 1442 | 4846 | 1346 | 1881 |
| 327 | W303.asm02.HP0_G0003280 | chrII | 1501 | 960 | 1629 | 1281 | 1174 | 1598 | 1510 | 1058 | 1836 | 3607 | 989 | 1710 | 4230 | 658 | 1183 | 3569 | 832 | 1485 |
| 328 | W303.asm02.HP0_G0003290 | chrII | 1508 | 245 | 377 | 1311 | 412 | 366 | 1355 | 267 | 407 | 2748 | 353 | 659 | 2158 | 294 | 544 | 1903 | 306 | 621 |
| 329 | W303.asm02.HP0_G0003300 | chrII | 2232 | 1294 | 2086 | 2176 | 1291 | 2136 | 2197 | 1285 | 2334 | 3204 | 1042 | 1858 | 3833 | 714 | 1326 | 3380 | 893 | 1574 |
| 330 | W303.asm02.HP0_G0003310 | chrII | 1787 | 644 | 542 | 1831 | 530 | 543 | 1721 | 664 | 443 | 1970 | 589 | 667 | 2327 | 454 | 638 | 2080 | 510 | 662 |
| 331 | W303.asm02.HP0_G0003320 | chrII | 7182 | 1711 | 1290 | 7049 | 1078 | 1316 | 6713 | 1478 | 1253 | 5118 | 1700 | 1782 | 6024 | 1537 | 2150 | 4207 | 1517 | 1703 |
| 332 | W303.asm02.HP0_G0003330 | chrII | 7031 | 883 | 1157 | 6817 | 1167 | 1210 | 6870 | 959 | 1501 | 9206 | 1133 | 2400 | 7378 | 975 | 2112 | 6901 | 1064 | 2107 |
| 333 | W303.asm02.HP0_G0003340 | chrII | 491 | 379 | 584 | 507 | 677 | 578 | 548 | 441 | 574 | 2634 | 422 | 673 | 2619 | 294 | 369 | 2383 | 358 | 492 |
| 334 | W303.asm02.HP0_G0003350 | chrII | 2792 | 1761 | 2059 | 2924 | 2641 | 1802 | 3119 | 2001 | 1946 | 15308 | 2344 | 2608 | 12713 | 1387 | 1513 | 13178 | 1902 | 2283 |
| 335 | W303.asm02.HP0_G0003360 | chrII | 9674 | 8693 | 10550 | 9169 | 9492 | 9679 | 9601 | 9159 | 11813 | 28856 | 8313 | 9768 | 29463 | 5672 | 7243 | 29825 | 7169 | 8771 |
| 336 | W303.asm02.HP0_G0003370 | chrII | 2009 | 807 | 776 | 1871 | 702 | 702 | 1846 | 653 | 706 | 2104 | 709 | 750 | 1791 | 391 | 559 | 1904 | 677 | 678 |
| 337 | W303.asm02.HP0_G0003380 | chrII | 11696 | 3529 | 4427 | 10911 | 4157 | 4200 | 11268 | 3653 | 5128 | 10739 | 3033 | 4166 | 14207 | 2542 | 3570 | 10628 | 2809 | 3764 |
| 338 | W303.asm02.HP0_G0003390 | chrII | 492 | 231 | 213 | 532 | 192 | 210 | 465 | 186 | 251 | 346 | 172 | 129 | 475 | 78 | 151 | 286 | 176 | 111 |
| 339 | W303.asm02.HP0_G0003400 | chrII | 243 | 82 | 76 | 235 | 78 | 76 | 216 | 76 | 72 | 322 | 80 | 101 | 276 | 44 | 44 | 263 | 61 | 93 |
| 340 | W303.asm02.HP0_G0003410 | chrII | 91 | 36 | 55 | 73 | 56 | 51 | 83 | 42 | 61 | 153 | 44 | 47 | 162 | 22 | 36 | 128 | 29 | 37 |
| 341 | W303.asm02.HP0_G0003420 | chrII | 14639 | 3795 | 3187 | 13555 | 2831 | 3449 | 14113 | 3619 | 3085 | 7975 | 3292 | 3758 | 10122 | 2893 | 3792 | 6855 | 3101 | 3411 |
| 342 | W303.asm02.HP0_G0003430 | chrII | 1699 | 320 | 270 | 1797 | 363 | 323 | 1493 | 298 | 377 | 2791 | 431 | 603 | 2226 | 235 | 449 | 2198 | 505 | 546 |
| 343 | W303.asm02.HP0_G0003440 | chrII | 8744 | 2407 | 2656 | 8460 | 2503 | 2497 | 8347 | 2447 | 3116 | 11896 | 3233 | 3794 | 12164 | 2637 | 3477 | 10922 | 2925 | 3518 |
| 344 | W303.asm02.HP0_G0003450 | chrII | 558 | 199 | 195 | 585 | 241 | 222 | 565 | 201 | 152 | 1923 | 267 | 339 | 1502 | 167 | 210 | 1337 | 277 | 283 |
| 345 | W303.asm02.HP0_G0003460 | chrII | 695 | 859 | 992 | 726 | 977 | 970 | 655 | 893 | 1009 | 3112 | 734 | 1004 | 3160 | 506 | 727 | 2819 | 764 | 857 |
| 346 | W303.asm02.HP0_G0003470 | chrII | 4071 | 3232 | 3829 | 3820 | 3969 | 3631 | 3992 | 3507 | 4336 | 11047 | 3020 | 3417 | 11185 | 2023 | 2548 | 11216 | 2569 | 2949 |
| 347 | W303.asm02.HP0_G0003480 | chrII | 945 | 1046 | 1045 | 841 | 1160 | 1014 | 1117 | 1078 | 920 | 6201 | 1248 | 1577 | 5373 | 868 | 793 | 5328 | 1032 | 1342 |
| 348 | W303.asm02.HP0_G0003490 | chrII | 1363 | 227 | 205 | 1398 | 239 | 227 | 1543 | 255 | 237 | 1854 | 331 | 452 | 1826 | 261 | 351 | 1526 | 339 | 399 |
| 349 | W303.asm02.HP0_G0003500 | chrII | 556 | 925 | 1886 | 531 | 989 | 2000 | 491 | 835 | 1265 | 2050 | 686 | 1602 | 1763 | 371 | 1079 | 1540 | 632 | 1198 |
| 350 | W303.asm02.HP0_G0003510 | chrII | 1980 | 3211 | 4306 | 1900 | 3932 | 4378 | 2190 | 3302 | 4462 | 19145 | 4063 | 6399 | 19261 | 3082 | 4754 | 16590 | 3290 | 5602 |
| 351 | W303.asm02.HP0_G0003520 | chrII | 4766 | 3332 | 3519 | 4523 | 2990 | 3358 | 4596 | 3443 | 3932 | 10380 | 3076 | 3589 | 11156 | 2086 | 2671 | 10720 | 2778 | 3223 |
| 352 | W303.asm02.HP0_G0003530 | chrII | 3874 | 2101 | 1422 | 4007 | 1509 | 1499 | 4018 | 1892 | 1275 | 5169 | 1693 | 1718 | 5974 | 1340 | 1404 | 4695 | 1507 | 1565 |
| 353 | W303.asm02.HP0_G0003540 | chrII | 817 | 644 | 350 | 796 | 334 | 381 | 658 | 542 | 399 | 2149 | 659 | 499 | 2367 | 444 | 494 | 1997 | 651 | 516 |
| 354 | W303.asm02.HP0_G0003550 | chrII | 13686 | 10608 | 8524 | 13109 | 8454 | 7770 | 14692 | 11015 | 8518 | 32170 | 12525 | 11099 | 30666 | 8843 | 8223 | 27768 | 10145 | 10052 |
| 355 | W303.asm02.HP0_G0003560 | chrII | 1154 | 1333 | 1904 | 834 | 1637 | 1943 | 859 | 1460 | 2437 | 3310 | 1236 | 1633 | 4285 | 808 | 1323 | 3655 | 1176 | 1330 |
| 356 | W303.asm02.HP0_G0003570 | chrII | 1010 | 858 | 809 | 949 | 799 | 758 | 972 | 840 | 760 | 3981 | 760 | 916 | 3460 | 438 | 629 | 3647 | 748 | 832 |
| 357 | W303.asm02.HP0_G0003580 | chrII | 2524 | 2501 | 2929 | 2403 | 2531 | 2962 | 2489 | 2473 | 3822 | 6778 | 2033 | 2453 | 7466 | 1356 | 1812 | 7610 | 1926 | 2160 |
| 358 | W303.asm02.HP0_G0003590 | chrII | 780 | 1252 | 842 | 892 | 858 | 841 | 723 | 1171 | 1002 | 2093 | 748 | 692 | 2386 | 429 | 526 | 2337 | 815 | 566 |
| 359 | W303.asm02.HP0_G0003600 | chrII | 5017 | 2525 | 2377 | 4519 | 2222 | 2289 | 4862 | 2565 | 2521 | 7831 | 2281 | 2659 | 8740 | 1538 | 1957 | 7665 | 2060 | 2393 |
| 360 | W303.asm02.HP0_G0003610 | chrII | 937 | 868 | 839 | 960 | 768 | 871 | 925 | 744 | 1155 | 2227 | 578 | 655 | 2953 | 354 | 463 | 2421 | 468 | 568 |
| 361 | W303.asm02.HP0_G0003620 | chrII | 744 | 852 | 779 | 727 | 639 | 743 | 670 | 780 | 843 | 2106 | 566 | 599 | 2317 | 304 | 435 | 2337 | 568 | 488 |
| 362 | W303.asm02.HP0_G0003630 | chrII | 13940 | 991 | 768 | 13731 | 1006 | 1379 | 11757 | 644 | 867 | 2135 | 1158 | 1284 | 1815 | 789 | 1363 | 1494 | 1285 | 1163 |
| 363 | W303.asm02.HP0_G0003640 | chrII | 3495 | 893 | 780 | 3557 | 676 | 904 | 3394 | 716 | 883 | 4858 | 1177 | 1694 | 3147 | 912 | 1264 | 3178 | 1093 | 1529 |
| 364 | W303.asm02.HP0_G0003650 | chrII | 2922 | 1946 | 2474 | 2873 | 1760 | 2450 | 2842 | 1908 | 2466 | 5397 | 1639 | 2572 | 5645 | 1109 | 1800 | 5612 | 1468 | 2156 |
| 365 | W303.asm02.HP0_G0003660 | chrII | 4092 | 689 | 1126 | 3779 | 1393 | 1354 | 4479 | 660 | 1344 | 10780 | 1328 | 2061 | 8112 | 856 | 1481 | 7006 | 1270 | 1822 |
| 366 | W303.asm02.HP0_G0003670 | chrII | 7041 | 2958 | 3038 | 6442 | 2598 | 3100 | 6593 | 3140 | 3241 | 6005 | 2532 | 2702 | 6674 | 1752 | 2368 | 5785 | 2283 | 2367 |
| 367 | W303.asm02.HP0_G0003680 | chrII | 419 | 923 | 731 | 306 | 694 | 626 | 413 | 849 | 770 | 2537 | 728 | 597 | 2491 | 519 | 542 | 2557 | 734 | 563 |
| 368 | W303.asm02.HP0_G0003690 | chrII | 1144 | 805 | 713 | 1019 | 588 | 760 | 1092 | 822 | 708 | 2356 | 709 | 902 | 2226 | 470 | 564 | 2338 | 671 | 700 |
| 369 | W303.asm02.HP0_G0003700 | chrII | 14841 | 1978 | 1530 | 14677 | 1241 | 1578 | 14829 | 1591 | 1780 | 11245 | 4098 | 4373 | 5857 | 2878 | 3335 | 6761 | 3856 | 3817 |
| 370 | W303.asm02.HP0_G0003710 | chrII | 5476 | 975 | 898 | 5449 | 899 | 930 | 5318 | 1026 | 957 | 6356 | 1416 | 1846 | 5511 | 1053 | 1387 | 5345 | 1290 | 1758 |

| | | | | | | | | | | | | | | | | | | | | |
|---|---|---|---|---|---|---|---|---|---|---|---|---|---|---|---|---|---|---|---|---|
| 371 | W303.asm02.HP0_G0003720 | chrII | 251 | 97 | 101 | 255 | 98 | 97 | 207 | 94 | 120 | 574 | 105 | 156 | 428 | 73 | 77 | 433 | 124 | 110 |
| 372 | W303.asm02.HP0_G0003730 | chrII | 9969 | 7846 | 10876 | 10100 | 10569 | 10759 | 11358 | 8004 | 13180 | 27100 | 7530 | 9209 | 27780 | 5757 | 7097 | 26195 | 6550 | 8322 |
| 373 | W303.asm02.HP0_G0003740 | chrII | 1508 | 194 | 170 | 1941 | 193 | 198 | 1618 | 135 | 218 | 1029 | 233 | 324 | 1430 | 162 | 251 | 637 | 224 | 253 |
| 374 | W303.asm02.HP0_G0003750 | chrII | 74 | 40 | 17 | 76 | 21 | 15 | 88 | 49 | 22 | 64 | 38 | 11 | 92 | 22 | 21 | 70 | 39 | 16 |
| 375 | W303.asm02.HP0_G0003760 | chrII | 1955 | 1042 | 666 | 1938 | 574 | 675 | 1848 | 626 | 713 | 1044 | 631 | 415 | 1639 | 404 | 520 | 984 | 766 | 364 |
| 376 | W303.asm02.HP0_G0003770 | chrII | 585 | 358 | 267 | 533 | 237 | 250 | 477 | 321 | 284 | 627 | 189 | 186 | 954 | 141 | 154 | 786 | 221 | 161 |
| 377 | W303.asm02.HP0_G0003780 | chrII | 3457 | 1420 | 1600 | 3230 | 1443 | 1617 | 2996 | 1507 | 1846 | 4177 | 1093 | 1428 | 5009 | 862 | 1193 | 4175 | 1090 | 1281 |
| 378 | W303.asm02.HP0_G0003790 | chrII | 509 | 267 | 222 | 488 | 244 | 221 | 539 | 265 | 233 | 909 | 173 | 196 | 793 | 124 | 132 | 920 | 183 | 147 |
| 379 | W303.asm02.HP0_G0003800 | chrII | 2757 | 2064 | 1803 | 2280 | 1681 | 1605 | 2620 | 1956 | 1956 | 8677 | 2581 | 2208 | 6402 | 1742 | 1615 | 7051 | 2305 | 1895 |
| 380 | W303.asm02.HP0_G0003810 | chrII | 981 | 730 | 758 | 907 | 661 | 826 | 863 | 790 | 904 | 1818 | 602 | 755 | 2161 | 455 | 542 | 1830 | 563 | 679 |
| 381 | W303.asm02.HP0_G0003820 | chrII | 12652 | 12438 | 15109 | 11207 | 14754 | 15332 | 12851 | 12621 | 17462 | 32318 | 11621 | 13496 | 33477 | 8209 | 10543 | 32149 | 9907 | 11851 |
| 382 | W303.asm02.HP0_G0003830 | chrII | 9313 | 7681 | 9897 | 9151 | 9412 | 9831 | 9796 | 7891 | 11508 | 20604 | 7482 | 8360 | 21728 | 5622 | 6891 | 20690 | 6614 | 7829 |
| 383 | W303.asm02.HP0_G0003840 | chrII | 3290 | 1544 | 1557 | 2890 | 1723 | 1491 | 2954 | 1632 | 1857 | 8163 | 1684 | 2189 | 7803 | 1334 | 1748 | 7518 | 1691 | 2030 |
| 384 | W303.asm02.HP0_G0003850 | chrII | 1980 | 970 | 760 | 2127 | 783 | 713 | 2103 | 1060 | 637 | 3104 | 967 | 993 | 2938 | 630 | 701 | 2870 | 942 | 875 |
| 385 | W303.asm02.HP0_G0003860 | chrII | 1015 | 529 | 516 | 947 | 434 | 596 | 923 | 515 | 761 | 1439 | 416 | 490 | 1607 | 287 | 375 | 1560 | 354 | 445 |
| 386 | W303.asm02.HP0_G0003870 | chrII | 401 | 737 | 465 | 301 | 574 | 486 | 379 | 644 | 468 | 2623 | 675 | 661 | 2144 | 464 | 409 | 2311 | 664 | 566 |
| 387 | W303.asm02.HP0_G0003880 | chrII | 42040 | 13511 | 12282 | 39211 | 11713 | 11852 | 42813 | 13131 | 12140 | 51098 | 18711 | 22126 | 49064 | 14373 | 17168 | 43307 | 16727 | 20478 |
| 388 | W303.asm02.HP0_G0003890 | chrII | 34 | 20 | 19 | 53 | 26 | 21 | 46 | 13 | 27 | 102 | 22 | 26 | 47 | 12 | 25 | 67 | 29 | 35 |
| 389 | W303.asm02.HP0_G0003900 | chrII | 924 | 334 | 354 | 711 | 324 | 408 | 803 | 304 | 441 | 1761 | 569 | 645 | 1316 | 407 | 537 | 1255 | 493 | 576 |
| 390 | W303.asm02.HP0_G0003910 | chrII | 5057 | 786 | 800 | 4570 | 835 | 996 | 4903 | 652 | 1054 | 7101 | 1629 | 2364 | 2820 | 1203 | 1472 | 3151 | 1446 | 2001 |
| 391 | W303.asm02.HP0_G0003920 | chrII | 7816 | 3420 | 2961 | 7331 | 2563 | 2808 | 7504 | 3189 | 2871 | 10815 | 3491 | 3678 | 10447 | 2227 | 3160 | 10538 | 3416 | 3352 |
| 392 | W303.asm02.HP0_G0003930 | chrII | 3226 | 711 | 736 | 3081 | 687 | 760 | 3082 | 624 | 895 | 5451 | 1197 | 1474 | 3266 | 757 | 1019 | 3648 | 1113 | 1350 |
| 393 | W303.asm02.HP0_G0003940 | chrII | 54 | 3 | 46 | 47 | 4 | 62 | 39 | 7 | 61 | 18 | 2 | 62 | 27 | 2 | 17 | 17 | 3 | 40 |
| 394 | W303.asm02.HP0_G0003950 | chrII | 1531 | 651 | 1937 | 1442 | 746 | 2240 | 1363 | 619 | 2694 | 2198 | 505 | 1732 | 3338 | 378 | 1290 | 2553 | 485 | 1527 |
| 395 | W303.asm02.HP0_G0003960 | chrII | 1364 | 729 | 728 | 1301 | 715 | 917 | 1441 | 609 | 934 | 7553 | 1399 | 1988 | 5495 | 1105 | 1458 | 4868 | 1254 | 1721 |
| 396 | W303.asm02.HP0_G0003970 | chrII | 3385 | 309 | 335 | 3702 | 435 | 321 | 3278 | 328 | 385 | 3589 | 508 | 516 | 5320 | 502 | 668 | 2115 | 461 | 462 |
| 397 | W303.asm02.HP0_G0003980 | chrII | 4070 | 440 | 389 | 3966 | 373 | 461 | 3929 | 386 | 528 | 2492 | 586 | 898 | 2271 | 575 | 753 | 1836 | 599 | 818 |
| 398 | W303.asm02.HP0_G0003990 | chrII | 5309 | 1780 | 1741 | 4823 | 1611 | 1782 | 5161 | 1880 | 1728 | 8481 | 2406 | 2925 | 7541 | 1765 | 2145 | 7361 | 2069 | 2667 |
| 399 | W303.asm02.HP0_G0004000 | chrII | 2821 | 1100 | 1321 | 3155 | 1146 | 1382 | 2642 | 1154 | 1550 | 7808 | 2077 | 2386 | 6727 | 1313 | 1809 | 6529 | 1975 | 2474 |
| 400 | W303.asm02.HP0_G0004010 | chrII | 4565 | 311 | 444 | 4586 | 404 | 495 | 4876 | 262 | 600 | 3345 | 677 | 1146 | 1974 | 712 | 910 | 1426 | 531 | 1017 |
| 401 | W303.asm02.HP0_G0004020 | chrII | 1221 | 790 | 955 | 1054 | 857 | 947 | 1191 | 814 | 1285 | 2512 | 659 | 742 | 3010 | 430 | 601 | 3119 | 626 | 669 |
| 402 | W303.asm02.HP0_G0004030 | chrII | 658 | 1033 | 718 | 808 | 772 | 740 | 666 | 855 | 1020 | 2200 | 584 | 508 | 2513 | 323 | 386 | 2497 | 687 | 458 |
| 403 | W303.asm02.HP0_G0004040 | chrII | 8572 | 971 | 988 | 8661 | 850 | 1121 | 8774 | 801 | 1285 | 7509 | 2355 | 2424 | 4142 | 1486 | 1897 | 4041 | 2105 | 2058 |
| 404 | W303.asm02.HP0_G0004050 | chrII | 1187 | 148 | 205 | 897 | 166 | 260 | 1155 | 119 | 264 | 722 | 194 | 394 | 260 | 161 | 184 | 236 | 163 | 329 |
| 405 | W303.asm02.HP0_G0004060 | chrII | 15879 | 443 | 294 | 14223 | 234 | 410 | 14452 | 269 | 425 | 1223 | 551 | 736 | 436 | 423 | 547 | 590 | 436 | 555 |
| 406 | W303.asm02.HP0_G0004070 | chrII | 2389 | 384 | 524 | 2527 | 527 | 572 | 2437 | 400 | 654 | 4414 | 767 | 1083 | 3131 | 518 | 839 | 3051 | 597 | 986 |
| 407 | W303.asm02.HP0_G0004080 | chrII | 1443 | 648 | 795 | 1392 | 761 | 694 | 1370 | 613 | 838 | 3621 | 620 | 971 | 3497 | 399 | 783 | 3515 | 643 | 847 |
| 408 | W303.asm02.HP0_G0004090 | chrII | 726 | 546 | 599 | 811 | 518 | 557 | 781 | 514 | 743 | 1627 | 364 | 410 | 1977 | 292 | 302 | 1714 | 361 | 392 |
| 409 | W303.asm02.HP0_G0004100 | chrII | 11142 | 1746 | 2147 | 9664 | 1917 | 2349 | 10671 | 1413 | 2906 | 8069 | 2480 | 4215 | 3295 | 2490 | 2755 | 2553 | 1814 | 3285 |
| 410 | W303.asm02.HP0_G0004110 | chrII | 1520 | 710 | 904 | 1391 | 762 | 982 | 1660 | 673 | 1125 | 5463 | 1457 | 1996 | 2596 | 986 | 1157 | 2923 | 1301 | 1758 |
| 411 | W303.asm02.HP0_G0004120 | chrII | 12969 | 5035 | 3959 | 11949 | 3841 | 4042 | 12928 | 5012 | 4018 | 20188 | 7465 | 7650 | 18072 | 5392 | 5667 | 17049 | 6269 | 6921 |
| 412 | W303.asm02.HP0_G0004130 | chrII | 6549 | 913 | 750 | 5764 | 699 | 887 | 6307 | 842 | 843 | 5535 | 1657 | 1796 | 5234 | 1327 | 1488 | 3546 | 1411 | 1561 |
| 413 | W303.asm02.HP0_G0004140 | chrII | 1621 | 131 | 188 | 1630 | 147 | 191 | 1566 | 106 | 221 | 1174 | 286 | 368 | 949 | 213 | 279 | 710 | 227 | 323 |
| 414 | W303.asm02.HP0_G0004150 | chrII | 1730 | 191 | 270 | 1740 | 258 | 300 | 1569 | 180 | 351 | 2520 | 418 | 744 | 1557 | 316 | 520 | 1404 | 362 | 638 |
| 415 | W303.asm02.HP0_G0004160 | chrII | 2377 | 847 | 737 | 2384 | 821 | 770 | 2528 | 892 | 773 | 5243 | 1159 | 1366 | 4006 | 759 | 911 | 4036 | 1048 | 1175 |
| 416 | W303.asm02.HP0_G0004170 | chrII | 1171 | 585 | 528 | 1100 | 547 | 494 | 1137 | 564 | 661 | 2044 | 430 | 577 | 2016 | 330 | 415 | 1992 | 497 | 447 |
| 417 | W303.asm02.HP0_G0004180 | chrII | 2400 | 647 | 704 | 2139 | 652 | 788 | 2210 | 713 | 855 | 6194 | 1318 | 1704 | 4192 | 1005 | 1281 | 4191 | 1120 | 1509 |
| 418 | W303.asm02.HP0_G0004190 | chrII | 2562 | 2093 | 1902 | 2630 | 1625 | 2189 | 2305 | 1950 | 2459 | 3355 | 1435 | 1435 | 4454 | 1108 | 1312 | 3280 | 1257 | 1172 |
| 419 | W303.asm02.HP0_G0004200 | chrII | 1338 | 485 | 496 | 788 | 483 | 569 | 820 | 490 | 640 | 677 | 412 | 351 | 1211 | 294 | 456 | 753 | 352 | 326 |
| 420 | W303.asm02.HP0_G0004210 | chrII | 1496 | 500 | 367 | 1273 | 408 | 440 | 1324 | 377 | 448 | 1807 | 579 | 683 | 1206 | 427 | 522 | 1215 | 554 | 578 |
| 421 | W303.asm02.HP0_G0004220 | chrII | 991 | 444 | 426 | 960 | 361 | 432 | 1046 | 402 | 438 | 2620 | 561 | 782 | 2316 | 395 | 534 | 2195 | 513 | 659 |
| 422 | W303.asm02.HP0_G0004230 | chrII | 340 | 522 | 675 | 263 | 633 | 668 | 308 | 535 | 833 | 1701 | 399 | 496 | 2054 | 250 | 410 | 1957 | 393 | 462 |
| 423 | W303.asm02.HP0_G0004240 | chrII | 10422 | 3689 | 2874 | 9471 | 2798 | 2975 | 10442 | 3518 | 2927 | 11644 | 3947 | 4006 | 12019 | 3019 | 3832 | 9963 | 3505 | 3670 |
| 424 | W303.asm02.HP0_G0004250 | chrII | 1817 | 425 | 591 | 1586 | 518 | 629 | 1771 | 415 | 717 | 3680 | 815 | 1162 | 2087 | 593 | 845 | 1998 | 717 | 994 |
| 425 | W303.asm02.HP0_G0004260 | chrII | 2273 | 1543 | 1264 | 2246 | 1380 | 1247 | 2205 | 1514 | 1353 | 5979 | 1395 | 1731 | 5816 | 965 | 1236 | 5701 | 1308 | 1526 |
| 426 | W303.asm02.HP0_G0004270 | chrII | 1985 | 483 | 551 | 1925 | 684 | 627 | 1820 | 474 | 600 | 4090 | 633 | 1031 | 3441 | 452 | 776 | 3209 | 728 | 893 |
| 427 | W303.asm02.HP0_G0004280 | chrII | 1406 | 700 | 1530 | 1181 | 1428 | 1301 | 1369 | 811 | 1588 | 14160 | 1152 | 2397 | 12897 | 904 | 1322 | 11117 | 978 | 1773 |
| 428 | W303.asm02.HP0_G0004290 | chrII | 1152 | 172 | 286 | 998 | 298 | 315 | 1211 | 165 | 372 | 2218 | 278 | 520 | 1568 | 234 | 385 | 1490 | 255 | 494 |
| 429 | W303.asm02.HP0_G0004300 | chrII | 175 | 163 | 230 | 141 | 160 | 217 | 178 | 145 | 270 | 526 | 143 | 249 | 709 | 79 | 141 | 452 | 142 | 179 |
| 430 | W303.asm02.HP0_G0004310 | chrII | 2676 | 279 | 272 | 2549 | 223 | 312 | 2599 | 230 | 316 | 1095 | 530 | 523 | 720 | 369 | 451 | 480 | 538 | 449 |
| 431 | W303.asm02.HP0_G0004320 | chrII | 1561 | 1696 | 1930 | 1690 | 1766 | 1837 | 1704 | 1651 | 2040 | 5393 | 1403 | 1586 | 5426 | 864 | 1066 | 5212 | 1269 | 1387 |
| 432 | W303.asm02.HP0_G0004330 | chrII | 1520 | 986 | 1193 | 1282 | 870 | 1139 | 1471 | 1107 | 1223 | 4639 | 1521 | 2053 | 3833 | 935 | 1367 | 3902 | 1384 | 1904 |
| 433 | W303.asm02.HP0_G0004340 | chrII | 498 | 520 | 798 | 384 | 606 | 827 | 380 | 551 | 1166 | 1491 | 447 | 488 | 1630 | 302 | 371 | 1632 | 389 | 445 |
| 434 | W303.asm02.HP0_G0004350 | chrII | 4767 | 566 | 851 | 4744 | 650 | 1082 | 5190 | 461 | 1229 | 8179 | 1156 | 2189 | 4287 | 925 | 1442 | 4056 | 988 | 1936 |
| 435 | W303.asm02.HP0_G0004360 | chrII | 3077 | 634 | 646 | 2919 | 561 | 650 | 3081 | 641 | 738 | 4329 | 934 | 1376 | 4352 | 736 | 908 | 3735 | 894 | 1179 |
| 436 | W303.asm02.HP0_G0004370 | chrII | 2830 | 1804 | 1400 | 3020 | 2014 | 1377 | 3469 | 1811 | 1276 | 12109 | 2185 | 2517 | 10644 | 1433 | 1248 | 10364 | 1803 | 2166 |
| 437 | W303.asm02.HP0_G0004380 | chrII | 1905 | 1321 | 1022 | 1461 | 1163 | 1072 | 2042 | 1263 | 1178 | 7759 | 2073 | 2341 | 4637 | 1477 | 1551 | 4939 | 1949 | 1966 |
| 438 | W303.asm02.HP0_G0004390 | chrII | 17106 | 8842 | 8059 | 16273 | 7992 | 7272 | 16699 | 9308 | 7310 | 23519 | 8955 | 9278 | 24839 | 6369 | 7099 | 23501 | 7441 | 8323 |
| 439 | W303.asm02.HP0_G0004400 | chrII | 213 | 269 | 114 | 262 | 169 | 123 | 225 | 220 | 112 | 375 | 148 | 99 | 320 | 63 | 50 | 376 | 181 | 93 |
| 440 | W303.asm02.HP0_G0004410 | chrII | 4584 | 1613 | 1173 | 4365 | 1193 | 1138 | 4160 | 1489 | 1265 | 4428 | 1510 | 1557 | 4464 | 1143 | 1284 | 4080 | 1432 | 1430 |
| 441 | W303.asm02.HP0_G0004420 | chrII | 1046 | 923 | 911 | 814 | 767 | 923 | 882 | 1019 | 1238 | 1531 | 731 | 631 | 1895 | 440 | 526 | 1680 | 661 | 550 |
| 442 | W303.asm02.HP0_G0004430 | chrII | 733 | 963 | 958 | 613 | 928 | 949 | 724 | 1017 | 1224 | 2542 | 725 | 723 | 2898 | 465 | 565 | 2648 | 624 | 684 |
| 443 | W303.asm02.HP0_G0004440 | chrII | 2919 | 429 | 380 | 3012 | 371 | 361 | 2661 | 369 | 469 | 2574 | 666 | 708 | 1708 | 421 | 578 | 1736 | 655 | 574 |
| 444 | W303.asm02.HP0_G0004450 | chrII | 672 | 312 | 421 | 639 | 390 | 432 | 625 | 295 | 547 | 1207 | 332 | 399 | 1235 | 210 | 332 | 1078 | 288 | 319 |
| 445 | W303.asm02.HP0_G0004460 | chrII | 3628 | 3332 | 3307 | 3232 | 2992 | 3371 | 3183 | 3399 | 4374 | 6905 | 2748 | 2509 | 8352 | 2162 | 2172 | 7351 | 2487 | 2202 |
| 446 | W303.asm02.HP0_G0004470 | chrII | 656 | 798 | 898 | 683 | 941 | 948 | 671 | 830 | 1146 | 2737 | 591 | 756 | 3426 | 399 | 509 | 2852 | 575 | 508 |
| 447 | W303.asm02.HP0_G0004480 | chrII | 587 | 471 | 476 | 527 | 381 | 488 | 538 | 530 | 652 | 1137 | 346 | 329 | 1680 | 241 | 305 | 1291 | 325 | 311 |
| 448 | W303.asm02.HP0_G0004490 | chrII | 720 | 770 | 921 | 769 | 846 | 871 | 789 | 854 | 936 | 3552 | 697 | 888 | 3887 | 489 | 629 | 3267 | 718 | 774 |
| 449 | W303.asm02.HP0_G0004500 | chrII | 2386 | 498 | 430 | 2407 | 396 | 522 | 2543 | 451 | 548 | 3918 | 868 | 1036 | 3289 | 586 | 858 | 3051 | 857 | 1041 |
| 450 | W303.asm02.HP0_G0004510 | chrII | 2209 | 1168 | 717 | 2121 | 787 | 813 | 2454 | 1158 | 787 | 3816 | 1404 | 1514 | 3281 | 913 | 974 | 3334 | 1197 | 1382 |
| 451 | W303.asm02.HP0_G0004520 | chrII | 1968 | 1072 | 1508 | 1750 | 1463 | 1460 | 1703 | 1107 | 2012 | 2845 | 884 | 1098 | 3623 | 774 | 959 | 2749 | 896 | 1040 |
| 452 | W303.asm02.HP0_G0004530 | chrII | 8722 | 3658 | 2094 | 7279 | 2453 | 2226 | 9268 | 3519 | 2328 | 13593 | 4451 | 3910 | 11591 | 3385 | 2729 | 11172 | 3816 | 3453 |
| 453 | W303.asm02.HP0_G0004540 | chrII | 785 | 637 | 606 | 717 | 506 | 621 | 701 | 682 | 613 | 1465 | 647 | 686 | 1598 | 446 | 485 | 1453 | 556 | 585 |
| 454 | W303.asm02.HP0_G0004550 | chrII | 2008 | 1633 | 1137 | 1734 | 1111 | 1143 | 1945 | 1499 | 1108 | 5314 | 1745 | 1680 | 4533 | 1248 | 1379 | 4447 | 1637 | 1535 |
| 455 | W303.asm02.HP0_G0004560 | chrII | 1367 | 970 | 858 | 1538 | 1102 | 920 | 1548 | 917 | 909 | 7702 | 1356 | 1799 | 6086 | 1085 | 889 | 5255 | 1171 | 1511 |
| 456 | W303.asm02.HP0_G0004570 | chrII | 2898 | 1332 | 1639 | 2379 | 1387 | 1565 | 2415 | 1483 | 2118 | 2797 | 1222 | 1263 | 3922 | 848 | 1277 | 2971 | 1044 | 1150 |
| 457 | W303.asm02.HP0_G0004580 | chrII | 2303 | 1591 | 1540 | 2100 | 1346 | 1441 | 2093 | 1569 | 1817 | 2505 | 1348 | 1269 | 3580 | 918 | 1200 | 2765 | 1225 | 1049 |
| 458 | W303.asm02.HP0_G0004590 | chrII | 1006 | 228 | 163 | 801 | 181 | 154 | 899 | 226 | 180 | 1085 | 324 | 261 | 850 | 189 | 214 | 767 | 307 | 217 |
| 459 | W303.asm02.HP0_G0004600 | chrII | 599 | 435 | 460 | 475 | 537 | 448 | 646 | 387 | 485 | 3548 | 515 | 735 | 2827 | 379 | 384 | 2711 | 443 | 625 |
| 460 | W303.asm02.HP0_G0004610 | chrII | 2628 | 1133 | 1113 | 2484 | 1034 | 1090 | 2413 | 1254 | 1233 | 3661 | 907 | 1092 | 4238 | 625 | 989 | 3619 | 829 | 1016 |
| 461 | W303.asm02.HP0_G0004620 | chrII | 5625 | 1155 | 826 | 5255 | 850 | 853 | 5302 | 1037 | 799 | 4002 | 1302 | 1452 | 3541 | 876 | 1076 | 3248 | 1190 | 1326 |
| 462 | W303.asm02.HP0_G0004630 | chrII | 1680 | 815 | 929 | 1535 | 832 | 813 | 1659 | 821 | 1038 | 3422 | 761 | 843 | 3236 | 503 | 687 | 3492 | 686 | 808 |
| 463 | W303.asm02.HP0_G0004640 | chrII | 3053 | 667 | 782 | 2993 | 902 | 838 | 3092 | 727 | 939 | 7608 | 1027 | 1789 | 5214 | 782 | 1222 | 5169 | 876 | 1624 |

| | | | | | | | | | | | | | | | | | | | | |
|---|---|---|---|---|---|---|---|---|---|---|---|---|---|---|---|---|---|---|---|---|
| 464 | W303.asm02.HP0_G0004650 | chrII | 2655 | 566 | 971 | 2407 | 811 | 967 | 2606 | 605 | 1226 | 5413 | 1287 | 1772 | 3406 | 944 | 1368 | 3219 | 1087 | 1546 |
| 465 | W303.asm02.HP0_G0004660 | chrII | 710 | 666 | 557 | 817 | 552 | 580 | 788 | 619 | 665 | 1546 | 515 | 542 | 1922 | 329 | 354 | 1447 | 421 | 424 |
| 466 | W303.asm02.HP0_G0004670 | chrII | 4431 | 1566 | 1418 | 4619 | 1321 | 1501 | 4321 | 1520 | 1213 | 6421 | 1264 | 2328 | 6420 | 980 | 1561 | 5577 | 1203 | 1932 |
| 467 | W303.asm02.HP0_G0004680 | chrII | 3397 | 361 | 298 | 3640 | 275 | 327 | 2965 | 319 | 371 | 1433 | 481 | 514 | 801 | 320 | 393 | 970 | 465 | 489 |
| 468 | W303.asm02.HP0_G0004690 | chrII | 6097 | 893 | 907 | 4235 | 904 | 871 | 5169 | 987 | 1062 | 6443 | 1201 | 1494 | 4728 | 853 | 1113 | 4942 | 1134 | 1314 |
| 469 | W303.asm02.HP0_G0004700 | chrII | 4263 | 1266 | 1808 | 3799 | 1593 | 1795 | 3827 | 1326 | 2213 | 3864 | 1072 | 1560 | 5422 | 873 | 1522 | 4147 | 1081 | 1372 |
| 470 | W303.asm02.HP0_G0004710 | chrII | 8817 | 4254 | 4046 | 8470 | 3638 | 3948 | 9122 | 4429 | 4514 | 20717 | 5700 | 5944 | 17090 | 4137 | 4784 | 17735 | 5053 | 5526 |
| 471 | W303.asm02.HP0_G0004720 | chrII | 2613 | 193 | 104 | 2158 | 131 | 120 | 2153 | 148 | 134 | 459 | 218 | 157 | 359 | 112 | 136 | 356 | 203 | 152 |
| 472 | W303.asm02.HP0_G0004730 | chrII | 815 | 110 | 120 | 942 | 120 | 82 | 682 | 101 | 128 | 187 | 69 | 63 | 325 | 44 | 91 | 218 | 59 | 58 |
| 473 | W303.asm02.HP0_G0004740 | chrII | 25046 | 8746 | 6236 | 24031 | 6181 | 6328 | 24358 | 9037 | 6527 | 28341 | 10767 | 9868 | 27184 | 7614 | 8197 | 24436 | 9120 | 8870 |
| 474 | W303.asm02.HP0_G0004750 | chrII | 4030 | 1416 | 1641 | 4411 | 1504 | 1525 | 3798 | 1356 | 1700 | 6638 | 1426 | 1772 | 7203 | 1105 | 1773 | 5711 | 1465 | 1540 |
| 475 | W303.asm02.HP0_G0004760 | chrII | 3287 | 1284 | 1344 | 3129 | 1328 | 1324 | 3208 | 1360 | 1569 | 5285 | 1119 | 1376 | 5934 | 846 | 1203 | 5438 | 975 | 1281 |
| 476 | W303.asm02.HP0_G0004770 | chrII | 3271 | 400 | 465 | 3053 | 459 | 558 | 3429 | 298 | 539 | 2778 | 604 | 952 | 1404 | 443 | 664 | 1321 | 503 | 800 |
| 477 | W303.asm02.HP0_G0004780 | chrII | 3821 | 1997 | 2021 | 3687 | 1836 | 1974 | 3392 | 1922 | 2507 | 5526 | 1532 | 1611 | 6391 | 937 | 1354 | 5246 | 1494 | 1459 |
| 478 | W303.asm02.HP0_G0004790 | chrII | 995 | 789 | 529 | 1096 | 638 | 513 | 1133 | 712 | 630 | 2116 | 639 | 599 | 2266 | 485 | 483 | 1925 | 617 | 585 |
| 479 | W303.asm02.HP0_G0004800 | chrII | 1330 | 987 | 867 | 1123 | 897 | 831 | 1128 | 950 | 924 | 2878 | 755 | 686 | 2742 | 434 | 500 | 2847 | 798 | 604 |
| 480 | W303.asm02.HP0_G0004810 | chrII | 862 | 217 | 229 | 744 | 263 | 257 | 864 | 254 | 297 | 2021 | 387 | 416 | 1709 | 315 | 416 | 1335 | 400 | 398 |
| 481 | W303.asm02.HP0_G0004820 | chrII | 1240 | 139 | 274 | 1195 | 290 | 339 | 1146 | 159 | 371 | 2218 | 274 | 650 | 2808 | 270 | 544 | 1274 | 215 | 566 |
| 482 | W303.asm02.HP0_G0004830 | chrII | 373 | 233 | 198 | 375 | 227 | 210 | 345 | 239 | 265 | 417 | 170 | 153 | 507 | 113 | 70 | 359 | 133 | 117 |
| 483 | W303.asm02.HP0_G0004840 | chrII | 2068 | 448 | 355 | 1834 | 458 | 373 | 2039 | 515 | 362 | 1732 | 477 | 314 | 1786 | 294 | 246 | 1645 | 400 | 327 |
| 484 | W303.asm02.HP0_G0004850 | chrII | 2334 | 223 | 120 | 2201 | 127 | 144 | 2174 | 115 | 128 | 1082 | 385 | 243 | 548 | 214 | 262 | 553 | 344 | 247 |
| 485 | W303.asm02.HP0_G0004860 | chrII | 9 | 6 | 1 | 7 | 0 | 2 | 11 | 2 | 0 | 22 | 8 | 0 | 14 | 1 | 0 | 8 | 6 | 1 |
| 486 | W303.asm02.HP0_G0004870 | chrII | 334 | 118 | 73 | 292 | 95 | 61 | 282 | 89 | 71 | 332 | 88 | 78 | 335 | 64 | 66 | 333 | 107 | 71 |
| 487 | W303.asm02.HP0_G0004880 | chrII | 74 | 63 | 38 | 78 | 54 | 61 | 67 | 35 | 29 | 92 | 18 | 28 | 71 | 10 | 14 | 84 | 22 | 23 |
| 488 | W303.asm02.HP0_G0004890 | chrII | 3517 | 1168 | 1162 | 2838 | 840 | 1300 | 3063 | 1090 | 1545 | 2951 | 850 | 1137 | 2980 | 570 | 813 | 3223 | 772 | 1098 |
| 489 | W303.asm02.HP0_G0004900 | chrII | 0 | 9 | 2 | 0 | 2 | 0 | 2 | 6 | 2 | 4 | 5 | 1 | 0 | 0 | 1 | 1 | 2 | 1 |
| 490 | W303.asm02.HP0_G0004920 | chrIII | 0 | 12 | 0 | 0 | 0 | 0 | 0 | 16 | 0 | 0 | 0 | 0 | 0 | 0 | 0 | 0 | 0 | 0 |
| 491 | W303.asm02.HP0_G0004930 | chrIII | 1 | 1 | 1 | 3 | 0 | 0 | 0 | 1 | 0 | 37 | 33 | 18 | 42 | 11 | 41 | 26 | 19 | 65 |
| 492 | W303.asm02.HP0_G0004940 | chrIII | 1 | 0 | 0 | 0 | 0 | 0 | 1 | 0 | 0 | 6 | 4 | 3 | 9 | 4 | 3 | 2 | 6 | 6 |
| 493 | W303.asm02.HP0_G0004950 | chrIII | 0 | 0 | 0 | 0 | 0 | 0 | 0 | 0 | 0 | 0 | 0 | 0 | 1 | 0 | 0 | 0 | 0 | 0 |
| 494 | W303.asm02.HP0_G0004960 | chrIII | 2 | 0 | 0 | 2 | 2 | 1 | 2 | 6 | 1 | 13 | 14 | 7 | 11 | 1 | 0 | 11 | 8 | 4 |
| 495 | W303.asm02.HP0_G0004970 | chrIII | 0 | 3 | 0 | 0 | 0 | 0 | 1 | 1 | 0 | 3 | 1 | 1 | 0 | 0 | 0 | 3 | 4 | 2 |
| 496 | W303.asm02.HP0_G0004980 | chrIII | 389 | 669 | 482 | 380 | 439 | 410 | 453 | 492 | 414 | 5481 | 749 | 693 | 3009 | 274 | 425 | 4171 | 748 | 611 |
| 497 | W303.asm02.HP0_G0004990 | chrIII | 555 | 564 | 618 | 531 | 699 | 600 | 627 | 600 | 655 | 8214 | 1453 | 2024 | 6616 | 958 | 1562 | 6550 | 1246 | 1980 |
| 498 | W303.asm02.HP0_G0005000 | chrIII | 2010 | 317 | 569 | 1998 | 627 | 550 | 2037 | 391 | 638 | 9129 | 1031 | 2319 | 7475 | 850 | 1570 | 6588 | 959 | 2030 |
| 499 | W303.asm02.HP0_G0005010 | chrIII | 3740 | 2337 | 2142 | 3527 | 2402 | 1969 | 4029 | 2439 | 2189 | 16428 | 4906 | 4523 | 16529 | 2952 | 2702 | 16238 | 3953 | 4062 |
| 500 | W303.asm02.HP0_G0005020 | chrIII | 395 | 613 | 717 | 276 | 871 | 789 | 389 | 653 | 748 | 3258 | 1116 | 1350 | 3658 | 765 | 915 | 3086 | 955 | 1125 |
| 501 | W303.asm02.HP0_G0005030 | chrIII | 2564 | 1907 | 1992 | 2257 | 1965 | 2026 | 2221 | 2078 | 2385 | 5476 | 3057 | 3092 | 6710 | 2287 | 2781 | 5562 | 2682 | 2639 |
| 502 | W303.asm02.HP0_G0005040 | chrIII | 8391 | 1533 | 1461 | 6980 | 1148 | 1664 | 8311 | 1215 | 1911 | 12971 | 4496 | 6071 | 5396 | 4001 | 4079 | 5554 | 3860 | 5078 |
| 503 | W303.asm02.HP0_G0005050 | chrIII | 840 | 537 | 587 | 656 | 586 | 583 | 684 | 546 | 748 | 2486 | 833 | 839 | 3071 | 478 | 713 | 2806 | 788 | 765 |
| 504 | W303.asm02.HP0_G0005060 | chrIII | 924 | 1433 | 1522 | 900 | 1292 | 1511 | 894 | 1223 | 1522 | 5564 | 1923 | 2228 | 6287 | 1123 | 1361 | 5752 | 2000 | 1914 |
| 505 | W303.asm02.HP0_G0005070 | chrIII | 2135 | 1650 | 2004 | 2401 | 2592 | 1839 | 2666 | 1873 | 2168 | 34452 | 5850 | 7962 | 27699 | 4119 | 4480 | 25540 | 4966 | 7012 |
| 506 | W303.asm02.HP0_G0005080 | chrIII | 7283 | 1953 | 1891 | 6813 | 1804 | 1757 | 6828 | 1922 | 1739 | 11698 | 3711 | 4022 | 12695 | 2738 | 3722 | 11124 | 3436 | 3876 |
| 507 | W303.asm02.HP0_G0005090 | chrIII | 2455 | 321 | 400 | 2576 | 355 | 496 | 2497 | 259 | 553 | 6524 | 1577 | 2064 | 4936 | 1020 | 1598 | 4335 | 1505 | 1814 |
| 508 | W303.asm02.HP0_G0005100 | chrIII | 9925 | 7207 | 6053 | 9532 | 5619 | 6302 | 9513 | 6161 | 5811 | 25487 | 9352 | 9563 | 24777 | 6631 | 7902 | 25797 | 8626 | 8796 |
| 509 | W303.asm02.HP0_G0005110 | chrIII | 13013 | 867 | 673 | 12138 | 659 | 711 | 12234 | 860 | 675 | 7249 | 2292 | 2006 | 9336 | 1943 | 1972 | 6151 | 2205 | 2074 |
| 510 | W303.asm02.HP0_G0005120 | chrIII | 314 | 79 | 43 | 283 | 43 | 62 | 279 | 72 | 77 | 75 | 88 | 71 | 443 | 134 | 141 | 74 | 107 | 49 |
| 511 | W303.asm02.HP0_G0005130 | chrIII | 653 | 78 | 101 | 538 | 99 | 115 | 506 | 78 | 120 | 316 | 124 | 116 | 654 | 78 | 107 | 337 | 130 | 103 |
| 512 | W303.asm02.HP0_G0005140 | chrIII | 1151 | 817 | 816 | 1048 | 668 | 817 | 984 | 784 | 813 | 4210 | 1267 | 1800 | 4316 | 843 | 1260 | 4114 | 1240 | 1688 |
| 513 | W303.asm02.HP0_G0005150 | chrIII | 6862 | 1515 | 1465 | 6042 | 1292 | 1532 | 6828 | 1420 | 1678 | 21912 | 6530 | 6473 | 13949 | 4944 | 4753 | 14404 | 5773 | 5784 |
| 514 | W303.asm02.HP0_G0005160 | chrIII | 1641 | 399 | 636 | 1600 | 281 | 805 | 1490 | 292 | 722 | 3131 | 1058 | 3112 | 1291 | 703 | 1528 | 1187 | 871 | 2566 |
| 515 | W303.asm02.HP0_G0005170 | chrIII | 20252 | 2710 | 5508 | 19326 | 2757 | 7289 | 21382 | 2182 | 6926 | 36329 | 10377 | 25102 | 24127 | 9088 | 18417 | 21854 | 9456 | 23277 |
| 516 | W303.asm02.HP0_G0005180 | chrIII | 40400 | 3328 | 2223 | 35453 | 1332 | 2817 | 36800 | 2323 | 2907 | 10176 | 10189 | 10087 | 8204 | 7902 | 9685 | 5608 | 10484 | 9013 |
| 517 | W303.asm02.HP0_G0005190 | chrIII | 4677 | 693 | 601 | 3999 | 595 | 700 | 4661 | 569 | 748 | 3378 | 1309 | 1645 | 2578 | 880 | 917 | 2394 | 1217 | 1327 |
| 518 | W303.asm02.HP0_G0005200 | chrIII | 2058 | 443 | 390 | 1608 | 340 | 401 | 1834 | 411 | 456 | 3813 | 1336 | 1451 | 4458 | 1071 | 1281 | 3077 | 1170 | 1206 |
| 519 | W303.asm02.HP0_G0005210 | chrIII | 324 | 45 | 74 | 286 | 84 | 98 | 426 | 40 | 119 | 1393 | 126 | 362 | 699 | 80 | 231 | 536 | 96 | 262 |
| 520 | W303.asm02.HP0_G0005220 | chrIII | 523 | 391 | 721 | 491 | 919 | 735 | 615 | 341 | 966 | 11138 | 1510 | 2033 | 6162 | 1369 | 1017 | 5153 | 1511 | 1568 |
| 521 | W303.asm02.HP0_G0005230 | chrIII | 5839 | 2525 | 2876 | 5340 | 2448 | 3045 | 4946 | 2492 | 3849 | 7742 | 4055 | 4689 | 10817 | 3069 | 4176 | 8257 | 3761 | 4170 |
| 522 | W303.asm02.HP0_G0005240 | chrIII | 4689 | 1588 | 817 | 4270 | 817 | 948 | 4302 | 1212 | 907 | 7023 | 3234 | 3507 | 5492 | 2355 | 2825 | 5569 | 3387 | 3081 |
| 523 | W303.asm02.HP0_G0005250 | chrIII | 2742 | 1177 | 1420 | 2289 | 1177 | 1399 | 2377 | 1278 | 1796 | 3818 | 2140 | 2201 | 6484 | 1726 | 2044 | 4084 | 2003 | 2007 |
| 524 | W303.asm02.HP0_G0005260 | chrIII | 2868 | 980 | 593 | 3131 | 602 | 564 | 2971 | 824 | 482 | 5944 | 1996 | 1989 | 6065 | 1373 | 1462 | 4794 | 1809 | 1852 |
| 525 | W303.asm02.HP0_G0005270 | chrIII | 2738 | 2403 | 3119 | 2702 | 3190 | 2895 | 2787 | 2448 | 3491 | 15925 | 3898 | 4886 | 16438 | 2528 | 3561 | 16265 | 3801 | 4379 |
| 526 | W303.asm02.HP0_G0005280 | chrIII | 5437 | 1900 | 1977 | 4404 | 1950 | 2152 | 4649 | 1608 | 2080 | 14106 | 5451 | 8616 | 8426 | 4344 | 5853 | 8362 | 4449 | 7411 |
| 527 | W303.asm02.HP0_G0005290 | chrIII | 1687 | 454 | 562 | 1613 | 501 | 581 | 1894 | 383 | 589 | 7269 | 1462 | 2545 | 5989 | 986 | 1532 | 5275 | 1332 | 2195 |
| 528 | W303.asm02.HP0_G0005300 | chrIII | 10394 | 2516 | 2289 | 10164 | 2213 | 2194 | 10069 | 2606 | 2533 | 20351 | 6615 | 7426 | 18340 | 4369 | 5875 | 17118 | 5844 | 6913 |
| 529 | W303.asm02.HP0_G0005310 | chrIII | 340 | 220 | 195 | 349 | 186 | 216 | 326 | 139 | 132 | 1442 | 502 | 528 | 928 | 187 | 245 | 1161 | 597 | 427 |
| 530 | W303.asm02.HP0_G0005320 | chrIII | 975 | 421 | 378 | 678 | 301 | 401 | 737 | 373 | 302 | 1247 | 728 | 546 | 1846 | 552 | 575 | 1193 | 685 | 528 |
| 531 | W303.asm02.HP0_G0005330 | chrIII | 81 | 108 | 147 | 75 | 74 | 124 | 79 | 102 | 137 | 443 | 321 | 346 | 443 | 166 | 194 | 444 | 255 | 255 |
| 532 | W303.asm02.HP0_G0005340 | chrIII | 3772 | 1268 | 2369 | 4149 | 2118 | 2561 | 3717 | 1140 | 2476 | 19971 | 4988 | 7778 | 16881 | 4524 | 6547 | 10564 | 4087 | 6138 |
| 533 | W303.asm02.HP0_G0005350 | chrIII | 2440 | 384 | 426 | 2376 | 484 | 494 | 2647 | 379 | 574 | 7975 | 1615 | 2262 | 5835 | 1190 | 1600 | 5570 | 1370 | 2074 |
| 534 | W303.asm02.HP0_G0005360 | chrIII | 547 | 165 | 224 | 454 | 208 | 245 | 491 | 206 | 291 | 626 | 183 | 293 | 542 | 120 | 209 | 570 | 201 | 299 |
| 535 | W303.asm02.HP0_G0005370 | chrIII | 820 | 996 | 1343 | 947 | 1329 | 1433 | 794 | 954 | 1345 | 4504 | 1203 | 1757 | 3245 | 615 | 1333 | 3800 | 1179 | 1401 |
| 536 | W303.asm02.HP0_G0005380 | chrIII | 5969 | 1300 | 1192 | 5801 | 1095 | 1329 | 5930 | 1225 | 1499 | 9140 | 2723 | 2983 | 7138 | 2203 | 2203 | 6096 | 2365 | 2749 |
| 537 | W303.asm02.HP0_G0005390 | chrIII | 1015 | 684 | 545 | 978 | 587 | 594 | 1083 | 640 | 536 | 2345 | 659 | 680 | 2343 | 379 | 489 | 2072 | 612 | 623 |
| 538 | W303.asm02.HP0_G0005400 | chrIII | 1929 | 424 | 472 | 1889 | 591 | 616 | 1964 | 369 | 598 | 5170 | 752 | 1300 | 2692 | 698 | 998 | 2735 | 669 | 1217 |
| 539 | W303.asm02.HP0_G0005410 | chrIII | 748 | 387 | 492 | 637 | 439 | 507 | 669 | 414 | 733 | 1211 | 340 | 311 | 2348 | 270 | 337 | 1378 | 306 | 275 |
| 540 | W303.asm02.HP0_G0005420 | chrIII | 10071 | 2214 | 1987 | 9689 | 2023 | 1986 | 9465 | 2048 | 2462 | 12285 | 4306 | 4076 | 6923 | 2906 | 2974 | 7612 | 3742 | 3582 |
| 541 | W303.asm02.HP0_G0005430 | chrIII | 1536 | 377 | 409 | 1494 | 313 | 411 | 1493 | 296 | 464 | 1872 | 701 | 883 | 862 | 495 | 543 | 909 | 678 | 734 |
| 542 | W303.asm02.HP0_G0005440 | chrIII | 9469 | 1702 | 1909 | 7470 | 1572 | 2313 | 9086 | 1297 | 2588 | 10299 | 2804 | 4008 | 2518 | 2241 | 2587 | 2411 | 2115 | 3431 |
| 543 | W303.asm02.HP0_G0005450 | chrIII | 2192 | 396 | 286 | 1753 | 278 | 350 | 2287 | 300 | 419 | 634 | 380 | 458 | 293 | 332 | 242 | 295 | 328 | 334 |
| 544 | W303.asm02.HP0_G0005460 | chrIII | 979 | 714 | 939 | 996 | 776 | 965 | 954 | 722 | 1215 | 2271 | 738 | 944 | 2604 | 522 | 713 | 2318 | 655 | 766 |
| 545 | W303.asm02.HP0_G0005470 | chrIII | 951 | 1027 | 637 | 958 | 745 | 665 | 940 | 1000 | 632 | 1901 | 793 | 647 | 1935 | 577 | 442 | 1731 | 714 | 566 |
| 546 | W303.asm02.HP0_G0005480 | chrIII | 1532 | 424 | 312 | 1495 | 272 | 327 | 1335 | 392 | 327 | 1744 | 519 | 494 | 1719 | 344 | 383 | 1490 | 555 | 478 |
| 547 | W303.asm02.HP0_G0005490 | chrIII | 954 | 910 | 1108 | 848 | 953 | 1005 | 927 | 938 | 1454 | 3539 | 769 | 803 | 3677 | 516 | 603 | 3949 | 697 | 652 |
| 548 | W303.asm02.HP0_G0005500 | chrIII | 2270 | 2160 | 2451 | 2253 | 1963 | 2403 | 2165 | 2257 | 2975 | 5489 | 1798 | 1924 | 5776 | 1170 | 1356 | 5864 | 1499 | 1629 |
| 549 | W303.asm02.HP0_G0005510 | chrIII | 109 | 67 | 48 | 119 | 45 | 39 | 128 | 84 | 59 | 187 | 73 | 51 | 147 | 46 | 30 | 198 | 75 | 41 |
| 550 | W303.asm02.HP0_G0005520 | chrIII | 193 | 128 | 72 | 172 | 77 | 63 | 158 | 122 | 87 | 195 | 111 | 70 | 183 | 64 | 32 | 216 | 89 | 51 |
| 551 | W303.asm02.HP0_G0005530 | chrIII | 5931 | 3588 | 2650 | 5852 | 2527 | 2718 | 5804 | 3530 | 2661 | 7729 | 3166 | 2992 | 7634 | 2225 | 2336 | 7206 | 2973 | 2836 |
| 552 | W303.asm02.HP0_G0005540 | chrIII | 3475 | 1730 | 1827 | 3289 | 1614 | 1908 | 2894 | 1650 | 2411 | 4079 | 1260 | 1435 | 5459 | 999 | 1226 | 4372 | 1315 | 1289 |
| 553 | W303.asm02.HP0_G0005550 | chrIII | 5050 | 3124 | 3395 | 4833 | 2953 | 3241 | 4546 | 3055 | 3713 | 7536 | 2796 | 3081 | 9810 | 2315 | 3258 | 7347 | 2543 | 2750 |
| 554 | W303.asm02.HP0_G0005560 | chrIII | 2062 | 4347 | 5128 | 2671 | 4400 | 4839 | 1625 | 4252 | 4867 | 5870 | 6172 | 5916 | 5916 | 2760 | 5113 | 6609 | 5747 | 5304 |
| 555 | W303.asm02.HP0_G0005570 | chrIII | 142 | 37 | 43 | 140 | 42 | 20 | 116 | 29 | 35 | 207 | 65 | 58 | 189 | 32 | 46 | 172 | 80 | 43 |
| 556 | W303.asm02.HP0_G0005580 | chrIII | 785 | 515 | 217 | 701 | 207 | 229 | 673 | 444 | 258 | 624 | 403 | 192 | 672 | 232 | 143 | 727 | 416 | 189 |

| | | | | | | | | | | | | | | | | | | | | |
|---|---|---|---|---|---|---|---|---|---|---|---|---|---|---|---|---|---|---|---|---|
| 557 | W303.asm02.HP0_G0005590 | chrIII | 1847 | 890 | 1275 | 2019 | 1340 | 1343 | 1744 | 958 | 1694 | 9761 | 1766 | 2535 | 6698 | 1054 | 1910 | 7461 | 1664 | 2271 |
| 558 | W303.asm02.HP0_G0005600 | chrIII | 9657 | 3939 | 4164 | 9371 | 3372 | 3939 | 9139 | 3843 | 4615 | 10273 | 3231 | 3522 | 10944 | 2185 | 2986 | 10426 | 2915 | 2999 |
| 559 | W303.asm02.HP0_G0005610 | chrIII | 286 | 66 | 57 | 390 | 66 | 56 | 336 | 62 | 67 | 285 | 96 | 79 | 384 | 65 | 78 | 229 | 112 | 54 |
| 560 | W303.asm02.HP0_G0005620 | chrIII | 5767 | 1436 | 1452 | 5752 | 1235 | 1525 | 5281 | 1406 | 1654 | 7258 | 2214 | 3004 | 4422 | 1578 | 2370 | 5030 | 2152 | 2843 |
| 561 | W303.asm02.HP0_G0005630 | chrIII | 74459 | 36409 | 43524 | 69704 | 37502 | 39459 | 76282 | 34254 | 42407 | 77328 | 36372 | 51452 | 71647 | 25491 | 41022 | 75109 | 33593 | 47966 |
| 562 | W303.asm02.HP0_G0005640 | chrIII | 2128 | 484 | 467 | 2011 | 545 | 507 | 1962 | 511 | 537 | 2351 | 533 | 625 | 2397 | 283 | 420 | 2203 | 533 | 552 |
| 563 | W303.asm02.HP0_G0005650 | chrIII | 902 | 582 | 612 | 939 | 573 | 637 | 905 | 611 | 788 | 2128 | 484 | 543 | 2255 | 361 | 461 | 2119 | 514 | 485 |
| 564 | W303.asm02.HP0_G0005660 | chrIII | 1432 | 1572 | 2143 | 1376 | 2397 | 2039 | 1726 | 1844 | 2656 | 7889 | 1656 | 1846 | 8350 | 963 | 1109 | 8107 | 1451 | 1515 |
| 565 | W303.asm02.HP0_G0005670 | chrIII | 10954 | 1409 | 1304 | 9507 | 1302 | 1463 | 10550 | 1298 | 1701 | 11171 | 3133 | 3206 | 5476 | 2305 | 2394 | 6370 | 2860 | 2980 |
| 566 | W303.asm02.HP0_G0005680 | chrIII | 843 | 1392 | 1664 | 734 | 1645 | 1660 | 779 | 1340 | 1891 | 2413 | 985 | 1031 | 2734 | 588 | 809 | 2607 | 911 | 970 |
| 567 | W303.asm02.HP0_G0005690 | chrIII | 1301 | 698 | 611 | 1258 | 607 | 565 | 1197 | 632 | 563 | 2548 | 666 | 724 | 2257 | 398 | 578 | 2353 | 645 | 642 |
| 568 | W303.asm02.HP0_G0005700 | chrIII | 356 | 316 | 153 | 267 | 152 | 151 | 304 | 314 | 188 | 375 | 224 | 134 | 501 | 111 | 110 | 478 | 190 | 99 |
| 569 | W303.asm02.HP0_G0005710 | chrIII | 886 | 471 | 585 | 642 | 551 | 567 | 728 | 551 | 808 | 1340 | 480 | 491 | 1790 | 296 | 452 | 1617 | 432 | 409 |
| 570 | W303.asm02.HP0_G0005720 | chrIII | 784 | 842 | 1050 | 578 | 893 | 963 | 628 | 863 | 1366 | 2598 | 681 | 657 | 3241 | 431 | 558 | 2977 | 650 | 551 |
| 571 | W303.asm02.HP0_G0005730 | chrIII | 13837 | 562 | 342 | 13213 | 1102 | 1216 | 12352 | 523 | 278 | 717 | 487 | 189 | 756 | 187 | 146 | 600 | 502 | 208 |
| 572 | W303.asm02.HP0_G0005740 | chrIII | 45 | 82 | 31 | 54 | 35 | 35 | 65 | 73 | 34 | 162 | 64 | 18 | 177 | 30 | 29 | 140 | 64 | 18 |
| 573 | W303.asm02.HP0_G0005750 | chrIII | 3282 | 1488 | 1421 | 3074 | 1343 | 1352 | 3028 | 1488 | 1435 | 5541 | 1565 | 1704 | 4516 | 970 | 1345 | 5305 | 1435 | 1665 |
| 574 | W303.asm02.HP0_G0005760 | chrIII | 3781 | 629 | 523 | 3999 | 447 | 584 | 3642 | 566 | 613 | 3260 | 731 | 1257 | 3453 | 682 | 991 | 2614 | 650 | 1076 |
| 575 | W303.asm02.HP0_G0005770 | chrIII | 144 | 77 | 39 | 135 | 52 | 52 | 109 | 71 | 58 | 204 | 97 | 70 | 207 | 61 | 67 | 133 | 78 | 49 |
| 576 | W303.asm02.HP0_G0005780 | chrIII | 2696 | 548 | 677 | 2718 | 648 | 717 | 2694 | 460 | 826 | 5819 | 1114 | 1575 | 3669 | 789 | 1018 | 3513 | 971 | 1404 |
| 577 | W303.asm02.HP0_G0005790 | chrIII | 741 | 783 | 958 | 728 | 947 | 914 | 654 | 787 | 1090 | 2950 | 705 | 896 | 3174 | 486 | 739 | 3055 | 697 | 862 |
| 578 | W303.asm02.HP0_G0005800 | chrIII | 586 | 622 | 610 | 593 | 666 | 604 | 604 | 614 | 693 | 4780 | 984 | 1013 | 3263 | 632 | 684 | 3383 | 847 | 932 |
| 579 | W303.asm02.HP0_G0005810 | chrIII | 4039 | 2835 | 3754 | 3632 | 2995 | 3783 | 3627 | 2924 | 5152 | 6605 | 2644 | 3121 | 7538 | 1696 | 2472 | 7530 | 2388 | 2751 |
| 580 | W303.asm02.HP0_G0005820 | chrIII | 9971 | 711 | 751 | 8907 | 633 | 915 | 9585 | 596 | 977 | 5592 | 1376 | 1772 | 1714 | 1047 | 1549 | 1564 | 1205 | 1541 |
| 581 | W303.asm02.HP0_G0005830 | chrIII | 12865 | 10222 | 11206 | 11670 | 10920 | 11452 | 12796 | 10204 | 12538 | 22057 | 9599 | 10379 | 22910 | 7265 | 8639 | 21856 | 8622 | 9460 |
| 582 | W303.asm02.HP0_G0005840 | chrIII | 3827 | 691 | 1027 | 3455 | 935 | 892 | 3396 | 702 | 1207 | 5118 | 644 | 1257 | 4773 | 486 | 1062 | 4774 | 692 | 1172 |
| 583 | W303.asm02.HP0_G0005850 | chrIII | 5062 | 437 | 655 | 5177 | 703 | 697 | 5030 | 386 | 794 | 6676 | 955 | 1645 | 3315 | 693 | 1162 | 3291 | 910 | 1453 |
| 584 | W303.asm02.HP0_G0005860 | chrIII | 4619 | 5040 | 4859 | 4415 | 4677 | 4353 | 4760 | 5113 | 4856 | 15161 | 4956 | 4767 | 15095 | 3374 | 3491 | 15373 | 4319 | 4133 |
| 585 | W303.asm02.HP0_G0005870 | chrIII | 2599 | 1865 | 2695 | 2521 | 2222 | 2757 | 2630 | 1916 | 2689 | 8464 | 1707 | 3082 | 8174 | 1093 | 1840 | 8067 | 1656 | 2560 |
| 586 | W303.asm02.HP0_G0005880 | chrIII | 3604 | 1940 | 1656 | 3497 | 1515 | 1582 | 3394 | 1888 | 1726 | 5974 | 1782 | 1655 | 7806 | 1446 | 1577 | 5651 | 1696 | 1662 |
| 587 | W303.asm02.HP0_G0005890 | chrIII | 3816 | 844 | 875 | 3541 | 961 | 843 | 3688 | 810 | 975 | 8279 | 1419 | 1879 | 5387 | 966 | 1302 | 6214 | 1342 | 1772 |
| 588 | W303.asm02.HP0_G0005900 | chrIII | 384 | 144 | 158 | 402 | 140 | 145 | 341 | 118 | 182 | 873 | 171 | 246 | 678 | 106 | 188 | 607 | 193 | 233 |
| 589 | W303.asm02.HP0_G0005910 | chrIII | 0 | 4 | 0 | 2 | 0 | 2 | 0 | 1 | 3 | 1 | 1 | 2 | 7 | 1 | 0 | 2 | 3 | 2 |
| 590 | W303.asm02.HP0_G0005920 | chrIII | 3224 | 810 | 979 | 3215 | 1072 | 1053 | 3187 | 875 | 1225 | 7203 | 1167 | 1883 | 4868 | 838 | 1530 | 4960 | 1110 | 1631 |
| 591 | W303.asm02.HP0_G0005930 | chrIII | 673 | 1474 | 1921 | 558 | 1972 | 1858 | 618 | 1544 | 2512 | 5434 | 1393 | 1375 | 7242 | 1051 | 1292 | 5771 | 1199 | 1355 |
| 592 | W303.asm02.HP0_G0005940 | chrIII | 1769 | 1218 | 1115 | 1434 | 1101 | 1059 | 1605 | 1198 | 1106 | 4108 | 1034 | 1028 | 4014 | 633 | 867 | 3967 | 1030 | 1023 |
| 593 | W303.asm02.HP0_G0005950 | chrIII | 590 | 190 | 207 | 508 | 198 | 213 | 593 | 133 | 261 | 1169 | 367 | 427 | 528 | 249 | 304 | 491 | 332 | 407 |
| 594 | W303.asm02.HP0_G0005960 | chrIII | 5488 | 829 | 1159 | 5039 | 1006 | 1049 | 5094 | 940 | 1314 | 4054 | 1436 | 1941 | 5409 | 1134 | 1770 | 3645 | 1384 | 1789 |
| 595 | W303.asm02.HP0_G0005970 | chrIII | 1695 | 1009 | 999 | 1584 | 1196 | 1033 | 1773 | 920 | 1340 | 5366 | 1999 | 1840 | 2693 | 1124 | 917 | 2996 | 1575 | 1635 |
| 596 | W303.asm02.HP0_G0005980 | chrIII | 2288 | 558 | 670 | 1726 | 528 | 672 | 2339 | 515 | 813 | 5025 | 1536 | 1371 | 1695 | 779 | 865 | 1895 | 1215 | 1279 |
| 597 | W303.asm02.HP0_G0005990 | chrIII | 4044 | 1835 | 1758 | 3514 | 1991 | 1609 | 4039 | 1963 | 2126 | 8087 | 3276 | 2643 | 7050 | 1936 | 2063 | 7195 | 2676 | 2538 |
| 598 | W303.asm02.HP0_G0006000 | chrIII | 6400 | 922 | 942 | 6121 | 809 | 1090 | 6908 | 693 | 1237 | 7991 | 2101 | 2556 | 3941 | 1631 | 1733 | 3197 | 1720 | 2254 |
| 599 | W303.asm02.HP0_G0006010 | chrIII | 13098 | 3128 | 3200 | 12111 | 2852 | 3351 | 13619 | 3120 | 3920 | 23683 | 6801 | 8384 | 20387 | 4788 | 5711 | 21152 | 5996 | 7439 |
| 600 | W303.asm02.HP0_G0006020 | chrIII | 633 | 553 | 769 | 586 | 863 | 692 | 629 | 548 | 807 | 4413 | 647 | 941 | 3774 | 358 | 531 | 4163 | 564 | 803 |
| 601 | W303.asm02.HP0_G0006030 | chrIII | 2666 | 1002 | 1524 | 2550 | 1755 | 1467 | 3106 | 1034 | 1944 | 14660 | 2468 | 3267 | 5694 | 1838 | 1486 | 5847 | 2175 | 2948 |
| 602 | W303.asm02.HP0_G0006040 | chrIII | 2532 | 1294 | 1216 | 1779 | 1253 | 982 | 2455 | 1423 | 1186 | 5263 | 2027 | 1773 | 6305 | 1343 | 1383 | 5273 | 1757 | 1588 |
| 603 | W303.asm02.HP0_G0006050 | chrIII | 2167 | 1045 | 1047 | 1737 | 1053 | 1029 | 1701 | 1113 | 1320 | 2659 | 1312 | 1166 | 3421 | 925 | 979 | 2746 | 1164 | 1015 |
| 604 | W303.asm02.HP0_G0006060 | chrIII | 2366 | 223 | 312 | 2093 | 219 | 385 | 2165 | 194 | 387 | 1160 | 530 | 645 | 889 | 387 | 474 | 765 | 483 | 567 |
| 605 | W303.asm02.HP0_G0006070 | chrIII | 316 | 73 | 113 | 246 | 110 | 120 | 307 | 86 | 137 | 639 | 176 | 246 | 580 | 121 | 158 | 504 | 148 | 187 |
| 606 | W303.asm02.HP0_G0006080 | chrIII | 463 | 354 | 405 | 420 | 389 | 480 | 495 | 238 | 463 | 2891 | 762 | 845 | 1351 | 432 | 614 | 1386 | 669 | 757 |
| 607 | W303.asm02.HP0_G0006090 | chrIII | 1089 | 221 | 252 | 1071 | 241 | 271 | 958 | 203 | 313 | 1860 | 340 | 480 | 1495 | 217 | 330 | 1501 | 316 | 414 |
| 608 | W303.asm02.HP0_G0006100 | chrIII | 4853 | 743 | 866 | 4759 | 747 | 961 | 5042 | 685 | 1063 | 9075 | 1755 | 2265 | 4989 | 1330 | 1662 | 4777 | 1522 | 2172 |
| 609 | W303.asm02.HP0_G0006110 | chrIII | 1090 | 207 | 220 | 972 | 180 | 289 | 1034 | 152 | 244 | 1347 | 580 | 600 | 407 | 401 | 379 | 455 | 561 | 497 |
| 610 | W303.asm02.HP0_G0006120 | chrIII | 5069 | 697 | 842 | 4680 | 634 | 982 | 4937 | 665 | 941 | 4241 | 1319 | 1743 | 3392 | 963 | 1384 | 3259 | 1218 | 1526 |
| 611 | W303.asm02.HP0_G0006130 | chrIII | 4132 | 1211 | 1597 | 3651 | 1542 | 1666 | 3484 | 1277 | 1945 | 3316 | 997 | 1382 | 4330 | 784 | 1314 | 3543 | 988 | 1302 |
| 612 | W303.asm02.HP0_G0006140 | chrIII | 317 | 417 | 585 | 314 | 690 | 675 | 390 | 385 | 728 | 5858 | 857 | 1245 | 3576 | 574 | 562 | 3758 | 803 | 1007 |
| 613 | W303.asm02.HP0_G0006150 | chrIII | 2207 | 356 | 473 | 2045 | 567 | 438 | 2272 | 396 | 495 | 4356 | 575 | 869 | 5180 | 596 | 805 | 2931 | 527 | 809 |
| 614 | W303.asm02.HP0_G0006160 | chrIII | 2730 | 425 | 385 | 2179 | 307 | 483 | 2570 | 313 | 467 | 2060 | 736 | 932 | 1252 | 507 | 631 | 1221 | 663 | 783 |
| 615 | W303.asm02.HP0_G0006170 | chrIII | 227 | 574 | 604 | 148 | 490 | 605 | 192 | 558 | 761 | 1145 | 421 | 427 | 1391 | 251 | 386 | 1214 | 389 | 391 |
| 616 | W303.asm02.HP0_G0006180 | chrIII | 174 | 214 | 271 | 100 | 224 | 284 | 115 | 188 | 329 | 481 | 118 | 180 | 658 | 102 | 192 | 536 | 149 | 141 |
| 617 | W303.asm02.HP0_G0006190 | chrIII | 1516 | 679 | 596 | 1485 | 519 | 597 | 1463 | 696 | 679 | 1547 | 627 | 568 | 1437 | 383 | 459 | 1494 | 600 | 544 |
| 618 | W303.asm02.HP0_G0006200 | chrIII | 12428 | 1186 | 1692 | 11409 | 1444 | 1735 | 12762 | 989 | 2312 | 3493 | 1608 | 2861 | 2293 | 1259 | 1524 | 2100 | 1331 | 2216 |
| 619 | W303.asm02.HP0_G0006210 | chrIII | 1282 | 422 | 457 | 1226 | 396 | 473 | 1171 | 345 | 579 | 1561 | 610 | 704 | 1082 | 451 | 527 | 936 | 570 | 567 |
| 620 | W303.asm02.HP0_G0006220 | chrIII | 1370 | 461 | 573 | 1386 | 562 | 528 | 1320 | 509 | 578 | 4226 | 520 | 1020 | 3888 | 382 | 669 | 3505 | 516 | 881 |
| 621 | W303.asm02.HP0_G0006230 | chrIII | 2223 | 1522 | 2062 | 2087 | 1882 | 2178 | 2165 | 1535 | 2552 | 4655 | 1265 | 1693 | 5694 | 938 | 1449 | 4741 | 1202 | 1462 |
| 622 | W303.asm02.HP0_G0006240 | chrIII | 1531 | 808 | 1126 | 1562 | 1009 | 1147 | 1398 | 845 | 1555 | 2215 | 852 | 839 | 3161 | 608 | 742 | 2398 | 712 | 723 |
| 623 | W303.asm02.HP0_G0006250 | chrIII | 9951 | 1158 | 1518 | 9067 | 1524 | 1832 | 10783 | 936 | 2097 | 4815 | 1565 | 3640 | 1914 | 1448 | 1903 | 1781 | 1113 | 2867 |
| 624 | W303.asm02.HP0_G0006260 | chrIII | 570 | 532 | 513 | 443 | 476 | 469 | 553 | 619 | 409 | 1138 | 494 | 476 | 1298 | 310 | 333 | 1068 | 419 | 410 |
| 625 | W303.asm02.HP0_G0006270 | chrIII | 1544 | 1046 | 1238 | 1414 | 1615 | 1274 | 1568 | 1286 | 1634 | 4954 | 1372 | 1143 | 4853 | 730 | 719 | 4426 | 1135 | 1095 |
| 626 | W303.asm02.HP0_G0006280 | chrIII | 13121 | 788 | 936 | 11824 | 716 | 1143 | 12962 | 541 | 1193 | 4447 | 1524 | 2316 | 1333 | 1465 | 1475 | 967 | 1110 | 1976 |
| 627 | W303.asm02.HP0_G0006290 | chrIII | 808 | 119 | 186 | 861 | 222 | 258 | 933 | 114 | 261 | 1666 | 247 | 512 | 556 | 174 | 255 | 520 | 178 | 385 |
| 628 | W303.asm02.HP0_G0006300 | chrIII | 1255 | 1178 | 1451 | 1212 | 1236 | 1418 | 1187 | 1172 | 2030 | 3307 | 1003 | 1183 | 3878 | 692 | 902 | 3836 | 952 | 1066 |
| 629 | W303.asm02.HP0_G0006310 | chrIII | 3556 | 104 | 96 | 3863 | 124 | 120 | 3373 | 93 | 129 | 617 | 141 | 193 | 400 | 119 | 157 | 324 | 124 | 161 |
| 630 | W303.asm02.HP0_G0006320 | chrIII | 1409 | 301 | 458 | 1303 | 473 | 465 | 1323 | 304 | 585 | 3055 | 556 | 887 | 1662 | 379 | 527 | 1723 | 447 | 811 |
| 631 | W303.asm02.HP0_G0006330 | chrIII | 5719 | 828 | 1458 | 5378 | 1634 | 1629 | 5344 | 822 | 1933 | 13828 | 1351 | 3386 | 7514 | 1126 | 2641 | 8405 | 1193 | 3047 |
| 632 | W303.asm02.HP0_G0006340 | chrIII | 2450 | 875 | 924 | 2309 | 828 | 865 | 2485 | 937 | 875 | 4399 | 944 | 1160 | 4171 | 669 | 953 | 4158 | 864 | 1116 |
| 633 | W303.asm02.HP0_G0006350 | chrIII | 1260 | 891 | 1131 | 1314 | 985 | 1142 | 1251 | 805 | 1474 | 3148 | 625 | 877 | 3550 | 367 | 611 | 3569 | 717 | 822 |
| 634 | W303.asm02.HP0_G0006360 | chrIII | 177 | 48 | 24 | 199 | 29 | 34 | 153 | 28 | 34 | 253 | 50 | 55 | 194 | 38 | 47 | 215 | 52 | 62 |
| 635 | W303.asm02.HP0_G0006370 | chrIII | 0 | 0 | 0 | 0 | 0 | 0 | 0 | 0 | 0 | 0 | 0 | 0 | 0 | 0 | 0 | 0 | 0 | 0 |
| 636 | W303.asm02.HP0_G0006380 | chrIII | 671 | 698 | 1708 | 753 | 922 | 1793 | 696 | 836 | 1851 | 3099 | 1762 | 3082 | 5871 | 1981 | 3028 | 2913 | 1287 | 1842 |
| 637 | W303.asm02.HP0_G0006390 | chrIII | 413 | 129 | 233 | 386 | 204 | 249 | 342 | 134 | 319 | 911 | 222 | 391 | 1474 | 202 | 339 | 900 | 212 | 314 |
| 638 | W303.asm02.HP0_G0006400 | chrIII | 680 | 174 | 299 | 602 | 220 | 239 | 627 | 208 | 282 | 2567 | 404 | 551 | 3997 | 462 | 637 | 2143 | 396 | 500 |
| 639 | W303.asm02.HP0_G0006410 | chrIII | 429 | 107 | 165 | 423 | 133 | 127 | 338 | 103 | 132 | 1570 | 282 | 283 | 2597 | 263 | 322 | 1398 | 235 | 286 |
| 640 | W303.asm02.HP0_G0006420 | chrIII | 500 | 583 | 857 | 479 | 611 | 757 | 444 | 486 | 827 | 1607 | 1143 | 1052 | 2424 | 1066 | 1245 | 1558 | 923 | 742 |
| 641 | W303.asm02.HP0_G0006430 | chrIII | 30 | 12 | 13 | 36 | 10 | 19 | 38 | 8 | 8 | 102 | 41 | 50 | 98 | 38 | 56 | 65 | 41 | 51 |
| 642 | W303.asm02.HP0_G0006440 | chrIII | 537 | 219 | 214 | 561 | 240 | 276 | 469 | 325 | 326 | 2132 | 602 | 607 | 1984 | 350 | 466 | 1723 | 558 | 578 |
| 643 | W303.asm02.HP0_G0006450 | chrIII | 777 | 296 | 443 | 680 | 467 | 394 | 702 | 328 | 465 | 6393 | 1052 | 1928 | 4477 | 736 | 1262 | 4694 | 980 | 1661 |
| 644 | W303.asm02.HP0_G0006460 | chrIII | 1330 | 1375 | 1897 | 1272 | 1555 | 1839 | 1171 | 1405 | 1900 | 5403 | 3453 | 3413 | 4825 | 2578 | 2878 | 4602 | 2739 | 2576 |
| 645 | W303.asm02.HP0_G0006470 | chrIII | 400 | 166 | 195 | 429 | 177 | 223 | 503 | 134 | 253 | 426 | 155 | 230 | 223 | 88 | 116 | 235 | 129 | 202 |
| 646 | W303.asm02.HP0_G0006480 | chrIV | 1747 | 857 | 898 | 1915 | 719 | 950 | 1876 | 874 | 987 | 3133 | 871 | 1159 | 3413 | 552 | 847 | 2925 | 726 | 1057 |
| 647 | W303.asm02.HP0_G0006490 | chrIV | 152 | 37 | 12 | 172 | 33 | 28 | 149 | 14 | 20 | 172 | 34 | 39 | 147 | 26 | 23 | 102 | 28 | 37 |
| 648 | W303.asm02.HP0_G0006500 | chrIV | 152 | 95 | 137 | 156 | 128 | 168 | 138 | 114 | 200 | 273 | 74 | 92 | 421 | 46 | 71 | 270 | 77 | 87 |
| 649 | W303.asm02.HP0_G0006510 | chrIV | 182 | 8 | 25 | 194 | 19 | 26 | 184 | 12 | 35 | 154 | 24 | 51 | 146 | 24 | 45 | 82 | 31 | 52 |

| | | | | | | | | | | | | | | | | | | | | |
|---|---|---|---|---|---|---|---|---|---|---|---|---|---|---|---|---|---|---|---|---|
| 650 | W303.asm02.HP0_G0006520 | chrIV | 16 | 4 | 5 | 16 | 1 | 4 | 13 | 2 | 3 | 10 | 5 | 5 | 8 | 1 | 2 | 3 | 3 | 2 |
| 651 | W303.asm02.HP0_G0006530 | chrIV | 368 | 125 | 116 | 324 | 105 | 100 | 317 | 132 | 124 | 234 | 84 | 87 | 332 | 62 | 57 | 227 | 86 | 78 |
| 652 | W303.asm02.HP0_G0006540 | chrIV | 748 | 694 | 717 | 642 | 630 | 691 | 663 | 601 | 625 | 788 | 1218 | 842 | 835 | 1200 | 981 | 629 | 964 | 535 |
| 653 | W303.asm02.HP0_G0006550 | chrIV | 159 | 48 | 42 | 144 | 34 | 38 | 119 | 52 | 33 | 210 | 57 | 55 | 234 | 55 | 57 | 190 | 66 | 44 |
| 654 | W303.asm02.HP0_G0006560 | chrIV | 1547 | 367 | 470 | 1664 | 486 | 456 | 1348 | 323 | 498 | 1805 | 246 | 261 | 2165 | 208 | 287 | 1991 | 225 | 293 |
| 655 | W303.asm02.HP0_G0006570 | chrIV | 386 | 303 | 607 | 364 | 634 | 588 | 407 | 302 | 570 | 4172 | 392 | 1048 | 3837 | 262 | 725 | 3509 | 391 | 927 |
| 656 | W303.asm02.HP0_G0006580 | chrIV | 727 | 137 | 219 | 600 | 158 | 222 | 653 | 153 | 328 | 1027 | 167 | 273 | 707 | 133 | 244 | 846 | 177 | 255 |
| 657 | W303.asm02.HP0_G0006590 | chrIV | 999 | 85 | 93 | 953 | 90 | 134 | 949 | 68 | 125 | 740 | 152 | 232 | 533 | 149 | 225 | 337 | 121 | 226 |
| 658 | W303.asm02.HP0_G0006600 | chrIV | 4053 | 1389 | 1367 | 3571 | 1223 | 1330 | 3838 | 1402 | 1326 | 5041 | 1364 | 1581 | 5592 | 1064 | 1406 | 4768 | 1180 | 1427 |
| 659 | W303.asm02.HP0_G0006610 | chrIV | 4470 | 2511 | 2093 | 3842 | 1892 | 2095 | 4389 | 2309 | 2174 | 7792 | 2562 | 3225 | 7185 | 1756 | 2283 | 7064 | 2324 | 2919 |
| 660 | W303.asm02.HP0_G0006620 | chrIV | 1793 | 2128 | 3069 | 1772 | 2368 | 3155 | 1740 | 2272 | 4035 | 6192 | 1747 | 2599 | 8780 | 1509 | 2090 | 7071 | 1641 | 2285 |
| 661 | W303.asm02.HP0_G0006630 | chrIV | 9710 | 585 | 512 | 11202 | 495 | 583 | 8957 | 524 | 684 | 3901 | 830 | 1109 | 2942 | 690 | 837 | 2429 | 836 | 883 |
| 662 | W303.asm02.HP0_G0006640 | chrIV | 2287 | 282 | 280 | 2139 | 264 | 279 | 1858 | 278 | 348 | 1892 | 439 | 555 | 2039 | 347 | 468 | 1565 | 396 | 531 |
| 663 | W303.asm02.HP0_G0006650 | chrIV | 2463 | 751 | 1260 | 2349 | 1264 | 1123 | 2529 | 794 | 1359 | 7891 | 1191 | 1927 | 6212 | 762 | 1389 | 6240 | 1109 | 1697 |
| 664 | W303.asm02.HP0_G0006660 | chrIV | 1936 | 571 | 308 | 1751 | 358 | 371 | 1761 | 391 | 374 | 1645 | 615 | 587 | 1543 | 463 | 568 | 1193 | 572 | 517 |
| 665 | W303.asm02.HP0_G0006670 | chrIV | 32349 | 8731 | 8553 | 30229 | 8690 | 8758 | 35997 | 8473 | 10178 | 47525 | 14214 | 19262 | 40191 | 11674 | 12974 | 38606 | 11647 | 16547 |
| 666 | W303.asm02.HP0_G0006680 | chrIV | 365 | 757 | 1251 | 311 | 1132 | 1144 | 461 | 900 | 1230 | 4472 | 980 | 1941 | 2435 | 506 | 1020 | 3957 | 836 | 1686 |
| 667 | W303.asm02.HP0_G0006690 | chrIV | 3492 | 1456 | 1302 | 2807 | 1246 | 1409 | 3309 | 1356 | 1411 | 8487 | 2238 | 3016 | 6492 | 1518 | 2243 | 6827 | 2063 | 2720 |
| 668 | W303.asm02.HP0_G0006700 | chrIV | 4102 | 1788 | 1373 | 3971 | 1387 | 1446 | 4035 | 1793 | 1349 | 7691 | 1914 | 2200 | 6299 | 1276 | 1513 | 6681 | 1712 | 2100 |
| 669 | W303.asm02.HP0_G0006710 | chrIV | 1274 | 360 | 496 | 1177 | 429 | 636 | 1355 | 300 | 651 | 4367 | 822 | 1515 | 1902 | 628 | 890 | 1871 | 730 | 1190 |
| 670 | W303.asm02.HP0_G0006720 | chrIV | 3003 | 251 | 295 | 3048 | 306 | 347 | 3025 | 219 | 394 | 2022 | 403 | 595 | 1135 | 289 | 414 | 1128 | 385 | 516 |
| 671 | W303.asm02.HP0_G0006730 | chrIV | 994 | 105 | 93 | 1233 | 87 | 91 | 965 | 106 | 139 | 306 | 225 | 195 | 304 | 131 | 121 | 227 | 239 | 161 |
| 672 | W303.asm02.HP0_G0006740 | chrIV | 771 | 140 | 203 | 758 | 175 | 230 | 741 | 103 | 234 | 1026 | 178 | 453 | 558 | 142 | 234 | 450 | 145 | 353 |
| 673 | W303.asm02.HP0_G0006750 | chrIV | 1086 | 1057 | 1188 | 1094 | 1112 | 1129 | 1116 | 1015 | 1563 | 3361 | 895 | 947 | 3909 | 631 | 759 | 3788 | 787 | 825 |
| 674 | W303.asm02.HP0_G0006760 | chrIV | 293 | 161 | 182 | 271 | 218 | 178 | 313 | 134 | 198 | 430 | 143 | 179 | 474 | 108 | 114 | 371 | 138 | 192 |
| 675 | W303.asm02.HP0_G0006770 | chrIV | 1213 | 955 | 1234 | 1124 | 1167 | 1118 | 1168 | 885 | 1434 | 4193 | 1043 | 1290 | 3928 | 712 | 995 | 3672 | 998 | 1102 |
| 676 | W303.asm02.HP0_G0006780 | chrIV | 1117 | 274 | 274 | 1132 | 325 | 279 | 950 | 311 | 253 | 1105 | 252 | 370 | 1207 | 168 | 285 | 1070 | 262 | 272 |
| 677 | W303.asm02.HP0_G0006790 | chrIV | 10640 | 1267 | 1302 | 9901 | 1297 | 1335 | 10098 | 1224 | 1390 | 6344 | 1591 | 2169 | 5860 | 1189 | 1786 | 5536 | 1643 | 1832 |
| 678 | W303.asm02.HP0_G0006800 | chrIV | 340 | 176 | 107 | 422 | 82 | 156 | 337 | 145 | 97 | 349 | 179 | 192 | 283 | 93 | 99 | 251 | 182 | 150 |
| 679 | W303.asm02.HP0_G0006810 | chrIV | 2571 | 2035 | 1717 | 2424 | 2311 | 1840 | 2706 | 2240 | 1862 | 7386 | 2321 | 2114 | 8325 | 1524 | 1530 | 6765 | 1837 | 1833 |
| 680 | W303.asm02.HP0_G0006820 | chrIV | 4577 | 3475 | 4851 | 4138 | 4211 | 4796 | 4437 | 3573 | 6693 | 10363 | 2928 | 3771 | 13228 | 2136 | 3048 | 11078 | 2543 | 3316 |
| 681 | W303.asm02.HP0_G0006830 | chrIV | 260 | 381 | 443 | 243 | 354 | 518 | 240 | 310 | 450 | 1396 | 260 | 550 | 1868 | 177 | 383 | 1143 | 247 | 481 |
| 682 | W303.asm02.HP0_G0006840 | chrIV | 354 | 197 | 142 | 330 | 113 | 127 | 305 | 155 | 139 | 409 | 160 | 157 | 435 | 108 | 88 | 339 | 160 | 136 |
| 683 | W303.asm02.HP0_G0006850 | chrIV | 261 | 480 | 347 | 234 | 425 | 419 | 292 | 445 | 453 | 2003 | 633 | 644 | 1345 | 462 | 382 | 1229 | 658 | 540 |
| 684 | W303.asm02.HP0_G0006860 | chrIV | 3601 | 4452 | 5363 | 3517 | 5285 | 5092 | 4002 | 4911 | 6474 | 12424 | 4375 | 4381 | 13660 | 2959 | 3267 | 13000 | 3704 | 4004 |
| 685 | W303.asm02.HP0_G0006870 | chrIV | 2646 | 736 | 607 | 2660 | 570 | 580 | 2763 | 736 | 586 | 3874 | 889 | 1177 | 2964 | 628 | 781 | 2813 | 765 | 952 |
| 686 | W303.asm02.HP0_G0006880 | chrIV | 1047 | 593 | 605 | 979 | 429 | 585 | 938 | 570 | 640 | 2308 | 651 | 647 | 2714 | 373 | 602 | 2303 | 620 | 638 |
| 687 | W303.asm02.HP0_G0006890 | chrIV | 1206 | 901 | 1483 | 995 | 1706 | 1402 | 1040 | 946 | 1637 | 6089 | 836 | 1406 | 5271 | 595 | 1050 | 5371 | 799 | 1215 |
| 688 | W303.asm02.HP0_G0006900 | chrIV | 13844 | 758 | 353 | 15315 | 286 | 347 | 13359 | 695 | 302 | 1032 | 740 | 482 | 1513 | 587 | 579 | 895 | 684 | 353 |
| 689 | W303.asm02.HP0_G0006910 | chrIV | 1869 | 393 | 555 | 1735 | 521 | 652 | 1900 | 362 | 691 | 3817 | 717 | 1179 | 1845 | 541 | 787 | 1842 | 680 | 1095 |
| 690 | W303.asm02.HP0_G0006920 | chrIV | 3876 | 1223 | 1144 | 3747 | 1090 | 1165 | 3818 | 1301 | 1311 | 4790 | 1341 | 1735 | 4687 | 1063 | 1264 | 4073 | 1203 | 1479 |
| 691 | W303.asm02.HP0_G0006930 | chrIV | 1445 | 1596 | 1349 | 1145 | 1566 | 1393 | 1507 | 1612 | 1233 | 6936 | 1614 | 1849 | 5882 | 969 | 1069 | 6041 | 1411 | 1649 |
| 692 | W303.asm02.HP0_G0006940 | chrIV | 720 | 771 | 893 | 664 | 681 | 915 | 578 | 817 | 1160 | 1733 | 577 | 667 | 2264 | 413 | 519 | 2061 | 597 | 527 |
| 693 | W303.asm02.HP0_G0006950 | chrIV | 932 | 204 | 183 | 931 | 179 | 228 | 790 | 175 | 221 | 965 | 258 | 255 | 696 | 168 | 222 | 648 | 215 | 219 |
| 694 | W303.asm02.HP0_G0006960 | chrIV | 4461 | 2505 | 2586 | 3872 | 2470 | 2620 | 4043 | 2342 | 2806 | 7771 | 2578 | 3621 | 7012 | 2239 | 3012 | 6105 | 2059 | 3081 |
| 695 | W303.asm02.HP0_G0006970 | chrIV | 1791 | 239 | 192 | 1711 | 203 | 246 | 1771 | 172 | 236 | 1480 | 450 | 478 | 898 | 284 | 319 | 831 | 399 | 449 |
| 696 | W303.asm02.HP0_G0006980 | chrIV | 15780 | 1235 | 1467 | 14247 | 1153 | 2167 | 16242 | 929 | 2114 | 2403 | 1239 | 2819 | 1590 | 1181 | 1267 | 1517 | 1048 | 1990 |
| 697 | W303.asm02.HP0_G0006990 | chrIV | 1688 | 111 | 122 | 1813 | 240 | 186 | 1527 | 103 | 149 | 1566 | 320 | 276 | 931 | 193 | 219 | 865 | 256 | 236 |
| 698 | W303.asm02.HP0_G0007000 | chrIV | 1876 | 1022 | 1301 | 1924 | 935 | 1573 | 1634 | 885 | 1209 | 3526 | 929 | 2104 | 3529 | 679 | 1196 | 3592 | 972 | 1785 |
| 699 | W303.asm02.HP0_G0007010 | chrIV | 6456 | 5561 | 8224 | 6618 | 6986 | 8295 | 6499 | 5596 | 10944 | 17036 | 5172 | 7295 | 20059 | 3868 | 5913 | 18876 | 4561 | 6678 |
| 700 | W303.asm02.HP0_G0007020 | chrIV | 6366 | 5547 | 6823 | 5757 | 6381 | 6691 | 5924 | 5458 | 8200 | 12527 | 5099 | 5593 | 13519 | 3942 | 4669 | 13170 | 4661 | 4943 |
| 701 | W303.asm02.HP0_G0007030 | chrIV | 6195 | 1692 | 1528 | 5722 | 1483 | 1552 | 5683 | 1708 | 1519 | 9833 | 1849 | 2589 | 8540 | 1273 | 2016 | 8583 | 1731 | 2356 |
| 702 | W303.asm02.HP0_G0007040 | chrIV | 1560 | 567 | 617 | 1601 | 630 | 696 | 1741 | 452 | 804 | 3726 | 762 | 1121 | 1846 | 579 | 690 | 2150 | 751 | 1028 |
| 703 | W303.asm02.HP0_G0007050 | chrIV | 3441 | 1099 | 1218 | 3075 | 1116 | 1159 | 3420 | 1151 | 1228 | 5595 | 1311 | 1679 | 5274 | 958 | 1424 | 4951 | 1170 | 1564 |
| 704 | W303.asm02.HP0_G0007060 | chrIV | 406 | 125 | 69 | 299 | 87 | 64 | 336 | 108 | 59 | 346 | 72 | 99 | 304 | 54 | 83 | 316 | 89 | 84 |
| 705 | W303.asm02.HP0_G0007070 | chrIV | 28996 | 3680 | 3594 | 25081 | 3429 | 4190 | 28511 | 3194 | 4754 | 28589 | 7969 | 10107 | 15297 | 6929 | 7502 | 14426 | 7082 | 8850 |
| 706 | W303.asm02.HP0_G0007080 | chrIV | 1879 | 329 | 185 | 2106 | 249 | 233 | 1676 | 284 | 182 | 978 | 281 | 282 | 986 | 199 | 204 | 893 | 345 | 228 |
| 707 | W303.asm02.HP0_G0007090 | chrIV | 1834 | 4505 | 3598 | 2192 | 3619 | 3269 | 2317 | 4421 | 3281 | 13322 | 5658 | 5345 | 9898 | 2921 | 3062 | 11809 | 4678 | 4660 |
| 708 | W303.asm02.HP0_G0007100 | chrIV | 1459 | 2459 | 2974 | 1225 | 2745 | 3064 | 1105 | 2584 | 3791 | 5112 | 2117 | 2189 | 7155 | 1666 | 2173 | 5433 | 1820 | 1623 |
| 709 | W303.asm02.HP0_G0007110 | chrIV | 1111 | 1183 | 1193 | 1163 | 1031 | 1105 | 972 | 980 | 1058 | 4587 | 1423 | 1466 | 3811 | 844 | 1203 | 3638 | 1373 | 1196 |
| 710 | W303.asm02.HP0_G0007120 | chrIV | 1505 | 1690 | 2318 | 1856 | 1733 | 2278 | 1646 | 1728 | 2178 | 5876 | 1590 | 2795 | 7551 | 1352 | 2229 | 5164 | 1342 | 2231 |
| 711 | W303.asm02.HP0_G0007130 | chrIV | 6122 | 2844 | 2621 | 5446 | 2242 | 2562 | 5728 | 2859 | 2738 | 8333 | 2772 | 3046 | 7511 | 1664 | 2077 | 8067 | 2653 | 2845 |
| 712 | W303.asm02.HP0_G0007140 | chrIV | 335 | 633 | 637 | 290 | 590 | 661 | 315 | 682 | 757 | 1352 | 489 | 405 | 1573 | 292 | 349 | 1371 | 446 | 373 |
| 713 | W303.asm02.HP0_G0007150 | chrIV | 1123 | 287 | 430 | 913 | 359 | 397 | 1088 | 326 | 427 | 2461 | 431 | 672 | 1930 | 295 | 490 | 2132 | 383 | 604 |
| 714 | W303.asm02.HP0_G0007160 | chrIV | 12538 | 1024 | 904 | 13304 | 874 | 1134 | 12653 | 839 | 1324 | 4875 | 1646 | 2508 | 3128 | 1209 | 1751 | 3139 | 1651 | 2292 |
| 715 | W303.asm02.HP0_G0007170 | chrIV | 2555 | 2059 | 1841 | 3280 | 1577 | 1759 | 3364 | 2059 | 1989 | 6316 | 3377 | 2612 | 6616 | 2157 | 2307 | 5857 | 3203 | 2449 |
| 716 | W303.asm02.HP0_G0007180 | chrIV | 8250 | 1462 | 1768 | 7402 | 1764 | 1795 | 7623 | 1253 | 2432 | 15771 | 2605 | 4238 | 5886 | 2233 | 2891 | 5852 | 2287 | 3508 |
| 717 | W303.asm02.HP0_G0007190 | chrIV | 1094 | 535 | 528 | 1127 | 570 | 544 | 1070 | 479 | 570 | 3572 | 908 | 1014 | 3185 | 609 | 672 | 3213 | 831 | 887 |
| 718 | W303.asm02.HP0_G0007200 | chrIV | 1898 | 315 | 234 | 1898 | 277 | 284 | 1583 | 278 | 272 | 402 | 232 | 241 | 702 | 170 | 279 | 393 | 258 | 183 |
| 719 | W303.asm02.HP0_G0007210 | chrIV | 2837 | 2334 | 1808 | 2456 | 1765 | 1801 | 2533 | 2053 | 2082 | 7414 | 3982 | 3682 | 5346 | 2613 | 2470 | 5764 | 3750 | 3063 |
| 720 | W303.asm02.HP0_G0007220 | chrIV | 1502 | 863 | 1194 | 1477 | 1440 | 1153 | 1644 | 1038 | 1417 | 10409 | 1680 | 2510 | 7268 | 1112 | 1383 | 7830 | 1580 | 2184 |
| 721 | W303.asm02.HP0_G0007230 | chrIV | 1402 | 1488 | 1811 | 1276 | 1852 | 1844 | 1410 | 1568 | 2188 | 4423 | 1218 | 1532 | 4603 | 760 | 1076 | 4301 | 1036 | 1251 |
| 722 | W303.asm02.HP0_G0007240 | chrIV | 1313 | 1111 | 1213 | 1088 | 1026 | 1199 | 1101 | 1218 | 1560 | 2790 | 895 | 968 | 3233 | 596 | 706 | 3041 | 817 | 832 |
| 723 | W303.asm02.HP0_G0007250 | chrIV | 1530 | 504 | 572 | 1467 | 483 | 648 | 1683 | 442 | 670 | 4524 | 899 | 1308 | 2332 | 667 | 825 | 2288 | 818 | 1231 |
| 724 | W303.asm02.HP0_G0007260 | chrIV | 7418 | 2915 | 2169 | 7844 | 2489 | 2201 | 7728 | 2871 | 2001 | 10743 | 2774 | 2836 | 10748 | 1806 | 2199 | 8605 | 2360 | 2597 |
| 725 | W303.asm02.HP0_G0007270 | chrIV | 717 | 433 | 551 | 545 | 455 | 536 | 581 | 426 | 760 | 990 | 364 | 430 | 1230 | 208 | 335 | 1249 | 322 | 359 |
| 726 | W303.asm02.HP0_G0007280 | chrIV | 3084 | 652 | 920 | 3557 | 775 | 828 | 2942 | 698 | 856 | 5321 | 665 | 1335 | 5282 | 476 | 1104 | 4820 | 592 | 1139 |
| 727 | W303.asm02.HP0_G0007290 | chrIV | 1175 | 440 | 407 | 1275 | 429 | 433 | 1144 | 411 | 426 | 3003 | 530 | 784 | 2488 | 348 | 537 | 2398 | 545 | 776 |
| 728 | W303.asm02.HP0_G0007300 | chrIV | 1101 | 1076 | 1543 | 1062 | 1418 | 1462 | 1101 | 1162 | 2092 | 3591 | 1108 | 1318 | 4150 | 760 | 992 | 4280 | 999 | 1168 |
| 729 | W303.asm02.HP0_G0007310 | chrIV | 461 | 92 | 127 | 451 | 123 | 147 | 499 | 65 | 199 | 1367 | 203 | 393 | 512 | 159 | 260 | 558 | 210 | 339 |
| 730 | W303.asm02.HP0_G0007320 | chrIV | 1214 | 746 | 785 | 1343 | 883 | 792 | 1234 | 701 | 779 | 3952 | 808 | 1228 | 3360 | 566 | 825 | 3066 | 708 | 1153 |
| 731 | W303.asm02.HP0_G0007330 | chrIV | 1499 | 529 | 607 | 1533 | 615 | 641 | 1308 | 504 | 789 | 1679 | 318 | 478 | 1685 | 223 | 393 | 1585 | 343 | 454 |
| 732 | W303.asm02.HP0_G0007340 | chrIV | 4862 | 2101 | 1858 | 5235 | 2881 | 1837 | 5417 | 2400 | 1852 | 16100 | 2937 | 2951 | 13651 | 1765 | 1668 | 13201 | 2305 | 2738 |
| 733 | W303.asm02.HP0_G0007350 | chrIV | 3211 | 780 | 911 | 3170 | 1026 | 917 | 3535 | 797 | 1047 | 8557 | 1643 | 1976 | 6470 | 1204 | 1350 | 5879 | 1431 | 1680 |
| 734 | W303.asm02.HP0_G0007360 | chrIV | 1859 | 261 | 339 | 1758 | 276 | 344 | 1627 | 278 | 382 | 1718 | 316 | 487 | 1363 | 216 | 408 | 1376 | 290 | 403 |
| 735 | W303.asm02.HP0_G0007370 | chrIV | 2231 | 1928 | 2105 | 2439 | 2680 | 1894 | 2710 | 2170 | 1901 | 16358 | 2453 | 2779 | 14252 | 1543 | 1533 | 14632 | 2140 | 2535 |
| 736 | W303.asm02.HP0_G0007380 | chrIV | 13657 | 5735 | 6464 | 14840 | 5581 | 6397 | 13953 | 5536 | 7037 | 17251 | 4827 | 5838 | 17786 | 3378 | 5023 | 17250 | 4240 | 5112 |
| 737 | W303.asm02.HP0_G0007390 | chrIV | 2337 | 325 | 550 | 2144 | 375 | 595 | 2238 | 351 | 697 | 2467 | 534 | 760 | 2181 | 327 | 548 | 2249 | 472 | 632 |
| 738 | W303.asm02.HP0_G0007400 | chrIV | 15575 | 2475 | 2839 | 14453 | 2433 | 3371 | 15682 | 2183 | 3641 | 22716 | 5528 | 7613 | 10748 | 4559 | 5163 | 10359 | 4881 | 6822 |
| 739 | W303.asm02.HP0_G0007410 | chrIV | 2298 | 1151 | 995 | 2136 | 935 | 1073 | 2230 | 968 | 1280 | 3914 | 1708 | 1714 | 2720 | 1316 | 1441 | 2338 | 1470 | 1611 |
| 740 | W303.asm02.HP0_G0007420 | chrIV | 8850 | 3662 | 2863 | 8403 | 3237 | 2854 | 9059 | 3555 | 2881 | 16998 | 4678 | 5602 | 14301 | 3663 | 3871 | 13895 | 3972 | 4960 |
| 741 | W303.asm02.HP0_G0007430 | chrIV | 1467 | 738 | 601 | 1463 | 568 | 597 | 1327 | 652 | 590 | 1995 | 511 | 653 | 1946 | 356 | 564 | 1962 | 542 | 558 |
| 742 | W303.asm02.HP0_G0007440 | chrIV | 1374 | 388 | 442 | 1249 | 482 | 558 | 1446 | 395 | 507 | 5875 | 1008 | 1477 | 3279 | 668 | 959 | 3831 | 861 | 1388 |

| | | | | | | | | | | | | | | | | | | | | |
|---|---|---|---|---|---|---|---|---|---|---|---|---|---|---|---|---|---|---|---|---|
| 743 | W303.asm02.HP0_G0007450 | chrIV | 7185 | 984 | 1467 | 6482 | 1403 | 1764 | 7051 | 744 | 1933 | 11632 | 1899 | 4168 | 3040 | 1620 | 2658 | 3101 | 1503 | 3446 |
| 744 | W303.asm02.HP0_G0007460 | chrIV | 1021 | 513 | 192 | 977 | 224 | 232 | 1034 | 458 | 190 | 892 | 355 | 248 | 989 | 275 | 169 | 694 | 396 | 203 |
| 745 | W303.asm02.HP0_G0007470 | chrIV | 3076 | 285 | 384 | 2758 | 391 | 471 | 3042 | 258 | 537 | 2378 | 592 | 861 | 1121 | 463 | 627 | 1114 | 507 | 811 |
| 746 | W303.asm02.HP0_G0007480 | chrIV | 6495 | 5571 | 7669 | 6138 | 5791 | 8098 | 6258 | 5422 | 10206 | 12431 | 4944 | 6699 | 16383 | 3768 | 5549 | 13256 | 4522 | 6078 |
| 747 | W303.asm02.HP0_G0007490 | chrIV | 4985 | 4886 | 5822 | 4882 | 5779 | 5850 | 4819 | 5010 | 7076 | 11449 | 4499 | 4659 | 12927 | 3351 | 4067 | 12368 | 4075 | 4158 |
| 748 | W303.asm02.HP0_G0007500 | chrIV | 3475 | 2175 | 2528 | 3399 | 2177 | 2436 | 3236 | 2200 | 3069 | 6234 | 1767 | 1938 | 7512 | 1222 | 1681 | 6638 | 1609 | 1747 |
| 749 | W303.asm02.HP0_G0007510 | chrIV | 6070 | 3516 | 3075 | 5784 | 2846 | 2972 | 6235 | 3476 | 3120 | 9209 | 3043 | 2943 | 10101 | 2037 | 2567 | 8836 | 2944 | 2558 |
| 750 | W303.asm02.HP0_G0007520 | chrIV | 875 | 1312 | 867 | 926 | 880 | 794 | 753 | 1141 | 720 | 3581 | 1113 | 981 | 3663 | 800 | 742 | 3199 | 1199 | 844 |
| 751 | W303.asm02.HP0_G0007530 | chrIV | 8235 | 1479 | 1490 | 8454 | 1479 | 1692 | 8197 | 1506 | 1795 | 9676 | 2049 | 3363 | 7196 | 1586 | 2555 | 7304 | 1964 | 2935 |
| 752 | W303.asm02.HP0_G0007540 | chrIV | 4154 | 5573 | 3399 | 4303 | 3575 | 3521 | 4631 | 4790 | 3031 | 12393 | 4392 | 4507 | 11023 | 2987 | 2848 | 10598 | 3787 | 3970 |
| 753 | W303.asm02.HP0_G0007550 | chrIV | 5506 | 2020 | 2036 | 4967 | 1576 | 2161 | 4531 | 2037 | 2630 | 2548 | 1702 | 1803 | 5181 | 1448 | 2098 | 2639 | 1634 | 1453 |
| 754 | W303.asm02.HP0_G0007560 | chrIV | 2141 | 1837 | 2279 | 1727 | 1995 | 2139 | 1857 | 1720 | 2708 | 4265 | 1733 | 1552 | 5475 | 1246 | 1313 | 4595 | 1526 | 1424 |
| 755 | W303.asm02.HP0_G0007570 | chrIV | 1085 | 388 | 201 | 1236 | 234 | 204 | 1129 | 347 | 230 | 2168 | 603 | 328 | 1365 | 335 | 185 | 1561 | 643 | 315 |
| 756 | W303.asm02.HP0_G0007580 | chrIV | 6250 | 1512 | 1603 | 6361 | 1159 | 1611 | 6482 | 1338 | 2058 | 8286 | 2614 | 3003 | 6402 | 1976 | 2277 | 5623 | 2420 | 2721 |
| 757 | W303.asm02.HP0_G0007590 | chrIV | 1049 | 596 | 629 | 1158 | 558 | 629 | 1165 | 613 | 570 | 2377 | 687 | 1059 | 2364 | 441 | 576 | 2455 | 669 | 853 |
| 758 | W303.asm02.HP0_G0007600 | chrIV | 37940 | 5403 | 5202 | 37361 | 4615 | 6019 | 37947 | 4562 | 6635 | 28955 | 9610 | 12421 | 17114 | 7971 | 9535 | 16190 | 8630 | 11158 |
| 759 | W303.asm02.HP0_G0007610 | chrIV | 6651 | 2765 | 4361 | 6159 | 2426 | 4632 | 6164 | 2752 | 4395 | 5559 | 2504 | 4689 | 6445 | 1592 | 3495 | 5729 | 2358 | 3884 |
| 760 | W303.asm02.HP0_G0007620 | chrIV | 23355 | 4542 | 5668 | 21843 | 2940 | 7086 | 22307 | 4179 | 5686 | 6525 | 4663 | 10425 | 8105 | 4067 | 7753 | 5990 | 4391 | 8593 |
| 761 | W303.asm02.HP0_G0007630 | chrIV | 1715 | 427 | 278 | 1718 | 202 | 313 | 1750 | 308 | 332 | 1610 | 639 | 588 | 1466 | 417 | 447 | 1191 | 701 | 528 |
| 762 | W303.asm02.HP0_G0007640 | chrIV | 4992 | 1980 | 2217 | 5228 | 2594 | 2249 | 5488 | 2002 | 2191 | 15781 | 2444 | 3678 | 14309 | 1709 | 2435 | 12651 | 2204 | 3183 |
| 763 | W303.asm02.HP0_G0007650 | chrIV | 1012 | 1388 | 1838 | 1002 | 1685 | 1843 | 1042 | 1464 | 2585 | 4756 | 1460 | 1548 | 5510 | 843 | 1004 | 5247 | 1192 | 1365 |
| 764 | W303.asm02.HP0_G0007660 | chrIV | 3461 | 1926 | 2303 | 3252 | 1934 | 2411 | 3124 | 2067 | 2799 | 5426 | 1819 | 2157 | 5623 | 1142 | 1650 | 5367 | 1561 | 1961 |
| 765 | W303.asm02.HP0_G0007670 | chrIV | 2037 | 1036 | 896 | 1715 | 852 | 906 | 1639 | 1051 | 997 | 2587 | 740 | 849 | 4864 | 695 | 864 | 2372 | 753 | 789 |
| 766 | W303.asm02.HP0_G0007680 | chrIV | 1857 | 956 | 1035 | 1797 | 1057 | 982 | 1988 | 1032 | 1204 | 9241 | 1399 | 2084 | 9555 | 1135 | 1660 | 6925 | 1293 | 1748 |
| 767 | W303.asm02.HP0_G0007690 | chrIV | 2458 | 1432 | 1322 | 2260 | 1412 | 1254 | 2402 | 1432 | 1220 | 7554 | 1457 | 1994 | 6344 | 993 | 1409 | 6681 | 1343 | 1802 |
| 768 | W303.asm02.HP0_G0007700 | chrIV | 3516 | 908 | 898 | 3741 | 915 | 893 | 3252 | 857 | 998 | 3322 | 739 | 977 | 3283 | 486 | 829 | 3087 | 722 | 928 |
| 769 | W303.asm02.HP0_G0007710 | chrIV | 56 | 20 | 38 | 49 | 25 | 36 | 46 | 24 | 48 | 49 | 21 | 65 | 53 | 15 | 26 | 40 | 27 | 54 |
| 770 | W303.asm02.HP0_G0007720 | chrIV | 3633 | 550 | 610 | 3710 | 550 | 551 | 3425 | 527 | 604 | 2907 | 705 | 855 | 2918 | 467 | 756 | 2602 | 744 | 774 |
| 771 | W303.asm02.HP0_G0007730 | chrIV | 3016 | 906 | 1318 | 3127 | 1592 | 1283 | 3531 | 1102 | 1662 | 14051 | 1698 | 2955 | 8663 | 1087 | 1601 | 10094 | 1426 | 2498 |
| 772 | W303.asm02.HP0_G0007740 | chrIV | 1538 | 1500 | 1872 | 1493 | 1907 | 1837 | 1641 | 1581 | 1671 | 6768 | 1411 | 2329 | 7127 | 1064 | 1476 | 6663 | 1351 | 1959 |
| 773 | W303.asm02.HP0_G0007750 | chrIV | 6222 | 1719 | 1579 | 5582 | 1187 | 1753 | 5344 | 1461 | 1635 | 2332 | 1245 | 1469 | 3513 | 1031 | 1598 | 2000 | 1346 | 1233 |
| 774 | W303.asm02.HP0_G0007760 | chrIV | 717 | 93 | 167 | 632 | 121 | 163 | 660 | 73 | 197 | 947 | 163 | 315 | 317 | 104 | 179 | 328 | 194 | 279 |
| 775 | W303.asm02.HP0_G0007770 | chrIV | 1595 | 949 | 797 | 1551 | 776 | 752 | 1635 | 921 | 837 | 2887 | 807 | 857 | 2823 | 540 | 656 | 2825 | 769 | 844 |
| 776 | W303.asm02.HP0_G0007780 | chrIV | 4209 | 966 | 1089 | 4083 | 953 | 1068 | 3834 | 928 | 1283 | 2495 | 661 | 800 | 3219 | 483 | 770 | 2518 | 644 | 663 |
| 777 | W303.asm02.HP0_G0007790 | chrIV | 4824 | 973 | 726 | 5196 | 731 | 719 | 4925 | 851 | 646 | 3585 | 733 | 1052 | 3368 | 556 | 744 | 3187 | 711 | 927 |
| 778 | W303.asm02.HP0_G0007800 | chrIV | 486 | 352 | 340 | 495 | 360 | 326 | 447 | 389 | 293 | 1340 | 272 | 379 | 1438 | 153 | 236 | 1354 | 253 | 325 |
| 779 | W303.asm02.HP0_G0007810 | chrIV | 1294 | 530 | 667 | 1389 | 587 | 655 | 1206 | 575 | 810 | 1961 | 355 | 515 | 2291 | 287 | 421 | 1903 | 332 | 472 |
| 780 | W303.asm02.HP0_G0007820 | chrIV | 4651 | 654 | 821 | 3983 | 691 | 886 | 4402 | 619 | 1094 | 5024 | 1149 | 1735 | 3590 | 854 | 1265 | 3581 | 1115 | 1507 |
| 781 | W303.asm02.HP0_G0007830 | chrIV | 1825 | 806 | 941 | 1715 | 1016 | 833 | 1903 | 855 | 948 | 6381 | 1076 | 1584 | 5666 | 738 | 1044 | 5340 | 895 | 1406 |
| 782 | W303.asm02.HP0_G0007840 | chrIV | 2160 | 1207 | 1005 | 2095 | 979 | 931 | 2109 | 1252 | 900 | 3049 | 1092 | 992 | 3315 | 690 | 754 | 3034 | 913 | 845 |
| 783 | W303.asm02.HP0_G0007850 | chrIV | 12076 | 6173 | 5734 | 11829 | 5104 | 6197 | 12288 | 6142 | 5360 | 15246 | 5509 | 6851 | 15357 | 3936 | 4831 | 14576 | 4900 | 6181 |
| 784 | W303.asm02.HP0_G0007860 | chrIV | 7207 | 2061 | 1543 | 7063 | 1648 | 1763 | 7014 | 2135 | 1526 | 5258 | 1688 | 1884 | 5388 | 1172 | 1609 | 4620 | 1489 | 1802 |
| 785 | W303.asm02.HP0_G0007870 | chrIV | 612 | 549 | 643 | 580 | 544 | 666 | 456 | 623 | 764 | 1739 | 399 | 492 | 1815 | 221 | 315 | 1955 | 430 | 406 |
| 786 | W303.asm02.HP0_G0007880 | chrIV | 13634 | 5206 | 4517 | 13832 | 4134 | 4617 | 13729 | 4952 | 4318 | 14416 | 4866 | 6113 | 14569 | 3504 | 4764 | 13371 | 4418 | 5528 |
| 787 | W303.asm02.HP0_G0007890 | chrIV | 5075 | 1598 | 2906 | 4496 | 1610 | 3436 | 5130 | 1456 | 3673 | 10286 | 2842 | 7345 | 4625 | 2446 | 3881 | 4365 | 2554 | 6327 |
| 788 | W303.asm02.HP0_G0007900 | chrIV | 1560 | 597 | 699 | 1407 | 587 | 680 | 1633 | 548 | 682 | 3959 | 823 | 1239 | 2949 | 532 | 742 | 3119 | 765 | 1124 |
| 789 | W303.asm02.HP0_G0007910 | chrIV | 2331 | 1663 | 2223 | 2213 | 1962 | 2277 | 2231 | 1809 | 2941 | 5822 | 1500 | 1801 | 6519 | 916 | 1525 | 5969 | 1324 | 1647 |
| 790 | W303.asm02.HP0_G0007920 | chrIV | 4283 | 710 | 567 | 4216 | 529 | 658 | 4119 | 655 | 567 | 2181 | 568 | 735 | 2111 | 367 | 637 | 1952 | 529 | 679 |
| 791 | W303.asm02.HP0_G0007930 | chrIV | 1571 | 1047 | 873 | 1371 | 878 | 834 | 1435 | 1023 | 771 | 3100 | 815 | 821 | 3140 | 491 | 616 | 3107 | 736 | 801 |
| 792 | W303.asm02.HP0_G0007940 | chrIV | 2188 | 853 | 963 | 2133 | 692 | 1022 | 2069 | 782 | 1057 | 3136 | 627 | 956 | 3357 | 461 | 746 | 3285 | 665 | 888 |
| 793 | W303.asm02.HP0_G0007950 | chrIV | 4384 | 1542 | 1625 | 4486 | 1726 | 1648 | 4359 | 1557 | 1669 | 6909 | 1483 | 1910 | 6886 | 995 | 1508 | 6110 | 1436 | 1627 |
| 794 | W303.asm02.HP0_G0007960 | chrIV | 752 | 303 | 204 | 740 | 192 | 171 | 703 | 233 | 219 | 1244 | 325 | 346 | 878 | 183 | 240 | 789 | 367 | 287 |
| 795 | W303.asm02.HP0_G0007970 | chrIV | 8259 | 1438 | 947 | 7330 | 916 | 1198 | 7968 | 1356 | 1062 | 5253 | 2059 | 2109 | 4955 | 1514 | 1771 | 4280 | 1896 | 1874 |
| 796 | W303.asm02.HP0_G0007980 | chrIV | 483 | 453 | 687 | 419 | 629 | 712 | 404 | 529 | 874 | 1594 | 378 | 548 | 1958 | 224 | 462 | 1789 | 337 | 394 |
| 797 | W303.asm02.HP0_G0007990 | chrIV | 585 | 137 | 39 | 621 | 38 | 53 | 541 | 99 | 47 | 156 | 132 | 57 | 132 | 135 | 60 | 105 | 139 | 65 |
| 798 | W303.asm02.HP0_G0008000 | chrIV | 11462 | 10251 | 9071 | 10266 | 8240 | 8588 | 11696 | 10591 | 8934 | 28671 | 9620 | 9898 | 28752 | 7069 | 7751 | 28569 | 9082 | 8946 |
| 799 | W303.asm02.HP0_G0008010 | chrIV | 10701 | 8307 | 11646 | 9484 | 10855 | 11125 | 10639 | 8456 | 13671 | 23559 | 7803 | 9351 | 25163 | 5761 | 7684 | 24134 | 6818 | 8331 |
| 800 | W303.asm02.HP0_G0008020 | chrIV | 13718 | 10319 | 9819 | 12628 | 10094 | 9848 | 13641 | 10435 | 9820 | 23652 | 9010 | 9271 | 23501 | 6919 | 7080 | 23034 | 8073 | 8386 |
| 801 | W303.asm02.HP0_G0008030 | chrIV | 8123 | 7006 | 8792 | 7378 | 8044 | 8829 | 7794 | 7338 | 10796 | 15099 | 6629 | 7490 | 17608 | 5009 | 6148 | 15894 | 5915 | 6497 |
| 802 | W303.asm02.HP0_G0008040 | chrIV | 1981 | 469 | 464 | 1741 | 418 | 454 | 2152 | 465 | 585 | 3201 | 805 | 971 | 1669 | 498 | 635 | 1710 | 766 | 877 |
| 803 | W303.asm02.HP0_G0008050 | chrIV | 2795 | 421 | 246 | 3354 | 265 | 225 | 2637 | 398 | 291 | 1148 | 371 | 256 | 1109 | 225 | 224 | 995 | 369 | 254 |
| 804 | W303.asm02.HP0_G0008060 | chrIV | 34 | 6 | 3 | 13 | 4 | 9 | 21 | 10 | 6 | 19 | 4 | 7 | 38 | 5 | 10 | 32 | 5 | 3 |
| 805 | W303.asm02.HP0_G0008070 | chrIV | 9868 | 3346 | 3050 | 9322 | 2809 | 3015 | 9560 | 3323 | 3248 | 9212 | 3282 | 3727 | 9327 | 2406 | 2996 | 8537 | 3037 | 3270 |
| 806 | W303.asm02.HP0_G0008080 | chrIV | 2942 | 477 | 461 | 2951 | 553 | 550 | 2695 | 493 | 605 | 4508 | 656 | 1002 | 3391 | 518 | 818 | 3477 | 723 | 892 |
| 807 | W303.asm02.HP0_G0008090 | chrIV | 3584 | 1155 | 827 | 3265 | 809 | 817 | 3139 | 1127 | 820 | 3394 | 1114 | 841 | 3532 | 694 | 851 | 3213 | 1067 | 825 |
| 808 | W303.asm02.HP0_G0008100 | chrIV | 6577 | 6213 | 8297 | 6577 | 7759 | 8261 | 6841 | 6168 | 10176 | 15995 | 6027 | 6954 | 18247 | 4179 | 5949 | 16816 | 5233 | 6134 |
| 809 | W303.asm02.HP0_G0008110 | chrIV | 3577 | 2189 | 2113 | 3578 | 2306 | 2219 | 3799 | 2388 | 2205 | 12012 | 2304 | 2601 | 10247 | 1475 | 1929 | 10520 | 2073 | 2356 |
| 810 | W303.asm02.HP0_G0008120 | chrIV | 961 | 1286 | 1766 | 850 | 1693 | 1502 | 972 | 1445 | 1867 | 8939 | 1466 | 1939 | 7482 | 812 | 1496 | 8856 | 1513 | 1840 |
| 811 | W303.asm02.HP0_G0008130 | chrIV | 8152 | 2783 | 2585 | 7690 | 2012 | 2483 | 7478 | 2843 | 3064 | 6288 | 2504 | 2663 | 9211 | 2064 | 2585 | 6614 | 2324 | 2265 |
| 812 | W303.asm02.HP0_G0008140 | chrIV | 6673 | 1122 | 1762 | 6397 | 1534 | 1806 | 6602 | 1068 | 1994 | 9321 | 1481 | 2302 | 8029 | 859 | 2025 | 8282 | 1404 | 2171 |
| 813 | W303.asm02.HP0_G0008150 | chrIV | 316 | 732 | 758 | 301 | 584 | 758 | 322 | 622 | 959 | 1496 | 491 | 527 | 1780 | 361 | 400 | 1731 | 535 | 424 |
| 814 | W303.asm02.HP0_G0008160 | chrIV | 3123 | 1808 | 2629 | 2522 | 2609 | 2651 | 2526 | 2105 | 3307 | 4695 | 1844 | 2169 | 5826 | 1500 | 2110 | 4936 | 1604 | 1688 |
| 815 | W303.asm02.HP0_G0008170 | chrIV | 3364 | 1382 | 1583 | 2621 | 1424 | 1746 | 3184 | 1483 | 1897 | 7517 | 1790 | 2703 | 7631 | 1545 | 2192 | 6330 | 1426 | 2248 |
| 816 | W303.asm02.HP0_G0008180 | chrIV | 3536 | 1832 | 1750 | 3621 | 1757 | 1682 | 3464 | 1973 | 1821 | 5475 | 1643 | 1584 | 6558 | 1130 | 1433 | 5611 | 1536 | 1545 |
| 817 | W303.asm02.HP0_G0008190 | chrIV | 2428 | 1288 | 1436 | 2048 | 1448 | 1330 | 2157 | 1327 | 1525 | 4727 | 1278 | 1436 | 4993 | 878 | 1169 | 4893 | 1139 | 1355 |
| 818 | W303.asm02.HP0_G0008200 | chrIV | 975 | 1319 | 2078 | 974 | 2492 | 1724 | 1214 | 1599 | 2078 | 13719 | 1674 | 2249 | 11616 | 1070 | 1186 | 12249 | 1492 | 1886 |
| 819 | W303.asm02.HP0_G0008210 | chrIV | 4728 | 2906 | 3701 | 4449 | 3464 | 3610 | 4557 | 2887 | 4150 | 8065 | 2707 | 3491 | 8944 | 2158 | 2933 | 8838 | 2567 | 3014 |
| 820 | W303.asm02.HP0_G0008220 | chrIV | 3169 | 1143 | 1313 | 3008 | 1719 | 1361 | 3562 | 1210 | 1611 | 15995 | 2356 | 3222 | 10231 | 1496 | 1658 | 10316 | 2025 | 2862 |
| 821 | W303.asm02.HP0_G0008230 | chrIV | 765 | 307 | 406 | 627 | 338 | 373 | 610 | 346 | 407 | 976 | 282 | 315 | 1073 | 202 | 270 | 940 | 258 | 260 |
| 822 | W303.asm02.HP0_G0008240 | chrIV | 2969 | 835 | 1176 | 2677 | 1261 | 1114 | 2957 | 863 | 1200 | 8730 | 906 | 1758 | 7382 | 626 | 1360 | 7311 | 877 | 1487 |
| 823 | W303.asm02.HP0_G0008250 | chrIV | 1743 | 270 | 233 | 1613 | 188 | 209 | 1549 | 261 | 244 | 987 | 268 | 323 | 1111 | 243 | 333 | 720 | 256 | 271 |
| 824 | W303.asm02.HP0_G0008260 | chrIV | 3523 | 413 | 538 | 3546 | 585 | 584 | 3300 | 455 | 741 | 4775 | 868 | 1231 | 2988 | 579 | 937 | 3165 | 793 | 1103 |
| 825 | W303.asm02.HP0_G0008270 | chrIV | 27996 | 19045 | 23080 | 26565 | 19841 | 21834 | 29444 | 20453 | 26383 | 49340 | 18596 | 21280 | 59148 | 15241 | 17404 | 50511 | 16418 | 19453 |
| 826 | W303.asm02.HP0_G0008280 | chrIV | 1655 | 856 | 667 | 1780 | 679 | 629 | 1660 | 824 | 626 | 2912 | 888 | 757 | 2560 | 534 | 602 | 2449 | 870 | 671 |
| 827 | W303.asm02.HP0_G0008290 | chrIV | 2870 | 1651 | 1663 | 3000 | 1596 | 1652 | 2965 | 1834 | 1916 | 5065 | 1789 | 1882 | 6467 | 1131 | 1581 | 4943 | 1460 | 1682 |
| 828 | W303.asm02.HP0_G0008300 | chrIV | 3848 | 727 | 733 | 3176 | 485 | 830 | 3841 | 677 | 872 | 3657 | 1374 | 1615 | 2664 | 936 | 1112 | 2795 | 1292 | 1510 |
| 829 | W303.asm02.HP0_G0008310 | chrIV | 2505 | 3719 | 4088 | 2510 | 4324 | 3981 | 2673 | 4120 | 4114 | 13749 | 3899 | 3990 | 13481 | 2468 | 2549 | 13431 | 3528 | 3371 |
| 830 | W303.asm02.HP0_G0008320 | chrIV | 368 | 433 | 570 | 362 | 470 | 542 | 371 | 455 | 621 | 1788 | 408 | 513 | 2396 | 338 | 373 | 1572 | 459 | 459 |
| 831 | W303.asm02.HP0_G0008330 | chrIV | 3587 | 275 | 572 | 3906 | 367 | 721 | 3563 | 232 | 863 | 3300 | 535 | 935 | 804 | 311 | 534 | 681 | 452 | 586 |
| 832 | W303.asm02.HP0_G0008340 | chrIV | 4816 | 1171 | 1254 | 5115 | 1069 | 1299 | 5015 | 951 | 1613 | 8060 | 2161 | 2157 | 3770 | 1530 | 1570 | 3792 | 2017 | 2033 |
| 833 | W303.asm02.HP0_G0008350 | chrIV | 12142 | 4644 | 3911 | 11541 | 3738 | 3687 | 11763 | 4795 | 3787 | 9555 | 4584 | 4165 | 11930 | 3566 | 4140 | 8900 | 4167 | 3816 |
| 834 | W303.asm02.HP0_G0008360 | chrIV | 2344 | 1289 | 1336 | 2287 | 1073 | 1359 | 2066 | 1366 | 1461 | 2814 | 952 | 1278 | 3084 | 735 | 1008 | 2650 | 1000 | 1150 |
| 835 | W303.asm02.HP0_G0008370 | chrIV | 3885 | 883 | 932 | 3761 | 923 | 909 | 3723 | 837 | 897 | 3348 | 717 | 952 | 4066 | 546 | 880 | 3179 | 754 | 817 |

| | | | | | | | | | | | | | | | | | | | | |
|---|---|---|---|---|---|---|---|---|---|---|---|---|---|---|---|---|---|---|---|---|
| 836 | W303.asm02.HP0_G0008380 | chrIV | 1729 | 1009 | 975 | 1693 | 888 | 1032 | 1651 | 902 | 1052 | 2674 | 691 | 986 | 2923 | 452 | 682 | 2569 | 707 | 839 |
| 837 | W303.asm02.HP0_G0008390 | chrIV | 820 | 1061 | 1195 | 855 | 1166 | 1107 | 727 | 1063 | 1283 | 3649 | 795 | 871 | 3827 | 524 | 722 | 3508 | 678 | 757 |
| 838 | W303.asm02.HP0_G0008400 | chrIV | 643 | 393 | 600 | 664 | 513 | 657 | 588 | 309 | 687 | 3032 | 523 | 854 | 2066 | 398 | 585 | 1967 | 447 | 699 |
| 839 | W303.asm02.HP0_G0008410 | chrIV | 9584 | 1599 | 1372 | 8492 | 1293 | 1454 | 9418 | 1648 | 1475 | 12663 | 2526 | 3321 | 9875 | 1823 | 2184 | 10263 | 2276 | 2857 |
| 840 | W303.asm02.HP0_G0008420 | chrIV | 769 | 381 | 516 | 1054 | 641 | 574 | 779 | 349 | 396 | 5554 | 268 | 345 | 3163 | 163 | 196 | 2657 | 256 | 322 |
| 841 | W303.asm02.HP0_G0008430 | chrIV | 1937 | 741 | 952 | 1971 | 1056 | 866 | 1970 | 843 | 939 | 4966 | 889 | 1029 | 4300 | 516 | 660 | 4671 | 774 | 962 |
| 842 | W303.asm02.HP0_G0008440 | chrIV | 1820 | 715 | 975 | 1765 | 911 | 938 | 1665 | 758 | 1076 | 6216 | 1268 | 1661 | 4677 | 824 | 1296 | 4506 | 1091 | 1522 |
| 843 | W303.asm02.HP0_G0008450 | chrIV | 1517 | 320 | 282 | 1546 | 291 | 264 | 1547 | 249 | 299 | 1972 | 478 | 563 | 1058 | 327 | 411 | 1188 | 480 | 482 |
| 844 | W303.asm02.HP0_G0008460 | chrIV | 2732 | 1028 | 1022 | 2792 | 1372 | 1071 | 3345 | 1082 | 1218 | 13771 | 2250 | 2820 | 8757 | 1390 | 1389 | 9091 | 1849 | 2495 |
| 845 | W303.asm02.HP0_G0008470 | chrIV | 1289 | 614 | 683 | 1299 | 709 | 603 | 1244 | 599 | 695 | 3863 | 735 | 986 | 3008 | 445 | 626 | 3544 | 679 | 897 |
| 846 | W303.asm02.HP0_G0008480 | chrIV | 9516 | 3792 | 3682 | 9087 | 3138 | 3497 | 9019 | 3732 | 3834 | 11982 | 4063 | 4788 | 13760 | 3039 | 3959 | 11648 | 3678 | 4173 |
| 847 | W303.asm02.HP0_G0008490 | chrIV | 2332 | 419 | 474 | 2356 | 503 | 492 | 2264 | 425 | 521 | 4021 | 707 | 1092 | 3097 | 483 | 872 | 3146 | 685 | 964 |
| 848 | W303.asm02.HP0_G0008500 | chrIV | 3755 | 1173 | 1574 | 3415 | 1111 | 1634 | 3346 | 1189 | 1557 | 3534 | 1029 | 1771 | 4200 | 787 | 1502 | 3396 | 930 | 1444 |
| 849 | W303.asm02.HP0_G0008510 | chrIV | 2607 | 425 | 501 | 2859 | 388 | 617 | 2439 | 360 | 663 | 3060 | 1163 | 1112 | 1393 | 744 | 889 | 1310 | 1082 | 904 |
| 850 | W303.asm02.HP0_G0008520 | chrIV | 629 | 176 | 85 | 736 | 91 | 91 | 579 | 118 | 100 | 312 | 156 | 192 | 351 | 106 | 98 | 280 | 209 | 185 |
| 851 | W303.asm02.HP0_G0008530 | chrIV | 6223 | 4568 | 1617 | 6252 | 952 | 1906 | 5939 | 2829 | 2035 | 9238 | 6104 | 4758 | 5477 | 3151 | 3334 | 5972 | 7914 | 3949 |
| 852 | W303.asm02.HP0_G0008540 | chrIV | 1977 | 272 | 272 | 2041 | 186 | 314 | 1663 | 246 | 282 | 442 | 279 | 370 | 404 | 162 | 292 | 447 | 349 | 354 |
| 853 | W303.asm02.HP0_G0008550 | chrIV | 6473 | 2544 | 1613 | 7531 | 1863 | 2005 | 6335 | 2142 | 1420 | 6871 | 3542 | 2801 | 6560 | 1902 | 2261 | 7130 | 3595 | 2520 |
| 854 | W303.asm02.HP0_G0008560 | chrIV | 10403 | 756 | 869 | 10514 | 857 | 996 | 9962 | 639 | 1098 | 4915 | 1257 | 2210 | 1907 | 1054 | 1469 | 1933 | 1123 | 1866 |
| 855 | W303.asm02.HP0_G0008570 | chrIV | 1116 | 895 | 1058 | 1112 | 867 | 1093 | 1027 | 984 | 1392 | 2696 | 754 | 816 | 3602 | 518 | 649 | 2887 | 609 | 717 |
| 856 | W303.asm02.HP0_G0008580 | chrIV | 1414 | 277 | 239 | 1430 | 268 | 248 | 1292 | 205 | 250 | 2185 | 432 | 526 | 1416 | 288 | 355 | 1468 | 457 | 431 |
| 857 | W303.asm02.HP0_G0008590 | chrIV | 7682 | 5336 | 5232 | 7518 | 4784 | 5055 | 7496 | 5376 | 6202 | 14737 | 4688 | 4435 | 15534 | 3163 | 3555 | 15548 | 4192 | 4008 |
| 858 | W303.asm02.HP0_G0008600 | chrIV | 9882 | 4047 | 3878 | 9295 | 4334 | 4087 | 10384 | 3595 | 4927 | 31656 | 9240 | 11730 | 16682 | 6088 | 7110 | 19155 | 7378 | 9831 |
| 859 | W303.asm02.HP0_G0008610 | chrIV | 2468 | 526 | 386 | 2646 | 482 | 467 | 2457 | 426 | 445 | 3485 | 905 | 982 | 2150 | 624 | 675 | 1851 | 920 | 824 |
| 860 | W303.asm02.HP0_G0008620 | chrIV | 575 | 416 | 311 | 566 | 288 | 359 | 614 | 417 | 384 | 1084 | 358 | 293 | 1264 | 225 | 237 | 1326 | 319 | 265 |
| 861 | W303.asm02.HP0_G0008630 | chrIV | 1320 | 1034 | 1272 | 1043 | 1031 | 1369 | 1168 | 1139 | 1778 | 3067 | 913 | 1007 | 3362 | 575 | 788 | 3407 | 849 | 932 |
| 862 | W303.asm02.HP0_G0008640 | chrIV | 1 | 1 | 1 | 2 | 0 | 1 | 3 | 6 | 0 | 3 | 3 | 1 | 2 | 5 | 1 | 3 | 7 | 1 |
| 863 | W303.asm02.HP0_G0008650 | chrIV | 14773 | 5666 | 4819 | 15092 | 4469 | 4621 | 15071 | 5242 | 4860 | 16085 | 5140 | 5652 | 15352 | 3739 | 4781 | 15718 | 4742 | 5273 |
| 864 | W303.asm02.HP0_G0008660 | chrIV | 1152 | 951 | 1049 | 1075 | 937 | 1059 | 1149 | 906 | 1207 | 3296 | 779 | 1003 | 3377 | 468 | 800 | 3435 | 750 | 878 |
| 865 | W303.asm02.HP0_G0008670 | chrIV | 2750 | 1257 | 1163 | 3072 | 1191 | 980 | 2881 | 1427 | 1106 | 5785 | 1492 | 1546 | 6490 | 971 | 1222 | 5746 | 1407 | 1465 |
| 866 | W303.asm02.HP0_G0008680 | chrIV | 4236 | 4072 | 5339 | 3901 | 4853 | 5142 | 3826 | 4300 | 6588 | 10082 | 3845 | 4503 | 11962 | 3101 | 3923 | 10456 | 3361 | 3678 |
| 867 | W303.asm02.HP0_G0008690 | chrIV | 1346 | 1097 | 1270 | 1356 | 1294 | 1238 | 1534 | 1024 | 1298 | 6490 | 1451 | 2126 | 5592 | 831 | 1425 | 5802 | 1277 | 1929 |
| 868 | W303.asm02.HP0_G0008700 | chrIV | 1862 | 1668 | 1536 | 2018 | 1458 | 1566 | 1752 | 1685 | 1863 | 4453 | 1463 | 1214 | 4559 | 884 | 889 | 4647 | 1337 | 1120 |
| 869 | W303.asm02.HP0_G0008710 | chrIV | 995 | 1420 | 1146 | 988 | 1089 | 1010 | 856 | 1180 | 1243 | 4303 | 1102 | 1029 | 4236 | 697 | 771 | 4493 | 1201 | 919 |
| 870 | W303.asm02.HP0_G0008720 | chrIV | 20616 | 1062 | 1021 | 18558 | 864 | 1199 | 18867 | 871 | 1228 | 6678 | 2539 | 2764 | 4982 | 2038 | 2678 | 3380 | 2442 | 2312 |
| 871 | W303.asm02.HP0_G0008730 | chrIV | 11124 | 7431 | 8507 | 10067 | 7890 | 8137 | 10852 | 7668 | 11295 | 21455 | 7677 | 7375 | 25057 | 5081 | 6133 | 22169 | 6708 | 6407 |
| 872 | W303.asm02.HP0_G0008740 | chrIV | 2385 | 1390 | 733 | 2577 | 693 | 736 | 2134 | 1272 | 677 | 2085 | 1298 | 758 | 2714 | 922 | 734 | 1975 | 1316 | 703 |
| 873 | W303.asm02.HP0_G0008750 | chrIV | 1912 | 1548 | 1118 | 1888 | 1100 | 1100 | 1839 | 1341 | 1188 | 4281 | 1486 | 1202 | 4317 | 843 | 998 | 4163 | 1493 | 1039 |
| 874 | W303.asm02.HP0_G0008760 | chrIV | 2877 | 1140 | 1400 | 2631 | 1194 | 1290 | 2672 | 1032 | 1457 | 3203 | 933 | 1235 | 3421 | 586 | 1022 | 3318 | 831 | 1032 |
| 875 | W303.asm02.HP0_G0008770 | chrIV | 2600 | 370 | 870 | 2898 | 827 | 1012 | 2473 | 339 | 1182 | 6595 | 821 | 1817 | 2201 | 558 | 965 | 2054 | 750 | 1323 |
| 876 | W303.asm02.HP0_G0008780 | chrIV | 574 | 440 | 397 | 392 | 440 | 413 | 446 | 390 | 466 | 1405 | 380 | 410 | 1440 | 267 | 288 | 1312 | 396 | 315 |
| 877 | W303.asm02.HP0_G0008790 | chrIV | 679 | 109 | 137 | 734 | 118 | 127 | 689 | 135 | 129 | 764 | 196 | 213 | 623 | 147 | 133 | 526 | 176 | 190 |
| 878 | W303.asm02.HP0_G0008800 | chrIV | 3893 | 2985 | 3217 | 4057 | 2982 | 3565 | 3815 | 2865 | 3639 | 31742 | 6329 | 7648 | 17028 | 5507 | 5968 | 20165 | 5548 | 6589 |
| 879 | W303.asm02.HP0_G0008810 | chrIV | 18722 | 6993 | 5138 | 17650 | 5382 | 5380 | 20484 | 6347 | 5756 | 23279 | 8016 | 8938 | 20929 | 6425 | 6596 | 19965 | 6876 | 7897 |
| 880 | W303.asm02.HP0_G0008820 | chrIV | 1186 | 811 | 803 | 951 | 685 | 760 | 1120 | 820 | 1075 | 2010 | 599 | 557 | 2403 | 409 | 507 | 2326 | 600 | 502 |
| 881 | W303.asm02.HP0_G0008830 | chrIV | 1266 | 369 | 717 | 1301 | 589 | 665 | 1115 | 454 | 780 | 2736 | 364 | 601 | 2810 | 235 | 466 | 2769 | 356 | 544 |
| 882 | W303.asm02.HP0_G0008840 | chrIV | 233 | 46 | 41 | 213 | 41 | 39 | 190 | 49 | 65 | 123 | 37 | 51 | 216 | 32 | 33 | 118 | 36 | 41 |
| 883 | W303.asm02.HP0_G0008850 | chrIV | 568 | 364 | 531 | 485 | 417 | 514 | 472 | 396 | 682 | 1115 | 304 | 337 | 1283 | 190 | 326 | 1195 | 272 | 252 |
| 884 | W303.asm02.HP0_G0008860 | chrIV | 3699 | 596 | 1045 | 3850 | 978 | 1144 | 3564 | 584 | 1174 | 9772 | 1350 | 2309 | 5936 | 961 | 1606 | 6078 | 1197 | 1970 |
| 885 | W303.asm02.HP0_G0008870 | chrIV | 1063 | 99 | 56 | 1088 | 69 | 67 | 871 | 111 | 67 | 274 | 127 | 72 | 466 | 108 | 100 | 325 | 145 | 82 |
| 886 | W303.asm02.HP0_G0008880 | chrIV | 728 | 318 | 364 | 499 | 343 | 383 | 659 | 361 | 422 | 1056 | 414 | 475 | 1155 | 321 | 283 | 803 | 328 | 368 |
| 887 | W303.asm02.HP0_G0008890 | chrIV | 715 | 847 | 1011 | 645 | 1182 | 914 | 705 | 925 | 1229 | 4139 | 868 | 990 | 4543 | 528 | 582 | 4112 | 781 | 741 |
| 888 | W303.asm02.HP0_G0008900 | chrIV | 555 | 727 | 901 | 573 | 1084 | 830 | 701 | 812 | 991 | 4569 | 694 | 785 | 4327 | 416 | 392 | 4194 | 699 | 674 |
| 889 | W303.asm02.HP0_G0008910 | chrIV | 817 | 500 | 494 | 692 | 394 | 492 | 686 | 478 | 521 | 829 | 360 | 361 | 992 | 276 | 342 | 827 | 320 | 334 |
| 890 | W303.asm02.HP0_G0008920 | chrIV | 18005 | 9780 | 9987 | 17904 | 9590 | 9433 | 18091 | 9917 | 10479 | 32645 | 10363 | 10310 | 31633 | 7025 | 7766 | 32645 | 9173 | 9185 |
| 891 | W303.asm02.HP0_G0008930 | chrIV | 10637 | 5832 | 6912 | 10400 | 6896 | 6206 | 10863 | 5893 | 6150 | 16286 | 5782 | 7491 | 16964 | 4816 | 5840 | 15560 | 5074 | 6668 |
| 892 | W303.asm02.HP0_G0008940 | chrIV | 2493 | 886 | 1206 | 2371 | 1102 | 1167 | 2504 | 922 | 1073 | 5971 | 1139 | 1970 | 5506 | 733 | 1333 | 5157 | 999 | 1741 |
| 893 | W303.asm02.HP0_G0008950 | chrIV | 4093 | 884 | 1286 | 4048 | 1002 | 1339 | 3722 | 934 | 1417 | 6166 | 1169 | 2052 | 4990 | 743 | 1648 | 5350 | 1151 | 1827 |
| 894 | W303.asm02.HP0_G0008960 | chrIV | 14328 | 1249 | 1464 | 13939 | 1545 | 1617 | 14289 | 1383 | 1743 | 13769 | 2898 | 3875 | 10812 | 2362 | 3342 | 8332 | 2604 | 3486 |
| 895 | W303.asm02.HP0_G0008970 | chrIV | 1049 | 372 | 332 | 1125 | 361 | 294 | 888 | 396 | 355 | 1570 | 363 | 376 | 1464 | 259 | 298 | 1483 | 362 | 342 |
| 896 | W303.asm02.HP0_G0008980 | chrIV | 1249 | 520 | 506 | 1012 | 475 | 575 | 947 | 464 | 808 | 1158 | 370 | 396 | 1609 | 253 | 375 | 1268 | 383 | 324 |
| 897 | W303.asm02.HP0_G0008990 | chrIV | 7644 | 3937 | 3753 | 7054 | 3380 | 3644 | 6989 | 3686 | 4420 | 7116 | 4094 | 3851 | 10292 | 3283 | 3747 | 7066 | 3618 | 3395 |
| 898 | W303.asm02.HP0_G0009000 | chrIV | 9476 | 28531 | 29687 | 9064 | 25852 | 28751 | 9328 | 28010 | 37429 | 53436 | 24788 | 22854 | 54016 | 18094 | 18168 | 53741 | 21351 | 19759 |
| 899 | W303.asm02.HP0_G0009010 | chrIV | 2336 | 666 | 789 | 2550 | 756 | 771 | 2430 | 725 | 808 | 5139 | 875 | 1243 | 3953 | 584 | 905 | 4265 | 773 | 1150 |
| 900 | W303.asm02.HP0_G0009020 | chrIV | 974 | 135 | 73 | 1035 | 65 | 74 | 817 | 132 | 101 | 107 | 89 | 49 | 456 | 85 | 78 | 121 | 82 | 36 |
| 901 | W303.asm02.HP0_G0009030 | chrIV | 8873 | 4922 | 5273 | 8862 | 5129 | 4780 | 8757 | 4962 | 5340 | 16613 | 5227 | 5611 | 16799 | 3614 | 4205 | 15394 | 4241 | 4916 |
| 902 | W303.asm02.HP0_G0009040 | chrIV | 6439 | 645 | 678 | 6137 | 633 | 824 | 6348 | 588 | 841 | 5448 | 1153 | 1577 | 4574 | 888 | 1105 | 4080 | 1074 | 1378 |
| 903 | W303.asm02.HP0_G0009050 | chrIV | 14337 | 6790 | 5849 | 13136 | 5874 | 5975 | 15075 | 6620 | 6785 | 37433 | 10553 | 12471 | 29112 | 8938 | 8122 | 28297 | 9243 | 10511 |
| 904 | W303.asm02.HP0_G0009060 | chrIV | 333 | 146 | 207 | 380 | 203 | 241 | 368 | 142 | 283 | 1293 | 264 | 414 | 705 | 201 | 280 | 676 | 183 | 356 |
| 905 | W303.asm02.HP0_G0009070 | chrIV | 0 | 0 | 0 | 0 | 0 | 0 | 0 | 0 | 0 | 0 | 0 | 0 | 0 | 0 | 0 | 0 | 0 | 0 |
| 906 | W303.asm02.HP0_G0009080 | chrIV | 1 | 0 | 0 | 0 | 0 | 1 | 0 | 0 | 0 | 1 | 0 | 0 | 0 | 1 | 0 | 1 | 1 | 0 |
| 907 | W303.asm02.HP0_G0009090 | chrIV | 32 | 15 | 12 | 39 | 14 | 11 | 40 | 21 | 6 | 92 | 24 | 20 | 47 | 18 | 11 | 71 | 16 | 20 |
| 908 | W303.asm02.HP0_G0009100 | chrIV | 5999 | 2241 | 2408 | 6080 | 2137 | 2363 | 5681 | 2181 | 2984 | 5447 | 1735 | 2186 | 7370 | 1387 | 2033 | 5485 | 1756 | 1994 |
| 909 | W303.asm02.HP0_G0009110 | chrIV | 395 | 155 | 134 | 378 | 128 | 142 | 417 | 160 | 194 | 521 | 149 | 109 | 376 | 53 | 62 | 607 | 118 | 85 |
| 910 | W303.asm02.HP0_G0009120 | chrIV | 1412 | 730 | 583 | 1784 | 599 | 681 | 1341 | 622 | 663 | 1460 | 612 | 565 | 1833 | 308 | 509 | 1246 | 558 | 535 |
| 911 | W303.asm02.HP0_G0009130 | chrIV | 7722 | 430 | 492 | 7052 | 475 | 578 | 7940 | 448 | 544 | 5001 | 892 | 1193 | 4318 | 721 | 852 | 3954 | 786 | 1021 |
| 912 | W303.asm02.HP0_G0009140 | chrIV | 710 | 973 | 1480 | 689 | 1436 | 1494 | 725 | 1058 | 2191 | 4100 | 958 | 1273 | 4876 | 633 | 888 | 4923 | 898 | 1125 |
| 913 | W303.asm02.HP0_G0009150 | chrIV | 3739 | 2585 | 3130 | 2547 | 2768 | 3046 | 3034 | 2382 | 3541 | 13408 | 4463 | 4937 | 8357 | 3148 | 3606 | 8574 | 3850 | 4140 |
| 914 | W303.asm02.HP0_G0009160 | chrIV | 6282 | 2833 | 2789 | 5184 | 2312 | 2652 | 5358 | 2672 | 3133 | 9018 | 3220 | 3736 | 8165 | 1936 | 2737 | 8347 | 2944 | 3416 |
| 915 | W303.asm02.HP0_G0009170 | chrIV | 2043 | 601 | 608 | 1958 | 719 | 587 | 1963 | 580 | 633 | 5689 | 1082 | 1299 | 4007 | 606 | 844 | 4181 | 931 | 1114 |
| 916 | W303.asm02.HP0_G0009180 | chrIV | 34727 | 21085 | 24345 | 32623 | 20318 | 23980 | 35547 | 20167 | 27699 | 40591 | 19987 | 24575 | 40082 | 14524 | 19754 | 39698 | 18707 | 22501 |
| 917 | W303.asm02.HP0_G0009190 | chrIV | 2997 | 1718 | 1814 | 2931 | 1665 | 1699 | 2992 | 1621 | 2080 | 5247 | 1407 | 1585 | 5059 | 848 | 1182 | 5699 | 1402 | 1495 |
| 918 | W303.asm02.HP0_G0009200 | chrIV | 2459 | 833 | 861 | 2658 | 944 | 946 | 2772 | 817 | 937 | 9514 | 1416 | 2028 | 7604 | 1019 | 1502 | 7249 | 1240 | 1786 |
| 919 | W303.asm02.HP0_G0009210 | chrIV | 7429 | 3015 | 3013 | 8215 | 3074 | 2814 | 7553 | 3163 | 2912 | 10991 | 3013 | 3072 | 12623 | 2039 | 2642 | 10635 | 2556 | 2849 |
| 920 | W303.asm02.HP0_G0009220 | chrIV | 27561 | 4894 | 4990 | 27268 | 4719 | 4719 | 27179 | 5345 | 5678 | 16002 | 5857 | 8286 | 25156 | 6828 | 8260 | 13607 | 5517 | 7039 |
| 921 | W303.asm02.HP0_G0009230 | chrIV | 4432 | 2064 | 2433 | 4046 | 2219 | 2480 | 4226 | 2168 | 3158 | 5870 | 1898 | 2164 | 7133 | 1220 | 1747 | 6007 | 1613 | 1901 |
| 922 | W303.asm02.HP0_G0009240 | chrIV | 2441 | 728 | 768 | 2375 | 814 | 785 | 2385 | 777 | 794 | 4104 | 862 | 1077 | 3829 | 527 | 782 | 3696 | 765 | 928 |
| 923 | W303.asm02.HP0_G0009250 | chrIV | 1816 | 595 | 499 | 1581 | 371 | 433 | 1457 | 563 | 587 | 1009 | 463 | 388 | 966 | 281 | 342 | 1105 | 514 | 385 |
| 924 | W303.asm02.HP0_G0009260 | chrIV | 1272 | 597 | 747 | 1278 | 653 | 770 | 1078 | 701 | 863 | 1430 | 630 | 732 | 2054 | 445 | 667 | 1426 | 513 | 643 |
| 925 | W303.asm02.HP0_G0009270 | chrIV | 4270 | 2070 | 2773 | 4373 | 3449 | 2590 | 4902 | 2386 | 2866 | 27008 | 3702 | 5555 | 22454 | 2470 | 3207 | 21806 | 2922 | 4834 |
| 926 | W303.asm02.HP0_G0009280 | chrIV | 5601 | 1297 | 1189 | 5609 | 1135 | 1127 | 5262 | 1236 | 1157 | 5838 | 1578 | 2039 | 5453 | 1160 | 1735 | 4781 | 1443 | 1855 |
| 927 | W303.asm02.HP0_G0009290 | chrIV | 5061 | 2546 | 2800 | 5220 | 2848 | 2619 | 5166 | 2945 | 2796 | 16024 | 3432 | 3699 | 16537 | 2616 | 3258 | 14700 | 3073 | 3638 |
| 928 | W303.asm02.HP0_G0009300 | chrIV | 3935 | 1350 | 1994 | 3457 | 1625 | 2013 | 3487 | 1375 | 2375 | 4494 | 1119 | 1604 | 5463 | 815 | 1357 | 4743 | 991 | 1313 |

| | | | | | | | | | | | | | | | | | | | | |
|---|---|---|---|---|---|---|---|---|---|---|---|---|---|---|---|---|---|---|---|---|
| 929 | W303.asm02.HP0_G0009310 | chrIV | 8557 | 8852 | 11513 | 7486 | 11010 | 11208 | 8836 | 8926 | 13167 | 22454 | 7914 | 9332 | 23368 | 5879 | 7791 | 21986 | 6996 | 8516 |
| 930 | W303.asm02.HP0_G0009320 | chrIV | 1065 | 588 | 748 | 1196 | 683 | 669 | 1030 | 564 | 875 | 2500 | 459 | 554 | 2460 | 328 | 453 | 2639 | 445 | 471 |
| 931 | W303.asm02.HP0_G0009330 | chrIV | 737 | 460 | 574 | 697 | 481 | 571 | 644 | 515 | 771 | 1488 | 390 | 456 | 1749 | 248 | 349 | 1645 | 357 | 388 |
| 932 | W303.asm02.HP0_G0009340 | chrIV | 1976 | 968 | 1114 | 1749 | 966 | 1157 | 1674 | 891 | 1317 | 2627 | 696 | 772 | 3045 | 466 | 701 | 3021 | 669 | 699 |
| 933 | W303.asm02.HP0_G0009350 | chrIV | 3541 | 1807 | 1618 | 3444 | 1609 | 1656 | 3477 | 1896 | 1942 | 5268 | 1319 | 1322 | 5983 | 930 | 1163 | 5435 | 1377 | 1199 |
| 934 | W303.asm02.HP0_G0009360 | chrIV | 2018 | 730 | 944 | 2277 | 633 | 892 | 1862 | 729 | 843 | 3092 | 827 | 1416 | 2693 | 441 | 937 | 3109 | 836 | 1132 |
| 935 | W303.asm02.HP0_G0009370 | chrIV | 1545 | 337 | 200 | 1471 | 170 | 256 | 1323 | 299 | 245 | 318 | 234 | 142 | 427 | 115 | 172 | 343 | 257 | 110 |
| 936 | W303.asm02.HP0_G0009380 | chrIV | 5741 | 3442 | 4372 | 5688 | 4022 | 4490 | 5524 | 3340 | 5195 | 9727 | 2792 | 3430 | 10692 | 1818 | 2922 | 10324 | 2535 | 3139 |
| 937 | W303.asm02.HP0_G0009390 | chrIV | 2176 | 1196 | 1262 | 2361 | 1139 | 1148 | 2040 | 1100 | 1323 | 8959 | 1765 | 2056 | 7957 | 1084 | 1577 | 7077 | 1717 | 1932 |
| 938 | W303.asm02.HP0_G0009400 | chrIV | 1281 | 924 | 775 | 1166 | 783 | 721 | 1134 | 910 | 853 | 2731 | 802 | 719 | 3535 | 612 | 644 | 2452 | 737 | 674 |
| 939 | W303.asm02.HP0_G0009410 | chrIV | 27882 | 1660 | 1220 | 27556 | 919 | 1422 | 25856 | 1166 | 1453 | 7475 | 3098 | 2536 | 5456 | 1961 | 2484 | 4400 | 3007 | 2185 |
| 940 | W303.asm02.HP0_G0009420 | chrIV | 668 | 790 | 1394 | 672 | 1406 | 1340 | 750 | 837 | 1508 | 4642 | 763 | 1214 | 4693 | 484 | 733 | 4808 | 769 | 1047 |
| 941 | W303.asm02.HP0_G0009430 | chrIV | 851 | 269 | 281 | 801 | 239 | 304 | 848 | 225 | 396 | 1250 | 279 | 438 | 1103 | 244 | 315 | 933 | 300 | 406 |
| 942 | W303.asm02.HP0_G0009440 | chrIV | 61593 | 11707 | 10065 | 61928 | 8842 | 11079 | 66309 | 10941 | 14546 | 59375 | 25891 | 22801 | 83230 | 35084 | 32062 | 38511 | 21138 | 21427 |
| 943 | W303.asm02.HP0_G0009450 | chrIV | 972 | 355 | 330 | 708 | 282 | 350 | 845 | 301 | 413 | 769 | 271 | 192 | 686 | 144 | 165 | 1022 | 269 | 198 |
| 944 | W303.asm02.HP0_G0009460 | chrIV | 2046 | 824 | 932 | 1873 | 786 | 997 | 1731 | 791 | 1184 | 2204 | 600 | 779 | 2577 | 426 | 661 | 2472 | 587 | 626 |
| 945 | W303.asm02.HP0_G0009470 | chrIV | 647 | 331 | 435 | 555 | 338 | 394 | 534 | 324 | 655 | 1253 | 309 | 359 | 1400 | 193 | 277 | 1490 | 281 | 366 |
| 946 | W303.asm02.HP0_G0009480 | chrIV | 4512 | 876 | 836 | 4546 | 912 | 831 | 4243 | 989 | 1058 | 6103 | 964 | 1089 | 4865 | 598 | 904 | 5315 | 916 | 1018 |
| 947 | W303.asm02.HP0_G0009490 | chrIV | 1909 | 503 | 605 | 1857 | 711 | 696 | 1884 | 489 | 753 | 5941 | 920 | 1397 | 4055 | 590 | 963 | 4064 | 802 | 1159 |
| 948 | W303.asm02.HP0_G0009500 | chrIV | 936 | 655 | 752 | 817 | 782 | 665 | 819 | 573 | 758 | 2318 | 466 | 555 | 2206 | 295 | 451 | 2332 | 481 | 491 |
| 949 | W303.asm02.HP0_G0009510 | chrIV | 2303 | 1528 | 1931 | 2240 | 2559 | 1787 | 2590 | 1751 | 2300 | 9648 | 1695 | 1801 | 9672 | 921 | 1059 | 9380 | 1359 | 1610 |
| 950 | W303.asm02.HP0_G0009520 | chrIV | 1707 | 972 | 1281 | 1669 | 1049 | 1287 | 1631 | 978 | 1783 | 2651 | 757 | 919 | 3514 | 508 | 764 | 2983 | 684 | 847 |
| 951 | W303.asm02.HP0_G0009530 | chrIV | 7340 | 635 | 467 | 7090 | 532 | 465 | 6741 | 594 | 473 | 3129 | 767 | 767 | 5474 | 850 | 1003 | 2397 | 733 | 692 |
| 952 | W303.asm02.HP0_G0009540 | chrIV | 3223 | 2648 | 4076 | 3058 | 3342 | 4179 | 3123 | 2774 | 5479 | 8068 | 2475 | 3429 | 10872 | 1833 | 3018 | 8953 | 2110 | 3098 |
| 953 | W303.asm02.HP0_G0009550 | chrIV | 1675 | 2108 | 2204 | 1517 | 2267 | 2145 | 1842 | 2139 | 2241 | 8244 | 2006 | 2431 | 7781 | 1167 | 1353 | 7601 | 1883 | 2103 |
| 954 | W303.asm02.HP0_G0009560 | chrIV | 1061 | 778 | 776 | 1115 | 873 | 750 | 1026 | 789 | 726 | 4032 | 739 | 919 | 3534 | 470 | 649 | 3478 | 739 | 842 |
| 955 | W303.asm02.HP0_G0009570 | chrIV | 2500 | 766 | 748 | 1895 | 701 | 734 | 2093 | 803 | 772 | 2741 | 726 | 767 | 2957 | 572 | 786 | 2724 | 583 | 758 |
| 956 | W303.asm02.HP0_G0009580 | chrIV | 667 | 974 | 1161 | 730 | 1102 | 1042 | 694 | 992 | 1104 | 3681 | 824 | 968 | 3775 | 552 | 794 | 3562 | 753 | 940 |
| 957 | W303.asm02.HP0_G0009590 | chrIV | 7681 | 4679 | 4576 | 7730 | 4738 | 4099 | 8520 | 4966 | 4166 | 21039 | 4778 | 5442 | 19463 | 3136 | 3466 | 19744 | 4292 | 4940 |
| 958 | W303.asm02.HP0_G0009600 | chrIV | 2691 | 1690 | 1661 | 2508 | 1458 | 1763 | 2552 | 1672 | 1790 | 3278 | 1277 | 1463 | 3218 | 888 | 1060 | 3219 | 1214 | 1308 |
| 959 | W303.asm02.HP0_G0009610 | chrIV | 2962 | 438 | 744 | 2415 | 710 | 844 | 2916 | 411 | 827 | 6714 | 986 | 2069 | 2706 | 790 | 1341 | 2915 | 831 | 1904 |
| 960 | W303.asm02.HP0_G0009620 | chrIV | 3567 | 286 | 430 | 4387 | 403 | 522 | 3328 | 310 | 548 | 4124 | 861 | 1031 | 2716 | 502 | 917 | 2721 | 720 | 911 |
| 961 | W303.asm02.HP0_G0009630 | chrIV | 1861 | 705 | 749 | 1874 | 833 | 770 | 2080 | 758 | 932 | 7031 | 1140 | 1673 | 4632 | 803 | 1053 | 4884 | 1072 | 1388 |
| 962 | W303.asm02.HP0_G0009640 | chrIV | 2929 | 2537 | 2866 | 2432 | 2266 | 2824 | 2729 | 2534 | 3410 | 6410 | 2175 | 2820 | 7200 | 1517 | 2037 | 6677 | 1976 | 2644 |
| 963 | W303.asm02.HP0_G0009650 | chrIV | 15104 | 5904 | 7706 | 15000 | 6913 | 7976 | 15310 | 6113 | 7777 | 20880 | 5692 | 8079 | 23441 | 4217 | 7253 | 19998 | 5153 | 7360 |
| 964 | W303.asm02.HP0_G0009660 | chrIV | 1754 | 849 | 959 | 1417 | 730 | 989 | 1440 | 823 | 1215 | 1929 | 581 | 755 | 2498 | 459 | 748 | 2092 | 548 | 670 |
| 965 | W303.asm02.HP0_G0009670 | chrIV | 1953 | 1631 | 2213 | 1916 | 2930 | 2008 | 2176 | 1791 | 2344 | 16464 | 2598 | 3641 | 13343 | 1643 | 1940 | 13869 | 2220 | 3205 |
| 966 | W303.asm02.HP0_G0009680 | chrIV | 2990 | 421 | 722 | 2709 | 614 | 771 | 2871 | 384 | 915 | 5419 | 847 | 1268 | 3421 | 538 | 967 | 3637 | 797 | 1134 |
| 967 | W303.asm02.HP0_G0009690 | chrIV | 834 | 192 | 233 | 962 | 221 | 235 | 830 | 165 | 274 | 2068 | 286 | 544 | 1431 | 224 | 363 | 1371 | 286 | 498 |
| 968 | W303.asm02.HP0_G0009700 | chrIV | 3018 | 992 | 843 | 2658 | 841 | 906 | 2805 | 1001 | 995 | 4125 | 1130 | 1251 | 3427 | 703 | 1034 | 3600 | 1102 | 1228 |
| 969 | W303.asm02.HP0_G0009710 | chrIV | 274 | 103 | 54 | 187 | 41 | 52 | 201 | 83 | 96 | 216 | 79 | 62 | 213 | 43 | 70 | 241 | 84 | 64 |
| 970 | W303.asm02.HP0_G0009720 | chrIV | 797 | 498 | 472 | 743 | 483 | 446 | 712 | 533 | 437 | 2211 | 533 | 557 | 2424 | 359 | 459 | 2013 | 457 | 482 |
| 971 | W303.asm02.HP0_G0009730 | chrIV | 977 | 731 | 670 | 854 | 747 | 687 | 1012 | 731 | 714 | 3730 | 708 | 890 | 3601 | 463 | 641 | 3636 | 629 | 836 |
| 972 | W303.asm02.HP0_G0009740 | chrIV | 1225 | 277 | 325 | 1092 | 333 | 341 | 1217 | 285 | 368 | 2398 | 537 | 754 | 1207 | 373 | 506 | 1311 | 510 | 649 |
| 973 | W303.asm02.HP0_G0009750 | chrIV | 1187 | 640 | 757 | 1051 | 868 | 657 | 1216 | 715 | 786 | 4882 | 848 | 1054 | 3517 | 522 | 650 | 4531 | 767 | 1006 |
| 974 | W303.asm02.HP0_G0009760 | chrIV | 2160 | 660 | 609 | 2005 | 567 | 564 | 2291 | 760 | 726 | 7254 | 1336 | 1269 | 4626 | 599 | 706 | 5641 | 1137 | 1143 |
| 975 | W303.asm02.HP0_G0009770 | chrIV | 1549 | 307 | 259 | 1544 | 229 | 284 | 1519 | 280 | 244 | 1407 | 426 | 507 | 1264 | 274 | 312 | 1188 | 362 | 437 |
| 976 | W303.asm02.HP0_G0009780 | chrIV | 1595 | 658 | 875 | 1138 | 727 | 828 | 1269 | 642 | 1133 | 1372 | 538 | 585 | 1934 | 418 | 491 | 1456 | 504 | 540 |
| 977 | W303.asm02.HP0_G0009790 | chrIV | 9460 | 1947 | 1695 | 8771 | 1879 | 1833 | 8975 | 2025 | 1782 | 5395 | 1582 | 1894 | 6742 | 1298 | 1732 | 4499 | 1434 | 1783 |
| 978 | W303.asm02.HP0_G0009800 | chrIV | 2693 | 986 | 1153 | 2554 | 895 | 1275 | 2691 | 1032 | 1034 | 5232 | 1095 | 2140 | 4690 | 769 | 1351 | 4745 | 1009 | 1877 |
| 979 | W303.asm02.HP0_G0009810 | chrIV | 1141 | 427 | 482 | 1105 | 482 | 465 | 1130 | 453 | 514 | 2288 | 399 | 561 | 2132 | 264 | 477 | 2427 | 392 | 501 |
| 980 | W303.asm02.HP0_G0009820 | chrIV | 2884 | 1577 | 1878 | 2428 | 1907 | 1713 | 2886 | 1811 | 2135 | 10614 | 1966 | 2252 | 8112 | 1318 | 1723 | 9113 | 1769 | 2220 |
| 981 | W303.asm02.HP0_G0009830 | chrIV | 881 | 61 | 47 | 946 | 26 | 39 | 781 | 39 | 40 | 82 | 35 | 19 | 64 | 16 | 17 | 54 | 39 | 17 |
| 982 | W303.asm02.HP0_G0009840 | chrIV | 2272 | 1202 | 1157 | 2516 | 1479 | 1152 | 2665 | 1283 | 1422 | 11781 | 2627 | 2587 | 7979 | 1529 | 1354 | 8592 | 2164 | 2168 |
| 983 | W303.asm02.HP0_G0009850 | chrIV | 1578 | 955 | 788 | 1387 | 1011 | 891 | 1460 | 1004 | 886 | 3064 | 822 | 935 | 3190 | 507 | 637 | 2876 | 711 | 850 |
| 984 | W303.asm02.HP0_G0009860 | chrIV | 5923 | 443 | 567 | 5140 | 476 | 748 | 5452 | 398 | 861 | 2264 | 662 | 1263 | 1287 | 598 | 963 | 934 | 534 | 1046 |
| 985 | W303.asm02.HP0_G0009870 | chrIV | 1185 | 332 | 397 | 1853 | 324 | 417 | 1053 | 279 | 410 | 2666 | 560 | 1022 | 2150 | 388 | 571 | 2056 | 477 | 860 |
| 986 | W303.asm02.HP0_G0009880 | chrIV | 901 | 171 | 186 | 944 | 147 | 176 | 862 | 190 | 181 | 1125 | 296 | 359 | 997 | 140 | 213 | 926 | 253 | 253 |
| 987 | W303.asm02.HP0_G0009890 | chrIV | 1335 | 238 | 218 | 1287 | 225 | 209 | 1164 | 223 | 295 | 1267 | 275 | 292 | 1099 | 181 | 249 | 1094 | 320 | 272 |
| 988 | W303.asm02.HP0_G0009900 | chrIV | 578 | 684 | 370 | 482 | 404 | 372 | 550 | 601 | 373 | 1209 | 480 | 366 | 1396 | 300 | 209 | 1227 | 463 | 272 |
| 989 | W303.asm02.HP0_G0009910 | chrIV | 13024 | 1923 | 2317 | 10940 | 2080 | 2544 | 12135 | 1583 | 3174 | 10512 | 3230 | 5846 | 3712 | 2957 | 3413 | 3530 | 2388 | 4725 |
| 990 | W303.asm02.HP0_G0009920 | chrIV | 2270 | 473 | 689 | 2182 | 592 | 688 | 2074 | 431 | 816 | 4765 | 828 | 1394 | 2772 | 619 | 871 | 2705 | 733 | 1252 |
| 991 | W303.asm02.HP0_G0009930 | chrIV | 24183 | 2394 | 1973 | 24730 | 1904 | 2044 | 23501 | 2120 | 2573 | 14628 | 4614 | 4879 | 8932 | 3611 | 3825 | 8673 | 4205 | 4227 |
| 992 | W303.asm02.HP0_G0009940 | chrIV | 1371 | 497 | 384 | 1339 | 369 | 432 | 1344 | 411 | 388 | 1448 | 442 | 516 | 1227 | 332 | 420 | 1241 | 384 | 494 |
| 993 | W303.asm02.HP0_G0009950 | chrIV | 1538 | 584 | 503 | 1459 | 479 | 491 | 1458 | 499 | 534 | 2079 | 490 | 623 | 2169 | 290 | 455 | 2014 | 497 | 567 |
| 994 | W303.asm02.HP0_G0009960 | chrIV | 1682 | 722 | 906 | 1957 | 724 | 809 | 1711 | 663 | 817 | 2338 | 721 | 1182 | 2155 | 453 | 812 | 2364 | 643 | 867 |
| 995 | W303.asm02.HP0_G0009970 | chrIV | 11091 | 7396 | 6149 | 11310 | 5777 | 5339 | 11254 | 7541 | 5362 | 10979 | 6517 | 5253 | 14769 | 5738 | 5329 | 9630 | 5494 | 4490 |
| 996 | W303.asm02.HP0_G0009980 | chrIV | 5520 | 2131 | 2590 | 5193 | 2310 | 2742 | 5641 | 2186 | 3252 | 13764 | 4642 | 5787 | 8948 | 3830 | 4550 | 8457 | 4002 | 4979 |
| 997 | W303.asm02.HP0_G0009990 | chrIV | 2718 | 953 | 1020 | 2684 | 1060 | 890 | 2562 | 1025 | 997 | 5469 | 958 | 1145 | 5037 | 602 | 905 | 5028 | 883 | 1065 |
| 998 | W303.asm02.HP0_G0010000 | chrIV | 2133 | 549 | 591 | 1939 | 627 | 610 | 2167 | 580 | 668 | 5601 | 805 | 1248 | 4220 | 498 | 853 | 4422 | 713 | 1134 |
| 999 | W303.asm02.HP0_G0010010 | chrIV | 1461 | 607 | 991 | 1161 | 844 | 977 | 1257 | 656 | 1271 | 2374 | 546 | 779 | 2665 | 339 | 687 | 2663 | 507 | 647 |
| 1000 | W303.asm02.HP0_G0010020 | chrIV | 1071 | 791 | 704 | 1091 | 787 | 769 | 1028 | 860 | 682 | 2565 | 746 | 838 | 2681 | 525 | 639 | 2359 | 611 | 768 |
| 1001 | W303.asm02.HP0_G0010030 | chrIV | 5117 | 655 | 920 | 5015 | 818 | 882 | 4821 | 706 | 1162 | 7214 | 903 | 1737 | 5164 | 736 | 1499 | 5141 | 815 | 1620 |
| 1002 | W303.asm02.HP0_G0010040 | chrIV | 636 | 567 | 547 | 712 | 564 | 569 | 565 | 504 | 635 | 2494 | 543 | 686 | 2443 | 309 | 525 | 2303 | 548 | 597 |
| 1003 | W303.asm02.HP0_G0010050 | chrIV | 2277 | 331 | 439 | 1966 | 421 | 505 | 2362 | 337 | 590 | 4743 | 760 | 1025 | 2349 | 549 | 714 | 2201 | 674 | 923 |
| 1004 | W303.asm02.HP0_G0010060 | chrIV | 939 | 1627 | 1890 | 917 | 1837 | 1713 | 1164 | 1789 | 1612 | 11399 | 1934 | 3104 | 10020 | 1330 | 2028 | 9565 | 1654 | 2863 |
| 1005 | W303.asm02.HP0_G0010070 | chrIV | 7808 | 1904 | 2011 | 7857 | 2262 | 1926 | 7946 | 2006 | 1756 | 9794 | 2171 | 2869 | 9090 | 1410 | 2112 | 8645 | 1814 | 2544 |
| 1006 | W303.asm02.HP0_G0010080 | chrIV | 4461 | 1425 | 997 | 4076 | 1173 | 935 | 4642 | 1458 | 1018 | 7660 | 1777 | 2059 | 7646 | 1444 | 1994 | 6832 | 1608 | 2007 |
| 1007 | W303.asm02.HP0_G0010090 | chrIV | 850 | 694 | 614 | 895 | 637 | 543 | 907 | 719 | 661 | 3262 | 834 | 859 | 2495 | 453 | 635 | 3215 | 737 | 838 |
| 1008 | W303.asm02.HP0_G0010100 | chrIV | 9539 | 2978 | 2264 | 8975 | 2317 | 2169 | 8717 | 2897 | 2094 | 10932 | 3752 | 3923 | 10997 | 2982 | 3356 | 9400 | 3230 | 3182 |
| 1009 | W303.asm02.HP0_G0010110 | chrIV | 5625 | 648 | 661 | 5885 | 751 | 907 | 5613 | 578 | 931 | 7223 | 1053 | 1983 | 3934 | 1017 | 1440 | 3930 | 903 | 1612 |
| 1010 | W303.asm02.HP0_G0010120 | chrIV | 4002 | 1716 | 1420 | 4411 | 1335 | 1517 | 3473 | 1574 | 1142 | 4841 | 1373 | 1383 | 4628 | 800 | 1020 | 4782 | 1300 | 1226 |
| 1011 | W303.asm02.HP0_G0010130 | chrIV | 3004 | 2475 | 2310 | 2999 | 2718 | 2402 | 3243 | 2516 | 2418 | 8966 | 2310 | 2449 | 8341 | 1461 | 1650 | 7994 | 1964 | 2219 |
| 1012 | W303.asm02.HP0_G0010140 | chrIV | 3769 | 1213 | 970 | 4028 | 969 | 1003 | 3901 | 1102 | 1094 | 6301 | 1549 | 1737 | 5383 | 1037 | 1286 | 4760 | 1399 | 1545 |
| 1013 | W303.asm02.HP0_G0010150 | chrIV | 20037 | 15110 | 16801 | 18732 | 15275 | 16906 | 19516 | 14126 | 16440 | 24460 | 12431 | 16273 | 27831 | 9424 | 12848 | 25294 | 11276 | 14266 |
| 1014 | W303.asm02.HP0_G0010160 | chrIV | 547 | 952 | 605 | 602 | 688 | 678 | 583 | 940 | 628 | 3317 | 1086 | 939 | 3543 | 736 | 680 | 2880 | 949 | 887 |
| 1015 | W303.asm02.HP0_G0010170 | chrIV | 7995 | 8730 | 8582 | 7350 | 8390 | 8342 | 7545 | 8442 | 7894 | 20761 | 7719 | 9855 | 19854 | 5757 | 7209 | 19407 | 6582 | 8538 |
| 1016 | W303.asm02.HP0_G0010180 | chrIV | 2126 | 524 | 687 | 2190 | 756 | 653 | 1963 | 544 | 762 | 6272 | 763 | 1299 | 4659 | 524 | 965 | 4801 | 689 | 1242 |
| 1017 | W303.asm02.HP0_G0010190 | chrIV | 2383 | 544 | 888 | 2353 | 711 | 851 | 2255 | 552 | 1059 | 3912 | 848 | 1512 | 3059 | 540 | 1151 | 3126 | 782 | 1370 |
| 1018 | W303.asm02.HP0_G0010200 | chrIV | 2473 | 1783 | 1316 | 2309 | 1688 | 1268 | 2541 | 1779 | 1271 | 8306 | 2041 | 2014 | 6732 | 1234 | 1043 | 7338 | 1792 | 1803 |
| 1019 | W303.asm02.HP0_G0010210 | chrIV | 2319 | 1319 | 1305 | 2298 | 1267 | 1265 | 2175 | 1362 | 1417 | 3879 | 1160 | 1231 | 4847 | 793 | 1043 | 3891 | 1029 | 1079 |
| 1020 | W303.asm02.HP0_G0010220 | chrIV | 2103 | 1084 | 980 | 1699 | 947 | 989 | 1616 | 1145 | 1086 | 2393 | 919 | 887 | 3134 | 635 | 773 | 2327 | 806 | 824 |
| 1021 | W303.asm02.HP0_G0010230 | chrIV | 4665 | 612 | 620 | 4314 | 610 | 581 | 4640 | 604 | 711 | 5922 | 1272 | 1537 | 4413 | 913 | 1087 | 4429 | 1123 | 1344 |

| | | | | | | | | | | | | | | | | | | | | |
|---|---|---|---|---|---|---|---|---|---|---|---|---|---|---|---|---|---|---|---|---|
| 1022 | W303.asm02.HP0_G0010240 | chrIV | 2061 | 2474 | 3217 | 2260 | 3383 | 2917 | 2348 | 2819 | 3176 | 11447 | 2391 | 2633 | 10860 | 1464 | 1676 | 11529 | 2019 | 2419 |
| 1023 | W303.asm02.HP0_G0010250 | chrIV | 3769 | 947 | 1230 | 3776 | 1191 | 1093 | 3675 | 1045 | 1194 | 6515 | 985 | 1475 | 5933 | 746 | 1195 | 6228 | 996 | 1392 |
| 1024 | W303.asm02.HP0_G0010260 | chrIV | 2650 | 1750 | 1920 | 2052 | 1777 | 1936 | 2353 | 1846 | 2564 | 4772 | 1526 | 1446 | 5584 | 979 | 1188 | 4698 | 1304 | 1344 |
| 1025 | W303.asm02.HP0_G0010270 | chrIV | 8696 | 3353 | 5173 | 8840 | 3876 | 5165 | 8129 | 3374 | 6476 | 11928 | 2592 | 3963 | 13266 | 1805 | 3209 | 12706 | 2469 | 3683 |
| 1026 | W303.asm02.HP0_G0010280 | chrIV | 3517 | 292 | 484 | 3576 | 356 | 590 | 3369 | 224 | 603 | 3590 | 695 | 1426 | 1667 | 572 | 1009 | 1632 | 633 | 1221 |
| 1027 | W303.asm02.HP0_G0010290 | chrIV | 101 | 12 | 4 | 81 | 4 | 6 | 94 | 17 | 5 | 67 | 14 | 24 | 40 | 11 | 10 | 47 | 27 | 12 |
| 1028 | W303.asm02.HP0_G0010300 | chrIV | 9027 | 1346 | 1636 | 8794 | 1730 | 1912 | 9092 | 1357 | 2096 | 9797 | 1840 | 3425 | 4733 | 1631 | 2637 | 4513 | 1574 | 3058 |
| 1029 | W303.asm02.HP0_G0010310 | chrIV | 33417 | 3278 | 1030 | 36385 | 1625 | 2315 | 31283 | 1833 | 728 | 2185 | 2761 | 1512 | 1580 | 1497 | 1739 | 1882 | 3066 | 1258 |
| 1030 | W303.asm02.HP0_G0010320 | chrIV | 10514 | 3924 | 3364 | 10280 | 3865 | 3461 | 11219 | 3808 | 3680 | 27738 | 6446 | 7933 | 22331 | 4829 | 5323 | 21004 | 5780 | 6734 |
| 1031 | W303.asm02.HP0_G0010330 | chrIV | 1105 | 786 | 396 | 799 | 419 | 414 | 908 | 694 | 458 | 1257 | 510 | 427 | 1223 | 307 | 305 | 1377 | 520 | 364 |
| 1032 | W303.asm02.HP0_G0010340 | chrIV | 9875 | 1610 | 1776 | 9880 | 1658 | 1933 | 9899 | 1569 | 2142 | 8896 | 2523 | 2980 | 6039 | 1766 | 2159 | 5717 | 2165 | 2599 |
| 1033 | W303.asm02.HP0_G0010350 | chrIV | 5578 | 2199 | 1423 | 5959 | 1550 | 1484 | 5566 | 2152 | 1482 | 6293 | 2336 | 2479 | 5709 | 1819 | 1882 | 5507 | 2276 | 2211 |
| 1034 | W303.asm02.HP0_G0010360 | chrIV | 3110 | 886 | 690 | 3208 | 722 | 762 | 3031 | 822 | 711 | 5548 | 1096 | 1319 | 4233 | 646 | 1061 | 4535 | 1075 | 1244 |
| 1035 | W303.asm02.HP0_G0010370 | chrIV | 5146 | 2843 | 2896 | 4663 | 2960 | 3056 | 4979 | 3022 | 3790 | 7193 | 2545 | 2305 | 8454 | 1794 | 2100 | 7556 | 2250 | 2228 |
| 1036 | W303.asm02.HP0_G0010380 | chrIV | 5964 | 2516 | 2453 | 5459 | 2272 | 2376 | 5389 | 2670 | 2768 | 5180 | 1984 | 2087 | 6868 | 1617 | 2148 | 5218 | 1800 | 1584 |
| 1037 | W303.asm02.HP0_G0010390 | chrIV | 239 | 307 | 261 | 245 | 263 | 263 | 235 | 318 | 408 | 713 | 244 | 212 | 1012 | 154 | 152 | 912 | 198 | 164 |
| 1038 | W303.asm02.HP0_G0010400 | chrIV | 394 | 549 | 730 | 388 | 706 | 713 | 377 | 563 | 886 | 2434 | 361 | 555 | 2641 | 295 | 378 | 2710 | 351 | 438 |
| 1039 | W303.asm02.HP0_G0010410 | chrIV | 668 | 364 | 536 | 722 | 599 | 468 | 791 | 350 | 544 | 4098 | 365 | 798 | 3801 | 254 | 590 | 3775 | 337 | 658 |
| 1040 | W303.asm02.HP0_G0010420 | chrIV | 1522 | 903 | 994 | 1636 | 1021 | 949 | 1477 | 882 | 1013 | 3527 | 709 | 917 | 3937 | 524 | 720 | 3376 | 656 | 817 |
| 1041 | W303.asm02.HP0_G0010430 | chrIV | 4409 | 1140 | 1065 | 3994 | 1077 | 1049 | 3883 | 1126 | 1153 | 6353 | 1429 | 1661 | 5629 | 1065 | 1457 | 5255 | 1317 | 1509 |
| 1042 | W303.asm02.HP0_G0010440 | chrIV | 105 | 48 | 32 | 92 | 25 | 34 | 75 | 44 | 47 | 80 | 29 | 29 | 125 | 25 | 15 | 101 | 39 | 22 |
| 1043 | W303.asm02.HP0_G0010450 | chrIV | 979 | 797 | 1044 | 861 | 864 | 1028 | 863 | 903 | 1360 | 2301 | 640 | 775 | 2836 | 420 | 537 | 2719 | 572 | 667 |
| 1044 | W303.asm02.HP0_G0010460 | chrIV | 901 | 1307 | 1686 | 1066 | 2077 | 1637 | 1140 | 1400 | 1573 | 7230 | 1032 | 1450 | 7627 | 785 | 875 | 6561 | 983 | 1164 |
| 1045 | W303.asm02.HP0_G0010470 | chrIV | 1053 | 412 | 514 | 836 | 474 | 509 | 899 | 403 | 633 | 1631 | 309 | 321 | 1725 | 189 | 214 | 1739 | 296 | 270 |
| 1046 | W303.asm02.HP0_G0010480 | chrIV | 4375 | 775 | 928 | 4442 | 962 | 1132 | 4296 | 761 | 1221 | 7474 | 1683 | 2285 | 4893 | 1047 | 1461 | 4321 | 1600 | 1898 |
| 1047 | W303.asm02.HP0_G0010490 | chrIV | 8412 | 2882 | 2874 | 7532 | 2702 | 3001 | 8776 | 2486 | 3733 | 17313 | 5671 | 6439 | 8323 | 3694 | 4016 | 9063 | 4882 | 5596 |
| 1048 | W303.asm02.HP0_G0010500 | chrIV | 4364 | 980 | 1236 | 4323 | 968 | 1419 | 4475 | 982 | 1531 | 7912 | 1600 | 2740 | 5909 | 1187 | 1807 | 5987 | 1436 | 2341 |
| 1049 | W303.asm02.HP0_G0010510 | chrIV | 6856 | 3767 | 3126 | 6944 | 3465 | 3015 | 7123 | 3691 | 3416 | 15044 | 4334 | 4627 | 13887 | 2977 | 3441 | 13532 | 3895 | 4079 |
| 1050 | W303.asm02.HP0_G0010520 | chrIV | 202 | 527 | 332 | 177 | 348 | 383 | 189 | 444 | 405 | 1206 | 407 | 497 | 1477 | 257 | 323 | 1343 | 431 | 500 |
| 1051 | W303.asm02.HP0_G0010530 | chrIV | 515 | 328 | 286 | 562 | 286 | 312 | 514 | 289 | 356 | 2223 | 494 | 651 | 1784 | 312 | 427 | 1532 | 430 | 582 |
| 1052 | W303.asm02.HP0_G0010540 | chrIV | 18102 | 4952 | 7088 | 18467 | 6951 | 6805 | 18012 | 5464 | 7627 | 24129 | 4530 | 7021 | 25889 | 3338 | 5461 | 23381 | 4416 | 6383 |
| 1053 | W303.asm02.HP0_G0010550 | chrIV | 4 | 6 | 0 | 3 | 1 | 3 | 5 | 9 | 4 | 9 | 9 | 3 | 7 | 4 | 1 | 9 | 11 | 1 |
| 1054 | W303.asm02.HP0_G0010560 | chrIV | 2356 | 711 | 521 | 2472 | 595 | 582 | 2531 | 675 | 570 | 4341 | 1046 | 1153 | 3316 | 608 | 709 | 3664 | 920 | 991 |
| 1055 | W303.asm02.HP0_G0010570 | chrIV | 3354 | 1522 | 1604 | 3179 | 1391 | 1617 | 3043 | 1527 | 1999 | 3859 | 1186 | 1207 | 4694 | 813 | 1112 | 4249 | 1079 | 1074 |
| 1056 | W303.asm02.HP0_G0010580 | chrIV | 2630 | 909 | 776 | 2688 | 797 | 802 | 2520 | 869 | 765 | 2953 | 711 | 875 | 3250 | 490 | 703 | 2921 | 735 | 793 |
| 1057 | W303.asm02.HP0_G0010590 | chrIV | 1477 | 735 | 984 | 1508 | 1023 | 841 | 1551 | 811 | 959 | 4494 | 753 | 914 | 4558 | 459 | 656 | 4263 | 660 | 792 |
| 1058 | W303.asm02.HP0_G0010600 | chrIV | 1886 | 434 | 442 | 2056 | 427 | 511 | 1842 | 417 | 564 | 3288 | 649 | 845 | 2610 | 490 | 703 | 2706 | 566 | 738 |
| 1059 | W303.asm02.HP0_G0010610 | chrIV | 608 | 590 | 696 | 542 | 600 | 725 | 447 | 593 | 958 | 1542 | 459 | 510 | 2011 | 290 | 421 | 1797 | 392 | 460 |
| 1060 | W303.asm02.HP0_G0010620 | chrIV | 2965 | 1151 | 934 | 2908 | 820 | 966 | 2562 | 1014 | 993 | 2875 | 760 | 819 | 2949 | 494 | 692 | 2968 | 869 | 659 |
| 1061 | W303.asm02.HP0_G0010630 | chrIV | 4185 | 706 | 638 | 3716 | 506 | 692 | 3643 | 744 | 774 | 2529 | 796 | 960 | 2569 | 553 | 826 | 2303 | 849 | 879 |
| 1062 | W303.asm02.HP0_G0010640 | chrIV | 934 | 554 | 547 | 954 | 504 | 489 | 935 | 540 | 558 | 3901 | 826 | 1048 | 3266 | 551 | 703 | 3069 | 741 | 908 |
| 1063 | W303.asm02.HP0_G0010650 | chrIV | 3748 | 1844 | 1892 | 3277 | 2075 | 1666 | 3855 | 1979 | 2130 | 11439 | 2544 | 2801 | 8646 | 1605 | 2070 | 10076 | 2325 | 2613 |
| 1064 | W303.asm02.HP0_G0010660 | chrIV | 1875 | 237 | 275 | 1924 | 259 | 359 | 1914 | 190 | 346 | 2590 | 546 | 688 | 930 | 357 | 507 | 940 | 403 | 569 |
| 1065 | W303.asm02.HP0_G0010670 | chrIV | 2846 | 635 | 950 | 3100 | 818 | 1096 | 2849 | 656 | 1184 | 6646 | 1231 | 2009 | 5115 | 920 | 1495 | 5018 | 1312 | 1867 |
| 1066 | W303.asm02.HP0_G0010680 | chrIV | 1817 | 1636 | 2692 | 1679 | 1950 | 2731 | 1481 | 1715 | 3075 | 4569 | 1260 | 2153 | 5397 | 1018 | 1710 | 4941 | 1175 | 1859 |
| 1067 | W303.asm02.HP0_G0010690 | chrIV | 1 | 3 | 1 | 2 | 0 | 1 | 3 | 1 | 2 | 2 | 0 | 4 | 3 | 2 | 0 | 5 | 1 | 2 |
| 1068 | W303.asm02.HP0_G0010700 | chrIV | 4659 | 2164 | 2074 | 4783 | 2518 | 2029 | 5023 | 2306 | 2242 | 13180 | 2466 | 3188 | 12211 | 1640 | 2364 | 12462 | 2298 | 2971 |
| 1069 | W303.asm02.HP0_G0010710 | chrIV | 8500 | 3328 | 3638 | 8043 | 3598 | 3477 | 8777 | 3327 | 3601 | 17479 | 3881 | 5373 | 15173 | 2603 | 3933 | 15924 | 3454 | 4859 |
| 1070 | W303.asm02.HP0_G0010720 | chrIV | 1465 | 162 | 196 | 1396 | 240 | 214 | 1433 | 143 | 207 | 2354 | 258 | 390 | 1684 | 169 | 313 | 1522 | 213 | 337 |
| 1071 | W303.asm02.HP0_G0010730 | chrIV | 6586 | 5108 | 3413 | 5720 | 3606 | 3668 | 6114 | 4120 | 3147 | 13917 | 4547 | 5767 | 12634 | 3962 | 5223 | 12533 | 4479 | 5197 |
| 1072 | W303.asm02.HP0_G0010740 | chrIV | 80 | 23 | 12 | 58 | 16 | 18 | 46 | 7 | 14 | 92 | 22 | 21 | 45 | 17 | 17 | 60 | 26 | 20 |
| 1073 | W303.asm02.HP0_G0010750 | chrIV | 4261 | 702 | 647 | 5536 | 759 | 757 | 3879 | 641 | 807 | 4180 | 1319 | 1513 | 3334 | 1036 | 1426 | 2705 | 1220 | 1383 |
| 1074 | W303.asm02.HP0_G0010760 | chrIV | 1317 | 451 | 632 | 1416 | 686 | 647 | 1425 | 441 | 732 | 6787 | 648 | 1417 | 4833 | 525 | 895 | 4902 | 646 | 1306 |
| 1075 | W303.asm02.HP0_G0010770 | chrIV | 285 | 81 | 80 | 294 | 62 | 72 | 278 | 57 | 74 | 206 | 67 | 88 | 196 | 32 | 35 | 220 | 64 | 74 |
| 1076 | W303.asm02.HP0_G0010780 | chrIV | 2375 | 641 | 737 | 2671 | 725 | 753 | 2144 | 566 | 820 | 5071 | 1008 | 1382 | 3474 | 645 | 1074 | 4118 | 1043 | 1266 |
| 1077 | W303.asm02.HP0_G0010790 | chrIV | 2312 | 687 | 693 | 2509 | 717 | 711 | 2471 | 689 | 745 | 5050 | 1173 | 1335 | 3458 | 763 | 909 | 3447 | 961 | 1274 |
| 1078 | W303.asm02.HP0_G0010800 | chrIV | 610 | 1516 | 1363 | 622 | 1303 | 1199 | 612 | 1335 | 1234 | 2693 | 1135 | 1202 | 1897 | 593 | 770 | 2952 | 1099 | 1086 |
| 1079 | W303.asm02.HP0_G0010810 | chrIV | 956 | 328 | 305 | 1006 | 292 | 244 | 944 | 343 | 291 | 1325 | 455 | 431 | 1684 | 311 | 315 | 1111 | 468 | 395 |
| 1080 | W303.asm02.HP0_G0010820 | chrIV | 5328 | 4578 | 7547 | 5227 | 7061 | 7661 | 5042 | 4709 | 7945 | 15617 | 3575 | 6293 | 16461 | 2704 | 5030 | 15375 | 3374 | 5412 |
| 1081 | W303.asm02.HP0_G0010830 | chrIV | 6895 | 4912 | 8050 | 6309 | 6811 | 7912 | 6634 | 4858 | 10415 | 14296 | 4038 | 5431 | 17193 | 2976 | 4312 | 14676 | 3791 | 4479 |
| 1082 | W303.asm02.HP0_G0010840 | chrIV | 8182 | 6335 | 8720 | 7195 | 7826 | 8188 | 8181 | 6502 | 9813 | 19601 | 6318 | 8157 | 19702 | 4170 | 6139 | 19974 | 5587 | 7318 |
| 1083 | W303.asm02.HP0_G0010850 | chrIV | 2151 | 428 | 634 | 2233 | 609 | 675 | 2157 | 469 | 760 | 5468 | 776 | 1366 | 4096 | 612 | 966 | 4073 | 690 | 1216 |
| 1084 | W303.asm02.HP0_G0010860 | chrIV | 3320 | 1026 | 748 | 3174 | 897 | 722 | 3607 | 1035 | 750 | 5691 | 1236 | 1308 | 5210 | 756 | 904 | 5329 | 1162 | 1120 |
| 1085 | W303.asm02.HP0_G0010870 | chrIV | 4455 | 769 | 536 | 4507 | 496 | 600 | 4029 | 608 | 588 | 3347 | 1051 | 1063 | 2333 | 692 | 810 | 2421 | 1143 | 1016 |
| 1086 | W303.asm02.HP0_G0010880 | chrIV | 2026 | 919 | 991 | 1776 | 765 | 1085 | 1782 | 937 | 1123 | 2051 | 910 | 1138 | 2466 | 679 | 981 | 2029 | 817 | 927 |
| 1087 | W303.asm02.HP0_G0010890 | chrIV | 3664 | 634 | 1106 | 3103 | 1103 | 1412 | 4045 | 571 | 1506 | 10318 | 1552 | 2779 | 4380 | 1145 | 1958 | 4209 | 1468 | 2266 |
| 1088 | W303.asm02.HP0_G0010900 | chrIV | 17143 | 5544 | 5364 | 17915 | 4900 | 5355 | 18538 | 5257 | 4474 | 19236 | 5187 | 6980 | 19605 | 4226 | 5171 | 17116 | 4477 | 6235 |
| 1089 | W303.asm02.HP0_G0010910 | chrIV | 4835 | 1489 | 1405 | 4917 | 1519 | 1514 | 5580 | 1261 | 1887 | 9701 | 2995 | 3386 | 4232 | 2017 | 1788 | 4280 | 2558 | 2749 |
| 1090 | W303.asm02.HP0_G0010920 | chrIV | 2127 | 1122 | 1363 | 2104 | 1114 | 1236 | 2014 | 1011 | 1576 | 4620 | 879 | 1069 | 4271 | 533 | 757 | 5267 | 873 | 935 |
| 1091 | W303.asm02.HP0_G0010930 | chrIV | 1215 | 880 | 1099 | 1085 | 893 | 1138 | 1102 | 884 | 1438 | 2082 | 711 | 835 | 2719 | 466 | 707 | 2250 | 678 | 713 |
| 1092 | W303.asm02.HP0_G0010940 | chrIV | 8775 | 1804 | 2143 | 8270 | 1874 | 2090 | 8013 | 1915 | 2452 | 7264 | 1882 | 2779 | 8058 | 1482 | 2318 | 7013 | 1815 | 2461 |
| 1093 | W303.asm02.HP0_G0010950 | chrIV | 12433 | 2938 | 3026 | 11825 | 2730 | 3179 | 12747 | 3015 | 3214 | 21073 | 4078 | 7391 | 18217 | 3280 | 5043 | 17062 | 3555 | 6533 |
| 1094 | W303.asm02.HP0_G0010960 | chrIV | 2505 | 386 | 443 | 2414 | 501 | 443 | 2491 | 406 | 472 | 3511 | 594 | 860 | 3088 | 444 | 567 | 2737 | 598 | 735 |
| 1095 | W303.asm02.HP0_G0010970 | chrIV | 879 | 679 | 652 | 813 | 742 | 603 | 917 | 657 | 593 | 2888 | 504 | 703 | 2819 | 352 | 420 | 2620 | 583 | 624 |
| 1096 | W303.asm02.HP0_G0010980 | chrIV | 1876 | 314 | 457 | 1659 | 430 | 431 | 1709 | 354 | 517 | 2409 | 493 | 679 | 2513 | 431 | 564 | 1765 | 452 | 630 |
| 1097 | W303.asm02.HP0_G0010990 | chrIV | 733 | 347 | 428 | 722 | 450 | 378 | 734 | 337 | 393 | 3455 | 471 | 681 | 2692 | 260 | 400 | 2677 | 469 | 549 |
| 1098 | W303.asm02.HP0_G0011000 | chrIV | 2834 | 800 | 670 | 3072 | 707 | 647 | 2720 | 778 | 685 | 4493 | 910 | 1120 | 4046 | 658 | 848 | 3725 | 920 | 1036 |
| 1099 | W303.asm02.HP0_G0011010 | chrIV | 3457 | 2822 | 2439 | 3259 | 2565 | 2316 | 3577 | 2804 | 2211 | 10760 | 2792 | 2962 | 10064 | 1924 | 2319 | 10108 | 2482 | 2643 |
| 1100 | W303.asm02.HP0_G0011020 | chrIV | 4053 | 630 | 403 | 4253 | 427 | 459 | 3874 | 586 | 384 | 1818 | 619 | 662 | 2271 | 527 | 495 | 1604 | 495 | 499 |
| 1101 | W303.asm02.HP0_G0011030 | chrIV | 249 | 31 | 19 | 191 | 29 | 38 | 210 | 34 | 29 | 89 | 38 | 12 | 85 | 32 | 32 | 42 | 26 | 27 |
| 1102 | W303.asm02.HP0_G0011040 | chrIV | 1992 | 128 | 115 | 2770 | 133 | 166 | 1826 | 85 | 160 | 1105 | 271 | 324 | 902 | 158 | 242 | 744 | 274 | 283 |
| 1103 | W303.asm02.HP0_G0011050 | chrIV | 1945 | 1159 | 1313 | 1673 | 1084 | 1293 | 1700 | 1098 | 1556 | 2099 | 919 | 856 | 2896 | 627 | 912 | 2410 | 818 | 857 |
| 1104 | W303.asm02.HP0_G0011060 | chrIV | 275 | 385 | 537 | 290 | 503 | 502 | 249 | 323 | 574 | 1918 | 283 | 388 | 2372 | 163 | 273 | 2149 | 311 | 343 |
| 1105 | W303.asm02.HP0_G0011070 | chrIV | 620 | 65 | 54 | 570 | 45 | 59 | 520 | 53 | 55 | 497 | 101 | 109 | 481 | 77 | 121 | 432 | 92 | 110 |
| 1106 | W303.asm02.HP0_G0011080 | chrIV | 10728 | 950 | 1149 | 10418 | 930 | 1156 | 10567 | 935 | 1316 | 8971 | 1826 | 2712 | 8444 | 1485 | 2301 | 6655 | 1603 | 2299 |
| 1107 | W303.asm02.HP0_G0011090 | chrIV | 466 | 284 | 179 | 410 | 194 | 165 | 430 | 255 | 173 | 721 | 252 | 248 | 665 | 157 | 137 | 654 | 259 | 220 |
| 1108 | W303.asm02.HP0_G0011100 | chrIV | 1497 | 458 | 455 | 959 | 445 | 446 | 1145 | 475 | 523 | 945 | 285 | 346 | 1139 | 214 | 291 | 1048 | 253 | 282 |
| 1109 | W303.asm02.HP0_G0011110 | chrIV | 1392 | 405 | 383 | 1338 | 352 | 358 | 1345 | 398 | 408 | 1847 | 428 | 513 | 1622 | 322 | 425 | 1865 | 396 | 455 |
| 1110 | W303.asm02.HP0_G0011120 | chrIV | 2293 | 341 | 386 | 2652 | 367 | 370 | 2150 | 323 | 339 | 1810 | 355 | 429 | 2082 | 259 | 463 | 1591 | 327 | 431 |
| 1111 | W303.asm02.HP0_G0011130 | chrIV | 1202 | 341 | 293 | 1197 | 294 | 259 | 1256 | 319 | 290 | 1834 | 416 | 378 | 1382 | 289 | 289 | 1310 | 353 | 307 |
| 1112 | W303.asm02.HP0_G0011140 | chrIV | 2617 | 1287 | 1113 | 2578 | 1152 | 1010 | 2633 | 1261 | 1029 | 5464 | 1254 | 1254 | 5243 | 883 | 939 | 4961 | 1088 | 1166 |
| 1113 | W303.asm02.HP0_G0011150 | chrIV | 30441 | 5284 | 3316 | 30314 | 3910 | 4632 | 26765 | 3714 | 2798 | 9898 | 4292 | 4721 | 9162 | 3292 | 4656 | 9262 | 4719 | 4279 |
| 1114 | W303.asm02.HP0_G0011160 | chrIV | 956 | 175 | 276 | 1077 | 231 | 285 | 861 | 152 | 246 | 1049 | 291 | 368 | 1545 | 285 | 395 | 876 | 263 | 347 |

| | | | | | | | | | | | | | | | | | | | | |
|---|---|---|---|---|---|---|---|---|---|---|---|---|---|---|---|---|---|---|---|---|
| 1115 | W303.asm02.HP0_G0011170 | chrIV | 1075 | 812 | 701 | 1127 | 663 | 659 | 1128 | 795 | 705 | 2392 | 903 | 843 | 2559 | 640 | 669 | 2241 | 771 | 830 |
| 1116 | W303.asm02.HP0_G0011180 | chrIV | 7775 | 1241 | 946 | 6306 | 954 | 997 | 7184 | 1270 | 894 | 6077 | 1392 | 1424 | 6977 | 1185 | 1278 | 4937 | 1223 | 1389 |
| 1117 | W303.asm02.HP0_G0011190 | chrIV | 3702 | 884 | 783 | 3739 | 733 | 821 | 3615 | 761 | 790 | 3118 | 1192 | 1322 | 2789 | 780 | 895 | 2572 | 1230 | 1120 |
| 1118 | W303.asm02.HP0_G0011200 | chrIV | 528 | 178 | 225 | 447 | 183 | 196 | 503 | 201 | 227 | 791 | 205 | 335 | 535 | 145 | 224 | 638 | 205 | 292 |
| 1119 | W303.asm02.HP0_G0011210 | chrIV | 3292 | 2151 | 2127 | 2985 | 1961 | 1990 | 3332 | 2218 | 2067 | 12808 | 3121 | 3365 | 10500 | 1802 | 2657 | 11876 | 2842 | 3844 |
| 1120 | W303.asm02.HP0_G0011220 | chrIV | 1594 | 546 | 702 | 1650 | 707 | 719 | 1506 | 511 | 713 | 2491 | 524 | 664 | 2527 | 320 | 550 | 2338 | 507 | 597 |
| 1121 | W303.asm02.HP0_G0011230 | chrIV | 2478 | 454 | 626 | 2446 | 611 | 750 | 2575 | 433 | 706 | 5681 | 942 | 1580 | 3445 | 696 | 1004 | 3392 | 780 | 1288 |
| 1122 | W303.asm02.HP0_G0011240 | chrIV | 2037 | 1518 | 1499 | 1882 | 1500 | 1371 | 2213 | 1516 | 1505 | 5525 | 1275 | 1353 | 5242 | 829 | 1056 | 5049 | 1162 | 1236 |
| 1123 | W303.asm02.HP0_G0011250 | chrIV | 2210 | 701 | 797 | 2046 | 795 | 740 | 2017 | 828 | 846 | 3110 | 783 | 977 | 3264 | 553 | 721 | 2996 | 711 | 820 |
| 1124 | W303.asm02.HP0_G0011260 | chrIV | 4603 | 2376 | 2521 | 4533 | 2232 | 2567 | 4003 | 2272 | 2924 | 8611 | 2609 | 2959 | 7214 | 1451 | 2207 | 7622 | 2506 | 2950 |
| 1125 | W303.asm02.HP0_G0011270 | chrIV | 833 | 95 | 82 | 650 | 77 | 73 | 692 | 87 | 88 | 581 | 183 | 155 | 379 | 83 | 114 | 433 | 192 | 142 |
| 1126 | W303.asm02.HP0_G0011280 | chrIV | 400 | 366 | 457 | 365 | 483 | 494 | 355 | 287 | 589 | 1982 | 388 | 508 | 2083 | 240 | 388 | 1627 | 399 | 472 |
| 1127 | W303.asm02.HP0_G0011290 | chrIV | 5165 | 3725 | 4415 | 4870 | 3939 | 4258 | 4574 | 3942 | 4921 | 7275 | 3403 | 3906 | 7793 | 2693 | 3604 | 7305 | 3141 | 3585 |
| 1128 | W303.asm02.HP0_G0011300 | chrIV | 5044 | 296 | 153 | 5538 | 232 | 198 | 4657 | 229 | 212 | 1953 | 520 | 398 | 1573 | 335 | 394 | 1153 | 479 | 310 |
| 1129 | W303.asm02.HP0_G0011310 | chrIV | 915 | 648 | 687 | 856 | 692 | 614 | 900 | 577 | 736 | 2885 | 632 | 752 | 2803 | 380 | 498 | 2943 | 562 | 643 |
| 1130 | W303.asm02.HP0_G0011320 | chrIV | 1460 | 1623 | 2427 | 1466 | 2356 | 2249 | 1477 | 1705 | 2850 | 7790 | 1421 | 2132 | 8175 | 896 | 1386 | 7359 | 1407 | 1810 |
| 1131 | W303.asm02.HP0_G0011330 | chrIV | 35 | 158 | 529 | 26 | 541 | 538 | 32 | 210 | 728 | 902 | 200 | 239 | 927 | 114 | 160 | 852 | 150 | 188 |
| 1132 | W303.asm02.HP0_G0011340 | chrIV | 1903 | 499 | 525 | 2133 | 483 | 513 | 1823 | 520 | 473 | 2213 | 443 | 624 | 2143 | 280 | 417 | 1999 | 388 | 565 |
| 1133 | W303.asm02.HP0_G0011350 | chrIV | 2134 | 489 | 720 | 2140 | 662 | 709 | 2101 | 526 | 918 | 5370 | 820 | 1530 | 3958 | 630 | 1102 | 3840 | 755 | 1387 |
| 1134 | W303.asm02.HP0_G0011360 | chrIV | 2516 | 1708 | 2008 | 2257 | 1618 | 2013 | 2404 | 1667 | 2283 | 5227 | 1388 | 1731 | 6337 | 990 | 1503 | 5209 | 1270 | 1628 |
| 1135 | W303.asm02.HP0_G0011370 | chrIV | 1184 | 479 | 435 | 1158 | 421 | 387 | 1227 | 467 | 434 | 1756 | 418 | 379 | 2072 | 208 | 257 | 1771 | 400 | 316 |
| 1136 | W303.asm02.HP0_G0011380 | chrIV | 550 | 280 | 317 | 413 | 310 | 371 | 434 | 278 | 521 | 844 | 214 | 253 | 1106 | 135 | 193 | 1010 | 200 | 208 |
| 1137 | W303.asm02.HP0_G0011390 | chrIV | 2775 | 755 | 572 | 2642 | 568 | 590 | 2473 | 774 | 519 | 2330 | 944 | 881 | 2511 | 691 | 806 | 2166 | 826 | 806 |
| 1138 | W303.asm02.HP0_G0011400 | chrIV | 1620 | 978 | 996 | 1462 | 951 | 975 | 1474 | 1010 | 1299 | 3006 | 850 | 786 | 3068 | 546 | 675 | 3146 | 755 | 742 |
| 1139 | W303.asm02.HP0_G0011410 | chrIV | 2784 | 853 | 908 | 2878 | 1009 | 878 | 2717 | 906 | 886 | 4183 | 872 | 1173 | 4107 | 528 | 813 | 3997 | 842 | 983 |
| 1140 | W303.asm02.HP0_G0011420 | chrIV | 1989 | 466 | 549 | 1732 | 624 | 555 | 1798 | 468 | 587 | 4707 | 738 | 1028 | 3828 | 502 | 760 | 3727 | 652 | 869 |
| 1141 | W303.asm02.HP0_G0011430 | chrIV | 5587 | 2289 | 2177 | 5873 | 2374 | 2093 | 6044 | 2444 | 2445 | 14786 | 2922 | 3578 | 11652 | 2001 | 2533 | 12813 | 2526 | 3291 |
| 1142 | W303.asm02.HP0_G0011440 | chrIV | 10539 | 872 | 1106 | 10069 | 1085 | 1385 | 10264 | 769 | 1432 | 6293 | 1439 | 2648 | 4912 | 1400 | 2467 | 3330 | 1203 | 2244 |
| 1143 | W303.asm02.HP0_G0011450 | chrIV | 9211 | 531 | 706 | 8752 | 552 | 732 | 8739 | 407 | 815 | 6526 | 1302 | 2002 | 3490 | 1197 | 1435 | 2471 | 1095 | 1627 |
| 1144 | W303.asm02.HP0_G0011460 | chrIV | 10161 | 1003 | 1130 | 9110 | 1000 | 1212 | 10249 | 920 | 1385 | 7208 | 1717 | 2308 | 4459 | 1456 | 1916 | 4260 | 1652 | 2064 |
| 1145 | W303.asm02.HP0_G0011470 | chrIV | 2449 | 568 | 570 | 2770 | 602 | 602 | 2483 | 589 | 665 | 3698 | 668 | 1288 | 3272 | 471 | 858 | 3071 | 641 | 1016 |
| 1146 | W303.asm02.HP0_G0011480 | chrIV | 8666 | 1391 | 1212 | 8520 | 1219 | 1219 | 8608 | 1374 | 1436 | 4907 | 1620 | 1890 | 4670 | 1257 | 1690 | 3872 | 1552 | 1734 |
| 1147 | W303.asm02.HP0_G0011490 | chrIV | 4738 | 1417 | 1526 | 4423 | 1606 | 1368 | 4944 | 1515 | 1504 | 8660 | 1982 | 2032 | 8737 | 1440 | 1510 | 7692 | 1725 | 1692 |
| 1148 | W303.asm02.HP0_G0011500 | chrIV | 4398 | 5802 | 6413 | 4296 | 5801 | 6279 | 4365 | 5972 | 7748 | 14204 | 5582 | 6008 | 15852 | 4531 | 4954 | 14186 | 4800 | 4813 |
| 1149 | W303.asm02.HP0_G0011510 | chrIV | 1177 | 1532 | 2075 | 1148 | 2595 | 1996 | 1450 | 1782 | 1841 | 12481 | 1630 | 2349 | 12039 | 1122 | 1331 | 10959 | 1404 | 2031 |
| 1150 | W303.asm02.HP0_G0011520 | chrIV | 2014 | 2569 | 2385 | 2040 | 2518 | 2160 | 2135 | 2567 | 2593 | 9624 | 2455 | 2241 | 8764 | 1337 | 1450 | 9130 | 2159 | 2161 |
| 1151 | W303.asm02.HP0_G0011530 | chrIV | 1662 | 442 | 657 | 1721 | 634 | 701 | 1495 | 402 | 710 | 5650 | 680 | 1390 | 3462 | 422 | 965 | 4000 | 611 | 1149 |
| 1152 | W303.asm02.HP0_G0011540 | chrIV | 880 | 814 | 1140 | 723 | 1070 | 1192 | 760 | 934 | 1615 | 3267 | 638 | 893 | 3948 | 479 | 734 | 3389 | 558 | 755 |
| 1153 | W303.asm02.HP0_G0011550 | chrIV | 1627 | 942 | 1186 | 1638 | 971 | 1056 | 1768 | 1035 | 1156 | 7162 | 1108 | 1888 | 5793 | 830 | 1294 | 6217 | 1065 | 1705 |
| 1154 | W303.asm02.HP0_G0011560 | chrIV | 3919 | 3900 | 3728 | 3341 | 3113 | 3781 | 3538 | 4093 | 3891 | 8977 | 3471 | 3754 | 8573 | 2456 | 2811 | 8609 | 3120 | 3334 |
| 1155 | W303.asm02.HP0_G0011570 | chrIV | 1578 | 842 | 815 | 1480 | 772 | 834 | 1494 | 871 | 1129 | 2305 | 727 | 708 | 2501 | 446 | 538 | 2374 | 687 | 696 |
| 1156 | W303.asm02.HP0_G0011580 | chrIV | 2942 | 1057 | 850 | 2956 | 768 | 876 | 2680 | 987 | 963 | 2994 | 1014 | 1115 | 2899 | 581 | 796 | 2910 | 928 | 896 |
| 1157 | W303.asm02.HP0_G0011590 | chrIV | 1865 | 1327 | 1135 | 1919 | 1079 | 1072 | 1874 | 1308 | 1290 | 4577 | 1162 | 1161 | 4367 | 719 | 937 | 4552 | 1078 | 1085 |
| 1158 | W303.asm02.HP0_G0011600 | chrIV | 1737 | 1074 | 1605 | 1579 | 1347 | 1511 | 1649 | 1169 | 2066 | 3774 | 850 | 1125 | 4822 | 594 | 932 | 4252 | 772 | 973 |
| 1159 | W303.asm02.HP0_G0011610 | chrIV | 1249 | 967 | 993 | 1331 | 866 | 973 | 1528 | 1056 | 977 | 6078 | 1508 | 1997 | 6322 | 951 | 1115 | 5445 | 1325 | 1694 |
| 1160 | W303.asm02.HP0_G0011620 | chrIV | 5078 | 558 | 788 | 5168 | 621 | 964 | 5324 | 514 | 942 | 6340 | 1118 | 2196 | 3034 | 899 | 1447 | 2596 | 906 | 2003 |
| 1161 | W303.asm02.HP0_G0011630 | chrIV | 2034 | 839 | 928 | 2169 | 890 | 849 | 2064 | 813 | 891 | 4222 | 884 | 1021 | 3988 | 550 | 799 | 3814 | 850 | 870 |
| 1162 | W303.asm02.HP0_G0011640 | chrIV | 2491 | 1368 | 1324 | 2678 | 1715 | 1358 | 2936 | 1489 | 1295 | 7334 | 1479 | 1800 | 6655 | 925 | 1082 | 6174 | 1261 | 1539 |
| 1163 | W303.asm02.HP0_G0011650 | chrIV | 1406 | 727 | 641 | 1702 | 747 | 823 | 1372 | 661 | 654 | 1919 | 615 | 630 | 2013 | 345 | 575 | 1549 | 588 | 561 |
| 1164 | W303.asm02.HP0_G0011660 | chrIV | 1570 | 193 | 254 | 1664 | 280 | 229 | 1447 | 177 | 242 | 2011 | 241 | 435 | 1629 | 173 | 376 | 1710 | 250 | 419 |
| 1165 | W303.asm02.HP0_G0011670 | chrIV | 718 | 354 | 538 | 727 | 443 | 498 | 717 | 374 | 706 | 1397 | 291 | 373 | 2053 | 240 | 359 | 1654 | 266 | 327 |
| 1166 | W303.asm02.HP0_G0011680 | chrIV | 7242 | 1202 | 1429 | 7540 | 1608 | 1535 | 6880 | 1336 | 1488 | 6701 | 1218 | 1845 | 7491 | 1113 | 1674 | 5697 | 1185 | 1700 |
| 1167 | W303.asm02.HP0_G0011690 | chrIV | 751 | 496 | 531 | 647 | 448 | 491 | 663 | 464 | 573 | 1348 | 333 | 429 | 1084 | 179 | 257 | 1497 | 358 | 377 |
| 1168 | W303.asm02.HP0_G0011700 | chrIV | 1152 | 668 | 715 | 1085 | 696 | 689 | 1098 | 659 | 825 | 2237 | 529 | 529 | 2980 | 332 | 510 | 2381 | 434 | 521 |
| 1169 | W303.asm02.HP0_G0011710 | chrIV | 1257 | 827 | 992 | 1153 | 706 | 897 | 1102 | 940 | 1141 | 2209 | 674 | 643 | 2255 | 426 | 632 | 2445 | 566 | 649 |
| 1170 | W303.asm02.HP0_G0011720 | chrIV | 2906 | 258 | 514 | 2851 | 305 | 627 | 2812 | 263 | 650 | 3348 | 566 | 1446 | 2150 | 404 | 873 | 1902 | 534 | 1229 |
| 1171 | W303.asm02.HP0_G0011730 | chrIV | 720 | 619 | 1040 | 411 | 793 | 1071 | 420 | 712 | 1353 | 1693 | 494 | 771 | 2559 | 348 | 742 | 2015 | 470 | 608 |
| 1172 | W303.asm02.HP0_G0011740 | chrIV | 2138 | 2831 | 3095 | 1844 | 2728 | 2672 | 2252 | 2862 | 3354 | 11123 | 2846 | 3125 | 10725 | 1725 | 1910 | 11752 | 2546 | 2808 |
| 1173 | W303.asm02.HP0_G0011750 | chrIV | 8690 | 1617 | 1385 | 8059 | 1467 | 1433 | 7889 | 1616 | 1327 | 5805 | 1604 | 2310 | 5719 | 1321 | 1829 | 5042 | 1514 | 2021 |
| 1174 | W303.asm02.HP0_G0011760 | chrIV | 2466 | 2646 | 3304 | 1844 | 3079 | 3208 | 1827 | 2735 | 4365 | 5613 | 2230 | 2307 | 7568 | 1762 | 2310 | 6255 | 1851 | 1913 |
| 1175 | W303.asm02.HP0_G0011770 | chrIV | 727 | 542 | 492 | 698 | 490 | 485 | 728 | 586 | 623 | 1661 | 469 | 458 | 1694 | 268 | 394 | 1780 | 439 | 419 |
| 1176 | W303.asm02.HP0_G0011780 | chrIV | 3455 | 2063 | 2451 | 3304 | 3085 | 2153 | 3946 | 2337 | 2515 | 20470 | 2725 | 3856 | 15987 | 1835 | 2239 | 17870 | 2272 | 3410 |
| 1177 | W303.asm02.HP0_G0011790 | chrIV | 1189 | 628 | 805 | 1079 | 890 | 795 | 1189 | 740 | 884 | 4660 | 725 | 1062 | 4234 | 502 | 921 | 4099 | 571 | 982 |
| 1178 | W303.asm02.HP0_G0011800 | chrIV | 2865 | 792 | 1142 | 2963 | 1144 | 1241 | 2936 | 801 | 1468 | 8073 | 1139 | 1663 | 5234 | 886 | 1352 | 4892 | 1033 | 1587 |
| 1179 | W303.asm02.HP0_G0011810 | chrIV | 8590 | 4081 | 4841 | 7757 | 4298 | 4548 | 8296 | 4427 | 5790 | 10604 | 4119 | 4189 | 12796 | 2628 | 3769 | 10863 | 3414 | 3729 |
| 1180 | W303.asm02.HP0_G0011820 | chrIV | 4171 | 1560 | 1448 | 4354 | 1406 | 1411 | 4227 | 1605 | 1403 | 4976 | 1386 | 1608 | 4786 | 864 | 1193 | 4765 | 1386 | 1294 |
| 1181 | W303.asm02.HP0_G0011830 | chrIV | 3616 | 654 | 518 | 3556 | 534 | 636 | 3414 | 584 | 601 | 3490 | 914 | 1090 | 2783 | 642 | 911 | 2676 | 906 | 958 |
| 1182 | W303.asm02.HP0_G0011840 | chrIV | 1155 | 1008 | 1781 | 1149 | 1578 | 1731 | 1100 | 1085 | 1972 | 5286 | 824 | 1354 | 5890 | 591 | 1065 | 5292 | 752 | 1179 |
| 1183 | W303.asm02.HP0_G0011850 | chrIV | 1644 | 443 | 645 | 1650 | 536 | 594 | 1571 | 483 | 660 | 2945 | 560 | 822 | 2705 | 332 | 597 | 2853 | 517 | 755 |
| 1184 | W303.asm02.HP0_G0011860 | chrIV | 2793 | 1304 | 1394 | 2648 | 1413 | 1330 | 2770 | 1315 | 1401 | 9061 | 1723 | 2201 | 7801 | 956 | 1544 | 8160 | 1654 | 2011 |
| 1185 | W303.asm02.HP0_G0011870 | chrIV | 2531 | 473 | 859 | 2641 | 883 | 907 | 2294 | 479 | 950 | 8852 | 814 | 2031 | 5760 | 536 | 1605 | 5751 | 708 | 1725 |
| 1186 | W303.asm02.HP0_G0011880 | chrIV | 7142 | 1141 | 1394 | 6682 | 1367 | 1505 | 6844 | 1300 | 1729 | 9743 | 1699 | 2804 | 7633 | 1474 | 2367 | 7615 | 1576 | 2652 |
| 1187 | W303.asm02.HP0_G0011890 | chrIV | 896 | 623 | 645 | 753 | 602 | 618 | 882 | 567 | 792 | 2129 | 417 | 497 | 2196 | 284 | 353 | 2368 | 410 | 452 |
| 1188 | W303.asm02.HP0_G0011900 | chrIV | 6627 | 2332 | 2501 | 6756 | 2407 | 2546 | 6396 | 2456 | 2905 | 6821 | 1966 | 2430 | 8120 | 1514 | 2092 | 6271 | 1779 | 2096 |
| 1189 | W303.asm02.HP0_G0011910 | chrIV | 1149 | 401 | 418 | 1252 | 482 | 413 | 1066 | 396 | 432 | 2791 | 612 | 654 | 2251 | 312 | 483 | 2145 | 626 | 591 |
| 1190 | W303.asm02.HP0_G0011920 | chrIV | 2132 | 1801 | 2091 | 1997 | 2162 | 2088 | 2060 | 1928 | 2213 | 5614 | 1594 | 1867 | 5683 | 992 | 1312 | 5530 | 1379 | 1517 |
| 1191 | W303.asm02.HP0_G0011930 | chrIV | 189 | 105 | 95 | 151 | 92 | 69 | 163 | 96 | 128 | 433 | 118 | 133 | 327 | 76 | 92 | 386 | 114 | 118 |
| 1192 | W303.asm02.HP0_G0011940 | chrIV | 12845 | 5281 | 3915 | 11731 | 4315 | 3414 | 13256 | 5624 | 3692 | 26542 | 7854 | 7324 | 21457 | 5368 | 5023 | 23280 | 6714 | 6757 |
| 1193 | W303.asm02.HP0_G0011950 | chrIV | 18786 | 11368 | 8297 | 17482 | 12918 | 7464 | 19094 | 10359 | 8534 | 53191 | 16395 | 11154 | 42644 | 9432 | 8489 | 43836 | 14558 | 10461 |
| 1194 | W303.asm02.HP0_G0011960 | chrIV | 8189 | 4057 | 5313 | 8511 | 5164 | 4977 | 8852 | 4331 | 5366 | 23170 | 4978 | 6878 | 22122 | 3124 | 5072 | 22033 | 4123 | 6406 |
| 1195 | W303.asm02.HP0_G0011970 | chrIV | 10520 | 1475 | 1643 | 10112 | 1612 | 1748 | 10405 | 1505 | 1925 | 8697 | 2306 | 3315 | 8115 | 1920 | 2789 | 6651 | 2183 | 3043 |
| 1196 | W303.asm02.HP0_G0011980 | chrIV | 4347 | 1048 | 1326 | 4061 | 1060 | 1317 | 4388 | 968 | 1451 | 9399 | 2235 | 3204 | 6730 | 1552 | 2149 | 6254 | 1970 | 2735 |
| 1197 | W303.asm02.HP0_G0011990 | chrIV | 9 | 4 | 3 | 11 | 5 | 7 | 17 | 5 | 2 | 29 | 9 | 5 | 25 | 4 | 6 | 15 | 4 | 4 |
| 1198 | W303.asm02.HP0_G0012000 | chrIV | 4866 | 1123 | 1153 | 4549 | 1266 | 1315 | 5243 | 1094 | 1319 | 9235 | 2414 | 2560 | 7875 | 1948 | 1874 | 6214 | 2135 | 2332 |
| 1199 | W303.asm02.HP0_G0012010 | chrIV | 1381 | 1070 | 1465 | 1327 | 1294 | 1279 | 1258 | 1141 | 1535 | 5258 | 1079 | 1445 | 7304 | 842 | 1256 | 4854 | 1177 | 1174 |
| 1200 | W303.asm02.HP0_G0012020 | chrIV | 1941 | 804 | 962 | 2046 | 1132 | 869 | 2070 | 785 | 1108 | 8249 | 1184 | 1730 | 7954 | 973 | 1467 | 6482 | 1181 | 1541 |
| 1201 | W303.asm02.HP0_G0012030 | chrIV | 1142 | 1292 | 1608 | 1115 | 1593 | 1480 | 1176 | 1409 | 1746 | 5820 | 1459 | 1576 | 5666 | 865 | 1176 | 5669 | 1245 | 1499 |
| 1202 | W303.asm02.HP0_G0012040 | chrIV | 13266 | 9435 | 10503 | 13568 | 9525 | 9789 | 13171 | 10083 | 11516 | 22894 | 10368 | 10546 | 25248 | 7083 | 9028 | 21936 | 9055 | 9260 |
| 1203 | W303.asm02.HP0_G0012050 | chrIV | 1395 | 1597 | 1350 | 1221 | 1337 | 1438 | 1379 | 1639 | 1588 | 7863 | 2227 | 2583 | 7080 | 1590 | 1961 | 6971 | 1896 | 2328 |
| 1204 | W303.asm02.HP0_G0012060 | chrIV | 1686 | 536 | 719 | 1715 | 733 | 715 | 1774 | 554 | 675 | 4095 | 587 | 978 | 3596 | 339 | 774 | 3601 | 501 | 878 |
| 1205 | W303.asm02.HP0_G0012070 | chrIV | 1484 | 744 | 1006 | 1399 | 835 | 1004 | 1282 | 810 | 1368 | 1878 | 711 | 789 | 2757 | 551 | 810 | 2327 | 583 | 773 |
| 1206 | W303.asm02.HP0_G0012080 | chrIV | 5472 | 849 | 699 | 5257 | 664 | 728 | 4915 | 794 | 675 | 2396 | 870 | 854 | 2725 | 590 | 895 | 2214 | 891 | 839 |
| 1207 | W303.asm02.HP0_G0012090 | chrIV | 2317 | 364 | 417 | 2104 | 404 | 489 | 2407 | 310 | 531 | 3418 | 593 | 1033 | 1183 | 463 | 675 | 1459 | 496 | 884 |

| | | | | | | | | | | | | | | | | | | | | |
|---|---|---|---|---|---|---|---|---|---|---|---|---|---|---|---|---|---|---|---|---|
| 1208 | W303.asm02.HP0_G0012100 | chrIV | 1364 | 2572 | 2613 | 1486 | 2772 | 2515 | 1585 | 2593 | 3351 | 9495 | 2380 | 2219 | 8907 | 1329 | 1202 | 10083 | 2129 | 2018 |
| 1209 | W303.asm02.HP0_G0012110 | chrIV | 2033 | 588 | 658 | 1942 | 599 | 757 | 2049 | 620 | 734 | 3746 | 791 | 1202 | 2903 | 523 | 816 | 2959 | 782 | 1110 |
| 1210 | W303.asm02.HP0_G0012120 | chrIV | 2009 | 402 | 526 | 2104 | 530 | 462 | 1957 | 445 | 491 | 3557 | 587 | 894 | 3350 | 364 | 596 | 2818 | 507 | 725 |
| 1211 | W303.asm02.HP0_G0012130 | chrIV | 2925 | 1854 | 2710 | 2760 | 2376 | 2671 | 2591 | 1969 | 3373 | 4578 | 1695 | 2215 | 5499 | 1172 | 2065 | 5127 | 1514 | 1942 |
| 1212 | W303.asm02.HP0_G0012140 | chrIV | 864 | 763 | 952 | 833 | 933 | 882 | 990 | 746 | 930 | 4417 | 729 | 1119 | 4273 | 536 | 769 | 3950 | 694 | 1008 |
| 1213 | W303.asm02.HP0_G0012150 | chrIV | 2561 | 1246 | 1201 | 2693 | 1632 | 1144 | 3086 | 1400 | 1235 | 12830 | 2122 | 2520 | 10064 | 1396 | 1378 | 10058 | 1824 | 2160 |
| 1214 | W303.asm02.HP0_G0012160 | chrIV | 1146 | 1079 | 1513 | 972 | 1344 | 1511 | 1117 | 1125 | 2105 | 4544 | 879 | 1086 | 4802 | 554 | 790 | 5306 | 811 | 1036 |
| 1215 | W303.asm02.HP0_G0012170 | chrIV | 4197 | 3642 | 2929 | 3743 | 2250 | 2846 | 3805 | 3348 | 3047 | 5668 | 2848 | 2654 | 6724 | 2105 | 2382 | 5567 | 2657 | 2371 |
| 1216 | W303.asm02.HP0_G0012180 | chrIV | 1751 | 501 | 623 | 1644 | 625 | 646 | 1521 | 601 | 852 | 2888 | 490 | 761 | 3078 | 321 | 578 | 2840 | 442 | 706 |
| 1217 | W303.asm02.HP0_G0012190 | chrIV | 827 | 999 | 990 | 753 | 1079 | 975 | 750 | 873 | 1079 | 3944 | 715 | 807 | 4078 | 456 | 596 | 4181 | 676 | 728 |
| 1218 | W303.asm02.HP0_G0012200 | chrIV | 1861 | 1050 | 1103 | 1736 | 969 | 1074 | 1802 | 1100 | 1247 | 3579 | 876 | 918 | 4034 | 592 | 816 | 3807 | 777 | 859 |
| 1219 | W303.asm02.HP0_G0012210 | chrIV | 5874 | 3168 | 4019 | 5294 | 3600 | 3743 | 6040 | 3402 | 4870 | 12689 | 2886 | 3262 | 14043 | 2013 | 2724 | 12955 | 2534 | 2930 |
| 1220 | W303.asm02.HP0_G0012220 | chrIV | 1230 | 1350 | 1828 | 995 | 1547 | 1802 | 1074 | 1263 | 2463 | 3983 | 972 | 1223 | 4871 | 651 | 1060 | 4506 | 919 | 1136 |
| 1221 | W303.asm02.HP0_G0012230 | chrIV | 140 | 141 | 33 | 106 | 52 | 76 | 96 | 90 | 45 | 45 | 39 | 15 | 81 | 27 | 16 | 56 | 50 | 13 |
| 1222 | W303.asm02.HP0_G0012240 | chrIV | 485 | 307 | 393 | 409 | 286 | 414 | 417 | 274 | 509 | 669 | 219 | 294 | 1055 | 161 | 265 | 814 | 205 | 252 |
| 1223 | W303.asm02.HP0_G0012250 | chrIV | 7913 | 1111 | 1063 | 7276 | 1128 | 1056 | 7572 | 1081 | 1114 | 5250 | 1355 | 1793 | 5582 | 1067 | 1650 | 4174 | 1286 | 1658 |
| 1224 | W303.asm02.HP0_G0012260 | chrIV | 3161 | 1021 | 998 | 2910 | 968 | 1143 | 2954 | 999 | 1101 | 4467 | 974 | 1358 | 4968 | 815 | 1081 | 3916 | 922 | 1272 |
| 1225 | W303.asm02.HP0_G0012270 | chrIV | 2440 | 2999 | 3929 | 2186 | 3396 | 3903 | 2012 | 2897 | 5401 | 7266 | 2367 | 3088 | 9124 | 2015 | 2650 | 8073 | 2224 | 2406 |
| 1226 | W303.asm02.HP0_G0012280 | chrIV | 1355 | 1170 | 1545 | 1232 | 1360 | 1539 | 1265 | 1273 | 1948 | 3493 | 1125 | 1253 | 4128 | 729 | 959 | 3735 | 1000 | 1188 |
| 1227 | W303.asm02.HP0_G0012290 | chrIV | 2909 | 621 | 629 | 2886 | 650 | 642 | 2846 | 630 | 694 | 4493 | 884 | 1034 | 4157 | 596 | 838 | 3489 | 807 | 962 |
| 1228 | W303.asm02.HP0_G0012300 | chrIV | 1649 | 334 | 402 | 1253 | 319 | 377 | 1328 | 348 | 562 | 532 | 224 | 220 | 1041 | 171 | 280 | 599 | 215 | 190 |
| 1229 | W303.asm02.HP0_G0012310 | chrIV | 1076 | 2103 | 1842 | 1037 | 1672 | 1903 | 1036 | 2068 | 1772 | 6383 | 1377 | 1673 | 4225 | 666 | 1151 | 5326 | 1480 | 1308 |
| 1230 | W303.asm02.HP0_G0012320 | chrIV | 7326 | 4729 | 6611 | 6168 | 5621 | 6345 | 6594 | 4514 | 7685 | 14371 | 4274 | 5071 | 15815 | 2745 | 4070 | 14187 | 3978 | 4454 |
| 1231 | W303.asm02.HP0_G0012330 | chrIV | 2133 | 773 | 809 | 1882 | 805 | 827 | 1772 | 776 | 1148 | 1931 | 658 | 678 | 2671 | 456 | 625 | 2212 | 599 | 632 |
| 1232 | W303.asm02.HP0_G0012340 | chrIV | 10277 | 9262 | 12102 | 8392 | 10796 | 11898 | 9193 | 9478 | 14680 | 20072 | 8802 | 10262 | 23303 | 6834 | 8582 | 21422 | 7944 | 8968 |
| 1233 | W303.asm02.HP0_G0012350 | chrIV | 985 | 757 | 605 | 831 | 468 | 591 | 849 | 695 | 724 | 1233 | 489 | 427 | 1624 | 361 | 347 | 1286 | 528 | 367 |
| 1234 | W303.asm02.HP0_G0012360 | chrIV | 745 | 650 | 389 | 744 | 612 | 399 | 797 | 657 | 545 | 4987 | 1151 | 677 | 2190 | 521 | 448 | 3373 | 1077 | 776 |
| 1235 | W303.asm02.HP0_G0012370 | chrIV | 26189 | 8560 | 7343 | 26429 | 7700 | 7224 | 29107 | 8182 | 8756 | 38505 | 13185 | 14195 | 30877 | 9920 | 10293 | 31705 | 11532 | 13441 |
| 1236 | W303.asm02.HP0_G0012380 | chrIV | 1378 | 595 | 460 | 1388 | 522 | 469 | 1372 | 625 | 439 | 2009 | 545 | 590 | 2146 | 395 | 440 | 1838 | 506 | 575 |
| 1237 | W303.asm02.HP0_G0012390 | chrIV | 2503 | 440 | 381 | 2204 | 374 | 361 | 2223 | 370 | 384 | 2331 | 633 | 695 | 1731 | 394 | 588 | 1677 | 557 | 543 |
| 1238 | W303.asm02.HP0_G0012400 | chrIV | 12377 | 1476 | 1284 | 11816 | 1171 | 1565 | 12208 | 1136 | 1868 | 6754 | 2691 | 3073 | 2658 | 2239 | 2146 | 2397 | 2178 | 2664 |
| 1239 | W303.asm02.HP0_G0012410 | chrIV | 1694 | 480 | 532 | 1779 | 609 | 621 | 1811 | 464 | 701 | 5332 | 776 | 1156 | 4186 | 550 | 779 | 4086 | 718 | 1001 |
| 1240 | W303.asm02.HP0_G0012420 | chrIV | 4123 | 1157 | 900 | 4444 | 986 | 821 | 4446 | 1157 | 877 | 6588 | 1379 | 1627 | 6016 | 976 | 1190 | 5829 | 1216 | 1520 |
| 1241 | W303.asm02.HP0_G0012430 | chrIV | 5036 | 1272 | 964 | 4063 | 804 | 1031 | 4175 | 1219 | 1059 | 1953 | 1004 | 923 | 2309 | 606 | 823 | 1807 | 966 | 788 |
| 1242 | W303.asm02.HP0_G0012440 | chrIV | 2743 | 1063 | 620 | 2414 | 721 | 694 | 2443 | 944 | 630 | 3143 | 944 | 947 | 2715 | 608 | 684 | 2673 | 1043 | 824 |
| 1243 | W303.asm02.HP0_G0012450 | chrIV | 2246 | 1850 | 1223 | 2341 | 1387 | 1357 | 2131 | 1819 | 1339 | 4361 | 1350 | 1223 | 4344 | 969 | 1055 | 4075 | 1488 | 1047 |
| 1244 | W303.asm02.HP0_G0012460 | chrIV | 12192 | 3889 | 2899 | 12753 | 2804 | 3139 | 12613 | 3568 | 2596 | 12563 | 3613 | 4817 | 11605 | 2768 | 3524 | 10721 | 3380 | 4235 |
| 1245 | W303.asm02.HP0_G0012470 | chrIV | 6455 | 2386 | 2343 | 6523 | 2768 | 2230 | 6973 | 2585 | 2617 | 17951 | 2985 | 3699 | 15310 | 2211 | 2702 | 14957 | 2585 | 3467 |
| 1246 | W303.asm02.HP0_G0012480 | chrIV | 1493 | 1521 | 1706 | 1455 | 1584 | 1678 | 1443 | 1536 | 2142 | 3859 | 1226 | 1293 | 4482 | 883 | 1006 | 3928 | 1105 | 1065 |
| 1247 | W303.asm02.HP0_G0012490 | chrIV | 2394 | 1554 | 1725 | 2578 | 1903 | 1858 | 2949 | 1524 | 2156 | 13926 | 2858 | 3606 | 10783 | 2082 | 1893 | 9588 | 2505 | 3087 |
| 1248 | W303.asm02.HP0_G0012500 | chrIV | 3497 | 3097 | 6553 | 3125 | 6606 | 5795 | 3558 | 3577 | 8147 | 12882 | 3149 | 4072 | 15248 | 2049 | 2883 | 12410 | 2423 | 3504 |
| 1249 | W303.asm02.HP0_G0012510 | chrIV | 1875 | 1356 | 2072 | 1600 | 1183 | 2084 | 1730 | 1369 | 2374 | 4447 | 1204 | 2417 | 4504 | 805 | 1451 | 4514 | 1134 | 2173 |
| 1250 | W303.asm02.HP0_G0012520 | chrIV | 335 | 115 | 83 | 307 | 75 | 84 | 323 | 87 | 91 | 479 | 127 | 120 | 325 | 76 | 63 | 311 | 136 | 96 |
| 1251 | W303.asm02.HP0_G0012530 | chrIV | 1171 | 429 | 341 | 1098 | 333 | 392 | 1167 | 395 | 411 | 821 | 342 | 268 | 989 | 250 | 210 | 734 | 361 | 263 |
| 1252 | W303.asm02.HP0_G0012540 | chrIV | 1614 | 1803 | 1430 | 1567 | 1510 | 1537 | 1600 | 1808 | 1515 | 5222 | 1643 | 1637 | 5568 | 1211 | 1331 | 4993 | 1420 | 1506 |
| 1253 | W303.asm02.HP0_G0012550 | chrIV | 5698 | 604 | 457 | 4989 | 411 | 611 | 5087 | 457 | 677 | 3029 | 1027 | 1052 | 1322 | 839 | 818 | 1202 | 997 | 888 |
| 1254 | W303.asm02.HP0_G0012560 | chrIV | 7394 | 369 | 461 | 7934 | 370 | 511 | 6883 | 319 | 534 | 1881 | 500 | 905 | 1042 | 383 | 619 | 1007 | 534 | 890 |
| 1255 | W303.asm02.HP0_G0012570 | chrIV | 2288 | 758 | 1037 | 2162 | 839 | 1185 | 2341 | 719 | 1273 | 6656 | 1051 | 1974 | 4661 | 798 | 1425 | 4873 | 972 | 1829 |
| 1256 | W303.asm02.HP0_G0012580 | chrIV | 2705 | 1728 | 1593 | 2131 | 1383 | 1502 | 2459 | 1830 | 1813 | 3888 | 1556 | 1463 | 3535 | 931 | 935 | 3539 | 1471 | 1262 |
| 1257 | W303.asm02.HP0_G0012590 | chrIV | 1550 | 615 | 537 | 1502 | 517 | 605 | 1480 | 552 | 726 | 3483 | 880 | 1015 | 1867 | 528 | 631 | 1923 | 885 | 895 |
| 1258 | W303.asm02.HP0_G0012600 | chrIV | 832 | 1432 | 1302 | 662 | 992 | 1244 | 784 | 1412 | 1636 | 2696 | 1019 | 890 | 2957 | 632 | 684 | 2942 | 1001 | 768 |
| 1259 | W303.asm02.HP0_G0012610 | chrIV | 2759 | 761 | 1086 | 2518 | 453 | 1200 | 2542 | 788 | 1297 | 3231 | 1248 | 2450 | 2469 | 939 | 1608 | 2458 | 1201 | 2152 |
| 1260 | W303.asm02.HP0_G0012620 | chrIV | 1657 | 1908 | 1607 | 1624 | 2044 | 1791 | 1827 | 2096 | 1734 | 7526 | 1981 | 2050 | 7117 | 1222 | 1235 | 6355 | 1689 | 1870 |
| 1261 | W303.asm02.HP0_G0012630 | chrIV | 675 | 633 | 541 | 628 | 495 | 539 | 683 | 641 | 675 | 2824 | 643 | 704 | 2278 | 454 | 498 | 2261 | 662 | 601 |
| 1262 | W303.asm02.HP0_G0012640 | chrIV | 2275 | 1019 | 717 | 2217 | 750 | 704 | 2151 | 943 | 797 | 3470 | 984 | 914 | 2880 | 598 | 739 | 2952 | 1013 | 826 |
| 1263 | W303.asm02.HP0_G0012650 | chrIV | 1983 | 816 | 633 | 1982 | 670 | 606 | 1956 | 795 | 692 | 4858 | 878 | 1107 | 4005 | 602 | 804 | 4189 | 992 | 942 |
| 1264 | W303.asm02.HP0_G0012660 | chrIV | 1366 | 923 | 632 | 1133 | 658 | 581 | 1287 | 994 | 619 | 3602 | 1012 | 930 | 3041 | 618 | 609 | 3227 | 928 | 862 |
| 1265 | W303.asm02.HP0_G0012670 | chrIV | 2549 | 238 | 520 | 2475 | 512 | 541 | 2751 | 184 | 679 | 3731 | 439 | 1092 | 1817 | 452 | 693 | 1038 | 397 | 936 |
| 1266 | W303.asm02.HP0_G0012680 | chrIV | 2482 | 477 | 510 | 2421 | 526 | 497 | 2299 | 466 | 551 | 2451 | 555 | 702 | 2255 | 416 | 582 | 2002 | 520 | 627 |
| 1267 | W303.asm02.HP0_G0012690 | chrIV | 1387 | 504 | 443 | 1329 | 452 | 508 | 1284 | 459 | 528 | 4121 | 849 | 972 | 3008 | 494 | 671 | 2914 | 782 | 904 |
| 1268 | W303.asm02.HP0_G0012700 | chrIV | 3108 | 769 | 575 | 3103 | 574 | 617 | 3006 | 732 | 516 | 2782 | 853 | 923 | 2814 | 556 | 813 | 2519 | 882 | 852 |
| 1269 | W303.asm02.HP0_G0012710 | chrIV | 1249 | 507 | 786 | 1043 | 711 | 766 | 1135 | 575 | 1121 | 1714 | 465 | 586 | 2441 | 334 | 611 | 1818 | 404 | 539 |
| 1270 | W303.asm02.HP0_G0012720 | chrIV | 2801 | 523 | 376 | 2936 | 340 | 390 | 2464 | 439 | 451 | 2581 | 642 | 659 | 2358 | 352 | 513 | 1983 | 632 | 571 |
| 1271 | W303.asm02.HP0_G0012730 | chrIV | 7042 | 4636 | 5140 | 7460 | 4345 | 5224 | 7245 | 4399 | 6525 | 12229 | 3160 | 3803 | 13100 | 2328 | 3224 | 13865 | 3095 | 3326 |
| 1272 | W303.asm02.HP0_G0012740 | chrIV | 1470 | 1433 | 1411 | 1194 | 1195 | 1299 | 1248 | 1407 | 1737 | 3683 | 1042 | 1004 | 3830 | 705 | 769 | 4170 | 979 | 935 |
| 1273 | W303.asm02.HP0_G0012750 | chrIV | 9539 | 5012 | 6079 | 9552 | 6169 | 5735 | 9599 | 5384 | 6534 | 16149 | 5054 | 5892 | 17030 | 3190 | 4327 | 15657 | 4201 | 5180 |
| 1274 | W303.asm02.HP0_G0012760 | chrIV | 12365 | 1315 | 1419 | 12396 | 1447 | 1558 | 11825 | 1197 | 1923 | 11129 | 2362 | 3668 | 7626 | 2043 | 2846 | 6827 | 2346 | 3334 |
| 1275 | W303.asm02.HP0_G0012770 | chrIV | 17795 | 2405 | 2527 | 18095 | 2284 | 2770 | 18640 | 1862 | 3313 | 11284 | 4326 | 5711 | 4393 | 3472 | 3011 | 3766 | 3216 | 4422 |
| 1276 | W303.asm02.HP0_G0012780 | chrIV | 1797 | 1929 | 2225 | 1789 | 1877 | 2352 | 1883 | 1906 | 2464 | 7040 | 1713 | 2536 | 8091 | 1270 | 1702 | 7020 | 1598 | 2293 |
| 1277 | W303.asm02.HP0_G0012790 | chrIV | 3721 | 789 | 589 | 3417 | 652 | 633 | 3310 | 725 | 580 | 2569 | 691 | 620 | 2885 | 527 | 663 | 2287 | 668 | 609 |
| 1278 | W303.asm02.HP0_G0012800 | chrIV | 3167 | 398 | 410 | 3591 | 322 | 477 | 3077 | 282 | 570 | 2276 | 710 | 776 | 1197 | 550 | 599 | 912 | 600 | 632 |
| 1279 | W303.asm02.HP0_G0012810 | chrIV | 418 | 100 | 121 | 458 | 131 | 99 | 409 | 119 | 138 | 861 | 192 | 184 | 798 | 86 | 153 | 688 | 171 | 178 |
| 1280 | W303.asm02.HP0_G0012820 | chrIV | 793 | 264 | 282 | 809 | 285 | 266 | 702 | 258 | 287 | 1594 | 362 | 422 | 1445 | 199 | 299 | 1353 | 350 | 361 |
| 1281 | W303.asm02.HP0_G0012830 | chrIV | 1148 | 456 | 425 | 1127 | 482 | 398 | 1164 | 473 | 439 | 2689 | 496 | 653 | 2326 | 326 | 383 | 2422 | 441 | 504 |
| 1282 | W303.asm02.HP0_G0012840 | chrIV | 1225 | 446 | 582 | 1137 | 641 | 589 | 1178 | 503 | 604 | 4291 | 601 | 1013 | 3685 | 373 | 672 | 3556 | 524 | 882 |
| 1283 | W303.asm02.HP0_G0012850 | chrIV | 2365 | 1075 | 1219 | 2192 | 1313 | 1144 | 2351 | 1115 | 1189 | 4723 | 1056 | 1232 | 5890 | 781 | 1004 | 4544 | 908 | 1123 |
| 1284 | W303.asm02.HP0_G0012860 | chrIV | 1860 | 253 | 342 | 1939 | 326 | 385 | 1776 | 206 | 405 | 2938 | 464 | 805 | 1560 | 425 | 630 | 1396 | 416 | 701 |
| 1285 | W303.asm02.HP0_G0012870 | chrIV | 2156 | 242 | 275 | 1990 | 330 | 327 | 2007 | 216 | 321 | 2727 | 431 | 624 | 1479 | 355 | 428 | 1685 | 438 | 580 |
| 1286 | W303.asm02.HP0_G0012880 | chrIV | 31 | 61 | 16 | 39 | 18 | 15 | 28 | 60 | 18 | 74 | 32 | 18 | 77 | 27 | 14 | 61 | 29 | 15 |
| 1287 | W303.asm02.HP0_G0012890 | chrIV | 5768 | 4417 | 6576 | 5154 | 5907 | 6399 | 5750 | 4278 | 8538 | 14657 | 3959 | 5087 | 16729 | 3000 | 4163 | 16477 | 3511 | 4498 |
| 1288 | W303.asm02.HP0_G0012900 | chrIV | 725 | 748 | 553 | 776 | 652 | 577 | 741 | 719 | 544 | 2559 | 613 | 736 | 2470 | 406 | 557 | 2424 | 739 | 647 |
| 1289 | W303.asm02.HP0_G0012910 | chrIV | 1479 | 1467 | 1887 | 1602 | 2176 | 1632 | 1725 | 1668 | 1944 | 8417 | 1447 | 1779 | 8045 | 940 | 1117 | 8640 | 1277 | 1592 |
| 1290 | W303.asm02.HP0_G0012920 | chrIV | 9324 | 8725 | 11597 | 8347 | 10489 | 11369 | 9237 | 8554 | 14325 | 23446 | 8037 | 9600 | 25890 | 5916 | 7615 | 24815 | 7123 | 8494 |
| 1291 | W303.asm02.HP0_G0012930 | chrIV | 440 | 862 | 665 | 306 | 778 | 684 | 406 | 907 | 637 | 2576 | 699 | 933 | 2925 | 527 | 740 | 2516 | 637 | 877 |
| 1292 | W303.asm02.HP0_G0012940 | chrIV | 8323 | 2791 | 2297 | 8048 | 2327 | 2233 | 8114 | 2916 | 2403 | 10455 | 3538 | 3301 | 8980 | 2300 | 2676 | 9289 | 3224 | 3126 |
| 1293 | W303.asm02.HP0_G0012950 | chrIV | 2835 | 187 | 200 | 2890 | 177 | 211 | 2569 | 174 | 239 | 585 | 179 | 173 | 742 | 113 | 177 | 562 | 137 | 172 |
| 1294 | W303.asm02.HP0_G0012960 | chrIV | 4317 | 3595 | 3860 | 3954 | 3761 | 3996 | 4214 | 3794 | 4411 | 10109 | 3635 | 3992 | 9166 | 2323 | 2910 | 9938 | 3125 | 3672 |
| 1295 | W303.asm02.HP0_G0012970 | chrIV | 3023 | 973 | 897 | 2844 | 803 | 934 | 2883 | 973 | 1062 | 6268 | 1742 | 1756 | 4803 | 1103 | 1355 | 5048 | 1557 | 1651 |
| 1296 | W303.asm02.HP0_G0012980 | chrIV | 10787 | 1143 | 1672 | 10298 | 1655 | 1773 | 10356 | 1254 | 2029 | 11375 | 1633 | 3354 | 7433 | 1387 | 2552 | 7053 | 1327 | 2859 |
| 1297 | W303.asm02.HP0_G0012990 | chrIV | 4107 | 1348 | 1483 | 4027 | 1494 | 1463 | 4037 | 1440 | 1422 | 7428 | 1324 | 1926 | 6181 | 901 | 1329 | 6546 | 1247 | 1662 |
| 1298 | W303.asm02.HP0_G0013000 | chrIV | 424 | 808 | 640 | 333 | 581 | 585 | 380 | 786 | 610 | 2167 | 632 | 639 | 2254 | 434 | 528 | 2029 | 628 | 561 |
| 1299 | W303.asm02.HP0_G0013010 | chrIV | 1766 | 914 | 717 | 1661 | 656 | 797 | 1697 | 855 | 658 | 2764 | 760 | 948 | 2823 | 543 | 642 | 2599 | 787 | 830 |
| 1300 | W303.asm02.HP0_G0013020 | chrIV | 569 | 403 | 275 | 320 | 236 | 298 | 314 | 401 | 345 | 417 | 258 | 260 | 632 | 167 | 239 | 495 | 280 | 204 |

| | | | | | | | | | | | | | | | | | | | | |
|---|---|---|---|---|---|---|---|---|---|---|---|---|---|---|---|---|---|---|---|---|
| 1301 | W303.asm02.HP0_G0013030 | chrIV | 3875 | 1416 | 1051 | 3798 | 1058 | 1133 | 3651 | 1382 | 1170 | 2533 | 1155 | 1201 | 2974 | 825 | 1033 | 2282 | 1163 | 1133 |
| 1302 | W303.asm02.HP0_G0013040 | chrIV | 4590 | 858 | 723 | 4614 | 661 | 923 | 4798 | 753 | 961 | 4306 | 1527 | 1667 | 2248 | 1021 | 1098 | 2441 | 1523 | 1505 |
| 1303 | W303.asm02.HP0_G0013050 | chrIV | 1915 | 294 | 356 | 1988 | 378 | 443 | 2160 | 214 | 481 | 3557 | 511 | 1023 | 1132 | 424 | 620 | 1074 | 459 | 857 |
| 1304 | W303.asm02.HP0_G0013060 | chrIV | 3485 | 1134 | 1141 | 3287 | 1451 | 1104 | 3746 | 1226 | 1260 | 9491 | 2006 | 2076 | 6888 | 1122 | 979 | 7713 | 1714 | 1718 |
| 1305 | W303.asm02.HP0_G0013070 | chrIV | 3227 | 382 | 543 | 3215 | 497 | 625 | 3189 | 339 | 734 | 5069 | 827 | 1253 | 2650 | 590 | 802 | 2713 | 696 | 1032 |
| 1306 | W303.asm02.HP0_G0013080 | chrIV | 2601 | 1598 | 1930 | 2623 | 1661 | 1990 | 2449 | 1624 | 2424 | 4533 | 1231 | 1540 | 5668 | 973 | 1241 | 4822 | 1121 | 1412 |
| 1307 | W303.asm02.HP0_G0013090 | chrIV | 1075 | 933 | 1216 | 924 | 1089 | 1271 | 964 | 1011 | 1642 | 2902 | 749 | 953 | 3631 | 537 | 734 | 3172 | 658 | 776 |
| 1308 | W303.asm02.HP0_G0013100 | chrIV | 3663 | 1043 | 661 | 3715 | 595 | 592 | 3453 | 951 | 692 | 3503 | 1167 | 1055 | 2960 | 760 | 834 | 2798 | 1325 | 960 |
| 1309 | W303.asm02.HP0_G0013110 | chrIV | 8611 | 6717 | 9336 | 8047 | 9265 | 9686 | 8776 | 6973 | 11642 | 21090 | 5837 | 7161 | 23058 | 4116 | 5698 | 21848 | 5239 | 6318 |
| 1310 | W303.asm02.HP0_G0013120 | chrIV | 1561 | 1473 | 1035 | 1468 | 1100 | 1131 | 1546 | 1422 | 1089 | 3688 | 1117 | 1205 | 3771 | 878 | 817 | 3202 | 1092 | 942 |
| 1311 | W303.asm02.HP0_G0013130 | chrIV | 2476 | 697 | 510 | 2637 | 622 | 517 | 2441 | 667 | 442 | 2795 | 658 | 752 | 2545 | 412 | 486 | 2336 | 621 | 644 |
| 1312 | W303.asm02.HP0_G0013140 | chrIV | 5302 | 472 | 524 | 5267 | 429 | 552 | 4789 | 413 | 602 | 3150 | 951 | 1092 | 1658 | 600 | 835 | 1515 | 931 | 906 |
| 1313 | W303.asm02.HP0_G0013150 | chrIV | 2325 | 550 | 438 | 1855 | 386 | 579 | 2083 | 429 | 535 | 2418 | 995 | 1207 | 1778 | 648 | 802 | 1756 | 932 | 1127 |
| 1314 | W303.asm02.HP0_G0013160 | chrIV | 8370 | 754 | 907 | 8269 | 695 | 940 | 8457 | 631 | 1019 | 6670 | 1502 | 1926 | 2406 | 1149 | 1422 | 2632 | 1288 | 1696 |
| 1315 | W303.asm02.HP0_G0013170 | chrIV | 466 | 397 | 418 | 472 | 488 | 451 | 465 | 423 | 553 | 1290 | 337 | 314 | 1349 | 206 | 221 | 1264 | 302 | 285 |
| 1316 | W303.asm02.HP0_G0013180 | chrIV | 3769 | 573 | 481 | 3614 | 534 | 545 | 3431 | 505 | 597 | 3106 | 838 | 922 | 2292 | 501 | 736 | 2351 | 807 | 892 |
| 1317 | W303.asm02.HP0_G0013190 | chrIV | 1164 | 620 | 443 | 1139 | 433 | 410 | 1129 | 586 | 416 | 1838 | 541 | 581 | 1742 | 370 | 416 | 1654 | 593 | 517 |
| 1318 | W303.asm02.HP0_G0013200 | chrIV | 2853 | 1842 | 2158 | 2733 | 1714 | 2170 | 2691 | 1736 | 2508 | 7990 | 3093 | 3348 | 5351 | 2040 | 2328 | 4849 | 2768 | 2674 |
| 1319 | W303.asm02.HP0_G0013210 | chrIV | 926 | 583 | 483 | 808 | 525 | 569 | 767 | 564 | 516 | 1485 | 489 | 561 | 1760 | 313 | 402 | 1350 | 419 | 520 |
| 1320 | W303.asm02.HP0_G0013220 | chrIV | 10218 | 4566 | 4113 | 9544 | 3965 | 4258 | 10082 | 4853 | 3977 | 13959 | 4396 | 4478 | 13946 | 3146 | 3398 | 12705 | 3675 | 3897 |
| 1321 | W303.asm02.HP0_G0013230 | chrIV | 2632 | 903 | 913 | 2779 | 844 | 956 | 2655 | 984 | 1007 | 3876 | 734 | 814 | 3620 | 431 | 695 | 3982 | 686 | 839 |
| 1322 | W303.asm02.HP0_G0013240 | chrIV | 3598 | 657 | 613 | 3841 | 700 | 639 | 3627 | 650 | 671 | 5614 | 1006 | 1229 | 4661 | 665 | 943 | 3802 | 946 | 1079 |
| 1323 | W303.asm02.HP0_G0013250 | chrIV | 4284 | 1760 | 1707 | 4399 | 1748 | 1718 | 4035 | 1724 | 2032 | 4385 | 1382 | 1529 | 5923 | 1109 | 1236 | 4389 | 1201 | 1309 |
| 1324 | W303.asm02.HP0_G0013260 | chrIV | 5850 | 4389 | 4837 | 5595 | 4422 | 4665 | 5746 | 4734 | 5674 | 9735 | 4102 | 4238 | 12716 | 3135 | 3731 | 9666 | 3568 | 3790 |
| 1325 | W303.asm02.HP0_G0013270 | chrIV | 894 | 239 | 267 | 776 | 254 | 255 | 854 | 185 | 357 | 1593 | 499 | 541 | 963 | 335 | 359 | 839 | 428 | 509 |
| 1326 | W303.asm02.HP0_G0013280 | chrIV | 803 | 565 | 459 | 684 | 523 | 458 | 819 | 529 | 409 | 2009 | 410 | 537 | 1995 | 296 | 327 | 1808 | 363 | 406 |
| 1327 | W303.asm02.HP0_G0013290 | chrIV | 2524 | 419 | 369 | 2166 | 433 | 385 | 2322 | 367 | 439 | 2609 | 792 | 836 | 1281 | 508 | 576 | 1364 | 739 | 790 |
| 1328 | W303.asm02.HP0_G0013300 | chrIV | 13 | 6 | 4 | 18 | 1 | 5 | 10 | 4 | 2 | 17 | 8 | 5 | 11 | 3 | 5 | 16 | 8 | 8 |
| 1329 | W303.asm02.HP0_G0013310 | chrIV | 837 | 1353 | 1769 | 907 | 2030 | 1492 | 921 | 1259 | 1680 | 8334 | 1549 | 1573 | 7223 | 945 | 1015 | 6988 | 1585 | 1284 |
| 1330 | W303.asm02.HP0_G0013320 | chrIV | 2882 | 795 | 1068 | 2352 | 984 | 1162 | 2405 | 795 | 1389 | 2513 | 595 | 900 | 3089 | 433 | 755 | 2577 | 607 | 790 |
| 1331 | W303.asm02.HP0_G0013330 | chrIV | 11464 | 2255 | 1896 | 11630 | 1947 | 1735 | 11149 | 2372 | 1932 | 8258 | 2532 | 2762 | 8564 | 1959 | 2324 | 7382 | 2552 | 2606 |
| 1332 | W303.asm02.HP0_G0013340 | chrIV | 2696 | 532 | 775 | 2782 | 775 | 742 | 2638 | 605 | 821 | 3899 | 583 | 878 | 3525 | 376 | 710 | 3863 | 505 | 854 |
| 1333 | W303.asm02.HP0_G0013350 | chrIV | 2370 | 2489 | 2792 | 2405 | 3424 | 2471 | 2864 | 2685 | 2402 | 19527 | 3071 | 3765 | 16294 | 1919 | 1946 | 16607 | 2606 | 3297 |
| 1334 | W303.asm02.HP0_G0013360 | chrIV | 12081 | 2670 | 2675 | 12130 | 2439 | 2575 | 13687 | 2884 | 2786 | 29138 | 9005 | 9466 | 20180 | 7339 | 7875 | 21028 | 7939 | 8635 |
| 1335 | W303.asm02.HP0_G0013370 | chrIV | 2321 | 652 | 407 | 2352 | 407 | 426 | 2347 | 590 | 392 | 3110 | 842 | 961 | 2793 | 607 | 633 | 2680 | 789 | 830 |
| 1336 | W303.asm02.HP0_G0013380 | chrIV | 1031 | 397 | 511 | 1051 | 500 | 472 | 933 | 471 | 463 | 2501 | 453 | 713 | 2154 | 272 | 517 | 2325 | 448 | 633 |
| 1337 | W303.asm02.HP0_G0013390 | chrIV | 5006 | 3620 | 4990 | 4238 | 4519 | 4862 | 4437 | 3843 | 5743 | 9666 | 3407 | 4060 | 11544 | 2331 | 3581 | 10390 | 2999 | 3431 |
| 1338 | W303.asm02.HP0_G0013400 | chrIV | 731 | 436 | 307 | 625 | 300 | 347 | 702 | 394 | 351 | 1307 | 428 | 455 | 1150 | 279 | 276 | 1206 | 435 | 383 |
| 1339 | W303.asm02.HP0_G0013410 | chrIV | 24292 | 3597 | 3659 | 21692 | 3241 | 3476 | 25279 | 3320 | 2949 | 17174 | 4994 | 6386 | 16542 | 4069 | 5264 | 15583 | 4242 | 5413 |
| 1340 | W303.asm02.HP0_G0013420 | chrIV | 590 | 818 | 1338 | 516 | 701 | 1555 | 633 | 777 | 1618 | 1851 | 508 | 1165 | 2564 | 367 | 783 | 2228 | 511 | 1029 |
| 1341 | W303.asm02.HP0_G0013430 | chrIV | 275 | 136 | 230 | 250 | 160 | 204 | 273 | 181 | 334 | 749 | 169 | 214 | 705 | 109 | 211 | 923 | 144 | 183 |
| 1342 | W303.asm02.HP0_G0013440 | chrIV | 3854 | 1267 | 1326 | 4027 | 1127 | 1226 | 3901 | 1268 | 1207 | 8124 | 1720 | 1934 | 8083 | 1371 | 1685 | 6816 | 1495 | 1649 |
| 1343 | W303.asm02.HP0_G0013450 | chrIV | 810 | 182 | 189 | 768 | 162 | 174 | 800 | 190 | 136 | 863 | 230 | 367 | 743 | 170 | 253 | 667 | 224 | 267 |
| 1344 | W303.asm02.HP0_G0013460 | chrIV | 4366 | 877 | 1205 | 4548 | 1195 | 1382 | 4979 | 943 | 1378 | 10208 | 1583 | 2839 | 8561 | 1247 | 1998 | 7345 | 1351 | 2589 |
| 1345 | W303.asm02.HP0_G0013470 | chrIV | 2495 | 3398 | 2888 | 2401 | 2777 | 3025 | 2725 | 3026 | 3365 | 22533 | 7891 | 7319 | 12219 | 5453 | 5106 | 13696 | 6788 | 6794 |
| 1346 | W303.asm02.HP0_G0013480 | chrIV | 3482 | 1378 | 1595 | 3153 | 1255 | 1675 | 3470 | 1389 | 1601 | 5430 | 1713 | 2330 | 4969 | 1349 | 1772 | 4921 | 1588 | 2116 |
| 1347 | W303.asm02.HP0_G0013490 | chrIV | 3373 | 1090 | 1383 | 2718 | 1192 | 1378 | 2983 | 1070 | 1875 | 2475 | 777 | 1061 | 3625 | 610 | 993 | 2632 | 828 | 943 |
| 1348 | W303.asm02.HP0_G0013500 | chrIV | 3773 | 2432 | 2589 | 3393 | 2049 | 2548 | 3266 | 2343 | 3112 | 4025 | 2017 | 1903 | 5206 | 1354 | 1897 | 4469 | 1898 | 1667 |
| 1349 | W303.asm02.HP0_G0013510 | chrIV | 1415 | 563 | 614 | 1325 | 721 | 556 | 1307 | 551 | 665 | 4681 | 816 | 1059 | 3524 | 504 | 643 | 3891 | 687 | 838 |
| 1350 | W303.asm02.HP0_G0013520 | chrIV | 2715 | 887 | 1585 | 3028 | 1507 | 1468 | 2702 | 988 | 1780 | 5378 | 685 | 1348 | 5384 | 499 | 1103 | 5418 | 680 | 1223 |
| 1351 | W303.asm02.HP0_G0013530 | chrIV | 25146 | 2407 | 1482 | 25041 | 1441 | 1543 | 23554 | 1739 | 1504 | 5290 | 2224 | 2072 | 6444 | 1788 | 2348 | 4454 | 2452 | 1972 |
| 1352 | W303.asm02.HP0_G0013540 | chrIV | 5002 | 1006 | 888 | 4431 | 665 | 1007 | 4838 | 845 | 1105 | 4579 | 1732 | 1909 | 2974 | 1259 | 1409 | 2511 | 1471 | 1762 |
| 1353 | W303.asm02.HP0_G0013550 | chrIV | 1922 | 775 | 1544 | 2062 | 750 | 1808 | 1923 | 779 | 1456 | 3087 | 740 | 2131 | 4067 | 566 | 1280 | 2862 | 634 | 1950 |
| 1354 | W303.asm02.HP0_G0013560 | chrIV | 2172 | 1419 | 2913 | 2053 | 1726 | 3037 | 2197 | 1547 | 3601 | 4368 | 1360 | 2845 | 5803 | 970 | 1919 | 4816 | 1191 | 2443 |
| 1355 | W303.asm02.HP0_G0013570 | chrIV | 829 | 405 | 484 | 839 | 475 | 489 | 804 | 438 | 497 | 3058 | 451 | 868 | 2552 | 259 | 562 | 2680 | 439 | 756 |
| 1356 | W303.asm02.HP0_G0013580 | chrIV | 323 | 123 | 99 | 307 | 139 | 94 | 300 | 134 | 119 | 466 | 98 | 96 | 444 | 64 | 67 | 504 | 103 | 88 |
| 1357 | W303.asm02.HP0_G0013590 | chrIV | 472 | 119 | 106 | 465 | 91 | 83 | 442 | 122 | 124 | 351 | 128 | 104 | 443 | 87 | 73 | 359 | 103 | 123 |
| 1358 | W303.asm02.HP0_G0013600 | chrIV | 2135 | 1575 | 1688 | 2073 | 1830 | 1627 | 2316 | 1680 | 1798 | 7134 | 1361 | 1580 | 8177 | 1025 | 1390 | 7038 | 1184 | 1450 |
| 1359 | W303.asm02.HP0_G0013610 | chrIV | 14777 | 10275 | 14608 | 14296 | 13363 | 14486 | 14162 | 10904 | 18230 | 22751 | 9903 | 13190 | 25858 | 7693 | 10807 | 23930 | 8993 | 11667 |
| 1360 | W303.asm02.HP0_G0013620 | chrIV | 845 | 527 | 502 | 853 | 461 | 474 | 726 | 552 | 613 | 1121 | 476 | 356 | 1695 | 382 | 347 | 1164 | 469 | 259 |
| 1361 | W303.asm02.HP0_G0013630 | chrIV | 1532 | 1459 | 1439 | 1438 | 1721 | 1457 | 1572 | 1455 | 1364 | 7109 | 1334 | 1565 | 6674 | 813 | 857 | 6229 | 1311 | 1213 |
| 1362 | W303.asm02.HP0_G0013640 | chrIV | 83 | 345 | 484 | 78 | 195 | 615 | 105 | 224 | 543 | 1399 | 438 | 1149 | 1445 | 270 | 710 | 1330 | 475 | 1116 |
| 1363 | W303.asm02.HP0_G0013650 | chrIV | 6310 | 5236 | 5863 | 5727 | 5591 | 5805 | 5744 | 5326 | 7077 | 11504 | 4215 | 4812 | 13349 | 3341 | 4112 | 11790 | 3837 | 3820 |
| 1364 | W303.asm02.HP0_G0013660 | chrIV | 4141 | 1330 | 797 | 3751 | 822 | 792 | 3899 | 1114 | 834 | 2507 | 1087 | 718 | 3046 | 693 | 753 | 2231 | 1017 | 682 |
| 1365 | W303.asm02.HP0_G0013670 | chrIV | 1904 | 2087 | 1813 | 1548 | 1771 | 1636 | 1615 | 1937 | 1938 | 6399 | 1653 | 1738 | 5324 | 1009 | 1302 | 6470 | 1644 | 1536 |
| 1366 | W303.asm02.HP0_G0013680 | chrIV | 1381 | 721 | 553 | 1282 | 573 | 554 | 1232 | 677 | 592 | 2317 | 466 | 536 | 2452 | 332 | 419 | 2247 | 519 | 461 |
| 1367 | W303.asm02.HP0_G0013690 | chrIV | 6161 | 3589 | 3604 | 5501 | 2598 | 3762 | 5330 | 3572 | 4158 | 3129 | 3670 | 3834 | 4625 | 3599 | 4466 | 3056 | 3179 | 2893 |
| 1368 | W303.asm02.HP0_G0013700 | chrIV | 187 | 39 | 42 | 192 | 44 | 62 | 193 | 25 | 65 | 293 | 65 | 105 | 118 | 31 | 60 | 126 | 80 | 130 |
| 1369 | W303.asm02.HP0_G0013710 | chrIV | 618 | 604 | 155 | 529 | 128 | 151 | 609 | 402 | 135 | 477 | 471 | 199 | 682 | 208 | 155 | 497 | 481 | 175 |
| 1370 | W303.asm02.HP0_G0013720 | chrIV | 1431 | 973 | 1048 | 1219 | 995 | 1068 | 1387 | 937 | 1098 | 3212 | 791 | 1039 | 3052 | 442 | 599 | 3024 | 733 | 908 |
| 1371 | W303.asm02.HP0_G0013730 | chrIV | 4297 | 942 | 892 | 4211 | 741 | 1044 | 4273 | 801 | 1099 | 5456 | 1497 | 1926 | 4349 | 1010 | 1316 | 4224 | 1459 | 1679 |
| 1372 | W303.asm02.HP0_G0013740 | chrIV | 807 | 353 | 570 | 890 | 322 | 583 | 748 | 333 | 690 | 936 | 199 | 425 | 1423 | 161 | 369 | 923 | 217 | 462 |
| 1373 | W303.asm02.HP0_G0013750 | chrIV | 1608 | 584 | 563 | 1412 | 578 | 582 | 1465 | 594 | 456 | 2302 | 516 | 695 | 2203 | 353 | 507 | 2138 | 448 | 622 |
| 1374 | W303.asm02.HP0_G0025560 | chrIX | 76 | 25 | 43 | 72 | 51 | 45 | 91 | 32 | 58 | 154 | 24 | 52 | 143 | 43 | 26 | 124 | 29 | 44 |
| 1375 | W303.asm02.HP0_G0025570 | chrIX | 0 | 0 | 0 | 0 | 0 | 0 | 0 | 0 | 0 | 0 | 0 | 0 | 0 | 0 | 0 | 0 | 0 | 0 |
| 1376 | W303.asm02.HP0_G0025580 | chrIX | 95 | 11 | 25 | 77 | 16 | 34 | 104 | 6 | 30 | 47 | 14 | 72 | 38 | 13 | 37 | 46 | 6 | 71 |
| 1377 | W303.asm02.HP0_G0025590 | chrIX | 460 | 33 | 90 | 426 | 49 | 134 | 484 | 40 | 125 | 325 | 67 | 306 | 264 | 55 | 226 | 203 | 61 | 332 |
| 1378 | W303.asm02.HP0_G0025600 | chrIX | 25332 | 594 | 1234 | 22815 | 1482 | 1411 | 25653 | 516 | 1596 | 15020 | 1400 | 4525 | 8916 | 1220 | 3709 | 6182 | 1159 | 3705 |
| 1379 | W303.asm02.HP0_G0025610 | chrIX | 325 | 269 | 365 | 343 | 338 | 334 | 261 | 322 | 382 | 714 | 362 | 377 | 914 | 250 | 315 | 709 | 341 | 286 |
| 1380 | W303.asm02.HP0_G0025620 | chrIX | 175 | 289 | 408 | 204 | 393 | 360 | 170 | 296 | 466 | 681 | 392 | 379 | 685 | 265 | 326 | 643 | 326 | 258 |
| 1381 | W303.asm02.HP0_G0025630 | chrIX | 847 | 270 | 275 | 864 | 249 | 257 | 816 | 230 | 323 | 1106 | 238 | 300 | 1054 | 145 | 178 | 1119 | 237 | 260 |
| 1382 | W303.asm02.HP0_G0025640 | chrIX | 350 | 117 | 153 | 291 | 141 | 151 | 308 | 116 | 186 | 355 | 127 | 136 | 303 | 83 | 115 | 237 | 126 | 108 |
| 1383 | W303.asm02.HP0_G0025650 | chrIX | 715 | 206 | 221 | 625 | 191 | 210 | 590 | 188 | 287 | 749 | 253 | 215 | 493 | 190 | 226 | 437 | 276 | 219 |
| 1384 | W303.asm02.HP0_G0025660 | chrIX | 125 | 31 | 35 | 86 | 36 | 29 | 92 | 28 | 41 | 150 | 37 | 43 | 142 | 24 | 41 | 131 | 34 | 39 |
| 1385 | W303.asm02.HP0_G0025670 | chrIX | 15305 | 968 | 747 | 18512 | 971 | 903 | 16136 | 944 | 859 | 6657 | 1371 | 1235 | 6421 | 973 | 1097 | 5645 | 1246 | 1238 |
| 1386 | W303.asm02.HP0_G0025680 | chrIX | 1414 | 342 | 494 | 1214 | 437 | 476 | 1342 | 423 | 561 | 1652 | 429 | 558 | 1782 | 275 | 468 | 1677 | 348 | 467 |
| 1387 | W303.asm02.HP0_G0025690 | chrIX | 516 | 77 | 56 | 560 | 54 | 78 | 518 | 62 | 91 | 385 | 104 | 139 | 245 | 63 | 100 | 206 | 100 | 136 |
| 1388 | W303.asm02.HP0_G0025700 | chrIX | 1000 | 195 | 327 | 965 | 357 | 331 | 980 | 232 | 485 | 1438 | 257 | 513 | 1095 | 190 | 333 | 1135 | 200 | 394 |
| 1389 | W303.asm02.HP0_G0025710 | chrIX | 659 | 847 | 1279 | 601 | 1463 | 1356 | 721 | 960 | 1796 | 3485 | 784 | 899 | 3964 | 483 | 744 | 3845 | 671 | 876 |
| 1390 | W303.asm02.HP0_G0025720 | chrIX | 7945 | 2003 | 2235 | 7710 | 1919 | 2096 | 7350 | 2191 | 2514 | 4794 | 1786 | 1985 | 6666 | 1296 | 1887 | 5144 | 1607 | 1809 |
| 1391 | W303.asm02.HP0_G0025730 | chrIX | 920 | 453 | 698 | 680 | 569 | 668 | 698 | 498 | 879 | 1013 | 382 | 463 | 1510 | 266 | 483 | 1214 | 366 | 393 |
| 1392 | W303.asm02.HP0_G0025740 | chrIX | 2170 | 253 | 413 | 2037 | 363 | 484 | 1983 | 250 | 521 | 1673 | 404 | 732 | 832 | 283 | 447 | 858 | 317 | 610 |
| 1393 | W303.asm02.HP0_G0025750 | chrIX | 1907 | 559 | 454 | 1924 | 424 | 564 | 1826 | 415 | 636 | 1262 | 785 | 861 | 537 | 653 | 573 | 522 | 641 | 624 |

| | | | | | | | | | | | | | | | | | | | | |
|---|---|---|---|---|---|---|---|---|---|---|---|---|---|---|---|---|---|---|---|---|
| 1394 | W303.asm02.HP0_G0025760 | chrIX | 7952 | 1393 | 1095 | 7351 | 879 | 1090 | 7446 | 1216 | 1256 | 3743 | 1790 | 1633 | 3867 | 1172 | 1622 | 2534 | 1473 | 1507 |
| 1395 | W303.asm02.HP0_G0025770 | chrIX | 1481 | 547 | 511 | 1143 | 405 | 547 | 1430 | 336 | 678 | 1714 | 723 | 830 | 702 | 524 | 560 | 658 | 653 | 730 |
| 1396 | W303.asm02.HP0_G0025780 | chrIX | 1802 | 482 | 405 | 1563 | 316 | 352 | 1591 | 394 | 395 | 1177 | 449 | 429 | 1220 | 345 | 350 | 1006 | 402 | 359 |
| 1397 | W303.asm02.HP0_G0025790 | chrIX | 1910 | 517 | 792 | 1724 | 729 | 830 | 1817 | 505 | 959 | 5437 | 927 | 1414 | 3882 | 655 | 1080 | 3433 | 763 | 1239 |
| 1398 | W303.asm02.HP0_G0025800 | chrIX | 464 | 66 | 175 | 555 | 169 | 149 | 452 | 52 | 180 | 1953 | 119 | 351 | 1133 | 93 | 167 | 1054 | 129 | 260 |
| 1399 | W303.asm02.HP0_G0025810 | chrIX | 4598 | 914 | 1120 | 4522 | 1256 | 1084 | 4636 | 1114 | 1032 | 11582 | 1360 | 2485 | 8335 | 926 | 1726 | 8985 | 1254 | 2131 |
| 1400 | W303.asm02.HP0_G0025820 | chrIX | 1735 | 609 | 816 | 1714 | 843 | 901 | 1759 | 626 | 981 | 7113 | 1037 | 1841 | 4855 | 770 | 1277 | 4721 | 869 | 1595 |
| 1401 | W303.asm02.HP0_G0025830 | chrIX | 753 | 168 | 187 | 785 | 192 | 222 | 764 | 161 | 224 | 1370 | 313 | 475 | 1031 | 226 | 348 | 863 | 267 | 371 |
| 1402 | W303.asm02.HP0_G0025840 | chrIX | 3228 | 1178 | 993 | 2927 | 993 | 1045 | 3358 | 1068 | 1218 | 5750 | 1539 | 1707 | 4465 | 1203 | 1281 | 4477 | 1461 | 1605 |
| 1403 | W303.asm02.HP0_G0025850 | chrIX | 1980 | 731 | 639 | 2016 | 775 | 583 | 1882 | 710 | 598 | 3624 | 807 | 955 | 3579 | 551 | 876 | 3313 | 809 | 928 |
| 1404 | W303.asm02.HP0_G0025860 | chrIX | 3786 | 772 | 985 | 3577 | 967 | 1080 | 3551 | 668 | 1317 | 6863 | 1667 | 2132 | 3369 | 1094 | 1291 | 3439 | 1593 | 1738 |
| 1405 | W303.asm02.HP0_G0025870 | chrIX | 9544 | 4288 | 3308 | 9436 | 3432 | 3423 | 9913 | 3933 | 2674 | 15243 | 4813 | 6497 | 12921 | 3592 | 4109 | 12639 | 4186 | 5492 |
| 1406 | W303.asm02.HP0_G0025880 | chrIX | 3799 | 921 | 1087 | 3825 | 919 | 1119 | 3959 | 988 | 1258 | 6935 | 1719 | 2574 | 7540 | 1431 | 1750 | 5306 | 1534 | 2232 |
| 1407 | W303.asm02.HP0_G0025890 | chrIX | 583 | 615 | 767 | 537 | 544 | 792 | 542 | 557 | 1041 | 1916 | 419 | 565 | 2158 | 282 | 408 | 2153 | 439 | 513 |
| 1408 | W303.asm02.HP0_G0025900 | chrIX | 4121 | 3446 | 3947 | 4211 | 3555 | 3780 | 4123 | 3790 | 5165 | 9481 | 3024 | 3343 | 11017 | 2110 | 2555 | 9986 | 2736 | 2922 |
| 1409 | W303.asm02.HP0_G0025910 | chrIX | 6173 | 713 | 627 | 5526 | 601 | 822 | 6035 | 571 | 925 | 6656 | 1365 | 1728 | 3491 | 1108 | 1173 | 3338 | 1343 | 1507 |
| 1410 | W303.asm02.HP0_G0025920 | chrIX | 13648 | 1263 | 568 | 13155 | 612 | 701 | 11873 | 1015 | 785 | 2123 | 1472 | 1186 | 1998 | 1134 | 1179 | 1432 | 1515 | 936 |
| 1411 | W303.asm02.HP0_G0025930 | chrIX | 4335 | 816 | 651 | 4436 | 506 | 685 | 4167 | 637 | 705 | 3378 | 1378 | 1229 | 2543 | 1014 | 965 | 2300 | 1416 | 1091 |
| 1412 | W303.asm02.HP0_G0025940 | chrIX | 19 | 13 | 5 | 18 | 1 | 0 | 22 | 9 | 5 | 19 | 16 | 8 | 10 | 11 | 3 | 12 | 14 | 3 |
| 1413 | W303.asm02.HP0_G0025950 | chrIX | 865 | 535 | 680 | 704 | 660 | 657 | 767 | 560 | 790 | 2150 | 569 | 744 | 2192 | 384 | 552 | 1974 | 569 | 704 |
| 1414 | W303.asm02.HP0_G0025960 | chrIX | 10260 | 6055 | 6577 | 9535 | 6916 | 6523 | 10247 | 6058 | 6293 | 21353 | 5780 | 6973 | 19480 | 4420 | 5067 | 19792 | 4848 | 6450 |
| 1415 | W303.asm02.HP0_G0025970 | chrIX | 463 | 118 | 74 | 487 | 111 | 115 | 447 | 112 | 79 | 426 | 137 | 130 | 427 | 103 | 91 | 357 | 152 | 110 |
| 1416 | W303.asm02.HP0_G0025980 | chrIX | 764 | 947 | 669 | 732 | 822 | 644 | 809 | 992 | 611 | 4998 | 1209 | 1246 | 4139 | 784 | 916 | 4546 | 1034 | 1228 |
| 1417 | W303.asm02.HP0_G0025990 | chrIX | 1659 | 422 | 840 | 1721 | 845 | 976 | 1644 | 448 | 1078 | 6457 | 937 | 1687 | 3912 | 519 | 1051 | 4210 | 895 | 1435 |
| 1418 | W303.asm02.HP0_G0026000 | chrIX | 3136 | 782 | 1071 | 2814 | 1055 | 1104 | 2813 | 775 | 1298 | 7950 | 962 | 2077 | 5891 | 726 | 1749 | 6413 | 919 | 1828 |
| 1419 | W303.asm02.HP0_G0026010 | chrIX | 1956 | 1235 | 1675 | 1833 | 1716 | 1637 | 2030 | 1341 | 1610 | 10077 | 1413 | 2350 | 8209 | 988 | 1675 | 8913 | 1276 | 2244 |
| 1420 | W303.asm02.HP0_G0026020 | chrIX | 1141 | 984 | 1746 | 1011 | 1939 | 1829 | 1173 | 1297 | 2318 | 5875 | 1174 | 1598 | 6391 | 626 | 941 | 6076 | 949 | 1380 |
| 1421 | W303.asm02.HP0_G0026030 | chrIX | 5661 | 1265 | 1446 | 5613 | 1436 | 1595 | 5841 | 1299 | 1580 | 12595 | 1827 | 3420 | 8502 | 1421 | 2464 | 8735 | 1580 | 3034 |
| 1422 | W303.asm02.HP0_G0026040 | chrIX | 15100 | 2584 | 1940 | 13633 | 1926 | 2201 | 14213 | 2309 | 2574 | 13159 | 4341 | 4257 | 7767 | 3636 | 3689 | 7600 | 3788 | 3528 |
| 1423 | W303.asm02.HP0_G0026050 | chrIX | 4924 | 2531 | 2732 | 4279 | 2333 | 2798 | 4600 | 2329 | 3445 | 5221 | 1772 | 2213 | 7755 | 1488 | 2213 | 5467 | 1605 | 1802 |
| 1424 | W303.asm02.HP0_G0026060 | chrIX | 10546 | 1162 | 1797 | 9768 | 1631 | 2082 | 11321 | 1125 | 2364 | 4419 | 2110 | 3310 | 2358 | 2105 | 2077 | 1503 | 1629 | 2913 |
| 1425 | W303.asm02.HP0_G0026070 | chrIX | 1403 | 199 | 186 | 1541 | 226 | 165 | 1416 | 179 | 214 | 1936 | 293 | 368 | 1796 | 212 | 247 | 1434 | 302 | 303 |
| 1426 | W303.asm02.HP0_G0026080 | chrIX | 2204 | 531 | 478 | 1694 | 373 | 457 | 1977 | 580 | 620 | 2220 | 756 | 731 | 2848 | 817 | 1046 | 1435 | 558 | 724 |
| 1427 | W303.asm02.HP0_G0026090 | chrIX | 812 | 371 | 426 | 742 | 382 | 432 | 738 | 346 | 472 | 1348 | 351 | 394 | 1242 | 193 | 290 | 1235 | 313 | 350 |
| 1428 | W303.asm02.HP0_G0026100 | chrIX | 1751 | 629 | 815 | 1513 | 861 | 749 | 1829 | 664 | 796 | 3052 | 538 | 544 | 2589 | 281 | 565 | 2700 | 500 | 462 |
| 1429 | W303.asm02.HP0_G0026110 | chrIX | 1464 | 1994 | 2789 | 1403 | 2757 | 2766 | 1492 | 2062 | 3342 | 7193 | 1778 | 2244 | 7496 | 1101 | 1804 | 7338 | 1524 | 2076 |
| 1430 | W303.asm02.HP0_G0026120 | chrIX | 6554 | 2007 | 1853 | 6422 | 1705 | 1581 | 6089 | 1832 | 1833 | 4617 | 1616 | 1808 | 9689 | 1879 | 2387 | 4178 | 1551 | 1763 |
| 1431 | W303.asm02.HP0_G0026130 | chrIX | 3150 | 2267 | 1834 | 2650 | 1674 | 1553 | 2919 | 2199 | 1733 | 4993 | 2043 | 2073 | 5245 | 1547 | 1727 | 4615 | 1735 | 1749 |
| 1432 | W303.asm02.HP0_G0026140 | chrIX | 5202 | 950 | 1153 | 4945 | 1222 | 1304 | 5298 | 931 | 1530 | 10498 | 1750 | 2434 | 6657 | 1398 | 1944 | 6312 | 1537 | 2140 |
| 1433 | W303.asm02.HP0_G0026150 | chrIX | 673 | 450 | 463 | 441 | 451 | 425 | 582 | 431 | 466 | 2416 | 521 | 517 | 2297 | 311 | 451 | 2110 | 501 | 510 |
| 1434 | W303.asm02.HP0_G0026160 | chrIX | 1239 | 161 | 144 | 1115 | 157 | 145 | 995 | 150 | 103 | 348 | 149 | 159 | 415 | 114 | 188 | 274 | 161 | 146 |
| 1435 | W303.asm02.HP0_G0026170 | chrIX | 6343 | 614 | 648 | 6164 | 840 | 900 | 6029 | 590 | 801 | 4810 | 832 | 1330 | 3328 | 684 | 1051 | 3045 | 806 | 1098 |
| 1436 | W303.asm02.HP0_G0026180 | chrIX | 4297 | 825 | 897 | 4052 | 731 | 965 | 3834 | 616 | 977 | 1565 | 564 | 647 | 2764 | 486 | 897 | 1422 | 557 | 552 |
| 1437 | W303.asm02.HP0_G0026190 | chrIX | 1344 | 1214 | 1787 | 1179 | 1823 | 1612 | 1375 | 1360 | 2065 | 6767 | 1170 | 1476 | 7027 | 720 | 891 | 6911 | 1059 | 1284 |
| 1438 | W303.asm02.HP0_G0026200 | chrIX | 12761 | 1619 | 2714 | 11157 | 1817 | 3328 | 12515 | 1314 | 3639 | 8458 | 2376 | 5981 | 3161 | 2376 | 3326 | 2163 | 1769 | 4876 |
| 1439 | W303.asm02.HP0_G0026210 | chrIX | 4286 | 475 | 898 | 3958 | 591 | 963 | 4197 | 438 | 1208 | 3181 | 1048 | 2081 | 2169 | 1031 | 1497 | 1304 | 1006 | 1806 |
| 1440 | W303.asm02.HP0_G0026220 | chrIX | 8781 | 685 | 450 | 8182 | 519 | 421 | 8027 | 773 | 515 | 3484 | 881 | 847 | 4826 | 911 | 959 | 2801 | 888 | 785 |
| 1441 | W303.asm02.HP0_G0026230 | chrIX | 2250 | 937 | 656 | 2262 | 623 | 720 | 2338 | 812 | 794 | 3907 | 1329 | 1294 | 3196 | 1014 | 1161 | 2788 | 1328 | 1335 |
| 1442 | W303.asm02.HP0_G0026240 | chrIX | 621 | 745 | 819 | 586 | 987 | 778 | 706 | 827 | 913 | 4994 | 767 | 945 | 4587 | 507 | 569 | 4379 | 736 | 822 |
| 1443 | W303.asm02.HP0_G0026250 | chrIX | 1487 | 1405 | 1642 | 1353 | 1826 | 1462 | 1556 | 1397 | 2041 | 6472 | 1245 | 1468 | 6446 | 752 | 944 | 6483 | 1111 | 1179 |
| 1444 | W303.asm02.HP0_G0026260 | chrIX | 322 | 220 | 339 | 258 | 247 | 364 | 244 | 229 | 397 | 639 | 169 | 297 | 688 | 101 | 202 | 691 | 162 | 233 |
| 1445 | W303.asm02.HP0_G0026270 | chrIX | 26 | 15 | 8 | 16 | 13 | 8 | 7 | 13 | 10 | 17 | 5 | 10 | 20 | 8 | 5 | 25 | 6 | 6 |
| 1446 | W303.asm02.HP0_G0026280 | chrIX | 1715 | 164 | 166 | 1791 | 174 | 236 | 1518 | 174 | 254 | 624 | 261 | 348 | 325 | 196 | 247 | 265 | 235 | 300 |
| 1447 | W303.asm02.HP0_G0026290 | chrIX | 727 | 101 | 135 | 691 | 98 | 131 | 746 | 96 | 192 | 395 | 257 | 217 | 281 | 141 | 134 | 176 | 189 | 194 |
| 1448 | W303.asm02.HP0_G0026300 | chrIX | 3942 | 954 | 1218 | 3627 | 1157 | 1231 | 3383 | 1042 | 1634 | 2983 | 858 | 993 | 3982 | 638 | 1020 | 2928 | 790 | 958 |
| 1449 | W303.asm02.HP0_G0026310 | chrIX | 4846 | 809 | 706 | 4964 | 599 | 718 | 4347 | 732 | 698 | 2086 | 555 | 609 | 2524 | 351 | 584 | 2300 | 568 | 566 |
| 1450 | W303.asm02.HP0_G0026320 | chrIX | 246 | 663 | 968 | 270 | 1156 | 884 | 313 | 718 | 1225 | 4682 | 649 | 847 | 4345 | 395 | 376 | 5195 | 585 | 671 |
| 1451 | W303.asm02.HP0_G0026330 | chrIX | 2582 | 329 | 401 | 2561 | 367 | 479 | 2502 | 336 | 463 | 3943 | 659 | 979 | 2837 | 441 | 686 | 2598 | 590 | 857 |
| 1452 | W303.asm02.HP0_G0026340 | chrIX | 4895 | 5881 | 5771 | 4998 | 5543 | 5512 | 5167 | 6190 | 6275 | 14418 | 5106 | 5518 | 13743 | 2979 | 3979 | 14703 | 4671 | 5172 |
| 1453 | W303.asm02.HP0_G0026350 | chrIX | 8474 | 1689 | 1504 | 8037 | 1478 | 1571 | 8205 | 1765 | 1416 | 5647 | 1639 | 2013 | 6181 | 1236 | 1731 | 4700 | 1375 | 1908 |
| 1454 | W303.asm02.HP0_G0026360 | chrIX | 530 | 239 | 276 | 488 | 285 | 266 | 550 | 211 | 267 | 2169 | 293 | 492 | 1612 | 215 | 300 | 1550 | 270 | 412 |
| 1455 | W303.asm02.HP0_G0026370 | chrIX | 1972 | 1309 | 1412 | 2119 | 1898 | 1283 | 2304 | 1437 | 1431 | 11926 | 1885 | 2267 | 9300 | 1225 | 1339 | 9302 | 1626 | 2154 |
| 1456 | W303.asm02.HP0_G0026380 | chrIX | 2707 | 829 | 1163 | 2499 | 1117 | 1183 | 2829 | 831 | 1411 | 7450 | 1520 | 1714 | 5371 | 1008 | 1258 | 5729 | 1413 | 1556 |
| 1457 | W303.asm02.HP0_G0026390 | chrIX | 537 | 524 | 330 | 496 | 322 | 343 | 483 | 475 | 415 | 982 | 421 | 252 | 1089 | 234 | 244 | 1086 | 419 | 244 |
| 1458 | W303.asm02.HP0_G0026400 | chrIX | 6834 | 1703 | 1807 | 6194 | 1555 | 1782 | 6229 | 1769 | 1839 | 7111 | 2435 | 2695 | 7612 | 1888 | 2531 | 6091 | 2174 | 2460 |
| 1459 | W303.asm02.HP0_G0026410 | chrIX | 2725 | 566 | 435 | 2317 | 404 | 407 | 2405 | 530 | 464 | 1096 | 458 | 393 | 1375 | 300 | 396 | 1060 | 487 | 337 |
| 1460 | W303.asm02.HP0_G0026420 | chrIX | 66 | 9 | 9 | 36 | 9 | 7 | 41 | 8 | 7 | 11 | 7 | 4 | 15 | 1 | 4 | 16 | 8 | 3 |
| 1461 | W303.asm02.HP0_G0026430 | chrIX | 3033 | 749 | 750 | 2567 | 720 | 769 | 2773 | 689 | 849 | 5877 | 1242 | 1547 | 3749 | 960 | 993 | 4101 | 1182 | 1365 |
| 1462 | W303.asm02.HP0_G0026440 | chrIX | 1803 | 194 | 203 | 1841 | 249 | 251 | 1648 | 162 | 255 | 2206 | 318 | 497 | 1571 | 205 | 322 | 1435 | 306 | 394 |
| 1463 | W303.asm02.HP0_G0026450 | chrIX | 2790 | 1134 | 1130 | 2313 | 993 | 1138 | 2720 | 1120 | 1049 | 4453 | 1092 | 1219 | 4166 | 738 | 893 | 4401 | 1097 | 1160 |
| 1464 | W303.asm02.HP0_G0026460 | chrIX | 23173 | 7851 | 6202 | 21898 | 6515 | 5827 | 23335 | 8031 | 6414 | 36691 | 11983 | 12352 | 29758 | 8242 | 8636 | 31555 | 10499 | 11366 |
| 1465 | W303.asm02.HP0_G0026470 | chrIX | 1732 | 701 | 393 | 1724 | 351 | 426 | 1470 | 551 | 393 | 1997 | 901 | 733 | 1502 | 557 | 684 | 1376 | 1040 | 658 |
| 1466 | W303.asm02.HP0_G0026480 | chrIX | 7503 | 3994 | 5019 | 7453 | 3905 | 5025 | 7438 | 4057 | 6402 | 11926 | 3502 | 4166 | 17409 | 2931 | 3573 | 12176 | 3102 | 3780 |
| 1467 | W303.asm02.HP0_G0026490 | chrIX | 24789 | 3667 | 3454 | 24073 | 3408 | 3801 | 25113 | 3424 | 3988 | 25493 | 6315 | 8168 | 18560 | 5241 | 7000 | 18010 | 5687 | 7426 |
| 1468 | W303.asm02.HP0_G0026500 | chrIX | 4573 | 2907 | 2860 | 3324 | 2340 | 2839 | 4087 | 2888 | 2423 | 8632 | 2476 | 3251 | 8457 | 1757 | 2318 | 8038 | 2251 | 2721 |
| 1469 | W303.asm02.HP0_G0026510 | chrIX | 1553 | 507 | 517 | 1585 | 506 | 483 | 1542 | 530 | 567 | 1497 | 333 | 401 | 1759 | 273 | 390 | 1459 | 310 | 388 |
| 1470 | W303.asm02.HP0_G0026520 | chrIX | 1219 | 189 | 254 | 1053 | 159 | 294 | 1059 | 161 | 251 | 568 | 155 | 258 | 745 | 130 | 176 | 554 | 169 | 219 |
| 1471 | W303.asm02.HP0_G0026530 | chrIX | 1925 | 653 | 788 | 1760 | 690 | 729 | 1587 | 654 | 838 | 2487 | 487 | 587 | 3016 | 317 | 539 | 2436 | 544 | 536 |
| 1472 | W303.asm02.HP0_G0026540 | chrIX | 11788 | 2295 | 2306 | 11349 | 2121 | 2190 | 11179 | 2250 | 2136 | 6968 | 2141 | 2992 | 8019 | 1747 | 2611 | 6398 | 2033 | 2785 |
| 1473 | W303.asm02.HP0_G0026550 | chrIX | 6926 | 6954 | 8863 | 6628 | 8707 | 8873 | 7061 | 7088 | 10449 | 20226 | 6292 | 7122 | 21944 | 4589 | 5646 | 20891 | 5278 | 6305 |
| 1474 | W303.asm02.HP0_G0026560 | chrIX | 4042 | 924 | 1061 | 4126 | 991 | 1056 | 3864 | 922 | 974 | 5841 | 1021 | 1548 | 5231 | 744 | 1207 | 5391 | 929 | 1482 |
| 1475 | W303.asm02.HP0_G0026570 | chrIX | 1248 | 273 | 306 | 1142 | 369 | 341 | 1207 | 258 | 369 | 2626 | 497 | 652 | 1744 | 368 | 504 | 1587 | 503 | 590 |
| 1476 | W303.asm02.HP0_G0026580 | chrIX | 989 | 48 | 87 | 942 | 76 | 111 | 990 | 44 | 101 | 208 | 75 | 106 | 157 | 60 | 68 | 109 | 50 | 72 |
| 1477 | W303.asm02.HP0_G0026590 | chrIX | 2331 | 843 | 904 | 2038 | 836 | 886 | 2022 | 825 | 1126 | 2140 | 809 | 854 | 3125 | 662 | 839 | 2047 | 713 | 721 |
| 1478 | W303.asm02.HP0_G0026600 | chrIX | 878 | 963 | 1168 | 758 | 1301 | 1034 | 959 | 1140 | 1509 | 3983 | 1177 | 1000 | 4172 | 693 | 625 | 4133 | 919 | 807 |
| 1479 | W303.asm02.HP0_G0026610 | chrIX | 3366 | 1876 | 1824 | 2964 | 1744 | 1829 | 3007 | 1905 | 2057 | 5943 | 1684 | 1706 | 7535 | 1298 | 1570 | 5437 | 1504 | 1483 |
| 1480 | W303.asm02.HP0_G0026620 | chrIX | 6219 | 2896 | 2943 | 5465 | 2569 | 2969 | 5768 | 2897 | 3371 | 5033 | 2490 | 2448 | 6739 | 2020 | 2523 | 5238 | 2176 | 2271 |
| 1481 | W303.asm02.HP0_G0026630 | chrIX | 897 | 241 | 197 | 1011 | 170 | 211 | 1031 | 187 | 221 | 1546 | 367 | 508 | 1435 | 290 | 279 | 1178 | 352 | 411 |
| 1482 | W303.asm02.HP0_G0026640 | chrIX | 720 | 212 | 147 | 645 | 166 | 140 | 619 | 163 | 185 | 677 | 266 | 200 | 621 | 173 | 141 | 498 | 260 | 167 |
| 1483 | W303.asm02.HP0_G0026650 | chrIX | 154 | 30 | 10 | 260 | 12 | 10 | 162 | 34 | 12 | 35 | 28 | 6 | 52 | 11 | 2 | 31 | 28 | 6 |
| 1484 | W303.asm02.HP0_G0026660 | chrIX | 1473 | 440 | 485 | 1574 | 431 | 507 | 1487 | 358 | 556 | 4861 | 908 | 999 | 3260 | 501 | 715 | 2822 | 1014 | 828 |
| 1485 | W303.asm02.HP0_G0026670 | chrIX | 1559 | 185 | 179 | 1657 | 132 | 166 | 1501 | 176 | 220 | 516 | 239 | 304 | 323 | 182 | 174 | 243 | 239 | 225 |
| 1486 | W303.asm02.HP0_G0026680 | chrIX | 20 | 10 | 14 | 33 | 11 | 14 | 20 | 25 | 15 | 53 | 20 | 9 | 48 | 19 | 9 | 37 | 19 | 10 |

| | | | | | | | | | | | | | | | | | | | | |
|---|---|---|---|---|---|---|---|---|---|---|---|---|---|---|---|---|---|---|---|---|
| 1487 | W303.asm02.HP0_G0026690 | chrIX | 4916 | 6932 | 9629 | 5101 | 8215 | 9450 | 5511 | 6776 | 9118 | 27389 | 6939 | 14363 | 22486 | 4196 | 8132 | 23636 | 6439 | 12501 |
| 1488 | W303.asm02.HP0_G0026700 | chrIX | 5547 | 5772 | 6874 | 4741 | 6588 | 6884 | 5281 | 5648 | 8283 | 13921 | 5077 | 5433 | 15615 | 3833 | 4408 | 14829 | 4796 | 4595 |
| 1489 | W303.asm02.HP0_G0026710 | chrIX | 5782 | 2671 | 2498 | 5139 | 2124 | 2466 | 5178 | 2693 | 2399 | 7871 | 2698 | 2883 | 7825 | 2147 | 2499 | 7361 | 2477 | 2622 |
| 1490 | W303.asm02.HP0_G0026720 | chrIX | 2613 | 961 | 533 | 2662 | 622 | 592 | 2430 | 921 | 498 | 3237 | 1144 | 1029 | 3228 | 829 | 893 | 2808 | 1097 | 843 |
| 1491 | W303.asm02.HP0_G0026730 | chrIX | 744 | 859 | 600 | 711 | 495 | 598 | 729 | 727 | 596 | 1490 | 462 | 493 | 1777 | 348 | 390 | 1657 | 555 | 435 |
| 1492 | W303.asm02.HP0_G0026740 | chrIX | 1818 | 1309 | 1280 | 2046 | 1166 | 1253 | 1790 | 1413 | 1429 | 8960 | 1789 | 2345 | 6686 | 1159 | 1556 | 7289 | 1577 | 2113 |
| 1493 | W303.asm02.HP0_G0026750 | chrIX | 1824 | 748 | 1290 | 1604 | 1492 | 1368 | 1806 | 745 | 1512 | 9568 | 1365 | 2311 | 5646 | 846 | 1443 | 5562 | 1210 | 2262 |
| 1494 | W303.asm02.HP0_G0026760 | chrIX | 1465 | 496 | 514 | 1307 | 547 | 604 | 1377 | 367 | 647 | 2800 | 874 | 1074 | 1160 | 568 | 642 | 1162 | 841 | 961 |
| 1495 | W303.asm02.HP0_G0026770 | chrIX | 4711 | 575 | 417 | 4554 | 346 | 410 | 3856 | 512 | 485 | 1538 | 420 | 409 | 1654 | 230 | 319 | 1434 | 453 | 390 |
| 1496 | W303.asm02.HP0_G0026780 | chrIX | 11825 | 4650 | 4272 | 10811 | 3836 | 4305 | 11993 | 4318 | 4688 | 17084 | 5654 | 6204 | 14583 | 3645 | 4517 | 14413 | 4892 | 5709 |
| 1497 | W303.asm02.HP0_G0026790 | chrIX | 848 | 357 | 321 | 811 | 229 | 380 | 747 | 220 | 368 | 936 | 415 | 587 | 296 | 273 | 326 | 269 | 440 | 403 |
| 1498 | W303.asm02.HP0_G0026800 | chrIX | 9221 | 2125 | 1554 | 8348 | 1561 | 1739 | 9283 | 2126 | 1649 | 8400 | 2859 | 3445 | 8401 | 2125 | 2569 | 7717 | 2681 | 3039 |
| 1499 | W303.asm02.HP0_G0026810 | chrIX | 274 | 1014 | 1363 | 189 | 1170 | 1375 | 266 | 1068 | 2057 | 2673 | 771 | 975 | 3134 | 481 | 649 | 2971 | 740 | 800 |
| 1500 | W303.asm02.HP0_G0026820 | chrIX | 4599 | 2043 | 1752 | 4054 | 1751 | 1631 | 4459 | 2040 | 1560 | 8253 | 2255 | 2447 | 7520 | 1572 | 1875 | 7336 | 2002 | 2399 |
| 1501 | W303.asm02.HP0_G0026830 | chrIX | 3892 | 1344 | 1390 | 4022 | 1451 | 1249 | 4028 | 1392 | 1436 | 12880 | 2196 | 2828 | 11243 | 1483 | 2085 | 10724 | 1985 | 2588 |
| 1502 | W303.asm02.HP0_G0026840 | chrIX | 392 | 199 | 174 | 378 | 138 | 176 | 302 | 166 | 199 | 772 | 177 | 239 | 513 | 74 | 146 | 547 | 202 | 184 |
| 1503 | W303.asm02.HP0_G0026850 | chrIX | 4974 | 460 | 560 | 4884 | 470 | 683 | 4862 | 388 | 782 | 4033 | 986 | 1147 | 2815 | 605 | 1048 | 2636 | 900 | 1106 |
| 1504 | W303.asm02.HP0_G0026860 | chrIX | 5076 | 2630 | 2587 | 5131 | 2667 | 2392 | 5037 | 2712 | 2894 | 9897 | 2376 | 2632 | 9670 | 1628 | 2096 | 9947 | 2265 | 2333 |
| 1505 | W303.asm02.HP0_G0026870 | chrIX | 9263 | 1841 | 1569 | 8728 | 1441 | 1605 | 9152 | 1953 | 1570 | 7854 | 2294 | 2671 | 8832 | 1745 | 2483 | 7215 | 2112 | 2359 |
| 1506 | W303.asm02.HP0_G0026880 | chrIX | 21159 | 1828 | 1616 | 20491 | 1251 | 1910 | 20737 | 1427 | 2192 | 8450 | 3254 | 3748 | 4874 | 2715 | 2930 | 4013 | 2765 | 3175 |
| 1507 | W303.asm02.HP0_G0026890 | chrIX | 20 | 7 | 8 | 25 | 7 | 2 | 17 | 10 | 9 | 23 | 8 | 3 | 22 | 6 | 6 | 14 | 8 | 8 |
| 1508 | W303.asm02.HP0_G0026900 | chrIX | 1351 | 277 | 518 | 1466 | 474 | 560 | 1226 | 293 | 613 | 4674 | 634 | 1083 | 3256 | 413 | 820 | 3395 | 608 | 958 |
| 1509 | W303.asm02.HP0_G0026910 | chrIX | 4174 | 1301 | 1785 | 4350 | 1542 | 1913 | 4375 | 1176 | 2253 | 13040 | 2379 | 3314 | 6232 | 1653 | 2277 | 6823 | 2030 | 2966 |
| 1510 | W303.asm02.HP0_G0026920 | chrIX | 880 | 407 | 245 | 824 | 197 | 239 | 779 | 373 | 315 | 792 | 250 | 167 | 1084 | 194 | 197 | 805 | 313 | 154 |
| 1511 | W303.asm02.HP0_G0026930 | chrIX | 371 | 137 | 136 | 366 | 120 | 137 | 359 | 119 | 138 | 575 | 150 | 142 | 621 | 79 | 88 | 408 | 171 | 129 |
| 1512 | W303.asm02.HP0_G0026940 | chrIX | 1034 | 815 | 944 | 974 | 769 | 944 | 1045 | 815 | 1356 | 2595 | 654 | 714 | 2853 | 404 | 564 | 2800 | 539 | 651 |
| 1513 | W303.asm02.HP0_G0026950 | chrIX | 2350 | 853 | 917 | 2415 | 913 | 907 | 2522 | 908 | 848 | 5933 | 961 | 1485 | 5618 | 678 | 1139 | 5370 | 846 | 1361 |
| 1514 | W303.asm02.HP0_G0026960 | chrIX | 649 | 156 | 165 | 611 | 181 | 202 | 628 | 164 | 206 | 508 | 202 | 191 | 495 | 147 | 171 | 315 | 172 | 170 |
| 1515 | W303.asm02.HP0_G0026970 | chrIX | 3202 | 626 | 588 | 2970 | 611 | 719 | 3131 | 577 | 757 | 2952 | 1009 | 1151 | 2730 | 1396 | 1214 | 1286 | 929 | 982 |
| 1516 | W303.asm02.HP0_G0026980 | chrIX | 12692 | 1884 | 1778 | 11418 | 1746 | 2200 | 12626 | 1459 | 2366 | 11542 | 3440 | 4218 | 5072 | 3014 | 2993 | 4599 | 3005 | 3667 |
| 1517 | W303.asm02.HP0_G0026990 | chrIX | 3951 | 2474 | 2281 | 3874 | 2369 | 2237 | 3896 | 2494 | 2335 | 8189 | 2330 | 2572 | 7939 | 1510 | 1904 | 7902 | 2119 | 2282 |
| 1518 | W303.asm02.HP0_G0027000 | chrIX | 1008 | 1009 | 1004 | 1031 | 1067 | 887 | 1002 | 1047 | 1037 | 4058 | 913 | 881 | 4265 | 605 | 607 | 3993 | 794 | 774 |
| 1519 | W303.asm02.HP0_G0027010 | chrIX | 783 | 909 | 768 | 804 | 1142 | 695 | 818 | 918 | 719 | 5841 | 1123 | 984 | 5276 | 611 | 491 | 5196 | 885 | 885 |
| 1520 | W303.asm02.HP0_G0027020 | chrIX | 23118 | 17023 | 18522 | 19723 | 18336 | 18264 | 24383 | 16652 | 19943 | 39768 | 16370 | 17949 | 39774 | 12786 | 14159 | 38782 | 14873 | 16423 |
| 1521 | W303.asm02.HP0_G0027030 | chrIX | 2478 | 453 | 443 | 2531 | 365 | 413 | 2249 | 423 | 452 | 2406 | 811 | 849 | 2012 | 500 | 769 | 1852 | 797 | 744 |
| 1522 | W303.asm02.HP0_G0027040 | chrIX | 1217 | 1069 | 1199 | 1025 | 1101 | 1166 | 1126 | 1171 | 1455 | 3742 | 1049 | 1143 | 3731 | 633 | 891 | 3564 | 960 | 955 |
| 1523 | W303.asm02.HP0_G0027050 | chrIX | 1483 | 3216 | 3486 | 1420 | 3007 | 3733 | 1793 | 2672 | 3006 | 8737 | 2271 | 3743 | 9495 | 1776 | 2680 | 7810 | 2234 | 3171 |
| 1524 | W303.asm02.HP0_G0027060 | chrIX | 628 | 443 | 475 | 541 | 401 | 451 | 515 | 419 | 668 | 892 | 389 | 417 | 801 | 221 | 314 | 1039 | 466 | 362 |
| 1525 | W303.asm02.HP0_G0027070 | chrIX | 1922 | 940 | 1123 | 1601 | 1157 | 1048 | 1747 | 1049 | 1199 | 6161 | 1300 | 1453 | 5254 | 740 | 1116 | 5286 | 1118 | 1358 |
| 1526 | W303.asm02.HP0_G0027080 | chrIX | 1969 | 338 | 374 | 1888 | 454 | 502 | 1932 | 303 | 382 | 2401 | 522 | 679 | 1784 | 389 | 521 | 1597 | 483 | 625 |
| 1527 | W303.asm02.HP0_G0027090 | chrIX | 62 | 9 | 16 | 41 | 5 | 8 | 42 | 14 | 8 | 32 | 11 | 7 | 21 | 8 | 7 | 18 | 11 | 6 |
| 1528 | W303.asm02.HP0_G0027100 | chrIX | 2762 | 110 | 128 | 2702 | 113 | 155 | 2851 | 81 | 129 | 1616 | 275 | 292 | 1202 | 196 | 264 | 876 | 212 | 247 |
| 1529 | W303.asm02.HP0_G0027110 | chrIX | 3452 | 652 | 557 | 3265 | 563 | 558 | 3389 | 620 | 554 | 3241 | 899 | 1179 | 3836 | 764 | 1028 | 3045 | 803 | 1001 |
| 1530 | W303.asm02.HP0_G0027120 | chrIX | 725 | 660 | 634 | 677 | 534 | 614 | 701 | 620 | 686 | 1318 | 513 | 553 | 1685 | 372 | 435 | 1266 | 462 | 447 |
| 1531 | W303.asm02.HP0_G0027130 | chrIX | 458 | 466 | 645 | 415 | 718 | 715 | 449 | 412 | 848 | 5132 | 914 | 1201 | 3352 | 526 | 754 | 3173 | 764 | 1114 |
| 1532 | W303.asm02.HP0_G0027140 | chrIX | 1397 | 1193 | 1580 | 1212 | 1509 | 1448 | 1319 | 1322 | 2122 | 4696 | 1339 | 1388 | 5529 | 930 | 1027 | 5395 | 1182 | 1311 |
| 1533 | W303.asm02.HP0_G0027150 | chrIX | 2429 | 1238 | 1064 | 2281 | 963 | 1006 | 2281 | 1136 | 1234 | 2375 | 919 | 751 | 2568 | 524 | 575 | 2343 | 911 | 676 |
| 1534 | W303.asm02.HP0_G0027160 | chrIX | 402 | 270 | 154 | 392 | 161 | 205 | 391 | 165 | 178 | 887 | 354 | 423 | 842 | 236 | 313 | 428 | 343 | 323 |
| 1535 | W303.asm02.HP0_G0027170 | chrIX | 4147 | 839 | 835 | 4356 | 791 | 836 | 4196 | 928 | 873 | 6495 | 1367 | 1647 | 5829 | 1061 | 1388 | 5161 | 1203 | 1602 |
| 1536 | W303.asm02.HP0_G0027180 | chrIX | 898 | 878 | 956 | 661 | 856 | 883 | 720 | 922 | 1132 | 2530 | 778 | 805 | 3068 | 566 | 674 | 2795 | 766 | 701 |
| 1537 | W303.asm02.HP0_G0027190 | chrIX | 526 | 733 | 590 | 461 | 522 | 503 | 497 | 728 | 600 | 2284 | 895 | 728 | 1648 | 512 | 424 | 2048 | 876 | 618 |
| 1538 | W303.asm02.HP0_G0027200 | chrIX | 1167 | 756 | 839 | 788 | 835 | 875 | 874 | 743 | 1085 | 1756 | 618 | 679 | 2141 | 389 | 619 | 1975 | 613 | 557 |
| 1539 | W303.asm02.HP0_G0027210 | chrIX | 2929 | 545 | 610 | 2753 | 569 | 690 | 2900 | 519 | 752 | 4497 | 1140 | 1462 | 2506 | 852 | 1055 | 2579 | 1076 | 1245 |
| 1540 | W303.asm02.HP0_G0027220 | chrIX | 1205 | 1162 | 1087 | 1300 | 741 | 1167 | 1272 | 1135 | 1070 | 2582 | 943 | 1269 | 2392 | 612 | 744 | 2630 | 965 | 1091 |
| 1541 | W303.asm02.HP0_G0027230 | chrIX | 3319 | 1442 | 887 | 3203 | 917 | 842 | 3252 | 1401 | 829 | 4398 | 1469 | 1386 | 4315 | 1055 | 910 | 3824 | 1358 | 1139 |
| 1542 | W303.asm02.HP0_G0027240 | chrIX | 3219 | 1046 | 956 | 3161 | 876 | 1056 | 3286 | 1158 | 993 | 6719 | 1785 | 2144 | 5307 | 1122 | 1370 | 5406 | 1736 | 1953 |
| 1543 | W303.asm02.HP0_G0027250 | chrIX | 11450 | 722 | 619 | 11019 | 524 | 712 | 11237 | 477 | 794 | 5377 | 1689 | 1755 | 1919 | 1230 | 1278 | 1773 | 1566 | 1592 |
| 1544 | W303.asm02.HP0_G0027260 | chrIX | 7495 | 2022 | 2399 | 7570 | 2401 | 2369 | 7506 | 2012 | 2704 | 10304 | 2076 | 2823 | 12590 | 1458 | 2521 | 9291 | 1795 | 2692 |
| 1545 | W303.asm02.HP0_G0027270 | chrIX | 513 | 505 | 435 | 460 | 447 | 464 | 449 | 477 | 702 | 1080 | 362 | 335 | 1137 | 197 | 237 | 1427 | 374 | 332 |
| 1546 | W303.asm02.HP0_G0027280 | chrIX | 16720 | 625 | 1078 | 14723 | 886 | 1380 | 16348 | 468 | 1360 | 1927 | 648 | 2312 | 1196 | 629 | 896 | 952 | 467 | 1424 |
| 1547 | W303.asm02.HP0_G0027290 | chrIX | 2085 | 175 | 195 | 1922 | 125 | 265 | 1970 | 144 | 239 | 1204 | 339 | 630 | 597 | 211 | 358 | 556 | 378 | 487 |
| 1548 | W303.asm02.HP0_G0027300 | chrIX | 1366 | 923 | 700 | 1400 | 745 | 660 | 1285 | 907 | 702 | 3905 | 1010 | 1067 | 3573 | 639 | 776 | 3362 | 865 | 983 |
| 1549 | W303.asm02.HP0_G0027310 | chrIX | 1078 | 549 | 445 | 1033 | 455 | 427 | 1042 | 517 | 357 | 1521 | 449 | 426 | 1908 | 353 | 338 | 1387 | 398 | 376 |
| 1550 | W303.asm02.HP0_G0027320 | chrIX | 1558 | 287 | 258 | 1631 | 305 | 314 | 1555 | 236 | 353 | 2331 | 492 | 733 | 1734 | 389 | 594 | 1600 | 562 | 648 |
| 1551 | W303.asm02.HP0_G0027330 | chrIX | 4427 | 2470 | 2448 | 4525 | 2394 | 2222 | 4287 | 2348 | 2621 | 7032 | 2088 | 2314 | 8268 | 1375 | 1861 | 7027 | 2068 | 2012 |
| 1552 | W303.asm02.HP0_G0027340 | chrIX | 1525 | 1518 | 1810 | 1231 | 2176 | 1661 | 1555 | 1538 | 2064 | 11827 | 2711 | 3313 | 9630 | 1937 | 2123 | 9208 | 2468 | 2902 |
| 1553 | W303.asm02.HP0_G0027350 | chrIX | 46 | 29 | 16 | 46 | 9 | 22 | 36 | 21 | 17 | 54 | 33 | 21 | 56 | 9 | 7 | 40 | 35 | 14 |
| 1554 | W303.asm02.HP0_G0027360 | chrIX | 571 | 49 | 34 | 538 | 39 | 45 | 455 | 20 | 60 | 163 | 79 | 73 | 71 | 67 | 64 | 88 | 78 | 74 |
| 1555 | W303.asm02.HP0_G0027370 | chrIX | 676 | 143 | 160 | 576 | 103 | 220 | 572 | 65 | 244 | 806 | 246 | 442 | 408 | 153 | 246 | 399 | 264 | 395 |
| 1556 | W303.asm02.HP0_G0027380 | chrIX | 938 | 229 | 262 | 839 | 215 | 279 | 842 | 177 | 270 | 723 | 291 | 420 | 691 | 189 | 364 | 400 | 242 | 351 |
| 1557 | W303.asm02.HP0_G0027390 | chrIX | 480 | 389 | 458 | 319 | 291 | 493 | 324 | 389 | 510 | 351 | 213 | 279 | 663 | 218 | 345 | 352 | 211 | 230 |
| 1558 | W303.asm02.HP0_G0027400 | chrIX | 737 | 111 | 165 | 698 | 158 | 201 | 568 | 89 | 181 | 992 | 193 | 363 | 991 | 117 | 254 | 759 | 187 | 315 |
| 1559 | W303.asm02.HP0_G0027410 | chrIX | 3548 | 64 | 59 | 2995 | 58 | 77 | 3717 | 42 | 81 | 297 | 46 | 111 | 174 | 55 | 84 | 127 | 46 | 89 |
| 1560 | W303.asm02.HP0_G0027420 | chrIX | 160 | 35 | 74 | 164 | 73 | 79 | 155 | 25 | 80 | 277 | 34 | 92 | 219 | 24 | 66 | 180 | 25 | 72 |
| 1561 | W303.asm02.HP0_G0027430 | chrIX | 152 | 16 | 41 | 136 | 33 | 22 | 116 | 12 | 43 | 236 | 21 | 67 | 140 | 17 | 51 | 101 | 29 | 49 |
| 1562 | W303.asm02.HP0_G0027440 | chrIX | 2626 | 744 | 753 | 2097 | 852 | 731 | 2472 | 773 | 744 | 3304 | 733 | 906 | 3537 | 536 | 701 | 2914 | 658 | 781 |
| 1563 | W303.asm02.HP0_G0027450 | chrIX | 8 | 4 | 5 | 10 | 0 | 3 | 8 | 3 | 9 | 11 | 4 | 5 | 7 | 1 | 5 | 7 | 3 | 5 |
| 1564 | W303.asm02.HP0_G0027460 | chrIX | 2482 | 1485 | 1588 | 2176 | 1203 | 1668 | 2298 | 1510 | 1846 | 3467 | 1235 | 1693 | 4223 | 954 | 1359 | 3721 | 1250 | 1542 |
| 1565 | W303.asm02.HP0_G0027470 | chrIX | 1932 | 325 | 422 | 1936 | 383 | 504 | 2168 | 250 | 553 | 4439 | 761 | 1173 | 2026 | 551 | 737 | 2230 | 642 | 1101 |
| 1566 | W303.asm02.HP0_G0027480 | chrIX | 2286 | 651 | 931 | 2317 | 819 | 976 | 2079 | 686 | 1211 | 2178 | 482 | 716 | 2987 | 394 | 624 | 2284 | 464 | 654 |
| 1567 | W303.asm02.HP0_G0027490 | chrIX | 1409 | 240 | 204 | 1450 | 249 | 243 | 1356 | 230 | 222 | 1582 | 299 | 490 | 1174 | 247 | 373 | 1018 | 263 | 464 |
| 1568 | W303.asm02.HP0_G0027500 | chrIX | 1854 | 954 | 759 | 1659 | 970 | 736 | 1938 | 937 | 705 | 6367 | 1494 | 1477 | 5137 | 892 | 757 | 5167 | 1184 | 1273 |
| 1569 | W303.asm02.HP0_G0027510 | chrIX | 459 | 71 | 109 | 451 | 96 | 105 | 479 | 62 | 89 | 353 | 146 | 170 | 289 | 89 | 101 | 263 | 113 | 144 |
| 1570 | W303.asm02.HP0_G0027520 | chrIX | 730 | 265 | 262 | 715 | 238 | 252 | 655 | 223 | 278 | 876 | 254 | 283 | 1615 | 313 | 323 | 702 | 263 | 270 |
| 1571 | W303.asm02.HP0_G0027530 | chrIX | 802 | 193 | 162 | 829 | 200 | 171 | 771 | 155 | 179 | 1158 | 283 | 291 | 946 | 190 | 210 | 727 | 269 | 205 |
| 1572 | W303.asm02.HP0_G0027540 | chrIX | 584 | 537 | 737 | 424 | 641 | 676 | 390 | 681 | 887 | 1771 | 487 | 618 | 1437 | 236 | 432 | 2084 | 473 | 515 |
| 1573 | W303.asm02.HP0_G0027550 | chrIX | 1241 | 636 | 564 | 1260 | 642 | 571 | 1250 | 557 | 605 | 2897 | 905 | 928 | 3156 | 599 | 741 | 2325 | 1029 | 761 |
| 1574 | W303.asm02.HP0_G0027560 | chrIX | 451 | 425 | 424 | 350 | 374 | 421 | 398 | 411 | 528 | 875 | 417 | 379 | 1489 | 324 | 407 | 809 | 335 | 316 |
| 1575 | W303.asm02.HP0_G0027570 | chrIX | 6693 | 940 | 1353 | 6421 | 1237 | 1450 | 6309 | 973 | 1710 | 12034 | 1990 | 3017 | 8689 | 1281 | 2111 | 8935 | 1849 | 2800 |
| 1576 | W303.asm02.HP0_G0027580 | chrIX | 3087 | 1235 | 1012 | 3147 | 924 | 1158 | 2932 | 1022 | 1225 | 4738 | 2036 | 2018 | 3194 | 1095 | 1425 | 3461 | 1748 | 1785 |
| 1577 | W303.asm02.HP0_G0027590 | chrIX | 2633 | 1198 | 1573 | 1924 | 1406 | 1642 | 2288 | 1239 | 1858 | 3606 | 1437 | 1996 | 3724 | 949 | 1503 | 3533 | 1316 | 1717 |
| 1578 | W303.asm02.HP0_G0027600 | chrIX | 2768 | 574 | 515 | 2360 | 451 | 516 | 2390 | 493 | 598 | 1759 | 770 | 873 | 1692 | 630 | 803 | 1375 | 714 | 788 |
| 1579 | W303.asm02.HP0_G0027610 | chrIX | 7227 | 2366 | 3004 | 6291 | 2450 | 3116 | 6667 | 2357 | 3604 | 5614 | 1986 | 2634 | 7401 | 1636 | 2407 | 5902 | 1845 | 2240 |

| | | | | | | | | | | | | | | | | | | | |
|---|---|---|---|---|---|---|---|---|---|---|---|---|---|---|---|---|---|---|---|
| 1580 | W303.asm02.HP0_G0027620 | chrIX | 4350 | 1190 | 1146 | 3789 | 1029 | 1165 | 3867 | 1104 | 1059 | 2765 | 1164 | 1421 | 4379 | 1042 | 1448 | 2609 | 1043 | 1172 |
| 1581 | W303.asm02.HP0_G0027630 | chrIX | 4087 | 396 | 415 | 3610 | 352 | 390 | 3644 | 427 | 406 | 1643 | 662 | 666 | 2765 | 634 | 818 | 1291 | 595 | 589 |
| 1582 | W303.asm02.HP0_G0027640 | chrIX | 102 | 11 | 3 | 90 | 6 | 11 | 114 | 6 | 15 | 31 | 8 | 12 | 99 | 19 | 25 | 17 | 11 | 15 |
| 1583 | W303.asm02.HP0_G0027650 | chrIX | 1000 | 594 | 624 | 1059 | 616 | 599 | 758 | 603 | 632 | 1668 | 465 | 479 | 2280 | 309 | 415 | 1517 | 438 | 429 |
| 1584 | W303.asm02.HP0_G0027660 | chrIX | 308 | 217 | 161 | 289 | 168 | 150 | 203 | 191 | 175 | 521 | 105 | 103 | 436 | 59 | 72 | 537 | 122 | 96 |
| 1585 | W303.asm02.HP0_G0027670 | chrIX | 37 | 46 | 29 | 30 | 35 | 24 | 14 | 44 | 36 | 127 | 28 | 16 | 62 | 21 | 10 | 107 | 27 | 24 |
| 1586 | W303.asm02.HP0_G0027680 | chrIX | 0 | 26 | 0 | 0 | 0 | 0 | 0 | 21 | 1 | 0 | 21 | 0 | 0 | 13 | 1 | 1 | 15 | 0 |
| 1587 | W303.asm02.HP0_ATP6 | chrMT | 0 | 2 | 1 | 0 | 1 | 0 | 0 | 0 | 6 | 1 | 0 | 4 | 0 | 2 | 17 | 1 | 0 | 0 |
| 1588 | W303.asm02.HP0_ATP8 | chrMT | 0 | 0 | 0 | 0 | 0 | 0 | 0 | 0 | 0 | 0 | 0 | 0 | 0 | 0 | 0 | 0 | 0 | 0 |
| 1589 | W303.asm02.HP0_ATP9 | chrMT | 0 | 4 | 10 | 0 | 8 | 17 | 0 | 2 | 26 | 8 | 1 | 5 | 13 | 5 | 39 | 9 | 5 | 6 |
| 1590 | W303.asm02.HP0_COB | chrMT | 0 | 5 | 5 | 0 | 5 | 13 | 0 | 4 | 20 | 12 | 1 | 7 | 18 | 2 | 41 | 18 | 4 | 9 |
| 1591 | W303.asm02.HP0_COX1 | chrMT | 0 | 7 | 9 | 0 | 10 | 15 | 1 | 1 | 23 | 50 | 2 | 5 | 27 | 3 | 50 | 77 | 2 | 5 |
| 1592 | W303.asm02.HP0_COX2 | chrMT | 0 | 1 | 2 | 0 | 3 | 12 | 0 | 0 | 8 | 10 | 1 | 3 | 10 | 0 | 21 | 8 | 6 | 4 |
| 1593 | W303.asm02.HP0_COX3 | chrMT | 0 | 8 | 6 | 0 | 2 | 8 | 0 | 1 | 7 | 10 | 2 | 4 | 12 | 2 | 8 | 11 | 6 | 5 |
| 1594 | W303.asm02.HP0_G0013760 | chrV | 1569 | 282 | 361 | 1374 | 376 | 444 | 1581 | 245 | 501 | 1340 | 329 | 656 | 706 | 261 | 415 | 622 | 287 | 529 |
| 1595 | W303.asm02.HP0_G0013770 | chrV | 1080 | 256 | 136 | 1045 | 131 | 150 | 772 | 209 | 127 | 874 | 449 | 293 | 725 | 256 | 273 | 759 | 432 | 274 |
| 1596 | W303.asm02.HP0_G0013780 | chrV | 510 | 645 | 450 | 326 | 440 | 471 | 459 | 588 | 499 | 848 | 373 | 237 | 1169 | 263 | 207 | 873 | 408 | 211 |
| 1597 | W303.asm02.HP0_G0013790 | chrV | 2197 | 5400 | 7366 | 1823 | 6265 | 7142 | 1879 | 5570 | 7187 | 16003 | 6332 | 7870 | 14836 | 3555 | 6599 | 15881 | 5660 | 7538 |
| 1598 | W303.asm02.HP0_G0013800 | chrV | 1353 | 38 | 30 | 1829 | 47 | 54 | 1473 | 25 | 29 | 126 | 40 | 44 | 252 | 62 | 44 | 91 | 38 | 36 |
| 1599 | W303.asm02.HP0_G0013810 | chrV | 216 | 12 | 18 | 274 | 22 | 21 | 233 | 14 | 29 | 254 | 21 | 42 | 143 | 17 | 26 | 103 | 15 | 39 |
| 1600 | W303.asm02.HP0_G0013820 | chrV | 1042 | 624 | 648 | 1012 | 640 | 682 | 1001 | 688 | 805 | 1686 | 491 | 537 | 2193 | 295 | 433 | 1725 | 458 | 455 |
| 1601 | W303.asm02.HP0_G0013830 | chrV | 744 | 765 | 700 | 716 | 716 | 728 | 772 | 778 | 676 | 1835 | 745 | 650 | 1994 | 537 | 494 | 1588 | 644 | 663 |
| 1602 | W303.asm02.HP0_G0013840 | chrV | 185 | 366 | 426 | 170 | 531 | 626 | 175 | 315 | 490 | 8778 | 1542 | 997 | 7217 | 529 | 1161 | 8544 | 1393 | 2164 |
| 1603 | W303.asm02.HP0_G0013850 | chrV | 269 | 333 | 480 | 274 | 355 | 419 | 270 | 317 | 472 | 1809 | 349 | 551 | 2052 | 239 | 481 | 1728 | 334 | 438 |
| 1604 | W303.asm02.HP0_G0013860 | chrV | 3609 | 787 | 1442 | 3731 | 1379 | 1410 | 3803 | 778 | 1575 | 9235 | 1567 | 2338 | 8988 | 1039 | 1832 | 6700 | 1427 | 1991 |
| 1605 | W303.asm02.HP0_G0013870 | chrV | 1433 | 489 | 775 | 1287 | 694 | 640 | 1290 | 518 | 705 | 3045 | 478 | 800 | 3238 | 303 | 594 | 2745 | 461 | 704 |
| 1606 | W303.asm02.HP0_G0013880 | chrV | 1529 | 478 | 720 | 1781 | 739 | 668 | 1605 | 507 | 736 | 4346 | 597 | 1032 | 3591 | 342 | 817 | 3931 | 579 | 925 |
| 1607 | W303.asm02.HP0_G0013890 | chrV | 29518 | 1300 | 1356 | 27840 | 1256 | 1842 | 27680 | 1033 | 1758 | 5623 | 2668 | 3124 | 2612 | 2718 | 2970 | 1491 | 2162 | 2762 |
| 1608 | W303.asm02.HP0_G0013900 | chrV | 888 | 311 | 525 | 625 | 454 | 528 | 679 | 356 | 630 | 724 | 284 | 384 | 1526 | 262 | 373 | 770 | 285 | 347 |
| 1609 | W303.asm02.HP0_G0013910 | chrV | 9652 | 2919 | 2516 | 7943 | 2221 | 2448 | 9832 | 2863 | 3199 | 12699 | 4188 | 4710 | 15325 | 4279 | 4439 | 9385 | 3793 | 4262 |
| 1610 | W303.asm02.HP0_G0013920 | chrV | 1330 | 302 | 235 | 1600 | 215 | 229 | 1307 | 253 | 253 | 574 | 194 | 158 | 699 | 135 | 121 | 647 | 198 | 130 |
| 1611 | W303.asm02.HP0_G0013930 | chrV | 2754 | 1295 | 1087 | 2767 | 1020 | 957 | 2536 | 1248 | 1073 | 4743 | 1350 | 1429 | 4453 | 846 | 1181 | 4284 | 1258 | 1256 |
| 1612 | W303.asm02.HP0_G0013940 | chrV | 4055 | 1878 | 1797 | 4060 | 2161 | 1740 | 4344 | 1980 | 1990 | 15762 | 2355 | 3124 | 12585 | 1668 | 2006 | 12962 | 2101 | 2824 |
| 1613 | W303.asm02.HP0_G0013950 | chrV | 10144 | 8916 | 10528 | 9883 | 10223 | 10368 | 10571 | 9005 | 12026 | 23851 | 8389 | 8633 | 24353 | 6018 | 6817 | 24036 | 7207 | 7868 |
| 1614 | W303.asm02.HP0_G0013960 | chrV | 3748 | 916 | 1082 | 3617 | 1086 | 1069 | 4061 | 982 | 942 | 8333 | 1175 | 1729 | 6248 | 830 | 1121 | 7148 | 1227 | 1552 |
| 1615 | W303.asm02.HP0_G0013970 | chrV | 4426 | 1972 | 1461 | 4151 | 1546 | 1533 | 4092 | 1872 | 1458 | 5968 | 1951 | 2027 | 5417 | 1319 | 1539 | 5068 | 1814 | 1688 |
| 1616 | W303.asm02.HP0_G0013980 | chrV | 5516 | 3271 | 4556 | 4628 | 4056 | 4665 | 5203 | 3324 | 5798 | 11796 | 3127 | 3941 | 12618 | 1996 | 3306 | 12511 | 2704 | 3579 |
| 1617 | W303.asm02.HP0_G0013990 | chrV | 10286 | 1514 | 1696 | 9833 | 1642 | 1659 | 9383 | 1537 | 2185 | 8460 | 2382 | 3527 | 7336 | 2113 | 2870 | 6048 | 2265 | 3030 |
| 1618 | W303.asm02.HP0_G0014000 | chrV | 299 | 76 | 71 | 275 | 77 | 69 | 315 | 89 | 69 | 278 | 104 | 78 | 417 | 76 | 53 | 237 | 99 | 64 |
| 1619 | W303.asm02.HP0_G0014010 | chrV | 134 | 498 | 469 | 122 | 448 | 419 | 155 | 563 | 541 | 1494 | 440 | 321 | 1696 | 293 | 204 | 1676 | 385 | 299 |
| 1620 | W303.asm02.HP0_G0014020 | chrV | 6739 | 916 | 804 | 5660 | 662 | 870 | 6279 | 740 | 1053 | 5159 | 1966 | 1911 | 3445 | 1458 | 1508 | 3222 | 1841 | 1620 |
| 1621 | W303.asm02.HP0_G0014030 | chrV | 4819 | 3066 | 4205 | 4835 | 3736 | 4634 | 5209 | 2321 | 5220 | 22216 | 7299 | 8878 | 11647 | 5022 | 5488 | 12664 | 6188 | 7909 |
| 1622 | W303.asm02.HP0_G0014040 | chrV | 58 | 8 | 34 | 52 | 15 | 30 | 47 | 18 | 29 | 63 | 27 | 27 | 44 | 13 | 20 | 32 | 25 | 28 |
| 1623 | W303.asm02.HP0_G0014050 | chrV | 1977 | 1387 | 1586 | 1624 | 1554 | 1510 | 1724 | 1361 | 1881 | 3388 | 1255 | 1048 | 3892 | 854 | 948 | 3599 | 1100 | 1018 |
| 1624 | W303.asm02.HP0_G0014060 | chrV | 4665 | 465 | 617 | 4426 | 617 | 744 | 4700 | 444 | 745 | 6004 | 1180 | 1573 | 3785 | 775 | 1215 | 3934 | 985 | 1524 |
| 1625 | W303.asm02.HP0_G0014070 | chrV | 3959 | 874 | 1289 | 3412 | 1143 | 1404 | 4410 | 830 | 1538 | 11561 | 1822 | 2725 | 6681 | 1470 | 1823 | 6634 | 1563 | 2470 |
| 1626 | W303.asm02.HP0_G0014080 | chrV | 1216 | 330 | 290 | 1134 | 255 | 318 | 1186 | 354 | 327 | 1048 | 371 | 459 | 1342 | 345 | 424 | 930 | 378 | 462 |
| 1627 | W303.asm02.HP0_G0014090 | chrV | 3728 | 3949 | 5256 | 4153 | 5368 | 4976 | 4355 | 4296 | 5427 | 36348 | 6529 | 9059 | 36146 | 5402 | 7259 | 30211 | 5369 | 8255 |
| 1628 | W303.asm02.HP0_G0014100 | chrV | 5631 | 896 | 812 | 5193 | 622 | 883 | 4783 | 658 | 974 | 1411 | 741 | 540 | 1890 | 359 | 639 | 1416 | 866 | 441 |
| 1629 | W303.asm02.HP0_G0014110 | chrV | 1991 | 856 | 815 | 1613 | 710 | 770 | 1777 | 744 | 865 | 3297 | 999 | 1199 | 3241 | 709 | 1016 | 2949 | 968 | 947 |
| 1630 | W303.asm02.HP0_G0014120 | chrV | 7757 | 2189 | 2129 | 7660 | 1966 | 2339 | 8211 | 1850 | 2616 | 13442 | 3687 | 4881 | 6975 | 2604 | 3430 | 7380 | 3430 | 4372 |
| 1631 | W303.asm02.HP0_G0014130 | chrV | 2278 | 1169 | 1164 | 1819 | 1114 | 1339 | 2452 | 909 | 1295 | 8838 | 2115 | 2600 | 5927 | 1672 | 1765 | 4645 | 1767 | 2247 |
| 1632 | W303.asm02.HP0_G0014140 | chrV | 84 | 25 | 15 | 83 | 10 | 20 | 80 | 36 | 8 | 102 | 44 | 22 | 124 | 28 | 12 | 70 | 22 | 13 |
| 1633 | W303.asm02.HP0_G0014150 | chrV | 4119 | 5097 | 5028 | 4334 | 4563 | 5775 | 4319 | 5383 | 5331 | 10759 | 5315 | 5650 | 10804 | 4075 | 4974 | 9951 | 4092 | 5544 |
| 1634 | W303.asm02.HP0_G0014160 | chrV | 1709 | 565 | 640 | 1608 | 597 | 727 | 1874 | 394 | 942 | 5265 | 1096 | 1445 | 1835 | 795 | 912 | 1654 | 935 | 1267 |
| 1635 | W303.asm02.HP0_G0014170 | chrV | 7800 | 1953 | 3272 | 6863 | 2111 | 3891 | 8672 | 1950 | 3820 | 23337 | 5095 | 10838 | 12017 | 3674 | 6108 | 12740 | 4286 | 9519 |
| 1636 | W303.asm02.HP0_G0014180 | chrV | 1244 | 415 | 449 | 1213 | 409 | 423 | 1157 | 461 | 473 | 1931 | 401 | 465 | 1833 | 266 | 374 | 1725 | 462 | 471 |
| 1637 | W303.asm02.HP0_G0014190 | chrV | 546 | 890 | 1107 | 448 | 1020 | 1004 | 504 | 913 | 1257 | 4224 | 761 | 999 | 4147 | 524 | 736 | 4395 | 790 | 981 |
| 1638 | W303.asm02.HP0_G0014200 | chrV | 318 | 297 | 203 | 258 | 211 | 186 | 304 | 279 | 261 | 553 | 233 | 154 | 737 | 181 | 116 | 565 | 240 | 126 |
| 1639 | W303.asm02.HP0_G0014210 | chrV | 8654 | 6272 | 7973 | 7927 | 7480 | 7798 | 8266 | 6469 | 9834 | 16610 | 5783 | 7157 | 18299 | 4392 | 5989 | 16801 | 5176 | 6569 |
| 1640 | W303.asm02.HP0_G0014220 | chrV | 2298 | 4355 | 5101 | 1847 | 4891 | 4903 | 2215 | 4299 | 6131 | 11130 | 3476 | 3485 | 13053 | 2582 | 2661 | 12340 | 3261 | 2981 |
| 1641 | W303.asm02.HP0_G0014230 | chrV | 1830 | 582 | 796 | 1784 | 700 | 745 | 1794 | 589 | 896 | 6253 | 1058 | 1484 | 4189 | 716 | 1196 | 4664 | 991 | 1400 |
| 1642 | W303.asm02.HP0_G0014240 | chrV | 5614 | 5342 | 5951 | 5232 | 5763 | 5355 | 5127 | 5422 | 6197 | 15038 | 4711 | 5523 | 14121 | 3334 | 4478 | 14232 | 4141 | 4275 |
| 1643 | W303.asm02.HP0_G0014250 | chrV | 590 | 252 | 258 | 593 | 281 | 261 | 654 | 254 | 300 | 1538 | 358 | 487 | 1088 | 217 | 341 | 1207 | 351 | 415 |
| 1644 | W303.asm02.HP0_G0014260 | chrV | 3603 | 1114 | 1359 | 3476 | 1390 | 1367 | 3843 | 1224 | 1782 | 12328 | 1693 | 2598 | 9157 | 1123 | 2030 | 10139 | 1510 | 2388 |
| 1645 | W303.asm02.HP0_G0014270 | chrV | 3284 | 1714 | 1358 | 3130 | 1580 | 1409 | 3141 | 1701 | 1442 | 7496 | 2340 | 2295 | 5866 | 1435 | 1731 | 6134 | 2008 | 2145 |
| 1646 | W303.asm02.HP0_G0014280 | chrV | 3633 | 1990 | 3419 | 3273 | 2820 | 3374 | 3385 | 2078 | 4604 | 6131 | 1727 | 2715 | 8030 | 1224 | 2421 | 6552 | 1632 | 2373 |
| 1647 | W303.asm02.HP0_G0014290 | chrV | 1621 | 241 | 327 | 1418 | 247 | 376 | 1460 | 165 | 451 | 1768 | 410 | 628 | 1022 | 311 | 417 | 878 | 424 | 551 |
| 1648 | W303.asm02.HP0_G0014300 | chrV | 304 | 62 | 126 | 335 | 74 | 94 | 291 | 49 | 127 | 622 | 115 | 250 | 401 | 66 | 106 | 335 | 107 | 192 |
| 1649 | W303.asm02.HP0_G0014310 | chrV | 1161 | 1052 | 1156 | 1113 | 1036 | 1187 | 1138 | 1055 | 1210 | 3318 | 801 | 966 | 3661 | 500 | 745 | 3145 | 830 | 908 |
| 1650 | W303.asm02.HP0_G0014320 | chrV | 2268 | 894 | 559 | 1976 | 555 | 621 | 2314 | 894 | 552 | 3580 | 1173 | 1202 | 3623 | 734 | 1007 | 3341 | 1089 | 1178 |
| 1651 | W303.asm02.HP0_G0014330 | chrV | 906 | 463 | 379 | 754 | 367 | 408 | 935 | 416 | 388 | 2125 | 717 | 699 | 1364 | 438 | 495 | 1464 | 606 | 641 |
| 1652 | W303.asm02.HP0_G0014340 | chrV | 5906 | 2166 | 1796 | 5796 | 1877 | 1660 | 5853 | 2105 | 1725 | 10461 | 2764 | 2906 | 8634 | 1818 | 2211 | 9145 | 2325 | 2593 |
| 1653 | W303.asm02.HP0_G0014350 | chrV | 5288 | 1725 | 1880 | 5374 | 1827 | 1774 | 5257 | 1853 | 2066 | 9709 | 2137 | 2560 | 9586 | 1576 | 2435 | 8941 | 1831 | 2555 |
| 1654 | W303.asm02.HP0_G0014360 | chrV | 3035 | 995 | 1053 | 2796 | 889 | 1128 | 2535 | 999 | 1365 | 2124 | 854 | 884 | 3398 | 543 | 794 | 2482 | 844 | 777 |
| 1655 | W303.asm02.HP0_G0014370 | chrV | 11024 | 361 | 230 | 12603 | 224 | 301 | 10767 | 311 | 258 | 1601 | 684 | 538 | 1054 | 519 | 701 | 940 | 624 | 488 |
| 1656 | W303.asm02.HP0_G0014380 | chrV | 19089 | 8623 | 11116 | 20541 | 10603 | 10832 | 21606 | 9836 | 11986 | 31918 | 9831 | 13765 | 32308 | 8579 | 11627 | 28368 | 8526 | 12110 |
| 1657 | W303.asm02.HP0_G0014390 | chrV | 123 | 40 | 40 | 101 | 32 | 22 | 132 | 34 | 42 | 157 | 35 | 27 | 175 | 25 | 29 | 131 | 42 | 30 |
| 1658 | W303.asm02.HP0_G0014400 | chrV | 2694 | 763 | 978 | 2795 | 782 | 1152 | 2702 | 723 | 1146 | 6399 | 1364 | 2177 | 3512 | 986 | 1830 | 3197 | 1156 | 1950 |
| 1659 | W303.asm02.HP0_G0014410 | chrV | 2831 | 1447 | 1597 | 2870 | 1426 | 1470 | 2534 | 1433 | 1603 | 5312 | 1429 | 1634 | 5626 | 961 | 1389 | 5090 | 1410 | 1431 |
| 1660 | W303.asm02.HP0_G0014420 | chrV | 2238 | 563 | 291 | 2035 | 287 | 306 | 2060 | 594 | 276 | 1380 | 556 | 391 | 1396 | 404 | 309 | 1208 | 589 | 316 |
| 1661 | W303.asm02.HP0_G0014430 | chrV | 1499 | 629 | 581 | 1325 | 564 | 585 | 1251 | 668 | 741 | 2183 | 553 | 490 | 2446 | 389 | 418 | 2440 | 595 | 476 |
| 1662 | W303.asm02.HP0_G0014440 | chrV | 1202 | 871 | 1191 | 950 | 1142 | 1159 | 1063 | 932 | 1633 | 2807 | 870 | 1061 | 3398 | 526 | 863 | 3402 | 826 | 851 |
| 1663 | W303.asm02.HP0_G0014450 | chrV | 8219 | 4183 | 4606 | 8020 | 3779 | 4726 | 8689 | 4272 | 5151 | 13054 | 3894 | 4717 | 14002 | 2646 | 3473 | 13429 | 3478 | 4231 |
| 1664 | W303.asm02.HP0_G0014460 | chrV | 4974 | 3126 | 3765 | 4685 | 3512 | 3784 | 4701 | 3383 | 4487 | 10000 | 3191 | 3435 | 11510 | 2113 | 2776 | 10222 | 2757 | 3246 |
| 1665 | W303.asm02.HP0_G0014470 | chrV | 4082 | 2427 | 3921 | 4294 | 3643 | 3467 | 4506 | 2894 | 3661 | 22384 | 4499 | 6150 | 19033 | 3151 | 4280 | 17985 | 3613 | 5084 |
| 1666 | W303.asm02.HP0_G0014480 | chrV | 3744 | 3591 | 2992 | 3802 | 3944 | 3022 | 3938 | 3903 | 3191 | 12012 | 3849 | 3488 | 12627 | 2575 | 2508 | 11244 | 3266 | 2969 |
| 1667 | W303.asm02.HP0_G0014490 | chrV | 5950 | 2116 | 2208 | 5374 | 1554 | 2454 | 6210 | 1989 | 2213 | 11145 | 3066 | 5318 | 9609 | 2310 | 3044 | 9615 | 2760 | 4755 |
| 1668 | W303.asm02.HP0_G0014500 | chrV | 6650 | 2410 | 1830 | 5975 | 1672 | 1891 | 6560 | 2188 | 2017 | 5786 | 2746 | 2828 | 5894 | 2260 | 2237 | 4931 | 2473 | 2491 |
| 1669 | W303.asm02.HP0_G0014510 | chrV | 2650 | 920 | 754 | 2299 | 740 | 817 | 2483 | 896 | 804 | 6147 | 1567 | 1595 | 4181 | 863 | 1053 | 4202 | 1361 | 1413 |
| 1670 | W303.asm02.HP0_G0014520 | chrV | 5875 | 3356 | 2948 | 6265 | 3679 | 2570 | 6885 | 3667 | 2928 | 19140 | 4900 | 4781 | 18862 | 3314 | 2781 | 17317 | 4129 | 4189 |
| 1671 | W303.asm02.HP0_G0014530 | chrV | 713 | 462 | 475 | 702 | 475 | 398 | 718 | 427 | 518 | 1990 | 435 | 415 | 1646 | 233 | 282 | 2147 | 472 | 374 |
| 1672 | W303.asm02.HP0_G0014540 | chrV | 1987 | 1807 | 2266 | 1974 | 2120 | 2238 | 2072 | 1924 | 2748 | 6945 | 1893 | 2124 | 7645 | 1175 | 1546 | 7518 | 1667 | 1825 |

| | | | | | | | | | | | | | | | | | | | | |
|---|---|---|---|---|---|---|---|---|---|---|---|---|---|---|---|---|---|---|---|---|
| 1673 | W303.asm02.HP0_G0014550 | chrV | 4402 | 522 | 814 | 4502 | 693 | 889 | 4281 | 497 | 860 | 7302 | 999 | 1810 | 5188 | 779 | 1280 | 4890 | 894 | 1593 |
| 1674 | W303.asm02.HP0_G0014560 | chrV | 5793 | 4330 | 5447 | 5076 | 4600 | 5468 | 5687 | 4372 | 7093 | 11190 | 4346 | 5157 | 12936 | 3177 | 4370 | 12051 | 3896 | 4582 |
| 1675 | W303.asm02.HP0_G0014570 | chrV | 2228 | 1067 | 1014 | 1811 | 924 | 984 | 2068 | 1033 | 914 | 2822 | 948 | 1005 | 3250 | 611 | 833 | 2853 | 891 | 870 |
| 1676 | W303.asm02.HP0_G0014580 | chrV | 1094 | 64 | 48 | 1319 | 87 | 101 | 1260 | 60 | 47 | 988 | 136 | 127 | 970 | 57 | 93 | 804 | 106 | 95 |
| 1677 | W303.asm02.HP0_G0014590 | chrV | 8077 | 3711 | 5088 | 8224 | 4238 | 5298 | 7844 | 4275 | 6134 | 10397 | 3423 | 4506 | 11203 | 2358 | 3656 | 10659 | 2871 | 3838 |
| 1678 | W303.asm02.HP0_G0014600 | chrV | 1429 | 338 | 371 | 1571 | 315 | 434 | 1308 | 269 | 427 | 3543 | 748 | 1266 | 1929 | 422 | 690 | 1969 | 805 | 1016 |
| 1679 | W303.asm02.HP0_G0014610 | chrV | 4285 | 400 | 431 | 4092 | 422 | 457 | 4060 | 369 | 511 | 3017 | 716 | 886 | 1959 | 557 | 759 | 1788 | 656 | 754 |
| 1680 | W303.asm02.HP0_G0014620 | chrV | 1693 | 273 | 293 | 1592 | 284 | 343 | 1529 | 331 | 375 | 1406 | 469 | 488 | 1434 | 296 | 341 | 1218 | 454 | 410 |
| 1681 | W303.asm02.HP0_G0014630 | chrV | 1878 | 1103 | 883 | 1628 | 1040 | 856 | 1827 | 1073 | 902 | 4678 | 1174 | 1203 | 4179 | 858 | 1050 | 4367 | 1077 | 1168 |
| 1682 | W303.asm02.HP0_G0014640 | chrV | 8198 | 1126 | 1438 | 7470 | 1191 | 1624 | 8162 | 1048 | 2017 | 7692 | 2397 | 3651 | 3707 | 1940 | 2210 | 3263 | 2278 | 3174 |
| 1683 | W303.asm02.HP0_G0014650 | chrV | 1306 | 546 | 595 | 1019 | 472 | 552 | 1148 | 555 | 685 | 1432 | 682 | 717 | 1862 | 448 | 673 | 1392 | 641 | 691 |
| 1684 | W303.asm02.HP0_G0014660 | chrV | 2378 | 873 | 665 | 1963 | 591 | 672 | 2477 | 758 | 793 | 3711 | 1580 | 1413 | 2635 | 1014 | 1122 | 2131 | 1402 | 1293 |
| 1685 | W303.asm02.HP0_G0014670 | chrV | 2321 | 1641 | 2110 | 1887 | 1756 | 2089 | 1924 | 1850 | 2600 | 3294 | 1522 | 1575 | 4100 | 1009 | 1518 | 3709 | 1352 | 1362 |
| 1686 | W303.asm02.HP0_G0014680 | chrV | 6312 | 1494 | 1150 | 6698 | 1030 | 1212 | 5672 | 1390 | 1081 | 4586 | 1904 | 1825 | 3944 | 1005 | 1413 | 3783 | 1888 | 1589 |
| 1687 | W303.asm02.HP0_G0014690 | chrV | 12162 | 5668 | 4999 | 12961 | 4848 | 4767 | 12497 | 5302 | 4945 | 17378 | 4944 | 5444 | 16097 | 3353 | 4694 | 17458 | 4680 | 4975 |
| 1688 | W303.asm02.HP0_G0014700 | chrV | 2737 | 1164 | 1013 | 2773 | 1106 | 1094 | 2717 | 1210 | 846 | 5607 | 1154 | 1300 | 4984 | 769 | 942 | 5027 | 1039 | 1206 |
| 1689 | W303.asm02.HP0_G0014710 | chrV | 5372 | 2917 | 2195 | 4393 | 2118 | 2227 | 5177 | 2642 | 2261 | 8707 | 3456 | 3784 | 8024 | 2719 | 3322 | 7323 | 3153 | 3338 |
| 1690 | W303.asm02.HP0_G0014720 | chrV | 992 | 423 | 419 | 949 | 467 | 394 | 973 | 387 | 438 | 2399 | 662 | 746 | 1789 | 517 | 578 | 1818 | 675 | 702 |
| 1691 | W303.asm02.HP0_G0014730 | chrV | 12982 | 3808 | 3576 | 12700 | 4040 | 3576 | 13884 | 3832 | 3799 | 23988 | 6095 | 7461 | 18875 | 4033 | 5040 | 18826 | 5178 | 6651 |
| 1692 | W303.asm02.HP0_G0014740 | chrV | 8364 | 3247 | 2719 | 8361 | 2225 | 2486 | 8333 | 3103 | 2972 | 10727 | 4291 | 3999 | 9896 | 2832 | 3458 | 10687 | 4029 | 3741 |
| 1693 | W303.asm02.HP0_G0014750 | chrV | 5847 | 636 | 817 | 5363 | 531 | 1015 | 5750 | 507 | 1016 | 4909 | 1228 | 2226 | 3140 | 875 | 1413 | 3062 | 1024 | 1891 |
| 1694 | W303.asm02.HP0_G0014760 | chrV | 605 | 138 | 167 | 615 | 196 | 185 | 596 | 111 | 194 | 1079 | 179 | 287 | 839 | 127 | 155 | 743 | 179 | 246 |
| 1695 | W303.asm02.HP0_G0014770 | chrV | 2053 | 1047 | 843 | 1881 | 927 | 868 | 1902 | 1164 | 964 | 3204 | 1043 | 900 | 3231 | 709 | 803 | 3192 | 921 | 838 |
| 1696 | W303.asm02.HP0_G0014780 | chrV | 3873 | 1853 | 2186 | 3794 | 2226 | 2329 | 3837 | 2230 | 2920 | 6756 | 1702 | 1831 | 8947 | 1193 | 1808 | 6906 | 1390 | 1749 |
| 1697 | W303.asm02.HP0_G0014790 | chrV | 4930 | 2654 | 2481 | 4435 | 2510 | 2500 | 4942 | 2712 | 2490 | 8446 | 3168 | 3304 | 8904 | 2326 | 3099 | 7985 | 2618 | 3110 |
| 1698 | W303.asm02.HP0_G0014800 | chrV | 1019 | 329 | 454 | 992 | 411 | 446 | 1083 | 280 | 496 | 3821 | 785 | 918 | 2165 | 596 | 824 | 2398 | 594 | 920 |
| 1699 | W303.asm02.HP0_G0014810 | chrV | 3091 | 126 | 182 | 3285 | 146 | 224 | 3093 | 126 | 203 | 1720 | 232 | 416 | 814 | 248 | 406 | 539 | 218 | 362 |
| 1700 | W303.asm02.HP0_G0014820 | chrV | 852 | 466 | 256 | 780 | 269 | 231 | 839 | 464 | 246 | 1326 | 598 | 451 | 1578 | 423 | 350 | 1181 | 549 | 413 |
| 1701 | W303.asm02.HP0_G0014830 | chrV | 1892 | 1099 | 570 | 1783 | 416 | 603 | 1470 | 914 | 537 | 1444 | 1021 | 755 | 1560 | 737 | 710 | 1339 | 1138 | 618 |
| 1702 | W303.asm02.HP0_G0014840 | chrV | 5941 | 5500 | 5103 | 5618 | 5580 | 4840 | 6375 | 5497 | 4859 | 29107 | 7435 | 8728 | 25611 | 5050 | 5596 | 25124 | 6680 | 7734 |
| 1703 | W303.asm02.HP0_G0014850 | chrV | 6843 | 2966 | 3913 | 5933 | 3103 | 3797 | 6308 | 2591 | 4549 | 5016 | 1714 | 1986 | 6065 | 1019 | 1699 | 5077 | 1785 | 1662 |
| 1704 | W303.asm02.HP0_G0014860 | chrV | 663 | 407 | 484 | 537 | 510 | 443 | 599 | 444 | 540 | 2150 | 330 | 447 | 1962 | 198 | 279 | 2117 | 355 | 397 |
| 1705 | W303.asm02.HP0_G0014870 | chrV | 201 | 318 | 126 | 185 | 125 | 120 | 231 | 216 | 144 | 374 | 208 | 165 | 384 | 113 | 147 | 397 | 206 | 139 |
| 1706 | W303.asm02.HP0_G0014880 | chrV | 51 | 120 | 39 | 33 | 40 | 40 | 32 | 98 | 39 | 93 | 81 | 44 | 109 | 42 | 50 | 76 | 89 | 38 |
| 1707 | W303.asm02.HP0_G0014890 | chrV | 1352 | 369 | 561 | 1332 | 433 | 592 | 1221 | 287 | 690 | 2953 | 732 | 1152 | 1065 | 458 | 774 | 1141 | 623 | 1043 |
| 1708 | W303.asm02.HP0_G0014900 | chrV | 1206 | 260 | 266 | 1086 | 286 | 246 | 1170 | 253 | 282 | 1912 | 437 | 539 | 1174 | 246 | 317 | 1273 | 409 | 460 |
| 1709 | W303.asm02.HP0_G0014910 | chrV | 6268 | 1441 | 1708 | 4284 | 1493 | 1705 | 5242 | 1340 | 2301 | 2493 | 823 | 773 | 3506 | 646 | 635 | 2942 | 669 | 556 |
| 1710 | W303.asm02.HP0_G0014920 | chrV | 36211 | 9667 | 9393 | 36908 | 9094 | 8294 | 40874 | 9182 | 9584 | 43637 | 13710 | 16036 | 39968 | 10863 | 12689 | 38795 | 12094 | 14535 |
| 1711 | W303.asm02.HP0_G0014930 | chrV | 7826 | 1824 | 2351 | 7174 | 1932 | 2267 | 7382 | 1537 | 2371 | 5896 | 1401 | 1715 | 6276 | 1010 | 1831 | 6249 | 1380 | 1771 |
| 1712 | W303.asm02.HP0_G0014940 | chrV | 333 | 123 | 152 | 367 | 144 | 109 | 344 | 122 | 153 | 707 | 111 | 115 | 758 | 79 | 109 | 687 | 117 | 110 |
| 1713 | W303.asm02.HP0_G0014950 | chrV | 6145 | 308 | 367 | 6078 | 475 | 405 | 6897 | 330 | 441 | 2540 | 504 | 673 | 3526 | 456 | 574 | 1907 | 461 | 566 |
| 1714 | W303.asm02.HP0_G0014960 | chrV | 436 | 411 | 402 | 318 | 391 | 477 | 317 | 393 | 420 | 895 | 306 | 277 | 1426 | 250 | 256 | 1016 | 260 | 202 |
| 1715 | W303.asm02.HP0_G0014970 | chrV | 1451 | 192 | 301 | 1789 | 289 | 345 | 1390 | 175 | 382 | 1842 | 347 | 627 | 687 | 244 | 344 | 659 | 289 | 514 |
| 1716 | W303.asm02.HP0_G0014980 | chrV | 6944 | 2608 | 1918 | 6681 | 2061 | 2004 | 6565 | 2517 | 1834 | 9108 | 2535 | 2877 | 8412 | 1886 | 2364 | 7857 | 2288 | 2618 |
| 1717 | W303.asm02.HP0_G0014990 | chrV | 1699 | 1751 | 2092 | 1532 | 1629 | 2031 | 1455 | 1733 | 2635 | 4487 | 1537 | 1526 | 6014 | 1097 | 1384 | 5170 | 1413 | 1381 |
| 1718 | W303.asm02.HP0_G0015000 | chrV | 1959 | 3276 | 3642 | 1951 | 4069 | 3697 | 2333 | 3400 | 3578 | 19195 | 3828 | 4949 | 15943 | 1972 | 2960 | 17361 | 3439 | 4568 |
| 1719 | W303.asm02.HP0_G0015010 | chrV | 3893 | 1692 | 2270 | 3776 | 1963 | 2438 | 3628 | 1649 | 3082 | 5353 | 1313 | 1945 | 6627 | 926 | 1762 | 5452 | 1173 | 1737 |
| 1720 | W303.asm02.HP0_G0015020 | chrV | 1101 | 728 | 666 | 1094 | 639 | 597 | 959 | 663 | 722 | 2519 | 682 | 742 | 2320 | 384 | 602 | 2580 | 659 | 667 |
| 1721 | W303.asm02.HP0_G0015030 | chrV | 3273 | 4364 | 4608 | 2541 | 4458 | 4578 | 2968 | 4240 | 3847 | 15288 | 3884 | 6236 | 13527 | 2880 | 4193 | 13506 | 3342 | 5306 |
| 1722 | W303.asm02.HP0_G0015040 | chrV | 11741 | 1373 | 1019 | 9927 | 801 | 1075 | 10343 | 1069 | 968 | 2946 | 1524 | 1393 | 3278 | 978 | 1628 | 2705 | 1599 | 1192 |
| 1723 | W303.asm02.HP0_G0015050 | chrV | 3494 | 243 | 150 | 3622 | 147 | 158 | 3058 | 187 | 151 | 599 | 281 | 208 | 554 | 205 | 197 | 471 | 238 | 157 |
| 1724 | W303.asm02.HP0_G0015060 | chrV | 3855 | 7190 | 8495 | 3190 | 7819 | 7870 | 3887 | 7693 | 8704 | 18085 | 6248 | 7946 | 19772 | 4709 | 6400 | 17454 | 4894 | 6899 |
| 1725 | W303.asm02.HP0_G0015070 | chrV | 1833 | 4541 | 7016 | 2046 | 7554 | 6366 | 1946 | 4740 | 7781 | 30745 | 6607 | 9725 | 23995 | 4604 | 6558 | 24610 | 5797 | 7862 |
| 1726 | W303.asm02.HP0_G0015080 | chrV | 6657 | 5179 | 6410 | 6047 | 6443 | 6401 | 6444 | 5146 | 7592 | 13275 | 4624 | 5104 | 14896 | 3474 | 4353 | 13830 | 4431 | 4527 |
| 1727 | W303.asm02.HP0_G0015090 | chrV | 4719 | 2036 | 1884 | 4041 | 1683 | 2043 | 4352 | 1923 | 1781 | 4284 | 1793 | 2249 | 5034 | 1419 | 1842 | 4035 | 1631 | 1956 |
| 1728 | W303.asm02.HP0_G0015100 | chrV | 1371 | 706 | 822 | 1087 | 653 | 838 | 1155 | 699 | 962 | 1832 | 586 | 703 | 2641 | 425 | 712 | 1754 | 568 | 642 |
| 1729 | W303.asm02.HP0_G0015110 | chrV | 3038 | 811 | 555 | 3170 | 545 | 580 | 2918 | 710 | 512 | 3248 | 770 | 1014 | 2772 | 574 | 747 | 2724 | 766 | 899 |
| 1730 | W303.asm02.HP0_G0015120 | chrV | 2450 | 832 | 1097 | 2045 | 864 | 1171 | 2219 | 712 | 1250 | 4913 | 1114 | 1998 | 3618 | 754 | 1421 | 3311 | 943 | 1784 |
| 1731 | W303.asm02.HP0_G0015130 | chrV | 1227 | 390 | 331 | 1341 | 410 | 313 | 1229 | 372 | 378 | 1635 | 415 | 474 | 1505 | 273 | 357 | 1135 | 374 | 397 |
| 1732 | W303.asm02.HP0_G0015140 | chrV | 2314 | 5893 | 4053 | 2009 | 3216 | 4215 | 2179 | 4473 | 4683 | 6324 | 4013 | 4362 | 9348 | 2250 | 3239 | 6153 | 4351 | 3724 |
| 1733 | W303.asm02.HP0_G0015150 | chrV | 5630 | 1045 | 1002 | 5118 | 867 | 1089 | 5255 | 1114 | 1086 | 4231 | 1287 | 1845 | 4848 | 1009 | 1566 | 3886 | 1112 | 1586 |
| 1734 | W303.asm02.HP0_G0015160 | chrV | 2242 | 733 | 921 | 2533 | 791 | 1086 | 2447 | 605 | 1022 | 6890 | 1496 | 2567 | 5560 | 1165 | 1792 | 4435 | 1298 | 2296 |
| 1735 | W303.asm02.HP0_G0015170 | chrV | 614 | 352 | 260 | 589 | 251 | 316 | 608 | 308 | 315 | 1942 | 621 | 636 | 1153 | 411 | 405 | 1219 | 661 | 554 |
| 1736 | W303.asm02.HP0_G0015180 | chrV | 925 | 353 | 200 | 952 | 198 | 177 | 779 | 227 | 262 | 666 | 237 | 170 | 770 | 125 | 155 | 666 | 283 | 151 |
| 1737 | W303.asm02.HP0_G0015190 | chrV | 13952 | 792 | 287 | 13813 | 397 | 429 | 11655 | 525 | 350 | 698 | 492 | 179 | 2083 | 454 | 540 | 777 | 468 | 220 |
| 1738 | W303.asm02.HP0_G0015200 | chrV | 2850 | 539 | 704 | 3169 | 621 | 848 | 2874 | 511 | 824 | 6152 | 1083 | 1980 | 5133 | 725 | 1336 | 4614 | 1059 | 1752 |
| 1739 | W303.asm02.HP0_G0015210 | chrV | 1357 | 2002 | 1812 | 1216 | 1647 | 1855 | 1286 | 1778 | 2153 | 16913 | 3661 | 3896 | 9266 | 2870 | 2535 | 11049 | 2987 | 3353 |
| 1740 | W303.asm02.HP0_G0015220 | chrV | 2494 | 1492 | 1980 | 2274 | 1710 | 2275 | 3105 | 1446 | 2537 | 16681 | 3697 | 5467 | 10028 | 2692 | 3262 | 9978 | 3214 | 5001 |
| 1741 | W303.asm02.HP0_G0015230 | chrV | 1457 | 757 | 913 | 1155 | 787 | 854 | 1276 | 777 | 1285 | 2210 | 670 | 773 | 2559 | 409 | 570 | 2730 | 587 | 689 |
| 1742 | W303.asm02.HP0_G0015240 | chrV | 4521 | 7073 | 9497 | 3940 | 8838 | 9642 | 4239 | 7069 | 11306 | 14869 | 6032 | 6760 | 18307 | 4556 | 6286 | 15957 | 5317 | 5931 |
| 1743 | W303.asm02.HP0_G0015250 | chrV | 608 | 1943 | 2244 | 704 | 2503 | 2340 | 616 | 1819 | 2637 | 13102 | 3461 | 4124 | 7768 | 2367 | 2344 | 7941 | 2906 | 3233 |
| 1744 | W303.asm02.HP0_G0015260 | chrV | 7986 | 6358 | 9426 | 6710 | 8494 | 9360 | 7739 | 6268 | 12218 | 17699 | 5774 | 7833 | 20439 | 4347 | 6212 | 19198 | 5296 | 6454 |
| 1745 | W303.asm02.HP0_G0015270 | chrV | 481 | 299 | 341 | 414 | 267 | 311 | 418 | 298 | 442 | 747 | 239 | 214 | 879 | 184 | 188 | 831 | 242 | 192 |
| 1746 | W303.asm02.HP0_G0015280 | chrV | 1036 | 323 | 489 | 1095 | 426 | 529 | 947 | 316 | 649 | 4954 | 641 | 1001 | 3179 | 457 | 590 | 3087 | 683 | 828 |
| 1747 | W303.asm02.HP0_G0015290 | chrV | 758 | 374 | 220 | 657 | 228 | 235 | 667 | 320 | 243 | 1247 | 432 | 394 | 1031 | 303 | 304 | 989 | 437 | 358 |
| 1748 | W303.asm02.HP0_G0015300 | chrV | 2595 | 1237 | 1417 | 2703 | 1447 | 1375 | 2614 | 1369 | 1410 | 7612 | 1469 | 1953 | 9661 | 1182 | 1541 | 7039 | 1434 | 1717 |
| 1749 | W303.asm02.HP0_G0015310 | chrV | 3269 | 750 | 684 | 3194 | 721 | 754 | 3129 | 698 | 751 | 4670 | 1040 | 1264 | 4235 | 663 | 938 | 4073 | 1030 | 1114 |
| 1750 | W303.asm02.HP0_G0015320 | chrV | 34 | 27 | 11 | 22 | 13 | 9 | 17 | 32 | 15 | 40 | 28 | 17 | 44 | 21 | 9 | 30 | 23 | 9 |
| 1751 | W303.asm02.HP0_G0015330 | chrV | 3869 | 493 | 271 | 4281 | 213 | 276 | 3417 | 410 | 300 | 925 | 613 | 453 | 1075 | 334 | 410 | 800 | 668 | 367 |
| 1752 | W303.asm02.HP0_G0015340 | chrV | 8732 | 1799 | 1509 | 8253 | 1430 | 1621 | 8336 | 1616 | 1548 | 10076 | 2301 | 2944 | 9075 | 1864 | 2246 | 8608 | 2152 | 2649 |
| 1753 | W303.asm02.HP0_G0015350 | chrV | 8014 | 3069 | 2626 | 6100 | 3015 | 2478 | 6910 | 3420 | 2358 | 5524 | 4041 | 4236 | 8016 | 3338 | 3793 | 5606 | 3543 | 3582 |
| 1754 | W303.asm02.HP0_G0015360 | chrV | 3906 | 1455 | 1375 | 3910 | 1814 | 1411 | 4204 | 1480 | 1350 | 11641 | 2169 | 2699 | 9884 | 1507 | 1608 | 8888 | 1935 | 2377 |
| 1755 | W303.asm02.HP0_G0015370 | chrV | 3677 | 656 | 659 | 3134 | 617 | 788 | 3344 | 637 | 846 | 4088 | 1269 | 1457 | 3345 | 882 | 1237 | 2973 | 1033 | 1351 |
| 1756 | W303.asm02.HP0_G0015380 | chrV | 91 | 24 | 19 | 59 | 12 | 20 | 58 | 27 | 20 | 83 | 31 | 40 | 72 | 22 | 18 | 57 | 48 | 25 |
| 1757 | W303.asm02.HP0_G0015390 | chrV | 36 | 13 | 11 | 40 | 10 | 10 | 37 | 13 | 14 | 51 | 14 | 18 | 78 | 7 | 11 | 40 | 17 | 6 |
| 1758 | W303.asm02.HP0_G0015400 | chrV | 7994 | 2328 | 2355 | 7024 | 2404 | 2554 | 8424 | 2414 | 2893 | 17814 | 4614 | 5521 | 13815 | 3425 | 3510 | 13895 | 3747 | 4936 |
| 1759 | W303.asm02.HP0_G0015410 | chrV | 5768 | 930 | 875 | 5207 | 896 | 896 | 5037 | 1021 | 881 | 3772 | 910 | 1195 | 3750 | 708 | 1072 | 3298 | 931 | 1090 |
| 1760 | W303.asm02.HP0_G0015420 | chrV | 2298 | 1641 | 2205 | 2220 | 1734 | 2353 | 2202 | 1736 | 2370 | 4586 | 1622 | 2572 | 5226 | 1329 | 1767 | 4421 | 1412 | 2255 |
| 1761 | W303.asm02.HP0_G0015430 | chrV | 7434 | 519 | 607 | 7771 | 662 | 764 | 7474 | 357 | 815 | 3251 | 720 | 1190 | 1252 | 586 | 688 | 1047 | 581 | 938 |
| 1762 | W303.asm02.HP0_G0015440 | chrV | 5458 | 886 | 835 | 4077 | 842 | 1102 | 5516 | 710 | 1166 | 6296 | 1576 | 2038 | 2002 | 1459 | 1518 | 2067 | 1374 | 1924 |
| 1763 | W303.asm02.HP0_G0015450 | chrV | 5953 | 4531 | 3258 | 5195 | 3347 | 3261 | 6093 | 4193 | 2992 | 13150 | 4658 | 4984 | 11884 | 3665 | 3453 | 11223 | 4000 | 4356 |
| 1764 | W303.asm02.HP0_G0015460 | chrV | 29721 | 4678 | 5655 | 19379 | 5274 | 5492 | 26003 | 4678 | 6124 | 19201 | 3836 | 6717 | 16961 | 4063 | 5665 | 14960 | 3433 | 4424 |
| 1765 | W303.asm02.HP0_G0015470 | chrV | 3 | 4 | 2 | 4 | 1 | 0 | 3 | 0 | 3 | 3 | 2 | 0 | 3 | 1 | 0 | 2 | 5 | 1 |

| | | | | | | | | | | | | | | | | | | | | |
|---|---|---|---|---|---|---|---|---|---|---|---|---|---|---|---|---|---|---|---|---|
| 1766 | W303.asm02.HP0_G0015480 | chrV | 1800 | 932 | 803 | 1652 | 759 | 822 | 1769 | 992 | 918 | 1979 | 759 | 663 | 2524 | 541 | 564 | 2168 | 638 | 604 |
| 1767 | W303.asm02.HP0_G0015490 | chrV | 2488 | 280 | 399 | 2387 | 428 | 455 | 2440 | 303 | 490 | 4229 | 515 | 934 | 2803 | 385 | 657 | 2579 | 458 | 804 |
| 1768 | W303.asm02.HP0_G0015500 | chrV | 1623 | 377 | 377 | 1454 | 357 | 406 | 1417 | 378 | 385 | 925 | 310 | 450 | 1021 | 207 | 328 | 749 | 302 | 388 |
| 1769 | W303.asm02.HP0_G0015510 | chrV | 6385 | 3994 | 3856 | 6217 | 3412 | 3833 | 6339 | 3617 | 3816 | 9363 | 3701 | 4208 | 10478 | 2740 | 3431 | 9058 | 3308 | 3774 |
| 1770 | W303.asm02.HP0_G0015520 | chrV | 2232 | 1373 | 652 | 2029 | 734 | 683 | 2204 | 1037 | 682 | 3936 | 1968 | 1416 | 3434 | 1472 | 1250 | 2693 | 1746 | 1299 |
| 1771 | W303.asm02.HP0_G0015530 | chrV | 538 | 128 | 112 | 579 | 120 | 91 | 499 | 165 | 157 | 756 | 269 | 219 | 496 | 164 | 144 | 543 | 288 | 207 |
| 1772 | W303.asm02.HP0_G0015540 | chrV | 2490 | 190 | 244 | 2631 | 255 | 274 | 2342 | 177 | 310 | 1829 | 332 | 419 | 1415 | 286 | 429 | 1306 | 323 | 392 |
| 1773 | W303.asm02.HP0_G0015550 | chrV | 3315 | 1582 | 1170 | 2867 | 1372 | 1013 | 2924 | 1475 | 1215 | 4715 | 1613 | 1446 | 4764 | 1158 | 1084 | 4205 | 1473 | 1322 |
| 1774 | W303.asm02.HP0_G0015560 | chrV | 5861 | 2187 | 1936 | 5411 | 2057 | 2051 | 5643 | 2370 | 1943 | 7369 | 2088 | 2457 | 7245 | 1307 | 1745 | 7089 | 1933 | 2258 |
| 1775 | W303.asm02.HP0_G0015570 | chrV | 2519 | 548 | 360 | 2424 | 361 | 372 | 2373 | 506 | 415 | 1477 | 571 | 443 | 1655 | 412 | 353 | 1385 | 560 | 425 |
| 1776 | W303.asm02.HP0_G0015580 | chrV | 9233 | 7021 | 9907 | 9600 | 9376 | 9358 | 9712 | 7071 | 12553 | 21401 | 6862 | 7930 | 22815 | 5183 | 6255 | 22414 | 5977 | 7213 |
| 1777 | W303.asm02.HP0_G0015590 | chrV | 9419 | 1848 | 994 | 10272 | 2165 | 2764 | 8047 | 992 | 686 | 2978 | 2555 | 1421 | 1589 | 1089 | 993 | 1876 | 2976 | 1131 |
| 1778 | W303.asm02.HP0_G0015600 | chrV | 927 | 304 | 353 | 1068 | 334 | 332 | 866 | 298 | 305 | 958 | 299 | 417 | 1102 | 203 | 258 | 908 | 262 | 332 |
| 1779 | W303.asm02.HP0_G0015610 | chrV | 4434 | 981 | 1249 | 4591 | 1277 | 1382 | 4523 | 981 | 1621 | 11334 | 1726 | 2796 | 6593 | 1334 | 2009 | 7009 | 1400 | 2510 |
| 1780 | W303.asm02.HP0_G0015620 | chrV | 92 | 44 | 61 | 77 | 59 | 53 | 89 | 57 | 76 | 168 | 39 | 47 | 248 | 20 | 30 | 147 | 41 | 42 |
| 1781 | W303.asm02.HP0_G0015630 | chrV | 2127 | 1306 | 1068 | 1940 | 1094 | 981 | 2166 | 1319 | 1098 | 6174 | 1625 | 1830 | 4946 | 1176 | 1311 | 5165 | 1601 | 1633 |
| 1782 | W303.asm02.HP0_G0015640 | chrV | 889 | 216 | 397 | 913 | 544 | 673 | 932 | 217 | 478 | 2405 | 462 | 821 | 1531 | 376 | 577 | 1249 | 423 | 725 |
| 1783 | W303.asm02.HP0_G0015650 | chrV | 4434 | 1846 | 1668 | 3695 | 2089 | 1788 | 4998 | 2005 | 2139 | 22192 | 4634 | 4835 | 15090 | 3140 | 2531 | 15807 | 3790 | 4585 |
| 1784 | W303.asm02.HP0_G0015660 | chrV | 1111 | 540 | 702 | 1256 | 730 | 741 | 1146 | 601 | 754 | 5308 | 653 | 1129 | 5737 | 541 | 1015 | 5008 | 529 | 1069 |
| 1785 | W303.asm02.HP0_G0015670 | chrV | 3870 | 2393 | 3159 | 4174 | 2958 | 3151 | 3814 | 2625 | 4182 | 7818 | 1941 | 2434 | 8797 | 1224 | 1998 | 8446 | 1680 | 2180 |
| 1786 | W303.asm02.HP0_G0015680 | chrV | 900 | 687 | 930 | 704 | 863 | 931 | 862 | 685 | 1121 | 6299 | 1092 | 1442 | 3864 | 667 | 1057 | 4753 | 950 | 1360 |
| 1787 | W303.asm02.HP0_G0015690 | chrV | 2622 | 435 | 448 | 2496 | 537 | 581 | 2589 | 350 | 628 | 3529 | 701 | 1113 | 1133 | 539 | 664 | 1170 | 575 | 927 |
| 1788 | W303.asm02.HP0_G0015700 | chrV | 1150 | 393 | 358 | 1060 | 358 | 343 | 1115 | 370 | 410 | 1632 | 417 | 435 | 1164 | 324 | 348 | 1271 | 460 | 342 |
| 1789 | W303.asm02.HP0_G0015710 | chrV | 1564 | 563 | 314 | 1490 | 351 | 320 | 1535 | 527 | 288 | 2010 | 614 | 501 | 1958 | 487 | 304 | 1458 | 588 | 382 |
| 1790 | W303.asm02.HP0_G0015720 | chrV | 7208 | 6685 | 8796 | 6669 | 8133 | 8696 | 7233 | 6704 | 11147 | 16397 | 6152 | 6664 | 18704 | 4577 | 5627 | 17595 | 5600 | 5931 |
| 1791 | W303.asm02.HP0_G0015730 | chrV | 1073 | 2067 | 2216 | 1030 | 1968 | 2094 | 1167 | 2200 | 2293 | 7231 | 1937 | 2205 | 9534 | 1471 | 1785 | 7123 | 1676 | 2204 |
| 1792 | W303.asm02.HP0_G0015740 | chrV | 2336 | 412 | 313 | 1783 | 260 | 409 | 2309 | 332 | 359 | 1682 | 619 | 751 | 960 | 521 | 678 | 729 | 573 | 700 |
| 1793 | W303.asm02.HP0_G0015750 | chrV | 8943 | 1754 | 1334 | 8408 | 1326 | 1553 | 9207 | 1706 | 1420 | 8762 | 3105 | 3164 | 8044 | 2280 | 2681 | 7584 | 2653 | 2885 |
| 1794 | W303.asm02.HP0_G0015760 | chrV | 433 | 123 | 97 | 464 | 88 | 76 | 362 | 95 | 110 | 181 | 84 | 78 | 308 | 60 | 94 | 181 | 98 | 62 |
| 1795 | W303.asm02.HP0_G0015770 | chrV | 6887 | 4184 | 3565 | 7149 | 3393 | 3582 | 7459 | 4207 | 3464 | 15745 | 4498 | 5812 | 13972 | 2964 | 3901 | 13813 | 3846 | 5214 |
| 1796 | W303.asm02.HP0_G0015780 | chrV | 1897 | 1064 | 1128 | 2147 | 1303 | 1155 | 2037 | 1123 | 1093 | 6948 | 1357 | 1682 | 5555 | 855 | 1220 | 5965 | 1247 | 1525 |
| 1797 | W303.asm02.HP0_G0015790 | chrV | 5445 | 4251 | 3637 | 5019 | 3694 | 3324 | 5447 | 3987 | 3773 | 12445 | 3001 | 3116 | 16275 | 2404 | 2892 | 13363 | 2720 | 2792 |
| 1798 | W303.asm02.HP0_G0015800 | chrV | 13771 | 1931 | 1875 | 12515 | 1856 | 2105 | 14073 | 1791 | 2472 | 16738 | 4113 | 4410 | 10153 | 3017 | 3445 | 9897 | 3519 | 3944 |
| 1799 | W303.asm02.HP0_G0015810 | chrV | 3013 | 3307 | 4463 | 2910 | 4581 | 4076 | 3270 | 3577 | 5048 | 12931 | 3227 | 3714 | 13196 | 2089 | 2348 | 12676 | 2817 | 3432 |
| 1800 | W303.asm02.HP0_G0015820 | chrV | 835 | 1620 | 2020 | 886 | 2295 | 2021 | 1041 | 1789 | 1950 | 8657 | 1705 | 2553 | 8399 | 1111 | 1360 | 7629 | 1498 | 2084 |
| 1801 | W303.asm02.HP0_G0015830 | chrV | 785 | 394 | 407 | 754 | 418 | 411 | 777 | 409 | 354 | 1373 | 374 | 510 | 1378 | 235 | 306 | 1253 | 344 | 449 |
| 1802 | W303.asm02.HP0_G0015840 | chrV | 3780 | 526 | 765 | 3231 | 786 | 883 | 3825 | 569 | 950 | 6604 | 1013 | 1727 | 3969 | 739 | 1293 | 3994 | 867 | 1601 |
| 1803 | W303.asm02.HP0_G0015850 | chrV | 1812 | 832 | 1172 | 1379 | 1092 | 1187 | 1663 | 845 | 1149 | 4821 | 907 | 1280 | 5120 | 615 | 1046 | 4235 | 964 | 1097 |
| 1804 | W303.asm02.HP0_G0015860 | chrV | 12407 | 8864 | 12314 | 12369 | 12277 | 12168 | 12651 | 9983 | 14385 | 26470 | 8997 | 10628 | 28423 | 6102 | 8914 | 26740 | 7356 | 9522 |
| 1805 | W303.asm02.HP0_G0015870 | chrV | 1566 | 374 | 650 | 1386 | 601 | 635 | 1562 | 426 | 784 | 4417 | 556 | 1174 | 3949 | 552 | 817 | 2377 | 475 | 976 |
| 1806 | W303.asm02.HP0_G0015880 | chrV | 1007 | 666 | 825 | 1016 | 687 | 779 | 868 | 663 | 909 | 2095 | 533 | 544 | 2657 | 337 | 496 | 2267 | 504 | 500 |
| 1807 | W303.asm02.HP0_G0015890 | chrV | 813 | 611 | 891 | 699 | 860 | 794 | 778 | 629 | 845 | 3884 | 708 | 900 | 3475 | 331 | 631 | 3631 | 617 | 803 |
| 1808 | W303.asm02.HP0_G0015900 | chrV | 15513 | 2126 | 2101 | 15042 | 1960 | 2145 | 14615 | 1955 | 2163 | 8523 | 2775 | 3680 | 6796 | 1825 | 3083 | 6812 | 2678 | 3247 |
| 1809 | W303.asm02.HP0_G0015910 | chrV | 2310 | 566 | 788 | 2569 | 661 | 855 | 2153 | 559 | 1032 | 1687 | 393 | 515 | 2204 | 354 | 517 | 1751 | 370 | 475 |
| 1810 | W303.asm02.HP0_G0015920 | chrV | 7266 | 1144 | 951 | 7293 | 883 | 895 | 6942 | 1071 | 977 | 4744 | 1419 | 1568 | 4576 | 1046 | 1339 | 4111 | 1290 | 1405 |
| 1811 | W303.asm02.HP0_G0015930 | chrV | 2845 | 2135 | 2711 | 2758 | 2391 | 2971 | 3028 | 2173 | 3088 | 15832 | 4433 | 4618 | 10800 | 2548 | 3498 | 11796 | 3919 | 5244 |
| 1812 | W303.asm02.HP0_G0015940 | chrV | 881 | 961 | 1622 | 685 | 1421 | 1693 | 788 | 991 | 2213 | 3336 | 964 | 1265 | 3891 | 567 | 1066 | 3566 | 817 | 1146 |
| 1813 | W303.asm02.HP0_G0015950 | chrV | 901 | 334 | 341 | 884 | 371 | 373 | 955 | 301 | 457 | 2519 | 708 | 765 | 1404 | 432 | 460 | 1334 | 610 | 637 |
| 1814 | W303.asm02.HP0_G0015960 | chrV | 4068 | 3338 | 3583 | 3857 | 3335 | 3395 | 4028 | 3385 | 3829 | 11819 | 3359 | 3814 | 11361 | 2295 | 2889 | 11372 | 3014 | 3522 |
| 1815 | W303.asm02.HP0_G0015970 | chrV | 1297 | 653 | 783 | 1261 | 770 | 767 | 1443 | 758 | 716 | 3561 | 629 | 862 | 3393 | 418 | 523 | 3349 | 568 | 791 |
| 1816 | W303.asm02.HP0_G0015980 | chrV | 7530 | 532 | 192 | 8139 | 729 | 808 | 6804 | 376 | 178 | 520 | 384 | 210 | 798 | 222 | 200 | 467 | 416 | 207 |
| 1817 | W303.asm02.HP0_G0015990 | chrV | 4546 | 779 | 1177 | 4310 | 1165 | 1280 | 4814 | 688 | 1649 | 7742 | 1620 | 2561 | 2737 | 1125 | 1380 | 2788 | 1235 | 2118 |
| 1818 | W303.asm02.HP0_G0016000 | chrV | 3801 | 863 | 1021 | 3302 | 852 | 1101 | 3666 | 722 | 1376 | 2789 | 1220 | 1563 | 1849 | 851 | 1122 | 1364 | 1091 | 1374 |
| 1819 | W303.asm02.HP0_G0016010 | chrV | 324 | 88 | 209 | 311 | 147 | 197 | 339 | 93 | 246 | 353 | 165 | 262 | 138 | 108 | 126 | 140 | 140 | 235 |
| 1820 | W303.asm02.HP0_G0016020 | chrV | 3988 | 809 | 1151 | 3557 | 977 | 1252 | 4031 | 749 | 1533 | 5190 | 1392 | 2116 | 2327 | 1082 | 1461 | 2041 | 1195 | 1859 |
| 1821 | W303.asm02.HP0_G0016030 | chrV | 6747 | 1468 | 2315 | 6791 | 2111 | 2439 | 6533 | 1524 | 2858 | 16305 | 1941 | 4333 | 11023 | 1542 | 3140 | 11001 | 1688 | 3780 |
| 1822 | W303.asm02.HP0_G0016040 | chrV | 5420 | 1478 | 1469 | 4730 | 1282 | 1598 | 5330 | 1434 | 1647 | 8559 | 2362 | 3653 | 6635 | 1728 | 2148 | 6516 | 2046 | 2988 |
| 1823 | W303.asm02.HP0_G0016050 | chrV | 2082 | 925 | 1158 | 2012 | 1102 | 1136 | 2072 | 1026 | 1080 | 5733 | 927 | 1377 | 5202 | 649 | 983 | 4993 | 870 | 1228 |
| 1824 | W303.asm02.HP0_G0016060 | chrV | 2631 | 372 | 457 | 3076 | 401 | 477 | 2550 | 339 | 515 | 2316 | 655 | 708 | 2088 | 382 | 489 | 1458 | 619 | 632 |
| 1825 | W303.asm02.HP0_G0016070 | chrV | 4615 | 2597 | 3017 | 4140 | 3113 | 3091 | 4238 | 2962 | 3695 | 6505 | 2521 | 2243 | 7453 | 1613 | 2168 | 6627 | 2132 | 2273 |
| 1826 | W303.asm02.HP0_G0016080 | chrV | 3801 | 672 | 626 | 3870 | 545 | 707 | 3807 | 597 | 686 | 3955 | 827 | 1525 | 3422 | 622 | 970 | 3031 | 694 | 1261 |
| 1827 | W303.asm02.HP0_G0016090 | chrV | 2854 | 251 | 258 | 2935 | 259 | 234 | 2631 | 219 | 259 | 1993 | 372 | 531 | 1243 | 231 | 366 | 1147 | 350 | 452 |
| 1828 | W303.asm02.HP0_G0016100 | chrV | 3497 | 1073 | 1032 | 3103 | 1047 | 1167 | 3021 | 1117 | 1048 | 2662 | 1090 | 1301 | 3300 | 897 | 1163 | 2427 | 986 | 1156 |
| 1829 | W303.asm02.HP0_G0016110 | chrV | 3996 | 406 | 507 | 3475 | 431 | 666 | 3859 | 332 | 657 | 6581 | 900 | 1569 | 1576 | 680 | 831 | 1908 | 732 | 1445 |
| 1830 | W303.asm02.HP0_G0016120 | chrV | 28727 | 4264 | 4234 | 25704 | 4058 | 4905 | 31703 | 3969 | 5319 | 34593 | 10366 | 11463 | 24670 | 8241 | 8245 | 24595 | 8626 | 10701 |
| 1831 | W303.asm02.HP0_G0016130 | chrV | 5200 | 984 | 1245 | 4765 | 1127 | 1240 | 5186 | 1032 | 1340 | 7760 | 1385 | 2559 | 5306 | 1164 | 1887 | 4639 | 1157 | 2187 |
| 1832 | W303.asm02.HP0_G0016140 | chrV | 2311 | 399 | 705 | 2678 | 641 | 703 | 2453 | 399 | 900 | 5758 | 734 | 1314 | 5249 | 678 | 1117 | 3351 | 625 | 1113 |
| 1833 | W303.asm02.HP0_G0016150 | chrV | 1127 | 1219 | 1180 | 1071 | 1137 | 1136 | 1211 | 1301 | 1274 | 5869 | 1199 | 1375 | 4940 | 752 | 850 | 5560 | 1177 | 1187 |
| 1834 | W303.asm02.HP0_G0016160 | chrV | 2570 | 317 | 452 | 2428 | 520 | 515 | 2513 | 266 | 586 | 4234 | 692 | 947 | 1802 | 511 | 564 | 1430 | 668 | 664 |
| 1835 | W303.asm02.HP0_G0016170 | chrV | 2235 | 921 | 911 | 2007 | 797 | 876 | 2090 | 908 | 1008 | 2252 | 765 | 757 | 2473 | 470 | 675 | 2295 | 693 | 708 |
| 1836 | W303.asm02.HP0_G0016180 | chrV | 3661 | 1044 | 1200 | 3233 | 1201 | 1139 | 3760 | 1080 | 1353 | 7745 | 1385 | 1766 | 6269 | 886 | 1254 | 6841 | 1184 | 1619 |
| 1837 | W303.asm02.HP0_G0016190 | chrV | 3359 | 667 | 892 | 3382 | 969 | 958 | 3256 | 664 | 1119 | 8952 | 1025 | 1895 | 5379 | 714 | 1282 | 5632 | 977 | 1538 |
| 1838 | W303.asm02.HP0_G0016200 | chrV | 816 | 308 | 418 | 843 | 331 | 473 | 927 | 290 | 515 | 2654 | 633 | 895 | 2046 | 438 | 598 | 2041 | 583 | 852 |
| 1839 | W303.asm02.HP0_G0016210 | chrV | 1392 | 1254 | 1754 | 1233 | 1636 | 1492 | 1390 | 1425 | 2182 | 4952 | 908 | 1508 | 4888 | 619 | 948 | 4718 | 861 | 1061 |
| 1840 | W303.asm02.HP0_G0016220 | chrV | 1249 | 1023 | 856 | 1213 | 796 | 954 | 1114 | 1027 | 879 | 2376 | 849 | 937 | 2658 | 496 | 870 | 2410 | 813 | 955 |
| 1841 | W303.asm02.HP0_G0016230 | chrV | 110 | 98 | 75 | 147 | 81 | 77 | 152 | 83 | 118 | 388 | 68 | 55 | 258 | 49 | 51 | 395 | 57 | 71 |
| 1842 | W303.asm02.HP0_G0016240 | chrV | 3577 | 589 | 681 | 3545 | 666 | 813 | 3474 | 488 | 746 | 6033 | 1125 | 1689 | 2704 | 788 | 969 | 2930 | 1038 | 1451 |
| 1843 | W303.asm02.HP0_G0016250 | chrV | 40700 | 11656 | 10570 | 40923 | 9042 | 11203 | 42405 | 11389 | 12761 | 41660 | 18046 | 21519 | 38675 | 14544 | 17631 | 35383 | 16146 | 19324 |
| 1844 | W303.asm02.HP0_G0016260 | chrV | 847 | 259 | 178 | 785 | 173 | 214 | 761 | 251 | 221 | 596 | 248 | 197 | 877 | 187 | 206 | 576 | 233 | 168 |
| 1845 | W303.asm02.HP0_G0016270 | chrV | 742 | 477 | 411 | 741 | 415 | 390 | 705 | 392 | 349 | 1861 | 405 | 523 | 1656 | 277 | 414 | 1792 | 384 | 452 |
| 1846 | W303.asm02.HP0_G0016280 | chrV | 376 | 208 | 176 | 279 | 180 | 140 | 285 | 186 | 133 | 697 | 191 | 207 | 695 | 137 | 155 | 651 | 204 | 189 |
| 1847 | W303.asm02.HP0_G0016290 | chrV | 9179 | 1902 | 1489 | 8121 | 1482 | 1471 | 8362 | 1859 | 1655 | 4513 | 1672 | 1783 | 5158 | 1199 | 1474 | 4131 | 1657 | 1488 |
| 1848 | W303.asm02.HP0_G0016300 | chrV | 1079 | 1142 | 965 | 933 | 853 | 993 | 1023 | 1076 | 1196 | 2456 | 844 | 824 | 2807 | 586 | 619 | 2645 | 811 | 684 |
| 1849 | W303.asm02.HP0_G0016310 | chrV | 742 | 127 | 208 | 863 | 219 | 194 | 657 | 145 | 249 | 1594 | 194 | 369 | 1146 | 141 | 293 | 1151 | 166 | 363 |
| 1850 | W303.asm02.HP0_G0016320 | chrV | 125 | 68 | 56 | 115 | 95 | 57 | 122 | 72 | 87 | 265 | 72 | 59 | 264 | 47 | 45 | 235 | 63 | 48 |
| 1851 | W303.asm02.HP0_G0016330 | chrV | 1392 | 825 | 832 | 1234 | 817 | 772 | 1197 | 852 | 976 | 3316 | 738 | 743 | 3139 | 517 | 597 | 3269 | 679 | 679 |
| 1852 | W303.asm02.HP0_G0016340 | chrV | 474 | 74 | 62 | 409 | 65 | 51 | 269 | 65 | 60 | 271 | 56 | 59 | 247 | 31 | 42 | 232 | 64 | 60 |
| 1853 | W303.asm02.HP0_G0016350 | chrV | 603 | 106 | 95 | 565 | 104 | 109 | 296 | 83 | 109 | 281 | 96 | 104 | 318 | 54 | 50 | 282 | 91 | 87 |
| 1854 | W303.asm02.HP0_G0016360 | chrV | 1313 | 342 | 278 | 972 | 248 | 306 | 601 | 255 | 293 | 520 | 191 | 166 | 501 | 108 | 111 | 518 | 216 | 162 |
| 1855 | W303.asm02.HP0_G0016370 | chrV | 72 | 15 | 17 | 50 | 23 | 17 | 65 | 20 | 14 | 33 | 17 | 34 | 16 | 16 | 6 | 11 | 20 | 21 |
| 1856 | W303.asm02.HP0_G0016380 | chrV | 0 | 0 | 0 | 0 | 0 | 0 | 0 | 0 | 0 | 0 | 0 | 0 | 0 | 0 | 0 | 0 | 0 | 0 |
| 1857 | W303.asm02.HP0_G0016390 | chrV | 0 | 0 | 0 | 0 | 0 | 0 | 0 | 0 | 0 | 0 | 0 | 0 | 0 | 0 | 0 | 0 | 0 | 0 |
| 1858 | W303.asm02.HP0_G0016400 | chrVI | 1045 | 370 | 409 | 1029 | 494 | 432 | 956 | 382 | 480 | 2973 | 560 | 682 | 2936 | 372 | 490 | 2033 | 476 | 566 |

| | | | | | | | | | | | | | | | | | | | | |
|---|---|---|---|---|---|---|---|---|---|---|---|---|---|---|---|---|---|---|---|---|
| 1859 | W303.asm02.HP0_G0016410 | chrVI | 3217 | 992 | 960 | 3070 | 1137 | 942 | 3286 | 969 | 1053 | 4833 | 992 | 1156 | 5626 | 774 | 1030 | 4374 | 957 | 1068 |
| 1860 | W303.asm02.HP0_G0016420 | chrVI | 3279 | 2506 | 2234 | 3106 | 2132 | 2146 | 3210 | 2379 | 2348 | 8140 | 2169 | 2386 | 9373 | 1593 | 1895 | 7897 | 1997 | 2229 |
| 1861 | W303.asm02.HP0_G0016430 | chrVI | 2619 | 1549 | 1248 | 2456 | 1306 | 1234 | 2643 | 1515 | 1351 | 7236 | 1853 | 2102 | 6489 | 1170 | 1579 | 6110 | 1956 | 1944 |
| 1862 | W303.asm02.HP0_G0016440 | chrVI | 1778 | 1179 | 1331 | 1473 | 1194 | 1472 | 1555 | 1225 | 1746 | 3183 | 916 | 1212 | 3543 | 632 | 899 | 3163 | 872 | 1031 |
| 1863 | W303.asm02.HP0_G0016450 | chrVI | 9388 | 8032 | 10534 | 8023 | 8469 | 10413 | 9557 | 8534 | 11911 | 21549 | 7309 | 10240 | 22862 | 5535 | 7333 | 22305 | 6708 | 8989 |
| 1864 | W303.asm02.HP0_G0016460 | chrVI | 3752 | 1086 | 913 | 3398 | 936 | 895 | 3455 | 1069 | 805 | 3264 | 985 | 971 | 3566 | 660 | 843 | 3112 | 891 | 922 |
| 1865 | W303.asm02.HP0_G0016470 | chrVI | 4785 | 460 | 301 | 4398 | 296 | 384 | 4596 | 345 | 347 | 2258 | 659 | 651 | 1466 | 432 | 501 | 1311 | 607 | 613 |
| 1866 | W303.asm02.HP0_G0016480 | chrVI | 177 | 138 | 65 | 221 | 54 | 70 | 183 | 111 | 68 | 252 | 113 | 85 | 327 | 67 | 63 | 235 | 143 | 84 |
| 1867 | W303.asm02.HP0_G0016490 | chrVI | 31 | 67 | 33 | 40 | 33 | 33 | 33 | 33 | 47 | 120 | 48 | 43 | 112 | 21 | 28 | 94 | 41 | 23 |
| 1868 | W303.asm02.HP0_G0016500 | chrVI | 1407 | 872 | 934 | 1317 | 884 | 918 | 1336 | 893 | 934 | 6444 | 1440 | 1601 | 5336 | 896 | 1149 | 5248 | 1276 | 1604 |
| 1869 | W303.asm02.HP0_G0016510 | chrVI | 408 | 120 | 137 | 418 | 131 | 121 | 350 | 109 | 145 | 501 | 170 | 185 | 757 | 104 | 125 | 393 | 166 | 131 |
| 1870 | W303.asm02.HP0_G0016520 | chrVI | 29100 | 9940 | 8963 | 28975 | 8443 | 8621 | 30408 | 9625 | 8509 | 32692 | 10443 | 14818 | 35544 | 10093 | 11288 | 28598 | 8685 | 12853 |
| 1871 | W303.asm02.HP0_G0016530 | chrVI | 10233 | 7387 | 7629 | 9672 | 6871 | 7739 | 10137 | 7370 | 9310 | 15430 | 6641 | 6749 | 17084 | 4504 | 5541 | 15618 | 5810 | 6135 |
| 1872 | W303.asm02.HP0_G0016540 | chrVI | 8267 | 736 | 1002 | 7619 | 1026 | 1196 | 7786 | 642 | 1387 | 9034 | 1339 | 2243 | 5433 | 1150 | 1919 | 5190 | 1179 | 2063 |
| 1873 | W303.asm02.HP0_G0016550 | chrVI | 1486 | 431 | 467 | 1321 | 396 | 547 | 1469 | 408 | 648 | 3151 | 779 | 1028 | 2533 | 487 | 814 | 2345 | 631 | 965 |
| 1874 | W303.asm02.HP0_G0016560 | chrVI | 1971 | 2670 | 3207 | 1628 | 2926 | 3228 | 1836 | 2755 | 3900 | 6910 | 1935 | 2308 | 8259 | 1664 | 2099 | 7353 | 1807 | 2130 |
| 1875 | W303.asm02.HP0_G0016570 | chrVI | 1211 | 379 | 365 | 1156 | 399 | 436 | 1089 | 375 | 423 | 3876 | 854 | 961 | 1830 | 486 | 654 | 1901 | 767 | 930 |
| 1876 | W303.asm02.HP0_G0016580 | chrVI | 3682 | 553 | 758 | 3431 | 631 | 710 | 3549 | 605 | 922 | 3624 | 640 | 1074 | 4276 | 720 | 1007 | 1873 | 598 | 965 |
| 1877 | W303.asm02.HP0_G0016590 | chrVI | 7115 | 2601 | 4353 | 7094 | 2854 | 4538 | 6588 | 2802 | 5558 | 10212 | 2746 | 5970 | 11355 | 2308 | 4238 | 8834 | 2862 | 5371 |
| 1878 | W303.asm02.HP0_G0016600 | chrVI | 790 | 370 | 308 | 668 | 298 | 329 | 657 | 359 | 398 | 779 | 486 | 478 | 663 | 366 | 376 | 507 | 489 | 349 |
| 1879 | W303.asm02.HP0_G0016610 | chrVI | 2495 | 784 | 526 | 2391 | 503 | 565 | 2468 | 733 | 585 | 2087 | 761 | 792 | 1812 | 522 | 572 | 1662 | 757 | 655 |
| 1880 | W303.asm02.HP0_G0016620 | chrVI | 2856 | 1378 | 1453 | 2427 | 1417 | 1457 | 2720 | 1322 | 1715 | 4112 | 1015 | 1250 | 4558 | 697 | 988 | 4408 | 986 | 1095 |
| 1881 | W303.asm02.HP0_G0016630 | chrVI | 1369 | 710 | 764 | 1361 | 847 | 718 | 1235 | 596 | 813 | 3921 | 608 | 665 | 3655 | 386 | 552 | 3514 | 679 | 599 |
| 1882 | W303.asm02.HP0_G0016640 | chrVI | 1953 | 8134 | 9959 | 2060 | 8949 | 9890 | 2304 | 7880 | 11013 | 24027 | 6631 | 6878 | 23979 | 4472 | 5752 | 23490 | 6126 | 6082 |
| 1883 | W303.asm02.HP0_G0016650 | chrVI | 1640 | 590 | 659 | 1294 | 647 | 656 | 1560 | 603 | 704 | 5518 | 889 | 1225 | 4786 | 675 | 1034 | 4298 | 788 | 1230 |
| 1884 | W303.asm02.HP0_G0016660 | chrVI | 228 | 253 | 285 | 245 | 287 | 283 | 214 | 215 | 327 | 1532 | 295 | 387 | 1066 | 155 | 245 | 1167 | 214 | 350 |
| 1885 | W303.asm02.HP0_G0016670 | chrVI | 1269 | 424 | 363 | 1407 | 412 | 365 | 1260 | 391 | 366 | 2478 | 504 | 509 | 2068 | 315 | 385 | 1971 | 535 | 454 |
| 1886 | W303.asm02.HP0_G0016680 | chrVI | 1218 | 518 | 463 | 1046 | 509 | 529 | 1183 | 498 | 657 | 5164 | 1063 | 1195 | 2391 | 717 | 567 | 2417 | 1100 | 1028 |
| 1887 | W303.asm02.HP0_G0016690 | chrVI | 11058 | 2526 | 2450 | 10114 | 2084 | 2941 | 12818 | 2017 | 3281 | 13902 | 4152 | 5801 | 6499 | 3012 | 3184 | 6517 | 3504 | 4649 |
| 1888 | W303.asm02.HP0_G0016700 | chrVI | 4166 | 631 | 781 | 3618 | 743 | 1031 | 3989 | 508 | 1013 | 3668 | 978 | 1557 | 1666 | 903 | 1161 | 1197 | 738 | 1248 |
| 1889 | W303.asm02.HP0_G0016710 | chrVI | 219 | 67 | 25 | 224 | 36 | 38 | 197 | 39 | 20 | 91 | 47 | 43 | 161 | 37 | 25 | 63 | 55 | 29 |
| 1890 | W303.asm02.HP0_G0016720 | chrVI | 12769 | 5098 | 3633 | 12502 | 3451 | 3625 | 12954 | 4970 | 4122 | 19011 | 7048 | 7546 | 17284 | 5785 | 6178 | 15619 | 6444 | 6658 |
| 1891 | W303.asm02.HP0_G0016730 | chrVI | 474 | 570 | 616 | 281 | 504 | 556 | 394 | 665 | 679 | 1429 | 379 | 464 | 1713 | 234 | 375 | 1609 | 404 | 341 |
| 1892 | W303.asm02.HP0_G0016740 | chrVI | 1642 | 989 | 900 | 1265 | 848 | 941 | 1301 | 964 | 944 | 2495 | 928 | 965 | 2724 | 662 | 867 | 2472 | 835 | 894 |
| 1893 | W303.asm02.HP0_G0016750 | chrVI | 8980 | 3464 | 2196 | 9248 | 2288 | 2732 | 8225 | 2287 | 2441 | 10198 | 4658 | 4172 | 5456 | 3035 | 3409 | 5949 | 5007 | 3696 |
| 1894 | W303.asm02.HP0_G0016760 | chrVI | 129 | 77 | 31 | 78 | 25 | 35 | 77 | 60 | 53 | 54 | 64 | 27 | 188 | 35 | 35 | 49 | 55 | 21 |
| 1895 | W303.asm02.HP0_G0016770 | chrVI | 7132 | 2682 | 2576 | 6887 | 2349 | 3072 | 5974 | 2952 | 4219 | 1654 | 1866 | 2870 | 7692 | 2526 | 3141 | 1731 | 2063 | 2351 |
| 1896 | W303.asm02.HP0_G0016780 | chrVI | 6185 | 866 | 780 | 6110 | 777 | 911 | 5956 | 781 | 838 | 6301 | 1279 | 1828 | 4784 | 873 | 1215 | 4440 | 1045 | 1579 |
| 1897 | W303.asm02.HP0_G0016790 | chrVI | 77 | 49 | 76 | 62 | 53 | 63 | 51 | 46 | 90 | 109 | 37 | 72 | 108 | 20 | 25 | 131 | 45 | 63 |
| 1898 | W303.asm02.HP0_G0016800 | chrVI | 278 | 20 | 30 | 335 | 20 | 27 | 278 | 26 | 28 | 80 | 42 | 38 | 73 | 23 | 20 | 58 | 26 | 32 |
| 1899 | W303.asm02.HP0_G0016810 | chrVI | 8750 | 2522 | 1955 | 7554 | 1927 | 2094 | 8030 | 2565 | 2283 | 8873 | 3685 | 3754 | 8272 | 2761 | 3326 | 7272 | 3249 | 3570 |
| 1900 | W303.asm02.HP0_G0016820 | chrVI | 2040 | 1035 | 1093 | 1854 | 1099 | 1086 | 1979 | 1014 | 1215 | 7797 | 1863 | 2214 | 5316 | 1231 | 1582 | 5582 | 1692 | 1987 |
| 1901 | W303.asm02.HP0_G0016830 | chrVI | 1899 | 620 | 896 | 2069 | 881 | 897 | 1970 | 657 | 791 | 5915 | 782 | 1585 | 5179 | 581 | 1184 | 5143 | 740 | 1485 |
| 1902 | W303.asm02.HP0_G0016840 | chrVI | 7679 | 962 | 1162 | 7726 | 1252 | 1331 | 7831 | 981 | 1568 | 11515 | 1707 | 3148 | 6307 | 1370 | 2204 | 6390 | 1425 | 2664 |
| 1903 | W303.asm02.HP0_G0016850 | chrVI | 7150 | 3486 | 4748 | 7055 | 4075 | 4460 | 7187 | 3603 | 5903 | 10298 | 3156 | 3782 | 12088 | 2239 | 3151 | 10895 | 2789 | 3308 |
| 1904 | W303.asm02.HP0_G0016860 | chrVI | 6464 | 3084 | 2707 | 6279 | 2934 | 2655 | 6778 | 3067 | 2787 | 15543 | 4212 | 4676 | 13878 | 3147 | 3858 | 13725 | 3528 | 4361 |
| 1905 | W303.asm02.HP0_G0016870 | chrVI | 516 | 196 | 146 | 463 | 173 | 171 | 407 | 199 | 199 | 753 | 161 | 155 | 638 | 110 | 85 | 718 | 172 | 152 |
| 1906 | W303.asm02.HP0_G0016880 | chrVI | 1796 | 1083 | 1003 | 1768 | 1184 | 960 | 2039 | 1024 | 1045 | 7751 | 1347 | 1610 | 6601 | 858 | 1010 | 6631 | 1343 | 1472 |
| 1907 | W303.asm02.HP0_G0016890 | chrVI | 1818 | 581 | 457 | 1764 | 492 | 489 | 1907 | 617 | 514 | 4261 | 958 | 993 | 3427 | 689 | 579 | 3581 | 896 | 904 |
| 1908 | W303.asm02.HP0_G0016900 | chrVI | 2162 | 1633 | 2127 | 2206 | 2231 | 2107 | 2212 | 1854 | 2959 | 7148 | 1765 | 1755 | 8916 | 1236 | 1302 | 7716 | 1510 | 1611 |
| 1909 | W303.asm02.HP0_G0016910 | chrVI | 5868 | 2803 | 2713 | 6049 | 2601 | 2555 | 6123 | 3000 | 2750 | 12666 | 3005 | 3227 | 11718 | 1953 | 2594 | 12026 | 2853 | 3003 |
| 1910 | W303.asm02.HP0_G0016920 | chrVI | 3117 | 939 | 1127 | 2926 | 871 | 1165 | 2727 | 958 | 1251 | 2355 | 843 | 1049 | 2953 | 605 | 983 | 2467 | 793 | 918 |
| 1911 | W303.asm02.HP0_G0016930 | chrVI | 7286 | 3987 | 4207 | 7343 | 3969 | 4176 | 7018 | 3864 | 4598 | 10633 | 3823 | 4255 | 10395 | 2405 | 3470 | 10118 | 3390 | 3935 |
| 1912 | W303.asm02.HP0_G0016940 | chrVI | 1526 | 1555 | 1374 | 1448 | 1395 | 1396 | 1511 | 1437 | 1496 | 5319 | 1097 | 1105 | 5439 | 708 | 904 | 5364 | 1154 | 1025 |
| 1913 | W303.asm02.HP0_G0016950 | chrVI | 8558 | 3486 | 2552 | 7612 | 2526 | 2552 | 7980 | 3371 | 2267 | 11642 | 3637 | 3752 | 10590 | 2698 | 3172 | 10226 | 3319 | 3392 |
| 1914 | W303.asm02.HP0_G0016960 | chrVI | 3146 | 820 | 488 | 2826 | 487 | 513 | 2772 | 676 | 441 | 2119 | 701 | 798 | 2230 | 451 | 537 | 1929 | 783 | 664 |
| 1915 | W303.asm02.HP0_G0016970 | chrVI | 1335 | 906 | 574 | 1382 | 516 | 567 | 1207 | 806 | 654 | 1564 | 581 | 387 | 1950 | 348 | 403 | 1615 | 636 | 405 |
| 1916 | W303.asm02.HP0_G0016980 | chrVI | 7056 | 2198 | 1695 | 6977 | 1872 | 1651 | 7216 | 2208 | 1588 | 13803 | 2793 | 3236 | 11345 | 2074 | 2283 | 11250 | 2570 | 3055 |
| 1917 | W303.asm02.HP0_G0016990 | chrVI | 9595 | 4593 | 3600 | 9533 | 3166 | 3515 | 8862 | 4265 | 3604 | 11342 | 4087 | 3887 | 10170 | 2744 | 3181 | 11345 | 3975 | 3566 |
| 1918 | W303.asm02.HP0_G0017000 | chrVI | 2356 | 1510 | 2158 | 2442 | 1954 | 2154 | 2168 | 1701 | 2934 | 4916 | 1505 | 1821 | 5838 | 1215 | 1511 | 5292 | 1365 | 1470 |
| 1919 | W303.asm02.HP0_G0017010 | chrVI | 108 | 121 | 103 | 119 | 96 | 104 | 119 | 108 | 108 | 330 | 112 | 119 | 283 | 58 | 68 | 277 | 150 | 87 |
| 1920 | W303.asm02.HP0_G0017020 | chrVI | 3545 | 763 | 769 | 3509 | 816 | 779 | 3683 | 764 | 886 | 6641 | 1295 | 1805 | 4926 | 792 | 1144 | 4926 | 1093 | 1547 |
| 1921 | W303.asm02.HP0_G0017030 | chrVI | 5808 | 1113 | 874 | 6318 | 854 | 910 | 5201 | 967 | 860 | 3953 | 1179 | 1203 | 4086 | 936 | 1097 | 3535 | 1115 | 1081 |
| 1922 | W303.asm02.HP0_G0017040 | chrVI | 13424 | 379 | 303 | 13204 | 285 | 331 | 12340 | 317 | 349 | 1321 | 558 | 517 | 1095 | 498 | 589 | 810 | 497 | 411 |
| 1923 | W303.asm02.HP0_G0017050 | chrVI | 8891 | 526 | 422 | 8520 | 423 | 588 | 8757 | 511 | 567 | 4876 | 843 | 1403 | 3102 | 784 | 1026 | 2612 | 710 | 1286 |
| 1924 | W303.asm02.HP0_G0017060 | chrVI | 4139 | 104 | 46 | 3796 | 47 | 58 | 3512 | 54 | 54 | 160 | 161 | 91 | 212 | 115 | 125 | 117 | 161 | 73 |
| 1925 | W303.asm02.HP0_G0017070 | chrVI | 5353 | 951 | 885 | 4731 | 788 | 866 | 5511 | 849 | 1001 | 3611 | 1903 | 1539 | 2484 | 1134 | 1162 | 2203 | 1629 | 1592 |
| 1926 | W303.asm02.HP0_G0017080 | chrVI | 4151 | 494 | 625 | 3740 | 606 | 674 | 3716 | 438 | 882 | 4843 | 721 | 1474 | 2392 | 570 | 1117 | 2404 | 599 | 1277 |
| 1927 | W303.asm02.HP0_G0017090 | chrVI | 1035 | 377 | 394 | 949 | 374 | 387 | 918 | 338 | 365 | 1346 | 315 | 349 | 1220 | 208 | 255 | 1216 | 349 | 265 |
| 1928 | W303.asm02.HP0_G0017100 | chrVI | 1062 | 288 | 305 | 933 | 262 | 408 | 1057 | 204 | 361 | 2056 | 509 | 828 | 1112 | 346 | 490 | 1110 | 443 | 671 |
| 1929 | W303.asm02.HP0_G0017110 | chrVI | 1530 | 131 | 167 | 1267 | 141 | 181 | 1531 | 112 | 196 | 794 | 247 | 388 | 318 | 158 | 215 | 301 | 182 | 281 |
| 1930 | W303.asm02.HP0_G0017120 | chrVI | 470 | 140 | 139 | 561 | 146 | 159 | 432 | 108 | 168 | 390 | 138 | 147 | 443 | 66 | 82 | 415 | 133 | 124 |
| 1931 | W303.asm02.HP0_G0017130 | chrVI | 6841 | 1941 | 1651 | 6276 | 1496 | 1691 | 6567 | 1758 | 1723 | 9323 | 2714 | 3068 | 7254 | 1745 | 2582 | 7374 | 2524 | 2980 |
| 1932 | W303.asm02.HP0_G0017140 | chrVI | 2447 | 1521 | 1337 | 2109 | 1172 | 1185 | 2347 | 1577 | 1444 | 4802 | 1595 | 1779 | 5263 | 1029 | 1438 | 4649 | 1503 | 1596 |
| 1933 | W303.asm02.HP0_G0017150 | chrVI | 342 | 129 | 214 | 316 | 163 | 332 | 349 | 133 | 274 | 344 | 160 | 279 | 358 | 99 | 166 | 349 | 164 | 261 |
| 1934 | W303.asm02.HP0_G0017160 | chrVI | 433 | 511 | 276 | 459 | 318 | 304 | 426 | 421 | 286 | 929 | 420 | 387 | 895 | 268 | 262 | 812 | 389 | 366 |
| 1935 | W303.asm02.HP0_G0017170 | chrVI | 1699 | 1336 | 1151 | 1657 | 1159 | 1159 | 1909 | 1260 | 1099 | 7842 | 2106 | 2423 | 6114 | 1422 | 1671 | 6362 | 2037 | 2224 |
| 1936 | W303.asm02.HP0_G0017180 | chrVI | 835 | 175 | 172 | 890 | 193 | 259 | 852 | 118 | 267 | 1636 | 418 | 542 | 539 | 251 | 373 | 579 | 405 | 494 |
| 1937 | W303.asm02.HP0_G0017190 | chrVI | 3430 | 779 | 808 | 2398 | 773 | 887 | 2924 | 801 | 972 | 3883 | 1156 | 1351 | 3134 | 1033 | 1202 | 2945 | 917 | 1216 |
| 1938 | W303.asm02.HP0_G0017200 | chrVI | 2420 | 559 | 680 | 2564 | 794 | 768 | 2486 | 606 | 826 | 5742 | 890 | 1524 | 4427 | 631 | 1057 | 4041 | 850 | 1330 |
| 1939 | W303.asm02.HP0_G0017210 | chrVI | 14744 | 10483 | 12235 | 14003 | 12240 | 11490 | 14915 | 10883 | 12577 | 34566 | 11376 | 13151 | 35300 | 8349 | 10554 | 34741 | 9988 | 12191 |
| 1940 | W303.asm02.HP0_G0017220 | chrVI | 28 | 62 | 6 | 41 | 18 | 20 | 25 | 44 | 10 | 119 | 87 | 9 | 94 | 45 | 10 | 112 | 88 | 10 |
| 1941 | W303.asm02.HP0_G0017230 | chrVI | 3129 | 1997 | 2601 | 2391 | 2473 | 2721 | 2563 | 2043 | 3260 | 5489 | 1747 | 2185 | 7032 | 1398 | 1934 | 6270 | 1654 | 1810 |
| 1942 | W303.asm02.HP0_G0017240 | chrVI | 204 | 183 | 217 | 201 | 204 | 246 | 220 | 177 | 216 | 343 | 149 | 146 | 454 | 103 | 109 | 287 | 162 | 124 |
| 1943 | W303.asm02.HP0_G0017250 | chrVI | 5140 | 4920 | 5323 | 4660 | 6090 | 5545 | 4527 | 5734 | 5319 | 13308 | 5076 | 5982 | 12681 | 3574 | 5020 | 11305 | 4458 | 5041 |
| 1944 | W303.asm02.HP0_G0017260 | chrVI | 1120 | 644 | 631 | 1024 | 621 | 714 | 1220 | 559 | 778 | 3317 | 1432 | 1408 | 1699 | 842 | 926 | 1838 | 1268 | 1261 |
| 1945 | W303.asm02.HP0_G0017270 | chrVI | 277 | 294 | 318 | 176 | 288 | 326 | 242 | 288 | 321 | 736 | 238 | 232 | 970 | 163 | 227 | 827 | 235 | 232 |
| 1946 | W303.asm02.HP0_G0017280 | chrVI | 217 | 284 | 212 | 204 | 189 | 206 | 192 | 248 | 183 | 480 | 210 | 211 | 449 | 106 | 100 | 402 | 163 | 169 |
| 1947 | W303.asm02.HP0_G0017290 | chrVI | 12703 | 2135 | 1477 | 11005 | 1490 | 1716 | 12416 | 1827 | 1871 | 9252 | 3434 | 3385 | 5030 | 2506 | 2311 | 5049 | 2963 | 2901 |
| 1948 | W303.asm02.HP0_G0017300 | chrVI | 1487 | 516 | 577 | 1447 | 591 | 638 | 1620 | 455 | 692 | 5561 | 1039 | 1483 | 2967 | 744 | 848 | 3273 | 1022 | 1271 |
| 1949 | W303.asm02.HP0_G0017310 | chrVI | 1724 | 1329 | 1391 | 1711 | 1321 | 1341 | 1684 | 1367 | 1291 | 4503 | 1184 | 1519 | 4370 | 808 | 1156 | 4267 | 1112 | 1350 |
| 1950 | W303.asm02.HP0_G0017320 | chrVI | 6078 | 831 | 1274 | 6243 | 1317 | 1305 | 5893 | 850 | 1452 | 11382 | 1506 | 2373 | 8421 | 933 | 1686 | 8845 | 1424 | 2011 |
| 1951 | W303.asm02.HP0_G0017330 | chrVI | 2420 | 1277 | 2585 | 2513 | 1354 | 2723 | 2417 | 1330 | 3222 | 4134 | 934 | 2267 | 4599 | 666 | 1530 | 4441 | 850 | 1955 |

| | | | | | | | | | | | | | | | | | | | | |
|---|---|---|---|---|---|---|---|---|---|---|---|---|---|---|---|---|---|---|---|---|
| 1952 | W303.asm02.HP0_G0017340 | chrVI | 843 | 847 | 1313 | 885 | 733 | 1476 | 759 | 782 | 1784 | 1595 | 544 | 1058 | 2316 | 412 | 852 | 1938 | 529 | 871 |
| 1953 | W303.asm02.HP0_G0017350 | chrVI | 1654 | 797 | 766 | 1648 | 743 | 765 | 1644 | 822 | 981 | 2210 | 622 | 580 | 2305 | 372 | 450 | 2347 | 566 | 526 |
| 1954 | W303.asm02.HP0_G0017360 | chrVI | 8540 | 3371 | 2280 | 7429 | 2022 | 2638 | 8533 | 2942 | 2717 | 13041 | 4372 | 4865 | 11241 | 3599 | 3726 | 9899 | 3960 | 4298 |
| 1955 | W303.asm02.HP0_G0017370 | chrVI | 2379 | 773 | 603 | 2394 | 589 | 638 | 2217 | 718 | 646 | 1703 | 525 | 775 | 2055 | 465 | 601 | 1501 | 540 | 644 |
| 1956 | W303.asm02.HP0_G0017380 | chrVI | 1155 | 675 | 530 | 1160 | 525 | 620 | 1182 | 697 | 514 | 1747 | 424 | 563 | 1975 | 265 | 405 | 1608 | 455 | 516 |
| 1957 | W303.asm02.HP0_G0017390 | chrVI | 4447 | 1224 | 1226 | 3796 | 975 | 1148 | 3949 | 1212 | 1251 | 3330 | 1167 | 1456 | 3258 | 807 | 997 | 3053 | 1130 | 1279 |
| 1958 | W303.asm02.HP0_G0017400 | chrVI | 1011 | 498 | 456 | 1042 | 458 | 437 | 1039 | 330 | 544 | 3609 | 750 | 935 | 2715 | 389 | 620 | 2694 | 800 | 818 |
| 1959 | W303.asm02.HP0_G0017410 | chrVI | 2603 | 1428 | 1806 | 2539 | 1647 | 1725 | 2323 | 1326 | 2447 | 3122 | 963 | 1203 | 4524 | 847 | 1331 | 3328 | 869 | 974 |
| 1960 | W303.asm02.HP0_G0017420 | chrVI | 7951 | 3092 | 3561 | 7786 | 3330 | 3480 | 7397 | 3149 | 3496 | 10740 | 2939 | 4368 | 11095 | 2326 | 3910 | 10212 | 2988 | 4125 |
| 1961 | W303.asm02.HP0_G0017430 | chrVI | 7509 | 1324 | 1308 | 6773 | 1030 | 1825 | 7475 | 962 | 1876 | 5655 | 1911 | 3676 | 2071 | 1516 | 1847 | 1965 | 1653 | 2879 |
| 1962 | W303.asm02.HP0_G0017440 | chrVI | 3730 | 2724 | 2730 | 3652 | 2342 | 2771 | 3627 | 2689 | 2955 | 7271 | 2245 | 2519 | 7675 | 1766 | 2100 | 6972 | 2047 | 2253 |
| 1963 | W303.asm02.HP0_G0017450 | chrVI | 58765 | 4905 | 2321 | 62153 | 3328 | 3194 | 55238 | 3651 | 2075 | 8894 | 4752 | 3171 | 15507 | 4768 | 4925 | 7438 | 4185 | 2932 |
| 1964 | W303.asm02.HP0_G0017460 | chrVI | 40 | 21 | 22 | 42 | 16 | 18 | 32 | 37 | 22 | 62 | 21 | 16 | 49 | 15 | 16 | 58 | 28 | 19 |
| 1965 | W303.asm02.HP0_G0017470 | chrVI | 30 | 5 | 9 | 26 | 4 | 8 | 27 | 6 | 6 | 34 | 10 | 6 | 23 | 2 | 7 | 26 | 1 | 7 |
| 1966 | W303.asm02.HP0_G0017480 | chrVI | 30 | 11 | 17 | 33 | 11 | 10 | 32 | 9 | 12 | 38 | 7 | 8 | 46 | 4 | 7 | 25 | 4 | 5 |
| 1967 | W303.asm02.HP0_G0017490 | chrVI | 3472 | 2081 | 2725 | 3156 | 2857 | 2606 | 3153 | 2336 | 3176 | 13950 | 3353 | 3609 | 10950 | 2303 | 2726 | 10746 | 2863 | 3211 |
| 1968 | W303.asm02.HP0_G0017500 | chrVI | 2 | 5 | 3 | 5 | 5 | 4 | 0 | 7 | 4 | 3 | 3 | 0 | 4 | 2 | 0 | 5 | 3 | 0 |
| 1969 | W303.asm02.HP0_G0017510 | chrVII | 0 | 15 | 0 | 0 | 0 | 0 | 0 | 25 | 0 | 0 | 13 | 0 | 0 | 4 | 0 | 0 | 10 | 0 |
| 1970 | W303.asm02.HP0_G0017520 | chrVII | 22 | 6 | 3 | 35 | 8 | 7 | 27 | 3 | 5 | 15 | 7 | 4 | 35 | 6 | 5 | 12 | 6 | 7 |
| 1971 | W303.asm02.HP0_G0017530 | chrVII | 27 | 2 | 0 | 31 | 2 | 8 | 23 | 2 | 3 | 12 | 3 | 2 | 12 | 1 | 4 | 11 | 4 | 5 |
| 1972 | W303.asm02.HP0_G0017540 | chrVII | 108 | 28 | 35 | 116 | 26 | 23 | 58 | 23 | 27 | 40 | 19 | 30 | 71 | 15 | 38 | 44 | 32 | 23 |
| 1973 | W303.asm02.HP0_G0017550 | chrVII | 255 | 160 | 138 | 214 | 139 | 166 | 182 | 153 | 207 | 351 | 107 | 95 | 505 | 86 | 63 | 356 | 97 | 87 |
| 1974 | W303.asm02.HP0_G0017560 | chrVII | 1780 | 1101 | 1241 | 1777 | 1147 | 1060 | 1819 | 1178 | 1261 | 6234 | 1313 | 1490 | 5337 | 1021 | 1248 | 5321 | 1177 | 1348 |
| 1975 | W303.asm02.HP0_G0017570 | chrVII | 1293 | 832 | 985 | 1273 | 916 | 959 | 1310 | 797 | 978 | 4746 | 918 | 1234 | 3896 | 592 | 807 | 3997 | 927 | 1070 |
| 1976 | W303.asm02.HP0_G0017580 | chrVII | 911 | 492 | 663 | 965 | 952 | 802 | 931 | 491 | 629 | 3715 | 726 | 681 | 3675 | 536 | 617 | 2776 | 668 | 569 |
| 1977 | W303.asm02.HP0_G0017590 | chrVII | 970 | 455 | 439 | 1023 | 481 | 450 | 1026 | 430 | 373 | 2101 | 431 | 643 | 1827 | 316 | 351 | 1655 | 383 | 517 |
| 1978 | W303.asm02.HP0_G0017600 | chrVII | 36737 | 13293 | 11342 | 38319 | 12275 | 11393 | 38610 | 13376 | 11329 | 48520 | 15367 | 15448 | 42897 | 10236 | 11891 | 44376 | 13564 | 15551 |
| 1979 | W303.asm02.HP0_G0017610 | chrVII | 6237 | 1859 | 1691 | 6221 | 1794 | 1658 | 6365 | 1954 | 1763 | 10013 | 2722 | 2968 | 8158 | 1782 | 2440 | 8364 | 2377 | 2692 |
| 1980 | W303.asm02.HP0_G0017620 | chrVII | 2049 | 298 | 368 | 1995 | 388 | 360 | 1993 | 324 | 417 | 2214 | 441 | 561 | 1997 | 283 | 453 | 1613 | 403 | 515 |
| 1981 | W303.asm02.HP0_G0017630 | chrVII | 1144 | 506 | 441 | 1081 | 353 | 473 | 958 | 462 | 465 | 1003 | 413 | 528 | 1138 | 274 | 450 | 954 | 340 | 435 |
| 1982 | W303.asm02.HP0_G0017640 | chrVII | 635 | 307 | 283 | 522 | 220 | 239 | 551 | 317 | 323 | 855 | 224 | 208 | 836 | 140 | 155 | 928 | 166 | 190 |
| 1983 | W303.asm02.HP0_G0017650 | chrVII | 6020 | 1400 | 1246 | 5420 | 1239 | 1383 | 5293 | 1386 | 1252 | 3612 | 1245 | 1500 | 4903 | 1084 | 1409 | 3412 | 1143 | 1283 |
| 1984 | W303.asm02.HP0_G0017660 | chrVII | 511 | 450 | 474 | 429 | 440 | 476 | 441 | 403 | 604 | 1483 | 393 | 411 | 1519 | 257 | 302 | 1565 | 347 | 321 |
| 1985 | W303.asm02.HP0_G0017670 | chrVII | 1491 | 1449 | 1679 | 1517 | 1695 | 1499 | 1607 | 1427 | 1825 | 6036 | 1231 | 1420 | 5967 | 717 | 975 | 6046 | 1176 | 1246 |
| 1986 | W303.asm02.HP0_G0017680 | chrVII | 19037 | 4135 | 3104 | 16547 | 3252 | 3259 | 19023 | 4088 | 3626 | 23997 | 6688 | 7763 | 18174 | 5277 | 5773 | 18496 | 6080 | 6991 |
| 1987 | W303.asm02.HP0_G0017690 | chrVII | 7446 | 3492 | 2476 | 7228 | 2665 | 2472 | 7395 | 3659 | 2426 | 13967 | 3831 | 3776 | 12155 | 2444 | 2992 | 11467 | 3338 | 3407 |
| 1988 | W303.asm02.HP0_G0017700 | chrVII | 963 | 750 | 836 | 894 | 795 | 774 | 876 | 718 | 1060 | 2358 | 595 | 632 | 2569 | 406 | 430 | 2522 | 599 | 583 |
| 1989 | W303.asm02.HP0_G0017710 | chrVII | 1392 | 1239 | 1621 | 1153 | 1461 | 1673 | 1292 | 1406 | 2207 | 4337 | 1090 | 1348 | 5884 | 818 | 1274 | 4568 | 931 | 1259 |
| 1990 | W303.asm02.HP0_G0017720 | chrVII | 2910 | 1259 | 1499 | 2803 | 1426 | 1417 | 2911 | 1363 | 1577 | 6433 | 1079 | 1489 | 6170 | 702 | 1071 | 6483 | 933 | 1273 |
| 1991 | W303.asm02.HP0_G0017730 | chrVII | 525 | 61 | 98 | 517 | 107 | 98 | 512 | 89 | 86 | 936 | 116 | 153 | 703 | 65 | 105 | 723 | 94 | 139 |
| 1992 | W303.asm02.HP0_G0017740 | chrVII | 5093 | 2162 | 2394 | 4437 | 2300 | 2220 | 5005 | 2461 | 2611 | 16171 | 2665 | 3683 | 13132 | 1970 | 2871 | 14757 | 2551 | 3391 |
| 1993 | W303.asm02.HP0_G0017750 | chrVII | 1809 | 551 | 364 | 1645 | 375 | 386 | 1663 | 562 | 381 | 1906 | 643 | 611 | 1822 | 406 | 446 | 1652 | 652 | 519 |
| 1994 | W303.asm02.HP0_G0017760 | chrVII | 3809 | 540 | 716 | 3323 | 608 | 762 | 3798 | 479 | 1005 | 3660 | 1026 | 1673 | 1739 | 794 | 952 | 1309 | 901 | 1345 |
| 1995 | W303.asm02.HP0_G0017770 | chrVII | 7064 | 1583 | 1685 | 4798 | 1347 | 1943 | 7192 | 1361 | 2249 | 10001 | 3880 | 4669 | 3306 | 2247 | 1923 | 3563 | 3123 | 3949 |
| 1996 | W303.asm02.HP0_G0017780 | chrVII | 3920 | 926 | 1099 | 3832 | 1140 | 1158 | 3668 | 992 | 1340 | 6965 | 1174 | 1754 | 6264 | 802 | 1445 | 6460 | 1069 | 1519 |
| 1997 | W303.asm02.HP0_G0017790 | chrVII | 2574 | 1552 | 1901 | 2374 | 1595 | 1749 | 2414 | 1563 | 2453 | 5717 | 1199 | 1503 | 5443 | 693 | 983 | 6180 | 1188 | 1279 |
| 1998 | W303.asm02.HP0_G0017800 | chrVII | 4426 | 2800 | 3058 | 4394 | 2554 | 2809 | 4298 | 2837 | 3798 | 6286 | 2397 | 2289 | 8801 | 1825 | 2075 | 6737 | 2143 | 2092 |
| 1999 | W303.asm02.HP0_G0017810 | chrVII | 170 | 106 | 52 | 122 | 47 | 39 | 135 | 84 | 90 | 159 | 76 | 57 | 300 | 52 | 37 | 227 | 97 | 28 |
| 2000 | W303.asm02.HP0_G0017820 | chrVII | 1474 | 479 | 361 | 1362 | 279 | 362 | 1308 | 494 | 410 | 1836 | 491 | 544 | 1663 | 302 | 466 | 1660 | 505 | 526 |
| 2001 | W303.asm02.HP0_G0017830 | chrVII | 5842 | 2824 | 1897 | 5709 | 1768 | 1946 | 6036 | 2518 | 1542 | 8878 | 2757 | 3086 | 7386 | 1834 | 2038 | 7591 | 2716 | 2780 |
| 2002 | W303.asm02.HP0_G0017840 | chrVII | 6685 | 695 | 543 | 6671 | 516 | 569 | 6234 | 616 | 648 | 3495 | 925 | 1022 | 3038 | 615 | 989 | 2845 | 882 | 928 |
| 2003 | W303.asm02.HP0_G0017850 | chrVII | 1364 | 1223 | 1541 | 1165 | 1265 | 1507 | 1231 | 1246 | 2080 | 2983 | 1006 | 1210 | 3603 | 720 | 919 | 3381 | 971 | 988 |
| 2004 | W303.asm02.HP0_G0017860 | chrVII | 1208 | 523 | 650 | 999 | 567 | 669 | 1055 | 589 | 792 | 1283 | 459 | 542 | 1666 | 322 | 459 | 1358 | 448 | 484 |
| 2005 | W303.asm02.HP0_G0017870 | chrVII | 164 | 90 | 64 | 134 | 45 | 52 | 165 | 82 | 42 | 384 | 74 | 63 | 251 | 70 | 56 | 284 | 86 | 61 |
| 2006 | W303.asm02.HP0_G0017880 | chrVII | 1296 | 3598 | 5106 | 1093 | 4257 | 4867 | 1337 | 3672 | 6049 | 11513 | 2877 | 4075 | 12243 | 2058 | 2985 | 11654 | 2683 | 3477 |
| 2007 | W303.asm02.HP0_G0017890 | chrVII | 1643 | 1742 | 1676 | 1466 | 1508 | 1747 | 1649 | 1803 | 1822 | 4298 | 1398 | 1461 | 4799 | 1149 | 1199 | 4189 | 1300 | 1345 |
| 2008 | W303.asm02.HP0_G0017900 | chrVII | 2564 | 1961 | 1364 | 2674 | 1290 | 1323 | 2650 | 1939 | 1442 | 4426 | 1447 | 1221 | 4688 | 885 | 904 | 4553 | 1372 | 1105 |
| 2009 | W303.asm02.HP0_G0017910 | chrVII | 923 | 427 | 159 | 900 | 155 | 162 | 735 | 444 | 153 | 1099 | 418 | 292 | 1128 | 276 | 203 | 1062 | 423 | 248 |
| 2010 | W303.asm02.HP0_G0017920 | chrVII | 3834 | 2943 | 1712 | 3456 | 1759 | 1832 | 3737 | 2721 | 1627 | 6134 | 2571 | 2431 | 5954 | 1901 | 1713 | 5321 | 2598 | 2203 |
| 2011 | W303.asm02.HP0_G0017930 | chrVII | 2302 | 2086 | 2284 | 1758 | 2097 | 2266 | 2017 | 2283 | 2999 | 4341 | 1930 | 1950 | 5612 | 1374 | 1653 | 4527 | 1743 | 1564 |
| 2012 | W303.asm02.HP0_G0017940 | chrVII | 6268 | 1519 | 917 | 5979 | 1025 | 931 | 6330 | 1457 | 897 | 5445 | 1471 | 1717 | 4887 | 1110 | 1220 | 4308 | 1437 | 1495 |
| 2013 | W303.asm02.HP0_G0017950 | chrVII | 863 | 248 | 281 | 800 | 260 | 337 | 905 | 186 | 365 | 2234 | 442 | 666 | 901 | 312 | 435 | 1015 | 386 | 567 |
| 2014 | W303.asm02.HP0_G0017960 | chrVII | 8021 | 1343 | 1960 | 7953 | 1855 | 2014 | 8552 | 1411 | 2551 | 8136 | 2451 | 3203 | 4300 | 2111 | 2020 | 4455 | 1935 | 2731 |
| 2015 | W303.asm02.HP0_G0017970 | chrVII | 1202 | 1925 | 1550 | 1234 | 1633 | 1522 | 1194 | 1887 | 1668 | 5818 | 1573 | 1528 | 5307 | 915 | 1113 | 5673 | 1389 | 1329 |
| 2016 | W303.asm02.HP0_G0017980 | chrVII | 993 | 173 | 154 | 917 | 167 | 168 | 900 | 143 | 160 | 1152 | 268 | 342 | 948 | 181 | 254 | 879 | 287 | 303 |
| 2017 | W303.asm02.HP0_G0017990 | chrVII | 1310 | 1204 | 1616 | 1150 | 1661 | 1489 | 1404 | 1252 | 1770 | 5201 | 1139 | 1471 | 5399 | 729 | 925 | 4803 | 1148 | 1282 |
| 2018 | W303.asm02.HP0_G0018000 | chrVII | 2612 | 1324 | 1234 | 2637 | 1174 | 1223 | 2583 | 1359 | 1309 | 3656 | 1294 | 1325 | 3693 | 800 | 951 | 3367 | 1144 | 1246 |
| 2019 | W303.asm02.HP0_G0018010 | chrVII | 470 | 174 | 343 | 395 | 382 | 399 | 426 | 122 | 466 | 4731 | 465 | 841 | 3073 | 229 | 499 | 2074 | 452 | 673 |
| 2020 | W303.asm02.HP0_G0018020 | chrVII | 3024 | 231 | 215 | 2888 | 176 | 268 | 2831 | 178 | 242 | 1132 | 367 | 501 | 680 | 264 | 354 | 608 | 325 | 405 |
| 2021 | W303.asm02.HP0_G0018030 | chrVII | 9122 | 1754 | 1743 | 8649 | 1825 | 2037 | 9211 | 1706 | 2150 | 15568 | 3241 | 4632 | 8958 | 2438 | 3315 | 9470 | 2784 | 4296 |
| 2022 | W303.asm02.HP0_G0018040 | chrVII | 16743 | 1505 | 2139 | 16770 | 2080 | 2248 | 16470 | 1709 | 2596 | 13516 | 2099 | 3877 | 11053 | 1874 | 3629 | 10523 | 2028 | 3713 |
| 2023 | W303.asm02.HP0_G0018050 | chrVII | 422 | 52 | 73 | 502 | 73 | 61 | 460 | 56 | 87 | 494 | 127 | 142 | 404 | 74 | 91 | 308 | 109 | 111 |
| 2024 | W303.asm02.HP0_G0018060 | chrVII | 808 | 383 | 422 | 689 | 380 | 382 | 696 | 492 | 505 | 1080 | 301 | 379 | 837 | 163 | 226 | 1248 | 298 | 346 |
| 2025 | W303.asm02.HP0_G0018070 | chrVII | 2820 | 871 | 788 | 2517 | 803 | 784 | 2888 | 882 | 842 | 6780 | 1737 | 1684 | 4205 | 1112 | 1244 | 4287 | 1577 | 1638 |
| 2026 | W303.asm02.HP0_G0018080 | chrVII | 12757 | 3168 | 2720 | 11111 | 2371 | 3154 | 12741 | 2865 | 3670 | 17973 | 6158 | 6692 | 11032 | 4922 | 4757 | 11036 | 5240 | 5907 |
| 2027 | W303.asm02.HP0_G0018090 | chrVII | 2319 | 461 | 561 | 1982 | 614 | 720 | 2363 | 471 | 820 | 5103 | 878 | 1488 | 2175 | 700 | 1045 | 2348 | 808 | 1281 |
| 2028 | W303.asm02.HP0_G0018100 | chrVII | 6545 | 5079 | 5294 | 6087 | 4096 | 5564 | 6613 | 5172 | 5704 | 10560 | 4476 | 5478 | 11803 | 3337 | 4063 | 10446 | 3932 | 4808 |
| 2029 | W303.asm02.HP0_G0018110 | chrVII | 3427 | 1902 | 1621 | 2635 | 1419 | 1480 | 2935 | 1899 | 1715 | 3438 | 1643 | 1384 | 4089 | 1167 | 1396 | 3531 | 1537 | 1233 |
| 2030 | W303.asm02.HP0_G0018120 | chrVII | 5044 | 410 | 502 | 4635 | 485 | 581 | 4841 | 347 | 645 | 4258 | 818 | 1510 | 1712 | 658 | 1096 | 1579 | 709 | 1321 |
| 2031 | W303.asm02.HP0_G0018130 | chrVII | 2600 | 1478 | 1543 | 2399 | 1349 | 1520 | 2401 | 1458 | 1714 | 3690 | 1156 | 1368 | 4151 | 824 | 1200 | 3585 | 1065 | 1195 |
| 2032 | W303.asm02.HP0_G0018140 | chrVII | 12273 | 1715 | 1926 | 10619 | 2036 | 2342 | 11759 | 1765 | 2956 | 15567 | 2907 | 5339 | 7880 | 2423 | 3726 | 7898 | 2503 | 4562 |
| 2033 | W303.asm02.HP0_G0018150 | chrVII | 464 | 220 | 432 | 412 | 276 | 489 | 381 | 245 | 641 | 732 | 189 | 399 | 1044 | 146 | 237 | 942 | 163 | 366 |
| 2034 | W303.asm02.HP0_G0018160 | chrVII | 986 | 380 | 437 | 1140 | 473 | 407 | 875 | 403 | 424 | 2083 | 320 | 469 | 2237 | 192 | 357 | 1983 | 300 | 446 |
| 2035 | W303.asm02.HP0_G0018170 | chrVII | 177 | 39 | 50 | 127 | 37 | 33 | 138 | 36 | 51 | 167 | 42 | 49 | 122 | 16 | 58 | 134 | 51 | 37 |
| 2036 | W303.asm02.HP0_G0018180 | chrVII | 2374 | 397 | 329 | 2401 | 358 | 335 | 2334 | 316 | 386 | 2578 | 635 | 711 | 2187 | 381 | 499 | 1967 | 637 | 609 |
| 2037 | W303.asm02.HP0_G0018190 | chrVII | 5587 | 3443 | 4410 | 5413 | 4034 | 4189 | 5241 | 3399 | 5294 | 7935 | 2900 | 3241 | 9473 | 2225 | 2897 | 8631 | 2631 | 2754 |
| 2038 | W303.asm02.HP0_G0018200 | chrVII | 3119 | 1681 | 1725 | 3701 | 1646 | 1672 | 3177 | 1579 | 1925 | 8051 | 1709 | 1890 | 8168 | 1161 | 1760 | 7479 | 1657 | 1775 |
| 2039 | W303.asm02.HP0_G0018210 | chrVII | 13323 | 8678 | 11409 | 12552 | 11194 | 11711 | 12753 | 9191 | 12280 | 21340 | 8101 | 10694 | 22101 | 6323 | 9078 | 20580 | 7514 | 9693 |
| 2040 | W303.asm02.HP0_G0018220 | chrVII | 2118 | 4072 | 4966 | 2072 | 4697 | 4910 | 2082 | 3912 | 6170 | 10000 | 3134 | 3661 | 10720 | 2420 | 3252 | 9759 | 2728 | 2934 |
| 2041 | W303.asm02.HP0_G0018230 | chrVII | 2220 | 1300 | 1202 | 1952 | 1206 | 1176 | 1914 | 1232 | 1371 | 6037 | 1543 | 1824 | 4798 | 1017 | 1377 | 4684 | 1439 | 1559 |
| 2042 | W303.asm02.HP0_G0018240 | chrVII | 1859 | 815 | 625 | 1850 | 631 | 655 | 1703 | 717 | 689 | 2039 | 584 | 603 | 2403 | 374 | 521 | 2047 | 574 | 550 |
| 2043 | W303.asm02.HP0_G0018250 | chrVII | 2965 | 229 | 302 | 1461 | 234 | 320 | 2074 | 212 | 280 | 398 | 130 | 194 | 553 | 102 | 154 | 395 | 132 | 121 |
| 2044 | W303.asm02.HP0_G0018260 | chrVII | 123 | 46 | 54 | 90 | 42 | 51 | 102 | 30 | 41 | 170 | 26 | 33 | 152 | 14 | 21 | 189 | 32 | 44 |

| | | | | | | | | | | | | | | | | | | | | |
|---|---|---|---|---|---|---|---|---|---|---|---|---|---|---|---|---|---|---|---|---|
| 2045 | W303.asm02.HP0_G0018270 | chrVII | 5843 | 1151 | 782 | 5716 | 813 | 913 | 5698 | 1011 | 862 | 5189 | 1596 | 1566 | 4247 | 1012 | 1362 | 4197 | 1441 | 1471 |
| 2046 | W303.asm02.HP0_G0018280 | chrVII | 1812 | 190 | 223 | 1773 | 190 | 268 | 1706 | 152 | 259 | 2007 | 292 | 463 | 1269 | 231 | 379 | 1329 | 297 | 410 |
| 2047 | W303.asm02.HP0_G0018290 | chrVII | 539 | 186 | 401 | 482 | 202 | 465 | 519 | 153 | 442 | 1230 | 259 | 981 | 1136 | 144 | 441 | 1057 | 220 | 768 |
| 2048 | W303.asm02.HP0_G0018300 | chrVII | 19 | 3 | 7 | 32 | 5 | 3 | 19 | 1 | 10 | 20 | 3 | 5 | 23 | 0 | 0 | 14 | 2 | 4 |
| 2049 | W303.asm02.HP0_G0018310 | chrVII | 2705 | 425 | 683 | 3152 | 751 | 747 | 2777 | 453 | 842 | 7814 | 912 | 1476 | 5794 | 504 | 962 | 5790 | 729 | 1265 |
| 2050 | W303.asm02.HP0_G0018320 | chrVII | 525 | 309 | 377 | 507 | 383 | 432 | 504 | 255 | 390 | 2584 | 393 | 649 | 2084 | 300 | 464 | 2167 | 385 | 584 |
| 2051 | W303.asm02.HP0_G0018330 | chrVII | 288 | 74 | 81 | 291 | 70 | 88 | 268 | 61 | 73 | 706 | 132 | 172 | 558 | 84 | 143 | 503 | 123 | 142 |
| 2052 | W303.asm02.HP0_G0018340 | chrVII | 1073 | 329 | 311 | 1015 | 298 | 353 | 1081 | 315 | 220 | 1235 | 316 | 483 | 1076 | 204 | 237 | 1009 | 281 | 336 |
| 2053 | W303.asm02.HP0_G0018350 | chrVII | 16489 | 1368 | 1835 | 15858 | 1500 | 2188 | 16704 | 1215 | 2453 | 16245 | 2912 | 5542 | 6687 | 2487 | 3773 | 7390 | 2436 | 4558 |
| 2054 | W303.asm02.HP0_G0018360 | chrVII | 2453 | 861 | 942 | 2093 | 1024 | 1098 | 2353 | 763 | 1241 | 5879 | 1485 | 2240 | 2829 | 1004 | 1403 | 3293 | 1410 | 1931 |
| 2055 | W303.asm02.HP0_G0018370 | chrVII | 2082 | 1090 | 956 | 2012 | 1280 | 871 | 2162 | 1266 | 1037 | 8360 | 1627 | 1785 | 6801 | 1037 | 905 | 6855 | 1348 | 1571 |
| 2056 | W303.asm02.HP0_G0018380 | chrVII | 128 | 59 | 56 | 137 | 43 | 40 | 117 | 76 | 73 | 364 | 69 | 58 | 285 | 42 | 33 | 327 | 63 | 53 |
| 2057 | W303.asm02.HP0_G0018390 | chrVII | 2200 | 1148 | 1085 | 2038 | 1185 | 953 | 2252 | 1255 | 993 | 6535 | 1439 | 1628 | 6077 | 913 | 979 | 5968 | 1320 | 1419 |
| 2058 | W303.asm02.HP0_G0018400 | chrVII | 4770 | 1027 | 1214 | 4338 | 1170 | 1336 | 4715 | 1102 | 1572 | 9577 | 1787 | 2570 | 7500 | 1396 | 2138 | 6883 | 1583 | 2430 |
| 2059 | W303.asm02.HP0_G0018410 | chrVII | 1445 | 308 | 275 | 1578 | 268 | 299 | 1295 | 348 | 278 | 1410 | 473 | 417 | 1044 | 312 | 258 | 1058 | 460 | 365 |
| 2060 | W303.asm02.HP0_G0018420 | chrVII | 2789 | 1065 | 902 | 2847 | 1012 | 1035 | 2783 | 1073 | 947 | 4377 | 1002 | 1202 | 4361 | 674 | 912 | 3675 | 846 | 1094 |
| 2061 | W303.asm02.HP0_G0018430 | chrVII | 2818 | 276 | 332 | 2547 | 433 | 418 | 2665 | 237 | 427 | 2723 | 483 | 686 | 1310 | 357 | 480 | 1337 | 436 | 623 |
| 2062 | W303.asm02.HP0_G0018440 | chrVII | 201 | 163 | 172 | 234 | 149 | 142 | 212 | 118 | 159 | 1769 | 248 | 248 | 1165 | 130 | 168 | 995 | 248 | 209 |
| 2063 | W303.asm02.HP0_G0018450 | chrVII | 7816 | 1760 | 1599 | 7475 | 1416 | 1536 | 7256 | 1778 | 1645 | 6393 | 2309 | 2275 | 5946 | 1691 | 1978 | 5432 | 2015 | 2182 |
| 2064 | W303.asm02.HP0_G0018460 | chrVII | 3391 | 1048 | 1065 | 3181 | 920 | 1036 | 3167 | 1066 | 983 | 5518 | 1299 | 1800 | 4481 | 845 | 1381 | 5007 | 1189 | 1736 |
| 2065 | W303.asm02.HP0_G0018470 | chrVII | 1181 | 856 | 1146 | 1108 | 983 | 1174 | 1058 | 923 | 1497 | 3745 | 694 | 902 | 4255 | 469 | 684 | 4240 | 639 | 775 |
| 2066 | W303.asm02.HP0_G0018480 | chrVII | 107 | 31 | 41 | 115 | 58 | 50 | 128 | 25 | 36 | 453 | 47 | 65 | 540 | 21 | 38 | 205 | 44 | 44 |
| 2067 | W303.asm02.HP0_G0018490 | chrVII | 7616 | 2366 | 2050 | 8223 | 1643 | 1982 | 8046 | 2578 | 2050 | 7788 | 2582 | 2967 | 12245 | 2351 | 2809 | 7100 | 2254 | 2933 |
| 2068 | W303.asm02.HP0_G0018500 | chrVII | 7068 | 452 | 626 | 6739 | 444 | 705 | 6661 | 443 | 722 | 1924 | 631 | 1447 | 1754 | 637 | 1025 | 1247 | 618 | 1258 |
| 2069 | W303.asm02.HP0_G0018510 | chrVII | 1613 | 824 | 769 | 1552 | 844 | 769 | 1609 | 819 | 757 | 4313 | 886 | 1139 | 3777 | 629 | 740 | 3712 | 733 | 994 |
| 2070 | W303.asm02.HP0_G0018520 | chrVII | 843 | 562 | 465 | 695 | 431 | 457 | 800 | 534 | 578 | 1377 | 448 | 375 | 1555 | 322 | 332 | 1462 | 409 | 360 |
| 2071 | W303.asm02.HP0_G0018530 | chrVII | 2209 | 605 | 458 | 2140 | 446 | 485 | 1968 | 613 | 461 | 2157 | 599 | 772 | 1937 | 403 | 488 | 1869 | 513 | 708 |
| 2072 | W303.asm02.HP0_G0018540 | chrVII | 3762 | 1010 | 1203 | 3678 | 1188 | 1152 | 3850 | 1156 | 1241 | 8697 | 1259 | 1916 | 7135 | 879 | 1533 | 7505 | 1181 | 1802 |
| 2073 | W303.asm02.HP0_G0018550 | chrVII | 8146 | 854 | 1351 | 8146 | 1265 | 1547 | 7848 | 905 | 1788 | 10083 | 1463 | 2850 | 5215 | 1179 | 2035 | 5027 | 1354 | 2466 |
| 2074 | W303.asm02.HP0_G0018560 | chrVII | 8518 | 2026 | 2310 | 7370 | 2075 | 2649 | 8749 | 1809 | 2925 | 14191 | 4413 | 5991 | 10163 | 3227 | 3934 | 10872 | 3702 | 5004 |
| 2075 | W303.asm02.HP0_G0018570 | chrVII | 10418 | 9294 | 11683 | 8753 | 11480 | 11625 | 10556 | 8965 | 14974 | 26381 | 8099 | 8924 | 28105 | 6063 | 7029 | 27918 | 7559 | 7747 |
| 2076 | W303.asm02.HP0_G0018580 | chrVII | 316 | 187 | 81 | 277 | 89 | 90 | 286 | 158 | 86 | 470 | 234 | 105 | 366 | 183 | 97 | 420 | 260 | 110 |
| 2077 | W303.asm02.HP0_G0018590 | chrVII | 2372 | 1308 | 1451 | 2219 | 1527 | 1416 | 2394 | 1364 | 1616 | 6143 | 1224 | 1470 | 5555 | 706 | 1058 | 6156 | 1067 | 1225 |
| 2078 | W303.asm02.HP0_G0018600 | chrVII | 599 | 274 | 360 | 559 | 348 | 342 | 643 | 290 | 344 | 2022 | 354 | 556 | 1688 | 208 | 269 | 1606 | 334 | 450 |
| 2079 | W303.asm02.HP0_G0018610 | chrVII | 5015 | 835 | 860 | 4622 | 930 | 796 | 4766 | 859 | 952 | 3415 | 869 | 1369 | 3658 | 632 | 903 | 2911 | 767 | 1110 |
| 2080 | W303.asm02.HP0_G0018620 | chrVII | 1390 | 1689 | 2011 | 1335 | 1718 | 1926 | 1350 | 1727 | 2166 | 6317 | 1409 | 1621 | 5853 | 754 | 1080 | 6335 | 1224 | 1469 |
| 2081 | W303.asm02.HP0_G0018630 | chrVII | 3756 | 361 | 606 | 3388 | 501 | 628 | 3347 | 368 | 722 | 4247 | 653 | 1062 | 2635 | 487 | 926 | 2625 | 649 | 922 |
| 2082 | W303.asm02.HP0_G0018640 | chrVII | 2382 | 776 | 1038 | 2291 | 937 | 921 | 2313 | 714 | 1194 | 8611 | 1678 | 2207 | 5351 | 1150 | 1578 | 5046 | 1403 | 2015 |
| 2083 | W303.asm02.HP0_G0018650 | chrVII | 1553 | 617 | 900 | 1577 | 950 | 896 | 1499 | 734 | 1044 | 8074 | 1227 | 1857 | 5420 | 861 | 1150 | 5820 | 1047 | 1730 |
| 2084 | W303.asm02.HP0_G0018660 | chrVII | 15 | 12 | 7 | 19 | 8 | 4 | 12 | 17 | 13 | 40 | 8 | 7 | 41 | 10 | 7 | 32 | 11 | 6 |
| 2085 | W303.asm02.HP0_G0018670 | chrVII | 12126 | 2034 | 2277 | 11213 | 1847 | 2693 | 11970 | 1880 | 2786 | 16313 | 3869 | 6671 | 9332 | 3105 | 4379 | 9369 | 3535 | 5986 |
| 2086 | W303.asm02.HP0_G0018680 | chrVII | 2628 | 771 | 625 | 2443 | 722 | 690 | 2327 | 732 | 732 | 2081 | 518 | 589 | 2243 | 358 | 484 | 2158 | 547 | 517 |
| 2087 | W303.asm02.HP0_G0018690 | chrVII | 9546 | 8396 | 9757 | 8276 | 9392 | 9784 | 9745 | 8277 | 10649 | 23072 | 7539 | 8241 | 22688 | 5599 | 6165 | 22800 | 6782 | 7328 |
| 2088 | W303.asm02.HP0_G0018700 | chrVII | 1507 | 537 | 271 | 1440 | 343 | 303 | 1487 | 490 | 327 | 2008 | 655 | 474 | 1410 | 409 | 396 | 1577 | 667 | 431 |
| 2089 | W303.asm02.HP0_G0018710 | chrVII | 4459 | 529 | 610 | 4280 | 748 | 740 | 4377 | 609 | 802 | 5965 | 1034 | 1636 | 4168 | 721 | 1249 | 3684 | 975 | 1433 |
| 2090 | W303.asm02.HP0_G0018720 | chrVII | 2976 | 282 | 500 | 2911 | 574 | 491 | 2582 | 325 | 543 | 4634 | 459 | 1065 | 3945 | 376 | 818 | 3415 | 467 | 930 |
| 2091 | W303.asm02.HP0_G0018730 | chrVII | 2792 | 871 | 726 | 2696 | 730 | 692 | 2686 | 861 | 758 | 4434 | 1070 | 1424 | 3499 | 824 | 977 | 3437 | 1031 | 1218 |
| 2092 | W303.asm02.HP0_G0018740 | chrVII | 8551 | 1847 | 1827 | 9081 | 1823 | 1880 | 8521 | 1803 | 1856 | 7916 | 1753 | 2810 | 8115 | 1332 | 2049 | 6885 | 1761 | 2444 |
| 2093 | W303.asm02.HP0_G0018750 | chrVII | 1433 | 385 | 274 | 1388 | 209 | 286 | 1330 | 344 | 334 | 1148 | 468 | 472 | 808 | 305 | 300 | 860 | 529 | 382 |
| 2094 | W303.asm02.HP0_G0018760 | chrVII | 343 | 571 | 788 | 348 | 348 | 864 | 291 | 515 | 835 | 1342 | 608 | 1158 | 1297 | 440 | 707 | 979 | 595 | 913 |
| 2095 | W303.asm02.HP0_G0018770 | chrVII | 2521 | 1346 | 2568 | 1973 | 1140 | 2696 | 2288 | 1318 | 3178 | 3446 | 1510 | 3753 | 4017 | 961 | 2363 | 2991 | 1420 | 3051 |
| 2096 | W303.asm02.HP0_G0018780 | chrVII | 4241 | 622 | 516 | 3287 | 452 | 556 | 3810 | 476 | 676 | 3982 | 940 | 1028 | 1964 | 704 | 878 | 2023 | 1007 | 851 |
| 2097 | W303.asm02.HP0_G0018790 | chrVII | 2185 | 1461 | 929 | 2195 | 1007 | 901 | 2075 | 1293 | 998 | 3884 | 1216 | 918 | 4383 | 843 | 781 | 3643 | 1204 | 884 |
| 2098 | W303.asm02.HP0_G0018800 | chrVII | 21026 | 22071 | 24745 | 19002 | 24409 | 23793 | 22503 | 21774 | 27794 | 58416 | 22780 | 24166 | 57939 | 16518 | 19319 | 57269 | 19571 | 21579 |
| 2099 | W303.asm02.HP0_G0018810 | chrVII | 8739 | 811 | 895 | 9072 | 979 | 1115 | 8844 | 613 | 1277 | 9773 | 1850 | 2439 | 4543 | 1536 | 1680 | 4012 | 1517 | 2015 |
| 2100 | W303.asm02.HP0_G0018820 | chrVII | 2809 | 313 | 390 | 2282 | 285 | 410 | 2175 | 305 | 501 | 402 | 264 | 332 | 992 | 268 | 392 | 436 | 218 | 319 |
| 2101 | W303.asm02.HP0_G0018830 | chrVII | 3218 | 1897 | 1850 | 3229 | 1873 | 1981 | 3893 | 2030 | 2178 | 18316 | 3875 | 4812 | 15003 | 2596 | 2443 | 14307 | 3586 | 4139 |
| 2102 | W303.asm02.HP0_G0018840 | chrVII | 3147 | 1033 | 895 | 2901 | 906 | 828 | 2858 | 924 | 867 | 5007 | 1119 | 1343 | 4294 | 770 | 974 | 4261 | 1298 | 1133 |
| 2103 | W303.asm02.HP0_G0018850 | chrVII | 952 | 2263 | 3324 | 921 | 2271 | 3564 | 981 | 1924 | 3391 | 3790 | 1298 | 2033 | 3817 | 832 | 1540 | 4107 | 1090 | 1731 |
| 2104 | W303.asm02.HP0_G0018860 | chrVII | 1787 | 856 | 596 | 1653 | 768 | 616 | 2038 | 737 | 659 | 4855 | 944 | 1086 | 4767 | 798 | 913 | 3884 | 909 | 986 |
| 2105 | W303.asm02.HP0_G0018870 | chrVII | 5038 | 2442 | 3058 | 5327 | 2807 | 2891 | 5172 | 2429 | 3628 | 9479 | 2101 | 2656 | 10498 | 1454 | 2259 | 9641 | 1996 | 2442 |
| 2106 | W303.asm02.HP0_G0018880 | chrVII | 1669 | 778 | 945 | 1445 | 774 | 983 | 1552 | 633 | 1224 | 4004 | 1852 | 1970 | 1650 | 1312 | 1198 | 1813 | 1604 | 1536 |
| 2107 | W303.asm02.HP0_G0018890 | chrVII | 911 | 471 | 483 | 890 | 611 | 531 | 833 | 485 | 489 | 3687 | 588 | 832 | 3075 | 383 | 449 | 3125 | 559 | 700 |
| 2108 | W303.asm02.HP0_G0018900 | chrVII | 3162 | 1018 | 849 | 3058 | 915 | 880 | 3154 | 910 | 909 | 6017 | 1290 | 1627 | 4764 | 838 | 1151 | 4888 | 1111 | 1494 |
| 2109 | W303.asm02.HP0_G0018910 | chrVII | 2751 | 1006 | 945 | 2719 | 1150 | 1011 | 3119 | 1093 | 1117 | 8868 | 1936 | 2149 | 6212 | 1311 | 1189 | 6676 | 1517 | 1900 |
| 2110 | W303.asm02.HP0_G0018920 | chrVII | 2339 | 590 | 568 | 2282 | 600 | 570 | 2193 | 580 | 692 | 3984 | 807 | 1060 | 3180 | 529 | 785 | 3207 | 722 | 892 |
| 2111 | W303.asm02.HP0_G0018930 | chrVII | 1802 | 653 | 621 | 1574 | 727 | 705 | 1582 | 766 | 690 | 1970 | 588 | 694 | 2018 | 404 | 543 | 1795 | 540 | 585 |
| 2112 | W303.asm02.HP0_G0018940 | chrVII | 6451 | 1885 | 1653 | 6378 | 1967 | 1781 | 6063 | 1846 | 1817 | 8540 | 2059 | 2598 | 7841 | 1444 | 1929 | 7042 | 1888 | 2401 |
| 2113 | W303.asm02.HP0_G0018950 | chrVII | 5059 | 3439 | 3128 | 4192 | 2996 | 3311 | 4639 | 3527 | 3550 | 8409 | 3322 | 3502 | 8567 | 2417 | 2679 | 7917 | 3097 | 3141 |
| 2114 | W303.asm02.HP0_G0018960 | chrVII | 17060 | 5519 | 4245 | 16247 | 4616 | 4412 | 17251 | 5598 | 3914 | 24922 | 7883 | 8777 | 24105 | 5998 | 6866 | 22410 | 6839 | 8094 |
| 2115 | W303.asm02.HP0_G0018970 | chrVII | 2806 | 651 | 620 | 2432 | 498 | 592 | 2545 | 605 | 646 | 2292 | 774 | 744 | 2193 | 469 | 645 | 2036 | 699 | 602 |
| 2116 | W303.asm02.HP0_G0018980 | chrVII | 11923 | 12415 | 14654 | 10380 | 13528 | 14245 | 11735 | 11929 | 18360 | 27907 | 10017 | 11204 | 29492 | 7540 | 9350 | 29148 | 9539 | 9746 |
| 2117 | W303.asm02.HP0_G0018990 | chrVII | 3324 | 1846 | 1585 | 3089 | 1348 | 1540 | 3304 | 1818 | 1590 | 6895 | 2133 | 2514 | 5920 | 1457 | 1739 | 6153 | 2030 | 2378 |
| 2118 | W303.asm02.HP0_G0019000 | chrVII | 1446 | 539 | 701 | 1449 | 718 | 748 | 1667 | 457 | 953 | 6007 | 1128 | 1649 | 2419 | 736 | 769 | 2615 | 944 | 1363 |
| 2119 | W303.asm02.HP0_G0019010 | chrVII | 2137 | 975 | 811 | 2187 | 824 | 822 | 2000 | 885 | 942 | 3454 | 872 | 835 | 3264 | 552 | 739 | 3328 | 914 | 828 |
| 2120 | W303.asm02.HP0_G0019020 | chrVII | 3002 | 1123 | 1171 | 2664 | 1315 | 1231 | 2932 | 1097 | 1519 | 9070 | 2177 | 2668 | 4667 | 1394 | 1659 | 5371 | 1971 | 2333 |
| 2121 | W303.asm02.HP0_G0019030 | chrVII | 2095 | 138 | 193 | 1620 | 141 | 163 | 1668 | 125 | 150 | 421 | 155 | 205 | 535 | 113 | 174 | 386 | 135 | 160 |
| 2122 | W303.asm02.HP0_G0019040 | chrVII | 4634 | 1327 | 1634 | 4961 | 1457 | 1535 | 4683 | 1418 | 1722 | 6330 | 1388 | 1764 | 6186 | 900 | 1374 | 6268 | 1184 | 1651 |
| 2123 | W303.asm02.HP0_G0019050 | chrVII | 2333 | 437 | 715 | 2348 | 700 | 742 | 2284 | 500 | 814 | 6934 | 945 | 1598 | 5277 | 766 | 1256 | 5300 | 834 | 1529 |
| 2124 | W303.asm02.HP0_G0019060 | chrVII | 2867 | 328 | 452 | 2745 | 448 | 489 | 2743 | 342 | 529 | 3202 | 598 | 921 | 1798 | 447 | 749 | 1975 | 573 | 850 |
| 2125 | W303.asm02.HP0_G0019070 | chrVII | 4682 | 982 | 1359 | 4633 | 1340 | 1379 | 4805 | 912 | 1798 | 10892 | 1660 | 2784 | 4633 | 1344 | 1857 | 4518 | 1404 | 2531 |
| 2126 | W303.asm02.HP0_G0019080 | chrVII | 1983 | 671 | 563 | 1887 | 588 | 666 | 1652 | 563 | 662 | 2592 | 780 | 1041 | 2031 | 530 | 774 | 2106 | 753 | 929 |
| 2127 | W303.asm02.HP0_G0019090 | chrVII | 1317 | 344 | 429 | 1325 | 393 | 431 | 1353 | 329 | 412 | 2057 | 436 | 637 | 1640 | 294 | 482 | 1629 | 391 | 560 |
| 2128 | W303.asm02.HP0_G0019100 | chrVII | 170 | 62 | 64 | 120 | 59 | 82 | 157 | 66 | 79 | 219 | 80 | 95 | 176 | 46 | 68 | 203 | 65 | 92 |
| 2129 | W303.asm02.HP0_G0019110 | chrVII | 3131 | 1074 | 1261 | 2900 | 1044 | 1226 | 2844 | 1138 | 1333 | 3344 | 1101 | 1382 | 4087 | 815 | 1223 | 3454 | 950 | 1328 |
| 2130 | W303.asm02.HP0_G0019120 | chrVII | 2949 | 852 | 999 | 2910 | 975 | 911 | 2905 | 920 | 1069 | 6027 | 866 | 1321 | 5256 | 678 | 932 | 5328 | 844 | 1141 |
| 2131 | W303.asm02.HP0_G0019130 | chrVII | 559 | 710 | 611 | 496 | 614 | 590 | 540 | 632 | 684 | 1878 | 587 | 592 | 1901 | 369 | 508 | 1840 | 567 | 523 |
| 2132 | W303.asm02.HP0_G0019140 | chrVII | 2997 | 1389 | 1600 | 2903 | 1257 | 1538 | 2753 | 1323 | 1682 | 5835 | 1395 | 2018 | 5388 | 979 | 1525 | 5475 | 1279 | 1859 |
| 2133 | W303.asm02.HP0_G0019150 | chrVII | 2940 | 416 | 434 | 2642 | 352 | 480 | 2635 | 343 | 536 | 4497 | 809 | 911 | 2147 | 496 | 655 | 2518 | 649 | 861 |
| 2134 | W303.asm02.HP0_G0019160 | chrVII | 6233 | 2107 | 2000 | 5738 | 1724 | 1892 | 5884 | 1994 | 1892 | 6441 | 1866 | 2088 | 6186 | 1087 | 1680 | 6168 | 1596 | 1810 |
| 2135 | W303.asm02.HP0_G0019170 | chrVII | 362 | 97 | 59 | 353 | 49 | 68 | 327 | 87 | 51 | 289 | 98 | 108 | 325 | 57 | 63 | 261 | 111 | 67 |
| 2136 | W303.asm02.HP0_G0019180 | chrVII | 2481 | 1206 | 1774 | 2456 | 1230 | 1774 | 2205 | 1118 | 2290 | 3648 | 847 | 1208 | 4867 | 564 | 982 | 4179 | 719 | 1009 |
| 2137 | W303.asm02.HP0_G0019190 | chrVII | 1061 | 717 | 927 | 1029 | 912 | 917 | 1015 | 865 | 1106 | 2482 | 653 | 750 | 2821 | 412 | 524 | 2531 | 591 | 659 |

| | | | | | | | | | | | | | | | | | | | |
|---|---|---|---|---|---|---|---|---|---|---|---|---|---|---|---|---|---|---|---|
| 2138 | W303.asm02.HP0_G0019200 | chrVII | 5272 | 2444 | 2833 | 5146 | 3514 | 2508 | 5645 | 2750 | 2400 | 19091 | 3696 | 4493 | 18031 | 2398 | 2641 | 16640 | 2904 | 3829 |
| 2139 | W303.asm02.HP0_G0019210 | chrVII | 3229 | 3771 | 3053 | 3112 | 3135 | 2572 | 3650 | 4095 | 2973 | 19023 | 5781 | 5377 | 16753 | 3832 | 4077 | 17113 | 4986 | 4626 |
| 2140 | W303.asm02.HP0_G0019220 | chrVII | 12213 | 11683 | 11273 | 11106 | 11173 | 11550 | 13275 | 11674 | 12847 | 26204 | 10320 | 10470 | 24554 | 7477 | 7827 | 24620 | 8800 | 9252 |
| 2141 | W303.asm02.HP0_G0019230 | chrVII | 406 | 308 | 305 | 503 | 283 | 262 | 388 | 252 | 264 | 1674 | 305 | 297 | 1777 | 164 | 255 | 1632 | 308 | 278 |
| 2142 | W303.asm02.HP0_G0019240 | chrVII | 7249 | 632 | 781 | 7192 | 655 | 1036 | 7123 | 539 | 939 | 5521 | 1157 | 2347 | 2885 | 875 | 1562 | 3142 | 1065 | 2061 |
| 2143 | W303.asm02.HP0_G0019250 | chrVII | 586 | 231 | 321 | 692 | 299 | 476 | 589 | 221 | 386 | 4341 | 733 | 1053 | 2390 | 407 | 677 | 2673 | 730 | 1086 |
| 2144 | W303.asm02.HP0_G0019260 | chrVII | 1843 | 1319 | 1813 | 1545 | 1752 | 1852 | 1844 | 1439 | 2380 | 4647 | 1067 | 1409 | 5227 | 656 | 1110 | 5022 | 972 | 1238 |
| 2145 | W303.asm02.HP0_G0019270 | chrVII | 8054 | 3217 | 4015 | 7338 | 3577 | 3800 | 7563 | 3207 | 4996 | 8309 | 2977 | 3694 | 10874 | 2262 | 3244 | 8545 | 2611 | 3222 |
| 2146 | W303.asm02.HP0_G0019280 | chrVII | 1576 | 1039 | 1017 | 1474 | 978 | 1007 | 1645 | 1151 | 917 | 4762 | 1136 | 1421 | 4042 | 713 | 956 | 4583 | 1061 | 1241 |
| 2147 | W303.asm02.HP0_G0019290 | chrVII | 3400 | 540 | 531 | 3434 | 590 | 578 | 3287 | 535 | 568 | 4586 | 985 | 1261 | 3587 | 554 | 965 | 3350 | 886 | 1162 |
| 2148 | W303.asm02.HP0_G0019300 | chrVII | 2175 | 982 | 859 | 2014 | 729 | 797 | 2228 | 871 | 784 | 4700 | 1108 | 1422 | 3964 | 728 | 1020 | 4361 | 1045 | 1271 |
| 2149 | W303.asm02.HP0_G0019310 | chrVII | 2447 | 608 | 562 | 2306 | 612 | 538 | 2495 | 573 | 595 | 3957 | 615 | 978 | 3998 | 449 | 646 | 3315 | 606 | 828 |
| 2150 | W303.asm02.HP0_G0019320 | chrVII | 1111 | 197 | 222 | 961 | 236 | 246 | 1169 | 146 | 315 | 1177 | 325 | 458 | 544 | 257 | 285 | 535 | 267 | 359 |
| 2151 | W303.asm02.HP0_G0019330 | chrVII | 4285 | 1646 | 2179 | 4419 | 1888 | 2947 | 4497 | 1402 | 2901 | 5850 | 2755 | 5577 | 2092 | 2451 | 3617 | 1672 | 2208 | 4737 |
| 2152 | W303.asm02.HP0_G0019340 | chrVII | 931 | 782 | 1014 | 835 | 868 | 1042 | 883 | 783 | 1231 | 2627 | 671 | 830 | 3157 | 433 | 565 | 3033 | 603 | 726 |
| 2153 | W303.asm02.HP0_G0019350 | chrVII | 1702 | 428 | 370 | 1609 | 416 | 408 | 1730 | 432 | 380 | 3327 | 664 | 926 | 2553 | 482 | 612 | 2508 | 568 | 804 |
| 2154 | W303.asm02.HP0_G0019360 | chrVII | 2119 | 888 | 850 | 2221 | 771 | 828 | 1832 | 886 | 886 | 2635 | 825 | 766 | 2663 | 484 | 695 | 2552 | 728 | 632 |
| 2155 | W303.asm02.HP0_G0019370 | chrVII | 1444 | 1459 | 1516 | 1460 | 1591 | 1632 | 1376 | 1372 | 1474 | 4722 | 1179 | 1518 | 5470 | 778 | 1148 | 4546 | 1201 | 1374 |
| 2156 | W303.asm02.HP0_G0019380 | chrVII | 249 | 569 | 498 | 232 | 442 | 497 | 209 | 516 | 520 | 1441 | 367 | 508 | 1608 | 242 | 356 | 1500 | 375 | 371 |
| 2157 | W303.asm02.HP0_G0019390 | chrVII | 2382 | 892 | 841 | 2171 | 782 | 985 | 2302 | 630 | 1009 | 7102 | 1624 | 2162 | 4632 | 1160 | 1261 | 4394 | 1568 | 1693 |
| 2158 | W303.asm02.HP0_G0019400 | chrVII | 15690 | 9183 | 9611 | 15401 | 11385 | 8047 | 17224 | 9236 | 9790 | 45750 | 12592 | 10815 | 40211 | 7416 | 6468 | 40177 | 11074 | 10085 |
| 2159 | W303.asm02.HP0_G0019410 | chrVII | 3388 | 2928 | 3520 | 3212 | 3096 | 3558 | 3176 | 3030 | 4872 | 8112 | 2401 | 2912 | 9655 | 1815 | 2587 | 9004 | 2130 | 2443 |
| 2160 | W303.asm02.HP0_G0019420 | chrVII | 2881 | 744 | 446 | 2825 | 363 | 458 | 2471 | 615 | 416 | 1540 | 648 | 576 | 1589 | 471 | 455 | 1315 | 734 | 541 |
| 2161 | W303.asm02.HP0_G0019430 | chrVII | 561 | 381 | 273 | 616 | 281 | 278 | 523 | 367 | 290 | 917 | 342 | 257 | 944 | 235 | 159 | 869 | 315 | 256 |
| 2162 | W303.asm02.HP0_G0019440 | chrVII | 1382 | 1214 | 1597 | 1134 | 1357 | 1650 | 1284 | 1276 | 1991 | 4748 | 1016 | 1439 | 4628 | 685 | 1045 | 4821 | 977 | 1362 |
| 2163 | W303.asm02.HP0_G0019450 | chrVII | 9627 | 938 | 1114 | 9408 | 1063 | 1221 | 9436 | 846 | 1374 | 10754 | 2122 | 2875 | 7616 | 1584 | 2124 | 5719 | 1659 | 2416 |
| 2164 | W303.asm02.HP0_G0019460 | chrVII | 14596 | 3933 | 2635 | 13767 | 2546 | 2985 | 14479 | 3600 | 2470 | 13442 | 4578 | 5796 | 10982 | 3717 | 4300 | 10730 | 4235 | 5110 |
| 2165 | W303.asm02.HP0_G0019470 | chrVII | 1411 | 706 | 898 | 1286 | 702 | 853 | 1283 | 790 | 1057 | 1708 | 629 | 758 | 2201 | 540 | 729 | 1695 | 565 | 725 |
| 2166 | W303.asm02.HP0_G0019480 | chrVII | 4096 | 279 | 226 | 4797 | 253 | 294 | 3784 | 233 | 239 | 1589 | 494 | 521 | 1482 | 318 | 403 | 1101 | 545 | 451 |
| 2167 | W303.asm02.HP0_G0019490 | chrVII | 2780 | 1014 | 851 | 2637 | 871 | 802 | 2638 | 944 | 852 | 4977 | 1271 | 1446 | 4314 | 798 | 1025 | 4098 | 1183 | 1295 |
| 2168 | W303.asm02.HP0_G0019500 | chrVII | 2690 | 2365 | 2071 | 2736 | 1880 | 2274 | 2846 | 2287 | 1929 | 7890 | 2281 | 3046 | 7651 | 1590 | 1843 | 7382 | 2025 | 2722 |
| 2169 | W303.asm02.HP0_G0019510 | chrVII | 1027 | 497 | 518 | 745 | 461 | 475 | 787 | 506 | 641 | 1460 | 455 | 390 | 1554 | 256 | 330 | 1507 | 430 | 387 |
| 2170 | W303.asm02.HP0_G0019520 | chrVII | 7937 | 2541 | 1693 | 7327 | 1789 | 1748 | 7432 | 2192 | 1853 | 9808 | 3366 | 3573 | 7730 | 2617 | 2880 | 7750 | 3141 | 3336 |
| 2171 | W303.asm02.HP0_G0019530 | chrVII | 1758 | 978 | 991 | 1482 | 1045 | 997 | 1765 | 994 | 912 | 4634 | 1036 | 1208 | 4817 | 675 | 929 | 4848 | 999 | 1305 |
| 2172 | W303.asm02.HP0_G0019540 | chrVII | 1363 | 908 | 1067 | 1431 | 829 | 1002 | 1246 | 839 | 823 | 4791 | 1242 | 2115 | 3804 | 564 | 1130 | 3876 | 1091 | 1646 |
| 2173 | W303.asm02.HP0_G0019550 | chrVII | 21747 | 6917 | 4150 | 22667 | 3310 | 4583 | 20570 | 5767 | 4085 | 7244 | 5084 | 4446 | 12519 | 4082 | 5230 | 7303 | 5310 | 4093 |
| 2174 | W303.asm02.HP0_G0019560 | chrVII | 4834 | 1149 | 917 | 4691 | 890 | 971 | 4575 | 1101 | 1104 | 6579 | 1729 | 1610 | 5309 | 1490 | 1591 | 4232 | 1728 | 1480 |
| 2175 | W303.asm02.HP0_G0019570 | chrVII | 1254 | 618 | 765 | 1387 | 754 | 852 | 1232 | 442 | 951 | 5354 | 1082 | 1337 | 3679 | 750 | 925 | 3167 | 1047 | 1043 |
| 2176 | W303.asm02.HP0_G0019580 | chrVII | 56 | 21 | 13 | 53 | 20 | 16 | 38 | 19 | 26 | 39 | 14 | 21 | 49 | 3 | 9 | 56 | 15 | 15 |
| 2177 | W303.asm02.HP0_G0019590 | chrVII | 1585 | 1339 | 1416 | 1479 | 1380 | 1439 | 1523 | 1172 | 1348 | 2259 | 897 | 937 | 2994 | 716 | 876 | 2408 | 808 | 858 |
| 2178 | W303.asm02.HP0_G0019600 | chrVII | 12582 | 8170 | 11323 | 11602 | 10900 | 11243 | 12526 | 8361 | 13651 | 26033 | 7575 | 9471 | 28538 | 5446 | 8061 | 26559 | 6771 | 8302 |
| 2179 | W303.asm02.HP0_G0019610 | chrVII | 8953 | 7359 | 9466 | 7401 | 8615 | 9227 | 8364 | 7514 | 11677 | 16687 | 6579 | 7564 | 18580 | 5244 | 6421 | 17675 | 6113 | 6473 |
| 2180 | W303.asm02.HP0_G0019620 | chrVII | 918 | 1029 | 1460 | 719 | 1780 | 1536 | 991 | 1300 | 2104 | 5371 | 1215 | 1160 | 5890 | 627 | 809 | 5681 | 1034 | 1009 |
| 2181 | W303.asm02.HP0_G0019630 | chrVII | 2387 | 1482 | 1819 | 2114 | 1770 | 2098 | 2711 | 1319 | 2325 | 12522 | 2759 | 3480 | 12495 | 2085 | 2829 | 9591 | 2409 | 3249 |
| 2182 | W303.asm02.HP0_G0019640 | chrVII | 5121 | 1133 | 1069 | 4477 | 983 | 1129 | 5011 | 1100 | 1199 | 8277 | 1946 | 2499 | 5394 | 1555 | 1706 | 5365 | 1794 | 2325 |
| 2183 | W303.asm02.HP0_G0019650 | chrVII | 7628 | 2339 | 1820 | 5903 | 1805 | 2122 | 7352 | 2163 | 2307 | 13080 | 4299 | 5062 | 7915 | 3482 | 3562 | 7872 | 3684 | 4356 |
| 2184 | W303.asm02.HP0_G0019660 | chrVII | 2101 | 464 | 443 | 2002 | 426 | 522 | 2066 | 381 | 560 | 2528 | 767 | 993 | 1534 | 459 | 704 | 1449 | 728 | 805 |
| 2185 | W303.asm02.HP0_G0019670 | chrVII | 2943 | 572 | 659 | 2642 | 668 | 826 | 2929 | 467 | 827 | 4202 | 1144 | 1572 | 1597 | 759 | 1026 | 1636 | 1046 | 1420 |
| 2186 | W303.asm02.HP0_G0019680 | chrVII | 6084 | 1597 | 2143 | 4897 | 1670 | 2310 | 5956 | 1392 | 2658 | 7232 | 2932 | 4212 | 2950 | 2281 | 2414 | 2584 | 2272 | 3545 |
| 2187 | W303.asm02.HP0_G0019690 | chrVII | 2810 | 1415 | 1125 | 2427 | 1166 | 1111 | 3204 | 1489 | 1354 | 6866 | 1963 | 1940 | 7115 | 1731 | 2082 | 6295 | 1853 | 1954 |
| 2188 | W303.asm02.HP0_G0019700 | chrVII | 2618 | 1655 | 1914 | 2357 | 1816 | 1921 | 2524 | 1742 | 2440 | 4967 | 1385 | 1539 | 5497 | 941 | 1250 | 5168 | 1205 | 1395 |
| 2189 | W303.asm02.HP0_G0019710 | chrVII | 4695 | 1792 | 1441 | 4404 | 1396 | 1391 | 4508 | 1815 | 1350 | 7508 | 2364 | 2534 | 7091 | 1623 | 1935 | 6253 | 1898 | 2359 |
| 2190 | W303.asm02.HP0_G0019720 | chrVII | 2126 | 492 | 395 | 2118 | 406 | 443 | 2094 | 486 | 404 | 1305 | 374 | 529 | 1322 | 321 | 362 | 1123 | 372 | 428 |
| 2191 | W303.asm02.HP0_G0019730 | chrVII | 1471 | 360 | 423 | 1493 | 342 | 423 | 1489 | 325 | 517 | 2390 | 547 | 691 | 1459 | 344 | 473 | 1569 | 575 | 678 |
| 2192 | W303.asm02.HP0_G0019740 | chrVII | 1959 | 1244 | 1438 | 1700 | 1473 | 1304 | 1909 | 1405 | 1682 | 8038 | 1290 | 1446 | 7383 | 788 | 1040 | 8506 | 1139 | 1419 |
| 2193 | W303.asm02.HP0_G0019750 | chrVII | 48 | 10 | 17 | 32 | 12 | 17 | 39 | 9 | 15 | 64 | 12 | 5 | 62 | 4 | 5 | 44 | 10 | 12 |
| 2194 | W303.asm02.HP0_G0019760 | chrVII | 2599 | 733 | 1157 | 2633 | 1234 | 1296 | 2604 | 659 | 1552 | 7690 | 1419 | 2311 | 2881 | 1019 | 1273 | 2750 | 1034 | 2067 |
| 2195 | W303.asm02.HP0_G0019770 | chrVII | 2547 | 689 | 829 | 2635 | 883 | 931 | 2489 | 726 | 920 | 6409 | 955 | 1745 | 4635 | 697 | 1279 | 4948 | 844 | 1551 |
| 2196 | W303.asm02.HP0_G0019780 | chrVII | 9415 | 5271 | 4488 | 8456 | 4212 | 4128 | 9100 | 5090 | 4597 | 17246 | 6279 | 6581 | 16090 | 4386 | 5076 | 15797 | 5607 | 6150 |
| 2197 | W303.asm02.HP0_G0019790 | chrVII | 10110 | 3998 | 4079 | 10369 | 3874 | 4116 | 10381 | 3871 | 3896 | 10661 | 3592 | 4747 | 10859 | 2842 | 3839 | 9894 | 3124 | 4322 |
| 2198 | W303.asm02.HP0_G0019800 | chrVII | 336 | 676 | 440 | 308 | 385 | 426 | 281 | 568 | 506 | 1028 | 480 | 312 | 1217 | 349 | 331 | 1129 | 462 | 315 |
| 2199 | W303.asm02.HP0_G0019810 | chrVII | 8035 | 9571 | 7450 | 8969 | 8527 | 8162 | 8646 | 9690 | 8540 | 44895 | 15565 | 19089 | 28266 | 9236 | 9883 | 36150 | 13478 | 15419 |
| 2200 | W303.asm02.HP0_G0019820 | chrVII | 39056 | 44791 | 35196 | 32679 | 35183 | 33628 | 37697 | 43719 | 40901 | 162369 | 68410 | 63312 | 124574 | 52887 | 47842 | 134079 | 60404 | 57817 |
| 2201 | W303.asm02.HP0_G0019830 | chrVII | 6983 | 549 | 847 | 7133 | 671 | 851 | 6220 | 547 | 1029 | 4347 | 1026 | 1566 | 2687 | 779 | 1415 | 2673 | 884 | 1337 |
| 2202 | W303.asm02.HP0_G0019840 | chrVII | 3350 | 1064 | 1203 | 3353 | 1058 | 1223 | 3168 | 1017 | 1481 | 4159 | 854 | 1047 | 4177 | 556 | 894 | 4091 | 916 | 1026 |
| 2203 | W303.asm02.HP0_G0019850 | chrVII | 2995 | 864 | 467 | 2797 | 506 | 490 | 2636 | 792 | 395 | 2579 | 919 | 812 | 2390 | 593 | 597 | 2301 | 870 | 734 |
| 2204 | W303.asm02.HP0_G0019860 | chrVII | 1155 | 383 | 360 | 1294 | 350 | 434 | 1118 | 271 | 457 | 2910 | 713 | 815 | 1760 | 406 | 540 | 1781 | 787 | 641 |
| 2205 | W303.asm02.HP0_G0019870 | chrVII | 1925 | 1089 | 824 | 1770 | 782 | 788 | 1756 | 1091 | 845 | 3064 | 997 | 765 | 3156 | 646 | 771 | 3036 | 962 | 764 |
| 2206 | W303.asm02.HP0_G0019880 | chrVII | 7739 | 3489 | 3197 | 7732 | 2803 | 2990 | 8009 | 3049 | 3252 | 9204 | 2963 | 2918 | 9022 | 1902 | 2329 | 9455 | 2970 | 2727 |
| 2207 | W303.asm02.HP0_G0019890 | chrVII | 2285 | 1398 | 1174 | 2231 | 1171 | 1094 | 2283 | 1550 | 1204 | 5208 | 1584 | 1643 | 4623 | 984 | 1064 | 5002 | 1462 | 1483 |
| 2208 | W303.asm02.HP0_G0019900 | chrVII | 3715 | 919 | 578 | 3700 | 504 | 580 | 3545 | 867 | 624 | 3950 | 1220 | 1186 | 3517 | 824 | 772 | 3209 | 1348 | 942 |
| 2209 | W303.asm02.HP0_G0019910 | chrVII | 1182 | 1014 | 767 | 1227 | 821 | 765 | 1195 | 979 | 699 | 4971 | 1210 | 1099 | 4260 | 692 | 747 | 4728 | 1313 | 1086 |
| 2210 | W303.asm02.HP0_G0019920 | chrVII | 854 | 748 | 595 | 734 | 607 | 568 | 804 | 749 | 628 | 3133 | 746 | 628 | 2920 | 463 | 533 | 3097 | 707 | 572 |
| 2211 | W303.asm02.HP0_G0019930 | chrVII | 6505 | 2188 | 1477 | 6534 | 1586 | 1518 | 6305 | 2057 | 1309 | 7267 | 2333 | 2310 | 6347 | 1443 | 1691 | 6269 | 2078 | 2072 |
| 2212 | W303.asm02.HP0_G0019940 | chrVII | 704 | 186 | 125 | 645 | 126 | 147 | 598 | 177 | 133 | 901 | 252 | 220 | 744 | 161 | 160 | 773 | 226 | 193 |
| 2213 | W303.asm02.HP0_G0019950 | chrVII | 3305 | 1619 | 1248 | 3469 | 1300 | 1280 | 3517 | 1544 | 1267 | 6573 | 2111 | 2009 | 5966 | 1467 | 1567 | 5587 | 1849 | 1694 |
| 2214 | W303.asm02.HP0_G0019960 | chrVII | 8453 | 1285 | 1727 | 7501 | 1128 | 2083 | 6883 | 1151 | 2165 | 1915 | 1123 | 1631 | 3300 | 882 | 1729 | 2078 | 1118 | 1241 |
| 2215 | W303.asm02.HP0_G0019970 | chrVII | 5088 | 806 | 829 | 5198 | 781 | 924 | 4877 | 761 | 1005 | 5916 | 1349 | 2171 | 3847 | 944 | 1431 | 4023 | 1286 | 1827 |
| 2216 | W303.asm02.HP0_G0019980 | chrVII | 1896 | 842 | 952 | 1659 | 666 | 880 | 1721 | 813 | 946 | 3986 | 1099 | 1405 | 3254 | 654 | 1031 | 3485 | 1012 | 1288 |
| 2217 | W303.asm02.HP0_G0019990 | chrVII | 1613 | 1139 | 862 | 1652 | 801 | 875 | 1664 | 958 | 960 | 4222 | 1254 | 1538 | 3611 | 878 | 992 | 3745 | 1239 | 1369 |
| 2218 | W303.asm02.HP0_G0020000 | chrVII | 4314 | 559 | 468 | 4089 | 514 | 532 | 4235 | 477 | 541 | 4537 | 1048 | 1175 | 2918 | 664 | 832 | 2785 | 991 | 1045 |
| 2219 | W303.asm02.HP0_G0020010 | chrVII | 1056 | 1012 | 1495 | 1016 | 1415 | 1867 | 1282 | 974 | 1906 | 9570 | 2005 | 3844 | 2640 | 1623 | 2299 | 2038 | 1499 | 3400 |
| 2220 | W303.asm02.HP0_G0020020 | chrVII | 1019 | 370 | 295 | 999 | 259 | 327 | 979 | 364 | 323 | 1542 | 477 | 486 | 1361 | 300 | 364 | 1366 | 459 | 395 |
| 2221 | W303.asm02.HP0_G0020030 | chrVII | 467 | 264 | 277 | 438 | 280 | 260 | 377 | 268 | 363 | 853 | 229 | 215 | 924 | 114 | 173 | 920 | 231 | 194 |
| 2222 | W303.asm02.HP0_G0020040 | chrVII | 2127 | 630 | 656 | 2133 | 637 | 629 | 2193 | 693 | 677 | 2973 | 742 | 959 | 3007 | 568 | 759 | 2759 | 768 | 828 |
| 2223 | W303.asm02.HP0_G0020050 | chrVII | 6915 | 1396 | 1028 | 7043 | 848 | 1128 | 6451 | 1093 | 1305 | 3926 | 1790 | 1968 | 3854 | 1340 | 1682 | 3002 | 1862 | 1770 |
| 2224 | W303.asm02.HP0_G0020060 | chrVII | 6401 | 3934 | 4947 | 5919 | 4284 | 4910 | 6139 | 4099 | 6059 | 9843 | 3523 | 3865 | 12046 | 2721 | 3577 | 10472 | 2969 | 3499 |
| 2225 | W303.asm02.HP0_G0020070 | chrVII | 6071 | 1292 | 882 | 5875 | 1062 | 967 | 5738 | 1087 | 901 | 3371 | 1170 | 1313 | 4266 | 949 | 1189 | 2517 | 1080 | 1097 |
| 2226 | W303.asm02.HP0_G0020080 | chrVII | 2340 | 237 | 120 | 2915 | 182 | 187 | 2211 | 132 | 189 | 1670 | 584 | 323 | 968 | 311 | 293 | 498 | 613 | 317 |
| 2227 | W303.asm02.HP0_G0020090 | chrVII | 1583 | 1349 | 1415 | 1307 | 1278 | 1332 | 1480 | 1275 | 1872 | 3546 | 1085 | 1007 | 4164 | 742 | 805 | 4121 | 1055 | 865 |
| 2228 | W303.asm02.HP0_G0020100 | chrVII | 5628 | 3540 | 3505 | 5159 | 2992 | 3572 | 5222 | 3572 | 3751 | 8863 | 2754 | 3011 | 9051 | 1888 | 2562 | 9098 | 2545 | 2677 |
| 2229 | W303.asm02.HP0_G0020110 | chrVII | 6355 | 5339 | 7030 | 6055 | 6431 | 7050 | 6504 | 5500 | 8629 | 14682 | 5196 | 6204 | 16317 | 3839 | 5258 | 15129 | 4702 | 5426 |
| 2230 | W303.asm02.HP0_G0020120 | chrVII | 4124 | 1420 | 2599 | 4003 | 1595 | 2827 | 3625 | 1395 | 3340 | 4344 | 1153 | 2435 | 5640 | 826 | 1739 | 4476 | 1221 | 2056 |

| | | | | | | | | | | | | | | | | | | | | |
|---|---|---|---|---|---|---|---|---|---|---|---|---|---|---|---|---|---|---|---|---|
| 2231 | W303.asm02.HP0_G0020130 | chrVII | 1397 | 1143 | 1170 | 1334 | 1155 | 1135 | 1300 | 1245 | 1557 | 3466 | 1164 | 1139 | 3514 | 688 | 815 | 3522 | 1090 | 1008 |
| 2232 | W303.asm02.HP0_G0020140 | chrVII | 333 | 781 | 831 | 363 | 817 | 800 | 366 | 728 | 1078 | 2320 | 555 | 520 | 2599 | 388 | 407 | 2691 | 493 | 465 |
| 2233 | W303.asm02.HP0_G0020150 | chrVII | 889 | 335 | 254 | 868 | 246 | 319 | 1005 | 280 | 300 | 1517 | 510 | 520 | 974 | 410 | 353 | 823 | 506 | 496 |
| 2234 | W303.asm02.HP0_G0020160 | chrVII | 7207 | 379 | 522 | 7198 | 624 | 642 | 7117 | 301 | 609 | 3080 | 613 | 1228 | 1360 | 624 | 727 | 1301 | 579 | 1097 |
| 2235 | W303.asm02.HP0_G0020170 | chrVII | 2954 | 1892 | 2074 | 2828 | 2237 | 2228 | 2856 | 2111 | 2271 | 6528 | 1445 | 1650 | 6798 | 966 | 1373 | 6225 | 1259 | 1474 |
| 2236 | W303.asm02.HP0_G0020180 | chrVII | 8067 | 6213 | 9332 | 7353 | 8350 | 9307 | 8090 | 6287 | 11945 | 18700 | 5835 | 7776 | 21608 | 4369 | 6464 | 19857 | 5320 | 6784 |
| 2237 | W303.asm02.HP0_G0020190 | chrVII | 245 | 567 | 555 | 240 | 648 | 564 | 234 | 272 | 706 | 3256 | 279 | 353 | 4032 | 158 | 256 | 3497 | 478 | 327 |
| 2238 | W303.asm02.HP0_G0020200 | chrVII | 237 | 195 | 197 | 156 | 217 | 200 | 211 | 198 | 285 | 900 | 211 | 212 | 818 | 97 | 167 | 788 | 200 | 200 |
| 2239 | W303.asm02.HP0_G0020210 | chrVII | 979 | 728 | 1005 | 896 | 900 | 856 | 766 | 751 | 1027 | 3723 | 749 | 785 | 3934 | 457 | 558 | 3596 | 709 | 718 |
| 2240 | W303.asm02.HP0_G0020220 | chrVII | 5793 | 4293 | 5749 | 5401 | 4215 | 5802 | 5402 | 4308 | 6289 | 9226 | 4019 | 5535 | 9610 | 2939 | 4061 | 10112 | 3886 | 4901 |
| 2241 | W303.asm02.HP0_G0020230 | chrVII | 1793 | 1378 | 1377 | 1686 | 1227 | 1412 | 1766 | 1379 | 1598 | 4574 | 1190 | 1383 | 4495 | 728 | 1024 | 4533 | 1135 | 1214 |
| 2242 | W303.asm02.HP0_G0020240 | chrVII | 1074 | 1107 | 1269 | 1256 | 1335 | 1086 | 1302 | 1120 | 1126 | 5889 | 1133 | 1515 | 6526 | 747 | 1147 | 5174 | 1096 | 1456 |
| 2243 | W303.asm02.HP0_G0020250 | chrVII | 1541 | 386 | 465 | 1317 | 419 | 497 | 1458 | 313 | 552 | 3396 | 682 | 992 | 2107 | 493 | 678 | 2207 | 599 | 861 |
| 2244 | W303.asm02.HP0_G0020260 | chrVII | 2044 | 823 | 807 | 2054 | 823 | 847 | 2043 | 781 | 973 | 2161 | 619 | 657 | 2785 | 439 | 555 | 2082 | 574 | 558 |
| 2245 | W303.asm02.HP0_G0020270 | chrVII | 2817 | 284 | 186 | 2988 | 145 | 167 | 2441 | 223 | 181 | 438 | 252 | 198 | 608 | 150 | 136 | 540 | 351 | 231 |
| 2246 | W303.asm02.HP0_G0020280 | chrVII | 5019 | 1732 | 1760 | 5021 | 1743 | 1809 | 4839 | 1675 | 2077 | 6308 | 1787 | 2331 | 8301 | 1626 | 2287 | 5620 | 1637 | 2024 |
| 2247 | W303.asm02.HP0_G0020290 | chrVII | 2020 | 1641 | 1763 | 2251 | 1620 | 1614 | 1863 | 1508 | 1992 | 5875 | 1631 | 1666 | 6538 | 1046 | 1415 | 5810 | 1457 | 1448 |
| 2248 | W303.asm02.HP0_G0020300 | chrVII | 1397 | 483 | 559 | 1404 | 549 | 552 | 1346 | 506 | 536 | 4417 | 728 | 1096 | 3835 | 391 | 804 | 3437 | 634 | 1003 |
| 2249 | W303.asm02.HP0_G0020310 | chrVII | 2944 | 1190 | 1270 | 3106 | 1051 | 1208 | 2902 | 1081 | 1275 | 4468 | 1150 | 1648 | 4558 | 864 | 1414 | 4237 | 1081 | 1470 |
| 2250 | W303.asm02.HP0_G0020320 | chrVII | 2819 | 1592 | 1102 | 2675 | 962 | 1093 | 2997 | 1142 | 1022 | 3696 | 966 | 1265 | 3354 | 562 | 817 | 3212 | 1013 | 1079 |
| 2251 | W303.asm02.HP0_G0020330 | chrVII | 136 | 133 | 136 | 99 | 94 | 139 | 127 | 92 | 151 | 422 | 116 | 84 | 401 | 57 | 78 | 438 | 149 | 66 |
| 2252 | W303.asm02.HP0_G0020340 | chrVII | 621 | 583 | 218 | 791 | 201 | 217 | 594 | 228 | 240 | 587 | 233 | 255 | 794 | 98 | 149 | 553 | 257 | 192 |
| 2253 | W303.asm02.HP0_G0020350 | chrVII | 649 | 173 | 166 | 625 | 162 | 170 | 505 | 152 | 214 | 404 | 128 | 143 | 446 | 81 | 131 | 396 | 137 | 101 |
| 2254 | W303.asm02.HP0_G0020360 | chrVII | 8484 | 1253 | 1616 | 7768 | 1468 | 1983 | 8955 | 1007 | 2210 | 17264 | 3045 | 4698 | 7260 | 2402 | 2924 | 7199 | 2589 | 4199 |
| 2255 | W303.asm02.HP0_G0020370 | chrVII | 2994 | 5413 | 5448 | 2044 | 4010 | 4941 | 2650 | 4944 | 5689 | 11173 | 6575 | 7805 | 8572 | 4997 | 5814 | 9114 | 5788 | 6442 |
| 2256 | W303.asm02.HP0_G0020380 | chrVII | 2003 | 332 | 517 | 2145 | 458 | 592 | 2100 | 345 | 618 | 4645 | 624 | 1278 | 2848 | 449 | 897 | 2906 | 579 | 1047 |
| 2257 | W303.asm02.HP0_G0020390 | chrVII | 1235 | 911 | 1134 | 1294 | 1017 | 1108 | 1185 | 964 | 1384 | 2853 | 708 | 814 | 2948 | 531 | 576 | 3235 | 716 | 670 |
| 2258 | W303.asm02.HP0_G0020400 | chrVII | 2485 | 519 | 470 | 2302 | 401 | 411 | 2522 | 475 | 478 | 2711 | 676 | 828 | 2432 | 436 | 570 | 2429 | 698 | 730 |
| 2259 | W303.asm02.HP0_G0020410 | chrVII | 336 | 94 | 56 | 336 | 40 | 41 | 286 | 76 | 54 | 155 | 73 | 51 | 224 | 47 | 38 | 161 | 92 | 40 |
| 2260 | W303.asm02.HP0_G0020420 | chrVII | 10599 | 7936 | 8376 | 11748 | 8349 | 8065 | 12943 | 7935 | 7265 | 28442 | 8177 | 10230 | 25994 | 5888 | 6941 | 26425 | 6950 | 9490 |
| 2261 | W303.asm02.HP0_G0020430 | chrVII | 12188 | 1590 | 1652 | 8965 | 1702 | 1825 | 12053 | 1305 | 2210 | 12136 | 3079 | 4582 | 4278 | 2461 | 2731 | 4371 | 2502 | 3749 |
| 2262 | W303.asm02.HP0_G0020440 | chrVII | 2482 | 823 | 1011 | 2216 | 784 | 1033 | 2290 | 800 | 1085 | 2516 | 636 | 949 | 3106 | 459 | 750 | 2533 | 640 | 862 |
| 2263 | W303.asm02.HP0_G0020450 | chrVII | 838 | 925 | 1445 | 737 | 1257 | 1427 | 738 | 956 | 1853 | 2269 | 761 | 1054 | 3048 | 485 | 942 | 2447 | 678 | 855 |
| 2264 | W303.asm02.HP0_G0020460 | chrVII | 893 | 1439 | 1460 | 729 | 1451 | 1257 | 731 | 1574 | 1395 | 5395 | 1560 | 1460 | 6003 | 977 | 1280 | 5337 | 1315 | 1464 |
| 2265 | W303.asm02.HP0_G0020470 | chrVII | 445 | 142 | 75 | 464 | 103 | 111 | 362 | 107 | 101 | 264 | 137 | 105 | 311 | 63 | 81 | 240 | 132 | 75 |
| 2266 | W303.asm02.HP0_G0020480 | chrVII | 408 | 106 | 99 | 425 | 98 | 92 | 389 | 66 | 81 | 290 | 127 | 136 | 425 | 94 | 80 | 264 | 118 | 88 |
| 2267 | W303.asm02.HP0_G0020490 | chrVII | 657 | 149 | 170 | 523 | 175 | 208 | 636 | 102 | 220 | 634 | 203 | 341 | 238 | 161 | 159 | 201 | 158 | 256 |
| 2268 | W303.asm02.HP0_G0020500 | chrVII | 4558 | 375 | 630 | 4733 | 479 | 666 | 4186 | 406 | 753 | 2327 | 554 | 1040 | 2924 | 437 | 1192 | 1965 | 459 | 942 |
| 2269 | W303.asm02.HP0_G0020510 | chrVII | 1040 | 650 | 890 | 1058 | 867 | 826 | 917 | 758 | 975 | 5060 | 702 | 960 | 4212 | 416 | 762 | 4806 | 704 | 862 |
| 2270 | W303.asm02.HP0_G0020520 | chrVII | 2512 | 768 | 665 | 2438 | 827 | 652 | 2527 | 791 | 781 | 3865 | 833 | 885 | 3538 | 504 | 734 | 3606 | 734 | 765 |
| 2271 | W303.asm02.HP0_G0020530 | chrVII | 623 | 986 | 984 | 663 | 908 | 969 | 591 | 1022 | 1177 | 2478 | 789 | 889 | 2552 | 525 | 682 | 2714 | 709 | 703 |
| 2272 | W303.asm02.HP0_G0020540 | chrVII | 3864 | 1035 | 1325 | 3583 | 1216 | 1279 | 3331 | 1127 | 1628 | 3313 | 829 | 1034 | 3912 | 549 | 946 | 3376 | 836 | 979 |
| 2273 | W303.asm02.HP0_G0020550 | chrVII | 2145 | 863 | 770 | 2230 | 789 | 859 | 2053 | 746 | 833 | 3541 | 777 | 1089 | 3447 | 627 | 780 | 2917 | 778 | 935 |
| 2274 | W303.asm02.HP0_G0020560 | chrVII | 1366 | 680 | 370 | 1280 | 532 | 518 | 1434 | 566 | 460 | 2758 | 742 | 953 | 1847 | 626 | 616 | 2075 | 688 | 857 |
| 2275 | W303.asm02.HP0_G0020570 | chrVII | 5499 | 2072 | 2271 | 5263 | 2262 | 2396 | 5429 | 1859 | 2293 | 9153 | 2062 | 3575 | 9925 | 1635 | 2116 | 7981 | 2226 | 2769 |
| 2276 | W303.asm02.HP0_G0020580 | chrVII | 1764 | 1349 | 1855 | 1484 | 1898 | 1844 | 1742 | 1405 | 2264 | 5557 | 1105 | 1415 | 5931 | 825 | 1000 | 5713 | 1082 | 1177 |
| 2277 | W303.asm02.HP0_G0020590 | chrVII | 8891 | 1801 | 1829 | 9001 | 1813 | 1744 | 8858 | 1779 | 1690 | 6070 | 1856 | 2709 | 6524 | 1503 | 1878 | 5443 | 1544 | 2240 |
| 2278 | W303.asm02.HP0_G0020600 | chrVII | 3968 | 1277 | 1535 | 4034 | 1689 | 1594 | 4429 | 1320 | 1827 | 12278 | 2192 | 2946 | 10138 | 1486 | 1971 | 9364 | 1951 | 2633 |
| 2279 | W303.asm02.HP0_G0020610 | chrVII | 7204 | 1662 | 1459 | 7070 | 1448 | 1547 | 6693 | 1645 | 1396 | 4950 | 1402 | 2119 | 5085 | 1249 | 1572 | 4192 | 1304 | 1751 |
| 2280 | W303.asm02.HP0_G0020620 | chrVII | 6074 | 6983 | 7795 | 6003 | 7631 | 7933 | 6159 | 6969 | 9428 | 19431 | 6715 | 6793 | 20205 | 5012 | 5290 | 19237 | 5942 | 6202 |
| 2281 | W303.asm02.HP0_G0020630 | chrVII | 21209 | 5378 | 4737 | 22459 | 3907 | 5177 | 21307 | 4611 | 6456 | 20657 | 7835 | 9406 | 15200 | 7211 | 7492 | 12430 | 6871 | 8015 |
| 2282 | W303.asm02.HP0_G0020640 | chrVII | 258 | 47 | 32 | 232 | 21 | 17 | 235 | 39 | 26 | 99 | 69 | 33 | 117 | 35 | 17 | 91 | 73 | 48 |
| 2283 | W303.asm02.HP0_G0020650 | chrVII | 8762 | 583 | 242 | 10016 | 186 | 238 | 9234 | 376 | 264 | 877 | 701 | 434 | 944 | 379 | 305 | 648 | 839 | 343 |
| 2284 | W303.asm02.HP0_G0020660 | chrVII | 3385 | 868 | 1269 | 3286 | 1202 | 1154 | 3490 | 998 | 1400 | 8713 | 1386 | 2216 | 8065 | 859 | 1543 | 7525 | 1140 | 1887 |
| 2285 | W303.asm02.HP0_G0020670 | chrVII | 2806 | 1543 | 2123 | 2923 | 2369 | 2026 | 3108 | 1640 | 2445 | 20298 | 2350 | 3701 | 15676 | 1652 | 2726 | 16595 | 1993 | 3398 |
| 2286 | W303.asm02.HP0_G0020680 | chrVII | 1360 | 620 | 519 | 1265 | 618 | 611 | 1368 | 587 | 549 | 3619 | 594 | 869 | 3019 | 398 | 546 | 3033 | 544 | 748 |
| 2287 | W303.asm02.HP0_G0020690 | chrVII | 3247 | 1051 | 750 | 3285 | 885 | 857 | 3187 | 971 | 924 | 5375 | 1423 | 1578 | 5537 | 1333 | 1539 | 3969 | 1273 | 1532 |
| 2288 | W303.asm02.HP0_G0020700 | chrVII | 1689 | 1069 | 1332 | 1658 | 1448 | 1286 | 1624 | 1190 | 1416 | 6092 | 1111 | 1359 | 5451 | 637 | 860 | 6057 | 1009 | 1197 |
| 2289 | W303.asm02.HP0_G0020710 | chrVII | 25158 | 6299 | 4932 | 22673 | 5004 | 5218 | 25459 | 5890 | 5939 | 45281 | 12347 | 13332 | 28648 | 9314 | 8716 | 30774 | 10866 | 12035 |
| 2290 | W303.asm02.HP0_G0020720 | chrVII | 2500 | 1502 | 1622 | 2255 | 1819 | 1630 | 2651 | 1681 | 1670 | 5057 | 1435 | 1462 | 5302 | 854 | 1010 | 4669 | 1250 | 1265 |
| 2291 | W303.asm02.HP0_G0020730 | chrVII | 835 | 178 | 176 | 812 | 184 | 160 | 846 | 174 | 227 | 1046 | 301 | 308 | 863 | 215 | 212 | 710 | 248 | 204 |
| 2292 | W303.asm02.HP0_G0020740 | chrVII | 8408 | 558 | 695 | 8172 | 727 | 887 | 9204 | 551 | 970 | 8800 | 1411 | 2015 | 7742 | 1529 | 1806 | 4457 | 1123 | 1711 |
| 2293 | W303.asm02.HP0_G0020750 | chrVII | 1121 | 527 | 848 | 1116 | 920 | 795 | 1019 | 561 | 836 | 5664 | 660 | 1273 | 4968 | 533 | 1097 | 5143 | 581 | 1189 |
| 2294 | W303.asm02.HP0_G0020760 | chrVII | 1651 | 666 | 618 | 1447 | 713 | 574 | 1595 | 737 | 613 | 2862 | 661 | 722 | 2769 | 487 | 662 | 2634 | 601 | 632 |
| 2295 | W303.asm02.HP0_G0020770 | chrVII | 5731 | 994 | 935 | 5578 | 955 | 969 | 5346 | 1071 | 847 | 6536 | 1390 | 1779 | 5533 | 1036 | 1475 | 5138 | 1236 | 1604 |
| 2296 | W303.asm02.HP0_G0020780 | chrVII | 3350 | 1001 | 1137 | 3431 | 990 | 1085 | 3020 | 1033 | 1310 | 3641 | 972 | 1313 | 3789 | 729 | 1109 | 3323 | 922 | 1121 |
| 2297 | W303.asm02.HP0_G0020790 | chrVII | 1328 | 823 | 719 | 1141 | 755 | 747 | 1107 | 797 | 786 | 2189 | 773 | 770 | 2783 | 573 | 707 | 2134 | 718 | 636 |
| 2298 | W303.asm02.HP0_G0020800 | chrVII | 6893 | 4702 | 4877 | 7191 | 6396 | 4166 | 7927 | 5416 | 5370 | 31775 | 6773 | 6499 | 27854 | 3784 | 3660 | 29807 | 5542 | 5888 |
| 2299 | W303.asm02.HP0_G0020810 | chrVII | 1219 | 806 | 608 | 1098 | 558 | 597 | 1175 | 786 | 612 | 2425 | 668 | 581 | 2520 | 467 | 518 | 2466 | 610 | 584 |
| 2300 | W303.asm02.HP0_G0020820 | chrVII | 1186 | 1467 | 1604 | 1000 | 1378 | 1572 | 996 | 1382 | 2152 | 2921 | 1288 | 1265 | 3770 | 887 | 1080 | 3387 | 1186 | 1129 |
| 2301 | W303.asm02.HP0_G0020830 | chrVII | 4518 | 5450 | 5538 | 4200 | 4978 | 5562 | 4346 | 5586 | 7493 | 12208 | 4207 | 4069 | 13768 | 3000 | 3139 | 13575 | 3846 | 3752 |
| 2302 | W303.asm02.HP0_G0020840 | chrVII | 608 | 1582 | 2323 | 528 | 2240 | 2209 | 666 | 1495 | 2508 | 5954 | 1449 | 1804 | 6240 | 1055 | 1687 | 6301 | 1228 | 1505 |
| 2303 | W303.asm02.HP0_G0020850 | chrVII | 254 | 196 | 199 | 234 | 205 | 182 | 235 | 213 | 205 | 914 | 224 | 220 | 632 | 128 | 148 | 885 | 187 | 220 |
| 2304 | W303.asm02.HP0_G0020860 | chrVII | 357 | 326 | 242 | 480 | 204 | 262 | 315 | 319 | 244 | 922 | 311 | 450 | 773 | 193 | 251 | 846 | 343 | 384 |
| 2305 | W303.asm02.HP0_G0020870 | chrVII | 6882 | 1614 | 1260 | 6569 | 1127 | 1211 | 6504 | 1563 | 1229 | 5170 | 1520 | 1434 | 5173 | 1019 | 1221 | 4956 | 1468 | 1295 |
| 2306 | W303.asm02.HP0_G0020880 | chrVII | 5652 | 854 | 795 | 5139 | 825 | 857 | 5141 | 783 | 783 | 3461 | 905 | 1370 | 3709 | 709 | 940 | 2842 | 933 | 1079 |
| 2307 | W303.asm02.HP0_G0020890 | chrVII | 2109 | 520 | 340 | 2316 | 383 | 380 | 2149 | 432 | 375 | 2856 | 740 | 835 | 2215 | 496 | 579 | 2475 | 669 | 806 |
| 2308 | W303.asm02.HP0_G0020900 | chrVII | 9645 | 2238 | 2459 | 9434 | 2365 | 2317 | 9247 | 2411 | 2733 | 24504 | 3576 | 5461 | 19037 | 2538 | 4036 | 19531 | 3195 | 5022 |
| 2309 | W303.asm02.HP0_G0020910 | chrVII | 1815 | 1300 | 1544 | 1588 | 1378 | 1648 | 1797 | 1356 | 1961 | 3892 | 959 | 1174 | 4105 | 615 | 834 | 4637 | 930 | 1104 |
| 2310 | W303.asm02.HP0_G0020920 | chrVII | 7761 | 6254 | 8838 | 7428 | 8424 | 8470 | 8152 | 6150 | 10826 | 19833 | 5978 | 7219 | 20826 | 4426 | 5618 | 20506 | 5102 | 6462 |
| 2311 | W303.asm02.HP0_G0020930 | chrVII | 5136 | 2344 | 2035 | 4826 | 2255 | 2016 | 4731 | 2266 | 2437 | 12426 | 3582 | 3719 | 9488 | 2400 | 2847 | 9517 | 3402 | 3480 |
| 2312 | W303.asm02.HP0_G0020940 | chrVII | 1541 | 1321 | 1268 | 1498 | 1250 | 1282 | 1521 | 1271 | 1526 | 3645 | 958 | 942 | 4367 | 654 | 778 | 4103 | 915 | 769 |
| 2313 | W303.asm02.HP0_G0020950 | chrVII | 1632 | 1004 | 930 | 1565 | 709 | 842 | 1562 | 827 | 819 | 2944 | 652 | 796 | 2834 | 397 | 542 | 3044 | 680 | 716 |
| 2314 | W303.asm02.HP0_G0020960 | chrVII | 32 | 37 | 24 | 34 | 21 | 28 | 26 | 46 | 19 | 117 | 37 | 27 | 68 | 23 | 20 | 101 | 44 | 24 |
| 2315 | W303.asm02.HP0_G0020970 | chrVII | 1387 | 532 | 589 | 1484 | 530 | 612 | 1398 | 435 | 709 | 2273 | 934 | 985 | 1039 | 736 | 699 | 1010 | 891 | 840 |
| 2316 | W303.asm02.HP0_G0020980 | chrVII | 2376 | 1887 | 2304 | 2276 | 2487 | 1939 | 2531 | 1951 | 2461 | 10767 | 1951 | 2065 | 9574 | 1147 | 1299 | 10593 | 1673 | 1804 |
| 2317 | W303.asm02.HP0_G0020990 | chrVII | 21870 | 7523 | 5567 | 22357 | 5671 | 5692 | 23718 | 7269 | 6309 | 32768 | 10336 | 11767 | 26758 | 7419 | 8217 | 26589 | 8776 | 10344 |
| 2318 | W303.asm02.HP0_G0021000 | chrVII | 2615 | 997 | 1206 | 2727 | 1232 | 1137 | 2473 | 1113 | 1371 | 6420 | 1152 | 1704 | 5712 | 862 | 1447 | 5585 | 1161 | 1393 |
| 2319 | W303.asm02.HP0_G0021010 | chrVII | 679 | 112 | 137 | 673 | 159 | 152 | 559 | 101 | 154 | 1592 | 231 | 253 | 1291 | 116 | 220 | 1184 | 236 | 234 |
| 2320 | W303.asm02.HP0_G0021020 | chrVII | 2358 | 363 | 325 | 1973 | 305 | 338 | 1949 | 332 | 349 | 1798 | 406 | 502 | 1512 | 248 | 459 | 1287 | 380 | 414 |
| 2321 | W303.asm02.HP0_G0021030 | chrVII | 2870 | 2266 | 2356 | 2917 | 2832 | 2055 | 3341 | 2535 | 2145 | 15379 | 2694 | 3238 | 13139 | 1705 | 1854 | 13700 | 2345 | 2746 |
| 2322 | W303.asm02.HP0_G0021040 | chrVII | 622 | 844 | 772 | 582 | 870 | 788 | 646 | 828 | 935 | 2417 | 700 | 646 | 2467 | 414 | 416 | 2415 | 647 | 570 |
| 2323 | W303.asm02.HP0_G0021050 | chrVII | 17852 | 1314 | 1316 | 17935 | 1228 | 1555 | 16929 | 1131 | 1553 | 6862 | 1947 | 2283 | 3835 | 1518 | 2241 | 3978 | 1764 | 2015 |

| | | | | | | | | | | | | | | | | | | | | |
|---|---|---|---|---|---|---|---|---|---|---|---|---|---|---|---|---|---|---|---|---|
| 2324 | W303.asm02.HP0_G0021060 | chrVII | 415 | 198 | 172 | 365 | 148 | 194 | 348 | 189 | 205 | 684 | 151 | 137 | 715 | 98 | 114 | 863 | 190 | 128 |
| 2325 | W303.asm02.HP0_G0021070 | chrVII | 13700 | 4306 | 5339 | 13620 | 4546 | 5081 | 13628 | 4170 | 6375 | 12392 | 3890 | 4572 | 13880 | 2745 | 3930 | 11896 | 3495 | 4018 |
| 2326 | W303.asm02.HP0_G0021080 | chrVII | 1197 | 326 | 255 | 1002 | 324 | 304 | 1062 | 338 | 289 | 1237 | 346 | 378 | 1126 | 253 | 319 | 1195 | 354 | 367 |
| 2327 | W303.asm02.HP0_G0021090 | chrVII | 1094 | 466 | 561 | 1100 | 588 | 598 | 1017 | 500 | 621 | 4459 | 756 | 1233 | 3166 | 418 | 791 | 3664 | 727 | 1067 |
| 2328 | W303.asm02.HP0_G0021100 | chrVII | 8939 | 5406 | 5615 | 8810 | 5051 | 5754 | 8528 | 5377 | 6646 | 11506 | 4541 | 4918 | 13345 | 3230 | 4213 | 11810 | 3880 | 4192 |
| 2329 | W303.asm02.HP0_G0021110 | chrVII | 11347 | 2984 | 2137 | 11883 | 2073 | 2064 | 11513 | 2928 | 2289 | 9982 | 3879 | 3453 | 11206 | 2924 | 3307 | 8841 | 3378 | 3224 |
| 2330 | W303.asm02.HP0_G0021120 | chrVII | 2947 | 226 | 605 | 2810 | 506 | 702 | 2572 | 258 | 866 | 2343 | 667 | 1325 | 1532 | 662 | 869 | 928 | 542 | 936 |
| 2331 | W303.asm02.HP0_G0021130 | chrVII | 1655 | 365 | 472 | 1725 | 501 | 462 | 1870 | 368 | 449 | 4609 | 553 | 908 | 3865 | 461 | 667 | 3687 | 525 | 854 |
| 2332 | W303.asm02.HP0_G0021140 | chrVII | 3125 | 498 | 440 | 3070 | 389 | 435 | 2650 | 503 | 553 | 2941 | 983 | 888 | 1880 | 701 | 760 | 1801 | 841 | 844 |
| 2333 | W303.asm02.HP0_G0021150 | chrVII | 16030 | 4633 | 1984 | 19970 | 3126 | 3512 | 13933 | 2535 | 1106 | 4208 | 3105 | 1408 | 3059 | 1146 | 1514 | 5228 | 3230 | 1333 |
| 2334 | W303.asm02.HP0_G0021160 | chrVII | 4429 | 611 | 664 | 4114 | 738 | 717 | 4047 | 591 | 779 | 4939 | 1150 | 1246 | 4332 | 797 | 1228 | 3341 | 1176 | 1188 |
| 2335 | W303.asm02.HP0_G0021170 | chrVII | 136 | 137 | 85 | 142 | 79 | 88 | 120 | 131 | 100 | 265 | 147 | 107 | 346 | 74 | 62 | 283 | 169 | 101 |
| 2336 | W303.asm02.HP0_G0021180 | chrVII | 1429 | 1532 | 1457 | 1470 | 2084 | 1367 | 1672 | 1791 | 1386 | 12534 | 2173 | 2307 | 10210 | 1296 | 1070 | 10420 | 1855 | 1987 |
| 2337 | W303.asm02.HP0_G0021190 | chrVII | 592 | 973 | 1058 | 674 | 1000 | 1304 | 561 | 825 | 957 | 3603 | 952 | 1091 | 3257 | 495 | 597 | 2517 | 823 | 973 |
| 2338 | W303.asm02.HP0_G0021200 | chrVII | 59 | 45 | 29 | 64 | 38 | 28 | 47 | 49 | 51 | 110 | 51 | 19 | 133 | 29 | 20 | 100 | 45 | 26 |
| 2339 | W303.asm02.HP0_G0021210 | chrVII | 4851 | 1867 | 1757 | 4696 | 1775 | 1698 | 4787 | 1833 | 1989 | 6706 | 1880 | 2041 | 6867 | 1279 | 1512 | 6157 | 1898 | 1875 |
| 2340 | W303.asm02.HP0_G0021220 | chrVII | 11103 | 6827 | 10404 | 10506 | 10622 | 10459 | 11136 | 7371 | 12680 | 26806 | 6721 | 8961 | 30236 | 5093 | 7723 | 28193 | 5873 | 8021 |
| 2341 | W303.asm02.HP0_G0021230 | chrVII | 4530 | 1139 | 856 | 4805 | 743 | 840 | 4407 | 1046 | 839 | 3452 | 1039 | 1057 | 4724 | 834 | 1178 | 3180 | 1026 | 1045 |
| 2342 | W303.asm02.HP0_G0021240 | chrVII | 7825 | 1864 | 2581 | 7714 | 2563 | 2411 | 7785 | 2067 | 2587 | 10960 | 1859 | 2601 | 11504 | 1339 | 1973 | 10279 | 1761 | 2294 |
| 2343 | W303.asm02.HP0_G0021250 | chrVII | 1236 | 1735 | 2114 | 1120 | 2020 | 2016 | 1325 | 1774 | 2277 | 6500 | 1431 | 1740 | 7028 | 902 | 1219 | 6447 | 1291 | 1598 |
| 2344 | W303.asm02.HP0_G0021260 | chrVII | 823 | 364 | 400 | 736 | 361 | 375 | 691 | 403 | 523 | 1079 | 315 | 314 | 2370 | 288 | 358 | 1120 | 248 | 307 |
| 2345 | W303.asm02.HP0_G0021270 | chrVII | 1076 | 544 | 580 | 857 | 503 | 609 | 989 | 506 | 633 | 1572 | 461 | 449 | 1723 | 275 | 380 | 1504 | 452 | 376 |
| 2346 | W303.asm02.HP0_G0021280 | chrVII | 17776 | 11003 | 9253 | 14340 | 9478 | 9024 | 17306 | 11048 | 8947 | 35427 | 12840 | 13479 | 29751 | 8586 | 9108 | 31758 | 11043 | 11306 |
| 2347 | W303.asm02.HP0_G0021290 | chrVII | 2797 | 378 | 317 | 2478 | 322 | 412 | 2789 | 365 | 468 | 1452 | 524 | 729 | 837 | 363 | 384 | 1139 | 470 | 543 |
| 2348 | W303.asm02.HP0_G0021300 | chrVII | 12008 | 4384 | 5678 | 11713 | 4819 | 5584 | 11937 | 4850 | 6420 | 23015 | 5190 | 7028 | 22741 | 4155 | 6192 | 24454 | 4504 | 6081 |
| 2349 | W303.asm02.HP0_G0021310 | chrVII | 1380 | 1638 | 2209 | 1425 | 2250 | 2166 | 1665 | 1774 | 2695 | 6148 | 1499 | 1762 | 7218 | 955 | 1127 | 6268 | 1319 | 1478 |
| 2350 | W303.asm02.HP0_G0021320 | chrVII | 2418 | 4633 | 7697 | 2391 | 9399 | 6569 | 3010 | 5286 | 8060 | 34642 | 7264 | 8356 | 33840 | 4145 | 4519 | 31595 | 5942 | 7008 |
| 2351 | W303.asm02.HP0_G0021330 | chrVII | 4060 | 875 | 881 | 4527 | 559 | 997 | 3619 | 566 | 812 | 2649 | 1075 | 1635 | 3814 | 776 | 1580 | 1631 | 993 | 1376 |
| 2352 | W303.asm02.HP0_G0021340 | chrVII | 17 | 14 | 10 | 10 | 8 | 10 | 11 | 12 | 6 | 19 | 18 | 6 | 34 | 9 | 4 | 13 | 14 | 6 |
| 2353 | W303.asm02.HP0_G0021350 | chrVII | 1130 | 931 | 1468 | 1250 | 971 | 1492 | 861 | 854 | 1798 | 1394 | 674 | 1073 | 2307 | 522 | 897 | 1318 | 653 | 858 |
| 2354 | W303.asm02.HP0_G0021360 | chrVII | 17988 | 2369 | 3102 | 17483 | 2814 | 3379 | 19004 | 1967 | 3934 | 28163 | 5889 | 8006 | 11694 | 4364 | 4737 | 11788 | 4869 | 6986 |
| 2355 | W303.asm02.HP0_G0021370 | chrVII | 2588 | 1477 | 1231 | 2370 | 1198 | 1415 | 2382 | 1542 | 1248 | 4639 | 1484 | 1632 | 4236 | 1000 | 1226 | 4399 | 1435 | 1515 |
| 2356 | W303.asm02.HP0_G0021380 | chrVII | 9335 | 2689 | 2235 | 8644 | 2168 | 2166 | 8919 | 2830 | 2307 | 8664 | 2731 | 2964 | 8601 | 2130 | 2366 | 7903 | 2702 | 2712 |
| 2357 | W303.asm02.HP0_G0021390 | chrVII | 1554 | 945 | 1065 | 1429 | 1124 | 960 | 1499 | 951 | 1143 | 5975 | 995 | 1144 | 8355 | 818 | 1023 | 5629 | 1010 | 1011 |
| 2358 | W303.asm02.HP0_G0021400 | chrVII | 8919 | 2921 | 3029 | 8648 | 2819 | 2981 | 8267 | 3106 | 2722 | 10914 | 3060 | 3955 | 12148 | 2414 | 3312 | 9950 | 2607 | 3514 |
| 2359 | W303.asm02.HP0_G0021410 | chrVII | 1951 | 1145 | 882 | 1671 | 724 | 955 | 1824 | 1051 | 1142 | 2421 | 839 | 714 | 3081 | 671 | 552 | 2647 | 888 | 628 |
| 2360 | W303.asm02.HP0_G0021420 | chrVII | 2045 | 956 | 548 | 1877 | 632 | 568 | 1978 | 824 | 621 | 3034 | 881 | 792 | 2567 | 482 | 528 | 2766 | 920 | 742 |
| 2361 | W303.asm02.HP0_G0021430 | chrVII | 2122 | 2314 | 2436 | 1802 | 2628 | 2374 | 1690 | 2591 | 3168 | 5897 | 2441 | 1853 | 7449 | 1480 | 1723 | 6371 | 1946 | 1726 |
| 2362 | W303.asm02.HP0_G0021440 | chrVII | 3890 | 721 | 807 | 3712 | 803 | 929 | 3801 | 685 | 957 | 4294 | 1075 | 1784 | 2615 | 846 | 1083 | 2781 | 894 | 1470 |
| 2363 | W303.asm02.HP0_G0021450 | chrVII | 3352 | 801 | 785 | 3294 | 699 | 779 | 3262 | 749 | 849 | 5008 | 1004 | 1452 | 4157 | 678 | 936 | 4107 | 991 | 1178 |
| 2364 | W303.asm02.HP0_G0021460 | chrVII | 1873 | 1461 | 1496 | 1705 | 1127 | 1390 | 1794 | 1483 | 1553 | 4304 | 1398 | 1662 | 4378 | 967 | 1172 | 4419 | 1417 | 1462 |
| 2365 | W303.asm02.HP0_G0021470 | chrVII | 1565 | 2395 | 1548 | 1366 | 1761 | 1411 | 1525 | 2386 | 1864 | 6938 | 2458 | 1892 | 6990 | 1624 | 1263 | 7152 | 2379 | 1723 |
| 2366 | W303.asm02.HP0_G0021480 | chrVII | 4280 | 651 | 622 | 3483 | 622 | 603 | 3409 | 648 | 779 | 2088 | 735 | 825 | 2558 | 593 | 811 | 1817 | 677 | 781 |
| 2367 | W303.asm02.HP0_G0021490 | chrVII | 10280 | 5438 | 6599 | 11055 | 6386 | 5824 | 12950 | 5228 | 6543 | 27369 | 5976 | 7903 | 22443 | 3873 | 5548 | 24103 | 5825 | 7172 |
| 2368 | W303.asm02.HP0_G0021500 | chrVII | 2518 | 569 | 369 | 2767 | 471 | 396 | 3162 | 545 | 430 | 5041 | 812 | 727 | 3276 | 497 | 517 | 4043 | 751 | 743 |
| 2369 | W303.asm02.HP0_G0021510 | chrVII | 16693 | 1854 | 1762 | 15213 | 1497 | 1931 | 16305 | 1573 | 2337 | 12154 | 3676 | 3949 | 5401 | 2931 | 2621 | 5455 | 3241 | 3539 |
| 2370 | W303.asm02.HP0_G0021520 | chrVII | 1246 | 535 | 376 | 1292 | 421 | 380 | 1293 | 488 | 291 | 1890 | 488 | 541 | 1763 | 343 | 448 | 1801 | 491 | 460 |
| 2371 | W303.asm02.HP0_G0021530 | chrVII | 16477 | 3994 | 3168 | 16431 | 3211 | 3277 | 17055 | 4047 | 2847 | 17566 | 4951 | 5421 | 17694 | 4005 | 4366 | 15292 | 4380 | 4578 |
| 2372 | W303.asm02.HP0_G0021540 | chrVII | 7 | 1 | 0 | 9 | 3 | 1 | 5 | 1 | 0 | 8 | 8 | 6 | 3 | 5 | 2 | 1 | 10 | 4 |
| 2373 | W303.asm02.HP0_G0021550 | chrVII | 2295 | 832 | 921 | 2115 | 846 | 993 | 2060 | 840 | 1111 | 3615 | 990 | 1329 | 4352 | 709 | 1075 | 3829 | 928 | 1175 |
| 2374 | W303.asm02.HP0_G0021560 | chrVII | 3102 | 4055 | 4278 | 2961 | 4109 | 4254 | 2815 | 4063 | 5644 | 7652 | 3378 | 2959 | 9021 | 2451 | 2587 | 7839 | 2780 | 2465 |
| 2375 | W303.asm02.HP0_G0021570 | chrVII | 8885 | 1028 | 1370 | 8988 | 1288 | 1379 | 8653 | 1078 | 1607 | 10972 | 1587 | 2889 | 8016 | 1275 | 2618 | 7712 | 1307 | 2608 |
| 2376 | W303.asm02.HP0_G0021580 | chrVII | 6599 | 3631 | 3307 | 5803 | 3300 | 3161 | 6663 | 3559 | 2995 | 16967 | 4401 | 5484 | 14263 | 3050 | 3655 | 14715 | 3793 | 4884 |
| 2377 | W303.asm02.HP0_G0021590 | chrVII | 8582 | 1432 | 1279 | 8341 | 1434 | 1346 | 8506 | 1411 | 1567 | 10046 | 2395 | 2719 | 6579 | 1654 | 2110 | 6820 | 2181 | 2562 |
| 2378 | W303.asm02.HP0_G0021600 | chrVII | 1059 | 1937 | 2414 | 1056 | 2754 | 2097 | 1047 | 2011 | 2629 | 10198 | 1749 | 2277 | 10324 | 1097 | 1475 | 10267 | 1583 | 2001 |
| 2379 | W303.asm02.HP0_G0021610 | chrVII | 478 | 177 | 290 | 427 | 323 | 374 | 364 | 164 | 333 | 1425 | 269 | 462 | 1170 | 150 | 316 | 1100 | 221 | 461 |
| 2380 | W303.asm02.HP0_G0021620 | chrVII | 10620 | 2242 | 2841 | 11398 | 2457 | 2981 | 11470 | 2416 | 3683 | 18412 | 4937 | 7169 | 20442 | 5550 | 5746 | 11317 | 4120 | 6263 |
| 2381 | W303.asm02.HP0_G0021630 | chrVII | 2639 | 1700 | 1753 | 2669 | 1803 | 1678 | 2705 | 1621 | 1973 | 14392 | 3300 | 3516 | 9557 | 2048 | 2050 | 9223 | 2631 | 2987 |
| 2382 | W303.asm02.HP0_G0021640 | chrVII | 96467 | 51490 | 47028 | 97985 | 43323 | 47397 | 102842 | 47577 | 50773 | 118374 | 71989 | 80322 | 108253 | 61373 | 66572 | 102943 | 64392 | 76361 |
| 2383 | W303.asm02.HP0_G0021650 | chrVII | 7043 | 1739 | 1454 | 7497 | 1456 | 1562 | 7123 | 1716 | 1438 | 8332 | 2277 | 2722 | 7739 | 1551 | 1996 | 7200 | 2023 | 2438 |
| 2384 | W303.asm02.HP0_G0021660 | chrVII | 7081 | 372 | 289 | 6536 | 257 | 367 | 6331 | 312 | 385 | 2161 | 686 | 686 | 1306 | 514 | 631 | 1170 | 721 | 602 |
| 2385 | W303.asm02.HP0_G0021670 | chrVII | 2303 | 1218 | 1114 | 2039 | 1293 | 1156 | 2220 | 1248 | 1087 | 4824 | 1108 | 1516 | 4632 | 805 | 1082 | 4361 | 1047 | 1310 |
| 2386 | W303.asm02.HP0_G0021680 | chrVII | 3178 | 578 | 525 | 3195 | 599 | 566 | 3134 | 537 | 492 | 3831 | 894 | 1160 | 3162 | 504 | 774 | 2993 | 768 | 1156 |
| 2387 | W303.asm02.HP0_G0021690 | chrVII | 1214 | 489 | 753 | 1194 | 582 | 865 | 1036 | 481 | 929 | 3862 | 1027 | 1231 | 2343 | 844 | 1027 | 2336 | 934 | 1058 |
| 2388 | W303.asm02.HP0_G0021700 | chrVII | 3107 | 1114 | 1670 | 3171 | 1738 | 1627 | 2952 | 1253 | 1727 | 11236 | 1537 | 2532 | 10110 | 1084 | 1869 | 10698 | 1456 | 2201 |
| 2389 | W303.asm02.HP0_G0021710 | chrVII | 2283 | 949 | 889 | 2090 | 825 | 872 | 1981 | 981 | 987 | 6238 | 1537 | 1801 | 3926 | 1008 | 1314 | 4619 | 1384 | 1735 |
| 2390 | W303.asm02.HP0_G0021720 | chrVII | 3158 | 1278 | 1219 | 3057 | 1474 | 1170 | 3344 | 1238 | 1544 | 13035 | 2583 | 3038 | 8045 | 1831 | 1772 | 8806 | 2246 | 2740 |
| 2391 | W303.asm02.HP0_G0021730 | chrVII | 2137 | 655 | 588 | 1914 | 548 | 651 | 1797 | 732 | 669 | 1493 | 527 | 520 | 1574 | 392 | 422 | 1514 | 525 | 543 |
| 2392 | W303.asm02.HP0_G0021740 | chrVII | 6676 | 1043 | 913 | 5976 | 816 | 983 | 5996 | 909 | 1107 | 6890 | 1732 | 1969 | 5882 | 1153 | 1560 | 5452 | 1635 | 1655 |
| 2393 | W303.asm02.HP0_G0021750 | chrVII | 674 | 607 | 638 | 526 | 570 | 591 | 625 | 585 | 745 | 1484 | 551 | 486 | 1482 | 382 | 470 | 1457 | 492 | 490 |
| 2394 | W303.asm02.HP0_G0021760 | chrVII | 13247 | 2366 | 2538 | 9907 | 2094 | 3220 | 13281 | 1688 | 3491 | 9243 | 3747 | 5434 | 2230 | 3301 | 3172 | 1990 | 2942 | 4478 |
| 2395 | W303.asm02.HP0_G0021770 | chrVII | 1797 | 582 | 376 | 1898 | 434 | 332 | 1669 | 513 | 357 | 1349 | 390 | 253 | 1602 | 284 | 311 | 1403 | 390 | 253 |
| 2396 | W303.asm02.HP0_G0021780 | chrVII | 688 | 402 | 448 | 572 | 359 | 374 | 558 | 378 | 611 | 998 | 271 | 287 | 1176 | 145 | 260 | 1122 | 286 | 231 |
| 2397 | W303.asm02.HP0_G0021790 | chrVII | 2980 | 2207 | 2171 | 2680 | 1865 | 2096 | 2665 | 2123 | 2415 | 4968 | 1750 | 1927 | 5702 | 1256 | 1491 | 4985 | 1586 | 1631 |
| 2398 | W303.asm02.HP0_G0021800 | chrVII | 1061 | 1315 | 1573 | 843 | 1505 | 1431 | 1119 | 1362 | 1619 | 5236 | 1174 | 1555 | 5500 | 686 | 1126 | 5233 | 1036 | 1403 |
| 2399 | W303.asm02.HP0_G0021810 | chrVII | 4560 | 2520 | 3889 | 3822 | 2800 | 3856 | 4010 | 2443 | 5079 | 4125 | 2482 | 2904 | 5481 | 1724 | 2581 | 4552 | 2148 | 2389 |
| 2400 | W303.asm02.HP0_G0021820 | chrVII | 4516 | 1571 | 1044 | 3486 | 1128 | 1187 | 4066 | 1230 | 1211 | 6487 | 2662 | 2226 | 2508 | 1899 | 1338 | 2934 | 2535 | 2043 |
| 2401 | W303.asm02.HP0_G0021830 | chrVII | 8781 | 3673 | 2221 | 10910 | 2792 | 2825 | 7457 | 2646 | 1928 | 15951 | 4439 | 4202 | 11012 | 2711 | 2700 | 13060 | 4267 | 3784 |
| 2402 | W303.asm02.HP0_G0021840 | chrVII | 1131 | 328 | 240 | 1202 | 293 | 295 | 1104 | 304 | 257 | 1615 | 297 | 417 | 1521 | 255 | 311 | 1301 | 300 | 332 |
| 2403 | W303.asm02.HP0_G0021850 | chrVII | 900 | 238 | 203 | 778 | 160 | 201 | 745 | 206 | 175 | 420 | 126 | 159 | 661 | 136 | 150 | 344 | 146 | 137 |
| 2404 | W303.asm02.HP0_G0021860 | chrVII | 16329 | 9597 | 9151 | 14956 | 10007 | 9755 | 17117 | 9998 | 8644 | 28137 | 9841 | 11759 | 26215 | 7359 | 8389 | 24465 | 7913 | 10432 |
| 2405 | W303.asm02.HP0_G0021870 | chrVII | 2085 | 1135 | 1575 | 1830 | 1341 | 1615 | 1725 | 1136 | 2041 | 3069 | 980 | 1257 | 4095 | 695 | 1110 | 3282 | 894 | 1137 |
| 2406 | W303.asm02.HP0_G0021880 | chrVII | 1887 | 884 | 898 | 1913 | 870 | 876 | 1888 | 976 | 1035 | 4843 | 1021 | 1053 | 4758 | 702 | 833 | 4606 | 910 | 965 |
| 2407 | W303.asm02.HP0_G0021890 | chrVII | 1062 | 310 | 551 | 1028 | 541 | 537 | 965 | 340 | 679 | 4682 | 553 | 1181 | 3365 | 475 | 894 | 2889 | 480 | 1051 |
| 2408 | W303.asm02.HP0_G0021900 | chrVII | 8095 | 1857 | 2101 | 7886 | 2031 | 2150 | 7891 | 1951 | 2702 | 17718 | 3683 | 4548 | 12538 | 2518 | 3532 | 14068 | 3390 | 4061 |
| 2409 | W303.asm02.HP0_G0021910 | chrVII | 5308 | 2003 | 2561 | 4991 | 2380 | 2510 | 4988 | 2167 | 3103 | 6467 | 1729 | 2411 | 7847 | 1385 | 1968 | 6337 | 1640 | 2108 |
| 2410 | W303.asm02.HP0_G0021920 | chrVII | 1260 | 294 | 294 | 1087 | 276 | 291 | 1175 | 262 | 345 | 1683 | 436 | 510 | 1616 | 291 | 396 | 1298 | 381 | 493 |
| 2411 | W303.asm02.HP0_G0021930 | chrVII | 1858 | 1100 | 900 | 1620 | 776 | 996 | 1576 | 1016 | 1116 | 2049 | 756 | 733 | 2460 | 469 | 727 | 2262 | 781 | 736 |
| 2412 | W303.asm02.HP0_G0021940 | chrVII | 1932 | 1005 | 701 | 2033 | 786 | 731 | 1871 | 875 | 729 | 2694 | 986 | 882 | 2864 | 672 | 808 | 2467 | 882 | 792 |
| 2413 | W303.asm02.HP0_G0021950 | chrVII | 671 | 119 | 130 | 587 | 125 | 136 | 764 | 94 | 162 | 1288 | 251 | 285 | 1340 | 194 | 245 | 844 | 195 | 209 |
| 2414 | W303.asm02.HP0_G0021960 | chrVII | 470 | 23 | 32 | 394 | 50 | 38 | 403 | 38 | 35 | 194 | 49 | 59 | 295 | 47 | 61 | 163 | 43 | 65 |
| 2415 | W303.asm02.HP0_G0021970 | chrVII | 1680 | 486 | 595 | 1627 | 473 | 620 | 1778 | 454 | 742 | 4831 | 1121 | 1295 | 3512 | 786 | 921 | 3671 | 931 | 1197 |
| 2416 | W303.asm02.HP0_G0021980 | chrVII | 4696 | 3365 | 3882 | 4944 | 4098 | 3622 | 5097 | 3598 | 3732 | 18371 | 4196 | 4787 | 17108 | 2858 | 3592 | 16051 | 3681 | 4301 |

| | | | | | | | | | | | | | | | | | | | | |
|---|---|---|---|---|---|---|---|---|---|---|---|---|---|---|---|---|---|---|---|---|
| 2417 | W303.asm02.HP0_G0021990 | chrVII | 494 | 498 | 579 | 447 | 531 | 627 | 454 | 483 | 800 | 1212 | 384 | 372 | 1948 | 293 | 399 | 1296 | 331 | 263 |
| 2418 | W303.asm02.HP0_G0022000 | chrVII | 14072 | 4495 | 4301 | 14471 | 3909 | 4384 | 14343 | 4300 | 4835 | 10835 | 4046 | 4607 | 12310 | 3364 | 4243 | 10030 | 3572 | 4481 |
| 2419 | W303.asm02.HP0_G0022010 | chrVII | 1554 | 1005 | 1341 | 1223 | 1061 | 1444 | 1237 | 1048 | 1770 | 3053 | 814 | 1126 | 3803 | 637 | 950 | 3254 | 773 | 1002 |
| 2420 | W303.asm02.HP0_G0022020 | chrVII | 3175 | 1047 | 1355 | 3232 | 1494 | 1445 | 3221 | 1090 | 1357 | 9541 | 1858 | 2839 | 8278 | 1437 | 2118 | 7292 | 1507 | 2454 |
| 2421 | W303.asm02.HP0_G0022030 | chrVII | 39611 | 26942 | 26795 | 43830 | 25998 | 26068 | 43739 | 25794 | 27314 | 54991 | 27572 | 25133 | 58092 | 19598 | 20195 | 54609 | 23032 | 22397 |
| 2422 | W303.asm02.HP0_G0022040 | chrVII | 4732 | 2170 | 2444 | 4224 | 2122 | 2387 | 4188 | 2217 | 2876 | 5282 | 1798 | 2101 | 6138 | 1254 | 1717 | 5248 | 1655 | 1754 |
| 2423 | W303.asm02.HP0_G0022050 | chrVII | 916 | 60 | 42 | 952 | 29 | 36 | 780 | 44 | 31 | 81 | 43 | 23 | 102 | 25 | 23 | 88 | 43 | 13 |
| 2424 | W303.asm02.HP0_G0022060 | chrVII | 5690 | 505 | 565 | 5611 | 452 | 541 | 5299 | 538 | 587 | 3700 | 807 | 1125 | 3329 | 482 | 941 | 3185 | 785 | 1057 |
| 2425 | W303.asm02.HP0_G0022070 | chrVII | 2975 | 242 | 327 | 2641 | 273 | 351 | 2673 | 248 | 338 | 2321 | 452 | 589 | 1356 | 264 | 478 | 1736 | 381 | 569 |
| 2426 | W303.asm02.HP0_G0022080 | chrVII | 675 | 412 | 439 | 524 | 435 | 443 | 564 | 368 | 488 | 2040 | 556 | 739 | 1454 | 413 | 443 | 1382 | 490 | 594 |
| 2427 | W303.asm02.HP0_G0022090 | chrVII | 42103 | 6495 | 6873 | 39573 | 6230 | 7453 | 43192 | 6098 | 8659 | 52207 | 12728 | 17352 | 36133 | 10516 | 12672 | 34206 | 11187 | 16140 |
| 2428 | W303.asm02.HP0_G0022100 | chrVII | 188 | 171 | 253 | 186 | 205 | 243 | 163 | 204 | 369 | 644 | 192 | 180 | 793 | 110 | 118 | 765 | 134 | 160 |
| 2429 | W303.asm02.HP0_G0022110 | chrVII | 4585 | 670 | 682 | 4439 | 562 | 760 | 4589 | 520 | 829 | 5915 | 1759 | 1614 | 3328 | 1211 | 1364 | 2984 | 1515 | 1472 |
| 2430 | W303.asm02.HP0_G0022120 | chrVII | 584 | 3150 | 323 | 673 | 348 | 333 | 501 | 2163 | 463 | 849 | 1589 | 224 | 1314 | 773 | 206 | 858 | 1957 | 184 |
| 2431 | W303.asm02.HP0_G0022130 | chrVII | 9940 | 4323 | 3834 | 10485 | 3315 | 3828 | 9408 | 4265 | 3951 | 9280 | 4068 | 5134 | 11870 | 3467 | 4577 | 8350 | 3546 | 4446 |
| 2432 | W303.asm02.HP0_G0022140 | chrVII | 2476 | 1478 | 1553 | 2410 | 2231 | 1433 | 2954 | 1609 | 1759 | 18013 | 2791 | 3229 | 13572 | 1744 | 1651 | 13385 | 2227 | 2755 |
| 2433 | W303.asm02.HP0_G0022150 | chrVII | 2320 | 788 | 713 | 2273 | 740 | 675 | 2410 | 769 | 632 | 3947 | 871 | 1089 | 3591 | 566 | 691 | 3348 | 818 | 1005 |
| 2434 | W303.asm02.HP0_G0022160 | chrVII | 1823 | 405 | 431 | 1724 | 395 | 370 | 1559 | 365 | 463 | 1809 | 396 | 490 | 1830 | 263 | 394 | 1703 | 333 | 457 |
| 2435 | W303.asm02.HP0_G0022170 | chrVII | 6084 | 343 | 172 | 5005 | 130 | 186 | 4921 | 246 | 148 | 313 | 301 | 251 | 438 | 172 | 238 | 256 | 314 | 197 |
| 2436 | W303.asm02.HP0_G0022180 | chrVII | 2587 | 38 | 69 | 3572 | 58 | 66 | 2310 | 30 | 71 | 692 | 72 | 107 | 324 | 60 | 62 | 201 | 80 | 74 |
| 2437 | W303.asm02.HP0_G0022190 | chrVII | 7724 | 1456 | 1441 | 8268 | 1606 | 1612 | 6749 | 1384 | 1440 | 8400 | 2045 | 2349 | 7001 | 1493 | 1781 | 6399 | 1826 | 2051 |
| 2438 | W303.asm02.HP0_G0022200 | chrVII | 563 | 643 | 570 | 565 | 694 | 611 | 675 | 625 | 605 | 2425 | 550 | 553 | 2469 | 349 | 326 | 2207 | 494 | 490 |
| 2439 | W303.asm02.HP0_G0022210 | chrVII | 2712 | 1400 | 1138 | 2429 | 1077 | 1080 | 2582 | 1264 | 1061 | 4543 | 1199 | 1237 | 4574 | 800 | 1008 | 4455 | 1112 | 1130 |
| 2440 | W303.asm02.HP0_G0022220 | chrVII | 8058 | 4485 | 4884 | 7741 | 4367 | 5097 | 7713 | 4374 | 5801 | 10497 | 3578 | 4167 | 11365 | 2567 | 3525 | 10473 | 3312 | 3778 |
| 2441 | W303.asm02.HP0_G0022230 | chrVII | 64308 | 7207 | 7608 | 63302 | 5750 | 7796 | 66364 | 6237 | 8525 | 19598 | 11280 | 14551 | 17778 | 10397 | 14825 | 13020 | 10717 | 13099 |
| 2442 | W303.asm02.HP0_G0022240 | chrVII | 6854 | 1185 | 1597 | 5867 | 1079 | 1592 | 6353 | 1107 | 1809 | 6266 | 1526 | 2677 | 6360 | 1076 | 2069 | 5155 | 1603 | 2480 |
| 2443 | W303.asm02.HP0_G0022250 | chrVII | 1561 | 424 | 217 | 1469 | 204 | 228 | 1528 | 317 | 213 | 651 | 475 | 247 | 760 | 212 | 181 | 510 | 445 | 252 |
| 2444 | W303.asm02.HP0_G0022260 | chrVII | 2845 | 1263 | 1389 | 2723 | 1307 | 1248 | 2638 | 1222 | 1366 | 5682 | 1377 | 1838 | 5831 | 1070 | 1403 | 5104 | 1304 | 1757 |
| 2445 | W303.asm02.HP0_G0022270 | chrVII | 4562 | 255 | 539 | 4358 | 431 | 622 | 4258 | 222 | 689 | 2825 | 477 | 1013 | 1443 | 354 | 622 | 1154 | 457 | 785 |
| 2446 | W303.asm02.HP0_G0022280 | chrVII | 1 | 1 | 0 | 0 | 0 | 0 | 1 | 0 | 0 | 2 | 1 | 0 | 2 | 1 | 1 | 0 | 1 | 2 |
| 2447 | W303.asm02.HP0_G0022290 | chrVII | 4792 | 1867 | 2148 | 3988 | 1657 | 1995 | 4283 | 1739 | 2560 | 12665 | 4001 | 4845 | 5444 | 1684 | 2444 | 7644 | 3615 | 4207 |
| 2448 | W303.asm02.HP0_G0022300 | chrVII | 4766 | 1368 | 1264 | 4649 | 1306 | 1276 | 4739 | 1478 | 1195 | 8414 | 1568 | 2056 | 7267 | 1099 | 1496 | 7095 | 1420 | 1895 |
| 2449 | W303.asm02.HP0_G0022310 | chrVII | 2775 | 1570 | 1474 | 2293 | 1470 | 1449 | 2549 | 1593 | 1695 | 4198 | 1312 | 1244 | 4272 | 811 | 1050 | 4249 | 1167 | 1149 |
| 2450 | W303.asm02.HP0_G0022320 | chrVII | 1292 | 442 | 522 | 1153 | 463 | 510 | 1147 | 497 | 564 | 2038 | 472 | 645 | 2147 | 272 | 420 | 1893 | 384 | 520 |
| 2451 | W303.asm02.HP0_G0022330 | chrVII | 8726 | 5278 | 4277 | 7296 | 4960 | 3933 | 8919 | 5410 | 4080 | 29806 | 6288 | 6345 | 24201 | 4195 | 4435 | 26639 | 5688 | 5744 |
| 2452 | W303.asm02.HP0_G0022340 | chrVII | 3204 | 1068 | 707 | 3234 | 796 | 706 | 3241 | 1000 | 803 | 6104 | 1536 | 1753 | 4324 | 1108 | 1280 | 4485 | 1396 | 1533 |
| 2453 | W303.asm02.HP0_G0022350 | chrVII | 4937 | 3129 | 3071 | 4698 | 2987 | 3060 | 4909 | 3248 | 3208 | 9877 | 3072 | 3255 | 9911 | 2161 | 2628 | 9193 | 2596 | 2972 |
| 2454 | W303.asm02.HP0_G0022360 | chrVII | 1895 | 221 | 183 | 1498 | 163 | 213 | 1321 | 168 | 183 | 936 | 414 | 380 | 742 | 317 | 318 | 590 | 408 | 389 |
| 2455 | W303.asm02.HP0_G0022370 | chrVII | 7825 | 690 | 784 | 7797 | 769 | 978 | 7630 | 597 | 1007 | 8254 | 1185 | 2407 | 5113 | 1040 | 1709 | 4828 | 1108 | 2043 |
| 2456 | W303.asm02.HP0_G0022380 | chrVII | 2480 | 546 | 904 | 2314 | 756 | 934 | 2397 | 538 | 1132 | 5830 | 770 | 1790 | 3413 | 511 | 1232 | 3675 | 639 | 1544 |
| 2457 | W303.asm02.HP0_G0022390 | chrVII | 811 | 843 | 1583 | 681 | 1699 | 1487 | 757 | 1044 | 2204 | 5066 | 827 | 1065 | 5454 | 498 | 704 | 5705 | 640 | 895 |
| 2458 | W303.asm02.HP0_G0022400 | chrVII | 21 | 14 | 2 | 18 | 5 | 9 | 15 | 3 | 5 | 19 | 6 | 4 | 22 | 1 | 2 | 22 | 11 | 3 |
| 2459 | W303.asm02.HP0_G0022410 | chrVII | 2352 | 609 | 877 | 2371 | 925 | 932 | 2270 | 589 | 891 | 6056 | 841 | 1706 | 4771 | 598 | 1148 | 4780 | 739 | 1454 |
| 2460 | W303.asm02.HP0_G0022420 | chrVII | 1732 | 1000 | 1370 | 1512 | 1154 | 1391 | 1409 | 1105 | 1734 | 3078 | 896 | 1216 | 3878 | 679 | 949 | 3452 | 807 | 1073 |
| 2461 | W303.asm02.HP0_G0022430 | chrVII | 2760 | 968 | 1042 | 2784 | 1094 | 988 | 2765 | 1046 | 1008 | 5242 | 1132 | 1321 | 4452 | 686 | 951 | 4634 | 917 | 1204 |
| 2462 | W303.asm02.HP0_G0022440 | chrVII | 2443 | 1243 | 1152 | 2076 | 1058 | 1133 | 2359 | 1163 | 1237 | 3942 | 1119 | 1446 | 4634 | 812 | 1245 | 3732 | 959 | 1337 |
| 2463 | W303.asm02.HP0_G0022450 | chrVII | 1142 | 530 | 507 | 1071 | 494 | 501 | 1069 | 523 | 461 | 2318 | 469 | 801 | 2626 | 346 | 504 | 2155 | 445 | 628 |
| 2464 | W303.asm02.HP0_G0022460 | chrVII | 15352 | 3821 | 4061 | 13340 | 3409 | 4464 | 15632 | 3672 | 5195 | 19852 | 7569 | 6632 | 16858 | 5522 | 5528 | 16460 | 6625 | 6421 |
| 2465 | W303.asm02.HP0_G0022470 | chrVII | 2918 | 933 | 891 | 2676 | 1414 | 955 | 2821 | 1145 | 986 | 6788 | 1324 | 1322 | 5591 | 912 | 796 | 5469 | 1045 | 1240 |
| 2466 | W303.asm02.HP0_G0022480 | chrVII | 4802 | 616 | 1126 | 4722 | 902 | 1198 | 4608 | 641 | 1313 | 8952 | 1117 | 2472 | 6478 | 939 | 2033 | 5774 | 1059 | 2212 |
| 2467 | W303.asm02.HP0_G0022490 | chrVII | 24072 | 17302 | 15117 | 22786 | 13458 | 14594 | 24309 | 16600 | 15903 | 29099 | 14325 | 13444 | 37879 | 13882 | 13383 | 26111 | 12910 | 12116 |
| 2468 | W303.asm02.HP0_G0022500 | chrVII | 1711 | 1124 | 1220 | 1514 | 1223 | 1135 | 1642 | 1150 | 1324 | 5186 | 984 | 1186 | 5356 | 769 | 748 | 4739 | 960 | 1014 |
| 2469 | W303.asm02.HP0_G0022510 | chrVII | 4603 | 4244 | 5172 | 4441 | 3433 | 5526 | 4179 | 3830 | 5762 | 8693 | 3001 | 4867 | 9863 | 2214 | 3561 | 9440 | 2822 | 4511 |
| 2470 | W303.asm02.HP0_G0022520 | chrVII | 18748 | 2916 | 2583 | 17543 | 2218 | 2865 | 19632 | 2572 | 3428 | 19775 | 6005 | 6839 | 16212 | 4557 | 5003 | 15498 | 5108 | 5792 |
| 2471 | W303.asm02.HP0_G0022530 | chrVII | 3265 | 2167 | 1961 | 3194 | 2173 | 1817 | 3466 | 2290 | 2176 | 6014 | 1758 | 1458 | 5767 | 1052 | 1035 | 5406 | 1428 | 1213 |
| 2472 | W303.asm02.HP0_G0022540 | chrVII | 1111 | 222 | 173 | 1031 | 180 | 171 | 964 | 209 | 192 | 921 | 248 | 282 | 1040 | 210 | 233 | 788 | 250 | 237 |
| 2473 | W303.asm02.HP0_G0022550 | chrVII | 979 | 288 | 199 | 888 | 239 | 183 | 869 | 290 | 185 | 1081 | 337 | 332 | 901 | 196 | 232 | 900 | 278 | 312 |
| 2474 | W303.asm02.HP0_G0022560 | chrVII | 1550 | 145 | 115 | 1633 | 137 | 113 | 1420 | 110 | 150 | 871 | 306 | 207 | 543 | 164 | 212 | 494 | 262 | 192 |
| 2475 | W303.asm02.HP0_G0022570 | chrVII | 2 | 3 | 1 | 5 | 1 | 1 | 5 | 3 | 1 | 6 | 1 | 0 | 7 | 0 | 2 | 5 | 2 | 1 |
| 2476 | W303.asm02.HP0_G0022580 | chrVII | 734 | 86 | 75 | 1097 | 62 | 60 | 805 | 76 | 53 | 285 | 79 | 75 | 420 | 73 | 58 | 255 | 75 | 59 |
| 2477 | W303.asm02.HP0_G0022590 | chrVII | 69 | 22 | 9 | 35 | 20 | 27 | 53 | 13 | 12 | 62 | 15 | 7 | 51 | 8 | 4 | 40 | 23 | 19 |
| 2478 | W303.asm02.HP0_G0022600 | chrVII | 8483 | 2601 | 1852 | 7495 | 1734 | 1902 | 7714 | 2578 | 1960 | 6295 | 2069 | 1847 | 7781 | 1714 | 1619 | 6232 | 1942 | 1683 |
| 2479 | W303.asm02.HP0_G0022610 | chrVII | 13 | 8 | 1 | 10 | 8 | 4 | 16 | 7 | 2 | 4 | 7 | 1 | 15 | 8 | 3 | 6 | 5 | 2 |
| 2480 | W303.asm02.HP0_G0022620 | chrVII | 1 | 0 | 0 | 0 | 0 | 0 | 0 | 0 | 0 | 0 | 1 | 0 | 0 | 0 | 0 | 0 | 0 | 0 |
| 2481 | W303.asm02.HP0_G0022630 | chrVIII | 1 | 6 | 4 | 7 | 1 | 2 | 4 | 4 | 7 | 12 | 4 | 7 | 1 | 1 | 1 | 3 | 2 | 5 |
| 2482 | W303.asm02.HP0_G0022640 | chrVIII | 10 | 15 | 6 | 8 | 10 | 7 | 21 | 13 | 7 | 13 | 11 | 5 | 3 | 1 | 1 | 14 | 9 | 4 |
| 2483 | W303.asm02.HP0_G0022650 | chrVIII | 2189 | 1519 | 1231 | 2214 | 969 | 1495 | 2747 | 1458 | 1159 | 2963 | 841 | 1185 | 2050 | 445 | 618 | 2652 | 863 | 1070 |
| 2484 | W303.asm02.HP0_G0022660 | chrVIII | 930 | 252 | 359 | 748 | 360 | 449 | 907 | 287 | 337 | 3791 | 676 | 547 | 2884 | 249 | 407 | 4101 | 569 | 1158 |
| 2485 | W303.asm02.HP0_G0022670 | chrVIII | 60 | 18 | 8 | 52 | 15 | 21 | 59 | 9 | 15 | 24 | 15 | 7 | 37 | 5 | 9 | 20 | 10 | 5 |
| 2486 | W303.asm02.HP0_G0022680 | chrVIII | 635 | 249 | 234 | 480 | 236 | 250 | 537 | 204 | 237 | 719 | 140 | 161 | 621 | 86 | 125 | 715 | 153 | 144 |
| 2487 | W303.asm02.HP0_G0022690 | chrVIII | 694 | 343 | 284 | 612 | 281 | 275 | 669 | 299 | 286 | 830 | 242 | 226 | 921 | 155 | 174 | 830 | 210 | 212 |
| 2488 | W303.asm02.HP0_G0022700 | chrVIII | 20 | 7 | 0 | 14 | 2 | 1 | 24 | 7 | 2 | 8 | 2 | 1 | 21 | 2 | 1 | 11 | 3 | 1 |
| 2489 | W303.asm02.HP0_G0022710 | chrVIII | 523 | 235 | 393 | 449 | 408 | 578 | 497 | 228 | 439 | 4164 | 1178 | 847 | 2528 | 430 | 798 | 3067 | 1040 | 1608 |
| 2490 | W303.asm02.HP0_G0022720 | chrVIII | 2349 | 1660 | 1620 | 2249 | 1743 | 1355 | 2603 | 1814 | 1590 | 8676 | 1801 | 1894 | 7820 | 1122 | 1166 | 8004 | 1451 | 1653 |
| 2491 | W303.asm02.HP0_G0022730 | chrVIII | 1687 | 474 | 636 | 1484 | 577 | 725 | 1642 | 503 | 780 | 3527 | 670 | 1120 | 2613 | 481 | 753 | 2634 | 561 | 985 |
| 2492 | W303.asm02.HP0_G0022740 | chrVIII | 49 | 14 | 4 | 42 | 6 | 12 | 52 | 11 | 7 | 30 | 9 | 14 | 38 | 8 | 6 | 25 | 12 | 8 |
| 2493 | W303.asm02.HP0_G0022750 | chrVIII | 1484 | 242 | 217 | 1093 | 216 | 269 | 1213 | 188 | 176 | 546 | 301 | 270 | 706 | 138 | 296 | 418 | 292 | 280 |
| 2494 | W303.asm02.HP0_G0022760 | chrVIII | 2612 | 371 | 666 | 2348 | 670 | 745 | 2316 | 406 | 773 | 6595 | 864 | 1159 | 7072 | 584 | 1330 | 6016 | 828 | 1541 |
| 2495 | W303.asm02.HP0_G0022770 | chrVIII | 25668 | 4462 | 3444 | 26038 | 3318 | 3732 | 25515 | 4433 | 3556 | 17515 | 6102 | 7425 | 16742 | 5075 | 5965 | 13878 | 5153 | 6396 |
| 2496 | W303.asm02.HP0_G0022780 | chrVIII | 5069 | 4505 | 4359 | 5373 | 4848 | 4430 | 5991 | 4171 | 4609 | 19655 | 5124 | 7520 | 15856 | 4273 | 4574 | 15955 | 3984 | 6205 |
| 2497 | W303.asm02.HP0_G0022790 | chrVIII | 1899 | 340 | 391 | 1666 | 389 | 509 | 1765 | 303 | 514 | 1562 | 500 | 648 | 704 | 409 | 510 | 685 | 445 | 550 |
| 2498 | W303.asm02.HP0_G0022800 | chrVIII | 3642 | 1167 | 941 | 3122 | 997 | 1019 | 3299 | 1198 | 912 | 3084 | 987 | 1101 | 3108 | 681 | 906 | 2769 | 866 | 998 |
| 2499 | W303.asm02.HP0_G0022810 | chrVIII | 6611 | 659 | 1080 | 6604 | 863 | 1260 | 6384 | 612 | 1466 | 6372 | 894 | 2003 | 3170 | 800 | 1779 | 3355 | 802 | 1681 |
| 2500 | W303.asm02.HP0_G0022820 | chrVIII | 627 | 709 | 824 | 628 | 948 | 862 | 642 | 653 | 974 | 5778 | 1004 | 1474 | 4861 | 723 | 986 | 4205 | 936 | 1248 |
| 2501 | W303.asm02.HP0_G0022830 | chrVIII | 803 | 213 | 270 | 672 | 198 | 318 | 813 | 139 | 305 | 499 | 237 | 403 | 136 | 181 | 251 | 161 | 165 | 295 |
| 2502 | W303.asm02.HP0_G0022840 | chrVIII | 2193 | 684 | 765 | 2125 | 831 | 811 | 2074 | 624 | 942 | 5302 | 1382 | 1716 | 3438 | 833 | 1168 | 3181 | 1284 | 1593 |
| 2503 | W303.asm02.HP0_G0022850 | chrVIII | 488 | 146 | 173 | 436 | 157 | 170 | 454 | 96 | 227 | 1011 | 240 | 258 | 549 | 198 | 218 | 439 | 221 | 228 |
| 2504 | W303.asm02.HP0_G0022860 | chrVIII | 4903 | 976 | 997 | 4448 | 1048 | 1098 | 4616 | 817 | 1252 | 3725 | 1198 | 1404 | 2829 | 859 | 1121 | 2521 | 1120 | 1235 |
| 2505 | W303.asm02.HP0_G0022870 | chrVIII | 1565 | 416 | 411 | 1279 | 412 | 393 | 1409 | 374 | 373 | 1964 | 756 | 592 | 1954 | 577 | 802 | 1573 | 656 | 569 |
| 2506 | W303.asm02.HP0_G0022880 | chrVIII | 1974 | 411 | 473 | 2102 | 434 | 580 | 2145 | 360 | 615 | 4343 | 861 | 1207 | 1788 | 580 | 750 | 1940 | 727 | 1064 |
| 2507 | W303.asm02.HP0_G0022890 | chrVIII | 413 | 160 | 115 | 350 | 121 | 110 | 351 | 183 | 147 | 739 | 219 | 199 | 785 | 128 | 130 | 757 | 233 | 187 |
| 2508 | W303.asm02.HP0_G0022900 | chrVIII | 15427 | 1971 | 874 | 16062 | 907 | 979 | 13404 | 1494 | 778 | 2596 | 1606 | 1070 | 3415 | 1006 | 1151 | 2391 | 1699 | 793 |
| 2509 | W303.asm02.HP0_G0022910 | chrVIII | 1825 | 646 | 554 | 1829 | 610 | 672 | 1749 | 591 | 618 | 4375 | 1345 | 1496 | 3220 | 865 | 1128 | 2976 | 1354 | 1386 |

| | | | | | | | | | | | | | | | | | | | | |
|---|---|---|---|---|---|---|---|---|---|---|---|---|---|---|---|---|---|---|---|---|
| 2510 | W303.asm02.HP0_G0022920 | chrVIII | 3856 | 286 | 293 | 3856 | 244 | 321 | 3763 | 241 | 371 | 2412 | 678 | 745 | 1563 | 499 | 613 | 1329 | 627 | 609 |
| 2511 | W303.asm02.HP0_G0022930 | chrVIII | 346 | 70 | 97 | 284 | 70 | 64 | 258 | 60 | 59 | 561 | 129 | 119 | 538 | 74 | 91 | 400 | 108 | 100 |
| 2512 | W303.asm02.HP0_G0022940 | chrVIII | 1588 | 888 | 981 | 1372 | 996 | 926 | 1430 | 835 | 1097 | 4553 | 955 | 1171 | 4250 | 576 | 915 | 4237 | 996 | 1078 |
| 2513 | W303.asm02.HP0_G0022950 | chrVIII | 531 | 282 | 140 | 398 | 160 | 133 | 444 | 258 | 173 | 802 | 454 | 265 | 624 | 271 | 234 | 691 | 467 | 246 |
| 2514 | W303.asm02.HP0_G0022960 | chrVIII | 16981 | 14333 | 18205 | 17090 | 17439 | 17972 | 17674 | 15016 | 22167 | 37598 | 14010 | 16375 | 40902 | 10487 | 13661 | 38976 | 12569 | 14615 |
| 2515 | W303.asm02.HP0_G0022970 | chrVIII | 2708 | 1064 | 779 | 2557 | 772 | 674 | 2547 | 1025 | 798 | 3269 | 822 | 749 | 3150 | 569 | 583 | 2965 | 829 | 698 |
| 2516 | W303.asm02.HP0_G0022980 | chrVIII | 992 | 1377 | 1526 | 1030 | 1550 | 1517 | 1007 | 1268 | 1767 | 5021 | 1109 | 1083 | 5639 | 645 | 653 | 5250 | 1053 | 927 |
| 2517 | W303.asm02.HP0_G0022990 | chrVIII | 478 | 395 | 270 | 470 | 290 | 302 | 440 | 350 | 349 | 1539 | 287 | 234 | 1345 | 205 | 163 | 1534 | 385 | 239 |
| 2518 | W303.asm02.HP0_G0023000 | chrVIII | 4265 | 3470 | 2959 | 3947 | 3268 | 2772 | 4306 | 3625 | 2998 | 11785 | 3904 | 3810 | 11794 | 2671 | 2653 | 11046 | 3228 | 3382 |
| 2519 | W303.asm02.HP0_G0023010 | chrVIII | 1556 | 313 | 244 | 1640 | 244 | 231 | 1508 | 277 | 268 | 1779 | 400 | 427 | 1534 | 282 | 427 | 1312 | 384 | 397 |
| 2520 | W303.asm02.HP0_G0023020 | chrVIII | 1809 | 457 | 350 | 1803 | 389 | 366 | 1700 | 481 | 384 | 2100 | 480 | 576 | 2014 | 364 | 438 | 1765 | 470 | 456 |
| 2521 | W303.asm02.HP0_G0023030 | chrVIII | 937 | 263 | 151 | 991 | 147 | 154 | 916 | 196 | 152 | 1769 | 312 | 266 | 555 | 254 | 201 | 706 | 353 | 262 |
| 2522 | W303.asm02.HP0_G0023040 | chrVIII | 2581 | 733 | 1093 | 2547 | 1010 | 1250 | 2814 | 613 | 1354 | 8115 | 1597 | 2592 | 2347 | 1170 | 1476 | 2475 | 1351 | 2107 |
| 2523 | W303.asm02.HP0_G0023050 | chrVIII | 329 | 322 | 320 | 312 | 259 | 272 | 336 | 289 | 354 | 602 | 258 | 235 | 668 | 174 | 186 | 586 | 226 | 228 |
| 2524 | W303.asm02.HP0_G0023060 | chrVIII | 13335 | 1455 | 1291 | 13077 | 1147 | 1495 | 14039 | 1241 | 1624 | 8022 | 2409 | 3148 | 7246 | 2032 | 2467 | 6131 | 2290 | 2780 |
| 2525 | W303.asm02.HP0_G0023070 | chrVIII | 3637 | 3149 | 2925 | 3318 | 2647 | 2674 | 3458 | 3070 | 3044 | 9561 | 2618 | 2419 | 9285 | 1793 | 2081 | 9471 | 2463 | 2186 |
| 2526 | W303.asm02.HP0_G0023080 | chrVIII | 7571 | 1622 | 675 | 7601 | 644 | 740 | 7469 | 1308 | 623 | 3736 | 1430 | 1267 | 3372 | 1185 | 947 | 2956 | 1374 | 1037 |
| 2527 | W303.asm02.HP0_G0023090 | chrVIII | 3030 | 3810 | 3894 | 2808 | 4050 | 4015 | 2807 | 3575 | 4440 | 9258 | 3185 | 3265 | 9432 | 2104 | 2468 | 9255 | 2985 | 2661 |
| 2528 | W303.asm02.HP0_G0023100 | chrVIII | 3427 | 2158 | 1427 | 3113 | 1420 | 1260 | 3152 | 2029 | 1322 | 5912 | 2135 | 1875 | 4728 | 1283 | 1370 | 5148 | 2187 | 1641 |
| 2529 | W303.asm02.HP0_G0023110 | chrVIII | 2237 | 2152 | 2510 | 2196 | 2242 | 2465 | 2236 | 2187 | 3163 | 4201 | 1818 | 1830 | 4306 | 1259 | 1539 | 4392 | 1605 | 1376 |
| 2530 | W303.asm02.HP0_G0023120 | chrVIII | 850 | 1017 | 539 | 876 | 528 | 543 | 684 | 825 | 560 | 1696 | 868 | 537 | 1740 | 549 | 433 | 1769 | 954 | 486 |
| 2531 | W303.asm02.HP0_G0023130 | chrVIII | 3715 | 1781 | 1470 | 3353 | 1518 | 1401 | 3718 | 1661 | 1572 | 6938 | 1750 | 1848 | 6197 | 1078 | 1291 | 6502 | 1723 | 1673 |
| 2532 | W303.asm02.HP0_G0023140 | chrVIII | 1524 | 878 | 657 | 1550 | 613 | 645 | 1417 | 815 | 686 | 2668 | 804 | 658 | 2608 | 504 | 565 | 2694 | 918 | 622 |
| 2533 | W303.asm02.HP0_G0023150 | chrVIII | 2535 | 3275 | 2748 | 2600 | 2945 | 2779 | 2794 | 2926 | 2629 | 10751 | 2789 | 2988 | 10343 | 1821 | 2271 | 10063 | 2731 | 2677 |
| 2534 | W303.asm02.HP0_G0023160 | chrVIII | 5795 | 3280 | 4905 | 5708 | 4204 | 4789 | 5342 | 3670 | 6225 | 8449 | 3357 | 4375 | 10738 | 2532 | 3804 | 9259 | 2915 | 3856 |
| 2535 | W303.asm02.HP0_G0023170 | chrVIII | 3803 | 839 | 551 | 3729 | 583 | 585 | 3662 | 696 | 630 | 4471 | 1564 | 1350 | 3638 | 1058 | 1074 | 2456 | 1728 | 1197 |
| 2536 | W303.asm02.HP0_G0023180 | chrVIII | 18214 | 6462 | 6220 | 18051 | 6416 | 5758 | 20423 | 6203 | 5379 | 38710 | 7873 | 9820 | 33475 | 6501 | 7456 | 32590 | 7024 | 8738 |
| 2537 | W303.asm02.HP0_G0023190 | chrVIII | 130 | 45 | 40 | 78 | 37 | 34 | 111 | 66 | 43 | 97 | 71 | 39 | 126 | 26 | 31 | 81 | 64 | 16 |
| 2538 | W303.asm02.HP0_G0023200 | chrVIII | 154 | 44 | 16 | 68 | 24 | 19 | 104 | 35 | 16 | 87 | 37 | 26 | 94 | 20 | 21 | 94 | 40 | 17 |
| 2539 | W303.asm02.HP0_G0023210 | chrVIII | 11866 | 3923 | 3076 | 10999 | 2844 | 2900 | 11041 | 3981 | 2977 | 8385 | 4972 | 4632 | 8958 | 3704 | 4046 | 7693 | 4683 | 3723 |
| 2540 | W303.asm02.HP0_G0023220 | chrVIII | 7981 | 1085 | 1144 | 7701 | 1244 | 1208 | 8201 | 957 | 1259 | 7064 | 1373 | 2137 | 5511 | 959 | 1738 | 5320 | 1376 | 1959 |
| 2541 | W303.asm02.HP0_G0023230 | chrVIII | 8473 | 6993 | 10153 | 8270 | 9794 | 10205 | 8735 | 7221 | 12754 | 20486 | 7051 | 8545 | 24062 | 5285 | 7075 | 22219 | 6309 | 7599 |
| 2542 | W303.asm02.HP0_G0023240 | chrVIII | 4139 | 510 | 318 | 3826 | 335 | 385 | 3814 | 431 | 367 | 2277 | 598 | 833 | 2126 | 504 | 669 | 1714 | 586 | 751 |
| 2543 | W303.asm02.HP0_G0023250 | chrVIII | 4057 | 1847 | 1955 | 4229 | 1836 | 1909 | 3837 | 1804 | 2205 | 5959 | 1563 | 1712 | 6215 | 1022 | 1495 | 6021 | 1491 | 1601 |
| 2544 | W303.asm02.HP0_G0023260 | chrVIII | 3442 | 2258 | 1830 | 3120 | 1875 | 1800 | 3290 | 2246 | 1868 | 6909 | 1985 | 1876 | 6850 | 1240 | 1483 | 6677 | 1745 | 1728 |
| 2545 | W303.asm02.HP0_G0023270 | chrVIII | 147 | 107 | 48 | 123 | 48 | 24 | 124 | 95 | 52 | 145 | 97 | 52 | 165 | 58 | 30 | 160 | 80 | 39 |
| 2546 | W303.asm02.HP0_G0023280 | chrVIII | 359 | 457 | 459 | 369 | 387 | 399 | 292 | 440 | 539 | 1951 | 380 | 432 | 1841 | 234 | 227 | 2046 | 397 | 408 |
| 2547 | W303.asm02.HP0_G0023290 | chrVIII | 10146 | 577 | 526 | 9278 | 384 | 575 | 9180 | 415 | 733 | 2435 | 949 | 1104 | 1236 | 711 | 962 | 1099 | 955 | 1027 |
| 2548 | W303.asm02.HP0_G0023300 | chrVIII | 2950 | 901 | 1068 | 2956 | 821 | 1121 | 2691 | 794 | 1155 | 3401 | 809 | 1304 | 3603 | 557 | 1042 | 3318 | 845 | 1190 |
| 2549 | W303.asm02.HP0_G0023310 | chrVIII | 5546 | 5715 | 4084 | 5481 | 4065 | 4200 | 5334 | 4876 | 3947 | 15419 | 6402 | 7687 | 12958 | 5412 | 6427 | 12298 | 5325 | 6693 |
| 2550 | W303.asm02.HP0_G0023320 | chrVIII | 17758 | 3465 | 2943 | 15688 | 2801 | 3668 | 18909 | 2756 | 3983 | 24628 | 8243 | 8922 | 14941 | 6507 | 6143 | 15224 | 6753 | 7570 |
| 2551 | W303.asm02.HP0_G0023330 | chrVIII | 13779 | 3275 | 2770 | 12017 | 3038 | 2766 | 14635 | 2985 | 3161 | 23572 | 5740 | 6161 | 16701 | 4557 | 4099 | 16266 | 5003 | 5615 |
| 2552 | W303.asm02.HP0_G0023340 | chrVIII | 5490 | 3860 | 5835 | 4275 | 5524 | 5900 | 4835 | 3932 | 6973 | 10313 | 3287 | 4711 | 12692 | 2378 | 4248 | 11761 | 3216 | 3847 |
| 2553 | W303.asm02.HP0_G0023350 | chrVIII | 618 | 282 | 171 | 556 | 128 | 120 | 513 | 229 | 187 | 480 | 149 | 116 | 945 | 160 | 139 | 448 | 175 | 74 |
| 2554 | W303.asm02.HP0_G0023360 | chrVIII | 3956 | 751 | 1106 | 4059 | 1206 | 1154 | 4151 | 911 | 1359 | 8539 | 939 | 1702 | 7524 | 821 | 1616 | 7110 | 806 | 1624 |
| 2555 | W303.asm02.HP0_G0023370 | chrVIII | 9888 | 1301 | 1416 | 9828 | 1318 | 1589 | 9649 | 1096 | 1824 | 9579 | 2521 | 3422 | 4825 | 2148 | 2260 | 4658 | 2251 | 2926 |
| 2556 | W303.asm02.HP0_G0023380 | chrVIII | 8835 | 1094 | 1384 | 7667 | 1178 | 1688 | 9256 | 894 | 1826 | 10966 | 2998 | 3773 | 3752 | 2073 | 2615 | 4509 | 2445 | 3356 |
| 2557 | W303.asm02.HP0_G0023390 | chrVIII | 5003 | 3388 | 3516 | 4303 | 3002 | 3253 | 4617 | 3308 | 4122 | 8562 | 3174 | 3149 | 9120 | 2251 | 2577 | 8842 | 2875 | 3005 |
| 2558 | W303.asm02.HP0_G0023400 | chrVIII | 17617 | 3524 | 3230 | 17903 | 3092 | 3454 | 17809 | 3154 | 3996 | 23374 | 5691 | 7429 | 15567 | 4421 | 5826 | 15011 | 5327 | 6949 |
| 2559 | W303.asm02.HP0_G0023410 | chrVIII | 7446 | 2152 | 1880 | 6852 | 1754 | 1844 | 6785 | 2100 | 2060 | 10781 | 2856 | 3064 | 9560 | 2392 | 2946 | 8788 | 2775 | 2702 |
| 2560 | W303.asm02.HP0_G0023420 | chrVIII | 1834 | 1953 | 1828 | 1766 | 1643 | 1844 | 1746 | 1907 | 2047 | 3666 | 1690 | 1765 | 4038 | 1247 | 1557 | 3596 | 1563 | 1508 |
| 2561 | W303.asm02.HP0_G0023430 | chrVIII | 12297 | 1398 | 1117 | 11260 | 1101 | 1265 | 12048 | 1366 | 1282 | 8247 | 1858 | 2826 | 10284 | 2083 | 2242 | 5495 | 1660 | 2436 |
| 2562 | W303.asm02.HP0_G0023440 | chrVIII | 601 | 437 | 366 | 512 | 454 | 387 | 573 | 373 | 377 | 3367 | 469 | 637 | 2837 | 365 | 384 | 2538 | 469 | 579 |
| 2563 | W303.asm02.HP0_G0023450 | chrVIII | 1127 | 726 | 988 | 923 | 893 | 967 | 1190 | 657 | 1227 | 5842 | 1465 | 1640 | 2694 | 1071 | 962 | 2434 | 1215 | 1453 |
| 2564 | W303.asm02.HP0_G0023460 | chrVIII | 315 | 239 | 136 | 258 | 156 | 189 | 335 | 255 | 204 | 1095 | 736 | 568 | 944 | 549 | 475 | 564 | 589 | 479 |
| 2565 | W303.asm02.HP0_G0023470 | chrVIII | 1113 | 332 | 235 | 1110 | 256 | 259 | 1010 | 276 | 249 | 1527 | 452 | 528 | 1435 | 286 | 329 | 1254 | 423 | 397 |
| 2566 | W303.asm02.HP0_G0023480 | chrVIII | 507 | 261 | 276 | 470 | 325 | 269 | 518 | 254 | 335 | 1432 | 236 | 301 | 1452 | 145 | 196 | 1382 | 240 | 223 |
| 2567 | W303.asm02.HP0_G0023490 | chrVIII | 599 | 579 | 512 | 482 | 577 | 519 | 612 | 555 | 441 | 2510 | 567 | 619 | 2603 | 342 | 422 | 2345 | 535 | 589 |
| 2568 | W303.asm02.HP0_G0023500 | chrVIII | 4089 | 2129 | 1749 | 4196 | 1583 | 2040 | 4043 | 1716 | 2204 | 10087 | 3256 | 3966 | 6894 | 2709 | 3100 | 6291 | 2954 | 3402 |
| 2569 | W303.asm02.HP0_G0023510 | chrVIII | 2808 | 717 | 699 | 2669 | 590 | 742 | 2359 | 800 | 729 | 2201 | 612 | 841 | 2359 | 481 | 633 | 2038 | 601 | 675 |
| 2570 | W303.asm02.HP0_G0023520 | chrVIII | 8764 | 2198 | 2430 | 8185 | 2178 | 2532 | 9007 | 1919 | 2740 | 13546 | 3465 | 4250 | 11151 | 2567 | 3225 | 9212 | 3031 | 4067 |
| 2571 | W303.asm02.HP0_G0023530 | chrVIII | 3520 | 2221 | 2838 | 2806 | 2657 | 2955 | 3138 | 2340 | 3545 | 5774 | 1841 | 2319 | 7167 | 1338 | 2116 | 6224 | 1744 | 1989 |
| 2572 | W303.asm02.HP0_G0023540 | chrVIII | 1345 | 344 | 264 | 1103 | 346 | 297 | 1320 | 269 | 352 | 2025 | 552 | 582 | 831 | 410 | 266 | 889 | 453 | 484 |
| 2573 | W303.asm02.HP0_G0023550 | chrVIII | 1149 | 451 | 671 | 791 | 577 | 637 | 1077 | 493 | 805 | 2225 | 574 | 769 | 2481 | 376 | 653 | 2315 | 501 | 688 |
| 2574 | W303.asm02.HP0_G0023560 | chrVIII | 7180 | 854 | 1088 | 5796 | 1024 | 1261 | 7535 | 642 | 1478 | 10295 | 2049 | 2437 | 2902 | 1596 | 1792 | 2802 | 1656 | 2245 |
| 2575 | W303.asm02.HP0_G0023570 | chrVIII | 5063 | 1562 | 1949 | 4409 | 1196 | 2087 | 4421 | 1556 | 2149 | 3781 | 1217 | 2180 | 5811 | 988 | 1738 | 3869 | 1065 | 1863 |
| 2576 | W303.asm02.HP0_G0023580 | chrVIII | 538 | 459 | 843 | 534 | 339 | 933 | 534 | 409 | 962 | 852 | 291 | 750 | 873 | 177 | 359 | 826 | 274 | 616 |
| 2577 | W303.asm02.HP0_G0023590 | chrVIII | 1957 | 995 | 1601 | 1842 | 1042 | 1465 | 1909 | 962 | 1536 | 6201 | 1243 | 2066 | 4988 | 764 | 1261 | 5349 | 1122 | 1804 |
| 2578 | W303.asm02.HP0_G0023600 | chrVIII | 705 | 835 | 761 | 512 | 810 | 809 | 638 | 798 | 908 | 3446 | 818 | 917 | 2929 | 534 | 664 | 3272 | 807 | 908 |
| 2579 | W303.asm02.HP0_G0023610 | chrVIII | 11220 | 2192 | 1838 | 9664 | 1915 | 1844 | 10428 | 2006 | 1954 | 12805 | 2625 | 2981 | 9308 | 2003 | 2435 | 9430 | 2397 | 2971 |
| 2580 | W303.asm02.HP0_G0023620 | chrVIII | 68 | 50 | 46 | 70 | 40 | 38 | 74 | 45 | 51 | 162 | 47 | 57 | 181 | 35 | 41 | 162 | 45 | 63 |
| 2581 | W303.asm02.HP0_G0023630 | chrVIII | 369 | 284 | 343 | 345 | 348 | 328 | 322 | 301 | 377 | 1607 | 519 | 460 | 1226 | 255 | 293 | 1245 | 421 | 363 |
| 2582 | W303.asm02.HP0_G0023640 | chrVIII | 3281 | 3759 | 3585 | 3267 | 3653 | 3370 | 3331 | 3747 | 3457 | 12447 | 3568 | 3632 | 15328 | 2746 | 3271 | 11973 | 3189 | 3345 |
| 2583 | W303.asm02.HP0_G0023650 | chrVIII | 1631 | 1155 | 1280 | 1657 | 1186 | 1202 | 1639 | 1165 | 1323 | 5827 | 1459 | 1566 | 5641 | 922 | 1162 | 5190 | 1393 | 1485 |
| 2584 | W303.asm02.HP0_G0023660 | chrVIII | 56 | 22 | 24 | 25 | 18 | 24 | 22 | 22 | 18 | 64 | 16 | 20 | 76 | 6 | 14 | 91 | 24 | 20 |
| 2585 | W303.asm02.HP0_G0023670 | chrVIII | 3359 | 3132 | 3962 | 3222 | 3649 | 4124 | 3189 | 3143 | 5246 | 7481 | 2287 | 3062 | 8548 | 1922 | 2743 | 7797 | 2057 | 2337 |
| 2586 | W303.asm02.HP0_G0023680 | chrVIII | 3232 | 3401 | 4820 | 3200 | 5010 | 4413 | 3420 | 3609 | 5893 | 14748 | 2857 | 3519 | 15726 | 1713 | 2418 | 15742 | 2401 | 3115 |
| 2587 | W303.asm02.HP0_G0023690 | chrVIII | 0 | 0 | 0 | 1 | 0 | 0 | 0 | 0 | 0 | 0 | 2 | 0 | 0 | 0 | 1 | 0 | 1 | 0 |
| 2588 | W303.asm02.HP0_G0023700 | chrVIII | 29 | 30 | 15 | 35 | 14 | 26 | 36 | 26 | 17 | 28 | 55 | 12 | 20 | 0 | 5 | 24 | 47 | 15 |
| 2589 | W303.asm02.HP0_G0023710 | chrVIII | 0 | 0 | 0 | 0 | 0 | 0 | 0 | 0 | 0 | 0 | 0 | 0 | 0 | 0 | 0 | 0 | 0 | 0 |
| 2590 | W303.asm02.HP0_G0023720 | chrVIII | 0 | 0 | 0 | 0 | 0 | 0 | 0 | 0 | 0 | 0 | 0 | 0 | 0 | 0 | 0 | 0 | 0 | 0 |
| 2591 | W303.asm02.HP0_G0023730 | chrVIII | 0 | 0 | 0 | 0 | 0 | 0 | 0 | 0 | 0 | 0 | 0 | 0 | 0 | 0 | 0 | 0 | 0 | 0 |
| 2592 | W303.asm02.HP0_G0023740 | chrVIII | 0 | 0 | 0 | 0 | 0 | 0 | 0 | 0 | 0 | 0 | 0 | 0 | 0 | 0 | 0 | 0 | 0 | 0 |
| 2593 | W303.asm02.HP0_G0023750 | chrVIII | 0 | 0 | 0 | 0 | 0 | 0 | 0 | 0 | 0 | 0 | 0 | 0 | 0 | 0 | 0 | 0 | 0 | 0 |
| 2594 | W303.asm02.HP0_G0023760 | chrVIII | 0 | 0 | 0 | 0 | 0 | 0 | 0 | 0 | 0 | 0 | 0 | 0 | 0 | 0 | 0 | 0 | 0 | 0 |
| 2595 | W303.asm02.HP0_G0023770 | chrVIII | 0 | 0 | 0 | 0 | 0 | 0 | 0 | 0 | 0 | 0 | 0 | 0 | 0 | 0 | 0 | 0 | 0 | 0 |
| 2596 | W303.asm02.HP0_G0023780 | chrVIII | 0 | 0 | 0 | 0 | 0 | 0 | 0 | 0 | 0 | 0 | 0 | 0 | 0 | 0 | 0 | 0 | 0 | 0 |
| 2597 | W303.asm02.HP0_G0023790 | chrVIII | 0 | 0 | 0 | 0 | 0 | 0 | 0 | 0 | 0 | 0 | 0 | 0 | 0 | 0 | 0 | 0 | 0 | 0 |
| 2598 | W303.asm02.HP0_G0023800 | chrVIII | 0 | 0 | 0 | 0 | 0 | 0 | 0 | 0 | 0 | 0 | 0 | 0 | 0 | 0 | 0 | 0 | 0 | 0 |
| 2599 | W303.asm02.HP0_G0023810 | chrVIII | 0 | 0 | 0 | 0 | 0 | 0 | 0 | 0 | 0 | 0 | 0 | 0 | 0 | 0 | 0 | 0 | 0 | 0 |
| 2600 | W303.asm02.HP0_G0023820 | chrVIII | 0 | 0 | 0 | 0 | 0 | 0 | 0 | 0 | 0 | 0 | 0 | 0 | 0 | 0 | 0 | 0 | 0 | 0 |
| 2601 | W303.asm02.HP0_G0023830 | chrVIII | 0 | 0 | 0 | 0 | 0 | 0 | 0 | 0 | 0 | 0 | 0 | 0 | 0 | 0 | 0 | 0 | 0 | 0 |
| 2602 | W303.asm02.HP0_G0023840 | chrVIII | 2004 | 951 | 821 | 2152 | 846 | 705 | 1910 | 993 | 679 | 5265 | 1132 | 1094 | 4532 | 652 | 789 | 4872 | 1051 | 1024 |

| | | | | | | | | | | | | | | | | | | | | |
|---|---|---|---|---|---|---|---|---|---|---|---|---|---|---|---|---|---|---|---|---|
| 2603 | W303.asm02.HP0_G0023850 | chrVIII | 4274 | 1400 | 1218 | 3887 | 1159 | 1265 | 4182 | 1373 | 1349 | 3964 | 1518 | 1467 | 3998 | 1060 | 1284 | 3366 | 1417 | 1314 |
| 2604 | W303.asm02.HP0_G0023860 | chrVIII | 843 | 690 | 410 | 830 | 417 | 413 | 852 | 627 | 356 | 2167 | 686 | 575 | 1826 | 442 | 373 | 1463 | 655 | 460 |
| 2605 | W303.asm02.HP0_G0023870 | chrVIII | 2407 | 1262 | 1093 | 2042 | 1050 | 1194 | 2073 | 1292 | 1474 | 2702 | 1014 | 1112 | 3349 | 818 | 902 | 2627 | 944 | 952 |
| 2606 | W303.asm02.HP0_G0023880 | chrVIII | 1285 | 840 | 931 | 1182 | 888 | 939 | 1172 | 882 | 1244 | 2769 | 729 | 738 | 3431 | 481 | 612 | 3071 | 600 | 706 |
| 2607 | W303.asm02.HP0_G0023890 | chrVIII | 399 | 461 | 275 | 346 | 269 | 272 | 399 | 403 | 274 | 1258 | 431 | 343 | 1114 | 241 | 256 | 1006 | 441 | 343 |
| 2608 | W303.asm02.HP0_G0023900 | chrVIII | 2375 | 1131 | 791 | 2063 | 891 | 805 | 2312 | 1080 | 681 | 3629 | 1011 | 1071 | 3580 | 727 | 666 | 3082 | 955 | 909 |
| 2609 | W303.asm02.HP0_G0023910 | chrVIII | 3621 | 2674 | 2724 | 3480 | 2506 | 2670 | 3591 | 2511 | 3316 | 8442 | 2070 | 2490 | 8510 | 1496 | 2070 | 8794 | 2015 | 2309 |
| 2610 | W303.asm02.HP0_G0023920 | chrVIII | 14612 | 8107 | 6669 | 13089 | 7243 | 6333 | 15387 | 8047 | 6656 | 27552 | 9293 | 10153 | 24850 | 6642 | 6533 | 25267 | 7965 | 8976 |
| 2611 | W303.asm02.HP0_G0023930 | chrVIII | 3923 | 2599 | 2055 | 3884 | 2257 | 1957 | 4356 | 2483 | 1965 | 10615 | 2672 | 2719 | 9480 | 1736 | 1850 | 9351 | 2421 | 2399 |
| 2612 | W303.asm02.HP0_G0023940 | chrVIII | 686 | 1679 | 1669 | 786 | 2407 | 1681 | 992 | 1831 | 1589 | 10332 | 1671 | 2088 | 9526 | 1093 | 1061 | 8757 | 1432 | 1722 |
| 2613 | W303.asm02.HP0_G0023950 | chrVIII | 1311 | 614 | 598 | 1303 | 487 | 671 | 1330 | 642 | 644 | 2593 | 702 | 855 | 2530 | 480 | 695 | 2480 | 669 | 743 |
| 2614 | W303.asm02.HP0_G0023960 | chrVIII | 5651 | 3894 | 4514 | 4694 | 4263 | 4155 | 5612 | 4032 | 4775 | 15423 | 3921 | 4148 | 14435 | 2235 | 2932 | 15361 | 3436 | 3810 |
| 2615 | W303.asm02.HP0_G0023970 | chrVIII | 1640 | 2036 | 1849 | 1471 | 2213 | 1755 | 1756 | 1977 | 2061 | 7433 | 1755 | 1764 | 7463 | 1088 | 1086 | 7146 | 1697 | 1470 |
| 2616 | W303.asm02.HP0_G0023980 | chrVIII | 1300 | 578 | 654 | 1218 | 781 | 831 | 1423 | 436 | 911 | 5705 | 1208 | 1521 | 3010 | 782 | 793 | 2617 | 1001 | 1361 |
| 2617 | W303.asm02.HP0_G0023990 | chrVIII | 2485 | 1852 | 2165 | 2340 | 2240 | 2154 | 2341 | 2077 | 2369 | 5267 | 1818 | 2152 | 6522 | 1344 | 1780 | 5245 | 1474 | 1922 |
| 2618 | W303.asm02.HP0_G0024000 | chrVIII | 5837 | 1887 | 1958 | 5357 | 1631 | 2097 | 6232 | 1472 | 2392 | 7664 | 3123 | 4169 | 3052 | 1998 | 2104 | 3418 | 2858 | 3576 |
| 2619 | W303.asm02.HP0_G0024010 | chrVIII | 2385 | 2135 | 3037 | 1783 | 2782 | 2818 | 2051 | 2240 | 3689 | 5536 | 2039 | 2363 | 6398 | 1428 | 1954 | 6315 | 1987 | 1991 |
| 2620 | W303.asm02.HP0_G0024020 | chrVIII | 7319 | 851 | 1203 | 6871 | 1008 | 1331 | 6965 | 866 | 1601 | 10246 | 1838 | 2730 | 5787 | 1366 | 1933 | 5446 | 1652 | 2444 |
| 2621 | W303.asm02.HP0_G0024030 | chrVIII | 6842 | 1151 | 1087 | 5729 | 988 | 1268 | 6675 | 1010 | 1525 | 8091 | 1987 | 2620 | 4283 | 1503 | 1857 | 4437 | 1980 | 2328 |
| 2622 | W303.asm02.HP0_G0024040 | chrVIII | 1700 | 313 | 266 | 1469 | 243 | 280 | 1426 | 294 | 302 | 975 | 314 | 316 | 1100 | 232 | 303 | 938 | 323 | 319 |
| 2623 | W303.asm02.HP0_G0024050 | chrVIII | 3248 | 1061 | 898 | 2863 | 831 | 882 | 2790 | 1010 | 978 | 4449 | 1225 | 1489 | 4090 | 855 | 1112 | 3741 | 1079 | 1414 |
| 2624 | W303.asm02.HP0_G0024060 | chrVIII | 6433 | 1298 | 1567 | 6352 | 1794 | 1619 | 6821 | 1461 | 1626 | 11072 | 1553 | 2616 | 9100 | 1113 | 2023 | 8960 | 1240 | 2363 |
| 2625 | W303.asm02.HP0_G0024070 | chrVIII | 2572 | 704 | 794 | 2254 | 679 | 709 | 2497 | 757 | 763 | 4350 | 913 | 1233 | 3666 | 572 | 864 | 3803 | 844 | 1177 |
| 2626 | W303.asm02.HP0_G0024080 | chrVIII | 1687 | 319 | 384 | 1745 | 379 | 416 | 1571 | 305 | 404 | 4555 | 675 | 946 | 2816 | 508 | 707 | 2853 | 523 | 878 |
| 2627 | W303.asm02.HP0_G0024090 | chrVIII | 25 | 10 | 10 | 13 | 8 | 14 | 14 | 5 | 6 | 15 | 4 | 4 | 30 | 3 | 1 | 14 | 6 | 2 |
| 2628 | W303.asm02.HP0_G0024100 | chrVIII | 3238 | 389 | 586 | 3168 | 472 | 636 | 2878 | 378 | 765 | 3287 | 814 | 1160 | 1803 | 585 | 902 | 1528 | 688 | 938 |
| 2629 | W303.asm02.HP0_G0024110 | chrVIII | 1593 | 1368 | 1613 | 1690 | 1773 | 1640 | 1693 | 1398 | 2297 | 5262 | 1363 | 1332 | 5989 | 799 | 876 | 5504 | 1130 | 1311 |
| 2630 | W303.asm02.HP0_G0024120 | chrVIII | 5240 | 1201 | 1221 | 5459 | 1222 | 1384 | 4820 | 1130 | 1344 | 10476 | 1958 | 2494 | 7647 | 1303 | 2299 | 8124 | 1819 | 2318 |
| 2631 | W303.asm02.HP0_G0024130 | chrVIII | 2586 | 619 | 513 | 2050 | 493 | 601 | 2538 | 556 | 574 | 3296 | 911 | 1207 | 2387 | 681 | 843 | 2369 | 825 | 1001 |
| 2632 | W303.asm02.HP0_G0024140 | chrVIII | 1006 | 461 | 717 | 787 | 649 | 813 | 967 | 495 | 918 | 6541 | 1338 | 1668 | 4237 | 784 | 1145 | 4358 | 1228 | 1509 |
| 2633 | W303.asm02.HP0_G0024150 | chrVIII | 563 | 1224 | 1673 | 540 | 1916 | 1505 | 684 | 1201 | 1798 | 5737 | 978 | 1273 | 6118 | 662 | 785 | 5895 | 909 | 1104 |
| 2634 | W303.asm02.HP0_G0024160 | chrVIII | 1874 | 715 | 650 | 1923 | 668 | 647 | 1846 | 649 | 696 | 4353 | 1113 | 1210 | 3808 | 906 | 1079 | 3176 | 1040 | 1159 |
| 2635 | W303.asm02.HP0_G0024170 | chrVIII | 25 | 21 | 5 | 23 | 12 | 4 | 25 | 20 | 14 | 43 | 33 | 9 | 53 | 9 | 5 | 54 | 21 | 8 |
| 2636 | W303.asm02.HP0_G0024180 | chrVIII | 6658 | 2113 | 584 | 6817 | 469 | 764 | 6019 | 1471 | 759 | 1026 | 1415 | 724 | 1368 | 655 | 674 | 998 | 1686 | 557 |
| 2637 | W303.asm02.HP0_G0024190 | chrVIII | 2140 | 1866 | 1913 | 2203 | 2305 | 1777 | 2363 | 1940 | 2081 | 7105 | 1878 | 1691 | 7131 | 1043 | 1024 | 7290 | 1686 | 1526 |
| 2638 | W303.asm02.HP0_G0024200 | chrVIII | 7677 | 4381 | 4756 | 7312 | 5469 | 4295 | 7595 | 4769 | 5473 | 18366 | 5776 | 5606 | 16990 | 3482 | 4250 | 17091 | 4616 | 4751 |
| 2639 | W303.asm02.HP0_G0024210 | chrVIII | 1665 | 953 | 683 | 1604 | 755 | 681 | 1523 | 826 | 575 | 2468 | 760 | 824 | 2579 | 461 | 558 | 2483 | 746 | 651 |
| 2640 | W303.asm02.HP0_G0024220 | chrVIII | 2331 | 628 | 732 | 2049 | 668 | 791 | 2214 | 515 | 951 | 5396 | 1157 | 1562 | 3670 | 789 | 1145 | 3234 | 1115 | 1342 |
| 2641 | W303.asm02.HP0_G0024230 | chrVIII | 13560 | 1637 | 828 | 13154 | 1640 | 899 | 12617 | 1563 | 1018 | 11534 | 3362 | 1400 | 9635 | 2293 | 1556 | 8285 | 2874 | 1226 |
| 2642 | W303.asm02.HP0_G0024240 | chrVIII | 84 | 28 | 26 | 47 | 22 | 20 | 55 | 14 | 27 | 163 | 42 | 48 | 128 | 18 | 21 | 106 | 50 | 37 |
| 2643 | W303.asm02.HP0_G0024250 | chrVIII | 231 | 3224 | 2927 | 191 | 2385 | 2536 | 242 | 2804 | 3813 | 17493 | 4801 | 6476 | 12912 | 2729 | 3043 | 12793 | 5090 | 5771 |
| 2644 | W303.asm02.HP0_G0024260 | chrVIII | 977 | 151 | 141 | 1034 | 152 | 168 | 927 | 137 | 173 | 898 | 281 | 266 | 711 | 130 | 186 | 643 | 303 | 222 |
| 2645 | W303.asm02.HP0_G0024270 | chrVIII | 10993 | 449 | 397 | 11530 | 340 | 447 | 9641 | 352 | 526 | 2652 | 726 | 881 | 2121 | 561 | 806 | 1588 | 712 | 697 |
| 2646 | W303.asm02.HP0_G0024280 | chrVIII | 5642 | 1069 | 2000 | 4861 | 1444 | 2308 | 5481 | 1093 | 2562 | 13526 | 2252 | 5375 | 9161 | 1897 | 3744 | 8890 | 2132 | 4798 |
| 2647 | W303.asm02.HP0_G0024290 | chrVIII | 6288 | 615 | 939 | 5512 | 995 | 1083 | 6018 | 630 | 1262 | 9179 | 998 | 2965 | 4107 | 944 | 2016 | 4391 | 808 | 2551 |
| 2648 | W303.asm02.HP0_G0024300 | chrVIII | 912 | 504 | 564 | 814 | 606 | 563 | 889 | 547 | 614 | 2135 | 526 | 640 | 2126 | 343 | 461 | 2155 | 466 | 572 |
| 2649 | W303.asm02.HP0_G0024310 | chrVIII | 1536 | 508 | 427 | 1384 | 413 | 434 | 1505 | 515 | 401 | 1736 | 419 | 506 | 1548 | 302 | 334 | 1638 | 413 | 432 |
| 2650 | W303.asm02.HP0_G0024320 | chrVIII | 2242 | 286 | 673 | 2280 | 505 | 782 | 2252 | 242 | 904 | 3559 | 595 | 1868 | 1486 | 505 | 1020 | 1248 | 483 | 1605 |
| 2651 | W303.asm02.HP0_G0024330 | chrVIII | 2873 | 917 | 1065 | 3133 | 1236 | 1086 | 2849 | 885 | 1173 | 7967 | 1291 | 1831 | 7100 | 1128 | 1544 | 6047 | 1220 | 1658 |
| 2652 | W303.asm02.HP0_G0024340 | chrVIII | 10598 | 3021 | 2280 | 8985 | 1975 | 2373 | 9449 | 2579 | 2226 | 5399 | 2196 | 2531 | 7652 | 1775 | 2529 | 5659 | 2201 | 2335 |
| 2653 | W303.asm02.HP0_G0024350 | chrVIII | 672 | 108 | 101 | 637 | 121 | 111 | 633 | 80 | 111 | 842 | 198 | 194 | 679 | 145 | 155 | 546 | 210 | 175 |
| 2654 | W303.asm02.HP0_G0024360 | chrVIII | 2397 | 870 | 980 | 2176 | 805 | 898 | 2134 | 873 | 1120 | 2332 | 815 | 996 | 2726 | 579 | 895 | 2116 | 784 | 846 |
| 2655 | W303.asm02.HP0_G0024370 | chrVIII | 5496 | 1338 | 1449 | 5331 | 1480 | 1474 | 6118 | 1198 | 1636 | 8603 | 1869 | 2459 | 6457 | 1374 | 1891 | 6343 | 1614 | 2223 |
| 2656 | W303.asm02.HP0_G0024380 | chrVIII | 10867 | 2094 | 1803 | 10127 | 1899 | 1918 | 10848 | 2014 | 1859 | 13503 | 3342 | 4362 | 10271 | 2472 | 3145 | 9812 | 2680 | 3910 |
| 2657 | W303.asm02.HP0_G0024390 | chrVIII | 1996 | 198 | 246 | 1966 | 255 | 274 | 1964 | 180 | 285 | 2117 | 326 | 615 | 1559 | 254 | 413 | 1487 | 300 | 519 |
| 2658 | W303.asm02.HP0_G0024400 | chrVIII | 2374 | 1243 | 1415 | 2183 | 1276 | 1513 | 2225 | 1169 | 1684 | 3834 | 1072 | 1331 | 4268 | 789 | 1067 | 3667 | 979 | 1255 |
| 2659 | W303.asm02.HP0_G0024410 | chrVIII | 1861 | 1326 | 883 | 1678 | 992 | 846 | 1812 | 1299 | 821 | 3927 | 1413 | 1473 | 3379 | 922 | 948 | 3186 | 1244 | 1180 |
| 2660 | W303.asm02.HP0_G0024420 | chrVIII | 3813 | 692 | 644 | 3070 | 512 | 714 | 3665 | 630 | 655 | 2726 | 936 | 1383 | 2435 | 792 | 976 | 1908 | 739 | 1118 |
| 2661 | W303.asm02.HP0_G0024430 | chrVIII | 6932 | 1254 | 1140 | 6238 | 984 | 1089 | 6562 | 1196 | 1260 | 5006 | 1566 | 1664 | 4686 | 1234 | 1510 | 3855 | 1435 | 1468 |
| 2662 | W303.asm02.HP0_G0024440 | chrVIII | 4528 | 675 | 573 | 3944 | 604 | 686 | 4015 | 701 | 671 | 4843 | 1025 | 1222 | 3275 | 702 | 863 | 3346 | 957 | 1112 |
| 2663 | W303.asm02.HP0_G0024450 | chrVIII | 2528 | 564 | 646 | 2296 | 563 | 652 | 2588 | 513 | 824 | 4833 | 1233 | 1371 | 1997 | 850 | 1020 | 2080 | 1132 | 1252 |
| 2664 | W303.asm02.HP0_G0024460 | chrVIII | 2770 | 643 | 494 | 2523 | 565 | 541 | 2406 | 652 | 548 | 1325 | 613 | 774 | 1759 | 526 | 623 | 1144 | 608 | 659 |
| 2665 | W303.asm02.HP0_G0024470 | chrVIII | 4170 | 1028 | 1059 | 3661 | 942 | 1160 | 4036 | 971 | 1262 | 6654 | 1538 | 2033 | 5002 | 1134 | 1359 | 4621 | 1416 | 1699 |
| 2666 | W303.asm02.HP0_G0024480 | chrVIII | 1538 | 560 | 658 | 1463 | 804 | 626 | 1449 | 614 | 610 | 2610 | 520 | 748 | 2713 | 367 | 594 | 2236 | 500 | 661 |
| 2667 | W303.asm02.HP0_G0024490 | chrVIII | 3369 | 417 | 566 | 3356 | 593 | 729 | 3538 | 419 | 758 | 4758 | 693 | 1370 | 2504 | 523 | 863 | 2287 | 634 | 1126 |
| 2668 | W303.asm02.HP0_G0024500 | chrVIII | 1031 | 421 | 680 | 909 | 677 | 634 | 932 | 479 | 716 | 4961 | 664 | 1069 | 3623 | 454 | 609 | 3909 | 629 | 1016 |
| 2669 | W303.asm02.HP0_G0024510 | chrVIII | 3557 | 2059 | 2403 | 3211 | 2271 | 2355 | 3300 | 2122 | 3126 | 6924 | 1794 | 2001 | 7752 | 1248 | 1824 | 7309 | 1554 | 1903 |
| 2670 | W303.asm02.HP0_G0024520 | chrVIII | 1357 | 1700 | 1447 | 1257 | 1444 | 1401 | 1326 | 1611 | 1649 | 3638 | 1339 | 1233 | 3928 | 936 | 798 | 3793 | 1308 | 998 |
| 2671 | W303.asm02.HP0_G0024530 | chrVIII | 1255 | 1189 | 1420 | 1339 | 1214 | 1499 | 1306 | 1368 | 1472 | 5101 | 1100 | 1589 | 5371 | 716 | 1262 | 4922 | 954 | 1556 |
| 2672 | W303.asm02.HP0_G0024540 | chrVIII | 793 | 333 | 244 | 852 | 266 | 226 | 787 | 316 | 296 | 1450 | 382 | 337 | 1281 | 252 | 246 | 1253 | 364 | 312 |
| 2673 | W303.asm02.HP0_G0024550 | chrVIII | 15 | 16 | 10 | 13 | 9 | 10 | 6 | 21 | 19 | 26 | 16 | 16 | 23 | 8 | 5 | 33 | 25 | 12 |
| 2674 | W303.asm02.HP0_G0024560 | chrVIII | 1053 | 991 | 797 | 1076 | 851 | 798 | 1116 | 935 | 734 | 2801 | 789 | 893 | 2704 | 497 | 583 | 2589 | 716 | 808 |
| 2675 | W303.asm02.HP0_G0024570 | chrVIII | 2969 | 3668 | 3549 | 3170 | 3682 | 3241 | 3227 | 3757 | 3282 | 15588 | 3665 | 4405 | 12139 | 1983 | 2818 | 14085 | 3148 | 4174 |
| 2676 | W303.asm02.HP0_G0024580 | chrVIII | 648 | 269 | 228 | 619 | 243 | 266 | 708 | 225 | 233 | 1713 | 360 | 544 | 1116 | 301 | 355 | 1258 | 378 | 503 |
| 2677 | W303.asm02.HP0_G0024590 | chrVIII | 1256 | 401 | 513 | 1058 | 433 | 466 | 1101 | 367 | 552 | 4506 | 818 | 888 | 2737 | 484 | 670 | 2614 | 790 | 831 |
| 2678 | W303.asm02.HP0_G0024600 | chrVIII | 5485 | 2338 | 1730 | 5657 | 1772 | 1761 | 5287 | 2186 | 1755 | 8358 | 2442 | 2580 | 7333 | 1760 | 1948 | 6992 | 2374 | 2320 |
| 2679 | W303.asm02.HP0_G0024610 | chrVIII | 1979 | 1226 | 1114 | 1709 | 933 | 1120 | 1699 | 1128 | 1325 | 2497 | 924 | 846 | 3059 | 665 | 816 | 2687 | 901 | 712 |
| 2680 | W303.asm02.HP0_G0024620 | chrVIII | 2785 | 2118 | 2588 | 2545 | 2278 | 2215 | 2743 | 2298 | 2728 | 7339 | 1870 | 2030 | 7908 | 1224 | 1562 | 7302 | 1676 | 1893 |
| 2681 | W303.asm02.HP0_G0024630 | chrVIII | 1283 | 527 | 647 | 1191 | 516 | 585 | 1102 | 568 | 696 | 2090 | 531 | 737 | 2321 | 444 | 563 | 1774 | 573 | 635 |
| 2682 | W303.asm02.HP0_G0024640 | chrVIII | 7245 | 1035 | 1306 | 7090 | 1275 | 1623 | 7654 | 923 | 1618 | 7151 | 1861 | 2938 | 2870 | 1403 | 1869 | 2405 | 1356 | 2423 |
| 2683 | W303.asm02.HP0_G0024650 | chrVIII | 129 | 704 | 1091 | 137 | 1075 | 1096 | 112 | 676 | 1287 | 972 | 518 | 503 | 1667 | 357 | 512 | 1125 | 398 | 350 |
| 2684 | W303.asm02.HP0_G0024660 | chrVIII | 601 | 1290 | 848 | 731 | 716 | 949 | 733 | 1018 | 1019 | 5057 | 1854 | 1311 | 2388 | 825 | 800 | 3111 | 2217 | 1165 |
| 2685 | W303.asm02.HP0_G0024670 | chrVIII | 3946 | 781 | 929 | 3712 | 786 | 975 | 3260 | 742 | 1199 | 1418 | 569 | 617 | 2499 | 480 | 692 | 1656 | 532 | 527 |
| 2686 | W303.asm02.HP0_G0024680 | chrVIII | 143 | 75 | 55 | 167 | 50 | 56 | 145 | 73 | 46 | 211 | 116 | 66 | 219 | 48 | 47 | 174 | 90 | 77 |
| 2687 | W303.asm02.HP0_G0024690 | chrVIII | 309 | 228 | 185 | 248 | 148 | 217 | 201 | 190 | 248 | 266 | 134 | 149 | 234 | 72 | 119 | 312 | 147 | 136 |
| 2688 | W303.asm02.HP0_G0024700 | chrVIII | 6952 | 4868 | 6758 | 6243 | 6210 | 6664 | 6741 | 4984 | 8406 | 13020 | 4377 | 5280 | 14990 | 3229 | 4522 | 14088 | 3929 | 4529 |
| 2689 | W303.asm02.HP0_G0024710 | chrVIII | 4707 | 2309 | 1971 | 4404 | 1606 | 1810 | 4479 | 2208 | 1992 | 6082 | 2097 | 2289 | 8617 | 2045 | 2083 | 5461 | 2126 | 2105 |
| 2690 | W303.asm02.HP0_G0024720 | chrVIII | 7540 | 5772 | 4480 | 7112 | 4942 | 4196 | 7895 | 6099 | 4046 | 18433 | 6098 | 5587 | 20631 | 4219 | 4510 | 16810 | 4818 | 4698 |
| 2691 | W303.asm02.HP0_G0024730 | chrVIII | 1267 | 1837 | 2702 | 1260 | 2353 | 2626 | 1324 | 1923 | 3406 | 5058 | 1664 | 2141 | 6524 | 1305 | 1648 | 5443 | 1467 | 1948 |
| 2692 | W303.asm02.HP0_G0024740 | chrVIII | 593 | 1072 | 1029 | 589 | 1018 | 1067 | 633 | 1002 | 1175 | 3321 | 717 | 779 | 3772 | 509 | 516 | 3449 | 736 | 692 |
| 2693 | W303.asm02.HP0_G0024750 | chrVIII | 4658 | 1804 | 1192 | 4858 | 1230 | 1105 | 4141 | 1903 | 1230 | 9231 | 2819 | 2359 | 6919 | 1677 | 1591 | 7666 | 2617 | 2171 |
| 2694 | W303.asm02.HP0_G0024760 | chrVIII | 4343 | 1420 | 1844 | 4104 | 1749 | 1779 | 3925 | 1444 | 2379 | 4652 | 1178 | 1738 | 5833 | 1038 | 1494 | 4556 | 1156 | 1545 |
| 2695 | W303.asm02.HP0_G0024770 | chrVIII | 1390 | 1265 | 1658 | 1412 | 2000 | 1559 | 1483 | 1382 | 1952 | 5648 | 1297 | 1482 | 6236 | 798 | 912 | 5867 | 1178 | 1224 |

| | | | | | | | | | | | | | | | | | | | | |
|---|---|---|---|---|---|---|---|---|---|---|---|---|---|---|---|---|---|---|---|---|
| 2696 | W303.asm02.HP0_G0024780 | chrVIII | 917 | 781 | 791 | 859 | 906 | 820 | 1162 | 801 | 802 | 6921 | 1130 | 1512 | 5646 | 784 | 932 | 5308 | 905 | 1377 |
| 2697 | W303.asm02.HP0_G0024790 | chrVIII | 958 | 369 | 386 | 908 | 347 | 365 | 850 | 345 | 398 | 2058 | 567 | 630 | 1437 | 337 | 422 | 1590 | 594 | 601 |
| 2698 | W303.asm02.HP0_G0024800 | chrVIII | 504 | 392 | 586 | 511 | 594 | 629 | 485 | 409 | 651 | 3085 | 515 | 862 | 2928 | 355 | 664 | 2785 | 477 | 748 |
| 2699 | W303.asm02.HP0_G0024810 | chrVIII | 3414 | 2443 | 2257 | 3150 | 2490 | 2402 | 3227 | 2576 | 2625 | 5832 | 2065 | 1846 | 7528 | 1512 | 1749 | 5961 | 1728 | 1565 |
| 2700 | W303.asm02.HP0_G0024820 | chrVIII | 98 | 159 | 137 | 85 | 136 | 128 | 98 | 155 | 188 | 340 | 94 | 110 | 390 | 100 | 82 | 400 | 97 | 97 |
| 2701 | W303.asm02.HP0_G0024830 | chrVIII | 1984 | 959 | 1122 | 2140 | 1178 | 1063 | 2365 | 1008 | 1231 | 7189 | 1283 | 1727 | 6291 | 789 | 1179 | 6500 | 1150 | 1561 |
| 2702 | W303.asm02.HP0_G0024840 | chrVIII | 2584 | 531 | 625 | 2290 | 576 | 646 | 2484 | 590 | 671 | 5177 | 866 | 1208 | 4049 | 640 | 963 | 4459 | 847 | 1108 |
| 2703 | W303.asm02.HP0_G0024850 | chrVIII | 1728 | 649 | 629 | 1883 | 635 | 675 | 1592 | 614 | 738 | 2140 | 467 | 431 | 2166 | 261 | 333 | 2390 | 411 | 390 |
| 2704 | W303.asm02.HP0_G0024860 | chrVIII | 3580 | 420 | 784 | 3563 | 638 | 891 | 3485 | 360 | 1008 | 5678 | 771 | 1422 | 2079 | 571 | 938 | 2220 | 694 | 1253 |
| 2705 | W303.asm02.HP0_G0024870 | chrVIII | 1713 | 291 | 288 | 1746 | 290 | 316 | 1595 | 273 | 278 | 1639 | 343 | 468 | 1429 | 256 | 346 | 1148 | 319 | 411 |
| 2706 | W303.asm02.HP0_G0024880 | chrVIII | 509 | 84 | 97 | 516 | 63 | 96 | 471 | 81 | 120 | 479 | 112 | 169 | 320 | 85 | 113 | 349 | 97 | 145 |
| 2707 | W303.asm02.HP0_G0024890 | chrVIII | 5355 | 920 | 812 | 5490 | 710 | 861 | 4924 | 820 | 896 | 4403 | 1561 | 1734 | 2738 | 1062 | 1393 | 2428 | 1347 | 1588 |
| 2708 | W303.asm02.HP0_G0024900 | chrVIII | 5848 | 5332 | 6417 | 5352 | 5882 | 6356 | 5658 | 5360 | 8221 | 10389 | 4511 | 5237 | 12839 | 3322 | 4434 | 11049 | 4048 | 4194 |
| 2709 | W303.asm02.HP0_G0024910 | chrVIII | 2575 | 3034 | 3285 | 2013 | 3151 | 3364 | 2415 | 3205 | 3489 | 9151 | 3169 | 3413 | 9822 | 2347 | 2984 | 8993 | 2662 | 3005 |
| 2710 | W303.asm02.HP0_G0024920 | chrVIII | 2012 | 252 | 335 | 2093 | 360 | 362 | 1937 | 234 | 393 | 3122 | 375 | 797 | 2024 | 329 | 614 | 1867 | 329 | 723 |
| 2711 | W303.asm02.HP0_G0024930 | chrVIII | 5606 | 561 | 931 | 5477 | 899 | 1129 | 5224 | 573 | 1216 | 3309 | 832 | 2255 | 2056 | 744 | 1318 | 2038 | 673 | 1828 |
| 2712 | W303.asm02.HP0_G0024940 | chrVIII | 1473 | 395 | 404 | 1400 | 441 | 420 | 1399 | 421 | 428 | 2373 | 573 | 771 | 1876 | 413 | 592 | 1961 | 553 | 747 |
| 2713 | W303.asm02.HP0_G0024950 | chrVIII | 307 | 503 | 680 | 230 | 611 | 630 | 273 | 529 | 826 | 1872 | 381 | 490 | 1906 | 221 | 334 | 2101 | 380 | 457 |
| 2714 | W303.asm02.HP0_G0024960 | chrVIII | 1057 | 369 | 412 | 881 | 428 | 399 | 961 | 317 | 511 | 2883 | 636 | 769 | 1951 | 370 | 532 | 1852 | 588 | 683 |
| 2715 | W303.asm02.HP0_G0024970 | chrVIII | 531 | 504 | 554 | 621 | 669 | 583 | 733 | 495 | 613 | 5901 | 1051 | 1370 | 4356 | 784 | 621 | 4335 | 924 | 1128 |
| 2716 | W303.asm02.HP0_G0024980 | chrVIII | 2717 | 3063 | 3305 | 2822 | 4095 | 2954 | 3217 | 3270 | 3605 | 21054 | 4354 | 4560 | 18305 | 2537 | 2519 | 18481 | 3810 | 3873 |
| 2717 | W303.asm02.HP0_G0024990 | chrVIII | 838 | 429 | 441 | 941 | 440 | 450 | 753 | 397 | 464 | 1866 | 455 | 554 | 2031 | 258 | 509 | 1882 | 454 | 438 |
| 2718 | W303.asm02.HP0_G0025000 | chrVIII | 780 | 322 | 351 | 774 | 431 | 357 | 886 | 395 | 399 | 3586 | 499 | 780 | 2807 | 333 | 564 | 3260 | 492 | 691 |
| 2719 | W303.asm02.HP0_G0025010 | chrVIII | 79 | 36 | 32 | 57 | 42 | 47 | 66 | 49 | 50 | 167 | 38 | 50 | 180 | 30 | 40 | 183 | 34 | 42 |
| 2720 | W303.asm02.HP0_G0025020 | chrVIII | 83894 | 35612 | 36421 | 79882 | 30072 | 38524 | 90084 | 30568 | 44463 | 113127 | 58998 | 70707 | 80790 | 46677 | 54981 | 85821 | 53837 | 67024 |
| 2721 | W303.asm02.HP0_G0025030 | chrVIII | 1304 | 437 | 475 | 1098 | 338 | 501 | 1220 | 367 | 559 | 2416 | 864 | 1073 | 1879 | 571 | 730 | 1872 | 802 | 965 |
| 2722 | W303.asm02.HP0_G0025040 | chrVIII | 100 | 21 | 17 | 66 | 14 | 25 | 80 | 34 | 23 | 139 | 43 | 52 | 89 | 33 | 49 | 121 | 40 | 43 |
| 2723 | W303.asm02.HP0_G0025050 | chrVIII | 1301 | 219 | 213 | 1112 | 196 | 259 | 1131 | 198 | 264 | 843 | 284 | 332 | 791 | 213 | 259 | 727 | 294 | 321 |
| 2724 | W303.asm02.HP0_G0025060 | chrVIII | 905 | 73 | 111 | 999 | 109 | 157 | 913 | 63 | 144 | 921 | 159 | 340 | 563 | 105 | 219 | 519 | 124 | 281 |
| 2725 | W303.asm02.HP0_G0025070 | chrVIII | 864 | 248 | 266 | 776 | 241 | 288 | 870 | 243 | 307 | 1776 | 423 | 636 | 1070 | 305 | 449 | 1323 | 413 | 593 |
| 2726 | W303.asm02.HP0_G0025080 | chrVIII | 12095 | 5479 | 6133 | 11086 | 5313 | 6339 | 11921 | 5435 | 6466 | 21450 | 10481 | 10831 | 17631 | 6657 | 7751 | 19346 | 9120 | 8894 |
| 2727 | W303.asm02.HP0_G0025090 | chrVIII | 511 | 425 | 231 | 488 | 204 | 234 | 493 | 428 | 246 | 556 | 300 | 176 | 688 | 250 | 155 | 497 | 329 | 147 |
| 2728 | W303.asm02.HP0_G0025100 | chrVIII | 1160 | 1475 | 1598 | 1107 | 1404 | 1572 | 1094 | 1522 | 1950 | 3941 | 1103 | 1145 | 4219 | 714 | 928 | 4273 | 1041 | 1025 |
| 2729 | W303.asm02.HP0_G0025110 | chrVIII | 1428 | 456 | 381 | 1464 | 338 | 352 | 1374 | 424 | 312 | 3073 | 704 | 714 | 2405 | 472 | 658 | 2351 | 658 | 655 |
| 2730 | W303.asm02.HP0_G0025120 | chrVIII | 16663 | 6097 | 6250 | 15524 | 5716 | 6888 | 17697 | 5994 | 7837 | 33075 | 11320 | 13479 | 24725 | 9612 | 11713 | 25642 | 10287 | 12672 |
| 2731 | W303.asm02.HP0_G0025130 | chrVIII | 231 | 256 | 381 | 250 | 285 | 328 | 223 | 241 | 501 | 932 | 183 | 220 | 898 | 81 | 137 | 925 | 178 | 197 |
| 2732 | W303.asm02.HP0_G0025140 | chrVIII | 24 | 23 | 18 | 22 | 13 | 17 | 15 | 25 | 26 | 49 | 13 | 6 | 44 | 15 | 9 | 57 | 21 | 8 |
| 2733 | W303.asm02.HP0_G0025150 | chrVIII | 2363 | 443 | 633 | 2264 | 630 | 723 | 2417 | 495 | 792 | 5529 | 846 | 1517 | 2880 | 645 | 1032 | 2796 | 702 | 1344 |
| 2734 | W303.asm02.HP0_G0025160 | chrVIII | 1263 | 1074 | 1011 | 1123 | 1156 | 927 | 1324 | 1101 | 1056 | 3606 | 1016 | 1065 | 3938 | 597 | 694 | 3485 | 824 | 878 |
| 2735 | W303.asm02.HP0_G0025170 | chrVIII | 3862 | 799 | 1113 | 3359 | 820 | 1320 | 4162 | 698 | 1363 | 6753 | 1793 | 2785 | 4267 | 1434 | 1918 | 3626 | 1472 | 2382 |
| 2736 | W303.asm02.HP0_G0025180 | chrVIII | 495 | 82 | 109 | 399 | 71 | 131 | 381 | 65 | 122 | 553 | 175 | 218 | 211 | 85 | 177 | 206 | 176 | 218 |
| 2737 | W303.asm02.HP0_G0025190 | chrVIII | 12335 | 5063 | 4050 | 12529 | 3954 | 4200 | 12796 | 4517 | 3714 | 15584 | 5241 | 5995 | 13094 | 3509 | 4484 | 13227 | 5076 | 5491 |
| 2738 | W303.asm02.HP0_G0025200 | chrVIII | 692 | 665 | 636 | 488 | 626 | 575 | 558 | 746 | 716 | 1599 | 535 | 504 | 1891 | 321 | 458 | 1542 | 543 | 434 |
| 2739 | W303.asm02.HP0_G0025210 | chrVIII | 1695 | 1229 | 1033 | 1653 | 964 | 1026 | 1704 | 1257 | 1016 | 2960 | 946 | 1054 | 3298 | 630 | 782 | 2846 | 904 | 931 |
| 2740 | W303.asm02.HP0_G0025220 | chrVIII | 9354 | 8660 | 7398 | 8873 | 7392 | 7813 | 9675 | 8862 | 8019 | 16145 | 8045 | 8122 | 17175 | 6107 | 6369 | 14499 | 6848 | 7191 |
| 2741 | W303.asm02.HP0_G0025230 | chrVIII | 2266 | 369 | 298 | 1986 | 338 | 346 | 2094 | 284 | 472 | 1738 | 624 | 735 | 817 | 401 | 519 | 787 | 530 | 630 |
| 2742 | W303.asm02.HP0_G0025240 | chrVIII | 3777 | 1034 | 993 | 3791 | 935 | 930 | 3519 | 965 | 1008 | 3687 | 1020 | 943 | 4473 | 603 | 895 | 3503 | 931 | 838 |
| 2743 | W303.asm02.HP0_G0025250 | chrVIII | 1123 | 2823 | 4098 | 1170 | 4245 | 3673 | 1341 | 2988 | 4780 | 15522 | 2485 | 3305 | 14761 | 1643 | 1989 | 16210 | 2376 | 2770 |
| 2744 | W303.asm02.HP0_G0025260 | chrVIII | 862 | 1463 | 1733 | 996 | 2363 | 1562 | 1146 | 1457 | 1792 | 12037 | 1522 | 2071 | 11221 | 997 | 1114 | 11046 | 1426 | 1804 |
| 2745 | W303.asm02.HP0_G0025270 | chrVIII | 2704 | 1650 | 1454 | 2350 | 1215 | 1415 | 2323 | 1587 | 1648 | 3276 | 1207 | 1153 | 3555 | 851 | 1047 | 3266 | 1204 | 1063 |
| 2746 | W303.asm02.HP0_G0025280 | chrVIII | 2632 | 1301 | 882 | 2492 | 724 | 987 | 2514 | 1105 | 987 | 3671 | 2113 | 1619 | 2680 | 1357 | 1353 | 2596 | 2040 | 1451 |
| 2747 | W303.asm02.HP0_G0025290 | chrVIII | 279 | 71 | 49 | 259 | 51 | 68 | 274 | 47 | 69 | 382 | 142 | 109 | 192 | 89 | 91 | 239 | 147 | 88 |
| 2748 | W303.asm02.HP0_G0025300 | chrVIII | 5804 | 1901 | 1557 | 5131 | 1474 | 1748 | 5762 | 1653 | 1641 | 6633 | 2424 | 2854 | 5399 | 1945 | 2233 | 5261 | 2175 | 2390 |
| 2749 | W303.asm02.HP0_G0025310 | chrVIII | 4272 | 2001 | 1971 | 4185 | 1855 | 1837 | 4277 | 1960 | 2121 | 8252 | 1976 | 2425 | 7303 | 1277 | 1876 | 7602 | 1894 | 2277 |
| 2750 | W303.asm02.HP0_G0025320 | chrVIII | 2877 | 302 | 224 | 2303 | 230 | 246 | 2477 | 279 | 243 | 1485 | 393 | 411 | 1003 | 305 | 350 | 980 | 415 | 354 |
| 2751 | W303.asm02.HP0_G0025330 | chrVIII | 15699 | 5814 | 6300 | 14227 | 6763 | 6282 | 16044 | 5703 | 6244 | 25871 | 7440 | 9583 | 23044 | 6077 | 6735 | 22319 | 6183 | 8629 |
| 2752 | W303.asm02.HP0_G0025340 | chrVIII | 1667 | 501 | 525 | 1548 | 459 | 582 | 1820 | 391 | 610 | 3762 | 1024 | 1066 | 1432 | 672 | 632 | 1480 | 825 | 873 |
| 2753 | W303.asm02.HP0_G0025350 | chrVIII | 4564 | 825 | 1137 | 4733 | 1062 | 1350 | 4851 | 741 | 1400 | 9779 | 2004 | 2990 | 4787 | 1614 | 2085 | 4523 | 1717 | 2714 |
| 2754 | W303.asm02.HP0_G0025360 | chrVIII | 1980 | 654 | 922 | 2063 | 956 | 1025 | 2140 | 694 | 1100 | 11263 | 1545 | 2251 | 9542 | 1111 | 1427 | 8466 | 1244 | 2015 |
| 2755 | W303.asm02.HP0_G0025370 | chrVIII | 1484 | 693 | 526 | 1546 | 526 | 453 | 1447 | 693 | 480 | 3240 | 717 | 674 | 2874 | 450 | 484 | 2795 | 670 | 616 |
| 2756 | W303.asm02.HP0_G0025380 | chrVIII | 12427 | 6609 | 6136 | 12808 | 5955 | 6596 | 13209 | 6210 | 7749 | 29429 | 12183 | 12906 | 18656 | 8848 | 8523 | 19521 | 10244 | 10908 |
| 2757 | W303.asm02.HP0_G0025390 | chrVIII | 8311 | 1431 | 1351 | 8225 | 1044 | 1256 | 7620 | 1516 | 1680 | 2962 | 1146 | 1170 | 8939 | 1725 | 1712 | 2831 | 969 | 1007 |
| 2758 | W303.asm02.HP0_G0025400 | chrVIII | 547 | 400 | 338 | 528 | 335 | 304 | 519 | 323 | 291 | 814 | 257 | 221 | 1108 | 170 | 215 | 709 | 229 | 184 |
| 2759 | W303.asm02.HP0_G0025410 | chrVIII | 1622 | 165 | 249 | 1340 | 233 | 339 | 1555 | 149 | 297 | 1502 | 281 | 508 | 685 | 218 | 325 | 652 | 202 | 503 |
| 2760 | W303.asm02.HP0_G0025420 | chrVIII | 149 | 57 | 74 | 122 | 65 | 73 | 119 | 45 | 92 | 186 | 46 | 52 | 284 | 31 | 42 | 264 | 47 | 49 |
| 2761 | W303.asm02.HP0_G0025430 | chrVIII | 63 | 42 | 52 | 60 | 52 | 63 | 59 | 51 | 98 | 150 | 33 | 33 | 184 | 17 | 20 | 145 | 22 | 25 |
| 2762 | W303.asm02.HP0_G0025440 | chrVIII | 63 | 11 | 23 | 48 | 20 | 17 | 51 | 7 | 17 | 61 | 14 | 22 | 66 | 8 | 11 | 52 | 8 | 13 |
| 2763 | W303.asm02.HP0_G0025450 | chrVIII | 52 | 5 | 9 | 58 | 3 | 4 | 58 | 3 | 4 | 71 | 11 | 32 | 74 | 10 | 18 | 44 | 7 | 23 |
| 2764 | W303.asm02.HP0_G0025460 | chrVIII | 166 | 4 | 11 | 127 | 14 | 16 | 158 | 5 | 15 | 200 | 26 | 61 | 140 | 26 | 32 | 75 | 19 | 53 |
| 2765 | W303.asm02.HP0_G0025470 | chrVIII | 60 | 5 | 7 | 37 | 8 | 9 | 58 | 2 | 6 | 127 | 15 | 32 | 98 | 34 | 28 | 54 | 9 | 47 |
| 2766 | W303.asm02.HP0_G0025480 | chrVIII | 137 | 7 | 10 | 131 | 26 | 15 | 154 | 4 | 22 | 335 | 35 | 101 | 268 | 62 | 79 | 128 | 26 | 84 |
| 2767 | W303.asm02.HP0_G0025490 | chrVIII | 532 | 43 | 37 | 496 | 44 | 58 | 567 | 33 | 73 | 1245 | 137 | 252 | 789 | 107 | 168 | 435 | 102 | 208 |
| 2768 | W303.asm02.HP0_G0025500 | chrVIII | 466 | 35 | 60 | 502 | 63 | 60 | 496 | 37 | 86 | 1082 | 162 | 261 | 713 | 100 | 163 | 431 | 109 | 237 |
| 2769 | W303.asm02.HP0_G0025510 | chrVIII | 16 | 28 | 24 | 27 | 19 | 26 | 7 | 22 | 37 | 83 | 34 | 28 | 91 | 18 | 36 | 78 | 35 | 45 |
| 2770 | W303.asm02.HP0_G0025520 | chrVIII | 27 | 22 | 26 | 13 | 20 | 33 | 8 | 23 | 19 | 100 | 57 | 67 | 114 | 31 | 33 | 103 | 55 | 60 |
| 2771 | W303.asm02.HP0_G0025530 | chrVIII | 661 | 5010 | 5117 | 614 | 5037 | 5128 | 324 | 5194 | 4371 | 11183 | 5667 | 4679 | 10031 | 3194 | 3261 | 10633 | 3906 | 3847 |
| 2772 | W303.asm02.HP0_G0025540 | chrVIII | 2533 | 1450 | 1725 | 2037 | 1262 | 1988 | 1107 | 1254 | 2058 | 9942 | 1652 | 3641 | 2521 | 575 | 1211 | 6353 | 1496 | 2797 |
| 2773 | W303.asm02.HP0_G0025550 | chrVIII | 4 | 6 | 0 | 5 | 1 | 5 | 2 | 11 | 1 | 5 | 2 | 3 | 1 | 5 | 1 | 4 | 2 | 2 |
| 2774 | W303.asm02.HP0_G0027690 | chrX | 83 | 47 | 48 | 88 | 28 | 47 | 99 | 33 | 65 | 171 | 20 | 59 | 170 | 32 | 43 | 129 | 34 | 53 |
| 2775 | W303.asm02.HP0_G0027700 | chrX | 0 | 0 | 0 | 0 | 0 | 0 | 0 | 0 | 0 | 0 | 0 | 0 | 0 | 0 | 0 | 0 | 0 | 0 |
| 2776 | W303.asm02.HP0_G0027710 | chrX | 423 | 74 | 76 | 454 | 85 | 112 | 419 | 61 | 108 | 274 | 77 | 106 | 244 | 60 | 60 | 184 | 64 | 101 |
| 2777 | W303.asm02.HP0_G0027720 | chrX | 962 | 514 | 532 | 918 | 508 | 564 | 839 | 380 | 508 | 1108 | 338 | 337 | 1647 | 214 | 286 | 1266 | 239 | 298 |
| 2778 | W303.asm02.HP0_G0027730 | chrX | 4738 | 4676 | 4673 | 4262 | 4540 | 4851 | 4477 | 4470 | 5419 | 9786 | 5049 | 4636 | 13917 | 4658 | 4352 | 9672 | 4370 | 4293 |
| 2779 | W303.asm02.HP0_G0027740 | chrX | 259 | 53 | 93 | 215 | 102 | 97 | 256 | 54 | 110 | 541 | 74 | 151 | 354 | 49 | 90 | 386 | 61 | 112 |
| 2780 | W303.asm02.HP0_G0027750 | chrX | 410 | 127 | 214 | 397 | 221 | 210 | 436 | 143 | 258 | 964 | 131 | 238 | 949 | 100 | 150 | 791 | 125 | 164 |
| 2781 | W303.asm02.HP0_G0027760 | chrX | 1022 | 280 | 356 | 997 | 418 | 485 | 789 | 214 | 410 | 906 | 234 | 393 | 707 | 125 | 268 | 661 | 249 | 331 |
| 2782 | W303.asm02.HP0_G0027770 | chrX | 1363 | 594 | 976 | 1032 | 761 | 1075 | 1328 | 558 | 1214 | 5115 | 1072 | 1325 | 2384 | 674 | 788 | 2595 | 841 | 998 |
| 2783 | W303.asm02.HP0_G0027780 | chrX | 1303 | 732 | 659 | 1180 | 551 | 696 | 1271 | 640 | 742 | 1900 | 817 | 850 | 1665 | 649 | 742 | 1358 | 705 | 637 |
| 2784 | W303.asm02.HP0_G0027790 | chrX | 960 | 594 | 765 | 967 | 841 | 717 | 949 | 625 | 726 | 4292 | 585 | 963 | 3822 | 430 | 658 | 3923 | 612 | 806 |
| 2785 | W303.asm02.HP0_G0027800 | chrX | 2718 | 1360 | 2241 | 2651 | 2365 | 2024 | 2804 | 1533 | 2405 | 8870 | 1489 | 2158 | 9886 | 1024 | 1474 | 8602 | 1179 | 1837 |
| 2786 | W303.asm02.HP0_G0027810 | chrX | 2952 | 926 | 1303 | 2911 | 1250 | 1264 | 2862 | 928 | 1549 | 8983 | 971 | 2307 | 7507 | 727 | 1666 | 8185 | 874 | 1928 |
| 2787 | W303.asm02.HP0_G0027820 | chrX | 853 | 306 | 370 | 861 | 349 | 370 | 812 | 292 | 328 | 1910 | 376 | 518 | 1821 | 204 | 348 | 1494 | 324 | 500 |
| 2788 | W303.asm02.HP0_G0027830 | chrX | 816 | 237 | 364 | 591 | 277 | 377 | 543 | 253 | 429 | 628 | 196 | 250 | 935 | 113 | 252 | 738 | 180 | 236 |

| | | | | | | | | | | | | | | | | | | | | |
|---|---|---|---|---|---|---|---|---|---|---|---|---|---|---|---|---|---|---|---|---|
| 2789 | W303.asm02.HP0_G0027840 | chrX | 2000 | 655 | 768 | 1959 | 796 | 689 | 1864 | 695 | 918 | 3838 | 597 | 770 | 3879 | 365 | 604 | 3909 | 526 | 668 |
| 2790 | W303.asm02.HP0_G0027850 | chrX | 1865 | 877 | 783 | 1800 | 923 | 809 | 1782 | 913 | 817 | 2866 | 708 | 844 | 3087 | 432 | 629 | 2704 | 645 | 770 |
| 2791 | W303.asm02.HP0_G0027860 | chrX | 1854 | 625 | 362 | 1773 | 386 | 373 | 2059 | 691 | 417 | 2832 | 861 | 757 | 2390 | 508 | 582 | 2531 | 810 | 774 |
| 2792 | W303.asm02.HP0_G0027870 | chrX | 5186 | 1578 | 1788 | 5695 | 1739 | 1847 | 6100 | 1464 | 2417 | 12780 | 2887 | 3770 | 7163 | 1747 | 2089 | 7104 | 2438 | 3180 |
| 2793 | W303.asm02.HP0_G0027880 | chrX | 477 | 261 | 192 | 437 | 190 | 236 | 407 | 270 | 268 | 422 | 232 | 179 | 591 | 184 | 187 | 408 | 252 | 169 |
| 2794 | W303.asm02.HP0_G0027890 | chrX | 1468 | 567 | 676 | 1435 | 716 | 715 | 1686 | 483 | 878 | 7827 | 1337 | 1520 | 3866 | 927 | 795 | 3663 | 1123 | 1312 |
| 2795 | W303.asm02.HP0_G0027900 | chrX | 2124 | 443 | 666 | 2018 | 662 | 687 | 2164 | 430 | 849 | 6461 | 937 | 1733 | 3432 | 692 | 1137 | 3528 | 857 | 1524 |
| 2796 | W303.asm02.HP0_G0027910 | chrX | 2010 | 2681 | 2483 | 1612 | 2104 | 2271 | 1787 | 2429 | 2947 | 4797 | 1812 | 1637 | 4647 | 1051 | 1380 | 5123 | 1742 | 1497 |
| 2797 | W303.asm02.HP0_G0027920 | chrX | 126 | 249 | 233 | 136 | 240 | 255 | 128 | 191 | 302 | 1525 | 277 | 438 | 1409 | 208 | 300 | 1092 | 341 | 375 |
| 2798 | W303.asm02.HP0_G0027930 | chrX | 511 | 685 | 621 | 540 | 595 | 560 | 582 | 675 | 667 | 2633 | 660 | 609 | 2242 | 368 | 453 | 2507 | 634 | 562 |
| 2799 | W303.asm02.HP0_G0027940 | chrX | 3004 | 1859 | 1369 | 2712 | 1345 | 1437 | 2721 | 1903 | 1419 | 5873 | 2008 | 2029 | 4904 | 1499 | 1475 | 5049 | 1923 | 1834 |
| 2800 | W303.asm02.HP0_G0027950 | chrX | 4233 | 3642 | 4273 | 3736 | 4065 | 4164 | 4089 | 3445 | 4793 | 7117 | 1860 | 1864 | 8981 | 1612 | 2022 | 7775 | 1657 | 1728 |
| 2801 | W303.asm02.HP0_G0027960 | chrX | 7701 | 7891 | 10826 | 6777 | 10135 | 10703 | 7789 | 7906 | 13350 | 20895 | 7288 | 8638 | 23019 | 5548 | 7078 | 22283 | 6541 | 7926 |
| 2802 | W303.asm02.HP0_G0027970 | chrX | 3205 | 2212 | 3077 | 2413 | 2897 | 3084 | 2670 | 2171 | 3790 | 6399 | 1762 | 2446 | 7940 | 1396 | 2086 | 7204 | 1797 | 1987 |
| 2803 | W303.asm02.HP0_G0027980 | chrX | 2212 | 733 | 635 | 1974 | 601 | 706 | 2264 | 741 | 709 | 4908 | 1136 | 1441 | 3597 | 843 | 874 | 3655 | 1072 | 1249 |
| 2804 | W303.asm02.HP0_G0027990 | chrX | 3120 | 1621 | 1695 | 2887 | 1515 | 1517 | 2978 | 1662 | 1858 | 9607 | 2072 | 2516 | 8277 | 1546 | 1908 | 7154 | 1951 | 2282 |
| 2805 | W303.asm02.HP0_G0028000 | chrX | 517 | 542 | 302 | 538 | 259 | 286 | 511 | 599 | 312 | 889 | 451 | 314 | 671 | 264 | 244 | 839 | 509 | 245 |
| 2806 | W303.asm02.HP0_G0028010 | chrX | 801 | 955 | 1159 | 657 | 1227 | 1203 | 704 | 1135 | 1531 | 2821 | 974 | 992 | 3427 | 636 | 836 | 2902 | 879 | 945 |
| 2807 | W303.asm02.HP0_G0028020 | chrX | 1126 | 1893 | 2040 | 1006 | 1813 | 1786 | 1141 | 1916 | 1946 | 7078 | 1773 | 1989 | 7623 | 1283 | 1625 | 7011 | 1746 | 1908 |
| 2808 | W303.asm02.HP0_G0028030 | chrX | 546 | 431 | 338 | 522 | 368 | 299 | 497 | 398 | 274 | 1773 | 445 | 486 | 1784 | 275 | 363 | 1507 | 444 | 504 |
| 2809 | W303.asm02.HP0_G0028040 | chrX | 2897 | 1357 | 1339 | 2950 | 1265 | 1332 | 2657 | 1466 | 1351 | 4451 | 1204 | 1515 | 5574 | 1023 | 1477 | 4019 | 1124 | 1381 |
| 2810 | W303.asm02.HP0_G0028050 | chrX | 1914 | 990 | 1210 | 1597 | 1150 | 1242 | 1666 | 1107 | 1641 | 2788 | 803 | 915 | 3426 | 589 | 837 | 3350 | 779 | 805 |
| 2811 | W303.asm02.HP0_G0028060 | chrX | 3342 | 2159 | 1913 | 3210 | 1820 | 1932 | 3265 | 2258 | 2188 | 5579 | 1890 | 1711 | 5790 | 1258 | 1204 | 5434 | 1648 | 1493 |
| 2812 | W303.asm02.HP0_G0028070 | chrX | 8769 | 6870 | 7422 | 9045 | 7482 | 7900 | 9567 | 6846 | 7903 | 19734 | 6709 | 7757 | 19098 | 4853 | 5648 | 18525 | 5467 | 6851 |
| 2813 | W303.asm02.HP0_G0028080 | chrX | 3744 | 941 | 773 | 4106 | 784 | 884 | 3966 | 793 | 994 | 6736 | 1646 | 1824 | 3859 | 1123 | 1259 | 4151 | 1660 | 1713 |
| 2814 | W303.asm02.HP0_G0028090 | chrX | 6141 | 3800 | 3127 | 5755 | 2942 | 2900 | 5709 | 4036 | 2949 | 11175 | 4121 | 3841 | 12698 | 2962 | 3045 | 10637 | 3799 | 3457 |
| 2815 | W303.asm02.HP0_G0028100 | chrX | 1242 | 1075 | 1417 | 1022 | 1102 | 1322 | 1134 | 1152 | 1541 | 2682 | 990 | 1100 | 3464 | 701 | 853 | 2892 | 948 | 907 |
| 2816 | W303.asm02.HP0_G0028110 | chrX | 2700 | 554 | 820 | 3221 | 666 | 923 | 2565 | 547 | 1108 | 3509 | 735 | 1030 | 2629 | 689 | 968 | 1990 | 673 | 994 |
| 2817 | W303.asm02.HP0_G0028120 | chrX | 9194 | 1118 | 1234 | 8966 | 1165 | 1131 | 8702 | 1097 | 1403 | 6022 | 1194 | 1424 | 9276 | 1142 | 1618 | 5007 | 1254 | 1367 |
| 2818 | W303.asm02.HP0_G0028130 | chrX | 351 | 269 | 139 | 275 | 117 | 142 | 284 | 193 | 102 | 307 | 160 | 112 | 436 | 127 | 59 | 268 | 196 | 85 |
| 2819 | W303.asm02.HP0_G0028140 | chrX | 982 | 940 | 770 | 1045 | 873 | 747 | 970 | 966 | 732 | 6788 | 1321 | 1652 | 5100 | 856 | 1194 | 5412 | 1366 | 1495 |
| 2820 | W303.asm02.HP0_G0028150 | chrX | 15190 | 9348 | 8648 | 14921 | 8103 | 9197 | 16255 | 9149 | 8263 | 23671 | 8765 | 10623 | 21556 | 5916 | 7883 | 20921 | 7866 | 9662 |
| 2821 | W303.asm02.HP0_G0028160 | chrX | 2858 | 2507 | 2730 | 2801 | 2647 | 2718 | 2534 | 2489 | 3197 | 5561 | 2192 | 2265 | 5491 | 1690 | 1915 | 5809 | 1937 | 1717 |
| 2822 | W303.asm02.HP0_G0028170 | chrX | 5607 | 640 | 613 | 5859 | 597 | 645 | 5009 | 538 | 697 | 5129 | 1115 | 1252 | 3341 | 789 | 997 | 3179 | 1154 | 1220 |
| 2823 | W303.asm02.HP0_G0028180 | chrX | 4929 | 490 | 283 | 4670 | 193 | 324 | 4452 | 425 | 271 | 1549 | 603 | 534 | 1531 | 466 | 632 | 1215 | 611 | 503 |
| 2824 | W303.asm02.HP0_G0028190 | chrX | 2010 | 451 | 266 | 1609 | 247 | 318 | 1587 | 348 | 298 | 1039 | 432 | 399 | 998 | 285 | 423 | 884 | 505 | 359 |
| 2825 | W303.asm02.HP0_G0028200 | chrX | 952 | 802 | 638 | 894 | 675 | 577 | 1024 | 703 | 581 | 3133 | 689 | 542 | 2889 | 337 | 366 | 2800 | 674 | 516 |
| 2826 | W303.asm02.HP0_G0028210 | chrX | 2949 | 408 | 284 | 2371 | 224 | 275 | 2229 | 426 | 362 | 451 | 354 | 265 | 1533 | 351 | 396 | 373 | 385 | 214 |
| 2827 | W303.asm02.HP0_G0028220 | chrX | 100 | 173 | 63 | 92 | 77 | 74 | 77 | 128 | 51 | 231 | 114 | 58 | 287 | 96 | 52 | 169 | 178 | 36 |
| 2828 | W303.asm02.HP0_G0028230 | chrX | 44285 | 7165 | 7033 | 45746 | 6622 | 8620 | 47153 | 5504 | 10395 | 37971 | 13917 | 15974 | 14567 | 14481 | 15536 | 9011 | 11032 | 14212 |
| 2829 | W303.asm02.HP0_G0028240 | chrX | 13788 | 2911 | 4476 | 12983 | 3680 | 4823 | 14283 | 2756 | 6130 | 19144 | 7232 | 9323 | 9096 | 5486 | 6410 | 7896 | 5265 | 8422 |
| 2830 | W303.asm02.HP0_G0028250 | chrX | 2239 | 1307 | 1484 | 2014 | 1650 | 1559 | 2362 | 1234 | 1722 | 11851 | 1587 | 2429 | 10786 | 1499 | 2015 | 8253 | 1430 | 2172 |
| 2831 | W303.asm02.HP0_G0028260 | chrX | 2040 | 1099 | 1197 | 1898 | 1195 | 1123 | 1934 | 1146 | 1315 | 5912 | 1211 | 1459 | 5470 | 801 | 1113 | 5127 | 1071 | 1330 |
| 2832 | W303.asm02.HP0_G0028270 | chrX | 6258 | 979 | 1059 | 6643 | 933 | 988 | 5717 | 991 | 1137 | 3748 | 885 | 1087 | 7507 | 862 | 1262 | 3486 | 875 | 889 |
| 2833 | W303.asm02.HP0_G0028280 | chrX | 5947 | 785 | 1338 | 5887 | 1117 | 1594 | 5627 | 772 | 1759 | 8640 | 1561 | 2838 | 5804 | 1178 | 2116 | 5774 | 1479 | 2520 |
| 2834 | W303.asm02.HP0_G0028290 | chrX | 3312 | 347 | 250 | 4167 | 251 | 337 | 3422 | 376 | 276 | 9527 | 3409 | 2969 | 5637 | 3033 | 3040 | 6524 | 2917 | 3009 |
| 2835 | W303.asm02.HP0_G0028300 | chrX | 7782 | 2427 | 2097 | 6615 | 2070 | 2296 | 6617 | 2389 | 2241 | 3941 | 2194 | 2103 | 4533 | 1552 | 1785 | 3965 | 2027 | 1949 |
| 2836 | W303.asm02.HP0_G0028310 | chrX | 1410 | 460 | 336 | 1465 | 354 | 407 | 1486 | 458 | 376 | 2584 | 591 | 741 | 2896 | 427 | 538 | 2217 | 611 | 674 |
| 2837 | W303.asm02.HP0_G0028320 | chrX | 1282 | 3172 | 3956 | 1299 | 4508 | 3838 | 1515 | 3524 | 5028 | 11681 | 2921 | 2835 | 12813 | 1962 | 2036 | 12232 | 2395 | 2592 |
| 2838 | W303.asm02.HP0_G0028330 | chrX | 407 | 734 | 733 | 465 | 622 | 683 | 485 | 693 | 771 | 1878 | 498 | 527 | 2068 | 293 | 363 | 1928 | 512 | 470 |
| 2839 | W303.asm02.HP0_G0028340 | chrX | 1292 | 892 | 656 | 1476 | 634 | 628 | 1280 | 904 | 673 | 2889 | 845 | 874 | 2865 | 560 | 660 | 2694 | 820 | 729 |
| 2840 | W303.asm02.HP0_G0028350 | chrX | 1890 | 2265 | 1880 | 1821 | 1995 | 1790 | 1887 | 2233 | 1906 | 7073 | 2101 | 1907 | 7001 | 1230 | 1411 | 7090 | 1923 | 1785 |
| 2841 | W303.asm02.HP0_G0028360 | chrX | 5015 | 1716 | 344 | 4804 | 736 | 905 | 3966 | 1454 | 307 | 355 | 1242 | 310 | 464 | 637 | 342 | 438 | 1285 | 336 |
| 2842 | W303.asm02.HP0_G0028370 | chrX | 7306 | 2961 | 3123 | 6511 | 3187 | 2856 | 6875 | 3367 | 3768 | 9558 | 3506 | 3535 | 11255 | 2416 | 3268 | 9010 | 2828 | 3309 |
| 2843 | W303.asm02.HP0_G0028380 | chrX | 11512 | 503 | 346 | 11422 | 368 | 511 | 10726 | 358 | 502 | 3171 | 1204 | 1188 | 1789 | 747 | 1075 | 1699 | 1121 | 1042 |
| 2844 | W303.asm02.HP0_G0028390 | chrX | 5722 | 2985 | 3268 | 4999 | 2920 | 3490 | 5183 | 3113 | 3942 | 8958 | 2652 | 3057 | 9418 | 1679 | 2478 | 8792 | 2265 | 2732 |
| 2845 | W303.asm02.HP0_G0028400 | chrX | 1957 | 749 | 644 | 2074 | 641 | 697 | 1918 | 726 | 749 | 3539 | 825 | 918 | 2791 | 482 | 656 | 2955 | 835 | 859 |
| 2846 | W303.asm02.HP0_G0028410 | chrX | 14230 | 8505 | 8232 | 15248 | 8207 | 8116 | 15581 | 8832 | 8323 | 23826 | 8569 | 8965 | 22257 | 6013 | 6674 | 22743 | 7472 | 8476 |
| 2847 | W303.asm02.HP0_G0028420 | chrX | 949 | 453 | 219 | 953 | 213 | 251 | 915 | 464 | 228 | 1375 | 394 | 400 | 1670 | 391 | 324 | 1135 | 476 | 299 |
| 2848 | W303.asm02.HP0_G0028430 | chrX | 6208 | 4194 | 5453 | 5989 | 5395 | 5633 | 5983 | 4323 | 6849 | 11086 | 3840 | 4584 | 13122 | 3131 | 3999 | 12223 | 3737 | 4164 |
| 2849 | W303.asm02.HP0_G0028440 | chrX | 3 | 4 | 1 | 7 | 1 | 1 | 2 | 4 | 0 | 3 | 1 | 0 | 7 | 3 | 0 | 2 | 4 | 3 |
| 2850 | W303.asm02.HP0_G0028450 | chrX | 1717 | 783 | 719 | 1514 | 627 | 630 | 1518 | 759 | 757 | 3429 | 912 | 849 | 2813 | 644 | 692 | 2790 | 900 | 774 |
| 2851 | W303.asm02.HP0_G0028460 | chrX | 689 | 911 | 685 | 833 | 583 | 761 | 600 | 795 | 676 | 844 | 930 | 752 | 1132 | 584 | 768 | 944 | 861 | 675 |
| 2852 | W303.asm02.HP0_G0028470 | chrX | 1999 | 1067 | 1064 | 1975 | 1144 | 1046 | 1950 | 1106 | 1010 | 5203 | 1248 | 1542 | 5201 | 886 | 1099 | 4502 | 1220 | 1285 |
| 2853 | W303.asm02.HP0_G0028480 | chrX | 916 | 401 | 292 | 845 | 270 | 280 | 800 | 384 | 285 | 1370 | 419 | 422 | 1451 | 326 | 372 | 1024 | 421 | 401 |
| 2854 | W303.asm02.HP0_G0028490 | chrX | 2712 | 723 | 861 | 2561 | 779 | 875 | 2361 | 754 | 919 | 2407 | 560 | 763 | 2845 | 430 | 649 | 2457 | 572 | 678 |
| 2855 | W303.asm02.HP0_G0028500 | chrX | 12310 | 2479 | 3142 | 11101 | 3379 | 3487 | 12881 | 2584 | 4114 | 22081 | 4559 | 9042 | 13983 | 4663 | 7062 | 11451 | 3830 | 7742 |
| 2856 | W303.asm02.HP0_G0028510 | chrX | 2612 | 685 | 973 | 2751 | 995 | 956 | 2551 | 708 | 1213 | 8016 | 1142 | 1799 | 5657 | 891 | 1167 | 4769 | 949 | 1553 |
| 2857 | W303.asm02.HP0_G0028520 | chrX | 3365 | 536 | 914 | 3211 | 747 | 1157 | 3417 | 480 | 1167 | 6600 | 1087 | 2058 | 4204 | 931 | 1530 | 3361 | 951 | 1885 |
| 2858 | W303.asm02.HP0_G0028530 | chrX | 1653 | 627 | 871 | 1437 | 764 | 857 | 1559 | 616 | 1047 | 1985 | 527 | 653 | 2339 | 400 | 525 | 1901 | 432 | 484 |
| 2859 | W303.asm02.HP0_G0028540 | chrX | 2193 | 147 | 126 | 1962 | 165 | 160 | 1984 | 116 | 184 | 2173 | 330 | 527 | 933 | 221 | 300 | 974 | 301 | 409 |
| 2860 | W303.asm02.HP0_G0028550 | chrX | 1497 | 628 | 446 | 1199 | 424 | 428 | 1413 | 546 | 483 | 1728 | 575 | 477 | 1708 | 367 | 397 | 1702 | 576 | 431 |
| 2861 | W303.asm02.HP0_G0028560 | chrX | 1635 | 1873 | 1474 | 1975 | 1825 | 1442 | 2017 | 1936 | 1489 | 7585 | 1932 | 1970 | 7010 | 1328 | 1136 | 6832 | 1761 | 1752 |
| 2862 | W303.asm02.HP0_G0028570 | chrX | 5613 | 2556 | 2686 | 5498 | 2538 | 2774 | 5920 | 2690 | 2674 | 8765 | 2517 | 3068 | 8875 | 1711 | 2228 | 8271 | 2060 | 2753 |
| 2863 | W303.asm02.HP0_G0028580 | chrX | 10947 | 3008 | 2374 | 10625 | 2559 | 2446 | 10571 | 3120 | 2438 | 11533 | 3672 | 3945 | 9859 | 2707 | 3202 | 9316 | 3242 | 3540 |
| 2864 | W303.asm02.HP0_G0028590 | chrX | 816 | 1730 | 2807 | 720 | 3288 | 2790 | 860 | 2169 | 3323 | 8716 | 1932 | 2356 | 9517 | 1199 | 1413 | 8570 | 1543 | 2065 |
| 2865 | W303.asm02.HP0_G0028600 | chrX | 4188 | 1637 | 1333 | 3441 | 1371 | 1367 | 4071 | 1811 | 1324 | 7400 | 2502 | 2678 | 6495 | 1571 | 1922 | 6563 | 2333 | 2488 |
| 2866 | W303.asm02.HP0_G0028610 | chrX | 507 | 219 | 189 | 433 | 217 | 205 | 463 | 202 | 244 | 974 | 348 | 298 | 806 | 299 | 242 | 709 | 354 | 344 |
| 2867 | W303.asm02.HP0_G0028620 | chrX | 1809 | 1781 | 1657 | 1819 | 1560 | 1781 | 1907 | 1814 | 1738 | 9177 | 2986 | 2835 | 7400 | 2130 | 2000 | 7571 | 2514 | 2741 |
| 2868 | W303.asm02.HP0_G0028630 | chrX | 929 | 567 | 525 | 870 | 567 | 535 | 914 | 494 | 489 | 1739 | 779 | 761 | 963 | 262 | 347 | 1241 | 750 | 705 |
| 2869 | W303.asm02.HP0_G0028640 | chrX | 1545 | 733 | 742 | 1366 | 770 | 756 | 1420 | 745 | 954 | 4286 | 1218 | 1391 | 3046 | 740 | 1122 | 2878 | 1107 | 1404 |
| 2870 | W303.asm02.HP0_G0028650 | chrX | 2821 | 1152 | 1031 | 2588 | 1051 | 1048 | 2691 | 1177 | 1072 | 6811 | 1661 | 1885 | 5237 | 1124 | 1444 | 5455 | 1522 | 1681 |
| 2871 | W303.asm02.HP0_G0028660 | chrX | 7583 | 5100 | 4753 | 6928 | 4857 | 4647 | 8057 | 5244 | 4896 | 23276 | 6553 | 7715 | 18895 | 4454 | 5430 | 20944 | 5614 | 6994 |
| 2872 | W303.asm02.HP0_G0028670 | chrX | 6342 | 1813 | 2560 | 6143 | 2953 | 2494 | 6817 | 1956 | 3079 | 29230 | 3424 | 5718 | 21324 | 2506 | 3696 | 20880 | 2933 | 5075 |
| 2873 | W303.asm02.HP0_G0028680 | chrX | 218 | 464 | 188 | 264 | 170 | 163 | 222 | 212 | 197 | 2195 | 535 | 423 | 1803 | 310 | 280 | 1116 | 677 | 395 |
| 2874 | W303.asm02.HP0_G0028690 | chrX | 396 | 124 | 104 | 379 | 104 | 118 | 411 | 120 | 112 | 749 | 151 | 175 | 660 | 120 | 147 | 547 | 197 | 183 |
| 2875 | W303.asm02.HP0_G0028700 | chrX | 681 | 263 | 181 | 803 | 213 | 201 | 729 | 304 | 219 | 934 | 307 | 247 | 801 | 193 | 193 | 877 | 238 | 235 |
| 2876 | W303.asm02.HP0_G0028710 | chrX | 5826 | 2573 | 2316 | 5512 | 2387 | 2408 | 5337 | 2865 | 2607 | 6813 | 2512 | 2586 | 7859 | 1836 | 2149 | 6448 | 2186 | 2434 |
| 2877 | W303.asm02.HP0_G0028720 | chrX | 1636 | 357 | 376 | 2051 | 473 | 334 | 1504 | 406 | 371 | 2598 | 528 | 550 | 2120 | 388 | 475 | 2176 | 414 | 425 |
| 2878 | W303.asm02.HP0_G0028730 | chrX | 1971 | 324 | 358 | 1957 | 396 | 433 | 1865 | 276 | 475 | 3183 | 548 | 792 | 1427 | 583 | 696 | 1597 | 447 | 691 |
| 2879 | W303.asm02.HP0_G0028740 | chrX | 6920 | 1172 | 1553 | 5672 | 1308 | 1750 | 6242 | 979 | 2052 | 5266 | 2447 | 2915 | 2059 | 1813 | 2127 | 2336 | 1887 | 2100 |
| 2880 | W303.asm02.HP0_G0028750 | chrX | 3012 | 330 | 432 | 2601 | 381 | 503 | 2847 | 322 | 535 | 2228 | 652 | 820 | 2577 | 514 | 703 | 1231 | 565 | 725 |
| 2881 | W303.asm02.HP0_G0028760 | chrX | 3615 | 560 | 591 | 3592 | 531 | 571 | 3691 | 525 | 732 | 3408 | 667 | 917 | 3582 | 741 | 768 | 2201 | 567 | 779 |

| | | | | | | | | | | | | | | | | | | | | |
|---|---|---|---|---|---|---|---|---|---|---|---|---|---|---|---|---|---|---|---|---|
| 2882 | W303.asm02.HP0_G0028770 | chrX | 1582 | 652 | 978 | 1589 | 1461 | 981 | 1692 | 675 | 1013 | 11654 | 1080 | 1864 | 9727 | 846 | 1162 | 8550 | 867 | 1802 |
| 2883 | W303.asm02.HP0_G0028780 | chrX | 2518 | 1640 | 1333 | 2215 | 1240 | 1288 | 2410 | 1919 | 1218 | 4659 | 1632 | 1852 | 4457 | 1080 | 1326 | 4498 | 1411 | 1726 |
| 2884 | W303.asm02.HP0_G0028790 | chrX | 3779 | 1074 | 1868 | 3426 | 1735 | 1876 | 3421 | 1076 | 2501 | 4163 | 846 | 1540 | 5052 | 708 | 1377 | 4246 | 875 | 1341 |
| 2885 | W303.asm02.HP0_G0028800 | chrX | 1388 | 213 | 313 | 1388 | 290 | 380 | 1402 | 229 | 413 | 2746 | 387 | 791 | 1403 | 311 | 510 | 1334 | 348 | 726 |
| 2886 | W303.asm02.HP0_G0028810 | chrX | 1214 | 427 | 564 | 1395 | 527 | 573 | 1084 | 483 | 635 | 3212 | 690 | 854 | 2545 | 519 | 715 | 2473 | 620 | 806 |
| 2887 | W303.asm02.HP0_G0028820 | chrX | 1236 | 781 | 857 | 1197 | 788 | 849 | 1188 | 764 | 885 | 4647 | 1246 | 1293 | 3636 | 884 | 993 | 3330 | 1093 | 1164 |
| 2888 | W303.asm02.HP0_G0028830 | chrX | 2107 | 353 | 489 | 2079 | 474 | 543 | 2305 | 338 | 564 | 3708 | 557 | 1093 | 2073 | 507 | 772 | 2155 | 487 | 984 |
| 2889 | W303.asm02.HP0_G0028840 | chrX | 1679 | 970 | 1715 | 1529 | 994 | 1616 | 1563 | 1049 | 1840 | 3782 | 810 | 1909 | 3913 | 586 | 1278 | 4150 | 765 | 1717 |
| 2890 | W303.asm02.HP0_G0028850 | chrX | 1178 | 541 | 805 | 1256 | 739 | 702 | 1281 | 607 | 747 | 4523 | 650 | 962 | 4084 | 372 | 644 | 4571 | 630 | 920 |
| 2891 | W303.asm02.HP0_G0028860 | chrX | 567 | 262 | 217 | 605 | 582 | 650 | 453 | 289 | 259 | 577 | 150 | 195 | 583 | 109 | 164 | 660 | 165 | 173 |
| 2892 | W303.asm02.HP0_G0028870 | chrX | 83 | 1076 | 1094 | 72 | 859 | 960 | 85 | 855 | 1028 | 7173 | 1997 | 1803 | 4644 | 1283 | 1181 | 6401 | 1648 | 1571 |
| 2893 | W303.asm02.HP0_G0028880 | chrX | 740 | 631 | 1094 | 759 | 1200 | 915 | 785 | 710 | 1040 | 6465 | 707 | 1197 | 5632 | 445 | 741 | 5842 | 588 | 1068 |
| 2894 | W303.asm02.HP0_G0028890 | chrX | 1722 | 1001 | 1428 | 1696 | 1139 | 1376 | 1750 | 1150 | 1334 | 4571 | 916 | 1327 | 4656 | 559 | 975 | 4968 | 808 | 1153 |
| 2895 | W303.asm02.HP0_G0028900 | chrX | 2178 | 140 | 308 | 2162 | 297 | 375 | 2112 | 126 | 428 | 3008 | 335 | 742 | 1528 | 235 | 476 | 1333 | 303 | 580 |
| 2896 | W303.asm02.HP0_G0028910 | chrX | 848 | 61 | 132 | 951 | 121 | 170 | 831 | 38 | 153 | 1118 | 144 | 273 | 445 | 82 | 215 | 429 | 110 | 253 |
| 2897 | W303.asm02.HP0_G0028920 | chrX | 7833 | 585 | 516 | 7926 | 565 | 662 | 7636 | 456 | 587 | 5436 | 1300 | 1340 | 4704 | 1428 | 1515 | 2882 | 1204 | 1299 |
| 2898 | W303.asm02.HP0_G0028930 | chrX | 5001 | 1182 | 1206 | 4642 | 1303 | 1319 | 5164 | 1206 | 1165 | 6742 | 1403 | 2320 | 5378 | 969 | 1515 | 5495 | 1205 | 2052 |
| 2899 | W303.asm02.HP0_G0028940 | chrX | 16359 | 2054 | 2441 | 14107 | 2176 | 2881 | 16728 | 1781 | 3102 | 25642 | 4542 | 6706 | 9272 | 3677 | 4509 | 9920 | 3704 | 6111 |
| 2900 | W303.asm02.HP0_G0028950 | chrX | 6181 | 622 | 805 | 4955 | 677 | 1118 | 6040 | 436 | 1016 | 2535 | 1174 | 1354 | 775 | 729 | 1207 | 791 | 866 | 1169 |
| 2901 | W303.asm02.HP0_G0028960 | chrX | 7142 | 998 | 907 | 5997 | 968 | 1209 | 7663 | 642 | 1234 | 2764 | 1196 | 1888 | 1327 | 966 | 947 | 1138 | 841 | 1375 |
| 2902 | W303.asm02.HP0_G0028970 | chrX | 345 | 81 | 79 | 249 | 75 | 104 | 319 | 64 | 86 | 260 | 85 | 121 | 171 | 65 | 82 | 198 | 63 | 96 |
| 2903 | W303.asm02.HP0_G0028980 | chrX | 4257 | 615 | 903 | 3948 | 881 | 1079 | 4211 | 535 | 1138 | 10594 | 1676 | 2788 | 3609 | 1185 | 1636 | 4087 | 1362 | 2465 |
| 2904 | W303.asm02.HP0_G0028990 | chrX | 2850 | 683 | 1155 | 3016 | 932 | 1299 | 3192 | 719 | 1260 | 6157 | 913 | 2004 | 5696 | 708 | 1316 | 5297 | 791 | 1767 |
| 2905 | W303.asm02.HP0_G0029000 | chrX | 1373 | 462 | 1468 | 1416 | 592 | 1655 | 1461 | 466 | 1601 | 4349 | 750 | 3531 | 3760 | 593 | 1948 | 3175 | 719 | 2820 |
| 2906 | W303.asm02.HP0_G0029010 | chrX | 1017 | 906 | 1144 | 830 | 1050 | 1152 | 902 | 984 | 1366 | 2504 | 705 | 813 | 3161 | 448 | 683 | 2825 | 580 | 723 |
| 2907 | W303.asm02.HP0_G0029020 | chrX | 1153 | 939 | 817 | 1039 | 914 | 732 | 1083 | 994 | 791 | 3686 | 885 | 982 | 3751 | 651 | 729 | 3405 | 812 | 881 |
| 2908 | W303.asm02.HP0_G0029030 | chrX | 2622 | 436 | 496 | 2419 | 421 | 541 | 2429 | 505 | 546 | 2230 | 571 | 716 | 2354 | 397 | 705 | 1832 | 510 | 641 |
| 2909 | W303.asm02.HP0_G0029040 | chrX | 897 | 919 | 1121 | 899 | 1193 | 1132 | 1108 | 831 | 1299 | 9499 | 1567 | 2214 | 5520 | 1182 | 1147 | 5722 | 1441 | 1871 |
| 2910 | W303.asm02.HP0_G0029050 | chrX | 5573 | 1627 | 1286 | 5238 | 1099 | 1295 | 5141 | 1563 | 1382 | 3474 | 1503 | 1595 | 4170 | 1130 | 1464 | 3117 | 1386 | 1531 |
| 2911 | W303.asm02.HP0_G0029060 | chrX | 9911 | 831 | 542 | 9813 | 509 | 623 | 9886 | 698 | 650 | 2247 | 925 | 1041 | 2858 | 892 | 1060 | 1771 | 888 | 913 |
| 2912 | W303.asm02.HP0_G0029070 | chrX | 1837 | 686 | 431 | 1696 | 489 | 544 | 1750 | 681 | 372 | 1382 | 595 | 614 | 1359 | 436 | 396 | 1141 | 484 | 520 |
| 2913 | W303.asm02.HP0_G0029080 | chrX | 7168 | 1449 | 1412 | 6098 | 1432 | 1453 | 6949 | 1501 | 1477 | 4993 | 1630 | 2122 | 5229 | 1231 | 1956 | 4154 | 1492 | 2011 |
| 2914 | W303.asm02.HP0_G0029090 | chrX | 4190 | 1686 | 2061 | 3317 | 1793 | 2117 | 3404 | 1801 | 2429 | 2707 | 1346 | 1690 | 3972 | 990 | 1624 | 2788 | 1299 | 1477 |
| 2915 | W303.asm02.HP0_G0029100 | chrX | 1347 | 717 | 876 | 1277 | 674 | 952 | 1460 | 676 | 1070 | 4620 | 1013 | 1750 | 2999 | 703 | 1078 | 2952 | 927 | 1551 |
| 2916 | W303.asm02.HP0_G0029110 | chrX | 2877 | 1817 | 1980 | 2731 | 2043 | 1899 | 2949 | 2015 | 1803 | 8670 | 1734 | 2174 | 7964 | 1175 | 1528 | 8211 | 1511 | 1996 |
| 2917 | W303.asm02.HP0_G0029120 | chrX | 5627 | 1186 | 1051 | 3753 | 754 | 1074 | 4757 | 1223 | 1009 | 3960 | 1194 | 1588 | 4835 | 1073 | 1460 | 3683 | 1064 | 1341 |
| 2918 | W303.asm02.HP0_G0029130 | chrX | 1110 | 614 | 484 | 904 | 419 | 490 | 1017 | 654 | 524 | 1310 | 501 | 490 | 1523 | 310 | 390 | 1446 | 442 | 392 |
| 2919 | W303.asm02.HP0_G0029140 | chrX | 421 | 297 | 333 | 437 | 307 | 301 | 418 | 294 | 371 | 2065 | 340 | 457 | 1828 | 198 | 292 | 1752 | 314 | 423 |
| 2920 | W303.asm02.HP0_G0029150 | chrX | 3325 | 761 | 645 | 3254 | 496 | 710 | 3018 | 619 | 640 | 2235 | 721 | 953 | 2253 | 443 | 822 | 2040 | 759 | 872 |
| 2921 | W303.asm02.HP0_G0029160 | chrX | 283 | 279 | 429 | 322 | 260 | 533 | 266 | 195 | 515 | 2111 | 478 | 990 | 1002 | 302 | 565 | 926 | 547 | 884 |
| 2922 | W303.asm02.HP0_G0029170 | chrX | 5197 | 1808 | 1501 | 4373 | 1393 | 1463 | 4642 | 1866 | 1489 | 4703 | 2110 | 2223 | 4746 | 1387 | 1885 | 4366 | 1917 | 2046 |
| 2923 | W303.asm02.HP0_G0029180 | chrX | 7232 | 1204 | 1014 | 7337 | 1064 | 1062 | 7612 | 1058 | 977 | 7490 | 1397 | 2301 | 6947 | 1105 | 1367 | 6055 | 1210 | 1830 |
| 2924 | W303.asm02.HP0_G0029190 | chrX | 4539 | 1664 | 1308 | 4360 | 981 | 1322 | 4437 | 1476 | 1241 | 3671 | 1284 | 1497 | 4033 | 849 | 1208 | 3667 | 1206 | 1302 |
| 2925 | W303.asm02.HP0_G0029200 | chrX | 61110 | 1700 | 2345 | 59815 | 1480 | 2836 | 63171 | 1469 | 1949 | 1102 | 2084 | 2576 | 1312 | 1257 | 2635 | 921 | 1775 | 2141 |
| 2926 | W303.asm02.HP0_G0029210 | chrX | 1839 | 603 | 601 | 1658 | 636 | 608 | 1677 | 566 | 657 | 3817 | 734 | 925 | 3430 | 630 | 849 | 3012 | 606 | 842 |
| 2927 | W303.asm02.HP0_G0029220 | chrX | 1741 | 1131 | 1157 | 1591 | 1662 | 1186 | 2087 | 1180 | 1456 | 16503 | 2445 | 3080 | 9860 | 1432 | 1500 | 10681 | 2081 | 2674 |
| 2928 | W303.asm02.HP0_G0029230 | chrX | 1722 | 1147 | 1265 | 1727 | 1173 | 1258 | 1692 | 1180 | 1490 | 3242 | 899 | 950 | 3388 | 585 | 737 | 3568 | 891 | 836 |
| 2929 | W303.asm02.HP0_G0029240 | chrX | 9282 | 2177 | 1905 | 9935 | 1686 | 1730 | 9341 | 2008 | 1934 | 3856 | 2118 | 1965 | 3591 | 1125 | 1591 | 3793 | 2003 | 1633 |
| 2930 | W303.asm02.HP0_G0029250 | chrX | 2308 | 625 | 735 | 2516 | 705 | 789 | 2252 | 631 | 624 | 4208 | 666 | 1230 | 3907 | 489 | 841 | 3559 | 638 | 1067 |
| 2931 | W303.asm02.HP0_G0029260 | chrX | 1803 | 943 | 1196 | 1733 | 1071 | 1086 | 1758 | 1057 | 1316 | 4105 | 876 | 1103 | 4347 | 628 | 871 | 4255 | 811 | 1042 |
| 2932 | W303.asm02.HP0_G0029270 | chrX | 702 | 140 | 188 | 658 | 174 | 180 | 658 | 150 | 227 | 430 | 163 | 174 | 369 | 123 | 123 | 276 | 157 | 151 |
| 2933 | W303.asm02.HP0_G0029280 | chrX | 3446 | 1120 | 838 | 3017 | 931 | 859 | 3264 | 1063 | 886 | 4952 | 1217 | 1259 | 4260 | 898 | 1035 | 4230 | 1119 | 1102 |
| 2934 | W303.asm02.HP0_G0029290 | chrX | 41 | 15 | 13 | 45 | 16 | 16 | 48 | 16 | 17 | 63 | 17 | 30 | 30 | 11 | 18 | 32 | 20 | 26 |
| 2935 | W303.asm02.HP0_G0029300 | chrX | 10698 | 1281 | 1806 | 10015 | 1639 | 2128 | 10153 | 1268 | 2171 | 11678 | 2527 | 4243 | 8659 | 2008 | 2890 | 8142 | 2080 | 3559 |
| 2936 | W303.asm02.HP0_G0029310 | chrX | 3058 | 1304 | 1675 | 3088 | 1848 | 1690 | 3132 | 1413 | 2033 | 14433 | 1701 | 3198 | 12022 | 1258 | 2268 | 12571 | 1522 | 2781 |
| 2937 | W303.asm02.HP0_G0029320 | chrX | 565 | 116 | 154 | 491 | 151 | 145 | 458 | 155 | 165 | 587 | 196 | 231 | 534 | 102 | 159 | 539 | 181 | 178 |
| 2938 | W303.asm02.HP0_G0029330 | chrX | 292 | 56 | 72 | 246 | 67 | 70 | 253 | 64 | 104 | 118 | 57 | 100 | 149 | 28 | 36 | 145 | 58 | 73 |
| 2939 | W303.asm02.HP0_G0029340 | chrX | 5164 | 1184 | 1551 | 5207 | 1271 | 1482 | 4691 | 1232 | 1608 | 5384 | 1228 | 1606 | 6257 | 759 | 1386 | 5393 | 1125 | 1442 |
| 2940 | W303.asm02.HP0_G0029350 | chrX | 968 | 482 | 827 | 854 | 428 | 962 | 898 | 445 | 984 | 1682 | 452 | 951 | 1729 | 288 | 546 | 1834 | 472 | 892 |
| 2941 | W303.asm02.HP0_G0029360 | chrX | 12236 | 5040 | 12498 | 12380 | 5168 | 14873 | 11986 | 4136 | 13805 | 32961 | 9309 | 28426 | 28424 | 8249 | 18713 | 26487 | 9040 | 24766 |
| 2942 | W303.asm02.HP0_G0029370 | chrX | 2256 | 652 | 835 | 2330 | 1218 | 789 | 2750 | 676 | 967 | 12845 | 1692 | 2039 | 8161 | 1139 | 891 | 8210 | 1323 | 1680 |
| 2943 | W303.asm02.HP0_G0029380 | chrX | 3064 | 1187 | 1790 | 3316 | 1210 | 1692 | 3126 | 1167 | 2014 | 4078 | 904 | 1525 | 4453 | 711 | 1193 | 4333 | 863 | 1342 |
| 2944 | W303.asm02.HP0_G0029390 | chrX | 1521 | 903 | 1212 | 1411 | 903 | 1169 | 1430 | 940 | 1503 | 2419 | 818 | 1069 | 3258 | 627 | 828 | 2524 | 700 | 862 |
| 2945 | W303.asm02.HP0_G0029400 | chrX | 3008 | 508 | 528 | 2957 | 520 | 578 | 2816 | 565 | 592 | 5499 | 922 | 1238 | 4459 | 670 | 1017 | 4114 | 809 | 1079 |
| 2946 | W303.asm02.HP0_G0029410 | chrX | 20 | 7 | 5 | 17 | 7 | 1 | 19 | 3 | 4 | 13 | 10 | 6 | 8 | 5 | 3 | 22 | 5 | 9 |
| 2947 | W303.asm02.HP0_G0029420 | chrX | 86 | 16 | 10 | 45 | 12 | 14 | 60 | 13 | 18 | 47 | 22 | 14 | 53 | 8 | 9 | 33 | 16 | 12 |
| 2948 | W303.asm02.HP0_G0029430 | chrX | 16569 | 2924 | 2694 | 15549 | 2614 | 2480 | 16749 | 2940 | 2485 | 12189 | 3477 | 4177 | 13535 | 3123 | 3357 | 10773 | 3088 | 3341 |
| 2949 | W303.asm02.HP0_G0029440 | chrX | 211 | 227 | 210 | 187 | 222 | 196 | 221 | 184 | 239 | 1753 | 303 | 403 | 1334 | 182 | 230 | 1308 | 293 | 347 |
| 2950 | W303.asm02.HP0_G0029450 | chrX | 1824 | 1256 | 1283 | 1721 | 1167 | 1274 | 1659 | 1247 | 1478 | 3362 | 964 | 1197 | 3671 | 737 | 922 | 3398 | 1050 | 1096 |
| 2951 | W303.asm02.HP0_G0029460 | chrX | 1234 | 521 | 476 | 1170 | 495 | 447 | 1121 | 510 | 446 | 1609 | 447 | 419 | 1860 | 296 | 334 | 1637 | 468 | 336 |
| 2952 | W303.asm02.HP0_G0029470 | chrX | 13144 | 703 | 773 | 12032 | 626 | 1006 | 12816 | 523 | 1003 | 892 | 519 | 855 | 1071 | 521 | 727 | 747 | 444 | 728 |
| 2953 | W303.asm02.HP0_G0029480 | chrX | 826 | 234 | 319 | 787 | 312 | 330 | 889 | 209 | 393 | 3442 | 591 | 727 | 1999 | 377 | 499 | 1885 | 558 | 689 |
| 2954 | W303.asm02.HP0_G0029490 | chrX | 7438 | 653 | 757 | 6503 | 723 | 752 | 6750 | 657 | 882 | 3623 | 850 | 1155 | 4096 | 629 | 1033 | 2864 | 820 | 986 |
| 2955 | W303.asm02.HP0_G0029500 | chrX | 12258 | 4277 | 3783 | 12009 | 3823 | 3514 | 12671 | 4313 | 3934 | 21414 | 5985 | 6818 | 17100 | 3885 | 4780 | 18582 | 5178 | 6158 |
| 2956 | W303.asm02.HP0_G0029510 | chrX | 2585 | 640 | 604 | 2532 | 660 | 574 | 2731 | 664 | 559 | 3266 | 665 | 790 | 3299 | 500 | 709 | 3079 | 695 | 766 |
| 2957 | W303.asm02.HP0_G0029520 | chrX | 7092 | 6222 | 5645 | 6868 | 5584 | 5381 | 7957 | 6154 | 5560 | 28282 | 9530 | 10605 | 25538 | 6681 | 8071 | 24127 | 8168 | 9568 |
| 2958 | W303.asm02.HP0_G0029530 | chrX | 165 | 716 | 809 | 140 | 810 | 868 | 149 | 766 | 1173 | 2616 | 684 | 698 | 2915 | 392 | 416 | 2799 | 611 | 577 |
| 2959 | W303.asm02.HP0_G0029540 | chrX | 1543 | 1388 | 1507 | 1386 | 1886 | 1448 | 1760 | 1681 | 1433 | 11697 | 1887 | 2206 | 9051 | 1107 | 1253 | 9978 | 1652 | 2020 |
| 2960 | W303.asm02.HP0_G0029550 | chrX | 9602 | 4914 | 4616 | 10011 | 4739 | 4151 | 10118 | 4955 | 4462 | 22417 | 6201 | 7109 | 19369 | 4099 | 5120 | 19680 | 5472 | 6285 |
| 2961 | W303.asm02.HP0_G0029560 | chrX | 564 | 856 | 840 | 552 | 809 | 845 | 566 | 812 | 1066 | 2252 | 636 | 650 | 2458 | 404 | 433 | 2541 | 760 | 563 |
| 2962 | W303.asm02.HP0_G0029570 | chrX | 8855 | 616 | 946 | 8308 | 911 | 1161 | 7984 | 686 | 1110 | 6681 | 1061 | 1959 | 4349 | 876 | 1964 | 4083 | 951 | 1703 |
| 2963 | W303.asm02.HP0_G0029580 | chrX | 1349 | 1262 | 1147 | 1178 | 1014 | 1115 | 1204 | 1335 | 1626 | 3356 | 1054 | 791 | 3694 | 684 | 641 | 3746 | 950 | 747 |
| 2964 | W303.asm02.HP0_G0029590 | chrX | 1021 | 617 | 755 | 932 | 571 | 725 | 820 | 599 | 965 | 1609 | 467 | 583 | 2032 | 321 | 475 | 1694 | 460 | 460 |
| 2965 | W303.asm02.HP0_G0029600 | chrX | 6873 | 3427 | 3514 | 6363 | 3088 | 3277 | 7125 | 3592 | 3417 | 15191 | 4847 | 5203 | 13330 | 3218 | 3932 | 13204 | 4391 | 4708 |
| 2966 | W303.asm02.HP0_G0029610 | chrX | 7146 | 3308 | 2949 | 7706 | 2647 | 2897 | 7159 | 3273 | 2821 | 7681 | 3208 | 3788 | 7895 | 2377 | 3042 | 7616 | 2952 | 3456 |
| 2967 | W303.asm02.HP0_G0029620 | chrX | 4302 | 1436 | 1223 | 3573 | 1000 | 1222 | 3925 | 1241 | 1396 | 7089 | 2581 | 2636 | 3934 | 1946 | 1914 | 3873 | 2363 | 2388 |
| 2968 | W303.asm02.HP0_G0029630 | chrX | 5129 | 1415 | 1244 | 4923 | 1677 | 1198 | 5567 | 1395 | 1469 | 13893 | 2759 | 2878 | 10023 | 1920 | 1525 | 10220 | 2414 | 2554 |
| 2969 | W303.asm02.HP0_G0029640 | chrX | 1691 | 1355 | 1424 | 1605 | 1513 | 1389 | 1832 | 1525 | 1450 | 6270 | 1292 | 1383 | 5580 | 764 | 908 | 6262 | 1156 | 1149 |
| 2970 | W303.asm02.HP0_G0029650 | chrX | 1305 | 298 | 310 | 1271 | 262 | 305 | 1333 | 287 | 363 | 2398 | 716 | 668 | 1562 | 490 | 483 | 1417 | 604 | 621 |
| 2971 | W303.asm02.HP0_G0029660 | chrX | 4486 | 1033 | 1172 | 4715 | 1207 | 1066 | 4509 | 1037 | 1038 | 7400 | 1332 | 1789 | 6781 | 954 | 1452 | 6918 | 1222 | 1743 |
| 2972 | W303.asm02.HP0_G0029670 | chrX | 79 | 11 | 9 | 43 | 13 | 25 | 68 | 23 | 15 | 28 | 14 | 8 | 20 | 11 | 6 | 35 | 10 | 15 |
| 2973 | W303.asm02.HP0_G0029680 | chrX | 4027 | 832 | 651 | 3737 | 647 | 716 | 3987 | 837 | 793 | 4954 | 1377 | 1505 | 3466 | 1091 | 1207 | 3381 | 1314 | 1365 |
| 2974 | W303.asm02.HP0_G0029690 | chrX | 7249 | 4375 | 5352 | 7547 | 5452 | 5036 | 7719 | 4794 | 4996 | 19222 | 4613 | 5571 | 19852 | 3069 | 4290 | 19042 | 3821 | 5000 |

| | | | | | | | | | | | | | | | | | | | | |
|---|---|---|---|---|---|---|---|---|---|---|---|---|---|---|---|---|---|---|---|---|
| 2975 | W303.asm02.HP0_G0029700 | chrX | 7518 | 811 | 555 | 7462 | 551 | 567 | 6930 | 769 | 589 | 2398 | 1042 | 727 | 3365 | 779 | 956 | 2078 | 1109 | 646 |
| 2976 | W303.asm02.HP0_G0029710 | chrX | 63794 | 21420 | 24820 | 65862 | 19880 | 24828 | 68143 | 21992 | 24907 | 60251 | 31508 | 39614 | 47453 | 18888 | 31001 | 52917 | 28309 | 39114 |
| 2977 | W303.asm02.HP0_G0029720 | chrX | 4702 | 862 | 830 | 3314 | 665 | 809 | 3980 | 824 | 825 | 2099 | 795 | 693 | 2524 | 676 | 749 | 2046 | 872 | 534 |
| 2978 | W303.asm02.HP0_G0029730 | chrX | 442 | 831 | 819 | 369 | 738 | 727 | 373 | 797 | 1028 | 1612 | 605 | 524 | 1961 | 353 | 370 | 1865 | 525 | 491 |
| 2979 | W303.asm02.HP0_G0029740 | chrX | 739 | 784 | 555 | 760 | 502 | 649 | 732 | 745 | 750 | 1645 | 592 | 478 | 1566 | 329 | 359 | 1873 | 628 | 384 |
| 2980 | W303.asm02.HP0_G0029750 | chrX | 1670 | 1276 | 960 | 1381 | 931 | 913 | 1426 | 1283 | 916 | 3758 | 1078 | 1069 | 3442 | 722 | 794 | 3370 | 1077 | 881 |
| 2981 | W303.asm02.HP0_G0029760 | chrX | 6133 | 2601 | 2129 | 5545 | 2261 | 2404 | 5907 | 2516 | 1962 | 7075 | 2448 | 2684 | 7021 | 1901 | 1988 | 6201 | 2192 | 2410 |
| 2982 | W303.asm02.HP0_G0029770 | chrX | 81 | 40 | 18 | 80 | 20 | 17 | 60 | 34 | 18 | 146 | 40 | 35 | 128 | 23 | 27 | 149 | 45 | 28 |
| 2983 | W303.asm02.HP0_G0029780 | chrX | 13 | 11 | 13 | 14 | 6 | 14 | 8 | 10 | 5 | 65 | 28 | 19 | 61 | 10 | 9 | 60 | 21 | 14 |
| 2984 | W303.asm02.HP0_G0029790 | chrX | 2830 | 1956 | 1689 | 2602 | 1619 | 1640 | 2874 | 2181 | 1696 | 6932 | 1978 | 2042 | 6542 | 1273 | 1631 | 6524 | 1799 | 1787 |
| 2985 | W303.asm02.HP0_G0029800 | chrX | 5648 | 5640 | 4539 | 5627 | 4282 | 5023 | 6104 | 4343 | 5938 | 27715 | 11649 | 11858 | 11392 | 7698 | 6643 | 13689 | 9937 | 9677 |
| 2986 | W303.asm02.HP0_G0029810 | chrX | 4316 | 1553 | 1199 | 3314 | 1280 | 1245 | 3912 | 1624 | 1308 | 4183 | 1585 | 1883 | 4546 | 1305 | 1711 | 3988 | 1554 | 1761 |
| 2987 | W303.asm02.HP0_G0029820 | chrX | 1612 | 666 | 416 | 1776 | 318 | 466 | 1682 | 542 | 461 | 1742 | 947 | 971 | 1541 | 633 | 651 | 1195 | 949 | 748 |
| 2988 | W303.asm02.HP0_G0029830 | chrX | 1041 | 215 | 119 | 862 | 95 | 118 | 850 | 153 | 113 | 585 | 198 | 203 | 618 | 149 | 190 | 491 | 228 | 187 |
| 2989 | W303.asm02.HP0_G0029840 | chrX | 404 | 495 | 631 | 339 | 411 | 712 | 283 | 569 | 905 | 852 | 351 | 486 | 1168 | 215 | 367 | 996 | 371 | 413 |
| 2990 | W303.asm02.HP0_G0029850 | chrX | 1895 | 1661 | 2134 | 1576 | 1762 | 1988 | 1890 | 1735 | 2806 | 5293 | 1418 | 1890 | 5817 | 889 | 1330 | 5239 | 1386 | 1660 |
| 2991 | W303.asm02.HP0_G0029860 | chrX | 3301 | 3278 | 3137 | 2898 | 2593 | 3126 | 2981 | 3132 | 3255 | 4351 | 2676 | 2824 | 4200 | 1385 | 2067 | 4585 | 2338 | 2475 |
| 2992 | W303.asm02.HP0_G0029870 | chrX | 1124 | 415 | 394 | 1003 | 456 | 486 | 1073 | 398 | 508 | 3365 | 609 | 965 | 2207 | 448 | 667 | 2342 | 575 | 892 |
| 2993 | W303.asm02.HP0_G0029880 | chrX | 1201 | 535 | 660 | 1181 | 802 | 667 | 1230 | 617 | 773 | 5250 | 648 | 1125 | 4627 | 459 | 902 | 4404 | 556 | 1047 |
| 2994 | W303.asm02.HP0_G0029890 | chrX | 1446 | 950 | 1553 | 1229 | 1448 | 1531 | 1524 | 1135 | 1827 | 5626 | 866 | 1440 | 6447 | 591 | 1057 | 5944 | 718 | 1191 |
| 2995 | W303.asm02.HP0_G0029900 | chrX | 3277 | 266 | 358 | 2849 | 361 | 425 | 2834 | 271 | 424 | 3448 | 528 | 1023 | 2230 | 416 | 841 | 2050 | 482 | 887 |
| 2996 | W303.asm02.HP0_G0029910 | chrX | 2509 | 548 | 788 | 2171 | 664 | 820 | 2272 | 560 | 1068 | 1625 | 520 | 624 | 2323 | 351 | 687 | 1586 | 442 | 617 |
| 2997 | W303.asm02.HP0_G0029920 | chrX | 2273 | 292 | 294 | 2186 | 375 | 357 | 2041 | 263 | 365 | 3110 | 482 | 789 | 2183 | 328 | 533 | 2179 | 415 | 656 |
| 2998 | W303.asm02.HP0_G0029930 | chrX | 1502 | 169 | 204 | 1760 | 158 | 186 | 1294 | 162 | 220 | 910 | 180 | 244 | 743 | 119 | 254 | 886 | 158 | 220 |
| 2999 | W303.asm02.HP0_G0029940 | chrX | 1953 | 303 | 434 | 1990 | 382 | 406 | 1839 | 335 | 520 | 2314 | 420 | 586 | 1881 | 289 | 512 | 1928 | 398 | 538 |
| 3000 | W303.asm02.HP0_G0029950 | chrX | 1159 | 512 | 644 | 956 | 612 | 706 | 1032 | 541 | 761 | 4812 | 1035 | 1496 | 3006 | 717 | 1051 | 3246 | 981 | 1340 |
| 3001 | W303.asm02.HP0_G0029960 | chrX | 1677 | 902 | 1234 | 1637 | 1310 | 1119 | 1848 | 1051 | 1347 | 12136 | 1459 | 2087 | 9919 | 910 | 1252 | 10178 | 1275 | 1920 |
| 3002 | W303.asm02.HP0_G0029970 | chrX | 5156 | 1744 | 1590 | 4499 | 1787 | 1492 | 5149 | 1844 | 1493 | 11061 | 2557 | 2872 | 8589 | 1640 | 2064 | 9226 | 2243 | 2682 |
| 3003 | W303.asm02.HP0_G0029980 | chrX | 1256 | 656 | 812 | 1175 | 754 | 714 | 1310 | 710 | 593 | 3679 | 672 | 994 | 3732 | 474 | 624 | 3372 | 620 | 929 |
| 3004 | W303.asm02.HP0_G0029990 | chrX | 3322 | 1618 | 1897 | 3130 | 1683 | 1889 | 2972 | 1759 | 1916 | 4415 | 1332 | 1651 | 5589 | 1093 | 1588 | 4406 | 1198 | 1524 |
| 3005 | W303.asm02.HP0_G0030000 | chrX | 51867 | 7564 | 8999 | 49617 | 7743 | 11437 | 52469 | 5278 | 11505 | 41666 | 13939 | 21293 | 20432 | 12391 | 16505 | 19765 | 12581 | 18830 |
| 3006 | W303.asm02.HP0_G0030010 | chrX | 1258 | 981 | 687 | 1208 | 761 | 1054 | 1225 | 580 | 840 | 4389 | 1459 | 1418 | 1596 | 885 | 899 | 2097 | 1479 | 1319 |
| 3007 | W303.asm02.HP0_G0030020 | chrX | 8507 | 5744 | 6242 | 8297 | 5779 | 5933 | 8276 | 6236 | 7043 | 11200 | 5109 | 6499 | 10155 | 3765 | 4523 | 10463 | 4475 | 5039 |
| 3008 | W303.asm02.HP0_G0030030 | chrX | 5499 | 935 | 978 | 5400 | 972 | 964 | 5348 | 1046 | 1068 | 5962 | 1549 | 1625 | 5249 | 1146 | 1470 | 4836 | 1429 | 1513 |
| 3009 | W303.asm02.HP0_G0030040 | chrX | 1630 | 946 | 971 | 1565 | 899 | 1019 | 1496 | 913 | 1056 | 2813 | 779 | 957 | 2708 | 503 | 702 | 2655 | 752 | 798 |
| 3010 | W303.asm02.HP0_G0030050 | chrX | 3205 | 1415 | 1207 | 2923 | 1213 | 1076 | 2857 | 1322 | 1353 | 5313 | 1566 | 1534 | 4894 | 1074 | 1359 | 4786 | 1457 | 1413 |
| 3011 | W303.asm02.HP0_G0030060 | chrX | 2634 | 666 | 712 | 2392 | 729 | 715 | 2421 | 673 | 687 | 2736 | 634 | 786 | 3094 | 485 | 679 | 2293 | 581 | 683 |
| 3012 | W303.asm02.HP0_G0030070 | chrX | 676 | 132 | 198 | 691 | 185 | 174 | 706 | 159 | 176 | 1885 | 198 | 314 | 1683 | 117 | 223 | 1554 | 182 | 307 |
| 3013 | W303.asm02.HP0_G0030080 | chrX | 597 | 579 | 2104 | 574 | 1257 | 1977 | 654 | 684 | 2318 | 5407 | 691 | 1960 | 7006 | 465 | 1272 | 5406 | 658 | 1536 |
| 3014 | W303.asm02.HP0_G0030090 | chrX | 411 | 573 | 664 | 434 | 566 | 748 | 418 | 519 | 776 | 1675 | 472 | 673 | 1615 | 287 | 366 | 1763 | 450 | 571 |
| 3015 | W303.asm02.HP0_G0030100 | chrX | 1374 | 835 | 651 | 1433 | 713 | 630 | 1249 | 776 | 624 | 2891 | 696 | 675 | 3023 | 418 | 521 | 2805 | 724 | 656 |
| 3016 | W303.asm02.HP0_G0030110 | chrX | 1174 | 1201 | 1127 | 1088 | 983 | 1178 | 1124 | 1176 | 1512 | 2835 | 902 | 835 | 3393 | 689 | 683 | 3125 | 860 | 766 |
| 3017 | W303.asm02.HP0_G0030120 | chrX | 886 | 1019 | 956 | 864 | 789 | 924 | 775 | 963 | 1225 | 2099 | 864 | 681 | 2297 | 559 | 599 | 2171 | 871 | 666 |
| 3018 | W303.asm02.HP0_G0030130 | chrX | 5874 | 651 | 633 | 4431 | 411 | 820 | 5171 | 486 | 705 | 1489 | 939 | 1188 | 700 | 652 | 664 | 619 | 841 | 887 |
| 3019 | W303.asm02.HP0_G0030140 | chrX | 2394 | 1087 | 782 | 2509 | 845 | 822 | 2406 | 991 | 898 | 5929 | 1990 | 1995 | 4531 | 1360 | 1483 | 4308 | 1667 | 1855 |
| 3020 | W303.asm02.HP0_G0030150 | chrX | 1842 | 447 | 458 | 1541 | 448 | 474 | 1567 | 462 | 492 | 2343 | 540 | 783 | 1921 | 449 | 641 | 1866 | 479 | 705 |
| 3021 | W303.asm02.HP0_G0030160 | chrX | 2867 | 839 | 904 | 3120 | 763 | 795 | 2769 | 846 | 791 | 4620 | 982 | 1276 | 3942 | 638 | 1031 | 3756 | 932 | 1125 |
| 3022 | W303.asm02.HP0_G0030170 | chrX | 0 | 2 | 0 | 0 | 0 | 0 | 0 | 1 | 0 | 0 | 0 | 0 | 0 | 1 | 0 | 0 | 1 | 0 |
| 3023 | W303.asm02.HP0_G0030180 | chrX | 180 | 385 | 721 | 165 | 743 | 706 | 172 | 471 | 891 | 1745 | 370 | 570 | 1773 | 203 | 353 | 1711 | 306 | 522 |
| 3024 | W303.asm02.HP0_G0030190 | chrX | 6922 | 2539 | 2722 | 6260 | 2672 | 2588 | 7187 | 2524 | 3129 | 16242 | 3952 | 5191 | 11931 | 2691 | 3523 | 12730 | 3574 | 4650 |
| 3025 | W303.asm02.HP0_G0030200 | chrX | 11998 | 2000 | 1728 | 10608 | 1386 | 1885 | 11264 | 1717 | 2032 | 12080 | 4075 | 4295 | 11370 | 3423 | 3761 | 9562 | 3674 | 3919 |
| 3026 | W303.asm02.HP0_G0030210 | chrX | 3593 | 616 | 931 | 3472 | 870 | 951 | 3208 | 596 | 1238 | 4809 | 712 | 1735 | 2834 | 550 | 1123 | 2958 | 661 | 1437 |
| 3027 | W303.asm02.HP0_G0030220 | chrX | 839 | 444 | 688 | 564 | 661 | 670 | 645 | 458 | 860 | 2018 | 380 | 504 | 2222 | 250 | 428 | 2110 | 355 | 444 |
| 3028 | W303.asm02.HP0_G0030230 | chrX | 3025 | 1705 | 1984 | 2749 | 1778 | 1889 | 2897 | 1786 | 2259 | 6330 | 1641 | 1757 | 6960 | 1077 | 1328 | 6494 | 1442 | 1611 |
| 3029 | W303.asm02.HP0_G0030240 | chrX | 1323 | 1802 | 1953 | 929 | 1875 | 1866 | 1193 | 1913 | 2272 | 5444 | 1575 | 1505 | 5528 | 966 | 1236 | 5571 | 1421 | 1392 |
| 3030 | W303.asm02.HP0_G0030250 | chrX | 5691 | 2652 | 2026 | 5306 | 2372 | 2024 | 5981 | 2705 | 2002 | 12548 | 3389 | 4239 | 10908 | 2375 | 2346 | 11099 | 2865 | 3529 |
| 3031 | W303.asm02.HP0_G0030260 | chrX | 3852 | 2522 | 1560 | 3457 | 1864 | 1501 | 3881 | 2407 | 1439 | 7984 | 2715 | 2521 | 7296 | 1895 | 1666 | 6745 | 2389 | 2122 |
| 3032 | W303.asm02.HP0_G0030270 | chrX | 9829 | 1223 | 1210 | 9278 | 921 | 1359 | 10059 | 1065 | 1440 | 8020 | 3777 | 4104 | 5617 | 3086 | 3808 | 5544 | 3338 | 3693 |
| 3033 | W303.asm02.HP0_G0030280 | chrX | 1526 | 863 | 786 | 1522 | 701 | 792 | 1568 | 906 | 735 | 2650 | 989 | 1191 | 2822 | 800 | 956 | 2445 | 940 | 1026 |
| 3034 | W303.asm02.HP0_G0030290 | chrX | 4438 | 2121 | 1849 | 4266 | 1730 | 1832 | 4741 | 2300 | 1790 | 9808 | 2695 | 2976 | 7762 | 1767 | 1973 | 8152 | 2598 | 2690 |
| 3035 | W303.asm02.HP0_G0030300 | chrX | 4475 | 3128 | 2606 | 4381 | 2365 | 2530 | 4629 | 3121 | 2484 | 8683 | 2740 | 3134 | 9342 | 1993 | 2513 | 7973 | 2502 | 2681 |
| 3036 | W303.asm02.HP0_G0030310 | chrX | 4289 | 3319 | 3722 | 3877 | 3262 | 4002 | 3940 | 3112 | 4962 | 16386 | 6735 | 8159 | 10073 | 5418 | 5531 | 9712 | 5628 | 6142 |
| 3037 | W303.asm02.HP0_G0030320 | chrX | 1240 | 441 | 484 | 1141 | 369 | 559 | 1180 | 437 | 584 | 873 | 633 | 929 | 668 | 346 | 461 | 588 | 627 | 817 |
| 3038 | W303.asm02.HP0_G0030330 | chrX | 4075 | 1470 | 1262 | 4074 | 1296 | 1278 | 3872 | 1430 | 1471 | 5887 | 1654 | 1980 | 6135 | 1286 | 1605 | 4939 | 1550 | 1615 |
| 3039 | W303.asm02.HP0_G0030340 | chrX | 1945 | 1209 | 1375 | 1569 | 1253 | 1370 | 1662 | 1281 | 1814 | 3280 | 1045 | 1030 | 4390 | 716 | 908 | 3614 | 925 | 910 |
| 3040 | W303.asm02.HP0_G0030350 | chrX | 1526 | 434 | 290 | 1570 | 243 | 309 | 1436 | 410 | 280 | 1480 | 512 | 509 | 1408 | 316 | 348 | 1285 | 545 | 411 |
| 3041 | W303.asm02.HP0_G0030360 | chrX | 3067 | 1949 | 2023 | 2826 | 1530 | 1942 | 2721 | 1937 | 2502 | 3013 | 1649 | 1517 | 3191 | 1223 | 1393 | 2761 | 1541 | 1387 |
| 3042 | W303.asm02.HP0_G0030370 | chrX | 911 | 874 | 716 | 711 | 619 | 768 | 680 | 873 | 961 | 1333 | 608 | 558 | 1638 | 385 | 508 | 1594 | 599 | 450 |
| 3043 | W303.asm02.HP0_G0030380 | chrX | 3910 | 3141 | 2826 | 4237 | 2420 | 2688 | 4140 | 3120 | 3419 | 7225 | 2469 | 2193 | 7891 | 1664 | 1769 | 7616 | 2380 | 2045 |
| 3044 | W303.asm02.HP0_G0030390 | chrX | 1349 | 629 | 610 | 1356 | 613 | 594 | 1302 | 669 | 590 | 4561 | 874 | 1263 | 3634 | 507 | 794 | 3640 | 882 | 1023 |
| 3045 | W303.asm02.HP0_G0030400 | chrX | 2848 | 623 | 892 | 3177 | 950 | 1037 | 2766 | 605 | 1056 | 7586 | 1061 | 2006 | 5845 | 753 | 1629 | 5787 | 896 | 1763 |
| 3046 | W303.asm02.HP0_G0030410 | chrX | 6388 | 403 | 683 | 6516 | 572 | 768 | 6169 | 386 | 804 | 5335 | 1102 | 1591 | 1759 | 794 | 1256 | 1612 | 893 | 1318 |
| 3047 | W303.asm02.HP0_G0030420 | chrX | 3330 | 533 | 650 | 3111 | 691 | 722 | 3360 | 486 | 822 | 3933 | 848 | 1674 | 1594 | 787 | 1065 | 1401 | 704 | 1340 |
| 3048 | W303.asm02.HP0_G0030430 | chrX | 2231 | 409 | 463 | 2295 | 380 | 518 | 2334 | 375 | 583 | 3625 | 851 | 1115 | 3003 | 589 | 763 | 2724 | 842 | 1000 |
| 3049 | W303.asm02.HP0_G0030440 | chrX | 784 | 118 | 93 | 798 | 71 | 103 | 643 | 100 | 91 | 869 | 139 | 142 | 474 | 71 | 101 | 709 | 139 | 111 |
| 3050 | W303.asm02.HP0_G0030450 | chrX | 4972 | 3867 | 5607 | 4641 | 5360 | 5591 | 4651 | 3985 | 7278 | 10492 | 3556 | 4392 | 12741 | 2509 | 3619 | 11368 | 3156 | 3654 |
| 3051 | W303.asm02.HP0_G0030460 | chrX | 215 | 86 | 33 | 199 | 50 | 42 | 162 | 69 | 52 | 117 | 94 | 46 | 232 | 81 | 36 | 118 | 123 | 30 |
| 3052 | W303.asm02.HP0_G0030470 | chrX | 5975 | 1118 | 1106 | 5874 | 883 | 1087 | 5481 | 1091 | 1276 | 2376 | 960 | 904 | 3394 | 689 | 817 | 2403 | 915 | 724 |
| 3053 | W303.asm02.HP0_G0030480 | chrX | 71 | 520 | 546 | 68 | 561 | 559 | 82 | 550 | 691 | 1961 | 409 | 440 | 2327 | 267 | 243 | 2135 | 435 | 394 |
| 3054 | W303.asm02.HP0_G0030490 | chrX | 1045 | 359 | 296 | 1097 | 254 | 258 | 1010 | 356 | 294 | 1918 | 469 | 458 | 1687 | 315 | 291 | 1555 | 467 | 419 |
| 3055 | W303.asm02.HP0_G0030500 | chrX | 1350 | 1096 | 827 | 1283 | 746 | 809 | 1172 | 1026 | 907 | 2018 | 889 | 855 | 2431 | 623 | 648 | 2161 | 883 | 754 |
| 3056 | W303.asm02.HP0_G0030510 | chrX | 3007 | 1521 | 1197 | 2755 | 1096 | 1154 | 2708 | 1471 | 1385 | 3585 | 1108 | 1022 | 3813 | 803 | 894 | 3742 | 1158 | 962 |
| 3057 | W303.asm02.HP0_G0030520 | chrX | 7435 | 1934 | 2528 | 6689 | 2374 | 2359 | 6828 | 2047 | 3192 | 6501 | 1701 | 2138 | 8227 | 1360 | 1836 | 6601 | 1730 | 1934 |
| 3058 | W303.asm02.HP0_G0030530 | chrX | 1499 | 1029 | 933 | 1256 | 706 | 933 | 1240 | 962 | 1181 | 2425 | 768 | 800 | 2922 | 518 | 581 | 2689 | 797 | 676 |
| 3059 | W303.asm02.HP0_G0030540 | chrX | 7895 | 750 | 569 | 7483 | 525 | 699 | 8023 | 600 | 787 | 3304 | 1345 | 1435 | 1578 | 1051 | 1044 | 1601 | 1223 | 1166 |
| 3060 | W303.asm02.HP0_G0030550 | chrX | 9704 | 8613 | 8898 | 8916 | 7771 | 9314 | 9310 | 8067 | 9956 | 13338 | 8651 | 9331 | 13175 | 6028 | 7133 | 12742 | 7750 | 7704 |
| 3061 | W303.asm02.HP0_G0030560 | chrX | 5813 | 5095 | 3483 | 4833 | 3573 | 3442 | 5824 | 4944 | 3693 | 15666 | 7358 | 7201 | 13822 | 4990 | 5168 | 14362 | 6716 | 6320 |
| 3062 | W303.asm02.HP0_G0030570 | chrX | 1445 | 253 | 488 | 1478 | 284 | 534 | 1368 | 234 | 557 | 2658 | 539 | 1233 | 1858 | 350 | 778 | 1685 | 420 | 1150 |
| 3063 | W303.asm02.HP0_G0030580 | chrX | 1502 | 266 | 355 | 1334 | 271 | 424 | 1478 | 286 | 449 | 1570 | 414 | 880 | 1228 | 307 | 604 | 1245 | 348 | 792 |
| 3064 | W303.asm02.HP0_G0030590 | chrX | 1456 | 372 | 566 | 1167 | 516 | 561 | 1077 | 457 | 806 | 1486 | 418 | 500 | 1778 | 291 | 505 | 1625 | 409 | 427 |
| 3065 | W303.asm02.HP0_G0030600 | chrX | 413 | 66 | 76 | 393 | 55 | 59 | 396 | 81 | 50 | 256 | 92 | 72 | 244 | 63 | 51 | 213 | 72 | 79 |
| 3066 | W303.asm02.HP0_G0030610 | chrX | 2478 | 3640 | 4586 | 2315 | 4232 | 4970 | 2398 | 3543 | 5755 | 19654 | 5561 | 10163 | 11065 | 4641 | 6389 | 10734 | 4383 | 9121 |
| 3067 | W303.asm02.HP0_G0030620 | chrX | 1113 | 444 | 487 | 938 | 478 | 490 | 941 | 444 | 551 | 2833 | 587 | 750 | 2041 | 392 | 632 | 2469 | 632 | 651 |

| | | | | | | | | | | | | | | | | | | | | |
|---|---|---|---|---|---|---|---|---|---|---|---|---|---|---|---|---|---|---|---|---|
| 3068 | W303.asm02.HP0_G0030630 | chrX | 1525 | 726 | 711 | 1475 | 889 | 676 | 1606 | 781 | 729 | 3610 | 801 | 991 | 3985 | 558 | 644 | 3366 | 673 | 787 |
| 3069 | W303.asm02.HP0_G0030640 | chrX | 218 | 599 | 466 | 205 | 572 | 461 | 222 | 570 | 426 | 1371 | 414 | 444 | 1847 | 310 | 314 | 1259 | 373 | 411 |
| 3070 | W303.asm02.HP0_G0030650 | chrX | 904 | 728 | 662 | 692 | 618 | 687 | 785 | 690 | 907 | 1371 | 576 | 513 | 1450 | 377 | 362 | 1467 | 566 | 490 |
| 3071 | W303.asm02.HP0_G0030660 | chrX | 5275 | 2073 | 1746 | 4946 | 1590 | 1648 | 5126 | 1970 | 1925 | 4804 | 2143 | 2483 | 5801 | 1711 | 1987 | 4358 | 2038 | 2238 |
| 3072 | W303.asm02.HP0_G0030670 | chrX | 718 | 142 | 47 | 729 | 55 | 58 | 631 | 98 | 41 | 238 | 140 | 80 | 286 | 66 | 42 | 212 | 112 | 58 |
| 3073 | W303.asm02.HP0_G0030680 | chrX | 734 | 359 | 273 | 619 | 225 | 315 | 707 | 273 | 305 | 1542 | 550 | 421 | 1358 | 315 | 346 | 1207 | 478 | 438 |
| 3074 | W303.asm02.HP0_G0030690 | chrX | 4864 | 2564 | 2013 | 4456 | 1866 | 2128 | 4589 | 2210 | 1637 | 11225 | 2546 | 3376 | 11000 | 2061 | 2728 | 10112 | 2499 | 3057 |
| 3075 | W303.asm02.HP0_G0030700 | chrX | 1705 | 1182 | 1719 | 1630 | 1248 | 1661 | 1665 | 1188 | 2293 | 3812 | 845 | 1184 | 4091 | 534 | 924 | 4571 | 732 | 1090 |
| 3076 | W303.asm02.HP0_G0030710 | chrX | 423 | 438 | 580 | 363 | 543 | 545 | 438 | 433 | 582 | 1846 | 456 | 542 | 1613 | 228 | 365 | 1635 | 412 | 496 |
| 3077 | W303.asm02.HP0_G0030720 | chrX | 162 | 33 | 25 | 126 | 13 | 22 | 148 | 28 | 16 | 197 | 48 | 66 | 137 | 30 | 41 | 165 | 50 | 25 |
| 3078 | W303.asm02.HP0_G0030730 | chrX | 23149 | 6216 | 6876 | 21574 | 6206 | 7897 | 21954 | 5689 | 9210 | 37642 | 13571 | 15738 | 30259 | 12165 | 12568 | 27403 | 10967 | 12337 |
| 3079 | W303.asm02.HP0_G0030740 | chrX | 3150 | 1261 | 1204 | 2957 | 1109 | 1292 | 2811 | 1260 | 1146 | 3809 | 1423 | 2034 | 3671 | 1245 | 1689 | 3277 | 1313 | 1603 |
| 3080 | W303.asm02.HP0_G0030750 | chrX | 25497 | 22308 | 26061 | 24161 | 25976 | 26627 | 26892 | 22514 | 28558 | 52987 | 21382 | 25031 | 53035 | 16266 | 19867 | 50320 | 18255 | 22595 |
| 3081 | W303.asm02.HP0_G0030760 | chrX | 753 | 1235 | 1029 | 645 | 1051 | 902 | 784 | 1238 | 1055 | 6142 | 1369 | 1153 | 5122 | 845 | 815 | 5535 | 1197 | 998 |
| 3082 | W303.asm02.HP0_G0030770 | chrX | 7201 | 1510 | 1141 | 6894 | 1120 | 1188 | 6603 | 1438 | 1230 | 6264 | 2057 | 2269 | 5973 | 1605 | 1982 | 5120 | 1869 | 2030 |
| 3083 | W303.asm02.HP0_G0030780 | chrX | 3209 | 847 | 825 | 2964 | 848 | 914 | 2871 | 732 | 973 | 6866 | 1682 | 1853 | 4630 | 1148 | 1404 | 4660 | 1555 | 1615 |
| 3084 | W303.asm02.HP0_G0030790 | chrX | 3299 | 231 | 303 | 2909 | 285 | 330 | 3066 | 247 | 344 | 1817 | 386 | 607 | 849 | 339 | 475 | 741 | 285 | 510 |
| 3085 | W303.asm02.HP0_G0030800 | chrX | 860 | 458 | 463 | 812 | 497 | 413 | 969 | 508 | 516 | 2024 | 564 | 598 | 2298 | 363 | 369 | 1706 | 460 | 535 |
| 3086 | W303.asm02.HP0_G0030810 | chrX | 3670 | 944 | 677 | 3472 | 797 | 698 | 3669 | 893 | 726 | 4312 | 1309 | 1258 | 3837 | 1043 | 1133 | 3299 | 1158 | 1206 |
| 3087 | W303.asm02.HP0_G0030820 | chrX | 1330 | 317 | 376 | 1260 | 369 | 420 | 1221 | 271 | 445 | 2009 | 404 | 592 | 1397 | 317 | 436 | 1322 | 363 | 498 |
| 3088 | W303.asm02.HP0_G0030830 | chrX | 2202 | 808 | 1189 | 1912 | 1123 | 1018 | 2192 | 916 | 1253 | 8752 | 1334 | 2024 | 6072 | 735 | 1292 | 7066 | 1145 | 1900 |
| 3089 | W303.asm02.HP0_G0030840 | chrX | 2154 | 930 | 1039 | 2113 | 1028 | 1030 | 2006 | 1053 | 1183 | 3935 | 1017 | 1279 | 4970 | 923 | 1175 | 3955 | 887 | 1132 |
| 3090 | W303.asm02.HP0_G0030850 | chrX | 3518 | 887 | 1042 | 3562 | 1096 | 1039 | 3576 | 927 | 1026 | 5903 | 1023 | 1598 | 5057 | 654 | 1068 | 4974 | 966 | 1352 |
| 3091 | W303.asm02.HP0_G0030860 | chrX | 566 | 342 | 370 | 534 | 369 | 310 | 572 | 344 | 384 | 1136 | 248 | 289 | 1238 | 189 | 227 | 1214 | 257 | 281 |
| 3092 | W303.asm02.HP0_G0030870 | chrX | 1472 | 760 | 969 | 1256 | 821 | 998 | 1296 | 724 | 1372 | 2120 | 596 | 856 | 2755 | 417 | 723 | 2527 | 582 | 698 |
| 3093 | W303.asm02.HP0_G0030880 | chrX | 770 | 710 | 516 | 733 | 591 | 571 | 787 | 627 | 543 | 2209 | 547 | 550 | 2207 | 344 | 350 | 2142 | 579 | 515 |
| 3094 | W303.asm02.HP0_G0030890 | chrX | 3767 | 963 | 1424 | 2392 | 1295 | 1508 | 2934 | 985 | 1795 | 6449 | 1268 | 2254 | 4805 | 1238 | 1948 | 4481 | 1180 | 1915 |
| 3095 | W303.asm02.HP0_G0030900 | chrX | 1739 | 262 | 376 | 1726 | 348 | 425 | 1590 | 312 | 448 | 3646 | 620 | 1135 | 1973 | 397 | 724 | 2055 | 581 | 1004 |
| 3096 | W303.asm02.HP0_G0030910 | chrX | 6224 | 5445 | 5924 | 5446 | 4771 | 5597 | 5917 | 5387 | 7104 | 13680 | 4931 | 5324 | 15242 | 3592 | 4105 | 14809 | 4508 | 4595 |
| 3097 | W303.asm02.HP0_G0030920 | chrX | 2354 | 570 | 902 | 2244 | 880 | 871 | 2297 | 611 | 908 | 5910 | 706 | 1675 | 5157 | 589 | 1165 | 4737 | 636 | 1472 |
| 3098 | W303.asm02.HP0_G0030930 | chrX | 805 | 621 | 989 | 654 | 799 | 1037 | 754 | 646 | 1217 | 2790 | 456 | 691 | 3186 | 300 | 493 | 3096 | 409 | 590 |
| 3099 | W303.asm02.HP0_G0030940 | chrX | 4648 | 1259 | 1249 | 4052 | 1145 | 1330 | 4162 | 1301 | 1366 | 4139 | 1143 | 1375 | 4968 | 914 | 1267 | 3986 | 1076 | 1242 |
| 3100 | W303.asm02.HP0_G0030950 | chrX | 5819 | 2363 | 2230 | 5141 | 1981 | 2264 | 6091 | 2194 | 2423 | 16538 | 4409 | 5090 | 11551 | 3401 | 3641 | 11561 | 3847 | 4574 |
| 3101 | W303.asm02.HP0_G0030960 | chrX | 1225 | 2436 | 2701 | 1229 | 2500 | 2640 | 1220 | 2505 | 2918 | 6165 | 2305 | 2445 | 7078 | 1629 | 1967 | 6085 | 1926 | 2023 |
| 3102 | W303.asm02.HP0_G0030970 | chrX | 19786 | 9128 | 8822 | 17518 | 9254 | 8708 | 20007 | 8733 | 8827 | 29716 | 10010 | 12483 | 26997 | 7923 | 9216 | 26104 | 8527 | 11052 |
| 3103 | W303.asm02.HP0_G0030980 | chrX | 6189 | 2055 | 2290 | 5877 | 1998 | 2729 | 6386 | 1907 | 2945 | 9286 | 3494 | 4630 | 4874 | 2376 | 3072 | 4964 | 3112 | 4197 |
| 3104 | W303.asm02.HP0_G0030990 | chrX | 1318 | 427 | 444 | 1182 | 412 | 440 | 1162 | 428 | 467 | 1255 | 413 | 458 | 1812 | 337 | 432 | 1088 | 356 | 386 |
| 3105 | W303.asm02.HP0_G0031000 | chrX | 139 | 39 | 24 | 146 | 85 | 62 | 140 | 17 | 20 | 65 | 22 | 26 | 76 | 11 | 15 | 52 | 13 | 13 |
| 3106 | W303.asm02.HP0_G0031010 | chrX | 772 | 71 | 152 | 720 | 106 | 200 | 845 | 49 | 212 | 599 | 135 | 542 | 261 | 105 | 254 | 260 | 88 | 414 |
| 3107 | W303.asm02.HP0_G0031020 | chrX | 566 | 227 | 289 | 574 | 317 | 272 | 565 | 266 | 269 | 1153 | 245 | 323 | 1147 | 198 | 245 | 955 | 229 | 284 |
| 3108 | W303.asm02.HP0_G0031030 | chrX | 430 | 131 | 206 | 428 | 218 | 188 | 447 | 154 | 180 | 880 | 181 | 252 | 709 | 92 | 152 | 665 | 131 | 254 |
| 3109 | W303.asm02.HP0_G0031040 | chrX | 490 | 75 | 73 | 346 | 93 | 87 | 410 | 65 | 98 | 500 | 150 | 199 | 250 | 135 | 121 | 199 | 137 | 191 |
| 3110 | W303.asm02.HP0_G0031050 | chrX | 548 | 15 | 26 | 461 | 34 | 26 | 412 | 14 | 34 | 144 | 37 | 60 | 71 | 42 | 43 | 69 | 38 | 50 |
| 3111 | W303.asm02.HP0_G0031060 | chrX | 1639 | 135 | 177 | 1301 | 186 | 187 | 1259 | 114 | 222 | 533 | 203 | 234 | 449 | 142 | 207 | 347 | 162 | 224 |
| 3112 | W303.asm02.HP0_G0031070 | chrX | 545 | 147 | 222 | 492 | 200 | 207 | 457 | 157 | 273 | 520 | 159 | 151 | 657 | 93 | 114 | 452 | 120 | 120 |
| 3113 | W303.asm02.HP0_G0031080 | chrX | 17 | 3 | 4 | 18 | 6 | 2 | 12 | 3 | 9 | 21 | 4 | 3 | 12 | 4 | 6 | 9 | 2 | 7 |
| 3114 | W303.asm02.HP0_G0031090 | chrX | 172 | 13 | 22 | 192 | 13 | 27 | 160 | 15 | 32 | 175 | 37 | 52 | 142 | 22 | 32 | 105 | 28 | 46 |
| 3115 | W303.asm02.HP0_G0031100 | chrX | 170 | 89 | 149 | 175 | 90 | 163 | 156 | 95 | 202 | 308 | 63 | 100 | 424 | 61 | 78 | 246 | 59 | 88 |
| 3116 | W303.asm02.HP0_G0031110 | chrX | 178 | 19 | 37 | 153 | 18 | 19 | 159 | 20 | 23 | 125 | 23 | 26 | 104 | 23 | 18 | 106 | 25 | 39 |
| 3117 | W303.asm02.HP0_G0031120 | chrXI | 1 | 12 | 0 | 0 | 0 | 0 | 0 | 24 | 0 | 0 | 12 | 0 | 0 | 6 | 0 | 2 | 11 | 0 |
| 3118 | W303.asm02.HP0_G0031130 | chrXI | 0 | 2 | 0 | 0 | 0 | 0 | 0 | 0 | 1 | 0 | 0 | 1 | 1 | 1 | 0 | 3 | 0 | 0 |
| 3119 | W303.asm02.HP0_G0031140 | chrXI | 1415 | 348 | 451 | 1438 | 459 | 415 | 1412 | 393 | 469 | 2728 | 521 | 685 | 2366 | 356 | 510 | 2279 | 479 | 538 |
| 3120 | W303.asm02.HP0_G0031150 | chrXI | 366 | 76 | 97 | 346 | 87 | 98 | 349 | 105 | 124 | 286 | 105 | 136 | 537 | 115 | 111 | 233 | 104 | 90 |
| 3121 | W303.asm02.HP0_G0031160 | chrXI | 452 | 127 | 127 | 346 | 182 | 149 | 396 | 104 | 159 | 1056 | 172 | 282 | 753 | 116 | 203 | 921 | 183 | 290 |
| 3122 | W303.asm02.HP0_G0031170 | chrXI | 736 | 504 | 565 | 577 | 489 | 565 | 654 | 513 | 715 | 2272 | 443 | 489 | 2136 | 291 | 369 | 2324 | 405 | 466 |
| 3123 | W303.asm02.HP0_G0031180 | chrXI | 1685 | 1102 | 1032 | 1804 | 906 | 906 | 1589 | 1043 | 1026 | 2660 | 1107 | 1171 | 3038 | 798 | 977 | 2393 | 939 | 976 |
| 3124 | W303.asm02.HP0_G0031190 | chrXI | 1052 | 218 | 264 | 1120 | 298 | 219 | 943 | 172 | 265 | 960 | 261 | 305 | 887 | 156 | 181 | 714 | 239 | 232 |
| 3125 | W303.asm02.HP0_G0031200 | chrXI | 2210 | 3003 | 2504 | 2366 | 3128 | 2365 | 2572 | 3082 | 2535 | 12631 | 3326 | 3027 | 11043 | 1999 | 1867 | 12066 | 2820 | 2835 |
| 3126 | W303.asm02.HP0_G0031210 | chrXI | 4249 | 499 | 604 | 3779 | 547 | 627 | 4118 | 467 | 751 | 3897 | 845 | 1590 | 1776 | 707 | 1007 | 1820 | 760 | 1394 |
| 3127 | W303.asm02.HP0_G0031220 | chrXI | 1949 | 1029 | 1293 | 1705 | 1199 | 1297 | 1855 | 1142 | 1553 | 3694 | 873 | 1023 | 3983 | 541 | 777 | 3811 | 790 | 934 |
| 3128 | W303.asm02.HP0_G0031230 | chrXI | 8345 | 1564 | 1417 | 8009 | 1472 | 1449 | 7862 | 1534 | 1539 | 8160 | 2239 | 2575 | 6561 | 1633 | 2283 | 6174 | 2022 | 2367 |
| 3129 | W303.asm02.HP0_G0031240 | chrXI | 9781 | 1944 | 1929 | 8659 | 2012 | 1934 | 9948 | 2016 | 2083 | 15876 | 3299 | 4391 | 13229 | 2831 | 3434 | 11691 | 2745 | 4007 |
| 3130 | W303.asm02.HP0_G0031250 | chrXI | 2880 | 1298 | 1221 | 2339 | 1070 | 1265 | 2779 | 1048 | 1462 | 8655 | 2476 | 3053 | 4341 | 1586 | 2038 | 5010 | 2074 | 2569 |
| 3131 | W303.asm02.HP0_G0031260 | chrXI | 17032 | 4148 | 3468 | 16713 | 3311 | 3521 | 16938 | 3747 | 4258 | 20598 | 6126 | 7173 | 14241 | 4616 | 5720 | 14450 | 5748 | 6708 |
| 3132 | W303.asm02.HP0_G0031270 | chrXI | 1888 | 2079 | 2714 | 1751 | 2668 | 2705 | 1945 | 2160 | 3265 | 19163 | 2837 | 4636 | 12443 | 2093 | 3340 | 14136 | 2639 | 4057 |
| 3133 | W303.asm02.HP0_G0031280 | chrXI | 1308 | 906 | 513 | 1284 | 499 | 544 | 1346 | 915 | 538 | 1564 | 753 | 589 | 1653 | 523 | 423 | 1455 | 704 | 529 |
| 3134 | W303.asm02.HP0_G0031290 | chrXI | 3405 | 1688 | 1399 | 2693 | 1310 | 1413 | 2952 | 1662 | 1590 | 5386 | 2186 | 2246 | 4306 | 1549 | 1643 | 4535 | 2064 | 2012 |
| 3135 | W303.asm02.HP0_G0031300 | chrXI | 1974 | 1265 | 1596 | 1754 | 1047 | 1674 | 1822 | 1178 | 1558 | 2971 | 966 | 1805 | 3269 | 715 | 1238 | 2952 | 901 | 1603 |
| 3136 | W303.asm02.HP0_G0031310 | chrXI | 3449 | 1400 | 2047 | 3483 | 1939 | 1827 | 3712 | 1583 | 2304 | 12787 | 1821 | 2724 | 11320 | 1353 | 1830 | 11752 | 1689 | 2524 |
| 3137 | W303.asm02.HP0_G0031320 | chrXI | 5707 | 499 | 630 | 5820 | 538 | 725 | 5568 | 330 | 787 | 2745 | 711 | 1593 | 794 | 589 | 654 | 724 | 565 | 1175 |
| 3138 | W303.asm02.HP0_G0031330 | chrXI | 2384 | 418 | 566 | 2255 | 574 | 671 | 2140 | 392 | 778 | 4065 | 635 | 1510 | 2285 | 558 | 916 | 2306 | 599 | 1244 |
| 3139 | W303.asm02.HP0_G0031340 | chrXI | 2059 | 628 | 640 | 1579 | 526 | 827 | 1910 | 604 | 749 | 2959 | 839 | 1476 | 2331 | 538 | 810 | 2411 | 890 | 1300 |
| 3140 | W303.asm02.HP0_G0031350 | chrXI | 925 | 357 | 337 | 901 | 343 | 405 | 971 | 265 | 421 | 767 | 328 | 413 | 451 | 268 | 221 | 455 | 327 | 315 |
| 3141 | W303.asm02.HP0_G0031360 | chrXI | 1918 | 279 | 358 | 2078 | 328 | 426 | 1849 | 246 | 391 | 2879 | 472 | 884 | 1921 | 384 | 603 | 1610 | 482 | 840 |
| 3142 | W303.asm02.HP0_G0031370 | chrXI | 5216 | 4321 | 5474 | 5127 | 4718 | 5666 | 5292 | 4375 | 7314 | 11789 | 4046 | 4520 | 14053 | 2831 | 3728 | 12438 | 3464 | 4064 |
| 3143 | W303.asm02.HP0_G0031380 | chrXI | 20244 | 3417 | 3137 | 20878 | 3064 | 3429 | 19808 | 3154 | 3264 | 12617 | 3414 | 4676 | 13028 | 2961 | 4166 | 11005 | 3578 | 4321 |
| 3144 | W303.asm02.HP0_G0031390 | chrXI | 2706 | 781 | 591 | 2361 | 631 | 633 | 2395 | 700 | 588 | 3212 | 777 | 1087 | 3292 | 597 | 778 | 2725 | 756 | 938 |
| 3145 | W303.asm02.HP0_G0031400 | chrXI | 4223 | 1627 | 1289 | 4258 | 1037 | 1272 | 3773 | 1382 | 1370 | 3504 | 1115 | 1036 | 4131 | 798 | 933 | 3851 | 1221 | 924 |
| 3146 | W303.asm02.HP0_G0031410 | chrXI | 7094 | 2503 | 2282 | 6350 | 2079 | 2334 | 6631 | 2311 | 2178 | 4688 | 1997 | 2857 | 5320 | 1934 | 2084 | 4083 | 1717 | 2407 |
| 3147 | W303.asm02.HP0_G0031420 | chrXI | 1316 | 666 | 690 | 1039 | 851 | 791 | 1599 | 551 | 1029 | 7153 | 1581 | 1785 | 3191 | 995 | 879 | 3696 | 1328 | 1571 |
| 3148 | W303.asm02.HP0_G0031430 | chrXI | 2348 | 2420 | 2351 | 2162 | 2214 | 2325 | 2340 | 2332 | 2758 | 5612 | 2193 | 2343 | 6530 | 1603 | 2075 | 5550 | 1949 | 2087 |
| 3149 | W303.asm02.HP0_G0031440 | chrXI | 1382 | 697 | 541 | 1405 | 441 | 519 | 1311 | 613 | 512 | 2185 | 576 | 687 | 2175 | 370 | 517 | 2038 | 565 | 585 |
| 3150 | W303.asm02.HP0_G0031450 | chrXI | 1745 | 521 | 503 | 1952 | 480 | 433 | 1563 | 473 | 544 | 2064 | 454 | 507 | 1776 | 260 | 374 | 1751 | 419 | 378 |
| 3151 | W303.asm02.HP0_G0031460 | chrXI | 1946 | 110 | 202 | 2034 | 140 | 190 | 1887 | 113 | 213 | 1051 | 187 | 313 | 975 | 178 | 279 | 741 | 175 | 269 |
| 3152 | W303.asm02.HP0_G0031470 | chrXI | 2524 | 1048 | 1260 | 2482 | 1233 | 1306 | 2282 | 998 | 1360 | 3864 | 820 | 1375 | 4124 | 621 | 1075 | 3516 | 839 | 1270 |
| 3153 | W303.asm02.HP0_G0031480 | chrXI | 5472 | 639 | 1288 | 5919 | 1216 | 1590 | 5739 | 619 | 1933 | 9972 | 1322 | 2639 | 10145 | 1289 | 2296 | 6851 | 1041 | 2323 |
| 3154 | W303.asm02.HP0_G0031490 | chrXI | 2939 | 932 | 1106 | 2844 | 1031 | 1082 | 3151 | 1089 | 1351 | 7999 | 1803 | 1916 | 5545 | 1208 | 1394 | 5525 | 1546 | 1709 |
| 3155 | W303.asm02.HP0_G0031500 | chrXI | 63 | 5 | 8 | 45 | 8 | 6 | 45 | 6 | 9 | 52 | 8 | 8 | 42 | 7 | 14 | 38 | 10 | 10 |
| 3156 | W303.asm02.HP0_G0031510 | chrXI | 1731 | 766 | 739 | 1617 | 704 | 678 | 1717 | 749 | 857 | 2940 | 912 | 850 | 3041 | 574 | 715 | 2175 | 727 | 696 |
| 3157 | W303.asm02.HP0_G0031520 | chrXI | 51515 | 6861 | 7849 | 47225 | 7975 | 7709 | 51852 | 6988 | 10104 | 62212 | 12582 | 18977 | 41644 | 10977 | 14674 | 39698 | 10681 | 16605 |
| 3158 | W303.asm02.HP0_G0031530 | chrXI | 6640 | 6295 | 5455 | 5859 | 5872 | 5151 | 7304 | 6770 | 5422 | 20707 | 7058 | 6437 | 19286 | 4651 | 4536 | 19064 | 6017 | 5818 |
| 3159 | W303.asm02.HP0_G0031540 | chrXI | 12347 | 7102 | 7797 | 11967 | 7824 | 8355 | 12902 | 7139 | 7789 | 19122 | 6762 | 8836 | 18030 | 5167 | 6271 | 16749 | 5542 | 7768 |
| 3160 | W303.asm02.HP0_G0031550 | chrXI | 3144 | 1124 | 1346 | 3373 | 1240 | 1343 | 2949 | 1065 | 1208 | 7215 | 1141 | 1900 | 5979 | 725 | 1378 | 6383 | 1055 | 1805 |

| | | | | | | | | | | | | | | | | | | | | |
|---|---|---|---|---|---|---|---|---|---|---|---|---|---|---|---|---|---|---|---|---|
| 3161 | W303.asm02.HP0_G0031560 | chrXI | 411 | 221 | 254 | 417 | 231 | 350 | 378 | 175 | 265 | 459 | 171 | 267 | 325 | 74 | 115 | 461 | 183 | 241 |
| 3162 | W303.asm02.HP0_G0031570 | chrXI | 2467 | 693 | 850 | 2235 | 1012 | 1000 | 2568 | 714 | 1107 | 7806 | 1254 | 1800 | 5895 | 930 | 1390 | 5799 | 1113 | 1657 |
| 3163 | W303.asm02.HP0_G0031580 | chrXI | 2087 | 881 | 912 | 2157 | 799 | 890 | 2183 | 826 | 976 | 5767 | 1408 | 1815 | 5041 | 956 | 1182 | 4702 | 1267 | 1584 |
| 3164 | W303.asm02.HP0_G0031590 | chrXI | 2211 | 1150 | 1516 | 2025 | 1209 | 1416 | 1948 | 1289 | 1630 | 5034 | 1339 | 1897 | 4770 | 799 | 1217 | 5032 | 1153 | 1590 |
| 3165 | W303.asm02.HP0_G0031600 | chrXI | 4082 | 431 | 602 | 3789 | 585 | 637 | 3754 | 382 | 764 | 5494 | 822 | 1484 | 3268 | 566 | 937 | 3339 | 729 | 1209 |
| 3166 | W303.asm02.HP0_G0031610 | chrXI | 4025 | 3440 | 4883 | 4449 | 5954 | 4851 | 4753 | 4182 | 4861 | 21180 | 4023 | 5090 | 21617 | 2237 | 3183 | 18891 | 3204 | 4608 |
| 3167 | W303.asm02.HP0_G0031620 | chrXI | 1874 | 306 | 315 | 2268 | 317 | 314 | 1779 | 248 | 394 | 2092 | 485 | 621 | 1253 | 314 | 467 | 1293 | 420 | 602 |
| 3168 | W303.asm02.HP0_G0031630 | chrXI | 6986 | 1663 | 1997 | 6556 | 1893 | 1996 | 6502 | 1729 | 2297 | 4986 | 1517 | 2191 | 6292 | 1350 | 1981 | 4756 | 1488 | 1996 |
| 3169 | W303.asm02.HP0_G0031640 | chrXI | 2627 | 238 | 277 | 2439 | 227 | 293 | 2214 | 203 | 384 | 1729 | 404 | 533 | 894 | 292 | 415 | 831 | 372 | 459 |
| 3170 | W303.asm02.HP0_G0031650 | chrXI | 3840 | 1181 | 1317 | 3625 | 1069 | 1349 | 3515 | 1176 | 1457 | 3010 | 904 | 1234 | 3725 | 725 | 1109 | 3135 | 903 | 1111 |
| 3171 | W303.asm02.HP0_G0031660 | chrXI | 125 | 228 | 161 | 125 | 151 | 164 | 118 | 209 | 159 | 809 | 235 | 195 | 804 | 151 | 118 | 831 | 269 | 199 |
| 3172 | W303.asm02.HP0_G0031670 | chrXI | 1843 | 1218 | 4980 | 1594 | 1579 | 4922 | 1827 | 1399 | 5009 | 9377 | 1481 | 8457 | 9190 | 1034 | 4410 | 9046 | 1458 | 7372 |
| 3173 | W303.asm02.HP0_G0031680 | chrXI | 19977 | 5923 | 5924 | 21976 | 5100 | 6855 | 22949 | 5035 | 8588 | 29573 | 10968 | 14261 | 31397 | 10853 | 12107 | 21121 | 9291 | 12102 |
| 3174 | W303.asm02.HP0_G0031690 | chrXI | 20553 | 419 | 450 | 20999 | 358 | 493 | 20107 | 430 | 773 | 1824 | 1077 | 959 | 4176 | 2949 | 2064 | 610 | 843 | 746 |
| 3175 | W303.asm02.HP0_G0031700 | chrXI | 578 | 156 | 192 | 418 | 140 | 167 | 425 | 147 | 234 | 320 | 143 | 146 | 761 | 121 | 141 | 303 | 119 | 106 |
| 3176 | W303.asm02.HP0_G0031710 | chrXI | 679 | 246 | 205 | 807 | 241 | 194 | 612 | 220 | 164 | 936 | 252 | 263 | 1004 | 133 | 220 | 887 | 233 | 203 |
| 3177 | W303.asm02.HP0_G0031720 | chrXI | 2032 | 825 | 775 | 1982 | 612 | 634 | 1916 | 848 | 787 | 3043 | 862 | 1114 | 9739 | 1725 | 1657 | 2595 | 763 | 981 |
| 3178 | W303.asm02.HP0_G0031730 | chrXI | 2521 | 2298 | 2602 | 2379 | 2513 | 2644 | 2253 | 2419 | 3035 | 6099 | 2243 | 2338 | 6916 | 1557 | 2079 | 6124 | 1870 | 2209 |
| 3179 | W303.asm02.HP0_G0031740 | chrXI | 1394 | 479 | 301 | 1383 | 283 | 313 | 1352 | 413 | 323 | 1460 | 464 | 411 | 1652 | 375 | 396 | 1364 | 448 | 371 |
| 3180 | W303.asm02.HP0_G0031750 | chrXI | 578 | 63 | 52 | 455 | 49 | 45 | 489 | 55 | 80 | 575 | 121 | 168 | 336 | 112 | 82 | 334 | 126 | 135 |
| 3181 | W303.asm02.HP0_G0031760 | chrXI | 12514 | 1502 | 1339 | 11674 | 1222 | 1536 | 12188 | 1385 | 1734 | 9842 | 2536 | 3335 | 7175 | 2280 | 2772 | 6087 | 2422 | 2938 |
| 3182 | W303.asm02.HP0_G0031770 | chrXI | 5434 | 4888 | 7391 | 4742 | 6959 | 7302 | 5177 | 5003 | 8833 | 14456 | 4296 | 5446 | 17089 | 2859 | 4407 | 15538 | 3861 | 4746 |
| 3183 | W303.asm02.HP0_G0031780 | chrXI | 3378 | 1207 | 1006 | 3054 | 1077 | 1088 | 3159 | 1076 | 1110 | 5907 | 1231 | 1493 | 5554 | 865 | 1213 | 4940 | 1270 | 1363 |
| 3184 | W303.asm02.HP0_G0031790 | chrXI | 727 | 1123 | 1468 | 604 | 1223 | 1416 | 661 | 1166 | 1725 | 3953 | 879 | 1079 | 4117 | 613 | 768 | 4052 | 775 | 970 |
| 3185 | W303.asm02.HP0_G0031800 | chrXI | 57255 | 20233 | 19980 | 56546 | 16136 | 20255 | 60238 | 19574 | 21826 | 49141 | 27564 | 31969 | 48478 | 20722 | 27340 | 46815 | 24878 | 30125 |
| 3186 | W303.asm02.HP0_G0031810 | chrXI | 7064 | 1026 | 650 | 6045 | 558 | 664 | 6137 | 857 | 690 | 2631 | 1295 | 1043 | 2823 | 943 | 998 | 2084 | 1355 | 961 |
| 3187 | W303.asm02.HP0_G0031820 | chrXI | 13234 | 6587 | 5779 | 12276 | 4654 | 5608 | 12504 | 6169 | 6378 | 11590 | 5869 | 5736 | 11256 | 3673 | 4593 | 11218 | 5375 | 4650 |
| 3188 | W303.asm02.HP0_G0031830 | chrXI | 762 | 800 | 622 | 852 | 577 | 576 | 769 | 694 | 655 | 2187 | 592 | 647 | 2221 | 327 | 370 | 2302 | 621 | 562 |
| 3189 | W303.asm02.HP0_G0031840 | chrXI | 11437 | 1893 | 1725 | 9510 | 1543 | 2129 | 10370 | 1379 | 2364 | 11945 | 3562 | 4390 | 6062 | 3207 | 3527 | 5561 | 3209 | 3021 |
| 3190 | W303.asm02.HP0_G0031850 | chrXI | 3064 | 1196 | 937 | 3121 | 903 | 1022 | 3121 | 1021 | 1187 | 6323 | 2221 | 2019 | 3796 | 1424 | 1406 | 3832 | 2023 | 1797 |
| 3191 | W303.asm02.HP0_G0031860 | chrXI | 15783 | 5575 | 4101 | 15636 | 4146 | 4150 | 15882 | 5196 | 3972 | 16661 | 6056 | 6632 | 14085 | 4299 | 5397 | 14358 | 5446 | 5985 |
| 3192 | W303.asm02.HP0_G0031870 | chrXI | 682 | 1266 | 1436 | 618 | 1375 | 1434 | 753 | 1368 | 1871 | 3738 | 1155 | 1183 | 4192 | 681 | 689 | 4148 | 1033 | 1024 |
| 3193 | W303.asm02.HP0_G0031880 | chrXI | 1870 | 1320 | 1071 | 1983 | 1595 | 979 | 2220 | 1418 | 1227 | 9667 | 1898 | 1792 | 8094 | 1221 | 928 | 7894 | 1695 | 1492 |
| 3194 | W303.asm02.HP0_G0031890 | chrXI | 11575 | 3436 | 3164 | 11647 | 2468 | 3208 | 11260 | 3232 | 3610 | 6859 | 2629 | 2779 | 9459 | 2185 | 2785 | 7594 | 2590 | 2530 |
| 3195 | W303.asm02.HP0_G0031900 | chrXI | 6028 | 3539 | 3423 | 5647 | 3619 | 3489 | 5454 | 3858 | 3527 | 6893 | 3056 | 3148 | 7533 | 2236 | 2701 | 6716 | 2660 | 2386 |
| 3196 | W303.asm02.HP0_G0031910 | chrXI | 4452 | 1686 | 1419 | 4226 | 1309 | 1503 | 4484 | 1665 | 1333 | 6517 | 1575 | 1594 | 6325 | 1068 | 1452 | 6172 | 1471 | 1499 |
| 3197 | W303.asm02.HP0_G0031920 | chrXI | 1699 | 678 | 772 | 1688 | 776 | 708 | 1575 | 611 | 854 | 4692 | 764 | 1002 | 3960 | 440 | 789 | 4452 | 836 | 916 |
| 3198 | W303.asm02.HP0_G0031930 | chrXI | 1043 | 1108 | 1339 | 901 | 1251 | 1418 | 882 | 1148 | 1822 | 2962 | 897 | 1087 | 3894 | 586 | 857 | 3407 | 849 | 922 |
| 3199 | W303.asm02.HP0_G0031940 | chrXI | 5106 | 1148 | 1269 | 4595 | 1140 | 1218 | 4471 | 1134 | 1343 | 3713 | 1024 | 1413 | 4457 | 852 | 1273 | 3605 | 1011 | 1312 |
| 3200 | W303.asm02.HP0_G0031950 | chrXI | 2779 | 982 | 1392 | 2471 | 1203 | 1362 | 2404 | 967 | 1928 | 3015 | 771 | 1247 | 4100 | 590 | 1044 | 3408 | 792 | 1075 |
| 3201 | W303.asm02.HP0_G0031960 | chrXI | 5318 | 1992 | 1768 | 4943 | 1672 | 1743 | 5317 | 1954 | 1624 | 8921 | 1997 | 2446 | 7469 | 1376 | 1820 | 7486 | 1823 | 2236 |
| 3202 | W303.asm02.HP0_G0031970 | chrXI | 1168 | 643 | 611 | 1137 | 648 | 638 | 1100 | 630 | 652 | 3889 | 699 | 979 | 3384 | 542 | 785 | 3308 | 647 | 871 |
| 3203 | W303.asm02.HP0_G0031980 | chrXI | 1177 | 449 | 401 | 1221 | 381 | 411 | 1065 | 421 | 479 | 1189 | 333 | 384 | 1477 | 203 | 403 | 1241 | 306 | 319 |
| 3204 | W303.asm02.HP0_G0031990 | chrXI | 530 | 225 | 210 | 506 | 227 | 196 | 456 | 223 | 180 | 1212 | 254 | 257 | 943 | 138 | 177 | 1049 | 220 | 214 |
| 3205 | W303.asm02.HP0_G0032000 | chrXI | 1788 | 1265 | 1207 | 1668 | 1111 | 1254 | 1704 | 1237 | 1371 | 3309 | 887 | 970 | 3539 | 615 | 770 | 3446 | 886 | 885 |
| 3206 | W303.asm02.HP0_G0032010 | chrXI | 3998 | 397 | 433 | 3663 | 367 | 546 | 3695 | 279 | 539 | 2525 | 592 | 996 | 1203 | 456 | 697 | 1041 | 576 | 929 |
| 3207 | W303.asm02.HP0_G0032020 | chrXI | 1901 | 1508 | 1710 | 1605 | 1514 | 1868 | 1786 | 1516 | 1760 | 5434 | 1521 | 2090 | 5094 | 928 | 1409 | 5197 | 1331 | 1900 |
| 3208 | W303.asm02.HP0_G0032030 | chrXI | 4243 | 2114 | 1805 | 3947 | 1853 | 1746 | 4319 | 2091 | 1684 | 11332 | 2651 | 2938 | 11580 | 1847 | 2011 | 10458 | 2344 | 2646 |
| 3209 | W303.asm02.HP0_G0032040 | chrXI | 8006 | 1914 | 1787 | 7165 | 1769 | 1697 | 7775 | 1883 | 1695 | 10950 | 2529 | 2565 | 11230 | 2049 | 2646 | 9153 | 2257 | 2455 |
| 3210 | W303.asm02.HP0_G0032050 | chrXI | 1586 | 821 | 782 | 1403 | 998 | 779 | 1627 | 856 | 853 | 6366 | 1263 | 1323 | 5055 | 729 | 825 | 5761 | 1220 | 1253 |
| 3211 | W303.asm02.HP0_G0032060 | chrXI | 2386 | 469 | 338 | 2127 | 256 | 378 | 2077 | 393 | 337 | 1862 | 649 | 767 | 1415 | 440 | 530 | 1470 | 661 | 687 |
| 3212 | W303.asm02.HP0_G0032070 | chrXI | 2886 | 1946 | 2195 | 2875 | 1862 | 2461 | 2778 | 2168 | 2816 | 5141 | 1754 | 1990 | 5991 | 1150 | 1575 | 5393 | 1568 | 1826 |
| 3213 | W303.asm02.HP0_G0032080 | chrXI | 3009 | 247 | 279 | 2886 | 205 | 283 | 2816 | 221 | 324 | 2326 | 533 | 662 | 1853 | 336 | 558 | 1647 | 453 | 551 |
| 3214 | W303.asm02.HP0_G0032090 | chrXI | 249 | 4978 | 5550 | 261 | 5884 | 5717 | 287 | 4789 | 5793 | 14215 | 4126 | 5286 | 11899 | 2476 | 3578 | 13899 | 3607 | 4361 |
| 3215 | W303.asm02.HP0_G0032100 | chrXI | 1866 | 919 | 1083 | 1646 | 852 | 1098 | 1821 | 917 | 1537 | 2812 | 737 | 868 | 3401 | 545 | 664 | 3105 | 719 | 792 |
| 3216 | W303.asm02.HP0_G0032110 | chrXI | 13947 | 6194 | 6942 | 13819 | 6211 | 7132 | 13838 | 6378 | 7403 | 16116 | 6005 | 7443 | 17333 | 4354 | 6293 | 16323 | 5088 | 6714 |
| 3217 | W303.asm02.HP0_G0032120 | chrXI | 1596 | 546 | 600 | 1623 | 598 | 640 | 1701 | 497 | 737 | 4568 | 748 | 1197 | 3844 | 635 | 842 | 3398 | 772 | 1064 |
| 3218 | W303.asm02.HP0_G0032130 | chrXI | 17 | 6 | 5 | 11 | 3 | 6 | 15 | 9 | 2 | 7 | 2 | 7 | 14 | 2 | 1 | 9 | 6 | 1 |
| 3219 | W303.asm02.HP0_G0032140 | chrXI | 2464 | 2347 | 2339 | 2507 | 2216 | 2311 | 2499 | 2280 | 2489 | 7646 | 2022 | 2446 | 7427 | 1229 | 1646 | 7190 | 1765 | 2152 |
| 3220 | W303.asm02.HP0_G0032150 | chrXI | 1377 | 1914 | 1347 | 1397 | 1480 | 1322 | 1521 | 1830 | 1421 | 6356 | 1973 | 1966 | 6218 | 1283 | 1274 | 6109 | 1787 | 1764 |
| 3221 | W303.asm02.HP0_G0032160 | chrXI | 1188 | 993 | 1031 | 1313 | 930 | 1152 | 1334 | 994 | 1289 | 8230 | 1874 | 2754 | 4928 | 1317 | 2094 | 5580 | 1710 | 2511 |
| 3222 | W303.asm02.HP0_G0032170 | chrXI | 1217 | 989 | 1154 | 1132 | 1191 | 1144 | 1186 | 937 | 1157 | 4960 | 1026 | 1250 | 5358 | 682 | 949 | 4885 | 915 | 1189 |
| 3223 | W303.asm02.HP0_G0032180 | chrXI | 6028 | 909 | 1365 | 7891 | 1265 | 1465 | 6068 | 893 | 1732 | 12188 | 2171 | 2740 | 10453 | 1430 | 2191 | 8585 | 1840 | 2152 |
| 3224 | W303.asm02.HP0_G0032190 | chrXI | 650 | 449 | 401 | 641 | 453 | 429 | 718 | 415 | 445 | 2021 | 523 | 658 | 1973 | 312 | 430 | 1940 | 450 | 507 |
| 3225 | W303.asm02.HP0_G0032200 | chrXI | 416 | 116 | 83 | 380 | 70 | 90 | 383 | 109 | 66 | 231 | 113 | 114 | 377 | 75 | 74 | 227 | 95 | 110 |
| 3226 | W303.asm02.HP0_G0032210 | chrXI | 373 | 1082 | 2001 | 387 | 1657 | 1897 | 436 | 1254 | 1951 | 7043 | 1229 | 2481 | 6225 | 610 | 1529 | 6801 | 1032 | 2376 |
| 3227 | W303.asm02.HP0_G0032220 | chrXI | 24786 | 3226 | 3873 | 25591 | 2737 | 4390 | 25953 | 3014 | 4984 | 23188 | 6607 | 10867 | 27853 | 7903 | 9346 | 16119 | 5869 | 9574 |
| 3228 | W303.asm02.HP0_G0032230 | chrXI | 8729 | 1123 | 1239 | 7319 | 878 | 1477 | 7883 | 981 | 1515 | 4177 | 1953 | 2472 | 3084 | 1400 | 2146 | 2912 | 1720 | 2056 |
| 3229 | W303.asm02.HP0_G0032240 | chrXI | 7104 | 711 | 1376 | 6652 | 1069 | 1482 | 7459 | 705 | 1619 | 7919 | 1215 | 3112 | 4753 | 1168 | 1984 | 4121 | 1085 | 2618 |
| 3230 | W303.asm02.HP0_G0032250 | chrXI | 8514 | 1270 | 1189 | 8448 | 974 | 1213 | 8157 | 1381 | 1211 | 7090 | 2058 | 2273 | 6281 | 1373 | 2059 | 5962 | 1821 | 2176 |
| 3231 | W303.asm02.HP0_G0032260 | chrXI | 1079 | 1371 | 1647 | 1064 | 2021 | 1599 | 1311 | 1556 | 2036 | 6728 | 1346 | 1426 | 6820 | 885 | 830 | 6462 | 1184 | 1292 |
| 3232 | W303.asm02.HP0_G0032270 | chrXI | 2326 | 572 | 447 | 2177 | 417 | 468 | 2199 | 443 | 520 | 2609 | 776 | 823 | 2214 | 607 | 629 | 1796 | 800 | 734 |
| 3233 | W303.asm02.HP0_G0032280 | chrXI | 22 | 14 | 3 | 25 | 2 | 3 | 35 | 16 | 7 | 19 | 19 | 6 | 17 | 11 | 5 | 11 | 19 | 5 |
| 3234 | W303.asm02.HP0_G0032290 | chrXI | 1707 | 946 | 1195 | 1611 | 1150 | 1116 | 1640 | 994 | 1273 | 1806 | 982 | 968 | 2210 | 657 | 926 | 1880 | 844 | 907 |
| 3235 | W303.asm02.HP0_G0032300 | chrXI | 6863 | 3980 | 2727 | 6575 | 2445 | 2475 | 6763 | 4212 | 2925 | 10695 | 3021 | 3905 | 23577 | 5245 | 5256 | 8906 | 2688 | 3646 |
| 3236 | W303.asm02.HP0_G0032310 | chrXI | 1207 | 220 | 190 | 1365 | 262 | 180 | 1311 | 209 | 161 | 1891 | 318 | 373 | 1466 | 234 | 284 | 1238 | 344 | 369 |
| 3237 | W303.asm02.HP0_G0032320 | chrXI | 5278 | 2643 | 2125 | 4631 | 2199 | 2271 | 5000 | 2569 | 2276 | 6956 | 2526 | 2691 | 6106 | 1571 | 2010 | 6254 | 2295 | 2440 |
| 3238 | W303.asm02.HP0_G0032330 | chrXI | 1741 | 572 | 543 | 1781 | 539 | 456 | 1661 | 516 | 524 | 2493 | 606 | 575 | 2586 | 393 | 501 | 2021 | 591 | 442 |
| 3239 | W303.asm02.HP0_G0032340 | chrXI | 1589 | 639 | 806 | 1578 | 799 | 757 | 1470 | 667 | 880 | 4644 | 706 | 1062 | 4247 | 446 | 906 | 4225 | 659 | 1007 |
| 3240 | W303.asm02.HP0_G0032350 | chrXI | 5316 | 791 | 486 | 5620 | 490 | 491 | 4955 | 661 | 492 | 1893 | 692 | 625 | 2318 | 509 | 669 | 1722 | 732 | 599 |
| 3241 | W303.asm02.HP0_G0032360 | chrXI | 1934 | 763 | 492 | 1750 | 476 | 520 | 1854 | 700 | 526 | 2530 | 715 | 602 | 2492 | 453 | 477 | 2499 | 695 | 552 |
| 3242 | W303.asm02.HP0_G0032370 | chrXI | 427 | 543 | 501 | 400 | 594 | 505 | 464 | 501 | 398 | 1957 | 436 | 550 | 2252 | 269 | 429 | 1851 | 434 | 546 |
| 3243 | W303.asm02.HP0_G0032380 | chrXI | 4109 | 687 | 666 | 4274 | 711 | 696 | 4058 | 621 | 726 | 5636 | 1149 | 1405 | 3930 | 694 | 1065 | 3708 | 1029 | 1260 |
| 3244 | W303.asm02.HP0_G0032390 | chrXI | 6907 | 1455 | 1736 | 6533 | 1692 | 1763 | 6411 | 1492 | 2108 | 5394 | 1507 | 2169 | 6325 | 1110 | 1945 | 4923 | 1451 | 1969 |
| 3245 | W303.asm02.HP0_G0032400 | chrXI | 150 | 102 | 83 | 128 | 70 | 105 | 136 | 102 | 90 | 157 | 120 | 76 | 295 | 103 | 70 | 117 | 108 | 59 |
| 3246 | W303.asm02.HP0_G0032410 | chrXI | 12602 | 2745 | 1832 | 11782 | 1840 | 1893 | 11798 | 2660 | 2100 | 11374 | 4323 | 3591 | 11124 | 4144 | 3668 | 8715 | 3656 | 2877 |
| 3247 | W303.asm02.HP0_G0032420 | chrXI | 1192 | 307 | 362 | 1017 | 342 | 344 | 1035 | 405 | 389 | 1728 | 493 | 600 | 1645 | 287 | 444 | 1651 | 427 | 602 |
| 3248 | W303.asm02.HP0_G0032430 | chrXI | 1902 | 1851 | 2430 | 1903 | 3048 | 2264 | 1976 | 2238 | 2311 | 12609 | 2026 | 2468 | 11994 | 1220 | 1559 | 11626 | 1695 | 2115 |
| 3249 | W303.asm02.HP0_G0032440 | chrXI | 18359 | 9254 | 7597 | 16721 | 8025 | 7542 | 19605 | 9222 | 7166 | 45610 | 13772 | 14585 | 38639 | 9902 | 11053 | 40473 | 12138 | 13267 |
| 3250 | W303.asm02.HP0_G0032450 | chrXI | 8461 | 6050 | 6967 | 7880 | 5923 | 6347 | 8543 | 6297 | 8406 | 17186 | 5383 | 5926 | 19079 | 3664 | 4875 | 18223 | 4751 | 5469 |
| 3251 | W303.asm02.HP0_G0032460 | chrXI | 4531 | 860 | 1015 | 4513 | 1127 | 972 | 4485 | 966 | 1036 | 5490 | 954 | 1201 | 6208 | 715 | 1070 | 4926 | 865 | 1046 |
| 3252 | W303.asm02.HP0_G0032470 | chrXI | 154 | 230 | 283 | 148 | 481 | 243 | 211 | 205 | 352 | 5351 | 512 | 661 | 3083 | 311 | 219 | 3441 | 445 | 602 |
| 3253 | W303.asm02.HP0_G0032480 | chrXI | 5196 | 4009 | 4648 | 4875 | 3956 | 4321 | 5240 | 4180 | 5302 | 13243 | 3665 | 3770 | 13751 | 2411 | 2881 | 14054 | 3286 | 3455 |

| | | | | | | | | | | | | | | | | | | | | |
|---|---|---|---|---|---|---|---|---|---|---|---|---|---|---|---|---|---|---|---|---|
| 3254 | W303.asm02.HP0_G0032490 | chrXI | 2318 | 1120 | 1137 | 2407 | 1146 | 1103 | 2402 | 1070 | 1285 | 4445 | 1017 | 1215 | 4740 | 596 | 838 | 4617 | 956 | 1070 |
| 3255 | W303.asm02.HP0_G0032500 | chrXI | 2356 | 868 | 1424 | 2292 | 1043 | 1333 | 2268 | 856 | 1398 | 5425 | 1030 | 2182 | 5196 | 674 | 1474 | 4758 | 1037 | 1840 |
| 3256 | W303.asm02.HP0_G0032510 | chrXI | 3351 | 1072 | 2539 | 3396 | 1063 | 2842 | 3427 | 1140 | 2792 | 7687 | 1488 | 5779 | 7548 | 1226 | 3192 | 6655 | 1404 | 4881 |
| 3257 | W303.asm02.HP0_G0032520 | chrXI | 890 | 341 | 457 | 914 | 545 | 429 | 858 | 391 | 429 | 4348 | 478 | 563 | 3258 | 302 | 393 | 3796 | 505 | 586 |
| 3258 | W303.asm02.HP0_G0032530 | chrXI | 659 | 356 | 415 | 636 | 352 | 412 | 632 | 347 | 398 | 1192 | 493 | 400 | 1141 | 321 | 304 | 1068 | 409 | 339 |
| 3259 | W303.asm02.HP0_G0032540 | chrXI | 572 | 403 | 378 | 527 | 394 | 387 | 625 | 406 | 377 | 1501 | 387 | 384 | 1440 | 283 | 285 | 1276 | 369 | 375 |
| 3260 | W303.asm02.HP0_G0032550 | chrXI | 699 | 859 | 820 | 713 | 776 | 802 | 716 | 899 | 895 | 2005 | 759 | 812 | 2322 | 679 | 634 | 1823 | 650 | 728 |
| 3261 | W303.asm02.HP0_G0032560 | chrXI | 342 | 1014 | 1933 | 213 | 1840 | 1926 | 304 | 1052 | 2376 | 3514 | 790 | 1264 | 3689 | 481 | 800 | 3857 | 815 | 1031 |
| 3262 | W303.asm02.HP0_G0032570 | chrXI | 3768 | 903 | 1325 | 3189 | 1246 | 1539 | 3956 | 799 | 1685 | 10476 | 2077 | 3683 | 3351 | 1531 | 1749 | 3695 | 1673 | 3292 |
| 3263 | W303.asm02.HP0_G0032580 | chrXI | 7413 | 3209 | 3321 | 6747 | 2814 | 3302 | 6555 | 3014 | 3916 | 7393 | 2992 | 3187 | 7836 | 1933 | 2894 | 7701 | 2829 | 2821 |
| 3264 | W303.asm02.HP0_G0032590 | chrXI | 297 | 122 | 88 | 151 | 43 | 83 | 184 | 109 | 108 | 145 | 64 | 59 | 157 | 50 | 55 | 186 | 92 | 54 |
| 3265 | W303.asm02.HP0_G0032600 | chrXI | 151 | 84 | 47 | 143 | 47 | 60 | 118 | 85 | 61 | 116 | 54 | 25 | 127 | 41 | 22 | 139 | 80 | 22 |
| 3266 | W303.asm02.HP0_G0032610 | chrXI | 9418 | 3472 | 3889 | 8992 | 3041 | 4030 | 8777 | 3536 | 4990 | 8149 | 2806 | 3154 | 13049 | 2545 | 3239 | 8390 | 2623 | 2822 |
| 3267 | W303.asm02.HP0_G0032620 | chrXI | 13858 | 1289 | 1410 | 13073 | 1291 | 1399 | 13545 | 1325 | 1679 | 9154 | 2179 | 2723 | 7018 | 1789 | 2361 | 6282 | 1904 | 2456 |
| 3268 | W303.asm02.HP0_G0032630 | chrXI | 2784 | 1101 | 1679 | 2811 | 1550 | 1579 | 2622 | 1261 | 1721 | 5410 | 1183 | 1680 | 6207 | 840 | 1384 | 5569 | 1001 | 1440 |
| 3269 | W303.asm02.HP0_G0032640 | chrXI | 4165 | 1559 | 1316 | 4352 | 1455 | 1100 | 4063 | 1644 | 1100 | 8026 | 1696 | 1807 | 11216 | 1462 | 1673 | 6333 | 1418 | 1354 |
| 3270 | W303.asm02.HP0_G0032650 | chrXI | 773 | 692 | 781 | 716 | 626 | 839 | 646 | 676 | 1115 | 1759 | 643 | 684 | 2345 | 388 | 553 | 1898 | 534 | 563 |
| 3271 | W303.asm02.HP0_G0032660 | chrXI | 93942 | 50677 | 48175 | 95991 | 44288 | 47098 | 101582 | 50209 | 48485 | 97013 | 55070 | 65760 | 91886 | 44756 | 51954 | 88207 | 48614 | 61257 |
| 3272 | W303.asm02.HP0_G0032670 | chrXI | 2275 | 536 | 367 | 2327 | 340 | 465 | 2195 | 416 | 464 | 3205 | 859 | 1032 | 1531 | 523 | 667 | 1731 | 879 | 909 |
| 3273 | W303.asm02.HP0_G0032680 | chrXI | 2348 | 1541 | 1699 | 1893 | 1443 | 1678 | 2080 | 1463 | 2173 | 3601 | 1279 | 1405 | 4228 | 853 | 1164 | 4004 | 1124 | 1219 |
| 3274 | W303.asm02.HP0_G0032690 | chrXI | 2361 | 2027 | 2731 | 2431 | 2478 | 2526 | 2468 | 2126 | 2862 | 12691 | 1996 | 3179 | 11382 | 1334 | 2362 | 12496 | 1951 | 2981 |
| 3275 | W303.asm02.HP0_G0032700 | chrXI | 14632 | 12912 | 15430 | 12935 | 12822 | 15094 | 14589 | 12691 | 19411 | 28120 | 12247 | 13042 | 32246 | 9045 | 11330 | 30532 | 10810 | 11315 |
| 3276 | W303.asm02.HP0_G0032710 | chrXI | 99 | 93 | 103 | 110 | 96 | 83 | 111 | 83 | 104 | 587 | 77 | 114 | 539 | 71 | 55 | 557 | 105 | 95 |
| 3277 | W303.asm02.HP0_G0032720 | chrXI | 14360 | 3176 | 2813 | 14386 | 3229 | 2840 | 14751 | 3678 | 3279 | 31656 | 6938 | 7610 | 29913 | 4604 | 6112 | 25233 | 5732 | 7108 |
| 3278 | W303.asm02.HP0_G0032730 | chrXI | 3407 | 1128 | 1557 | 3096 | 1260 | 1544 | 3053 | 1105 | 2268 | 2462 | 937 | 1372 | 4166 | 790 | 1235 | 2733 | 889 | 1071 |
| 3279 | W303.asm02.HP0_G0032740 | chrXI | 941 | 1365 | 980 | 858 | 980 | 898 | 903 | 1138 | 1113 | 3246 | 1375 | 1240 | 3062 | 892 | 979 | 2857 | 1455 | 1117 |
| 3280 | W303.asm02.HP0_G0032750 | chrXI | 2044 | 867 | 1121 | 2119 | 1060 | 1140 | 1877 | 746 | 1212 | 4778 | 1126 | 1415 | 3468 | 729 | 1073 | 3154 | 1039 | 1274 |
| 3281 | W303.asm02.HP0_G0032760 | chrXI | 1137 | 113 | 177 | 1046 | 173 | 175 | 1101 | 97 | 201 | 900 | 211 | 307 | 754 | 128 | 228 | 514 | 217 | 283 |
| 3282 | W303.asm02.HP0_G0032770 | chrXI | 1147 | 857 | 861 | 1193 | 864 | 879 | 1169 | 843 | 1002 | 2582 | 643 | 701 | 3153 | 447 | 552 | 2381 | 566 | 629 |
| 3283 | W303.asm02.HP0_G0032780 | chrXI | 570 | 319 | 421 | 592 | 419 | 459 | 513 | 265 | 421 | 2249 | 415 | 873 | 1845 | 285 | 606 | 1645 | 494 | 769 |
| 3284 | W303.asm02.HP0_G0032790 | chrXI | 2916 | 1272 | 966 | 2440 | 977 | 916 | 2908 | 1335 | 992 | 4570 | 1243 | 1243 | 4241 | 717 | 901 | 4533 | 1102 | 1120 |
| 3285 | W303.asm02.HP0_G0032800 | chrXI | 6174 | 1550 | 1850 | 5873 | 1711 | 1673 | 6094 | 1720 | 2050 | 10212 | 2860 | 3253 | 9323 | 2051 | 2528 | 8018 | 2440 | 3015 |
| 3286 | W303.asm02.HP0_G0032810 | chrXI | 2391 | 1341 | 1121 | 2272 | 1236 | 1068 | 2402 | 1247 | 1092 | 5043 | 1422 | 1443 | 4987 | 859 | 1002 | 4566 | 1261 | 1322 |
| 3287 | W303.asm02.HP0_G0032820 | chrXI | 139 | 29 | 40 | 82 | 30 | 37 | 92 | 17 | 52 | 169 | 48 | 59 | 68 | 31 | 65 | 66 | 30 | 52 |
| 3288 | W303.asm02.HP0_G0032830 | chrXI | 1108 | 312 | 559 | 1208 | 373 | 692 | 1033 | 251 | 785 | 2423 | 518 | 981 | 1122 | 321 | 878 | 824 | 429 | 972 |
| 3289 | W303.asm02.HP0_G0032840 | chrXI | 1444 | 399 | 343 | 1334 | 294 | 318 | 1466 | 442 | 302 | 2011 | 423 | 685 | 2089 | 326 | 405 | 1954 | 427 | 500 |
| 3290 | W303.asm02.HP0_G0032850 | chrXI | 2513 | 1153 | 1114 | 2123 | 1047 | 1019 | 2101 | 1177 | 1229 | 3224 | 1096 | 966 | 3575 | 662 | 960 | 3329 | 952 | 980 |
| 3291 | W303.asm02.HP0_G0032860 | chrXI | 3116 | 2385 | 2738 | 2974 | 2464 | 2560 | 2844 | 2415 | 3412 | 6081 | 1967 | 2172 | 6903 | 1196 | 1778 | 6357 | 1840 | 1861 |
| 3292 | W303.asm02.HP0_G0032870 | chrXI | 9271 | 2565 | 2325 | 8836 | 2297 | 2165 | 9052 | 2625 | 2170 | 11975 | 3226 | 3471 | 13704 | 2425 | 3210 | 10954 | 2944 | 3348 |
| 3293 | W303.asm02.HP0_G0032880 | chrXI | 2584 | 218 | 431 | 2447 | 345 | 458 | 2299 | 200 | 443 | 2465 | 453 | 978 | 831 | 308 | 620 | 770 | 382 | 755 |
| 3294 | W303.asm02.HP0_G0032890 | chrXI | 126 | 48 | 33 | 100 | 26 | 34 | 104 | 37 | 32 | 71 | 45 | 16 | 110 | 27 | 35 | 88 | 39 | 16 |
| 3295 | W303.asm02.HP0_G0032900 | chrXI | 17460 | 4049 | 2581 | 16616 | 2554 | 2727 | 15516 | 3641 | 2405 | 10030 | 3891 | 3766 | 9416 | 2937 | 3684 | 8711 | 3563 | 3393 |
| 3296 | W303.asm02.HP0_G0032910 | chrXI | 2475 | 296 | 447 | 2442 | 316 | 510 | 2313 | 306 | 494 | 3155 | 730 | 1069 | 1836 | 494 | 720 | 1948 | 671 | 953 |
| 3297 | W303.asm02.HP0_G0032920 | chrXI | 4197 | 2659 | 3256 | 3647 | 2872 | 3028 | 3872 | 2790 | 4074 | 7592 | 2292 | 2534 | 8196 | 1576 | 2200 | 8169 | 2059 | 2384 |
| 3298 | W303.asm02.HP0_G0032930 | chrXI | 1006 | 672 | 677 | 1103 | 818 | 662 | 1034 | 720 | 832 | 5076 | 833 | 1054 | 4623 | 528 | 749 | 4557 | 773 | 990 |
| 3299 | W303.asm02.HP0_G0032940 | chrXI | 2564 | 247 | 374 | 2692 | 371 | 508 | 2647 | 213 | 483 | 1763 | 394 | 789 | 506 | 285 | 498 | 375 | 252 | 721 |
| 3300 | W303.asm02.HP0_G0032950 | chrXI | 1797 | 152 | 174 | 2033 | 185 | 212 | 2011 | 115 | 245 | 1395 | 301 | 509 | 285 | 240 | 201 | 241 | 212 | 436 |
| 3301 | W303.asm02.HP0_G0032960 | chrXI | 2231 | 677 | 924 | 1794 | 1044 | 1002 | 2339 | 629 | 1237 | 7253 | 1925 | 2323 | 1770 | 993 | 1005 | 2109 | 1465 | 1872 |
| 3302 | W303.asm02.HP0_G0032970 | chrXI | 3014 | 2744 | 2549 | 2966 | 2500 | 2529 | 3262 | 2627 | 2649 | 9491 | 2601 | 2570 | 10272 | 1614 | 2072 | 9256 | 2468 | 2353 |
| 3303 | W303.asm02.HP0_G0032980 | chrXI | 2586 | 1893 | 1710 | 2545 | 1602 | 1492 | 2687 | 1813 | 1821 | 7622 | 2014 | 2346 | 7392 | 1430 | 1643 | 6956 | 1860 | 2102 |
| 3304 | W303.asm02.HP0_G0032990 | chrXI | 2537 | 312 | 210 | 2682 | 189 | 188 | 2303 | 269 | 275 | 506 | 200 | 163 | 706 | 140 | 196 | 540 | 241 | 154 |
| 3305 | W303.asm02.HP0_G0033000 | chrXI | 3783 | 536 | 609 | 3600 | 547 | 700 | 3739 | 430 | 781 | 5289 | 1127 | 1372 | 2097 | 787 | 1015 | 2737 | 985 | 1284 |
| 3306 | W303.asm02.HP0_G0033010 | chrXI | 1840 | 1150 | 1423 | 1502 | 1162 | 1460 | 1857 | 1244 | 1541 | 4214 | 1085 | 1637 | 4014 | 741 | 1175 | 3952 | 1055 | 1454 |
| 3307 | W303.asm02.HP0_G0033020 | chrXI | 629 | 258 | 264 | 352 | 256 | 261 | 452 | 244 | 192 | 833 | 202 | 341 | 905 | 133 | 265 | 926 | 255 | 275 |
| 3308 | W303.asm02.HP0_G0033030 | chrXI | 1848 | 766 | 550 | 1906 | 578 | 680 | 1846 | 709 | 558 | 2034 | 634 | 685 | 2033 | 378 | 561 | 1720 | 595 | 663 |
| 3309 | W303.asm02.HP0_G0033040 | chrXI | 1449 | 490 | 570 | 1447 | 548 | 677 | 1477 | 457 | 684 | 4069 | 737 | 1467 | 2878 | 580 | 965 | 2898 | 692 | 1239 |
| 3310 | W303.asm02.HP0_G0033050 | chrXI | 3194 | 1834 | 1756 | 3049 | 2125 | 1607 | 3160 | 1950 | 2068 | 10547 | 2327 | 2313 | 8413 | 1285 | 1506 | 9743 | 2162 | 2116 |
| 3311 | W303.asm02.HP0_G0033060 | chrXI | 3071 | 538 | 751 | 3515 | 653 | 799 | 2930 | 426 | 957 | 5602 | 1139 | 1595 | 2901 | 709 | 1134 | 2940 | 1184 | 1394 |
| 3312 | W303.asm02.HP0_G0033070 | chrXI | 3051 | 1663 | 1496 | 2360 | 1291 | 1311 | 2779 | 1718 | 1457 | 5039 | 1631 | 1977 | 5148 | 1146 | 1405 | 4776 | 1625 | 1796 |
| 3313 | W303.asm02.HP0_G0033080 | chrXI | 622 | 814 | 877 | 332 | 812 | 779 | 362 | 823 | 1051 | 1899 | 755 | 734 | 2350 | 497 | 731 | 1979 | 659 | 528 |
| 3314 | W303.asm02.HP0_G0033090 | chrXI | 980 | 926 | 641 | 997 | 638 | 667 | 966 | 916 | 650 | 2994 | 777 | 856 | 2745 | 576 | 567 | 2496 | 722 | 718 |
| 3315 | W303.asm02.HP0_G0033100 | chrXI | 1744 | 579 | 490 | 1757 | 491 | 451 | 1727 | 520 | 457 | 2941 | 589 | 706 | 2488 | 349 | 542 | 2633 | 567 | 612 |
| 3316 | W303.asm02.HP0_G0033110 | chrXI | 5109 | 4926 | 5959 | 4779 | 5437 | 6165 | 4767 | 5192 | 6916 | 10899 | 4412 | 5359 | 11774 | 3347 | 4324 | 10898 | 3710 | 4347 |
| 3317 | W303.asm02.HP0_G0033120 | chrXI | 2932 | 1348 | 1723 | 3068 | 1713 | 1658 | 3051 | 1399 | 1663 | 8553 | 1396 | 2093 | 7760 | 1004 | 1689 | 7774 | 1346 | 1935 |
| 3318 | W303.asm02.HP0_G0033130 | chrXI | 1824 | 790 | 1147 | 1881 | 1421 | 1062 | 2100 | 832 | 1474 | 14901 | 1320 | 2465 | 9200 | 930 | 1558 | 10800 | 1268 | 2134 |
| 3319 | W303.asm02.HP0_G0033140 | chrXI | 6688 | 3178 | 3870 | 6545 | 3242 | 3893 | 6540 | 3103 | 4813 | 7639 | 2899 | 3403 | 10246 | 2363 | 3262 | 8049 | 2535 | 3123 |
| 3320 | W303.asm02.HP0_G0033150 | chrXI | 5296 | 1044 | 934 | 5443 | 903 | 1024 | 5054 | 978 | 818 | 5120 | 1107 | 1471 | 5353 | 804 | 1092 | 4319 | 1041 | 1346 |
| 3321 | W303.asm02.HP0_G0033160 | chrXI | 1290 | 793 | 773 | 1217 | 695 | 781 | 1247 | 695 | 1012 | 2295 | 517 | 568 | 2388 | 304 | 397 | 2534 | 567 | 518 |
| 3322 | W303.asm02.HP0_G0033170 | chrXI | 9531 | 1064 | 1404 | 9335 | 1417 | 1504 | 9205 | 1064 | 1582 | 10483 | 1903 | 2843 | 7246 | 1440 | 2286 | 6979 | 1719 | 2464 |
| 3323 | W303.asm02.HP0_G0033180 | chrXI | 3213 | 3814 | 5533 | 2998 | 5700 | 5201 | 3586 | 4064 | 6778 | 14551 | 3668 | 4196 | 14983 | 2361 | 2840 | 15217 | 3026 | 3688 |
| 3324 | W303.asm02.HP0_G0033190 | chrXI | 3239 | 2066 | 2034 | 2726 | 1868 | 1951 | 3210 | 1994 | 2256 | 6871 | 1620 | 1433 | 7179 | 1039 | 1117 | 7358 | 1504 | 1365 |
| 3325 | W303.asm02.HP0_G0033200 | chrXI | 8696 | 3250 | 2794 | 8702 | 2547 | 2874 | 8415 | 3274 | 2968 | 7741 | 2703 | 2720 | 9002 | 2061 | 2610 | 7480 | 2528 | 2655 |
| 3326 | W303.asm02.HP0_G0033210 | chrXI | 1226 | 667 | 762 | 1104 | 669 | 739 | 1133 | 668 | 1005 | 1790 | 536 | 568 | 2238 | 380 | 497 | 1983 | 484 | 489 |
| 3327 | W303.asm02.HP0_G0033220 | chrXI | 6519 | 5323 | 6826 | 5367 | 6617 | 6962 | 5890 | 5043 | 8319 | 12522 | 4516 | 5721 | 13588 | 3286 | 4688 | 13104 | 4292 | 4771 |
| 3328 | W303.asm02.HP0_G0033230 | chrXI | 4159 | 1162 | 587 | 3845 | 750 | 580 | 3842 | 1008 | 564 | 4303 | 1257 | 873 | 3508 | 795 | 650 | 3544 | 1313 | 810 |
| 3329 | W303.asm02.HP0_G0033240 | chrXI | 2012 | 1039 | 1037 | 2024 | 1058 | 1076 | 2137 | 1029 | 1208 | 9270 | 2088 | 2135 | 7255 | 1426 | 1529 | 6983 | 1986 | 2067 |
| 3330 | W303.asm02.HP0_G0033250 | chrXI | 3106 | 1394 | 1524 | 2528 | 1386 | 1520 | 2636 | 1361 | 1888 | 3489 | 1122 | 1353 | 4260 | 865 | 1194 | 3670 | 1181 | 1226 |
| 3331 | W303.asm02.HP0_G0033260 | chrXI | 3710 | 1797 | 1395 | 3683 | 1450 | 1443 | 3574 | 1854 | 1483 | 4521 | 1658 | 1492 | 5001 | 1059 | 1216 | 4122 | 1481 | 1423 |
| 3332 | W303.asm02.HP0_G0033270 | chrXI | 2201 | 1468 | 1689 | 1425 | 1284 | 1673 | 1542 | 1535 | 2210 | 1954 | 1023 | 768 | 2682 | 853 | 764 | 2084 | 935 | 548 |
| 3333 | W303.asm02.HP0_G0033280 | chrXI | 14174 | 3172 | 2420 | 14220 | 2201 | 2360 | 14166 | 3105 | 2516 | 18294 | 5003 | 5808 | 15366 | 3644 | 4604 | 15149 | 4697 | 5420 |
| 3334 | W303.asm02.HP0_G0033290 | chrXI | 3559 | 776 | 581 | 3410 | 626 | 678 | 3644 | 660 | 750 | 5536 | 1310 | 1408 | 3256 | 897 | 1077 | 3471 | 1257 | 1305 |
| 3335 | W303.asm02.HP0_G0033300 | chrXI | 3668 | 1483 | 1126 | 3468 | 1048 | 1144 | 3497 | 1503 | 989 | 5191 | 1571 | 1785 | 6188 | 1260 | 1653 | 4731 | 1524 | 1641 |
| 3336 | W303.asm02.HP0_G0033310 | chrXI | 746 | 698 | 485 | 711 | 445 | 514 | 647 | 679 | 693 | 1844 | 552 | 461 | 1924 | 402 | 447 | 2108 | 581 | 379 |
| 3337 | W303.asm02.HP0_G0033320 | chrXI | 775 | 373 | 258 | 678 | 248 | 259 | 674 | 336 | 256 | 950 | 344 | 287 | 906 | 218 | 214 | 781 | 431 | 255 |
| 3338 | W303.asm02.HP0_G0033330 | chrXI | 7969 | 2587 | 2757 | 7614 | 2718 | 2722 | 7448 | 2699 | 3053 | 7524 | 2388 | 2747 | 9092 | 1801 | 2406 | 7264 | 2318 | 2598 |
| 3339 | W303.asm02.HP0_G0033340 | chrXI | 1741 | 1085 | 1201 | 1529 | 1136 | 1187 | 1574 | 1040 | 1397 | 3307 | 959 | 1034 | 3943 | 645 | 883 | 3477 | 898 | 916 |
| 3340 | W303.asm02.HP0_G0033350 | chrXI | 3075 | 1814 | 1687 | 3378 | 1629 | 1655 | 3096 | 1773 | 1551 | 8180 | 1847 | 2119 | 7226 | 1015 | 1408 | 7994 | 1711 | 1950 |
| 3341 | W303.asm02.HP0_G0033360 | chrXI | 2381 | 579 | 481 | 2481 | 436 | 515 | 2320 | 535 | 500 | 2055 | 627 | 634 | 2123 | 427 | 502 | 1749 | 614 | 568 |
| 3342 | W303.asm02.HP0_G0033370 | chrXI | 972 | 293 | 297 | 913 | 268 | 274 | 1101 | 253 | 304 | 2798 | 554 | 739 | 1853 | 403 | 572 | 2144 | 540 | 674 |
| 3343 | W303.asm02.HP0_G0033380 | chrXI | 4174 | 1874 | 2093 | 4301 | 1338 | 2342 | 3999 | 1638 | 2411 | 3162 | 1211 | 1661 | 3481 | 777 | 1353 | 3644 | 1240 | 1342 |
| 3344 | W303.asm02.HP0_G0033390 | chrXI | 6673 | 1849 | 2023 | 6498 | 1888 | 2352 | 7614 | 1762 | 2489 | 11303 | 5125 | 6571 | 7629 | 3057 | 3374 | 6059 | 3957 | 5642 |
| 3345 | W303.asm02.HP0_G0033400 | chrXI | 6029 | 3150 | 3336 | 6013 | 2932 | 3216 | 5848 | 3196 | 3910 | 8374 | 2825 | 2874 | 9084 | 1911 | 2531 | 8928 | 2487 | 2725 |
| 3346 | W303.asm02.HP0_G0033410 | chrXI | 795 | 286 | 325 | 744 | 323 | 300 | 819 | 314 | 310 | 1248 | 338 | 369 | 1259 | 189 | 306 | 1151 | 292 | 325 |

| | | | | | | | | | | | | | | | | | | | | |
|---|---|---|---|---|---|---|---|---|---|---|---|---|---|---|---|---|---|---|---|---|
| 3347 | W303.asm02.HP0_G0033420 | chrXI | 6832 | 1896 | 2041 | 6972 | 2110 | 2085 | 6559 | 1929 | 2102 | 10300 | 2199 | 3032 | 9761 | 1741 | 2476 | 8845 | 1988 | 2490 |
| 3348 | W303.asm02.HP0_G0033430 | chrXI | 1408 | 485 | 520 | 1481 | 523 | 470 | 1279 | 430 | 557 | 2216 | 479 | 522 | 2152 | 274 | 412 | 2135 | 535 | 439 |
| 3349 | W303.asm02.HP0_G0033440 | chrXI | 13844 | 1834 | 1485 | 13688 | 1643 | 1454 | 13961 | 1845 | 1480 | 10862 | 2715 | 2728 | 9224 | 1876 | 2327 | 8503 | 2405 | 2532 |
| 3350 | W303.asm02.HP0_G0033450 | chrXI | 1162 | 216 | 173 | 1083 | 220 | 217 | 1108 | 207 | 217 | 1373 | 284 | 391 | 1030 | 200 | 237 | 929 | 257 | 332 |
| 3351 | W303.asm02.HP0_G0033460 | chrXI | 1355 | 558 | 539 | 1257 | 489 | 498 | 1185 | 530 | 459 | 1895 | 501 | 604 | 1960 | 340 | 497 | 1775 | 455 | 634 |
| 3352 | W303.asm02.HP0_G0033470 | chrXI | 2767 | 554 | 704 | 2815 | 697 | 806 | 2871 | 478 | 791 | 6167 | 1107 | 1533 | 4627 | 863 | 1279 | 4121 | 1023 | 1409 |
| 3353 | W303.asm02.HP0_G0033480 | chrXI | 1841 | 908 | 870 | 1737 | 954 | 930 | 1774 | 896 | 873 | 2796 | 771 | 935 | 2652 | 431 | 677 | 2770 | 653 | 828 |
| 3354 | W303.asm02.HP0_G0033490 | chrXI | 1986 | 774 | 828 | 2034 | 904 | 796 | 2053 | 813 | 877 | 4294 | 754 | 1012 | 3864 | 501 | 771 | 3531 | 731 | 933 |
| 3355 | W303.asm02.HP0_G0033500 | chrXI | 482 | 493 | 668 | 483 | 938 | 687 | 560 | 497 | 765 | 8920 | 956 | 1535 | 5560 | 727 | 714 | 5378 | 921 | 1314 |
| 3356 | W303.asm02.HP0_G0033510 | chrXI | 850 | 1062 | 866 | 918 | 1088 | 952 | 944 | 1001 | 854 | 5278 | 1368 | 1415 | 4562 | 838 | 816 | 4349 | 1156 | 1243 |
| 3357 | W303.asm02.HP0_G0033520 | chrXI | 3211 | 2060 | 2074 | 2989 | 2022 | 1813 | 3163 | 2158 | 2303 | 7797 | 2212 | 2176 | 7704 | 1305 | 1676 | 7756 | 1911 | 1938 |
| 3358 | W303.asm02.HP0_G0033530 | chrXI | 1729 | 519 | 618 | 1753 | 549 | 604 | 1734 | 591 | 673 | 4648 | 1033 | 1445 | 3606 | 598 | 977 | 3827 | 925 | 1295 |
| 3359 | W303.asm02.HP0_G0033540 | chrXI | 4052 | 991 | 1301 | 4014 | 1234 | 1439 | 4130 | 1024 | 1584 | 11317 | 1908 | 2998 | 7626 | 1431 | 2216 | 7417 | 1635 | 2699 |
| 3360 | W303.asm02.HP0_G0033550 | chrXI | 2041 | 860 | 782 | 2285 | 853 | 805 | 2090 | 917 | 770 | 4597 | 906 | 1122 | 4324 | 601 | 917 | 4331 | 936 | 1039 |
| 3361 | W303.asm02.HP0_G0033560 | chrXI | 1434 | 1425 | 1637 | 1453 | 1477 | 1643 | 1343 | 1334 | 2137 | 4199 | 1095 | 1256 | 4254 | 675 | 898 | 4776 | 1068 | 1070 |
| 3362 | W303.asm02.HP0_G0033570 | chrXI | 3636 | 498 | 793 | 3876 | 741 | 881 | 3394 | 470 | 1051 | 6073 | 787 | 1745 | 3425 | 602 | 1266 | 3321 | 707 | 1422 |
| 3363 | W303.asm02.HP0_G0033580 | chrXI | 30 | 2 | 0 | 26 | 2 | 3 | 27 | 1 | 1 | 15 | 2 | 7 | 18 | 1 | 1 | 7 | 2 | 5 |
| 3364 | W303.asm02.HP0_G0033590 | chrXI | 662 | 151 | 167 | 658 | 167 | 203 | 668 | 184 | 136 | 387 | 187 | 206 | 747 | 137 | 218 | 342 | 157 | 180 |
| 3365 | W303.asm02.HP0_G0033600 | chrXI | 4106 | 1940 | 2549 | 4252 | 2432 | 2573 | 4068 | 1964 | 2951 | 6499 | 1632 | 2264 | 7985 | 1211 | 1919 | 5975 | 1426 | 1919 |
| 3366 | W303.asm02.HP0_G0033610 | chrXI | 1228 | 427 | 433 | 1092 | 540 | 498 | 1202 | 426 | 538 | 4170 | 703 | 967 | 2749 | 466 | 590 | 2779 | 625 | 778 |
| 3367 | W303.asm02.HP0_G0033620 | chrXI | 1451 | 717 | 720 | 1298 | 681 | 748 | 1273 | 721 | 833 | 2193 | 658 | 653 | 2499 | 399 | 548 | 2224 | 627 | 656 |
| 3368 | W303.asm02.HP0_G0033630 | chrXI | 2698 | 1591 | 1051 | 2415 | 1062 | 1056 | 2860 | 1407 | 1183 | 5741 | 2036 | 1843 | 4265 | 1447 | 1390 | 4338 | 1941 | 1805 |
| 3369 | W303.asm02.HP0_G0033640 | chrXI | 3742 | 2674 | 3811 | 4326 | 3489 | 4171 | 3882 | 2528 | 4448 | 15848 | 5399 | 5775 | 16299 | 6017 | 8124 | 8932 | 4447 | 5016 |
| 3370 | W303.asm02.HP0_G0033650 | chrXI | 58 | 20 | 45 | 61 | 38 | 41 | 68 | 20 | 52 | 107 | 20 | 46 | 75 | 22 | 28 | 55 | 33 | 25 |
| 3371 | W303.asm02.HP0_G0033660 | chrXI | 21857 | 5737 | 6429 | 20759 | 5436 | 7312 | 22737 | 4734 | 8296 | 13735 | 7640 | 10414 | 5795 | 5122 | 5438 | 6429 | 6287 | 7868 |
| 3372 | W303.asm02.HP0_G0033670 | chrXI | 5888 | 2865 | 2511 | 5289 | 2689 | 2456 | 6532 | 2982 | 2839 | 11833 | 3915 | 4207 | 11867 | 2824 | 3252 | 10327 | 3194 | 3822 |
| 3373 | W303.asm02.HP0_G0033680 | chrXI | 793 | 521 | 711 | 694 | 676 | 617 | 832 | 557 | 666 | 3884 | 848 | 999 | 4027 | 724 | 646 | 2735 | 736 | 887 |
| 3374 | W303.asm02.HP0_G0033690 | chrXI | 237 | 70 | 65 | 186 | 93 | 89 | 231 | 79 | 92 | 550 | 144 | 176 | 411 | 109 | 85 | 396 | 129 | 159 |
| 3375 | W303.asm02.HP0_G0033700 | chrXI | 15112 | 1846 | 1977 | 14492 | 1549 | 2099 | 15203 | 1680 | 2265 | 6752 | 2724 | 3642 | 5853 | 2242 | 3055 | 4502 | 2668 | 3428 |
| 3376 | W303.asm02.HP0_G0033710 | chrXI | 12533 | 1393 | 1237 | 10176 | 1108 | 1492 | 12238 | 946 | 1742 | 8025 | 2896 | 3384 | 2746 | 2151 | 2194 | 2620 | 2338 | 2831 |
| 3377 | W303.asm02.HP0_G0033720 | chrXI | 2982 | 477 | 452 | 2695 | 430 | 482 | 2607 | 491 | 478 | 1057 | 532 | 588 | 1300 | 432 | 492 | 735 | 455 | 523 |
| 3378 | W303.asm02.HP0_G0033730 | chrXI | 764 | 148 | 269 | 728 | 169 | 337 | 771 | 106 | 352 | 955 | 261 | 673 | 435 | 231 | 334 | 378 | 213 | 548 |
| 3379 | W303.asm02.HP0_G0033740 | chrXI | 806 | 719 | 836 | 823 | 728 | 751 | 720 | 583 | 855 | 2824 | 607 | 685 | 2772 | 364 | 598 | 2698 | 588 | 656 |
| 3380 | W303.asm02.HP0_G0033750 | chrXI | 1410 | 808 | 823 | 1662 | 680 | 831 | 1316 | 717 | 841 | 3050 | 998 | 969 | 3065 | 538 | 830 | 2969 | 894 | 974 |
| 3381 | W303.asm02.HP0_G0033760 | chrXI | 704 | 233 | 293 | 583 | 290 | 332 | 605 | 258 | 297 | 898 | 257 | 401 | 852 | 219 | 259 | 861 | 262 | 320 |
| 3382 | W303.asm02.HP0_G0033770 | chrXI | 2738 | 470 | 627 | 2364 | 694 | 662 | 2678 | 410 | 797 | 4701 | 575 | 1478 | 3128 | 551 | 1137 | 2995 | 567 | 1301 |
| 3383 | W303.asm02.HP0_G0033780 | chrXI | 531 | 677 | 730 | 624 | 695 | 671 | 497 | 663 | 701 | 2153 | 536 | 645 | 2055 | 324 | 470 | 2196 | 525 | 581 |
| 3384 | W303.asm02.HP0_G0033790 | chrXI | 1707 | 902 | 808 | 2157 | 978 | 857 | 2355 | 901 | 879 | 8197 | 1369 | 1818 | 6705 | 1060 | 959 | 6252 | 1162 | 1505 |
| 3385 | W303.asm02.HP0_G0033800 | chrXI | 3606 | 3811 | 5267 | 2881 | 4696 | 5348 | 3436 | 3619 | 6809 | 10095 | 3341 | 4075 | 11078 | 2459 | 3220 | 11272 | 3145 | 3716 |
| 3386 | W303.asm02.HP0_G0033810 | chrXI | 3555 | 354 | 232 | 3409 | 232 | 232 | 3147 | 327 | 274 | 1082 | 442 | 284 | 1119 | 222 | 316 | 942 | 380 | 272 |
| 3387 | W303.asm02.HP0_G0033820 | chrXI | 11643 | 8205 | 7581 | 11659 | 7338 | 7544 | 12533 | 8450 | 8046 | 21093 | 8199 | 8557 | 20618 | 5923 | 6580 | 20116 | 7089 | 7830 |
| 3388 | W303.asm02.HP0_G0033830 | chrXI | 735 | 889 | 725 | 565 | 956 | 705 | 630 | 952 | 730 | 3904 | 797 | 846 | 3338 | 457 | 431 | 3976 | 716 | 723 |
| 3389 | W303.asm02.HP0_G0033840 | chrXI | 3315 | 730 | 858 | 2952 | 743 | 756 | 3185 | 765 | 909 | 2288 | 678 | 814 | 5390 | 796 | 854 | 1974 | 646 | 622 |
| 3390 | W303.asm02.HP0_G0033850 | chrXI | 2173 | 1126 | 1079 | 2100 | 1193 | 1111 | 1984 | 1151 | 1064 | 4261 | 1020 | 1328 | 4012 | 704 | 992 | 3874 | 978 | 1116 |
| 3391 | W303.asm02.HP0_G0033860 | chrXI | 1124 | 500 | 525 | 1051 | 737 | 605 | 1132 | 548 | 617 | 3948 | 571 | 885 | 3004 | 361 | 540 | 3336 | 527 | 775 |
| 3392 | W303.asm02.HP0_G0033870 | chrXI | 1711 | 681 | 634 | 1793 | 716 | 698 | 1753 | 693 | 669 | 4253 | 787 | 1042 | 3528 | 516 | 752 | 3690 | 762 | 980 |
| 3393 | W303.asm02.HP0_G0033880 | chrXI | 3915 | 1581 | 1455 | 3719 | 1242 | 1466 | 3657 | 1544 | 1602 | 3584 | 1306 | 1595 | 3789 | 1010 | 1219 | 3334 | 1158 | 1333 |
| 3394 | W303.asm02.HP0_G0033890 | chrXI | 9656 | 1726 | 1240 | 8529 | 1035 | 1521 | 8933 | 1519 | 1580 | 6081 | 2365 | 2687 | 4044 | 1678 | 1894 | 4186 | 2364 | 2415 |
| 3395 | W303.asm02.HP0_G0033900 | chrXI | 5498 | 530 | 565 | 5363 | 457 | 622 | 5014 | 545 | 738 | 3231 | 992 | 1080 | 2271 | 697 | 1026 | 2376 | 946 | 1033 |
| 3396 | W303.asm02.HP0_G0033910 | chrXI | 2423 | 2801 | 2653 | 2439 | 2188 | 2754 | 2494 | 2701 | 3269 | 5073 | 2098 | 2139 | 6421 | 1441 | 1686 | 5521 | 2017 | 1835 |
| 3397 | W303.asm02.HP0_G0033920 | chrXI | 3422 | 595 | 408 | 2054 | 442 | 455 | 2664 | 460 | 464 | 1572 | 732 | 524 | 1094 | 497 | 521 | 1034 | 773 | 466 |
| 3398 | W303.asm02.HP0_G0033930 | chrXI | 2405 | 982 | 813 | 1885 | 778 | 773 | 2144 | 924 | 830 | 3577 | 985 | 1029 | 3472 | 658 | 851 | 3326 | 980 | 851 |
| 3399 | W303.asm02.HP0_G0033940 | chrXI | 4513 | 7437 | 8441 | 4907 | 8195 | 8178 | 4595 | 7894 | 8903 | 16353 | 6950 | 6965 | 16181 | 4176 | 5422 | 16131 | 6536 | 5903 |
| 3400 | W303.asm02.HP0_G0033950 | chrXI | 3831 | 515 | 698 | 3603 | 777 | 917 | 3874 | 476 | 915 | 4885 | 842 | 1086 | 2083 | 661 | 775 | 1863 | 719 | 954 |
| 3401 | W303.asm02.HP0_G0033960 | chrXI | 178 | 48 | 63 | 146 | 84 | 86 | 153 | 56 | 72 | 109 | 46 | 44 | 147 | 27 | 22 | 120 | 53 | 26 |
| 3402 | W303.asm02.HP0_G0033970 | chrXI | 2936 | 1429 | 1652 | 2658 | 1372 | 1624 | 2545 | 1430 | 2250 | 3757 | 1327 | 1346 | 4727 | 882 | 1122 | 4233 | 1140 | 1169 |
| 3403 | W303.asm02.HP0_G0033980 | chrXI | 3738 | 896 | 497 | 4533 | 1063 | 549 | 3881 | 765 | 513 | 4450 | 641 | 342 | 4743 | 405 | 287 | 2869 | 625 | 270 |
| 3404 | W303.asm02.HP0_G0033990 | chrXI | 4606 | 447 | 363 | 4461 | 314 | 492 | 4447 | 398 | 448 | 1172 | 822 | 790 | 1875 | 908 | 982 | 706 | 732 | 649 |
| 3405 | W303.asm02.HP0_G0034000 | chrXI | 801 | 253 | 212 | 763 | 188 | 283 | 858 | 230 | 298 | 1418 | 439 | 491 | 1426 | 365 | 368 | 1053 | 358 | 422 |
| 3406 | W303.asm02.HP0_G0034010 | chrXI | 2235 | 534 | 561 | 2224 | 644 | 564 | 2129 | 567 | 582 | 3194 | 538 | 688 | 2825 | 368 | 546 | 2861 | 533 | 621 |
| 3407 | W303.asm02.HP0_G0034020 | chrXI | 1179 | 1201 | 1383 | 1068 | 1644 | 1185 | 1337 | 1358 | 1517 | 10124 | 1527 | 1940 | 9708 | 1171 | 1311 | 8681 | 1271 | 1721 |
| 3408 | W303.asm02.HP0_G0034030 | chrXI | 1388 | 740 | 807 | 896 | 701 | 748 | 1044 | 774 | 938 | 2401 | 594 | 619 | 2644 | 411 | 469 | 2554 | 547 | 537 |
| 3409 | W303.asm02.HP0_G0034040 | chrXI | 1807 | 2761 | 2989 | 1983 | 3707 | 2835 | 2151 | 3175 | 3057 | 13309 | 3015 | 3265 | 11932 | 1849 | 1955 | 12281 | 2439 | 2919 |
| 3410 | W303.asm02.HP0_G0034050 | chrXI | 4670 | 1608 | 1720 | 4608 | 1956 | 1522 | 4567 | 1725 | 1585 | 11327 | 2017 | 2594 | 9828 | 1288 | 1901 | 10365 | 1835 | 2292 |
| 3411 | W303.asm02.HP0_G0034060 | chrXI | 552 | 422 | 647 | 506 | 582 | 648 | 480 | 442 | 830 | 1638 | 289 | 450 | 1793 | 175 | 392 | 1844 | 304 | 389 |
| 3412 | W303.asm02.HP0_G0034070 | chrXI | 3557 | 1566 | 1377 | 3290 | 1444 | 1361 | 3669 | 1648 | 1499 | 6949 | 1637 | 1823 | 5942 | 973 | 1313 | 6345 | 1475 | 1710 |
| 3413 | W303.asm02.HP0_G0034080 | chrXI | 25 | 11 | 15 | 22 | 15 | 12 | 22 | 24 | 12 | 62 | 25 | 19 | 55 | 8 | 17 | 42 | 20 | 11 |
| 3414 | W303.asm02.HP0_G0034090 | chrXI | 4417 | 1261 | 1942 | 4079 | 1640 | 2038 | 4162 | 1306 | 2682 | 4113 | 1082 | 1627 | 5626 | 892 | 1489 | 4214 | 1015 | 1538 |
| 3415 | W303.asm02.HP0_G0034100 | chrXI | 1307 | 404 | 505 | 1234 | 527 | 514 | 1235 | 427 | 572 | 4358 | 603 | 938 | 3322 | 429 | 663 | 3119 | 581 | 832 |
| 3416 | W303.asm02.HP0_G0034110 | chrXI | 1308 | 887 | 691 | 1445 | 717 | 682 | 1282 | 758 | 748 | 2814 | 716 | 809 | 2779 | 441 | 548 | 2861 | 810 | 663 |
| 3417 | W303.asm02.HP0_G0034120 | chrXI | 2339 | 1198 | 1265 | 2399 | 1063 | 1196 | 2231 | 1263 | 1331 | 4283 | 1149 | 1319 | 4179 | 820 | 1057 | 4533 | 1067 | 1196 |
| 3418 | W303.asm02.HP0_G0034130 | chrXI | 4463 | 675 | 556 | 4185 | 515 | 677 | 4234 | 549 | 682 | 4405 | 978 | 1318 | 1826 | 679 | 776 | 2138 | 957 | 1058 |
| 3419 | W303.asm02.HP0_G0034140 | chrXI | 2896 | 729 | 551 | 2871 | 609 | 607 | 3106 | 727 | 672 | 4960 | 1089 | 1088 | 4482 | 825 | 976 | 3711 | 925 | 1011 |
| 3420 | W303.asm02.HP0_G0034150 | chrXI | 6345 | 618 | 584 | 6593 | 448 | 640 | 6239 | 521 | 817 | 3015 | 1023 | 1179 | 5603 | 1487 | 1650 | 1579 | 922 | 1032 |
| 3421 | W303.asm02.HP0_G0034160 | chrXI | 2156 | 1946 | 1992 | 2178 | 2424 | 1857 | 2539 | 2105 | 2370 | 16399 | 3394 | 3048 | 13018 | 2255 | 1953 | 12730 | 2856 | 2809 |
| 3422 | W303.asm02.HP0_G0034170 | chrXI | 2727 | 785 | 2195 | 3535 | 1932 | 2608 | 2446 | 788 | 2664 | 10317 | 1341 | 3294 | 7383 | 1102 | 3016 | 5545 | 1049 | 2683 |
| 3423 | W303.asm02.HP0_G0034180 | chrXI | 9000 | 1105 | 1264 | 9123 | 1348 | 1248 | 8837 | 1155 | 1218 | 11844 | 1263 | 2725 | 9815 | 1046 | 2056 | 9872 | 1061 | 2535 |
| 3424 | W303.asm02.HP0_G0034190 | chrXI | 269 | 343 | 566 | 234 | 479 | 551 | 214 | 296 | 690 | 1287 | 325 | 464 | 1370 | 208 | 333 | 1253 | 254 | 416 |
| 3425 | W303.asm02.HP0_G0034200 | chrXI | 4291 | 525 | 700 | 4636 | 654 | 809 | 4116 | 532 | 747 | 5359 | 891 | 1598 | 4590 | 663 | 1214 | 3705 | 833 | 1354 |
| 3426 | W303.asm02.HP0_G0034210 | chrXI | 805 | 224 | 244 | 730 | 204 | 221 | 801 | 204 | 233 | 599 | 197 | 189 | 859 | 144 | 138 | 576 | 173 | 147 |
| 3427 | W303.asm02.HP0_G0034220 | chrXI | 3886 | 240 | 285 | 3626 | 352 | 292 | 3642 | 270 | 351 | 2825 | 555 | 560 | 2613 | 414 | 571 | 2051 | 483 | 490 |
| 3428 | W303.asm02.HP0_G0034230 | chrXI | 406 | 371 | 415 | 426 | 593 | 417 | 423 | 364 | 420 | 5112 | 581 | 859 | 3932 | 472 | 454 | 3646 | 540 | 690 |
| 3429 | W303.asm02.HP0_G0034240 | chrXI | 3062 | 1486 | 1511 | 2952 | 1489 | 1427 | 3076 | 1520 | 1695 | 5741 | 1493 | 1674 | 6237 | 1061 | 1362 | 5342 | 1348 | 1509 |
| 3430 | W303.asm02.HP0_G0034250 | chrXI | 837 | 316 | 411 | 769 | 327 | 365 | 728 | 372 | 362 | 1398 | 400 | 523 | 1370 | 253 | 337 | 1450 | 375 | 503 |
| 3431 | W303.asm02.HP0_G0034260 | chrXI | 1459 | 72 | 102 | 1179 | 88 | 98 | 1261 | 51 | 106 | 761 | 110 | 228 | 318 | 109 | 174 | 258 | 114 | 215 |
| 3432 | W303.asm02.HP0_G0034270 | chrXI | 887 | 89 | 183 | 672 | 178 | 178 | 748 | 116 | 200 | 1057 | 130 | 308 | 698 | 107 | 233 | 806 | 137 | 318 |
| 3433 | W303.asm02.HP0_G0034280 | chrXI | 300 | 44 | 64 | 235 | 79 | 83 | 308 | 61 | 93 | 312 | 65 | 123 | 201 | 45 | 95 | 272 | 40 | 107 |
| 3434 | W303.asm02.HP0_G0034290 | chrXI | 387 | 93 | 141 | 487 | 128 | 139 | 453 | 75 | 140 | 975 | 140 | 253 | 751 | 84 | 135 | 654 | 131 | 191 |
| 3435 | W303.asm02.HP0_G0034300 | chrXI | 924 | 342 | 320 | 1011 | 323 | 328 | 739 | 276 | 302 | 979 | 268 | 299 | 681 | 108 | 154 | 826 | 249 | 273 |
| 3436 | W303.asm02.HP0_G0034310 | chrXII | 30 | 9 | 12 | 61 | 13 | 10 | 55 | 3 | 19 | 61 | 20 | 31 | 18 | 21 | 13 | 25 | 20 | 28 |
| 3437 | W303.asm02.HP0_G0034320 | chrXII | 2 | 0 | 0 | 0 | 0 | 1 | 2 | 0 | 0 | 1 | 0 | 0 | 1 | 0 | 0 | 0 | 1 | 0 |
| 3438 | W303.asm02.HP0_G0034330 | chrXII | 235 | 33 | 26 | 256 | 36 | 46 | 223 | 22 | 37 | 125 | 44 | 75 | 49 | 36 | 44 | 55 | 42 | 75 |
| 3439 | W303.asm02.HP0_G0034340 | chrXII | 0 | 6 | 0 | 0 | 0 | 0 | 0 | 8 | 0 | 0 | 3 | 0 | 0 | 3 | 0 | 0 | 2 | 0 |

| | | | | | | | | | | | | | | | | | | | | |
|---|---|---|---|---|---|---|---|---|---|---|---|---|---|---|---|---|---|---|---|---|
| 3440 | W303.asm02.HP0_G0034350 | chrXII | 5 | 3 | 1 | 8 | 0 | 2 | 3 | 0 | 0 | 1 | 1 | 1 | 1 | 0 | 0 | 1 | 1 | 0 |
| 3441 | W303.asm02.HP0_G0034360 | chrXII | 587 | 274 | 147 | 469 | 163 | 151 | 550 | 223 | 117 | 845 | 250 | 181 | 719 | 141 | 147 | 682 | 224 | 175 |
| 3442 | W303.asm02.HP0_G0034370 | chrXII | 3454 | 384 | 335 | 2152 | 312 | 349 | 2706 | 398 | 370 | 908 | 348 | 305 | 1104 | 362 | 313 | 703 | 292 | 260 |
| 3443 | W303.asm02.HP0_G0034380 | chrXII | 2345 | 479 | 408 | 1227 | 427 | 373 | 1760 | 442 | 379 | 1621 | 475 | 434 | 1099 | 422 | 374 | 1042 | 406 | 335 |
| 3444 | W303.asm02.HP0_G0034390 | chrXII | 2710 | 2056 | 2689 | 2553 | 2065 | 2665 | 2435 | 2058 | 2733 | 1128 | 2914 | 2433 | 1745 | 1990 | 2276 | 952 | 2301 | 1761 |
| 3445 | W303.asm02.HP0_G0034400 | chrXII | 200 | 62 | 71 | 217 | 56 | 79 | 183 | 66 | 91 | 380 | 93 | 119 | 306 | 70 | 81 | 253 | 104 | 97 |
| 3446 | W303.asm02.HP0_G0034410 | chrXII | 1137 | 297 | 322 | 901 | 246 | 353 | 1024 | 241 | 423 | 1760 | 423 | 693 | 906 | 252 | 421 | 1003 | 395 | 606 |
| 3447 | W303.asm02.HP0_G0034420 | chrXII | 556 | 248 | 222 | 492 | 204 | 221 | 477 | 244 | 232 | 969 | 233 | 234 | 902 | 150 | 185 | 843 | 229 | 205 |
| 3448 | W303.asm02.HP0_G0034430 | chrXII | 335 | 154 | 139 | 331 | 186 | 208 | 332 | 129 | 149 | 543 | 134 | 192 | 500 | 115 | 117 | 467 | 147 | 180 |
| 3449 | W303.asm02.HP0_G0034440 | chrXII | 1240 | 325 | 482 | 1172 | 465 | 436 | 964 | 265 | 493 | 1200 | 296 | 321 | 1034 | 179 | 300 | 1021 | 269 | 282 |
| 3450 | W303.asm02.HP0_G0034450 | chrXII | 1893 | 541 | 771 | 1683 | 812 | 598 | 1636 | 522 | 763 | 4199 | 528 | 765 | 3727 | 349 | 639 | 3693 | 518 | 717 |
| 3451 | W303.asm02.HP0_G0034460 | chrXII | 404 | 404 | 471 | 362 | 674 | 521 | 409 | 250 | 307 | 788 | 347 | 269 | 501 | 125 | 187 | 614 | 323 | 248 |
| 3452 | W303.asm02.HP0_G0034470 | chrXII | 1883 | 549 | 803 | 1847 | 898 | 837 | 1858 | 617 | 798 | 5668 | 1067 | 1242 | 5842 | 615 | 1011 | 5162 | 921 | 1382 |
| 3453 | W303.asm02.HP0_G0034480 | chrXII | 5102 | 3390 | 4415 | 4174 | 3549 | 4308 | 4470 | 3263 | 5481 | 7588 | 2864 | 3635 | 10098 | 2097 | 3048 | 8959 | 2752 | 3130 |
| 3454 | W303.asm02.HP0_G0034490 | chrXII | 783 | 620 | 750 | 621 | 613 | 800 | 601 | 577 | 1055 | 1578 | 492 | 488 | 2023 | 306 | 471 | 1841 | 450 | 460 |
| 3455 | W303.asm02.HP0_G0034500 | chrXII | 5664 | 1450 | 2049 | 4981 | 2009 | 2164 | 5394 | 1641 | 2738 | 15944 | 2936 | 4367 | 9525 | 2172 | 3049 | 10153 | 2612 | 3919 |
| 3456 | W303.asm02.HP0_G0034510 | chrXII | 147 | 82 | 64 | 139 | 47 | 68 | 124 | 85 | 66 | 108 | 74 | 30 | 148 | 31 | 30 | 110 | 54 | 26 |
| 3457 | W303.asm02.HP0_G0034520 | chrXII | 26636 | 14986 | 13759 | 24760 | 14709 | 14125 | 28005 | 15198 | 13170 | 38756 | 15115 | 16049 | 37427 | 11706 | 12692 | 36622 | 12701 | 14887 |
| 3458 | W303.asm02.HP0_G0034530 | chrXII | 3160 | 870 | 964 | 2928 | 1006 | 1028 | 3109 | 844 | 1292 | 6674 | 1572 | 1630 | 3386 | 1091 | 1261 | 3674 | 1317 | 1541 |
| 3459 | W303.asm02.HP0_G0034540 | chrXII | 644 | 125 | 168 | 540 | 152 | 140 | 516 | 155 | 228 | 505 | 115 | 126 | 585 | 63 | 78 | 653 | 126 | 114 |
| 3460 | W303.asm02.HP0_G0034550 | chrXII | 8121 | 2828 | 2086 | 8124 | 2507 | 2094 | 8102 | 3047 | 1991 | 9046 | 3178 | 3570 | 8083 | 2712 | 2773 | 7248 | 2696 | 2555 |
| 3461 | W303.asm02.HP0_G0034560 | chrXII | 9519 | 655 | 803 | 9711 | 688 | 915 | 8856 | 628 | 985 | 5690 | 973 | 2191 | 3288 | 953 | 1813 | 2980 | 848 | 1859 |
| 3462 | W303.asm02.HP0_G0034570 | chrXII | 712 | 203 | 207 | 742 | 214 | 221 | 639 | 159 | 306 | 2010 | 328 | 462 | 1041 | 239 | 302 | 1033 | 341 | 360 |
| 3463 | W303.asm02.HP0_G0034580 | chrXII | 3703 | 1071 | 886 | 3379 | 1007 | 887 | 3598 | 1053 | 841 | 5652 | 1180 | 1501 | 5098 | 844 | 1088 | 4657 | 1115 | 1230 |
| 3464 | W303.asm02.HP0_G0034590 | chrXII | 442 | 256 | 496 | 429 | 408 | 520 | 520 | 291 | 558 | 5011 | 563 | 1249 | 3804 | 408 | 655 | 3521 | 469 | 1124 |
| 3465 | W303.asm02.HP0_G0034600 | chrXII | 789 | 814 | 1044 | 809 | 1216 | 918 | 992 | 854 | 1116 | 10829 | 1543 | 2109 | 7386 | 1035 | 1136 | 7676 | 1342 | 1865 |
| 3466 | W303.asm02.HP0_G0034610 | chrXII | 387 | 322 | 330 | 336 | 332 | 372 | 380 | 330 | 445 | 1094 | 269 | 335 | 1194 | 156 | 207 | 1211 | 245 | 276 |
| 3467 | W303.asm02.HP0_G0034620 | chrXII | 1569 | 558 | 627 | 1685 | 721 | 618 | 1579 | 619 | 601 | 4756 | 631 | 936 | 4289 | 452 | 702 | 4297 | 528 | 839 |
| 3468 | W303.asm02.HP0_G0034630 | chrXII | 1804 | 718 | 987 | 1782 | 807 | 980 | 1781 | 727 | 1084 | 7606 | 1602 | 1970 | 3586 | 1193 | 1382 | 4237 | 1418 | 1852 |
| 3469 | W303.asm02.HP0_G0034640 | chrXII | 10122 | 1029 | 892 | 9239 | 854 | 1205 | 9209 | 799 | 1174 | 6465 | 2017 | 2459 | 2834 | 1448 | 1881 | 3005 | 1863 | 2396 |
| 3470 | W303.asm02.HP0_G0034650 | chrXII | 981 | 1012 | 1104 | 990 | 1096 | 1205 | 918 | 901 | 1430 | 8904 | 1887 | 1914 | 3913 | 1376 | 1395 | 3696 | 1618 | 1718 |
| 3471 | W303.asm02.HP0_G0034660 | chrXII | 2643 | 2118 | 1568 | 2525 | 1708 | 1582 | 2630 | 2024 | 1621 | 4846 | 1866 | 1795 | 4770 | 1246 | 1240 | 4386 | 1798 | 1422 |
| 3472 | W303.asm02.HP0_G0034670 | chrXII | 77571 | 6557 | 4690 | 73866 | 5683 | 7072 | 69332 | 4368 | 4960 | 23421 | 8772 | 8255 | 14209 | 5744 | 8385 | 17880 | 9306 | 7116 |
| 3473 | W303.asm02.HP0_G0034680 | chrXII | 302 | 89 | 92 | 252 | 62 | 73 | 292 | 78 | 72 | 201 | 105 | 74 | 263 | 58 | 68 | 174 | 84 | 80 |
| 3474 | W303.asm02.HP0_G0034690 | chrXII | 17373 | 10633 | 12769 | 16215 | 11250 | 14502 | 17646 | 7880 | 14201 | 70126 | 20328 | 28313 | 43139 | 16008 | 20627 | 49512 | 18199 | 26722 |
| 3475 | W303.asm02.HP0_G0034700 | chrXII | 5275 | 795 | 855 | 4605 | 676 | 886 | 4923 | 741 | 992 | 2891 | 1065 | 1069 | 2157 | 716 | 738 | 2009 | 967 | 944 |
| 3476 | W303.asm02.HP0_G0034710 | chrXII | 1446 | 453 | 329 | 1302 | 341 | 381 | 1541 | 358 | 434 | 2643 | 950 | 831 | 1112 | 656 | 533 | 1007 | 776 | 786 |
| 3477 | W303.asm02.HP0_G0034720 | chrXII | 3419 | 586 | 825 | 3224 | 827 | 1040 | 3765 | 573 | 1110 | 7120 | 1344 | 2264 | 2768 | 1054 | 1379 | 2791 | 1030 | 2013 |
| 3478 | W303.asm02.HP0_G0034730 | chrXII | 2447 | 586 | 678 | 3051 | 630 | 830 | 2450 | 444 | 883 | 3860 | 1357 | 1578 | 2389 | 999 | 1173 | 1732 | 1145 | 1210 |
| 3479 | W303.asm02.HP0_G0034740 | chrXII | 1107 | 188 | 232 | 1257 | 286 | 204 | 1011 | 200 | 226 | 1308 | 295 | 290 | 1246 | 211 | 211 | 1030 | 222 | 261 |
| 3480 | W303.asm02.HP0_G0034750 | chrXII | 22258 | 2624 | 1892 | 19587 | 1917 | 2411 | 22292 | 1977 | 2509 | 20743 | 4968 | 5510 | 12641 | 4055 | 4071 | 12623 | 4599 | 4985 |
| 3481 | W303.asm02.HP0_G0034760 | chrXII | 0 | 0 | 0 | 0 | 0 | 0 | 0 | 0 | 0 | 0 | 0 | 0 | 0 | 0 | 0 | 0 | 0 | 0 |
| 3482 | W303.asm02.HP0_G0034770 | chrXII | 2056 | 432 | 804 | 2160 | 464 | 804 | 1948 | 385 | 711 | 2853 | 478 | 1170 | 2270 | 337 | 741 | 2376 | 485 | 1034 |
| 3483 | W303.asm02.HP0_G0034780 | chrXII | 5970 | 863 | 1164 | 5451 | 1124 | 1133 | 5471 | 989 | 1446 | 8105 | 1372 | 1939 | 6287 | 1139 | 1742 | 5760 | 1320 | 1761 |
| 3484 | W303.asm02.HP0_G0034790 | chrXII | 1500 | 1259 | 1429 | 1317 | 1278 | 1492 | 1383 | 1272 | 1998 | 3069 | 1103 | 1172 | 3547 | 784 | 969 | 3324 | 909 | 1091 |
| 3485 | W303.asm02.HP0_G0034800 | chrXII | 4 | 8 | 11 | 5 | 20 | 15 | 8 | 9 | 20 | 16 | 3 | 9 | 25 | 4 | 16 | 14 | 4 | 7 |
| 3486 | W303.asm02.HP0_G0034810 | chrXII | 1477 | 120 | 285 | 1639 | 300 | 315 | 1411 | 122 | 387 | 919 | 141 | 338 | 532 | 110 | 228 | 396 | 126 | 280 |
| 3487 | W303.asm02.HP0_G0034820 | chrXII | 1230 | 455 | 598 | 1154 | 603 | 616 | 1457 | 377 | 723 | 4519 | 774 | 1015 | 4499 | 635 | 741 | 2758 | 712 | 881 |
| 3488 | W303.asm02.HP0_G0034830 | chrXII | 793 | 1957 | 2427 | 778 | 2735 | 2079 | 856 | 1996 | 2311 | 10434 | 1796 | 2372 | 9948 | 1176 | 1436 | 9549 | 1682 | 1937 |
| 3489 | W303.asm02.HP0_G0034840 | chrXII | 593 | 160 | 254 | 660 | 289 | 343 | 625 | 139 | 354 | 2845 | 363 | 628 | 2066 | 205 | 477 | 1848 | 349 | 515 |
| 3490 | W303.asm02.HP0_G0034850 | chrXII | 2462 | 1065 | 1392 | 1917 | 1137 | 1368 | 1898 | 1118 | 1818 | 2091 | 892 | 1149 | 3064 | 708 | 1070 | 2322 | 905 | 983 |
| 3491 | W303.asm02.HP0_G0034860 | chrXII | 2308 | 1268 | 1097 | 2416 | 1392 | 1167 | 2827 | 1072 | 1385 | 13820 | 2747 | 2816 | 6654 | 1837 | 1245 | 7143 | 2205 | 2403 |
| 3492 | W303.asm02.HP0_G0034870 | chrXII | 1335 | 418 | 529 | 1212 | 505 | 518 | 1281 | 429 | 532 | 2374 | 406 | 554 | 2181 | 244 | 476 | 2375 | 332 | 491 |
| 3493 | W303.asm02.HP0_G0034880 | chrXII | 1459 | 402 | 356 | 1392 | 333 | 328 | 1258 | 421 | 298 | 2458 | 485 | 542 | 2104 | 341 | 455 | 1964 | 465 | 516 |
| 3494 | W303.asm02.HP0_G0034890 | chrXII | 1237 | 332 | 358 | 1068 | 342 | 336 | 1169 | 331 | 424 | 2003 | 419 | 574 | 1679 | 335 | 502 | 1817 | 419 | 563 |
| 3495 | W303.asm02.HP0_G0034900 | chrXII | 755 | 622 | 1097 | 738 | 1027 | 985 | 822 | 662 | 1234 | 4033 | 569 | 971 | 4265 | 348 | 653 | 4246 | 515 | 806 |
| 3496 | W303.asm02.HP0_G0034910 | chrXII | 2076 | 416 | 442 | 2043 | 506 | 513 | 2038 | 394 | 496 | 3928 | 665 | 985 | 2854 | 434 | 747 | 3128 | 617 | 886 |
| 3497 | W303.asm02.HP0_G0034920 | chrXII | 1168 | 583 | 415 | 1026 | 365 | 430 | 1141 | 453 | 445 | 2249 | 595 | 604 | 1711 | 382 | 448 | 1862 | 616 | 549 |
| 3498 | W303.asm02.HP0_G0034930 | chrXII | 3737 | 847 | 912 | 3490 | 842 | 963 | 3545 | 794 | 1130 | 5974 | 1514 | 1988 | 3707 | 1113 | 1542 | 3913 | 1392 | 1698 |
| 3499 | W303.asm02.HP0_G0034940 | chrXII | 2767 | 798 | 642 | 2757 | 546 | 617 | 2453 | 797 | 621 | 3591 | 1187 | 1073 | 2665 | 751 | 916 | 2904 | 1127 | 970 |
| 3500 | W303.asm02.HP0_G0034950 | chrXII | 2386 | 1518 | 1581 | 2434 | 1848 | 1290 | 2604 | 1533 | 1516 | 12633 | 2038 | 2233 | 10531 | 1236 | 1317 | 11002 | 1954 | 1990 |
| 3501 | W303.asm02.HP0_G0034960 | chrXII | 1746 | 1632 | 2019 | 1729 | 2219 | 1993 | 1888 | 1822 | 2568 | 7728 | 1595 | 1832 | 8034 | 1010 | 1098 | 8090 | 1409 | 1560 |
| 3502 | W303.asm02.HP0_G0034970 | chrXII | 516 | 242 | 209 | 527 | 236 | 191 | 502 | 205 | 168 | 776 | 210 | 232 | 862 | 161 | 168 | 680 | 260 | 190 |
| 3503 | W303.asm02.HP0_G0034980 | chrXII | 2612 | 796 | 603 | 2432 | 600 | 663 | 2632 | 710 | 622 | 4834 | 1379 | 1413 | 3117 | 811 | 891 | 3489 | 1200 | 1244 |
| 3504 | W303.asm02.HP0_G0034990 | chrXII | 2068 | 389 | 411 | 2466 | 344 | 467 | 2003 | 319 | 470 | 3465 | 879 | 1005 | 2337 | 596 | 846 | 1944 | 807 | 863 |
| 3505 | W303.asm02.HP0_G0035000 | chrXII | 1194 | 1150 | 1261 | 964 | 1285 | 1203 | 1124 | 1159 | 1368 | 3933 | 1115 | 1134 | 4325 | 698 | 761 | 3788 | 1049 | 992 |
| 3506 | W303.asm02.HP0_G0035010 | chrXII | 3977 | 1831 | 2450 | 3505 | 2391 | 2253 | 3660 | 1847 | 2984 | 6908 | 1661 | 2372 | 8037 | 1423 | 1940 | 7018 | 1581 | 2139 |
| 3507 | W303.asm02.HP0_G0035020 | chrXII | 2040 | 2309 | 2760 | 2131 | 3239 | 2663 | 2324 | 2657 | 2978 | 12241 | 2555 | 3208 | 11739 | 1770 | 1885 | 11308 | 2184 | 2657 |
| 3508 | W303.asm02.HP0_G0035030 | chrXII | 493 | 229 | 239 | 514 | 195 | 246 | 470 | 231 | 293 | 509 | 176 | 134 | 593 | 131 | 117 | 592 | 202 | 123 |
| 3509 | W303.asm02.HP0_G0035040 | chrXII | 837 | 562 | 717 | 787 | 646 | 680 | 726 | 646 | 748 | 1794 | 587 | 653 | 2066 | 339 | 523 | 1954 | 539 | 640 |
| 3510 | W303.asm02.HP0_G0035050 | chrXII | 28 | 14 | 6 | 24 | 5 | 9 | 22 | 12 | 9 | 15 | 10 | 2 | 7 | 4 | 6 | 15 | 6 | 1 |
| 3511 | W303.asm02.HP0_G0035060 | chrXII | 62 | 10 | 8 | 47 | 8 | 10 | 37 | 12 | 16 | 20 | 3 | 8 | 29 | 12 | 7 | 28 | 4 | 8 |
| 3512 | W303.asm02.HP0_G0035070 | chrXII | 562 | 292 | 438 | 531 | 464 | 362 | 615 | 246 | 474 | 3715 | 441 | 707 | 2932 | 252 | 409 | 2963 | 387 | 582 |
| 3513 | W303.asm02.HP0_G0035080 | chrXII | 1393 | 406 | 382 | 1373 | 437 | 395 | 1476 | 417 | 426 | 3118 | 588 | 734 | 2450 | 371 | 432 | 2354 | 476 | 662 |
| 3514 | W303.asm02.HP0_G0035090 | chrXII | 1519 | 1008 | 1183 | 1347 | 1125 | 1242 | 1385 | 1003 | 1489 | 2390 | 743 | 888 | 2783 | 467 | 635 | 2504 | 733 | 739 |
| 3515 | W303.asm02.HP0_G0035100 | chrXII | 3203 | 2202 | 2326 | 3075 | 2608 | 2127 | 3364 | 2495 | 2447 | 9254 | 2442 | 2451 | 9801 | 1497 | 1720 | 9542 | 2148 | 2271 |
| 3516 | W303.asm02.HP0_G0035110 | chrXII | 1086 | 1486 | 1451 | 953 | 1504 | 1466 | 999 | 1427 | 1468 | 4645 | 1277 | 1600 | 4510 | 803 | 1082 | 4390 | 1177 | 1412 |
| 3517 | W303.asm02.HP0_G0035120 | chrXII | 2036 | 832 | 999 | 2162 | 975 | 1008 | 2290 | 793 | 1144 | 7291 | 1617 | 2043 | 6902 | 1112 | 1562 | 4867 | 1415 | 1748 |
| 3518 | W303.asm02.HP0_G0035130 | chrXII | 2747 | 1081 | 1184 | 2524 | 1226 | 1137 | 2795 | 1174 | 1401 | 7541 | 1655 | 1820 | 6210 | 1063 | 1334 | 6216 | 1450 | 1717 |
| 3519 | W303.asm02.HP0_G0035140 | chrXII | 1596 | 1016 | 1064 | 1378 | 925 | 1081 | 1567 | 1066 | 1334 | 2687 | 861 | 755 | 3281 | 536 | 688 | 2822 | 704 | 721 |
| 3520 | W303.asm02.HP0_G0035150 | chrXII | 1198 | 1403 | 1768 | 1205 | 1850 | 1771 | 1242 | 1586 | 2122 | 5387 | 1147 | 1351 | 5515 | 770 | 1021 | 5423 | 1142 | 1251 |
| 3521 | W303.asm02.HP0_G0035160 | chrXII | 2538 | 1494 | 1595 | 2515 | 1947 | 1370 | 2573 | 1465 | 1547 | 7556 | 2058 | 2119 | 6284 | 1209 | 1631 | 6277 | 1972 | 1977 |
| 3522 | W303.asm02.HP0_G0035170 | chrXII | 2328 | 281 | 523 | 2337 | 503 | 518 | 2271 | 266 | 667 | 4036 | 369 | 1063 | 2499 | 290 | 764 | 2707 | 340 | 867 |
| 3523 | W303.asm02.HP0_G0035180 | chrXII | 7385 | 3022 | 2378 | 7255 | 2421 | 2514 | 6872 | 3042 | 2469 | 7319 | 2479 | 2399 | 7894 | 1676 | 2178 | 6769 | 2181 | 2388 |
| 3524 | W303.asm02.HP0_G0035190 | chrXII | 1684 | 796 | 621 | 1861 | 658 | 658 | 1764 | 695 | 636 | 3244 | 660 | 1013 | 2949 | 504 | 687 | 2756 | 696 | 868 |
| 3525 | W303.asm02.HP0_G0035200 | chrXII | 8175 | 4067 | 3636 | 7298 | 3375 | 3931 | 8393 | 3687 | 4311 | 17323 | 6451 | 7905 | 13054 | 4607 | 5683 | 13421 | 5799 | 7417 |
| 3526 | W303.asm02.HP0_G0035210 | chrXII | 16951 | 3779 | 3319 | 15612 | 2987 | 3638 | 16761 | 3516 | 3783 | 18031 | 6024 | 7551 | 14221 | 4621 | 5933 | 13285 | 5491 | 6923 |
| 3527 | W303.asm02.HP0_G0035220 | chrXII | 19903 | 20179 | 16344 | 16891 | 16424 | 16349 | 20279 | 19737 | 18244 | 38528 | 19374 | 16560 | 38317 | 14797 | 13402 | 38653 | 17010 | 15310 |
| 3528 | W303.asm02.HP0_G0035230 | chrXII | 1433 | 380 | 300 | 1338 | 268 | 248 | 1276 | 317 | 321 | 927 | 239 | 194 | 1228 | 157 | 186 | 998 | 263 | 160 |
| 3529 | W303.asm02.HP0_G0035240 | chrXII | 2044 | 447 | 518 | 1845 | 554 | 581 | 1967 | 448 | 688 | 4570 | 671 | 1126 | 3158 | 487 | 762 | 2942 | 649 | 928 |
| 3530 | W303.asm02.HP0_G0035250 | chrXII | 2144 | 1612 | 1884 | 2307 | 1776 | 1758 | 2399 | 1652 | 2123 | 6887 | 1389 | 1643 | 6964 | 824 | 1214 | 7266 | 1325 | 1509 |
| 3531 | W303.asm02.HP0_G0035260 | chrXII | 1422 | 1067 | 1255 | 1411 | 1158 | 1254 | 1420 | 1215 | 1318 | 6780 | 1620 | 1394 | 5928 | 871 | 1396 | 7256 | 1431 | 1727 |
| 3532 | W303.asm02.HP0_G0035270 | chrXII | 1357 | 224 | 249 | 1197 | 242 | 232 | 1260 | 215 | 249 | 2175 | 385 | 448 | 1860 | 283 | 342 | 1601 | 378 | 420 |

| | | | | | | | | | | | | | | | | | | | | |
|---|---|---|---|---|---|---|---|---|---|---|---|---|---|---|---|---|---|---|---|---|
| 3533 | W303.asm02.HP0_G0035280 | chrXII | 1083 | 755 | 837 | 1049 | 777 | 826 | 1042 | 797 | 1060 | 2323 | 556 | 650 | 2473 | 393 | 492 | 2764 | 561 | 591 |
| 3534 | W303.asm02.HP0_G0035290 | chrXII | 228 | 68 | 46 | 215 | 53 | 65 | 232 | 62 | 52 | 283 | 104 | 81 | 266 | 80 | 65 | 182 | 89 | 69 |
| 3535 | W303.asm02.HP0_G0035300 | chrXII | 2665 | 2628 | 3269 | 2532 | 2891 | 3139 | 2512 | 2646 | 4353 | 5902 | 2281 | 2750 | 6499 | 1698 | 2183 | 6557 | 2052 | 2160 |
| 3536 | W303.asm02.HP0_G0035310 | chrXII | 2894 | 639 | 745 | 2605 | 735 | 777 | 2620 | 579 | 858 | 5424 | 930 | 1386 | 3845 | 646 | 1081 | 4067 | 808 | 1295 |
| 3537 | W303.asm02.HP0_G0035320 | chrXII | 989 | 601 | 485 | 994 | 455 | 419 | 991 | 646 | 445 | 2075 | 677 | 651 | 2915 | 519 | 416 | 1711 | 583 | 535 |
| 3538 | W303.asm02.HP0_G0035330 | chrXII | 188 | 601 | 441 | 152 | 410 | 447 | 158 | 647 | 469 | 1169 | 495 | 716 | 2220 | 425 | 337 | 965 | 399 | 506 |
| 3539 | W303.asm02.HP0_G0035340 | chrXII | 2924 | 4004 | 4671 | 2595 | 3921 | 4506 | 2766 | 3936 | 5960 | 7434 | 3523 | 3802 | 8990 | 2438 | 3347 | 8352 | 3259 | 3231 |
| 3540 | W303.asm02.HP0_G0035350 | chrXII | 68808 | 21433 | 22026 | 64711 | 19283 | 26384 | 76942 | 16864 | 29412 | 47175 | 27688 | 39458 | 25837 | 28767 | 23380 | 23726 | 23415 | 33097 |
| 3541 | W303.asm02.HP0_G0035360 | chrXII | 3926 | 593 | 516 | 3924 | 560 | 583 | 4075 | 490 | 573 | 5460 | 1025 | 1543 | 2760 | 786 | 1022 | 2875 | 858 | 1350 |
| 3542 | W303.asm02.HP0_G0035370 | chrXII | 476 | 241 | 367 | 351 | 295 | 343 | 402 | 301 | 515 | 2146 | 326 | 448 | 2138 | 261 | 397 | 2106 | 295 | 480 |
| 3543 | W303.asm02.HP0_G0035380 | chrXII | 744 | 169 | 254 | 726 | 192 | 252 | 699 | 189 | 234 | 2453 | 460 | 605 | 1411 | 269 | 448 | 1535 | 452 | 628 |
| 3544 | W303.asm02.HP0_G0035390 | chrXII | 17164 | 10433 | 13543 | 17211 | 13757 | 13665 | 19296 | 11262 | 13206 | 38377 | 11317 | 14839 | 37605 | 8117 | 11095 | 35309 | 8924 | 13800 |
| 3545 | W303.asm02.HP0_G0035400 | chrXII | 2273 | 1849 | 1675 | 2088 | 1502 | 1650 | 2114 | 1749 | 2022 | 5037 | 1358 | 1574 | 6415 | 1075 | 1322 | 5596 | 1369 | 1334 |
| 3546 | W303.asm02.HP0_G0035410 | chrXII | 2326 | 1342 | 1833 | 1658 | 1501 | 1927 | 2034 | 1306 | 1990 | 3344 | 1104 | 1620 | 3894 | 668 | 1076 | 3386 | 943 | 1450 |
| 3547 | W303.asm02.HP0_G0035420 | chrXII | 830 | 1320 | 2153 | 915 | 2047 | 2187 | 910 | 1279 | 3071 | 6131 | 1102 | 1533 | 7564 | 720 | 1047 | 6524 | 994 | 1309 |
| 3548 | W303.asm02.HP0_G0035430 | chrXII | 1602 | 876 | 931 | 1603 | 918 | 934 | 1553 | 1002 | 871 | 4119 | 1001 | 1338 | 4393 | 760 | 864 | 3558 | 922 | 1145 |
| 3549 | W303.asm02.HP0_G0035440 | chrXII | 177 | 88 | 66 | 150 | 53 | 68 | 151 | 78 | 72 | 219 | 72 | 65 | 230 | 43 | 46 | 247 | 80 | 69 |
| 3550 | W303.asm02.HP0_G0035450 | chrXII | 1106 | 177 | 210 | 1197 | 178 | 164 | 861 | 154 | 200 | 421 | 149 | 167 | 569 | 115 | 145 | 369 | 158 | 125 |
| 3551 | W303.asm02.HP0_G0035460 | chrXII | 701 | 522 | 572 | 648 | 523 | 535 | 667 | 390 | 698 | 2093 | 831 | 999 | 817 | 480 | 484 | 880 | 779 | 860 |
| 3552 | W303.asm02.HP0_G0035470 | chrXII | 8277 | 8227 | 7627 | 9302 | 8212 | 7186 | 10249 | 8143 | 7334 | 27188 | 9098 | 9027 | 22256 | 5168 | 6373 | 26133 | 7719 | 8758 |
| 3553 | W303.asm02.HP0_G0035480 | chrXII | 2144 | 829 | 680 | 1993 | 794 | 731 | 2172 | 883 | 780 | 5773 | 1304 | 1464 | 5169 | 989 | 1095 | 4537 | 1182 | 1394 |
| 3554 | W303.asm02.HP0_G0035490 | chrXII | 1988 | 655 | 849 | 1381 | 578 | 997 | 1844 | 579 | 1202 | 4879 | 1315 | 1839 | 2706 | 1042 | 1039 | 2785 | 1159 | 1375 |
| 3555 | W303.asm02.HP0_G0035500 | chrXII | 3447 | 918 | 772 | 3068 | 801 | 700 | 3564 | 903 | 627 | 3139 | 808 | 1227 | 3427 | 651 | 800 | 2860 | 703 | 974 |
| 3556 | W303.asm02.HP0_G0035510 | chrXII | 17656 | 3536 | 3100 | 16372 | 3096 | 3279 | 18582 | 3289 | 3957 | 21609 | 6129 | 7050 | 15367 | 4664 | 4769 | 15950 | 5361 | 6166 |
| 3557 | W303.asm02.HP0_G0035520 | chrXII | 6466 | 5512 | 7044 | 6018 | 6616 | 7242 | 6583 | 5581 | 7759 | 14256 | 4824 | 6181 | 15519 | 3881 | 4934 | 14635 | 4360 | 5613 |
| 3558 | W303.asm02.HP0_G0035530 | chrXII | 239 | 377 | 365 | 245 | 496 | 353 | 267 | 427 | 409 | 3299 | 548 | 563 | 3163 | 311 | 271 | 3130 | 550 | 465 |
| 3559 | W303.asm02.HP0_G0035540 | chrXII | 3748 | 2443 | 2515 | 3291 | 2209 | 2524 | 3276 | 2410 | 2545 | 5884 | 1776 | 1899 | 6680 | 1170 | 1688 | 6260 | 1619 | 1617 |
| 3560 | W303.asm02.HP0_G0035550 | chrXII | 1981 | 1367 | 1677 | 1760 | 1463 | 1690 | 1800 | 1533 | 2191 | 4551 | 1341 | 1391 | 5033 | 863 | 1138 | 4748 | 1129 | 1220 |
| 3561 | W303.asm02.HP0_G0035560 | chrXII | 2738 | 1498 | 1958 | 2655 | 1562 | 1890 | 2664 | 1551 | 2298 | 4466 | 1101 | 1443 | 5172 | 885 | 1328 | 4890 | 1006 | 1292 |
| 3562 | W303.asm02.HP0_G0035570 | chrXII | 1599 | 718 | 1039 | 1508 | 1096 | 839 | 1457 | 755 | 1034 | 5945 | 794 | 1161 | 5828 | 552 | 893 | 5109 | 754 | 1024 |
| 3563 | W303.asm02.HP0_G0035580 | chrXII | 823 | 594 | 990 | 750 | 1240 | 954 | 866 | 772 | 1500 | 3889 | 644 | 722 | 4580 | 370 | 601 | 3527 | 486 | 667 |
| 3564 | W303.asm02.HP0_G0035590 | chrXII | 22073 | 2206 | 2106 | 21864 | 2063 | 2191 | 21447 | 2213 | 2703 | 13920 | 3765 | 4756 | 13546 | 3231 | 3986 | 9816 | 3421 | 4516 |
| 3565 | W303.asm02.HP0_G0035600 | chrXII | 1623 | 276 | 248 | 1429 | 221 | 292 | 1450 | 222 | 312 | 697 | 248 | 293 | 644 | 164 | 208 | 616 | 268 | 284 |
| 3566 | W303.asm02.HP0_G0035610 | chrXII | 2970 | 624 | 942 | 2773 | 844 | 1007 | 2914 | 693 | 998 | 7009 | 1129 | 1777 | 4940 | 833 | 1308 | 5196 | 1047 | 1553 |
| 3567 | W303.asm02.HP0_G0035620 | chrXII | 2539 | 517 | 463 | 2155 | 451 | 603 | 2486 | 420 | 652 | 2566 | 766 | 1045 | 1294 | 633 | 647 | 1189 | 706 | 917 |
| 3568 | W303.asm02.HP0_G0035630 | chrXII | 292 | 519 | 978 | 314 | 1177 | 1035 | 337 | 650 | 944 | 2766 | 411 | 754 | 3169 | 372 | 468 | 2558 | 397 | 697 |
| 3569 | W303.asm02.HP0_G0035640 | chrXII | 1793 | 1611 | 1723 | 1790 | 1994 | 1676 | 1948 | 1636 | 2037 | 6832 | 2245 | 2117 | 5907 | 1430 | 1285 | 5938 | 1949 | 1805 |
| 3570 | W303.asm02.HP0_G0035650 | chrXII | 24693 | 22866 | 22505 | 22905 | 21597 | 22224 | 25558 | 22151 | 26036 | 42776 | 21242 | 20402 | 43332 | 16423 | 16705 | 42455 | 19406 | 18315 |
| 3571 | W303.asm02.HP0_G0035660 | chrXII | 4497 | 658 | 506 | 3968 | 506 | 611 | 4276 | 563 | 652 | 3097 | 1120 | 1212 | 1829 | 756 | 831 | 1824 | 1068 | 1064 |
| 3572 | W303.asm02.HP0_G0035670 | chrXII | 1501 | 1096 | 1360 | 1415 | 910 | 1293 | 1273 | 1038 | 1594 | 3156 | 868 | 1145 | 3483 | 600 | 844 | 3279 | 835 | 1067 |
| 3573 | W303.asm02.HP0_G0035680 | chrXII | 5359 | 972 | 1042 | 4864 | 782 | 1065 | 4872 | 734 | 1334 | 3824 | 1379 | 1817 | 2143 | 936 | 1339 | 2145 | 1276 | 1445 |
| 3574 | W303.asm02.HP0_G0035690 | chrXII | 1918 | 512 | 381 | 1606 | 298 | 390 | 1460 | 456 | 465 | 778 | 336 | 314 | 1117 | 225 | 288 | 656 | 360 | 321 |
| 3575 | W303.asm02.HP0_G0035700 | chrXII | 309 | 102 | 86 | 269 | 106 | 110 | 295 | 98 | 110 | 604 | 103 | 124 | 457 | 77 | 75 | 459 | 131 | 130 |
| 3576 | W303.asm02.HP0_G0035710 | chrXII | 1403 | 498 | 569 | 1506 | 502 | 516 | 1354 | 457 | 593 | 2085 | 561 | 618 | 2027 | 263 | 453 | 2178 | 564 | 609 |
| 3577 | W303.asm02.HP0_G0035720 | chrXII | 3203 | 2817 | 2815 | 3010 | 2985 | 2542 | 3475 | 3189 | 2843 | 16196 | 3767 | 3787 | 14005 | 2336 | 2786 | 15305 | 3059 | 3621 |
| 3578 | W303.asm02.HP0_G0035730 | chrXII | 664 | 944 | 1469 | 628 | 1768 | 1446 | 668 | 1084 | 1528 | 8170 | 1324 | 2214 | 6952 | 836 | 1674 | 6596 | 1153 | 2080 |
| 3579 | W303.asm02.HP0_G0035740 | chrXII | 2157 | 822 | 733 | 1985 | 675 | 717 | 2078 | 837 | 641 | 3349 | 783 | 883 | 3096 | 556 | 681 | 2939 | 722 | 775 |
| 3580 | W303.asm02.HP0_G0035750 | chrXII | 1956 | 323 | 532 | 1802 | 556 | 604 | 1983 | 320 | 577 | 5531 | 633 | 1278 | 3798 | 466 | 814 | 3617 | 516 | 1196 |
| 3581 | W303.asm02.HP0_G0035760 | chrXII | 2343 | 356 | 574 | 2371 | 551 | 601 | 2187 | 387 | 704 | 3844 | 499 | 1304 | 2468 | 406 | 860 | 2640 | 457 | 1057 |
| 3582 | W303.asm02.HP0_G0035770 | chrXII | 1673 | 866 | 1109 | 1487 | 891 | 1056 | 1618 | 968 | 1082 | 5556 | 1180 | 1665 | 4618 | 873 | 1263 | 4699 | 1065 | 1586 |
| 3583 | W303.asm02.HP0_G0035780 | chrXII | 7411 | 849 | 811 | 6499 | 786 | 897 | 7584 | 691 | 1074 | 6159 | 1459 | 1861 | 3573 | 1170 | 1277 | 3478 | 1257 | 1553 |
| 3584 | W303.asm02.HP0_G0035790 | chrXII | 3836 | 477 | 580 | 4008 | 630 | 692 | 3748 | 397 | 794 | 3452 | 798 | 1140 | 2081 | 576 | 872 | 1931 | 688 | 950 |
| 3585 | W303.asm02.HP0_G0035800 | chrXII | 810 | 155 | 198 | 746 | 200 | 193 | 723 | 164 | 249 | 1319 | 265 | 360 | 860 | 181 | 238 | 856 | 262 | 327 |
| 3586 | W303.asm02.HP0_G0035810 | chrXII | 667 | 189 | 270 | 642 | 228 | 296 | 669 | 172 | 290 | 1235 | 269 | 474 | 982 | 186 | 330 | 845 | 179 | 384 |
| 3587 | W303.asm02.HP0_G0035820 | chrXII | 3737 | 1980 | 2069 | 3802 | 1866 | 2019 | 3508 | 1891 | 2634 | 4474 | 1480 | 1635 | 5925 | 1156 | 1484 | 4944 | 1397 | 1500 |
| 3588 | W303.asm02.HP0_G0035830 | chrXII | 843 | 197 | 216 | 869 | 233 | 261 | 799 | 139 | 283 | 618 | 225 | 346 | 301 | 155 | 143 | 258 | 221 | 274 |
| 3589 | W303.asm02.HP0_G0035840 | chrXII | 2110 | 417 | 537 | 2088 | 341 | 721 | 2305 | 294 | 765 | 3131 | 844 | 1635 | 1050 | 557 | 820 | 1039 | 683 | 1381 |
| 3590 | W303.asm02.HP0_G0035850 | chrXII | 3149 | 470 | 687 | 3009 | 691 | 723 | 3236 | 406 | 902 | 5458 | 998 | 1613 | 2108 | 775 | 944 | 2081 | 860 | 1363 |
| 3591 | W303.asm02.HP0_G0035860 | chrXII | 1825 | 823 | 917 | 1706 | 803 | 896 | 1654 | 843 | 961 | 2674 | 724 | 795 | 2606 | 468 | 633 | 2682 | 694 | 712 |
| 3592 | W303.asm02.HP0_G0035870 | chrXII | 1310 | 318 | 350 | 1344 | 366 | 355 | 1221 | 352 | 387 | 2361 | 443 | 634 | 2216 | 338 | 441 | 1887 | 456 | 545 |
| 3593 | W303.asm02.HP0_G0035880 | chrXII | 1693 | 572 | 706 | 1487 | 439 | 815 | 1933 | 501 | 783 | 2534 | 854 | 1434 | 2605 | 516 | 787 | 2142 | 809 | 1212 |
| 3594 | W303.asm02.HP0_G0035890 | chrXII | 335 | 424 | 479 | 232 | 434 | 468 | 287 | 482 | 602 | 859 | 392 | 427 | 1011 | 245 | 318 | 917 | 400 | 342 |
| 3595 | W303.asm02.HP0_G0035900 | chrXII | 3617 | 4241 | 4315 | 3058 | 3514 | 4230 | 3610 | 4008 | 4439 | 12285 | 4272 | 5462 | 11708 | 2791 | 3901 | 11824 | 3962 | 5221 |
| 3596 | W303.asm02.HP0_G0035910 | chrXII | 1694 | 675 | 413 | 1482 | 348 | 412 | 1582 | 637 | 368 | 1066 | 506 | 387 | 1118 | 326 | 359 | 1043 | 473 | 345 |
| 3597 | W303.asm02.HP0_G0035920 | chrXII | 635 | 723 | 578 | 623 | 595 | 508 | 734 | 718 | 533 | 3102 | 802 | 899 | 2541 | 506 | 657 | 2911 | 719 | 807 |
| 3598 | W303.asm02.HP0_G0035930 | chrXII | 805 | 995 | 1647 | 796 | 898 | 1809 | 776 | 1081 | 2052 | 2132 | 816 | 1598 | 2733 | 641 | 1079 | 2236 | 803 | 1345 |
| 3599 | W303.asm02.HP0_G0035940 | chrXII | 2386 | 711 | 651 | 2114 | 572 | 685 | 2233 | 683 | 695 | 2654 | 827 | 942 | 2551 | 460 | 599 | 2344 | 705 | 813 |
| 3600 | W303.asm02.HP0_G0035950 | chrXII | 1104 | 951 | 1522 | 1101 | 1515 | 1513 | 954 | 871 | 1783 | 4903 | 846 | 1263 | 5199 | 487 | 846 | 5159 | 887 | 1047 |
| 3601 | W303.asm02.HP0_G0035960 | chrXII | 545 | 1463 | 1314 | 610 | 1019 | 1353 | 493 | 1263 | 1190 | 2049 | 2396 | 2401 | 1439 | 1026 | 1902 | 2106 | 2371 | 2163 |
| 3602 | W303.asm02.HP0_G0035970 | chrXII | 16115 | 16633 | 16097 | 15075 | 14677 | 16771 | 14804 | 16120 | 15526 | 27492 | 17048 | 20085 | 30754 | 13881 | 17727 | 28054 | 15900 | 18497 |
| 3603 | W303.asm02.HP0_G0035980 | chrXII | 27015 | 12042 | 12589 | 25612 | 11485 | 12369 | 27127 | 13469 | 14286 | 31880 | 17857 | 19625 | 36648 | 15290 | 18047 | 31955 | 16065 | 17934 |
| 3604 | W303.asm02.HP0_G0035990 | chrXII | 76 | 31 | 18 | 82 | 20 | 13 | 86 | 34 | 18 | 74 | 98 | 26 | 80 | 35 | 19 | 59 | 93 | 24 |
| 3605 | W303.asm02.HP0_G0036000 | chrXII | 11475 | 3880 | 4240 | 10841 | 3562 | 4034 | 11125 | 4005 | 4338 | 15866 | 4603 | 6021 | 15331 | 3335 | 4818 | 14829 | 4157 | 5501 |
| 3606 | W303.asm02.HP0_G0036010 | chrXII | 2941 | 817 | 1113 | 2660 | 1114 | 1133 | 2986 | 858 | 1403 | 10318 | 1893 | 2462 | 6187 | 1180 | 1848 | 6689 | 1653 | 2345 |
| 3607 | W303.asm02.HP0_G0036020 | chrXII | 3101 | 319 | 406 | 3424 | 436 | 445 | 3152 | 307 | 459 | 4569 | 660 | 1166 | 3071 | 450 | 778 | 3087 | 567 | 985 |
| 3608 | W303.asm02.HP0_G0036030 | chrXII | 1843 | 197 | 389 | 1804 | 376 | 500 | 1793 | 143 | 528 | 748 | 226 | 543 | 526 | 153 | 241 | 451 | 198 | 347 |
| 3609 | W303.asm02.HP0_G0036040 | chrXII | 3092 | 450 | 533 | 2923 | 576 | 530 | 2904 | 436 | 537 | 4664 | 888 | 1176 | 3532 | 562 | 792 | 3437 | 807 | 1029 |
| 3610 | W303.asm02.HP0_G0036050 | chrXII | 3194 | 1165 | 1135 | 2794 | 984 | 1156 | 2891 | 1116 | 1247 | 3573 | 1118 | 1162 | 3857 | 760 | 1035 | 3323 | 943 | 1089 |
| 3611 | W303.asm02.HP0_G0036060 | chrXII | 1846 | 415 | 575 | 1580 | 467 | 589 | 1430 | 393 | 566 | 1703 | 322 | 542 | 1852 | 216 | 453 | 1872 | 337 | 435 |
| 3612 | W303.asm02.HP0_G0036070 | chrXII | 4831 | 987 | 1323 | 5959 | 870 | 1406 | 4325 | 918 | 1495 | 6146 | 1724 | 2414 | 5540 | 1400 | 1976 | 4118 | 1612 | 2137 |
| 3613 | W303.asm02.HP0_G0036080 | chrXII | 3256 | 396 | 608 | 2872 | 447 | 709 | 3247 | 387 | 804 | 1973 | 664 | 1111 | 1604 | 669 | 686 | 970 | 516 | 864 |
| 3614 | W303.asm02.HP0_G0036090 | chrXII | 287 | 34 | 43 | 300 | 52 | 46 | 295 | 44 | 67 | 217 | 60 | 63 | 230 | 41 | 53 | 176 | 48 | 51 |
| 3615 | W303.asm02.HP0_G0036100 | chrXII | 150 | 46 | 44 | 129 | 46 | 55 | 154 | 24 | 48 | 397 | 63 | 87 | 256 | 40 | 81 | 265 | 40 | 86 |
| 3616 | W303.asm02.HP0_G0036110 | chrXII | 754 | 781 | 1037 | 737 | 654 | 1129 | 750 | 734 | 1075 | 2579 | 787 | 1337 | 2465 | 519 | 892 | 2763 | 729 | 1277 |
| 3617 | W303.asm02.HP0_G0036120 | chrXII | 2045 | 447 | 609 | 2033 | 569 | 596 | 1897 | 390 | 595 | 4201 | 586 | 852 | 3132 | 365 | 612 | 3740 | 554 | 814 |
| 3618 | W303.asm02.HP0_G0036130 | chrXII | 1596 | 439 | 444 | 1508 | 487 | 458 | 1472 | 468 | 431 | 1614 | 451 | 530 | 1974 | 332 | 427 | 1590 | 389 | 479 |
| 3619 | W303.asm02.HP0_G0036140 | chrXII | 2092 | 1302 | 1873 | 1879 | 2278 | 1749 | 2225 | 1333 | 2303 | 18363 | 2353 | 3699 | 12286 | 1656 | 2217 | 13456 | 2198 | 3254 |
| 3620 | W303.asm02.HP0_G0036150 | chrXII | 2436 | 1825 | 1876 | 2195 | 1819 | 1755 | 2410 | 1867 | 1932 | 7040 | 2255 | 2235 | 5545 | 1434 | 1717 | 5788 | 2086 | 1882 |
| 3621 | W303.asm02.HP0_G0036160 | chrXII | 1870 | 223 | 277 | 1682 | 242 | 348 | 1983 | 168 | 383 | 2195 | 395 | 865 | 544 | 393 | 599 | 579 | 315 | 708 |
| 3622 | W303.asm02.HP0_G0036170 | chrXII | 1268 | 322 | 327 | 1322 | 326 | 307 | 1170 | 341 | 308 | 1774 | 389 | 551 | 1613 | 258 | 466 | 1591 | 361 | 435 |
| 3623 | W303.asm02.HP0_G0036180 | chrXII | 3370 | 296 | 274 | 3616 | 272 | 375 | 3369 | 238 | 391 | 3158 | 685 | 1136 | 1329 | 567 | 820 | 1294 | 595 | 906 |
| 3624 | W303.asm02.HP0_G0036190 | chrXII | 871 | 145 | 180 | 747 | 175 | 205 | 850 | 173 | 210 | 1058 | 233 | 352 | 712 | 168 | 246 | 592 | 180 | 279 |
| 3625 | W303.asm02.HP0_G0036200 | chrXII | 1452 | 458 | 406 | 1507 | 436 | 390 | 1448 | 481 | 388 | 2784 | 541 | 660 | 2504 | 371 | 468 | 2251 | 539 | 602 |

| | | | | | | | | | | | | | | | | | | | | |
|---|---|---|---|---|---|---|---|---|---|---|---|---|---|---|---|---|---|---|---|---|
| 3626 | W303.asm02.HP0_G0036210 | chrXII | 208 | 163 | 231 | 261 | 212 | 332 | 191 | 161 | 167 | 2458 | 409 | 251 | 3170 | 91 | 316 | 3557 | 361 | 778 |
| 3627 | W303.asm02.HP0_G0036220 | chrXII | 885 | 302 | 242 | 610 | 193 | 276 | 741 | 247 | 318 | 1433 | 412 | 448 | 1037 | 257 | 384 | 1205 | 421 | 493 |
| 3628 | W303.asm02.HP0_G0036230 | chrXII | 4012 | 901 | 1299 | 3893 | 1330 | 1269 | 3983 | 1053 | 1517 | 10340 | 1922 | 2403 | 8842 | 1422 | 1928 | 6903 | 1630 | 2183 |
| 3629 | W303.asm02.HP0_G0036240 | chrXII | 1807 | 185 | 200 | 1770 | 190 | 224 | 1755 | 133 | 291 | 482 | 196 | 346 | 271 | 162 | 150 | 268 | 168 | 269 |
| 3630 | W303.asm02.HP0_G0036250 | chrXII | 1449 | 527 | 503 | 1148 | 519 | 524 | 1273 | 558 | 670 | 2045 | 774 | 655 | 1951 | 430 | 528 | 1612 | 593 | 636 |
| 3631 | W303.asm02.HP0_G0036260 | chrXII | 878 | 244 | 225 | 1127 | 231 | 247 | 854 | 257 | 231 | 1305 | 300 | 290 | 1574 | 321 | 344 | 798 | 292 | 294 |
| 3632 | W303.asm02.HP0_G0036270 | chrXII | 1516 | 545 | 641 | 1458 | 702 | 635 | 1578 | 539 | 748 | 6003 | 938 | 1316 | 4094 | 577 | 838 | 4573 | 920 | 1152 |
| 3633 | W303.asm02.HP0_G0036280 | chrXII | 1884 | 417 | 542 | 1663 | 500 | 518 | 1846 | 385 | 697 | 3753 | 570 | 907 | 2928 | 450 | 765 | 2549 | 567 | 823 |
| 3634 | W303.asm02.HP0_G0036290 | chrXII | 906 | 874 | 876 | 692 | 876 | 887 | 829 | 873 | 1091 | 2249 | 676 | 604 | 2793 | 475 | 505 | 2391 | 589 | 559 |
| 3635 | W303.asm02.HP0_G0036300 | chrXII | 365 | 329 | 373 | 367 | 342 | 302 | 380 | 305 | 451 | 1163 | 272 | 273 | 1342 | 194 | 228 | 1272 | 247 | 233 |
| 3636 | W303.asm02.HP0_G0036310 | chrXII | 0 | 0 | 0 | 0 | 0 | 0 | 0 | 0 | 0 | 0 | 0 | 0 | 0 | 0 | 0 | 0 | 0 | 0 |
| 3637 | W303.asm02.HP0_G0036320 | chrXII | 0 | 0 | 0 | 0 | 0 | 0 | 0 | 0 | 0 | 0 | 0 | 0 | 0 | 0 | 0 | 0 | 0 | 0 |
| 3638 | W303.asm02.HP0_G0036330 | chrXII | 269 | 43 | 64 | 260 | 70 | 61 | 238 | 58 | 85 | 503 | 62 | 91 | 387 | 49 | 73 | 431 | 52 | 95 |
| 3639 | W303.asm02.HP0_G0036340 | chrXII | 0 | 0 | 0 | 0 | 0 | 0 | 0 | 0 | 0 | 0 | 0 | 0 | 0 | 0 | 0 | 0 | 0 | 0 |
| 3640 | W303.asm02.HP0_G0036350 | chrXII | 0 | 0 | 0 | 0 | 0 | 0 | 0 | 0 | 0 | 0 | 1 | 0 | 0 | 0 | 0 | 0 | 1 | 0 |
| 3641 | W303.asm02.HP0_G0036360 | chrXII | 0 | 0 | 0 | 0 | 0 | 0 | 0 | 0 | 0 | 0 | 0 | 0 | 0 | 0 | 0 | 0 | 0 | 0 |
| 3642 | W303.asm02.HP0_G0036370 | chrXII | 0 | 0 | 0 | 0 | 0 | 0 | 0 | 0 | 0 | 0 | 0 | 0 | 0 | 0 | 0 | 0 | 0 | 0 |
| 3643 | W303.asm02.HP0_G0036380 | chrXII | 0 | 0 | 0 | 0 | 0 | 0 | 0 | 0 | 0 | 0 | 0 | 0 | 0 | 0 | 0 | 0 | 0 | 0 |
| 3644 | W303.asm02.HP0_G0036390 | chrXII | 0 | 0 | 0 | 0 | 0 | 0 | 0 | 0 | 0 | 0 | 0 | 0 | 0 | 0 | 0 | 0 | 0 | 0 |
| 3645 | W303.asm02.HP0_G0036400 | chrXII | 0 | 0 | 0 | 0 | 0 | 0 | 0 | 0 | 0 | 0 | 0 | 0 | 0 | 0 | 0 | 0 | 0 | 0 |
| 3646 | W303.asm02.HP0_G0036410 | chrXII | 0 | 0 | 0 | 0 | 0 | 0 | 0 | 0 | 0 | 0 | 0 | 0 | 0 | 0 | 0 | 0 | 0 | 0 |
| 3647 | W303.asm02.HP0_G0036420 | chrXII | 0 | 0 | 0 | 0 | 0 | 0 | 0 | 0 | 0 | 0 | 0 | 0 | 0 | 0 | 0 | 0 | 0 | 0 |
| 3648 | W303.asm02.HP0_G0036430 | chrXII | 0 | 0 | 0 | 0 | 0 | 0 | 0 | 0 | 0 | 0 | 0 | 0 | 0 | 0 | 0 | 0 | 0 | 0 |
| 3649 | W303.asm02.HP0_G0036440 | chrXII | 973 | 104 | 101 | 922 | 106 | 110 | 951 | 132 | 93 | 1056 | 263 | 296 | 1188 | 241 | 251 | 876 | 209 | 275 |
| 3650 | W303.asm02.HP0_G0036450 | chrXII | 643 | 169 | 128 | 675 | 143 | 140 | 602 | 205 | 138 | 393 | 128 | 119 | 615 | 133 | 115 | 344 | 118 | 103 |
| 3651 | W303.asm02.HP0_G0036460 | chrXII | 1254 | 416 | 322 | 1166 | 295 | 292 | 1104 | 374 | 334 | 1330 | 635 | 485 | 1014 | 388 | 478 | 1208 | 550 | 442 |
| 3652 | W303.asm02.HP0_G0036470 | chrXII | 12921 | 7371 | 5711 | 13027 | 5558 | 6040 | 14368 | 6183 | 6318 | 35552 | 10800 | 13012 | 25429 | 7839 | 8776 | 26451 | 9860 | 11811 |
| 3653 | W303.asm02.HP0_G0036480 | chrXII | 701 | 380 | 588 | 550 | 550 | 576 | 668 | 433 | 511 | 1317 | 367 | 547 | 1654 | 290 | 387 | 1380 | 301 | 442 |
| 3654 | W303.asm02.HP0_G0036490 | chrXII | 3205 | 1135 | 935 | 2953 | 928 | 1022 | 3203 | 864 | 869 | 2828 | 1231 | 1307 | 2705 | 936 | 1038 | 2223 | 1259 | 1228 |
| 3655 | W303.asm02.HP0_G0036500 | chrXII | 587 | 131 | 168 | 490 | 117 | 147 | 516 | 78 | 143 | 403 | 145 | 197 | 189 | 105 | 124 | 174 | 150 | 177 |
| 3656 | W303.asm02.HP0_G0036510 | chrXII | 4 | 1 | 5 | 3 | 1 | 3 | 5 | 5 | 8 | 8 | 2 | 0 | 1 | 5 | 4 | 4 | 8 | 3 |
| 3657 | W303.asm02.HP0_G0036520 | chrXII | 7570 | 2450 | 1673 | 7649 | 1654 | 1515 | 7485 | 2373 | 1774 | 9629 | 3308 | 3302 | 8357 | 2467 | 2535 | 8027 | 3246 | 2950 |
| 3658 | W303.asm02.HP0_G0036530 | chrXII | 207 | 404 | 169 | 157 | 113 | 145 | 170 | 302 | 150 | 346 | 185 | 104 | 403 | 161 | 113 | 316 | 268 | 135 |
| 3659 | W303.asm02.HP0_G0036540 | chrXII | 1484 | 578 | 461 | 1160 | 394 | 406 | 1140 | 476 | 504 | 1230 | 358 | 312 | 1230 | 190 | 266 | 1196 | 440 | 268 |
| 3660 | W303.asm02.HP0_G0036550 | chrXII | 4185 | 1904 | 2360 | 3950 | 1760 | 2309 | 3843 | 1877 | 2907 | 7464 | 1419 | 1829 | 7618 | 928 | 1447 | 7960 | 1414 | 1488 |
| 3661 | W303.asm02.HP0_G0036560 | chrXII | 20152 | 17779 | 22940 | 19153 | 21775 | 22604 | 20131 | 18756 | 28754 | 42690 | 17511 | 18760 | 48436 | 13053 | 16237 | 44474 | 14789 | 16894 |
| 3662 | W303.asm02.HP0_G0036570 | chrXII | 1403 | 2917 | 1875 | 1344 | 2234 | 1948 | 1466 | 2797 | 2182 | 7791 | 2790 | 2205 | 7878 | 2004 | 1866 | 7435 | 2688 | 1833 |
| 3663 | W303.asm02.HP0_G0036580 | chrXII | 1309 | 1204 | 1210 | 1188 | 1004 | 1261 | 1264 | 1200 | 1542 | 2337 | 992 | 1030 | 2836 | 687 | 781 | 2419 | 917 | 908 |
| 3664 | W303.asm02.HP0_G0036590 | chrXII | 7007 | 2505 | 2064 | 6920 | 2376 | 1945 | 7945 | 2592 | 1993 | 9763 | 2741 | 2577 | 10035 | 1985 | 1820 | 9223 | 2337 | 2425 |
| 3665 | W303.asm02.HP0_G0036600 | chrXII | 2071 | 444 | 314 | 2032 | 324 | 280 | 1934 | 373 | 380 | 2317 | 764 | 738 | 1711 | 640 | 571 | 1391 | 805 | 642 |
| 3666 | W303.asm02.HP0_G0036610 | chrXII | 650 | 284 | 176 | 540 | 160 | 154 | 583 | 240 | 195 | 650 | 341 | 309 | 695 | 301 | 243 | 469 | 400 | 251 |
| 3667 | W303.asm02.HP0_G0036620 | chrXII | 10318 | 4231 | 3996 | 9294 | 4671 | 3992 | 10542 | 4540 | 4480 | 26092 | 6814 | 8126 | 22235 | 4728 | 5732 | 21758 | 5753 | 7421 |
| 3668 | W303.asm02.HP0_G0036630 | chrXII | 1327 | 112 | 174 | 1294 | 168 | 195 | 1219 | 134 | 257 | 659 | 150 | 286 | 552 | 114 | 188 | 466 | 125 | 228 |
| 3669 | W303.asm02.HP0_G0036640 | chrXII | 4880 | 510 | 403 | 4953 | 285 | 386 | 4256 | 420 | 468 | 1483 | 762 | 520 | 1351 | 455 | 569 | 1015 | 717 | 477 |
| 3670 | W303.asm02.HP0_G0036650 | chrXII | 8542 | 1048 | 616 | 6919 | 443 | 639 | 6901 | 842 | 640 | 1164 | 816 | 768 | 1683 | 702 | 829 | 898 | 865 | 730 |
| 3671 | W303.asm02.HP0_G0036660 | chrXII | 9647 | 3449 | 2700 | 7346 | 2604 | 2884 | 8892 | 3371 | 3157 | 8981 | 4289 | 4054 | 8713 | 3404 | 3413 | 7715 | 3727 | 3728 |
| 3672 | W303.asm02.HP0_G0036670 | chrXII | 6817 | 1174 | 1297 | 3843 | 1221 | 1484 | 6114 | 968 | 1551 | 9152 | 2209 | 2795 | 7406 | 1753 | 1755 | 6928 | 1793 | 2043 |
| 3673 | W303.asm02.HP0_G0036680 | chrXII | 3013 | 1673 | 1994 | 3165 | 1897 | 1930 | 2919 | 1863 | 2192 | 5991 | 1674 | 2064 | 6948 | 1105 | 1675 | 6066 | 1487 | 1844 |
| 3674 | W303.asm02.HP0_G0036690 | chrXII | 4266 | 786 | 873 | 4427 | 924 | 869 | 4160 | 780 | 971 | 6891 | 1149 | 1728 | 5674 | 762 | 1369 | 5682 | 1062 | 1579 |
| 3675 | W303.asm02.HP0_G0036700 | chrXII | 2063 | 946 | 1325 | 1918 | 1055 | 1258 | 2216 | 1045 | 1134 | 1635 | 427 | 651 | 1644 | 306 | 448 | 1654 | 446 | 566 |
| 3676 | W303.asm02.HP0_G0036710 | chrXII | 4718 | 3445 | 4882 | 4114 | 4527 | 4921 | 4280 | 3386 | 5882 | 10432 | 3060 | 3928 | 11213 | 2353 | 3243 | 10860 | 2838 | 3363 |
| 3677 | W303.asm02.HP0_G0036720 | chrXII | 1969 | 1625 | 1867 | 1873 | 2004 | 1741 | 2170 | 1778 | 1686 | 6963 | 1765 | 2436 | 6463 | 1268 | 1511 | 6545 | 1531 | 2142 |
| 3678 | W303.asm02.HP0_G0036730 | chrXII | 505 | 99 | 117 | 511 | 90 | 142 | 474 | 60 | 103 | 887 | 143 | 322 | 299 | 116 | 202 | 259 | 120 | 254 |
| 3679 | W303.asm02.HP0_G0036740 | chrXII | 2724 | 718 | 943 | 2215 | 771 | 1101 | 2730 | 624 | 1140 | 6494 | 1620 | 2477 | 3141 | 1192 | 1452 | 3290 | 1295 | 2128 |
| 3680 | W303.asm02.HP0_G0036750 | chrXII | 2791 | 708 | 758 | 2763 | 777 | 767 | 2505 | 747 | 858 | 5059 | 1170 | 1525 | 3360 | 835 | 1177 | 3781 | 1037 | 1407 |
| 3681 | W303.asm02.HP0_G0036760 | chrXII | 3263 | 1268 | 1179 | 2844 | 1164 | 1412 | 3354 | 1176 | 1438 | 10758 | 2330 | 2617 | 7528 | 1865 | 1990 | 7409 | 1935 | 2357 |
| 3682 | W303.asm02.HP0_G0036770 | chrXII | 1110 | 918 | 703 | 1282 | 833 | 706 | 1123 | 933 | 752 | 3681 | 1072 | 1148 | 2945 | 725 | 816 | 3107 | 959 | 1028 |
| 3683 | W303.asm02.HP0_G0036780 | chrXII | 7619 | 2807 | 2200 | 6713 | 2124 | 2385 | 7511 | 2742 | 2607 | 14295 | 4680 | 4868 | 11727 | 3197 | 3522 | 11929 | 4044 | 4471 |
| 3684 | W303.asm02.HP0_G0036790 | chrXII | 3750 | 557 | 489 | 3308 | 481 | 460 | 3316 | 572 | 551 | 2292 | 695 | 746 | 1623 | 479 | 598 | 1646 | 613 | 607 |
| 3685 | W303.asm02.HP0_G0036800 | chrXII | 5919 | 2443 | 1929 | 6110 | 1720 | 1912 | 5686 | 2622 | 2341 | 7222 | 2932 | 3618 | 12963 | 3189 | 3521 | 6127 | 2613 | 3007 |
| 3686 | W303.asm02.HP0_G0036810 | chrXII | 4514 | 2932 | 2873 | 4268 | 2451 | 2615 | 4426 | 2836 | 3033 | 8182 | 2256 | 2519 | 8868 | 1623 | 1800 | 8307 | 2218 | 2258 |
| 3687 | W303.asm02.HP0_G0036820 | chrXII | 3256 | 2123 | 2169 | 3486 | 2816 | 1986 | 3770 | 2261 | 2130 | 19123 | 3277 | 3788 | 16319 | 2257 | 2278 | 14901 | 2812 | 3421 |
| 3688 | W303.asm02.HP0_G0036830 | chrXII | 11359 | 8514 | 9788 | 11284 | 9805 | 8999 | 12126 | 9313 | 11249 | 34597 | 8566 | 9133 | 32795 | 5360 | 6046 | 35413 | 7624 | 7996 |
| 3689 | W303.asm02.HP0_G0036840 | chrXII | 3668 | 1170 | 1176 | 3738 | 951 | 1206 | 3916 | 1223 | 1146 | 3784 | 977 | 1778 | 3696 | 812 | 1144 | 3348 | 926 | 1527 |
| 3690 | W303.asm02.HP0_G0036850 | chrXII | 1602 | 1386 | 1854 | 1523 | 1631 | 1878 | 1628 | 1615 | 2255 | 4559 | 1241 | 1634 | 5528 | 884 | 1264 | 4418 | 1109 | 1431 |
| 3691 | W303.asm02.HP0_G0036860 | chrXII | 3816 | 1025 | 780 | 3510 | 661 | 699 | 3411 | 974 | 781 | 2796 | 962 | 875 | 3383 | 713 | 816 | 2756 | 918 | 824 |
| 3692 | W303.asm02.HP0_G0036870 | chrXII | 14419 | 1119 | 1220 | 13027 | 967 | 1501 | 14254 | 786 | 1691 | 4470 | 1336 | 2247 | 2181 | 1281 | 1475 | 1830 | 1210 | 1858 |
| 3693 | W303.asm02.HP0_G0036880 | chrXII | 3913 | 1394 | 1603 | 3364 | 1612 | 1591 | 3185 | 1467 | 1952 | 2906 | 1182 | 1352 | 3868 | 978 | 1300 | 2639 | 1164 | 1219 |
| 3694 | W303.asm02.HP0_G0036890 | chrXII | 3679 | 1413 | 1236 | 3918 | 1029 | 1322 | 4041 | 1124 | 1101 | 3133 | 1434 | 1523 | 2153 | 494 | 847 | 3223 | 1651 | 1627 |
| 3695 | W303.asm02.HP0_G0036900 | chrXII | 12093 | 645 | 620 | 10969 | 645 | 805 | 11553 | 537 | 756 | 6723 | 1434 | 1982 | 4775 | 1075 | 1670 | 3936 | 1176 | 1681 |
| 3696 | W303.asm02.HP0_G0036910 | chrXII | 2821 | 459 | 600 | 2657 | 417 | 735 | 2809 | 428 | 707 | 3601 | 854 | 1606 | 2115 | 662 | 995 | 2059 | 737 | 1434 |
| 3697 | W303.asm02.HP0_G0036920 | chrXII | 6427 | 3481 | 3903 | 5982 | 3214 | 3707 | 6460 | 3609 | 3760 | 13121 | 3733 | 5012 | 12885 | 2634 | 3718 | 12338 | 3286 | 4580 |
| 3698 | W303.asm02.HP0_G0036930 | chrXII | 3813 | 653 | 462 | 3164 | 513 | 539 | 3591 | 596 | 506 | 2967 | 877 | 965 | 2373 | 751 | 840 | 2061 | 815 | 828 |
| 3699 | W303.asm02.HP0_G0036940 | chrXII | 362 | 559 | 350 | 340 | 463 | 383 | 375 | 498 | 268 | 2044 | 531 | 519 | 2015 | 371 | 406 | 1838 | 486 | 513 |
| 3700 | W303.asm02.HP0_G0036950 | chrXII | 663 | 566 | 602 | 679 | 555 | 567 | 579 | 550 | 727 | 1468 | 441 | 478 | 1838 | 278 | 361 | 1578 | 386 | 409 |
| 3701 | W303.asm02.HP0_G0036960 | chrXII | 987 | 866 | 675 | 920 | 696 | 627 | 1111 | 835 | 664 | 3784 | 1099 | 1172 | 3167 | 644 | 796 | 3379 | 993 | 1090 |
| 3702 | W303.asm02.HP0_G0036970 | chrXII | 202 | 399 | 458 | 167 | 405 | 451 | 194 | 316 | 501 | 2662 | 605 | 575 | 1377 | 276 | 460 | 2015 | 528 | 667 |
| 3703 | W303.asm02.HP0_G0036980 | chrXII | 1249 | 9154 | 12835 | 1337 | 12161 | 11646 | 1249 | 10391 | 13907 | 50311 | 13882 | 13039 | 35877 | 6654 | 9723 | 50064 | 11993 | 15452 |
| 3704 | W303.asm02.HP0_G0036990 | chrXII | 924 | 747 | 644 | 977 | 737 | 670 | 1011 | 781 | 650 | 2957 | 747 | 858 | 2632 | 447 | 544 | 2693 | 731 | 700 |
| 3705 | W303.asm02.HP0_G0037000 | chrXII | 11076 | 10930 | 10534 | 11518 | 10227 | 11400 | 10690 | 8837 | 10040 | 19814 | 6645 | 8496 | 21878 | 5218 | 7294 | 20745 | 6454 | 7526 |
| 3706 | W303.asm02.HP0_G0037010 | chrXII | 1292 | 450 | 417 | 1000 | 365 | 479 | 1114 | 456 | 508 | 824 | 366 | 345 | 1556 | 267 | 515 | 790 | 329 | 324 |
| 3707 | W303.asm02.HP0_G0037020 | chrXII | 2373 | 508 | 614 | 2600 | 508 | 727 | 2428 | 389 | 726 | 2763 | 1002 | 1219 | 991 | 692 | 819 | 814 | 681 | 985 |
| 3708 | W303.asm02.HP0_G0037030 | chrXII | 2597 | 1790 | 1356 | 2371 | 1438 | 1239 | 2345 | 1772 | 1524 | 5587 | 1785 | 1905 | 5141 | 1210 | 1473 | 5312 | 1590 | 1473 |
| 3709 | W303.asm02.HP0_G0037040 | chrXII | 2832 | 2148 | 2592 | 2537 | 2607 | 2510 | 2877 | 2499 | 3182 | 8854 | 2236 | 2166 | 9679 | 1378 | 1441 | 9106 | 1868 | 1881 |
| 3710 | W303.asm02.HP0_G0037050 | chrXII | 1947 | 1395 | 1640 | 2002 | 1945 | 1521 | 2365 | 1449 | 1792 | 13750 | 2158 | 3016 | 10565 | 1428 | 1574 | 10719 | 1888 | 2363 |
| 3711 | W303.asm02.HP0_G0037060 | chrXII | 983 | 416 | 834 | 942 | 1055 | 961 | 1065 | 409 | 1013 | 10194 | 911 | 1748 | 6265 | 626 | 917 | 5436 | 958 | 1520 |
| 3712 | W303.asm02.HP0_G0037070 | chrXII | 1336 | 1351 | 1473 | 1457 | 1530 | 1447 | 1361 | 1326 | 1341 | 6384 | 1455 | 2042 | 5991 | 994 | 1425 | 5511 | 1318 | 1729 |
| 3713 | W303.asm02.HP0_G0037080 | chrXII | 3169 | 2056 | 2382 | 2972 | 2214 | 2387 | 2942 | 1777 | 2621 | 7639 | 1772 | 2097 | 8774 | 1031 | 1721 | 8050 | 2068 | 1857 |
| 3714 | W303.asm02.HP0_G0037090 | chrXII | 842 | 646 | 970 | 809 | 923 | 837 | 793 | 590 | 871 | 4380 | 670 | 945 | 4357 | 395 | 645 | 4272 | 677 | 762 |
| 3715 | W303.asm02.HP0_G0037100 | chrXII | 945 | 545 | 712 | 831 | 670 | 752 | 869 | 526 | 1002 | 2379 | 485 | 669 | 2610 | 320 | 450 | 2607 | 448 | 565 |
| 3716 | W303.asm02.HP0_G0037110 | chrXII | 2317 | 307 | 337 | 2131 | 363 | 373 | 2265 | 249 | 442 | 3458 | 687 | 735 | 1860 | 473 | 568 | 1897 | 636 | 635 |
| 3717 | W303.asm02.HP0_G0037120 | chrXII | 5390 | 2972 | 3086 | 5849 | 2982 | 3104 | 5900 | 3184 | 3352 | 9291 | 3000 | 3484 | 9964 | 2138 | 2714 | 9173 | 2657 | 3134 |
| 3718 | W303.asm02.HP0_G0037130 | chrXII | 2143 | 764 | 611 | 2274 | 510 | 714 | 2287 | 546 | 653 | 4374 | 1429 | 1846 | 1267 | 619 | 782 | 2436 | 1405 | 1713 |

| | | | | | | | | | | | | | | | | | | | | |
|---|---|---|---|---|---|---|---|---|---|---|---|---|---|---|---|---|---|---|---|---|
| 3719 | W303.asm02.HP0_G0037140 | chrXII | 807 | 416 | 460 | 816 | 414 | 398 | 807 | 379 | 416 | 2221 | 464 | 570 | 1893 | 305 | 416 | 2020 | 388 | 508 |
| 3720 | W303.asm02.HP0_G0037150 | chrXII | 322 | 83 | 87 | 340 | 104 | 108 | 349 | 54 | 123 | 366 | 109 | 187 | 154 | 90 | 97 | 130 | 102 | 166 |
| 3721 | W303.asm02.HP0_G0037160 | chrXII | 46 | 15 | 16 | 48 | 21 | 29 | 51 | 9 | 30 | 52 | 17 | 31 | 48 | 13 | 27 | 28 | 19 | 34 |
| 3722 | W303.asm02.HP0_G0037170 | chrXII | 1061 | 355 | 501 | 967 | 507 | 555 | 1030 | 287 | 564 | 1968 | 550 | 898 | 1075 | 427 | 540 | 911 | 515 | 804 |
| 3723 | W303.asm02.HP0_G0037180 | chrXII | 1431 | 358 | 428 | 1403 | 469 | 445 | 1212 | 358 | 463 | 2653 | 433 | 705 | 2471 | 259 | 614 | 2522 | 457 | 613 |
| 3724 | W303.asm02.HP0_G0037190 | chrXII | 4562 | 728 | 770 | 4321 | 706 | 719 | 4229 | 679 | 836 | 3871 | 858 | 1210 | 3627 | 690 | 908 | 3346 | 828 | 1149 |
| 3725 | W303.asm02.HP0_G0037200 | chrXII | 3392 | 410 | 496 | 3268 | 477 | 560 | 3330 | 420 | 703 | 4160 | 699 | 1035 | 2808 | 480 | 792 | 2993 | 615 | 905 |
| 3726 | W303.asm02.HP0_G0037210 | chrXII | 5489 | 969 | 994 | 5139 | 829 | 1067 | 5298 | 908 | 1139 | 5851 | 1491 | 2183 | 5127 | 1141 | 1696 | 4711 | 1392 | 1916 |
| 3727 | W303.asm02.HP0_G0037220 | chrXII | 479 | 332 | 497 | 422 | 380 | 452 | 485 | 355 | 569 | 1428 | 263 | 367 | 1355 | 173 | 241 | 1564 | 282 | 310 |
| 3728 | W303.asm02.HP0_G0037230 | chrXII | 2246 | 1340 | 1770 | 2240 | 1707 | 1791 | 2368 | 1530 | 1850 | 5590 | 1315 | 1685 | 5873 | 787 | 1158 | 5477 | 1052 | 1544 |
| 3729 | W303.asm02.HP0_G0037240 | chrXII | 5517 | 1163 | 1069 | 4950 | 1113 | 1186 | 5821 | 1120 | 1352 | 10052 | 2278 | 2447 | 8864 | 1447 | 1857 | 9340 | 2018 | 2276 |
| 3730 | W303.asm02.HP0_G0037250 | chrXII | 461 | 113 | 131 | 365 | 113 | 144 | 416 | 100 | 158 | 880 | 185 | 215 | 910 | 167 | 181 | 777 | 184 | 164 |
| 3731 | W303.asm02.HP0_G0037260 | chrXII | 751 | 799 | 700 | 695 | 651 | 665 | 722 | 783 | 722 | 2070 | 749 | 678 | 2426 | 518 | 624 | 1885 | 705 | 654 |
| 3732 | W303.asm02.HP0_G0037270 | chrXII | 4246 | 720 | 971 | 3962 | 1025 | 1047 | 3978 | 694 | 1180 | 6542 | 992 | 1852 | 4684 | 818 | 1533 | 4663 | 852 | 1799 |
| 3733 | W303.asm02.HP0_G0037280 | chrXII | 7402 | 878 | 1060 | 7641 | 874 | 1228 | 7052 | 778 | 1195 | 7588 | 1787 | 2551 | 4224 | 1490 | 1810 | 3954 | 1619 | 2094 |
| 3734 | W303.asm02.HP0_G0037290 | chrXII | 81980 | 22481 | 19321 | 75844 | 21455 | 19930 | 89925 | 22749 | 23464 | 136108 | 40318 | 48222 | 108329 | 32973 | 31052 | 105732 | 35253 | 43767 |
| 3735 | W303.asm02.HP0_G0037300 | chrXII | 4504 | 2469 | 2978 | 4412 | 2397 | 2908 | 4176 | 2409 | 3609 | 6903 | 2253 | 2488 | 8628 | 1676 | 2109 | 7493 | 2100 | 2212 |
| 3736 | W303.asm02.HP0_G0037310 | chrXII | 1793 | 1038 | 699 | 1772 | 488 | 746 | 1521 | 794 | 844 | 1037 | 650 | 508 | 1450 | 413 | 522 | 1186 | 712 | 436 |
| 3737 | W303.asm02.HP0_G0037320 | chrXII | 6537 | 1227 | 1110 | 6185 | 1164 | 1177 | 6609 | 1215 | 1255 | 7263 | 1956 | 2500 | 5365 | 1442 | 1860 | 5228 | 1719 | 2207 |
| 3738 | W303.asm02.HP0_G0037330 | chrXII | 1938 | 815 | 677 | 1988 | 672 | 669 | 1896 | 685 | 925 | 1565 | 601 | 510 | 2132 | 395 | 524 | 1625 | 613 | 508 |
| 3739 | W303.asm02.HP0_G0037340 | chrXII | 3415 | 540 | 614 | 3043 | 629 | 654 | 3250 | 522 | 709 | 3869 | 715 | 1110 | 1661 | 541 | 844 | 1693 | 659 | 964 |
| 3740 | W303.asm02.HP0_G0037350 | chrXII | 19626 | 5172 | 4075 | 19974 | 3963 | 4265 | 18612 | 4821 | 4134 | 12056 | 4889 | 4836 | 12627 | 3292 | 3971 | 10874 | 4268 | 4201 |
| 3741 | W303.asm02.HP0_G0037360 | chrXII | 26424 | 942 | 783 | 26235 | 694 | 916 | 23589 | 768 | 984 | 3656 | 1670 | 1667 | 4067 | 2046 | 2161 | 2054 | 1526 | 1439 |
| 3742 | W303.asm02.HP0_G0037370 | chrXII | 39374 | 9647 | 7495 | 38854 | 7383 | 9210 | 39163 | 6963 | 9170 | 38121 | 13997 | 18743 | 25735 | 14029 | 15754 | 22567 | 13500 | 17677 |
| 3743 | W303.asm02.HP0_G0037380 | chrXII | 2780 | 1026 | 763 | 2895 | 767 | 836 | 2660 | 977 | 724 | 4312 | 1301 | 1483 | 4184 | 935 | 1140 | 3569 | 1306 | 1409 |
| 3744 | W303.asm02.HP0_G0037390 | chrXII | 1712 | 1335 | 1922 | 1228 | 1621 | 1848 | 1384 | 1379 | 2424 | 4128 | 1356 | 1698 | 4659 | 884 | 1371 | 4605 | 1215 | 1350 |
| 3745 | W303.asm02.HP0_G0037400 | chrXII | 1064 | 199 | 155 | 943 | 187 | 148 | 1024 | 193 | 155 | 1757 | 297 | 293 | 1230 | 189 | 207 | 1221 | 284 | 326 |
| 3746 | W303.asm02.HP0_G0037410 | chrXII | 2687 | 4875 | 6285 | 2631 | 5432 | 5891 | 2884 | 5074 | 7341 | 12660 | 4383 | 4298 | 14640 | 3662 | 3529 | 13446 | 3749 | 3976 |
| 3747 | W303.asm02.HP0_G0037420 | chrXII | 2301 | 909 | 858 | 1973 | 819 | 791 | 2158 | 930 | 978 | 2686 | 623 | 540 | 3286 | 492 | 564 | 3025 | 612 | 528 |
| 3748 | W303.asm02.HP0_G0037430 | chrXII | 1391 | 901 | 1068 | 1236 | 878 | 983 | 1241 | 915 | 1231 | 3938 | 832 | 1240 | 3435 | 540 | 888 | 3965 | 872 | 1081 |
| 3749 | W303.asm02.HP0_G0037440 | chrXII | 773 | 468 | 342 | 779 | 331 | 320 | 659 | 381 | 347 | 1069 | 336 | 285 | 1167 | 181 | 227 | 1098 | 358 | 283 |
| 3750 | W303.asm02.HP0_G0037450 | chrXII | 3002 | 1429 | 1379 | 2834 | 1432 | 1468 | 2933 | 1493 | 1392 | 5619 | 1392 | 1727 | 6135 | 1071 | 1335 | 4748 | 1143 | 1656 |
| 3751 | W303.asm02.HP0_G0037460 | chrXII | 7404 | 1513 | 1191 | 6875 | 955 | 1082 | 6790 | 1336 | 1129 | 4258 | 1525 | 1434 | 5352 | 1165 | 1603 | 4136 | 1578 | 1425 |
| 3752 | W303.asm02.HP0_G0037470 | chrXII | 1954 | 342 | 372 | 1978 | 354 | 387 | 1683 | 321 | 422 | 1284 | 313 | 383 | 1438 | 245 | 325 | 1124 | 301 | 332 |
| 3753 | W303.asm02.HP0_G0037480 | chrXII | 1368 | 785 | 863 | 1280 | 1062 | 958 | 1439 | 876 | 794 | 5605 | 798 | 1100 | 5253 | 572 | 833 | 4925 | 736 | 1054 |
| 3754 | W303.asm02.HP0_G0037490 | chrXII | 937 | 253 | 226 | 850 | 217 | 235 | 828 | 217 | 206 | 1535 | 272 | 381 | 1310 | 253 | 293 | 1234 | 269 | 314 |
| 3755 | W303.asm02.HP0_G0037500 | chrXII | 2416 | 1639 | 1780 | 2477 | 1629 | 1480 | 2466 | 1711 | 1873 | 8248 | 1489 | 1848 | 8754 | 1186 | 1646 | 7988 | 1438 | 1651 |
| 3756 | W303.asm02.HP0_G0037510 | chrXII | 473 | 481 | 832 | 536 | 771 | 868 | 539 | 510 | 1046 | 2429 | 444 | 636 | 2476 | 286 | 493 | 2501 | 356 | 602 |
| 3757 | W303.asm02.HP0_G0037520 | chrXII | 5470 | 3587 | 4746 | 5831 | 5521 | 4223 | 6185 | 4143 | 5289 | 19899 | 3699 | 3924 | 20150 | 2376 | 2509 | 19519 | 3127 | 3552 |
| 3758 | W303.asm02.HP0_G0037530 | chrXII | 1770 | 493 | 545 | 1629 | 604 | 583 | 1809 | 487 | 636 | 5177 | 791 | 1168 | 3878 | 548 | 851 | 4075 | 748 | 1164 |
| 3759 | W303.asm02.HP0_G0037540 | chrXII | 4306 | 845 | 1378 | 4403 | 1377 | 1432 | 4434 | 997 | 1629 | 7756 | 1027 | 2144 | 5502 | 592 | 1600 | 5012 | 865 | 1901 |
| 3760 | W303.asm02.HP0_G0037550 | chrXII | 778 | 215 | 177 | 669 | 169 | 173 | 692 | 220 | 209 | 376 | 184 | 139 | 474 | 89 | 115 | 396 | 154 | 104 |
| 3761 | W303.asm02.HP0_G0037560 | chrXII | 1147 | 552 | 533 | 988 | 457 | 595 | 931 | 532 | 691 | 1570 | 415 | 436 | 1730 | 242 | 358 | 1727 | 333 | 356 |
| 3762 | W303.asm02.HP0_G0037570 | chrXII | 794 | 439 | 366 | 884 | 317 | 350 | 743 | 435 | 388 | 939 | 309 | 261 | 1121 | 220 | 249 | 915 | 335 | 223 |
| 3763 | W303.asm02.HP0_G0037580 | chrXII | 1521 | 1437 | 1836 | 1240 | 1660 | 1625 | 1338 | 1466 | 2067 | 6082 | 1250 | 1483 | 6121 | 899 | 1204 | 6229 | 1272 | 1462 |
| 3764 | W303.asm02.HP0_G0037590 | chrXII | 92 | 22 | 14 | 51 | 8 | 9 | 58 | 18 | 12 | 36 | 15 | 26 | 46 | 12 | 27 | 39 | 14 | 8 |
| 3765 | W303.asm02.HP0_G0037600 | chrXII | 30088 | 10482 | 6440 | 29783 | 6521 | 6191 | 32238 | 10204 | 6489 | 35130 | 13197 | 12188 | 38546 | 10429 | 9196 | 30057 | 10357 | 9967 |
| 3766 | W303.asm02.HP0_G0037610 | chrXII | 1786 | 1736 | 2000 | 1809 | 1970 | 1829 | 1862 | 1833 | 2205 | 7017 | 1546 | 1748 | 6715 | 1022 | 1250 | 7231 | 1362 | 1559 |
| 3767 | W303.asm02.HP0_G0037620 | chrXII | 1385 | 701 | 495 | 1286 | 538 | 499 | 1304 | 686 | 521 | 2058 | 647 | 613 | 2154 | 422 | 494 | 1950 | 622 | 573 |
| 3768 | W303.asm02.HP0_G0037630 | chrXII | 1997 | 522 | 455 | 2033 | 431 | 447 | 1900 | 418 | 487 | 2651 | 601 | 775 | 2689 | 396 | 676 | 2141 | 590 | 746 |
| 3769 | W303.asm02.HP0_G0037640 | chrXII | 1863 | 860 | 893 | 1902 | 655 | 982 | 1717 | 745 | 1213 | 1939 | 572 | 659 | 2381 | 353 | 572 | 2171 | 529 | 553 |
| 3770 | W303.asm02.HP0_G0037650 | chrXII | 2402 | 2046 | 2014 | 2433 | 2068 | 1958 | 2667 | 2024 | 1912 | 9853 | 2016 | 2201 | 9569 | 1221 | 1605 | 9827 | 1794 | 2162 |
| 3771 | W303.asm02.HP0_G0037660 | chrXII | 2393 | 1768 | 3092 | 1955 | 2177 | 3277 | 2229 | 1949 | 3697 | 6406 | 1595 | 2780 | 7272 | 1090 | 2035 | 6742 | 1493 | 2435 |
| 3772 | W303.asm02.HP0_G0037670 | chrXII | 12612 | 8032 | 7635 | 11314 | 7753 | 7836 | 12967 | 8029 | 7211 | 24938 | 8255 | 9707 | 24233 | 6062 | 7233 | 24155 | 7425 | 8690 |
| 3773 | W303.asm02.HP0_G0037680 | chrXII | 4014 | 4495 | 4808 | 3712 | 4417 | 4858 | 3786 | 4620 | 5535 | 10216 | 3869 | 4204 | 11627 | 3129 | 3671 | 10005 | 3416 | 3283 |
| 3774 | W303.asm02.HP0_G0037690 | chrXII | 872 | 375 | 628 | 868 | 585 | 616 | 784 | 340 | 810 | 1136 | 408 | 308 | 1507 | 252 | 381 | 1039 | 440 | 223 |
| 3775 | W303.asm02.HP0_G0037700 | chrXII | 2360 | 889 | 881 | 2193 | 931 | 922 | 2217 | 978 | 906 | 2882 | 727 | 738 | 3245 | 495 | 584 | 2778 | 721 | 669 |
| 3776 | W303.asm02.HP0_G0037710 | chrXII | 6812 | 536 | 775 | 6400 | 530 | 832 | 6783 | 482 | 962 | 3399 | 1326 | 1690 | 1878 | 1290 | 1521 | 1432 | 1162 | 1365 |
| 3777 | W303.asm02.HP0_G0037720 | chrXII | 7712 | 7696 | 11324 | 6543 | 9571 | 10024 | 8602 | 8706 | 12554 | 38358 | 10836 | 15635 | 50307 | 11739 | 14643 | 31815 | 8899 | 13721 |
| 3778 | W303.asm02.HP0_G0037730 | chrXII | 2470 | 2791 | 3492 | 2040 | 2748 | 3425 | 2514 | 2695 | 4533 | 9139 | 2557 | 3050 | 8839 | 1482 | 2094 | 9873 | 2306 | 2852 |
| 3779 | W303.asm02.HP0_G0037740 | chrXII | 20197 | 6890 | 7131 | 12839 | 5294 | 7294 | 17152 | 7109 | 7071 | 9708 | 4966 | 8338 | 12889 | 5756 | 6045 | 8641 | 4083 | 5980 |
| 3780 | W303.asm02.HP0_G0037750 | chrXII | 23457 | 7405 | 5698 | 21390 | 6391 | 5945 | 22534 | 6909 | 6922 | 43196 | 14814 | 18015 | 39833 | 13841 | 14361 | 31257 | 13492 | 16419 |
| 3781 | W303.asm02.HP0_G0037760 | chrXII | 4259 | 997 | 1386 | 3931 | 1414 | 1370 | 4020 | 1169 | 1700 | 11472 | 1664 | 2743 | 7833 | 1158 | 1989 | 8465 | 1479 | 2416 |
| 3782 | W303.asm02.HP0_G0037770 | chrXII | 1169 | 458 | 497 | 899 | 441 | 447 | 1037 | 389 | 653 | 1297 | 360 | 407 | 1365 | 173 | 336 | 1447 | 300 | 350 |
| 3783 | W303.asm02.HP0_G0037780 | chrXII | 201 | 92 | 122 | 225 | 95 | 91 | 176 | 106 | 107 | 230 | 112 | 115 | 285 | 51 | 71 | 208 | 91 | 97 |
| 3784 | W303.asm02.HP0_G0037790 | chrXII | 109 | 73 | 85 | 83 | 61 | 79 | 79 | 68 | 90 | 79 | 37 | 55 | 105 | 14 | 11 | 102 | 39 | 40 |
| 3785 | W303.asm02.HP0_G0037800 | chrXII | 25 | 6 | 6 | 23 | 8 | 5 | 14 | 6 | 6 | 23 | 12 | 6 | 25 | 3 | 5 | 21 | 16 | 5 |
| 3786 | W303.asm02.HP0_G0037810 | chrXII | 5390 | 1343 | 1998 | 5671 | 1935 | 1878 | 5201 | 1604 | 2060 | 9105 | 1500 | 2103 | 8957 | 899 | 1597 | 8633 | 1341 | 1927 |
| 3787 | W303.asm02.HP0_G0037820 | chrXII | 6677 | 928 | 1143 | 6451 | 1099 | 1176 | 6503 | 863 | 1353 | 10481 | 1797 | 2738 | 7440 | 1335 | 2015 | 6890 | 1493 | 2451 |
| 3788 | W303.asm02.HP0_G0037830 | chrXII | 698 | 273 | 190 | 744 | 229 | 204 | 719 | 252 | 202 | 735 | 324 | 216 | 843 | 196 | 216 | 679 | 326 | 185 |
| 3789 | W303.asm02.HP0_G0037840 | chrXII | 6246 | 2062 | 3058 | 5974 | 2877 | 3234 | 5812 | 2175 | 4031 | 6753 | 1604 | 2372 | 7868 | 1214 | 1975 | 6595 | 1548 | 2089 |
| 3790 | W303.asm02.HP0_G0037850 | chrXII | 34 | 27 | 46 | 19 | 44 | 49 | 31 | 17 | 53 | 185 | 39 | 67 | 148 | 22 | 61 | 136 | 44 | 63 |
| 3791 | W303.asm02.HP0_G0037860 | chrXII | 282 | 152 | 180 | 223 | 203 | 219 | 277 | 167 | 213 | 1181 | 318 | 411 | 911 | 235 | 309 | 843 | 266 | 390 |
| 3792 | W303.asm02.HP0_G0037870 | chrXII | 8194 | 2182 | 2145 | 8067 | 2005 | 2108 | 8040 | 2230 | 2291 | 14238 | 3677 | 4166 | 11464 | 2707 | 3333 | 11321 | 3369 | 3840 |
| 3793 | W303.asm02.HP0_G0037880 | chrXII | 1459 | 426 | 445 | 1397 | 454 | 503 | 1458 | 347 | 570 | 2530 | 653 | 856 | 1945 | 467 | 603 | 1775 | 594 | 766 |
| 3794 | W303.asm02.HP0_G0037890 | chrXII | 11 | 1 | 1 | 9 | 2 | 4 | 19 | 6 | 8 | 25 | 2 | 1 | 25 | 2 | 0 | 27 | 2 | 1 |
| 3795 | W303.asm02.HP0_G0037900 | chrXII | 453 | 374 | 485 | 449 | 479 | 492 | 458 | 368 | 548 | 1622 | 350 | 471 | 1538 | 183 | 312 | 1606 | 294 | 384 |
| 3796 | W303.asm02.HP0_G0037910 | chrXII | 2258 | 322 | 469 | 2005 | 444 | 493 | 2072 | 311 | 593 | 3752 | 590 | 929 | 2100 | 453 | 659 | 2089 | 585 | 862 |
| 3797 | W303.asm02.HP0_G0037920 | chrXII | 1068 | 246 | 418 | 1015 | 492 | 380 | 1009 | 247 | 476 | 3772 | 371 | 794 | 3217 | 298 | 592 | 2958 | 310 | 749 |
| 3798 | W303.asm02.HP0_G0037930 | chrXII | 1485 | 794 | 632 | 1286 | 649 | 640 | 1369 | 804 | 626 | 4231 | 1001 | 1286 | 3531 | 693 | 868 | 3649 | 897 | 1190 |
| 3799 | W303.asm02.HP0_G0037940 | chrXII | 891 | 567 | 582 | 939 | 621 | 641 | 999 | 586 | 516 | 2340 | 460 | 740 | 2315 | 297 | 419 | 2057 | 410 | 604 |
| 3800 | W303.asm02.HP0_G0037950 | chrXII | 5102 | 398 | 427 | 4522 | 354 | 502 | 4606 | 352 | 503 | 2941 | 726 | 929 | 1579 | 540 | 776 | 1519 | 686 | 826 |
| 3801 | W303.asm02.HP0_G0037960 | chrXII | 6722 | 6416 | 8407 | 6147 | 7345 | 8253 | 6462 | 6377 | 10442 | 15198 | 5801 | 6747 | 17677 | 4412 | 6019 | 17207 | 5089 | 5944 |
| 3802 | W303.asm02.HP0_G0037970 | chrXII | 2001 | 644 | 600 | 1986 | 564 | 610 | 1953 | 679 | 564 | 1724 | 601 | 768 | 2158 | 487 | 635 | 1614 | 603 | 676 |
| 3803 | W303.asm02.HP0_G0037980 | chrXII | 9310 | 695 | 319 | 9447 | 1101 | 1284 | 7700 | 520 | 442 | 442 | 510 | 251 | 1329 | 471 | 480 | 573 | 446 | 270 |
| 3804 | W303.asm02.HP0_G0037990 | chrXII | 1644 | 1102 | 978 | 1563 | 877 | 981 | 1629 | 1011 | 997 | 5990 | 1266 | 1745 | 5263 | 833 | 1219 | 5209 | 1205 | 1559 |
| 3805 | W303.asm02.HP0_G0038000 | chrXII | 273 | 155 | 102 | 219 | 81 | 88 | 241 | 180 | 91 | 346 | 106 | 96 | 354 | 82 | 78 | 331 | 106 | 72 |
| 3806 | W303.asm02.HP0_G0038010 | chrXII | 14885 | 2004 | 1529 | 15208 | 1531 | 1730 | 15670 | 1859 | 1812 | 13125 | 3449 | 3892 | 11261 | 2773 | 3036 | 9327 | 3105 | 3348 |
| 3807 | W303.asm02.HP0_G0038020 | chrXII | 2264 | 899 | 943 | 2593 | 1027 | 833 | 2331 | 950 | 902 | 6879 | 1177 | 1557 | 7375 | 780 | 1182 | 5440 | 1089 | 1433 |
| 3808 | W303.asm02.HP0_G0038030 | chrXII | 3775 | 4284 | 6044 | 3264 | 5834 | 5872 | 3424 | 4336 | 7817 | 13697 | 3979 | 4982 | 15888 | 2968 | 3935 | 14807 | 3591 | 4204 |
| 3809 | W303.asm02.HP0_G0038040 | chrXII | 7812 | 853 | 1092 | 7646 | 957 | 1293 | 8334 | 711 | 1500 | 11811 | 2248 | 2936 | 5660 | 1617 | 1940 | 5962 | 1936 | 2550 |
| 3810 | W303.asm02.HP0_G0038050 | chrXII | 1577 | 768 | 990 | 1605 | 1309 | 870 | 1778 | 887 | 1032 | 9530 | 1084 | 1525 | 7046 | 690 | 854 | 7300 | 1004 | 1389 |
| 3811 | W303.asm02.HP0_G0038060 | chrXII | 5472 | 377 | 388 | 5144 | 393 | 551 | 5575 | 276 | 577 | 309 | 289 | 398 | 540 | 272 | 369 | 313 | 249 | 332 |

| | | | | | | | | | | | | | | | | | | | | |
|---|---|---|---|---|---|---|---|---|---|---|---|---|---|---|---|---|---|---|---|---|
| 3812 | W303.asm02.HP0_G0038070 | chrXII | 27604 | 9323 | 8226 | 25673 | 8445 | 9178 | 29162 | 9739 | 8970 | 40765 | 14472 | 16504 | 39773 | 11875 | 12573 | 38758 | 12833 | 15233 |
| 3813 | W303.asm02.HP0_G0038080 | chrXII | 346 | 159 | 190 | 318 | 191 | 202 | 330 | 162 | 234 | 768 | 127 | 152 | 912 | 113 | 106 | 730 | 135 | 129 |
| 3814 | W303.asm02.HP0_G0038090 | chrXII | 34419 | 5493 | 7535 | 32494 | 7152 | 7591 | 34692 | 5759 | 8923 | 50049 | 11036 | 16779 | 37281 | 11049 | 13654 | 27122 | 9286 | 14619 |
| 3815 | W303.asm02.HP0_G0038100 | chrXII | 370 | 56 | 56 | 263 | 50 | 62 | 291 | 54 | 89 | 166 | 49 | 111 | 132 | 45 | 115 | 123 | 41 | 54 |
| 3816 | W303.asm02.HP0_G0038110 | chrXII | 1009 | 221 | 128 | 778 | 120 | 131 | 954 | 206 | 152 | 356 | 172 | 154 | 327 | 144 | 97 | 263 | 177 | 119 |
| 3817 | W303.asm02.HP0_G0038120 | chrXII | 7122 | 5583 | 6977 | 6540 | 6391 | 6776 | 7017 | 5622 | 8198 | 15796 | 5018 | 5639 | 17437 | 3678 | 4499 | 16521 | 4495 | 5062 |
| 3818 | W303.asm02.HP0_G0038130 | chrXII | 6717 | 1598 | 1344 | 6885 | 1184 | 1370 | 6200 | 1549 | 1389 | 3616 | 1288 | 1286 | 4539 | 970 | 1551 | 3818 | 1355 | 1124 |
| 3819 | W303.asm02.HP0_G0038140 | chrXII | 79 | 148 | 120 | 64 | 107 | 143 | 78 | 123 | 191 | 260 | 133 | 123 | 459 | 90 | 128 | 277 | 141 | 119 |
| 3820 | W303.asm02.HP0_G0038150 | chrXII | 3410 | 1500 | 1570 | 3703 | 1638 | 1700 | 3437 | 1612 | 1710 | 12448 | 2374 | 3143 | 9464 | 1659 | 2161 | 8936 | 2108 | 2920 |
| 3821 | W303.asm02.HP0_G0038160 | chrXII | 1453 | 1293 | 1401 | 1258 | 1141 | 1525 | 1337 | 1202 | 1627 | 5513 | 1997 | 2151 | 3912 | 1178 | 1532 | 4176 | 1775 | 1975 |
| 3822 | W303.asm02.HP0_G0038170 | chrXII | 211 | 120 | 77 | 186 | 44 | 76 | 191 | 99 | 90 | 116 | 86 | 68 | 159 | 60 | 46 | 105 | 93 | 59 |
| 3823 | W303.asm02.HP0_G0038180 | chrXII | 6849 | 1436 | 1216 | 7307 | 912 | 1264 | 6414 | 1343 | 1052 | 3923 | 1415 | 1909 | 3852 | 877 | 1158 | 3401 | 1317 | 1553 |
| 3824 | W303.asm02.HP0_G0038190 | chrXII | 3543 | 2850 | 2657 | 3043 | 2441 | 2651 | 3301 | 2789 | 2803 | 6528 | 2462 | 2677 | 6754 | 1650 | 1959 | 6383 | 2252 | 2438 |
| 3825 | W303.asm02.HP0_G0038200 | chrXII | 1174 | 489 | 360 | 994 | 283 | 384 | 1132 | 440 | 412 | 2503 | 642 | 918 | 1536 | 444 | 583 | 1861 | 645 | 786 |
| 3826 | W303.asm02.HP0_G0038210 | chrXII | 737 | 487 | 447 | 710 | 481 | 505 | 750 | 459 | 552 | 3393 | 846 | 878 | 2877 | 593 | 722 | 2312 | 675 | 794 |
| 3827 | W303.asm02.HP0_G0038220 | chrXII | 19477 | 6334 | 5342 | 17341 | 4350 | 5696 | 19767 | 6371 | 5429 | 23293 | 9421 | 12207 | 24130 | 7277 | 8942 | 22136 | 8330 | 11254 |
| 3828 | W303.asm02.HP0_G0038230 | chrXII | 2861 | 5465 | 7240 | 2459 | 6260 | 8644 | 3189 | 4776 | 9166 | 33712 | 10916 | 15273 | 26733 | 8013 | 9830 | 25429 | 9204 | 12780 |
| 3829 | W303.asm02.HP0_G0038240 | chrXII | 2496 | 539 | 559 | 2315 | 454 | 561 | 2221 | 622 | 572 | 1669 | 718 | 937 | 2344 | 567 | 844 | 1568 | 651 | 729 |
| 3830 | W303.asm02.HP0_G0038250 | chrXII | 631 | 164 | 235 | 657 | 215 | 306 | 717 | 166 | 308 | 1715 | 310 | 602 | 1092 | 216 | 434 | 925 | 266 | 546 |
| 3831 | W303.asm02.HP0_G0038260 | chrXII | 9208 | 2504 | 2472 | 6549 | 2371 | 2138 | 8478 | 2467 | 2350 | 11875 | 2944 | 3486 | 9866 | 1818 | 2270 | 10617 | 2583 | 3067 |
| 3832 | W303.asm02.HP0_G0038270 | chrXII | 1719 | 605 | 531 | 1698 | 541 | 540 | 1878 | 622 | 628 | 3372 | 758 | 878 | 2699 | 558 | 645 | 2712 | 722 | 773 |
| 3833 | W303.asm02.HP0_G0038280 | chrXII | 1186 | 381 | 477 | 899 | 400 | 494 | 849 | 396 | 656 | 1027 | 310 | 385 | 1520 | 261 | 373 | 1154 | 287 | 318 |
| 3834 | W303.asm02.HP0_G0038290 | chrXII | 3555 | 703 | 608 | 3568 | 609 | 611 | 3209 | 580 | 678 | 4268 | 968 | 1196 | 3777 | 658 | 858 | 3230 | 891 | 1047 |
| 3835 | W303.asm02.HP0_G0038300 | chrXII | 918 | 1172 | 924 | 921 | 780 | 959 | 825 | 1056 | 1204 | 2581 | 670 | 650 | 2924 | 457 | 521 | 3029 | 712 | 602 |
| 3836 | W303.asm02.HP0_G0038310 | chrXII | 417 | 763 | 983 | 359 | 1079 | 931 | 411 | 852 | 1155 | 3023 | 688 | 835 | 3111 | 414 | 506 | 3286 | 581 | 679 |
| 3837 | W303.asm02.HP0_G0038320 | chrXII | 562 | 484 | 449 | 321 | 338 | 435 | 457 | 512 | 564 | 687 | 303 | 327 | 917 | 201 | 228 | 742 | 310 | 269 |
| 3838 | W303.asm02.HP0_G0038330 | chrXII | 3464 | 3508 | 3984 | 3317 | 3628 | 3938 | 3710 | 3409 | 5238 | 8098 | 2861 | 2960 | 9530 | 2154 | 2850 | 9279 | 2618 | 2674 |
| 3839 | W303.asm02.HP0_G0038340 | chrXII | 1633 | 637 | 501 | 1740 | 592 | 490 | 1568 | 614 | 559 | 3211 | 713 | 896 | 2398 | 392 | 590 | 2815 | 732 | 753 |
| 3840 | W303.asm02.HP0_G0038350 | chrXII | 3090 | 946 | 829 | 3165 | 862 | 877 | 2809 | 959 | 815 | 5455 | 1188 | 1503 | 4261 | 786 | 1114 | 4449 | 1142 | 1388 |
| 3841 | W303.asm02.HP0_G0038360 | chrXII | 6016 | 2157 | 2391 | 5823 | 2120 | 2509 | 5723 | 2222 | 2407 | 5805 | 2073 | 2324 | 6766 | 1624 | 2239 | 5789 | 1750 | 2214 |
| 3842 | W303.asm02.HP0_G0038370 | chrXII | 3039 | 434 | 674 | 3059 | 617 | 663 | 2972 | 438 | 778 | 5329 | 782 | 1470 | 3276 | 596 | 997 | 2975 | 705 | 1260 |
| 3843 | W303.asm02.HP0_G0038380 | chrXII | 5059 | 6231 | 7859 | 4774 | 7564 | 7344 | 5609 | 6958 | 9017 | 21141 | 6011 | 6512 | 22211 | 3966 | 4937 | 21971 | 4839 | 5909 |
| 3844 | W303.asm02.HP0_G0038390 | chrXII | 2383 | 565 | 777 | 2279 | 676 | 820 | 2473 | 647 | 890 | 5019 | 904 | 1656 | 3672 | 572 | 1083 | 3984 | 866 | 1362 |
| 3845 | W303.asm02.HP0_G0038400 | chrXII | 3593 | 1214 | 1228 | 3204 | 1063 | 1333 | 3474 | 1169 | 1486 | 4520 | 1854 | 1856 | 3587 | 1148 | 1493 | 3479 | 1645 | 1731 |
| 3846 | W303.asm02.HP0_G0038410 | chrXII | 285 | 561 | 336 | 179 | 312 | 372 | 243 | 554 | 351 | 771 | 327 | 230 | 1061 | 257 | 257 | 839 | 404 | 237 |
| 3847 | W303.asm02.HP0_G0038420 | chrXII | 399 | 170 | 70 | 368 | 76 | 86 | 432 | 159 | 79 | 329 | 135 | 104 | 350 | 90 | 74 | 277 | 135 | 88 |
| 3848 | W303.asm02.HP0_G0038430 | chrXII | 6812 | 1634 | 2887 | 6126 | 1651 | 3544 | 6886 | 1302 | 3805 | 13424 | 3594 | 6663 | 5255 | 2980 | 4269 | 5295 | 3078 | 6149 |
| 3849 | W303.asm02.HP0_G0038440 | chrXII | 3618 | 454 | 692 | 3259 | 497 | 805 | 3688 | 376 | 832 | 5075 | 961 | 1697 | 1257 | 660 | 1077 | 1346 | 782 | 1484 |
| 3850 | W303.asm02.HP0_G0038450 | chrXII | 1240 | 462 | 379 | 1223 | 415 | 411 | 1282 | 411 | 459 | 2079 | 490 | 587 | 1911 | 311 | 447 | 1931 | 449 | 474 |
| 3851 | W303.asm02.HP0_G0038460 | chrXII | 8092 | 498 | 706 | 7235 | 611 | 850 | 7660 | 404 | 882 | 4751 | 1015 | 1815 | 2505 | 1026 | 1176 | 1853 | 978 | 1548 |
| 3852 | W303.asm02.HP0_G0038470 | chrXII | 1684 | 361 | 498 | 1772 | 534 | 517 | 1698 | 408 | 561 | 3458 | 540 | 927 | 2946 | 368 | 663 | 2871 | 478 | 871 |
| 3853 | W303.asm02.HP0_G0038480 | chrXII | 3600 | 1316 | 1403 | 3306 | 1691 | 1383 | 3911 | 1344 | 1692 | 14582 | 2685 | 3312 | 9031 | 1808 | 2096 | 9476 | 2329 | 3050 |
| 3854 | W303.asm02.HP0_G0038490 | chrXII | 67 | 13 | 24 | 70 | 35 | 24 | 89 | 17 | 24 | 402 | 51 | 56 | 294 | 36 | 25 | 234 | 52 | 59 |
| 3855 | W303.asm02.HP0_G0038500 | chrXII | 2565 | 565 | 622 | 2345 | 672 | 663 | 2401 | 547 | 682 | 4794 | 772 | 1254 | 3919 | 625 | 938 | 3658 | 708 | 1149 |
| 3856 | W303.asm02.HP0_G0038510 | chrXII | 5225 | 2686 | 2548 | 5415 | 2374 | 2387 | 5238 | 2690 | 2491 | 7618 | 2173 | 2560 | 7281 | 1398 | 1936 | 7199 | 2003 | 2196 |
| 3857 | W303.asm02.HP0_G0038520 | chrXII | 2661 | 1999 | 2493 | 2254 | 2367 | 2542 | 2323 | 2064 | 2854 | 4752 | 1757 | 2035 | 5339 | 1151 | 1774 | 5039 | 1594 | 1762 |
| 3858 | W303.asm02.HP0_G0038530 | chrXII | 6306 | 1488 | 1066 | 5847 | 1189 | 1147 | 6258 | 1575 | 1152 | 10313 | 2255 | 2243 | 7598 | 1666 | 1657 | 7664 | 2044 | 1964 |
| 3859 | W303.asm02.HP0_G0038540 | chrXII | 1334 | 517 | 533 | 906 | 497 | 525 | 1049 | 511 | 510 | 1169 | 396 | 489 | 1668 | 350 | 398 | 1137 | 425 | 447 |
| 3860 | W303.asm02.HP0_G0038550 | chrXII | 18591 | 2983 | 3086 | 17503 | 2805 | 3449 | 18864 | 2748 | 3968 | 18963 | 6496 | 6947 | 23115 | 6236 | 7507 | 15082 | 5533 | 6327 |
| 3861 | W303.asm02.HP0_G0038560 | chrXII | 1542 | 715 | 479 | 1725 | 514 | 444 | 1423 | 609 | 456 | 2779 | 631 | 608 | 2479 | 430 | 515 | 2345 | 645 | 493 |
| 3862 | W303.asm02.HP0_G0038570 | chrXII | 887 | 132 | 131 | 704 | 129 | 141 | 746 | 112 | 163 | 947 | 267 | 252 | 347 | 160 | 154 | 376 | 195 | 201 |
| 3863 | W303.asm02.HP0_G0038580 | chrXII | 1123 | 183 | 217 | 995 | 216 | 218 | 1189 | 167 | 267 | 1228 | 396 | 455 | 515 | 249 | 307 | 506 | 283 | 399 |
| 3864 | W303.asm02.HP0_G0038590 | chrXII | 2803 | 3133 | 4116 | 2608 | 3709 | 4058 | 2512 | 3172 | 5107 | 6910 | 2645 | 3158 | 7968 | 1956 | 2666 | 7279 | 2272 | 2406 |
| 3865 | W303.asm02.HP0_G0038600 | chrXII | 886 | 651 | 479 | 781 | 430 | 483 | 872 | 611 | 556 | 3204 | 749 | 856 | 2608 | 470 | 642 | 2751 | 750 | 763 |
| 3866 | W303.asm02.HP0_G0038610 | chrXII | 2433 | 660 | 730 | 2026 | 837 | 736 | 2457 | 663 | 1043 | 7990 | 1457 | 1860 | 4770 | 916 | 1086 | 4722 | 1272 | 1554 |
| 3867 | W303.asm02.HP0_G0038620 | chrXII | 5926 | 706 | 1033 | 5723 | 895 | 1117 | 5635 | 667 | 1257 | 7726 | 1259 | 2438 | 3721 | 942 | 1528 | 3727 | 1130 | 2134 |
| 3868 | W303.asm02.HP0_G0038630 | chrXII | 2403 | 1004 | 1035 | 2545 | 907 | 1147 | 2182 | 851 | 1334 | 5760 | 1539 | 2328 | 1788 | 1200 | 1500 | 2153 | 1507 | 2060 |
| 3869 | W303.asm02.HP0_G0038640 | chrXII | 3048 | 622 | 819 | 2958 | 846 | 864 | 3391 | 567 | 976 | 7800 | 1550 | 1925 | 3523 | 1004 | 892 | 3627 | 1269 | 1660 |
| 3870 | W303.asm02.HP0_G0038650 | chrXII | 1818 | 636 | 918 | 2051 | 900 | 945 | 1914 | 605 | 1170 | 7188 | 1305 | 1815 | 4225 | 768 | 1299 | 4182 | 1205 | 1632 |
| 3871 | W303.asm02.HP0_G0038660 | chrXII | 437 | 625 | 626 | 399 | 538 | 635 | 418 | 691 | 661 | 1809 | 494 | 498 | 1994 | 341 | 419 | 1695 | 460 | 442 |
| 3872 | W303.asm02.HP0_G0038670 | chrXII | 880 | 917 | 671 | 898 | 697 | 597 | 889 | 849 | 700 | 3554 | 998 | 996 | 3132 | 617 | 699 | 3027 | 969 | 924 |
| 3873 | W303.asm02.HP0_G0038680 | chrXII | 6017 | 5501 | 6699 | 6205 | 6806 | 6557 | 6350 | 5669 | 8582 | 16087 | 5073 | 5126 | 17752 | 3448 | 4167 | 16312 | 4242 | 4688 |
| 3874 | W303.asm02.HP0_G0038690 | chrXII | 13 | 14 | 6 | 9 | 10 | 10 | 8 | 14 | 7 | 17 | 18 | 7 | 20 | 9 | 5 | 19 | 21 | 3 |
| 3875 | W303.asm02.HP0_G0038700 | chrXII | 559 | 1120 | 1043 | 537 | 1292 | 1035 | 619 | 1107 | 1027 | 4389 | 857 | 1058 | 4468 | 636 | 714 | 4263 | 904 | 892 |
| 3876 | W303.asm02.HP0_G0038710 | chrXII | 244 | 330 | 376 | 302 | 254 | 364 | 248 | 329 | 493 | 584 | 255 | 251 | 727 | 139 | 195 | 704 | 224 | 199 |
| 3877 | W303.asm02.HP0_G0038720 | chrXII | 2669 | 1255 | 1353 | 2865 | 1737 | 1406 | 3241 | 1217 | 1660 | 15260 | 2358 | 3235 | 8780 | 1639 | 1668 | 9082 | 1998 | 2855 |
| 3878 | W303.asm02.HP0_G0038730 | chrXII | 8583 | 1048 | 1229 | 7963 | 1249 | 1431 | 8334 | 972 | 1480 | 10075 | 1960 | 3140 | 5438 | 1616 | 2010 | 5276 | 1688 | 2574 |
| 3879 | W303.asm02.HP0_G0038740 | chrXII | 675 | 393 | 439 | 538 | 438 | 382 | 535 | 397 | 598 | 1233 | 290 | 324 | 1325 | 201 | 229 | 1315 | 337 | 311 |
| 3880 | W303.asm02.HP0_G0038750 | chrXII | 1891 | 960 | 901 | 1975 | 946 | 1004 | 1958 | 1093 | 891 | 3660 | 854 | 1011 | 3836 | 650 | 663 | 3428 | 750 | 898 |
| 3881 | W303.asm02.HP0_G0038760 | chrXII | 52 | 9 | 7 | 31 | 6 | 4 | 37 | 6 | 1 | 30 | 4 | 2 | 20 | 3 | 5 | 39 | 3 | 2 |
| 3882 | W303.asm02.HP0_G0038770 | chrXII | 3147 | 1519 | 1602 | 3076 | 1773 | 1675 | 3631 | 1663 | 2171 | 21752 | 3771 | 3251 | 15358 | 2599 | 2298 | 13786 | 3148 | 3155 |
| 3883 | W303.asm02.HP0_G0038780 | chrXII | 4389 | 1506 | 1641 | 4775 | 1554 | 1619 | 3981 | 1540 | 1816 | 4175 | 1272 | 1539 | 6406 | 1115 | 1515 | 3865 | 1193 | 1376 |
| 3884 | W303.asm02.HP0_G0038790 | chrXII | 59 | 7 | 8 | 60 | 7 | 11 | 46 | 6 | 9 | 41 | 22 | 12 | 35 | 18 | 12 | 26 | 12 | 12 |
| 3885 | W303.asm02.HP0_G0038800 | chrXII | 1466 | 326 | 300 | 1468 | 311 | 324 | 1458 | 327 | 333 | 2115 | 447 | 576 | 1785 | 386 | 450 | 1706 | 467 | 477 |
| 3886 | W303.asm02.HP0_G0038810 | chrXII | 2945 | 1729 | 1767 | 3111 | 1600 | 1704 | 2975 | 1773 | 1820 | 5908 | 1606 | 1921 | 5751 | 928 | 1317 | 5986 | 1356 | 1677 |
| 3887 | W303.asm02.HP0_G0038820 | chrXII | 2339 | 458 | 800 | 2328 | 768 | 846 | 2452 | 473 | 958 | 8316 | 805 | 1918 | 5140 | 587 | 1220 | 5387 | 687 | 1662 |
| 3888 | W303.asm02.HP0_G0038830 | chrXII | 3490 | 1997 | 1796 | 2940 | 1912 | 1660 | 3115 | 2154 | 1939 | 6867 | 1841 | 1796 | 6874 | 1171 | 1311 | 6781 | 1651 | 1659 |
| 3889 | W303.asm02.HP0_G0038840 | chrXII | 3771 | 2362 | 2888 | 3760 | 2441 | 2915 | 3563 | 2288 | 3750 | 6548 | 1998 | 2278 | 7359 | 1429 | 1985 | 6943 | 1786 | 2097 |
| 3890 | W303.asm02.HP0_G0038850 | chrXII | 2465 | 537 | 535 | 2382 | 570 | 629 | 2365 | 463 | 745 | 4798 | 663 | 1367 | 2835 | 584 | 992 | 2891 | 699 | 1163 |
| 3891 | W303.asm02.HP0_G0038860 | chrXII | 1152 | 414 | 1140 | 1143 | 502 | 1385 | 1040 | 390 | 1549 | 1391 | 299 | 1019 | 1936 | 238 | 646 | 1690 | 290 | 888 |
| 3892 | W303.asm02.HP0_G0038870 | chrXII | 842 | 296 | 354 | 826 | 377 | 392 | 793 | 283 | 421 | 2179 | 371 | 744 | 1668 | 236 | 447 | 1941 | 381 | 663 |
| 3893 | W303.asm02.HP0_G0038880 | chrXII | 1733 | 208 | 328 | 1832 | 287 | 360 | 1603 | 203 | 381 | 1341 | 225 | 617 | 924 | 195 | 392 | 900 | 185 | 461 |
| 3894 | W303.asm02.HP0_G0038890 | chrXII | 553 | 219 | 271 | 490 | 235 | 277 | 607 | 252 | 252 | 1512 | 313 | 505 | 1171 | 268 | 366 | 1174 | 347 | 451 |
| 3895 | W303.asm02.HP0_G0038900 | chrXII | 2748 | 774 | 926 | 2559 | 831 | 953 | 2814 | 772 | 1021 | 6021 | 1311 | 1882 | 4591 | 848 | 1311 | 4666 | 1156 | 1677 |
| 3896 | W303.asm02.HP0_G0038910 | chrXII | 13370 | 1749 | 2016 | 12185 | 1392 | 2462 | 13984 | 1493 | 2477 | 12756 | 3391 | 6015 | 8046 | 2706 | 3527 | 7240 | 3144 | 5139 |
| 3897 | W303.asm02.HP0_G0038920 | chrXII | 4846 | 659 | 789 | 4721 | 800 | 912 | 5042 | 540 | 1102 | 7669 | 1229 | 2066 | 3740 | 991 | 1411 | 3370 | 886 | 1769 |
| 3898 | W303.asm02.HP0_G0038930 | chrXII | 1950 | 438 | 624 | 1865 | 518 | 605 | 1840 | 456 | 633 | 2105 | 477 | 696 | 2313 | 281 | 519 | 1838 | 442 | 530 |
| 3899 | W303.asm02.HP0_G0038940 | chrXII | 3634 | 3580 | 3562 | 3403 | 2971 | 3580 | 3877 | 3608 | 3849 | 23043 | 5867 | 9127 | 14325 | 3954 | 5021 | 17837 | 5105 | 7410 |
| 3900 | W303.asm02.HP0_G0038950 | chrXII | 2782 | 675 | 707 | 2465 | 793 | 633 | 2607 | 706 | 653 | 3695 | 676 | 800 | 3452 | 454 | 622 | 3620 | 670 | 780 |
| 3901 | W303.asm02.HP0_G0038960 | chrXII | 1171 | 1826 | 2655 | 1113 | 2890 | 2533 | 1246 | 2073 | 3442 | 8184 | 2000 | 2022 | 8858 | 1091 | 1367 | 8462 | 1576 | 1794 |
| 3902 | W303.asm02.HP0_G0038970 | chrXII | 2251 | 595 | 832 | 1895 | 729 | 866 | 2138 | 606 | 927 | 5976 | 1086 | 1718 | 3988 | 713 | 1265 | 4247 | 1091 | 1489 |
| 3903 | W303.asm02.HP0_G0038980 | chrXII | 768 | 704 | 1005 | 580 | 1089 | 1062 | 640 | 768 | 1176 | 2570 | 593 | 769 | 3383 | 428 | 819 | 2586 | 506 | 648 |
| 3904 | W303.asm02.HP0_G0038990 | chrXII | 336 | 118 | 84 | 311 | 108 | 121 | 331 | 89 | 94 | 246 | 132 | 137 | 95 | 118 | 141 | 102 | 129 | 147 |

| | | | | | | | | | | | | | | | | | | | |
|---|---|---|---|---|---|---|---|---|---|---|---|---|---|---|---|---|---|---|---|
| 3905 | W303.asm02.HP0_G0039000 | chrXII | 2240 | 1826 | 2100 | 1856 | 1985 | 2154 | 1914 | 1999 | 2919 | 4107 | 1850 | 1623 | 5287 | 1258 | 1367 | 4405 | 1489 | 1471 |
| 3906 | W303.asm02.HP0_G0039010 | chrXII | 10029 | 449 | 505 | 8716 | 430 | 639 | 9249 | 343 | 671 | 3073 | 788 | 1219 | 875 | 661 | 745 | 714 | 620 | 945 |
| 3907 | W303.asm02.HP0_G0039020 | chrXII | 3675 | 565 | 596 | 3674 | 586 | 549 | 3607 | 599 | 602 | 4805 | 887 | 1252 | 2846 | 661 | 902 | 3347 | 779 | 1098 |
| 3908 | W303.asm02.HP0_G0039030 | chrXII | 24292 | 15345 | 20481 | 23792 | 18624 | 19887 | 25687 | 15793 | 25555 | 45553 | 14234 | 16454 | 48571 | 10552 | 13319 | 48259 | 12340 | 15083 |
| 3909 | W303.asm02.HP0_G0039040 | chrXII | 2679 | 471 | 565 | 2756 | 671 | 596 | 2701 | 493 | 719 | 4914 | 711 | 1118 | 4717 | 610 | 874 | 3643 | 725 | 1013 |
| 3910 | W303.asm02.HP0_G0039050 | chrXII | 1526 | 1026 | 1069 | 1426 | 1140 | 1094 | 1454 | 1031 | 1155 | 4632 | 1019 | 1076 | 6515 | 853 | 1051 | 4115 | 911 | 959 |
| 3911 | W303.asm02.HP0_G0039060 | chrXII | 18 | 12 | 10 | 20 | 2 | 7 | 22 | 11 | 19 | 28 | 12 | 7 | 40 | 8 | 5 | 33 | 13 | 13 |
| 3912 | W303.asm02.HP0_G0039070 | chrXII | 1084 | 835 | 589 | 1088 | 548 | 549 | 1083 | 839 | 643 | 2226 | 642 | 608 | 2104 | 432 | 412 | 2380 | 649 | 535 |
| 3913 | W303.asm02.HP0_G0039080 | chrXII | 11400 | 6329 | 5561 | 10602 | 5101 | 5627 | 11694 | 6061 | 6266 | 14549 | 4652 | 4669 | 15527 | 3573 | 3957 | 14807 | 4381 | 4223 |
| 3914 | W303.asm02.HP0_G0039090 | chrXII | 6078 | 3537 | 5472 | 5820 | 5454 | 5612 | 6940 | 3551 | 5605 | 16911 | 2867 | 4991 | 17261 | 2481 | 3695 | 16035 | 2627 | 4663 |
| 3915 | W303.asm02.HP0_G0039100 | chrXII | 5686 | 1739 | 1612 | 5479 | 1905 | 1606 | 6096 | 1823 | 1754 | 12637 | 2534 | 2756 | 13023 | 1900 | 2078 | 10650 | 2032 | 2502 |
| 3916 | W303.asm02.HP0_G0039110 | chrXII | 5023 | 833 | 913 | 4637 | 838 | 890 | 4901 | 809 | 948 | 6164 | 1268 | 1634 | 5066 | 932 | 1322 | 4814 | 1044 | 1582 |
| 3917 | W303.asm02.HP0_G0039120 | chrXII | 2970 | 348 | 407 | 2752 | 508 | 460 | 3058 | 355 | 490 | 4869 | 768 | 1079 | 2842 | 576 | 707 | 2859 | 643 | 985 |
| 3918 | W303.asm02.HP0_G0039130 | chrXII | 1178 | 1945 | 2344 | 1117 | 2459 | 2324 | 1215 | 1747 | 2012 | 9884 | 1439 | 1963 | 7830 | 776 | 1281 | 9633 | 1356 | 1770 |
| 3919 | W303.asm02.HP0_G0039140 | chrXII | 523 | 426 | 368 | 470 | 335 | 299 | 490 | 387 | 360 | 1470 | 277 | 334 | 1462 | 196 | 216 | 1393 | 254 | 256 |
| 3920 | W303.asm02.HP0_G0039150 | chrXII | 6906 | 854 | 1228 | 6820 | 1114 | 1394 | 6344 | 967 | 1648 | 7005 | 918 | 1942 | 5878 | 831 | 1759 | 5871 | 896 | 1717 |
| 3921 | W303.asm02.HP0_G0039160 | chrXII | 1004 | 953 | 1060 | 1091 | 1066 | 989 | 817 | 894 | 1174 | 3137 | 735 | 867 | 3499 | 500 | 684 | 3463 | 775 | 787 |
| 3922 | W303.asm02.HP0_G0039170 | chrXII | 488 | 314 | 368 | 446 | 348 | 358 | 423 | 306 | 457 | 847 | 177 | 232 | 949 | 128 | 214 | 943 | 194 | 186 |
| 3923 | W303.asm02.HP0_G0039180 | chrXII | 995 | 383 | 355 | 971 | 322 | 357 | 978 | 392 | 335 | 1297 | 353 | 572 | 1471 | 216 | 340 | 1270 | 367 | 493 |
| 3924 | W303.asm02.HP0_G0039190 | chrXII | 893 | 1338 | 1437 | 808 | 1273 | 1468 | 894 | 1327 | 1459 | 4467 | 1143 | 1302 | 4516 | 653 | 1110 | 4435 | 994 | 1269 |
| 3925 | W303.asm02.HP0_G0039200 | chrXII | 562 | 921 | 1140 | 520 | 783 | 926 | 460 | 807 | 1147 | 1195 | 962 | 687 | 1525 | 860 | 825 | 1306 | 775 | 540 |
| 3926 | W303.asm02.HP0_G0039210 | chrXII | 10 | 5 | 12 | 32 | 5 | 12 | 14 | 13 | 10 | 18 | 9 | 14 | 15 | 7 | 13 | 10 | 8 | 12 |
| 3927 | W303.asm02.HP0_G0039220 | chrXII | 32 | 16 | 19 | 37 | 19 | 27 | 37 | 10 | 12 | 26 | 13 | 7 | 21 | 11 | 15 | 19 | 13 | 18 |
| 3928 | W303.asm02.HP0_G0039230 | chrXII | 0 | 0 | 0 | 1 | 0 | 0 | 0 | 0 | 1 | 0 | 0 | 0 | 0 | 0 | 0 | 0 | 0 | 0 |
| 3929 | W303.asm02.HP0_G0039240 | chrXIII | 3486 | 1118 | 968 | 2935 | 803 | 985 | 2881 | 1211 | 1028 | 3632 | 1153 | 991 | 3316 | 567 | 689 | 3612 | 1138 | 976 |
| 3930 | W303.asm02.HP0_G0039250 | chrXIII | 3802 | 3048 | 2954 | 3553 | 2293 | 2587 | 3306 | 3072 | 3131 | 3494 | 3678 | 2646 | 5954 | 2705 | 2950 | 3460 | 3273 | 2099 |
| 3931 | W303.asm02.HP0_G0039260 | chrXIII | 4566 | 4014 | 11520 | 4481 | 4234 | 12902 | 4650 | 3628 | 11574 | 10905 | 3584 | 13238 | 12764 | 2309 | 8229 | 11282 | 3760 | 11033 |
| 3932 | W303.asm02.HP0_G0039270 | chrXIII | 4025 | 987 | 917 | 3589 | 891 | 998 | 3449 | 905 | 915 | 1882 | 747 | 1035 | 2495 | 642 | 847 | 1826 | 726 | 935 |
| 3933 | W303.asm02.HP0_G0039280 | chrXIII | 17017 | 1145 | 556 | 15901 | 520 | 696 | 15036 | 925 | 697 | 1556 | 1236 | 1027 | 2161 | 875 | 1000 | 1266 | 1405 | 861 |
| 3934 | W303.asm02.HP0_G0039290 | chrXIII | 2431 | 1046 | 1079 | 2171 | 928 | 1142 | 2483 | 1009 | 1185 | 6772 | 1644 | 2356 | 5284 | 983 | 1565 | 5301 | 1430 | 2051 |
| 3935 | W303.asm02.HP0_G0039300 | chrXIII | 9379 | 6606 | 4940 | 8033 | 4602 | 4992 | 10352 | 5678 | 4754 | 25399 | 9029 | 9633 | 19485 | 5586 | 5824 | 21115 | 7777 | 8714 |
| 3936 | W303.asm02.HP0_G0039310 | chrXIII | 3039 | 1807 | 2203 | 2659 | 2090 | 2245 | 3105 | 1855 | 2192 | 8060 | 1809 | 2481 | 7733 | 1286 | 1580 | 7346 | 1662 | 2259 |
| 3937 | W303.asm02.HP0_G0039320 | chrXIII | 6762 | 1621 | 1543 | 6385 | 1412 | 1780 | 7089 | 1474 | 2065 | 11073 | 2913 | 4133 | 7893 | 2075 | 2766 | 7952 | 2498 | 3802 |
| 3938 | W303.asm02.HP0_G0039330 | chrXIII | 1661 | 12223 | 16345 | 1729 | 16893 | 15212 | 1847 | 11947 | 14877 | 42744 | 14374 | 18315 | 49339 | 12678 | 13669 | 36243 | 11402 | 15695 |
| 3939 | W303.asm02.HP0_G0039340 | chrXIII | 29 | 66 | 42 | 13 | 30 | 25 | 25 | 59 | 48 | 74 | 47 | 33 | 111 | 25 | 20 | 107 | 48 | 26 |
| 3940 | W303.asm02.HP0_G0039350 | chrXIII | 2550 | 1387 | 1441 | 2199 | 1336 | 1308 | 2401 | 1395 | 1498 | 4450 | 1390 | 1485 | 4668 | 907 | 1215 | 4430 | 1372 | 1299 |
| 3941 | W303.asm02.HP0_G0039360 | chrXIII | 10049 | 2316 | 1647 | 10803 | 1729 | 1984 | 9604 | 1981 | 2015 | 8758 | 4150 | 3749 | 4884 | 3152 | 3050 | 4671 | 3715 | 2906 |
| 3942 | W303.asm02.HP0_G0039370 | chrXIII | 4129 | 1502 | 880 | 3641 | 1014 | 734 | 4042 | 1390 | 787 | 3614 | 1308 | 1078 | 4339 | 1051 | 900 | 3357 | 1357 | 930 |
| 3943 | W303.asm02.HP0_G0039380 | chrXIII | 1649 | 76 | 101 | 1876 | 76 | 97 | 1461 | 61 | 113 | 582 | 106 | 192 | 492 | 58 | 129 | 283 | 99 | 136 |
| 3944 | W303.asm02.HP0_G0039390 | chrXIII | 14971 | 760 | 910 | 15113 | 827 | 1150 | 14401 | 618 | 1198 | 7424 | 1528 | 2649 | 4863 | 1500 | 2455 | 3369 | 1357 | 2273 |
| 3945 | W303.asm02.HP0_G0039400 | chrXIII | 696 | 2028 | 2893 | 797 | 2042 | 2487 | 690 | 2241 | 3227 | 3504 | 2914 | 3054 | 2804 | 1475 | 2249 | 3284 | 2387 | 2298 |
| 3946 | W303.asm02.HP0_G0039410 | chrXIII | 1565 | 1277 | 1267 | 1498 | 1098 | 1264 | 1434 | 1261 | 1461 | 7178 | 1732 | 2206 | 4534 | 1060 | 1435 | 5424 | 1615 | 2077 |
| 3947 | W303.asm02.HP0_G0039420 | chrXIII | 2088 | 360 | 413 | 2066 | 373 | 444 | 2277 | 267 | 523 | 3785 | 853 | 1019 | 2366 | 646 | 620 | 2103 | 789 | 915 |
| 3948 | W303.asm02.HP0_G0039430 | chrXIII | 638 | 942 | 982 | 583 | 1140 | 890 | 674 | 1023 | 1044 | 4637 | 1153 | 1206 | 4845 | 828 | 802 | 3836 | 967 | 1030 |
| 3949 | W303.asm02.HP0_G0039440 | chrXIII | 1477 | 326 | 333 | 1540 | 327 | 362 | 1541 | 306 | 389 | 2069 | 452 | 658 | 2042 | 360 | 444 | 1647 | 419 | 572 |
| 3950 | W303.asm02.HP0_G0039450 | chrXIII | 2119 | 606 | 922 | 2055 | 947 | 951 | 2129 | 658 | 1149 | 8669 | 1066 | 1785 | 7637 | 877 | 1460 | 6654 | 982 | 1633 |
| 3951 | W303.asm02.HP0_G0039460 | chrXIII | 12520 | 4232 | 3906 | 11470 | 3437 | 3846 | 11947 | 4288 | 4397 | 9682 | 3990 | 4231 | 11267 | 3329 | 3649 | 9362 | 3756 | 3860 |
| 3952 | W303.asm02.HP0_G0039470 | chrXIII | 1041 | 219 | 215 | 1067 | 212 | 222 | 1153 | 176 | 241 | 2851 | 522 | 674 | 1667 | 393 | 413 | 1511 | 473 | 614 |
| 3953 | W303.asm02.HP0_G0039480 | chrXIII | 379 | 776 | 1183 | 321 | 1056 | 1200 | 335 | 858 | 1638 | 2934 | 749 | 1057 | 3374 | 455 | 596 | 3261 | 656 | 865 |
| 3954 | W303.asm02.HP0_G0039490 | chrXIII | 941 | 689 | 700 | 828 | 614 | 615 | 883 | 661 | 732 | 2105 | 522 | 555 | 2208 | 343 | 440 | 2156 | 555 | 470 |
| 3955 | W303.asm02.HP0_G0039500 | chrXIII | 6530 | 4793 | 6285 | 6336 | 6187 | 6034 | 7102 | 5536 | 6929 | 18034 | 5406 | 6526 | 18428 | 3687 | 5098 | 17205 | 4341 | 5593 |
| 3956 | W303.asm02.HP0_G0039510 | chrXIII | 4976 | 2268 | 2800 | 4244 | 2594 | 2698 | 4678 | 2372 | 2969 | 7972 | 2119 | 2628 | 8363 | 1331 | 2070 | 8172 | 1805 | 2318 |
| 3957 | W303.asm02.HP0_G0039520 | chrXIII | 1260 | 536 | 634 | 1202 | 662 | 593 | 1225 | 642 | 604 | 4723 | 740 | 1052 | 3644 | 629 | 780 | 3975 | 684 | 1013 |
| 3958 | W303.asm02.HP0_G0039530 | chrXIII | 4291 | 1201 | 1702 | 4100 | 1687 | 1695 | 4119 | 1439 | 2073 | 11061 | 1367 | 2638 | 9094 | 1002 | 1938 | 9905 | 1233 | 2325 |
| 3959 | W303.asm02.HP0_G0039540 | chrXIII | 1480 | 616 | 571 | 1315 | 474 | 688 | 1652 | 470 | 783 | 2613 | 959 | 1250 | 1145 | 705 | 768 | 1156 | 970 | 1111 |
| 3960 | W303.asm02.HP0_G0039550 | chrXIII | 693 | 449 | 547 | 644 | 472 | 542 | 587 | 456 | 728 | 1122 | 364 | 392 | 1490 | 264 | 346 | 1158 | 399 | 404 |
| 3961 | W303.asm02.HP0_G0039560 | chrXIII | 27364 | 746 | 626 | 21097 | 467 | 798 | 22195 | 532 | 804 | 1741 | 927 | 1078 | 1016 | 712 | 1170 | 995 | 753 | 816 |
| 3962 | W303.asm02.HP0_G0039570 | chrXIII | 2152 | 403 | 499 | 2198 | 545 | 492 | 2109 | 403 | 600 | 4219 | 713 | 1125 | 2435 | 535 | 697 | 2528 | 683 | 1018 |
| 3963 | W303.asm02.HP0_G0039580 | chrXIII | 1205 | 1180 | 1149 | 1105 | 1317 | 1267 | 1210 | 1338 | 1509 | 3469 | 1127 | 1084 | 4498 | 737 | 1016 | 3409 | 927 | 1071 |
| 3964 | W303.asm02.HP0_G0039590 | chrXIII | 2160 | 1342 | 2233 | 2080 | 1883 | 2152 | 2071 | 1500 | 2331 | 6339 | 1168 | 2118 | 6818 | 745 | 1423 | 6295 | 1083 | 1826 |
| 3965 | W303.asm02.HP0_G0039600 | chrXIII | 1123 | 363 | 454 | 1160 | 436 | 425 | 1041 | 330 | 411 | 1572 | 319 | 525 | 1523 | 245 | 360 | 1478 | 260 | 401 |
| 3966 | W303.asm02.HP0_G0039610 | chrXIII | 2442 | 1344 | 1646 | 2201 | 1602 | 1769 | 2273 | 1411 | 2047 | 4645 | 1241 | 1567 | 5206 | 765 | 1191 | 4536 | 1147 | 1306 |
| 3967 | W303.asm02.HP0_G0039620 | chrXIII | 3097 | 1162 | 1172 | 3460 | 1563 | 1176 | 3827 | 1182 | 1238 | 16371 | 2258 | 3315 | 10862 | 1566 | 1611 | 9849 | 1905 | 2747 |
| 3968 | W303.asm02.HP0_G0039630 | chrXIII | 8875 | 3450 | 3786 | 8319 | 3252 | 3492 | 8377 | 3597 | 3605 | 9257 | 3346 | 4222 | 10173 | 2531 | 3528 | 9384 | 3013 | 3717 |
| 3969 | W303.asm02.HP0_G0039640 | chrXIII | 2551 | 1188 | 2362 | 3078 | 2305 | 2294 | 2556 | 1408 | 2827 | 13866 | 2045 | 2996 | 12478 | 1697 | 2704 | 12542 | 1684 | 2282 |
| 3970 | W303.asm02.HP0_G0039650 | chrXIII | 42 | 11 | 11 | 26 | 11 | 10 | 29 | 13 | 7 | 71 | 15 | 16 | 78 | 8 | 6 | 70 | 10 | 8 |
| 3971 | W303.asm02.HP0_G0039660 | chrXIII | 2087 | 1334 | 1376 | 1934 | 1432 | 1511 | 2250 | 1318 | 1438 | 7732 | 1614 | 2260 | 7801 | 1127 | 1525 | 6310 | 1396 | 1860 |
| 3972 | W303.asm02.HP0_G0039670 | chrXIII | 359 | 364 | 257 | 354 | 325 | 204 | 329 | 346 | 264 | 1275 | 386 | 336 | 1320 | 295 | 287 | 911 | 389 | 236 |
| 3973 | W303.asm02.HP0_G0039680 | chrXIII | 7477 | 1334 | 1251 | 6185 | 1183 | 1418 | 6854 | 1238 | 1543 | 10199 | 2692 | 3143 | 6224 | 2081 | 2291 | 6588 | 2490 | 2784 |
| 3974 | W303.asm02.HP0_G0039690 | chrXIII | 11833 | 1960 | 1604 | 11308 | 1467 | 1789 | 12436 | 1567 | 2035 | 11661 | 3558 | 4070 | 6422 | 2958 | 3056 | 6629 | 3190 | 3669 |
| 3975 | W303.asm02.HP0_G0039700 | chrXIII | 293 | 101 | 76 | 255 | 77 | 75 | 258 | 93 | 67 | 282 | 138 | 91 | 197 | 95 | 56 | 213 | 146 | 84 |
| 3976 | W303.asm02.HP0_G0039710 | chrXIII | 1304 | 320 | 427 | 1097 | 471 | 517 | 1223 | 246 | 560 | 3904 | 640 | 1026 | 1331 | 502 | 507 | 1271 | 505 | 731 |
| 3977 | W303.asm02.HP0_G0039720 | chrXIII | 1284 | 1395 | 1950 | 1065 | 1827 | 1991 | 1071 | 1520 | 2496 | 3669 | 1367 | 1724 | 4705 | 1065 | 1678 | 4098 | 1242 | 1435 |
| 3978 | W303.asm02.HP0_G0039730 | chrXIII | 1528 | 512 | 803 | 1568 | 828 | 892 | 1606 | 606 | 1057 | 7537 | 918 | 1657 | 4334 | 659 | 1151 | 4586 | 871 | 1481 |
| 3979 | W303.asm02.HP0_G0039740 | chrXIII | 711 | 1392 | 1188 | 826 | 1526 | 1139 | 931 | 1458 | 1168 | 7301 | 1562 | 1737 | 6422 | 1047 | 967 | 6365 | 1322 | 1496 |
| 3980 | W303.asm02.HP0_G0039750 | chrXIII | 3716 | 1506 | 1812 | 3484 | 1549 | 1851 | 3290 | 1646 | 2375 | 4756 | 1598 | 1894 | 5915 | 1189 | 1676 | 4989 | 1464 | 1774 |
| 3981 | W303.asm02.HP0_G0039760 | chrXIII | 6169 | 3396 | 3678 | 5109 | 3143 | 3545 | 5618 | 3293 | 4567 | 7746 | 2837 | 3294 | 9518 | 2326 | 2704 | 7771 | 2777 | 2669 |
| 3982 | W303.asm02.HP0_G0039770 | chrXIII | 820 | 1144 | 1438 | 666 | 1236 | 1437 | 785 | 1187 | 1821 | 3689 | 951 | 1024 | 4074 | 599 | 831 | 3923 | 768 | 961 |
| 3983 | W303.asm02.HP0_G0039780 | chrXIII | 1845 | 365 | 452 | 1767 | 380 | 467 | 1823 | 361 | 517 | 3191 | 628 | 1051 | 2522 | 450 | 751 | 2337 | 590 | 843 |
| 3984 | W303.asm02.HP0_G0039790 | chrXIII | 4984 | 1872 | 1742 | 4909 | 1794 | 1940 | 5899 | 1558 | 1889 | 18898 | 3379 | 4197 | 11746 | 2167 | 3036 | 12307 | 2949 | 3897 |
| 3985 | W303.asm02.HP0_G0039800 | chrXIII | 13351 | 3270 | 2341 | 11273 | 2461 | 2538 | 12773 | 3282 | 1900 | 14129 | 3826 | 5085 | 12356 | 2993 | 3383 | 11683 | 3385 | 4106 |
| 3986 | W303.asm02.HP0_G0039810 | chrXIII | 7012 | 6509 | 7645 | 6520 | 7521 | 8162 | 7467 | 6367 | 8623 | 19081 | 5390 | 6259 | 19202 | 4121 | 5104 | 19002 | 5019 | 5909 |
| 3987 | W303.asm02.HP0_G0039820 | chrXIII | 18270 | 1929 | 2182 | 16620 | 2080 | 2695 | 17591 | 1889 | 2523 | 17072 | 3192 | 4422 | 12036 | 2728 | 3984 | 11143 | 2766 | 4135 |
| 3988 | W303.asm02.HP0_G0039830 | chrXIII | 2764 | 966 | 1231 | 2622 | 1128 | 1112 | 2781 | 1112 | 1058 | 5871 | 1009 | 1550 | 7171 | 763 | 1268 | 5386 | 946 | 1482 |
| 3989 | W303.asm02.HP0_G0039840 | chrXIII | 7309 | 1286 | 898 | 6479 | 767 | 1018 | 6623 | 1048 | 1177 | 6510 | 2040 | 2487 | 3991 | 1663 | 1772 | 3783 | 2135 | 2101 |
| 3990 | W303.asm02.HP0_G0039850 | chrXIII | 7700 | 2209 | 2172 | 7263 | 1964 | 2042 | 7375 | 2165 | 2384 | 12050 | 2977 | 3888 | 9529 | 2070 | 2955 | 9947 | 2629 | 3517 |
| 3991 | W303.asm02.HP0_G0039860 | chrXIII | 1832 | 651 | 776 | 1778 | 690 | 698 | 1727 | 645 | 759 | 2639 | 626 | 736 | 2899 | 408 | 664 | 2421 | 573 | 666 |
| 3992 | W303.asm02.HP0_G0039870 | chrXIII | 2253 | 2007 | 2185 | 2298 | 1897 | 2268 | 2275 | 2032 | 2988 | 5645 | 1510 | 1647 | 7004 | 1105 | 1270 | 6241 | 1382 | 1504 |
| 3993 | W303.asm02.HP0_G0039880 | chrXIII | 294 | 174 | 110 | 263 | 135 | 128 | 278 | 136 | 99 | 540 | 138 | 133 | 730 | 91 | 86 | 435 | 124 | 111 |
| 3994 | W303.asm02.HP0_G0039890 | chrXIII | 2107 | 487 | 594 | 2133 | 717 | 638 | 2239 | 525 | 686 | 6305 | 881 | 1233 | 3979 | 617 | 897 | 4400 | 754 | 1138 |
| 3995 | W303.asm02.HP0_G0039900 | chrXIII | 2080 | 957 | 707 | 1968 | 794 | 653 | 1993 | 1003 | 731 | 2702 | 971 | 837 | 3326 | 743 | 803 | 2821 | 857 | 785 |
| 3996 | W303.asm02.HP0_G0039910 | chrXIII | 15833 | 12298 | 16001 | 14860 | 14782 | 15861 | 16679 | 12584 | 18580 | 38528 | 11936 | 14328 | 40359 | 8960 | 11239 | 39730 | 10413 | 12986 |
| 3997 | W303.asm02.HP0_G0039920 | chrXIII | 2581 | 1289 | 1332 | 2338 | 1437 | 1366 | 2383 | 1288 | 1336 | 5106 | 1044 | 1308 | 5709 | 662 | 1060 | 4683 | 943 | 1085 |

| | | | | | | | | | | | | | | | | | | | | |
|---|---|---|---|---|---|---|---|---|---|---|---|---|---|---|---|---|---|---|---|---|
| 3998 | W303.asm02.HP0_G0039930 | chrXIII | 2468 | 700 | 647 | 2102 | 758 | 689 | 2369 | 666 | 661 | 5201 | 930 | 1203 | 4170 | 688 | 852 | 3938 | 813 | 1063 |
| 3999 | W303.asm02.HP0_G0039940 | chrXIII | 315 | 509 | 673 | 263 | 739 | 600 | 332 | 539 | 637 | 3179 | 646 | 828 | 3329 | 356 | 480 | 3036 | 513 | 693 |
| 4000 | W303.asm02.HP0_G0039950 | chrXIII | 1561 | 605 | 805 | 1223 | 678 | 894 | 1481 | 551 | 1018 | 5795 | 1069 | 1792 | 2236 | 670 | 1210 | 2930 | 1067 | 1681 |
| 4001 | W303.asm02.HP0_G0039960 | chrXIII | 226 | 124 | 133 | 149 | 124 | 152 | 153 | 129 | 160 | 212 | 102 | 98 | 355 | 56 | 72 | 291 | 100 | 90 |
| 4002 | W303.asm02.HP0_G0039970 | chrXIII | 5353 | 6046 | 5315 | 5206 | 5207 | 5362 | 5345 | 6047 | 5445 | 10422 | 4984 | 4879 | 11301 | 3751 | 4496 | 10317 | 4495 | 4511 |
| 4003 | W303.asm02.HP0_G0039980 | chrXIII | 7684 | 783 | 791 | 7383 | 679 | 962 | 7453 | 584 | 1080 | 5020 | 1571 | 1897 | 3311 | 1396 | 1677 | 2165 | 1356 | 1727 |
| 4004 | W303.asm02.HP0_G0039990 | chrXIII | 4373 | 3484 | 4417 | 3426 | 4407 | 4189 | 4775 | 3532 | 4821 | 29860 | 6001 | 7898 | 18806 | 3568 | 4303 | 24138 | 5420 | 6696 |
| 4005 | W303.asm02.HP0_G0040000 | chrXIII | 1284 | 1084 | 1578 | 999 | 1209 | 1525 | 1067 | 1144 | 2167 | 3517 | 876 | 1193 | 4400 | 629 | 932 | 3965 | 736 | 1141 |
| 4006 | W303.asm02.HP0_G0040010 | chrXIII | 6309 | 519 | 359 | 7600 | 299 | 415 | 5943 | 427 | 416 | 1159 | 727 | 734 | 1050 | 544 | 598 | 850 | 776 | 556 |
| 4007 | W303.asm02.HP0_G0040020 | chrXIII | 116 | 81 | 42 | 100 | 28 | 37 | 87 | 67 | 38 | 84 | 66 | 35 | 67 | 40 | 23 | 81 | 76 | 33 |
| 4008 | W303.asm02.HP0_G0040030 | chrXIII | 1011 | 1217 | 947 | 996 | 1082 | 1005 | 865 | 1321 | 1184 | 2689 | 972 | 732 | 3151 | 520 | 632 | 2858 | 982 | 692 |
| 4009 | W303.asm02.HP0_G0040040 | chrXIII | 2399 | 3406 | 2588 | 1967 | 2512 | 2485 | 2096 | 3564 | 2653 | 8261 | 3925 | 3585 | 6080 | 2363 | 2790 | 7093 | 3525 | 3281 |
| 4010 | W303.asm02.HP0_G0040050 | chrXIII | 2378 | 896 | 796 | 2033 | 750 | 765 | 2293 | 837 | 794 | 4698 | 1225 | 1383 | 4345 | 1020 | 1076 | 3914 | 1208 | 1156 |
| 4011 | W303.asm02.HP0_G0040060 | chrXIII | 1745 | 632 | 638 | 1651 | 600 | 617 | 1510 | 598 | 721 | 1933 | 489 | 623 | 2121 | 331 | 518 | 1919 | 474 | 503 |
| 4012 | W303.asm02.HP0_G0040070 | chrXIII | 2938 | 427 | 731 | 2920 | 690 | 729 | 2930 | 416 | 862 | 6390 | 845 | 1683 | 3613 | 632 | 1099 | 3445 | 692 | 1474 |
| 4013 | W303.asm02.HP0_G0040080 | chrXIII | 12786 | 4245 | 3771 | 12488 | 3483 | 3496 | 12669 | 4331 | 3941 | 16221 | 5681 | 5636 | 13455 | 3903 | 4855 | 13847 | 4824 | 5118 |
| 4014 | W303.asm02.HP0_G0040090 | chrXIII | 27 | 4 | 10 | 14 | 8 | 10 | 13 | 4 | 6 | 41 | 14 | 10 | 28 | 4 | 7 | 28 | 10 | 4 |
| 4015 | W303.asm02.HP0_G0040100 | chrXIII | 1056 | 962 | 1475 | 932 | 1255 | 1320 | 1058 | 1016 | 1525 | 6705 | 1069 | 1522 | 5862 | 591 | 997 | 6782 | 1055 | 1355 |
| 4016 | W303.asm02.HP0_G0040110 | chrXIII | 169 | 368 | 908 | 160 | 1059 | 834 | 184 | 436 | 958 | 5059 | 490 | 1040 | 4703 | 349 | 556 | 4470 | 522 | 755 |
| 4017 | W303.asm02.HP0_G0040120 | chrXIII | 2074 | 180 | 136 | 2054 | 171 | 175 | 1937 | 165 | 166 | 900 | 236 | 281 | 862 | 160 | 188 | 709 | 229 | 244 |
| 4018 | W303.asm02.HP0_G0040130 | chrXIII | 1222 | 981 | 831 | 1228 | 791 | 745 | 1059 | 894 | 853 | 2004 | 668 | 674 | 2357 | 394 | 482 | 2254 | 736 | 584 |
| 4019 | W303.asm02.HP0_G0040140 | chrXIII | 1171 | 1062 | 1132 | 1294 | 1212 | 1118 | 1155 | 1089 | 1145 | 6147 | 1595 | 1652 | 5497 | 970 | 1283 | 5075 | 1443 | 1524 |
| 4020 | W303.asm02.HP0_G0040150 | chrXIII | 356 | 110 | 93 | 369 | 107 | 111 | 344 | 94 | 119 | 1016 | 222 | 236 | 761 | 148 | 163 | 734 | 200 | 205 |
| 4021 | W303.asm02.HP0_G0040160 | chrXIII | 1063 | 626 | 1062 | 917 | 949 | 1041 | 986 | 685 | 1382 | 2364 | 549 | 757 | 3013 | 380 | 588 | 2783 | 461 | 667 |
| 4022 | W303.asm02.HP0_G0040170 | chrXIII | 5997 | 856 | 1015 | 5350 | 1045 | 1075 | 5675 | 821 | 1358 | 9505 | 1800 | 2277 | 5428 | 1395 | 1678 | 5156 | 1551 | 1991 |
| 4023 | W303.asm02.HP0_G0040180 | chrXIII | 5540 | 583 | 960 | 5194 | 835 | 1148 | 5567 | 553 | 1251 | 5866 | 1158 | 2026 | 4510 | 832 | 1613 | 4179 | 1048 | 1816 |
| 4024 | W303.asm02.HP0_G0040190 | chrXIII | 2733 | 773 | 725 | 2718 | 711 | 790 | 2669 | 612 | 808 | 4969 | 1111 | 1491 | 3582 | 843 | 1142 | 3367 | 1000 | 1294 |
| 4025 | W303.asm02.HP0_G0040200 | chrXIII | 1688 | 730 | 956 | 1682 | 863 | 994 | 1754 | 625 | 1049 | 7288 | 1556 | 2113 | 4065 | 1116 | 1339 | 3968 | 1270 | 1933 |
| 4026 | W303.asm02.HP0_G0040210 | chrXIII | 4930 | 1414 | 1785 | 4537 | 1403 | 1766 | 4309 | 1507 | 2471 | 4081 | 1269 | 1669 | 5414 | 1003 | 1488 | 4181 | 1191 | 1501 |
| 4027 | W303.asm02.HP0_G0040220 | chrXIII | 2432 | 454 | 641 | 2369 | 355 | 763 | 2464 | 372 | 802 | 3617 | 806 | 1684 | 1952 | 560 | 996 | 1905 | 771 | 1446 |
| 4028 | W303.asm02.HP0_G0040230 | chrXIII | 18132 | 13182 | 16853 | 16986 | 12160 | 17983 | 17712 | 12996 | 18687 | 21709 | 10966 | 16381 | 23242 | 8090 | 12026 | 20922 | 9628 | 14133 |
| 4029 | W303.asm02.HP0_G0040240 | chrXIII | 381 | 372 | 383 | 343 | 300 | 438 | 417 | 258 | 475 | 1693 | 731 | 907 | 1154 | 512 | 634 | 1127 | 628 | 898 |
| 4030 | W303.asm02.HP0_G0040250 | chrXIII | 11478 | 8488 | 13024 | 11155 | 12385 | 12508 | 11856 | 8866 | 16220 | 28203 | 8531 | 10685 | 30859 | 6212 | 8992 | 29665 | 7391 | 9396 |
| 4031 | W303.asm02.HP0_G0040260 | chrXIII | 6252 | 973 | 771 | 6390 | 733 | 914 | 6058 | 817 | 1040 | 4099 | 1392 | 1681 | 2606 | 1137 | 1228 | 2324 | 1307 | 1484 |
| 4032 | W303.asm02.HP0_G0040270 | chrXIII | 6931 | 4660 | 7066 | 5866 | 6410 | 6782 | 6566 | 4773 | 8514 | 14969 | 4225 | 5780 | 17024 | 3211 | 4991 | 16306 | 3968 | 4936 |
| 4033 | W303.asm02.HP0_G0040280 | chrXIII | 1397 | 1166 | 1157 | 1273 | 1197 | 1008 | 1515 | 1092 | 1096 | 4841 | 1035 | 935 | 4682 | 635 | 708 | 4363 | 1022 | 884 |
| 4034 | W303.asm02.HP0_G0040290 | chrXIII | 3627 | 3651 | 4588 | 2907 | 4333 | 4445 | 3513 | 3726 | 6345 | 10048 | 3264 | 3572 | 11187 | 1984 | 2604 | 10721 | 2750 | 3039 |
| 4035 | W303.asm02.HP0_G0040300 | chrXIII | 1197 | 926 | 1007 | 1015 | 846 | 934 | 1103 | 865 | 1068 | 3337 | 811 | 882 | 3554 | 500 | 698 | 3532 | 732 | 816 |
| 4036 | W303.asm02.HP0_G0040310 | chrXIII | 2032 | 485 | 491 | 2109 | 469 | 498 | 1900 | 406 | 571 | 3244 | 745 | 984 | 1984 | 491 | 705 | 1929 | 802 | 857 |
| 4037 | W303.asm02.HP0_G0040320 | chrXIII | 2235 | 1574 | 1908 | 2053 | 1655 | 1785 | 2069 | 1563 | 2079 | 5167 | 1294 | 1408 | 5608 | 902 | 1022 | 5600 | 1236 | 1376 |
| 4038 | W303.asm02.HP0_G0040330 | chrXIII | 488 | 1288 | 1721 | 413 | 1965 | 1679 | 481 | 1420 | 1716 | 7400 | 1117 | 1472 | 7294 | 732 | 1002 | 6835 | 1042 | 1256 |
| 4039 | W303.asm02.HP0_G0040340 | chrXIII | 1015 | 267 | 373 | 1034 | 369 | 476 | 1007 | 246 | 510 | 2664 | 442 | 779 | 1388 | 330 | 564 | 1514 | 389 | 681 |
| 4040 | W303.asm02.HP0_G0040350 | chrXIII | 5273 | 1777 | 1712 | 5222 | 1698 | 1495 | 5367 | 1773 | 1773 | 10291 | 2489 | 2509 | 14497 | 1864 | 2174 | 9253 | 2351 | 2305 |
| 4041 | W303.asm02.HP0_G0040360 | chrXIII | 2209 | 909 | 748 | 1978 | 800 | 767 | 2111 | 841 | 698 | 3754 | 885 | 1173 | 4173 | 683 | 772 | 3217 | 848 | 950 |
| 4042 | W303.asm02.HP0_G0040370 | chrXIII | 1544 | 547 | 416 | 1433 | 432 | 435 | 1623 | 523 | 526 | 2770 | 859 | 898 | 1858 | 564 | 468 | 1913 | 807 | 735 |
| 4043 | W303.asm02.HP0_G0040380 | chrXIII | 3712 | 423 | 485 | 3281 | 371 | 610 | 3599 | 322 | 556 | 4073 | 962 | 1267 | 2231 | 751 | 935 | 2066 | 848 | 1119 |
| 4044 | W303.asm02.HP0_G0040390 | chrXIII | 5324 | 4468 | 5344 | 4685 | 4188 | 5465 | 5282 | 4534 | 6224 | 10405 | 3870 | 4782 | 11996 | 3062 | 3809 | 10109 | 3433 | 4427 |
| 4045 | W303.asm02.HP0_G0040400 | chrXIII | 717 | 628 | 670 | 526 | 528 | 677 | 559 | 611 | 755 | 1777 | 508 | 606 | 1974 | 341 | 432 | 1905 | 478 | 454 |
| 4046 | W303.asm02.HP0_G0040410 | chrXIII | 9352 | 1465 | 1581 | 8753 | 1576 | 1660 | 9187 | 1235 | 2250 | 9169 | 3132 | 3815 | 4083 | 2181 | 2175 | 4172 | 2626 | 3391 |
| 4047 | W303.asm02.HP0_G0040420 | chrXIII | 2488 | 869 | 1513 | 2211 | 1407 | 1501 | 2099 | 995 | 1919 | 2873 | 779 | 1291 | 3892 | 590 | 1199 | 3051 | 703 | 1127 |
| 4048 | W303.asm02.HP0_G0040430 | chrXIII | 10181 | 5450 | 4527 | 10544 | 4289 | 4595 | 11738 | 4346 | 5116 | 21297 | 6495 | 6924 | 13041 | 5159 | 4956 | 14482 | 6149 | 6408 |
| 4049 | W303.asm02.HP0_G0040440 | chrXIII | 10609 | 2763 | 2644 | 10869 | 2832 | 2771 | 10877 | 2757 | 2844 | 13162 | 3492 | 4342 | 11902 | 2428 | 3631 | 11004 | 3120 | 3921 |
| 4050 | W303.asm02.HP0_G0040450 | chrXIII | 919 | 348 | 444 | 1219 | 501 | 496 | 945 | 312 | 503 | 4793 | 723 | 1023 | 3472 | 409 | 651 | 3372 | 698 | 856 |
| 4051 | W303.asm02.HP0_G0040460 | chrXIII | 853 | 1027 | 969 | 890 | 894 | 941 | 924 | 916 | 1039 | 3931 | 955 | 925 | 4278 | 548 | 542 | 3794 | 984 | 796 |
| 4052 | W303.asm02.HP0_G0040470 | chrXIII | 11242 | 3225 | 2420 | 10583 | 2099 | 2459 | 10436 | 2876 | 2320 | 8286 | 2976 | 3100 | 8455 | 2127 | 2566 | 7434 | 2935 | 2983 |
| 4053 | W303.asm02.HP0_G0040480 | chrXIII | 2329 | 848 | 735 | 2301 | 677 | 783 | 2132 | 820 | 782 | 3845 | 947 | 996 | 3360 | 522 | 766 | 3688 | 1041 | 883 |
| 4054 | W303.asm02.HP0_G0040490 | chrXIII | 2653 | 2046 | 2618 | 2282 | 2161 | 2619 | 2417 | 1910 | 3354 | 5934 | 1710 | 2280 | 6454 | 1220 | 1746 | 6390 | 1734 | 1997 |
| 4055 | W303.asm02.HP0_G0040500 | chrXIII | 1422 | 964 | 855 | 1391 | 935 | 819 | 1474 | 1016 | 876 | 7263 | 1656 | 1879 | 6043 | 1197 | 1593 | 6367 | 1482 | 1799 |
| 4056 | W303.asm02.HP0_G0040510 | chrXIII | 68 | 69 | 39 | 68 | 39 | 34 | 72 | 66 | 42 | 567 | 96 | 109 | 354 | 86 | 60 | 511 | 95 | 95 |
| 4057 | W303.asm02.HP0_G0040520 | chrXIII | 6586 | 5310 | 4972 | 6188 | 4718 | 4674 | 6099 | 5764 | 5290 | 13997 | 7104 | 7666 | 14118 | 4911 | 6005 | 13781 | 6643 | 6597 |
| 4058 | W303.asm02.HP0_G0040530 | chrXIII | 414 | 715 | 777 | 365 | 775 | 769 | 381 | 754 | 984 | 1920 | 639 | 621 | 2579 | 459 | 572 | 2228 | 564 | 552 |
| 4059 | W303.asm02.HP0_G0040540 | chrXIII | 5113 | 981 | 765 | 5072 | 799 | 844 | 4955 | 923 | 928 | 5357 | 1473 | 1602 | 4314 | 885 | 1241 | 4586 | 1287 | 1503 |
| 4060 | W303.asm02.HP0_G0040550 | chrXIII | 3772 | 1575 | 1219 | 3687 | 1369 | 1318 | 3777 | 1667 | 1143 | 6159 | 1793 | 2265 | 5735 | 1250 | 1483 | 5366 | 1620 | 1808 |
| 4061 | W303.asm02.HP0_G0040560 | chrXIII | 12237 | 1501 | 1508 | 10891 | 1338 | 1812 | 12526 | 1297 | 1902 | 16129 | 4106 | 4296 | 10783 | 3839 | 3765 | 8581 | 3290 | 3730 |
| 4062 | W303.asm02.HP0_G0040570 | chrXIII | 3217 | 1670 | 1904 | 3011 | 1587 | 1785 | 3149 | 1627 | 1925 | 3497 | 1192 | 1353 | 3985 | 830 | 1125 | 3441 | 1146 | 1200 |
| 4063 | W303.asm02.HP0_G0040580 | chrXIII | 885 | 1812 | 2537 | 834 | 2329 | 2338 | 870 | 1830 | 2509 | 9860 | 1961 | 2493 | 11627 | 1477 | 2088 | 9472 | 1781 | 2285 |
| 4064 | W303.asm02.HP0_G0040590 | chrXIII | 13431 | 4316 | 3085 | 19413 | 4316 | 2983 | 13776 | 4152 | 3376 | 31963 | 6413 | 5350 | 30526 | 4554 | 3761 | 24562 | 5111 | 4841 |
| 4065 | W303.asm02.HP0_G0040600 | chrXIII | 21484 | 3050 | 3207 | 21975 | 3322 | 3524 | 21790 | 3059 | 3639 | 31610 | 5737 | 8520 | 22336 | 4834 | 6450 | 21907 | 4758 | 7486 |
| 4066 | W303.asm02.HP0_G0040610 | chrXIII | 452 | 709 | 1106 | 349 | 718 | 1190 | 369 | 727 | 1274 | 2896 | 693 | 1400 | 2742 | 434 | 969 | 3097 | 671 | 1318 |
| 4067 | W303.asm02.HP0_G0040620 | chrXIII | 1242 | 1609 | 1827 | 1258 | 2271 | 1675 | 1557 | 1720 | 1665 | 10314 | 1854 | 2241 | 9754 | 1159 | 1409 | 8709 | 1642 | 1959 |
| 4068 | W303.asm02.HP0_G0040630 | chrXIII | 1921 | 5521 | 5233 | 2416 | 4815 | 4738 | 2930 | 4958 | 4573 | 18366 | 4422 | 4944 | 14381 | 2891 | 3738 | 16783 | 4034 | 4400 |
| 4069 | W303.asm02.HP0_G0040640 | chrXIII | 2406 | 339 | 585 | 2753 | 593 | 648 | 2666 | 293 | 727 | 6343 | 990 | 1461 | 1904 | 700 | 914 | 1964 | 757 | 1279 |
| 4070 | W303.asm02.HP0_G0040650 | chrXIII | 269 | 86 | 70 | 231 | 74 | 59 | 217 | 64 | 69 | 204 | 71 | 57 | 233 | 42 | 56 | 139 | 57 | 49 |
| 4071 | W303.asm02.HP0_G0040660 | chrXIII | 115 | 54 | 78 | 121 | 91 | 72 | 117 | 55 | 92 | 322 | 61 | 108 | 248 | 29 | 59 | 333 | 44 | 99 |
| 4072 | W303.asm02.HP0_G0040670 | chrXIII | 993 | 417 | 487 | 1039 | 499 | 467 | 1032 | 382 | 435 | 3761 | 705 | 944 | 2664 | 471 | 646 | 2762 | 577 | 869 |
| 4073 | W303.asm02.HP0_G0040680 | chrXIII | 2919 | 676 | 814 | 2805 | 752 | 766 | 2671 | 683 | 837 | 2620 | 767 | 870 | 3002 | 458 | 641 | 2474 | 682 | 735 |
| 4074 | W303.asm02.HP0_G0040690 | chrXIII | 1457 | 967 | 786 | 1674 | 911 | 851 | 1258 | 872 | 792 | 3059 | 828 | 839 | 3154 | 468 | 573 | 2954 | 755 | 715 |
| 4075 | W303.asm02.HP0_G0040700 | chrXIII | 3111 | 1468 | 1962 | 2788 | 1537 | 2085 | 2678 | 1466 | 2428 | 3774 | 1121 | 1635 | 4567 | 769 | 1299 | 4164 | 1033 | 1374 |
| 4076 | W303.asm02.HP0_G0040710 | chrXIII | 1497 | 661 | 596 | 1400 | 612 | 655 | 1363 | 614 | 710 | 2037 | 501 | 499 | 2155 | 297 | 377 | 1901 | 575 | 398 |
| 4077 | W303.asm02.HP0_G0040720 | chrXIII | 12816 | 2612 | 2864 | 12674 | 2804 | 2704 | 12438 | 2706 | 3071 | 9261 | 2440 | 3126 | 11525 | 1999 | 2740 | 8947 | 2323 | 2845 |
| 4078 | W303.asm02.HP0_G0040730 | chrXIII | 463 | 306 | 272 | 462 | 261 | 304 | 369 | 309 | 373 | 744 | 214 | 201 | 777 | 122 | 186 | 750 | 204 | 202 |
| 4079 | W303.asm02.HP0_G0040740 | chrXIII | 1288 | 537 | 474 | 1251 | 487 | 537 | 1188 | 469 | 493 | 2876 | 778 | 1032 | 1869 | 519 | 662 | 2217 | 701 | 873 |
| 4080 | W303.asm02.HP0_G0040750 | chrXIII | 7801 | 1913 | 1571 | 7604 | 1393 | 1564 | 7446 | 1782 | 1645 | 7688 | 2427 | 2711 | 6582 | 1684 | 2158 | 6714 | 2327 | 2629 |
| 4081 | W303.asm02.HP0_G0040760 | chrXIII | 1884 | 1098 | 1237 | 1913 | 1104 | 1322 | 1911 | 1039 | 1639 | 3020 | 786 | 979 | 3371 | 484 | 714 | 3170 | 751 | 840 |
| 4082 | W303.asm02.HP0_G0040770 | chrXIII | 4974 | 1225 | 1505 | 5227 | 1285 | 1465 | 4580 | 1193 | 1608 | 5326 | 1107 | 1554 | 5456 | 690 | 1139 | 5460 | 1020 | 1343 |
| 4083 | W303.asm02.HP0_G0040780 | chrXIII | 1105 | 299 | 253 | 1268 | 260 | 224 | 941 | 263 | 262 | 1272 | 339 | 348 | 1235 | 210 | 308 | 1142 | 389 | 313 |
| 4084 | W303.asm02.HP0_G0040790 | chrXIII | 10885 | 1261 | 1121 | 10378 | 1107 | 1176 | 10048 | 1214 | 1410 | 7782 | 2290 | 2253 | 4802 | 1387 | 2031 | 4561 | 1969 | 2048 |
| 4085 | W303.asm02.HP0_G0040800 | chrXIII | 2 | 2 | 0 | 4 | 1 | 0 | 2 | 0 | 0 | 4 | 1 | 3 | 1 | 2 | 0 | 3 | 4 | 0 |
| 4086 | W303.asm02.HP0_G0040810 | chrXIII | 1983 | 534 | 473 | 1805 | 532 | 547 | 2164 | 515 | 621 | 4611 | 847 | 1118 | 3218 | 672 | 970 | 3392 | 739 | 1013 |
| 4087 | W303.asm02.HP0_G0040820 | chrXIII | 25 | 2 | 1 | 10 | 2 | 3 | 6 | 1 | 4 | 11 | 3 | 1 | 7 | 1 | 1 | 16 | 3 | 1 |
| 4088 | W303.asm02.HP0_G0040830 | chrXIII | 4251 | 2088 | 2253 | 4322 | 2132 | 2182 | 4321 | 2196 | 2045 | 8117 | 1909 | 2342 | 7440 | 1165 | 1733 | 8070 | 1677 | 2121 |
| 4089 | W303.asm02.HP0_G0040840 | chrXIII | 253 | 199 | 164 | 330 | 212 | 175 | 210 | 170 | 183 | 628 | 160 | 146 | 714 | 100 | 106 | 537 | 180 | 158 |
| 4090 | W303.asm02.HP0_G0040850 | chrXIII | 894 | 360 | 447 | 821 | 383 | 399 | 724 | 327 | 470 | 1115 | 283 | 320 | 1164 | 200 | 262 | 1112 | 249 | 253 |

| | | | | | | | | | | | | | | | | | | | | |
|---|---|---|---|---|---|---|---|---|---|---|---|---|---|---|---|---|---|---|---|---|
| 4091 | W303.asm02.HP0_G0040860 | chrXIII | 770 | 372 | 473 | 899 | 528 | 477 | 775 | 330 | 421 | 2526 | 423 | 653 | 2429 | 250 | 449 | 2196 | 401 | 561 |
| 4092 | W303.asm02.HP0_G0040870 | chrXIII | 1895 | 760 | 977 | 1621 | 794 | 1029 | 2002 | 695 | 1046 | 8116 | 1514 | 2416 | 5430 | 907 | 1569 | 5616 | 1453 | 2110 |
| 4093 | W303.asm02.HP0_G0040880 | chrXIII | 4092 | 4533 | 4984 | 3772 | 4786 | 4842 | 3862 | 4601 | 5304 | 9668 | 4377 | 4509 | 10385 | 2885 | 4107 | 9943 | 3911 | 4090 |
| 4094 | W303.asm02.HP0_G0040890 | chrXIII | 7137 | 2890 | 3066 | 7349 | 2752 | 3006 | 6819 | 3197 | 3489 | 8831 | 2514 | 2320 | 11171 | 1747 | 2007 | 8660 | 2231 | 2011 |
| 4095 | W303.asm02.HP0_G0040900 | chrXIII | 585 | 221 | 341 | 528 | 101 | 483 | 471 | 181 | 515 | 227 | 121 | 320 | 365 | 112 | 205 | 260 | 154 | 271 |
| 4096 | W303.asm02.HP0_G0040910 | chrXIII | 906 | 995 | 656 | 902 | 648 | 677 | 798 | 966 | 681 | 2359 | 866 | 793 | 2862 | 597 | 625 | 2248 | 855 | 728 |
| 4097 | W303.asm02.HP0_G0040920 | chrXIII | 646 | 537 | 402 | 642 | 440 | 397 | 757 | 551 | 466 | 2170 | 662 | 570 | 2310 | 445 | 493 | 1702 | 661 | 521 |
| 4098 | W303.asm02.HP0_G0040930 | chrXIII | 2572 | 708 | 585 | 2680 | 710 | 686 | 2378 | 633 | 586 | 4585 | 933 | 1437 | 3549 | 589 | 989 | 3402 | 808 | 1210 |
| 4099 | W303.asm02.HP0_G0040940 | chrXIII | 6767 | 1023 | 1671 | 5615 | 1383 | 1795 | 6513 | 914 | 2150 | 8712 | 2044 | 3667 | 3322 | 1480 | 2073 | 3291 | 1814 | 3173 |
| 4100 | W303.asm02.HP0_G0040950 | chrXIII | 1056 | 680 | 519 | 913 | 522 | 484 | 1041 | 692 | 520 | 2351 | 689 | 782 | 2122 | 434 | 584 | 2147 | 652 | 705 |
| 4101 | W303.asm02.HP0_G0040960 | chrXIII | 3556 | 1279 | 1384 | 3726 | 1732 | 1443 | 4474 | 1243 | 1687 | 18473 | 3320 | 3684 | 10691 | 2176 | 1773 | 11113 | 2784 | 3250 |
| 4102 | W303.asm02.HP0_G0040970 | chrXIII | 659 | 460 | 408 | 635 | 356 | 392 | 632 | 434 | 453 | 1034 | 355 | 270 | 1185 | 202 | 265 | 1104 | 316 | 261 |
| 4103 | W303.asm02.HP0_G0040980 | chrXIII | 1452 | 216 | 228 | 1503 | 218 | 259 | 1285 | 194 | 291 | 1028 | 265 | 337 | 1014 | 147 | 373 | 890 | 279 | 331 |
| 4104 | W303.asm02.HP0_G0040990 | chrXIII | 5609 | 1115 | 1657 | 5121 | 1285 | 1869 | 5366 | 1189 | 2079 | 11543 | 2309 | 4445 | 8612 | 1926 | 3260 | 7561 | 2122 | 4239 |
| 4105 | W303.asm02.HP0_G0041000 | chrXIII | 611 | 293 | 302 | 664 | 263 | 309 | 578 | 248 | 301 | 1047 | 261 | 372 | 1384 | 203 | 285 | 922 | 236 | 307 |
| 4106 | W303.asm02.HP0_G0041010 | chrXIII | 473 | 273 | 209 | 425 | 235 | 217 | 377 | 220 | 246 | 715 | 323 | 279 | 648 | 275 | 297 | 552 | 295 | 244 |
| 4107 | W303.asm02.HP0_G0041020 | chrXIII | 837 | 1462 | 1567 | 796 | 1559 | 2288 | 929 | 1259 | 1785 | 22235 | 5813 | 3806 | 11384 | 1802 | 3115 | 17262 | 5464 | 6604 |
| 4108 | W303.asm02.HP0_G0041030 | chrXIII | 595 | 474 | 597 | 497 | 570 | 591 | 488 | 470 | 689 | 1122 | 324 | 394 | 1332 | 197 | 310 | 1227 | 332 | 315 |
| 4109 | W303.asm02.HP0_G0041040 | chrXIII | 1573 | 1393 | 1142 | 1492 | 1087 | 1193 | 1447 | 1401 | 1390 | 2875 | 983 | 988 | 3048 | 678 | 841 | 3094 | 945 | 864 |
| 4110 | W303.asm02.HP0_G0041050 | chrXIII | 6108 | 2160 | 2023 | 5819 | 2017 | 2026 | 6115 | 2200 | 2183 | 9391 | 1967 | 2069 | 8636 | 1110 | 1530 | 9975 | 1887 | 1914 |
| 4111 | W303.asm02.HP0_G0041060 | chrXIII | 2854 | 4070 | 3155 | 2633 | 3073 | 2962 | 2663 | 4032 | 3140 | 9869 | 3480 | 3242 | 9453 | 2416 | 2342 | 9261 | 2942 | 3020 |
| 4112 | W303.asm02.HP0_G0041070 | chrXIII | 258 | 122 | 111 | 280 | 128 | 109 | 275 | 143 | 104 | 896 | 217 | 197 | 641 | 143 | 136 | 608 | 173 | 166 |
| 4113 | W303.asm02.HP0_G0041080 | chrXIII | 3147 | 1296 | 1198 | 3026 | 1254 | 1267 | 2914 | 1224 | 1116 | 5465 | 1180 | 1478 | 4872 | 789 | 1211 | 4558 | 1112 | 1320 |
| 4114 | W303.asm02.HP0_G0041090 | chrXIII | 434 | 509 | 573 | 433 | 498 | 564 | 438 | 459 | 589 | 1608 | 331 | 357 | 1881 | 183 | 320 | 1703 | 386 | 319 |
| 4115 | W303.asm02.HP0_G0041100 | chrXIII | 1804 | 495 | 666 | 1878 | 619 | 583 | 1781 | 565 | 677 | 3857 | 605 | 869 | 3534 | 392 | 633 | 3312 | 547 | 742 |
| 4116 | W303.asm02.HP0_G0041110 | chrXIII | 3953 | 592 | 550 | 3729 | 467 | 588 | 3893 | 581 | 503 | 4029 | 882 | 1129 | 3852 | 620 | 852 | 3347 | 762 | 1039 |
| 4117 | W303.asm02.HP0_G0041120 | chrXIII | 1208 | 361 | 234 | 1094 | 207 | 256 | 1080 | 273 | 260 | 1091 | 446 | 405 | 851 | 276 | 336 | 774 | 457 | 376 |
| 4118 | W303.asm02.HP0_G0041130 | chrXIII | 173 | 68 | 73 | 119 | 68 | 84 | 137 | 58 | 89 | 429 | 99 | 84 | 236 | 59 | 58 | 246 | 102 | 94 |
| 4119 | W303.asm02.HP0_G0041140 | chrXIII | 738 | 627 | 741 | 1000 | 681 | 836 | 790 | 539 | 898 | 3587 | 880 | 1337 | 1899 | 504 | 720 | 1643 | 805 | 1135 |
| 4120 | W303.asm02.HP0_G0041150 | chrXIII | 5705 | 3652 | 3675 | 5686 | 3103 | 3906 | 5530 | 3652 | 4121 | 7313 | 2593 | 3283 | 8035 | 2063 | 2505 | 7577 | 2449 | 2820 |
| 4121 | W303.asm02.HP0_G0041160 | chrXIII | 824 | 980 | 930 | 754 | 770 | 881 | 778 | 991 | 1069 | 2021 | 781 | 791 | 2159 | 446 | 535 | 2166 | 710 | 643 |
| 4122 | W303.asm02.HP0_G0041170 | chrXIII | 2039 | 1448 | 1721 | 1755 | 1479 | 1691 | 1796 | 1497 | 2322 | 3610 | 1369 | 1354 | 4572 | 883 | 1138 | 3980 | 1201 | 1266 |
| 4123 | W303.asm02.HP0_G0041180 | chrXIII | 2558 | 777 | 1006 | 2619 | 935 | 799 | 2647 | 853 | 891 | 5512 | 771 | 1189 | 5417 | 532 | 877 | 5176 | 830 | 960 |
| 4124 | W303.asm02.HP0_G0041190 | chrXIII | 2139 | 650 | 766 | 2210 | 775 | 770 | 2329 | 697 | 836 | 5450 | 876 | 1420 | 4194 | 616 | 987 | 4666 | 750 | 1312 |
| 4125 | W303.asm02.HP0_G0041200 | chrXIII | 1254 | 725 | 976 | 1202 | 858 | 982 | 1229 | 735 | 1273 | 2731 | 559 | 734 | 3366 | 367 | 509 | 3110 | 556 | 608 |
| 4126 | W303.asm02.HP0_G0041210 | chrXIII | 1182 | 517 | 686 | 1081 | 575 | 764 | 1214 | 492 | 727 | 3590 | 721 | 1474 | 2933 | 616 | 879 | 2843 | 695 | 1175 |
| 4127 | W303.asm02.HP0_G0041220 | chrXIII | 2976 | 1842 | 2041 | 3218 | 1918 | 2009 | 3352 | 1842 | 2130 | 8671 | 1970 | 2816 | 8141 | 1605 | 2033 | 7728 | 1817 | 2684 |
| 4128 | W303.asm02.HP0_G0041230 | chrXIII | 4332 | 940 | 1165 | 3758 | 1344 | 1262 | 4457 | 881 | 1402 | 13258 | 1931 | 3033 | 7961 | 1497 | 1975 | 8141 | 1678 | 2761 |
| 4129 | W303.asm02.HP0_G0041240 | chrXIII | 6523 | 297 | 210 | 7731 | 229 | 205 | 6456 | 272 | 263 | 1084 | 277 | 290 | 1350 | 134 | 214 | 918 | 243 | 243 |
| 4130 | W303.asm02.HP0_G0041250 | chrXIII | 14137 | 4036 | 3822 | 13020 | 3554 | 3699 | 14154 | 4352 | 4191 | 16059 | 5410 | 6078 | 14796 | 4138 | 5191 | 13909 | 4871 | 5500 |
| 4131 | W303.asm02.HP0_G0041260 | chrXIII | 1175 | 100 | 105 | 1346 | 61 | 112 | 870 | 81 | 130 | 388 | 175 | 195 | 229 | 96 | 199 | 184 | 169 | 211 |
| 4132 | W303.asm02.HP0_G0041270 | chrXIII | 6618 | 469 | 792 | 6356 | 598 | 868 | 6449 | 414 | 850 | 5096 | 967 | 1732 | 2310 | 751 | 1205 | 2119 | 769 | 1520 |
| 4133 | W303.asm02.HP0_G0041280 | chrXIII | 1822 | 422 | 374 | 1588 | 348 | 384 | 1699 | 350 | 427 | 1129 | 388 | 420 | 1101 | 276 | 382 | 1016 | 357 | 424 |
| 4134 | W303.asm02.HP0_G0041290 | chrXIII | 1952 | 752 | 951 | 1918 | 934 | 897 | 1877 | 815 | 1154 | 5636 | 1266 | 1541 | 3962 | 881 | 1166 | 4143 | 1105 | 1342 |
| 4135 | W303.asm02.HP0_G0041300 | chrXIII | 15866 | 1123 | 1550 | 15795 | 1538 | 1758 | 15939 | 984 | 2174 | 11616 | 2401 | 3752 | 8872 | 2115 | 2936 | 7004 | 2027 | 3386 |
| 4136 | W303.asm02.HP0_G0041310 | chrXIII | 3021 | 1308 | 1359 | 2544 | 894 | 1684 | 2551 | 1149 | 1681 | 1065 | 1206 | 1382 | 1615 | 832 | 1152 | 1033 | 1142 | 887 |
| 4137 | W303.asm02.HP0_G0041320 | chrXIII | 2551 | 1192 | 1052 | 2157 | 1239 | 1067 | 2264 | 1209 | 986 | 7512 | 1516 | 1923 | 6263 | 935 | 1458 | 6277 | 1309 | 1827 |
| 4138 | W303.asm02.HP0_G0041330 | chrXIII | 5521 | 1773 | 1412 | 5070 | 1296 | 1463 | 5238 | 1583 | 1638 | 7774 | 2219 | 2883 | 6077 | 1641 | 2296 | 5962 | 2224 | 2620 |
| 4139 | W303.asm02.HP0_G0041340 | chrXIII | 743 | 961 | 1065 | 682 | 1173 | 1079 | 818 | 1022 | 1281 | 8031 | 1323 | 1863 | 7350 | 952 | 1168 | 6948 | 1285 | 1657 |
| 4140 | W303.asm02.HP0_G0041350 | chrXIII | 188 | 74 | 136 | 187 | 140 | 149 | 161 | 109 | 189 | 1188 | 157 | 239 | 860 | 105 | 156 | 916 | 127 | 217 |
| 4141 | W303.asm02.HP0_G0041360 | chrXIII | 299 | 106 | 157 | 292 | 151 | 155 | 274 | 148 | 204 | 688 | 144 | 265 | 564 | 113 | 152 | 482 | 120 | 229 |
| 4142 | W303.asm02.HP0_G0041370 | chrXIII | 1352 | 396 | 393 | 1312 | 405 | 368 | 1363 | 430 | 432 | 1267 | 433 | 583 | 1276 | 362 | 406 | 886 | 328 | 531 |
| 4143 | W303.asm02.HP0_G0041380 | chrXIII | 2243 | 1011 | 1045 | 2216 | 884 | 1112 | 2126 | 934 | 1094 | 2668 | 739 | 1004 | 2576 | 496 | 797 | 2678 | 744 | 878 |
| 4144 | W303.asm02.HP0_G0041390 | chrXIII | 4749 | 1198 | 1044 | 4483 | 1078 | 1067 | 4628 | 1200 | 1223 | 5793 | 1098 | 1564 | 5480 | 827 | 1267 | 5182 | 1120 | 1376 |
| 4145 | W303.asm02.HP0_G0041400 | chrXIII | 8393 | 2890 | 2506 | 8198 | 2236 | 2376 | 8629 | 2960 | 2434 | 8656 | 3025 | 3250 | 8939 | 2184 | 2400 | 8391 | 2607 | 2953 |
| 4146 | W303.asm02.HP0_G0041410 | chrXIII | 1368 | 501 | 746 | 1542 | 807 | 758 | 1345 | 517 | 839 | 5176 | 577 | 1000 | 5100 | 389 | 843 | 4590 | 535 | 914 |
| 4147 | W303.asm02.HP0_G0041420 | chrXIII | 153 | 43 | 59 | 178 | 63 | 78 | 153 | 61 | 88 | 156 | 39 | 35 | 230 | 37 | 29 | 155 | 30 | 38 |
| 4148 | W303.asm02.HP0_G0041430 | chrXIII | 1600 | 395 | 457 | 1776 | 486 | 527 | 1547 | 365 | 563 | 5755 | 761 | 1177 | 3748 | 506 | 845 | 3747 | 669 | 1117 |
| 4149 | W303.asm02.HP0_G0041440 | chrXIII | 14678 | 445 | 387 | 14519 | 393 | 387 | 13080 | 452 | 483 | 2983 | 1165 | 920 | 3490 | 1262 | 1173 | 1387 | 980 | 796 |
| 4150 | W303.asm02.HP0_G0041450 | chrXIII | 28141 | 537 | 442 | 25929 | 455 | 579 | 24944 | 478 | 487 | 1778 | 740 | 786 | 2424 | 677 | 1074 | 1308 | 790 | 705 |
| 4151 | W303.asm02.HP0_G0041460 | chrXIII | 52 | 15 | 14 | 51 | 17 | 25 | 57 | 12 | 9 | 97 | 22 | 45 | 56 | 11 | 15 | 51 | 14 | 24 |
| 4152 | W303.asm02.HP0_G0041470 | chrXIII | 621 | 153 | 195 | 637 | 185 | 216 | 614 | 137 | 218 | 1707 | 266 | 528 | 1062 | 222 | 312 | 1012 | 235 | 406 |
| 4153 | W303.asm02.HP0_G0041480 | chrXIII | 1273 | 25 | 36 | 1398 | 71 | 71 | 962 | 40 | 42 | 77 | 18 | 28 | 102 | 18 | 19 | 94 | 45 | 28 |
| 4154 | W303.asm02.HP0_G0041490 | chrXIII | 9908 | 6730 | 6518 | 10414 | 7118 | 7032 | 10624 | 6084 | 7891 | 33874 | 11690 | 12631 | 21036 | 8921 | 8760 | 21060 | 9808 | 10814 |
| 4155 | W303.asm02.HP0_G0041500 | chrXIII | 5957 | 729 | 865 | 5099 | 675 | 1019 | 5555 | 566 | 1143 | 2282 | 737 | 1473 | 1708 | 622 | 863 | 1529 | 646 | 1218 |
| 4156 | W303.asm02.HP0_G0041510 | chrXIII | 10689 | 2448 | 2316 | 10369 | 1875 | 2096 | 10295 | 2366 | 2274 | 5881 | 1874 | 1853 | 6679 | 1248 | 1676 | 5964 | 1739 | 1585 |
| 4157 | W303.asm02.HP0_G0041520 | chrXIII | 1150 | 441 | 307 | 1291 | 308 | 330 | 1100 | 385 | 298 | 1843 | 420 | 551 | 1679 | 265 | 371 | 1495 | 571 | 420 |
| 4158 | W303.asm02.HP0_G0041530 | chrXIII | 743 | 554 | 812 | 646 | 550 | 913 | 573 | 550 | 959 | 1551 | 430 | 770 | 1975 | 299 | 550 | 1666 | 406 | 645 |
| 4159 | W303.asm02.HP0_G0041540 | chrXIII | 1516 | 730 | 611 | 1407 | 623 | 577 | 1489 | 714 | 601 | 3109 | 711 | 942 | 2895 | 515 | 637 | 2800 | 783 | 834 |
| 4160 | W303.asm02.HP0_G0041550 | chrXIII | 4477 | 617 | 448 | 4118 | 489 | 496 | 3917 | 626 | 508 | 1452 | 601 | 483 | 1746 | 430 | 462 | 1448 | 565 | 464 |
| 4161 | W303.asm02.HP0_G0041560 | chrXIII | 3460 | 993 | 1075 | 3513 | 1005 | 1061 | 3273 | 1086 | 1007 | 4762 | 1001 | 1217 | 4653 | 677 | 929 | 4405 | 916 | 1102 |
| 4162 | W303.asm02.HP0_G0041570 | chrXIII | 16631 | 17471 | 18176 | 13787 | 17517 | 17466 | 17491 | 17391 | 20462 | 43141 | 16677 | 17193 | 43600 | 12654 | 14053 | 42002 | 15307 | 15932 |
| 4163 | W303.asm02.HP0_G0041580 | chrXIII | 513 | 391 | 368 | 542 | 337 | 324 | 514 | 403 | 303 | 1735 | 342 | 383 | 1761 | 251 | 320 | 1710 | 371 | 394 |
| 4164 | W303.asm02.HP0_G0041590 | chrXIII | 440 | 640 | 384 | 443 | 337 | 390 | 433 | 568 | 374 | 1080 | 356 | 267 | 1403 | 286 | 215 | 1169 | 416 | 244 |
| 4165 | W303.asm02.HP0_G0041600 | chrXIII | 1935 | 902 | 1044 | 1991 | 840 | 957 | 1914 | 978 | 1114 | 5404 | 1323 | 1451 | 5249 | 873 | 1282 | 5094 | 1050 | 1332 |
| 4166 | W303.asm02.HP0_G0041610 | chrXIII | 3557 | 566 | 642 | 1798 | 470 | 601 | 3256 | 500 | 665 | 1364 | 515 | 568 | 1675 | 366 | 399 | 1325 | 476 | 484 |
| 4167 | W303.asm02.HP0_G0041620 | chrXIII | 1535 | 1571 | 1044 | 1274 | 974 | 1139 | 1343 | 1405 | 1199 | 2572 | 1160 | 1026 | 2818 | 778 | 830 | 2693 | 1098 | 923 |
| 4168 | W303.asm02.HP0_G0041630 | chrXIII | 6734 | 3259 | 3820 | 6474 | 3394 | 3970 | 6482 | 3086 | 4279 | 8600 | 3442 | 5366 | 12702 | 3591 | 5078 | 7718 | 2962 | 4435 |
| 4169 | W303.asm02.HP0_G0041640 | chrXIII | 1360 | 1522 | 1646 | 1320 | 1497 | 1737 | 1425 | 1557 | 2001 | 4793 | 1514 | 1772 | 5059 | 976 | 1191 | 4812 | 1342 | 1531 |
| 4170 | W303.asm02.HP0_G0041650 | chrXIII | 6362 | 378 | 684 | 6622 | 606 | 734 | 6831 | 361 | 873 | 5618 | 956 | 1549 | 3638 | 1013 | 1370 | 2151 | 825 | 1284 |
| 4171 | W303.asm02.HP0_G0041660 | chrXIII | 7705 | 2901 | 3887 | 7807 | 3864 | 3597 | 7906 | 3159 | 4019 | 18620 | 3031 | 4408 | 16508 | 2186 | 3598 | 17643 | 2788 | 4124 |
| 4172 | W303.asm02.HP0_G0041670 | chrXIII | 389 | 364 | 558 | 409 | 520 | 497 | 382 | 352 | 584 | 2000 | 328 | 489 | 2183 | 226 | 347 | 2125 | 346 | 433 |
| 4173 | W303.asm02.HP0_G0041680 | chrXIII | 742 | 690 | 829 | 699 | 776 | 788 | 790 | 729 | 1032 | 2494 | 540 | 648 | 2924 | 385 | 455 | 2668 | 507 | 537 |
| 4174 | W303.asm02.HP0_G0041690 | chrXIII | 2442 | 545 | 647 | 2565 | 827 | 650 | 2963 | 509 | 852 | 9618 | 1294 | 1736 | 4028 | 904 | 796 | 4082 | 1060 | 1438 |
| 4175 | W303.asm02.HP0_G0041700 | chrXIII | 2286 | 759 | 1091 | 2247 | 1084 | 1142 | 2330 | 832 | 1180 | 9461 | 1323 | 2458 | 5178 | 1060 | 1612 | 6002 | 1124 | 2189 |
| 4176 | W303.asm02.HP0_G0041710 | chrXIII | 2627 | 1623 | 1922 | 2335 | 1658 | 1939 | 2577 | 1791 | 2239 | 5443 | 1355 | 1711 | 5164 | 890 | 1270 | 5764 | 1199 | 1529 |
| 4177 | W303.asm02.HP0_G0041720 | chrXIII | 3721 | 2877 | 2787 | 3729 | 3783 | 2679 | 4246 | 3095 | 2730 | 20732 | 4338 | 5067 | 16710 | 2798 | 2769 | 17488 | 3820 | 4465 |
| 4178 | W303.asm02.HP0_G0041730 | chrXIII | 1185 | 900 | 1145 | 1089 | 1102 | 1069 | 1151 | 967 | 1379 | 3543 | 755 | 953 | 3794 | 496 | 697 | 3953 | 783 | 808 |
| 4179 | W303.asm02.HP0_G0041740 | chrXIII | 2225 | 1467 | 1504 | 2263 | 1348 | 1519 | 2337 | 1430 | 1588 | 4523 | 1142 | 1247 | 4343 | 713 | 901 | 4603 | 1131 | 1124 |
| 4180 | W303.asm02.HP0_G0041750 | chrXIII | 5124 | 1091 | 810 | 4969 | 784 | 913 | 4671 | 972 | 880 | 3218 | 1246 | 1297 | 3539 | 854 | 1154 | 2744 | 1135 | 1170 |
| 4181 | W303.asm02.HP0_G0041760 | chrXIII | 1556 | 292 | 304 | 1592 | 296 | 358 | 1584 | 222 | 404 | 2410 | 491 | 801 | 1092 | 319 | 534 | 797 | 379 | 565 |
| 4182 | W303.asm02.HP0_G0041770 | chrXIII | 1104 | 299 | 251 | 1002 | 422 | 442 | 1010 | 306 | 302 | 1783 | 398 | 498 | 1477 | 261 | 365 | 1349 | 415 | 473 |
| 4183 | W303.asm02.HP0_G0041780 | chrXIII | 911 | 443 | 567 | 742 | 582 | 519 | 799 | 509 | 618 | 1897 | 601 | 601 | 2086 | 334 | 424 | 1887 | 466 | 551 |

| | | | | | | | | | | | | | | | | | | | | |
|---|---|---|---|---|---|---|---|---|---|---|---|---|---|---|---|---|---|---|---|---|
| 4184 | W303.asm02.HP0_G0041790 | chrXIII | 3809 | 685 | 885 | 3820 | 760 | 871 | 3576 | 711 | 864 | 3278 | 774 | 1283 | 3622 | 594 | 1049 | 2925 | 671 | 1092 |
| 4185 | W303.asm02.HP0_G0041800 | chrXIII | 3520 | 349 | 268 | 3578 | 256 | 338 | 3194 | 242 | 288 | 2773 | 876 | 754 | 2012 | 497 | 579 | 1800 | 796 | 674 |
| 4186 | W303.asm02.HP0_G0041810 | chrXIII | 15305 | 11807 | 12220 | 14141 | 12283 | 12968 | 16003 | 11810 | 13379 | 28515 | 11112 | 12494 | 27796 | 8045 | 9303 | 26374 | 9251 | 10983 |
| 4187 | W303.asm02.HP0_G0041820 | chrXIII | 8995 | 8539 | 10500 | 8438 | 9708 | 10496 | 9264 | 8324 | 14015 | 21526 | 7391 | 8262 | 23874 | 5484 | 6724 | 22816 | 6726 | 7291 |
| 4188 | W303.asm02.HP0_G0041830 | chrXIII | 539 | 1037 | 683 | 538 | 717 | 685 | 593 | 976 | 700 | 2350 | 846 | 758 | 2657 | 629 | 623 | 2418 | 793 | 707 |
| 4189 | W303.asm02.HP0_G0041840 | chrXIII | 15302 | 7335 | 6979 | 16434 | 6832 | 7160 | 15718 | 7367 | 8911 | 38876 | 14504 | 14645 | 31101 | 11962 | 10659 | 27736 | 12403 | 12513 |
| 4190 | W303.asm02.HP0_G0041850 | chrXIII | 16065 | 7278 | 5835 | 17099 | 6236 | 5786 | 17736 | 7198 | 5592 | 21474 | 8182 | 8577 | 20456 | 5869 | 5974 | 19154 | 6936 | 7486 |
| 4191 | W303.asm02.HP0_G0041860 | chrXIII | 543 | 383 | 321 | 519 | 383 | 296 | 580 | 346 | 261 | 1553 | 334 | 390 | 1354 | 186 | 237 | 1484 | 352 | 333 |
| 4192 | W303.asm02.HP0_G0041870 | chrXIII | 985 | 869 | 725 | 943 | 645 | 759 | 962 | 827 | 769 | 1780 | 607 | 705 | 1853 | 432 | 579 | 1671 | 565 | 645 |
| 4193 | W303.asm02.HP0_G0041880 | chrXIII | 3833 | 2675 | 2813 | 3710 | 2277 | 2958 | 3849 | 2709 | 2784 | 7334 | 2369 | 3368 | 7661 | 1618 | 2275 | 7007 | 2115 | 2919 |
| 4194 | W303.asm02.HP0_G0041890 | chrXIII | 1457 | 906 | 1162 | 1256 | 1066 | 1124 | 1365 | 945 | 1577 | 2285 | 838 | 872 | 3195 | 574 | 821 | 2495 | 710 | 824 |
| 4195 | W303.asm02.HP0_G0041900 | chrXIII | 4815 | 1454 | 1029 | 4547 | 993 | 1024 | 4370 | 1458 | 1074 | 4481 | 1434 | 1566 | 4949 | 1048 | 1371 | 4169 | 1427 | 1420 |
| 4196 | W303.asm02.HP0_G0041910 | chrXIII | 2683 | 664 | 649 | 2596 | 530 | 776 | 2697 | 525 | 793 | 4718 | 1215 | 1729 | 2933 | 877 | 1073 | 3157 | 1110 | 1454 |
| 4197 | W303.asm02.HP0_G0041920 | chrXIII | 1280 | 317 | 279 | 1170 | 270 | 310 | 1142 | 341 | 367 | 1845 | 390 | 481 | 1425 | 260 | 368 | 1487 | 367 | 459 |
| 4198 | W303.asm02.HP0_G0041930 | chrXIII | 1138 | 804 | 542 | 1062 | 559 | 492 | 965 | 689 | 563 | 2667 | 754 | 712 | 2503 | 465 | 628 | 2251 | 724 | 678 |
| 4199 | W303.asm02.HP0_G0041940 | chrXIII | 1071 | 424 | 274 | 995 | 251 | 263 | 932 | 467 | 263 | 1032 | 331 | 302 | 936 | 218 | 183 | 851 | 351 | 233 |
| 4200 | W303.asm02.HP0_G0041950 | chrXIII | 3598 | 1372 | 1290 | 3461 | 1209 | 1309 | 3178 | 1298 | 1669 | 3322 | 1010 | 1114 | 4426 | 707 | 936 | 3395 | 1039 | 1034 |
| 4201 | W303.asm02.HP0_G0041960 | chrXIII | 4662 | 1287 | 1895 | 4490 | 1575 | 1899 | 4394 | 1372 | 2603 | 4397 | 1091 | 1577 | 5714 | 859 | 1317 | 4443 | 1048 | 1329 |
| 4202 | W303.asm02.HP0_G0041970 | chrXIII | 737 | 183 | 154 | 769 | 142 | 147 | 719 | 168 | 113 | 502 | 156 | 140 | 611 | 89 | 132 | 487 | 152 | 118 |
| 4203 | W303.asm02.HP0_G0041980 | chrXIII | 3726 | 385 | 351 | 3648 | 319 | 395 | 3184 | 283 | 435 | 2286 | 458 | 809 | 1600 | 358 | 580 | 1666 | 454 | 595 |
| 4204 | W303.asm02.HP0_G0041990 | chrXIII | 2605 | 1612 | 1806 | 2798 | 1520 | 1807 | 2621 | 1488 | 1943 | 5207 | 1421 | 1666 | 5349 | 868 | 1284 | 5309 | 1359 | 1504 |
| 4205 | W303.asm02.HP0_G0042000 | chrXIII | 1844 | 472 | 667 | 1736 | 609 | 750 | 1641 | 481 | 836 | 5186 | 862 | 1470 | 2772 | 625 | 1158 | 2997 | 707 | 1307 |
| 4206 | W303.asm02.HP0_G0042010 | chrXIII | 338 | 207 | 289 | 285 | 227 | 289 | 319 | 206 | 282 | 1507 | 267 | 549 | 1590 | 216 | 429 | 1387 | 250 | 524 |
| 4207 | W303.asm02.HP0_G0042020 | chrXIII | 1169 | 194 | 314 | 1223 | 229 | 344 | 1058 | 177 | 370 | 2518 | 410 | 737 | 2007 | 289 | 502 | 1567 | 333 | 664 |
| 4208 | W303.asm02.HP0_G0042030 | chrXIII | 3084 | 411 | 464 | 3103 | 402 | 521 | 2852 | 371 | 578 | 3570 | 676 | 956 | 1724 | 491 | 638 | 1782 | 622 | 809 |
| 4209 | W303.asm02.HP0_G0042040 | chrXIII | 3806 | 661 | 801 | 3506 | 817 | 773 | 3710 | 704 | 802 | 4637 | 1011 | 1330 | 4849 | 805 | 1154 | 3593 | 896 | 1270 |
| 4210 | W303.asm02.HP0_G0042050 | chrXIII | 2133 | 463 | 576 | 2222 | 655 | 533 | 2139 | 483 | 535 | 4194 | 554 | 904 | 3553 | 376 | 709 | 3415 | 485 | 810 |
| 4211 | W303.asm02.HP0_G0042060 | chrXIII | 1580 | 484 | 527 | 1530 | 539 | 487 | 1512 | 522 | 539 | 3562 | 592 | 837 | 2867 | 403 | 648 | 3034 | 541 | 746 |
| 4212 | W303.asm02.HP0_G0042070 | chrXIII | 5599 | 689 | 313 | 6955 | 241 | 288 | 5504 | 575 | 306 | 1009 | 711 | 367 | 1354 | 422 | 355 | 1073 | 878 | 354 |
| 4213 | W303.asm02.HP0_G0042080 | chrXIII | 1954 | 532 | 383 | 1826 | 347 | 412 | 1734 | 511 | 391 | 1458 | 499 | 591 | 1964 | 423 | 549 | 1181 | 465 | 491 |
| 4214 | W303.asm02.HP0_G0042090 | chrXIII | 2766 | 661 | 711 | 2556 | 781 | 810 | 2706 | 563 | 927 | 6171 | 1262 | 1419 | 4578 | 826 | 1128 | 4459 | 1257 | 1189 |
| 4215 | W303.asm02.HP0_G0042100 | chrXIII | 1513 | 379 | 370 | 1437 | 412 | 400 | 1536 | 316 | 436 | 3105 | 573 | 836 | 2284 | 391 | 475 | 2154 | 519 | 759 |
| 4216 | W303.asm02.HP0_G0042110 | chrXIII | 20817 | 5700 | 8232 | 17196 | 5276 | 8847 | 18833 | 6208 | 9317 | 10127 | 5783 | 7385 | 13938 | 3376 | 6113 | 9348 | 4938 | 5986 |
| 4217 | W303.asm02.HP0_G0042120 | chrXIII | 2093 | 284 | 296 | 1572 | 274 | 340 | 1563 | 272 | 394 | 553 | 278 | 252 | 845 | 200 | 283 | 524 | 259 | 219 |
| 4218 | W303.asm02.HP0_G0042130 | chrXIII | 170 | 84 | 56 | 135 | 39 | 43 | 111 | 64 | 84 | 79 | 47 | 28 | 103 | 31 | 19 | 96 | 52 | 37 |
| 4219 | W303.asm02.HP0_G0042140 | chrXIII | 25 | 6 | 10 | 21 | 5 | 11 | 10 | 6 | 8 | 24 | 4 | 6 | 37 | 1 | 4 | 27 | 7 | 2 |
| 4220 | W303.asm02.HP0_G0042150 | chrXIII | 2208 | 377 | 583 | 2010 | 614 | 567 | 2095 | 407 | 705 | 4768 | 647 | 1207 | 3310 | 501 | 825 | 3219 | 577 | 1112 |
| 4221 | W303.asm02.HP0_G0042160 | chrXIII | 1004 | 574 | 645 | 905 | 746 | 591 | 973 | 600 | 646 | 4208 | 827 | 809 | 3940 | 448 | 657 | 3698 | 752 | 709 |
| 4222 | W303.asm02.HP0_G0042170 | chrXIII | 4093 | 1289 | 1640 | 3650 | 1431 | 1491 | 3638 | 1275 | 1978 | 4451 | 1213 | 1330 | 5204 | 728 | 1124 | 4903 | 1133 | 1245 |
| 4223 | W303.asm02.HP0_G0042180 | chrXIII | 903 | 337 | 406 | 891 | 454 | 429 | 811 | 325 | 435 | 3418 | 451 | 745 | 2922 | 336 | 500 | 2578 | 442 | 710 |
| 4224 | W303.asm02.HP0_G0042190 | chrXIII | 2482 | 691 | 639 | 1998 | 509 | 617 | 2073 | 756 | 653 | 1635 | 532 | 599 | 2392 | 396 | 476 | 1732 | 533 | 544 |
| 4225 | W303.asm02.HP0_G0042200 | chrXIII | 4391 | 368 | 363 | 4229 | 296 | 364 | 4101 | 347 | 422 | 2002 | 566 | 590 | 1970 | 440 | 589 | 1262 | 526 | 533 |
| 4226 | W303.asm02.HP0_G0042210 | chrXIII | 1987 | 127 | 175 | 1775 | 206 | 228 | 1803 | 121 | 244 | 1490 | 325 | 382 | 518 | 209 | 253 | 445 | 278 | 352 |
| 4227 | W303.asm02.HP0_G0042220 | chrXIII | 7399 | 1634 | 1489 | 7078 | 1339 | 1542 | 7727 | 1536 | 1713 | 10020 | 3032 | 3414 | 7307 | 2197 | 2585 | 7116 | 2711 | 3279 |
| 4228 | W303.asm02.HP0_G0042230 | chrXIII | 1486 | 882 | 2333 | 1277 | 977 | 2628 | 1379 | 880 | 2842 | 2448 | 679 | 2250 | 4550 | 682 | 1520 | 2814 | 726 | 1925 |
| 4229 | W303.asm02.HP0_G0042240 | chrXIII | 810 | 812 | 1133 | 862 | 1243 | 1074 | 931 | 851 | 1168 | 5455 | 736 | 1150 | 5776 | 601 | 758 | 5532 | 725 | 1078 |
| 4230 | W303.asm02.HP0_G0042250 | chrXIII | 38008 | 30477 | 20212 | 37827 | 20508 | 19752 | 38396 | 25887 | 15689 | 84138 | 29348 | 34881 | 70061 | 24388 | 26761 | 76296 | 27323 | 32043 |
| 4231 | W303.asm02.HP0_G0042260 | chrXIII | 992 | 568 | 686 | 977 | 565 | 692 | 1032 | 554 | 690 | 2301 | 506 | 761 | 2344 | 326 | 489 | 2257 | 540 | 655 |
| 4232 | W303.asm02.HP0_G0042270 | chrXIII | 6554 | 2173 | 2350 | 6713 | 2087 | 2285 | 6390 | 2371 | 2792 | 6065 | 2028 | 2136 | 7141 | 1393 | 1872 | 6170 | 1956 | 2010 |
| 4233 | W303.asm02.HP0_G0042280 | chrXIII | 1108 | 249 | 341 | 874 | 252 | 377 | 1021 | 239 | 421 | 2452 | 644 | 857 | 1487 | 511 | 492 | 1305 | 562 | 633 |
| 4234 | W303.asm02.HP0_G0042290 | chrXIII | 1462 | 284 | 424 | 1389 | 429 | 447 | 1327 | 257 | 481 | 3481 | 465 | 991 | 2170 | 382 | 730 | 2254 | 411 | 989 |
| 4235 | W303.asm02.HP0_G0042300 | chrXIII | 3554 | 1655 | 1982 | 3678 | 1628 | 2177 | 3530 | 1593 | 2034 | 5380 | 1649 | 2733 | 6295 | 1181 | 1766 | 5395 | 1517 | 2359 |
| 4236 | W303.asm02.HP0_G0042310 | chrXIII | 3478 | 273 | 470 | 3039 | 408 | 490 | 3616 | 261 | 664 | 4852 | 738 | 1103 | 4559 | 620 | 977 | 3049 | 627 | 959 |
| 4237 | W303.asm02.HP0_G0042320 | chrXIII | 5917 | 1816 | 1594 | 6018 | 1475 | 1574 | 5502 | 1752 | 1695 | 4902 | 1415 | 1679 | 5579 | 1154 | 1386 | 4964 | 1479 | 1503 |
| 4238 | W303.asm02.HP0_G0042330 | chrXIII | 3754 | 2675 | 3822 | 3356 | 3609 | 3729 | 3572 | 2725 | 4830 | 8068 | 2076 | 2727 | 9623 | 1565 | 2350 | 8867 | 2113 | 2300 |
| 4239 | W303.asm02.HP0_G0042340 | chrXIII | 256 | 139 | 110 | 274 | 92 | 108 | 249 | 105 | 102 | 287 | 105 | 114 | 366 | 93 | 133 | 229 | 108 | 128 |
| 4240 | W303.asm02.HP0_G0042350 | chrXIII | 720 | 934 | 1451 | 821 | 1159 | 1376 | 675 | 806 | 1682 | 1693 | 695 | 886 | 2053 | 351 | 700 | 1498 | 535 | 707 |
| 4241 | W303.asm02.HP0_G0042360 | chrXIII | 9597 | 542 | 505 | 8700 | 520 | 570 | 8381 | 484 | 737 | 2919 | 978 | 1074 | 1837 | 920 | 1052 | 1505 | 1020 | 947 |
| 4242 | W303.asm02.HP0_G0042370 | chrXIII | 3725 | 1509 | 1412 | 3598 | 1345 | 1406 | 3612 | 1467 | 1386 | 3966 | 1238 | 1344 | 4934 | 994 | 1227 | 3571 | 1082 | 1237 |
| 4243 | W303.asm02.HP0_G0042380 | chrXIII | 600 | 431 | 382 | 569 | 409 | 375 | 661 | 436 | 307 | 2262 | 535 | 580 | 1830 | 327 | 548 | 2056 | 480 | 612 |
| 4244 | W303.asm02.HP0_G0042390 | chrXIII | 790 | 720 | 1146 | 955 | 1235 | 1270 | 884 | 748 | 1084 | 5609 | 1079 | 1990 | 4417 | 685 | 1230 | 4058 | 848 | 1738 |
| 4245 | W303.asm02.HP0_G0042400 | chrXIII | 5339 | 3011 | 2848 | 5209 | 2608 | 2863 | 5131 | 2990 | 2756 | 8300 | 2735 | 2903 | 10437 | 2200 | 2487 | 7890 | 2402 | 2651 |
| 4246 | W303.asm02.HP0_G0042410 | chrXIII | 1705 | 552 | 345 | 1777 | 381 | 386 | 1684 | 422 | 355 | 1979 | 482 | 610 | 1714 | 307 | 365 | 1536 | 536 | 515 |
| 4247 | W303.asm02.HP0_G0042420 | chrXIII | 5837 | 5683 | 6058 | 5509 | 5642 | 6389 | 6299 | 5171 | 5953 | 14862 | 4904 | 6094 | 15542 | 3571 | 4880 | 13696 | 4350 | 5921 |
| 4248 | W303.asm02.HP0_G0042430 | chrXIII | 18913 | 5504 | 5912 | 17491 | 5500 | 5306 | 18614 | 5646 | 5968 | 19482 | 5486 | 6668 | 21011 | 4151 | 5598 | 18565 | 4963 | 6186 |
| 4249 | W303.asm02.HP0_G0042440 | chrXIII | 519 | 193 | 146 | 582 | 117 | 124 | 517 | 156 | 138 | 1018 | 215 | 293 | 853 | 113 | 220 | 735 | 208 | 242 |
| 4250 | W303.asm02.HP0_G0042450 | chrXIII | 34663 | 4336 | 5499 | 32762 | 4993 | 6059 | 36517 | 4018 | 7236 | 44572 | 10129 | 13601 | 29014 | 7484 | 10077 | 28609 | 9082 | 12585 |
| 4251 | W303.asm02.HP0_G0042460 | chrXIII | 1070 | 105 | 57 | 1160 | 81 | 82 | 1014 | 83 | 88 | 278 | 136 | 85 | 247 | 71 | 89 | 229 | 100 | 104 |
| 4252 | W303.asm02.HP0_G0042470 | chrXIII | 1674 | 272 | 512 | 1535 | 510 | 549 | 1616 | 265 | 630 | 3051 | 575 | 1175 | 1288 | 457 | 698 | 1326 | 453 | 971 |
| 4253 | W303.asm02.HP0_G0042480 | chrXIII | 5542 | 1906 | 1867 | 5670 | 1588 | 1673 | 5679 | 1904 | 1959 | 9849 | 2429 | 2942 | 7907 | 1494 | 2189 | 8137 | 2248 | 2953 |
| 4254 | W303.asm02.HP0_G0042490 | chrXIII | 1016 | 1042 | 1149 | 900 | 1017 | 1130 | 926 | 1044 | 1260 | 3941 | 970 | 1067 | 3543 | 531 | 759 | 4234 | 900 | 991 |
| 4255 | W303.asm02.HP0_G0042500 | chrXIII | 4080 | 995 | 725 | 3788 | 984 | 1091 | 3500 | 868 | 929 | 3462 | 941 | 975 | 2788 | 535 | 847 | 3158 | 922 | 880 |
| 4256 | W303.asm02.HP0_G0042510 | chrXIII | 1132 | 950 | 1068 | 1114 | 1144 | 1016 | 1108 | 896 | 1132 | 4577 | 806 | 1042 | 4170 | 536 | 716 | 4285 | 765 | 931 |
| 4257 | W303.asm02.HP0_G0042520 | chrXIII | 2913 | 1799 | 2234 | 2371 | 2164 | 2054 | 2875 | 2090 | 2492 | 10537 | 1899 | 2082 | 10227 | 1206 | 1437 | 10654 | 1595 | 1890 |
| 4258 | W303.asm02.HP0_G0042530 | chrXIII | 2027 | 181 | 164 | 2177 | 209 | 241 | 2010 | 156 | 240 | 2055 | 423 | 528 | 1329 | 271 | 313 | 1155 | 341 | 443 |
| 4259 | W303.asm02.HP0_G0042540 | chrXIII | 2035 | 720 | 1169 | 2103 | 625 | 1450 | 2036 | 652 | 1448 | 4570 | 1441 | 3364 | 2371 | 1118 | 1759 | 2149 | 1317 | 2896 |
| 4260 | W303.asm02.HP0_G0042550 | chrXIII | 2879 | 2675 | 2063 | 2302 | 2131 | 1966 | 2729 | 2631 | 1833 | 8424 | 2023 | 2014 | 7978 | 1785 | 1844 | 7656 | 1855 | 1999 |
| 4261 | W303.asm02.HP0_G0042560 | chrXIII | 2456 | 724 | 1223 | 2725 | 1256 | 1175 | 2404 | 712 | 1544 | 8123 | 1404 | 2061 | 4778 | 1014 | 1533 | 4450 | 1288 | 1645 |
| 4262 | W303.asm02.HP0_G0042570 | chrXIII | 7958 | 3480 | 3295 | 7151 | 3616 | 3585 | 9131 | 3212 | 4477 | 30006 | 8908 | 8605 | 15884 | 6012 | 4407 | 18602 | 7790 | 7424 |
| 4263 | W303.asm02.HP0_G0042580 | chrXIII | 3119 | 696 | 1121 | 2905 | 953 | 1087 | 2842 | 761 | 1237 | 6068 | 907 | 1674 | 5474 | 612 | 1345 | 5355 | 780 | 1511 |
| 4264 | W303.asm02.HP0_G0042590 | chrXIII | 7785 | 333 | 423 | 7838 | 413 | 617 | 7557 | 280 | 563 | 4744 | 629 | 1673 | 2582 | 489 | 1032 | 2509 | 557 | 1417 |
| 4265 | W303.asm02.HP0_G0042600 | chrXIII | 3269 | 2685 | 1973 | 3074 | 2031 | 1952 | 3427 | 2493 | 1946 | 8580 | 2781 | 2902 | 7001 | 1839 | 2049 | 7695 | 2462 | 2753 |
| 4266 | W303.asm02.HP0_G0042610 | chrXIII | 2437 | 2141 | 2030 | 2501 | 2054 | 1928 | 2655 | 2264 | 1987 | 7611 | 1889 | 2150 | 8105 | 1198 | 1446 | 7282 | 1716 | 1820 |
| 4267 | W303.asm02.HP0_G0042620 | chrXIII | 2720 | 728 | 539 | 2213 | 546 | 559 | 2533 | 633 | 556 | 2160 | 649 | 681 | 2148 | 438 | 627 | 2129 | 632 | 581 |
| 4268 | W303.asm02.HP0_G0042630 | chrXIII | 940 | 892 | 1163 | 912 | 1104 | 1089 | 875 | 863 | 1361 | 5092 | 778 | 826 | 5016 | 448 | 596 | 5228 | 717 | 774 |
| 4269 | W303.asm02.HP0_G0042640 | chrXIII | 1203 | 366 | 347 | 1299 | 343 | 333 | 1172 | 305 | 320 | 2784 | 492 | 625 | 2121 | 377 | 409 | 1960 | 509 | 545 |
| 4270 | W303.asm02.HP0_G0042650 | chrXIII | 1927 | 790 | 1136 | 1779 | 1072 | 1122 | 1787 | 839 | 1450 | 2539 | 672 | 1028 | 2994 | 483 | 800 | 2562 | 696 | 923 |
| 4271 | W303.asm02.HP0_G0042660 | chrXIII | 8945 | 4957 | 4862 | 8647 | 4323 | 4836 | 8725 | 5132 | 5284 | 11711 | 4789 | 5433 | 14553 | 3569 | 4468 | 10727 | 4292 | 4858 |
| 4272 | W303.asm02.HP0_G0042670 | chrXIII | 2072 | 222 | 291 | 1997 | 344 | 395 | 2031 | 217 | 377 | 3130 | 447 | 859 | 2076 | 309 | 573 | 1810 | 378 | 639 |
| 4273 | W303.asm02.HP0_G0042680 | chrXIII | 2067 | 733 | 925 | 2007 | 870 | 900 | 1919 | 781 | 1154 | 3113 | 602 | 817 | 3370 | 464 | 688 | 3213 | 545 | 781 |
| 4274 | W303.asm02.HP0_G0042690 | chrXIII | 6297 | 1616 | 1627 | 5947 | 2267 | 1840 | 6885 | 1603 | 2047 | 22723 | 2997 | 4667 | 14136 | 2063 | 2648 | 14811 | 2433 | 4427 |
| 4275 | W303.asm02.HP0_G0042700 | chrXIII | 5060 | 4858 | 5829 | 5191 | 5592 | 5885 | 5317 | 4937 | 7743 | 12327 | 4289 | 4384 | 14425 | 3125 | 3648 | 13457 | 3624 | 4087 |
| 4276 | W303.asm02.HP0_G0042710 | chrXIII | 8 | 13 | 2 | 6 | 6 | 3 | 2 | 6 | 7 | 27 | 4 | 2 | 26 | 2 | 0 | 35 | 5 | 2 |

| | | | | | | | | | | | | | | | | | | | | |
|---|---|---|---|---|---|---|---|---|---|---|---|---|---|---|---|---|---|---|---|---|
| 4277 | W303.asm02.HP0_G0042720 | chrXIII | 1881 | 546 | 528 | 1782 | 559 | 540 | 1712 | 605 | 593 | 4546 | 722 | 1061 | 3470 | 562 | 809 | 3607 | 695 | 934 |
| 4278 | W303.asm02.HP0_G0042730 | chrXIII | 585 | 135 | 109 | 625 | 103 | 111 | 504 | 155 | 131 | 471 | 122 | 137 | 381 | 63 | 92 | 447 | 126 | 123 |
| 4279 | W303.asm02.HP0_G0042740 | chrXIII | 1284 | 619 | 810 | 1168 | 686 | 774 | 1192 | 581 | 904 | 2520 | 530 | 572 | 2596 | 321 | 443 | 2603 | 443 | 494 |
| 4280 | W303.asm02.HP0_G0042750 | chrXIII | 1573 | 916 | 1252 | 1382 | 944 | 1319 | 1521 | 908 | 1422 | 3328 | 669 | 1131 | 3416 | 418 | 731 | 3490 | 683 | 997 |
| 4281 | W303.asm02.HP0_G0042760 | chrXIII | 5491 | 4139 | 3573 | 4906 | 3831 | 3649 | 5545 | 4262 | 3717 | 14704 | 4125 | 4466 | 14193 | 2686 | 3455 | 13977 | 3653 | 4002 |
| 4282 | W303.asm02.HP0_G0042770 | chrXIII | 1831 | 1934 | 1527 | 1623 | 1472 | 1605 | 1731 | 1924 | 1754 | 3750 | 1721 | 1560 | 4119 | 1200 | 1249 | 3516 | 1526 | 1446 |
| 4283 | W303.asm02.HP0_G0042780 | chrXIII | 7739 | 1159 | 1051 | 7365 | 991 | 1187 | 7499 | 1071 | 1462 | 8052 | 1925 | 2592 | 5895 | 1558 | 2020 | 5712 | 1820 | 2154 |
| 4284 | W303.asm02.HP0_G0042790 | chrXIII | 4289 | 1441 | 1550 | 4219 | 1505 | 1428 | 4059 | 1498 | 1743 | 7809 | 2111 | 2543 | 9714 | 2249 | 2350 | 6196 | 1906 | 2284 |
| 4285 | W303.asm02.HP0_G0042800 | chrXIII | 1477 | 1226 | 1644 | 1474 | 1965 | 1392 | 1750 | 1327 | 1906 | 8477 | 1396 | 1418 | 8405 | 846 | 812 | 8173 | 1149 | 1185 |
| 4286 | W303.asm02.HP0_G0042810 | chrXIII | 1054 | 548 | 320 | 965 | 371 | 357 | 938 | 480 | 304 | 2034 | 539 | 567 | 1584 | 326 | 345 | 1564 | 505 | 526 |
| 4287 | W303.asm02.HP0_G0042820 | chrXIII | 1331 | 3159 | 3412 | 1033 | 3402 | 3187 | 1314 | 3413 | 3658 | 12876 | 3890 | 4613 | 10859 | 2592 | 3640 | 11827 | 3359 | 4068 |
| 4288 | W303.asm02.HP0_G0042830 | chrXIII | 10806 | 8252 | 11927 | 10540 | 11138 | 11926 | 11501 | 8528 | 14924 | 25866 | 7757 | 9737 | 29089 | 5623 | 7777 | 27098 | 6589 | 8687 |
| 4289 | W303.asm02.HP0_G0042840 | chrXIII | 6585 | 4365 | 4310 | 6405 | 4651 | 3961 | 6819 | 4492 | 4287 | 19874 | 4820 | 4809 | 18768 | 3296 | 3523 | 18869 | 4376 | 4326 |
| 4290 | W303.asm02.HP0_G0042850 | chrXIII | 91 | 36 | 30 | 76 | 51 | 63 | 97 | 19 | 22 | 117 | 56 | 40 | 54 | 17 | 28 | 69 | 54 | 34 |
| 4291 | W303.asm02.HP0_G0042860 | chrXIII | 2502 | 908 | 1001 | 1927 | 879 | 988 | 2007 | 833 | 1138 | 2076 | 816 | 842 | 2271 | 544 | 717 | 2057 | 704 | 774 |
| 4292 | W303.asm02.HP0_G0042870 | chrXIII | 11002 | 1913 | 2170 | 9328 | 2182 | 2343 | 11370 | 1477 | 2913 | 20637 | 4855 | 4548 | 7590 | 3455 | 2705 | 8229 | 4104 | 3918 |
| 4293 | W303.asm02.HP0_G0042880 | chrXIII | 3006 | 1085 | 1389 | 2856 | 1260 | 1359 | 2818 | 1142 | 1534 | 8872 | 1127 | 1999 | 7292 | 771 | 1368 | 8501 | 1105 | 1833 |
| 4294 | W303.asm02.HP0_G0042890 | chrXIII | 12343 | 506 | 358 | 10252 | 271 | 575 | 11509 | 403 | 548 | 1371 | 849 | 1088 | 1074 | 820 | 1025 | 670 | 853 | 883 |
| 4295 | W303.asm02.HP0_G0042900 | chrXIII | 282 | 108 | 112 | 287 | 103 | 142 | 222 | 102 | 128 | 318 | 116 | 112 | 335 | 68 | 86 | 298 | 123 | 147 |
| 4296 | W303.asm02.HP0_G0042910 | chrXIII | 8988 | 4901 | 5320 | 9151 | 4058 | 5697 | 8330 | 4354 | 7129 | 4427 | 3841 | 4166 | 7035 | 2985 | 3838 | 4880 | 3627 | 3361 |
| 4297 | W303.asm02.HP0_G0042920 | chrXIII | 1176 | 263 | 383 | 1020 | 258 | 355 | 1135 | 298 | 378 | 1120 | 306 | 358 | 1134 | 191 | 303 | 937 | 288 | 306 |
| 4298 | W303.asm02.HP0_G0042930 | chrXIII | 1920 | 401 | 359 | 1677 | 313 | 339 | 1638 | 358 | 412 | 1855 | 585 | 574 | 1194 | 377 | 419 | 1296 | 528 | 512 |
| 4299 | W303.asm02.HP0_G0042940 | chrXIII | 3784 | 1417 | 1403 | 3777 | 1446 | 1421 | 3587 | 1404 | 1582 | 4080 | 1271 | 1266 | 5088 | 880 | 1150 | 4291 | 995 | 1174 |
| 4300 | W303.asm02.HP0_G0042950 | chrXIII | 1370 | 1471 | 1827 | 1078 | 1579 | 1855 | 1162 | 1432 | 2450 | 3005 | 1153 | 1459 | 3502 | 874 | 1245 | 3562 | 1140 | 1039 |
| 4301 | W303.asm02.HP0_G0042960 | chrXIII | 999 | 431 | 819 | 881 | 763 | 732 | 928 | 563 | 909 | 4201 | 583 | 1093 | 4064 | 417 | 914 | 3787 | 605 | 966 |
| 4302 | W303.asm02.HP0_G0042970 | chrXIII | 3263 | 1071 | 1088 | 2688 | 948 | 1010 | 2965 | 1164 | 1107 | 4473 | 1175 | 1340 | 4366 | 856 | 1207 | 4537 | 1125 | 1320 |
| 4303 | W303.asm02.HP0_G0042980 | chrXIII | 2146 | 1017 | 1259 | 2107 | 1553 | 1050 | 2317 | 1004 | 1471 | 11352 | 1179 | 1847 | 9820 | 832 | 1298 | 9987 | 1061 | 1566 |
| 4304 | W303.asm02.HP0_G0042990 | chrXIII | 8140 | 5655 | 7311 | 8438 | 6857 | 6911 | 8200 | 6153 | 9456 | 18205 | 5988 | 6629 | 20448 | 4174 | 5477 | 20075 | 5041 | 5905 |
| 4305 | W303.asm02.HP0_G0043000 | chrXIII | 10328 | 1047 | 1234 | 8583 | 977 | 1412 | 9531 | 930 | 1439 | 8682 | 1986 | 2233 | 6986 | 1538 | 2236 | 6046 | 1826 | 2014 |
| 4306 | W303.asm02.HP0_G0043010 | chrXIII | 2465 | 440 | 289 | 1962 | 241 | 298 | 2295 | 347 | 274 | 1163 | 478 | 497 | 1326 | 340 | 467 | 1100 | 524 | 427 |
| 4307 | W303.asm02.HP0_G0043020 | chrXIII | 1763 | 1107 | 1038 | 1567 | 1033 | 1013 | 1545 | 1073 | 1290 | 3288 | 927 | 893 | 3632 | 556 | 744 | 3145 | 915 | 776 |
| 4308 | W303.asm02.HP0_G0043030 | chrXIII | 2501 | 1535 | 1789 | 2222 | 1521 | 1665 | 2360 | 1580 | 1937 | 4773 | 1394 | 1704 | 6009 | 1002 | 1473 | 5035 | 1328 | 1693 |
| 4309 | W303.asm02.HP0_G0043040 | chrXIII | 1297 | 602 | 615 | 1132 | 624 | 569 | 1209 | 538 | 586 | 2948 | 832 | 841 | 2341 | 411 | 535 | 2219 | 802 | 679 |
| 4310 | W303.asm02.HP0_G0043050 | chrXIII | 3906 | 1141 | 1830 | 3920 | 1851 | 1793 | 3826 | 1251 | 2127 | 15609 | 3098 | 3620 | 9426 | 1634 | 2214 | 9973 | 2430 | 3299 |
| 4311 | W303.asm02.HP0_G0043060 | chrXIII | 6487 | 1920 | 2153 | 6252 | 1983 | 2090 | 6283 | 2188 | 2392 | 7077 | 1920 | 2403 | 7669 | 1379 | 2082 | 7027 | 1824 | 2132 |
| 4312 | W303.asm02.HP0_G0043070 | chrXIII | 693 | 447 | 577 | 679 | 618 | 545 | 645 | 442 | 575 | 2655 | 461 | 611 | 2772 | 270 | 404 | 2464 | 491 | 488 |
| 4313 | W303.asm02.HP0_G0043080 | chrXIII | 1339 | 903 | 833 | 983 | 1041 | 802 | 1162 | 1058 | 1058 | 3699 | 985 | 754 | 3996 | 552 | 572 | 3713 | 792 | 745 |
| 4314 | W303.asm02.HP0_G0043090 | chrXIII | 1670 | 1305 | 1410 | 1543 | 1339 | 1302 | 1612 | 1248 | 1410 | 4824 | 1200 | 1268 | 4784 | 835 | 1024 | 4762 | 1172 | 1177 |
| 4315 | W303.asm02.HP0_G0043100 | chrXIII | 2143 | 403 | 277 | 1846 | 251 | 271 | 1824 | 361 | 321 | 701 | 344 | 283 | 821 | 257 | 269 | 730 | 388 | 312 |
| 4316 | W303.asm02.HP0_G0043110 | chrXIII | 6185 | 1663 | 1303 | 5676 | 1207 | 1332 | 6451 | 1527 | 1384 | 9200 | 2870 | 2901 | 7194 | 2027 | 2385 | 6880 | 2539 | 2674 |
| 4317 | W303.asm02.HP0_G0043120 | chrXIII | 1936 | 135 | 244 | 1805 | 220 | 279 | 1788 | 99 | 303 | 1182 | 245 | 506 | 411 | 150 | 248 | 376 | 183 | 419 |
| 4318 | W303.asm02.HP0_G0043130 | chrXIII | 529 | 653 | 666 | 503 | 549 | 651 | 513 | 715 | 730 | 2078 | 515 | 631 | 2007 | 284 | 385 | 1917 | 448 | 553 |
| 4319 | W303.asm02.HP0_G0043140 | chrXIII | 2535 | 437 | 570 | 2472 | 599 | 716 | 2533 | 437 | 752 | 4967 | 921 | 1430 | 2707 | 693 | 1065 | 2442 | 798 | 1336 |
| 4320 | W303.asm02.HP0_G0043150 | chrXIII | 11654 | 2021 | 1668 | 11256 | 1405 | 1966 | 12477 | 1620 | 2160 | 9561 | 4233 | 4877 | 3759 | 3197 | 3366 | 3378 | 3399 | 4215 |
| 4321 | W303.asm02.HP0_G0043160 | chrXIII | 3720 | 559 | 626 | 2863 | 514 | 877 | 3923 | 443 | 920 | 5145 | 1233 | 1878 | 1263 | 889 | 924 | 1342 | 990 | 1442 |
| 4322 | W303.asm02.HP0_G0043170 | chrXIII | 3705 | 504 | 458 | 3555 | 428 | 417 | 3398 | 459 | 512 | 3389 | 684 | 974 | 2519 | 569 | 708 | 2195 | 602 | 802 |
| 4323 | W303.asm02.HP0_G0043180 | chrXIII | 843 | 98 | 112 | 658 | 114 | 145 | 756 | 81 | 160 | 985 | 182 | 372 | 411 | 153 | 264 | 449 | 158 | 355 |
| 4324 | W303.asm02.HP0_G0043190 | chrXIII | 1837 | 190 | 196 | 1984 | 212 | 227 | 1555 | 158 | 219 | 975 | 264 | 370 | 922 | 183 | 291 | 714 | 222 | 276 |
| 4325 | W303.asm02.HP0_G0043200 | chrXIII | 1820 | 528 | 486 | 1605 | 478 | 485 | 1514 | 566 | 494 | 1956 | 444 | 486 | 2212 | 312 | 429 | 1875 | 407 | 460 |
| 4326 | W303.asm02.HP0_G0043210 | chrXIII | 1371 | 993 | 1225 | 1244 | 1081 | 1124 | 1247 | 983 | 1342 | 4629 | 916 | 1185 | 4670 | 614 | 960 | 4461 | 800 | 1020 |
| 4327 | W303.asm02.HP0_G0043220 | chrXIII | 1593 | 558 | 735 | 1668 | 664 | 548 | 1574 | 596 | 722 | 3102 | 564 | 651 | 2933 | 367 | 465 | 2931 | 502 | 577 |
| 4328 | W303.asm02.HP0_G0043230 | chrXIII | 2522 | 736 | 840 | 2721 | 753 | 787 | 2232 | 732 | 1049 | 3020 | 583 | 681 | 3554 | 401 | 632 | 3278 | 648 | 611 |
| 4329 | W303.asm02.HP0_G0043240 | chrXIII | 2854 | 747 | 678 | 2412 | 688 | 674 | 2646 | 636 | 702 | 4425 | 940 | 1225 | 3420 | 600 | 760 | 3673 | 888 | 1067 |
| 4330 | W303.asm02.HP0_G0043250 | chrXIII | 2499 | 926 | 1384 | 1970 | 1246 | 1315 | 1990 | 908 | 1793 | 2818 | 821 | 1114 | 3621 | 567 | 1009 | 3015 | 731 | 976 |
| 4331 | W303.asm02.HP0_G0043260 | chrXIII | 1110 | 356 | 525 | 993 | 522 | 548 | 1081 | 384 | 588 | 2919 | 455 | 836 | 2340 | 321 | 642 | 2322 | 437 | 760 |
| 4332 | W303.asm02.HP0_G0043270 | chrXIII | 1339 | 323 | 571 | 1217 | 531 | 494 | 1104 | 347 | 548 | 3038 | 375 | 717 | 2787 | 260 | 500 | 2709 | 394 | 656 |
| 4333 | W303.asm02.HP0_G0043280 | chrXIII | 1589 | 392 | 607 | 1643 | 688 | 593 | 1572 | 448 | 613 | 5138 | 693 | 1100 | 4161 | 448 | 737 | 3987 | 639 | 932 |
| 4334 | W303.asm02.HP0_G0043290 | chrXIII | 3703 | 1532 | 1631 | 3269 | 1614 | 1516 | 3463 | 1648 | 1763 | 5307 | 1528 | 1733 | 6138 | 1074 | 1480 | 5023 | 1324 | 1579 |
| 4335 | W303.asm02.HP0_G0043300 | chrXIII | 4163 | 1690 | 1777 | 3860 | 2153 | 1746 | 4700 | 1725 | 2035 | 17777 | 3136 | 4034 | 12841 | 1973 | 2001 | 14097 | 2603 | 3428 |
| 4336 | W303.asm02.HP0_G0043310 | chrXIII | 2702 | 245 | 257 | 3408 | 236 | 286 | 2583 | 203 | 315 | 2024 | 383 | 647 | 2108 | 187 | 558 | 1150 | 417 | 550 |
| 4337 | W303.asm02.HP0_G0043320 | chrXIII | 456 | 622 | 1078 | 381 | 913 | 963 | 429 | 631 | 1357 | 2303 | 548 | 764 | 2659 | 361 | 533 | 2664 | 482 | 633 |
| 4338 | W303.asm02.HP0_G0043330 | chrXIII | 1802 | 553 | 504 | 1736 | 415 | 423 | 1637 | 464 | 494 | 2194 | 457 | 661 | 2369 | 375 | 502 | 1982 | 533 | 541 |
| 4339 | W303.asm02.HP0_G0043340 | chrXIII | 1722 | 886 | 789 | 1627 | 878 | 730 | 1746 | 909 | 819 | 2804 | 732 | 647 | 3165 | 486 | 535 | 2616 | 716 | 637 |
| 4340 | W303.asm02.HP0_G0043350 | chrXIII | 8595 | 3934 | 4279 | 8177 | 4041 | 4103 | 8685 | 4098 | 4837 | 8374 | 3265 | 3434 | 11705 | 2796 | 3183 | 8322 | 2903 | 3075 |
| 4341 | W303.asm02.HP0_G0043360 | chrXIII | 7123 | 1255 | 1437 | 6053 | 1129 | 1716 | 6951 | 1023 | 1797 | 10289 | 3079 | 3738 | 4263 | 2426 | 2409 | 3655 | 2520 | 3087 |
| 4342 | W303.asm02.HP0_G0043370 | chrXIII | 31583 | 6996 | 5427 | 28214 | 4583 | 5287 | 30082 | 6909 | 5666 | 20452 | 9604 | 9737 | 22070 | 7370 | 8112 | 17222 | 8616 | 8763 |
| 4343 | W303.asm02.HP0_G0043380 | chrXIII | 2574 | 1531 | 1530 | 2285 | 1368 | 1510 | 2314 | 1490 | 1791 | 3180 | 1276 | 1217 | 3578 | 886 | 989 | 3427 | 1145 | 1049 |
| 4344 | W303.asm02.HP0_G0043390 | chrXIII | 1451 | 832 | 993 | 1382 | 874 | 1013 | 1364 | 816 | 1262 | 2577 | 637 | 756 | 3405 | 396 | 616 | 2993 | 574 | 672 |
| 4345 | W303.asm02.HP0_G0043400 | chrXIII | 1518 | 1122 | 1320 | 1132 | 1229 | 1161 | 1552 | 1222 | 1522 | 8805 | 1958 | 2572 | 6646 | 1305 | 1675 | 7236 | 1808 | 2377 |
| 4346 | W303.asm02.HP0_G0043410 | chrXIII | 3469 | 616 | 689 | 2904 | 697 | 804 | 3651 | 550 | 945 | 5340 | 1334 | 1631 | 2703 | 896 | 1045 | 2799 | 1075 | 1499 |
| 4347 | W303.asm02.HP0_G0043420 | chrXIII | 5813 | 1020 | 1246 | 5614 | 1067 | 1410 | 5422 | 944 | 1541 | 7351 | 1894 | 2740 | 4078 | 1451 | 2048 | 4148 | 1758 | 2469 |
| 4348 | W303.asm02.HP0_G0043430 | chrXIII | 309 | 122 | 153 | 410 | 187 | 192 | 430 | 109 | 219 | 1051 | 192 | 270 | 761 | 110 | 172 | 675 | 238 | 240 |
| 4349 | W303.asm02.HP0_G0043440 | chrXIII | 6272 | 628 | 759 | 6294 | 683 | 833 | 5780 | 629 | 987 | 5859 | 972 | 1501 | 4529 | 857 | 1317 | 3760 | 841 | 1294 |
| 4350 | W303.asm02.HP0_G0043450 | chrXIII | 18670 | 4426 | 4307 | 19348 | 4080 | 4376 | 21006 | 5020 | 4796 | 22589 | 8948 | 9359 | 18184 | 5382 | 5496 | 16582 | 6638 | 7973 |
| 4351 | W303.asm02.HP0_G0043460 | chrXIII | 1258 | 222 | 301 | 1149 | 278 | 306 | 1238 | 217 | 325 | 1550 | 260 | 441 | 1422 | 223 | 274 | 1036 | 174 | 369 |
| 4352 | W303.asm02.HP0_G0043470 | chrXIII | 28361 | 15856 | 16607 | 30007 | 16783 | 15056 | 31041 | 17606 | 15766 | 60951 | 21328 | 24218 | 58142 | 14256 | 17109 | 56071 | 17632 | 21894 |
| 4353 | W303.asm02.HP0_G0043480 | chrXIII | 7112 | 2108 | 2291 | 6510 | 2329 | 2354 | 7772 | 2113 | 2824 | 21667 | 3979 | 5412 | 15948 | 2940 | 3697 | 15307 | 3533 | 5010 |
| 4354 | W303.asm02.HP0_G0043490 | chrXIII | 9417 | 5306 | 7313 | 9314 | 7339 | 6967 | 10168 | 5761 | 7244 | 33592 | 5918 | 7881 | 30317 | 4263 | 5537 | 31140 | 5221 | 7165 |
| 4355 | W303.asm02.HP0_G0043500 | chrXIII | 1371 | 492 | 553 | 1559 | 578 | 537 | 1747 | 504 | 600 | 4444 | 952 | 1067 | 4456 | 695 | 632 | 3474 | 737 | 935 |
| 4356 | W303.asm02.HP0_G0043510 | chrXIII | 8627 | 2233 | 1974 | 8653 | 1736 | 2028 | 7883 | 2210 | 1952 | 5258 | 2065 | 2368 | 6183 | 1431 | 1922 | 4881 | 1885 | 2190 |
| 4357 | W303.asm02.HP0_G0043520 | chrXIII | 1396 | 1037 | 1067 | 1377 | 1160 | 1071 | 1404 | 1138 | 1133 | 4333 | 1087 | 1091 | 4104 | 632 | 662 | 4045 | 907 | 993 |
| 4358 | W303.asm02.HP0_G0043530 | chrXIII | 2218 | 639 | 713 | 2120 | 663 | 620 | 1992 | 615 | 657 | 4528 | 951 | 1075 | 3397 | 583 | 867 | 3811 | 908 | 1099 |
| 4359 | W303.asm02.HP0_G0043540 | chrXIII | 9558 | 3199 | 2960 | 8779 | 2535 | 2968 | 8503 | 2902 | 3053 | 9645 | 3621 | 4559 | 8619 | 2880 | 3666 | 8730 | 3227 | 4064 |
| 4360 | W303.asm02.HP0_G0043550 | chrXIII | 7095 | 2028 | 1976 | 6326 | 1514 | 2062 | 6578 | 1939 | 1713 | 6245 | 2171 | 3656 | 10335 | 2240 | 3154 | 5381 | 2087 | 3173 |
| 4361 | W303.asm02.HP0_G0043560 | chrXIII | 739 | 134 | 161 | 775 | 138 | 137 | 630 | 105 | 140 | 401 | 79 | 109 | 496 | 61 | 81 | 383 | 96 | 122 |
| 4362 | W303.asm02.HP0_G0043570 | chrXIII | 3737 | 221 | 323 | 2811 | 332 | 434 | 3208 | 218 | 394 | 2072 | 343 | 713 | 1283 | 235 | 463 | 1029 | 260 | 628 |
| 4363 | W303.asm02.HP0_G0043580 | chrXIII | 13818 | 6282 | 6347 | 12563 | 5872 | 5837 | 13882 | 6518 | 6615 | 16460 | 9257 | 9232 | 15893 | 6989 | 7393 | 13334 | 7618 | 7076 |
| 4364 | W303.asm02.HP0_G0043590 | chrXIII | 4545 | 1744 | 1710 | 4180 | 1787 | 1615 | 4437 | 1857 | 1894 | 13431 | 2803 | 2315 | 13062 | 1815 | 2411 | 11990 | 2384 | 2754 |
| 4365 | W303.asm02.HP0_G0043600 | chrXIII | 34 | 10 | 7 | 21 | 14 | 5 | 41 | 11 | 14 | 29 | 10 | 8 | 15 | 8 | 4 | 31 | 7 | 15 |
| 4366 | W303.asm02.HP0_G0043620 | chrXIV | 0 | 0 | 0 | 0 | 0 | 0 | 0 | 0 | 0 | 0 | 0 | 0 | 0 | 0 | 0 | 0 | 0 | 0 |
| 4367 | W303.asm02.HP0_G0043630 | chrXIV | 8 | 10 | 0 | 3 | 0 | 1 | 4 | 14 | 0 | 0 | 8 | 0 | 3 | 3 | 0 | 1 | 6 | 1 |
| 4368 | W303.asm02.HP0_G0043640 | chrXIV | 3772 | 1322 | 1296 | 3218 | 1081 | 1599 | 3347 | 1299 | 1743 | 2940 | 968 | 1170 | 3495 | 664 | 827 | 3133 | 848 | 1139 |
| 4369 | W303.asm02.HP0_G0043650 | chrXIV | 1350 | 186 | 207 | 1261 | 243 | 215 | 1250 | 219 | 198 | 648 | 219 | 254 | 692 | 188 | 210 | 547 | 234 | 286 |

| | | | | | | | | | | | | | | | | | | | | |
|---|---|---|---|---|---|---|---|---|---|---|---|---|---|---|---|---|---|---|---|---|
| 4370 | W303.asm02.HP0_G0043660 | chrXIV | 678 | 538 | 630 | 638 | 560 | 674 | 596 | 585 | 815 | 1359 | 455 | 577 | 1496 | 267 | 352 | 1338 | 418 | 524 |
| 4371 | W303.asm02.HP0_G0043670 | chrXIV | 2012 | 822 | 779 | 1836 | 808 | 829 | 1839 | 845 | 827 | 1715 | 849 | 1133 | 1617 | 539 | 668 | 1382 | 758 | 886 |
| 4372 | W303.asm02.HP0_G0043680 | chrXIV | 582 | 234 | 154 | 588 | 149 | 202 | 573 | 188 | 152 | 276 | 149 | 178 | 400 | 95 | 110 | 231 | 190 | 187 |
| 4373 | W303.asm02.HP0_G0043690 | chrXIV | 1095 | 668 | 615 | 1046 | 563 | 635 | 1041 | 635 | 689 | 1754 | 861 | 734 | 1597 | 668 | 598 | 1420 | 809 | 545 |
| 4374 | W303.asm02.HP0_G0043700 | chrXIV | 3780 | 1294 | 979 | 3756 | 1076 | 1067 | 3656 | 1206 | 1062 | 6256 | 1861 | 1968 | 4319 | 1187 | 1527 | 4600 | 1634 | 1852 |
| 4375 | W303.asm02.HP0_G0043710 | chrXIV | 2287 | 381 | 610 | 2358 | 509 | 666 | 2322 | 354 | 748 | 4698 | 773 | 1456 | 2663 | 537 | 1022 | 2732 | 603 | 1277 |
| 4376 | W303.asm02.HP0_G0043720 | chrXIV | 166 | 43 | 82 | 170 | 78 | 77 | 169 | 43 | 102 | 601 | 105 | 167 | 426 | 70 | 128 | 365 | 81 | 152 |
| 4377 | W303.asm02.HP0_G0043730 | chrXIV | 16091 | 3330 | 3861 | 17138 | 3805 | 4733 | 17827 | 2722 | 5613 | 31333 | 6923 | 9588 | 21057 | 6138 | 7205 | 16825 | 5430 | 8379 |
| 4378 | W303.asm02.HP0_G0043740 | chrXIV | 452 | 779 | 758 | 407 | 743 | 790 | 405 | 728 | 844 | 2185 | 544 | 622 | 2227 | 356 | 439 | 2183 | 530 | 514 |
| 4379 | W303.asm02.HP0_G0043750 | chrXIV | 1214 | 613 | 661 | 1139 | 628 | 663 | 1164 | 665 | 709 | 2786 | 604 | 775 | 2808 | 371 | 562 | 2642 | 588 | 677 |
| 4380 | W303.asm02.HP0_G0043760 | chrXIV | 28 | 13 | 9 | 8 | 2 | 3 | 14 | 15 | 3 | 16 | 5 | 6 | 35 | 1 | 2 | 20 | 4 | 2 |
| 4381 | W303.asm02.HP0_G0043770 | chrXIV | 1970 | 1123 | 1130 | 1784 | 1145 | 1096 | 1830 | 1216 | 1267 | 4721 | 1104 | 1270 | 4800 | 751 | 978 | 4460 | 982 | 1128 |
| 4382 | W303.asm02.HP0_G0043780 | chrXIV | 4467 | 923 | 1006 | 4173 | 705 | 1108 | 4355 | 738 | 1237 | 3451 | 1331 | 1660 | 1805 | 945 | 1112 | 1540 | 1249 | 1515 |
| 4383 | W303.asm02.HP0_G0043790 | chrXIV | 3963 | 511 | 509 | 3601 | 456 | 623 | 3960 | 381 | 674 | 3909 | 1064 | 1228 | 1509 | 817 | 968 | 1438 | 885 | 1142 |
| 4384 | W303.asm02.HP0_G0043800 | chrXIV | 828 | 592 | 624 | 622 | 548 | 580 | 720 | 609 | 604 | 1965 | 544 | 727 | 1905 | 357 | 510 | 1961 | 503 | 590 |
| 4385 | W303.asm02.HP0_G0043810 | chrXIV | 53 | 17 | 16 | 71 | 23 | 11 | 43 | 14 | 22 | 38 | 24 | 14 | 56 | 10 | 13 | 28 | 15 | 13 |
| 4386 | W303.asm02.HP0_G0043820 | chrXIV | 2534 | 321 | 321 | 2516 | 292 | 407 | 2518 | 242 | 410 | 2267 | 480 | 801 | 1286 | 413 | 536 | 1163 | 493 | 650 |
| 4387 | W303.asm02.HP0_G0043830 | chrXIV | 984 | 744 | 617 | 901 | 608 | 557 | 1013 | 776 | 516 | 2246 | 575 | 766 | 2103 | 418 | 483 | 1982 | 547 | 615 |
| 4388 | W303.asm02.HP0_G0043840 | chrXIV | 10918 | 1810 | 1998 | 10379 | 1910 | 1821 | 10803 | 1908 | 2166 | 7873 | 1722 | 2353 | 9701 | 1326 | 2047 | 7624 | 1609 | 2123 |
| 4389 | W303.asm02.HP0_G0043850 | chrXIV | 963 | 231 | 190 | 899 | 193 | 205 | 872 | 255 | 184 | 959 | 267 | 307 | 1096 | 232 | 234 | 826 | 253 | 280 |
| 4390 | W303.asm02.HP0_G0043860 | chrXIV | 1377 | 880 | 1026 | 1591 | 1228 | 1225 | 1887 | 888 | 1318 | 11509 | 1928 | 2687 | 6531 | 1405 | 1471 | 6384 | 1622 | 2387 |
| 4391 | W303.asm02.HP0_G0043870 | chrXIV | 1928 | 2319 | 1962 | 1780 | 1808 | 1844 | 1919 | 2152 | 2050 | 4460 | 1838 | 1754 | 4869 | 1220 | 1294 | 4640 | 1580 | 1474 |
| 4392 | W303.asm02.HP0_G0043880 | chrXIV | 613 | 881 | 794 | 732 | 930 | 716 | 590 | 855 | 642 | 3824 | 772 | 952 | 3515 | 538 | 644 | 3302 | 773 | 825 |
| 4393 | W303.asm02.HP0_G0043890 | chrXIV | 1872 | 445 | 424 | 1568 | 397 | 468 | 1712 | 409 | 434 | 1552 | 427 | 662 | 1606 | 353 | 486 | 1323 | 396 | 579 |
| 4394 | W303.asm02.HP0_G0043900 | chrXIV | 535 | 277 | 246 | 497 | 211 | 274 | 573 | 171 | 289 | 1560 | 468 | 562 | 687 | 338 | 320 | 650 | 336 | 496 |
| 4395 | W303.asm02.HP0_G0043910 | chrXIV | 4170 | 1973 | 1185 | 4402 | 1757 | 1343 | 4889 | 1933 | 1248 | 13227 | 3031 | 3091 | 10279 | 1988 | 1677 | 9626 | 2398 | 2725 |
| 4396 | W303.asm02.HP0_G0043920 | chrXIV | 9531 | 3483 | 3155 | 9615 | 2896 | 2913 | 10001 | 3473 | 3216 | 14432 | 3988 | 4540 | 13316 | 3073 | 3909 | 12576 | 3603 | 4109 |
| 4397 | W303.asm02.HP0_G0043930 | chrXIV | 5510 | 2298 | 2365 | 4969 | 2139 | 2262 | 4950 | 2405 | 3009 | 5788 | 2191 | 2438 | 7367 | 1701 | 2298 | 5832 | 1980 | 2202 |
| 4398 | W303.asm02.HP0_G0043940 | chrXIV | 4738 | 923 | 484 | 4752 | 425 | 555 | 4364 | 868 | 592 | 1839 | 1162 | 986 | 2220 | 840 | 921 | 1658 | 1205 | 928 |
| 4399 | W303.asm02.HP0_G0043950 | chrXIV | 731 | 492 | 251 | 728 | 271 | 299 | 749 | 391 | 320 | 1268 | 445 | 532 | 960 | 248 | 340 | 1022 | 466 | 436 |
| 4400 | W303.asm02.HP0_G0043960 | chrXIV | 9322 | 8613 | 11238 | 8123 | 9936 | 10716 | 8873 | 8415 | 13381 | 20925 | 7924 | 8801 | 23084 | 5859 | 7252 | 22139 | 7211 | 7679 |
| 4401 | W303.asm02.HP0_G0043970 | chrXIV | 4435 | 3203 | 3315 | 3711 | 3450 | 3369 | 4628 | 3125 | 3436 | 11726 | 2632 | 3709 | 10435 | 1778 | 2826 | 11407 | 2443 | 3485 |
| 4402 | W303.asm02.HP0_G0043980 | chrXIV | 1644 | 625 | 1037 | 1391 | 840 | 1202 | 1499 | 596 | 1262 | 4730 | 1644 | 2377 | 3294 | 1115 | 1569 | 2998 | 1339 | 2181 |
| 4403 | W303.asm02.HP0_G0043990 | chrXIV | 310 | 308 | 366 | 301 | 463 | 379 | 350 | 326 | 357 | 4030 | 488 | 741 | 2733 | 350 | 367 | 2792 | 425 | 622 |
| 4404 | W303.asm02.HP0_G0044000 | chrXIV | 1033 | 391 | 473 | 972 | 455 | 629 | 1133 | 292 | 625 | 4716 | 840 | 1202 | 1597 | 618 | 713 | 1585 | 648 | 1039 |
| 4405 | W303.asm02.HP0_G0044010 | chrXIV | 3206 | 585 | 954 | 3054 | 824 | 1018 | 2842 | 616 | 1170 | 7280 | 902 | 1988 | 5249 | 667 | 1559 | 5305 | 810 | 1786 |
| 4406 | W303.asm02.HP0_G0044020 | chrXIV | 1873 | 332 | 426 | 2156 | 478 | 476 | 1728 | 292 | 566 | 3640 | 642 | 939 | 2398 | 441 | 696 | 2285 | 575 | 854 |
| 4407 | W303.asm02.HP0_G0044030 | chrXIV | 4186 | 1327 | 1125 | 3947 | 1168 | 1164 | 4154 | 1313 | 1148 | 6088 | 1630 | 1869 | 7387 | 1458 | 1791 | 4637 | 1426 | 1751 |
| 4408 | W303.asm02.HP0_G0044040 | chrXIV | 2516 | 403 | 387 | 2449 | 381 | 460 | 2316 | 348 | 489 | 2725 | 661 | 915 | 2353 | 659 | 781 | 1605 | 654 | 816 |
| 4409 | W303.asm02.HP0_G0044050 | chrXIV | 2514 | 1561 | 1404 | 2391 | 1515 | 1421 | 2452 | 1554 | 1445 | 6462 | 1751 | 1828 | 5758 | 1010 | 1159 | 5679 | 1341 | 1668 |
| 4410 | W303.asm02.HP0_G0044060 | chrXIV | 703 | 448 | 499 | 692 | 478 | 488 | 729 | 450 | 517 | 3522 | 663 | 873 | 2609 | 482 | 632 | 2627 | 550 | 803 |
| 4411 | W303.asm02.HP0_G0044070 | chrXIV | 2750 | 1614 | 1181 | 2369 | 1289 | 1196 | 2805 | 1567 | 1184 | 5492 | 1569 | 1815 | 5142 | 1174 | 1372 | 4855 | 1478 | 1593 |
| 4412 | W303.asm02.HP0_G0044080 | chrXIV | 185 | 519 | 1034 | 119 | 845 | 1075 | 221 | 588 | 1095 | 3073 | 816 | 1836 | 3028 | 565 | 1246 | 2394 | 771 | 1663 |
| 4413 | W303.asm02.HP0_G0044090 | chrXIV | 3392 | 990 | 734 | 3731 | 834 | 869 | 3511 | 1051 | 826 | 6116 | 1485 | 1601 | 6147 | 1084 | 1224 | 5161 | 1415 | 1394 |
| 4414 | W303.asm02.HP0_G0044100 | chrXIV | 12885 | 3799 | 4549 | 12164 | 4170 | 4552 | 13206 | 4247 | 4579 | 23965 | 4748 | 7657 | 22985 | 3920 | 5970 | 21791 | 4297 | 6945 |
| 4415 | W303.asm02.HP0_G0044110 | chrXIV | 2009 | 704 | 847 | 2028 | 652 | 775 | 1977 | 740 | 871 | 3075 | 799 | 1121 | 2501 | 518 | 780 | 2645 | 800 | 996 |
| 4416 | W303.asm02.HP0_G0044120 | chrXIV | 3157 | 2382 | 2253 | 2976 | 2175 | 2330 | 3019 | 2375 | 2612 | 6423 | 2100 | 2455 | 7163 | 1491 | 2050 | 6201 | 2003 | 2251 |
| 4417 | W303.asm02.HP0_G0044130 | chrXIV | 1383 | 579 | 755 | 1314 | 671 | 892 | 1610 | 456 | 946 | 3798 | 1282 | 1632 | 926 | 839 | 958 | 848 | 1003 | 1406 |
| 4418 | W303.asm02.HP0_G0044140 | chrXIV | 631 | 353 | 289 | 608 | 343 | 262 | 678 | 307 | 271 | 1936 | 471 | 425 | 1777 | 329 | 310 | 1530 | 485 | 387 |
| 4419 | W303.asm02.HP0_G0044150 | chrXIV | 3381 | 4132 | 4916 | 2974 | 4249 | 5039 | 3229 | 3734 | 6025 | 9973 | 3007 | 3962 | 11628 | 2377 | 3313 | 10541 | 2774 | 3696 |
| 4420 | W303.asm02.HP0_G0044160 | chrXIV | 3074 | 1850 | 1643 | 2583 | 1449 | 1539 | 3016 | 1752 | 1740 | 8543 | 2364 | 2444 | 6375 | 1742 | 1871 | 6902 | 2176 | 2245 |
| 4421 | W303.asm02.HP0_G0044170 | chrXIV | 369 | 341 | 194 | 345 | 199 | 203 | 364 | 278 | 179 | 868 | 381 | 259 | 635 | 187 | 170 | 739 | 400 | 281 |
| 4422 | W303.asm02.HP0_G0044180 | chrXIV | 611 | 103 | 174 | 588 | 158 | 221 | 546 | 91 | 213 | 1000 | 198 | 441 | 318 | 132 | 212 | 322 | 147 | 386 |
| 4423 | W303.asm02.HP0_G0044190 | chrXIV | 82 | 29 | 24 | 56 | 36 | 36 | 79 | 29 | 42 | 22 | 22 | 14 | 104 | 24 | 33 | 22 | 13 | 20 |
| 4424 | W303.asm02.HP0_G0044200 | chrXIV | 1533 | 481 | 396 | 1052 | 414 | 472 | 1282 | 415 | 478 | 1691 | 599 | 638 | 1176 | 534 | 585 | 1103 | 512 | 562 |
| 4425 | W303.asm02.HP0_G0044210 | chrXIV | 321 | 433 | 651 | 314 | 582 | 646 | 308 | 437 | 597 | 2809 | 523 | 820 | 2549 | 341 | 504 | 2249 | 480 | 667 |
| 4426 | W303.asm02.HP0_G0044220 | chrXIV | 7139 | 1316 | 1044 | 6289 | 845 | 1089 | 6233 | 1198 | 991 | 1991 | 1141 | 1267 | 3278 | 1146 | 1528 | 1854 | 1089 | 1050 |
| 4427 | W303.asm02.HP0_G0044230 | chrXIV | 763 | 325 | 432 | 716 | 478 | 431 | 785 | 411 | 497 | 3825 | 518 | 903 | 2844 | 358 | 666 | 2843 | 454 | 916 |
| 4428 | W303.asm02.HP0_G0044240 | chrXIV | 2896 | 928 | 1028 | 2967 | 1135 | 987 | 2973 | 1018 | 965 | 6528 | 1107 | 1599 | 6059 | 748 | 1178 | 5867 | 905 | 1468 |
| 4429 | W303.asm02.HP0_G0044250 | chrXIV | 4487 | 415 | 713 | 4481 | 656 | 827 | 4474 | 373 | 937 | 2482 | 523 | 1337 | 1819 | 429 | 762 | 1782 | 434 | 1006 |
| 4430 | W303.asm02.HP0_G0044260 | chrXIV | 532 | 69 | 53 | 421 | 86 | 59 | 416 | 73 | 67 | 309 | 117 | 86 | 379 | 58 | 78 | 246 | 89 | 85 |
| 4431 | W303.asm02.HP0_G0044270 | chrXIV | 60 | 14 | 10 | 57 | 13 | 14 | 56 | 14 | 17 | 52 | 13 | 19 | 40 | 8 | 7 | 46 | 12 | 15 |
| 4432 | W303.asm02.HP0_G0044280 | chrXIV | 4559 | 2397 | 2456 | 4380 | 2601 | 2686 | 4528 | 2352 | 2791 | 18741 | 4770 | 5101 | 11379 | 3003 | 3608 | 11785 | 4151 | 4597 |
| 4433 | W303.asm02.HP0_G0044290 | chrXIV | 5944 | 742 | 786 | 6142 | 803 | 936 | 5946 | 717 | 931 | 8153 | 1417 | 2308 | 5247 | 1062 | 1671 | 5359 | 1215 | 2104 |
| 4434 | W303.asm02.HP0_G0044300 | chrXIV | 2580 | 1391 | 1625 | 2492 | 1381 | 1581 | 2411 | 1470 | 1958 | 3996 | 1048 | 1323 | 4822 | 721 | 1079 | 4267 | 965 | 1081 |
| 4435 | W303.asm02.HP0_G0044310 | chrXIV | 2842 | 498 | 444 | 2569 | 419 | 399 | 2799 | 568 | 465 | 2944 | 592 | 645 | 2694 | 431 | 515 | 2579 | 515 | 578 |
| 4436 | W303.asm02.HP0_G0044320 | chrXIV | 3498 | 1757 | 1928 | 3505 | 1813 | 1734 | 3687 | 1860 | 1721 | 6165 | 1811 | 2119 | 6584 | 1056 | 1593 | 6212 | 1564 | 1876 |
| 4437 | W303.asm02.HP0_G0044330 | chrXIV | 2928 | 590 | 673 | 2669 | 739 | 713 | 2832 | 630 | 948 | 5798 | 841 | 1664 | 3028 | 735 | 1219 | 3172 | 727 | 1620 |
| 4438 | W303.asm02.HP0_G0044340 | chrXIV | 1611 | 1035 | 1317 | 1444 | 1167 | 1285 | 1592 | 1128 | 1480 | 3519 | 794 | 888 | 3826 | 561 | 689 | 3594 | 716 | 750 |
| 4439 | W303.asm02.HP0_G0044350 | chrXIV | 1047 | 956 | 824 | 1057 | 745 | 746 | 1029 | 1013 | 853 | 2519 | 919 | 842 | 2541 | 604 | 648 | 2664 | 927 | 784 |
| 4440 | W303.asm02.HP0_G0044360 | chrXIV | 357 | 340 | 492 | 274 | 409 | 514 | 290 | 329 | 676 | 1115 | 345 | 403 | 1435 | 237 | 346 | 1346 | 332 | 374 |
| 4441 | W303.asm02.HP0_G0044370 | chrXIV | 2879 | 1266 | 1722 | 2711 | 1574 | 1521 | 2720 | 1337 | 1794 | 6270 | 1184 | 1643 | 6080 | 739 | 1232 | 6418 | 1086 | 1533 |
| 4442 | W303.asm02.HP0_G0044380 | chrXIV | 2831 | 501 | 858 | 2898 | 740 | 986 | 2695 | 530 | 1071 | 4976 | 785 | 1746 | 3471 | 608 | 1332 | 4104 | 780 | 1524 |
| 4443 | W303.asm02.HP0_G0044390 | chrXIV | 3363 | 1449 | 1397 | 3411 | 1528 | 1453 | 3856 | 1573 | 1685 | 12149 | 2594 | 2812 | 9227 | 1648 | 1907 | 10024 | 2194 | 2651 |
| 4444 | W303.asm02.HP0_G0044400 | chrXIV | 1005 | 640 | 372 | 863 | 412 | 332 | 894 | 568 | 361 | 2776 | 679 | 497 | 2767 | 497 | 383 | 2163 | 839 | 438 |
| 4445 | W303.asm02.HP0_G0044410 | chrXIV | 1269 | 1337 | 1113 | 1333 | 1072 | 1036 | 1421 | 1305 | 1311 | 3543 | 995 | 896 | 3677 | 605 | 699 | 3594 | 1048 | 741 |
| 4446 | W303.asm02.HP0_G0044420 | chrXIV | 8684 | 1945 | 1802 | 8082 | 1834 | 1968 | 8176 | 2157 | 1976 | 6956 | 1689 | 2277 | 7816 | 1398 | 1989 | 6044 | 1601 | 1972 |
| 4447 | W303.asm02.HP0_G0044430 | chrXIV | 7230 | 946 | 535 | 6294 | 604 | 647 | 7446 | 765 | 671 | 7467 | 2162 | 1597 | 2738 | 1498 | 1102 | 3498 | 2055 | 1483 |
| 4448 | W303.asm02.HP0_G0044440 | chrXIV | 2228 | 440 | 656 | 2292 | 790 | 712 | 2319 | 466 | 850 | 4496 | 483 | 837 | 4234 | 369 | 700 | 3840 | 470 | 753 |
| 4449 | W303.asm02.HP0_G0044450 | chrXIV | 813 | 436 | 821 | 696 | 673 | 820 | 752 | 478 | 885 | 1810 | 409 | 671 | 1837 | 226 | 468 | 1821 | 361 | 577 |
| 4450 | W303.asm02.HP0_G0044460 | chrXIV | 4389 | 2967 | 3586 | 4381 | 4104 | 3196 | 5066 | 3386 | 3717 | 17138 | 3448 | 4136 | 15557 | 2154 | 2546 | 15992 | 2893 | 3682 |
| 4451 | W303.asm02.HP0_G0044470 | chrXIV | 8895 | 2156 | 1948 | 8826 | 2091 | 2076 | 9181 | 2067 | 2562 | 16525 | 4050 | 4973 | 9914 | 2948 | 2851 | 10181 | 3292 | 4243 |
| 4452 | W303.asm02.HP0_G0044480 | chrXIV | 3015 | 1508 | 1326 | 2951 | 1310 | 1251 | 3038 | 1579 | 1447 | 5818 | 1738 | 1827 | 6057 | 1255 | 1569 | 5238 | 1565 | 1636 |
| 4453 | W303.asm02.HP0_G0044490 | chrXIV | 4145 | 4153 | 5691 | 3659 | 5159 | 5575 | 3892 | 4146 | 7239 | 10003 | 3771 | 4553 | 12210 | 2752 | 3790 | 10817 | 3381 | 3913 |
| 4454 | W303.asm02.HP0_G0044500 | chrXIV | 7755 | 1315 | 1458 | 6730 | 1253 | 1849 | 8032 | 1104 | 1795 | 9459 | 2698 | 3657 | 4665 | 2027 | 2655 | 3915 | 2318 | 3234 |
| 4455 | W303.asm02.HP0_G0044510 | chrXIV | 2293 | 329 | 473 | 2211 | 434 | 446 | 1854 | 296 | 578 | 2701 | 517 | 893 | 1751 | 386 | 670 | 1564 | 446 | 675 |
| 4456 | W303.asm02.HP0_G0044520 | chrXIV | 21678 | 3306 | 3533 | 18849 | 2478 | 4322 | 22101 | 2392 | 4725 | 2136 | 2331 | 4194 | 1928 | 2522 | 2732 | 1445 | 1997 | 3008 |
| 4457 | W303.asm02.HP0_G0044530 | chrXIV | 2508 | 561 | 662 | 2029 | 657 | 671 | 2580 | 474 | 793 | 4207 | 1367 | 1232 | 2728 | 855 | 815 | 2203 | 1139 | 1092 |
| 4458 | W303.asm02.HP0_G0044540 | chrXIV | 7328 | 1194 | 944 | 6534 | 902 | 1089 | 7389 | 1164 | 1123 | 7393 | 2135 | 2563 | 7396 | 1945 | 2143 | 5966 | 1760 | 2150 |
| 4459 | W303.asm02.HP0_G0044550 | chrXIV | 1740 | 796 | 1009 | 1423 | 930 | 1145 | 1682 | 750 | 1127 | 7571 | 1582 | 2151 | 3893 | 1204 | 1383 | 3945 | 1359 | 1925 |
| 4460 | W303.asm02.HP0_G0044560 | chrXIV | 1373 | 246 | 223 | 1188 | 182 | 254 | 1232 | 177 | 272 | 1230 | 360 | 425 | 781 | 241 | 315 | 733 | 359 | 421 |
| 4461 | W303.asm02.HP0_G0044570 | chrXIV | 3435 | 638 | 858 | 3132 | 822 | 845 | 3283 | 722 | 901 | 6388 | 1006 | 1549 | 5073 | 763 | 1168 | 5123 | 873 | 1427 |
| 4462 | W303.asm02.HP0_G0044580 | chrXIV | 1112 | 286 | 604 | 1112 | 614 | 681 | 1060 | 282 | 562 | 2747 | 468 | 844 | 2265 | 222 | 374 | 2103 | 428 | 636 |

| | | | | | | | | | | | | | | | | | | | | |
|---|---|---|---|---|---|---|---|---|---|---|---|---|---|---|---|---|---|---|---|---|
| 4463 | W303.asm02.HP0_G0044590 | chrXIV | 2688 | 276 | 468 | 2512 | 436 | 603 | 2767 | 273 | 548 | 4127 | 559 | 1093 | 2680 | 410 | 666 | 2534 | 432 | 945 |
| 4464 | W303.asm02.HP0_G0044600 | chrXIV | 1678 | 1799 | 1628 | 1534 | 1886 | 1550 | 1785 | 1776 | 1686 | 5801 | 1476 | 1599 | 5576 | 996 | 1191 | 5399 | 1418 | 1446 |
| 4465 | W303.asm02.HP0_G0044610 | chrXIV | 2315 | 2666 | 3618 | 2281 | 3257 | 3555 | 2567 | 2621 | 4070 | 10528 | 2614 | 3223 | 10488 | 1385 | 2563 | 11026 | 2459 | 3054 |
| 4466 | W303.asm02.HP0_G0044620 | chrXIV | 1150 | 199 | 164 | 1006 | 133 | 177 | 1019 | 141 | 187 | 1459 | 418 | 438 | 1146 | 238 | 279 | 1050 | 389 | 403 |
| 4467 | W303.asm02.HP0_G0044630 | chrXIV | 6285 | 2559 | 2665 | 6887 | 2643 | 2632 | 6642 | 2715 | 2835 | 10009 | 2524 | 3199 | 11175 | 1935 | 2496 | 8952 | 2247 | 2845 |
| 4468 | W303.asm02.HP0_G0044640 | chrXIV | 2248 | 1173 | 782 | 2312 | 1074 | 860 | 2464 | 1370 | 842 | 5772 | 1331 | 1234 | 5764 | 928 | 796 | 5058 | 1180 | 1108 |
| 4469 | W303.asm02.HP0_G0044650 | chrXIV | 1005 | 956 | 1186 | 1031 | 1094 | 1089 | 931 | 888 | 1153 | 3300 | 735 | 919 | 3378 | 445 | 650 | 3352 | 718 | 771 |
| 4470 | W303.asm02.HP0_G0044660 | chrXIV | 2373 | 523 | 538 | 2343 | 562 | 558 | 2121 | 513 | 587 | 3833 | 661 | 1072 | 3454 | 469 | 795 | 2939 | 649 | 931 |
| 4471 | W303.asm02.HP0_G0044670 | chrXIV | 1165 | 284 | 256 | 1174 | 253 | 262 | 1089 | 263 | 310 | 1360 | 346 | 470 | 1308 | 254 | 423 | 974 | 386 | 377 |
| 4472 | W303.asm02.HP0_G0044680 | chrXIV | 934 | 666 | 770 | 941 | 649 | 791 | 949 | 743 | 1077 | 2054 | 539 | 771 | 2102 | 363 | 559 | 2485 | 519 | 710 |
| 4473 | W303.asm02.HP0_G0044690 | chrXIV | 2356 | 508 | 860 | 2123 | 957 | 911 | 2453 | 549 | 1098 | 7045 | 993 | 1908 | 3190 | 712 | 1008 | 3435 | 911 | 1594 |
| 4474 | W303.asm02.HP0_G0044700 | chrXIV | 3173 | 1441 | 1652 | 2796 | 1540 | 1533 | 2677 | 1353 | 1939 | 4928 | 1899 | 2201 | 4067 | 1343 | 1637 | 3914 | 1772 | 1865 |
| 4475 | W303.asm02.HP0_G0044710 | chrXIV | 3860 | 2233 | 1953 | 3821 | 1810 | 1691 | 3951 | 2263 | 1837 | 10642 | 2658 | 2728 | 9074 | 1726 | 1997 | 10094 | 2374 | 2471 |
| 4476 | W303.asm02.HP0_G0044720 | chrXIV | 592 | 489 | 751 | 581 | 594 | 677 | 652 | 506 | 676 | 3439 | 672 | 1058 | 3070 | 445 | 653 | 2807 | 578 | 941 |
| 4477 | W303.asm02.HP0_G0044730 | chrXIV | 4579 | 1629 | 2175 | 4542 | 1944 | 2239 | 4410 | 1629 | 2299 | 6062 | 1512 | 2005 | 6431 | 1072 | 1664 | 5813 | 1377 | 1869 |
| 4478 | W303.asm02.HP0_G0044740 | chrXIV | 2099 | 787 | 826 | 2030 | 720 | 859 | 2171 | 691 | 979 | 7391 | 1655 | 2038 | 4386 | 1056 | 1311 | 4599 | 1470 | 1833 |
| 4479 | W303.asm02.HP0_G0044750 | chrXIV | 4311 | 994 | 604 | 4003 | 762 | 692 | 3911 | 896 | 624 | 3919 | 1068 | 1168 | 3463 | 752 | 893 | 2946 | 996 | 1052 |
| 4480 | W303.asm02.HP0_G0044760 | chrXIV | 996 | 766 | 643 | 1188 | 483 | 655 | 990 | 640 | 884 | 1356 | 511 | 501 | 1584 | 280 | 391 | 1555 | 501 | 466 |
| 4481 | W303.asm02.HP0_G0044770 | chrXIV | 1439 | 1165 | 1181 | 1329 | 1196 | 1262 | 1337 | 1147 | 1249 | 3654 | 905 | 1119 | 4260 | 704 | 834 | 3153 | 860 | 970 |
| 4482 | W303.asm02.HP0_G0044780 | chrXIV | 7143 | 1914 | 2042 | 6270 | 1677 | 2124 | 6586 | 2053 | 2215 | 8720 | 2402 | 3608 | 8501 | 1589 | 2663 | 7969 | 2349 | 3150 |
| 4483 | W303.asm02.HP0_G0044790 | chrXIV | 438 | 354 | 496 | 372 | 369 | 477 | 362 | 365 | 644 | 1224 | 243 | 318 | 1198 | 162 | 263 | 1425 | 228 | 236 |
| 4484 | W303.asm02.HP0_G0044800 | chrXIV | 126 | 130 | 88 | 78 | 85 | 75 | 106 | 126 | 89 | 463 | 124 | 97 | 323 | 80 | 66 | 494 | 130 | 89 |
| 4485 | W303.asm02.HP0_G0044810 | chrXIV | 29831 | 11324 | 8935 | 26679 | 9658 | 8751 | 33054 | 10546 | 8691 | 43772 | 14081 | 18521 | 37858 | 12329 | 12175 | 35635 | 11249 | 16046 |
| 4486 | W303.asm02.HP0_G0044820 | chrXIV | 10396 | 4432 | 4762 | 9902 | 4288 | 4458 | 9578 | 4222 | 5245 | 7520 | 4544 | 4608 | 12532 | 4052 | 4569 | 7609 | 4015 | 4361 |
| 4487 | W303.asm02.HP0_G0044830 | chrXIV | 1207 | 1544 | 1456 | 1208 | 1776 | 1322 | 1358 | 1576 | 1519 | 8739 | 1796 | 2056 | 8051 | 1189 | 1184 | 7945 | 1652 | 1803 |
| 4488 | W303.asm02.HP0_G0044840 | chrXIV | 2051 | 1638 | 1865 | 1923 | 1727 | 1795 | 2146 | 1719 | 1916 | 6015 | 1315 | 1630 | 6093 | 807 | 1213 | 6009 | 1156 | 1462 |
| 4489 | W303.asm02.HP0_G0044850 | chrXIV | 211 | 43 | 63 | 189 | 59 | 56 | 189 | 44 | 70 | 327 | 78 | 120 | 151 | 42 | 64 | 157 | 60 | 109 |
| 4490 | W303.asm02.HP0_G0044860 | chrXIV | 839 | 161 | 172 | 887 | 166 | 197 | 729 | 160 | 252 | 506 | 222 | 341 | 391 | 149 | 215 | 299 | 226 | 273 |
| 4491 | W303.asm02.HP0_G0044870 | chrXIV | 1066 | 578 | 1110 | 1138 | 1143 | 961 | 1032 | 590 | 1269 | 9811 | 900 | 1796 | 6532 | 633 | 1015 | 6895 | 952 | 1367 |
| 4492 | W303.asm02.HP0_G0044880 | chrXIV | 6174 | 738 | 701 | 5727 | 582 | 719 | 5704 | 717 | 799 | 1817 | 699 | 727 | 2811 | 536 | 890 | 1662 | 654 | 692 |
| 4493 | W303.asm02.HP0_G0044890 | chrXIV | 3016 | 616 | 760 | 3114 | 792 | 807 | 2964 | 479 | 960 | 5641 | 1206 | 1735 | 2809 | 763 | 1084 | 3129 | 1183 | 1483 |
| 4494 | W303.asm02.HP0_G0044900 | chrXIV | 2907 | 509 | 891 | 3159 | 795 | 1000 | 2961 | 560 | 1017 | 10334 | 1373 | 2653 | 6537 | 956 | 1831 | 6212 | 1215 | 2469 |
| 4495 | W303.asm02.HP0_G0044910 | chrXIV | 444 | 175 | 177 | 392 | 155 | 159 | 367 | 182 | 248 | 432 | 124 | 131 | 582 | 108 | 105 | 454 | 102 | 89 |
| 4496 | W303.asm02.HP0_G0044920 | chrXIV | 1421 | 136 | 191 | 1529 | 208 | 228 | 1415 | 144 | 220 | 472 | 145 | 187 | 481 | 95 | 112 | 433 | 127 | 151 |
| 4497 | W303.asm02.HP0_G0044930 | chrXIV | 3095 | 79 | 53 | 3502 | 83 | 83 | 2658 | 73 | 82 | 343 | 127 | 118 | 201 | 73 | 78 | 216 | 120 | 117 |
| 4498 | W303.asm02.HP0_G0044940 | chrXIV | 2382 | 439 | 373 | 2126 | 325 | 381 | 2156 | 428 | 378 | 2271 | 523 | 759 | 2062 | 368 | 468 | 1991 | 469 | 683 |
| 4499 | W303.asm02.HP0_G0044950 | chrXIV | 10919 | 1686 | 2174 | 11728 | 2009 | 2103 | 10431 | 1535 | 2999 | 16509 | 2989 | 4409 | 13019 | 2664 | 3532 | 10434 | 2645 | 3856 |
| 4500 | W303.asm02.HP0_G0044960 | chrXIV | 1896 | 828 | 906 | 1255 | 1032 | 903 | 1591 | 845 | 845 | 3060 | 638 | 736 | 3757 | 476 | 574 | 2757 | 590 | 633 |
| 4501 | W303.asm02.HP0_G0044970 | chrXIV | 8597 | 1885 | 2677 | 8034 | 2081 | 2934 | 8544 | 1624 | 3337 | 18149 | 4350 | 5228 | 9377 | 3878 | 4099 | 8783 | 3777 | 4740 |
| 4502 | W303.asm02.HP0_G0044980 | chrXIV | 7832 | 3322 | 2987 | 7610 | 2906 | 2778 | 7948 | 3368 | 2858 | 18489 | 4351 | 5247 | 16413 | 3067 | 3612 | 16502 | 3899 | 4684 |
| 4503 | W303.asm02.HP0_G0044990 | chrXIV | 1492 | 762 | 1077 | 1392 | 1062 | 999 | 1477 | 885 | 985 | 4221 | 701 | 1102 | 4345 | 471 | 797 | 3791 | 531 | 906 |
| 4504 | W303.asm02.HP0_G0045000 | chrXIV | 678 | 126 | 179 | 629 | 201 | 220 | 679 | 120 | 189 | 1395 | 260 | 425 | 839 | 204 | 254 | 747 | 223 | 333 |
| 4505 | W303.asm02.HP0_G0045010 | chrXIV | 3389 | 957 | 1363 | 3387 | 1655 | 1550 | 3836 | 919 | 1683 | 14251 | 1986 | 3060 | 9971 | 1733 | 1942 | 8819 | 1740 | 2578 |
| 4506 | W303.asm02.HP0_G0045020 | chrXIV | 5613 | 1346 | 1665 | 5005 | 1629 | 1619 | 4964 | 1467 | 1875 | 5113 | 1400 | 2219 | 6706 | 1202 | 1846 | 4955 | 1312 | 2144 |
| 4507 | W303.asm02.HP0_G0045030 | chrXIV | 3298 | 1129 | 1062 | 3559 | 1003 | 1178 | 3047 | 1047 | 1101 | 5972 | 1322 | 1964 | 6050 | 1028 | 1629 | 5046 | 1134 | 1721 |
| 4508 | W303.asm02.HP0_G0045040 | chrXIV | 1436 | 1639 | 2133 | 1322 | 2774 | 1931 | 1697 | 1890 | 2171 | 13006 | 1785 | 2299 | 11623 | 1102 | 1287 | 11774 | 1517 | 2062 |
| 4509 | W303.asm02.HP0_G0045050 | chrXIV | 4815 | 1132 | 1255 | 4813 | 1224 | 1179 | 4722 | 1279 | 1230 | 5406 | 1272 | 1431 | 5026 | 800 | 1089 | 5074 | 1208 | 1379 |
| 4510 | W303.asm02.HP0_G0045060 | chrXIV | 3763 | 3472 | 3008 | 3558 | 2600 | 3312 | 3378 | 3576 | 3152 | 9885 | 3456 | 4374 | 10790 | 2509 | 3628 | 8760 | 3015 | 3890 |
| 4511 | W303.asm02.HP0_G0045070 | chrXIV | 238 | 104 | 108 | 134 | 76 | 85 | 156 | 115 | 76 | 286 | 92 | 100 | 410 | 56 | 123 | 336 | 79 | 84 |
| 4512 | W303.asm02.HP0_G0045080 | chrXIV | 16074 | 19208 | 19682 | 14115 | 18659 | 19297 | 16324 | 18238 | 23938 | 41542 | 17173 | 17005 | 43812 | 12686 | 13444 | 41857 | 15438 | 14668 |
| 4513 | W303.asm02.HP0_G0045090 | chrXIV | 7501 | 2427 | 2407 | 7391 | 2246 | 2357 | 6878 | 2433 | 2590 | 8584 | 2377 | 3045 | 9209 | 1896 | 2629 | 8038 | 2360 | 2672 |
| 4514 | W303.asm02.HP0_G0045100 | chrXIV | 4205 | 716 | 724 | 4307 | 657 | 820 | 4536 | 612 | 864 | 6161 | 1502 | 1895 | 5727 | 1329 | 1574 | 3958 | 1281 | 1474 |
| 4515 | W303.asm02.HP0_G0045110 | chrXIV | 3760 | 1982 | 1771 | 3739 | 2397 | 1617 | 4217 | 2157 | 1666 | 11978 | 2526 | 2660 | 11340 | 1599 | 1432 | 10509 | 2067 | 2237 |
| 4516 | W303.asm02.HP0_G0045120 | chrXIV | 10178 | 1258 | 989 | 10526 | 801 | 1058 | 9675 | 1001 | 962 | 3621 | 1213 | 1481 | 4068 | 989 | 1454 | 2938 | 1114 | 1301 |
| 4517 | W303.asm02.HP0_G0045130 | chrXIV | 1242 | 332 | 585 | 1215 | 532 | 628 | 1176 | 311 | 704 | 3696 | 555 | 1585 | 2052 | 494 | 998 | 2024 | 520 | 1314 |
| 4518 | W303.asm02.HP0_G0045140 | chrXIV | 3765 | 923 | 783 | 3659 | 723 | 760 | 3837 | 751 | 885 | 7386 | 2266 | 2633 | 4072 | 1628 | 1996 | 4955 | 2086 | 2268 |
| 4519 | W303.asm02.HP0_G0045150 | chrXIV | 2985 | 1499 | 1130 | 2673 | 1182 | 1189 | 2714 | 1398 | 1048 | 4341 | 1553 | 1840 | 4635 | 1306 | 1499 | 3789 | 1467 | 1592 |
| 4520 | W303.asm02.HP0_G0045160 | chrXIV | 3150 | 496 | 646 | 3398 | 601 | 650 | 2967 | 506 | 809 | 5604 | 895 | 1361 | 4416 | 666 | 1124 | 3894 | 836 | 1213 |
| 4521 | W303.asm02.HP0_G0045170 | chrXIV | 2768 | 1403 | 948 | 2403 | 1094 | 991 | 2652 | 1305 | 882 | 5625 | 1626 | 1747 | 5010 | 1076 | 1279 | 4737 | 1441 | 1568 |
| 4522 | W303.asm02.HP0_G0045180 | chrXIV | 1265 | 544 | 512 | 1280 | 582 | 547 | 1199 | 575 | 484 | 3138 | 810 | 768 | 3356 | 531 | 577 | 2721 | 646 | 727 |
| 4523 | W303.asm02.HP0_G0045190 | chrXIV | 612 | 729 | 1138 | 586 | 1221 | 1077 | 682 | 744 | 1257 | 4291 | 731 | 1048 | 4698 | 454 | 682 | 4299 | 632 | 948 |
| 4524 | W303.asm02.HP0_G0045200 | chrXIV | 4470 | 700 | 692 | 4184 | 904 | 779 | 4570 | 678 | 950 | 9736 | 1450 | 1923 | 5391 | 1048 | 1204 | 5556 | 1356 | 1809 |
| 4525 | W303.asm02.HP0_G0045210 | chrXIV | 94 | 374 | 517 | 103 | 591 | 528 | 105 | 374 | 737 | 1736 | 282 | 343 | 2169 | 185 | 193 | 1915 | 255 | 315 |
| 4526 | W303.asm02.HP0_G0045220 | chrXIV | 5048 | 3818 | 5409 | 4207 | 4964 | 5279 | 4776 | 3798 | 6657 | 10678 | 3348 | 4161 | 12156 | 2478 | 3351 | 11595 | 3213 | 3424 |
| 4527 | W303.asm02.HP0_G0045230 | chrXIV | 4091 | 731 | 871 | 3857 | 700 | 1047 | 4082 | 585 | 1013 | 6991 | 1249 | 2339 | 3835 | 1056 | 1508 | 3463 | 1238 | 2019 |
| 4528 | W303.asm02.HP0_G0045240 | chrXIV | 33653 | 11339 | 7370 | 28902 | 7538 | 7674 | 30953 | 10488 | 5980 | 10787 | 9271 | 8889 | 7644 | 5905 | 6917 | 9231 | 8018 | 7346 |
| 4529 | W303.asm02.HP0_G0045250 | chrXIV | 2297 | 907 | 637 | 2316 | 595 | 675 | 2062 | 808 | 612 | 2160 | 801 | 693 | 2459 | 567 | 661 | 2101 | 821 | 679 |
| 4530 | W303.asm02.HP0_G0045260 | chrXIV | 1195 | 584 | 315 | 1093 | 360 | 300 | 1170 | 594 | 299 | 1978 | 766 | 606 | 1737 | 446 | 460 | 1812 | 630 | 567 |
| 4531 | W303.asm02.HP0_G0045270 | chrXIV | 3359 | 1700 | 1511 | 3463 | 1311 | 1535 | 3229 | 1710 | 1898 | 3448 | 1453 | 1452 | 4018 | 925 | 1142 | 3453 | 1377 | 1226 |
| 4532 | W303.asm02.HP0_G0045280 | chrXIV | 3188 | 2954 | 3029 | 3462 | 2657 | 2881 | 2871 | 2694 | 3351 | 7580 | 2859 | 2858 | 7165 | 1731 | 2433 | 7239 | 2670 | 2483 |
| 4533 | W303.asm02.HP0_G0045290 | chrXIV | 4143 | 1612 | 2374 | 4031 | 2016 | 2455 | 3909 | 1502 | 2688 | 5717 | 1156 | 1704 | 6173 | 899 | 1653 | 5807 | 1042 | 1495 |
| 4534 | W303.asm02.HP0_G0045300 | chrXIV | 8613 | 1831 | 1888 | 8473 | 1811 | 1888 | 9267 | 1751 | 2233 | 16301 | 3715 | 4281 | 10949 | 2676 | 3042 | 10729 | 3381 | 3833 |
| 4535 | W303.asm02.HP0_G0045310 | chrXIV | 1982 | 2161 | 2251 | 1572 | 2207 | 2231 | 1788 | 2383 | 2848 | 5648 | 2111 | 1802 | 6158 | 1279 | 1403 | 5744 | 1749 | 1688 |
| 4536 | W303.asm02.HP0_G0045320 | chrXIV | 1066 | 388 | 277 | 1130 | 250 | 346 | 1214 | 293 | 360 | 2361 | 650 | 671 | 1442 | 397 | 488 | 1558 | 598 | 542 |
| 4537 | W303.asm02.HP0_G0045330 | chrXIV | 1709 | 1272 | 1609 | 1911 | 1900 | 1641 | 1949 | 1547 | 1857 | 7927 | 1654 | 1889 | 7513 | 999 | 1260 | 7394 | 1362 | 1915 |
| 4538 | W303.asm02.HP0_G0045340 | chrXIV | 3025 | 2356 | 2913 | 2581 | 2551 | 2955 | 2864 | 2402 | 3757 | 5751 | 2344 | 2602 | 6744 | 1486 | 2037 | 6078 | 2036 | 2224 |
| 4539 | W303.asm02.HP0_G0045350 | chrXIV | 1640 | 681 | 703 | 1484 | 568 | 688 | 1486 | 686 | 753 | 1895 | 744 | 835 | 1903 | 464 | 641 | 1820 | 708 | 702 |
| 4540 | W303.asm02.HP0_G0045360 | chrXIV | 866 | 850 | 1154 | 776 | 893 | 1253 | 846 | 899 | 1687 | 2499 | 655 | 949 | 2787 | 386 | 689 | 2819 | 587 | 789 |
| 4541 | W303.asm02.HP0_G0045370 | chrXIV | 40 | 47 | 36 | 23 | 34 | 36 | 28 | 32 | 37 | 88 | 25 | 29 | 99 | 23 | 32 | 106 | 29 | 20 |
| 4542 | W303.asm02.HP0_G0045380 | chrXIV | 335 | 261 | 191 | 311 | 171 | 224 | 317 | 243 | 261 | 613 | 176 | 165 | 732 | 127 | 169 | 700 | 217 | 145 |
| 4543 | W303.asm02.HP0_G0045390 | chrXIV | 4159 | 375 | 205 | 4649 | 240 | 198 | 3827 | 303 | 204 | 1588 | 420 | 225 | 1243 | 222 | 175 | 907 | 461 | 249 |
| 4544 | W303.asm02.HP0_G0045400 | chrXIV | 628 | 798 | 898 | 682 | 855 | 959 | 763 | 701 | 969 | 3644 | 1288 | 1135 | 4797 | 1333 | 1520 | 2161 | 1111 | 970 |
| 4545 | W303.asm02.HP0_G0045410 | chrXIV | 1104 | 1097 | 1167 | 981 | 1709 | 1029 | 1096 | 1179 | 1205 | 9772 | 1652 | 1580 | 6835 | 820 | 783 | 8814 | 1393 | 1403 |
| 4546 | W303.asm02.HP0_G0045420 | chrXIV | 4271 | 624 | 946 | 4109 | 902 | 956 | 3844 | 641 | 1017 | 7691 | 918 | 2040 | 5655 | 694 | 1465 | 5739 | 781 | 1734 |
| 4547 | W303.asm02.HP0_G0045440 | chrXIV | 10947 | 1268 | 1360 | 9157 | 1105 | 1638 | 9925 | 971 | 1845 | 7718 | 2071 | 2980 | 2992 | 1921 | 2258 | 2537 | 1766 | 2664 |
| 4548 | W303.asm02.HP0_G0045450 | chrXIV | 15069 | 3839 | 4220 | 15391 | 3797 | 4206 | 15132 | 4117 | 4416 | 12710 | 3666 | 4393 | 14317 | 2588 | 3468 | 12125 | 3247 | 4053 |
| 4549 | W303.asm02.HP0_G0045460 | chrXIV | 3518 | 1063 | 809 | 3306 | 914 | 807 | 3356 | 1083 | 664 | 4161 | 1018 | 1153 | 3898 | 611 | 930 | 3334 | 918 | 1089 |
| 4550 | W303.asm02.HP0_G0045470 | chrXIV | 8315 | 4364 | 5109 | 8131 | 4200 | 4751 | 8116 | 4313 | 5876 | 10459 | 4355 | 5076 | 11933 | 3246 | 4252 | 10906 | 4054 | 4560 |
| 4551 | W303.asm02.HP0_G0045480 | chrXIV | 11570 | 7619 | 7754 | 11229 | 5762 | 8737 | 10886 | 6126 | 9654 | 11446 | 16212 | 16854 | 5723 | 13880 | 13321 | 5375 | 13062 | 13011 |
| 4552 | W303.asm02.HP0_G0045490 | chrXIV | 659 | 591 | 405 | 594 | 405 | 465 | 526 | 598 | 534 | 997 | 422 | 369 | 1189 | 293 | 277 | 1070 | 428 | 291 |
| 4553 | W303.asm02.HP0_G0045500 | chrXIV | 1918 | 587 | 808 | 1915 | 1023 | 963 | 2415 | 640 | 1190 | 13379 | 1611 | 2570 | 6345 | 1071 | 1257 | 6857 | 1334 | 2322 |
| 4554 | W303.asm02.HP0_G0045510 | chrXIV | 6608 | 3104 | 4290 | 6516 | 4109 | 4501 | 6329 | 3219 | 5482 | 10643 | 2713 | 4169 | 12329 | 2054 | 3522 | 10794 | 2400 | 3833 |
| 4555 | W303.asm02.HP0_G0045520 | chrXIV | 3033 | 2381 | 2269 | 3311 | 1973 | 2161 | 3172 | 2428 | 2608 | 8168 | 2431 | 2709 | 8884 | 1848 | 2347 | 8726 | 2299 | 2566 |

| | | | | | | | | | | | | | | | | | | | | |
|---|---|---|---|---|---|---|---|---|---|---|---|---|---|---|---|---|---|---|---|---|
| 4556 | W303.asm02.HP0_G0045530 | chrXIV | 521 | 770 | 878 | 431 | 790 | 860 | 458 | 867 | 1118 | 2627 | 581 | 662 | 3085 | 420 | 526 | 3049 | 562 | 626 |
| 4557 | W303.asm02.HP0_G0045540 | chrXIV | 273 | 170 | 115 | 238 | 102 | 92 | 245 | 129 | 115 | 356 | 133 | 134 | 306 | 74 | 95 | 371 | 141 | 110 |
| 4558 | W303.asm02.HP0_G0045550 | chrXIV | 3086 | 359 | 478 | 3164 | 517 | 515 | 2770 | 393 | 524 | 4387 | 588 | 858 | 3565 | 392 | 755 | 3405 | 589 | 813 |
| 4559 | W303.asm02.HP0_G0045560 | chrXIV | 826 | 443 | 620 | 872 | 616 | 547 | 907 | 498 | 665 | 3441 | 545 | 776 | 2766 | 354 | 672 | 3182 | 533 | 758 |
| 4560 | W303.asm02.HP0_G0045570 | chrXIV | 1427 | 447 | 459 | 1469 | 543 | 566 | 1422 | 371 | 577 | 2817 | 1041 | 836 | 1402 | 587 | 637 | 1221 | 927 | 753 |
| 4561 | W303.asm02.HP0_G0045580 | chrXIV | 1059 | 331 | 613 | 1107 | 731 | 792 | 1295 | 278 | 886 | 5687 | 1009 | 1365 | 1658 | 636 | 707 | 1502 | 804 | 1062 |
| 4562 | W303.asm02.HP0_G0045590 | chrXIV | 4810 | 1737 | 1942 | 4531 | 2096 | 1922 | 4989 | 1825 | 2108 | 15061 | 2889 | 3609 | 12396 | 2125 | 2550 | 11030 | 2517 | 3147 |
| 4563 | W303.asm02.HP0_G0045600 | chrXIV | 2721 | 781 | 1215 | 2234 | 1028 | 1112 | 2220 | 796 | 1521 | 2170 | 569 | 921 | 3010 | 436 | 838 | 2514 | 534 | 752 |
| 4564 | W303.asm02.HP0_G0045610 | chrXIV | 21883 | 4152 | 4072 | 22306 | 4238 | 4180 | 22577 | 4135 | 3970 | 21932 | 4581 | 6444 | 21194 | 3606 | 4739 | 18815 | 3832 | 5765 |
| 4565 | W303.asm02.HP0_G0045620 | chrXIV | 1639 | 1603 | 2220 | 1862 | 2391 | 2077 | 1913 | 1693 | 2792 | 7436 | 1276 | 1715 | 7622 | 867 | 1015 | 7792 | 1215 | 1429 |
| 4566 | W303.asm02.HP0_G0045630 | chrXIV | 8078 | 1632 | 1884 | 8264 | 1699 | 1862 | 7981 | 1678 | 2107 | 14189 | 2795 | 3718 | 10483 | 1577 | 2922 | 10629 | 2562 | 3415 |
| 4567 | W303.asm02.HP0_G0045640 | chrXIV | 278 | 109 | 88 | 221 | 81 | 72 | 254 | 87 | 93 | 324 | 98 | 88 | 222 | 85 | 101 | 216 | 119 | 89 |
| 4568 | W303.asm02.HP0_G0045650 | chrXIV | 683 | 246 | 187 | 643 | 205 | 262 | 696 | 190 | 224 | 1472 | 451 | 534 | 492 | 327 | 340 | 633 | 443 | 492 |
| 4569 | W303.asm02.HP0_G0045660 | chrXIV | 4660 | 743 | 583 | 4311 | 524 | 605 | 4218 | 726 | 774 | 3155 | 1220 | 1133 | 2837 | 952 | 1137 | 1910 | 1212 | 1074 |
| 4570 | W303.asm02.HP0_G0045670 | chrXIV | 1343 | 2248 | 3118 | 1137 | 3172 | 3037 | 1330 | 2371 | 3781 | 7947 | 1893 | 2571 | 9503 | 1316 | 1755 | 8299 | 1711 | 2152 |
| 4571 | W303.asm02.HP0_G0045680 | chrXIV | 1769 | 1762 | 3417 | 1382 | 4331 | 2949 | 2055 | 1895 | 4285 | 27282 | 4937 | 5034 | 19335 | 2679 | 1829 | 19472 | 4269 | 3945 |
| 4572 | W303.asm02.HP0_G0045690 | chrXIV | 2747 | 2963 | 5928 | 2998 | 3868 | 5987 | 3289 | 2970 | 7344 | 8094 | 1950 | 5191 | 7660 | 1175 | 2836 | 8645 | 1816 | 4350 |
| 4573 | W303.asm02.HP0_G0045700 | chrXIV | 2491 | 2649 | 3746 | 2586 | 3801 | 3569 | 2872 | 2969 | 4570 | 11860 | 2888 | 3061 | 12295 | 1715 | 1936 | 12381 | 2344 | 2623 |
| 4574 | W303.asm02.HP0_G0045710 | chrXIV | 758 | 566 | 578 | 625 | 431 | 595 | 745 | 498 | 692 | 3155 | 763 | 1091 | 2142 | 545 | 728 | 2416 | 793 | 950 |
| 4575 | W303.asm02.HP0_G0045720 | chrXIV | 921 | 1015 | 1168 | 971 | 951 | 1236 | 906 | 995 | 1297 | 2952 | 834 | 1026 | 3331 | 476 | 691 | 3069 | 808 | 926 |
| 4576 | W303.asm02.HP0_G0045730 | chrXIV | 1083 | 193 | 287 | 948 | 258 | 286 | 966 | 169 | 318 | 2259 | 288 | 615 | 991 | 261 | 419 | 868 | 299 | 486 |
| 4577 | W303.asm02.HP0_G0045740 | chrXIV | 9469 | 4492 | 5151 | 9235 | 4552 | 5897 | 9810 | 4060 | 6710 | 22301 | 7758 | 10674 | 19673 | 6628 | 7885 | 16798 | 6496 | 8866 |
| 4578 | W303.asm02.HP0_G0045750 | chrXIV | 1526 | 597 | 870 | 1800 | 739 | 1030 | 1622 | 499 | 1137 | 5846 | 1442 | 2072 | 2701 | 911 | 1271 | 2650 | 1248 | 1777 |
| 4579 | W303.asm02.HP0_G0045760 | chrXIV | 1526 | 430 | 649 | 1548 | 726 | 741 | 1761 | 448 | 766 | 6254 | 790 | 1626 | 3568 | 605 | 1074 | 3722 | 635 | 1469 |
| 4580 | W303.asm02.HP0_G0045770 | chrXIV | 3030 | 1060 | 1148 | 3325 | 1102 | 1135 | 3280 | 1116 | 1227 | 8878 | 1922 | 2314 | 7607 | 1427 | 1735 | 6915 | 1786 | 2101 |
| 4581 | W303.asm02.HP0_G0045780 | chrXIV | 3105 | 2369 | 2639 | 3036 | 2334 | 2612 | 2689 | 2255 | 3331 | 5455 | 1823 | 2223 | 5781 | 1265 | 1957 | 5335 | 1787 | 1839 |
| 4582 | W303.asm02.HP0_G0045790 | chrXIV | 1857 | 766 | 560 | 1950 | 598 | 531 | 1852 | 658 | 533 | 2960 | 805 | 958 | 2728 | 567 | 747 | 2577 | 716 | 858 |
| 4583 | W303.asm02.HP0_G0045800 | chrXIV | 10430 | 4420 | 4143 | 10564 | 3798 | 4266 | 9649 | 4012 | 3897 | 12456 | 3861 | 5226 | 12887 | 2614 | 4321 | 10865 | 3497 | 4583 |
| 4584 | W303.asm02.HP0_G0045810 | chrXIV | 4882 | 796 | 509 | 4693 | 479 | 522 | 4614 | 667 | 580 | 2875 | 963 | 874 | 2063 | 673 | 752 | 2063 | 945 | 815 |
| 4585 | W303.asm02.HP0_G0045820 | chrXIV | 8112 | 4422 | 4804 | 7686 | 5013 | 5185 | 8635 | 4341 | 4854 | 15314 | 3585 | 4301 | 15934 | 2791 | 3380 | 15308 | 3366 | 4049 |
| 4586 | W303.asm02.HP0_G0045830 | chrXIV | 534 | 260 | 377 | 521 | 452 | 365 | 493 | 295 | 356 | 3371 | 520 | 649 | 2134 | 287 | 302 | 2095 | 433 | 567 |
| 4587 | W303.asm02.HP0_G0045840 | chrXIV | 3323 | 594 | 733 | 3089 | 724 | 811 | 3449 | 571 | 865 | 5288 | 1244 | 1535 | 3633 | 856 | 1167 | 3631 | 1138 | 1462 |
| 4588 | W303.asm02.HP0_G0045850 | chrXIV | 1147 | 38 | 27 | 1294 | 23 | 27 | 865 | 31 | 19 | 73 | 43 | 25 | 62 | 24 | 22 | 46 | 38 | 31 |
| 4589 | W303.asm02.HP0_G0045860 | chrXIV | 833 | 133 | 117 | 791 | 88 | 132 | 724 | 133 | 120 | 515 | 147 | 217 | 450 | 114 | 179 | 409 | 142 | 168 |
| 4590 | W303.asm02.HP0_G0045870 | chrXIV | 9818 | 673 | 821 | 9082 | 839 | 1012 | 9756 | 612 | 963 | 10009 | 1519 | 2438 | 5999 | 1155 | 1760 | 5416 | 1310 | 2012 |
| 4591 | W303.asm02.HP0_G0045880 | chrXIV | 732 | 1106 | 1116 | 590 | 1056 | 983 | 720 | 1055 | 1236 | 3960 | 1050 | 1211 | 4050 | 736 | 995 | 4144 | 927 | 1124 |
| 4592 | W303.asm02.HP0_G0045890 | chrXIV | 4024 | 775 | 983 | 4203 | 1181 | 1029 | 4353 | 869 | 1096 | 8282 | 1125 | 1930 | 5953 | 855 | 1342 | 6448 | 997 | 1654 |
| 4593 | W303.asm02.HP0_G0045900 | chrXIV | 5767 | 1997 | 2115 | 5175 | 2112 | 2097 | 5950 | 2042 | 2230 | 15537 | 2855 | 3289 | 11104 | 1639 | 2188 | 12903 | 2435 | 2977 |
| 4594 | W303.asm02.HP0_G0045910 | chrXIV | 503 | 324 | 392 | 452 | 392 | 381 | 453 | 382 | 438 | 1015 | 354 | 313 | 1163 | 203 | 256 | 1019 | 282 | 330 |
| 4595 | W303.asm02.HP0_G0045920 | chrXIV | 5934 | 1855 | 2245 | 6278 | 1996 | 2111 | 6213 | 2123 | 2451 | 15958 | 2557 | 3885 | 14483 | 1887 | 2996 | 13832 | 2362 | 3676 |
| 4596 | W303.asm02.HP0_G0045930 | chrXIV | 5806 | 2787 | 3387 | 5932 | 2851 | 3385 | 5604 | 2709 | 4349 | 8721 | 2165 | 2394 | 9913 | 1467 | 2137 | 9785 | 1994 | 2093 |
| 4597 | W303.asm02.HP0_G0045940 | chrXIV | 2656 | 1317 | 1137 | 2584 | 1422 | 1273 | 2605 | 1225 | 1248 | 6892 | 1585 | 1862 | 5586 | 1055 | 1362 | 5724 | 1520 | 1661 |
| 4598 | W303.asm02.HP0_G0045950 | chrXIV | 4403 | 2021 | 2933 | 4026 | 2480 | 2952 | 4057 | 2058 | 3747 | 5866 | 1744 | 2469 | 7538 | 1248 | 2177 | 6186 | 1569 | 2228 |
| 4599 | W303.asm02.HP0_G0045960 | chrXIV | 1640 | 669 | 505 | 1609 | 496 | 503 | 1679 | 595 | 483 | 3484 | 810 | 960 | 3516 | 571 | 744 | 3001 | 787 | 847 |
| 4600 | W303.asm02.HP0_G0045970 | chrXIV | 12953 | 7961 | 7936 | 12654 | 7029 | 8654 | 13165 | 8506 | 8006 | 18410 | 7564 | 9117 | 20830 | 5444 | 7101 | 16422 | 6349 | 8368 |
| 4601 | W303.asm02.HP0_G0045980 | chrXIV | 1663 | 829 | 844 | 1732 | 626 | 968 | 1727 | 531 | 1096 | 3154 | 1137 | 1765 | 1056 | 1016 | 868 | 739 | 958 | 1345 |
| 4602 | W303.asm02.HP0_G0045990 | chrXIV | 4147 | 1339 | 404 | 5396 | 597 | 719 | 3275 | 719 | 421 | 1813 | 1466 | 690 | 1011 | 612 | 547 | 1237 | 1823 | 738 |
| 4603 | W303.asm02.HP0_G0046000 | chrXIV | 3648 | 781 | 805 | 3932 | 787 | 746 | 3938 | 780 | 824 | 5941 | 1143 | 1398 | 5563 | 760 | 1110 | 4830 | 1090 | 1285 |
| 4604 | W303.asm02.HP0_G0046010 | chrXIV | 1503 | 1749 | 1810 | 1367 | 2269 | 1690 | 1659 | 1921 | 2054 | 7531 | 1742 | 1974 | 7182 | 1008 | 1142 | 7218 | 1570 | 1590 |
| 4605 | W303.asm02.HP0_G0046020 | chrXIV | 6253 | 748 | 788 | 6740 | 655 | 847 | 6865 | 603 | 946 | 6112 | 1472 | 1674 | 6283 | 1210 | 1604 | 5220 | 1432 | 1505 |
| 4606 | W303.asm02.HP0_G0046030 | chrXIV | 5595 | 1157 | 907 | 5394 | 896 | 906 | 5366 | 1031 | 860 | 5465 | 1375 | 1749 | 6110 | 1112 | 1469 | 4514 | 1278 | 1590 |
| 4607 | W303.asm02.HP0_G0046040 | chrXIV | 610 | 327 | 198 | 546 | 183 | 266 | 603 | 253 | 251 | 1310 | 437 | 372 | 895 | 273 | 270 | 921 | 434 | 319 |
| 4608 | W303.asm02.HP0_G0046050 | chrXIV | 18357 | 2641 | 2350 | 16052 | 2053 | 2751 | 17968 | 2078 | 3046 | 14678 | 5504 | 6501 | 8322 | 4389 | 4530 | 7667 | 4670 | 5668 |
| 4609 | W303.asm02.HP0_G0046060 | chrXIV | 2479 | 1104 | 1788 | 2079 | 1553 | 1785 | 2027 | 1124 | 2340 | 3609 | 864 | 1478 | 4723 | 637 | 1402 | 4215 | 844 | 1242 |
| 4610 | W303.asm02.HP0_G0046070 | chrXIV | 14540 | 12483 | 14260 | 14340 | 13999 | 14156 | 15620 | 12719 | 14883 | 31350 | 12042 | 13383 | 31973 | 8690 | 10426 | 30594 | 10278 | 12067 |
| 4611 | W303.asm02.HP0_G0046080 | chrXIV | 2511 | 429 | 618 | 2323 | 650 | 668 | 2601 | 350 | 801 | 3767 | 807 | 1068 | 1553 | 579 | 758 | 1464 | 593 | 939 |
| 4612 | W303.asm02.HP0_G0046090 | chrXIV | 4272 | 7184 | 9369 | 3571 | 8183 | 9157 | 4256 | 7219 | 11702 | 18794 | 6245 | 7619 | 21363 | 4678 | 6210 | 19534 | 5342 | 6939 |
| 4613 | W303.asm02.HP0_G0046100 | chrXIV | 2488 | 1620 | 2165 | 2129 | 2230 | 2349 | 2966 | 1734 | 2812 | 13332 | 2613 | 3974 | 12438 | 1896 | 3174 | 10094 | 2197 | 3708 |
| 4614 | W303.asm02.HP0_G0046110 | chrXIV | 143 | 414 | 1709 | 130 | 2016 | 1514 | 152 | 429 | 2219 | 11513 | 853 | 2139 | 7319 | 301 | 859 | 8827 | 1227 | 1368 |
| 4615 | W303.asm02.HP0_G0046120 | chrXIV | 10689 | 7249 | 6906 | 11047 | 6446 | 7425 | 11253 | 5669 | 7578 | 38160 | 11398 | 14115 | 27544 | 8554 | 10024 | 31090 | 10533 | 12216 |
| 4616 | W303.asm02.HP0_G0046130 | chrXIV | 1653 | 777 | 422 | 1579 | 401 | 480 | 1462 | 769 | 396 | 1500 | 836 | 632 | 1201 | 625 | 443 | 1257 | 897 | 585 |
| 4617 | W303.asm02.HP0_G0046140 | chrXIV | 1125 | 1249 | 1346 | 1259 | 1325 | 1318 | 1396 | 1218 | 1388 | 7550 | 1567 | 2341 | 6008 | 988 | 1183 | 6653 | 1278 | 1961 |
| 4618 | W303.asm02.HP0_G0046150 | chrXIV | 9055 | 1951 | 1769 | 8454 | 2415 | 1971 | 9938 | 2029 | 1999 | 23095 | 4211 | 4824 | 18301 | 2833 | 2804 | 16914 | 3320 | 4432 |
| 4619 | W303.asm02.HP0_G0046160 | chrXIV | 3977 | 957 | 908 | 3991 | 972 | 961 | 4225 | 1026 | 1160 | 7550 | 1526 | 1937 | 6512 | 1106 | 1472 | 5972 | 1398 | 1742 |
| 4620 | W303.asm02.HP0_G0046170 | chrXIV | 8207 | 1067 | 1053 | 7807 | 961 | 1163 | 8243 | 1091 | 1303 | 5717 | 1811 | 1980 | 9529 | 2196 | 2586 | 4257 | 1416 | 1775 |
| 4621 | W303.asm02.HP0_G0046180 | chrXIV | 2877 | 1432 | 1098 | 2543 | 1112 | 1130 | 2683 | 1404 | 1170 | 3871 | 1330 | 1109 | 4874 | 1091 | 1141 | 3665 | 1123 | 1054 |
| 4622 | W303.asm02.HP0_G0046190 | chrXIV | 16413 | 10633 | 8674 | 15660 | 8477 | 8734 | 16461 | 11068 | 8590 | 18746 | 10557 | 12894 | 20247 | 9608 | 10096 | 16482 | 9111 | 9860 |
| 4623 | W303.asm02.HP0_G0046200 | chrXIV | 0 | 0 | 0 | 0 | 0 | 0 | 0 | 0 | 0 | 0 | 0 | 0 | 0 | 0 | 0 | 0 | 0 | 0 |
| 4624 | W303.asm02.HP0_G0046210 | chrXIV | 863 | 158 | 337 | 855 | 266 | 404 | 944 | 155 | 409 | 1673 | 252 | 683 | 1004 | 187 | 447 | 948 | 221 | 651 |
| 4625 | W303.asm02.HP0_G0046220 | chrXIV | 1230 | 542 | 426 | 1172 | 492 | 403 | 1284 | 466 | 513 | 3649 | 776 | 801 | 3713 | 667 | 641 | 2491 | 767 | 706 |
| 4626 | W303.asm02.HP0_G0046230 | chrXIV | 4713 | 4024 | 4709 | 4498 | 4494 | 4447 | 4619 | 4101 | 5316 | 10223 | 3535 | 4732 | 9509 | 2755 | 4010 | 9262 | 3178 | 3708 |
| 4627 | W303.asm02.HP0_G0046240 | chrXIV | 2103 | 1238 | 1135 | 2261 | 1054 | 1184 | 2008 | 1165 | 1307 | 3637 | 881 | 957 | 3720 | 544 | 760 | 4037 | 875 | 807 |
| 4628 | W303.asm02.HP0_G0046250 | chrXIV | 1622 | 1542 | 1760 | 1647 | 1791 | 1759 | 1795 | 1558 | 1941 | 5069 | 1222 | 1468 | 5394 | 741 | 1034 | 4837 | 1113 | 1231 |
| 4629 | W303.asm02.HP0_G0046260 | chrXIV | 3127 | 890 | 1451 | 2955 | 994 | 1420 | 2994 | 966 | 1558 | 7839 | 1347 | 2977 | 6686 | 869 | 1922 | 6466 | 1188 | 2810 |
| 4630 | W303.asm02.HP0_G0046270 | chrXIV | 2342 | 1293 | 1086 | 2332 | 1042 | 1063 | 2370 | 1258 | 1117 | 6217 | 1722 | 1552 | 4877 | 1087 | 1116 | 5594 | 1572 | 1405 |
| 4631 | W303.asm02.HP0_G0046280 | chrXIV | 791 | 272 | 416 | 812 | 377 | 478 | 819 | 218 | 550 | 3028 | 520 | 832 | 1630 | 370 | 588 | 1721 | 547 | 726 |
| 4632 | W303.asm02.HP0_G0046290 | chrXIV | 2045 | 969 | 688 | 1784 | 779 | 724 | 1917 | 980 | 727 | 2443 | 980 | 878 | 3018 | 828 | 757 | 2086 | 870 | 722 |
| 4633 | W303.asm02.HP0_G0046300 | chrXIV | 6231 | 763 | 667 | 5352 | 601 | 775 | 5546 | 650 | 932 | 5790 | 1541 | 1763 | 3785 | 1180 | 1355 | 3953 | 1495 | 1595 |
| 4634 | W303.asm02.HP0_G0046310 | chrXIV | 2387 | 2205 | 2625 | 2591 | 2076 | 2535 | 2446 | 2161 | 3158 | 5976 | 2188 | 2428 | 6868 | 1652 | 2098 | 6262 | 1842 | 2299 |
| 4635 | W303.asm02.HP0_G0046320 | chrXIV | 62 | 15 | 15 | 33 | 6 | 12 | 44 | 8 | 10 | 13 | 12 | 10 | 14 | 5 | 5 | 22 | 25 | 8 |
| 4636 | W303.asm02.HP0_G0046330 | chrXIV | 1793 | 188 | 199 | 1734 | 214 | 229 | 1702 | 112 | 227 | 1779 | 394 | 396 | 1491 | 257 | 284 | 1231 | 367 | 312 |
| 4637 | W303.asm02.HP0_G0046340 | chrXIV | 2748 | 1286 | 1538 | 2867 | 1313 | 1430 | 2660 | 1388 | 1585 | 7176 | 1377 | 1906 | 5823 | 871 | 1417 | 6543 | 1336 | 1715 |
| 4638 | W303.asm02.HP0_G0046350 | chrXIV | 3286 | 1637 | 1635 | 3101 | 1509 | 1581 | 3109 | 1601 | 1804 | 6588 | 1683 | 2078 | 7591 | 1318 | 1690 | 5952 | 1548 | 1831 |
| 4639 | W303.asm02.HP0_G0046360 | chrXIV | 4757 | 597 | 607 | 4526 | 568 | 643 | 4804 | 529 | 719 | 4693 | 1121 | 1377 | 3308 | 802 | 1064 | 3184 | 1044 | 1294 |
| 4640 | W303.asm02.HP0_G0046370 | chrXIV | 1482 | 798 | 1128 | 1446 | 947 | 1038 | 1485 | 874 | 1491 | 3320 | 676 | 846 | 3424 | 460 | 638 | 3681 | 615 | 751 |
| 4641 | W303.asm02.HP0_G0046380 | chrXIV | 7276 | 5967 | 4546 | 6484 | 4469 | 4078 | 6498 | 6000 | 4153 | 14950 | 7576 | 6462 | 16900 | 5421 | 5876 | 14472 | 7527 | 6521 |
| 4642 | W303.asm02.HP0_G0046390 | chrXIV | 782 | 10874 | 9174 | 765 | 7234 | 9626 | 782 | 11386 | 9506 | 9063 | 9374 | 11269 | 16154 | 7934 | 9869 | 10330 | 8016 | 10399 |
| 4643 | W303.asm02.HP0_G0046400 | chrXIV | 1759 | 1353 | 956 | 1659 | 965 | 871 | 1800 | 1308 | 944 | 4941 | 1322 | 1294 | 4755 | 928 | 843 | 4681 | 1152 | 1139 |
| 4644 | W303.asm02.HP0_G0046410 | chrXIV | 172 | 23 | 27 | 160 | 28 | 28 | 181 | 16 | 32 | 124 | 22 | 41 | 97 | 21 | 39 | 74 | 20 | 25 |
| 4645 | W303.asm02.HP0_G0046420 | chrXIV | 90 | 25 | 32 | 112 | 25 | 23 | 117 | 26 | 43 | 269 | 52 | 61 | 225 | 34 | 51 | 137 | 14 | 59 |
| 4646 | W303.asm02.HP0_G0046430 | chrXIV | 2694 | 1588 | 1757 | 2673 | 1636 | 1564 | 2569 | 1598 | 1992 | 5385 | 1333 | 1484 | 7211 | 911 | 1307 | 5437 | 1232 | 1276 |
| 4647 | W303.asm02.HP0_G0046440 | chrXIV | 9357 | 6084 | 8959 | 9417 | 7539 | 8446 | 9308 | 6133 | 9642 | 18556 | 5937 | 8835 | 19080 | 4584 | 7244 | 18217 | 5307 | 7952 |
| 4648 | W303.asm02.HP0_G0046450 | chrXIV | 5667 | 5727 | 9342 | 5682 | 7881 | 9013 | 5324 | 6085 | 11397 | 14450 | 5149 | 7292 | 16459 | 3568 | 6033 | 15448 | 4287 | 6389 |

| | | | | | | | | | | | | | | | | | | | | |
|---|---|---|---|---|---|---|---|---|---|---|---|---|---|---|---|---|---|---|---|---|
| 4649 | W303.asm02.HP0_G0046460 | chrXIV | 1172 | 347 | 256 | 1021 | 311 | 283 | 1150 | 335 | 318 | 2482 | 582 | 584 | 1735 | 397 | 442 | 1917 | 542 | 556 |
| 4650 | W303.asm02.HP0_G0046470 | chrXIV | 4001 | 562 | 640 | 4512 | 692 | 682 | 4052 | 566 | 595 | 4405 | 841 | 1054 | 4341 | 528 | 868 | 3981 | 804 | 960 |
| 4651 | W303.asm02.HP0_G0046480 | chrXIV | 4794 | 1224 | 1443 | 4734 | 1123 | 1357 | 4429 | 1289 | 1454 | 5873 | 1301 | 2101 | 5735 | 985 | 1610 | 5521 | 1267 | 1952 |
| 4652 | W303.asm02.HP0_G0046490 | chrXIV | 1403 | 339 | 346 | 1524 | 293 | 312 | 1253 | 356 | 272 | 1997 | 482 | 525 | 1708 | 296 | 394 | 1693 | 431 | 439 |
| 4653 | W303.asm02.HP0_G0046500 | chrXIV | 1399 | 1262 | 1777 | 1182 | 1378 | 1643 | 1277 | 1292 | 2308 | 3056 | 1029 | 1357 | 4080 | 770 | 1111 | 3358 | 958 | 1187 |
| 4654 | W303.asm02.HP0_G0046510 | chrXIV | 831 | 283 | 433 | 556 | 518 | 402 | 728 | 240 | 468 | 1549 | 256 | 255 | 1716 | 166 | 181 | 1388 | 242 | 184 |
| 4655 | W303.asm02.HP0_G0046520 | chrXIV | 1695 | 264 | 449 | 1722 | 514 | 452 | 1987 | 250 | 581 | 5675 | 725 | 1087 | 2802 | 483 | 535 | 2736 | 577 | 973 |
| 4656 | W303.asm02.HP0_G0046530 | chrXIV | 1930 | 1178 | 1184 | 1978 | 1511 | 1209 | 2157 | 1153 | 1172 | 7131 | 1167 | 1558 | 6833 | 716 | 915 | 6275 | 964 | 1376 |
| 4657 | W303.asm02.HP0_G0046540 | chrXIV | 2225 | 490 | 706 | 2388 | 735 | 648 | 2266 | 502 | 761 | 6004 | 949 | 1414 | 3895 | 653 | 1045 | 3897 | 870 | 1315 |
| 4658 | W303.asm02.HP0_G0046550 | chrXIV | 1834 | 360 | 421 | 1970 | 376 | 456 | 1778 | 326 | 470 | 3049 | 614 | 895 | 2134 | 404 | 667 | 2057 | 537 | 758 |
| 4659 | W303.asm02.HP0_G0046560 | chrXIV | 289 | 33 | 45 | 348 | 30 | 44 | 237 | 30 | 60 | 229 | 37 | 62 | 160 | 34 | 55 | 140 | 31 | 58 |
| 4660 | W303.asm02.HP0_G0046570 | chrXIV | 218 | 39 | 53 | 215 | 40 | 47 | 243 | 30 | 44 | 170 | 35 | 54 | 166 | 14 | 33 | 109 | 18 | 37 |
| 4661 | W303.asm02.HP0_G0046580 | chrXIV | 19656 | 4498 | 4369 | 19225 | 4275 | 4118 | 19887 | 4739 | 4409 | 24690 | 6856 | 8104 | 21677 | 4664 | 6265 | 21274 | 6233 | 7438 |
| 4662 | W303.asm02.HP0_G0046590 | chrXIV | 5029 | 1609 | 2821 | 5037 | 2103 | 3136 | 4672 | 1730 | 3807 | 4546 | 1735 | 2937 | 7837 | 1620 | 2618 | 4783 | 1534 | 2349 |
| 4663 | W303.asm02.HP0_G0046600 | chrXIV | 1234 | 99 | 132 | 1148 | 122 | 136 | 1076 | 88 | 139 | 785 | 145 | 281 | 450 | 113 | 176 | 407 | 167 | 218 |
| 4664 | W303.asm02.HP0_G0046610 | chrXIV | 2254 | 822 | 577 | 2276 | 611 | 718 | 2018 | 770 | 646 | 1529 | 577 | 403 | 2300 | 438 | 449 | 1555 | 636 | 416 |
| 4665 | W303.asm02.HP0_G0046620 | chrXIV | 1576 | 1034 | 709 | 1417 | 597 | 777 | 1457 | 927 | 805 | 2035 | 714 | 660 | 2351 | 512 | 570 | 2079 | 825 | 514 |
| 4666 | W303.asm02.HP0_G0046630 | chrXIV | 5909 | 3175 | 3913 | 5167 | 3115 | 3876 | 5752 | 3244 | 4606 | 9419 | 2897 | 3741 | 10172 | 2176 | 2960 | 10168 | 2685 | 3360 |
| 4667 | W303.asm02.HP0_G0046640 | chrXIV | 884 | 457 | 306 | 1054 | 288 | 277 | 830 | 451 | 356 | 974 | 385 | 219 | 1008 | 236 | 177 | 868 | 396 | 206 |
| 4668 | W303.asm02.HP0_G0046650 | chrXIV | 2664 | 1485 | 1084 | 2432 | 946 | 1055 | 2518 | 1435 | 1114 | 4869 | 1575 | 1301 | 5244 | 953 | 1027 | 5049 | 1569 | 1221 |
| 4669 | W303.asm02.HP0_G0046660 | chrXIV | 15815 | 5572 | 4047 | 15889 | 3743 | 4787 | 13517 | 4345 | 4205 | 17562 | 6763 | 6299 | 9664 | 4183 | 5502 | 14300 | 6978 | 5977 |
| 4670 | W303.asm02.HP0_G0046670 | chrXIV | 3509 | 3416 | 1908 | 3522 | 2422 | 2573 | 3094 | 3007 | 1797 | 5587 | 2700 | 1877 | 5132 | 1687 | 1608 | 5588 | 2852 | 1795 |
| 4671 | W303.asm02.HP0_G0046680 | chrXIV | 11445 | 3394 | 2770 | 10862 | 2668 | 2871 | 10935 | 3411 | 3037 | 9449 | 3303 | 3316 | 10466 | 2563 | 3010 | 9179 | 2939 | 3137 |
| 4672 | W303.asm02.HP0_G0046690 | chrXIV | 7770 | 2624 | 1793 | 7204 | 2061 | 1774 | 7474 | 2486 | 1888 | 10624 | 3604 | 3354 | 8724 | 2401 | 2522 | 8278 | 3343 | 3241 |
| 4673 | W303.asm02.HP0_G0046700 | chrXIV | 2492 | 1044 | 866 | 2152 | 779 | 835 | 2223 | 1024 | 1001 | 3368 | 1155 | 1078 | 3423 | 854 | 972 | 3028 | 1211 | 958 |
| 4674 | W303.asm02.HP0_G0046710 | chrXIV | 4779 | 4902 | 4803 | 5263 | 5542 | 4646 | 5581 | 5263 | 5559 | 16587 | 4959 | 4374 | 16588 | 3056 | 2798 | 16392 | 4343 | 3994 |
| 4675 | W303.asm02.HP0_G0046720 | chrXIV | 1351 | 1933 | 1710 | 1490 | 1597 | 1516 | 1401 | 1816 | 1984 | 5516 | 1519 | 1446 | 5357 | 986 | 898 | 6156 | 1603 | 1217 |
| 4676 | W303.asm02.HP0_G0046730 | chrXIV | 12901 | 6973 | 4054 | 12344 | 4562 | 4067 | 11180 | 6567 | 4357 | 16287 | 8614 | 5890 | 16267 | 6593 | 6149 | 15136 | 8031 | 5089 |
| 4677 | W303.asm02.HP0_G0046740 | chrXIV | 1132 | 393 | 180 | 1056 | 167 | 167 | 929 | 328 | 192 | 661 | 395 | 269 | 708 | 347 | 240 | 525 | 478 | 227 |
| 4678 | W303.asm02.HP0_G0046750 | chrXIV | 3197 | 1401 | 865 | 2955 | 1082 | 820 | 3325 | 1430 | 802 | 5249 | 1740 | 1253 | 4920 | 1048 | 752 | 4993 | 1492 | 1116 |
| 4679 | W303.asm02.HP0_G0046760 | chrXIV | 515 | 500 | 385 | 506 | 361 | 371 | 495 | 522 | 433 | 1166 | 405 | 273 | 1353 | 249 | 207 | 1337 | 434 | 251 |
| 4680 | W303.asm02.HP0_G0046770 | chrXIV | 6525 | 521 | 451 | 6530 | 419 | 547 | 6523 | 441 | 539 | 4357 | 1157 | 1296 | 4647 | 897 | 1132 | 3350 | 1069 | 1124 |
| 4681 | W303.asm02.HP0_G0046780 | chrXIV | 3756 | 1002 | 873 | 3886 | 736 | 835 | 3412 | 964 | 932 | 2630 | 957 | 925 | 2980 | 577 | 761 | 2497 | 1002 | 857 |
| 4682 | W303.asm02.HP0_G0046790 | chrXIV | 2192 | 457 | 590 | 1666 | 569 | 613 | 1915 | 490 | 705 | 5785 | 1144 | 1604 | 3521 | 799 | 1211 | 3607 | 1067 | 1558 |
| 4683 | W303.asm02.HP0_G0046800 | chrXIV | 376 | 609 | 586 | 349 | 789 | 605 | 370 | 783 | 541 | 2410 | 626 | 648 | 2662 | 496 | 597 | 2275 | 548 | 587 |
| 4684 | W303.asm02.HP0_G0046810 | chrXIV | 1314 | 516 | 464 | 1199 | 469 | 465 | 1096 | 523 | 488 | 1202 | 384 | 380 | 1389 | 266 | 285 | 1250 | 373 | 294 |
| 4685 | W303.asm02.HP0_G0046820 | chrXIV | 2236 | 484 | 417 | 2221 | 460 | 448 | 2256 | 427 | 490 | 4457 | 979 | 1099 | 2431 | 691 | 641 | 2361 | 1018 | 956 |
| 4686 | W303.asm02.HP0_G0046830 | chrXIV | 1057 | 1454 | 1375 | 1082 | 1734 | 1163 | 1266 | 1481 | 1444 | 8144 | 1544 | 1385 | 8014 | 1003 | 846 | 7752 | 1359 | 1259 |
| 4687 | W303.asm02.HP0_G0046840 | chrXIV | 3400 | 697 | 899 | 2962 | 796 | 1064 | 3603 | 621 | 1087 | 7482 | 1516 | 2138 | 2464 | 1174 | 1581 | 2274 | 1224 | 1961 |
| 4688 | W303.asm02.HP0_G0046850 | chrXIV | 826 | 229 | 145 | 1027 | 165 | 183 | 795 | 179 | 215 | 949 | 394 | 297 | 629 | 226 | 273 | 557 | 415 | 263 |
| 4689 | W303.asm02.HP0_G0046860 | chrXIV | 1085 | 492 | 530 | 1052 | 574 | 587 | 1258 | 493 | 638 | 4842 | 1263 | 1356 | 2693 | 718 | 706 | 2534 | 1057 | 1080 |
| 4690 | W303.asm02.HP0_G0046870 | chrXIV | 35831 | 5842 | 5599 | 32139 | 5965 | 5874 | 35929 | 5375 | 7317 | 38727 | 9775 | 16010 | 17790 | 9485 | 11029 | 16692 | 8192 | 14434 |
| 4691 | W303.asm02.HP0_G0046880 | chrXIV | 10925 | 2090 | 1884 | 10195 | 1721 | 2170 | 10925 | 2035 | 2566 | 9816 | 3998 | 4051 | 5616 | 3434 | 3113 | 4516 | 3381 | 3838 |
| 4692 | W303.asm02.HP0_G0046890 | chrXIV | 2070 | 901 | 1014 | 1878 | 1065 | 1152 | 2069 | 837 | 1287 | 6238 | 1589 | 1975 | 4596 | 1115 | 1378 | 4613 | 1333 | 1694 |
| 4693 | W303.asm02.HP0_G0046900 | chrXIV | 5573 | 1454 | 1490 | 5403 | 1261 | 1642 | 5524 | 1071 | 1652 | 7525 | 1584 | 2338 | 3507 | 1119 | 1641 | 4514 | 1676 | 2183 |
| 4694 | W303.asm02.HP0_G0046910 | chrXIV | 14 | 4 | 3 | 10 | 6 | 5 | 16 | 2 | 13 | 27 | 2 | 10 | 17 | 1 | 2 | 16 | 0 | 8 |
| 4695 | W303.asm02.HP0_G0046920 | chrXIV | 763 | 642 | 739 | 672 | 828 | 712 | 641 | 689 | 927 | 2755 | 592 | 833 | 2656 | 384 | 602 | 2355 | 531 | 716 |
| 4696 | W303.asm02.HP0_G0046930 | chrXIV | 7182 | 3326 | 2715 | 6738 | 2696 | 2598 | 7354 | 3352 | 2342 | 12291 | 3211 | 3465 | 11218 | 2360 | 2373 | 11380 | 2762 | 2971 |
| 4697 | W303.asm02.HP0_G0046940 | chrXIV | 4440 | 1321 | 1956 | 4008 | 1742 | 2157 | 3955 | 1472 | 2398 | 4213 | 1214 | 1926 | 5426 | 918 | 1507 | 4282 | 1159 | 1655 |
| 4698 | W303.asm02.HP0_G0046950 | chrXIV | 2833 | 593 | 509 | 2703 | 498 | 538 | 2853 | 489 | 522 | 3294 | 664 | 960 | 2626 | 484 | 720 | 2600 | 701 | 909 |
| 4699 | W303.asm02.HP0_G0046960 | chrXIV | 1289 | 721 | 865 | 1153 | 1037 | 878 | 1223 | 840 | 1134 | 3295 | 762 | 788 | 3708 | 466 | 605 | 3252 | 639 | 693 |
| 4700 | W303.asm02.HP0_G0046970 | chrXIV | 2379 | 1395 | 1796 | 2105 | 1433 | 1953 | 2361 | 1389 | 1746 | 5487 | 1304 | 2554 | 5573 | 907 | 1552 | 4945 | 1145 | 2227 |
| 4701 | W303.asm02.HP0_G0046980 | chrXIV | 1544 | 460 | 667 | 1283 | 518 | 683 | 1507 | 423 | 871 | 3256 | 748 | 1104 | 1936 | 592 | 687 | 1818 | 757 | 943 |
| 4702 | W303.asm02.HP0_G0046990 | chrXIV | 1394 | 1406 | 1363 | 1290 | 1403 | 1336 | 1283 | 1434 | 1337 | 5053 | 1218 | 1279 | 4688 | 792 | 926 | 4856 | 1066 | 1145 |
| 4703 | W303.asm02.HP0_G0047000 | chrXIV | 2231 | 892 | 680 | 2105 | 827 | 760 | 2199 | 865 | 726 | 4119 | 1087 | 1192 | 3258 | 735 | 837 | 3271 | 986 | 1056 |
| 4704 | W303.asm02.HP0_G0047010 | chrXIV | 2479 | 966 | 1440 | 1999 | 1044 | 1510 | 2178 | 1098 | 1576 | 6735 | 1485 | 2791 | 5245 | 1098 | 2058 | 5210 | 1299 | 2396 |
| 4705 | W303.asm02.HP0_G0047020 | chrXIV | 2313 | 345 | 654 | 2287 | 638 | 698 | 2076 | 367 | 927 | 4050 | 571 | 1260 | 1811 | 424 | 887 | 1986 | 510 | 1098 |
| 4706 | W303.asm02.HP0_G0047030 | chrXIV | 1124 | 825 | 833 | 1136 | 815 | 832 | 1103 | 816 | 877 | 2709 | 735 | 833 | 2807 | 481 | 685 | 2405 | 678 | 743 |
| 4707 | W303.asm02.HP0_G0047040 | chrXIV | 918 | 527 | 614 | 725 | 479 | 591 | 784 | 502 | 786 | 1440 | 468 | 421 | 1879 | 276 | 398 | 1795 | 395 | 437 |
| 4708 | W303.asm02.HP0_G0047050 | chrXIV | 5394 | 757 | 774 | 5228 | 706 | 921 | 5166 | 597 | 1020 | 5756 | 1448 | 1903 | 2894 | 1153 | 1216 | 2579 | 1405 | 1758 |
| 4709 | W303.asm02.HP0_G0047060 | chrXIV | 2578 | 589 | 260 | 2559 | 222 | 261 | 2142 | 377 | 264 | 1563 | 705 | 511 | 1438 | 329 | 423 | 1401 | 786 | 469 |
| 4710 | W303.asm02.HP0_G0047070 | chrXIV | 7651 | 246 | 219 | 6510 | 193 | 255 | 5997 | 226 | 270 | 273 | 179 | 170 | 725 | 174 | 224 | 373 | 187 | 151 |
| 4711 | W303.asm02.HP0_G0047080 | chrXIV | 8100 | 4343 | 3604 | 7605 | 3172 | 3463 | 7718 | 4228 | 3650 | 9542 | 3625 | 3317 | 11363 | 2727 | 3184 | 9350 | 3581 | 3058 |
| 4712 | W303.asm02.HP0_G0047090 | chrXIV | 5976 | 1920 | 2069 | 4846 | 1636 | 2075 | 4960 | 2031 | 2509 | 4205 | 1902 | 2250 | 5319 | 1434 | 2219 | 4210 | 1741 | 1808 |
| 4713 | W303.asm02.HP0_G0047100 | chrXIV | 2772 | 1008 | 1147 | 2312 | 989 | 1181 | 2217 | 962 | 1310 | 1878 | 788 | 1060 | 2522 | 579 | 958 | 1876 | 761 | 949 |
| 4714 | W303.asm02.HP0_G0047110 | chrXIV | 607 | 825 | 735 | 585 | 832 | 738 | 676 | 778 | 800 | 7007 | 1199 | 1521 | 5085 | 918 | 867 | 5060 | 1151 | 1282 |
| 4715 | W303.asm02.HP0_G0047120 | chrXIV | 1070 | 526 | 612 | 913 | 497 | 691 | 1014 | 464 | 650 | 3548 | 684 | 1130 | 3103 | 419 | 774 | 2896 | 700 | 1010 |
| 4716 | W303.asm02.HP0_G0047130 | chrXIV | 2521 | 735 | 786 | 2410 | 705 | 837 | 2441 | 754 | 829 | 2143 | 606 | 960 | 2658 | 507 | 754 | 1931 | 657 | 909 |
| 4717 | W303.asm02.HP0_G0047140 | chrXIV | 1398 | 786 | 692 | 1264 | 563 | 665 | 1371 | 640 | 886 | 2983 | 1214 | 1251 | 2098 | 909 | 923 | 1829 | 995 | 1087 |
| 4718 | W303.asm02.HP0_G0047150 | chrXIV | 8068 | 2900 | 2349 | 7800 | 2470 | 2464 | 8903 | 2411 | 2248 | 9465 | 2622 | 3576 | 7386 | 2041 | 2235 | 7517 | 2415 | 3202 |
| 4719 | W303.asm02.HP0_G0047160 | chrXIV | 1038 | 203 | 285 | 902 | 326 | 375 | 1237 | 141 | 301 | 2807 | 437 | 552 | 1492 | 235 | 361 | 1411 | 497 | 546 |
| 4720 | W303.asm02.HP0_G0047170 | chrXIV | 2825 | 460 | 370 | 2951 | 353 | 383 | 2506 | 331 | 371 | 2242 | 445 | 603 | 1964 | 384 | 523 | 1645 | 528 | 529 |
| 4721 | W303.asm02.HP0_G0047180 | chrXIV | 1243 | 1354 | 971 | 1221 | 1038 | 998 | 1374 | 1124 | 810 | 3300 | 1052 | 1238 | 3386 | 782 | 708 | 2876 | 852 | 1018 |
| 4722 | W303.asm02.HP0_G0047190 | chrXIV | 2287 | 255 | 291 | 2430 | 246 | 338 | 2084 | 213 | 389 | 1199 | 426 | 474 | 562 | 283 | 368 | 464 | 432 | 410 |
| 4723 | W303.asm02.HP0_G0047200 | chrXIV | 665 | 625 | 478 | 757 | 509 | 480 | 717 | 562 | 447 | 2194 | 585 | 633 | 1986 | 363 | 389 | 1854 | 562 | 525 |
| 4724 | W303.asm02.HP0_G0047210 | chrXIV | 2178 | 664 | 422 | 2087 | 490 | 485 | 2051 | 650 | 434 | 1519 | 587 | 621 | 1357 | 367 | 364 | 1327 | 544 | 480 |
| 4725 | W303.asm02.HP0_G0047220 | chrXIV | 1637 | 4079 | 3771 | 1814 | 3320 | 3588 | 2046 | 3999 | 3913 | 9748 | 3200 | 3186 | 9228 | 1706 | 2247 | 9694 | 2833 | 2948 |
| 4726 | W303.asm02.HP0_G0047230 | chrXIV | 6751 | 1131 | 1185 | 6613 | 1241 | 1272 | 7527 | 1111 | 1395 | 12517 | 1908 | 2858 | 9299 | 1616 | 2103 | 8427 | 1631 | 2706 |
| 4727 | W303.asm02.HP0_G0047240 | chrXIV | 1710 | 1127 | 1082 | 1736 | 1081 | 1112 | 1575 | 1023 | 1213 | 6857 | 1548 | 2169 | 4721 | 989 | 1554 | 4754 | 1445 | 1826 |
| 4728 | W303.asm02.HP0_G0047250 | chrXIV | 1526 | 1787 | 3649 | 1634 | 4182 | 3095 | 1867 | 1946 | 4152 | 17198 | 2025 | 3569 | 15847 | 1423 | 2151 | 15260 | 1926 | 2911 |
| 4729 | W303.asm02.HP0_G0047260 | chrXIV | 1606 | 1105 | 815 | 1501 | 1021 | 928 | 1841 | 1136 | 809 | 4831 | 958 | 1127 | 4468 | 717 | 700 | 4252 | 821 | 967 |
| 4730 | W303.asm02.HP0_G0047270 | chrXIV | 4108 | 1035 | 895 | 3495 | 969 | 889 | 4053 | 1026 | 1028 | 6495 | 1891 | 1849 | 4864 | 1362 | 1474 | 4498 | 1639 | 1627 |
| 4731 | W303.asm02.HP0_G0047280 | chrXIV | 635 | 123 | 120 | 541 | 132 | 129 | 622 | 132 | 127 | 934 | 234 | 268 | 872 | 166 | 242 | 705 | 209 | 292 |
| 4732 | W303.asm02.HP0_G0047290 | chrXIV | 620 | 220 | 169 | 538 | 185 | 159 | 570 | 224 | 159 | 552 | 163 | 149 | 713 | 198 | 154 | 447 | 145 | 123 |
| 4733 | W303.asm02.HP0_G0047300 | chrXIV | 414 | 198 | 181 | 359 | 167 | 178 | 403 | 198 | 191 | 678 | 203 | 221 | 962 | 186 | 178 | 503 | 186 | 192 |
| 4734 | W303.asm02.HP0_G0047310 | chrXIV | 730 | 136 | 169 | 676 | 112 | 190 | 718 | 99 | 138 | 1522 | 198 | 300 | 748 | 105 | 197 | 1185 | 200 | 336 |
| 4735 | W303.asm02.HP0_G0047320 | chrXIV | 547 | 1217 | 1479 | 429 | 956 | 1575 | 488 | 1117 | 1911 | 2847 | 785 | 1037 | 3691 | 482 | 729 | 3082 | 703 | 890 |
| 4736 | W303.asm02.HP0_G0047330 | chrXIV | 266 | 144 | 216 | 293 | 178 | 230 | 319 | 168 | 230 | 1097 | 203 | 362 | 934 | 111 | 247 | 911 | 187 | 258 |
| 4737 | W303.asm02.HP0_G0047340 | chrXIV | 304 | 52 | 93 | 318 | 93 | 91 | 330 | 53 | 114 | 1185 | 108 | 255 | 717 | 80 | 154 | 797 | 84 | 207 |
| 4738 | W303.asm02.HP0_G0047350 | chrXIV | 391 | 65 | 90 | 377 | 96 | 92 | 328 | 87 | 112 | 588 | 115 | 175 | 439 | 72 | 117 | 346 | 86 | 146 |
| 4739 | W303.asm02.HP0_G0047360 | chrXIV | 1511 | 324 | 541 | 1293 | 444 | 468 | 1310 | 390 | 712 | 2419 | 451 | 869 | 4648 | 922 | 1223 | 1541 | 415 | 799 |
| 4740 | W303.asm02.HP0_G0047370 | chrXIV | 961 | 229 | 366 | 949 | 305 | 342 | 883 | 222 | 469 | 3312 | 331 | 754 | 5685 | 717 | 813 | 1943 | 329 | 554 |
| 4741 | W303.asm02.HP0_G0047380 | chrXIV | 7930 | 1696 | 2289 | 7808 | 2226 | 2609 | 8779 | 1441 | 2914 | 15437 | 2575 | 4410 | 13461 | 2545 | 3791 | 9510 | 2231 | 4010 |

| | | | | | | | | | | | | | | | | | | | | |
|---|---|---|---|---|---|---|---|---|---|---|---|---|---|---|---|---|---|---|---|---|
| 4742 | W303.asm02.HP0_G0047390 | chrXIV | 2082 | 952 | 852 | 1716 | 889 | 863 | 1825 | 574 | 563 | 1447 | 444 | 532 | 1262 | 274 | 434 | 1329 | 447 | 525 |
| 4743 | W303.asm02.HP0_G0047400 | chrXIV | 53 | 192 | 419 | 152 | 295 | 409 | 41 | 227 | 377 | 1860 | 211 | 550 | 1295 | 106 | 301 | 1177 | 208 | 510 |
| 4744 | W303.asm02.HP0_G0047410 | chrXIV | 28 | 93 | 254 | 45 | 172 | 298 | 31 | 91 | 255 | 660 | 101 | 302 | 456 | 36 | 159 | 390 | 95 | 248 |
| 4745 | W303.asm02.HP0_G0047420 | chrXIV | 10 | 7 | 0 | 4 | 2 | 0 | 3 | 10 | 1 | 0 | 9 | 1 | 0 | 11 | 1 | 0 | 10 | 0 |
| 4746 | W303.asm02.HP0_G0047440 | chrXV | 2 | 13 | 1 | 0 | 0 | 0 | 1 | 18 | 0 | 0 | 9 | 0 | 0 | 11 | 0 | 0 | 12 | 0 |
| 4747 | W303.asm02.HP0_G0047450 | chrXV | 171 | 114 | 87 | 127 | 98 | 79 | 118 | 111 | 79 | 103 | 73 | 68 | 120 | 36 | 54 | 116 | 98 | 46 |
| 4748 | W303.asm02.HP0_G0047460 | chrXV | 153 | 105 | 65 | 150 | 42 | 60 | 134 | 93 | 73 | 92 | 48 | 61 | 118 | 38 | 43 | 104 | 60 | 48 |
| 4749 | W303.asm02.HP0_G0047470 | chrXV | 1035 | 593 | 602 | 955 | 521 | 681 | 963 | 560 | 644 | 2170 | 660 | 849 | 1834 | 376 | 585 | 1814 | 637 | 771 |
| 4750 | W303.asm02.HP0_G0047480 | chrXV | 572 | 90 | 87 | 420 | 72 | 94 | 489 | 99 | 97 | 202 | 96 | 88 | 319 | 69 | 73 | 182 | 85 | 94 |
| 4751 | W303.asm02.HP0_G0047490 | chrXV | 199 | 49 | 43 | 132 | 34 | 52 | 144 | 47 | 45 | 78 | 30 | 38 | 95 | 39 | 33 | 72 | 40 | 28 |
| 4752 | W303.asm02.HP0_G0047500 | chrXV | 3 | 7 | 2 | 4 | 2 | 3 | 3 | 2 | 0 | 0 | 3 | 2 | 2 | 1 | 0 | 3 | 1 | 0 |
| 4753 | W303.asm02.HP0_G0047510 | chrXV | 3 | 1 | 0 | 3 | 0 | 0 | 2 | 0 | 0 | 0 | 1 | 1 | 0 | 1 | 1 | 1 | 0 | 0 |
| 4754 | W303.asm02.HP0_G0047520 | chrXV | 426 | 235 | 245 | 314 | 183 | 270 | 361 | 235 | 310 | 485 | 150 | 220 | 691 | 137 | 197 | 507 | 134 | 167 |
| 4755 | W303.asm02.HP0_G0047530 | chrXV | 486 | 237 | 231 | 346 | 155 | 224 | 387 | 227 | 229 | 436 | 166 | 163 | 638 | 120 | 162 | 576 | 137 | 133 |
| 4756 | W303.asm02.HP0_G0047540 | chrXV | 990 | 582 | 564 | 1021 | 491 | 576 | 1012 | 554 | 818 | 1304 | 412 | 427 | 2770 | 500 | 465 | 1340 | 389 | 404 |
| 4757 | W303.asm02.HP0_G0047550 | chrXV | 1482 | 459 | 603 | 1338 | 614 | 624 | 1339 | 468 | 643 | 5170 | 893 | 745 | 3413 | 370 | 721 | 5335 | 831 | 1320 |
| 4758 | W303.asm02.HP0_G0047560 | chrXV | 974 | 151 | 191 | 858 | 151 | 143 | 835 | 159 | 248 | 465 | 135 | 119 | 646 | 118 | 97 | 401 | 99 | 121 |
| 4759 | W303.asm02.HP0_G0047570 | chrXV | 503 | 76 | 170 | 464 | 91 | 187 | 521 | 87 | 213 | 469 | 85 | 346 | 405 | 79 | 191 | 297 | 92 | 316 |
| 4760 | W303.asm02.HP0_G0047580 | chrXV | 24548 | 812 | 929 | 24870 | 912 | 1062 | 25452 | 608 | 1327 | 7176 | 1614 | 3863 | 3799 | 1589 | 2607 | 2238 | 1228 | 2880 |
| 4761 | W303.asm02.HP0_G0047590 | chrXV | 1146 | 385 | 348 | 826 | 301 | 304 | 993 | 368 | 368 | 1120 | 294 | 297 | 1179 | 210 | 200 | 1169 | 265 | 211 |
| 4762 | W303.asm02.HP0_G0047600 | chrXV | 586 | 24 | 26 | 523 | 29 | 42 | 512 | 25 | 36 | 252 | 69 | 74 | 192 | 49 | 74 | 153 | 62 | 64 |
| 4763 | W303.asm02.HP0_G0047610 | chrXV | 1629 | 69 | 62 | 1449 | 69 | 98 | 1397 | 56 | 89 | 595 | 143 | 171 | 485 | 102 | 147 | 311 | 137 | 152 |
| 4764 | W303.asm02.HP0_G0047620 | chrXV | 981 | 41 | 47 | 897 | 46 | 44 | 807 | 39 | 56 | 280 | 74 | 92 | 197 | 64 | 74 | 155 | 79 | 78 |
| 4765 | W303.asm02.HP0_G0047630 | chrXV | 2258 | 2845 | 2510 | 2281 | 2264 | 2749 | 2021 | 3110 | 3046 | 13353 | 5562 | 5343 | 14914 | 4763 | 5294 | 10799 | 4804 | 5645 |
| 4766 | W303.asm02.HP0_G0047640 | chrXV | 1143 | 5067 | 3789 | 1180 | 2643 | 3574 | 1163 | 4159 | 3814 | 5351 | 5027 | 4658 | 5679 | 3039 | 3290 | 4793 | 5164 | 3552 |
| 4767 | W303.asm02.HP0_G0047650 | chrXV | 1402 | 1409 | 1722 | 1382 | 1753 | 1653 | 1338 | 1392 | 2349 | 4855 | 1165 | 1261 | 5835 | 753 | 1091 | 5332 | 1069 | 1179 |
| 4768 | W303.asm02.HP0_G0047660 | chrXV | 1533 | 405 | 508 | 1483 | 399 | 505 | 1447 | 415 | 538 | 3475 | 645 | 1083 | 2699 | 419 | 696 | 2613 | 552 | 927 |
| 4769 | W303.asm02.HP0_G0047670 | chrXV | 2477 | 1233 | 1122 | 2351 | 1166 | 1176 | 2594 | 1221 | 1082 | 2876 | 1066 | 1204 | 3359 | 785 | 876 | 2759 | 929 | 1001 |
| 4770 | W303.asm02.HP0_G0047680 | chrXV | 496 | 424 | 568 | 410 | 483 | 608 | 492 | 474 | 716 | 1653 | 334 | 466 | 1976 | 261 | 331 | 1735 | 347 | 428 |
| 4771 | W303.asm02.HP0_G0047690 | chrXV | 7792 | 2009 | 2180 | 7842 | 2357 | 2198 | 7968 | 2065 | 2214 | 13921 | 2496 | 3359 | 11377 | 1570 | 2560 | 11821 | 2188 | 3180 |
| 4772 | W303.asm02.HP0_G0047700 | chrXV | 1010 | 450 | 430 | 1146 | 582 | 503 | 1254 | 449 | 575 | 5279 | 812 | 1017 | 4028 | 503 | 566 | 3896 | 622 | 828 |
| 4773 | W303.asm02.HP0_G0047710 | chrXV | 3902 | 1795 | 1950 | 3563 | 1520 | 1946 | 3818 | 1891 | 1926 | 4308 | 1831 | 1989 | 4854 | 1184 | 1618 | 4359 | 1494 | 1832 |
| 4774 | W303.asm02.HP0_G0047720 | chrXV | 874 | 1434 | 1435 | 852 | 1530 | 1413 | 1022 | 1495 | 1537 | 4800 | 1310 | 1464 | 4872 | 930 | 939 | 4599 | 1197 | 1270 |
| 4775 | W303.asm02.HP0_G0047730 | chrXV | 134 | 96 | 124 | 168 | 157 | 142 | 192 | 85 | 196 | 2699 | 245 | 305 | 1601 | 125 | 150 | 1623 | 198 | 305 |
| 4776 | W303.asm02.HP0_G0047740 | chrXV | 9830 | 6998 | 9599 | 8996 | 8766 | 9335 | 9611 | 7329 | 12144 | 21284 | 6803 | 8025 | 23402 | 4625 | 6510 | 22179 | 5782 | 7224 |
| 4777 | W303.asm02.HP0_G0047750 | chrXV | 1683 | 239 | 297 | 1507 | 336 | 357 | 1670 | 200 | 419 | 2783 | 369 | 714 | 1528 | 329 | 536 | 1341 | 372 | 631 |
| 4778 | W303.asm02.HP0_G0047760 | chrXV | 1028 | 935 | 934 | 913 | 870 | 876 | 1026 | 1011 | 987 | 4327 | 1115 | 1227 | 3902 | 718 | 895 | 4318 | 986 | 1060 |
| 4779 | W303.asm02.HP0_G0047770 | chrXV | 2798 | 1048 | 1464 | 3204 | 1630 | 1363 | 2955 | 946 | 1563 | 12952 | 1666 | 2313 | 11398 | 1073 | 1758 | 10302 | 1674 | 1979 |
| 4780 | W303.asm02.HP0_G0047780 | chrXV | 1891 | 398 | 293 | 1660 | 214 | 345 | 1939 | 273 | 353 | 1547 | 627 | 625 | 578 | 427 | 406 | 619 | 565 | 565 |
| 4781 | W303.asm02.HP0_G0047790 | chrXV | 1557 | 1961 | 1791 | 1554 | 1700 | 1898 | 1475 | 1895 | 2154 | 3570 | 1638 | 1390 | 4354 | 1106 | 1192 | 3676 | 1505 | 1332 |
| 4782 | W303.asm02.HP0_G0047800 | chrXV | 357 | 117 | 82 | 310 | 88 | 94 | 319 | 90 | 87 | 295 | 117 | 104 | 309 | 52 | 51 | 323 | 124 | 82 |
| 4783 | W303.asm02.HP0_G0047810 | chrXV | 122 | 24 | 10 | 79 | 11 | 16 | 104 | 11 | 4 | 28 | 10 | 11 | 35 | 5 | 5 | 27 | 18 | 10 |
| 4784 | W303.asm02.HP0_G0047820 | chrXV | 2933 | 917 | 1269 | 2900 | 1425 | 1236 | 3010 | 889 | 1407 | 10254 | 2179 | 2446 | 6095 | 1526 | 1650 | 6015 | 1802 | 2253 |
| 4785 | W303.asm02.HP0_G0047830 | chrXV | 4101 | 2181 | 2651 | 3821 | 2090 | 2617 | 3750 | 1972 | 3285 | 5448 | 1728 | 1950 | 7102 | 1356 | 1925 | 5842 | 1593 | 1884 |
| 4786 | W303.asm02.HP0_G0047840 | chrXV | 345 | 796 | 460 | 354 | 603 | 509 | 355 | 705 | 528 | 1807 | 472 | 392 | 1959 | 345 | 257 | 1732 | 558 | 358 |
| 4787 | W303.asm02.HP0_G0047850 | chrXV | 9226 | 9453 | 12523 | 7975 | 11342 | 12148 | 9114 | 9166 | 15357 | 23271 | 8170 | 9901 | 26820 | 6327 | 8497 | 25156 | 7485 | 8874 |
| 4788 | W303.asm02.HP0_G0047860 | chrXV | 1424 | 698 | 351 | 1761 | 426 | 387 | 1307 | 501 | 364 | 1962 | 841 | 883 | 2047 | 551 | 732 | 1332 | 916 | 746 |
| 4789 | W303.asm02.HP0_G0047870 | chrXV | 1459 | 735 | 565 | 1146 | 726 | 600 | 1340 | 711 | 670 | 4589 | 920 | 966 | 3245 | 590 | 628 | 3451 | 930 | 861 |
| 4790 | W303.asm02.HP0_G0047880 | chrXV | 190 | 1437 | 1835 | 210 | 2054 | 1639 | 280 | 1510 | 1971 | 9010 | 1745 | 2121 | 8641 | 1005 | 1069 | 8419 | 1462 | 1759 |
| 4791 | W303.asm02.HP0_G0047890 | chrXV | 22260 | 3730 | 2816 | 21968 | 2873 | 2718 | 22975 | 4117 | 2886 | 22115 | 5762 | 5226 | 17934 | 3709 | 4360 | 19826 | 4771 | 5021 |
| 4792 | W303.asm02.HP0_G0047900 | chrXV | 2786 | 1454 | 1752 | 2583 | 1418 | 1625 | 2600 | 1496 | 1835 | 6600 | 1737 | 2704 | 8244 | 1433 | 1941 | 5823 | 1626 | 2218 |
| 4793 | W303.asm02.HP0_G0047910 | chrXV | 5680 | 5421 | 6937 | 5071 | 6302 | 6803 | 5422 | 5501 | 8833 | 13753 | 4826 | 5413 | 16857 | 3597 | 4615 | 15299 | 4440 | 4515 |
| 4794 | W303.asm02.HP0_G0047920 | chrXV | 10800 | 10949 | 10885 | 9304 | 10894 | 11065 | 11120 | 10716 | 11561 | 26481 | 9657 | 10674 | 24706 | 7718 | 8086 | 24660 | 8859 | 9724 |
| 4795 | W303.asm02.HP0_G0047930 | chrXV | 1339 | 1318 | 1359 | 1314 | 1137 | 1244 | 1264 | 1296 | 1526 | 3341 | 2560 | 2202 | 2491 | 1901 | 1849 | 2454 | 2232 | 1810 |
| 4796 | W303.asm02.HP0_G0047940 | chrXV | 1839 | 431 | 652 | 1826 | 476 | 687 | 1535 | 466 | 766 | 1814 | 410 | 797 | 2089 | 304 | 608 | 1880 | 431 | 645 |
| 4797 | W303.asm02.HP0_G0047950 | chrXV | 2015 | 847 | 694 | 2086 | 733 | 772 | 1994 | 838 | 685 | 2937 | 858 | 1089 | 2355 | 522 | 749 | 2746 | 767 | 1002 |
| 4798 | W303.asm02.HP0_G0047960 | chrXV | 2374 | 672 | 688 | 1838 | 733 | 660 | 2345 | 620 | 735 | 5698 | 1020 | 1229 | 3742 | 596 | 813 | 4589 | 939 | 1083 |
| 4799 | W303.asm02.HP0_G0047970 | chrXV | 278 | 122 | 81 | 227 | 81 | 101 | 262 | 100 | 98 | 291 | 109 | 87 | 274 | 61 | 65 | 268 | 130 | 88 |
| 4800 | W303.asm02.HP0_G0047980 | chrXV | 4670 | 758 | 763 | 5197 | 724 | 744 | 4877 | 771 | 718 | 4163 | 985 | 1138 | 5529 | 705 | 1112 | 3511 | 856 | 1139 |
| 4801 | W303.asm02.HP0_G0047990 | chrXV | 2694 | 950 | 757 | 2486 | 863 | 831 | 2779 | 887 | 818 | 4003 | 1365 | 1601 | 4457 | 1264 | 1395 | 2799 | 1182 | 1457 |
| 4802 | W303.asm02.HP0_G0048000 | chrXV | 3396 | 474 | 375 | 3174 | 322 | 399 | 3452 | 409 | 441 | 2333 | 758 | 695 | 1934 | 524 | 595 | 1877 | 642 | 625 |
| 4803 | W303.asm02.HP0_G0048010 | chrXV | 1552 | 407 | 399 | 1180 | 373 | 423 | 1259 | 343 | 485 | 1093 | 521 | 442 | 839 | 366 | 394 | 821 | 455 | 374 |
| 4804 | W303.asm02.HP0_G0048020 | chrXV | 10303 | 7330 | 11048 | 10254 | 10130 | 10728 | 9795 | 7843 | 14021 | 19872 | 6697 | 9335 | 23279 | 5032 | 7433 | 21206 | 6003 | 7960 |
| 4805 | W303.asm02.HP0_G0048030 | chrXV | 2752 | 999 | 704 | 2739 | 631 | 779 | 2542 | 1013 | 953 | 2039 | 911 | 749 | 2500 | 679 | 713 | 1986 | 918 | 688 |
| 4806 | W303.asm02.HP0_G0048040 | chrXV | 1067 | 578 | 649 | 1025 | 468 | 723 | 1006 | 657 | 723 | 2106 | 565 | 766 | 2512 | 441 | 594 | 2228 | 533 | 685 |
| 4807 | W303.asm02.HP0_G0048050 | chrXV | 536 | 206 | 240 | 447 | 213 | 243 | 503 | 126 | 309 | 691 | 262 | 399 | 458 | 154 | 226 | 326 | 218 | 335 |
| 4808 | W303.asm02.HP0_G0048060 | chrXV | 637 | 209 | 301 | 622 | 296 | 256 | 605 | 194 | 257 | 836 | 202 | 458 | 714 | 156 | 292 | 687 | 188 | 404 |
| 4809 | W303.asm02.HP0_G0048070 | chrXV | 2864 | 1793 | 2157 | 2408 | 1976 | 2017 | 2506 | 2004 | 2378 | 9326 | 2219 | 2446 | 9438 | 1562 | 1909 | 8540 | 1959 | 2119 |
| 4810 | W303.asm02.HP0_G0048080 | chrXV | 1123 | 780 | 1027 | 1162 | 926 | 966 | 1204 | 887 | 1361 | 2838 | 684 | 729 | 3253 | 422 | 557 | 3023 | 655 | 714 |
| 4811 | W303.asm02.HP0_G0048090 | chrXV | 1969 | 660 | 744 | 1754 | 829 | 613 | 1684 | 699 | 874 | 3482 | 893 | 639 | 3688 | 388 | 401 | 3948 | 849 | 627 |
| 4812 | W303.asm02.HP0_G0048100 | chrXV | 3636 | 389 | 558 | 3675 | 502 | 574 | 3434 | 345 | 709 | 4110 | 741 | 1126 | 3135 | 676 | 949 | 2698 | 647 | 973 |
| 4813 | W303.asm02.HP0_G0048110 | chrXV | 8439 | 1425 | 1570 | 7603 | 1546 | 1633 | 8583 | 1503 | 1971 | 14679 | 2583 | 3460 | 10496 | 1928 | 2530 | 10076 | 2207 | 3133 |
| 4814 | W303.asm02.HP0_G0048120 | chrXV | 171 | 94 | 76 | 105 | 95 | 89 | 146 | 93 | 123 | 300 | 53 | 67 | 413 | 39 | 28 | 336 | 55 | 27 |
| 4815 | W303.asm02.HP0_G0048130 | chrXV | 8056 | 3909 | 3232 | 7357 | 3524 | 3119 | 8394 | 4156 | 3164 | 17265 | 4523 | 4940 | 14706 | 2864 | 3510 | 15262 | 3846 | 4372 |
| 4816 | W303.asm02.HP0_G0048140 | chrXV | 4807 | 907 | 504 | 4198 | 557 | 589 | 4220 | 793 | 588 | 2080 | 726 | 644 | 2394 | 524 | 618 | 1944 | 754 | 607 |
| 4817 | W303.asm02.HP0_G0048150 | chrXV | 588 | 237 | 319 | 539 | 280 | 313 | 509 | 227 | 366 | 2251 | 423 | 600 | 1372 | 315 | 393 | 1516 | 411 | 546 |
| 4818 | W303.asm02.HP0_G0048160 | chrXV | 959 | 748 | 729 | 819 | 686 | 759 | 947 | 746 | 707 | 2839 | 649 | 910 | 2931 | 519 | 653 | 2713 | 631 | 787 |
| 4819 | W303.asm02.HP0_G0048170 | chrXV | 3435 | 1276 | 1074 | 3214 | 1190 | 1167 | 3640 | 1355 | 957 | 4637 | 1276 | 1463 | 4683 | 945 | 977 | 3989 | 1069 | 1327 |
| 4820 | W303.asm02.HP0_G0048180 | chrXV | 1758 | 2055 | 2092 | 1667 | 2057 | 1963 | 1807 | 2284 | 2127 | 7229 | 2158 | 2138 | 6817 | 1377 | 1372 | 6800 | 1838 | 1957 |
| 4821 | W303.asm02.HP0_G0048190 | chrXV | 219 | 208 | 265 | 228 | 220 | 217 | 191 | 178 | 302 | 549 | 145 | 170 | 851 | 95 | 145 | 619 | 146 | 139 |
| 4822 | W303.asm02.HP0_G0048200 | chrXV | 1638 | 693 | 937 | 1495 | 834 | 1005 | 1703 | 720 | 1080 | 5978 | 1024 | 1798 | 5355 | 887 | 1386 | 4769 | 1036 | 1670 |
| 4823 | W303.asm02.HP0_G0048210 | chrXV | 1245 | 517 | 756 | 1366 | 778 | 745 | 1195 | 589 | 748 | 4205 | 605 | 1123 | 3700 | 354 | 730 | 3775 | 589 | 970 |
| 4824 | W303.asm02.HP0_G0048220 | chrXV | 2411 | 1392 | 1486 | 2264 | 1201 | 1660 | 2374 | 1487 | 1679 | 4052 | 1144 | 1599 | 4179 | 842 | 1068 | 4009 | 1009 | 1491 |
| 4825 | W303.asm02.HP0_G0048230 | chrXV | 6360 | 564 | 586 | 5754 | 564 | 658 | 5862 | 521 | 725 | 5104 | 1118 | 1325 | 3670 | 883 | 1200 | 3195 | 988 | 1093 |
| 4826 | W303.asm02.HP0_G0048240 | chrXV | 887 | 379 | 595 | 613 | 495 | 637 | 710 | 382 | 813 | 1056 | 327 | 469 | 1707 | 245 | 426 | 1280 | 342 | 362 |
| 4827 | W303.asm02.HP0_G0048250 | chrXV | 69642 | 27927 | 28672 | 72337 | 24057 | 27957 | 73793 | 26667 | 34690 | 78457 | 43184 | 46979 | 69984 | 33568 | 41529 | 69044 | 39379 | 46288 |
| 4828 | W303.asm02.HP0_G0048260 | chrXV | 143 | 32 | 12 | 122 | 13 | 9 | 145 | 15 | 11 | 59 | 36 | 27 | 54 | 24 | 21 | 43 | 44 | 20 |
| 4829 | W303.asm02.HP0_G0048270 | chrXV | 385 | 58 | 118 | 375 | 54 | 139 | 330 | 39 | 172 | 396 | 118 | 341 | 357 | 96 | 204 | 232 | 127 | 241 |
| 4830 | W303.asm02.HP0_G0048280 | chrXV | 2132 | 531 | 436 | 2115 | 369 | 444 | 1736 | 510 | 485 | 925 | 483 | 505 | 1321 | 360 | 515 | 921 | 442 | 470 |
| 4831 | W303.asm02.HP0_G0048290 | chrXV | 4249 | 935 | 1011 | 4044 | 954 | 1068 | 4217 | 950 | 897 | 2995 | 1002 | 1388 | 3869 | 784 | 1268 | 2663 | 897 | 1205 |
| 4832 | W303.asm02.HP0_G0048300 | chrXV | 10466 | 782 | 1098 | 12097 | 1267 | 1497 | 9553 | 731 | 1597 | 8505 | 1142 | 2131 | 4771 | 844 | 1576 | 4873 | 976 | 1747 |
| 4833 | W303.asm02.HP0_G0048310 | chrXV | 471 | 552 | 803 | 529 | 1112 | 746 | 612 | 651 | 824 | 5691 | 734 | 1165 | 5338 | 394 | 586 | 5194 | 553 | 935 |
| 4834 | W303.asm02.HP0_G0048320 | chrXV | 980 | 231 | 371 | 1075 | 396 | 382 | 954 | 250 | 437 | 3717 | 476 | 801 | 2848 | 320 | 554 | 2622 | 409 | 778 |

| | | | | | | | | | | | | | | | | | | | | |
|---|---|---|---|---|---|---|---|---|---|---|---|---|---|---|---|---|---|---|---|---|
| 4835 | W303.asm02.HP0_G0048330 | chrXV | 1086 | 1137 | 1919 | 683 | 1705 | 1852 | 741 | 1152 | 2484 | 3750 | 946 | 1613 | 5217 | 795 | 1433 | 4362 | 868 | 1077 |
| 4836 | W303.asm02.HP0_G0048340 | chrXV | 2818 | 3548 | 5025 | 3097 | 5278 | 4677 | 3302 | 4139 | 6151 | 15575 | 3828 | 4341 | 16045 | 2430 | 2852 | 16874 | 3296 | 3934 |
| 4837 | W303.asm02.HP0_G0048350 | chrXV | 929 | 979 | 1345 | 884 | 1375 | 1176 | 969 | 1170 | 1404 | 7763 | 1014 | 1501 | 6557 | 699 | 1170 | 7785 | 981 | 1430 |
| 4838 | W303.asm02.HP0_G0048360 | chrXV | 1465 | 380 | 505 | 1345 | 545 | 512 | 1362 | 412 | 642 | 3303 | 507 | 882 | 2126 | 356 | 597 | 2331 | 485 | 810 |
| 4839 | W303.asm02.HP0_G0048370 | chrXV | 2385 | 620 | 431 | 2147 | 405 | 510 | 2024 | 517 | 530 | 2250 | 638 | 801 | 2101 | 453 | 694 | 1957 | 637 | 722 |
| 4840 | W303.asm02.HP0_G0048380 | chrXV | 1646 | 409 | 279 | 1575 | 297 | 329 | 1720 | 296 | 296 | 2010 | 453 | 596 | 1661 | 306 | 419 | 1518 | 419 | 537 |
| 4841 | W303.asm02.HP0_G0048390 | chrXV | 4822 | 1376 | 1666 | 4371 | 1362 | 1696 | 4104 | 1420 | 2225 | 2794 | 1093 | 1204 | 4204 | 832 | 1235 | 2794 | 1009 | 1163 |
| 4842 | W303.asm02.HP0_G0048400 | chrXV | 3540 | 1436 | 1138 | 3236 | 1353 | 1181 | 3565 | 1368 | 1239 | 6953 | 1973 | 2146 | 6290 | 1408 | 1734 | 5478 | 1623 | 1955 |
| 4843 | W303.asm02.HP0_G0048410 | chrXV | 660 | 501 | 504 | 641 | 566 | 441 | 667 | 513 | 526 | 2017 | 415 | 506 | 2043 | 324 | 485 | 2099 | 336 | 469 |
| 4844 | W303.asm02.HP0_G0048420 | chrXV | 3479 | 992 | 1118 | 3335 | 1152 | 1172 | 3138 | 1002 | 1275 | 7837 | 1444 | 2051 | 5498 | 1015 | 1582 | 5894 | 1366 | 1759 |
| 4845 | W303.asm02.HP0_G0048430 | chrXV | 1378 | 221 | 310 | 1320 | 259 | 275 | 1141 | 171 | 366 | 2089 | 473 | 557 | 1436 | 345 | 457 | 1280 | 438 | 507 |
| 4846 | W303.asm02.HP0_G0048440 | chrXV | 1674 | 588 | 398 | 1493 | 382 | 425 | 1577 | 414 | 415 | 3157 | 776 | 936 | 1747 | 446 | 568 | 2095 | 751 | 773 |
| 4847 | W303.asm02.HP0_G0048450 | chrXV | 1144 | 315 | 325 | 1176 | 301 | 332 | 1043 | 324 | 400 | 1845 | 363 | 510 | 1493 | 300 | 370 | 1557 | 392 | 414 |
| 4848 | W303.asm02.HP0_G0048460 | chrXV | 4596 | 3019 | 4531 | 3769 | 3625 | 4349 | 4463 | 3095 | 5366 | 9000 | 2441 | 3439 | 10180 | 1587 | 2895 | 9382 | 1984 | 3100 |
| 4849 | W303.asm02.HP0_G0048470 | chrXV | 1638 | 424 | 556 | 1546 | 556 | 552 | 1636 | 431 | 662 | 3617 | 731 | 1019 | 2695 | 463 | 813 | 2673 | 658 | 935 |
| 4850 | W303.asm02.HP0_G0048480 | chrXV | 5229 | 2212 | 1803 | 4549 | 1889 | 1750 | 4900 | 2159 | 1769 | 7747 | 2318 | 2330 | 7441 | 1742 | 2126 | 7037 | 2169 | 2192 |
| 4851 | W303.asm02.HP0_G0048490 | chrXV | 6043 | 2752 | 2674 | 5183 | 2844 | 2428 | 6329 | 2934 | 3009 | 13484 | 3716 | 4252 | 12294 | 2485 | 2976 | 11657 | 3357 | 3723 |
| 4852 | W303.asm02.HP0_G0048500 | chrXV | 5249 | 738 | 809 | 4492 | 692 | 973 | 5162 | 648 | 935 | 3766 | 1362 | 1717 | 1975 | 1079 | 1317 | 1572 | 1209 | 1500 |
| 4853 | W303.asm02.HP0_G0048510 | chrXV | 8040 | 1735 | 1783 | 8014 | 1502 | 1817 | 8425 | 1284 | 2283 | 6065 | 2449 | 3202 | 2344 | 1415 | 1584 | 2741 | 2191 | 2808 |
| 4854 | W303.asm02.HP0_G0048520 | chrXV | 906 | 2219 | 2098 | 672 | 1785 | 2350 | 890 | 1775 | 2481 | 14026 | 4549 | 5396 | 7883 | 3224 | 3107 | 10131 | 3413 | 4138 |
| 4855 | W303.asm02.HP0_G0048530 | chrXV | 5013 | 1001 | 1142 | 4469 | 869 | 1392 | 5198 | 909 | 1591 | 8203 | 1794 | 3201 | 4978 | 1499 | 2082 | 4891 | 1726 | 2849 |
| 4856 | W303.asm02.HP0_G0048540 | chrXV | 1120 | 586 | 654 | 999 | 673 | 596 | 1094 | 678 | 679 | 3483 | 697 | 755 | 3089 | 342 | 509 | 3390 | 675 | 736 |
| 4857 | W303.asm02.HP0_G0048550 | chrXV | 871 | 772 | 711 | 757 | 507 | 694 | 687 | 646 | 626 | 2339 | 653 | 1034 | 2442 | 377 | 694 | 2068 | 730 | 888 |
| 4858 | W303.asm02.HP0_G0048560 | chrXV | 1312 | 1168 | 956 | 1513 | 1083 | 1005 | 1489 | 1070 | 1072 | 4259 | 1120 | 1034 | 5820 | 670 | 631 | 3866 | 1046 | 908 |
| 4859 | W303.asm02.HP0_G0048570 | chrXV | 3287 | 811 | 653 | 3382 | 710 | 627 | 2970 | 803 | 731 | 2722 | 787 | 694 | 3083 | 530 | 683 | 2524 | 789 | 670 |
| 4860 | W303.asm02.HP0_G0048580 | chrXV | 4086 | 1096 | 780 | 4407 | 707 | 1043 | 3438 | 934 | 873 | 529 | 765 | 778 | 689 | 370 | 517 | 503 | 807 | 621 |
| 4861 | W303.asm02.HP0_G0048590 | chrXV | 3173 | 1128 | 890 | 2829 | 930 | 948 | 3019 | 1045 | 869 | 6020 | 1408 | 1879 | 5122 | 1025 | 1223 | 4592 | 1282 | 1513 |
| 4862 | W303.asm02.HP0_G0048600 | chrXV | 8672 | 602 | 917 | 8443 | 874 | 1190 | 9239 | 491 | 1190 | 10956 | 1986 | 2780 | 3431 | 1392 | 1989 | 3701 | 1593 | 2440 |
| 4863 | W303.asm02.HP0_G0048610 | chrXV | 6424 | 2791 | 2409 | 5794 | 2314 | 2357 | 6272 | 3029 | 2521 | 11879 | 3833 | 4204 | 10551 | 2828 | 3355 | 10656 | 3370 | 3743 |
| 4864 | W303.asm02.HP0_G0048620 | chrXV | 2105 | 804 | 1013 | 2137 | 727 | 1105 | 1899 | 753 | 1240 | 1846 | 582 | 806 | 2708 | 477 | 871 | 1955 | 625 | 650 |
| 4865 | W303.asm02.HP0_G0048630 | chrXV | 1045 | 98 | 96 | 921 | 66 | 84 | 864 | 77 | 102 | 233 | 126 | 118 | 354 | 86 | 97 | 178 | 124 | 121 |
| 4866 | W303.asm02.HP0_G0048640 | chrXV | 3860 | 370 | 716 | 3608 | 724 | 743 | 3836 | 513 | 895 | 6366 | 782 | 1319 | 6166 | 620 | 900 | 4773 | 678 | 1127 |
| 4867 | W303.asm02.HP0_G0048650 | chrXV | 1140 | 600 | 670 | 1211 | 639 | 588 | 1128 | 571 | 740 | 2158 | 478 | 545 | 2864 | 349 | 414 | 2223 | 463 | 529 |
| 4868 | W303.asm02.HP0_G0048660 | chrXV | 1471 | 592 | 420 | 1484 | 429 | 417 | 1438 | 510 | 418 | 2003 | 591 | 653 | 1650 | 380 | 451 | 1731 | 550 | 564 |
| 4869 | W303.asm02.HP0_G0048670 | chrXV | 2236 | 1294 | 1758 | 2244 | 1434 | 1624 | 2210 | 1285 | 1974 | 4970 | 1022 | 1449 | 5146 | 676 | 1134 | 5128 | 962 | 1214 |
| 4870 | W303.asm02.HP0_G0048680 | chrXV | 3444 | 2438 | 2219 | 3557 | 2860 | 2173 | 3889 | 2578 | 2183 | 14835 | 2672 | 3144 | 13303 | 1851 | 1875 | 13065 | 2436 | 2684 |
| 4871 | W303.asm02.HP0_G0048690 | chrXV | 13507 | 7848 | 8436 | 12983 | 8765 | 8851 | 13604 | 7889 | 8244 | 21761 | 7735 | 10325 | 21390 | 6616 | 7831 | 20195 | 6822 | 9498 |
| 4872 | W303.asm02.HP0_G0048700 | chrXV | 8160 | 7544 | 9050 | 6860 | 8377 | 8964 | 7442 | 7330 | 11248 | 15023 | 6564 | 7186 | 17614 | 4899 | 5968 | 16057 | 5917 | 6245 |
| 4873 | W303.asm02.HP0_G0048710 | chrXV | 5701 | 3640 | 3687 | 5610 | 3454 | 3964 | 5576 | 3618 | 3761 | 9271 | 3294 | 4272 | 9354 | 2489 | 3457 | 8399 | 3060 | 3639 |
| 4874 | W303.asm02.HP0_G0048720 | chrXV | 2303 | 777 | 787 | 2229 | 961 | 978 | 2447 | 639 | 848 | 6706 | 1658 | 1928 | 4928 | 989 | 1316 | 4704 | 1588 | 1753 |
| 4875 | W303.asm02.HP0_G0048730 | chrXV | 1209 | 500 | 684 | 1336 | 629 | 696 | 1279 | 482 | 737 | 3991 | 661 | 1186 | 3170 | 386 | 770 | 3516 | 676 | 1063 |
| 4876 | W303.asm02.HP0_G0048740 | chrXV | 3308 | 935 | 999 | 2903 | 905 | 941 | 3003 | 832 | 1151 | 4122 | 810 | 1300 | 4125 | 612 | 956 | 3668 | 840 | 1095 |
| 4877 | W303.asm02.HP0_G0048750 | chrXV | 3607 | 1880 | 779 | 3989 | 784 | 901 | 3109 | 1471 | 765 | 1544 | 1158 | 648 | 1694 | 685 | 611 | 1780 | 1196 | 561 |
| 4878 | W303.asm02.HP0_G0048760 | chrXV | 2166 | 1345 | 4995 | 2392 | 1495 | 5775 | 2316 | 1351 | 4884 | 5297 | 1142 | 5935 | 6917 | 899 | 3416 | 5255 | 1130 | 5062 |
| 4879 | W303.asm02.HP0_G0048770 | chrXV | 16271 | 2758 | 2705 | 16420 | 2087 | 2933 | 17469 | 2532 | 3082 | 14638 | 5665 | 5892 | 13024 | 4882 | 4962 | 10165 | 4699 | 5063 |
| 4880 | W303.asm02.HP0_G0048780 | chrXV | 805 | 266 | 329 | 787 | 386 | 382 | 964 | 242 | 415 | 2089 | 444 | 661 | 1474 | 327 | 448 | 1405 | 371 | 565 |
| 4881 | W303.asm02.HP0_G0048790 | chrXV | 1175 | 284 | 311 | 1118 | 335 | 290 | 1343 | 286 | 297 | 3079 | 622 | 816 | 2000 | 477 | 519 | 2004 | 543 | 581 |
| 4882 | W303.asm02.HP0_G0048800 | chrXV | 10960 | 1049 | 1084 | 11647 | 995 | 1449 | 11308 | 1011 | 1434 | 8859 | 2217 | 3387 | 7732 | 1777 | 2098 | 6596 | 1880 | 2783 |
| 4883 | W303.asm02.HP0_G0048810 | chrXV | 771 | 670 | 597 | 666 | 507 | 637 | 651 | 694 | 741 | 1498 | 723 | 604 | 1546 | 452 | 478 | 1446 | 638 | 559 |
| 4884 | W303.asm02.HP0_G0048820 | chrXV | 1292 | 233 | 286 | 1418 | 244 | 323 | 1212 | 211 | 318 | 1984 | 402 | 632 | 1268 | 237 | 353 | 1358 | 392 | 525 |
| 4885 | W303.asm02.HP0_G0048830 | chrXV | 47 | 11 | 14 | 46 | 11 | 9 | 32 | 10 | 15 | 48 | 18 | 23 | 39 | 9 | 7 | 35 | 17 | 16 |
| 4886 | W303.asm02.HP0_G0048840 | chrXV | 2745 | 302 | 483 | 2825 | 470 | 553 | 2497 | 270 | 601 | 3945 | 606 | 1340 | 3275 | 470 | 949 | 2707 | 668 | 1145 |
| 4887 | W303.asm02.HP0_G0048850 | chrXV | 3463 | 1620 | 1364 | 3226 | 1559 | 1372 | 3630 | 1824 | 1480 | 7422 | 2030 | 1844 | 7002 | 1257 | 1099 | 6686 | 1845 | 1718 |
| 4888 | W303.asm02.HP0_G0048860 | chrXV | 3100 | 732 | 1046 | 2898 | 1042 | 1165 | 3401 | 616 | 1332 | 11125 | 1834 | 2769 | 4735 | 1281 | 1619 | 4956 | 1637 | 2388 |
| 4889 | W303.asm02.HP0_G0048870 | chrXV | 1372 | 957 | 1661 | 1275 | 1478 | 1651 | 1445 | 871 | 2031 | 9372 | 2117 | 3346 | 4691 | 1465 | 2147 | 4737 | 1919 | 2965 |
| 4890 | W303.asm02.HP0_G0048880 | chrXV | 0 | 2 | 1 | 2 | 2 | 0 | 0 | 0 | 0 | 0 | 2 | 0 | 3 | 3 | 0 | 2 | 1 | 0 |
| 4891 | W303.asm02.HP0_G0048890 | chrXV | 2777 | 1135 | 1915 | 3194 | 1334 | 2033 | 3183 | 1351 | 2007 | 9728 | 2047 | 4776 | 11316 | 1944 | 3167 | 7495 | 1777 | 3844 |
| 4892 | W303.asm02.HP0_G0048900 | chrXV | 3968 | 1545 | 1415 | 4016 | 1488 | 1415 | 3826 | 1489 | 1474 | 4971 | 1454 | 1413 | 4886 | 894 | 1079 | 4693 | 1259 | 1346 |
| 4893 | W303.asm02.HP0_G0048910 | chrXV | 1160 | 609 | 515 | 1189 | 463 | 477 | 1123 | 591 | 601 | 2529 | 719 | 717 | 2285 | 388 | 507 | 2406 | 661 | 706 |
| 4894 | W303.asm02.HP0_G0048920 | chrXV | 11989 | 1080 | 978 | 15092 | 746 | 884 | 10705 | 868 | 857 | 3088 | 980 | 1030 | 3681 | 666 | 1014 | 2929 | 986 | 1009 |
| 4895 | W303.asm02.HP0_G0048930 | chrXV | 492 | 129 | 127 | 558 | 145 | 120 | 520 | 104 | 103 | 673 | 132 | 171 | 484 | 72 | 131 | 558 | 155 | 143 |
| 4896 | W303.asm02.HP0_G0048940 | chrXV | 1763 | 463 | 382 | 2233 | 393 | 356 | 1426 | 438 | 428 | 1313 | 301 | 242 | 1387 | 181 | 167 | 1349 | 300 | 212 |
| 4897 | W303.asm02.HP0_G0048950 | chrXV | 1591 | 914 | 1564 | 1774 | 783 | 1595 | 1468 | 949 | 1426 | 4614 | 1037 | 2279 | 4152 | 707 | 1528 | 4399 | 1007 | 2011 |
| 4898 | W303.asm02.HP0_G0048960 | chrXV | 1546 | 1427 | 1561 | 1495 | 1365 | 1553 | 1571 | 1363 | 1844 | 3869 | 1191 | 1425 | 3699 | 755 | 1105 | 3897 | 1021 | 1236 |
| 4899 | W303.asm02.HP0_G0048970 | chrXV | 1226 | 505 | 516 | 1030 | 490 | 614 | 1156 | 398 | 699 | 3086 | 918 | 1181 | 1403 | 615 | 671 | 1665 | 757 | 1008 |
| 4900 | W303.asm02.HP0_G0048980 | chrXV | 925 | 1505 | 1796 | 975 | 1942 | 1579 | 1081 | 1451 | 1829 | 7714 | 1543 | 2066 | 7595 | 1071 | 1253 | 6787 | 1464 | 1771 |
| 4901 | W303.asm02.HP0_G0048990 | chrXV | 1649 | 727 | 513 | 1510 | 502 | 501 | 1484 | 663 | 577 | 1905 | 647 | 504 | 2033 | 440 | 411 | 1832 | 616 | 463 |
| 4902 | W303.asm02.HP0_G0049000 | chrXV | 1746 | 834 | 945 | 1439 | 810 | 885 | 1495 | 728 | 1089 | 2149 | 636 | 746 | 2722 | 444 | 609 | 2404 | 586 | 594 |
| 4903 | W303.asm02.HP0_G0049010 | chrXV | 1493 | 814 | 1258 | 1739 | 945 | 1127 | 1800 | 886 | 1066 | 6029 | 1224 | 2447 | 6511 | 848 | 1307 | 5137 | 971 | 1981 |
| 4904 | W303.asm02.HP0_G0049020 | chrXV | 2787 | 1021 | 970 | 2816 | 1017 | 1172 | 3007 | 893 | 1199 | 8019 | 1614 | 2462 | 5182 | 1130 | 1610 | 5342 | 1631 | 2245 |
| 4905 | W303.asm02.HP0_G0049030 | chrXV | 1854 | 910 | 906 | 1652 | 894 | 918 | 1736 | 992 | 854 | 2265 | 1017 | 1198 | 2261 | 704 | 818 | 2050 | 858 | 1056 |
| 4906 | W303.asm02.HP0_G0049040 | chrXV | 7014 | 874 | 1125 | 6762 | 1091 | 1362 | 6976 | 775 | 1515 | 4954 | 1006 | 2522 | 2243 | 947 | 1695 | 2289 | 803 | 2089 |
| 4907 | W303.asm02.HP0_G0049050 | chrXV | 1293 | 969 | 824 | 1126 | 716 | 808 | 1132 | 868 | 923 | 2942 | 736 | 832 | 3044 | 547 | 650 | 2901 | 773 | 792 |
| 4908 | W303.asm02.HP0_G0049060 | chrXV | 3140 | 1966 | 1830 | 2642 | 1894 | 1677 | 2971 | 1851 | 2326 | 5551 | 1567 | 1173 | 6792 | 1059 | 1017 | 5870 | 1547 | 995 |
| 4909 | W303.asm02.HP0_G0049070 | chrXV | 945 | 1056 | 577 | 886 | 595 | 541 | 953 | 1037 | 603 | 2288 | 899 | 601 | 2262 | 544 | 490 | 2455 | 844 | 598 |
| 4910 | W303.asm02.HP0_G0049080 | chrXV | 4471 | 1012 | 887 | 4059 | 967 | 918 | 4849 | 940 | 1059 | 9278 | 2026 | 2210 | 6253 | 1298 | 1272 | 7011 | 1862 | 1855 |
| 4911 | W303.asm02.HP0_G0049090 | chrXV | 4133 | 2092 | 2728 | 3622 | 1866 | 2547 | 4163 | 2169 | 2756 | 8539 | 2298 | 3578 | 8321 | 1517 | 2433 | 8245 | 2202 | 3244 |
| 4912 | W303.asm02.HP0_G0049100 | chrXV | 1615 | 942 | 599 | 1392 | 549 | 565 | 1400 | 900 | 732 | 1773 | 845 | 591 | 2646 | 766 | 536 | 1545 | 897 | 524 |
| 4913 | W303.asm02.HP0_G0049110 | chrXV | 1133 | 507 | 525 | 1105 | 675 | 510 | 1220 | 541 | 643 | 4908 | 1080 | 1137 | 3743 | 744 | 636 | 3819 | 921 | 1023 |
| 4914 | W303.asm02.HP0_G0049120 | chrXV | 2566 | 795 | 492 | 2576 | 464 | 541 | 2240 | 707 | 529 | 2834 | 1019 | 730 | 2321 | 565 | 539 | 2204 | 1054 | 659 |
| 4915 | W303.asm02.HP0_G0049130 | chrXV | 2069 | 1592 | 1335 | 1919 | 1583 | 1243 | 1896 | 1624 | 1550 | 5276 | 1730 | 1501 | 4635 | 928 | 1109 | 5059 | 1719 | 1459 |
| 4916 | W303.asm02.HP0_G0049140 | chrXV | 22211 | 10540 | 8772 | 22696 | 8805 | 9255 | 22636 | 9466 | 7409 | 25481 | 9933 | 11222 | 24640 | 7303 | 9168 | 23260 | 8694 | 10492 |
| 4917 | W303.asm02.HP0_G0049150 | chrXV | 4329 | 1528 | 1420 | 4574 | 1288 | 1426 | 4332 | 1430 | 1522 | 9782 | 2902 | 2640 | 7535 | 1908 | 2183 | 7001 | 2472 | 2360 |
| 4918 | W303.asm02.HP0_G0049160 | chrXV | 217 | 11 | 8 | 123 | 16 | 7 | 182 | 17 | 14 | 255 | 25 | 20 | 129 | 10 | 26 | 170 | 22 | 14 |
| 4919 | W303.asm02.HP0_G0049170 | chrXV | 743 | 25 | 34 | 612 | 60 | 50 | 789 | 25 | 49 | 674 | 100 | 125 | 183 | 62 | 85 | 169 | 87 | 94 |
| 4920 | W303.asm02.HP0_G0049180 | chrXV | 690 | 64 | 70 | 627 | 85 | 67 | 714 | 87 | 75 | 478 | 135 | 140 | 509 | 79 | 86 | 414 | 98 | 117 |
| 4921 | W303.asm02.HP0_G0049190 | chrXV | 596 | 272 | 462 | 647 | 376 | 489 | 546 | 301 | 558 | 2647 | 359 | 708 | 1889 | 195 | 429 | 2071 | 351 | 652 |
| 4922 | W303.asm02.HP0_G0049200 | chrXV | 38 | 18 | 20 | 39 | 14 | 25 | 28 | 22 | 19 | 43 | 17 | 20 | 46 | 9 | 6 | 65 | 12 | 11 |
| 4923 | W303.asm02.HP0_G0049210 | chrXV | 351 | 175 | 201 | 310 | 174 | 168 | 294 | 167 | 216 | 839 | 178 | 176 | 849 | 102 | 173 | 827 | 166 | 174 |
| 4924 | W303.asm02.HP0_G0049220 | chrXV | 9688 | 990 | 1357 | 9967 | 1266 | 1506 | 10014 | 1059 | 1858 | 11869 | 2262 | 3026 | 10192 | 1867 | 2716 | 8671 | 1971 | 2812 |
| 4925 | W303.asm02.HP0_G0049230 | chrXV | 1944 | 1277 | 1548 | 1965 | 1400 | 1443 | 1980 | 1340 | 1882 | 4854 | 1175 | 1267 | 5392 | 749 | 1059 | 4901 | 998 | 1067 |
| 4926 | W303.asm02.HP0_G0049240 | chrXV | 2456 | 545 | 696 | 2535 | 736 | 713 | 2313 | 539 | 709 | 5492 | 691 | 1198 | 5002 | 518 | 860 | 4481 | 668 | 1046 |
| 4927 | W303.asm02.HP0_G0049250 | chrXV | 1925 | 455 | 502 | 1582 | 493 | 548 | 1944 | 385 | 562 | 3122 | 649 | 834 | 2757 | 361 | 762 | 2459 | 629 | 763 |

| | | | | | | | | | | | | | | | | | | | |
|---|---|---|---|---|---|---|---|---|---|---|---|---|---|---|---|---|---|---|---|
| 4928 | W303.asm02.HP0_G0049260 | chrXV | 2260 | 527 | 424 | 2642 | 424 | 359 | 1968 | 478 | 432 | 1353 | 456 | 325 | 1394 | 241 | 247 | 1451 | 420 | 314 |
| 4929 | W303.asm02.HP0_G0049270 | chrXV | 6670 | 5813 | 7148 | 5831 | 6209 | 6881 | 5765 | 5360 | 8848 | 9925 | 4282 | 5711 | 13687 | 3888 | 5335 | 10707 | 4251 | 5104 |
| 4930 | W303.asm02.HP0_G0049280 | chrXV | 1897 | 938 | 948 | 1642 | 788 | 1041 | 1447 | 823 | 1363 | 1633 | 680 | 865 | 2103 | 426 | 707 | 1922 | 681 | 657 |
| 4931 | W303.asm02.HP0_G0049290 | chrXV | 3196 | 2691 | 3037 | 2868 | 3154 | 2823 | 3136 | 2959 | 3984 | 10443 | 2842 | 2685 | 10680 | 1726 | 1811 | 10824 | 2521 | 2456 |
| 4932 | W303.asm02.HP0_G0049300 | chrXV | 4918 | 1187 | 1099 | 5208 | 1053 | 1151 | 4832 | 976 | 1200 | 6964 | 1709 | 1897 | 3676 | 1084 | 1415 | 3616 | 1609 | 1652 |
| 4933 | W303.asm02.HP0_G0049310 | chrXV | 990 | 782 | 472 | 875 | 446 | 535 | 1103 | 639 | 587 | 2721 | 1188 | 1179 | 2046 | 881 | 1082 | 2137 | 1227 | 1155 |
| 4934 | W303.asm02.HP0_G0049320 | chrXV | 1155 | 1142 | 638 | 930 | 702 | 661 | 1118 | 965 | 685 | 2790 | 909 | 744 | 2546 | 618 | 604 | 2581 | 939 | 681 |
| 4935 | W303.asm02.HP0_G0049330 | chrXV | 24323 | 9012 | 5577 | 25676 | 6078 | 6454 | 24147 | 6615 | 5311 | 28961 | 9050 | 11141 | 20882 | 7197 | 9229 | 23866 | 8831 | 9968 |
| 4936 | W303.asm02.HP0_G0049340 | chrXV | 1657 | 686 | 373 | 2069 | 340 | 305 | 1615 | 539 | 398 | 1706 | 611 | 429 | 2138 | 333 | 476 | 1608 | 567 | 442 |
| 4937 | W303.asm02.HP0_G0049350 | chrXV | 71 | 43 | 25 | 56 | 24 | 22 | 58 | 30 | 20 | 128 | 38 | 18 | 122 | 19 | 16 | 118 | 25 | 16 |
| 4938 | W303.asm02.HP0_G0049360 | chrXV | 1778 | 310 | 424 | 1693 | 427 | 455 | 1577 | 321 | 504 | 4166 | 697 | 945 | 3307 | 459 | 699 | 3152 | 578 | 846 |
| 4939 | W303.asm02.HP0_G0049370 | chrXV | 801 | 141 | 129 | 696 | 135 | 158 | 619 | 129 | 176 | 370 | 124 | 164 | 378 | 95 | 134 | 326 | 151 | 143 |
| 4940 | W303.asm02.HP0_G0049380 | chrXV | 1299 | 126 | 225 | 1838 | 215 | 214 | 1222 | 139 | 249 | 1303 | 173 | 287 | 1360 | 116 | 251 | 783 | 161 | 207 |
| 4941 | W303.asm02.HP0_G0049390 | chrXV | 126 | 10 | 11 | 114 | 15 | 15 | 113 | 12 | 11 | 65 | 20 | 22 | 44 | 11 | 17 | 61 | 17 | 22 |
| 4942 | W303.asm02.HP0_G0049400 | chrXV | 2006 | 511 | 575 | 2013 | 624 | 610 | 2067 | 493 | 576 | 4019 | 653 | 908 | 3362 | 485 | 715 | 3118 | 606 | 859 |
| 4943 | W303.asm02.HP0_G0049410 | chrXV | 1684 | 479 | 588 | 1610 | 639 | 533 | 1718 | 469 | 655 | 4206 | 700 | 874 | 3421 | 477 | 707 | 3280 | 561 | 838 |
| 4944 | W303.asm02.HP0_G0049420 | chrXV | 82 | 10 | 7 | 66 | 13 | 20 | 87 | 6 | 21 | 66 | 15 | 38 | 56 | 14 | 25 | 53 | 13 | 28 |
| 4945 | W303.asm02.HP0_G0049430 | chrXV | 4455 | 360 | 476 | 4790 | 428 | 587 | 4173 | 329 | 572 | 3336 | 523 | 993 | 3496 | 513 | 879 | 2238 | 494 | 896 |
| 4946 | W303.asm02.HP0_G0049440 | chrXV | 6373 | 1498 | 1291 | 6150 | 1221 | 1463 | 5600 | 1283 | 1446 | 3276 | 1039 | 1132 | 4660 | 811 | 1187 | 3147 | 995 | 1005 |
| 4947 | W303.asm02.HP0_G0049450 | chrXV | 3290 | 892 | 879 | 3117 | 664 | 893 | 3008 | 817 | 788 | 3018 | 688 | 1173 | 2846 | 472 | 704 | 2791 | 765 | 1006 |
| 4948 | W303.asm02.HP0_G0049460 | chrXV | 1004 | 451 | 467 | 915 | 553 | 475 | 1021 | 442 | 611 | 3971 | 654 | 869 | 2793 | 444 | 668 | 2948 | 618 | 877 |
| 4949 | W303.asm02.HP0_G0049470 | chrXV | 4853 | 2713 | 3235 | 4791 | 3051 | 3135 | 4967 | 2843 | 3604 | 9710 | 2491 | 2979 | 9900 | 1766 | 2448 | 10052 | 2286 | 2646 |
| 4950 | W303.asm02.HP0_G0049480 | chrXV | 2841 | 784 | 746 | 2581 | 682 | 646 | 2598 | 849 | 862 | 1960 | 715 | 642 | 2423 | 479 | 568 | 2236 | 666 | 648 |
| 4951 | W303.asm02.HP0_G0049490 | chrXV | 9835 | 1203 | 846 | 9388 | 818 | 1077 | 9638 | 1072 | 1035 | 7600 | 2394 | 2611 | 6671 | 1722 | 2091 | 5528 | 2209 | 2340 |
| 4952 | W303.asm02.HP0_G0049500 | chrXV | 4039 | 1063 | 1247 | 3608 | 1145 | 1300 | 4021 | 1016 | 1422 | 9383 | 1958 | 2335 | 7070 | 1347 | 1911 | 6539 | 1835 | 1986 |
| 4953 | W303.asm02.HP0_G0049510 | chrXV | 250 | 492 | 350 | 374 | 328 | 391 | 312 | 396 | 470 | 1144 | 283 | 239 | 1261 | 189 | 227 | 1270 | 366 | 270 |
| 4954 | W303.asm02.HP0_G0049520 | chrXV | 1679 | 1113 | 1482 | 1344 | 1305 | 1478 | 1435 | 1167 | 2013 | 2627 | 1054 | 1288 | 3356 | 772 | 1133 | 2957 | 942 | 1031 |
| 4955 | W303.asm02.HP0_G0049530 | chrXV | 5157 | 2962 | 2649 | 4958 | 2552 | 2450 | 5147 | 3120 | 2402 | 12521 | 3427 | 3813 | 10876 | 2186 | 2581 | 11631 | 3017 | 3363 |
| 4956 | W303.asm02.HP0_G0049540 | chrXV | 1623 | 530 | 651 | 1796 | 730 | 696 | 1714 | 379 | 688 | 5233 | 697 | 1194 | 4216 | 368 | 813 | 3850 | 788 | 1037 |
| 4957 | W303.asm02.HP0_G0049550 | chrXV | 3930 | 777 | 1130 | 3875 | 1082 | 1215 | 4257 | 708 | 1279 | 9249 | 1764 | 2620 | 4044 | 1337 | 1498 | 4139 | 1483 | 2163 |
| 4958 | W303.asm02.HP0_G0049560 | chrXV | 1704 | 276 | 212 | 1763 | 264 | 207 | 1646 | 216 | 283 | 1953 | 422 | 437 | 1490 | 276 | 371 | 1443 | 402 | 439 |
| 4959 | W303.asm02.HP0_G0049570 | chrXV | 7108 | 3404 | 3987 | 6848 | 4165 | 3893 | 7590 | 3310 | 4060 | 14042 | 2735 | 2915 | 15525 | 1829 | 2462 | 13641 | 2377 | 2571 |
| 4960 | W303.asm02.HP0_G0049580 | chrXV | 7682 | 2775 | 3530 | 8343 | 3118 | 3644 | 7264 | 2265 | 4028 | 6990 | 2039 | 2438 | 8641 | 1235 | 2252 | 6808 | 2049 | 2072 |
| 4961 | W303.asm02.HP0_G0049590 | chrXV | 3420 | 393 | 319 | 3542 | 339 | 440 | 3139 | 336 | 455 | 2271 | 806 | 943 | 1349 | 425 | 777 | 1219 | 913 | 897 |
| 4962 | W303.asm02.HP0_G0049600 | chrXV | 3069 | 1823 | 1472 | 3112 | 2054 | 1417 | 3299 | 1940 | 1322 | 9707 | 2176 | 2237 | 9541 | 1435 | 1362 | 8141 | 1810 | 2018 |
| 4963 | W303.asm02.HP0_G0049610 | chrXV | 2235 | 639 | 508 | 2132 | 498 | 540 | 2201 | 566 | 560 | 3567 | 814 | 1012 | 2927 | 615 | 694 | 3125 | 835 | 892 |
| 4964 | W303.asm02.HP0_G0049620 | chrXV | 2775 | 924 | 939 | 2952 | 878 | 882 | 2575 | 959 | 872 | 5247 | 1200 | 1452 | 4829 | 785 | 1080 | 4876 | 1050 | 1260 |
| 4965 | W303.asm02.HP0_G0049630 | chrXV | 4317 | 810 | 882 | 4376 | 867 | 922 | 3928 | 778 | 908 | 3963 | 981 | 1404 | 2697 | 639 | 1077 | 3256 | 959 | 1317 |
| 4966 | W303.asm02.HP0_G0049640 | chrXV | 1209 | 163 | 309 | 1381 | 307 | 314 | 1172 | 170 | 311 | 1847 | 297 | 554 | 1517 | 205 | 372 | 1377 | 263 | 451 |
| 4967 | W303.asm02.HP0_G0049650 | chrXV | 8639 | 2555 | 2480 | 8535 | 2374 | 2366 | 8721 | 2543 | 2683 | 12393 | 3202 | 3622 | 12203 | 2305 | 3044 | 11357 | 2879 | 3409 |
| 4968 | W303.asm02.HP0_G0049660 | chrXV | 634 | 590 | 446 | 616 | 373 | 419 | 598 | 428 | 451 | 1351 | 475 | 489 | 1457 | 319 | 416 | 1041 | 590 | 364 |
| 4969 | W303.asm02.HP0_G0049670 | chrXV | 47097 | 14361 | 11533 | 42941 | 11717 | 12909 | 50296 | 12401 | 14485 | 59511 | 24149 | 25702 | 45641 | 20673 | 18653 | 46458 | 20804 | 22190 |
| 4970 | W303.asm02.HP0_G0049680 | chrXV | 474 | 895 | 547 | 442 | 559 | 586 | 496 | 834 | 622 | 1969 | 801 | 540 | 1838 | 446 | 457 | 1911 | 689 | 474 |
| 4971 | W303.asm02.HP0_G0049690 | chrXV | 11737 | 2799 | 2683 | 11964 | 2450 | 2963 | 12227 | 2808 | 3474 | 18156 | 6045 | 5748 | 13683 | 4709 | 4605 | 13536 | 4927 | 4458 |
| 4972 | W303.asm02.HP0_G0049700 | chrXV | 1797 | 235 | 328 | 1919 | 327 | 385 | 1688 | 188 | 451 | 2210 | 610 | 628 | 938 | 458 | 489 | 779 | 480 | 563 |
| 4973 | W303.asm02.HP0_G0049710 | chrXV | 2294 | 1376 | 1826 | 1970 | 1502 | 1648 | 2221 | 1500 | 1822 | 6724 | 1409 | 1896 | 7124 | 934 | 1396 | 6996 | 1191 | 1646 |
| 4974 | W303.asm02.HP0_G0049720 | chrXV | 1591 | 212 | 258 | 1628 | 180 | 288 | 1695 | 234 | 267 | 2319 | 490 | 756 | 1541 | 325 | 495 | 1424 | 482 | 693 |
| 4975 | W303.asm02.HP0_G0049730 | chrXV | 2230 | 229 | 404 | 2396 | 292 | 410 | 2194 | 215 | 470 | 2872 | 642 | 894 | 1456 | 419 | 680 | 1532 | 611 | 786 |
| 4976 | W303.asm02.HP0_G0049740 | chrXV | 1290 | 413 | 616 | 1232 | 500 | 566 | 1302 | 386 | 699 | 4847 | 1047 | 1416 | 2248 | 555 | 746 | 2705 | 946 | 1213 |
| 4977 | W303.asm02.HP0_G0049750 | chrXV | 303 | 376 | 347 | 294 | 444 | 359 | 340 | 393 | 308 | 2352 | 431 | 558 | 2228 | 278 | 452 | 2112 | 394 | 542 |
| 4978 | W303.asm02.HP0_G0049760 | chrXV | 1425 | 733 | 709 | 1260 | 533 | 702 | 1389 | 691 | 778 | 3702 | 1049 | 1102 | 2785 | 679 | 806 | 2904 | 981 | 1134 |
| 4979 | W303.asm02.HP0_G0049770 | chrXV | 1129 | 1080 | 1268 | 1151 | 1077 | 1277 | 1069 | 1022 | 1696 | 3287 | 823 | 1043 | 3771 | 517 | 778 | 3777 | 826 | 870 |
| 4980 | W303.asm02.HP0_G0049780 | chrXV | 1453 | 779 | 801 | 1524 | 731 | 776 | 1557 | 819 | 745 | 3990 | 829 | 1067 | 3567 | 527 | 708 | 3837 | 882 | 973 |
| 4981 | W303.asm02.HP0_G0049790 | chrXV | 510 | 193 | 100 | 470 | 125 | 142 | 490 | 163 | 90 | 712 | 184 | 242 | 524 | 130 | 137 | 511 | 202 | 212 |
| 4982 | W303.asm02.HP0_G0049800 | chrXV | 807 | 1518 | 1696 | 757 | 1826 | 1793 | 885 | 1633 | 1654 | 6091 | 1291 | 1793 | 5925 | 973 | 1074 | 5501 | 1330 | 1672 |
| 4983 | W303.asm02.HP0_G0049810 | chrXV | 226 | 840 | 1081 | 174 | 927 | 1001 | 204 | 819 | 1323 | 3003 | 543 | 704 | 4524 | 434 | 628 | 3281 | 588 | 680 |
| 4984 | W303.asm02.HP0_G0049820 | chrXV | 1991 | 647 | 981 | 1874 | 862 | 896 | 1917 | 710 | 1081 | 4687 | 646 | 950 | 5729 | 479 | 859 | 4742 | 634 | 907 |
| 4985 | W303.asm02.HP0_G0049830 | chrXV | 2662 | 325 | 347 | 2445 | 338 | 429 | 2659 | 286 | 432 | 2955 | 731 | 828 | 1470 | 484 | 645 | 1525 | 673 | 764 |
| 4986 | W303.asm02.HP0_G0049840 | chrXV | 608 | 104 | 136 | 579 | 131 | 156 | 567 | 80 | 168 | 314 | 153 | 233 | 160 | 121 | 112 | 153 | 103 | 185 |
| 4987 | W303.asm02.HP0_G0049850 | chrXV | 559 | 141 | 240 | 457 | 211 | 254 | 597 | 121 | 294 | 525 | 220 | 349 | 240 | 160 | 223 | 216 | 163 | 303 |
| 4988 | W303.asm02.HP0_G0049860 | chrXV | 3035 | 1255 | 1756 | 2925 | 1602 | 1680 | 3042 | 1427 | 1587 | 7526 | 1410 | 2466 | 7810 | 1045 | 1778 | 6763 | 1237 | 2123 |
| 4989 | W303.asm02.HP0_G0049870 | chrXV | 8811 | 1247 | 1728 | 8014 | 1496 | 1768 | 8241 | 1242 | 2010 | 10379 | 1991 | 2984 | 6960 | 1623 | 2714 | 6316 | 1671 | 2693 |
| 4990 | W303.asm02.HP0_G0049880 | chrXV | 1850 | 1080 | 1017 | 1734 | 851 | 899 | 1758 | 1090 | 870 | 4381 | 1150 | 1315 | 3992 | 750 | 1041 | 3871 | 1114 | 1176 |
| 4991 | W303.asm02.HP0_G0049890 | chrXV | 6412 | 3060 | 2512 | 6433 | 2416 | 2514 | 5886 | 2907 | 2751 | 8384 | 2691 | 2395 | 9331 | 1743 | 2183 | 7928 | 2545 | 2263 |
| 4992 | W303.asm02.HP0_G0049900 | chrXV | 4977 | 808 | 766 | 4679 | 699 | 865 | 5006 | 769 | 996 | 6201 | 1399 | 1621 | 4823 | 939 | 1295 | 4337 | 1235 | 1434 |
| 4993 | W303.asm02.HP0_G0049910 | chrXV | 3102 | 949 | 757 | 3082 | 809 | 894 | 3633 | 783 | 983 | 6880 | 1884 | 2090 | 4418 | 1421 | 1208 | 3970 | 1627 | 1738 |
| 4994 | W303.asm02.HP0_G0049920 | chrXV | 1265 | 320 | 349 | 1224 | 304 | 369 | 1199 | 332 | 427 | 3460 | 705 | 838 | 2247 | 414 | 513 | 2277 | 744 | 702 |
| 4995 | W303.asm02.HP0_G0049930 | chrXV | 1989 | 800 | 951 | 1951 | 919 | 898 | 1895 | 837 | 1059 | 6001 | 840 | 1428 | 4786 | 665 | 1119 | 5248 | 842 | 1402 |
| 4996 | W303.asm02.HP0_G0049940 | chrXV | 1124 | 1020 | 955 | 1126 | 924 | 905 | 1027 | 1026 | 1113 | 3241 | 860 | 814 | 3452 | 514 | 662 | 3406 | 807 | 755 |
| 4997 | W303.asm02.HP0_G0049950 | chrXV | 946 | 2146 | 2976 | 838 | 3296 | 2917 | 1126 | 2238 | 3653 | 9522 | 1881 | 2111 | 9406 | 1142 | 1231 | 10063 | 1673 | 1793 |
| 4998 | W303.asm02.HP0_G0049960 | chrXV | 8344 | 6856 | 9711 | 6812 | 8683 | 9566 | 8325 | 6472 | 12525 | 21889 | 5876 | 7699 | 24713 | 4503 | 6130 | 23730 | 5630 | 6433 |
| 4999 | W303.asm02.HP0_G0049970 | chrXV | 1565 | 794 | 721 | 1382 | 603 | 715 | 1365 | 823 | 824 | 1936 | 673 | 659 | 2356 | 475 | 547 | 1915 | 556 | 584 |
| 5000 | W303.asm02.HP0_G0049980 | chrXV | 2927 | 795 | 900 | 2904 | 933 | 1071 | 3026 | 680 | 1181 | 7477 | 1450 | 2314 | 2778 | 1102 | 1155 | 2539 | 1291 | 1898 |
| 5001 | W303.asm02.HP0_G0049990 | chrXV | 8510 | 3322 | 5124 | 8164 | 3542 | 5425 | 8527 | 3505 | 5245 | 11313 | 3150 | 5382 | 12424 | 2087 | 3769 | 10975 | 2620 | 4782 |
| 5002 | W303.asm02.HP0_G0050000 | chrXV | 349 | 91 | 53 | 319 | 53 | 68 | 252 | 81 | 37 | 167 | 115 | 55 | 200 | 68 | 62 | 146 | 119 | 53 |
| 5003 | W303.asm02.HP0_G0050010 | chrXV | 1437 | 1899 | 2242 | 1632 | 2516 | 2063 | 1740 | 1891 | 2383 | 9258 | 2016 | 2024 | 8232 | 1174 | 1244 | 8441 | 1662 | 1721 |
| 5004 | W303.asm02.HP0_G0050020 | chrXV | 1575 | 1546 | 2363 | 1431 | 1654 | 2482 | 1463 | 1595 | 3251 | 3968 | 1166 | 1912 | 4907 | 835 | 1409 | 4457 | 1038 | 1675 |
| 5005 | W303.asm02.HP0_G0050030 | chrXV | 2080 | 656 | 865 | 1964 | 832 | 796 | 1896 | 685 | 896 | 4026 | 808 | 1160 | 5353 | 707 | 992 | 3440 | 765 | 994 |
| 5006 | W303.asm02.HP0_G0050040 | chrXV | 1390 | 879 | 1002 | 1410 | 1126 | 1048 | 1368 | 907 | 1065 | 3833 | 780 | 868 | 4015 | 433 | 633 | 3563 | 630 | 763 |
| 5007 | W303.asm02.HP0_G0050050 | chrXV | 196 | 334 | 579 | 182 | 526 | 498 | 198 | 279 | 522 | 2346 | 319 | 436 | 1985 | 180 | 220 | 1667 | 331 | 308 |
| 5008 | W303.asm02.HP0_G0050060 | chrXV | 2665 | 2734 | 2595 | 2090 | 2865 | 2592 | 2795 | 2782 | 2936 | 16895 | 4118 | 4991 | 11185 | 2536 | 2848 | 13015 | 3388 | 4208 |
| 5009 | W303.asm02.HP0_G0050070 | chrXV | 5677 | 780 | 932 | 4990 | 835 | 942 | 5231 | 684 | 1265 | 5997 | 1236 | 1763 | 4002 | 936 | 1244 | 3836 | 1146 | 1468 |
| 5010 | W303.asm02.HP0_G0050080 | chrXV | 2273 | 642 | 548 | 2145 | 560 | 619 | 2248 | 600 | 569 | 3147 | 746 | 1030 | 2718 | 568 | 736 | 2493 | 733 | 965 |
| 5011 | W303.asm02.HP0_G0050090 | chrXV | 2004 | 718 | 862 | 2091 | 765 | 833 | 1894 | 782 | 1122 | 2316 | 554 | 723 | 2975 | 451 | 603 | 2389 | 510 | 683 |
| 5012 | W303.asm02.HP0_G0050100 | chrXV | 3489 | 385 | 380 | 3155 | 342 | 424 | 3366 | 299 | 494 | 3649 | 702 | 968 | 2141 | 596 | 703 | 2051 | 593 | 838 |
| 5013 | W303.asm02.HP0_G0050110 | chrXV | 3820 | 370 | 380 | 4054 | 386 | 436 | 3822 | 355 | 533 | 3027 | 750 | 882 | 1719 | 543 | 690 | 1570 | 724 | 780 |
| 5014 | W303.asm02.HP0_G0050120 | chrXV | 610 | 337 | 332 | 525 | 295 | 355 | 518 | 374 | 328 | 921 | 315 | 349 | 1092 | 180 | 229 | 1091 | 294 | 312 |
| 5015 | W303.asm02.HP0_G0050130 | chrXV | 2747 | 1512 | 1999 | 2563 | 1687 | 1914 | 2582 | 1587 | 2419 | 4838 | 1238 | 1535 | 5500 | 926 | 1152 | 5279 | 1142 | 1395 |
| 5016 | W303.asm02.HP0_G0050140 | chrXV | 2142 | 726 | 1161 | 2074 | 1236 | 1232 | 2268 | 732 | 1537 | 9405 | 1424 | 2640 | 3712 | 965 | 1346 | 3734 | 1260 | 2238 |
| 5017 | W303.asm02.HP0_G0050150 | chrXV | 8868 | 4066 | 4065 | 9376 | 3680 | 3830 | 8999 | 3816 | 4209 | 13924 | 4022 | 4957 | 14064 | 2720 | 3978 | 13610 | 3558 | 4474 |
| 5018 | W303.asm02.HP0_G0050160 | chrXV | 3180 | 1416 | 1459 | 3190 | 1570 | 1386 | 3302 | 1508 | 1635 | 7800 | 1612 | 1866 | 6518 | 1068 | 1344 | 7190 | 1403 | 1731 |
| 5019 | W303.asm02.HP0_G0050170 | chrXV | 1644 | 636 | 783 | 1768 | 1083 | 761 | 1856 | 664 | 858 | 6544 | 924 | 1087 | 5752 | 554 | 606 | 5369 | 733 | 890 |
| 5020 | W303.asm02.HP0_G0050180 | chrXV | 9443 | 1209 | 1231 | 8840 | 1083 | 1525 | 8141 | 1083 | 1320 | 1824 | 1042 | 1272 | 2837 | 1087 | 1433 | 1588 | 949 | 975 |

| | | | | | | | | | | | | | | | | | | | | |
|---|---|---|---|---|---|---|---|---|---|---|---|---|---|---|---|---|---|---|---|---|
| 5021 | W303.asm02.HP0_G0050190 | chrXV | 7117 | 5987 | 7885 | 7086 | 6663 | 7838 | 7036 | 6174 | 10024 | 13453 | 5772 | 6596 | 16854 | 4026 | 5564 | 14296 | 5023 | 6035 |
| 5022 | W303.asm02.HP0_G0050200 | chrXV | 6972 | 2111 | 1579 | 7082 | 1800 | 1758 | 7184 | 2172 | 1440 | 9513 | 2214 | 2708 | 7992 | 1389 | 1998 | 7164 | 1991 | 2508 |
| 5023 | W303.asm02.HP0_G0050210 | chrXV | 7798 | 1294 | 1441 | 7956 | 1443 | 1409 | 7461 | 1336 | 1658 | 8612 | 1379 | 2241 | 8017 | 1140 | 1961 | 7213 | 1366 | 1979 |
| 5024 | W303.asm02.HP0_G0050220 | chrXV | 2989 | 908 | 1250 | 2868 | 1104 | 1293 | 2630 | 873 | 1615 | 2805 | 696 | 948 | 3580 | 475 | 869 | 2848 | 709 | 832 |
| 5025 | W303.asm02.HP0_G0050230 | chrXV | 1354 | 309 | 416 | 1148 | 326 | 373 | 1243 | 312 | 488 | 2032 | 423 | 572 | 1747 | 356 | 455 | 1396 | 457 | 543 |
| 5026 | W303.asm02.HP0_G0050240 | chrXV | 5691 | 591 | 753 | 5666 | 719 | 865 | 5981 | 589 | 947 | 7795 | 1213 | 1853 | 6000 | 1159 | 1548 | 4145 | 1068 | 1646 |
| 5027 | W303.asm02.HP0_G0050250 | chrXV | 2449 | 379 | 415 | 1725 | 357 | 379 | 2116 | 264 | 377 | 1639 | 495 | 585 | 1444 | 295 | 458 | 1310 | 493 | 483 |
| 5028 | W303.asm02.HP0_G0050260 | chrXV | 483 | 350 | 381 | 487 | 445 | 385 | 526 | 402 | 465 | 3073 | 526 | 796 | 2396 | 379 | 536 | 2653 | 492 | 765 |
| 5029 | W303.asm02.HP0_G0050270 | chrXV | 813 | 1500 | 1448 | 707 | 1128 | 1288 | 771 | 1455 | 1659 | 2774 | 1037 | 1073 | 2734 | 643 | 852 | 3067 | 929 | 842 |
| 5030 | W303.asm02.HP0_G0050280 | chrXV | 3820 | 1131 | 1021 | 3334 | 998 | 1036 | 3273 | 1282 | 1054 | 3115 | 1108 | 1141 | 3419 | 806 | 1011 | 2981 | 982 | 1039 |
| 5031 | W303.asm02.HP0_G0050290 | chrXV | 6413 | 751 | 549 | 6215 | 577 | 582 | 6198 | 638 | 461 | 4008 | 835 | 1129 | 3903 | 651 | 912 | 3324 | 690 | 959 |
| 5032 | W303.asm02.HP0_G0050300 | chrXV | 17317 | 4537 | 3876 | 16346 | 3584 | 3764 | 19133 | 4264 | 4782 | 22344 | 7863 | 8720 | 17869 | 6125 | 6162 | 17871 | 6931 | 8150 |
| 5033 | W303.asm02.HP0_G0050310 | chrXV | 2535 | 132 | 73 | 2693 | 83 | 94 | 2414 | 113 | 81 | 293 | 137 | 63 | 622 | 100 | 98 | 312 | 129 | 87 |
| 5034 | W303.asm02.HP0_G0050320 | chrXV | 10654 | 5813 | 5292 | 9341 | 5366 | 5193 | 10034 | 5905 | 4920 | 17340 | 7556 | 8043 | 19545 | 6133 | 6931 | 15239 | 6794 | 7430 |
| 5035 | W303.asm02.HP0_G0050330 | chrXV | 688 | 342 | 655 | 821 | 486 | 586 | 655 | 316 | 619 | 1889 | 428 | 547 | 2402 | 262 | 586 | 1625 | 415 | 476 |
| 5036 | W303.asm02.HP0_G0050340 | chrXV | 3367 | 848 | 902 | 2616 | 887 | 928 | 3271 | 758 | 1065 | 4437 | 1317 | 1372 | 3056 | 741 | 978 | 3085 | 1222 | 1282 |
| 5037 | W303.asm02.HP0_G0050350 | chrXV | 957 | 198 | 362 | 1075 | 338 | 468 | 1017 | 152 | 503 | 1966 | 358 | 774 | 630 | 234 | 365 | 487 | 287 | 620 |
| 5038 | W303.asm02.HP0_G0050360 | chrXV | 3251 | 639 | 590 | 3616 | 597 | 696 | 3283 | 566 | 779 | 6654 | 1169 | 1731 | 4536 | 836 | 1066 | 4387 | 1048 | 1454 |
| 5039 | W303.asm02.HP0_G0050370 | chrXV | 8507 | 4457 | 4099 | 8844 | 3854 | 4042 | 8647 | 4518 | 4134 | 13076 | 5035 | 5912 | 15205 | 4152 | 5145 | 11815 | 4338 | 5144 |
| 5040 | W303.asm02.HP0_G0050380 | chrXV | 851 | 1163 | 1828 | 715 | 1635 | 1609 | 825 | 1167 | 2143 | 4640 | 840 | 1086 | 5737 | 591 | 870 | 5315 | 768 | 1002 |
| 5041 | W303.asm02.HP0_G0050390 | chrXV | 555 | 400 | 469 | 555 | 546 | 459 | 622 | 448 | 360 | 3199 | 525 | 747 | 2499 | 369 | 477 | 2580 | 455 | 658 |
| 5042 | W303.asm02.HP0_G0050400 | chrXV | 2702 | 2750 | 2452 | 2638 | 3230 | 2279 | 2950 | 2916 | 2590 | 11538 | 3538 | 3233 | 10147 | 2277 | 2039 | 10062 | 2787 | 3072 |
| 5043 | W303.asm02.HP0_G0050410 | chrXV | 1341 | 1093 | 1513 | 1582 | 1465 | 1380 | 1230 | 1004 | 1582 | 6058 | 1047 | 1416 | 5548 | 625 | 1018 | 6070 | 1103 | 1200 |
| 5044 | W303.asm02.HP0_G0050420 | chrXV | 688 | 582 | 679 | 543 | 536 | 643 | 618 | 628 | 850 | 1481 | 506 | 467 | 1749 | 356 | 403 | 1585 | 453 | 443 |
| 5045 | W303.asm02.HP0_G0050430 | chrXV | 1102 | 583 | 620 | 1181 | 606 | 557 | 1260 | 630 | 613 | 3884 | 715 | 872 | 3283 | 423 | 584 | 3527 | 601 | 786 |
| 5046 | W303.asm02.HP0_G0050440 | chrXV | 3215 | 1733 | 2537 | 2928 | 2154 | 2534 | 3076 | 1711 | 3370 | 4511 | 1388 | 2045 | 5848 | 1070 | 1746 | 4657 | 1238 | 1688 |
| 5047 | W303.asm02.HP0_G0050450 | chrXV | 9001 | 2462 | 2798 | 8327 | 2971 | 3059 | 9291 | 2553 | 3357 | 23958 | 4338 | 6312 | 16066 | 3377 | 4672 | 16016 | 3858 | 5779 |
| 5048 | W303.asm02.HP0_G0050460 | chrXV | 1011 | 332 | 338 | 894 | 279 | 386 | 885 | 382 | 466 | 901 | 299 | 356 | 1079 | 207 | 293 | 1030 | 312 | 284 |
| 5049 | W303.asm02.HP0_G0050470 | chrXV | 12902 | 1771 | 1922 | 12467 | 1699 | 2079 | 12799 | 1601 | 2415 | 20050 | 4511 | 6977 | 11651 | 4011 | 5242 | 8630 | 4929 | 8543 |
| 5050 | W303.asm02.HP0_G0050480 | chrXV | 835 | 871 | 1087 | 954 | 1241 | 1012 | 985 | 930 | 1050 | 5739 | 861 | 1161 | 4912 | 479 | 663 | 5156 | 811 | 898 |
| 5051 | W303.asm02.HP0_G0050490 | chrXV | 4915 | 828 | 734 | 4345 | 669 | 733 | 4558 | 750 | 776 | 3816 | 1088 | 1250 | 3630 | 766 | 1113 | 2926 | 1000 | 1093 |
| 5052 | W303.asm02.HP0_G0050500 | chrXV | 1001 | 189 | 228 | 1028 | 226 | 259 | 1054 | 149 | 307 | 2529 | 392 | 621 | 1612 | 201 | 372 | 1554 | 380 | 483 |
| 5053 | W303.asm02.HP0_G0050510 | chrXV | 5367 | 2649 | 2355 | 4344 | 2104 | 2407 | 5390 | 2526 | 2335 | 7750 | 2585 | 3167 | 7667 | 1986 | 2638 | 7184 | 2380 | 2910 |
| 5054 | W303.asm02.HP0_G0050520 | chrXV | 6227 | 2381 | 2690 | 6525 | 2641 | 2694 | 6274 | 2455 | 3173 | 8338 | 1915 | 2700 | 10020 | 1485 | 2419 | 8240 | 1847 | 2497 |
| 5055 | W303.asm02.HP0_G0050530 | chrXV | 962 | 1020 | 1182 | 867 | 1199 | 1197 | 899 | 1063 | 1560 | 3194 | 938 | 987 | 3332 | 628 | 777 | 3309 | 731 | 896 |
| 5056 | W303.asm02.HP0_G0050540 | chrXV | 2435 | 440 | 528 | 2376 | 484 | 529 | 2464 | 378 | 634 | 6223 | 888 | 1313 | 3892 | 625 | 1039 | 3964 | 836 | 1221 |
| 5057 | W303.asm02.HP0_G0050550 | chrXV | 1122 | 1070 | 637 | 1148 | 445 | 647 | 1049 | 675 | 728 | 2571 | 1165 | 996 | 2008 | 807 | 973 | 1375 | 1281 | 789 |
| 5058 | W303.asm02.HP0_G0050560 | chrXV | 2027 | 626 | 644 | 2292 | 602 | 596 | 2177 | 625 | 759 | 4188 | 950 | 1079 | 3390 | 761 | 927 | 3415 | 863 | 939 |
| 5059 | W303.asm02.HP0_G0050570 | chrXV | 2646 | 1334 | 1158 | 2411 | 1077 | 1261 | 2518 | 1366 | 1006 | 2880 | 1260 | 1456 | 3183 | 975 | 1158 | 2554 | 1087 | 1148 |
| 5060 | W303.asm02.HP0_G0050580 | chrXV | 3732 | 2120 | 2212 | 3373 | 1888 | 2084 | 3472 | 2093 | 2688 | 6623 | 1740 | 1792 | 6929 | 1169 | 1480 | 7317 | 1638 | 1620 |
| 5061 | W303.asm02.HP0_G0050590 | chrXV | 8525 | 2762 | 2707 | 8428 | 2586 | 2449 | 9001 | 2924 | 2887 | 12751 | 2903 | 3165 | 11854 | 2081 | 2446 | 12288 | 2668 | 2942 |
| 5062 | W303.asm02.HP0_G0050600 | chrXV | 1788 | 946 | 1059 | 1567 | 1060 | 1030 | 1670 | 844 | 968 | 4776 | 745 | 933 | 4794 | 497 | 716 | 4685 | 702 | 816 |
| 5063 | W303.asm02.HP0_G0050610 | chrXV | 12587 | 3744 | 3574 | 12313 | 3664 | 3192 | 12884 | 3886 | 4008 | 26175 | 5530 | 5797 | 21446 | 3614 | 3949 | 22380 | 5261 | 5174 |
| 5064 | W303.asm02.HP0_G0050620 | chrXV | 3205 | 1172 | 980 | 3020 | 1021 | 1019 | 3351 | 1079 | 942 | 7121 | 1495 | 2008 | 5192 | 1051 | 1317 | 5846 | 1425 | 1843 |
| 5065 | W303.asm02.HP0_G0050630 | chrXV | 1384 | 281 | 704 | 1503 | 528 | 698 | 1268 | 299 | 803 | 3791 | 540 | 1362 | 2790 | 339 | 936 | 2974 | 608 | 1203 |
| 5066 | W303.asm02.HP0_G0050640 | chrXV | 9261 | 387 | 245 | 7925 | 201 | 269 | 7238 | 381 | 285 | 560 | 433 | 426 | 995 | 358 | 565 | 386 | 467 | 400 |
| 5067 | W303.asm02.HP0_G0050650 | chrXV | 1711 | 1193 | 977 | 1541 | 881 | 1025 | 1673 | 1126 | 952 | 2976 | 883 | 1107 | 2944 | 616 | 689 | 2808 | 829 | 905 |
| 5068 | W303.asm02.HP0_G0050660 | chrXV | 7772 | 2196 | 1651 | 7159 | 1787 | 1693 | 7926 | 2363 | 1575 | 10519 | 2591 | 2588 | 9604 | 1777 | 2182 | 8955 | 2176 | 2450 |
| 5069 | W303.asm02.HP0_G0050670 | chrXV | 8164 | 1900 | 1864 | 8263 | 1299 | 1760 | 7930 | 1985 | 1852 | 7449 | 2238 | 2783 | 7634 | 1514 | 2090 | 6858 | 1943 | 2417 |
| 5070 | W303.asm02.HP0_G0050680 | chrXV | 687 | 269 | 275 | 650 | 286 | 273 | 598 | 241 | 281 | 747 | 160 | 213 | 1111 | 141 | 177 | 682 | 177 | 190 |
| 5071 | W303.asm02.HP0_G0050690 | chrXV | 7670 | 211 | 265 | 8695 | 308 | 208 | 7671 | 220 | 314 | 1689 | 358 | 350 | 1683 | 220 | 304 | 1325 | 268 | 331 |
| 5072 | W303.asm02.HP0_G0050700 | chrXV | 1261 | 863 | 946 | 1129 | 852 | 860 | 1188 | 849 | 1192 | 2223 | 689 | 665 | 2777 | 502 | 537 | 2268 | 550 | 557 |
| 5073 | W303.asm02.HP0_G0050710 | chrXV | 183 | 175 | 161 | 191 | 137 | 161 | 169 | 182 | 182 | 685 | 183 | 185 | 747 | 119 | 162 | 640 | 182 | 189 |
| 5074 | W303.asm02.HP0_G0050720 | chrXV | 2151 | 261 | 255 | 1829 | 250 | 312 | 2016 | 166 | 354 | 356 | 199 | 298 | 418 | 153 | 230 | 293 | 199 | 273 |
| 5075 | W303.asm02.HP0_G0050730 | chrXV | 17 | 15 | 15 | 9 | 12 | 12 | 25 | 23 | 17 | 23 | 9 | 7 | 27 | 9 | 5 | 18 | 12 | 6 |
| 5076 | W303.asm02.HP0_G0050740 | chrXV | 3039 | 3616 | 3040 | 2476 | 2950 | 2843 | 2908 | 3498 | 2901 | 9891 | 3499 | 3428 | 8796 | 2116 | 2335 | 9502 | 3084 | 3222 |
| 5077 | W303.asm02.HP0_G0050750 | chrXV | 3001 | 1014 | 786 | 2518 | 593 | 893 | 2561 | 991 | 779 | 1513 | 785 | 1038 | 2363 | 736 | 923 | 1472 | 839 | 871 |
| 5078 | W303.asm02.HP0_G0050760 | chrXV | 91 | 101 | 21 | 76 | 37 | 42 | 71 | 65 | 22 | 134 | 58 | 22 | 130 | 31 | 20 | 130 | 58 | 21 |
| 5079 | W303.asm02.HP0_G0050770 | chrXV | 21629 | 5634 | 5224 | 21552 | 5108 | 5407 | 21522 | 5668 | 5686 | 24194 | 7163 | 9515 | 24580 | 6296 | 8401 | 20509 | 6422 | 8503 |
| 5080 | W303.asm02.HP0_G0050780 | chrXV | 1368 | 257 | 447 | 1329 | 381 | 514 | 1558 | 255 | 545 | 4386 | 672 | 1169 | 2352 | 508 | 761 | 2096 | 547 | 1042 |
| 5081 | W303.asm02.HP0_G0050790 | chrXV | 1658 | 785 | 879 | 1516 | 770 | 1005 | 1564 | 852 | 939 | 2354 | 708 | 1098 | 2599 | 536 | 845 | 2244 | 648 | 955 |
| 5082 | W303.asm02.HP0_G0050800 | chrXV | 423 | 151 | 153 | 445 | 155 | 183 | 463 | 171 | 170 | 666 | 176 | 254 | 579 | 120 | 144 | 558 | 179 | 237 |
| 5083 | W303.asm02.HP0_G0050810 | chrXV | 3332 | 481 | 647 | 3311 | 682 | 721 | 3313 | 449 | 760 | 6580 | 915 | 1726 | 3749 | 691 | 1150 | 3869 | 793 | 1564 |
| 5084 | W303.asm02.HP0_G0050820 | chrXV | 764 | 121 | 216 | 826 | 153 | 175 | 830 | 91 | 180 | 860 | 185 | 293 | 1036 | 201 | 273 | 521 | 169 | 264 |
| 5085 | W303.asm02.HP0_G0050830 | chrXV | 1477 | 727 | 841 | 1430 | 727 | 961 | 1383 | 812 | 1198 | 2147 | 458 | 787 | 3087 | 354 | 593 | 2713 | 574 | 750 |
| 5086 | W303.asm02.HP0_G0050840 | chrXV | 4698 | 1662 | 1618 | 4646 | 1715 | 1758 | 4654 | 1717 | 1623 | 7199 | 1801 | 2546 | 6842 | 1351 | 2105 | 6326 | 1633 | 2463 |
| 5087 | W303.asm02.HP0_G0050850 | chrXV | 1616 | 778 | 1359 | 1750 | 1459 | 1378 | 1847 | 982 | 1476 | 6060 | 801 | 1303 | 6847 | 576 | 1015 | 5963 | 683 | 1177 |
| 5088 | W303.asm02.HP0_G0050860 | chrXV | 2811 | 1534 | 1435 | 2706 | 1433 | 1222 | 2814 | 1524 | 1545 | 7045 | 1682 | 1918 | 5609 | 1170 | 1424 | 6140 | 1690 | 1686 |
| 5089 | W303.asm02.HP0_G0050870 | chrXV | 7621 | 1462 | 1355 | 7219 | 1244 | 1537 | 7641 | 1251 | 1918 | 6737 | 2469 | 2599 | 2914 | 1892 | 1672 | 2939 | 2009 | 2384 |
| 5090 | W303.asm02.HP0_G0050880 | chrXV | 15323 | 5405 | 8019 | 15431 | 5753 | 8459 | 16301 | 5446 | 7998 | 22800 | 5227 | 10563 | 23019 | 4321 | 6983 | 19838 | 4554 | 9340 |
| 5091 | W303.asm02.HP0_G0050890 | chrXV | 1568 | 1067 | 1094 | 1552 | 1135 | 1211 | 1616 | 1007 | 1191 | 5624 | 1329 | 1874 | 4864 | 1053 | 1448 | 4253 | 1148 | 1634 |
| 5092 | W303.asm02.HP0_G0050900 | chrXV | 819 | 726 | 574 | 600 | 580 | 585 | 737 | 717 | 536 | 1258 | 546 | 516 | 1176 | 434 | 457 | 1031 | 543 | 484 |
| 5093 | W303.asm02.HP0_G0050910 | chrXV | 9071 | 1501 | 3089 | 8702 | 2871 | 2985 | 8483 | 1419 | 3837 | 20503 | 3122 | 5290 | 15047 | 1850 | 3251 | 14014 | 3213 | 4379 |
| 5094 | W303.asm02.HP0_G0050920 | chrXV | 3669 | 497 | 634 | 3654 | 651 | 588 | 3851 | 513 | 662 | 4867 | 789 | 1217 | 3896 | 610 | 838 | 3519 | 725 | 1018 |
| 5095 | W303.asm02.HP0_G0050930 | chrXV | 5261 | 3204 | 4470 | 5440 | 4883 | 3932 | 6070 | 3543 | 4463 | 22644 | 4052 | 5064 | 20368 | 2319 | 3038 | 21117 | 3424 | 4414 |
| 5096 | W303.asm02.HP0_G0050940 | chrXV | 4777 | 1224 | 1202 | 4628 | 1415 | 1154 | 4979 | 1252 | 1343 | 14333 | 2123 | 2849 | 10772 | 1609 | 1897 | 10253 | 1884 | 2530 |
| 5097 | W303.asm02.HP0_G0050950 | chrXV | 3046 | 238 | 479 | 3295 | 450 | 588 | 3010 | 236 | 593 | 3646 | 554 | 901 | 4832 | 506 | 982 | 2307 | 583 | 833 |
| 5098 | W303.asm02.HP0_G0050960 | chrXV | 10203 | 3980 | 3432 | 9828 | 3677 | 3394 | 10415 | 4053 | 3097 | 16582 | 4135 | 4882 | 14252 | 2644 | 3495 | 15582 | 3604 | 4296 |
| 5099 | W303.asm02.HP0_G0050970 | chrXV | 1890 | 1465 | 2224 | 1679 | 1974 | 2093 | 1835 | 1554 | 2921 | 5159 | 1374 | 1625 | 6212 | 949 | 1383 | 5912 | 1060 | 1422 |
| 5100 | W303.asm02.HP0_G0050980 | chrXV | 4329 | 777 | 885 | 3870 | 988 | 946 | 4144 | 772 | 1011 | 6786 | 1094 | 1576 | 5340 | 791 | 1261 | 5350 | 984 | 1464 |
| 5101 | W303.asm02.HP0_G0050990 | chrXV | 7607 | 2387 | 2108 | 7311 | 2314 | 2127 | 7666 | 2351 | 1874 | 11523 | 2743 | 3264 | 10680 | 1958 | 2512 | 9908 | 2432 | 2977 |
| 5102 | W303.asm02.HP0_G0051000 | chrXV | 906 | 585 | 1010 | 905 | 964 | 972 | 853 | 588 | 1206 | 3021 | 503 | 845 | 3442 | 349 | 617 | 3134 | 456 | 763 |
| 5103 | W303.asm02.HP0_G0051010 | chrXV | 36 | 25 | 16 | 29 | 8 | 15 | 26 | 22 | 22 | 38 | 36 | 9 | 50 | 13 | 7 | 38 | 30 | 18 |
| 5104 | W303.asm02.HP0_G0051020 | chrXV | 5238 | 1739 | 2085 | 5239 | 1777 | 2174 | 4923 | 1788 | 2809 | 3968 | 1501 | 1586 | 5368 | 1143 | 1637 | 4135 | 1327 | 1430 |
| 5105 | W303.asm02.HP0_G0051030 | chrXV | 2276 | 1234 | 1440 | 2378 | 1619 | 1464 | 2413 | 1238 | 1329 | 7705 | 1239 | 1888 | 6993 | 847 | 1338 | 6654 | 1055 | 1650 |
| 5106 | W303.asm02.HP0_G0051040 | chrXV | 3680 | 968 | 985 | 3359 | 981 | 974 | 3789 | 931 | 1178 | 9825 | 1915 | 2256 | 5880 | 1355 | 1590 | 6076 | 1589 | 2037 |
| 5107 | W303.asm02.HP0_G0051050 | chrXV | 3165 | 448 | 556 | 2887 | 499 | 502 | 2879 | 443 | 562 | 3956 | 804 | 1044 | 2475 | 471 | 770 | 2789 | 740 | 982 |
| 5108 | W303.asm02.HP0_G0051060 | chrXV | 4890 | 1705 | 1201 | 5445 | 960 | 1224 | 4528 | 1407 | 1481 | 2342 | 1060 | 873 | 3233 | 648 | 871 | 2514 | 1156 | 760 |
| 5109 | W303.asm02.HP0_G0051070 | chrXV | 3491 | 642 | 639 | 3443 | 575 | 737 | 3407 | 532 | 747 | 3224 | 1042 | 1262 | 2846 | 766 | 982 | 2619 | 969 | 1067 |
| 5110 | W303.asm02.HP0_G0051080 | chrXV | 3154 | 3028 | 2809 | 3078 | 2552 | 2412 | 3167 | 3040 | 2909 | 8054 | 3041 | 2800 | 9136 | 2213 | 2371 | 7804 | 2554 | 2728 |
| 5111 | W303.asm02.HP0_G0051090 | chrXV | 2246 | 842 | 389 | 2238 | 336 | 366 | 2027 | 691 | 438 | 1384 | 797 | 569 | 1184 | 542 | 481 | 1104 | 924 | 525 |
| 5112 | W303.asm02.HP0_G0051100 | chrXV | 2751 | 2023 | 2383 | 2264 | 2526 | 2380 | 2401 | 2304 | 2391 | 6212 | 1615 | 2564 | 6656 | 1346 | 1734 | 5618 | 1440 | 2144 |
| 5113 | W303.asm02.HP0_G0051110 | chrXV | 223 | 659 | 654 | 172 | 538 | 725 | 188 | 683 | 693 | 1985 | 1153 | 1033 | 1488 | 544 | 722 | 1693 | 1028 | 947 |

| | | | | | | | | | | | | | | | | | | | | |
|---|---|---|---|---|---|---|---|---|---|---|---|---|---|---|---|---|---|---|---|---|
| 5114 | W303.asm02.HP0_G0051120 | chrXV | 9979 | 680 | 882 | 8940 | 716 | 937 | 9850 | 577 | 1115 | 4399 | 1248 | 1762 | 2585 | 1121 | 1753 | 2157 | 1065 | 1555 |
| 5115 | W303.asm02.HP0_G0051130 | chrXV | 1573 | 450 | 394 | 1649 | 315 | 371 | 1539 | 425 | 384 | 1483 | 432 | 447 | 1628 | 298 | 405 | 1375 | 401 | 387 |
| 5116 | W303.asm02.HP0_G0051140 | chrXV | 920 | 301 | 326 | 887 | 307 | 346 | 912 | 283 | 414 | 1439 | 311 | 331 | 1180 | 230 | 279 | 1116 | 324 | 302 |
| 5117 | W303.asm02.HP0_G0051150 | chrXV | 24076 | 3357 | 2474 | 21320 | 2475 | 2992 | 23358 | 2917 | 2671 | 12153 | 4023 | 4603 | 12233 | 3805 | 4647 | 9343 | 3744 | 4153 |
| 5118 | W303.asm02.HP0_G0051160 | chrXV | 2372 | 356 | 417 | 2044 | 396 | 466 | 2364 | 271 | 559 | 2250 | 577 | 727 | 805 | 361 | 507 | 899 | 478 | 607 |
| 5119 | W303.asm02.HP0_G0051170 | chrXV | 10632 | 3268 | 5889 | 10888 | 5090 | 6009 | 10609 | 3528 | 7479 | 14917 | 2791 | 5016 | 17346 | 1933 | 4328 | 15495 | 2476 | 4658 |
| 5120 | W303.asm02.HP0_G0051180 | chrXV | 501 | 357 | 479 | 553 | 469 | 540 | 559 | 328 | 587 | 4289 | 772 | 1157 | 2291 | 561 | 723 | 2234 | 651 | 970 |
| 5121 | W303.asm02.HP0_G0051190 | chrXV | 4500 | 2965 | 4995 | 4135 | 4794 | 5033 | 4228 | 3182 | 6155 | 9699 | 2670 | 3800 | 11265 | 2033 | 3235 | 10480 | 2404 | 3456 |
| 5122 | W303.asm02.HP0_G0051200 | chrXV | 422 | 450 | 616 | 323 | 420 | 708 | 356 | 451 | 653 | 1524 | 462 | 996 | 1451 | 327 | 515 | 1397 | 450 | 820 |
| 5123 | W303.asm02.HP0_G0051210 | chrXV | 84 | 129 | 44 | 90 | 54 | 43 | 77 | 103 | 58 | 233 | 135 | 54 | 263 | 64 | 33 | 233 | 105 | 40 |
| 5124 | W303.asm02.HP0_G0051220 | chrXV | 1930 | 1298 | 1056 | 1812 | 1003 | 1154 | 1829 | 1428 | 1180 | 3289 | 1081 | 906 | 4659 | 830 | 802 | 3257 | 1053 | 837 |
| 5125 | W303.asm02.HP0_G0051230 | chrXV | 2455 | 350 | 343 | 2477 | 303 | 324 | 2488 | 315 | 303 | 3632 | 787 | 1151 | 2228 | 563 | 592 | 2404 | 687 | 1057 |
| 5126 | W303.asm02.HP0_G0051240 | chrXV | 5309 | 862 | 933 | 4932 | 766 | 1130 | 5168 | 878 | 1029 | 6244 | 1568 | 2610 | 4098 | 1129 | 1674 | 4266 | 1310 | 2317 |
| 5127 | W303.asm02.HP0_G0051250 | chrXV | 2841 | 1939 | 2046 | 2566 | 2078 | 1953 | 2915 | 2003 | 2192 | 8269 | 1822 | 2197 | 7820 | 1258 | 1698 | 7673 | 1643 | 1969 |
| 5128 | W303.asm02.HP0_G0051260 | chrXV | 249 | 75 | 65 | 208 | 64 | 87 | 258 | 76 | 91 | 453 | 79 | 95 | 338 | 55 | 77 | 448 | 67 | 114 |
| 5129 | W303.asm02.HP0_G0051270 | chrXV | 5219 | 1688 | 2050 | 4754 | 2241 | 2027 | 5712 | 1936 | 2122 | 13017 | 2380 | 3346 | 11257 | 1452 | 2113 | 11294 | 1882 | 3043 |
| 5130 | W303.asm02.HP0_G0051280 | chrXV | 1414 | 1002 | 1028 | 1625 | 1091 | 1006 | 1442 | 944 | 946 | 3995 | 799 | 1145 | 3680 | 541 | 826 | 3691 | 829 | 951 |
| 5131 | W303.asm02.HP0_G0051290 | chrXV | 2438 | 1135 | 1089 | 2459 | 975 | 1096 | 2265 | 1073 | 1236 | 3150 | 885 | 942 | 3696 | 570 | 760 | 3135 | 910 | 894 |
| 5132 | W303.asm02.HP0_G0051300 | chrXV | 1911 | 1785 | 1429 | 1545 | 1505 | 1384 | 2018 | 1874 | 1703 | 6575 | 2247 | 2096 | 5222 | 1434 | 1477 | 5476 | 1982 | 2067 |
| 5133 | W303.asm02.HP0_G0051310 | chrXV | 14116 | 3206 | 5129 | 14455 | 4220 | 5307 | 15276 | 3103 | 6237 | 29108 | 7921 | 10657 | 26635 | 6465 | 9018 | 21666 | 6472 | 10018 |
| 5134 | W303.asm02.HP0_G0051320 | chrXV | 1880 | 701 | 820 | 1911 | 752 | 690 | 1719 | 776 | 863 | 3150 | 596 | 658 | 3150 | 321 | 579 | 3325 | 570 | 641 |
| 5135 | W303.asm02.HP0_G0051330 | chrXV | 1893 | 628 | 555 | 1787 | 491 | 671 | 1826 | 577 | 575 | 2801 | 687 | 1152 | 2602 | 514 | 777 | 2353 | 654 | 1007 |
| 5136 | W303.asm02.HP0_G0051340 | chrXV | 3358 | 1331 | 902 | 2771 | 940 | 872 | 3158 | 1334 | 954 | 4167 | 1376 | 1290 | 4182 | 918 | 984 | 4116 | 1243 | 1110 |
| 5137 | W303.asm02.HP0_G0051350 | chrXV | 1849 | 1193 | 1219 | 1562 | 1434 | 1149 | 1738 | 1358 | 1594 | 4232 | 1241 | 1017 | 4373 | 744 | 649 | 4404 | 1037 | 932 |
| 5138 | W303.asm02.HP0_G0051360 | chrXV | 700 | 1169 | 1490 | 616 | 1398 | 1493 | 640 | 1298 | 1927 | 4064 | 988 | 1140 | 4748 | 736 | 963 | 4273 | 888 | 1012 |
| 5139 | W303.asm02.HP0_G0051370 | chrXV | 7593 | 3610 | 6183 | 7164 | 4383 | 5824 | 7513 | 4190 | 6698 | 19835 | 3890 | 7035 | 20386 | 2755 | 4967 | 19783 | 3482 | 6376 |
| 5140 | W303.asm02.HP0_G0051380 | chrXV | 108 | 29 | 9 | 112 | 8 | 12 | 85 | 17 | 15 | 24 | 12 | 6 | 30 | 8 | 10 | 13 | 18 | 7 |
| 5141 | W303.asm02.HP0_G0051390 | chrXV | 2670 | 766 | 851 | 2692 | 834 | 796 | 2465 | 773 | 936 | 5659 | 1135 | 1444 | 4542 | 767 | 1212 | 4395 | 1101 | 1330 |
| 5142 | W303.asm02.HP0_G0051400 | chrXV | 2230 | 992 | 1032 | 1838 | 1050 | 1060 | 1761 | 1151 | 1208 | 2542 | 819 | 778 | 2729 | 499 | 700 | 2735 | 737 | 770 |
| 5143 | W303.asm02.HP0_G0051410 | chrXV | 622 | 328 | 354 | 609 | 347 | 330 | 599 | 331 | 492 | 1082 | 231 | 266 | 1224 | 147 | 165 | 1103 | 250 | 211 |
| 5144 | W303.asm02.HP0_G0051420 | chrXV | 12077 | 5124 | 3962 | 11879 | 3921 | 3961 | 11766 | 4881 | 3653 | 13550 | 4784 | 5386 | 12873 | 3657 | 4405 | 12198 | 4436 | 4794 |
| 5145 | W303.asm02.HP0_G0051430 | chrXV | 5533 | 1627 | 1857 | 5673 | 1902 | 1971 | 6429 | 1723 | 2582 | 13632 | 2855 | 3480 | 9334 | 1959 | 2431 | 9946 | 2378 | 3175 |
| 5146 | W303.asm02.HP0_G0051440 | chrXV | 7012 | 4406 | 6030 | 7268 | 4882 | 6068 | 6696 | 4207 | 8002 | 11785 | 3344 | 4038 | 13914 | 2365 | 3312 | 13213 | 3111 | 3508 |
| 5147 | W303.asm02.HP0_G0051450 | chrXV | 673 | 1619 | 920 | 643 | 1058 | 913 | 697 | 1462 | 988 | 3812 | 1226 | 960 | 3820 | 795 | 643 | 3759 | 1273 | 860 |
| 5148 | W303.asm02.HP0_G0051460 | chrXV | 2260 | 1053 | 832 | 2541 | 981 | 763 | 2351 | 947 | 797 | 4475 | 1021 | 1160 | 4685 | 857 | 969 | 3883 | 946 | 1068 |
| 5149 | W303.asm02.HP0_G0051470 | chrXV | 514 | 367 | 362 | 503 | 361 | 356 | 533 | 346 | 419 | 1217 | 351 | 439 | 1401 | 265 | 365 | 1182 | 299 | 392 |
| 5150 | W303.asm02.HP0_G0051480 | chrXV | 1044 | 833 | 1090 | 821 | 1022 | 1211 | 894 | 887 | 1562 | 2663 | 694 | 893 | 2978 | 474 | 725 | 3153 | 666 | 765 |
| 5151 | W303.asm02.HP0_G0051490 | chrXV | 1249 | 804 | 800 | 1194 | 801 | 752 | 1103 | 788 | 777 | 3148 | 680 | 875 | 3183 | 512 | 584 | 3119 | 647 | 735 |
| 5152 | W303.asm02.HP0_G0051500 | chrXV | 7426 | 575 | 758 | 6840 | 791 | 1033 | 6964 | 482 | 946 | 5067 | 1036 | 1480 | 1632 | 704 | 938 | 1501 | 884 | 1096 |
| 5153 | W303.asm02.HP0_G0051510 | chrXV | 175 | 26 | 10 | 137 | 14 | 20 | 124 | 14 | 13 | 27 | 19 | 5 | 24 | 12 | 11 | 31 | 12 | 11 |
| 5154 | W303.asm02.HP0_G0051520 | chrXV | 271 | 283 | 360 | 261 | 327 | 314 | 281 | 257 | 297 | 2106 | 292 | 556 | 1713 | 201 | 330 | 1429 | 283 | 484 |
| 5155 | W303.asm02.HP0_G0051530 | chrXV | 14515 | 7030 | 5611 | 12788 | 5736 | 5483 | 14306 | 7128 | 5786 | 31919 | 8861 | 8780 | 26292 | 6178 | 7004 | 27387 | 7977 | 8212 |
| 5156 | W303.asm02.HP0_G0051540 | chrXV | 2300 | 1311 | 1350 | 1977 | 1448 | 1608 | 2570 | 1314 | 1731 | 8774 | 2506 | 3025 | 5942 | 1609 | 1905 | 6670 | 2273 | 2670 |
| 5157 | W303.asm02.HP0_G0051550 | chrXV | 1446 | 2078 | 1740 | 1628 | 2094 | 1510 | 1969 | 2282 | 1737 | 11181 | 2837 | 2569 | 9319 | 1683 | 1271 | 9550 | 2445 | 2292 |
| 5158 | W303.asm02.HP0_G0051560 | chrXV | 2125 | 730 | 588 | 2107 | 233 | 654 | 1895 | 506 | 666 | 1646 | 1359 | 1163 | 770 | 608 | 1065 | 773 | 1121 | 1001 |
| 5159 | W303.asm02.HP0_G0051570 | chrXV | 1933 | 800 | 859 | 2011 | 847 | 793 | 1877 | 839 | 873 | 3892 | 870 | 1195 | 3402 | 467 | 803 | 3539 | 778 | 1135 |
| 5160 | W303.asm02.HP0_G0051580 | chrXV | 3912 | 1173 | 874 | 3961 | 891 | 943 | 3779 | 1121 | 831 | 4090 | 1078 | 1256 | 3692 | 775 | 1070 | 3772 | 1115 | 1100 |
| 5161 | W303.asm02.HP0_G0051590 | chrXV | 4246 | 2993 | 4192 | 4083 | 3707 | 4184 | 4436 | 3089 | 5561 | 11056 | 2822 | 3577 | 12616 | 1836 | 2741 | 11717 | 2408 | 3196 |
| 5162 | W303.asm02.HP0_G0051600 | chrXV | 1832 | 747 | 540 | 1692 | 464 | 490 | 1622 | 735 | 520 | 1749 | 556 | 477 | 1843 | 433 | 402 | 1824 | 521 | 397 |
| 5163 | W303.asm02.HP0_G0051610 | chrXV | 3505 | 1907 | 1384 | 3112 | 1274 | 1251 | 3034 | 1832 | 1422 | 5103 | 1553 | 1437 | 5421 | 880 | 1097 | 4995 | 1637 | 1252 |
| 5164 | W303.asm02.HP0_G0051620 | chrXV | 3503 | 2564 | 2823 | 3481 | 2775 | 2722 | 3555 | 2666 | 3327 | 8391 | 2203 | 2236 | 9549 | 1425 | 1820 | 8410 | 2102 | 1934 |
| 5165 | W303.asm02.HP0_G0051630 | chrXV | 2902 | 1953 | 2071 | 2308 | 2071 | 1982 | 2864 | 1956 | 2630 | 5787 | 1521 | 1617 | 6769 | 1006 | 1226 | 6102 | 1512 | 1392 |
| 5166 | W303.asm02.HP0_G0051640 | chrXV | 789 | 556 | 415 | 722 | 357 | 459 | 743 | 503 | 425 | 1288 | 502 | 500 | 1280 | 330 | 344 | 1318 | 451 | 463 |
| 5167 | W303.asm02.HP0_G0051650 | chrXV | 5274 | 2800 | 2717 | 4861 | 2521 | 2973 | 4730 | 2518 | 3665 | 4390 | 2371 | 2149 | 6150 | 1741 | 2241 | 4740 | 2265 | 2114 |
| 5168 | W303.asm02.HP0_G0051660 | chrXV | 4975 | 2439 | 2051 | 4405 | 2065 | 2250 | 4463 | 2457 | 2360 | 4975 | 2093 | 2031 | 5934 | 1698 | 1836 | 4869 | 1856 | 1846 |
| 5169 | W303.asm02.HP0_G0051670 | chrXV | 371 | 754 | 931 | 353 | 1042 | 951 | 412 | 746 | 1065 | 4014 | 672 | 808 | 3953 | 446 | 445 | 3698 | 610 | 645 |
| 5170 | W303.asm02.HP0_G0051680 | chrXV | 2194 | 1029 | 2196 | 2054 | 791 | 2634 | 1980 | 1011 | 2290 | 2803 | 827 | 2688 | 3142 | 603 | 1554 | 3010 | 802 | 2297 |
| 5171 | W303.asm02.HP0_G0051690 | chrXV | 2764 | 410 | 583 | 2729 | 383 | 658 | 2570 | 400 | 742 | 837 | 445 | 670 | 1103 | 278 | 620 | 839 | 427 | 579 |
| 5172 | W303.asm02.HP0_G0051700 | chrXV | 7412 | 605 | 799 | 7360 | 846 | 951 | 7609 | 572 | 1009 | 5592 | 1027 | 2163 | 2582 | 821 | 1418 | 2600 | 868 | 1763 |
| 5173 | W303.asm02.HP0_G0051710 | chrXV | 2363 | 339 | 533 | 1877 | 533 | 568 | 2100 | 378 | 709 | 4241 | 716 | 1169 | 1934 | 560 | 875 | 2023 | 627 | 1039 |
| 5174 | W303.asm02.HP0_G0051720 | chrXV | 1798 | 825 | 492 | 1403 | 469 | 526 | 1570 | 815 | 474 | 1565 | 556 | 441 | 1719 | 383 | 411 | 1530 | 596 | 455 |
| 5175 | W303.asm02.HP0_G0051730 | chrXV | 8051 | 6180 | 9252 | 7533 | 8659 | 9137 | 8238 | 6312 | 11263 | 17722 | 5856 | 7400 | 20115 | 4388 | 6283 | 18807 | 5154 | 6615 |
| 5176 | W303.asm02.HP0_G0051740 | chrXV | 7 | 19 | 3 | 9 | 3 | 4 | 5 | 13 | 7 | 16 | 12 | 4 | 22 | 6 | 5 | 20 | 4 | 1 |
| 5177 | W303.asm02.HP0_G0051750 | chrXV | 1128 | 1620 | 2234 | 1284 | 2714 | 2205 | 1343 | 1801 | 2485 | 8783 | 1733 | 2156 | 8298 | 1015 | 1296 | 8154 | 1488 | 2038 |
| 5178 | W303.asm02.HP0_G0051760 | chrXV | 931 | 903 | 714 | 920 | 769 | 690 | 962 | 919 | 749 | 2378 | 714 | 641 | 2374 | 478 | 472 | 2169 | 671 | 555 |
| 5179 | W303.asm02.HP0_G0051770 | chrXV | 1282 | 439 | 534 | 1199 | 473 | 570 | 1132 | 478 | 654 | 3483 | 540 | 900 | 2779 | 424 | 674 | 2830 | 548 | 819 |
| 5180 | W303.asm02.HP0_G0051780 | chrXV | 1326 | 900 | 1110 | 1306 | 1081 | 1091 | 1345 | 901 | 1268 | 3203 | 706 | 949 | 3556 | 510 | 722 | 3315 | 690 | 795 |
| 5181 | W303.asm02.HP0_G0051790 | chrXV | 185 | 300 | 139 | 234 | 148 | 128 | 166 | 241 | 142 | 737 | 257 | 167 | 498 | 123 | 106 | 749 | 338 | 130 |
| 5182 | W303.asm02.HP0_G0051800 | chrXV | 8904 | 10701 | 7467 | 8661 | 7016 | 7333 | 7763 | 9469 | 7509 | 16096 | 8541 | 5758 | 14443 | 5374 | 5330 | 17360 | 8243 | 5520 |
| 5183 | W303.asm02.HP0_G0051810 | chrXV | 2242 | 1081 | 925 | 2252 | 875 | 979 | 2269 | 1144 | 1051 | 6864 | 1700 | 1854 | 6354 | 1186 | 1446 | 5700 | 1756 | 1744 |
| 5184 | W303.asm02.HP0_G0051820 | chrXV | 812 | 379 | 397 | 875 | 384 | 482 | 870 | 346 | 441 | 2921 | 618 | 800 | 2204 | 522 | 637 | 1840 | 554 | 719 |
| 5185 | W303.asm02.HP0_G0051830 | chrXV | 2025 | 2938 | 3179 | 1815 | 3301 | 3369 | 1878 | 3151 | 3483 | 18766 | 5617 | 6439 | 18180 | 4587 | 4936 | 16829 | 5321 | 6047 |
| 5186 | W303.asm02.HP0_G0051840 | chrXV | 4268 | 639 | 879 | 4313 | 895 | 945 | 4182 | 621 | 1111 | 7827 | 1075 | 1960 | 5451 | 825 | 1323 | 5369 | 1015 | 1770 |
| 5187 | W303.asm02.HP0_G0051850 | chrXV | 568 | 535 | 619 | 363 | 568 | 624 | 428 | 575 | 806 | 1415 | 415 | 380 | 1703 | 273 | 352 | 1628 | 396 | 316 |
| 5188 | W303.asm02.HP0_G0051860 | chrXV | 1381 | 1069 | 1215 | 1268 | 1253 | 1198 | 1295 | 1203 | 1474 | 2876 | 826 | 861 | 3180 | 594 | 744 | 2977 | 684 | 734 |
| 5189 | W303.asm02.HP0_G0051870 | chrXV | 2561 | 573 | 1062 | 2395 | 1368 | 1099 | 2549 | 673 | 1259 | 8769 | 871 | 1273 | 13242 | 1028 | 1281 | 5155 | 831 | 1082 |
| 5190 | W303.asm02.HP0_G0051880 | chrXV | 2402 | 1126 | 1244 | 2179 | 1350 | 1269 | 2197 | 1124 | 1315 | 5051 | 1003 | 1332 | 4720 | 666 | 1000 | 4414 | 885 | 1257 |
| 5191 | W303.asm02.HP0_G0051890 | chrXV | 1525 | 177 | 263 | 1481 | 266 | 256 | 1471 | 178 | 283 | 3022 | 360 | 583 | 2063 | 255 | 379 | 2034 | 320 | 477 |
| 5192 | W303.asm02.HP0_G0051900 | chrXV | 11183 | 2970 | 2910 | 11171 | 3190 | 2899 | 12024 | 2845 | 3517 | 27457 | 6635 | 8088 | 16141 | 4972 | 4964 | 17427 | 5455 | 6767 |
| 5193 | W303.asm02.HP0_G0051910 | chrXV | 2234 | 1465 | 1700 | 2160 | 1448 | 1891 | 2344 | 1552 | 1726 | 5436 | 1434 | 1998 | 5081 | 1003 | 1445 | 5184 | 1296 | 1805 |
| 5194 | W303.asm02.HP0_G0051920 | chrXV | 12179 | 9352 | 13463 | 11628 | 12581 | 13104 | 12524 | 9642 | 17360 | 28434 | 8830 | 10959 | 31771 | 6381 | 8892 | 30068 | 7698 | 9789 |
| 5195 | W303.asm02.HP0_G0051930 | chrXV | 225 | 323 | 569 | 232 | 490 | 559 | 194 | 384 | 635 | 1029 | 255 | 385 | 327 | 61 | 133 | 1079 | 215 | 279 |
| 5196 | W303.asm02.HP0_G0051940 | chrXV | 551 | 3803 | 4567 | 401 | 4600 | 4339 | 654 | 4285 | 5065 | 10096 | 2797 | 2991 | 7124 | 1244 | 1848 | 10764 | 2329 | 2635 |
| 5197 | W303.asm02.HP0_G0051950 | chrXV | 2283 | 1481 | 1456 | 2075 | 1285 | 1434 | 2383 | 1479 | 1447 | 8225 | 2152 | 2036 | 8112 | 1229 | 1864 | 7992 | 2075 | 2394 |
| 5198 | W303.asm02.HP0_G0051960 | chrXV | 19 | 18 | 19 | 28 | 27 | 23 | 32 | 21 | 29 | 153 | 40 | 30 | 142 | 25 | 23 | 99 | 34 | 28 |
| 5199 | W303.asm02.HP0_G0051970 | chrXV | 18729 | 3733 | 3130 | 18115 | 2635 | 3304 | 17937 | 3594 | 3897 | 16970 | 5370 | 6953 | 15691 | 4493 | 6304 | 12721 | 5293 | 6438 |
| 5200 | W303.asm02.HP0_G0051980 | chrXV | 42 | 48 | 32 | 40 | 20 | 19 | 45 | 35 | 34 | 84 | 30 | 19 | 115 | 21 | 12 | 89 | 41 | 21 |
| 5201 | W303.asm02.HP0_G0051990 | chrXV | 1126 | 745 | 844 | 1334 | 798 | 792 | 1110 | 763 | 996 | 3010 | 606 | 707 | 2825 | 393 | 489 | 2994 | 522 | 586 |
| 5202 | W303.asm02.HP0_G0052000 | chrXV | 2022 | 1888 | 2349 | 2021 | 2066 | 2289 | 2123 | 2050 | 2746 | 6810 | 1577 | 1755 | 7527 | 1028 | 1413 | 7550 | 1383 | 1738 |
| 5203 | W303.asm02.HP0_G0052010 | chrXV | 2095 | 352 | 627 | 1769 | 358 | 864 | 2167 | 328 | 782 | 3226 | 685 | 1752 | 1923 | 482 | 1075 | 1802 | 624 | 1618 |
| 5204 | W303.asm02.HP0_G0052020 | chrXV | 1964 | 469 | 669 | 1949 | 634 | 797 | 1958 | 404 | 802 | 5945 | 1096 | 1742 | 2827 | 784 | 1170 | 2638 | 949 | 1604 |
| 5205 | W303.asm02.HP0_G0052030 | chrXV | 7906 | 2058 | 1491 | 7098 | 1496 | 1679 | 7993 | 1924 | 1832 | 11145 | 3279 | 3771 | 7814 | 2327 | 2575 | 8480 | 2983 | 3260 |
| 5206 | W303.asm02.HP0_G0052040 | chrXV | 703 | 210 | 454 | 721 | 366 | 510 | 665 | 214 | 604 | 3086 | 442 | 964 | 2066 | 307 | 672 | 1940 | 398 | 963 |

| | | | | | | | | | | | | | | | | | | | | |
|---|---|---|---|---|---|---|---|---|---|---|---|---|---|---|---|---|---|---|---|---|
| 5207 | W303.asm02.HP0_G0052050 | chrXV | 12015 | 3855 | 3209 | 11876 | 3256 | 3442 | 12578 | 4118 | 3769 | 25384 | 5450 | 6903 | 18644 | 4359 | 5132 | 19207 | 5073 | 6520 |
| 5208 | W303.asm02.HP0_G0052060 | chrXV | 2233 | 1523 | 2466 | 1907 | 2323 | 2354 | 1931 | 1552 | 3101 | 5307 | 1405 | 1916 | 7143 | 980 | 1636 | 5713 | 1222 | 1567 |
| 5209 | W303.asm02.HP0_G0052070 | chrXV | 1062 | 182 | 372 | 1027 | 360 | 321 | 1001 | 229 | 427 | 2436 | 370 | 576 | 1922 | 238 | 364 | 1732 | 331 | 516 |
| 5210 | W303.asm02.HP0_G0052080 | chrXV | 7395 | 417 | 560 | 6474 | 476 | 631 | 7510 | 358 | 685 | 5081 | 985 | 1315 | 2710 | 827 | 1043 | 2020 | 810 | 1185 |
| 5211 | W303.asm02.HP0_G0052090 | chrXV | 1557 | 369 | 576 | 1570 | 582 | 667 | 1523 | 385 | 724 | 4808 | 761 | 1309 | 2099 | 552 | 852 | 2212 | 674 | 1169 |
| 5212 | W303.asm02.HP0_G0052100 | chrXV | 11908 | 3025 | 2175 | 10781 | 2157 | 2558 | 12188 | 2769 | 2196 | 9499 | 3084 | 3875 | 9594 | 2681 | 3053 | 8535 | 2551 | 3337 |
| 5213 | W303.asm02.HP0_G0052110 | chrXV | 0 | 0 | 0 | 1 | 0 | 0 | 0 | 0 | 0 | 0 | 1 | 0 | 0 | 0 | 0 | 0 | 0 | 0 |
| 5214 | W303.asm02.HP0_G0052120 | chrXV | 564 | 340 | 469 | 538 | 458 | 448 | 535 | 363 | 466 | 1974 | 311 | 445 | 2069 | 193 | 370 | 1860 | 305 | 425 |
| 5215 | W303.asm02.HP0_G0052130 | chrXV | 11359 | 6755 | 6171 | 10020 | 6815 | 5499 | 11793 | 7267 | 6582 | 36486 | 8813 | 9048 | 31731 | 6021 | 6281 | 31840 | 7753 | 8056 |
| 5216 | W303.asm02.HP0_G0052140 | chrXV | 1864 | 674 | 881 | 1799 | 707 | 940 | 1753 | 714 | 994 | 5435 | 944 | 1773 | 3954 | 738 | 1208 | 4041 | 888 | 1544 |
| 5217 | W303.asm02.HP0_G0052150 | chrXV | 522 | 269 | 503 | 519 | 557 | 548 | 616 | 240 | 659 | 3663 | 488 | 979 | 1834 | 399 | 488 | 1602 | 403 | 780 |
| 5218 | W303.asm02.HP0_G0052160 | chrXV | 603 | 291 | 1135 | 608 | 1373 | 1148 | 533 | 230 | 1285 | 3203 | 420 | 1007 | 4178 | 149 | 455 | 3277 | 567 | 575 |
| 5219 | W303.asm02.HP0_G0052170 | chrXV | 332 | 176 | 177 | 241 | 168 | 184 | 284 | 147 | 193 | 399 | 128 | 127 | 500 | 74 | 70 | 411 | 140 | 123 |
| 5220 | W303.asm02.HP0_G0052180 | chrXV | 1179 | 1897 | 1928 | 1157 | 2467 | 2017 | 1258 | 2024 | 1908 | 8336 | 1816 | 2228 | 7881 | 1222 | 1231 | 7784 | 1609 | 1943 |
| 5221 | W303.asm02.HP0_G0052190 | chrXV | 8351 | 3060 | 2876 | 7886 | 4020 | 2891 | 9236 | 3262 | 3566 | 35050 | 5286 | 7148 | 23669 | 3761 | 4284 | 25314 | 4604 | 6630 |
| 5222 | W303.asm02.HP0_G0052200 | chrXV | 2085 | 1767 | 1971 | 2229 | 2344 | 1989 | 2245 | 1934 | 1893 | 9455 | 1675 | 2073 | 9030 | 1180 | 1364 | 8380 | 1318 | 1855 |
| 5223 | W303.asm02.HP0_G0052210 | chrXV | 112 | 28 | 33 | 110 | 53 | 40 | 112 | 29 | 35 | 217 | 54 | 62 | 236 | 44 | 49 | 158 | 54 | 51 |
| 5224 | W303.asm02.HP0_G0052220 | chrXV | 3763 | 979 | 4062 | 5262 | 2810 | 3683 | 3525 | 1069 | 3646 | 10499 | 1290 | 4920 | 10537 | 570 | 2731 | 9931 | 1196 | 3902 |
| 5225 | W303.asm02.HP0_G0052230 | chrXV | 1973 | 870 | 1037 | 1860 | 1070 | 945 | 2012 | 1060 | 1146 | 8617 | 1400 | 1883 | 6595 | 1008 | 1246 | 6957 | 1224 | 1728 |
| 5226 | W303.asm02.HP0_G0052240 | chrXV | 2642 | 261 | 411 | 3178 | 235 | 540 | 2529 | 232 | 508 | 1274 | 618 | 1064 | 643 | 499 | 844 | 412 | 478 | 927 |
| 5227 | W303.asm02.HP0_G0052250 | chrXV | 495 | 663 | 510 | 461 | 362 | 510 | 492 | 502 | 699 | 938 | 1130 | 1012 | 370 | 563 | 594 | 384 | 989 | 727 |
| 5228 | W303.asm02.HP0_G0052260 | chrXV | 1067 | 257 | 262 | 955 | 235 | 287 | 1089 | 233 | 322 | 2488 | 368 | 596 | 1670 | 224 | 404 | 1759 | 425 | 502 |
| 5229 | W303.asm02.HP0_G0052270 | chrXV | 1380 | 494 | 598 | 1403 | 603 | 575 | 1337 | 489 | 621 | 2971 | 532 | 810 | 2540 | 379 | 585 | 2625 | 503 | 692 |
| 5230 | W303.asm02.HP0_G0052280 | chrXV | 112 | 35 | 47 | 99 | 51 | 60 | 116 | 35 | 37 | 273 | 43 | 47 | 220 | 27 | 29 | 284 | 54 | 39 |
| 5231 | W303.asm02.HP0_G0052290 | chrXV | 1948 | 624 | 597 | 2345 | 575 | 637 | 1932 | 619 | 636 | 1986 | 548 | 573 | 2232 | 354 | 492 | 2029 | 436 | 477 |
| 5232 | W303.asm02.HP0_G0052300 | chrXV | 2043 | 896 | 848 | 2064 | 923 | 828 | 2027 | 965 | 822 | 5003 | 978 | 1175 | 5371 | 696 | 1014 | 4464 | 937 | 1090 |
| 5233 | W303.asm02.HP0_G0052310 | chrXV | 13325 | 2494 | 2974 | 13701 | 2797 | 2816 | 13233 | 2654 | 3016 | 12063 | 2540 | 3924 | 12761 | 1951 | 3165 | 11487 | 2660 | 3482 |
| 5234 | W303.asm02.HP0_G0052320 | chrXV | 4267 | 1962 | 2557 | 4433 | 2565 | 2397 | 4623 | 2345 | 2474 | 15511 | 2565 | 3424 | 15298 | 2184 | 3108 | 14578 | 2284 | 3236 |
| 5235 | W303.asm02.HP0_G0052330 | chrXV | 4813 | 1097 | 1130 | 4539 | 1074 | 1176 | 4722 | 1024 | 1428 | 6208 | 1808 | 2352 | 2605 | 1353 | 1428 | 3237 | 1728 | 1900 |
| 5236 | W303.asm02.HP0_G0052340 | chrXV | 1356 | 988 | 970 | 1071 | 813 | 1004 | 1192 | 914 | 1284 | 2172 | 685 | 664 | 2494 | 496 | 574 | 2404 | 702 | 635 |
| 5237 | W303.asm02.HP0_G0052350 | chrXV | 3374 | 515 | 394 | 2910 | 383 | 384 | 3195 | 434 | 434 | 1959 | 678 | 707 | 1538 | 538 | 614 | 1289 | 711 | 667 |
| 5238 | W303.asm02.HP0_G0052360 | chrXV | 317 | 281 | 647 | 324 | 711 | 737 | 392 | 272 | 838 | 6314 | 639 | 1351 | 3924 | 470 | 764 | 3609 | 651 | 990 |
| 5239 | W303.asm02.HP0_G0052370 | chrXV | 868 | 1895 | 3910 | 919 | 3761 | 3452 | 861 | 1739 | 4393 | 11867 | 1719 | 2783 | 12589 | 998 | 1858 | 11737 | 1976 | 2063 |
| 5240 | W303.asm02.HP0_G0052380 | chrXV | 15601 | 6307 | 5558 | 14299 | 5896 | 5194 | 16636 | 6354 | 5792 | 32802 | 8420 | 9488 | 27650 | 5887 | 6662 | 28283 | 7617 | 8558 |
| 5241 | W303.asm02.HP0_G0052390 | chrXV | 10325 | 5942 | 6271 | 10051 | 5340 | 6143 | 10365 | 5686 | 7281 | 12994 | 5043 | 5264 | 14508 | 3562 | 4545 | 12546 | 4458 | 4721 |
| 5242 | W303.asm02.HP0_G0052400 | chrXV | 2472 | 311 | 396 | 2675 | 374 | 403 | 2425 | 278 | 447 | 2978 | 584 | 841 | 2197 | 388 | 658 | 1892 | 511 | 677 |
| 5243 | W303.asm02.HP0_G0052410 | chrXV | 486 | 239 | 228 | 459 | 207 | 206 | 480 | 229 | 211 | 831 | 240 | 222 | 749 | 118 | 163 | 846 | 207 | 188 |
| 5244 | W303.asm02.HP0_G0052420 | chrXV | 996 | 914 | 818 | 896 | 662 | 826 | 886 | 879 | 886 | 2015 | 648 | 615 | 2629 | 480 | 570 | 2234 | 586 | 570 |
| 5245 | W303.asm02.HP0_G0052430 | chrXV | 1341 | 930 | 893 | 1322 | 779 | 893 | 1334 | 915 | 1086 | 2403 | 606 | 678 | 2807 | 447 | 484 | 2619 | 694 | 561 |
| 5246 | W303.asm02.HP0_G0052440 | chrXV | 9946 | 10291 | 13617 | 8685 | 12393 | 13206 | 9885 | 9893 | 17332 | 25406 | 8942 | 10840 | 28320 | 6696 | 9020 | 27571 | 8085 | 9644 |
| 5247 | W303.asm02.HP0_G0052450 | chrXV | 8840 | 1554 | 1720 | 8346 | 1723 | 1846 | 9539 | 1465 | 2130 | 13192 | 3222 | 3999 | 7970 | 2628 | 3129 | 7411 | 2759 | 3706 |
| 5248 | W303.asm02.HP0_G0052460 | chrXV | 1247 | 183 | 274 | 1190 | 248 | 315 | 1108 | 148 | 338 | 2377 | 369 | 869 | 941 | 272 | 390 | 948 | 302 | 679 |
| 5249 | W303.asm02.HP0_G0052470 | chrXV | 2547 | 313 | 484 | 2737 | 431 | 519 | 2687 | 280 | 626 | 4101 | 675 | 1102 | 3163 | 460 | 806 | 3059 | 701 | 1009 |
| 5250 | W303.asm02.HP0_G0052480 | chrXV | 1388 | 214 | 283 | 1242 | 272 | 323 | 1424 | 174 | 350 | 2671 | 526 | 787 | 982 | 367 | 486 | 1021 | 480 | 698 |
| 5251 | W303.asm02.HP0_G0052490 | chrXV | 9239 | 2101 | 1183 | 11528 | 1384 | 1457 | 8885 | 1631 | 1608 | 4583 | 2300 | 2025 | 6592 | 3519 | 3349 | 2734 | 2327 | 1809 |
| 5252 | W303.asm02.HP0_G0052500 | chrXV | 7095 | 9330 | 7041 | 7690 | 7949 | 7603 | 7899 | 8996 | 8749 | 43413 | 17471 | 16861 | 35199 | 15497 | 14160 | 32437 | 15178 | 14954 |
| 5253 | W303.asm02.HP0_G0052510 | chrXV | 49 | 18 | 16 | 41 | 9 | 6 | 32 | 11 | 17 | 35 | 20 | 11 | 39 | 12 | 6 | 38 | 24 | 4 |
| 5254 | W303.asm02.HP0_G0052520 | chrXV | 647 | 40 | 51 | 744 | 61 | 69 | 602 | 29 | 75 | 400 | 78 | 102 | 403 | 45 | 75 | 354 | 58 | 75 |
| 5255 | W303.asm02.HP0_G0052530 | chrXV | 1752 | 425 | 554 | 1517 | 562 | 453 | 1758 | 532 | 516 | 5507 | 980 | 891 | 3641 | 414 | 637 | 4888 | 797 | 875 |
| 5256 | W303.asm02.HP0_G0052540 | chrXV | 415 | 83 | 133 | 398 | 125 | 113 | 411 | 88 | 152 | 955 | 202 | 261 | 484 | 139 | 155 | 489 | 170 | 251 |
| 5257 | W303.asm02.HP0_G0052550 | chrXV | 696 | 198 | 213 | 604 | 240 | 268 | 702 | 134 | 258 | 1061 | 313 | 494 | 654 | 220 | 316 | 609 | 272 | 400 |
| 5258 | W303.asm02.HP0_G0052560 | chrXV | 578 | 163 | 210 | 543 | 205 | 213 | 514 | 142 | 241 | 2035 | 291 | 346 | 1615 | 170 | 286 | 2011 | 306 | 448 |
| 5259 | W303.asm02.HP0_G0052570 | chrXV | 28 | 25 | 16 | 35 | 12 | 19 | 28 | 24 | 22 | 90 | 18 | 29 | 71 | 5 | 8 | 92 | 30 | 33 |
| 5260 | W303.asm02.HP0_G0052580 | chrXV | 528 | 172 | 195 | 515 | 310 | 345 | 533 | 171 | 186 | 4701 | 941 | 389 | 5242 | 309 | 582 | 6133 | 833 | 2500 |
| 5261 | W303.asm02.HP0_G0052590 | chrXV | 951 | 115 | 172 | 936 | 265 | 336 | 897 | 101 | 155 | 4585 | 855 | 457 | 4491 | 234 | 550 | 5408 | 632 | 2759 |
| 5262 | W303.asm02.HP0_G0052600 | chrXV | 1479 | 105 | 145 | 1525 | 138 | 163 | 1465 | 82 | 172 | 1835 | 273 | 469 | 812 | 160 | 309 | 945 | 266 | 445 |
| 5263 | W303.asm02.HP0_G0052610 | chrXV | 3649 | 587 | 697 | 4866 | 678 | 641 | 3241 | 569 | 765 | 2167 | 445 | 581 | 2508 | 269 | 415 | 2091 | 469 | 541 |
| 5264 | W303.asm02.HP0_G0052620 | chrXV | 5214 | 442 | 575 | 5230 | 456 | 501 | 4426 | 455 | 492 | 2104 | 503 | 682 | 2505 | 350 | 758 | 1780 | 452 | 623 |
| 5265 | W303.asm02.HP0_G0052630 | chrXV | 924 | 47 | 59 | 735 | 46 | 70 | 830 | 56 | 69 | 248 | 62 | 93 | 235 | 63 | 122 | 225 | 64 | 104 |
| 5266 | W303.asm02.HP0_G0052640 | chrXV | 127 | 9 | 12 | 117 | 10 | 9 | 129 | 8 | 17 | 66 | 15 | 16 | 79 | 14 | 17 | 45 | 18 | 18 |
| 5267 | W303.asm02.HP0_G0052650 | chrXV | 2236 | 295 | 358 | 1889 | 391 | 358 | 1881 | 295 | 434 | 1298 | 411 | 492 | 1221 | 321 | 389 | 856 | 356 | 496 |
| 5268 | W303.asm02.HP0_G0052660 | chrXVI | 0 | 0 | 0 | 0 | 0 | 0 | 0 | 0 | 0 | 0 | 0 | 0 | 0 | 0 | 0 | 0 | 0 | 0 |
| 5269 | W303.asm02.HP0_G0052670 | chrXVI | 54 | 9 | 4 | 41 | 6 | 1 | 55 | 10 | 2 | 10 | 16 | 8 | 2 | 8 | 3 | 4 | 8 | 10 |
| 5270 | W303.asm02.HP0_G0052680 | chrXVI | 50 | 5 | 5 | 32 | 4 | 5 | 41 | 8 | 4 | 11 | 6 | 5 | 11 | 5 | 1 | 5 | 8 | 3 |
| 5271 | W303.asm02.HP0_G0052690 | chrXVI | 29 | 0 | 1 | 15 | 3 | 3 | 27 | 2 | 2 | 4 | 2 | 3 | 9 | 2 | 1 | 2 | 0 | 0 |
| 5272 | W303.asm02.HP0_G0052700 | chrXVI | 666 | 42 | 58 | 616 | 57 | 86 | 507 | 47 | 61 | 108 | 57 | 78 | 113 | 71 | 71 | 61 | 48 | 63 |
| 5273 | W303.asm02.HP0_G0052710 | chrXVI | 60 | 8 | 5 | 85 | 11 | 10 | 37 | 5 | 9 | 21 | 4 | 7 | 18 | 1 | 0 | 13 | 4 | 4 |
| 5274 | W303.asm02.HP0_G0052720 | chrXVI | 631 | 183 | 123 | 665 | 127 | 130 | 533 | 149 | 169 | 246 | 132 | 118 | 393 | 97 | 89 | 211 | 123 | 102 |
| 5275 | W303.asm02.HP0_G0052730 | chrXVI | 160 | 336 | 550 | 155 | 519 | 535 | 158 | 410 | 633 | 2016 | 274 | 409 | 1984 | 190 | 290 | 2144 | 301 | 398 |
| 5276 | W303.asm02.HP0_G0052740 | chrXVI | 1063 | 131 | 192 | 885 | 148 | 160 | 899 | 117 | 168 | 398 | 135 | 183 | 431 | 100 | 124 | 224 | 132 | 163 |
| 5277 | W303.asm02.HP0_G0052750 | chrXVI | 118 | 17 | 13 | 81 | 19 | 8 | 99 | 10 | 11 | 162 | 31 | 43 | 90 | 22 | 32 | 102 | 31 | 55 |
| 5278 | W303.asm02.HP0_G0052760 | chrXVI | 3249 | 1923 | 2052 | 1740 | 1597 | 1815 | 2396 | 1606 | 2108 | 7545 | 2989 | 3249 | 4814 | 2475 | 2578 | 4584 | 2594 | 2546 |
| 5279 | W303.asm02.HP0_G0052770 | chrXVI | 5469 | 5836 | 8639 | 4882 | 6904 | 8141 | 6116 | 6027 | 11039 | 18167 | 5269 | 7171 | 19699 | 3270 | 5103 | 19182 | 4481 | 6125 |
| 5280 | W303.asm02.HP0_G0052780 | chrXVI | 235 | 185 | 114 | 186 | 118 | 132 | 216 | 131 | 128 | 741 | 214 | 218 | 571 | 162 | 168 | 581 | 236 | 193 |
| 5281 | W303.asm02.HP0_G0052790 | chrXVI | 2736 | 2435 | 3075 | 2116 | 2732 | 3120 | 2028 | 2485 | 3805 | 4816 | 2130 | 2544 | 6186 | 1630 | 2420 | 5471 | 1926 | 2065 |
| 5282 | W303.asm02.HP0_G0052800 | chrXVI | 3160 | 831 | 920 | 2802 | 947 | 1043 | 2929 | 746 | 1053 | 5281 | 1472 | 1722 | 3178 | 996 | 1142 | 3288 | 1311 | 1484 |
| 5283 | W303.asm02.HP0_G0052810 | chrXVI | 409 | 107 | 173 | 388 | 150 | 191 | 430 | 104 | 232 | 1024 | 196 | 354 | 542 | 146 | 290 | 618 | 185 | 349 |
| 5284 | W303.asm02.HP0_G0052820 | chrXVI | 1610 | 623 | 1114 | 1737 | 1001 | 1036 | 1547 | 709 | 1046 | 5357 | 929 | 1461 | 4573 | 575 | 1063 | 4489 | 800 | 1280 |
| 5285 | W303.asm02.HP0_G0052830 | chrXVI | 451 | 581 | 725 | 419 | 728 | 774 | 424 | 577 | 837 | 1571 | 455 | 534 | 2109 | 258 | 430 | 1828 | 360 | 483 |
| 5286 | W303.asm02.HP0_G0052840 | chrXVI | 2455 | 1733 | 1789 | 2442 | 2260 | 1631 | 2642 | 1872 | 1899 | 9332 | 2130 | 2239 | 9204 | 1283 | 1343 | 9196 | 1815 | 2087 |
| 5287 | W303.asm02.HP0_G0052850 | chrXVI | 719 | 2269 | 2457 | 822 | 2392 | 2629 | 732 | 2082 | 1934 | 7817 | 3153 | 3103 | 5451 | 1516 | 2774 | 6961 | 2973 | 3051 |
| 5288 | W303.asm02.HP0_G0052860 | chrXVI | 365 | 229 | 206 | 279 | 187 | 174 | 343 | 197 | 230 | 712 | 223 | 205 | 690 | 159 | 215 | 579 | 228 | 232 |
| 5289 | W303.asm02.HP0_G0052870 | chrXVI | 2385 | 1178 | 777 | 2261 | 1069 | 891 | 2666 | 1143 | 815 | 9357 | 1850 | 1883 | 5899 | 1168 | 1087 | 6730 | 1448 | 1782 |
| 5290 | W303.asm02.HP0_G0052880 | chrXVI | 8411 | 3338 | 2430 | 7847 | 2539 | 2584 | 8337 | 3250 | 2703 | 12890 | 4846 | 5632 | 12830 | 4230 | 4344 | 9859 | 4152 | 4788 |
| 5291 | W303.asm02.HP0_G0052890 | chrXVI | 11641 | 3183 | 3623 | 11928 | 3172 | 3421 | 11569 | 3118 | 3469 | 11631 | 2821 | 3546 | 13012 | 1907 | 3013 | 11868 | 2794 | 3286 |
| 5292 | W303.asm02.HP0_G0052900 | chrXVI | 3711 | 823 | 636 | 3856 | 659 | 624 | 3601 | 704 | 673 | 4439 | 949 | 1098 | 3951 | 642 | 887 | 3956 | 935 | 1017 |
| 5293 | W303.asm02.HP0_G0052910 | chrXVI | 534 | 216 | 148 | 469 | 169 | 184 | 462 | 149 | 180 | 716 | 252 | 307 | 837 | 190 | 213 | 607 | 274 | 263 |
| 5294 | W303.asm02.HP0_G0052920 | chrXVI | 719 | 212 | 264 | 668 | 193 | 227 | 628 | 216 | 278 | 861 | 235 | 194 | 882 | 150 | 162 | 866 | 219 | 206 |
| 5295 | W303.asm02.HP0_G0052930 | chrXVI | 2300 | 1105 | 1569 | 2359 | 1616 | 1727 | 2655 | 1333 | 1997 | 13607 | 2644 | 4439 | 13616 | 2058 | 2869 | 10412 | 2202 | 3982 |
| 5296 | W303.asm02.HP0_G0052940 | chrXVI | 721 | 418 | 462 | 700 | 482 | 447 | 678 | 410 | 446 | 2336 | 473 | 730 | 2097 | 286 | 468 | 1986 | 480 | 610 |
| 5297 | W303.asm02.HP0_G0052950 | chrXVI | 885 | 1045 | 1161 | 823 | 1264 | 1067 | 979 | 1150 | 1314 | 6137 | 1109 | 1326 | 5879 | 665 | 1025 | 6179 | 929 | 1310 |
| 5298 | W303.asm02.HP0_G0052960 | chrXVI | 695 | 537 | 412 | 705 | 422 | 380 | 742 | 487 | 340 | 2397 | 587 | 666 | 2071 | 401 | 522 | 2324 | 484 | 614 |
| 5299 | W303.asm02.HP0_G0052970 | chrXVI | 1510 | 602 | 620 | 1263 | 533 | 643 | 1434 | 576 | 706 | 3192 | 935 | 1202 | 3192 | 691 | 886 | 2636 | 862 | 1111 |

| | | | | | | | | | | | | | | | | | | | | |
|---|---|---|---|---|---|---|---|---|---|---|---|---|---|---|---|---|---|---|---|---|
| 5300 | W303.asm02.HP0_G0052980 | chrXVI | 3386 | 1039 | 809 | 3641 | 948 | 1009 | 2833 | 942 | 601 | 897 | 715 | 666 | 1264 | 522 | 541 | 898 | 630 | 624 |
| 5301 | W303.asm02.HP0_G0052990 | chrXVI | 9979 | 8235 | 10115 | 9473 | 9904 | 10232 | 9718 | 8468 | 10829 | 20049 | 8556 | 10035 | 21106 | 6402 | 8093 | 19669 | 7285 | 8900 |
| 5302 | W303.asm02.HP0_G0053000 | chrXVI | 5245 | 509 | 388 | 5010 | 412 | 474 | 4891 | 487 | 478 | 3050 | 745 | 879 | 2049 | 587 | 663 | 1877 | 713 | 800 |
| 5303 | W303.asm02.HP0_G0053010 | chrXVI | 1296 | 326 | 302 | 1238 | 352 | 323 | 1085 | 312 | 336 | 2041 | 445 | 494 | 1331 | 260 | 360 | 1594 | 407 | 443 |
| 5304 | W303.asm02.HP0_G0053020 | chrXVI | 8040 | 593 | 370 | 7452 | 290 | 492 | 7233 | 376 | 487 | 1964 | 1160 | 855 | 728 | 594 | 724 | 734 | 1093 | 797 |
| 5305 | W303.asm02.HP0_G0053030 | chrXVI | 2986 | 1038 | 926 | 2631 | 959 | 1064 | 2821 | 924 | 1123 | 5002 | 1479 | 1639 | 3940 | 1435 | 1352 | 3229 | 1364 | 1454 |
| 5306 | W303.asm02.HP0_G0053040 | chrXVI | 1544 | 898 | 623 | 1275 | 661 | 547 | 1632 | 866 | 610 | 3613 | 900 | 716 | 3075 | 594 | 514 | 3197 | 946 | 558 |
| 5307 | W303.asm02.HP0_G0053050 | chrXVI | 2096 | 1411 | 1759 | 1933 | 1213 | 1731 | 2083 | 1379 | 1903 | 5655 | 1446 | 2316 | 5217 | 880 | 1468 | 5116 | 1414 | 2033 |
| 5308 | W303.asm02.HP0_G0053060 | chrXVI | 6120 | 4691 | 4640 | 6314 | 4456 | 4342 | 6344 | 4731 | 4648 | 16032 | 3992 | 4485 | 15819 | 2585 | 3055 | 15733 | 3515 | 3958 |
| 5309 | W303.asm02.HP0_G0053070 | chrXVI | 4455 | 1407 | 1489 | 4171 | 1501 | 1482 | 4385 | 1441 | 1659 | 9749 | 1761 | 2524 | 10637 | 1375 | 2298 | 8895 | 1551 | 2316 |
| 5310 | W303.asm02.HP0_G0053080 | chrXVI | 645 | 478 | 436 | 552 | 399 | 422 | 620 | 549 | 610 | 1587 | 448 | 373 | 2011 | 301 | 306 | 1866 | 400 | 353 |
| 5311 | W303.asm02.HP0_G0053090 | chrXVI | 37638 | 11263 | 6762 | 36056 | 6997 | 8403 | 33516 | 7258 | 5143 | 24988 | 10600 | 12051 | 15947 | 7347 | 10676 | 22495 | 11228 | 10564 |
| 5312 | W303.asm02.HP0_G0053100 | chrXVI | 2424 | 2774 | 2397 | 2677 | 2669 | 2731 | 2355 | 2776 | 2429 | 7497 | 2436 | 2310 | 7515 | 1551 | 1477 | 7413 | 2178 | 2144 |
| 5313 | W303.asm02.HP0_G0053110 | chrXVI | 7071 | 5744 | 6642 | 6927 | 6660 | 6625 | 6914 | 6269 | 7204 | 17514 | 5756 | 6305 | 17276 | 3869 | 4673 | 16754 | 4803 | 5656 |
| 5314 | W303.asm02.HP0_G0053120 | chrXVI | 1475 | 450 | 360 | 1475 | 360 | 377 | 1460 | 447 | 378 | 2057 | 452 | 496 | 1931 | 284 | 406 | 1805 | 455 | 462 |
| 5315 | W303.asm02.HP0_G0053130 | chrXVI | 2754 | 1289 | 1100 | 2617 | 1074 | 985 | 2814 | 1242 | 1143 | 5612 | 1628 | 1622 | 4909 | 1091 | 1241 | 4927 | 1503 | 1448 |
| 5316 | W303.asm02.HP0_G0053140 | chrXVI | 3476 | 1929 | 1875 | 2870 | 1737 | 1753 | 3079 | 2046 | 2132 | 4995 | 1813 | 1621 | 5320 | 1246 | 1441 | 5106 | 1624 | 1461 |
| 5317 | W303.asm02.HP0_G0053150 | chrXVI | 837 | 393 | 327 | 845 | 388 | 294 | 843 | 315 | 306 | 1498 | 394 | 430 | 1496 | 287 | 302 | 1211 | 425 | 404 |
| 5318 | W303.asm02.HP0_G0053160 | chrXVI | 10285 | 2898 | 2141 | 10083 | 2435 | 2203 | 9710 | 2773 | 2079 | 8554 | 2937 | 3230 | 9427 | 2622 | 2790 | 6995 | 2529 | 2753 |
| 5319 | W303.asm02.HP0_G0053170 | chrXVI | 68145 | 5413 | 5037 | 65542 | 4694 | 5937 | 70810 | 4654 | 6829 | 38093 | 10951 | 16380 | 16776 | 11432 | 11024 | 12547 | 8311 | 13263 |
| 5320 | W303.asm02.HP0_G0053180 | chrXVI | 4367 | 285 | 168 | 4375 | 168 | 159 | 4099 | 264 | 219 | 756 | 350 | 229 | 1029 | 246 | 332 | 572 | 348 | 249 |
| 5321 | W303.asm02.HP0_G0053190 | chrXVI | 809 | 810 | 625 | 871 | 644 | 644 | 761 | 740 | 624 | 1928 | 732 | 756 | 1751 | 447 | 426 | 1577 | 740 | 622 |
| 5322 | W303.asm02.HP0_G0053200 | chrXVI | 1983 | 982 | 857 | 2007 | 822 | 784 | 1693 | 910 | 841 | 5334 | 1096 | 1374 | 4364 | 590 | 991 | 4974 | 1133 | 1217 |
| 5323 | W303.asm02.HP0_G0053210 | chrXVI | 1068 | 1937 | 2677 | 1030 | 2358 | 2515 | 1165 | 2100 | 3254 | 6073 | 1470 | 1672 | 7277 | 926 | 1325 | 6636 | 1277 | 1476 |
| 5324 | W303.asm02.HP0_G0053220 | chrXVI | 4354 | 2087 | 2763 | 3832 | 2811 | 2893 | 4940 | 1920 | 3482 | 29970 | 4754 | 5848 | 16752 | 3305 | 3351 | 15474 | 4119 | 5081 |
| 5325 | W303.asm02.HP0_G0053230 | chrXVI | 6308 | 3859 | 4023 | 5744 | 3513 | 4077 | 5824 | 3919 | 5308 | 8653 | 3429 | 3555 | 10235 | 2405 | 3115 | 9619 | 3153 | 3226 |
| 5326 | W303.asm02.HP0_G0053240 | chrXVI | 1869 | 723 | 692 | 1807 | 749 | 720 | 1787 | 669 | 743 | 3480 | 821 | 950 | 2878 | 459 | 737 | 2952 | 781 | 881 |
| 5327 | W303.asm02.HP0_G0053250 | chrXVI | 464 | 182 | 117 | 314 | 115 | 145 | 401 | 165 | 169 | 268 | 149 | 164 | 318 | 119 | 112 | 263 | 219 | 153 |
| 5328 | W303.asm02.HP0_G0053260 | chrXVI | 2088 | 281 | 221 | 1673 | 176 | 262 | 1809 | 253 | 278 | 1034 | 499 | 525 | 773 | 436 | 436 | 549 | 505 | 516 |
| 5329 | W303.asm02.HP0_G0053270 | chrXVI | 1816 | 844 | 1325 | 1732 | 1114 | 1364 | 1678 | 744 | 1596 | 7283 | 1565 | 2602 | 5745 | 1282 | 1735 | 3737 | 1439 | 2090 |
| 5330 | W303.asm02.HP0_G0053280 | chrXVI | 7995 | 5357 | 7153 | 6525 | 6433 | 6982 | 8370 | 5210 | 9017 | 16506 | 5181 | 6100 | 17019 | 3631 | 4573 | 16770 | 4482 | 5196 |
| 5331 | W303.asm02.HP0_G0053290 | chrXVI | 977 | 277 | 231 | 1188 | 236 | 253 | 886 | 236 | 234 | 1412 | 280 | 348 | 1288 | 157 | 297 | 1105 | 363 | 317 |
| 5332 | W303.asm02.HP0_G0053300 | chrXVI | 3530 | 3440 | 5253 | 3204 | 4020 | 5200 | 3494 | 3491 | 6655 | 9150 | 2739 | 4307 | 10202 | 1950 | 3111 | 9996 | 2488 | 3838 |
| 5333 | W303.asm02.HP0_G0053310 | chrXVI | 3909 | 1003 | 1206 | 3661 | 1571 | 1226 | 4273 | 1029 | 1581 | 15491 | 2241 | 2996 | 8382 | 1522 | 1720 | 8244 | 1840 | 2629 |
| 5334 | W303.asm02.HP0_G0053320 | chrXVI | 1590 | 747 | 926 | 1440 | 914 | 902 | 1518 | 826 | 1047 | 3559 | 657 | 878 | 3530 | 433 | 589 | 3628 | 600 | 786 |
| 5335 | W303.asm02.HP0_G0053330 | chrXVI | 5920 | 1798 | 1718 | 5878 | 1557 | 1768 | 5612 | 1808 | 1775 | 6086 | 1585 | 2312 | 6590 | 1215 | 1756 | 5529 | 1530 | 1967 |
| 5336 | W303.asm02.HP0_G0053340 | chrXVI | 1337 | 845 | 862 | 1253 | 787 | 839 | 1265 | 783 | 880 | 2804 | 876 | 911 | 2835 | 548 | 792 | 2701 | 795 | 856 |
| 5337 | W303.asm02.HP0_G0053350 | chrXVI | 1120 | 1031 | 1229 | 1023 | 1172 | 1200 | 1043 | 1059 | 1279 | 2815 | 769 | 934 | 3630 | 488 | 724 | 2832 | 691 | 869 |
| 5338 | W303.asm02.HP0_G0053360 | chrXVI | 1809 | 2145 | 1740 | 1695 | 2008 | 1665 | 2016 | 2185 | 1882 | 11436 | 2962 | 3069 | 8973 | 1965 | 1889 | 8980 | 2508 | 2596 |
| 5339 | W303.asm02.HP0_G0053370 | chrXVI | 1153 | 1693 | 2599 | 1052 | 2658 | 2353 | 1173 | 1876 | 3421 | 7275 | 1762 | 1999 | 7481 | 1021 | 1326 | 8244 | 1488 | 1776 |
| 5340 | W303.asm02.HP0_G0053380 | chrXVI | 8015 | 4176 | 4672 | 8414 | 4616 | 4182 | 8265 | 4465 | 4686 | 17363 | 3749 | 4540 | 17277 | 2681 | 3573 | 17382 | 3520 | 4266 |
| 5341 | W303.asm02.HP0_G0053390 | chrXVI | 219 | 140 | 130 | 219 | 143 | 131 | 240 | 147 | 106 | 727 | 171 | 266 | 716 | 104 | 162 | 690 | 143 | 205 |
| 5342 | W303.asm02.HP0_G0053400 | chrXVI | 1699 | 390 | 434 | 1398 | 390 | 437 | 1766 | 322 | 556 | 3414 | 602 | 863 | 2503 | 434 | 616 | 2532 | 605 | 838 |
| 5343 | W303.asm02.HP0_G0053410 | chrXVI | 4311 | 2022 | 1765 | 4325 | 2116 | 1582 | 4725 | 2081 | 2039 | 16690 | 3590 | 3349 | 11343 | 2183 | 1990 | 12192 | 2936 | 2986 |
| 5344 | W303.asm02.HP0_G0053420 | chrXVI | 3572 | 2529 | 1937 | 3163 | 1741 | 1891 | 3269 | 2312 | 2025 | 4522 | 1900 | 1786 | 4655 | 1459 | 1555 | 4150 | 1926 | 1600 |
| 5345 | W303.asm02.HP0_G0053430 | chrXVI | 206 | 51 | 31 | 145 | 28 | 42 | 142 | 37 | 41 | 258 | 58 | 65 | 229 | 52 | 45 | 223 | 65 | 59 |
| 5346 | W303.asm02.HP0_G0053440 | chrXVI | 5136 | 1368 | 1425 | 5083 | 1389 | 1628 | 5397 | 1203 | 1641 | 10095 | 2598 | 3317 | 7543 | 1818 | 2171 | 6629 | 2315 | 2730 |
| 5347 | W303.asm02.HP0_G0053450 | chrXVI | 1969 | 341 | 580 | 1959 | 509 | 599 | 1851 | 366 | 604 | 3881 | 565 | 1020 | 3797 | 451 | 722 | 3301 | 584 | 812 |
| 5348 | W303.asm02.HP0_G0053460 | chrXVI | 304 | 42 | 45 | 304 | 62 | 56 | 262 | 36 | 45 | 362 | 50 | 47 | 280 | 34 | 31 | 351 | 44 | 44 |
| 5349 | W303.asm02.HP0_G0053470 | chrXVI | 31 | 6 | 11 | 31 | 15 | 11 | 39 | 10 | 14 | 106 | 12 | 7 | 94 | 5 | 9 | 85 | 17 | 11 |
| 5350 | W303.asm02.HP0_G0053480 | chrXVI | 3264 | 1651 | 1573 | 3021 | 1607 | 1516 | 3132 | 1726 | 1526 | 5119 | 1394 | 1528 | 4891 | 970 | 1112 | 4991 | 1266 | 1376 |
| 5351 | W303.asm02.HP0_G0053490 | chrXVI | 3712 | 4483 | 4410 | 3452 | 4820 | 4696 | 4058 | 4622 | 4882 | 12153 | 3913 | 4058 | 14567 | 3213 | 3540 | 12053 | 3590 | 4034 |
| 5352 | W303.asm02.HP0_G0053500 | chrXVI | 4146 | 917 | 680 | 3740 | 628 | 658 | 3592 | 826 | 767 | 2262 | 756 | 698 | 2566 | 486 | 622 | 2226 | 683 | 550 |
| 5353 | W303.asm02.HP0_G0053510 | chrXVI | 5556 | 1367 | 1500 | 5269 | 1653 | 1532 | 5353 | 1480 | 1675 | 9164 | 1531 | 2214 | 7779 | 1024 | 1751 | 8334 | 1416 | 1947 |
| 5354 | W303.asm02.HP0_G0053520 | chrXVI | 966 | 221 | 326 | 1022 | 312 | 341 | 903 | 257 | 326 | 2118 | 328 | 509 | 1759 | 221 | 389 | 1551 | 271 | 451 |
| 5355 | W303.asm02.HP0_G0053530 | chrXVI | 1461 | 616 | 552 | 1531 | 728 | 497 | 1548 | 744 | 591 | 4249 | 1021 | 903 | 3493 | 635 | 553 | 3649 | 804 | 798 |
| 5356 | W303.asm02.HP0_G0053540 | chrXVI | 194 | 91 | 87 | 166 | 99 | 86 | 199 | 120 | 90 | 399 | 96 | 91 | 334 | 75 | 96 | 419 | 102 | 80 |
| 5357 | W303.asm02.HP0_G0053550 | chrXVI | 1475 | 539 | 717 | 1305 | 489 | 719 | 1386 | 546 | 804 | 1752 | 422 | 600 | 2159 | 288 | 556 | 1862 | 382 | 533 |
| 5358 | W303.asm02.HP0_G0053560 | chrXVI | 7632 | 940 | 975 | 7371 | 930 | 1253 | 7860 | 736 | 1348 | 4365 | 1597 | 2189 | 1668 | 1242 | 957 | 1471 | 1168 | 1713 |
| 5359 | W303.asm02.HP0_G0053570 | chrXVI | 890 | 872 | 823 | 825 | 804 | 806 | 806 | 819 | 1126 | 1558 | 684 | 547 | 2034 | 477 | 463 | 1537 | 621 | 449 |
| 5360 | W303.asm02.HP0_G0053580 | chrXVI | 363 | 81 | 97 | 270 | 107 | 94 | 376 | 78 | 93 | 591 | 84 | 120 | 498 | 40 | 60 | 466 | 84 | 86 |
| 5361 | W303.asm02.HP0_G0053590 | chrXVI | 3859 | 1029 | 812 | 3071 | 748 | 782 | 3525 | 1170 | 861 | 4177 | 1367 | 1248 | 4776 | 1015 | 1112 | 3921 | 1141 | 1166 |
| 5362 | W303.asm02.HP0_G0053600 | chrXVI | 38 | 22 | 18 | 40 | 22 | 32 | 41 | 21 | 20 | 175 | 76 | 57 | 206 | 58 | 60 | 194 | 58 | 58 |
| 5363 | W303.asm02.HP0_G0053610 | chrXVI | 4037 | 277 | 213 | 4239 | 206 | 191 | 3593 | 262 | 236 | 519 | 247 | 185 | 790 | 183 | 200 | 452 | 179 | 166 |
| 5364 | W303.asm02.HP0_G0053620 | chrXVI | 3874 | 794 | 1134 | 3402 | 908 | 1169 | 3938 | 855 | 1303 | 7338 | 1560 | 2413 | 5354 | 1096 | 1789 | 4851 | 1323 | 2196 |
| 5365 | W303.asm02.HP0_G0053630 | chrXVI | 1348 | 701 | 876 | 1106 | 753 | 875 | 991 | 770 | 932 | 2122 | 657 | 1019 | 2507 | 519 | 812 | 1934 | 612 | 854 |
| 5366 | W303.asm02.HP0_G0053640 | chrXVI | 2104 | 1044 | 1437 | 2009 | 1672 | 1268 | 2417 | 1066 | 1621 | 13884 | 1556 | 2167 | 10321 | 996 | 1307 | 11199 | 1319 | 1935 |
| 5367 | W303.asm02.HP0_G0053650 | chrXVI | 6522 | 1230 | 1265 | 6620 | 1291 | 1185 | 6733 | 1213 | 1371 | 8131 | 2077 | 2171 | 6684 | 1448 | 1755 | 5792 | 1810 | 1929 |
| 5368 | W303.asm02.HP0_G0053660 | chrXVI | 2477 | 313 | 323 | 2454 | 323 | 422 | 2385 | 286 | 390 | 3550 | 617 | 1051 | 1637 | 483 | 631 | 1809 | 571 | 996 |
| 5369 | W303.asm02.HP0_G0053670 | chrXVI | 6394 | 3116 | 3245 | 6784 | 3146 | 3154 | 6572 | 3263 | 3283 | 14167 | 3860 | 4805 | 13677 | 2726 | 3434 | 12245 | 3356 | 4187 |
| 5370 | W303.asm02.HP0_G0053680 | chrXVI | 3272 | 2440 | 2633 | 3023 | 2324 | 2628 | 2998 | 2293 | 3264 | 5855 | 1881 | 2050 | 6824 | 1366 | 1639 | 6413 | 1737 | 1788 |
| 5371 | W303.asm02.HP0_G0053690 | chrXVI | 2267 | 954 | 994 | 2235 | 856 | 1079 | 1992 | 944 | 1206 | 5100 | 1897 | 2038 | 4156 | 1215 | 1503 | 3822 | 1535 | 1716 |
| 5372 | W303.asm02.HP0_G0053700 | chrXVI | 3780 | 761 | 847 | 3357 | 713 | 789 | 3605 | 742 | 954 | 5646 | 1486 | 1647 | 5182 | 1287 | 1457 | 3652 | 1295 | 1487 |
| 5373 | W303.asm02.HP0_G0053710 | chrXVI | 2919 | 777 | 883 | 2654 | 952 | 814 | 2818 | 798 | 845 | 4117 | 716 | 1025 | 5478 | 659 | 887 | 3554 | 645 | 943 |
| 5374 | W303.asm02.HP0_G0053720 | chrXVI | 981 | 364 | 686 | 946 | 618 | 590 | 924 | 401 | 742 | 3797 | 432 | 803 | 3215 | 319 | 633 | 3362 | 354 | 733 |
| 5375 | W303.asm02.HP0_G0053730 | chrXVI | 7406 | 1518 | 1750 | 7504 | 1706 | 1825 | 7288 | 1528 | 1907 | 6255 | 1424 | 2190 | 6970 | 1140 | 1813 | 6037 | 1430 | 1976 |
| 5376 | W303.asm02.HP0_G0053740 | chrXVI | 2844 | 826 | 1051 | 2825 | 976 | 1022 | 2719 | 800 | 1066 | 5421 | 1141 | 1800 | 5720 | 838 | 1413 | 4512 | 1054 | 1629 |
| 5377 | W303.asm02.HP0_G0053750 | chrXVI | 1689 | 455 | 667 | 1674 | 446 | 639 | 1476 | 454 | 756 | 513 | 1236 | 884 | 813 | 1387 | 1270 | 533 | 913 | 576 |
| 5378 | W303.asm02.HP0_G0053760 | chrXVI | 4668 | 1559 | 1882 | 4360 | 1796 | 2208 | 3871 | 1575 | 2183 | 4183 | 1370 | 1696 | 6040 | 1223 | 1866 | 4252 | 1279 | 1592 |
| 5379 | W303.asm02.HP0_G0053770 | chrXVI | 5138 | 1318 | 1207 | 5091 | 1218 | 1297 | 5210 | 1339 | 1360 | 10108 | 2464 | 2751 | 7570 | 1592 | 1936 | 7463 | 2103 | 2560 |
| 5380 | W303.asm02.HP0_G0053780 | chrXVI | 1659 | 571 | 691 | 1731 | 665 | 713 | 1491 | 522 | 722 | 2155 | 453 | 622 | 2600 | 289 | 536 | 2256 | 401 | 554 |
| 5381 | W303.asm02.HP0_G0053790 | chrXVI | 1214 | 236 | 468 | 1123 | 473 | 444 | 1092 | 243 | 530 | 3021 | 289 | 652 | 3027 | 173 | 540 | 2709 | 258 | 591 |
| 5382 | W303.asm02.HP0_G0053800 | chrXVI | 1383 | 175 | 189 | 1479 | 146 | 183 | 1238 | 133 | 172 | 825 | 210 | 221 | 1081 | 160 | 229 | 706 | 185 | 171 |
| 5383 | W303.asm02.HP0_G0053810 | chrXVI | 1771 | 231 | 231 | 1694 | 262 | 206 | 1612 | 187 | 206 | 1581 | 329 | 406 | 1264 | 248 | 348 | 1166 | 301 | 411 |
| 5384 | W303.asm02.HP0_G0053820 | chrXVI | 6124 | 857 | 1422 | 6034 | 1129 | 1434 | 7057 | 1085 | 1442 | 7509 | 1931 | 3054 | 9198 | 1530 | 2028 | 6298 | 1410 | 2518 |
| 5385 | W303.asm02.HP0_G0053830 | chrXVI | 967 | 574 | 594 | 1051 | 604 | 612 | 924 | 509 | 647 | 3082 | 559 | 817 | 2974 | 414 | 562 | 2509 | 524 | 653 |
| 5386 | W303.asm02.HP0_G0053840 | chrXVI | 1593 | 1071 | 1158 | 1662 | 973 | 1090 | 1583 | 1147 | 1054 | 4559 | 1006 | 1372 | 4167 | 611 | 963 | 4445 | 985 | 1203 |
| 5387 | W303.asm02.HP0_G0053850 | chrXVI | 13676 | 4465 | 4109 | 12857 | 4623 | 4300 | 14153 | 4560 | 4668 | 30690 | 6140 | 7642 | 22789 | 4437 | 5332 | 23368 | 5260 | 7016 |
| 5388 | W303.asm02.HP0_G0053860 | chrXVI | 3061 | 1166 | 1165 | 3233 | 1204 | 1241 | 3227 | 1102 | 1058 | 4465 | 1050 | 1423 | 4477 | 678 | 918 | 3920 | 1080 | 1254 |
| 5389 | W303.asm02.HP0_G0053870 | chrXVI | 2291 | 759 | 757 | 2600 | 857 | 867 | 2659 | 728 | 927 | 6280 | 1174 | 1559 | 5577 | 930 | 1293 | 5193 | 1013 | 1560 |
| 5390 | W303.asm02.HP0_G0053880 | chrXVI | 891 | 527 | 669 | 905 | 783 | 640 | 877 | 553 | 757 | 3789 | 701 | 933 | 3387 | 515 | 619 | 3290 | 628 | 715 |
| 5391 | W303.asm02.HP0_G0053890 | chrXVI | 1345 | 249 | 238 | 1395 | 228 | 290 | 1194 | 258 | 265 | 719 | 232 | 228 | 717 | 132 | 170 | 698 | 205 | 209 |
| 5392 | W303.asm02.HP0_G0053900 | chrXVI | 1616 | 344 | 298 | 1539 | 346 | 367 | 1749 | 294 | 401 | 2719 | 669 | 864 | 1460 | 579 | 624 | 1407 | 574 | 720 |

| | | | | | | | | | | | | | | | | | | | | |
|---|---|---|---|---|---|---|---|---|---|---|---|---|---|---|---|---|---|---|---|---|
| 5393 | W303.asm02.HP0_G0053910 | chrXVI | 17023 | 3017 | 2893 | 16351 | 2563 | 3195 | 16224 | 2982 | 3399 | 13076 | 4980 | 6324 | 14812 | 4365 | 6132 | 10342 | 4564 | 5838 |
| 5394 | W303.asm02.HP0_G0053920 | chrXVI | 633 | 150 | 237 | 564 | 226 | 265 | 654 | 119 | 273 | 1736 | 312 | 580 | 718 | 218 | 402 | 826 | 253 | 521 |
| 5395 | W303.asm02.HP0_G0053930 | chrXVI | 2246 | 610 | 458 | 1904 | 393 | 441 | 2017 | 561 | 472 | 1730 | 533 | 585 | 1951 | 386 | 477 | 1806 | 497 | 534 |
| 5396 | W303.asm02.HP0_G0053940 | chrXVI | 1968 | 497 | 503 | 1967 | 716 | 686 | 1857 | 376 | 621 | 2439 | 725 | 922 | 1146 | 532 | 598 | 1310 | 703 | 770 |
| 5397 | W303.asm02.HP0_G0053950 | chrXVI | 2651 | 519 | 607 | 2707 | 629 | 634 | 2534 | 490 | 679 | 4004 | 817 | 1346 | 3193 | 588 | 1021 | 3135 | 812 | 1273 |
| 5398 | W303.asm02.HP0_G0053960 | chrXVI | 1402 | 661 | 801 | 1530 | 769 | 739 | 1189 | 689 | 875 | 1832 | 475 | 460 | 1866 | 291 | 342 | 2050 | 449 | 438 |
| 5399 | W303.asm02.HP0_G0053970 | chrXVI | 330 | 281 | 264 | 336 | 268 | 286 | 349 | 280 | 315 | 820 | 250 | 173 | 769 | 135 | 159 | 887 | 231 | 166 |
| 5400 | W303.asm02.HP0_G0053980 | chrXVI | 254 | 6 | 5 | 298 | 15 | 6 | 188 | 9 | 6 | 42 | 21 | 8 | 57 | 14 | 9 | 42 | 10 | 7 |
| 5401 | W303.asm02.HP0_G0053990 | chrXVI | 1488 | 486 | 711 | 1528 | 491 | 632 | 1268 | 513 | 741 | 1889 | 458 | 513 | 2033 | 258 | 427 | 1796 | 441 | 432 |
| 5402 | W303.asm02.HP0_G0054000 | chrXVI | 2286 | 2284 | 3548 | 2678 | 3777 | 3212 | 2809 | 2677 | 4204 | 13433 | 2210 | 2787 | 13570 | 1491 | 1744 | 13679 | 1901 | 2254 |
| 5403 | W303.asm02.HP0_G0054010 | chrXVI | 14093 | 4888 | 4570 | 14492 | 4528 | 4312 | 15498 | 5504 | 4354 | 26134 | 6682 | 6965 | 25910 | 4613 | 5680 | 25109 | 5747 | 6690 |
| 5404 | W303.asm02.HP0_G0054020 | chrXVI | 605 | 581 | 612 | 486 | 585 | 585 | 532 | 647 | 801 | 1121 | 473 | 479 | 1364 | 317 | 420 | 1295 | 437 | 416 |
| 5405 | W303.asm02.HP0_G0054030 | chrXVI | 6876 | 5709 | 8043 | 6233 | 7456 | 7941 | 6455 | 5917 | 9675 | 15003 | 5430 | 6310 | 16591 | 3962 | 5474 | 16314 | 4957 | 5727 |
| 5406 | W303.asm02.HP0_G0054040 | chrXVI | 2093 | 490 | 673 | 2152 | 681 | 805 | 2262 | 402 | 864 | 6399 | 1127 | 1720 | 5248 | 1031 | 1492 | 4317 | 940 | 1685 |
| 5407 | W303.asm02.HP0_G0054050 | chrXVI | 3857 | 565 | 530 | 3332 | 585 | 563 | 3723 | 608 | 674 | 4461 | 1012 | 1127 | 3183 | 738 | 916 | 3002 | 984 | 992 |
| 5408 | W303.asm02.HP0_G0054060 | chrXVI | 2288 | 1086 | 929 | 2006 | 926 | 847 | 2121 | 1017 | 905 | 4922 | 1154 | 1348 | 3947 | 691 | 987 | 4289 | 1085 | 1196 |
| 5409 | W303.asm02.HP0_G0054070 | chrXVI | 1215 | 587 | 710 | 1267 | 689 | 729 | 1334 | 607 | 759 | 2675 | 619 | 774 | 2829 | 383 | 582 | 2566 | 528 | 698 |
| 5410 | W303.asm02.HP0_G0054080 | chrXVI | 1595 | 642 | 1427 | 1711 | 1087 | 1495 | 1545 | 678 | 1908 | 9159 | 1505 | 2929 | 7241 | 1000 | 2023 | 5916 | 1359 | 2779 |
| 5411 | W303.asm02.HP0_G0054090 | chrXVI | 2256 | 3561 | 3420 | 2110 | 3096 | 3036 | 1974 | 3873 | 3348 | 10298 | 5012 | 4951 | 12825 | 3530 | 4230 | 9760 | 4424 | 4758 |
| 5412 | W303.asm02.HP0_G0054100 | chrXVI | 1760 | 462 | 369 | 1698 | 452 | 415 | 1673 | 501 | 448 | 1751 | 771 | 725 | 1768 | 640 | 619 | 1309 | 714 | 602 |
| 5413 | W303.asm02.HP0_G0054110 | chrXVI | 1360 | 330 | 186 | 1216 | 305 | 252 | 1237 | 236 | 196 | 1505 | 426 | 354 | 1184 | 270 | 282 | 1101 | 353 | 369 |
| 5414 | W303.asm02.HP0_G0054120 | chrXVI | 1738 | 1011 | 1212 | 1661 | 1200 | 1261 | 1557 | 971 | 1328 | 3460 | 808 | 1324 | 4110 | 590 | 1025 | 3232 | 775 | 1103 |
| 5415 | W303.asm02.HP0_G0054130 | chrXVI | 32637 | 15483 | 11895 | 30584 | 12656 | 11652 | 34103 | 14847 | 11685 | 46705 | 18453 | 19842 | 40915 | 15105 | 14077 | 39864 | 15817 | 17497 |
| 5416 | W303.asm02.HP0_G0054140 | chrXVI | 35 | 37 | 4 | 30 | 9 | 8 | 42 | 32 | 16 | 74 | 46 | 5 | 56 | 20 | 4 | 90 | 46 | 7 |
| 5417 | W303.asm02.HP0_G0054150 | chrXVI | 5755 | 2015 | 1798 | 5548 | 1775 | 2028 | 5622 | 2049 | 1632 | 6876 | 1879 | 2600 | 6038 | 1348 | 1833 | 5970 | 1624 | 2364 |
| 5418 | W303.asm02.HP0_G0054160 | chrXVI | 778 | 323 | 482 | 696 | 475 | 530 | 802 | 287 | 635 | 3232 | 576 | 927 | 1638 | 388 | 651 | 1647 | 520 | 836 |
| 5419 | W303.asm02.HP0_G0054170 | chrXVI | 6964 | 1054 | 1089 | 6557 | 973 | 1041 | 6731 | 1075 | 1185 | 5809 | 1917 | 2308 | 4399 | 1375 | 2039 | 3949 | 1564 | 1988 |
| 5420 | W303.asm02.HP0_G0054180 | chrXVI | 2656 | 1751 | 2115 | 2734 | 2563 | 1938 | 3172 | 1869 | 2333 | 19703 | 2698 | 3916 | 15269 | 1893 | 1953 | 14797 | 2399 | 3380 |
| 5421 | W303.asm02.HP0_G0054190 | chrXVI | 3416 | 1127 | 1182 | 3200 | 1186 | 1110 | 3478 | 1199 | 1474 | 9616 | 1521 | 1802 | 6967 | 1047 | 1439 | 7856 | 1424 | 1770 |
| 5422 | W303.asm02.HP0_G0054200 | chrXVI | 567 | 287 | 248 | 583 | 230 | 247 | 603 | 292 | 201 | 745 | 239 | 259 | 821 | 161 | 218 | 762 | 179 | 223 |
| 5423 | W303.asm02.HP0_G0054210 | chrXVI | 5142 | 998 | 1002 | 4884 | 760 | 997 | 4343 | 917 | 1194 | 1668 | 605 | 595 | 2802 | 488 | 727 | 1743 | 622 | 565 |
| 5424 | W303.asm02.HP0_G0054220 | chrXVI | 1119 | 662 | 774 | 1040 | 770 | 689 | 1099 | 647 | 805 | 4073 | 736 | 863 | 3974 | 434 | 625 | 4120 | 692 | 745 |
| 5425 | W303.asm02.HP0_G0054230 | chrXVI | 34 | 21 | 9 | 29 | 13 | 19 | 35 | 19 | 16 | 62 | 25 | 11 | 78 | 14 | 13 | 54 | 9 | 16 |
| 5426 | W303.asm02.HP0_G0054240 | chrXVI | 1487 | 630 | 597 | 1490 | 594 | 613 | 1490 | 643 | 613 | 2592 | 610 | 780 | 2512 | 425 | 609 | 2411 | 479 | 744 |
| 5427 | W303.asm02.HP0_G0054250 | chrXVI | 98 | 38 | 37 | 61 | 28 | 42 | 85 | 32 | 39 | 126 | 31 | 54 | 127 | 30 | 44 | 110 | 35 | 49 |
| 5428 | W303.asm02.HP0_G0054260 | chrXVI | 1186 | 139 | 157 | 1198 | 133 | 146 | 987 | 128 | 135 | 638 | 233 | 167 | 407 | 152 | 148 | 385 | 173 | 151 |
| 5429 | W303.asm02.HP0_G0054270 | chrXVI | 12007 | 3305 | 3851 | 11706 | 3770 | 3741 | 11582 | 3590 | 4341 | 11710 | 3172 | 3957 | 13830 | 2440 | 3519 | 11459 | 2813 | 3676 |
| 5430 | W303.asm02.HP0_G0054280 | chrXVI | 5319 | 5806 | 6917 | 4977 | 6842 | 6411 | 5374 | 5737 | 8086 | 17053 | 4710 | 5452 | 17728 | 3060 | 3967 | 18356 | 4411 | 4933 |
| 5431 | W303.asm02.HP0_G0054290 | chrXVI | 1363 | 814 | 616 | 1337 | 711 | 693 | 1293 | 714 | 682 | 4284 | 1118 | 1359 | 3533 | 850 | 1292 | 3244 | 967 | 1260 |
| 5432 | W303.asm02.HP0_G0054300 | chrXVI | 2682 | 385 | 719 | 2709 | 650 | 758 | 2568 | 393 | 790 | 6165 | 764 | 1540 | 3607 | 572 | 1152 | 3649 | 665 | 1335 |
| 5433 | W303.asm02.HP0_G0054310 | chrXVI | 672 | 301 | 161 | 605 | 165 | 169 | 605 | 238 | 149 | 753 | 245 | 226 | 716 | 162 | 179 | 678 | 258 | 202 |
| 5434 | W303.asm02.HP0_G0054320 | chrXVI | 1984 | 2452 | 2186 | 1603 | 2016 | 2130 | 1823 | 2443 | 2407 | 5609 | 1834 | 1837 | 6050 | 1042 | 1230 | 6355 | 1643 | 1447 |
| 5435 | W303.asm02.HP0_G0054330 | chrXVI | 1185 | 717 | 499 | 1467 | 692 | 587 | 1127 | 692 | 683 | 4811 | 1222 | 1215 | 3597 | 868 | 826 | 3623 | 1116 | 1160 |
| 5436 | W303.asm02.HP0_G0054340 | chrXVI | 6169 | 567 | 759 | 6066 | 651 | 858 | 5942 | 479 | 1008 | 5082 | 1066 | 1955 | 5283 | 1332 | 1731 | 2570 | 950 | 1703 |
| 5437 | W303.asm02.HP0_G0054350 | chrXVI | 1240 | 500 | 480 | 1322 | 453 | 477 | 1188 | 524 | 513 | 2072 | 471 | 623 | 2037 | 326 | 522 | 1851 | 474 | 518 |
| 5438 | W303.asm02.HP0_G0054360 | chrXVI | 294 | 556 | 536 | 256 | 554 | 610 | 284 | 541 | 681 | 1363 | 390 | 469 | 1573 | 273 | 367 | 1307 | 391 | 436 |
| 5439 | W303.asm02.HP0_G0054370 | chrXVI | 704 | 967 | 865 | 691 | 932 | 925 | 683 | 1008 | 888 | 2702 | 884 | 898 | 2808 | 552 | 624 | 2619 | 805 | 730 |
| 5440 | W303.asm02.HP0_G0054380 | chrXVI | 24742 | 8657 | 8768 | 24526 | 8814 | 10164 | 25231 | 6745 | 9589 | 46224 | 13386 | 20101 | 34130 | 9075 | 12716 | 36317 | 12050 | 17769 |
| 5441 | W303.asm02.HP0_G0054390 | chrXVI | 7968 | 831 | 1576 | 7721 | 1562 | 1834 | 8047 | 743 | 1988 | 14118 | 1864 | 3533 | 9214 | 1209 | 2603 | 8280 | 1629 | 3163 |
| 5442 | W303.asm02.HP0_G0054400 | chrXVI | 5927 | 580 | 579 | 5722 | 561 | 665 | 5558 | 472 | 692 | 5060 | 1074 | 1616 | 3434 | 834 | 1062 | 3439 | 985 | 1243 |
| 5443 | W303.asm02.HP0_G0054410 | chrXVI | 3823 | 631 | 653 | 3702 | 553 | 616 | 3713 | 621 | 663 | 3864 | 849 | 1124 | 3230 | 604 | 827 | 2984 | 790 | 1054 |
| 5444 | W303.asm02.HP0_G0054420 | chrXVI | 2070 | 596 | 637 | 2260 | 607 | 715 | 2242 | 555 | 841 | 5442 | 1224 | 1517 | 3399 | 818 | 927 | 3517 | 1134 | 1389 |
| 5445 | W303.asm02.HP0_G0054430 | chrXVI | 3521 | 561 | 397 | 3504 | 345 | 396 | 3170 | 419 | 424 | 2475 | 620 | 766 | 1880 | 413 | 605 | 1810 | 685 | 637 |
| 5446 | W303.asm02.HP0_G0054440 | chrXVI | 1617 | 757 | 789 | 1603 | 611 | 788 | 1462 | 713 | 964 | 2153 | 554 | 668 | 2562 | 391 | 559 | 2331 | 546 | 597 |
| 5447 | W303.asm02.HP0_G0054450 | chrXVI | 3024 | 1482 | 1376 | 2524 | 1241 | 1388 | 2664 | 1514 | 1484 | 3729 | 1395 | 1496 | 3907 | 912 | 1203 | 3737 | 1214 | 1333 |
| 5448 | W303.asm02.HP0_G0054460 | chrXVI | 3685 | 1266 | 1396 | 3574 | 1361 | 1244 | 3292 | 1348 | 1513 | 5688 | 1355 | 1706 | 6443 | 919 | 1422 | 5214 | 1340 | 1569 |
| 5449 | W303.asm02.HP0_G0054470 | chrXVI | 510 | 229 | 275 | 388 | 301 | 292 | 461 | 279 | 338 | 681 | 148 | 204 | 1031 | 120 | 196 | 822 | 172 | 186 |
| 5450 | W303.asm02.HP0_G0054480 | chrXVI | 2040 | 1011 | 931 | 1838 | 866 | 868 | 1713 | 902 | 1066 | 2841 | 724 | 918 | 3151 | 477 | 697 | 2939 | 758 | 750 |
| 5451 | W303.asm02.HP0_G0054490 | chrXVI | 898 | 1068 | 915 | 797 | 1020 | 1047 | 828 | 991 | 1060 | 3075 | 814 | 708 | 2905 | 483 | 546 | 2728 | 836 | 605 |
| 5452 | W303.asm02.HP0_G0054500 | chrXVI | 4098 | 3277 | 5554 | 3653 | 3742 | 5911 | 4016 | 3555 | 6351 | 8829 | 2895 | 5271 | 10133 | 1958 | 3628 | 8717 | 2577 | 4574 |
| 5453 | W303.asm02.HP0_G0054510 | chrXVI | 3671 | 2354 | 3573 | 3894 | 4069 | 3494 | 4739 | 2679 | 3803 | 25582 | 3385 | 6833 | 22130 | 2392 | 3348 | 20016 | 3042 | 5794 |
| 5454 | W303.asm02.HP0_G0054520 | chrXVI | 496 | 682 | 591 | 507 | 535 | 529 | 468 | 638 | 556 | 1927 | 764 | 676 | 1822 | 516 | 521 | 1638 | 656 | 547 |
| 5455 | W303.asm02.HP0_G0054530 | chrXVI | 8056 | 3487 | 2872 | 7697 | 2515 | 2653 | 7878 | 3214 | 3274 | 10745 | 4833 | 4919 | 8533 | 3436 | 4028 | 8867 | 4301 | 4095 |
| 5456 | W303.asm02.HP0_G0054540 | chrXVI | 13404 | 10923 | 13052 | 14052 | 12959 | 12689 | 15576 | 10887 | 16436 | 32016 | 10091 | 10543 | 32970 | 7384 | 8234 | 31885 | 8752 | 9678 |
| 5457 | W303.asm02.HP0_G0054550 | chrXVI | 4424 | 398 | 653 | 4568 | 620 | 719 | 4536 | 365 | 803 | 4363 | 625 | 1266 | 1615 | 609 | 824 | 1155 | 558 | 1021 |
| 5458 | W303.asm02.HP0_G0054560 | chrXVI | 3399 | 706 | 784 | 3455 | 652 | 739 | 3171 | 688 | 793 | 2373 | 792 | 1024 | 6412 | 982 | 1027 | 1941 | 679 | 801 |
| 5459 | W303.asm02.HP0_G0054570 | chrXVI | 7198 | 1273 | 1366 | 6255 | 988 | 1369 | 6101 | 1093 | 1541 | 3540 | 1105 | 1500 | 4148 | 826 | 1568 | 3454 | 1186 | 1354 |
| 5460 | W303.asm02.HP0_G0054580 | chrXVI | 2581 | 1368 | 921 | 2544 | 1286 | 883 | 2880 | 1338 | 1028 | 10122 | 2285 | 2231 | 6433 | 1448 | 1258 | 7743 | 2121 | 1991 |
| 5461 | W303.asm02.HP0_G0054590 | chrXVI | 5372 | 328 | 704 | 4979 | 554 | 955 | 5420 | 308 | 888 | 2118 | 505 | 2260 | 1178 | 527 | 1088 | 1013 | 345 | 1687 |
| 5462 | W303.asm02.HP0_G0054600 | chrXVI | 6378 | 683 | 637 | 5988 | 615 | 744 | 6086 | 569 | 897 | 8037 | 1438 | 1789 | 4402 | 990 | 1352 | 4762 | 1235 | 1564 |
| 5463 | W303.asm02.HP0_G0054610 | chrXVI | 1412 | 460 | 510 | 1338 | 570 | 494 | 1312 | 493 | 471 | 3244 | 589 | 783 | 2672 | 409 | 555 | 2697 | 505 | 718 |
| 5464 | W303.asm02.HP0_G0054620 | chrXVI | 1932 | 425 | 782 | 1811 | 827 | 860 | 1960 | 370 | 958 | 7245 | 724 | 1931 | 3686 | 589 | 1166 | 3525 | 676 | 1540 |
| 5465 | W303.asm02.HP0_G0054630 | chrXVI | 1881 | 4420 | 4564 | 1543 | 4529 | 4930 | 1901 | 4365 | 4619 | 12448 | 4030 | 5992 | 12885 | 3415 | 4295 | 12246 | 3537 | 5599 |
| 5466 | W303.asm02.HP0_G0054640 | chrXVI | 7359 | 6617 | 9431 | 6403 | 8997 | 9259 | 7599 | 6762 | 11255 | 19717 | 6316 | 7737 | 21451 | 4716 | 6391 | 19897 | 5666 | 7040 |
| 5467 | W303.asm02.HP0_G0054650 | chrXVI | 6200 | 6386 | 7647 | 5515 | 6774 | 7442 | 5728 | 6462 | 9694 | 15707 | 5751 | 6944 | 17398 | 4656 | 5954 | 15447 | 5024 | 5419 |
| 5468 | W303.asm02.HP0_G0054660 | chrXVI | 609 | 474 | 360 | 697 | 334 | 383 | 669 | 426 | 326 | 1192 | 407 | 402 | 1148 | 228 | 288 | 1085 | 386 | 361 |
| 5469 | W303.asm02.HP0_G0054670 | chrXVI | 1617 | 402 | 337 | 1566 | 306 | 319 | 1494 | 403 | 379 | 1694 | 406 | 439 | 1318 | 259 | 320 | 1369 | 365 | 391 |
| 5470 | W303.asm02.HP0_G0054680 | chrXVI | 4570 | 1013 | 1139 | 4316 | 1104 | 1158 | 4628 | 938 | 1376 | 7258 | 1816 | 2610 | 4587 | 1120 | 2133 | 5168 | 1677 | 2552 |
| 5471 | W303.asm02.HP0_G0054690 | chrXVI | 3402 | 694 | 606 | 3449 | 595 | 591 | 3279 | 550 | 555 | 4623 | 962 | 1121 | 3703 | 561 | 811 | 3742 | 859 | 979 |
| 5472 | W303.asm02.HP0_G0054700 | chrXVI | 1113 | 429 | 320 | 928 | 375 | 347 | 1082 | 372 | 313 | 2376 | 551 | 652 | 1575 | 400 | 510 | 1860 | 534 | 609 |
| 5473 | W303.asm02.HP0_G0054710 | chrXVI | 868 | 706 | 797 | 834 | 689 | 793 | 904 | 756 | 1012 | 1980 | 552 | 558 | 2694 | 458 | 540 | 2125 | 527 | 541 |
| 5474 | W303.asm02.HP0_G0054720 | chrXVI | 4242 | 1049 | 1424 | 4141 | 1277 | 1333 | 4147 | 1122 | 1472 | 6474 | 1112 | 1465 | 8590 | 965 | 1310 | 5940 | 997 | 1220 |
| 5475 | W303.asm02.HP0_G0054730 | chrXVI | 493 | 748 | 857 | 487 | 790 | 856 | 463 | 693 | 923 | 2884 | 608 | 708 | 3006 | 339 | 510 | 2982 | 590 | 678 |
| 5476 | W303.asm02.HP0_G0054740 | chrXVI | 423 | 167 | 128 | 392 | 166 | 159 | 436 | 168 | 124 | 1182 | 274 | 292 | 1025 | 183 | 149 | 1014 | 265 | 238 |
| 5477 | W303.asm02.HP0_G0054750 | chrXVI | 1730 | 361 | 287 | 1467 | 280 | 370 | 1527 | 350 | 361 | 1361 | 471 | 602 | 2025 | 515 | 551 | 1093 | 475 | 563 |
| 5478 | W303.asm02.HP0_G0054760 | chrXVI | 2279 | 555 | 627 | 2415 | 528 | 622 | 2235 | 451 | 689 | 3854 | 960 | 1175 | 4247 | 640 | 976 | 3392 | 891 | 1065 |
| 5479 | W303.asm02.HP0_G0054770 | chrXVI | 1348 | 786 | 847 | 1280 | 689 | 720 | 1288 | 777 | 935 | 2065 | 639 | 595 | 2162 | 375 | 538 | 2389 | 540 | 530 |
| 5480 | W303.asm02.HP0_G0054780 | chrXVI | 2088 | 1507 | 1350 | 1944 | 1327 | 1304 | 1993 | 1340 | 1590 | 4261 | 979 | 1011 | 4572 | 615 | 809 | 4135 | 1037 | 892 |
| 5481 | W303.asm02.HP0_G0054790 | chrXVI | 10017 | 1311 | 1304 | 9096 | 1282 | 1668 | 9657 | 1184 | 1848 | 11322 | 2540 | 3323 | 7336 | 1940 | 2367 | 6787 | 2247 | 2972 |
| 5482 | W303.asm02.HP0_G0054800 | chrXVI | 1990 | 12524 | 10273 | 3077 | 8852 | 10913 | 2129 | 11521 | 11177 | 38100 | 14239 | 17390 | 36001 | 11420 | 14565 | 32332 | 13499 | 15827 |
| 5483 | W303.asm02.HP0_G0054810 | chrXVI | 1250 | 458 | 440 | 1305 | 453 | 419 | 1112 | 424 | 533 | 1707 | 434 | 449 | 1671 | 233 | 333 | 1886 | 472 | 370 |
| 5484 | W303.asm02.HP0_G0054820 | chrXVI | 4048 | 1611 | 1610 | 4017 | 1472 | 1563 | 4017 | 1559 | 1975 | 4297 | 1579 | 1944 | 5251 | 1220 | 1542 | 4398 | 1429 | 1691 |
| 5485 | W303.asm02.HP0_G0054830 | chrXVI | 1578 | 654 | 962 | 1633 | 903 | 1017 | 1571 | 641 | 1156 | 8663 | 1093 | 2102 | 6447 | 995 | 1550 | 5560 | 1044 | 1855 |

| | | | | | | | | | | | | | | | | | | | | |
|---|---|---|---|---|---|---|---|---|---|---|---|---|---|---|---|---|---|---|---|---|
| 5486 | W303.asm02.HP0_G0054840 | chrXVI | 6449 | 847 | 751 | 8570 | 896 | 684 | 5963 | 877 | 705 | 6571 | 1258 | 1104 | 7815 | 1031 | 1047 | 5081 | 1077 | 921 |
| 5487 | W303.asm02.HP0_G0054850 | chrXVI | 977 | 77 | 85 | 811 | 113 | 106 | 767 | 58 | 105 | 306 | 81 | 123 | 303 | 53 | 100 | 179 | 88 | 102 |
| 5488 | W303.asm02.HP0_G0054860 | chrXVI | 2846 | 442 | 568 | 2555 | 402 | 710 | 2582 | 323 | 677 | 2166 | 700 | 1198 | 1134 | 522 | 688 | 1060 | 636 | 1019 |
| 5489 | W303.asm02.HP0_G0054870 | chrXVI | 1379 | 188 | 107 | 1078 | 124 | 128 | 1126 | 165 | 122 | 345 | 167 | 95 | 520 | 104 | 130 | 328 | 177 | 105 |
| 5490 | W303.asm02.HP0_G0054880 | chrXVI | 6155 | 2643 | 2706 | 6344 | 2476 | 2488 | 6286 | 2547 | 2861 | 9666 | 2555 | 2838 | 11675 | 2078 | 2377 | 9205 | 2375 | 2621 |
| 5491 | W303.asm02.HP0_G0054890 | chrXVI | 781 | 505 | 767 | 762 | 716 | 771 | 771 | 472 | 1040 | 2661 | 387 | 591 | 3895 | 400 | 438 | 2676 | 450 | 474 |
| 5492 | W303.asm02.HP0_G0054900 | chrXVI | 868 | 768 | 1021 | 751 | 898 | 991 | 828 | 739 | 1267 | 2434 | 594 | 725 | 2971 | 347 | 652 | 2712 | 563 | 667 |
| 5493 | W303.asm02.HP0_G0054910 | chrXVI | 3567 | 3596 | 4885 | 3203 | 4579 | 4608 | 3654 | 3853 | 5545 | 15811 | 3296 | 4615 | 16861 | 2194 | 3636 | 15849 | 2777 | 4265 |
| 5494 | W303.asm02.HP0_G0054920 | chrXVI | 5465 | 1024 | 1156 | 5412 | 1198 | 1389 | 5602 | 833 | 1551 | 9053 | 2380 | 2628 | 4694 | 1670 | 2003 | 3868 | 2113 | 2249 |
| 5495 | W303.asm02.HP0_G0054930 | chrXVI | 13538 | 2172 | 1750 | 11369 | 1633 | 2147 | 12666 | 1694 | 2322 | 6117 | 2438 | 3179 | 3398 | 2335 | 2157 | 2969 | 2406 | 2614 |
| 5496 | W303.asm02.HP0_G0054940 | chrXVI | 575 | 764 | 740 | 540 | 702 | 729 | 585 | 754 | 753 | 2273 | 688 | 547 | 2569 | 420 | 434 | 2183 | 598 | 517 |
| 5497 | W303.asm02.HP0_G0054950 | chrXVI | 679 | 419 | 535 | 649 | 516 | 561 | 587 | 454 | 751 | 1300 | 349 | 487 | 1534 | 251 | 360 | 1354 | 328 | 409 |
| 5498 | W303.asm02.HP0_G0054960 | chrXVI | 2823 | 897 | 952 | 2733 | 1014 | 1037 | 2768 | 903 | 978 | 5513 | 1063 | 1386 | 4723 | 673 | 1135 | 4851 | 1044 | 1335 |
| 5499 | W303.asm02.HP0_G0054970 | chrXVI | 2425 | 1039 | 1509 | 2297 | 2187 | 1482 | 2767 | 1243 | 1626 | 15354 | 1929 | 2964 | 12206 | 1206 | 1556 | 11143 | 1552 | 2566 |
| 5500 | W303.asm02.HP0_G0054980 | chrXVI | 1052 | 443 | 522 | 1047 | 505 | 506 | 1007 | 376 | 607 | 3713 | 841 | 995 | 2727 | 484 | 686 | 2599 | 799 | 949 |
| 5501 | W303.asm02.HP0_G0054990 | chrXVI | 762 | 235 | 312 | 606 | 242 | 309 | 715 | 191 | 399 | 957 | 212 | 299 | 1075 | 141 | 214 | 1021 | 210 | 266 |
| 5502 | W303.asm02.HP0_G0055000 | chrXVI | 6464 | 436 | 576 | 5931 | 530 | 633 | 6039 | 448 | 684 | 5099 | 926 | 1466 | 3219 | 735 | 1017 | 2928 | 816 | 1285 |
| 5503 | W303.asm02.HP0_G0055010 | chrXVI | 644 | 610 | 560 | 585 | 555 | 554 | 603 | 606 | 625 | 1705 | 482 | 509 | 1706 | 325 | 359 | 1667 | 513 | 485 |
| 5504 | W303.asm02.HP0_G0055020 | chrXVI | 11 | 14 | 6 | 8 | 3 | 6 | 5 | 2 | 1 | 22 | 8 | 2 | 14 | 3 | 4 | 28 | 7 | 2 |
| 5505 | W303.asm02.HP0_G0055030 | chrXVI | 802 | 835 | 863 | 757 | 843 | 884 | 675 | 911 | 1125 | 2311 | 635 | 647 | 2458 | 356 | 425 | 2234 | 603 | 522 |
| 5506 | W303.asm02.HP0_G0055040 | chrXVI | 7403 | 7087 | 8352 | 6820 | 7729 | 8173 | 6964 | 7376 | 9849 | 15953 | 6447 | 6821 | 18920 | 5074 | 6036 | 16527 | 5868 | 6266 |
| 5507 | W303.asm02.HP0_G0055050 | chrXVI | 350 | 153 | 126 | 343 | 152 | 131 | 377 | 140 | 198 | 1238 | 306 | 259 | 458 | 184 | 115 | 560 | 303 | 222 |
| 5508 | W303.asm02.HP0_G0055060 | chrXVI | 759 | 595 | 592 | 792 | 528 | 569 | 789 | 529 | 701 | 1524 | 407 | 443 | 1588 | 278 | 349 | 1576 | 336 | 448 |
| 5509 | W303.asm02.HP0_G0055070 | chrXVI | 21 | 27 | 19 | 21 | 12 | 11 | 15 | 24 | 32 | 39 | 23 | 13 | 58 | 11 | 7 | 42 | 11 | 9 |
| 5510 | W303.asm02.HP0_G0055080 | chrXVI | 3969 | 802 | 1227 | 3975 | 1048 | 1380 | 4179 | 706 | 1444 | 11736 | 1912 | 2880 | 8964 | 1600 | 2340 | 8385 | 1703 | 2703 |
| 5511 | W303.asm02.HP0_G0055090 | chrXVI | 4670 | 1081 | 868 | 4353 | 796 | 854 | 4561 | 984 | 892 | 3858 | 1108 | 1465 | 3607 | 888 | 1279 | 3401 | 1062 | 1288 |
| 5512 | W303.asm02.HP0_G0055100 | chrXVI | 489 | 306 | 423 | 468 | 437 | 441 | 646 | 268 | 496 | 4987 | 532 | 1056 | 2344 | 350 | 503 | 2680 | 556 | 868 |
| 5513 | W303.asm02.HP0_G0055110 | chrXVI | 3187 | 549 | 610 | 3049 | 597 | 642 | 3343 | 600 | 719 | 5732 | 1034 | 1496 | 4510 | 869 | 1030 | 3972 | 1006 | 1371 |
| 5514 | W303.asm02.HP0_G0055120 | chrXVI | 12474 | 9815 | 9503 | 12084 | 7941 | 9216 | 13435 | 9049 | 10062 | 21522 | 8799 | 9978 | 20417 | 5611 | 7322 | 21301 | 7962 | 9203 |
| 5515 | W303.asm02.HP0_G0055130 | chrXVI | 155 | 62 | 52 | 148 | 20 | 36 | 173 | 61 | 33 | 110 | 51 | 23 | 86 | 37 | 21 | 88 | 49 | 33 |
| 5516 | W303.asm02.HP0_G0055140 | chrXVI | 636 | 1019 | 1732 | 714 | 2207 | 1615 | 583 | 1142 | 1646 | 5668 | 1138 | 1800 | 5015 | 610 | 1356 | 4503 | 968 | 1394 |
| 5517 | W303.asm02.HP0_G0055150 | chrXVI | 25 | 19 | 10 | 14 | 21 | 16 | 6 | 13 | 17 | 56 | 17 | 10 | 77 | 2 | 14 | 43 | 22 | 7 |
| 5518 | W303.asm02.HP0_G0055160 | chrXVI | 687 | 1243 | 977 | 547 | 1144 | 983 | 697 | 1276 | 1214 | 2325 | 1092 | 845 | 3226 | 829 | 800 | 2325 | 910 | 757 |
| 5519 | W303.asm02.HP0_G0055170 | chrXVI | 3588 | 819 | 998 | 3427 | 906 | 1101 | 3514 | 724 | 1198 | 7159 | 1677 | 2108 | 3794 | 1214 | 1483 | 3526 | 1422 | 1975 |
| 5520 | W303.asm02.HP0_G0055180 | chrXVI | 3698 | 591 | 1053 | 3656 | 832 | 1132 | 3422 | 569 | 1219 | 6245 | 990 | 2002 | 4551 | 691 | 1336 | 4688 | 781 | 1677 |
| 5521 | W303.asm02.HP0_G0055190 | chrXVI | 8 | 1 | 0 | 5 | 1 | 3 | 6 | 5 | 1 | 6 | 1 | 1 | 3 | 0 | 1 | 4 | 2 | 3 |
| 5522 | W303.asm02.HP0_G0055200 | chrXVI | 3387 | 629 | 976 | 3613 | 1004 | 927 | 3345 | 640 | 1157 | 5458 | 781 | 1283 | 5218 | 547 | 938 | 5235 | 676 | 1109 |
| 5523 | W303.asm02.HP0_G0055210 | chrXVI | 4497 | 6338 | 8721 | 4282 | 7714 | 8265 | 4825 | 6780 | 8087 | 27610 | 7643 | 10992 | 27186 | 5128 | 8109 | 26734 | 6133 | 9281 |
| 5524 | W303.asm02.HP0_G0055220 | chrXVI | 802 | 192 | 292 | 596 | 231 | 336 | 718 | 161 | 328 | 1469 | 447 | 701 | 797 | 332 | 448 | 756 | 389 | 524 |
| 5525 | W303.asm02.HP0_G0055230 | chrXVI | 674 | 150 | 240 | 506 | 230 | 307 | 616 | 136 | 300 | 913 | 292 | 501 | 527 | 232 | 359 | 457 | 259 | 446 |
| 5526 | W303.asm02.HP0_G0055240 | chrXVI | 2129 | 296 | 444 | 1985 | 375 | 455 | 2395 | 229 | 558 | 4250 | 520 | 1108 | 2384 | 323 | 734 | 2634 | 385 | 965 |
| 5527 | W303.asm02.HP0_G0055250 | chrXVI | 4704 | 584 | 658 | 4257 | 587 | 703 | 4422 | 470 | 797 | 3278 | 964 | 1111 | 2015 | 880 | 998 | 1680 | 780 | 1000 |
| 5528 | W303.asm02.HP0_G0055260 | chrXVI | 3263 | 217 | 234 | 3643 | 214 | 290 | 2976 | 174 | 318 | 1300 | 405 | 482 | 517 | 229 | 329 | 530 | 303 | 360 |
| 5529 | W303.asm02.HP0_G0055270 | chrXVI | 5927 | 2245 | 2376 | 4850 | 2013 | 2420 | 4983 | 2337 | 2979 | 3695 | 1792 | 1947 | 5348 | 1397 | 1760 | 4146 | 1762 | 1740 |
| 5530 | W303.asm02.HP0_G0055280 | chrXVI | 2362 | 1939 | 2440 | 2433 | 3277 | 2266 | 2720 | 2138 | 2772 | 30943 | 3850 | 5545 | 25259 | 2391 | 2999 | 24268 | 3088 | 4936 |
| 5531 | W303.asm02.HP0_G0055290 | chrXVI | 1092 | 1140 | 1176 | 1154 | 1199 | 1178 | 1162 | 1107 | 1149 | 3605 | 960 | 1148 | 3680 | 610 | 811 | 3599 | 808 | 974 |
| 5532 | W303.asm02.HP0_G0055300 | chrXVI | 5065 | 2645 | 4206 | 4458 | 2926 | 4388 | 4891 | 2684 | 5657 | 8252 | 2297 | 3586 | 10006 | 1652 | 2799 | 9493 | 2080 | 3053 |
| 5533 | W303.asm02.HP0_G0055310 | chrXVI | 9330 | 1131 | 870 | 8892 | 920 | 935 | 9686 | 1085 | 1022 | 9651 | 1869 | 2228 | 5731 | 1539 | 1586 | 6026 | 1684 | 2123 |
| 5534 | W303.asm02.HP0_G0055320 | chrXVI | 2667 | 520 | 722 | 2429 | 653 | 647 | 2660 | 534 | 792 | 4164 | 760 | 1198 | 2999 | 545 | 902 | 3191 | 748 | 1025 |
| 5535 | W303.asm02.HP0_G0055330 | chrXVI | 1258 | 736 | 732 | 1318 | 695 | 665 | 1180 | 697 | 733 | 3128 | 681 | 793 | 3090 | 440 | 610 | 3163 | 672 | 715 |
| 5536 | W303.asm02.HP0_G0055340 | chrXVI | 2964 | 678 | 782 | 2742 | 692 | 770 | 2607 | 697 | 833 | 4864 | 1060 | 1423 | 3998 | 832 | 1221 | 3451 | 926 | 1256 |
| 5537 | W303.asm02.HP0_G0055350 | chrXVI | 977 | 218 | 239 | 1052 | 453 | 477 | 824 | 194 | 289 | 1452 | 364 | 409 | 1281 | 226 | 305 | 1099 | 337 | 357 |
| 5538 | W303.asm02.HP0_G0055360 | chrXVI | 16248 | 3747 | 3099 | 16501 | 2276 | 3587 | 15374 | 2829 | 4098 | 13348 | 6242 | 6363 | 6359 | 4549 | 5433 | 6751 | 5987 | 5251 |
| 5539 | W303.asm02.HP0_G0055370 | chrXVI | 2038 | 919 | 463 | 1621 | 417 | 503 | 1707 | 885 | 525 | 1874 | 864 | 663 | 1895 | 613 | 607 | 1811 | 901 | 524 |
| 5540 | W303.asm02.HP0_G0055380 | chrXVI | 1626 | 1556 | 938 | 1482 | 856 | 973 | 1559 | 1471 | 1043 | 2629 | 1369 | 854 | 3039 | 987 | 706 | 2703 | 1268 | 818 |
| 5541 | W303.asm02.HP0_G0055390 | chrXVI | 690 | 2190 | 1425 | 612 | 1170 | 1490 | 666 | 2031 | 1754 | 3624 | 1575 | 1215 | 4020 | 1145 | 876 | 3960 | 1690 | 1117 |
| 5542 | W303.asm02.HP0_G0055400 | chrXVI | 460 | 409 | 194 | 444 | 206 | 185 | 456 | 378 | 222 | 814 | 485 | 250 | 896 | 365 | 220 | 668 | 512 | 220 |
| 5543 | W303.asm02.HP0_G0055410 | chrXVI | 287 | 698 | 516 | 313 | 547 | 458 | 313 | 645 | 417 | 1960 | 1276 | 786 | 2589 | 991 | 892 | 1739 | 1218 | 836 |
| 5544 | W303.asm02.HP0_G0055420 | chrXVI | 2011 | 660 | 671 | 1940 | 574 | 639 | 1735 | 675 | 740 | 2935 | 936 | 893 | 3026 | 601 | 716 | 2543 | 850 | 707 |
| 5545 | W303.asm02.HP0_G0055430 | chrXVI | 3547 | 2697 | 2616 | 3267 | 2494 | 2396 | 3193 | 2722 | 3117 | 8413 | 2702 | 2849 | 9524 | 2038 | 2394 | 8195 | 2505 | 2472 |
| 5546 | W303.asm02.HP0_G0055440 | chrXVI | 1837 | 512 | 356 | 1814 | 367 | 328 | 1759 | 474 | 307 | 1512 | 479 | 426 | 3795 | 644 | 500 | 1382 | 463 | 382 |
| 5547 | W303.asm02.HP0_G0055450 | chrXVI | 2173 | 651 | 851 | 2096 | 773 | 892 | 1918 | 672 | 1142 | 2181 | 651 | 838 | 3094 | 422 | 645 | 2315 | 551 | 690 |
| 5548 | W303.asm02.HP0_G0055460 | chrXVI | 551 | 67 | 80 | 566 | 86 | 71 | 548 | 64 | 80 | 368 | 94 | 115 | 447 | 54 | 64 | 304 | 89 | 86 |
| 5549 | W303.asm02.HP0_G0055470 | chrXVI | 4320 | 552 | 841 | 4613 | 865 | 838 | 4680 | 517 | 953 | 7049 | 1152 | 1571 | 4939 | 698 | 1157 | 4881 | 1053 | 1340 |
| 5550 | W303.asm02.HP0_G0055480 | chrXVI | 730 | 642 | 988 | 669 | 894 | 946 | 700 | 600 | 843 | 3539 | 435 | 746 | 3901 | 377 | 575 | 3101 | 467 | 646 |
| 5551 | W303.asm02.HP0_G0055490 | chrXVI | 5699 | 2059 | 2244 | 5257 | 2820 | 2373 | 6632 | 1994 | 2915 | 29172 | 5163 | 6498 | 14869 | 3522 | 3348 | 16290 | 4085 | 5755 |
| 5552 | W303.asm02.HP0_G0055500 | chrXVI | 1020 | 1676 | 2013 | 978 | 1882 | 2026 | 854 | 1490 | 2608 | 3767 | 1284 | 1687 | 4154 | 931 | 1344 | 4306 | 1194 | 1293 |
| 5553 | W303.asm02.HP0_G0055510 | chrXVI | 3258 | 963 | 841 | 2758 | 820 | 769 | 3110 | 877 | 970 | 3183 | 937 | 1031 | 3384 | 729 | 913 | 2728 | 921 | 890 |
| 5554 | W303.asm02.HP0_G0055520 | chrXVI | 2112 | 176 | 162 | 2421 | 191 | 153 | 2048 | 140 | 161 | 1515 | 248 | 233 | 1027 | 151 | 194 | 1007 | 223 | 219 |
| 5555 | W303.asm02.HP0_G0055530 | chrXVI | 1345 | 98 | 125 | 1139 | 100 | 108 | 1200 | 97 | 159 | 250 | 117 | 101 | 364 | 51 | 73 | 240 | 84 | 83 |
| 5556 | W303.asm02.HP0_G0055540 | chrXVI | 1815 | 3550 | 3206 | 1647 | 3467 | 2870 | 1742 | 3577 | 3178 | 11149 | 3548 | 3643 | 10675 | 2411 | 2558 | 10752 | 3100 | 3152 |
| 5557 | W303.asm02.HP0_G0055550 | chrXVI | 43 | 27 | 34 | 18 | 29 | 42 | 23 | 23 | 39 | 94 | 22 | 37 | 90 | 7 | 31 | 96 | 20 | 16 |
| 5558 | W303.asm02.HP0_G0055560 | chrXVI | 579 | 500 | 609 | 448 | 599 | 602 | 487 | 522 | 643 | 1538 | 456 | 482 | 1607 | 290 | 320 | 1739 | 381 | 419 |
| 5559 | W303.asm02.HP0_G0055570 | chrXVI | 1686 | 506 | 568 | 1840 | 626 | 584 | 1995 | 566 | 426 | 4837 | 705 | 1098 | 4308 | 542 | 734 | 3629 | 561 | 984 |
| 5560 | W303.asm02.HP0_G0055580 | chrXVI | 2532 | 497 | 708 | 2311 | 581 | 848 | 2683 | 361 | 880 | 4138 | 825 | 1715 | 1857 | 618 | 981 | 1766 | 614 | 1363 |
| 5561 | W303.asm02.HP0_G0055590 | chrXVI | 1643 | 1584 | 2335 | 1342 | 2184 | 2300 | 1258 | 1727 | 3062 | 4583 | 1404 | 1842 | 5517 | 1131 | 1659 | 5183 | 1211 | 1390 |
| 5562 | W303.asm02.HP0_G0055600 | chrXVI | 3405 | 917 | 1120 | 3146 | 1061 | 1006 | 3158 | 1049 | 1144 | 4833 | 984 | 1352 | 4450 | 796 | 1148 | 4645 | 895 | 1172 |
| 5563 | W303.asm02.HP0_G0055610 | chrXVI | 2690 | 451 | 759 | 2463 | 689 | 852 | 2695 | 402 | 919 | 5674 | 758 | 1673 | 3513 | 541 | 1081 | 3456 | 714 | 1370 |
| 5564 | W303.asm02.HP0_G0055620 | chrXVI | 5111 | 925 | 822 | 4541 | 812 | 903 | 4869 | 777 | 1019 | 4801 | 1328 | 1516 | 3091 | 995 | 1176 | 3070 | 1294 | 1395 |
| 5565 | W303.asm02.HP0_G0055630 | chrXVI | 8346 | 1380 | 1588 | 8044 | 1288 | 1789 | 8219 | 1045 | 2001 | 10286 | 2775 | 3627 | 5089 | 2542 | 2633 | 4851 | 2362 | 3284 |
| 5566 | W303.asm02.HP0_G0055640 | chrXVI | 2728 | 1125 | 1160 | 2748 | 1041 | 1271 | 2715 | 1060 | 1184 | 3766 | 771 | 1235 | 3747 | 514 | 858 | 3642 | 783 | 1022 |
| 5567 | W303.asm02.HP0_G0055650 | chrXVI | 1889 | 237 | 259 | 1805 | 222 | 281 | 1650 | 181 | 336 | 1781 | 445 | 552 | 1014 | 286 | 461 | 1024 | 411 | 483 |
| 5568 | W303.asm02.HP0_G0055660 | chrXVI | 434 | 386 | 361 | 535 | 378 | 342 | 481 | 319 | 431 | 1260 | 242 | 242 | 1201 | 156 | 209 | 1236 | 222 | 222 |
| 5569 | W303.asm02.HP0_G0055670 | chrXVI | 5669 | 4498 | 5377 | 5507 | 4105 | 5519 | 5582 | 4181 | 7241 | 9929 | 3803 | 4324 | 11963 | 2554 | 3443 | 10920 | 3340 | 3722 |
| 5570 | W303.asm02.HP0_G0055680 | chrXVI | 7280 | 2445 | 2412 | 6639 | 2312 | 2362 | 7269 | 2611 | 2330 | 12326 | 2622 | 3147 | 10919 | 1766 | 2513 | 11274 | 2403 | 2942 |
| 5571 | W303.asm02.HP0_G0055690 | chrXVI | 3421 | 187 | 198 | 4214 | 202 | 212 | 3071 | 154 | 215 | 1487 | 300 | 334 | 1093 | 191 | 243 | 1057 | 320 | 255 |
| 5572 | W303.asm02.HP0_G0055700 | chrXVI | 604 | 364 | 425 | 548 | 501 | 381 | 624 | 366 | 443 | 3241 | 542 | 756 | 2605 | 386 | 494 | 2603 | 505 | 701 |
| 5573 | W303.asm02.HP0_G0055710 | chrXVI | 1648 | 375 | 441 | 1582 | 461 | 498 | 1548 | 350 | 543 | 4449 | 857 | 1153 | 2428 | 530 | 752 | 2459 | 721 | 1116 |
| 5574 | W303.asm02.HP0_G0055720 | chrXVI | 6950 | 5230 | 5087 | 5928 | 5249 | 4741 | 6981 | 5210 | 5120 | 20573 | 6070 | 6755 | 18035 | 3898 | 4532 | 18528 | 5402 | 6136 |
| 5575 | W303.asm02.HP0_G0055730 | chrXVI | 4269 | 1301 | 1083 | 3938 | 1065 | 1200 | 4553 | 1095 | 1214 | 7935 | 2476 | 2418 | 4574 | 1720 | 1871 | 4625 | 2167 | 2326 |
| 5576 | W303.asm02.HP0_G0055740 | chrXVI | 4746 | 3412 | 5491 | 5201 | 6156 | 5119 | 4701 | 3443 | 5547 | 25489 | 4779 | 6962 | 23605 | 3235 | 4523 | 19130 | 3951 | 5362 |
| 5577 | W303.asm02.HP0_G0055750 | chrXVI | 15148 | 3836 | 2811 | 13160 | 2855 | 2894 | 15449 | 3557 | 2948 | 14906 | 4883 | 5516 | 14027 | 3726 | 4533 | 12559 | 4468 | 4882 |
| 5578 | W303.asm02.HP0_G0055760 | chrXVI | 2605 | 4364 | 6285 | 2491 | 5370 | 6860 | 2087 | 4327 | 6603 | 7339 | 4692 | 5570 | 9143 | 2837 | 5587 | 7481 | 4658 | 5232 |

| | | | | | | | | | | | | | | | | | | | | |
|---|---|---|---|---|---|---|---|---|---|---|---|---|---|---|---|---|---|---|---|---|
| 5579 | W303.asm02.HP0_G0055770 | chrXVI | 3162 | 908 | 975 | 2754 | 905 | 928 | 2888 | 953 | 984 | 2799 | 1062 | 1062 | 3120 | 738 | 939 | 2686 | 965 | 937 |
| 5580 | W303.asm02.HP0_G0055780 | chrXVI | 1869 | 297 | 293 | 1790 | 230 | 398 | 1731 | 249 | 412 | 1559 | 533 | 644 | 1269 | 344 | 414 | 1118 | 520 | 552 |
| 5581 | W303.asm02.HP0_G0055790 | chrXVI | 9165 | 4933 | 4685 | 9220 | 4685 | 4152 | 9411 | 5069 | 5524 | 22673 | 7276 | 7251 | 17467 | 4580 | 5397 | 18970 | 6275 | 6393 |
| 5582 | W303.asm02.HP0_G0055800 | chrXVI | 2221 | 258 | 311 | 2649 | 356 | 336 | 2118 | 252 | 387 | 3099 | 509 | 653 | 2007 | 276 | 526 | 2187 | 435 | 597 |
| 5583 | W303.asm02.HP0_G0055810 | chrXVI | 6487 | 5134 | 6603 | 5570 | 6381 | 6657 | 5952 | 5330 | 7466 | 11130 | 4679 | 5405 | 13547 | 3664 | 4929 | 11927 | 4371 | 4740 |
| 5584 | W303.asm02.HP0_G0055820 | chrXVI | 2084 | 794 | 697 | 2096 | 722 | 693 | 2032 | 767 | 815 | 3589 | 740 | 899 | 2969 | 441 | 761 | 3361 | 712 | 870 |
| 5585 | W303.asm02.HP0_G0055830 | chrXVI | 194 | 630 | 699 | 161 | 588 | 717 | 159 | 644 | 947 | 1387 | 449 | 448 | 1867 | 289 | 406 | 1718 | 461 | 383 |
| 5586 | W303.asm02.HP0_G0055840 | chrXVI | 7717 | 1160 | 1000 | 7905 | 954 | 1058 | 7230 | 1118 | 1067 | 3708 | 1025 | 1390 | 4859 | 889 | 1239 | 3225 | 995 | 1222 |
| 5587 | W303.asm02.HP0_G0055850 | chrXVI | 910 | 1004 | 991 | 859 | 1010 | 923 | 1045 | 1073 | 861 | 5126 | 1350 | 1602 | 3769 | 816 | 942 | 4475 | 1277 | 1475 |
| 5588 | W303.asm02.HP0_G0055860 | chrXVI | 1378 | 265 | 301 | 1467 | 373 | 358 | 1405 | 306 | 415 | 2913 | 473 | 739 | 1965 | 328 | 477 | 2143 | 465 | 687 |
| 5589 | W303.asm02.HP0_G0055870 | chrXVI | 1294 | 969 | 1221 | 1090 | 1112 | 1213 | 1358 | 1062 | 1537 | 3067 | 924 | 923 | 3484 | 574 | 808 | 3070 | 858 | 859 |
| 5590 | W303.asm02.HP0_G0055880 | chrXVI | 2339 | 1803 | 2955 | 2079 | 2619 | 2886 | 2158 | 1953 | 3250 | 4896 | 1459 | 2298 | 5291 | 934 | 1935 | 4800 | 1208 | 1977 |
| 5591 | W303.asm02.HP0_G0055890 | chrXVI | 299 | 170 | 129 | 284 | 109 | 146 | 330 | 107 | 188 | 690 | 277 | 335 | 198 | 178 | 172 | 232 | 257 | 250 |
| 5592 | W303.asm02.HP0_G0055900 | chrXVI | 2338 | 682 | 839 | 2524 | 743 | 783 | 2251 | 647 | 848 | 5345 | 713 | 1362 | 4346 | 440 | 964 | 4746 | 727 | 1235 |
| 5593 | W303.asm02.HP0_G0055910 | chrXVI | 1668 | 1165 | 1384 | 1572 | 1295 | 1328 | 1620 | 1162 | 1530 | 4403 | 963 | 1113 | 5282 | 624 | 924 | 4619 | 936 | 1025 |
| 5594 | W303.asm02.HP0_G0055920 | chrXVI | 1740 | 1296 | 963 | 1588 | 967 | 954 | 1727 | 1177 | 934 | 3372 | 995 | 1002 | 3210 | 607 | 763 | 3081 | 952 | 924 |
| 5595 | W303.asm02.HP0_G0055930 | chrXVI | 1887 | 1086 | 1071 | 1636 | 1006 | 900 | 1915 | 1150 | 1168 | 5231 | 1459 | 1521 | 4520 | 957 | 1118 | 4677 | 1354 | 1367 |
| 5596 | W303.asm02.HP0_G0055940 | chrXVI | 925 | 2172 | 2342 | 970 | 2118 | 2269 | 968 | 2164 | 2612 | 6395 | 1738 | 1927 | 6868 | 1145 | 1232 | 6831 | 1689 | 1664 |
| 5597 | W303.asm02.HP0_G0055950 | chrXVI | 169 | 487 | 316 | 173 | 249 | 316 | 170 | 427 | 269 | 795 | 372 | 305 | 811 | 213 | 219 | 700 | 436 | 245 |
| 5598 | W303.asm02.HP0_G0055960 | chrXVI | 3477 | 2430 | 2721 | 3035 | 2639 | 2607 | 3200 | 2597 | 3245 | 7716 | 2731 | 2741 | 7859 | 1769 | 2234 | 8022 | 2394 | 2587 |
| 5599 | W303.asm02.HP0_G0055970 | chrXVI | 4627 | 1090 | 826 | 4343 | 733 | 915 | 4886 | 1079 | 911 | 3947 | 1319 | 1466 | 4261 | 991 | 1125 | 3675 | 1025 | 1267 |
| 5600 | W303.asm02.HP0_G0055980 | chrXVI | 930 | 610 | 802 | 1067 | 606 | 883 | 991 | 461 | 994 | 7898 | 1734 | 2135 | 4381 | 920 | 1465 | 4625 | 1800 | 1805 |
| 5601 | W303.asm02.HP0_G0055990 | chrXVI | 1023 | 472 | 348 | 867 | 327 | 293 | 897 | 447 | 392 | 980 | 421 | 336 | 1229 | 263 | 276 | 1018 | 415 | 293 |
| 5602 | W303.asm02.HP0_G0056000 | chrXVI | 2326 | 1221 | 1124 | 2336 | 989 | 1236 | 2076 | 1324 | 1222 | 2450 | 1066 | 1168 | 2685 | 755 | 1088 | 2472 | 1018 | 1038 |
| 5603 | W303.asm02.HP0_G0056010 | chrXVI | 552 | 776 | 526 | 554 | 443 | 498 | 572 | 652 | 558 | 1951 | 692 | 656 | 1587 | 341 | 442 | 2031 | 759 | 590 |
| 5604 | W303.asm02.HP0_G0056020 | chrXVI | 5238 | 4608 | 4379 | 4627 | 4052 | 4188 | 5090 | 4558 | 5304 | 11945 | 4243 | 4284 | 13100 | 3205 | 3646 | 12517 | 4098 | 3994 |
| 5605 | W303.asm02.HP0_G0056030 | chrXVI | 1915 | 1062 | 1447 | 1906 | 1349 | 1208 | 1955 | 1051 | 1368 | 5965 | 1001 | 1293 | 6191 | 657 | 1050 | 5949 | 912 | 1121 |
| 5606 | W303.asm02.HP0_G0056040 | chrXVI | 493 | 704 | 608 | 482 | 591 | 528 | 450 | 726 | 596 | 2280 | 692 | 536 | 1928 | 352 | 348 | 2342 | 643 | 468 |
| 5607 | W303.asm02.HP0_G0056050 | chrXVI | 6503 | 1749 | 1541 | 6911 | 1640 | 1580 | 6662 | 1839 | 1392 | 10145 | 1951 | 2743 | 9664 | 1483 | 1901 | 8957 | 1707 | 2436 |
| 5608 | W303.asm02.HP0_G0056060 | chrXVI | 2754 | 1876 | 2666 | 2721 | 2045 | 2605 | 2667 | 1945 | 3620 | 5980 | 1710 | 2277 | 7045 | 1105 | 1711 | 7166 | 1602 | 2045 |
| 5609 | W303.asm02.HP0_G0056070 | chrXVI | 14081 | 5536 | 7403 | 12218 | 6674 | 7794 | 14821 | 5479 | 8931 | 46672 | 12165 | 16996 | 36397 | 9955 | 13183 | 34456 | 10960 | 15235 |
| 5610 | W303.asm02.HP0_G0056080 | chrXVI | 5 | 4 | 3 | 8 | 1 | 0 | 3 | 3 | 2 | 4 | 4 | 1 | 7 | 1 | 0 | 4 | 0 | 0 |
| 5611 | W303.asm02.HP0_G0056090 | chrXVI | 4396 | 1369 | 908 | 4168 | 869 | 976 | 4497 | 1418 | 928 | 5498 | 1862 | 1649 | 5418 | 1455 | 1304 | 4453 | 1739 | 1494 |
| 5612 | W303.asm02.HP0_G0056100 | chrXVI | 1598 | 390 | 214 | 1389 | 162 | 200 | 1445 | 426 | 214 | 627 | 366 | 199 | 1900 | 432 | 253 | 545 | 352 | 189 |
| 5613 | W303.asm02.HP0_G0056110 | chrXVI | 1931 | 685 | 643 | 1727 | 415 | 679 | 1751 | 577 | 648 | 2100 | 685 | 1147 | 1973 | 475 | 841 | 1682 | 766 | 988 |
| 5614 | W303.asm02.HP0_G0056120 | chrXVI | 17399 | 9179 | 7575 | 17725 | 8064 | 7584 | 19888 | 8624 | 8931 | 31434 | 14878 | 15701 | 24840 | 12556 | 10938 | 25233 | 13103 | 14633 |
| 5615 | W303.asm02.HP0_G0056130 | chrXVI | 3629 | 516 | 387 | 3344 | 388 | 397 | 3423 | 408 | 426 | 2704 | 632 | 760 | 1963 | 485 | 588 | 1903 | 607 | 661 |
| 5616 | W303.asm02.HP0_G0056140 | chrXVI | 619 | 616 | 742 | 623 | 704 | 793 | 632 | 694 | 1063 | 2130 | 555 | 521 | 2696 | 415 | 432 | 2247 | 470 | 494 |
| 5617 | W303.asm02.HP0_G0056150 | chrXVI | 347 | 534 | 535 | 338 | 589 | 461 | 312 | 505 | 463 | 2498 | 607 | 629 | 2628 | 340 | 483 | 2285 | 496 | 578 |
| 5618 | W303.asm02.HP0_G0056160 | chrXVI | 1746 | 267 | 322 | 1648 | 373 | 337 | 1565 | 312 | 354 | 2874 | 494 | 618 | 2761 | 324 | 414 | 2513 | 439 | 521 |
| 5619 | W303.asm02.HP0_G0056170 | chrXVI | 1644 | 874 | 744 | 1627 | 802 | 732 | 1638 | 824 | 820 | 2806 | 743 | 823 | 2756 | 515 | 638 | 2745 | 644 | 755 |
| 5620 | W303.asm02.HP0_G0056180 | chrXVI | 5007 | 2725 | 3018 | 4876 | 2801 | 2819 | 4955 | 2933 | 3467 | 9703 | 2693 | 3024 | 10292 | 1785 | 2422 | 9574 | 2418 | 2942 |
| 5621 | W303.asm02.HP0_G0056190 | chrXVI | 7287 | 2482 | 2229 | 6844 | 2160 | 2076 | 7325 | 2397 | 2654 | 12373 | 3787 | 3628 | 8579 | 2472 | 2804 | 9333 | 3391 | 3361 |
| 5622 | W303.asm02.HP0_G0056200 | chrXVI | 4790 | 676 | 831 | 4910 | 813 | 875 | 4845 | 697 | 983 | 7585 | 1240 | 1803 | 5011 | 1003 | 1227 | 4919 | 1085 | 1613 |
| 5623 | W303.asm02.HP0_G0056210 | chrXVI | 5839 | 621 | 639 | 5194 | 518 | 678 | 5353 | 574 | 726 | 5529 | 1239 | 1571 | 3666 | 955 | 1140 | 3564 | 1049 | 1342 |
| 5624 | W303.asm02.HP0_G0056220 | chrXVI | 1774 | 390 | 400 | 1715 | 360 | 371 | 1490 | 395 | 392 | 1504 | 459 | 528 | 1440 | 359 | 395 | 1102 | 406 | 433 |
| 5625 | W303.asm02.HP0_G0056230 | chrXVI | 374 | 363 | 463 | 294 | 381 | 448 | 298 | 344 | 657 | 984 | 322 | 355 | 1175 | 200 | 245 | 1050 | 310 | 275 |
| 5626 | W303.asm02.HP0_G0056240 | chrXVI | 1264 | 240 | 380 | 1244 | 344 | 421 | 1233 | 237 | 488 | 2660 | 416 | 746 | 1176 | 328 | 525 | 1095 | 324 | 601 |
| 5627 | W303.asm02.HP0_G0056250 | chrXVI | 151 | 14 | 17 | 107 | 12 | 8 | 121 | 11 | 20 | 141 | 23 | 22 | 59 | 19 | 11 | 50 | 19 | 23 |
| 5628 | W303.asm02.HP0_G0056260 | chrXVI | 2902 | 745 | 1266 | 3130 | 1385 | 1194 | 2847 | 865 | 1225 | 8834 | 969 | 1607 | 8119 | 657 | 1226 | 7934 | 918 | 1482 |
| 5629 | W303.asm02.HP0_G0056270 | chrXVI | 1930 | 1097 | 928 | 1798 | 835 | 989 | 1712 | 1006 | 1239 | 1778 | 794 | 732 | 2390 | 493 | 662 | 1985 | 758 | 609 |
| 5630 | W303.asm02.HP0_G0056280 | chrXVI | 3858 | 1872 | 2108 | 3178 | 1827 | 2164 | 3290 | 1898 | 2761 | 4522 | 1668 | 1855 | 5291 | 1295 | 1702 | 4504 | 1584 | 1665 |
| 5631 | W303.asm02.HP0_G0056290 | chrXVI | 1451 | 1787 | 1167 | 1312 | 1017 | 1225 | 1242 | 1661 | 1482 | 2320 | 1152 | 779 | 2712 | 744 | 615 | 2721 | 1315 | 682 |
| 5632 | W303.asm02.HP0_G0056300 | chrXVI | 9464 | 7157 | 9270 | 8928 | 8554 | 8986 | 9418 | 7197 | 11970 | 19338 | 6600 | 7485 | 21253 | 5021 | 5955 | 20331 | 5838 | 6877 |
| 5633 | W303.asm02.HP0_G0056310 | chrXVI | 9967 | 6541 | 6200 | 9095 | 5369 | 6054 | 9101 | 6388 | 6817 | 12935 | 5095 | 5227 | 13790 | 3599 | 4665 | 13089 | 4681 | 4811 |
| 5634 | W303.asm02.HP0_G0056320 | chrXVI | 1231 | 389 | 453 | 1213 | 534 | 461 | 1266 | 367 | 500 | 4769 | 680 | 1063 | 3603 | 507 | 789 | 3250 | 661 | 1046 |
| 5635 | W303.asm02.HP0_G0056330 | chrXVI | 2042 | 1306 | 1245 | 1656 | 1267 | 1281 | 1875 | 1382 | 1331 | 8017 | 1576 | 1741 | 7253 | 1083 | 1447 | 7131 | 1406 | 1712 |
| 5636 | W303.asm02.HP0_G0056340 | chrXVI | 1425 | 570 | 323 | 1292 | 311 | 339 | 1299 | 429 | 299 | 2108 | 546 | 485 | 1877 | 369 | 405 | 1992 | 550 | 485 |
| 5637 | W303.asm02.HP0_G0056350 | chrXVI | 1676 | 923 | 1093 | 1649 | 880 | 1076 | 1478 | 853 | 1306 | 2069 | 639 | 813 | 2323 | 386 | 639 | 2419 | 614 | 684 |
| 5638 | W303.asm02.HP0_G0056360 | chrXVI | 10545 | 6041 | 6441 | 10705 | 5742 | 6428 | 10567 | 5751 | 7345 | 16116 | 4787 | 5558 | 17096 | 3389 | 4951 | 16444 | 4323 | 5017 |
| 5639 | W303.asm02.HP0_G0056370 | chrXVI | 1856 | 1127 | 1379 | 1847 | 1090 | 1448 | 1675 | 1044 | 1940 | 3195 | 819 | 1037 | 3672 | 551 | 814 | 3354 | 846 | 859 |
| 5640 | W303.asm02.HP0_G0056380 | chrXVI | 4740 | 4057 | 3669 | 4712 | 4090 | 3429 | 5179 | 4346 | 3540 | 16316 | 4633 | 4615 | 15438 | 3122 | 2913 | 15488 | 3944 | 4201 |
| 5641 | W303.asm02.HP0_G0056390 | chrXVI | 1052 | 662 | 498 | 1104 | 486 | 498 | 1016 | 614 | 401 | 2566 | 708 | 743 | 2013 | 447 | 593 | 2121 | 689 | 648 |
| 5642 | W303.asm02.HP0_G0056400 | chrXVI | 2285 | 672 | 915 | 2436 | 1280 | 962 | 2664 | 681 | 1153 | 12031 | 1362 | 2266 | 8097 | 1005 | 1125 | 7492 | 1101 | 1883 |
| 5643 | W303.asm02.HP0_G0056410 | chrXVI | 6723 | 3154 | 3467 | 5954 | 3192 | 3300 | 6588 | 3120 | 4131 | 8335 | 2887 | 3113 | 8697 | 2086 | 2558 | 7966 | 2437 | 2778 |
| 5644 | W303.asm02.HP0_G0056420 | chrXVI | 3463 | 1060 | 1305 | 2976 | 1116 | 1321 | 3537 | 1075 | 1489 | 7001 | 1672 | 2045 | 5891 | 1269 | 1710 | 5130 | 1428 | 1907 |
| 5645 | W303.asm02.HP0_G0056430 | chrXVI | 3273 | 542 | 673 | 3176 | 611 | 794 | 3024 | 489 | 810 | 4936 | 1057 | 1460 | 2837 | 880 | 1207 | 2446 | 945 | 1275 |
| 5646 | W303.asm02.HP0_G0056440 | chrXVI | 466 | 48 | 150 | 652 | 95 | 135 | 398 | 33 | 159 | 976 | 42 | 274 | 775 | 29 | 172 | 769 | 43 | 236 |
| 5647 | W303.asm02.HP0_G0056450 | chrXVI | 2926 | 377 | 828 | 2644 | 695 | 875 | 2555 | 377 | 1128 | 5401 | 585 | 2218 | 3597 | 477 | 1477 | 3644 | 509 | 1863 |
| 5648 | W303.asm02.HP0_G0056460 | chrXVI | 2526 | 2821 | 3346 | 2297 | 3266 | 3035 | 2680 | 3088 | 3814 | 11453 | 3004 | 3395 | 10688 | 1827 | 2280 | 11430 | 2614 | 2909 |
| 5649 | W303.asm02.HP0_G0056470 | chrXVI | 592 | 1384 | 1527 | 533 | 1613 | 1385 | 578 | 1524 | 1649 | 8173 | 1589 | 1596 | 7574 | 1041 | 1329 | 8188 | 1331 | 1569 |
| 5650 | W303.asm02.HP0_G0056480 | chrXVI | 567 | 799 | 839 | 506 | 948 | 760 | 516 | 822 | 809 | 2872 | 744 | 749 | 3163 | 453 | 549 | 3001 | 690 | 699 |
| 5651 | W303.asm02.HP0_G0056490 | chrXVI | 480 | 207 | 196 | 415 | 177 | 167 | 457 | 169 | 204 | 1232 | 386 | 327 | 904 | 158 | 223 | 869 | 383 | 263 |
| 5652 | W303.asm02.HP0_G0056500 | chrXVI | 3532 | 787 | 1146 | 3464 | 1099 | 1158 | 3332 | 791 | 1167 | 7530 | 1199 | 1773 | 5401 | 748 | 1384 | 6167 | 1215 | 1712 |
| 5653 | W303.asm02.HP0_G0056510 | chrXVI | 11278 | 9203 | 10496 | 12487 | 10218 | 10696 | 11192 | 9817 | 10118 | 41587 | 13098 | 14836 | 40207 | 9680 | 14008 | 37730 | 10817 | 16129 |
| 5654 | W303.asm02.HP0_G0056520 | chrXVI | 4847 | 1715 | 2281 | 5366 | 2369 | 2424 | 5188 | 1837 | 2193 | 11045 | 1997 | 3563 | 10556 | 1444 | 2229 | 9747 | 1807 | 2930 |
| 5655 | W303.asm02.HP0_G0056530 | chrXVI | 4556 | 819 | 562 | 3685 | 515 | 614 | 4059 | 671 | 675 | 2928 | 1046 | 1085 | 2085 | 695 | 803 | 2109 | 1054 | 969 |
| 5656 | W303.asm02.HP0_G0056540 | chrXVI | 980 | 1521 | 1211 | 916 | 1121 | 1190 | 968 | 1394 | 1388 | 4282 | 1208 | 1249 | 3898 | 739 | 860 | 4050 | 1164 | 1087 |
| 5657 | W303.asm02.HP0_G0056550 | chrXVI | 4619 | 959 | 969 | 4819 | 762 | 1143 | 5175 | 750 | 1278 | 5478 | 1596 | 1967 | 3471 | 1145 | 1377 | 3429 | 1332 | 1819 |
| 5658 | W303.asm02.HP0_G0056560 | chrXVI | 1696 | 1822 | 937 | 1663 | 1024 | 942 | 1582 | 1812 | 945 | 2608 | 1201 | 806 | 3092 | 874 | 687 | 2580 | 1377 | 717 |
| 5659 | W303.asm02.HP0_G0056570 | chrXVI | 5826 | 6934 | 8043 | 4907 | 7778 | 7908 | 5639 | 6637 | 10133 | 17239 | 6226 | 6619 | 18051 | 4588 | 5217 | 17800 | 5503 | 5839 |
| 5660 | W303.asm02.HP0_G0056580 | chrXVI | 6653 | 2711 | 2096 | 6840 | 2420 | 2231 | 6778 | 2777 | 2006 | 9082 | 2638 | 2880 | 8893 | 1864 | 2100 | 7966 | 2390 | 2458 |
| 5661 | W303.asm02.HP0_G0056590 | chrXVI | 3178 | 1201 | 1674 | 2642 | 1459 | 1612 | 2658 | 1241 | 1759 | 3157 | 1029 | 1631 | 3668 | 817 | 1379 | 3215 | 996 | 1366 |
| 5662 | W303.asm02.HP0_G0056600 | chrXVI | 1468 | 1141 | 939 | 1427 | 905 | 1004 | 1348 | 1158 | 1118 | 2071 | 789 | 755 | 2433 | 506 | 549 | 2130 | 799 | 645 |
| 5663 | W303.asm02.HP0_G0056610 | chrXVI | 2444 | 646 | 641 | 2417 | 645 | 707 | 2335 | 603 | 777 | 4749 | 975 | 1307 | 3102 | 680 | 923 | 2890 | 944 | 1152 |
| 5664 | W303.asm02.HP0_G0056620 | chrXVI | 2456 | 1434 | 1688 | 2435 | 2303 | 1632 | 2794 | 1540 | 1567 | 15739 | 1839 | 3090 | 12057 | 1308 | 1823 | 11927 | 1663 | 2805 |
| 5665 | W303.asm02.HP0_G0056630 | chrXVI | 1305 | 714 | 1498 | 1347 | 1478 | 1463 | 1261 | 757 | 1660 | 8211 | 1575 | 2044 | 6867 | 1154 | 1845 | 5608 | 1337 | 1766 |
| 5666 | W303.asm02.HP0_G0056640 | chrXVI | 2202 | 1575 | 1709 | 2350 | 1502 | 1617 | 2079 | 1549 | 1785 | 5805 | 1528 | 2034 | 5447 | 1033 | 1350 | 5465 | 1434 | 1790 |
| 5667 | W303.asm02.HP0_G0056650 | chrXVI | 4011 | 682 | 806 | 3778 | 683 | 764 | 3674 | 689 | 953 | 4217 | 978 | 1329 | 3698 | 687 | 1026 | 3367 | 977 | 1159 |
| 5668 | W303.asm02.HP0_G0056660 | chrXVI | 868 | 524 | 645 | 964 | 704 | 591 | 883 | 585 | 604 | 3537 | 656 | 830 | 3152 | 359 | 703 | 3601 | 517 | 718 |
| 5669 | W303.asm02.HP0_G0056670 | chrXVI | 1404 | 2216 | 2551 | 1402 | 2878 | 2358 | 1618 | 2549 | 2904 | 8688 | 2338 | 2192 | 9775 | 1332 | 1476 | 8907 | 1874 | 1992 |
| 5670 | W303.asm02.HP0_G0056680 | chrXVI | 1929 | 1059 | 1008 | 2192 | 1136 | 974 | 2271 | 1163 | 1107 | 9012 | 1660 | 2054 | 7448 | 1125 | 1062 | 7760 | 1403 | 1733 |
| 5671 | W303.asm02.HP0_G0056690 | chrXVI | 4703 | 6275 | 8388 | 4659 | 9354 | 8771 | 5304 | 7000 | 9648 | 42076 | 9769 | 15091 | 37176 | 7248 | 11028 | 35313 | 7614 | 13064 |

| | | | | | | | | | | | | | | | | | | | | |
|---|---|---|---|---|---|---|---|---|---|---|---|---|---|---|---|---|---|---|---|---|
| 5672 | W303.asm02.HP0_G0056700 | chrXVI | 769 | 58 | 56 | 796 | 48 | 59 | 526 | 61 | 94 | 208 | 67 | 65 | 200 | 28 | 47 | 229 | 88 | 64 |
| 5673 | W303.asm02.HP0_G0056710 | chrXVI | 2118 | 1151 | 1309 | 1980 | 1138 | 1277 | 1976 | 1155 | 1575 | 4087 | 1180 | 1312 | 3971 | 719 | 957 | 4098 | 1078 | 1278 |
| 5674 | W303.asm02.HP0_G0056720 | chrXVI | 12198 | 2554 | 1976 | 12880 | 1835 | 2077 | 12508 | 2461 | 1849 | 9726 | 2669 | 3789 | 11094 | 2571 | 2853 | 8107 | 2408 | 3214 |
| 5675 | W303.asm02.HP0_G0056730 | chrXVI | 6566 | 12220 | 9852 | 6859 | 8593 | 10004 | 6297 | 12037 | 11928 | 20486 | 8782 | 7469 | 20562 | 6173 | 6896 | 20361 | 7882 | 6603 |
| 5676 | W303.asm02.HP0_G0056740 | chrXVI | 1149 | 413 | 314 | 1198 | 245 | 380 | 1004 | 329 | 328 | 311 | 276 | 238 | 371 | 138 | 187 | 306 | 335 | 229 |
| 5677 | W303.asm02.HP0_G0056750 | chrXVI | 1040 | 193 | 192 | 1000 | 253 | 226 | 923 | 207 | 258 | 1378 | 365 | 467 | 736 | 212 | 238 | 798 | 376 | 355 |
| 5678 | W303.asm02.HP0_G0056760 | chrXVI | 366 | 67 | 48 | 360 | 61 | 82 | 258 | 35 | 62 | 409 | 81 | 115 | 234 | 46 | 57 | 288 | 71 | 103 |
| 5679 | W303.asm02.HP0_G0056770 | chrXVI | 5786 | 318 | 276 | 6464 | 362 | 488 | 4978 | 215 | 331 | 2414 | 522 | 635 | 1893 | 366 | 595 | 1608 | 516 | 563 |
| 5680 | W303.asm02.HP0_G0056780 | chrXVI | 1246 | 859 | 737 | 1372 | 749 | 659 | 1129 | 832 | 646 | 3620 | 910 | 968 | 3257 | 616 | 841 | 3307 | 927 | 871 |
| 5681 | W303.asm02.HP0_G0056790 | chrXVI | 2644 | 697 | 821 | 2239 | 678 | 908 | 2258 | 593 | 1008 | 5443 | 1631 | 1852 | 2476 | 1087 | 1329 | 2682 | 1261 | 1605 |
| 5682 | W303.asm02.HP0_G0056800 | chrXVI | 362 | 96 | 198 | 295 | 150 | 211 | 289 | 87 | 243 | 534 | 190 | 278 | 352 | 75 | 165 | 334 | 184 | 236 |
| 5683 | W303.asm02.HP0_G0056810 | chrXVI | 1259 | 2231 | 862 | 1110 | 934 | 1082 | 996 | 1608 | 621 | 2364 | 1510 | 846 | 2504 | 902 | 857 | 2763 | 1635 | 812 |
| 5684 | W303.asm02.HP0_G0056820 | chrXVI | 7829 | 2207 | 2276 | 7627 | 2091 | 2264 | 8548 | 2246 | 2693 | 16456 | 4619 | 4938 | 11988 | 3571 | 3594 | 10313 | 3893 | 4501 |
| 5685 | W303.asm02.HP0_G0056830 | chrXVI | 48497 | 649 | 569 | 49189 | 593 | 699 | 44108 | 629 | 775 | 2686 | 847 | 1029 | 7310 | 1932 | 2659 | 1962 | 826 | 905 |
| 5686 | W303.asm02.HP0_G0056840 | chrXVI | 3563 | 631 | 708 | 3821 | 688 | 785 | 3737 | 544 | 854 | 6258 | 1228 | 1770 | 4286 | 948 | 1235 | 3811 | 1048 | 1552 |
| 5687 | W303.asm02.HP0_G0056850 | chrXVI | 2279 | 447 | 461 | 2137 | 464 | 463 | 2116 | 467 | 478 | 2972 | 579 | 799 | 2554 | 463 | 560 | 2526 | 530 | 748 |
| 5688 | W303.asm02.HP0_G0056860 | chrXVI | 17203 | 4726 | 4032 | 16734 | 4452 | 3747 | 17658 | 4717 | 4443 | 27297 | 7551 | 8159 | 21223 | 5211 | 5610 | 21853 | 6607 | 7268 |
| 5689 | W303.asm02.HP0_G0056870 | chrXVI | 1443 | 478 | 571 | 1427 | 638 | 543 | 1536 | 443 | 624 | 5981 | 657 | 1261 | 4413 | 456 | 878 | 4507 | 586 | 1145 |
| 5690 | W303.asm02.HP0_G0056880 | chrXVI | 13532 | 6202 | 6673 | 13839 | 6232 | 6692 | 13397 | 6328 | 7286 | 15481 | 5470 | 6720 | 20131 | 4662 | 6680 | 15127 | 4808 | 6117 |
| 5691 | W303.asm02.HP0_G0056890 | chrXVI | 1631 | 808 | 1240 | 1512 | 964 | 1254 | 1419 | 782 | 1879 | 2408 | 584 | 1134 | 3176 | 489 | 894 | 2675 | 630 | 925 |
| 5692 | W303.asm02.HP0_G0056900 | chrXVI | 1306 | 1213 | 1396 | 1011 | 1178 | 1493 | 1152 | 1222 | 1590 | 1494 | 829 | 770 | 2572 | 813 | 823 | 1424 | 715 | 617 |
| 5693 | W303.asm02.HP0_G0056910 | chrXVI | 479 | 168 | 177 | 413 | 153 | 149 | 397 | 183 | 131 | 877 | 206 | 239 | 860 | 148 | 179 | 741 | 205 | 195 |
| 5694 | W303.asm02.HP0_G0056920 | chrXVI | 3628 | 1937 | 1764 | 3739 | 1870 | 1639 | 3830 | 1927 | 1729 | 10212 | 2376 | 2611 | 8484 | 1401 | 1785 | 8872 | 2166 | 2325 |
| 5695 | W303.asm02.HP0_G0056930 | chrXVI | 2783 | 1575 | 2060 | 2163 | 1785 | 2036 | 2515 | 1594 | 2613 | 4088 | 1367 | 1639 | 4860 | 916 | 1504 | 4505 | 1317 | 1356 |
| 5696 | W303.asm02.HP0_G0056940 | chrXVI | 1102 | 355 | 277 | 977 | 250 | 333 | 1014 | 254 | 348 | 2279 | 533 | 623 | 1024 | 343 | 413 | 1086 | 599 | 517 |
| 5697 | W303.asm02.HP0_G0056950 | chrXVI | 2269 | 961 | 733 | 2139 | 706 | 719 | 1940 | 1028 | 723 | 2300 | 681 | 858 | 2503 | 525 | 653 | 2437 | 687 | 766 |
| 5698 | W303.asm02.HP0_G0056960 | chrXVI | 5803 | 1713 | 1762 | 5607 | 1769 | 1751 | 5748 | 1678 | 2083 | 8238 | 2298 | 2718 | 7442 | 1607 | 2262 | 6675 | 1902 | 2607 |
| 5699 | W303.asm02.HP0_G0056970 | chrXVI | 668 | 178 | 237 | 763 | 174 | 263 | 682 | 156 | 306 | 1160 | 346 | 518 | 687 | 244 | 327 | 598 | 296 | 468 |
| 5700 | W303.asm02.HP0_G0056980 | chrXVI | 714 | 446 | 543 | 689 | 502 | 614 | 640 | 358 | 604 | 3085 | 705 | 985 | 1980 | 390 | 627 | 2049 | 656 | 874 |
| 5701 | W303.asm02.HP0_G0056990 | chrXVI | 1860 | 1498 | 1016 | 1749 | 1034 | 968 | 1923 | 1452 | 1093 | 3336 | 1355 | 1172 | 3141 | 854 | 946 | 3177 | 1343 | 1110 |
| 5702 | W303.asm02.HP0_G0057000 | chrXVI | 2059 | 736 | 471 | 2187 | 506 | 523 | 2106 | 640 | 541 | 2485 | 783 | 916 | 2100 | 562 | 719 | 1876 | 763 | 870 |
| 5703 | W303.asm02.HP0_G0057010 | chrXVI | 2468 | 833 | 710 | 2232 | 753 | 715 | 2252 | 795 | 696 | 3877 | 759 | 989 | 3354 | 481 | 693 | 3592 | 772 | 858 |
| 5704 | W303.asm02.HP0_G0057020 | chrXVI | 2221 | 1416 | 1304 | 2179 | 1236 | 1352 | 2046 | 1321 | 1592 | 4057 | 1090 | 1123 | 4320 | 741 | 888 | 4131 | 1006 | 1017 |
| 5705 | W303.asm02.HP0_G0057030 | chrXVI | 10797 | 2536 | 3924 | 10260 | 2815 | 4736 | 12100 | 2176 | 5218 | 20736 | 5084 | 8112 | 14006 | 3720 | 5742 | 13224 | 4440 | 7318 |
| 5706 | W303.asm02.HP0_G0057040 | chrXVI | 595 | 773 | 657 | 496 | 708 | 738 | 497 | 838 | 786 | 1162 | 633 | 582 | 1277 | 409 | 418 | 1141 | 554 | 481 |
| 5707 | W303.asm02.HP0_G0057050 | chrXVI | 11268 | 2185 | 1954 | 10589 | 1456 | 2224 | 11724 | 2271 | 2125 | 7182 | 2964 | 4290 | 7386 | 2374 | 3115 | 5933 | 2654 | 3800 |
| 5708 | W303.asm02.HP0_G0057060 | chrXVI | 19782 | 407 | 360 | 19269 | 360 | 453 | 17627 | 316 | 475 | 1278 | 525 | 808 | 981 | 612 | 762 | 650 | 422 | 700 |
| 5709 | W303.asm02.HP0_G0057070 | chrXVI | 3797 | 469 | 507 | 3629 | 518 | 549 | 3842 | 468 | 581 | 4153 | 858 | 1173 | 3541 | 673 | 1106 | 2875 | 698 | 1060 |
| 5710 | W303.asm02.HP0_G0057080 | chrXVI | 258 | 797 | 811 | 234 | 868 | 777 | 315 | 836 | 865 | 3076 | 560 | 681 | 3205 | 365 | 523 | 3019 | 590 | 637 |
| 5711 | W303.asm02.HP0_G0057090 | chrXVI | 3933 | 3766 | 5137 | 3689 | 4997 | 4964 | 3917 | 4090 | 6481 | 11819 | 3458 | 4030 | 13741 | 2474 | 3233 | 12646 | 3047 | 3624 |
| 5712 | W303.asm02.HP0_G0057100 | chrXVI | 1289 | 1116 | 1358 | 965 | 1168 | 1369 | 1083 | 1156 | 1683 | 3008 | 850 | 939 | 3950 | 679 | 766 | 3386 | 855 | 836 |
| 5713 | W303.asm02.HP0_G0057110 | chrXVI | 2562 | 626 | 956 | 2522 | 944 | 966 | 2625 | 636 | 1252 | 7582 | 952 | 1787 | 4797 | 674 | 1314 | 5175 | 858 | 1663 |
| 5714 | W303.asm02.HP0_G0057120 | chrXVI | 1010 | 410 | 647 | 865 | 685 | 744 | 1226 | 385 | 846 | 8039 | 1057 | 1541 | 4492 | 801 | 926 | 4338 | 882 | 1253 |
| 5715 | W303.asm02.HP0_G0057130 | chrXVI | 11917 | 3846 | 2672 | 11448 | 2832 | 2923 | 11535 | 3576 | 2954 | 14240 | 4810 | 5495 | 11770 | 4291 | 4384 | 10984 | 4226 | 4251 |
| 5716 | W303.asm02.HP0_G0057140 | chrXVI | 451 | 60 | 95 | 478 | 106 | 122 | 449 | 67 | 114 | 297 | 131 | 108 | 172 | 63 | 92 | 163 | 86 | 109 |
| 5717 | W303.asm02.HP0_G0057150 | chrXVI | 279 | 32 | 65 | 290 | 55 | 53 | 248 | 41 | 72 | 171 | 49 | 60 | 223 | 48 | 65 | 108 | 30 | 41 |
| 5718 | W303.asm02.HP0_G0057160 | chrXVI | 1544 | 640 | 927 | 1537 | 705 | 864 | 1363 | 619 | 891 | 3182 | 673 | 1010 | 3544 | 640 | 1283 | 2595 | 618 | 865 |
| 5719 | W303.asm02.HP0_G0057170 | chrXVI | 607 | 151 | 241 | 548 | 203 | 221 | 553 | 185 | 251 | 545 | 138 | 147 | 991 | 133 | 194 | 429 | 121 | 158 |
| 5720 | W303.asm02.HP0_G0057180 | chrXVI | 656 | 316 | 300 | 541 | 311 | 310 | 636 | 307 | 296 | 1070 | 221 | 247 | 1226 | 154 | 173 | 996 | 226 | 246 |
| 5721 | W303.asm02.HP0_G0057190 | chrXVI | 1435 | 1357 | 1210 | 1394 | 1053 | 1097 | 1420 | 1349 | 1349 | 4851 | 1197 | 1174 | 5327 | 824 | 977 | 4906 | 1126 | 1137 |
| 5722 | W303.asm02.HP0_G0057200 | chrXVI | 1468 | 1446 | 937 | 1458 | 952 | 943 | 1448 | 1310 | 984 | 2970 | 1079 | 855 | 3362 | 634 | 663 | 2938 | 1124 | 772 |
| 5723 | W303.asm02.HP0_G0057210 | chrXVI | 174 | 281 | 159 | 154 | 130 | 170 | 138 | 249 | 201 | 261 | 157 | 98 | 424 | 110 | 85 | 310 | 201 | 78 |
| 5724 | W303.asm02.HP0_G0057220 | chrXVI | 319 | 313 | 209 | 327 | 195 | 200 | 290 | 290 | 204 | 722 | 263 | 196 | 627 | 146 | 161 | 616 | 304 | 196 |
| 5725 | W303.asm02.HP0_G0057230 | chrXVI | 12 | 16 | 5 | 6 | 6 | 0 | 15 | 29 | 3 | 28 | 18 | 6 | 8 | 4 | 1 | 16 | 14 | 5 |
| 5726 | W303.asm02.HP0_G0057240 | chrXVI | 118 | 41 | 40 | 154 | 41 | 53 | 125 | 22 | 58 | 106 | 43 | 86 | 46 | 43 | 25 | 53 | 32 | 54 |

**Km_counts**

| | locus_tag | protein_id | chr | Km-V-1 | Km-V-1_NaCl | Km-V-1_TM | Km-V-2 | Km-V-2_NaCl | Km-V-2_TM | Km-V-3 | Km-V-3_NaCl | Km-V-3_TM |
|---|---|---|---|---|---|---|---|---|---|---|---|---|
| 1 | FIM1_1 | QGN13365.1 | CP015054.1 | 0 | 0 | 0 | 0 | 0 | 0 | 0 | 0 | 0 |
| 2 | FIM1_2 | QGN13366.1 | CP015054.1 | 0 | 0 | 0 | 0 | 0 | 0 | 0 | 0 | 0 |
| 3 | FIM1_3 | QGN13367.1 | CP015054.1 | 10467 | 2637 | 2698 | 8031 | 1720 | 2308 | 10421 | 2837 | 2403 |
| 4 | FIM1_5 | QGN13368.1 | CP015054.1 | 1991 | 307 | 621 | 1580 | 201 | 488 | 1749 | 346 | 455 |
| 5 | FIM1_6 | QGN13369.1 | CP015054.1 | 823 | 247 | 338 | 635 | 186 | 355 | 998 | 265 | 316 |
| 6 | FIM1_7 | QGN13370.1 | CP015054.1 | 178 | 45 | 32 | 135 | 39 | 35 | 140 | 47 | 31 |
| 7 | FIM1_8 | QGN13371.1 | CP015054.1 | 1201 | 244 | 320 | 918 | 112 | 301 | 1846 | 242 | 224 |
| 8 | FIM1_9 | QGN13372.1 | CP015054.1 | 7356 | 4437 | 1025 | 5928 | 1913 | 897 | 9953 | 4267 | 774 |
| 9 | FIM1_10 | QGN13373.1 | CP015054.1 | 9530 | 2250 | 1021 | 8067 | 724 | 755 | 17186 | 2212 | 618 |
| 10 | FIM1_11 | QGN13374.1 | CP015054.1 | 857 | 207 | 186 | 656 | 155 | 178 | 940 | 262 | 165 |
| 11 | FIM1_12 | QGN13375.1 | CP015054.1 | 3052 | 938 | 523 | 2836 | 351 | 495 | 4762 | 495 | 410 |
| 12 | FIM1_13 | QGN13376.1 | CP015054.1 | 53355 | 31454 | 29179 | 42768 | 17221 | 28075 | 55188 | 31401 | 29597 |
| 13 | FIM1_14 | QGN13377.1 | CP015054.1 | 34 | 14 | 15 | 35 | 10 | 10 | 20 | 15 | 11 |
| 14 | FIM1_15 | QGN13378.1 | CP015054.1 | 18355 | 13270 | 12821 | 16287 | 8124 | 13522 | 16930 | 13516 | 12877 |
| 15 | FIM1_16 | QGN13379.1 | CP015054.1 | 16629 | 8964 | 11937 | 12690 | 5378 | 11862 | 14770 | 9817 | 11901 |
| 16 | FIM1_17 | QGN13380.1 | CP015054.1 | 27151 | 18793 | 17889 | 23718 | 11178 | 17714 | 26010 | 19539 | 18302 |
| 17 | FIM1_18 | QGN13381.1 | CP015054.1 | 3586 | 2649 | 1709 | 2808 | 779 | 1445 | 2596 | 2022 | 1644 |
| 18 | FIM1_19 | QGN13382.1 | CP015054.1 | 2666 | 354 | 973 | 1996 | 268 | 908 | 2947 | 391 | 771 |
| 19 | FIM1_20 | QGN13383.1 | CP015054.1 | 6182 | 2073 | 3203 | 5071 | 1517 | 3066 | 6297 | 2207 | 3037 |
| 20 | FIM1_21 | QGN13384.1 | CP015054.1 | 1093 | 328 | 524 | 745 | 236 | 552 | 998 | 391 | 449 |
| 21 | FIM1_22 | QGN13385.1 | CP015054.1 | 3859 | 1025 | 1282 | 2939 | 715 | 1170 | 3978 | 1058 | 1080 |
| 22 | FIM1_23 | QGN13386.1 | CP015054.1 | 511 | 107 | 171 | 452 | 79 | 180 | 571 | 107 | 172 |
| 23 | FIM1_24 | QGN13387.1 | CP015054.1 | 2115 | 1302 | 793 | 1712 | 1010 | 862 | 800 | 1379 | 784 |
| 24 | FIM1_25 | QGN13388.1 | CP015054.1 | 703 | 328 | 423 | 591 | 265 | 446 | 366 | 347 | 371 |
| 25 | FIM1_26 | QGN13389.1 | CP015054.1 | 465 | 90 | 249 | 354 | 95 | 244 | 430 | 114 | 217 |
| 26 | FIM1_27 | QGN13390.1 | CP015054.1 | 1514 | 347 | 637 | 1272 | 288 | 661 | 1052 | 337 | 457 |
| 27 | FIM1_28 | QGN13391.1 | CP015054.1 | 28833 | 9341 | 11292 | 24353 | 7400 | 10327 | 27566 | 10181 | 10368 |
| 28 | FIM1_29 | QGN13392.1 | CP015054.1 | 8643 | 1044 | 2889 | 6816 | 616 | 2727 | 11042 | 1125 | 2490 |
| 29 | FIM1_30 | QGN13393.1 | CP015054.1 | 3287 | 1350 | 1838 | 2703 | 969 | 1784 | 3115 | 1394 | 1631 |
| 30 | FIM1_31 | QGN13394.1 | CP015054.1 | 14449 | 3081 | 3669 | 9947 | 2232 | 3436 | 10463 | 3249 | 3203 |
| 31 | FIM1_32 | QGN13395.1 | CP015054.1 | 4942 | 1577 | 2703 | 4286 | 1108 | 2729 | 5621 | 1721 | 2415 |
| 32 | FIM1_33 | QGN13396.1 | CP015054.1 | 1602 | 497 | 758 | 1400 | 385 | 673 | 1385 | 519 | 576 |
| 33 | FIM1_34 | QGN13397.1 | CP015054.1 | 4053 | 1211 | 1408 | 2998 | 822 | 1339 | 3851 | 1341 | 1160 |
| 34 | FIM1_35 | QGN13398.1 | CP015054.1 | 95 | 17 | 24 | 87 | 15 | 34 | 104 | 21 | 31 |
| 35 | FIM1_36 | QGN13399.1 | CP015054.1 | 10021 | 4721 | 7517 | 9302 | 5379 | 6795 | 1634 | 5027 | 7474 |
| 36 | FIM1_37 | QGN13400.1 | CP015054.1 | 1360 | 651 | 543 | 1197 | 558 | 470 | 1340 | 728 | 479 |
| 37 | FIM1_38 | QGN13401.1 | CP015054.1 | 1657 | 1433 | 783 | 1315 | 1171 | 799 | 852 | 1604 | 909 |
| 38 | FIM1_39 | QGN13402.1 | CP015054.1 | 3955 | 1563 | 1322 | 2901 | 1175 | 1209 | 1477 | 1684 | 1215 |
| 39 | FIM1_40 | QGN13403.1 | CP015054.1 | 1637 | 367 | 2147 | 1307 | 300 | 2014 | 2098 | 405 | 1767 |
| 40 | FIM1_41 | QGN13404.1 | CP015054.1 | 2454 | 731 | 749 | 1971 | 533 | 765 | 2215 | 779 | 712 |
| 41 | FIM1_42 | QGN13405.1 | CP015054.1 | 1588 | 844 | 866 | 1546 | 741 | 884 | 1361 | 989 | 881 |
| 42 | FIM1_43 | QGN13406.1 | CP015054.1 | 5928 | 1652 | 1899 | 4896 | 1214 | 1789 | 5473 | 1797 | 1587 |
| 43 | FIM1_44 | QGN13407.1 | CP015054.1 | 280 | 110 | 255 | 251 | 86 | 247 | 289 | 127 | 201 |
| 44 | FIM1_45 | QGN13408.1 | CP015054.1 | 1449 | 340 | 1050 | 1219 | 271 | 1106 | 1605 | 346 | 977 |
| 45 | FIM1_46 | QGN13409.1 | CP015054.1 | 9303 | 3061 | 2736 | 7507 | 1842 | 2525 | 12196 | 3207 | 2531 |
| 46 | FIM1_47 | QGN13410.1 | CP015054.1 | 6343 | 995 | 1085 | 4591 | 709 | 1017 | 5215 | 1055 | 969 |

| | | | | | | | | | | | | |
|---|---|---|---|---|---|---|---|---|---|---|---|---|
| 47 | FIM1_48 | QGN13411.1 | CP015054.1 | 4111 | 1474 | 3002 | 3276 | 986 | 2818 | 3880 | 1445 | 2573 |
| 48 | FIM1_49 | QGN13412.1 | CP015054.1 | 12049 | 14791 | 6964 | 8763 | 9696 | 6335 | 3225 | 15582 | 6736 |
| 49 | FIM1_50 | QGN13413.1 | CP015054.1 | 5142 | 2181 | 3701 | 3647 | 1232 | 3840 | 3250 | 2508 | 3607 |
| 50 | FIM1_51 | QGN13414.1 | CP015054.1 | 5739 | 2066 | 2583 | 4658 | 1786 | 2640 | 4019 | 2282 | 2405 |
| 51 | FIM1_52 | QGN13415.1 | CP015054.1 | 6015 | 2659 | 3220 | 4836 | 1633 | 2724 | 6882 | 2700 | 2840 |
| 52 | FIM1_53 | QGN13416.1 | CP015054.1 | 51058 | 33063 | 28924 | 46643 | 26688 | 27398 | 49317 | 33487 | 28591 |
| 53 | FIM1_54 | QGN13417.1 | CP015054.1 | 14892 | 1350 | 3187 | 9865 | 766 | 2896 | 13430 | 1293 | 2898 |
| 54 | FIM1_55 | QGN13418.1 | CP015054.1 | 514 | 107 | 1087 | 451 | 55 | 949 | 1351 | 71 | 881 |
| 55 | FIM1_56 | QGN13419.1 | CP015054.1 | 45 | 7 | 5 | 34 | 11 | 23 | 113 | 13 | 11 |
| 56 | FIM1_57 | QGN13420.1 | CP015054.1 | 688 | 285 | 306 | 569 | 181 | 318 | 1107 | 282 | 277 |
| 57 | FIM1_58 | QGN13421.1 | CP015054.1 | 6196 | 3927 | 3227 | 4721 | 2838 | 2962 | 3157 | 4318 | 2990 |
| 58 | FIM1_59 | QGN13422.1 | CP015054.1 | 21472 | 2860 | 3938 | 15788 | 2057 | 3837 | 22798 | 3086 | 3606 |
| 59 | FIM1_60 | QGN13423.1 | CP015054.1 | 5408 | 1752 | 1714 | 4213 | 1270 | 1713 | 3649 | 1960 | 1580 |
| 60 | FIM1_61 | QGN13424.1 | CP015054.1 | 5121 | 580 | 1247 | 3917 | 504 | 1095 | 5486 | 615 | 978 |
| 61 | FIM1_62 | QGN13425.1 | CP015054.1 | 2265 | 932 | 968 | 1649 | 810 | 1109 | 1260 | 1074 | 992 |
| 62 | FIM1_63 | QGN13426.1 | CP015054.1 | 132 | 32 | 71 | 110 | 16 | 86 | 117 | 36 | 53 |
| 63 | FIM1_64 | QGN13427.1 | CP015054.1 | 3435 | 1239 | 1966 | 3272 | 963 | 1825 | 2796 | 1326 | 1640 |
| 64 | FIM1_65 | QGN13428.1 | CP015054.1 | 857 | 118 | 210 | 737 | 97 | 149 | 752 | 107 | 140 |
| 65 | FIM1_66 | QGN13429.1 | CP015054.1 | 8376 | 4670 | 3815 | 7064 | 3394 | 3626 | 4848 | 5035 | 3835 |
| 66 | FIM1_67 | QGN13430.1 | CP015054.1 | 1636 | 614 | 264 | 1094 | 389 | 252 | 1354 | 791 | 219 |
| 67 | FIM1_68 | QGN13431.1 | CP015054.1 | 25206 | 14858 | 14435 | 23276 | 12276 | 13564 | 24096 | 14923 | 14097 |
| 68 | FIM1_69 | QGN13432.1 | CP015054.1 | 21724 | 14167 | 12722 | 20629 | 11815 | 12584 | 20915 | 14643 | 12358 |
| 69 | FIM1_70 | QGN13433.1 | CP015054.1 | 2464 | 1589 | 906 | 1799 | 887 | 833 | 1238 | 1709 | 839 |
| 70 | FIM1_71 | QGN13434.1 | CP015054.1 | 1682 | 449 | 586 | 1351 | 349 | 627 | 1165 | 483 | 591 |
| 71 | FIM1_72 | QGN13435.1 | CP015054.1 | 1174 | 441 | 474 | 957 | 290 | 511 | 1128 | 489 | 440 |
| 72 | FIM1_73 | QGN13436.1 | CP015054.1 | 27045 | 11556 | 11287 | 24060 | 8720 | 10227 | 26685 | 12214 | 10674 |
| 73 | FIM1_74 | QGN13437.1 | CP015054.1 | 1782 | 575 | 669 | 1439 | 495 | 621 | 1517 | 679 | 663 |
| 74 | FIM1_75 | QGN13438.1 | CP015054.1 | 40911 | 17196 | 19454 | 35872 | 9490 | 17333 | 65110 | 17550 | 17690 |
| 75 | FIM1_76 | QGN13439.1 | CP015054.1 | 1233 | 173 | 243 | 1037 | 123 | 235 | 1799 | 192 | 205 |
| 76 | FIM1_77 | QGN13440.1 | CP015054.1 | 2216 | 551 | 523 | 1863 | 423 | 504 | 3083 | 616 | 524 |
| 77 | FIM1_78 | QGN13441.1 | CP015054.1 | 9180 | 1892 | 2733 | 6997 | 1443 | 2482 | 9941 | 2055 | 2498 |
| 78 | FIM1_79 | QGN13442.1 | CP015054.1 | 12737 | 522 | 898 | 9115 | 475 | 853 | 27733 | 594 | 853 |
| 79 | FIM1_80 | QGN13443.1 | CP015054.1 | 468 | 27 | 68 | 373 | 25 | 66 | 647 | 35 | 51 |
| 80 | FIM1_81 | QGN13444.1 | CP015054.1 | 5551 | 660 | 972 | 4057 | 567 | 934 | 6621 | 690 | 849 |
| 81 | FIM1_82 | QGN13445.1 | CP015054.1 | 5407 | 1404 | 1891 | 3990 | 1117 | 1603 | 4251 | 1619 | 1400 |
| 82 | FIM1_83 | QGN13446.1 | CP015054.1 | 758 | 167 | 333 | 638 | 173 | 329 | 889 | 215 | 339 |
| 83 | FIM1_84 | QGN13447.1 | CP015054.1 | 2078 | 194 | 867 | 1586 | 183 | 749 | 2694 | 250 | 622 |
| 84 | FIM1_85 | QGN13448.1 | CP015054.1 | 1666 | 354 | 1031 | 1266 | 252 | 989 | 1656 | 395 | 904 |
| 85 | FIM1_86 | QGN13449.1 | CP015054.1 | 992 | 238 | 1304 | 861 | 165 | 1201 | 1049 | 218 | 1084 |
| 86 | FIM1_87 | QGN13450.1 | CP015054.1 | 1164 | 512 | 529 | 784 | 261 | 437 | 694 | 561 | 408 |
| 87 | FIM1_88 | QGN13451.1 | CP015054.1 | 1969 | 257 | 719 | 1708 | 180 | 653 | 2385 | 250 | 577 |
| 88 | FIM1_89 | QGN13452.1 | CP015054.1 | 2623 | 168 | 495 | 1893 | 80 | 357 | 4544 | 175 | 265 |
| 89 | FIM1_90 | QGN13453.1 | CP015054.1 | 8953 | 2337 | 2437 | 7066 | 1736 | 2196 | 7507 | 2527 | 1954 |
| 90 | FIM1_91 | QGN13454.1 | CP015054.1 | 1889 | 4903 | 877 | 1240 | 2874 | 811 | 705 | 5107 | 968 |
| 91 | FIM1_92 | QGN13455.1 | CP015054.1 | 833 | 132 | 357 | 575 | 100 | 327 | 912 | 160 | 278 |
| 92 | FIM1_93 | QGN13456.1 | CP015054.1 | 708 | 112 | 286 | 478 | 77 | 269 | 524 | 104 | 242 |
| 93 | FIM1_94 | QGN13457.1 | CP015054.1 | 826 | 78 | 376 | 709 | 71 | 367 | 1072 | 74 | 274 |
| 94 | FIM1_95 | QGN13458.1 | CP015054.1 | 17169 | 3953 | 4153 | 13658 | 2692 | 3838 | 15418 | 4424 | 3780 |

| | | | | | | | | | | | | |
|---|---|---|---|---|---|---|---|---|---|---|---|---|
| 95 | FIM1_96 | QGN13459.1 | CP015054.1 | 1769 | 82 | 298 | 1389 | 76 | 246 | 3311 | 91 | 240 |
| 96 | FIM1_97 | QGN13460.1 | CP015054.1 | 6621 | 784 | 1844 | 4059 | 560 | 1315 | 6483 | 718 | 1900 |
| 97 | FIM1_98 | QGN13461.1 | CP015054.1 | 328 | 79 | 159 | 274 | 79 | 170 | 239 | 85 | 156 |
| 98 | FIM1_99 | QGN13462.1 | CP015054.1 | 1182 | 236 | 444 | 968 | 173 | 461 | 1071 | 237 | 380 |
| 99 | FIM1_100 | QGN13463.1 | CP015054.1 | 9712 | 2621 | 3073 | 8556 | 1889 | 2829 | 10026 | 2936 | 2636 |
| 100 | FIM1_101 | QGN13464.1 | CP015054.1 | 4846 | 494 | 726 | 3731 | 351 | 695 | 6159 | 521 | 587 |
| 101 | FIM1_102 | QGN13465.1 | CP015054.1 | 533 | 96 | 181 | 430 | 81 | 173 | 457 | 109 | 158 |
| 102 | FIM1_103 | QGN13466.1 | CP015054.1 | 3758 | 1169 | 1674 | 3007 | 838 | 1608 | 3485 | 1259 | 1403 |
| 103 | FIM1_104 | QGN13467.1 | CP015054.1 | 1861 | 295 | 630 | 1393 | 243 | 561 | 1679 | 328 | 497 |
| 104 | FIM1_105 | QGN13468.1 | CP015054.1 | 7581 | 3367 | 3085 | 6948 | 3248 | 2944 | 5523 | 3838 | 2883 |
| 105 | FIM1_107 | QGN13469.1 | CP015054.1 | 2132 | 638 | 877 | 1722 | 439 | 847 | 1762 | 624 | 815 |
| 106 | FIM1_108 | QGN13470.1 | CP015054.1 | 2961 | 1865 | 1808 | 2779 | 1456 | 1911 | 2099 | 1991 | 1671 |
| 107 | FIM1_109 | QGN13471.1 | CP015054.1 | 17295 | 5054 | 5027 | 14593 | 2746 | 2712 | 21754 | 5348 | 4670 |
| 108 | FIM1_110 | QGN13472.1 | CP015054.1 | 3428 | 693 | 822 | 2438 | 437 | 635 | 3739 | 777 | 705 |
| 109 | FIM1_111 | QGN13473.1 | CP015054.1 | 12595 | 2723 | 6825 | 9579 | 2073 | 6482 | 13853 | 2910 | 6253 |
| 110 | FIM1_112 | QGN13474.1 | CP015054.1 | 1731 | 814 | 805 | 1274 | 642 | 771 | 897 | 904 | 767 |
| 111 | FIM1_113 | QGN13475.1 | CP015054.1 | 4534 | 468 | 822 | 3449 | 333 | 819 | 5256 | 489 | 734 |
| 112 | FIM1_114 | QGN13476.1 | CP015054.1 | 2160 | 566 | 880 | 1726 | 425 | 895 | 1980 | 690 | 750 |
| 113 | FIM1_115 | QGN13477.1 | CP015054.1 | 4905 | 349 | 807 | 3727 | 317 | 717 | 3629 | 400 | 656 |
| 114 | FIM1_116 | QGN13478.1 | CP015054.1 | 1148 | 192 | 322 | 918 | 190 | 311 | 792 | 210 | 341 |
| 115 | FIM1_117 | QGN13479.1 | CP015054.1 | 20782 | 10299 | 9707 | 17984 | 8133 | 9052 | 17996 | 10841 | 9154 |
| 116 | FIM1_118 | QGN13480.1 | CP015054.1 | 345 | 60 | 169 | 230 | 52 | 155 | 520 | 50 | 139 |
| 117 | FIM1_119 | QGN13481.1 | CP015054.1 | 12559 | 7723 | 6052 | 11742 | 7402 | 5713 | 8024 | 8180 | 5802 |
| 118 | FIM1_120 | QGN13482.1 | CP015054.1 | 2595 | 1062 | 1191 | 2002 | 792 | 1129 | 1994 | 1183 | 1048 |
| 119 | FIM1_121 | QGN13483.1 | CP015054.1 | 6595 | 4375 | 2626 | 4574 | 3096 | 2554 | 2117 | 4737 | 2608 |
| 120 | FIM1_122 | QGN13484.1 | CP015054.1 | 8146 | 2465 | 2804 | 6073 | 1857 | 2617 | 6005 | 2688 | 2544 |
| 121 | FIM1_123 | QGN13485.1 | CP015054.1 | 5091 | 4376 | 4176 | 4544 | 2634 | 4145 | 6092 | 4306 | 4160 |
| 122 | FIM1_124 | QGN13486.1 | CP015054.1 | 2696 | 572 | 674 | 2101 | 341 | 654 | 2681 | 624 | 594 |
| 123 | FIM1_125 | QGN13487.1 | CP015054.1 | 1147 | 204 | 353 | 855 | 99 | 255 | 968 | 181 | 251 |
| 124 | FIM1_126 | QGN13488.1 | CP015054.1 | 3124 | 733 | 1195 | 2591 | 605 | 1067 | 3054 | 858 | 1092 |
| 125 | FIM1_127 | QGN13489.1 | CP015054.1 | 977 | 331 | 662 | 976 | 210 | 613 | 1411 | 394 | 546 |
| 126 | FIM1_128 | QGN13490.1 | CP015054.1 | 30211 | 13456 | 14706 | 24098 | 9521 | 13777 | 25907 | 14204 | 14617 |
| 127 | FIM1_129 | QGN13491.1 | CP015054.1 | 2373 | 866 | 826 | 1723 | 623 | 791 | 1549 | 941 | 791 |
| 128 | FIM1_130 | QGN13492.1 | CP015054.1 | 17067 | 5499 | 10142 | 14058 | 3274 | 9238 | 22935 | 5653 | 9143 |
| 129 | FIM1_131 | QGN13493.1 | CP015054.1 | 8857 | 1844 | 1833 | 6975 | 1283 | 1641 | 7955 | 1886 | 1561 |
| 130 | FIM1_132 | QGN13494.1 | CP015054.1 | 33135 | 21788 | 19004 | 31663 | 17430 | 17162 | 32594 | 22040 | 18527 |
| 131 | FIM1_133 | QGN13495.1 | CP015054.1 | 14852 | 4340 | 3870 | 11412 | 3335 | 3373 | 11470 | 4336 | 3262 |
| 132 | FIM1_134 | QGN13496.1 | CP015054.1 | 10291 | 3500 | 4158 | 8573 | 2043 | 3917 | 11731 | 3725 | 3683 |
| 133 | FIM1_135 | QGN13497.1 | CP015054.1 | 2966 | 1020 | 1270 | 2901 | 756 | 1200 | 3290 | 1017 | 1190 |
| 134 | FIM1_136 | QGN13498.1 | CP015054.1 | 598 | 35 | 78 | 527 | 19 | 63 | 1201 | 41 | 69 |
| 135 | FIM1_137 | QGN13499.1 | CP015054.1 | 2417 | 123 | 382 | 2049 | 85 | 364 | 5190 | 142 | 378 |
| 136 | FIM1_138 | QGN13500.1 | CP015054.1 | 14599 | 11596 | 10549 | 13861 | 9941 | 10218 | 14183 | 11926 | 10182 |
| 137 | FIM1_140 | QGN13501.1 | CP015054.1 | 26080 | 17116 | 16001 | 25391 | 14578 | 15434 | 25088 | 18017 | 15476 |
| 138 | FIM1_141 | QGN13502.1 | CP015054.1 | 1945 | 506 | 655 | 1451 | 316 | 679 | 2429 | 514 | 607 |
| 139 | FIM1_142 | QGN13503.1 | CP015054.1 | 24106 | 10091 | 10964 | 20647 | 7109 | 10466 | 24824 | 10565 | 10325 |
| 140 | FIM1_143 | QGN13504.1 | CP015054.1 | 677 | 130 | 132 | 485 | 69 | 100 | 1101 | 154 | 102 |
| 141 | FIM1_144 | QGN13505.1 | CP015054.1 | 1186 | 459 | 693 | 947 | 262 | 672 | 1049 | 564 | 679 |
| 142 | FIM1_145 | QGN13506.1 | CP015054.1 | 40145 | 14415 | 10917 | 32895 | 11592 | 10167 | 30794 | 14853 | 10324 |

| | | | | | | | | | | | | |
|---|---|---|---|---|---|---|---|---|---|---|---|---|
| 143 | FIM1_146 | QGN13507.1 | CP015054.1 | 6544 | 1858 | 1941 | 5605 | 1304 | 1750 | 6045 | 2022 | 1712 |
| 144 | FIM1_147 | QGN13508.1 | CP015054.1 | 5661 | 1184 | 1745 | 4097 | 914 | 1643 | 4612 | 1236 | 1537 |
| 145 | FIM1_148 | QGN13509.1 | CP015054.1 | 2288 | 480 | 710 | 1699 | 360 | 669 | 2316 | 481 | 635 |
| 146 | FIM1_149 | QGN13510.1 | CP015054.1 | 1099 | 361 | 586 | 867 | 255 | 540 | 895 | 410 | 472 |
| 147 | FIM1_150 | QGN13511.1 | CP015054.1 | 2641 | 509 | 1088 | 2020 | 391 | 1009 | 2077 | 552 | 904 |
| 148 | FIM1_151 | QGN13512.1 | CP015054.1 | 2217 | 429 | 763 | 1855 | 351 | 714 | 2639 | 494 | 619 |
| 149 | FIM1_153 | QGN13513.1 | CP015054.1 | 775 | 384 | 734 | 543 | 318 | 686 | 371 | 412 | 585 |
| 150 | FIM1_154 | QGN13514.1 | CP015054.1 | 7085 | 865 | 1659 | 5135 | 711 | 1621 | 8785 | 944 | 1474 |
| 151 | FIM1_155 | QGN13515.1 | CP015054.1 | 10817 | 2286 | 3078 | 8637 | 1795 | 2844 | 10355 | 2504 | 3058 |
| 152 | FIM1_156 | QGN13516.1 | CP015054.1 | 3166 | 620 | 952 | 2306 | 437 | 834 | 4440 | 689 | 865 |
| 153 | FIM1_157 | QGN13517.1 | CP015054.1 | 5028 | 528 | 1240 | 4755 | 270 | 1067 | 12666 | 533 | 1058 |
| 154 | FIM1_158 | QGN13518.1 | CP015054.1 | 13138 | 4929 | 6749 | 11588 | 3147 | 6072 | 16431 | 5299 | 6120 |
| 155 | FIM1_159 | QGN13519.1 | CP015054.1 | 16281 | 4783 | 5948 | 13357 | 3342 | 5440 | 15326 | 4941 | 5385 |
| 156 | FIM1_160 | QGN13520.1 | CP015054.1 | 13920 | 4113 | 5229 | 11136 | 3028 | 4738 | 13877 | 4493 | 4614 |
| 157 | FIM1_161 | QGN13521.1 | CP015054.1 | 2324 | 350 | 498 | 1846 | 187 | 503 | 2732 | 380 | 519 |
| 158 | FIM1_162 | QGN13522.1 | CP015054.1 | 3312 | 579 | 735 | 2738 | 543 | 688 | 2284 | 698 | 721 |
| 159 | FIM1_163 | QGN13523.1 | CP015054.1 | 677 | 140 | 287 | 511 | 93 | 221 | 1022 | 160 | 235 |
| 160 | FIM1_164 | QGN13524.1 | CP015054.1 | 2915 | 510 | 706 | 2152 | 395 | 683 | 2755 | 554 | 665 |
| 161 | FIM1_165 | QGN13525.1 | CP015054.1 | 822 | 449 | 508 | 783 | 346 | 530 | 717 | 511 | 463 |
| 162 | FIM1_166 | QGN13526.1 | CP015054.1 | 1905 | 380 | 795 | 1708 | 363 | 775 | 1111 | 401 | 615 |
| 163 | FIM1_167 | QGN13527.1 | CP015054.1 | 1110 | 207 | 360 | 929 | 142 | 295 | 1222 | 246 | 256 |
| 164 | FIM1_168 | QGN13528.1 | CP015054.1 | 2777 | 886 | 1253 | 2064 | 694 | 1180 | 2112 | 918 | 1006 |
| 165 | FIM1_169 | QGN13529.1 | CP015054.1 | 6079 | 783 | 1486 | 4531 | 629 | 1413 | 6658 | 844 | 1175 |
| 166 | FIM1_170 | QGN13530.1 | CP015054.1 | 24794 | 9289 | 9783 | 20527 | 6905 | 9131 | 24648 | 10070 | 9181 |
| 167 | FIM1_171 | QGN13531.1 | CP015054.1 | 2840 | 1559 | 1658 | 2331 | 1200 | 1697 | 1987 | 1731 | 1551 |
| 168 | FIM1_172 | QGN13532.1 | CP015054.1 | 1398 | 417 | 743 | 1188 | 313 | 683 | 1521 | 453 | 593 |
| 169 | FIM1_173 | QGN13533.1 | CP015054.1 | 1539 | 765 | 805 | 1375 | 526 | 709 | 1220 | 780 | 651 |
| 170 | FIM1_174 | QGN13534.1 | CP015054.1 | 662 | 172 | 366 | 681 | 136 | 320 | 623 | 193 | 292 |
| 171 | FIM1_175 | QGN13535.1 | CP015054.1 | 173 | 43 | 98 | 176 | 50 | 76 | 139 | 34 | 59 |
| 172 | FIM1_176 | QGN13536.1 | CP015054.1 | 6287 | 1458 | 1710 | 4327 | 1173 | 1714 | 3652 | 1633 | 1580 |
| 173 | FIM1_177 | QGN13537.1 | CP015054.1 | 11102 | 7959 | 7316 | 10936 | 6625 | 7457 | 8790 | 8340 | 7378 |
| 174 | FIM1_178 | QGN13538.1 | CP015054.1 | 1770 | 535 | 860 | 1310 | 393 | 884 | 1779 | 594 | 767 |
| 175 | FIM1_179 | QGN13539.1 | CP015054.1 | 6052 | 1752 | 1968 | 4749 | 1296 | 1772 | 5937 | 1730 | 1716 |
| 176 | FIM1_180 | QGN13540.1 | CP015054.1 | 4626 | 1034 | 1378 | 4071 | 790 | 1320 | 4627 | 1145 | 1202 |
| 177 | FIM1_181 | QGN13541.1 | CP015054.1 | 22461 | 10845 | 9932 | 19451 | 8355 | 9204 | 21567 | 11223 | 9213 |
| 178 | FIM1_182 | QGN13542.1 | CP015054.1 | 15628 | 4082 | 4268 | 11789 | 3694 | 3741 | 11945 | 4306 | 4021 |
| 179 | FIM1_183 | QGN13543.1 | CP015054.1 | 144 | 34 | 87 | 105 | 36 | 76 | 155 | 61 | 75 |
| 180 | FIM1_184 | QGN13544.1 | CP015054.1 | 10867 | 3408 | 3878 | 7677 | 2352 | 3520 | 8994 | 3539 | 3519 |
| 181 | FIM1_185 | QGN13545.1 | CP015054.1 | 1886 | 398 | 353 | 1297 | 340 | 327 | 2110 | 425 | 373 |
| 182 | FIM1_186 | QGN13546.1 | CP015054.1 | 1055 | 122 | 294 | 769 | 63 | 205 | 1184 | 117 | 169 |
| 183 | FIM1_187 | QGN13547.1 | CP015054.1 | 15553 | 6902 | 7247 | 12272 | 4308 | 6571 | 7895 | 6559 | 6701 |
| 184 | FIM1_189 | QGN13548.1 | CP015054.1 | 3438 | 1128 | 1069 | 2628 | 818 | 1028 | 2539 | 1200 | 937 |
| 185 | FIM1_190 | QGN13549.1 | CP015054.1 | 252 | 125 | 171 | 196 | 86 | 151 | 312 | 104 | 172 |
| 186 | FIM1_191 | QGN13550.1 | CP015054.1 | 2182 | 181 | 481 | 1743 | 172 | 431 | 2424 | 250 | 385 |
| 187 | FIM1_192 | QGN13551.1 | CP015054.1 | 16431 | 1614 | 3138 | 12348 | 1113 | 2899 | 18387 | 1708 | 2953 |
| 188 | FIM1_193 | QGN13552.1 | CP015054.1 | 1222 | 6 | 16 | 739 | 11 | 12 | 872 | 12 | 14 |
| 189 | FIM1_194 | QGN13553.1 | CP015054.1 | 9518 | 1574 | 2217 | 6426 | 1286 | 2028 | 6928 | 1697 | 1835 |
| 190 | FIM1_195 | QGN13554.1 | CP015054.1 | 1969 | 844 | 1267 | 1691 | 690 | 1178 | 1335 | 903 | 1017 |

| | | | | | | | | | | | |
|---|---|---|---|---|---|---|---|---|---|---|---|
| 191 | FIM1_196 | QGN13555.1 | CP015054.1 | 5833 | 2532 | 2471 | 5513 | 1974 | 2350 | 4802 | 2749 | 2270 |
| 192 | FIM1_197 | QGN13556.1 | CP015054.1 | 4665 | 1522 | 1450 | 4175 | 682 | 1377 | 8267 | 1551 | 1182 |
| 193 | FIM1_198 | QGN13557.1 | CP015054.1 | 13757 | 4028 | 2833 | 10414 | 2781 | 2492 | 13473 | 4329 | 2499 |
| 194 | FIM1_199 | QGN13558.1 | CP015054.1 | 3268 | 614 | 1056 | 2714 | 366 | 972 | 2681 | 581 | 814 |
| 195 | FIM1_200 | QGN13559.1 | CP015054.1 | 5088 | 1989 | 2040 | 3818 | 1598 | 1881 | 4046 | 2219 | 1811 |
| 196 | FIM1_201 | QGN13560.1 | CP015054.1 | 5209 | 1272 | 1928 | 4269 | 1020 | 1835 | 3899 | 1450 | 1657 |
| 197 | FIM1_202 | QGN13561.1 | CP015054.1 | 26 | 7 | 10 | 22 | 0 | 14 | 18 | 4 | 11 |
| 198 | FIM1_203 | QGN13562.1 | CP015054.1 | 2425 | 534 | 1348 | 2014 | 411 | 1298 | 2494 | 566 | 1033 |
| 199 | FIM1_204 | QGN13563.1 | CP015054.1 | 323 | 58 | 119 | 262 | 40 | 126 | 454 | 42 | 116 |
| 200 | FIM1_205 | QGN13564.1 | CP015054.1 | 5494 | 1402 | 1227 | 4445 | 1015 | 1129 | 1773 | 1524 | 1060 |
| 201 | FIM1_206 | QGN13565.1 | CP015054.1 | 7303 | 1944 | 2646 | 5673 | 1285 | 2500 | 7658 | 2143 | 2373 |
| 202 | FIM1_207 | QGN13566.1 | CP015054.1 | 1084 | 423 | 852 | 946 | 290 | 735 | 1265 | 482 | 770 |
| 203 | FIM1_208 | QGN13567.1 | CP015054.1 | 5366 | 2017 | 3030 | 4759 | 1226 | 2796 | 6586 | 1989 | 2746 |
| 204 | FIM1_209 | QGN13568.1 | CP015054.1 | 15236 | 2113 | 3079 | 11491 | 1707 | 2862 | 14712 | 2468 | 2776 |
| 205 | FIM1_210 | QGN13569.1 | CP015054.1 | 5712 | 2231 | 2639 | 5155 | 1910 | 2638 | 5320 | 2416 | 2401 |
| 206 | FIM1_211 | QGN13570.1 | CP015054.1 | 3649 | 601 | 1327 | 2934 | 468 | 1272 | 4111 | 613 | 1031 |
| 207 | FIM1_212 | QGN13571.1 | CP015054.1 | 4284 | 582 | 1197 | 3165 | 480 | 1060 | 4434 | 648 | 1034 |
| 208 | FIM1_213 | QGN13572.1 | CP015054.1 | 3246 | 1035 | 1143 | 2376 | 723 | 1175 | 2724 | 1059 | 1069 |
| 209 | FIM1_214 | QGN13573.1 | CP015054.1 | 587 | 66 | 329 | 490 | 69 | 339 | 699 | 78 | 230 |
| 210 | FIM1_215 | QGN13574.1 | CP015054.1 | 2295 | 385 | 615 | 1817 | 334 | 608 | 2149 | 426 | 570 |
| 211 | FIM1_216 | QGN13575.1 | CP015054.1 | 12675 | 2548 | 3040 | 10749 | 1854 | 2766 | 13380 | 2550 | 2908 |
| 212 | FIM1_217 | QGN13576.1 | CP015054.1 | 5192 | 1852 | 2300 | 5025 | 1444 | 2146 | 5336 | 2125 | 2118 |
| 213 | FIM1_218 | QGN13577.1 | CP015054.1 | 5122 | 211 | 762 | 3508 | 166 | 695 | 5881 | 262 | 685 |
| 214 | FIM1_219 | QGN13578.1 | CP015054.1 | 3251 | 377 | 819 | 2687 | 323 | 811 | 3896 | 403 | 785 |
| 215 | FIM1_220 | QGN13579.1 | CP015054.1 | 719 | 190 | 421 | 598 | 155 | 354 | 823 | 218 | 318 |
| 216 | FIM1_221 | QGN13580.1 | CP015054.1 | 2014 | 854 | 1083 | 1672 | 597 | 1145 | 1998 | 884 | 958 |
| 217 | FIM1_222 | QGN13581.1 | CP015054.1 | 5256 | 905 | 1426 | 4061 | 633 | 1355 | 5576 | 995 | 1199 |
| 218 | FIM1_224 | QGN13582.1 | CP015054.1 | 3582 | 1621 | 1933 | 2761 | 1254 | 1765 | 2252 | 1769 | 1719 |
| 219 | FIM1_225 | QGN13583.1 | CP015054.1 | 3148 | 449 | 1203 | 2806 | 353 | 1129 | 3500 | 390 | 1074 |
| 220 | FIM1_226 | QGN13584.1 | CP015054.1 | 708 | 86 | 412 | 681 | 59 | 370 | 1517 | 96 | 328 |
| 221 | FIM1_227 | QGN13585.1 | CP015054.1 | 7779 | 1081 | 1908 | 6748 | 668 | 1705 | 8936 | 1074 | 1673 |
| 222 | FIM1_228 | QGN13586.1 | CP015054.1 | 165 | 34 | 130 | 116 | 33 | 143 | 119 | 61 | 123 |
| 223 | FIM1_229 | QGN13587.1 | CP015054.1 | 19716 | 5385 | 9606 | 16067 | 3620 | 8849 | 20195 | 5973 | 8546 |
| 224 | FIM1_230 | QGN13588.1 | CP015054.1 | 3615 | 1089 | 1836 | 3123 | 716 | 1864 | 4586 | 1146 | 1637 |
| 225 | FIM1_231 | QGN13589.1 | CP015054.1 | 193 | 29 | 210 | 152 | 27 | 196 | 267 | 41 | 220 |
| 226 | FIM1_232 | QGN13590.1 | CP015054.1 | 1 | 1 | 0 | 0 | 2 | 0 | 1 | 1 | 1 |
| 227 | FIM1_233 | QGN13591.1 | CP015054.1 | 13093 | 3943 | 4426 | 10906 | 2826 | 4028 | 10983 | 4254 | 3926 |
| 228 | FIM1_234 | QGN13592.1 | CP015054.1 | 5035 | 1118 | 1712 | 3880 | 806 | 1444 | 4574 | 1203 | 1370 |
| 229 | FIM1_235 | QGN13593.1 | CP015054.1 | 1817 | 292 | 804 | 1422 | 228 | 812 | 2249 | 272 | 653 |
| 230 | FIM1_236 | QGN13594.1 | CP015054.1 | 7708 | 1163 | 2155 | 6104 | 886 | 2004 | 9398 | 1319 | 1800 |
| 231 | FIM1_237 | QGN13595.1 | CP015054.1 | 944 | 172 | 699 | 881 | 125 | 570 | 1876 | 186 | 555 |
| 232 | FIM1_238 | QGN13596.1 | CP015054.1 | 972 | 152 | 492 | 975 | 126 | 447 | 1095 | 179 | 385 |
| 233 | FIM1_239 | QGN13597.1 | CP015054.1 | 166 | 25 | 94 | 111 | 22 | 87 | 219 | 42 | 62 |
| 234 | FIM1_240 | QGN13598.1 | CP015054.1 | 1814 | 134 | 485 | 1410 | 127 | 448 | 2830 | 160 | 422 |
| 235 | FIM1_241 | QGN13599.1 | CP015054.1 | 8705 | 2817 | 1886 | 6801 | 1968 | 1863 | 10170 | 2860 | 1762 |
| 236 | FIM1_242 | QGN13600.1 | CP015054.1 | 120 | 26 | 66 | 78 | 19 | 60 | 103 | 35 | 40 |
| 237 | FIM1_243 | QGN13601.1 | CP015054.1 | 3880 | 853 | 995 | 3017 | 579 | 951 | 3818 | 873 | 886 |
| 238 | FIM1_244 | QGN13602.1 | CP015054.1 | 38 | 3 | 11 | 41 | 3 | 4 | 22 | 4 | 0 |

| | | | | | | | | | | | | |
|---|---|---|---|---|---|---|---|---|---|---|---|---|
| 239 | FIM1_245 | QGN13603.1 | CP015054.1 | 7646 | 5785 | 4004 | 6730 | 4393 | 3667 | 3016 | 6177 | 3819 |
| 240 | FIM1_246 | QGN13604.1 | CP015054.1 | 6853 | 1388 | 1939 | 5248 | 831 | 1685 | 7307 | 1440 | 1597 |
| 241 | FIM1_247 | QGN13605.1 | CP015054.1 | 7061 | 175 | 1143 | 5465 | 95 | 984 | 6473 | 214 | 973 |
| 242 | FIM1_248 | QGN13606.1 | CP015054.1 | 2494 | 868 | 787 | 1962 | 526 | 673 | 2583 | 839 | 658 |
| 243 | FIM1_249 | QGN13607.1 | CP015054.1 | 1466 | 1504 | 817 | 932 | 959 | 712 | 241 | 1672 | 678 |
| 244 | FIM1_250 | QGN13608.1 | CP015054.1 | 1739 | 423 | 651 | 1090 | 339 | 558 | 1278 | 471 | 628 |
| 245 | FIM1_251 | QGN13609.1 | CP015054.1 | 1128 | 222 | 588 | 815 | 219 | 565 | 839 | 282 | 467 |
| 246 | FIM1_252 | QGN13610.1 | CP015054.1 | 5124 | 1236 | 1862 | 4476 | 1057 | 1682 | 4273 | 1356 | 1566 |
| 247 | FIM1_253 | QGN13611.1 | CP015054.1 | 10398 | 3699 | 3669 | 9312 | 2863 | 3319 | 10194 | 3972 | 3307 |
| 248 | FIM1_254 | QGN13612.1 | CP015054.1 | 1213 | 319 | 636 | 1096 | 247 | 584 | 917 | 355 | 450 |
| 249 | FIM1_255 | QGN13613.1 | CP015054.1 | 8678 | 2318 | 3990 | 7753 | 1390 | 3359 | 11772 | 2418 | 3467 |
| 250 | FIM1_256 | QGN13614.1 | CP015054.1 | 7223 | 3230 | 2514 | 6109 | 1863 | 2230 | 4771 | 3497 | 2484 |
| 251 | FIM1_257 | QGN13615.1 | CP015054.1 | 2852 | 619 | 887 | 2271 | 458 | 748 | 1830 | 618 | 737 |
| 252 | FIM1_258 | QGN13616.1 | CP015054.1 | 3885 | 604 | 1184 | 2933 | 505 | 1092 | 3622 | 666 | 980 |
| 253 | FIM1_259 | QGN13617.1 | CP015054.1 | 5909 | 2572 | 2719 | 5214 | 1998 | 2685 | 4848 | 2876 | 2577 |
| 254 | FIM1_260 | QGN13618.1 | CP015054.1 | 1148 | 333 | 320 | 819 | 232 | 338 | 1392 | 321 | 267 |
| 255 | FIM1_261 | QGN13619.1 | CP015054.1 | 4285 | 825 | 1425 | 3195 | 643 | 1377 | 4234 | 857 | 1210 |
| 256 | FIM1_262 | QGN13620.1 | CP015054.1 | 2106 | 1538 | 912 | 1599 | 1068 | 891 | 994 | 1590 | 913 |
| 257 | FIM1_263 | QGN13621.1 | CP015054.1 | 8694 | 2706 | 2401 | 7283 | 1698 | 2097 | 9995 | 3013 | 1901 |
| 258 | FIM1_264 | QGN13622.1 | CP015054.1 | 2901 | 632 | 826 | 2266 | 484 | 754 | 2660 | 706 | 718 |
| 259 | FIM1_265 | QGN13623.1 | CP015054.1 | 5152 | 865 | 1386 | 3982 | 659 | 1287 | 4008 | 895 | 1121 |
| 260 | FIM1_266 | QGN13624.1 | CP015054.1 | 4173 | 534 | 789 | 3116 | 415 | 690 | 2998 | 585 | 661 |
| 261 | FIM1_267 | QGN13625.1 | CP015054.1 | 12822 | 4834 | 4222 | 9814 | 3364 | 3932 | 9227 | 5207 | 3816 |
| 262 | FIM1_268 | QGN13626.1 | CP015054.1 | 11577 | 3546 | 5390 | 9550 | 1933 | 4940 | 16416 | 3579 | 4754 |
| 263 | FIM1_269 | QGN13627.1 | CP015054.1 | 3574 | 852 | 1285 | 2786 | 648 | 1144 | 2172 | 884 | 1035 |
| 264 | FIM1_270 | QGN13628.1 | CP015054.1 | 187 | 76 | 163 | 195 | 54 | 162 | 191 | 60 | 137 |
| 265 | FIM1_271 | QGN13629.1 | CP015054.1 | 8870 | 1803 | 2399 | 6497 | 1422 | 2228 | 9470 | 1984 | 2202 |
| 266 | FIM1_272 | QGN13630.1 | CP015054.1 | 2039 | 430 | 642 | 1611 | 345 | 514 | 1738 | 467 | 504 |
| 267 | FIM1_273 | QGN13631.1 | CP015054.1 | 1331 | 103 | 570 | 1147 | 71 | 484 | 1349 | 115 | 304 |
| 268 | FIM1_274 | QGN13632.1 | CP015054.1 | 3217 | 1776 | 1456 | 2316 | 1321 | 1338 | 1395 | 1961 | 1351 |
| 269 | FIM1_275 | QGN13633.1 | CP015054.1 | 2705 | 1124 | 1524 | 2513 | 935 | 1563 | 2217 | 1189 | 1392 |
| 270 | FIM1_276 | QGN13634.1 | CP015054.1 | 3565 | 522 | 884 | 2581 | 386 | 858 | 3032 | 543 | 701 |
| 271 | FIM1_277 | QGN13635.1 | CP015054.1 | 1017 | 131 | 292 | 758 | 114 | 269 | 882 | 166 | 228 |
| 272 | FIM1_278 | QGN13636.1 | CP015054.1 | 1342 | 470 | 696 | 1087 | 304 | 594 | 1070 | 463 | 596 |
| 273 | FIM1_279 | QGN13637.1 | CP015054.1 | 5722 | 3018 | 2947 | 5079 | 2303 | 2811 | 6628 | 3085 | 2818 |
| 274 | FIM1_280 | QGN13638.1 | CP015054.1 | 63 | 20 | 16 | 35 | 15 | 14 | 27 | 16 | 12 |
| 275 | FIM1_281 | QGN13639.1 | CP015054.1 | 9724 | 2037 | 2062 | 7406 | 1495 | 1839 | 9786 | 2142 | 1805 |
| 276 | FIM1_282 | QGN13640.1 | CP015054.1 | 1534 | 208 | 272 | 1170 | 175 | 252 | 1808 | 214 | 229 |
| 277 | FIM1_283 | QGN13641.1 | CP015054.1 | 2939 | 507 | 311 | 1702 | 322 | 297 | 2099 | 503 | 305 |
| 278 | FIM1_284 | QGN13642.1 | CP015054.1 | 2372 | 1123 | 827 | 1881 | 644 | 765 | 1886 | 1188 | 725 |
| 279 | FIM1_285 | QGN13643.1 | CP015054.1 | 5621 | 1488 | 1418 | 4309 | 990 | 1200 | 4851 | 1550 | 1194 |
| 280 | FIM1_286 | QGN13644.1 | CP015054.1 | 11634 | 1968 | 2557 | 9084 | 1501 | 2481 | 13946 | 2082 | 2614 |
| 281 | FIM1_287 | QGN13645.1 | CP015054.1 | 1857 | 410 | 569 | 1556 | 361 | 526 | 1663 | 522 | 513 |
| 282 | FIM1_288 | QGN13646.1 | CP015054.1 | 243 | 93 | 112 | 256 | 55 | 137 | 215 | 101 | 110 |
| 283 | FIM1_290 | QGN13647.1 | CP015054.1 | 1063 | 342 | 337 | 987 | 275 | 279 | 887 | 349 | 314 |
| 284 | FIM1_291 | QGN13648.1 | CP015054.1 | 3995 | 901 | 1167 | 3276 | 643 | 1044 | 5723 | 931 | 1002 |
| 285 | FIM1_292 | QGN13649.1 | CP015054.1 | 2594 | 297 | 273 | 2012 | 183 | 238 | 6254 | 326 | 253 |
| 286 | FIM1_293 | QGN13650.1 | CP015054.1 | 9604 | 1600 | 1970 | 7583 | 1317 | 1780 | 12259 | 1833 | 1774 |

| | | | | | | | | | | | | |
|---|---|---|---|---|---|---|---|---|---|---|---|---|
| 287 | FIM1_294 | QGN13651.1 | CP015054.1 | 2842 | 104 | 126 | 1465 | 86 | 140 | 740 | 113 | 128 |
| 288 | FIM1_295 | QGN13652.1 | CP015054.1 | 124 | 21 | 25 | 76 | 9 | 16 | 100 | 17 | 24 |
| 289 | FIM1_296 | QGN13653.1 | CP015054.1 | 233 | 55 | 44 | 127 | 82 | 61 | 193 | 34 | 54 |
| 290 | FIM1_297 | QGN13654.1 | CP015054.1 | 112 | 11 | 20 | 94 | 9 | 17 | 111 | 11 | 10 |
| 291 | FIM1_298 | QGN13655.1 | CP015054.1 | 211 | 12 | 27 | 158 | 15 | 32 | 132 | 15 | 24 |
| 292 | FIM1_299 | QGN13656.1 | CP015054.1 | 2130 | 367 | 590 | 1445 | 269 | 505 | 1899 | 390 | 512 |
| 293 | FIM1_300 | QGN13657.1 | CP015054.1 | 3196 | 2587 | 1412 | 2201 | 1603 | 1229 | 442 | 2700 | 1268 |
| 294 | FIM1_301 | QGN13658.1 | CP015054.1 | 671 | 95 | 229 | 475 | 59 | 209 | 399 | 86 | 160 |
| 295 | FIM1_302 | QGN13659.1 | CP015054.1 | 2066 | 287 | 722 | 1621 | 218 | 635 | 1372 | 348 | 612 |
| 296 | FIM1_303 | QGN13660.1 | CP015054.1 | 52014 | 29714 | 23724 | 49567 | 17949 | 20404 | 84015 | 30479 | 21494 |
| 297 | FIM1_304 | QGN13661.1 | CP015054.1 | 4005 | 228 | 491 | 3115 | 153 | 406 | 6818 | 241 | 392 |
| 298 | FIM1_305 | QGN13662.1 | CP015054.1 | 1582 | 63 | 173 | 1338 | 30 | 157 | 4627 | 58 | 169 |
| 299 | FIM1_306 | QGN13663.1 | CP015054.1 | 389 | 85 | 105 | 272 | 36 | 84 | 1473 | 86 | 70 |
| 300 | FIM1_307 | QGN13664.1 | CP015054.1 | 2733 | 806 | 1225 | 1820 | 579 | 1053 | 1734 | 852 | 933 |
| 301 | FIM1_308 | QGN13665.1 | CP015054.1 | 4896 | 2295 | 1958 | 3741 | 1865 | 1845 | 3046 | 2654 | 1832 |
| 302 | FIM1_309 | QGN13666.1 | CP015054.1 | 10427 | 3530 | 5028 | 8292 | 2589 | 4628 | 8559 | 3767 | 4615 |
| 303 | FIM1_310 | QGN13667.1 | CP015054.1 | 10114 | 4645 | 3103 | 7887 | 3136 | 3041 | 5943 | 4903 | 2932 |
| 304 | FIM1_311 | QGN13668.1 | CP015054.1 | 8030 | 2308 | 2473 | 6419 | 1646 | 2183 | 7029 | 2506 | 1964 |
| 305 | FIM1_312 | QGN13669.1 | CP015054.1 | 7715 | 2065 | 3603 | 6292 | 1197 | 3285 | 10502 | 2101 | 3207 |
| 306 | FIM1_313 | QGN13670.1 | CP015054.1 | 7416 | 2035 | 2343 | 6323 | 1746 | 2363 | 5137 | 2233 | 2139 |
| 307 | FIM1_314 | QGN13671.1 | CP015054.1 | 5331 | 788 | 3019 | 4467 | 709 | 2554 | 4719 | 882 | 1932 |
| 308 | FIM1_315 | QGN13672.1 | CP015054.1 | 2212 | 253 | 574 | 1747 | 183 | 514 | 1832 | 274 | 497 |
| 309 | FIM1_316 | QGN13673.1 | CP015054.1 | 1608 | 1325 | 956 | 1242 | 917 | 747 | 360 | 1489 | 859 |
| 310 | FIM1_317 | QGN13674.1 | CP015054.1 | 107 | 19 | 137 | 120 | 8 | 123 | 192 | 13 | 99 |
| 311 | FIM1_318 | QGN13675.1 | CP015054.1 | 19655 | 7360 | 6434 | 16285 | 5428 | 5679 | 17770 | 7792 | 5885 |
| 312 | FIM1_319 | QGN13676.1 | CP015054.1 | 816 | 178 | 223 | 672 | 135 | 217 | 679 | 160 | 208 |
| 313 | FIM1_320 | QGN13677.1 | CP015054.1 | 4215 | 1095 | 1201 | 3513 | 476 | 1088 | 6673 | 903 | 1168 |
| 314 | FIM1_321 | QGN13678.1 | CP015054.1 | 109 | 142 | 96 | 100 | 42 | 55 | 53 | 129 | 112 |
| 315 | FIM1_322 | QGN13679.1 | CP015054.1 | 28580 | 14380 | 13106 | 25394 | 11572 | 12397 | 27907 | 14656 | 12462 |
| 316 | FIM1_323 | QGN13680.1 | CP015054.1 | 1365 | 208 | 561 | 1031 | 101 | 509 | 1352 | 141 | 562 |
| 317 | FIM1_324 | QGN13681.1 | CP015054.1 | 7511 | 4757 | 4641 | 6130 | 1616 | 3957 | 8586 | 3813 | 4654 |
| 318 | FIM1_325 | QGN13682.1 | CP015054.1 | 52 | 11 | 16 | 40 | 7 | 12 | 16 | 7 | 16 |
| 319 | FIM1_326 | QGN13683.1 | CP015054.1 | 8033 | 162 | 116 | 3305 | 93 | 150 | 26716 | 133 | 82 |
| 320 | FIM1_327 | QGN13684.1 | CP015054.1 | 3740 | 739 | 1096 | 3043 | 552 | 996 | 4071 | 706 | 988 |
| 321 | FIM1_328 | QGN13685.1 | CP015054.1 | 530 | 110 | 97 | 371 | 70 | 123 | 711 | 112 | 117 |
| 322 | FIM1_329 | QGN13686.1 | CP015054.1 | 3179 | 207 | 617 | 2355 | 154 | 592 | 3449 | 226 | 501 |
| 323 | FIM1_330 | QGN13687.1 | CP015054.1 | 21688 | 8174 | 8139 | 19507 | 5278 | 7267 | 21877 | 8729 | 7867 |
| 324 | FIM1_331 | QGN13688.1 | CP015054.1 | 994 | 180 | 286 | 854 | 103 | 255 | 1107 | 220 | 255 |
| 325 | FIM1_332 | QGN13689.1 | CP015054.1 | 2958 | 532 | 973 | 2746 | 350 | 900 | 3486 | 624 | 770 |
| 326 | FIM1_333 | QGN13690.1 | CP015054.1 | 3713 | 476 | 797 | 3030 | 334 | 764 | 4156 | 469 | 713 |
| 327 | FIM1_334 | QGN13691.1 | CP015054.1 | 5025 | 1519 | 1727 | 3948 | 1157 | 1500 | 3646 | 1680 | 1421 |
| 328 | FIM1_335 | QGN13692.1 | CP015054.1 | 2052 | 1337 | 1054 | 1703 | 928 | 934 | 509 | 1440 | 963 |
| 329 | FIM1_336 | QGN13693.1 | CP015054.1 | 2160 | 152 | 499 | 1891 | 102 | 414 | 3382 | 152 | 407 |
| 330 | FIM1_337 | QGN13694.1 | CP015054.1 | 7542 | 2056 | 3467 | 7255 | 1397 | 3180 | 8885 | 2189 | 3159 |
| 331 | FIM1_338 | QGN13695.1 | CP015054.1 | 6613 | 2101 | 1114 | 4802 | 1425 | 1082 | 4163 | 2182 | 1117 |
| 332 | FIM1_339 | QGN13696.1 | CP015054.1 | 8986 | 3842 | 3092 | 6802 | 1633 | 2731 | 9570 | 3844 | 2361 |
| 333 | FIM1_340 | QGN13697.1 | CP015054.1 | 1025 | 151 | 274 | 773 | 104 | 261 | 1017 | 168 | 253 |
| 334 | FIM1_341 | QGN13698.1 | CP015054.1 | 12256 | 4007 | 3465 | 9580 | 3060 | 3285 | 8777 | 4378 | 3274 |

| | | | | | | | | | | | |
|---|---|---|---|---|---|---|---|---|---|---|---|
| 335 | FIM1_342 | QGN13699.1 | CP015054.1 | 2455 | 743 | 1036 | 2290 | 540 | 890 | 2295 | 798 | 851 |
| 336 | FIM1_343 | QGN13700.1 | CP015054.1 | 8637 | 1397 | 1985 | 6691 | 978 | 1871 | 7882 | 1471 | 1939 |
| 337 | FIM1_344 | QGN13701.1 | CP015054.1 | 3286 | 1049 | 1587 | 2937 | 732 | 1496 | 3771 | 1136 | 1457 |
| 338 | FIM1_345 | QGN13702.1 | CP015054.1 | 7223 | 3203 | 885 | 6797 | 1851 | 794 | 15277 | 3402 | 795 |
| 339 | FIM1_346 | QGN13703.1 | CP015054.1 | 16817 | 4521 | 4071 | 14254 | 3197 | 3790 | 19589 | 4880 | 3684 |
| 340 | FIM1_347 | QGN13704.1 | CP015054.1 | 3341 | 1202 | 1732 | 2958 | 927 | 1543 | 3240 | 1351 | 1579 |
| 341 | FIM1_348 | QGN13705.1 | CP015054.1 | 3090 | 2955 | 1577 | 2486 | 2270 | 1299 | 907 | 3263 | 1495 |
| 342 | FIM1_349 | QGN13706.1 | CP015054.1 | 8131 | 1420 | 2266 | 6374 | 1173 | 2120 | 6914 | 1507 | 2002 |
| 343 | FIM1_350 | QGN13707.1 | CP015054.1 | 1786 | 231 | 518 | 1276 | 186 | 476 | 1644 | 266 | 453 |
| 344 | FIM1_351 | QGN13708.1 | CP015054.1 | 5350 | 1033 | 1530 | 4271 | 760 | 1295 | 6163 | 1117 | 1294 |
| 345 | FIM1_352 | QGN13709.1 | CP015054.1 | 22509 | 2990 | 5085 | 17795 | 2203 | 4734 | 26753 | 3340 | 4912 |
| 346 | FIM1_354 | QGN13710.1 | CP015054.1 | 2027 | 99 | 559 | 1775 | 61 | 532 | 1829 | 106 | 467 |
| 347 | FIM1_355 | QGN13711.1 | CP015054.1 | 5708 | 1721 | 2809 | 5280 | 1238 | 2381 | 3984 | 1917 | 2318 |
| 348 | FIM1_356 | QGN13712.1 | CP015054.1 | 6350 | 383 | 594 | 4409 | 413 | 541 | 5938 | 469 | 569 |
| 349 | FIM1_357 | QGN13713.1 | CP015054.1 | 15693 | 1906 | 3946 | 13615 | 1349 | 3273 | 20368 | 2153 | 3347 |
| 350 | FIM1_358 | QGN13714.1 | CP015054.1 | 11098 | 8339 | 4560 | 8937 | 6293 | 3777 | 4170 | 9221 | 4223 |
| 351 | FIM1_359 | QGN13715.1 | CP015054.1 | 21063 | 6248 | 6409 | 16904 | 4398 | 5751 | 22125 | 6987 | 5723 |
| 352 | FIM1_360 | QGN13716.1 | CP015054.1 | 3527 | 915 | 1128 | 3346 | 769 | 1055 | 2989 | 980 | 1005 |
| 353 | FIM1_361 | QGN13717.1 | CP015054.1 | 1804 | 307 | 442 | 1488 | 258 | 423 | 1225 | 346 | 492 |
| 354 | FIM1_362 | QGN13718.1 | CP015054.1 | 8592 | 1286 | 2342 | 7061 | 992 | 2074 | 10749 | 1421 | 1949 |
| 355 | FIM1_363 | QGN13719.1 | CP015054.1 | 3498 | 767 | 1190 | 2491 | 565 | 1076 | 2483 | 838 | 1031 |
| 356 | FIM1_364 | QGN13720.1 | CP015054.1 | 6702 | 1658 | 2635 | 5149 | 1236 | 2420 | 7119 | 1779 | 2262 |
| 357 | FIM1_365 | QGN13721.1 | CP015054.1 | 3482 | 1318 | 1333 | 2670 | 926 | 1275 | 1840 | 1410 | 1190 |
| 358 | FIM1_366 | QGN13722.1 | CP015054.1 | 5980 | 908 | 1800 | 4327 | 757 | 1652 | 5116 | 937 | 1477 |
| 359 | FIM1_367 | QGN13723.1 | CP015054.1 | 9404 | 2909 | 1860 | 7888 | 2021 | 1652 | 6973 | 3143 | 1704 |
| 360 | FIM1_368 | QGN13724.1 | CP015054.1 | 1571 | 343 | 372 | 1136 | 238 | 326 | 1072 | 324 | 338 |
| 361 | FIM1_369 | QGN13725.1 | CP015054.1 | 82 | 24 | 41 | 65 | 20 | 39 | 56 | 17 | 26 |
| 362 | FIM1_370 | QGN13726.1 | CP015054.1 | 1806 | 358 | 977 | 1516 | 328 | 756 | 1720 | 465 | 704 |
| 363 | FIM1_371 | QGN13727.1 | CP015054.1 | 581 | 38 | 78 | 472 | 42 | 76 | 358 | 59 | 78 |
| 364 | FIM1_372 | QGN13728.1 | CP015054.1 | 36542 | 13535 | 13594 | 32431 | 8448 | 12074 | 41988 | 13570 | 13375 |
| 365 | FIM1_373 | QGN13729.1 | CP015054.1 | 5081 | 820 | 1333 | 4359 | 612 | 1122 | 4941 | 899 | 1192 |
| 366 | FIM1_374 | QGN13730.1 | CP015054.1 | 3770 | 1021 | 1298 | 3107 | 748 | 1187 | 4119 | 1072 | 1147 |
| 367 | FIM1_375 | QGN13731.1 | CP015054.1 | 5632 | 818 | 1363 | 4947 | 665 | 1208 | 6250 | 930 | 1345 |
| 368 | FIM1_376 | QGN13732.1 | CP015054.1 | 1376 | 209 | 870 | 1137 | 103 | 695 | 2095 | 192 | 700 |
| 369 | FIM1_377 | QGN13733.1 | CP015054.1 | 1397 | 305 | 988 | 1328 | 188 | 883 | 1474 | 313 | 844 |
| 370 | FIM1_378 | QGN13734.1 | CP015054.1 | 76990 | 50059 | 37391 | 72595 | 31379 | 32741 | 107682 | 49759 | 35055 |
| 371 | FIM1_379 | QGN13735.1 | CP015054.1 | 2358 | 347 | 432 | 1968 | 266 | 403 | 2730 | 380 | 405 |
| 372 | FIM1_380 | QGN13736.1 | CP015054.1 | 11001 | 1063 | 1866 | 8744 | 792 | 1788 | 12235 | 1189 | 1741 |
| 373 | FIM1_381 | QGN13737.1 | CP015054.1 | 10528 | 2868 | 2504 | 7911 | 2122 | 2316 | 7928 | 3204 | 2263 |
| 374 | FIM1_382 | QGN13738.1 | CP015054.1 | 8 | 2 | 1 | 5 | 2 | 0 | 14 | 1 | 1 |
| 375 | FIM1_383 | QGN13739.1 | CP015054.1 | 8292 | 3147 | 1652 | 7912 | 853 | 1514 | 25405 | 1824 | 1265 |
| 376 | FIM1_384 | QGN13740.1 | CP015054.1 | 94777 | 48979 | 39704 | 82767 | 38573 | 34986 | 88829 | 51091 | 38427 |
| 377 | FIM1_385 | QGN13741.1 | CP015054.1 | 3285 | 380 | 707 | 3146 | 348 | 614 | 2513 | 455 | 565 |
| 378 | FIM1_386 | QGN13742.1 | CP015054.1 | 12128 | 826 | 2805 | 10220 | 769 | 2475 | 9266 | 934 | 2379 |
| 379 | FIM1_387 | QGN13743.1 | CP015054.1 | 1851 | 513 | 873 | 1468 | 387 | 942 | 1746 | 529 | 819 |
| 380 | FIM1_388 | QGN13744.1 | CP015054.1 | 2137 | 265 | 578 | 1875 | 216 | 500 | 2283 | 325 | 518 |
| 381 | FIM1_389 | QGN13745.1 | CP015054.1 | 8811 | 2093 | 3112 | 7838 | 1409 | 2834 | 11370 | 2331 | 2646 |
| 382 | FIM1_390 | QGN13746.1 | CP015054.1 | 872 | 156 | 448 | 627 | 151 | 432 | 871 | 202 | 290 |

| | | | | | | | | | | | |
|---|---|---|---|---|---|---|---|---|---|---|---|
| 383 | FIM1_391 | QGN13747.1 | CP015054.1 | 3547 | 666 | 1357 | 2846 | 526 | 1187 | 4078 | 818 | 1162 |
| 384 | FIM1_392 | QGN13748.1 | CP015054.1 | 6871 | 3009 | 3781 | 6032 | 2165 | 3577 | 7125 | 3202 | 3514 |
| 385 | FIM1_393 | QGN13749.1 | CP015054.1 | 7365 | 1392 | 1348 | 5430 | 994 | 1329 | 6491 | 1546 | 1242 |
| 386 | FIM1_394 | QGN13750.1 | CP015054.1 | 909 | 157 | 188 | 618 | 124 | 126 | 650 | 152 | 121 |
| 387 | FIM1_395 | QGN13751.1 | CP015054.1 | 4653 | 693 | 1079 | 3438 | 540 | 1029 | 5109 | 843 | 986 |
| 388 | FIM1_396 | QGN13752.1 | CP015054.1 | 4850 | 2196 | 1904 | 4415 | 1723 | 1756 | 4073 | 2321 | 1806 |
| 389 | FIM1_397 | QGN13753.1 | CP015054.1 | 128 | 27 | 32 | 132 | 21 | 33 | 95 | 22 | 28 |
| 390 | FIM1_398 | QGN13754.1 | CP015054.1 | 5122 | 1179 | 1956 | 3972 | 988 | 1917 | 5754 | 1316 | 1742 |
| 391 | FIM1_399 | QGN13755.1 | CP015054.1 | 3685 | 1843 | 3079 | 3058 | 1286 | 2957 | 4071 | 1868 | 2895 |
| 392 | FIM1_401 | QGN13756.1 | CP015054.1 | 3379 | 883 | 1380 | 2603 | 700 | 1231 | 2736 | 1037 | 1143 |
| 393 | FIM1_402 | QGN13757.1 | CP015054.1 | 1096 | 91 | 202 | 942 | 76 | 196 | 2016 | 101 | 154 |
| 394 | FIM1_403 | QGN13758.1 | CP015054.1 | 2558 | 325 | 551 | 2217 | 190 | 499 | 3987 | 339 | 530 |
| 395 | FIM1_404 | QGN13759.1 | CP015054.1 | 1299 | 293 | 396 | 1182 | 160 | 367 | 2351 | 299 | 365 |
| 396 | FIM1_405 | QGN13760.1 | CP015054.1 | 21503 | 6794 | 6070 | 18594 | 5148 | 5468 | 19213 | 7264 | 5477 |
| 397 | FIM1_406 | QGN13761.1 | CP015054.1 | 26379 | 8410 | 8058 | 22600 | 6530 | 7214 | 24014 | 8918 | 7487 |
| 398 | FIM1_407 | QGN13762.1 | CP015054.1 | 8905 | 1663 | 2666 | 7528 | 1060 | 2391 | 10381 | 1927 | 2434 |
| 399 | FIM1_408 | QGN13763.1 | CP015054.1 | 4445 | 918 | 1319 | 4025 | 687 | 1164 | 5045 | 1022 | 1155 |
| 400 | FIM1_409 | QGN13764.1 | CP015054.1 | 12083 | 2997 | 4314 | 9976 | 1814 | 3686 | 16004 | 3111 | 3738 |
| 401 | FIM1_410 | QGN13765.1 | CP015054.1 | 400 | 52 | 169 | 294 | 33 | 159 | 538 | 66 | 134 |
| 402 | FIM1_411 | QGN13766.1 | CP015054.1 | 6742 | 882 | 1154 | 5254 | 286 | 1036 | 12246 | 938 | 995 |
| 403 | FIM1_412 | QGN13767.1 | CP015054.1 | 1927 | 413 | 488 | 1840 | 395 | 409 | 2177 | 496 | 393 |
| 404 | FIM1_413 | QGN13768.1 | CP015054.1 | 11630 | 3074 | 2856 | 9983 | 2560 | 2389 | 12217 | 3512 | 2506 |
| 405 | FIM1_414 | QGN13769.1 | CP015054.1 | 2008 | 137 | 412 | 1525 | 153 | 358 | 2201 | 165 | 335 |
| 406 | FIM1_415 | QGN13770.1 | CP015054.1 | 1299 | 195 | 330 | 1052 | 135 | 298 | 1565 | 199 | 307 |
| 407 | FIM1_416 | QGN13771.1 | CP015054.1 | 3773 | 622 | 1031 | 2773 | 590 | 930 | 3903 | 752 | 1018 |
| 408 | FIM1_417 | QGN13772.1 | CP015054.1 | 12220 | 3359 | 5087 | 10187 | 1701 | 4640 | 16656 | 3491 | 4185 |
| 409 | FIM1_418 | QGN13773.1 | CP015054.1 | 1631 | 476 | 1126 | 737 | 229 | 995 | 927 | 402 | 903 |
| 410 | FIM1_419 | QGN13774.1 | CP015054.1 | 3074 | 401 | 822 | 2291 | 288 | 771 | 2668 | 388 | 655 |
| 411 | FIM1_420 | QGN13775.1 | CP015054.1 | 1937 | 494 | 664 | 1381 | 371 | 639 | 1261 | 514 | 605 |
| 412 | FIM1_421 | QGN13776.1 | CP015054.1 | 3195 | 699 | 771 | 2312 | 552 | 723 | 2898 | 725 | 788 |
| 413 | FIM1_422 | QGN13777.1 | CP015054.1 | 1937 | 223 | 560 | 1649 | 187 | 474 | 2115 | 247 | 436 |
| 414 | FIM1_423 | QGN13778.1 | CP015054.1 | 329 | 47 | 42 | 216 | 11 | 39 | 434 | 32 | 24 |
| 415 | FIM1_425 | QGN13779.1 | CP015054.1 | 5568 | 5011 | 2635 | 4361 | 3743 | 2406 | 1783 | 5406 | 2599 |
| 416 | FIM1_426 | QGN13780.1 | CP015054.1 | 13356 | 3314 | 3496 | 9951 | 2412 | 3294 | 12369 | 3645 | 3183 |
| 417 | FIM1_427 | QGN13781.1 | CP015054.1 | 647 | 224 | 331 | 522 | 172 | 297 | 672 | 233 | 296 |
| 418 | FIM1_428 | QGN13782.1 | CP015054.1 | 867 | 297 | 275 | 706 | 228 | 264 | 910 | 278 | 232 |
| 419 | FIM1_429 | QGN13783.1 | CP015054.1 | 2683 | 1111 | 1333 | 2663 | 746 | 1205 | 2724 | 1168 | 1201 |
| 420 | FIM1_430 | QGN13784.1 | CP015054.1 | 2771 | 708 | 858 | 2132 | 486 | 794 | 2699 | 775 | 718 |
| 421 | FIM1_431 | QGN13785.1 | CP015054.1 | 1938 | 409 | 528 | 1421 | 273 | 446 | 1997 | 398 | 449 |
| 422 | FIM1_432 | QGN13786.1 | CP015054.1 | 368 | 170 | 281 | 300 | 137 | 238 | 186 | 174 | 211 |
| 423 | FIM1_433 | QGN13787.1 | CP015054.1 | 3862 | 1624 | 1744 | 3215 | 1228 | 1531 | 2614 | 1704 | 1563 |
| 424 | FIM1_434 | QGN13788.1 | CP015054.1 | 3138 | 652 | 892 | 2469 | 496 | 814 | 2874 | 679 | 810 |
| 425 | FIM1_435 | QGN13789.1 | CP015054.1 | 2051 | 679 | 755 | 1767 | 493 | 696 | 1774 | 746 | 714 |
| 426 | FIM1_436 | QGN13790.1 | CP015054.1 | 810 | 271 | 434 | 543 | 184 | 361 | 373 | 292 | 325 |
| 427 | FIM1_437 | QGN13791.1 | CP015054.1 | 1068 | 220 | 545 | 1086 | 187 | 515 | 1157 | 209 | 439 |
| 428 | FIM1_438 | QGN13792.1 | CP015054.1 | 120 | 7 | 34 | 72 | 7 | 17 | 176 | 4 | 16 |
| 429 | FIM1_439 | QGN13793.1 | CP015054.1 | 5114 | 2478 | 1542 | 3624 | 1558 | 1422 | 1466 | 2622 | 1380 |
| 430 | FIM1_440 | QGN13794.1 | CP015054.1 | 1292 | 345 | 693 | 1124 | 275 | 610 | 1687 | 356 | 500 |

| | | | | | | | | | | | | |
|---|---|---|---|---|---|---|---|---|---|---|---|---|
| 431 | FIM1_441 | QGN13795.1 | CP015054.1 | 2681 | 484 | 631 | 2114 | 347 | 602 | 3301 | 519 | 545 |
| 432 | FIM1_442 | QGN13796.1 | CP015054.1 | 4061 | 461 | 1139 | 3329 | 293 | 1058 | 4849 | 471 | 937 |
| 433 | FIM1_443 | QGN13797.1 | CP015054.1 | 812 | 68 | 173 | 605 | 40 | 142 | 990 | 60 | 137 |
| 434 | FIM1_444 | QGN13798.1 | CP015054.1 | 2642 | 844 | 1260 | 2231 | 567 | 1199 | 2577 | 999 | 1128 |
| 435 | FIM1_445 | QGN13799.1 | CP015054.1 | 5548 | 932 | 1839 | 4344 | 693 | 1638 | 5769 | 956 | 1441 |
| 436 | FIM1_446 | QGN13800.1 | CP015054.1 | 503 | 131 | 461 | 315 | 99 | 388 | 433 | 127 | 287 |
| 437 | FIM1_447 | QGN13801.1 | CP015054.1 | 3456 | 513 | 742 | 2547 | 378 | 639 | 3321 | 575 | 621 |
| 438 | FIM1_448 | QGN13802.1 | CP015054.1 | 7749 | 1421 | 2290 | 6305 | 1087 | 2143 | 9186 | 1491 | 1921 |
| 439 | FIM1_449 | QGN13803.1 | CP015054.1 | 13732 | 2400 | 4185 | 10485 | 1826 | 3738 | 15280 | 2422 | 3376 |
| 440 | FIM1_450 | QGN13804.1 | CP015054.1 | 146 | 74 | 122 | 130 | 59 | 89 | 55 | 73 | 98 |
| 441 | FIM1_451 | QGN13805.1 | CP015054.1 | 6549 | 1744 | 1627 | 4620 | 1260 | 1412 | 3223 | 1839 | 1383 |
| 442 | FIM1_452 | QGN13806.1 | CP015054.1 | 3315 | 391 | 779 | 2770 | 251 | 639 | 2891 | 434 | 619 |
| 443 | FIM1_453 | QGN13807.1 | CP015054.1 | 4070 | 2023 | 3411 | 3729 | 1647 | 2947 | 3620 | 2249 | 2854 |
| 444 | FIM1_454 | QGN13808.1 | CP015054.1 | 1527 | 125 | 244 | 1062 | 81 | 215 | 2066 | 129 | 210 |
| 445 | FIM1_455 | QGN13809.1 | CP015054.1 | 2685 | 879 | 1129 | 2804 | 623 | 1069 | 2779 | 960 | 998 |
| 446 | FIM1_456 | QGN13810.1 | CP015054.1 | 2949 | 746 | 980 | 2391 | 557 | 925 | 3654 | 808 | 897 |
| 447 | FIM1_457 | QGN13811.1 | CP015054.1 | 23190 | 13864 | 13281 | 22156 | 11321 | 12425 | 22529 | 14113 | 13022 |
| 448 | FIM1_458 | QGN13812.1 | CP015054.1 | 31087 | 20423 | 18351 | 30048 | 17227 | 17762 | 31117 | 21195 | 17784 |
| 449 | FIM1_459 | QGN13813.1 | CP015054.1 | 3338 | 1195 | 1154 | 2596 | 1049 | 1011 | 2683 | 1237 | 1054 |
| 450 | FIM1_460 | QGN13814.1 | CP015054.1 | 741 | 283 | 399 | 539 | 208 | 376 | 565 | 286 | 321 |
| 451 | FIM1_461 | QGN13815.1 | CP015054.1 | 2480 | 379 | 824 | 1934 | 304 | 649 | 2599 | 433 | 641 |
| 452 | FIM1_462 | QGN13816.1 | CP015054.1 | 1792 | 430 | 654 | 1490 | 314 | 567 | 1496 | 452 | 521 |
| 453 | FIM1_463 | QGN13817.1 | CP015054.1 | 3423 | 2309 | 2389 | 2448 | 1603 | 2090 | 1141 | 2537 | 2353 |
| 454 | FIM1_464 | QGN13818.1 | CP015054.1 | 16331 | 3454 | 4869 | 13165 | 2155 | 4616 | 20335 | 3665 | 4280 |
| 455 | FIM1_465 | QGN13819.1 | CP015054.1 | 41 | 10 | 17 | 45 | 7 | 31 | 32 | 17 | 22 |
| 456 | FIM1_466 | QGN13820.1 | CP015054.1 | 20614 | 7572 | 5846 | 16303 | 5713 | 5443 | 16814 | 8210 | 5528 |
| 457 | FIM1_467 | QGN13821.1 | CP015054.1 | 4607 | 2536 | 1709 | 4078 | 1941 | 1564 | 2199 | 2757 | 1579 |
| 458 | FIM1_468 | QGN13822.1 | CP015054.1 | 148 | 35 | 69 | 130 | 32 | 52 | 101 | 32 | 46 |
| 459 | FIM1_469 | QGN13823.1 | CP015054.1 | 1393 | 474 | 855 | 1279 | 391 | 755 | 904 | 521 | 645 |
| 460 | FIM1_470 | QGN13824.1 | CP015054.1 | 4900 | 552 | 1176 | 4316 | 357 | 1020 | 5490 | 556 | 1042 |
| 461 | FIM1_471 | QGN13825.1 | CP015054.1 | 3795 | 389 | 821 | 3276 | 261 | 755 | 6071 | 424 | 676 |
| 462 | FIM1_472 | QGN13826.1 | CP015054.1 | 1245 | 353 | 309 | 958 | 280 | 285 | 497 | 366 | 271 |
| 463 | FIM1_473 | QGN13827.1 | CP015054.1 | 5802 | 767 | 1315 | 4449 | 603 | 1113 | 6534 | 785 | 1114 |
| 464 | FIM1_474 | QGN13828.1 | CP015054.1 | 2886 | 1300 | 1558 | 2694 | 992 | 1434 | 2157 | 1463 | 1316 |
| 465 | FIM1_475 | QGN13829.1 | CP015054.1 | 2175 | 164 | 659 | 1560 | 166 | 594 | 2913 | 193 | 516 |
| 466 | FIM1_476 | QGN13830.1 | CP015054.1 | 15250 | 8488 | 8097 | 13875 | 6646 | 7634 | 14920 | 8718 | 7877 |
| 467 | FIM1_477 | QGN13831.1 | CP015054.1 | 16872 | 7467 | 7444 | 15027 | 5209 | 6884 | 17752 | 8063 | 7149 |
| 468 | FIM1_478 | QGN13832.1 | CP015054.1 | 12397 | 5426 | 2828 | 9343 | 3634 | 2464 | 3265 | 5570 | 2482 |
| 469 | FIM1_479 | QGN13833.1 | CP015054.1 | 5512 | 925 | 1287 | 4052 | 655 | 1140 | 4675 | 902 | 1150 |
| 470 | FIM1_480 | QGN13834.1 | CP015054.1 | 1793 | 458 | 1142 | 1529 | 372 | 1064 | 1675 | 511 | 1022 |
| 471 | FIM1_481 | QGN13835.1 | CP015054.1 | 4532 | 412 | 936 | 3480 | 312 | 858 | 5423 | 431 | 706 |
| 472 | FIM1_482 | QGN13836.1 | CP015054.1 | 4212 | 417 | 1033 | 3152 | 346 | 851 | 4531 | 394 | 816 |
| 473 | FIM1_483 | QGN13837.1 | CP015054.1 | 5826 | 1466 | 1721 | 4437 | 1073 | 1649 | 4534 | 1607 | 1692 |
| 474 | FIM1_484 | QGN13838.1 | CP015054.1 | 60 | 5 | 15 | 40 | 3 | 8 | 35 | 7 | 7 |
| 475 | FIM1_485 | QGN13839.1 | CP015054.1 | 283 | 44 | 113 | 249 | 39 | 123 | 314 | 45 | 103 |
| 476 | FIM1_486 | QGN13840.1 | CP015054.1 | 3809 | 265 | 1039 | 3391 | 223 | 986 | 5197 | 314 | 736 |
| 477 | FIM1_487 | QGN13841.1 | CP015054.1 | 1185 | 488 | 724 | 1424 | 353 | 652 | 1304 | 510 | 576 |
| 478 | FIM1_488 | QGN13842.1 | CP015054.1 | 9749 | 2984 | 5473 | 8705 | 2048 | 4682 | 9596 | 3052 | 4893 |

| | | | | | | | | | | | | |
|---|---|---|---|---|---|---|---|---|---|---|---|---|
| 479 | FIM1_489 | QGN13843.1 | CP015054.1 | 17355 | 2588 | 2642 | 12621 | 1845 | 2501 | 15565 | 2615 | 2535 |
| 480 | FIM1_490 | QGN13844.1 | CP015054.1 | 6731 | 773 | 1526 | 5257 | 569 | 1355 | 8677 | 835 | 1269 |
| 481 | FIM1_491 | QGN13845.1 | CP015054.1 | 21114 | 13529 | 10982 | 19221 | 10064 | 10023 | 22164 | 12946 | 10810 |
| 482 | FIM1_492 | QGN13846.1 | CP015054.1 | 5305 | 1185 | 1811 | 5193 | 937 | 1595 | 6200 | 1322 | 1483 |
| 483 | FIM1_493 | QGN13847.1 | CP015054.1 | 2314 | 995 | 1204 | 2035 | 809 | 1149 | 1742 | 1106 | 1029 |
| 484 | FIM1_494 | QGN13848.1 | CP015054.1 | 3958 | 1741 | 2004 | 3074 | 1247 | 1709 | 2498 | 1825 | 1617 |
| 485 | FIM1_495 | QGN13849.1 | CP015054.1 | 6819 | 2112 | 3738 | 6355 | 1577 | 3330 | 6559 | 2391 | 3149 |
| 486 | FIM1_496 | QGN13850.1 | CP015054.1 | 12714 | 4043 | 4603 | 10638 | 2892 | 4134 | 13668 | 4181 | 4237 |
| 487 | FIM1_497 | QGN13851.1 | CP015054.1 | 3587 | 764 | 1681 | 2917 | 586 | 1639 | 3425 | 795 | 1460 |
| 488 | FIM1_498 | QGN13852.1 | CP015054.1 | 1164 | 392 | 604 | 1259 | 364 | 574 | 926 | 459 | 567 |
| 489 | FIM1_499 | QGN13853.1 | CP015054.1 | 6097 | 2702 | 3295 | 5839 | 2002 | 3122 | 6104 | 2827 | 3013 |
| 490 | FIM1_500 | QGN13854.1 | CP015054.1 | 740 | 239 | 274 | 629 | 208 | 246 | 724 | 257 | 253 |
| 491 | FIM1_501 | QGN13855.1 | CP015054.1 | 1372 | 490 | 520 | 1214 | 413 | 454 | 1379 | 538 | 500 |
| 492 | FIM1_502 | QGN13856.1 | CP015054.1 | 5209 | 2042 | 1870 | 4979 | 1397 | 1678 | 4588 | 2177 | 1661 |
| 493 | FIM1_503 | QGN13857.1 | CP015054.1 | 3615 | 268 | 727 | 2895 | 264 | 622 | 4689 | 340 | 717 |
| 494 | FIM1_504 | QGN13858.1 | CP015054.1 | 3134 | 535 | 1000 | 2370 | 322 | 917 | 3176 | 576 | 859 |
| 495 | FIM1_505 | QGN13859.1 | CP015054.1 | 2440 | 974 | 835 | 1906 | 647 | 795 | 1218 | 1030 | 825 |
| 496 | FIM1_506 | QGN13860.1 | CP015054.1 | 6046 | 3847 | 2530 | 5494 | 3270 | 2051 | 2479 | 4068 | 2161 |
| 497 | FIM1_507 | QGN13861.1 | CP015054.1 | 3114 | 892 | 1613 | 2920 | 764 | 1453 | 2493 | 956 | 1387 |
| 498 | FIM1_508 | QGN13862.1 | CP015054.1 | 3270 | 431 | 785 | 2707 | 313 | 728 | 3694 | 455 | 684 |
| 499 | FIM1_509 | QGN13863.1 | CP015054.1 | 3075 | 258 | 542 | 2436 | 152 | 449 | 5541 | 303 | 415 |
| 500 | FIM1_510 | QGN13864.1 | CP015054.1 | 1814 | 599 | 760 | 1718 | 389 | 731 | 1983 | 633 | 720 |
| 501 | FIM1_511 | QGN13865.1 | CP015054.1 | 2588 | 860 | 1405 | 2407 | 352 | 1219 | 3946 | 902 | 1154 |
| 502 | FIM1_512 | QGN13866.1 | CP015054.1 | 450 | 87 | 166 | 402 | 90 | 183 | 546 | 119 | 152 |
| 503 | FIM1_513 | QGN13867.1 | CP015054.1 | 6199 | 2735 | 2750 | 5729 | 2135 | 2473 | 4847 | 2932 | 2431 |
| 504 | FIM1_514 | QGN13868.1 | CP015054.1 | 5681 | 711 | 1687 | 4573 | 622 | 1470 | 7399 | 813 | 1376 |
| 505 | FIM1_515 | QGN13869.1 | CP015054.1 | 4426 | 1071 | 1623 | 3779 | 833 | 1381 | 3837 | 1098 | 1353 |
| 506 | FIM1_516 | QGN13870.1 | CP015054.1 | 3496 | 742 | 1243 | 2838 | 579 | 1065 | 3328 | 789 | 988 |
| 507 | FIM1_517 | QGN13871.1 | CP015054.1 | 2771 | 1557 | 1049 | 2234 | 1213 | 958 | 856 | 1707 | 979 |
| 508 | FIM1_518 | QGN13872.1 | CP015054.1 | 51486 | 22968 | 21304 | 45346 | 17609 | 19387 | 51177 | 24062 | 19941 |
| 509 | FIM1_519 | QGN13873.1 | CP015054.1 | 1429 | 61 | 220 | 995 | 47 | 175 | 3149 | 54 | 124 |
| 510 | FIM1_521 | QGN13874.1 | CP015054.1 | 19 | 6 | 5 | 17 | 5 | 8 | 15 | 5 | 8 |
| 511 | FIM1_522 | QGN13875.1 | CP015054.1 | 6420 | 2625 | 2137 | 4724 | 2121 | 1891 | 2220 | 2964 | 1850 |
| 512 | FIM1_523 | QGN13876.1 | CP015054.1 | 851 | 416 | 673 | 797 | 354 | 598 | 506 | 508 | 538 |
| 513 | FIM1_524 | QGN13877.1 | CP015054.1 | 6378 | 791 | 1545 | 4992 | 636 | 1344 | 5720 | 917 | 1222 |
| 514 | FIM1_525 | QGN13878.1 | CP015054.1 | 4241 | 862 | 1802 | 3212 | 757 | 1530 | 3939 | 920 | 1490 |
| 515 | FIM1_526 | QGN13879.1 | CP015054.1 | 343 | 29 | 120 | 319 | 41 | 133 | 445 | 42 | 75 |
| 516 | FIM1_527 | QGN13880.1 | CP015054.1 | 48522 | 11032 | 6034 | 39364 | 5243 | 5342 | 84063 | 11090 | 4985 |
| 517 | FIM1_528 | QGN13881.1 | CP015054.1 | 12760 | 2138 | 2500 | 10508 | 1240 | 2335 | 19468 | 2278 | 2512 |
| 518 | FIM1_529 | QGN13882.1 | CP015054.1 | 60 | 19 | 17 | 51 | 10 | 11 | 79 | 17 | 12 |
| 519 | FIM1_530 | QGN13883.1 | CP015054.1 | 6356 | 2564 | 2978 | 6005 | 2237 | 2738 | 5065 | 2840 | 2837 |
| 520 | FIM1_531 | QGN13884.1 | CP015054.1 | 22538 | 9336 | 7650 | 20408 | 5829 | 6986 | 28515 | 10159 | 7611 |
| 521 | FIM1_532 | QGN13885.1 | CP015054.1 | 10923 | 6078 | 2583 | 8248 | 3654 | 2333 | 2580 | 6311 | 2402 |
| 522 | FIM1_533 | QGN13886.1 | CP015054.1 | 4406 | 649 | 864 | 3422 | 433 | 777 | 3591 | 722 | 728 |
| 523 | FIM1_534 | QGN13887.1 | CP015054.1 | 2899 | 500 | 916 | 2623 | 327 | 775 | 2934 | 570 | 768 |
| 524 | FIM1_535 | QGN13888.1 | CP015054.1 | 5013 | 1144 | 1268 | 4301 | 670 | 1125 | 6639 | 1184 | 1162 |
| 525 | FIM1_536 | QGN13889.1 | CP015054.1 | 4264 | 812 | 781 | 2462 | 447 | 674 | 2265 | 797 | 693 |
| 526 | FIM1_537 | QGN13890.1 | CP015054.1 | 5279 | 373 | 774 | 4121 | 308 | 684 | 5217 | 434 | 686 |

| | | | | | | | | | | | | |
|---|---|---|---|---|---|---|---|---|---|---|---|---|
| 527 | FIM1_538 | QGN13891.1 | CP015054.1 | 2661 | 632 | 997 | 2293 | 471 | 979 | 2531 | 644 | 917 |
| 528 | FIM1_539 | QGN13892.1 | CP015054.1 | 1194 | 229 | 373 | 899 | 214 | 384 | 1075 | 274 | 380 |
| 529 | FIM1_540 | QGN13893.1 | CP015054.1 | 7932 | 1241 | 1998 | 6111 | 1050 | 1894 | 6676 | 1391 | 1910 |
| 530 | FIM1_541 | QGN13894.1 | CP015054.1 | 1124 | 330 | 511 | 929 | 269 | 476 | 1206 | 353 | 430 |
| 531 | FIM1_542 | QGN13895.1 | CP015054.1 | 1670 | 540 | 848 | 1281 | 368 | 738 | 1391 | 634 | 652 |
| 532 | FIM1_543 | QGN13896.1 | CP015054.1 | 2297 | 362 | 425 | 1771 | 259 | 367 | 2075 | 323 | 371 |
| 533 | FIM1_544 | QGN13897.1 | CP015054.1 | 5784 | 1531 | 1585 | 5305 | 1259 | 1383 | 5233 | 1634 | 1311 |
| 534 | FIM1_545 | QGN13898.1 | CP015054.1 | 2367 | 612 | 3099 | 1980 | 464 | 2858 | 3001 | 634 | 2845 |
| 535 | FIM1_546 | QGN13899.1 | CP015054.1 | 10037 | 4050 | 2677 | 7885 | 2975 | 2596 | 10915 | 4450 | 2510 |
| 536 | FIM1_547 | QGN13900.1 | CP015054.1 | 7736 | 1039 | 1844 | 5778 | 874 | 1648 | 7279 | 1206 | 1504 |
| 537 | FIM1_549 | QGN13901.1 | CP015054.1 | 4385 | 881 | 728 | 3417 | 530 | 694 | 3920 | 896 | 737 |
| 538 | FIM1_550 | QGN13902.1 | CP015054.1 | 26327 | 13395 | 9939 | 21328 | 9929 | 8839 | 19255 | 13654 | 9397 |
| 539 | FIM1_551 | QGN13903.1 | CP015054.1 | 1054 | 380 | 603 | 849 | 303 | 573 | 698 | 465 | 554 |
| 540 | FIM1_552 | QGN13904.1 | CP015054.1 | 2749 | 668 | 1076 | 2197 | 502 | 1030 | 2456 | 749 | 920 |
| 541 | FIM1_553 | QGN13905.1 | CP015054.1 | 4829 | 775 | 1537 | 3530 | 574 | 1455 | 4857 | 803 | 1314 |
| 542 | FIM1_554 | QGN13906.1 | CP015054.1 | 3563 | 1225 | 1730 | 3023 | 929 | 1744 | 4126 | 1386 | 1676 |
| 543 | FIM1_555 | QGN13907.1 | CP015054.1 | 1169 | 283 | 638 | 812 | 238 | 554 | 916 | 341 | 518 |
| 544 | FIM1_556 | QGN13908.1 | CP015054.1 | 1829 | 899 | 953 | 1417 | 589 | 897 | 1852 | 931 | 809 |
| 545 | FIM1_557 | QGN13909.1 | CP015054.1 | 2230 | 827 | 730 | 1814 | 580 | 699 | 2514 | 935 | 662 |
| 546 | FIM1_558 | QGN13910.1 | CP015054.1 | 2267 | 1243 | 1155 | 2102 | 891 | 1032 | 1544 | 1301 | 1007 |
| 547 | FIM1_559 | QGN13911.1 | CP015054.1 | 1866 | 525 | 759 | 1492 | 371 | 637 | 1286 | 540 | 584 |
| 548 | FIM1_560 | QGN13912.1 | CP015054.1 | 3999 | 1185 | 2855 | 3376 | 925 | 2631 | 2914 | 1281 | 2410 |
| 549 | FIM1_561 | QGN13913.1 | CP015054.1 | 2855 | 521 | 719 | 2099 | 394 | 638 | 3564 | 500 | 560 |
| 550 | FIM1_562 | QGN13914.1 | CP015054.1 | 4932 | 1034 | 1520 | 4130 | 697 | 1415 | 6061 | 1140 | 1311 |
| 551 | FIM1_563 | QGN13915.1 | CP015054.1 | 270 | 121 | 159 | 258 | 75 | 131 | 400 | 114 | 111 |
| 552 | FIM1_564 | QGN13916.1 | CP015054.1 | 2677 | 1344 | 1513 | 2453 | 999 | 1400 | 2128 | 1476 | 1302 |
| 553 | FIM1_565 | QGN13917.1 | CP015054.1 | 3762 | 1719 | 1257 | 2493 | 1215 | 1119 | 1896 | 1877 | 1099 |
| 554 | FIM1_566 | QGN13918.1 | CP015054.1 | 1388 | 723 | 734 | 1115 | 572 | 723 | 937 | 797 | 699 |
| 555 | FIM1_567 | QGN13919.1 | CP015054.1 | 3152 | 1169 | 1545 | 2622 | 867 | 1427 | 2312 | 1209 | 1364 |
| 556 | FIM1_568 | QGN13920.1 | CP015054.1 | 7473 | 2420 | 2181 | 6075 | 957 | 2144 | 5846 | 2964 | 2494 |
| 557 | FIM1_569 | QGN13921.1 | CP015054.1 | 4503 | 1755 | 1858 | 3404 | 928 | 2005 | 3927 | 2099 | 1876 |
| 558 | FIM1_570 | QGN13922.1 | CP015054.1 | 1302 | 2073 | 959 | 1086 | 1417 | 809 | 1496 | 2179 | 896 |
| 559 | FIM1_571 | QGN13923.1 | CP015054.1 | 7092 | 816 | 2356 | 6059 | 489 | 2312 | 7503 | 950 | 2071 |
| 560 | FIM1_572 | QGN13924.1 | CP015054.1 | 1137 | 147 | 905 | 1020 | 96 | 797 | 619 | 169 | 788 |
| 561 | FIM1_573 | QGN13925.1 | CP015054.1 | 9915 | 3917 | 3082 | 8013 | 2505 | 3111 | 10414 | 4094 | 2762 |
| 562 | FIM1_574 | QGN13926.1 | CP015054.1 | 6535 | 4541 | 2652 | 4495 | 2823 | 2408 | 1585 | 4708 | 2476 |
| 563 | FIM1_575 | QGN13927.1 | CP015054.1 | 4699 | 2328 | 2207 | 3966 | 1542 | 2018 | 3777 | 2317 | 2014 |
| 564 | FIM1_576 | QGN13928.1 | CP015054.1 | 13816 | 4893 | 4394 | 10708 | 3988 | 4209 | 10674 | 5254 | 4278 |
| 565 | FIM1_577 | QGN13929.1 | CP015054.1 | 1371 | 547 | 730 | 931 | 472 | 723 | 1083 | 602 | 639 |
| 566 | FIM1_578 | QGN13930.1 | CP015054.1 | 8857 | 2750 | 4671 | 7010 | 1656 | 4160 | 11150 | 2733 | 4102 |
| 567 | FIM1_579 | QGN13931.1 | CP015054.1 | 3200 | 409 | 765 | 2234 | 345 | 757 | 3159 | 487 | 650 |
| 568 | FIM1_580 | QGN13932.1 | CP015054.1 | 574 | 125 | 238 | 533 | 103 | 253 | 502 | 127 | 205 |
| 569 | FIM1_581 | QGN13933.1 | CP015054.1 | 2579 | 501 | 688 | 1884 | 358 | 591 | 1630 | 550 | 554 |
| 570 | FIM1_582 | QGN13934.1 | CP015054.1 | 4495 | 4650 | 2511 | 3500 | 3360 | 2387 | 930 | 4955 | 2325 |
| 571 | FIM1_583 | QGN13935.1 | CP015054.1 | 530 | 190 | 250 | 392 | 172 | 253 | 313 | 200 | 231 |
| 572 | FIM1_584 | QGN13936.1 | CP015054.1 | 1969 | 422 | 927 | 1588 | 360 | 936 | 1774 | 473 | 767 |
| 573 | FIM1_585 | QGN13937.1 | CP015054.1 | 27463 | 7493 | 8565 | 23108 | 6472 | 8024 | 19498 | 8243 | 8176 |
| 574 | FIM1_586 | QGN13938.1 | CP015054.1 | 5353 | 2180 | 1327 | 3454 | 1409 | 1280 | 3617 | 2207 | 1293 |

| | | | | | | | | | | | | |
|---|---|---|---|---|---|---|---|---|---|---|---|---|
| 575 | FIM1_587 | QGN13939.1 | CP015054.1 | 1229 | 457 | 645 | 859 | 427 | 600 | 1092 | 441 | 587 |
| 576 | FIM1_588 | QGN13940.1 | CP015054.1 | 1092 | 601 | 576 | 837 | 495 | 613 | 792 | 643 | 555 |
| 577 | FIM1_589 | QGN13941.1 | CP015054.1 | 2587 | 526 | 781 | 2125 | 383 | 713 | 2866 | 552 | 695 |
| 578 | FIM1_590 | QGN13942.1 | CP015054.1 | 2663 | 436 | 643 | 1973 | 323 | 616 | 2819 | 485 | 574 |
| 579 | FIM1_591 | QGN13943.1 | CP015054.1 | 1751 | 462 | 584 | 1182 | 315 | 494 | 1935 | 456 | 450 |
| 580 | FIM1_592 | QGN13944.1 | CP015054.1 | 16881 | 3681 | 5256 | 13308 | 2979 | 5016 | 15157 | 4132 | 4797 |
| 581 | FIM1_593 | QGN13945.1 | CP015054.1 | 2651 | 723 | 1169 | 2390 | 657 | 1035 | 1902 | 828 | 898 |
| 582 | FIM1_594 | QGN13946.1 | CP015054.1 | 3909 | 1691 | 1519 | 2940 | 1512 | 1402 | 3001 | 1964 | 1507 |
| 583 | FIM1_595 | QGN13947.1 | CP015054.1 | 4607 | 2134 | 1683 | 3436 | 1615 | 1618 | 2598 | 2301 | 1898 |
| 584 | FIM1_596 | QGN13948.1 | CP015054.1 | 7035 | 2626 | 3691 | 6162 | 2076 | 3500 | 5317 | 2803 | 3297 |
| 585 | FIM1_597 | QGN13949.1 | CP015054.1 | 4319 | 1088 | 1183 | 3487 | 873 | 1052 | 4770 | 1100 | 1096 |
| 586 | FIM1_598 | QGN13950.1 | CP015054.1 | 1785 | 806 | 916 | 1536 | 624 | 842 | 1247 | 849 | 731 |
| 587 | FIM1_599 | QGN13951.1 | CP015054.1 | 1690 | 318 | 546 | 1257 | 254 | 553 | 1274 | 360 | 449 |
| 588 | FIM1_600 | QGN13952.1 | CP015054.1 | 3669 | 834 | 1565 | 3066 | 459 | 1496 | 5263 | 833 | 1311 |
| 589 | FIM1_601 | QGN13953.1 | CP015054.1 | 4829 | 3775 | 2256 | 3604 | 2317 | 1936 | 1132 | 3953 | 1882 |
| 590 | FIM1_602 | QGN13954.1 | CP015054.1 | 5899 | 1359 | 1766 | 4632 | 935 | 1526 | 4872 | 1396 | 1408 |
| 591 | FIM1_603 | QGN13955.1 | CP015054.1 | 20170 | 2325 | 5061 | 19033 | 1779 | 4451 | 20735 | 2435 | 4254 |
| 592 | FIM1_604 | QGN13956.1 | CP015054.1 | 3796 | 1603 | 1958 | 3725 | 1318 | 1729 | 3720 | 1748 | 1657 |
| 593 | FIM1_605 | QGN13957.1 | CP015054.1 | 788 | 226 | 524 | 767 | 154 | 522 | 1148 | 300 | 480 |
| 594 | FIM1_606 | QGN13958.1 | CP015054.1 | 2402 | 299 | 1627 | 2068 | 238 | 1516 | 3532 | 292 | 1223 |
| 595 | FIM1_607 | QGN13959.1 | CP015054.1 | 1202 | 983 | 504 | 661 | 574 | 478 | 391 | 1093 | 445 |
| 596 | FIM1_608 | QGN13960.1 | CP015054.1 | 1465 | 360 | 537 | 1223 | 243 | 537 | 1197 | 359 | 510 |
| 597 | FIM1_609 | QGN13961.1 | CP015054.1 | 26054 | 4119 | 5208 | 20032 | 3152 | 5004 | 33667 | 4324 | 4820 |
| 598 | FIM1_610 | QGN13962.1 | CP015054.1 | 1168 | 255 | 557 | 1022 | 187 | 461 | 1186 | 306 | 459 |
| 599 | FIM1_611 | QGN13963.1 | CP015054.1 | 786 | 69 | 136 | 673 | 43 | 141 | 1154 | 69 | 99 |
| 600 | FIM1_612 | QGN13964.1 | CP015054.1 | 5595 | 736 | 1308 | 4207 | 590 | 1278 | 6519 | 828 | 1130 |
| 601 | FIM1_613 | QGN13965.1 | CP015054.1 | 4059 | 721 | 1216 | 3054 | 502 | 1101 | 3750 | 720 | 1033 |
| 602 | FIM1_615 | QGN13966.1 | CP015054.1 | 3254 | 353 | 750 | 2445 | 219 | 649 | 3943 | 346 | 604 |
| 603 | FIM1_616 | QGN13967.1 | CP015054.1 | 4288 | 1793 | 2127 | 3384 | 1057 | 1966 | 5073 | 1822 | 1900 |
| 604 | FIM1_617 | QGN13968.1 | CP015054.1 | 662 | 357 | 391 | 554 | 288 | 369 | 252 | 410 | 326 |
| 605 | FIM1_618 | QGN13969.1 | CP015054.1 | 7948 | 751 | 907 | 5680 | 517 | 888 | 8163 | 832 | 868 |
| 606 | FIM1_619 | QGN13970.1 | CP015054.1 | 29021 | 8555 | 9947 | 23788 | 5266 | 8792 | 35402 | 9020 | 8673 |
| 607 | FIM1_620 | QGN13971.1 | CP015054.1 | 2481 | 415 | 938 | 2184 | 339 | 841 | 2305 | 430 | 788 |
| 608 | FIM1_621 | QGN13972.1 | CP015054.1 | 1797 | 205 | 1035 | 1480 | 192 | 951 | 1299 | 229 | 895 |
| 609 | FIM1_622 | QGN13973.1 | CP015054.1 | 466 | 126 | 388 | 350 | 118 | 377 | 402 | 154 | 327 |
| 610 | FIM1_623 | QGN13974.1 | CP015054.1 | 5423 | 2616 | 2526 | 4584 | 1823 | 2528 | 5612 | 2575 | 2316 |
| 611 | FIM1_624 | QGN13975.1 | CP015054.1 | 4 | 2 | 0 | 4 | 0 | 1 | 3 | 1 | 2 |
| 612 | FIM1_625 | QGN13976.1 | CP015054.1 | 2758 | 269 | 184 | 2541 | 134 | 219 | 7542 | 315 | 142 |
| 613 | FIM1_626 | QGN13977.1 | CP015054.1 | 4037 | 2542 | 2837 | 3932 | 1161 | 2835 | 7727 | 2612 | 2658 |
| 614 | FIM1_627 | QGN13978.1 | CP015054.1 | 3633 | 342 | 797 | 2630 | 274 | 665 | 3352 | 392 | 711 |
| 615 | FIM1_628 | QGN13979.1 | CP015054.1 | 6371 | 1074 | 2989 | 5791 | 567 | 2636 | 7972 | 1124 | 2618 |
| 616 | FIM1_629 | QGN13980.1 | CP015054.1 | 7215 | 2361 | 1827 | 5834 | 1485 | 1620 | 6137 | 2725 | 1906 |
| 617 | FIM1_630 | QGN13981.1 | CP015054.1 | 2543 | 458 | 824 | 2040 | 282 | 696 | 1589 | 467 | 652 |
| 618 | FIM1_631 | QGN13982.1 | CP015054.1 | 8681 | 1097 | 1372 | 6553 | 862 | 1340 | 8826 | 1250 | 1335 |
| 619 | FIM1_632 | QGN13983.1 | CP015054.1 | 917 | 109 | 263 | 698 | 101 | 228 | 1418 | 111 | 171 |
| 620 | FIM1_633 | QGN13984.1 | CP015054.1 | 10977 | 5121 | 4888 | 9453 | 3673 | 4272 | 9168 | 5540 | 4642 |
| 621 | FIM1_634 | QGN13985.1 | CP015054.1 | 1242 | 345 | 422 | 1013 | 238 | 362 | 1032 | 364 | 383 |
| 622 | FIM1_635 | QGN13986.1 | CP015054.1 | 6787 | 3057 | 3345 | 6154 | 2478 | 3116 | 6146 | 3228 | 3007 |

| | | | | | | | | | | | | |
|---|---|---|---|---|---|---|---|---|---|---|---|---|
| 623 | FIM1_636 | QGN13987.1 | CP015054.1 | 1701 | 311 | 452 | 1319 | 236 | 435 | 1755 | 341 | 354 |
| 624 | FIM1_637 | QGN13988.1 | CP015054.1 | 13188 | 7797 | 7334 | 12983 | 6773 | 7179 | 11953 | 8213 | 6851 |
| 625 | FIM1_638 | QGN13989.1 | CP015054.1 | 4084 | 945 | 1399 | 3026 | 725 | 1243 | 2993 | 1099 | 1236 |
| 626 | FIM1_639 | QGN13990.1 | CP015054.1 | 170 | 39 | 79 | 171 | 26 | 55 | 124 | 48 | 52 |
| 627 | FIM1_640 | QGN13991.1 | CP015054.1 | 3366 | 806 | 998 | 2546 | 583 | 973 | 2956 | 813 | 786 |
| 628 | FIM1_641 | QGN13992.1 | CP015054.1 | 1392 | 198 | 532 | 1126 | 174 | 454 | 1459 | 242 | 391 |
| 629 | FIM1_642 | QGN13993.1 | CP015054.1 | 7858 | 2859 | 2944 | 6156 | 1329 | 2692 | 9307 | 2833 | 2387 |
| 630 | FIM1_644 | QGN13994.1 | CP015054.1 | 2923 | 1832 | 1490 | 2180 | 1352 | 1527 | 1234 | 2127 | 1420 |
| 631 | FIM1_645 | QGN13995.1 | CP015054.1 | 5643 | 583 | 871 | 4465 | 448 | 752 | 6296 | 651 | 793 |
| 632 | FIM1_646 | QGN13996.1 | CP015054.1 | 8781 | 2368 | 2247 | 6832 | 1846 | 2038 | 10586 | 2771 | 1965 |
| 633 | FIM1_647 | QGN13997.1 | CP015054.1 | 4169 | 1549 | 1267 | 3394 | 1290 | 1215 | 2231 | 1733 | 1120 |
| 634 | FIM1_648 | QGN13998.1 | CP015054.1 | 436 | 59 | 132 | 338 | 47 | 140 | 441 | 89 | 122 |
| 635 | FIM1_650 | QGN13999.1 | CP015054.1 | 1726 | 243 | 516 | 1581 | 183 | 435 | 2179 | 245 | 429 |
| 636 | FIM1_651 | QGN14000.1 | CP015054.1 | 4037 | 411 | 752 | 2888 | 350 | 719 | 4831 | 403 | 697 |
| 637 | FIM1_652 | QGN14001.1 | CP015054.1 | 6096 | 1749 | 2018 | 4432 | 1475 | 1897 | 5121 | 1883 | 1841 |
| 638 | FIM1_653 | QGN14002.1 | CP015054.1 | 5960 | 5318 | 2800 | 3973 | 3387 | 2515 | 1329 | 5741 | 2586 |
| 639 | FIM1_654 | QGN14003.1 | CP015054.1 | 1389 | 294 | 570 | 1133 | 227 | 525 | 1361 | 312 | 490 |
| 640 | FIM1_655 | QGN14004.1 | CP015054.1 | 483 | 198 | 359 | 352 | 121 | 417 | 619 | 217 | 361 |
| 641 | FIM1_656 | QGN14005.1 | CP015054.1 | 3740 | 1434 | 1561 | 3046 | 1039 | 1441 | 3237 | 1577 | 1429 |
| 642 | FIM1_657 | QGN14006.1 | CP015054.1 | 2600 | 385 | 749 | 1864 | 344 | 704 | 2556 | 446 | 686 |
| 643 | FIM1_658 | QGN14007.1 | CP015054.1 | 2043 | 217 | 650 | 1611 | 180 | 547 | 2512 | 255 | 492 |
| 644 | FIM1_659 | QGN14008.1 | CP015054.1 | 4854 | 1593 | 2577 | 4506 | 1168 | 2250 | 4896 | 1703 | 2160 |
| 645 | FIM1_660 | QGN14009.1 | CP015054.1 | 1923 | 313 | 802 | 1597 | 228 | 725 | 1950 | 333 | 636 |
| 646 | FIM1_661 | QGN14010.1 | CP015054.1 | 3054 | 489 | 723 | 2141 | 305 | 596 | 2889 | 471 | 466 |
| 647 | FIM1_662 | QGN14011.1 | CP015054.1 | 24147 | 15038 | 11305 | 20347 | 12692 | 9498 | 14588 | 15667 | 10271 |
| 648 | FIM1_663 | QGN14012.1 | CP015054.1 | 4072 | 327 | 501 | 2034 | 194 | 413 | 3922 | 386 | 439 |
| 649 | FIM1_664 | QGN14013.1 | CP015054.1 | 4922 | 2094 | 2575 | 4628 | 1416 | 2365 | 5228 | 2327 | 2372 |
| 650 | FIM1_665 | QGN14014.1 | CP015054.1 | 1328 | 364 | 639 | 988 | 266 | 529 | 1529 | 395 | 482 |
| 651 | FIM1_666 | QGN14015.1 | CP015054.1 | 696 | 85 | 219 | 610 | 72 | 194 | 829 | 89 | 158 |
| 652 | FIM1_667 | QGN14016.1 | CP015054.1 | 1312 | 102 | 196 | 795 | 91 | 230 | 1528 | 97 | 160 |
| 653 | FIM1_668 | QGN14017.1 | CP015054.1 | 12143 | 4053 | 3028 | 9498 | 2645 | 2846 | 8480 | 4275 | 2842 |
| 654 | FIM1_669 | QGN14018.1 | CP015054.1 | 5004 | 856 | 1655 | 3875 | 645 | 1518 | 5379 | 861 | 1356 |
| 655 | FIM1_670 | QGN14019.1 | CP015054.1 | 8165 | 3538 | 2232 | 5678 | 762 | 1900 | 10979 | 3274 | 1620 |
| 656 | FIM1_671 | QGN14020.1 | CP015054.1 | 1679 | 360 | 538 | 1295 | 345 | 495 | 1423 | 412 | 477 |
| 657 | FIM1_672 | QGN14021.1 | CP015054.1 | 2521 | 203 | 543 | 1752 | 225 | 537 | 2601 | 263 | 478 |
| 658 | FIM1_673 | QGN14022.1 | CP015054.1 | 1406 | 59 | 152 | 676 | 58 | 139 | 995 | 93 | 116 |
| 659 | FIM1_674 | QGN14023.1 | CP015054.1 | 15668 | 2287 | 4278 | 11581 | 1594 | 3915 | 21943 | 2428 | 3844 |
| 660 | FIM1_675 | QGN14024.1 | CP015054.1 | 4284 | 795 | 1406 | 3459 | 573 | 1224 | 4014 | 889 | 1244 |
| 661 | FIM1_676 | QGN14025.1 | CP015054.1 | 7762 | 1447 | 3953 | 5693 | 1012 | 3668 | 7450 | 1480 | 3479 |
| 662 | FIM1_677 | QGN14026.1 | CP015054.1 | 4389 | 693 | 1201 | 3901 | 408 | 1036 | 7193 | 825 | 931 |
| 663 | FIM1_678 | QGN14027.1 | CP015054.1 | 10055 | 1661 | 1994 | 8612 | 1113 | 1740 | 11968 | 1770 | 1810 |
| 664 | FIM1_679 | QGN14028.1 | CP015054.1 | 9196 | 2048 | 2112 | 6468 | 1253 | 2039 | 9083 | 2204 | 1864 |
| 665 | FIM1_680 | QGN14029.1 | CP015054.1 | 8026 | 2221 | 6025 | 6346 | 1456 | 5316 | 18906 | 2369 | 4617 |
| 666 | FIM1_681 | QGN14030.1 | CP015054.1 | 6842 | 1276 | 1200 | 5520 | 863 | 1085 | 6946 | 1319 | 1238 |
| 667 | FIM1_682 | QGN14031.1 | CP015054.1 | 5748 | 349 | 1097 | 4245 | 315 | 893 | 6284 | 360 | 869 |
| 668 | FIM1_683 | QGN14032.1 | CP015054.1 | 11329 | 2510 | 2731 | 9100 | 1888 | 2359 | 10351 | 2694 | 2309 |
| 669 | FIM1_684 | QGN14033.1 | CP015054.1 | 6175 | 775 | 1412 | 4766 | 541 | 1304 | 6927 | 759 | 1067 |
| 670 | FIM1_685 | QGN14034.1 | CP015054.1 | 1536 | 192 | 448 | 1336 | 168 | 397 | 2253 | 206 | 384 |

| | | | | | | | | | | | | |
|---|---|---|---|---|---|---|---|---|---|---|---|---|
| 671 | FIM1_686 | QGN14035.1 | CP015054.1 | 7590 | 2724 | 3241 | 6810 | 1848 | 2984 | 9136 | 2833 | 2840 |
| 672 | FIM1_687 | QGN14036.1 | CP015054.1 | 2443 | 553 | 799 | 1775 | 462 | 757 | 2616 | 635 | 753 |
| 673 | FIM1_688 | QGN14037.1 | CP015054.1 | 1979 | 508 | 1136 | 1618 | 371 | 1122 | 2419 | 613 | 970 |
| 674 | FIM1_689 | QGN14038.1 | CP015054.1 | 230 | 115 | 103 | 201 | 96 | 97 | 123 | 127 | 102 |
| 675 | FIM1_690 | QGN14039.1 | CP015054.1 | 4132 | 1118 | 1114 | 2953 | 783 | 1065 | 2894 | 1220 | 972 |
| 676 | FIM1_691 | QGN14040.1 | CP015054.1 | 3511 | 1472 | 1395 | 2827 | 1119 | 1372 | 2336 | 1610 | 1282 |
| 677 | FIM1_692 | QGN14041.1 | CP015054.1 | 2743 | 887 | 1199 | 2240 | 617 | 1070 | 2723 | 1000 | 1041 |
| 678 | FIM1_693 | QGN14042.1 | CP015054.1 | 3380 | 767 | 1010 | 2518 | 547 | 962 | 3742 | 768 | 906 |
| 679 | FIM1_694 | QGN14043.1 | CP015054.1 | 1462 | 206 | 469 | 1100 | 175 | 515 | 1573 | 241 | 411 |
| 680 | FIM1_695 | QGN14044.1 | CP015054.1 | 3108 | 532 | 795 | 2367 | 411 | 819 | 2699 | 561 | 675 |
| 681 | FIM1_696 | QGN14045.1 | CP015054.1 | 3663 | 1083 | 1166 | 3067 | 751 | 1245 | 3501 | 1217 | 1099 |
| 682 | FIM1_697 | QGN14046.1 | CP015054.1 | 2337 | 442 | 730 | 1936 | 286 | 690 | 2713 | 446 | 634 |
| 683 | FIM1_698 | QGN14047.1 | CP015054.1 | 2047 | 667 | 637 | 1998 | 602 | 600 | 1444 | 831 | 549 |
| 684 | FIM1_699 | QGN14048.1 | CP015054.1 | 10447 | 1865 | 2519 | 7835 | 1487 | 2384 | 8360 | 2028 | 2055 |
| 685 | FIM1_700 | QGN14049.1 | CP015054.1 | 8745 | 6123 | 5666 | 7658 | 4582 | 5205 | 9190 | 6713 | 5025 |
| 686 | FIM1_701 | QGN14050.1 | CP015054.1 | 2806 | 1919 | 1553 | 2255 | 1319 | 1549 | 1313 | 2037 | 1411 |
| 687 | FIM1_702 | QGN14051.1 | CP015054.1 | 2156 | 943 | 1001 | 1922 | 624 | 937 | 2303 | 996 | 864 |
| 688 | FIM1_703 | QGN14052.1 | CP015054.1 | 8118 | 2305 | 2180 | 6099 | 1960 | 2049 | 5179 | 2599 | 2185 |
| 689 | FIM1_704 | QGN14053.1 | CP015054.1 | 27277 | 5470 | 4840 | 22937 | 2626 | 4339 | 20512 | 5784 | 3889 |
| 690 | FIM1_705 | QGN14054.1 | CP015054.1 | 5058 | 872 | 947 | 3255 | 657 | 836 | 2704 | 1108 | 903 |
| 691 | FIM1_706 | QGN14055.1 | CP015054.1 | 4855 | 1475 | 1735 | 3622 | 1225 | 1744 | 4119 | 1632 | 1702 |
| 692 | FIM1_707 | QGN14056.1 | CP015054.1 | 2846 | 706 | 774 | 2231 | 624 | 723 | 2362 | 773 | 698 |
| 693 | FIM1_708 | QGN14057.1 | CP015054.1 | 1642 | 370 | 596 | 1170 | 272 | 571 | 1298 | 374 | 495 |
| 694 | FIM1_709 | QGN14058.1 | CP015054.1 | 3104 | 693 | 1066 | 2442 | 620 | 1100 | 2938 | 786 | 1007 |
| 695 | FIM1_710 | QGN14059.1 | CP015054.1 | 1448 | 322 | 581 | 1091 | 248 | 607 | 1888 | 354 | 545 |
| 696 | FIM1_711 | QGN14060.1 | CP015054.1 | 937 | 270 | 445 | 921 | 237 | 482 | 820 | 304 | 440 |
| 697 | FIM1_712 | QGN14061.1 | CP015054.1 | 101 | 13 | 37 | 67 | 14 | 26 | 49 | 14 | 21 |
| 698 | FIM1_713 | QGN14062.1 | CP015054.1 | 2645 | 499 | 1110 | 1880 | 827 | 1245 | 1507 | 590 | 1202 |
| 699 | FIM1_714 | QGN14063.1 | CP015054.1 | 1843 | 624 | 532 | 1551 | 468 | 518 | 2299 | 596 | 564 |
| 700 | FIM1_715 | QGN14064.1 | CP015054.1 | 2793 | 433 | 2002 | 1841 | 347 | 1860 | 2453 | 524 | 1837 |
| 701 | FIM1_716 | QGN14065.1 | CP015054.1 | 0 | 0 | 0 | 0 | 0 | 0 | 0 | 0 | 0 |
| 702 | FIM1_717 | QGN14066.1 | CP015054.1 | 0 | 1 | 0 | 0 | 0 | 0 | 0 | 0 | 1 |
| 703 | FIM1_718 | QGN14067.1 | CP015054.1 | 8234 | 2470 | 4302 | 6021 | 1571 | 4247 | 10550 | 2471 | 4020 |
| 704 | FIM1_719 | QGN14068.1 | CP015054.1 | 10449 | 6569 | 6395 | 9644 | 5489 | 6783 | 9332 | 6750 | 5911 |
| 705 | FIM1_720 | QGN14069.1 | CP015054.1 | 1792 | 318 | 470 | 1217 | 279 | 415 | 1557 | 349 | 392 |
| 706 | FIM1_721 | QGN14070.1 | CP015054.1 | 4807 | 1505 | 1790 | 3959 | 1388 | 1913 | 3648 | 1706 | 1609 |
| 707 | FIM1_722 | QGN14071.1 | CP015054.1 | 11385 | 4018 | 3936 | 8945 | 3018 | 3974 | 9122 | 4166 | 3628 |
| 708 | FIM1_723 | QGN14072.1 | CP015054.1 | 3009 | 418 | 628 | 2414 | 342 | 614 | 2489 | 424 | 580 |
| 709 | FIM1_724 | QGN14073.1 | CP015054.1 | 771 | 361 | 286 | 586 | 278 | 357 | 410 | 347 | 289 |
| 710 | FIM1_725 | QGN14074.1 | CP015054.1 | 9682 | 7301 | 4039 | 7880 | 5764 | 4104 | 4557 | 7590 | 3909 |
| 711 | FIM1_726 | QGN14075.1 | CP015054.1 | 2187 | 630 | 723 | 1798 | 465 | 799 | 1695 | 690 | 612 |
| 712 | FIM1_727 | QGN14076.1 | CP015054.1 | 2778 | 543 | 1118 | 2166 | 437 | 1238 | 3152 | 693 | 1022 |
| 713 | FIM1_728 | QGN14077.1 | CP015054.1 | 6000 | 2075 | 1678 | 4523 | 1922 | 1715 | 3518 | 2365 | 1594 |
| 714 | FIM1_729 | QGN14078.1 | CP015054.1 | 5633 | 2084 | 2049 | 4165 | 1561 | 2117 | 3967 | 2271 | 1871 |
| 715 | FIM1_730 | QGN14079.1 | CP015054.1 | 4450 | 1237 | 1677 | 3587 | 966 | 1625 | 3720 | 1334 | 1493 |
| 716 | FIM1_731 | QGN14080.1 | CP015054.1 | 1799 | 216 | 433 | 1331 | 194 | 370 | 1607 | 256 | 316 |
| 717 | FIM1_732 | QGN14081.1 | CP015054.1 | 13623 | 8078 | 7458 | 12885 | 6992 | 7895 | 11785 | 8550 | 7240 |
| 718 | FIM1_733 | QGN14082.1 | CP015054.1 | 4650 | 1897 | 1592 | 3296 | 1019 | 1674 | 2971 | 1866 | 1439 |

| | | | | | | | | | | | | |
|---|---|---|---|---|---|---|---|---|---|---|---|---|
| 719 | FIM1_734 | QGN14083.1 | CP015054.1 | 4723 | 1228 | 2487 | 3881 | 951 | 2417 | 4325 | 1308 | 2156 |
| 720 | FIM1_735 | QGN14084.1 | CP015054.1 | 2049 | 297 | 649 | 1809 | 212 | 659 | 2212 | 292 | 493 |
| 721 | FIM1_736 | QGN14085.1 | CP015054.1 | 7216 | 1320 | 2087 | 5559 | 1165 | 2192 | 8449 | 1474 | 2149 |
| 722 | FIM1_737 | QGN14086.1 | CP015054.1 | 5861 | 1504 | 1957 | 4652 | 1093 | 2112 | 5833 | 1636 | 1798 |
| 723 | FIM1_738 | QGN14087.1 | CP015054.1 | 2176 | 1218 | 1775 | 2078 | 859 | 1906 | 2149 | 1262 | 1612 |
| 724 | FIM1_739 | QGN14088.1 | CP015054.1 | 833 | 246 | 481 | 690 | 222 | 495 | 650 | 291 | 378 |
| 725 | FIM1_740 | QGN14089.1 | CP015054.1 | 1818 | 471 | 747 | 1409 | 435 | 789 | 1492 | 589 | 641 |
| 726 | FIM1_741 | QGN14090.1 | CP015054.1 | 3452 | 1502 | 1457 | 2768 | 1262 | 1536 | 3180 | 1710 | 1337 |
| 727 | FIM1_742 | QGN14091.1 | CP015054.1 | 6129 | 1266 | 1746 | 5329 | 1066 | 1753 | 5180 | 1472 | 1612 |
| 728 | FIM1_743 | QGN14092.1 | CP015054.1 | 3905 | 1520 | 1823 | 3209 | 1194 | 1851 | 4066 | 1619 | 1579 |
| 729 | FIM1_744 | QGN14093.1 | CP015054.1 | 887 | 182 | 577 | 871 | 155 | 585 | 896 | 194 | 472 |
| 730 | FIM1_745 | QGN14094.1 | CP015054.1 | 1679 | 214 | 401 | 1466 | 160 | 395 | 1986 | 199 | 365 |
| 731 | FIM1_746 | QGN14095.1 | CP015054.1 | 3720 | 855 | 864 | 3043 | 550 | 904 | 5336 | 917 | 799 |
| 732 | FIM1_747 | QGN14096.1 | CP015054.1 | 2098 | 553 | 1329 | 1850 | 359 | 1368 | 2810 | 611 | 1080 |
| 733 | FIM1_748 | QGN14097.1 | CP015054.1 | 2429 | 455 | 732 | 1898 | 403 | 658 | 2601 | 511 | 669 |
| 734 | FIM1_749 | QGN14098.1 | CP015054.1 | 2248 | 333 | 1086 | 1901 | 231 | 1031 | 3257 | 368 | 807 |
| 735 | FIM1_750 | QGN14099.1 | CP015054.1 | 3480 | 762 | 1204 | 2518 | 653 | 1219 | 2716 | 808 | 937 |
| 736 | FIM1_751 | QGN14100.1 | CP015054.1 | 9903 | 2149 | 3360 | 7819 | 1651 | 3330 | 10597 | 2202 | 2976 |
| 737 | FIM1_752 | QGN14101.1 | CP015054.1 | 12625 | 4759 | 3439 | 10081 | 3278 | 3402 | 12446 | 5096 | 2926 |
| 738 | FIM1_753 | QGN14102.1 | CP015054.1 | 8167 | 2450 | 1365 | 7403 | 1207 | 1252 | 9491 | 2448 | 1110 |
| 739 | FIM1_754 | QGN14103.1 | CP015054.1 | 8999 | 823 | 1744 | 7147 | 733 | 1713 | 9435 | 995 | 1558 |
| 740 | FIM1_755 | QGN14104.1 | CP015054.1 | 5970 | 1390 | 2517 | 4621 | 1054 | 2416 | 5750 | 1462 | 2062 |
| 741 | FIM1_756 | QGN14105.1 | CP015054.1 | 1274 | 418 | 708 | 949 | 331 | 790 | 1255 | 451 | 689 |
| 742 | FIM1_757 | QGN14106.1 | CP015054.1 | 4542 | 1397 | 1931 | 3866 | 1109 | 2087 | 3379 | 1532 | 1874 |
| 743 | FIM1_758 | QGN14107.1 | CP015054.1 | 1027 | 147 | 345 | 912 | 130 | 321 | 1189 | 150 | 243 |
| 744 | FIM1_759 | QGN14108.1 | CP015054.1 | 1398 | 384 | 596 | 1171 | 391 | 650 | 1218 | 473 | 556 |
| 745 | FIM1_760 | QGN14109.1 | CP015054.1 | 7813 | 1435 | 2047 | 5923 | 1181 | 2111 | 7145 | 1663 | 1860 |
| 746 | FIM1_761 | QGN14110.1 | CP015054.1 | 6496 | 1330 | 3109 | 6239 | 1056 | 2921 | 6619 | 1506 | 2473 |
| 747 | FIM1_762 | QGN14111.1 | CP015054.1 | 1142 | 820 | 1140 | 1106 | 725 | 1137 | 958 | 811 | 1034 |
| 748 | FIM1_763 | QGN14112.1 | CP015054.1 | 1910 | 361 | 632 | 1546 | 335 | 584 | 1871 | 426 | 553 |
| 749 | FIM1_764 | QGN14113.1 | CP015054.1 | 5618 | 1590 | 1890 | 4904 | 1481 | 1828 | 4781 | 1717 | 1661 |
| 750 | FIM1_765 | QGN14114.1 | CP015054.1 | 869 | 234 | 328 | 756 | 210 | 315 | 763 | 295 | 284 |
| 751 | FIM1_767 | QGN14115.1 | CP015054.1 | 2503 | 381 | 400 | 1965 | 258 | 395 | 7495 | 389 | 371 |
| 752 | FIM1_768 | QGN14116.1 | CP015054.1 | 909 | 149 | 346 | 711 | 120 | 359 | 1013 | 159 | 258 |
| 753 | FIM1_769 | QGN14117.1 | CP015054.1 | 6694 | 1569 | 2005 | 5559 | 1300 | 2008 | 7008 | 1783 | 1821 |
| 754 | FIM1_770 | QGN14118.1 | CP015054.1 | 7763 | 2543 | 2602 | 5633 | 1659 | 2549 | 5543 | 2682 | 2376 |
| 755 | FIM1_771 | QGN14119.1 | CP015054.1 | 46 | 11 | 9 | 35 | 9 | 7 | 15 | 9 | 8 |
| 756 | FIM1_772 | QGN14120.1 | CP015054.1 | 119 | 32 | 101 | 98 | 22 | 93 | 95 | 39 | 109 |
| 757 | FIM1_773 | QGN14121.1 | CP015054.1 | 4298 | 2488 | 1960 | 3619 | 1889 | 1940 | 1958 | 2611 | 1705 |
| 758 | FIM1_774 | QGN14122.1 | CP015054.1 | 2422 | 485 | 619 | 1833 | 423 | 682 | 2239 | 521 | 525 |
| 759 | FIM1_776 | QGN14123.1 | CP015054.1 | 6181 | 1514 | 1902 | 5488 | 1174 | 1976 | 6539 | 1702 | 1776 |
| 760 | FIM1_777 | QGN14124.1 | CP015054.1 | 7122 | 1369 | 2259 | 5527 | 1097 | 2310 | 6952 | 1386 | 1906 |
| 761 | FIM1_778 | QGN14125.1 | CP015054.1 | 1915 | 280 | 600 | 1605 | 194 | 552 | 2786 | 297 | 475 |
| 762 | FIM1_779 | QGN14126.1 | CP015054.1 | 30017 | 16479 | 16122 | 27064 | 11661 | 14922 | 24360 | 17299 | 14490 |
| 763 | FIM1_780 | QGN14127.1 | CP015054.1 | 1469 | 320 | 267 | 1156 | 156 | 304 | 1279 | 315 | 257 |
| 764 | FIM1_781 | QGN14128.1 | CP015054.1 | 761 | 142 | 257 | 689 | 115 | 236 | 1634 | 139 | 193 |
| 765 | FIM1_782 | QGN14129.1 | CP015054.1 | 6741 | 255 | 1084 | 4783 | 263 | 1077 | 9668 | 318 | 1022 |
| 766 | FIM1_783 | QGN14130.1 | CP015054.1 | 2284 | 384 | 753 | 2290 | 325 | 781 | 3290 | 396 | 748 |

| | | | | | | | | | | | | |
|---|---|---|---|---|---|---|---|---|---|---|---|---|
| 767 | FIM1_784 | QGN14131.1 | CP015054.1 | 1410 | 90 | 296 | 1322 | 96 | 272 | 2179 | 115 | 223 |
| 768 | FIM1_785 | QGN14132.1 | CP015054.1 | 10099 | 1986 | 3706 | 8919 | 1810 | 3666 | 7817 | 2267 | 3103 |
| 769 | FIM1_786 | QGN14133.1 | CP015054.1 | 5019 | 1409 | 2084 | 4554 | 1258 | 2098 | 4515 | 1537 | 1728 |
| 770 | FIM1_787 | QGN14134.1 | CP015054.1 | 13856 | 2888 | 3049 | 12444 | 3361 | 3012 | 16327 | 3209 | 2583 |
| 771 | FIM1_788 | QGN14135.1 | CP015054.1 | 544 | 124 | 106 | 458 | 90 | 137 | 536 | 116 | 136 |
| 772 | FIM1_789 | QGN14136.1 | CP015054.1 | 1675 | 663 | 1257 | 1606 | 369 | 1294 | 2023 | 615 | 1117 |
| 773 | FIM1_790 | QGN14137.1 | CP015054.1 | 1284 | 461 | 171 | 1016 | 349 | 162 | 1939 | 546 | 121 |
| 774 | FIM1_791 | QGN14138.1 | CP015054.1 | 13233 | 2780 | 1843 | 10384 | 2079 | 1807 | 17488 | 3109 | 1573 |
| 775 | FIM1_792 | QGN14139.1 | CP015054.1 | 2899 | 738 | 948 | 2258 | 547 | 948 | 2863 | 735 | 867 |
| 776 | FIM1_793 | QGN14140.1 | CP015054.1 | 10248 | 2646 | 2694 | 8468 | 1617 | 2752 | 11850 | 2833 | 2453 |
| 777 | FIM1_794 | QGN14141.1 | CP015054.1 | 2528 | 652 | 791 | 2182 | 495 | 739 | 2426 | 672 | 631 |
| 778 | FIM1_795 | QGN14142.1 | CP015054.1 | 5675 | 2418 | 2347 | 4791 | 1971 | 2555 | 3554 | 2594 | 2107 |
| 779 | FIM1_796 | QGN14143.1 | CP015054.1 | 3711 | 574 | 1426 | 2677 | 438 | 1440 | 3131 | 594 | 1068 |
| 780 | FIM1_797 | QGN14144.1 | CP015054.1 | 559 | 233 | 359 | 507 | 191 | 435 | 374 | 262 | 327 |
| 781 | FIM1_798 | QGN14145.1 | CP015054.1 | 1293 | 413 | 636 | 1195 | 330 | 694 | 1286 | 424 | 581 |
| 782 | FIM1_799 | QGN14146.1 | CP015054.1 | 3228 | 857 | 1036 | 2589 | 316 | 1017 | 7216 | 856 | 793 |
| 783 | FIM1_800 | QGN14147.1 | CP015054.1 | 3838 | 2325 | 2721 | 3975 | 1780 | 2765 | 4727 | 2397 | 2413 |
| 784 | FIM1_801 | QGN14148.1 | CP015054.1 | 7636 | 1940 | 657 | 6588 | 1392 | 611 | 12679 | 2216 | 577 |
| 785 | FIM1_802 | QGN14149.1 | CP015054.1 | 3080 | 768 | 706 | 2382 | 529 | 699 | 1590 | 738 | 555 |
| 786 | FIM1_803 | QGN14150.1 | CP015054.1 | 3272 | 578 | 954 | 2633 | 491 | 992 | 3360 | 654 | 836 |
| 787 | FIM1_804 | QGN14151.1 | CP015054.1 | 18715 | 5853 | 4022 | 13282 | 3999 | 3875 | 9328 | 6171 | 3568 |
| 788 | FIM1_805 | QGN14152.1 | CP015054.1 | 2439 | 319 | 681 | 2174 | 225 | 735 | 3225 | 378 | 541 |
| 789 | FIM1_806 | QGN14153.1 | CP015054.1 | 4071 | 184 | 704 | 3257 | 159 | 684 | 9929 | 213 | 557 |
| 790 | FIM1_807 | QGN14154.1 | CP015054.1 | 493 | 148 | 213 | 383 | 129 | 202 | 546 | 131 | 194 |
| 791 | FIM1_808 | QGN14155.1 | CP015054.1 | 3158 | 685 | 1294 | 2426 | 547 | 1293 | 3544 | 773 | 1073 |
| 792 | FIM1_809 | QGN14156.1 | CP015054.1 | 21948 | 9423 | 28172 | 19427 | 6186 | 27740 | 27246 | 9527 | 27516 |
| 793 | FIM1_810 | QGN14157.1 | CP015054.1 | 3703 | 916 | 998 | 2931 | 718 | 1173 | 3802 | 959 | 933 |
| 794 | FIM1_811 | QGN14158.1 | CP015054.1 | 6762 | 1449 | 1863 | 5410 | 1008 | 1864 | 8227 | 1492 | 1516 |
| 795 | FIM1_812 | QGN14159.1 | CP015054.1 | 2530 | 1461 | 2157 | 2129 | 933 | 2140 | 2408 | 1474 | 1823 |
| 796 | FIM1_813 | QGN14160.1 | CP015054.1 | 2260 | 487 | 560 | 1573 | 315 | 534 | 2009 | 485 | 430 |
| 797 | FIM1_814 | QGN14161.1 | CP015054.1 | 195 | 35 | 27 | 118 | 19 | 25 | 113 | 24 | 18 |
| 798 | FIM1_815 | QGN14162.1 | CP015054.1 | 546 | 51 | 48 | 415 | 52 | 64 | 248 | 72 | 53 |
| 799 | FIM1_816 | QGN14163.1 | CP015054.1 | 590 | 280 | 274 | 644 | 179 | 311 | 933 | 279 | 235 |
| 800 | FIM1_817 | QGN14164.1 | CP015054.1 | 3104 | 1306 | 1054 | 2092 | 922 | 1023 | 1999 | 1396 | 931 |
| 801 | FIM1_818 | QGN14165.1 | CP015054.1 | 251 | 173 | 360 | 192 | 176 | 340 | 132 | 193 | 320 |
| 802 | FIM1_819 | QGN14166.1 | CP015054.1 | 3322 | 745 | 1213 | 2784 | 605 | 1181 | 3206 | 740 | 960 |
| 803 | FIM1_820 | QGN14167.1 | CP015054.1 | 13202 | 4989 | 4471 | 11773 | 2777 | 4310 | 20946 | 5524 | 3901 |
| 804 | FIM1_821 | QGN14168.1 | CP015054.1 | 14414 | 7468 | 6775 | 13139 | 5967 | 6883 | 12407 | 7508 | 6671 |
| 805 | FIM1_822 | QGN14169.1 | CP015054.1 | 2056 | 591 | 901 | 1666 | 527 | 924 | 1746 | 660 | 909 |
| 806 | FIM1_823 | QGN14170.1 | CP015054.1 | 1813 | 471 | 1218 | 1361 | 426 | 1203 | 1658 | 561 | 961 |
| 807 | FIM1_824 | QGN14171.1 | CP015054.1 | 7524 | 7589 | 3370 | 5010 | 4802 | 3260 | 1389 | 7874 | 3056 |
| 808 | FIM1_825 | QGN14172.1 | CP015054.1 | 5787 | 2381 | 2993 | 4387 | 1775 | 2890 | 4556 | 2638 | 2393 |
| 809 | FIM1_826 | QGN14173.1 | CP015054.1 | 144 | 61 | 103 | 106 | 39 | 82 | 301 | 55 | 49 |
| 810 | FIM1_827 | QGN14174.1 | CP015054.1 | 30 | 4 | 6 | 18 | 7 | 15 | 24 | 3 | 8 |
| 811 | FIM1_828 | QGN14175.1 | CP015054.1 | 1930 | 760 | 1738 | 1280 | 581 | 1773 | 1705 | 849 | 1339 |
| 812 | FIM1_829 | QGN14176.1 | CP015054.1 | 582 | 271 | 573 | 485 | 216 | 605 | 486 | 296 | 566 |
| 813 | FIM1_830 | QGN14177.1 | CP015054.1 | 5523 | 1523 | 2436 | 4929 | 899 | 2441 | 7109 | 1590 | 2153 |
| 814 | FIM1_831 | QGN14178.1 | CP015054.1 | 5893 | 3880 | 3946 | 5924 | 3129 | 3877 | 4683 | 3994 | 3767 |

| | | | | | | | | | | | | |
|---|---|---|---|---|---|---|---|---|---|---|---|---|
| 815 | FIM1_832 | QGN14179.1 | CP015054.1 | 1566 | 712 | 680 | 1163 | 450 | 709 | 908 | 684 | 567 |
| 816 | FIM1_833 | QGN14180.1 | CP015054.1 | 7392 | 4585 | 3115 | 5817 | 3455 | 3056 | 3246 | 4973 | 2975 |
| 817 | FIM1_834 | QGN14181.1 | CP015054.1 | 3772 | 716 | 1164 | 3088 | 587 | 1169 | 3930 | 859 | 1062 |
| 818 | FIM1_835 | QGN14182.1 | CP015054.1 | 2979 | 538 | 772 | 2146 | 442 | 803 | 2943 | 606 | 533 |
| 819 | FIM1_836 | QGN14183.1 | CP015054.1 | 2199 | 460 | 274 | 1738 | 309 | 326 | 5559 | 559 | 190 |
| 820 | FIM1_837 | QGN14184.1 | CP015054.1 | 419 | 187 | 276 | 280 | 179 | 333 | 745 | 214 | 262 |
| 821 | FIM1_838 | QGN14185.1 | CP015054.1 | 381 | 182 | 260 | 271 | 172 | 335 | 621 | 220 | 253 |
| 822 | FIM1_839 | QGN14186.1 | CP015054.1 | 108 | 31 | 74 | 69 | 16 | 84 | 174 | 39 | 67 |
| 823 | FIM1_840 | QGN14187.1 | CP015054.1 | 1047 | 311 | 463 | 740 | 188 | 507 | 1389 | 242 | 437 |
| 824 | FIM1_841 | QGN14188.1 | CP015054.1 | 306 | 40 | 43 | 286 | 18 | 32 | 402 | 47 | 36 |
| 825 | FIM1_842 | QGN14189.1 | CP015054.1 | 655 | 124 | 114 | 591 | 60 | 104 | 588 | 132 | 116 |
| 826 | FIM1_843 | QGN14190.1 | CP015054.1 | 453 | 126 | 165 | 432 | 42 | 115 | 453 | 125 | 205 |
| 827 | FIM1_844 | QGN14191.1 | CP015054.1 | 308 | 22 | 68 | 244 | 23 | 64 | 332 | 37 | 70 |
| 828 | FIM1_846 | QGN14192.1 | CP015054.1 | 124 | 31 | 32 | 123 | 30 | 52 | 193 | 30 | 34 |
| 829 | FIM1_847 | QGN14193.1 | CP015054.1 | 45 | 7 | 7 | 42 | 7 | 10 | 55 | 5 | 8 |
| 830 | FIM1_850 | QGN14194.1 | CP015055.1 | 1506 | 346 | 541 | 1305 | 199 | 474 | 1658 | 368 | 492 |
| 831 | FIM1_851 | QGN14195.1 | CP015055.1 | 921 | 148 | 313 | 766 | 87 | 234 | 805 | 130 | 250 |
| 832 | FIM1_852 | QGN14196.1 | CP015055.1 | 88 | 45 | 61 | 65 | 32 | 60 | 114 | 57 | 48 |
| 833 | FIM1_853 | QGN14197.1 | CP015055.1 | 146 | 77 | 53 | 165 | 58 | 47 | 155 | 86 | 44 |
| 834 | FIM1_854 | QGN14198.1 | CP015055.1 | 2299 | 859 | 911 | 1951 | 561 | 817 | 2324 | 974 | 697 |
| 835 | FIM1_855 | QGN14199.1 | CP015055.1 | 500 | 165 | 218 | 464 | 131 | 195 | 546 | 195 | 220 |
| 836 | FIM1_856 | QGN14200.1 | CP015055.1 | 743 | 168 | 241 | 656 | 103 | 228 | 724 | 183 | 236 |
| 837 | FIM1_857 | QGN14201.1 | CP015055.1 | 518 | 103 | 122 | 356 | 66 | 87 | 548 | 125 | 101 |
| 838 | FIM1_858 | QGN14202.1 | CP015055.1 | 442 | 213 | 313 | 361 | 107 | 183 | 421 | 285 | 497 |
| 839 | FIM1_859 | QGN14203.1 | CP015055.1 | 847 | 227 | 374 | 679 | 156 | 365 | 839 | 233 | 385 |
| 840 | FIM1_860 | QGN14204.1 | CP015055.1 | 1102 | 159 | 469 | 925 | 161 | 432 | 1168 | 177 | 367 |
| 841 | FIM1_861 | QGN14205.1 | CP015055.1 | 1918 | 192 | 774 | 1662 | 203 | 659 | 1360 | 254 | 468 |
| 842 | FIM1_862 | QGN14206.1 | CP015055.1 | 406 | 76 | 117 | 364 | 60 | 112 | 384 | 76 | 95 |
| 843 | FIM1_863 | QGN14207.1 | CP015055.1 | 71 | 8 | 53 | 57 | 8 | 44 | 86 | 11 | 45 |
| 844 | FIM1_864 | QGN14208.1 | CP015055.1 | 1565 | 223 | 666 | 1134 | 213 | 570 | 2257 | 252 | 566 |
| 845 | FIM1_865 | QGN14209.1 | CP015055.1 | 1413 | 543 | 893 | 1416 | 278 | 822 | 2109 | 623 | 754 |
| 846 | FIM1_866 | QGN14210.1 | CP015055.1 | 1246 | 214 | 444 | 1036 | 115 | 364 | 1890 | 253 | 374 |
| 847 | FIM1_867 | QGN14211.1 | CP015055.1 | 4290 | 627 | 973 | 3083 | 474 | 902 | 4570 | 709 | 817 |
| 848 | FIM1_868 | QGN14212.1 | CP015055.1 | 2298 | 449 | 748 | 2007 | 292 | 713 | 2656 | 523 | 616 |
| 849 | FIM1_869 | QGN14213.1 | CP015055.1 | 1214 | 507 | 709 | 989 | 305 | 627 | 1241 | 504 | 598 |
| 850 | FIM1_870 | QGN14214.1 | CP015055.1 | 2918 | 1390 | 1281 | 1812 | 732 | 1027 | 1619 | 1486 | 939 |
| 851 | FIM1_871 | QGN14215.1 | CP015055.1 | 1586 | 758 | 1151 | 1356 | 590 | 1113 | 1497 | 840 | 1032 |
| 852 | FIM1_872 | QGN14216.1 | CP015055.1 | 5459 | 2017 | 2312 | 4827 | 1490 | 2252 | 4573 | 2092 | 2122 |
| 853 | FIM1_873 | QGN14217.1 | CP015055.1 | 2107 | 471 | 687 | 1620 | 317 | 610 | 2516 | 526 | 660 |
| 854 | FIM1_874 | QGN14218.1 | CP015055.1 | 16729 | 1704 | 2538 | 13015 | 1025 | 2279 | 27850 | 1944 | 2172 |
| 855 | FIM1_875 | QGN14219.1 | CP015055.1 | 1324 | 225 | 311 | 1088 | 151 | 246 | 1295 | 213 | 255 |
| 856 | FIM1_876 | QGN14220.1 | CP015055.1 | 7795 | 2173 | 2461 | 6298 | 1351 | 2161 | 6407 | 2346 | 1903 |
| 857 | FIM1_877 | QGN14221.1 | CP015055.1 | 1897 | 897 | 1246 | 1763 | 697 | 1107 | 2009 | 990 | 1067 |
| 858 | FIM1_878 | QGN14222.1 | CP015055.1 | 7088 | 2664 | 3189 | 6436 | 1872 | 2995 | 7973 | 3013 | 2946 |
| 859 | FIM1_879 | QGN14223.1 | CP015055.1 | 40 | 18 | 9 | 20 | 19 | 13 | 32 | 11 | 7 |
| 860 | FIM1_880 | QGN14224.1 | CP015055.1 | 3999 | 422 | 1120 | 2785 | 266 | 990 | 4819 | 444 | 981 |
| 861 | FIM1_881 | QGN14225.1 | CP015055.1 | 3693 | 927 | 1230 | 3049 | 672 | 1159 | 3748 | 1019 | 1073 |
| 862 | FIM1_882 | QGN14226.1 | CP015055.1 | 1210 | 228 | 323 | 934 | 200 | 390 | 1232 | 241 | 349 |

| | | | | | | | | | | | | |
|---|---|---|---|---|---|---|---|---|---|---|---|---|
| 863 | FIM1_883 | QGN14227.1 | CP015055.1 | 4216 | 877 | 1469 | 3348 | 519 | 1362 | 5622 | 870 | 1194 |
| 864 | FIM1_884 | QGN14228.1 | CP015055.1 | 753 | 95 | 295 | 567 | 56 | 286 | 1223 | 84 | 222 |
| 865 | FIM1_885 | QGN14229.1 | CP015055.1 | 4278 | 1567 | 1704 | 3679 | 1105 | 1497 | 4463 | 1653 | 1480 |
| 866 | FIM1_886 | QGN14230.1 | CP015055.1 | 12484 | 2528 | 3776 | 9977 | 1551 | 3430 | 14303 | 2530 | 3243 |
| 867 | FIM1_887 | QGN14231.1 | CP015055.1 | 1000 | 310 | 784 | 754 | 231 | 715 | 1008 | 345 | 614 |
| 868 | FIM1_888 | QGN14232.1 | CP015055.1 | 3373 | 740 | 1494 | 2483 | 532 | 1382 | 3900 | 795 | 1276 |
| 869 | FIM1_889 | QGN14233.1 | CP015055.1 | 2504 | 558 | 898 | 2055 | 462 | 807 | 2372 | 638 | 704 |
| 870 | FIM1_890 | QGN14234.1 | CP015055.1 | 517 | 274 | 307 | 528 | 215 | 281 | 432 | 291 | 236 |
| 871 | FIM1_891 | QGN14235.1 | CP015055.1 | 10541 | 1339 | 2919 | 8652 | 1020 | 2707 | 14053 | 1448 | 2420 |
| 872 | FIM1_892 | QGN14236.1 | CP015055.1 | 231 | 38 | 130 | 196 | 29 | 163 | 477 | 33 | 134 |
| 873 | FIM1_893 | QGN14237.1 | CP015055.1 | 105 | 13 | 14 | 66 | 14 | 9 | 165 | 18 | 10 |
| 874 | FIM1_894 | QGN14238.1 | CP015055.1 | 17737 | 645 | 1387 | 15169 | 501 | 1576 | 26209 | 694 | 1268 |
| 875 | FIM1_895 | QGN14239.1 | CP015055.1 | 2508 | 345 | 583 | 2130 | 288 | 570 | 2199 | 413 | 531 |
| 876 | FIM1_896 | QGN14240.1 | CP015055.1 | 325 | 74 | 157 | 267 | 61 | 134 | 336 | 88 | 106 |
| 877 | FIM1_897 | QGN14241.1 | CP015055.1 | 1580 | 343 | 602 | 1223 | 269 | 525 | 1333 | 375 | 480 |
| 878 | FIM1_898 | QGN14242.1 | CP015055.1 | 15942 | 3091 | 4812 | 13102 | 1976 | 4312 | 17514 | 3239 | 4318 |
| 879 | FIM1_899 | QGN14243.1 | CP015055.1 | 1275 | 579 | 600 | 782 | 412 | 609 | 769 | 601 | 521 |
| 880 | FIM1_900 | QGN14244.1 | CP015055.1 | 5149 | 1025 | 1663 | 3648 | 704 | 1486 | 6171 | 1045 | 1396 |
| 881 | FIM1_901 | QGN14245.1 | CP015055.1 | 9537 | 2396 | 2752 | 7365 | 1630 | 2651 | 10770 | 2759 | 2465 |
| 882 | FIM1_902 | QGN14246.1 | CP015055.1 | 1275 | 268 | 510 | 1126 | 239 | 466 | 1086 | 305 | 392 |
| 883 | FIM1_903 | QGN14247.1 | CP015055.1 | 2582 | 1917 | 1275 | 2302 | 1683 | 1148 | 823 | 2222 | 1117 |
| 884 | FIM1_904 | QGN14248.1 | CP015055.1 | 3127 | 1395 | 1063 | 1713 | 858 | 953 | 816 | 1394 | 897 |
| 885 | FIM1_905 | QGN14249.1 | CP015055.1 | 2668 | 315 | 595 | 2135 | 212 | 545 | 2948 | 312 | 430 |
| 886 | FIM1_906 | QGN14250.1 | CP015055.1 | 17647 | 11247 | 11369 | 16663 | 9616 | 10970 | 15781 | 11971 | 10633 |
| 887 | FIM1_907 | QGN14251.1 | CP015055.1 | 30142 | 16344 | 14770 | 28627 | 13518 | 13281 | 29214 | 16883 | 14164 |
| 888 | FIM1_908 | QGN14252.1 | CP015055.1 | 25 | 0 | 4 | 23 | 1 | 9 | 20 | 4 | 6 |
| 889 | FIM1_909 | QGN14253.1 | CP015055.1 | 1963 | 828 | 1084 | 1660 | 579 | 990 | 1145 | 962 | 909 |
| 890 | FIM1_910 | QGN14254.1 | CP015055.1 | 1112 | 45 | 173 | 848 | 25 | 158 | 1928 | 50 | 142 |
| 891 | FIM1_912 | QGN14255.1 | CP015055.1 | 240 | 9 | 60 | 169 | 11 | 44 | 354 | 14 | 31 |
| 892 | FIM1_913 | QGN14256.1 | CP015055.1 | 7731 | 953 | 1751 | 6021 | 1082 | 1632 | 7640 | 1065 | 1263 |
| 893 | FIM1_914 | QGN14257.1 | CP015055.1 | 1730 | 302 | 598 | 1265 | 208 | 484 | 1539 | 299 | 397 |
| 894 | FIM1_915 | QGN14258.1 | CP015055.1 | 13184 | 3004 | 3263 | 10487 | 2502 | 3052 | 9820 | 3406 | 2895 |
| 895 | FIM1_916 | QGN14259.1 | CP015055.1 | 4676 | 652 | 906 | 3647 | 389 | 822 | 5505 | 728 | 814 |
| 896 | FIM1_917 | QGN14260.1 | CP015055.1 | 27818 | 15174 | 4841 | 21197 | 9609 | 4114 | 28434 | 15736 | 4510 |
| 897 | FIM1_918 | QGN14261.1 | CP015055.1 | 765 | 148 | 259 | 724 | 126 | 244 | 766 | 166 | 234 |
| 898 | FIM1_919 | QGN14262.1 | CP015055.1 | 1412 | 117 | 225 | 1054 | 101 | 217 | 1382 | 145 | 232 |
| 899 | FIM1_920 | QGN14263.1 | CP015055.1 | 5986 | 1081 | 1591 | 5626 | 835 | 1427 | 5384 | 1304 | 1461 |
| 900 | FIM1_921 | QGN14264.1 | CP015055.1 | 4492 | 1388 | 2639 | 4107 | 1021 | 2322 | 5165 | 1570 | 2309 |
| 901 | FIM1_922 | QGN14265.1 | CP015055.1 | 1138 | 181 | 566 | 973 | 155 | 502 | 1022 | 189 | 410 |
| 902 | FIM1_923 | QGN14266.1 | CP015055.1 | 1145 | 273 | 332 | 885 | 192 | 295 | 1091 | 283 | 316 |
| 903 | FIM1_924 | QGN14267.1 | CP015055.1 | 1930 | 266 | 632 | 1969 | 260 | 575 | 2295 | 307 | 528 |
| 904 | FIM1_925 | QGN14268.1 | CP015055.1 | 567 | 79 | 219 | 543 | 42 | 170 | 644 | 54 | 114 |
| 905 | FIM1_926 | QGN14269.1 | CP015055.1 | 13248 | 937 | 1446 | 11742 | 620 | 1375 | 24706 | 908 | 1236 |
| 906 | FIM1_927 | QGN14270.1 | CP015055.1 | 13469 | 6120 | 5817 | 11882 | 4817 | 5338 | 12320 | 6645 | 5324 |
| 907 | FIM1_928 | QGN14271.1 | CP015055.1 | 1584 | 596 | 1056 | 1417 | 482 | 880 | 1315 | 717 | 873 |
| 908 | FIM1_929 | QGN14272.1 | CP015055.1 | 5425 | 878 | 1272 | 4514 | 567 | 1083 | 5529 | 901 | 1063 |
| 909 | FIM1_930 | QGN14273.1 | CP015055.1 | 7379 | 958 | 1515 | 5581 | 639 | 1312 | 8974 | 1028 | 1240 |
| 910 | FIM1_931 | QGN14274.1 | CP015055.1 | 3141 | 1266 | 1533 | 2537 | 992 | 1360 | 2610 | 1353 | 1306 |

| | | | | | | | | | | | | |
|---|---|---|---|---|---|---|---|---|---|---|---|---|
| 911 | FIM1_932 | QGN14275.1 | CP015055.1 | 1475 | 274 | 466 | 1225 | 167 | 408 | 1752 | 251 | 356 |
| 912 | FIM1_933 | QGN14276.1 | CP015055.1 | 6814 | 1003 | 1792 | 5045 | 716 | 1605 | 7557 | 1096 | 1451 |
| 913 | FIM1_934 | QGN14277.1 | CP015055.1 | 3270 | 680 | 1417 | 2645 | 528 | 1174 | 2504 | 757 | 1077 |
| 914 | FIM1_935 | QGN14278.1 | CP015055.1 | 4373 | 710 | 689 | 3734 | 340 | 643 | 6713 | 723 | 605 |
| 915 | FIM1_936 | QGN14279.1 | CP015055.1 | 27329 | 5030 | 7970 | 21874 | 3229 | 7345 | 32396 | 5181 | 7025 |
| 916 | FIM1_937 | QGN14280.1 | CP015055.1 | 2927 | 1645 | 2394 | 2584 | 1158 | 2158 | 3488 | 1735 | 2010 |
| 917 | FIM1_938 | QGN14281.1 | CP015055.1 | 9732 | 2724 | 2637 | 8053 | 1804 | 2340 | 9437 | 2911 | 2298 |
| 918 | FIM1_939 | QGN14282.1 | CP015055.1 | 4300 | 1462 | 1852 | 3346 | 995 | 1553 | 3248 | 1554 | 1545 |
| 919 | FIM1_940 | QGN14283.1 | CP015055.1 | 8268 | 1974 | 2141 | 6556 | 1670 | 1897 | 8902 | 2219 | 1957 |
| 920 | FIM1_941 | QGN14284.1 | CP015055.1 | 1089 | 473 | 355 | 915 | 198 | 295 | 1652 | 568 | 323 |
| 921 | FIM1_942 | QGN14285.1 | CP015055.1 | 5483 | 2152 | 728 | 3264 | 1464 | 673 | 2455 | 2247 | 582 |
| 922 | FIM1_943 | QGN14286.1 | CP015055.1 | 13235 | 7755 | 7553 | 13009 | 7129 | 7489 | 12321 | 8330 | 7188 |
| 923 | FIM1_944 | QGN14287.1 | CP015055.1 | 2302 | 1348 | 1280 | 1443 | 912 | 1143 | 750 | 1491 | 1140 |
| 924 | FIM1_945 | QGN14288.1 | CP015055.1 | 6122 | 1285 | 1550 | 5061 | 945 | 1513 | 6576 | 1466 | 1384 |
| 925 | FIM1_946 | QGN14289.1 | CP015055.1 | 156 | 28 | 39 | 131 | 18 | 38 | 153 | 21 | 40 |
| 926 | FIM1_947 | QGN14290.1 | CP015055.1 | 14507 | 3898 | 4192 | 10927 | 3219 | 3975 | 11694 | 4376 | 3841 |
| 927 | FIM1_948 | QGN14291.1 | CP015055.1 | 21958 | 4060 | 4180 | 15975 | 3052 | 3974 | 20326 | 4264 | 3726 |
| 928 | FIM1_949 | QGN14292.1 | CP015055.1 | 7268 | 302 | 1074 | 6018 | 284 | 989 | 9278 | 301 | 954 |
| 929 | FIM1_950 | QGN14293.1 | CP015055.1 | 1622 | 261 | 418 | 1344 | 128 | 345 | 1177 | 264 | 400 |
| 930 | FIM1_951 | QGN14294.1 | CP015055.1 | 39656 | 18398 | 12872 | 37303 | 11223 | 11227 | 63864 | 18157 | 11687 |
| 931 | FIM1_952 | QGN14295.1 | CP015055.1 | 70 | 16 | 21 | 64 | 13 | 11 | 131 | 19 | 16 |
| 932 | FIM1_953 | QGN14296.1 | CP015055.1 | 10807 | 1996 | 3027 | 10543 | 1361 | 2801 | 13181 | 2146 | 2757 |
| 933 | FIM1_954 | QGN14297.1 | CP015055.1 | 5786 | 988 | 1637 | 4110 | 1022 | 1478 | 2971 | 1099 | 1631 |
| 934 | FIM1_955 | QGN14298.1 | CP015055.1 | 8547 | 629 | 1043 | 6503 | 507 | 860 | 6962 | 673 | 788 |
| 935 | FIM1_956 | QGN14299.1 | CP015055.1 | 55 | 0 | 4 | 49 | 1 | 1 | 66 | 1 | 5 |
| 936 | FIM1_957 | QGN14300.1 | CP015055.1 | 28037 | 4107 | 10715 | 25013 | 6299 | 10967 | 29538 | 4526 | 10210 |
| 937 | FIM1_958 | QGN14301.1 | CP015055.1 | 6530 | 3358 | 3281 | 6091 | 1748 | 2937 | 10405 | 3341 | 2965 |
| 938 | FIM1_959 | QGN14302.1 | CP015055.1 | 3094 | 2436 | 878 | 2120 | 1200 | 767 | 1466 | 2659 | 793 |
| 939 | FIM1_960 | QGN14303.1 | CP015055.1 | 5252 | 2217 | 2064 | 4312 | 1528 | 1892 | 3602 | 2352 | 1689 |
| 940 | FIM1_961 | QGN14304.1 | CP015055.1 | 10488 | 2968 | 4771 | 8740 | 2301 | 4510 | 10320 | 3222 | 4250 |
| 941 | FIM1_962 | QGN14305.1 | CP015055.1 | 5725 | 2132 | 2429 | 4072 | 1606 | 2235 | 5675 | 2409 | 2169 |
| 942 | FIM1_963 | QGN14306.1 | CP015055.1 | 2120 | 399 | 625 | 1772 | 293 | 595 | 2462 | 429 | 511 |
| 943 | FIM1_964 | QGN14307.1 | CP015055.1 | 425 | 140 | 289 | 391 | 86 | 229 | 412 | 147 | 201 |
| 944 | FIM1_965 | QGN14308.1 | CP015055.1 | 3092 | 433 | 817 | 2202 | 364 | 688 | 2816 | 511 | 736 |
| 945 | FIM1_966 | QGN14309.1 | CP015055.1 | 7374 | 2501 | 4237 | 5703 | 1720 | 3875 | 8614 | 2610 | 3751 |
| 946 | FIM1_967 | QGN14310.1 | CP015055.1 | 2502 | 666 | 976 | 1773 | 522 | 892 | 1963 | 731 | 913 |
| 947 | FIM1_968 | QGN14311.1 | CP015055.1 | 1633 | 416 | 594 | 1334 | 293 | 514 | 1579 | 444 | 476 |
| 948 | FIM1_969 | QGN14312.1 | CP015055.1 | 1578 | 215 | 266 | 943 | 149 | 263 | 1175 | 202 | 249 |
| 949 | FIM1_970 | QGN14313.1 | CP015055.1 | 2791 | 520 | 798 | 1866 | 417 | 805 | 2740 | 539 | 675 |
| 950 | FIM1_971 | QGN14314.1 | CP015055.1 | 404 | 46 | 115 | 279 | 44 | 107 | 292 | 70 | 88 |
| 951 | FIM1_972 | QGN14315.1 | CP015055.1 | 273 | 119 | 145 | 282 | 118 | 148 | 168 | 109 | 127 |
| 952 | FIM1_973 | QGN14316.1 | CP015055.1 | 4786 | 559 | 938 | 3607 | 442 | 864 | 4770 | 639 | 751 |
| 953 | FIM1_974 | QGN14317.1 | CP015055.1 | 3364 | 701 | 970 | 2386 | 620 | 797 | 3434 | 729 | 810 |
| 954 | FIM1_975 | QGN14318.1 | CP015055.1 | 7790 | 1307 | 3188 | 5990 | 895 | 2776 | 8292 | 1361 | 2697 |
| 955 | FIM1_976 | QGN14319.1 | CP015055.1 | 463 | 22 | 43 | 391 | 12 | 31 | 332 | 38 | 21 |
| 956 | FIM1_977 | QGN14320.1 | CP015055.1 | 8992 | 3659 | 2622 | 6405 | 2149 | 2402 | 4729 | 3732 | 2600 |
| 957 | FIM1_978 | QGN14321.1 | CP015055.1 | 10201 | 6275 | 3110 | 8068 | 3812 | 2887 | 2341 | 6313 | 3103 |
| 958 | FIM1_979 | QGN14322.1 | CP015055.1 | 4084 | 2977 | 1623 | 3795 | 2373 | 1428 | 1291 | 3197 | 1517 |

| | | | | | | | | | | | | |
|---|---|---|---|---|---|---|---|---|---|---|---|---|
| 959 | FIM1_980 | QGN14323.1 | CP015055.1 | 9841 | 1306 | 1749 | 7826 | 1328 | 1605 | 10201 | 1283 | 1544 |
| 960 | FIM1_981 | QGN14324.1 | CP015055.1 | 3639 | 770 | 535 | 3184 | 221 | 499 | 8720 | 645 | 400 |
| 961 | FIM1_982 | QGN14325.1 | CP015055.1 | 2196 | 865 | 1075 | 2319 | 426 | 1004 | 3308 | 935 | 925 |
| 962 | FIM1_983 | QGN14326.1 | CP015055.1 | 21494 | 9228 | 10887 | 19609 | 5765 | 9589 | 26155 | 9592 | 10098 |
| 963 | FIM1_984 | QGN14327.1 | CP015055.1 | 32 | 10 | 11 | 32 | 7 | 10 | 298 | 12 | 15 |
| 964 | FIM1_988 | QGN14328.1 | CP015055.1 | 5056 | 1654 | 2042 | 4486 | 920 | 1796 | 7139 | 1735 | 1810 |
| 965 | FIM1_989 | QGN14329.1 | CP015055.1 | 304 | 5 | 24 | 189 | 7 | 26 | 397 | 4 | 14 |
| 966 | FIM1_990 | QGN14330.1 | CP015055.1 | 221 | 9 | 16 | 157 | 5 | 18 | 300 | 5 | 9 |
| 967 | FIM1_991 | QGN14331.1 | CP015055.1 | 3946 | 825 | 1152 | 3056 | 485 | 1043 | 4229 | 813 | 1080 |
| 968 | FIM1_992 | QGN14332.1 | CP015055.1 | 8139 | 1260 | 1903 | 6129 | 864 | 1701 | 8188 | 1224 | 1728 |
| 969 | FIM1_993 | QGN14333.1 | CP015055.1 | 2719 | 1265 | 1302 | 1929 | 877 | 1175 | 1110 | 1314 | 1274 |
| 970 | FIM1_994 | QGN14334.1 | CP015055.1 | 4537 | 1961 | 1763 | 3775 | 1385 | 1654 | 2472 | 2149 | 1689 |
| 971 | FIM1_995 | QGN14335.1 | CP015055.1 | 3754 | 2605 | 1685 | 3040 | 1777 | 1450 | 1316 | 2800 | 1587 |
| 972 | FIM1_996 | QGN14336.1 | CP015055.1 | 4449 | 4103 | 2578 | 3150 | 2544 | 2197 | 836 | 4416 | 2317 |
| 973 | FIM1_997 | QGN14337.1 | CP015055.1 | 777 | 59 | 111 | 388 | 51 | 85 | 456 | 68 | 98 |
| 974 | FIM1_998 | QGN14338.1 | CP015055.1 | 3848 | 557 | 941 | 2854 | 488 | 857 | 3068 | 573 | 812 |
| 975 | FIM1_999 | QGN14339.1 | CP015055.1 | 2316 | 1396 | 1245 | 1907 | 1036 | 1100 | 1594 | 1417 | 1117 |
| 976 | FIM1_1000 | QGN14340.1 | CP015055.1 | 7409 | 181 | 491 | 5074 | 145 | 442 | 5850 | 158 | 443 |
| 977 | FIM1_1001 | QGN14341.1 | CP015055.1 | 802 | 199 | 283 | 684 | 157 | 274 | 619 | 244 | 287 |
| 978 | FIM1_1002 | QGN14342.1 | CP015055.1 | 979 | 215 | 295 | 794 | 161 | 276 | 855 | 210 | 263 |
| 979 | FIM1_1003 | QGN14343.1 | CP015055.1 | 7612 | 4861 | 2856 | 5763 | 2888 | 2507 | 2636 | 5200 | 2528 |
| 980 | FIM1_1004 | QGN14344.1 | CP015055.1 | 7615 | 1669 | 3204 | 6470 | 1012 | 2829 | 9193 | 1806 | 3007 |
| 981 | FIM1_1005 | QGN14345.1 | CP015055.1 | 3904 | 732 | 1005 | 2863 | 575 | 925 | 3887 | 834 | 896 |
| 982 | FIM1_1006 | QGN14346.1 | CP015055.1 | 12624 | 2153 | 3115 | 9597 | 1438 | 2768 | 13348 | 2244 | 2662 |
| 983 | FIM1_1007 | QGN14347.1 | CP015055.1 | 1458 | 325 | 730 | 1448 | 236 | 759 | 1485 | 383 | 643 |
| 984 | FIM1_1008 | QGN14348.1 | CP015055.1 | 1035 | 114 | 265 | 939 | 96 | 223 | 1038 | 137 | 235 |
| 985 | FIM1_1009 | QGN14349.1 | CP015055.1 | 2055 | 406 | 629 | 1611 | 290 | 566 | 1381 | 431 | 516 |
| 986 | FIM1_1010 | QGN14350.1 | CP015055.1 | 14116 | 1039 | 1621 | 10822 | 775 | 1508 | 16406 | 1040 | 1353 |
| 987 | FIM1_1011 | QGN14351.1 | CP015055.1 | 25619 | 6903 | 11623 | 21044 | 4670 | 10349 | 26119 | 6843 | 10984 |
| 988 | FIM1_1012 | QGN14352.1 | CP015055.1 | 5970 | 1523 | 1674 | 5274 | 616 | 1475 | 12645 | 1566 | 1449 |
| 989 | FIM1_1013 | QGN14353.1 | CP015055.1 | 6938 | 1558 | 3218 | 5915 | 1199 | 2961 | 7985 | 1680 | 2696 |
| 990 | FIM1_1014 | QGN14354.1 | CP015055.1 | 5539 | 676 | 740 | 5085 | 262 | 373 | 7610 | 652 | 766 |
| 991 | FIM1_1015 | QGN14355.1 | CP015055.1 | 4777 | 2534 | 1891 | 4211 | 1798 | 1595 | 1515 | 2676 | 1602 |
| 992 | FIM1_1016 | QGN14356.1 | CP015055.1 | 152 | 137 | 167 | 91 | 90 | 152 | 109 | 139 | 178 |
| 993 | FIM1_1017 | QGN14357.1 | CP015055.1 | 7343 | 4763 | 3197 | 5758 | 3165 | 2950 | 4858 | 5312 | 3120 |
| 994 | FIM1_1018 | QGN14358.1 | CP015055.1 | 2441 | 941 | 1207 | 1982 | 781 | 1106 | 2024 | 1093 | 1105 |
| 995 | FIM1_1019 | QGN14359.1 | CP015055.1 | 3401 | 476 | 661 | 2230 | 356 | 716 | 2821 | 545 | 589 |
| 996 | FIM1_1020 | QGN14360.1 | CP015055.1 | 432 | 40 | 111 | 336 | 19 | 135 | 710 | 33 | 112 |
| 997 | FIM1_1021 | QGN14361.1 | CP015055.1 | 3220 | 109 | 238 | 1909 | 70 | 222 | 5021 | 97 | 192 |
| 998 | FIM1_1022 | QGN14362.1 | CP015055.1 | 16916 | 6530 | 6061 | 15406 | 4575 | 5410 | 18772 | 6638 | 5396 |
| 999 | FIM1_1023 | QGN14363.1 | CP015055.1 | 31 | 11 | 6 | 41 | 7 | 1 | 43 | 6 | 5 |
| 1000 | FIM1_1024 | QGN14364.1 | CP015055.1 | 7510 | 662 | 1085 | 5725 | 542 | 977 | 11720 | 684 | 1007 |
| 1001 | FIM1_1025 | QGN14365.1 | CP015055.1 | 8824 | 4693 | 4698 | 8615 | 3172 | 4464 | 6891 | 4885 | 3888 |
| 1002 | FIM1_1026 | QGN14366.1 | CP015055.1 | 11422 | 5657 | 6414 | 11320 | 4026 | 5927 | 9502 | 5847 | 5481 |
| 1003 | FIM1_1027 | QGN14367.1 | CP015055.1 | 2341 | 421 | 987 | 1833 | 292 | 911 | 3483 | 485 | 854 |
| 1004 | FIM1_1028 | QGN14368.1 | CP015055.1 | 804 | 65 | 134 | 454 | 51 | 95 | 786 | 66 | 86 |
| 1005 | FIM1_1029 | QGN14369.1 | CP015055.1 | 23189 | 11419 | 10676 | 20410 | 7645 | 9835 | 25996 | 12098 | 10544 |
| 1006 | FIM1_1030 | QGN14370.1 | CP015055.1 | 9106 | 6663 | 3101 | 6434 | 4462 | 2790 | 2088 | 7169 | 2937 |

| | | | | | | | | | | | | |
|---|---|---|---|---|---|---|---|---|---|---|---|---|
| 1007 | FIM1_1031 | QGN14371.1 | CP015055.1 | 139 | 46 | 59 | 114 | 35 | 53 | 173 | 41 | 53 |
| 1008 | FIM1_1032 | QGN14372.1 | CP015055.1 | 1516 | 277 | 612 | 1282 | 209 | 571 | 1642 | 285 | 514 |
| 1009 | FIM1_1033 | QGN14373.1 | CP015055.1 | 307 | 120 | 173 | 252 | 112 | 201 | 203 | 129 | 147 |
| 1010 | FIM1_1034 | QGN14374.1 | CP015055.1 | 2517 | 280 | 603 | 2173 | 234 | 579 | 2432 | 294 | 482 |
| 1011 | FIM1_1035 | QGN14375.1 | CP015055.1 | 538 | 64 | 172 | 510 | 75 | 166 | 440 | 86 | 143 |
| 1012 | FIM1_1036 | QGN14376.1 | CP015055.1 | 895 | 305 | 590 | 657 | 309 | 603 | 907 | 343 | 579 |
| 1013 | FIM1_1037 | QGN14377.1 | CP015055.1 | 3802 | 766 | 1235 | 2950 | 590 | 1120 | 3656 | 883 | 985 |
| 1014 | FIM1_1038 | QGN14378.1 | CP015055.1 | 7741 | 2613 | 3303 | 6547 | 1880 | 2952 | 6885 | 2973 | 2889 |
| 1015 | FIM1_1039 | QGN14379.1 | CP015055.1 | 9009 | 2263 | 1947 | 7274 | 1569 | 1758 | 10800 | 2473 | 1781 |
| 1016 | FIM1_1040 | QGN14380.1 | CP015055.1 | 5749 | 795 | 1349 | 4725 | 587 | 1226 | 7537 | 886 | 1198 |
| 1017 | FIM1_1041 | QGN14381.1 | CP015055.1 | 7001 | 1876 | 1412 | 6166 | 1212 | 1173 | 6927 | 2266 | 1312 |
| 1018 | FIM1_1042 | QGN14382.1 | CP015055.1 | 238 | 66 | 53 | 187 | 60 | 48 | 194 | 61 | 56 |
| 1019 | FIM1_1043 | QGN14383.1 | CP015055.1 | 916 | 113 | 169 | 615 | 85 | 149 | 363 | 87 | 185 |
| 1020 | FIM1_1044 | QGN14384.1 | CP015055.1 | 8148 | 1153 | 1588 | 6427 | 861 | 1409 | 9005 | 1248 | 1406 |
| 1021 | FIM1_1045 | QGN14385.1 | CP015055.1 | 882 | 218 | 475 | 864 | 168 | 459 | 1024 | 261 | 398 |
| 1022 | FIM1_1046 | QGN14386.1 | CP015055.1 | 2555 | 341 | 824 | 1995 | 236 | 783 | 3082 | 347 | 634 |
| 1023 | FIM1_1047 | QGN14387.1 | CP015055.1 | 4262 | 795 | 1292 | 3360 | 594 | 1263 | 3906 | 835 | 1100 |
| 1024 | FIM1_1048 | QGN14388.1 | CP015055.1 | 4835 | 1112 | 1575 | 3825 | 903 | 1515 | 4192 | 1251 | 1332 |
| 1025 | FIM1_1049 | QGN14389.1 | CP015055.1 | 4484 | 4089 | 2128 | 2864 | 2434 | 1867 | 979 | 4313 | 1763 |
| 1026 | FIM1_1050 | QGN14390.1 | CP015055.1 | 247 | 14 | 102 | 170 | 15 | 125 | 216 | 26 | 146 |
| 1027 | FIM1_1051 | QGN14391.1 | CP015055.1 | 1030 | 323 | 586 | 945 | 158 | 507 | 1481 | 317 | 499 |
| 1028 | FIM1_1052 | QGN14392.1 | CP015055.1 | 1611 | 230 | 1617 | 1578 | 157 | 1595 | 1851 | 242 | 1503 |
| 1029 | FIM1_1053 | QGN14393.1 | CP015055.1 | 241 | 53 | 89 | 218 | 54 | 92 | 254 | 52 | 79 |
| 1030 | FIM1_1054 | QGN14394.1 | CP015055.1 | 5871 | 2044 | 1718 | 4921 | 1499 | 1686 | 4349 | 2110 | 1666 |
| 1031 | FIM1_1055 | QGN14395.1 | CP015055.1 | 353 | 62 | 243 | 304 | 55 | 199 | 333 | 82 | 188 |
| 1032 | FIM1_1056 | QGN14396.1 | CP015055.1 | 1246 | 433 | 498 | 847 | 274 | 445 | 669 | 438 | 427 |
| 1033 | FIM1_1057 | QGN14397.1 | CP015055.1 | 5490 | 756 | 1141 | 4222 | 540 | 1025 | 5693 | 833 | 1034 |
| 1034 | FIM1_1058 | QGN14398.1 | CP015055.1 | 4880 | 1802 | 2938 | 4076 | 1145 | 2488 | 4092 | 1974 | 2589 |
| 1035 | FIM1_1059 | QGN14399.1 | CP015055.1 | 26 | 5 | 11 | 14 | 6 | 23 | 23 | 3 | 9 |
| 1036 | FIM1_1060 | QGN14400.1 | CP015055.1 | 10152 | 5908 | 3369 | 7682 | 3992 | 2986 | 4639 | 6426 | 3024 |
| 1037 | FIM1_1061 | QGN14401.1 | CP015055.1 | 2011 | 646 | 638 | 1424 | 355 | 541 | 1696 | 688 | 526 |
| 1038 | FIM1_1062 | QGN14402.1 | CP015055.1 | 2248 | 476 | 654 | 1598 | 383 | 631 | 2025 | 520 | 573 |
| 1039 | FIM1_1063 | QGN14403.1 | CP015055.1 | 1760 | 267 | 665 | 1598 | 218 | 644 | 2301 | 281 | 477 |
| 1040 | FIM1_1064 | QGN14404.1 | CP015055.1 | 2619 | 578 | 1086 | 2222 | 337 | 939 | 4556 | 594 | 858 |
| 1041 | FIM1_1065 | QGN14405.1 | CP015055.1 | 3217 | 733 | 1061 | 2521 | 608 | 971 | 3883 | 804 | 918 |
| 1042 | FIM1_1066 | QGN14406.1 | CP015055.1 | 896 | 312 | 479 | 732 | 215 | 409 | 808 | 352 | 354 |
| 1043 | FIM1_1067 | QGN14407.1 | CP015055.1 | 1197 | 217 | 402 | 950 | 144 | 375 | 1222 | 227 | 317 |
| 1044 | FIM1_1068 | QGN14408.1 | CP015055.1 | 984 | 228 | 143 | 658 | 154 | 129 | 341 | 288 | 112 |
| 1045 | FIM1_1069 | QGN14409.1 | CP015055.1 | 1816 | 163 | 370 | 1515 | 100 | 318 | 1961 | 160 | 350 |
| 1046 | FIM1_1070 | QGN14410.1 | CP015055.1 | 3792 | 419 | 1127 | 2960 | 352 | 1074 | 4382 | 436 | 976 |
| 1047 | FIM1_1071 | QGN14411.1 | CP015055.1 | 4236 | 1063 | 1256 | 3381 | 720 | 1158 | 3753 | 1135 | 1169 |
| 1048 | FIM1_1072 | QGN14412.1 | CP015055.1 | 3782 | 1846 | 828 | 2168 | 963 | 768 | 1732 | 1812 | 805 |
| 1049 | FIM1_1074 | QGN14413.1 | CP015055.1 | 2472 | 959 | 1110 | 2236 | 591 | 976 | 2396 | 992 | 941 |
| 1050 | FIM1_1075 | QGN14414.1 | CP015055.1 | 8121 | 2325 | 2213 | 7070 | 1697 | 1939 | 6721 | 2505 | 2045 |
| 1051 | FIM1_1076 | QGN14415.1 | CP015055.1 | 1893 | 220 | 424 | 1473 | 178 | 428 | 2187 | 221 | 376 |
| 1052 | FIM1_1077 | QGN14416.1 | CP015055.1 | 331 | 100 | 69 | 265 | 94 | 90 | 241 | 120 | 98 |
| 1053 | FIM1_1078 | QGN14417.1 | CP015055.1 | 882 | 187 | 277 | 751 | 156 | 215 | 555 | 242 | 235 |
| 1054 | FIM1_1079 | QGN14418.1 | CP015055.1 | 1788 | 143 | 294 | 1244 | 149 | 285 | 2238 | 168 | 327 |

| | | | | | | | | | | | | |
|---|---|---|---|---|---|---|---|---|---|---|---|---|
| 1055 | FIM1_1080 | QGN14419.1 | CP015055.1 | 3087 | 604 | 898 | 2411 | 457 | 873 | 3251 | 705 | 972 |
| 1056 | FIM1_1081 | QGN14420.1 | CP015055.1 | 6842 | 711 | 1324 | 6021 | 394 | 1130 | 10067 | 806 | 1189 |
| 1057 | FIM1_1082 | QGN14421.1 | CP015055.1 | 4380 | 865 | 688 | 3279 | 644 | 567 | 4239 | 905 | 639 |
| 1058 | FIM1_1083 | QGN14422.1 | CP015055.1 | 4753 | 766 | 1239 | 4047 | 586 | 1219 | 4667 | 770 | 1053 |
| 1059 | FIM1_1084 | QGN14423.1 | CP015055.1 | 2290 | 362 | 662 | 1807 | 298 | 571 | 2397 | 354 | 509 |
| 1060 | FIM1_1085 | QGN14424.1 | CP015055.1 | 1146 | 287 | 551 | 880 | 260 | 529 | 1054 | 349 | 462 |
| 1061 | FIM1_1086 | QGN14425.1 | CP015055.1 | 1141 | 220 | 795 | 944 | 166 | 731 | 1336 | 221 | 558 |
| 1062 | FIM1_1087 | QGN14426.1 | CP015055.1 | 2776 | 420 | 702 | 1990 | 333 | 564 | 2066 | 457 | 467 |
| 1063 | FIM1_1088 | QGN14427.1 | CP015055.1 | 5947 | 1595 | 1979 | 4609 | 1197 | 1801 | 5240 | 1750 | 1654 |
| 1064 | FIM1_1089 | QGN14428.1 | CP015055.1 | 209 | 67 | 127 | 189 | 69 | 123 | 160 | 68 | 108 |
| 1065 | FIM1_1090 | QGN14429.1 | CP015055.1 | 10200 | 4011 | 3879 | 8887 | 2638 | 3536 | 12525 | 4383 | 3444 |
| 1066 | FIM1_1091 | QGN14430.1 | CP015055.1 | 4929 | 3122 | 1871 | 3530 | 2037 | 1639 | 1714 | 3299 | 1621 |
| 1067 | FIM1_1092 | QGN14431.1 | CP015055.1 | 3096 | 609 | 949 | 2258 | 405 | 884 | 2476 | 663 | 754 |
| 1068 | FIM1_1093 | QGN14432.1 | CP015055.1 | 2276 | 437 | 919 | 1805 | 338 | 839 | 2969 | 506 | 754 |
| 1069 | FIM1_1094 | QGN14433.1 | CP015055.1 | 4568 | 1217 | 1287 | 3818 | 930 | 1188 | 4139 | 1339 | 1159 |
| 1070 | FIM1_1095 | QGN14434.1 | CP015055.1 | 4267 | 687 | 970 | 3319 | 496 | 895 | 4036 | 679 | 809 |
| 1071 | FIM1_1096 | QGN14435.1 | CP015055.1 | 956 | 310 | 359 | 1004 | 212 | 382 | 1041 | 300 | 299 |
| 1072 | FIM1_1097 | QGN14436.1 | CP015055.1 | 25079 | 8044 | 5191 | 19633 | 4437 | 4494 | 35126 | 8354 | 4115 |
| 1073 | FIM1_1098 | QGN14437.1 | CP015055.1 | 5615 | 2698 | 3100 | 5563 | 2122 | 2724 | 5315 | 2952 | 2644 |
| 1074 | FIM1_1099 | QGN14438.1 | CP015055.1 | 3098 | 255 | 741 | 2520 | 225 | 635 | 3807 | 276 | 549 |
| 1075 | FIM1_1100 | QGN14439.1 | CP015055.1 | 17425 | 6958 | 6412 | 15280 | 5371 | 5678 | 16841 | 7600 | 6064 |
| 1076 | FIM1_1101 | QGN14440.1 | CP015055.1 | 225 | 34 | 58 | 175 | 27 | 68 | 250 | 39 | 55 |
| 1077 | FIM1_1102 | QGN14441.1 | CP015055.1 | 2308 | 867 | 1255 | 2194 | 688 | 1154 | 2033 | 969 | 1161 |
| 1078 | FIM1_1103 | QGN14442.1 | CP015055.1 | 2147 | 728 | 948 | 1865 | 568 | 834 | 830 | 795 | 873 |
| 1079 | FIM1_1104 | QGN14443.1 | CP015055.1 | 4269 | 973 | 1578 | 3818 | 721 | 1311 | 3574 | 1082 | 1355 |
| 1080 | FIM1_1105 | QGN14444.1 | CP015055.1 | 8215 | 5671 | 3853 | 7083 | 3661 | 3476 | 9221 | 5753 | 3573 |
| 1081 | FIM1_1106 | QGN14445.1 | CP015055.1 | 17287 | 4574 | 6096 | 14569 | 3304 | 5645 | 16920 | 4805 | 5393 |
| 1082 | FIM1_1107 | QGN14446.1 | CP015055.1 | 2074 | 716 | 1037 | 1854 | 521 | 909 | 1470 | 726 | 878 |
| 1083 | FIM1_1108 | QGN14447.1 | CP015055.1 | 3060 | 2466 | 1359 | 2577 | 2491 | 1214 | 1196 | 2636 | 1279 |
| 1084 | FIM1_1109 | QGN14448.1 | CP015055.1 | 14935 | 2922 | 3757 | 11647 | 1773 | 3395 | 19935 | 3145 | 3143 |
| 1085 | FIM1_1110 | QGN14449.1 | CP015055.1 | 2431 | 943 | 1252 | 2212 | 722 | 1234 | 2459 | 998 | 1112 |
| 1086 | FIM1_1111 | QGN14450.1 | CP015055.1 | 2149 | 598 | 1120 | 1559 | 468 | 1023 | 2488 | 663 | 954 |
| 1087 | FIM1_1112 | QGN14451.1 | CP015055.1 | 11745 | 2337 | 2615 | 8893 | 1631 | 2386 | 10877 | 2453 | 2314 |
| 1088 | FIM1_1113 | QGN14452.1 | CP015055.1 | 6655 | 1061 | 1100 | 5426 | 773 | 1082 | 5747 | 1125 | 1149 |
| 1089 | FIM1_1114 | QGN14453.1 | CP015055.1 | 2744 | 352 | 230 | 1819 | 188 | 220 | 10336 | 325 | 180 |
| 1090 | FIM1_1115 | QGN14454.1 | CP015055.1 | 1354 | 109 | 189 | 926 | 106 | 152 | 1388 | 111 | 169 |
| 1091 | FIM1_1116 | QGN14455.1 | CP015055.1 | 18557 | 4091 | 6597 | 15851 | 3409 | 6325 | 16260 | 4597 | 6208 |
| 1092 | FIM1_1117 | QGN14456.1 | CP015055.1 | 4477 | 582 | 2369 | 3498 | 548 | 2193 | 4837 | 624 | 1723 |
| 1093 | FIM1_1118 | QGN14457.1 | CP015055.1 | 46 | 13 | 17 | 28 | 6 | 13 | 28 | 20 | 12 |
| 1094 | FIM1_1119 | QGN14458.1 | CP015055.1 | 8965 | 1994 | 2259 | 6808 | 1202 | 1983 | 9812 | 2046 | 1991 |
| 1095 | FIM1_1120 | QGN14459.1 | CP015055.1 | 114 | 13 | 23 | 75 | 6 | 24 | 84 | 12 | 18 |
| 1096 | FIM1_1121 | QGN14460.1 | CP015055.1 | 25983 | 9233 | 10986 | 20910 | 5331 | 10297 | 36921 | 9715 | 10206 |
| 1097 | FIM1_1122 | QGN14461.1 | CP015055.1 | 15900 | 3865 | 4568 | 13092 | 2917 | 4356 | 16732 | 4185 | 4019 |
| 1098 | FIM1_1123 | QGN14462.1 | CP015055.1 | 981 | 220 | 452 | 885 | 170 | 336 | 975 | 272 | 330 |
| 1099 | FIM1_1124 | QGN14463.1 | CP015055.1 | 12224 | 1514 | 2660 | 10055 | 1140 | 2412 | 11888 | 1710 | 2170 |
| 1100 | FIM1_1125 | QGN14464.1 | CP015055.1 | 36978 | 21797 | 20100 | 35491 | 17628 | 18455 | 35139 | 22428 | 19003 |
| 1101 | FIM1_1126 | QGN14465.1 | CP015055.1 | 1469 | 534 | 511 | 1334 | 250 | 446 | 2543 | 529 | 403 |
| 1102 | FIM1_1127 | QGN14466.1 | CP015055.1 | 1209 | 112 | 385 | 882 | 92 | 358 | 1607 | 119 | 320 |

| | | | | | | | | | | | | |
|---|---|---|---|---|---|---|---|---|---|---|---|---|
| 1103 | FIM1_1128 | QGN14467.1 | CP015055.1 | 231 | 66 | 143 | 220 | 46 | 136 | 402 | 65 | 142 |
| 1104 | FIM1_1129 | QGN14468.1 | CP015055.1 | 7454 | 1999 | 2749 | 5866 | 1443 | 2495 | 7346 | 2153 | 2428 |
| 1105 | FIM1_1130 | QGN14469.1 | CP015055.1 | 1296 | 280 | 473 | 981 | 196 | 378 | 878 | 281 | 374 |
| 1106 | FIM1_1131 | QGN14470.1 | CP015055.1 | 3373 | 851 | 1283 | 2715 | 574 | 1079 | 3077 | 937 | 1032 |
| 1107 | FIM1_1132 | QGN14471.1 | CP015055.1 | 3525 | 266 | 450 | 2416 | 129 | 349 | 3216 | 279 | 306 |
| 1108 | FIM1_1133 | QGN14472.1 | CP015055.1 | 2978 | 438 | 516 | 2345 | 336 | 450 | 3162 | 522 | 420 |
| 1109 | FIM1_1134 | QGN14473.1 | CP015055.1 | 1141 | 214 | 300 | 797 | 109 | 270 | 986 | 203 | 269 |
| 1110 | FIM1_1135 | QGN14474.1 | CP015055.1 | 28807 | 14248 | 11128 | 23534 | 9616 | 10090 | 31432 | 14859 | 11060 |
| 1111 | FIM1_1136 | QGN14475.1 | CP015055.1 | 2229 | 425 | 983 | 1816 | 334 | 968 | 2796 | 445 | 948 |
| 1112 | FIM1_1137 | QGN14476.1 | CP015055.1 | 11679 | 1696 | 3727 | 9422 | 1166 | 3411 | 9675 | 1547 | 3853 |
| 1113 | FIM1_1138 | QGN14477.1 | CP015055.1 | 1075 | 211 | 526 | 843 | 149 | 533 | 1255 | 199 | 455 |
| 1114 | FIM1_1139 | QGN14478.1 | CP015055.1 | 21788 | 5383 | 7590 | 17987 | 4198 | 6877 | 21349 | 5899 | 6960 |
| 1115 | FIM1_1140 | QGN14479.1 | CP015055.1 | 4506 | 1726 | 2923 | 3356 | 805 | 2633 | 3794 | 2076 | 2889 |
| 1116 | FIM1_1141 | QGN14480.1 | CP015055.1 | 5025 | 2028 | 3105 | 3703 | 1055 | 3058 | 4319 | 2577 | 3275 |
| 1117 | FIM1_1142 | QGN14481.1 | CP015055.1 | 2647 | 526 | 753 | 2234 | 397 | 662 | 2263 | 565 | 625 |
| 1118 | FIM1_1143 | QGN14482.1 | CP015055.1 | 4364 | 1227 | 1475 | 3576 | 941 | 1391 | 3499 | 1289 | 1293 |
| 1119 | FIM1_1144 | QGN14483.1 | CP015055.1 | 1920 | 558 | 866 | 1701 | 513 | 851 | 1305 | 650 | 717 |
| 1120 | FIM1_1145 | QGN14484.1 | CP015055.1 | 23505 | 9432 | 7001 | 19891 | 7447 | 6483 | 18952 | 9861 | 6545 |
| 1121 | FIM1_1146 | QGN14485.1 | CP015055.1 | 1231 | 327 | 582 | 1020 | 198 | 535 | 1771 | 351 | 540 |
| 1122 | FIM1_1147 | QGN14486.1 | CP015055.1 | 7753 | 3529 | 2672 | 6679 | 2498 | 2472 | 6405 | 3815 | 2432 |
| 1123 | FIM1_1148 | QGN14487.1 | CP015055.1 | 1420 | 486 | 618 | 1261 | 315 | 572 | 928 | 472 | 508 |
| 1124 | FIM1_1149 | QGN14488.1 | CP015055.1 | 595 | 194 | 243 | 513 | 149 | 220 | 459 | 218 | 231 |
| 1125 | FIM1_1150 | QGN14489.1 | CP015055.1 | 1657 | 648 | 813 | 1446 | 547 | 866 | 1460 | 754 | 750 |
| 1126 | FIM1_1151 | QGN14490.1 | CP015055.1 | 12316 | 4438 | 4412 | 10215 | 3027 | 3995 | 15100 | 5029 | 3978 |
| 1127 | FIM1_1152 | QGN14491.1 | CP015055.1 | 3158 | 433 | 617 | 2433 | 331 | 586 | 3938 | 479 | 602 |
| 1128 | FIM1_1153 | QGN14492.1 | CP015055.1 | 289 | 17 | 70 | 199 | 25 | 64 | 245 | 20 | 58 |
| 1129 | FIM1_1154 | QGN14493.1 | CP015055.1 | 3099 | 673 | 1367 | 2464 | 557 | 1248 | 2398 | 775 | 1183 |
| 1130 | FIM1_1155 | QGN14494.1 | CP015055.1 | 310 | 57 | 77 | 216 | 54 | 53 | 243 | 51 | 64 |
| 1131 | FIM1_1156 | QGN14495.1 | CP015055.1 | 17770 | 8522 | 7895 | 16091 | 6498 | 7338 | 17859 | 8705 | 7281 |
| 1132 | FIM1_1157 | QGN14496.1 | CP015055.1 | 8438 | 1399 | 2574 | 7043 | 1113 | 2242 | 9830 | 1648 | 1909 |
| 1133 | FIM1_1158 | QGN14497.1 | CP015055.1 | 3761 | 139 | 1216 | 2732 | 132 | 1092 | 4310 | 158 | 870 |
| 1134 | FIM1_1159 | QGN14498.1 | CP015055.1 | 4797 | 1415 | 1743 | 4003 | 990 | 1524 | 3944 | 1536 | 1569 |
| 1135 | FIM1_1160 | QGN14499.1 | CP015055.1 | 22563 | 10838 | 7704 | 18674 | 7236 | 6955 | 24526 | 10923 | 7103 |
| 1136 | FIM1_1161 | QGN14500.1 | CP015055.1 | 279 | 51 | 50 | 163 | 30 | 40 | 320 | 54 | 35 |
| 1137 | FIM1_1162 | QGN14501.1 | CP015055.1 | 22167 | 3419 | 7574 | 15609 | 1299 | 6288 | 27037 | 3126 | 4174 |
| 1138 | FIM1_1163 | QGN14502.1 | CP015055.1 | 3252 | 741 | 768 | 2330 | 563 | 758 | 3038 | 723 | 732 |
| 1139 | FIM1_1164 | QGN14503.1 | CP015055.1 | 97 | 23 | 31 | 73 | 24 | 44 | 42 | 24 | 29 |
| 1140 | FIM1_1165 | QGN14504.1 | CP015055.1 | 1947 | 652 | 837 | 1563 | 510 | 875 | 1778 | 753 | 719 |
| 1141 | FIM1_1166 | QGN14505.1 | CP015055.1 | 11512 | 3946 | 2840 | 8814 | 2821 | 2587 | 14928 | 4152 | 2573 |
| 1142 | FIM1_1167 | QGN14506.1 | CP015055.1 | 2969 | 548 | 735 | 2519 | 321 | 688 | 3710 | 532 | 641 |
| 1143 | FIM1_1168 | QGN14507.1 | CP015055.1 | 5074 | 1055 | 1453 | 3478 | 801 | 1224 | 4308 | 1206 | 1200 |
| 1144 | FIM1_1169 | QGN14508.1 | CP015055.1 | 1299 | 728 | 844 | 945 | 448 | 810 | 1816 | 812 | 747 |
| 1145 | FIM1_1170 | QGN14509.1 | CP015055.1 | 2401 | 915 | 943 | 2031 | 565 | 932 | 2152 | 919 | 859 |
| 1146 | FIM1_1171 | QGN14510.1 | CP015055.1 | 178 | 62 | 32 | 111 | 48 | 36 | 133 | 66 | 27 |
| 1147 | FIM1_1172 | QGN14511.1 | CP015055.1 | 5982 | 2443 | 2476 | 4713 | 1520 | 2116 | 5870 | 2600 | 2143 |
| 1148 | FIM1_1173 | QGN14512.1 | CP015055.1 | 884 | 443 | 594 | 492 | 339 | 544 | 611 | 424 | 608 |
| 1149 | FIM1_1174 | QGN14513.1 | CP015055.1 | 2091 | 229 | 242 | 1238 | 168 | 248 | 1762 | 231 | 281 |
| 1150 | FIM1_1175 | QGN14514.1 | CP015055.1 | 380 | 85 | 144 | 302 | 67 | 135 | 261 | 88 | 112 |

| | | | | | | | | | | | | |
|---|---|---|---|---|---|---|---|---|---|---|---|---|
| 1151 | FIM1_1177 | QGN14515.1 | CP015055.1 | 3964 | 1743 | 2045 | 3575 | 1269 | 1884 | 3727 | 1814 | 1784 |
| 1152 | FIM1_1178 | QGN14516.1 | CP015055.1 | 27852 | 16973 | 11004 | 22164 | 13151 | 10135 | 15861 | 17935 | 10541 |
| 1153 | FIM1_1179 | QGN14517.1 | CP015055.1 | 2493 | 569 | 792 | 1805 | 421 | 783 | 1911 | 580 | 693 |
| 1154 | FIM1_1180 | QGN14518.1 | CP015055.1 | 2108 | 867 | 803 | 1422 | 704 | 783 | 1910 | 916 | 787 |
| 1155 | FIM1_1181 | QGN14519.1 | CP015055.1 | 5033 | 1560 | 1954 | 4221 | 746 | 1722 | 5382 | 1651 | 1744 |
| 1156 | FIM1_1182 | QGN14520.1 | CP015055.1 | 381 | 90 | 45 | 299 | 57 | 54 | 294 | 109 | 39 |
| 1157 | FIM1_1183 | QGN14521.1 | CP015055.1 | 2042 | 1708 | 508 | 1306 | 980 | 431 | 1572 | 1767 | 435 |
| 1158 | FIM1_1184 | QGN14522.1 | CP015055.1 | 1337 | 414 | 664 | 977 | 298 | 523 | 809 | 414 | 454 |
| 1159 | FIM1_1185 | QGN14523.1 | CP015055.1 | 16 | 10 | 8 | 9 | 3 | 6 | 6 | 9 | 2 |
| 1160 | FIM1_1186 | QGN14524.1 | CP015055.1 | 2139 | 372 | 647 | 1664 | 265 | 550 | 1919 | 441 | 482 |
| 1161 | FIM1_1187 | QGN14525.1 | CP015055.1 | 16436 | 4889 | 4898 | 12848 | 3686 | 4409 | 15588 | 5109 | 4328 |
| 1162 | FIM1_1188 | QGN14526.1 | CP015055.1 | 2617 | 247 | 344 | 1949 | 184 | 297 | 2587 | 275 | 275 |
| 1163 | FIM1_1189 | QGN14527.1 | CP015055.1 | 3687 | 274 | 725 | 2711 | 217 | 638 | 5439 | 325 | 607 |
| 1164 | FIM1_1190 | QGN14528.1 | CP015055.1 | 1289 | 92 | 222 | 964 | 61 | 177 | 1566 | 111 | 192 |
| 1165 | FIM1_1191 | QGN14529.1 | CP015055.1 | 94 | 16 | 26 | 71 | 27 | 36 | 60 | 18 | 20 |
| 1166 | FIM1_1192 | QGN14530.1 | CP015055.1 | 2007 | 723 | 884 | 1703 | 640 | 870 | 1159 | 847 | 824 |
| 1167 | FIM1_1193 | QGN14531.1 | CP015055.1 | 12319 | 4206 | 4199 | 9646 | 3162 | 3910 | 11105 | 4415 | 3833 |
| 1168 | FIM1_1194 | QGN14532.1 | CP015055.1 | 6533 | 3113 | 2468 | 4798 | 2217 | 2340 | 4587 | 3266 | 2262 |
| 1169 | FIM1_1195 | QGN14533.1 | CP015055.1 | 506 | 121 | 361 | 389 | 95 | 361 | 350 | 138 | 314 |
| 1170 | FIM1_1196 | QGN14534.1 | CP015055.1 | 26551 | 16580 | 14697 | 24649 | 13859 | 14092 | 25954 | 17224 | 14273 |
| 1171 | FIM1_1197 | QGN14535.1 | CP015055.1 | 7333 | 2497 | 4424 | 6299 | 1575 | 3954 | 8465 | 2549 | 3709 |
| 1172 | FIM1_1198 | QGN14536.1 | CP015055.1 | 5084 | 1140 | 1099 | 3357 | 666 | 1052 | 4174 | 1131 | 997 |
| 1173 | FIM1_1199 | QGN14537.1 | CP015055.1 | 4419 | 886 | 1165 | 3374 | 702 | 1093 | 4708 | 907 | 1080 |
| 1174 | FIM1_1200 | QGN14538.1 | CP015055.1 | 13486 | 3635 | 3172 | 11976 | 2131 | 2886 | 18549 | 3721 | 2792 |
| 1175 | FIM1_1201 | QGN14539.1 | CP015055.1 | 464 | 116 | 203 | 387 | 91 | 190 | 473 | 122 | 170 |
| 1176 | FIM1_1202 | QGN14540.1 | CP015055.1 | 846 | 171 | 308 | 720 | 158 | 296 | 1042 | 186 | 271 |
| 1177 | FIM1_1203 | QGN14541.1 | CP015055.1 | 1741 | 462 | 1051 | 1912 | 213 | 896 | 3299 | 506 | 851 |
| 1178 | FIM1_1204 | QGN14542.1 | CP015055.1 | 12613 | 1779 | 2936 | 10577 | 1345 | 2734 | 16130 | 1843 | 2625 |
| 1179 | FIM1_1205 | QGN14543.1 | CP015055.1 | 2053 | 369 | 740 | 1732 | 292 | 680 | 2123 | 425 | 637 |
| 1180 | FIM1_1206 | QGN14544.1 | CP015055.1 | 1938 | 442 | 610 | 1453 | 355 | 632 | 1937 | 536 | 693 |
| 1181 | FIM1_1207 | QGN14545.1 | CP015055.1 | 6488 | 945 | 1639 | 4850 | 754 | 1465 | 5371 | 1085 | 1405 |
| 1182 | FIM1_1208 | QGN14546.1 | CP015055.1 | 4029 | 806 | 1343 | 3110 | 755 | 1302 | 3560 | 936 | 1280 |
| 1183 | FIM1_1209 | QGN14547.1 | CP015055.1 | 2868 | 738 | 982 | 2107 | 513 | 872 | 2326 | 831 | 828 |
| 1184 | FIM1_1210 | QGN14548.1 | CP015055.1 | 634 | 118 | 388 | 555 | 103 | 441 | 677 | 158 | 338 |
| 1185 | FIM1_1211 | QGN14549.1 | CP015055.1 | 2938 | 553 | 1632 | 2366 | 352 | 1576 | 4206 | 570 | 1481 |
| 1186 | FIM1_1212 | QGN14550.1 | CP015055.1 | 20083 | 11405 | 7850 | 16190 | 7345 | 7316 | 17046 | 11363 | 7095 |
| 1187 | FIM1_1213 | QGN14551.1 | CP015055.1 | 14005 | 7958 | 7855 | 13374 | 6734 | 7362 | 14008 | 8159 | 7270 |
| 1188 | FIM1_1214 | QGN14552.1 | CP015055.1 | 29654 | 18646 | 16184 | 26832 | 14588 | 14444 | 31363 | 18789 | 15811 |
| 1189 | FIM1_1215 | QGN14553.1 | CP015055.1 | 2688 | 546 | 797 | 2075 | 388 | 752 | 2668 | 559 | 677 |
| 1190 | FIM1_1216 | QGN14554.1 | CP015055.1 | 3565 | 575 | 689 | 2451 | 418 | 657 | 3804 | 634 | 672 |
| 1191 | FIM1_1217 | QGN14555.1 | CP015055.1 | 6844 | 5790 | 3037 | 4676 | 3604 | 2644 | 1656 | 5850 | 2858 |
| 1192 | FIM1_1218 | QGN14556.1 | CP015055.1 | 774 | 118 | 546 | 523 | 109 | 484 | 706 | 117 | 427 |
| 1193 | FIM1_1219 | QGN14557.1 | CP015055.1 | 3942 | 963 | 1000 | 2921 | 647 | 998 | 2934 | 1099 | 818 |
| 1194 | FIM1_1220 | QGN14558.1 | CP015055.1 | 1610 | 334 | 418 | 1189 | 255 | 384 | 1564 | 399 | 383 |
| 1195 | FIM1_1221 | QGN14559.1 | CP015055.1 | 1988 | 488 | 708 | 1594 | 393 | 698 | 1745 | 526 | 603 |
| 1196 | FIM1_1222 | QGN14560.1 | CP015055.1 | 15089 | 3144 | 4940 | 12445 | 2333 | 4639 | 14705 | 3302 | 4153 |
| 1197 | FIM1_1223 | QGN14561.1 | CP015055.1 | 1449 | 112 | 269 | 1077 | 104 | 269 | 1642 | 123 | 221 |
| 1198 | FIM1_1224 | QGN14562.1 | CP015055.1 | 523 | 145 | 260 | 390 | 120 | 267 | 618 | 150 | 203 |

| | | | | | | | | | | | | |
|---|---|---|---|---|---|---|---|---|---|---|---|---|
| 1199 | FIM1_1225 | QGN14563.1 | CP015055.1 | 8455 | 1562 | 1597 | 5961 | 1062 | 1536 | 7574 | 1667 | 1449 |
| 1200 | FIM1_1226 | QGN14564.1 | CP015055.1 | 13 | 1 | 5 | 14 | 1 | 5 | 19 | 3 | 4 |
| 1201 | FIM1_1227 | QGN14565.1 | CP015055.1 | 18007 | 2226 | 3042 | 13036 | 1743 | 2819 | 18752 | 2439 | 2666 |
| 1202 | FIM1_1228 | QGN14566.1 | CP015055.1 | 3827 | 793 | 1154 | 3088 | 566 | 1081 | 3457 | 809 | 948 |
| 1203 | FIM1_1229 | QGN14567.1 | CP015055.1 | 23 | 0 | 6 | 15 | 1 | 7 | 11 | 1 | 3 |
| 1204 | FIM1_1230 | QGN14568.1 | CP015055.1 | 4842 | 2124 | 1554 | 3376 | 1488 | 1559 | 2727 | 2150 | 1558 |
| 1205 | FIM1_1231 | QGN14569.1 | CP015055.1 | 9443 | 3420 | 2872 | 6961 | 2445 | 2762 | 6123 | 3625 | 2655 |
| 1206 | FIM1_1232 | QGN14570.1 | CP015055.1 | 5308 | 3148 | 2230 | 3847 | 1898 | 2005 | 4297 | 3322 | 2148 |
| 1207 | FIM1_1233 | QGN14571.1 | CP015055.1 | 1549 | 686 | 1254 | 1636 | 599 | 1171 | 1339 | 735 | 1132 |
| 1208 | FIM1_1234 | QGN14572.1 | CP015055.1 | 18915 | 8159 | 7975 | 16465 | 6182 | 7365 | 14502 | 8750 | 7156 |
| 1209 | FIM1_1235 | QGN14573.1 | CP015055.1 | 3877 | 1332 | 1652 | 3526 | 759 | 1496 | 5748 | 1439 | 1382 |
| 1210 | FIM1_1236 | QGN14574.1 | CP015055.1 | 760 | 263 | 307 | 641 | 167 | 298 | 870 | 319 | 281 |
| 1211 | FIM1_1237 | QGN14575.1 | CP015055.1 | 12 | 2 | 6 | 13 | 3 | 8 | 8 | 6 | 2 |
| 1212 | FIM1_1238 | QGN14576.1 | CP015055.1 | 8 | 10 | 13 | 11 | 1 | 13 | 3 | 9 | 16 |
| 1213 | FIM1_1239 | QGN14577.1 | CP015055.1 | 4462 | 1705 | 2513 | 3848 | 1125 | 2390 | 5458 | 1751 | 2293 |
| 1214 | FIM1_1240 | QGN14578.1 | CP015055.1 | 648 | 145 | 285 | 568 | 112 | 265 | 498 | 135 | 222 |
| 1215 | FIM1_1241 | QGN14579.1 | CP015055.1 | 4 | 0 | 1 | 5 | 0 | 0 | 4 | 0 | 1 |
| 1216 | FIM1_1242 | QGN14580.1 | CP015055.1 | 2184 | 428 | 780 | 1603 | 304 | 686 | 1629 | 461 | 545 |
| 1217 | FIM1_1243 | QGN14581.1 | CP015055.1 | 8906 | 2306 | 2991 | 6735 | 1926 | 2722 | 8422 | 2559 | 2584 |
| 1218 | FIM1_1244 | QGN14582.1 | CP015055.1 | 1700 | 385 | 865 | 1230 | 349 | 787 | 1674 | 410 | 693 |
| 1219 | FIM1_1245 | QGN14583.1 | CP015055.1 | 3806 | 668 | 970 | 2631 | 513 | 991 | 3965 | 768 | 792 |
| 1220 | FIM1_1246 | QGN14584.1 | CP015055.1 | 1441 | 337 | 539 | 1156 | 249 | 476 | 1295 | 426 | 464 |
| 1221 | FIM1_1247 | QGN14585.1 | CP015055.1 | 1529 | 242 | 821 | 1241 | 192 | 761 | 1948 | 275 | 651 |
| 1222 | FIM1_1248 | QGN14586.1 | CP015055.1 | 2862 | 1113 | 1515 | 2721 | 909 | 1443 | 2332 | 1242 | 1387 |
| 1223 | FIM1_1249 | QGN14587.1 | CP015055.1 | 14422 | 4164 | 2578 | 12828 | 2669 | 2430 | 15338 | 4601 | 2784 |
| 1224 | FIM1_1250 | QGN14588.1 | CP015055.1 | 12192 | 809 | 1406 | 10244 | 739 | 1206 | 8631 | 800 | 1131 |
| 1225 | FIM1_1251 | QGN14589.1 | CP015055.1 | 10472 | 2508 | 3086 | 9239 | 1614 | 2831 | 13422 | 2542 | 2593 |
| 1226 | FIM1_1252 | QGN14590.1 | CP015055.1 | 3567 | 1533 | 1530 | 3323 | 1121 | 1418 | 2457 | 1637 | 1290 |
| 1227 | FIM1_1253 | QGN14591.1 | CP015055.1 | 1120 | 184 | 389 | 867 | 145 | 337 | 1059 | 212 | 318 |
| 1228 | FIM1_1254 | QGN14592.1 | CP015055.1 | 2361 | 958 | 1064 | 2030 | 776 | 1072 | 1311 | 1044 | 974 |
| 1229 | FIM1_1255 | QGN14593.1 | CP015055.1 | 4616 | 2290 | 1772 | 3178 | 1648 | 1628 | 1144 | 2478 | 1534 |
| 1230 | FIM1_1256 | QGN14594.1 | CP015055.1 | 3847 | 488 | 1064 | 2947 | 390 | 1005 | 3784 | 558 | 912 |
| 1231 | FIM1_1257 | QGN14595.1 | CP015055.1 | 5687 | 961 | 1492 | 4246 | 761 | 1402 | 5804 | 1007 | 1253 |
| 1232 | FIM1_1258 | QGN14596.1 | CP015055.1 | 1741 | 617 | 1241 | 1374 | 388 | 1210 | 1767 | 648 | 1097 |
| 1233 | FIM1_1259 | QGN14597.1 | CP015055.1 | 793 | 134 | 309 | 615 | 74 | 325 | 961 | 132 | 265 |
| 1234 | FIM1_1261 | QGN14598.1 | CP015055.1 | 4782 | 2063 | 1848 | 3937 | 1777 | 1842 | 3328 | 2284 | 1743 |
| 1235 | FIM1_1262 | QGN14599.1 | CP015055.1 | 2273 | 1107 | 3179 | 1912 | 725 | 3014 | 3359 | 1134 | 2971 |
| 1236 | FIM1_1263 | QGN14600.1 | CP015055.1 | 4574 | 1379 | 1814 | 4197 | 1002 | 1672 | 4208 | 1511 | 1676 |
| 1237 | FIM1_1264 | QGN14601.1 | CP015055.1 | 4583 | 2266 | 2524 | 4602 | 1747 | 2295 | 5030 | 2362 | 2287 |
| 1238 | FIM1_1265 | QGN14602.1 | CP015055.1 | 15072 | 3171 | 4567 | 12240 | 2190 | 3872 | 16239 | 3438 | 3845 |
| 1239 | FIM1_1266 | QGN14603.1 | CP015055.1 | 1758 | 745 | 785 | 1308 | 492 | 665 | 846 | 862 | 645 |
| 1240 | FIM1_1267 | QGN14604.1 | CP015055.1 | 7786 | 5829 | 3077 | 5834 | 3896 | 2673 | 2015 | 6151 | 2656 |
| 1241 | FIM1_1268 | QGN14605.1 | CP015055.1 | 1695 | 409 | 775 | 1467 | 312 | 732 | 1283 | 403 | 591 |
| 1242 | FIM1_1269 | QGN14606.1 | CP015055.1 | 6919 | 5432 | 2522 | 4675 | 3352 | 2207 | 1632 | 5663 | 2295 |
| 1243 | FIM1_1270 | QGN14607.1 | CP015055.1 | 4765 | 2823 | 2458 | 3631 | 2078 | 2162 | 3086 | 3046 | 2184 |
| 1244 | FIM1_1271 | QGN14608.1 | CP015055.1 | 9222 | 2006 | 2851 | 7288 | 1299 | 2595 | 10704 | 2093 | 2495 |
| 1245 | FIM1_1272 | QGN14609.1 | CP015055.1 | 17317 | 6909 | 6215 | 14487 | 4795 | 5612 | 14619 | 7467 | 5449 |
| 1246 | FIM1_1273 | QGN14610.1 | CP015055.1 | 1686 | 251 | 598 | 1396 | 202 | 496 | 1784 | 255 | 450 |

| | | | | | | | | | | | | |
|---|---|---|---|---|---|---|---|---|---|---|---|---|
| 1247 | FIM1_1274 | QGN14611.1 | CP015055.1 | 344 | 27 | 83 | 300 | 30 | 68 | 263 | 34 | 60 |
| 1248 | FIM1_1275 | QGN14612.1 | CP015055.1 | 6767 | 1274 | 2667 | 6716 | 1068 | 2550 | 7578 | 1507 | 2215 |
| 1249 | FIM1_1276 | QGN14613.1 | CP015055.1 | 7466 | 1823 | 2996 | 5986 | 1297 | 2668 | 7817 | 1851 | 2564 |
| 1250 | FIM1_1277 | QGN14614.1 | CP015055.1 | 7966 | 957 | 2063 | 5949 | 661 | 1879 | 11681 | 1134 | 1720 |
| 1251 | FIM1_1279 | QGN14615.1 | CP015055.1 | 2269 | 450 | 1019 | 1857 | 352 | 1028 | 2625 | 494 | 891 |
| 1252 | FIM1_1280 | QGN14616.1 | CP015055.1 | 7487 | 3344 | 8993 | 6028 | 2192 | 8340 | 9174 | 3484 | 8503 |
| 1253 | FIM1_1281 | QGN14617.1 | CP015055.1 | 4997 | 2025 | 2021 | 4028 | 1404 | 1887 | 3431 | 2188 | 1721 |
| 1254 | FIM1_1282 | QGN14618.1 | CP015055.1 | 2656 | 377 | 563 | 2184 | 272 | 513 | 2841 | 406 | 462 |
| 1255 | FIM1_1283 | QGN14619.1 | CP015055.1 | 5733 | 1734 | 3659 | 4335 | 778 | 2146 | 8607 | 1294 | 2552 |
| 1256 | FIM1_1284 | QGN14620.1 | CP015055.1 | 7635 | 388 | 422 | 3693 | 279 | 373 | 3772 | 468 | 382 |
| 1257 | FIM1_1285 | QGN14621.1 | CP015055.1 | 12071 | 3232 | 3332 | 9138 | 2317 | 3075 | 13130 | 3425 | 3132 |
| 1258 | FIM1_1286 | QGN14622.1 | CP015055.1 | 733 | 320 | 506 | 692 | 288 | 525 | 676 | 352 | 466 |
| 1259 | FIM1_1287 | QGN14623.1 | CP015055.1 | 3941 | 643 | 1398 | 3305 | 420 | 1295 | 4245 | 637 | 1191 |
| 1260 | FIM1_1288 | QGN14624.1 | CP015055.1 | 2096 | 386 | 449 | 1756 | 288 | 470 | 1657 | 461 | 441 |
| 1261 | FIM1_1289 | QGN14625.1 | CP015055.1 | 4819 | 2010 | 1924 | 3835 | 1388 | 1704 | 3018 | 2093 | 1718 |
| 1262 | FIM1_1290 | QGN14626.1 | CP015055.1 | 11979 | 2081 | 3936 | 9229 | 1677 | 3568 | 11657 | 2283 | 3461 |
| 1263 | FIM1_1291 | QGN14627.1 | CP015055.1 | 1571 | 792 | 990 | 1133 | 658 | 907 | 1522 | 899 | 926 |
| 1264 | FIM1_1292 | QGN14628.1 | CP015055.1 | 440 | 195 | 192 | 324 | 159 | 211 | 354 | 197 | 191 |
| 1265 | FIM1_1293 | QGN14629.1 | CP015055.1 | 4524 | 3555 | 2044 | 3098 | 2157 | 1786 | 1786 | 3654 | 1933 |
| 1266 | FIM1_1294 | QGN14630.1 | CP015055.1 | 2394 | 1134 | 900 | 1645 | 883 | 893 | 613 | 1218 | 913 |
| 1267 | FIM1_1295 | QGN14631.1 | CP015055.1 | 9383 | 1540 | 2723 | 8195 | 1489 | 2497 | 9778 | 1702 | 2433 |
| 1268 | FIM1_1296 | QGN14632.1 | CP015055.1 | 4442 | 197 | 719 | 3285 | 158 | 635 | 4559 | 207 | 618 |
| 1269 | FIM1_1297 | QGN14633.1 | CP015055.1 | 2158 | 585 | 544 | 1654 | 281 | 539 | 5519 | 491 | 424 |
| 1270 | FIM1_1298 | QGN14634.1 | CP015055.1 | 6763 | 839 | 3033 | 5285 | 581 | 2780 | 9820 | 1013 | 2625 |
| 1271 | FIM1_1299 | QGN14635.1 | CP015055.1 | 2255 | 550 | 903 | 1846 | 450 | 903 | 2255 | 721 | 758 |
| 1272 | FIM1_1300 | QGN14636.1 | CP015055.1 | 3939 | 1241 | 1073 | 2755 | 805 | 967 | 2625 | 1354 | 967 |
| 1273 | FIM1_1301 | QGN14637.1 | CP015055.1 | 815 | 151 | 311 | 647 | 128 | 262 | 768 | 158 | 265 |
| 1274 | FIM1_1302 | QGN14638.1 | CP015055.1 | 1825 | 474 | 560 | 1223 | 343 | 542 | 1620 | 474 | 532 |
| 1275 | FIM1_1303 | QGN14639.1 | CP015055.1 | 3315 | 2576 | 1666 | 2669 | 1874 | 1477 | 1627 | 2724 | 1409 |
| 1276 | FIM1_1304 | QGN14640.1 | CP015055.1 | 5532 | 1058 | 1149 | 3963 | 737 | 1206 | 5863 | 1123 | 1147 |
| 1277 | FIM1_1305 | QGN14641.1 | CP015055.1 | 11303 | 2827 | 3212 | 8639 | 1585 | 3045 | 17541 | 3038 | 2724 |
| 1278 | FIM1_1306 | QGN14642.1 | CP015055.1 | 206 | 49 | 97 | 200 | 24 | 86 | 185 | 59 | 92 |
| 1279 | FIM1_1307 | QGN14643.1 | CP015055.1 | 2742 | 650 | 1039 | 2133 | 515 | 1015 | 3157 | 715 | 851 |
| 1280 | FIM1_1308 | QGN14644.1 | CP015055.1 | 2509 | 579 | 704 | 1891 | 446 | 644 | 1995 | 649 | 619 |
| 1281 | FIM1_1309 | QGN14645.1 | CP015055.1 | 2838 | 658 | 949 | 2536 | 338 | 894 | 3193 | 680 | 880 |
| 1282 | FIM1_1310 | QGN14646.1 | CP015055.1 | 5302 | 1880 | 1844 | 4282 | 1278 | 1668 | 4828 | 2018 | 1667 |
| 1283 | FIM1_1311 | QGN14647.1 | CP015055.1 | 6146 | 2403 | 434 | 3678 | 1400 | 380 | 8423 | 2730 | 363 |
| 1284 | FIM1_1312 | QGN14648.1 | CP015055.1 | 3638 | 1115 | 1492 | 2943 | 582 | 1304 | 4884 | 1127 | 1174 |
| 1285 | FIM1_1313 | QGN14649.1 | CP015055.1 | 57047 | 26905 | 34698 | 49591 | 20576 | 31355 | 59984 | 27542 | 33628 |
| 1286 | FIM1_1314 | QGN14650.1 | CP015055.1 | 2554 | 661 | 646 | 2156 | 604 | 601 | 1417 | 749 | 623 |
| 1287 | FIM1_1315 | QGN14651.1 | CP015055.1 | 5836 | 2977 | 2893 | 5421 | 2333 | 3011 | 4907 | 3244 | 2695 |
| 1288 | FIM1_1316 | QGN14652.1 | CP015055.1 | 2249 | 324 | 947 | 1694 | 220 | 792 | 1410 | 351 | 564 |
| 1289 | FIM1_1317 | QGN14653.1 | CP015055.1 | 1431 | 422 | 537 | 1268 | 316 | 493 | 1070 | 479 | 549 |
| 1290 | FIM1_1318 | QGN14654.1 | CP015055.1 | 2161 | 830 | 1428 | 2015 | 659 | 1317 | 2359 | 921 | 1208 |
| 1291 | FIM1_1319 | QGN14655.1 | CP015055.1 | 16135 | 2081 | 2961 | 13082 | 1270 | 2833 | 18545 | 1951 | 2478 |
| 1292 | FIM1_1320 | QGN14656.1 | CP015055.1 | 39072 | 9186 | 11136 | 32512 | 8192 | 10520 | 28727 | 9878 | 10585 |
| 1293 | FIM1_1321 | QGN14657.1 | CP015055.1 | 1906 | 434 | 653 | 1560 | 343 | 593 | 1850 | 455 | 573 |
| 1294 | FIM1_1322 | QGN14658.1 | CP015055.1 | 16576 | 5826 | 7266 | 14526 | 3408 | 6918 | 17693 | 5986 | 6763 |

| | | | | | | | | | | | | |
|---|---|---|---|---|---|---|---|---|---|---|---|---|
| 1295 | FIM1_1323 | QGN14659.1 | CP015055.1 | 1549 | 173 | 344 | 1332 | 113 | 296 | 2701 | 176 | 243 |
| 1296 | FIM1_1324 | QGN14660.1 | CP015055.1 | 1581 | 1370 | 916 | 1346 | 986 | 903 | 570 | 1385 | 860 |
| 1297 | FIM1_1325 | QGN14661.1 | CP015055.1 | 4382 | 733 | 1236 | 3600 | 530 | 1131 | 6998 | 808 | 1007 |
| 1298 | FIM1_1326 | QGN14662.1 | CP015055.1 | 2398 | 531 | 924 | 1875 | 386 | 811 | 2975 | 561 | 696 |
| 1299 | FIM1_1327 | QGN14663.1 | CP015055.1 | 5988 | 793 | 1583 | 4422 | 613 | 1524 | 7149 | 854 | 1435 |
| 1300 | FIM1_1328 | QGN14664.1 | CP015055.1 | 5453 | 719 | 1350 | 4009 | 482 | 1278 | 5647 | 774 | 1216 |
| 1301 | FIM1_1329 | QGN14665.1 | CP015055.1 | 6185 | 380 | 518 | 3710 | 238 | 514 | 7236 | 357 | 436 |
| 1302 | FIM1_1331 | QGN14666.1 | CP015055.1 | 548 | 56 | 168 | 340 | 34 | 137 | 782 | 52 | 137 |
| 1303 | FIM1_1332 | QGN14667.1 | CP015055.1 | 1136 | 265 | 543 | 898 | 211 | 617 | 1314 | 265 | 540 |
| 1304 | FIM1_1333 | QGN14668.1 | CP015055.1 | 14638 | 8554 | 5857 | 12071 | 6723 | 5292 | 6718 | 8928 | 5224 |
| 1305 | FIM1_1334 | QGN14669.1 | CP015055.1 | 6225 | 1424 | 2946 | 4942 | 1161 | 2687 | 7505 | 1611 | 2970 |
| 1306 | FIM1_1335 | QGN14670.1 | CP015055.1 | 1983 | 601 | 918 | 1635 | 424 | 867 | 2273 | 597 | 823 |
| 1307 | FIM1_1336 | QGN14671.1 | CP015055.1 | 8606 | 2020 | 822 | 5703 | 996 | 702 | 6073 | 2205 | 750 |
| 1308 | FIM1_1337 | QGN14672.1 | CP015055.1 | 8621 | 6040 | 2757 | 6169 | 4005 | 2301 | 2611 | 6193 | 2496 |
| 1309 | FIM1_1338 | QGN14673.1 | CP015055.1 | 1715 | 170 | 548 | 1421 | 151 | 475 | 2899 | 219 | 416 |
| 1310 | FIM1_1339 | QGN14674.1 | CP015055.1 | 18732 | 3507 | 3930 | 15314 | 2559 | 3462 | 13942 | 3802 | 3441 |
| 1311 | FIM1_1340 | QGN14675.1 | CP015055.1 | 6781 | 1392 | 2051 | 6219 | 1127 | 1812 | 5884 | 1597 | 1674 |
| 1312 | FIM1_1341 | QGN14676.1 | CP015055.1 | 1660 | 341 | 764 | 1452 | 289 | 750 | 1501 | 440 | 630 |
| 1313 | FIM1_1342 | QGN14677.1 | CP015055.1 | 6666 | 1706 | 2069 | 5413 | 1400 | 1889 | 5862 | 1978 | 1965 |
| 1314 | FIM1_1344 | QGN14678.1 | CP015055.1 | 1302 | 153 | 266 | 940 | 92 | 254 | 1611 | 153 | 235 |
| 1315 | FIM1_1345 | QGN14679.1 | CP015055.1 | 3951 | 946 | 993 | 3161 | 662 | 903 | 3190 | 1070 | 790 |
| 1316 | FIM1_1346 | QGN14680.1 | CP015055.1 | 13420 | 7603 | 6625 | 12371 | 5625 | 5981 | 12992 | 7845 | 6256 |
| 1317 | FIM1_1347 | QGN14681.1 | CP015055.1 | 23587 | 15627 | 14943 | 22737 | 13693 | 14711 | 23642 | 16264 | 14523 |
| 1318 | FIM1_1348 | QGN14682.1 | CP015055.1 | 24097 | 9400 | 8565 | 19284 | 7137 | 8290 | 23759 | 10040 | 8200 |
| 1319 | FIM1_1349 | QGN14683.1 | CP015055.1 | 2699 | 526 | 1213 | 2133 | 475 | 1124 | 2657 | 582 | 1044 |
| 1320 | FIM1_1350 | QGN14684.1 | CP015055.1 | 14210 | 7201 | 7680 | 13443 | 6301 | 7391 | 12183 | 7681 | 7374 |
| 1321 | FIM1_1352 | QGN14685.1 | CP015055.1 | 3410 | 1384 | 2273 | 3050 | 1026 | 2121 | 3703 | 1449 | 2170 |
| 1322 | FIM1_1353 | QGN14686.1 | CP015055.1 | 17182 | 4956 | 9736 | 14115 | 3467 | 8983 | 19980 | 5259 | 8689 |
| 1323 | FIM1_1354 | QGN14687.1 | CP015055.1 | 2112 | 650 | 949 | 1746 | 508 | 950 | 2118 | 675 | 855 |
| 1324 | FIM1_1355 | QGN14688.1 | CP015055.1 | 5586 | 987 | 1284 | 4020 | 744 | 1150 | 5378 | 1040 | 1104 |
| 1325 | FIM1_1356 | QGN14689.1 | CP015055.1 | 1660 | 364 | 659 | 1382 | 299 | 620 | 1492 | 401 | 552 |
| 1326 | FIM1_1357 | QGN14690.1 | CP015055.1 | 2341 | 569 | 948 | 1938 | 427 | 868 | 2656 | 672 | 769 |
| 1327 | FIM1_1358 | QGN14691.1 | CP015055.1 | 3183 | 1050 | 1216 | 2868 | 725 | 1077 | 2530 | 1118 | 1023 |
| 1328 | FIM1_1359 | QGN14692.1 | CP015055.1 | 1615 | 253 | 435 | 1389 | 209 | 391 | 1786 | 306 | 395 |
| 1329 | FIM1_1360 | QGN14693.1 | CP015055.1 | 12937 | 1072 | 1529 | 10510 | 1004 | 1343 | 13968 | 1321 | 1176 |
| 1330 | FIM1_1361 | QGN14694.1 | CP015055.1 | 429 | 120 | 214 | 394 | 61 | 198 | 439 | 130 | 157 |
| 1331 | FIM1_1362 | QGN14695.1 | CP015055.1 | 701 | 248 | 361 | 615 | 153 | 384 | 922 | 273 | 388 |
| 1332 | FIM1_1363 | QGN14696.1 | CP015055.1 | 6480 | 5123 | 2695 | 4580 | 3417 | 2566 | 1978 | 5520 | 2554 |
| 1333 | FIM1_1364 | QGN14697.1 | CP015055.1 | 353 | 241 | 361 | 236 | 178 | 301 | 133 | 206 | 289 |
| 1334 | FIM1_1365 | QGN14698.1 | CP015055.1 | 7054 | 1903 | 2236 | 5352 | 1139 | 2071 | 8089 | 1896 | 1971 |
| 1335 | FIM1_1366 | QGN14699.1 | CP015055.1 | 2981 | 1155 | 1136 | 2296 | 854 | 1126 | 2031 | 1268 | 995 |
| 1336 | FIM1_1367 | QGN14700.1 | CP015055.1 | 3128 | 334 | 816 | 2654 | 250 | 736 | 2941 | 388 | 702 |
| 1337 | FIM1_1368 | QGN14701.1 | CP015055.1 | 13849 | 4032 | 5302 | 11469 | 2493 | 4851 | 15735 | 4230 | 4760 |
| 1338 | FIM1_1369 | QGN14702.1 | CP015055.1 | 7073 | 4797 | 1548 | 5544 | 2246 | 1343 | 9195 | 4809 | 1435 |
| 1339 | FIM1_1370 | QGN14703.1 | CP015055.1 | 86 | 39 | 5 | 89 | 23 | 14 | 68 | 40 | 16 |
| 1340 | FIM1_1371 | QGN14704.1 | CP015055.1 | 3413 | 727 | 1392 | 2912 | 510 | 1403 | 2795 | 705 | 1209 |
| 1341 | FIM1_1372 | QGN14705.1 | CP015055.1 | 298 | 87 | 368 | 285 | 69 | 356 | 283 | 100 | 328 |
| 1342 | FIM1_1373 | QGN14706.1 | CP015055.1 | 6925 | 1666 | 2607 | 5796 | 1008 | 2337 | 8689 | 1752 | 2190 |

| | | | | | | | | | | | | |
|---|---|---|---|---|---|---|---|---|---|---|---|---|
| 1343 | FIM1_1374 | QGN14707.1 | CP015055.1 | 4292 | 1344 | 1982 | 3317 | 1033 | 1778 | 4145 | 1401 | 1780 |
| 1344 | FIM1_1375 | QGN14708.1 | CP015055.1 | 6771 | 750 | 1368 | 4772 | 493 | 1074 | 6670 | 792 | 1043 |
| 1345 | FIM1_1376 | QGN14709.1 | CP015055.1 | 168 | 34 | 43 | 145 | 26 | 55 | 218 | 36 | 49 |
| 1346 | FIM1_1377 | QGN14710.1 | CP015055.1 | 14464 | 2582 | 2790 | 10443 | 1991 | 2603 | 9660 | 2745 | 2309 |
| 1347 | FIM1_1378 | QGN14711.1 | CP015055.1 | 476 | 192 | 215 | 344 | 200 | 273 | 581 | 178 | 258 |
| 1348 | FIM1_1379 | QGN14712.1 | CP015055.1 | 923 | 64 | 232 | 786 | 40 | 177 | 1455 | 76 | 185 |
| 1349 | FIM1_1380 | QGN14713.1 | CP015055.1 | 6680 | 894 | 1460 | 5204 | 608 | 1260 | 11932 | 953 | 1112 |
| 1350 | FIM1_1382 | QGN14714.1 | CP015055.1 | 11487 | 2760 | 3392 | 8717 | 2118 | 3173 | 11233 | 2890 | 2940 |
| 1351 | FIM1_1383 | QGN14715.1 | CP015055.1 | 6082 | 1533 | 2329 | 4868 | 1055 | 2138 | 7373 | 1678 | 2095 |
| 1352 | FIM1_1384 | QGN14716.1 | CP015055.1 | 1975 | 891 | 959 | 1910 | 721 | 922 | 1981 | 1037 | 838 |
| 1353 | FIM1_1385 | QGN14717.1 | CP015055.1 | 10701 | 3138 | 3605 | 8947 | 1578 | 3394 | 18673 | 3106 | 3173 |
| 1354 | FIM1_1386 | QGN14718.1 | CP015055.1 | 1945 | 333 | 563 | 1656 | 252 | 510 | 1758 | 357 | 500 |
| 1355 | FIM1_1387 | QGN14719.1 | CP015055.1 | 7284 | 2127 | 2667 | 5816 | 1633 | 2485 | 6632 | 2350 | 2335 |
| 1356 | FIM1_1388 | QGN14720.1 | CP015055.1 | 1006 | 146 | 315 | 831 | 118 | 319 | 974 | 164 | 295 |
| 1357 | FIM1_1389 | QGN14721.1 | CP015055.1 | 914 | 237 | 652 | 684 | 207 | 609 | 1004 | 248 | 498 |
| 1358 | FIM1_1390 | QGN14722.1 | CP015055.1 | 9879 | 1372 | 2292 | 7276 | 1135 | 2159 | 9082 | 1528 | 2024 |
| 1359 | FIM1_1391 | QGN14723.1 | CP015055.1 | 11012 | 1606 | 4452 | 9381 | 951 | 4043 | 13315 | 1560 | 3961 |
| 1360 | FIM1_1392 | QGN14724.1 | CP015055.1 | 2382 | 383 | 1435 | 2152 | 172 | 1203 | 2997 | 400 | 1178 |
| 1361 | FIM1_1393 | QGN14725.1 | CP015055.1 | 1880 | 628 | 1186 | 2015 | 293 | 1054 | 2564 | 669 | 1054 |
| 1362 | FIM1_1394 | QGN14726.1 | CP015055.1 | 801 | 46 | 399 | 653 | 25 | 364 | 927 | 64 | 253 |
| 1363 | FIM1_1395 | QGN14727.1 | CP015055.1 | 2956 | 305 | 859 | 2347 | 279 | 760 | 2913 | 340 | 644 |
| 1364 | FIM1_1396 | QGN14728.1 | CP015055.1 | 4600 | 923 | 1183 | 3571 | 574 | 1048 | 4308 | 950 | 1036 |
| 1365 | FIM1_1397 | QGN14729.1 | CP015055.1 | 9010 | 2872 | 4506 | 8102 | 1696 | 3998 | 9909 | 3041 | 3973 |
| 1366 | FIM1_1399 | QGN14730.1 | CP015055.1 | 5677 | 1534 | 2312 | 4975 | 1126 | 2024 | 6518 | 1694 | 1925 |
| 1367 | FIM1_1400 | QGN14731.1 | CP015055.1 | 8851 | 1173 | 1547 | 7588 | 772 | 1389 | 8141 | 1200 | 1314 |
| 1368 | FIM1_1401 | QGN14732.1 | CP015055.1 | 15668 | 2472 | 2914 | 12712 | 2050 | 2433 | 8995 | 2741 | 2480 |
| 1369 | FIM1_1402 | QGN14733.1 | CP015055.1 | 2841 | 539 | 682 | 2161 | 414 | 635 | 3011 | 597 | 610 |
| 1370 | FIM1_1403 | QGN14734.1 | CP015055.1 | 1429 | 431 | 569 | 1266 | 260 | 531 | 2334 | 454 | 421 |
| 1371 | FIM1_1404 | QGN14735.1 | CP015055.1 | 698 | 196 | 365 | 714 | 306 | 504 | 894 | 363 | 420 |
| 1372 | FIM1_1405 | QGN14736.1 | CP015055.1 | 1832 | 727 | 1124 | 1465 | 299 | 969 | 1166 | 765 | 863 |
| 1373 | FIM1_1406 | QGN14737.1 | CP015055.1 | 19620 | 4125 | 5541 | 14337 | 2493 | 5127 | 14654 | 4214 | 5037 |
| 1374 | FIM1_1407 | QGN14738.1 | CP015055.1 | 4868 | 785 | 1261 | 3496 | 620 | 1153 | 3741 | 819 | 987 |
| 1375 | FIM1_1408 | QGN14739.1 | CP015055.1 | 14510 | 6149 | 4221 | 11216 | 3853 | 3957 | 18066 | 6402 | 3799 |
| 1376 | FIM1_1409 | QGN14740.1 | CP015055.1 | 4045 | 2420 | 1819 | 3233 | 1698 | 1681 | 1667 | 2588 | 1856 |
| 1377 | FIM1_1410 | QGN14741.1 | CP015055.1 | 8307 | 3205 | 3776 | 8013 | 2370 | 3503 | 8205 | 3424 | 3487 |
| 1378 | FIM1_1411 | QGN14742.1 | CP015055.1 | 3060 | 572 | 1159 | 2227 | 452 | 1092 | 3561 | 620 | 1125 |
| 1379 | FIM1_1412 | QGN14743.1 | CP015055.1 | 17423 | 3385 | 3101 | 13492 | 2153 | 2582 | 20860 | 3568 | 2916 |
| 1380 | FIM1_1413 | QGN14744.1 | CP015055.1 | 2976 | 302 | 514 | 2240 | 172 | 460 | 3225 | 295 | 462 |
| 1381 | FIM1_1414 | QGN14745.1 | CP015055.1 | 1042 | 171 | 273 | 823 | 132 | 266 | 999 | 229 | 273 |
| 1382 | FIM1_1415 | QGN14746.1 | CP015055.1 | 4544 | 797 | 1677 | 4010 | 640 | 1526 | 4171 | 899 | 1416 |
| 1383 | FIM1_1416 | QGN14747.1 | CP015055.1 | 2042 | 412 | 1350 | 1686 | 313 | 1229 | 2298 | 410 | 1061 |
| 1384 | FIM1_1418 | QGN14748.1 | CP015055.1 | 2468 | 759 | 976 | 1891 | 482 | 961 | 2200 | 764 | 846 |
| 1385 | FIM1_1419 | QGN14749.1 | CP015055.1 | 10 | 1 | 2 | 13 | 1 | 5 | 5 | 1 | 0 |
| 1386 | FIM1_1420 | QGN14750.1 | CP015055.1 | 3397 | 1036 | 1288 | 2960 | 672 | 1272 | 3084 | 1102 | 1151 |
| 1387 | FIM1_1421 | QGN14751.1 | CP015055.1 | 2060 | 255 | 522 | 1352 | 187 | 532 | 1905 | 270 | 427 |
| 1388 | FIM1_1422 | QGN14752.1 | CP015055.1 | 1245 | 196 | 594 | 1105 | 183 | 515 | 1069 | 256 | 440 |
| 1389 | FIM1_1423 | QGN14753.1 | CP015055.1 | 3082 | 388 | 698 | 2245 | 325 | 689 | 2772 | 464 | 574 |
| 1390 | FIM1_1424 | QGN14754.1 | CP015055.1 | 3572 | 667 | 1147 | 2869 | 514 | 1082 | 4067 | 780 | 1051 |

| | | | | | | | | | | | | |
|---|---|---|---|---|---|---|---|---|---|---|---|---|
| 1391 | FIM1_1425 | QGN14755.1 | CP015055.1 | 3438 | 543 | 1046 | 3035 | 274 | 838 | 4540 | 621 | 856 |
| 1392 | FIM1_1426 | QGN14756.1 | CP015055.1 | 15063 | 3315 | 2340 | 12456 | 2019 | 2401 | 18239 | 3420 | 2530 |
| 1393 | FIM1_1427 | QGN14757.1 | CP015055.1 | 217 | 26 | 38 | 191 | 20 | 23 | 123 | 18 | 36 |
| 1394 | FIM1_1428 | QGN14758.1 | CP015055.1 | 6603 | 1002 | 2048 | 4879 | 848 | 2070 | 7076 | 1147 | 1711 |
| 1395 | FIM1_1429 | QGN14759.1 | CP015055.1 | 7865 | 1394 | 1795 | 5760 | 928 | 1639 | 9046 | 1481 | 1644 |
| 1396 | FIM1_1430 | QGN14760.1 | CP015055.1 | 4124 | 433 | 1120 | 3309 | 302 | 948 | 5233 | 479 | 898 |
| 1397 | FIM1_1431 | QGN14761.1 | CP015055.1 | 7774 | 846 | 1964 | 5557 | 413 | 1560 | 8018 | 832 | 1674 |
| 1398 | FIM1_1432 | QGN14762.1 | CP015055.1 | 27495 | 10531 | 10769 | 22495 | 8587 | 10369 | 21897 | 10963 | 10516 |
| 1399 | FIM1_1433 | QGN14763.1 | CP015055.1 | 4020 | 1302 | 1672 | 3139 | 865 | 1484 | 4869 | 1357 | 1457 |
| 1400 | FIM1_1434 | QGN14764.1 | CP015055.1 | 7751 | 1206 | 1663 | 5791 | 781 | 1431 | 7137 | 1356 | 1363 |
| 1401 | FIM1_1435 | QGN14765.1 | CP015055.1 | 13800 | 3577 | 4395 | 12121 | 2189 | 3919 | 17729 | 3814 | 3900 |
| 1402 | FIM1_1436 | QGN14766.1 | CP015055.1 | 1124 | 194 | 485 | 842 | 175 | 391 | 1322 | 232 | 361 |
| 1403 | FIM1_1437 | QGN14767.1 | CP015055.1 | 741 | 206 | 327 | 686 | 160 | 299 | 804 | 237 | 279 |
| 1404 | FIM1_1438 | QGN14768.1 | CP015055.1 | 131722 | 61544 | 52237 | 118272 | 47788 | 46869 | 135986 | 62725 | 50082 |
| 1405 | FIM1_1439 | QGN14769.1 | CP015055.1 | 4961 | 471 | 844 | 3733 | 311 | 777 | 6142 | 520 | 644 |
| 1406 | FIM1_1440 | QGN14770.1 | CP015055.1 | 843 | 209 | 321 | 625 | 144 | 283 | 710 | 204 | 284 |
| 1407 | FIM1_1441 | QGN14771.1 | CP015055.1 | 1067 | 106 | 107 | 708 | 103 | 101 | 1037 | 104 | 126 |
| 1408 | FIM1_1442 | QGN14772.1 | CP015055.1 | 816 | 38 | 38 | 539 | 31 | 46 | 1996 | 47 | 36 |
| 1409 | FIM1_1443 | QGN14773.1 | CP015055.1 | 1186 | 357 | 452 | 1019 | 302 | 392 | 941 | 373 | 341 |
| 1410 | FIM1_1444 | QGN14774.1 | CP015055.1 | 15365 | 12390 | 11494 | 15369 | 10603 | 10845 | 14983 | 12825 | 10987 |
| 1411 | FIM1_1445 | QGN14775.1 | CP015055.1 | 2072 | 1977 | 807 | 1412 | 1022 | 702 | 622 | 2096 | 762 |
| 1412 | FIM1_1446 | QGN14776.1 | CP015055.1 | 1486 | 242 | 644 | 1288 | 201 | 556 | 1135 | 241 | 488 |
| 1413 | FIM1_1447 | QGN14777.1 | CP015055.1 | 1233 | 662 | 801 | 1266 | 619 | 852 | 830 | 740 | 702 |
| 1414 | FIM1_1448 | QGN14778.1 | CP015055.1 | 990 | 450 | 513 | 900 | 329 | 501 | 831 | 433 | 435 |
| 1415 | FIM1_1449 | QGN14779.1 | CP015055.1 | 3620 | 2415 | 2199 | 3004 | 1507 | 1885 | 4190 | 2568 | 1916 |
| 1416 | FIM1_1450 | QGN14780.1 | CP015055.1 | 3909 | 601 | 646 | 2728 | 461 | 635 | 3092 | 641 | 683 |
| 1417 | FIM1_1451 | QGN14781.1 | CP015055.1 | 8787 | 3012 | 9040 | 8028 | 3951 | 8813 | 3392 | 3664 | 9003 |
| 1418 | FIM1_1452 | QGN14782.1 | CP015055.1 | 8 | 6 | 8 | 9 | 7 | 2 | 2 | 8 | 9 |
| 1419 | FIM1_1453 | QGN14783.1 | CP015055.1 | 10804 | 2892 | 3425 | 7313 | 1136 | 3199 | 12562 | 2735 | 2768 |
| 1420 | FIM1_1454 | QGN14784.1 | CP015055.1 | 3272 | 485 | 756 | 2340 | 348 | 739 | 3009 | 544 | 671 |
| 1421 | FIM1_1455 | QGN14785.1 | CP015055.1 | 1844 | 1055 | 1335 | 1448 | 817 | 1274 | 741 | 1108 | 1298 |
| 1422 | FIM1_1456 | QGN14786.1 | CP015055.1 | 1349 | 446 | 459 | 1223 | 358 | 484 | 1116 | 483 | 387 |
| 1423 | FIM1_1457 | QGN14787.1 | CP015055.1 | 10415 | 1398 | 2836 | 7325 | 1111 | 2612 | 8714 | 1399 | 2535 |
| 1424 | FIM1_1458 | QGN14788.1 | CP015055.1 | 2970 | 911 | 1272 | 2730 | 722 | 1243 | 2503 | 984 | 1061 |
| 1425 | FIM1_1460 | QGN14789.1 | CP015055.1 | 2617 | 656 | 1126 | 2083 | 490 | 1062 | 2544 | 676 | 972 |
| 1426 | FIM1_1461 | QGN14790.1 | CP015055.1 | 545 | 478 | 1321 | 623 | 389 | 1284 | 1152 | 508 | 1271 |
| 1427 | FIM1_1462 | QGN14791.1 | CP015055.1 | 3412 | 1020 | 1298 | 2700 | 810 | 1216 | 2289 | 1158 | 1210 |
| 1428 | FIM1_1463 | QGN14792.1 | CP015055.1 | 512 | 170 | 196 | 469 | 148 | 231 | 419 | 181 | 177 |
| 1429 | FIM1_1464 | QGN14793.1 | CP015055.1 | 64 | 6 | 12 | 46 | 5 | 14 | 123 | 4 | 12 |
| 1430 | FIM1_1465 | QGN14794.1 | CP015055.1 | 3597 | 586 | 872 | 2919 | 503 | 828 | 3890 | 625 | 823 |
| 1431 | FIM1_1466 | QGN14795.1 | CP015055.1 | 2597 | 580 | 1176 | 2126 | 486 | 1138 | 2677 | 648 | 965 |
| 1432 | FIM1_1467 | QGN14796.1 | CP015055.1 | 7900 | 1076 | 2014 | 5929 | 763 | 1996 | 9615 | 1119 | 1833 |
| 1433 | FIM1_1468 | QGN14797.1 | CP015055.1 | 7683 | 2197 | 2836 | 6379 | 1471 | 2679 | 8363 | 2261 | 2418 |
| 1434 | FIM1_1469 | QGN14798.1 | CP015055.1 | 278 | 43 | 316 | 152 | 25 | 258 | 199 | 38 | 234 |
| 1435 | FIM1_1470 | QGN14799.1 | CP015055.1 | 6720 | 1273 | 1841 | 5145 | 856 | 1671 | 5331 | 1288 | 1481 |
| 1436 | FIM1_1471 | QGN14800.1 | CP015055.1 | 5081 | 3449 | 4183 | 4856 | 2563 | 4272 | 5474 | 3595 | 4057 |
| 1437 | FIM1_1472 | QGN14801.1 | CP015055.1 | 942 | 579 | 1448 | 975 | 406 | 1383 | 1029 | 596 | 1319 |
| 1438 | FIM1_1473 | QGN14802.1 | CP015055.1 | 2773 | 1026 | 739 | 2232 | 762 | 826 | 2425 | 1105 | 754 |

| | | | | | | | | | | | |
|---|---|---|---|---|---|---|---|---|---|---|---|
| 1439 | FIM1_1474 | QGN14803.1 | CP015055.1 | 6857 | 1938 | 2225 | 5512 | 1357 | 2130 | 6928 | 2058 | 2069 |
| 1440 | FIM1_1475 | QGN14804.1 | CP015055.1 | 4178 | 787 | 969 | 3016 | 681 | 912 | 3802 | 890 | 918 |
| 1441 | FIM1_1476 | QGN14805.1 | CP015055.1 | 634 | 58 | 53 | 571 | 36 | 49 | 1557 | 52 | 45 |
| 1442 | FIM1_1477 | QGN14806.1 | CP015055.1 | 6342 | 968 | 1831 | 5161 | 735 | 1672 | 5723 | 972 | 1466 |
| 1443 | FIM1_1478 | QGN14807.1 | CP015055.1 | 6580 | 510 | 1305 | 5007 | 554 | 1236 | 5334 | 522 | 1051 |
| 1444 | FIM1_1479 | QGN14808.1 | CP015055.1 | 1720 | 275 | 774 | 1297 | 231 | 707 | 1844 | 343 | 670 |
| 1445 | FIM1_1480 | QGN14809.1 | CP015055.1 | 1549 | 834 | 867 | 1201 | 672 | 848 | 568 | 888 | 710 |
| 1446 | FIM1_1481 | QGN14810.1 | CP015055.1 | 688 | 264 | 406 | 505 | 217 | 445 | 593 | 253 | 405 |
| 1447 | FIM1_1482 | QGN14811.1 | CP015055.1 | 23347 | 3996 | 4453 | 19325 | 2952 | 4209 | 19976 | 4533 | 4035 |
| 1448 | FIM1_1483 | QGN14812.1 | CP015055.1 | 1096 | 255 | 379 | 958 | 181 | 383 | 685 | 264 | 286 |
| 1449 | FIM1_1484 | QGN14813.1 | CP015055.1 | 2108 | 287 | 776 | 1602 | 235 | 673 | 2125 | 278 | 686 |
| 1450 | FIM1_1485 | QGN14814.1 | CP015055.1 | 7108 | 1937 | 3235 | 5646 | 1477 | 2820 | 6327 | 2100 | 2691 |
| 1451 | FIM1_1486 | QGN14815.1 | CP015055.1 | 2529 | 1272 | 2713 | 2005 | 718 | 2710 | 3236 | 1205 | 2611 |
| 1452 | FIM1_1487 | QGN14816.1 | CP015055.1 | 2822 | 2769 | 1513 | 1909 | 1683 | 1443 | 757 | 2918 | 1398 |
| 1453 | FIM1_1488 | QGN14817.1 | CP015055.1 | 14772 | 3626 | 4365 | 11587 | 2876 | 4166 | 12755 | 4109 | 4227 |
| 1454 | FIM1_1489 | QGN14818.1 | CP015055.1 | 787 | 350 | 470 | 717 | 255 | 434 | 829 | 372 | 384 |
| 1455 | FIM1_1490 | QGN14819.1 | CP015055.1 | 994 | 235 | 550 | 902 | 160 | 524 | 1457 | 263 | 494 |
| 1456 | FIM1_1491 | QGN14820.1 | CP015055.1 | 2682 | 575 | 959 | 1924 | 484 | 907 | 2208 | 655 | 831 |
| 1457 | FIM1_1492 | QGN14821.1 | CP015055.1 | 3110 | 917 | 1716 | 2565 | 640 | 1625 | 3050 | 995 | 1394 |
| 1458 | FIM1_1493 | QGN14822.1 | CP015055.1 | 1352 | 186 | 676 | 1195 | 136 | 601 | 2093 | 187 | 534 |
| 1459 | FIM1_1494 | QGN14823.1 | CP015055.1 | 425 | 91 | 224 | 339 | 75 | 208 | 346 | 94 | 202 |
| 1460 | FIM1_1495 | QGN14824.1 | CP015055.1 | 204 | 28 | 67 | 172 | 30 | 60 | 198 | 33 | 53 |
| 1461 | FIM1_1496 | QGN14825.1 | CP015055.1 | 2584 | 604 | 696 | 1942 | 461 | 641 | 2344 | 637 | 611 |
| 1462 | FIM1_1497 | QGN14826.1 | CP015055.1 | 1869 | 315 | 526 | 1297 | 243 | 491 | 1894 | 390 | 433 |
| 1463 | FIM1_1498 | QGN14827.1 | CP015055.1 | 1281 | 346 | 427 | 820 | 304 | 448 | 1099 | 369 | 430 |
| 1464 | FIM1_1499 | QGN14828.1 | CP015055.1 | 5133 | 1180 | 2627 | 4413 | 849 | 2503 | 4981 | 1248 | 2384 |
| 1465 | FIM1_1500 | QGN14829.1 | CP015055.1 | 4481 | 736 | 1491 | 3669 | 673 | 1344 | 4390 | 757 | 1234 |
| 1466 | FIM1_1501 | QGN14830.1 | CP015055.1 | 3018 | 811 | 1462 | 2355 | 638 | 1281 | 2330 | 852 | 1212 |
| 1467 | FIM1_1502 | QGN14831.1 | CP015055.1 | 12 | 2 | 1 | 16 | 3 | 2 | 12 | 3 | 2 |
| 1468 | FIM1_1503 | QGN14832.1 | CP015055.1 | 6048 | 1414 | 2268 | 5336 | 1154 | 2054 | 5349 | 1701 | 1951 |
| 1469 | FIM1_1504 | QGN14833.1 | CP015055.1 | 2717 | 489 | 778 | 2114 | 365 | 736 | 2493 | 500 | 649 |
| 1470 | FIM1_1505 | QGN14834.1 | CP015055.1 | 1525 | 158 | 435 | 1073 | 124 | 471 | 1900 | 151 | 376 |
| 1471 | FIM1_1506 | QGN14835.1 | CP015055.1 | 389 | 46 | 123 | 365 | 41 | 130 | 267 | 38 | 84 |
| 1472 | FIM1_1507 | QGN14836.1 | CP015055.1 | 8938 | 3256 | 5325 | 7415 | 2050 | 4990 | 10607 | 3555 | 4900 |
| 1473 | FIM1_1508 | QGN14837.1 | CP015055.1 | 4593 | 1000 | 1531 | 3703 | 712 | 1542 | 4592 | 1115 | 1507 |
| 1474 | FIM1_1509 | QGN14838.1 | CP015055.1 | 3083 | 1366 | 1360 | 2638 | 1038 | 1408 | 2136 | 1392 | 1276 |
| 1475 | FIM1_1510 | QGN14839.1 | CP015055.1 | 2356 | 1041 | 1130 | 1699 | 674 | 1112 | 1837 | 1024 | 1084 |
| 1476 | FIM1_1511 | QGN14840.1 | CP015055.1 | 454 | 102 | 181 | 392 | 89 | 193 | 426 | 133 | 181 |
| 1477 | FIM1_1512 | QGN14841.1 | CP015055.1 | 1318 | 431 | 528 | 1155 | 317 | 515 | 1201 | 512 | 488 |
| 1478 | FIM1_1513 | QGN14842.1 | CP015055.1 | 8886 | 2696 | 4792 | 7604 | 1873 | 4467 | 10874 | 2992 | 4133 |
| 1479 | FIM1_1514 | QGN14843.1 | CP015055.1 | 21304 | 6658 | 11919 | 17031 | 4465 | 10912 | 28199 | 6951 | 11142 |
| 1480 | FIM1_1515 | QGN14844.1 | CP015055.1 | 872 | 90 | 251 | 602 | 66 | 231 | 994 | 88 | 205 |
| 1481 | FIM1_1516 | QGN14845.1 | CP015055.1 | 165 | 60 | 53 | 110 | 114 | 40 | 508 | 46 | 31 |
| 1482 | FIM1_1518 | QGN14846.1 | CP015055.1 | 2644 | 1450 | 1318 | 2242 | 1033 | 1286 | 1802 | 1473 | 1148 |
| 1483 | FIM1_1519 | QGN14847.1 | CP015055.1 | 1151 | 236 | 651 | 893 | 230 | 536 | 839 | 288 | 520 |
| 1484 | FIM1_1520 | QGN14848.1 | CP015055.1 | 8869 | 2178 | 4119 | 7781 | 1427 | 3808 | 11163 | 2387 | 3571 |
| 1485 | FIM1_1521 | QGN14849.1 | CP015055.1 | 9052 | 5707 | 2934 | 6031 | 3440 | 2583 | 1586 | 5732 | 2724 |
| 1486 | FIM1_1523 | QGN14850.1 | CP015055.1 | 939 | 181 | 339 | 858 | 155 | 312 | 967 | 203 | 266 |

| | | | | | | | | | | | | |
|---|---|---|---|---|---|---|---|---|---|---|---|---|
| 1487 | FIM1_1524 | QGN14851.1 | CP015055.1 | 2057 | 734 | 1030 | 1707 | 637 | 1078 | 1368 | 889 | 994 |
| 1488 | FIM1_1525 | QGN14852.1 | CP015055.1 | 4067 | 948 | 1593 | 3331 | 655 | 1601 | 4146 | 1048 | 1421 |
| 1489 | FIM1_1526 | QGN14853.1 | CP015055.1 | 8073 | 2286 | 10124 | 6288 | 1379 | 9087 | 10776 | 2196 | 9297 |
| 1490 | FIM1_1527 | QGN14854.1 | CP015055.1 | 1243 | 284 | 470 | 1077 | 230 | 392 | 1203 | 278 | 430 |
| 1491 | FIM1_1528 | QGN14855.1 | CP015055.1 | 1963 | 581 | 672 | 1584 | 451 | 603 | 692 | 641 | 545 |
| 1492 | FIM1_1529 | QGN14856.1 | CP015055.1 | 3453 | 257 | 1115 | 2647 | 272 | 996 | 4081 | 277 | 871 |
| 1493 | FIM1_1530 | QGN14857.1 | CP015055.1 | 9433 | 1064 | 1809 | 7123 | 798 | 1604 | 9633 | 1183 | 1377 |
| 1494 | FIM1_1531 | QGN14858.1 | CP015055.1 | 2023 | 229 | 867 | 1884 | 149 | 678 | 2826 | 256 | 632 |
| 1495 | FIM1_1532 | QGN14859.1 | CP015055.1 | 1318 | 314 | 743 | 1158 | 190 | 718 | 2508 | 337 | 729 |
| 1496 | FIM1_1533 | QGN14860.1 | CP015055.1 | 1471 | 489 | 629 | 1298 | 385 | 609 | 1358 | 535 | 554 |
| 1497 | FIM1_1534 | QGN14861.1 | CP015055.1 | 1117 | 228 | 385 | 865 | 175 | 345 | 864 | 269 | 332 |
| 1498 | FIM1_1535 | QGN14862.1 | CP015055.1 | 1749 | 468 | 739 | 1415 | 370 | 665 | 1552 | 495 | 618 |
| 1499 | FIM1_1536 | QGN14863.1 | CP015055.1 | 2043 | 510 | 548 | 1609 | 382 | 511 | 1596 | 524 | 469 |
| 1500 | FIM1_1537 | QGN14864.1 | CP015055.1 | 5784 | 1488 | 1685 | 4603 | 1149 | 1429 | 4289 | 1632 | 1416 |
| 1501 | FIM1_1538 | QGN14865.1 | CP015055.1 | 6765 | 940 | 1639 | 5151 | 782 | 1512 | 6917 | 997 | 1387 |
| 1502 | FIM1_1539 | QGN14866.1 | CP015055.1 | 0 | 1 | 1 | 0 | 0 | 2 | 2 | 2 | 1 |
| 1503 | FIM1_1540 | QGN14867.1 | CP015055.1 | 2149 | 352 | 853 | 1567 | 307 | 829 | 2557 | 365 | 663 |
| 1504 | FIM1_1541 | QGN14868.1 | CP015055.1 | 8064 | 1363 | 1576 | 6871 | 970 | 1426 | 10217 | 1479 | 1349 |
| 1505 | FIM1_1542 | QGN14869.1 | CP015055.1 | 7835 | 6036 | 3025 | 5058 | 3795 | 2707 | 2309 | 6353 | 2737 |
| 1506 | FIM1_1543 | QGN14870.1 | CP015055.1 | 6155 | 4813 | 2625 | 3898 | 3105 | 2385 | 1130 | 5070 | 2404 |
| 1507 | FIM1_1544 | QGN14871.1 | CP015055.1 | 150 | 95 | 251 | 129 | 47 | 245 | 125 | 100 | 150 |
| 1508 | FIM1_1545 | QGN14872.1 | CP015055.1 | 2282 | 466 | 933 | 1870 | 474 | 920 | 2252 | 527 | 893 |
| 1509 | FIM1_1546 | QGN14873.1 | CP015055.1 | 474 | 125 | 337 | 363 | 120 | 301 | 360 | 151 | 258 |
| 1510 | FIM1_1547 | QGN14874.1 | CP015055.1 | 1725 | 503 | 862 | 1341 | 430 | 863 | 1423 | 510 | 727 |
| 1511 | FIM1_1548 | QGN14875.1 | CP015055.1 | 2786 | 1431 | 1326 | 2278 | 1122 | 1365 | 1272 | 1585 | 1273 |
| 1512 | FIM1_1549 | QGN14876.1 | CP015055.1 | 4154 | 972 | 2494 | 3283 | 801 | 2347 | 4353 | 1180 | 2216 |
| 1513 | FIM1_1550 | QGN14877.1 | CP015055.1 | 2673 | 782 | 1846 | 1764 | 560 | 1805 | 2069 | 818 | 1717 |
| 1514 | FIM1_1551 | QGN14878.1 | CP015055.1 | 22684 | 7796 | 34043 | 18434 | 5000 | 30023 | 28528 | 8178 | 32975 |
| 1515 | FIM1_1552 | QGN14879.1 | CP015055.1 | 649 | 445 | 220 | 507 | 273 | 224 | 344 | 432 | 215 |
| 1516 | FIM1_1553 | QGN14880.1 | CP015055.1 | 6015 | 5463 | 2628 | 3795 | 3425 | 2477 | 1088 | 5768 | 2516 |
| 1517 | FIM1_1554 | QGN14881.1 | CP015055.1 | 2783 | 828 | 780 | 2223 | 565 | 769 | 1767 | 944 | 709 |
| 1518 | FIM1_1556 | QGN14882.1 | CP015055.1 | 766 | 347 | 416 | 530 | 218 | 397 | 574 | 349 | 361 |
| 1519 | FIM1_1557 | QGN14883.1 | CP015055.1 | 498 | 238 | 307 | 431 | 215 | 353 | 341 | 274 | 312 |
| 1520 | FIM1_1558 | QGN14884.1 | CP015055.1 | 6828 | 1116 | 1859 | 4979 | 845 | 1715 | 7383 | 1184 | 1570 |
| 1521 | FIM1_1559 | QGN14885.1 | CP015055.1 | 14399 | 4831 | 5572 | 11908 | 3155 | 4827 | 14075 | 4957 | 4858 |
| 1522 | FIM1_1560 | QGN14886.1 | CP015055.1 | 3457 | 553 | 1833 | 2537 | 434 | 1700 | 3285 | 570 | 1438 |
| 1523 | FIM1_1561 | QGN14887.1 | CP015055.1 | 5331 | 648 | 1439 | 3669 | 532 | 1334 | 4619 | 719 | 1345 |
| 1524 | FIM1_1562 | QGN14888.1 | CP015055.1 | 991 | 372 | 474 | 913 | 299 | 522 | 551 | 393 | 452 |
| 1525 | FIM1_1563 | QGN14889.1 | CP015055.1 | 3602 | 808 | 790 | 3016 | 671 | 686 | 3364 | 963 | 635 |
| 1526 | FIM1_1564 | QGN14890.1 | CP015055.1 | 725 | 15 | 31 | 401 | 15 | 34 | 1094 | 20 | 35 |
| 1527 | FIM1_1565 | QGN14891.1 | CP015055.1 | 33 | 7 | 5 | 18 | 5 | 4 | 28 | 6 | 8 |
| 1528 | FIM1_1566 | QGN14892.1 | CP015055.1 | 8109 | 1675 | 1820 | 6299 | 1070 | 1588 | 9468 | 1824 | 1517 |
| 1529 | FIM1_1567 | QGN14893.1 | CP015055.1 | 5277 | 658 | 633 | 4315 | 450 | 586 | 6177 | 673 | 490 |
| 1530 | FIM1_1568 | QGN14894.1 | CP015055.1 | 17615 | 5196 | 6644 | 14171 | 3237 | 5916 | 20540 | 5376 | 5749 |
| 1531 | FIM1_1569 | QGN14895.1 | CP015055.1 | 514 | 30 | 198 | 503 | 21 | 143 | 891 | 34 | 129 |
| 1532 | FIM1_1570 | QGN14896.1 | CP015055.1 | 3248 | 467 | 1032 | 2399 | 349 | 879 | 3222 | 493 | 789 |
| 1533 | FIM1_1571 | QGN14897.1 | CP015055.1 | 867 | 256 | 331 | 536 | 178 | 321 | 644 | 241 | 274 |
| 1534 | FIM1_1572 | QGN14898.1 | CP015055.1 | 1272 | 276 | 460 | 1015 | 219 | 413 | 1244 | 271 | 380 |

| | | | | | | | | | | | | |
|---|---|---|---|---|---|---|---|---|---|---|---|---|
| 1535 | FIM1_1573 | QGN14899.1 | CP015055.1 | 2558 | 832 | 899 | 2127 | 665 | 795 | 2235 | 854 | 829 |
| 1536 | FIM1_1574 | QGN14900.1 | CP015055.1 | 8491 | 2094 | 2645 | 7211 | 1638 | 2303 | 8771 | 2381 | 2356 |
| 1537 | FIM1_1575 | QGN14901.1 | CP015055.1 | 2597 | 320 | 836 | 2089 | 235 | 730 | 2781 | 358 | 679 |
| 1538 | FIM1_1576 | QGN14902.1 | CP015055.1 | 54 | 15 | 41 | 77 | 10 | 34 | 104 | 14 | 34 |
| 1539 | FIM1_1577 | QGN14903.1 | CP015055.1 | 1183 | 218 | 770 | 1067 | 205 | 741 | 1585 | 241 | 721 |
| 1540 | FIM1_1578 | QGN14904.1 | CP015055.1 | 22885 | 9122 | 8089 | 19632 | 7199 | 7190 | 18635 | 9475 | 7482 |
| 1541 | FIM1_1579 | QGN14905.1 | CP015055.1 | 3264 | 2568 | 1366 | 2389 | 1834 | 1233 | 978 | 2693 | 1379 |
| 1542 | FIM1_1580 | QGN14906.1 | CP015055.1 | 3383 | 1844 | 1133 | 2247 | 1051 | 1136 | 1523 | 2028 | 1166 |
| 1543 | FIM1_1581 | QGN14907.1 | CP015055.1 | 9386 | 4102 | 3511 | 7822 | 2889 | 3386 | 10522 | 4451 | 3192 |
| 1544 | FIM1_1582 | QGN14908.1 | CP015055.1 | 131 | 21 | 70 | 106 | 18 | 55 | 273 | 26 | 60 |
| 1545 | FIM1_1583 | QGN14909.1 | CP015055.1 | 2603 | 427 | 648 | 2047 | 316 | 641 | 2756 | 449 | 595 |
| 1546 | FIM1_1584 | QGN14910.1 | CP015055.1 | 14437 | 4479 | 4141 | 12047 | 2737 | 3791 | 17168 | 4524 | 3819 |
| 1547 | FIM1_1585 | QGN14911.1 | CP015055.1 | 1426 | 381 | 819 | 1528 | 244 | 762 | 2050 | 412 | 783 |
| 1548 | FIM1_1586 | QGN14912.1 | CP015055.1 | 2993 | 529 | 976 | 2538 | 308 | 846 | 7406 | 630 | 859 |
| 1549 | FIM1_1587 | QGN14913.1 | CP015055.1 | 18369 | 6148 | 6388 | 15895 | 4709 | 5945 | 18830 | 6590 | 5985 |
| 1550 | FIM1_1588 | QGN14914.1 | CP015055.1 | 17568 | 6474 | 5219 | 13542 | 4640 | 4641 | 15111 | 6960 | 4722 |
| 1551 | FIM1_1589 | QGN14915.1 | CP015055.1 | 322 | 126 | 147 | 242 | 97 | 140 | 260 | 147 | 161 |
| 1552 | FIM1_1590 | QGN14916.1 | CP015055.1 | 1368 | 709 | 883 | 1070 | 574 | 877 | 774 | 745 | 743 |
| 1553 | FIM1_1591 | QGN14917.1 | CP015055.1 | 331 | 122 | 151 | 247 | 100 | 166 | 167 | 109 | 117 |
| 1554 | FIM1_1592 | QGN14918.1 | CP015055.1 | 388 | 72 | 249 | 274 | 58 | 223 | 323 | 75 | 173 |
| 1555 | FIM1_1593 | QGN14919.1 | CP015055.1 | 14927 | 901 | 1912 | 11900 | 434 | 1718 | 25194 | 894 | 1549 |
| 1556 | FIM1_1594 | QGN14920.1 | CP015055.1 | 6627 | 3560 | 2363 | 4522 | 2083 | 2061 | 2145 | 3873 | 1791 |
| 1557 | FIM1_1595 | QGN14921.1 | CP015055.1 | 2634 | 1266 | 1351 | 2382 | 935 | 1307 | 1588 | 1336 | 1188 |
| 1558 | FIM1_1596 | QGN14922.1 | CP015055.1 | 5846 | 579 | 1412 | 4760 | 458 | 1319 | 6682 | 628 | 1297 |
| 1559 | FIM1_1597 | QGN14923.1 | CP015055.1 | 3173 | 2606 | 1174 | 2439 | 1718 | 1092 | 816 | 2753 | 1211 |
| 1560 | FIM1_1598 | QGN14924.1 | CP015055.1 | 5887 | 4121 | 2317 | 4128 | 2639 | 2029 | 1175 | 4589 | 1945 |
| 1561 | FIM1_1599 | QGN14925.1 | CP015055.1 | 6652 | 1935 | 2440 | 5071 | 1218 | 2204 | 6095 | 2188 | 1983 |
| 1562 | FIM1_1601 | QGN14926.1 | CP015055.1 | 37438 | 22696 | 20746 | 36121 | 18937 | 19297 | 36043 | 23146 | 20196 |
| 1563 | FIM1_1602 | QGN14927.1 | CP015055.1 | 3257 | 1190 | 2000 | 2960 | 1009 | 1890 | 2976 | 1365 | 1706 |
| 1564 | FIM1_1603 | QGN14928.1 | CP015055.1 | 10856 | 4008 | 3823 | 9147 | 2842 | 3546 | 7865 | 4276 | 3422 |
| 1565 | FIM1_1604 | QGN14929.1 | CP015055.1 | 8183 | 1247 | 1962 | 5574 | 658 | 1683 | 5615 | 1277 | 1780 |
| 1566 | FIM1_1605 | QGN14930.1 | CP015055.1 | 5352 | 4844 | 2080 | 3749 | 3256 | 1803 | 1050 | 4861 | 1850 |
| 1567 | FIM1_1606 | QGN14931.1 | CP015055.1 | 18792 | 5578 | 6732 | 15508 | 3891 | 6369 | 18304 | 5880 | 6455 |
| 1568 | FIM1_1607 | QGN14932.1 | CP015055.1 | 3575 | 614 | 1285 | 2675 | 537 | 1162 | 3036 | 668 | 1068 |
| 1569 | FIM1_1608 | QGN14933.1 | CP015055.1 | 7493 | 2239 | 2451 | 5328 | 1627 | 2265 | 4733 | 2374 | 2094 |
| 1570 | FIM1_1609 | QGN14934.1 | CP015055.1 | 3151 | 981 | 1237 | 2787 | 785 | 1064 | 1833 | 1079 | 1070 |
| 1571 | FIM1_1610 | QGN14935.1 | CP015055.1 | 4058 | 1652 | 1811 | 3550 | 1018 | 1764 | 4002 | 1671 | 1589 |
| 1572 | FIM1_1611 | QGN14936.1 | CP015055.1 | 4825 | 1039 | 1993 | 3334 | 880 | 1678 | 3419 | 1150 | 1542 |
| 1573 | FIM1_1612 | QGN14937.1 | CP015055.1 | 2957 | 1604 | 1353 | 2597 | 1262 | 1287 | 1824 | 1680 | 1265 |
| 1574 | FIM1_1613 | QGN14938.1 | CP015055.1 | 4580 | 816 | 1114 | 3454 | 526 | 1020 | 4194 | 766 | 886 |
| 1575 | FIM1_1614 | QGN14939.1 | CP015055.1 | 961 | 200 | 617 | 741 | 179 | 578 | 1064 | 193 | 470 |
| 1576 | FIM1_1615 | QGN14940.1 | CP015055.1 | 6619 | 2290 | 2647 | 5747 | 1705 | 2612 | 8226 | 2427 | 2425 |
| 1577 | FIM1_1616 | QGN14941.1 | CP015055.1 | 2490 | 561 | 767 | 1898 | 411 | 673 | 1591 | 565 | 696 |
| 1578 | FIM1_1617 | QGN14942.1 | CP015055.1 | 7505 | 3864 | 2568 | 5601 | 2550 | 2469 | 6036 | 4259 | 2512 |
| 1579 | FIM1_1618 | QGN14943.1 | CP015055.1 | 462 | 453 | 222 | 327 | 294 | 205 | 219 | 443 | 206 |
| 1580 | FIM1_1619 | QGN14944.1 | CP015055.1 | 2339 | 1481 | 1416 | 1677 | 962 | 1185 | 1639 | 1526 | 1187 |
| 1581 | FIM1_1621 | QGN14945.1 | CP015055.1 | 10480 | 2057 | 2927 | 7524 | 1628 | 2743 | 9680 | 2267 | 2584 |
| 1582 | FIM1_1622 | QGN14946.1 | CP015055.1 | 1458 | 612 | 395 | 1176 | 388 | 377 | 2218 | 636 | 425 |

| | | | | | | | | | | | |
|---|---|---|---|---|---|---|---|---|---|---|---|
| 1583 | FIM1_1623 | QGN14947.1 | CP015055.1 | 1003 | 212 | 245 | 789 | 195 | 257 | 990 | 269 | 270 |
| 1584 | FIM1_1624 | QGN14948.1 | CP015055.1 | 712 | 448 | 593 | 580 | 325 | 626 | 590 | 487 | 530 |
| 1585 | FIM1_1625 | QGN14949.1 | CP015055.1 | 2610 | 1014 | 1593 | 2240 | 638 | 1507 | 3117 | 947 | 1459 |
| 1586 | FIM1_1627 | QGN14950.1 | CP015055.1 | 4290 | 1693 | 2414 | 2942 | 1051 | 2192 | 2594 | 1724 | 2056 |
| 1587 | FIM1_1628 | QGN14951.1 | CP015055.1 | 648 | 189 | 385 | 460 | 202 | 205 | 1476 | 138 | 238 |
| 1588 | FIM1_1629 | QGN14952.1 | CP015055.1 | 781 | 205 | 332 | 643 | 184 | 340 | 701 | 200 | 312 |
| 1589 | FIM1_1630 | QGN14953.1 | CP015055.1 | 9492 | 1311 | 2371 | 7153 | 991 | 2041 | 10533 | 1383 | 1946 |
| 1590 | FIM1_1631 | QGN14954.1 | CP015055.1 | 1714 | 432 | 634 | 1261 | 186 | 571 | 2108 | 466 | 539 |
| 1591 | FIM1_1632 | QGN14955.1 | CP015055.1 | 6186 | 757 | 1130 | 4603 | 627 | 1034 | 5018 | 851 | 1091 |
| 1592 | FIM1_1633 | QGN14956.1 | CP015055.1 | 744 | 1357 | 1064 | 647 | 623 | 865 | 214 | 1377 | 851 |
| 1593 | FIM1_1634 | QGN14957.1 | CP015055.1 | 5993 | 1362 | 1842 | 5395 | 1075 | 1832 | 5945 | 1494 | 1766 |
| 1594 | FIM1_1635 | QGN14958.1 | CP015055.1 | 15335 | 4098 | 5362 | 13001 | 2774 | 5031 | 17165 | 4329 | 4847 |
| 1595 | FIM1_1636 | QGN14959.1 | CP015055.1 | 1692 | 527 | 903 | 1614 | 374 | 834 | 1765 | 607 | 810 |
| 1596 | FIM1_1637 | QGN14960.1 | CP015055.1 | 529 | 137 | 228 | 470 | 92 | 229 | 391 | 163 | 222 |
| 1597 | FIM1_1638 | QGN14961.1 | CP015055.1 | 89 | 18 | 17 | 53 | 19 | 27 | 159 | 17 | 20 |
| 1598 | FIM1_1640 | QGN14962.1 | CP015055.1 | 6810 | 1496 | 2133 | 4818 | 1162 | 1998 | 5862 | 1491 | 1987 |
| 1599 | FIM1_1641 | QGN14963.1 | CP015055.1 | 10755 | 3401 | 4434 | 8957 | 2539 | 4135 | 9287 | 3600 | 4056 |
| 1600 | FIM1_1642 | QGN14964.1 | CP015055.1 | 5743 | 703 | 1759 | 4656 | 490 | 1506 | 6377 | 740 | 1430 |
| 1601 | FIM1_1643 | QGN14965.1 | CP015055.1 | 3765 | 878 | 1357 | 3081 | 608 | 1238 | 5047 | 941 | 1195 |
| 1602 | FIM1_1644 | QGN14966.1 | CP015055.1 | 1711 | 306 | 587 | 1293 | 256 | 584 | 1351 | 324 | 483 |
| 1603 | FIM1_1645 | QGN14967.1 | CP015055.1 | 1720 | 334 | 487 | 1384 | 284 | 462 | 1322 | 341 | 432 |
| 1604 | FIM1_1646 | QGN14968.1 | CP015055.1 | 11446 | 803 | 2201 | 9378 | 730 | 1998 | 13552 | 839 | 1794 |
| 1605 | FIM1_1647 | QGN14969.1 | CP015055.1 | 201 | 63 | 82 | 145 | 70 | 72 | 140 | 54 | 61 |
| 1606 | FIM1_1648 | QGN14970.1 | CP015055.1 | 5203 | 2322 | 3397 | 4070 | 2270 | 3166 | 3923 | 2450 | 3200 |
| 1607 | FIM1_1649 | QGN14971.1 | CP015055.1 | 4960 | 2985 | 2804 | 4278 | 2386 | 2688 | 2849 | 3274 | 2582 |
| 1608 | FIM1_1650 | QGN14972.1 | CP015055.1 | 2159 | 916 | 894 | 1798 | 440 | 844 | 3815 | 916 | 749 |
| 1609 | FIM1_1651 | QGN14973.1 | CP015055.1 | 753 | 133 | 450 | 699 | 113 | 466 | 1178 | 119 | 399 |
| 1610 | FIM1_1652 | QGN14974.1 | CP015055.1 | 6643 | 2083 | 2887 | 5388 | 1336 | 2606 | 5347 | 2302 | 2676 |
| 1611 | FIM1_1653 | QGN14975.1 | CP015055.1 | 2056 | 244 | 424 | 1718 | 206 | 369 | 1703 | 262 | 413 |
| 1612 | FIM1_1654 | QGN14976.1 | CP015055.1 | 1701 | 86 | 422 | 1254 | 118 | 369 | 1581 | 98 | 364 |
| 1613 | FIM1_1655 | QGN14977.1 | CP015055.1 | 3695 | 856 | 1182 | 3014 | 608 | 1071 | 3327 | 906 | 970 |
| 1614 | FIM1_1656 | QGN14978.1 | CP015055.1 | 4830 | 201 | 1620 | 4428 | 131 | 1428 | 4665 | 218 | 1144 |
| 1615 | FIM1_1657 | QGN14979.1 | CP015055.1 | 61514 | 30840 | 26580 | 55827 | 24171 | 23364 | 60147 | 31230 | 25468 |
| 1616 | FIM1_1658 | QGN14980.1 | CP015055.1 | 2041 | 707 | 898 | 1411 | 583 | 859 | 1480 | 858 | 889 |
| 1617 | FIM1_1659 | QGN14981.1 | CP015055.1 | 9984 | 1701 | 2455 | 7697 | 1179 | 2342 | 10366 | 1780 | 2197 |
| 1618 | FIM1_1660 | QGN14982.1 | CP015055.1 | 5333 | 1396 | 2454 | 3817 | 1047 | 2346 | 4343 | 1558 | 2083 |
| 1619 | FIM1_1661 | QGN14983.1 | CP015055.1 | 3035 | 1356 | 1496 | 2378 | 927 | 1614 | 2667 | 1497 | 1356 |
| 1620 | FIM1_1664 | QGN14984.1 | CP015055.1 | 4604 | 1494 | 2005 | 4122 | 1141 | 1908 | 4173 | 1567 | 1811 |
| 1621 | FIM1_1665 | QGN14985.1 | CP015055.1 | 905 | 208 | 218 | 643 | 130 | 241 | 886 | 208 | 237 |
| 1622 | FIM1_1666 | QGN14986.1 | CP015055.1 | 2084 | 586 | 796 | 1699 | 430 | 697 | 2008 | 650 | 710 |
| 1623 | FIM1_1667 | QGN14987.1 | CP015055.1 | 2247 | 3635 | 1000 | 1244 | 1872 | 963 | 782 | 3677 | 1060 |
| 1624 | FIM1_1669 | QGN14988.1 | CP015055.1 | 5355 | 1631 | 1521 | 3599 | 1064 | 1460 | 4786 | 1782 | 1451 |
| 1625 | FIM1_1670 | QGN14989.1 | CP015055.1 | 2860 | 1774 | 1613 | 2739 | 1560 | 1570 | 1729 | 2029 | 1375 |
| 1626 | FIM1_1671 | QGN14990.1 | CP015055.1 | 10029 | 7953 | 4155 | 6724 | 5149 | 3916 | 2561 | 8428 | 4069 |
| 1627 | FIM1_1672 | QGN14991.1 | CP015055.1 | 10127 | 3075 | 3755 | 8314 | 1807 | 3516 | 10721 | 3204 | 3511 |
| 1628 | FIM1_1673 | QGN14992.1 | CP015055.1 | 10534 | 2521 | 2646 | 8150 | 1896 | 2428 | 10908 | 2742 | 2414 |
| 1629 | FIM1_1674 | QGN14993.1 | CP015055.1 | 4099 | 506 | 2023 | 2692 | 328 | 1853 | 3428 | 568 | 1946 |
| 1630 | FIM1_1675 | QGN14994.1 | CP015055.1 | 29 | 18 | 24 | 22 | 4 | 14 | 28 | 19 | 14 |

| | | | | | | | | | | | | |
|---|---|---|---|---|---|---|---|---|---|---|---|---|
| 1631 | FIM1_1676 | QGN14995.1 | CP015055.1 | 1415 | 337 | 572 | 1139 | 256 | 496 | 1343 | 367 | 463 |
| 1632 | FIM1_1677 | QGN14996.1 | CP015055.1 | 1110 | 215 | 472 | 923 | 114 | 472 | 3815 | 219 | 341 |
| 1633 | FIM1_1678 | QGN14997.1 | CP015055.1 | 504 | 86 | 138 | 427 | 45 | 161 | 798 | 94 | 126 |
| 1634 | FIM1_1679 | QGN14998.1 | CP015055.1 | 1942 | 425 | 404 | 1242 | 173 | 399 | 2711 | 411 | 410 |
| 1635 | FIM1_1680 | QGN14999.1 | CP015055.1 | 1056 | 594 | 348 | 837 | 338 | 375 | 920 | 595 | 436 |
| 1636 | FIM1_1681 | QGN15000.1 | CP015055.1 | 110 | 10 | 45 | 68 | 10 | 49 | 121 | 21 | 44 |
| 1637 | FIM1_1682 | QGN15001.1 | CP015055.1 | 2188 | 644 | 644 | 1687 | 468 | 644 | 1640 | 771 | 694 |
| 1638 | FIM1_1683 | QGN15002.1 | CP015055.1 | 8481 | 3921 | 3070 | 6988 | 2210 | 2881 | 10421 | 4320 | 2689 |
| 1639 | FIM1_1684 | QGN15003.1 | CP015055.1 | 1776 | 678 | 2458 | 1267 | 460 | 2417 | 1818 | 714 | 2541 |
| 1640 | FIM1_1685 | QGN15004.1 | CP015055.1 | 3170 | 1224 | 1927 | 2586 | 778 | 1742 | 3358 | 1274 | 1627 |
| 1641 | FIM1_1686 | QGN15005.1 | CP015055.1 | 2110 | 615 | 903 | 1654 | 481 | 917 | 2202 | 730 | 860 |
| 1642 | FIM1_1687 | QGN15006.1 | CP015055.1 | 3912 | 755 | 3180 | 3143 | 484 | 3124 | 4175 | 843 | 3157 |
| 1643 | FIM1_1688 | QGN15007.1 | CP015055.1 | 358 | 163 | 127 | 273 | 117 | 144 | 512 | 168 | 115 |
| 1644 | FIM1_1689 | QGN15008.1 | CP015055.1 | 817 | 229 | 142 | 630 | 202 | 138 | 562 | 193 | 148 |
| 1645 | FIM1_1690 | QGN15009.1 | CP015055.1 | 10703 | 1671 | 3390 | 8249 | 1355 | 3174 | 9544 | 1843 | 3074 |
| 1646 | FIM1_1691 | QGN15010.1 | CP015055.1 | 5921 | 679 | 351 | 4678 | 363 | 320 | 18118 | 807 | 254 |
| 1647 | FIM1_1692 | QGN15011.1 | CP015055.1 | 3487 | 394 | 412 | 2466 | 302 | 385 | 5420 | 421 | 330 |
| 1648 | FIM1_1693 | QGN15012.1 | CP015055.1 | 1268 | 249 | 380 | 947 | 127 | 391 | 1903 | 258 | 332 |
| 1649 | FIM1_1694 | QGN15013.1 | CP015055.1 | 142 | 41 | 45 | 102 | 23 | 41 | 212 | 31 | 49 |
| 1650 | FIM1_1695 | QGN15014.1 | CP015055.1 | 604 | 64 | 131 | 473 | 40 | 93 | 1034 | 72 | 123 |
| 1651 | FIM1_1697 | QGN15015.1 | CP015055.1 | 468 | 105 | 115 | 354 | 80 | 91 | 495 | 160 | 102 |
| 1652 | FIM1_1698 | QGN15016.1 | CP015055.1 | 3356 | 1995 | 1631 | 2669 | 1760 | 1822 | 3608 | 2327 | 1743 |
| 1653 | FIM1_1700 | QGN15017.1 | CP015056.1 | 0 | 0 | 0 | 0 | 0 | 0 | 0 | 0 | 0 |
| 1654 | FIM1_1701 | QGN15018.1 | CP015056.1 | 967 | 421 | 339 | 875 | 257 | 393 | 940 | 409 | 288 |
| 1655 | FIM1_1703 | QGN15019.1 | CP015056.1 | 269 | 39 | 64 | 183 | 19 | 89 | 435 | 38 | 49 |
| 1656 | FIM1_1704 | QGN15020.1 | CP015056.1 | 44 | 14 | 11 | 28 | 11 | 10 | 56 | 8 | 9 |
| 1657 | FIM1_1705 | QGN15021.1 | CP015056.1 | 44 | 16 | 19 | 41 | 10 | 11 | 60 | 16 | 14 |
| 1658 | FIM1_1706 | QGN15022.1 | CP015056.1 | 0 | 0 | 0 | 0 | 0 | 0 | 2 | 0 | 0 |
| 1659 | FIM1_1707 | QGN15023.1 | CP015056.1 | 1258 | 249 | 258 | 883 | 189 | 221 | 1338 | 296 | 255 |
| 1660 | FIM1_1708 | QGN15024.1 | CP015056.1 | 9643 | 3665 | 341 | 5140 | 2034 | 306 | 8245 | 3920 | 343 |
| 1661 | FIM1_1709 | QGN15025.1 | CP015056.1 | 2973 | 537 | 716 | 2390 | 441 | 636 | 2580 | 593 | 650 |
| 1662 | FIM1_1710 | QGN15026.1 | CP015056.1 | 526 | 172 | 237 | 396 | 142 | 219 | 521 | 174 | 220 |
| 1663 | FIM1_1711 | QGN15027.1 | CP015056.1 | 2516 | 329 | 1094 | 2020 | 125 | 993 | 7694 | 310 | 891 |
| 1664 | FIM1_1712 | QGN15028.1 | CP015056.1 | 3405 | 5730 | 1861 | 2848 | 1355 | 1528 | 2227 | 4962 | 1742 |
| 1665 | FIM1_1713 | QGN15029.1 | CP015056.1 | 125 | 50 | 46 | 118 | 34 | 36 | 154 | 65 | 28 |
| 1666 | FIM1_1714 | QGN15030.1 | CP015056.1 | 608 | 116 | 130 | 470 | 85 | 137 | 628 | 113 | 128 |
| 1667 | FIM1_1715 | QGN15031.1 | CP015056.1 | 651 | 68 | 241 | 429 | 46 | 204 | 723 | 56 | 182 |
| 1668 | FIM1_1716 | QGN15032.1 | CP015056.1 | 7314 | 3416 | 4954 | 6153 | 1415 | 4699 | 12113 | 2955 | 4550 |
| 1669 | FIM1_1717 | QGN15033.1 | CP015056.1 | 563 | 92 | 150 | 461 | 88 | 134 | 567 | 112 | 138 |
| 1670 | FIM1_1718 | QGN15034.1 | CP015056.1 | 414 | 109 | 194 | 319 | 88 | 189 | 388 | 110 | 134 |
| 1671 | FIM1_1719 | QGN15035.1 | CP015056.1 | 677 | 134 | 186 | 465 | 98 | 196 | 716 | 125 | 169 |
| 1672 | FIM1_1720 | QGN15036.1 | CP015056.1 | 6 | 1 | 3 | 8 | 0 | 7 | 6 | 6 | 2 |
| 1673 | FIM1_1721 | QGN15037.1 | CP015056.1 | 80 | 20 | 15 | 60 | 14 | 20 | 41 | 26 | 22 |
| 1674 | FIM1_1722 | QGN15038.1 | CP015056.1 | 0 | 0 | 0 | 0 | 0 | 0 | 0 | 0 | 0 |
| 1675 | FIM1_1723 | QGN15039.1 | CP015056.1 | 3 | 3 | 0 | 4 | 0 | 3 | 11 | 1 | 2 |
| 1676 | FIM1_1724 | QGN15040.1 | CP015056.1 | 26542 | 14160 | 3008 | 23906 | 6021 | 2717 | 40404 | 13244 | 2195 |
| 1677 | FIM1_1725 | QGN15041.1 | CP015056.1 | 31558 | 18593 | 15185 | 27713 | 13405 | 13772 | 27941 | 18693 | 15210 |
| 1678 | FIM1_1727 | QGN15042.1 | CP015056.1 | 6094 | 721 | 2210 | 5102 | 471 | 1901 | 7830 | 745 | 1935 |

| | | | | | | | | | | | | |
|---|---|---|---|---|---|---|---|---|---|---|---|---|
| 1679 | FIM1_1728 | QGN15043.1 | CP015056.1 | 14717 | 4340 | 5306 | 12224 | 2556 | 4970 | 18687 | 4393 | 4586 |
| 1680 | FIM1_1729 | QGN15044.1 | CP015056.1 | 8765 | 2186 | 3843 | 7029 | 1398 | 3310 | 9010 | 2437 | 3310 |
| 1681 | FIM1_1730 | QGN15045.1 | CP015056.1 | 12393 | 3212 | 4325 | 10278 | 2314 | 3923 | 13114 | 3712 | 3893 |
| 1682 | FIM1_1731 | QGN15046.1 | CP015056.1 | 10501 | 3338 | 3803 | 8793 | 2450 | 3329 | 10842 | 3447 | 3287 |
| 1683 | FIM1_1732 | QGN15047.1 | CP015056.1 | 2382 | 804 | 980 | 2231 | 607 | 910 | 2399 | 842 | 869 |
| 1684 | FIM1_1733 | QGN15048.1 | CP015056.1 | 23832 | 4416 | 8031 | 19517 | 2752 | 7494 | 36472 | 4494 | 7357 |
| 1685 | FIM1_1734 | QGN15049.1 | CP015056.1 | 1375 | 309 | 519 | 998 | 270 | 474 | 1807 | 336 | 458 |
| 1686 | FIM1_1735 | QGN15050.1 | CP015056.1 | 3678 | 1590 | 1855 | 3390 | 1109 | 1697 | 4154 | 1600 | 1782 |
| 1687 | FIM1_1736 | QGN15051.1 | CP015056.1 | 1744 | 380 | 650 | 1404 | 266 | 647 | 1145 | 409 | 525 |
| 1688 | FIM1_1737 | QGN15052.1 | CP015056.1 | 1386 | 114 | 522 | 1026 | 101 | 443 | 2033 | 103 | 391 |
| 1689 | FIM1_1738 | QGN15053.1 | CP015056.1 | 2814 | 693 | 1061 | 2244 | 474 | 946 | 3235 | 735 | 894 |
| 1690 | FIM1_1739 | QGN15054.1 | CP015056.1 | 5579 | 3016 | 2637 | 4482 | 2189 | 2535 | 3355 | 3171 | 2362 |
| 1691 | FIM1_1740 | QGN15055.1 | CP015056.1 | 2102 | 505 | 906 | 1797 | 329 | 892 | 2207 | 526 | 808 |
| 1692 | FIM1_1741 | QGN15056.1 | CP015056.1 | 7906 | 4726 | 2976 | 6737 | 3672 | 2611 | 6166 | 5213 | 2875 |
| 1693 | FIM1_1742 | QGN15057.1 | CP015056.1 | 2771 | 1055 | 881 | 2394 | 892 | 888 | 1783 | 1210 | 816 |
| 1694 | FIM1_1743 | QGN15058.1 | CP015056.1 | 11975 | 3694 | 3138 | 9087 | 2818 | 2894 | 7568 | 3926 | 3335 |
| 1695 | FIM1_1744 | QGN15059.1 | CP015056.1 | 12056 | 1668 | 2107 | 8747 | 1184 | 1939 | 11465 | 1974 | 1881 |
| 1696 | FIM1_1745 | QGN15060.1 | CP015056.1 | 3777 | 899 | 1294 | 3107 | 723 | 1173 | 3426 | 1047 | 1075 |
| 1697 | FIM1_1746 | QGN15061.1 | CP015056.1 | 1047 | 344 | 452 | 1034 | 303 | 413 | 650 | 353 | 398 |
| 1698 | FIM1_1747 | QGN15062.1 | CP015056.1 | 10783 | 1687 | 2633 | 8267 | 1290 | 2475 | 10178 | 1841 | 2336 |
| 1699 | FIM1_1748 | QGN15063.1 | CP015056.1 | 463 | 63 | 48 | 467 | 30 | 62 | 862 | 71 | 51 |
| 1700 | FIM1_1749 | QGN15064.1 | CP015056.1 | 10785 | 2947 | 3611 | 9477 | 1845 | 3164 | 13376 | 3036 | 2914 |
| 1701 | FIM1_1750 | QGN15065.1 | CP015056.1 | 12393 | 3590 | 4652 | 10251 | 2589 | 3969 | 11755 | 3941 | 3938 |
| 1702 | FIM1_1751 | QGN15066.1 | CP015056.1 | 2548 | 1540 | 1007 | 2138 | 1338 | 877 | 812 | 1878 | 908 |
| 1703 | FIM1_1752 | QGN15067.1 | CP015056.1 | 3407 | 746 | 824 | 2079 | 444 | 748 | 2752 | 775 | 825 |
| 1704 | FIM1_1753 | QGN15068.1 | CP015056.1 | 1994 | 317 | 766 | 1592 | 230 | 665 | 2259 | 315 | 607 |
| 1705 | FIM1_1754 | QGN15069.1 | CP015056.1 | 2456 | 227 | 246 | 1754 | 184 | 238 | 2385 | 257 | 213 |
| 1706 | FIM1_1755 | QGN15070.1 | CP015056.1 | 6029 | 1506 | 1595 | 5369 | 1132 | 1390 | 6257 | 1757 | 1407 |
| 1707 | FIM1_1756 | QGN15071.1 | CP015056.1 | 5456 | 767 | 954 | 4966 | 561 | 889 | 5717 | 805 | 944 |
| 1708 | FIM1_1757 | QGN15072.1 | CP015056.1 | 3734 | 1009 | 1295 | 2554 | 701 | 1187 | 3864 | 1074 | 1058 |
| 1709 | FIM1_1758 | QGN15073.1 | CP015056.1 | 552 | 244 | 339 | 408 | 196 | 304 | 398 | 308 | 296 |
| 1710 | FIM1_1759 | QGN15074.1 | CP015056.1 | 1417 | 207 | 318 | 1089 | 171 | 282 | 1453 | 237 | 297 |
| 1711 | FIM1_1760 | QGN15075.1 | CP015056.1 | 16895 | 2437 | 3161 | 13596 | 1876 | 2831 | 15373 | 2702 | 2738 |
| 1712 | FIM1_1761 | QGN15076.1 | CP015056.1 | 11104 | 1361 | 2141 | 8898 | 1041 | 1913 | 12797 | 1469 | 2026 |
| 1713 | FIM1_1762 | QGN15077.1 | CP015056.1 | 9474 | 7425 | 3978 | 8358 | 5663 | 3473 | 3099 | 7830 | 3622 |
| 1714 | FIM1_1763 | QGN15078.1 | CP015056.1 | 163 | 58 | 144 | 140 | 70 | 100 | 67 | 65 | 89 |
| 1715 | FIM1_1764 | QGN15079.1 | CP015056.1 | 9240 | 3496 | 3349 | 7332 | 2617 | 3000 | 6136 | 3764 | 2925 |
| 1716 | FIM1_1765 | QGN15080.1 | CP015056.1 | 5061 | 1086 | 1598 | 4093 | 802 | 1326 | 6987 | 1250 | 1352 |
| 1717 | FIM1_1766 | QGN15081.1 | CP015056.1 | 6762 | 953 | 1533 | 4801 | 520 | 1192 | 7309 | 970 | 1184 |
| 1718 | FIM1_1767 | QGN15082.1 | CP015056.1 | 3654 | 1081 | 1574 | 3211 | 630 | 1548 | 4820 | 1071 | 1387 |
| 1719 | FIM1_1769 | QGN15083.1 | CP015056.1 | 80 | 10 | 13 | 63 | 6 | 20 | 12 | 9 | 11 |
| 1720 | FIM1_1770 | QGN15084.1 | CP015056.1 | 4654 | 2918 | 2646 | 4563 | 2408 | 2262 | 4671 | 2906 | 2460 |
| 1721 | FIM1_1771 | QGN15085.1 | CP015056.1 | 3389 | 1703 | 1052 | 2358 | 994 | 939 | 2773 | 1848 | 902 |
| 1722 | FIM1_1772 | QGN15086.1 | CP015056.1 | 2851 | 1438 | 1455 | 2209 | 1085 | 1468 | 1829 | 1557 | 1375 |
| 1723 | FIM1_1774 | QGN15087.1 | CP015056.1 | 1521 | 530 | 798 | 1223 | 347 | 806 | 1848 | 534 | 757 |
| 1724 | FIM1_1775 | QGN15088.1 | CP015056.1 | 15214 | 4720 | 6042 | 12203 | 2760 | 5521 | 19620 | 4824 | 5403 |
| 1725 | FIM1_1776 | QGN15089.1 | CP015056.1 | 9479 | 1878 | 2337 | 7297 | 1340 | 2171 | 10469 | 2171 | 2040 |
| 1726 | FIM1_1777 | QGN15090.1 | CP015056.1 | 1606 | 1796 | 2127 | 1582 | 986 | 1867 | 2041 | 2104 | 1511 |

| | | | | | | | | | | | | |
|---|---|---|---|---|---|---|---|---|---|---|---|---|
| 1727 | FIM1_1778 | QGN15091.1 | CP015056.1 | 9860 | 1603 | 2275 | 7613 | 1261 | 2142 | 10920 | 1826 | 1950 |
| 1728 | FIM1_1779 | QGN15092.1 | CP015056.1 | 13943 | 4298 | 4574 | 10068 | 1678 | 4057 | 16288 | 4118 | 3147 |
| 1729 | FIM1_1780 | QGN15093.1 | CP015056.1 | 1933 | 631 | 717 | 1677 | 483 | 725 | 2059 | 751 | 730 |
| 1730 | FIM1_1781 | QGN15094.1 | CP015056.1 | 1845 | 270 | 375 | 1386 | 177 | 413 | 1802 | 290 | 364 |
| 1731 | FIM1_1782 | QGN15095.1 | CP015056.1 | 2882 | 501 | 901 | 2348 | 332 | 774 | 3345 | 545 | 732 |
| 1732 | FIM1_1783 | QGN15096.1 | CP015056.1 | 879 | 78 | 499 | 761 | 56 | 476 | 1007 | 95 | 375 |
| 1733 | FIM1_1784 | QGN15097.1 | CP015056.1 | 6186 | 879 | 2081 | 4278 | 614 | 1797 | 5450 | 930 | 1634 |
| 1734 | FIM1_1785 | QGN15098.1 | CP015056.1 | 1926 | 316 | 856 | 1420 | 282 | 885 | 2452 | 368 | 738 |
| 1735 | FIM1_1786 | QGN15099.1 | CP015056.1 | 134 | 30 | 41 | 79 | 27 | 36 | 113 | 27 | 40 |
| 1736 | FIM1_1787 | QGN15100.1 | CP015056.1 | 3963 | 3616 | 1961 | 3256 | 2364 | 1697 | 1113 | 3713 | 1788 |
| 1737 | FIM1_1788 | QGN15101.1 | CP015056.1 | 1364 | 451 | 517 | 1001 | 318 | 492 | 939 | 464 | 472 |
| 1738 | FIM1_1789 | QGN15102.1 | CP015056.1 | 4921 | 524 | 1300 | 3643 | 411 | 1197 | 5315 | 595 | 1281 |
| 1739 | FIM1_1790 | QGN15103.1 | CP015056.1 | 1824 | 349 | 688 | 1595 | 184 | 731 | 2409 | 352 | 516 |
| 1740 | FIM1_1791 | QGN15104.1 | CP015056.1 | 1766 | 225 | 372 | 1352 | 145 | 386 | 1732 | 238 | 368 |
| 1741 | FIM1_1792 | QGN15105.1 | CP015056.1 | 16992 | 4895 | 5499 | 14403 | 3348 | 5151 | 20656 | 5009 | 5360 |
| 1742 | FIM1_1793 | QGN15106.1 | CP015056.1 | 4563 | 885 | 1454 | 3473 | 897 | 1269 | 3811 | 985 | 1319 |
| 1743 | FIM1_1794 | QGN15107.1 | CP015056.1 | 2246 | 175 | 734 | 1792 | 151 | 630 | 2851 | 218 | 581 |
| 1744 | FIM1_1795 | QGN15108.1 | CP015056.1 | 2777 | 311 | 742 | 2300 | 274 | 664 | 2696 | 346 | 671 |
| 1745 | FIM1_1796 | QGN15109.1 | CP015056.1 | 133 | 38 | 73 | 102 | 40 | 63 | 134 | 42 | 65 |
| 1746 | FIM1_1797 | QGN15110.1 | CP015056.1 | 4273 | 843 | 1517 | 4042 | 650 | 1330 | 3334 | 896 | 1300 |
| 1747 | FIM1_1798 | QGN15111.1 | CP015056.1 | 7836 | 802 | 1189 | 6157 | 546 | 1086 | 9381 | 863 | 1126 |
| 1748 | FIM1_1799 | QGN15112.1 | CP015056.1 | 3503 | 1255 | 1576 | 3522 | 998 | 1374 | 3014 | 1486 | 1267 |
| 1749 | FIM1_1800 | QGN15113.1 | CP015056.1 | 2413 | 141 | 481 | 1824 | 172 | 457 | 2351 | 243 | 415 |
| 1750 | FIM1_1801 | QGN15114.1 | CP015056.1 | 2402 | 3120 | 1567 | 1746 | 1789 | 1366 | 691 | 3445 | 1513 |
| 1751 | FIM1_1802 | QGN15115.1 | CP015056.1 | 13323 | 1635 | 1902 | 11427 | 1065 | 1772 | 15737 | 1724 | 1872 |
| 1752 | FIM1_1803 | QGN15116.1 | CP015056.1 | 9622 | 1254 | 1891 | 8044 | 659 | 1689 | 15146 | 1326 | 1746 |
| 1753 | FIM1_1804 | QGN15117.1 | CP015056.1 | 2277 | 294 | 478 | 1909 | 205 | 436 | 2331 | 365 | 410 |
| 1754 | FIM1_1805 | QGN15118.1 | CP015056.1 | 113 | 12 | 30 | 89 | 14 | 14 | 75 | 10 | 23 |
| 1755 | FIM1_1806 | QGN15119.1 | CP015056.1 | 3897 | 693 | 913 | 3244 | 470 | 878 | 4913 | 766 | 832 |
| 1756 | FIM1_1807 | QGN15120.1 | CP015056.1 | 609 | 80 | 73 | 500 | 57 | 60 | 702 | 79 | 76 |
| 1757 | FIM1_1808 | QGN15121.1 | CP015056.1 | 12110 | 935 | 1675 | 9592 | 753 | 1683 | 11045 | 1002 | 1686 |
| 1758 | FIM1_1809 | QGN15122.1 | CP015056.1 | 16306 | 1947 | 2749 | 12743 | 1211 | 2569 | 20407 | 1935 | 2859 |
| 1759 | FIM1_1810 | QGN15123.1 | CP015056.1 | 4910 | 826 | 1327 | 4140 | 558 | 1217 | 5583 | 939 | 1195 |
| 1760 | FIM1_1811 | QGN15124.1 | CP015056.1 | 3410 | 985 | 1326 | 3003 | 725 | 1190 | 3062 | 978 | 1160 |
| 1761 | FIM1_1812 | QGN15125.1 | CP015056.1 | 1905 | 197 | 609 | 1555 | 156 | 547 | 2168 | 227 | 501 |
| 1762 | FIM1_1813 | QGN15126.1 | CP015056.1 | 1347 | 286 | 453 | 1283 | 237 | 405 | 1273 | 281 | 343 |
| 1763 | FIM1_1814 | QGN15127.1 | CP015056.1 | 2428 | 438 | 919 | 2021 | 333 | 738 | 2345 | 495 | 719 |
| 1764 | FIM1_1815 | QGN15128.1 | CP015056.1 | 1861 | 277 | 390 | 1696 | 197 | 348 | 2409 | 269 | 378 |
| 1765 | FIM1_1816 | QGN15129.1 | CP015056.1 | 7239 | 1384 | 1919 | 5609 | 1084 | 1712 | 6297 | 1526 | 1585 |
| 1766 | FIM1_1817 | QGN15130.1 | CP015056.1 | 5846 | 1069 | 1513 | 4565 | 690 | 1380 | 5784 | 1150 | 1278 |
| 1767 | FIM1_1818 | QGN15131.1 | CP015056.1 | 46159 | 25999 | 20792 | 41380 | 19640 | 19202 | 46467 | 25530 | 20399 |
| 1768 | FIM1_1819 | QGN15132.1 | CP015056.1 | 8116 | 1706 | 2746 | 6475 | 1197 | 2433 | 8702 | 1717 | 2353 |
| 1769 | FIM1_1820 | QGN15133.1 | CP015056.1 | 1113 | 384 | 394 | 990 | 295 | 458 | 734 | 407 | 318 |
| 1770 | FIM1_1821 | QGN15134.1 | CP015056.1 | 3458 | 1520 | 1621 | 3062 | 1182 | 1515 | 2501 | 1722 | 1309 |
| 1771 | FIM1_1822 | QGN15135.1 | CP015056.1 | 1448 | 205 | 467 | 1248 | 147 | 400 | 1205 | 242 | 355 |
| 1772 | FIM1_1823 | QGN15136.1 | CP015056.1 | 317 | 152 | 197 | 521 | 182 | 229 | 215 | 177 | 205 |
| 1773 | FIM1_1824 | QGN15137.1 | CP015056.1 | 3606 | 603 | 1006 | 2973 | 497 | 909 | 3117 | 638 | 787 |
| 1774 | FIM1_1825 | QGN15138.1 | CP015056.1 | 7719 | 3191 | 4066 | 6806 | 2940 | 3961 | 6020 | 3329 | 3947 |

| | | | | | | | | | | | | |
|---|---|---|---|---|---|---|---|---|---|---|---|---|
| 1775 | FIM1_1826 | QGN15139.1 | CP015056.1 | 21391 | 3405 | 6127 | 17592 | 1792 | 5736 | 27066 | 3618 | 5065 |
| 1776 | FIM1_1827 | QGN15140.1 | CP015056.1 | 10521 | 3493 | 4776 | 9688 | 2784 | 3996 | 10651 | 3553 | 4354 |
| 1777 | FIM1_1828 | QGN15141.1 | CP015056.1 | 12087 | 1937 | 3238 | 9644 | 1284 | 3013 | 13482 | 2108 | 2714 |
| 1778 | FIM1_1829 | QGN15142.1 | CP015056.1 | 590 | 152 | 313 | 541 | 93 | 315 | 734 | 160 | 243 |
| 1779 | FIM1_1830 | QGN15143.1 | CP015056.1 | 5404 | 1091 | 1570 | 4285 | 921 | 1322 | 4303 | 1201 | 1337 |
| 1780 | FIM1_1831 | QGN15144.1 | CP015056.1 | 1028 | 482 | 522 | 705 | 360 | 476 | 762 | 508 | 420 |
| 1781 | FIM1_1832 | QGN15145.1 | CP015056.1 | 957 | 176 | 382 | 865 | 111 | 375 | 1211 | 230 | 336 |
| 1782 | FIM1_1833 | QGN15146.1 | CP015056.1 | 1833 | 465 | 721 | 1508 | 336 | 688 | 1793 | 433 | 628 |
| 1783 | FIM1_1834 | QGN15147.1 | CP015056.1 | 2932 | 272 | 706 | 2207 | 198 | 615 | 4644 | 290 | 570 |
| 1784 | FIM1_1835 | QGN15148.1 | CP015056.1 | 2005 | 272 | 718 | 1633 | 218 | 648 | 2149 | 282 | 581 |
| 1785 | FIM1_1836 | QGN15149.1 | CP015056.1 | 6843 | 1542 | 2188 | 5464 | 1251 | 2001 | 5316 | 1705 | 1746 |
| 1786 | FIM1_1837 | QGN15150.1 | CP015056.1 | 14507 | 5681 | 4132 | 11447 | 4738 | 3766 | 7875 | 6179 | 3982 |
| 1787 | FIM1_1838 | QGN15151.1 | CP015056.1 | 394 | 122 | 140 | 350 | 117 | 110 | 331 | 144 | 141 |
| 1788 | FIM1_1839 | QGN15152.1 | CP015056.1 | 5510 | 1911 | 3150 | 5217 | 1434 | 2983 | 5243 | 2021 | 2809 |
| 1789 | FIM1_1840 | QGN15153.1 | CP015056.1 | 19528 | 6156 | 5777 | 16360 | 4436 | 5173 | 19678 | 6436 | 5493 |
| 1790 | FIM1_1841 | QGN15154.1 | CP015056.1 | 2275 | 468 | 910 | 2064 | 342 | 887 | 2392 | 506 | 757 |
| 1791 | FIM1_1842 | QGN15155.1 | CP015056.1 | 4197 | 390 | 992 | 3034 | 307 | 935 | 4351 | 426 | 820 |
| 1792 | FIM1_1843 | QGN15156.1 | CP015056.1 | 26479 | 7028 | 6808 | 21104 | 5451 | 6336 | 21483 | 7941 | 6240 |
| 1793 | FIM1_1844 | QGN15157.1 | CP015056.1 | 135 | 50 | 49 | 134 | 47 | 62 | 136 | 54 | 55 |
| 1794 | FIM1_1845 | QGN15158.1 | CP015056.1 | 1394 | 350 | 533 | 1126 | 325 | 471 | 1065 | 384 | 450 |
| 1795 | FIM1_1846 | QGN15159.1 | CP015056.1 | 10223 | 3483 | 3071 | 8578 | 2335 | 2714 | 7063 | 3499 | 2664 |
| 1796 | FIM1_1847 | QGN15160.1 | CP015056.1 | 12157 | 2325 | 2751 | 10105 | 1234 | 2614 | 14082 | 2670 | 2463 |
| 1797 | FIM1_1848 | QGN15161.1 | CP015056.1 | 16511 | 5498 | 6209 | 14927 | 3604 | 6052 | 18634 | 5965 | 5877 |
| 1798 | FIM1_1849 | QGN15162.1 | CP015056.1 | 3211 | 1905 | 4654 | 3474 | 1111 | 4288 | 6950 | 2103 | 4372 |
| 1799 | FIM1_1850 | QGN15163.1 | CP015056.1 | 1830 | 397 | 718 | 1487 | 253 | 601 | 2408 | 460 | 579 |
| 1800 | FIM1_1851 | QGN15164.1 | CP015056.1 | 6119 | 2785 | 3602 | 5481 | 1854 | 3141 | 6239 | 2973 | 3315 |
| 1801 | FIM1_1853 | QGN15165.1 | CP015056.1 | 118049 | 50147 | 45321 | 105978 | 39156 | 41322 | 110233 | 50576 | 42922 |
| 1802 | FIM1_1854 | QGN15166.1 | CP015056.1 | 19198 | 5506 | 4184 | 16304 | 3666 | 3765 | 20785 | 5698 | 3925 |
| 1803 | FIM1_1855 | QGN15167.1 | CP015056.1 | 3264 | 446 | 580 | 2710 | 202 | 494 | 3591 | 411 | 456 |
| 1804 | FIM1_1856 | QGN15168.1 | CP015056.1 | 5612 | 749 | 1557 | 4649 | 556 | 1311 | 6654 | 845 | 1222 |
| 1805 | FIM1_1857 | QGN15169.1 | CP015056.1 | 24193 | 4452 | 5179 | 18219 | 3573 | 4630 | 20857 | 4814 | 4687 |
| 1806 | FIM1_1858 | QGN15170.1 | CP015056.1 | 2331 | 779 | 1004 | 2019 | 515 | 964 | 2984 | 809 | 874 |
| 1807 | FIM1_1859 | QGN15171.1 | CP015056.1 | 3465 | 1668 | 1799 | 2807 | 1289 | 1729 | 2018 | 1827 | 1509 |
| 1808 | FIM1_1860 | QGN15172.1 | CP015056.1 | 6045 | 580 | 1203 | 4307 | 440 | 1144 | 7372 | 597 | 1042 |
| 1809 | FIM1_1861 | QGN15173.1 | CP015056.1 | 1367 | 390 | 473 | 1087 | 419 | 461 | 1335 | 460 | 432 |
| 1810 | FIM1_1862 | QGN15174.1 | CP015056.1 | 3050 | 562 | 713 | 2444 | 467 | 652 | 2689 | 590 | 667 |
| 1811 | FIM1_1863 | QGN15175.1 | CP015056.1 | 1125 | 436 | 561 | 885 | 337 | 617 | 1072 | 431 | 576 |
| 1812 | FIM1_1864 | QGN15176.1 | CP015056.1 | 2784 | 646 | 948 | 2092 | 471 | 921 | 3489 | 669 | 864 |
| 1813 | FIM1_1865 | QGN15177.1 | CP015056.1 | 2768 | 644 | 892 | 1954 | 470 | 754 | 1873 | 705 | 719 |
| 1814 | FIM1_1866 | QGN15178.1 | CP015056.1 | 942 | 134 | 169 | 617 | 94 | 162 | 821 | 146 | 184 |
| 1815 | FIM1_1867 | QGN15179.1 | CP015056.1 | 922 | 120 | 139 | 656 | 43 | 146 | 1063 | 116 | 123 |
| 1816 | FIM1_1868 | QGN15180.1 | CP015056.1 | 4280 | 1339 | 1530 | 3608 | 842 | 1423 | 4387 | 1453 | 1447 |
| 1817 | FIM1_1869 | QGN15181.1 | CP015056.1 | 4252 | 1839 | 1848 | 3266 | 1352 | 1760 | 2899 | 1891 | 1748 |
| 1818 | FIM1_1870 | QGN15182.1 | CP015056.1 | 2967 | 609 | 1168 | 2230 | 473 | 1127 | 3004 | 675 | 1030 |
| 1819 | FIM1_1871 | QGN15183.1 | CP015056.1 | 5644 | 734 | 1232 | 4460 | 499 | 1113 | 7508 | 811 | 973 |
| 1820 | FIM1_1872 | QGN15184.1 | CP015056.1 | 3428 | 450 | 618 | 2425 | 281 | 536 | 2677 | 514 | 521 |
| 1821 | FIM1_1873 | QGN15185.1 | CP015056.1 | 2750 | 518 | 915 | 2097 | 403 | 950 | 2364 | 622 | 783 |
| 1822 | FIM1_1874 | QGN15186.1 | CP015056.1 | 7387 | 302 | 1599 | 5560 | 203 | 1419 | 6332 | 351 | 1190 |

| | | | | | | | | | | | | |
|---|---|---|---|---|---|---|---|---|---|---|---|---|
| 1823 | FIM1_1875 | QGN15187.1 | CP015056.1 | 226 | 9 | 53 | 173 | 4 | 57 | 200 | 6 | 40 |
| 1824 | FIM1_1876 | QGN15188.1 | CP015056.1 | 2575 | 467 | 519 | 2179 | 369 | 453 | 857 | 468 | 395 |
| 1825 | FIM1_1877 | QGN15189.1 | CP015056.1 | 2382 | 1862 | 913 | 1502 | 1028 | 691 | 428 | 1839 | 758 |
| 1826 | FIM1_1878 | QGN15190.1 | CP015056.1 | 11298 | 1335 | 2083 | 8679 | 806 | 1979 | 17607 | 1556 | 1827 |
| 1827 | FIM1_1879 | QGN15191.1 | CP015056.1 | 1051 | 231 | 395 | 967 | 175 | 398 | 1178 | 280 | 336 |
| 1828 | FIM1_1880 | QGN15192.1 | CP015056.1 | 6646 | 2528 | 2965 | 5698 | 1556 | 2682 | 9162 | 2576 | 2688 |
| 1829 | FIM1_1881 | QGN15193.1 | CP015056.1 | 2403 | 404 | 597 | 1811 | 326 | 574 | 2411 | 444 | 563 |
| 1830 | FIM1_1882 | QGN15194.1 | CP015056.1 | 10146 | 1736 | 2777 | 8653 | 1490 | 2403 | 10120 | 1987 | 2380 |
| 1831 | FIM1_1883 | QGN15195.1 | CP015056.1 | 688 | 61 | 102 | 457 | 56 | 94 | 627 | 57 | 84 |
| 1832 | FIM1_1884 | QGN15196.1 | CP015056.1 | 11526 | 2734 | 4045 | 9754 | 1612 | 3802 | 13866 | 2794 | 3617 |
| 1833 | FIM1_1885 | QGN15197.1 | CP015056.1 | 3233 | 940 | 1594 | 2552 | 611 | 1557 | 2617 | 977 | 1486 |
| 1834 | FIM1_1886 | QGN15198.1 | CP015056.1 | 2178 | 342 | 915 | 1547 | 292 | 803 | 2093 | 400 | 777 |
| 1835 | FIM1_1887 | QGN15199.1 | CP015056.1 | 11477 | 4772 | 5333 | 10189 | 3572 | 4967 | 10109 | 5042 | 4927 |
| 1836 | FIM1_1888 | QGN15200.1 | CP015056.1 | 4912 | 1646 | 2524 | 4383 | 1265 | 2392 | 4868 | 1784 | 2097 |
| 1837 | FIM1_1889 | QGN15201.1 | CP015056.1 | 4589 | 1211 | 1321 | 3413 | 765 | 1173 | 3195 | 1254 | 1164 |
| 1838 | FIM1_1890 | QGN15202.1 | CP015056.1 | 1825 | 2074 | 818 | 925 | 1062 | 744 | 273 | 2157 | 820 |
| 1839 | FIM1_1891 | QGN15203.1 | CP015056.1 | 6867 | 362 | 1414 | 5158 | 221 | 1207 | 6337 | 355 | 1112 |
| 1840 | FIM1_1892 | QGN15204.1 | CP015056.1 | 1734 | 577 | 881 | 1428 | 371 | 834 | 1602 | 537 | 813 |
| 1841 | FIM1_1893 | QGN15205.1 | CP015056.1 | 3706 | 1358 | 1379 | 3284 | 1063 | 1166 | 2967 | 1608 | 1305 |
| 1842 | FIM1_1894 | QGN15206.1 | CP015056.1 | 4853 | 602 | 1699 | 3356 | 473 | 1444 | 5129 | 653 | 1334 |
| 1843 | FIM1_1895 | QGN15207.1 | CP015056.1 | 2578 | 809 | 1104 | 2093 | 573 | 1014 | 2911 | 887 | 879 |
| 1844 | FIM1_1896 | QGN15208.1 | CP015056.1 | 2515 | 724 | 1092 | 1788 | 492 | 1041 | 2247 | 843 | 966 |
| 1845 | FIM1_1897 | QGN15209.1 | CP015056.1 | 3756 | 1481 | 1742 | 3052 | 1095 | 1585 | 2777 | 1645 | 1456 |
| 1846 | FIM1_1898 | QGN15210.1 | CP015056.1 | 4192 | 1084 | 1588 | 3625 | 923 | 1447 | 3893 | 1273 | 1301 |
| 1847 | FIM1_1899 | QGN15211.1 | CP015056.1 | 15110 | 6423 | 7029 | 14466 | 4200 | 6316 | 24957 | 6782 | 6097 |
| 1848 | FIM1_1900 | QGN15212.1 | CP015056.1 | 1050 | 249 | 397 | 734 | 167 | 378 | 802 | 260 | 341 |
| 1849 | FIM1_1901 | QGN15213.1 | CP015056.1 | 13123 | 2656 | 3513 | 10298 | 1771 | 3172 | 15379 | 2832 | 3135 |
| 1850 | FIM1_1902 | QGN15214.1 | CP015056.1 | 2047 | 681 | 673 | 1491 | 518 | 664 | 1186 | 747 | 628 |
| 1851 | FIM1_1903 | QGN15215.1 | CP015056.1 | 2626 | 669 | 579 | 1905 | 368 | 484 | 2210 | 668 | 503 |
| 1852 | FIM1_1904 | QGN15216.1 | CP015056.1 | 9142 | 5466 | 2592 | 7300 | 3345 | 2348 | 11698 | 5911 | 2307 |
| 1853 | FIM1_1905 | QGN15217.1 | CP015056.1 | 2431 | 640 | 974 | 1992 | 571 | 857 | 1821 | 747 | 815 |
| 1854 | FIM1_1907 | QGN15218.1 | CP015056.1 | 2080 | 939 | 1029 | 1934 | 725 | 1055 | 1620 | 1044 | 981 |
| 1855 | FIM1_1908 | QGN15219.1 | CP015056.1 | 6454 | 3973 | 3782 | 5731 | 3067 | 3657 | 6446 | 4009 | 3768 |
| 1856 | FIM1_1909 | QGN15220.1 | CP015056.1 | 13517 | 1900 | 3074 | 11210 | 1507 | 2924 | 16052 | 2167 | 2837 |
| 1857 | FIM1_1910 | QGN15221.1 | CP015056.1 | 3335 | 517 | 1573 | 2478 | 339 | 1570 | 3498 | 535 | 1400 |
| 1858 | FIM1_1911 | QGN15222.1 | CP015056.1 | 16027 | 6501 | 6300 | 13515 | 4861 | 5655 | 16059 | 6959 | 5834 |
| 1859 | FIM1_1912 | QGN15223.1 | CP015056.1 | 10228 | 2016 | 2510 | 7787 | 1528 | 2290 | 10881 | 2313 | 2157 |
| 1860 | FIM1_1913 | QGN15224.1 | CP015056.1 | 3531 | 3313 | 2163 | 2947 | 2497 | 2032 | 1294 | 3622 | 2120 |
| 1861 | FIM1_1914 | QGN15225.1 | CP015056.1 | 641 | 395 | 538 | 687 | 286 | 549 | 607 | 428 | 446 |
| 1862 | FIM1_1915 | QGN15226.1 | CP015056.1 | 9562 | 1454 | 1787 | 7057 | 1043 | 1737 | 6397 | 1609 | 1604 |
| 1863 | FIM1_1916 | QGN15227.1 | CP015056.1 | 3815 | 588 | 1238 | 2629 | 490 | 1004 | 1855 | 628 | 890 |
| 1864 | FIM1_1917 | QGN15228.1 | CP015056.1 | 1467 | 998 | 892 | 1166 | 691 | 832 | 539 | 1122 | 766 |
| 1865 | FIM1_1918 | QGN15229.1 | CP015056.1 | 1645 | 597 | 833 | 1633 | 405 | 790 | 1363 | 625 | 718 |
| 1866 | FIM1_1919 | QGN15230.1 | CP015056.1 | 4036 | 986 | 882 | 3009 | 737 | 850 | 4567 | 1034 | 847 |
| 1867 | FIM1_1920 | QGN15231.1 | CP015056.1 | 3408 | 4010 | 1629 | 2310 | 2528 | 1603 | 750 | 4274 | 1669 |
| 1868 | FIM1_1921 | QGN15232.1 | CP015056.1 | 3142 | 1589 | 2738 | 2215 | 1301 | 2505 | 2787 | 1761 | 2135 |
| 1869 | FIM1_1922 | QGN15233.1 | CP015056.1 | 5637 | 1494 | 2102 | 4188 | 1095 | 1846 | 4769 | 1567 | 1716 |
| 1870 | FIM1_1923 | QGN15234.1 | CP015056.1 | 5542 | 2707 | 2213 | 4951 | 1954 | 1898 | 4693 | 3037 | 1978 |

| | | | | | | | | | | | | |
|---|---|---|---|---|---|---|---|---|---|---|---|---|
| 1871 | FIM1_1924 | QGN15235.1 | CP015056.1 | 1603 | 858 | 990 | 1645 | 698 | 944 | 1298 | 1008 | 867 |
| 1872 | FIM1_1925 | QGN15236.1 | CP015056.1 | 5302 | 1124 | 1858 | 4008 | 853 | 1707 | 4169 | 1236 | 1660 |
| 1873 | FIM1_1926 | QGN15237.1 | CP015056.1 | 1233 | 349 | 502 | 1022 | 252 | 483 | 1205 | 371 | 402 |
| 1874 | FIM1_1927 | QGN15238.1 | CP015056.1 | 7643 | 5489 | 2824 | 5190 | 4034 | 2494 | 1968 | 6032 | 2668 |
| 1875 | FIM1_1928 | QGN15239.1 | CP015056.1 | 4903 | 808 | 1469 | 3595 | 721 | 1386 | 3964 | 967 | 1227 |
| 1876 | FIM1_1929 | QGN15240.1 | CP015056.1 | 2239 | 261 | 470 | 1583 | 213 | 463 | 1746 | 305 | 397 |
| 1877 | FIM1_1930 | QGN15241.1 | CP015056.1 | 1634 | 412 | 538 | 1451 | 410 | 584 | 1203 | 443 | 604 |
| 1878 | FIM1_1931 | QGN15242.1 | CP015056.1 | 16156 | 7622 | 13115 | 14155 | 5497 | 12164 | 18233 | 8104 | 12308 |
| 1879 | FIM1_1932 | QGN15243.1 | CP015056.1 | 1173 | 250 | 524 | 1027 | 205 | 442 | 1538 | 278 | 431 |
| 1880 | FIM1_1933 | QGN15244.1 | CP015056.1 | 15194 | 4019 | 4472 | 11325 | 2647 | 3949 | 14977 | 4242 | 3927 |
| 1881 | FIM1_1934 | QGN15245.1 | CP015056.1 | 1726 | 137 | 331 | 1513 | 111 | 281 | 1772 | 115 | 267 |
| 1882 | FIM1_1935 | QGN15246.1 | CP015056.1 | 8287 | 463 | 809 | 5403 | 371 | 779 | 6208 | 516 | 700 |
| 1883 | FIM1_1936 | QGN15247.1 | CP015056.1 | 16008 | 4786 | 4199 | 12718 | 3344 | 3915 | 14538 | 5228 | 3807 |
| 1884 | FIM1_1937 | QGN15248.1 | CP015056.1 | 4148 | 2330 | 2379 | 3763 | 1666 | 2249 | 4952 | 2477 | 2094 |
| 1885 | FIM1_1938 | QGN15249.1 | CP015056.1 | 1946 | 1058 | 1182 | 1879 | 859 | 1248 | 1477 | 1152 | 1041 |
| 1886 | FIM1_1939 | QGN15250.1 | CP015056.1 | 398 | 266 | 212 | 380 | 154 | 184 | 672 | 295 | 168 |
| 1887 | FIM1_1940 | QGN15251.1 | CP015056.1 | 8715 | 1694 | 2992 | 6707 | 1436 | 2658 | 7404 | 1830 | 2546 |
| 1888 | FIM1_1941 | QGN15252.1 | CP015056.1 | 5276 | 803 | 1166 | 4300 | 482 | 1092 | 7801 | 839 | 1040 |
| 1889 | FIM1_1942 | QGN15253.1 | CP015056.1 | 2354 | 439 | 708 | 2106 | 254 | 705 | 3764 | 421 | 615 |
| 1890 | FIM1_1943 | QGN15254.1 | CP015056.1 | 2707 | 724 | 947 | 1999 | 607 | 871 | 2533 | 821 | 960 |
| 1891 | FIM1_1944 | QGN15255.1 | CP015056.1 | 1587 | 293 | 447 | 1224 | 199 | 475 | 1499 | 290 | 397 |
| 1892 | FIM1_1945 | QGN15256.1 | CP015056.1 | 1673 | 451 | 668 | 1290 | 304 | 619 | 1794 | 493 | 620 |
| 1893 | FIM1_1946 | QGN15257.1 | CP015056.1 | 7455 | 907 | 1638 | 5829 | 828 | 1426 | 8792 | 965 | 1498 |
| 1894 | FIM1_1947 | QGN15258.1 | CP015056.1 | 2075 | 635 | 984 | 1699 | 508 | 935 | 2000 | 713 | 809 |
| 1895 | FIM1_1948 | QGN15259.1 | CP015056.1 | 5916 | 3328 | 1492 | 4821 | 2066 | 1377 | 5678 | 3451 | 1455 |
| 1896 | FIM1_1949 | QGN15260.1 | CP015056.1 | 753 | 240 | 424 | 551 | 195 | 409 | 667 | 316 | 340 |
| 1897 | FIM1_1950 | QGN15261.1 | CP015056.1 | 2673 | 480 | 1076 | 2243 | 383 | 970 | 2849 | 558 | 912 |
| 1898 | FIM1_1951 | QGN15262.1 | CP015056.1 | 2286 | 259 | 1346 | 2231 | 145 | 1146 | 3219 | 288 | 1121 |
| 1899 | FIM1_1952 | QGN15263.1 | CP015056.1 | 6516 | 2365 | 1762 | 5407 | 1418 | 1512 | 5858 | 2323 | 1504 |
| 1900 | FIM1_1953 | QGN15264.1 | CP015056.1 | 10980 | 1499 | 1887 | 8715 | 1086 | 1756 | 12495 | 1582 | 1659 |
| 1901 | FIM1_1954 | QGN15265.1 | CP015056.1 | 4666 | 3235 | 1642 | 3345 | 2139 | 1425 | 989 | 3758 | 1433 |
| 1902 | FIM1_1955 | QGN15266.1 | CP015056.1 | 2149 | 307 | 429 | 1585 | 247 | 428 | 2370 | 330 | 379 |
| 1903 | FIM1_1956 | QGN15267.1 | CP015056.1 | 1918 | 520 | 550 | 1210 | 475 | 526 | 1860 | 532 | 497 |
| 1904 | FIM1_1957 | QGN15268.1 | CP015056.1 | 2962 | 852 | 1026 | 2475 | 719 | 1063 | 2953 | 968 | 962 |
| 1905 | FIM1_1958 | QGN15269.1 | CP015056.1 | 3398 | 283 | 943 | 2784 | 198 | 838 | 3950 | 312 | 770 |
| 1906 | FIM1_1959 | QGN15270.1 | CP015056.1 | 2231 | 610 | 616 | 2056 | 451 | 582 | 2066 | 640 | 601 |
| 1907 | FIM1_1960 | QGN15271.1 | CP015056.1 | 7181 | 2027 | 2397 | 6202 | 1519 | 2114 | 6892 | 2163 | 2099 |
| 1908 | FIM1_1961 | QGN15272.1 | CP015056.1 | 17330 | 356 | 903 | 12099 | 443 | 794 | 15639 | 422 | 546 |
| 1909 | FIM1_1962 | QGN15273.1 | CP015056.1 | 74265 | 35908 | 31634 | 69985 | 27683 | 28516 | 67467 | 36231 | 30222 |
| 1910 | FIM1_1963 | QGN15274.1 | CP015056.1 | 1522 | 292 | 507 | 1437 | 274 | 459 | 1195 | 344 | 465 |
| 1911 | FIM1_1964 | QGN15275.1 | CP015056.1 | 29159 | 11944 | 9779 | 23955 | 8554 | 8629 | 25908 | 12356 | 9436 |
| 1912 | FIM1_1965 | QGN15276.1 | CP015056.1 | 4770 | 722 | 925 | 3863 | 520 | 850 | 4472 | 793 | 826 |
| 1913 | FIM1_1966 | QGN15277.1 | CP015056.1 | 3727 | 848 | 1125 | 2860 | 629 | 1086 | 4349 | 874 | 994 |
| 1914 | FIM1_1967 | QGN15278.1 | CP015056.1 | 7419 | 974 | 1706 | 5989 | 681 | 1532 | 9411 | 1012 | 1416 |
| 1915 | FIM1_1968 | QGN15279.1 | CP015056.1 | 7655 | 2451 | 1869 | 5356 | 1743 | 1742 | 3445 | 2769 | 1584 |
| 1916 | FIM1_1969 | QGN15280.1 | CP015056.1 | 0 | 1 | 1 | 1 | 0 | 0 | 1 | 0 | 0 |
| 1917 | FIM1_1970 | QGN15281.1 | CP015056.1 | 873 | 314 | 522 | 849 | 298 | 479 | 749 | 360 | 506 |
| 1918 | FIM1_1971 | QGN15282.1 | CP015056.1 | 872 | 239 | 400 | 890 | 157 | 382 | 815 | 241 | 361 |

| | | | | | | | | | | | | |
|---|---|---|---|---|---|---|---|---|---|---|---|---|
| 1919 | FIM1_1972 | QGN15283.1 | CP015056.1 | 5926 | 647 | 2109 | 5199 | 260 | 1734 | 16863 | 689 | 1663 |
| 1920 | FIM1_1973 | QGN15284.1 | CP015056.1 | 20740 | 7671 | 5598 | 16272 | 4923 | 4863 | 18998 | 8186 | 5400 |
| 1921 | FIM1_1974 | QGN15285.1 | CP015056.1 | 2 | 0 | 1 | 1 | 1 | 2 | 2 | 1 | 3 |
| 1922 | FIM1_1975 | QGN15286.1 | CP015056.1 | 2794 | 729 | 1252 | 2486 | 479 | 1149 | 2602 | 767 | 1192 |
| 1923 | FIM1_1976 | QGN15287.1 | CP015056.1 | 10668 | 1837 | 2046 | 8109 | 1412 | 1797 | 11029 | 1983 | 1745 |
| 1924 | FIM1_1977 | QGN15288.1 | CP015056.1 | 2808 | 587 | 723 | 2274 | 368 | 653 | 2524 | 598 | 608 |
| 1925 | FIM1_1978 | QGN15289.1 | CP015056.1 | 7291 | 2316 | 2065 | 6129 | 1538 | 1881 | 7848 | 2487 | 1986 |
| 1926 | FIM1_1979 | QGN15290.1 | CP015056.1 | 2562 | 321 | 716 | 2130 | 281 | 654 | 2614 | 359 | 539 |
| 1927 | FIM1_1980 | QGN15291.1 | CP015056.1 | 19911 | 4611 | 8174 | 18456 | 4163 | 7278 | 14787 | 5107 | 7044 |
| 1928 | FIM1_1981 | QGN15292.1 | CP015056.1 | 14624 | 8965 | 9016 | 13630 | 7857 | 8670 | 11256 | 9129 | 8831 |
| 1929 | FIM1_1982 | QGN15293.1 | CP015056.1 | 240 | 9 | 14 | 171 | 9 | 14 | 505 | 9 | 6 |
| 1930 | FIM1_1983 | QGN15294.1 | CP015056.1 | 11874 | 4209 | 3591 | 10195 | 2879 | 3413 | 12259 | 4429 | 3305 |
| 1931 | FIM1_1984 | QGN15295.1 | CP015056.1 | 1220 | 394 | 465 | 861 | 304 | 487 | 1162 | 412 | 376 |
| 1932 | FIM1_1985 | QGN15296.1 | CP015056.1 | 8820 | 1951 | 2479 | 6547 | 1480 | 2251 | 6114 | 2108 | 2169 |
| 1933 | FIM1_1986 | QGN15297.1 | CP015056.1 | 5332 | 1362 | 1762 | 4136 | 1088 | 1610 | 4932 | 1496 | 1459 |
| 1934 | FIM1_1987 | QGN15298.1 | CP015056.1 | 2153 | 629 | 730 | 1801 | 425 | 681 | 1841 | 694 | 600 |
| 1935 | FIM1_1988 | QGN15299.1 | CP015056.1 | 6998 | 1474 | 2416 | 6257 | 1252 | 2347 | 7459 | 1755 | 2094 |
| 1936 | FIM1_1989 | QGN15300.1 | CP015056.1 | 4301 | 1384 | 568 | 2773 | 738 | 520 | 4005 | 1429 | 631 |
| 1937 | FIM1_1990 | QGN15301.1 | CP015056.1 | 6020 | 2070 | 1525 | 4496 | 1452 | 1505 | 4734 | 2147 | 1445 |
| 1938 | FIM1_1991 | QGN15302.1 | CP015056.1 | 7964 | 1145 | 1874 | 5358 | 949 | 1722 | 6206 | 1176 | 1644 |
| 1939 | FIM1_1992 | QGN15303.1 | CP015056.1 | 14408 | 2831 | 3990 | 11504 | 2220 | 3559 | 13822 | 3107 | 3406 |
| 1940 | FIM1_1993 | QGN15304.1 | CP015056.1 | 4300 | 553 | 1558 | 3988 | 330 | 1584 | 5935 | 622 | 1351 |
| 1941 | FIM1_1994 | QGN15305.1 | CP015056.1 | 38380 | 16131 | 11654 | 32240 | 12126 | 10623 | 32308 | 17144 | 10804 |
| 1942 | FIM1_1995 | QGN15306.1 | CP015056.1 | 12955 | 2288 | 2760 | 10132 | 1660 | 2461 | 12325 | 2418 | 2435 |
| 1943 | FIM1_1996 | QGN15307.1 | CP015056.1 | 15562 | 9123 | 6682 | 12256 | 6376 | 6095 | 8475 | 9455 | 6108 |
| 1944 | FIM1_1997 | QGN15308.1 | CP015056.1 | 3139 | 1075 | 1281 | 2319 | 784 | 1172 | 3490 | 1212 | 1169 |
| 1945 | FIM1_1998 | QGN15309.1 | CP015056.1 | 4027 | 950 | 1267 | 3037 | 752 | 1185 | 3659 | 1040 | 1175 |
| 1946 | FIM1_1999 | QGN15310.1 | CP015056.1 | 1696 | 264 | 169 | 1430 | 149 | 170 | 2843 | 263 | 200 |
| 1947 | FIM1_2000 | QGN15311.1 | CP015056.1 | 12070 | 4775 | 4849 | 11150 | 3673 | 4696 | 10036 | 5066 | 4576 |
| 1948 | FIM1_2001 | QGN15312.1 | CP015056.1 | 6895 | 4528 | 2383 | 5165 | 2836 | 2083 | 2323 | 4721 | 2089 |
| 1949 | FIM1_2002 | QGN15313.1 | CP015056.1 | 7814 | 3355 | 2639 | 6028 | 2229 | 2390 | 5526 | 3562 | 2268 |
| 1950 | FIM1_2003 | QGN15314.1 | CP015056.1 | 12152 | 3702 | 4900 | 9506 | 2314 | 4677 | 10807 | 3995 | 4347 |
| 1951 | FIM1_2004 | QGN15315.1 | CP015056.1 | 1469 | 244 | 397 | 1089 | 261 | 363 | 1069 | 275 | 297 |
| 1952 | FIM1_2005 | QGN15316.1 | CP015056.1 | 82613 | 24925 | 27526 | 68294 | 22887 | 26735 | 68640 | 25741 | 26590 |
| 1953 | FIM1_2006 | QGN15317.1 | CP015056.1 | 4175 | 489 | 898 | 3156 | 409 | 821 | 3369 | 618 | 733 |
| 1954 | FIM1_2007 | QGN15318.1 | CP015056.1 | 1574 | 479 | 521 | 1101 | 365 | 509 | 1228 | 437 | 455 |
| 1955 | FIM1_2008 | QGN15319.1 | CP015056.1 | 1698 | 266 | 544 | 1252 | 222 | 428 | 1537 | 279 | 425 |
| 1956 | FIM1_2009 | QGN15320.1 | CP015056.1 | 584 | 132 | 225 | 514 | 114 | 233 | 1748 | 122 | 203 |
| 1957 | FIM1_2010 | QGN15321.1 | CP015056.1 | 238 | 19 | 31 | 184 | 16 | 20 | 144 | 33 | 21 |
| 1958 | FIM1_2011 | QGN15322.1 | CP015056.1 | 8574 | 1971 | 2106 | 6159 | 1505 | 1889 | 8494 | 2035 | 1831 |
| 1959 | FIM1_2012 | QGN15323.1 | CP015056.1 | 9099 | 2118 | 1894 | 6401 | 1280 | 1624 | 8480 | 2268 | 1506 |
| 1960 | FIM1_2013 | QGN15324.1 | CP015056.1 | 3030 | 246 | 896 | 2280 | 201 | 850 | 4602 | 252 | 798 |
| 1961 | FIM1_2014 | QGN15325.1 | CP015056.1 | 4622 | 708 | 1529 | 3974 | 553 | 1345 | 5585 | 750 | 1285 |
| 1962 | FIM1_2015 | QGN15326.1 | CP015056.1 | 24 | 7 | 14 | 22 | 5 | 18 | 17 | 10 | 22 |
| 1963 | FIM1_2016 | QGN15327.1 | CP015056.1 | 7125 | 544 | 2195 | 5685 | 473 | 2084 | 6865 | 588 | 1791 |
| 1964 | FIM1_2017 | QGN15328.1 | CP015056.1 | 4385 | 431 | 772 | 2854 | 405 | 732 | 2954 | 487 | 748 |
| 1965 | FIM1_2018 | QGN15329.1 | CP015056.1 | 263 | 24 | 94 | 240 | 31 | 111 | 387 | 32 | 75 |
| 1966 | FIM1_2019 | QGN15330.1 | CP015056.1 | 3624 | 833 | 1277 | 2438 | 585 | 1206 | 3604 | 858 | 1069 |

| | | | | | | | | | | | | |
|---|---|---|---|---|---|---|---|---|---|---|---|---|
| 1967 | FIM1_2020 | QGN15331.1 | CP015056.1 | 10245 | 1396 | 2310 | 8029 | 1303 | 2242 | 11471 | 1408 | 2152 |
| 1968 | FIM1_2021 | QGN15332.1 | CP015056.1 | 10926 | 3173 | 3424 | 8192 | 2300 | 3118 | 7600 | 3269 | 3019 |
| 1969 | FIM1_2022 | QGN15333.1 | CP015056.1 | 2902 | 1082 | 1488 | 2559 | 824 | 1432 | 2380 | 1182 | 1311 |
| 1970 | FIM1_2023 | QGN15334.1 | CP015056.1 | 8675 | 661 | 1837 | 7458 | 614 | 1616 | 8371 | 746 | 1471 |
| 1971 | FIM1_2024 | QGN15335.1 | CP015056.1 | 1998 | 84 | 158 | 1251 | 52 | 157 | 2756 | 62 | 97 |
| 1972 | FIM1_2025 | QGN15336.1 | CP015056.1 | 458 | 58 | 166 | 357 | 68 | 192 | 382 | 78 | 154 |
| 1973 | FIM1_2026 | QGN15337.1 | CP015056.1 | 3216 | 652 | 1108 | 2238 | 517 | 1063 | 2894 | 681 | 971 |
| 1974 | FIM1_2027 | QGN15338.1 | CP015056.1 | 1615 | 613 | 945 | 1364 | 495 | 860 | 1694 | 611 | 787 |
| 1975 | FIM1_2028 | QGN15339.1 | CP015056.1 | 4675 | 1801 | 1660 | 4086 | 1570 | 1595 | 2508 | 2012 | 1555 |
| 1976 | FIM1_2029 | QGN15340.1 | CP015056.1 | 4100 | 866 | 2394 | 3589 | 614 | 2213 | 4670 | 887 | 2064 |
| 1977 | FIM1_2030 | QGN15341.1 | CP015056.1 | 3088 | 219 | 369 | 2112 | 165 | 348 | 4812 | 237 | 305 |
| 1978 | FIM1_2031 | QGN15342.1 | CP015056.1 | 8538 | 253 | 1037 | 6171 | 274 | 889 | 10219 | 273 | 548 |
| 1979 | FIM1_2032 | QGN15343.1 | CP015056.1 | 151 | 15 | 13 | 106 | 10 | 5 | 225 | 11 | 5 |
| 1980 | FIM1_2033 | QGN15344.1 | CP015056.1 | 7622 | 760 | 1273 | 5765 | 608 | 1118 | 7751 | 886 | 1079 |
| 1981 | FIM1_2034 | QGN15345.1 | CP015056.1 | 4974 | 1261 | 1747 | 4240 | 858 | 1607 | 6309 | 1393 | 1429 |
| 1982 | FIM1_2035 | QGN15346.1 | CP015056.1 | 362 | 129 | 166 | 283 | 93 | 171 | 424 | 124 | 173 |
| 1983 | FIM1_2037 | QGN15347.1 | CP015056.1 | 472 | 120 | 407 | 386 | 104 | 375 | 788 | 136 | 316 |
| 1984 | FIM1_2038 | QGN15348.1 | CP015056.1 | 8466 | 1153 | 1886 | 5706 | 1142 | 1750 | 6357 | 1214 | 1606 |
| 1985 | FIM1_2039 | QGN15349.1 | CP015056.1 | 981 | 726 | 444 | 815 | 419 | 443 | 607 | 700 | 440 |
| 1986 | FIM1_2040 | QGN15350.1 | CP015056.1 | 734 | 545 | 275 | 637 | 394 | 289 | 549 | 544 | 301 |
| 1987 | FIM1_2041 | QGN15351.1 | CP015056.1 | 10748 | 10059 | 1461 | 7929 | 8524 | 1376 | 6812 | 11231 | 1229 |
| 1988 | FIM1_2042 | QGN15352.1 | CP015056.1 | 7782 | 2804 | 3004 | 6130 | 2106 | 2746 | 7692 | 2882 | 2727 |
| 1989 | FIM1_2043 | QGN15353.1 | CP015056.1 | 4453 | 2964 | 1812 | 3876 | 2288 | 1687 | 2377 | 3294 | 1718 |
| 1990 | FIM1_2044 | QGN15354.1 | CP015056.1 | 2215 | 372 | 760 | 1597 | 308 | 649 | 2552 | 371 | 594 |
| 1991 | FIM1_2045 | QGN15355.1 | CP015056.1 | 779 | 162 | 290 | 586 | 147 | 268 | 818 | 166 | 307 |
| 1992 | FIM1_2046 | QGN15356.1 | CP015056.1 | 454 | 130 | 256 | 378 | 112 | 255 | 793 | 141 | 233 |
| 1993 | FIM1_2047 | QGN15357.1 | CP015056.1 | 2020 | 411 | 543 | 1659 | 209 | 520 | 3123 | 400 | 455 |
| 1994 | FIM1_2048 | QGN15358.1 | CP015056.1 | 3377 | 1432 | 1662 | 3086 | 1031 | 1680 | 4073 | 1671 | 1556 |
| 1995 | FIM1_2049 | QGN15359.1 | CP015056.1 | 783 | 279 | 416 | 577 | 217 | 366 | 693 | 308 | 357 |
| 1996 | FIM1_2050 | QGN15360.1 | CP015056.1 | 6308 | 2692 | 2941 | 5483 | 1941 | 2801 | 6774 | 2873 | 2533 |
| 1997 | FIM1_2051 | QGN15361.1 | CP015056.1 | 4779 | 1487 | 1881 | 3997 | 955 | 1705 | 5459 | 1627 | 1590 |
| 1998 | FIM1_2052 | QGN15362.1 | CP015056.1 | 5025 | 1806 | 4174 | 4336 | 1130 | 3450 | 5806 | 1902 | 2664 |
| 1999 | FIM1_2053 | QGN15363.1 | CP015056.1 | 15371 | 6130 | 6453 | 13360 | 4254 | 5820 | 14925 | 6354 | 5856 |
| 2000 | FIM1_2054 | QGN15364.1 | CP015056.1 | 2238 | 841 | 1976 | 2190 | 553 | 1726 | 3059 | 773 | 1735 |
| 2001 | FIM1_2055 | QGN15365.1 | CP015056.1 | 3570 | 471 | 869 | 2859 | 590 | 699 | 2673 | 625 | 734 |
| 2002 | FIM1_2056 | QGN15366.1 | CP015056.1 | 3203 | 417 | 906 | 2909 | 502 | 736 | 2442 | 537 | 754 |
| 2003 | FIM1_2057 | QGN15367.1 | CP015056.1 | 1886 | 395 | 590 | 1514 | 336 | 639 | 1831 | 359 | 531 |
| 2004 | FIM1_2058 | QGN15368.1 | CP015056.1 | 1696 | 229 | 363 | 1343 | 182 | 402 | 1512 | 221 | 326 |
| 2005 | FIM1_2059 | QGN15369.1 | CP015056.1 | 1628 | 564 | 457 | 1255 | 529 | 442 | 1855 | 622 | 527 |
| 2006 | FIM1_2060 | QGN15370.1 | CP015056.1 | 5693 | 3405 | 3849 | 4634 | 1600 | 3582 | 9915 | 2967 | 3623 |
| 2007 | FIM1_2061 | QGN15371.1 | CP015056.1 | 4141 | 1430 | 1611 | 3666 | 1135 | 1622 | 3799 | 1517 | 1425 |
| 2008 | FIM1_2062 | QGN15372.1 | CP015056.1 | 4009 | 427 | 750 | 2943 | 328 | 589 | 6159 | 435 | 562 |
| 2009 | FIM1_2063 | QGN15373.1 | CP015056.1 | 11255 | 1261 | 2236 | 9056 | 994 | 2257 | 17188 | 1355 | 2007 |
| 2010 | FIM1_2064 | QGN15374.1 | CP015056.1 | 4716 | 143 | 484 | 3582 | 109 | 374 | 8435 | 160 | 448 |
| 2011 | FIM1_2065 | QGN15375.1 | CP015056.1 | 219 | 40 | 33 | 187 | 24 | 35 | 291 | 35 | 39 |
| 2012 | FIM1_2066 | QGN15376.1 | CP015056.1 | 6091 | 568 | 365 | 4262 | 236 | 356 | 13961 | 577 | 329 |
| 2013 | FIM1_2067 | QGN15377.1 | CP015056.1 | 6491 | 1129 | 2518 | 5758 | 589 | 2171 | 6479 | 1145 | 1901 |
| 2014 | FIM1_2068 | QGN15378.1 | CP015056.1 | 4564 | 844 | 959 | 3344 | 546 | 848 | 3939 | 900 | 879 |

| | | | | | | | | | | | | |
|---|---|---|---|---|---|---|---|---|---|---|---|---|
| 2015 | FIM1_2069 | QGN15379.1 | CP015056.1 | 4930 | 457 | 1121 | 3871 | 364 | 1002 | 5727 | 511 | 902 |
| 2016 | FIM1_2070 | QGN15380.1 | CP015056.1 | 149 | 31 | 67 | 151 | 36 | 56 | 92 | 58 | 72 |
| 2017 | FIM1_2071 | QGN15381.1 | CP015056.1 | 5214 | 1354 | 1712 | 4104 | 979 | 1655 | 5432 | 1499 | 1591 |
| 2018 | FIM1_2072 | QGN15382.1 | CP015056.1 | 2916 | 1109 | 1361 | 2561 | 887 | 1333 | 2206 | 1221 | 1160 |
| 2019 | FIM1_2073 | QGN15383.1 | CP015056.1 | 6097 | 3223 | 2998 | 5503 | 2802 | 2825 | 2773 | 3509 | 2536 |
| 2020 | FIM1_2074 | QGN15384.1 | CP015056.1 | 1819 | 568 | 706 | 1492 | 382 | 655 | 1814 | 601 | 620 |
| 2021 | FIM1_2075 | QGN15385.1 | CP015056.1 | 10989 | 3450 | 6456 | 9111 | 2252 | 5813 | 12657 | 3725 | 5991 |
| 2022 | FIM1_2076 | QGN15386.1 | CP015056.1 | 3276 | 760 | 1415 | 2773 | 523 | 1294 | 3364 | 847 | 1235 |
| 2023 | FIM1_2077 | QGN15387.1 | CP015056.1 | 7451 | 2081 | 2118 | 6082 | 1397 | 1954 | 7778 | 2292 | 1668 |
| 2024 | FIM1_2078 | QGN15388.1 | CP015056.1 | 4500 | 793 | 1428 | 4296 | 618 | 1159 | 4123 | 826 | 1180 |
| 2025 | FIM1_2079 | QGN15389.1 | CP015056.1 | 2040 | 116 | 4478 | 1874 | 119 | 3664 | 3240 | 141 | 2662 |
| 2026 | FIM1_2080 | QGN15390.1 | CP015056.1 | 169 | 16 | 29 | 126 | 15 | 11 | 134 | 27 | 20 |
| 2027 | FIM1_2081 | QGN15391.1 | CP015056.1 | 17980 | 16777 | 7918 | 14887 | 12503 | 6878 | 6812 | 18072 | 7476 |
| 2028 | FIM1_2082 | QGN15392.1 | CP015056.1 | 26 | 13 | 8 | 28 | 13 | 13 | 21 | 14 | 6 |
| 2029 | FIM1_2083 | QGN15393.1 | CP015056.1 | 2567 | 338 | 803 | 1949 | 242 | 637 | 2689 | 373 | 493 |
| 2030 | FIM1_2084 | QGN15394.1 | CP015056.1 | 12506 | 5370 | 6373 | 10165 | 3643 | 5784 | 12029 | 5664 | 5488 |
| 2031 | FIM1_2085 | QGN15395.1 | CP015056.1 | 6846 | 1578 | 2474 | 5448 | 1077 | 2354 | 7295 | 1691 | 2210 |
| 2032 | FIM1_2086 | QGN15396.1 | CP015056.1 | 7864 | 1837 | 3331 | 6952 | 1322 | 3097 | 7682 | 2120 | 2989 |
| 2033 | FIM1_2087 | QGN15397.1 | CP015056.1 | 9912 | 2955 | 3911 | 7661 | 2057 | 3642 | 8834 | 3014 | 3782 |
| 2034 | FIM1_2088 | QGN15398.1 | CP015056.1 | 4789 | 1643 | 1147 | 3355 | 1004 | 1083 | 3099 | 1768 | 1049 |
| 2035 | FIM1_2089 | QGN15399.1 | CP015056.1 | 672 | 178 | 260 | 623 | 146 | 276 | 703 | 218 | 238 |
| 2036 | FIM1_2090 | QGN15400.1 | CP015056.1 | 2307 | 498 | 661 | 1753 | 424 | 632 | 2099 | 539 | 590 |
| 2037 | FIM1_2091 | QGN15401.1 | CP015056.1 | 1759 | 1100 | 1055 | 1474 | 816 | 1051 | 1369 | 1189 | 937 |
| 2038 | FIM1_2092 | QGN15402.1 | CP015056.1 | 1069 | 659 | 607 | 712 | 505 | 623 | 272 | 733 | 545 |
| 2039 | FIM1_2093 | QGN15403.1 | CP015056.1 | 6440 | 5050 | 2531 | 4139 | 3095 | 2221 | 1454 | 5225 | 2378 |
| 2040 | FIM1_2094 | QGN15404.1 | CP015056.1 | 14784 | 8315 | 6668 | 12202 | 6425 | 6377 | 9545 | 8894 | 6176 |
| 2041 | FIM1_2095 | QGN15405.1 | CP015056.1 | 3027 | 208 | 746 | 2520 | 188 | 691 | 5159 | 230 | 588 |
| 2042 | FIM1_2096 | QGN15406.1 | CP015056.1 | 4970 | 293 | 597 | 4579 | 274 | 490 | 8702 | 273 | 394 |
| 2043 | FIM1_2097 | QGN15407.1 | CP015056.1 | 1582 | 638 | 1370 | 1300 | 450 | 1268 | 1443 | 720 | 1162 |
| 2044 | FIM1_2098 | QGN15408.1 | CP015056.1 | 993 | 3 | 157 | 634 | 6 | 120 | 2709 | 6 | 116 |
| 2045 | FIM1_2099 | QGN15409.1 | CP015056.1 | 26 | 12 | 15 | 22 | 15 | 16 | 11 | 12 | 11 |
| 2046 | FIM1_2100 | QGN15410.1 | CP015056.1 | 3607 | 1876 | 1612 | 2892 | 1375 | 1458 | 1862 | 2058 | 1420 |
| 2047 | FIM1_2101 | QGN15411.1 | CP015056.1 | 1914 | 608 | 687 | 1559 | 382 | 743 | 1289 | 631 | 662 |
| 2048 | FIM1_2102 | QGN15412.1 | CP015056.1 | 2690 | 571 | 971 | 2183 | 449 | 883 | 2758 | 635 | 887 |
| 2049 | FIM1_2103 | QGN15413.1 | CP015056.1 | 17309 | 3770 | 4641 | 14999 | 1814 | 4348 | 27084 | 3921 | 4126 |
| 2050 | FIM1_2104 | QGN15414.1 | CP015056.1 | 1071 | 590 | 558 | 1022 | 451 | 594 | 840 | 671 | 561 |
| 2051 | FIM1_2105 | QGN15415.1 | CP015056.1 | 905 | 311 | 493 | 695 | 266 | 458 | 636 | 327 | 439 |
| 2052 | FIM1_2106 | QGN15416.1 | CP015056.1 | 2356 | 277 | 790 | 1809 | 202 | 842 | 3552 | 300 | 643 |
| 2053 | FIM1_2107 | QGN15417.1 | CP015056.1 | 2618 | 573 | 692 | 2053 | 403 | 599 | 2041 | 605 | 561 |
| 2054 | FIM1_2108 | QGN15418.1 | CP015056.1 | 1657 | 276 | 618 | 1142 | 262 | 563 | 1397 | 310 | 515 |
| 2055 | FIM1_2109 | QGN15419.1 | CP015056.1 | 386 | 191 | 197 | 382 | 158 | 199 | 307 | 194 | 143 |
| 2056 | FIM1_2110 | QGN15420.1 | CP015056.1 | 1161 | 289 | 325 | 846 | 234 | 358 | 825 | 284 | 306 |
| 2057 | FIM1_2111 | QGN15421.1 | CP015056.1 | 141 | 27 | 39 | 112 | 21 | 36 | 137 | 28 | 41 |
| 2058 | FIM1_2112 | QGN15422.1 | CP015056.1 | 2299 | 345 | 876 | 1817 | 277 | 752 | 3093 | 371 | 685 |
| 2059 | FIM1_2113 | QGN15423.1 | CP015056.1 | 4208 | 659 | 1228 | 3334 | 531 | 1237 | 4176 | 780 | 1207 |
| 2060 | FIM1_2115 | QGN15424.1 | CP015056.1 | 2439 | 876 | 1056 | 2006 | 651 | 996 | 2199 | 902 | 976 |
| 2061 | FIM1_2116 | QGN15425.1 | CP015056.1 | 3248 | 554 | 970 | 2598 | 484 | 899 | 3241 | 584 | 930 |
| 2062 | FIM1_2117 | QGN15426.1 | CP015056.1 | 6512 | 1788 | 1951 | 5247 | 1414 | 1769 | 6170 | 1891 | 1762 |

| | | | | | | | | | | | | |
|---|---|---|---|---|---|---|---|---|---|---|---|---|
| 2063 | FIM1_2118 | QGN15427.1 | CP015056.1 | 4420 | 906 | 1165 | 3587 | 736 | 1092 | 4130 | 987 | 1041 |
| 2064 | FIM1_2119 | QGN15428.1 | CP015056.1 | 1323 | 988 | 810 | 888 | 617 | 743 | 310 | 1092 | 697 |
| 2065 | FIM1_2120 | QGN15429.1 | CP015056.1 | 2559 | 1174 | 1712 | 2359 | 872 | 1634 | 2765 | 1278 | 1555 |
| 2066 | FIM1_2121 | QGN15430.1 | CP015056.1 | 9038 | 2417 | 2862 | 7340 | 1803 | 2571 | 10843 | 2558 | 2604 |
| 2067 | FIM1_2122 | QGN15431.1 | CP015056.1 | 2790 | 898 | 1141 | 2176 | 609 | 1142 | 2766 | 1011 | 1026 |
| 2068 | FIM1_2123 | QGN15432.1 | CP015056.1 | 25528 | 4733 | 6689 | 19362 | 3403 | 6033 | 27081 | 5179 | 6079 |
| 2069 | FIM1_2124 | QGN15433.1 | CP015056.1 | 7375 | 617 | 1854 | 5561 | 519 | 1621 | 11200 | 606 | 1410 |
| 2070 | FIM1_2125 | QGN15434.1 | CP015056.1 | 3938 | 653 | 992 | 2814 | 479 | 818 | 4291 | 677 | 882 |
| 2071 | FIM1_2126 | QGN15435.1 | CP015056.1 | 2056 | 499 | 839 | 1595 | 305 | 832 | 2923 | 518 | 712 |
| 2072 | FIM1_2127 | QGN15436.1 | CP015056.1 | 4218 | 624 | 951 | 3445 | 461 | 839 | 4972 | 598 | 717 |
| 2073 | FIM1_2128 | QGN15437.1 | CP015056.1 | 1408 | 413 | 651 | 1156 | 404 | 645 | 1528 | 404 | 645 |
| 2074 | FIM1_2129 | QGN15438.1 | CP015056.1 | 5768 | 2754 | 2534 | 4552 | 2061 | 2518 | 4157 | 3144 | 2474 |
| 2075 | FIM1_2130 | QGN15439.1 | CP015056.1 | 6931 | 1733 | 1490 | 6144 | 1110 | 1417 | 16285 | 1776 | 1339 |
| 2076 | FIM1_2131 | QGN15440.1 | CP015056.1 | 19411 | 9638 | 9624 | 17419 | 6784 | 8851 | 19935 | 9728 | 9370 |
| 2077 | FIM1_2132 | QGN15441.1 | CP015056.1 | 3943 | 3713 | 1721 | 2380 | 2222 | 1569 | 648 | 3936 | 1514 |
| 2078 | FIM1_2133 | QGN15442.1 | CP015056.1 | 2396 | 900 | 851 | 1894 | 741 | 826 | 2235 | 1043 | 760 |
| 2079 | FIM1_2134 | QGN15443.1 | CP015056.1 | 1787 | 535 | 926 | 1515 | 434 | 882 | 1458 | 598 | 791 |
| 2080 | FIM1_2135 | QGN15444.1 | CP015056.1 | 5369 | 1994 | 2037 | 4350 | 1413 | 1901 | 4438 | 2194 | 1744 |
| 2081 | FIM1_2136 | QGN15445.1 | CP015056.1 | 4201 | 1186 | 1839 | 3117 | 939 | 1801 | 3742 | 1247 | 1626 |
| 2082 | FIM1_2137 | QGN15446.1 | CP015056.1 | 1293 | 650 | 810 | 1142 | 554 | 787 | 1071 | 689 | 734 |
| 2083 | FIM1_2138 | QGN15447.1 | CP015056.1 | 2948 | 710 | 950 | 2296 | 547 | 898 | 2178 | 778 | 825 |
| 2084 | FIM1_2139 | QGN15448.1 | CP015056.1 | 3209 | 2034 | 2609 | 2922 | 1340 | 2619 | 3946 | 2249 | 2388 |
| 2085 | FIM1_2140 | QGN15449.1 | CP015056.1 | 3268 | 502 | 931 | 2770 | 376 | 846 | 3697 | 577 | 721 |
| 2086 | FIM1_2141 | QGN15450.1 | CP015056.1 | 2839 | 1083 | 1050 | 2209 | 783 | 1197 | 2024 | 1190 | 1109 |
| 2087 | FIM1_2142 | QGN15451.1 | CP015056.1 | 447 | 83 | 184 | 325 | 52 | 148 | 531 | 89 | 135 |
| 2088 | FIM1_2143 | QGN15452.1 | CP015056.1 | 2011 | 560 | 828 | 1580 | 435 | 752 | 1527 | 570 | 746 |
| 2089 | FIM1_2144 | QGN15453.1 | CP015056.1 | 3652 | 767 | 1153 | 2663 | 552 | 1255 | 3361 | 837 | 1026 |
| 2090 | FIM1_2145 | QGN15454.1 | CP015056.1 | 1943 | 414 | 720 | 1699 | 274 | 659 | 3009 | 431 | 577 |
| 2091 | FIM1_2146 | QGN15455.1 | CP015056.1 | 19732 | 6688 | 12913 | 18417 | 5016 | 11643 | 23245 | 7111 | 11482 |
| 2092 | FIM1_2147 | QGN15456.1 | CP015056.1 | 1756 | 576 | 591 | 1265 | 550 | 528 | 1309 | 576 | 535 |
| 2093 | FIM1_2148 | QGN15457.1 | CP015056.1 | 1174 | 297 | 533 | 902 | 275 | 518 | 1259 | 327 | 417 |
| 2094 | FIM1_2149 | QGN15458.1 | CP015056.1 | 2487 | 611 | 845 | 1921 | 395 | 822 | 2001 | 696 | 771 |
| 2095 | FIM1_2150 | QGN15459.1 | CP015056.1 | 2462 | 684 | 1332 | 2295 | 361 | 1286 | 3730 | 737 | 1084 |
| 2096 | FIM1_2151 | QGN15460.1 | CP015056.1 | 2053 | 1277 | 624 | 1527 | 745 | 589 | 2068 | 1325 | 591 |
| 2097 | FIM1_2152 | QGN15461.1 | CP015056.1 | 655 | 178 | 157 | 560 | 151 | 140 | 836 | 224 | 148 |
| 2098 | FIM1_2153 | QGN15462.1 | CP015056.1 | 1197 | 182 | 380 | 865 | 132 | 421 | 1059 | 196 | 435 |
| 2099 | FIM1_2154 | QGN15463.1 | CP015056.1 | 1685 | 1923 | 1024 | 1248 | 1347 | 1072 | 267 | 1981 | 870 |
| 2100 | FIM1_2155 | QGN15464.1 | CP015056.1 | 2642 | 460 | 747 | 1910 | 356 | 645 | 2082 | 488 | 570 |
| 2101 | FIM1_2156 | QGN15465.1 | CP015056.1 | 1917 | 967 | 1190 | 1736 | 772 | 1233 | 1345 | 1019 | 1012 |
| 2102 | FIM1_2157 | QGN15466.1 | CP015056.1 | 8488 | 2014 | 4792 | 6513 | 1381 | 4408 | 7870 | 2078 | 3969 |
| 2103 | FIM1_2158 | QGN15467.1 | CP015056.1 | 7950 | 2649 | 2232 | 6084 | 1951 | 2078 | 6225 | 2879 | 2150 |
| 2104 | FIM1_2159 | QGN15468.1 | CP015056.1 | 220 | 42 | 58 | 191 | 26 | 58 | 286 | 51 | 48 |
| 2105 | FIM1_2160 | QGN15469.1 | CP015056.1 | 3218 | 1028 | 1487 | 3059 | 786 | 1433 | 3877 | 1033 | 1399 |
| 2106 | FIM1_2161 | QGN15470.1 | CP015056.1 | 3629 | 714 | 1455 | 2872 | 595 | 1260 | 3635 | 790 | 1246 |
| 2107 | FIM1_2162 | QGN15471.1 | CP015056.1 | 1404 | 550 | 557 | 1011 | 417 | 533 | 1088 | 643 | 563 |
| 2108 | FIM1_2163 | QGN15472.1 | CP015056.1 | 1460 | 355 | 709 | 1261 | 306 | 642 | 2136 | 391 | 625 |
| 2109 | FIM1_2164 | QGN15473.1 | CP015056.1 | 1842 | 432 | 736 | 1631 | 395 | 667 | 1606 | 494 | 627 |
| 2110 | FIM1_2165 | QGN15474.1 | CP015056.1 | 7196 | 2241 | 2248 | 5773 | 1641 | 1912 | 4902 | 2519 | 1861 |

| | | | | | | | | | | | | |
|---|---|---|---|---|---|---|---|---|---|---|---|---|
| 2111 | FIM1_2166 | QGN15475.1 | CP015056.1 | 892 | 253 | 335 | 619 | 151 | 324 | 882 | 235 | 310 |
| 2112 | FIM1_2167 | QGN15476.1 | CP015056.1 | 11577 | 2926 | 4619 | 9759 | 1813 | 4232 | 13929 | 3037 | 4051 |
| 2113 | FIM1_2168 | QGN15477.1 | CP015056.1 | 9816 | 3893 | 3981 | 7631 | 2909 | 3524 | 8225 | 4254 | 3570 |
| 2114 | FIM1_2169 | QGN15478.1 | CP015056.1 | 7380 | 1784 | 2253 | 5677 | 1283 | 1949 | 5863 | 2073 | 1836 |
| 2115 | FIM1_2170 | QGN15479.1 | CP015056.1 | 7787 | 1733 | 3480 | 6665 | 1060 | 3083 | 9983 | 1868 | 2790 |
| 2116 | FIM1_2171 | QGN15480.1 | CP015056.1 | 2856 | 724 | 1330 | 2336 | 542 | 1135 | 1996 | 813 | 1137 |
| 2117 | FIM1_2172 | QGN15481.1 | CP015056.1 | 6403 | 1257 | 1332 | 5637 | 1056 | 1230 | 6750 | 1482 | 1182 |
| 2118 | FIM1_2173 | QGN15482.1 | CP015056.1 | 1534 | 533 | 854 | 1263 | 359 | 948 | 1274 | 615 | 736 |
| 2119 | FIM1_2174 | QGN15483.1 | CP015056.1 | 2660 | 628 | 864 | 2185 | 356 | 894 | 4469 | 735 | 807 |
| 2120 | FIM1_2175 | QGN15484.1 | CP015056.1 | 11093 | 2664 | 3030 | 8762 | 2025 | 2738 | 10335 | 2826 | 2692 |
| 2121 | FIM1_2176 | QGN15485.1 | CP015056.1 | 1630 | 169 | 1090 | 1283 | 139 | 1023 | 2279 | 201 | 866 |
| 2122 | FIM1_2177 | QGN15486.1 | CP015056.1 | 190 | 44 | 46 | 127 | 20 | 37 | 189 | 51 | 31 |
| 2123 | FIM1_2178 | QGN15487.1 | CP015056.1 | 9548 | 2808 | 4975 | 7285 | 1573 | 4607 | 10705 | 2650 | 3963 |
| 2124 | FIM1_2179 | QGN15488.1 | CP015056.1 | 21275 | 5077 | 8682 | 17477 | 3467 | 7801 | 26012 | 5540 | 7886 |
| 2125 | FIM1_2180 | QGN15489.1 | CP015056.1 | 5993 | 4083 | 2252 | 4345 | 2604 | 2039 | 1349 | 4306 | 2138 |
| 2126 | FIM1_2181 | QGN15490.1 | CP015056.1 | 7819 | 2644 | 2368 | 6753 | 1906 | 2007 | 5639 | 2958 | 2251 |
| 2127 | FIM1_2182 | QGN15491.1 | CP015056.1 | 1659 | 1226 | 557 | 1472 | 319 | 528 | 3724 | 501 | 460 |
| 2128 | FIM1_2183 | QGN15492.1 | CP015056.1 | 8129 | 1465 | 1873 | 6140 | 1082 | 1774 | 9252 | 1592 | 1577 |
| 2129 | FIM1_2184 | QGN15493.1 | CP015056.1 | 8450 | 1768 | 2607 | 7362 | 1270 | 2382 | 9615 | 1921 | 2390 |
| 2130 | FIM1_2185 | QGN15494.1 | CP015056.1 | 8684 | 1618 | 1800 | 6821 | 875 | 1630 | 13344 | 1873 | 1477 |
| 2131 | FIM1_2186 | QGN15495.1 | CP015056.1 | 3086 | 1263 | 1435 | 2652 | 850 | 1469 | 2280 | 1318 | 1277 |
| 2132 | FIM1_2187 | QGN15496.1 | CP015056.1 | 3884 | 2068 | 1423 | 2961 | 1395 | 1256 | 1544 | 2145 | 1287 |
| 2133 | FIM1_2188 | QGN15497.1 | CP015056.1 | 10217 | 2951 | 3506 | 7948 | 2215 | 3300 | 10855 | 3235 | 3234 |
| 2134 | FIM1_2189 | QGN15498.1 | CP015056.1 | 1602 | 868 | 872 | 1322 | 637 | 874 | 951 | 936 | 765 |
| 2135 | FIM1_2190 | QGN15499.1 | CP015056.1 | 7243 | 1288 | 1822 | 5747 | 937 | 1613 | 6830 | 1352 | 1493 |
| 2136 | FIM1_2191 | QGN15500.1 | CP015056.1 | 17226 | 6463 | 6939 | 14094 | 4176 | 6714 | 16225 | 6838 | 6557 |
| 2137 | FIM1_2192 | QGN15501.1 | CP015056.1 | 1839 | 718 | 809 | 1182 | 473 | 761 | 1101 | 772 | 713 |
| 2138 | FIM1_2193 | QGN15502.1 | CP015056.1 | 7765 | 6800 | 3128 | 5595 | 4268 | 2791 | 1931 | 7036 | 2897 |
| 2139 | FIM1_2194 | QGN15503.1 | CP015056.1 | 1381 | 689 | 589 | 1013 | 443 | 571 | 520 | 841 | 479 |
| 2140 | FIM1_2195 | QGN15504.1 | CP015056.1 | 1688 | 558 | 642 | 1431 | 427 | 598 | 1582 | 572 | 512 |
| 2141 | FIM1_2196 | QGN15505.1 | CP015056.1 | 1121 | 202 | 267 | 852 | 124 | 216 | 848 | 206 | 222 |
| 2142 | FIM1_2197 | QGN15506.1 | CP015056.1 | 3537 | 963 | 1117 | 2984 | 739 | 1066 | 3119 | 1086 | 1126 |
| 2143 | FIM1_2198 | QGN15507.1 | CP015056.1 | 5106 | 1459 | 1465 | 4198 | 1184 | 1316 | 5480 | 1285 | 1729 |
| 2144 | FIM1_2199 | QGN15508.1 | CP015056.1 | 343 | 145 | 80 | 284 | 56 | 63 | 224 | 138 | 58 |
| 2145 | FIM1_2200 | QGN15509.1 | CP015056.1 | 2686 | 1060 | 1293 | 2619 | 656 | 1186 | 2856 | 1148 | 1075 |
| 2146 | FIM1_2201 | QGN15510.1 | CP015056.1 | 2909 | 790 | 784 | 2083 | 546 | 733 | 2449 | 831 | 678 |
| 2147 | FIM1_2202 | QGN15511.1 | CP015056.1 | 11221 | 1950 | 4413 | 8707 | 1463 | 4210 | 12337 | 2064 | 4241 |
| 2148 | FIM1_2203 | QGN15512.1 | CP015056.1 | 2279 | 312 | 937 | 1761 | 248 | 796 | 2818 | 319 | 801 |
| 2149 | FIM1_2204 | QGN15513.1 | CP015056.1 | 2693 | 248 | 923 | 2303 | 247 | 869 | 3181 | 315 | 758 |
| 2150 | FIM1_2205 | QGN15514.1 | CP015056.1 | 2376 | 524 | 829 | 1543 | 358 | 766 | 1703 | 546 | 711 |
| 2151 | FIM1_2206 | QGN15515.1 | CP015056.1 | 1857 | 1754 | 848 | 1552 | 1202 | 779 | 415 | 1806 | 849 |
| 2152 | FIM1_2207 | QGN15516.1 | CP015056.1 | 2655 | 180 | 394 | 2108 | 138 | 342 | 2380 | 233 | 329 |
| 2153 | FIM1_2208 | QGN15517.1 | CP015056.1 | 2 | 0 | 0 | 7 | 0 | 0 | 2 | 1 | 0 |
| 2154 | FIM1_2209 | QGN15518.1 | CP015056.1 | 4540 | 908 | 1363 | 3329 | 1017 | 1348 | 4306 | 1052 | 1307 |
| 2155 | FIM1_2210 | QGN15519.1 | CP015056.1 | 9058 | 7479 | 3379 | 6422 | 4974 | 2968 | 2850 | 7669 | 3141 |
| 2156 | FIM1_2211 | QGN15520.1 | CP015056.1 | 7969 | 3618 | 3867 | 7054 | 2902 | 3530 | 6802 | 3822 | 3534 |
| 2157 | FIM1_2212 | QGN15521.1 | CP015056.1 | 2081 | 418 | 1100 | 1535 | 319 | 985 | 2352 | 433 | 935 |
| 2158 | FIM1_2213 | QGN15522.1 | CP015056.1 | 2025 | 683 | 750 | 1560 | 444 | 715 | 1439 | 687 | 665 |

| | | | | | | | | | | | | |
|---|---|---|---|---|---|---|---|---|---|---|---|---|
| 2159 | FIM1_2214 | QGN15523.1 | CP015056.1 | 1576 | 351 | 665 | 1208 | 257 | 541 | 1160 | 422 | 508 |
| 2160 | FIM1_2215 | QGN15524.1 | CP015056.1 | 10818 | 6660 | 5016 | 8741 | 5541 | 4883 | 5516 | 7077 | 4818 |
| 2161 | FIM1_2216 | QGN15525.1 | CP015056.1 | 2400 | 631 | 894 | 1730 | 484 | 922 | 1635 | 623 | 771 |
| 2162 | FIM1_2217 | QGN15526.1 | CP015056.1 | 3960 | 356 | 662 | 2972 | 306 | 589 | 4739 | 434 | 517 |
| 2163 | FIM1_2218 | QGN15527.1 | CP015056.1 | 16047 | 5288 | 8386 | 13264 | 3612 | 7764 | 18489 | 5531 | 7677 |
| 2164 | FIM1_2219 | QGN15528.1 | CP015056.1 | 5955 | 2408 | 2263 | 4751 | 1767 | 2076 | 4133 | 2519 | 2054 |
| 2165 | FIM1_2220 | QGN15529.1 | CP015056.1 | 4289 | 915 | 1378 | 3286 | 670 | 1281 | 4183 | 974 | 1140 |
| 2166 | FIM1_2221 | QGN15530.1 | CP015056.1 | 1878 | 502 | 755 | 1600 | 331 | 749 | 1940 | 538 | 762 |
| 2167 | FIM1_2222 | QGN15531.1 | CP015056.1 | 1246 | 255 | 537 | 1033 | 206 | 522 | 2028 | 258 | 458 |
| 2168 | FIM1_2223 | QGN15532.1 | CP015056.1 | 3431 | 637 | 870 | 2443 | 464 | 838 | 2896 | 667 | 732 |
| 2169 | FIM1_2224 | QGN15533.1 | CP015056.1 | 6866 | 3678 | 3117 | 5577 | 2933 | 3042 | 4160 | 4004 | 3078 |
| 2170 | FIM1_2225 | QGN15534.1 | CP015056.1 | 6588 | 654 | 848 | 4751 | 543 | 738 | 5986 | 893 | 746 |
| 2171 | FIM1_2226 | QGN15535.1 | CP015056.1 | 19945 | 8847 | 8381 | 17190 | 6025 | 7747 | 21853 | 9132 | 7851 |
| 2172 | FIM1_2227 | QGN15536.1 | CP015056.1 | 7413 | 2395 | 1353 | 5546 | 1403 | 1154 | 4524 | 2710 | 1204 |
| 2173 | FIM1_2228 | QGN15537.1 | CP015056.1 | 3707 | 628 | 834 | 2851 | 439 | 803 | 3457 | 654 | 744 |
| 2174 | FIM1_2229 | QGN15538.1 | CP015056.1 | 2725 | 698 | 699 | 2602 | 555 | 581 | 2147 | 786 | 669 |
| 2175 | FIM1_2230 | QGN15539.1 | CP015056.1 | 1477 | 196 | 248 | 1185 | 93 | 235 | 1854 | 139 | 202 |
| 2176 | FIM1_2231 | QGN15540.1 | CP015056.1 | 2598 | 527 | 1003 | 1759 | 407 | 960 | 2441 | 504 | 826 |
| 2177 | FIM1_2232 | QGN15541.1 | CP015056.1 | 2667 | 271 | 235 | 1437 | 203 | 253 | 1768 | 253 | 220 |
| 2178 | FIM1_2233 | QGN15542.1 | CP015056.1 | 2934 | 1612 | 1532 | 2406 | 1071 | 1475 | 1815 | 1744 | 1312 |
| 2179 | FIM1_2234 | QGN15543.1 | CP015056.1 | 2076 | 207 | 483 | 1600 | 161 | 451 | 1939 | 228 | 392 |
| 2180 | FIM1_2235 | QGN15544.1 | CP015056.1 | 1105 | 404 | 516 | 993 | 308 | 551 | 656 | 399 | 481 |
| 2181 | FIM1_2236 | QGN15545.1 | CP015056.1 | 2301 | 658 | 1389 | 1497 | 498 | 1268 | 1649 | 760 | 1113 |
| 2182 | FIM1_2237 | QGN15546.1 | CP015056.1 | 96 | 9 | 22 | 69 | 9 | 20 | 91 | 14 | 13 |
| 2183 | FIM1_2238 | QGN15547.1 | CP015056.1 | 1656 | 414 | 646 | 1386 | 287 | 647 | 2430 | 437 | 605 |
| 2184 | FIM1_2239 | QGN15548.1 | CP015056.1 | 1196 | 269 | 883 | 928 | 247 | 855 | 952 | 319 | 727 |
| 2185 | FIM1_2240 | QGN15549.1 | CP015056.1 | 1412 | 291 | 713 | 1253 | 232 | 762 | 1346 | 379 | 610 |
| 2186 | FIM1_2241 | QGN15550.1 | CP015056.1 | 6452 | 1472 | 2604 | 5886 | 1127 | 2277 | 7303 | 1722 | 2019 |
| 2187 | FIM1_2242 | QGN15551.1 | CP015056.1 | 32 | 4 | 7 | 19 | 2 | 11 | 25 | 3 | 6 |
| 2188 | FIM1_2243 | QGN15552.1 | CP015056.1 | 2755 | 859 | 993 | 2297 | 702 | 921 | 1725 | 891 | 839 |
| 2189 | FIM1_2244 | QGN15553.1 | CP015056.1 | 14653 | 3783 | 3919 | 11311 | 2942 | 3525 | 11295 | 4192 | 3518 |
| 2190 | FIM1_2245 | QGN15554.1 | CP015056.1 | 3633 | 1266 | 1048 | 3148 | 647 | 978 | 5314 | 1425 | 974 |
| 2191 | FIM1_2246 | QGN15555.1 | CP015056.1 | 25814 | 10404 | 10389 | 21317 | 7973 | 9718 | 20513 | 11010 | 9704 |
| 2192 | FIM1_2247 | QGN15556.1 | CP015056.1 | 2655 | 695 | 1022 | 1947 | 482 | 963 | 2832 | 736 | 785 |
| 2193 | FIM1_2248 | QGN15557.1 | CP015056.1 | 3517 | 4279 | 2083 | 2382 | 2690 | 1814 | 795 | 4669 | 1988 |
| 2194 | FIM1_2249 | QGN15558.1 | CP015056.1 | 9789 | 6060 | 3372 | 6356 | 3739 | 3110 | 2575 | 6370 | 2969 |
| 2195 | FIM1_2250 | QGN15559.1 | CP015056.1 | 264 | 584 | 359 | 199 | 332 | 339 | 104 | 582 | 303 |
| 2196 | FIM1_2251 | QGN15560.1 | CP015056.1 | 1851 | 336 | 406 | 1415 | 225 | 371 | 1515 | 332 | 363 |
| 2197 | FIM1_2252 | QGN15561.1 | CP015056.1 | 9006 | 2190 | 3168 | 7750 | 1350 | 2991 | 11184 | 2225 | 2791 |
| 2198 | FIM1_2253 | QGN15562.1 | CP015056.1 | 45586 | 14721 | 12134 | 35934 | 11013 | 11130 | 39095 | 15658 | 11319 |
| 2199 | FIM1_2254 | QGN15563.1 | CP015056.1 | 4201 | 1711 | 2527 | 3492 | 1219 | 2410 | 4835 | 1871 | 2278 |
| 2200 | FIM1_2255 | QGN15564.1 | CP015056.1 | 26828 | 12285 | 11830 | 22693 | 8455 | 10681 | 29245 | 12355 | 11266 |
| 2201 | FIM1_2256 | QGN15565.1 | CP015056.1 | 1107 | 504 | 1486 | 964 | 308 | 1168 | 1816 | 529 | 1221 |
| 2202 | FIM1_2257 | QGN15566.1 | CP015056.1 | 2509 | 320 | 910 | 1930 | 89 | 682 | 7764 | 257 | 661 |
| 2203 | FIM1_2258 | QGN15567.1 | CP015056.1 | 4493 | 1449 | 1252 | 3845 | 1526 | 1277 | 4276 | 1702 | 1370 |
| 2204 | FIM1_2259 | QGN15568.1 | CP015056.1 | 769 | 197 | 389 | 657 | 189 | 399 | 760 | 277 | 307 |
| 2205 | FIM1_2260 | QGN15569.1 | CP015056.1 | 4034 | 1090 | 1574 | 3012 | 777 | 1515 | 4469 | 1233 | 1346 |
| 2206 | FIM1_2261 | QGN15570.1 | CP015056.1 | 6903 | 2567 | 2726 | 5128 | 1919 | 2602 | 5835 | 2815 | 2594 |

| | | | | | | | | | | | | |
|---|---|---|---|---|---|---|---|---|---|---|---|---|
| 2207 | FIM1_2262 | QGN15571.1 | CP015056.1 | 4030 | 1552 | 2682 | 3729 | 990 | 2568 | 5921 | 1676 | 2405 |
| 2208 | FIM1_2263 | QGN15572.1 | CP015056.1 | 6489 | 2855 | 4731 | 5924 | 1868 | 4740 | 7684 | 3104 | 4552 |
| 2209 | FIM1_2264 | QGN15573.1 | CP015056.1 | 3388 | 867 | 1113 | 2881 | 669 | 1069 | 3870 | 988 | 1015 |
| 2210 | FIM1_2265 | QGN15574.1 | CP015056.1 | 3310 | 706 | 842 | 2623 | 488 | 753 | 3199 | 684 | 788 |
| 2211 | FIM1_2266 | QGN15575.1 | CP015056.1 | 3224 | 1140 | 1411 | 3010 | 799 | 1294 | 2829 | 1371 | 1181 |
| 2212 | FIM1_2267 | QGN15576.1 | CP015056.1 | 3912 | 1118 | 1313 | 3141 | 923 | 1170 | 5464 | 1165 | 1325 |
| 2213 | FIM1_2268 | QGN15577.1 | CP015056.1 | 3053 | 1064 | 1191 | 2060 | 816 | 1209 | 2414 | 1129 | 1050 |
| 2214 | FIM1_2270 | QGN15578.1 | CP015056.1 | 1674 | 710 | 817 | 1298 | 488 | 730 | 772 | 814 | 626 |
| 2215 | FIM1_2271 | QGN15579.1 | CP015056.1 | 3635 | 896 | 1026 | 2653 | 835 | 1073 | 3256 | 938 | 1057 |
| 2216 | FIM1_2272 | QGN15580.1 | CP015056.1 | 1625 | 203 | 1023 | 1423 | 119 | 897 | 1881 | 179 | 827 |
| 2217 | FIM1_2273 | QGN15581.1 | CP015056.1 | 322 | 49 | 170 | 270 | 30 | 149 | 387 | 64 | 141 |
| 2218 | FIM1_2274 | QGN15582.1 | CP015056.1 | 4753 | 691 | 2759 | 4139 | 597 | 2127 | 9560 | 750 | 2044 |
| 2219 | FIM1_2275 | QGN15583.1 | CP015056.1 | 3684 | 1399 | 1580 | 3247 | 752 | 1450 | 5569 | 1426 | 1407 |
| 2220 | FIM1_2276 | QGN15584.1 | CP015056.1 | 2590 | 542 | 721 | 2096 | 449 | 610 | 2211 | 581 | 561 |
| 2221 | FIM1_2277 | QGN15585.1 | CP015056.1 | 20 | 6 | 4 | 10 | 5 | 4 | 8 | 8 | 3 |
| 2222 | FIM1_2278 | QGN15586.1 | CP015056.1 | 5911 | 1115 | 1491 | 4338 | 738 | 1341 | 5663 | 1227 | 1349 |
| 2223 | FIM1_2279 | QGN15587.1 | CP015056.1 | 928 | 174 | 185 | 698 | 151 | 188 | 1062 | 170 | 168 |
| 2224 | FIM1_2280 | QGN15588.1 | CP015056.1 | 4857 | 1196 | 2028 | 4404 | 839 | 2010 | 5704 | 1318 | 1889 |
| 2225 | FIM1_2281 | QGN15589.1 | CP015056.1 | 2726 | 248 | 551 | 1991 | 272 | 460 | 2011 | 272 | 499 |
| 2226 | FIM1_2282 | QGN15590.1 | CP015056.1 | 877 | 63 | 299 | 811 | 50 | 276 | 1149 | 89 | 220 |
| 2227 | FIM1_2283 | QGN15591.1 | CP015056.1 | 2029 | 431 | 1807 | 2006 | 342 | 1480 | 2070 | 475 | 1241 |
| 2228 | FIM1_2284 | QGN15592.1 | CP015056.1 | 9084 | 1129 | 2255 | 6750 | 831 | 1990 | 9798 | 1259 | 1792 |
| 2229 | FIM1_2285 | QGN15593.1 | CP015056.1 | 10392 | 2712 | 4170 | 8921 | 1406 | 3786 | 11407 | 2570 | 3329 |
| 2230 | FIM1_2286 | QGN15594.1 | CP015056.1 | 1806 | 639 | 820 | 1601 | 458 | 834 | 2137 | 685 | 735 |
| 2231 | FIM1_2287 | QGN15595.1 | CP015056.1 | 2309 | 394 | 742 | 1977 | 329 | 736 | 2776 | 447 | 568 |
| 2232 | FIM1_2288 | QGN15596.1 | CP015056.1 | 4768 | 1412 | 1604 | 3354 | 1101 | 1497 | 3331 | 1595 | 1508 |
| 2233 | FIM1_2289 | QGN15597.1 | CP015056.1 | 5648 | 3398 | 3005 | 3505 | 2347 | 2868 | 3586 | 3536 | 2733 |
| 2234 | FIM1_2290 | QGN15598.1 | CP015056.1 | 8317 | 2180 | 1553 | 5490 | 1113 | 1402 | 7941 | 2191 | 1416 |
| 2235 | FIM1_2291 | QGN15599.1 | CP015056.1 | 6202 | 3678 | 2963 | 5712 | 1762 | 2713 | 7117 | 3763 | 2766 |
| 2236 | FIM1_2292 | QGN15600.1 | CP015056.1 | 1617 | 610 | 760 | 1404 | 501 | 824 | 1292 | 742 | 769 |
| 2237 | FIM1_2293 | QGN15601.1 | CP015056.1 | 1074 | 374 | 656 | 1002 | 309 | 609 | 652 | 438 | 539 |
| 2238 | FIM1_2294 | QGN15602.1 | CP015056.1 | 22404 | 8302 | 8143 | 18763 | 6399 | 7057 | 18298 | 9068 | 7397 |
| 2239 | FIM1_2295 | QGN15603.1 | CP015056.1 | 1564 | 452 | 690 | 1172 | 402 | 624 | 1543 | 545 | 635 |
| 2240 | FIM1_2297 | QGN15604.1 | CP015056.1 | 2775 | 682 | 1045 | 2204 | 616 | 933 | 2372 | 836 | 896 |
| 2241 | FIM1_2298 | QGN15605.1 | CP015056.1 | 6798 | 674 | 1258 | 5436 | 654 | 1245 | 9861 | 723 | 1119 |
| 2242 | FIM1_2299 | QGN15606.1 | CP015056.1 | 4035 | 792 | 1445 | 3267 | 685 | 1312 | 3880 | 915 | 1415 |
| 2243 | FIM1_2300 | QGN15607.1 | CP015056.1 | 1278 | 196 | 433 | 1055 | 206 | 490 | 1753 | 251 | 421 |
| 2244 | FIM1_2301 | QGN15608.1 | CP015056.1 | 4503 | 2243 | 2489 | 4243 | 1427 | 2348 | 5301 | 2286 | 2223 |
| 2245 | FIM1_2302 | QGN15609.1 | CP015056.1 | 5221 | 1700 | 4044 | 3626 | 1296 | 3607 | 4280 | 1727 | 3733 |
| 2246 | FIM1_2303 | QGN15610.1 | CP015056.1 | 1254 | 146 | 225 | 867 | 122 | 185 | 1404 | 169 | 170 |
| 2247 | FIM1_2304 | QGN15611.1 | CP015056.1 | 884 | 103 | 412 | 701 | 65 | 405 | 1085 | 97 | 304 |
| 2248 | FIM1_2305 | QGN15612.1 | CP015056.1 | 2165 | 322 | 628 | 1759 | 283 | 617 | 2259 | 382 | 495 |
| 2249 | FIM1_2306 | QGN15613.1 | CP015056.1 | 878 | 32 | 196 | 752 | 26 | 178 | 1199 | 39 | 153 |
| 2250 | FIM1_2307 | QGN15614.1 | CP015056.1 | 5938 | 2234 | 1876 | 4961 | 1613 | 1865 | 4207 | 2416 | 1791 |
| 2251 | FIM1_2308 | QGN15615.1 | CP015056.1 | 3496 | 1906 | 1648 | 2810 | 1453 | 1515 | 1707 | 2099 | 1430 |
| 2252 | FIM1_2309 | QGN15616.1 | CP015056.1 | 575 | 83 | 217 | 449 | 57 | 226 | 1341 | 95 | 192 |
| 2253 | FIM1_2310 | QGN15617.1 | CP015056.1 | 573 | 332 | 457 | 520 | 182 | 372 | 473 | 329 | 314 |
| 2254 | FIM1_2311 | QGN15618.1 | CP015056.1 | 6761 | 955 | 1496 | 4776 | 570 | 1320 | 5661 | 1006 | 1352 |

| | | | | | | | | | | | | |
|---|---|---|---|---|---|---|---|---|---|---|---|---|
| 2255 | FIM1_2312 | QGN15619.1 | CP015056.1 | 11105 | 3618 | 3525 | 8333 | 2542 | 3422 | 12044 | 3832 | 3258 |
| 2256 | FIM1_2313 | QGN15620.1 | CP015056.1 | 3183 | 1678 | 1078 | 2157 | 797 | 929 | 2106 | 1668 | 921 |
| 2257 | FIM1_2314 | QGN15621.1 | CP015056.1 | 2268 | 414 | 901 | 1986 | 393 | 898 | 2228 | 483 | 734 |
| 2258 | FIM1_2315 | QGN15622.1 | CP015056.1 | 4673 | 261 | 728 | 3361 | 246 | 566 | 4993 | 329 | 361 |
| 2259 | FIM1_2316 | QGN15623.1 | CP015056.1 | 3048 | 491 | 711 | 2627 | 337 | 724 | 3999 | 528 | 703 |
| 2260 | FIM1_2317 | QGN15624.1 | CP015056.1 | 9900 | 1261 | 1732 | 7929 | 974 | 1585 | 9260 | 1334 | 1394 |
| 2261 | FIM1_2318 | QGN15625.1 | CP015056.1 | 15296 | 14079 | 8582 | 14250 | 7770 | 7085 | 11731 | 14420 | 8292 |
| 2262 | FIM1_2319 | QGN15626.1 | CP015056.1 | 761 | 79 | 139 | 592 | 68 | 127 | 952 | 80 | 126 |
| 2263 | FIM1_2320 | QGN15627.1 | CP015056.1 | 335 | 179 | 187 | 253 | 138 | 180 | 312 | 175 | 150 |
| 2264 | FIM1_2321 | QGN15628.1 | CP015056.1 | 4219 | 1182 | 1454 | 3062 | 886 | 1375 | 3557 | 1219 | 1235 |
| 2265 | FIM1_2322 | QGN15629.1 | CP015056.1 | 1519 | 405 | 903 | 1214 | 334 | 845 | 1629 | 437 | 717 |
| 2266 | FIM1_2323 | QGN15630.1 | CP015056.1 | 7261 | 2583 | 2390 | 5966 | 2002 | 2289 | 5289 | 2873 | 2156 |
| 2267 | FIM1_2324 | QGN15631.1 | CP015056.1 | 14932 | 2765 | 3509 | 11379 | 2337 | 3160 | 11102 | 2908 | 3057 |
| 2268 | FIM1_2325 | QGN15632.1 | CP015056.1 | 94 | 14 | 32 | 80 | 13 | 41 | 116 | 17 | 29 |
| 2269 | FIM1_2326 | QGN15633.1 | CP015056.1 | 1043 | 237 | 329 | 889 | 192 | 371 | 964 | 206 | 322 |
| 2270 | FIM1_2327 | QGN15634.1 | CP015056.1 | 8288 | 1799 | 2600 | 7141 | 1296 | 2429 | 9827 | 2033 | 2237 |
| 2271 | FIM1_2328 | QGN15635.1 | CP015056.1 | 22653 | 6657 | 6276 | 17817 | 4680 | 5838 | 20524 | 6828 | 5519 |
| 2272 | FIM1_2329 | QGN15636.1 | CP015056.1 | 1266 | 220 | 267 | 896 | 150 | 260 | 802 | 252 | 247 |
| 2273 | FIM1_2330 | QGN15637.1 | CP015056.1 | 1032 | 377 | 365 | 641 | 273 | 331 | 705 | 376 | 363 |
| 2274 | FIM1_2331 | QGN15638.1 | CP015056.1 | 5230 | 2253 | 2453 | 5117 | 1291 | 2271 | 6474 | 2510 | 2145 |
| 2275 | FIM1_2332 | QGN15639.1 | CP015056.1 | 14083 | 8248 | 8947 | 11236 | 4169 | 7959 | 18380 | 8859 | 8024 |
| 2276 | FIM1_2333 | QGN15640.1 | CP015056.1 | 1072 | 234 | 347 | 738 | 174 | 318 | 932 | 273 | 315 |
| 2277 | FIM1_2334 | QGN15641.1 | CP015056.1 | 1580 | 246 | 407 | 1283 | 180 | 401 | 2628 | 270 | 370 |
| 2278 | FIM1_2335 | QGN15642.1 | CP015056.1 | 15300 | 4419 | 5728 | 14853 | 3081 | 5164 | 17890 | 4583 | 5172 |
| 2279 | FIM1_2336 | QGN15643.1 | CP015056.1 | 7888 | 473 | 884 | 7105 | 342 | 907 | 13016 | 478 | 725 |
| 2280 | FIM1_2337 | QGN15644.1 | CP015056.1 | 700 | 113 | 117 | 542 | 94 | 137 | 873 | 122 | 132 |
| 2281 | FIM1_2338 | QGN15645.1 | CP015056.1 | 4599 | 1014 | 1247 | 3368 | 897 | 1130 | 3950 | 1164 | 1059 |
| 2282 | FIM1_2339 | QGN15646.1 | CP015056.1 | 2700 | 202 | 504 | 1883 | 173 | 494 | 2939 | 243 | 445 |
| 2283 | FIM1_2340 | QGN15647.1 | CP015056.1 | 2131 | 532 | 1623 | 1768 | 478 | 1469 | 1954 | 563 | 1243 |
| 2284 | FIM1_2341 | QGN15648.1 | CP015056.1 | 1468 | 247 | 353 | 1299 | 205 | 313 | 2049 | 251 | 265 |
| 2285 | FIM1_2342 | QGN15649.1 | CP015056.1 | 3095 | 1137 | 1525 | 2732 | 764 | 1481 | 3475 | 1308 | 1485 |
| 2286 | FIM1_2343 | QGN15650.1 | CP015056.1 | 89 | 11 | 25 | 79 | 8 | 26 | 201 | 13 | 25 |
| 2287 | FIM1_2344 | QGN15651.1 | CP015056.1 | 1969 | 146 | 355 | 1632 | 74 | 326 | 4106 | 172 | 266 |
| 2288 | FIM1_2345 | QGN15652.1 | CP015056.1 | 2600 | 332 | 478 | 1962 | 301 | 425 | 3842 | 393 | 442 |
| 2289 | FIM1_2346 | QGN15653.1 | CP015056.1 | 9775 | 2809 | 3150 | 7571 | 2177 | 3041 | 8121 | 2934 | 3045 |
| 2290 | FIM1_2347 | QGN15654.1 | CP015056.1 | 1250 | 389 | 618 | 1141 | 351 | 585 | 841 | 461 | 454 |
| 2291 | FIM1_2348 | QGN15655.1 | CP015056.1 | 6906 | 2416 | 2424 | 5014 | 1750 | 2408 | 5157 | 2564 | 2197 |
| 2292 | FIM1_2349 | QGN15656.1 | CP015056.1 | 6857 | 963 | 1892 | 4931 | 795 | 1781 | 7558 | 1034 | 1755 |
| 2293 | FIM1_2350 | QGN15657.1 | CP015056.1 | 5423 | 1884 | 2893 | 4715 | 1173 | 2688 | 6689 | 1938 | 2507 |
| 2294 | FIM1_2351 | QGN15658.1 | CP015056.1 | 2771 | 645 | 987 | 2209 | 397 | 973 | 3230 | 684 | 709 |
| 2295 | FIM1_2352 | QGN15659.1 | CP015056.1 | 4541 | 582 | 1008 | 3507 | 439 | 833 | 4653 | 552 | 663 |
| 2296 | FIM1_2353 | QGN15660.1 | CP015056.1 | 1828 | 535 | 790 | 1495 | 443 | 809 | 1635 | 487 | 753 |
| 2297 | FIM1_2354 | QGN15661.1 | CP015056.1 | 1677 | 255 | 678 | 1246 | 174 | 548 | 1444 | 236 | 558 |
| 2298 | FIM1_2355 | QGN15662.1 | CP015056.1 | 402 | 95 | 124 | 285 | 92 | 117 | 265 | 106 | 110 |
| 2299 | FIM1_2356 | QGN15663.1 | CP015056.1 | 1502 | 127 | 353 | 1089 | 93 | 273 | 2283 | 167 | 290 |
| 2300 | FIM1_2357 | QGN15664.1 | CP015056.1 | 5400 | 2481 | 1740 | 3870 | 1831 | 1565 | 2504 | 2720 | 1562 |
| 2301 | FIM1_2358 | QGN15665.1 | CP015056.1 | 1474 | 679 | 979 | 1138 | 512 | 978 | 1333 | 622 | 911 |
| 2302 | FIM1_2359 | QGN15666.1 | CP015056.1 | 8922 | 3873 | 3727 | 7549 | 2821 | 3457 | 6610 | 4245 | 3394 |

| | | | | | | | | | | | | |
|---|---|---|---|---|---|---|---|---|---|---|---|---|
| 2303 | FIM1_2360 | QGN15667.1 | CP015056.1 | 5653 | 1370 | 1344 | 4649 | 678 | 1290 | 9029 | 1391 | 1152 |
| 2304 | FIM1_2361 | QGN15668.1 | CP015056.1 | 6808 | 2124 | 2941 | 5777 | 1505 | 2678 | 7660 | 2310 | 2590 |
| 2305 | FIM1_2362 | QGN15669.1 | CP015056.1 | 3451 | 1125 | 1233 | 2670 | 834 | 1177 | 2771 | 1209 | 1086 |
| 2306 | FIM1_2363 | QGN15670.1 | CP015056.1 | 6173 | 1375 | 1670 | 4894 | 1154 | 1529 | 5251 | 1426 | 1474 |
| 2307 | FIM1_2364 | QGN15671.1 | CP015056.1 | 8203 | 1639 | 2144 | 6090 | 1308 | 1949 | 7261 | 1708 | 1853 |
| 2308 | FIM1_2365 | QGN15672.1 | CP015056.1 | 29524 | 19908 | 17729 | 27552 | 16493 | 16927 | 27663 | 20593 | 17408 |
| 2309 | FIM1_2366 | QGN15673.1 | CP015056.1 | 1525 | 267 | 572 | 1278 | 185 | 503 | 1803 | 251 | 432 |
| 2310 | FIM1_2367 | QGN15674.1 | CP015056.1 | 1647 | 357 | 433 | 1121 | 285 | 387 | 834 | 398 | 405 |
| 2311 | FIM1_2368 | QGN15675.1 | CP015056.1 | 9922 | 1317 | 2138 | 6505 | 896 | 1964 | 6799 | 1336 | 1950 |
| 2312 | FIM1_2369 | QGN15676.1 | CP015056.1 | 2556 | 495 | 1121 | 1948 | 454 | 994 | 2946 | 601 | 985 |
| 2313 | FIM1_2370 | QGN15677.1 | CP015056.1 | 732 | 248 | 367 | 623 | 228 | 359 | 222 | 183 | 380 |
| 2314 | FIM1_2371 | QGN15678.1 | CP015056.1 | 20701 | 1528 | 2288 | 15451 | 1010 | 2179 | 26467 | 1476 | 2167 |
| 2315 | FIM1_2372 | QGN15679.1 | CP015056.1 | 13930 | 4166 | 4192 | 12520 | 3209 | 3879 | 12875 | 4519 | 3419 |
| 2316 | FIM1_2373 | QGN15680.1 | CP015056.1 | 747 | 31 | 657 | 619 | 32 | 718 | 2005 | 57 | 633 |
| 2317 | FIM1_2374 | QGN15681.1 | CP015056.1 | 3426 | 543 | 837 | 2736 | 411 | 818 | 3154 | 621 | 687 |
| 2318 | FIM1_2375 | QGN15682.1 | CP015056.1 | 5386 | 1139 | 1694 | 4498 | 838 | 1488 | 5452 | 1223 | 1468 |
| 2319 | FIM1_2376 | QGN15683.1 | CP015056.1 | 3689 | 790 | 1562 | 3110 | 587 | 1290 | 3044 | 880 | 1202 |
| 2320 | FIM1_2377 | QGN15684.1 | CP015056.1 | 24566 | 8515 | 6415 | 19342 | 6325 | 5954 | 23246 | 9105 | 6219 |
| 2321 | FIM1_2378 | QGN15685.1 | CP015056.1 | 1981 | 353 | 615 | 1428 | 309 | 579 | 2306 | 343 | 554 |
| 2322 | FIM1_2379 | QGN15686.1 | CP015056.1 | 6753 | 1097 | 1133 | 4674 | 954 | 1049 | 4470 | 1179 | 994 |
| 2323 | FIM1_2380 | QGN15687.1 | CP015056.1 | 1344 | 558 | 650 | 1266 | 428 | 725 | 987 | 574 | 654 |
| 2324 | FIM1_2381 | QGN15688.1 | CP015056.1 | 3562 | 1234 | 1603 | 2780 | 879 | 1481 | 3283 | 1306 | 1337 |
| 2325 | FIM1_2382 | QGN15689.1 | CP015056.1 | 1850 | 374 | 528 | 1498 | 239 | 472 | 1775 | 432 | 440 |
| 2326 | FIM1_2383 | QGN15690.1 | CP015056.1 | 3898 | 351 | 620 | 2540 | 247 | 514 | 3922 | 367 | 461 |
| 2327 | FIM1_2384 | QGN15691.1 | CP015056.1 | 16795 | 9898 | 9444 | 13914 | 7702 | 8462 | 12205 | 11053 | 8597 |
| 2328 | FIM1_2385 | QGN15692.1 | CP015056.1 | 2118 | 526 | 936 | 1794 | 379 | 876 | 2264 | 568 | 782 |
| 2329 | FIM1_2386 | QGN15693.1 | CP015056.1 | 1507 | 184 | 747 | 1352 | 163 | 757 | 2456 | 180 | 676 |
| 2330 | FIM1_2387 | QGN15694.1 | CP015056.1 | 1142 | 254 | 403 | 875 | 199 | 392 | 1244 | 274 | 348 |
| 2331 | FIM1_2388 | QGN15695.1 | CP015056.1 | 8689 | 4526 | 5632 | 7392 | 3387 | 5248 | 8538 | 4778 | 5213 |
| 2332 | FIM1_2389 | QGN15696.1 | CP015056.1 | 7016 | 1522 | 1578 | 4435 | 815 | 1339 | 5583 | 1634 | 1394 |
| 2333 | FIM1_2390 | QGN15697.1 | CP015056.1 | 3201 | 503 | 805 | 2428 | 386 | 875 | 3538 | 573 | 710 |
| 2334 | FIM1_2391 | QGN15698.1 | CP015056.1 | 2281 | 939 | 931 | 1827 | 575 | 896 | 1524 | 1010 | 811 |
| 2335 | FIM1_2392 | QGN15699.1 | CP015056.1 | 1852 | 659 | 1064 | 1694 | 523 | 1019 | 1109 | 780 | 914 |
| 2336 | FIM1_2393 | QGN15700.1 | CP015056.1 | 376 | 207 | 463 | 381 | 161 | 518 | 247 | 226 | 426 |
| 2337 | FIM1_2394 | QGN15701.1 | CP015056.1 | 3983 | 140 | 723 | 3041 | 130 | 671 | 6744 | 145 | 553 |
| 2338 | FIM1_2395 | QGN15702.1 | CP015056.1 | 33731 | 20541 | 15202 | 30582 | 15455 | 13984 | 28458 | 20336 | 14001 |
| 2339 | FIM1_2396 | QGN15703.1 | CP015056.1 | 6710 | 1641 | 2001 | 5041 | 1154 | 1902 | 7873 | 1778 | 1907 |
| 2340 | FIM1_2397 | QGN15704.1 | CP015056.1 | 9969 | 1277 | 2502 | 7715 | 1125 | 2418 | 10631 | 1431 | 2226 |
| 2341 | FIM1_2398 | QGN15705.1 | CP015056.1 | 2144 | 878 | 2537 | 1778 | 604 | 2681 | 2069 | 939 | 2428 |
| 2342 | FIM1_2399 | QGN15706.1 | CP015056.1 | 960 | 266 | 1155 | 705 | 210 | 1203 | 770 | 363 | 1036 |
| 2343 | FIM1_2400 | QGN15707.1 | CP015056.1 | 955 | 168 | 462 | 708 | 164 | 481 | 1043 | 206 | 376 |
| 2344 | FIM1_2401 | QGN15708.1 | CP015056.1 | 1484 | 422 | 633 | 1319 | 311 | 624 | 1618 | 461 | 539 |
| 2345 | FIM1_2402 | QGN15709.1 | CP015056.1 | 3061 | 1227 | 1502 | 2532 | 920 | 1508 | 2437 | 1350 | 1353 |
| 2346 | FIM1_2403 | QGN15710.1 | CP015056.1 | 5325 | 1161 | 1624 | 4106 | 919 | 1673 | 5887 | 1299 | 1540 |
| 2347 | FIM1_2404 | QGN15711.1 | CP015056.1 | 13728 | 3998 | 4217 | 11965 | 3391 | 3587 | 13952 | 4270 | 3600 |
| 2348 | FIM1_2405 | QGN15712.1 | CP015056.1 | 3981 | 1326 | 1595 | 3452 | 1020 | 1554 | 3383 | 1505 | 1409 |
| 2349 | FIM1_2406 | QGN15713.1 | CP015056.1 | 27874 | 6805 | 6158 | 20480 | 2508 | 5315 | 31900 | 6679 | 4839 |
| 2350 | FIM1_2407 | QGN15714.1 | CP015056.1 | 2755 | 1691 | 1157 | 1849 | 999 | 990 | 1431 | 1837 | 1005 |

| | | | | | | | | | | | |
|---|---|---|---|---|---|---|---|---|---|---|---|
| 2351 | FIM1_2408 | QGN15715.1 | CP015056.1 | 903 | 115 | 266 | 637 | 89 | 252 | 734 | 130 | 209 |
| 2352 | FIM1_2409 | QGN15716.1 | CP015056.1 | 1210 | 288 | 524 | 985 | 247 | 495 | 882 | 308 | 451 |
| 2353 | FIM1_2410 | QGN15717.1 | CP015056.1 | 11195 | 4422 | 4810 | 9194 | 3411 | 4621 | 9171 | 4824 | 4766 |
| 2354 | FIM1_2411 | QGN15718.1 | CP015056.1 | 1861 | 432 | 543 | 1455 | 321 | 592 | 1784 | 431 | 481 |
| 2355 | FIM1_2412 | QGN15719.1 | CP015056.1 | 1281 | 423 | 521 | 1026 | 355 | 532 | 942 | 473 | 500 |
| 2356 | FIM1_2413 | QGN15720.1 | CP015056.1 | 6734 | 1438 | 1918 | 5036 | 978 | 1813 | 6010 | 1514 | 1726 |
| 2357 | FIM1_2414 | QGN15721.1 | CP015056.1 | 3756 | 790 | 1214 | 3363 | 569 | 1185 | 3816 | 875 | 1073 |
| 2358 | FIM1_2415 | QGN15722.1 | CP015056.1 | 2156 | 699 | 715 | 1745 | 514 | 710 | 2149 | 730 | 646 |
| 2359 | FIM1_2416 | QGN15723.1 | CP015056.1 | 9422 | 2622 | 3883 | 7192 | 2216 | 3574 | 7573 | 2997 | 3360 |
| 2360 | FIM1_2417 | QGN15724.1 | CP015056.1 | 3188 | 541 | 1089 | 2444 | 392 | 961 | 2708 | 560 | 853 |
| 2361 | FIM1_2418 | QGN15725.1 | CP015056.1 | 33226 | 13024 | 12307 | 27480 | 9862 | 11149 | 34342 | 13974 | 11318 |
| 2362 | FIM1_2419 | QGN15726.1 | CP015056.1 | 1863 | 535 | 408 | 1381 | 403 | 455 | 2965 | 585 | 477 |
| 2363 | FIM1_2420 | QGN15727.1 | CP015056.1 | 4290 | 1297 | 1299 | 2930 | 792 | 1237 | 2999 | 1331 | 1178 |
| 2364 | FIM1_2422 | QGN15728.1 | CP015056.1 | 14847 | 2444 | 3496 | 11274 | 2030 | 3363 | 15550 | 2686 | 3176 |
| 2365 | FIM1_2423 | QGN15729.1 | CP015056.1 | 2558 | 286 | 516 | 1728 | 242 | 481 | 1971 | 313 | 426 |
| 2366 | FIM1_2424 | QGN15730.1 | CP015056.1 | 3125 | 572 | 1004 | 2516 | 384 | 888 | 5459 | 594 | 882 |
| 2367 | FIM1_2425 | QGN15731.1 | CP015056.1 | 1016 | 367 | 588 | 998 | 285 | 592 | 730 | 409 | 464 |
| 2368 | FIM1_2426 | QGN15732.1 | CP015056.1 | 9080 | 1199 | 2679 | 7006 | 993 | 2391 | 11639 | 1289 | 2159 |
| 2369 | FIM1_2427 | QGN15733.1 | CP015056.1 | 487 | 140 | 220 | 483 | 112 | 242 | 462 | 162 | 150 |
| 2370 | FIM1_2428 | QGN15734.1 | CP015056.1 | 1183 | 113 | 302 | 920 | 82 | 292 | 1493 | 95 | 256 |
| 2371 | FIM1_2429 | QGN15735.1 | CP015056.1 | 3299 | 655 | 838 | 2623 | 544 | 828 | 3004 | 657 | 846 |
| 2372 | FIM1_2430 | QGN15736.1 | CP015056.1 | 4616 | 707 | 1416 | 3604 | 575 | 1320 | 4441 | 763 | 1221 |
| 2373 | FIM1_2431 | QGN15737.1 | CP015056.1 | 4513 | 985 | 2258 | 3673 | 665 | 2130 | 3905 | 1133 | 1972 |
| 2374 | FIM1_2432 | QGN15738.1 | CP015056.1 | 4066 | 368 | 1051 | 2931 | 265 | 923 | 5608 | 365 | 823 |
| 2375 | FIM1_2433 | QGN15739.1 | CP015056.1 | 32757 | 31460 | 25187 | 25680 | 14068 | 23863 | 36414 | 30182 | 25349 |
| 2376 | FIM1_2434 | QGN15740.1 | CP015056.1 | 9104 | 4638 | 4777 | 7178 | 1695 | 4719 | 10323 | 4359 | 4147 |
| 2377 | FIM1_2435 | QGN15741.1 | CP015056.1 | 4401 | 908 | 1398 | 3939 | 722 | 1311 | 5061 | 1078 | 1297 |
| 2378 | FIM1_2436 | QGN15742.1 | CP015056.1 | 888 | 180 | 215 | 616 | 119 | 222 | 1138 | 199 | 245 |
| 2379 | FIM1_2437 | QGN15743.1 | CP015056.1 | 548 | 28 | 88 | 422 | 23 | 81 | 622 | 38 | 59 |
| 2380 | FIM1_2438 | QGN15744.1 | CP015056.1 | 4747 | 812 | 1022 | 3652 | 621 | 974 | 5433 | 821 | 983 |
| 2381 | FIM1_2439 | QGN15745.1 | CP015056.1 | 1429 | 168 | 276 | 1073 | 157 | 254 | 1563 | 196 | 218 |
| 2382 | FIM1_2440 | QGN15746.1 | CP015056.1 | 363 | 98 | 205 | 261 | 34 | 126 | 211 | 45 | 122 |
| 2383 | FIM1_2441 | QGN15747.1 | CP015057.1 | 0 | 0 | 0 | 0 | 0 | 0 | 0 | 0 | 0 |
| 2384 | FIM1_2444 | QGN15748.1 | CP015057.1 | 29998 | 20445 | 15525 | 20920 | 15466 | 16880 | 41803 | 22665 | 17355 |
| 2385 | FIM1_2445 | QGN15749.1 | CP015057.1 | 2480 | 587 | 575 | 2209 | 540 | 615 | 2222 | 647 | 596 |
| 2386 | FIM1_2446 | QGN15750.1 | CP015057.1 | 546 | 166 | 200 | 474 | 108 | 195 | 632 | 230 | 197 |
| 2387 | FIM1_2447 | QGN15751.1 | CP015057.1 | 1062 | 264 | 482 | 830 | 149 | 420 | 1010 | 279 | 423 |
| 2388 | FIM1_2448 | QGN15752.1 | CP015057.1 | 82 | 10 | 32 | 57 | 8 | 40 | 137 | 15 | 31 |
| 2389 | FIM1_2449 | QGN15753.1 | CP015057.1 | 89 | 27 | 28 | 90 | 16 | 35 | 268 | 31 | 29 |
| 2390 | FIM1_2450 | QGN15754.1 | CP015057.1 | 3086 | 215 | 389 | 2681 | 141 | 329 | 5259 | 193 | 253 |
| 2391 | FIM1_2451 | QGN15755.1 | CP015057.1 | 3628 | 607 | 565 | 3278 | 166 | 494 | 7761 | 619 | 334 |
| 2392 | FIM1_2452 | QGN15756.1 | CP015057.1 | 196 | 20 | 39 | 180 | 26 | 36 | 170 | 31 | 27 |
| 2393 | FIM1_2453 | QGN15757.1 | CP015057.1 | 625 | 84 | 168 | 542 | 57 | 158 | 878 | 104 | 148 |
| 2394 | FIM1_2454 | QGN15758.1 | CP015057.1 | 1322 | 92 | 254 | 1034 | 71 | 204 | 1970 | 102 | 151 |
| 2395 | FIM1_2455 | QGN15759.1 | CP015057.1 | 7216 | 329 | 721 | 5300 | 172 | 639 | 9810 | 303 | 581 |
| 2396 | FIM1_2456 | QGN15760.1 | CP015057.1 | 601 | 56 | 263 | 442 | 53 | 246 | 851 | 71 | 182 |
| 2397 | FIM1_2457 | QGN15761.1 | CP015057.1 | 1862 | 166 | 338 | 1387 | 114 | 316 | 1960 | 155 | 221 |
| 2398 | FIM1_2458 | QGN15762.1 | CP015057.1 | 4849 | 1086 | 1459 | 4540 | 515 | 1297 | 7332 | 1118 | 1258 |

| | | | | | | | | | | | |
|---|---|---|---|---|---|---|---|---|---|---|---|
| 2399 | FIM1_2459 | QGN15763.1 | CP015057.1 | 3772 | 2343 | 1551 | 2955 | 1618 | 1383 | 1313 | 2473 | 1474 |
| 2400 | FIM1_2460 | QGN15764.1 | CP015057.1 | 5929 | 5518 | 3055 | 4533 | 3407 | 2836 | 1441 | 5606 | 2732 |
| 2401 | FIM1_2461 | QGN15765.1 | CP015057.1 | 10545 | 1956 | 2496 | 7925 | 1431 | 2112 | 9431 | 2184 | 2024 |
| 2402 | FIM1_2462 | QGN15766.1 | CP015057.1 | 13615 | 4235 | 4183 | 10370 | 3157 | 3687 | 12979 | 4616 | 3912 |
| 2403 | FIM1_2463 | QGN15767.1 | CP015057.1 | 3430 | 460 | 732 | 2662 | 313 | 619 | 2866 | 490 | 605 |
| 2404 | FIM1_2464 | QGN15768.1 | CP015057.1 | 8149 | 777 | 1891 | 5544 | 852 | 1592 | 7358 | 1064 | 1551 |
| 2405 | FIM1_2465 | QGN15769.1 | CP015057.1 | 1025 | 272 | 771 | 810 | 222 | 685 | 922 | 277 | 584 |
| 2406 | FIM1_2466 | QGN15770.1 | CP015057.1 | 3537 | 225 | 718 | 2772 | 170 | 639 | 5037 | 236 | 573 |
| 2407 | FIM1_2467 | QGN15771.1 | CP015057.1 | 1287 | 223 | 285 | 1123 | 136 | 232 | 1149 | 240 | 289 |
| 2408 | FIM1_2468 | QGN15772.1 | CP015057.1 | 5414 | 1385 | 2128 | 4524 | 1174 | 2193 | 5356 | 1546 | 1866 |
| 2409 | FIM1_2469 | QGN15773.1 | CP015057.1 | 4727 | 1030 | 1833 | 4039 | 793 | 1642 | 4506 | 1131 | 1478 |
| 2410 | FIM1_2470 | QGN15774.1 | CP015057.1 | 20858 | 4093 | 6684 | 16260 | 3184 | 6074 | 20224 | 4513 | 5714 |
| 2411 | FIM1_2471 | QGN15775.1 | CP015057.1 | 2065 | 606 | 1144 | 1847 | 503 | 1193 | 2177 | 755 | 1104 |
| 2412 | FIM1_2472 | QGN15776.1 | CP015057.1 | 5752 | 860 | 1637 | 4395 | 701 | 1451 | 7051 | 865 | 1471 |
| 2413 | FIM1_2473 | QGN15777.1 | CP015057.1 | 2890 | 2046 | 1242 | 2074 | 1395 | 1146 | 1639 | 2124 | 1025 |
| 2414 | FIM1_2474 | QGN15778.1 | CP015057.1 | 8463 | 2622 | 2459 | 7116 | 1869 | 2417 | 10131 | 2941 | 2250 |
| 2415 | FIM1_2475 | QGN15779.1 | CP015057.1 | 3647 | 483 | 1291 | 2765 | 428 | 1231 | 3852 | 491 | 1094 |
| 2416 | FIM1_2476 | QGN15780.1 | CP015057.1 | 14836 | 9159 | 8919 | 14560 | 6858 | 7789 | 15176 | 9009 | 8596 |
| 2417 | FIM1_2477 | QGN15781.1 | CP015057.1 | 5514 | 1297 | 1806 | 4345 | 1148 | 1574 | 4596 | 1419 | 1500 |
| 2418 | FIM1_2478 | QGN15782.1 | CP015057.1 | 1926 | 792 | 1094 | 1550 | 612 | 1006 | 1475 | 808 | 1007 |
| 2419 | FIM1_2479 | QGN15783.1 | CP015057.1 | 1890 | 471 | 697 | 1559 | 404 | 684 | 1597 | 517 | 635 |
| 2420 | FIM1_2480 | QGN15784.1 | CP015057.1 | 5639 | 1233 | 1599 | 4374 | 868 | 1552 | 6164 | 1297 | 1504 |
| 2421 | FIM1_2481 | QGN15785.1 | CP015057.1 | 3814 | 648 | 1038 | 2885 | 529 | 911 | 3425 | 700 | 1016 |
| 2422 | FIM1_2482 | QGN15786.1 | CP015057.1 | 3685 | 901 | 1314 | 2694 | 658 | 1165 | 3666 | 920 | 1076 |
| 2423 | FIM1_2483 | QGN15787.1 | CP015057.1 | 3934 | 588 | 1066 | 2973 | 468 | 992 | 3844 | 632 | 835 |
| 2424 | FIM1_2484 | QGN15788.1 | CP015057.1 | 1111 | 170 | 288 | 881 | 136 | 276 | 955 | 197 | 261 |
| 2425 | FIM1_2485 | QGN15789.1 | CP015057.1 | 4224 | 506 | 1256 | 3336 | 371 | 1120 | 5867 | 577 | 971 |
| 2426 | FIM1_2486 | QGN15790.1 | CP015057.1 | 15472 | 11691 | 10478 | 15270 | 10157 | 10610 | 14897 | 11982 | 10383 |
| 2427 | FIM1_2487 | QGN15791.1 | CP015057.1 | 6702 | 1527 | 2073 | 5253 | 1175 | 1897 | 7107 | 1630 | 1917 |
| 2428 | FIM1_2488 | QGN15792.1 | CP015057.1 | 986 | 196 | 331 | 872 | 162 | 323 | 911 | 233 | 285 |
| 2429 | FIM1_2489 | QGN15793.1 | CP015057.1 | 1124 | 251 | 715 | 1043 | 200 | 644 | 1021 | 265 | 602 |
| 2430 | FIM1_2490 | QGN15794.1 | CP015057.1 | 3043 | 686 | 1134 | 3129 | 494 | 1076 | 4098 | 793 | 985 |
| 2431 | FIM1_2491 | QGN15795.1 | CP015057.1 | 814 | 40 | 97 | 776 | 77 | 93 | 2234 | 50 | 61 |
| 2432 | FIM1_2492 | QGN15796.1 | CP015057.1 | 6890 | 2016 | 2034 | 5960 | 1576 | 1825 | 6578 | 2184 | 1782 |
| 2433 | FIM1_2493 | QGN15797.1 | CP015057.1 | 4406 | 1094 | 1453 | 3935 | 828 | 1341 | 4037 | 1251 | 1306 |
| 2434 | FIM1_2494 | QGN15798.1 | CP015057.1 | 3496 | 444 | 823 | 2587 | 369 | 733 | 2707 | 465 | 628 |
| 2435 | FIM1_2495 | QGN15799.1 | CP015057.1 | 9904 | 2872 | 4895 | 7905 | 2264 | 4433 | 9938 | 3062 | 4354 |
| 2436 | FIM1_2496 | QGN15800.1 | CP015057.1 | 2889 | 236 | 1027 | 2520 | 259 | 975 | 3127 | 308 | 809 |
| 2437 | FIM1_2497 | QGN15801.1 | CP015057.1 | 1330 | 347 | 376 | 1004 | 301 | 344 | 1252 | 374 | 394 |
| 2438 | FIM1_2498 | QGN15802.1 | CP015057.1 | 202 | 67 | 78 | 159 | 72 | 94 | 257 | 79 | 105 |
| 2439 | FIM1_2499 | QGN15803.1 | CP015057.1 | 3874 | 746 | 943 | 2792 | 427 | 859 | 3368 | 823 | 881 |
| 2440 | FIM1_2500 | QGN15804.1 | CP015057.1 | 4888 | 223 | 684 | 3826 | 174 | 638 | 6859 | 225 | 623 |
| 2441 | FIM1_2501 | QGN15805.1 | CP015057.1 | 16 | 0 | 4 | 9 | 1 | 4 | 35 | 0 | 5 |
| 2442 | FIM1_2502 | QGN15806.1 | CP015057.1 | 5584 | 1423 | 3053 | 4814 | 888 | 2663 | 6377 | 1585 | 2876 |
| 2443 | FIM1_2503 | QGN15807.1 | CP015057.1 | 1348 | 374 | 633 | 1050 | 262 | 516 | 771 | 413 | 475 |
| 2444 | FIM1_2504 | QGN15808.1 | CP015057.1 | 2257 | 386 | 738 | 1744 | 314 | 680 | 2425 | 415 | 723 |
| 2445 | FIM1_2505 | QGN15809.1 | CP015057.1 | 954 | 379 | 568 | 715 | 273 | 548 | 1128 | 350 | 547 |
| 2446 | FIM1_2506 | QGN15810.1 | CP015057.1 | 1802 | 110 | 350 | 1461 | 71 | 343 | 3130 | 106 | 282 |

| | | | | | | | | | | | | |
|---|---|---|---|---|---|---|---|---|---|---|---|---|
| 2447 | FIM1_2507 | QGN15811.1 | CP015057.1 | 2031 | 706 | 738 | 1442 | 476 | 695 | 1962 | 749 | 731 |
| 2448 | FIM1_2508 | QGN15812.1 | CP015057.1 | 65 | 10 | 22 | 61 | 10 | 16 | 34 | 12 | 10 |
| 2449 | FIM1_2509 | QGN15813.1 | CP015057.1 | 17 | 4 | 16 | 11 | 5 | 11 | 10 | 8 | 10 |
| 2450 | FIM1_2510 | QGN15814.1 | CP015057.1 | 497 | 198 | 327 | 377 | 124 | 235 | 332 | 229 | 225 |
| 2451 | FIM1_2511 | QGN15815.1 | CP015057.1 | 12879 | 7682 | 3858 | 9727 | 5375 | 3412 | 6364 | 8008 | 3662 |
| 2452 | FIM1_2512 | QGN15816.1 | CP015057.1 | 2714 | 523 | 1146 | 2124 | 436 | 1064 | 3216 | 568 | 1076 |
| 2453 | FIM1_2513 | QGN15817.1 | CP015057.1 | 10711 | 6094 | 5379 | 9968 | 4610 | 5238 | 9852 | 6335 | 5204 |
| 2454 | FIM1_2514 | QGN15818.1 | CP015057.1 | 1412 | 173 | 371 | 1116 | 143 | 363 | 1839 | 220 | 299 |
| 2455 | FIM1_2515 | QGN15819.1 | CP015057.1 | 297 | 31 | 70 | 235 | 24 | 64 | 226 | 45 | 51 |
| 2456 | FIM1_2516 | QGN15820.1 | CP015057.1 | 11130 | 2936 | 3111 | 8167 | 2223 | 2894 | 9032 | 3255 | 2950 |
| 2457 | FIM1_2517 | QGN15821.1 | CP015057.1 | 3649 | 997 | 1209 | 3133 | 520 | 1068 | 4968 | 1124 | 1051 |
| 2458 | FIM1_2518 | QGN15822.1 | CP015057.1 | 5402 | 1377 | 1826 | 3959 | 1041 | 1694 | 4225 | 1505 | 1586 |
| 2459 | FIM1_2519 | QGN15823.1 | CP015057.1 | 7284 | 4285 | 2139 | 4811 | 2615 | 1927 | 1211 | 4369 | 1892 |
| 2460 | FIM1_2520 | QGN15824.1 | CP015057.1 | 1725 | 582 | 587 | 1320 | 396 | 543 | 945 | 707 | 506 |
| 2461 | FIM1_2521 | QGN15825.1 | CP015057.1 | 1576 | 627 | 720 | 1158 | 432 | 645 | 1051 | 651 | 633 |
| 2462 | FIM1_2522 | QGN15826.1 | CP015057.1 | 919 | 360 | 448 | 715 | 229 | 354 | 1026 | 361 | 348 |
| 2463 | FIM1_2523 | QGN15827.1 | CP015057.1 | 12563 | 3487 | 3772 | 9995 | 2441 | 3501 | 12775 | 3589 | 3383 |
| 2464 | FIM1_2524 | QGN15828.1 | CP015057.1 | 8715 | 1114 | 1220 | 5655 | 676 | 1206 | 7621 | 1244 | 1242 |
| 2465 | FIM1_2525 | QGN15829.1 | CP015057.1 | 1217 | 453 | 744 | 1129 | 298 | 738 | 1076 | 416 | 664 |
| 2466 | FIM1_2526 | QGN15830.1 | CP015057.1 | 5695 | 2323 | 6967 | 5985 | 1757 | 5820 | 4982 | 2447 | 5493 |
| 2467 | FIM1_2527 | QGN15831.1 | CP015057.1 | 351 | 75 | 394 | 287 | 67 | 310 | 317 | 73 | 371 |
| 2468 | FIM1_2528 | QGN15832.1 | CP015057.1 | 45246 | 21341 | 18809 | 41362 | 17800 | 17786 | 44875 | 21653 | 18770 |
| 2469 | FIM1_2529 | QGN15833.1 | CP015057.1 | 24725 | 6764 | 6464 | 19006 | 2331 | 5798 | 36158 | 6230 | 5181 |
| 2470 | FIM1_2530 | QGN15834.1 | CP015057.1 | 5601 | 398 | 893 | 4524 | 304 | 819 | 7564 | 388 | 696 |
| 2471 | FIM1_2531 | QGN15835.1 | CP015057.1 | 9766 | 2389 | 2580 | 8413 | 1671 | 2151 | 8379 | 2687 | 2217 |
| 2472 | FIM1_2532 | QGN15836.1 | CP015057.1 | 746 | 200 | 650 | 1014 | 293 | 745 | 450 | 202 | 537 |
| 2473 | FIM1_2533 | QGN15837.1 | CP015057.1 | 6337 | 1388 | 2755 | 5548 | 1050 | 2477 | 6472 | 1545 | 2445 |
| 2474 | FIM1_2534 | QGN15838.1 | CP015057.1 | 5402 | 1009 | 1627 | 4323 | 718 | 1565 | 6325 | 1120 | 1379 |
| 2475 | FIM1_2535 | QGN15839.1 | CP015057.1 | 1087 | 276 | 349 | 949 | 246 | 339 | 898 | 313 | 344 |
| 2476 | FIM1_2536 | QGN15840.1 | CP015057.1 | 857 | 276 | 260 | 706 | 197 | 242 | 814 | 276 | 346 |
| 2477 | FIM1_2537 | QGN15841.1 | CP015057.1 | 2089 | 615 | 537 | 1727 | 711 | 435 | 3926 | 666 | 522 |
| 2478 | FIM1_2538 | QGN15842.1 | CP015057.1 | 1754 | 520 | 737 | 1365 | 346 | 585 | 2001 | 480 | 646 |
| 2479 | FIM1_2539 | QGN15843.1 | CP015057.1 | 1732 | 464 | 554 | 1685 | 323 | 468 | 1502 | 487 | 471 |
| 2480 | FIM1_2540 | QGN15844.1 | CP015057.1 | 10993 | 1898 | 2659 | 8792 | 1389 | 2476 | 13038 | 2079 | 2427 |
| 2481 | FIM1_2541 | QGN15845.1 | CP015057.1 | 2934 | 987 | 1128 | 2708 | 813 | 992 | 2198 | 1099 | 918 |
| 2482 | FIM1_2542 | QGN15846.1 | CP015057.1 | 619 | 287 | 405 | 723 | 225 | 368 | 458 | 323 | 344 |
| 2483 | FIM1_2543 | QGN15847.1 | CP015057.1 | 4303 | 606 | 995 | 3609 | 457 | 910 | 4553 | 680 | 786 |
| 2484 | FIM1_2544 | QGN15848.1 | CP015057.1 | 2662 | 571 | 817 | 1954 | 345 | 713 | 2259 | 567 | 651 |
| 2485 | FIM1_2546 | QGN15849.1 | CP015057.1 | 2735 | 723 | 1257 | 2794 | 484 | 1175 | 2548 | 664 | 1084 |
| 2486 | FIM1_2547 | QGN15850.1 | CP015057.1 | 8606 | 606 | 1377 | 6246 | 740 | 1217 | 9911 | 686 | 1399 |
| 2487 | FIM1_2548 | QGN15851.1 | CP015057.1 | 15857 | 3961 | 3152 | 12175 | 4254 | 2861 | 13508 | 4021 | 3080 |
| 2488 | FIM1_2549 | QGN15852.1 | CP015057.1 | 19867 | 2886 | 4622 | 16831 | 2097 | 4225 | 25005 | 3127 | 3989 |
| 2489 | FIM1_2550 | QGN15853.1 | CP015057.1 | 3677 | 630 | 1608 | 3095 | 845 | 1486 | 2575 | 771 | 1397 |
| 2490 | FIM1_2551 | QGN15854.1 | CP015057.1 | 27 | 3 | 3 | 20 | 2 | 5 | 63 | 1 | 5 |
| 2491 | FIM1_2552 | QGN15855.1 | CP015057.1 | 5078 | 1076 | 964 | 4092 | 655 | 960 | 13194 | 1226 | 913 |
| 2492 | FIM1_2553 | QGN15856.1 | CP015057.1 | 2113 | 217 | 426 | 1414 | 156 | 368 | 2134 | 225 | 366 |
| 2493 | FIM1_2554 | QGN15857.1 | CP015057.1 | 5178 | 330 | 1247 | 3732 | 408 | 1140 | 4375 | 437 | 1140 |
| 2494 | FIM1_2555 | QGN15858.1 | CP015057.1 | 20912 | 16168 | 1132 | 16796 | 5742 | 927 | 57484 | 15881 | 813 |

| | | | | | | | | | | | | |
|---|---|---|---|---|---|---|---|---|---|---|---|---|
| 2495 | FIM1_2556 | QGN15859.1 | CP015057.1 | 1432 | 49 | 64 | 923 | 55 | 58 | 2991 | 48 | 83 |
| 2496 | FIM1_2557 | QGN15860.1 | CP015057.1 | 27073 | 10772 | 8192 | 22081 | 5981 | 7946 | 37882 | 9086 | 7728 |
| 2497 | FIM1_2558 | QGN15861.1 | CP015057.1 | 1766 | 1310 | 1008 | 1501 | 991 | 971 | 669 | 1475 | 975 |
| 2498 | FIM1_2559 | QGN15862.1 | CP015057.1 | 1626 | 167 | 367 | 1466 | 145 | 286 | 3172 | 185 | 259 |
| 2499 | FIM1_2560 | QGN15863.1 | CP015057.1 | 7794 | 2571 | 2812 | 6668 | 1470 | 2619 | 7987 | 2926 | 2566 |
| 2500 | FIM1_2561 | QGN15864.1 | CP015057.1 | 919 | 155 | 253 | 758 | 105 | 227 | 1728 | 159 | 227 |
| 2501 | FIM1_2562 | QGN15865.1 | CP015057.1 | 9655 | 1238 | 2156 | 7565 | 903 | 2044 | 12210 | 1390 | 1873 |
| 2502 | FIM1_2563 | QGN15866.1 | CP015057.1 | 8571 | 2231 | 2270 | 6953 | 1572 | 2019 | 8650 | 2464 | 1965 |
| 2503 | FIM1_2564 | QGN15867.1 | CP015057.1 | 3744 | 524 | 1055 | 3460 | 428 | 870 | 5206 | 589 | 669 |
| 2504 | FIM1_2565 | QGN15868.1 | CP015057.1 | 1938 | 619 | 1047 | 1895 | 505 | 958 | 2111 | 666 | 834 |
| 2505 | FIM1_2566 | QGN15869.1 | CP015057.1 | 10380 | 957 | 1878 | 7942 | 704 | 1701 | 13167 | 931 | 1482 |
| 2506 | FIM1_2567 | QGN15870.1 | CP015057.1 | 3181 | 702 | 802 | 2327 | 475 | 724 | 2556 | 746 | 621 |
| 2507 | FIM1_2568 | QGN15871.1 | CP015057.1 | 1775 | 1090 | 1034 | 1627 | 886 | 1008 | 949 | 1188 | 991 |
| 2508 | FIM1_2570 | QGN15872.1 | CP015057.1 | 1024 | 111 | 350 | 1003 | 110 | 251 | 763 | 155 | 244 |
| 2509 | FIM1_2571 | QGN15873.1 | CP015057.1 | 736 | 451 | 302 | 445 | 221 | 293 | 368 | 489 | 275 |
| 2510 | FIM1_2572 | QGN15874.1 | CP015057.1 | 390 | 284 | 218 | 308 | 160 | 206 | 221 | 307 | 179 |
| 2511 | FIM1_2573 | QGN15875.1 | CP015057.1 | 1066 | 200 | 505 | 923 | 152 | 475 | 1174 | 181 | 389 |
| 2512 | FIM1_2574 | QGN15876.1 | CP015057.1 | 4129 | 1386 | 1528 | 3614 | 1075 | 1420 | 3242 | 1624 | 1367 |
| 2513 | FIM1_2575 | QGN15877.1 | CP015057.1 | 2401 | 722 | 810 | 2003 | 535 | 747 | 1735 | 787 | 757 |
| 2514 | FIM1_2576 | QGN15878.1 | CP015057.1 | 35305 | 13382 | 12041 | 29987 | 10141 | 11168 | 35746 | 13971 | 10982 |
| 2515 | FIM1_2577 | QGN15879.1 | CP015057.1 | 14717 | 3854 | 3903 | 11508 | 3103 | 3513 | 15322 | 4127 | 2966 |
| 2516 | FIM1_2578 | QGN15880.1 | CP015057.1 | 200 | 53 | 49 | 176 | 41 | 44 | 161 | 43 | 48 |
| 2517 | FIM1_2579 | QGN15881.1 | CP015057.1 | 783 | 159 | 132 | 713 | 88 | 128 | 838 | 149 | 111 |
| 2518 | FIM1_2580 | QGN15882.1 | CP015057.1 | 3734 | 331 | 544 | 3409 | 217 | 498 | 7132 | 356 | 437 |
| 2519 | FIM1_2581 | QGN15883.1 | CP015057.1 | 32008 | 21049 | 16005 | 30561 | 13463 | 14238 | 49918 | 21575 | 14715 |
| 2520 | FIM1_2582 | QGN15884.1 | CP015057.1 | 1057 | 194 | 646 | 1044 | 192 | 587 | 1543 | 256 | 544 |
| 2521 | FIM1_2583 | QGN15885.1 | CP015057.1 | 414 | 260 | 560 | 381 | 175 | 585 | 433 | 287 | 496 |
| 2522 | FIM1_2584 | QGN15886.1 | CP015057.1 | 226 | 17 | 40 | 236 | 50 | 76 | 115 | 59 | 44 |
| 2523 | FIM1_2585 | QGN15887.1 | CP015057.1 | 9553 | 3426 | 3633 | 7240 | 2572 | 3414 | 8800 | 3579 | 3408 |
| 2524 | FIM1_2586 | QGN15888.1 | CP015057.1 | 3133 | 778 | 907 | 2264 | 555 | 947 | 2314 | 838 | 904 |
| 2525 | FIM1_2587 | QGN15889.1 | CP015057.1 | 12266 | 3176 | 3056 | 9249 | 2618 | 2752 | 10550 | 3372 | 2788 |
| 2526 | FIM1_2588 | QGN15890.1 | CP015057.1 | 8726 | 1828 | 8690 | 6852 | 1246 | 8209 | 8836 | 1977 | 9756 |
| 2527 | FIM1_2589 | QGN15891.1 | CP015057.1 | 6 | 0 | 4 | 4 | 3 | 6 | 3 | 1 | 6 |
| 2528 | FIM1_2590 | QGN15892.1 | CP015057.1 | 5538 | 269 | 1190 | 3880 | 134 | 1041 | 7521 | 242 | 1001 |
| 2529 | FIM1_2591 | QGN15893.1 | CP015057.1 | 2584 | 662 | 738 | 1865 | 479 | 658 | 1532 | 710 | 663 |
| 2530 | FIM1_2592 | QGN15894.1 | CP015057.1 | 5264 | 993 | 1412 | 4224 | 799 | 1381 | 4525 | 1086 | 1253 |
| 2531 | FIM1_2593 | QGN15895.1 | CP015057.1 | 5296 | 636 | 1312 | 4026 | 435 | 1139 | 5530 | 680 | 992 |
| 2532 | FIM1_2594 | QGN15896.1 | CP015057.1 | 4803 | 1309 | 2025 | 3994 | 892 | 1802 | 5009 | 1388 | 1740 |
| 2533 | FIM1_2595 | QGN15897.1 | CP015057.1 | 3283 | 936 | 1150 | 2661 | 700 | 1013 | 2464 | 1090 | 998 |
| 2534 | FIM1_2596 | QGN15898.1 | CP015057.1 | 6062 | 881 | 1655 | 5042 | 805 | 1526 | 7263 | 1009 | 1348 |
| 2535 | FIM1_2597 | QGN15899.1 | CP015057.1 | 2812 | 859 | 1067 | 1928 | 519 | 899 | 1878 | 842 | 752 |
| 2536 | FIM1_2598 | QGN15900.1 | CP015057.1 | 1879 | 455 | 1073 | 1318 | 416 | 1022 | 1666 | 505 | 947 |
| 2537 | FIM1_2599 | QGN15901.1 | CP015057.1 | 2320 | 873 | 904 | 1734 | 612 | 848 | 977 | 881 | 762 |
| 2538 | FIM1_2600 | QGN15902.1 | CP015057.1 | 2731 | 1116 | 1128 | 2269 | 903 | 1141 | 1599 | 1177 | 1021 |
| 2539 | FIM1_2601 | QGN15903.1 | CP015057.1 | 5853 | 1860 | 1815 | 4569 | 1399 | 1673 | 4048 | 1959 | 1603 |
| 2540 | FIM1_2602 | QGN15904.1 | CP015057.1 | 6615 | 1229 | 1542 | 4826 | 894 | 1458 | 6583 | 1341 | 1433 |
| 2541 | FIM1_2603 | QGN15905.1 | CP015057.1 | 1179 | 113 | 273 | 864 | 83 | 238 | 1182 | 141 | 255 |
| 2542 | FIM1_2604 | QGN15906.1 | CP015057.1 | 1234 | 257 | 539 | 940 | 212 | 531 | 1388 | 266 | 457 |

| | | | | | | | | | | | | |
|---|---|---|---|---|---|---|---|---|---|---|---|---|
| 2543 | FIM1_2605 | QGN15907.1 | CP015057.1 | 5 | 0 | 0 | 6 | 0 | 0 | 8 | 0 | 0 |
| 2544 | FIM1_2606 | QGN15908.1 | CP015057.1 | 62 | 28 | 15 | 37 | 23 | 18 | 62 | 23 | 22 |
| 2545 | FIM1_2607 | QGN15909.1 | CP015057.1 | 7751 | 2970 | 2539 | 5777 | 2421 | 2242 | 3996 | 3341 | 2344 |
| 2546 | FIM1_2608 | QGN15910.1 | CP015057.1 | 2219 | 535 | 737 | 1802 | 408 | 646 | 1890 | 584 | 631 |
| 2547 | FIM1_2609 | QGN15911.1 | CP015057.1 | 1207 | 174 | 430 | 1052 | 142 | 388 | 1229 | 201 | 371 |
| 2548 | FIM1_2610 | QGN15912.1 | CP015057.1 | 3110 | 462 | 809 | 2634 | 420 | 807 | 2609 | 492 | 746 |
| 2549 | FIM1_2611 | QGN15913.1 | CP015057.1 | 6091 | 816 | 2238 | 4485 | 642 | 1889 | 2812 | 867 | 1767 |
| 2550 | FIM1_2612 | QGN15914.1 | CP015057.1 | 11946 | 1820 | 2410 | 10572 | 1377 | 2163 | 13931 | 1902 | 2224 |
| 2551 | FIM1_2613 | QGN15915.1 | CP015057.1 | 13171 | 1475 | 2351 | 10376 | 1142 | 2347 | 18473 | 1556 | 2138 |
| 2552 | FIM1_2614 | QGN15916.1 | CP015057.1 | 1589 | 261 | 856 | 1456 | 219 | 769 | 1540 | 273 | 647 |
| 2553 | FIM1_2615 | QGN15917.1 | CP015057.1 | 126 | 32 | 87 | 99 | 44 | 63 | 124 | 47 | 56 |
| 2554 | FIM1_2616 | QGN15918.1 | CP015057.1 | 1981 | 383 | 571 | 1528 | 353 | 522 | 2107 | 458 | 523 |
| 2555 | FIM1_2617 | QGN15919.1 | CP015057.1 | 2535 | 677 | 1143 | 1818 | 453 | 1056 | 2171 | 710 | 887 |
| 2556 | FIM1_2618 | QGN15920.1 | CP015057.1 | 889 | 180 | 387 | 803 | 166 | 397 | 821 | 217 | 388 |
| 2557 | FIM1_2619 | QGN15921.1 | CP015057.1 | 556 | 107 | 261 | 521 | 94 | 295 | 551 | 112 | 246 |
| 2558 | FIM1_2620 | QGN15922.1 | CP015057.1 | 815 | 418 | 615 | 638 | 303 | 593 | 893 | 393 | 569 |
| 2559 | FIM1_2621 | QGN15923.1 | CP015057.1 | 26115 | 6410 | 6179 | 20746 | 4466 | 5872 | 28064 | 6706 | 5703 |
| 2560 | FIM1_2622 | QGN15924.1 | CP015057.1 | 1716 | 263 | 665 | 1214 | 241 | 618 | 2011 | 294 | 570 |
| 2561 | FIM1_2623 | QGN15925.1 | CP015057.1 | 3631 | 1224 | 1532 | 2871 | 946 | 1484 | 4115 | 1379 | 1358 |
| 2562 | FIM1_2624 | QGN15926.1 | CP015057.1 | 17533 | 4930 | 5897 | 14686 | 3035 | 5302 | 22372 | 5135 | 5166 |
| 2563 | FIM1_2625 | QGN15927.1 | CP015057.1 | 1619 | 184 | 553 | 1303 | 164 | 490 | 1458 | 198 | 423 |
| 2564 | FIM1_2626 | QGN15928.1 | CP015057.1 | 4330 | 891 | 1803 | 3509 | 611 | 1742 | 4929 | 969 | 1641 |
| 2565 | FIM1_2627 | QGN15929.1 | CP015057.1 | 1147 | 384 | 1551 | 955 | 353 | 1546 | 916 | 398 | 1402 |
| 2566 | FIM1_2628 | QGN15930.1 | CP015057.1 | 1757 | 603 | 916 | 1511 | 430 | 810 | 1456 | 599 | 661 |
| 2567 | FIM1_2629 | QGN15931.1 | CP015057.1 | 33083 | 6419 | 5075 | 24675 | 7780 | 4512 | 16964 | 7734 | 4853 |
| 2568 | FIM1_2630 | QGN15932.1 | CP015057.1 | 870 | 486 | 588 | 594 | 285 | 522 | 589 | 558 | 464 |
| 2569 | FIM1_2631 | QGN15933.1 | CP015057.1 | 729 | 114 | 161 | 517 | 89 | 142 | 782 | 130 | 118 |
| 2570 | FIM1_2632 | QGN15934.1 | CP015057.1 | 1439 | 416 | 722 | 1018 | 268 | 680 | 1303 | 404 | 521 |
| 2571 | FIM1_2633 | QGN15935.1 | CP015057.1 | 3546 | 380 | 631 | 2636 | 337 | 591 | 3583 | 393 | 506 |
| 2572 | FIM1_2634 | QGN15936.1 | CP015057.1 | 8668 | 3925 | 5888 | 9350 | 2047 | 5475 | 14106 | 4110 | 5311 |
| 2573 | FIM1_2635 | QGN15937.1 | CP015057.1 | 44136 | 31471 | 28293 | 39588 | 25533 | 24863 | 53542 | 31421 | 27722 |
| 2574 | FIM1_2636 | QGN15938.1 | CP015057.1 | 12501 | 2448 | 3243 | 9464 | 1907 | 2984 | 12122 | 2636 | 3029 |
| 2575 | FIM1_2637 | QGN15939.1 | CP015057.1 | 7707 | 2081 | 5179 | 6414 | 1506 | 4805 | 9276 | 2138 | 4663 |
| 2576 | FIM1_2638 | QGN15940.1 | CP015057.1 | 4142 | 943 | 1240 | 3098 | 712 | 1133 | 3146 | 1108 | 1166 |
| 2577 | FIM1_2639 | QGN15941.1 | CP015057.1 | 7782 | 1438 | 3319 | 6289 | 1400 | 2973 | 10535 | 1687 | 3167 |
| 2578 | FIM1_2640 | QGN15942.1 | CP015057.1 | 7697 | 1345 | 2524 | 6456 | 1151 | 2254 | 9545 | 1461 | 2257 |
| 2579 | FIM1_2641 | QGN15943.1 | CP015057.1 | 1960 | 457 | 939 | 1693 | 442 | 817 | 1728 | 505 | 767 |
| 2580 | FIM1_2642 | QGN15944.1 | CP015057.1 | 1195 | 176 | 301 | 992 | 126 | 265 | 1447 | 169 | 247 |
| 2581 | FIM1_2643 | QGN15945.1 | CP015057.1 | 6529 | 875 | 2062 | 5441 | 902 | 1789 | 8020 | 940 | 1711 |
| 2582 | FIM1_2644 | QGN15946.1 | CP015057.1 | 3651 | 909 | 1854 | 3290 | 671 | 1552 | 3553 | 923 | 1490 |
| 2583 | FIM1_2645 | QGN15947.1 | CP015057.1 | 1466 | 376 | 606 | 1332 | 273 | 543 | 1673 | 398 | 499 |
| 2584 | FIM1_2646 | QGN15948.1 | CP015057.1 | 6046 | 1658 | 2405 | 5166 | 1089 | 2264 | 6945 | 1778 | 2184 |
| 2585 | FIM1_2647 | QGN15949.1 | CP015057.1 | 9099 | 1765 | 2900 | 7189 | 1195 | 2671 | 10514 | 1800 | 2456 |
| 2586 | FIM1_2648 | QGN15950.1 | CP015057.1 | 1034 | 170 | 227 | 816 | 135 | 248 | 1092 | 186 | 212 |
| 2587 | FIM1_2649 | QGN15951.1 | CP015057.1 | 2119 | 681 | 651 | 1696 | 513 | 658 | 1768 | 714 | 653 |
| 2588 | FIM1_2650 | QGN15952.1 | CP015057.1 | 17219 | 14142 | 13123 | 16769 | 11803 | 11755 | 16754 | 14033 | 12692 |
| 2589 | FIM1_2651 | QGN15953.1 | CP015057.1 | 1126 | 222 | 236 | 831 | 170 | 278 | 1144 | 241 | 229 |
| 2590 | FIM1_2652 | QGN15954.1 | CP015057.1 | 19382 | 6577 | 6131 | 16365 | 5288 | 5850 | 17222 | 6996 | 5942 |

| | | | | | | | | | | | | |
|---|---|---|---|---|---|---|---|---|---|---|---|---|
| 2591 | FIM1_2653 | QGN15955.1 | CP015057.1 | 3356 | 1026 | 1517 | 2780 | 761 | 1428 | 3003 | 1083 | 1363 |
| 2592 | FIM1_2654 | QGN15956.1 | CP015057.1 | 5840 | 1340 | 1976 | 5048 | 774 | 1665 | 6738 | 1397 | 1620 |
| 2593 | FIM1_2655 | QGN15957.1 | CP015057.1 | 5503 | 1097 | 1323 | 4131 | 773 | 1322 | 4478 | 1121 | 1179 |
| 2594 | FIM1_2656 | QGN15958.1 | CP015057.1 | 1843 | 106 | 164 | 1428 | 33 | 153 | 2137 | 96 | 105 |
| 2595 | FIM1_2657 | QGN15959.1 | CP015057.1 | 2114 | 620 | 753 | 1812 | 455 | 732 | 2140 | 561 | 642 |
| 2596 | FIM1_2658 | QGN15960.1 | CP015057.1 | 1115 | 3 | 34 | 883 | 2 | 21 | 2666 | 4 | 21 |
| 2597 | FIM1_2659 | QGN15961.1 | CP015057.1 | 5299 | 863 | 1374 | 3457 | 673 | 1204 | 4417 | 970 | 1136 |
| 2598 | FIM1_2660 | QGN15962.1 | CP015057.1 | 1970 | 826 | 1975 | 1670 | 785 | 1762 | 1441 | 804 | 1919 |
| 2599 | FIM1_2661 | QGN15963.1 | CP015057.1 | 6075 | 505 | 1135 | 4636 | 466 | 1029 | 10309 | 609 | 992 |
| 2600 | FIM1_2662 | QGN15964.1 | CP015057.1 | 20070 | 3469 | 4111 | 15853 | 2827 | 3862 | 15928 | 3862 | 3778 |
| 2601 | FIM1_2663 | QGN15965.1 | CP015057.1 | 621 | 359 | 365 | 418 | 286 | 359 | 244 | 393 | 390 |
| 2602 | FIM1_2664 | QGN15966.1 | CP015057.1 | 5541 | 1297 | 2434 | 4145 | 1059 | 2285 | 5370 | 1400 | 2145 |
| 2603 | FIM1_2665 | QGN15967.1 | CP015057.1 | 1021 | 61 | 219 | 982 | 58 | 215 | 2272 | 68 | 194 |
| 2604 | FIM1_2666 | QGN15968.1 | CP015057.1 | 4543 | 348 | 505 | 3701 | 197 | 414 | 6933 | 354 | 407 |
| 2605 | FIM1_2667 | QGN15969.1 | CP015057.1 | 9851 | 2046 | 2595 | 7714 | 1469 | 2429 | 10182 | 2133 | 2232 |
| 2606 | FIM1_2668 | QGN15970.1 | CP015057.1 | 13494 | 1966 | 2952 | 9597 | 1376 | 2570 | 10887 | 2205 | 2327 |
| 2607 | FIM1_2669 | QGN15971.1 | CP015057.1 | 122 | 30 | 34 | 101 | 16 | 31 | 195 | 14 | 38 |
| 2608 | FIM1_2671 | QGN15972.1 | CP015057.1 | 42 | 14 | 38 | 34 | 30 | 41 | 49 | 14 | 34 |
| 2609 | FIM1_2672 | QGN15973.1 | CP015057.1 | 2633 | 492 | 766 | 1973 | 374 | 761 | 2103 | 513 | 689 |
| 2610 | FIM1_2674 | QGN15974.1 | CP015057.1 | 14156 | 2497 | 5726 | 10115 | 1526 | 5065 | 8874 | 2507 | 5027 |
| 2611 | FIM1_2675 | QGN15975.1 | CP015057.1 | 35588 | 1086 | 147 | 31698 | 182 | 74 | 78194 | 772 | 81 |
| 2612 | FIM1_2676 | QGN15976.1 | CP015057.1 | 60108 | 33831 | 10282 | 57085 | 13353 | 7343 | 97482 | 33465 | 7760 |
| 2613 | FIM1_2677 | QGN15977.1 | CP015057.1 | 9147 | 3390 | 3335 | 7509 | 2545 | 2882 | 6202 | 3591 | 2951 |
| 2614 | FIM1_2678 | QGN15978.1 | CP015057.1 | 1147 | 364 | 509 | 1116 | 301 | 487 | 929 | 407 | 456 |
| 2615 | FIM1_2679 | QGN15979.1 | CP015057.1 | 6034 | 478 | 1088 | 4775 | 404 | 923 | 6947 | 541 | 831 |
| 2616 | FIM1_2680 | QGN15980.1 | CP015057.1 | 8688 | 4711 | 3273 | 7097 | 3656 | 2987 | 5477 | 5123 | 2904 |
| 2617 | FIM1_2681 | QGN15981.1 | CP015057.1 | 8247 | 3417 | 1933 | 7944 | 1758 | 1685 | 15815 | 3329 | 1675 |
| 2618 | FIM1_2682 | QGN15982.1 | CP015057.1 | 2933 | 347 | 372 | 2215 | 238 | 351 | 4719 | 390 | 385 |
| 2619 | FIM1_2683 | QGN15983.1 | CP015057.1 | 3651 | 1010 | 1992 | 3100 | 498 | 1713 | 3938 | 1050 | 1856 |
| 2620 | FIM1_2684 | QGN15984.1 | CP015057.1 | 9629 | 7577 | 3906 | 6950 | 4980 | 3541 | 3252 | 8353 | 3700 |
| 2621 | FIM1_2685 | QGN15985.1 | CP015057.1 | 1909 | 467 | 864 | 1569 | 470 | 870 | 1946 | 548 | 727 |
| 2622 | FIM1_2686 | QGN15986.1 | CP015057.1 | 1173 | 43 | 128 | 964 | 51 | 124 | 1558 | 43 | 104 |
| 2623 | FIM1_2687 | QGN15987.1 | CP015057.1 | 3570 | 300 | 622 | 2731 | 227 | 577 | 4719 | 329 | 497 |
| 2624 | FIM1_2688 | QGN15988.1 | CP015057.1 | 7015 | 1315 | 1392 | 5493 | 928 | 1278 | 5729 | 1425 | 1359 |
| 2625 | FIM1_2689 | QGN15989.1 | CP015057.1 | 3646 | 926 | 1671 | 3259 | 663 | 1639 | 4113 | 997 | 1464 |
| 2626 | FIM1_2690 | QGN15990.1 | CP015057.1 | 12465 | 5287 | 6002 | 10278 | 3564 | 5568 | 12310 | 5504 | 5429 |
| 2627 | FIM1_2691 | QGN15991.1 | CP015057.1 | 3931 | 910 | 1191 | 2867 | 705 | 1105 | 3456 | 946 | 1145 |
| 2628 | FIM1_2692 | QGN15992.1 | CP015057.1 | 1745 | 390 | 567 | 1325 | 344 | 524 | 1450 | 432 | 506 |
| 2629 | FIM1_2693 | QGN15993.1 | CP015057.1 | 5031 | 1592 | 2054 | 4266 | 1254 | 2078 | 4445 | 1751 | 1831 |
| 2630 | FIM1_2694 | QGN15994.1 | CP015057.1 | 1420 | 375 | 390 | 1044 | 300 | 384 | 1316 | 384 | 386 |
| 2631 | FIM1_2695 | QGN15995.1 | CP015057.1 | 5065 | 2605 | 2338 | 4257 | 1932 | 2219 | 3476 | 2873 | 2171 |
| 2632 | FIM1_2696 | QGN15996.1 | CP015057.1 | 12296 | 8032 | 5598 | 10674 | 6503 | 4782 | 6066 | 8409 | 4983 |
| 2633 | FIM1_2697 | QGN15997.1 | CP015057.1 | 683 | 329 | 434 | 516 | 271 | 377 | 260 | 389 | 374 |
| 2634 | FIM1_2698 | QGN15998.1 | CP015057.1 | 7603 | 1743 | 3052 | 6001 | 1303 | 2861 | 8801 | 1974 | 2908 |
| 2635 | FIM1_2699 | QGN15999.1 | CP015057.1 | 3628 | 3194 | 1673 | 2617 | 1976 | 1560 | 1287 | 3487 | 1456 |
| 2636 | FIM1_2700 | QGN16000.1 | CP015057.1 | 1614 | 240 | 382 | 1243 | 159 | 315 | 1940 | 301 | 357 |
| 2637 | FIM1_2701 | QGN16001.1 | CP015057.1 | 6929 | 898 | 1958 | 5193 | 928 | 1707 | 5218 | 894 | 1653 |
| 2638 | FIM1_2702 | QGN16002.1 | CP015057.1 | 483 | 122 | 157 | 350 | 78 | 129 | 529 | 89 | 123 |

| | | | | | | | | | | | | |
|---|---|---|---|---|---|---|---|---|---|---|---|---|
| 2639 | FIM1_2703 | QGN16003.1 | CP015057.1 | 106 | 31 | 87 | 59 | 24 | 79 | 289 | 21 | 74 |
| 2640 | FIM1_2704 | QGN16004.1 | CP015057.1 | 4676 | 942 | 1212 | 3507 | 659 | 1166 | 3473 | 1004 | 1024 |
| 2641 | FIM1_2705 | QGN16005.1 | CP015057.1 | 5717 | 4686 | 1934 | 3974 | 2096 | 1825 | 4343 | 4729 | 1798 |
| 2642 | FIM1_2706 | QGN16006.1 | CP015057.1 | 47 | 16 | 10 | 37 | 6 | 7 | 41 | 16 | 10 |
| 2643 | FIM1_2707 | QGN16007.1 | CP015057.1 | 1 | 1 | 0 | 1 | 2 | 0 | 0 | 1 | 3 |
| 2644 | FIM1_2708 | QGN16008.1 | CP015057.1 | 11048 | 8722 | 4122 | 8245 | 6000 | 3697 | 3175 | 9088 | 3828 |
| 2645 | FIM1_2709 | QGN16009.1 | CP015057.1 | 6199 | 2629 | 2790 | 5249 | 1917 | 2722 | 5677 | 2814 | 2666 |
| 2646 | FIM1_2710 | QGN16010.1 | CP015057.1 | 13390 | 8748 | 7825 | 12606 | 6437 | 7762 | 14004 | 9051 | 7596 |
| 2647 | FIM1_2711 | QGN16011.1 | CP015057.1 | 1015 | 207 | 240 | 859 | 173 | 223 | 1051 | 233 | 221 |
| 2648 | FIM1_2712 | QGN16012.1 | CP015057.1 | 6684 | 2511 | 2583 | 5956 | 2170 | 2516 | 4903 | 2693 | 2433 |
| 2649 | FIM1_2713 | QGN16013.1 | CP015057.1 | 5008 | 1062 | 1691 | 4028 | 839 | 1658 | 4005 | 1151 | 1567 |
| 2650 | FIM1_2714 | QGN16014.1 | CP015057.1 | 3888 | 750 | 1334 | 2896 | 531 | 1247 | 4014 | 799 | 1186 |
| 2651 | FIM1_2715 | QGN16015.1 | CP015057.1 | 6982 | 2644 | 2641 | 5530 | 1989 | 2325 | 5413 | 2792 | 2558 |
| 2652 | FIM1_2716 | QGN16016.1 | CP015057.1 | 12068 | 6730 | 5655 | 9107 | 4273 | 5190 | 16039 | 7894 | 5090 |
| 2653 | FIM1_2717 | QGN16017.1 | CP015057.1 | 34 | 8 | 13 | 27 | 5 | 14 | 8 | 6 | 13 |
| 2654 | FIM1_2718 | QGN16018.1 | CP015057.1 | 19072 | 2903 | 3849 | 14908 | 2406 | 3509 | 17108 | 3486 | 3268 |
| 2655 | FIM1_2719 | QGN16019.1 | CP015057.1 | 5080 | 1788 | 2067 | 3447 | 1526 | 1956 | 5611 | 1919 | 1812 |
| 2656 | FIM1_2720 | QGN16020.1 | CP015057.1 | 9089 | 1663 | 3921 | 6944 | 1262 | 3689 | 9619 | 1812 | 3610 |
| 2657 | FIM1_2721 | QGN16021.1 | CP015057.1 | 2237 | 905 | 521 | 1467 | 397 | 448 | 1915 | 827 | 496 |
| 2658 | FIM1_2722 | QGN16022.1 | CP015057.1 | 4180 | 2371 | 2373 | 3380 | 1709 | 2393 | 2783 | 2557 | 2144 |
| 2659 | FIM1_2723 | QGN16023.1 | CP015057.1 | 1319 | 328 | 871 | 1129 | 236 | 836 | 1217 | 340 | 900 |
| 2660 | FIM1_2724 | QGN16024.1 | CP015057.1 | 5639 | 668 | 2247 | 3970 | 368 | 1954 | 5432 | 709 | 1827 |
| 2661 | FIM1_2725 | QGN16025.1 | CP015057.1 | 3 | 0 | 3 | 2 | 0 | 1 | 1 | 0 | 0 |
| 2662 | FIM1_2726 | QGN16026.1 | CP015057.1 | 2262 | 229 | 753 | 1904 | 123 | 764 | 3392 | 240 | 687 |
| 2663 | FIM1_2727 | QGN16027.1 | CP015057.1 | 302 | 32 | 79 | 178 | 31 | 78 | 239 | 24 | 58 |
| 2664 | FIM1_2728 | QGN16028.1 | CP015057.1 | 1270 | 80 | 422 | 846 | 94 | 403 | 1409 | 80 | 344 |
| 2665 | FIM1_2729 | QGN16029.1 | CP015057.1 | 2701 | 48 | 502 | 2265 | 9 | 443 | 3283 | 40 | 192 |
| 2666 | FIM1_2730 | QGN16030.1 | CP015057.1 | 291 | 18 | 66 | 232 | 16 | 52 | 506 | 13 | 38 |
| 2667 | FIM1_2731 | QGN16031.1 | CP015057.1 | 13742 | 3023 | 3338 | 10964 | 2138 | 3023 | 12832 | 3297 | 3040 |
| 2668 | FIM1_2732 | QGN16032.1 | CP015057.1 | 24310 | 6171 | 7125 | 19623 | 4390 | 6660 | 24899 | 6793 | 6534 |
| 2669 | FIM1_2733 | QGN16033.1 | CP015057.1 | 868 | 215 | 301 | 676 | 157 | 252 | 715 | 274 | 232 |
| 2670 | FIM1_2734 | QGN16034.1 | CP015057.1 | 540 | 143 | 210 | 407 | 106 | 237 | 526 | 176 | 198 |
| 2671 | FIM1_2735 | QGN16035.1 | CP015057.1 | 1409 | 471 | 442 | 939 | 362 | 457 | 1474 | 520 | 498 |
| 2672 | FIM1_2736 | QGN16036.1 | CP015057.1 | 857 | 389 | 622 | 707 | 303 | 634 | 606 | 387 | 499 |
| 2673 | FIM1_2737 | QGN16037.1 | CP015057.1 | 5344 | 2081 | 2087 | 4244 | 1514 | 1875 | 3449 | 2181 | 1844 |
| 2674 | FIM1_2738 | QGN16038.1 | CP015057.1 | 1236 | 180 | 508 | 1012 | 178 | 461 | 1605 | 198 | 521 |
| 2675 | FIM1_2739 | QGN16039.1 | CP015057.1 | 5138 | 631 | 2270 | 4494 | 425 | 2138 | 6389 | 698 | 1993 |
| 2676 | FIM1_2740 | QGN16040.1 | CP015057.1 | 12135 | 4998 | 6092 | 9663 | 3388 | 5235 | 11696 | 5197 | 5107 |
| 2677 | FIM1_2741 | QGN16041.1 | CP015057.1 | 1790 | 627 | 866 | 1394 | 512 | 735 | 1512 | 680 | 751 |
| 2678 | FIM1_2742 | QGN16042.1 | CP015057.1 | 16715 | 4407 | 6461 | 15106 | 2014 | 6136 | 23137 | 4655 | 5503 |
| 2679 | FIM1_2743 | QGN16043.1 | CP015057.1 | 2324 | 821 | 681 | 1620 | 649 | 626 | 863 | 856 | 694 |
| 2680 | FIM1_2744 | QGN16044.1 | CP015057.1 | 12819 | 3391 | 8386 | 11753 | 2462 | 7198 | 11684 | 3669 | 7071 |
| 2681 | FIM1_2745 | QGN16045.1 | CP015057.1 | 2195 | 1164 | 1437 | 1956 | 1001 | 1528 | 1429 | 1236 | 1280 |
| 2682 | FIM1_2746 | QGN16046.1 | CP015057.1 | 2462 | 675 | 1493 | 2107 | 497 | 1564 | 2473 | 718 | 1308 |
| 2683 | FIM1_2747 | QGN16047.1 | CP015057.1 | 6117 | 2563 | 2372 | 5361 | 2144 | 2262 | 4315 | 2725 | 2327 |
| 2684 | FIM1_2748 | QGN16048.1 | CP015057.1 | 1451 | 217 | 434 | 1140 | 196 | 410 | 1275 | 251 | 393 |
| 2685 | FIM1_2749 | QGN16049.1 | CP015057.1 | 3272 | 202 | 547 | 2376 | 214 | 549 | 3918 | 235 | 537 |
| 2686 | FIM1_2750 | QGN16050.1 | CP015057.1 | 243 | 50 | 58 | 155 | 51 | 59 | 295 | 57 | 59 |

| | | | | | | | | | | | |
|---|---|---|---|---|---|---|---|---|---|---|---|
| 2687 | FIM1_2751 | QGN16051.1 | CP015057.1 | 925 | 458 | 400 | 829 | 266 | 351 | 1155 | 487 | 355 |
| 2688 | FIM1_2752 | QGN16052.1 | CP015057.1 | 3528 | 698 | 1206 | 2839 | 557 | 1068 | 3931 | 805 | 977 |
| 2689 | FIM1_2753 | QGN16053.1 | CP015057.1 | 4125 | 1089 | 838 | 3241 | 646 | 769 | 5245 | 1231 | 804 |
| 2690 | FIM1_2754 | QGN16054.1 | CP015057.1 | 9095 | 1446 | 1349 | 6932 | 956 | 1347 | 17201 | 1574 | 1283 |
| 2691 | FIM1_2755 | QGN16055.1 | CP015057.1 | 2462 | 626 | 899 | 1908 | 446 | 884 | 2271 | 654 | 845 |
| 2692 | FIM1_2756 | QGN16056.1 | CP015057.1 | 1982 | 622 | 808 | 1727 | 499 | 809 | 1376 | 680 | 762 |
| 2693 | FIM1_2757 | QGN16057.1 | CP015057.1 | 11172 | 8829 | 4138 | 7909 | 5813 | 3753 | 2935 | 9346 | 3956 |
| 2694 | FIM1_2758 | QGN16058.1 | CP015057.1 | 1687 | 444 | 548 | 1484 | 298 | 532 | 1114 | 481 | 483 |
| 2695 | FIM1_2759 | QGN16059.1 | CP015057.1 | 178 | 176 | 154 | 162 | 96 | 142 | 146 | 194 | 131 |
| 2696 | FIM1_2760 | QGN16060.1 | CP015057.1 | 26598 | 6376 | 5737 | 20806 | 4456 | 5392 | 26396 | 6913 | 5694 |
| 2697 | FIM1_2761 | QGN16061.1 | CP015057.1 | 4876 | 849 | 1278 | 3758 | 689 | 1199 | 4331 | 951 | 1121 |
| 2698 | FIM1_2762 | QGN16062.1 | CP015057.1 | 7101 | 1801 | 2237 | 5788 | 1207 | 1966 | 7778 | 2039 | 1972 |
| 2699 | FIM1_2763 | QGN16063.1 | CP015057.1 | 2627 | 151 | 15 | 2480 | 25 | 20 | 4350 | 129 | 8 |
| 2700 | FIM1_2764 | QGN16064.1 | CP015057.1 | 2212 | 524 | 1818 | 2029 | 249 | 1689 | 3125 | 495 | 1618 |
| 2701 | FIM1_2765 | QGN16065.1 | CP015057.1 | 1634 | 911 | 1522 | 1409 | 562 | 1321 | 1295 | 989 | 1241 |
| 2702 | FIM1_2766 | QGN16066.1 | CP015057.1 | 7587 | 2943 | 2696 | 6317 | 1959 | 2351 | 8268 | 3139 | 2400 |
| 2703 | FIM1_2767 | QGN16067.1 | CP015057.1 | 6354 | 838 | 2279 | 4954 | 746 | 2203 | 7234 | 993 | 1945 |
| 2704 | FIM1_2768 | QGN16068.1 | CP015057.1 | 2789 | 609 | 972 | 2584 | 329 | 979 | 3812 | 605 | 906 |
| 2705 | FIM1_2769 | QGN16069.1 | CP015057.1 | 3610 | 1303 | 2484 | 3173 | 953 | 2189 | 3553 | 1385 | 2088 |
| 2706 | FIM1_2770 | QGN16070.1 | CP015057.1 | 4151 | 370 | 1240 | 2894 | 291 | 1139 | 5970 | 373 | 925 |
| 2707 | FIM1_2771 | QGN16071.1 | CP015057.1 | 1098 | 280 | 575 | 795 | 219 | 473 | 1019 | 289 | 476 |
| 2708 | FIM1_2772 | QGN16072.1 | CP015057.1 | 6894 | 770 | 1038 | 5090 | 619 | 916 | 7858 | 821 | 837 |
| 2709 | FIM1_2773 | QGN16073.1 | CP015057.1 | 937 | 228 | 459 | 858 | 195 | 366 | 849 | 255 | 355 |
| 2710 | FIM1_2774 | QGN16074.1 | CP015057.1 | 2925 | 614 | 628 | 2240 | 290 | 516 | 2212 | 607 | 448 |
| 2711 | FIM1_2775 | QGN16075.1 | CP015057.1 | 10235 | 3134 | 3936 | 7872 | 2345 | 3573 | 8578 | 3419 | 3529 |
| 2712 | FIM1_2776 | QGN16076.1 | CP015057.1 | 2587 | 363 | 658 | 1872 | 286 | 627 | 1701 | 404 | 520 |
| 2713 | FIM1_2777 | QGN16077.1 | CP015057.1 | 1916 | 203 | 741 | 1535 | 210 | 635 | 2516 | 276 | 561 |
| 2714 | FIM1_2778 | QGN16078.1 | CP015057.1 | 1494 | 324 | 562 | 1205 | 294 | 536 | 1394 | 391 | 429 |
| 2715 | FIM1_2779 | QGN16079.1 | CP015057.1 | 2021 | 836 | 761 | 1576 | 521 | 777 | 1599 | 844 | 806 |
| 2716 | FIM1_2780 | QGN16080.1 | CP015057.1 | 487 | 47 | 113 | 449 | 40 | 119 | 906 | 58 | 172 |
| 2717 | FIM1_2781 | QGN16081.1 | CP015057.1 | 7014 | 459 | 1095 | 5705 | 267 | 915 | 8941 | 485 | 792 |
| 2718 | FIM1_2782 | QGN16082.1 | CP015057.1 | 2369 | 1296 | 1425 | 1928 | 961 | 1443 | 1559 | 1474 | 1209 |
| 2719 | FIM1_2783 | QGN16083.1 | CP015057.1 | 12209 | 9659 | 4709 | 8377 | 5966 | 4225 | 2862 | 10494 | 4272 |
| 2720 | FIM1_2784 | QGN16084.1 | CP015057.1 | 2509 | 425 | 783 | 1885 | 338 | 769 | 2323 | 488 | 738 |
| 2721 | FIM1_2785 | QGN16085.1 | CP015057.1 | 5287 | 1629 | 2755 | 4138 | 1246 | 2779 | 4756 | 1776 | 2442 |
| 2722 | FIM1_2786 | QGN16086.1 | CP015057.1 | 7619 | 1457 | 2513 | 5754 | 1056 | 2397 | 7042 | 1630 | 2253 |
| 2723 | FIM1_2787 | QGN16087.1 | CP015057.1 | 23461 | 4024 | 4614 | 18297 | 3145 | 4321 | 22718 | 4583 | 4558 |
| 2724 | FIM1_2788 | QGN16088.1 | CP015057.1 | 969 | 87 | 327 | 817 | 54 | 287 | 1552 | 95 | 219 |
| 2725 | FIM1_2789 | QGN16089.1 | CP015057.1 | 418 | 143 | 303 | 410 | 113 | 330 | 385 | 153 | 258 |
| 2726 | FIM1_2790 | QGN16090.1 | CP015057.1 | 2304 | 434 | 880 | 1697 | 335 | 850 | 3018 | 459 | 690 |
| 2727 | FIM1_2791 | QGN16091.1 | CP015057.1 | 744 | 79 | 134 | 466 | 58 | 105 | 577 | 98 | 98 |
| 2728 | FIM1_2792 | QGN16092.1 | CP015057.1 | 1098 | 385 | 786 | 944 | 298 | 792 | 1231 | 443 | 740 |
| 2729 | FIM1_2793 | QGN16093.1 | CP015057.1 | 4931 | 978 | 1121 | 4320 | 726 | 1077 | 5172 | 1058 | 968 |
| 2730 | FIM1_2794 | QGN16094.1 | CP015057.1 | 5013 | 1613 | 1864 | 4361 | 1000 | 1810 | 4767 | 1681 | 1760 |
| 2731 | FIM1_2795 | QGN16095.1 | CP015057.1 | 2089 | 426 | 531 | 1635 | 297 | 445 | 1713 | 427 | 411 |
| 2732 | FIM1_2796 | QGN16096.1 | CP015057.1 | 851 | 105 | 137 | 661 | 100 | 130 | 2641 | 101 | 124 |
| 2733 | FIM1_2797 | QGN16097.1 | CP015057.1 | 142 | 6 | 46 | 134 | 19 | 57 | 134 | 24 | 45 |
| 2734 | FIM1_2798 | QGN16098.1 | CP015057.1 | 2406 | 1188 | 1925 | 1650 | 777 | 1747 | 2267 | 1243 | 1743 |

| | | | | | | | | | | | | |
|---|---|---|---|---|---|---|---|---|---|---|---|---|
| 2735 | FIM1_2800 | QGN16099.1 | CP015057.1 | 8111 | 3361 | 2253 | 6471 | 2031 | 2089 | 7476 | 3799 | 1999 |
| 2736 | FIM1_2801 | QGN16100.1 | CP015057.1 | 8376 | 3414 | 3443 | 7268 | 2434 | 3348 | 8713 | 3439 | 3192 |
| 2737 | FIM1_2802 | QGN16101.1 | CP015057.1 | 2063 | 490 | 639 | 1615 | 348 | 622 | 2483 | 540 | 612 |
| 2738 | FIM1_2803 | QGN16102.1 | CP015057.1 | 3885 | 803 | 974 | 2806 | 620 | 966 | 4160 | 801 | 914 |
| 2739 | FIM1_2804 | QGN16103.1 | CP015057.1 | 5725 | 1385 | 1721 | 4280 | 1062 | 1617 | 5333 | 1539 | 1547 |
| 2740 | FIM1_2805 | QGN16104.1 | CP015057.1 | 3275 | 1212 | 1507 | 2614 | 967 | 1440 | 2658 | 1332 | 1296 |
| 2741 | FIM1_2806 | QGN16105.1 | CP015057.1 | 702 | 109 | 191 | 527 | 86 | 190 | 539 | 93 | 153 |
| 2742 | FIM1_2807 | QGN16106.1 | CP015057.1 | 2237 | 671 | 875 | 2002 | 473 | 883 | 2187 | 717 | 777 |
| 2743 | FIM1_2808 | QGN16107.1 | CP015057.1 | 3637 | 997 | 1290 | 3438 | 743 | 1292 | 3771 | 1089 | 1194 |
| 2744 | FIM1_2809 | QGN16108.1 | CP015057.1 | 535 | 42 | 143 | 402 | 36 | 121 | 869 | 56 | 114 |
| 2745 | FIM1_2810 | QGN16109.1 | CP015057.1 | 171 | 146 | 173 | 140 | 157 | 242 | 156 | 136 | 132 |
| 2746 | FIM1_2811 | QGN16110.1 | CP015057.1 | 1221 | 286 | 429 | 1027 | 212 | 368 | 882 | 345 | 361 |
| 2747 | FIM1_2812 | QGN16111.1 | CP015057.1 | 5403 | 1750 | 2213 | 4533 | 1145 | 2093 | 6909 | 1883 | 1848 |
| 2748 | FIM1_2813 | QGN16112.1 | CP015057.1 | 2419 | 424 | 820 | 2197 | 297 | 773 | 3020 | 476 | 680 |
| 2749 | FIM1_2814 | QGN16113.1 | CP015057.1 | 5284 | 924 | 1453 | 4180 | 784 | 1310 | 6690 | 1044 | 1174 |
| 2750 | FIM1_2815 | QGN16114.1 | CP015057.1 | 359 | 213 | 348 | 306 | 184 | 329 | 306 | 223 | 300 |
| 2751 | FIM1_2816 | QGN16115.1 | CP015057.1 | 592 | 242 | 351 | 543 | 208 | 312 | 488 | 258 | 326 |
| 2752 | FIM1_2817 | QGN16116.1 | CP015057.1 | 828 | 327 | 379 | 750 | 315 | 420 | 669 | 390 | 414 |
| 2753 | FIM1_2818 | QGN16117.1 | CP015057.1 | 1446 | 178 | 346 | 1091 | 144 | 322 | 1137 | 194 | 292 |
| 2754 | FIM1_2819 | QGN16118.1 | CP015057.1 | 3708 | 1302 | 1624 | 2872 | 868 | 1555 | 3126 | 1418 | 1591 |
| 2755 | FIM1_2820 | QGN16119.1 | CP015057.1 | 20919 | 904 | 1809 | 18414 | 725 | 1763 | 30392 | 851 | 1258 |
| 2756 | FIM1_2821 | QGN16120.1 | CP015057.1 | 6117 | 913 | 1666 | 4591 | 660 | 1573 | 6791 | 968 | 1349 |
| 2757 | FIM1_2822 | QGN16121.1 | CP015057.1 | 2268 | 446 | 898 | 2158 | 239 | 951 | 2605 | 464 | 714 |
| 2758 | FIM1_2823 | QGN16122.1 | CP015057.1 | 7202 | 2749 | 2871 | 6330 | 2055 | 2669 | 7165 | 2862 | 2718 |
| 2759 | FIM1_2824 | QGN16123.1 | CP015057.1 | 4057 | 737 | 869 | 2936 | 593 | 878 | 4044 | 807 | 807 |
| 2760 | FIM1_2825 | QGN16124.1 | CP015057.1 | 2682 | 546 | 997 | 2078 | 472 | 939 | 2863 | 552 | 837 |
| 2761 | FIM1_2826 | QGN16125.1 | CP015057.1 | 3589 | 281 | 825 | 2267 | 210 | 732 | 3243 | 352 | 680 |
| 2762 | FIM1_2827 | QGN16126.1 | CP015057.1 | 6562 | 4523 | 2681 | 4972 | 2922 | 2509 | 2631 | 4881 | 2494 |
| 2763 | FIM1_2828 | QGN16127.1 | CP015057.1 | 2470 | 272 | 664 | 1852 | 191 | 654 | 2564 | 296 | 539 |
| 2764 | FIM1_2829 | QGN16128.1 | CP015057.1 | 8218 | 1977 | 2403 | 6072 | 1387 | 2120 | 5537 | 2176 | 1982 |
| 2765 | FIM1_2830 | QGN16129.1 | CP015057.1 | 7078 | 1277 | 1231 | 5402 | 1136 | 1148 | 6516 | 1428 | 1493 |
| 2766 | FIM1_2831 | QGN16130.1 | CP015057.1 | 382 | 88 | 100 | 354 | 109 | 82 | 1465 | 57 | 54 |
| 2767 | FIM1_2832 | QGN16131.1 | CP015057.1 | 6211 | 783 | 1127 | 4803 | 566 | 1005 | 7140 | 894 | 874 |
| 2768 | FIM1_2833 | QGN16132.1 | CP015057.1 | 4163 | 2529 | 1601 | 2846 | 1590 | 1544 | 1027 | 2574 | 1375 |
| 2769 | FIM1_2834 | QGN16133.1 | CP015057.1 | 11095 | 551 | 1154 | 8613 | 367 | 1072 | 11791 | 602 | 1107 |
| 2770 | FIM1_2835 | QGN16134.1 | CP015057.1 | 4236 | 634 | 1005 | 3649 | 532 | 960 | 4255 | 723 | 843 |
| 2771 | FIM1_2836 | QGN16135.1 | CP015057.1 | 3185 | 413 | 732 | 2184 | 347 | 685 | 2950 | 512 | 591 |
| 2772 | FIM1_2837 | QGN16136.1 | CP015057.1 | 2577 | 418 | 1086 | 2040 | 233 | 980 | 3145 | 396 | 965 |
| 2773 | FIM1_2838 | QGN16137.1 | CP015057.1 | 7419 | 1953 | 1741 | 5863 | 1349 | 1700 | 7842 | 2047 | 1479 |
| 2774 | FIM1_2839 | QGN16138.1 | CP015057.1 | 2261 | 295 | 536 | 1752 | 219 | 517 | 2578 | 276 | 487 |
| 2775 | FIM1_2840 | QGN16139.1 | CP015057.1 | 329 | 46 | 78 | 240 | 33 | 78 | 305 | 43 | 64 |
| 2776 | FIM1_2841 | QGN16140.1 | CP015057.1 | 2797 | 751 | 836 | 2196 | 579 | 823 | 1927 | 827 | 767 |
| 2777 | FIM1_2842 | QGN16141.1 | CP015057.1 | 6100 | 2913 | 2634 | 4704 | 2106 | 2505 | 4018 | 3182 | 2290 |
| 2778 | FIM1_2843 | QGN16142.1 | CP015057.1 | 536 | 85 | 229 | 421 | 74 | 220 | 523 | 101 | 201 |
| 2779 | FIM1_2844 | QGN16143.1 | CP015057.1 | 987 | 205 | 534 | 841 | 181 | 505 | 1008 | 256 | 396 |
| 2780 | FIM1_2845 | QGN16144.1 | CP015057.1 | 78334 | 48790 | 31121 | 73184 | 31809 | 27224 | 111232 | 48870 | 27449 |
| 2781 | FIM1_2846 | QGN16145.1 | CP015057.1 | 2809 | 1354 | 908 | 2266 | 885 | 770 | 1465 | 1474 | 873 |
| 2782 | FIM1_2847 | QGN16146.1 | CP015057.1 | 15750 | 182 | 2022 | 12348 | 516 | 2015 | 14583 | 228 | 1404 |

| | | | | | | | | | | | | |
|---|---|---|---|---|---|---|---|---|---|---|---|---|
| 2783 | FIM1_2848 | QGN16147.1 | CP015057.1 | 1377 | 386 | 499 | 1163 | 256 | 436 | 1365 | 381 | 383 |
| 2784 | FIM1_2849 | QGN16148.1 | CP015057.1 | 23640 | 14287 | 13210 | 22284 | 12088 | 12530 | 22283 | 15004 | 12433 |
| 2785 | FIM1_2850 | QGN16149.1 | CP015057.1 | 3849 | 1636 | 1982 | 3253 | 1390 | 1993 | 2583 | 1935 | 1658 |
| 2786 | FIM1_2851 | QGN16150.1 | CP015057.1 | 8242 | 1314 | 2121 | 6505 | 886 | 1926 | 10097 | 1418 | 1776 |
| 2787 | FIM1_2852 | QGN16151.1 | CP015057.1 | 1085 | 427 | 778 | 947 | 295 | 790 | 1354 | 471 | 687 |
| 2788 | FIM1_2853 | QGN16152.1 | CP015057.1 | 7416 | 5773 | 2646 | 5210 | 4005 | 2500 | 1974 | 6293 | 2624 |
| 2789 | FIM1_2854 | QGN16153.1 | CP015057.1 | 3301 | 1291 | 1546 | 3064 | 814 | 1579 | 3821 | 1325 | 1398 |
| 2790 | FIM1_2855 | QGN16154.1 | CP015057.1 | 245 | 39 | 113 | 183 | 33 | 100 | 272 | 48 | 94 |
| 2791 | FIM1_2856 | QGN16155.1 | CP015057.1 | 2061 | 187 | 482 | 1547 | 171 | 418 | 2554 | 204 | 383 |
| 2792 | FIM1_2857 | QGN16156.1 | CP015057.1 | 1398 | 739 | 681 | 1075 | 503 | 612 | 854 | 808 | 590 |
| 2793 | FIM1_2858 | QGN16157.1 | CP015057.1 | 3180 | 775 | 898 | 2578 | 512 | 785 | 3128 | 883 | 728 |
| 2794 | FIM1_2859 | QGN16158.1 | CP015057.1 | 6128 | 2943 | 3879 | 6356 | 1390 | 3695 | 7550 | 3081 | 3509 |
| 2795 | FIM1_2860 | QGN16159.1 | CP015057.1 | 21213 | 10911 | 7036 | 18407 | 6137 | 4389 | 13120 | 11090 | 7360 |
| 2796 | FIM1_2862 | QGN16160.1 | CP015057.1 | 313 | 122 | 227 | 318 | 102 | 241 | 236 | 137 | 185 |
| 2797 | FIM1_2863 | QGN16161.1 | CP015057.1 | 4804 | 804 | 1240 | 3085 | 437 | 1156 | 3822 | 846 | 1057 |
| 2798 | FIM1_2864 | QGN16162.1 | CP015057.1 | 5143 | 3558 | 2615 | 4041 | 2182 | 2554 | 3142 | 3820 | 2326 |
| 2799 | FIM1_2865 | QGN16163.1 | CP015057.1 | 1890 | 913 | 1029 | 1306 | 686 | 989 | 952 | 1041 | 879 |
| 2800 | FIM1_2866 | QGN16164.1 | CP015057.1 | 1686 | 267 | 500 | 1356 | 180 | 512 | 2375 | 277 | 472 |
| 2801 | FIM1_2867 | QGN16165.1 | CP015057.1 | 3517 | 714 | 1366 | 2755 | 617 | 1352 | 2424 | 790 | 1166 |
| 2802 | FIM1_2868 | QGN16166.1 | CP015057.1 | 198 | 8 | 36 | 134 | 8 | 33 | 185 | 14 | 28 |
| 2803 | FIM1_2869 | QGN16167.1 | CP015057.1 | 576 | 13 | 92 | 416 | 9 | 70 | 822 | 13 | 56 |
| 2804 | FIM1_2870 | QGN16168.1 | CP015057.1 | 12222 | 3994 | 4000 | 9077 | 2744 | 3858 | 9541 | 4073 | 3767 |
| 2805 | FIM1_2871 | QGN16169.1 | CP015057.1 | 2819 | 622 | 817 | 2183 | 484 | 768 | 2249 | 713 | 730 |
| 2806 | FIM1_2872 | QGN16170.1 | CP015057.1 | 13159 | 895 | 1246 | 10419 | 780 | 1129 | 14659 | 888 | 1068 |
| 2807 | FIM1_2873 | QGN16171.1 | CP015057.1 | 1241 | 292 | 244 | 1166 | 243 | 195 | 1532 | 301 | 225 |
| 2808 | FIM1_2874 | QGN16172.1 | CP015057.1 | 13544 | 2679 | 3390 | 11398 | 2068 | 2977 | 16517 | 2781 | 2969 |
| 2809 | FIM1_2875 | QGN16173.1 | CP015057.1 | 4721 | 2428 | 1574 | 4240 | 1563 | 1469 | 5725 | 2576 | 1486 |
| 2810 | FIM1_2876 | QGN16174.1 | CP015057.1 | 5408 | 1386 | 1909 | 4805 | 997 | 1779 | 4712 | 1634 | 1628 |
| 2811 | FIM1_2877 | QGN16175.1 | CP015057.1 | 4978 | 5596 | 2714 | 3705 | 3539 | 2493 | 1374 | 5903 | 2495 |
| 2812 | FIM1_2878 | QGN16176.1 | CP015057.1 | 3289 | 621 | 857 | 2244 | 490 | 888 | 3143 | 690 | 797 |
| 2813 | FIM1_2879 | QGN16177.1 | CP015057.1 | 4471 | 738 | 785 | 3324 | 601 | 722 | 6575 | 836 | 659 |
| 2814 | FIM1_2880 | QGN16178.1 | CP015057.1 | 3001 | 815 | 829 | 2410 | 682 | 753 | 2882 | 911 | 827 |
| 2815 | FIM1_2881 | QGN16179.1 | CP015057.1 | 2273 | 956 | 995 | 1783 | 799 | 993 | 1270 | 1083 | 964 |
| 2816 | FIM1_2882 | QGN16180.1 | CP015057.1 | 4475 | 824 | 914 | 3236 | 569 | 935 | 5527 | 846 | 892 |
| 2817 | FIM1_2883 | QGN16181.1 | CP015057.1 | 6414 | 1216 | 1857 | 5080 | 953 | 1776 | 6888 | 1462 | 1628 |
| 2818 | FIM1_2884 | QGN16182.1 | CP015057.1 | 2000 | 455 | 598 | 1434 | 302 | 537 | 1574 | 451 | 431 |
| 2819 | FIM1_2885 | QGN16183.1 | CP015057.1 | 2275 | 2443 | 1299 | 1564 | 1418 | 1216 | 441 | 2532 | 1163 |
| 2820 | FIM1_2886 | QGN16184.1 | CP015057.1 | 1043 | 1351 | 664 | 635 | 865 | 661 | 196 | 1528 | 621 |
| 2821 | FIM1_2887 | QGN16185.1 | CP015057.1 | 6950 | 6432 | 3484 | 4877 | 4315 | 3155 | 2077 | 7075 | 3296 |
| 2822 | FIM1_2888 | QGN16186.1 | CP015057.1 | 742 | 77 | 340 | 625 | 68 | 321 | 1047 | 92 | 317 |
| 2823 | FIM1_2889 | QGN16187.1 | CP015057.1 | 5657 | 1213 | 1803 | 4256 | 1046 | 1682 | 4382 | 1360 | 1587 |
| 2824 | FIM1_2890 | QGN16188.1 | CP015057.1 | 7888 | 2694 | 4334 | 6816 | 1678 | 3799 | 9769 | 2831 | 3695 |
| 2825 | FIM1_2891 | QGN16189.1 | CP015057.1 | 9414 | 2933 | 2600 | 7883 | 1950 | 2403 | 11232 | 3072 | 2265 |
| 2826 | FIM1_2892 | QGN16190.1 | CP015057.1 | 953 | 103 | 140 | 819 | 86 | 169 | 966 | 132 | 136 |
| 2827 | FIM1_2893 | QGN16191.1 | CP015057.1 | 1708 | 287 | 454 | 1522 | 231 | 363 | 1384 | 314 | 436 |
| 2828 | FIM1_2894 | QGN16192.1 | CP015057.1 | 3456 | 417 | 663 | 2486 | 388 | 663 | 4122 | 444 | 580 |
| 2829 | FIM1_2895 | QGN16193.1 | CP015057.1 | 5965 | 2009 | 2903 | 5054 | 1544 | 2773 | 6985 | 2270 | 2513 |
| 2830 | FIM1_2896 | QGN16194.1 | CP015057.1 | 4025 | 1088 | 1129 | 2959 | 751 | 998 | 4509 | 1106 | 928 |

| | | | | | | | | | | | | |
|---|---|---|---|---|---|---|---|---|---|---|---|---|
| 2831 | FIM1_2897 | QGN16195.1 | CP015057.1 | 17602 | 4305 | 3257 | 14661 | 2806 | 2960 | 22224 | 4546 | 2766 |
| 2832 | FIM1_2898 | QGN16196.1 | CP015057.1 | 26478 | 16163 | 14210 | 24805 | 13058 | 13705 | 26470 | 16419 | 14128 |
| 2833 | FIM1_2899 | QGN16197.1 | CP015057.1 | 21256 | 12772 | 5608 | 16701 | 10381 | 5024 | 9562 | 13387 | 5576 |
| 2834 | FIM1_2900 | QGN16198.1 | CP015057.1 | 15531 | 7731 | 2717 | 11999 | 4561 | 2481 | 16024 | 8329 | 2563 |
| 2835 | FIM1_2901 | QGN16199.1 | CP015057.1 | 3882 | 1366 | 1323 | 2751 | 868 | 1287 | 3340 | 1362 | 1255 |
| 2836 | FIM1_2903 | QGN16200.1 | CP015057.1 | 7709 | 1174 | 1855 | 6119 | 735 | 1685 | 10307 | 1309 | 1481 |
| 2837 | FIM1_2904 | QGN16201.1 | CP015057.1 | 5103 | 1032 | 1164 | 4448 | 727 | 1021 | 5937 | 1020 | 1073 |
| 2838 | FIM1_2905 | QGN16202.1 | CP015057.1 | 3671 | 530 | 674 | 2947 | 346 | 634 | 3731 | 547 | 569 |
| 2839 | FIM1_2906 | QGN16203.1 | CP015057.1 | 493 | 71 | 181 | 386 | 77 | 154 | 291 | 81 | 133 |
| 2840 | FIM1_2907 | QGN16204.1 | CP015057.1 | 2131 | 745 | 988 | 1664 | 629 | 899 | 1125 | 808 | 902 |
| 2841 | FIM1_2908 | QGN16205.1 | CP015057.1 | 7459 | 2032 | 2112 | 5729 | 1417 | 2063 | 5132 | 2153 | 2185 |
| 2842 | FIM1_2909 | QGN16206.1 | CP015057.1 | 3480 | 902 | 875 | 2213 | 626 | 800 | 2715 | 1004 | 881 |
| 2843 | FIM1_2911 | QGN16207.1 | CP015057.1 | 778 | 165 | 387 | 662 | 103 | 398 | 1248 | 170 | 363 |
| 2844 | FIM1_2912 | QGN16208.1 | CP015057.1 | 10695 | 1855 | 1950 | 7956 | 1180 | 1734 | 11850 | 1859 | 1662 |
| 2845 | FIM1_2913 | QGN16209.1 | CP015057.1 | 69 | 13 | 12 | 23 | 8 | 9 | 71 | 18 | 3 |
| 2846 | FIM1_2914 | QGN16210.1 | CP015057.1 | 6042 | 2029 | 1736 | 4863 | 1431 | 1637 | 2252 | 2146 | 1706 |
| 2847 | FIM1_2915 | QGN16211.1 | CP015057.1 | 1375 | 521 | 1968 | 1418 | 328 | 1830 | 1959 | 486 | 1748 |
| 2848 | FIM1_2916 | QGN16212.1 | CP015057.1 | 10652 | 3201 | 5599 | 8496 | 2587 | 5315 | 9758 | 3384 | 5795 |
| 2849 | FIM1_2917 | QGN16213.1 | CP015057.1 | 4104 | 594 | 1036 | 3058 | 402 | 1005 | 4808 | 626 | 913 |
| 2850 | FIM1_2918 | QGN16214.1 | CP015057.1 | 1947 | 676 | 593 | 1573 | 459 | 622 | 1996 | 681 | 614 |
| 2851 | FIM1_2919 | QGN16215.1 | CP015057.1 | 9448 | 1794 | 2553 | 7020 | 1473 | 2448 | 8859 | 2023 | 2347 |
| 2852 | FIM1_2920 | QGN16216.1 | CP015057.1 | 637 | 277 | 323 | 571 | 190 | 295 | 305 | 289 | 230 |
| 2853 | FIM1_2921 | QGN16217.1 | CP015057.1 | 5864 | 3106 | 2245 | 4845 | 2444 | 2098 | 2096 | 3345 | 2147 |
| 2854 | FIM1_2922 | QGN16218.1 | CP015057.1 | 3209 | 323 | 912 | 2404 | 251 | 849 | 3525 | 375 | 714 |
| 2855 | FIM1_2923 | QGN16219.1 | CP015057.1 | 7500 | 4008 | 3200 | 6164 | 2650 | 2972 | 8014 | 3863 | 3211 |
| 2856 | FIM1_2924 | QGN16220.1 | CP015057.1 | 10954 | 6147 | 5159 | 10155 | 4606 | 5030 | 10050 | 6127 | 5113 |
| 2857 | FIM1_2925 | QGN16221.1 | CP015057.1 | 160 | 16 | 35 | 118 | 25 | 34 | 183 | 28 | 28 |
| 2858 | FIM1_2926 | QGN16222.1 | CP015057.1 | 5645 | 1520 | 2058 | 4727 | 924 | 1860 | 5586 | 1714 | 1801 |
| 2859 | FIM1_2927 | QGN16223.1 | CP015057.1 | 11123 | 3827 | 5499 | 9733 | 2622 | 5081 | 11501 | 4266 | 4795 |
| 2860 | FIM1_2928 | QGN16224.1 | CP015057.1 | 4422 | 422 | 456 | 3549 | 273 | 375 | 9874 | 438 | 370 |
| 2861 | FIM1_2929 | QGN16225.1 | CP015057.1 | 6711 | 561 | 1381 | 5056 | 446 | 1114 | 9837 | 572 | 1113 |
| 2862 | FIM1_2930 | QGN16226.1 | CP015057.1 | 527 | 365 | 326 | 328 | 213 | 253 | 130 | 397 | 255 |
| 2863 | FIM1_2931 | QGN16227.1 | CP015057.1 | 5583 | 3329 | 2268 | 4501 | 2507 | 2233 | 3689 | 3460 | 2241 |
| 2864 | FIM1_2932 | QGN16228.1 | CP015057.1 | 1910 | 563 | 800 | 1641 | 347 | 766 | 2107 | 615 | 773 |
| 2865 | FIM1_2933 | QGN16229.1 | CP015057.1 | 9380 | 1614 | 2246 | 7310 | 1303 | 2092 | 8791 | 1744 | 2028 |
| 2866 | FIM1_2934 | QGN16230.1 | CP015057.1 | 9303 | 4023 | 2494 | 6230 | 2539 | 2335 | 5306 | 4240 | 2217 |
| 2867 | FIM1_2935 | QGN16231.1 | CP015057.1 | 31495 | 13198 | 12030 | 27384 | 9643 | 10736 | 28620 | 13626 | 11719 |
| 2868 | FIM1_2936 | QGN16232.1 | CP015057.1 | 13479 | 5209 | 3584 | 9753 | 3134 | 3126 | 14023 | 5674 | 2901 |
| 2869 | FIM1_2937 | QGN16233.1 | CP015057.1 | 2435 | 722 | 692 | 1896 | 487 | 615 | 1927 | 737 | 724 |
| 2870 | FIM1_2939 | QGN16234.1 | CP015057.1 | 1644 | 141 | 470 | 1323 | 125 | 435 | 2220 | 158 | 376 |
| 2871 | FIM1_2940 | QGN16235.1 | CP015057.1 | 10 | 2 | 5 | 21 | 1 | 3 | 10 | 4 | 5 |
| 2872 | FIM1_2941 | QGN16236.1 | CP015057.1 | 2794 | 1154 | 1452 | 2687 | 876 | 1377 | 2280 | 1271 | 1319 |
| 2873 | FIM1_2942 | QGN16237.1 | CP015057.1 | 13417 | 3826 | 9316 | 10734 | 2695 | 8536 | 14591 | 3949 | 8475 |
| 2874 | FIM1_2943 | QGN16238.1 | CP015057.1 | 3268 | 681 | 1063 | 2680 | 520 | 994 | 2871 | 718 | 934 |
| 2875 | FIM1_2944 | QGN16239.1 | CP015057.1 | 1977 | 754 | 817 | 1414 | 615 | 809 | 1437 | 843 | 717 |
| 2876 | FIM1_2945 | QGN16240.1 | CP015057.1 | 1431 | 605 | 731 | 1155 | 485 | 687 | 805 | 671 | 633 |
| 2877 | FIM1_2946 | QGN16241.1 | CP015057.1 | 6172 | 792 | 1694 | 4686 | 675 | 1525 | 6757 | 912 | 1386 |
| 2878 | FIM1_2947 | QGN16242.1 | CP015057.1 | 6602 | 3075 | 3298 | 5580 | 1896 | 3145 | 7219 | 3331 | 3131 |

| | | | | | | | | | | | | |
|---|---|---|---|---|---|---|---|---|---|---|---|---|
| 2879 | FIM1_2948 | QGN16243.1 | CP015057.1 | 3250 | 445 | 1093 | 2376 | 421 | 1014 | 3293 | 465 | 948 |
| 2880 | FIM1_2949 | QGN16244.1 | CP015057.1 | 7618 | 1734 | 3311 | 6678 | 1171 | 3063 | 9012 | 1895 | 2777 |
| 2881 | FIM1_2950 | QGN16245.1 | CP015057.1 | 367 | 28 | 68 | 317 | 34 | 71 | 495 | 41 | 46 |
| 2882 | FIM1_2951 | QGN16246.1 | CP015057.1 | 6658 | 1550 | 2031 | 5514 | 1155 | 1914 | 6940 | 1830 | 1839 |
| 2883 | FIM1_2952 | QGN16247.1 | CP015057.1 | 2098 | 419 | 802 | 1725 | 291 | 708 | 2863 | 438 | 632 |
| 2884 | FIM1_2953 | QGN16248.1 | CP015057.1 | 4837 | 1003 | 1402 | 3775 | 666 | 1375 | 5877 | 1130 | 1323 |
| 2885 | FIM1_2954 | QGN16249.1 | CP015057.1 | 1971 | 373 | 524 | 1440 | 291 | 520 | 1575 | 398 | 477 |
| 2886 | FIM1_2955 | QGN16250.1 | CP015057.1 | 1806 | 918 | 985 | 1632 | 726 | 1067 | 1315 | 1035 | 917 |
| 2887 | FIM1_2956 | QGN16251.1 | CP015057.1 | 863 | 230 | 270 | 684 | 208 | 300 | 545 | 260 | 289 |
| 2888 | FIM1_2957 | QGN16252.1 | CP015057.1 | 8141 | 5541 | 2797 | 5627 | 3266 | 2617 | 2839 | 5835 | 2553 |
| 2889 | FIM1_2958 | QGN16253.1 | CP015057.1 | 5686 | 1247 | 1542 | 3955 | 938 | 1501 | 6307 | 1387 | 1326 |
| 2890 | FIM1_2959 | QGN16254.1 | CP015057.1 | 7912 | 4834 | 2996 | 7133 | 4679 | 2979 | 5092 | 5372 | 3025 |
| 2891 | FIM1_2960 | QGN16255.1 | CP015057.1 | 7080 | 4495 | 2307 | 5126 | 2896 | 2174 | 2396 | 4466 | 2053 |
| 2892 | FIM1_2961 | QGN16256.1 | CP015057.1 | 8911 | 5268 | 3030 | 6378 | 3657 | 2801 | 4277 | 5588 | 3009 |
| 2893 | FIM1_2962 | QGN16257.1 | CP015057.1 | 1190 | 204 | 228 | 902 | 128 | 235 | 1353 | 176 | 186 |
| 2894 | FIM1_2963 | QGN16258.1 | CP015057.1 | 931 | 274 | 372 | 767 | 200 | 340 | 888 | 260 | 317 |
| 2895 | FIM1_2964 | QGN16259.1 | CP015057.1 | 11740 | 6472 | 6565 | 11648 | 4975 | 6099 | 10866 | 6842 | 6078 |
| 2896 | FIM1_2965 | QGN16260.1 | CP015057.1 | 2893 | 938 | 1173 | 2258 | 693 | 1099 | 2906 | 1076 | 1083 |
| 2897 | FIM1_2966 | QGN16261.1 | CP015057.1 | 840 | 238 | 323 | 834 | 188 | 313 | 774 | 248 | 282 |
| 2898 | FIM1_2967 | QGN16262.1 | CP015057.1 | 4307 | 868 | 1322 | 3272 | 646 | 1188 | 3713 | 903 | 1097 |
| 2899 | FIM1_2968 | QGN16263.1 | CP015057.1 | 5562 | 762 | 1920 | 4302 | 621 | 1698 | 5301 | 831 | 1645 |
| 2900 | FIM1_2969 | QGN16264.1 | CP015057.1 | 9017 | 5047 | 5032 | 8507 | 4068 | 5037 | 8858 | 5164 | 4939 |
| 2901 | FIM1_2970 | QGN16265.1 | CP015057.1 | 2412 | 402 | 521 | 2048 | 332 | 452 | 2697 | 446 | 397 |
| 2902 | FIM1_2971 | QGN16266.1 | CP015057.1 | 839 | 687 | 534 | 707 | 540 | 483 | 297 | 793 | 477 |
| 2903 | FIM1_2972 | QGN16267.1 | CP015057.1 | 3288 | 338 | 581 | 2723 | 265 | 569 | 2571 | 381 | 487 |
| 2904 | FIM1_2973 | QGN16268.1 | CP015057.1 | 3059 | 88 | 314 | 2938 | 74 | 285 | 3472 | 114 | 192 |
| 2905 | FIM1_2974 | QGN16269.1 | CP015057.1 | 7409 | 1168 | 1698 | 5147 | 863 | 1524 | 6220 | 1265 | 1403 |
| 2906 | FIM1_2975 | QGN16270.1 | CP015057.1 | 5605 | 3547 | 2033 | 4202 | 2262 | 1930 | 2933 | 3760 | 1948 |
| 2907 | FIM1_2976 | QGN16271.1 | CP015057.1 | 1454 | 421 | 485 | 1151 | 316 | 445 | 1383 | 501 | 454 |
| 2908 | FIM1_2977 | QGN16272.1 | CP015057.1 | 1673 | 343 | 778 | 1166 | 230 | 688 | 2157 | 315 | 670 |
| 2909 | FIM1_2978 | QGN16273.1 | CP015057.1 | 4938 | 2518 | 2663 | 4735 | 1681 | 2530 | 5176 | 2760 | 2379 |
| 2910 | FIM1_2979 | QGN16274.1 | CP015057.1 | 616 | 100 | 151 | 498 | 86 | 162 | 555 | 124 | 135 |
| 2911 | FIM1_2980 | QGN16275.1 | CP015057.1 | 2742 | 355 | 791 | 2123 | 260 | 751 | 2662 | 375 | 639 |
| 2912 | FIM1_2981 | QGN16276.1 | CP015057.1 | 9628 | 6852 | 3723 | 7342 | 4701 | 3302 | 3655 | 7482 | 3585 |
| 2913 | FIM1_2982 | QGN16277.1 | CP015057.1 | 9558 | 2061 | 3122 | 7671 | 1351 | 2814 | 11494 | 2314 | 2562 |
| 2914 | FIM1_2983 | QGN16278.1 | CP015057.1 | 10227 | 6839 | 5454 | 8005 | 5081 | 5062 | 5642 | 7120 | 5169 |
| 2915 | FIM1_2984 | QGN16279.1 | CP015057.1 | 586 | 645 | 613 | 423 | 465 | 583 | 175 | 735 | 559 |
| 2916 | FIM1_2985 | QGN16280.1 | CP015057.1 | 30629 | 12855 | 17411 | 27009 | 6909 | 15799 | 35813 | 13867 | 15957 |
| 2917 | FIM1_2986 | QGN16281.1 | CP015057.1 | 2738 | 495 | 922 | 2141 | 410 | 832 | 2472 | 558 | 763 |
| 2918 | FIM1_2987 | QGN16282.1 | CP015057.1 | 5281 | 1277 | 1604 | 4585 | 1071 | 1488 | 4762 | 1403 | 1538 |
| 2919 | FIM1_2988 | QGN16283.1 | CP015057.1 | 3546 | 522 | 899 | 2854 | 415 | 805 | 3804 | 588 | 599 |
| 2920 | FIM1_2989 | QGN16284.1 | CP015057.1 | 1525 | 913 | 818 | 1368 | 810 | 821 | 868 | 1040 | 823 |
| 2921 | FIM1_2990 | QGN16285.1 | CP015057.1 | 32157 | 10214 | 4038 | 25733 | 7621 | 3405 | 30147 | 11206 | 3568 |
| 2922 | FIM1_2991 | QGN16286.1 | CP015057.1 | 387 | 50 | 578 | 366 | 27 | 483 | 289 | 54 | 491 |
| 2923 | FIM1_2992 | QGN16287.1 | CP015057.1 | 202 | 24 | 80 | 141 | 17 | 62 | 118 | 23 | 74 |
| 2924 | FIM1_2993 | QGN16288.1 | CP015057.1 | 2632 | 489 | 925 | 2081 | 440 | 860 | 2713 | 491 | 807 |
| 2925 | FIM1_2994 | QGN16289.1 | CP015057.1 | 2492 | 378 | 859 | 2055 | 355 | 855 | 2580 | 412 | 736 |
| 2926 | FIM1_2995 | QGN16290.1 | CP015057.1 | 927 | 27 | 273 | 692 | 34 | 238 | 1201 | 30 | 196 |

| | | | | | | | | | | | | |
|---|---|---|---|---|---|---|---|---|---|---|---|---|
| 2927 | FIM1_2996 | QGN16291.1 | CP015057.1 | 1225 | 89 | 454 | 1179 | 23 | 390 | 3900 | 97 | 307 |
| 2928 | FIM1_2997 | QGN16292.1 | CP015057.1 | 5064 | 843 | 1280 | 4000 | 632 | 1214 | 6763 | 979 | 1161 |
| 2929 | FIM1_2998 | QGN16293.1 | CP015057.1 | 7006 | 1838 | 3772 | 5894 | 1154 | 3403 | 8448 | 1902 | 3384 |
| 2930 | FIM1_2999 | QGN16294.1 | CP015057.1 | 2333 | 386 | 907 | 1846 | 273 | 783 | 2694 | 424 | 767 |
| 2931 | FIM1_3000 | QGN16295.1 | CP015057.1 | 3693 | 1895 | 981 | 3138 | 1046 | 902 | 4059 | 1982 | 875 |
| 2932 | FIM1_3001 | QGN16296.1 | CP015057.1 | 3008 | 778 | 1123 | 2161 | 604 | 1016 | 1935 | 925 | 984 |
| 2933 | FIM1_3002 | QGN16297.1 | CP015057.1 | 846 | 173 | 409 | 594 | 133 | 379 | 996 | 187 | 337 |
| 2934 | FIM1_3003 | QGN16298.1 | CP015057.1 | 2062 | 772 | 1054 | 1437 | 622 | 1032 | 683 | 924 | 788 |
| 2935 | FIM1_3004 | QGN16299.1 | CP015057.1 | 14642 | 3946 | 5068 | 11583 | 2958 | 4814 | 13344 | 4325 | 4525 |
| 2936 | FIM1_3005 | QGN16300.1 | CP015057.1 | 2295 | 2628 | 1865 | 1857 | 2170 | 1782 | 629 | 2966 | 1708 |
| 2937 | FIM1_3006 | QGN16301.1 | CP015057.1 | 13099 | 4961 | 3621 | 10634 | 3816 | 3304 | 8139 | 5301 | 3220 |
| 2938 | FIM1_3007 | QGN16302.1 | CP015057.1 | 8379 | 3429 | 3244 | 6049 | 2371 | 3055 | 5866 | 3706 | 2897 |
| 2939 | FIM1_3008 | QGN16303.1 | CP015057.1 | 5361 | 1256 | 2032 | 4163 | 975 | 1782 | 4968 | 1357 | 1645 |
| 2940 | FIM1_3009 | QGN16304.1 | CP015057.1 | 4626 | 831 | 1511 | 3412 | 608 | 1378 | 4021 | 854 | 1333 |
| 2941 | FIM1_3010 | QGN16305.1 | CP015057.1 | 1345 | 344 | 662 | 1082 | 280 | 629 | 1085 | 418 | 499 |
| 2942 | FIM1_3011 | QGN16306.1 | CP015057.1 | 1913 | 319 | 414 | 1499 | 257 | 417 | 2028 | 338 | 333 |
| 2943 | FIM1_3012 | QGN16307.1 | CP015057.1 | 5188 | 2400 | 2072 | 3940 | 1557 | 1955 | 3815 | 2478 | 1741 |
| 2944 | FIM1_3013 | QGN16308.1 | CP015057.1 | 6979 | 1343 | 2130 | 5316 | 1010 | 2004 | 6266 | 1417 | 1980 |
| 2945 | FIM1_3014 | QGN16309.1 | CP015057.1 | 11897 | 8782 | 3966 | 8463 | 6215 | 3520 | 3903 | 9504 | 3913 |
| 2946 | FIM1_3015 | QGN16310.1 | CP015057.1 | 6392 | 4078 | 4523 | 5767 | 2896 | 4356 | 6409 | 4340 | 4253 |
| 2947 | FIM1_3016 | QGN16311.1 | CP015057.1 | 3220 | 263 | 683 | 2225 | 213 | 581 | 2224 | 281 | 456 |
| 2948 | FIM1_3017 | QGN16312.1 | CP015057.1 | 39385 | 21845 | 18908 | 36772 | 16962 | 17747 | 40238 | 21352 | 18492 |
| 2949 | FIM1_3018 | QGN16313.1 | CP015057.1 | 800 | 462 | 498 | 625 | 346 | 505 | 468 | 499 | 466 |
| 2950 | FIM1_3019 | QGN16314.1 | CP015057.1 | 10648 | 2860 | 5227 | 9006 | 2143 | 4846 | 11181 | 2970 | 4597 |
| 2951 | FIM1_3020 | QGN16315.1 | CP015057.1 | 465 | 59 | 68 | 373 | 53 | 62 | 294 | 68 | 49 |
| 2952 | FIM1_3021 | QGN16316.1 | CP015057.1 | 6959 | 3544 | 2671 | 5101 | 2214 | 2469 | 6024 | 3870 | 2529 |
| 2953 | FIM1_3022 | QGN16317.1 | CP015057.1 | 488 | 198 | 147 | 356 | 142 | 158 | 332 | 221 | 131 |
| 2954 | FIM1_3023 | QGN16318.1 | CP015057.1 | 1949 | 104 | 499 | 1631 | 93 | 423 | 3468 | 123 | 418 |
| 2955 | FIM1_3024 | QGN16319.1 | CP015057.1 | 140 | 23 | 28 | 89 | 12 | 38 | 120 | 30 | 18 |
| 2956 | FIM1_3025 | QGN16320.1 | CP015057.1 | 3425 | 419 | 861 | 2596 | 309 | 752 | 3479 | 435 | 696 |
| 2957 | FIM1_3026 | QGN16321.1 | CP015057.1 | 7640 | 1639 | 1795 | 5480 | 1268 | 1605 | 6349 | 1690 | 1638 |
| 2958 | FIM1_3027 | QGN16322.1 | CP015057.1 | 12 | 0 | 3 | 14 | 6 | 11 | 11 | 9 | 5 |
| 2959 | FIM1_3028 | QGN16323.1 | CP015057.1 | 15839 | 2728 | 3438 | 12440 | 2359 | 3225 | 12750 | 3034 | 3115 |
| 2960 | FIM1_3029 | QGN16324.1 | CP015057.1 | 3794 | 701 | 1350 | 3029 | 531 | 1253 | 4921 | 755 | 1216 |
| 2961 | FIM1_3030 | QGN16325.1 | CP015057.1 | 6512 | 1870 | 4100 | 5101 | 1526 | 3741 | 7142 | 2018 | 3513 |
| 2962 | FIM1_3031 | QGN16326.1 | CP015057.1 | 2858 | 566 | 1209 | 2457 | 453 | 1154 | 4172 | 599 | 1014 |
| 2963 | FIM1_3032 | QGN16327.1 | CP015057.1 | 5433 | 2054 | 2564 | 5075 | 1552 | 2344 | 5217 | 2252 | 2346 |
| 2964 | FIM1_3033 | QGN16328.1 | CP015057.1 | 3027 | 656 | 1291 | 2684 | 517 | 1092 | 2224 | 758 | 1029 |
| 2965 | FIM1_3034 | QGN16329.1 | CP015057.1 | 8946 | 636 | 2793 | 7848 | 522 | 2387 | 13522 | 722 | 2533 |
| 2966 | FIM1_3035 | QGN16330.1 | CP015057.1 | 17624 | 9297 | 10792 | 16497 | 5756 | 9811 | 24253 | 9692 | 9785 |
| 2967 | FIM1_3036 | QGN16331.1 | CP015057.1 | 10606 | 2095 | 2484 | 9026 | 1430 | 2136 | 13066 | 2303 | 2147 |
| 2968 | FIM1_3037 | QGN16332.1 | CP015057.1 | 1393 | 449 | 1260 | 1619 | 386 | 1137 | 1704 | 531 | 1129 |
| 2969 | FIM1_3038 | QGN16333.1 | CP015057.1 | 24678 | 17160 | 15462 | 23299 | 14492 | 14756 | 24111 | 17838 | 15298 |
| 2970 | FIM1_3039 | QGN16334.1 | CP015057.1 | 5407 | 2794 | 2602 | 4484 | 2007 | 2341 | 3342 | 3127 | 2243 |
| 2971 | FIM1_3040 | QGN16335.1 | CP015057.1 | 3644 | 3841 | 2115 | 2364 | 2389 | 2120 | 794 | 4022 | 1867 |
| 2972 | FIM1_3041 | QGN16336.1 | CP015057.1 | 3625 | 995 | 1520 | 2797 | 761 | 1448 | 3657 | 1025 | 1299 |
| 2973 | FIM1_3043 | QGN16337.1 | CP015057.1 | 589 | 138 | 121 | 445 | 90 | 152 | 533 | 132 | 134 |
| 2974 | FIM1_3044 | QGN16338.1 | CP015057.1 | 5461 | 327 | 305 | 4962 | 155 | 311 | 25189 | 316 | 264 |

| | | | | | | | | | | | | |
|---|---|---|---|---|---|---|---|---|---|---|---|---|
| 2975 | FIM1_3045 | QGN16339.1 | CP015057.1 | 1771 | 184 | 399 | 1215 | 136 | 398 | 2617 | 186 | 382 |
| 2976 | FIM1_3046 | QGN16340.1 | CP015057.1 | 6637 | 700 | 1325 | 5000 | 489 | 1299 | 7334 | 761 | 1236 |
| 2977 | FIM1_3047 | QGN16341.1 | CP015057.1 | 1826 | 540 | 738 | 1335 | 378 | 724 | 1443 | 545 | 666 |
| 2978 | FIM1_3048 | QGN16342.1 | CP015057.1 | 4229 | 3279 | 2666 | 3915 | 2706 | 2494 | 2357 | 3543 | 2422 |
| 2979 | FIM1_3049 | QGN16343.1 | CP015057.1 | 1964 | 480 | 1003 | 1388 | 355 | 1000 | 1737 | 487 | 836 |
| 2980 | FIM1_3050 | QGN16344.1 | CP015057.1 | 1496 | 592 | 769 | 1246 | 392 | 735 | 1358 | 640 | 613 |
| 2981 | FIM1_3051 | QGN16345.1 | CP015057.1 | 2213 | 267 | 931 | 1765 | 209 | 858 | 3325 | 273 | 856 |
| 2982 | FIM1_3052 | QGN16346.1 | CP015057.1 | 2253 | 423 | 742 | 1838 | 307 | 706 | 1951 | 458 | 625 |
| 2983 | FIM1_3053 | QGN16347.1 | CP015057.1 | 3277 | 1144 | 1879 | 2649 | 884 | 1645 | 3067 | 1198 | 1708 |
| 2984 | FIM1_3054 | QGN16348.1 | CP015057.1 | 4434 | 978 | 852 | 2980 | 537 | 762 | 4437 | 1083 | 590 |
| 2985 | FIM1_3055 | QGN16349.1 | CP015057.1 | 12 | 2 | 4 | 13 | 2 | 4 | 6 | 5 | 7 |
| 2986 | FIM1_3056 | QGN16350.1 | CP015057.1 | 13207 | 3428 | 3543 | 9935 | 2386 | 3159 | 12603 | 3903 | 3032 |
| 2987 | FIM1_3057 | QGN16351.1 | CP015057.1 | 698 | 246 | 452 | 607 | 215 | 487 | 488 | 237 | 416 |
| 2988 | FIM1_3058 | QGN16352.1 | CP015057.1 | 8827 | 1392 | 2488 | 6540 | 1052 | 2133 | 8585 | 1431 | 2119 |
| 2989 | FIM1_3059 | QGN16353.1 | CP015057.1 | 2180 | 330 | 773 | 1724 | 233 | 777 | 2474 | 375 | 664 |
| 2990 | FIM1_3060 | QGN16354.1 | CP015057.1 | 1724 | 363 | 631 | 1344 | 298 | 642 | 1200 | 443 | 514 |
| 2991 | FIM1_3061 | QGN16355.1 | CP015057.1 | 9547 | 2096 | 3964 | 7216 | 1362 | 3504 | 10189 | 2140 | 3364 |
| 2992 | FIM1_3062 | QGN16356.1 | CP015057.1 | 3099 | 1307 | 1680 | 2458 | 918 | 1477 | 1512 | 1448 | 1499 |
| 2993 | FIM1_3063 | QGN16357.1 | CP015057.1 | 2948 | 400 | 997 | 2155 | 300 | 968 | 3215 | 360 | 923 |
| 2994 | FIM1_3064 | QGN16358.1 | CP015057.1 | 15055 | 6083 | 5392 | 12958 | 4204 | 4655 | 13296 | 6619 | 4557 |
| 2995 | FIM1_3065 | QGN16359.1 | CP015057.1 | 6082 | 1777 | 2257 | 4722 | 1238 | 2115 | 5121 | 1932 | 1935 |
| 2996 | FIM1_3066 | QGN16360.1 | CP015057.1 | 8387 | 3385 | 4824 | 7333 | 2462 | 4693 | 8429 | 3716 | 4466 |
| 2997 | FIM1_3067 | QGN16361.1 | CP015057.1 | 1420 | 514 | 521 | 1224 | 393 | 507 | 1283 | 571 | 473 |
| 2998 | FIM1_3068 | QGN16362.1 | CP015057.1 | 17154 | 3623 | 4326 | 13360 | 2762 | 3852 | 15535 | 3965 | 4015 |
| 2999 | FIM1_3069 | QGN16363.1 | CP015057.1 | 1987 | 983 | 1069 | 1717 | 728 | 956 | 1223 | 1081 | 866 |
| 3000 | FIM1_3070 | QGN16364.1 | CP015057.1 | 15231 | 6727 | 6313 | 13503 | 5149 | 5533 | 11371 | 7246 | 5446 |
| 3001 | FIM1_3071 | QGN16365.1 | CP015057.1 | 6086 | 1899 | 3269 | 5158 | 1224 | 3177 | 7323 | 1960 | 2887 |
| 3002 | FIM1_3072 | QGN16366.1 | CP015057.1 | 3375 | 526 | 1913 | 2509 | 388 | 1683 | 5804 | 571 | 1625 |
| 3003 | FIM1_3074 | QGN16367.1 | CP015057.1 | 2137 | 569 | 810 | 1649 | 381 | 775 | 2042 | 577 | 718 |
| 3004 | FIM1_3075 | QGN16368.1 | CP015057.1 | 2691 | 1337 | 961 | 1793 | 879 | 896 | 1552 | 1464 | 836 |
| 3005 | FIM1_3076 | QGN16369.1 | CP015057.1 | 3438 | 588 | 779 | 2330 | 478 | 756 | 3006 | 662 | 758 |
| 3006 | FIM1_3077 | QGN16370.1 | CP015057.1 | 6387 | 1569 | 1613 | 4472 | 1280 | 1492 | 6005 | 1668 | 1427 |
| 3007 | FIM1_3078 | QGN16371.1 | CP015057.1 | 1715 | 572 | 739 | 1280 | 378 | 666 | 1771 | 591 | 637 |
| 3008 | FIM1_3079 | QGN16372.1 | CP015057.1 | 3250 | 1963 | 1874 | 2514 | 1406 | 1861 | 2361 | 2247 | 1708 |
| 3009 | FIM1_3080 | QGN16373.1 | CP015057.1 | 3959 | 1111 | 1100 | 3070 | 796 | 997 | 3243 | 1216 | 918 |
| 3010 | FIM1_3081 | QGN16374.1 | CP015057.1 | 6981 | 862 | 1024 | 4624 | 889 | 1029 | 5249 | 961 | 1036 |
| 3011 | FIM1_3082 | QGN16375.1 | CP015057.1 | 5568 | 975 | 1419 | 4106 | 776 | 1230 | 5707 | 1063 | 1236 |
| 3012 | FIM1_3083 | QGN16376.1 | CP015057.1 | 7541 | 7944 | 3612 | 5489 | 4978 | 3236 | 2051 | 8395 | 3325 |
| 3013 | FIM1_3085 | QGN16377.1 | CP015057.1 | 88 | 24 | 19 | 67 | 19 | 28 | 97 | 30 | 13 |
| 3014 | FIM1_3086 | QGN16378.1 | CP015057.1 | 3645 | 911 | 435 | 2398 | 474 | 438 | 3954 | 995 | 363 |
| 3015 | FIM1_3087 | QGN16379.1 | CP015057.1 | 13228 | 8798 | 4985 | 9615 | 5586 | 4470 | 5951 | 9364 | 4444 |
| 3016 | FIM1_3088 | QGN16380.1 | CP015057.1 | 887 | 121 | 444 | 695 | 84 | 435 | 1463 | 134 | 359 |
| 3017 | FIM1_3089 | QGN16381.1 | CP015057.1 | 1290 | 281 | 388 | 1074 | 208 | 432 | 1682 | 318 | 415 |
| 3018 | FIM1_3090 | QGN16382.1 | CP015057.1 | 1030 | 364 | 180 | 708 | 230 | 224 | 810 | 368 | 195 |
| 3019 | FIM1_3092 | QGN16383.1 | CP015057.1 | 1315 | 351 | 230 | 893 | 158 | 230 | 1301 | 397 | 231 |
| 3020 | FIM1_3097 | QGN16384.1 | CP015057.1 | 3 | 0 | 0 | 1 | 2 | 1 | 4 | 4 | 0 |
| 3021 | FIM1_3098 | QGN16385.1 | CP015058.1 | 2303 | 629 | 1660 | 1969 | 410 | 1802 | 3355 | 656 | 1580 |
| 3022 | FIM1_3099 | QGN16386.1 | CP015058.1 | 4259 | 1694 | 1796 | 3684 | 859 | 1718 | 6288 | 1859 | 1510 |

| | | | | | | | | | | | | |
|---|---|---|---|---|---|---|---|---|---|---|---|---|
| 3023 | FIM1_3100 | QGN16387.1 | CP015058.1 | 1734 | 541 | 765 | 1370 | 349 | 773 | 2309 | 563 | 696 |
| 3024 | FIM1_3101 | QGN16388.1 | CP015058.1 | 1988 | 651 | 904 | 1568 | 431 | 907 | 2579 | 707 | 776 |
| 3025 | FIM1_3102 | QGN16389.1 | CP015058.1 | 20557 | 16394 | 15009 | 20499 | 14006 | 14440 | 19438 | 17063 | 14942 |
| 3026 | FIM1_3103 | QGN16390.1 | CP015058.1 | 320 | 125 | 120 | 285 | 83 | 117 | 278 | 143 | 107 |
| 3027 | FIM1_3104 | QGN16391.1 | CP015058.1 | 5406 | 963 | 1648 | 4126 | 791 | 1530 | 6046 | 1100 | 1449 |
| 3028 | FIM1_3105 | QGN16392.1 | CP015058.1 | 27 | 12 | 6 | 17 | 11 | 12 | 16 | 11 | 13 |
| 3029 | FIM1_3106 | QGN16393.1 | CP015058.1 | 3905 | 721 | 1240 | 3283 | 565 | 1194 | 4056 | 808 | 1190 |
| 3030 | FIM1_3107 | QGN16394.1 | CP015058.1 | 981 | 292 | 484 | 782 | 195 | 533 | 1457 | 178 | 414 |
| 3031 | FIM1_3108 | QGN16395.1 | CP015058.1 | 5245 | 948 | 1180 | 3694 | 587 | 1081 | 7293 | 971 | 1054 |
| 3032 | FIM1_3109 | QGN16396.1 | CP015058.1 | 5086 | 4225 | 2883 | 4394 | 2941 | 2850 | 2419 | 4575 | 2609 |
| 3033 | FIM1_3110 | QGN16397.1 | CP015058.1 | 7709 | 2829 | 573 | 5876 | 620 | 482 | 23494 | 2743 | 444 |
| 3034 | FIM1_3111 | QGN16398.1 | CP015058.1 | 2998 | 620 | 349 | 2539 | 223 | 347 | 8570 | 582 | 289 |
| 3035 | FIM1_3112 | QGN16399.1 | CP015058.1 | 1467 | 430 | 823 | 1119 | 274 | 745 | 1059 | 425 | 637 |
| 3036 | FIM1_3113 | QGN16400.1 | CP015058.1 | 9207 | 578 | 1640 | 7723 | 509 | 1387 | 7168 | 616 | 1389 |
| 3037 | FIM1_3114 | QGN16401.1 | CP015058.1 | 3128 | 1264 | 1312 | 2576 | 509 | 1203 | 3461 | 1265 | 1249 |
| 3038 | FIM1_3115 | QGN16402.1 | CP015058.1 | 670 | 81 | 267 | 559 | 63 | 217 | 705 | 84 | 166 |
| 3039 | FIM1_3116 | QGN16403.1 | CP015058.1 | 5411 | 849 | 1266 | 3847 | 511 | 1223 | 6764 | 930 | 1153 |
| 3040 | FIM1_3117 | QGN16404.1 | CP015058.1 | 18 | 11 | 17 | 29 | 6 | 15 | 27 | 10 | 15 |
| 3041 | FIM1_3118 | QGN16405.1 | CP015058.1 | 12062 | 1768 | 4407 | 10085 | 1722 | 4467 | 14230 | 1837 | 3552 |
| 3042 | FIM1_3119 | QGN16406.1 | CP015058.1 | 2170 | 515 | 841 | 1778 | 357 | 762 | 2788 | 567 | 754 |
| 3043 | FIM1_3120 | QGN16407.1 | CP015058.1 | 6957 | 1892 | 2421 | 5645 | 1392 | 2339 | 7292 | 2066 | 2262 |
| 3044 | FIM1_3121 | QGN16408.1 | CP015058.1 | 3321 | 579 | 912 | 2620 | 509 | 861 | 3906 | 622 | 783 |
| 3045 | FIM1_3122 | QGN16409.1 | CP015058.1 | 4663 | 862 | 2437 | 4055 | 574 | 2334 | 3468 | 957 | 2009 |
| 3046 | FIM1_3123 | QGN16410.1 | CP015058.1 | 6541 | 978 | 2370 | 5139 | 744 | 2315 | 7720 | 1052 | 2232 |
| 3047 | FIM1_3124 | QGN16411.1 | CP015058.1 | 12527 | 3132 | 3347 | 10126 | 2265 | 3076 | 11226 | 3372 | 2859 |
| 3048 | FIM1_3125 | QGN16412.1 | CP015058.1 | 2187 | 319 | 422 | 1482 | 280 | 413 | 1975 | 339 | 443 |
| 3049 | FIM1_3126 | QGN16413.1 | CP015058.1 | 7628 | 1150 | 1907 | 6370 | 940 | 1648 | 9356 | 1272 | 1727 |
| 3050 | FIM1_3127 | QGN16414.1 | CP015058.1 | 4546 | 891 | 1812 | 3867 | 609 | 1603 | 5653 | 948 | 1443 |
| 3051 | FIM1_3128 | QGN16415.1 | CP015058.1 | 19760 | 7880 | 10991 | 16339 | 5557 | 9988 | 20694 | 8128 | 10212 |
| 3052 | FIM1_3129 | QGN16416.1 | CP015058.1 | 8786 | 3261 | 3294 | 7037 | 2517 | 2973 | 7735 | 3393 | 3121 |
| 3053 | FIM1_3130 | QGN16417.1 | CP015058.1 | 2371 | 1013 | 1185 | 1663 | 735 | 1078 | 1003 | 1154 | 924 |
| 3054 | FIM1_3131 | QGN16418.1 | CP015058.1 | 2474 | 554 | 867 | 2050 | 375 | 822 | 2281 | 569 | 819 |
| 3055 | FIM1_3132 | QGN16419.1 | CP015058.1 | 6818 | 3099 | 2974 | 5353 | 2101 | 2678 | 5121 | 3280 | 2464 |
| 3056 | FIM1_3133 | QGN16420.1 | CP015058.1 | 1930 | 378 | 303 | 1308 | 275 | 342 | 1888 | 360 | 366 |
| 3057 | FIM1_3134 | QGN16421.1 | CP015058.1 | 32742 | 10386 | 9672 | 27115 | 6203 | 8418 | 29748 | 11317 | 8018 |
| 3058 | FIM1_3135 | QGN16422.1 | CP015058.1 | 2323 | 769 | 1023 | 1786 | 539 | 962 | 2180 | 829 | 941 |
| 3059 | FIM1_3136 | QGN16423.1 | CP015058.1 | 8870 | 5748 | 3320 | 6791 | 4213 | 3115 | 4515 | 6247 | 3242 |
| 3060 | FIM1_3137 | QGN16424.1 | CP015058.1 | 55 | 13 | 21 | 66 | 4 | 34 | 59 | 20 | 8 |
| 3061 | FIM1_3138 | QGN16425.1 | CP015058.1 | 12398 | 8601 | 4687 | 8880 | 5851 | 4335 | 3837 | 9204 | 4413 |
| 3062 | FIM1_3139 | QGN16426.1 | CP015058.1 | 1347 | 753 | 883 | 884 | 524 | 810 | 781 | 880 | 710 |
| 3063 | FIM1_3140 | QGN16427.1 | CP015058.1 | 1100 | 233 | 399 | 810 | 180 | 395 | 1232 | 288 | 348 |
| 3064 | FIM1_3141 | QGN16428.1 | CP015058.1 | 1847 | 862 | 1091 | 1267 | 597 | 1069 | 1061 | 876 | 1008 |
| 3065 | FIM1_3142 | QGN16429.1 | CP015058.1 | 849 | 102 | 170 | 633 | 63 | 151 | 1527 | 87 | 124 |
| 3066 | FIM1_3143 | QGN16430.1 | CP015058.1 | 10248 | 5819 | 4845 | 8940 | 3941 | 4354 | 9193 | 6003 | 4552 |
| 3067 | FIM1_3144 | QGN16431.1 | CP015058.1 | 590 | 144 | 411 | 472 | 147 | 349 | 516 | 171 | 309 |
| 3068 | FIM1_3145 | QGN16432.1 | CP015058.1 | 3622 | 877 | 1236 | 2745 | 662 | 1163 | 2382 | 913 | 1106 |
| 3069 | FIM1_3146 | QGN16433.1 | CP015058.1 | 1663 | 386 | 504 | 1190 | 294 | 482 | 1651 | 373 | 432 |
| 3070 | FIM1_3147 | QGN16434.1 | CP015058.1 | 3554 | 341 | 919 | 2642 | 231 | 826 | 5014 | 381 | 662 |

| | | | | | | | | | | | | |
|---|---|---|---|---|---|---|---|---|---|---|---|---|
| 3071 | FIM1_3148 | QGN16435.1 | CP015058.1 | 1976 | 311 | 985 | 1455 | 241 | 1004 | 1939 | 313 | 755 |
| 3072 | FIM1_3149 | QGN16436.1 | CP015058.1 | 2448 | 601 | 948 | 1820 | 431 | 961 | 2385 | 599 | 917 |
| 3073 | FIM1_3150 | QGN16437.1 | CP015058.1 | 8605 | 7388 | 3777 | 6211 | 5159 | 3440 | 2861 | 8197 | 3614 |
| 3074 | FIM1_3151 | QGN16438.1 | CP015058.1 | 1533 | 652 | 495 | 1096 | 580 | 427 | 1873 | 682 | 455 |
| 3075 | FIM1_3152 | QGN16439.1 | CP015058.1 | 545 | 142 | 203 | 461 | 132 | 201 | 506 | 159 | 161 |
| 3076 | FIM1_3153 | QGN16440.1 | CP015058.1 | 1695 | 360 | 628 | 1348 | 246 | 612 | 1876 | 355 | 589 |
| 3077 | FIM1_3154 | QGN16441.1 | CP015058.1 | 5244 | 2037 | 2052 | 3654 | 1413 | 1886 | 2133 | 2149 | 1838 |
| 3078 | FIM1_3155 | QGN16442.1 | CP015058.1 | 7533 | 1684 | 1667 | 6072 | 1124 | 1721 | 10177 | 1723 | 1684 |
| 3079 | FIM1_3156 | QGN16443.1 | CP015058.1 | 6341 | 2787 | 3132 | 5836 | 1904 | 3265 | 7671 | 2896 | 2948 |
| 3080 | FIM1_3157 | QGN16444.1 | CP015058.1 | 1471 | 285 | 1742 | 1506 | 112 | 1646 | 3374 | 337 | 1685 |
| 3081 | FIM1_3158 | QGN16445.1 | CP015058.1 | 970 | 168 | 296 | 909 | 119 | 260 | 993 | 178 | 182 |
| 3082 | FIM1_3159 | QGN16446.1 | CP015058.1 | 676 | 192 | 329 | 594 | 183 | 301 | 681 | 173 | 315 |
| 3083 | FIM1_3160 | QGN16447.1 | CP015058.1 | 1194 | 104 | 272 | 840 | 74 | 248 | 1523 | 117 | 232 |
| 3084 | FIM1_3161 | QGN16448.1 | CP015058.1 | 5051 | 715 | 392 | 3527 | 393 | 380 | 4244 | 1008 | 440 |
| 3085 | FIM1_3162 | QGN16449.1 | CP015058.1 | 4699 | 1063 | 1469 | 3865 | 490 | 1335 | 5222 | 1326 | 1204 |
| 3086 | FIM1_3163 | QGN16450.1 | CP015058.1 | 1050 | 415 | 796 | 922 | 322 | 778 | 576 | 445 | 704 |
| 3087 | FIM1_3164 | QGN16451.1 | CP015058.1 | 8782 | 1894 | 2032 | 6754 | 1374 | 2008 | 8550 | 2004 | 1886 |
| 3088 | FIM1_3165 | QGN16452.1 | CP015058.1 | 1564 | 662 | 768 | 1290 | 499 | 671 | 1037 | 674 | 595 |
| 3089 | FIM1_3166 | QGN16453.1 | CP015058.1 | 2961 | 739 | 1048 | 2095 | 551 | 899 | 2847 | 826 | 915 |
| 3090 | FIM1_3167 | QGN16454.1 | CP015058.1 | 7014 | 5188 | 2604 | 4645 | 3090 | 2255 | 1698 | 5402 | 2177 |
| 3091 | FIM1_3168 | QGN16455.1 | CP015058.1 | 2839 | 686 | 1159 | 2573 | 479 | 1019 | 2308 | 694 | 940 |
| 3092 | FIM1_3169 | QGN16456.1 | CP015058.1 | 2400 | 743 | 1043 | 2096 | 558 | 929 | 1777 | 813 | 863 |
| 3093 | FIM1_3170 | QGN16457.1 | CP015058.1 | 217 | 21 | 50 | 159 | 25 | 36 | 205 | 39 | 30 |
| 3094 | FIM1_3171 | QGN16458.1 | CP015058.1 | 3106 | 1555 | 1557 | 2576 | 1093 | 1450 | 1909 | 1643 | 1381 |
| 3095 | FIM1_3172 | QGN16459.1 | CP015058.1 | 3401 | 1000 | 1669 | 2914 | 581 | 1478 | 3468 | 1040 | 1405 |
| 3096 | FIM1_3173 | QGN16460.1 | CP015058.1 | 5583 | 775 | 1022 | 4047 | 520 | 1012 | 6367 | 850 | 898 |
| 3097 | FIM1_3174 | QGN16461.1 | CP015058.1 | 1422 | 376 | 429 | 1276 | 292 | 402 | 1912 | 472 | 388 |
| 3098 | FIM1_3175 | QGN16462.1 | CP015058.1 | 3313 | 844 | 1257 | 3068 | 660 | 1134 | 3356 | 880 | 1127 |
| 3099 | FIM1_3176 | QGN16463.1 | CP015058.1 | 1428 | 228 | 313 | 1267 | 163 | 291 | 1686 | 222 | 282 |
| 3100 | FIM1_3177 | QGN16464.1 | CP015058.1 | 6635 | 1534 | 1888 | 4826 | 1213 | 1897 | 5357 | 1638 | 1614 |
| 3101 | FIM1_3178 | QGN16465.1 | CP015058.1 | 3367 | 1569 | 1748 | 3169 | 976 | 1714 | 4442 | 1674 | 1618 |
| 3102 | FIM1_3179 | QGN16466.1 | CP015058.1 | 1302 | 119 | 261 | 983 | 99 | 220 | 3134 | 152 | 207 |
| 3103 | FIM1_3180 | QGN16467.1 | CP015058.1 | 4583 | 635 | 1055 | 3506 | 435 | 964 | 5483 | 739 | 943 |
| 3104 | FIM1_3181 | QGN16468.1 | CP015058.1 | 5862 | 4284 | 2301 | 3979 | 2610 | 2094 | 1797 | 4450 | 2130 |
| 3105 | FIM1_3182 | QGN16469.1 | CP015058.1 | 3381 | 1581 | 1589 | 2475 | 1147 | 1546 | 2399 | 1684 | 1449 |
| 3106 | FIM1_3183 | QGN16470.1 | CP015058.1 | 17401 | 5440 | 5017 | 14327 | 3458 | 4379 | 18791 | 5772 | 4570 |
| 3107 | FIM1_3184 | QGN16471.1 | CP015058.1 | 1261 | 561 | 640 | 1233 | 401 | 616 | 912 | 562 | 536 |
| 3108 | FIM1_3185 | QGN16472.1 | CP015058.1 | 2256 | 1172 | 961 | 1956 | 800 | 842 | 1135 | 1169 | 840 |
| 3109 | FIM1_3186 | QGN16473.1 | CP015058.1 | 10221 | 1921 | 4108 | 7504 | 1475 | 3848 | 11283 | 2063 | 3658 |
| 3110 | FIM1_3187 | QGN16474.1 | CP015058.1 | 1220 | 414 | 590 | 792 | 352 | 576 | 810 | 492 | 601 |
| 3111 | FIM1_3188 | QGN16475.1 | CP015058.1 | 1701 | 430 | 570 | 1182 | 336 | 617 | 1500 | 439 | 539 |
| 3112 | FIM1_3189 | QGN16476.1 | CP015058.1 | 952 | 223 | 402 | 858 | 219 | 455 | 693 | 266 | 316 |
| 3113 | FIM1_3190 | QGN16477.1 | CP015058.1 | 1636 | 350 | 601 | 1300 | 293 | 562 | 1129 | 441 | 493 |
| 3114 | FIM1_3191 | QGN16478.1 | CP015058.1 | 1893 | 714 | 868 | 1385 | 493 | 811 | 1247 | 687 | 686 |
| 3115 | FIM1_3192 | QGN16479.1 | CP015058.1 | 2632 | 392 | 620 | 1999 | 331 | 536 | 1867 | 407 | 550 |
| 3116 | FIM1_3193 | QGN16480.1 | CP015058.1 | 116 | 43 | 27 | 124 | 45 | 27 | 76 | 39 | 20 |
| 3117 | FIM1_3194 | QGN16481.1 | CP015058.1 | 7693 | 1567 | 1974 | 5970 | 1006 | 1821 | 6207 | 1761 | 1695 |
| 3118 | FIM1_3195 | QGN16482.1 | CP015058.1 | 5222 | 3967 | 2429 | 4228 | 3017 | 2280 | 2004 | 4194 | 2277 |

| | | | | | | | | | | | | |
|---|---|---|---|---|---|---|---|---|---|---|---|---|
| 3119 | FIM1_3196 | QGN16483.1 | CP015058.1 | 990 | 288 | 416 | 884 | 229 | 398 | 883 | 268 | 379 |
| 3120 | FIM1_3197 | QGN16484.1 | CP015058.1 | 4957 | 584 | 467 | 2875 | 254 | 426 | 5249 | 574 | 424 |
| 3121 | FIM1_3198 | QGN16485.1 | CP015058.1 | 1853 | 256 | 434 | 1416 | 167 | 409 | 1603 | 304 | 438 |
| 3122 | FIM1_3199 | QGN16486.1 | CP015058.1 | 5775 | 1720 | 1781 | 4325 | 1189 | 1679 | 6332 | 1806 | 1726 |
| 3123 | FIM1_3200 | QGN16487.1 | CP015058.1 | 19965 | 8284 | 8097 | 17058 | 6466 | 7319 | 18690 | 8994 | 7542 |
| 3124 | FIM1_3201 | QGN16488.1 | CP015058.1 | 1045 | 349 | 541 | 894 | 299 | 538 | 702 | 337 | 435 |
| 3125 | FIM1_3202 | QGN16489.1 | CP015058.1 | 1700 | 243 | 506 | 1212 | 225 | 440 | 1371 | 256 | 356 |
| 3126 | FIM1_3203 | QGN16490.1 | CP015058.1 | 6365 | 684 | 1142 | 4869 | 519 | 1031 | 7309 | 757 | 899 |
| 3127 | FIM1_3204 | QGN16491.1 | CP015058.1 | 1997 | 480 | 755 | 1511 | 306 | 640 | 1857 | 492 | 602 |
| 3128 | FIM1_3205 | QGN16492.1 | CP015058.1 | 10151 | 3586 | 4255 | 9084 | 2259 | 3708 | 11165 | 3655 | 3688 |
| 3129 | FIM1_3206 | QGN16493.1 | CP015058.1 | 3882 | 1137 | 1592 | 3137 | 938 | 1445 | 3354 | 1229 | 1404 |
| 3130 | FIM1_3207 | QGN16494.1 | CP015058.1 | 3445 | 405 | 920 | 2542 | 268 | 782 | 4139 | 481 | 759 |
| 3131 | FIM1_3208 | QGN16495.1 | CP015058.1 | 5616 | 2399 | 3388 | 4627 | 1651 | 3151 | 6209 | 2526 | 3157 |
| 3132 | FIM1_3209 | QGN16496.1 | CP015058.1 | 17128 | 4991 | 6781 | 13849 | 3784 | 6483 | 17342 | 5394 | 6212 |
| 3133 | FIM1_3210 | QGN16497.1 | CP015058.1 | 415 | 180 | 1171 | 376 | 144 | 987 | 549 | 198 | 759 |
| 3134 | FIM1_3211 | QGN16498.1 | CP015058.1 | 889 | 206 | 465 | 776 | 149 | 428 | 902 | 233 | 373 |
| 3135 | FIM1_3212 | QGN16499.1 | CP015058.1 | 6407 | 2435 | 2756 | 5508 | 1766 | 2623 | 6009 | 2630 | 2644 |
| 3136 | FIM1_3213 | QGN16500.1 | CP015058.1 | 10169 | 2370 | 3231 | 7837 | 1764 | 2842 | 9607 | 2488 | 2837 |
| 3137 | FIM1_3214 | QGN16501.1 | CP015058.1 | 5429 | 5064 | 2736 | 3653 | 3441 | 2597 | 752 | 5455 | 2574 |
| 3138 | FIM1_3215 | QGN16502.1 | CP015058.1 | 9613 | 1644 | 1755 | 7292 | 1259 | 1573 | 8978 | 1793 | 1620 |
| 3139 | FIM1_3216 | QGN16503.1 | CP015058.1 | 144 | 16 | 80 | 98 | 21 | 64 | 233 | 26 | 69 |
| 3140 | FIM1_3217 | QGN16504.1 | CP015058.1 | 6590 | 1770 | 1610 | 4819 | 1092 | 1567 | 6767 | 1867 | 1560 |
| 3141 | FIM1_3218 | QGN16505.1 | CP015058.1 | 272 | 79 | 327 | 225 | 78 | 231 | 568 | 69 | 252 |
| 3142 | FIM1_3219 | QGN16506.1 | CP015058.1 | 6078 | 1102 | 3058 | 5121 | 855 | 2717 | 7769 | 1198 | 2898 |
| 3143 | FIM1_3220 | QGN16507.1 | CP015058.1 | 5934 | 1553 | 751 | 4741 | 747 | 696 | 9752 | 1577 | 664 |
| 3144 | FIM1_3221 | QGN16508.1 | CP015058.1 | 15741 | 5214 | 5867 | 13323 | 3553 | 5404 | 15895 | 5679 | 5156 |
| 3145 | FIM1_3222 | QGN16509.1 | CP015058.1 | 7409 | 2164 | 3460 | 6409 | 1523 | 3068 | 7889 | 2157 | 2985 |
| 3146 | FIM1_3223 | QGN16510.1 | CP015058.1 | 13448 | 4385 | 4206 | 10923 | 3352 | 3830 | 11312 | 4831 | 3896 |
| 3147 | FIM1_3224 | QGN16511.1 | CP015058.1 | 8687 | 735 | 1678 | 6243 | 653 | 1482 | 8490 | 753 | 1318 |
| 3148 | FIM1_3225 | QGN16512.1 | CP015058.1 | 147 | 52 | 140 | 123 | 46 | 125 | 117 | 64 | 92 |
| 3149 | FIM1_3226 | QGN16513.1 | CP015058.1 | 2460 | 392 | 862 | 2117 | 318 | 674 | 2622 | 479 | 659 |
| 3150 | FIM1_3227 | QGN16514.1 | CP015058.1 | 6144 | 1930 | 1947 | 5008 | 1644 | 1776 | 4089 | 2049 | 1832 |
| 3151 | FIM1_3228 | QGN16515.1 | CP015058.1 | 3307 | 997 | 1721 | 2850 | 773 | 1727 | 3032 | 1102 | 1545 |
| 3152 | FIM1_3229 | QGN16516.1 | CP015058.1 | 6160 | 3517 | 3496 | 5722 | 2772 | 3138 | 5701 | 3551 | 3043 |
| 3153 | FIM1_3230 | QGN16517.1 | CP015058.1 | 8474 | 3505 | 3927 | 7612 | 2655 | 3752 | 8499 | 3719 | 3617 |
| 3154 | FIM1_3231 | QGN16518.1 | CP015058.1 | 10262 | 4804 | 4086 | 8445 | 3055 | 3668 | 8903 | 4961 | 3856 |
| 3155 | FIM1_3232 | QGN16519.1 | CP015058.1 | 1178 | 115 | 106 | 961 | 91 | 90 | 1271 | 135 | 101 |
| 3156 | FIM1_3233 | QGN16520.1 | CP015058.1 | 1398 | 222 | 2439 | 1201 | 199 | 2298 | 1957 | 240 | 2342 |
| 3157 | FIM1_3234 | QGN16521.1 | CP015058.1 | 6470 | 5953 | 2925 | 4651 | 3790 | 2601 | 1433 | 6421 | 2816 |
| 3158 | FIM1_3235 | QGN16522.1 | CP015058.1 | 3150 | 616 | 1421 | 2655 | 517 | 1291 | 3285 | 621 | 1155 |
| 3159 | FIM1_3236 | QGN16523.1 | CP015058.1 | 34282 | 9188 | 6875 | 27354 | 6539 | 6380 | 26499 | 9653 | 6640 |
| 3160 | FIM1_3237 | QGN16524.1 | CP015058.1 | 12627 | 2826 | 4824 | 10980 | 1816 | 4217 | 14418 | 3018 | 3921 |
| 3161 | FIM1_3238 | QGN16525.1 | CP015058.1 | 928 | 195 | 435 | 929 | 192 | 413 | 968 | 227 | 368 |
| 3162 | FIM1_3239 | QGN16526.1 | CP015058.1 | 6604 | 1845 | 2280 | 6088 | 1351 | 1931 | 7389 | 1995 | 1930 |
| 3163 | FIM1_3240 | QGN16527.1 | CP015058.1 | 6428 | 2031 | 2076 | 5530 | 1676 | 1850 | 5655 | 2286 | 1854 |
| 3164 | FIM1_3241 | QGN16528.1 | CP015058.1 | 2514 | 452 | 1133 | 2388 | 248 | 948 | 3908 | 451 | 884 |
| 3165 | FIM1_3242 | QGN16529.1 | CP015058.1 | 2310 | 352 | 572 | 1714 | 276 | 573 | 2975 | 416 | 535 |
| 3166 | FIM1_3243 | QGN16530.1 | CP015058.1 | 5679 | 1750 | 1933 | 4392 | 1545 | 1927 | 4298 | 1926 | 1949 |

| | | | | | | | | | | | | |
|---|---|---|---|---|---|---|---|---|---|---|---|---|
| 3167 | FIM1_3244 | QGN16531.1 | CP015058.1 | 1454 | 553 | 1446 | 990 | 381 | 1355 | 1091 | 612 | 1151 |
| 3168 | FIM1_3245 | QGN16532.1 | CP015058.1 | 4093 | 584 | 1224 | 3082 | 430 | 1035 | 3939 | 687 | 1004 |
| 3169 | FIM1_3246 | QGN16533.1 | CP015058.1 | 4082 | 389 | 659 | 3013 | 379 | 631 | 4796 | 450 | 536 |
| 3170 | FIM1_3247 | QGN16534.1 | CP015058.1 | 22033 | 3551 | 4568 | 16999 | 2532 | 4238 | 25073 | 3970 | 4025 |
| 3171 | FIM1_3248 | QGN16535.1 | CP015058.1 | 660 | 48 | 200 | 647 | 49 | 185 | 776 | 55 | 181 |
| 3172 | FIM1_3249 | QGN16536.1 | CP015058.1 | 6087 | 2057 | 3336 | 5569 | 1279 | 2905 | 7451 | 2143 | 2915 |
| 3173 | FIM1_3250 | QGN16537.1 | CP015058.1 | 1403 | 564 | 798 | 1262 | 442 | 782 | 1136 | 576 | 726 |
| 3174 | FIM1_3251 | QGN16538.1 | CP015058.1 | 2378 | 468 | 549 | 1634 | 360 | 531 | 2397 | 484 | 497 |
| 3175 | FIM1_3252 | QGN16539.1 | CP015058.1 | 99 | 12 | 37 | 75 | 18 | 34 | 122 | 14 | 35 |
| 3176 | FIM1_3253 | QGN16540.1 | CP015058.1 | 6669 | 570 | 3777 | 6031 | 365 | 3087 | 16446 | 553 | 2947 |
| 3177 | FIM1_3254 | QGN16541.1 | CP015058.1 | 11680 | 3427 | 4138 | 9597 | 2332 | 3784 | 12496 | 3707 | 3600 |
| 3178 | FIM1_3255 | QGN16542.1 | CP015058.1 | 2239 | 434 | 601 | 1840 | 333 | 585 | 2392 | 444 | 570 |
| 3179 | FIM1_3256 | QGN16543.1 | CP015058.1 | 69 | 29 | 32 | 53 | 31 | 24 | 52 | 19 | 28 |
| 3180 | FIM1_3258 | QGN16544.1 | CP015058.1 | 166 | 195 | 306 | 96 | 162 | 334 | 153 | 236 | 290 |
| 3181 | FIM1_3259 | QGN16545.1 | CP015058.1 | 383 | 263 | 541 | 235 | 196 | 525 | 327 | 235 | 470 |
| 3182 | FIM1_3260 | QGN16546.1 | CP015058.1 | 5309 | 884 | 1041 | 3865 | 755 | 938 | 4527 | 937 | 985 |
| 3183 | FIM1_3261 | QGN16547.1 | CP015058.1 | 1581 | 326 | 570 | 1428 | 272 | 480 | 1190 | 361 | 484 |
| 3184 | FIM1_3262 | QGN16548.1 | CP015058.1 | 47453 | 30702 | 25444 | 42869 | 24924 | 24452 | 47267 | 30966 | 25781 |
| 3185 | FIM1_3263 | QGN16549.1 | CP015058.1 | 5474 | 1249 | 1777 | 4068 | 877 | 1659 | 4775 | 1325 | 1472 |
| 3186 | FIM1_3264 | QGN16550.1 | CP015058.1 | 3437 | 1161 | 1513 | 2582 | 853 | 1468 | 2760 | 1161 | 1278 |
| 3187 | FIM1_3265 | QGN16551.1 | CP015058.1 | 3561 | 891 | 985 | 2794 | 660 | 965 | 3415 | 1016 | 1024 |
| 3188 | FIM1_3266 | QGN16552.1 | CP015058.1 | 1249 | 569 | 594 | 994 | 382 | 550 | 664 | 654 | 527 |
| 3189 | FIM1_3267 | QGN16553.1 | CP015058.1 | 8271 | 3174 | 3120 | 6507 | 2352 | 2977 | 5318 | 3226 | 2751 |
| 3190 | FIM1_3268 | QGN16554.1 | CP015058.1 | 1543 | 652 | 766 | 1439 | 547 | 794 | 1077 | 736 | 696 |
| 3191 | FIM1_3269 | QGN16555.1 | CP015058.1 | 1405 | 249 | 420 | 1043 | 212 | 398 | 1768 | 250 | 458 |
| 3192 | FIM1_3270 | QGN16556.1 | CP015058.1 | 12700 | 2306 | 2875 | 10839 | 1095 | 3049 | 22396 | 2085 | 2237 |
| 3193 | FIM1_3271 | QGN16557.1 | CP015058.1 | 6860 | 2314 | 3347 | 5127 | 1917 | 3107 | 6228 | 2330 | 2854 |
| 3194 | FIM1_3272 | QGN16558.1 | CP015058.1 | 2924 | 428 | 1163 | 2093 | 360 | 1011 | 3225 | 464 | 868 |
| 3195 | FIM1_3273 | QGN16559.1 | CP015058.1 | 2591 | 2051 | 1734 | 2394 | 1834 | 1591 | 1013 | 2226 | 1601 |
| 3196 | FIM1_3274 | QGN16560.1 | CP015058.1 | 6851 | 1331 | 2545 | 5546 | 943 | 2247 | 6854 | 1376 | 2238 |
| 3197 | FIM1_3275 | QGN16561.1 | CP015058.1 | 7174 | 3639 | 3053 | 5879 | 2400 | 2654 | 5706 | 3669 | 2726 |
| 3198 | FIM1_3276 | QGN16562.1 | CP015058.1 | 2253 | 91 | 393 | 1877 | 72 | 353 | 3039 | 123 | 252 |
| 3199 | FIM1_3277 | QGN16563.1 | CP015058.1 | 5551 | 1744 | 1879 | 4222 | 1367 | 1912 | 4410 | 1949 | 2006 |
| 3200 | FIM1_3278 | QGN16564.1 | CP015058.1 | 18690 | 4757 | 5757 | 14154 | 3703 | 5341 | 18711 | 5403 | 5129 |
| 3201 | FIM1_3279 | QGN16565.1 | CP015058.1 | 8414 | 3244 | 3039 | 7549 | 2293 | 2770 | 5987 | 3496 | 2760 |
| 3202 | FIM1_3280 | QGN16566.1 | CP015058.1 | 4630 | 1088 | 2079 | 3694 | 741 | 1890 | 5464 | 1127 | 1731 |
| 3203 | FIM1_3281 | QGN16567.1 | CP015058.1 | 5183 | 3016 | 5140 | 5078 | 2201 | 4846 | 6077 | 3185 | 4628 |
| 3204 | FIM1_3282 | QGN16568.1 | CP015058.1 | 18865 | 4455 | 4568 | 14057 | 3080 | 4311 | 18875 | 4610 | 4295 |
| 3205 | FIM1_3283 | QGN16569.1 | CP015058.1 | 1309 | 416 | 599 | 1113 | 376 | 609 | 1527 | 491 | 509 |
| 3206 | FIM1_3284 | QGN16570.1 | CP015058.1 | 4182 | 366 | 1280 | 3013 | 408 | 1217 | 4181 | 398 | 1031 |
| 3207 | FIM1_3286 | QGN16571.1 | CP015058.1 | 6178 | 1053 | 1362 | 4791 | 880 | 1298 | 4646 | 1122 | 1191 |
| 3208 | FIM1_3287 | QGN16572.1 | CP015058.1 | 1208 | 325 | 332 | 917 | 346 | 343 | 721 | 371 | 330 |
| 3209 | FIM1_3288 | QGN16573.1 | CP015058.1 | 48566 | 20298 | 23370 | 39835 | 14373 | 21032 | 48456 | 22132 | 22113 |
| 3210 | FIM1_3289 | QGN16574.1 | CP015058.1 | 43 | 13 | 19 | 32 | 4 | 22 | 36 | 6 | 16 |
| 3211 | FIM1_3290 | QGN16575.1 | CP015058.1 | 4517 | 873 | 1507 | 4138 | 695 | 1434 | 4706 | 971 | 1427 |
| 3212 | FIM1_3291 | QGN16576.1 | CP015058.1 | 74 | 6 | 7 | 59 | 3 | 10 | 29 | 8 | 6 |
| 3213 | FIM1_3292 | QGN16577.1 | CP015058.1 | 49577 | 27568 | 25975 | 46132 | 22507 | 23944 | 46457 | 28310 | 24752 |
| 3214 | FIM1_3293 | QGN16578.1 | CP015058.1 | 685 | 344 | 341 | 668 | 299 | 325 | 407 | 377 | 308 |

| | | | | | | | | | | | | |
|---|---|---|---|---|---|---|---|---|---|---|---|---|
| 3215 | FIM1_3294 | QGN16579.1 | CP015058.1 | 704 | 276 | 222 | 463 | 232 | 206 | 525 | 303 | 214 |
| 3216 | FIM1_3295 | QGN16580.1 | CP015058.1 | 7766 | 538 | 931 | 5481 | 372 | 767 | 9036 | 539 | 833 |
| 3217 | FIM1_3297 | QGN16581.1 | CP015058.1 | 4405 | 1638 | 2163 | 3220 | 1012 | 1904 | 3785 | 1736 | 1796 |
| 3218 | FIM1_3298 | QGN16582.1 | CP015058.1 | 8391 | 4855 | 2773 | 5535 | 3144 | 2596 | 2777 | 5106 | 2616 |
| 3219 | FIM1_3299 | QGN16583.1 | CP015058.1 | 18171 | 3826 | 3946 | 13124 | 2725 | 3625 | 16680 | 3994 | 3602 |
| 3220 | FIM1_3300 | QGN16584.1 | CP015058.1 | 15026 | 11518 | 11035 | 15163 | 10025 | 10820 | 13921 | 12037 | 10617 |
| 3221 | FIM1_3301 | QGN16585.1 | CP015058.1 | 3443 | 2021 | 1455 | 2948 | 1273 | 1425 | 4122 | 2114 | 1376 |
| 3222 | FIM1_3302 | QGN16586.1 | CP015058.1 | 624 | 996 | 537 | 452 | 651 | 574 | 123 | 1103 | 551 |
| 3223 | FIM1_3303 | QGN16587.1 | CP015058.1 | 1028 | 410 | 463 | 778 | 315 | 539 | 1092 | 420 | 463 |
| 3224 | FIM1_3304 | QGN16588.1 | CP015058.1 | 21580 | 15461 | 7658 | 16407 | 10559 | 6977 | 8076 | 16091 | 7401 |
| 3225 | FIM1_3305 | QGN16589.1 | CP015058.1 | 27663 | 5474 | 10553 | 21663 | 3315 | 8928 | 33432 | 5503 | 9262 |
| 3226 | FIM1_3306 | QGN16590.1 | CP015058.1 | 810 | 195 | 378 | 778 | 171 | 332 | 509 | 202 | 338 |
| 3227 | FIM1_3307 | QGN16591.1 | CP015058.1 | 2714 | 576 | 771 | 2499 | 419 | 704 | 3954 | 616 | 687 |
| 3228 | FIM1_3308 | QGN16592.1 | CP015058.1 | 1975 | 251 | 270 | 1252 | 186 | 283 | 2416 | 290 | 274 |
| 3229 | FIM1_3309 | QGN16593.1 | CP015058.1 | 12415 | 3661 | 3564 | 9951 | 2777 | 3167 | 9344 | 3847 | 3290 |
| 3230 | FIM1_3310 | QGN16594.1 | CP015058.1 | 4503 | 883 | 1443 | 3643 | 699 | 1329 | 4734 | 963 | 1274 |
| 3231 | FIM1_3311 | QGN16595.1 | CP015058.1 | 11993 | 5080 | 5470 | 10334 | 3720 | 4842 | 11355 | 5457 | 4933 |
| 3232 | FIM1_3312 | QGN16596.1 | CP015058.1 | 15711 | 1077 | 1777 | 12049 | 992 | 1600 | 16751 | 1189 | 1613 |
| 3233 | FIM1_3313 | QGN16597.1 | CP015058.1 | 2865 | 127 | 385 | 2140 | 87 | 324 | 3817 | 155 | 285 |
| 3234 | FIM1_3314 | QGN16598.1 | CP015058.1 | 1607 | 293 | 511 | 1367 | 253 | 438 | 1485 | 308 | 405 |
| 3235 | FIM1_3315 | QGN16599.1 | CP015058.1 | 1443 | 513 | 646 | 1364 | 343 | 574 | 1186 | 530 | 561 |
| 3236 | FIM1_3316 | QGN16600.1 | CP015058.1 | 4355 | 1714 | 1363 | 3501 | 1462 | 1265 | 3637 | 1884 | 1233 |
| 3237 | FIM1_3317 | QGN16601.1 | CP015058.1 | 234 | 68 | 183 | 183 | 47 | 165 | 235 | 51 | 134 |
| 3238 | FIM1_3318 | QGN16602.1 | CP015058.1 | 1076 | 1258 | 734 | 898 | 853 | 702 | 299 | 1337 | 669 |
| 3239 | FIM1_3319 | QGN16603.1 | CP015058.1 | 3669 | 1170 | 1345 | 2561 | 950 | 1167 | 2329 | 1360 | 1190 |
| 3240 | FIM1_3320 | QGN16604.1 | CP015058.1 | 7101 | 3882 | 1580 | 5604 | 981 | 1310 | 10705 | 3988 | 1278 |
| 3241 | FIM1_3321 | QGN16605.1 | CP015058.1 | 7041 | 3763 | 1245 | 4742 | 2301 | 1133 | 2596 | 4024 | 1063 |
| 3242 | FIM1_3322 | QGN16606.1 | CP015058.1 | 2297 | 238 | 307 | 1628 | 254 | 286 | 1803 | 284 | 279 |
| 3243 | FIM1_3323 | QGN16607.1 | CP015058.1 | 2388 | 365 | 1246 | 2006 | 329 | 1118 | 3452 | 444 | 878 |
| 3244 | FIM1_3324 | QGN16608.1 | CP015058.1 | 5160 | 905 | 1404 | 3852 | 574 | 1259 | 7852 | 945 | 1292 |
| 3245 | FIM1_3325 | QGN16609.1 | CP015058.1 | 4 | 0 | 11 | 1 | 4 | 11 | 3 | 0 | 5 |
| 3246 | FIM1_3326 | QGN16610.1 | CP015058.1 | 30310 | 9981 | 8662 | 25507 | 6870 | 7908 | 29930 | 10347 | 8051 |
| 3247 | FIM1_3327 | QGN16611.1 | CP015058.1 | 4161 | 1145 | 1006 | 2791 | 828 | 973 | 3897 | 1217 | 975 |
| 3248 | FIM1_3328 | QGN16612.1 | CP015058.1 | 646 | 123 | 184 | 571 | 99 | 196 | 723 | 134 | 148 |
| 3249 | FIM1_3329 | QGN16613.1 | CP015058.1 | 508 | 297 | 177 | 395 | 204 | 181 | 441 | 289 | 140 |
| 3250 | FIM1_3331 | QGN16614.1 | CP015058.1 | 613 | 259 | 370 | 583 | 215 | 369 | 661 | 285 | 367 |
| 3251 | FIM1_3332 | QGN16615.1 | CP015058.1 | 1141 | 443 | 598 | 951 | 246 | 592 | 1041 | 474 | 545 |
| 3252 | FIM1_3333 | QGN16616.1 | CP015058.1 | 5909 | 5749 | 3287 | 4375 | 3035 | 2875 | 2663 | 5902 | 2577 |
| 3253 | FIM1_3334 | QGN16617.1 | CP015058.1 | 73575 | 33231 | 29449 | 69055 | 25965 | 27027 | 68299 | 33531 | 28627 |
| 3254 | FIM1_3335 | QGN16618.1 | CP015058.1 | 3571 | 1430 | 1259 | 2563 | 944 | 1058 | 1601 | 1545 | 1072 |
| 3255 | FIM1_3336 | QGN16619.1 | CP015058.1 | 1996 | 962 | 1015 | 1740 | 789 | 1042 | 1096 | 1072 | 850 |
| 3256 | FIM1_3337 | QGN16620.1 | CP015058.1 | 5769 | 704 | 2636 | 4997 | 514 | 2438 | 5488 | 816 | 2201 |
| 3257 | FIM1_3338 | QGN16621.1 | CP015058.1 | 29 | 5 | 19 | 30 | 9 | 14 | 20 | 8 | 18 |
| 3258 | FIM1_3339 | QGN16622.1 | CP015058.1 | 2494 | 592 | 1133 | 2063 | 440 | 1082 | 2713 | 619 | 1114 |
| 3259 | FIM1_3340 | QGN16623.1 | CP015058.1 | 6623 | 2061 | 8664 | 5442 | 1210 | 8025 | 7798 | 2222 | 7749 |
| 3260 | FIM1_3341 | QGN16624.1 | CP015058.1 | 3381 | 644 | 1097 | 2547 | 555 | 1054 | 3403 | 637 | 1003 |
| 3261 | FIM1_3342 | QGN16625.1 | CP015058.1 | 378 | 74 | 137 | 319 | 49 | 123 | 515 | 44 | 121 |
| 3262 | FIM1_3343 | QGN16626.1 | CP015058.1 | 26139 | 6209 | 9055 | 21826 | 4167 | 8152 | 22181 | 6843 | 8152 |

| | | | | | | | | | | | | |
|---|---|---|---|---|---|---|---|---|---|---|---|---|
| 3263 | FIM1_3344 | QGN16627.1 | CP015058.1 | 56384 | 9922 | 21539 | 41327 | 5044 | 18282 | 57352 | 9210 | 19419 |
| 3264 | FIM1_3345 | QGN16628.1 | CP015058.1 | 10742 | 5190 | 8476 | 9385 | 2595 | 7864 | 11970 | 4938 | 7697 |
| 3265 | FIM1_3346 | QGN16629.1 | CP015058.1 | 691 | 145 | 205 | 495 | 72 | 165 | 586 | 144 | 168 |
| 3266 | FIM1_3347 | QGN16630.1 | CP015058.1 | 7775 | 641 | 868 | 5732 | 364 | 779 | 8276 | 642 | 741 |
| 3267 | FIM1_3348 | QGN16631.1 | CP015058.1 | 4076 | 139 | 587 | 2911 | 196 | 456 | 4503 | 175 | 437 |
| 3268 | FIM1_3349 | QGN16632.1 | CP015058.1 | 5560 | 858 | 1447 | 4512 | 557 | 1313 | 7815 | 920 | 1303 |
| 3269 | FIM1_3350 | QGN16633.1 | CP015058.1 | 23709 | 9191 | 5775 | 19746 | 4338 | 4709 | 27431 | 8930 | 4433 |
| 3270 | FIM1_3351 | QGN16634.1 | CP015058.1 | 4763 | 799 | 1761 | 4081 | 567 | 1741 | 5496 | 869 | 1491 |
| 3271 | FIM1_3353 | QGN16635.1 | CP015058.1 | 12223 | 8410 | 7663 | 11160 | 7085 | 7699 | 11875 | 8828 | 7443 |
| 3272 | FIM1_3355 | QGN16636.1 | CP015058.1 | 3020 | 929 | 1189 | 2555 | 631 | 1148 | 3620 | 1035 | 1054 |
| 3273 | FIM1_3356 | QGN16637.1 | CP015058.1 | 4676 | 354 | 1027 | 3724 | 278 | 980 | 6754 | 407 | 910 |
| 3274 | FIM1_3357 | QGN16638.1 | CP015058.1 | 778 | 295 | 415 | 637 | 208 | 402 | 1152 | 290 | 344 |
| 3275 | FIM1_3358 | QGN16639.1 | CP015058.1 | 9 | 5 | 6 | 9 | 7 | 12 | 12 | 3 | 12 |
| 3276 | FIM1_3359 | QGN16640.1 | CP015058.1 | 13069 | 4535 | 4342 | 10981 | 3593 | 4175 | 11944 | 4872 | 3865 |
| 3277 | FIM1_3360 | QGN16641.1 | CP015058.1 | 7345 | 1812 | 3709 | 6105 | 1347 | 3650 | 8050 | 1944 | 3315 |
| 3278 | FIM1_3361 | QGN16642.1 | CP015058.1 | 15425 | 6588 | 5516 | 12137 | 4710 | 5067 | 15730 | 6866 | 5212 |
| 3279 | FIM1_3362 | QGN16643.1 | CP015058.1 | 12591 | 1632 | 2119 | 9104 | 1134 | 1914 | 15172 | 1762 | 1884 |
| 3280 | FIM1_3363 | QGN16644.1 | CP015058.1 | 3010 | 1082 | 1303 | 2613 | 669 | 1251 | 3732 | 1051 | 1102 |
| 3281 | FIM1_3364 | QGN16645.1 | CP015058.1 | 5949 | 827 | 1692 | 4561 | 639 | 1541 | 7081 | 971 | 1357 |
| 3282 | FIM1_3365 | QGN16646.1 | CP015058.1 | 840 | 171 | 340 | 538 | 145 | 338 | 536 | 213 | 271 |
| 3283 | FIM1_3366 | QGN16647.1 | CP015058.1 | 2879 | 78 | 383 | 2014 | 45 | 293 | 2727 | 70 | 261 |
| 3284 | FIM1_3367 | QGN16648.1 | CP015058.1 | 1753 | 315 | 649 | 1170 | 302 | 677 | 1859 | 296 | 556 |
| 3285 | FIM1_3368 | QGN16649.1 | CP015058.1 | 789 | 837 | 508 | 698 | 467 | 451 | 927 | 696 | 494 |
| 3286 | FIM1_3369 | QGN16650.1 | CP015058.1 | 2501 | 903 | 810 | 1992 | 529 | 741 | 2113 | 942 | 747 |
| 3287 | FIM1_3370 | QGN16651.1 | CP015058.1 | 1145 | 570 | 959 | 1110 | 362 | 955 | 1174 | 692 | 840 |
| 3288 | FIM1_3371 | QGN16652.1 | CP015058.1 | 1154 | 657 | 873 | 954 | 523 | 811 | 854 | 653 | 742 |
| 3289 | FIM1_3372 | QGN16653.1 | CP015058.1 | 2186 | 419 | 768 | 1718 | 321 | 691 | 1823 | 429 | 613 |
| 3290 | FIM1_3373 | QGN16654.1 | CP015058.1 | 1176 | 1099 | 657 | 927 | 818 | 632 | 278 | 1235 | 586 |
| 3291 | FIM1_3374 | QGN16655.1 | CP015058.1 | 18244 | 4663 | 3173 | 13203 | 1785 | 2918 | 19858 | 4469 | 2865 |
| 3292 | FIM1_3375 | QGN16656.1 | CP015058.1 | 17063 | 8197 | 4466 | 14401 | 5383 | 3970 | 21240 | 8408 | 4016 |
| 3293 | FIM1_3376 | QGN16657.1 | CP015058.1 | 85 | 12 | 13 | 63 | 3 | 14 | 42 | 15 | 10 |
| 3294 | FIM1_3377 | QGN16658.1 | CP015058.1 | 3904 | 1786 | 1163 | 2890 | 1192 | 1113 | 1868 | 1886 | 1135 |
| 3295 | FIM1_3378 | QGN16659.1 | CP015058.1 | 61 | 16 | 13 | 46 | 12 | 17 | 35 | 8 | 22 |
| 3296 | FIM1_3379 | QGN16660.1 | CP015058.1 | 1542 | 304 | 713 | 1167 | 176 | 687 | 1004 | 278 | 611 |
| 3297 | FIM1_3380 | QGN16661.1 | CP015058.1 | 1155 | 221 | 466 | 840 | 104 | 416 | 812 | 233 | 340 |
| 3298 | FIM1_3381 | QGN16662.1 | CP015058.1 | 23702 | 19280 | 16510 | 21545 | 15671 | 15338 | 25454 | 19306 | 16626 |
| 3299 | FIM1_3382 | QGN16663.1 | CP015058.1 | 1030 | 301 | 323 | 961 | 172 | 331 | 968 | 346 | 298 |
| 3300 | FIM1_3383 | QGN16664.1 | CP015058.1 | 7268 | 1546 | 1815 | 5875 | 1056 | 1746 | 7105 | 1669 | 1742 |
| 3301 | FIM1_3384 | QGN16665.1 | CP015058.1 | 17116 | 15341 | 6780 | 12631 | 11000 | 6153 | 4547 | 17126 | 6684 |
| 3302 | FIM1_3385 | QGN16666.1 | CP015058.1 | 239 | 45 | 62 | 193 | 57 | 81 | 124 | 52 | 60 |
| 3303 | FIM1_3386 | QGN16667.1 | CP015058.1 | 665 | 823 | 741 | 426 | 524 | 775 | 235 | 903 | 618 |
| 3304 | FIM1_3387 | QGN16668.1 | CP015058.1 | 7673 | 972 | 1588 | 6006 | 729 | 1399 | 6956 | 1055 | 1277 |
| 3305 | FIM1_3388 | QGN16669.1 | CP015058.1 | 8212 | 2169 | 2328 | 5795 | 1384 | 2389 | 9686 | 2279 | 2353 |
| 3306 | FIM1_3389 | QGN16670.1 | CP015058.1 | 1801 | 837 | 1238 | 1634 | 269 | 1202 | 3486 | 915 | 1158 |
| 3307 | FIM1_3390 | QGN16671.1 | CP015058.1 | 11808 | 2772 | 2733 | 9560 | 1518 | 2530 | 11730 | 3034 | 2438 |
| 3308 | FIM1_3391 | QGN16672.1 | CP015058.1 | 5147 | 1295 | 1111 | 4105 | 1069 | 945 | 3566 | 1274 | 1009 |
| 3309 | FIM1_3392 | QGN16673.1 | CP015058.1 | 2109 | 381 | 1026 | 1688 | 332 | 822 | 2549 | 410 | 735 |
| 3310 | FIM1_3393 | QGN16674.1 | CP015058.1 | 121 | 85 | 105 | 118 | 89 | 104 | 409 | 57 | 91 |

| | | | | | | | | | | | | |
|---|---|---|---|---|---|---|---|---|---|---|---|---|
| 3311 | FIM1_3394 | QGN16675.1 | CP015058.1 | 8386 | 1080 | 2150 | 6776 | 660 | 1863 | 7463 | 1090 | 1820 |
| 3312 | FIM1_3395 | QGN16676.1 | CP015058.1 | 7524 | 1949 | 1921 | 6705 | 1518 | 1837 | 6326 | 2053 | 2080 |
| 3313 | FIM1_3396 | QGN16677.1 | CP015058.1 | 2230 | 576 | 864 | 2095 | 518 | 821 | 1670 | 639 | 799 |
| 3314 | FIM1_3397 | QGN16678.1 | CP015058.1 | 25175 | 3395 | 7813 | 21670 | 3381 | 7333 | 26044 | 3531 | 7473 |
| 3315 | FIM1_3398 | QGN16679.1 | CP015058.1 | 14 | 11 | 6 | 13 | 4 | 9 | 8 | 5 | 8 |
| 3316 | FIM1_3399 | QGN16680.1 | CP015058.1 | 3028 | 467 | 988 | 2528 | 409 | 870 | 3511 | 487 | 808 |
| 3317 | FIM1_3400 | QGN16681.1 | CP015058.1 | 4730 | 796 | 1290 | 4063 | 655 | 1183 | 4168 | 867 | 1047 |
| 3318 | FIM1_3401 | QGN16682.1 | CP015058.1 | 5662 | 885 | 1076 | 4656 | 615 | 958 | 4638 | 1038 | 832 |
| 3319 | FIM1_3402 | QGN16683.1 | CP015058.1 | 7483 | 3481 | 3426 | 5957 | 2195 | 3163 | 6118 | 3658 | 3037 |
| 3320 | FIM1_3403 | QGN16684.1 | CP015058.1 | 13011 | 7229 | 3439 | 9122 | 4727 | 2974 | 4037 | 7543 | 2998 |
| 3321 | FIM1_3404 | QGN16685.1 | CP015058.1 | 66 | 11 | 17 | 50 | 11 | 17 | 138 | 10 | 13 |
| 3322 | FIM1_3405 | QGN16686.1 | CP015058.1 | 23322 | 11939 | 7662 | 18522 | 9211 | 6795 | 13660 | 12594 | 7340 |
| 3323 | FIM1_3407 | QGN16687.1 | CP015058.1 | 2941 | 1050 | 1266 | 2760 | 792 | 1164 | 2489 | 1132 | 1123 |
| 3324 | FIM1_3408 | QGN16688.1 | CP015058.1 | 3588 | 552 | 886 | 2919 | 445 | 860 | 4336 | 618 | 749 |
| 3325 | FIM1_3409 | QGN16689.1 | CP015058.1 | 16378 | 8531 | 8793 | 16181 | 5444 | 8125 | 19040 | 8465 | 8500 |
| 3326 | FIM1_3410 | QGN16690.1 | CP015058.1 | 4914 | 1353 | 1303 | 3301 | 934 | 1144 | 1774 | 1524 | 1071 |
| 3327 | FIM1_3411 | QGN16691.1 | CP015058.1 | 3633 | 1814 | 1624 | 2987 | 1365 | 1360 | 1587 | 1949 | 1511 |
| 3328 | FIM1_3412 | QGN16692.1 | CP015058.1 | 3173 | 956 | 1306 | 2763 | 680 | 1188 | 2972 | 961 | 1166 |
| 3329 | FIM1_3413 | QGN16693.1 | CP015058.1 | 6913 | 2302 | 2409 | 5089 | 1607 | 2152 | 5044 | 2445 | 2090 |
| 3330 | FIM1_3414 | QGN16694.1 | CP015058.1 | 3603 | 1629 | 2911 | 3211 | 1274 | 2655 | 2745 | 1799 | 2489 |
| 3331 | FIM1_3415 | QGN16695.1 | CP015058.1 | 4791 | 794 | 837 | 3841 | 506 | 771 | 4867 | 879 | 839 |
| 3332 | FIM1_3416 | QGN16696.1 | CP015058.1 | 1940 | 301 | 389 | 1552 | 193 | 348 | 2363 | 335 | 381 |
| 3333 | FIM1_3417 | QGN16697.1 | CP015058.1 | 21947 | 14365 | 13548 | 21059 | 11735 | 12381 | 21215 | 14597 | 12842 |
| 3334 | FIM1_3418 | QGN16698.1 | CP015058.1 | 3943 | 2447 | 2284 | 3286 | 1825 | 2044 | 2772 | 2657 | 1963 |
| 3335 | FIM1_3419 | QGN16699.1 | CP015058.1 | 9333 | 1950 | 2333 | 7402 | 1341 | 2144 | 10745 | 2024 | 2003 |
| 3336 | FIM1_3420 | QGN16700.1 | CP015058.1 | 252 | 18 | 80 | 213 | 14 | 62 | 683 | 17 | 46 |
| 3337 | FIM1_3421 | QGN16701.1 | CP015058.1 | 302 | 77 | 110 | 218 | 39 | 101 | 382 | 82 | 90 |
| 3338 | FIM1_3422 | QGN16702.1 | CP015058.1 | 374 | 49 | 77 | 250 | 30 | 41 | 722 | 49 | 57 |
| 3339 | FIM1_3423 | QGN16703.1 | CP015058.1 | 2880 | 96 | 300 | 1867 | 102 | 273 | 3083 | 110 | 272 |
| 3340 | FIM1_3424 | QGN16704.1 | CP015058.1 | 2816 | 499 | 646 | 2543 | 337 | 628 | 3844 | 514 | 631 |
| 3341 | FIM1_3425 | QGN16705.1 | CP015058.1 | 2819 | 326 | 938 | 1987 | 170 | 759 | 2731 | 408 | 353 |
| 3342 | FIM1_3426 | QGN16706.1 | CP015058.1 | 20836 | 3623 | 1491 | 13111 | 691 | 1144 | 28959 | 2971 | 1093 |
| 3343 | FIM1_3427 | QGN16707.1 | CP015058.1 | 7088 | 549 | 965 | 5381 | 405 | 835 | 9678 | 567 | 841 |
| 3344 | FIM1_3428 | QGN16708.1 | CP015058.1 | 358 | 27 | 45 | 265 | 23 | 55 | 319 | 31 | 37 |
| 3345 | FIM1_3429 | QGN16709.1 | CP015058.1 | 6596 | 2729 | 2742 | 5515 | 1932 | 2487 | 4913 | 2958 | 2474 |
| 3346 | FIM1_3430 | QGN16710.1 | CP015058.1 | 3353 | 1378 | 1298 | 3019 | 878 | 1201 | 2969 | 1394 | 1070 |
| 3347 | FIM1_3431 | QGN16711.1 | CP015058.1 | 778 | 429 | 167 | 604 | 198 | 122 | 912 | 404 | 142 |
| 3348 | FIM1_3432 | QGN16712.1 | CP015058.1 | 2084 | 628 | 701 | 1707 | 442 | 684 | 1999 | 592 | 605 |
| 3349 | FIM1_3433 | QGN16713.1 | CP015058.1 | 13240 | 2749 | 3283 | 10145 | 1933 | 3049 | 11512 | 2689 | 2776 |
| 3350 | FIM1_3434 | QGN16714.1 | CP015058.1 | 2115 | 269 | 806 | 1641 | 217 | 666 | 2399 | 340 | 639 |
| 3351 | FIM1_3435 | QGN16715.1 | CP015058.1 | 11572 | 2465 | 2181 | 8645 | 1962 | 1946 | 8259 | 2549 | 1970 |
| 3352 | FIM1_3437 | QGN16716.1 | CP015058.1 | 38086 | 10212 | 17090 | 30436 | 7546 | 16062 | 45812 | 10536 | 16395 |
| 3353 | FIM1_3438 | QGN16717.1 | CP015058.1 | 6465 | 1588 | 2576 | 6138 | 1295 | 2212 | 4358 | 1884 | 2532 |
| 3354 | FIM1_3439 | QGN16718.1 | CP015058.1 | 7276 | 707 | 2126 | 5225 | 260 | 1793 | 12220 | 677 | 1030 |
| 3355 | FIM1_3440 | QGN16719.1 | CP015058.1 | 8546 | 2312 | 2429 | 6695 | 1429 | 2184 | 10397 | 2456 | 2076 |
| 3356 | FIM1_3441 | QGN16720.1 | CP015058.1 | 1995 | 279 | 732 | 1828 | 203 | 658 | 2984 | 323 | 580 |
| 3357 | FIM1_3442 | QGN16721.1 | CP015058.1 | 16133 | 3133 | 4083 | 12572 | 2902 | 3751 | 15651 | 3326 | 3685 |
| 3358 | FIM1_3443 | QGN16722.1 | CP015058.1 | 110 | 2 | 81 | 71 | 3 | 55 | 48 | 4 | 54 |

| | | | | | | | | | | | | |
|---|---|---|---|---|---|---|---|---|---|---|---|---|
| 3359 | FIM1_3444 | QGN16723.1 | CP015058.1 | 8765 | 3804 | 3624 | 7326 | 2672 | 2990 | 5902 | 4032 | 3220 |
| 3360 | FIM1_3445 | QGN16724.1 | CP015058.1 | 6359 | 2918 | 3047 | 6119 | 2407 | 2933 | 5005 | 3151 | 2818 |
| 3361 | FIM1_3446 | QGN16725.1 | CP015058.1 | 3753 | 821 | 1212 | 3185 | 647 | 1157 | 3709 | 902 | 1135 |
| 3362 | FIM1_3447 | QGN16726.1 | CP015058.1 | 6143 | 1813 | 2453 | 4461 | 1544 | 2320 | 3985 | 2069 | 2503 |
| 3363 | FIM1_3448 | QGN16727.1 | CP015058.1 | 2725 | 343 | 616 | 2083 | 281 | 597 | 2752 | 347 | 513 |
| 3364 | FIM1_3449 | QGN16728.1 | CP015058.1 | 1683 | 482 | 698 | 1313 | 402 | 677 | 1941 | 544 | 703 |
| 3365 | FIM1_3450 | QGN16729.1 | CP015058.1 | 8804 | 6871 | 3397 | 6330 | 4323 | 3079 | 2368 | 7325 | 3203 |
| 3366 | FIM1_3451 | QGN16730.1 | CP015058.1 | 0 | 0 | 0 | 0 | 0 | 0 | 0 | 0 | 0 |
| 3367 | FIM1_3452 | QGN16731.1 | CP015058.1 | 5291 | 895 | 1557 | 3642 | 695 | 1302 | 4472 | 971 | 1329 |
| 3368 | FIM1_3453 | QGN16732.1 | CP015058.1 | 59 | 1 | 0 | 36 | 2 | 0 | 31 | 0 | 1 |
| 3369 | FIM1_3454 | QGN16733.1 | CP015058.1 | 54978 | 1148 | 1439 | 38111 | 1833 | 1191 | 57679 | 1404 | 784 |
| 3370 | FIM1_3455 | QGN16734.1 | CP015058.1 | 3402 | 32 | 56 | 2267 | 43 | 45 | 3028 | 44 | 28 |
| 3371 | FIM1_3456 | QGN16735.1 | CP015058.1 | 361 | 22 | 31 | 249 | 16 | 11 | 390 | 28 | 19 |
| 3372 | FIM1_3457 | QGN16736.1 | CP015058.1 | 3921 | 283 | 411 | 3296 | 342 | 375 | 5280 | 349 | 371 |
| 3373 | FIM1_3458 | QGN16737.1 | CP015058.1 | 46168 | 3300 | 3781 | 35892 | 3097 | 3296 | 63915 | 3738 | 2901 |
| 3374 | FIM1_3460 | QGN16738.1 | CP015058.1 | 16093 | 5326 | 4454 | 13832 | 3640 | 4174 | 18557 | 5589 | 4144 |
| 3375 | FIM1_3461 | QGN16739.1 | CP015058.1 | 4309 | 985 | 1429 | 3397 | 721 | 1241 | 3414 | 1090 | 1096 |
| 3376 | FIM1_3462 | QGN16740.1 | CP015058.1 | 2459 | 415 | 790 | 1833 | 306 | 727 | 2605 | 414 | 670 |
| 3377 | FIM1_3463 | QGN16741.1 | CP015058.1 | 16525 | 2033 | 2021 | 11936 | 1100 | 1895 | 22637 | 1984 | 2007 |
| 3378 | FIM1_3464 | QGN16742.1 | CP015058.1 | 764 | 176 | 267 | 602 | 143 | 220 | 503 | 181 | 212 |
| 3379 | FIM1_3465 | QGN16743.1 | CP015058.1 | 592 | 49 | 162 | 504 | 28 | 166 | 852 | 48 | 148 |
| 3380 | FIM1_3466 | QGN16744.1 | CP015058.1 | 8365 | 1861 | 2600 | 7156 | 1406 | 2299 | 8078 | 1933 | 2167 |
| 3381 | FIM1_3467 | QGN16745.1 | CP015058.1 | 3654 | 388 | 703 | 2908 | 208 | 644 | 5558 | 403 | 592 |
| 3382 | FIM1_3468 | QGN16746.1 | CP015058.1 | 4227 | 2139 | 2270 | 4278 | 1852 | 2107 | 3812 | 2287 | 2069 |
| 3383 | FIM1_3469 | QGN16747.1 | CP015058.1 | 9076 | 938 | 1759 | 7200 | 750 | 1673 | 10895 | 1024 | 1685 |
| 3384 | FIM1_3470 | QGN16748.1 | CP015058.1 | 2979 | 465 | 631 | 2262 | 368 | 601 | 3012 | 486 | 588 |
| 3385 | FIM1_3471 | QGN16749.1 | CP015058.1 | 4107 | 2502 | 2304 | 3621 | 2046 | 2179 | 3077 | 2581 | 2079 |
| 3386 | FIM1_3472 | QGN16750.1 | CP015058.1 | 2142 | 294 | 617 | 1662 | 209 | 585 | 1766 | 344 | 490 |
| 3387 | FIM1_3473 | QGN16751.1 | CP015058.1 | 8611 | 5051 | 2572 | 5844 | 3328 | 2209 | 2075 | 5244 | 2167 |
| 3388 | FIM1_3474 | QGN16752.1 | CP015058.1 | 5734 | 72 | 345 | 4306 | 60 | 285 | 3255 | 74 | 214 |
| 3389 | FIM1_3475 | QGN16753.1 | CP015058.1 | 9204 | 2254 | 2767 | 7984 | 1799 | 2547 | 9223 | 2426 | 2546 |
| 3390 | FIM1_3476 | QGN16754.1 | CP015058.1 | 4498 | 747 | 1275 | 3502 | 604 | 1230 | 5142 | 732 | 1049 |
| 3391 | FIM1_3477 | QGN16755.1 | CP015058.1 | 4596 | 1244 | 1440 | 4205 | 819 | 1355 | 4062 | 1343 | 1316 |
| 3392 | FIM1_3478 | QGN16756.1 | CP015058.1 | 614 | 102 | 264 | 431 | 94 | 218 | 334 | 109 | 240 |
| 3393 | FIM1_3479 | QGN16757.1 | CP015058.1 | 13826 | 4862 | 7669 | 11742 | 3563 | 7075 | 14832 | 5368 | 6893 |
| 3394 | FIM1_3480 | QGN16758.1 | CP015058.1 | 4061 | 4367 | 2443 | 3125 | 3318 | 2192 | 923 | 4610 | 2297 |
| 3395 | FIM1_3481 | QGN16759.1 | CP015058.1 | 10243 | 2427 | 2737 | 8110 | 1763 | 2567 | 9245 | 2677 | 2576 |
| 3396 | FIM1_3482 | QGN16760.1 | CP015058.1 | 3873 | 1501 | 4433 | 3298 | 945 | 4217 | 4535 | 1557 | 4011 |
| 3397 | FIM1_3483 | QGN16761.1 | CP015058.1 | 2026 | 428 | 808 | 1684 | 351 | 792 | 1914 | 491 | 792 |
| 3398 | FIM1_3484 | QGN16762.1 | CP015058.1 | 3744 | 591 | 1396 | 2933 | 416 | 1302 | 4554 | 709 | 1278 |
| 3399 | FIM1_3485 | QGN16763.1 | CP015058.1 | 661 | 307 | 505 | 708 | 208 | 493 | 491 | 312 | 441 |
| 3400 | FIM1_3486 | QGN16764.1 | CP015058.1 | 6646 | 2138 | 3565 | 5491 | 1430 | 3291 | 7239 | 2240 | 3147 |
| 3401 | FIM1_3487 | QGN16765.1 | CP015058.1 | 1573 | 535 | 851 | 1509 | 464 | 952 | 1048 | 635 | 725 |
| 3402 | FIM1_3488 | QGN16766.1 | CP015058.1 | 1338 | 207 | 336 | 1087 | 161 | 347 | 1336 | 238 | 296 |
| 3403 | FIM1_3489 | QGN16767.1 | CP015058.1 | 1672 | 542 | 591 | 1479 | 334 | 525 | 1437 | 565 | 522 |
| 3404 | FIM1_3490 | QGN16768.1 | CP015058.1 | 2397 | 764 | 1018 | 2311 | 619 | 926 | 1991 | 816 | 872 |
| 3405 | FIM1_3491 | QGN16769.1 | CP015058.1 | 5149 | 1011 | 1754 | 3662 | 863 | 1605 | 5900 | 1066 | 1335 |
| 3406 | FIM1_3492 | QGN16770.1 | CP015058.1 | 2366 | 234 | 838 | 1815 | 240 | 694 | 1808 | 309 | 611 |

| | | | | | | | | | | | | |
|---|---|---|---|---|---|---|---|---|---|---|---|---|
| 3407 | FIM1_3493 | QGN16771.1 | CP015058.1 | 675 | 67 | 127 | 455 | 59 | 96 | 256 | 76 | 87 |
| 3408 | FIM1_3494 | QGN16772.1 | CP015058.1 | 22791 | 9171 | 7902 | 20123 | 7626 | 7257 | 23388 | 9805 | 7853 |
| 3409 | FIM1_3495 | QGN16773.1 | CP015058.1 | 1058 | 96 | 128 | 786 | 51 | 99 | 1322 | 97 | 102 |
| 3410 | FIM1_3496 | QGN16774.1 | CP015058.1 | 1342 | 499 | 510 | 1240 | 460 | 473 | 801 | 576 | 432 |
| 3411 | FIM1_3497 | QGN16775.1 | CP015058.1 | 6520 | 293 | 893 | 4798 | 289 | 821 | 8963 | 278 | 713 |
| 3412 | FIM1_3498 | QGN16776.1 | CP015058.1 | 346 | 89 | 150 | 261 | 100 | 157 | 156 | 127 | 142 |
| 3413 | FIM1_3499 | QGN16777.1 | CP015058.1 | 7549 | 1470 | 2632 | 6098 | 1170 | 2387 | 8447 | 1604 | 2183 |
| 3414 | FIM1_3500 | QGN16778.1 | CP015058.1 | 3990 | 1308 | 1712 | 3693 | 708 | 1532 | 4240 | 1311 | 1486 |
| 3415 | FIM1_3501 | QGN16779.1 | CP015058.1 | 9765 | 989 | 2080 | 8372 | 940 | 1833 | 10215 | 1099 | 1679 |
| 3416 | FIM1_3503 | QGN16780.1 | CP015058.1 | 1772 | 191 | 367 | 1355 | 75 | 289 | 2158 | 209 | 297 |
| 3417 | FIM1_3504 | QGN16781.1 | CP015058.1 | 985 | 728 | 464 | 658 | 256 | 347 | 539 | 738 | 389 |
| 3418 | FIM1_3505 | QGN16782.1 | CP015058.1 | 7021 | 3283 | 2961 | 6021 | 2478 | 2769 | 6286 | 3518 | 2710 |
| 3419 | FIM1_3506 | QGN16783.1 | CP015058.1 | 1861 | 423 | 652 | 1635 | 314 | 632 | 1576 | 437 | 599 |
| 3420 | FIM1_3507 | QGN16784.1 | CP015058.1 | 3407 | 1092 | 1877 | 2751 | 936 | 1539 | 2731 | 1254 | 1479 |
| 3421 | FIM1_3508 | QGN16785.1 | CP015058.1 | 2368 | 785 | 849 | 1570 | 552 | 770 | 1441 | 821 | 708 |
| 3422 | FIM1_3509 | QGN16786.1 | CP015058.1 | 110 | 7 | 7 | 56 | 6 | 9 | 78 | 6 | 9 |
| 3423 | FIM1_3510 | QGN16787.1 | CP015058.1 | 11 | 1 | 0 | 8 | 0 | 0 | 34 | 1 | 0 |
| 3424 | FIM1_3511 | QGN16788.1 | CP015058.1 | 9298 | 3269 | 4545 | 8185 | 2290 | 4294 | 9687 | 3399 | 4237 |
| 3425 | FIM1_3512 | QGN16789.1 | CP015058.1 | 13321 | 2524 | 3813 | 11236 | 1513 | 3517 | 16905 | 2592 | 3393 |
| 3426 | FIM1_3513 | QGN16790.1 | CP015058.1 | 1709 | 388 | 627 | 1434 | 262 | 585 | 2683 | 371 | 505 |
| 3427 | FIM1_3514 | QGN16791.1 | CP015058.1 | 10281 | 1660 | 2032 | 7933 | 1169 | 1826 | 10778 | 1700 | 1716 |
| 3428 | FIM1_3515 | QGN16792.1 | CP015058.1 | 1 | 2 | 0 | 5 | 2 | 0 | 1 | 0 | 0 |
| 3429 | FIM1_3516 | QGN16793.1 | CP015058.1 | 24511 | 7825 | 6084 | 20023 | 5653 | 5497 | 13676 | 8197 | 5862 |
| 3430 | FIM1_3517 | QGN16794.1 | CP015058.1 | 1132 | 220 | 517 | 1050 | 157 | 401 | 1218 | 262 | 387 |
| 3431 | FIM1_3518 | QGN16795.1 | CP015058.1 | 8113 | 3921 | 2325 | 6389 | 2805 | 2063 | 3850 | 4249 | 2149 |
| 3432 | FIM1_3519 | QGN16796.1 | CP015058.1 | 2197 | 533 | 675 | 1777 | 408 | 715 | 1556 | 577 | 739 |
| 3433 | FIM1_3520 | QGN16797.1 | CP015058.1 | 5294 | 647 | 899 | 4314 | 480 | 883 | 5388 | 696 | 934 |
| 3434 | FIM1_3522 | QGN16798.1 | CP015058.1 | 8104 | 5392 | 2600 | 6266 | 3262 | 2244 | 1934 | 5435 | 2452 |
| 3435 | FIM1_3523 | QGN16799.1 | CP015058.1 | 22831 | 6694 | 5918 | 18234 | 4737 | 5580 | 22061 | 7194 | 5295 |
| 3436 | FIM1_3524 | QGN16800.1 | CP015058.1 | 3824 | 1628 | 1501 | 3342 | 1332 | 1366 | 2380 | 1815 | 1422 |
| 3437 | FIM1_3525 | QGN16801.1 | CP015058.1 | 1671 | 310 | 481 | 1286 | 236 | 439 | 982 | 299 | 390 |
| 3438 | FIM1_3526 | QGN16802.1 | CP015058.1 | 5895 | 2040 | 2368 | 5446 | 1524 | 2225 | 6272 | 2352 | 2171 |
| 3439 | FIM1_3527 | QGN16803.1 | CP015058.1 | 4133 | 934 | 1688 | 3423 | 722 | 1522 | 4662 | 1036 | 1479 |
| 3440 | FIM1_3528 | QGN16804.1 | CP015058.1 | 10018 | 1433 | 1464 | 7293 | 810 | 1419 | 16698 | 1513 | 1413 |
| 3441 | FIM1_3529 | QGN16805.1 | CP015058.1 | 1560 | 418 | 590 | 1044 | 322 | 519 | 895 | 454 | 531 |
| 3442 | FIM1_3530 | QGN16806.1 | CP015058.1 | 820 | 252 | 392 | 650 | 178 | 367 | 458 | 264 | 367 |
| 3443 | FIM1_3531 | QGN16807.1 | CP015058.1 | 3534 | 691 | 1161 | 2818 | 601 | 1056 | 3481 | 754 | 1028 |
| 3444 | FIM1_3532 | QGN16808.1 | CP015058.1 | 1997 | 629 | 794 | 1501 | 553 | 807 | 1473 | 746 | 810 |
| 3445 | FIM1_3533 | QGN16809.1 | CP015058.1 | 1793 | 200 | 360 | 1249 | 135 | 352 | 1421 | 217 | 283 |
| 3446 | FIM1_3534 | QGN16810.1 | CP015058.1 | 6634 | 1506 | 2239 | 4953 | 1063 | 2141 | 7516 | 1626 | 2198 |
| 3447 | FIM1_3535 | QGN16811.1 | CP015058.1 | 3275 | 1121 | 1498 | 2765 | 800 | 1302 | 3042 | 1282 | 1235 |
| 3448 | FIM1_3536 | QGN16812.1 | CP015058.1 | 875 | 292 | 404 | 731 | 229 | 424 | 721 | 331 | 373 |
| 3449 | FIM1_3537 | QGN16813.1 | CP015058.1 | 14733 | 11050 | 6845 | 10763 | 8479 | 6022 | 4862 | 11608 | 6512 |
| 3450 | FIM1_3538 | QGN16814.1 | CP015058.1 | 2277 | 447 | 898 | 1712 | 373 | 832 | 2498 | 534 | 736 |
| 3451 | FIM1_3539 | QGN16815.1 | CP015058.1 | 916 | 393 | 543 | 874 | 268 | 528 | 1046 | 429 | 476 |
| 3452 | FIM1_3540 | QGN16816.1 | CP015058.1 | 6505 | 2271 | 2613 | 5623 | 1513 | 2269 | 7760 | 2437 | 2221 |
| 3453 | FIM1_3541 | QGN16817.1 | CP015058.1 | 365 | 381 | 210 | 221 | 168 | 184 | 362 | 473 | 205 |
| 3454 | FIM1_3542 | QGN16818.1 | CP015058.1 | 1245 | 216 | 324 | 843 | 195 | 297 | 1311 | 222 | 321 |

| | | | | | | | | | | | | |
|---|---|---|---|---|---|---|---|---|---|---|---|---|
| 3455 | FIM1_3543 | QGN16819.1 | CP015058.1 | 41 | 2 | 18 | 19 | 3 | 15 | 56 | 5 | 19 |
| 3456 | FIM1_3544 | QGN16820.1 | CP015058.1 | 6588 | 2125 | 2430 | 5189 | 1660 | 2329 | 5591 | 2259 | 2386 |
| 3457 | FIM1_3545 | QGN16821.1 | CP015058.1 | 617 | 24 | 33 | 370 | 15 | 31 | 837 | 39 | 27 |
| 3458 | FIM1_3546 | QGN16822.1 | CP015058.1 | 5301 | 1623 | 1054 | 3525 | 1132 | 945 | 3898 | 1596 | 886 |
| 3459 | FIM1_3547 | QGN16823.1 | CP015058.1 | 844 | 772 | 1131 | 868 | 585 | 1069 | 1228 | 877 | 1006 |
| 3460 | FIM1_3548 | QGN16824.1 | CP015058.1 | 1647 | 371 | 429 | 1669 | 255 | 374 | 1746 | 407 | 310 |
| 3461 | FIM1_3549 | QGN16825.1 | CP015058.1 | 461 | 82 | 120 | 361 | 79 | 130 | 329 | 98 | 111 |
| 3462 | FIM1_3550 | QGN16826.1 | CP015058.1 | 2173 | 462 | 1607 | 1793 | 486 | 1552 | 1874 | 468 | 1253 |
| 3463 | FIM1_3551 | QGN16827.1 | CP015058.1 | 4777 | 1226 | 1498 | 5007 | 941 | 1534 | 3968 | 1303 | 1416 |
| 3464 | FIM1_3552 | QGN16828.1 | CP015058.1 | 1066 | 295 | 548 | 982 | 281 | 551 | 1096 | 384 | 488 |
| 3465 | FIM1_3553 | QGN16829.1 | CP015058.1 | 1471 | 278 | 397 | 1170 | 247 | 389 | 1405 | 317 | 385 |
| 3466 | FIM1_3554 | QGN16830.1 | CP015058.1 | 949 | 367 | 657 | 873 | 279 | 658 | 767 | 375 | 627 |
| 3467 | FIM1_3555 | QGN16831.1 | CP015058.1 | 1469 | 251 | 587 | 1157 | 200 | 533 | 1285 | 262 | 463 |
| 3468 | FIM1_3556 | QGN16832.1 | CP015058.1 | 6388 | 1622 | 1811 | 5154 | 1012 | 1615 | 6084 | 1662 | 1521 |
| 3469 | FIM1_3557 | QGN16833.1 | CP015058.1 | 7824 | 5827 | 3551 | 6114 | 4121 | 3020 | 3134 | 6464 | 3248 |
| 3470 | FIM1_3558 | QGN16834.1 | CP015058.1 | 2849 | 786 | 976 | 2403 | 459 | 922 | 3438 | 761 | 949 |
| 3471 | FIM1_3559 | QGN16835.1 | CP015058.1 | 533 | 106 | 216 | 497 | 76 | 169 | 554 | 128 | 149 |
| 3472 | FIM1_3560 | QGN16836.1 | CP015058.1 | 3128 | 769 | 985 | 2774 | 582 | 899 | 3625 | 794 | 905 |
| 3473 | FIM1_3561 | QGN16837.1 | CP015058.1 | 2957 | 166 | 536 | 2170 | 128 | 503 | 6082 | 164 | 458 |
| 3474 | FIM1_3562 | QGN16838.1 | CP015058.1 | 4947 | 1088 | 989 | 3787 | 699 | 992 | 6080 | 1163 | 936 |
| 3475 | FIM1_3563 | QGN16839.1 | CP015058.1 | 4179 | 1334 | 1561 | 2960 | 1053 | 1458 | 4108 | 1411 | 1357 |
| 3476 | FIM1_3565 | QGN16840.1 | CP015058.1 | 885 | 61 | 80 | 725 | 51 | 67 | 310 | 64 | 46 |
| 3477 | FIM1_3566 | QGN16841.1 | CP015058.1 | 728 | 126 | 172 | 564 | 94 | 160 | 365 | 119 | 112 |
| 3478 | FIM1_3567 | QGN16842.1 | CP015058.1 | 16926 | 6703 | 6355 | 13697 | 5186 | 5762 | 14122 | 6924 | 6149 |
| 3479 | FIM1_3568 | QGN16843.1 | CP015058.1 | 16609 | 2026 | 4710 | 13661 | 1324 | 4467 | 15466 | 2191 | 4068 |
| 3480 | FIM1_3569 | QGN16844.1 | CP015058.1 | 3674 | 2940 | 1483 | 3017 | 1819 | 1261 | 676 | 3079 | 1436 |
| 3481 | FIM1_3570 | QGN16845.1 | CP015058.1 | 2592 | 1070 | 1368 | 2116 | 730 | 1207 | 1072 | 1140 | 1107 |
| 3482 | FIM1_3571 | QGN16846.1 | CP015058.1 | 9510 | 3056 | 3480 | 8235 | 2435 | 3156 | 7469 | 3450 | 3117 |
| 3483 | FIM1_3572 | QGN16847.1 | CP015058.1 | 1080 | 275 | 530 | 1029 | 230 | 477 | 1090 | 327 | 412 |
| 3484 | FIM1_3573 | QGN16848.1 | CP015058.1 | 10336 | 3782 | 3833 | 7792 | 2848 | 3499 | 7362 | 3975 | 3402 |
| 3485 | FIM1_3574 | QGN16849.1 | CP015058.1 | 14527 | 1157 | 1499 | 12287 | 674 | 1445 | 27751 | 1176 | 1267 |
| 3486 | FIM1_3575 | QGN16850.1 | CP015058.1 | 4722 | 964 | 909 | 4315 | 746 | 734 | 6961 | 975 | 747 |
| 3487 | FIM1_3576 | QGN16851.1 | CP015058.1 | 8026 | 2250 | 3206 | 6749 | 1407 | 2982 | 8302 | 2382 | 2753 |
| 3488 | FIM1_3577 | QGN16852.1 | CP015058.1 | 1003 | 196 | 602 | 753 | 168 | 495 | 1075 | 222 | 429 |
| 3489 | FIM1_3579 | QGN16853.1 | CP015058.1 | 8383 | 1384 | 1881 | 6407 | 1054 | 1770 | 7647 | 1543 | 1712 |
| 3490 | FIM1_3580 | QGN16854.1 | CP015058.1 | 7942 | 7488 | 3895 | 5713 | 4920 | 3527 | 1924 | 8185 | 3778 |
| 3491 | FIM1_3581 | QGN16855.1 | CP015058.1 | 8487 | 1726 | 5654 | 3279 | 1005 | 6456 | 8426 | 1842 | 5207 |
| 3492 | FIM1_3582 | QGN16856.1 | CP015058.1 | 6551 | 1421 | 1550 | 5730 | 1057 | 1304 | 6793 | 1480 | 1178 |
| 3493 | FIM1_3583 | QGN16857.1 | CP015058.1 | 8968 | 1645 | 1622 | 6791 | 1350 | 1431 | 8011 | 1736 | 1514 |
| 3494 | FIM1_3584 | QGN16858.1 | CP015058.1 | 1192 | 403 | 639 | 1066 | 410 | 596 | 985 | 431 | 610 |
| 3495 | FIM1_3585 | QGN16859.1 | CP015058.1 | 3434 | 432 | 1121 | 2727 | 472 | 1154 | 2699 | 443 | 1090 |
| 3496 | FIM1_3586 | QGN16860.1 | CP015058.1 | 11732 | 2379 | 2190 | 9337 | 1892 | 2312 | 10242 | 2533 | 2249 |
| 3497 | FIM1_3587 | QGN16861.1 | CP015058.1 | 557 | 39 | 56 | 413 | 66 | 60 | 270 | 56 | 41 |
| 3498 | FIM1_3588 | QGN16862.1 | CP015058.1 | 3082 | 638 | 962 | 2093 | 477 | 855 | 2816 | 717 | 830 |
| 3499 | FIM1_3589 | QGN16863.1 | CP015058.1 | 14688 | 1994 | 2272 | 11303 | 1281 | 2036 | 17039 | 2179 | 1875 |
| 3500 | FIM1_3590 | QGN16864.1 | CP015058.1 | 13 | 0 | 1 | 24 | 0 | 0 | 7 | 1 | 1 |
| 3501 | FIM1_3591 | QGN16865.1 | CP015058.1 | 1412 | 141 | 519 | 1097 | 120 | 456 | 2306 | 157 | 415 |
| 3502 | FIM1_3592 | QGN16866.1 | CP015058.1 | 640 | 1951 | 2804 | 323 | 734 | 2268 | 259 | 2019 | 2242 |

| | | | | | | | | | | | | |
|---|---|---|---|---|---|---|---|---|---|---|---|---|
| 3503 | FIM1_3593 | QGN16867.1 | CP015058.1 | 3 | 0 | 0 | 2 | 1 | 0 | 0 | 1 | 0 |
| 3504 | FIM1_3594 | QGN16868.1 | CP015058.1 | 9192 | 2756 | 2657 | 8160 | 1921 | 2371 | 7626 | 2895 | 2427 |
| 3505 | FIM1_3595 | QGN16869.1 | CP015058.1 | 12489 | 3197 | 5875 | 10046 | 2149 | 4942 | 13776 | 3238 | 5075 |
| 3506 | FIM1_3596 | QGN16870.1 | CP015058.1 | 140 | 17 | 29 | 162 | 21 | 38 | 78 | 25 | 17 |
| 3507 | FIM1_3597 | QGN16871.1 | CP015058.1 | 14805 | 9449 | 10572 | 14673 | 6995 | 9517 | 15939 | 9588 | 9859 |
| 3508 | FIM1_3598 | QGN16872.1 | CP015058.1 | 5597 | 3123 | 1755 | 4525 | 2273 | 1526 | 2566 | 3309 | 1623 |
| 3509 | FIM1_3599 | QGN16873.1 | CP015058.1 | 15711 | 5041 | 4566 | 12816 | 3378 | 4237 | 16137 | 5247 | 4210 |
| 3510 | FIM1_3600 | QGN16874.1 | CP015058.1 | 839 | 225 | 421 | 639 | 214 | 435 | 529 | 279 | 341 |
| 3511 | FIM1_3601 | QGN16875.1 | CP015058.1 | 7846 | 1243 | 1868 | 5710 | 957 | 1744 | 7424 | 1389 | 1608 |
| 3512 | FIM1_3603 | QGN16876.1 | CP015058.1 | 1010 | 329 | 587 | 661 | 285 | 561 | 1078 | 340 | 486 |
| 3513 | FIM1_3604 | QGN16877.1 | CP015058.1 | 7266 | 4738 | 3015 | 5651 | 3308 | 2827 | 3706 | 5027 | 2823 |
| 3514 | FIM1_3605 | QGN16878.1 | CP015058.1 | 3388 | 627 | 1188 | 2791 | 446 | 1065 | 3324 | 629 | 980 |
| 3515 | FIM1_3606 | QGN16879.1 | CP015058.1 | 1563 | 196 | 634 | 1064 | 175 | 619 | 1331 | 217 | 523 |
| 3516 | FIM1_3607 | QGN16880.1 | CP015058.1 | 3224 | 1363 | 1221 | 2336 | 914 | 1126 | 1993 | 1462 | 1057 |
| 3517 | FIM1_3608 | QGN16881.1 | CP015058.1 | 1828 | 336 | 489 | 1223 | 227 | 407 | 1403 | 376 | 431 |
| 3518 | FIM1_3609 | QGN16882.1 | CP015058.1 | 3297 | 310 | 978 | 2441 | 401 | 857 | 5680 | 366 | 736 |
| 3519 | FIM1_3610 | QGN16883.1 | CP015058.1 | 3609 | 956 | 1789 | 2989 | 734 | 1687 | 2516 | 967 | 1548 |
| 3520 | FIM1_3611 | QGN16884.1 | CP015058.1 | 2547 | 531 | 747 | 1826 | 421 | 737 | 2367 | 570 | 659 |
| 3521 | FIM1_3612 | QGN16885.1 | CP015058.1 | 4189 | 846 | 1252 | 3271 | 666 | 1261 | 4795 | 970 | 1256 |
| 3522 | FIM1_3613 | QGN16886.1 | CP015058.1 | 2279 | 244 | 1259 | 1812 | 163 | 1185 | 3256 | 289 | 1100 |
| 3523 | FIM1_3614 | QGN16887.1 | CP015058.1 | 304 | 179 | 369 | 252 | 133 | 392 | 155 | 200 | 318 |
| 3524 | FIM1_3615 | QGN16888.1 | CP015058.1 | 2642 | 1517 | 1127 | 2090 | 1154 | 1122 | 1355 | 1762 | 973 |
| 3525 | FIM1_3616 | QGN16889.1 | CP015058.1 | 7866 | 3814 | 2790 | 5958 | 2891 | 2566 | 2632 | 4196 | 2548 |
| 3526 | FIM1_3617 | QGN16890.1 | CP015058.1 | 371 | 183 | 190 | 338 | 205 | 186 | 179 | 227 | 180 |
| 3527 | FIM1_3618 | QGN16891.1 | CP015058.1 | 6847 | 1167 | 2655 | 5302 | 979 | 2510 | 8143 | 1304 | 2507 |
| 3528 | FIM1_3619 | QGN16892.1 | CP015058.1 | 671 | 185 | 325 | 602 | 172 | 312 | 397 | 205 | 269 |
| 3529 | FIM1_3620 | QGN16893.1 | CP015058.1 | 4325 | 549 | 1207 | 3457 | 437 | 1133 | 5912 | 647 | 986 |
| 3530 | FIM1_3621 | QGN16894.1 | CP015058.1 | 3265 | 822 | 1429 | 2784 | 678 | 1358 | 2884 | 901 | 1294 |
| 3531 | FIM1_3622 | QGN16895.1 | CP015058.1 | 6372 | 1523 | 2093 | 4931 | 1089 | 1945 | 7197 | 1611 | 1781 |
| 3532 | FIM1_3623 | QGN16896.1 | CP015058.1 | 683 | 41 | 148 | 477 | 35 | 127 | 3189 | 43 | 120 |
| 3533 | FIM1_3624 | QGN16897.1 | CP015058.1 | 1098 | 36 | 383 | 965 | 24 | 354 | 2738 | 32 | 296 |
| 3534 | FIM1_3625 | QGN16898.1 | CP015058.1 | 42769 | 19518 | 16461 | 38331 | 14568 | 14892 | 44264 | 20209 | 15406 |
| 3535 | FIM1_3626 | QGN16899.1 | CP015058.1 | 433 | 62 | 237 | 381 | 51 | 198 | 452 | 73 | 190 |
| 3536 | FIM1_3627 | QGN16900.1 | CP015058.1 | 2613 | 448 | 426 | 1887 | 299 | 379 | 3438 | 517 | 401 |
| 3537 | FIM1_3628 | QGN16901.1 | CP015058.1 | 29087 | 10226 | 10190 | 24050 | 6197 | 9337 | 31090 | 11109 | 9979 |
| 3538 | FIM1_3629 | QGN16902.1 | CP015058.1 | 15004 | 2816 | 4358 | 12732 | 1976 | 4034 | 16064 | 3022 | 3778 |
| 3539 | FIM1_3630 | QGN16903.1 | CP015058.1 | 730 | 69 | 126 | 583 | 54 | 145 | 1230 | 66 | 118 |
| 3540 | FIM1_3631 | QGN16904.1 | CP015058.1 | 1125 | 335 | 462 | 936 | 253 | 496 | 1020 | 317 | 426 |
| 3541 | FIM1_3632 | QGN16905.1 | CP015058.1 | 21296 | 7235 | 8050 | 17359 | 5422 | 7464 | 16216 | 7907 | 7081 |
| 3542 | FIM1_3633 | QGN16906.1 | CP015058.1 | 4856 | 242 | 1135 | 4300 | 221 | 1252 | 13166 | 261 | 1064 |
| 3543 | FIM1_3634 | QGN16907.1 | CP015058.1 | 6880 | 1867 | 2627 | 6006 | 1468 | 2568 | 6124 | 2060 | 2379 |
| 3544 | FIM1_3635 | QGN16908.1 | CP015058.1 | 7334 | 3079 | 2896 | 5995 | 2404 | 2856 | 5733 | 3434 | 2720 |
| 3545 | FIM1_3636 | QGN16909.1 | CP015058.1 | 6291 | 1055 | 1653 | 4628 | 840 | 1501 | 5985 | 1151 | 1341 |
| 3546 | FIM1_3637 | QGN16910.1 | CP015058.1 | 4731 | 1453 | 2107 | 3484 | 1025 | 1939 | 4123 | 1548 | 1740 |
| 3547 | FIM1_3638 | QGN16911.1 | CP015058.1 | 11043 | 3333 | 4986 | 9464 | 3545 | 5058 | 6884 | 3684 | 4985 |
| 3548 | FIM1_3639 | QGN16912.1 | CP015058.1 | 534 | 152 | 237 | 392 | 120 | 255 | 623 | 169 | 200 |
| 3549 | FIM1_3640 | QGN16913.1 | CP015058.1 | 1662 | 455 | 651 | 1276 | 405 | 657 | 1364 | 497 | 605 |
| 3550 | FIM1_3641 | QGN16914.1 | CP015058.1 | 1682 | 506 | 577 | 1166 | 364 | 546 | 1430 | 578 | 517 |

| | | | | | | | | | | | | |
|---|---|---|---|---|---|---|---|---|---|---|---|---|
| 3551 | FIM1_3642 | QGN16915.1 | CP015058.1 | 4794 | 864 | 1706 | 3149 | 588 | 1617 | 4872 | 1000 | 1502 |
| 3552 | FIM1_3643 | QGN16916.1 | CP015058.1 | 2142 | 468 | 833 | 1640 | 407 | 805 | 2032 | 506 | 737 |
| 3553 | FIM1_3644 | QGN16917.1 | CP015058.1 | 4495 | 1157 | 1650 | 3544 | 888 | 1526 | 4935 | 1244 | 1383 |
| 3554 | FIM1_3645 | QGN16918.1 | CP015058.1 | 5782 | 1172 | 1862 | 4546 | 969 | 1718 | 5046 | 1316 | 1577 |
| 3555 | FIM1_3646 | QGN16919.1 | CP015058.1 | 2297 | 728 | 756 | 1815 | 564 | 782 | 1236 | 785 | 739 |
| 3556 | FIM1_3647 | QGN16920.1 | CP015058.1 | 29969 | 22480 | 20407 | 28248 | 18984 | 19730 | 29629 | 23124 | 20263 |
| 3557 | FIM1_3648 | QGN16921.1 | CP015058.1 | 3457 | 2897 | 1439 | 2094 | 1691 | 1285 | 593 | 3001 | 1263 |
| 3558 | FIM1_3649 | QGN16922.1 | CP015058.1 | 2258 | 1100 | 1067 | 1705 | 588 | 946 | 1410 | 1098 | 831 |
| 3559 | FIM1_3650 | QGN16923.1 | CP015058.1 | 2895 | 796 | 832 | 2274 | 616 | 814 | 2330 | 903 | 763 |
| 3560 | FIM1_3651 | QGN16924.1 | CP015058.1 | 2245 | 752 | 754 | 1520 | 587 | 770 | 1938 | 801 | 818 |
| 3561 | FIM1_3652 | QGN16925.1 | CP015058.1 | 328 | 147 | 163 | 328 | 56 | 177 | 747 | 139 | 152 |
| 3562 | FIM1_3653 | QGN16926.1 | CP015058.1 | 9636 | 4376 | 3451 | 7767 | 3231 | 3128 | 8435 | 4705 | 3134 |
| 3563 | FIM1_3654 | QGN16927.1 | CP015058.1 | 12137 | 4937 | 5472 | 9853 | 3546 | 5263 | 11747 | 5124 | 5052 |
| 3564 | FIM1_3655 | QGN16928.1 | CP015058.1 | 7411 | 2092 | 3153 | 5662 | 1538 | 2969 | 7326 | 2328 | 2862 |
| 3565 | FIM1_3656 | QGN16929.1 | CP015058.1 | 8876 | 3586 | 3969 | 7294 | 2618 | 3767 | 7628 | 3901 | 3492 |
| 3566 | FIM1_3657 | QGN16930.1 | CP015058.1 | 1712 | 275 | 445 | 1291 | 199 | 369 | 1772 | 354 | 280 |
| 3567 | FIM1_3658 | QGN16931.1 | CP015058.1 | 2364 | 2017 | 1316 | 1777 | 1380 | 1228 | 1084 | 2085 | 1119 |
| 3568 | FIM1_3659 | QGN16932.1 | CP015058.1 | 4253 | 848 | 1248 | 3256 | 567 | 1080 | 3798 | 928 | 1049 |
| 3569 | FIM1_3660 | QGN16933.1 | CP015058.1 | 2049 | 805 | 789 | 1641 | 722 | 826 | 1788 | 937 | 795 |
| 3570 | FIM1_3661 | QGN16934.1 | CP015058.1 | 3184 | 994 | 1462 | 2440 | 808 | 1336 | 3390 | 1038 | 1354 |
| 3571 | FIM1_3662 | QGN16935.1 | CP015058.1 | 2605 | 349 | 827 | 1896 | 315 | 760 | 2886 | 402 | 607 |
| 3572 | FIM1_3663 | QGN16936.1 | CP015058.1 | 5474 | 3540 | 2947 | 3361 | 1931 | 2700 | 1899 | 3976 | 2084 |
| 3573 | FIM1_3664 | QGN16937.1 | CP015058.1 | 2313 | 1173 | 813 | 1488 | 805 | 739 | 1461 | 1236 | 874 |
| 3574 | FIM1_3665 | QGN16938.1 | CP015058.1 | 1870 | 357 | 838 | 1353 | 324 | 855 | 1395 | 385 | 634 |
| 3575 | FIM1_3666 | QGN16939.1 | CP015058.1 | 2881 | 791 | 919 | 2010 | 609 | 901 | 1880 | 839 | 842 |
| 3576 | FIM1_3667 | QGN16940.1 | CP015058.1 | 3556 | 918 | 1504 | 2607 | 810 | 1388 | 2681 | 1029 | 1354 |
| 3577 | FIM1_3668 | QGN16941.1 | CP015058.1 | 2635 | 795 | 942 | 1974 | 548 | 844 | 1795 | 759 | 812 |
| 3578 | FIM1_3669 | QGN16942.1 | CP015058.1 | 2357 | 332 | 718 | 1935 | 326 | 615 | 2415 | 377 | 589 |
| 3579 | FIM1_3670 | QGN16943.1 | CP015058.1 | 1190 | 248 | 476 | 1140 | 208 | 488 | 1084 | 263 | 431 |
| 3580 | FIM1_3671 | QGN16944.1 | CP015058.1 | 859 | 165 | 330 | 622 | 170 | 361 | 1007 | 220 | 327 |
| 3581 | FIM1_3672 | QGN16945.1 | CP015058.1 | 3777 | 911 | 1211 | 3447 | 618 | 1005 | 4119 | 964 | 956 |
| 3582 | FIM1_3673 | QGN16946.1 | CP015058.1 | 3980 | 1923 | 1331 | 2431 | 1179 | 1298 | 1867 | 2057 | 1161 |
| 3583 | FIM1_3674 | QGN16947.1 | CP015058.1 | 128 | 31 | 34 | 83 | 18 | 22 | 96 | 29 | 26 |
| 3584 | FIM1_3675 | QGN16948.1 | CP015058.1 | 6917 | 2468 | 3003 | 5840 | 1931 | 2883 | 6069 | 2659 | 2691 |
| 3585 | FIM1_3676 | QGN16949.1 | CP015058.1 | 344 | 65 | 67 | 221 | 54 | 54 | 271 | 63 | 55 |
| 3586 | FIM1_3677 | QGN16950.1 | CP015058.1 | 2915 | 737 | 1037 | 2118 | 539 | 975 | 2867 | 781 | 867 |
| 3587 | FIM1_3678 | QGN16951.1 | CP015058.1 | 3404 | 2766 | 1727 | 2617 | 1944 | 1592 | 1009 | 2962 | 1597 |
| 3588 | FIM1_3679 | QGN16952.1 | CP015058.1 | 103048 | 51342 | 42962 | 92885 | 39955 | 38303 | 99144 | 51607 | 40794 |
| 3589 | FIM1_3680 | QGN16953.1 | CP015058.1 | 360 | 141 | 82 | 256 | 98 | 93 | 179 | 142 | 90 |
| 3590 | FIM1_3681 | QGN16954.1 | CP015058.1 | 2013 | 501 | 748 | 1550 | 371 | 692 | 1917 | 552 | 599 |
| 3591 | FIM1_3682 | QGN16955.1 | CP015058.1 | 4457 | 1867 | 1149 | 3259 | 1258 | 1112 | 4734 | 2066 | 1227 |
| 3592 | FIM1_3683 | QGN16956.1 | CP015058.1 | 1477 | 754 | 999 | 1475 | 589 | 1016 | 1275 | 846 | 801 |
| 3593 | FIM1_3684 | QGN16957.1 | CP015058.1 | 1278 | 281 | 366 | 843 | 183 | 334 | 1352 | 311 | 322 |
| 3594 | FIM1_3685 | QGN16958.1 | CP015058.1 | 1458 | 285 | 600 | 1115 | 213 | 612 | 1813 | 309 | 476 |
| 3595 | FIM1_3686 | QGN16959.1 | CP015058.1 | 1401 | 863 | 699 | 912 | 455 | 738 | 449 | 868 | 598 |
| 3596 | FIM1_3687 | QGN16960.1 | CP015058.1 | 968 | 221 | 458 | 817 | 181 | 471 | 817 | 255 | 411 |
| 3597 | FIM1_3688 | QGN16961.1 | CP015058.1 | 2345 | 452 | 965 | 1496 | 358 | 852 | 2714 | 497 | 751 |
| 3598 | FIM1_3689 | QGN16962.1 | CP015058.1 | 3962 | 654 | 1198 | 3063 | 504 | 1061 | 3408 | 719 | 953 |

| | | | | | | | | | | | | |
|---|---|---|---|---|---|---|---|---|---|---|---|---|
| 3599 | FIM1_3690 | QGN16963.1 | CP015058.1 | 1727 | 178 | 738 | 1316 | 111 | 618 | 2244 | 167 | 523 |
| 3600 | FIM1_3691 | QGN16964.1 | CP015058.1 | 6904 | 1787 | 2431 | 5321 | 1407 | 2341 | 6935 | 2017 | 2082 |
| 3601 | FIM1_3692 | QGN16965.1 | CP015058.1 | 3379 | 1457 | 1460 | 2589 | 1008 | 1402 | 2906 | 1579 | 1150 |
| 3602 | FIM1_3693 | QGN16966.1 | CP015058.1 | 3975 | 810 | 1301 | 3130 | 550 | 1324 | 4108 | 789 | 1136 |
| 3603 | FIM1_3694 | QGN16967.1 | CP015058.1 | 1856 | 539 | 648 | 1440 | 450 | 628 | 2041 | 645 | 652 |
| 3604 | FIM1_3695 | QGN16968.1 | CP015058.1 | 9473 | 1892 | 2626 | 8218 | 1191 | 2458 | 13092 | 1974 | 2007 |
| 3605 | FIM1_3696 | QGN16969.1 | CP015058.1 | 5834 | 1844 | 2399 | 4686 | 1438 | 2269 | 6618 | 1997 | 2151 |
| 3606 | FIM1_3697 | QGN16970.1 | CP015058.1 | 16648 | 8113 | 8278 | 13138 | 3595 | 7741 | 20749 | 7977 | 7203 |
| 3607 | FIM1_3698 | QGN16971.1 | CP015058.1 | 6461 | 2042 | 1862 | 4475 | 1318 | 1703 | 5819 | 2166 | 1621 |
| 3608 | FIM1_3699 | QGN16972.1 | CP015058.1 | 714 | 70 | 109 | 529 | 46 | 102 | 2082 | 59 | 79 |
| 3609 | FIM1_3700 | QGN16973.1 | CP015058.1 | 1621 | 115 | 326 | 1251 | 82 | 408 | 3488 | 117 | 331 |
| 3610 | FIM1_3701 | QGN16974.1 | CP015058.1 | 1173 | 478 | 610 | 974 | 471 | 645 | 796 | 526 | 567 |
| 3611 | FIM1_3702 | QGN16975.1 | CP015058.1 | 912 | 119 | 293 | 789 | 73 | 276 | 1663 | 146 | 214 |
| 3612 | FIM1_3703 | QGN16976.1 | CP015058.1 | 6281 | 1033 | 1609 | 4513 | 816 | 1520 | 5912 | 1146 | 1372 |
| 3613 | FIM1_3704 | QGN16977.1 | CP015058.1 | 1572 | 403 | 812 | 1074 | 402 | 916 | 1578 | 460 | 732 |
| 3614 | FIM1_3705 | QGN16978.1 | CP015058.1 | 4874 | 1317 | 1960 | 3658 | 995 | 1902 | 3934 | 1364 | 1754 |
| 3615 | FIM1_3706 | QGN16979.1 | CP015058.1 | 8213 | 2966 | 3335 | 6457 | 2120 | 2925 | 6950 | 3122 | 2969 |
| 3616 | FIM1_3707 | QGN16980.1 | CP015058.1 | 1236 | 314 | 670 | 989 | 189 | 595 | 977 | 297 | 532 |
| 3617 | FIM1_3708 | QGN16981.1 | CP015058.1 | 10882 | 2846 | 3937 | 8932 | 1851 | 3710 | 11024 | 2891 | 3624 |
| 3618 | FIM1_3709 | QGN16982.1 | CP015058.1 | 3146 | 1327 | 1546 | 2564 | 1091 | 1461 | 1708 | 1490 | 1307 |
| 3619 | FIM1_3710 | QGN16983.1 | CP015058.1 | 5944 | 964 | 2057 | 4722 | 716 | 2071 | 6463 | 973 | 1767 |
| 3620 | FIM1_3711 | QGN16984.1 | CP015058.1 | 7394 | 1168 | 1949 | 5397 | 919 | 1951 | 7125 | 1297 | 1715 |
| 3621 | FIM1_3712 | QGN16985.1 | CP015058.1 | 5363 | 1774 | 2687 | 3914 | 1010 | 2569 | 6803 | 1804 | 2324 |
| 3622 | FIM1_3713 | QGN16986.1 | CP015058.1 | 6905 | 1301 | 2040 | 4905 | 966 | 2003 | 6667 | 1388 | 1948 |
| 3623 | FIM1_3714 | QGN16987.1 | CP015058.1 | 1173 | 247 | 406 | 1111 | 91 | 348 | 1659 | 252 | 347 |
| 3624 | FIM1_3715 | QGN16988.1 | CP015058.1 | 821 | 147 | 263 | 636 | 117 | 231 | 642 | 151 | 236 |
| 3625 | FIM1_3716 | QGN16989.1 | CP015058.1 | 8678 | 2054 | 4384 | 6826 | 1585 | 4156 | 8560 | 2213 | 4117 |
| 3626 | FIM1_3717 | QGN16990.1 | CP015058.1 | 12492 | 4865 | 5970 | 10255 | 2479 | 5758 | 17564 | 4518 | 5494 |
| 3627 | FIM1_3718 | QGN16991.1 | CP015058.1 | 2025 | 653 | 1197 | 1548 | 529 | 1158 | 2005 | 720 | 1166 |
| 3628 | FIM1_3719 | QGN16992.1 | CP015058.1 | 2064 | 319 | 735 | 1584 | 235 | 800 | 3135 | 335 | 824 |
| 3629 | FIM1_3720 | QGN16993.1 | CP015058.1 | 2527 | 633 | 856 | 2120 | 411 | 923 | 3565 | 713 | 847 |
| 3630 | FIM1_3721 | QGN16994.1 | CP015058.1 | 21 | 4 | 6 | 15 | 1 | 10 | 31 | 5 | 10 |
| 3631 | FIM1_3722 | QGN16995.1 | CP015058.1 | 174 | 45 | 67 | 157 | 50 | 84 | 277 | 52 | 69 |
| 3632 | FIM1_3723 | QGN16996.1 | CP015058.1 | 863 | 114 | 204 | 815 | 73 | 163 | 1450 | 120 | 172 |
| 3633 | FIM1_3725 | QGN16997.1 | CP015058.1 | 257 | 80 | 59 | 175 | 45 | 62 | 391 | 55 | 61 |
| 3634 | FIM1_3726 | QGN16998.1 | CP015058.1 | 0 | 0 | 0 | 0 | 0 | 0 | 0 | 0 | 0 |
| 3635 | FIM1_3727 | QGN16999.1 | CP015058.1 | 533 | 320 | 352 | 710 | 277 | 378 | 488 | 321 | 323 |
| 3636 | FIM1_3728 | QGN17000.1 | CP015058.1 | 0 | 0 | 0 | 0 | 0 | 1 | 4 | 0 | 0 |
| 3637 | FIM1_3730 | QGN17001.1 | CP015059.1 | 2 | 3 | 1 | 0 | 3 | 3 | 5 | 3 | 6 |
| 3638 | FIM1_3731 | QGN17002.1 | CP015059.1 | 28 | 152 | 122 | 11 | 81 | 112 | 33 | 152 | 115 |
| 3639 | FIM1_3732 | QGN17003.1 | CP015059.1 | 387 | 182 | 241 | 378 | 191 | 228 | 281 | 198 | 241 |
| 3640 | FIM1_3733 | QGN17004.1 | CP015059.1 | 2820 | 686 | 1218 | 1920 | 448 | 1049 | 2140 | 684 | 937 |
| 3641 | FIM1_3734 | QGN17005.1 | CP015059.1 | 1425 | 391 | 1312 | 1021 | 331 | 1332 | 1292 | 448 | 1047 |
| 3642 | FIM1_3735 | QGN17006.1 | CP015059.1 | 4572 | 915 | 1719 | 3423 | 673 | 1625 | 4228 | 1009 | 1478 |
| 3643 | FIM1_3736 | QGN17007.1 | CP015059.1 | 3902 | 1077 | 1046 | 2547 | 849 | 1001 | 2592 | 1238 | 978 |
| 3644 | FIM1_3737 | QGN17008.1 | CP015059.1 | 2297 | 61 | 232 | 1894 | 40 | 178 | 5648 | 55 | 142 |
| 3645 | FIM1_3738 | QGN17009.1 | CP015059.1 | 1430 | 152 | 216 | 1134 | 110 | 197 | 1993 | 160 | 183 |
| 3646 | FIM1_3739 | QGN17010.1 | CP015059.1 | 2748 | 452 | 1000 | 2592 | 247 | 906 | 5066 | 532 | 851 |

| | | | | | | | | | | | | |
|---|---|---|---|---|---|---|---|---|---|---|---|---|
| 3647 | FIM1_3740 | QGN17011.1 | CP015059.1 | 3450 | 2031 | 1739 | 3373 | 1200 | 1534 | 4510 | 1951 | 1444 |
| 3648 | FIM1_3741 | QGN17012.1 | CP015059.1 | 10549 | 974 | 2310 | 8054 | 791 | 2004 | 14240 | 1008 | 1761 |
| 3649 | FIM1_3742 | QGN17013.1 | CP015059.1 | 6556 | 1027 | 2135 | 4944 | 777 | 1915 | 7142 | 1062 | 1808 |
| 3650 | FIM1_3743 | QGN17014.1 | CP015059.1 | 9813 | 2480 | 2904 | 7570 | 1777 | 2545 | 9237 | 2619 | 2465 |
| 3651 | FIM1_3744 | QGN17015.1 | CP015059.1 | 4846 | 719 | 1611 | 3744 | 572 | 1393 | 4251 | 786 | 1390 |
| 3652 | FIM1_3745 | QGN17016.1 | CP015059.1 | 22918 | 11810 | 10275 | 19745 | 8389 | 9070 | 23637 | 12271 | 9748 |
| 3653 | FIM1_3746 | QGN17017.1 | CP015059.1 | 4424 | 1140 | 1607 | 4063 | 790 | 1495 | 5799 | 1218 | 1304 |
| 3654 | FIM1_3747 | QGN17018.1 | CP015059.1 | 42155 | 12970 | 11612 | 33318 | 10781 | 11176 | 39928 | 14224 | 11343 |
| 3655 | FIM1_3748 | QGN17019.1 | CP015059.1 | 10905 | 8199 | 7728 | 9726 | 5795 | 7776 | 9227 | 8876 | 7518 |
| 3656 | FIM1_3749 | QGN17020.1 | CP015059.1 | 10547 | 3061 | 4126 | 8071 | 2228 | 3845 | 9331 | 3213 | 3622 |
| 3657 | FIM1_3750 | QGN17021.1 | CP015059.1 | 2135 | 1706 | 1000 | 1436 | 1201 | 980 | 524 | 1821 | 931 |
| 3658 | FIM1_3751 | QGN17022.1 | CP015059.1 | 4980 | 1075 | 1769 | 3641 | 857 | 1654 | 4515 | 1185 | 1463 |
| 3659 | FIM1_3752 | QGN17023.1 | CP015059.1 | 7344 | 1105 | 2024 | 5524 | 906 | 1864 | 6830 | 1113 | 1724 |
| 3660 | FIM1_3753 | QGN17024.1 | CP015059.1 | 8011 | 2430 | 1607 | 6240 | 1283 | 1649 | 9006 | 2431 | 1554 |
| 3661 | FIM1_3754 | QGN17025.1 | CP015059.1 | 8421 | 2426 | 1609 | 6313 | 1556 | 1499 | 5622 | 2516 | 1516 |
| 3662 | FIM1_3755 | QGN17026.1 | CP015059.1 | 374 | 222 | 208 | 317 | 127 | 200 | 219 | 215 | 179 |
| 3663 | FIM1_3756 | QGN17027.1 | CP015059.1 | 35620 | 22604 | 19941 | 33551 | 18329 | 18915 | 34419 | 23063 | 19588 |
| 3664 | FIM1_3757 | QGN17028.1 | CP015059.1 | 1763 | 356 | 610 | 1353 | 293 | 611 | 1384 | 402 | 495 |
| 3665 | FIM1_3758 | QGN17029.1 | CP015059.1 | 2144 | 1218 | 1555 | 2143 | 1142 | 1617 | 1448 | 1449 | 1271 |
| 3666 | FIM1_3759 | QGN17030.1 | CP015059.1 | 20055 | 8344 | 8088 | 17207 | 5691 | 7799 | 22518 | 8847 | 7631 |
| 3667 | FIM1_3760 | QGN17031.1 | CP015059.1 | 1700 | 373 | 591 | 1496 | 266 | 512 | 1573 | 408 | 481 |
| 3668 | FIM1_3761 | QGN17032.1 | CP015059.1 | 2263 | 455 | 603 | 1737 | 323 | 573 | 1945 | 502 | 526 |
| 3669 | FIM1_3762 | QGN17033.1 | CP015059.1 | 5695 | 1331 | 1656 | 5088 | 904 | 1467 | 5818 | 1501 | 1415 |
| 3670 | FIM1_3763 | QGN17034.1 | CP015059.1 | 9035 | 2132 | 3311 | 6790 | 1835 | 2970 | 6662 | 2317 | 3073 |
| 3671 | FIM1_3764 | QGN17035.1 | CP015059.1 | 820 | 234 | 681 | 635 | 176 | 620 | 956 | 260 | 583 |
| 3672 | FIM1_3765 | QGN17036.1 | CP015059.1 | 23470 | 5728 | 9420 | 18805 | 4073 | 8469 | 26995 | 6325 | 8682 |
| 3673 | FIM1_3766 | QGN17037.1 | CP015059.1 | 279 | 47 | 50 | 235 | 23 | 57 | 284 | 50 | 51 |
| 3674 | FIM1_3767 | QGN17038.1 | CP015059.1 | 4715 | 1334 | 1836 | 3677 | 782 | 1753 | 5346 | 1349 | 1616 |
| 3675 | FIM1_3768 | QGN17039.1 | CP015059.1 | 32712 | 24428 | 21559 | 26712 | 17620 | 20703 | 21867 | 25413 | 21810 |
| 3676 | FIM1_3769 | QGN17040.1 | CP015059.1 | 5944 | 1646 | 1675 | 4886 | 1222 | 1490 | 5231 | 1810 | 1415 |
| 3677 | FIM1_3770 | QGN17041.1 | CP015059.1 | 15907 | 5239 | 6219 | 12687 | 3463 | 5565 | 15455 | 5594 | 5268 |
| 3678 | FIM1_3771 | QGN17042.1 | CP015059.1 | 3975 | 982 | 1197 | 3141 | 724 | 1100 | 2780 | 1031 | 1079 |
| 3679 | FIM1_3772 | QGN17043.1 | CP015059.1 | 5728 | 1494 | 1888 | 4342 | 808 | 1787 | 7181 | 1577 | 1700 |
| 3680 | FIM1_3773 | QGN17044.1 | CP015059.1 | 23419 | 6857 | 8583 | 17107 | 5057 | 7630 | 22696 | 7789 | 7495 |
| 3681 | FIM1_3774 | QGN17045.1 | CP015059.1 | 8409 | 1733 | 4658 | 7001 | 1346 | 4360 | 10047 | 1945 | 4271 |
| 3682 | FIM1_3775 | QGN17046.1 | CP015059.1 | 628 | 118 | 253 | 468 | 77 | 206 | 576 | 130 | 206 |
| 3683 | FIM1_3776 | QGN17047.1 | CP015059.1 | 911 | 118 | 587 | 789 | 93 | 540 | 909 | 121 | 527 |
| 3684 | FIM1_3777 | QGN17048.1 | CP015059.1 | 25603 | 6044 | 10499 | 19514 | 3903 | 9542 | 29818 | 6259 | 9583 |
| 3685 | FIM1_3778 | QGN17049.1 | CP015059.1 | 1538 | 185 | 201 | 1003 | 149 | 164 | 1197 | 197 | 171 |
| 3686 | FIM1_3779 | QGN17050.1 | CP015059.1 | 2450 | 862 | 1457 | 2007 | 659 | 1302 | 2227 | 936 | 1185 |
| 3687 | FIM1_3780 | QGN17051.1 | CP015059.1 | 16481 | 9104 | 9958 | 14306 | 6353 | 9259 | 17934 | 9021 | 9649 |
| 3688 | FIM1_3782 | QGN17052.1 | CP015059.1 | 9431 | 1464 | 2788 | 7185 | 1046 | 2541 | 10917 | 1587 | 2364 |
| 3689 | FIM1_3783 | QGN17053.1 | CP015059.1 | 5466 | 623 | 1390 | 3709 | 373 | 1204 | 5582 | 636 | 1033 |
| 3690 | FIM1_3784 | QGN17054.1 | CP015059.1 | 5038 | 2016 | 3960 | 4649 | 1316 | 3787 | 6008 | 2066 | 3575 |
| 3691 | FIM1_3785 | QGN17055.1 | CP015059.1 | 672 | 167 | 411 | 590 | 82 | 398 | 804 | 181 | 296 |
| 3692 | FIM1_3786 | QGN17056.1 | CP015059.1 | 9478 | 2559 | 3970 | 8103 | 1850 | 3565 | 10204 | 2622 | 3287 |
| 3693 | FIM1_3787 | QGN17057.1 | CP015059.1 | 271 | 49 | 47 | 190 | 36 | 60 | 186 | 51 | 47 |
| 3694 | FIM1_3788 | QGN17058.1 | CP015059.1 | 12021 | 4661 | 5969 | 10195 | 2737 | 5515 | 14678 | 4944 | 5320 |

| | | | | | | | | | | | | |
|---|---|---|---|---|---|---|---|---|---|---|---|---|
| 3695 | FIM1_3789 | QGN17059.1 | CP015059.1 | 1902 | 404 | 597 | 1473 | 293 | 569 | 1996 | 411 | 463 |
| 3696 | FIM1_3790 | QGN17060.1 | CP015059.1 | 1160 | 193 | 333 | 798 | 171 | 364 | 1155 | 219 | 299 |
| 3697 | FIM1_3791 | QGN17061.1 | CP015059.1 | 8979 | 2113 | 2670 | 6895 | 1573 | 2369 | 9933 | 2159 | 2532 |
| 3698 | FIM1_3792 | QGN17062.1 | CP015059.1 | 9495 | 1844 | 2306 | 7518 | 983 | 2224 | 9818 | 2140 | 2235 |
| 3699 | FIM1_3793 | QGN17063.1 | CP015059.1 | 8805 | 4475 | 5177 | 8131 | 2946 | 5384 | 9868 | 4954 | 4891 |
| 3700 | FIM1_3794 | QGN17064.1 | CP015059.1 | 14080 | 9895 | 8869 | 13747 | 8156 | 8044 | 14075 | 10086 | 8443 |
| 3701 | FIM1_3795 | QGN17065.1 | CP015059.1 | 3878 | 595 | 676 | 2804 | 295 | 553 | 2980 | 581 | 515 |
| 3702 | FIM1_3796 | QGN17066.1 | CP015059.1 | 744 | 223 | 261 | 504 | 156 | 251 | 672 | 229 | 276 |
| 3703 | FIM1_3797 | QGN17067.1 | CP015059.1 | 10327 | 2747 | 3579 | 9559 | 1809 | 3175 | 10794 | 3046 | 2989 |
| 3704 | FIM1_3798 | QGN17068.1 | CP015059.1 | 5802 | 1050 | 2069 | 5760 | 761 | 1865 | 4539 | 1217 | 1601 |
| 3705 | FIM1_3799 | QGN17069.1 | CP015059.1 | 26337 | 8037 | 14169 | 22553 | 4661 | 12834 | 37669 | 8234 | 12705 |
| 3706 | FIM1_3801 | QGN17070.1 | CP015059.1 | 7327 | 1687 | 1984 | 5672 | 910 | 1927 | 7517 | 1789 | 1587 |
| 3707 | FIM1_3802 | QGN17071.1 | CP015059.1 | 3242 | 1676 | 1657 | 2482 | 679 | 1550 | 6895 | 1465 | 1432 |
| 3708 | FIM1_3803 | QGN17072.1 | CP015059.1 | 545 | 173 | 297 | 563 | 168 | 252 | 505 | 174 | 223 |
| 3709 | FIM1_3804 | QGN17073.1 | CP015059.1 | 538 | 49 | 296 | 464 | 42 | 274 | 756 | 57 | 209 |
| 3710 | FIM1_3805 | QGN17074.1 | CP015059.1 | 1059 | 189 | 282 | 913 | 147 | 300 | 1401 | 215 | 234 |
| 3711 | FIM1_3806 | QGN17075.1 | CP015059.1 | 1219 | 777 | 972 | 955 | 564 | 836 | 422 | 824 | 736 |
| 3712 | FIM1_3807 | QGN17076.1 | CP015059.1 | 26875 | 9122 | 8391 | 22654 | 6921 | 7713 | 25661 | 9697 | 7773 |
| 3713 | FIM1_3808 | QGN17077.1 | CP015059.1 | 1721 | 234 | 413 | 1471 | 152 | 395 | 1819 | 266 | 349 |
| 3714 | FIM1_3809 | QGN17078.1 | CP015059.1 | 2810 | 2989 | 1648 | 2123 | 2038 | 1459 | 600 | 3193 | 1508 |
| 3715 | FIM1_3810 | QGN17079.1 | CP015059.1 | 3679 | 1695 | 1760 | 2965 | 1315 | 1733 | 2305 | 1782 | 1605 |
| 3716 | FIM1_3811 | QGN17080.1 | CP015059.1 | 11857 | 4070 | 2230 | 10235 | 2368 | 2062 | 18179 | 4431 | 2060 |
| 3717 | FIM1_3812 | QGN17081.1 | CP015059.1 | 71 | 6 | 9 | 63 | 7 | 9 | 78 | 11 | 10 |
| 3718 | FIM1_3813 | QGN17082.1 | CP015059.1 | 8685 | 3865 | 3434 | 7426 | 2702 | 2975 | 7202 | 4020 | 2998 |
| 3719 | FIM1_3814 | QGN17083.1 | CP015059.1 | 58 | 15 | 19 | 73 | 9 | 19 | 34 | 11 | 18 |
| 3720 | FIM1_3815 | QGN17084.1 | CP015059.1 | 33172 | 8308 | 14363 | 28248 | 5408 | 13174 | 45561 | 9259 | 12659 |
| 3721 | FIM1_3816 | QGN17085.1 | CP015059.1 | 4036 | 2456 | 2123 | 3010 | 1761 | 1941 | 1977 | 2714 | 1909 |
| 3722 | FIM1_3817 | QGN17086.1 | CP015059.1 | 6248 | 862 | 4662 | 4674 | 688 | 4286 | 6157 | 845 | 3598 |
| 3723 | FIM1_3818 | QGN17087.1 | CP015059.1 | 336 | 40 | 58 | 245 | 40 | 47 | 405 | 46 | 50 |
| 3724 | FIM1_3819 | QGN17088.1 | CP015059.1 | 7518 | 949 | 1773 | 6284 | 553 | 1661 | 10104 | 997 | 1489 |
| 3725 | FIM1_3820 | QGN17089.1 | CP015059.1 | 3109 | 2103 | 1083 | 2028 | 1707 | 1122 | 1411 | 2365 | 1079 |
| 3726 | FIM1_3821 | QGN17090.1 | CP015059.1 | 2133 | 742 | 833 | 1419 | 541 | 767 | 1289 | 817 | 717 |
| 3727 | FIM1_3822 | QGN17091.1 | CP015059.1 | 14109 | 3394 | 3843 | 12688 | 2235 | 3350 | 16563 | 3458 | 3274 |
| 3728 | FIM1_3823 | QGN17092.1 | CP015059.1 | 418 | 126 | 75 | 383 | 100 | 71 | 385 | 141 | 52 |
| 3729 | FIM1_3824 | QGN17093.1 | CP015059.1 | 63362 | 15473 | 9438 | 51375 | 7978 | 8113 | 109851 | 15634 | 7776 |
| 3730 | FIM1_3825 | QGN17094.1 | CP015059.1 | 272 | 80 | 307 | 208 | 88 | 332 | 419 | 81 | 200 |
| 3731 | FIM1_3826 | QGN17095.1 | CP015059.1 | 15479 | 5342 | 4842 | 12988 | 3913 | 4523 | 13448 | 5619 | 4477 |
| 3732 | FIM1_3827 | QGN17096.1 | CP015059.1 | 946 | 265 | 460 | 920 | 271 | 424 | 890 | 313 | 373 |
| 3733 | FIM1_3828 | QGN17097.1 | CP015059.1 | 5161 | 2758 | 2544 | 4628 | 1966 | 2471 | 4764 | 2846 | 2239 |
| 3734 | FIM1_3829 | QGN17098.1 | CP015059.1 | 3317 | 1660 | 2356 | 3073 | 1354 | 2250 | 2696 | 1803 | 2142 |
| 3735 | FIM1_3830 | QGN17099.1 | CP015059.1 | 9836 | 1734 | 2544 | 8138 | 1063 | 2345 | 12005 | 1840 | 2298 |
| 3736 | FIM1_3831 | QGN17100.1 | CP015059.1 | 2100 | 133 | 596 | 1786 | 107 | 490 | 3359 | 141 | 505 |
| 3737 | FIM1_3832 | QGN17101.1 | CP015059.1 | 1454 | 557 | 862 | 1334 | 409 | 788 | 1007 | 593 | 694 |
| 3738 | FIM1_3833 | QGN17102.1 | CP015059.1 | 402 | 16 | 55 | 182 | 22 | 60 | 166 | 26 | 37 |
| 3739 | FIM1_3834 | QGN17103.1 | CP015059.1 | 7483 | 2104 | 2503 | 5948 | 1358 | 2228 | 8701 | 2178 | 2280 |
| 3740 | FIM1_3835 | QGN17104.1 | CP015059.1 | 1360 | 202 | 229 | 1154 | 162 | 207 | 1413 | 223 | 284 |
| 3741 | FIM1_3836 | QGN17105.1 | CP015059.1 | 9773 | 3727 | 3405 | 8296 | 3205 | 3394 | 7682 | 4183 | 3231 |
| 3742 | FIM1_3837 | QGN17106.1 | CP015059.1 | 4274 | 1571 | 2128 | 3388 | 1124 | 1961 | 3954 | 1694 | 1969 |

| | | | | | | | | | | | | |
|---|---|---|---|---|---|---|---|---|---|---|---|---|
| 3743 | FIM1_3838 | QGN17107.1 | CP015059.1 | 6723 | 1346 | 1630 | 5079 | 976 | 1482 | 6077 | 1399 | 1318 |
| 3744 | FIM1_3839 | QGN17108.1 | CP015059.1 | 9047 | 720 | 1741 | 6397 | 848 | 1862 | 5258 | 857 | 1912 |
| 3745 | FIM1_3840 | QGN17109.1 | CP015059.1 | 1621 | 1011 | 773 | 1279 | 675 | 629 | 1515 | 1048 | 674 |
| 3746 | FIM1_3841 | QGN17110.1 | CP015059.1 | 9445 | 1297 | 1874 | 7313 | 958 | 1760 | 8195 | 1438 | 1647 |
| 3747 | FIM1_3842 | QGN17111.1 | CP015059.1 | 622 | 92 | 136 | 474 | 76 | 108 | 598 | 108 | 90 |
| 3748 | FIM1_3843 | QGN17112.1 | CP015059.1 | 26837 | 11669 | 10019 | 22861 | 8909 | 8987 | 22259 | 12256 | 9011 |
| 3749 | FIM1_3844 | QGN17113.1 | CP015059.1 | 3339 | 1157 | 898 | 1956 | 648 | 755 | 1470 | 1128 | 818 |
| 3750 | FIM1_3845 | QGN17114.1 | CP015059.1 | 1221 | 1352 | 654 | 824 | 763 | 672 | 439 | 1438 | 675 |
| 3751 | FIM1_3846 | QGN17115.1 | CP015059.1 | 2167 | 837 | 588 | 1550 | 593 | 599 | 1670 | 864 | 594 |
| 3752 | FIM1_3847 | QGN17116.1 | CP015059.1 | 57644 | 33859 | 27320 | 49648 | 25966 | 25235 | 59854 | 33294 | 27005 |
| 3753 | FIM1_3848 | QGN17117.1 | CP015059.1 | 2294 | 477 | 595 | 1576 | 396 | 650 | 1095 | 495 | 585 |
| 3754 | FIM1_3849 | QGN17118.1 | CP015059.1 | 61836 | 29856 | 26966 | 54698 | 21847 | 24385 | 61164 | 30405 | 26040 |
| 3755 | FIM1_3850 | QGN17119.1 | CP015059.1 | 7163 | 1567 | 2134 | 6871 | 916 | 1933 | 8357 | 1663 | 1971 |
| 3756 | FIM1_3851 | QGN17120.1 | CP015059.1 | 3925 | 506 | 716 | 3168 | 461 | 652 | 3804 | 585 | 561 |
| 3757 | FIM1_3852 | QGN17121.1 | CP015059.1 | 1799 | 987 | 958 | 1617 | 799 | 1025 | 1573 | 1113 | 831 |
| 3758 | FIM1_3853 | QGN17122.1 | CP015059.1 | 5168 | 1015 | 1476 | 3779 | 818 | 1336 | 4426 | 1050 | 1182 |
| 3759 | FIM1_3854 | QGN17123.1 | CP015059.1 | 11061 | 3434 | 5253 | 9071 | 2184 | 4938 | 14666 | 3730 | 4641 |
| 3760 | FIM1_3855 | QGN17124.1 | CP015059.1 | 8026 | 2115 | 3232 | 6146 | 1434 | 3112 | 9341 | 2246 | 2839 |
| 3761 | FIM1_3856 | QGN17125.1 | CP015059.1 | 1721 | 417 | 1744 | 1551 | 312 | 1635 | 2460 | 450 | 1555 |
| 3762 | FIM1_3857 | QGN17126.1 | CP015059.1 | 13418 | 6006 | 6315 | 11607 | 4593 | 5714 | 13233 | 6271 | 5834 |
| 3763 | FIM1_3858 | QGN17127.1 | CP015059.1 | 2345 | 466 | 1021 | 1854 | 369 | 992 | 3015 | 536 | 873 |
| 3764 | FIM1_3859 | QGN17128.1 | CP015059.1 | 54 | 8 | 29 | 41 | 10 | 24 | 38 | 17 | 19 |
| 3765 | FIM1_3860 | QGN17129.1 | CP015059.1 | 6811 | 1203 | 2002 | 5587 | 995 | 1698 | 7736 | 1267 | 1577 |
| 3766 | FIM1_3861 | QGN17130.1 | CP015059.1 | 2047 | 314 | 724 | 1667 | 236 | 620 | 2520 | 337 | 622 |
| 3767 | FIM1_3862 | QGN17131.1 | CP015059.1 | 8881 | 3093 | 4252 | 8257 | 1157 | 4036 | 14426 | 2482 | 3546 |
| 3768 | FIM1_3863 | QGN17132.1 | CP015059.1 | 17061 | 4255 | 4553 | 12815 | 3472 | 4155 | 16460 | 4805 | 4482 |
| 3769 | FIM1_3864 | QGN17133.1 | CP015059.1 | 2246 | 13 | 248 | 1689 | 2 | 207 | 1267 | 17 | 100 |
| 3770 | FIM1_3865 | QGN17134.1 | CP015059.1 | 3343 | 60 | 1056 | 2889 | 32 | 747 | 2023 | 70 | 528 |
| 3771 | FIM1_3866 | QGN17135.1 | CP015059.1 | 6555 | 590 | 1610 | 4884 | 515 | 1342 | 4719 | 653 | 982 |
| 3772 | FIM1_3867 | QGN17136.1 | CP015059.1 | 3611 | 1152 | 1231 | 3196 | 908 | 1227 | 3093 | 1179 | 1137 |
| 3773 | FIM1_3868 | QGN17137.1 | CP015059.1 | 1033 | 125 | 302 | 927 | 107 | 290 | 1388 | 119 | 259 |
| 3774 | FIM1_3869 | QGN17138.1 | CP015059.1 | 33373 | 14018 | 12305 | 28640 | 9528 | 11754 | 35668 | 14429 | 11919 |
| 3775 | FIM1_3870 | QGN17139.1 | CP015059.1 | 2391 | 613 | 1583 | 1900 | 558 | 1640 | 3482 | 643 | 1282 |
| 3776 | FIM1_3871 | QGN17140.1 | CP015059.1 | 5387 | 2891 | 2265 | 4076 | 2004 | 2097 | 1979 | 3201 | 2073 |
| 3777 | FIM1_3872 | QGN17141.1 | CP015059.1 | 12672 | 4143 | 3000 | 9998 | 2883 | 2829 | 12846 | 4232 | 2850 |
| 3778 | FIM1_3873 | QGN17142.1 | CP015059.1 | 3689 | 528 | 2135 | 3045 | 426 | 1925 | 4350 | 545 | 1746 |
| 3779 | FIM1_3874 | QGN17143.1 | CP015059.1 | 1228 | 147 | 375 | 951 | 104 | 392 | 1610 | 159 | 334 |
| 3780 | FIM1_3875 | QGN17144.1 | CP015059.1 | 841 | 94 | 253 | 658 | 67 | 239 | 818 | 82 | 262 |
| 3781 | FIM1_3876 | QGN17145.1 | CP015059.1 | 232 | 15 | 76 | 236 | 22 | 69 | 325 | 7 | 69 |
| 3782 | FIM1_3877 | QGN17146.1 | CP015059.1 | 850 | 17 | 66 | 698 | 9 | 57 | 998 | 13 | 58 |
| 3783 | FIM1_3878 | QGN17147.1 | CP015059.1 | 142 | 3 | 12 | 96 | 7 | 17 | 57 | 4 | 10 |
| 3784 | FIM1_3879 | QGN17148.1 | CP015059.1 | 69064 | 6802 | 10859 | 56559 | 4945 | 9813 | 107349 | 7315 | 9548 |
| 3785 | FIM1_3880 | QGN17149.1 | CP015059.1 | 5449 | 1892 | 2925 | 4975 | 1200 | 2996 | 6903 | 1952 | 2650 |
| 3786 | FIM1_3881 | QGN17150.1 | CP015059.1 | 15062 | 4018 | 7330 | 11902 | 2875 | 6720 | 16063 | 4431 | 6701 |
| 3787 | FIM1_3882 | QGN17151.1 | CP015059.1 | 123 | 20 | 61 | 83 | 8 | 57 | 173 | 22 | 37 |
| 3788 | FIM1_3883 | QGN17152.1 | CP015059.1 | 901 | 111 | 436 | 819 | 81 | 383 | 1277 | 119 | 318 |
| 3789 | FIM1_3884 | QGN17153.1 | CP015059.1 | 1041 | 160 | 365 | 797 | 138 | 356 | 1954 | 199 | 307 |
| 3790 | FIM1_3885 | QGN17154.1 | CP015059.1 | 7147 | 1282 | 2491 | 5771 | 1058 | 2146 | 6399 | 1416 | 2159 |

| | | | | | | | | | | | | |
|---|---|---|---|---|---|---|---|---|---|---|---|---|
| 3791 | FIM1_3886 | QGN17155.1 | CP015059.1 | 6454 | 2412 | 3682 | 5861 | 1480 | 3443 | 8421 | 2485 | 3271 |
| 3792 | FIM1_3887 | QGN17156.1 | CP015059.1 | 780 | 146 | 486 | 600 | 126 | 376 | 837 | 188 | 362 |
| 3793 | FIM1_3888 | QGN17157.1 | CP015059.1 | 4901 | 2096 | 2872 | 4772 | 1718 | 2621 | 4545 | 2228 | 2548 |
| 3794 | FIM1_3889 | QGN17158.1 | CP015059.1 | 2331 | 452 | 1321 | 2099 | 362 | 1148 | 2669 | 453 | 1050 |
| 3795 | FIM1_3890 | QGN17159.1 | CP015059.1 | 4658 | 1754 | 1922 | 4705 | 984 | 1757 | 6508 | 1808 | 1822 |
| 3796 | FIM1_3891 | QGN17160.1 | CP015059.1 | 1114 | 89 | 237 | 893 | 72 | 190 | 1960 | 88 | 212 |
| 3797 | FIM1_3892 | QGN17161.1 | CP015059.1 | 11495 | 356 | 1249 | 8174 | 406 | 1183 | 9984 | 405 | 1096 |
| 3798 | FIM1_3893 | QGN17162.1 | CP015059.1 | 498 | 84 | 146 | 320 | 72 | 94 | 388 | 103 | 99 |
| 3799 | FIM1_3894 | QGN17163.1 | CP015059.1 | 2261 | 321 | 597 | 1825 | 234 | 602 | 2857 | 354 | 525 |
| 3800 | FIM1_3895 | QGN17164.1 | CP015059.1 | 368 | 79 | 150 | 375 | 76 | 135 | 327 | 119 | 103 |
| 3801 | FIM1_3896 | QGN17165.1 | CP015059.1 | 3672 | 526 | 1061 | 2900 | 410 | 1018 | 4464 | 583 | 918 |
| 3802 | FIM1_3897 | QGN17166.1 | CP015059.1 | 1098 | 440 | 452 | 769 | 255 | 417 | 760 | 439 | 388 |
| 3803 | FIM1_3898 | QGN17167.1 | CP015059.1 | 8507 | 7778 | 3551 | 5852 | 4763 | 3120 | 1817 | 8033 | 3345 |
| 3804 | FIM1_3899 | QGN17168.1 | CP015059.1 | 3414 | 473 | 802 | 2389 | 366 | 796 | 3476 | 484 | 660 |
| 3805 | FIM1_3900 | QGN17169.1 | CP015059.1 | 756 | 295 | 272 | 612 | 210 | 272 | 714 | 290 | 264 |
| 3806 | FIM1_3901 | QGN17170.1 | CP015059.1 | 4468 | 1050 | 1256 | 3342 | 751 | 1144 | 3992 | 1145 | 1014 |
| 3807 | FIM1_3902 | QGN17171.1 | CP015059.1 | 2648 | 375 | 456 | 1869 | 232 | 352 | 2523 | 420 | 359 |
| 3808 | FIM1_3903 | QGN17172.1 | CP015059.1 | 1188 | 329 | 454 | 975 | 222 | 400 | 860 | 343 | 345 |
| 3809 | FIM1_3904 | QGN17173.1 | CP015059.1 | 12392 | 7141 | 6872 | 11477 | 5651 | 6707 | 12698 | 7721 | 6564 |
| 3810 | FIM1_3905 | QGN17174.1 | CP015059.1 | 29647 | 4535 | 6421 | 22975 | 2726 | 5788 | 25965 | 4694 | 5399 |
| 3811 | FIM1_3906 | QGN17175.1 | CP015059.1 | 1742 | 257 | 1121 | 1409 | 216 | 1014 | 972 | 288 | 915 |
| 3812 | FIM1_3907 | QGN17176.1 | CP015059.1 | 3710 | 659 | 2145 | 3214 | 486 | 1950 | 4235 | 741 | 1902 |
| 3813 | FIM1_3908 | QGN17177.1 | CP015059.1 | 4703 | 955 | 1447 | 3650 | 711 | 1368 | 3905 | 1037 | 1257 |
| 3814 | FIM1_3909 | QGN17178.1 | CP015059.1 | 1718 | 1334 | 1097 | 1153 | 960 | 1003 | 516 | 1393 | 1031 |
| 3815 | FIM1_3910 | QGN17179.1 | CP015059.1 | 13523 | 5332 | 5359 | 12421 | 3704 | 4831 | 14616 | 5870 | 4795 |
| 3816 | FIM1_3911 | QGN17180.1 | CP015059.1 | 424 | 55 | 163 | 391 | 31 | 185 | 409 | 48 | 110 |
| 3817 | FIM1_3912 | QGN17181.1 | CP015059.1 | 23438 | 1182 | 1469 | 19473 | 659 | 1188 | 29489 | 1268 | 1091 |
| 3818 | FIM1_3913 | QGN17182.1 | CP015059.1 | 3169 | 322 | 319 | 2560 | 213 | 293 | 3216 | 373 | 253 |
| 3819 | FIM1_3914 | QGN17183.1 | CP015059.1 | 1458 | 279 | 371 | 1256 | 221 | 306 | 1665 | 301 | 328 |
| 3820 | FIM1_3915 | QGN17184.1 | CP015059.1 | 8121 | 2028 | 2093 | 6458 | 1674 | 1837 | 6540 | 2398 | 1830 |
| 3821 | FIM1_3916 | QGN17185.1 | CP015059.1 | 2558 | 431 | 666 | 1981 | 369 | 611 | 2859 | 452 | 644 |
| 3822 | FIM1_3917 | QGN17186.1 | CP015059.1 | 6249 | 1129 | 1521 | 4707 | 948 | 1402 | 5621 | 1296 | 1408 |
| 3823 | FIM1_3918 | QGN17187.1 | CP015059.1 | 1761 | 624 | 1575 | 1202 | 470 | 1378 | 725 | 653 | 1283 |
| 3824 | FIM1_3919 | QGN17188.1 | CP015059.1 | 724 | 66 | 232 | 565 | 67 | 217 | 746 | 66 | 200 |
| 3825 | FIM1_3920 | QGN17189.1 | CP015059.1 | 479 | 98 | 182 | 378 | 82 | 198 | 584 | 121 | 189 |
| 3826 | FIM1_3921 | QGN17190.1 | CP015059.1 | 8165 | 1659 | 2247 | 5975 | 1187 | 2222 | 8977 | 1831 | 1985 |
| 3827 | FIM1_3922 | QGN17191.1 | CP015059.1 | 957 | 222 | 492 | 714 | 208 | 483 | 692 | 255 | 419 |
| 3828 | FIM1_3923 | QGN17192.1 | CP015059.1 | 4580 | 2422 | 2388 | 4360 | 1867 | 2299 | 4203 | 2664 | 2221 |
| 3829 | FIM1_3924 | QGN17193.1 | CP015059.1 | 824 | 1313 | 482 | 476 | 732 | 411 | 131 | 1244 | 451 |
| 3830 | FIM1_3925 | QGN17194.1 | CP015059.1 | 433 | 27 | 150 | 412 | 20 | 123 | 848 | 30 | 82 |
| 3831 | FIM1_3926 | QGN17195.1 | CP015059.1 | 5780 | 1002 | 1239 | 4644 | 554 | 1153 | 7828 | 982 | 1110 |
| 3832 | FIM1_3927 | QGN17196.1 | CP015059.1 | 441 | 95 | 377 | 303 | 49 | 320 | 357 | 88 | 372 |
| 3833 | FIM1_3928 | QGN17197.1 | CP015059.1 | 9206 | 1605 | 1646 | 6161 | 1363 | 1438 | 7118 | 1754 | 1521 |
| 3834 | FIM1_3929 | QGN17198.1 | CP015059.1 | 2524 | 1198 | 1423 | 2129 | 986 | 1442 | 1444 | 1295 | 1257 |
| 3835 | FIM1_3930 | QGN17199.1 | CP015059.1 | 2645 | 637 | 917 | 1962 | 541 | 897 | 2250 | 768 | 818 |
| 3836 | FIM1_3931 | QGN17200.1 | CP015059.1 | 0 | 0 | 1 | 6 | 5 | 4 | 3 | 1 | 6 |
| 3837 | FIM1_3932 | QGN17201.1 | CP015059.1 | 3751 | 1030 | 2111 | 2619 | 868 | 1993 | 3368 | 1169 | 1847 |
| 3838 | FIM1_3933 | QGN17202.1 | CP015059.1 | 980 | 291 | 565 | 936 | 236 | 518 | 846 | 330 | 475 |

| | | | | | | | | | | | | |
|---|---|---|---|---|---|---|---|---|---|---|---|---|
| 3839 | FIM1_3934 | QGN17203.1 | CP015059.1 | 3112 | 588 | 659 | 2376 | 478 | 692 | 3328 | 692 | 596 |
| 3840 | FIM1_3935 | QGN17204.1 | CP015059.1 | 3160 | 1047 | 1157 | 2636 | 622 | 1135 | 4377 | 1073 | 1013 |
| 3841 | FIM1_3936 | QGN17205.1 | CP015059.1 | 2801 | 361 | 733 | 2162 | 298 | 665 | 2476 | 397 | 573 |
| 3842 | FIM1_3937 | QGN17206.1 | CP015059.1 | 107 | 4 | 29 | 98 | 9 | 29 | 34 | 5 | 29 |
| 3843 | FIM1_3938 | QGN17207.1 | CP015059.1 | 275 | 46 | 59 | 256 | 66 | 69 | 206 | 75 | 41 |
| 3844 | FIM1_3939 | QGN17208.1 | CP015059.1 | 1670 | 397 | 228 | 1375 | 166 | 213 | 3281 | 424 | 191 |
| 3845 | FIM1_3940 | QGN17209.1 | CP015059.1 | 13570 | 2030 | 1891 | 9744 | 1277 | 1775 | 9279 | 2113 | 1772 |
| 3846 | FIM1_3941 | QGN17210.1 | CP015059.1 | 320 | 16 | 25 | 251 | 15 | 13 | 224 | 16 | 23 |
| 3847 | FIM1_3942 | QGN17211.1 | CP015059.1 | 1736 | 285 | 619 | 1422 | 229 | 597 | 1586 | 307 | 540 |
| 3848 | FIM1_3943 | QGN17212.1 | CP015059.1 | 14541 | 3514 | 3527 | 11007 | 2689 | 3477 | 12356 | 3976 | 3343 |
| 3849 | FIM1_3944 | QGN17213.1 | CP015059.1 | 1910 | 548 | 927 | 1323 | 466 | 832 | 1500 | 607 | 854 |
| 3850 | FIM1_3945 | QGN17214.1 | CP015059.1 | 1348 | 737 | 827 | 1169 | 653 | 896 | 607 | 843 | 706 |
| 3851 | FIM1_3946 | QGN17215.1 | CP015059.1 | 2676 | 1418 | 1197 | 1919 | 1083 | 1118 | 1203 | 1531 | 1185 |
| 3852 | FIM1_3947 | QGN17216.1 | CP015059.1 | 4544 | 677 | 1184 | 3078 | 528 | 1081 | 3319 | 775 | 1112 |
| 3853 | FIM1_3948 | QGN17217.1 | CP015059.1 | 5472 | 748 | 1436 | 3961 | 535 | 1395 | 5716 | 771 | 1164 |
| 3854 | FIM1_3949 | QGN17218.1 | CP015059.1 | 4756 | 646 | 1268 | 3429 | 477 | 1259 | 5125 | 714 | 1023 |
| 3855 | FIM1_3950 | QGN17219.1 | CP015059.1 | 3656 | 736 | 1226 | 3186 | 575 | 1115 | 3116 | 751 | 1060 |
| 3856 | FIM1_3951 | QGN17220.1 | CP015059.1 | 5578 | 1185 | 1612 | 4670 | 850 | 1486 | 6118 | 1282 | 1407 |
| 3857 | FIM1_3952 | QGN17221.1 | CP015059.1 | 8026 | 1032 | 1312 | 5632 | 717 | 1196 | 7057 | 1046 | 1161 |
| 3858 | FIM1_3953 | QGN17222.1 | CP015059.1 | 4424 | 976 | 1890 | 3974 | 712 | 1662 | 4240 | 874 | 1703 |
| 3859 | FIM1_3954 | QGN17223.1 | CP015059.1 | 1615 | 363 | 490 | 1088 | 231 | 443 | 1667 | 479 | 428 |
| 3860 | FIM1_3955 | QGN17224.1 | CP015059.1 | 3040 | 281 | 647 | 2295 | 245 | 540 | 2094 | 289 | 481 |
| 3861 | FIM1_3956 | QGN17225.1 | CP015059.1 | 4627 | 4271 | 1996 | 3167 | 2963 | 1873 | 1745 | 4507 | 2129 |
| 3862 | FIM1_3957 | QGN17226.1 | CP015059.1 | 9362 | 1578 | 2110 | 6933 | 1084 | 1831 | 8727 | 1610 | 1754 |
| 3863 | FIM1_3958 | QGN17227.1 | CP015059.1 | 7963 | 6058 | 2292 | 5875 | 3292 | 2049 | 5374 | 6341 | 2031 |
| 3864 | FIM1_3959 | QGN17228.1 | CP015059.1 | 2273 | 360 | 618 | 1797 | 253 | 580 | 2581 | 391 | 437 |
| 3865 | FIM1_3960 | QGN17229.1 | CP015059.1 | 530 | 230 | 142 | 456 | 185 | 132 | 328 | 222 | 161 |
| 3866 | FIM1_3961 | QGN17230.1 | CP015059.1 | 2942 | 492 | 851 | 2080 | 482 | 699 | 2600 | 500 | 702 |
| 3867 | FIM1_3962 | QGN17231.1 | CP015059.1 | 2055 | 835 | 1036 | 1989 | 535 | 970 | 2348 | 835 | 951 |
| 3868 | FIM1_3963 | QGN17232.1 | CP015059.1 | 5203 | 899 | 1243 | 4249 | 667 | 1099 | 4229 | 1052 | 1132 |
| 3869 | FIM1_3964 | QGN17233.1 | CP015059.1 | 10079 | 1909 | 2027 | 7941 | 1175 | 1792 | 8712 | 2105 | 1533 |
| 3870 | FIM1_3965 | QGN17234.1 | CP015059.1 | 1296 | 321 | 531 | 1175 | 262 | 500 | 920 | 359 | 464 |
| 3871 | FIM1_3966 | QGN17235.1 | CP015059.1 | 3426 | 197 | 395 | 2722 | 52 | 353 | 3323 | 174 | 280 |
| 3872 | FIM1_3967 | QGN17236.1 | CP015059.1 | 2758 | 1029 | 1391 | 2474 | 859 | 1297 | 2267 | 1169 | 1256 |
| 3873 | FIM1_3968 | QGN17237.1 | CP015059.1 | 4396 | 843 | 1142 | 3040 | 601 | 1183 | 3083 | 866 | 993 |
| 3874 | FIM1_3969 | QGN17238.1 | CP015059.1 | 6727 | 1376 | 2053 | 5692 | 1039 | 1851 | 7978 | 1606 | 1750 |
| 3875 | FIM1_3970 | QGN17239.1 | CP015059.1 | 3564 | 603 | 653 | 2574 | 537 | 605 | 2003 | 712 | 630 |
| 3876 | FIM1_3971 | QGN17240.1 | CP015059.1 | 1490 | 141 | 382 | 1007 | 109 | 358 | 1240 | 148 | 340 |
| 3877 | FIM1_3972 | QGN17241.1 | CP015059.1 | 2036 | 184 | 521 | 1548 | 135 | 441 | 2048 | 195 | 396 |
| 3878 | FIM1_3973 | QGN17242.1 | CP015059.1 | 11082 | 6171 | 4943 | 9596 | 5027 | 4627 | 8006 | 6574 | 4678 |
| 3879 | FIM1_3974 | QGN17243.1 | CP015059.1 | 2318 | 838 | 1040 | 2030 | 475 | 1043 | 3113 | 848 | 966 |
| 3880 | FIM1_3975 | QGN17244.1 | CP015059.1 | 3086 | 292 | 629 | 2426 | 210 | 600 | 4126 | 290 | 486 |
| 3881 | FIM1_3976 | QGN17245.1 | CP015059.1 | 20083 | 5266 | 6030 | 17279 | 2888 | 5447 | 30905 | 5543 | 5284 |
| 3882 | FIM1_3977 | QGN17246.1 | CP015059.1 | 4853 | 66 | 274 | 2954 | 70 | 227 | 5413 | 54 | 219 |
| 3883 | FIM1_3978 | QGN17247.1 | CP015059.1 | 2885 | 899 | 948 | 2315 | 711 | 869 | 1605 | 959 | 925 |
| 3884 | FIM1_3979 | QGN17248.1 | CP015059.1 | 1696 | 450 | 653 | 1199 | 350 | 649 | 1241 | 517 | 592 |
| 3885 | FIM1_3980 | QGN17249.1 | CP015059.1 | 22859 | 16363 | 14305 | 21308 | 13157 | 13923 | 21344 | 16418 | 14360 |
| 3886 | FIM1_3981 | QGN17250.1 | CP015059.1 | 1110 | 99 | 133 | 971 | 40 | 126 | 2943 | 74 | 106 |

| | | | | | | | | | | | | |
|---|---|---|---|---|---|---|---|---|---|---|---|---|
| 3887 | FIM1_3982 | QGN17251.1 | CP015059.1 | 7635 | 1894 | 2690 | 6108 | 1316 | 2390 | 8871 | 1980 | 2360 |
| 3888 | FIM1_3983 | QGN17252.1 | CP015059.1 | 4291 | 1070 | 1570 | 3529 | 907 | 1492 | 3204 | 1182 | 1317 |
| 3889 | FIM1_3984 | QGN17253.1 | CP015059.1 | 568 | 145 | 238 | 501 | 126 | 254 | 465 | 173 | 234 |
| 3890 | FIM1_3985 | QGN17254.1 | CP015059.1 | 1752 | 326 | 662 | 1428 | 287 | 587 | 1941 | 355 | 574 |
| 3891 | FIM1_3986 | QGN17255.1 | CP015059.1 | 5191 | 1039 | 1624 | 4178 | 861 | 1388 | 3311 | 1181 | 1439 |
| 3892 | FIM1_3987 | QGN17256.1 | CP015059.1 | 2075 | 462 | 703 | 1765 | 339 | 613 | 1711 | 495 | 587 |
| 3893 | FIM1_3988 | QGN17257.1 | CP015059.1 | 2497 | 417 | 977 | 1997 | 342 | 993 | 2432 | 419 | 900 |
| 3894 | FIM1_3989 | QGN17258.1 | CP015059.1 | 30458 | 7381 | 9134 | 22641 | 5720 | 8652 | 26962 | 8211 | 8338 |
| 3895 | FIM1_3990 | QGN17259.1 | CP015059.1 | 5133 | 510 | 1494 | 4182 | 405 | 1409 | 8203 | 530 | 1346 |
| 3896 | FIM1_3991 | QGN17260.1 | CP015059.1 | 3127 | 1872 | 1245 | 2438 | 1151 | 1018 | 1136 | 1849 | 1092 |
| 3897 | FIM1_3992 | QGN17261.1 | CP015059.1 | 1638 | 446 | 664 | 1342 | 330 | 611 | 1677 | 467 | 549 |
| 3898 | FIM1_3993 | QGN17262.1 | CP015059.1 | 19994 | 7414 | 6566 | 16636 | 5452 | 5964 | 19424 | 7901 | 6130 |
| 3899 | FIM1_3994 | QGN17263.1 | CP015059.1 | 207 | 35 | 37 | 141 | 24 | 52 | 161 | 23 | 25 |
| 3900 | FIM1_3995 | QGN17264.1 | CP015059.1 | 578 | 83 | 107 | 470 | 89 | 104 | 493 | 104 | 107 |
| 3901 | FIM1_3996 | QGN17265.1 | CP015059.1 | 4422 | 1764 | 1293 | 2819 | 1081 | 1076 | 1495 | 1825 | 1170 |
| 3902 | FIM1_3997 | QGN17266.1 | CP015059.1 | 6095 | 1332 | 1968 | 4293 | 920 | 1837 | 5538 | 1410 | 1595 |
| 3903 | FIM1_3998 | QGN17267.1 | CP015059.1 | 2883 | 566 | 1208 | 2019 | 452 | 1067 | 3301 | 602 | 1026 |
| 3904 | FIM1_3999 | QGN17268.1 | CP015059.1 | 228 | 139 | 226 | 210 | 142 | 225 | 195 | 159 | 267 |
| 3905 | FIM1_4000 | QGN17269.1 | CP015059.1 | 1993 | 634 | 986 | 1729 | 590 | 875 | 1421 | 778 | 786 |
| 3906 | FIM1_4001 | QGN17270.1 | CP015059.1 | 271 | 53 | 112 | 232 | 66 | 93 | 190 | 87 | 85 |
| 3907 | FIM1_4002 | QGN17271.1 | CP015059.1 | 10080 | 4257 | 4076 | 8924 | 3206 | 3688 | 9219 | 4470 | 4000 |
| 3908 | FIM1_4003 | QGN17272.1 | CP015059.1 | 6691 | 2883 | 4043 | 6456 | 2172 | 3735 | 7616 | 3081 | 3661 |
| 3909 | FIM1_4004 | QGN17273.1 | CP015059.1 | 2035 | 752 | 1182 | 1838 | 589 | 1169 | 2162 | 802 | 1132 |
| 3910 | FIM1_4005 | QGN17274.1 | CP015059.1 | 7288 | 1420 | 3519 | 5897 | 942 | 3146 | 9485 | 1587 | 2936 |
| 3911 | FIM1_4006 | QGN17275.1 | CP015059.1 | 5628 | 1875 | 2087 | 4197 | 1218 | 1919 | 5043 | 1980 | 1887 |
| 3912 | FIM1_4007 | QGN17276.1 | CP015059.1 | 4194 | 1024 | 1523 | 3431 | 753 | 1370 | 3319 | 1048 | 1332 |
| 3913 | FIM1_4008 | QGN17277.1 | CP015059.1 | 1069 | 588 | 271 | 517 | 319 | 261 | 768 | 617 | 286 |
| 3914 | FIM1_4009 | QGN17278.1 | CP015059.1 | 5880 | 3486 | 2108 | 4616 | 2016 | 1959 | 4233 | 3945 | 2060 |
| 3915 | FIM1_4010 | QGN17279.1 | CP015059.1 | 5041 | 955 | 1618 | 3792 | 727 | 1544 | 5404 | 1085 | 1399 |
| 3916 | FIM1_4011 | QGN17280.1 | CP015059.1 | 4711 | 1013 | 1737 | 3581 | 755 | 1648 | 4588 | 1113 | 1568 |
| 3917 | FIM1_4012 | QGN17281.1 | CP015059.1 | 5962 | 1764 | 1987 | 4612 | 1351 | 1862 | 4121 | 1873 | 1692 |
| 3918 | FIM1_4013 | QGN17282.1 | CP015059.1 | 4847 | 1412 | 1566 | 4029 | 1048 | 1441 | 3853 | 1475 | 1304 |
| 3919 | FIM1_4014 | QGN17283.1 | CP015059.1 | 533 | 201 | 394 | 391 | 130 | 404 | 499 | 206 | 328 |
| 3920 | FIM1_4015 | QGN17284.1 | CP015059.1 | 5226 | 1388 | 3343 | 3695 | 1099 | 2999 | 5716 | 1466 | 2552 |
| 3921 | FIM1_4016 | QGN17285.1 | CP015059.1 | 6423 | 1717 | 2200 | 5314 | 990 | 2125 | 8945 | 1899 | 2048 |
| 3922 | FIM1_4017 | QGN17286.1 | CP015059.1 | 856 | 388 | 325 | 591 | 249 | 323 | 415 | 385 | 293 |
| 3923 | FIM1_4018 | QGN17287.1 | CP015059.1 | 8018 | 6152 | 3010 | 5676 | 4172 | 2765 | 1683 | 6797 | 2978 |
| 3924 | FIM1_4019 | QGN17288.1 | CP015059.1 | 2300 | 421 | 621 | 1490 | 304 | 629 | 1696 | 471 | 519 |
| 3925 | FIM1_4020 | QGN17289.1 | CP015059.1 | 7669 | 3279 | 2459 | 6279 | 2425 | 2344 | 6929 | 3438 | 2349 |
| 3926 | FIM1_4021 | QGN17290.1 | CP015059.1 | 2492 | 598 | 1023 | 1881 | 485 | 984 | 1557 | 654 | 879 |
| 3927 | FIM1_4022 | QGN17291.1 | CP015059.1 | 50 | 24 | 24 | 40 | 18 | 24 | 30 | 25 | 24 |
| 3928 | FIM1_4023 | QGN17292.1 | CP015059.1 | 16581 | 5138 | 5239 | 14428 | 3734 | 4868 | 17046 | 5548 | 4864 |
| 3929 | FIM1_4024 | QGN17293.1 | CP015059.1 | 4617 | 2354 | 2182 | 3683 | 993 | 2080 | 7899 | 1908 | 2066 |
| 3930 | FIM1_4025 | QGN17294.1 | CP015059.1 | 416 | 123 | 308 | 378 | 92 | 293 | 397 | 111 | 291 |
| 3931 | FIM1_4026 | QGN17295.1 | CP015059.1 | 396 | 108 | 168 | 379 | 85 | 163 | 424 | 93 | 72 |
| 3932 | FIM1_4027 | QGN17296.1 | CP015059.1 | 464 | 172 | 300 | 357 | 91 | 244 | 992 | 109 | 258 |
| 3933 | FIM1_4028 | QGN17297.1 | CP015059.1 | 2574 | 1003 | 987 | 1989 | 782 | 950 | 2406 | 1109 | 955 |
| 3934 | FIM1_4029 | QGN17298.1 | CP015059.1 | 527 | 69 | 59 | 339 | 47 | 70 | 485 | 95 | 48 |

| | | | | | | | | | | | | |
|---|---|---|---|---|---|---|---|---|---|---|---|---|
| 3935 | FIM1_4030 | QGN17299.1 | CP015059.1 | 11181 | 8419 | 4041 | 7931 | 6021 | 3523 | 3543 | 9165 | 3747 |
| 3936 | FIM1_4031 | QGN17300.1 | CP015059.1 | 5801 | 2681 | 2916 | 5126 | 2021 | 2796 | 4535 | 3038 | 2611 |
| 3937 | FIM1_4033 | QGN17301.1 | CP015059.1 | 1757 | 479 | 769 | 1214 | 332 | 733 | 1197 | 510 | 641 |
| 3938 | FIM1_4034 | QGN17302.1 | CP015059.1 | 936 | 645 | 419 | 774 | 431 | 353 | 620 | 716 | 380 |
| 3939 | FIM1_4035 | QGN17303.1 | CP015059.1 | 634 | 489 | 282 | 541 | 344 | 314 | 323 | 533 | 348 |
| 3940 | FIM1_4036 | QGN17304.1 | CP015059.1 | 4138 | 1431 | 1477 | 3462 | 978 | 1544 | 3532 | 1459 | 1321 |
| 3941 | FIM1_4037 | QGN17305.1 | CP015059.1 | 3709 | 678 | 1190 | 2646 | 529 | 1096 | 3268 | 790 | 907 |
| 3942 | FIM1_4038 | QGN17306.1 | CP015059.1 | 1762 | 469 | 587 | 1326 | 353 | 625 | 1911 | 578 | 580 |
| 3943 | FIM1_4039 | QGN17307.1 | CP015059.1 | 2198 | 1078 | 644 | 1523 | 780 | 587 | 1995 | 1135 | 619 |
| 3944 | FIM1_4040 | QGN17308.1 | CP015059.1 | 23 | 3 | 4 | 13 | 5 | 2 | 18 | 7 | 4 |
| 3945 | FIM1_4041 | QGN17309.1 | CP015059.1 | 2043 | 1404 | 845 | 1381 | 905 | 752 | 788 | 1572 | 780 |
| 3946 | FIM1_4042 | QGN17310.1 | CP015059.1 | 4408 | 1096 | 1174 | 3225 | 851 | 1086 | 3735 | 1170 | 1050 |
| 3947 | FIM1_4043 | QGN17311.1 | CP015059.1 | 4607 | 963 | 1278 | 3574 | 721 | 1233 | 3936 | 1036 | 1234 |
| 3948 | FIM1_4044 | QGN17312.1 | CP015059.1 | 1042 | 220 | 249 | 758 | 153 | 208 | 681 | 247 | 203 |
| 3949 | FIM1_4045 | QGN17313.1 | CP015059.1 | 10529 | 4576 | 3606 | 8180 | 3120 | 3450 | 5434 | 4871 | 3493 |
| 3950 | FIM1_4046 | QGN17314.1 | CP015059.1 | 2235 | 971 | 1166 | 2028 | 764 | 1227 | 1544 | 1048 | 986 |
| 3951 | FIM1_4047 | QGN17315.1 | CP015059.1 | 24506 | 6790 | 6382 | 18002 | 4811 | 5726 | 21152 | 7200 | 5794 |
| 3952 | FIM1_4048 | QGN17316.1 | CP015059.1 | 1031 | 454 | 389 | 813 | 272 | 348 | 466 | 418 | 353 |
| 3953 | FIM1_4049 | QGN17317.1 | CP015059.1 | 1749 | 1986 | 1313 | 1392 | 1427 | 1187 | 370 | 2016 | 1240 |
| 3954 | FIM1_4050 | QGN17318.1 | CP015059.1 | 13365 | 2625 | 3165 | 10014 | 1868 | 2880 | 12726 | 2699 | 2809 |
| 3955 | FIM1_4051 | QGN17319.1 | CP015059.1 | 13489 | 4040 | 10806 | 12472 | 2487 | 9430 | 10698 | 4005 | 8625 |
| 3956 | FIM1_4052 | QGN17320.1 | CP015059.1 | 303 | 46 | 39 | 257 | 41 | 42 | 246 | 48 | 42 |
| 3957 | FIM1_4053 | QGN17321.1 | CP015059.1 | 4185 | 226 | 547 | 3740 | 130 | 551 | 5267 | 248 | 478 |
| 3958 | FIM1_4054 | QGN17322.1 | CP015059.1 | 9896 | 1427 | 2423 | 7636 | 1146 | 2083 | 10809 | 1541 | 1985 |
| 3959 | FIM1_4055 | QGN17323.1 | CP015059.1 | 4415 | 760 | 657 | 3510 | 381 | 618 | 7606 | 834 | 627 |
| 3960 | FIM1_4056 | QGN17324.1 | CP015059.1 | 5631 | 693 | 1083 | 4479 | 507 | 915 | 7536 | 675 | 901 |
| 3961 | FIM1_4057 | QGN17325.1 | CP015059.1 | 2832 | 647 | 992 | 2462 | 472 | 883 | 3079 | 706 | 830 |
| 3962 | FIM1_4058 | QGN17326.1 | CP015059.1 | 829 | 229 | 365 | 743 | 157 | 380 | 850 | 278 | 309 |
| 3963 | FIM1_4059 | QGN17327.1 | CP015059.1 | 14186 | 3784 | 2691 | 11208 | 2349 | 2629 | 17057 | 4017 | 2669 |
| 3964 | FIM1_4060 | QGN17328.1 | CP015059.1 | 3564 | 2827 | 1833 | 3470 | 2485 | 1700 | 1632 | 3247 | 1631 |
| 3965 | FIM1_4061 | QGN17329.1 | CP015059.1 | 8371 | 9868 | 4665 | 6149 | 5100 | 3916 | 2297 | 10387 | 3575 |
| 3966 | FIM1_4062 | QGN17330.1 | CP015059.1 | 1760 | 2521 | 1266 | 1512 | 1306 | 1041 | 393 | 2691 | 948 |
| 3967 | FIM1_4063 | QGN17331.1 | CP015059.1 | 7025 | 1213 | 1866 | 5583 | 843 | 1761 | 7607 | 1346 | 1597 |
| 3968 | FIM1_4064 | QGN17332.1 | CP015059.1 | 8104 | 5353 | 5143 | 7455 | 4103 | 4879 | 5101 | 5489 | 4713 |
| 3969 | FIM1_4065 | QGN17333.1 | CP015059.1 | 8865 | 6248 | 3388 | 7139 | 4376 | 2704 | 2382 | 6863 | 3104 |
| 3970 | FIM1_4066 | QGN17334.1 | CP015059.1 | 8971 | 2139 | 2267 | 7745 | 1341 | 2063 | 11407 | 2372 | 1947 |
| 3971 | FIM1_4067 | QGN17335.1 | CP015059.1 | 2807 | 577 | 680 | 2476 | 446 | 638 | 3290 | 572 | 601 |
| 3972 | FIM1_4068 | QGN17336.1 | CP015059.1 | 762 | 283 | 302 | 660 | 187 | 291 | 613 | 275 | 279 |
| 3973 | FIM1_4069 | QGN17337.1 | CP015059.1 | 11616 | 1512 | 2034 | 8454 | 1153 | 1838 | 11369 | 1636 | 1716 |
| 3974 | FIM1_4070 | QGN17338.1 | CP015059.1 | 9967 | 1399 | 2368 | 7908 | 780 | 2047 | 7910 | 1493 | 1934 |
| 3975 | FIM1_4071 | QGN17339.1 | CP015059.1 | 349 | 64 | 53 | 266 | 42 | 42 | 275 | 63 | 44 |
| 3976 | FIM1_4072 | QGN17340.1 | CP015059.1 | 4790 | 805 | 1475 | 3664 | 659 | 1372 | 3861 | 835 | 1251 |
| 3977 | FIM1_4073 | QGN17341.1 | CP015059.1 | 2116 | 545 | 937 | 1584 | 361 | 850 | 1747 | 643 | 853 |
| 3978 | FIM1_4074 | QGN17342.1 | CP015059.1 | 4041 | 1160 | 1054 | 3495 | 459 | 974 | 5726 | 1276 | 913 |
| 3979 | FIM1_4075 | QGN17343.1 | CP015059.1 | 4063 | 134 | 687 | 3954 | 92 | 595 | 4904 | 166 | 389 |
| 3980 | FIM1_4076 | QGN17344.1 | CP015059.1 | 63 | 7 | 10 | 46 | 7 | 13 | 44 | 8 | 7 |
| 3981 | FIM1_4077 | QGN17345.1 | CP015059.1 | 83116 | 42167 | 24213 | 70927 | 25481 | 21069 | 121026 | 42697 | 21529 |
| 3982 | FIM1_4078 | QGN17346.1 | CP015059.1 | 1572 | 132 | 156 | 1147 | 76 | 135 | 1419 | 118 | 126 |

| | | | | | | | | | | | | |
|---|---|---|---|---|---|---|---|---|---|---|---|---|
| 3983 | FIM1_4079 | QGN17347.1 | CP015059.1 | 58 | 3 | 5 | 56 | 3 | 4 | 148 | 10 | 1 |
| 3984 | FIM1_4080 | QGN17348.1 | CP015059.1 | 22640 | 1374 | 1844 | 19797 | 619 | 1398 | 24243 | 1483 | 1220 |
| 3985 | FIM1_4081 | QGN17349.1 | CP015059.1 | 57 | 12 | 13 | 54 | 11 | 8 | 82 | 24 | 13 |
| 3986 | FIM1_4082 | QGN17350.1 | CP015059.1 | 215 | 57 | 14 | 145 | 24 | 11 | 109 | 54 | 12 |
| 3987 | FIM1_4083 | QGN17351.1 | CP015059.1 | 1070 | 494 | 514 | 828 | 326 | 424 | 596 | 518 | 466 |
| 3988 | FIM1_4084 | QGN17352.1 | CP015059.1 | 7489 | 3259 | 3202 | 6184 | 2378 | 2765 | 4639 | 3408 | 2788 |
| 3989 | FIM1_4085 | QGN17353.1 | CP015059.1 | 27868 | 9170 | 8511 | 22253 | 6650 | 7480 | 22920 | 9790 | 7673 |
| 3990 | FIM1_4086 | QGN17354.1 | CP015059.1 | 21849 | 16113 | 8531 | 16277 | 11424 | 7783 | 9346 | 17737 | 8335 |
| 3991 | FIM1_4087 | QGN17355.1 | CP015059.1 | 899 | 178 | 253 | 758 | 181 | 232 | 493 | 215 | 166 |
| 3992 | FIM1_4088 | QGN17356.1 | CP015059.1 | 10179 | 1697 | 1757 | 7611 | 1051 | 1581 | 9625 | 1650 | 1531 |
| 3993 | FIM1_4089 | QGN17357.1 | CP015059.1 | 632 | 99 | 125 | 514 | 56 | 116 | 600 | 91 | 122 |
| 3994 | FIM1_4090 | QGN17358.1 | CP015059.1 | 682 | 106 | 165 | 512 | 81 | 147 | 629 | 132 | 110 |
| 3995 | FIM1_4091 | QGN17359.1 | CP015059.1 | 4143 | 4602 | 2096 | 2915 | 2941 | 1849 | 749 | 4899 | 2022 |
| 3996 | FIM1_4092 | QGN17360.1 | CP015059.1 | 2599 | 667 | 1132 | 2152 | 483 | 1057 | 2894 | 741 | 926 |
| 3997 | FIM1_4093 | QGN17361.1 | CP015059.1 | 7857 | 687 | 1202 | 4912 | 482 | 1056 | 7459 | 793 | 965 |
| 3998 | FIM1_4094 | QGN17362.1 | CP015059.1 | 15338 | 6754 | 2383 | 12898 | 4898 | 1909 | 14872 | 7706 | 1828 |
| 3999 | FIM1_4095 | QGN17363.1 | CP015059.1 | 2871 | 1464 | 1565 | 2723 | 1075 | 1532 | 2464 | 1546 | 1443 |
| 4000 | FIM1_4096 | QGN17364.1 | CP015059.1 | 2436 | 1267 | 1993 | 2477 | 1096 | 1847 | 2279 | 1445 | 1741 |
| 4001 | FIM1_4097 | QGN17365.1 | CP015059.1 | 3870 | 2501 | 1347 | 3423 | 1800 | 1259 | 947 | 2676 | 1370 |
| 4002 | FIM1_4098 | QGN17366.1 | CP015059.1 | 3301 | 969 | 2231 | 2959 | 773 | 1889 | 2401 | 1046 | 1946 |
| 4003 | FIM1_4099 | QGN17367.1 | CP015059.1 | 662 | 169 | 500 | 625 | 139 | 436 | 505 | 194 | 400 |
| 4004 | FIM1_4100 | QGN17368.1 | CP015059.1 | 1208 | 161 | 273 | 1139 | 123 | 282 | 1274 | 176 | 230 |
| 4005 | FIM1_4101 | QGN17369.1 | CP015059.1 | 2559 | 352 | 695 | 1881 | 249 | 607 | 2880 | 324 | 568 |
| 4006 | FIM1_4102 | QGN17370.1 | CP015059.1 | 6327 | 753 | 1822 | 5034 | 573 | 1598 | 7396 | 826 | 1524 |
| 4007 | FIM1_4103 | QGN17371.1 | CP015059.1 | 5666 | 1455 | 8121 | 4464 | 1026 | 7579 | 7299 | 1600 | 7662 |
| 4008 | FIM1_4104 | QGN17372.1 | CP015059.1 | 3340 | 1265 | 1349 | 2334 | 760 | 1193 | 2139 | 1470 | 1303 |
| 4009 | FIM1_4105 | QGN17373.1 | CP015059.1 | 335 | 166 | 208 | 251 | 138 | 201 | 415 | 189 | 235 |
| 4010 | FIM1_4106 | QGN17374.1 | CP015059.1 | 22371 | 11493 | 3265 | 18827 | 7463 | 2679 | 15453 | 11822 | 3249 |
| 4011 | FIM1_4107 | QGN17375.1 | CP015059.1 | 2866 | 884 | 1041 | 2284 | 653 | 973 | 2061 | 959 | 881 |
| 4012 | FIM1_4108 | QGN17376.1 | CP015059.1 | 5319 | 2278 | 2247 | 4311 | 1708 | 2070 | 4666 | 2423 | 2018 |
| 4013 | FIM1_4109 | QGN17377.1 | CP015059.1 | 690 | 190 | 363 | 618 | 158 | 314 | 470 | 199 | 272 |
| 4014 | FIM1_4110 | QGN17378.1 | CP015059.1 | 2376 | 768 | 855 | 1794 | 529 | 667 | 1819 | 806 | 761 |
| 4015 | FIM1_4111 | QGN17379.1 | CP015059.1 | 6465 | 1710 | 2135 | 5215 | 1278 | 1969 | 6001 | 2016 | 1908 |
| 4016 | FIM1_4112 | QGN17380.1 | CP015059.1 | 8462 | 1964 | 1754 | 6707 | 854 | 1635 | 9804 | 2060 | 1725 |
| 4017 | FIM1_4113 | QGN17381.1 | CP015059.1 | 4302 | 540 | 1361 | 2826 | 627 | 1020 | 3272 | 500 | 1053 |
| 4018 | FIM1_4114 | QGN17382.1 | CP015059.1 | 4582 | 3196 | 1276 | 3481 | 2083 | 1178 | 1269 | 3461 | 1429 |
| 4019 | FIM1_4115 | QGN17383.1 | CP015059.1 | 5063 | 761 | 1037 | 3699 | 512 | 949 | 3088 | 717 | 856 |
| 4020 | FIM1_4116 | QGN17384.1 | CP015059.1 | 868 | 237 | 254 | 726 | 192 | 225 | 424 | 241 | 237 |
| 4021 | FIM1_4117 | QGN17385.1 | CP015059.1 | 475 | 139 | 143 | 425 | 96 | 144 | 226 | 148 | 131 |
| 4022 | FIM1_4118 | QGN17386.1 | CP015059.1 | 6398 | 1325 | 1745 | 5557 | 805 | 1614 | 8721 | 1441 | 1532 |
| 4023 | FIM1_4119 | QGN17387.1 | CP015059.1 | 4506 | 1658 | 1382 | 4088 | 1315 | 1265 | 4444 | 1618 | 1340 |
| 4024 | FIM1_4120 | QGN17388.1 | CP015059.1 | 6739 | 3551 | 2432 | 5362 | 2529 | 2088 | 1124 | 3844 | 2087 |
| 4025 | FIM1_4121 | QGN17389.1 | CP015059.1 | 2112 | 945 | 927 | 1276 | 622 | 841 | 651 | 1030 | 795 |
| 4026 | FIM1_4122 | QGN17390.1 | CP015059.1 | 2024 | 287 | 568 | 1611 | 195 | 602 | 2204 | 281 | 496 |
| 4027 | FIM1_4123 | QGN17391.1 | CP015059.1 | 3352 | 498 | 872 | 3033 | 379 | 818 | 3395 | 537 | 788 |
| 4028 | FIM1_4124 | QGN17392.1 | CP015059.1 | 191 | 37 | 58 | 177 | 19 | 69 | 187 | 31 | 52 |
| 4029 | FIM1_4125 | QGN17393.1 | CP015059.1 | 22213 | 4179 | 5673 | 19916 | 3294 | 5480 | 29430 | 4390 | 5146 |
| 4030 | FIM1_4126 | QGN17394.1 | CP015059.1 | 993 | 233 | 329 | 738 | 173 | 262 | 1142 | 250 | 249 |

| | | | | | | | | | | | | |
|---|---|---|---|---|---|---|---|---|---|---|---|---|
| 4031 | FIM1_4127 | QGN17395.1 | CP015059.1 | 198 | 16 | 44 | 146 | 19 | 57 | 135 | 14 | 43 |
| 4032 | FIM1_4128 | QGN17396.1 | CP015059.1 | 3174 | 1170 | 1107 | 2311 | 878 | 1063 | 2028 | 1228 | 1007 |
| 4033 | FIM1_4129 | QGN17397.1 | CP015059.1 | 3367 | 2074 | 2248 | 3150 | 1734 | 2346 | 2341 | 2256 | 2129 |
| 4034 | FIM1_4130 | QGN17398.1 | CP015059.1 | 8709 | 939 | 1605 | 6724 | 821 | 1318 | 6741 | 1067 | 1182 |
| 4035 | FIM1_4131 | QGN17399.1 | CP015059.1 | 28 | 3 | 10 | 24 | 2 | 12 | 22 | 1 | 6 |
| 4036 | FIM1_4132 | QGN17400.1 | CP015059.1 | 621 | 86 | 224 | 487 | 54 | 198 | 582 | 100 | 168 |
| 4037 | FIM1_4133 | QGN17401.1 | CP015059.1 | 1707 | 257 | 770 | 1562 | 194 | 699 | 2534 | 284 | 574 |
| 4038 | FIM1_4134 | QGN17402.1 | CP015059.1 | 3162 | 927 | 1426 | 2576 | 675 | 1293 | 3639 | 997 | 1250 |
| 4039 | FIM1_4135 | QGN17403.1 | CP015059.1 | 4347 | 640 | 827 | 3302 | 491 | 743 | 4226 | 621 | 706 |
| 4040 | FIM1_4136 | QGN17404.1 | CP015059.1 | 4856 | 1571 | 2083 | 4554 | 1018 | 1908 | 5603 | 1641 | 1841 |
| 4041 | FIM1_4137 | QGN17405.1 | CP015059.1 | 10573 | 2151 | 2699 | 7544 | 1661 | 2476 | 9388 | 2399 | 2410 |
| 4042 | FIM1_4138 | QGN17406.1 | CP015059.1 | 3821 | 1063 | 1147 | 2994 | 722 | 1025 | 3389 | 1126 | 1017 |
| 4043 | FIM1_4139 | QGN17407.1 | CP015059.1 | 6343 | 4549 | 2537 | 4739 | 3345 | 2560 | 2356 | 5151 | 2533 |
| 4044 | FIM1_4140 | QGN17408.1 | CP015059.1 | 3305 | 774 | 1068 | 2566 | 588 | 967 | 2180 | 807 | 848 |
| 4045 | FIM1_4141 | QGN17409.1 | CP015059.1 | 15270 | 7656 | 7720 | 13776 | 5658 | 7347 | 14461 | 7893 | 7461 |
| 4046 | FIM1_4142 | QGN17410.1 | CP015059.1 | 2735 | 452 | 786 | 2246 | 358 | 709 | 2744 | 537 | 613 |
| 4047 | FIM1_4143 | QGN17411.1 | CP015059.1 | 2058 | 634 | 879 | 1751 | 491 | 774 | 1521 | 650 | 668 |
| 4048 | FIM1_4144 | QGN17412.1 | CP015059.1 | 607 | 169 | 396 | 618 | 151 | 417 | 468 | 206 | 307 |
| 4049 | FIM1_4145 | QGN17413.1 | CP015059.1 | 1292 | 147 | 484 | 1158 | 134 | 435 | 1103 | 180 | 371 |
| 4050 | FIM1_4146 | QGN17414.1 | CP015059.1 | 2789 | 2163 | 1451 | 2082 | 1507 | 1271 | 355 | 2345 | 1191 |
| 4051 | FIM1_4147 | QGN17415.1 | CP015059.1 | 2422 | 472 | 788 | 2053 | 312 | 703 | 2626 | 497 | 682 |
| 4052 | FIM1_4148 | QGN17416.1 | CP015059.1 | 14399 | 1748 | 2426 | 11047 | 1276 | 2252 | 15005 | 1876 | 2232 |
| 4053 | FIM1_4149 | QGN17417.1 | CP015059.1 | 18108 | 8700 | 11390 | 16071 | 6352 | 10563 | 19327 | 9344 | 11574 |
| 4054 | FIM1_4150 | QGN17418.1 | CP015059.1 | 5480 | 1337 | 2079 | 4803 | 1101 | 1785 | 5374 | 1536 | 1771 |
| 4055 | FIM1_4151 | QGN17419.1 | CP015059.1 | 11415 | 1290 | 2108 | 9069 | 948 | 1983 | 13645 | 1435 | 1896 |
| 4056 | FIM1_4152 | QGN17420.1 | CP015059.1 | 2670 | 2835 | 1302 | 1581 | 1858 | 1131 | 465 | 3126 | 1213 |
| 4057 | FIM1_4153 | QGN17421.1 | CP015059.1 | 6150 | 900 | 1380 | 4561 | 537 | 1212 | 6732 | 965 | 1132 |
| 4058 | FIM1_4154 | QGN17422.1 | CP015059.1 | 1797 | 304 | 659 | 1504 | 193 | 582 | 2241 | 315 | 530 |
| 4059 | FIM1_4155 | QGN17423.1 | CP015059.1 | 5501 | 3249 | 3246 | 5665 | 2723 | 3398 | 3896 | 3537 | 3128 |
| 4060 | FIM1_4156 | QGN17424.1 | CP015059.1 | 7214 | 5029 | 2873 | 5304 | 3408 | 2467 | 2080 | 5231 | 2629 |
| 4061 | FIM1_4157 | QGN17425.1 | CP015059.1 | 7237 | 2598 | 2065 | 5921 | 1541 | 1974 | 6485 | 2759 | 2045 |
| 4062 | FIM1_4158 | QGN17426.1 | CP015059.1 | 13765 | 3378 | 3462 | 10931 | 2499 | 3094 | 11451 | 3609 | 3015 |
| 4063 | FIM1_4159 | QGN17427.1 | CP015059.1 | 2898 | 710 | 1043 | 2574 | 517 | 916 | 3169 | 762 | 899 |
| 4064 | FIM1_4160 | QGN17428.1 | CP015059.1 | 764 | 193 | 325 | 748 | 151 | 314 | 663 | 231 | 298 |
| 4065 | FIM1_4161 | QGN17429.1 | CP015059.1 | 28257 | 13176 | 13842 | 26387 | 9237 | 12484 | 31104 | 13484 | 12782 |
| 4066 | FIM1_4162 | QGN17430.1 | CP015059.1 | 2701 | 827 | 1668 | 2361 | 490 | 1507 | 3032 | 950 | 1367 |
| 4067 | FIM1_4163 | QGN17431.1 | CP015059.1 | 11394 | 6239 | 6688 | 10833 | 4389 | 6079 | 10207 | 6398 | 6111 |
| 4068 | FIM1_4164 | QGN17432.1 | CP015059.1 | 144 | 23 | 67 | 95 | 20 | 57 | 180 | 27 | 48 |
| 4069 | FIM1_4165 | QGN17433.1 | CP015059.1 | 519 | 407 | 409 | 465 | 231 | 369 | 547 | 355 | 355 |
| 4070 | FIM1_4166 | QGN17434.1 | CP015059.1 | 9961 | 779 | 1106 | 6739 | 395 | 970 | 11895 | 799 | 885 |
| 4071 | FIM1_4167 | QGN17435.1 | CP015059.1 | 4194 | 484 | 1030 | 2976 | 383 | 982 | 4958 | 551 | 908 |
| 4072 | FIM1_4169 | QGN17436.1 | CP015059.1 | 2348 | 360 | 694 | 1751 | 239 | 599 | 2420 | 404 | 533 |
| 4073 | FIM1_4170 | QGN17437.1 | CP015059.1 | 1505 | 199 | 823 | 1236 | 179 | 682 | 1145 | 208 | 636 |
| 4074 | FIM1_4171 | QGN17438.1 | CP015059.1 | 49 | 10 | 104 | 65 | 18 | 124 | 50 | 4 | 75 |
| 4075 | FIM1_4172 | QGN17439.1 | CP015059.1 | 3586 | 475 | 858 | 2986 | 342 | 723 | 4268 | 509 | 742 |
| 4076 | FIM1_4173 | QGN17440.1 | CP015059.1 | 2250 | 533 | 962 | 1695 | 385 | 805 | 2121 | 581 | 786 |
| 4077 | FIM1_4174 | QGN17441.1 | CP015059.1 | 13844 | 7283 | 3762 | 10439 | 5733 | 3423 | 4782 | 8149 | 3688 |
| 4078 | FIM1_4175 | QGN17442.1 | CP015059.1 | 4126 | 863 | 1589 | 3422 | 651 | 1541 | 3922 | 1019 | 1476 |

| | | | | | | | | | | | | |
|---|---|---|---|---|---|---|---|---|---|---|---|---|
| 4079 | FIM1_4176 | QGN17443.1 | CP015059.1 | 3458 | 620 | 1319 | 2972 | 514 | 1156 | 3443 | 674 | 1117 |
| 4080 | FIM1_4177 | QGN17444.1 | CP015059.1 | 8944 | 6299 | 6736 | 8933 | 4784 | 6498 | 7030 | 6458 | 6242 |
| 4081 | FIM1_4178 | QGN17445.1 | CP015059.1 | 8676 | 5770 | 6420 | 8767 | 4316 | 5913 | 6211 | 5920 | 5775 |
| 4082 | FIM1_4180 | QGN17446.1 | CP015059.1 | 56 | 19 | 20 | 52 | 10 | 20 | 26 | 13 | 12 |
| 4083 | FIM1_4181 | QGN17447.1 | CP015059.1 | 8476 | 1899 | 2208 | 6898 | 1150 | 1899 | 12936 | 1992 | 1956 |
| 4084 | FIM1_4182 | QGN17448.1 | CP015059.1 | 17121 | 7528 | 7172 | 15048 | 5923 | 6594 | 16260 | 7933 | 6640 |
| 4085 | FIM1_4183 | QGN17449.1 | CP015059.1 | 3937 | 1023 | 1409 | 3239 | 745 | 1326 | 2867 | 1168 | 1277 |
| 4086 | FIM1_4184 | QGN17450.1 | CP015059.1 | 6801 | 1637 | 1848 | 5073 | 1095 | 1617 | 5264 | 1758 | 1560 |
| 4087 | FIM1_4185 | QGN17451.1 | CP015059.1 | 1539 | 477 | 684 | 1159 | 375 | 612 | 1427 | 486 | 549 |
| 4088 | FIM1_4186 | QGN17452.1 | CP015059.1 | 5155 | 592 | 2917 | 4347 | 641 | 2671 | 5454 | 644 | 2801 |
| 4089 | FIM1_4187 | QGN17453.1 | CP015059.1 | 2841 | 385 | 613 | 2163 | 272 | 560 | 2753 | 424 | 507 |
| 4090 | FIM1_4188 | QGN17454.1 | CP015059.1 | 4244 | 1080 | 1031 | 3300 | 798 | 883 | 3061 | 1152 | 936 |
| 4091 | FIM1_4189 | QGN17455.1 | CP015059.1 | 9413 | 1490 | 2529 | 7029 | 1250 | 2602 | 12491 | 1626 | 2529 |
| 4092 | FIM1_4190 | QGN17456.1 | CP015059.1 | 3571 | 1137 | 2050 | 2933 | 830 | 1912 | 3455 | 1181 | 1793 |
| 4093 | FIM1_4191 | QGN17457.1 | CP015059.1 | 6191 | 1243 | 1945 | 4904 | 786 | 1669 | 5693 | 1299 | 1504 |
| 4094 | FIM1_4192 | QGN17458.1 | CP015059.1 | 1457 | 1114 | 819 | 949 | 711 | 817 | 599 | 1185 | 798 |
| 4095 | FIM1_4193 | QGN17459.1 | CP015059.1 | 655 | 126 | 197 | 475 | 104 | 220 | 886 | 152 | 218 |
| 4096 | FIM1_4194 | QGN17460.1 | CP015059.1 | 3346 | 474 | 1022 | 2555 | 373 | 851 | 4385 | 604 | 865 |
| 4097 | FIM1_4195 | QGN17461.1 | CP015059.1 | 5139 | 1523 | 2104 | 4560 | 1080 | 1853 | 5567 | 1606 | 1756 |
| 4098 | FIM1_4196 | QGN17462.1 | CP015059.1 | 11738 | 1347 | 2466 | 9016 | 971 | 2225 | 15260 | 1401 | 2199 |
| 4099 | FIM1_4197 | QGN17463.1 | CP015059.1 | 3042 | 254 | 1299 | 2487 | 353 | 1182 | 5557 | 355 | 1131 |
| 4100 | FIM1_4198 | QGN17464.1 | CP015059.1 | 6067 | 574 | 1536 | 4662 | 499 | 1356 | 8433 | 584 | 1228 |
| 4101 | FIM1_4199 | QGN17465.1 | CP015059.1 | 1112 | 186 | 236 | 1019 | 140 | 210 | 1800 | 165 | 212 |
| 4102 | FIM1_4201 | QGN17466.1 | CP015059.1 | 1532 | 230 | 532 | 1187 | 199 | 452 | 1363 | 257 | 406 |
| 4103 | FIM1_4203 | QGN17467.1 | CP015059.1 | 577 | 172 | 282 | 543 | 134 | 263 | 660 | 178 | 243 |
| 4104 | FIM1_4204 | QGN17468.1 | CP015059.1 | 2416 | 364 | 898 | 1853 | 302 | 837 | 2430 | 388 | 687 |
| 4105 | FIM1_4205 | QGN17469.1 | CP015059.1 | 18838 | 2023 | 1062 | 13015 | 1112 | 875 | 22366 | 2095 | 953 |
| 4106 | FIM1_4206 | QGN17470.1 | CP015059.1 | 4497 | 1417 | 1310 | 4100 | 819 | 1116 | 3807 | 1490 | 1104 |
| 4107 | FIM1_4207 | QGN17471.1 | CP015059.1 | 5870 | 1101 | 2413 | 4892 | 824 | 2001 | 5820 | 1252 | 1675 |
| 4108 | FIM1_4208 | QGN17472.1 | CP015059.1 | 4523 | 449 | 975 | 3096 | 279 | 824 | 5055 | 434 | 699 |
| 4109 | FIM1_4209 | QGN17473.1 | CP015059.1 | 2011 | 243 | 583 | 1610 | 184 | 483 | 2446 | 254 | 449 |
| 4110 | FIM1_4210 | QGN17474.1 | CP015059.1 | 5873 | 1585 | 1876 | 4940 | 1043 | 1713 | 5118 | 1641 | 1563 |
| 4111 | FIM1_4211 | QGN17475.1 | CP015059.1 | 1604 | 607 | 644 | 1501 | 496 | 540 | 869 | 751 | 563 |
| 4112 | FIM1_4212 | QGN17476.1 | CP015059.1 | 10617 | 2138 | 2724 | 9006 | 1552 | 2385 | 9983 | 2341 | 2323 |
| 4113 | FIM1_4213 | QGN17477.1 | CP015059.1 | 1140 | 221 | 346 | 1008 | 161 | 359 | 1353 | 234 | 306 |
| 4114 | FIM1_4214 | QGN17478.1 | CP015059.1 | 2368 | 748 | 991 | 2209 | 482 | 917 | 2628 | 828 | 828 |
| 4115 | FIM1_4215 | QGN17479.1 | CP015059.1 | 17929 | 4368 | 4149 | 13904 | 1729 | 3552 | 24843 | 4306 | 3325 |
| 4116 | FIM1_4216 | QGN17480.1 | CP015059.1 | 1902 | 285 | 532 | 1498 | 188 | 509 | 3499 | 369 | 524 |
| 4117 | FIM1_4217 | QGN17481.1 | CP015059.1 | 2552 | 246 | 705 | 2196 | 139 | 629 | 4351 | 250 | 586 |
| 4118 | FIM1_4218 | QGN17482.1 | CP015059.1 | 97 | 18 | 38 | 91 | 11 | 31 | 123 | 14 | 32 |
| 4119 | FIM1_4219 | QGN17483.1 | CP015059.1 | 1095 | 285 | 408 | 1130 | 268 | 396 | 763 | 277 | 371 |
| 4120 | FIM1_4220 | QGN17484.1 | CP015059.1 | 1788 | 485 | 674 | 1521 | 425 | 616 | 1415 | 583 | 609 |
| 4121 | FIM1_4221 | QGN17485.1 | CP015059.1 | 138 | 111 | 259 | 127 | 77 | 234 | 179 | 121 | 229 |
| 4122 | FIM1_4222 | QGN17486.1 | CP015059.1 | 4446 | 982 | 1472 | 3590 | 668 | 1296 | 4444 | 1032 | 1293 |
| 4123 | FIM1_4223 | QGN17487.1 | CP015059.1 | 6095 | 1886 | 2413 | 5587 | 1136 | 2116 | 6188 | 2024 | 2222 |
| 4124 | FIM1_4224 | QGN17488.1 | CP015059.1 | 5737 | 4416 | 5198 | 6040 | 3498 | 4960 | 5695 | 4662 | 4775 |
| 4125 | FIM1_4225 | QGN17489.1 | CP015059.1 | 3545 | 389 | 527 | 2470 | 262 | 464 | 3495 | 367 | 564 |
| 4126 | FIM1_4226 | QGN17490.1 | CP015059.1 | 91877 | 49810 | 17474 | 84475 | 25491 | 15116 | 151192 | 49673 | 14037 |

| | | | | | | | | | | | | |
|---|---|---|---|---|---|---|---|---|---|---|---|---|
| 4127 | FIM1_4227 | QGN17491.1 | CP015059.1 | 23677 | 585 | 1634 | 19116 | 1058 | 1515 | 24206 | 755 | 1405 |
| 4128 | FIM1_4228 | QGN17492.1 | CP015059.1 | 55 | 5 | 12 | 54 | 8 | 11 | 44 | 14 | 12 |
| 4129 | FIM1_4229 | QGN17493.1 | CP015059.1 | 4828 | 183 | 333 | 2445 | 177 | 289 | 1902 | 178 | 310 |
| 4130 | FIM1_4230 | QGN17494.1 | CP015059.1 | 6553 | 2224 | 1974 | 5988 | 1727 | 1705 | 3641 | 2485 | 1931 |
| 4131 | FIM1_4231 | QGN17495.1 | CP015059.1 | 17264 | 5852 | 4453 | 14539 | 4355 | 3968 | 10861 | 6231 | 4165 |
| 4132 | FIM1_4232 | QGN17496.1 | CP015059.1 | 1195 | 279 | 416 | 964 | 181 | 361 | 1115 | 268 | 384 |
| 4133 | FIM1_4233 | QGN17497.1 | CP015059.1 | 3111 | 386 | 704 | 2592 | 287 | 542 | 2537 | 387 | 605 |
| 4134 | FIM1_4234 | QGN17498.1 | CP015059.1 | 2056 | 946 | 1005 | 1688 | 778 | 1007 | 1419 | 1045 | 912 |
| 4135 | FIM1_4235 | QGN17499.1 | CP015059.1 | 2202 | 609 | 735 | 1563 | 424 | 756 | 1931 | 692 | 753 |
| 4136 | FIM1_4236 | QGN17500.1 | CP015059.1 | 7370 | 1367 | 2162 | 5261 | 1094 | 2049 | 7074 | 1412 | 1988 |
| 4137 | FIM1_4237 | QGN17501.1 | CP015059.1 | 19717 | 5989 | 5700 | 15552 | 4701 | 5532 | 19644 | 6570 | 5759 |
| 4138 | FIM1_4238 | QGN17502.1 | CP015059.1 | 5855 | 2574 | 2444 | 5160 | 2029 | 2336 | 4387 | 3090 | 2287 |
| 4139 | FIM1_4239 | QGN17503.1 | CP015059.1 | 7184 | 2227 | 1975 | 5353 | 1451 | 1826 | 4747 | 2424 | 1772 |
| 4140 | FIM1_4240 | QGN17504.1 | CP015059.1 | 2003 | 1799 | 1187 | 1578 | 1196 | 1047 | 670 | 1901 | 1034 |
| 4141 | FIM1_4241 | QGN17505.1 | CP015059.1 | 8910 | 1627 | 2014 | 6754 | 1137 | 1844 | 7584 | 1824 | 1710 |
| 4142 | FIM1_4242 | QGN17506.1 | CP015059.1 | 1054 | 86 | 176 | 893 | 57 | 130 | 814 | 76 | 124 |
| 4143 | FIM1_4243 | QGN17507.1 | CP015059.1 | 1172 | 550 | 695 | 991 | 345 | 635 | 1067 | 553 | 603 |
| 4144 | FIM1_4244 | QGN17508.1 | CP015059.1 | 2011 | 191 | 828 | 1786 | 137 | 656 | 2812 | 218 | 618 |
| 4145 | FIM1_4245 | QGN17509.1 | CP015059.1 | 142 | 4 | 18 | 134 | 4 | 20 | 172 | 8 | 10 |
| 4146 | FIM1_4246 | QGN17510.1 | CP015059.1 | 1894 | 232 | 485 | 1411 | 208 | 473 | 2123 | 275 | 398 |
| 4147 | FIM1_4247 | QGN17511.1 | CP015059.1 | 1844 | 327 | 312 | 1280 | 233 | 293 | 1624 | 334 | 268 |
| 4148 | FIM1_4248 | QGN17512.1 | CP015059.1 | 1958 | 759 | 1044 | 1624 | 584 | 1051 | 1810 | 864 | 857 |
| 4149 | FIM1_4249 | QGN17513.1 | CP015059.1 | 18623 | 4284 | 4241 | 15188 | 2737 | 3704 | 20980 | 4505 | 3538 |
| 4150 | FIM1_4250 | QGN17514.1 | CP015059.1 | 3239 | 883 | 674 | 2736 | 637 | 649 | 3144 | 927 | 629 |
| 4151 | FIM1_4251 | QGN17515.1 | CP015059.1 | 6574 | 2256 | 3109 | 5466 | 1418 | 2811 | 7032 | 2330 | 2512 |
| 4152 | FIM1_4252 | QGN17516.1 | CP015059.1 | 3026 | 479 | 1035 | 2786 | 419 | 926 | 3233 | 564 | 923 |
| 4153 | FIM1_4253 | QGN17517.1 | CP015059.1 | 10776 | 1489 | 2793 | 9818 | 926 | 2606 | 19857 | 1501 | 2148 |
| 4154 | FIM1_4254 | QGN17518.1 | CP015059.1 | 3015 | 694 | 853 | 2364 | 465 | 775 | 4137 | 793 | 771 |
| 4155 | FIM1_4255 | QGN17519.1 | CP015059.1 | 4597 | 1169 | 1671 | 4374 | 1101 | 1578 | 3542 | 1272 | 1422 |
| 4156 | FIM1_4256 | QGN17520.1 | CP015059.1 | 1106 | 2 | 7 | 1214 | 1 | 6 | 1842 | 1 | 2 |
| 4157 | FIM1_4257 | QGN17521.1 | CP015059.1 | 18744 | 1198 | 2323 | 15932 | 746 | 1971 | 22810 | 1229 | 1788 |
| 4158 | FIM1_4258 | QGN17522.1 | CP015059.1 | 83 | 26 | 28 | 73 | 32 | 34 | 92 | 28 | 36 |
| 4159 | FIM1_4259 | QGN17523.1 | CP015059.1 | 8459 | 1549 | 2984 | 6784 | 1021 | 2626 | 10150 | 1691 | 2590 |
| 4160 | FIM1_4260 | QGN17524.1 | CP015059.1 | 3334 | 346 | 762 | 2458 | 295 | 660 | 2742 | 399 | 664 |
| 4161 | FIM1_4261 | QGN17525.1 | CP015059.1 | 2397 | 661 | 642 | 1790 | 355 | 451 | 2380 | 760 | 398 |
| 4162 | FIM1_4262 | QGN17526.1 | CP015059.1 | 8552 | 3224 | 2445 | 6820 | 1816 | 2056 | 7503 | 3444 | 2112 |
| 4163 | FIM1_4263 | QGN17527.1 | CP015059.1 | 1434 | 202 | 475 | 1146 | 169 | 444 | 1669 | 194 | 389 |
| 4164 | FIM1_4264 | QGN17528.1 | CP015059.1 | 0 | 0 | 0 | 0 | 0 | 0 | 0 | 0 | 0 |
| 4165 | FIM1_4265 | QGN17529.1 | CP015059.1 | 1944 | 724 | 842 | 1406 | 495 | 795 | 1283 | 770 | 701 |
| 4166 | FIM1_4266 | QGN17530.1 | CP015059.1 | 32681 | 9770 | 9788 | 26890 | 7488 | 8774 | 24216 | 10421 | 9202 |
| 4167 | FIM1_4267 | QGN17531.1 | CP015059.1 | 3003 | 1230 | 1249 | 2292 | 881 | 1146 | 1918 | 1314 | 1118 |
| 4168 | FIM1_4268 | QGN17532.1 | CP015059.1 | 12942 | 2814 | 2937 | 9539 | 2002 | 2634 | 9066 | 3003 | 2566 |
| 4169 | FIM1_4269 | QGN17533.1 | CP015059.1 | 1039 | 344 | 314 | 894 | 251 | 297 | 965 | 361 | 302 |
| 4170 | FIM1_4270 | QGN17534.1 | CP015059.1 | 6421 | 1254 | 1705 | 5178 | 882 | 1490 | 9120 | 1355 | 1365 |
| 4171 | FIM1_4271 | QGN17535.1 | CP015059.1 | 2146 | 526 | 594 | 1882 | 445 | 601 | 2624 | 620 | 622 |
| 4172 | FIM1_4272 | QGN17536.1 | CP015059.1 | 1210 | 404 | 529 | 1103 | 200 | 462 | 2289 | 483 | 465 |
| 4173 | FIM1_4273 | QGN17537.1 | CP015059.1 | 2958 | 2939 | 1502 | 2000 | 1768 | 1374 | 707 | 3125 | 1302 |
| 4174 | FIM1_4274 | QGN17538.1 | CP015059.1 | 1286 | 148 | 209 | 964 | 98 | 173 | 1261 | 125 | 141 |

| | | | | | | | | | | | | |
|---|---|---|---|---|---|---|---|---|---|---|---|---|
| 4175 | FIM1_4275 | QGN17539.1 | CP015059.1 | 45 | 9 | 12 | 29 | 10 | 19 | 40 | 7 | 7 |
| 4176 | FIM1_4276 | QGN17540.1 | CP015059.1 | 4772 | 1651 | 2146 | 4121 | 1265 | 2055 | 4116 | 1732 | 1859 |
| 4177 | FIM1_4277 | QGN17541.1 | CP015059.1 | 404 | 66 | 196 | 353 | 42 | 179 | 486 | 53 | 170 |
| 4178 | FIM1_4278 | QGN17542.1 | CP015059.1 | 7570 | 2008 | 2704 | 6345 | 1339 | 2431 | 8500 | 2100 | 2448 |
| 4179 | FIM1_4279 | QGN17543.1 | CP015059.1 | 4527 | 4597 | 2786 | 3615 | 3686 | 2516 | 1328 | 4963 | 2553 |
| 4180 | FIM1_4280 | QGN17544.1 | CP015059.1 | 4768 | 1839 | 3021 | 4708 | 1087 | 2686 | 6105 | 1972 | 2587 |
| 4181 | FIM1_4281 | QGN17545.1 | CP015059.1 | 14523 | 4058 | 4459 | 11845 | 2840 | 3995 | 13294 | 4152 | 3964 |
| 4182 | FIM1_4282 | QGN17546.1 | CP015059.1 | 5552 | 1352 | 532 | 5422 | 483 | 454 | 12602 | 1515 | 361 |
| 4183 | FIM1_4283 | QGN17547.1 | CP015059.1 | 145 | 23 | 28 | 97 | 11 | 33 | 200 | 16 | 14 |
| 4184 | FIM1_4284 | QGN17548.1 | CP015059.1 | 10026 | 3095 | 6393 | 9105 | 1854 | 5780 | 8852 | 3425 | 5496 |
| 4185 | FIM1_4285 | QGN17549.1 | CP015059.1 | 5372 | 1790 | 2293 | 4807 | 1186 | 2128 | 6138 | 1944 | 2017 |
| 4186 | FIM1_4286 | QGN17550.1 | CP015059.1 | 7544 | 2400 | 2366 | 5719 | 1345 | 2020 | 5624 | 2593 | 1951 |
| 4187 | FIM1_4287 | QGN17551.1 | CP015059.1 | 38 | 6 | 14 | 32 | 5 | 10 | 38 | 7 | 5 |
| 4188 | FIM1_4288 | QGN17552.1 | CP015059.1 | 4258 | 1065 | 1119 | 3532 | 679 | 972 | 4017 | 1224 | 916 |
| 4189 | FIM1_4289 | QGN17553.1 | CP015059.1 | 5576 | 1378 | 2088 | 4939 | 1015 | 1897 | 5489 | 1580 | 1847 |
| 4190 | FIM1_4290 | QGN17554.1 | CP015059.1 | 3208 | 912 | 1145 | 2534 | 608 | 1117 | 3433 | 927 | 1017 |
| 4191 | FIM1_4291 | QGN17555.1 | CP015059.1 | 1887 | 582 | 749 | 1464 | 470 | 649 | 1476 | 662 | 677 |
| 4192 | FIM1_4292 | QGN17556.1 | CP015059.1 | 1761 | 799 | 847 | 1228 | 549 | 807 | 1334 | 821 | 813 |
| 4193 | FIM1_4293 | QGN17557.1 | CP015059.1 | 1441 | 436 | 603 | 1181 | 355 | 544 | 1316 | 491 | 496 |
| 4194 | FIM1_4294 | QGN17558.1 | CP015059.1 | 2490 | 401 | 907 | 1940 | 328 | 858 | 3317 | 442 | 790 |
| 4195 | FIM1_4295 | QGN17559.1 | CP015059.1 | 2150 | 482 | 990 | 1731 | 406 | 902 | 1965 | 526 | 792 |
| 4196 | FIM1_4296 | QGN17560.1 | CP015059.1 | 2203 | 758 | 980 | 1816 | 523 | 922 | 2392 | 781 | 884 |
| 4197 | FIM1_4297 | QGN17561.1 | CP015059.1 | 3206 | 216 | 615 | 2186 | 129 | 552 | 4022 | 244 | 522 |
| 4198 | FIM1_4298 | QGN17562.1 | CP015059.1 | 1114 | 153 | 201 | 843 | 114 | 203 | 1066 | 197 | 219 |
| 4199 | FIM1_4299 | QGN17563.1 | CP015059.1 | 1201 | 290 | 261 | 854 | 204 | 208 | 1303 | 276 | 229 |
| 4200 | FIM1_4300 | QGN17564.1 | CP015059.1 | 715 | 135 | 166 | 559 | 111 | 154 | 714 | 132 | 163 |
| 4201 | FIM1_4301 | QGN17565.1 | CP015059.1 | 18 | 6 | 4 | 15 | 4 | 3 | 33 | 9 | 8 |
| 4202 | FIM1_4303 | QGN17566.1 | CP015059.1 | 119 | 97 | 41 | 74 | 72 | 54 | 108 | 83 | 50 |
| 4203 | FIM1_4304 | QGN17567.1 | CP015059.1 | 150 | 7 | 26 | 81 | 2 | 15 | 237 | 11 | 16 |
| 4204 | FIM1_4306 | QGN17568.1 | CP015059.1 | 0 | 0 | 0 | 0 | 0 | 0 | 0 | 0 | 0 |
| 4205 | FIM1_4307 | QGN17569.1 | CP015059.1 | 835 | 271 | 357 | 774 | 233 | 317 | 728 | 346 | 286 |
| 4206 | FIM1_4308 | QGN17570.1 | CP015059.1 | 269 | 121 | 104 | 206 | 62 | 97 | 240 | 148 | 94 |
| 4207 | FIM1_4314 | QGN17571.1 | CP015059.1 | 1124 | 882 | 504 | 980 | 513 | 477 | 793 | 1131 | 490 |
| 4208 | FIM1_4315 | QGN17572.1 | CP015059.1 | 0 | 0 | 0 | 0 | 0 | 0 | 0 | 0 | 0 |
| 4209 | FIM1_4779 | QGN17573.1 | CP015060.1 | 0 | 2 | 0 | 0 | 0 | 1 | 3 | 0 | 1 |
| 4210 | FIM1_4780 | QGN17574.1 | CP015060.1 | 214 | 165 | 40 | 161 | 69 | 54 | 138 | 231 | 48 |
| 4211 | FIM1_4781 | QGN17575.1 | CP015060.1 | 161 | 48 | 24 | 114 | 22 | 23 | 211 | 45 | 25 |
| 4212 | FIM1_4782 | QGN17576.1 | CP015060.1 | 1141 | 108 | 216 | 833 | 78 | 211 | 1132 | 114 | 204 |
| 4213 | FIM1_4783 | QGN17577.1 | CP015060.1 | 3344 | 360 | 2207 | 2755 | 193 | 1939 | 3577 | 362 | 1742 |
| 4214 | FIM1_4785 | QGN17578.1 | CP015060.1 | 3036 | 680 | 585 | 2391 | 445 | 628 | 3333 | 735 | 645 |
| 4215 | FIM1_4786 | QGN17579.1 | CP015060.1 | 12812 | 3694 | 761 | 8911 | 2917 | 738 | 22561 | 4614 | 815 |
| 4216 | FIM1_4787 | QGN17580.1 | CP015060.1 | 642 | 150 | 106 | 460 | 129 | 98 | 599 | 174 | 96 |
| 4217 | FIM1_4788 | QGN17581.1 | CP015060.1 | 1223 | 91 | 175 | 782 | 76 | 187 | 3961 | 107 | 157 |
| 4218 | FIM1_4789 | QGN17582.1 | CP015060.1 | 1312 | 76 | 187 | 812 | 197 | 233 | 484 | 95 | 204 |
| 4219 | FIM1_4790 | QGN17583.1 | CP015060.1 | 1441 | 466 | 550 | 1240 | 295 | 514 | 1636 | 523 | 472 |
| 4220 | FIM1_4791 | QGN17584.1 | CP015060.1 | 4612 | 2534 | 1214 | 4204 | 2213 | 1187 | 2123 | 2773 | 1296 |
| 4221 | FIM1_4792 | QGN17585.1 | CP015060.1 | 2039 | 263 | 435 | 1486 | 163 | 363 | 1491 | 297 | 313 |
| 4222 | FIM1_4793 | QGN17586.1 | CP015060.1 | 8523 | 976 | 1432 | 6671 | 657 | 1269 | 9896 | 1035 | 1237 |

| | | | | | | | | | | | | |
|---|---|---|---|---|---|---|---|---|---|---|---|---|
| 4223 | FIM1_4794 | QGN17587.1 | CP015060.1 | 11137 | 780 | 1780 | 9511 | 526 | 1524 | 17245 | 808 | 1345 |
| 4224 | FIM1_4795 | QGN17588.1 | CP015060.1 | 28418 | 2426 | 4705 | 22023 | 3103 | 4260 | 31169 | 2717 | 4169 |
| 4225 | FIM1_4796 | QGN17589.1 | CP015060.1 | 5053 | 790 | 1774 | 4165 | 636 | 1515 | 4639 | 942 | 1387 |
| 4226 | FIM1_4797 | QGN17590.1 | CP015060.1 | 7932 | 6073 | 3302 | 6950 | 4381 | 2835 | 2613 | 6508 | 3032 |
| 4227 | FIM1_4798 | QGN17591.1 | CP015060.1 | 1122 | 631 | 1046 | 1095 | 489 | 827 | 985 | 744 | 893 |
| 4228 | FIM1_4799 | QGN17592.1 | CP015060.1 | 2315 | 526 | 928 | 1932 | 423 | 812 | 1795 | 541 | 856 |
| 4229 | FIM1_4800 | QGN17593.1 | CP015060.1 | 35 | 8 | 11 | 24 | 13 | 12 | 18 | 10 | 6 |
| 4230 | FIM1_4801 | QGN17594.1 | CP015060.1 | 768 | 268 | 259 | 640 | 266 | 242 | 575 | 288 | 213 |
| 4231 | FIM1_4802 | QGN17595.1 | CP015060.1 | 2469 | 495 | 697 | 1894 | 380 | 612 | 2365 | 568 | 582 |
| 4232 | FIM1_4803 | QGN17596.1 | CP015060.1 | 5028 | 1041 | 1652 | 4208 | 768 | 1456 | 4402 | 1189 | 1528 |
| 4233 | FIM1_4804 | QGN17597.1 | CP015060.1 | 8483 | 5175 | 4877 | 8109 | 3825 | 4738 | 8781 | 5255 | 4944 |
| 4234 | FIM1_4805 | QGN17598.1 | CP015060.1 | 7577 | 6160 | 3730 | 5548 | 4183 | 3341 | 4697 | 6752 | 3516 |
| 4235 | FIM1_4806 | QGN17599.1 | CP015060.1 | 5700 | 1474 | 1761 | 4488 | 999 | 1589 | 5719 | 1544 | 1580 |
| 4236 | FIM1_4807 | QGN17600.1 | CP015060.1 | 7478 | 1936 | 2364 | 6816 | 1616 | 2159 | 5171 | 2238 | 2196 |
| 4237 | FIM1_4808 | QGN17601.1 | CP015060.1 | 7101 | 1128 | 1918 | 5437 | 748 | 1785 | 8423 | 1192 | 1682 |
| 4238 | FIM1_4809 | QGN17602.1 | CP015060.1 | 1244 | 466 | 593 | 1102 | 351 | 526 | 1291 | 532 | 517 |
| 4239 | FIM1_4810 | QGN17603.1 | CP015060.1 | 26461 | 8622 | 10557 | 22773 | 6712 | 10017 | 26358 | 9194 | 9991 |
| 4240 | FIM1_4811 | QGN17604.1 | CP015060.1 | 4680 | 1087 | 1498 | 3976 | 535 | 1250 | 9266 | 1100 | 1236 |
| 4241 | FIM1_4812 | QGN17605.1 | CP015060.1 | 726 | 88 | 222 | 569 | 66 | 234 | 600 | 100 | 180 |
| 4242 | FIM1_4813 | QGN17606.1 | CP015060.1 | 3556 | 2658 | 1222 | 2676 | 1737 | 982 | 1091 | 2725 | 1122 |
| 4243 | FIM1_4814 | QGN17607.1 | CP015060.1 | 2910 | 810 | 1018 | 2230 | 534 | 903 | 3036 | 830 | 849 |
| 4244 | FIM1_4815 | QGN17608.1 | CP015060.1 | 2585 | 393 | 745 | 1923 | 297 | 743 | 2593 | 383 | 590 |
| 4245 | FIM1_4816 | QGN17609.1 | CP015060.1 | 2863 | 441 | 1053 | 1999 | 404 | 968 | 2733 | 498 | 974 |
| 4246 | FIM1_4817 | QGN17610.1 | CP015060.1 | 3229 | 1276 | 1388 | 2616 | 923 | 1267 | 1906 | 1306 | 1282 |
| 4247 | FIM1_4818 | QGN17611.1 | CP015060.1 | 2663 | 433 | 819 | 2080 | 343 | 808 | 2927 | 474 | 795 |
| 4248 | FIM1_4819 | QGN17612.1 | CP015060.1 | 9100 | 3472 | 8545 | 7928 | 2213 | 7688 | 10214 | 3557 | 7739 |
| 4249 | FIM1_4820 | QGN17613.1 | CP015060.1 | 4045 | 985 | 1382 | 3096 | 719 | 1354 | 5027 | 1049 | 1202 |
| 4250 | FIM1_4821 | QGN17614.1 | CP015060.1 | 1964 | 583 | 951 | 1550 | 437 | 926 | 2173 | 655 | 832 |
| 4251 | FIM1_4822 | QGN17615.1 | CP015060.1 | 3018 | 689 | 1738 | 2848 | 538 | 1670 | 3997 | 764 | 1485 |
| 4252 | FIM1_4823 | QGN17616.1 | CP015060.1 | 3004 | 597 | 1303 | 2236 | 456 | 1194 | 3049 | 645 | 954 |
| 4253 | FIM1_4824 | QGN17617.1 | CP015060.1 | 6976 | 3891 | 2667 | 4917 | 2560 | 2383 | 2929 | 4028 | 2273 |
| 4254 | FIM1_4825 | QGN17618.1 | CP015060.1 | 936 | 124 | 277 | 814 | 60 | 252 | 1453 | 137 | 180 |
| 4255 | FIM1_4826 | QGN17619.1 | CP015060.1 | 3161 | 728 | 841 | 2163 | 533 | 798 | 2681 | 739 | 735 |
| 4256 | FIM1_4827 | QGN17620.1 | CP015060.1 | 3727 | 975 | 1175 | 2650 | 790 | 1117 | 2582 | 1067 | 1024 |
| 4257 | FIM1_4828 | QGN17621.1 | CP015060.1 | 1901 | 465 | 802 | 1841 | 325 | 713 | 2176 | 586 | 696 |
| 4258 | FIM1_4829 | QGN17622.1 | CP015060.1 | 10759 | 576 | 1756 | 8194 | 345 | 1612 | 14974 | 577 | 1610 |
| 4259 | FIM1_4830 | QGN17623.1 | CP015060.1 | 9714 | 659 | 2277 | 6908 | 500 | 2136 | 10998 | 692 | 1947 |
| 4260 | FIM1_4831 | QGN17624.1 | CP015060.1 | 3158 | 1502 | 1872 | 2468 | 1189 | 1783 | 3203 | 1632 | 1554 |
| 4261 | FIM1_4832 | QGN17625.1 | CP015060.1 | 7966 | 1919 | 3430 | 6680 | 1455 | 3072 | 8103 | 2126 | 3163 |
| 4262 | FIM1_4833 | QGN17626.1 | CP015060.1 | 11843 | 3023 | 3497 | 9740 | 2194 | 3010 | 10852 | 3604 | 3096 |
| 4263 | FIM1_4834 | QGN17627.1 | CP015060.1 | 5915 | 751 | 895 | 4511 | 383 | 876 | 8093 | 844 | 892 |
| 4264 | FIM1_4835 | QGN17628.1 | CP015060.1 | 5411 | 3639 | 3325 | 5438 | 3024 | 3173 | 4842 | 3819 | 3077 |
| 4265 | FIM1_4836 | QGN17629.1 | CP015060.1 | 643 | 271 | 749 | 477 | 236 | 649 | 1035 | 246 | 646 |
| 4266 | FIM1_4837 | QGN17630.1 | CP015060.1 | 23 | 18 | 70 | 21 | 13 | 62 | 27 | 13 | 57 |
| 4267 | FIM1_4838 | QGN17631.1 | CP015060.1 | 47 | 8 | 7 | 41 | 3 | 6 | 49 | 5 | 3 |
| 4268 | FIM1_4839 | QGN17632.1 | CP015060.1 | 2492 | 173 | 588 | 1915 | 154 | 479 | 2823 | 215 | 504 |
| 4269 | FIM1_4840 | QGN17633.1 | CP015060.1 | 18336 | 8075 | 7542 | 16177 | 6325 | 6559 | 15206 | 8413 | 6950 |
| 4270 | FIM1_4841 | QGN17634.1 | CP015060.1 | 614 | 60 | 134 | 577 | 52 | 128 | 748 | 71 | 112 |

| | | | | | | | | | | | |
|---|---|---|---|---|---|---|---|---|---|---|---|
| 4271 | FIM1_4842 | QGN17635.1 | CP015060.1 | 91417 | 58702 | 52341 | 85723 | 42335 | 45678 | 113113 | 58758 | 48554 |
| 4272 | FIM1_4843 | QGN17636.1 | CP015060.1 | 30 | 10 | 4 | 12 | 1 | 5 | 79 | 7 | 1 |
| 4273 | FIM1_4844 | QGN17637.1 | CP015060.1 | 647 | 220 | 291 | 654 | 58 | 208 | 2848 | 87 | 196 |
| 4274 | FIM1_4845 | QGN17638.1 | CP015060.1 | 1542 | 297 | 359 | 1258 | 150 | 366 | 2872 | 330 | 316 |
| 4275 | FIM1_4846 | QGN17639.1 | CP015060.1 | 2609 | 114 | 606 | 2120 | 101 | 480 | 2188 | 136 | 458 |
| 4276 | FIM1_4847 | QGN17640.1 | CP015060.1 | 4594 | 215 | 411 | 3320 | 167 | 366 | 4516 | 243 | 387 |
| 4277 | FIM1_4848 | QGN17641.1 | CP015060.1 | 12828 | 1658 | 1900 | 10569 | 1082 | 1651 | 16601 | 1810 | 1559 |
| 4278 | FIM1_4849 | QGN17642.1 | CP015060.1 | 14112 | 3359 | 2553 | 11315 | 1955 | 2380 | 16605 | 3745 | 2812 |
| 4279 | FIM1_4850 | QGN17643.1 | CP015060.1 | 1675 | 73 | 343 | 1254 | 70 | 273 | 2332 | 56 | 255 |
| 4280 | FIM1_4851 | QGN17644.1 | CP015060.1 | 1433 | 485 | 783 | 1422 | 353 | 693 | 1582 | 504 | 593 |
| 4281 | FIM1_4852 | QGN17645.1 | CP015060.1 | 770 | 170 | 512 | 588 | 174 | 453 | 673 | 194 | 438 |
| 4282 | FIM1_4853 | QGN17646.1 | CP015060.1 | 1831 | 190 | 277 | 1357 | 145 | 263 | 2377 | 231 | 255 |
| 4283 | FIM1_4854 | QGN17647.1 | CP015060.1 | 11991 | 2739 | 2176 | 9569 | 2079 | 1945 | 9238 | 3103 | 2199 |
| 4284 | FIM1_4855 | QGN17648.1 | CP015060.1 | 4851 | 670 | 754 | 3800 | 461 | 645 | 3645 | 731 | 752 |
| 4285 | FIM1_4856 | QGN17649.1 | CP015060.1 | 230 | 19 | 102 | 200 | 18 | 105 | 327 | 34 | 74 |
| 4286 | FIM1_4857 | QGN17650.1 | CP015060.1 | 155 | 29 | 24 | 134 | 19 | 15 | 187 | 33 | 30 |
| 4287 | FIM1_4858 | QGN17651.1 | CP015060.1 | 2230 | 2738 | 1696 | 1537 | 1861 | 1566 | 558 | 2822 | 1564 |
| 4288 | FIM1_4859 | QGN17652.1 | CP015060.1 | 6230 | 864 | 1374 | 4723 | 675 | 1156 | 5944 | 948 | 1094 |
| 4289 | FIM1_4860 | QGN17653.1 | CP015060.1 | 1899 | 276 | 571 | 1602 | 235 | 515 | 2501 | 283 | 464 |
| 4290 | FIM1_4861 | QGN17654.1 | CP015060.1 | 1898 | 484 | 797 | 1791 | 376 | 683 | 1980 | 569 | 677 |
| 4291 | FIM1_4862 | QGN17655.1 | CP015060.1 | 3013 | 1453 | 1393 | 2588 | 1104 | 1355 | 2072 | 1526 | 1170 |
| 4292 | FIM1_4863 | QGN17656.1 | CP015060.1 | 8719 | 7745 | 3986 | 6973 | 5429 | 3566 | 2531 | 8110 | 3887 |
| 4293 | FIM1_4864 | QGN17657.1 | CP015060.1 | 115 | 43 | 67 | 92 | 29 | 67 | 51 | 38 | 53 |
| 4294 | FIM1_4865 | QGN17658.1 | CP015060.1 | 2599 | 410 | 636 | 2116 | 275 | 561 | 2176 | 411 | 530 |
| 4295 | FIM1_4866 | QGN17659.1 | CP015060.1 | 5054 | 355 | 1688 | 3897 | 229 | 1490 | 8038 | 380 | 1487 |
| 4296 | FIM1_4867 | QGN17660.1 | CP015060.1 | 5484 | 1714 | 1343 | 4450 | 1401 | 1312 | 3034 | 1756 | 1302 |
| 4297 | FIM1_4868 | QGN17661.1 | CP015060.1 | 2624 | 988 | 1188 | 2536 | 783 | 1193 | 2349 | 1068 | 982 |
| 4298 | FIM1_4869 | QGN17662.1 | CP015060.1 | 2566 | 1525 | 2174 | 2539 | 1206 | 2216 | 2611 | 1627 | 2133 |
| 4299 | FIM1_4870 | QGN17663.1 | CP015060.1 | 1933 | 537 | 542 | 1336 | 364 | 502 | 969 | 637 | 484 |
| 4300 | FIM1_4871 | QGN17664.1 | CP015060.1 | 1389 | 259 | 271 | 1099 | 182 | 276 | 2021 | 248 | 271 |
| 4301 | FIM1_4872 | QGN17665.1 | CP015060.1 | 1489 | 103 | 178 | 1145 | 101 | 153 | 1555 | 119 | 150 |
| 4302 | FIM1_4873 | QGN17666.1 | CP015060.1 | 2665 | 1527 | 1249 | 2138 | 1036 | 1146 | 1404 | 1627 | 1089 |
| 4303 | FIM1_4874 | QGN17667.1 | CP015060.1 | 7399 | 2604 | 2287 | 5478 | 2136 | 2203 | 3300 | 2961 | 2241 |
| 4304 | FIM1_4875 | QGN17668.1 | CP015060.1 | 4686 | 307 | 1234 | 3856 | 178 | 1195 | 8830 | 311 | 1101 |
| 4305 | FIM1_4876 | QGN17669.1 | CP015060.1 | 75 | 18 | 56 | 87 | 16 | 63 | 124 | 27 | 62 |
| 4306 | FIM1_4877 | QGN17670.1 | CP015060.1 | 5051 | 507 | 742 | 3421 | 366 | 640 | 5139 | 530 | 629 |
| 4307 | FIM1_4878 | QGN17671.1 | CP015060.1 | 31680 | 19506 | 17238 | 28614 | 16196 | 16407 | 30454 | 19976 | 16786 |
| 4308 | FIM1_4879 | QGN17672.1 | CP015060.1 | 6615 | 1966 | 2324 | 5018 | 1423 | 2135 | 5379 | 2142 | 1910 |
| 4309 | FIM1_4880 | QGN17673.1 | CP015060.1 | 12531 | 4199 | 3880 | 10114 | 2983 | 3510 | 11305 | 4347 | 3616 |
| 4310 | FIM1_4881 | QGN17674.1 | CP015060.1 | 8219 | 1481 | 1798 | 6210 | 1143 | 1504 | 6922 | 1565 | 1486 |
| 4311 | FIM1_4882 | QGN17675.1 | CP015060.1 | 1506 | 18 | 15 | 530 | 31 | 36 | 321 | 14 | 13 |
| 4312 | FIM1_4883 | QGN17676.1 | CP015060.1 | 11729 | 2007 | 2795 | 8877 | 1443 | 2680 | 11956 | 2072 | 2483 |
| 4313 | FIM1_4884 | QGN17677.1 | CP015060.1 | 6483 | 1374 | 2080 | 4785 | 958 | 1989 | 6434 | 1549 | 1870 |
| 4314 | FIM1_4885 | QGN17678.1 | CP015060.1 | 2941 | 430 | 832 | 2503 | 292 | 813 | 3311 | 420 | 735 |
| 4315 | FIM1_4886 | QGN17679.1 | CP015060.1 | 610 | 135 | 254 | 527 | 75 | 255 | 556 | 131 | 205 |
| 4316 | FIM1_4887 | QGN17680.1 | CP015060.1 | 884 | 221 | 437 | 691 | 154 | 418 | 721 | 229 | 375 |
| 4317 | FIM1_4888 | QGN17681.1 | CP015060.1 | 2433 | 345 | 933 | 2342 | 282 | 794 | 2771 | 398 | 805 |
| 4318 | FIM1_4889 | QGN17682.1 | CP015060.1 | 940 | 152 | 572 | 919 | 96 | 559 | 1596 | 175 | 434 |

| | | | | | | | | | | | | |
|---|---|---|---|---|---|---|---|---|---|---|---|---|
| 4319 | FIM1_4890 | QGN17683.1 | CP015060.1 | 1579 | 344 | 565 | 1234 | 235 | 567 | 1643 | 326 | 461 |
| 4320 | FIM1_4891 | QGN17684.1 | CP015060.1 | 2405 | 741 | 869 | 2083 | 624 | 774 | 1899 | 841 | 708 |
| 4321 | FIM1_4892 | QGN17685.1 | CP015060.1 | 17293 | 3950 | 3544 | 14263 | 2798 | 3200 | 16225 | 4162 | 3322 |
| 4322 | FIM1_4893 | QGN17686.1 | CP015060.1 | 31616 | 8643 | 7412 | 26097 | 6364 | 6762 | 27247 | 9217 | 7147 |
| 4323 | FIM1_4894 | QGN17687.1 | CP015060.1 | 3572 | 693 | 1462 | 2965 | 531 | 1395 | 3589 | 784 | 1272 |
| 4324 | FIM1_4895 | QGN17688.1 | CP015060.1 | 6257 | 1119 | 1534 | 5209 | 853 | 1310 | 4705 | 1189 | 1331 |
| 4325 | FIM1_4896 | QGN17689.1 | CP015060.1 | 6970 | 494 | 1149 | 5499 | 386 | 994 | 6541 | 550 | 834 |
| 4326 | FIM1_4897 | QGN17690.1 | CP015060.1 | 4096 | 566 | 854 | 3252 | 428 | 784 | 3931 | 580 | 824 |
| 4327 | FIM1_4898 | QGN17691.1 | CP015060.1 | 3011 | 505 | 894 | 2595 | 381 | 853 | 3453 | 563 | 680 |
| 4328 | FIM1_4899 | QGN17692.1 | CP015060.1 | 9500 | 6940 | 7819 | 9114 | 5129 | 7439 | 7857 | 7498 | 7070 |
| 4329 | FIM1_4900 | QGN17693.1 | CP015060.1 | 8219 | 4165 | 4731 | 8442 | 3029 | 4257 | 6288 | 4417 | 4190 |
| 4330 | FIM1_4901 | QGN17694.1 | CP015060.1 | 883 | 202 | 381 | 1043 | 122 | 353 | 1225 | 258 | 313 |
| 4331 | FIM1_4902 | QGN17695.1 | CP015060.1 | 3089 | 427 | 814 | 2732 | 367 | 733 | 2538 | 499 | 714 |
| 4332 | FIM1_4903 | QGN17696.1 | CP015060.1 | 3634 | 491 | 883 | 3114 | 434 | 765 | 4271 | 557 | 740 |
| 4333 | FIM1_4904 | QGN17697.1 | CP015060.1 | 1623 | 744 | 1796 | 1507 | 644 | 1383 | 1003 | 838 | 1336 |
| 4334 | FIM1_4905 | QGN17698.1 | CP015060.1 | 2072 | 174 | 459 | 1685 | 141 | 402 | 2125 | 172 | 344 |
| 4335 | FIM1_4906 | QGN17699.1 | CP015060.1 | 4042 | 1015 | 1430 | 3252 | 687 | 1134 | 3154 | 958 | 1053 |
| 4336 | FIM1_4907 | QGN17700.1 | CP015060.1 | 4957 | 2801 | 1500 | 3445 | 1738 | 1336 | 1231 | 2894 | 1370 |
| 4337 | FIM1_4908 | QGN17701.1 | CP015060.1 | 981 | 348 | 552 | 946 | 308 | 601 | 718 | 405 | 458 |
| 4338 | FIM1_4909 | QGN17702.1 | CP015060.1 | 5335 | 2564 | 2366 | 4519 | 1928 | 2237 | 3738 | 2805 | 2167 |
| 4339 | FIM1_4910 | QGN17703.1 | CP015060.1 | 2336 | 428 | 663 | 1793 | 313 | 581 | 1314 | 464 | 461 |
| 4340 | FIM1_4911 | QGN17704.1 | CP015060.1 | 4211 | 803 | 1185 | 3069 | 635 | 1165 | 2019 | 902 | 1006 |
| 4341 | FIM1_4912 | QGN17705.1 | CP015060.1 | 2275 | 743 | 906 | 1675 | 398 | 825 | 2708 | 889 | 855 |
| 4342 | FIM1_4913 | QGN17706.1 | CP015060.1 | 2739 | 2853 | 1238 | 1801 | 1771 | 1087 | 966 | 2951 | 1197 |
| 4343 | FIM1_4914 | QGN17707.1 | CP015060.1 | 5526 | 2509 | 1804 | 4190 | 1497 | 1636 | 5593 | 2721 | 1584 |
| 4344 | FIM1_4915 | QGN17708.1 | CP015060.1 | 8003 | 1019 | 1693 | 5934 | 702 | 1517 | 9466 | 1126 | 1431 |
| 4345 | FIM1_4916 | QGN17709.1 | CP015060.1 | 1375 | 2075 | 1195 | 892 | 1181 | 1045 | 210 | 2225 | 1206 |
| 4346 | FIM1_4917 | QGN17710.1 | CP015060.1 | 4962 | 854 | 1225 | 4082 | 673 | 1104 | 4721 | 907 | 1052 |
| 4347 | FIM1_4918 | QGN17711.1 | CP015060.1 | 431 | 143 | 124 | 334 | 97 | 122 | 386 | 156 | 146 |
| 4348 | FIM1_4919 | QGN17712.1 | CP015060.1 | 2627 | 1384 | 1198 | 2086 | 1227 | 1110 | 1068 | 1630 | 1140 |
| 4349 | FIM1_4920 | QGN17713.1 | CP015060.1 | 1422 | 86 | 346 | 1075 | 66 | 348 | 2285 | 74 | 280 |
| 4350 | FIM1_4921 | QGN17714.1 | CP015060.1 | 19366 | 9022 | 8450 | 15296 | 7238 | 8123 | 14797 | 9417 | 8316 |
| 4351 | FIM1_4922 | QGN17715.1 | CP015060.1 | 2345 | 907 | 934 | 1870 | 648 | 880 | 1372 | 961 | 863 |
| 4352 | FIM1_4923 | QGN17716.1 | CP015060.1 | 1696 | 489 | 491 | 1377 | 387 | 500 | 1943 | 537 | 511 |
| 4353 | FIM1_4924 | QGN17717.1 | CP015060.1 | 1899 | 590 | 2686 | 1423 | 992 | 2483 | 2428 | 683 | 2373 |
| 4354 | FIM1_4925 | QGN17718.1 | CP015060.1 | 594 | 144 | 93 | 410 | 113 | 112 | 575 | 173 | 126 |
| 4355 | FIM1_4926 | QGN17719.1 | CP015060.1 | 4045 | 823 | 1230 | 3025 | 569 | 1112 | 3617 | 878 | 1078 |
| 4356 | FIM1_4927 | QGN17720.1 | CP015060.1 | 1983 | 1635 | 1149 | 1331 | 1067 | 1055 | 488 | 1793 | 989 |
| 4357 | FIM1_4928 | QGN17721.1 | CP015060.1 | 1435 | 1432 | 907 | 829 | 851 | 895 | 302 | 1519 | 822 |
| 4358 | FIM1_4929 | QGN17722.1 | CP015060.1 | 1488 | 332 | 830 | 1116 | 237 | 779 | 1430 | 353 | 741 |
| 4359 | FIM1_4931 | QGN17723.1 | CP015060.1 | 11113 | 3210 | 3227 | 9802 | 2303 | 2855 | 11001 | 3476 | 2737 |
| 4360 | FIM1_4932 | QGN17724.1 | CP015060.1 | 20074 | 2822 | 2905 | 15039 | 1922 | 2660 | 24477 | 3045 | 2500 |
| 4361 | FIM1_4933 | QGN17725.1 | CP015060.1 | 4018 | 1006 | 1517 | 3013 | 767 | 1433 | 4112 | 1040 | 1295 |
| 4362 | FIM1_4935 | QGN17726.1 | CP015060.1 | 5263 | 511 | 824 | 4215 | 342 | 667 | 8758 | 552 | 641 |
| 4363 | FIM1_4936 | QGN17727.1 | CP015060.1 | 4914 | 565 | 1198 | 4017 | 351 | 1055 | 5803 | 585 | 995 |
| 4364 | FIM1_4937 | QGN17728.1 | CP015060.1 | 21072 | 4029 | 7786 | 16978 | 2691 | 7234 | 24853 | 4240 | 6867 |
| 4365 | FIM1_4938 | QGN17729.1 | CP015060.1 | 3111 | 933 | 883 | 2429 | 762 | 850 | 1841 | 1124 | 924 |
| 4366 | FIM1_4939 | QGN17730.1 | CP015060.1 | 1817 | 577 | 1056 | 1363 | 468 | 1045 | 1519 | 673 | 925 |

| | | | | | | | | | | | |
|---|---|---|---|---|---|---|---|---|---|---|---|
| 4367 | FIM1_4940 | QGN17731.1 | CP015060.1 | 2335 | 1876 | 1243 | 1691 | 1247 | 1142 | 747 | 2050 | 1120 |
| 4368 | FIM1_4941 | QGN17732.1 | CP015060.1 | 16646 | 8780 | 7222 | 14865 | 6346 | 6358 | 14997 | 8950 | 6722 |
| 4369 | FIM1_4942 | QGN17733.1 | CP015060.1 | 5522 | 763 | 1002 | 3906 | 620 | 934 | 3911 | 799 | 946 |
| 4370 | FIM1_4943 | QGN17734.1 | CP015060.1 | 6974 | 1002 | 2417 | 4834 | 600 | 2187 | 6309 | 1037 | 2167 |
| 4371 | FIM1_4944 | QGN17735.1 | CP015060.1 | 5681 | 618 | 758 | 4006 | 362 | 740 | 4876 | 620 | 706 |
| 4372 | FIM1_4945 | QGN17736.1 | CP015060.1 | 19490 | 5349 | 6583 | 16755 | 4015 | 6071 | 21157 | 5616 | 5984 |
| 4373 | FIM1_4946 | QGN17737.1 | CP015060.1 | 11258 | 2320 | 2383 | 8580 | 1812 | 2273 | 9430 | 2540 | 2200 |
| 4374 | FIM1_4947 | QGN17738.1 | CP015060.1 | 4174 | 902 | 1715 | 3225 | 677 | 1569 | 4562 | 1008 | 1419 |
| 4375 | FIM1_4948 | QGN17739.1 | CP015060.1 | 1355 | 227 | 445 | 1170 | 164 | 414 | 1328 | 223 | 405 |
| 4376 | FIM1_4949 | QGN17740.1 | CP015060.1 | 3508 | 775 | 1681 | 2808 | 587 | 1448 | 3442 | 791 | 1315 |
| 4377 | FIM1_4950 | QGN17741.1 | CP015060.1 | 5305 | 1847 | 3511 | 4368 | 1393 | 3110 | 4956 | 1876 | 3029 |
| 4378 | FIM1_4951 | QGN17742.1 | CP015060.1 | 9319 | 1359 | 1987 | 7201 | 1018 | 1926 | 9252 | 1412 | 1671 |
| 4379 | FIM1_4952 | QGN17743.1 | CP015060.1 | 10882 | 9738 | 5479 | 8047 | 6785 | 4820 | 3010 | 10682 | 5005 |
| 4380 | FIM1_4953 | QGN17744.1 | CP015060.1 | 3737 | 3770 | 1620 | 2354 | 2295 | 1451 | 773 | 3881 | 1459 |
| 4381 | FIM1_4954 | QGN17745.1 | CP015060.1 | 320 | 153 | 284 | 239 | 151 | 248 | 89 | 197 | 244 |
| 4382 | FIM1_4955 | QGN17746.1 | CP015060.1 | 19058 | 9828 | 8406 | 14828 | 7539 | 7505 | 14695 | 10516 | 7842 |
| 4383 | FIM1_4956 | QGN17747.1 | CP015060.1 | 3392 | 1270 | 632 | 2538 | 613 | 543 | 5616 | 1350 | 564 |
| 4384 | FIM1_4957 | QGN17748.1 | CP015060.1 | 1107 | 645 | 319 | 961 | 319 | 255 | 1120 | 672 | 291 |
| 4385 | FIM1_4958 | QGN17749.1 | CP015060.1 | 614 | 22 | 158 | 657 | 24 | 156 | 1749 | 19 | 123 |
| 4386 | FIM1_4959 | QGN17750.1 | CP015060.1 | 4 | 9 | 9 | 2 | 21 | 35 | 5 | 6 | 3 |
| 4387 | FIM1_4960 | QGN17751.1 | CP015060.1 | 9336 | 2896 | 3359 | 7954 | 2165 | 3218 | 7702 | 3291 | 3344 |
| 4388 | FIM1_4961 | QGN17752.1 | CP015060.1 | 7736 | 3083 | 2252 | 6096 | 1768 | 2147 | 5325 | 3512 | 2227 |
| 4389 | FIM1_4962 | QGN17753.1 | CP015060.1 | 1439 | 803 | 571 | 1235 | 495 | 575 | 1387 | 892 | 468 |
| 4390 | FIM1_4963 | QGN17754.1 | CP015060.1 | 1237 | 389 | 525 | 1037 | 288 | 501 | 1474 | 427 | 439 |
| 4391 | FIM1_4964 | QGN17755.1 | CP015060.1 | 449 | 47 | 53 | 327 | 26 | 53 | 388 | 47 | 42 |
| 4392 | FIM1_4965 | QGN17756.1 | CP015060.1 | 19096 | 6748 | 6171 | 15463 | 5377 | 5692 | 14521 | 7016 | 5794 |
| 4393 | FIM1_4966 | QGN17757.1 | CP015060.1 | 6023 | 2058 | 3814 | 4936 | 1333 | 3496 | 6831 | 2119 | 3385 |
| 4394 | FIM1_4967 | QGN17758.1 | CP015060.1 | 4880 | 929 | 1362 | 3823 | 542 | 1352 | 6256 | 935 | 1204 |
| 4395 | FIM1_4968 | QGN17759.1 | CP015060.1 | 3527 | 1845 | 1715 | 2482 | 1270 | 1372 | 917 | 1860 | 1441 |
| 4396 | FIM1_4969 | QGN17760.1 | CP015060.1 | 22239 | 9776 | 11427 | 19524 | 6617 | 10081 | 21904 | 9828 | 11042 |
| 4397 | FIM1_4970 | QGN17761.1 | CP015060.1 | 36273 | 13081 | 11925 | 28301 | 9324 | 10812 | 29067 | 13757 | 10558 |
| 4398 | FIM1_4971 | QGN17762.1 | CP015060.1 | 21129 | 2582 | 4409 | 16594 | 1920 | 3825 | 23532 | 2766 | 3894 |
| 4399 | FIM1_4972 | QGN17763.1 | CP015060.1 | 8089 | 896 | 1544 | 6974 | 669 | 1351 | 8194 | 899 | 1348 |
| 4400 | FIM1_4973 | QGN17764.1 | CP015060.1 | 5419 | 847 | 1837 | 4421 | 562 | 1737 | 3922 | 796 | 1751 |
| 4401 | FIM1_4974 | QGN17765.1 | CP015060.1 | 1019 | 255 | 442 | 825 | 207 | 417 | 1196 | 332 | 419 |
| 4402 | FIM1_4975 | QGN17766.1 | CP015060.1 | 35 | 3 | 34 | 17 | 3 | 39 | 35 | 2 | 29 |
| 4403 | FIM1_4976 | QGN17767.1 | CP015060.1 | 23464 | 6349 | 6570 | 18837 | 4764 | 6218 | 23071 | 6738 | 5999 |
| 4404 | FIM1_4977 | QGN17768.1 | CP015060.1 | 2426 | 2375 | 1199 | 1971 | 1748 | 1078 | 371 | 2623 | 1183 |
| 4405 | FIM1_4978 | QGN17769.1 | CP015060.1 | 1495 | 379 | 1103 | 1246 | 258 | 835 | 1139 | 361 | 860 |
| 4406 | FIM1_4979 | QGN17770.1 | CP015060.1 | 37459 | 13325 | 14284 | 30464 | 9031 | 12570 | 36538 | 13333 | 13145 |
| 4407 | FIM1_4980 | QGN17771.1 | CP015060.1 | 1743 | 161 | 560 | 1148 | 128 | 489 | 1272 | 176 | 473 |
| 4408 | FIM1_4981 | QGN17772.1 | CP015060.1 | 31158 | 14794 | 13035 | 28265 | 10166 | 11320 | 28967 | 14894 | 12018 |
| 4409 | FIM1_4982 | QGN17773.1 | CP015060.1 | 2729 | 585 | 857 | 2006 | 452 | 853 | 2629 | 618 | 746 |
| 4410 | FIM1_4983 | QGN17774.1 | CP015060.1 | 2418 | 508 | 831 | 2047 | 369 | 759 | 2333 | 526 | 749 |
| 4411 | FIM1_4984 | QGN17775.1 | CP015060.1 | 5824 | 317 | 1047 | 4718 | 275 | 898 | 9084 | 396 | 882 |
| 4412 | FIM1_4985 | QGN17776.1 | CP015060.1 | 8309 | 1326 | 2241 | 6804 | 953 | 2107 | 10431 | 1437 | 1862 |
| 4413 | FIM1_4986 | QGN17777.1 | CP015060.1 | 11272 | 2560 | 3709 | 9317 | 1855 | 3534 | 14194 | 2737 | 3346 |
| 4414 | FIM1_4987 | QGN17778.1 | CP015060.1 | 4464 | 1232 | 3013 | 3847 | 853 | 2725 | 4900 | 1246 | 2664 |

| | | | | | | | | | | | | |
|---|---|---|---|---|---|---|---|---|---|---|---|---|
| 4415 | FIM1_4988 | QGN17779.1 | CP015060.1 | 375 | 81 | 132 | 356 | 54 | 140 | 417 | 82 | 133 |
| 4416 | FIM1_4989 | QGN17780.1 | CP015060.1 | 23434 | 6301 | 4028 | 19067 | 4283 | 3870 | 28668 | 6745 | 3508 |
| 4417 | FIM1_4990 | QGN17781.1 | CP015060.1 | 5 | 1 | 1 | 8 | 0 | 3 | 2 | 0 | 2 |
| 4418 | FIM1_4991 | QGN17782.1 | CP015060.1 | 10927 | 2477 | 3177 | 9173 | 1809 | 2913 | 11055 | 2892 | 2770 |
| 4419 | FIM1_4992 | QGN17783.1 | CP015060.1 | 1907 | 602 | 1349 | 1664 | 432 | 1183 | 2289 | 664 | 1083 |
| 4420 | FIM1_4993 | QGN17784.1 | CP015060.1 | 9506 | 1636 | 2006 | 6953 | 1351 | 1862 | 7885 | 1772 | 1820 |
| 4421 | FIM1_4994 | QGN17785.1 | CP015060.1 | 2 | 0 | 1 | 1 | 1 | 0 | 2 | 1 | 0 |
| 4422 | FIM1_4995 | QGN17786.1 | CP015060.1 | 3277 | 1488 | 921 | 2586 | 918 | 799 | 5329 | 1601 | 830 |
| 4423 | FIM1_4996 | QGN17787.1 | CP015060.1 | 1607 | 227 | 374 | 1238 | 152 | 397 | 2183 | 211 | 355 |
| 4424 | FIM1_4997 | QGN17788.1 | CP015060.1 | 3481 | 809 | 1542 | 2836 | 644 | 1478 | 3939 | 828 | 1294 |
| 4425 | FIM1_4998 | QGN17789.1 | CP015060.1 | 6855 | 1396 | 2043 | 5676 | 806 | 1753 | 10506 | 1616 | 1693 |
| 4426 | FIM1_4999 | QGN17790.1 | CP015060.1 | 7899 | 3510 | 2962 | 6203 | 2676 | 2878 | 4439 | 3889 | 2831 |
| 4427 | FIM1_5000 | QGN17791.1 | CP015060.1 | 2549 | 288 | 327 | 2228 | 197 | 322 | 3257 | 297 | 334 |
| 4428 | FIM1_5001 | QGN17792.1 | CP015060.1 | 3336 | 502 | 485 | 2574 | 236 | 399 | 6372 | 548 | 338 |
| 4429 | FIM1_5002 | QGN17793.1 | CP015060.1 | 7242 | 794 | 1491 | 5289 | 614 | 1353 | 8841 | 832 | 1197 |
| 4430 | FIM1_5003 | QGN17794.1 | CP015060.1 | 2820 | 543 | 1231 | 2314 | 490 | 1014 | 2041 | 531 | 973 |
| 4431 | FIM1_5004 | QGN17795.1 | CP015060.1 | 6463 | 2922 | 2365 | 5125 | 2082 | 2196 | 2873 | 3085 | 2253 |
| 4432 | FIM1_5005 | QGN17796.1 | CP015060.1 | 32806 | 20567 | 17732 | 29801 | 16951 | 17462 | 33411 | 21056 | 17845 |
| 4433 | FIM1_5006 | QGN17797.1 | CP015060.1 | 427 | 126 | 273 | 294 | 79 | 267 | 566 | 89 | 216 |
| 4434 | FIM1_5007 | QGN17798.1 | CP015060.1 | 12090 | 1989 | 1774 | 9144 | 1358 | 1672 | 10566 | 2080 | 1693 |
| 4435 | FIM1_5008 | QGN17799.1 | CP015060.1 | 11959 | 3040 | 3464 | 10251 | 2134 | 2978 | 11527 | 3276 | 2916 |
| 4436 | FIM1_5009 | QGN17800.1 | CP015060.1 | 123 | 9 | 44 | 106 | 15 | 30 | 102 | 13 | 38 |
| 4437 | FIM1_5010 | QGN17801.1 | CP015060.1 | 3131 | 605 | 896 | 2343 | 482 | 791 | 3013 | 645 | 812 |
| 4438 | FIM1_5011 | QGN17802.1 | CP015060.1 | 412 | 112 | 186 | 339 | 104 | 148 | 311 | 128 | 129 |
| 4439 | FIM1_5012 | QGN17803.1 | CP015060.1 | 1160 | 555 | 1164 | 1155 | 418 | 1168 | 1001 | 641 | 980 |
| 4440 | FIM1_5013 | QGN17804.1 | CP015060.1 | 3291 | 638 | 1145 | 2746 | 520 | 1100 | 2950 | 731 | 962 |
| 4441 | FIM1_5014 | QGN17805.1 | CP015060.1 | 4298 | 1031 | 2938 | 3751 | 748 | 2612 | 3438 | 1120 | 2221 |
| 4442 | FIM1_5015 | QGN17806.1 | CP015060.1 | 7138 | 1934 | 2527 | 6437 | 1567 | 2419 | 5537 | 2003 | 2315 |
| 4443 | FIM1_5016 | QGN17807.1 | CP015060.1 | 3025 | 256 | 473 | 2588 | 135 | 395 | 4282 | 225 | 341 |
| 4444 | FIM1_5017 | QGN17808.1 | CP015060.1 | 8865 | 1660 | 2232 | 7687 | 1166 | 2009 | 7664 | 1796 | 1903 |
| 4445 | FIM1_5018 | QGN17809.1 | CP015060.1 | 875 | 175 | 340 | 799 | 146 | 361 | 1074 | 207 | 315 |
| 4446 | FIM1_5019 | QGN17810.1 | CP015060.1 | 5042 | 736 | 824 | 3830 | 512 | 711 | 4618 | 810 | 751 |
| 4447 | FIM1_5020 | QGN17811.1 | CP015060.1 | 5146 | 680 | 771 | 3315 | 423 | 697 | 4129 | 740 | 677 |
| 4448 | FIM1_5021 | QGN17812.1 | CP015060.1 | 2369 | 188 | 544 | 1819 | 163 | 479 | 2948 | 220 | 431 |
| 4449 | FIM1_5022 | QGN17813.1 | CP015060.1 | 8746 | 1432 | 1998 | 6655 | 1021 | 1739 | 10173 | 1447 | 1683 |
| 4450 | FIM1_5023 | QGN17814.1 | CP015060.1 | 4341 | 1612 | 1595 | 3588 | 1338 | 1459 | 3621 | 1785 | 1453 |
| 4451 | FIM1_5024 | QGN17815.1 | CP015060.1 | 7589 | 3133 | 2690 | 5678 | 2233 | 2607 | 5052 | 3430 | 2540 |
| 4452 | FIM1_5025 | QGN17816.1 | CP015060.1 | 8738 | 1276 | 2638 | 7652 | 1662 | 2523 | 8833 | 1558 | 2225 |
| 4453 | FIM1_5026 | QGN17817.1 | CP015060.1 | 6265 | 2093 | 1040 | 4514 | 1114 | 949 | 6749 | 2223 | 927 |
| 4454 | FIM1_5027 | QGN17818.1 | CP015060.1 | 6857 | 4198 | 2322 | 5178 | 3044 | 2264 | 3433 | 4359 | 2628 |
| 4455 | FIM1_5028 | QGN17819.1 | CP015060.1 | 11044 | 2526 | 2864 | 8134 | 1791 | 2550 | 10228 | 2772 | 2518 |
| 4456 | FIM1_5029 | QGN17820.1 | CP015060.1 | 7401 | 1656 | 2517 | 5506 | 1185 | 2462 | 6971 | 1766 | 2167 |
| 4457 | FIM1_5030 | QGN17821.1 | CP015060.1 | 6220 | 2365 | 4130 | 5307 | 1658 | 3861 | 5317 | 2511 | 3672 |
| 4458 | FIM1_5031 | QGN17822.1 | CP015060.1 | 9849 | 6540 | 3484 | 7758 | 4885 | 3229 | 6629 | 6828 | 3676 |
| 4459 | FIM1_5032 | QGN17823.1 | CP015060.1 | 3722 | 3558 | 1560 | 2582 | 1976 | 1424 | 903 | 3684 | 1474 |
| 4460 | FIM1_5033 | QGN17824.1 | CP015060.1 | 5011 | 927 | 1851 | 4016 | 695 | 1517 | 4857 | 1019 | 1491 |
| 4461 | FIM1_5034 | QGN17825.1 | CP015060.1 | 6595 | 2217 | 6075 | 5841 | 1276 | 5487 | 9099 | 2408 | 5434 |
| 4462 | FIM1_5035 | QGN17826.1 | CP015060.1 | 3309 | 672 | 998 | 2588 | 504 | 798 | 2519 | 665 | 832 |

| | | | | | | | | | | | | |
|---|---|---|---|---|---|---|---|---|---|---|---|---|
| 4463 | FIM1_5036 | QGN17827.1 | CP015060.1 | 2155 | 908 | 1234 | 2158 | 796 | 1142 | 1771 | 1079 | 1028 |
| 4464 | FIM1_5037 | QGN17828.1 | CP015060.1 | 3661 | 814 | 1217 | 3018 | 565 | 1125 | 4002 | 870 | 960 |
| 4465 | FIM1_5038 | QGN17829.1 | CP015060.1 | 8809 | 2554 | 2866 | 7027 | 1942 | 2743 | 7884 | 2745 | 2723 |
| 4466 | FIM1_5039 | QGN17830.1 | CP015060.1 | 4447 | 746 | 1285 | 3663 | 527 | 1170 | 5729 | 681 | 1073 |
| 4467 | FIM1_5040 | QGN17831.1 | CP015060.1 | 25908 | 15504 | 13840 | 24200 | 13223 | 13403 | 22784 | 16110 | 13343 |
| 4468 | FIM1_5041 | QGN17832.1 | CP015060.1 | 1363 | 281 | 373 | 1111 | 151 | 313 | 1553 | 321 | 281 |
| 4469 | FIM1_5042 | QGN17833.1 | CP015060.1 | 57258 | 15196 | 14612 | 43268 | 9987 | 13149 | 58109 | 15479 | 13220 |
| 4470 | FIM1_5043 | QGN17834.1 | CP015060.1 | 2050 | 553 | 881 | 2027 | 434 | 751 | 1820 | 641 | 711 |
| 4471 | FIM1_5044 | QGN17835.1 | CP015060.1 | 774 | 133 | 277 | 636 | 128 | 277 | 702 | 168 | 207 |
| 4472 | FIM1_5045 | QGN17836.1 | CP015060.1 | 7079 | 976 | 3079 | 6896 | 944 | 2676 | 5570 | 973 | 2454 |
| 4473 | FIM1_5046 | QGN17837.1 | CP015060.1 | 16246 | 1694 | 1827 | 13648 | 1173 | 1692 | 23073 | 1803 | 1508 |
| 4474 | FIM1_5047 | QGN17838.1 | CP015060.1 | 4210 | 1693 | 1070 | 3201 | 1382 | 989 | 3578 | 1706 | 1014 |
| 4475 | FIM1_5048 | QGN17839.1 | CP015060.1 | 956 | 337 | 765 | 831 | 258 | 779 | 883 | 392 | 677 |
| 4476 | FIM1_5049 | QGN17840.1 | CP015060.1 | 5928 | 1616 | 2123 | 4763 | 1193 | 1912 | 5860 | 1775 | 1877 |
| 4477 | FIM1_5050 | QGN17841.1 | CP015060.1 | 7982 | 899 | 1593 | 6265 | 708 | 1415 | 9743 | 919 | 1350 |
| 4478 | FIM1_5051 | QGN17842.1 | CP015060.1 | 91 | 5 | 17 | 105 | 7 | 17 | 70 | 11 | 8 |
| 4479 | FIM1_5052 | QGN17843.1 | CP015060.1 | 1254 | 671 | 737 | 1093 | 427 | 666 | 914 | 684 | 702 |
| 4480 | FIM1_5053 | QGN17844.1 | CP015060.1 | 2813 | 1055 | 1133 | 2491 | 769 | 1101 | 2017 | 1146 | 1061 |
| 4481 | FIM1_5054 | QGN17845.1 | CP015060.1 | 1326 | 352 | 652 | 987 | 310 | 597 | 1153 | 480 | 522 |
| 4482 | FIM1_5055 | QGN17846.1 | CP015060.1 | 8285 | 2716 | 3012 | 6850 | 2140 | 2796 | 5970 | 3053 | 2647 |
| 4483 | FIM1_5056 | QGN17847.1 | CP015060.1 | 3964 | 616 | 1178 | 2943 | 422 | 1243 | 4428 | 716 | 1094 |
| 4484 | FIM1_5057 | QGN17848.1 | CP015060.1 | 9038 | 9659 | 4366 | 6120 | 6363 | 4008 | 1774 | 10303 | 4175 |
| 4485 | FIM1_5058 | QGN17849.1 | CP015060.1 | 3251 | 1183 | 2931 | 2205 | 1209 | 2845 | 2294 | 1286 | 2433 |
| 4486 | FIM1_5059 | QGN17850.1 | CP015060.1 | 1971 | 406 | 634 | 1484 | 306 | 570 | 1647 | 463 | 555 |
| 4487 | FIM1_5060 | QGN17851.1 | CP015060.1 | 29 | 6 | 8 | 22 | 5 | 2 | 24 | 10 | 4 |
| 4488 | FIM1_5061 | QGN17852.1 | CP015060.1 | 2513 | 1866 | 1648 | 1970 | 1211 | 1580 | 2416 | 1963 | 1540 |
| 4489 | FIM1_5062 | QGN17853.1 | CP015060.1 | 1739 | 246 | 611 | 1500 | 152 | 555 | 2025 | 250 | 479 |
| 4490 | FIM1_5063 | QGN17854.1 | CP015060.1 | 3503 | 1508 | 1651 | 3082 | 1197 | 1577 | 3047 | 1614 | 1550 |
| 4491 | FIM1_5064 | QGN17855.1 | CP015060.1 | 1886 | 594 | 717 | 1547 | 370 | 622 | 1138 | 662 | 524 |
| 4492 | FIM1_5065 | QGN17856.1 | CP015060.1 | 13426 | 2832 | 3008 | 9525 | 1995 | 2645 | 10882 | 2871 | 2677 |
| 4493 | FIM1_5066 | QGN17857.1 | CP015060.1 | 1298 | 230 | 417 | 1076 | 204 | 360 | 1421 | 289 | 344 |
| 4494 | FIM1_5067 | QGN17858.1 | CP015060.1 | 8538 | 2829 | 2740 | 7122 | 2216 | 2600 | 5699 | 3179 | 2498 |
| 4495 | FIM1_5068 | QGN17859.1 | CP015060.1 | 1862 | 670 | 965 | 1584 | 518 | 894 | 1778 | 688 | 918 |
| 4496 | FIM1_5069 | QGN17860.1 | CP015060.1 | 3883 | 107 | 113 | 3160 | 36 | 108 | 6277 | 99 | 70 |
| 4497 | FIM1_5070 | QGN17861.1 | CP015060.1 | 129 | 28 | 76 | 104 | 14 | 73 | 95 | 30 | 75 |
| 4498 | FIM1_5071 | QGN17862.1 | CP015060.1 | 8799 | 4900 | 7763 | 8012 | 3348 | 6889 | 11142 | 5009 | 7080 |
| 4499 | FIM1_5072 | QGN17863.1 | CP015060.1 | 402 | 34 | 170 | 306 | 28 | 166 | 384 | 56 | 134 |
| 4500 | FIM1_5073 | QGN17864.1 | CP015060.1 | 1226 | 291 | 639 | 1191 | 265 | 597 | 826 | 327 | 575 |
| 4501 | FIM1_5074 | QGN17865.1 | CP015060.1 | 1415 | 342 | 505 | 1301 | 262 | 462 | 1706 | 336 | 453 |
| 4502 | FIM1_5075 | QGN17866.1 | CP015060.1 | 6649 | 375 | 763 | 5180 | 288 | 694 | 8037 | 409 | 691 |
| 4503 | FIM1_5076 | QGN17867.1 | CP015060.1 | 5187 | 2869 | 3790 | 5302 | 2071 | 3502 | 5375 | 3032 | 3504 |
| 4504 | FIM1_5077 | QGN17868.1 | CP015060.1 | 6558 | 1156 | 1960 | 5933 | 889 | 1796 | 7457 | 1227 | 1718 |
| 4505 | FIM1_5078 | QGN17869.1 | CP015060.1 | 1444 | 194 | 399 | 1062 | 133 | 320 | 1626 | 202 | 347 |
| 4506 | FIM1_5079 | QGN17870.1 | CP015060.1 | 12024 | 4453 | 3939 | 11001 | 3676 | 3772 | 10503 | 5001 | 4304 |
| 4507 | FIM1_5080 | QGN17871.1 | CP015060.1 | 3056 | 569 | 970 | 2220 | 451 | 903 | 2642 | 639 | 813 |
| 4508 | FIM1_5081 | QGN17872.1 | CP015060.1 | 3745 | 1046 | 1416 | 3160 | 700 | 1319 | 4324 | 1080 | 1215 |
| 4509 | FIM1_5082 | QGN17873.1 | CP015060.1 | 928 | 483 | 507 | 888 | 354 | 522 | 918 | 475 | 508 |
| 4510 | FIM1_5083 | QGN17874.1 | CP015060.1 | 14561 | 4022 | 4873 | 11302 | 2707 | 4461 | 12889 | 4409 | 4365 |

| | | | | | | | | | | | | |
|---|---|---|---|---|---|---|---|---|---|---|---|---|
| 4511 | FIM1_5084 | QGN17875.1 | CP015060.1 | 1346 | 806 | 777 | 912 | 451 | 700 | 538 | 855 | 703 |
| 4512 | FIM1_5085 | QGN17876.1 | CP015060.1 | 8572 | 814 | 1461 | 5798 | 719 | 1258 | 7779 | 982 | 1270 |
| 4513 | FIM1_5086 | QGN17877.1 | CP015060.1 | 8835 | 1751 | 2487 | 6749 | 1319 | 2212 | 7989 | 1815 | 1979 |
| 4514 | FIM1_5087 | QGN17878.1 | CP015060.1 | 1979 | 774 | 1991 | 1796 | 588 | 2030 | 2162 | 922 | 1849 |
| 4515 | FIM1_5088 | QGN17879.1 | CP015060.1 | 24551 | 7857 | 9122 | 20698 | 4973 | 8648 | 29408 | 8121 | 8190 |
| 4516 | FIM1_5089 | QGN17880.1 | CP015060.1 | 4382 | 1479 | 2736 | 3572 | 1164 | 2494 | 3729 | 1604 | 2403 |
| 4517 | FIM1_5090 | QGN17881.1 | CP015060.1 | 1845 | 143 | 311 | 1436 | 91 | 276 | 2238 | 186 | 309 |
| 4518 | FIM1_5091 | QGN17882.1 | CP015060.1 | 932 | 158 | 576 | 951 | 100 | 435 | 833 | 164 | 469 |
| 4519 | FIM1_5092 | QGN17883.1 | CP015060.1 | 11 | 6 | 5 | 10 | 3 | 5 | 6 | 1 | 2 |
| 4520 | FIM1_5093 | QGN17884.1 | CP015060.1 | 3163 | 678 | 1362 | 2321 | 407 | 1135 | 3249 | 730 | 1067 |
| 4521 | FIM1_5094 | QGN17885.1 | CP015060.1 | 143061 | 89594 | 80554 | 127375 | 70435 | 72535 | 155398 | 88268 | 79584 |
| 4522 | FIM1_5095 | QGN17886.1 | CP015060.1 | 1049 | 353 | 516 | 915 | 263 | 439 | 758 | 411 | 439 |
| 4523 | FIM1_5096 | QGN17887.1 | CP015060.1 | 2746 | 1439 | 1072 | 2153 | 899 | 1055 | 1706 | 1576 | 971 |
| 4524 | FIM1_5097 | QGN17888.1 | CP015060.1 | 26537 | 8486 | 7478 | 22309 | 6363 | 6796 | 26455 | 8715 | 6885 |
| 4525 | FIM1_5098 | QGN17889.1 | CP015060.1 | 1053 | 171 | 227 | 804 | 130 | 179 | 1409 | 181 | 198 |
| 4526 | FIM1_5099 | QGN17890.1 | CP015060.1 | 3060 | 1563 | 1649 | 2296 | 1211 | 1650 | 2000 | 1742 | 1492 |
| 4527 | FIM1_5100 | QGN17891.1 | CP015060.1 | 3105 | 566 | 950 | 2406 | 444 | 847 | 3070 | 669 | 792 |
| 4528 | FIM1_5101 | QGN17892.1 | CP015060.1 | 2962 | 465 | 922 | 2429 | 374 | 897 | 3312 | 516 | 764 |
| 4529 | FIM1_5102 | QGN17893.1 | CP015060.1 | 10429 | 1412 | 1632 | 9091 | 928 | 1339 | 17684 | 1461 | 1293 |
| 4530 | FIM1_5103 | QGN17894.1 | CP015060.1 | 888 | 263 | 489 | 883 | 217 | 452 | 1020 | 291 | 395 |
| 4531 | FIM1_5104 | QGN17895.1 | CP015060.1 | 1704 | 458 | 613 | 1541 | 353 | 560 | 1500 | 489 | 526 |
| 4532 | FIM1_5105 | QGN17896.1 | CP015060.1 | 5187 | 1855 | 2055 | 4703 | 1393 | 1914 | 5145 | 2068 | 1847 |
| 4533 | FIM1_5106 | QGN17897.1 | CP015060.1 | 3354 | 265 | 713 | 2762 | 167 | 657 | 4776 | 279 | 562 |
| 4534 | FIM1_5107 | QGN17898.1 | CP015060.1 | 7670 | 4077 | 3495 | 6822 | 3029 | 3151 | 7560 | 4203 | 3353 |
| 4535 | FIM1_5108 | QGN17899.1 | CP015060.1 | 2606 | 717 | 939 | 2232 | 529 | 728 | 1759 | 789 | 746 |
| 4536 | FIM1_5109 | QGN17900.1 | CP015060.1 | 2448 | 904 | 811 | 1663 | 691 | 715 | 1713 | 973 | 804 |
| 4537 | FIM1_5110 | QGN17901.1 | CP015060.1 | 4765 | 849 | 1150 | 3737 | 624 | 1109 | 4971 | 883 | 985 |
| 4538 | FIM1_5111 | QGN17902.1 | CP015060.1 | 5153 | 1291 | 1527 | 3826 | 1092 | 1443 | 4021 | 1458 | 1426 |
| 4539 | FIM1_5112 | QGN17903.1 | CP015060.1 | 2115 | 552 | 817 | 1732 | 430 | 729 | 2262 | 577 | 643 |
| 4540 | FIM1_5113 | QGN17904.1 | CP015060.1 | 377 | 128 | 122 | 307 | 84 | 138 | 338 | 140 | 113 |
| 4541 | FIM1_5114 | QGN17905.1 | CP015060.1 | 928 | 497 | 412 | 591 | 288 | 385 | 481 | 520 | 340 |
| 4542 | FIM1_5115 | QGN17906.1 | CP015060.1 | 2829 | 1734 | 1365 | 2504 | 1209 | 1239 | 2038 | 1862 | 1292 |
| 4543 | FIM1_5116 | QGN17907.1 | CP015060.1 | 2957 | 1117 | 1266 | 2432 | 780 | 1260 | 2590 | 1197 | 1211 |
| 4544 | FIM1_5117 | QGN17908.1 | CP015060.1 | 1876 | 348 | 477 | 1354 | 266 | 501 | 1238 | 390 | 487 |
| 4545 | FIM1_5118 | QGN17909.1 | CP015060.1 | 4487 | 2439 | 2812 | 4320 | 1750 | 2699 | 4270 | 2648 | 2624 |
| 4546 | FIM1_5119 | QGN17910.1 | CP015060.1 | 2536 | 764 | 1436 | 2323 | 558 | 1300 | 2703 | 811 | 1180 |
| 4547 | FIM1_5120 | QGN17911.1 | CP015060.1 | 4980 | 805 | 1122 | 3510 | 456 | 1011 | 4824 | 874 | 966 |
| 4548 | FIM1_5121 | QGN17912.1 | CP015060.1 | 2614 | 1578 | 2658 | 2630 | 1324 | 2602 | 2444 | 1684 | 2346 |
| 4549 | FIM1_5122 | QGN17913.1 | CP015060.1 | 21024 | 2426 | 3120 | 16163 | 1572 | 2609 | 23439 | 2739 | 2531 |
| 4550 | FIM1_5123 | QGN17914.1 | CP015060.1 | 7699 | 4370 | 4665 | 6726 | 2841 | 4146 | 4988 | 3402 | 4369 |
| 4551 | FIM1_5124 | QGN17915.1 | CP015060.1 | 2122 | 605 | 819 | 1620 | 529 | 787 | 2403 | 703 | 847 |
| 4552 | FIM1_5125 | QGN17916.1 | CP015060.1 | 4919 | 2572 | 2883 | 2975 | 1812 | 2511 | 2204 | 2774 | 2450 |
| 4553 | FIM1_5126 | QGN17917.1 | CP015060.1 | 4587 | 1138 | 1387 | 3827 | 816 | 1226 | 4367 | 1194 | 1183 |
| 4554 | FIM1_5127 | QGN17918.1 | CP015060.1 | 12738 | 3521 | 3801 | 10560 | 1967 | 3286 | 16706 | 3638 | 3250 |
| 4555 | FIM1_5128 | QGN17919.1 | CP015060.1 | 9567 | 2202 | 3098 | 7345 | 1402 | 2753 | 10265 | 2497 | 2649 |
| 4556 | FIM1_5129 | QGN17920.1 | CP015060.1 | 3628 | 960 | 1102 | 3120 | 738 | 926 | 2514 | 1044 | 940 |
| 4557 | FIM1_5130 | QGN17921.1 | CP015060.1 | 6029 | 1317 | 2414 | 5152 | 757 | 2164 | 8370 | 1462 | 2087 |
| 4558 | FIM1_5131 | QGN17922.1 | CP015060.1 | 8787 | 606 | 2321 | 6120 | 192 | 2092 | 9871 | 599 | 1588 |

| | | | | | | | | | | | | |
|---|---|---|---|---|---|---|---|---|---|---|---|---|
| 4559 | FIM1_5132 | QGN17923.1 | CP015060.1 | 686 | 125 | 181 | 566 | 93 | 159 | 661 | 150 | 144 |
| 4560 | FIM1_5133 | QGN17924.1 | CP015060.1 | 5061 | 632 | 1394 | 4117 | 488 | 1276 | 5471 | 728 | 1201 |
| 4561 | FIM1_5134 | QGN17925.1 | CP015060.1 | 867 | 466 | 599 | 706 | 367 | 567 | 591 | 583 | 570 |
| 4562 | FIM1_5135 | QGN17926.1 | CP015060.1 | 2630 | 1006 | 1238 | 2414 | 809 | 1196 | 2109 | 1096 | 1137 |
| 4563 | FIM1_5136 | QGN17927.1 | CP015060.1 | 4549 | 1697 | 2715 | 4052 | 1172 | 2558 | 5394 | 1788 | 2332 |
| 4564 | FIM1_5137 | QGN17928.1 | CP015060.1 | 2379 | 580 | 1106 | 1860 | 459 | 994 | 2315 | 595 | 922 |
| 4565 | FIM1_5138 | QGN17929.1 | CP015060.1 | 5522 | 1677 | 2084 | 5729 | 1187 | 1834 | 5920 | 1789 | 1735 |
| 4566 | FIM1_5139 | QGN17930.1 | CP015060.1 | 9267 | 1870 | 3506 | 8427 | 1384 | 3082 | 10431 | 2078 | 3163 |
| 4567 | FIM1_5140 | QGN17931.1 | CP015060.1 | 7007 | 2382 | 2634 | 6034 | 1162 | 2439 | 7348 | 2563 | 2347 |
| 4568 | FIM1_5141 | QGN17932.1 | CP015060.1 | 12348 | 1114 | 1680 | 9168 | 1093 | 1528 | 10430 | 1282 | 1525 |
| 4569 | FIM1_5142 | QGN17933.1 | CP015060.1 | 5138 | 784 | 1550 | 3869 | 641 | 1440 | 6681 | 753 | 1231 |
| 4570 | FIM1_5143 | QGN17934.1 | CP015060.1 | 5890 | 1660 | 2045 | 4714 | 1297 | 1816 | 5939 | 1779 | 1851 |
| 4571 | FIM1_5144 | QGN17935.1 | CP015060.1 | 4688 | 760 | 1852 | 3980 | 702 | 1769 | 3950 | 859 | 1566 |
| 4572 | FIM1_5145 | QGN17936.1 | CP015060.1 | 3853 | 1395 | 2419 | 3156 | 811 | 2395 | 5311 | 1512 | 2393 |
| 4573 | FIM1_5146 | QGN17937.1 | CP015060.1 | 1105 | 800 | 494 | 938 | 465 | 479 | 547 | 814 | 452 |
| 4574 | FIM1_5147 | QGN17938.1 | CP015060.1 | 43358 | 29665 | 26233 | 40348 | 24203 | 24279 | 43549 | 30096 | 25532 |
| 4575 | FIM1_5148 | QGN17939.1 | CP015060.1 | 706 | 133 | 402 | 550 | 103 | 343 | 435 | 126 | 424 |
| 4576 | FIM1_5149 | QGN17940.1 | CP015060.1 | 4455 | 412 | 3049 | 3965 | 577 | 2946 | 11722 | 579 | 3346 |
| 4577 | FIM1_5150 | QGN17941.1 | CP015060.1 | 28185 | 8348 | 7591 | 23943 | 6431 | 7245 | 27800 | 9141 | 7321 |
| 4578 | FIM1_5151 | QGN17942.1 | CP015060.1 | 550 | 145 | 182 | 480 | 117 | 174 | 591 | 147 | 194 |
| 4579 | FIM1_5152 | QGN17943.1 | CP015060.1 | 384 | 64 | 322 | 294 | 60 | 292 | 849 | 69 | 240 |
| 4580 | FIM1_5153 | QGN17944.1 | CP015060.1 | 17576 | 4649 | 4695 | 13412 | 3688 | 4558 | 15596 | 5056 | 4379 |
| 4581 | FIM1_5154 | QGN17945.1 | CP015060.1 | 15 | 4 | 5 | 8 | 2 | 4 | 4 | 2 | 2 |
| 4582 | FIM1_5155 | QGN17946.1 | CP015060.1 | 2518 | 569 | 995 | 1749 | 486 | 1009 | 1824 | 621 | 792 |
| 4583 | FIM1_5156 | QGN17947.1 | CP015060.1 | 2220 | 749 | 719 | 1882 | 598 | 756 | 1668 | 873 | 677 |
| 4584 | FIM1_5157 | QGN17948.1 | CP015060.1 | 5364 | 1771 | 2118 | 4428 | 1353 | 1986 | 4725 | 1897 | 1930 |
| 4585 | FIM1_5158 | QGN17949.1 | CP015060.1 | 4542 | 593 | 1029 | 3361 | 469 | 958 | 5117 | 597 | 974 |
| 4586 | FIM1_5159 | QGN17950.1 | CP015060.1 | 2424 | 515 | 854 | 1849 | 374 | 796 | 1797 | 544 | 661 |
| 4587 | FIM1_5160 | QGN17951.1 | CP015060.1 | 24845 | 15780 | 8944 | 19978 | 13178 | 7988 | 12475 | 16827 | 9017 |
| 4588 | FIM1_5161 | QGN17952.1 | CP015060.1 | 7880 | 985 | 1323 | 5952 | 756 | 1282 | 7817 | 1202 | 1270 |
| 4589 | FIM1_5162 | QGN17953.1 | CP015060.1 | 5696 | 2053 | 2351 | 4873 | 1503 | 2228 | 5393 | 2177 | 2062 |
| 4590 | FIM1_5163 | QGN17954.1 | CP015060.1 | 8477 | 1893 | 3377 | 6820 | 1328 | 3054 | 6919 | 2016 | 2987 |
| 4591 | FIM1_5164 | QGN17955.1 | CP015060.1 | 606 | 148 | 162 | 584 | 107 | 129 | 594 | 134 | 146 |
| 4592 | FIM1_5165 | QGN17956.1 | CP015060.1 | 4591 | 497 | 877 | 3511 | 444 | 753 | 4196 | 563 | 693 |
| 4593 | FIM1_5166 | QGN17957.1 | CP015060.1 | 8708 | 1915 | 2683 | 6583 | 1421 | 2555 | 9227 | 1980 | 2446 |
| 4594 | FIM1_5167 | QGN17958.1 | CP015060.1 | 1939 | 347 | 643 | 1387 | 279 | 615 | 1746 | 349 | 499 |
| 4595 | FIM1_5168 | QGN17959.1 | CP015060.1 | 2369 | 299 | 659 | 1797 | 215 | 628 | 2479 | 281 | 509 |
| 4596 | FIM1_5169 | QGN17960.1 | CP015060.1 | 366 | 148 | 222 | 360 | 123 | 219 | 371 | 152 | 217 |
| 4597 | FIM1_5170 | QGN17961.1 | CP015060.1 | 5572 | 1030 | 1701 | 4416 | 783 | 1694 | 6176 | 1083 | 1601 |
| 4598 | FIM1_5171 | QGN17962.1 | CP015060.1 | 2496 | 559 | 728 | 2183 | 409 | 613 | 2887 | 627 | 606 |
| 4599 | FIM1_5172 | QGN17963.1 | CP015060.1 | 12653 | 7042 | 6012 | 11270 | 5646 | 5690 | 11322 | 7457 | 5701 |
| 4600 | FIM1_5173 | QGN17964.1 | CP015060.1 | 4259 | 848 | 1302 | 3342 | 684 | 1163 | 5053 | 921 | 1144 |
| 4601 | FIM1_5174 | QGN17965.1 | CP015060.1 | 10938 | 2697 | 3739 | 7867 | 1749 | 3342 | 9714 | 2751 | 3419 |
| 4602 | FIM1_5175 | QGN17966.1 | CP015060.1 | 6955 | 2171 | 2632 | 5888 | 1266 | 2530 | 6874 | 2394 | 2597 |
| 4603 | FIM1_5176 | QGN17967.1 | CP015060.1 | 618 | 770 | 410 | 438 | 451 | 361 | 467 | 739 | 373 |
| 4604 | FIM1_5177 | QGN17968.1 | CP015060.1 | 2126 | 2208 | 1262 | 1420 | 1287 | 1166 | 412 | 2409 | 1271 |
| 4605 | FIM1_5178 | QGN17969.1 | CP015060.1 | 2190 | 887 | 941 | 1772 | 643 | 862 | 2062 | 1017 | 837 |
| 4606 | FIM1_5179 | QGN17970.1 | CP015060.1 | 7902 | 6321 | 3436 | 5820 | 4464 | 3210 | 2198 | 6825 | 3353 |

| | | | | | | | | | | | | |
|---|---|---|---|---|---|---|---|---|---|---|---|---|
| 4607 | FIM1_5180 | QGN17971.1 | CP015060.1 | 1999 | 450 | 591 | 1594 | 327 | 621 | 1885 | 498 | 548 |
| 4608 | FIM1_5181 | QGN17972.1 | CP015060.1 | 10029 | 3277 | 4727 | 8473 | 1967 | 4449 | 12176 | 3339 | 4223 |
| 4609 | FIM1_5182 | QGN17973.1 | CP015060.1 | 1922 | 721 | 1353 | 1904 | 617 | 1251 | 1814 | 827 | 1156 |
| 4610 | FIM1_5183 | QGN17974.1 | CP015060.1 | 10909 | 4380 | 4164 | 8512 | 3686 | 3945 | 6887 | 5056 | 3961 |
| 4611 | FIM1_5184 | QGN17975.1 | CP015060.1 | 1757 | 563 | 608 | 1253 | 460 | 611 | 1433 | 614 | 526 |
| 4612 | FIM1_5185 | QGN17976.1 | CP015060.1 | 6983 | 2757 | 3000 | 5751 | 1894 | 2872 | 8165 | 3017 | 2622 |
| 4613 | FIM1_5186 | QGN17977.1 | CP015060.1 | 3790 | 889 | 1703 | 3230 | 669 | 1635 | 4235 | 1087 | 1530 |
| 4614 | FIM1_5187 | QGN17978.1 | CP015060.1 | 7140 | 668 | 1486 | 5403 | 417 | 1423 | 10623 | 732 | 1308 |
| 4615 | FIM1_5188 | QGN17979.1 | CP015060.1 | 17431 | 5180 | 6943 | 14459 | 5466 | 6993 | 11758 | 5569 | 6711 |
| 4616 | FIM1_5189 | QGN17980.1 | CP015060.1 | 419 | 134 | 260 | 416 | 108 | 264 | 447 | 170 | 221 |
| 4617 | FIM1_5190 | QGN17981.1 | CP015060.1 | 2555 | 611 | 1090 | 2145 | 454 | 1039 | 2588 | 668 | 897 |
| 4618 | FIM1_5191 | QGN17982.1 | CP015060.1 | 3644 | 938 | 939 | 2458 | 558 | 890 | 3182 | 1108 | 757 |
| 4619 | FIM1_5192 | QGN17983.1 | CP015060.1 | 7521 | 306 | 332 | 3461 | 319 | 310 | 1667 | 293 | 253 |
| 4620 | FIM1_5193 | QGN17984.1 | CP015060.1 | 32029 | 14392 | 14420 | 29620 | 12092 | 13199 | 30018 | 14810 | 13643 |
| 4621 | FIM1_5194 | QGN17985.1 | CP015060.1 | 6687 | 3138 | 2881 | 5324 | 2044 | 2762 | 9469 | 3316 | 2752 |
| 4622 | FIM1_5195 | QGN17986.1 | CP015060.1 | 16587 | 3539 | 3546 | 13261 | 2463 | 3228 | 13047 | 3744 | 3087 |
| 4623 | FIM1_5196 | QGN17987.1 | CP015060.1 | 506 | 105 | 270 | 385 | 87 | 240 | 634 | 112 | 193 |
| 4624 | FIM1_5197 | QGN17988.1 | CP015060.1 | 935 | 319 | 322 | 810 | 227 | 292 | 831 | 320 | 304 |
| 4625 | FIM1_5199 | QGN17989.1 | CP015060.1 | 8299 | 2441 | 2367 | 6394 | 1834 | 2276 | 7772 | 2606 | 2229 |
| 4626 | FIM1_5200 | QGN17990.1 | CP015060.1 | 2290 | 103 | 132 | 1770 | 107 | 129 | 1947 | 128 | 150 |
| 4627 | FIM1_5201 | QGN17991.1 | CP015060.1 | 2592 | 222 | 326 | 1774 | 178 | 308 | 1857 | 268 | 293 |
| 4628 | FIM1_5202 | QGN17992.1 | CP015060.1 | 1446 | 389 | 587 | 1100 | 301 | 547 | 1478 | 403 | 484 |
| 4629 | FIM1_5203 | QGN17993.1 | CP015060.1 | 2082 | 425 | 607 | 1674 | 330 | 535 | 1994 | 471 | 536 |
| 4630 | FIM1_5204 | QGN17994.1 | CP015060.1 | 1935 | 247 | 448 | 1604 | 173 | 409 | 1274 | 249 | 327 |
| 4631 | FIM1_5205 | QGN17995.1 | CP015060.1 | 4234 | 1217 | 1263 | 3537 | 835 | 1218 | 3617 | 1233 | 1204 |
| 4632 | FIM1_5206 | QGN17996.1 | CP015060.1 | 2679 | 660 | 711 | 1924 | 463 | 790 | 2892 | 722 | 712 |
| 4633 | FIM1_5207 | QGN17997.1 | CP015060.1 | 635 | 103 | 284 | 485 | 94 | 252 | 708 | 107 | 172 |
| 4634 | FIM1_5208 | QGN17998.1 | CP015060.1 | 20 | 12 | 11 | 19 | 12 | 10 | 41 | 12 | 14 |
| 4635 | FIM1_5209 | QGN17999.1 | CP015060.1 | 112 | 24 | 30 | 80 | 13 | 36 | 139 | 26 | 26 |
| 4636 | FIM1_4316 | QGN18000.1 | CP015061.1 | 6465 | 4311 | 3409 | 5083 | 3131 | 3386 | 7012 | 4217 | 3534 |
| 4637 | FIM1_4317 | QGN18001.1 | CP015061.1 | 0 | 0 | 0 | 0 | 0 | 0 | 0 | 0 | 0 |
| 4638 | FIM1_4318 | QGN18002.1 | CP015061.1 | 651 | 56 | 172 | 546 | 43 | 160 | 960 | 84 | 147 |
| 4639 | FIM1_4319 | QGN18003.1 | CP015061.1 | 1196 | 398 | 481 | 816 | 277 | 500 | 1310 | 437 | 455 |
| 4640 | FIM1_4320 | QGN18004.1 | CP015061.1 | 11181 | 2584 | 3000 | 8494 | 1811 | 2793 | 13519 | 2799 | 2643 |
| 4641 | FIM1_4321 | QGN18005.1 | CP015061.1 | 3057 | 802 | 1886 | 2721 | 631 | 1760 | 2607 | 869 | 1418 |
| 4642 | FIM1_4322 | QGN18006.1 | CP015061.1 | 3137 | 1482 | 1162 | 2350 | 875 | 1090 | 2293 | 1523 | 1023 |
| 4643 | FIM1_4323 | QGN18007.1 | CP015061.1 | 41084 | 13642 | 12202 | 33294 | 9565 | 11029 | 44532 | 14734 | 11177 |
| 4644 | FIM1_4324 | QGN18008.1 | CP015061.1 | 89 | 12 | 27 | 46 | 9 | 36 | 79 | 12 | 24 |
| 4645 | FIM1_4325 | QGN18009.1 | CP015061.1 | 4959 | 1113 | 1686 | 3644 | 804 | 1609 | 4662 | 1086 | 1473 |
| 4646 | FIM1_4326 | QGN18010.1 | CP015061.1 | 2985 | 931 | 1218 | 2363 | 748 | 1105 | 1697 | 1021 | 1042 |
| 4647 | FIM1_4327 | QGN18011.1 | CP015061.1 | 7229 | 1712 | 1470 | 5624 | 1326 | 1465 | 6154 | 2000 | 1559 |
| 4648 | FIM1_4328 | QGN18012.1 | CP015061.1 | 1578 | 833 | 692 | 1212 | 547 | 692 | 725 | 785 | 647 |
| 4649 | FIM1_4329 | QGN18013.1 | CP015061.1 | 1573 | 227 | 920 | 1119 | 162 | 749 | 1111 | 221 | 595 |
| 4650 | FIM1_4330 | QGN18014.1 | CP015061.1 | 8538 | 1434 | 1731 | 6374 | 1221 | 1540 | 7735 | 1512 | 1507 |
| 4651 | FIM1_4331 | QGN18015.1 | CP015061.1 | 8507 | 5779 | 2100 | 6932 | 3297 | 1813 | 14643 | 5727 | 1789 |
| 4652 | FIM1_4332 | QGN18016.1 | CP015061.1 | 4341 | 1180 | 1419 | 3223 | 810 | 1341 | 4439 | 1209 | 1210 |
| 4653 | FIM1_4333 | QGN18017.1 | CP015061.1 | 5671 | 2082 | 2551 | 4630 | 1437 | 2423 | 5956 | 2286 | 2156 |
| 4654 | FIM1_4334 | QGN18018.1 | CP015061.1 | 1561 | 471 | 767 | 1469 | 386 | 774 | 1672 | 529 | 726 |

| | | | | | | | | | | | | |
|---|---|---|---|---|---|---|---|---|---|---|---|---|
| 4655 | FIM1_4335 | QGN18019.1 | CP015061.1 | 5747 | 1235 | 2739 | 4345 | 836 | 2554 | 6201 | 1230 | 2277 |
| 4656 | FIM1_4336 | QGN18020.1 | CP015061.1 | 2179 | 408 | 884 | 1537 | 420 | 990 | 1669 | 492 | 795 |
| 4657 | FIM1_4337 | QGN18021.1 | CP015061.1 | 3363 | 960 | 1395 | 2699 | 739 | 1283 | 2786 | 1045 | 1174 |
| 4658 | FIM1_4338 | QGN18022.1 | CP015061.1 | 10936 | 3170 | 3657 | 8864 | 2146 | 3497 | 10175 | 3536 | 3246 |
| 4659 | FIM1_4339 | QGN18023.1 | CP015061.1 | 863 | 143 | 595 | 610 | 134 | 574 | 742 | 155 | 437 |
| 4660 | FIM1_4340 | QGN18024.1 | CP015061.1 | 11439 | 7094 | 14095 | 10594 | 5324 | 13387 | 11534 | 7359 | 13866 |
| 4661 | FIM1_4341 | QGN18025.1 | CP015061.1 | 630 | 232 | 342 | 496 | 155 | 367 | 524 | 261 | 395 |
| 4662 | FIM1_4342 | QGN18026.1 | CP015061.1 | 964 | 313 | 588 | 724 | 254 | 629 | 906 | 394 | 600 |
| 4663 | FIM1_4343 | QGN18027.1 | CP015061.1 | 38070 | 25208 | 22693 | 35952 | 20714 | 20918 | 37578 | 25441 | 21807 |
| 4664 | FIM1_4344 | QGN18028.1 | CP015061.1 | 3436 | 3418 | 2023 | 2441 | 2223 | 1972 | 1248 | 3553 | 1780 |
| 4665 | FIM1_4345 | QGN18029.1 | CP015061.1 | 720 | 273 | 618 | 683 | 176 | 599 | 789 | 265 | 616 |
| 4666 | FIM1_4346 | QGN18030.1 | CP015061.1 | 1417 | 1890 | 1144 | 1036 | 1416 | 1194 | 206 | 2008 | 1160 |
| 4667 | FIM1_4347 | QGN18031.1 | CP015061.1 | 4618 | 710 | 1870 | 3528 | 489 | 1778 | 5431 | 791 | 1615 |
| 4668 | FIM1_4348 | QGN18032.1 | CP015061.1 | 7229 | 1355 | 1991 | 5319 | 1076 | 1927 | 6851 | 1513 | 1643 |
| 4669 | FIM1_4349 | QGN18033.1 | CP015061.1 | 1077 | 35 | 61 | 335 | 18 | 74 | 374 | 29 | 44 |
| 4670 | FIM1_4350 | QGN18034.1 | CP015061.1 | 7509 | 1783 | 1720 | 5668 | 1126 | 1489 | 7683 | 1921 | 1507 |
| 4671 | FIM1_4351 | QGN18035.1 | CP015061.1 | 11921 | 4037 | 3896 | 9093 | 2994 | 3568 | 9110 | 4438 | 3545 |
| 4672 | FIM1_4352 | QGN18036.1 | CP015061.1 | 2103 | 1224 | 1102 | 1559 | 930 | 1154 | 1024 | 1169 | 931 |
| 4673 | FIM1_4353 | QGN18037.1 | CP015061.1 | 6246 | 1975 | 1765 | 4402 | 1419 | 1719 | 4012 | 2120 | 1607 |
| 4674 | FIM1_4354 | QGN18038.1 | CP015061.1 | 2122 | 760 | 1891 | 1739 | 512 | 1863 | 2731 | 823 | 1700 |
| 4675 | FIM1_4355 | QGN18039.1 | CP015061.1 | 77 | 4 | 30 | 56 | 9 | 16 | 52 | 10 | 20 |
| 4676 | FIM1_4356 | QGN18040.1 | CP015061.1 | 38184 | 20958 | 22431 | 33174 | 14742 | 20797 | 43125 | 21557 | 21666 |
| 4677 | FIM1_4357 | QGN18041.1 | CP015061.1 | 3094 | 818 | 2121 | 2628 | 616 | 2069 | 3107 | 848 | 1786 |
| 4678 | FIM1_4358 | QGN18042.1 | CP015061.1 | 7060 | 3003 | 4193 | 6192 | 2057 | 4034 | 7957 | 3166 | 4088 |
| 4679 | FIM1_4359 | QGN18043.1 | CP015061.1 | 3837 | 1269 | 1988 | 3335 | 986 | 1909 | 4569 | 1334 | 1864 |
| 4680 | FIM1_4360 | QGN18044.1 | CP015061.1 | 20496 | 6661 | 6362 | 16205 | 4965 | 5780 | 20535 | 6873 | 5907 |
| 4681 | FIM1_4361 | QGN18045.1 | CP015061.1 | 6458 | 4005 | 2033 | 4098 | 2203 | 1945 | 2862 | 4141 | 1878 |
| 4682 | FIM1_4363 | QGN18046.1 | CP015061.1 | 1642 | 1334 | 756 | 1158 | 748 | 689 | 629 | 1468 | 608 |
| 4683 | FIM1_4364 | QGN18047.1 | CP015061.1 | 670 | 227 | 420 | 566 | 191 | 468 | 564 | 252 | 376 |
| 4684 | FIM1_4365 | QGN18048.1 | CP015061.1 | 2121 | 833 | 1102 | 1694 | 630 | 1082 | 1475 | 832 | 976 |
| 4685 | FIM1_4366 | QGN18049.1 | CP015061.1 | 20998 | 6934 | 8182 | 15523 | 4854 | 7265 | 20351 | 7312 | 7516 |
| 4686 | FIM1_4367 | QGN18050.1 | CP015061.1 | 4094 | 1282 | 1403 | 3211 | 830 | 1337 | 3698 | 1329 | 1324 |
| 4687 | FIM1_4368 | QGN18051.1 | CP015061.1 | 3225 | 1091 | 1380 | 2878 | 869 | 1391 | 2697 | 1114 | 1261 |
| 4688 | FIM1_4369 | QGN18052.1 | CP015061.1 | 4748 | 4470 | 2252 | 3232 | 3040 | 2118 | 1339 | 4866 | 2176 |
| 4689 | FIM1_4370 | QGN18053.1 | CP015061.1 | 1279 | 1166 | 670 | 930 | 554 | 720 | 1006 | 1159 | 585 |
| 4690 | FIM1_4371 | QGN18054.1 | CP015061.1 | 7681 | 2345 | 2989 | 5648 | 1409 | 2776 | 7202 | 2357 | 2625 |
| 4691 | FIM1_4372 | QGN18055.1 | CP015061.1 | 4376 | 1986 | 2268 | 4117 | 1961 | 2276 | 3852 | 2274 | 1980 |
| 4692 | FIM1_4373 | QGN18056.1 | CP015061.1 | 19249 | 10140 | 6048 | 14493 | 7408 | 5220 | 8197 | 10795 | 5400 |
| 4693 | FIM1_4374 | QGN18057.1 | CP015061.1 | 2981 | 651 | 1102 | 2316 | 452 | 1026 | 2544 | 600 | 837 |
| 4694 | FIM1_4376 | QGN18058.1 | CP015061.1 | 568 | 180 | 198 | 439 | 143 | 201 | 415 | 204 | 209 |
| 4695 | FIM1_4377 | QGN18059.1 | CP015061.1 | 15610 | 5536 | 4053 | 12472 | 4489 | 3840 | 8305 | 6175 | 3964 |
| 4696 | FIM1_4378 | QGN18060.1 | CP015061.1 | 198 | 142 | 176 | 197 | 90 | 211 | 147 | 152 | 210 |
| 4697 | FIM1_4379 | QGN18061.1 | CP015061.1 | 8185 | 2009 | 4063 | 5879 | 1322 | 4050 | 8813 | 2249 | 3750 |
| 4698 | FIM1_4380 | QGN18062.1 | CP015061.1 | 4671 | 925 | 1213 | 3449 | 728 | 1061 | 4681 | 1039 | 1005 |
| 4699 | FIM1_4381 | QGN18063.1 | CP015061.1 | 3809 | 1192 | 1250 | 2793 | 907 | 1163 | 2701 | 1316 | 1150 |
| 4700 | FIM1_4382 | QGN18064.1 | CP015061.1 | 2051 | 591 | 733 | 1588 | 492 | 796 | 1990 | 690 | 775 |
| 4701 | FIM1_4383 | QGN18065.1 | CP015061.1 | 13314 | 4716 | 4931 | 11383 | 3461 | 4500 | 12488 | 5007 | 4485 |
| 4702 | FIM1_4384 | QGN18066.1 | CP015061.1 | 2776 | 1040 | 960 | 2057 | 431 | 861 | 3953 | 1154 | 829 |

| | | | | | | | | | | | | |
|---|---|---|---|---|---|---|---|---|---|---|---|---|
| 4703 | FIM1_4385 | QGN18067.1 | CP015061.1 | 5915 | 879 | 1530 | 4472 | 654 | 1374 | 6410 | 1002 | 1242 |
| 4704 | FIM1_4386 | QGN18068.1 | CP015061.1 | 27240 | 13628 | 15711 | 25005 | 9437 | 13827 | 29510 | 14157 | 14471 |
| 4705 | FIM1_4387 | QGN18069.1 | CP015061.1 | 64 | 21 | 21 | 55 | 21 | 25 | 44 | 25 | 27 |
| 4706 | FIM1_4388 | QGN18070.1 | CP015061.1 | 6629 | 4309 | 1218 | 5102 | 3737 | 1066 | 3302 | 4503 | 1183 |
| 4707 | FIM1_4389 | QGN18071.1 | CP015061.1 | 796 | 114 | 196 | 748 | 73 | 167 | 951 | 101 | 211 |
| 4708 | FIM1_4390 | QGN18072.1 | CP015061.1 | 279 | 45 | 136 | 204 | 35 | 98 | 203 | 58 | 92 |
| 4709 | FIM1_4391 | QGN18073.1 | CP015061.1 | 19544 | 12924 | 8275 | 15533 | 10419 | 7470 | 10479 | 13591 | 8060 |
| 4710 | FIM1_4392 | QGN18074.1 | CP015061.1 | 6163 | 1617 | 1864 | 6276 | 1531 | 1607 | 6621 | 1606 | 1667 |
| 4711 | FIM1_4393 | QGN18075.1 | CP015061.1 | 290 | 47 | 103 | 290 | 19 | 85 | 438 | 58 | 89 |
| 4712 | FIM1_4394 | QGN18076.1 | CP015061.1 | 1565 | 293 | 557 | 1308 | 264 | 487 | 1467 | 353 | 428 |
| 4713 | FIM1_4396 | QGN18077.1 | CP015061.1 | 5400 | 2095 | 3453 | 4660 | 1309 | 3260 | 6271 | 2272 | 3089 |
| 4714 | FIM1_4397 | QGN18078.1 | CP015061.1 | 3983 | 443 | 1111 | 3856 | 348 | 1143 | 4694 | 464 | 1011 |
| 4715 | FIM1_4398 | QGN18079.1 | CP015061.1 | 1040 | 45 | 80 | 1035 | 52 | 82 | 1387 | 25 | 65 |
| 4716 | FIM1_4399 | QGN18080.1 | CP015061.1 | 34937 | 13091 | 11493 | 28093 | 9292 | 10524 | 32077 | 13713 | 11019 |
| 4717 | FIM1_4400 | QGN18081.1 | CP015061.1 | 2988 | 591 | 1060 | 2598 | 481 | 990 | 2683 | 678 | 875 |
| 4718 | FIM1_4401 | QGN18082.1 | CP015061.1 | 5455 | 2345 | 2207 | 4444 | 1596 | 1994 | 3517 | 2359 | 1950 |
| 4719 | FIM1_4402 | QGN18083.1 | CP015061.1 | 286 | 156 | 151 | 253 | 142 | 116 | 204 | 161 | 119 |
| 4720 | FIM1_4403 | QGN18084.1 | CP015061.1 | 20968 | 3093 | 3286 | 16117 | 1798 | 3113 | 29589 | 3289 | 3151 |
| 4721 | FIM1_4404 | QGN18085.1 | CP015061.1 | 1107 | 207 | 409 | 835 | 202 | 414 | 824 | 235 | 327 |
| 4722 | FIM1_4405 | QGN18086.1 | CP015061.1 | 1382 | 314 | 541 | 1079 | 265 | 518 | 1287 | 316 | 466 |
| 4723 | FIM1_4406 | QGN18087.1 | CP015061.1 | 3246 | 431 | 970 | 2657 | 375 | 827 | 3591 | 475 | 785 |
| 4724 | FIM1_4407 | QGN18088.1 | CP015061.1 | 160 | 16 | 40 | 123 | 11 | 30 | 177 | 27 | 36 |
| 4725 | FIM1_4408 | QGN18089.1 | CP015061.1 | 1481 | 80 | 417 | 1041 | 53 | 388 | 1611 | 103 | 414 |
| 4726 | FIM1_4409 | QGN18090.1 | CP015061.1 | 5811 | 10860 | 2858 | 5104 | 3648 | 2449 | 2284 | 11722 | 3273 |
| 4727 | FIM1_4410 | QGN18091.1 | CP015061.1 | 1918 | 438 | 816 | 1676 | 277 | 685 | 2448 | 447 | 708 |
| 4728 | FIM1_4412 | QGN18092.1 | CP015061.1 | 838 | 118 | 146 | 562 | 60 | 108 | 964 | 126 | 132 |
| 4729 | FIM1_4413 | QGN18093.1 | CP015061.1 | 3479 | 822 | 1392 | 2536 | 703 | 1260 | 2825 | 969 | 1167 |
| 4730 | FIM1_4414 | QGN18094.1 | CP015061.1 | 11028 | 2743 | 2796 | 7909 | 2273 | 2600 | 9135 | 2969 | 2480 |
| 4731 | FIM1_4415 | QGN18095.1 | CP015061.1 | 15556 | 1274 | 2690 | 11828 | 963 | 2438 | 17341 | 1357 | 2307 |
| 4732 | FIM1_4416 | QGN18096.1 | CP015061.1 | 5768 | 842 | 1566 | 4629 | 665 | 1443 | 7843 | 905 | 1427 |
| 4733 | FIM1_4417 | QGN18097.1 | CP015061.1 | 4245 | 806 | 1081 | 2867 | 650 | 952 | 3898 | 872 | 886 |
| 4734 | FIM1_4418 | QGN18098.1 | CP015061.1 | 13924 | 5371 | 5066 | 11406 | 4279 | 4634 | 10263 | 5923 | 4471 |
| 4735 | FIM1_4419 | QGN18099.1 | CP015061.1 | 2487 | 643 | 1169 | 2071 | 420 | 1020 | 2514 | 649 | 925 |
| 4736 | FIM1_4420 | QGN18100.1 | CP015061.1 | 3044 | 487 | 1072 | 2608 | 325 | 1016 | 3278 | 495 | 887 |
| 4737 | FIM1_4421 | QGN18101.1 | CP015061.1 | 1536 | 446 | 786 | 1310 | 342 | 681 | 1403 | 451 | 666 |
| 4738 | FIM1_4422 | QGN18102.1 | CP015061.1 | 22 | 8 | 10 | 11 | 5 | 6 | 34 | 8 | 13 |
| 4739 | FIM1_4423 | QGN18103.1 | CP015061.1 | 3510 | 1233 | 1698 | 3557 | 1078 | 1562 | 2807 | 1253 | 1364 |
| 4740 | FIM1_4424 | QGN18104.1 | CP015061.1 | 2990 | 112 | 787 | 2050 | 312 | 836 | 2388 | 116 | 600 |
| 4741 | FIM1_4425 | QGN18105.1 | CP015061.1 | 1924 | 440 | 527 | 1436 | 366 | 485 | 1768 | 537 | 491 |
| 4742 | FIM1_4426 | QGN18106.1 | CP015061.1 | 1863 | 454 | 576 | 1584 | 387 | 565 | 1737 | 545 | 482 |
| 4743 | FIM1_4427 | QGN18107.1 | CP015061.1 | 11649 | 3521 | 4359 | 9715 | 2394 | 4444 | 14180 | 3732 | 3882 |
| 4744 | FIM1_4428 | QGN18108.1 | CP015061.1 | 22194 | 2838 | 6391 | 19394 | 3226 | 5959 | 27961 | 2994 | 4692 |
| 4745 | FIM1_4429 | QGN18109.1 | CP015061.1 | 6991 | 1524 | 1671 | 5788 | 1116 | 1434 | 6442 | 1599 | 1453 |
| 4746 | FIM1_4430 | QGN18110.1 | CP015061.1 | 564 | 281 | 324 | 485 | 241 | 315 | 530 | 292 | 304 |
| 4747 | FIM1_4431 | QGN18111.1 | CP015061.1 | 7344 | 1052 | 1477 | 6016 | 775 | 1353 | 8759 | 1215 | 1257 |
| 4748 | FIM1_4432 | QGN18112.1 | CP015061.1 | 14386 | 4975 | 4957 | 12383 | 3738 | 4406 | 12892 | 5231 | 4363 |
| 4749 | FIM1_4433 | QGN18113.1 | CP015061.1 | 1229 | 391 | 595 | 958 | 303 | 586 | 980 | 485 | 465 |
| 4750 | FIM1_4434 | QGN18114.1 | CP015061.1 | 4960 | 1106 | 1723 | 3963 | 795 | 1621 | 4703 | 1186 | 1552 |

| | | | | | | | | | | | | |
|---|---|---|---|---|---|---|---|---|---|---|---|---|
| 4751 | FIM1_4435 | QGN18115.1 | CP015061.1 | 13010 | 2307 | 3365 | 9774 | 1744 | 3008 | 14567 | 2580 | 2907 |
| 4752 | FIM1_4436 | QGN18116.1 | CP015061.1 | 4621 | 838 | 1531 | 3610 | 615 | 1261 | 4877 | 895 | 1228 |
| 4753 | FIM1_4437 | QGN18117.1 | CP015061.1 | 1508 | 265 | 575 | 1074 | 243 | 586 | 1370 | 285 | 556 |
| 4754 | FIM1_4438 | QGN18118.1 | CP015061.1 | 4735 | 471 | 1117 | 3362 | 417 | 896 | 5388 | 458 | 910 |
| 4755 | FIM1_4439 | QGN18119.1 | CP015061.1 | 2924 | 990 | 1318 | 2649 | 756 | 1303 | 2666 | 956 | 1236 |
| 4756 | FIM1_4440 | QGN18120.1 | CP015061.1 | 3660 | 328 | 1374 | 3255 | 224 | 1274 | 6219 | 351 | 1208 |
| 4757 | FIM1_4441 | QGN18121.1 | CP015061.1 | 11682 | 1834 | 3050 | 9275 | 1356 | 2880 | 14991 | 1938 | 2892 |
| 4758 | FIM1_4443 | QGN18122.1 | CP015061.1 | 1142 | 94 | 246 | 807 | 88 | 241 | 2350 | 79 | 253 |
| 4759 | FIM1_4444 | QGN18123.1 | CP015061.1 | 57457 | 28896 | 25589 | 50707 | 20684 | 23128 | 56848 | 29561 | 24712 |
| 4760 | FIM1_4445 | QGN18124.1 | CP015061.1 | 3926 | 774 | 1247 | 2872 | 666 | 1129 | 2763 | 872 | 1068 |
| 4761 | FIM1_4446 | QGN18125.1 | CP015061.1 | 11294 | 3153 | 4923 | 9592 | 2067 | 4375 | 13275 | 3287 | 4300 |
| 4762 | FIM1_4447 | QGN18126.1 | CP015061.1 | 4792 | 948 | 1349 | 3599 | 728 | 1200 | 4885 | 1005 | 1219 |
| 4763 | FIM1_4448 | QGN18127.1 | CP015061.1 | 1051 | 155 | 197 | 784 | 151 | 202 | 976 | 145 | 182 |
| 4764 | FIM1_4449 | QGN18128.1 | CP015061.1 | 2335 | 365 | 534 | 2076 | 131 | 471 | 3807 | 399 | 455 |
| 4765 | FIM1_4450 | QGN18129.1 | CP015061.1 | 7950 | 1904 | 3634 | 6604 | 1497 | 3408 | 7712 | 2085 | 3356 |
| 4766 | FIM1_4451 | QGN18130.1 | CP015061.1 | 4206 | 740 | 2552 | 3348 | 652 | 2208 | 6619 | 787 | 1998 |
| 4767 | FIM1_4452 | QGN18131.1 | CP015061.1 | 15517 | 9317 | 9207 | 15070 | 8130 | 9101 | 14113 | 9930 | 8653 |
| 4768 | FIM1_4454 | QGN18132.1 | CP015061.1 | 14438 | 3066 | 3125 | 11995 | 2218 | 2692 | 14358 | 3313 | 2716 |
| 4769 | FIM1_4455 | QGN18133.1 | CP015061.1 | 2498 | 1239 | 1195 | 2105 | 906 | 1170 | 1668 | 1222 | 1133 |
| 4770 | FIM1_4456 | QGN18134.1 | CP015061.1 | 3255 | 495 | 1468 | 2776 | 389 | 1290 | 4990 | 638 | 1207 |
| 4771 | FIM1_4457 | QGN18135.1 | CP015061.1 | 3733 | 1231 | 1899 | 3331 | 1030 | 1732 | 3872 | 1333 | 1573 |
| 4772 | FIM1_4458 | QGN18136.1 | CP015061.1 | 3411 | 743 | 1469 | 2683 | 638 | 1422 | 2590 | 827 | 1374 |
| 4773 | FIM1_4460 | QGN18137.1 | CP015061.1 | 800 | 304 | 665 | 561 | 153 | 562 | 955 | 256 | 467 |
| 4774 | FIM1_4461 | QGN18138.1 | CP015061.1 | 248 | 162 | 256 | 141 | 90 | 301 | 333 | 155 | 216 |
| 4775 | FIM1_4462 | QGN18139.1 | CP015061.1 | 2831 | 741 | 908 | 1939 | 550 | 906 | 2592 | 743 | 806 |
| 4776 | FIM1_4463 | QGN18140.1 | CP015061.1 | 6861 | 437 | 817 | 4918 | 284 | 721 | 6138 | 418 | 563 |
| 4777 | FIM1_4464 | QGN18141.1 | CP015061.1 | 2802 | 456 | 830 | 2265 | 330 | 752 | 2508 | 481 | 743 |
| 4778 | FIM1_4465 | QGN18142.1 | CP015061.1 | 2194 | 763 | 1277 | 2102 | 607 | 1352 | 2328 | 839 | 1069 |
| 4779 | FIM1_4466 | QGN18143.1 | CP015061.1 | 3282 | 744 | 1117 | 2647 | 431 | 1106 | 3373 | 820 | 1057 |
| 4780 | FIM1_4467 | QGN18144.1 | CP015061.1 | 6349 | 2374 | 2422 | 5534 | 1622 | 2290 | 5659 | 2627 | 2125 |
| 4781 | FIM1_4468 | QGN18145.1 | CP015061.1 | 574 | 112 | 254 | 455 | 113 | 255 | 580 | 126 | 239 |
| 4782 | FIM1_4469 | QGN18146.1 | CP015061.1 | 6393 | 3723 | 2551 | 4832 | 2736 | 2409 | 2338 | 3866 | 2516 |
| 4783 | FIM1_4470 | QGN18147.1 | CP015061.1 | 10594 | 2945 | 2371 | 8313 | 1845 | 2420 | 6250 | 3151 | 2374 |
| 4784 | FIM1_4471 | QGN18148.1 | CP015061.1 | 1230 | 586 | 820 | 1263 | 383 | 760 | 1674 | 608 | 731 |
| 4785 | FIM1_4472 | QGN18149.1 | CP015061.1 | 7099 | 1227 | 2060 | 5570 | 907 | 1890 | 7171 | 1237 | 1829 |
| 4786 | FIM1_4473 | QGN18150.1 | CP015061.1 | 1953 | 439 | 675 | 1578 | 366 | 599 | 1706 | 446 | 526 |
| 4787 | FIM1_4474 | QGN18151.1 | CP015061.1 | 24071 | 9224 | 10435 | 22705 | 6598 | 9555 | 29655 | 9998 | 9953 |
| 4788 | FIM1_4475 | QGN18152.1 | CP015061.1 | 1903 | 296 | 1114 | 1805 | 240 | 1051 | 1310 | 313 | 980 |
| 4789 | FIM1_4476 | QGN18153.1 | CP015061.1 | 2322 | 982 | 1523 | 2104 | 782 | 1401 | 1177 | 1128 | 1378 |
| 4790 | FIM1_4477 | QGN18154.1 | CP015061.1 | 5814 | 1289 | 2127 | 5089 | 959 | 2027 | 5157 | 1369 | 1864 |
| 4791 | FIM1_4478 | QGN18155.1 | CP015061.1 | 4180 | 1100 | 1690 | 3638 | 752 | 1737 | 4911 | 1187 | 1578 |
| 4792 | FIM1_4479 | QGN18156.1 | CP015061.1 | 6687 | 1719 | 1804 | 6027 | 1087 | 1644 | 7531 | 1869 | 1680 |
| 4793 | FIM1_4480 | QGN18157.1 | CP015061.1 | 1204 | 236 | 439 | 943 | 168 | 387 | 953 | 281 | 395 |
| 4794 | FIM1_4481 | QGN18158.1 | CP015061.1 | 6384 | 1071 | 1922 | 4743 | 860 | 1788 | 6029 | 1174 | 1529 |
| 4795 | FIM1_4482 | QGN18159.1 | CP015061.1 | 7013 | 19 | 43 | 5010 | 22 | 34 | 13560 | 9 | 22 |
| 4796 | FIM1_4483 | QGN18160.1 | CP015061.1 | 42 | 2 | 9 | 33 | 1 | 3 | 38 | 3 | 6 |
| 4797 | FIM1_4484 | QGN18161.1 | CP015061.1 | 28 | 20 | 11 | 24 | 15 | 31 | 20 | 5 | 7 |
| 4798 | FIM1_4485 | QGN18162.1 | CP015061.1 | 18 | 11 | 4 | 15 | 12 | 6 | 13 | 13 | 7 |

| | | | | | | | | | | | | |
|---|---|---|---|---|---|---|---|---|---|---|---|---|
| 4799 | FIM1_4486 | QGN18163.1 | CP015061.1 | 7521 | 1399 | 891 | 5083 | 765 | 854 | 4375 | 1457 | 1062 |
| 4800 | FIM1_4487 | QGN18164.1 | CP015061.1 | 1207 | 87 | 181 | 787 | 59 | 154 | 2013 | 93 | 147 |
| 4801 | FIM1_4488 | QGN18165.1 | CP015061.1 | 53 | 26 | 21 | 55 | 13 | 25 | 71 | 17 | 32 |
| 4802 | FIM1_4489 | QGN18166.1 | CP015061.1 | 3299 | 826 | 1121 | 2538 | 593 | 1000 | 3780 | 878 | 947 |
| 4803 | FIM1_4490 | QGN18167.1 | CP015061.1 | 22600 | 4555 | 7213 | 21213 | 2095 | 6158 | 45393 | 4786 | 5916 |
| 4804 | FIM1_4491 | QGN18168.1 | CP015061.1 | 3794 | 591 | 665 | 2842 | 343 | 645 | 3692 | 640 | 582 |
| 4805 | FIM1_4492 | QGN18169.1 | CP015061.1 | 1453 | 732 | 627 | 1156 | 476 | 621 | 1315 | 738 | 551 |
| 4806 | FIM1_4493 | QGN18170.1 | CP015061.1 | 29926 | 5034 | 5711 | 22667 | 2640 | 4560 | 28616 | 5241 | 2491 |
| 4807 | FIM1_4494 | QGN18171.1 | CP015061.1 | 102 | 52 | 19 | 86 | 35 | 25 | 84 | 80 | 29 |
| 4808 | FIM1_4495 | QGN18172.1 | CP015061.1 | 3161 | 654 | 845 | 2767 | 422 | 729 | 4828 | 695 | 614 |
| 4809 | FIM1_4496 | QGN18173.1 | CP015061.1 | 1670 | 414 | 565 | 1388 | 307 | 487 | 1678 | 506 | 470 |
| 4810 | FIM1_4497 | QGN18174.1 | CP015061.1 | 7396 | 3635 | 2501 | 6960 | 2421 | 2456 | 8702 | 4067 | 2366 |
| 4811 | FIM1_4498 | QGN18175.1 | CP015061.1 | 29757 | 18952 | 17091 | 27421 | 15828 | 16465 | 28637 | 19552 | 16488 |
| 4812 | FIM1_4499 | QGN18176.1 | CP015061.1 | 7002 | 1461 | 3766 | 5210 | 1077 | 3556 | 8649 | 1645 | 3320 |
| 4813 | FIM1_4500 | QGN18177.1 | CP015061.1 | 8777 | 1607 | 2309 | 7522 | 1180 | 2037 | 10539 | 1730 | 1978 |
| 4814 | FIM1_4501 | QGN18178.1 | CP015061.1 | 2781 | 1197 | 1792 | 1845 | 820 | 1666 | 1229 | 1211 | 1600 |
| 4815 | FIM1_4502 | QGN18179.1 | CP015061.1 | 475 | 112 | 218 | 459 | 120 | 214 | 305 | 128 | 170 |
| 4816 | FIM1_4503 | QGN18180.1 | CP015061.1 | 1722 | 315 | 715 | 1310 | 278 | 616 | 1742 | 326 | 611 |
| 4817 | FIM1_4504 | QGN18181.1 | CP015061.1 | 4506 | 513 | 855 | 3360 | 380 | 778 | 4164 | 569 | 757 |
| 4818 | FIM1_4505 | QGN18182.1 | CP015061.1 | 5291 | 3815 | 2108 | 3603 | 2397 | 1879 | 976 | 4119 | 1905 |
| 4819 | FIM1_4506 | QGN18183.1 | CP015061.1 | 2408 | 1290 | 1250 | 2050 | 992 | 1249 | 1417 | 1323 | 1193 |
| 4820 | FIM1_4507 | QGN18184.1 | CP015061.1 | 5745 | 1803 | 2270 | 4427 | 1387 | 2118 | 5475 | 1971 | 2075 |
| 4821 | FIM1_4508 | QGN18185.1 | CP015061.1 | 2564 | 683 | 1107 | 2113 | 498 | 1006 | 1909 | 698 | 897 |
| 4822 | FIM1_4509 | QGN18186.1 | CP015061.1 | 4598 | 1849 | 2864 | 3867 | 1232 | 2586 | 4211 | 1885 | 2425 |
| 4823 | FIM1_4510 | QGN18187.1 | CP015061.1 | 9744 | 4999 | 4205 | 8373 | 3756 | 3792 | 7998 | 5271 | 3858 |
| 4824 | FIM1_4512 | QGN18188.1 | CP015061.1 | 1961 | 361 | 882 | 1446 | 290 | 849 | 2009 | 413 | 807 |
| 4825 | FIM1_4513 | QGN18189.1 | CP015061.1 | 360 | 155 | 237 | 276 | 124 | 233 | 262 | 179 | 231 |
| 4826 | FIM1_4514 | QGN18190.1 | CP015061.1 | 625 | 129 | 348 | 515 | 109 | 333 | 629 | 137 | 257 |
| 4827 | FIM1_4515 | QGN18191.1 | CP015061.1 | 3045 | 571 | 1236 | 2520 | 394 | 1152 | 3533 | 593 | 1015 |
| 4828 | FIM1_4516 | QGN18192.1 | CP015061.1 | 1074 | 985 | 585 | 671 | 638 | 488 | 177 | 1090 | 557 |
| 4829 | FIM1_4517 | QGN18193.1 | CP015061.1 | 3775 | 288 | 1035 | 2660 | 206 | 845 | 6556 | 318 | 828 |
| 4830 | FIM1_4518 | QGN18194.1 | CP015061.1 | 2085 | 349 | 650 | 1704 | 352 | 690 | 1879 | 400 | 622 |
| 4831 | FIM1_4519 | QGN18195.1 | CP015061.1 | 2627 | 768 | 889 | 2248 | 577 | 853 | 2591 | 754 | 833 |
| 4832 | FIM1_4520 | QGN18196.1 | CP015061.1 | 4952 | 1600 | 1488 | 3967 | 1331 | 1350 | 3706 | 1775 | 1455 |
| 4833 | FIM1_4521 | QGN18197.1 | CP015061.1 | 8078 | 1315 | 2150 | 6280 | 944 | 2015 | 8231 | 1392 | 1857 |
| 4834 | FIM1_4522 | QGN18198.1 | CP015061.1 | 2466 | 117 | 304 | 1692 | 67 | 260 | 2077 | 135 | 209 |
| 4835 | FIM1_4523 | QGN18199.1 | CP015061.1 | 8595 | 2618 | 2701 | 7001 | 1819 | 2371 | 7924 | 2684 | 2185 |
| 4836 | FIM1_4524 | QGN18200.1 | CP015061.1 | 2542 | 1390 | 1638 | 2436 | 900 | 1536 | 2736 | 1479 | 1438 |
| 4837 | FIM1_4525 | QGN18201.1 | CP015061.1 | 623 | 155 | 242 | 598 | 100 | 258 | 608 | 158 | 241 |
| 4838 | FIM1_4526 | QGN18202.1 | CP015061.1 | 13434 | 5142 | 6292 | 11506 | 3276 | 5310 | 13912 | 5434 | 5528 |
| 4839 | FIM1_4527 | QGN18203.1 | CP015061.1 | 43802 | 12058 | 17510 | 38695 | 9271 | 16137 | 44504 | 14034 | 17449 |
| 4840 | FIM1_4528 | QGN18204.1 | CP015061.1 | 638 | 41 | 133 | 378 | 22 | 120 | 604 | 45 | 98 |
| 4841 | FIM1_4529 | QGN18205.1 | CP015061.1 | 1100 | 132 | 304 | 871 | 73 | 297 | 2920 | 114 | 280 |
| 4842 | FIM1_4530 | QGN18206.1 | CP015061.1 | 7295 | 1391 | 2163 | 5422 | 992 | 1992 | 7328 | 1427 | 1765 |
| 4843 | FIM1_4531 | QGN18207.1 | CP015061.1 | 4448 | 1788 | 4892 | 3943 | 1308 | 4452 | 4473 | 1948 | 4304 |
| 4844 | FIM1_4532 | QGN18208.1 | CP015061.1 | 2134 | 1067 | 1622 | 2187 | 745 | 1550 | 2737 | 1173 | 1481 |
| 4845 | FIM1_4533 | QGN18209.1 | CP015061.1 | 16090 | 12222 | 7115 | 12520 | 7769 | 6390 | 6647 | 13066 | 6867 |
| 4846 | FIM1_4534 | QGN18210.1 | CP015061.1 | 1214 | 292 | 828 | 979 | 265 | 780 | 1121 | 341 | 658 |

| | | | | | | | | | | | | |
|---|---|---|---|---|---|---|---|---|---|---|---|---|
| 4847 | FIM1_4535 | QGN18211.1 | CP015061.1 | 2895 | 1512 | 1500 | 2785 | 1213 | 1568 | 2239 | 1703 | 1439 |
| 4848 | FIM1_4536 | QGN18212.1 | CP015061.1 | 6614 | 1223 | 2057 | 4972 | 478 | 1822 | 8790 | 1204 | 1790 |
| 4849 | FIM1_4537 | QGN18213.1 | CP015061.1 | 2362 | 654 | 755 | 1625 | 545 | 678 | 1785 | 728 | 728 |
| 4850 | FIM1_4538 | QGN18214.1 | CP015061.1 | 1032 | 209 | 562 | 736 | 154 | 313 | 1482 | 166 | 328 |
| 4851 | FIM1_4539 | QGN18215.1 | CP015061.1 | 2149 | 659 | 912 | 1659 | 561 | 851 | 2262 | 677 | 790 |
| 4852 | FIM1_4540 | QGN18216.1 | CP015061.1 | 719 | 247 | 730 | 495 | 227 | 709 | 792 | 266 | 504 |
| 4853 | FIM1_4542 | QGN18217.1 | CP015061.1 | 10501 | 2800 | 4173 | 8165 | 1660 | 3936 | 12663 | 2837 | 3474 |
| 4854 | FIM1_4543 | QGN18218.1 | CP015061.1 | 6583 | 6227 | 2907 | 4285 | 3719 | 2579 | 1223 | 6539 | 2673 |
| 4855 | FIM1_4544 | QGN18219.1 | CP015061.1 | 2032 | 164 | 566 | 1624 | 125 | 475 | 2867 | 176 | 454 |
| 4856 | FIM1_4545 | QGN18220.1 | CP015061.1 | 5229 | 2764 | 1731 | 3581 | 1525 | 1515 | 1934 | 2940 | 1582 |
| 4857 | FIM1_4546 | QGN18221.1 | CP015061.1 | 9742 | 6963 | 3474 | 6351 | 4387 | 2946 | 2557 | 7380 | 3234 |
| 4858 | FIM1_4547 | QGN18222.1 | CP015061.1 | 116 | 84 | 97 | 98 | 85 | 66 | 39 | 90 | 83 |
| 4859 | FIM1_4548 | QGN18223.1 | CP015061.1 | 2788 | 702 | 1134 | 2089 | 589 | 1009 | 2906 | 764 | 988 |
| 4860 | FIM1_4549 | QGN18224.1 | CP015061.1 | 6802 | 543 | 2389 | 4540 | 347 | 2241 | 6322 | 507 | 2066 |
| 4861 | FIM1_4550 | QGN18225.1 | CP015061.1 | 193 | 29 | 47 | 108 | 24 | 42 | 142 | 41 | 32 |
| 4862 | FIM1_4551 | QGN18226.1 | CP015061.1 | 269 | 64 | 110 | 278 | 49 | 108 | 269 | 57 | 104 |
| 4863 | FIM1_4552 | QGN18227.1 | CP015061.1 | 2923 | 655 | 964 | 2529 | 439 | 850 | 2873 | 702 | 859 |
| 4864 | FIM1_4553 | QGN18228.1 | CP015061.1 | 1328 | 354 | 459 | 1246 | 323 | 405 | 1034 | 373 | 417 |
| 4865 | FIM1_4554 | QGN18229.1 | CP015061.1 | 7417 | 1004 | 2440 | 5765 | 737 | 2068 | 7139 | 1073 | 1913 |
| 4866 | FIM1_4555 | QGN18230.1 | CP015061.1 | 16169 | 4278 | 4270 | 12089 | 3102 | 3718 | 15372 | 4733 | 3891 |
| 4867 | FIM1_4556 | QGN18231.1 | CP015061.1 | 6028 | 2139 | 2330 | 5181 | 1183 | 2287 | 7368 | 2121 | 2074 |
| 4868 | FIM1_4557 | QGN18232.1 | CP015061.1 | 8866 | 3493 | 4481 | 8045 | 2485 | 4011 | 9098 | 3689 | 4018 |
| 4869 | FIM1_4558 | QGN18233.1 | CP015061.1 | 3953 | 488 | 1801 | 3088 | 312 | 1689 | 5802 | 495 | 1602 |
| 4870 | FIM1_4559 | QGN18234.1 | CP015061.1 | 2045 | 274 | 1216 | 1632 | 218 | 1160 | 2763 | 299 | 1117 |
| 4871 | FIM1_4560 | QGN18235.1 | CP015061.1 | 272 | 39 | 178 | 231 | 40 | 194 | 277 | 45 | 127 |
| 4872 | FIM1_4561 | QGN18236.1 | CP015061.1 | 14449 | 2378 | 3274 | 11244 | 1622 | 3029 | 15988 | 2463 | 2685 |
| 4873 | FIM1_4562 | QGN18237.1 | CP015061.1 | 2732 | 334 | 212 | 1189 | 139 | 195 | 2032 | 294 | 214 |
| 4874 | FIM1_4563 | QGN18238.1 | CP015061.1 | 3710 | 2336 | 1938 | 3437 | 2054 | 1985 | 2123 | 2563 | 1654 |
| 4875 | FIM1_4564 | QGN18239.1 | CP015061.1 | 345 | 127 | 444 | 370 | 133 | 449 | 276 | 150 | 337 |
| 4876 | FIM1_4565 | QGN18240.1 | CP015061.1 | 16347 | 1935 | 4414 | 12619 | 1347 | 4004 | 19200 | 2102 | 4013 |
| 4877 | FIM1_4566 | QGN18241.1 | CP015061.1 | 39221 | 4640 | 14499 | 32358 | 1187 | 13150 | 27540 | 4698 | 9951 |
| 4878 | FIM1_4567 | QGN18242.1 | CP015061.1 | 13198 | 4985 | 4647 | 10864 | 3601 | 4418 | 13273 | 5035 | 4363 |
| 4879 | FIM1_4568 | QGN18243.1 | CP015061.1 | 13755 | 3621 | 4177 | 11445 | 2642 | 3815 | 13771 | 3972 | 3819 |
| 4880 | FIM1_4569 | QGN18244.1 | CP015061.1 | 3904 | 1681 | 2243 | 3141 | 1205 | 2261 | 3964 | 1779 | 1984 |
| 4881 | FIM1_4570 | QGN18245.1 | CP015061.1 | 4250 | 764 | 1169 | 3085 | 596 | 1021 | 4652 | 774 | 1044 |
| 4882 | FIM1_4571 | QGN18246.1 | CP015061.1 | 2548 | 751 | 1051 | 2033 | 607 | 957 | 1865 | 824 | 901 |
| 4883 | FIM1_4572 | QGN18247.1 | CP015061.1 | 16329 | 4881 | 3544 | 13084 | 3683 | 3381 | 11459 | 5526 | 3454 |
| 4884 | FIM1_4573 | QGN18248.1 | CP015061.1 | 8013 | 648 | 1647 | 6484 | 541 | 1543 | 8684 | 757 | 1305 |
| 4885 | FIM1_4574 | QGN18249.1 | CP015061.1 | 1766 | 331 | 609 | 1293 | 244 | 580 | 1455 | 342 | 485 |
| 4886 | FIM1_4575 | QGN18250.1 | CP015061.1 | 12669 | 6843 | 7536 | 11369 | 4875 | 6575 | 13249 | 7056 | 6721 |
| 4887 | FIM1_4576 | QGN18251.1 | CP015061.1 | 3014 | 1296 | 1511 | 2337 | 1043 | 1530 | 2244 | 1393 | 1360 |
| 4888 | FIM1_4577 | QGN18252.1 | CP015061.1 | 1243 | 205 | 423 | 1186 | 113 | 449 | 1481 | 220 | 416 |
| 4889 | FIM1_4578 | QGN18253.1 | CP015061.1 | 879 | 303 | 352 | 724 | 174 | 324 | 826 | 309 | 311 |
| 4890 | FIM1_4579 | QGN18254.1 | CP015061.1 | 1090 | 267 | 304 | 831 | 165 | 221 | 990 | 294 | 265 |
| 4891 | FIM1_4580 | QGN18255.1 | CP015061.1 | 4765 | 3920 | 2481 | 3953 | 2646 | 2184 | 2613 | 4015 | 2340 |
| 4892 | FIM1_4582 | QGN18256.1 | CP015061.1 | 6360 | 994 | 1264 | 4635 | 788 | 1203 | 7351 | 1145 | 1194 |
| 4893 | FIM1_4583 | QGN18257.1 | CP015061.1 | 5657 | 935 | 1233 | 4049 | 627 | 1161 | 5159 | 929 | 1014 |
| 4894 | FIM1_4584 | QGN18258.1 | CP015061.1 | 4400 | 1344 | 1579 | 3569 | 804 | 1548 | 5519 | 1287 | 1288 |

| | | | | | | | | | | | | |
|---|---|---|---|---|---|---|---|---|---|---|---|---|
| 4895 | FIM1_4585 | QGN18259.1 | CP015061.1 | 7428 | 1462 | 1845 | 5714 | 881 | 1662 | 8437 | 1476 | 1551 |
| 4896 | FIM1_4586 | QGN18260.1 | CP015061.1 | 36831 | 12150 | 13041 | 30315 | 8245 | 11990 | 33859 | 12648 | 12214 |
| 4897 | FIM1_4587 | QGN18261.1 | CP015061.1 | 4668 | 3120 | 1687 | 3206 | 2137 | 1516 | 989 | 3386 | 1435 |
| 4898 | FIM1_4588 | QGN18262.1 | CP015061.1 | 5148 | 2175 | 3693 | 4124 | 1513 | 3451 | 4138 | 2397 | 3237 |
| 4899 | FIM1_4589 | QGN18263.1 | CP015061.1 | 3347 | 3582 | 1971 | 2397 | 2251 | 1883 | 1182 | 3745 | 1705 |
| 4900 | FIM1_4590 | QGN18264.1 | CP015061.1 | 1099 | 390 | 539 | 746 | 265 | 468 | 1057 | 474 | 442 |
| 4901 | FIM1_4591 | QGN18265.1 | CP015061.1 | 4816 | 2063 | 2230 | 3810 | 1596 | 2072 | 3480 | 2296 | 2117 |
| 4902 | FIM1_4592 | QGN18266.1 | CP015061.1 | 2239 | 985 | 935 | 1766 | 718 | 877 | 2029 | 1028 | 873 |
| 4903 | FIM1_4593 | QGN18267.1 | CP015061.1 | 1394 | 275 | 798 | 1146 | 214 | 769 | 1770 | 323 | 600 |
| 4904 | FIM1_4594 | QGN18268.1 | CP015061.1 | 3811 | 208 | 533 | 3795 | 243 | 441 | 5459 | 220 | 412 |
| 4905 | FIM1_4595 | QGN18269.1 | CP015061.1 | 11987 | 3651 | 3308 | 9513 | 2647 | 3007 | 8882 | 3846 | 3240 |
| 4906 | FIM1_4596 | QGN18270.1 | CP015061.1 | 5656 | 5176 | 2634 | 3756 | 3104 | 2365 | 2320 | 5505 | 2490 |
| 4907 | FIM1_4597 | QGN18271.1 | CP015061.1 | 4996 | 1356 | 1925 | 4181 | 916 | 1752 | 5688 | 1546 | 1584 |
| 4908 | FIM1_4598 | QGN18272.1 | CP015061.1 | 1014 | 248 | 477 | 779 | 202 | 437 | 1096 | 254 | 367 |
| 4909 | FIM1_4599 | QGN18273.1 | CP015061.1 | 549 | 192 | 229 | 468 | 129 | 205 | 364 | 202 | 181 |
| 4910 | FIM1_4600 | QGN18274.1 | CP015061.1 | 11890 | 3062 | 2932 | 9096 | 2176 | 2661 | 9587 | 3317 | 2680 |
| 4911 | FIM1_4601 | QGN18275.1 | CP015061.1 | 2075 | 428 | 786 | 1801 | 329 | 723 | 1759 | 460 | 658 |
| 4912 | FIM1_4602 | QGN18276.1 | CP015061.1 | 660 | 322 | 501 | 703 | 282 | 499 | 363 | 400 | 384 |
| 4913 | FIM1_4603 | QGN18277.1 | CP015061.1 | 9946 | 2912 | 3587 | 8491 | 2201 | 3093 | 8191 | 3018 | 3235 |
| 4914 | FIM1_4604 | QGN18278.1 | CP015061.1 | 374 | 49 | 167 | 351 | 46 | 158 | 254 | 76 | 101 |
| 4915 | FIM1_4605 | QGN18279.1 | CP015061.1 | 2328 | 799 | 921 | 1886 | 558 | 860 | 2041 | 863 | 818 |
| 4916 | FIM1_4606 | QGN18280.1 | CP015061.1 | 1341 | 581 | 663 | 1036 | 383 | 619 | 1170 | 595 | 574 |
| 4917 | FIM1_4607 | QGN18281.1 | CP015061.1 | 5425 | 1620 | 2041 | 4694 | 1160 | 1834 | 5915 | 1825 | 1757 |
| 4918 | FIM1_4608 | QGN18282.1 | CP015061.1 | 940 | 94 | 363 | 792 | 77 | 280 | 1321 | 84 | 257 |
| 4919 | FIM1_4609 | QGN18283.1 | CP015061.1 | 8451 | 2819 | 3305 | 6282 | 2345 | 2879 | 3456 | 2907 | 2771 |
| 4920 | FIM1_4610 | QGN18284.1 | CP015061.1 | 93 | 1 | 4 | 37 | 3 | 7 | 40 | 2 | 3 |
| 4921 | FIM1_4611 | QGN18285.1 | CP015061.1 | 3 | 0 | 0 | 2 | 0 | 0 | 1 | 0 | 0 |
| 4922 | FIM1_4612 | QGN18286.1 | CP015061.1 | 21704 | 4836 | 6440 | 16830 | 3642 | 5969 | 19921 | 5138 | 5814 |
| 4923 | FIM1_4613 | QGN18287.1 | CP015061.1 | 19046 | 12471 | 12059 | 16358 | 9407 | 11188 | 18532 | 13190 | 11570 |
| 4924 | FIM1_4614 | QGN18288.1 | CP015061.1 | 2085 | 361 | 476 | 1809 | 249 | 399 | 1992 | 387 | 371 |
| 4925 | FIM1_4615 | QGN18289.1 | CP015061.1 | 3886 | 944 | 1161 | 3263 | 628 | 1073 | 4351 | 903 | 1079 |
| 4926 | FIM1_4616 | QGN18290.1 | CP015061.1 | 8685 | 3024 | 2732 | 7061 | 2090 | 2491 | 6265 | 3093 | 2353 |
| 4927 | FIM1_4617 | QGN18291.1 | CP015061.1 | 11838 | 1692 | 3350 | 8905 | 1361 | 2988 | 13249 | 1972 | 2971 |
| 4928 | FIM1_4618 | QGN18292.1 | CP015061.1 | 11184 | 2572 | 4004 | 7901 | 1814 | 3664 | 10026 | 2766 | 3499 |
| 4929 | FIM1_4619 | QGN18293.1 | CP015061.1 | 1965 | 523 | 878 | 1573 | 437 | 818 | 1943 | 615 | 736 |
| 4930 | FIM1_4620 | QGN18294.1 | CP015061.1 | 2364 | 441 | 644 | 2020 | 409 | 583 | 1616 | 594 | 551 |
| 4931 | FIM1_4621 | QGN18295.1 | CP015061.1 | 508 | 93 | 112 | 350 | 88 | 98 | 369 | 114 | 124 |
| 4932 | FIM1_4622 | QGN18296.1 | CP015061.1 | 4767 | 409 | 901 | 3941 | 312 | 835 | 6169 | 416 | 690 |
| 4933 | FIM1_4623 | QGN18297.1 | CP015061.1 | 2996 | 940 | 1384 | 2333 | 635 | 1282 | 2435 | 954 | 1210 |
| 4934 | FIM1_4624 | QGN18298.1 | CP015061.1 | 3072 | 1337 | 2448 | 2717 | 838 | 2490 | 3454 | 1384 | 2160 |
| 4935 | FIM1_4625 | QGN18299.1 | CP015061.1 | 3354 | 614 | 837 | 2607 | 441 | 748 | 3157 | 678 | 678 |
| 4936 | FIM1_4626 | QGN18300.1 | CP015061.1 | 1174 | 151 | 306 | 865 | 125 | 268 | 2170 | 210 | 254 |
| 4937 | FIM1_4627 | QGN18301.1 | CP015061.1 | 6974 | 1176 | 2293 | 5449 | 751 | 2051 | 7985 | 1253 | 1965 |
| 4938 | FIM1_4628 | QGN18302.1 | CP015061.1 | 528 | 86 | 318 | 530 | 81 | 236 | 827 | 90 | 254 |
| 4939 | FIM1_4629 | QGN18303.1 | CP015061.1 | 1658 | 349 | 630 | 1372 | 296 | 632 | 1408 | 372 | 571 |
| 4940 | FIM1_4630 | QGN18304.1 | CP015061.1 | 3402 | 498 | 1063 | 3181 | 395 | 971 | 3709 | 501 | 886 |
| 4941 | FIM1_4631 | QGN18305.1 | CP015061.1 | 4591 | 1146 | 2052 | 4072 | 698 | 1847 | 5675 | 1163 | 1826 |
| 4942 | FIM1_4632 | QGN18306.1 | CP015061.1 | 7833 | 1356 | 1718 | 5518 | 873 | 1546 | 7197 | 1426 | 1489 |

| | | | | | | | | | | | | |
|---|---|---|---|---|---|---|---|---|---|---|---|---|
| 4943 | FIM1_4633 | QGN18307.1 | CP015061.1 | 153 | 32 | 51 | 120 | 11 | 54 | 121 | 21 | 40 |
| 4944 | FIM1_4634 | QGN18308.1 | CP015061.1 | 10317 | 1087 | 3070 | 7439 | 528 | 2811 | 9422 | 1041 | 2711 |
| 4945 | FIM1_4635 | QGN18309.1 | CP015061.1 | 3341 | 497 | 959 | 2499 | 360 | 924 | 4357 | 537 | 952 |
| 4946 | FIM1_4636 | QGN18310.1 | CP015061.1 | 1151 | 340 | 425 | 1003 | 270 | 413 | 1254 | 334 | 436 |
| 4947 | FIM1_4637 | QGN18311.1 | CP015061.1 | 2242 | 615 | 782 | 1994 | 382 | 895 | 2875 | 660 | 756 |
| 4948 | FIM1_4638 | QGN18312.1 | CP015061.1 | 3369 | 602 | 1097 | 2733 | 480 | 1044 | 2893 | 714 | 825 |
| 4949 | FIM1_4639 | QGN18313.1 | CP015061.1 | 9038 | 1153 | 1804 | 6931 | 873 | 1693 | 10550 | 1272 | 1534 |
| 4950 | FIM1_4640 | QGN18314.1 | CP015061.1 | 2126 | 629 | 839 | 1780 | 470 | 799 | 1382 | 713 | 671 |
| 4951 | FIM1_4641 | QGN18315.1 | CP015061.1 | 12218 | 5358 | 5478 | 10886 | 3994 | 5060 | 11385 | 5660 | 4771 |
| 4952 | FIM1_4642 | QGN18316.1 | CP015061.1 | 8568 | 2329 | 2957 | 8249 | 1639 | 2768 | 8809 | 2666 | 2515 |
| 4953 | FIM1_4643 | QGN18317.1 | CP015061.1 | 768 | 367 | 496 | 767 | 264 | 420 | 747 | 424 | 382 |
| 4954 | FIM1_4644 | QGN18318.1 | CP015061.1 | 1814 | 387 | 798 | 1477 | 327 | 752 | 2182 | 429 | 730 |
| 4955 | FIM1_4645 | QGN18319.1 | CP015061.1 | 2867 | 848 | 1057 | 2087 | 635 | 985 | 1912 | 970 | 948 |
| 4956 | FIM1_4646 | QGN18320.1 | CP015061.1 | 3213 | 1225 | 1506 | 2379 | 854 | 1348 | 2222 | 1318 | 1285 |
| 4957 | FIM1_4647 | QGN18321.1 | CP015061.1 | 2950 | 1574 | 1790 | 2444 | 1184 | 1722 | 1997 | 1709 | 1631 |
| 4958 | FIM1_4648 | QGN18322.1 | CP015061.1 | 3974 | 1098 | 1514 | 3345 | 811 | 1402 | 3773 | 1243 | 1295 |
| 4959 | FIM1_4649 | QGN18323.1 | CP015061.1 | 7418 | 1489 | 2165 | 5750 | 1067 | 1961 | 9246 | 1602 | 1822 |
| 4960 | FIM1_4650 | QGN18324.1 | CP015061.1 | 3279 | 1165 | 1394 | 2766 | 851 | 1291 | 2757 | 1208 | 1233 |
| 4961 | FIM1_4651 | QGN18325.1 | CP015061.1 | 11836 | 3620 | 2445 | 8940 | 2291 | 2429 | 14781 | 3815 | 2540 |
| 4962 | FIM1_4652 | QGN18326.1 | CP015061.1 | 3880 | 1124 | 1440 | 3350 | 811 | 1345 | 3421 | 1272 | 1176 |
| 4963 | FIM1_4653 | QGN18327.1 | CP015061.1 | 3299 | 696 | 988 | 2688 | 531 | 887 | 3968 | 779 | 847 |
| 4964 | FIM1_4654 | QGN18328.1 | CP015061.1 | 1376 | 630 | 875 | 1384 | 348 | 787 | 1849 | 732 | 773 |
| 4965 | FIM1_4655 | QGN18329.1 | CP015061.1 | 6373 | 816 | 1390 | 5317 | 680 | 1307 | 6298 | 831 | 1173 |
| 4966 | FIM1_4656 | QGN18330.1 | CP015061.1 | 9503 | 1658 | 2173 | 7778 | 1066 | 2062 | 9667 | 1747 | 2070 |
| 4967 | FIM1_4657 | QGN18331.1 | CP015061.1 | 2484 | 728 | 829 | 2005 | 500 | 670 | 2727 | 839 | 737 |
| 4968 | FIM1_4658 | QGN18332.1 | CP015061.1 | 39524 | 22134 | 17827 | 35438 | 16925 | 15859 | 29997 | 22491 | 18359 |
| 4969 | FIM1_4659 | QGN18333.1 | CP015061.1 | 50 | 4 | 4 | 32 | 5 | 0 | 31 | 1 | 2 |
| 4970 | FIM1_4660 | QGN18334.1 | CP015061.1 | 520 | 50 | 62 | 588 | 34 | 37 | 620 | 60 | 26 |
| 4971 | FIM1_4661 | QGN18335.1 | CP015061.1 | 708 | 379 | 162 | 765 | 267 | 132 | 615 | 423 | 150 |
| 4972 | FIM1_4662 | QGN18336.1 | CP015061.1 | 3528 | 4343 | 2424 | 2509 | 3069 | 2061 | 887 | 4782 | 2060 |
| 4973 | FIM1_4664 | QGN18337.1 | CP015061.1 | 65 | 14 | 34 | 60 | 8 | 38 | 51 | 16 | 31 |
| 4974 | FIM1_4665 | QGN18338.1 | CP015061.1 | 11039 | 4875 | 2603 | 8349 | 3227 | 2197 | 8108 | 5192 | 2357 |
| 4975 | FIM1_4666 | QGN18339.1 | CP015061.1 | 2339 | 272 | 1211 | 1775 | 235 | 1176 | 4365 | 287 | 1108 |
| 4976 | FIM1_4667 | QGN18340.1 | CP015061.1 | 14246 | 7562 | 3488 | 10203 | 5078 | 3354 | 3913 | 7616 | 3171 |
| 4977 | FIM1_4668 | QGN18341.1 | CP015061.1 | 2304 | 417 | 298 | 1719 | 429 | 244 | 4246 | 485 | 275 |
| 4978 | FIM1_4669 | QGN18342.1 | CP015061.1 | 186 | 18 | 39 | 146 | 6 | 24 | 132 | 19 | 23 |
| 4979 | FIM1_4670 | QGN18343.1 | CP015061.1 | 9410 | 1424 | 2027 | 7039 | 1170 | 1928 | 8627 | 1565 | 1910 |
| 4980 | FIM1_4671 | QGN18344.1 | CP015061.1 | 3711 | 494 | 964 | 2929 | 361 | 933 | 5165 | 564 | 845 |
| 4981 | FIM1_4672 | QGN18345.1 | CP015061.1 | 2302 | 648 | 861 | 1861 | 460 | 776 | 2595 | 657 | 705 |
| 4982 | FIM1_4673 | QGN18346.1 | CP015061.1 | 1472 | 188 | 327 | 1152 | 122 | 297 | 1231 | 202 | 281 |
| 4983 | FIM1_4674 | QGN18347.1 | CP015061.1 | 642 | 139 | 220 | 520 | 111 | 196 | 552 | 124 | 149 |
| 4984 | FIM1_4675 | QGN18348.1 | CP015061.1 | 10304 | 3897 | 3682 | 8908 | 3071 | 3165 | 8924 | 4097 | 3203 |
| 4985 | FIM1_4676 | QGN18349.1 | CP015061.1 | 6793 | 1748 | 1798 | 5028 | 1129 | 1588 | 5409 | 1779 | 1594 |
| 4986 | FIM1_4677 | QGN18350.1 | CP015061.1 | 2 | 1 | 0 | 0 | 0 | 1 | 0 | 0 | 0 |
| 4987 | FIM1_4678 | QGN18351.1 | CP015061.1 | 6890 | 2156 | 1736 | 5477 | 1623 | 1502 | 3040 | 2459 | 1567 |
| 4988 | FIM1_4679 | QGN18352.1 | CP015061.1 | 3632 | 1254 | 1033 | 2761 | 834 | 929 | 5152 | 1320 | 969 |
| 4989 | FIM1_4680 | QGN18353.1 | CP015061.1 | 1810 | 462 | 543 | 1486 | 297 | 493 | 2062 | 450 | 434 |
| 4990 | FIM1_4681 | QGN18354.1 | CP015061.1 | 6882 | 2552 | 1653 | 5616 | 1421 | 1380 | 9347 | 2662 | 1250 |

| | | | | | | | | | | | | |
|---|---|---|---|---|---|---|---|---|---|---|---|---|
| 4991 | FIM1_4682 | QGN18355.1 | CP015061.1 | 15630 | 4900 | 5468 | 13017 | 3734 | 4803 | 13481 | 5392 | 4851 |
| 4992 | FIM1_4683 | QGN18356.1 | CP015061.1 | 48 | 116 | 202 | 36 | 98 | 164 | 15 | 109 | 179 |
| 4993 | FIM1_4684 | QGN18357.1 | CP015061.1 | 4337 | 1377 | 1450 | 3178 | 1036 | 1286 | 2288 | 1493 | 1156 |
| 4994 | FIM1_4685 | QGN18358.1 | CP015061.1 | 5927 | 465 | 1326 | 5024 | 827 | 1072 | 5974 | 508 | 936 |
| 4995 | FIM1_4686 | QGN18359.1 | CP015061.1 | 1829 | 54 | 73 | 1671 | 15 | 63 | 3932 | 48 | 67 |
| 4996 | FIM1_4687 | QGN18360.1 | CP015061.1 | 15 | 4 | 5 | 10 | 6 | 6 | 7 | 7 | 4 |
| 4997 | FIM1_4688 | QGN18361.1 | CP015061.1 | 2774 | 1035 | 906 | 2350 | 535 | 797 | 2064 | 1069 | 902 |
| 4998 | FIM1_4689 | QGN18362.1 | CP015061.1 | 3212 | 751 | 1009 | 2274 | 598 | 931 | 3091 | 821 | 893 |
| 4999 | FIM1_4690 | QGN18363.1 | CP015061.1 | 8064 | 1953 | 2811 | 6487 | 1543 | 2428 | 8421 | 2133 | 2300 |
| 5000 | FIM1_4691 | QGN18364.1 | CP015061.1 | 4803 | 2012 | 717 | 3728 | 1376 | 747 | 1839 | 2233 | 930 |
| 5001 | FIM1_4692 | QGN18365.1 | CP015061.1 | 2493 | 1138 | 979 | 2521 | 996 | 963 | 2353 | 1249 | 781 |
| 5002 | FIM1_4693 | QGN18366.1 | CP015061.1 | 1104 | 201 | 255 | 919 | 128 | 207 | 1355 | 223 | 211 |
| 5003 | FIM1_4694 | QGN18367.1 | CP015061.1 | 2824 | 336 | 693 | 2413 | 294 | 651 | 3534 | 435 | 583 |
| 5004 | FIM1_4695 | QGN18368.1 | CP015061.1 | 2959 | 456 | 508 | 2228 | 326 | 498 | 2259 | 526 | 552 |
| 5005 | FIM1_4696 | QGN18369.1 | CP015061.1 | 20 | 4 | 5 | 24 | 2 | 3 | 22 | 5 | 4 |
| 5006 | FIM1_4697 | QGN18370.1 | CP015061.1 | 4262 | 2340 | 930 | 3560 | 1438 | 850 | 1488 | 2419 | 963 |
| 5007 | FIM1_4698 | QGN18371.1 | CP015061.1 | 3681 | 693 | 1177 | 3330 | 528 | 1032 | 3703 | 745 | 959 |
| 5008 | FIM1_4699 | QGN18372.1 | CP015061.1 | 4015 | 553 | 676 | 3141 | 380 | 687 | 4233 | 626 | 684 |
| 5009 | FIM1_4700 | QGN18373.1 | CP015061.1 | 65 | 16 | 33 | 91 | 17 | 23 | 49 | 28 | 14 |
| 5010 | FIM1_4701 | QGN18374.1 | CP015061.1 | 570 | 593 | 1068 | 709 | 369 | 997 | 993 | 694 | 992 |
| 5011 | FIM1_4702 | QGN18375.1 | CP015061.1 | 7004 | 1397 | 2017 | 5437 | 1111 | 1842 | 7004 | 1497 | 1848 |
| 5012 | FIM1_4703 | QGN18376.1 | CP015061.1 | 23232 | 5177 | 8756 | 18689 | 3504 | 8086 | 26533 | 5324 | 7845 |
| 5013 | FIM1_4704 | QGN18377.1 | CP015061.1 | 3169 | 1070 | 1240 | 2652 | 634 | 1116 | 4292 | 1063 | 1160 |
| 5014 | FIM1_4705 | QGN18378.1 | CP015061.1 | 3426 | 966 | 1103 | 2767 | 723 | 958 | 3232 | 1079 | 995 |
| 5015 | FIM1_4706 | QGN18379.1 | CP015061.1 | 2959 | 924 | 1149 | 2828 | 721 | 1031 | 2666 | 999 | 990 |
| 5016 | FIM1_4707 | QGN18380.1 | CP015061.1 | 1500 | 323 | 656 | 1419 | 258 | 552 | 1327 | 367 | 570 |
| 5017 | FIM1_4708 | QGN18381.1 | CP015061.1 | 3536 | 921 | 1012 | 2807 | 683 | 898 | 1648 | 1025 | 918 |
| 5018 | FIM1_4709 | QGN18382.1 | CP015061.1 | 7724 | 1418 | 1307 | 6022 | 1071 | 1083 | 7275 | 1500 | 1137 |
| 5019 | FIM1_4710 | QGN18383.1 | CP015061.1 | 3837 | 673 | 1124 | 2912 | 702 | 1128 | 2811 | 761 | 1130 |
| 5020 | FIM1_4711 | QGN18384.1 | CP015061.1 | 7787 | 1203 | 1866 | 6059 | 819 | 1777 | 8589 | 1240 | 1791 |
| 5021 | FIM1_4712 | QGN18385.1 | CP015061.1 | 2140 | 302 | 427 | 1628 | 227 | 420 | 1886 | 333 | 421 |
| 5022 | FIM1_4713 | QGN18386.1 | CP015061.1 | 6880 | 1352 | 2049 | 5007 | 1030 | 1779 | 6712 | 1469 | 1779 |
| 5023 | FIM1_4714 | QGN18387.1 | CP015061.1 | 4881 | 815 | 1152 | 4272 | 735 | 1172 | 6191 | 1029 | 1201 |
| 5024 | FIM1_4715 | QGN18388.1 | CP015061.1 | 13931 | 3109 | 3672 | 10972 | 2233 | 3344 | 16770 | 3308 | 3237 |
| 5025 | FIM1_4716 | QGN18389.1 | CP015061.1 | 15294 | 2195 | 2688 | 10182 | 1395 | 2368 | 13973 | 2398 | 2375 |
| 5026 | FIM1_4717 | QGN18390.1 | CP015061.1 | 6548 | 6133 | 4448 | 5701 | 4381 | 3714 | 2195 | 6755 | 4220 |
| 5027 | FIM1_4718 | QGN18391.1 | CP015061.1 | 3467 | 2819 | 1523 | 2265 | 1747 | 1298 | 536 | 2977 | 1456 |
| 5028 | FIM1_4719 | QGN18392.1 | CP015061.1 | 985 | 287 | 301 | 830 | 212 | 295 | 626 | 313 | 299 |
| 5029 | FIM1_4720 | QGN18393.1 | CP015061.1 | 6287 | 4068 | 2077 | 4696 | 2500 | 1779 | 1941 | 4340 | 1852 |
| 5030 | FIM1_4721 | QGN18394.1 | CP015061.1 | 1553 | 468 | 1324 | 1477 | 287 | 1274 | 1744 | 460 | 1228 |
| 5031 | FIM1_4722 | QGN18395.1 | CP015061.1 | 4314 | 3000 | 1306 | 3088 | 1783 | 1243 | 954 | 3159 | 1492 |
| 5032 | FIM1_4723 | QGN18396.1 | CP015061.1 | 17 | 5 | 11 | 18 | 3 | 11 | 35 | 3 | 6 |
| 5033 | FIM1_4724 | QGN18397.1 | CP015061.1 | 1101 | 320 | 352 | 993 | 320 | 359 | 872 | 357 | 340 |
| 5034 | FIM1_4725 | QGN18398.1 | CP015061.1 | 414 | 119 | 185 | 409 | 125 | 135 | 507 | 148 | 151 |
| 5035 | FIM1_4726 | QGN18399.1 | CP015061.1 | 100 | 40 | 60 | 87 | 35 | 44 | 106 | 54 | 44 |
| 5036 | FIM1_4727 | QGN18400.1 | CP015061.1 | 559 | 251 | 345 | 483 | 188 | 288 | 634 | 282 | 293 |
| 5037 | FIM1_4728 | QGN18401.1 | CP015061.1 | 289 | 57 | 115 | 289 | 19 | 88 | 619 | 44 | 83 |
| 5038 | FIM1_4729 | QGN18402.1 | CP015061.1 | 1664 | 399 | 694 | 1512 | 299 | 575 | 1971 | 424 | 572 |

| | | | | | | | | | | | | |
|---|---|---|---|---|---|---|---|---|---|---|---|---|
| 5039 | FIM1_4730 | QGN18403.1 | CP015061.1 | 632 | 401 | 309 | 626 | 415 | 265 | 216 | 391 | 313 |
| 5040 | FIM1_4731 | QGN18404.1 | CP015061.1 | 2978 | 1831 | 1149 | 2570 | 1252 | 962 | 1144 | 1925 | 1047 |
| 5041 | FIM1_4732 | QGN18405.1 | CP015061.1 | 8714 | 1838 | 3259 | 7861 | 1130 | 2997 | 10776 | 1902 | 2821 |
| 5042 | FIM1_4733 | QGN18406.1 | CP015061.1 | 3953 | 671 | 925 | 3310 | 459 | 907 | 4523 | 661 | 941 |
| 5043 | FIM1_4734 | QGN18407.1 | CP015061.1 | 1429 | 200 | 245 | 1071 | 225 | 211 | 1604 | 246 | 224 |
| 5044 | FIM1_4736 | QGN18408.1 | CP015061.1 | 926 | 181 | 170 | 708 | 161 | 155 | 1108 | 167 | 153 |
| 5045 | FIM1_4737 | QGN18409.1 | CP015061.1 | 2453 | 330 | 426 | 1742 | 208 | 383 | 2667 | 375 | 328 |
| 5046 | FIM1_4738 | QGN18410.1 | CP015061.1 | 784 | 45 | 128 | 629 | 47 | 141 | 358 | 68 | 91 |
| 5047 | FIM1_4739 | QGN18411.1 | CP015061.1 | 1829 | 354 | 323 | 1442 | 209 | 309 | 1434 | 383 | 349 |
| 5048 | FIM1_4740 | QGN18412.1 | CP015061.1 | 6503 | 322 | 1462 | 5131 | 286 | 1186 | 9007 | 388 | 1002 |
| 5049 | FIM1_4741 | QGN18413.1 | CP015061.1 | 1810 | 174 | 521 | 1563 | 125 | 458 | 2411 | 204 | 418 |
| 5050 | FIM1_4742 | QGN18414.1 | CP015061.1 | 1304 | 134 | 198 | 998 | 84 | 184 | 1811 | 153 | 189 |
| 5051 | FIM1_4743 | QGN18415.1 | CP015061.1 | 154 | 11 | 16 | 131 | 10 | 18 | 295 | 8 | 8 |
| 5052 | FIM1_4744 | QGN18416.1 | CP015061.1 | 6002 | 2304 | 2512 | 5763 | 1467 | 2214 | 6272 | 2495 | 2318 |
| 5053 | FIM1_4745 | QGN18417.1 | CP015061.1 | 1836 | 377 | 969 | 1479 | 326 | 847 | 2253 | 385 | 769 |
| 5054 | FIM1_4746 | QGN18418.1 | CP015061.1 | 6448 | 2705 | 2057 | 5160 | 1999 | 1706 | 5202 | 2768 | 1779 |
| 5055 | FIM1_4747 | QGN18419.1 | CP015061.1 | 1517 | 441 | 610 | 1137 | 351 | 537 | 1313 | 537 | 575 |
| 5056 | FIM1_4748 | QGN18420.1 | CP015061.1 | 9113 | 2347 | 4251 | 7678 | 1576 | 3810 | 10188 | 2589 | 3651 |
| 5057 | FIM1_4749 | QGN18421.1 | CP015061.1 | 955 | 173 | 378 | 887 | 151 | 345 | 1215 | 207 | 325 |
| 5058 | FIM1_4750 | QGN18422.1 | CP015061.1 | 13613 | 3483 | 3710 | 10588 | 2707 | 3388 | 13177 | 3783 | 3372 |
| 5059 | FIM1_4751 | QGN18423.1 | CP015061.1 | 573 | 162 | 145 | 443 | 126 | 169 | 299 | 162 | 125 |
| 5060 | FIM1_4752 | QGN18424.1 | CP015061.1 | 9 | 3 | 3 | 2 | 2 | 1 | 5 | 0 | 2 |
| 5061 | FIM1_4753 | QGN18425.1 | CP015061.1 | 4178 | 3335 | 1564 | 2458 | 2078 | 1571 | 1292 | 3373 | 1531 |
| 5062 | FIM1_4754 | QGN18426.1 | CP015061.1 | 4334 | 1514 | 1826 | 3525 | 1060 | 1771 | 4395 | 1659 | 1578 |
| 5063 | FIM1_4755 | QGN18427.1 | CP015061.1 | 985 | 199 | 420 | 897 | 156 | 432 | 949 | 210 | 367 |
| 5064 | FIM1_4756 | QGN18428.1 | CP015061.1 | 4961 | 1877 | 1462 | 3567 | 1336 | 1319 | 2617 | 1985 | 1208 |
| 5065 | FIM1_4757 | QGN18429.1 | CP015061.1 | 2619 | 2005 | 1660 | 1913 | 1239 | 1571 | 842 | 2160 | 1397 |
| 5066 | FIM1_4758 | QGN18430.1 | CP015061.1 | 2518 | 227 | 476 | 2029 | 158 | 385 | 3130 | 246 | 399 |
| 5067 | FIM1_4759 | QGN18431.1 | CP015061.1 | 12900 | 5244 | 4609 | 11593 | 3182 | 4086 | 18223 | 5701 | 4109 |
| 5068 | FIM1_4760 | QGN18432.1 | CP015061.1 | 14423 | 3120 | 3106 | 11628 | 2031 | 2823 | 12401 | 3851 | 3126 |
| 5069 | FIM1_4761 | QGN18433.1 | CP015061.1 | 810 | 249 | 346 | 698 | 148 | 287 | 899 | 259 | 261 |
| 5070 | FIM1_4762 | QGN18434.1 | CP015061.1 | 447 | 76 | 90 | 345 | 63 | 90 | 367 | 73 | 88 |
| 5071 | FIM1_4763 | QGN18435.1 | CP015061.1 | 581 | 116 | 126 | 444 | 89 | 90 | 473 | 131 | 129 |
| 5072 | FIM1_4764 | QGN18436.1 | CP015061.1 | 771 | 199 | 250 | 565 | 181 | 244 | 783 | 209 | 274 |
| 5073 | FIM1_4765 | QGN18437.1 | CP015061.1 | 474 | 123 | 279 | 386 | 56 | 268 | 815 | 96 | 193 |
| 5074 | FIM1_4766 | QGN18438.1 | CP015061.1 | 320 | 102 | 155 | 248 | 71 | 151 | 416 | 137 | 137 |
| 5075 | FIM1_4767 | QGN18439.1 | CP015061.1 | 1522 | 502 | 587 | 1054 | 289 | 592 | 1722 | 548 | 522 |
| 5076 | FIM1_4769 | QGN18440.1 | CP015061.1 | 211 | 81 | 64 | 154 | 47 | 77 | 272 | 89 | 65 |
| 5077 | FIM1_4770 | QGN18441.1 | CP015061.1 | 712 | 515 | 349 | 450 | 399 | 440 | 1087 | 577 | 426 |
| 5078 | FIM1_4771 | QGN18442.1 | CP015061.1 | 2208 | 531 | 484 | 1764 | 470 | 506 | 2046 | 631 | 464 |
| 5079 | FIM1_4772 | QGN18443.1 | CP015061.1 | 172 | 19 | 45 | 141 | 6 | 40 | 242 | 14 | 33 |
| 5080 | FIM1_4773 | QGN18444.1 | CP015061.1 | 2354 | 1242 | 3776 | 2096 | 586 | 3404 | 2321 | 1164 | 3301 |
| 5081 | FIM1_4776 | QGN18445.1 | CP015061.1 | 0 | 0 | 0 | 0 | 0 | 0 | 0 | 0 | 0 |

**KS_counts**

| | Name | locus_tag | chr | KS-1-1 | KS-1-1_NaCl | KS-1-1_TM | KS-1-2 | KS-1-2_NaCl | KS-1-2_TM | KS-1-3 | KS-1-3_NaCl | KS-1-3_TM | KS-3-1 | KS-3-1_NaCl | KS-3-1_TM | KS-3-2 | KS-3-2_NaCl | KS-3-2_TM | KS-3-3 | KS-3-3_NaCl | KS-3-3_TM |
|---|---|---|---|---|---|---|---|---|---|---|---|---|---|---|---|---|---|---|---|---|---|
| 1 | W303.asm02.HP0_G0000010.mRNA.1.CDS.1 | NA | chrI | 0 | 0 | 0 | 0 | 0 | 0 | 0 | 0 | 0 | 0 | 0 | 0 | 0 | 0 | 0 | 0 | 0 | 0 |
| 2 | W303.asm02.HP0_G0000020.mRNA.1.CDS.1 | NA | chrI | 37 | 3 | 4 | 119 | 3 | 4 | 30 | 2 | 4 | 0 | 0 | 0 | 0 | 0 | 0 | 0 | 0 | 0 |
| 3 | W303.asm02.HP0_G0000030.mRNA.1.CDS.1 | NA | chrI | 22 | 2 | 2 | 42 | 4 | 0 | 15 | 6 | 3 | 0 | 0 | 0 | 0 | 0 | 0 | 0 | 0 | 0 |
| 4 | W303.asm02.HP0_G0000040.mRNA.1.CDS.1 | NA | chrI | 2106 | 1354 | 1215 | 8219 | 2513 | 1396 | 3809 | 1393 | 1651 | 0 | 0 | 0 | 0 | 0 | 0 | 0 | 0 | 0 |
| 5 | W303.asm02.HP0_G0000050.mRNA.1.CDS.1 | NA | chrI | 2328 | 89 | 89 | 1480 | 67 | 81 | 773 | 54 | 101 | 0 | 0 | 0 | 0 | 0 | 0 | 0 | 0 | 0 |
| 6 | W303.asm02.HP0_G0000060.mRNA.1.CDS.1 | NA | chrI | 3172 | 81 | 87 | 2646 | 115 | 104 | 1526 | 91 | 115 | 0 | 0 | 0 | 0 | 0 | 0 | 0 | 0 | 0 |
| 7 | W303.asm02.HP0_G0000070.mRNA.1.CDS.1 | NA | chrI | 4779 | 262 | 308 | 4881 | 358 | 326 | 3646 | 283 | 408 | 0 | 0 | 0 | 1 | 0 | 0 | 0 | 0 | 0 |
| 8 | W303.asm02.HP0_G0000080.mRNA.1.CDS.1 | NA | chrI | 705 | 354 | 176 | 1065 | 282 | 170 | 635 | 205 | 249 | 0 | 0 | 0 | 0 | 0 | 0 | 0 | 0 | 0 |
| 9 | W303.asm02.HP0_G0000090.mRNA.1.CDS.1 | NA | chrI | 1112 | 432 | 327 | 1520 | 316 | 285 | 1347 | 279 | 473 | 0 | 0 | 0 | 0 | 0 | 0 | 0 | 0 | 0 |
| 10 | W303.asm02.HP0_G0000100.mRNA.1.CDS.1 | NA | chrI | 29122 | 5396 | 6723 | 60996 | 7616 | 5815 | 56039 | 7294 | 8883 | 1 | 0 | 0 | 0 | 0 | 0 | 0 | 1 | 0 |
| 11 | W303.asm02.HP0_G0000110.mRNA.1.CDS.1 | NA | chrI | 16656 | 3771 | 2503 | 18952 | 3441 | 2310 | 15361 | 3113 | 3315 | 3 | 0 | 0 | 0 | 0 | 0 | 0 | 0 | 0 |
| 12 | W303.asm02.HP0_G0000120.mRNA.1.CDS.1 | NA | chrI | 2454 | 507 | 680 | 5176 | 388 | 481 | 3174 | 446 | 866 | 0 | 0 | 0 | 0 | 0 | 0 | 1 | 1 | 0 |
| 13 | W303.asm02.HP0_G0000130.mRNA.1.CDS.1 | NA | chrI | 18242 | 1927 | 1598 | 26183 | 1982 | 1555 | 21994 | 1699 | 2176 | 0 | 1 | 0 | 0 | 0 | 0 | 0 | 0 | 0 |
| 14 | W303.asm02.HP0_G0000140.mRNA.1.CDS.1 | NA | chrI | 1957 | 2034 | 1217 | 1788 | 1548 | 1269 | 1849 | 1425 | 1729 | 2 | 0 | 0 | 2 | 0 | 0 | 0 | 0 | 0 |
| 15 | W303.asm02.HP0_G0000150.mRNA.1.CDS.1 | NA | chrI | 6766 | 1341 | 1003 | 10469 | 1371 | 1041 | 9100 | 1249 | 1422 | 1 | 0 | 0 | 0 | 0 | 0 | 0 | 0 | 0 |
| 16 | W303.asm02.HP0_G0000160.mRNA.1.CDS.1 | NA | chrI | 4610 | 1240 | 1623 | 9872 | 1730 | 1378 | 6720 | 1342 | 2106 | 1 | 0 | 0 | 2 | 0 | 0 | 0 | 0 | 0 |
| 17 | W303.asm02.HP0_G0000170.mRNA.1.CDS.1 | NA | chrI | 903 | 322 | 547 | 1340 | 266 | 453 | 961 | 265 | 800 | 0 | 0 | 0 | 1 | 0 | 0 | 0 | 0 | 0 |
| 18 | W303.asm02.HP0_G0000180.mRNA.1.CDS.1 | NA | chrI | 1718 | 423 | 497 | 2980 | 395 | 456 | 2268 | 401 | 639 | 0 | 0 | 0 | 0 | 0 | 0 | 0 | 0 | 0 |
| 19 | W303.asm02.HP0_G0000190.mRNA.1.CDS.1 | NA | chrI | 18844 | 2650 | 6023 | 28581 | 4358 | 5883 | 24534 | 3749 | 8112 | 1 | 1 | 0 | 0 | 0 | 0 | 0 | 0 | 0 |
| 20 | W303.asm02.HP0_G0000200.mRNA.1.CDS.1 | NA | chrI | 1258 | 719 | 899 | 5003 | 921 | 779 | 1858 | 769 | 1052 | 0 | 0 | 0 | 0 | 0 | 0 | 0 | 0 | 0 |
| 21 | W303.asm02.HP0_G0000210.mRNA.1.CDS.1 | NA | chrI | 2049 | 524 | 389 | 3090 | 422 | 429 | 2578 | 394 | 593 | 1 | 0 | 0 | 2 | 0 | 1 | 0 | 0 | 0 |
| 22 | W303.asm02.HP0_G0000220.mRNA.1.CDS.1 | NA | chrI | 3976 | 370 | 690 | 5787 | 743 | 569 | 4302 | 736 | 895 | 0 | 0 | 0 | 0 | 0 | 0 | 0 | 0 | 0 |
| 23 | W303.asm02.HP0_G0000230.mRNA.1.CDS.1 | NA | chrI | 2134 | 746 | 728 | 3202 | 1061 | 756 | 2548 | 765 | 916 | 0 | 0 | 0 | 0 | 0 | 0 | 0 | 0 | 0 |
| 24 | W303.asm02.HP0_G0000240.mRNA.1.CDS.1 | NA | chrI | 328 | 454 | 455 | 968 | 466 | 446 | 579 | 348 | 500 | 0 | 0 | 0 | 0 | 0 | 0 | 0 | 0 | 0 |
| 25 | W303.asm02.HP0_G0000250.mRNA.1.CDS.1 | NA | chrI | 956 | 782 | 589 | 1362 | 565 | 541 | 985 | 523 | 767 | 0 | 0 | 0 | 0 | 0 | 0 | 0 | 0 | 0 |
| 26 | W303.asm02.HP0_G0000260.mRNA.1.CDS.1 | NA | chrI | 198 | 222 | 234 | 556 | 164 | 203 | 399 | 166 | 333 | 0 | 0 | 0 | 0 | 0 | 0 | 0 | 0 | 0 |
| 27 | W303.asm02.HP0_G0000270.mRNA.1.CDS.1 | NA | chrI | 5665 | 1194 | 1308 | 5758 | 908 | 1063 | 5659 | 1262 | 1827 | 0 | 0 | 0 | 0 | 0 | 0 | 0 | 1 | 0 |
| 28 | W303.asm02.HP0_G0000280.mRNA.1.CDS.1 | NA | chrI | 13408 | 3183 | 2427 | 13819 | 2599 | 2038 | 13616 | 2928 | 3115 | 1 | 1 | 0 | 0 | 0 | 0 | 0 | 0 | 0 |
| 29 | W303.asm02.HP0_G0000290.mRNA.1.CDS.1 | NA | chrI | 4022 | 1106 | 1920 | 6431 | 1391 | 1800 | 5025 | 1292 | 2433 | 0 | 0 | 0 | 0 | 0 | 0 | 0 | 0 | 0 |
| 30 | W303.asm02.HP0_G0000300.mRNA.1.CDS.1 | NA | chrI | 5521 | 1328 | 1627 | 7376 | 1984 | 1445 | 6690 | 1738 | 2128 | 0 | 0 | 0 | 0 | 0 | 0 | 0 | 0 | 0 |
| 31 | W303.asm02.HP0_G0000310.mRNA.1.CDS.1 | NA | chrI | 4181 | 1066 | 672 | 2691 | 810 | 566 | 3462 | 784 | 944 | 0 | 1 | 0 | 0 | 0 | 0 | 0 | 0 | 0 |
| 32 | W303.asm02.HP0_G0000320.mRNA.1.CDS.1 | NA | chrI | 53823 | 6553 | 10795 | 69137 | 8057 | 10460 | 63942 | 5192 | 12887 | 18 | 17 | 1 | 25 | 1 | 2 | 0 | 0 | 0 |
| 33 | W303.asm02.HP0_G0000330.mRNA.1.CDS.1 | NA | chrI | 221 | 234 | 195 | 981 | 273 | 179 | 486 | 189 | 263 | 0 | 0 | 0 | 0 | 0 | 0 | 0 | 0 | 0 |
| 34 | W303.asm02.HP0_G0000340.mRNA.1.CDS.1 | NA | chrI | 3162 | 3303 | 2674 | 3240 | 3071 | 2350 | 3503 | 2921 | 3444 | 3 | 1 | 1 | 1 | 0 | 0 | 0 | 0 | 0 |
| 35 | W303.asm02.HP0_G0000350.mRNA.1.CDS.1 | NA | chrI | 10985 | 5951 | 5817 | 17759 | 7163 | 5377 | 15765 | 6937 | 7729 | 3 | 2 | 0 | 2 | 0 | 1 | 0 | 0 | 0 |
| 36 | W303.asm02.HP0_G0000360.mRNA.1.CDS.1 | NA | chrI | 163 | 243 | 257 | 310 | 262 | 351 | 172 | 264 | 456 | 0 | 0 | 0 | 0 | 0 | 0 | 0 | 0 | 0 |
| 37 | W303.asm02.HP0_G0000370.mRNA.1.CDS.1 | NA | chrI | 916 | 536 | 618 | 1867 | 632 | 706 | 1149 | 594 | 967 | 0 | 0 | 0 | 0 | 0 | 0 | 0 | 1 | 0 |
| 38 | W303.asm02.HP0_G0000380.mRNA.1.CDS.1 | NA | chrI | 753 | 714 | 395 | 1319 | 664 | 495 | 1004 | 522 | 646 | 1 | 1 | 0 | 0 | 0 | 0 | 0 | 0 | 0 |
| 39 | W303.asm02.HP0_G0000390.mRNA.1.CDS.1 | NA | chrI | 1967 | 567 | 569 | 2911 | 611 | 585 | 2526 | 601 | 821 | 1 | 0 | 0 | 0 | 0 | 0 | 0 | 1 | 0 |
| 40 | W303.asm02.HP0_G0000400.mRNA.1.CDS.1 | NA | chrI | 3459 | 452 | 708 | 4583 | 706 | 663 | 3669 | 722 | 1006 | 1 | 0 | 0 | 0 | 0 | 0 | 0 | 0 | 0 |
| 41 | W303.asm02.HP0_G0000410.mRNA.1.CDS.1 | NA | chrI | 8 | 5 | 8 | 97 | 17 | 9 | 19 | 3 | 5 | 0 | 0 | 0 | 0 | 0 | 0 | 0 | 0 | 0 |
| 42 | W303.asm02.HP0_G0000410.mRNA.1.CDS.2 | NA | chrI | 301 | 195 | 194 | 752 | 165 | 227 | 460 | 105 | 334 | 1 | 0 | 0 | 0 | 0 | 0 | 0 | 0 | 0 |
| 43 | W303.asm02.HP0_G0000420.mRNA.1.CDS.1 | NA | chrI | 9113 | 2107 | 2264 | 11855 | 2802 | 2168 | 10557 | 3098 | 3486 | 0 | 0 | 0 | 3 | 0 | 0 | 0 | 0 | 0 |
| 44 | W303.asm02.HP0_G0000430.mRNA.1.CDS.1 | NA | chrI | 1274 | 919 | 617 | 2173 | 657 | 512 | 1574 | 786 | 929 | 0 | 0 | 0 | 0 | 0 | 0 | 0 | 0 | 0 |
| 45 | W303.asm02.HP0_G0000440.mRNA.1.CDS.1 | NA | chrI | 902 | 213 | 223 | 1353 | 238 | 229 | 1030 | 209 | 347 | 0 | 0 | 0 | 0 | 0 | 0 | 0 | 0 | 0 |
| 46 | W303.asm02.HP0_G0000450.mRNA.1.CDS.1 | NA | chrI | 1850 | 1090 | 1269 | 4434 | 1444 | 1157 | 3085 | 1498 | 1999 | 0 | 0 | 0 | 2 | 0 | 0 | 0 | 0 | 0 |
| 47 | W303.asm02.HP0_G0000460.mRNA.1.CDS.1 | NA | chrI | 6741 | 5737 | 3841 | 7545 | 5246 | 4357 | 8580 | 4962 | 6111 | 1 | 0 | 0 | 2 | 0 | 0 | 0 | 1 | 0 |
| 48 | W303.asm02.HP0_G0000470.mRNA.1.CDS.1 | NA | chrI | 2504 | 818 | 849 | 4591 | 1308 | 1118 | 2839 | 1002 | 1379 | 0 | 0 | 0 | 0 | 0 | 0 | 0 | 0 | 0 |
| 49 | W303.asm02.HP0_G0000480.mRNA.1.CDS.1 | NA | chrI | 6011 | 4273 | 7695 | 11433 | 6738 | 8230 | 8394 | 6125 | 11245 | 2 | 0 | 0 | 3 | 0 | 0 | 0 | 0 | 0 |
| 50 | W303.asm02.HP0_G0000490.mRNA.1.CDS.1 | NA | chrI | 5998 | 2369 | 1778 | 7322 | 3301 | 2082 | 5487 | 2626 | 2627 | 0 | 0 | 0 | 0 | 0 | 0 | 0 | 0 | 0 |
| 51 | W303.asm02.HP0_G0000500.mRNA.1.CDS.1 | NA | chrI | 21890 | 1656 | 2418 | 35338 | 2608 | 2073 | 32999 | 2752 | 3807 | 0 | 0 | 0 | 0 | 0 | 0 | 0 | 0 | 0 |
| 52 | W303.asm02.HP0_G0000510.mRNA.1.CDS.1 | NA | chrI | 957 | 781 | 331 | 1273 | 770 | 353 | 1243 | 622 | 509 | 1 | 0 | 0 | 0 | 0 | 0 | 0 | 0 | 0 |
| 53 | W303.asm02.HP0_G0000520.mRNA.1.CDS.1 | NA | chrI | 6791 | 796 | 1330 | 8352 | 1321 | 1240 | 9362 | 1161 | 1983 | 2 | 0 | 0 | 0 | 0 | 0 | 0 | 0 | 0 |
| 54 | W303.asm02.HP0_G0000530.mRNA.1.CDS.1 | NA | chrI | 2956 | 2402 | 2877 | 6576 | 3162 | 3142 | 4804 | 2641 | 4401 | 0 | 0 | 0 | 0 | 0 | 0 | 0 | 1 | 0 |
| 55 | W303.asm02.HP0_G0000540.mRNA.1.CDS.1 | NA | chrI | 2893 | 696 | 1124 | 6155 | 1382 | 1312 | 3170 | 1065 | 1697 | 1 | 0 | 0 | 0 | 0 | 0 | 0 | 0 | 0 |
| 56 | W303.asm02.HP0_G0000550.mRNA.1.CDS.1 | NA | chrI | 2854 | 713 | 1041 | 6034 | 1117 | 1105 | 3628 | 972 | 1567 | 2 | 0 | 0 | 0 | 0 | 0 | 0 | 0 | 0 |
| 57 | W303.asm02.HP0_G0000560.mRNA.1.CDS.1 | NA | chrI | 5442 | 1315 | 885 | 7284 | 1547 | 1006 | 5324 | 1464 | 1292 | 0 | 0 | 0 | 0 | 0 | 0 | 0 | 0 | 0 |
| 58 | W303.asm02.HP0_G0000570.mRNA.1.CDS.1 | NA | chrI | 2448 | 1129 | 1298 | 3446 | 1157 | 1187 | 3268 | 1165 | 1960 | 1 | 0 | 0 | 1 | 0 | 0 | 0 | 0 | 0 |
| 59 | W303.asm02.HP0_G0000580.mRNA.1.CDS.1 | NA | chrI | 5732 | 1539 | 1500 | 7507 | 1521 | 1448 | 7576 | 1768 | 2272 | 0 | 0 | 0 | 0 | 0 | 0 | 0 | 1 | 0 |
| 60 | W303.asm02.HP0_G0000590.mRNA.1.CDS.1 | NA | chrI | 15505 | 6268 | 5521 | 31774 | 6172 | 6219 | 29775 | 5427 | 8311 | 3 | 1 | 0 | 3 | 0 | 1 | 0 | 0 | 0 |
| 61 | W303.asm02.HP0_G0000600.mRNA.1.CDS.1 | NA | chrI | 1911 | 460 | 466 | 2192 | 532 | 536 | 1944 | 600 | 795 | 0 | 0 | 0 | 0 | 0 | 0 | 0 | 0 | 0 |
| 62 | W303.asm02.HP0_G0000610.mRNA.1.CDS.1 | NA | chrI | 1664 | 799 | 992 | 4082 | 1298 | 1133 | 2686 | 1110 | 1531 | 0 | 0 | 0 | 0 | 0 | 0 | 0 | 0 | 0 |
| 63 | W303.asm02.HP0_G0000620.mRNA.1.CDS.1 | NA | chrI | 901 | 619 | 388 | 1741 | 563 | 449 | 1070 | 461 | 573 | 0 | 0 | 0 | 0 | 0 | 1 | 0 | 0 | 0 |
| 64 | W303.asm02.HP0_G0000630.mRNA.1.CDS.1 | NA | chrI | 2342 | 695 | 842 | 4440 | 843 | 893 | 3181 | 714 | 1302 | 0 | 0 | 0 | 0 | 0 | 0 | 0 | 0 | 0 |
| 65 | W303.asm02.HP0_G0000640.mRNA.1.CDS.1 | NA | chrI | 2310 | 1749 | 2108 | 3228 | 1446 | 2219 | 2861 | 1325 | 3300 | 2 | 1 | 0 | 0 | 0 | 0 | 0 | 0 | 0 |
| 66 | W303.asm02.HP0_G0000650.mRNA.1.CDS.1 | NA | chrI | 3471 | 8608 | 7337 | 6489 | 6500 | 6172 | 5768 | 7951 | 9505 | 30 | 10 | 1 | 23 | 0 | 2 | 0 | 0 | 3 |
| 67 | W303.asm02.HP0_G0000660.mRNA.1.CDS.1 | NA | chrI | 77 | 49 | 26 | 155 | 60 | 30 | 101 | 60 | 37 | 0 | 0 | 0 | 0 | 0 | 0 | 0 | 0 | 0 |
| 68 | W303.asm02.HP0_G0000660.mRNA.1.CDS.2 | NA | chrI | 5957 | 8083 | 4577 | 5764 | 4603 | 3869 | 6848 | 5857 | 6387 | 14 | 7 | 1 | 6 | 1 | 1 | 1 | 2 | 0 |
| 69 | W303.asm02.HP0_G0000670.mRNA.1.CDS.1 | NA | chrI | 6024 | 2983 | 2362 | 10426 | 3224 | 2496 | 6854 | 2973 | 3631 | 0 | 0 | 0 | 0 | 0 | 0 | 0 | 0 | 0 |
| 70 | W303.asm02.HP0_G0000680.mRNA.1.CDS.1 | NA | chrI | 57 | 21 | 26 | 100 | 29 | 12 | 81 | 23 | 16 | 0 | 0 | 0 | 0 | 0 | 0 | 0 | 0 | 0 |
| 71 | W303.asm02.HP0_G0000680.mRNA.1.CDS.2 | NA | chrI | 4491 | 830 | 1033 | 8029 | 1222 | 1010 | 6110 | 940 | 1442 | 1 | 1 | 0 | 0 | 0 | 0 | 0 | 0 | 0 |
| 72 | W303.asm02.HP0_G0000690.mRNA.1.CDS.1 | NA | chrI | 1 | 1 | 1 | 3 | 2 | 1 | 3 | 1 | 1 | 0 | 0 | 0 | 0 | 0 | 0 | 0 | 0 | 0 |
| 73 | W303.asm02.HP0_G0000690.mRNA.1.CDS.2 | NA | chrI | 0 | 0 | 0 | 0 | 0 | 0 | 0 | 0 | 0 | 0 | 0 | 0 | 0 | 0 | 0 | 0 | 0 | 0 |
| 74 | W303.asm02.HP0_G0000700.mRNA.1.CDS.1 | NA | chrI | 3510 | 2360 | 1673 | 4634 | 2648 | 1789 | 3826 | 2203 | 2443 | 1 | 0 | 0 | 1 | 0 | 0 | 0 | 0 | 0 |
| 75 | W303.asm02.HP0_G0000710.mRNA.1.CDS.1 | NA | chrI | 3127 | 3887 | 3393 | 5392 | 2763 | 3445 | 3976 | 2535 | 4412 | 2 | 1 | 0 | 0 | 0 | 0 | 0 | 1 | 0 |
| 76 | W303.asm02.HP0_G0000720.mRNA.1.CDS.1 | NA | chrI | 2402 | 1357 | 2059 | 3998 | 1301 | 1760 | 2709 | 1251 | 2590 | 0 | 0 | 0 | 1 | 0 | 0 | 0 | 0 | 0 |
| 77 | W303.asm02.HP0_G0000730.mRNA.1.CDS.1 | NA | chrI | 1101 | 1425 | 2039 | 3741 | 1484 | 1707 | 2328 | 1294 | 2587 | 1 | 0 | 1 | 0 | 0 | 0 | 0 | 1 | 0 |
| 78 | W303.asm02.HP0_G0000740.mRNA.1.CDS.1 | NA | chrI | 276 | 673 | 536 | 962 | 565 | 512 | 550 | 481 | 678 | 2 | 1 | 0 | 0 | 0 | 0 | 0 | 0 | 0 |
| 79 | W303.asm02.HP0_G0000750.mRNA.1.CDS.1 | NA | chrI | 2784 | 1020 | 1298 | 4806 | 1039 | 1065 | 3362 | 1155 | 1784 | 0 | 1 | 0 | 2 | 0 | 1 | 0 | 0 | 0 |
| 80 | W303.asm02.HP0_G0000760.mRNA.1.CDS.1 | NA | chrI | 422 | 255 | 318 | 1853 | 212 | 282 | 1027 | 234 | 453 | 0 | 0 | 0 | 0 | 0 | 0 | 0 | 0 | 0 |
| 81 | W303.asm02.HP0_G0000770.mRNA.1.CDS.1 | NA | chrI | 980 | 1211 | 694 | 1789 | 1183 | 645 | 1225 | 1185 | 977 | 0 | 0 | 0 | 0 | 0 | 0 | 0 | 0 | 0 |
| 82 | W303.asm02.HP0_G0000780.mRNA.1.CDS.1 | NA | chrI | 4673 | 3665 | 3520 | 9934 | 3796 | 3382 | 7222 | 3162 | 4441 | 0 | 1 | 0 | 0 | 0 | 0 | 0 | 1 | 0 |

| | | | | | | | | | | | | | | | | | | | | | |
|---|---|---|---|---|---|---|---|---|---|---|---|---|---|---|---|---|---|---|---|---|---|
| 83 | W303.asm02.HP0_G0000790.mRNA.1.CDS.1 | NA | chrI | 920 | 200 | 278 | 2193 | 446 | 231 | 1560 | 330 | 343 | 0 | 0 | 0 | 0 | 0 | 0 | 0 | 0 | 0 |
| 84 | W303.asm02.HP0_G0000790.mRNA.1.CDS.2 | NA | chrI | 604 | 69 | 130 | 1685 | 233 | 181 | 945 | 181 | 136 | 0 | 0 | 0 | 0 | 0 | 0 | 0 | 0 | 0 |
| 85 | W303.asm02.HP0_G0000800.mRNA.1.CDS.1 | NA | chrI | 2629 | 635 | 347 | 7148 | 516 | 349 | 3646 | 400 | 444 | 0 | 0 | 0 | 0 | 0 | 0 | 0 | 0 | 0 |
| 86 | W303.asm02.HP0_G0000810.mRNA.1.CDS.1 | NA | chrI | 2359 | 1282 | 1116 | 4632 | 1004 | 1065 | 3479 | 972 | 1527 | 2 | 1 | 0 | 0 | 0 | 0 | 0 | 0 | 0 |
| 87 | W303.asm02.HP0_G0000820.mRNA.1.CDS.1 | NA | chrI | 2595 | 292 | 271 | 2918 | 314 | 254 | 2863 | 215 | 301 | 0 | 0 | 0 | 0 | 0 | 0 | 0 | 0 | 0 |
| 88 | W303.asm02.HP0_G0000830.mRNA.1.CDS.1 | NA | chrI | 1496 | 208 | 190 | 2324 | 353 | 225 | 1369 | 145 | 269 | 0 | 0 | 0 | 0 | 0 | 0 | 0 | 0 | 0 |
| 89 | W303.asm02.HP0_G0000830.mRNA.1.CDS.2 | NA | chrI | 5721 | 1363 | 917 | 10400 | 1239 | 827 | 8017 | 880 | 1147 | 0 | 0 | 0 | 0 | 0 | 0 | 0 | 0 | 0 |
| 90 | W303.asm02.HP0_G0000830.mRNA.1.CDS.3 | NA | chrI | 948 | 881 | 582 | 2464 | 631 | 548 | 1543 | 565 | 750 | 0 | 0 | 0 | 0 | 0 | 0 | 0 | 0 | 0 |
| 91 | W303.asm02.HP0_G0000840.mRNA.1.CDS.1 | NA | chrI | 3775 | 342 | 462 | 4047 | 562 | 387 | 2627 | 498 | 576 | 0 | 0 | 0 | 0 | 0 | 0 | 0 | 1 | 0 |
| 92 | W303.asm02.HP0_G0000850.mRNA.1.CDS.1 | NA | chrI | 16785 | 1394 | 1655 | 21445 | 1838 | 1525 | 19342 | 1476 | 2322 | 0 | 0 | 0 | 0 | 0 | 0 | 0 | 0 | 0 |
| 93 | W303.asm02.HP0_G0004920.mRNA.1.CDS.1 | NA | chrIII | 0 | 0 | 0 | 0 | 0 | 0 | 0 | 0 | 0 | 0 | 0 | 0 | 0 | 0 | 0 | 0 | 0 | 0 |
| 94 | W303.asm02.HP0_G0004930.mRNA.1.CDS.1 | NA | chrIII | 0 | 2 | 0 | 0 | 16 | 0 | 0 | 0 | 0 | 14884 | 2464 | 5915 | 15569 | 3801 | 4763 | 15721 | 3476 | 3772 |
| 95 | W303.asm02.HP0_G0004940.mRNA.1.CDS.1 | NA | chrIII | 0 | 0 | 0 | 0 | 0 | 0 | 0 | 0 | 0 | 209 | 38 | 66 | 182 | 53 | 59 | 140 | 64 | 39 |
| 96 | W303.asm02.HP0_G0004940.mRNA.1.CDS.2 | NA | chrIII | 0 | 5 | 0 | 0 | 19 | 0 | 0 | 0 | 1 | 13461 | 2757 | 5989 | 13635 | 4766 | 5327 | 12355 | 3022 | 3925 |
| 97 | W303.asm02.HP0_G0004950.mRNA.1.CDS.1 | NA | chrIII | 0 | 0 | 0 | 0 | 0 | 0 | 0 | 0 | 0 | 243 | 35 | 45 | 220 | 24 | 36 | 248 | 27 | 27 |
| 98 | W303.asm02.HP0_G0004960.mRNA.1.CDS.1 | NA | chrIII | 0 | 1 | 0 | 0 | 3 | 0 | 0 | 0 | 0 | 1653 | 272 | 470 | 1487 | 276 | 405 | 1752 | 280 | 307 |
| 99 | W303.asm02.HP0_G0004970.mRNA.1.CDS.1 | NA | chrIII | 0 | 0 | 0 | 0 | 1 | 0 | 0 | 0 | 0 | 1241 | 161 | 233 | 870 | 135 | 270 | 1120 | 159 | 157 |
| 100 | W303.asm02.HP0_G0004970.mRNA.1.CDS.2 | NA | chrIII | 0 | 0 | 0 | 0 | 0 | 0 | 0 | 0 | 0 | 248 | 19 | 46 | 131 | 18 | 37 | 209 | 21 | 19 |
| 101 | W303.asm02.HP0_G0004980.mRNA.1.CDS.1 | NA | chrIII | 0 | 1 | 0 | 0 | 2 | 0 | 0 | 0 | 0 | 3358 | 652 | 934 | 3624 | 861 | 765 | 4042 | 659 | 577 |
| 102 | W303.asm02.HP0_G0004990.mRNA.1.CDS.1 | NA | chrIII | 0 | 0 | 0 | 0 | 3 | 0 | 0 | 0 | 0 | 3113 | 741 | 724 | 3577 | 883 | 770 | 3133 | 776 | 533 |
| 103 | W303.asm02.HP0_G0005000.mRNA.1.CDS.1 | NA | chrIII | 0 | 3 | 0 | 1 | 13 | 0 | 0 | 0 | 0 | 11457 | 2219 | 2467 | 11540 | 1948 | 2174 | 11157 | 1894 | 1559 |
| 104 | W303.asm02.HP0_G0005010.mRNA.1.CDS.1 | NA | chrIII | 0 | 6 | 0 | 0 | 23 | 1 | 0 | 0 | 0 | 18007 | 4675 | 5544 | 16284 | 3990 | 5222 | 17728 | 4365 | 3632 |
| 105 | W303.asm02.HP0_G0005020.mRNA.1.CDS.1 | NA | chrIII | 0 | 0 | 1 | 0 | 1 | 0 | 0 | 0 | 0 | 1469 | 445 | 502 | 1014 | 352 | 520 | 1169 | 591 | 276 |
| 106 | W303.asm02.HP0_G0005030.mRNA.1.CDS.1 | NA | chrIII | 0 | 1 | 0 | 0 | 2 | 1 | 0 | 0 | 0 | 1145 | 243 | 302 | 939 | 204 | 305 | 1080 | 203 | 186 |
| 107 | W303.asm02.HP0_G0005040.mRNA.1.CDS.1 | NA | chrIII | 0 | 0 | 0 | 1 | 4 | 0 | 0 | 0 | 0 | 2151 | 408 | 1639 | 3815 | 762 | 1384 | 3008 | 477 | 1004 |
| 108 | W303.asm02.HP0_G0005050.mRNA.1.CDS.1 | NA | chrIII | 0 | 0 | 0 | 0 | 5 | 0 | 0 | 0 | 0 | 3090 | 520 | 642 | 2254 | 599 | 623 | 2818 | 598 | 399 |
| 109 | W303.asm02.HP0_G0005060.mRNA.1.CDS.1 | NA | chrIII | 0 | 2 | 0 | 0 | 1 | 0 | 0 | 1 | 0 | 5488 | 819 | 1233 | 4201 | 841 | 1234 | 5301 | 847 | 779 |
| 110 | W303.asm02.HP0_G0005070.mRNA.1.CDS.1 | NA | chrIII | 0 | 5 | 0 | 1 | 23 | 0 | 0 | 0 | 0 | 17323 | 4651 | 4941 | 19173 | 4552 | 4565 | 17400 | 4185 | 3248 |
| 111 | W303.asm02.HP0_G0005080.mRNA.1.CDS.1 | NA | chrIII | 0 | 0 | 1 | 1 | 7 | 1 | 0 | 0 | 0 | 5212 | 713 | 1375 | 4978 | 755 | 1308 | 5042 | 694 | 870 |
| 112 | W303.asm02.HP0_G0005090.mRNA.1.CDS.1 | NA | chrIII | 0 | 0 | 0 | 0 | 6 | 0 | 0 | 0 | 0 | 2741 | 553 | 1035 | 3528 | 469 | 908 | 2906 | 447 | 637 |
| 113 | W303.asm02.HP0_G0005100.mRNA.1.CDS.1 | NA | chrIII | 0 | 11 | 1 | 2 | 46 | 0 | 0 | 1 | 1 | 22262 | 5465 | 9122 | 19827 | 5187 | 8170 | 23184 | 5274 | 5697 |
| 114 | W303.asm02.HP0_G0005110.mRNA.1.CDS.1 | NA | chrIII | 0 | 1 | 0 | 2 | 13 | 1 | 0 | 0 | 0 | 7456 | 1571 | 2582 | 6721 | 1471 | 2593 | 6886 | 1302 | 1635 |
| 115 | W303.asm02.HP0_G0005120.mRNA.1.CDS.1 | NA | chrIII | 0 | 1 | 0 | 0 | 5 | 0 | 0 | 0 | 0 | 2352 | 417 | 554 | 1619 | 398 | 553 | 2026 | 448 | 362 |
| 116 | W303.asm02.HP0_G0005130.mRNA.1.CDS.1 | NA | chrIII | 0 | 2 | 0 | 0 | 28 | 0 | 0 | 0 | 0 | 21597 | 4328 | 6911 | 19217 | 3948 | 5802 | 20971 | 4091 | 4370 |
| 117 | W303.asm02.HP0_G0005140.mRNA.1.CDS.1 | NA | chrIII | 0 | 0 | 0 | 0 | 3 | 0 | 0 | 0 | 0 | 1456 | 272 | 690 | 1283 | 272 | 610 | 1435 | 288 | 433 |
| 118 | W303.asm02.HP0_G0005150.mRNA.1.CDS.1 | NA | chrIII | 0 | 0 | 0 | 1 | 15 | 0 | 0 | 0 | 0 | 9224 | 2373 | 2786 | 12716 | 2452 | 2359 | 10693 | 2245 | 1669 |
| 119 | W303.asm02.HP0_G0005160.mRNA.1.CDS.1 | NA | chrIII | 0 | 0 | 0 | 0 | 6 | 0 | 0 | 0 | 0 | 1222 | 656 | 1993 | 4361 | 673 | 2000 | 2656 | 679 | 1396 |
| 120 | W303.asm02.HP0_G0005170.mRNA.1.CDS.1 | NA | chrIII | 0 | 11 | 1 | 4 | 32 | 4 | 1 | 2 | 1 | 15244 | 6061 | 13242 | 24662 | 6607 | 13043 | 20971 | 5550 | 9176 |
| 121 | W303.asm02.HP0_G0005180.mRNA.1.CDS.1 | NA | chrIII | 4 | 0 | 0 | 7 | 6 | 0 | 0 | 0 | 2 | 1873 | 1053 | 1615 | 4083 | 1054 | 1659 | 2602 | 957 | 1042 |
| 122 | W303.asm02.HP0_G0005190.mRNA.1.CDS.1 | NA | chrIII | 0 | 1 | 0 | 0 | 1 | 0 | 0 | 0 | 0 | 2819 | 734 | 1257 | 4307 | 831 | 1339 | 3524 | 733 | 829 |
| 123 | W303.asm02.HP0_G0005200.mRNA.1.CDS.1 | NA | chrIII | 0 | 1 | 0 | 0 | 9 | 0 | 0 | 0 | 0 | 4880 | 918 | 1790 | 4851 | 846 | 1722 | 4757 | 808 | 1149 |
| 124 | W303.asm02.HP0_G0005210.mRNA.1.CDS.1 | NA | chrIII | 0 | 0 | 0 | 0 | 5 | 0 | 0 | 0 | 0 | 972 | 456 | 696 | 2232 | 482 | 712 | 1527 | 436 | 497 |
| 125 | W303.asm02.HP0_G0005220.mRNA.1.CDS.1 | NA | chrIII | 0 | 1 | 0 | 0 | 26 | 0 | 0 | 0 | 0 | 8801 | 4134 | 4681 | 15479 | 3581 | 4462 | 11740 | 3413 | 3388 |
| 126 | W303.asm02.HP0_G0005230.mRNA.1.CDS.1 | NA | chrIII | 0 | 5 | 0 | 0 | 4 | 0 | 0 | 0 | 0 | 6037 | 1003 | 1786 | 4587 | 1293 | 1644 | 5680 | 1166 | 1121 |
| 127 | W303.asm02.HP0_G0005240.mRNA.1.CDS.1 | NA | chrIII | 0 | 1 | 0 | 0 | 5 | 0 | 0 | 0 | 0 | 3566 | 897 | 1884 | 5154 | 1040 | 1825 | 3873 | 877 | 1197 |
| 128 | W303.asm02.HP0_G0005250.mRNA.1.CDS.1 | NA | chrIII | 0 | 2 | 0 | 0 | 0 | 3 | 0 | 0 | 0 | 1721 | 207 | 570 | 1292 | 374 | 660 | 1414 | 245 | 341 |
| 129 | W303.asm02.HP0_G0005260.mRNA.1.CDS.1 | NA | chrIII | 0 | 4 | 0 | 0 | 4 | 1 | 0 | 0 | 1 | 3833 | 592 | 1089 | 4285 | 588 | 1122 | 3746 | 541 | 750 |
| 130 | W303.asm02.HP0_G0005270.mRNA.1.CDS.1 | NA | chrIII | 0 | 1 | 0 | 0 | 23 | 3 | 0 | 0 | 1 | 18256 | 4098 | 4027 | 15121 | 3726 | 3798 | 17388 | 3765 | 2699 |
| 131 | W303.asm02.HP0_G0005280.mRNA.1.CDS.1 | NA | chrIII | 0 | 1 | 0 | 1 | 7 | 0 | 0 | 0 | 0 | 5497 | 1185 | 3180 | 7425 | 1281 | 2999 | 5601 | 1046 | 2104 |
| 132 | W303.asm02.HP0_G0005290.mRNA.1.CDS.1 | NA | chrIII | 0 | 0 | 0 | 0 | 1 | 0 | 0 | 0 | 0 | 2985 | 774 | 1453 | 4170 | 861 | 1394 | 3331 | 805 | 978 |
| 133 | W303.asm02.HP0_G0005300.mRNA.1.CDS.1 | NA | chrIII | 0 | 2 | 0 | 1 | 9 | 1 | 0 | 0 | 0 | 5370 | 1417 | 2296 | 6940 | 1532 | 2414 | 5863 | 1290 | 1573 |
| 134 | W303.asm02.HP0_G0005310.mRNA.1.CDS.1 | NA | chrIII | 0 | 0 | 0 | 0 | 1 | 1 | 0 | 0 | 0 | 1032 | 118 | 481 | 1139 | 125 | 456 | 1050 | 104 | 302 |
| 135 | W303.asm02.HP0_G0005320.mRNA.1.CDS.1 | NA | chrIII | 0 | 1 | 0 | 0 | 6 | 0 | 0 | 0 | 0 | 3989 | 495 | 1284 | 2947 | 522 | 966 | 3421 | 573 | 717 |
| 136 | W303.asm02.HP0_G0005330.mRNA.1.CDS.1 | NA | chrIII | 0 | 13 | 0 | 0 | 21 | 0 | 0 | 0 | 0 | 23396 | 3767 | 10581 | 17051 | 3189 | 8187 | 22902 | 6103 | 7159 |
| 137 | W303.asm02.HP0_G0005340.mRNA.1.CDS.1 | NA | chrIII | 1 | 0 | 0 | 1 | 12 | 2 | 0 | 0 | 0 | 7826 | 2187 | 2779 | 11370 | 1874 | 2721 | 8910 | 1887 | 1780 |
| 138 | W303.asm02.HP0_G0005350.mRNA.1.CDS.1 | NA | chrIII | 0 | 1 | 0 | 1 | 2 | 0 | 0 | 0 | 0 | 3540 | 735 | 832 | 4864 | 836 | 799 | 4099 | 678 | 582 |
| 139 | W303.asm02.HP0_G0005360.mRNA.1.CDS.1 | NA | chrIII | 0 | 0 | 0 | 0 | 2 | 0 | 0 | 0 | 0 | 1460 | 206 | 435 | 1142 | 257 | 439 | 1299 | 236 | 262 |
| 140 | W303.asm02.HP0_G0005370.mRNA.1.CDS.1 | NA | chrIII | 0 | 0 | 0 | 0 | 1 | 0 | 0 | 0 | 0 | 747 | 154 | 254 | 816 | 158 | 291 | 591 | 166 | 162 |
| 141 | W303.asm02.HP0_G0005370.mRNA.1.CDS.2 | NA | chrIII | 0 | 1 | 0 | 0 | 1 | 0 | 0 | 0 | 0 | 1951 | 359 | 613 | 2190 | 355 | 619 | 1842 | 318 | 360 |
| 142 | W303.asm02.HP0_G0005370.mRNA.1.CDS.3 | NA | chrIII | 0 | 0 | 0 | 0 | 0 | 0 | 0 | 0 | 0 | 325 | 72 | 103 | 316 | 65 | 98 | 272 | 73 | 66 |
| 143 | W303.asm02.HP0_G0005380.mRNA.1.CDS.1 | NA | chrIII | 0 | 1 | 1 | 0 | 9 | 1 | 0 | 0 | 0 | 5445 | 1060 | 2036 | 6565 | 1229 | 1644 | 5922 | 1252 | 1313 |
| 144 | W303.asm02.HP0_G0005390.mRNA.1.CDS.1 | NA | chrIII | 0 | 0 | 0 | 0 | 4 | 0 | 0 | 0 | 0 | 3387 | 504 | 789 | 3220 | 465 | 721 | 3130 | 441 | 557 |
| 145 | W303.asm02.HP0_G0005400.mRNA.1.CDS.1 | NA | chrIII | 0 | 2 | 0 | 1 | 15 | 0 | 0 | 0 | 0 | 10066 | 1844 | 2151 | 10824 | 2408 | 2047 | 9941 | 2071 | 1416 |
| 146 | W303.asm02.HP0_G0005410.mRNA.1.CDS.1 | NA | chrIII | 0 | 0 | 0 | 0 | 1 | 0 | 0 | 0 | 0 | 50 | 6 | 18 | 35 | 3 | 16 | 30 | 5 | 8 |
| 147 | W303.asm02.HP0_G0005410.mRNA.1.CDS.2 | NA | chrIII | 0 | 4 | 0 | 0 | 2 | 0 | 0 | 0 | 0 | 2312 | 293 | 713 | 1614 | 490 | 631 | 2036 | 367 | 431 |
| 148 | W303.asm02.HP0_G0005420.mRNA.1.CDS.1 | NA | chrIII | 1 | 2 | 0 | 2 | 17 | 1 | 0 | 0 | 0 | 8914 | 3491 | 4036 | 13766 | 3277 | 3960 | 10064 | 3254 | 2793 |
| 149 | W303.asm02.HP0_G0005430.mRNA.1.CDS.1 | NA | chrIII | 0 | 1 | 0 | 0 | 4 | 0 | 0 | 0 | 0 | 1106 | 559 | 843 | 2665 | 578 | 821 | 1615 | 597 | 585 |
| 150 | W303.asm02.HP0_G0005440.mRNA.1.CDS.1 | NA | chrIII | 0 | 4 | 1 | 2 | 22 | 1 | 0 | 0 | 0 | 3453 | 3238 | 3711 | 13072 | 3749 | 4364 | 8757 | 3433 | 2621 |
| 151 | W303.asm02.HP0_G0005450.mRNA.1.CDS.1 | NA | chrIII | 0 | 0 | 0 | 0 | 0 | 0 | 0 | 0 | 0 | 698 | 445 | 737 | 1666 | 510 | 694 | 847 | 477 | 473 |
| 152 | W303.asm02.HP0_G0005460.mRNA.1.CDS.1 | NA | chrIII | 0 | 0 | 0 | 0 | 0 | 0 | 0 | 0 | 0 | 147 | 26 | 27 | 134 | 35 | 26 | 103 | 50 | 29 |
| 153 | W303.asm02.HP0_G0005460.mRNA.1.CDS.2 | NA | chrIII | 0 | 0 | 0 | 0 | 0 | 0 | 0 | 0 | 0 | 61 | 13 | 17 | 50 | 11 | 13 | 35 | 13 | 8 |
| 154 | W303.asm02.HP0_G0005460.mRNA.1.CDS.3 | NA | chrIII | 0 | 1 | 0 | 0 | 1 | 1 | 0 | 0 | 0 | 1310 | 232 | 336 | 964 | 229 | 284 | 1163 | 258 | 207 |
| 155 | W303.asm02.HP0_G0005470.mRNA.1.CDS.1 | NA | chrIII | 0 | 3 | 0 | 0 | 7 | 0 | 0 | 2 | 1 | 4675 | 936 | 1787 | 4382 | 834 | 1930 | 4603 | 797 | 1212 |
| 156 | W303.asm02.HP0_G0005480.mRNA.1.CDS.1 | NA | chrIII | 0 | 2 | 0 | 0 | 4 | 0 | 0 | 0 | 0 | 4116 | 870 | 1372 | 4601 | 877 | 1243 | 4035 | 819 | 839 |
| 157 | W303.asm02.HP0_G0005490.mRNA.1.CDS.1 | NA | chrIII | 0 | 0 | 0 | 0 | 0 | 0 | 0 | 0 | 0 | 24 | 2 | 5 | 19 | 2 | 3 | 27 | 4 | 4 |
| 158 | W303.asm02.HP0_G0005490.mRNA.1.CDS.2 | NA | chrIII | 0 | 1 | 0 | 1 | 7 | 0 | 0 | 0 | 0 | 4528 | 808 | 942 | 3258 | 685 | 918 | 4379 | 741 | 553 |
| 159 | W303.asm02.HP0_G0005500.mRNA.1.CDS.1 | NA | chrIII | 0 | 1 | 0 | 0 | 4 | 1 | 0 | 0 | 0 | 4196 | 687 | 835 | 3151 | 689 | 893 | 4347 | 636 | 550 |
| 160 | W303.asm02.HP0_G0005510.mRNA.1.CDS.1 | NA | chrIII | 0 | 0 | 0 | 0 | 2 | 0 | 0 | 0 | 0 | 1127 | 130 | 184 | 733 | 119 | 158 | 1103 | 118 | 120 |
| 161 | W303.asm02.HP0_G0005520.mRNA.1.CDS.1 | NA | chrIII | 0 | 4 | 0 | 0 | 3 | 0 | 0 | 0 | 0 | 3272 | 716 | 812 | 2572 | 664 | 794 | 3264 | 806 | 546 |
| 162 | W303.asm02.HP0_G0005530.mRNA.1.CDS.1 | NA | chrIII | 0 | 2 | 0 | 1 | 7 | 0 | 0 | 0 | 0 | 7758 | 1618 | 2429 | 7179 | 1859 | 2190 | 7748 | 1641 | 1541 |
| 163 | W303.asm02.HP0_G0005540.mRNA.1.CDS.1 | NA | chrIII | 0 | 1 | 0 | 1 | 6 | 0 | 0 | 0 | 0 | 5688 | 679 | 1001 | 3907 | 829 | 969 | 5158 | 806 | 597 |
| 164 | W303.asm02.HP0_G0005550.mRNA.1.CDS.1 | NA | chrIII | 0 | 1 | 0 | 0 | 5 | 1 | 0 | 0 | 0 | 4938 | 841 | 1111 | 4121 | 945 | 1159 | 4500 | 766 | 808 |
| 165 | W303.asm02.HP0_G0005560.mRNA.1.CDS.1 | NA | chrIII | 0 | 2 | 0 | 0 | 1 | 0 | 0 | 0 | 0 | 2615 | 299 | 737 | 2440 | 251 | 566 | 2669 | 273 | 418 |
| 166 | W303.asm02.HP0_G0005570.mRNA.1.CDS.1 | NA | chrIII | 0 | 0 | 0 | 0 | 0 | 0 | 0 | 0 | 0 | 125 | 16 | 19 | 118 | 14 | 27 | 171 | 12 | 13 |

| | | | | | | | | | | | | | | | | | | | | | |
|---|---|---|---|---|---|---|---|---|---|---|---|---|---|---|---|---|---|---|---|---|---|
| 167 | W303.asm02.HP0_G0005570.mRNA.1.CDS.2 | NA | chrIII | 0 | 0 | 0 | 0 | 0 | 0 | 0 | 0 | 0 | 521 | 66 | 99 | 462 | 64 | 97 | 435 | 58 | 71 |
| 168 | W303.asm02.HP0_G0005580.mRNA.1.CDS.1 | NA | chrIII | 0 | 0 | 0 | 0 | 3 | 0 | 0 | 0 | 0 | 1568 | 352 | 424 | 1257 | 279 | 397 | 1599 | 336 | 245 |
| 169 | W303.asm02.HP0_G0005590.mRNA.1.CDS.1 | NA | chrIII | 0 | 2 | 0 | 0 | 16 | 0 | 0 | 0 | 0 | 15625 | 3423 | 3530 | 15993 | 2786 | 3197 | 15699 | 2678 | 2231 |
| 170 | W303.asm02.HP0_G0005600.mRNA.1.CDS.1 | NA | chrIII | 0 | 2 | 0 | 2 | 6 | 1 | 0 | 0 | 0 | 7186 | 1045 | 1819 | 5782 | 939 | 1643 | 6860 | 1086 | 1128 |
| 171 | W303.asm02.HP0_G0005610.mRNA.1.CDS.1 | NA | chrIII | 0 | 0 | 0 | 0 | 2 | 0 | 0 | 0 | 0 | 1383 | 464 | 436 | 1852 | 439 | 447 | 1480 | 426 | 282 |
| 172 | W303.asm02.HP0_G0005620.mRNA.1.CDS.1 | NA | chrIII | 0 | 0 | 0 | 0 | 20 | 0 | 0 | 0 | 0 | 10631 | 2553 | 5110 | 13090 | 2406 | 4819 | 11348 | 2370 | 3341 |
| 173 | W303.asm02.HP0_G0005630.mRNA.1.CDS.1 | NA | chrIII | 9 | 22 | 4 | 34 | 62 | 12 | 3 | 2 | 2 | 25724 | 5953 | 13011 | 24135 | 8082 | 13038 | 24951 | 6547 | 8006 |
| 174 | W303.asm02.HP0_G0005640.mRNA.1.CDS.1 | NA | chrIII | 0 | 1 | 0 | 0 | 6 | 0 | 0 | 0 | 0 | 6740 | 1039 | 1610 | 5752 | 838 | 1544 | 6517 | 796 | 980 |
| 175 | W303.asm02.HP0_G0005650.mRNA.1.CDS.1 | NA | chrIII | 0 | 0 | 0 | 0 | 0 | 0 | 0 | 0 | 0 | 3530 | 453 | 789 | 2630 | 458 | 640 | 3552 | 394 | 464 |
| 176 | W303.asm02.HP0_G0005660.mRNA.1.CDS.1 | NA | chrIII | 1 | 6 | 1 | 0 | 25 | 0 | 0 | 0 | 0 | 13564 | 2893 | 2652 | 9859 | 2866 | 2776 | 13704 | 2983 | 1711 |
| 177 | W303.asm02.HP0_G0005670.mRNA.1.CDS.1 | NA | chrIII | 1 | 0 | 1 | 2 | 4 | 0 | 0 | 0 | 0 | 6664 | 1489 | 1866 | 8807 | 1674 | 1858 | 6994 | 1355 | 1248 |
| 178 | W303.asm02.HP0_G0005680.mRNA.1.CDS.1 | NA | chrIII | 0 | 1 | 0 | 0 | 2 | 0 | 0 | 2 | 0 | 3134 | 492 | 542 | 2176 | 500 | 544 | 2850 | 570 | 343 |
| 179 | W303.asm02.HP0_G0005690.mRNA.1.CDS.1 | NA | chrIII | 0 | 0 | 0 | 0 | 0 | 0 | 0 | 0 | 0 | 0 | 0 | 0 | 0 | 0 | 0 | 0 | 0 | 0 |
| 180 | W303.asm02.HP0_G0005700.mRNA.1.CDS.1 | NA | chrIII | 0 | 0 | 0 | 0 | 0 | 0 | 0 | 0 | 0 | 0 | 0 | 0 | 0 | 0 | 0 | 0 | 0 | 0 |
| 181 | W303.asm02.HP0_G0005710.mRNA.1.CDS.1 | NA | chrIII | 0 | 0 | 0 | 0 | 0 | 0 | 0 | 0 | 0 | 0 | 0 | 0 | 0 | 0 | 0 | 0 | 0 | 0 |
| 182 | W303.asm02.HP0_G0005720.mRNA.1.CDS.1 | NA | chrIII | 0 | 0 | 1 | 0 | 0 | 0 | 0 | 0 | 0 | 0 | 0 | 0 | 2 | 0 | 0 | 0 | 0 | 0 |
| 183 | W303.asm02.HP0_G0005730.mRNA.1.CDS.1 | NA | chrIII | 1 | 0 | 0 | 1 | 0 | 0 | 0 | 0 | 0 | 0 | 0 | 0 | 0 | 0 | 0 | 0 | 0 | 0 |
| 184 | W303.asm02.HP0_G0005740.mRNA.1.CDS.1 | NA | chrIII | 0 | 0 | 0 | 0 | 0 | 0 | 0 | 0 | 0 | 0 | 0 | 0 | 0 | 0 | 0 | 0 | 0 | 0 |
| 185 | W303.asm02.HP0_G0005750.mRNA.1.CDS.1 | NA | chrIII | 0 | 0 | 0 | 0 | 0 | 0 | 0 | 0 | 0 | 2 | 0 | 0 | 0 | 0 | 0 | 0 | 0 | 0 |
| 186 | W303.asm02.HP0_G0005760.mRNA.1.CDS.1 | NA | chrIII | 0 | 0 | 0 | 0 | 0 | 1 | 0 | 0 | 0 | 0 | 0 | 0 | 0 | 0 | 0 | 0 | 1 | 0 |
| 187 | W303.asm02.HP0_G0005770.mRNA.1.CDS.1 | NA | chrIII | 0 | 0 | 0 | 0 | 0 | 0 | 0 | 0 | 0 | 0 | 0 | 0 | 0 | 0 | 0 | 0 | 0 | 0 |
| 188 | W303.asm02.HP0_G0005780.mRNA.1.CDS.1 | NA | chrIII | 0 | 0 | 1 | 1 | 0 | 0 | 0 | 0 | 0 | 0 | 0 | 0 | 0 | 0 | 0 | 0 | 0 | 0 |
| 189 | W303.asm02.HP0_G0005790.mRNA.1.CDS.1 | NA | chrIII | 0 | 0 | 0 | 0 | 0 | 0 | 0 | 0 | 0 | 0 | 0 | 0 | 0 | 0 | 0 | 0 | 0 | 0 |
| 190 | W303.asm02.HP0_G0005800.mRNA.1.CDS.1 | NA | chrIII | 0 | 1 | 0 | 0 | 10 | 0 | 0 | 0 | 0 | 5704 | 1141 | 1388 | 6012 | 907 | 1323 | 5811 | 899 | 903 |
| 191 | W303.asm02.HP0_G0005810.mRNA.1.CDS.1 | NA | chrIII | 0 | 1 | 0 | 0 | 5 | 0 | 0 | 0 | 0 | 2025 | 349 | 726 | 1419 | 369 | 698 | 1889 | 369 | 489 |
| 192 | W303.asm02.HP0_G0005810.mRNA.1.CDS.2 | NA | chrIII | 0 | 0 | 0 | 0 | 3 | 0 | 0 | 0 | 0 | 1681 | 242 | 514 | 1100 | 209 | 541 | 1494 | 231 | 304 |
| 193 | W303.asm02.HP0_G0005820.mRNA.1.CDS.1 | NA | chrIII | 1 | 1 | 0 | 0 | 6 | 0 | 0 | 0 | 0 | 4350 | 1126 | 2013 | 9045 | 1394 | 1945 | 6977 | 1001 | 1304 |
| 194 | W303.asm02.HP0_G0005830.mRNA.1.CDS.1 | NA | chrIII | 0 | 0 | 0 | 0 | 1 | 0 | 0 | 0 | 0 | 93 | 18 | 18 | 42 | 10 | 13 | 69 | 14 | 10 |
| 195 | W303.asm02.HP0_G0005830.mRNA.1.CDS.2 | NA | chrIII | 2 | 21 | 3 | 5 | 87 | 2 | 0 | 0 | 0 | 18492 | 6930 | 6528 | 16413 | 5958 | 6777 | 18382 | 6306 | 4624 |
| 196 | W303.asm02.HP0_G0005840.mRNA.1.CDS.1 | NA | chrIII | 0 | 3 | 0 | 0 | 6 | 0 | 0 | 0 | 0 | 8936 | 1363 | 3150 | 7789 | 1036 | 2691 | 8505 | 1127 | 1975 |
| 197 | W303.asm02.HP0_G0005850.mRNA.1.CDS.1 | NA | chrIII | 1 | 4 | 0 | 0 | 15 | 0 | 0 | 0 | 0 | 7764 | 1887 | 2661 | 11911 | 1904 | 2389 | 8904 | 1770 | 1861 |
| 198 | W303.asm02.HP0_G0005860.mRNA.1.CDS.1 | NA | chrIII | 0 | 3 | 0 | 1 | 19 | 1 | 0 | 0 | 1 | 12348 | 3226 | 3152 | 9896 | 2848 | 2928 | 11403 | 2848 | 1789 |
| 199 | W303.asm02.HP0_G0005870.mRNA.1.CDS.1 | NA | chrIII | 0 | 3 | 0 | 0 | 17 | 1 | 0 | 0 | 0 | 9213 | 2289 | 3433 | 8678 | 2538 | 3300 | 9223 | 2484 | 2399 |
| 200 | W303.asm02.HP0_G0005880.mRNA.1.CDS.1 | NA | chrIII | 0 | 3 | 0 | 0 | 19 | 1 | 0 | 0 | 0 | 10510 | 2396 | 4129 | 9332 | 2075 | 4191 | 9968 | 2217 | 2761 |
| 201 | W303.asm02.HP0_G0005890.mRNA.1.CDS.1 | NA | chrIII | 0 | 1 | 0 | 1 | 3 | 0 | 0 | 2 | 0 | 4730 | 744 | 1044 | 5523 | 602 | 958 | 4609 | 595 | 651 |
| 202 | W303.asm02.HP0_G0005900.mRNA.1.CDS.1 | NA | chrIII | 0 | 0 | 0 | 0 | 2 | 0 | 0 | 0 | 0 | 1257 | 239 | 359 | 1439 | 260 | 302 | 1330 | 229 | 235 |
| 203 | W303.asm02.HP0_G0005910.mRNA.1.CDS.1 | NA | chrIII | 0 | 0 | 0 | 0 | 0 | 0 | 0 | 0 | 0 | 0 | 0 | 0 | 0 | 0 | 0 | 0 | 0 | 0 |
| 204 | W303.asm02.HP0_G0005910.mRNA.1.CDS.2 | NA | chrIII | 0 | 0 | 0 | 0 | 0 | 0 | 0 | 0 | 0 | 0 | 0 | 0 | 0 | 0 | 0 | 0 | 0 | 0 |
| 205 | W303.asm02.HP0_G0005910.mRNA.1.CDS.3 | NA | chrIII | 0 | 0 | 0 | 0 | 0 | 0 | 0 | 0 | 0 | 5 | 0 | 2 | 3 | 0 | 1 | 5 | 1 | 0 |
| 206 | W303.asm02.HP0_G0005920.mRNA.1.CDS.1 | NA | chrIII | 0 | 2 | 0 | 0 | 9 | 0 | 0 | 0 | 0 | 7140 | 1364 | 2089 | 8964 | 1410 | 1956 | 7678 | 1211 | 1430 |
| 207 | W303.asm02.HP0_G0005930.mRNA.1.CDS.1 | NA | chrIII | 0 | 0 | 0 | 0 | 1 | 0 | 0 | 0 | 0 | 2077 | 401 | 262 | 1425 | 304 | 314 | 2017 | 375 | 191 |
| 208 | W303.asm02.HP0_G0005940.mRNA.1.CDS.1 | NA | chrIII | 0 | 1 | 0 | 0 | 4 | 1 | 0 | 0 | 0 | 1238 | 391 | 330 | 1247 | 350 | 323 | 1153 | 364 | 192 |
| 209 | W303.asm02.HP0_G0005950.mRNA.1.CDS.1 | NA | chrIII | 0 | 0 | 0 | 0 | 0 | 0 | 0 | 0 | 0 | 2 | 1 | 2 | 6 | 1 | 1 | 3 | 1 | 2 |
| 210 | W303.asm02.HP0_G0005950.mRNA.1.CDS.2 | NA | chrIII | 0 | 1 | 0 | 0 | 3 | 0 | 0 | 0 | 0 | 3525 | 1462 | 1985 | 6102 | 1486 | 2005 | 4582 | 1389 | 1296 |
| 211 | W303.asm02.HP0_G0005960.mRNA.1.CDS.1 | NA | chrIII | 0 | 0 | 0 | 0 | 0 | 1 | 0 | 0 | 0 | 2042 | 464 | 815 | 1944 | 568 | 908 | 1837 | 577 | 553 |
| 212 | W303.asm02.HP0_G0005970.mRNA.1.CDS.1 | NA | chrIII | 0 | 2 | 0 | 0 | 7 | 0 | 0 | 0 | 0 | 3104 | 1510 | 1493 | 5151 | 1499 | 1824 | 3638 | 1417 | 1078 |
| 213 | W303.asm02.HP0_G0005980.mRNA.1.CDS.1 | NA | chrIII | 0 | 1 | 0 | 0 | 4 | 0 | 1 | 0 | 0 | 3621 | 2154 | 2702 | 9894 | 2369 | 2804 | 6802 | 2053 | 1772 |
| 214 | W303.asm02.HP0_G0005990.mRNA.1.CDS.1 | NA | chrIII | 1 | 3 | 0 | 2 | 10 | 0 | 0 | 0 | 0 | 7457 | 2597 | 2686 | 7620 | 2250 | 3107 | 7132 | 2145 | 1884 |
| 215 | W303.asm02.HP0_G0006000.mRNA.1.CDS.1 | NA | chrIII | 0 | 2 | 0 | 0 | 13 | 1 | 0 | 0 | 0 | 4574 | 2095 | 2886 | 8811 | 2080 | 2857 | 6536 | 1942 | 1879 |
| 216 | W303.asm02.HP0_G0006010.mRNA.1.CDS.1 | NA | chrIII | 1 | 9 | 0 | 1 | 44 | 0 | 0 | 0 | 0 | 24620 | 6280 | 8786 | 27166 | 6716 | 8241 | 24342 | 5657 | 6047 |
| 217 | W303.asm02.HP0_G0006020.mRNA.1.CDS.1 | NA | chrIII | 0 | 0 | 0 | 0 | 3 | 0 | 0 | 0 | 0 | 6735 | 918 | 1127 | 5358 | 892 | 970 | 5935 | 854 | 760 |
| 218 | W303.asm02.HP0_G0006030.mRNA.1.CDS.1 | NA | chrIII | 0 | 3 | 0 | 0 | 26 | 0 | 0 | 0 | 0 | 9496 | 4062 | 5109 | 20116 | 3800 | 5013 | 14064 | 3580 | 3463 |
| 219 | W303.asm02.HP0_G0006040.mRNA.1.CDS.1 | NA | chrIII | 0 | 3 | 0 | 1 | 21 | 0 | 0 | 0 | 0 | 7984 | 2604 | 2382 | 7037 | 2247 | 2883 | 7128 | 2339 | 1708 |
| 220 | W303.asm02.HP0_G0006050.mRNA.1.CDS.1 | NA | chrIII | 0 | 0 | 0 | 0 | 4 | 0 | 0 | 0 | 0 | 2811 | 769 | 1130 | 2591 | 715 | 1327 | 2543 | 717 | 799 |
| 221 | W303.asm02.HP0_G0006060.mRNA.1.CDS.1 | NA | chrIII | 0 | 0 | 0 | 0 | 1 | 0 | 0 | 0 | 0 | 519 | 512 | 599 | 1775 | 682 | 619 | 851 | 783 | 436 |
| 222 | W303.asm02.HP0_G0006070.mRNA.1.CDS.1 | NA | chrIII | 1 | 0 | 0 | 0 | 0 | 0 | 0 | 0 | 0 | 1297 | 436 | 359 | 1333 | 359 | 316 | 1306 | 393 | 201 |
| 223 | W303.asm02.HP0_G0006080.mRNA.1.CDS.1 | NA | chrIII | 0 | 0 | 0 | 0 | 4 | 0 | 0 | 0 | 0 | 2773 | 941 | 1182 | 4341 | 881 | 1400 | 3591 | 757 | 754 |
| 224 | W303.asm02.HP0_G0006090.mRNA.1.CDS.1 | NA | chrIII | 0 | 1 | 0 | 0 | 2 | 0 | 0 | 0 | 0 | 2932 | 485 | 806 | 3466 | 456 | 677 | 3084 | 436 | 558 |
| 225 | W303.asm02.HP0_G0006100.mRNA.1.CDS.1 | NA | chrIII | 0 | 4 | 0 | 1 | 5 | 0 | 0 | 0 | 0 | 7276 | 1629 | 2974 | 10964 | 1663 | 2756 | 8764 | 1362 | 1907 |
| 226 | W303.asm02.HP0_G0006110.mRNA.1.CDS.1 | NA | chrIII | 0 | 0 | 0 | 0 | 6 | 0 | 0 | 0 | 0 | 1650 | 785 | 1474 | 6848 | 900 | 1352 | 3809 | 707 | 972 |
| 227 | W303.asm02.HP0_G0006120.mRNA.1.CDS.1 | NA | chrIII | 0 | 1 | 0 | 1 | 3 | 0 | 1 | 0 | 0 | 1448 | 573 | 1227 | 4334 | 583 | 1234 | 2619 | 431 | 857 |
| 228 | W303.asm02.HP0_G0006130.mRNA.1.CDS.1 | NA | chrIII | 0 | 0 | 0 | 0 | 3 | 0 | 0 | 0 | 0 | 2781 | 436 | 553 | 2255 | 449 | 505 | 2326 | 478 | 353 |
| 229 | W303.asm02.HP0_G0006140.mRNA.1.CDS.1 | NA | chrIII | 0 | 2 | 0 | 0 | 10 | 0 | 0 | 0 | 0 | 4524 | 1716 | 1798 | 6437 | 1632 | 1737 | 5077 | 1596 | 1190 |
| 230 | W303.asm02.HP0_G0006150.mRNA.1.CDS.1 | NA | chrIII | 0 | 5 | 0 | 0 | 22 | 0 | 0 | 0 | 0 | 12796 | 2368 | 5946 | 14183 | 1910 | 5667 | 12494 | 1891 | 3735 |
| 231 | W303.asm02.HP0_G0006160.mRNA.1.CDS.1 | NA | chrIII | 0 | 1 | 0 | 0 | 5 | 0 | 0 | 0 | 0 | 1534 | 713 | 1814 | 3724 | 803 | 2016 | 2521 | 559 | 1301 |
| 232 | W303.asm02.HP0_G0006170.mRNA.1.CDS.1 | NA | chrIII | 0 | 0 | 0 | 0 | 0 | 0 | 0 | 0 | 0 | 967 | 136 | 256 | 817 | 109 | 296 | 1026 | 98 | 149 |
| 233 | W303.asm02.HP0_G0006180.mRNA.1.CDS.1 | NA | chrIII | 0 | 0 | 0 | 0 | 0 | 0 | 0 | 0 | 0 | 552 | 61 | 151 | 369 | 86 | 131 | 412 | 78 | 73 |
| 234 | W303.asm02.HP0_G0006190.mRNA.1.CDS.1 | NA | chrIII | 0 | 0 | 0 | 0 | 1 | 0 | 0 | 0 | 0 | 2348 | 414 | 563 | 2181 | 447 | 748 | 2238 | 342 | 481 |
| 235 | W303.asm02.HP0_G0006200.mRNA.1.CDS.1 | NA | chrIII | 0 | 0 | 0 | 2 | 6 | 1 | 0 | 0 | 0 | 2383 | 1525 | 2281 | 6447 | 1792 | 2005 | 3647 | 1986 | 1468 |
| 236 | W303.asm02.HP0_G0006210.mRNA.1.CDS.1 | NA | chrIII | 0 | 1 | 0 | 0 | 6 | 0 | 0 | 0 | 0 | 4260 | 1347 | 1826 | 5799 | 1531 | 1943 | 4592 | 1304 | 1296 |
| 237 | W303.asm02.HP0_G0006220.mRNA.1.CDS.1 | NA | chrIII | 0 | 3 | 0 | 0 | 12 | 0 | 0 | 0 | 0 | 12519 | 1968 | 2755 | 11765 | 1877 | 2525 | 12086 | 1747 | 1760 |
| 238 | W303.asm02.HP0_G0006230.mRNA.1.CDS.1 | NA | chrIII | 0 | 3 | 1 | 0 | 4 | 0 | 0 | 0 | 0 | 4032 | 629 | 1321 | 3176 | 665 | 1238 | 3705 | 625 | 777 |
| 239 | W303.asm02.HP0_G0006240.mRNA.1.CDS.1 | NA | chrIII | 0 | 0 | 0 | 0 | 2 | 0 | 0 | 0 | 0 | 1983 | 293 | 411 | 1342 | 277 | 388 | 1780 | 253 | 246 |
| 240 | W303.asm02.HP0_G0006250.mRNA.1.CDS.1 | NA | chrIII | 0 | 2 | 0 | 0 | 21 | 3 | 0 | 0 | 1 | 2965 | 2845 | 4834 | 16755 | 2534 | 4660 | 6304 | 2906 | 3109 |
| 241 | W303.asm02.HP0_G0006260.mRNA.1.CDS.1 | NA | chrIII | 0 | 0 | 0 | 0 | 4 | 0 | 0 | 0 | 0 | 1837 | 443 | 402 | 1631 | 355 | 427 | 1706 | 298 | 258 |
| 242 | W303.asm02.HP0_G0006270.mRNA.1.CDS.1 | NA | chrIII | 0 | 2 | 0 | 0 | 17 | 0 | 0 | 0 | 0 | 5568 | 1894 | 1633 | 4523 | 1653 | 1732 | 5271 | 1934 | 1115 |
| 243 | W303.asm02.HP0_G0006280.mRNA.1.CDS.1 | NA | chrIII | 1 | 4 | 0 | 1 | 4 | 0 | 0 | 0 | 0 | 1734 | 1716 | 3348 | 11219 | 2441 | 3129 | 5132 | 2054 | 2178 |
| 244 | W303.asm02.HP0_G0006290.mRNA.1.CDS.1 | NA | chrIII | 0 | 1 | 0 | 0 | 4 | 0 | 0 | 0 | 0 | 2237 | 668 | 1058 | 3919 | 613 | 1055 | 2862 | 676 | 679 |
| 245 | W303.asm02.HP0_G0006300.mRNA.1.CDS.1 | NA | chrIII | 0 | 7 | 1 | 0 | 8 | 0 | 0 | 0 | 0 | 6531 | 1437 | 1850 | 4907 | 1213 | 1722 | 6429 | 1323 | 1172 |
| 246 | W303.asm02.HP0_G0006310.mRNA.1.CDS.1 | NA | chrIII | 0 | 2 | 0 | 0 | 0 | 0 | 0 | 0 | 0 | 2812 | 510 | 720 | 2956 | 414 | 679 | 2896 | 363 | 476 |
| 247 | W303.asm02.HP0_G0006320.mRNA.1.CDS.1 | NA | chrIII | 0 | 0 | 0 | 0 | 0 | 0 | 0 | 0 | 0 | 19 | 2 | 0 | 19 | 6 | 2 | 27 | 2 | 3 |
| 248 | W303.asm02.HP0_G0006320.mRNA.1.CDS.2 | NA | chrIII | 0 | 1 | 0 | 0 | 2 | 0 | 0 | 0 | 0 | 4139 | 752 | 1207 | 4125 | 691 | 1203 | 4191 | 676 | 723 |
| 249 | W303.asm02.HP0_G0006330.mRNA.1.CDS.1 | NA | chrIII | 0 | 5 | 0 | 0 | 20 | 1 | 0 | 0 | 0 | 13421 | 2761 | 4716 | 18538 | 2866 | 4362 | 15699 | 2763 | 3235 |
| 250 | W303.asm02.HP0_G0006340.mRNA.1.CDS.1 | NA | chrIII | 0 | 3 | 0 | 1 | 10 | 0 | 0 | 0 | 0 | 6938 | 1029 | 1769 | 6010 | 1053 | 1633 | 6708 | 953 | 1141 |

| | | | | | | | | | | | | | | | | | | | | | |
|---|---|---|---|---|---|---|---|---|---|---|---|---|---|---|---|---|---|---|---|---|---|
| 251 | W303.asm02.HP0_G0006350.mRNA.1.CDS.1 | NA | chrIII | 0 | 0 | 0 | 0 | 3 | 0 | 0 | 0 | 0 | 3260 | 455 | 782 | 2302 | 327 | 674 | 3057 | 359 | 471 |
| 252 | W303.asm02.HP0_G0006360.mRNA.1.CDS.1 | NA | chrIII | 0 | 0 | 0 | 0 | 0 | 0 | 0 | 0 | 0 | 433 | 93 | 132 | 409 | 71 | 139 | 425 | 74 | 81 |
| 253 | W303.asm02.HP0_G0006370.mRNA.1.CDS.1 | NA | chrIII | 0 | 0 | 0 | 0 | 0 | 0 | 0 | 0 | 0 | 0 | 0 | 0 | 0 | 0 | 0 | 0 | 0 | 0 |
| 254 | W303.asm02.HP0_G0006370.mRNA.1.CDS.2 | NA | chrIII | 0 | 0 | 0 | 0 | 0 | 0 | 0 | 0 | 0 | 0 | 0 | 0 | 0 | 0 | 0 | 0 | 0 | 0 |
| 255 | W303.asm02.HP0_G0006370.mRNA.1.CDS.3 | NA | chrIII | 0 | 0 | 0 | 0 | 0 | 0 | 0 | 0 | 0 | 121 | 13 | 15 | 79 | 24 | 15 | 114 | 9 | 14 |
| 256 | W303.asm02.HP0_G0006380.mRNA.1.CDS.1 | NA | chrIII | 0 | 1 | 0 | 0 | 5 | 0 | 0 | 0 | 0 | 7387 | 735 | 1505 | 6303 | 1972 | 1397 | 6431 | 1063 | 977 |
| 257 | W303.asm02.HP0_G0006390.mRNA.1.CDS.1 | NA | chrIII | 0 | 1 | 0 | 0 | 2 | 0 | 0 | 0 | 0 | 1183 | 104 | 607 | 1011 | 122 | 573 | 1096 | 99 | 427 |
| 258 | W303.asm02.HP0_G0006390.mRNA.1.CDS.2 | NA | chrIII | 0 | 0 | 0 | 0 | 0 | 0 | 0 | 0 | 0 | 805 | 80 | 170 | 555 | 88 | 130 | 703 | 122 | 103 |
| 259 | W303.asm02.HP0_G0006400.mRNA.1.CDS.1 | NA | chrIII | 0 | 1 | 0 | 1 | 0 | 0 | 0 | 0 | 0 | 5022 | 606 | 2614 | 5092 | 660 | 2428 | 5006 | 487 | 1734 |
| 260 | W303.asm02.HP0_G0006410.mRNA.1.CDS.1 | NA | chrIII | 0 | 0 | 0 | 0 | 4 | 0 | 0 | 0 | 0 | 3595 | 332 | 1405 | 3294 | 349 | 1199 | 3416 | 248 | 872 |
| 261 | W303.asm02.HP0_G0006420.mRNA.1.CDS.1 | NA | chrIII | 0 | 6 | 0 | 0 | 21 | 0 | 0 | 1 | 0 | 9620 | 2404 | 2631 | 7890 | 2279 | 2566 | 8580 | 2448 | 1657 |
| 262 | W303.asm02.HP0_G0006430.mRNA.1.CDS.1 | NA | chrIII | 0 | 0 | 0 | 0 | 1 | 0 | 0 | 0 | 0 | 836 | 266 | 205 | 845 | 234 | 266 | 778 | 219 | 142 |
| 263 | W303.asm02.HP0_G0006440.mRNA.1.CDS.1 | NA | chrIII | 0 | 0 | 1 | 0 | 6 | 0 | 0 | 0 | 0 | 3834 | 789 | 916 | 3695 | 730 | 962 | 3511 | 617 | 673 |
| 264 | W303.asm02.HP0_G0006450.mRNA.1.CDS.1 | NA | chrIII | 0 | 0 | 0 | 0 | 3 | 0 | 0 | 0 | 0 | 5010 | 809 | 1088 | 4883 | 668 | 1012 | 4724 | 778 | 671 |
| 265 | W303.asm02.HP0_G0006460.mRNA.1.CDS.1 | NA | chrIII | 0 | 2 | 0 | 0 | 6 | 0 | 0 | 0 | 1 | 4171 | 931 | 1138 | 4052 | 809 | 1148 | 3925 | 801 | 752 |
| 266 | W303.asm02.HP0_G0006470.mRNA.1.CDS.1 | NA | chrIII | 0 | 0 | 0 | 0 | 0 | 0 | 0 | 0 | 0 | 1 | 0 | 0 | 0 | 0 | 0 | 0 | 0 | 0 |
| 267 | W303.asm02.HP0_G0006470.mRNA.1.CDS.2 | NA | chrIII | 1 | 0 | 0 | 1 | 0 | 3 | 1 | 1 | 0 | 1 | 0 | 1 | 3 | 0 | 2 | 0 | 0 | 0 |
| 268 | QGN13365.1 | FIM1_1 | CP015054. | 0 | 0 | 0 | 0 | 0 | 0 | 0 | 0 | 0 | 0 | 0 | 0 | 0 | 0 | 0 | 0 | 0 | 0 |
| 269 | QGN13366.1 | FIM1_2 | CP015054. | 1 | 0 | 0 | 0 | 0 | 0 | 0 | 0 | 0 | 0 | 0 | 0 | 0 | 0 | 0 | 0 | 0 | 0 |
| 270 | QGN13367.1 | FIM1_3 | CP015054. | 2669 | 2018 | 1383 | 4157 | 1426 | 941 | 3033 | 1480 | 1572 | 6246 | 2355 | 2771 | 10035 | 2407 | 2635 | 7934 | 2031 | 1911 |
| 271 | QGN13368.1 | FIM1_5 | CP015054. | 1616 | 70 | 370 | 2375 | 316 | 322 | 1506 | 327 | 498 | 1241 | 327 | 591 | 2313 | 299 | 538 | 1559 | 298 | 385 |
| 272 | QGN13369.1 | FIM1_6 | CP015054. | 863 | 133 | 257 | 866 | 227 | 211 | 784 | 259 | 334 | 577 | 274 | 426 | 1316 | 268 | 393 | 1004 | 299 | 251 |
| 273 | QGN13370.1 | FIM1_7 | CP015054. | 209 | 23 | 11 | 384 | 27 | 11 | 209 | 21 | 16 | 272 | 35 | 25 | 168 | 22 | 16 | 216 | 32 | 9 |
| 274 | QGN13371.1 | FIM1_8 | CP015054. | 648 | 96 | 146 | 1357 | 156 | 140 | 613 | 75 | 217 | 1810 | 194 | 294 | 1427 | 224 | 223 | 1691 | 184 | 173 |
| 275 | QGN13372.1 | FIM1_9 | CP015054. | 1878 | 823 | 664 | 12550 | 1155 | 610 | 4933 | 620 | 892 | 9823 | 2475 | 928 | 10534 | 4137 | 1016 | 10104 | 2087 | 718 |
| 276 | QGN13373.1 | FIM1_10 | CP015054. | 22256 | 472 | 668 | 22251 | 291 | 455 | 20841 | 370 | 928 | 2647 | 261 | 432 | 2859 | 901 | 491 | 2568 | 479 | 327 |
| 277 | QGN13374.1 | FIM1_11 | CP015054. | 1383 | 176 | 103 | 1248 | 95 | 65 | 1161 | 124 | 147 | 1370 | 174 | 217 | 998 | 215 | 227 | 1198 | 193 | 158 |
| 278 | QGN13375.1 | FIM1_12 | CP015054. | 10649 | 300 | 694 | 22688 | 352 | 508 | 13024 | 273 | 937 | 1629 | 193 | 389 | 1303 | 338 | 223 | 1526 | 337 | 237 |
| 279 | QGN13376.1 | FIM1_13 | CP015054. | 1737 | 33795 | 32038 | 21650 | 31923 | 25217 | 35209 | 40982 | 34974 | 84661 | 21666 | 31011 | 82615 | 25998 | 33016 | 89173 | 25115 | 24314 |
| 280 | QGN13377.1 | FIM1_14 | CP015054. | 13 | 17 | 9 | 55 | 25 | 8 | 23 | 7 | 10 | 76 | 8 | 7 | 45 | 8 | 11 | 65 | 10 | 6 |
| 281 | QGN13378.1 | FIM1_15 | CP015054. | 210 | 18231 | 12541 | 5750 | 13482 | 11003 | 9229 | 15220 | 14532 | 38090 | 9248 | 13277 | 30554 | 10257 | 12847 | 39529 | 10969 | 9631 |
| 282 | QGN13379.1 | FIM1_16 | CP015054. | 2295 | 12468 | 14923 | 8420 | 11909 | 11383 | 9982 | 17005 | 16971 | 44866 | 7429 | 13137 | 39815 | 8120 | 13717 | 46187 | 8677 | 9081 |
| 283 | QGN13380.1 | FIM1_17 | CP015054. | 479 | 23923 | 18558 | 4114 | 19423 | 15343 | 11655 | 22714 | 21131 | 56143 | 13531 | 18083 | 46813 | 16027 | 19692 | 56142 | 15528 | 13607 |
| 284 | QGN13381.1 | FIM1_18 | CP015054. | 2000 | 870 | 1334 | 3126 | 1927 | 1382 | 2004 | 1911 | 2072 | 9488 | 1541 | 1673 | 8465 | 1399 | 1293 | 8411 | 2159 | 1115 |
| 285 | QGN13382.1 | FIM1_19 | CP015054. | 1568 | 384 | 720 | 2208 | 425 | 639 | 1808 | 436 | 1000 | 3943 | 381 | 1043 | 3902 | 335 | 945 | 3904 | 288 | 652 |
| 286 | QGN13383.1 | FIM1_20 | CP015054. | 3305 | 2300 | 2256 | 4738 | 2083 | 1866 | 3686 | 1759 | 2881 | 11853 | 2172 | 3702 | 10211 | 1901 | 3672 | 11610 | 1852 | 2647 |
| 287 | QGN13384.1 | FIM1_21 | CP015054. | 1025 | 248 | 401 | 904 | 252 | 343 | 925 | 283 | 507 | 2948 | 325 | 613 | 2290 | 308 | 554 | 3079 | 258 | 370 |
| 288 | QGN13385.1 | FIM1_22 | CP015054. | 2074 | 914 | 889 | 2603 | 1008 | 772 | 2412 | 959 | 1153 | 5118 | 929 | 1508 | 4917 | 937 | 1355 | 5058 | 794 | 894 |
| 289 | QGN13386.1 | FIM1_23 | CP015054. | 388 | 154 | 118 | 631 | 145 | 137 | 432 | 92 | 167 | 1106 | 100 | 210 | 793 | 119 | 232 | 1124 | 102 | 135 |
| 290 | QGN13387.1 | FIM1_24 | CP015054. | 1096 | 1266 | 379 | 520 | 1228 | 480 | 781 | 844 | 562 | 4090 | 1174 | 810 | 3378 | 877 | 862 | 4137 | 1176 | 560 |
| 291 | QGN13388.1 | FIM1_25 | CP015054. | 758 | 424 | 284 | 417 | 261 | 248 | 690 | 262 | 363 | 1773 | 299 | 428 | 1228 | 247 | 449 | 1590 | 314 | 272 |
| 292 | QGN13389.1 | FIM1_26 | CP015054. | 750 | 176 | 258 | 369 | 150 | 227 | 541 | 196 | 323 | 1246 | 146 | 313 | 892 | 108 | 282 | 1167 | 122 | 167 |
| 293 | QGN13390.1 | FIM1_27 | CP015054. | 589 | 543 | 493 | 988 | 402 | 403 | 681 | 384 | 627 | 3001 | 352 | 623 | 2482 | 310 | 612 | 2874 | 342 | 402 |
| 294 | QGN13391.1 | FIM1_28 | CP015054. | 9405 | 10227 | 7219 | 9719 | 7322 | 5953 | 9345 | 6604 | 9068 | 10833 | 9360 | 11238 | 28758 | 8218 | 11039 | 20838 | 8489 | 8096 |
| 295 | QGN13392.1 | FIM1_29 | CP015054. | 5174 | 1152 | 2215 | 9017 | 1196 | 1898 | 6278 | 1461 | 2902 | 3314 | 819 | 3185 | 7459 | 973 | 2859 | 4788 | 931 | 2133 |
| 296 | QGN13393.1 | FIM1_30 | CP015054. | 3117 | 1641 | 1370 | 3067 | 1261 | 1234 | 2972 | 1075 | 1749 | 8468 | 1213 | 2171 | 6162 | 1117 | 1876 | 8171 | 1099 | 1361 |
| 297 | QGN13394.1 | FIM1_31 | CP015054. | 6291 | 2144 | 2460 | 8424 | 2552 | 2230 | 6645 | 2244 | 3163 | 11802 | 2520 | 3490 | 14706 | 2264 | 3441 | 12882 | 2275 | 2495 |
| 298 | QGN13395.1 | FIM1_32 | CP015054. | 2515 | 2320 | 2086 | 4108 | 1901 | 1992 | 3055 | 1467 | 2836 | 9105 | 1591 | 2984 | 7494 | 1669 | 3173 | 8734 | 1481 | 2113 |
| 299 | QGN13396.1 | FIM1_33 | CP015054. | 473 | 598 | 498 | 1185 | 472 | 451 | 653 | 394 | 683 | 2934 | 426 | 687 | 2202 | 386 | 663 | 2690 | 424 | 420 |
| 300 | QGN13397.1 | FIM1_34 | CP015054. | 1419 | 919 | 950 | 1775 | 1063 | 845 | 1715 | 940 | 1268 | 6504 | 1064 | 1544 | 6432 | 1158 | 1317 | 6359 | 1129 | 1018 |
| 301 | QGN13398.1 | FIM1_35 | CP015054. | 305 | 16 | 23 | 181 | 12 | 12 | 152 | 14 | 24 | 93 | 18 | 47 | 99 | 5 | 36 | 100 | 20 | 17 |
| 302 | QGN13399.1 | FIM1_36 | CP015054. | 125 | 3547 | 3468 | 651 | 6235 | 4061 | 138 | 3631 | 4840 | 18723 | 5947 | 6442 | 19271 | 3040 | 6177 | 19234 | 4377 | 4294 |
| 303 | QGN13400.1 | FIM1_37 | CP015054. | 296 | 623 | 391 | 623 | 648 | 377 | 453 | 481 | 519 | 2401 | 574 | 555 | 2219 | 414 | 564 | 2206 | 446 | 380 |
| 304 | QGN13401.1 | FIM1_38 | CP015054. | 543 | 1080 | 463 | 785 | 1103 | 426 | 605 | 885 | 543 | 4723 | 1230 | 921 | 3688 | 960 | 834 | 4353 | 1140 | 590 |
| 305 | QGN13402.1 | FIM1_39 | CP015054. | 2225 | 1379 | 740 | 1345 | 1639 | 856 | 1781 | 1160 | 1034 | 4014 | 1220 | 1136 | 4734 | 1147 | 1165 | 4189 | 1207 | 771 |
| 306 | QGN13403.1 | FIM1_40 | CP015054. | 2040 | 560 | 1694 | 2083 | 456 | 1398 | 2023 | 594 | 2012 | 3905 | 420 | 2338 | 3185 | 364 | 2209 | 3842 | 364 | 1555 |
| 307 | QGN13404.1 | FIM1_41 | CP015054. | 1271 | 853 | 555 | 1791 | 655 | 489 | 1550 | 572 | 719 | 4175 | 612 | 738 | 3042 | 576 | 737 | 4032 | 650 | 464 |
| 308 | QGN13405.1 | FIM1_42 | CP015054. | 568 | 1377 | 629 | 608 | 934 | 565 | 414 | 835 | 851 | 3530 | 936 | 1010 | 2685 | 833 | 1016 | 3201 | 856 | 617 |
| 309 | QGN13406.1 | FIM1_43 | CP015054. | 1124 | 1838 | 1359 | 1743 | 1866 | 1158 | 1424 | 1586 | 1749 | 9195 | 1703 | 2204 | 8949 | 1803 | 1989 | 8470 | 1613 | 1480 |
| 310 | QGN13407.1 | FIM1_44 | CP015054. | 125 | 205 | 179 | 261 | 132 | 185 | 135 | 122 | 272 | 853 | 130 | 310 | 532 | 123 | 289 | 811 | 118 | 166 |
| 311 | QGN13408.1 | FIM1_45 | CP015054. | 805 | 497 | 855 | 1312 | 403 | 747 | 868 | 411 | 1187 | 3401 | 313 | 1380 | 2582 | 381 | 1246 | 3287 | 376 | 843 |
| 312 | QGN13409.1 | FIM1_46 | CP015054. | 6502 | 2514 | 1861 | 11433 | 2501 | 1680 | 8045 | 1914 | 2386 | 12761 | 2237 | 2971 | 11923 | 2597 | 2653 | 12667 | 2148 | 1897 |
| 313 | QGN13410.1 | FIM1_47 | CP015054. | 4952 | 762 | 724 | 5061 | 1025 | 664 | 5769 | 970 | 981 | 2258 | 371 | 529 | 2919 | 382 | 457 | 2489 | 344 | 371 |
| 314 | QGN13411.1 | FIM1_48 | CP015054. | 2576 | 1939 | 2456 | 3545 | 1774 | 2164 | 3137 | 1644 | 3268 | 6924 | 1201 | 2957 | 6441 | 1165 | 2739 | 7102 | 1070 | 1874 |
| 315 | QGN13412.1 | FIM1_49 | CP015054. | 11274 | 6066 | 2996 | 5286 | 7011 | 3533 | 9235 | 5555 | 3988 | 20654 | 9568 | 6467 | 21640 | 9189 | 5842 | 20723 | 10566 | 4625 |
| 316 | QGN13413.1 | FIM1_50 | CP015054. | 1817 | 4975 | 6537 | 2105 | 5794 | 4927 | 3163 | 9299 | 7120 | 12299 | 1783 | 4487 | 15021 | 2051 | 4337 | 13353 | 2474 | 3051 |
| 317 | QGN13414.1 | FIM1_51 | CP015054. | 7047 | 1938 | 2184 | 6525 | 3000 | 1813 | 5472 | 2205 | 2674 | 12250 | 2526 | 2851 | 10168 | 2109 | 2735 | 11935 | 1952 | 1784 |
| 318 | QGN13415.1 | FIM1_52 | CP015054. | 2599 | 2152 | 2385 | 5288 | 2261 | 2057 | 3965 | 1973 | 3021 | 10062 | 1784 | 3409 | 8744 | 1935 | 3017 | 8993 | 1859 | 2228 |
| 319 | QGN13416.1 | FIM1_53 | CP015054. | 25988 | 28234 | 18862 | 26575 | 23033 | 17846 | 21970 | 19652 | 21840 | 74816 | 28606 | 27879 | 66984 | 25495 | 28468 | 77160 | 24988 | 21135 |
| 320 | QGN13417.1 | FIM1_54 | CP015054. | 10781 | 1074 | 1741 | 12283 | 1612 | 1436 | 10560 | 1621 | 2340 | 2499 | 1020 | 2839 | 7915 | 974 | 2740 | 4265 | 872 | 1909 |
| 321 | QGN13418.1 | FIM1_55 | CP015054. | 1207 | 752 | 1915 | 1872 | 244 | 982 | 785 | 483 | 2119 | 1871 | 181 | 1350 | 2114 | 284 | 1207 | 2079 | 157 | 666 |
| 322 | QGN13419.1 | FIM1_56 | CP015054. | 1273 | 13 | 30 | 1391 | 24 | 24 | 983 | 16 | 31 | 84 | 16 | 34 | 109 | 11 | 24 | 135 | 12 | 20 |
| 323 | QGN13420.1 | FIM1_57 | CP015054. | 260 | 308 | 321 | 1210 | 300 | 264 | 454 | 283 | 446 | 1146 | 206 | 413 | 1307 | 266 | 348 | 1125 | 228 | 266 |
| 324 | QGN13421.1 | FIM1_58 | CP015054. | 4710 | 2385 | 1860 | 2966 | 3157 | 1715 | 4517 | 2489 | 2336 | 9372 | 2953 | 3088 | 8473 | 2736 | 2885 | 9112 | 2805 | 2128 |
| 325 | QGN13422.1 | FIM1_59 | CP015054. | 10743 | 1762 | 2543 | 15780 | 2388 | 1913 | 13980 | 2305 | 3315 | 2842 | 1931 | 4178 | 15077 | 2187 | 3597 | 6114 | 2184 | 2813 |
| 326 | QGN13423.1 | FIM1_60 | CP015054. | 1194 | 1999 | 1158 | 1651 | 1798 | 1172 | 1386 | 1424 | 1544 | 6964 | 1483 | 1866 | 6942 | 1372 | 1475 | 6883 | 1405 | 1152 |
| 327 | QGN13424.1 | FIM1_61 | CP015054. | 2997 | 715 | 860 | 4025 | 769 | 791 | 2975 | 688 | 1136 | 7029 | 606 | 1304 | 6502 | 539 | 1222 | 6227 | 531 | 972 |
| 328 | QGN13425.1 | FIM1_62 | CP015054. | 1046 | 835 | 648 | 1075 | 986 | 582 | 1080 | 696 | 866 | 5818 | 994 | 1234 | 4559 | 815 | 1117 | 5787 | 838 | 770 |
| 329 | QGN13426.1 | FIM1_63 | CP015054. | 47 | 22 | 48 | 231 | 41 | 35 | 119 | 24 | 56 | 184 | 22 | 68 | 172 | 33 | 60 | 175 | 21 | 41 |
| 330 | QGN13427.1 | FIM1_64 | CP015054. | 1658 | 2051 | 1691 | 2247 | 1863 | 1592 | 1746 | 1608 | 2219 | 5384 | 1150 | 2112 | 5498 | 1059 | 2080 | 5217 | 999 | 1382 |
| 331 | QGN13428.1 | FIM1_65 | CP015054. | 666 | 133 | 184 | 862 | 125 | 132 | 713 | 149 | 201 | 820 | 98 | 179 | 719 | 106 | 180 | 666 | 102 | 123 |
| 332 | QGN13429.1 | FIM1_66 | CP015054. | 4679 | 2736 | 2753 | 4137 | 3130 | 2413 | 4544 | 2385 | 3528 | 10199 | 3705 | 4360 | 15550 | 3423 | 3996 | 13081 | 3860 | 3031 |
| 333 | QGN13430.1 | FIM1_67 | CP015054. | 1060 | 159 | 143 | 1239 | 272 | 150 | 1065 | 164 | 230 | 1678 | 297 | 255 | 1398 | 377 | 340 | 1539 | 454 | 183 |
| 334 | QGN13431.1 | FIM1_68 | CP015054. | 12826 | 16282 | 8499 | 9681 | 10727 | 7756 | 7463 | 8840 | 10446 | 31804 | 12643 | 13513 | 30742 | 11404 | 13296 | 32983 | 11107 | 10127 |

| | | | | | | | | | | | | | | | | | | | | | |
|---|---|---|---|---|---|---|---|---|---|---|---|---|---|---|---|---|---|---|---|---|---|
| 335 | QGN13432.1 | FIM1_69 | CP015054. | 10645 | 14679 | 8606 | 10751 | 10768 | 8078 | 8973 | 9089 | 9839 | 29860 | 12368 | 12470 | 27108 | 10757 | 12193 | 31012 | 10832 | 9091 |
| 336 | QGN13433.1 | FIM1_70 | CP015054. | 2527 | 1025 | 463 | 1904 | 1128 | 514 | 2272 | 777 | 649 | 3858 | 980 | 802 | 3632 | 1051 | 719 | 3780 | 1124 | 544 |
| 337 | QGN13434.1 | FIM1_71 | CP015054. | 996 | 633 | 480 | 1459 | 509 | 484 | 1116 | 531 | 652 | 2835 | 400 | 646 | 1956 | 374 | 584 | 2594 | 378 | 414 |
| 338 | QGN13435.1 | FIM1_72 | CP015054. | 641 | 638 | 328 | 789 | 421 | 293 | 654 | 408 | 489 | 2569 | 430 | 462 | 1807 | 334 | 477 | 2776 | 369 | 298 |
| 339 | QGN13436.1 | FIM1_73 | CP015054. | 6657 | 11335 | 7909 | 11401 | 10788 | 7090 | 9191 | 9558 | 9611 | 31792 | 11133 | 12106 | 32872 | 10881 | 11958 | 31196 | 9870 | 8865 |
| 340 | QGN13437.1 | FIM1_74 | CP015054. | 461 | 655 | 535 | 546 | 629 | 512 | 402 | 558 | 695 | 2412 | 579 | 645 | 2281 | 477 | 618 | 2391 | 543 | 416 |
| 341 | QGN13438.1 | FIM1_75 | CP015054. | 73961 | 9893 | 14681 | 75972 | 10162 | 14053 | 68882 | 8833 | 17684 | 34455 | 9675 | 20882 | 39834 | 12581 | 20816 | 34792 | 10327 | 14449 |
| 342 | QGN13439.1 | FIM1_76 | CP015054. | 926 | 152 | 185 | 1220 | 228 | 151 | 904 | 231 | 239 | 674 | 146 | 211 | 1016 | 176 | 179 | 744 | 154 | 127 |
| 343 | QGN13440.1 | FIM1_77 | CP015054. | 1904 | 607 | 430 | 2285 | 680 | 385 | 1953 | 587 | 542 | 2399 | 491 | 573 | 2496 | 512 | 492 | 2454 | 482 | 343 |
| 344 | QGN13441.1 | FIM1_78 | CP015054. | 4875 | 1616 | 2114 | 5546 | 2358 | 1829 | 5351 | 2267 | 2631 | 6511 | 2006 | 2876 | 10705 | 1856 | 2841 | 8309 | 1746 | 2055 |
| 345 | QGN13442.1 | FIM1_79 | CP015054. | 20289 | 531 | 694 | 22010 | 647 | 600 | 22990 | 687 | 995 | 5361 | 497 | 895 | 7127 | 469 | 734 | 5567 | 461 | 565 |
| 346 | QGN13443.1 | FIM1_80 | CP015054. | 427 | 53 | 52 | 644 | 58 | 29 | 472 | 51 | 64 | 175 | 41 | 89 | 330 | 29 | 65 | 234 | 33 | 50 |
| 347 | QGN13444.1 | FIM1_81 | CP015054. | 3475 | 603 | 841 | 5298 | 894 | 706 | 3159 | 721 | 962 | 3063 | 732 | 1158 | 5044 | 671 | 964 | 3929 | 632 | 742 |
| 348 | QGN13445.1 | FIM1_82 | CP015054. | 2543 | 943 | 1210 | 2374 | 1239 | 1115 | 2388 | 1058 | 1587 | 7064 | 1356 | 1707 | 7579 | 1347 | 1576 | 6993 | 1266 | 1197 |
| 349 | QGN13446.1 | FIM1_83 | CP015054. | 646 | 335 | 249 | 520 | 225 | 193 | 459 | 241 | 355 | 1391 | 230 | 441 | 1299 | 216 | 410 | 1378 | 191 | 242 |
| 350 | QGN13447.1 | FIM1_84 | CP015054. | 1635 | 489 | 725 | 1362 | 386 | 611 | 1322 | 498 | 958 | 2632 | 335 | 970 | 2666 | 285 | 897 | 2711 | 246 | 588 |
| 351 | QGN13448.1 | FIM1_85 | CP015054. | 793 | 375 | 652 | 1235 | 392 | 612 | 845 | 317 | 900 | 2748 | 385 | 1104 | 2395 | 326 | 1064 | 2602 | 335 | 754 |
| 352 | QGN13449.1 | FIM1_86 | CP015054. | 1452 | 470 | 1098 | 1528 | 400 | 1024 | 1222 | 407 | 1538 | 2148 | 243 | 1329 | 1635 | 196 | 1338 | 1928 | 229 | 826 |
| 353 | QGN13450.1 | FIM1_87 | CP015054. | 1382 | 322 | 252 | 949 | 355 | 281 | 1341 | 234 | 396 | 1183 | 276 | 405 | 1216 | 364 | 360 | 1233 | 353 | 298 |
| 354 | QGN13451.1 | FIM1_88 | CP015054. | 1746 | 424 | 528 | 3625 | 591 | 606 | 2121 | 383 | 694 | 2073 | 269 | 828 | 2516 | 349 | 711 | 2097 | 235 | 508 |
| 355 | QGN13452.1 | FIM1_89 | CP015054. | 3453 | 70 | 261 | 2325 | 89 | 374 | 3074 | 59 | 466 | 2160 | 44 | 139 | 2022 | 206 | 173 | 1729 | 77 | 75 |
| 356 | QGN13453.1 | FIM1_90 | CP015054. | 2049 | 1737 | 1765 | 2197 | 2285 | 1461 | 2090 | 1888 | 2204 | 10339 | 2181 | 2099 | 11041 | 1951 | 2002 | 10171 | 1941 | 1440 |
| 357 | QGN13454.1 | FIM1_91 | CP015054. | 1714 | 1091 | 284 | 1141 | 1185 | 384 | 1659 | 768 | 382 | 5226 | 2050 | 1077 | 3690 | 2584 | 918 | 4594 | 3059 | 669 |
| 358 | QGN13455.1 | FIM1_92 | CP015054. | 472 | 120 | 280 | 513 | 184 | 198 | 383 | 187 | 356 | 995 | 170 | 349 | 1254 | 163 | 296 | 1092 | 121 | 222 |
| 359 | QGN13456.1 | FIM1_93 | CP015054. | 410 | 185 | 225 | 429 | 126 | 190 | 381 | 157 | 303 | 772 | 113 | 247 | 858 | 117 | 241 | 770 | 107 | 189 |
| 360 | QGN13457.1 | FIM1_94 | CP015054. | 673 | 172 | 315 | 933 | 154 | 270 | 578 | 170 | 428 | 1534 | 100 | 445 | 1338 | 88 | 361 | 1533 | 101 | 245 |
| 361 | QGN13458.1 | FIM1_95 | CP015054. | 5133 | 4441 | 3038 | 5117 | 4251 | 2637 | 4794 | 4328 | 4041 | 11931 | 3633 | 4664 | 18631 | 4046 | 4315 | 13788 | 3466 | 3157 |
| 362 | QGN13459.1 | FIM1_96 | CP015054. | 6168 | 178 | 287 | 3756 | 202 | 169 | 4585 | 191 | 320 | 759 | 115 | 469 | 1853 | 114 | 401 | 993 | 81 | 313 |
| 363 | QGN13460.1 | FIM1_97 | CP015054. | 18727 | 3693 | 4436 | 5281 | 5128 | 5168 | 6855 | 4918 | 5806 | 6763 | 1825 | 2313 | 9484 | 1480 | 1947 | 8245 | 1268 | 1924 |
| 364 | QGN13461.1 | FIM1_98 | CP015054. | 384 | 206 | 146 | 218 | 100 | 114 | 221 | 138 | 184 | 815 | 140 | 227 | 602 | 95 | 233 | 726 | 104 | 121 |
| 365 | QGN13462.1 | FIM1_99 | CP015054. | 440 | 454 | 369 | 325 | 341 | 279 | 354 | 383 | 486 | 761 | 279 | 522 | 1663 | 252 | 532 | 1126 | 251 | 351 |
| 366 | QGN13463.1 | FIM1_100 | CP015054. | 4112 | 4084 | 2644 | 5245 | 3183 | 2175 | 4541 | 2984 | 3327 | 4791 | 2455 | 2875 | 9356 | 2587 | 3092 | 6918 | 2377 | 2152 |
| 367 | QGN13464.1 | FIM1_101 | CP015054. | 10128 | 682 | 584 | 7462 | 510 | 498 | 8651 | 496 | 845 | 1667 | 463 | 700 | 3285 | 429 | 775 | 2014 | 501 | 532 |
| 368 | QGN13465.1 | FIM1_102 | CP015054. | 419 | 130 | 133 | 666 | 95 | 134 | 455 | 107 | 214 | 1050 | 106 | 214 | 720 | 69 | 185 | 1065 | 94 | 129 |
| 369 | QGN13466.1 | FIM1_103 | CP015054. | 1306 | 899 | 1030 | 1921 | 1122 | 994 | 1638 | 856 | 1333 | 8934 | 1064 | 1854 | 6822 | 1135 | 1673 | 8325 | 1022 | 1191 |
| 370 | QGN13467.1 | FIM1_104 | CP015054. | 738 | 382 | 437 | 982 | 342 | 383 | 882 | 315 | 590 | 2560 | 296 | 700 | 2620 | 288 | 641 | 2598 | 275 | 450 |
| 371 | QGN13468.1 | FIM1_105 | CP015054. | 6032 | 3310 | 1359 | 5159 | 2977 | 1781 | 5816 | 2017 | 1914 | 11010 | 3331 | 2787 | 9279 | 2395 | 2670 | 10375 | 2893 | 1917 |
| 372 | QGN13469.1 | FIM1_107 | CP015054. | 646 | 773 | 628 | 1134 | 581 | 598 | 943 | 506 | 867 | 3876 | 569 | 983 | 3022 | 504 | 982 | 3515 | 488 | 636 |
| 373 | QGN13470.1 | FIM1_108 | CP015054. | 1012 | 2419 | 1388 | 988 | 1972 | 1244 | 1034 | 1603 | 1702 | 7715 | 1696 | 1758 | 5377 | 1480 | 1759 | 7596 | 1633 | 1109 |
| 374 | QGN13471.1 | FIM1_109 | CP015054. | 1741 | 1038 | 741 | 2212 | 728 | 719 | 2067 | 726 | 1030 | 3016 | 711 | 994 | 3443 | 735 | 992 | 2984 | 682 | 669 |
| 375 | QGN13472.1 | FIM1_110 | CP015054. | 1787 | 278 | 254 | 1427 | 298 | 267 | 1700 | 275 | 384 | 2342 | 272 | 437 | 1998 | 238 | 375 | 2270 | 238 | 283 |
| 376 | QGN13473.1 | FIM1_111 | CP015054. | 5520 | 2177 | 4862 | 7137 | 2958 | 4109 | 6066 | 2785 | 6261 | 20502 | 3212 | 8033 | 20595 | 2939 | 7124 | 20563 | 2520 | 5554 |
| 377 | QGN13474.1 | FIM1_112 | CP015054. | 815 | 661 | 434 | 614 | 678 | 436 | 786 | 585 | 567 | 3811 | 768 | 783 | 3125 | 628 | 708 | 3683 | 681 | 548 |
| 378 | QGN13475.1 | FIM1_113 | CP015054. | 1782 | 403 | 732 | 3033 | 541 | 568 | 2348 | 487 | 972 | 4475 | 469 | 983 | 5352 | 453 | 834 | 4618 | 383 | 671 |
| 379 | QGN13476.1 | FIM1_114 | CP015054. | 1459 | 741 | 659 | 1255 | 603 | 537 | 1187 | 496 | 824 | 4272 | 559 | 915 | 3332 | 568 | 884 | 4122 | 566 | 603 |
| 380 | QGN13477.1 | FIM1_115 | CP015054. | 2766 | 127 | 525 | 4645 | 487 | 457 | 3182 | 368 | 733 | 3860 | 313 | 820 | 4268 | 300 | 663 | 3846 | 254 | 559 |
| 381 | QGN13478.1 | FIM1_116 | CP015054. | 788 | 325 | 300 | 1269 | 263 | 274 | 727 | 224 | 439 | 1208 | 237 | 405 | 1241 | 190 | 400 | 1184 | 229 | 314 |
| 382 | QGN13479.1 | FIM1_117 | CP015054. | 3468 | 5818 | 6564 | 9834 | 8910 | 5410 | 7395 | 7290 | 7592 | 34502 | 9880 | 10390 | 31021 | 9639 | 9888 | 34338 | 8759 | 7222 |
| 383 | QGN13480.1 | FIM1_118 | CP015054. | 456 | 54 | 193 | 758 | 73 | 156 | 422 | 69 | 292 | 287 | 61 | 302 | 505 | 51 | 215 | 386 | 45 | 150 |
| 384 | QGN13481.1 | FIM1_119 | CP015054. | 4862 | 7652 | 3417 | 3788 | 5605 | 3851 | 4167 | 4451 | 4434 | 18515 | 7763 | 5362 | 15656 | 5433 | 5808 | 17254 | 6355 | 4037 |
| 385 | QGN13482.1 | FIM1_120 | CP015054. | 1706 | 1013 | 1021 | 1907 | 1176 | 833 | 1846 | 1044 | 1237 | 2523 | 873 | 1171 | 4208 | 860 | 1136 | 3137 | 873 | 820 |
| 386 | QGN13483.1 | FIM1_121 | CP015054. | 6680 | 1567 | 1207 | 3406 | 2448 | 1270 | 6289 | 1864 | 1532 | 12795 | 2818 | 2499 | 10125 | 2765 | 2122 | 11923 | 3082 | 1775 |
| 387 | QGN13484.1 | FIM1_122 | CP015054. | 3407 | 2245 | 2032 | 4406 | 2278 | 1763 | 4332 | 1871 | 2592 | 12977 | 2269 | 3137 | 11486 | 2133 | 2944 | 12490 | 2088 | 2093 |
| 388 | QGN13485.1 | FIM1_123 | CP015054. | 3450 | 4349 | 3663 | 4678 | 3662 | 2938 | 3617 | 3120 | 4285 | 14930 | 3510 | 4983 | 11977 | 3666 | 4538 | 15165 | 3130 | 3190 |
| 389 | QGN13486.1 | FIM1_124 | CP015054. | 562 | 560 | 444 | 528 | 525 | 355 | 364 | 468 | 585 | 1832 | 399 | 515 | 2521 | 484 | 497 | 2122 | 411 | 351 |
| 390 | QGN13487.1 | FIM1_125 | CP015054. | 4000 | 486 | 458 | 5569 | 592 | 497 | 3148 | 450 | 682 | 350 | 113 | 213 | 604 | 117 | 241 | 461 | 116 | 151 |
| 391 | QGN13488.1 | FIM1_126 | CP015054. | 968 | 1271 | 958 | 1107 | 998 | 812 | 875 | 986 | 1149 | 4698 | 903 | 1128 | 4676 | 809 | 1227 | 4410 | 665 | 750 |
| 392 | QGN13489.1 | FIM1_127 | CP015054. | 1030 | 799 | 741 | 2291 | 571 | 575 | 1378 | 495 | 998 | 2538 | 319 | 827 | 1955 | 418 | 752 | 2236 | 326 | 502 |
| 393 | QGN13490.1 | FIM1_128 | CP015054. | 9564 | 12278 | 11656 | 9148 | 12460 | 8752 | 8877 | 11294 | 13373 | 26045 | 13165 | 17402 | 45425 | 12231 | 15159 | 35706 | 11562 | 11145 |
| 394 | QGN13491.1 | FIM1_129 | CP015054. | 2773 | 887 | 491 | 1615 | 852 | 502 | 2656 | 804 | 627 | 1193 | 692 | 716 | 2553 | 633 | 725 | 1723 | 652 | 498 |
| 395 | QGN13492.1 | FIM1_130 | CP015054. | 17359 | 5149 | 9090 | 22523 | 6079 | 7247 | 18033 | 5811 | 10897 | 18355 | 4376 | 11347 | 25285 | 5295 | 10294 | 20930 | 4386 | 7415 |
| 396 | QGN13493.1 | FIM1_131 | CP015054. | 1795 | 1399 | 1218 | 3358 | 1922 | 1119 | 2460 | 1614 | 1594 | 3529 | 1544 | 1472 | 7000 | 1563 | 1309 | 4796 | 1436 | 1017 |
| 397 | QGN13494.1 | FIM1_132 | CP015054. | 13492 | 17448 | 13121 | 15042 | 15553 | 10971 | 13805 | 14190 | 14886 | 40382 | 19981 | 18468 | 42978 | 17564 | 18955 | 41737 | 16375 | 14657 |
| 398 | QGN13495.1 | FIM1_133 | CP015054. | 5543 | 4095 | 3163 | 6812 | 5277 | 2797 | 5800 | 4859 | 4007 | 7334 | 3213 | 3252 | 14895 | 2994 | 3074 | 10262 | 2501 | 2189 |
| 399 | QGN13496.1 | FIM1_134 | CP015054. | 4057 | 2415 | 2873 | 6732 | 2836 | 2362 | 5700 | 2745 | 3601 | 9148 | 2724 | 4282 | 12583 | 2791 | 3974 | 10006 | 2625 | 2808 |
| 400 | QGN13497.1 | FIM1_135 | CP015054. | 1212 | 1108 | 932 | 1359 | 926 | 807 | 1198 | 890 | 1270 | 4938 | 799 | 1105 | 4307 | 819 | 1087 | 4783 | 682 | 756 |
| 401 | QGN13498.1 | FIM1_136 | CP015054. | 2645 | 46 | 64 | 1410 | 36 | 57 | 2398 | 52 | 73 | 150 | 32 | 62 | 230 | 30 | 61 | 187 | 12 | 47 |
| 402 | QGN13499.1 | FIM1_137 | CP015054. | 4536 | 253 | 456 | 5797 | 327 | 363 | 3850 | 262 | 609 | 1289 | 141 | 681 | 2327 | 154 | 651 | 1589 | 117 | 432 |
| 403 | QGN13500.1 | FIM1_138 | CP015054. | 9135 | 11711 | 6498 | 6808 | 8215 | 5619 | 7088 | 7267 | 7680 | 24087 | 9584 | 9787 | 19333 | 8177 | 9771 | 22897 | 8574 | 7122 |
| 404 | QGN13501.1 | FIM1_140 | CP015054. | 13753 | 16924 | 10565 | 13254 | 12882 | 10088 | 11101 | 10871 | 12424 | 36995 | 15503 | 14677 | 32170 | 13323 | 15407 | 37175 | 13762 | 11163 |
| 405 | QGN13502.1 | FIM1_141 | CP015054. | 291 | 537 | 556 | 440 | 652 | 403 | 231 | 630 | 709 | 2081 | 453 | 578 | 2435 | 495 | 513 | 2326 | 433 | 357 |
| 406 | QGN13503.1 | FIM1_142 | CP015054. | 8185 | 10273 | 7508 | 11140 | 9061 | 6543 | 8613 | 7829 | 9464 | 30705 | 8812 | 11685 | 32105 | 8959 | 11009 | 30891 | 8315 | 8054 |
| 407 | QGN13504.1 | FIM1_143 | CP015054. | 1590 | 81 | 117 | 1009 | 111 | 92 | 1123 | 116 | 128 | 366 | 71 | 131 | 711 | 125 | 86 | 471 | 102 | 65 |
| 408 | QGN13505.1 | FIM1_144 | CP015054. | 787 | 281 | 525 | 2850 | 1140 | 753 | 1012 | 296 | 628 | 3764 | 450 | 1117 | 3680 | 622 | 1053 | 3629 | 569 | 752 |
| 409 | QGN13506.1 | FIM1_145 | CP015054. | 13784 | 7568 | 6080 | 28244 | 8513 | 5840 | 25723 | 7188 | 8071 | 14305 | 10767 | 10381 | 31623 | 9254 | 10207 | 23109 | 9987 | 7720 |
| 410 | QGN13507.1 | FIM1_146 | CP015054. | 4023 | 2245 | 1383 | 4265 | 2006 | 1357 | 3973 | 1751 | 1921 | 4663 | 1540 | 2053 | 6862 | 1565 | 2055 | 5144 | 1590 | 1356 |
| 411 | QGN13508.1 | FIM1_147 | CP015054. | 3036 | 1060 | 1301 | 3239 | 1373 | 1146 | 2926 | 1196 | 1638 | 5957 | 1168 | 2069 | 7554 | 1091 | 1831 | 6573 | 1022 | 1364 |
| 412 | QGN13509.1 | FIM1_148 | CP015054. | 1072 | 481 | 484 | 1209 | 505 | 396 | 1338 | 468 | 637 | 2870 | 459 | 678 | 2872 | 415 | 578 | 2696 | 358 | 446 |
| 413 | QGN13510.1 | FIM1_149 | CP015054. | 371 | 440 | 452 | 389 | 331 | 362 | 376 | 397 | 535 | 2411 | 374 | 583 | 1941 | 352 | 561 | 2186 | 325 | 374 |
| 414 | QGN13511.1 | FIM1_150 | CP015054. | 1359 | 555 | 702 | 1653 | 589 | 663 | 1491 | 635 | 909 | 4038 | 501 | 982 | 3870 | 429 | 1018 | 3732 | 421 | 678 |
| 415 | QGN13512.1 | FIM1_151 | CP015054. | 1762 | 574 | 578 | 2110 | 445 | 483 | 1674 | 413 | 739 | 4068 | 425 | 947 | 3261 | 437 | 902 | 4235 | 435 | 559 |
| 416 | QGN13513.1 | FIM1_153 | CP015054. | 398 | 328 | 455 | 287 | 360 | 383 | 327 | 326 | 502 | 2493 | 376 | 635 | 2113 | 330 | 545 | 2516 | 356 | 397 |
| 417 | QGN13514.1 | FIM1_154 | CP015054. | 6728 | 1029 | 1228 | 5192 | 1017 | 922 | 5356 | 1134 | 1770 | 1273 | 852 | 1835 | 5942 | 733 | 1707 | 2803 | 766 | 1246 |
| 418 | QGN13515.1 | FIM1_155 | CP015054. | 5999 | 2548 | 2528 | 5228 | 2233 | 1795 | 5355 | 2564 | 3272 | 2798 | 2038 | 3479 | 8743 | 1877 | 3485 | 4722 | 1969 | 2440 |

| | | | | | | | | | | | | | | | | | | | | | |
|---|---|---|---|---|---|---|---|---|---|---|---|---|---|---|---|---|---|---|---|---|---|
| 419 | QGN13516.1 | FIM1_156 | CP015054. | 5034 | 1449 | 1024 | 4830 | 1147 | 845 | 4469 | 1027 | 1483 | 2482 | 633 | 897 | 3247 | 574 | 1011 | 2847 | 549 | 628 |
| 420 | QGN13517.1 | FIM1_157 | CP015054. | 12898 | 1556 | 1363 | 10143 | 934 | 898 | 10158 | 1133 | 1856 | 1504 | 472 | 1367 | 4804 | 956 | 1319 | 3105 | 563 | 928 |
| 421 | QGN13518.1 | FIM1_158 | CP015054. | 3882 | 5834 | 5617 | 6616 | 5054 | 5020 | 4492 | 4815 | 7275 | 17038 | 4359 | 7480 | 17626 | 4773 | 6523 | 16200 | 4027 | 5148 |
| 422 | QGN13519.1 | FIM1_159 | CP015054. | 6558 | 4536 | 4254 | 7613 | 4565 | 3665 | 6776 | 4250 | 5430 | 16295 | 4218 | 6168 | 18504 | 3972 | 5988 | 16857 | 3591 | 4221 |
| 423 | QGN13520.1 | FIM1_160 | CP015054. | 4800 | 4512 | 3627 | 7438 | 4038 | 3338 | 5814 | 3674 | 4669 | 14439 | 3579 | 5650 | 15763 | 3589 | 5179 | 14425 | 3334 | 3694 |
| 424 | QGN13521.1 | FIM1_161 | CP015054. | 1525 | 304 | 532 | 2616 | 304 | 381 | 1384 | 324 | 639 | 1011 | 219 | 625 | 2060 | 286 | 606 | 1404 | 240 | 422 |
| 425 | QGN13522.1 | FIM1_162 | CP015054. | 2139 | 760 | 809 | 2337 | 1174 | 702 | 1644 | 720 | 1077 | 5487 | 939 | 941 | 6438 | 788 | 812 | 5127 | 703 | 677 |
| 426 | QGN13523.1 | FIM1_163 | CP015054. | 2272 | 183 | 196 | 3302 | 108 | 126 | 3197 | 145 | 223 | 1730 | 167 | 431 | 1691 | 412 | 382 | 1535 | 237 | 266 |
| 427 | QGN13524.1 | FIM1_164 | CP015054. | 2389 | 438 | 675 | 2380 | 470 | 498 | 2686 | 362 | 752 | 1777 | 400 | 770 | 2039 | 396 | 710 | 1862 | 333 | 454 |
| 428 | QGN13525.1 | FIM1_165 | CP015054. | 247 | 531 | 382 | 335 | 443 | 368 | 268 | 351 | 532 | 2786 | 431 | 533 | 1981 | 399 | 498 | 2578 | 431 | 324 |
| 429 | QGN13526.1 | FIM1_166 | CP015054. | 680 | 351 | 387 | 996 | 422 | 415 | 1105 | 356 | 575 | 3962 | 417 | 593 | 3185 | 323 | 586 | 4042 | 356 | 430 |
| 430 | QGN13527.1 | FIM1_167 | CP015054. | 3110 | 131 | 191 | 2100 | 147 | 223 | 1908 | 108 | 296 | 2144 | 177 | 438 | 1699 | 175 | 385 | 1947 | 177 | 253 |
| 431 | QGN13528.1 | FIM1_168 | CP015054. | 1345 | 713 | 779 | 1334 | 787 | 737 | 1330 | 706 | 1129 | 6093 | 865 | 1247 | 4759 | 693 | 1154 | 5817 | 812 | 843 |
| 432 | QGN13529.1 | FIM1_169 | CP015054. | 11132 | 834 | 1204 | 8573 | 912 | 935 | 10717 | 904 | 1494 | 5805 | 770 | 1547 | 5718 | 624 | 1399 | 5661 | 674 | 894 |
| 433 | QGN13530.1 | FIM1_170 | CP015054. | 13646 | 7161 | 5811 | 16489 | 7297 | 5304 | 13520 | 6036 | 7015 | 27455 | 8525 | 11037 | 30409 | 7984 | 10456 | 28005 | 8163 | 7397 |
| 434 | QGN13531.1 | FIM1_171 | CP015054. | 644 | 1659 | 1122 | 788 | 1348 | 1113 | 793 | 1080 | 1526 | 6979 | 1313 | 1740 | 5569 | 1242 | 1598 | 6799 | 1383 | 1156 |
| 435 | QGN13532.1 | FIM1_172 | CP015054. | 1004 | 615 | 494 | 1058 | 439 | 510 | 878 | 330 | 706 | 2258 | 393 | 699 | 1878 | 355 | 735 | 2030 | 392 | 469 |
| 436 | QGN13533.1 | FIM1_173 | CP015054. | 1383 | 543 | 529 | 1303 | 621 | 449 | 1228 | 521 | 656 | 2960 | 668 | 813 | 2364 | 575 | 826 | 2715 | 667 | 539 |
| 437 | QGN13534.1 | FIM1_174 | CP015054. | 13 | 219 | 225 | 254 | 264 | 181 | 116 | 186 | 296 | 1479 | 243 | 349 | 1153 | 232 | 333 | 1351 | 207 | 232 |
| 438 | QGN13535.1 | FIM1_175 | CP015054. | 13 | 89 | 51 | 53 | 76 | 60 | 20 | 49 | 88 | 392 | 68 | 94 | 274 | 65 | 90 | 312 | 60 | 72 |
| 439 | QGN13536.1 | FIM1_176 | CP015054. | 1807 | 1373 | 1050 | 2378 | 1729 | 937 | 2008 | 1319 | 1333 | 9377 | 1436 | 1815 | 8096 | 1319 | 1699 | 9189 | 1342 | 1294 |
| 440 | QGN13537.1 | FIM1_177 | CP015054. | 524 | 8730 | 4711 | 1856 | 6347 | 4455 | 1075 | 5351 | 5761 | 22270 | 7833 | 8391 | 17826 | 6709 | 7962 | 21075 | 7099 | 5617 |
| 441 | QGN13538.1 | FIM1_178 | CP015054. | 958 | 501 | 667 | 1133 | 564 | 587 | 951 | 517 | 835 | 3356 | 527 | 955 | 2921 | 546 | 868 | 3398 | 502 | 653 |
| 442 | QGN13539.1 | FIM1_179 | CP015054. | 4842 | 1216 | 1481 | 4296 | 1587 | 1203 | 3989 | 1379 | 1781 | 10951 | 1843 | 2103 | 9617 | 1540 | 2181 | 10635 | 1342 | 1518 |
| 443 | QGN13540.1 | FIM1_180 | CP015054. | 4902 | 1142 | 1011 | 3738 | 1069 | 970 | 4120 | 1011 | 1407 | 4691 | 891 | 1394 | 5440 | 917 | 1281 | 4665 | 866 | 925 |
| 444 | QGN13541.1 | FIM1_181 | CP015054. | 11855 | 9184 | 6129 | 13635 | 9199 | 6062 | 11692 | 7337 | 7835 | 32248 | 10045 | 10794 | 31815 | 8960 | 10181 | 32318 | 8975 | 7417 |
| 445 | QGN13542.1 | FIM1_182 | CP015054. | 1023 | 3429 | 4161 | 3745 | 5254 | 3549 | 2858 | 4559 | 5299 | 21400 | 4060 | 5269 | 21274 | 3284 | 4485 | 19895 | 3099 | 3438 |
| 446 | QGN13543.1 | FIM1_183 | CP015054. | 66 | 43 | 62 | 218 | 59 | 42 | 66 | 38 | 74 | 393 | 54 | 107 | 323 | 52 | 89 | 373 | 45 | 78 |
| 447 | QGN13544.1 | FIM1_184 | CP015054. | 4964 | 1826 | 2835 | 6072 | 2812 | 2410 | 5514 | 2535 | 3613 | 8247 | 2724 | 4642 | 14085 | 2585 | 4056 | 10766 | 2599 | 3071 |
| 448 | QGN13545.1 | FIM1_185 | CP015054. | 14283 | 228 | 335 | 9335 | 430 | 245 | 10560 | 344 | 393 | 1185 | 358 | 454 | 1856 | 314 | 375 | 1347 | 305 | 292 |
| 449 | QGN13546.1 | FIM1_186 | CP015054. | 1578 | 238 | 254 | 1194 | 147 | 170 | 1335 | 198 | 300 | 348 | 113 | 239 | 1064 | 140 | 237 | 554 | 105 | 162 |
| 450 | QGN13547.1 | FIM1_187 | CP015054. | 7170 | 3753 | 5185 | 16157 | 8424 | 6778 | 7213 | 5202 | 6378 | 10232 | 4136 | 6186 | 18867 | 3769 | 5086 | 13932 | 5255 | 4272 |
| 451 | QGN13548.1 | FIM1_189 | CP015054. | 1589 | 1153 | 804 | 2091 | 1069 | 758 | 1333 | 963 | 1087 | 3781 | 977 | 994 | 3721 | 860 | 1107 | 3537 | 804 | 690 |
| 452 | QGN13549.1 | FIM1_190 | CP015054. | 29 | 106 | 132 | 77 | 144 | 124 | 20 | 104 | 168 | 592 | 106 | 216 | 588 | 119 | 171 | 614 | 104 | 116 |
| 453 | QGN13550.1 | FIM1_191 | CP015054. | 1721 | 510 | 458 | 2248 | 529 | 363 | 1902 | 561 | 607 | 1385 | 261 | 559 | 2135 | 242 | 476 | 1507 | 202 | 346 |
| 454 | QGN13551.1 | FIM1_192 | CP015054. | 12174 | 1031 | 2401 | 13352 | 2007 | 1794 | 10854 | 2100 | 3079 | 1291 | 1075 | 3262 | 8677 | 1135 | 2914 | 3275 | 1221 | 2244 |
| 455 | QGN13552.1 | FIM1_193 | CP015054. | 10436 | 34 | 22 | 3823 | 15 | 11 | 6626 | 28 | 25 | 37 | 9 | 34 | 92 | 15 | 29 | 61 | 6 | 14 |
| 456 | QGN13553.1 | FIM1_194 | CP015054. | 5090 | 1560 | 1630 | 3791 | 2078 | 1244 | 3788 | 2202 | 2134 | 3174 | 1494 | 2044 | 8011 | 1304 | 1926 | 5098 | 1240 | 1362 |
| 457 | QGN13554.1 | FIM1_195 | CP015054. | 1117 | 1327 | 905 | 1051 | 977 | 870 | 1041 | 849 | 1274 | 4745 | 878 | 1030 | 3491 | 644 | 1172 | 4212 | 671 | 768 |
| 458 | QGN13555.1 | FIM1_196 | CP015054. | 1462 | 3698 | 1767 | 1590 | 2562 | 1637 | 1508 | 2367 | 2309 | 6977 | 2279 | 2269 | 7568 | 2172 | 2277 | 6714 | 2187 | 1681 |
| 459 | QGN13556.1 | FIM1_197 | CP015054. | 3326 | 1176 | 1437 | 5596 | 1033 | 1054 | 3417 | 1022 | 1871 | 6788 | 831 | 1790 | 6728 | 1568 | 1600 | 6312 | 1026 | 1078 |
| 460 | QGN13557.1 | FIM1_198 | CP015054. | 14930 | 2610 | 2035 | 12521 | 3029 | 1790 | 13336 | 2684 | 2606 | 5823 | 2848 | 2705 | 11918 | 3043 | 2677 | 7615 | 2962 | 1894 |
| 461 | QGN13558.1 | FIM1_199 | CP015054. | 2191 | 822 | 780 | 3198 | 629 | 615 | 2530 | 745 | 950 | 1116 | 447 | 879 | 2661 | 421 | 859 | 1762 | 438 | 626 |
| 462 | QGN13559.1 | FIM1_200 | CP015054. | 1925 | 912 | 1369 | 1978 | 1765 | 1089 | 2295 | 1450 | 1612 | 10659 | 1906 | 2485 | 11257 | 1769 | 2119 | 10688 | 1685 | 1592 |
| 463 | QGN13560.1 | FIM1_201 | CP015054. | 2279 | 1589 | 1361 | 2856 | 1226 | 1227 | 2224 | 1218 | 1950 | 8356 | 1240 | 1758 | 7064 | 1018 | 1743 | 7805 | 1033 | 1217 |
| 464 | QGN13561.1 | FIM1_202 | CP015054. | 22 | 5 | 9 | 31 | 6 | 5 | 19 | 12 | 8 | 77 | 8 | 22 | 59 | 7 | 16 | 75 | 6 | 8 |
| 465 | QGN13562.1 | FIM1_203 | CP015054. | 1337 | 715 | 891 | 1539 | 538 | 760 | 1425 | 531 | 1120 | 5906 | 562 | 1620 | 4590 | 571 | 1461 | 6001 | 495 | 1018 |
| 466 | QGN13563.1 | FIM1_204 | CP015054. | 295 | 87 | 113 | 427 | 127 | 130 | 167 | 93 | 157 | 533 | 67 | 137 | 542 | 57 | 121 | 545 | 62 | 80 |
| 467 | QGN13564.1 | FIM1_205 | CP015054. | 2587 | 1409 | 750 | 1644 | 1506 | 746 | 2175 | 1345 | 1024 | 1505 | 1155 | 1005 | 3820 | 1014 | 955 | 2179 | 1105 | 733 |
| 468 | QGN13565.1 | FIM1_206 | CP015054. | 2460 | 2367 | 1860 | 4178 | 2244 | 1717 | 3235 | 1880 | 2458 | 9129 | 1618 | 2499 | 8597 | 1807 | 2489 | 9062 | 1537 | 1680 |
| 469 | QGN13566.1 | FIM1_207 | CP015054. | 417 | 869 | 599 | 640 | 499 | 568 | 495 | 461 | 821 | 2708 | 502 | 1046 | 2463 | 527 | 1029 | 2417 | 463 | 681 |
| 470 | QGN13567.1 | FIM1_208 | CP015054. | 2572 | 2092 | 2418 | 4741 | 2131 | 1929 | 3668 | 1978 | 2834 | 9999 | 1861 | 3602 | 9445 | 2196 | 3433 | 9443 | 1780 | 2421 |
| 471 | QGN13568.1 | FIM1_209 | CP015054. | 8231 | 2159 | 2328 | 12639 | 2729 | 2000 | 12101 | 2349 | 3102 | 9196 | 2265 | 3607 | 14959 | 2267 | 3170 | 11404 | 1925 | 2415 |
| 472 | QGN13569.1 | FIM1_210 | CP015054. | 1104 | 2886 | 1747 | 2093 | 2217 | 1820 | 1561 | 1882 | 2385 | 8243 | 2195 | 2900 | 7620 | 2051 | 2775 | 8028 | 2240 | 2012 |
| 473 | QGN13570.1 | FIM1_211 | CP015054. | 2051 | 707 | 908 | 2981 | 669 | 824 | 2100 | 622 | 1262 | 7055 | 628 | 1653 | 5369 | 571 | 1395 | 6732 | 584 | 966 |
| 474 | QGN13571.1 | FIM1_212 | CP015054. | 1045 | 848 | 847 | 2290 | 772 | 703 | 1503 | 801 | 1156 | 6135 | 724 | 1377 | 5360 | 669 | 1313 | 5546 | 557 | 889 |
| 475 | QGN13572.1 | FIM1_213 | CP015054. | 871 | 927 | 919 | 922 | 979 | 818 | 1032 | 836 | 1112 | 6156 | 990 | 1411 | 5648 | 929 | 1232 | 5793 | 958 | 872 |
| 476 | QGN13573.1 | FIM1_214 | CP015054. | 472 | 167 | 300 | 549 | 146 | 256 | 297 | 181 | 390 | 2274 | 114 | 437 | 1613 | 86 | 386 | 2212 | 101 | 271 |
| 477 | QGN13574.1 | FIM1_215 | CP015054. | 1250 | 398 | 452 | 1362 | 461 | 374 | 1095 | 423 | 585 | 3040 | 399 | 717 | 2867 | 370 | 698 | 2832 | 398 | 512 |
| 478 | QGN13575.1 | FIM1_216 | CP015054. | 2009 | 3117 | 2420 | 5797 | 3494 | 1974 | 3806 | 3159 | 2950 | 7951 | 2350 | 3411 | 12251 | 2308 | 2981 | 9093 | 2053 | 2282 |
| 479 | QGN13576.1 | FIM1_217 | CP015054. | 2095 | 3056 | 1804 | 2873 | 1965 | 1508 | 2189 | 1702 | 2340 | 6988 | 1864 | 2281 | 6784 | 1821 | 2465 | 6264 | 1638 | 1633 |
| 480 | QGN13577.1 | FIM1_218 | CP015054. | 16401 | 527 | 760 | 10040 | 513 | 498 | 12400 | 626 | 983 | 1225 | 436 | 1386 | 4329 | 659 | 1127 | 2438 | 352 | 884 |
| 481 | QGN13578.1 | FIM1_219 | CP015054. | 1840 | 426 | 728 | 2252 | 418 | 593 | 2089 | 419 | 935 | 3616 | 502 | 1023 | 4109 | 473 | 916 | 3627 | 358 | 682 |
| 482 | QGN13579.1 | FIM1_220 | CP015054. | 1419 | 293 | 245 | 1217 | 203 | 250 | 1342 | 202 | 369 | 1199 | 197 | 371 | 1085 | 153 | 327 | 1318 | 134 | 221 |
| 483 | QGN13580.1 | FIM1_221 | CP015054. | 874 | 1023 | 860 | 1211 | 691 | 591 | 1095 | 622 | 1015 | 4891 | 830 | 1284 | 3667 | 751 | 1134 | 4850 | 787 | 775 |
| 484 | QGN13581.1 | FIM1_222 | CP015054. | 4339 | 808 | 873 | 4494 | 851 | 763 | 3927 | 744 | 1160 | 5475 | 988 | 1873 | 6353 | 1048 | 1598 | 5722 | 962 | 1258 |
| 485 | QGN13582.1 | FIM1_224 | CP015054. | 867 | 1682 | 1389 | 859 | 1413 | 1339 | 771 | 1282 | 1743 | 7200 | 1606 | 1798 | 5930 | 1385 | 1936 | 6656 | 1457 | 1240 |
| 486 | QGN13583.1 | FIM1_225 | CP015054. | 1172 | 1166 | 1015 | 1166 | 868 | 907 | 1110 | 926 | 1319 | 2195 | 466 | 1267 | 2846 | 476 | 1119 | 2130 | 487 | 803 |
| 487 | QGN13584.1 | FIM1_226 | CP015054. | 893 | 229 | 488 | 1207 | 118 | 331 | 1022 | 185 | 687 | 747 | 85 | 426 | 774 | 133 | 436 | 656 | 92 | 278 |
| 488 | QGN13585.1 | FIM1_227 | CP015054. | 3517 | 990 | 1162 | 3293 | 860 | 808 | 3391 | 962 | 1506 | 971 | 840 | 1411 | 1997 | 1089 | 1656 | 1544 | 971 | 1068 |
| 489 | QGN13586.1 | FIM1_228 | CP015054. | 89 | 77 | 89 | 115 | 70 | 69 | 88 | 73 | 111 | 185 | 44 | 158 | 186 | 51 | 149 | 167 | 47 | 99 |
| 490 | QGN13587.1 | FIM1_229 | CP015054. | 3412 | 4283 | 6452 | 7060 | 4003 | 5574 | 4953 | 3245 | 8277 | 16197 | 4120 | 9602 | 21751 | 4689 | 9185 | 17951 | 4164 | 6626 |
| 491 | QGN13588.1 | FIM1_230 | CP015054. | 3071 | 1431 | 1340 | 3216 | 1127 | 1191 | 2763 | 1189 | 1780 | 5914 | 1051 | 2086 | 5799 | 1054 | 2092 | 6158 | 1018 | 1323 |
| 492 | QGN13589.1 | FIM1_231 | CP015054. | 348 | 126 | 305 | 217 | 128 | 264 | 104 | 212 | 366 | 449 | 59 | 206 | 580 | 29 | 171 | 516 | 29 | 120 |
| 493 | QGN13590.1 | FIM1_232 | CP015054. | 2 | 2 | 2 | 1 | 1 | 2 | 0 | 0 | 0 | 2 | 1 | 0 | 3 | 1 | 1 | 1 | 0 | 0 |
| 494 | QGN13591.1 | FIM1_233 | CP015054. | 1253 | 3711 | 3035 | 3099 | 3654 | 2483 | 2048 | 3067 | 3845 | 11711 | 3531 | 4453 | 13885 | 3627 | 4059 | 11567 | 3400 | 3042 |
| 495 | QGN13592.1 | FIM1_234 | CP015054. | 2865 | 1097 | 1293 | 3209 | 1139 | 993 | 2931 | 1190 | 1530 | 6227 | 1098 | 1762 | 6489 | 1004 | 1533 | 6087 | 991 | 1143 |
| 496 | QGN13593.1 | FIM1_235 | CP015054. | 1305 | 307 | 642 | 1642 | 310 | 491 | 1313 | 360 | 795 | 4591 | 269 | 926 | 3609 | 251 | 778 | 4443 | 282 | 598 |
| 497 | QGN13594.1 | FIM1_236 | CP015054. | 12813 | 1189 | 1500 | 12302 | 1237 | 1237 | 12617 | 1161 | 2006 | 3739 | 1072 | 2205 | 6854 | 1291 | 2114 | 4744 | 1110 | 1472 |
| 498 | QGN13595.1 | FIM1_237 | CP015054. | 605 | 165 | 504 | 1026 | 224 | 386 | 670 | 362 | 630 | 628 | 168 | 730 | 1936 | 144 | 601 | 998 | 166 | 518 |
| 499 | QGN13596.1 | FIM1_238 | CP015054. | 409 | 375 | 409 | 648 | 228 | 330 | 460 | 300 | 518 | 1499 | 201 | 450 | 1471 | 199 | 473 | 1258 | 169 | 340 |
| 500 | QGN13597.1 | FIM1_239 | CP015054. | 133 | 90 | 98 | 210 | 63 | 63 | 114 | 99 | 101 | 270 | 53 | 79 | 342 | 40 | 88 | 277 | 47 | 57 |
| 501 | QGN13598.1 | FIM1_240 | CP015054. | 1127 | 311 | 513 | 1261 | 259 | 327 | 851 | 353 | 636 | 1830 | 207 | 611 | 2546 | 279 | 558 | 1866 | 163 | 402 |
| 502 | QGN13599.1 | FIM1_241 | CP015054. | 6506 | 2814 | 1878 | 7446 | 2864 | 1553 | 6931 | 2581 | 2378 | 11689 | 2538 | 2353 | 12919 | 2562 | 2183 | 12022 | 2229 | 1577 |

| | | | | | | | | | | | | | | | | | | | | | |
|---|---|---|---|---|---|---|---|---|---|---|---|---|---|---|---|---|---|---|---|---|---|
| 503 | QGN13600.1 | FIM1_242 | CP015054. | 40 | 41 | 41 | 86 | 39 | 38 | 35 | 27 | 65 | 252 | 33 | 70 | 230 | 29 | 62 | 226 | 41 | 42 |
| 504 | QGN13601.1 | FIM1_243 | CP015054. | 1106 | 971 | 698 | 1590 | 923 | 672 | 1048 | 862 | 1035 | 3636 | 773 | 853 | 4136 | 764 | 823 | 3583 | 777 | 534 |
| 505 | QGN13602.1 | FIM1_244 | CP015054. | 70 | 7 | 4 | 105 | 16 | 7 | 64 | 6 | 4 | 25 | 4 | 4 | 28 | 4 | 4 | 18 | 5 | 4 |
| 506 | QGN13603.1 | FIM1_245 | CP015054. | 4673 | 4590 | 2263 | 2824 | 3761 | 2261 | 4008 | 3544 | 3022 | 12082 | 4989 | 3656 | 10916 | 3607 | 3852 | 11532 | 4702 | 2662 |
| 507 | QGN13604.1 | FIM1_246 | CP015054. | 5918 | 1274 | 1451 | 6872 | 1269 | 1270 | 5853 | 1205 | 1893 | 4804 | 1005 | 2043 | 6711 | 1229 | 1944 | 5060 | 959 | 1287 |
| 508 | QGN13605.1 | FIM1_247 | CP015054. | 3113 | 484 | 1188 | 2843 | 246 | 737 | 3959 | 422 | 1315 | 3873 | 221 | 1643 | 5183 | 363 | 1322 | 3914 | 198 | 1037 |
| 509 | QGN13606.1 | FIM1_248 | CP015054. | 2671 | 805 | 643 | 3700 | 728 | 506 | 2390 | 613 | 702 | 3102 | 633 | 760 | 2908 | 761 | 699 | 2892 | 734 | 551 |
| 510 | QGN13607.1 | FIM1_249 | CP015054. | 1221 | 836 | 345 | 676 | 653 | 377 | 1093 | 560 | 487 | 2681 | 817 | 545 | 2051 | 913 | 599 | 2425 | 1082 | 404 |
| 511 | QGN13608.1 | FIM1_250 | CP015054. | 959 | 441 | 553 | 1118 | 582 | 461 | 1005 | 547 | 653 | 4095 | 494 | 846 | 3411 | 401 | 737 | 3784 | 420 | 570 |
| 512 | QGN13609.1 | FIM1_251 | CP015054. | 251 | 328 | 367 | 321 | 325 | 338 | 331 | 351 | 504 | 2608 | 324 | 584 | 2309 | 244 | 520 | 2354 | 280 | 401 |
| 513 | QGN13610.1 | FIM1_252 | CP015054. | 2570 | 1586 | 1397 | 2743 | 1578 | 1305 | 2440 | 1357 | 1868 | 5966 | 1387 | 1836 | 6149 | 1190 | 1846 | 5595 | 1182 | 1244 |
| 514 | QGN13611.1 | FIM1_253 | CP015054. | 8387 | 3629 | 2501 | 7661 | 2539 | 2104 | 7245 | 2580 | 3203 | 12866 | 3561 | 4017 | 13288 | 3305 | 4059 | 12525 | 3282 | 2736 |
| 515 | QGN13612.1 | FIM1_254 | CP015054. | 1224 | 534 | 396 | 1386 | 340 | 400 | 1158 | 523 | 563 | 1937 | 364 | 559 | 1971 | 299 | 562 | 1774 | 321 | 381 |
| 516 | QGN13613.1 | FIM1_255 | CP015054. | 3683 | 3438 | 3222 | 6469 | 2726 | 2415 | 4952 | 2735 | 3912 | 6484 | 2143 | 3940 | 10679 | 2336 | 4265 | 8469 | 1874 | 2726 |
| 517 | QGN13614.1 | FIM1_256 | CP015054. | 3116 | 2147 | 1885 | 2927 | 2635 | 1471 | 2800 | 2501 | 2508 | 3628 | 2057 | 2004 | 6942 | 2279 | 2504 | 4502 | 1968 | 1371 |
| 518 | QGN13615.1 | FIM1_257 | CP015054. | 2072 | 783 | 568 | 2430 | 784 | 572 | 1897 | 739 | 819 | 2361 | 489 | 768 | 2561 | 481 | 737 | 2320 | 516 | 484 |
| 519 | QGN13616.1 | FIM1_258 | CP015054. | 3225 | 580 | 847 | 3184 | 781 | 720 | 2944 | 750 | 1108 | 3094 | 684 | 1378 | 4505 | 580 | 1182 | 3426 | 575 | 916 |
| 520 | QGN13617.1 | FIM1_259 | CP015054. | 1588 | 2093 | 1740 | 2867 | 2080 | 1642 | 2324 | 1627 | 2264 | 14025 | 2443 | 3228 | 10571 | 2308 | 2743 | 13294 | 2401 | 1959 |
| 521 | QGN13618.1 | FIM1_260 | CP015054. | 511 | 539 | 439 | 881 | 410 | 329 | 757 | 445 | 531 | 2231 | 305 | 397 | 1847 | 287 | 416 | 2243 | 247 | 240 |
| 522 | QGN13619.1 | FIM1_261 | CP015054. | 1673 | 656 | 1037 | 1331 | 875 | 804 | 1467 | 884 | 1297 | 1967 | 906 | 1571 | 5787 | 803 | 1351 | 3846 | 782 | 1110 |
| 523 | QGN13620.1 | FIM1_262 | CP015054. | 1541 | 1149 | 501 | 1270 | 810 | 533 | 1490 | 712 | 686 | 3619 | 1116 | 895 | 3171 | 916 | 938 | 3811 | 1126 | 593 |
| 524 | QGN13621.1 | FIM1_263 | CP015054. | 26142 | 1971 | 1584 | 22498 | 2319 | 1625 | 18951 | 1963 | 2299 | 7905 | 1897 | 2029 | 9238 | 2113 | 1931 | 8526 | 1845 | 1433 |
| 525 | QGN13622.1 | FIM1_264 | CP015054. | 2274 | 606 | 582 | 2230 | 539 | 504 | 2147 | 588 | 812 | 1908 | 586 | 790 | 3028 | 612 | 741 | 2200 | 553 | 559 |
| 526 | QGN13623.1 | FIM1_265 | CP015054. | 1731 | 672 | 891 | 1959 | 766 | 738 | 1853 | 768 | 1236 | 5343 | 791 | 1302 | 6002 | 805 | 1191 | 5431 | 743 | 944 |
| 527 | QGN13624.1 | FIM1_266 | CP015054. | 1765 | 589 | 517 | 2287 | 671 | 469 | 1701 | 685 | 698 | 1940 | 479 | 815 | 3455 | 459 | 711 | 2158 | 477 | 527 |
| 528 | QGN13625.1 | FIM1_267 | CP015054. | 2052 | 4055 | 2838 | 2677 | 4551 | 2651 | 2442 | 3960 | 3686 | 14259 | 3995 | 4532 | 17988 | 3944 | 3974 | 14428 | 3995 | 2821 |
| 529 | QGN13626.1 | FIM1_268 | CP015054. | 9552 | 2656 | 3987 | 14734 | 2849 | 3241 | 11465 | 2689 | 5008 | 9374 | 2592 | 6689 | 14157 | 3276 | 6055 | 10891 | 2822 | 4427 |
| 530 | QGN13627.1 | FIM1_269 | CP015054. | 2009 | 1099 | 886 | 2333 | 955 | 746 | 2112 | 868 | 1119 | 3936 | 792 | 1239 | 3995 | 783 | 1206 | 3709 | 822 | 874 |
| 531 | QGN13628.1 | FIM1_270 | CP015054. | 80 | 150 | 109 | 106 | 81 | 145 | 67 | 82 | 208 | 477 | 61 | 128 | 423 | 53 | 152 | 445 | 61 | 82 |
| 532 | QGN13629.1 | FIM1_271 | CP015054. | 3652 | 1492 | 1569 | 5283 | 1692 | 1272 | 4607 | 1428 | 2090 | 5576 | 1679 | 2627 | 10940 | 1586 | 2445 | 8316 | 1476 | 1831 |
| 533 | QGN13630.1 | FIM1_272 | CP015054. | 1169 | 541 | 483 | 1118 | 398 | 412 | 1080 | 424 | 603 | 1143 | 407 | 621 | 2152 | 408 | 585 | 1482 | 422 | 411 |
| 534 | QGN13631.1 | FIM1_273 | CP015054. | 561 | 157 | 256 | 1525 | 176 | 225 | 981 | 157 | 306 | 1234 | 121 | 768 | 1867 | 106 | 631 | 1337 | 106 | 520 |
| 535 | QGN13632.1 | FIM1_274 | CP015054. | 1845 | 1481 | 682 | 1105 | 1253 | 751 | 1618 | 982 | 967 | 5640 | 1564 | 1299 | 5163 | 1343 | 1475 | 5391 | 1522 | 881 |
| 536 | QGN13633.1 | FIM1_275 | CP015054. | 1320 | 1688 | 988 | 1925 | 1172 | 1040 | 1429 | 945 | 1301 | 5470 | 1057 | 1599 | 3749 | 947 | 1546 | 4730 | 1109 | 1030 |
| 537 | QGN13634.1 | FIM1_276 | CP015054. | 668 | 706 | 623 | 782 | 627 | 527 | 666 | 651 | 812 | 1806 | 457 | 918 | 3403 | 522 | 764 | 2301 | 485 | 579 |
| 538 | QGN13635.1 | FIM1_277 | CP015054. | 691 | 168 | 220 | 688 | 160 | 168 | 638 | 174 | 298 | 853 | 173 | 259 | 945 | 113 | 287 | 821 | 153 | 196 |
| 539 | QGN13636.1 | FIM1_278 | CP015054. | 743 | 534 | 492 | 523 | 375 | 426 | 633 | 360 | 661 | 2100 | 352 | 544 | 1817 | 395 | 628 | 1865 | 361 | 406 |
| 540 | QGN13637.1 | FIM1_279 | CP015054. | 4011 | 3239 | 2342 | 3921 | 2843 | 1878 | 3390 | 2525 | 2632 | 9428 | 2795 | 2822 | 8320 | 2601 | 3025 | 10011 | 2133 | 2224 |
| 541 | QGN13638.1 | FIM1_280 | CP015054. | 26 | 33 | 16 | 27 | 42 | 33 | 17 | 22 | 12 | 104 | 20 | 20 | 64 | 17 | 22 | 99 | 11 | 14 |
| 542 | QGN13639.1 | FIM1_281 | CP015054. | 1966 | 1714 | 2251 | 3886 | 2979 | 1824 | 3269 | 2741 | 2908 | 7288 | 2276 | 1943 | 9177 | 2577 | 2102 | 8018 | 1970 | 1501 |
| 543 | QGN13640.1 | FIM1_282 | CP015054. | 1604 | 207 | 231 | 1407 | 212 | 155 | 1326 | 266 | 291 | 122 | 115 | 264 | 929 | 145 | 213 | 289 | 138 | 162 |
| 544 | QGN13641.1 | FIM1_283 | CP015054. | 28772 | 668 | 329 | 11707 | 309 | 206 | 12924 | 573 | 449 | 300 | 327 | 350 | 920 | 386 | 379 | 461 | 383 | 231 |
| 545 | QGN13642.1 | FIM1_284 | CP015054. | 2887 | 763 | 522 | 2175 | 625 | 466 | 2455 | 688 | 667 | 2515 | 534 | 780 | 2570 | 683 | 764 | 2029 | 720 | 556 |
| 546 | QGN13643.1 | FIM1_285 | CP015054. | 941 | 1126 | 873 | 1329 | 891 | 692 | 984 | 1005 | 1264 | 695 | 790 | 1134 | 4447 | 956 | 1031 | 1591 | 929 | 753 |
| 547 | QGN13644.1 | FIM1_286 | CP015054. | 8225 | 1308 | 2165 | 8468 | 2236 | 1563 | 7795 | 2216 | 2759 | 4214 | 1691 | 3028 | 8211 | 1709 | 2821 | 5681 | 1584 | 2078 |
| 548 | QGN13645.1 | FIM1_287 | CP015054. | 838 | 640 | 435 | 1187 | 461 | 379 | 1119 | 478 | 572 | 1826 | 380 | 564 | 1858 | 376 | 595 | 1822 | 385 | 363 |
| 549 | QGN13646.1 | FIM1_288 | CP015054. | 114 | 122 | 96 | 204 | 99 | 91 | 189 | 111 | 111 | 375 | 96 | 121 | 357 | 72 | 110 | 327 | 92 | 54 |
| 550 | QGN13647.1 | FIM1_290 | CP015054. | 465 | 201 | 280 | 481 | 364 | 179 | 406 | 309 | 275 | 934 | 299 | 306 | 1237 | 246 | 292 | 1090 | 261 | 202 |
| 551 | QGN13648.1 | FIM1_291 | CP015054. | 4274 | 1681 | 1086 | 4778 | 982 | 905 | 3555 | 983 | 1541 | 3660 | 881 | 1200 | 4221 | 952 | 1327 | 3889 | 753 | 836 |
| 552 | QGN13649.1 | FIM1_292 | CP015054. | 9218 | 221 | 792 | 9311 | 318 | 414 | 6568 | 306 | 860 | 1296 | 299 | 989 | 2341 | 574 | 842 | 1737 | 304 | 583 |
| 553 | QGN13650.1 | FIM1_293 | CP015054. | 20434 | 1595 | 1704 | 16913 | 1941 | 1336 | 17599 | 1656 | 2137 | 5894 | 1443 | 2150 | 8923 | 1312 | 1911 | 7054 | 1224 | 1388 |
| 554 | QGN13651.1 | FIM1_294 | CP015054. | 39744 | 123 | 76 | 22772 | 297 | 99 | 21767 | 109 | 82 | 125 | 77 | 142 | 440 | 83 | 113 | 220 | 74 | 80 |
| 555 | QGN13652.1 | FIM1_295 | CP015054. | 623 | 9 | 18 | 303 | 15 | 14 | 392 | 21 | 22 | 34 | 13 | 21 | 104 | 12 | 15 | 59 | 10 | 12 |
| 556 | QGN13653.1 | FIM1_296 | CP015054. | 787 | 28 | 48 | 605 | 35 | 22 | 428 | 33 | 46 | 96 | 30 | 65 | 211 | 22 | 40 | 133 | 27 | 34 |
| 557 | QGN13654.1 | FIM1_297 | CP015054. | 242 | 11 | 13 | 125 | 14 | 10 | 121 | 15 | 21 | 43 | 15 | 12 | 104 | 6 | 21 | 44 | 9 | 15 |
| 558 | QGN13655.1 | FIM1_298 | CP015054. | 229 | 12 | 29 | 259 | 12 | 18 | 149 | 16 | 30 | 45 | 16 | 25 | 78 | 11 | 22 | 57 | 15 | 19 |
| 559 | QGN13656.1 | FIM1_299 | CP015054. | 1910 | 244 | 424 | 1139 | 301 | 269 | 1831 | 394 | 552 | 497 | 337 | 553 | 1568 | 324 | 551 | 855 | 260 | 411 |
| 560 | QGN13657.1 | FIM1_300 | CP015054. | 2390 | 1402 | 495 | 1229 | 1451 | 544 | 1947 | 1243 | 724 | 3858 | 1457 | 1015 | 4089 | 1597 | 991 | 4028 | 1784 | 754 |
| 561 | QGN13658.1 | FIM1_301 | CP015054. | 385 | 58 | 150 | 570 | 156 | 114 | 376 | 126 | 190 | 277 | 71 | 176 | 568 | 97 | 194 | 376 | 61 | 138 |
| 562 | QGN13659.1 | FIM1_302 | CP015054. | 2118 | 542 | 542 | 2273 | 495 | 515 | 1696 | 503 | 749 | 872 | 275 | 679 | 1696 | 290 | 703 | 970 | 305 | 491 |
| 563 | QGN13660.1 | FIM1_303 | CP015054. | 102144 | 18393 | 18701 | 90636 | 15827 | 16045 | 90692 | 13717 | 22259 | 46528 | 16899 | 23603 | 50831 | 21455 | 24062 | 48526 | 17180 | 16724 |
| 564 | QGN13661.1 | FIM1_304 | CP015054. | 3799 | 273 | 365 | 4103 | 247 | 264 | 3606 | 286 | 464 | 250 | 182 | 397 | 512 | 281 | 472 | 379 | 243 | 346 |
| 565 | QGN13662.1 | FIM1_305 | CP015054. | 7270 | 176 | 351 | 6986 | 128 | 171 | 6428 | 160 | 375 | 1648 | 76 | 422 | 1628 | 199 | 412 | 1571 | 65 | 250 |
| 566 | QGN13663.1 | FIM1_306 | CP015054. | 1552 | 96 | 180 | 3187 | 27 | 74 | 2937 | 115 | 202 | 665 | 45 | 202 | 613 | 103 | 190 | 532 | 80 | 128 |
| 567 | QGN13664.1 | FIM1_307 | CP015054. | 400 | 579 | 818 | 414 | 792 | 710 | 381 | 730 | 1010 | 4601 | 799 | 1109 | 4727 | 744 | 1003 | 4376 | 699 | 716 |
| 568 | QGN13665.1 | FIM1_308 | CP015054. | 2809 | 1875 | 1068 | 2203 | 1954 | 1102 | 2803 | 1583 | 1366 | 8020 | 1906 | 2096 | 7494 | 1679 | 1886 | 7422 | 1958 | 1399 |
| 569 | QGN13666.1 | FIM1_309 | CP015054. | 4374 | 2487 | 3156 | 4842 | 3237 | 2861 | 4800 | 2808 | 4188 | 13640 | 3039 | 5319 | 13687 | 2707 | 4805 | 13624 | 2701 | 3479 |
| 570 | QGN13667.1 | FIM1_310 | CP015054. | 7493 | 2251 | 1715 | 5450 | 2881 | 1710 | 6177 | 2221 | 2326 | 10426 | 3194 | 2929 | 12296 | 3070 | 2710 | 10132 | 3280 | 2111 |
| 571 | QGN13668.1 | FIM1_311 | CP015054. | 17018 | 1704 | 1971 | 14561 | 2208 | 1659 | 14990 | 2211 | 2623 | 10524 | 1941 | 2057 | 12100 | 1827 | 1954 | 10978 | 1764 | 1439 |
| 572 | QGN13669.1 | FIM1_312 | CP015054. | 2230 | 2857 | 3115 | 6088 | 2285 | 2430 | 3834 | 2206 | 3711 | 12289 | 1641 | 4175 | 11404 | 1987 | 3971 | 11676 | 1590 | 2790 |
| 573 | QGN13670.1 | FIM1_313 | CP015054. | 839 | 2489 | 1578 | 2108 | 2275 | 1491 | 1672 | 1839 | 2073 | 7207 | 1946 | 2461 | 7853 | 1672 | 2487 | 7518 | 1868 | 1648 |
| 574 | QGN13671.1 | FIM1_314 | CP015054. | 1468 | 1092 | 2189 | 2030 | 1380 | 1886 | 1311 | 1532 | 2979 | 3757 | 1006 | 2333 | 9474 | 657 | 2301 | 6378 | 733 | 1653 |
| 575 | QGN13672.1 | FIM1_315 | CP015054. | 1661 | 431 | 417 | 1683 | 356 | 406 | 1475 | 371 | 593 | 2400 | 231 | 553 | 2394 | 182 | 510 | 2271 | 212 | 378 |
| 576 | QGN13673.1 | FIM1_316 | CP015054. | 1609 | 1091 | 442 | 404 | 1062 | 438 | 1190 | 974 | 578 | 2522 | 951 | 763 | 2591 | 838 | 765 | 2487 | 942 | 547 |
| 577 | QGN13674.1 | FIM1_317 | CP015054. | 159 | 55 | 143 | 174 | 19 | 107 | 163 | 59 | 202 | 475 | 12 | 127 | 344 | 12 | 121 | 465 | 21 | 66 |
| 578 | QGN13675.1 | FIM1_318 | CP015054. | 7079 | 6127 | 3696 | 7540 | 5874 | 3192 | 7000 | 5184 | 4599 | 18590 | 6338 | 6834 | 23327 | 6146 | 6286 | 19285 | 5906 | 4694 |
| 579 | QGN13676.1 | FIM1_319 | CP015054. | 700 | 268 | 185 | 534 | 211 | 170 | 791 | 210 | 266 | 747 | 169 | 243 | 846 | 133 | 201 | 775 | 164 | 150 |
| 580 | QGN13677.1 | FIM1_320 | CP015054. | 4005 | 973 | 1184 | 10667 | 865 | 833 | 5972 | 1001 | 1503 | 4171 | 574 | 1207 | 4626 | 859 | 1026 | 4302 | 821 | 832 |
| 581 | QGN13678.1 | FIM1_321 | CP015054. | 70 | 84 | 106 | 127 | 53 | 75 | 65 | 129 | 139 | 331 | 50 | 80 | 232 | 52 | 57 | 282 | 90 | 82 |
| 582 | QGN13679.1 | FIM1_322 | CP015054. | 14377 | 13922 | 8577 | 14007 | 10914 | 7542 | 12342 | 9398 | 10070 | 25314 | 12808 | 11665 | 28817 | 11601 | 12016 | 25530 | 10884 | 9204 |
| 583 | QGN13680.1 | FIM1_323 | CP015054. | 825 | 704 | 630 | 813 | 397 | 449 | 380 | 674 | 927 | 1717 | 181 | 438 | 1571 | 208 | 420 | 1580 | 206 | 345 |
| 584 | QGN13681.1 | FIM1_324 | CP015054. | 6295 | 8444 | 5850 | 10361 | 8675 | 5229 | 5970 | 9703 | 8226 | 4775 | 2545 | 3486 | 10538 | 2305 | 2754 | 7413 | 3343 | 2499 |
| 585 | QGN13682.1 | FIM1_325 | CP015054. | 57 | 19 | 7 | 41 | 12 | 13 | 43 | 21 | 17 | 9 | 8 | 8 | 20 | 7 | 10 | 14 | 13 | 8 |
| 586 | QGN13683.1 | FIM1_326 | CP015054. | 69902 | 118 | 72 | 50550 | 125 | 82 | 59740 | 163 | 131 | 193 | 40 | 81 | 413 | 80 | 62 | 279 | 48 | 55 |

| | | | | | | | | | | | | | | | | | | | | | |
|---|---|---|---|---|---|---|---|---|---|---|---|---|---|---|---|---|---|---|---|---|---|
| 587 | QGN13684.1 | FIM1_327 | CP015054. | 2595 | 1107 | 720 | 2217 | 818 | 689 | 2378 | 779 | 1001 | 3040 | 599 | 1134 | 3354 | 539 | 1063 | 2988 | 600 | 757 |
| 588 | QGN13685.1 | FIM1_328 | CP015054. | 465 | 118 | 108 | 1893 | 115 | 82 | 591 | 122 | 126 | 167 | 73 | 100 | 320 | 93 | 119 | 172 | 53 | 60 |
| 589 | QGN13686.1 | FIM1_329 | CP015054. | 1409 | 247 | 446 | 1811 | 251 | 335 | 1437 | 438 | 593 | 859 | 195 | 672 | 2732 | 246 | 523 | 1532 | 210 | 467 |
| 590 | QGN13687.1 | FIM1_330 | CP015054. | 17856 | 10971 | 5683 | 13790 | 7550 | 4782 | 15021 | 7939 | 7260 | 13270 | 8869 | 9376 | 24612 | 9448 | 10609 | 17731 | 8505 | 7086 |
| 591 | QGN13688.1 | FIM1_331 | CP015054. | 3868 | 456 | 266 | 1640 | 222 | 189 | 2095 | 314 | 312 | 1040 | 253 | 307 | 1145 | 303 | 398 | 1070 | 231 | 191 |
| 592 | QGN13689.1 | FIM1_332 | CP015054. | 2149 | 1251 | 754 | 2816 | 786 | 718 | 2222 | 894 | 947 | 1373 | 602 | 1032 | 2806 | 726 | 1093 | 1815 | 644 | 748 |
| 593 | QGN13690.1 | FIM1_333 | CP015054. | 3356 | 739 | 613 | 3187 | 582 | 481 | 2786 | 595 | 744 | 1316 | 517 | 990 | 2757 | 526 | 865 | 1665 | 501 | 608 |
| 594 | QGN13691.1 | FIM1_334 | CP015054. | 1945 | 749 | 1220 | 1927 | 1379 | 983 | 1861 | 1249 | 1481 | 5141 | 1491 | 1590 | 6710 | 1512 | 1577 | 5334 | 1331 | 1086 |
| 595 | QGN13692.1 | FIM1_335 | CP015054. | 1426 | 1259 | 534 | 593 | 1158 | 600 | 1124 | 1010 | 765 | 2833 | 1085 | 834 | 2722 | 944 | 951 | 2479 | 1037 | 588 |
| 596 | QGN13693.1 | FIM1_336 | CP015054. | 2825 | 239 | 457 | 2873 | 271 | 378 | 2544 | 309 | 585 | 900 | 157 | 530 | 2040 | 199 | 442 | 1240 | 145 | 364 |
| 597 | QGN13694.1 | FIM1_337 | CP015054. | 4282 | 3755 | 2902 | 7356 | 2751 | 2470 | 5062 | 2853 | 3648 | 4519 | 2029 | 3727 | 7475 | 2172 | 4023 | 5266 | 1872 | 2717 |
| 598 | QGN13695.1 | FIM1_338 | CP015054. | 6186 | 1495 | 606 | 4358 | 1598 | 575 | 6405 | 1303 | 741 | 1023 | 1432 | 1223 | 5088 | 1421 | 1093 | 2076 | 1740 | 764 |
| 599 | QGN13696.1 | FIM1_339 | CP015054. | 7836 | 1919 | 1934 | 11596 | 1826 | 1734 | 9423 | 1753 | 2480 | 11210 | 2491 | 2729 | 10811 | 3837 | 2944 | 10369 | 2391 | 1950 |
| 600 | QGN13697.1 | FIM1_340 | CP015054. | 267 | 20 | 121 | 395 | 147 | 127 | 281 | 172 | 216 | 1050 | 158 | 275 | 1480 | 151 | 223 | 1168 | 134 | 156 |
| 601 | QGN13698.1 | FIM1_341 | CP015054. | 5412 | 3077 | 2121 | 4509 | 3280 | 1902 | 5610 | 3220 | 2908 | 12108 | 3612 | 3365 | 15310 | 3116 | 3256 | 13138 | 3350 | 2421 |
| 602 | QGN13699.1 | FIM1_342 | CP015054. | 1004 | 1002 | 664 | 1934 | 697 | 610 | 1478 | 679 | 916 | 3724 | 698 | 998 | 3330 | 706 | 1062 | 3124 | 684 | 690 |
| 603 | QGN13700.1 | FIM1_343 | CP015054. | 4223 | 1355 | 1617 | 4988 | 1415 | 1216 | 3845 | 1482 | 2181 | 1620 | 1168 | 2287 | 6625 | 1199 | 2114 | 3032 | 1152 | 1609 |
| 604 | QGN13701.1 | FIM1_344 | CP015054. | 3386 | 1897 | 1226 | 3278 | 1310 | 1144 | 2673 | 1218 | 1617 | 4176 | 1051 | 1625 | 4040 | 895 | 1687 | 4195 | 912 | 1120 |
| 605 | QGN13702.1 | FIM1_345 | CP015054. | 7725 | 3209 | 1290 | 15339 | 3504 | 975 | 8751 | 2461 | 1506 | 7510 | 2854 | 1320 | 13408 | 3980 | 1210 | 9588 | 3179 | 848 |
| 606 | QGN13703.1 | FIM1_346 | CP015054. | 12666 | 5401 | 3572 | 13642 | 5686 | 2965 | 12613 | 5319 | 4555 | 6803 | 4314 | 4423 | 16316 | 4235 | 4780 | 11415 | 3883 | 3044 |
| 607 | QGN13704.1 | FIM1_347 | CP015054. | 5072 | 1606 | 1514 | 3846 | 1450 | 1268 | 4179 | 1562 | 2003 | 3097 | 1079 | 1841 | 3598 | 877 | 2010 | 3163 | 995 | 1141 |
| 608 | QGN13705.1 | FIM1_348 | CP015054. | 2279 | 2048 | 615 | 902 | 1657 | 782 | 2110 | 1471 | 852 | 4277 | 2050 | 1174 | 4387 | 1473 | 1335 | 4154 | 2083 | 882 |
| 609 | QGN13706.1 | FIM1_349 | CP015054. | 3184 | 1449 | 1533 | 4999 | 1501 | 1222 | 4362 | 1316 | 2073 | 5690 | 1340 | 2480 | 9157 | 1358 | 2152 | 6943 | 1257 | 1685 |
| 610 | QGN13707.1 | FIM1_350 | CP015054. | 641 | 305 | 328 | 1045 | 289 | 323 | 772 | 289 | 481 | 2589 | 258 | 532 | 2333 | 223 | 458 | 2456 | 226 | 350 |
| 611 | QGN13708.1 | FIM1_351 | CP015054. | 2687 | 1051 | 1034 | 3421 | 1122 | 921 | 2927 | 993 | 1356 | 5679 | 1019 | 1645 | 6055 | 1118 | 1448 | 5633 | 1013 | 1038 |
| 612 | QGN13709.1 | FIM1_352 | CP015054. | 19306 | 2987 | 3741 | 10772 | 1729 | 1910 | 18353 | 3725 | 4767 | 3799 | 2647 | 5657 | 15822 | 2699 | 5203 | 7894 | 2600 | 3649 |
| 613 | QGN13710.1 | FIM1_354 | CP015054. | 1659 | 203 | 272 | 1726 | 124 | 228 | 1664 | 169 | 411 | 369 | 76 | 650 | 1826 | 89 | 554 | 522 | 75 | 380 |
| 614 | QGN13711.1 | FIM1_355 | CP015054. | 2061 | 2456 | 2044 | 2423 | 2003 | 1768 | 1920 | 1828 | 2722 | 7115 | 1962 | 2766 | 7674 | 1967 | 2862 | 6534 | 1552 | 1894 |
| 615 | QGN13712.1 | FIM1_356 | CP015054. | 19368 | 802 | 683 | 9272 | 645 | 433 | 17977 | 981 | 823 | 3810 | 482 | 762 | 4637 | 340 | 749 | 3607 | 378 | 493 |
| 616 | QGN13713.1 | FIM1_357 | CP015054. | 4633 | 1224 | 2594 | 9324 | 2118 | 2080 | 6987 | 1621 | 3477 | 2137 | 1495 | 5261 | 10186 | 1871 | 5271 | 4628 | 1585 | 3642 |
| 617 | QGN13714.1 | FIM1_358 | CP015054. | 8015 | 5428 | 2044 | 3554 | 5334 | 2236 | 6558 | 4830 | 2678 | 4938 | 6015 | 3980 | 13303 | 4653 | 3968 | 8460 | 6070 | 2963 |
| 618 | QGN13715.1 | FIM1_359 | CP015054. | 12132 | 5086 | 4013 | 13152 | 5242 | 3601 | 11909 | 4924 | 5081 | 18029 | 5267 | 6492 | 20942 | 5470 | 6031 | 17428 | 4990 | 4331 |
| 619 | QGN13716.1 | FIM1_360 | CP015054. | 1957 | 1200 | 854 | 2267 | 993 | 738 | 1820 | 891 | 1147 | 3486 | 871 | 1171 | 3944 | 836 | 1117 | 3317 | 857 | 766 |
| 620 | QGN13717.1 | FIM1_361 | CP015054. | 1090 | 466 | 319 | 937 | 392 | 268 | 981 | 443 | 453 | 799 | 336 | 506 | 1631 | 308 | 476 | 1031 | 347 | 354 |
| 621 | QGN13718.1 | FIM1_362 | CP015054. | 2716 | 1337 | 1674 | 5285 | 1505 | 1370 | 3423 | 1460 | 2173 | 3725 | 1416 | 2406 | 9134 | 1437 | 2304 | 6240 | 1216 | 1626 |
| 622 | QGN13719.1 | FIM1_363 | CP015054. | 2537 | 480 | 725 | 2043 | 715 | 690 | 2069 | 681 | 992 | 2682 | 722 | 1268 | 3924 | 692 | 1114 | 2871 | 629 | 832 |
| 623 | QGN13720.1 | FIM1_364 | CP015054. | 3609 | 1646 | 2138 | 4798 | 1924 | 1891 | 3981 | 1758 | 2683 | 7861 | 1532 | 3146 | 8819 | 1530 | 2714 | 7830 | 1396 | 1976 |
| 624 | QGN13721.1 | FIM1_365 | CP015054. | 2242 | 1295 | 766 | 1576 | 1030 | 763 | 2046 | 968 | 1070 | 5068 | 1138 | 1168 | 4251 | 963 | 1162 | 4789 | 1047 | 811 |
| 625 | QGN13722.1 | FIM1_366 | CP015054. | 4188 | 714 | 1389 | 3493 | 858 | 1090 | 3953 | 942 | 1766 | 4372 | 963 | 2150 | 8753 | 953 | 1773 | 5641 | 941 | 1458 |
| 626 | QGN13723.1 | FIM1_367 | CP015054. | 26551 | 3130 | 1677 | 32165 | 7608 | 3502 | 21725 | 2640 | 2063 | 2763 | 1422 | 1038 | 4362 | 1233 | 1031 | 2795 | 1447 | 717 |
| 627 | QGN13724.1 | FIM1_368 | CP015054. | 2041 | 242 | 213 | 1419 | 253 | 153 | 1938 | 236 | 274 | 225 | 230 | 408 | 1251 | 260 | 341 | 490 | 295 | 256 |
| 628 | QGN13725.1 | FIM1_369 | CP015054. | 79 | 31 | 28 | 62 | 19 | 21 | 76 | 20 | 42 | 8 | 22 | 27 | 90 | 23 | 35 | 25 | 17 | 22 |
| 629 | QGN13726.1 | FIM1_370 | CP015054. | 740 | 562 | 700 | 665 | 479 | 517 | 691 | 504 | 896 | 1692 | 409 | 875 | 2593 | 339 | 807 | 1987 | 344 | 604 |
| 630 | QGN13727.1 | FIM1_371 | CP015054. | 346 | 59 | 36 | 279 | 61 | 44 | 164 | 63 | 65 | 37 | 36 | 71 | 116 | 46 | 70 | 83 | 42 | 47 |
| 631 | QGN13728.1 | FIM1_372 | CP015054. | 28746 | 11350 | 9298 | 36899 | 9412 | 6678 | 27738 | 10973 | 11615 | 8081 | 8342 | 13935 | 27717 | 9113 | 12747 | 15946 | 9638 | 9338 |
| 632 | QGN13729.1 | FIM1_373 | CP015054. | 3354 | 743 | 927 | 3332 | 739 | 634 | 3651 | 995 | 1158 | 1074 | 689 | 1060 | 2635 | 716 | 1236 | 1619 | 747 | 866 |
| 633 | QGN13730.1 | FIM1_374 | CP015054. | 4914 | 2390 | 1308 | 4475 | 1549 | 1166 | 4746 | 1621 | 1834 | 2417 | 956 | 1312 | 3615 | 860 | 1401 | 2942 | 852 | 941 |
| 634 | QGN13731.1 | FIM1_375 | CP015054. | 5074 | 1135 | 1037 | 5025 | 1004 | 729 | 4549 | 1148 | 1317 | 854 | 717 | 1454 | 2808 | 652 | 1429 | 1241 | 816 | 1013 |
| 635 | QGN13732.1 | FIM1_376 | CP015054. | 723 | 526 | 907 | 1862 | 268 | 555 | 1708 | 499 | 1145 | 992 | 217 | 942 | 1593 | 390 | 1214 | 1398 | 254 | 698 |
| 636 | QGN13733.1 | FIM1_377 | CP015054. | 727 | 716 | 941 | 950 | 410 | 689 | 1229 | 614 | 1237 | 927 | 346 | 879 | 1662 | 470 | 1151 | 1275 | 415 | 622 |
| 637 | QGN13734.1 | FIM1_378 | CP015054. | 132679 | 26886 | 27954 | 126707 | 23668 | 23341 | 116924 | 22392 | 32556 | 41207 | 25507 | 35320 | 69752 | 31687 | 35203 | 53480 | 27380 | 26386 |
| 638 | QGN13735.1 | FIM1_379 | CP015054. | 943 | 104 | 293 | 1759 | 357 | 204 | 1540 | 438 | 318 | 385 | 285 | 472 | 1392 | 301 | 389 | 636 | 320 | 296 |
| 639 | QGN13736.1 | FIM1_380 | CP015054. | 4339 | 758 | 1321 | 5737 | 1420 | 1044 | 3723 | 1445 | 1857 | 1858 | 843 | 1863 | 7203 | 944 | 1743 | 3369 | 860 | 1340 |
| 640 | QGN13737.1 | FIM1_381 | CP015054. | 3193 | 1261 | 1670 | 3440 | 1633 | 1226 | 3294 | 1573 | 2148 | 1809 | 1960 | 2366 | 10175 | 1910 | 2262 | 5037 | 2013 | 1674 |
| 641 | QGN13738.1 | FIM1_382 | CP015054. | 11 | 4 | 0 | 50 | 2 | 1 | 31 | 1 | 1 | 6 | 3 | 1 | 6 | 0 | 0 | 4 | 1 | 0 |
| 642 | QGN13739.1 | FIM1_383 | CP015054. | 37747 | 703 | 1381 | 56982 | 556 | 1220 | 44205 | 834 | 1558 | 3696 | 572 | 1348 | 3970 | 992 | 779 | 3464 | 1011 | 762 |
| 643 | QGN13740.1 | FIM1_384 | CP015054. | 32941 | 18565 | 28334 | 57995 | 49679 | 23856 | 47249 | 42518 | 32283 | 88190 | 37814 | 32933 | 121933 | 36883 | 29825 | 102079 | 32798 | 23166 |
| 644 | QGN13741.1 | FIM1_385 | CP015054. | 1448 | 459 | 378 | 2250 | 332 | 319 | 2028 | 356 | 556 | 845 | 286 | 403 | 1465 | 212 | 522 | 992 | 269 | 297 |
| 645 | QGN13742.1 | FIM1_386 | CP015054. | 2632 | 925 | 2131 | 8531 | 1725 | 1886 | 5406 | 1350 | 2966 | 3168 | 1115 | 3557 | 8940 | 1003 | 2966 | 4959 | 852 | 2151 |
| 646 | QGN13743.1 | FIM1_387 | CP015054. | 605 | 636 | 652 | 1085 | 486 | 551 | 703 | 460 | 788 | 4268 | 544 | 978 | 3150 | 409 | 913 | 4160 | 414 | 701 |
| 647 | QGN13744.1 | FIM1_388 | CP015054. | 985 | 355 | 383 | 1772 | 305 | 354 | 1181 | 394 | 542 | 1194 | 317 | 531 | 1806 | 320 | 565 | 1155 | 303 | 397 |
| 648 | QGN13745.1 | FIM1_389 | CP015054. | 2467 | 1951 | 2424 | 5322 | 2311 | 1945 | 3528 | 2043 | 3295 | 5623 | 1955 | 3040 | 9184 | 2053 | 2807 | 6708 | 1856 | 1981 |
| 649 | QGN13746.1 | FIM1_390 | CP015054. | 681 | 251 | 384 | 385 | 224 | 247 | 497 | 306 | 387 | 883 | 203 | 442 | 1495 | 146 | 359 | 1087 | 165 | 306 |
| 650 | QGN13747.1 | FIM1_391 | CP015054. | 1138 | 768 | 892 | 2243 | 808 | 754 | 1513 | 778 | 1139 | 3260 | 816 | 1325 | 3752 | 767 | 1403 | 3217 | 678 | 906 |
| 651 | QGN13748.1 | FIM1_392 | CP015054. | 2556 | 3741 | 2965 | 4534 | 2932 | 2532 | 3462 | 2469 | 3606 | 13060 | 2694 | 4094 | 10420 | 2624 | 3879 | 12623 | 2511 | 2614 |
| 652 | QGN13749.1 | FIM1_393 | CP015054. | 3226 | 1370 | 952 | 3591 | 1235 | 844 | 3239 | 1313 | 1299 | 4299 | 1156 | 1465 | 6137 | 1217 | 1276 | 4170 | 1102 | 966 |
| 653 | QGN13750.1 | FIM1_394 | CP015054. | 352 | 64 | 117 | 407 | 166 | 62 | 342 | 171 | 127 | 646 | 124 | 154 | 755 | 171 | 110 | 567 | 129 | 100 |
| 654 | QGN13751.1 | FIM1_395 | CP015054. | 1668 | 583 | 918 | 2625 | 795 | 662 | 2138 | 782 | 1145 | 3620 | 726 | 1317 | 6143 | 733 | 1197 | 4459 | 583 | 918 |
| 655 | QGN13752.1 | FIM1_396 | CP015054. | 1299 | 2176 | 1028 | 1436 | 1914 | 944 | 1066 | 1447 | 1316 | 8768 | 2018 | 1978 | 7096 | 1688 | 2029 | 8637 | 1739 | 1375 |
| 656 | QGN13753.1 | FIM1_397 | CP015054. | 53 | 30 | 22 | 142 | 48 | 19 | 70 | 27 | 20 | 194 | 24 | 32 | 136 | 19 | 34 | 225 | 14 | 18 |
| 657 | QGN13754.1 | FIM1_398 | CP015054. | 2183 | 1331 | 1287 | 3429 | 1406 | 1261 | 2956 | 1051 | 1808 | 9813 | 1227 | 2228 | 7815 | 1151 | 2017 | 9720 | 1011 | 1377 |
| 658 | QGN13755.1 | FIM1_399 | CP015054. | 1224 | 2190 | 2568 | 2385 | 1852 | 2205 | 1822 | 1596 | 3185 | 8721 | 1603 | 3555 | 6815 | 1452 | 3198 | 8576 | 1401 | 2317 |
| 659 | QGN13756.1 | FIM1_401 | CP015054. | 1147 | 565 | 940 | 1441 | 891 | 646 | 1419 | 808 | 1093 | 6638 | 902 | 1601 | 6054 | 864 | 1345 | 6052 | 786 | 1009 |
| 660 | QGN13757.1 | FIM1_402 | CP015054. | 1608 | 165 | 215 | 2325 | 131 | 147 | 2108 | 143 | 301 | 1824 | 102 | 237 | 1432 | 115 | 222 | 1540 | 90 | 170 |
| 661 | QGN13758.1 | FIM1_403 | CP015054. | 4296 | 485 | 428 | 4726 | 349 | 301 | 4584 | 437 | 604 | 2165 | 327 | 580 | 2632 | 484 | 668 | 2138 | 342 | 483 |
| 662 | QGN13759.1 | FIM1_404 | CP015054. | 1029 | 290 | 296 | 1906 | 192 | 208 | 1561 | 289 | 370 | 1800 | 229 | 484 | 1715 | 401 | 420 | 1697 | 254 | 299 |
| 663 | QGN13760.1 | FIM1_405 | CP015054. | 6559 | 5527 | 3655 | 8283 | 6015 | 3140 | 6855 | 5546 | 4478 | 15694 | 6013 | 6399 | 23877 | 5925 | 5875 | 17726 | 5485 | 4466 |
| 664 | QGN13761.1 | FIM1_406 | CP015054. | 10128 | 7682 | 4822 | 12946 | 7413 | 4284 | 10857 | 6412 | 6007 | 18949 | 8107 | 8583 | 27525 | 7516 | 8649 | 21123 | 7228 | 5979 |
| 665 | QGN13762.1 | FIM1_407 | CP015054. | 5055 | 2714 | 2186 | 4894 | 1617 | 1371 | 3498 | 2478 | 2917 | 10641 | 2264 | 3368 | 13599 | 3109 | 3717 | 10463 | 2151 | 2463 |
| 666 | QGN13763.1 | FIM1_408 | CP015054. | 4131 | 1009 | 1049 | 4661 | 974 | 843 | 4208 | 1108 | 1296 | 3524 | 1008 | 1540 | 4668 | 1043 | 1494 | 3508 | 967 | 1008 |
| 667 | QGN13764.1 | FIM1_409 | CP015054. | 6537 | 3265 | 3349 | 7710 | 3014 | 2452 | 6851 | 2811 | 4076 | 10431 | 2390 | 4329 | 14513 | 3132 | 4357 | 12121 | 2320 | 2931 |
| 668 | QGN13765.1 | FIM1_410 | CP015054. | 744 | 125 | 164 | 823 | 89 | 117 | 481 | 126 | 218 | 365 | 51 | 218 | 605 | 80 | 180 | 482 | 56 | 145 |
| 669 | QGN13766.1 | FIM1_411 | CP015054. | 7975 | 804 | 1282 | 12806 | 613 | 732 | 11898 | 980 | 1616 | 1693 | 451 | 1425 | 4761 | 935 | 1119 | 2929 | 643 | 827 |
| 670 | QGN13767.1 | FIM1_412 | CP015054. | 1835 | 497 | 368 | 2178 | 564 | 419 | 2122 | 397 | 517 | 1970 | 381 | 384 | 1919 | 377 | 430 | 1819 | 390 | 249 |

| | | | | | | | | | | | | | | | | | | | | | |
|---|---|---|---|---|---|---|---|---|---|---|---|---|---|---|---|---|---|---|---|---|---|
| 671 | QGN13768.1 | FIM1_413 | CP015054. | 6839 | 5324 | 3733 | 10179 | 5269 | 3018 | 8565 | 5001 | 4616 | 10581 | 3232 | 2938 | 13569 | 2967 | 3093 | 11487 | 2418 | 2024 |
| 672 | QGN13769.1 | FIM1_414 | CP015054. | 8699 | 271 | 289 | 3044 | 287 | 207 | 5602 | 324 | 384 | 541 | 226 | 450 | 1300 | 169 | 452 | 744 | 175 | 328 |
| 673 | QGN13770.1 | FIM1_415 | CP015054. | 523 | 264 | 330 | 1030 | 226 | 234 | 676 | 281 | 424 | 1118 | 154 | 418 | 1475 | 238 | 402 | 1254 | 204 | 260 |
| 674 | QGN13771.1 | FIM1_416 | CP015054. | 4039 | 633 | 770 | 4288 | 757 | 650 | 3526 | 658 | 946 | 2508 | 772 | 1426 | 3988 | 711 | 1265 | 2828 | 709 | 893 |
| 675 | QGN13772.1 | FIM1_417 | CP015054. | 18124 | 4528 | 5075 | 24389 | 3725 | 4299 | 18477 | 3527 | 6506 | 11960 | 1873 | 4830 | 12713 | 2889 | 4378 | 11554 | 2116 | 3215 |
| 676 | QGN13773.1 | FIM1_418 | CP015054. | 2837 | 314 | 701 | 2922 | 449 | 711 | 2384 | 362 | 929 | 1321 | 284 | 1219 | 1557 | 374 | 1151 | 1461 | 295 | 806 |
| 677 | QGN13774.1 | FIM1_419 | CP015054. | 1733 | 352 | 597 | 2029 | 486 | 520 | 1328 | 426 | 786 | 1664 | 393 | 817 | 2989 | 374 | 793 | 2061 | 340 | 550 |
| 678 | QGN13775.1 | FIM1_420 | CP015054. | 1290 | 411 | 451 | 1065 | 476 | 418 | 1192 | 439 | 598 | 2656 | 456 | 727 | 2582 | 395 | 634 | 2893 | 383 | 448 |
| 679 | QGN13776.1 | FIM1_421 | CP015054. | 2627 | 506 | 629 | 3083 | 690 | 553 | 2678 | 541 | 770 | 4480 | 590 | 1028 | 3843 | 606 | 972 | 4334 | 637 | 690 |
| 680 | QGN13777.1 | FIM1_422 | CP015054. | 1382 | 436 | 451 | 1521 | 268 | 382 | 1263 | 377 | 648 | 1741 | 234 | 552 | 2386 | 204 | 566 | 1923 | 212 | 374 |
| 681 | QGN13778.1 | FIM1_423 | CP015054. | 259 | 19 | 29 | 1018 | 15 | 26 | 532 | 12 | 30 | 203 | 22 | 23 | 221 | 83 | 32 | 192 | 10 | 28 |
| 682 | QGN13779.1 | FIM1_425 | CP015054. | 3058 | 3248 | 1170 | 1375 | 2684 | 1392 | 2385 | 2188 | 1564 | 11168 | 3537 | 2203 | 9131 | 2691 | 2116 | 10243 | 3697 | 1496 |
| 683 | QGN13780.1 | FIM1_426 | CP015054. | 5496 | 2098 | 2337 | 7308 | 3108 | 2014 | 6391 | 2693 | 2950 | 10383 | 2886 | 4021 | 16594 | 2806 | 3548 | 12269 | 2830 | 2637 |
| 684 | QGN13781.1 | FIM1_427 | CP015054. | 254 | 271 | 234 | 507 | 257 | 230 | 414 | 223 | 314 | 1292 | 231 | 335 | 992 | 212 | 331 | 1205 | 205 | 210 |
| 685 | QGN13782.1 | FIM1_428 | CP015054. | 115 | 238 | 206 | 255 | 279 | 163 | 110 | 283 | 262 | 715 | 285 | 329 | 1353 | 262 | 301 | 883 | 271 | 231 |
| 686 | QGN13783.1 | FIM1_429 | CP015054. | 493 | 1689 | 914 | 679 | 1314 | 784 | 495 | 1050 | 1303 | 3253 | 1066 | 1168 | 3688 | 1098 | 1112 | 3136 | 998 | 718 |
| 687 | QGN13784.1 | FIM1_430 | CP015054. | 1076 | 915 | 726 | 1515 | 650 | 647 | 1333 | 679 | 1012 | 3357 | 580 | 855 | 3178 | 567 | 787 | 3323 | 525 | 541 |
| 688 | QGN13785.1 | FIM1_431 | CP015054. | 1002 | 149 | 325 | 1338 | 444 | 284 | 1250 | 270 | 403 | 2057 | 327 | 524 | 2371 | 382 | 452 | 2158 | 318 | 328 |
| 689 | QGN13786.1 | FIM1_432 | CP015054. | 308 | 211 | 140 | 178 | 138 | 155 | 299 | 139 | 211 | 778 | 175 | 169 | 576 | 125 | 194 | 621 | 195 | 113 |
| 690 | QGN13787.1 | FIM1_433 | CP015054. | 992 | 1968 | 1171 | 907 | 1463 | 1090 | 759 | 1457 | 1542 | 5981 | 1489 | 1835 | 6369 | 1382 | 1745 | 6064 | 1484 | 1145 |
| 691 | QGN13788.1 | FIM1_434 | CP015054. | 1295 | 479 | 536 | 1572 | 748 | 477 | 1480 | 634 | 649 | 2702 | 627 | 923 | 4534 | 633 | 856 | 3206 | 618 | 650 |
| 692 | QGN13789.1 | FIM1_435 | CP015054. | 1381 | 826 | 564 | 1762 | 690 | 533 | 1750 | 623 | 714 | 3496 | 665 | 954 | 2986 | 709 | 957 | 3124 | 617 | 615 |
| 693 | QGN13790.1 | FIM1_436 | CP015054. | 819 | 253 | 224 | 530 | 209 | 204 | 841 | 205 | 307 | 1545 | 214 | 287 | 1200 | 220 | 318 | 1555 | 247 | 226 |
| 694 | QGN13791.1 | FIM1_437 | CP015054. | 618 | 389 | 525 | 1335 | 319 | 406 | 926 | 363 | 642 | 1805 | 260 | 541 | 1668 | 239 | 484 | 1539 | 246 | 331 |
| 695 | QGN13792.1 | FIM1_438 | CP015054. | 50 | 10 | 9 | 177 | 16 | 11 | 90 | 13 | 23 | 18 | 7 | 16 | 67 | 4 | 22 | 42 | 7 | 17 |
| 696 | QGN13793.1 | FIM1_439 | CP015054. | 3004 | 1477 | 668 | 1634 | 1308 | 669 | 2723 | 1311 | 963 | 3469 | 1408 | 1104 | 4393 | 1643 | 1117 | 3295 | 1741 | 802 |
| 697 | QGN13794.1 | FIM1_440 | CP015054. | 2247 | 780 | 709 | 2291 | 581 | 549 | 2308 | 600 | 913 | 2087 | 368 | 657 | 1928 | 403 | 668 | 1785 | 346 | 454 |
| 698 | QGN13795.1 | FIM1_441 | CP015054. | 3326 | 385 | 470 | 3747 | 521 | 362 | 3636 | 459 | 605 | 2511 | 481 | 674 | 2464 | 555 | 565 | 2481 | 440 | 449 |
| 699 | QGN13796.1 | FIM1_442 | CP015054. | 3484 | 397 | 902 | 3383 | 542 | 751 | 2774 | 564 | 1219 | 1615 | 434 | 1305 | 3902 | 432 | 1154 | 2416 | 435 | 824 |
| 700 | QGN13797.1 | FIM1_443 | CP015054. | 497 | 19 | 129 | 605 | 76 | 70 | 465 | 88 | 126 | 209 | 48 | 192 | 547 | 52 | 137 | 289 | 52 | 109 |
| 701 | QGN13798.1 | FIM1_444 | CP015054. | 1798 | 1178 | 884 | 2500 | 676 | 826 | 1954 | 676 | 1143 | 3919 | 735 | 1261 | 3300 | 723 | 1279 | 3589 | 747 | 873 |
| 702 | QGN13799.1 | FIM1_445 | CP015054. | 3940 | 970 | 1243 | 4325 | 1012 | 936 | 4442 | 1135 | 1508 | 3246 | 957 | 1932 | 5973 | 902 | 1769 | 4139 | 841 | 1238 |
| 703 | QGN13800.1 | FIM1_446 | CP015054. | 334 | 151 | 303 | 293 | 143 | 237 | 292 | 219 | 352 | 1455 | 160 | 396 | 1323 | 103 | 401 | 1424 | 131 | 327 |
| 704 | QGN13801.1 | FIM1_447 | CP015054. | 1600 | 489 | 468 | 1851 | 502 | 408 | 1752 | 584 | 617 | 2775 | 474 | 824 | 3424 | 495 | 667 | 2799 | 447 | 504 |
| 705 | QGN13802.1 | FIM1_448 | CP015054. | 2695 | 1052 | 1767 | 5118 | 1284 | 1383 | 4012 | 1172 | 2204 | 7503 | 1429 | 2668 | 10177 | 1425 | 2449 | 8220 | 1242 | 1776 |
| 706 | QGN13803.1 | FIM1_449 | CP015054. | 5967 | 1912 | 2945 | 5588 | 2566 | 2261 | 4521 | 2765 | 3620 | 6453 | 2723 | 4399 | 17243 | 2540 | 4005 | 11150 | 2504 | 2982 |
| 707 | QGN13804.1 | FIM1_450 | CP015054. | 54 | 133 | 53 | 62 | 80 | 45 | 46 | 72 | 91 | 531 | 168 | 169 | 538 | 140 | 190 | 476 | 143 | 106 |
| 708 | QGN13805.1 | FIM1_451 | CP015054. | 5657 | 1456 | 810 | 3414 | 1481 | 749 | 4953 | 1587 | 1168 | 1935 | 1235 | 1272 | 4899 | 1319 | 1206 | 2773 | 1341 | 954 |
| 709 | QGN13806.1 | FIM1_452 | CP015054. | 1142 | 368 | 617 | 1028 | 410 | 443 | 902 | 551 | 733 | 1881 | 399 | 605 | 3314 | 330 | 639 | 2260 | 303 | 398 |
| 710 | QGN13807.1 | FIM1_453 | CP015054. | 949 | 2193 | 2702 | 1121 | 1760 | 2258 | 686 | 1489 | 3595 | 3698 | 2162 | 2729 | 6901 | 1832 | 3559 | 5911 | 1879 | 2091 |
| 711 | QGN13808.1 | FIM1_454 | CP015054. | 348 | 67 | 190 | 754 | 132 | 121 | 488 | 193 | 229 | 116 | 72 | 260 | 688 | 90 | 183 | 239 | 108 | 136 |
| 712 | QGN13809.1 | FIM1_455 | CP015054. | 517 | 1660 | 944 | 1548 | 860 | 812 | 1015 | 965 | 1252 | 1748 | 781 | 1072 | 2861 | 884 | 1170 | 2188 | 813 | 781 |
| 713 | QGN13810.1 | FIM1_456 | CP015054. | 5049 | 1302 | 1403 | 3686 | 1121 | 885 | 3111 | 1429 | 1785 | 1124 | 666 | 997 | 3014 | 699 | 1070 | 1699 | 625 | 641 |
| 714 | QGN13811.1 | FIM1_457 | CP015054. | 14022 | 14547 | 8067 | 13019 | 9987 | 7352 | 10803 | 8637 | 9624 | 29348 | 11837 | 12739 | 28532 | 10845 | 12567 | 30399 | 10404 | 9469 |
| 715 | QGN13812.1 | FIM1_458 | CP015054. | 17383 | 19267 | 12133 | 17364 | 15123 | 11620 | 14446 | 13004 | 14508 | 41994 | 18214 | 17717 | 38065 | 15918 | 17995 | 43249 | 15940 | 13401 |
| 716 | QGN13813.1 | FIM1_459 | CP015054. | 1027 | 1253 | 1272 | 855 | 1448 | 1163 | 474 | 1653 | 1649 | 2859 | 1131 | 1312 | 5198 | 1010 | 1228 | 3574 | 1018 | 991 |
| 717 | QGN13814.1 | FIM1_460 | CP015054. | 1037 | 220 | 212 | 654 | 222 | 198 | 909 | 294 | 296 | 1504 | 239 | 355 | 1373 | 193 | 351 | 1528 | 261 | 241 |
| 718 | QGN13815.1 | FIM1_461 | CP015054. | 1942 | 535 | 667 | 1950 | 516 | 508 | 1661 | 508 | 841 | 1413 | 383 | 815 | 2628 | 334 | 790 | 1753 | 342 | 526 |
| 719 | QGN13816.1 | FIM1_462 | CP015054. | 1039 | 542 | 503 | 1139 | 498 | 395 | 987 | 533 | 621 | 1759 | 438 | 638 | 2023 | 440 | 697 | 1708 | 390 | 475 |
| 720 | QGN13817.1 | FIM1_463 | CP015054. | 975 | 1482 | 1399 | 558 | 1632 | 1380 | 865 | 1459 | 1752 | 4671 | 1740 | 2417 | 5438 | 1451 | 2194 | 4319 | 1659 | 1596 |
| 721 | QGN13818.1 | FIM1_464 | CP015054. | 4064 | 3868 | 3781 | 7605 | 3619 | 2843 | 6430 | 4228 | 4558 | 9397 | 3209 | 5536 | 16944 | 3310 | 4500 | 11662 | 3047 | 3435 |
| 722 | QGN13819.1 | FIM1_465 | CP015054. | 9 | 39 | 12 | 37 | 41 | 23 | 14 | 22 | 29 | 70 | 19 | 24 | 53 | 14 | 33 | 41 | 28 | 19 |
| 723 | QGN13820.1 | FIM1_466 | CP015054. | 5202 | 5418 | 3434 | 4491 | 5235 | 2979 | 5178 | 4830 | 4289 | 13910 | 6458 | 5746 | 22323 | 5938 | 5412 | 15550 | 6110 | 4157 |
| 724 | QGN13821.1 | FIM1_467 | CP015054. | 2987 | 2782 | 1007 | 1946 | 2122 | 1158 | 2768 | 1833 | 1441 | 5968 | 2065 | 1446 | 5047 | 1593 | 1592 | 4980 | 1987 | 1136 |
| 725 | QGN13822.1 | FIM1_468 | CP015054. | 390 | 72 | 46 | 421 | 121 | 80 | 395 | 67 | 74 | 114 | 50 | 109 | 175 | 47 | 103 | 128 | 54 | 67 |
| 726 | QGN13823.1 | FIM1_469 | CP015054. | 2163 | 918 | 572 | 1079 | 583 | 549 | 1502 | 630 | 784 | 1525 | 457 | 632 | 1797 | 347 | 829 | 1532 | 432 | 505 |
| 727 | QGN13824.1 | FIM1_470 | CP015054. | 5226 | 789 | 1044 | 5769 | 806 | 720 | 4998 | 1073 | 1352 | 826 | 478 | 1203 | 2725 | 536 | 1228 | 1561 | 498 | 802 |
| 728 | QGN13825.1 | FIM1_471 | CP015054. | 1864 | 210 | 616 | 3126 | 452 | 428 | 1999 | 576 | 730 | 681 | 308 | 752 | 2813 | 346 | 760 | 1535 | 374 | 489 |
| 729 | QGN13826.1 | FIM1_472 | CP015054. | 1427 | 274 | 252 | 1556 | 372 | 188 | 794 | 392 | 269 | 554 | 360 | 355 | 1229 | 261 | 386 | 902 | 334 | 220 |
| 730 | QGN13827.1 | FIM1_473 | CP015054. | 2761 | 787 | 935 | 3728 | 922 | 825 | 3077 | 945 | 1284 | 2061 | 807 | 1472 | 6075 | 695 | 1375 | 3695 | 641 | 987 |
| 731 | QGN13828.1 | FIM1_474 | CP015054. | 723 | 1584 | 1003 | 798 | 1309 | 906 | 737 | 1152 | 1376 | 6832 | 1259 | 1495 | 5319 | 1155 | 1552 | 5997 | 1206 | 1034 |
| 732 | QGN13829.1 | FIM1_475 | CP015054. | 2355 | 244 | 546 | 1670 | 281 | 403 | 1895 | 447 | 664 | 1565 | 240 | 607 | 3468 | 194 | 540 | 2360 | 189 | 462 |
| 733 | QGN13830.1 | FIM1_476 | CP015054. | 3069 | 7886 | 5872 | 7487 | 7988 | 5200 | 6164 | 6815 | 6900 | 25518 | 8999 | 9368 | 23112 | 8621 | 9487 | 25177 | 7732 | 7170 |
| 734 | QGN13831.1 | FIM1_477 | CP015054. | 13592 | 8284 | 4734 | 14710 | 5806 | 4290 | 12601 | 5206 | 5910 | 19027 | 6380 | 7560 | 19938 | 6149 | 7837 | 18186 | 5571 | 5323 |
| 735 | QGN13832.1 | FIM1_478 | CP015054. | 6434 | 3149 | 1088 | 3086 | 2652 | 1227 | 5748 | 2724 | 1579 | 5799 | 3096 | 2198 | 9249 | 2872 | 2039 | 6027 | 3814 | 1530 |
| 736 | QGN13833.1 | FIM1_479 | CP015054. | 2284 | 769 | 879 | 2909 | 881 | 680 | 2529 | 863 | 1190 | 1146 | 648 | 1379 | 4352 | 686 | 1134 | 2076 | 746 | 949 |
| 737 | QGN13834.1 | FIM1_480 | CP015054. | 728 | 759 | 1416 | 1221 | 847 | 1027 | 958 | 939 | 1733 | 3717 | 531 | 1448 | 3206 | 443 | 1209 | 3456 | 436 | 867 |
| 738 | QGN13835.1 | FIM1_481 | CP015054. | 4729 | 407 | 807 | 4964 | 552 | 661 | 4635 | 577 | 1006 | 4630 | 488 | 1012 | 5128 | 423 | 954 | 4801 | 385 | 661 |
| 739 | QGN13836.1 | FIM1_482 | CP015054. | 3222 | 491 | 675 | 3310 | 675 | 514 | 3207 | 652 | 944 | 2871 | 476 | 1128 | 5211 | 410 | 1010 | 3468 | 424 | 702 |
| 740 | QGN13837.1 | FIM1_483 | CP015054. | 1881 | 1189 | 1208 | 2467 | 1359 | 1065 | 2317 | 1303 | 1645 | 7081 | 1376 | 1801 | 8053 | 1118 | 1769 | 7271 | 1183 | 1305 |
| 741 | QGN13838.1 | FIM1_484 | CP015054. | 9 | 8 | 13 | 49 | 13 | 13 | 17 | 17 | 15 | 28 | 2 | 2 | 29 | 3 | 0 | 25 | 3 | 1 |
| 742 | QGN13839.1 | FIM1_485 | CP015054. | 399 | 81 | 94 | 472 | 109 | 79 | 274 | 83 | 109 | 114 | 45 | 77 | 228 | 35 | 69 | 137 | 23 | 54 |
| 743 | QGN13840.1 | FIM1_486 | CP015054. | 1836 | 392 | 771 | 2966 | 520 | 546 | 1682 | 537 | 905 | 2504 | 371 | 763 | 3792 | 448 | 734 | 2851 | 300 | 537 |
| 744 | QGN13841.1 | FIM1_487 | CP015054. | 300 | 790 | 482 | 641 | 473 | 526 | 319 | 371 | 693 | 1543 | 482 | 588 | 1470 | 420 | 652 | 1456 | 464 | 388 |
| 745 | QGN13842.1 | FIM1_488 | CP015054. | 3966 | 4534 | 4417 | 7852 | 4396 | 4222 | 2968 | 2835 | 5676 | 8438 | 2847 | 5685 | 10933 | 2962 | 5618 | 9486 | 2830 | 3759 |
| 746 | QGN13843.1 | FIM1_489 | CP015054. | 5461 | 1928 | 1730 | 6277 | 2534 | 1261 | 5590 | 2580 | 2197 | 2628 | 2080 | 2896 | 11710 | 2060 | 2359 | 5250 | 2135 | 1832 |
| 747 | QGN13844.1 | FIM1_490 | CP015054. | 3061 | 959 | 1028 | 4563 | 1006 | 910 | 3553 | 1048 | 1474 | 4944 | 718 | 1931 | 6882 | 835 | 1414 | 5265 | 699 | 1152 |
| 748 | QGN13845.1 | FIM1_491 | CP015054. | 11256 | 11527 | 7365 | 11599 | 9707 | 6095 | 9707 | 8805 | 8403 | 19302 | 10376 | 10192 | 23211 | 9628 | 10041 | 22016 | 9109 | 7886 |
| 749 | QGN13846.1 | FIM1_492 | CP015054. | 3445 | 2225 | 1995 | 4598 | 1854 | 1651 | 3450 | 2033 | 2511 | 5638 | 1219 | 1825 | 6120 | 1387 | 1955 | 5007 | 1150 | 1230 |
| 750 | QGN13847.1 | FIM1_493 | CP015054. | 1169 | 1178 | 772 | 1135 | 941 | 828 | 1076 | 842 | 1100 | 4471 | 1034 | 1263 | 3405 | 902 | 1260 | 4034 | 980 | 835 |
| 751 | QGN13848.1 | FIM1_494 | CP015054. | 542 | 1579 | 1141 | 748 | 1657 | 1143 | 650 | 1500 | 1427 | 6765 | 1555 | 1826 | 6744 | 1470 | 1714 | 6532 | 1522 | 1206 |
| 752 | QGN13849.1 | FIM1_495 | CP015054. | 1199 | 1351 | 2577 | 1434 | 1816 | 2042 | 987 | 1605 | 3157 | 10555 | 2422 | 3892 | 11723 | 2256 | 3810 | 11488 | 2229 | 2424 |
| 753 | QGN13850.1 | FIM1_496 | CP015054. | 4220 | 4568 | 3628 | 5734 | 4349 | 3196 | 4466 | 4071 | 4657 | 13156 | 3601 | 5449 | 17098 | 3552 | 4837 | 14310 | 3330 | 3503 |
| 754 | QGN13851.1 | FIM1_497 | CP015054. | 2802 | 1326 | 1663 | 2752 | 1390 | 1442 | 2032 | 1525 | 2115 | 856 | 642 | 1659 | 3099 | 610 | 1519 | 1636 | 601 | 1150 |

| | | | | | | | | | | | | | | | | | | | | | |
|---|---|---|---|---|---|---|---|---|---|---|---|---|---|---|---|---|---|---|---|---|---|
| 755 | QGN13852.1 | FIM1_498 | CP015054. | 750 | 620 | 439 | 705 | 433 | 411 | 659 | 377 | 585 | 1477 | 410 | 607 | 1411 | 394 | 664 | 1291 | 434 | 437 |
| 756 | QGN13853.1 | FIM1_499 | CP015054. | 2358 | 2800 | 2132 | 4197 | 2420 | 2204 | 2943 | 2182 | 2798 | 6359 | 2314 | 3137 | 7054 | 2318 | 3138 | 6641 | 2438 | 2125 |
| 757 | QGN13854.1 | FIM1_500 | CP015054. | 520 | 403 | 179 | 577 | 279 | 187 | 484 | 305 | 275 | 530 | 294 | 264 | 882 | 230 | 355 | 552 | 246 | 201 |
| 758 | QGN13855.1 | FIM1_501 | CP015054. | 922 | 372 | 331 | 769 | 597 | 251 | 729 | 601 | 451 | 914 | 571 | 582 | 1734 | 483 | 561 | 1147 | 537 | 372 |
| 759 | QGN13856.1 | FIM1_502 | CP015054. | 2097 | 2847 | 1506 | 3310 | 1772 | 1421 | 2876 | 1909 | 2185 | 6119 | 1691 | 1866 | 6476 | 1741 | 2050 | 6031 | 1658 | 1253 |
| 760 | QGN13857.1 | FIM1_503 | CP015054. | 2957 | 297 | 730 | 2673 | 466 | 523 | 2222 | 509 | 826 | 2647 | 312 | 887 | 3348 | 275 | 804 | 2702 | 220 | 540 |
| 761 | QGN13858.1 | FIM1_504 | CP015054. | 5927 | 499 | 772 | 5336 | 542 | 686 | 5986 | 509 | 1018 | 2532 | 442 | 1088 | 2924 | 482 | 891 | 2492 | 451 | 676 |
| 762 | QGN13859.1 | FIM1_505 | CP015054. | 1889 | 1057 | 446 | 1764 | 936 | 500 | 2480 | 897 | 637 | 1654 | 637 | 700 | 2535 | 710 | 670 | 1774 | 729 | 472 |
| 763 | QGN13860.1 | FIM1_506 | CP015054. | 3353 | 3455 | 1239 | 1600 | 2474 | 1212 | 2487 | 2262 | 1727 | 6738 | 3484 | 1847 | 7639 | 2476 | 2180 | 6354 | 3114 | 1432 |
| 764 | QGN13861.1 | FIM1_507 | CP015054. | 2225 | 1408 | 1231 | 1831 | 1064 | 1153 | 1644 | 1187 | 1730 | 3990 | 875 | 1367 | 4411 | 741 | 1501 | 3644 | 762 | 923 |
| 765 | QGN13862.1 | FIM1_508 | CP015054. | 3995 | 607 | 547 | 4160 | 464 | 556 | 3176 | 730 | 819 | 1718 | 370 | 821 | 2498 | 362 | 712 | 1742 | 356 | 510 |
| 766 | QGN13863.1 | FIM1_509 | CP015054. | 4640 | 384 | 562 | 4493 | 345 | 379 | 4334 | 391 | 699 | 1424 | 244 | 673 | 2731 | 411 | 649 | 1874 | 257 | 431 |
| 767 | QGN13864.1 | FIM1_510 | CP015054. | 621 | 794 | 516 | 1089 | 562 | 492 | 922 | 678 | 703 | 3318 | 500 | 723 | 2792 | 488 | 731 | 2840 | 446 | 511 |
| 768 | QGN13865.1 | FIM1_511 | CP015054. | 5932 | 937 | 1304 | 5817 | 1034 | 1257 | 4871 | 841 | 1663 | 1767 | 468 | 1359 | 2742 | 874 | 1463 | 1962 | 590 | 944 |
| 769 | QGN13866.1 | FIM1_512 | CP015054. | 1513 | 360 | 200 | 1350 | 141 | 165 | 1480 | 277 | 255 | 566 | 133 | 291 | 610 | 95 | 310 | 524 | 132 | 181 |
| 770 | QGN13867.1 | FIM1_513 | CP015054. | 2319 | 3475 | 1962 | 1993 | 2784 | 1711 | 1885 | 2331 | 2474 | 7467 | 2735 | 2618 | 9018 | 2342 | 2859 | 7806 | 2314 | 1852 |
| 771 | QGN13868.1 | FIM1_514 | CP015054. | 4245 | 909 | 1513 | 3674 | 1170 | 1097 | 3461 | 1327 | 1845 | 3694 | 853 | 1852 | 7186 | 742 | 1832 | 4870 | 708 | 1272 |
| 772 | QGN13869.1 | FIM1_515 | CP015054. | 2434 | 1440 | 1189 | 2874 | 1233 | 951 | 2484 | 1339 | 1522 | 4167 | 1159 | 1627 | 5037 | 1192 | 1623 | 4108 | 1074 | 1153 |
| 773 | QGN13870.1 | FIM1_516 | CP015054. | 1955 | 1405 | 900 | 2477 | 912 | 757 | 2181 | 943 | 1147 | 3510 | 821 | 1309 | 3839 | 683 | 1210 | 3250 | 655 | 852 |
| 774 | QGN13871.1 | FIM1_517 | CP015054. | 1827 | 1358 | 489 | 887 | 1204 | 558 | 1439 | 1104 | 635 | 2820 | 1189 | 920 | 3661 | 1005 | 868 | 2977 | 1312 | 647 |
| 775 | QGN13872.1 | FIM1_518 | CP015054. | 21902 | 23682 | 13079 | 23794 | 18411 | 12004 | 20375 | 17681 | 16283 | 46343 | 19637 | 21174 | 53274 | 17333 | 20280 | 47444 | 17268 | 14980 |
| 776 | QGN13873.1 | FIM1_519 | CP015054. | 7225 | 108 | 198 | 3921 | 101 | 139 | 4346 | 142 | 304 | 132 | 41 | 154 | 699 | 55 | 163 | 394 | 36 | 106 |
| 777 | QGN13874.1 | FIM1_521 | CP015054. | 25 | 10 | 2 | 63 | 14 | 4 | 15 | 5 | 3 | 23 | 3 | 4 | 21 | 1 | 5 | 23 | 7 | 2 |
| 778 | QGN13875.1 | FIM1_522 | CP015054. | 2689 | 1985 | 1215 | 1308 | 2225 | 1215 | 2204 | 1930 | 1684 | 7954 | 2305 | 1804 | 7933 | 1930 | 1763 | 7469 | 1990 | 1260 |
| 779 | QGN13876.1 | FIM1_523 | CP015054. | 235 | 597 | 485 | 189 | 447 | 478 | 182 | 387 | 697 | 2735 | 502 | 603 | 2059 | 418 | 623 | 2588 | 427 | 397 |
| 780 | QGN13877.1 | FIM1_524 | CP015054. | 1265 | 930 | 1043 | 2411 | 846 | 966 | 1877 | 718 | 1414 | 6306 | 965 | 1807 | 7560 | 1055 | 1757 | 6648 | 937 | 1381 |
| 781 | QGN13878.1 | FIM1_525 | CP015054. | 2892 | 1346 | 1546 | 3169 | 1336 | 1331 | 3154 | 1310 | 2083 | 4013 | 871 | 1510 | 4789 | 735 | 1593 | 4232 | 730 | 1058 |
| 782 | QGN13879.1 | FIM1_526 | CP015054. | 685 | 25 | 27 | 428 | 55 | 37 | 346 | 11 | 53 | 912 | 70 | 70 | 688 | 35 | 72 | 1063 | 34 | 43 |
| 783 | QGN13880.1 | FIM1_527 | CP015054. | 74341 | 3925 | 4327 | 83214 | 3814 | 3108 | 86397 | 4220 | 5469 | 3652 | 3814 | 5205 | 23559 | 8284 | 4452 | 10880 | 5931 | 3638 |
| 784 | QGN13881.1 | FIM1_528 | CP015054. | 11078 | 1809 | 1801 | 14519 | 1885 | 1255 | 12255 | 2395 | 2333 | 1262 | 1220 | 2943 | 6755 | 1664 | 2456 | 2216 | 1569 | 1591 |
| 785 | QGN13882.1 | FIM1_529 | CP015054. | 77 | 39 | 16 | 60 | 26 | 14 | 55 | 21 | 18 | 60 | 8 | 12 | 79 | 15 | 12 | 61 | 14 | 11 |
| 786 | QGN13883.1 | FIM1_530 | CP015054. | 672 | 5949 | 2373 | 1147 | 3561 | 1901 | 935 | 3826 | 3213 | 11345 | 3607 | 2810 | 10526 | 2793 | 2866 | 10217 | 2748 | 2023 |
| 787 | QGN13884.1 | FIM1_531 | CP015054. | 3349 | 12579 | 6406 | 9669 | 8494 | 4392 | 6396 | 10497 | 8303 | 7155 | 7234 | 9439 | 26791 | 9117 | 8515 | 14947 | 7828 | 6532 |
| 788 | QGN13885.1 | FIM1_532 | CP015054. | 11041 | 4014 | 1074 | 4942 | 3433 | 1128 | 10277 | 3650 | 1590 | 1432 | 3043 | 1794 | 7041 | 3045 | 1771 | 3215 | 4038 | 1377 |
| 789 | QGN13886.1 | FIM1_533 | CP015054. | 2793 | 865 | 690 | 2705 | 751 | 541 | 2564 | 687 | 897 | 2189 | 573 | 888 | 3321 | 563 | 788 | 2393 | 574 | 593 |
| 790 | QGN13887.1 | FIM1_534 | CP015054. | 1360 | 1013 | 861 | 2108 | 690 | 693 | 1622 | 723 | 1102 | 3491 | 548 | 879 | 3670 | 667 | 1053 | 3282 | 492 | 624 |
| 791 | QGN13888.1 | FIM1_535 | CP015054. | 1577 | 1427 | 945 | 3148 | 854 | 673 | 3278 | 966 | 1324 | 645 | 653 | 1416 | 4422 | 937 | 1330 | 1905 | 944 | 893 |
| 792 | QGN13889.1 | FIM1_536 | CP015054. | 7079 | 294 | 425 | 4300 | 480 | 407 | 5111 | 446 | 630 | 406 | 322 | 341 | 1447 | 404 | 376 | 918 | 400 | 249 |
| 793 | QGN13890.1 | FIM1_537 | CP015054. | 2387 | 344 | 698 | 3017 | 588 | 459 | 2017 | 596 | 940 | 736 | 330 | 817 | 3502 | 373 | 621 | 1630 | 262 | 522 |
| 794 | QGN13891.1 | FIM1_538 | CP015054. | 431 | 632 | 690 | 942 | 676 | 601 | 600 | 657 | 985 | 2209 | 690 | 1175 | 3844 | 732 | 981 | 2847 | 647 | 760 |
| 795 | QGN13892.1 | FIM1_539 | CP015054. | 577 | 237 | 257 | 609 | 246 | 229 | 510 | 241 | 349 | 2093 | 252 | 415 | 1684 | 181 | 405 | 2133 | 230 | 260 |
| 796 | QGN13893.1 | FIM1_540 | CP015054. | 4012 | 1029 | 1390 | 4403 | 1273 | 1238 | 4409 | 1170 | 1868 | 4377 | 1382 | 2260 | 6190 | 1189 | 2176 | 4887 | 1275 | 1614 |
| 797 | QGN13894.1 | FIM1_541 | CP015054. | 493 | 371 | 367 | 517 | 362 | 307 | 476 | 333 | 479 | 2179 | 362 | 617 | 2096 | 338 | 555 | 2139 | 337 | 387 |
| 798 | QGN13895.1 | FIM1_542 | CP015054. | 489 | 664 | 582 | 756 | 519 | 523 | 603 | 524 | 719 | 3640 | 522 | 790 | 3037 | 481 | 756 | 3769 | 516 | 493 |
| 799 | QGN13896.1 | FIM1_543 | CP015054. | 1342 | 229 | 282 | 1572 | 311 | 221 | 1413 | 358 | 342 | 573 | 260 | 455 | 1657 | 280 | 325 | 902 | 255 | 310 |
| 800 | QGN13897.1 | FIM1_544 | CP015054. | 1797 | 2356 | 881 | 2064 | 1452 | 838 | 1842 | 1209 | 1267 | 4551 | 1482 | 1291 | 5483 | 1254 | 1306 | 4580 | 1265 | 929 |
| 801 | QGN13898.1 | FIM1_545 | CP015054. | 368 | 1192 | 2486 | 873 | 919 | 2029 | 448 | 1037 | 3113 | 2382 | 581 | 3406 | 3645 | 574 | 3065 | 3050 | 520 | 2290 |
| 802 | QGN13899.1 | FIM1_546 | CP015054. | 14528 | 2408 | 2313 | 15270 | 3589 | 2035 | 14249 | 2795 | 2862 | 15345 | 3028 | 3078 | 15288 | 3252 | 2970 | 15391 | 2876 | 1999 |
| 803 | QGN13900.1 | FIM1_547 | CP015054. | 5870 | 864 | 1375 | 4737 | 1289 | 1038 | 4423 | 1296 | 1688 | 4096 | 1052 | 1890 | 7804 | 941 | 1802 | 5131 | 895 | 1264 |
| 804 | QGN13901.1 | FIM1_549 | CP015054. | 2237 | 863 | 474 | 2619 | 649 | 423 | 2568 | 739 | 655 | 568 | 487 | 651 | 1317 | 595 | 693 | 736 | 586 | 439 |
| 805 | QGN13902.1 | FIM1_550 | CP015054. | 13151 | 7302 | 4939 | 18909 | 8807 | 5441 | 12422 | 6740 | 6257 | 22941 | 9740 | 9221 | 30195 | 9250 | 8785 | 24423 | 8835 | 6634 |
| 806 | QGN13903.1 | FIM1_551 | CP015054. | 103 | 484 | 380 | 259 | 432 | 380 | 204 | 383 | 561 | 2333 | 319 | 644 | 1882 | 331 | 542 | 2154 | 307 | 427 |
| 807 | QGN13904.1 | FIM1_552 | CP015054. | 1103 | 496 | 730 | 1665 | 661 | 633 | 1496 | 597 | 904 | 5983 | 684 | 1073 | 5181 | 602 | 1021 | 5636 | 578 | 762 |
| 808 | QGN13905.1 | FIM1_553 | CP015054. | 2915 | 509 | 1133 | 3008 | 746 | 866 | 2966 | 618 | 1416 | 5876 | 691 | 1561 | 6412 | 714 | 1436 | 5867 | 691 | 1105 |
| 809 | QGN13906.1 | FIM1_554 | CP015054. | 1688 | 1231 | 1274 | 2278 | 1250 | 1122 | 2067 | 1016 | 1556 | 7292 | 1288 | 1942 | 6201 | 1117 | 1787 | 7148 | 1062 | 1173 |
| 810 | QGN13907.1 | FIM1_555 | CP015054. | 642 | 345 | 457 | 558 | 328 | 364 | 539 | 311 | 553 | 2992 | 308 | 567 | 2319 | 215 | 573 | 2933 | 308 | 370 |
| 811 | QGN13908.1 | FIM1_556 | CP015054. | 1272 | 996 | 927 | 2313 | 947 | 765 | 1486 | 864 | 1213 | 4422 | 723 | 1081 | 3519 | 754 | 1005 | 4193 | 694 | 642 |
| 812 | QGN13909.1 | FIM1_557 | CP015054. | 1831 | 837 | 576 | 2060 | 626 | 484 | 1849 | 621 | 681 | 2307 | 532 | 760 | 2568 | 691 | 710 | 2306 | 671 | 487 |
| 813 | QGN13910.1 | FIM1_558 | CP015054. | 332 | 1102 | 663 | 454 | 939 | 695 | 492 | 832 | 868 | 4547 | 1173 | 1129 | 3975 | 1026 | 1298 | 4023 | 1210 | 839 |
| 814 | QGN13911.1 | FIM1_559 | CP015054. | 891 | 388 | 465 | 933 | 476 | 429 | 859 | 443 | 658 | 3289 | 521 | 775 | 3119 | 514 | 779 | 3067 | 415 | 594 |
| 815 | QGN13912.1 | FIM1_560 | CP015054. | 991 | 1271 | 1705 | 1486 | 1221 | 1729 | 1379 | 1062 | 2354 | 7787 | 1165 | 3118 | 6431 | 999 | 2888 | 7631 | 1076 | 2119 |
| 816 | QGN13913.1 | FIM1_561 | CP015054. | 3560 | 340 | 437 | 2666 | 415 | 353 | 3408 | 400 | 539 | 2903 | 411 | 672 | 3222 | 410 | 583 | 2847 | 384 | 452 |
| 817 | QGN13914.1 | FIM1_562 | CP015054. | 7114 | 1230 | 908 | 7297 | 1029 | 869 | 6614 | 904 | 1250 | 3465 | 856 | 1589 | 4555 | 917 | 1417 | 3984 | 868 | 1014 |
| 818 | QGN13915.1 | FIM1_563 | CP015054. | 1057 | 124 | 132 | 642 | 49 | 80 | 1036 | 84 | 159 | 830 | 99 | 181 | 606 | 91 | 190 | 750 | 98 | 113 |
| 819 | QGN13916.1 | FIM1_564 | CP015054. | 476 | 1546 | 874 | 798 | 1010 | 895 | 843 | 800 | 1210 | 5624 | 1254 | 1493 | 4420 | 1097 | 1549 | 5338 | 1196 | 1004 |
| 820 | QGN13917.1 | FIM1_565 | CP015054. | 1232 | 1048 | 594 | 769 | 928 | 676 | 1225 | 805 | 941 | 6025 | 1229 | 1024 | 5470 | 1095 | 1009 | 5819 | 1284 | 733 |
| 821 | QGN13918.1 | FIM1_566 | CP015054. | 413 | 826 | 557 | 408 | 634 | 508 | 394 | 542 | 681 | 3258 | 574 | 712 | 2427 | 515 | 727 | 3079 | 569 | 472 |
| 822 | QGN13919.1 | FIM1_567 | CP015054. | 1376 | 1314 | 1126 | 1718 | 1069 | 1010 | 1445 | 1002 | 1469 | 7161 | 1105 | 1510 | 5928 | 934 | 1460 | 7123 | 877 | 1013 |
| 823 | QGN13920.1 | FIM1_568 | CP015054. | 3006 | 2478 | 4992 | 1978 | 1282 | 2406 | 5521 | 6092 | 5749 | 6670 | 877 | 1676 | 7669 | 1446 | 1850 | 7247 | 1543 | 1246 |
| 824 | QGN13921.1 | FIM1_569 | CP015054. | 874 | 2173 | 3112 | 1705 | 1539 | 1924 | 2861 | 3272 | 3368 | 8414 | 866 | 1788 | 7001 | 1241 | 2032 | 8229 | 1527 | 1311 |
| 825 | QGN13922.1 | FIM1_570 | CP015054. | 894 | 509 | 457 | 1623 | 980 | 520 | 1202 | 590 | 562 | 3079 | 1156 | 892 | 3078 | 864 | 704 | 3150 | 1122 | 534 |
| 826 | QGN13923.1 | FIM1_571 | CP015054. | 1801 | 818 | 1183 | 11645 | 1255 | 1479 | 5530 | 595 | 1518 | 5086 | 390 | 1579 | 4631 | 431 | 1483 | 4732 | 369 | 1013 |
| 827 | QGN13924.1 | FIM1_572 | CP015054. | 262 | 213 | 560 | 1716 | 715 | 681 | 547 | 255 | 649 | 887 | 63 | 492 | 706 | 45 | 482 | 829 | 70 | 305 |
| 828 | QGN13925.1 | FIM1_573 | CP015054. | 7598 | 3872 | 2100 | 11872 | 3049 | 2139 | 9618 | 2287 | 2872 | 11437 | 2906 | 2859 | 9606 | 3274 | 2905 | 10636 | 2898 | 1980 |
| 829 | QGN13926.1 | FIM1_574 | CP015054. | 3921 | 2466 | 1243 | 1285 | 2641 | 1428 | 3128 | 2239 | 1758 | 7114 | 3117 | 2291 | 8907 | 3026 | 2162 | 7411 | 3290 | 1580 |
| 830 | QGN13927.1 | FIM1_575 | CP015054. | 3992 | 2255 | 1468 | 5033 | 2194 | 1432 | 4621 | 1500 | 1972 | 7349 | 1983 | 2151 | 6638 | 1843 | 2417 | 7158 | 1879 | 1535 |
| 831 | QGN13928.1 | FIM1_576 | CP015054. | 4981 | 4241 | 3181 | 7021 | 4625 | 2804 | 6027 | 3860 | 4089 | 19063 | 4688 | 4924 | 18484 | 3914 | 4550 | 18331 | 3896 | 3311 |
| 832 | QGN13929.1 | FIM1_577 | CP015054. | 513 | 783 | 778 | 853 | 982 | 648 | 711 | 864 | 1024 | 3123 | 661 | 826 | 2618 | 528 | 774 | 3254 | 579 | 521 |
| 833 | QGN13930.1 | FIM1_578 | CP015054. | 3420 | 3032 | 3449 | 6290 | 2679 | 2898 | 4610 | 2650 | 4225 | 13638 | 2375 | 5133 | 13347 | 2657 | 4730 | 13901 | 2212 | 3336 |
| 834 | QGN13931.1 | FIM1_579 | CP015054. | 648 | 361 | 513 | 1128 | 502 | 410 | 907 | 384 | 620 | 3567 | 434 | 907 | 3897 | 395 | 731 | 3909 | 390 | 555 |
| 835 | QGN13932.1 | FIM1_580 | CP015054. | 443 | 199 | 183 | 474 | 144 | 144 | 512 | 142 | 275 | 1023 | 131 | 237 | 960 | 118 | 221 | 1009 | 110 | 141 |
| 836 | QGN13933.1 | FIM1_581 | CP015054. | 1699 | 386 | 434 | 1213 | 499 | 388 | 1443 | 447 | 591 | 4039 | 467 | 700 | 3677 | 480 | 513 | 3903 | 409 | 485 |
| 837 | QGN13934.1 | FIM1_582 | CP015054. | 2949 | 2606 | 1080 | 1214 | 2454 | 1215 | 2414 | 2141 | 1549 | 7886 | 3365 | 1966 | 7136 | 2789 | 2112 | 7920 | 3562 | 1344 |
| 838 | QGN13935.1 | FIM1_583 | CP015054. | 236 | 224 | 174 | 331 | 169 | 176 | 242 | 147 | 237 | 1111 | 161 | 199 | 741 | 95 | 173 | 1049 | 152 | 140 |

| | | | | | | | | | | | | | | | | | | | | | |
|---|---|---|---|---|---|---|---|---|---|---|---|---|---|---|---|---|---|---|---|---|---|
| 839 | QGN13936.1 | FIM1_584 | CP015054. | 918 | 592 | 617 | 1160 | 608 | 594 | 1033 | 531 | 809 | 4750 | 476 | 978 | 3455 | 379 | 846 | 4497 | 390 | 618 |
| 840 | QGN13937.1 | FIM1_585 | CP015054. | 2074 | 7741 | 5706 | 5741 | 9602 | 5021 | 3398 | 7739 | 7071 | 25466 | 8597 | 9118 | 33743 | 7984 | 8578 | 28190 | 6787 | 6275 |
| 841 | QGN13938.1 | FIM1_586 | CP015054. | 5804 | 775 | 720 | 3481 | 1304 | 721 | 6124 | 1001 | 836 | 4299 | 1287 | 1230 | 5587 | 1600 | 1113 | 4679 | 1514 | 868 |
| 842 | QGN13939.1 | FIM1_587 | CP015054. | 504 | 818 | 694 | 1072 | 587 | 584 | 524 | 592 | 872 | 2788 | 496 | 629 | 2304 | 301 | 602 | 2871 | 346 | 393 |
| 843 | QGN13940.1 | FIM1_588 | CP015054. | 239 | 716 | 420 | 300 | 502 | 417 | 293 | 454 | 564 | 2932 | 574 | 623 | 2081 | 430 | 506 | 2712 | 482 | 377 |
| 844 | QGN13941.1 | FIM1_589 | CP015054. | 1558 | 473 | 471 | 2473 | 579 | 414 | 1790 | 427 | 614 | 4139 | 546 | 804 | 3733 | 542 | 823 | 4022 | 526 | 560 |
| 845 | QGN13942.1 | FIM1_590 | CP015054. | 2286 | 433 | 443 | 2705 | 393 | 392 | 2492 | 378 | 572 | 3977 | 409 | 758 | 3209 | 354 | 661 | 3616 | 381 | 478 |
| 846 | QGN13943.1 | FIM1_591 | CP015054. | 1749 | 509 | 569 | 2324 | 508 | 437 | 2059 | 525 | 719 | 3386 | 369 | 592 | 2685 | 323 | 501 | 3131 | 372 | 380 |
| 847 | QGN13944.1 | FIM1_592 | CP015054. | 7585 | 3150 | 3978 | 9406 | 3675 | 3392 | 8736 | 3325 | 5401 | 9746 | 3445 | 5340 | 20621 | 3196 | 5057 | 13847 | 3210 | 3842 |
| 848 | QGN13945.1 | FIM1_593 | CP015054. | 1109 | 949 | 784 | 1390 | 681 | 760 | 1075 | 509 | 1054 | 4665 | 735 | 1052 | 3754 | 653 | 1030 | 4376 | 723 | 674 |
| 849 | QGN13946.1 | FIM1_594 | CP015054. | 1171 | 1005 | 1584 | 1650 | 1838 | 1224 | 877 | 1811 | 1930 | 7908 | 1634 | 1686 | 8636 | 1314 | 1504 | 8680 | 1400 | 1180 |
| 850 | QGN13947.1 | FIM1_595 | CP015054. | 1578 | 1588 | 1310 | 2204 | 1727 | 1131 | 2497 | 1331 | 1568 | 6413 | 1809 | 2187 | 7086 | 1703 | 1832 | 6703 | 1741 | 1422 |
| 851 | QGN13948.1 | FIM1_596 | CP015054. | 828 | 3493 | 2667 | 2006 | 3371 | 2477 | 1291 | 2731 | 3295 | 9136 | 3289 | 4079 | 10554 | 3051 | 4216 | 8680 | 3007 | 2773 |
| 852 | QGN13949.1 | FIM1_597 | CP015054. | 6037 | 1144 | 1040 | 6145 | 1014 | 920 | 6036 | 1073 | 1435 | 6354 | 963 | 1048 | 4871 | 762 | 1095 | 5798 | 778 | 730 |
| 853 | QGN13950.1 | FIM1_598 | CP015054. | 572 | 735 | 536 | 634 | 562 | 497 | 715 | 599 | 749 | 3752 | 729 | 909 | 2769 | 630 | 865 | 3361 | 664 | 596 |
| 854 | QGN13951.1 | FIM1_599 | CP015054. | 845 | 407 | 371 | 1027 | 324 | 360 | 962 | 367 | 514 | 2477 | 319 | 594 | 2215 | 298 | 568 | 2400 | 323 | 377 |
| 855 | QGN13952.1 | FIM1_600 | CP015054. | 1520 | 850 | 1284 | 3358 | 951 | 1071 | 2010 | 862 | 1637 | 6243 | 827 | 2195 | 6413 | 1230 | 1928 | 6293 | 904 | 1463 |
| 856 | QGN13953.1 | FIM1_601 | CP015054. | 3742 | 2217 | 1044 | 1599 | 2236 | 1147 | 3025 | 1885 | 1409 | 6719 | 2659 | 1765 | 6493 | 2551 | 1699 | 6149 | 2715 | 1333 |
| 857 | QGN13954.1 | FIM1_602 | CP015054. | 4627 | 1277 | 1280 | 4481 | 1424 | 1182 | 4221 | 1239 | 1590 | 7455 | 1187 | 1887 | 7183 | 1204 | 1650 | 6938 | 1087 | 1270 |
| 858 | QGN13955.1 | FIM1_603 | CP015054. | 10623 | 4412 | 5898 | 20755 | 4142 | 4567 | 13337 | 5051 | 7354 | 18858 | 2660 | 7513 | 20134 | 2790 | 6999 | 18398 | 2395 | 4848 |
| 859 | QGN13956.1 | FIM1_604 | CP015054. | 1293 | 1873 | 1384 | 2103 | 1552 | 1212 | 1380 | 1328 | 1864 | 8456 | 2013 | 2104 | 8060 | 1668 | 2380 | 8263 | 1532 | 1494 |
| 860 | QGN13957.1 | FIM1_605 | CP015054. | 135 | 351 | 508 | 208 | 303 | 439 | 111 | 252 | 719 | 2185 | 245 | 684 | 1707 | 333 | 665 | 2041 | 236 | 391 |
| 861 | QGN13958.1 | FIM1_606 | CP015054. | 2087 | 521 | 1587 | 3333 | 544 | 1246 | 2207 | 578 | 2055 | 4987 | 325 | 1797 | 3910 | 300 | 1523 | 4611 | 287 | 1162 |
| 862 | QGN13959.1 | FIM1_607 | CP015054. | 1062 | 470 | 219 | 614 | 581 | 268 | 1029 | 322 | 303 | 2035 | 660 | 450 | 1684 | 810 | 441 | 1930 | 684 | 321 |
| 863 | QGN13960.1 | FIM1_608 | CP015054. | 1054 | 502 | 414 | 950 | 339 | 402 | 1021 | 317 | 606 | 1803 | 343 | 501 | 1724 | 276 | 507 | 1804 | 276 | 304 |
| 864 | QGN13961.1 | FIM1_609 | CP015054. | 25933 | 4404 | 3629 | 21433 | 4403 | 2836 | 13880 | 4345 | 4700 | 8551 | 4063 | 5481 | 22618 | 4403 | 4353 | 14938 | 3331 | 3744 |
| 865 | QGN13962.1 | FIM1_610 | CP015054. | 775 | 390 | 396 | 773 | 273 | 359 | 728 | 303 | 545 | 2375 | 287 | 541 | 2037 | 238 | 569 | 2305 | 232 | 409 |
| 866 | QGN13963.1 | FIM1_611 | CP015054. | 368 | 103 | 139 | 695 | 96 | 112 | 419 | 106 | 158 | 497 | 63 | 149 | 638 | 78 | 139 | 475 | 51 | 103 |
| 867 | QGN13964.1 | FIM1_612 | CP015054. | 5092 | 803 | 1072 | 5730 | 840 | 886 | 5333 | 796 | 1446 | 5635 | 758 | 1507 | 6694 | 759 | 1333 | 5947 | 675 | 957 |
| 868 | QGN13965.1 | FIM1_613 | CP015054. | 1670 | 819 | 809 | 2755 | 723 | 748 | 2135 | 751 | 1186 | 3846 | 635 | 1261 | 4686 | 586 | 1193 | 4023 | 565 | 808 |
| 869 | QGN13966.1 | FIM1_615 | CP015054. | 2291 | 530 | 698 | 3494 | 479 | 545 | 2478 | 548 | 887 | 3650 | 351 | 760 | 3473 | 397 | 746 | 3220 | 301 | 560 |
| 870 | QGN13967.1 | FIM1_616 | CP015054. | 2074 | 1943 | 1577 | 2898 | 1567 | 1319 | 2597 | 1499 | 1985 | 9042 | 1687 | 2552 | 8018 | 1745 | 2560 | 9163 | 1476 | 1626 |
| 871 | QGN13968.1 | FIM1_617 | CP015054. | 251 | 468 | 257 | 1093 | 455 | 317 | 308 | 338 | 351 | 1646 | 357 | 369 | 1253 | 276 | 390 | 1404 | 299 | 226 |
| 872 | QGN13969.1 | FIM1_618 | CP015054. | 5368 | 622 | 673 | 4938 | 696 | 554 | 4082 | 863 | 962 | 583 | 444 | 907 | 1743 | 523 | 865 | 752 | 543 | 567 |
| 873 | QGN13970.1 | FIM1_619 | CP015054. | 42499 | 6652 | 9446 | 51914 | 8548 | 7859 | 56355 | 9055 | 11847 | 15847 | 6505 | 8748 | 30929 | 7137 | 8863 | 22667 | 5470 | 6369 |
| 874 | QGN13971.1 | FIM1_620 | CP015054. | 179 | 773 | 636 | 1121 | 713 | 616 | 601 | 713 | 987 | 2501 | 465 | 1010 | 3079 | 558 | 937 | 2583 | 443 | 633 |
| 875 | QGN13972.1 | FIM1_621 | CP015054. | 536 | 435 | 490 | 571 | 330 | 641 | 327 | 296 | 869 | 319 | 303 | 777 | 1129 | 171 | 983 | 671 | 243 | 636 |
| 876 | QGN13973.1 | FIM1_622 | CP015054. | 340 | 278 | 282 | 330 | 197 | 231 | 271 | 222 | 336 | 826 | 165 | 388 | 738 | 134 | 402 | 754 | 156 | 257 |
| 877 | QGN13974.1 | FIM1_623 | CP015054. | 2194 | 2678 | 2313 | 3254 | 2644 | 1931 | 2678 | 2247 | 2780 | 11809 | 2314 | 2719 | 9123 | 2191 | 2621 | 12783 | 2171 | 1896 |
| 878 | QGN13975.1 | FIM1_624 | CP015054. | 8 | 3 | 0 | 51 | 4 | 6 | 4 | 2 | 4 | 18 | 3 | 2 | 11 | 1 | 2 | 19 | 4 | 3 |
| 879 | QGN13976.1 | FIM1_625 | CP015054. | 19044 | 171 | 177 | 15935 | 130 | 137 | 17003 | 157 | 237 | 1429 | 117 | 197 | 1300 | 281 | 166 | 1158 | 175 | 145 |
| 880 | QGN13977.1 | FIM1_626 | CP015054. | 11789 | 1922 | 2641 | 15720 | 1370 | 2240 | 11173 | 1166 | 3149 | 7616 | 951 | 2927 | 6177 | 1705 | 2578 | 6579 | 1517 | 1762 |
| 881 | QGN13978.1 | FIM1_627 | CP015054. | 4851 | 505 | 565 | 3802 | 411 | 466 | 4762 | 418 | 731 | 2582 | 462 | 779 | 3496 | 397 | 772 | 2619 | 401 | 672 |
| 882 | QGN13979.1 | FIM1_628 | CP015054. | 5431 | 1940 | 2621 | 9253 | 1992 | 2293 | 5301 | 1241 | 3262 | 3689 | 891 | 3407 | 6637 | 1101 | 3569 | 4994 | 886 | 2394 |
| 883 | QGN13980.1 | FIM1_629 | CP015054. | 5130 | 1281 | 1589 | 4863 | 2556 | 1413 | 4735 | 1571 | 2260 | 2679 | 1648 | 2097 | 5183 | 1884 | 1855 | 3707 | 1846 | 1387 |
| 884 | QGN13981.1 | FIM1_630 | CP015054. | 1423 | 460 | 436 | 1788 | 359 | 446 | 1846 | 375 | 629 | 2035 | 371 | 771 | 2308 | 407 | 768 | 1939 | 389 | 491 |
| 885 | QGN13982.1 | FIM1_631 | CP015054. | 5669 | 797 | 969 | 5515 | 1072 | 825 | 5217 | 1043 | 1393 | 1394 | 945 | 1614 | 6898 | 1048 | 1413 | 3138 | 975 | 1136 |
| 886 | QGN13983.1 | FIM1_632 | CP015054. | 883 | 218 | 263 | 1100 | 218 | 213 | 671 | 240 | 303 | 768 | 162 | 334 | 1165 | 119 | 301 | 879 | 107 | 186 |
| 887 | QGN13984.1 | FIM1_633 | CP015054. | 1503 | 5694 | 3542 | 2421 | 4871 | 2867 | 2138 | 4271 | 4300 | 14968 | 4765 | 5351 | 16024 | 4696 | 5078 | 15045 | 4162 | 3719 |
| 888 | QGN13985.1 | FIM1_634 | CP015054. | 1133 | 328 | 280 | 1091 | 275 | 236 | 1109 | 258 | 363 | 1765 | 276 | 425 | 1465 | 262 | 431 | 1691 | 246 | 285 |
| 889 | QGN13986.1 | FIM1_635 | CP015054. | 2394 | 3778 | 2235 | 3148 | 2837 | 1991 | 2691 | 2459 | 2747 | 10200 | 3112 | 3378 | 9411 | 2875 | 3543 | 9673 | 2974 | 2403 |
| 890 | QGN13987.1 | FIM1_636 | CP015054. | 1002 | 367 | 237 | 1116 | 254 | 215 | 991 | 315 | 339 | 1350 | 241 | 381 | 1614 | 257 | 361 | 1396 | 262 | 246 |
| 891 | QGN13988.1 | FIM1_637 | CP015054. | 6540 | 8635 | 4683 | 6943 | 6176 | 4559 | 6337 | 5993 | 6004 | 19755 | 7411 | 8071 | 18546 | 6190 | 8000 | 20053 | 6900 | 5375 |
| 892 | QGN13989.1 | FIM1_638 | CP015054. | 2413 | 932 | 864 | 2526 | 908 | 819 | 2360 | 817 | 1148 | 5899 | 784 | 1397 | 4898 | 749 | 1276 | 6053 | 789 | 863 |
| 893 | QGN13990.1 | FIM1_639 | CP015054. | 190 | 48 | 40 | 433 | 41 | 47 | 239 | 45 | 54 | 333 | 25 | 47 | 228 | 30 | 47 | 346 | 24 | 21 |
| 894 | QGN13991.1 | FIM1_640 | CP015054. | 1001 | 692 | 713 | 1194 | 851 | 665 | 1056 | 744 | 922 | 4593 | 743 | 1110 | 4868 | 769 | 909 | 4337 | 644 | 729 |
| 895 | QGN13992.1 | FIM1_641 | CP015054. | 607 | 268 | 352 | 1195 | 317 | 375 | 686 | 315 | 503 | 2143 | 238 | 460 | 1961 | 215 | 436 | 1998 | 231 | 283 |
| 896 | QGN13993.1 | FIM1_642 | CP015054. | 3274 | 2103 | 1880 | 10100 | 1862 | 1564 | 6838 | 1852 | 2509 | 8972 | 1777 | 2762 | 10037 | 2883 | 2793 | 8757 | 1931 | 1934 |
| 897 | QGN13994.1 | FIM1_644 | CP015054. | 1840 | 1592 | 847 | 1380 | 1360 | 837 | 1617 | 1144 | 1058 | 5953 | 1363 | 1460 | 4491 | 1135 | 1346 | 6003 | 1456 | 961 |
| 898 | QGN13995.1 | FIM1_645 | CP015054. | 2054 | 467 | 650 | 4088 | 618 | 533 | 2567 | 684 | 903 | 489 | 284 | 562 | 1084 | 409 | 594 | 746 | 384 | 404 |
| 899 | QGN13996.1 | FIM1_646 | CP015054. | 4012 | 1784 | 1983 | 5607 | 2802 | 1617 | 5226 | 2068 | 2608 | 4702 | 1794 | 1180 | 6926 | 1708 | 1286 | 5484 | 1680 | 795 |
| 900 | QGN13997.1 | FIM1_647 | CP015054. | 2007 | 1216 | 647 | 1358 | 1013 | 629 | 1894 | 961 | 858 | 6186 | 1372 | 1140 | 5004 | 978 | 1109 | 5272 | 1279 | 751 |
| 901 | QGN13998.1 | FIM1_648 | CP015054. | 379 | 144 | 125 | 450 | 112 | 105 | 333 | 118 | 187 | 697 | 85 | 168 | 606 | 68 | 175 | 685 | 78 | 111 |
| 902 | QGN13999.1 | FIM1_650 | CP015054. | 1850 | 596 | 386 | 1867 | 393 | 377 | 1471 | 401 | 543 | 1682 | 310 | 591 | 1798 | 294 | 626 | 1550 | 240 | 419 |
| 903 | QGN14000.1 | FIM1_651 | CP015054. | 8708 | 432 | 574 | 5460 | 506 | 435 | 6763 | 490 | 768 | 1179 | 480 | 957 | 3926 | 398 | 857 | 2051 | 414 | 639 |
| 904 | QGN14001.1 | FIM1_652 | CP015054. | 3458 | 1663 | 1475 | 4161 | 1755 | 1273 | 3033 | 1370 | 1795 | 8845 | 1905 | 2331 | 8260 | 1495 | 2184 | 8703 | 1571 | 1598 |
| 905 | QGN14002.1 | FIM1_653 | CP015054. | 5738 | 2504 | 1040 | 2855 | 2850 | 1285 | 5491 | 2297 | 1580 | 9756 | 3212 | 2316 | 9210 | 3484 | 2235 | 9746 | 3603 | 1739 |
| 906 | QGN14003.1 | FIM1_654 | CP015054. | 888 | 350 | 435 | 1211 | 314 | 362 | 809 | 355 | 634 | 3120 | 282 | 609 | 2465 | 243 | 566 | 3009 | 268 | 380 |
| 907 | QGN14004.1 | FIM1_655 | CP015054. | 805 | 493 | 554 | 836 | 431 | 494 | 710 | 451 | 760 | 1729 | 223 | 483 | 1223 | 174 | 455 | 1781 | 198 | 273 |
| 908 | QGN14005.1 | FIM1_656 | CP015054. | 1210 | 1540 | 989 | 1895 | 1206 | 911 | 1588 | 961 | 1367 | 6985 | 1342 | 1699 | 5789 | 1249 | 1531 | 6705 | 1179 | 1088 |
| 909 | QGN14006.1 | FIM1_657 | CP015054. | 941 | 345 | 701 | 1365 | 513 | 552 | 1019 | 522 | 884 | 1701 | 452 | 952 | 3330 | 420 | 850 | 2392 | 386 | 657 |
| 910 | QGN14007.1 | FIM1_658 | CP015054. | 1757 | 336 | 528 | 1660 | 313 | 406 | 1408 | 449 | 617 | 1318 | 314 | 745 | 2400 | 212 | 660 | 1627 | 238 | 503 |
| 911 | QGN14008.1 | FIM1_659 | CP015054. | 3135 | 2180 | 1803 | 3253 | 1495 | 1541 | 2622 | 1401 | 2382 | 5011 | 1679 | 2474 | 5936 | 1575 | 2842 | 4988 | 1362 | 1827 |
| 912 | QGN14009.1 | FIM1_660 | CP015054. | 2452 | 634 | 917 | 3508 | 647 | 828 | 2274 | 530 | 1187 | 2225 | 337 | 818 | 2339 | 351 | 779 | 2172 | 263 | 531 |
| 913 | QGN14010.1 | FIM1_661 | CP015054. | 2628 | 388 | 525 | 2646 | 504 | 403 | 2397 | 476 | 622 | 1301 | 344 | 695 | 3153 | 443 | 598 | 1864 | 409 | 523 |
| 914 | QGN14011.1 | FIM1_662 | CP015054. | 10667 | 10751 | 5757 | 7399 | 11198 | 5568 | 9696 | 10392 | 7272 | 25229 | 14183 | 9864 | 33120 | 10667 | 9969 | 28768 | 11969 | 7130 |
| 915 | QGN14012.1 | FIM1_663 | CP015054. | 1645 | 112 | 395 | 2863 | 715 | 335 | 1478 | 432 | 456 | 1258 | 271 | 528 | 2277 | 273 | 605 | 1380 | 263 | 335 |
| 916 | QGN14013.1 | FIM1_664 | CP015054. | 2430 | 2814 | 2473 | 5296 | 2749 | 2246 | 3343 | 2370 | 3113 | 9145 | 1734 | 2624 | 8519 | 1867 | 2375 | 8523 | 1646 | 1595 |
| 917 | QGN14014.1 | FIM1_665 | CP015054. | 464 | 395 | 474 | 550 | 387 | 370 | 382 | 312 | 582 | 2347 | 418 | 795 | 2397 | 363 | 711 | 2267 | 358 | 497 |
| 918 | QGN14015.1 | FIM1_666 | CP015054. | 251 | 47 | 145 | 680 | 84 | 133 | 451 | 138 | 201 | 881 | 95 | 196 | 831 | 75 | 191 | 819 | 87 | 147 |
| 919 | QGN14016.1 | FIM1_667 | CP015054. | 211 | 172 | 160 | 439 | 164 | 125 | 243 | 109 | 205 | 743 | 136 | 251 | 869 | 109 | 231 | 768 | 103 | 161 |
| 920 | QGN14017.1 | FIM1_668 | CP015054. | 4728 | 1898 | 1798 | 4993 | 2924 | 1674 | 5145 | 2372 | 2403 | 9510 | 2671 | 2904 | 12539 | 2803 | 2516 | 10432 | 2888 | 1870 |
| 921 | QGN14018.1 | FIM1_669 | CP015054. | 1125 | 782 | 1135 | 1454 | 739 | 894 | 1261 | 725 | 1456 | 599 | 445 | 1018 | 2686 | 452 | 930 | 1219 | 403 | 676 |
| 922 | QGN14019.1 | FIM1_670 | CP015054. | 4791 | 1039 | 1332 | 17525 | 975 | 1189 | 13167 | 793 | 1867 | 5199 | 749 | 1125 | 6430 | 2881 | 1427 | 5303 | 878 | 975 |

| | | | | | | | | | | | | | | | | | | | | | |
|---|---|---|---|---|---|---|---|---|---|---|---|---|---|---|---|---|---|---|---|---|---|
| 923 | QGN14020.1 | FIM1_671 | CP015054. | 529 | 613 | 406 | 690 | 507 | 367 | 474 | 498 | 521 | 2087 | 490 | 488 | 1999 | 325 | 517 | 1892 | 328 | 354 |
| 924 | QGN14021.1 | FIM1_672 | CP015054. | 1319 | 267 | 440 | 1368 | 330 | 369 | 944 | 344 | 575 | 2104 | 338 | 622 | 3071 | 287 | 556 | 2300 | 241 | 465 |
| 925 | QGN14022.1 | FIM1_673 | CP015054. | 2220 | 258 | 185 | 3416 | 97 | 124 | 1047 | 223 | 236 | 476 | 80 | 195 | 620 | 71 | 125 | 461 | 77 | 119 |
| 926 | QGN14023.1 | FIM1_674 | CP015054. | 10681 | 2544 | 3996 | 12636 | 2997 | 3077 | 9888 | 2798 | 5068 | 11584 | 2191 | 5604 | 17348 | 2629 | 4622 | 13020 | 2022 | 3516 |
| 927 | QGN14024.1 | FIM1_675 | CP015054. | 2503 | 1154 | 1042 | 2094 | 880 | 915 | 2367 | 1031 | 1500 | 2466 | 740 | 1551 | 3739 | 699 | 1599 | 2980 | 707 | 1033 |
| 928 | QGN14025.1 | FIM1_676 | CP015054. | 5588 | 1314 | 3328 | 6438 | 1629 | 2529 | 5137 | 2119 | 4185 | 2544 | 1370 | 4204 | 9051 | 1048 | 4015 | 4760 | 1181 | 3017 |
| 929 | QGN14026.1 | FIM1_677 | CP015054. | 10200 | 520 | 999 | 9583 | 514 | 848 | 11062 | 513 | 1221 | 4397 | 519 | 1276 | 4163 | 673 | 1313 | 4138 | 560 | 837 |
| 930 | QGN14027.1 | FIM1_678 | CP015054. | 4566 | 1791 | 1729 | 6476 | 1884 | 1686 | 4700 | 1583 | 2219 | 4253 | 1310 | 2278 | 8054 | 1885 | 1782 | 5316 | 1411 | 1511 |
| 931 | QGN14028.1 | FIM1_679 | CP015054. | 3551 | 1629 | 1736 | 2881 | 763 | 893 | 1975 | 1667 | 2136 | 6774 | 2337 | 2456 | 10975 | 3307 | 3324 | 6957 | 2422 | 2014 |
| 932 | QGN14029.1 | FIM1_680 | CP015054. | 20012 | 2580 | 3830 | 13409 | 2628 | 3502 | 15697 | 4974 | 5581 | 10875 | 2878 | 4302 | 10562 | 2371 | 4597 | 10946 | 2449 | 2750 |
| 933 | QGN14030.1 | FIM1_681 | CP015054. | 6422 | 1399 | 1039 | 6336 | 1891 | 852 | 6305 | 1686 | 1397 | 603 | 793 | 1291 | 4889 | 1048 | 1094 | 1955 | 981 | 828 |
| 934 | QGN14031.1 | FIM1_682 | CP015054. | 6983 | 593 | 770 | 4132 | 604 | 574 | 4203 | 854 | 1163 | 842 | 420 | 989 | 3570 | 328 | 838 | 1556 | 388 | 632 |
| 935 | QGN14032.1 | FIM1_683 | CP015054. | 9209 | 2557 | 1733 | 7800 | 2406 | 1582 | 8336 | 2152 | 2277 | 4942 | 2191 | 2113 | 8868 | 1984 | 2203 | 6366 | 1889 | 1538 |
| 936 | QGN14033.1 | FIM1_684 | CP015054. | 4820 | 756 | 966 | 5451 | 805 | 797 | 5187 | 887 | 1203 | 4034 | 708 | 1268 | 5575 | 644 | 1200 | 4361 | 604 | 932 |
| 937 | QGN14034.1 | FIM1_685 | CP015054. | 964 | 350 | 404 | 1739 | 296 | 345 | 1187 | 257 | 512 | 1934 | 239 | 588 | 2141 | 238 | 539 | 1945 | 226 | 367 |
| 938 | QGN14035.1 | FIM1_686 | CP015054. | 6087 | 2585 | 2307 | 7350 | 2645 | 2105 | 6128 | 2243 | 2963 | 10487 | 2348 | 3579 | 10316 | 2610 | 3389 | 10008 | 2247 | 2354 |
| 939 | QGN14036.1 | FIM1_687 | CP015054. | 2062 | 575 | 710 | 1692 | 627 | 558 | 1888 | 691 | 827 | 3797 | 703 | 981 | 4132 | 539 | 932 | 4060 | 555 | 637 |
| 940 | QGN14037.1 | FIM1_688 | CP015054. | 1153 | 461 | 826 | 1459 | 589 | 721 | 1252 | 516 | 1042 | 3919 | 484 | 1287 | 3476 | 448 | 1155 | 3830 | 437 | 861 |
| 941 | QGN14038.1 | FIM1_689 | CP015054. | 272 | 183 | 82 | 186 | 125 | 73 | 120 | 87 | 120 | 473 | 129 | 119 | 375 | 85 | 108 | 429 | 94 | 63 |
| 942 | QGN14039.1 | FIM1_690 | CP015054. | 2207 | 991 | 885 | 2234 | 1186 | 822 | 2267 | 1013 | 1279 | 6634 | 886 | 1001 | 5599 | 837 | 1019 | 6614 | 831 | 715 |
| 943 | QGN14040.1 | FIM1_691 | CP015054. | 1455 | 1196 | 744 | 1291 | 1002 | 699 | 1327 | 840 | 1037 | 5697 | 1269 | 1511 | 5148 | 1170 | 1340 | 5733 | 1258 | 971 |
| 944 | QGN14041.1 | FIM1_692 | CP015054. | 859 | 926 | 696 | 1552 | 837 | 699 | 1123 | 671 | 945 | 5101 | 720 | 1103 | 3890 | 737 | 1019 | 5238 | 722 | 749 |
| 945 | QGN14042.1 | FIM1_693 | CP015054. | 1667 | 601 | 674 | 2306 | 658 | 612 | 1888 | 598 | 909 | 6211 | 676 | 1165 | 5193 | 572 | 1118 | 6052 | 605 | 782 |
| 946 | QGN14043.1 | FIM1_694 | CP015054. | 512 | 229 | 378 | 795 | 324 | 314 | 544 | 305 | 504 | 2796 | 245 | 615 | 2721 | 239 | 479 | 2773 | 232 | 400 |
| 947 | QGN14044.1 | FIM1_695 | CP015054. | 2038 | 502 | 593 | 1940 | 539 | 541 | 2024 | 534 | 779 | 3131 | 553 | 872 | 3883 | 522 | 795 | 3279 | 509 | 632 |
| 948 | QGN14045.1 | FIM1_696 | CP015054. | 1360 | 1021 | 900 | 2024 | 922 | 757 | 1734 | 804 | 1145 | 7284 | 1021 | 1482 | 5647 | 1000 | 1287 | 7169 | 1041 | 886 |
| 949 | QGN14046.1 | FIM1_697 | CP015054. | 1862 | 626 | 546 | 2438 | 508 | 462 | 2076 | 487 | 741 | 3583 | 380 | 736 | 2819 | 417 | 759 | 3412 | 325 | 486 |
| 950 | QGN14047.1 | FIM1_698 | CP015054. | 222 | 719 | 470 | 225 | 795 | 458 | 172 | 612 | 731 | 2262 | 748 | 452 | 2695 | 698 | 444 | 2254 | 699 | 357 |
| 951 | QGN14048.1 | FIM1_699 | CP015054. | 5400 | 1404 | 1854 | 4620 | 1995 | 1463 | 4356 | 2035 | 2413 | 3426 | 1870 | 2550 | 11520 | 1679 | 2189 | 6413 | 1739 | 1748 |
| 952 | QGN14049.1 | FIM1_700 | CP015054. | 6297 | 4718 | 4394 | 6612 | 5661 | 3902 | 6315 | 4457 | 5543 | 21596 | 5762 | 6038 | 18909 | 5319 | 6069 | 21396 | 5324 | 4108 |
| 953 | QGN14050.1 | FIM1_701 | CP015054. | 2100 | 1529 | 759 | 1209 | 1422 | 766 | 1735 | 1102 | 1025 | 5414 | 1410 | 1409 | 4694 | 1216 | 1262 | 5409 | 1500 | 915 |
| 954 | QGN14051.1 | FIM1_702 | CP015054. | 1185 | 921 | 683 | 1671 | 678 | 596 | 1362 | 655 | 867 | 4803 | 788 | 1199 | 3836 | 809 | 997 | 4926 | 765 | 699 |
| 955 | QGN14052.1 | FIM1_703 | CP015054. | 2459 | 1693 | 1867 | 2569 | 2354 | 1511 | 3010 | 2082 | 2430 | 7206 | 1979 | 2487 | 10953 | 1782 | 2173 | 9486 | 1731 | 1649 |
| 956 | QGN14053.1 | FIM1_704 | CP015054. | 13492 | 2249 | 3220 | 12625 | 4949 | 2905 | 12649 | 3738 | 4354 | 6298 | 2928 | 3503 | 17831 | 3137 | 3255 | 10459 | 3314 | 2451 |
| 957 | QGN14054.1 | FIM1_705 | CP015054. | 2237 | 790 | 723 | 1343 | 687 | 519 | 1497 | 949 | 998 | 1510 | 861 | 862 | 5166 | 1006 | 979 | 2940 | 851 | 634 |
| 958 | QGN14055.1 | FIM1_706 | CP015054. | 1352 | 986 | 1356 | 1948 | 1464 | 1194 | 1284 | 1083 | 1853 | 8549 | 1686 | 2548 | 10000 | 1464 | 2153 | 9609 | 1431 | 1666 |
| 959 | QGN14056.1 | FIM1_707 | CP015054. | 755 | 649 | 581 | 950 | 723 | 540 | 712 | 613 | 784 | 3895 | 702 | 898 | 3831 | 590 | 743 | 3957 | 629 | 537 |
| 960 | QGN14057.1 | FIM1_708 | CP015054. | 1052 | 382 | 380 | 1076 | 333 | 331 | 1095 | 328 | 516 | 3553 | 318 | 655 | 2586 | 287 | 555 | 3535 | 306 | 424 |
| 961 | QGN14058.1 | FIM1_709 | CP015054. | 1514 | 618 | 823 | 1944 | 831 | 712 | 1557 | 696 | 1041 | 6275 | 722 | 1336 | 5139 | 635 | 1196 | 6079 | 643 | 831 |
| 962 | QGN14059.1 | FIM1_710 | CP015054. | 1037 | 350 | 451 | 1437 | 387 | 343 | 970 | 440 | 567 | 1935 | 344 | 761 | 1820 | 372 | 627 | 1982 | 320 | 443 |
| 963 | QGN14060.1 | FIM1_711 | CP015054. | 263 | 456 | 334 | 393 | 268 | 309 | 289 | 229 | 440 | 1614 | 224 | 440 | 1225 | 219 | 434 | 1541 | 243 | 295 |
| 964 | QGN14061.1 | FIM1_712 | CP015054. | 5 | 8 | 8 | 29 | 3 | 10 | 22 | 12 | 18 | 125 | 15 | 12 | 62 | 7 | 9 | 82 | 13 | 3 |
| 965 | QGN14062.1 | FIM1_713 | CP015054. | 2254 | 1763 | 1029 | 1535 | 1149 | 934 | 939 | 1131 | 1564 | 5305 | 1278 | 1362 | 4844 | 556 | 1257 | 5539 | 835 | 966 |
| 966 | QGN14063.1 | FIM1_714 | CP015054. | 4095 | 494 | 515 | 2851 | 531 | 371 | 3657 | 523 | 648 | 2613 | 457 | 737 | 2312 | 436 | 609 | 2562 | 494 | 387 |
| 967 | QGN14064.1 | FIM1_715 | CP015054. | 1028 | 163 | 1321 | 1344 | 521 | 1038 | 1278 | 474 | 1691 | 5593 | 485 | 2373 | 4567 | 420 | 1931 | 5043 | 386 | 1554 |
| 968 | QGN14065.1 | FIM1_716 | CP015054. | 0 | 0 | 0 | 0 | 0 | 0 | 0 | 0 | 0 | 0 | 0 | 0 | 0 | 0 | 0 | 0 | 0 | 0 |
| 969 | QGN14066.1 | FIM1_717 | CP015054. | 1 | 0 | 0 | 0 | 0 | 0 | 0 | 0 | 0 | 2 | 0 | 0 | 4 | 0 | 1 | 2 | 0 | 1 |
| 970 | QGN14067.1 | FIM1_718 | CP015054. | 4926 | 1328 | 3225 | 8053 | 2430 | 2523 | 5991 | 2276 | 3807 | 10290 | 1951 | 5171 | 11348 | 2159 | 4733 | 10263 | 1748 | 3359 |
| 971 | QGN14068.1 | FIM1_719 | CP015054. | 5518 | 5734 | 3882 | 5675 | 4660 | 3901 | 4885 | 3527 | 4844 | 22120 | 6213 | 6945 | 17460 | 5307 | 6599 | 22573 | 5654 | 4810 |
| 972 | QGN14069.1 | FIM1_720 | CP015054. | 1493 | 225 | 321 | 1570 | 377 | 248 | 1644 | 273 | 389 | 2242 | 283 | 404 | 1818 | 286 | 461 | 2056 | 286 | 298 |
| 973 | QGN14070.1 | FIM1_721 | CP015054. | 3249 | 1904 | 1202 | 3214 | 1491 | 1149 | 3326 | 1234 | 1681 | 7707 | 1382 | 1849 | 6437 | 1230 | 1696 | 7434 | 1231 | 1219 |
| 974 | QGN14071.1 | FIM1_722 | CP015054. | 4919 | 2758 | 2625 | 5729 | 3239 | 2512 | 5627 | 2491 | 3469 | 16805 | 2940 | 4106 | 15750 | 2966 | 3465 | 16059 | 2844 | 2617 |
| 975 | QGN14072.1 | FIM1_723 | CP015054. | 2080 | 422 | 456 | 2586 | 637 | 404 | 2554 | 394 | 502 | 3405 | 377 | 596 | 3450 | 352 | 529 | 3408 | 338 | 399 |
| 976 | QGN14073.1 | FIM1_724 | CP015054. | 365 | 295 | 203 | 362 | 270 | 177 | 419 | 172 | 255 | 1910 | 275 | 304 | 1309 | 253 | 278 | 1881 | 275 | 204 |
| 977 | QGN14074.1 | FIM1_725 | CP015054. | 6386 | 4453 | 2094 | 4078 | 4694 | 2225 | 5699 | 3593 | 2621 | 17615 | 5483 | 4196 | 15058 | 4559 | 3764 | 17290 | 5625 | 2751 |
| 978 | QGN14075.1 | FIM1_726 | CP015054. | 1546 | 492 | 485 | 1577 | 521 | 418 | 1582 | 459 | 604 | 4018 | 545 | 758 | 3041 | 470 | 672 | 3932 | 478 | 484 |
| 979 | QGN14076.1 | FIM1_727 | CP015054. | 1099 | 467 | 837 | 1766 | 578 | 692 | 1225 | 483 | 1112 | 7357 | 588 | 1463 | 5547 | 602 | 1125 | 6725 | 531 | 904 |
| 980 | QGN14077.1 | FIM1_728 | CP015054. | 2812 | 1654 | 1362 | 2663 | 2140 | 1292 | 1824 | 1904 | 1767 | 5434 | 1859 | 1856 | 7631 | 1302 | 1671 | 6475 | 1547 | 1217 |
| 981 | QGN14078.1 | FIM1_729 | CP015054. | 712 | 1948 | 1363 | 1014 | 1791 | 1267 | 817 | 1403 | 1816 | 11350 | 1740 | 2141 | 9144 | 1687 | 1910 | 11317 | 1618 | 1357 |
| 982 | QGN14079.1 | FIM1_730 | CP015054. | 2026 | 1171 | 1009 | 2681 | 1213 | 972 | 2531 | 1070 | 1347 | 7962 | 1120 | 1776 | 6606 | 1021 | 1534 | 7860 | 1030 | 1176 |
| 983 | QGN14080.1 | FIM1_731 | CP015054. | 839 | 213 | 302 | 984 | 279 | 200 | 780 | 259 | 350 | 1581 | 200 | 378 | 1701 | 210 | 307 | 1694 | 188 | 235 |
| 984 | QGN14081.1 | FIM1_732 | CP015054. | 6865 | 8861 | 5034 | 6544 | 6749 | 4922 | 5875 | 5767 | 6307 | 24922 | 7981 | 8224 | 20074 | 6851 | 8026 | 23266 | 7207 | 5845 |
| 985 | QGN14082.1 | FIM1_733 | CP015054. | 1660 | 1069 | 906 | 5757 | 1257 | 1035 | 3567 | 943 | 1257 | 8751 | 1370 | 1539 | 7355 | 1778 | 1516 | 8210 | 1308 | 1080 |
| 986 | QGN14083.1 | FIM1_734 | CP015054. | 2366 | 1365 | 1867 | 2752 | 1292 | 1620 | 2614 | 1336 | 2374 | 6359 | 1031 | 2357 | 6030 | 994 | 2268 | 5905 | 943 | 1550 |
| 987 | QGN14084.1 | FIM1_735 | CP015054. | 1376 | 448 | 519 | 1612 | 427 | 448 | 1284 | 487 | 703 | 2105 | 212 | 526 | 2143 | 228 | 494 | 1937 | 159 | 319 |
| 988 | QGN14085.1 | FIM1_736 | CP015054. | 3769 | 1063 | 1504 | 5044 | 1543 | 1224 | 3481 | 1278 | 1883 | 11118 | 1506 | 3351 | 11613 | 1391 | 2927 | 11160 | 1271 | 2093 |
| 989 | QGN14086.1 | FIM1_737 | CP015054. | 3149 | 1334 | 1416 | 4344 | 1336 | 1353 | 3791 | 1249 | 1895 | 8715 | 1166 | 2140 | 7715 | 1171 | 1916 | 8643 | 1204 | 1350 |
| 990 | QGN14087.1 | FIM1_738 | CP015054. | 1125 | 1628 | 1469 | 1909 | 1166 | 1486 | 1643 | 1241 | 1804 | 6040 | 1089 | 2097 | 4215 | 1178 | 1897 | 5155 | 1477 | 1368 |
| 991 | QGN14088.1 | FIM1_739 | CP015054. | 570 | 350 | 280 | 1177 | 331 | 349 | 874 | 224 | 391 | 2069 | 250 | 501 | 1348 | 227 | 457 | 1895 | 260 | 267 |
| 992 | QGN14089.1 | FIM1_740 | CP015054. | 1068 | 555 | 488 | 1342 | 455 | 403 | 1183 | 425 | 637 | 3778 | 501 | 756 | 2677 | 471 | 763 | 3586 | 485 | 465 |
| 993 | QGN14090.1 | FIM1_741 | CP015054. | 2233 | 1209 | 1042 | 2186 | 1378 | 863 | 1966 | 1160 | 1265 | 7128 | 1478 | 1833 | 5751 | 1372 | 1679 | 6899 | 1370 | 1104 |
| 994 | QGN14091.1 | FIM1_742 | CP015054. | 1194 | 1042 | 1253 | 1960 | 1399 | 1097 | 1419 | 1072 | 1647 | 5607 | 1479 | 1648 | 8552 | 1416 | 1536 | 6874 | 1344 | 1155 |
| 995 | QGN14092.1 | FIM1_743 | CP015054. | 3007 | 1664 | 1633 | 3756 | 1724 | 1551 | 3024 | 1431 | 2166 | 8129 | 1409 | 1962 | 6431 | 1261 | 1888 | 7328 | 1256 | 1280 |
| 996 | QGN14093.1 | FIM1_744 | CP015054. | 477 | 323 | 354 | 722 | 197 | 327 | 494 | 257 | 553 | 2103 | 202 | 560 | 1532 | 156 | 560 | 2005 | 171 | 351 |
| 997 | QGN14094.1 | FIM1_745 | CP015054. | 2075 | 272 | 332 | 2295 | 267 | 322 | 1938 | 236 | 441 | 1846 | 180 | 409 | 1562 | 173 | 407 | 1786 | 144 | 294 |
| 998 | QGN14095.1 | FIM1_746 | CP015054. | 14661 | 700 | 731 | 11538 | 833 | 579 | 11648 | 602 | 928 | 2931 | 647 | 1084 | 4228 | 762 | 1058 | 3428 | 634 | 719 |
| 999 | QGN14096.1 | FIM1_747 | CP015054. | 950 | 852 | 1020 | 1791 | 619 | 795 | 1159 | 613 | 1196 | 4495 | 526 | 1515 | 3919 | 564 | 1279 | 4177 | 479 | 1008 |
| 1000 | QGN14097.1 | FIM1_748 | CP015054. | 894 | 137 | 504 | 1085 | 476 | 407 | 750 | 486 | 608 | 2739 | 478 | 771 | 3557 | 468 | 720 | 2853 | 413 | 538 |
| 1001 | QGN14098.1 | FIM1_749 | CP015054. | 2697 | 556 | 869 | 3458 | 441 | 800 | 2932 | 478 | 1148 | 4377 | 350 | 1104 | 3485 | 400 | 1072 | 4082 | 315 | 738 |
| 1002 | QGN14099.1 | FIM1_750 | CP015054. | 2822 | 934 | 946 | 2771 | 962 | 798 | 2648 | 939 | 1161 | 4819 | 820 | 1427 | 4567 | 673 | 1200 | 4495 | 625 | 990 |
| 1003 | QGN14100.1 | FIM1_751 | CP015054. | 3383 | 1862 | 2424 | 4905 | 2172 | 2129 | 3882 | 2008 | 3161 | 10512 | 1969 | 3769 | 12723 | 2005 | 3451 | 11052 | 1783 | 2622 |
| 1004 | QGN14101.1 | FIM1_752 | CP015054. | 16431 | 3130 | 2668 | 15327 | 4032 | 2348 | 13800 | 3991 | 3342 | 13145 | 3554 | 2927 | 14028 | 3534 | 2973 | 12389 | 3438 | 1955 |
| 1005 | QGN14102.1 | FIM1_753 | CP015054. | 1851 | 676 | 770 | 3221 | 892 | 730 | 2119 | 785 | 1149 | 6314 | 1524 | 1070 | 6219 | 1829 | 1255 | 6343 | 1657 | 819 |
| 1006 | QGN14103.1 | FIM1_754 | CP015054. | 4879 | 1082 | 1354 | 6399 | 1100 | 1011 | 4931 | 1180 | 1761 | 2244 | 880 | 1887 | 6082 | 813 | 1668 | 3389 | 857 | 1262 |

| | | | | | | | | | | | | | | | | | | | | | |
|---|---|---|---|---|---|---|---|---|---|---|---|---|---|---|---|---|---|---|---|---|---|
| 1007 | QGN14104.1 | FIM1_755 | CP015054. | 2272 | 1180 | 1756 | 3020 | 1350 | 1699 | 2534 | 1322 | 2335 | 8520 | 1559 | 2969 | 9691 | 1451 | 2764 | 8896 | 1444 | 1961 |
| 1008 | QGN14105.1 | FIM1_756 | CP015054. | 1412 | 451 | 753 | 1759 | 512 | 554 | 1442 | 451 | 888 | 3110 | 399 | 862 | 2421 | 429 | 798 | 3149 | 385 | 495 |
| 1009 | QGN14106.1 | FIM1_757 | CP015054. | 974 | 1370 | 1326 | 1285 | 1280 | 1199 | 1275 | 1020 | 1773 | 8111 | 1400 | 1925 | 6700 | 1329 | 1724 | 7783 | 1294 | 1299 |
| 1010 | QGN14107.1 | FIM1_758 | CP015054. | 697 | 228 | 291 | 908 | 240 | 263 | 669 | 241 | 390 | 1502 | 139 | 305 | 1505 | 162 | 251 | 1512 | 120 | 167 |
| 1011 | QGN14108.1 | FIM1_759 | CP015054. | 746 | 522 | 549 | 958 | 545 | 498 | 889 | 484 | 727 | 3481 | 407 | 690 | 2969 | 358 | 569 | 3118 | 383 | 427 |
| 1012 | QGN14109.1 | FIM1_760 | CP015054. | 2722 | 962 | 1461 | 3501 | 1492 | 1127 | 2872 | 1299 | 1788 | 5131 | 1575 | 2268 | 10410 | 1508 | 2122 | 7225 | 1362 | 1580 |
| 1013 | QGN14110.1 | FIM1_761 | CP015054. | 2272 | 1739 | 1971 | 6193 | 1630 | 1837 | 3540 | 1476 | 2543 | 7230 | 1585 | 3118 | 7876 | 1699 | 3201 | 6983 | 1388 | 2137 |
| 1014 | QGN14111.1 | FIM1_762 | CP015054. | 362 | 1037 | 765 | 408 | 816 | 893 | 324 | 652 | 1019 | 3634 | 906 | 1436 | 2865 | 800 | 1428 | 3345 | 768 | 903 |
| 1015 | QGN14112.1 | FIM1_763 | CP015054. | 1892 | 467 | 645 | 1826 | 462 | 465 | 1640 | 627 | 870 | 2116 | 373 | 612 | 2374 | 291 | 555 | 2233 | 324 | 376 |
| 1016 | QGN14113.1 | FIM1_764 | CP015054. | 3385 | 1660 | 1592 | 3775 | 2031 | 1476 | 3179 | 1746 | 2013 | 6454 | 1895 | 2054 | 7260 | 1589 | 2134 | 6647 | 1494 | 1406 |
| 1017 | QGN14114.1 | FIM1_765 | CP015054. | 762 | 132 | 248 | 405 | 266 | 186 | 459 | 252 | 261 | 515 | 272 | 263 | 1070 | 239 | 323 | 643 | 230 | 243 |
| 1018 | QGN14115.1 | FIM1_767 | CP015054. | 6298 | 346 | 625 | 10106 | 419 | 377 | 6984 | 418 | 775 | 1084 | 282 | 587 | 2260 | 490 | 453 | 1453 | 325 | 330 |
| 1019 | QGN14116.1 | FIM1_768 | CP015054. | 656 | 263 | 338 | 1059 | 156 | 241 | 812 | 230 | 441 | 913 | 161 | 341 | 904 | 166 | 345 | 929 | 152 | 243 |
| 1020 | QGN14117.1 | FIM1_769 | CP015054. | 2494 | 1513 | 1525 | 3171 | 1829 | 1334 | 2705 | 1561 | 1949 | 9711 | 1759 | 2569 | 10128 | 1921 | 2240 | 9684 | 1613 | 1560 |
| 1021 | QGN14118.1 | FIM1_770 | CP015054. | 5705 | 1504 | 1953 | 4377 | 2054 | 1825 | 5179 | 1778 | 2714 | 945 | 749 | 1026 | 2935 | 969 | 937 | 1682 | 904 | 712 |
| 1022 | QGN14119.1 | FIM1_771 | CP015054. | 18 | 5 | 9 | 14 | 12 | 10 | 12 | 6 | 7 | 15 | 3 | 2 | 13 | 2 | 2 | 15 | 0 | 0 |
| 1023 | QGN14120.1 | FIM1_772 | CP015054. | 55 | 39 | 51 | 171 | 70 | 52 | 33 | 52 | 59 | 23 | 7 | 33 | 42 | 3 | 18 | 29 | 5 | 16 |
| 1024 | QGN14121.1 | FIM1_773 | CP015054. | 1983 | 2191 | 1029 | 1516 | 1751 | 1082 | 1789 | 1475 | 1403 | 6987 | 1807 | 1633 | 6164 | 1522 | 1610 | 6648 | 1813 | 1069 |
| 1025 | QGN14122.1 | FIM1_774 | CP015054. | 819 | 399 | 477 | 1441 | 544 | 419 | 1162 | 443 | 651 | 2058 | 463 | 766 | 2511 | 508 | 668 | 2204 | 438 | 533 |
| 1026 | QGN14123.1 | FIM1_776 | CP015054. | 1192 | 1376 | 1433 | 2180 | 1632 | 1228 | 1362 | 1229 | 1991 | 4110 | 1325 | 1617 | 6530 | 1309 | 1535 | 5390 | 1369 | 1147 |
| 1027 | QGN14124.1 | FIM1_777 | CP015054. | 2482 | 1227 | 1784 | 2873 | 1740 | 1472 | 2296 | 1665 | 2147 | 6761 | 1381 | 2325 | 8717 | 1262 | 2016 | 7089 | 1173 | 1523 |
| 1028 | QGN14125.1 | FIM1_778 | CP015054. | 1157 | 566 | 509 | 1586 | 357 | 411 | 1114 | 388 | 648 | 1581 | 294 | 541 | 2248 | 319 | 555 | 1854 | 273 | 376 |
| 1029 | QGN14126.1 | FIM1_779 | CP015054. | 6134 | 14693 | 10623 | 7188 | 13023 | 9912 | 5958 | 12165 | 13656 | 19959 | 10580 | 8545 | 28000 | 9866 | 8793 | 20596 | 9640 | 6607 |
| 1030 | QGN14127.1 | FIM1_780 | CP015054. | 2086 | 484 | 267 | 1146 | 347 | 227 | 1729 | 373 | 402 | 226 | 149 | 168 | 556 | 177 | 212 | 299 | 190 | 142 |
| 1031 | QGN14128.1 | FIM1_781 | CP015054. | 3102 | 384 | 321 | 2043 | 242 | 187 | 2216 | 299 | 417 | 1134 | 123 | 241 | 1091 | 135 | 230 | 971 | 112 | 142 |
| 1032 | QGN14129.1 | FIM1_782 | CP015054. | 5587 | 574 | 1393 | 4081 | 887 | 951 | 4148 | 916 | 1713 | 2998 | 472 | 1695 | 6588 | 469 | 1636 | 4583 | 289 | 1117 |
| 1033 | QGN14130.1 | FIM1_783 | CP015054. | 2413 | 902 | 703 | 2121 | 905 | 580 | 2116 | 927 | 957 | 668 | 397 | 722 | 1428 | 384 | 772 | 933 | 381 | 506 |
| 1034 | QGN14131.1 | FIM1_784 | CP015054. | 2468 | 230 | 356 | 2287 | 278 | 300 | 1667 | 315 | 475 | 491 | 150 | 349 | 1182 | 145 | 285 | 770 | 114 | 234 |
| 1035 | QGN14132.1 | FIM1_785 | CP015054. | 2527 | 1734 | 2253 | 3374 | 2231 | 1976 | 2783 | 2070 | 3175 | 6175 | 2153 | 2920 | 11348 | 1918 | 2648 | 8322 | 2084 | 2087 |
| 1036 | QGN14133.1 | FIM1_786 | CP015054. | 3030 | 2407 | 1676 | 3504 | 1763 | 1577 | 2873 | 1657 | 2242 | 6295 | 1473 | 2027 | 5676 | 1295 | 1983 | 5728 | 1330 | 1295 |
| 1037 | QGN14134.1 | FIM1_787 | CP015054. | 17320 | 2382 | 2184 | 15818 | 3839 | 2257 | 12508 | 2824 | 3002 | 10315 | 2918 | 2204 | 10686 | 2083 | 2442 | 10608 | 2219 | 1507 |
| 1038 | QGN14135.1 | FIM1_788 | CP015054. | 312 | 112 | 89 | 1120 | 160 | 96 | 369 | 128 | 129 | 553 | 98 | 138 | 617 | 79 | 144 | 593 | 93 | 102 |
| 1039 | QGN14136.1 | FIM1_789 | CP015054. | 437 | 782 | 923 | 1141 | 567 | 873 | 603 | 440 | 1142 | 3011 | 517 | 1165 | 2578 | 548 | 1147 | 2771 | 464 | 753 |
| 1040 | QGN14137.1 | FIM1_790 | CP015054. | 374 | 138 | 110 | 787 | 115 | 89 | 729 | 124 | 185 | 648 | 238 | 180 | 906 | 395 | 176 | 677 | 289 | 136 |
| 1041 | QGN14138.1 | FIM1_791 | CP015054. | 4306 | 2144 | 1889 | 12516 | 2798 | 1706 | 7657 | 2157 | 2458 | 6258 | 1400 | 1141 | 7591 | 1806 | 1071 | 5884 | 1434 | 733 |
| 1042 | QGN14139.1 | FIM1_792 | CP015054. | 967 | 620 | 681 | 1473 | 615 | 562 | 888 | 511 | 875 | 4860 | 660 | 1060 | 4068 | 602 | 977 | 4604 | 630 | 699 |
| 1043 | QGN14140.1 | FIM1_793 | CP015054. | 10038 | 2017 | 1943 | 10764 | 1906 | 1355 | 9139 | 2088 | 2481 | 10489 | 2087 | 2831 | 13076 | 3039 | 2749 | 10835 | 2248 | 1919 |
| 1044 | QGN14141.1 | FIM1_794 | CP015054. | 3562 | 666 | 477 | 3354 | 563 | 459 | 3467 | 459 | 666 | 2310 | 475 | 683 | 2376 | 527 | 701 | 2272 | 570 | 471 |
| 1045 | QGN14142.1 | FIM1_795 | CP015054. | 2572 | 1929 | 1459 | 2731 | 1988 | 1527 | 2562 | 1604 | 2003 | 11041 | 2187 | 2373 | 8426 | 1995 | 2250 | 10622 | 2049 | 1648 |
| 1046 | QGN14143.1 | FIM1_796 | CP015054. | 2955 | 677 | 806 | 3018 | 561 | 778 | 2816 | 539 | 1183 | 3707 | 571 | 1283 | 3942 | 501 | 1284 | 3399 | 502 | 832 |
| 1047 | QGN14144.1 | FIM1_797 | CP015054. | 174 | 315 | 237 | 182 | 187 | 211 | 173 | 187 | 338 | 1481 | 168 | 342 | 931 | 203 | 354 | 1225 | 224 | 200 |
| 1048 | QGN14145.1 | FIM1_798 | CP015054. | 368 | 481 | 459 | 842 | 370 | 391 | 676 | 316 | 625 | 2667 | 362 | 683 | 2033 | 360 | 658 | 2380 | 377 | 391 |
| 1049 | QGN14146.1 | FIM1_799 | CP015054. | 6098 | 797 | 1171 | 8993 | 701 | 858 | 6140 | 625 | 1420 | 4697 | 406 | 1403 | 4963 | 992 | 1483 | 4727 | 582 | 860 |
| 1050 | QGN14147.1 | FIM1_800 | CP015054. | 2909 | 2389 | 2029 | 5163 | 1674 | 1958 | 3615 | 1514 | 2500 | 6568 | 1703 | 2615 | 5105 | 1759 | 2486 | 5962 | 1898 | 1735 |
| 1051 | QGN14148.1 | FIM1_801 | CP015054. | 3712 | 1739 | 571 | 9851 | 1906 | 469 | 6919 | 1266 | 822 | 1091 | 1010 | 301 | 3412 | 1365 | 307 | 2519 | 1167 | 219 |
| 1052 | QGN14149.1 | FIM1_802 | CP015054. | 1575 | 694 | 387 | 1121 | 646 | 380 | 1515 | 680 | 552 | 1649 | 505 | 533 | 2498 | 512 | 447 | 1782 | 569 | 346 |
| 1053 | QGN14150.1 | FIM1_803 | CP015054. | 1497 | 690 | 713 | 2032 | 731 | 683 | 1426 | 769 | 983 | 4283 | 623 | 992 | 4052 | 599 | 990 | 4019 | 582 | 686 |
| 1054 | QGN14151.1 | FIM1_804 | CP015054. | 7771 | 3128 | 1958 | 5262 | 3994 | 1854 | 6940 | 4084 | 2723 | 7458 | 3789 | 3133 | 13369 | 3857 | 2990 | 8159 | 4071 | 2186 |
| 1055 | QGN14152.1 | FIM1_805 | CP015054. | 3069 | 659 | 775 | 4742 | 696 | 632 | 3196 | 651 | 1014 | 3359 | 311 | 753 | 3172 | 433 | 690 | 3064 | 320 | 512 |
| 1056 | QGN14153.1 | FIM1_806 | CP015054. | 6656 | 268 | 839 | 8320 | 350 | 602 | 6284 | 371 | 1011 | 4262 | 221 | 928 | 4400 | 403 | 797 | 3849 | 220 | 598 |
| 1057 | QGN14154.1 | FIM1_807 | CP015054. | 6241 | 144 | 157 | 4128 | 155 | 143 | 6296 | 178 | 237 | 691 | 123 | 227 | 722 | 119 | 180 | 684 | 123 | 134 |
| 1058 | QGN14155.1 | FIM1_808 | CP015054. | 2573 | 950 | 960 | 2534 | 718 | 931 | 2705 | 844 | 1324 | 1601 | 624 | 1106 | 3167 | 538 | 1170 | 2321 | 528 | 835 |
| 1059 | QGN14156.1 | FIM1_809 | CP015054. | 13449 | 13140 | 23325 | 25586 | 12684 | 20926 | 16182 | 12289 | 28252 | 32038 | 7841 | 33442 | 34925 | 8899 | 31120 | 33729 | 7858 | 23246 |
| 1060 | QGN14157.1 | FIM1_810 | CP015054. | 1972 | 930 | 771 | 1983 | 933 | 724 | 1760 | 778 | 1043 | 6137 | 1022 | 1269 | 5414 | 918 | 1133 | 5965 | 870 | 758 |
| 1061 | QGN14158.1 | FIM1_811 | CP015054. | 1905 | 1217 | 1812 | 4667 | 1583 | 1435 | 3212 | 1491 | 2216 | 8387 | 1146 | 2112 | 9121 | 1372 | 1764 | 8258 | 1066 | 1387 |
| 1062 | QGN14159.1 | FIM1_812 | CP015054. | 1225 | 1583 | 1529 | 1935 | 1222 | 1379 | 1463 | 1074 | 1959 | 7580 | 1402 | 2444 | 5288 | 1311 | 2288 | 7073 | 1391 | 1523 |
| 1063 | QGN14160.1 | FIM1_813 | CP015054. | 1914 | 323 | 371 | 1807 | 362 | 309 | 2113 | 306 | 463 | 2973 | 321 | 506 | 2723 | 370 | 477 | 2869 | 343 | 350 |
| 1064 | QGN14161.1 | FIM1_814 | CP015054. | 174 | 33 | 18 | 403 | 55 | 37 | 255 | 29 | 22 | 170 | 17 | 36 | 156 | 18 | 37 | 174 | 26 | 23 |
| 1065 | QGN14162.1 | FIM1_815 | CP015054. | 117 | 71 | 57 | 297 | 86 | 63 | 245 | 69 | 66 | 352 | 48 | 61 | 264 | 35 | 41 | 351 | 44 | 33 |
| 1066 | QGN14163.1 | FIM1_816 | CP015054. | 594 | 321 | 261 | 1596 | 307 | 236 | 839 | 207 | 361 | 1110 | 197 | 246 | 903 | 198 | 259 | 1208 | 224 | 153 |
| 1067 | QGN14164.1 | FIM1_817 | CP015054. | 5185 | 1066 | 609 | 3078 | 1017 | 611 | 4529 | 786 | 791 | 3437 | 877 | 1013 | 3505 | 1005 | 845 | 3360 | 987 | 697 |
| 1068 | QGN14165.1 | FIM1_818 | CP015054. | 173 | 250 | 265 | 184 | 236 | 233 | 149 | 232 | 338 | 1806 | 214 | 330 | 1192 | 170 | 314 | 1824 | 193 | 242 |
| 1069 | QGN14166.1 | FIM1_819 | CP015054. | 3213 | 829 | 790 | 2389 | 806 | 740 | 2415 | 825 | 1021 | 3440 | 710 | 952 | 3777 | 494 | 974 | 3539 | 540 | 622 |
| 1070 | QGN14167.1 | FIM1_820 | CP015054. | 21224 | 3255 | 3090 | 28944 | 3215 | 2840 | 24560 | 2531 | 3994 | 14632 | 2845 | 4443 | 13547 | 3568 | 4471 | 14340 | 2980 | 3013 |
| 1071 | QGN14168.1 | FIM1_821 | CP015054. | 2770 | 6429 | 4749 | 6530 | 6190 | 4282 | 5187 | 5056 | 5698 | 22765 | 7103 | 7587 | 20388 | 6652 | 7354 | 22227 | 6027 | 5531 |
| 1072 | QGN14169.1 | FIM1_822 | CP015054. | 1449 | 781 | 869 | 1345 | 633 | 739 | 1163 | 558 | 1080 | 883 | 599 | 985 | 2073 | 548 | 952 | 1544 | 576 | 697 |
| 1073 | QGN14170.1 | FIM1_823 | CP015054. | 958 | 649 | 949 | 1212 | 704 | 856 | 1010 | 768 | 1255 | 3774 | 532 | 1197 | 3141 | 502 | 1140 | 3504 | 449 | 803 |
| 1074 | QGN14171.1 | FIM1_824 | CP015054. | 6523 | 2566 | 1542 | 2629 | 3115 | 1670 | 5690 | 2522 | 2085 | 12855 | 4039 | 2784 | 12198 | 4329 | 2533 | 12266 | 4712 | 1911 |
| 1075 | QGN14172.1 | FIM1_825 | CP015054. | 3432 | 1963 | 1951 | 4720 | 2657 | 1870 | 4006 | 1796 | 2377 | 10862 | 2588 | 3934 | 10443 | 2288 | 3193 | 10525 | 2369 | 2659 |
| 1076 | QGN14173.1 | FIM1_826 | CP015054. | 105 | 40 | 49 | 393 | 41 | 40 | 190 | 46 | 60 | 359 | 62 | 83 | 332 | 47 | 81 | 336 | 62 | 62 |
| 1077 | QGN14174.1 | FIM1_827 | CP015054. | 5 | 10 | 5 | 27 | 5 | 3 | 23 | 14 | 6 | 80 | 8 | 10 | 53 | 5 | 12 | 62 | 6 | 10 |
| 1078 | QGN14175.1 | FIM1_828 | CP015054. | 2826 | 852 | 1376 | 2507 | 741 | 1148 | 2118 | 1047 | 1723 | 4004 | 637 | 1500 | 3429 | 595 | 1317 | 4237 | 612 | 868 |
| 1079 | QGN14176.1 | FIM1_829 | CP015054. | 278 | 342 | 415 | 475 | 334 | 328 | 359 | 264 | 488 | 1976 | 243 | 721 | 1413 | 205 | 601 | 1987 | 226 | 461 |
| 1080 | QGN14177.1 | FIM1_830 | CP015054. | 1379 | 1484 | 2457 | 8534 | 2052 | 2438 | 2733 | 1427 | 3125 | 9481 | 1356 | 3241 | 9436 | 2085 | 2326 | 9225 | 1491 | 2088 |
| 1081 | QGN14178.1 | FIM1_831 | CP015054. | 1512 | 3183 | 2746 | 2714 | 3171 | 2534 | 2112 | 2626 | 3341 | 12622 | 3682 | 4053 | 10698 | 3488 | 4169 | 11489 | 3142 | 2961 |
| 1082 | QGN14179.1 | FIM1_832 | CP015054. | 1107 | 409 | 385 | 1038 | 407 | 419 | 1107 | 278 | 516 | 3132 | 470 | 614 | 2229 | 532 | 597 | 3009 | 570 | 417 |
| 1083 | QGN14180.1 | FIM1_833 | CP015054. | 5666 | 2761 | 1463 | 3127 | 3019 | 1731 | 4918 | 2092 | 2044 | 9348 | 3421 | 2995 | 9744 | 2892 | 2924 | 9509 | 3543 | 2073 |
| 1084 | QGN14181.1 | FIM1_834 | CP015054. | 591 | 757 | 880 | 1142 | 836 | 748 | 785 | 667 | 1179 | 3778 | 736 | 1090 | 4514 | 766 | 965 | 4089 | 662 | 734 |
| 1085 | QGN14182.1 | FIM1_835 | CP015054. | 1920 | 471 | 538 | 1456 | 526 | 509 | 1483 | 475 | 746 | 4808 | 464 | 633 | 4277 | 425 | 586 | 4612 | 455 | 427 |
| 1086 | QGN14183.1 | FIM1_836 | CP015054. | 3067 | 340 | 322 | 2603 | 357 | 216 | 2689 | 302 | 460 | 5238 | 958 | 1107 | 4860 | 1205 | 814 | 5798 | 908 | 619 |
| 1087 | QGN14184.1 | FIM1_837 | CP015054. | 533 | 194 | 213 | 725 | 278 | 195 | 463 | 147 | 296 | 27 | 3 | 21 | 20 | 6 | 12 | 25 | 2 | 22 |
| 1088 | QGN14185.1 | FIM1_838 | CP015054. | 405 | 201 | 146 | 403 | 161 | 124 | 432 | 134 | 229 | 1 | 0 | 0 | 2 | 0 | 0 | 5 | 0 | 1 |
| 1089 | QGN14186.1 | FIM1_839 | CP015054. | 123 | 47 | 43 | 199 | 44 | 38 | 149 | 38 | 59 | 7 | 1 | 3 | 5 | 0 | 2 | 8 | 0 | 0 |
| 1090 | QGN14187.1 | FIM1_840 | CP015054. | 676 | 346 | 380 | 989 | 252 | 291 | 967 | 250 | 478 | 1134 | 229 | 542 | 1230 | 210 | 518 | 1217 | 182 | 425 |

| | | | | | | | | | | | | | | | | | | | | | |
|---|---|---|---|---|---|---|---|---|---|---|---|---|---|---|---|---|---|---|---|---|---|
| 1091 | QGN14188.1 | FIM1_841 | CP015054. | 608 | 107 | 101 | 388 | 36 | 50 | 837 | 99 | 107 | 138 | 32 | 44 | 185 | 37 | 28 | 151 | 37 | 34 |
| 1092 | QGN14189.1 | FIM1_842 | CP015054. | 261 | 296 | 291 | 702 | 274 | 194 | 571 | 354 | 425 | 553 | 99 | 111 | 428 | 153 | 124 | 441 | 100 | 87 |
| 1093 | QGN14190.1 | FIM1_843 | CP015054. | 1116 | 1766 | 3081 | 1357 | 367 | 1519 | 1844 | 2913 | 4753 | 969 | 89 | 264 | 688 | 104 | 158 | 820 | 118 | 187 |
| 1094 | QGN14191.1 | FIM1_844 | CP015054. | 537 | 85 | 98 | 361 | 72 | 55 | 416 | 102 | 103 | 134 | 44 | 79 | 247 | 44 | 63 | 182 | 45 | 48 |
| 1095 | QGN14192.1 | FIM1_846 | CP015054. | 559 | 76 | 68 | 341 | 44 | 37 | 479 | 53 | 84 | 68 | 18 | 56 | 104 | 27 | 44 | 71 | 27 | 28 |
| 1096 | QGN14193.1 | FIM1_847 | CP015054. | 87 | 7 | 8 | 95 | 3 | 3 | 78 | 6 | 13 | 24 | 5 | 13 | 34 | 5 | 5 | 24 | 6 | 5 |
| 1097 | QGN14194.1 | FIM1_850 | CP015055. | 1158 | 463 | 421 | 5534 | 584 | 385 | 2406 | 407 | 498 | 591 | 257 | 466 | 1449 | 318 | 427 | 921 | 218 | 321 |
| 1098 | QGN14195.1 | FIM1_851 | CP015055. | 761 | 54 | 199 | 1330 | 168 | 162 | 691 | 160 | 220 | 701 | 100 | 280 | 1170 | 113 | 230 | 792 | 112 | 230 |
| 1099 | QGN14196.1 | FIM1_852 | CP015055. | 62 | 53 | 40 | 96 | 43 | 40 | 62 | 34 | 54 | 112 | 49 | 59 | 200 | 49 | 73 | 180 | 40 | 51 |
| 1100 | QGN14197.1 | FIM1_853 | CP015055. | 361 | 39 | 35 | 416 | 62 | 37 | 233 | 26 | 42 | 465 | 62 | 68 | 348 | 76 | 66 | 372 | 74 | 35 |
| 1101 | QGN14198.1 | FIM1_854 | CP015055. | 3810 | 306 | 458 | 9236 | 755 | 500 | 4450 | 272 | 542 | 7726 | 1060 | 1111 | 7055 | 1364 | 1011 | 7601 | 959 | 832 |
| 1102 | QGN14199.1 | FIM1_855 | CP015055. | 392 | 116 | 158 | 692 | 155 | 120 | 269 | 160 | 203 | 1136 | 221 | 255 | 959 | 228 | 251 | 1190 | 174 | 154 |
| 1103 | QGN14200.1 | FIM1_856 | CP015055. | 329 | 191 | 183 | 725 | 253 | 154 | 644 | 198 | 255 | 1037 | 166 | 223 | 1011 | 197 | 225 | 1008 | 141 | 146 |
| 1104 | QGN14201.1 | FIM1_857 | CP015055. | 607 | 153 | 108 | 614 | 102 | 90 | 564 | 134 | 156 | 565 | 94 | 143 | 630 | 106 | 100 | 563 | 110 | 84 |
| 1105 | QGN14202.1 | FIM1_858 | CP015055. | 446 | 213 | 496 | 561 | 374 | 405 | 358 | 388 | 791 | 4267 | 437 | 951 | 4948 | 446 | 602 | 4089 | 543 | 691 |
| 1106 | QGN14203.1 | FIM1_859 | CP015055. | 1233 | 256 | 302 | 1184 | 281 | 302 | 1076 | 243 | 345 | 979 | 266 | 564 | 1183 | 288 | 499 | 979 | 249 | 327 |
| 1107 | QGN14204.1 | FIM1_860 | CP015055. | 1123 | 391 | 505 | 1350 | 318 | 393 | 1174 | 335 | 727 | 1874 | 188 | 481 | 1768 | 187 | 434 | 1850 | 166 | 302 |
| 1108 | QGN14205.1 | FIM1_861 | CP015055. | 863 | 452 | 503 | 802 | 371 | 434 | 823 | 411 | 655 | 3385 | 312 | 730 | 3605 | 257 | 632 | 3203 | 252 | 529 |
| 1109 | QGN14206.1 | FIM1_862 | CP015055. | 292 | 72 | 71 | 668 | 74 | 71 | 469 | 58 | 89 | 651 | 84 | 116 | 567 | 118 | 106 | 529 | 78 | 93 |
| 1110 | QGN14207.1 | FIM1_863 | CP015055. | 27 | 5 | 27 | 176 | 11 | 23 | 69 | 9 | 36 | 18 | 8 | 57 | 42 | 10 | 47 | 30 | 7 | 25 |
| 1111 | QGN14208.1 | FIM1_864 | CP015055. | 769 | 442 | 439 | 1398 | 363 | 379 | 866 | 354 | 609 | 352 | 246 | 697 | 1421 | 239 | 656 | 807 | 209 | 441 |
| 1112 | QGN14209.1 | FIM1_865 | CP015055. | 1647 | 678 | 773 | 2547 | 316 | 613 | 1778 | 383 | 1051 | 2817 | 399 | 1057 | 2542 | 574 | 999 | 2740 | 522 | 639 |
| 1113 | QGN14210.1 | FIM1_866 | CP015055. | 759 | 237 | 433 | 1319 | 256 | 272 | 990 | 286 | 482 | 1859 | 232 | 568 | 1808 | 280 | 497 | 1762 | 206 | 361 |
| 1114 | QGN14211.1 | FIM1_867 | CP015055. | 4843 | 656 | 702 | 4581 | 527 | 554 | 4754 | 565 | 947 | 1429 | 525 | 1075 | 4065 | 563 | 850 | 2241 | 575 | 699 |
| 1115 | QGN14212.1 | FIM1_868 | CP015055. | 1061 | 511 | 520 | 2079 | 482 | 431 | 1372 | 370 | 716 | 4371 | 453 | 892 | 3360 | 551 | 821 | 4316 | 435 | 540 |
| 1116 | QGN14213.1 | FIM1_869 | CP015055. | 416 | 516 | 478 | 677 | 445 | 397 | 465 | 437 | 569 | 2917 | 369 | 685 | 2417 | 444 | 606 | 2685 | 380 | 436 |
| 1117 | QGN14214.1 | FIM1_870 | CP015055. | 3264 | 645 | 556 | 1990 | 769 | 573 | 3159 | 670 | 782 | 4226 | 916 | 817 | 3351 | 1175 | 839 | 3927 | 926 | 638 |
| 1118 | QGN14215.1 | FIM1_871 | CP015055. | 801 | 806 | 726 | 1003 | 714 | 694 | 783 | 679 | 943 | 3958 | 796 | 1078 | 3299 | 716 | 1014 | 3892 | 759 | 683 |
| 1119 | QGN14216.1 | FIM1_872 | CP015055. | 3608 | 2146 | 1503 | 3665 | 1954 | 1380 | 3689 | 1658 | 1990 | 10925 | 1988 | 2543 | 9123 | 1749 | 2569 | 10342 | 1760 | 1833 |
| 1120 | QGN14217.1 | FIM1_873 | CP015055. | 3982 | 436 | 495 | 3109 | 434 | 396 | 3890 | 424 | 657 | 3608 | 461 | 875 | 2667 | 529 | 738 | 3293 | 474 | 509 |
| 1121 | QGN14218.1 | FIM1_874 | CP015055. | 24935 | 954 | 2184 | 28248 | 1429 | 1663 | 29129 | 1516 | 2775 | 5363 | 1013 | 2925 | 11064 | 1436 | 2547 | 7020 | 1040 | 1961 |
| 1122 | QGN14219.1 | FIM1_875 | CP015055. | 1100 | 275 | 203 | 1002 | 183 | 157 | 1155 | 213 | 249 | 330 | 162 | 268 | 841 | 160 | 331 | 416 | 184 | 182 |
| 1123 | QGN14220.1 | FIM1_876 | CP015055. | 9091 | 1380 | 1632 | 5987 | 1860 | 1295 | 7152 | 1898 | 2194 | 1910 | 1440 | 2314 | 7951 | 1417 | 2120 | 4228 | 1461 | 1457 |
| 1124 | QGN14221.1 | FIM1_877 | CP015055. | 848 | 1148 | 766 | 974 | 702 | 689 | 820 | 726 | 1054 | 4777 | 1018 | 1310 | 3669 | 832 | 1369 | 4717 | 856 | 874 |
| 1125 | QGN14222.1 | FIM1_878 | CP015055. | 3624 | 2929 | 2262 | 5303 | 2243 | 2001 | 3892 | 1932 | 2895 | 11972 | 2504 | 3504 | 10205 | 2573 | 3679 | 11190 | 2326 | 2504 |
| 1126 | QGN14223.1 | FIM1_879 | CP015055. | 69 | 31 | 19 | 184 | 21 | 17 | 75 | 22 | 26 | 92 | 10 | 15 | 65 | 8 | 18 | 74 | 13 | 10 |
| 1127 | QGN14224.1 | FIM1_880 | CP015055. | 2917 | 430 | 762 | 3284 | 437 | 614 | 2618 | 423 | 1030 | 4531 | 430 | 1351 | 5101 | 414 | 1131 | 4743 | 378 | 881 |
| 1128 | QGN14225.1 | FIM1_881 | CP015055. | 2388 | 1009 | 843 | 3488 | 842 | 772 | 2921 | 792 | 1093 | 5881 | 893 | 1413 | 5041 | 961 | 1437 | 5523 | 845 | 1048 |
| 1129 | QGN14226.1 | FIM1_882 | CP015055. | 1028 | 345 | 305 | 1327 | 262 | 245 | 1176 | 293 | 376 | 2228 | 259 | 502 | 1744 | 219 | 440 | 2336 | 230 | 296 |
| 1130 | QGN14227.1 | FIM1_883 | CP015055. | 1328 | 1003 | 1092 | 3843 | 857 | 863 | 2281 | 784 | 1280 | 5366 | 712 | 1645 | 5269 | 788 | 1537 | 5392 | 663 | 1060 |
| 1131 | QGN14228.1 | FIM1_884 | CP015055. | 264 | 188 | 281 | 747 | 120 | 194 | 392 | 118 | 302 | 1895 | 128 | 465 | 1521 | 130 | 348 | 1941 | 108 | 278 |
| 1132 | QGN14229.1 | FIM1_885 | CP015055. | 3794 | 1191 | 1264 | 4005 | 1260 | 1103 | 3892 | 1000 | 1516 | 6791 | 1182 | 1724 | 6501 | 1451 | 1519 | 5916 | 1257 | 1046 |
| 1133 | QGN14230.1 | FIM1_886 | CP015055. | 3379 | 2217 | 2964 | 6541 | 2610 | 2280 | 4246 | 2422 | 3637 | 4814 | 2274 | 4628 | 14819 | 2728 | 3901 | 8657 | 2381 | 3057 |
| 1134 | QGN14231.1 | FIM1_887 | CP015055. | 589 | 326 | 555 | 461 | 384 | 491 | 480 | 408 | 640 | 1251 | 342 | 730 | 1764 | 318 | 834 | 1606 | 295 | 531 |
| 1135 | QGN14232.1 | FIM1_888 | CP015055. | 1376 | 937 | 1543 | 2903 | 1012 | 1194 | 2019 | 1036 | 1958 | 3736 | 700 | 1492 | 4141 | 670 | 1401 | 3908 | 617 | 1018 |
| 1136 | QGN14233.1 | FIM1_889 | CP015055. | 957 | 716 | 508 | 1524 | 604 | 493 | 1275 | 621 | 706 | 3384 | 668 | 983 | 3391 | 577 | 864 | 3281 | 573 | 581 |
| 1137 | QGN14234.1 | FIM1_890 | CP015055. | 237 | 199 | 180 | 345 | 240 | 137 | 284 | 257 | 225 | 1539 | 276 | 299 | 1181 | 217 | 281 | 1429 | 263 | 190 |
| 1138 | QGN14235.1 | FIM1_891 | CP015055. | 8777 | 1913 | 2777 | 15523 | 2867 | 2371 | 11214 | 2058 | 3551 | 13803 | 1590 | 3707 | 15819 | 1933 | 3457 | 13570 | 1359 | 2518 |
| 1139 | QGN14236.1 | FIM1_892 | CP015055. | 330 | 140 | 175 | 651 | 72 | 76 | 380 | 119 | 212 | 261 | 91 | 178 | 541 | 67 | 248 | 377 | 73 | 139 |
| 1140 | QGN14237.1 | FIM1_893 | CP015055. | 87 | 15 | 13 | 233 | 19 | 7 | 111 | 13 | 10 | 47 | 5 | 12 | 79 | 11 | 12 | 55 | 7 | 7 |
| 1141 | QGN14238.1 | FIM1_894 | CP015055. | 12480 | 893 | 938 | 26130 | 1699 | 718 | 17301 | 1255 | 1209 | 4616 | 1132 | 2136 | 14442 | 1605 | 1965 | 7826 | 829 | 1497 |
| 1142 | QGN14239.1 | FIM1_895 | CP015055. | 911 | 593 | 359 | 1248 | 416 | 316 | 1213 | 496 | 500 | 1579 | 453 | 642 | 2328 | 378 | 596 | 1684 | 427 | 446 |
| 1143 | QGN14240.1 | FIM1_896 | CP015055. | 193 | 94 | 101 | 271 | 77 | 75 | 279 | 125 | 113 | 333 | 89 | 117 | 427 | 76 | 131 | 366 | 69 | 74 |
| 1144 | QGN14241.1 | FIM1_897 | CP015055. | 805 | 554 | 434 | 595 | 300 | 311 | 568 | 372 | 597 | 758 | 333 | 535 | 1896 | 330 | 550 | 1366 | 287 | 373 |
| 1145 | QGN14242.1 | FIM1_898 | CP015055. | 4370 | 3194 | 3621 | 6598 | 3075 | 2541 | 4595 | 3500 | 4536 | 7332 | 3033 | 5564 | 18586 | 3801 | 5617 | 11704 | 2709 | 3866 |
| 1146 | QGN14243.1 | FIM1_899 | CP015055. | 1291 | 342 | 311 | 782 | 449 | 336 | 1175 | 362 | 420 | 2175 | 528 | 648 | 2125 | 573 | 545 | 2141 | 542 | 476 |
| 1147 | QGN14244.1 | FIM1_900 | CP015055. | 1350 | 1060 | 1233 | 3611 | 1295 | 1083 | 2362 | 1137 | 1609 | 6631 | 1090 | 2073 | 7647 | 1162 | 1848 | 6856 | 1012 | 1299 |
| 1148 | QGN14245.1 | FIM1_901 | CP015055. | 6722 | 1838 | 2739 | 8571 | 2712 | 2211 | 9156 | 2288 | 3479 | 6199 | 1878 | 1692 | 7173 | 1893 | 1710 | 6513 | 1845 | 1209 |
| 1149 | QGN14246.1 | FIM1_902 | CP015055. | 543 | 170 | 317 | 648 | 247 | 285 | 609 | 262 | 412 | 1952 | 245 | 497 | 2196 | 219 | 473 | 1898 | 249 | 333 |
| 1150 | QGN14247.1 | FIM1_903 | CP015055. | 1538 | 1858 | 566 | 578 | 1427 | 720 | 1145 | 1154 | 853 | 4586 | 1689 | 974 | 3445 | 1147 | 1187 | 4045 | 1801 | 694 |
| 1151 | QGN14248.1 | FIM1_904 | CP015055. | 2466 | 609 | 579 | 1277 | 1047 | 600 | 1898 | 745 | 761 | 3005 | 910 | 941 | 4215 | 1061 | 746 | 3458 | 1020 | 587 |
| 1152 | QGN14249.1 | FIM1_905 | CP015055. | 3369 | 244 | 470 | 4053 | 422 | 402 | 3874 | 381 | 642 | 2696 | 214 | 655 | 2942 | 279 | 557 | 2767 | 244 | 409 |
| 1153 | QGN14250.1 | FIM1_906 | CP015055. | 8782 | 14118 | 7302 | 8331 | 9977 | 6969 | 6820 | 8712 | 8590 | 23780 | 10307 | 10274 | 21262 | 9134 | 10897 | 21979 | 9416 | 7729 |
| 1154 | QGN14251.1 | FIM1_907 | CP015055. | 15915 | 16339 | 9215 | 15655 | 12279 | 8392 | 13151 | 10288 | 10758 | 29957 | 13949 | 14104 | 34202 | 12512 | 13858 | 31569 | 12452 | 10697 |
| 1155 | QGN14252.1 | FIM1_908 | CP015055. | 48 | 9 | 6 | 55 | 4 | 3 | 24 | 12 | 5 | 11 | 3 | 4 | 14 | 2 | 2 | 14 | 2 | 5 |
| 1156 | QGN14253.1 | FIM1_909 | CP015055. | 397 | 1175 | 746 | 358 | 885 | 698 | 356 | 907 | 1005 | 3689 | 722 | 965 | 3548 | 752 | 897 | 3437 | 758 | 627 |
| 1157 | QGN14254.1 | FIM1_910 | CP015055. | 792 | 147 | 176 | 1264 | 105 | 133 | 774 | 117 | 196 | 621 | 70 | 204 | 876 | 76 | 181 | 713 | 45 | 129 |
| 1158 | QGN14255.1 | FIM1_912 | CP015055. | 1136 | 36 | 33 | 650 | 26 | 32 | 682 | 26 | 46 | 187 | 23 | 66 | 239 | 21 | 76 | 203 | 22 | 55 |
| 1159 | QGN14256.1 | FIM1_913 | CP015055. | 6171 | 903 | 644 | 4220 | 1260 | 661 | 4935 | 1196 | 934 | 2526 | 1173 | 1532 | 7720 | 899 | 1508 | 4587 | 999 | 1140 |
| 1160 | QGN14257.1 | FIM1_914 | CP015055. | 2047 | 263 | 322 | 1915 | 294 | 290 | 2122 | 313 | 435 | 1198 | 251 | 438 | 1746 | 246 | 409 | 1321 | 217 | 321 |
| 1161 | QGN14258.1 | FIM1_915 | CP015055. | 4389 | 1950 | 2296 | 5694 | 2680 | 1971 | 5107 | 2284 | 3037 | 12762 | 3133 | 3602 | 17299 | 2595 | 3337 | 13865 | 2738 | 2490 |
| 1162 | QGN14259.1 | FIM1_916 | CP015055. | 1737 | 939 | 712 | 3709 | 857 | 628 | 2452 | 866 | 930 | 2483 | 528 | 955 | 3613 | 641 | 893 | 2648 | 545 | 656 |
| 1163 | QGN14260.1 | FIM1_917 | CP015055. | 31058 | 5293 | 4274 | 26483 | 7011 | 3318 | 27761 | 7010 | 5379 | 4065 | 6666 | 5337 | 23111 | 8063 | 4788 | 10663 | 9133 | 3651 |
| 1164 | QGN14261.1 | FIM1_918 | CP015055. | 189 | 242 | 172 | 212 | 165 | 142 | 225 | 307 | 211 | 154 | 108 | 227 | 344 | 87 | 172 | 179 | 167 | 107 |
| 1165 | QGN14262.1 | FIM1_919 | CP015055. | 1154 | 58 | 151 | 999 | 154 | 122 | 1089 | 223 | 206 | 211 | 104 | 201 | 426 | 98 | 187 | 307 | 132 | 149 |
| 1166 | QGN14263.1 | FIM1_920 | CP015055. | 4679 | 1799 | 1180 | 3630 | 1554 | 948 | 3762 | 1677 | 1629 | 1516 | 969 | 1151 | 3435 | 922 | 1419 | 2032 | 971 | 929 |
| 1167 | QGN14264.1 | FIM1_921 | CP015055. | 5784 | 2683 | 2209 | 5056 | 1611 | 1883 | 4950 | 1630 | 2846 | 5803 | 1353 | 2538 | 5588 | 1323 | 3092 | 5634 | 1269 | 1864 |
| 1168 | QGN14265.1 | FIM1_922 | CP015055. | 1260 | 440 | 415 | 965 | 221 | 358 | 1056 | 388 | 588 | 895 | 187 | 430 | 921 | 182 | 515 | 815 | 194 | 325 |
| 1169 | QGN14266.1 | FIM1_923 | CP015055. | 841 | 268 | 225 | 1117 | 243 | 172 | 1189 | 264 | 285 | 412 | 212 | 383 | 1092 | 234 | 365 | 635 | 251 | 224 |
| 1170 | QGN14267.1 | FIM1_924 | CP015055. | 2399 | 1009 | 767 | 3694 | 805 | 712 | 2249 | 863 | 1085 | 1666 | 373 | 618 | 2300 | 334 | 647 | 1719 | 304 | 401 |
| 1171 | QGN14268.1 | FIM1_925 | CP015055. | 632 | 215 | 231 | 838 | 114 | 161 | 564 | 157 | 279 | 609 | 70 | 157 | 634 | 73 | 158 | 565 | 78 | 88 |
| 1172 | QGN14269.1 | FIM1_926 | CP015055. | 43673 | 1131 | 1450 | 36399 | 1164 | 1083 | 35549 | 1167 | 2035 | 718 | 626 | 1682 | 4560 | 897 | 1576 | 1541 | 729 | 1076 |
| 1173 | QGN14270.1 | FIM1_927 | CP015055. | 6030 | 6109 | 3554 | 6111 | 5156 | 3322 | 5523 | 4524 | 4485 | 16719 | 6352 | 5958 | 17706 | 5575 | 6153 | 16679 | 5640 | 4323 |
| 1174 | QGN14271.1 | FIM1_928 | CP015055. | 429 | 1384 | 1013 | 823 | 936 | 862 | 621 | 1038 | 1272 | 1431 | 622 | 902 | 2026 | 469 | 979 | 1686 | 524 | 638 |

| | | | | | | | | | | | | | | | | | | | | | |
|---|---|---|---|---|---|---|---|---|---|---|---|---|---|---|---|---|---|---|---|---|---|
| 1175 | QGN14272.1 | FIM1_929 | CP015055. | 3493 | 714 | 899 | 3553 | 757 | 682 | 3470 | 934 | 1158 | 1090 | 668 | 1199 | 3979 | 691 | 1184 | 2032 | 691 | 801 |
| 1176 | QGN14273.1 | FIM1_930 | CP015055. | 3561 | 877 | 1014 | 3832 | 1037 | 796 | 3564 | 1123 | 1277 | 2134 | 791 | 1408 | 5300 | 827 | 1300 | 3179 | 840 | 994 |
| 1177 | QGN14274.1 | FIM1_931 | CP015055. | 1238 | 1815 | 1146 | 1052 | 1384 | 981 | 1247 | 1312 | 1428 | 5173 | 1328 | 1600 | 4630 | 1047 | 1451 | 4626 | 1090 | 1019 |
| 1178 | QGN14275.1 | FIM1_932 | CP015055. | 641 | 250 | 357 | 666 | 280 | 249 | 670 | 286 | 393 | 1838 | 254 | 533 | 2077 | 250 | 431 | 1832 | 274 | 316 |
| 1179 | QGN14276.1 | FIM1_933 | CP015055. | 4046 | 1148 | 1366 | 4778 | 1201 | 1202 | 4448 | 1267 | 1779 | 6624 | 996 | 2009 | 8260 | 974 | 1746 | 6946 | 901 | 1360 |
| 1180 | QGN14277.1 | FIM1_934 | CP015055. | 1995 | 1186 | 885 | 1915 | 876 | 883 | 1625 | 950 | 1195 | 3602 | 875 | 1202 | 4230 | 657 | 1230 | 3489 | 749 | 816 |
| 1181 | QGN14278.1 | FIM1_935 | CP015055. | 740 | 737 | 885 | 4557 | 911 | 621 | 2206 | 882 | 1096 | 1682 | 470 | 812 | 4587 | 826 | 671 | 2725 | 525 | 538 |
| 1182 | QGN14279.1 | FIM1_936 | CP015055. | 9005 | 4542 | 5826 | 20349 | 5669 | 4524 | 13402 | 5542 | 7127 | 7082 | 4087 | 8636 | 24768 | 4587 | 7607 | 13519 | 4153 | 5808 |
| 1183 | QGN14280.1 | FIM1_937 | CP015055. | 3287 | 2819 | 2237 | 2842 | 2300 | 1861 | 2621 | 2013 | 2816 | 7157 | 1744 | 2432 | 6514 | 1804 | 2628 | 6305 | 1504 | 1677 |
| 1184 | QGN14281.1 | FIM1_938 | CP015055. | 6528 | 3276 | 1758 | 6155 | 1893 | 1262 | 5965 | 2308 | 2281 | 3546 | 2377 | 2900 | 9462 | 2529 | 2911 | 5718 | 2313 | 1955 |
| 1185 | QGN14282.1 | FIM1_939 | CP015055. | 1345 | 1269 | 1152 | 1586 | 1304 | 954 | 1395 | 1324 | 1365 | 6369 | 1336 | 1710 | 6921 | 1383 | 1731 | 6048 | 1228 | 1203 |
| 1186 | QGN14283.1 | FIM1_940 | CP015055. | 1610 | 1796 | 1693 | 4259 | 1937 | 1385 | 3112 | 1614 | 2119 | 9274 | 2251 | 2843 | 13642 | 2126 | 2454 | 10303 | 2004 | 1961 |
| 1187 | QGN14284.1 | FIM1_941 | CP015055. | 655 | 306 | 341 | 705 | 232 | 253 | 575 | 335 | 461 | 1490 | 224 | 403 | 1904 | 384 | 324 | 1528 | 417 | 332 |
| 1188 | QGN14285.1 | FIM1_942 | CP015055. | 10579 | 704 | 315 | 6976 | 831 | 308 | 9505 | 568 | 380 | 2445 | 768 | 447 | 2620 | 1133 | 420 | 2260 | 1043 | 303 |
| 1189 | QGN14286.1 | FIM1_943 | CP015055. | 7166 | 7682 | 5074 | 6758 | 6471 | 5378 | 5762 | 5273 | 5887 | 20794 | 7410 | 7711 | 17226 | 6524 | 7644 | 20422 | 7188 | 5657 |
| 1190 | QGN14287.1 | FIM1_944 | CP015055. | 1125 | 876 | 708 | 555 | 1158 | 662 | 862 | 979 | 909 | 2439 | 1020 | 1156 | 3778 | 1176 | 1012 | 2853 | 1079 | 787 |
| 1191 | QGN14288.1 | FIM1_945 | CP015055. | 5322 | 2293 | 1368 | 6221 | 1683 | 1339 | 6109 | 1543 | 1910 | 6844 | 1135 | 1541 | 6314 | 1072 | 1558 | 6529 | 1015 | 1078 |
| 1192 | QGN14289.1 | FIM1_946 | CP015055. | 63 | 13 | 39 | 85 | 26 | 23 | 80 | 34 | 46 | 99 | 24 | 41 | 127 | 28 | 39 | 99 | 28 | 27 |
| 1193 | QGN14290.1 | FIM1_947 | CP015055. | 4441 | 2122 | 3040 | 6959 | 3235 | 2482 | 6182 | 2737 | 3589 | 22058 | 3805 | 4949 | 21311 | 3314 | 4087 | 20711 | 3406 | 3148 |
| 1194 | QGN14291.1 | FIM1_948 | CP015055. | 10261 | 2617 | 2813 | 10867 | 3390 | 2468 | 9955 | 3027 | 3666 | 12316 | 3353 | 4187 | 19380 | 3036 | 3761 | 14766 | 2974 | 2933 |
| 1195 | QGN14292.1 | FIM1_949 | CP015055. | 2363 | 572 | 962 | 3446 | 730 | 669 | 2307 | 702 | 1359 | 1739 | 499 | 1426 | 4847 | 674 | 1195 | 2380 | 373 | 894 |
| 1196 | QGN14293.1 | FIM1_950 | CP015055. | 1130 | 406 | 427 | 921 | 273 | 265 | 854 | 346 | 581 | 335 | 183 | 348 | 840 | 290 | 400 | 537 | 223 | 253 |
| 1197 | QGN14294.1 | FIM1_951 | CP015055. | 71262 | 12865 | 9958 | 67189 | 10912 | 8576 | 65784 | 10482 | 12353 | 28437 | 9704 | 12533 | 34919 | 13353 | 12061 | 29374 | 10854 | 9396 |
| 1198 | QGN14295.1 | FIM1_952 | CP015055. | 331 | 32 | 12 | 205 | 18 | 13 | 180 | 14 | 19 | 24 | 30 | 26 | 58 | 23 | 25 | 42 | 10 | 20 |
| 1199 | QGN14296.1 | FIM1_953 | CP015055. | 8995 | 4047 | 2219 | 10795 | 2912 | 2097 | 8208 | 2881 | 3055 | 5129 | 1794 | 3261 | 8286 | 2235 | 3160 | 5391 | 1840 | 2081 |
| 1200 | QGN14297.1 | FIM1_954 | CP015055. | 1380 | 1256 | 1309 | 1327 | 1973 | 1062 | 1236 | 2019 | 1747 | 3655 | 1373 | 1784 | 7896 | 1026 | 1547 | 5103 | 1058 | 1106 |
| 1201 | QGN14298.1 | FIM1_955 | CP015055. | 13068 | 1016 | 1031 | 9094 | 654 | 646 | 11498 | 926 | 1241 | 772 | 461 | 748 | 3827 | 448 | 654 | 1770 | 472 | 537 |
| 1202 | QGN14299.1 | FIM1_956 | CP015055. | 58 | 3 | 4 | 117 | 2 | 4 | 110 | 7 | 9 | 37 | 1 | 6 | 62 | 1 | 1 | 70 | 1 | 3 |
| 1203 | QGN14300.1 | FIM1_957 | CP015055. | 19414 | 8559 | 6672 | 8860 | 9622 | 6677 | 5582 | 8099 | 8634 | 28364 | 10620 | 12822 | 40546 | 7120 | 12702 | 33027 | 6355 | 9167 |
| 1204 | QGN14301.1 | FIM1_958 | CP015055. | 8567 | 2908 | 3078 | 11815 | 1793 | 2089 | 10101 | 2384 | 3897 | 10619 | 2063 | 3558 | 9738 | 2733 | 3588 | 10118 | 2234 | 2352 |
| 1205 | QGN14302.1 | FIM1_959 | CP015055. | 5644 | 1044 | 379 | 3269 | 925 | 402 | 6037 | 765 | 537 | 2220 | 1025 | 706 | 2903 | 1514 | 774 | 2253 | 1433 | 573 |
| 1206 | QGN14303.1 | FIM1_960 | CP015055. | 1514 | 2402 | 1358 | 1422 | 1991 | 1297 | 1377 | 1734 | 1702 | 7747 | 1838 | 1867 | 6932 | 1713 | 1834 | 7257 | 1801 | 1279 |
| 1207 | QGN14304.1 | FIM1_961 | CP015055. | 3202 | 3288 | 3344 | 5240 | 3465 | 2965 | 3762 | 3058 | 4356 | 16341 | 3076 | 5196 | 14651 | 2922 | 4742 | 16575 | 2693 | 3446 |
| 1208 | QGN14305.1 | FIM1_962 | CP015055. | 2748 | 1608 | 1685 | 2356 | 1847 | 1391 | 2221 | 1651 | 1985 | 11098 | 2086 | 2810 | 10725 | 1777 | 2530 | 10662 | 1757 | 1939 |
| 1209 | QGN14306.1 | FIM1_963 | CP015055. | 549 | 468 | 454 | 1205 | 413 | 349 | 592 | 427 | 607 | 2526 | 411 | 666 | 2554 | 390 | 635 | 2536 | 407 | 426 |
| 1210 | QGN14307.1 | FIM1_964 | CP015055. | 145 | 116 | 196 | 197 | 136 | 187 | 168 | 144 | 215 | 881 | 143 | 221 | 749 | 152 | 242 | 880 | 136 | 136 |
| 1211 | QGN14308.1 | FIM1_965 | CP015055. | 2118 | 501 | 555 | 2231 | 513 | 480 | 2060 | 452 | 731 | 4773 | 565 | 1089 | 4376 | 437 | 892 | 4623 | 449 | 708 |
| 1212 | QGN14309.1 | FIM1_966 | CP015055. | 1839 | 2537 | 3340 | 4574 | 2697 | 2967 | 3335 | 2463 | 4090 | 12117 | 1990 | 4849 | 11424 | 2259 | 4119 | 12184 | 2097 | 3127 |
| 1213 | QGN14310.1 | FIM1_967 | CP015055. | 1689 | 560 | 565 | 1302 | 578 | 487 | 1666 | 527 | 755 | 3310 | 615 | 1110 | 3210 | 560 | 948 | 3389 | 574 | 743 |
| 1214 | QGN14311.1 | FIM1_968 | CP015055. | 3523 | 481 | 387 | 2277 | 381 | 433 | 2396 | 364 | 548 | 1741 | 319 | 570 | 1696 | 349 | 623 | 1549 | 298 | 381 |
| 1215 | QGN14312.1 | FIM1_969 | CP015055. | 1998 | 160 | 158 | 2599 | 194 | 129 | 1403 | 237 | 198 | 1273 | 189 | 286 | 1269 | 137 | 265 | 1160 | 162 | 181 |
| 1216 | QGN14313.1 | FIM1_970 | CP015055. | 1440 | 423 | 627 | 1510 | 526 | 513 | 1778 | 528 | 772 | 3826 | 551 | 934 | 4113 | 504 | 808 | 3685 | 457 | 634 |
| 1217 | QGN14314.1 | FIM1_971 | CP015055. | 401 | 69 | 71 | 378 | 43 | 58 | 408 | 58 | 98 | 654 | 56 | 98 | 446 | 52 | 83 | 643 | 45 | 54 |
| 1218 | QGN14315.1 | FIM1_972 | CP015055. | 131 | 158 | 98 | 110 | 131 | 100 | 101 | 107 | 157 | 311 | 124 | 149 | 399 | 90 | 160 | 339 | 109 | 130 |
| 1219 | QGN14316.1 | FIM1_973 | CP015055. | 3204 | 458 | 636 | 3256 | 602 | 551 | 3131 | 511 | 840 | 3375 | 453 | 933 | 4315 | 498 | 780 | 3607 | 470 | 555 |
| 1220 | QGN14317.1 | FIM1_974 | CP015055. | 2454 | 876 | 733 | 1504 | 733 | 647 | 1503 | 758 | 1036 | 2819 | 527 | 695 | 2923 | 362 | 685 | 2748 | 402 | 498 |
| 1221 | QGN14318.1 | FIM1_975 | CP015055. | 3060 | 1374 | 2494 | 3960 | 1790 | 1950 | 3716 | 1737 | 3145 | 1900 | 869 | 2923 | 5264 | 971 | 2562 | 3028 | 870 | 1972 |
| 1222 | QGN14319.1 | FIM1_976 | CP015055. | 754 | 36 | 26 | 698 | 29 | 15 | 690 | 37 | 29 | 37 | 32 | 41 | 156 | 34 | 32 | 85 | 23 | 26 |
| 1223 | QGN14320.1 | FIM1_977 | CP015055. | 7462 | 2311 | 1623 | 3864 | 2778 | 1303 | 6200 | 2947 | 2293 | 1603 | 2285 | 2701 | 8189 | 2865 | 2575 | 3451 | 2778 | 1840 |
| 1224 | QGN14321.1 | FIM1_978 | CP015055. | 10002 | 3613 | 1476 | 3727 | 3676 | 1314 | 8300 | 3797 | 2071 | 1181 | 3073 | 2514 | 6557 | 3345 | 2399 | 2384 | 4279 | 1806 |
| 1225 | QGN14322.1 | FIM1_979 | CP015055. | 2654 | 2609 | 740 | 1100 | 1886 | 758 | 2302 | 1995 | 974 | 2685 | 2124 | 1245 | 4118 | 1574 | 1459 | 2824 | 2054 | 920 |
| 1226 | QGN14323.1 | FIM1_980 | CP015055. | 3777 | 2417 | 1353 | 5472 | 2155 | 1168 | 3520 | 2097 | 1816 | 2039 | 1687 | 1621 | 6657 | 1594 | 1961 | 3347 | 1119 | 1149 |
| 1227 | QGN14324.1 | FIM1_981 | CP015055. | 3713 | 858 | 814 | 12450 | 469 | 579 | 6403 | 575 | 1134 | 1358 | 377 | 786 | 2601 | 583 | 688 | 1939 | 439 | 501 |
| 1228 | QGN14325.1 | FIM1_982 | CP015055. | 4091 | 1664 | 1268 | 5770 | 1021 | 1005 | 4099 | 1094 | 1626 | 3249 | 575 | 1160 | 2946 | 854 | 1303 | 2833 | 692 | 785 |
| 1229 | QGN14326.1 | FIM1_983 | CP015055. | 14946 | 8764 | 8903 | 20331 | 8494 | 7040 | 15783 | 8304 | 10857 | 21895 | 8739 | 12589 | 32103 | 10095 | 12865 | 27649 | 8131 | 9163 |
| 1230 | QGN14327.1 | FIM1_984 | CP015055. | 473 | 20 | 44 | 1279 | 7 | 18 | 1018 | 43 | 52 | 49 | 10 | 16 | 65 | 18 | 31 | 66 | 10 | 25 |
| 1231 | QGN14328.1 | FIM1_988 | CP015055. | 4534 | 2490 | 2014 | 5625 | 2051 | 1713 | 4803 | 1943 | 2639 | 6569 | 1254 | 2094 | 6628 | 1638 | 2329 | 6506 | 1361 | 1382 |
| 1232 | QGN14329.1 | FIM1_989 | CP015055. | 312 | 21 | 30 | 527 | 18 | 31 | 385 | 23 | 51 | 267 | 14 | 28 | 261 | 13 | 22 | 269 | 15 | 15 |
| 1233 | QGN14330.1 | FIM1_990 | CP015055. | 239 | 12 | 27 | 347 | 14 | 20 | 282 | 16 | 34 | 152 | 3 | 20 | 133 | 4 | 25 | 154 | 5 | 16 |
| 1234 | QGN14331.1 | FIM1_991 | CP015055. | 2265 | 604 | 999 | 3469 | 666 | 705 | 2323 | 705 | 1347 | 1036 | 619 | 1466 | 3576 | 713 | 1369 | 1876 | 652 | 930 |
| 1235 | QGN14332.1 | FIM1_992 | CP015055. | 4110 | 1342 | 1472 | 4279 | 1288 | 1166 | 3820 | 1509 | 1925 | 1971 | 1037 | 2143 | 6598 | 1040 | 1963 | 3603 | 1052 | 1458 |
| 1236 | QGN14333.1 | FIM1_993 | CP015055. | 700 | 820 | 805 | 379 | 1298 | 729 | 665 | 1187 | 1083 | 2617 | 1099 | 1431 | 4721 | 1108 | 1275 | 3390 | 1074 | 932 |
| 1237 | QGN14334.1 | FIM1_994 | CP015055. | 2851 | 2103 | 1104 | 1827 | 1562 | 1096 | 2495 | 1418 | 1470 | 5119 | 1759 | 1702 | 5385 | 1504 | 1778 | 4717 | 1572 | 1282 |
| 1238 | QGN14335.1 | FIM1_995 | CP015055. | 3822 | 2070 | 799 | 1811 | 1751 | 816 | 3086 | 1615 | 968 | 4383 | 2030 | 1524 | 5234 | 1814 | 1475 | 4144 | 1976 | 1037 |
| 1239 | QGN14336.1 | FIM1_996 | CP015055. | 4161 | 1894 | 1138 | 1332 | 2343 | 1246 | 3576 | 2102 | 1405 | 3003 | 2533 | 2383 | 6378 | 2520 | 2194 | 4303 | 2813 | 1681 |
| 1240 | QGN14337.1 | FIM1_997 | CP015055. | 2329 | 82 | 60 | 1005 | 56 | 48 | 1257 | 68 | 78 | 513 | 92 | 158 | 557 | 66 | 142 | 495 | 68 | 99 |
| 1241 | QGN14338.1 | FIM1_998 | CP015055. | 1814 | 779 | 614 | 2294 | 818 | 598 | 2099 | 955 | 850 | 2561 | 625 | 894 | 3893 | 585 | 910 | 2703 | 538 | 676 |
| 1242 | QGN14339.1 | FIM1_999 | CP015055. | 686 | 1813 | 882 | 744 | 1367 | 866 | 684 | 1143 | 1112 | 3616 | 1067 | 1165 | 3434 | 1048 | 1197 | 3155 | 1159 | 783 |
| 1243 | QGN14340.1 | FIM1_1000 | CP015055. | 4825 | 176 | 397 | 4188 | 282 | 291 | 4569 | 302 | 541 | 515 | 182 | 525 | 2914 | 130 | 430 | 1297 | 140 | 363 |
| 1244 | QGN14341.1 | FIM1_1001 | CP015055. | 179 | 270 | 226 | 187 | 283 | 185 | 183 | 234 | 280 | 1139 | 215 | 279 | 1084 | 231 | 245 | 1114 | 226 | 203 |
| 1245 | QGN14342.1 | FIM1_1002 | CP015055. | 404 | 290 | 202 | 276 | 228 | 212 | 324 | 213 | 311 | 948 | 204 | 276 | 1058 | 193 | 266 | 990 | 205 | 147 |
| 1246 | QGN14343.1 | FIM1_1003 | CP015055. | 6915 | 2635 | 1287 | 3532 | 2857 | 1272 | 5372 | 2588 | 1669 | 4372 | 2987 | 2259 | 6885 | 3184 | 2288 | 4849 | 3282 | 1702 |
| 1247 | QGN14344.1 | FIM1_1004 | CP015055. | 4635 | 1909 | 2642 | 7570 | 1551 | 2013 | 5588 | 1525 | 3373 | 2148 | 1181 | 3618 | 6623 | 1505 | 3497 | 3637 | 1307 | 2540 |
| 1248 | QGN14345.1 | FIM1_1005 | CP015055. | 2310 | 796 | 765 | 2257 | 743 | 623 | 2111 | 747 | 943 | 1228 | 795 | 913 | 2861 | 741 | 954 | 1866 | 695 | 661 |
| 1249 | QGN14346.1 | FIM1_1006 | CP015055. | 4264 | 1899 | 2307 | 4507 | 2405 | 2017 | 3738 | 2139 | 2949 | 5057 | 2031 | 3474 | 12430 | 2325 | 3209 | 7531 | 2082 | 2200 |
| 1250 | QGN14347.1 | FIM1_1007 | CP015055. | 878 | 399 | 472 | 1164 | 412 | 342 | 984 | 437 | 582 | 1962 | 376 | 674 | 2499 | 399 | 678 | 2077 | 366 | 451 |
| 1251 | QGN14348.1 | FIM1_1008 | CP015055. | 884 | 237 | 236 | 843 | 126 | 139 | 1010 | 176 | 257 | 1550 | 172 | 291 | 1347 | 141 | 321 | 1360 | 135 | 180 |
| 1252 | QGN14349.1 | FIM1_1009 | CP015055. | 1334 | 502 | 470 | 1312 | 418 | 397 | 1648 | 460 | 613 | 2450 | 421 | 638 | 2530 | 460 | 635 | 2213 | 341 | 453 |
| 1253 | QGN14350.1 | FIM1_1010 | CP015055. | 6640 | 1031 | 1399 | 8786 | 1398 | 1075 | 6753 | 1481 | 1913 | 1918 | 867 | 1936 | 9200 | 1184 | 1557 | 4308 | 944 | 1279 |
| 1254 | QGN14351.1 | FIM1_1011 | CP015055. | 8276 | 5463 | 7913 | 10303 | 6936 | 6852 | 8424 | 6562 | 10182 | 8008 | 6675 | 15479 | 28321 | 6596 | 14507 | 15991 | 6214 | 10486 |
| 1255 | QGN14352.1 | FIM1_1012 | CP015055. | 6374 | 1084 | 1725 | 15779 | 818 | 1033 | 10099 | 1210 | 2224 | 1175 | 776 | 2247 | 5160 | 1606 | 2166 | 2307 | 1162 | 1482 |
| 1256 | QGN14353.1 | FIM1_1013 | CP015055. | 2059 | 1386 | 2150 | 1654 | 1810 | 1745 | 1156 | 1757 | 2912 | 3590 | 1751 | 3523 | 10966 | 1697 | 3069 | 6178 | 1656 | 2298 |
| 1257 | QGN14354.1 | FIM1_1014 | CP015055. | 2395 | 3291 | 1380 | 1163 | 2158 | 995 | 578 | 3380 | 1786 | 4669 | 1944 | 1502 | 8465 | 2541 | 793 | 7083 | 1700 | 1271 |
| 1258 | QGN14355.1 | FIM1_1015 | CP015055. | 3951 | 2137 | 858 | 1857 | 1647 | 869 | 3046 | 1624 | 1173 | 3322 | 1765 | 1283 | 4422 | 1570 | 1437 | 3082 | 1914 | 894 |

| | | | | | | | | | | | | | | | | | | | | | |
|---|---|---|---|---|---|---|---|---|---|---|---|---|---|---|---|---|---|---|---|---|---|
| 1259 | QGN14356.1 | FIM1_1016 | CP015055. | 326 | 296 | 191 | 136 | 176 | 159 | 131 | 200 | 207 | 347 | 120 | 200 | 379 | 146 | 204 | 372 | 125 | 112 |
| 1260 | QGN14357.1 | FIM1_1017 | CP015055. | 1547 | 4406 | 2048 | 1494 | 3102 | 1811 | 1477 | 2871 | 2632 | 13075 | 3725 | 3299 | 12446 | 3848 | 3326 | 12087 | 3838 | 2185 |
| 1261 | QGN14358.1 | FIM1_1018 | CP015055. | 596 | 756 | 901 | 757 | 908 | 753 | 543 | 828 | 1062 | 4794 | 1124 | 1339 | 4868 | 934 | 1378 | 4425 | 914 | 914 |
| 1262 | QGN14359.1 | FIM1_1019 | CP015055. | 2058 | 248 | 500 | 2033 | 395 | 372 | 2190 | 426 | 637 | 1963 | 425 | 630 | 3463 | 408 | 583 | 2460 | 414 | 452 |
| 1263 | QGN14360.1 | FIM1_1020 | CP015055. | 57 | 80 | 173 | 158 | 68 | 94 | 114 | 88 | 187 | 201 | 50 | 140 | 739 | 50 | 145 | 368 | 41 | 125 |
| 1264 | QGN14361.1 | FIM1_1021 | CP015055. | 4852 | 31 | 199 | 5320 | 181 | 254 | 2979 | 97 | 260 | 999 | 68 | 244 | 1365 | 71 | 198 | 992 | 57 | 145 |
| 1265 | QGN14362.1 | FIM1_1022 | CP015055. | 14219 | 8283 | 4143 | 12246 | 5958 | 3503 | 11648 | 5726 | 5036 | 15392 | 4960 | 6054 | 16655 | 5165 | 5932 | 15283 | 4837 | 4250 |
| 1266 | QGN14363.1 | FIM1_1023 | CP015055. | 25 | 5 | 3 | 47 | 13 | 4 | 16 | 7 | 5 | 23 | 10 | 18 | 40 | 14 | 13 | 28 | 10 | 3 |
| 1267 | QGN14364.1 | FIM1_1024 | CP015055. | 5585 | 918 | 1077 | 5950 | 1054 | 696 | 4220 | 1296 | 1464 | 1585 | 605 | 1468 | 4712 | 755 | 1305 | 2435 | 660 | 911 |
| 1268 | QGN14365.1 | FIM1_1025 | CP015055. | 8469 | 4892 | 3856 | 10763 | 5155 | 4825 | 7778 | 3515 | 5041 | 10621 | 2811 | 3710 | 9193 | 2524 | 3809 | 10118 | 2856 | 2491 |
| 1269 | QGN14366.1 | FIM1_1026 | CP015055. | 12369 | 6883 | 5200 | 9969 | 4048 | 4467 | 11126 | 4650 | 6812 | 14294 | 4436 | 5663 | 12583 | 3960 | 5929 | 12423 | 4090 | 4219 |
| 1270 | QGN14367.1 | FIM1_1027 | CP015055. | 3130 | 318 | 714 | 3646 | 413 | 521 | 2724 | 433 | 894 | 1298 | 402 | 1272 | 2836 | 466 | 1024 | 1965 | 422 | 823 |
| 1271 | QGN14368.1 | FIM1_1028 | CP015055. | 1279 | 72 | 115 | 1093 | 90 | 82 | 1198 | 112 | 162 | 254 | 60 | 118 | 452 | 50 | 110 | 361 | 77 | 86 |
| 1272 | QGN14369.1 | FIM1_1029 | CP015055. | 3780 | 7240 | 8011 | 10823 | 10608 | 5754 | 7236 | 9887 | 9220 | 15498 | 11396 | 14174 | 38210 | 12176 | 12128 | 26851 | 11022 | 9711 |
| 1273 | QGN14370.1 | FIM1_1030 | CP015055. | 5967 | 3610 | 1280 | 2042 | 3459 | 1341 | 4767 | 3209 | 1744 | 4140 | 4267 | 2383 | 9199 | 3874 | 2347 | 5985 | 4777 | 1902 |
| 1274 | QGN14371.1 | FIM1_1031 | CP015055. | 234 | 68 | 51 | 185 | 64 | 51 | 79 | 47 | 74 | 225 | 49 | 83 | 277 | 47 | 70 | 206 | 47 | 68 |
| 1275 | QGN14372.1 | FIM1_1032 | CP015055. | 1849 | 431 | 674 | 2282 | 459 | 569 | 1616 | 521 | 926 | 2654 | 285 | 618 | 2600 | 252 | 539 | 2404 | 213 | 417 |
| 1276 | QGN14373.1 | FIM1_1033 | CP015055. | 74 | 192 | 140 | 132 | 164 | 139 | 71 | 129 | 194 | 1279 | 164 | 239 | 778 | 149 | 212 | 1220 | 155 | 116 |
| 1277 | QGN14374.1 | FIM1_1034 | CP015055. | 1046 | 470 | 457 | 1428 | 331 | 379 | 1107 | 363 | 579 | 1600 | 241 | 576 | 2027 | 244 | 546 | 1645 | 224 | 407 |
| 1278 | QGN14375.1 | FIM1_1035 | CP015055. | 259 | 57 | 118 | 363 | 85 | 84 | 241 | 90 | 149 | 505 | 73 | 166 | 579 | 64 | 123 | 499 | 84 | 100 |
| 1279 | QGN14376.1 | FIM1_1036 | CP015055. | 122 | 326 | 468 | 178 | 410 | 356 | 108 | 362 | 583 | 2424 | 395 | 823 | 2457 | 304 | 693 | 2641 | 340 | 493 |
| 1280 | QGN14377.1 | FIM1_1037 | CP015055. | 1430 | 702 | 773 | 1956 | 747 | 698 | 1466 | 712 | 967 | 4175 | 728 | 1264 | 4489 | 745 | 1131 | 4160 | 607 | 829 |
| 1281 | QGN14378.1 | FIM1_1038 | CP015055. | 2310 | 2542 | 2255 | 2794 | 2440 | 1960 | 1904 | 2218 | 2781 | 10159 | 2502 | 3151 | 10372 | 2218 | 3128 | 9482 | 2134 | 2083 |
| 1282 | QGN14379.1 | FIM1_1039 | CP015055. | 6129 | 1685 | 1832 | 9174 | 2493 | 1498 | 7314 | 1972 | 2370 | 9953 | 1714 | 1984 | 10407 | 1817 | 1823 | 9729 | 1703 | 1279 |
| 1283 | QGN14380.1 | FIM1_1040 | CP015055. | 3696 | 811 | 1242 | 5503 | 964 | 1098 | 3685 | 864 | 1745 | 1599 | 781 | 1386 | 4895 | 799 | 1384 | 2858 | 684 | 887 |
| 1284 | QGN14381.1 | FIM1_1041 | CP015055. | 1944 | 1278 | 1179 | 6277 | 2653 | 1212 | 5057 | 2119 | 1810 | 1564 | 889 | 507 | 4357 | 1097 | 517 | 2733 | 993 | 356 |
| 1285 | QGN14382.1 | FIM1_1042 | CP015055. | 129 | 85 | 41 | 169 | 65 | 46 | 125 | 47 | 65 | 146 | 45 | 37 | 142 | 36 | 68 | 150 | 32 | 30 |
| 1286 | QGN14383.1 | FIM1_1043 | CP015055. | 334 | 121 | 163 | 455 | 128 | 158 | 186 | 112 | 194 | 391 | 91 | 118 | 466 | 67 | 121 | 434 | 69 | 85 |
| 1287 | QGN14384.1 | FIM1_1044 | CP015055. | 4390 | 557 | 1061 | 4665 | 1329 | 879 | 3925 | 1164 | 1323 | 2645 | 1201 | 1570 | 6930 | 1194 | 1418 | 4331 | 1091 | 1026 |
| 1288 | QGN14385.1 | FIM1_1045 | CP015055. | 1050 | 435 | 382 | 1022 | 274 | 317 | 1010 | 250 | 470 | 1231 | 270 | 448 | 1316 | 206 | 441 | 1069 | 207 | 296 |
| 1289 | QGN14386.1 | FIM1_1046 | CP015055. | 917 | 428 | 596 | 1616 | 399 | 503 | 1137 | 433 | 778 | 2717 | 378 | 808 | 3366 | 373 | 701 | 2749 | 308 | 583 |
| 1290 | QGN14387.1 | FIM1_1047 | CP015055. | 2663 | 731 | 931 | 3034 | 826 | 847 | 2508 | 803 | 1269 | 5783 | 754 | 1439 | 5381 | 709 | 1247 | 5313 | 687 | 960 |
| 1291 | QGN14388.1 | FIM1_1048 | CP015055. | 1197 | 1004 | 1063 | 2213 | 1097 | 871 | 1736 | 976 | 1322 | 6428 | 1073 | 1665 | 7312 | 1049 | 1490 | 6331 | 968 | 1101 |
| 1292 | QGN14389.1 | FIM1_1049 | CP015055. | 3487 | 1997 | 853 | 1251 | 1904 | 874 | 2581 | 1462 | 1174 | 6052 | 2216 | 1641 | 6359 | 2420 | 1443 | 6297 | 2627 | 1144 |
| 1293 | QGN14390.1 | FIM1_1050 | CP015055. | 353 | 54 | 119 | 319 | 65 | 131 | 140 | 41 | 170 | 99 | 33 | 147 | 227 | 17 | 125 | 161 | 25 | 71 |
| 1294 | QGN14391.1 | FIM1_1051 | CP015055. | 701 | 485 | 508 | 1177 | 351 | 474 | 1056 | 348 | 753 | 1478 | 248 | 588 | 1770 | 346 | 576 | 1443 | 276 | 405 |
| 1295 | QGN14392.1 | FIM1_1052 | CP015055. | 590 | 460 | 1516 | 1354 | 392 | 1465 | 509 | 378 | 1777 | 2809 | 269 | 1627 | 1959 | 283 | 1630 | 2221 | 203 | 1028 |
| 1296 | QGN14393.1 | FIM1_1053 | CP015055. | 72 | 42 | 61 | 71 | 48 | 66 | 40 | 80 | 88 | 281 | 43 | 93 | 309 | 37 | 93 | 227 | 44 | 68 |
| 1297 | QGN14394.1 | FIM1_1054 | CP015055. | 1550 | 1896 | 1173 | 1758 | 1884 | 1114 | 1684 | 1610 | 1593 | 5338 | 1740 | 1889 | 7331 | 1705 | 1702 | 5642 | 1681 | 1191 |
| 1298 | QGN14395.1 | FIM1_1055 | CP015055. | 363 | 154 | 186 | 336 | 100 | 165 | 337 | 111 | 231 | 926 | 92 | 185 | 681 | 52 | 222 | 862 | 71 | 159 |
| 1299 | QGN14396.1 | FIM1_1056 | CP015055. | 813 | 370 | 290 | 802 | 349 | 259 | 942 | 360 | 373 | 2388 | 351 | 500 | 2179 | 352 | 442 | 2376 | 354 | 362 |
| 1300 | QGN14397.1 | FIM1_1057 | CP015055. | 2208 | 720 | 833 | 2876 | 818 | 688 | 2495 | 859 | 1095 | 3336 | 745 | 1184 | 5264 | 759 | 1133 | 3867 | 683 | 776 |
| 1301 | QGN14398.1 | FIM1_1058 | CP015055. | 1408 | 1479 | 1878 | 2473 | 1499 | 1686 | 1946 | 1467 | 2494 | 6805 | 1666 | 3016 | 7514 | 1715 | 2806 | 6268 | 1636 | 2030 |
| 1302 | QGN14399.1 | FIM1_1059 | CP015055. | 86 | 3 | 3 | 112 | 6 | 6 | 151 | 6 | 11 | 77 | 12 | 45 | 88 | 2 | 8 | 153 | 17 | 16 |
| 1303 | QGN14400.1 | FIM1_1060 | CP015055. | 5263 | 2563 | 1406 | 3607 | 3183 | 1580 | 5489 | 2623 | 1954 | 13493 | 4168 | 2857 | 13460 | 3662 | 2615 | 12867 | 4399 | 2076 |
| 1304 | QGN14401.1 | FIM1_1061 | CP015055. | 1876 | 370 | 358 | 1822 | 471 | 344 | 1915 | 351 | 497 | 2446 | 432 | 655 | 2252 | 683 | 610 | 2250 | 496 | 419 |
| 1305 | QGN14402.1 | FIM1_1062 | CP015055. | 2086 | 386 | 503 | 2024 | 423 | 434 | 2134 | 378 | 615 | 3541 | 480 | 705 | 2933 | 425 | 651 | 3462 | 401 | 438 |
| 1306 | QGN14403.1 | FIM1_1063 | CP015055. | 2031 | 338 | 518 | 2741 | 290 | 448 | 2731 | 298 | 715 | 3440 | 270 | 711 | 2556 | 280 | 676 | 3191 | 261 | 440 |
| 1307 | QGN14404.1 | FIM1_1064 | CP015055. | 5008 | 541 | 863 | 5651 | 585 | 671 | 4990 | 615 | 1069 | 2348 | 380 | 921 | 2436 | 485 | 902 | 2286 | 431 | 583 |
| 1308 | QGN14405.1 | FIM1_1065 | CP015055. | 1628 | 760 | 863 | 1697 | 940 | 646 | 1668 | 787 | 1172 | 4150 | 747 | 962 | 4320 | 790 | 953 | 3964 | 699 | 669 |
| 1309 | QGN14406.1 | FIM1_1066 | CP015055. | 402 | 334 | 263 | 460 | 269 | 215 | 412 | 226 | 362 | 1981 | 282 | 436 | 1595 | 245 | 396 | 1991 | 245 | 281 |
| 1310 | QGN14407.1 | FIM1_1067 | CP015055. | 437 | 188 | 258 | 607 | 190 | 233 | 568 | 202 | 361 | 2139 | 192 | 354 | 1660 | 157 | 333 | 2082 | 228 | 230 |
| 1311 | QGN14408.1 | FIM1_1068 | CP015055. | 533 | 166 | 166 | 708 | 199 | 127 | 392 | 201 | 215 | 272 | 114 | 98 | 489 | 103 | 81 | 327 | 120 | 63 |
| 1312 | QGN14409.1 | FIM1_1069 | CP015055. | 994 | 27 | 278 | 1427 | 154 | 180 | 1049 | 148 | 314 | 916 | 153 | 432 | 1659 | 143 | 320 | 946 | 130 | 285 |
| 1313 | QGN14410.1 | FIM1_1070 | CP015055. | 2681 | 532 | 930 | 3302 | 529 | 737 | 2895 | 494 | 1199 | 2472 | 444 | 1272 | 3838 | 407 | 1100 | 3014 | 355 | 848 |
| 1314 | QGN14411.1 | FIM1_1071 | CP015055. | 1531 | 948 | 1004 | 1864 | 1157 | 854 | 1598 | 991 | 1309 | 5165 | 969 | 1392 | 5392 | 949 | 1286 | 4599 | 860 | 904 |
| 1315 | QGN14412.1 | FIM1_1072 | CP015055. | 6596 | 615 | 385 | 3559 | 996 | 346 | 5529 | 927 | 518 | 397 | 879 | 726 | 2064 | 1446 | 634 | 782 | 1222 | 508 |
| 1316 | QGN14413.1 | FIM1_1074 | CP015055. | 1094 | 1213 | 793 | 1295 | 807 | 763 | 999 | 792 | 1124 | 4381 | 774 | 1107 | 3699 | 808 | 1076 | 3906 | 710 | 696 |
| 1317 | QGN14414.1 | FIM1_1075 | CP015055. | 6462 | 1948 | 1753 | 5597 | 2513 | 1459 | 5949 | 2234 | 2320 | 5011 | 2171 | 2388 | 8495 | 1906 | 2385 | 6148 | 1880 | 1585 |
| 1318 | QGN14415.1 | FIM1_1076 | CP015055. | 399 | 226 | 304 | 514 | 255 | 209 | 301 | 282 | 405 | 287 | 285 | 502 | 971 | 226 | 414 | 494 | 241 | 284 |
| 1319 | QGN14416.1 | FIM1_1077 | CP015055. | 299 | 151 | 52 | 222 | 110 | 50 | 154 | 89 | 77 | 58 | 115 | 105 | 280 | 86 | 98 | 123 | 73 | 60 |
| 1320 | QGN14417.1 | FIM1_1078 | CP015055. | 495 | 222 | 144 | 371 | 186 | 137 | 458 | 184 | 215 | 959 | 193 | 203 | 990 | 165 | 228 | 940 | 201 | 142 |
| 1321 | QGN14418.1 | FIM1_1079 | CP015055. | 568 | 471 | 232 | 892 | 323 | 184 | 455 | 306 | 305 | 585 | 197 | 240 | 945 | 177 | 202 | 666 | 159 | 148 |
| 1322 | QGN14419.1 | FIM1_1080 | CP015055. | 841 | 722 | 765 | 1471 | 1106 | 464 | 1421 | 1081 | 906 | 1365 | 819 | 908 | 4243 | 733 | 782 | 2387 | 612 | 620 |
| 1323 | QGN14420.1 | FIM1_1081 | CP015055. | 785 | 875 | 669 | 3118 | 774 | 513 | 1935 | 849 | 912 | 1170 | 557 | 764 | 3337 | 627 | 650 | 2208 | 529 | 553 |
| 1324 | QGN14421.1 | FIM1_1082 | CP015055. | 1280 | 997 | 656 | 2590 | 829 | 550 | 1401 | 878 | 818 | 1134 | 680 | 667 | 3393 | 640 | 667 | 2012 | 626 | 504 |
| 1325 | QGN14422.1 | FIM1_1083 | CP015055. | 2559 | 1220 | 1216 | 4007 | 1081 | 967 | 2672 | 1193 | 1461 | 4471 | 725 | 1163 | 4900 | 608 | 1244 | 4076 | 584 | 775 |
| 1326 | QGN14423.1 | FIM1_1084 | CP015055. | 1418 | 646 | 698 | 2674 | 621 | 511 | 1486 | 718 | 864 | 632 | 138 | 237 | 1186 | 100 | 236 | 833 | 113 | 165 |
| 1327 | QGN14424.1 | FIM1_1085 | CP015055. | 1079 | 356 | 417 | 1138 | 324 | 296 | 988 | 311 | 485 | 2398 | 322 | 495 | 2067 | 265 | 548 | 2368 | 293 | 394 |
| 1328 | QGN14425.1 | FIM1_1086 | CP015055. | 380 | 331 | 622 | 677 | 343 | 476 | 477 | 475 | 852 | 2992 | 236 | 751 | 3107 | 220 | 704 | 2863 | 195 | 484 |
| 1329 | QGN14426.1 | FIM1_1087 | CP015055. | 998 | 321 | 402 | 1674 | 445 | 358 | 1436 | 365 | 492 | 2963 | 386 | 612 | 3266 | 381 | 584 | 2913 | 368 | 460 |
| 1330 | QGN14427.1 | FIM1_1088 | CP015055. | 2875 | 1421 | 1347 | 4071 | 1439 | 1152 | 3273 | 1379 | 1700 | 8702 | 1480 | 2028 | 7608 | 1246 | 1932 | 8115 | 1363 | 1302 |
| 1331 | QGN14428.1 | FIM1_1089 | CP015055. | 70 | 143 | 117 | 53 | 84 | 92 | 61 | 89 | 134 | 587 | 94 | 108 | 390 | 87 | 118 | 482 | 65 | 77 |
| 1332 | QGN14429.1 | FIM1_1090 | CP015055. | 7155 | 3587 | 2674 | 11977 | 3024 | 2408 | 8987 | 2569 | 3313 | 11577 | 3433 | 4239 | 13527 | 3558 | 4585 | 12447 | 3213 | 3249 |
| 1333 | QGN14430.1 | FIM1_1091 | CP015055. | 3619 | 1946 | 783 | 2149 | 1873 | 858 | 3077 | 1741 | 1090 | 4253 | 2015 | 1542 | 5654 | 1952 | 1433 | 4487 | 2273 | 1101 |
| 1334 | QGN14431.1 | FIM1_1092 | CP015055. | 1584 | 538 | 583 | 1876 | 484 | 465 | 1887 | 543 | 724 | 3035 | 546 | 817 | 3631 | 539 | 803 | 3061 | 466 | 604 |
| 1335 | QGN14432.1 | FIM1_1093 | CP015055. | 3426 | 634 | 1100 | 2876 | 802 | 775 | 3036 | 782 | 1296 | 2444 | 485 | 1121 | 3857 | 428 | 1054 | 3042 | 428 | 723 |
| 1336 | QGN14433.1 | FIM1_1094 | CP015055. | 2562 | 995 | 847 | 2718 | 988 | 740 | 2476 | 858 | 1077 | 4787 | 1144 | 1570 | 5811 | 1155 | 1403 | 4973 | 1174 | 1047 |
| 1337 | QGN14434.1 | FIM1_1095 | CP015055. | 1680 | 558 | 668 | 2543 | 721 | 593 | 2170 | 699 | 859 | 3764 | 622 | 1057 | 4882 | 598 | 941 | 3979 | 537 | 745 |
| 1338 | QGN14435.1 | FIM1_1096 | CP015055. | 853 | 330 | 231 | 1253 | 209 | 216 | 900 | 192 | 324 | 1350 | 216 | 352 | 1109 | 257 | 390 | 1082 | 292 | 202 |
| 1339 | QGN14436.1 | FIM1_1097 | CP015055. | 23523 | 2491 | 3129 | 25724 | 3996 | 2651 | 28855 | 3588 | 3890 | 16298 | 4084 | 4839 | 22347 | 6297 | 4251 | 17513 | 4849 | 3381 |
| 1340 | QGN14437.1 | FIM1_1098 | CP015055. | 1436 | 3901 | 2150 | 3382 | 2763 | 2029 | 2628 | 2566 | 2848 | 9740 | 2867 | 3449 | 8558 | 2551 | 3626 | 8743 | 2732 | 2507 |
| 1341 | QGN14438.1 | FIM1_1099 | CP015055. | 2140 | 432 | 516 | 2857 | 412 | 451 | 2196 | 500 | 689 | 1146 | 309 | 685 | 2219 | 280 | 667 | 1462 | 285 | 534 |
| 1342 | QGN14439.1 | FIM1_1100 | CP015055. | 8393 | 6811 | 3698 | 9669 | 5119 | 3469 | 9181 | 4798 | 4816 | 21255 | 6479 | 6336 | 20962 | 5933 | 6544 | 19862 | 5650 | 4503 |

| | | | | | | | | | | | | | | | | | | | | |
|---|---|---|---|---|---|---|---|---|---|---|---|---|---|---|---|---|---|---|---|---|
| 1343 | QGN14440.1 | FIM1_1101 | CP015055. | 84 | 27 | 27 | 778 | 111 | 75 | 178 | 24 | 39 | 65 | 20 | 40 | 132 | 21 | 46 | 114 | 15 | 34 |
| 1344 | QGN14441.1 | FIM1_1102 | CP015055. | 974 | 1116 | 766 | 973 | 942 | 722 | 767 | 727 | 1007 | 3691 | 960 | 1180 | 2973 | 803 | 1229 | 3209 | 795 | 831 |
| 1345 | QGN14442.1 | FIM1_1103 | CP015055. | 865 | 1112 | 684 | 485 | 809 | 607 | 724 | 867 | 882 | 2075 | 672 | 904 | 2913 | 611 | 869 | 2124 | 608 | 585 |
| 1346 | QGN14443.1 | FIM1_1104 | CP015055. | 882 | 1332 | 1057 | 1481 | 1232 | 886 | 1006 | 1310 | 1373 | 1466 | 834 | 1456 | 3585 | 921 | 1503 | 2104 | 882 | 987 |
| 1347 | QGN14444.1 | FIM1_1105 | CP015055. | 5415 | 2975 | 2256 | 5945 | 3375 | 1956 | 4970 | 2736 | 2739 | 10501 | 4154 | 3799 | 13143 | 4144 | 3734 | 11538 | 3773 | 2649 |
| 1348 | QGN14445.1 | FIM1_1106 | CP015055. | 5266 | 3590 | 4436 | 8074 | 4997 | 3463 | 5785 | 4892 | 5518 | 12343 | 5022 | 6518 | 25485 | 4664 | 6445 | 19424 | 3965 | 4410 |
| 1349 | QGN14446.1 | FIM1_1107 | CP015055. | 707 | 999 | 604 | 1436 | 794 | 585 | 994 | 704 | 817 | 2071 | 666 | 864 | 2335 | 669 | 981 | 1912 | 694 | 641 |
| 1350 | QGN14447.1 | FIM1_1108 | CP015055. | 1136 | 1817 | 466 | 508 | 1590 | 699 | 912 | 1197 | 687 | 4609 | 1868 | 954 | 4184 | 1199 | 1038 | 4249 | 1701 | 619 |
| 1351 | QGN14448.1 | FIM1_1109 | CP015055. | 10104 | 2375 | 2575 | 15127 | 2204 | 2126 | 11687 | 1933 | 3296 | 7134 | 2286 | 3962 | 13419 | 2889 | 3913 | 9619 | 2456 | 2652 |
| 1352 | QGN14449.1 | FIM1_1110 | CP015055. | 1355 | 1147 | 737 | 1946 | 843 | 737 | 1648 | 643 | 958 | 5361 | 897 | 1303 | 4030 | 780 | 1239 | 5174 | 947 | 822 |
| 1353 | QGN14450.1 | FIM1_1111 | CP015055. | 1163 | 1108 | 1092 | 1420 | 1020 | 957 | 1202 | 978 | 1453 | 1614 | 640 | 1121 | 2748 | 583 | 1098 | 2182 | 543 | 683 |
| 1354 | QGN14451.1 | FIM1_1112 | CP015055. | 16283 | 1701 | 2033 | 13888 | 2245 | 1685 | 14802 | 2269 | 2454 | 7804 | 2030 | 2535 | 9711 | 1764 | 2598 | 8150 | 1671 | 1804 |
| 1355 | QGN14452.1 | FIM1_1113 | CP015055. | 2067 | 1058 | 741 | 2577 | 1034 | 649 | 2256 | 1121 | 1104 | 534 | 732 | 1062 | 2599 | 727 | 909 | 1056 | 790 | 723 |
| 1356 | QGN14453.1 | FIM1_1114 | CP015055. | 24834 | 374 | 414 | 28369 | 233 | 220 | 24066 | 474 | 473 | 472 | 216 | 383 | 1268 | 335 | 283 | 687 | 232 | 244 |
| 1357 | QGN14454.1 | FIM1_1115 | CP015055. | 2296 | 98 | 147 | 3065 | 170 | 115 | 1413 | 174 | 163 | 202 | 93 | 178 | 501 | 114 | 154 | 316 | 114 | 122 |
| 1358 | QGN14455.1 | FIM1_1116 | CP015055. | 6114 | 5892 | 3874 | 4271 | 4725 | 3094 | 3688 | 5184 | 4963 | 12002 | 4829 | 6846 | 19177 | 4695 | 6519 | 14082 | 4432 | 4958 |
| 1359 | QGN14456.1 | FIM1_1117 | CP015055. | 4215 | 1168 | 1589 | 3447 | 730 | 1376 | 3417 | 790 | 2237 | 3556 | 781 | 2497 | 5384 | 606 | 2367 | 4030 | 619 | 1727 |
| 1360 | QGN14457.1 | FIM1_1118 | CP015055. | 104 | 16 | 7 | 104 | 8 | 4 | 87 | 11 | 9 | 62 | 7 | 15 | 59 | 5 | 12 | 68 | 6 | 7 |
| 1361 | QGN14458.1 | FIM1_1119 | CP015055. | 4036 | 1885 | 1782 | 7548 | 2451 | 1557 | 5979 | 2315 | 2391 | 6165 | 1423 | 2272 | 8189 | 1707 | 2205 | 6630 | 1407 | 1527 |
| 1362 | QGN14459.1 | FIM1_1120 | CP015055. | 183 | 14 | 20 | 268 | 15 | 21 | 204 | 21 | 33 | 163 | 19 | 40 | 135 | 10 | 31 | 159 | 14 | 13 |
| 1363 | QGN14460.1 | FIM1_1121 | CP015055. | 20820 | 4927 | 8429 | 30995 | 6994 | 6732 | 25983 | 6130 | 10484 | 34505 | 7065 | 13726 | 40600 | 8873 | 12130 | 35519 | 7503 | 9197 |
| 1364 | QGN14461.1 | FIM1_1122 | CP015055. | 5768 | 3404 | 3065 | 8686 | 3605 | 2628 | 6541 | 2983 | 4000 | 13547 | 3491 | 4395 | 17378 | 3432 | 3965 | 14358 | 3473 | 2985 |
| 1365 | QGN14462.1 | FIM1_1123 | CP015055. | 1243 | 351 | 261 | 958 | 296 | 252 | 1233 | 331 | 358 | 926 | 216 | 327 | 1279 | 253 | 344 | 957 | 233 | 242 |
| 1366 | QGN14463.1 | FIM1_1124 | CP015055. | 5343 | 2079 | 1840 | 8487 | 2083 | 1728 | 6259 | 2701 | 2416 | 5983 | 1571 | 2509 | 10298 | 1572 | 2482 | 7595 | 1446 | 1826 |
| 1367 | QGN14464.1 | FIM1_1125 | CP015055. | 19643 | 23902 | 12160 | 18003 | 16143 | 11197 | 15351 | 15004 | 14546 | 32720 | 19246 | 17645 | 40126 | 17295 | 18515 | 35345 | 16254 | 13587 |
| 1368 | QGN14465.1 | FIM1_1126 | CP015055. | 995 | 574 | 665 | 3698 | 385 | 438 | 2409 | 456 | 851 | 1603 | 239 | 529 | 1691 | 452 | 508 | 1588 | 403 | 282 |
| 1369 | QGN14466.1 | FIM1_1127 | CP015055. | 1317 | 94 | 348 | 1183 | 216 | 248 | 895 | 273 | 443 | 808 | 147 | 412 | 1473 | 118 | 387 | 1107 | 116 | 271 |
| 1370 | QGN14467.1 | FIM1_1128 | CP015055. | 761 | 189 | 159 | 330 | 122 | 138 | 322 | 127 | 223 | 315 | 115 | 235 | 579 | 77 | 213 | 436 | 89 | 146 |
| 1371 | QGN14468.1 | FIM1_1129 | CP015055. | 2742 | 1607 | 2141 | 4484 | 2351 | 1785 | 3838 | 1713 | 2690 | 12125 | 1794 | 3323 | 11813 | 1748 | 3002 | 11566 | 1561 | 2138 |
| 1372 | QGN14469.1 | FIM1_1130 | CP015055. | 963 | 358 | 291 | 698 | 278 | 246 | 713 | 277 | 337 | 1601 | 240 | 360 | 1321 | 198 | 322 | 1356 | 208 | 220 |
| 1373 | QGN14470.1 | FIM1_1131 | CP015055. | 1345 | 735 | 842 | 1158 | 830 | 706 | 1013 | 882 | 1028 | 5852 | 875 | 1261 | 5717 | 864 | 1202 | 4894 | 731 | 874 |
| 1374 | QGN14471.1 | FIM1_1132 | CP015055. | 8166 | 141 | 373 | 6156 | 163 | 295 | 6389 | 186 | 525 | 921 | 181 | 652 | 2072 | 239 | 622 | 1158 | 206 | 453 |
| 1375 | QGN14472.1 | FIM1_1133 | CP015055. | 2446 | 281 | 255 | 2097 | 320 | 245 | 1517 | 260 | 369 | 294 | 266 | 453 | 1085 | 308 | 535 | 496 | 274 | 307 |
| 1376 | QGN14473.1 | FIM1_1134 | CP015055. | 615 | 218 | 223 | 419 | 192 | 134 | 703 | 314 | 284 | 146 | 135 | 240 | 395 | 191 | 230 | 245 | 202 | 183 |
| 1377 | QGN14474.1 | FIM1_1135 | CP015055. | 3855 | 4953 | 7388 | 11697 | 9994 | 5271 | 9364 | 9601 | 8112 | 35051 | 13646 | 14245 | 45194 | 13579 | 12184 | 37369 | 12212 | 9656 |
| 1378 | QGN14475.1 | FIM1_1136 | CP015055. | 2098 | 813 | 1032 | 1818 | 903 | 895 | 1934 | 1015 | 1342 | 1695 | 432 | 1083 | 2769 | 420 | 1037 | 2098 | 388 | 646 |
| 1379 | QGN14476.1 | FIM1_1137 | CP015055. | 5495 | 2482 | 3315 | 8950 | 4821 | 2843 | 4448 | 3915 | 3931 | 2999 | 1479 | 3414 | 10360 | 1452 | 2959 | 5758 | 1712 | 2285 |
| 1380 | QGN14477.1 | FIM1_1138 | CP015055. | 1049 | 435 | 589 | 1520 | 373 | 467 | 1102 | 334 | 762 | 2368 | 199 | 582 | 1886 | 189 | 582 | 2237 | 167 | 426 |
| 1381 | QGN14478.1 | FIM1_1139 | CP015055. | 7145 | 7886 | 7082 | 9102 | 7375 | 5487 | 6717 | 7229 | 8137 | 19073 | 6158 | 10198 | 29939 | 6164 | 8773 | 23526 | 5732 | 6341 |
| 1382 | QGN14479.1 | FIM1_1140 | CP015055. | 453 | 1081 | 4018 | 581 | 2641 | 2632 | 1676 | 4849 | 4232 | 9077 | 1305 | 4658 | 13963 | 1973 | 3989 | 10405 | 1988 | 3162 |
| 1383 | QGN14480.1 | FIM1_1141 | CP015055. | 325 | 3103 | 4567 | 587 | 3202 | 3035 | 1328 | 5415 | 4798 | 7830 | 1541 | 4773 | 13506 | 2489 | 4529 | 10295 | 2384 | 3292 |
| 1384 | QGN14481.1 | FIM1_1142 | CP015055. | 1466 | 784 | 556 | 1615 | 583 | 504 | 1443 | 578 | 805 | 1799 | 456 | 700 | 2511 | 458 | 750 | 1846 | 422 | 510 |
| 1385 | QGN14482.1 | FIM1_1143 | CP015055. | 2268 | 1344 | 935 | 2225 | 1236 | 869 | 2149 | 1115 | 1377 | 4314 | 1096 | 1542 | 5997 | 1034 | 1379 | 4975 | 1040 | 999 |
| 1386 | QGN14483.1 | FIM1_1144 | CP015055. | 170 | 725 | 536 | 464 | 503 | 506 | 326 | 451 | 709 | 4359 | 647 | 872 | 2951 | 502 | 832 | 3881 | 591 | 547 |
| 1387 | QGN14484.1 | FIM1_1145 | CP015055. | 15616 | 6440 | 3838 | 15117 | 6920 | 3952 | 13980 | 5886 | 5014 | 19707 | 7837 | 7018 | 23269 | 6890 | 6701 | 19257 | 7389 | 4924 |
| 1388 | QGN14485.1 | FIM1_1146 | CP015055. | 769 | 456 | 542 | 730 | 353 | 431 | 697 | 443 | 649 | 2215 | 260 | 680 | 2419 | 347 | 675 | 2194 | 294 | 438 |
| 1389 | QGN14486.1 | FIM1_1147 | CP015055. | 5277 | 2262 | 1561 | 4995 | 2436 | 1470 | 4301 | 1857 | 2012 | 11710 | 2896 | 2808 | 10512 | 2816 | 2979 | 10511 | 2972 | 1947 |
| 1390 | QGN14487.1 | FIM1_1148 | CP015055. | 1631 | 531 | 334 | 1306 | 427 | 339 | 1392 | 367 | 510 | 1716 | 364 | 672 | 1750 | 345 | 663 | 1624 | 395 | 445 |
| 1391 | QGN14488.1 | FIM1_1149 | CP015055. | 342 | 305 | 206 | 406 | 173 | 165 | 287 | 178 | 239 | 1241 | 178 | 227 | 963 | 145 | 231 | 1174 | 144 | 152 |
| 1392 | QGN14489.1 | FIM1_1150 | CP015055. | 813 | 932 | 539 | 838 | 660 | 536 | 759 | 547 | 773 | 3756 | 665 | 858 | 2540 | 586 | 827 | 3488 | 611 | 543 |
| 1393 | QGN14490.1 | FIM1_1151 | CP015055. | 7575 | 4035 | 3531 | 16861 | 4742 | 3062 | 11000 | 4232 | 4227 | 18281 | 3948 | 5305 | 17779 | 4277 | 5123 | 16867 | 3772 | 3607 |
| 1394 | QGN14491.1 | FIM1_1152 | CP015055. | 1734 | 510 | 513 | 2370 | 411 | 335 | 1937 | 448 | 591 | 531 | 365 | 684 | 1928 | 381 | 645 | 1067 | 349 | 462 |
| 1395 | QGN14492.1 | FIM1_1153 | CP015055. | 141 | 18 | 47 | 202 | 36 | 34 | 109 | 23 | 38 | 200 | 43 | 117 | 306 | 34 | 103 | 262 | 32 | 65 |
| 1396 | QGN14493.1 | FIM1_1154 | CP015055. | 1934 | 846 | 1050 | 1481 | 840 | 930 | 1572 | 848 | 1354 | 4736 | 737 | 1462 | 4856 | 648 | 1438 | 4617 | 620 | 985 |
| 1397 | QGN14494.1 | FIM1_1155 | CP015055. | 97 | 48 | 50 | 475 | 110 | 60 | 121 | 53 | 72 | 176 | 49 | 86 | 287 | 43 | 61 | 239 | 46 | 61 |
| 1398 | QGN14495.1 | FIM1_1156 | CP015055. | 7691 | 8163 | 5324 | 12439 | 6498 | 4726 | 11039 | 5456 | 6525 | 27861 | 8884 | 8325 | 24321 | 8237 | 8886 | 26123 | 7437 | 6278 |
| 1399 | QGN14496.1 | FIM1_1157 | CP015055. | 4170 | 1777 | 1990 | 5350 | 1840 | 1673 | 3699 | 1724 | 2511 | 11928 | 1651 | 2698 | 11183 | 1438 | 2468 | 11552 | 1287 | 1748 |
| 1400 | QGN14497.1 | FIM1_1158 | CP015055. | 3360 | 233 | 944 | 3262 | 266 | 575 | 2778 | 397 | 1093 | 896 | 203 | 1480 | 3504 | 173 | 1270 | 1589 | 181 | 977 |
| 1401 | QGN14498.1 | FIM1_1159 | CP015055. | 1969 | 1320 | 1064 | 2195 | 1297 | 1057 | 1823 | 1132 | 1450 | 6021 | 1107 | 1365 | 5601 | 1144 | 1366 | 5568 | 1123 | 944 |
| 1402 | QGN14499.1 | FIM1_1160 | CP015055. | 24870 | 9679 | 7672 | 30249 | 10356 | 5515 | 22587 | 8557 | 9178 | 30637 | 11672 | 8911 | 38100 | 12118 | 9295 | 34743 | 10271 | 6308 |
| 1403 | QGN14500.1 | FIM1_1161 | CP015055. | 422 | 61 | 48 | 1208 | 24 | 31 | 425 | 84 | 62 | 237 | 47 | 59 | 280 | 82 | 87 | 243 | 34 | 46 |
| 1404 | QGN14501.1 | FIM1_1162 | CP015055. | 9964 | 1415 | 4019 | 29217 | 2031 | 3897 | 19469 | 1500 | 5525 | 15788 | 2586 | 4648 | 21066 | 6372 | 5704 | 16997 | 2084 | 3919 |
| 1405 | QGN14502.1 | FIM1_1163 | CP015055. | 2476 | 646 | 559 | 2529 | 663 | 590 | 2517 | 558 | 823 | 4593 | 641 | 956 | 4162 | 603 | 818 | 4679 | 613 | 606 |
| 1406 | QGN14503.1 | FIM1_1164 | CP015055. | 49 | 24 | 19 | 63 | 24 | 27 | 60 | 21 | 28 | 145 | 17 | 31 | 97 | 30 | 40 | 109 | 27 | 22 |
| 1407 | QGN14504.1 | FIM1_1165 | CP015055. | 399 | 743 | 569 | 885 | 618 | 486 | 687 | 519 | 730 | 3894 | 635 | 847 | 2918 | 557 | 857 | 3816 | 575 | 584 |
| 1408 | QGN14505.1 | FIM1_1166 | CP015055. | 32839 | 3649 | 2411 | 22654 | 4418 | 2449 | 22366 | 3571 | 3186 | 13380 | 3443 | 3030 | 14601 | 3180 | 3009 | 13435 | 2807 | 2164 |
| 1409 | QGN14506.1 | FIM1_1167 | CP015055. | 1627 | 646 | 579 | 2762 | 446 | 465 | 1924 | 512 | 671 | 1918 | 367 | 614 | 2535 | 432 | 641 | 1902 | 360 | 439 |
| 1410 | QGN14507.1 | FIM1_1168 | CP015055. | 2100 | 729 | 916 | 2798 | 1096 | 809 | 2350 | 990 | 1219 | 4311 | 958 | 1214 | 5685 | 891 | 1203 | 4248 | 861 | 887 |
| 1411 | QGN14508.1 | FIM1_1169 | CP015055. | 347 | 508 | 690 | 598 | 610 | 534 | 368 | 461 | 827 | 4250 | 689 | 1150 | 3697 | 769 | 1038 | 4092 | 612 | 706 |
| 1412 | QGN14509.1 | FIM1_1170 | CP015055. | 377 | 904 | 721 | 700 | 717 | 605 | 533 | 633 | 910 | 4075 | 744 | 919 | 3240 | 768 | 1072 | 3919 | 666 | 669 |
| 1413 | QGN14510.1 | FIM1_1171 | CP015055. | 124 | 27 | 31 | 167 | 53 | 24 | 111 | 46 | 37 | 48 | 30 | 29 | 83 | 26 | 18 | 67 | 22 | 19 |
| 1414 | QGN14511.1 | FIM1_1172 | CP015055. | 3999 | 2056 | 2117 | 5149 | 2301 | 1739 | 4327 | 1925 | 2556 | 8527 | 1682 | 2305 | 8383 | 1962 | 2053 | 8178 | 1655 | 1468 |
| 1415 | QGN14512.1 | FIM1_1173 | CP015055. | 228 | 335 | 573 | 369 | 563 | 460 | 251 | 572 | 737 | 2499 | 465 | 515 | 2251 | 297 | 498 | 2766 | 335 | 322 |
| 1416 | QGN14513.1 | FIM1_1174 | CP015055. | 1787 | 173 | 294 | 3663 | 415 | 282 | 1740 | 361 | 367 | 1088 | 211 | 329 | 1610 | 190 | 310 | 1153 | 225 | 206 |
| 1417 | QGN14514.1 | FIM1_1175 | CP015055. | 253 | 78 | 105 | 611 | 145 | 146 | 468 | 94 | 136 | 352 | 62 | 108 | 300 | 42 | 104 | 343 | 66 | 62 |
| 1418 | QGN14515.1 | FIM1_1177 | CP015055. | 2966 | 1246 | 1585 | 3418 | 1357 | 1429 | 2861 | 1201 | 2117 | 8578 | 1426 | 2100 | 6973 | 1413 | 1957 | 8191 | 1269 | 1357 |
| 1419 | QGN14516.1 | FIM1_1178 | CP015055. | 13753 | 10983 | 5792 | 11962 | 10258 | 6026 | 13710 | 8918 | 7520 | 34515 | 12656 | 9957 | 35663 | 10694 | 9415 | 34649 | 12004 | 7222 |
| 1420 | QGN14517.1 | FIM1_1179 | CP015055. | 1368 | 467 | 503 | 1836 | 529 | 504 | 1603 | 371 | 656 | 4244 | 493 | 946 | 4021 | 496 | 826 | 4185 | 475 | 595 |
| 1421 | QGN14518.1 | FIM1_1180 | CP015055. | 3820 | 988 | 1185 | 2539 | 1145 | 955 | 3152 | 1081 | 1507 | 2429 | 779 | 1065 | 2027 | 685 | 978 | 2310 | 678 | 605 |
| 1422 | QGN14519.1 | FIM1_1181 | CP015055. | 3377 | 1859 | 1739 | 3488 | 1256 | 1238 | 2461 | 1524 | 2199 | 4602 | 1155 | 2233 | 6408 | 1947 | 2310 | 4443 | 1236 | 1573 |
| 1423 | QGN14520.1 | FIM1_1182 | CP015055. | 382 | 37 | 41 | 437 | 60 | 44 | 361 | 40 | 51 | 168 | 42 | 53 | 246 | 65 | 51 | 139 | 74 | 43 |
| 1424 | QGN14521.1 | FIM1_1183 | CP015055. | 6620 | 701 | 490 | 4775 | 1045 | 525 | 4777 | 748 | 625 | 1537 | 917 | 400 | 2291 | 1039 | 400 | 1752 | 1107 | 310 |
| 1425 | QGN14522.1 | FIM1_1184 | CP015055. | 498 | 476 | 360 | 560 | 400 | 383 | 684 | 343 | 557 | 2167 | 384 | 547 | 1946 | 367 | 524 | 2087 | 394 | 363 |
| 1426 | QGN14523.1 | FIM1_1185 | CP015055. | 3 | 3 | 2 | 6 | 7 | 1 | 1 | 1 | 4 | 29 | 4 | 6 | 26 | 2 | 3 | 32 | 4 | 5 |

| | | | | | | | | | | | | | | | | | | | | | |
|---|---|---|---|---|---|---|---|---|---|---|---|---|---|---|---|---|---|---|---|---|---|
| 1427 | QGN14524.1 | FIM1_1186 | CP015055. | 1678 | 400 | 391 | 1782 | 368 | 375 | 1719 | 375 | 501 | 2333 | 364 | 584 | 2543 | 335 | 497 | 2290 | 386 | 395 |
| 1428 | QGN14525.1 | FIM1_1187 | CP015055. | 5755 | 3932 | 3406 | 7566 | 4467 | 2828 | 6810 | 3828 | 4418 | 20861 | 4345 | 5425 | 21890 | 4179 | 4686 | 20966 | 4066 | 3670 |
| 1429 | QGN14526.1 | FIM1_1188 | CP015055. | 17135 | 224 | 219 | 7512 | 305 | 175 | 15199 | 276 | 265 | 972 | 211 | 373 | 1571 | 220 | 319 | 1140 | 231 | 236 |
| 1430 | QGN14527.1 | FIM1_1189 | CP015055. | 18128 | 318 | 610 | 8944 | 559 | 458 | 11616 | 522 | 748 | 1377 | 317 | 909 | 2855 | 472 | 830 | 1856 | 319 | 585 |
| 1431 | QGN14528.1 | FIM1_1190 | CP015055. | 13340 | 167 | 263 | 5276 | 142 | 182 | 9154 | 158 | 326 | 604 | 75 | 261 | 834 | 160 | 247 | 652 | 90 | 185 |
| 1432 | QGN14529.1 | FIM1_1191 | CP015055. | 898 | 23 | 15 | 342 | 26 | 15 | 202 | 26 | 15 | 360 | 59 | 139 | 524 | 21 | 20 | 593 | 38 | 41 |
| 1433 | QGN14530.1 | FIM1_1192 | CP015055. | 739 | 1172 | 633 | 600 | 832 | 654 | 540 | 711 | 905 | 3549 | 785 | 905 | 3286 | 684 | 846 | 3104 | 760 | 537 |
| 1434 | QGN14531.1 | FIM1_1193 | CP015055. | 6153 | 3177 | 2577 | 6586 | 3930 | 2395 | 6061 | 3054 | 3332 | 13186 | 3554 | 4234 | 14745 | 3332 | 4208 | 13697 | 3368 | 2929 |
| 1435 | QGN14532.1 | FIM1_1194 | CP015055. | 1078 | 2740 | 1742 | 1449 | 2670 | 1772 | 1209 | 2010 | 2298 | 12023 | 2467 | 2537 | 10449 | 2270 | 2300 | 11760 | 2423 | 1620 |
| 1436 | QGN14533.1 | FIM1_1195 | CP015055. | 188 | 298 | 346 | 229 | 183 | 270 | 153 | 245 | 407 | 1127 | 154 | 306 | 908 | 119 | 341 | 1030 | 125 | 195 |
| 1437 | QGN14534.1 | FIM1_1196 | CP015055. | 13794 | 15617 | 9043 | 13743 | 12170 | 8782 | 11265 | 10277 | 10729 | 36593 | 13785 | 14204 | 31702 | 12378 | 13625 | 36149 | 12575 | 10248 |
| 1438 | QGN14535.1 | FIM1_1197 | CP015055. | 2994 | 3340 | 3451 | 5426 | 2896 | 3170 | 3788 | 2660 | 4404 | 13502 | 2361 | 4474 | 11813 | 2371 | 4712 | 12083 | 2123 | 3077 |
| 1439 | QGN14536.1 | FIM1_1198 | CP015055. | 5060 | 564 | 636 | 3576 | 818 | 595 | 4948 | 734 | 934 | 1879 | 744 | 1106 | 3973 | 1018 | 943 | 2492 | 836 | 728 |
| 1440 | QGN14537.1 | FIM1_1199 | CP015055. | 1577 | 890 | 796 | 2698 | 980 | 682 | 1763 | 837 | 992 | 4272 | 766 | 1265 | 4734 | 637 | 1120 | 4495 | 658 | 815 |
| 1441 | QGN14538.1 | FIM1_1200 | CP015055. | 6801 | 2223 | 2815 | 10723 | 3185 | 1948 | 8195 | 3007 | 3395 | 4624 | 2514 | 3393 | 12284 | 3461 | 2891 | 7812 | 3016 | 2153 |
| 1442 | QGN14539.1 | FIM1_1201 | CP015055. | 355 | 226 | 141 | 325 | 138 | 120 | 290 | 152 | 225 | 272 | 113 | 209 | 483 | 121 | 208 | 325 | 102 | 131 |
| 1443 | QGN14540.1 | FIM1_1202 | CP015055. | 811 | 190 | 253 | 708 | 239 | 190 | 674 | 243 | 318 | 261 | 158 | 276 | 833 | 165 | 279 | 412 | 180 | 174 |
| 1444 | QGN14541.1 | FIM1_1203 | CP015055. | 1229 | 791 | 1446 | 3331 | 522 | 1186 | 2443 | 554 | 1911 | 2889 | 318 | 1126 | 2579 | 759 | 1339 | 2465 | 428 | 760 |
| 1445 | QGN14542.1 | FIM1_1204 | CP015055. | 14000 | 2627 | 2515 | 13806 | 2505 | 2149 | 12565 | 2630 | 3366 | 4937 | 1922 | 3382 | 11254 | 1808 | 3250 | 7282 | 1609 | 2438 |
| 1446 | QGN14543.1 | FIM1_1205 | CP015055. | 1300 | 976 | 781 | 1890 | 645 | 614 | 1710 | 720 | 995 | 1534 | 384 | 627 | 2291 | 345 | 661 | 1731 | 362 | 402 |
| 1447 | QGN14544.1 | FIM1_1206 | CP015055. | 1059 | 401 | 499 | 1605 | 475 | 490 | 1421 | 406 | 664 | 2547 | 522 | 707 | 2429 | 423 | 676 | 2490 | 388 | 478 |
| 1448 | QGN14545.1 | FIM1_1207 | CP015055. | 3931 | 710 | 1058 | 3890 | 1040 | 883 | 3629 | 957 | 1414 | 5644 | 989 | 1743 | 7359 | 897 | 1604 | 5846 | 841 | 1227 |
| 1449 | QGN14546.1 | FIM1_1208 | CP015055. | 1316 | 873 | 1158 | 1464 | 1018 | 936 | 1099 | 917 | 1484 | 8615 | 1097 | 1665 | 7667 | 788 | 1463 | 8309 | 844 | 1098 |
| 1450 | QGN14547.1 | FIM1_1209 | CP015055. | 948 | 678 | 643 | 935 | 683 | 572 | 871 | 739 | 874 | 3167 | 598 | 980 | 4224 | 636 | 816 | 3416 | 644 | 623 |
| 1451 | QGN14548.1 | FIM1_1210 | CP015055. | 274 | 567 | 427 | 206 | 244 | 320 | 208 | 334 | 494 | 1363 | 211 | 509 | 1268 | 144 | 429 | 1310 | 150 | 286 |
| 1452 | QGN14549.1 | FIM1_1211 | CP015055. | 1399 | 965 | 1255 | 2474 | 670 | 1075 | 1866 | 764 | 1651 | 3885 | 482 | 2064 | 4131 | 525 | 1793 | 4103 | 442 | 1382 |
| 1453 | QGN14550.1 | FIM1_1212 | CP015055. | 11120 | 6554 | 4663 | 18326 | 6221 | 4681 | 14774 | 5265 | 5897 | 18323 | 8519 | 7206 | 23161 | 7877 | 7313 | 20061 | 8768 | 5278 |
| 1454 | QGN14551.1 | FIM1_1213 | CP015055. | 8545 | 8785 | 5189 | 6934 | 6193 | 4661 | 6915 | 5725 | 6319 | 17116 | 7681 | 6937 | 16483 | 6510 | 7213 | 16348 | 6460 | 5579 |
| 1455 | QGN14552.1 | FIM1_1214 | CP015055. | 14405 | 14186 | 10458 | 13984 | 13782 | 9098 | 12285 | 11609 | 11998 | 39094 | 16369 | 15515 | 39794 | 14393 | 15796 | 42171 | 13289 | 12003 |
| 1456 | QGN14553.1 | FIM1_1215 | CP015055. | 1834 | 711 | 609 | 1719 | 588 | 522 | 1629 | 541 | 826 | 2162 | 521 | 776 | 2965 | 497 | 734 | 2265 | 515 | 553 |
| 1457 | QGN14554.1 | FIM1_1216 | CP015055. | 1863 | 337 | 508 | 2066 | 582 | 411 | 1789 | 554 | 612 | 2191 | 463 | 827 | 3397 | 542 | 651 | 2406 | 466 | 463 |
| 1458 | QGN14555.1 | FIM1_1217 | CP015055. | 5578 | 2818 | 1224 | 2712 | 3028 | 1277 | 5265 | 2642 | 1610 | 7339 | 3456 | 2469 | 8997 | 3292 | 2410 | 7528 | 4110 | 1723 |
| 1459 | QGN14556.1 | FIM1_1218 | CP015055. | 737 | 193 | 373 | 878 | 155 | 308 | 863 | 217 | 464 | 2028 | 137 | 509 | 1541 | 96 | 505 | 2107 | 131 | 358 |
| 1460 | QGN14557.1 | FIM1_1219 | CP015055. | 2797 | 781 | 723 | 2559 | 784 | 661 | 3005 | 758 | 960 | 4127 | 840 | 928 | 4636 | 799 | 1020 | 4399 | 770 | 663 |
| 1461 | QGN14558.1 | FIM1_1220 | CP015055. | 1504 | 128 | 330 | 1824 | 371 | 292 | 1644 | 327 | 416 | 2120 | 389 | 563 | 2489 | 342 | 478 | 2452 | 286 | 364 |
| 1462 | QGN14559.1 | FIM1_1221 | CP015055. | 2307 | 655 | 506 | 2108 | 558 | 454 | 2261 | 554 | 690 | 2821 | 538 | 834 | 2996 | 465 | 767 | 2790 | 484 | 510 |
| 1463 | QGN14560.1 | FIM1_1222 | CP015055. | 8398 | 2946 | 3659 | 10380 | 3258 | 3021 | 8678 | 3340 | 4824 | 18745 | 3109 | 5381 | 19461 | 2958 | 4884 | 18457 | 2716 | 3647 |
| 1464 | QGN14561.1 | FIM1_1223 | CP015055. | 650 | 159 | 221 | 1249 | 132 | 168 | 691 | 156 | 315 | 1259 | 118 | 347 | 1437 | 146 | 293 | 1379 | 124 | 188 |
| 1465 | QGN14562.1 | FIM1_1224 | CP015055. | 406 | 132 | 218 | 557 | 174 | 186 | 429 | 174 | 295 | 1194 | 156 | 295 | 995 | 130 | 261 | 1208 | 130 | 162 |
| 1466 | QGN14563.1 | FIM1_1225 | CP015055. | 6180 | 674 | 1049 | 7511 | 1306 | 1053 | 6434 | 894 | 1468 | 6414 | 1216 | 1757 | 9188 | 1460 | 1399 | 7294 | 1169 | 1144 |
| 1467 | QGN14564.1 | FIM1_1226 | CP015055. | 18 | 3 | 3 | 36 | 3 | 3 | 7 | 2 | 1 | 15 | 1 | 4 | 16 | 4 | 6 | 15 | 3 | 1 |
| 1468 | QGN14565.1 | FIM1_1227 | CP015055. | 12767 | 2067 | 2347 | 13392 | 2554 | 2005 | 11546 | 2384 | 3257 | 4152 | 1891 | 2844 | 12528 | 1929 | 2938 | 7581 | 1812 | 2129 |
| 1469 | QGN14566.1 | FIM1_1228 | CP015055. | 1294 | 883 | 876 | 3131 | 902 | 771 | 1798 | 902 | 1113 | 4004 | 576 | 1073 | 4245 | 662 | 920 | 3973 | 611 | 734 |
| 1470 | QGN14567.1 | FIM1_1229 | CP015055. | 14 | 7 | 0 | 36 | 4 | 1 | 11 | 3 | 2 | 17 | 4 | 5 | 19 | 3 | 3 | 20 | 2 | 4 |
| 1471 | QGN14568.1 | FIM1_1230 | CP015055. | 2170 | 1018 | 1039 | 1932 | 1466 | 839 | 2160 | 1112 | 1288 | 9166 | 1636 | 1888 | 8013 | 1576 | 1616 | 8864 | 1579 | 1182 |
| 1472 | QGN14569.1 | FIM1_1231 | CP015055. | 6560 | 1439 | 1865 | 5089 | 2434 | 1664 | 6660 | 2137 | 2497 | 9278 | 2765 | 3133 | 12366 | 2577 | 2800 | 10586 | 2603 | 2214 |
| 1473 | QGN14570.1 | FIM1_1232 | CP015055. | 3590 | 2250 | 1849 | 3055 | 2916 | 1443 | 3378 | 3715 | 2366 | 9260 | 2742 | 2341 | 12521 | 2366 | 2111 | 10386 | 2674 | 1728 |
| 1474 | QGN14571.1 | FIM1_1233 | CP015055. | 1003 | 1547 | 721 | 560 | 984 | 690 | 659 | 801 | 958 | 4154 | 953 | 1199 | 3397 | 860 | 1213 | 3931 | 930 | 796 |
| 1475 | QGN14572.1 | FIM1_1234 | CP015055. | 6380 | 7334 | 5178 | 7509 | 6064 | 4815 | 6467 | 5368 | 6632 | 25004 | 7238 | 7401 | 23944 | 6334 | 7571 | 24764 | 6464 | 5033 |
| 1476 | QGN14573.1 | FIM1_1235 | CP015055. | 5847 | 1217 | 1173 | 6509 | 991 | 1024 | 6359 | 1021 | 1511 | 4451 | 944 | 1615 | 4754 | 1142 | 1639 | 4271 | 973 | 1064 |
| 1477 | QGN14574.1 | FIM1_1236 | CP015055. | 960 | 162 | 220 | 838 | 286 | 206 | 668 | 192 | 238 | 675 | 287 | 424 | 1056 | 303 | 402 | 829 | 253 | 251 |
| 1478 | QGN14575.1 | FIM1_1237 | CP015055. | 367 | 9 | 12 | 120 | 4 | 5 | 234 | 10 | 14 | 9 | 4 | 3 | 14 | 1 | 8 | 21 | 6 | 7 |
| 1479 | QGN14576.1 | FIM1_1238 | CP015055. | 12 | 4 | 2 | 25 | 23 | 6 | 5 | 2 | 5 | 40 | 4 | 8 | 42 | 8 | 6 | 33 | 8 | 8 |
| 1480 | QGN14577.1 | FIM1_1239 | CP015055. | 1347 | 3151 | 2227 | 3917 | 2503 | 2109 | 2744 | 2192 | 2903 | 8103 | 1469 | 2507 | 6958 | 1422 | 2510 | 7774 | 1373 | 1771 |
| 1481 | QGN14578.1 | FIM1_1240 | CP015055. | 565 | 249 | 186 | 646 | 144 | 178 | 647 | 164 | 237 | 963 | 116 | 224 | 873 | 85 | 238 | 794 | 124 | 155 |
| 1482 | QGN14579.1 | FIM1_1241 | CP015055. | 21 | 0 | 0 | 85 | 4 | 4 | 21 | 3 | 1 | 8 | 2 | 3 | 7 | 2 | 2 | 9 | 1 | 1 |
| 1483 | QGN14580.1 | FIM1_1242 | CP015055. | 1834 | 549 | 454 | 1364 | 435 | 438 | 1443 | 449 | 569 | 2852 | 389 | 643 | 2712 | 375 | 617 | 2524 | 347 | 446 |
| 1484 | QGN14581.1 | FIM1_1243 | CP015055. | 4609 | 2155 | 2024 | 5034 | 2312 | 1869 | 4513 | 2036 | 2627 | 11541 | 2271 | 3142 | 11307 | 1938 | 2979 | 11869 | 1938 | 2057 |
| 1485 | QGN14582.1 | FIM1_1244 | CP015055. | 992 | 524 | 754 | 1410 | 603 | 617 | 1176 | 568 | 949 | 3903 | 474 | 925 | 3435 | 353 | 903 | 3858 | 333 | 607 |
| 1486 | QGN14583.1 | FIM1_1245 | CP015055. | 2340 | 480 | 677 | 2645 | 795 | 535 | 2504 | 683 | 811 | 5356 | 742 | 1127 | 4641 | 647 | 974 | 5091 | 612 | 720 |
| 1487 | QGN14584.1 | FIM1_1246 | CP015055. | 426 | 449 | 401 | 668 | 339 | 378 | 546 | 396 | 558 | 1816 | 348 | 578 | 2096 | 303 | 550 | 1824 | 289 | 386 |
| 1488 | QGN14585.1 | FIM1_1247 | CP015055. | 3138 | 598 | 826 | 2728 | 498 | 638 | 2782 | 604 | 1076 | 1726 | 335 | 885 | 2326 | 261 | 833 | 1920 | 238 | 517 |
| 1489 | QGN14586.1 | FIM1_1248 | CP015055. | 1402 | 1874 | 1100 | 1285 | 1190 | 1038 | 1336 | 1071 | 1506 | 3285 | 1081 | 1276 | 3524 | 1000 | 1412 | 3251 | 975 | 980 |
| 1490 | QGN14587.1 | FIM1_1249 | CP015055. | 4619 | 2950 | 2413 | 9478 | 3084 | 1905 | 9941 | 3821 | 3364 | 2059 | 2203 | 2466 | 8905 | 2836 | 2135 | 3618 | 2877 | 1591 |
| 1491 | QGN14588.1 | FIM1_1250 | CP015055. | 11458 | 782 | 990 | 13561 | 931 | 902 | 14650 | 928 | 1415 | 3103 | 628 | 1172 | 6115 | 573 | 1156 | 4593 | 474 | 851 |
| 1492 | QGN14589.1 | FIM1_1251 | CP015055. | 8003 | 2994 | 2376 | 10089 | 2293 | 2140 | 8599 | 2067 | 3078 | 8160 | 1859 | 3235 | 10350 | 2307 | 3075 | 8426 | 1965 | 2176 |
| 1493 | QGN14590.1 | FIM1_1252 | CP015055. | 1585 | 2191 | 1155 | 1274 | 1632 | 1090 | 1444 | 1521 | 1470 | 5023 | 1361 | 1381 | 4586 | 1236 | 1426 | 4645 | 1304 | 900 |
| 1494 | QGN14591.1 | FIM1_1253 | CP015055. | 362 | 296 | 335 | 505 | 330 | 310 | 389 | 299 | 419 | 1198 | 218 | 382 | 1689 | 212 | 344 | 1297 | 165 | 240 |
| 1495 | QGN14592.1 | FIM1_1254 | CP015055. | 980 | 1249 | 717 | 926 | 920 | 737 | 910 | 746 | 971 | 5556 | 954 | 1145 | 3830 | 812 | 1055 | 4988 | 936 | 726 |
| 1496 | QGN14593.1 | FIM1_1255 | CP015055. | 2745 | 1407 | 845 | 1293 | 1733 | 853 | 2593 | 1476 | 1117 | 8087 | 1904 | 1560 | 6943 | 1641 | 1448 | 7192 | 1911 | 1059 |
| 1497 | QGN14594.1 | FIM1_1256 | CP015055. | 2115 | 387 | 790 | 2671 | 593 | 634 | 2190 | 493 | 975 | 6993 | 598 | 1334 | 6118 | 472 | 1024 | 6745 | 489 | 866 |
| 1498 | QGN14595.1 | FIM1_1257 | CP015055. | 4547 | 908 | 1258 | 5074 | 1352 | 1105 | 4178 | 1083 | 1621 | 3167 | 939 | 1792 | 5704 | 1000 | 1597 | 4134 | 834 | 1121 |
| 1499 | QGN14596.1 | FIM1_1258 | CP015055. | 1234 | 869 | 1039 | 911 | 719 | 867 | 984 | 732 | 1306 | 4351 | 731 | 1196 | 3825 | 651 | 1241 | 4367 | 578 | 801 |
| 1500 | QGN14597.1 | FIM1_1259 | CP015055. | 1113 | 136 | 341 | 741 | 198 | 264 | 605 | 234 | 433 | 1120 | 118 | 317 | 1221 | 110 | 324 | 1095 | 95 | 198 |
| 1501 | QGN14598.1 | FIM1_1261 | CP015055. | 842 | 2041 | 1091 | 1798 | 1853 | 1095 | 1623 | 1455 | 1353 | 8523 | 2153 | 2013 | 7205 | 1678 | 1951 | 8767 | 1911 | 1373 |
| 1502 | QGN14599.1 | FIM1_1262 | CP015055. | 811 | 1865 | 3659 | 2331 | 1673 | 2816 | 1763 | 1681 | 4397 | 5431 | 1059 | 3440 | 4599 | 1002 | 3274 | 5371 | 854 | 2125 |
| 1503 | QGN14600.1 | FIM1_1263 | CP015055. | 2773 | 2305 | 1562 | 3991 | 1502 | 1277 | 3387 | 1751 | 1952 | 1944 | 1201 | 1869 | 4139 | 1132 | 1955 | 2635 | 1208 | 1278 |
| 1504 | QGN14601.1 | FIM1_1264 | CP015055. | 1157 | 2936 | 2238 | 2545 | 3342 | 2603 | 1224 | 1972 | 2784 | 9254 | 2134 | 2546 | 7775 | 1976 | 2198 | 8640 | 1831 | 1688 |
| 1505 | QGN14602.1 | FIM1_1265 | CP015055. | 9424 | 2280 | 2986 | 11725 | 2920 | 2475 | 10982 | 2493 | 3696 | 6534 | 2767 | 4997 | 16406 | 3108 | 4437 | 10384 | 2815 | 3358 |
| 1506 | QGN14603.1 | FIM1_1266 | CP015055. | 723 | 609 | 394 | 529 | 552 | 377 | 746 | 512 | 543 | 2482 | 673 | 650 | 2505 | 604 | 718 | 2540 | 642 | 497 |
| 1507 | QGN14604.1 | FIM1_1267 | CP015055. | 5673 | 2743 | 1297 | 2620 | 3582 | 1468 | 4720 | 3040 | 1736 | 8055 | 3957 | 2734 | 10759 | 3709 | 2554 | 9237 | 4268 | 1977 |
| 1508 | QGN14605.1 | FIM1_1268 | CP015055. | 1081 | 582 | 554 | 1547 | 538 | 508 | 984 | 415 | 729 | 3015 | 430 | 781 | 2450 | 328 | 781 | 2888 | 380 | 521 |
| 1509 | QGN14606.1 | FIM1_1269 | CP015055. | 6061 | 2442 | 1058 | 2658 | 2736 | 1203 | 5550 | 2356 | 1533 | 8497 | 3467 | 2148 | 9332 | 3113 | 2014 | 8436 | 3752 | 1546 |
| 1510 | QGN14607.1 | FIM1_1270 | CP015055. | 2612 | 2015 | 1758 | 2342 | 2203 | 1509 | 3031 | 2112 | 2170 | 6265 | 2299 | 2651 | 7685 | 2094 | 2395 | 6203 | 2186 | 1802 |

| | | | | | | | | | | | | | | | | | | | | | |
|---|---|---|---|---|---|---|---|---|---|---|---|---|---|---|---|---|---|---|---|---|---|
| 1511 | QGN14608.1 | FIM1_1271 | CP015055. | 2910 | 2085 | 2069 | 3919 | 1733 | 1458 | 2660 | 1853 | 2498 | 13894 | 2188 | 3051 | 13334 | 2543 | 3293 | 13057 | 1741 | 2179 |
| 1512 | QGN14609.1 | FIM1_1272 | CP015055. | 18153 | 5543 | 3630 | 14994 | 4557 | 3144 | 14068 | 4495 | 4628 | 15729 | 5300 | 5904 | 17842 | 5354 | 6012 | 15044 | 5244 | 4241 |
| 1513 | QGN14610.1 | FIM1_1273 | CP015055. | 1269 | 286 | 390 | 776 | 236 | 379 | 785 | 302 | 574 | 1217 | 255 | 505 | 1919 | 242 | 467 | 1625 | 251 | 389 |
| 1514 | QGN14611.1 | FIM1_1274 | CP015055. | 230 | 22 | 60 | 181 | 36 | 47 | 170 | 64 | 53 | 136 | 30 | 60 | 244 | 25 | 76 | 163 | 30 | 39 |
| 1515 | QGN14612.1 | FIM1_1275 | CP015055. | 2168 | 1897 | 2269 | 2029 | 2031 | 1909 | 1552 | 2127 | 3080 | 6216 | 1601 | 2734 | 10064 | 1393 | 2692 | 7968 | 1433 | 1790 |
| 1516 | QGN14613.1 | FIM1_1276 | CP015055. | 6019 | 2210 | 2249 | 6984 | 2127 | 2104 | 6039 | 1893 | 3028 | 7813 | 1590 | 3208 | 8817 | 1521 | 3132 | 7832 | 1424 | 2100 |
| 1517 | QGN14614.1 | FIM1_1277 | CP015055. | 5532 | 728 | 1429 | 8294 | 1176 | 1201 | 6215 | 1059 | 1870 | 5721 | 847 | 2547 | 8533 | 1035 | 2234 | 6914 | 808 | 1573 |
| 1518 | QGN14615.1 | FIM1_1279 | CP015055. | 1888 | 601 | 970 | 2529 | 807 | 859 | 1572 | 684 | 1298 | 3987 | 548 | 1105 | 3583 | 586 | 1000 | 3802 | 442 | 675 |
| 1519 | QGN14616.1 | FIM1_1280 | CP015055. | 3916 | 4113 | 7926 | 6616 | 4067 | 6921 | 5072 | 3913 | 9661 | 17932 | 3300 | 11005 | 15612 | 3470 | 10375 | 17044 | 3010 | 7439 |
| 1520 | QGN14617.1 | FIM1_1281 | CP015055. | 1065 | 2003 | 1314 | 1066 | 1664 | 1307 | 1018 | 1445 | 1876 | 6901 | 1630 | 1787 | 6974 | 1516 | 1660 | 6710 | 1560 | 1189 |
| 1521 | QGN14618.1 | FIM1_1282 | CP015055. | 1868 | 136 | 438 | 2394 | 451 | 299 | 1914 | 499 | 516 | 909 | 313 | 599 | 2463 | 361 | 543 | 1413 | 343 | 361 |
| 1522 | QGN14619.1 | FIM1_1283 | CP015055. | 10001 | 1161 | 2627 | 21360 | 3083 | 3669 | 14004 | 1608 | 3341 | 7386 | 1495 | 4072 | 8408 | 1866 | 4305 | 7610 | 1541 | 2808 |
| 1523 | QGN14620.1 | FIM1_1284 | CP015055. | 5044 | 230 | 391 | 3515 | 299 | 291 | 2943 | 300 | 460 | 1088 | 327 | 567 | 2097 | 366 | 608 | 1425 | 335 | 378 |
| 1524 | QGN14621.1 | FIM1_1285 | CP015055. | 10343 | 1705 | 2114 | 10067 | 2501 | 1867 | 9541 | 2126 | 2832 | 14734 | 2788 | 4151 | 16304 | 2892 | 3651 | 15771 | 2632 | 2659 |
| 1525 | QGN14622.1 | FIM1_1286 | CP015055. | 226 | 346 | 284 | 306 | 225 | 248 | 288 | 230 | 371 | 1734 | 318 | 452 | 1277 | 201 | 424 | 1757 | 259 | 268 |
| 1526 | QGN14623.1 | FIM1_1287 | CP015055. | 455 | 798 | 1066 | 1784 | 823 | 944 | 1163 | 833 | 1397 | 5197 | 614 | 1560 | 5550 | 800 | 1418 | 5048 | 597 | 1020 |
| 1527 | QGN14624.1 | FIM1_1288 | CP015055. | 635 | 648 | 341 | 486 | 533 | 334 | 415 | 473 | 481 | 1589 | 379 | 482 | 1936 | 417 | 451 | 1613 | 370 | 300 |
| 1528 | QGN14625.1 | FIM1_1289 | CP015055. | 590 | 1996 | 1349 | 754 | 1922 | 1215 | 690 | 1575 | 1691 | 7494 | 1707 | 2050 | 7985 | 1676 | 1814 | 7408 | 1596 | 1348 |
| 1529 | QGN14626.1 | FIM1_1290 | CP015055. | 5356 | 1910 | 2900 | 10199 | 2743 | 2845 | 7319 | 2021 | 3795 | 11636 | 2344 | 4237 | 14633 | 2260 | 3844 | 12633 | 2028 | 2994 |
| 1530 | QGN14627.1 | FIM1_1291 | CP015055. | 180 | 624 | 864 | 452 | 799 | 739 | 212 | 733 | 1154 | 4666 | 828 | 1069 | 3801 | 663 | 1001 | 4454 | 639 | 650 |
| 1531 | QGN14628.1 | FIM1_1292 | CP015055. | 268 | 167 | 90 | 245 | 140 | 102 | 235 | 110 | 138 | 1430 | 163 | 236 | 902 | 179 | 190 | 1335 | 152 | 135 |
| 1532 | QGN14629.1 | FIM1_1293 | CP015055. | 3517 | 1459 | 924 | 2244 | 1776 | 945 | 3595 | 1370 | 1267 | 4837 | 1982 | 1901 | 6580 | 2222 | 1800 | 5304 | 2304 | 1421 |
| 1533 | QGN14630.1 | FIM1_1294 | CP015055. | 1259 | 754 | 458 | 674 | 1090 | 447 | 1368 | 832 | 601 | 4345 | 981 | 938 | 3925 | 860 | 887 | 4330 | 1010 | 616 |
| 1534 | QGN14631.1 | FIM1_1295 | CP015055. | 10531 | 2366 | 2133 | 8428 | 2288 | 1877 | 8325 | 2467 | 2821 | 4203 | 1524 | 3155 | 6447 | 1325 | 3002 | 4587 | 1262 | 1974 |
| 1535 | QGN14632.1 | FIM1_1296 | CP015055. | 4965 | 264 | 598 | 3223 | 502 | 427 | 3461 | 484 | 784 | 276 | 142 | 797 | 2122 | 156 | 644 | 701 | 189 | 489 |
| 1536 | QGN14633.1 | FIM1_1297 | CP015055. | 9908 | 556 | 1094 | 18492 | 746 | 818 | 9493 | 567 | 1247 | 5735 | 547 | 1563 | 4476 | 887 | 1112 | 5641 | 857 | 885 |
| 1537 | QGN14634.1 | FIM1_1298 | CP015055. | 2286 | 998 | 2400 | 7495 | 1177 | 1849 | 4823 | 997 | 2934 | 9070 | 946 | 3636 | 10266 | 1137 | 3181 | 9402 | 882 | 2408 |
| 1538 | QGN14635.1 | FIM1_1299 | CP015055. | 1348 | 500 | 655 | 1508 | 515 | 560 | 1155 | 550 | 822 | 4565 | 537 | 957 | 4045 | 465 | 848 | 4104 | 526 | 577 |
| 1539 | QGN14636.1 | FIM1_1300 | CP015055. | 2839 | 534 | 643 | 2625 | 930 | 595 | 2760 | 790 | 850 | 5630 | 936 | 1126 | 5137 | 960 | 1028 | 5245 | 963 | 822 |
| 1540 | QGN14637.1 | FIM1_1301 | CP015055. | 420 | 232 | 271 | 784 | 212 | 211 | 597 | 196 | 320 | 1648 | 165 | 315 | 1290 | 139 | 297 | 1460 | 140 | 185 |
| 1541 | QGN14638.1 | FIM1_1302 | CP015055. | 1701 | 349 | 336 | 1791 | 423 | 321 | 1663 | 321 | 445 | 3396 | 357 | 683 | 2684 | 346 | 566 | 3137 | 366 | 466 |
| 1542 | QGN14639.1 | FIM1_1303 | CP015055. | 2711 | 1710 | 691 | 1953 | 1335 | 788 | 2535 | 1043 | 1011 | 5860 | 1805 | 1527 | 4904 | 1452 | 1428 | 5935 | 1917 | 962 |
| 1543 | QGN14640.1 | FIM1_1304 | CP015055. | 4670 | 795 | 912 | 3878 | 834 | 563 | 4164 | 919 | 1122 | 1031 | 743 | 1382 | 4900 | 757 | 1144 | 2452 | 788 | 895 |
| 1544 | QGN14641.1 | FIM1_1305 | CP015055. | 9263 | 2232 | 2482 | 13193 | 2604 | 1921 | 10674 | 2348 | 3050 | 11397 | 2163 | 3774 | 13722 | 2878 | 3423 | 12039 | 2327 | 2525 |
| 1545 | QGN14642.1 | FIM1_1306 | CP015055. | 317 | 81 | 62 | 333 | 48 | 64 | 314 | 42 | 106 | 698 | 46 | 103 | 407 | 41 | 92 | 576 | 48 | 74 |
| 1546 | QGN14643.1 | FIM1_1307 | CP015055. | 625 | 523 | 688 | 1115 | 649 | 580 | 639 | 537 | 885 | 4960 | 649 | 1081 | 4893 | 622 | 952 | 5403 | 582 | 682 |
| 1547 | QGN14644.1 | FIM1_1308 | CP015055. | 1916 | 495 | 451 | 1628 | 587 | 418 | 1912 | 402 | 567 | 3202 | 538 | 696 | 2867 | 498 | 660 | 2998 | 484 | 474 |
| 1548 | QGN14645.1 | FIM1_1309 | CP015055. | 1030 | 839 | 748 | 1888 | 607 | 658 | 1505 | 640 | 1071 | 3116 | 573 | 957 | 3037 | 811 | 1056 | 2591 | 625 | 659 |
| 1549 | QGN14646.1 | FIM1_1310 | CP015055. | 1291 | 1691 | 1146 | 2369 | 1534 | 965 | 1912 | 1367 | 1438 | 6221 | 1472 | 1761 | 7055 | 1536 | 1689 | 6164 | 1458 | 1224 |
| 1550 | QGN14647.1 | FIM1_1311 | CP015055. | 9130 | 265 | 316 | 14869 | 293 | 272 | 13591 | 195 | 428 | 1780 | 527 | 725 | 3351 | 1064 | 626 | 1882 | 1121 | 475 |
| 1551 | QGN14648.1 | FIM1_1312 | CP015055. | 2456 | 1446 | 1548 | 6363 | 2249 | 1674 | 3333 | 1350 | 1994 | 3763 | 688 | 1508 | 4990 | 1053 | 1336 | 3915 | 778 | 978 |
| 1552 | QGN14649.1 | FIM1_1313 | CP015055. | 9385 | 21780 | 26077 | 20660 | 24288 | 21632 | 14408 | 20423 | 29689 | 54354 | 25511 | 38126 | 82962 | 25134 | 33574 | 68232 | 22018 | 26293 |
| 1553 | QGN14650.1 | FIM1_1314 | CP015055. | 132 | 1120 | 539 | 336 | 828 | 459 | 279 | 824 | 631 | 3699 | 675 | 632 | 3258 | 576 | 687 | 3194 | 520 | 432 |
| 1554 | QGN14651.1 | FIM1_1315 | CP015055. | 2055 | 3284 | 1972 | 3932 | 2374 | 1833 | 3356 | 1995 | 2512 | 11248 | 2928 | 3008 | 8279 | 2738 | 3073 | 11356 | 2730 | 2190 |
| 1555 | QGN14652.1 | FIM1_1316 | CP015055. | 272 | 785 | 923 | 579 | 1109 | 859 | 357 | 882 | 1299 | 2823 | 627 | 640 | 3978 | 741 | 728 | 3162 | 401 | 466 |
| 1556 | QGN14653.1 | FIM1_1317 | CP015055. | 272 | 815 | 447 | 243 | 642 | 428 | 327 | 569 | 600 | 972 | 441 | 610 | 2087 | 477 | 493 | 1326 | 428 | 369 |
| 1557 | QGN14654.1 | FIM1_1318 | CP015055. | 1354 | 1465 | 1143 | 1198 | 982 | 974 | 1260 | 884 | 1496 | 5143 | 893 | 1430 | 4104 | 882 | 1398 | 4670 | 797 | 956 |
| 1558 | QGN14655.1 | FIM1_1319 | CP015055. | 4617 | 2230 | 1989 | 9496 | 2321 | 1647 | 6413 | 2616 | 2730 | 3503 | 1605 | 3138 | 10335 | 2096 | 2614 | 5426 | 1684 | 2138 |
| 1559 | QGN14656.1 | FIM1_1320 | CP015055. | 9839 | 6228 | 7607 | 12352 | 10881 | 6130 | 11948 | 10326 | 9085 | 18095 | 11547 | 12029 | 43117 | 11189 | 11721 | 31202 | 9221 | 7932 |
| 1560 | QGN14657.1 | FIM1_1321 | CP015055. | 1075 | 464 | 499 | 2042 | 491 | 400 | 1282 | 407 | 592 | 3006 | 490 | 739 | 2830 | 466 | 692 | 2725 | 428 | 473 |
| 1561 | QGN14658.1 | FIM1_1322 | CP015055. | 2397 | 5887 | 5402 | 9176 | 5089 | 4465 | 5806 | 4997 | 6694 | 22327 | 5058 | 9026 | 23671 | 6665 | 8316 | 22142 | 5628 | 6163 |
| 1562 | QGN14659.1 | FIM1_1323 | CP015055. | 5537 | 293 | 371 | 6135 | 235 | 250 | 5420 | 240 | 388 | 2190 | 199 | 442 | 1885 | 225 | 362 | 1884 | 152 | 264 |
| 1563 | QGN14660.1 | FIM1_1324 | CP015055. | 1143 | 1169 | 524 | 620 | 880 | 567 | 1026 | 775 | 720 | 4302 | 979 | 892 | 2941 | 854 | 850 | 4149 | 912 | 562 |
| 1564 | QGN14661.1 | FIM1_1325 | CP015055. | 7055 | 933 | 1546 | 6582 | 1270 | 1234 | 6963 | 1249 | 2020 | 5039 | 695 | 1369 | 6617 | 706 | 1191 | 5518 | 619 | 896 |
| 1565 | QGN14662.1 | FIM1_1326 | CP015055. | 2926 | 641 | 667 | 3210 | 555 | 558 | 3044 | 514 | 905 | 3224 | 451 | 879 | 3100 | 463 | 843 | 3201 | 376 | 565 |
| 1566 | QGN14663.1 | FIM1_1327 | CP015055. | 8617 | 746 | 1057 | 7802 | 934 | 946 | 7742 | 766 | 1455 | 6157 | 917 | 1986 | 7595 | 977 | 1749 | 6471 | 858 | 1368 |
| 1567 | QGN14664.1 | FIM1_1328 | CP015055. | 3346 | 750 | 1044 | 3837 | 801 | 846 | 3201 | 766 | 1386 | 5025 | 714 | 1455 | 5201 | 656 | 1387 | 4957 | 541 | 926 |
| 1568 | QGN14665.1 | FIM1_1329 | CP015055. | 21655 | 417 | 500 | 10163 | 523 | 411 | 11154 | 410 | 718 | 1771 | 303 | 714 | 2981 | 388 | 586 | 2166 | 314 | 422 |
| 1569 | QGN14666.1 | FIM1_1331 | CP015055. | 204 | 15 | 100 | 320 | 72 | 85 | 206 | 73 | 144 | 622 | 47 | 164 | 822 | 60 | 135 | 635 | 39 | 127 |
| 1570 | QGN14667.1 | FIM1_1332 | CP015055. | 864 | 316 | 511 | 725 | 320 | 418 | 589 | 320 | 690 | 1436 | 276 | 700 | 1974 | 259 | 634 | 1713 | 221 | 472 |
| 1571 | QGN14668.1 | FIM1_1333 | CP015055. | 7428 | 8084 | 2769 | 4573 | 5653 | 3002 | 7080 | 5296 | 4053 | 17879 | 7471 | 4755 | 19207 | 5520 | 5079 | 16525 | 6715 | 3665 |
| 1572 | QGN14669.1 | FIM1_1334 | CP015055. | 2893 | 2824 | 2684 | 3419 | 2296 | 1847 | 4301 | 2809 | 3062 | 3815 | 1926 | 4254 | 9701 | 1534 | 3308 | 6331 | 1739 | 2684 |
| 1573 | QGN14670.1 | FIM1_1335 | CP015055. | 1905 | 944 | 733 | 1103 | 706 | 614 | 1608 | 766 | 953 | 1534 | 567 | 1033 | 2523 | 554 | 1018 | 1858 | 477 | 709 |
| 1574 | QGN14671.1 | FIM1_1336 | CP015055. | 4994 | 326 | 388 | 4477 | 549 | 328 | 6451 | 492 | 567 | 670 | 698 | 547 | 2169 | 1216 | 573 | 1163 | 1129 | 428 |
| 1575 | QGN14672.1 | FIM1_1337 | CP015055. | 5859 | 2936 | 1009 | 2208 | 2838 | 1146 | 4874 | 2413 | 1447 | 6268 | 3288 | 1975 | 8802 | 2777 | 1782 | 7131 | 4021 | 1381 |
| 1576 | QGN14673.1 | FIM1_1338 | CP015055. | 4500 | 437 | 549 | 3463 | 326 | 363 | 3636 | 380 | 701 | 3200 | 260 | 584 | 3023 | 337 | 628 | 2819 | 199 | 384 |
| 1577 | QGN14674.1 | FIM1_1339 | CP015055. | 6808 | 2773 | 2721 | 10026 | 4313 | 2426 | 7383 | 3964 | 3623 | 7467 | 3500 | 3981 | 17529 | 3433 | 3961 | 11363 | 3114 | 2704 |
| 1578 | QGN14675.1 | FIM1_1340 | CP015055. | 1711 | 2170 | 1469 | 2620 | 1788 | 1261 | 2114 | 1787 | 1890 | 4924 | 1462 | 1715 | 7246 | 1229 | 1925 | 5053 | 1189 | 1273 |
| 1579 | QGN14676.1 | FIM1_1341 | CP015055. | 215 | 497 | 566 | 809 | 453 | 502 | 375 | 458 | 836 | 2920 | 408 | 669 | 2276 | 344 | 674 | 2588 | 333 | 473 |
| 1580 | QGN14677.1 | FIM1_1342 | CP015055. | 1347 | 1197 | 1563 | 2617 | 1708 | 1319 | 1866 | 1468 | 1992 | 5732 | 1676 | 2332 | 9623 | 1583 | 2047 | 7602 | 1498 | 1555 |
| 1581 | QGN14678.1 | FIM1_1344 | CP015055. | 1678 | 181 | 226 | 1984 | 199 | 183 | 1935 | 207 | 299 | 1238 | 154 | 281 | 1331 | 147 | 290 | 1208 | 124 | 173 |
| 1582 | QGN14679.1 | FIM1_1345 | CP015055. | 3807 | 1117 | 976 | 6232 | 1546 | 942 | 4587 | 1116 | 1263 | 3806 | 663 | 701 | 3787 | 722 | 715 | 3480 | 542 | 498 |
| 1583 | QGN14680.1 | FIM1_1346 | CP015055. | 4986 | 6299 | 4501 | 8659 | 6211 | 3887 | 7844 | 5127 | 5487 | 23098 | 6638 | 6981 | 20622 | 6579 | 6960 | 22786 | 6114 | 4922 |
| 1584 | QGN14681.1 | FIM1_1347 | CP015055. | 13499 | 16154 | 10475 | 13267 | 12043 | 10368 | 11589 | 10336 | 12089 | 34666 | 14494 | 14559 | 30650 | 12818 | 14932 | 36153 | 13262 | 11179 |
| 1585 | QGN14682.1 | FIM1_1348 | CP015055. | 8522 | 7033 | 5520 | 12144 | 7770 | 5126 | 10248 | 6227 | 6867 | 34190 | 7681 | 9069 | 30418 | 7154 | 8151 | 33150 | 7303 | 5796 |
| 1586 | QGN14683.1 | FIM1_1349 | CP015055. | 1439 | 659 | 940 | 1589 | 790 | 790 | 1227 | 761 | 1219 | 5286 | 736 | 1451 | 5829 | 562 | 1342 | 5215 | 572 | 935 |
| 1587 | QGN14684.1 | FIM1_1350 | CP015055. | 9728 | 7419 | 4627 | 8092 | 5206 | 4453 | 7221 | 4039 | 5850 | 22635 | 7086 | 7869 | 18723 | 6062 | 7744 | 22591 | 6091 | 5788 |
| 1588 | QGN14685.1 | FIM1_1352 | CP015055. | 2131 | 1926 | 2302 | 3145 | 1853 | 1996 | 2292 | 1672 | 2667 | 6901 | 1467 | 2613 | 6186 | 1172 | 2500 | 6601 | 1165 | 1747 |
| 1589 | QGN14686.1 | FIM1_1353 | CP015055. | 6346 | 4153 | 6854 | 10539 | 4767 | 5932 | 7495 | 4339 | 8569 | 21827 | 4445 | 10505 | 22405 | 4330 | 9556 | 21764 | 3977 | 6952 |
| 1590 | QGN14687.1 | FIM1_1354 | CP015055. | 764 | 781 | 684 | 1624 | 630 | 584 | 1196 | 620 | 896 | 4714 | 665 | 1119 | 3591 | 571 | 1016 | 4318 | 600 | 677 |
| 1591 | QGN14688.1 | FIM1_1355 | CP015055. | 3264 | 980 | 892 | 3590 | 953 | 838 | 3450 | 816 | 1218 | 4961 | 846 | 1455 | 5963 | 784 | 1274 | 5051 | 777 | 895 |
| 1592 | QGN14689.1 | FIM1_1356 | CP015055. | 958 | 425 | 463 | 1267 | 460 | 408 | 894 | 425 | 596 | 3484 | 324 | 709 | 2739 | 338 | 644 | 3294 | 355 | 469 |
| 1593 | QGN14690.1 | FIM1_1357 | CP015055. | 805 | 771 | 793 | 1797 | 668 | 683 | 1275 | 604 | 925 | 3559 | 572 | 960 | 3144 | 632 | 1050 | 3356 | 513 | 638 |
| 1594 | QGN14691.1 | FIM1_1358 | CP015055. | 374 | 906 | 802 | 633 | 1194 | 716 | 390 | 937 | 1025 | 3537 | 1132 | 1216 | 4533 | 1066 | 1111 | 4064 | 1071 | 774 |

| | | | | | | | | | | | | | | | | | | | | | |
|---|---|---|---|---|---|---|---|---|---|---|---|---|---|---|---|---|---|---|---|---|---|
| 1595 | QGN14692.1 | FIM1_1359 | CP015055. | 769 | 386 | 401 | 940 | 318 | 264 | 622 | 373 | 464 | 820 | 240 | 424 | 1433 | 255 | 384 | 1038 | 248 | 293 |
| 1596 | QGN14693.1 | FIM1_1360 | CP015055. | 3431 | 884 | 1042 | 8136 | 1732 | 903 | 4798 | 1088 | 1401 | 6751 | 1842 | 1551 | 14645 | 2011 | 1549 | 10151 | 1278 | 1114 |
| 1597 | QGN14694.1 | FIM1_1361 | CP015055. | 1096 | 103 | 118 | 1272 | 122 | 136 | 933 | 71 | 189 | 1120 | 113 | 265 | 891 | 172 | 237 | 975 | 145 | 175 |
| 1598 | QGN14695.1 | FIM1_1362 | CP015055. | 1424 | 328 | 329 | 1763 | 205 | 279 | 1167 | 226 | 400 | 1504 | 237 | 496 | 1227 | 253 | 461 | 1405 | 227 | 311 |
| 1599 | QGN14696.1 | FIM1_1363 | CP015055. | 5337 | 2347 | 1309 | 2353 | 2870 | 1357 | 4756 | 2296 | 1765 | 12964 | 3398 | 2569 | 10109 | 3227 | 2315 | 12380 | 3747 | 1736 |
| 1600 | QGN14697.1 | FIM1_1364 | CP015055. | 310 | 223 | 197 | 227 | 193 | 199 | 313 | 187 | 244 | 1438 | 163 | 319 | 929 | 161 | 288 | 1404 | 173 | 187 |
| 1601 | QGN14698.1 | FIM1_1365 | CP015055. | 2809 | 1371 | 1429 | 6078 | 1522 | 1389 | 4206 | 1043 | 2016 | 7323 | 1190 | 2421 | 7643 | 1553 | 2151 | 7393 | 1282 | 1429 |
| 1602 | QGN14699.1 | FIM1_1366 | CP015055. | 1124 | 1223 | 757 | 1135 | 1011 | 811 | 1091 | 764 | 1052 | 5678 | 976 | 1127 | 4491 | 958 | 1103 | 5477 | 1062 | 779 |
| 1603 | QGN14700.1 | FIM1_1367 | CP015055. | 1834 | 528 | 680 | 2730 | 520 | 578 | 2147 | 508 | 970 | 3796 | 386 | 873 | 3625 | 421 | 791 | 3649 | 307 | 613 |
| 1604 | QGN14701.1 | FIM1_1368 | CP015055. | 7541 | 3104 | 3382 | 11480 | 3238 | 3030 | 10127 | 2784 | 4367 | 15992 | 3147 | 6170 | 15903 | 3472 | 5970 | 16535 | 3189 | 4240 |
| 1605 | QGN14702.1 | FIM1_1369 | CP015055. | 17700 | 1151 | 769 | 16542 | 1639 | 869 | 18119 | 1074 | 1028 | 7423 | 1726 | 1375 | 6507 | 2901 | 1318 | 6722 | 2387 | 983 |
| 1606 | QGN14703.1 | FIM1_1370 | CP015055. | 114 | 21 | 4 | 173 | 19 | 11 | 125 | 11 | 10 | 86 | 14 | 13 | 48 | 31 | 9 | 59 | 25 | 9 |
| 1607 | QGN14704.1 | FIM1_1371 | CP015055. | 1633 | 1135 | 913 | 2047 | 878 | 976 | 1702 | 865 | 1294 | 2174 | 528 | 1142 | 2582 | 562 | 1152 | 2251 | 603 | 715 |
| 1608 | QGN14705.1 | FIM1_1372 | CP015055. | 257 | 325 | 606 | 159 | 204 | 455 | 159 | 239 | 716 | 362 | 77 | 321 | 348 | 62 | 293 | 350 | 60 | 176 |
| 1609 | QGN14706.1 | FIM1_1373 | CP015055. | 5408 | 1403 | 1877 | 7697 | 1617 | 1636 | 5885 | 1331 | 2234 | 4170 | 679 | 1434 | 4571 | 892 | 1293 | 3842 | 775 | 933 |
| 1610 | QGN14707.1 | FIM1_1374 | CP015055. | 1733 | 1699 | 1634 | 2694 | 1880 | 1422 | 2335 | 1653 | 2108 | 8512 | 1350 | 2237 | 7207 | 1265 | 1969 | 8361 | 1119 | 1367 |
| 1611 | QGN14708.1 | FIM1_1375 | CP015055. | 5620 | 397 | 814 | 5518 | 564 | 521 | 5406 | 639 | 942 | 944 | 543 | 1528 | 5304 | 600 | 1336 | 2094 | 584 | 1059 |
| 1612 | QGN14709.1 | FIM1_1376 | CP015055. | 237 | 77 | 66 | 302 | 59 | 41 | 336 | 73 | 85 | 303 | 46 | 111 | 354 | 35 | 75 | 356 | 42 | 68 |
| 1613 | QGN14710.1 | FIM1_1377 | CP015055. | 7827 | 2068 | 1723 | 8980 | 2447 | 1689 | 8820 | 2204 | 2450 | 10433 | 2371 | 2690 | 13076 | 1998 | 2435 | 10736 | 2100 | 1841 |
| 1614 | QGN14711.1 | FIM1_1378 | CP015055. | 511 | 255 | 191 | 524 | 259 | 180 | 479 | 186 | 255 | 1072 | 245 | 333 | 966 | 182 | 308 | 1130 | 210 | 196 |
| 1615 | QGN14712.1 | FIM1_1379 | CP015055. | 1073 | 31 | 149 | 1264 | 64 | 113 | 1237 | 86 | 168 | 614 | 66 | 233 | 841 | 79 | 214 | 635 | 43 | 165 |
| 1616 | QGN14713.1 | FIM1_1380 | CP015055. | 9684 | 531 | 1071 | 13797 | 871 | 776 | 13049 | 1045 | 1275 | 2289 | 733 | 1292 | 6243 | 749 | 1199 | 4020 | 716 | 883 |
| 1617 | QGN14714.1 | FIM1_1382 | CP015055. | 5960 | 2559 | 2204 | 5842 | 2759 | 1991 | 5949 | 2525 | 3081 | 12483 | 2442 | 3509 | 13108 | 2229 | 3320 | 13304 | 2225 | 2452 |
| 1618 | QGN14715.1 | FIM1_1383 | CP015055. | 2951 | 2237 | 1634 | 4875 | 1690 | 1405 | 3849 | 1561 | 2156 | 7518 | 1376 | 2555 | 7924 | 1531 | 2479 | 7835 | 1273 | 1717 |
| 1619 | QGN14716.1 | FIM1_1384 | CP015055. | 2046 | 1085 | 676 | 2041 | 870 | 682 | 1708 | 630 | 815 | 4468 | 796 | 913 | 3204 | 726 | 929 | 4078 | 858 | 619 |
| 1620 | QGN14717.1 | FIM1_1385 | CP015055. | 14974 | 2347 | 3405 | 18652 | 2388 | 2560 | 14863 | 2319 | 4191 | 9002 | 1916 | 4561 | 12880 | 2771 | 4119 | 10457 | 2097 | 2913 |
| 1621 | QGN14718.1 | FIM1_1386 | CP015055. | 546 | 346 | 401 | 642 | 441 | 313 | 531 | 431 | 550 | 1199 | 341 | 408 | 2140 | 341 | 416 | 1432 | 354 | 302 |
| 1622 | QGN14719.1 | FIM1_1387 | CP015055. | 3439 | 1938 | 1840 | 3767 | 1992 | 1602 | 3541 | 1886 | 2301 | 9493 | 1926 | 2726 | 9735 | 1906 | 2387 | 9209 | 1804 | 1734 |
| 1623 | QGN14720.1 | FIM1_1388 | CP015055. | 806 | 214 | 248 | 1045 | 206 | 222 | 839 | 171 | 323 | 1552 | 151 | 404 | 1328 | 144 | 337 | 1530 | 162 | 244 |
| 1624 | QGN14721.1 | FIM1_1389 | CP015055. | 798 | 482 | 572 | 874 | 409 | 514 | 663 | 414 | 726 | 2257 | 317 | 727 | 2002 | 241 | 673 | 2202 | 249 | 427 |
| 1625 | QGN14722.1 | FIM1_1390 | CP015055. | 6207 | 1051 | 1743 | 7207 | 1356 | 1370 | 6928 | 1395 | 2146 | 3195 | 1418 | 2481 | 9560 | 1281 | 2348 | 5776 | 1242 | 1674 |
| 1626 | QGN14723.1 | FIM1_1391 | CP015055. | 8080 | 2181 | 2635 | 8425 | 1835 | 1912 | 7556 | 2194 | 3576 | 882 | 1148 | 5292 | 7699 | 1265 | 4706 | 2587 | 1257 | 3418 |
| 1627 | QGN14724.1 | FIM1_1392 | CP015055. | 2991 | 631 | 907 | 3392 | 567 | 772 | 2781 | 517 | 1173 | 1142 | 386 | 994 | 2228 | 549 | 1164 | 1549 | 503 | 710 |
| 1628 | QGN14725.1 | FIM1_1393 | CP015055. | 1124 | 1077 | 914 | 1559 | 625 | 739 | 1090 | 519 | 1200 | 2486 | 479 | 1119 | 2564 | 722 | 1177 | 2332 | 555 | 771 |
| 1629 | QGN14726.1 | FIM1_1394 | CP015055. | 3340 | 180 | 579 | 4208 | 117 | 352 | 4267 | 275 | 694 | 348 | 83 | 510 | 869 | 104 | 510 | 539 | 69 | 316 |
| 1630 | QGN14727.1 | FIM1_1395 | CP015055. | 1200 | 275 | 503 | 1419 | 383 | 404 | 1277 | 362 | 686 | 834 | 305 | 808 | 3436 | 244 | 693 | 1900 | 324 | 560 |
| 1631 | QGN14728.1 | FIM1_1396 | CP015055. | 846 | 847 | 773 | 1895 | 935 | 661 | 1461 | 872 | 975 | 2685 | 818 | 1320 | 4940 | 962 | 1321 | 3426 | 755 | 874 |
| 1632 | QGN14729.1 | FIM1_1397 | CP015055. | 3064 | 3842 | 3559 | 5925 | 2937 | 3326 | 4038 | 2610 | 4557 | 9496 | 2814 | 4739 | 11054 | 3141 | 5374 | 9665 | 2505 | 3447 |
| 1633 | QGN14730.1 | FIM1_1399 | CP015055. | 10020 | 1890 | 1603 | 8316 | 1529 | 1360 | 8883 | 1670 | 2190 | 2020 | 1339 | 2641 | 6173 | 1635 | 2734 | 3589 | 1569 | 1758 |
| 1634 | QGN14731.1 | FIM1_1400 | CP015055. | 2186 | 1230 | 932 | 5232 | 1408 | 877 | 3586 | 1069 | 1344 | 2029 | 912 | 1350 | 5731 | 1074 | 1223 | 3455 | 913 | 949 |
| 1635 | QGN14732.1 | FIM1_1401 | CP015055. | 9964 | 499 | 1090 | 14277 | 1602 | 1249 | 16603 | 830 | 1472 | 5330 | 1254 | 1914 | 9956 | 1818 | 1727 | 6704 | 1391 | 1449 |
| 1636 | QGN14733.1 | FIM1_1402 | CP015055. | 1570 | 391 | 497 | 1904 | 491 | 381 | 1918 | 464 | 559 | 2828 | 403 | 657 | 3190 | 489 | 610 | 2934 | 462 | 472 |
| 1637 | QGN14734.1 | FIM1_1403 | CP015055. | 1895 | 421 | 464 | 2764 | 419 | 427 | 2333 | 374 | 633 | 1640 | 330 | 690 | 2149 | 414 | 563 | 1800 | 322 | 403 |
| 1638 | QGN14735.1 | FIM1_1404 | CP015055. | 377 | 2297 | 1521 | 388 | 798 | 706 | 630 | 1786 | 1451 | 995 | 557 | 1214 | 1224 | 318 | 712 | 1240 | 439 | 490 |
| 1639 | QGN14736.1 | FIM1_1405 | CP015055. | 362 | 489 | 824 | 813 | 640 | 858 | 473 | 428 | 1020 | 2454 | 367 | 720 | 2595 | 478 | 759 | 2405 | 377 | 480 |
| 1640 | QGN14737.1 | FIM1_1406 | CP015055. | 15923 | 2052 | 3671 | 15807 | 3521 | 3035 | 14392 | 2996 | 4786 | 4705 | 2777 | 6063 | 16230 | 3195 | 5535 | 9445 | 2931 | 4175 |
| 1641 | QGN14738.1 | FIM1_1407 | CP015055. | 2366 | 561 | 729 | 2607 | 713 | 661 | 2344 | 657 | 999 | 3703 | 867 | 1273 | 5516 | 771 | 1117 | 4163 | 733 | 904 |
| 1642 | QGN14739.1 | FIM1_1408 | CP015055. | 40247 | 3690 | 3221 | 30263 | 4291 | 2904 | 31582 | 3820 | 4003 | 16122 | 4366 | 4641 | 18593 | 4792 | 4207 | 16346 | 4137 | 3077 |
| 1643 | QGN14740.1 | FIM1_1409 | CP015055. | 5439 | 1674 | 892 | 2163 | 1545 | 872 | 4895 | 1622 | 1243 | 804 | 1377 | 1796 | 3376 | 1221 | 1611 | 1641 | 1726 | 1177 |
| 1644 | QGN14741.1 | FIM1_1410 | CP015055. | 2289 | 4127 | 2971 | 5040 | 3035 | 2771 | 4010 | 2877 | 3721 | 6666 | 2725 | 3306 | 7968 | 2559 | 3384 | 6709 | 2667 | 2344 |
| 1645 | QGN14742.1 | FIM1_1411 | CP015055. | 3765 | 581 | 806 | 3002 | 938 | 597 | 3208 | 847 | 1060 | 516 | 551 | 1298 | 2859 | 447 | 1151 | 1242 | 528 | 847 |
| 1646 | QGN14743.1 | FIM1_1412 | CP015055. | 16708 | 1923 | 2443 | 14314 | 2887 | 1784 | 15137 | 2834 | 3141 | 4696 | 2504 | 3531 | 12728 | 2984 | 3259 | 7509 | 2349 | 2237 |
| 1647 | QGN14744.1 | FIM1_1413 | CP015055. | 1012 | 99 | 426 | 1714 | 334 | 299 | 1541 | 403 | 560 | 985 | 213 | 488 | 2201 | 255 | 435 | 1321 | 241 | 357 |
| 1648 | QGN14745.1 | FIM1_1414 | CP015055. | 621 | 134 | 196 | 603 | 208 | 171 | 659 | 206 | 240 | 291 | 129 | 316 | 630 | 134 | 273 | 411 | 165 | 208 |
| 1649 | QGN14746.1 | FIM1_1415 | CP015055. | 2320 | 1255 | 1253 | 2387 | 1037 | 1146 | 2261 | 949 | 1711 | 4719 | 751 | 1489 | 4808 | 582 | 1427 | 4345 | 627 | 974 |
| 1650 | QGN14747.1 | FIM1_1416 | CP015055. | 1197 | 510 | 853 | 1766 | 507 | 907 | 1457 | 477 | 1274 | 2752 | 423 | 1268 | 2456 | 387 | 1223 | 2781 | 365 | 888 |
| 1651 | QGN14748.1 | FIM1_1418 | CP015055. | 507 | 865 | 665 | 788 | 847 | 688 | 546 | 760 | 905 | 3323 | 543 | 1062 | 3681 | 604 | 881 | 3479 | 561 | 618 |
| 1652 | QGN14749.1 | FIM1_1419 | CP015055. | 8 | 3 | 4 | 13 | 6 | 3 | 5 | 13 | 4 | 9 | 1 | 6 | 12 | 4 | 4 | 11 | 4 | 3 |
| 1653 | QGN14750.1 | FIM1_1420 | CP015055. | 2474 | 1540 | 815 | 2304 | 1043 | 811 | 2368 | 1063 | 1192 | 2966 | 826 | 1177 | 3365 | 775 | 1217 | 3213 | 782 | 745 |
| 1654 | QGN14751.1 | FIM1_1421 | CP015055. | 1001 | 201 | 392 | 1628 | 314 | 337 | 1336 | 397 | 538 | 523 | 242 | 519 | 1585 | 227 | 454 | 955 | 219 | 383 |
| 1655 | QGN14752.1 | FIM1_1422 | CP015055. | 864 | 468 | 393 | 1075 | 299 | 353 | 908 | 357 | 527 | 472 | 211 | 408 | 1207 | 211 | 449 | 743 | 209 | 284 |
| 1656 | QGN14753.1 | FIM1_1423 | CP015055. | 785 | 434 | 482 | 973 | 466 | 428 | 905 | 525 | 675 | 2606 | 457 | 677 | 2996 | 506 | 585 | 2783 | 440 | 431 |
| 1657 | QGN14754.1 | FIM1_1424 | CP015055. | 4635 | 915 | 839 | 4538 | 806 | 714 | 4181 | 786 | 1067 | 3282 | 728 | 1271 | 3715 | 676 | 1243 | 3164 | 685 | 782 |
| 1658 | QGN14755.1 | FIM1_1425 | CP015055. | 4314 | 769 | 799 | 5808 | 585 | 686 | 5753 | 664 | 1062 | 2726 | 447 | 880 | 3015 | 589 | 1147 | 2327 | 476 | 646 |
| 1659 | QGN14756.1 | FIM1_1426 | CP015055. | 1914 | 3659 | 2024 | 6647 | 2401 | 1363 | 4655 | 3472 | 2798 | 1531 | 1966 | 3163 | 7758 | 2727 | 2504 | 2462 | 2530 | 1929 |
| 1660 | QGN14757.1 | FIM1_1427 | CP015055. | 50 | 35 | 19 | 78 | 37 | 17 | 51 | 11 | 16 | 97 | 19 | 24 | 109 | 17 | 28 | 90 | 12 | 17 |
| 1661 | QGN14758.1 | FIM1_1428 | CP015055. | 3336 | 1016 | 1512 | 2669 | 1106 | 1224 | 3240 | 1170 | 2000 | 4334 | 1206 | 2208 | 8327 | 1002 | 1982 | 6433 | 1025 | 1552 |
| 1662 | QGN14759.1 | FIM1_1429 | CP015055. | 4382 | 872 | 1382 | 4411 | 1321 | 1052 | 4316 | 1246 | 1754 | 4151 | 1211 | 1873 | 8600 | 1240 | 1746 | 5865 | 1046 | 1225 |
| 1663 | QGN14760.1 | FIM1_1430 | CP015055. | 1728 | 809 | 849 | 2566 | 601 | 632 | 1951 | 727 | 1105 | 1432 | 480 | 1073 | 3233 | 457 | 1109 | 2212 | 387 | 751 |
| 1664 | QGN14761.1 | FIM1_1431 | CP015055. | 8472 | 672 | 1225 | 9027 | 1113 | 1050 | 10107 | 955 | 1632 | 2555 | 626 | 1880 | 5030 | 741 | 1910 | 3628 | 555 | 1252 |
| 1665 | QGN14762.1 | FIM1_1432 | CP015055. | 3720 | 11123 | 7856 | 7064 | 10642 | 6597 | 5876 | 9559 | 9886 | 38503 | 12814 | 13086 | 41261 | 11047 | 13274 | 38715 | 10310 | 8940 |
| 1666 | QGN14763.1 | FIM1_1433 | CP015055. | 3821 | 1078 | 1183 | 4254 | 1158 | 1006 | 4487 | 1076 | 1500 | 6321 | 1144 | 1757 | 5671 | 1067 | 1644 | 6041 | 943 | 1160 |
| 1667 | QGN14764.1 | FIM1_1434 | CP015055. | 5123 | 544 | 1064 | 4944 | 1019 | 766 | 5037 | 1070 | 1312 | 1555 | 898 | 1816 | 5281 | 1014 | 1554 | 2573 | 941 | 1129 |
| 1668 | QGN14765.1 | FIM1_1435 | CP015055. | 14103 | 4249 | 3233 | 15531 | 3448 | 2823 | 13838 | 3269 | 4413 | 5725 | 3112 | 4972 | 13257 | 3514 | 5294 | 8619 | 2830 | 3437 |
| 1669 | QGN14766.1 | FIM1_1436 | CP015055. | 2102 | 297 | 420 | 1659 | 327 | 346 | 1682 | 401 | 549 | 1187 | 275 | 550 | 1612 | 215 | 479 | 1327 | 209 | 298 |
| 1670 | QGN14767.1 | FIM1_1437 | CP015055. | 279 | 253 | 214 | 353 | 245 | 228 | 236 | 216 | 290 | 1061 | 218 | 352 | 1133 | 211 | 288 | 1163 | 207 | 244 |
| 1671 | QGN14768.1 | FIM1_1438 | CP015055. | 75039 | 41678 | 30942 | 76416 | 43109 | 28561 | 62959 | 37969 | 37235 | 85409 | 49485 | 50223 | 122409 | 46512 | 47498 | 95923 | 44408 | 37100 |
| 1672 | QGN14769.1 | FIM1_1439 | CP015055. | 4931 | 563 | 594 | 5152 | 544 | 529 | 5140 | 561 | 820 | 1602 | 391 | 775 | 2559 | 447 | 720 | 1585 | 396 | 539 |
| 1673 | QGN14770.1 | FIM1_1440 | CP015055. | 1144 | 347 | 278 | 1104 | 266 | 217 | 1164 | 234 | 365 | 1343 | 171 | 349 | 1163 | 170 | 295 | 1208 | 160 | 202 |
| 1674 | QGN14771.1 | FIM1_1441 | CP015055. | 3222 | 70 | 95 | 1334 | 147 | 72 | 2249 | 115 | 116 | 187 | 101 | 107 | 548 | 102 | 101 | 335 | 96 | 65 |
| 1675 | QGN14772.1 | FIM1_1442 | CP015055. | 7338 | 18 | 34 | 6236 | 43 | 28 | 6869 | 48 | 45 | 140 | 29 | 51 | 220 | 19 | 48 | 172 | 24 | 35 |
| 1676 | QGN14773.1 | FIM1_1443 | CP015055. | 624 | 440 | 273 | 853 | 298 | 225 | 879 | 364 | 317 | 1851 | 275 | 375 | 1463 | 253 | 377 | 1627 | 269 | 312 |
| 1677 | QGN14774.1 | FIM1_1444 | CP015055. | 9608 | 13083 | 7144 | 8457 | 9720 | 6707 | 7075 | 8321 | 8493 | 25801 | 10563 | 10935 | 21715 | 9447 | 11051 | 24744 | 9504 | 8199 |
| 1678 | QGN14775.1 | FIM1_1445 | CP015055. | 1540 | 896 | 365 | 798 | 1037 | 426 | 1185 | 659 | 494 | 3433 | 1064 | 671 | 3053 | 1451 | 634 | 3139 | 1445 | 492 |

| | | | | | | | | | | | | | | | | | | | | | |
|---|---|---|---|---|---|---|---|---|---|---|---|---|---|---|---|---|---|---|---|---|---|
| 1679 | QGN14776.1 | FIM1_1446 | CP015055. | 683 | 553 | 628 | 1110 | 396 | 561 | 621 | 468 | 843 | 2840 | 257 | 602 | 2347 | 227 | 594 | 2542 | 203 | 355 |
| 1680 | QGN14777.1 | FIM1_1447 | CP015055. | 248 | 745 | 490 | 508 | 543 | 494 | 456 | 430 | 601 | 2514 | 597 | 775 | 1710 | 511 | 767 | 2164 | 663 | 476 |
| 1681 | QGN14778.1 | FIM1_1448 | CP015055. | 271 | 537 | 365 | 470 | 319 | 336 | 237 | 282 | 460 | 2237 | 312 | 503 | 1572 | 328 | 431 | 1969 | 333 | 266 |
| 1682 | QGN14779.1 | FIM1_1449 | CP015055. | 1758 | 1731 | 1758 | 2793 | 2284 | 1394 | 1893 | 2120 | 2112 | 6264 | 2057 | 2336 | 6760 | 2232 | 2078 | 6444 | 1801 | 1579 |
| 1683 | QGN14780.1 | FIM1_1450 | CP015055. | 1978 | 423 | 502 | 2283 | 513 | 373 | 2234 | 503 | 589 | 1815 | 394 | 850 | 2942 | 376 | 602 | 2123 | 479 | 507 |
| 1684 | QGN14781.1 | FIM1_1451 | CP015055. | 1611 | 9178 | 8164 | 1477 | 7674 | 6380 | 1385 | 7007 | 10042 | 21497 | 7547 | 10681 | 22918 | 4843 | 9364 | 23086 | 4449 | 7342 |
| 1685 | QGN14782.1 | FIM1_1452 | CP015055. | 3 | 8 | 5 | 2 | 10 | 5 | 3 | 5 | 8 | 13 | 2 | 4 | 10 | 1 | 7 | 12 | 6 | 1 |
| 1686 | QGN14783.1 | FIM1_1453 | CP015055. | 3801 | 1916 | 2319 | 13119 | 1895 | 1980 | 9496 | 1770 | 3254 | 10619 | 1873 | 2734 | 11879 | 3312 | 2992 | 10466 | 1540 | 2013 |
| 1687 | QGN14784.1 | FIM1_1454 | CP015055. | 4902 | 560 | 604 | 3732 | 549 | 498 | 3553 | 633 | 807 | 3968 | 490 | 806 | 3462 | 356 | 703 | 3985 | 395 | 474 |
| 1688 | QGN14785.1 | FIM1_1455 | CP015055. | 1061 | 753 | 889 | 1576 | 954 | 902 | 1668 | 774 | 1129 | 2250 | 722 | 1190 | 2385 | 677 | 1043 | 2315 | 680 | 816 |
| 1689 | QGN14786.1 | FIM1_1456 | CP015055. | 1082 | 395 | 288 | 1290 | 314 | 289 | 1051 | 237 | 383 | 2222 | 276 | 387 | 1538 | 262 | 397 | 1896 | 314 | 249 |
| 1690 | QGN14787.1 | FIM1_1457 | CP015055. | 6408 | 1350 | 1913 | 6190 | 1672 | 1630 | 5560 | 1541 | 2473 | 4814 | 1517 | 3305 | 10209 | 1237 | 2847 | 6629 | 1225 | 2152 |
| 1691 | QGN14788.1 | FIM1_1458 | CP015055. | 1653 | 1319 | 870 | 1939 | 932 | 893 | 1616 | 867 | 1270 | 4622 | 928 | 1257 | 3668 | 809 | 1190 | 4071 | 904 | 868 |
| 1692 | QGN14789.1 | FIM1_1460 | CP015055. | 604 | 915 | 770 | 959 | 884 | 700 | 606 | 795 | 1039 | 3561 | 645 | 1198 | 4155 | 622 | 963 | 3876 | 593 | 733 |
| 1693 | QGN14790.1 | FIM1_1461 | CP015055. | 13 | 723 | 918 | 78 | 616 | 971 | 21 | 500 | 1167 | 3880 | 607 | 1512 | 2960 | 519 | 1435 | 3829 | 457 | 922 |
| 1694 | QGN14791.1 | FIM1_1462 | CP015055. | 867 | 1228 | 938 | 914 | 1257 | 853 | 839 | 1068 | 1198 | 5940 | 1015 | 1405 | 5348 | 910 | 1314 | 5627 | 893 | 922 |
| 1695 | QGN14792.1 | FIM1_1463 | CP015055. | 247 | 194 | 118 | 367 | 135 | 122 | 263 | 139 | 172 | 1094 | 141 | 224 | 726 | 132 | 193 | 912 | 179 | 124 |
| 1696 | QGN14793.1 | FIM1_1464 | CP015055. | 26 | 6 | 9 | 210 | 14 | 15 | 36 | 5 | 16 | 114 | 11 | 27 | 112 | 9 | 21 | 107 | 5 | 16 |
| 1697 | QGN14794.1 | FIM1_1465 | CP015055. | 4594 | 817 | 716 | 4113 | 711 | 611 | 4542 | 641 | 860 | 4712 | 604 | 1026 | 3951 | 546 | 1011 | 4192 | 515 | 737 |
| 1698 | QGN14795.1 | FIM1_1466 | CP015055. | 1888 | 812 | 949 | 1811 | 607 | 787 | 1664 | 626 | 1259 | 4771 | 604 | 1205 | 4014 | 515 | 1234 | 4453 | 514 | 755 |
| 1699 | QGN14796.1 | FIM1_1467 | CP015055. | 3431 | 810 | 1603 | 5595 | 1252 | 1302 | 3724 | 1172 | 2098 | 6615 | 936 | 2596 | 9614 | 1085 | 2141 | 7799 | 935 | 1561 |
| 1700 | QGN14797.1 | FIM1_1468 | CP015055. | 3716 | 2252 | 1983 | 5860 | 1954 | 1731 | 4635 | 1688 | 2648 | 11258 | 1892 | 3116 | 10448 | 2151 | 2923 | 11112 | 1760 | 2014 |
| 1701 | QGN14798.1 | FIM1_1469 | CP015055. | 158 | 86 | 213 | 310 | 69 | 158 | 341 | 93 | 309 | 558 | 46 | 440 | 502 | 44 | 357 | 510 | 37 | 262 |
| 1702 | QGN14799.1 | FIM1_1470 | CP015055. | 2571 | 1168 | 1038 | 2874 | 1134 | 973 | 2576 | 1168 | 1542 | 4471 | 1011 | 1636 | 5807 | 1008 | 1529 | 4743 | 929 | 1058 |
| 1703 | QGN14800.1 | FIM1_1471 | CP015055. | 3436 | 3639 | 2944 | 4855 | 2810 | 2574 | 3422 | 2485 | 3648 | 13262 | 3480 | 5058 | 10327 | 3251 | 4976 | 13275 | 3250 | 3441 |
| 1704 | QGN14801.1 | FIM1_1472 | CP015055. | 1008 | 777 | 1014 | 1101 | 543 | 874 | 795 | 464 | 1155 | 3137 | 553 | 1492 | 2237 | 574 | 1389 | 2751 | 656 | 1017 |
| 1705 | QGN14802.1 | FIM1_1473 | CP015055. | 3969 | 508 | 444 | 4554 | 689 | 394 | 4475 | 457 | 490 | 4494 | 717 | 888 | 3421 | 669 | 839 | 4120 | 752 | 587 |
| 1706 | QGN14803.1 | FIM1_1474 | CP015055. | 1242 | 1511 | 1586 | 2912 | 1838 | 1401 | 2269 | 1369 | 2035 | 10666 | 1633 | 2519 | 9645 | 1731 | 2209 | 10309 | 1548 | 1613 |
| 1707 | QGN14804.1 | FIM1_1475 | CP015055. | 2795 | 674 | 692 | 3083 | 905 | 592 | 2540 | 757 | 895 | 6248 | 880 | 1245 | 5487 | 829 | 1187 | 5891 | 753 | 797 |
| 1708 | QGN14805.1 | FIM1_1476 | CP015055. | 15010 | 45 | 51 | 15577 | 51 | 58 | 10524 | 45 | 79 | 502 | 51 | 82 | 439 | 81 | 103 | 469 | 66 | 64 |
| 1709 | QGN14806.1 | FIM1_1477 | CP015055. | 3989 | 1363 | 1281 | 4566 | 1246 | 1118 | 3933 | 1348 | 1875 | 1086 | 721 | 1608 | 5119 | 690 | 1466 | 2194 | 733 | 1044 |
| 1710 | QGN14807.1 | FIM1_1478 | CP015055. | 1505 | 910 | 1595 | 4004 | 1765 | 1525 | 1655 | 1475 | 2164 | 6423 | 724 | 1157 | 7276 | 453 | 931 | 6478 | 470 | 784 |
| 1711 | QGN14808.1 | FIM1_1479 | CP015055. | 924 | 346 | 674 | 1075 | 392 | 562 | 996 | 381 | 858 | 4354 | 334 | 1042 | 3825 | 324 | 885 | 3946 | 278 | 704 |
| 1712 | QGN14809.1 | FIM1_1480 | CP015055. | 644 | 802 | 507 | 571 | 730 | 513 | 579 | 582 | 652 | 3398 | 762 | 840 | 2735 | 553 | 710 | 3109 | 689 | 475 |
| 1713 | QGN14810.1 | FIM1_1481 | CP015055. | 615 | 507 | 413 | 868 | 366 | 366 | 480 | 354 | 554 | 2356 | 301 | 554 | 1597 | 216 | 442 | 2347 | 255 | 306 |
| 1714 | QGN14811.1 | FIM1_1482 | CP015055. | 11658 | 4326 | 3786 | 20454 | 6045 | 3662 | 20314 | 4982 | 5449 | 10621 | 2785 | 2155 | 12994 | 2634 | 2224 | 10306 | 2623 | 1451 |
| 1715 | QGN14812.1 | FIM1_1483 | CP015055. | 214 | 382 | 270 | 478 | 229 | 239 | 378 | 230 | 343 | 1888 | 216 | 299 | 1575 | 196 | 337 | 1744 | 228 | 201 |
| 1716 | QGN14813.1 | FIM1_1484 | CP015055. | 596 | 320 | 629 | 1009 | 400 | 504 | 672 | 452 | 817 | 3009 | 391 | 844 | 3390 | 287 | 751 | 3141 | 277 | 527 |
| 1717 | QGN14814.1 | FIM1_1485 | CP015055. | 2845 | 1907 | 2257 | 2925 | 2027 | 1965 | 2800 | 2107 | 3036 | 11121 | 2081 | 3120 | 10375 | 1765 | 3180 | 10803 | 1643 | 2219 |
| 1718 | QGN14815.1 | FIM1_1486 | CP015055. | 977 | 1255 | 1965 | 2208 | 809 | 1662 | 1348 | 794 | 2390 | 6074 | 825 | 3050 | 4316 | 877 | 2776 | 6009 | 958 | 2077 |
| 1719 | QGN14816.1 | FIM1_1487 | CP015055. | 2056 | 1127 | 605 | 1233 | 1305 | 668 | 2237 | 1042 | 862 | 6699 | 1733 | 1329 | 5546 | 1840 | 1266 | 6685 | 1920 | 937 |
| 1720 | QGN14817.1 | FIM1_1488 | CP015055. | 5473 | 2733 | 2904 | 5327 | 3249 | 2692 | 5098 | 2671 | 3984 | 15753 | 3567 | 4657 | 19184 | 3439 | 4313 | 16035 | 3307 | 3225 |
| 1721 | QGN14818.1 | FIM1_1489 | CP015055. | 186 | 525 | 339 | 238 | 373 | 297 | 151 | 289 | 477 | 1972 | 414 | 490 | 1545 | 368 | 485 | 1930 | 349 | 297 |
| 1722 | QGN14819.1 | FIM1_1490 | CP015055. | 257 | 462 | 429 | 462 | 403 | 392 | 289 | 363 | 599 | 2039 | 293 | 602 | 1829 | 393 | 629 | 1941 | 311 | 335 |
| 1723 | QGN14820.1 | FIM1_1491 | CP015055. | 1270 | 751 | 764 | 1402 | 726 | 656 | 1384 | 703 | 1013 | 2075 | 599 | 1039 | 2889 | 523 | 925 | 2330 | 517 | 701 |
| 1724 | QGN14821.1 | FIM1_1492 | CP015055. | 1827 | 1041 | 1103 | 2183 | 821 | 1157 | 1845 | 735 | 1560 | 5555 | 802 | 1751 | 4141 | 714 | 1658 | 5602 | 764 | 1082 |
| 1725 | QGN14822.1 | FIM1_1493 | CP015055. | 1065 | 344 | 597 | 1247 | 241 | 485 | 847 | 316 | 781 | 2290 | 285 | 791 | 2038 | 276 | 789 | 2160 | 210 | 508 |
| 1726 | QGN14823.1 | FIM1_1494 | CP015055. | 344 | 278 | 234 | 291 | 153 | 175 | 360 | 169 | 267 | 474 | 114 | 216 | 605 | 108 | 227 | 524 | 95 | 183 |
| 1727 | QGN14824.1 | FIM1_1495 | CP015055. | 140 | 9 | 39 | 165 | 52 | 38 | 179 | 59 | 67 | 236 | 36 | 57 | 307 | 29 | 70 | 237 | 29 | 36 |
| 1728 | QGN14825.1 | FIM1_1496 | CP015055. | 1939 | 558 | 441 | 1559 | 540 | 455 | 1931 | 538 | 604 | 2791 | 491 | 722 | 2675 | 544 | 617 | 2734 | 486 | 419 |
| 1729 | QGN14826.1 | FIM1_1497 | CP015055. | 731 | 225 | 391 | 788 | 246 | 371 | 698 | 313 | 550 | 1560 | 302 | 452 | 2047 | 269 | 413 | 1688 | 229 | 270 |
| 1730 | QGN14827.1 | FIM1_1498 | CP015055. | 505 | 395 | 509 | 614 | 391 | 325 | 617 | 458 | 556 | 2787 | 410 | 582 | 2174 | 291 | 505 | 2735 | 305 | 359 |
| 1731 | QGN14828.1 | FIM1_1499 | CP015055. | 809 | 1446 | 1936 | 1526 | 1509 | 1751 | 827 | 1338 | 2593 | 1680 | 854 | 2244 | 4570 | 936 | 2105 | 2709 | 840 | 1654 |
| 1732 | QGN14829.1 | FIM1_1500 | CP015055. | 2866 | 744 | 1077 | 3204 | 793 | 920 | 2808 | 889 | 1506 | 3190 | 829 | 1552 | 5811 | 684 | 1331 | 4054 | 700 | 1036 |
| 1733 | QGN14830.1 | FIM1_1501 | CP015055. | 1214 | 1037 | 1046 | 845 | 994 | 826 | 915 | 918 | 1278 | 6335 | 937 | 1283 | 5999 | 714 | 1302 | 6402 | 656 | 919 |
| 1734 | QGN14831.1 | FIM1_1502 | CP015055. | 7 | 4 | 3 | 8 | 2 | 6 | 8 | 8 | 6 | 11 | 5 | 2 | 14 | 0 | 3 | 20 | 5 | 2 |
| 1735 | QGN14832.1 | FIM1_1503 | CP015055. | 1248 | 1238 | 1413 | 1709 | 1570 | 1268 | 1518 | 1365 | 1872 | 5516 | 1575 | 1894 | 7367 | 1448 | 1711 | 6252 | 1389 | 1247 |
| 1736 | QGN14833.1 | FIM1_1504 | CP015055. | 1210 | 520 | 519 | 1604 | 455 | 481 | 1409 | 488 | 718 | 4221 | 513 | 847 | 3798 | 410 | 801 | 4042 | 407 | 556 |
| 1737 | QGN14834.1 | FIM1_1505 | CP015055. | 1275 | 239 | 416 | 1173 | 253 | 353 | 1046 | 260 | 549 | 2057 | 183 | 589 | 2255 | 163 | 463 | 2189 | 145 | 355 |
| 1738 | QGN14835.1 | FIM1_1506 | CP015055. | 333 | 121 | 91 | 385 | 51 | 113 | 318 | 46 | 172 | 830 | 54 | 124 | 570 | 49 | 110 | 696 | 52 | 84 |
| 1739 | QGN14836.1 | FIM1_1507 | CP015055. | 3229 | 3505 | 4114 | 6150 | 3075 | 3584 | 4319 | 2687 | 4956 | 15336 | 2755 | 6057 | 13167 | 2931 | 5491 | 15150 | 2581 | 3895 |
| 1740 | QGN14837.1 | FIM1_1508 | CP015055. | 1647 | 1341 | 968 | 3091 | 1166 | 938 | 2199 | 971 | 1370 | 5498 | 881 | 1638 | 5834 | 989 | 1591 | 5804 | 878 | 1082 |
| 1741 | QGN14838.1 | FIM1_1509 | CP015055. | 1087 | 1522 | 917 | 1184 | 1282 | 872 | 1021 | 1071 | 1209 | 6891 | 1324 | 1516 | 5325 | 1175 | 1389 | 6755 | 1316 | 996 |
| 1742 | QGN14839.1 | FIM1_1510 | CP015055. | 862 | 1012 | 794 | 1046 | 916 | 781 | 961 | 779 | 1022 | 6331 | 974 | 1388 | 5010 | 856 | 1155 | 6595 | 922 | 837 |
| 1743 | QGN14840.1 | FIM1_1511 | CP015055. | 269 | 169 | 157 | 420 | 118 | 129 | 308 | 126 | 201 | 1140 | 103 | 212 | 731 | 115 | 180 | 1151 | 121 | 134 |
| 1744 | QGN14841.1 | FIM1_1512 | CP015055. | 632 | 595 | 386 | 637 | 395 | 315 | 620 | 424 | 522 | 2644 | 455 | 567 | 2053 | 362 | 603 | 2546 | 438 | 363 |
| 1745 | QGN14842.1 | FIM1_1513 | CP015055. | 4709 | 3132 | 3647 | 6741 | 3205 | 3224 | 4863 | 3002 | 4490 | 14678 | 2787 | 5221 | 14190 | 2914 | 5171 | 14278 | 2562 | 3508 |
| 1746 | QGN14843.1 | FIM1_1514 | CP015055. | 15858 | 7161 | 11035 | 23860 | 9352 | 9514 | 18688 | 8358 | 13288 | 18311 | 5457 | 13429 | 26057 | 5566 | 12227 | 21278 | 5084 | 8862 |
| 1747 | QGN14844.1 | FIM1_1515 | CP015055. | 1530 | 207 | 283 | 1986 | 212 | 231 | 1220 | 246 | 319 | 616 | 129 | 336 | 966 | 82 | 280 | 701 | 94 | 217 |
| 1748 | QGN14845.1 | FIM1_1516 | CP015055. | 1058 | 156 | 172 | 1522 | 79 | 97 | 1571 | 183 | 244 | 325 | 39 | 77 | 328 | 47 | 54 | 362 | 45 | 41 |
| 1749 | QGN14846.1 | FIM1_1518 | CP015055. | 1417 | 1323 | 889 | 1271 | 1138 | 819 | 1353 | 957 | 1161 | 5604 | 1263 | 1273 | 4407 | 1184 | 1300 | 5366 | 1221 | 876 |
| 1750 | QGN14847.1 | FIM1_1519 | CP015055. | 948 | 476 | 456 | 1062 | 368 | 439 | 995 | 353 | 590 | 1947 | 302 | 574 | 1488 | 250 | 617 | 1709 | 286 | 401 |
| 1751 | QGN14848.1 | FIM1_1520 | CP015055. | 3584 | 2709 | 3315 | 7887 | 2198 | 2834 | 5038 | 2215 | 4054 | 13291 | 2272 | 4506 | 12505 | 2344 | 4675 | 12765 | 1915 | 2989 |
| 1752 | QGN14849.1 | FIM1_1521 | CP015055. | 6175 | 2330 | 1202 | 2775 | 3190 | 1292 | 5509 | 2746 | 1699 | 8311 | 3571 | 2505 | 11383 | 3721 | 2222 | 8829 | 4106 | 1744 |
| 1753 | QGN14850.1 | FIM1_1523 | CP015055. | 461 | 147 | 220 | 704 | 230 | 203 | 532 | 230 | 281 | 1141 | 253 | 438 | 1192 | 231 | 364 | 1047 | 239 | 248 |
| 1754 | QGN14851.1 | FIM1_1524 | CP015055. | 654 | 820 | 729 | 888 | 814 | 656 | 821 | 640 | 918 | 4159 | 798 | 1197 | 3576 | 663 | 1095 | 3967 | 750 | 740 |
| 1755 | QGN14852.1 | FIM1_1525 | CP015055. | 1832 | 843 | 1104 | 2651 | 969 | 954 | 2076 | 797 | 1386 | 5573 | 941 | 2088 | 5489 | 967 | 1825 | 5593 | 874 | 1254 |
| 1756 | QGN14853.1 | FIM1_1526 | CP015055. | 9294 | 3129 | 10786 | 11098 | 2803 | 7670 | 10292 | 3490 | 11763 | 10759 | 1916 | 15967 | 11921 | 2138 | 12862 | 11177 | 1894 | 9850 |
| 1757 | QGN14854.1 | FIM1_1527 | CP015055. | 925 | 408 | 372 | 974 | 300 | 330 | 989 | 299 | 468 | 1829 | 293 | 457 | 1667 | 288 | 480 | 1679 | 289 | 310 |
| 1758 | QGN14855.1 | FIM1_1528 | CP015055. | 1385 | 433 | 320 | 588 | 527 | 298 | 985 | 528 | 420 | 454 | 468 | 585 | 1649 | 494 | 540 | 850 | 533 | 361 |
| 1759 | QGN14856.1 | FIM1_1529 | CP015055. | 7376 | 715 | 867 | 6127 | 678 | 786 | 6520 | 735 | 1164 | 3887 | 371 | 1284 | 4483 | 283 | 1202 | 3650 | 321 | 858 |
| 1760 | QGN14857.1 | FIM1_1530 | CP015055. | 8935 | 595 | 1192 | 8453 | 1030 | 953 | 8375 | 1067 | 1523 | 7134 | 991 | 1781 | 8912 | 962 | 1690 | 7579 | 821 | 1210 |
| 1761 | QGN14858.1 | FIM1_1531 | CP015055. | 496 | 626 | 876 | 759 | 342 | 629 | 438 | 579 | 1236 | 1265 | 290 | 823 | 2096 | 294 | 921 | 1701 | 231 | 600 |
| 1762 | QGN14859.1 | FIM1_1532 | CP015055. | 130 | 690 | 832 | 364 | 450 | 605 | 184 | 546 | 1138 | 2373 | 380 | 943 | 2645 | 553 | 949 | 2403 | 352 | 608 |

| | | | | | | | | | | | | | | | | | | | | | |
|---|---|---|---|---|---|---|---|---|---|---|---|---|---|---|---|---|---|---|---|---|---|
| 1763 | QGN14860.1 | FIM1_1533 | CP015055. | 414 | 427 | 524 | 621 | 497 | 382 | 426 | 437 | 607 | 3522 | 511 | 738 | 2817 | 493 | 605 | 3382 | 508 | 460 |
| 1764 | QGN14861.1 | FIM1_1534 | CP015055. | 427 | 147 | 241 | 509 | 237 | 206 | 476 | 207 | 293 | 1576 | 213 | 384 | 1829 | 211 | 339 | 1625 | 196 | 286 |
| 1765 | QGN14862.1 | FIM1_1535 | CP015055. | 752 | 641 | 491 | 884 | 546 | 400 | 768 | 489 | 624 | 3066 | 491 | 800 | 2927 | 433 | 695 | 2980 | 476 | 546 |
| 1766 | QGN14863.1 | FIM1_1536 | CP015055. | 1547 | 590 | 363 | 1280 | 501 | 357 | 1555 | 498 | 509 | 1820 | 409 | 579 | 2239 | 456 | 459 | 1702 | 437 | 329 |
| 1767 | QGN14864.1 | FIM1_1537 | CP015055. | 1333 | 1661 | 1263 | 2054 | 1574 | 992 | 1756 | 1605 | 1546 | 4610 | 1472 | 1618 | 6605 | 1306 | 1527 | 5167 | 1262 | 1126 |
| 1768 | QGN14865.1 | FIM1_1538 | CP015055. | 3078 | 900 | 1091 | 3708 | 1156 | 926 | 3000 | 934 | 1383 | 4516 | 1104 | 1965 | 7815 | 934 | 1787 | 5714 | 868 | 1286 |
| 1769 | QGN14866.1 | FIM1_1539 | CP015055. | 4 | 0 | 0 | 3 | 0 | 0 | 0 | 0 | 0 | 2 | 0 | 3 | 5 | 0 | 2 | 1 | 0 | 1 |
| 1770 | QGN14867.1 | FIM1_1540 | CP015055. | 1490 | 476 | 927 | 1414 | 658 | 700 | 1362 | 728 | 1078 | 3350 | 511 | 968 | 4232 | 362 | 810 | 3699 | 394 | 642 |
| 1771 | QGN14868.1 | FIM1_1541 | CP015055. | 6836 | 1267 | 1433 | 10396 | 1708 | 1238 | 7831 | 1495 | 1945 | 2421 | 1145 | 1598 | 6786 | 1280 | 1595 | 4198 | 1142 | 1143 |
| 1772 | QGN14869.1 | FIM1_1542 | CP015055. | 6855 | 2393 | 1364 | 3554 | 3146 | 1374 | 6358 | 2095 | 1741 | 12917 | 3881 | 2693 | 10800 | 3915 | 2289 | 12591 | 4224 | 1853 |
| 1773 | QGN14870.1 | FIM1_1543 | CP015055. | 4418 | 2077 | 1135 | 2031 | 2773 | 1202 | 3967 | 2176 | 1575 | 10847 | 3218 | 2167 | 9238 | 3207 | 2092 | 10566 | 3397 | 1543 |
| 1774 | QGN14871.1 | FIM1_1544 | CP015055. | 28 | 114 | 168 | 49 | 116 | 181 | 27 | 143 | 240 | 949 | 109 | 259 | 750 | 105 | 237 | 900 | 129 | 169 |
| 1775 | QGN14872.1 | FIM1_1545 | CP015055. | 373 | 800 | 779 | 567 | 712 | 595 | 503 | 796 | 1034 | 7081 | 778 | 1265 | 5756 | 573 | 1075 | 6782 | 544 | 804 |
| 1776 | QGN14873.1 | FIM1_1546 | CP015055. | 256 | 249 | 241 | 327 | 214 | 193 | 224 | 219 | 311 | 1502 | 166 | 343 | 1109 | 112 | 298 | 1399 | 140 | 204 |
| 1777 | QGN14874.1 | FIM1_1547 | CP015055. | 1064 | 800 | 826 | 1701 | 886 | 733 | 1335 | 741 | 1173 | 3974 | 507 | 905 | 2850 | 386 | 806 | 3484 | 391 | 525 |
| 1778 | QGN14875.1 | FIM1_1548 | CP015055. | 1335 | 1299 | 738 | 1096 | 1138 | 839 | 1334 | 918 | 1034 | 5258 | 1221 | 1327 | 4362 | 1085 | 1294 | 5175 | 1227 | 892 |
| 1779 | QGN14876.1 | FIM1_1549 | CP015055. | 907 | 945 | 1650 | 1828 | 1173 | 1456 | 1034 | 1103 | 2135 | 5725 | 949 | 2751 | 5930 | 831 | 2468 | 5570 | 818 | 1746 |
| 1780 | QGN14877.1 | FIM1_1550 | CP015055. | 1347 | 695 | 1966 | 927 | 999 | 1430 | 1092 | 1154 | 2307 | 1589 | 792 | 1975 | 2877 | 615 | 1850 | 2269 | 643 | 1216 |
| 1781 | QGN14878.1 | FIM1_1551 | CP015055. | 2756 | 7270 | 29636 | 9395 | 11701 | 23104 | 5246 | 11640 | 33042 | 22531 | 8052 | 38919 | 39759 | 8604 | 36213 | 32312 | 7107 | 27601 |
| 1782 | QGN14879.1 | FIM1_1552 | CP015055. | 538 | 305 | 158 | 564 | 315 | 176 | 480 | 181 | 182 | 1849 | 320 | 259 | 1302 | 346 | 237 | 1740 | 401 | 184 |
| 1783 | QGN14880.1 | FIM1_1553 | CP015055. | 3600 | 2462 | 1085 | 1666 | 2948 | 1200 | 3101 | 2191 | 1367 | 10300 | 3234 | 2226 | 9269 | 3352 | 2058 | 9863 | 3616 | 1629 |
| 1784 | QGN14881.1 | FIM1_1554 | CP015055. | 1396 | 604 | 486 | 1339 | 668 | 445 | 1413 | 511 | 593 | 4569 | 733 | 845 | 3533 | 684 | 723 | 4464 | 693 | 540 |
| 1785 | QGN14882.1 | FIM1_1556 | CP015055. | 150 | 130 | 255 | 274 | 285 | 174 | 221 | 193 | 272 | 2150 | 330 | 475 | 1679 | 305 | 409 | 2047 | 277 | 270 |
| 1786 | QGN14883.1 | FIM1_1557 | CP015055. | 149 | 257 | 226 | 255 | 287 | 214 | 192 | 198 | 286 | 1835 | 278 | 428 | 1267 | 291 | 370 | 1714 | 258 | 294 |
| 1787 | QGN14884.1 | FIM1_1558 | CP015055. | 2266 | 905 | 1365 | 3626 | 1311 | 1106 | 2680 | 1065 | 1719 | 8948 | 1087 | 2125 | 9056 | 1165 | 1765 | 8765 | 990 | 1362 |
| 1788 | QGN14885.1 | FIM1_1559 | CP015055. | 3913 | 4350 | 3576 | 6721 | 4090 | 3052 | 5117 | 3841 | 4677 | 5690 | 3688 | 5263 | 15664 | 3826 | 5055 | 9912 | 3558 | 3738 |
| 1789 | QGN14886.1 | FIM1_1560 | CP015055. | 1775 | 494 | 1273 | 1654 | 625 | 1021 | 1876 | 661 | 1652 | 4175 | 593 | 1777 | 5421 | 536 | 1563 | 4561 | 503 | 1189 |
| 1790 | QGN14887.1 | FIM1_1561 | CP015055. | 6101 | 862 | 1279 | 3826 | 906 | 1077 | 4969 | 1005 | 1723 | 7020 | 953 | 1779 | 7889 | 690 | 1603 | 7334 | 778 | 1240 |
| 1791 | QGN14888.1 | FIM1_1562 | CP015055. | 537 | 510 | 340 | 920 | 343 | 359 | 684 | 350 | 511 | 2589 | 320 | 480 | 1638 | 276 | 465 | 2360 | 325 | 290 |
| 1792 | QGN14889.1 | FIM1_1563 | CP015055. | 867 | 840 | 586 | 1251 | 1055 | 522 | 891 | 863 | 822 | 3091 | 903 | 684 | 3839 | 745 | 643 | 3280 | 736 | 504 |
| 1793 | QGN14890.1 | FIM1_1564 | CP015055. | 3092 | 6 | 41 | 3816 | 43 | 27 | 4053 | 29 | 49 | 203 | 15 | 57 | 292 | 15 | 51 | 244 | 14 | 22 |
| 1794 | QGN14891.1 | FIM1_1565 | CP015055. | 43 | 9 | 4 | 94 | 4 | 2 | 56 | 5 | 10 | 34 | 2 | 4 | 34 | 1 | 6 | 25 | 4 | 3 |
| 1795 | QGN14892.1 | FIM1_1566 | CP015055. | 10181 | 623 | 1129 | 8218 | 1310 | 903 | 9016 | 1086 | 1433 | 4776 | 1374 | 1650 | 7342 | 1663 | 1675 | 5905 | 1323 | 1193 |
| 1796 | QGN14893.1 | FIM1_1567 | CP015055. | 1901 | 550 | 487 | 4441 | 570 | 400 | 4334 | 528 | 655 | 1126 | 523 | 555 | 2923 | 728 | 532 | 1746 | 514 | 383 |
| 1797 | QGN14894.1 | FIM1_1568 | CP015055. | 7671 | 4427 | 4489 | 9687 | 4409 | 3777 | 7938 | 3912 | 5811 | 18926 | 4299 | 7136 | 22942 | 5337 | 6573 | 19541 | 4158 | 4742 |
| 1798 | QGN14895.1 | FIM1_1569 | CP015055. | 472 | 76 | 157 | 1018 | 77 | 118 | 413 | 97 | 230 | 477 | 39 | 181 | 741 | 39 | 173 | 504 | 36 | 117 |
| 1799 | QGN14896.1 | FIM1_1570 | CP015055. | 2453 | 617 | 644 | 2019 | 579 | 586 | 2155 | 661 | 890 | 3028 | 477 | 891 | 3811 | 350 | 857 | 3264 | 413 | 657 |
| 1800 | QGN14897.1 | FIM1_1571 | CP015055. | 319 | 253 | 215 | 435 | 259 | 233 | 386 | 192 | 334 | 1514 | 227 | 350 | 1161 | 213 | 384 | 1522 | 237 | 210 |
| 1801 | QGN14898.1 | FIM1_1572 | CP015055. | 1928 | 362 | 289 | 1610 | 255 | 258 | 1643 | 254 | 369 | 2023 | 268 | 420 | 1664 | 209 | 356 | 1994 | 243 | 288 |
| 1802 | QGN14899.1 | FIM1_1573 | CP015055. | 1517 | 968 | 730 | 1507 | 881 | 640 | 1421 | 899 | 978 | 4253 | 840 | 968 | 4159 | 686 | 939 | 4020 | 650 | 641 |
| 1803 | QGN14900.1 | FIM1_1574 | CP015055. | 4758 | 2470 | 1920 | 5574 | 2093 | 1736 | 4750 | 1908 | 2511 | 10378 | 1964 | 2654 | 10239 | 1846 | 2518 | 9724 | 1550 | 1768 |
| 1804 | QGN14901.1 | FIM1_1575 | CP015055. | 1014 | 499 | 736 | 1496 | 516 | 512 | 1212 | 550 | 917 | 2744 | 412 | 933 | 3393 | 371 | 792 | 2650 | 278 | 561 |
| 1805 | QGN14902.1 | FIM1_1576 | CP015055. | 45 | 36 | 40 | 84 | 29 | 36 | 70 | 24 | 46 | 56 | 17 | 43 | 81 | 19 | 42 | 70 | 20 | 25 |
| 1806 | QGN14903.1 | FIM1_1577 | CP015055. | 1247 | 648 | 703 | 1089 | 496 | 704 | 916 | 430 | 956 | 990 | 275 | 706 | 1247 | 260 | 717 | 1075 | 213 | 469 |
| 1807 | QGN14904.1 | FIM1_1578 | CP015055. | 10175 | 9267 | 4822 | 10304 | 7949 | 4405 | 9516 | 7264 | 6339 | 19279 | 8268 | 7134 | 23339 | 7498 | 7964 | 20299 | 7898 | 5243 |
| 1808 | QGN14905.1 | FIM1_1579 | CP015055. | 1835 | 2104 | 642 | 797 | 1733 | 762 | 1556 | 1415 | 882 | 3574 | 1812 | 1185 | 4136 | 1524 | 1232 | 3513 | 1819 | 836 |
| 1809 | QGN14906.1 | FIM1_1580 | CP015055. | 3918 | 966 | 697 | 3179 | 1214 | 688 | 3682 | 870 | 865 | 6488 | 1230 | 1341 | 5356 | 1480 | 1151 | 5959 | 1443 | 954 |
| 1810 | QGN14907.1 | FIM1_1581 | CP015055. | 9896 | 2949 | 2375 | 9488 | 3162 | 2124 | 9028 | 2826 | 3047 | 15522 | 3339 | 3884 | 14025 | 3298 | 3430 | 16248 | 3215 | 2500 |
| 1811 | QGN14908.1 | FIM1_1582 | CP015055. | 536 | 98 | 118 | 439 | 43 | 59 | 432 | 79 | 136 | 306 | 29 | 112 | 246 | 39 | 110 | 276 | 25 | 56 |
| 1812 | QGN14909.1 | FIM1_1583 | CP015055. | 2772 | 672 | 554 | 2085 | 607 | 442 | 2356 | 760 | 713 | 1795 | 403 | 715 | 2262 | 376 | 635 | 1944 | 351 | 463 |
| 1813 | QGN14910.1 | FIM1_1584 | CP015055. | 5382 | 3698 | 3213 | 5839 | 4662 | 2890 | 4187 | 4102 | 4289 | 10651 | 3485 | 4594 | 17846 | 4010 | 3904 | 12920 | 3407 | 2898 |
| 1814 | QGN14911.1 | FIM1_1585 | CP015055. | 3158 | 776 | 657 | 3247 | 557 | 622 | 3288 | 503 | 862 | 1705 | 294 | 773 | 1558 | 369 | 796 | 1537 | 293 | 555 |
| 1815 | QGN14912.1 | FIM1_1586 | CP015055. | 12729 | 732 | 832 | 15219 | 595 | 639 | 16256 | 584 | 1072 | 1862 | 483 | 979 | 2453 | 754 | 1039 | 1845 | 491 | 731 |
| 1816 | QGN14913.1 | FIM1_1587 | CP015055. | 13208 | 5030 | 4916 | 11704 | 5908 | 3996 | 11331 | 5610 | 5810 | 13573 | 6038 | 6437 | 21177 | 6057 | 6413 | 17209 | 5011 | 4869 |
| 1817 | QGN14914.1 | FIM1_1588 | CP015055. | 7355 | 4179 | 3483 | 7244 | 5259 | 2726 | 7470 | 4645 | 4202 | 7887 | 5089 | 5498 | 21066 | 4738 | 5057 | 14848 | 4878 | 3944 |
| 1818 | QGN14915.1 | FIM1_1589 | CP015055. | 101 | 120 | 128 | 100 | 133 | 114 | 116 | 128 | 181 | 624 | 128 | 166 | 517 | 123 | 161 | 574 | 133 | 96 |
| 1819 | QGN14916.1 | FIM1_1590 | CP015055. | 620 | 1018 | 549 | 337 | 782 | 577 | 586 | 705 | 781 | 3281 | 698 | 806 | 2825 | 561 | 809 | 2723 | 645 | 538 |
| 1820 | QGN14917.1 | FIM1_1591 | CP015055. | 136 | 188 | 134 | 152 | 117 | 128 | 88 | 97 | 211 | 975 | 188 | 203 | 745 | 107 | 191 | 912 | 133 | 121 |
| 1821 | QGN14918.1 | FIM1_1592 | CP015055. | 148 | 60 | 195 | 222 | 110 | 143 | 159 | 122 | 226 | 1350 | 90 | 259 | 1036 | 68 | 250 | 1182 | 69 | 164 |
| 1822 | QGN14919.1 | FIM1_1593 | CP015055. | 4611 | 1435 | 2299 | 22982 | 2089 | 1934 | 7104 | 1927 | 2899 | 11727 | 1183 | 2555 | 15715 | 1716 | 2077 | 13827 | 754 | 1761 |
| 1823 | QGN14920.1 | FIM1_1594 | CP015055. | 5675 | 1382 | 1127 | 2717 | 1805 | 1047 | 4331 | 1680 | 1463 | 5971 | 1850 | 1713 | 6709 | 2240 | 1616 | 6006 | 2133 | 1142 |
| 1824 | QGN14921.1 | FIM1_1595 | CP015055. | 731 | 1473 | 980 | 708 | 1172 | 987 | 652 | 913 | 1300 | 4803 | 1028 | 1407 | 4190 | 1015 | 1257 | 4263 | 1127 | 942 |
| 1825 | QGN14922.1 | FIM1_1596 | CP015055. | 3261 | 935 | 1479 | 3372 | 885 | 1182 | 2757 | 950 | 1964 | 1733 | 634 | 1461 | 5769 | 626 | 1389 | 3562 | 530 | 1028 |
| 1826 | QGN14923.1 | FIM1_1597 | CP015055. | 2099 | 1738 | 494 | 920 | 1400 | 507 | 1904 | 1209 | 673 | 2704 | 1452 | 892 | 3336 | 1327 | 899 | 2768 | 1812 | 616 |
| 1827 | QGN14924.1 | FIM1_1598 | CP015055. | 4502 | 2285 | 1055 | 2024 | 2578 | 1170 | 4082 | 2086 | 1335 | 7764 | 2688 | 1768 | 7351 | 2815 | 1813 | 7847 | 2926 | 1259 |
| 1828 | QGN14925.1 | FIM1_1599 | CP015055. | 4079 | 1649 | 1633 | 4645 | 1672 | 1345 | 4796 | 1389 | 2039 | 8204 | 1418 | 2394 | 7793 | 1669 | 2332 | 7830 | 1619 | 1642 |
| 1829 | QGN14926.1 | FIM1_1601 | CP015055. | 19884 | 22435 | 13369 | 16684 | 16566 | 12581 | 14849 | 14594 | 16111 | 44201 | 19545 | 19387 | 42946 | 17655 | 19022 | 44148 | 17680 | 14110 |
| 1830 | QGN14927.1 | FIM1_1602 | CP015055. | 2491 | 2911 | 2127 | 2506 | 2445 | 2053 | 2489 | 2156 | 2721 | 5437 | 1259 | 1779 | 5144 | 1241 | 1877 | 5035 | 1172 | 1194 |
| 1831 | QGN14928.1 | FIM1_1603 | CP015055. | 5348 | 4069 | 2569 | 4640 | 3632 | 2416 | 4601 | 3142 | 3402 | 14515 | 3318 | 3728 | 13978 | 3180 | 3423 | 14611 | 3181 | 2547 |
| 1832 | QGN14929.1 | FIM1_1604 | CP015055. | 8089 | 780 | 1321 | 6901 | 978 | 1235 | 7131 | 784 | 1604 | 3592 | 769 | 2010 | 5067 | 876 | 2003 | 3819 | 763 | 1297 |
| 1833 | QGN14930.1 | FIM1_1605 | CP015055. | 3857 | 2360 | 790 | 1679 | 2121 | 809 | 3383 | 1956 | 1064 | 5272 | 2522 | 1644 | 6834 | 2362 | 1515 | 5787 | 3254 | 1137 |
| 1834 | QGN14931.1 | FIM1_1606 | CP015055. | 5080 | 5333 | 4537 | 10540 | 5734 | 3917 | 7872 | 5023 | 5863 | 15462 | 5714 | 7761 | 23682 | 5691 | 7072 | 17791 | 5224 | 5253 |
| 1835 | QGN14932.1 | FIM1_1607 | CP015055. | 1099 | 510 | 959 | 1771 | 630 | 735 | 1161 | 569 | 1198 | 4232 | 753 | 1576 | 5814 | 654 | 1373 | 4632 | 618 | 995 |
| 1836 | QGN14933.1 | FIM1_1608 | CP015055. | 2020 | 1852 | 1617 | 1955 | 2261 | 1552 | 1946 | 2048 | 2178 | 5740 | 1982 | 2893 | 9582 | 1716 | 2238 | 7065 | 1751 | 1775 |
| 1837 | QGN14934.1 | FIM1_1609 | CP015055. | 1045 | 1499 | 775 | 781 | 1224 | 747 | 831 | 1260 | 979 | 2823 | 997 | 1055 | 3514 | 821 | 1172 | 2834 | 896 | 734 |
| 1838 | QGN14935.1 | FIM1_1610 | CP015055. | 782 | 1871 | 1314 | 2610 | 1392 | 1103 | 1738 | 1353 | 1636 | 7775 | 1360 | 2122 | 6265 | 1574 | 2072 | 7184 | 1578 | 1374 |
| 1839 | QGN14936.1 | FIM1_1611 | CP015055. | 2509 | 1275 | 1445 | 1643 | 1314 | 1145 | 2104 | 1551 | 1772 | 4692 | 1098 | 1767 | 6888 | 778 | 1577 | 5419 | 842 | 1194 |
| 1840 | QGN14937.1 | FIM1_1612 | CP015055. | 1813 | 1731 | 800 | 993 | 1170 | 831 | 1386 | 995 | 1122 | 3156 | 1398 | 1177 | 3221 | 1015 | 1316 | 3500 | 1266 | 786 |
| 1841 | QGN14938.1 | FIM1_1613 | CP015055. | 2801 | 871 | 664 | 2801 | 855 | 612 | 2546 | 949 | 899 | 2869 | 753 | 1122 | 4149 | 760 | 1042 | 2971 | 720 | 783 |
| 1842 | QGN14939.1 | FIM1_1614 | CP015055. | 939 | 316 | 527 | 955 | 283 | 376 | 704 | 371 | 590 | 2028 | 230 | 621 | 1968 | 179 | 564 | 1961 | 190 | 401 |
| 1843 | QGN14940.1 | FIM1_1615 | CP015055. | 4830 | 2444 | 2049 | 6590 | 2427 | 1752 | 6219 | 1826 | 2642 | 7313 | 2036 | 2132 | 7051 | 1938 | 2115 | 7797 | 1924 | 1447 |
| 1844 | QGN14941.1 | FIM1_1616 | CP015055. | 1888 | 596 | 483 | 1534 | 535 | 433 | 1727 | 471 | 680 | 2420 | 448 | 674 | 2579 | 423 | 716 | 2288 | 435 | 477 |
| 1845 | QGN14942.1 | FIM1_1617 | CP015055. | 6237 | 2232 | 1624 | 5763 | 2332 | 1382 | 6248 | 1874 | 2033 | 8193 | 2452 | 2769 | 9678 | 2719 | 2470 | 8255 | 2811 | 1887 |
| 1846 | QGN14943.1 | FIM1_1618 | CP015055. | 744 | 250 | 100 | 466 | 252 | 88 | 613 | 175 | 110 | 998 | 296 | 199 | 693 | 226 | 218 | 896 | 338 | 130 |

| | | | | | | | | | | | | | | | | | | | | | |
|---|---|---|---|---|---|---|---|---|---|---|---|---|---|---|---|---|---|---|---|---|---|
| 1847 | QGN14944.1 | FIM1_1619 | CP015055. | 2426 | 1644 | 1139 | 2152 | 1599 | 1160 | 1955 | 1378 | 1552 | 3644 | 1123 | 1307 | 3299 | 1196 | 1326 | 3473 | 1111 | 840 |
| 1848 | QGN14945.1 | FIM1_1621 | CP015055. | 5532 | 1745 | 2125 | 6099 | 2499 | 1903 | 4898 | 2221 | 2795 | 10232 | 1927 | 3211 | 11783 | 1873 | 3066 | 9507 | 1740 | 2134 |
| 1849 | QGN14946.1 | FIM1_1622 | CP015055. | 4485 | 489 | 440 | 4246 | 547 | 360 | 3581 | 516 | 618 | 1941 | 371 | 437 | 1767 | 430 | 474 | 1839 | 409 | 290 |
| 1850 | QGN14947.1 | FIM1_1623 | CP015055. | 682 | 214 | 215 | 995 | 277 | 183 | 536 | 221 | 260 | 620 | 251 | 368 | 1139 | 244 | 324 | 769 | 228 | 219 |
| 1851 | QGN14948.1 | FIM1_1624 | CP015055. | 353 | 824 | 579 | 550 | 564 | 569 | 597 | 638 | 849 | 2406 | 413 | 586 | 1770 | 383 | 580 | 2356 | 376 | 384 |
| 1852 | QGN14949.1 | FIM1_1625 | CP015055. | 4319 | 1433 | 1298 | 3697 | 964 | 1167 | 3858 | 929 | 1708 | 5649 | 838 | 1859 | 4427 | 864 | 1796 | 5585 | 793 | 1179 |
| 1853 | QGN14950.1 | FIM1_1627 | CP015055. | 653 | 1838 | 1646 | 609 | 1668 | 1315 | 667 | 1632 | 2130 | 3086 | 1460 | 2452 | 6265 | 1437 | 2119 | 4527 | 1438 | 1555 |
| 1854 | QGN14951.1 | FIM1_1628 | CP015055. | 841 | 835 | 1217 | 3369 | 265 | 703 | 2933 | 735 | 1531 | 790 | 116 | 249 | 771 | 142 | 211 | 865 | 183 | 143 |
| 1855 | QGN14952.1 | FIM1_1629 | CP015055. | 456 | 271 | 245 | 399 | 258 | 222 | 288 | 238 | 307 | 1409 | 242 | 407 | 1362 | 183 | 365 | 1481 | 197 | 279 |
| 1856 | QGN14953.1 | FIM1_1630 | CP015055. | 8126 | 1449 | 1711 | 10263 | 1692 | 1541 | 8581 | 1691 | 2300 | 4756 | 1276 | 2328 | 7880 | 1162 | 2034 | 6019 | 1091 | 1573 |
| 1857 | QGN14954.1 | FIM1_1631 | CP015055. | 983 | 530 | 513 | 1663 | 426 | 413 | 1447 | 474 | 642 | 2973 | 298 | 637 | 2442 | 491 | 656 | 2482 | 373 | 402 |
| 1858 | QGN14955.1 | FIM1_1632 | CP015055. | 5982 | 707 | 1015 | 4391 | 1079 | 697 | 4538 | 1287 | 1249 | 830 | 672 | 1065 | 4588 | 573 | 1058 | 1841 | 578 | 721 |
| 1859 | QGN14956.1 | FIM1_1633 | CP015055. | 733 | 1174 | 927 | 300 | 1018 | 800 | 329 | 1306 | 1288 | 4078 | 996 | 991 | 3198 | 938 | 937 | 3785 | 882 | 686 |
| 1860 | QGN14957.1 | FIM1_1634 | CP015055. | 710 | 1474 | 1277 | 1640 | 1557 | 1202 | 1190 | 1223 | 1786 | 7085 | 1710 | 1844 | 7836 | 1461 | 1679 | 7329 | 1376 | 1261 |
| 1861 | QGN14958.1 | FIM1_1635 | CP015055. | 8844 | 3935 | 3537 | 12947 | 4427 | 3237 | 9818 | 3558 | 4477 | 15002 | 3500 | 5856 | 16737 | 3708 | 5508 | 15438 | 3176 | 3985 |
| 1862 | QGN14959.1 | FIM1_1636 | CP015055. | 662 | 900 | 686 | 787 | 541 | 642 | 652 | 521 | 987 | 1515 | 491 | 783 | 1594 | 487 | 944 | 1560 | 470 | 524 |
| 1863 | QGN14960.1 | FIM1_1637 | CP015055. | 337 | 270 | 160 | 300 | 177 | 185 | 379 | 156 | 227 | 399 | 132 | 213 | 505 | 136 | 258 | 428 | 138 | 149 |
| 1864 | QGN14961.1 | FIM1_1638 | CP015055. | 598 | 22 | 27 | 645 | 29 | 24 | 455 | 29 | 43 | 68 | 15 | 46 | 113 | 12 | 26 | 76 | 14 | 21 |
| 1865 | QGN14962.1 | FIM1_1640 | CP015055. | 3797 | 1268 | 1607 | 3825 | 1911 | 1444 | 3279 | 1532 | 2205 | 7541 | 1691 | 2864 | 10001 | 1423 | 2512 | 8114 | 1422 | 1860 |
| 1866 | QGN14963.1 | FIM1_1641 | CP015055. | 2686 | 4306 | 2990 | 2959 | 3297 | 2765 | 2542 | 2959 | 3849 | 12987 | 3165 | 4939 | 13773 | 2885 | 4482 | 13703 | 2918 | 3291 |
| 1867 | QGN14964.1 | FIM1_1642 | CP015055. | 3562 | 936 | 1391 | 4098 | 912 | 1042 | 3240 | 1041 | 1817 | 4664 | 772 | 1942 | 5958 | 779 | 1706 | 4675 | 599 | 1236 |
| 1868 | QGN14965.1 | FIM1_1643 | CP015055. | 2405 | 1012 | 966 | 4053 | 905 | 875 | 3095 | 816 | 1273 | 5180 | 794 | 1505 | 4983 | 811 | 1415 | 5148 | 714 | 976 |
| 1869 | QGN14966.1 | FIM1_1644 | CP015055. | 1101 | 397 | 378 | 1253 | 368 | 351 | 1194 | 303 | 494 | 3079 | 321 | 546 | 2508 | 276 | 502 | 2924 | 269 | 396 |
| 1870 | QGN14967.1 | FIM1_1645 | CP015055. | 1109 | 394 | 344 | 1506 | 343 | 356 | 1440 | 375 | 563 | 2630 | 356 | 561 | 2013 | 319 | 545 | 2339 | 320 | 343 |
| 1871 | QGN14968.1 | FIM1_1646 | CP015055. | 13250 | 1466 | 1705 | 15445 | 1656 | 1696 | 12976 | 1398 | 2511 | 4100 | 977 | 2031 | 9101 | 761 | 1773 | 6141 | 819 | 1427 |
| 1872 | QGN14969.1 | FIM1_1647 | CP015055. | 57 | 62 | 28 | 103 | 81 | 40 | 51 | 40 | 40 | 265 | 80 | 44 | 241 | 54 | 58 | 250 | 47 | 34 |
| 1873 | QGN14970.1 | FIM1_1648 | CP015055. | 4238 | 3434 | 2904 | 3118 | 3141 | 2577 | 3062 | 3161 | 3826 | 10918 | 3442 | 3431 | 10310 | 2001 | 3430 | 10506 | 2583 | 2379 |
| 1874 | QGN14971.1 | FIM1_1649 | CP015055. | 2342 | 2723 | 1667 | 1775 | 2057 | 1693 | 2219 | 1729 | 2196 | 7927 | 2383 | 2415 | 6487 | 1779 | 2412 | 7488 | 2220 | 1704 |
| 1875 | QGN14972.1 | FIM1_1650 | CP015055. | 1295 | 710 | 1136 | 3274 | 479 | 711 | 2285 | 520 | 1416 | 4454 | 520 | 1222 | 3518 | 837 | 1064 | 4235 | 739 | 735 |
| 1876 | QGN14973.1 | FIM1_1651 | CP015055. | 932 | 288 | 423 | 1168 | 226 | 373 | 1110 | 283 | 553 | 2114 | 166 | 476 | 1649 | 100 | 451 | 1971 | 136 | 353 |
| 1877 | QGN14974.1 | FIM1_1652 | CP015055. | 1351 | 2540 | 2019 | 1551 | 1985 | 1664 | 1026 | 2164 | 2580 | 12246 | 2072 | 2703 | 10696 | 2522 | 3034 | 11380 | 1649 | 1906 |
| 1878 | QGN14975.1 | FIM1_1653 | CP015055. | 1586 | 388 | 303 | 1475 | 336 | 289 | 1541 | 262 | 429 | 2545 | 239 | 473 | 2169 | 171 | 429 | 2233 | 222 | 285 |
| 1879 | QGN14976.1 | FIM1_1654 | CP015055. | 739 | 268 | 405 | 500 | 316 | 311 | 372 | 370 | 499 | 1365 | 220 | 599 | 1986 | 174 | 595 | 1450 | 152 | 433 |
| 1880 | QGN14977.1 | FIM1_1655 | CP015055. | 2836 | 1277 | 848 | 2425 | 983 | 833 | 2605 | 887 | 1251 | 4831 | 847 | 1145 | 4826 | 776 | 1111 | 4888 | 746 | 721 |
| 1881 | QGN14978.1 | FIM1_1656 | CP015055. | 3052 | 345 | 1047 | 3304 | 396 | 910 | 2914 | 460 | 1439 | 3168 | 232 | 1999 | 4964 | 261 | 2012 | 3996 | 188 | 1288 |
| 1882 | QGN14979.1 | FIM1_1657 | CP015055. | 28292 | 28953 | 15937 | 27127 | 23390 | 14938 | 24192 | 21800 | 19476 | 45973 | 24363 | 23942 | 59666 | 22937 | 24472 | 50299 | 22362 | 18487 |
| 1883 | QGN14980.1 | FIM1_1658 | CP015055. | 1220 | 676 | 773 | 1074 | 974 | 715 | 1221 | 860 | 989 | 3306 | 759 | 977 | 2928 | 598 | 929 | 3150 | 631 | 586 |
| 1884 | QGN14981.1 | FIM1_1659 | CP015055. | 4706 | 1285 | 1696 | 6432 | 1685 | 1475 | 6255 | 1493 | 2243 | 7210 | 1440 | 2735 | 9934 | 1621 | 2328 | 7751 | 1439 | 1834 |
| 1885 | QGN14982.1 | FIM1_1660 | CP015055. | 2018 | 1067 | 1460 | 2596 | 1172 | 1310 | 2366 | 983 | 1937 | 10703 | 1245 | 2565 | 8263 | 1167 | 2349 | 10432 | 1169 | 1711 |
| 1886 | QGN14983.1 | FIM1_1661 | CP015055. | 628 | 1302 | 932 | 975 | 1044 | 895 | 831 | 908 | 1238 | 7599 | 1199 | 1771 | 5715 | 1036 | 1619 | 7798 | 1142 | 1022 |
| 1887 | QGN14984.1 | FIM1_1664 | CP015055. | 3210 | 1621 | 1432 | 4018 | 1371 | 1322 | 3266 | 1269 | 1828 | 6149 | 1352 | 2049 | 5591 | 1346 | 1964 | 5594 | 1371 | 1380 |
| 1888 | QGN14985.1 | FIM1_1665 | CP015055. | 596 | 67 | 153 | 821 | 202 | 135 | 609 | 172 | 222 | 1293 | 176 | 307 | 1292 | 201 | 239 | 1446 | 157 | 194 |
| 1889 | QGN14986.1 | FIM1_1666 | CP015055. | 1360 | 530 | 569 | 1610 | 686 | 447 | 1203 | 516 | 739 | 3034 | 557 | 924 | 3177 | 587 | 846 | 2998 | 527 | 606 |
| 1890 | QGN14987.1 | FIM1_1667 | CP015055. | 4519 | 1113 | 423 | 2452 | 1255 | 471 | 4344 | 762 | 527 | 4502 | 1586 | 1080 | 3561 | 1969 | 927 | 4388 | 2327 | 767 |
| 1891 | QGN14988.1 | FIM1_1669 | CP015055. | 4247 | 989 | 1100 | 3712 | 1456 | 996 | 4516 | 1081 | 1432 | 6733 | 1246 | 1786 | 7164 | 1252 | 1508 | 6740 | 1182 | 1102 |
| 1892 | QGN14989.1 | FIM1_1670 | CP015055. | 1666 | 2003 | 951 | 1414 | 1479 | 1157 | 1435 | 1138 | 1363 | 6021 | 1824 | 1507 | 4527 | 1388 | 1531 | 5547 | 1747 | 1020 |
| 1893 | QGN14990.1 | FIM1_1671 | CP015055. | 6671 | 2917 | 1882 | 3835 | 4532 | 2070 | 6097 | 3193 | 2471 | 15822 | 5266 | 3811 | 14529 | 5670 | 3486 | 15540 | 5586 | 2654 |
| 1894 | QGN14991.1 | FIM1_1672 | CP015055. | 5479 | 2189 | 2532 | 9844 | 3297 | 2588 | 6739 | 2033 | 2979 | 11857 | 2226 | 3980 | 12132 | 2634 | 3503 | 11590 | 2200 | 2616 |
| 1895 | QGN14992.1 | FIM1_1673 | CP015055. | 5436 | 1598 | 1802 | 6828 | 2275 | 1474 | 5899 | 1826 | 2277 | 11745 | 2355 | 2951 | 13148 | 2426 | 2587 | 12298 | 2144 | 1972 |
| 1896 | QGN14993.1 | FIM1_1674 | CP015055. | 3145 | 417 | 1201 | 4207 | 626 | 1091 | 2982 | 414 | 1611 | 3828 | 640 | 2908 | 5528 | 586 | 2454 | 4817 | 607 | 1821 |
| 1897 | QGN14994.1 | FIM1_1675 | CP015055. | 13 | 17 | 12 | 53 | 18 | 8 | 22 | 15 | 11 | 39 | 10 | 22 | 41 | 9 | 27 | 37 | 8 | 9 |
| 1898 | QGN14995.1 | FIM1_1676 | CP015055. | 791 | 295 | 356 | 1009 | 371 | 327 | 658 | 263 | 447 | 2090 | 307 | 622 | 1920 | 357 | 555 | 1993 | 374 | 347 |
| 1899 | QGN14996.1 | FIM1_1677 | CP015055. | 693 | 189 | 607 | 3757 | 188 | 472 | 1619 | 217 | 755 | 2068 | 168 | 630 | 1866 | 280 | 557 | 1995 | 162 | 397 |
| 1900 | QGN14997.1 | FIM1_1678 | CP015055. | 539 | 70 | 134 | 1239 | 80 | 95 | 551 | 70 | 160 | 722 | 69 | 181 | 716 | 119 | 179 | 681 | 79 | 105 |
| 1901 | QGN14998.1 | FIM1_1679 | CP015055. | 1047 | 286 | 418 | 1856 | 242 | 282 | 1284 | 283 | 544 | 2727 | 248 | 551 | 2373 | 484 | 484 | 2659 | 292 | 366 |
| 1902 | QGN14999.1 | FIM1_1680 | CP015055. | 772 | 260 | 284 | 606 | 419 | 260 | 724 | 302 | 376 | 1146 | 377 | 389 | 1067 | 464 | 331 | 1064 | 430 | 204 |
| 1903 | QGN15000.1 | FIM1_1681 | CP015055. | 70 | 17 | 39 | 272 | 34 | 32 | 110 | 19 | 63 | 270 | 14 | 80 | 193 | 27 | 85 | 342 | 23 | 51 |
| 1904 | QGN15001.1 | FIM1_1682 | CP015055. | 1829 | 465 | 462 | 1685 | 713 | 383 | 1613 | 642 | 582 | 1316 | 571 | 750 | 2857 | 565 | 707 | 1648 | 604 | 478 |
| 1905 | QGN15002.1 | FIM1_1683 | CP015055. | 3166 | 2880 | 2785 | 8212 | 2881 | 2358 | 5686 | 2481 | 3607 | 16705 | 2694 | 3514 | 14999 | 3433 | 3406 | 16273 | 2788 | 2351 |
| 1906 | QGN15003.1 | FIM1_1684 | CP015055. | 939 | 704 | 1928 | 1198 | 837 | 1724 | 994 | 617 | 2351 | 5409 | 622 | 3595 | 4452 | 599 | 3102 | 5380 | 557 | 2163 |
| 1907 | QGN15004.1 | FIM1_1685 | CP015055. | 5587 | 1358 | 2127 | 6390 | 1604 | 1758 | 6123 | 1469 | 2713 | 5550 | 852 | 2244 | 5069 | 1080 | 2079 | 5866 | 898 | 1335 |
| 1908 | QGN15005.1 | FIM1_1686 | CP015055. | 1046 | 841 | 646 | 1756 | 686 | 670 | 1502 | 544 | 987 | 3795 | 587 | 1101 | 3220 | 620 | 1033 | 3727 | 541 | 713 |
| 1909 | QGN15006.1 | FIM1_1687 | CP015055. | 2786 | 1086 | 2972 | 4363 | 1198 | 2484 | 3235 | 1071 | 3532 | 6987 | 724 | 4800 | 6226 | 791 | 4181 | 6506 | 608 | 2854 |
| 1910 | QGN15007.1 | FIM1_1688 | CP015055. | 910 | 376 | 383 | 934 | 157 | 195 | 1284 | 312 | 497 | 835 | 122 | 195 | 687 | 142 | 173 | 811 | 102 | 116 |
| 1911 | QGN15008.1 | FIM1_1689 | CP015055. | 740 | 204 | 118 | 945 | 231 | 112 | 483 | 171 | 147 | 511 | 216 | 143 | 685 | 137 | 182 | 577 | 149 | 112 |
| 1912 | QGN15009.1 | FIM1_1690 | CP015055. | 4454 | 2354 | 2918 | 5027 | 2745 | 2480 | 3786 | 2380 | 3807 | 13988 | 1811 | 3788 | 14710 | 1654 | 3742 | 13618 | 1318 | 2611 |
| 1913 | QGN15010.1 | FIM1_1691 | CP015055. | 14769 | 125 | 91 | 34726 | 384 | 134 | 16217 | 64 | 101 | 2918 | 355 | 367 | 3384 | 483 | 359 | 2679 | 300 | 255 |
| 1914 | QGN15011.1 | FIM1_1692 | CP015055. | 7654 | 280 | 348 | 10249 | 406 | 269 | 5689 | 248 | 392 | 2981 | 408 | 593 | 2846 | 374 | 508 | 2842 | 346 | 364 |
| 1915 | QGN15012.1 | FIM1_1693 | CP015055. | 4414 | 476 | 855 | 2677 | 230 | 417 | 5265 | 622 | 1053 | 697 | 182 | 447 | 1077 | 235 | 404 | 787 | 171 | 269 |
| 1916 | QGN15013.1 | FIM1_1694 | CP015055. | 182 | 18 | 28 | 200 | 27 | 20 | 171 | 27 | 45 | 44 | 19 | 38 | 143 | 25 | 35 | 77 | 21 | 33 |
| 1917 | QGN15014.1 | FIM1_1695 | CP015055. | 935 | 119 | 121 | 867 | 83 | 61 | 832 | 128 | 160 | 46 | 39 | 100 | 277 | 42 | 97 | 137 | 60 | 78 |
| 1918 | QGN15015.1 | FIM1_1697 | CP015055. | 371 | 188 | 132 | 1256 | 98 | 82 | 638 | 135 | 155 | 453 | 121 | 125 | 521 | 131 | 128 | 401 | 93 | 70 |
| 1919 | QGN15016.1 | FIM1_1698 | CP015055. | 3059 | 687 | 966 | 4648 | 2124 | 778 | 3764 | 1248 | 1109 | 1691 | 1772 | 1784 | 4261 | 1693 | 1680 | 2536 | 1850 | 1351 |
| 1920 | QGN15017.1 | FIM1_1700 | CP015056. | 0 | 0 | 0 | 0 | 0 | 0 | 0 | 0 | 0 | 0 | 0 | 0 | 0 | 0 | 0 | 0 | 0 | 0 |
| 1921 | QGN15018.1 | FIM1_1701 | CP015056. | 962 | 242 | 247 | 1766 | 427 | 280 | 1007 | 215 | 347 | 2431 | 338 | 422 | 1851 | 428 | 363 | 2477 | 329 | 304 |
| 1922 | QGN15019.1 | FIM1_1703 | CP015056. | 79 | 33 | 48 | 245 | 44 | 38 | 142 | 32 | 69 | 365 | 33 | 92 | 372 | 23 | 61 | 426 | 25 | 65 |
| 1923 | QGN15020.1 | FIM1_1704 | CP015056. | 45 | 8 | 4 | 88 | 14 | 3 | 67 | 4 | 3 | 50 | 15 | 13 | 56 | 9 | 10 | 50 | 9 | 9 |
| 1924 | QGN15021.1 | FIM1_1705 | CP015056. | 36 | 4 | 19 | 59 | 9 | 8 | 57 | 11 | 10 | 89 | 15 | 13 | 97 | 10 | 22 | 146 | 6 | 10 |
| 1925 | QGN15022.1 | FIM1_1706 | CP015056. | 0 | 0 | 0 | 1 | 1 | 1 | 0 | 1 | 0 | 1 | 1 | 0 | 1 | 0 | 0 | 1 | 0 | 1 |
| 1926 | QGN15023.1 | FIM1_1707 | CP015056. | 828 | 125 | 187 | 839 | 272 | 117 | 768 | 204 | 182 | 1042 | 263 | 237 | 1348 | 237 | 225 | 1219 | 218 | 193 |
| 1927 | QGN15024.1 | FIM1_1708 | CP015056. | 32480 | 126 | 237 | 15375 | 602 | 155 | 18048 | 419 | 289 | 365 | 560 | 364 | 1880 | 869 | 315 | 550 | 1539 | 268 |
| 1928 | QGN15025.1 | FIM1_1709 | CP015056. | 10328 | 615 | 511 | 3227 | 688 | 420 | 3970 | 679 | 640 | 2515 | 571 | 756 | 3646 | 619 | 679 | 2841 | 485 | 533 |
| 1929 | QGN15026.1 | FIM1_1710 | CP015056. | 5024 | 179 | 187 | 1446 | 283 | 182 | 1660 | 188 | 231 | 732 | 186 | 253 | 771 | 171 | 248 | 678 | 138 | 158 |
| 1930 | QGN15027.1 | FIM1_1711 | CP015056. | 8229 | 557 | 1503 | 17340 | 302 | 801 | 6530 | 637 | 1848 | 2138 | 177 | 1122 | 2701 | 373 | 913 | 2131 | 217 | 606 |

| | | | | | | | | | | | | | | | | | | | | | |
|---|---|---|---|---|---|---|---|---|---|---|---|---|---|---|---|---|---|---|---|---|---|
| 1931 | QGN15028.1 | FIM1_1712 | CP015056. | 2675 | 3048 | 2185 | 5024 | 2448 | 1635 | 1773 | 4030 | 3374 | 3302 | 1297 | 990 | 4297 | 1866 | 737 | 3413 | 2132 | 707 |
| 1932 | QGN15029.1 | FIM1_1713 | CP015056. | 592 | 50 | 42 | 358 | 63 | 38 | 545 | 31 | 47 | 184 | 38 | 49 | 193 | 31 | 56 | 168 | 34 | 42 |
| 1933 | QGN15030.1 | FIM1_1714 | CP015056. | 2787 | 112 | 79 | 1751 | 118 | 73 | 2574 | 90 | 114 | 653 | 108 | 152 | 700 | 110 | 142 | 611 | 112 | 109 |
| 1934 | QGN15031.1 | FIM1_1715 | CP015056. | 229 | 89 | 174 | 712 | 84 | 166 | 367 | 89 | 222 | 583 | 60 | 246 | 700 | 71 | 208 | 610 | 56 | 140 |
| 1935 | QGN15032.1 | FIM1_1716 | CP015056. | 3313 | 3044 | 4668 | 10934 | 2599 | 3682 | 5867 | 2324 | 5331 | 10774 | 1927 | 5954 | 11452 | 3068 | 5095 | 10621 | 2612 | 3734 |
| 1936 | QGN15033.1 | FIM1_1717 | CP015056. | 289 | 106 | 93 | 486 | 102 | 66 | 318 | 119 | 108 | 558 | 96 | 182 | 660 | 81 | 139 | 600 | 84 | 113 |
| 1937 | QGN15034.1 | FIM1_1718 | CP015056. | 279 | 98 | 104 | 394 | 85 | 58 | 303 | 67 | 123 | 696 | 89 | 183 | 640 | 96 | 174 | 716 | 94 | 125 |
| 1938 | QGN15035.1 | FIM1_1719 | CP015056. | 448 | 138 | 146 | 906 | 148 | 115 | 667 | 133 | 155 | 584 | 118 | 171 | 637 | 111 | 147 | 575 | 83 | 113 |
| 1939 | QGN15036.1 | FIM1_1720 | CP015056. | 6 | 6 | 5 | 24 | 2 | 1 | 14 | 2 | 5 | 19 | 6 | 4 | 16 | 1 | 3 | 26 | 2 | 3 |
| 1940 | QGN15037.1 | FIM1_1721 | CP015056. | 13 | 35 | 17 | 46 | 18 | 13 | 17 | 14 | 6 | 289 | 35 | 27 | 206 | 37 | 31 | 331 | 22 | 22 |
| 1941 | QGN15038.1 | FIM1_1722 | CP015056. | 0 | 0 | 0 | 0 | 0 | 0 | 0 | 0 | 0 | 0 | 0 | 0 | 0 | 0 | 0 | 0 | 0 | 0 |
| 1942 | QGN15039.1 | FIM1_1723 | CP015056. | 10 | 1 | 1 | 45 | 3 | 2 | 9 | 1 | 1 | 8 | 1 | 1 | 8 | 2 | 1 | 10 | 0 | 0 |
| 1943 | QGN15040.1 | FIM1_1724 | CP015056. | 36939 | 3620 | 2021 | 47279 | 3433 | 1729 | 44629 | 3376 | 3030 | 25132 | 4753 | 2544 | 24540 | 12867 | 2931 | 21832 | 7211 | 1997 |
| 1944 | QGN15041.1 | FIM1_1725 | CP015056. | 2850 | 10821 | 10678 | 10010 | 14938 | 9016 | 6522 | 13162 | 12607 | 47805 | 18134 | 18356 | 48761 | 17561 | 17136 | 48290 | 15832 | 12971 |
| 1945 | QGN15042.1 | FIM1_1727 | CP015056. | 2091 | 1052 | 1887 | 4499 | 1049 | 1367 | 2784 | 1263 | 2303 | 2439 | 731 | 2456 | 6164 | 775 | 2162 | 3679 | 691 | 1632 |
| 1946 | QGN15043.1 | FIM1_1728 | CP015056. | 7201 | 3193 | 4211 | 11628 | 3816 | 3465 | 8469 | 3698 | 5319 | 11777 | 3449 | 6353 | 18252 | 4215 | 5822 | 13311 | 3320 | 4286 |
| 1947 | QGN15044.1 | FIM1_1729 | CP015056. | 2979 | 2051 | 2823 | 4650 | 1877 | 2215 | 3567 | 2101 | 3567 | 7026 | 2081 | 3869 | 10492 | 2345 | 4185 | 7356 | 1939 | 2853 |
| 1948 | QGN15045.1 | FIM1_1730 | CP015056. | 4372 | 3750 | 2854 | 6506 | 3587 | 2596 | 5050 | 3302 | 3789 | 11507 | 3255 | 4631 | 13103 | 3067 | 4624 | 11276 | 2828 | 3167 |
| 1949 | QGN15046.1 | FIM1_1731 | CP015056. | 3464 | 3544 | 2917 | 5432 | 3532 | 2774 | 4241 | 3308 | 3885 | 12695 | 3328 | 4519 | 14725 | 3288 | 4354 | 12617 | 2999 | 3023 |
| 1950 | QGN15047.1 | FIM1_1732 | CP015056. | 1811 | 1427 | 694 | 1952 | 794 | 641 | 1707 | 790 | 881 | 1634 | 660 | 924 | 2465 | 547 | 958 | 1871 | 606 | 628 |
| 1951 | QGN15048.1 | FIM1_1733 | CP015056. | 7412 | 3776 | 6210 | 20268 | 5343 | 4584 | 11413 | 4637 | 7489 | 6450 | 3554 | 10027 | 24019 | 4397 | 8279 | 13105 | 3767 | 6593 |
| 1952 | QGN15049.1 | FIM1_1734 | CP015056. | 359 | 280 | 366 | 922 | 348 | 295 | 365 | 293 | 452 | 1869 | 392 | 561 | 2460 | 261 | 499 | 2016 | 242 | 376 |
| 1953 | QGN15050.1 | FIM1_1735 | CP015056. | 2250 | 2730 | 1642 | 4692 | 2109 | 1590 | 2580 | 1696 | 2089 | 6840 | 1579 | 2107 | 6283 | 1563 | 1871 | 6306 | 1386 | 1414 |
| 1954 | QGN15051.1 | FIM1_1736 | CP015056. | 1286 | 378 | 416 | 1029 | 376 | 406 | 1309 | 351 | 584 | 2690 | 332 | 576 | 2192 | 298 | 561 | 2460 | 305 | 393 |
| 1955 | QGN15052.1 | FIM1_1737 | CP015056. | 761 | 292 | 471 | 1193 | 239 | 385 | 847 | 283 | 635 | 1048 | 131 | 572 | 1715 | 131 | 540 | 1205 | 121 | 365 |
| 1956 | QGN15053.1 | FIM1_1738 | CP015056. | 2060 | 1259 | 1189 | 4395 | 1100 | 951 | 3328 | 1136 | 1508 | 3884 | 675 | 1107 | 3675 | 619 | 1027 | 3688 | 535 | 724 |
| 1957 | QGN15054.1 | FIM1_1739 | CP015056. | 3381 | 2201 | 1308 | 2779 | 2109 | 1429 | 3067 | 1977 | 1793 | 8617 | 2508 | 2599 | 8396 | 2060 | 2568 | 7863 | 2408 | 1769 |
| 1958 | QGN15055.1 | FIM1_1740 | CP015056. | 1421 | 798 | 751 | 1888 | 533 | 624 | 1480 | 616 | 993 | 2792 | 477 | 972 | 2373 | 396 | 953 | 2491 | 405 | 616 |
| 1959 | QGN15056.1 | FIM1_1741 | CP015056. | 17620 | 3913 | 1592 | 10692 | 3065 | 1745 | 12491 | 2729 | 2179 | 6684 | 3392 | 2653 | 7770 | 2800 | 2821 | 6302 | 3171 | 1975 |
| 1960 | QGN15057.1 | FIM1_1742 | CP015056. | 2083 | 1166 | 433 | 2055 | 852 | 469 | 2275 | 870 | 616 | 2497 | 1039 | 892 | 3094 | 827 | 979 | 2304 | 986 | 623 |
| 1961 | QGN15058.1 | FIM1_1743 | CP015056. | 2441 | 1754 | 2079 | 2521 | 3358 | 1753 | 2190 | 2830 | 2569 | 3710 | 2321 | 2757 | 11171 | 2123 | 2584 | 6349 | 2255 | 1828 |
| 1962 | QGN15059.1 | FIM1_1744 | CP015056. | 6862 | 1457 | 1401 | 6969 | 1500 | 1046 | 6821 | 1600 | 1885 | 1434 | 1259 | 1906 | 7751 | 1401 | 1849 | 3348 | 1385 | 1250 |
| 1963 | QGN15060.1 | FIM1_1745 | CP015056. | 1265 | 1358 | 868 | 1873 | 1024 | 797 | 1460 | 1034 | 1165 | 3658 | 885 | 1226 | 4007 | 757 | 1252 | 3412 | 850 | 822 |
| 1964 | QGN15061.1 | FIM1_1746 | CP015056. | 545 | 454 | 270 | 628 | 377 | 315 | 564 | 344 | 381 | 2067 | 345 | 490 | 1553 | 300 | 434 | 1695 | 337 | 321 |
| 1965 | QGN15062.1 | FIM1_1747 | CP015056. | 6504 | 1975 | 2019 | 6641 | 1761 | 1683 | 6300 | 1633 | 2700 | 4363 | 1858 | 3043 | 11295 | 1932 | 2780 | 6742 | 1599 | 2111 |
| 1966 | QGN15063.1 | FIM1_1748 | CP015056. | 1918 | 30 | 23 | 3385 | 28 | 25 | 2441 | 38 | 33 | 41 | 17 | 22 | 95 | 15 | 23 | 73 | 16 | 16 |
| 1967 | QGN15064.1 | FIM1_1749 | CP015056. | 9027 | 3580 | 2662 | 12020 | 3208 | 2173 | 9907 | 3202 | 3383 | 5497 | 2489 | 3533 | 11786 | 2935 | 3678 | 7752 | 2584 | 2535 |
| 1968 | QGN15065.1 | FIM1_1750 | CP015056. | 3175 | 3020 | 3084 | 5064 | 3254 | 2606 | 3486 | 3105 | 3941 | 2661 | 3315 | 4606 | 13145 | 3251 | 4627 | 7218 | 3072 | 3158 |
| 1969 | QGN15066.1 | FIM1_1751 | CP015056. | 1888 | 1428 | 408 | 673 | 883 | 406 | 1482 | 923 | 622 | 928 | 1178 | 784 | 2723 | 900 | 855 | 1604 | 1381 | 555 |
| 1970 | QGN15067.1 | FIM1_1752 | CP015056. | 4299 | 483 | 636 | 3513 | 711 | 494 | 3971 | 782 | 870 | 540 | 484 | 986 | 2668 | 593 | 914 | 1107 | 510 | 578 |
| 1971 | QGN15068.1 | FIM1_1753 | CP015056. | 935 | 530 | 702 | 1471 | 464 | 562 | 1137 | 574 | 818 | 2472 | 321 | 885 | 2613 | 308 | 900 | 2302 | 263 | 672 |
| 1972 | QGN15069.1 | FIM1_1754 | CP015056. | 3033 | 324 | 197 | 3067 | 438 | 155 | 2054 | 468 | 287 | 314 | 201 | 256 | 1037 | 179 | 192 | 482 | 187 | 162 |
| 1973 | QGN15070.1 | FIM1_1755 | CP015056. | 6738 | 2175 | 1193 | 6044 | 1384 | 872 | 6479 | 1615 | 1573 | 797 | 1129 | 1466 | 4727 | 1265 | 1505 | 1641 | 1272 | 1056 |
| 1974 | QGN15071.1 | FIM1_1756 | CP015056. | 1036 | 1237 | 797 | 2781 | 1048 | 700 | 1738 | 1158 | 1139 | 985 | 673 | 916 | 2540 | 610 | 908 | 1468 | 663 | 668 |
| 1975 | QGN15072.1 | FIM1_1757 | CP015056. | 7671 | 1481 | 926 | 3707 | 838 | 694 | 5620 | 981 | 1341 | 1636 | 746 | 1201 | 3880 | 772 | 1255 | 2886 | 727 | 903 |
| 1976 | QGN15073.1 | FIM1_1758 | CP015056. | 310 | 395 | 217 | 138 | 323 | 189 | 230 | 296 | 301 | 376 | 257 | 295 | 908 | 230 | 351 | 641 | 221 | 222 |
| 1977 | QGN15074.1 | FIM1_1759 | CP015056. | 1199 | 264 | 243 | 477 | 182 | 166 | 888 | 275 | 312 | 187 | 197 | 253 | 429 | 185 | 238 | 289 | 203 | 180 |
| 1978 | QGN15075.1 | FIM1_1760 | CP015056. | 8416 | 1837 | 2260 | 8526 | 2282 | 1854 | 8143 | 2354 | 3051 | 2943 | 2123 | 3105 | 12018 | 2037 | 2947 | 5632 | 2041 | 2019 |
| 1979 | QGN15076.1 | FIM1_1761 | CP015056. | 8599 | 1196 | 1531 | 9525 | 1620 | 1209 | 7909 | 1746 | 2116 | 1549 | 1057 | 2512 | 7189 | 1250 | 2175 | 3073 | 1216 | 1700 |
| 1980 | QGN15077.1 | FIM1_1762 | CP015056. | 5590 | 4931 | 1567 | 3049 | 4092 | 1808 | 5010 | 4036 | 2200 | 8181 | 5059 | 3105 | 11003 | 3893 | 3252 | 8354 | 5152 | 2447 |
| 1981 | QGN15078.1 | FIM1_1763 | CP015056. | 61 | 189 | 83 | 35 | 80 | 81 | 49 | 103 | 114 | 246 | 73 | 80 | 231 | 59 | 101 | 216 | 66 | 70 |
| 1982 | QGN15079.1 | FIM1_1764 | CP015056. | 1840 | 3164 | 2294 | 2148 | 3151 | 2063 | 1858 | 2854 | 2911 | 9266 | 3113 | 3291 | 12170 | 3006 | 3048 | 9234 | 3009 | 2301 |
| 1983 | QGN15080.1 | FIM1_1765 | CP015056. | 8484 | 1303 | 1493 | 8107 | 1338 | 1248 | 9304 | 1170 | 1827 | 4072 | 954 | 1376 | 4450 | 864 | 1378 | 4110 | 777 | 996 |
| 1984 | QGN15081.1 | FIM1_1766 | CP015056. | 7162 | 825 | 960 | 7238 | 963 | 720 | 6799 | 1098 | 1232 | 889 | 603 | 1350 | 4446 | 819 | 1299 | 2088 | 681 | 944 |
| 1985 | QGN15082.1 | FIM1_1767 | CP015056. | 2216 | 1368 | 1195 | 3418 | 1019 | 1064 | 2693 | 933 | 1687 | 5283 | 834 | 1940 | 5362 | 1039 | 1753 | 5348 | 824 | 1170 |
| 1986 | QGN15083.1 | FIM1_1769 | CP015056. | 40 | 15 | 8 | 33 | 32 | 15 | 16 | 12 | 14 | 131 | 7 | 12 | 86 | 7 | 18 | 97 | 15 | 11 |
| 1987 | QGN15084.1 | FIM1_1770 | CP015056. | 2523 | 2513 | 1840 | 2473 | 2272 | 1703 | 2281 | 1991 | 2093 | 5976 | 2657 | 2499 | 5638 | 2476 | 2488 | 5808 | 2339 | 1895 |
| 1988 | QGN15085.1 | FIM1_1771 | CP015056. | 5148 | 781 | 794 | 4310 | 1209 | 830 | 4555 | 862 | 1013 | 2975 | 1199 | 1002 | 3762 | 1358 | 987 | 3284 | 1146 | 712 |
| 1989 | QGN15086.1 | FIM1_1772 | CP015056. | 858 | 1860 | 1018 | 909 | 1417 | 980 | 928 | 1147 | 1356 | 6959 | 1352 | 1475 | 4855 | 1213 | 1405 | 6271 | 1354 | 1037 |
| 1990 | QGN15087.1 | FIM1_1774 | CP015056. | 1362 | 792 | 686 | 1680 | 534 | 567 | 1445 | 507 | 867 | 3124 | 510 | 960 | 2786 | 544 | 1006 | 3319 | 480 | 632 |
| 1991 | QGN15088.1 | FIM1_1775 | CP015056. | 5232 | 3889 | 4631 | 9799 | 4618 | 3935 | 6580 | 4084 | 5675 | 17571 | 3959 | 7058 | 22131 | 4708 | 6688 | 18705 | 3587 | 4712 |
| 1992 | QGN15089.1 | FIM1_1776 | CP015056. | 2984 | 1242 | 1562 | 4236 | 1918 | 1306 | 4037 | 1499 | 2077 | 5491 | 1687 | 2428 | 10838 | 1876 | 2257 | 7773 | 1628 | 1638 |
| 1993 | QGN15090.1 | FIM1_1777 | CP015056. | 3903 | 2419 | 2906 | 7358 | 2693 | 2604 | 4729 | 2941 | 4030 | 11735 | 2429 | 2052 | 10067 | 3155 | 2308 | 11536 | 2478 | 1491 |
| 1994 | QGN15091.1 | FIM1_1778 | CP015056. | 5485 | 1353 | 1608 | 6260 | 1510 | 1315 | 5195 | 1671 | 2102 | 4293 | 1474 | 2228 | 9347 | 1445 | 2073 | 6250 | 1330 | 1668 |
| 1995 | QGN15092.1 | FIM1_1779 | CP015056. | 6944 | 2082 | 2522 | 16750 | 2387 | 2488 | 12665 | 1792 | 3651 | 19961 | 3155 | 3553 | 18206 | 5446 | 3866 | 19535 | 2651 | 2732 |
| 1996 | QGN15093.1 | FIM1_1780 | CP015056. | 1569 | 763 | 511 | 1732 | 595 | 452 | 1745 | 461 | 655 | 3869 | 645 | 1012 | 3219 | 571 | 887 | 3780 | 531 | 661 |
| 1997 | QGN15094.1 | FIM1_1781 | CP015056. | 1765 | 178 | 292 | 2127 | 340 | 273 | 1942 | 283 | 394 | 2624 | 265 | 528 | 2366 | 307 | 435 | 2486 | 261 | 375 |
| 1998 | QGN15095.1 | FIM1_1782 | CP015056. | 2955 | 571 | 684 | 3310 | 594 | 570 | 3060 | 569 | 881 | 3759 | 443 | 921 | 3947 | 533 | 903 | 3490 | 459 | 653 |
| 1999 | QGN15096.1 | FIM1_1783 | CP015056. | 1104 | 190 | 343 | 1427 | 208 | 338 | 963 | 133 | 474 | 1527 | 138 | 595 | 1121 | 123 | 641 | 1286 | 135 | 337 |
| 2000 | QGN15097.1 | FIM1_1784 | CP015056. | 4821 | 783 | 1326 | 4872 | 1024 | 1053 | 4921 | 1013 | 1730 | 4066 | 830 | 2398 | 7125 | 864 | 1980 | 5179 | 784 | 1564 |
| 2001 | QGN15098.1 | FIM1_1785 | CP015056. | 738 | 377 | 737 | 1193 | 411 | 515 | 808 | 484 | 889 | 4544 | 401 | 1129 | 4076 | 341 | 904 | 4427 | 353 | 678 |
| 2002 | QGN15099.1 | FIM1_1786 | CP015056. | 99 | 39 | 32 | 112 | 45 | 37 | 65 | 36 | 30 | 114 | 37 | 63 | 203 | 24 | 60 | 143 | 28 | 42 |
| 2003 | QGN15100.1 | FIM1_1787 | CP015056. | 3068 | 2746 | 954 | 1492 | 1997 | 1087 | 2434 | 1744 | 1250 | 6468 | 2586 | 1715 | 6485 | 2076 | 1835 | 6171 | 2676 | 1250 |
| 2004 | QGN15101.1 | FIM1_1788 | CP015056. | 658 | 449 | 317 | 575 | 319 | 270 | 646 | 285 | 421 | 1938 | 366 | 401 | 1745 | 308 | 440 | 1940 | 334 | 259 |
| 2005 | QGN15102.1 | FIM1_1789 | CP015056. | 2640 | 545 | 917 | 3057 | 606 | 619 | 2970 | 680 | 1168 | 1014 | 523 | 1542 | 4470 | 502 | 1330 | 2022 | 531 | 1005 |
| 2006 | QGN15103.1 | FIM1_1790 | CP015056. | 1785 | 825 | 787 | 3093 | 514 | 737 | 1845 | 545 | 1065 | 1499 | 264 | 555 | 1690 | 329 | 570 | 1565 | 257 | 367 |
| 2007 | QGN15104.1 | FIM1_1791 | CP015056. | 617 | 223 | 214 | 747 | 183 | 166 | 603 | 193 | 294 | 162 | 90 | 385 | 244 | 144 | 328 | 218 | 145 | 251 |
| 2008 | QGN15105.1 | FIM1_1792 | CP015056. | 4207 | 3358 | 4600 | 8081 | 5015 | 3230 | 5612 | 4622 | 5340 | 6224 | 4478 | 7159 | 21217 | 5024 | 6064 | 11545 | 4349 | 4341 |
| 2009 | QGN15106.1 | FIM1_1793 | CP015056. | 2730 | 1455 | 1183 | 2607 | 1403 | 918 | 2493 | 1358 | 1491 | 5040 | 1382 | 1912 | 6414 | 1033 | 1856 | 5189 | 891 | 1293 |
| 2010 | QGN15107.1 | FIM1_1794 | CP015056. | 1673 | 74 | 629 | 1309 | 300 | 426 | 1373 | 424 | 789 | 836 | 234 | 774 | 3175 | 239 | 689 | 1699 | 189 | 486 |
| 2011 | QGN15108.1 | FIM1_1795 | CP015056. | 1556 | 598 | 517 | 1742 | 596 | 453 | 1558 | 508 | 740 | 2331 | 364 | 667 | 2553 | 335 | 686 | 2349 | 362 | 439 |
| 2012 | QGN15109.1 | FIM1_1796 | CP015056. | 68 | 98 | 70 | 102 | 74 | 49 | 53 | 84 | 67 | 136 | 66 | 105 | 321 | 39 | 95 | 212 | 56 | 54 |
| 2013 | QGN15110.1 | FIM1_1797 | CP015056. | 2973 | 1187 | 943 | 1222 | 1421 | 774 | 1195 | 1256 | 1216 | 4270 | 1078 | 1598 | 6007 | 1172 | 1535 | 4729 | 913 | 931 |
| 2014 | QGN15111.1 | FIM1_1798 | CP015056. | 9085 | 790 | 982 | 7429 | 924 | 709 | 7616 | 1055 | 1226 | 952 | 656 | 1338 | 4517 | 754 | 1249 | 1814 | 683 | 933 |

| | | | | | | | | | | | | | | | | | | | | | |
|---|---|---|---|---|---|---|---|---|---|---|---|---|---|---|---|---|---|---|---|---|---|
| 2015 | QGN15112.1 | FIM1_1799 | CP015056. | 2433 | 1713 | 1015 | 2794 | 1234 | 981 | 2483 | 1163 | 1422 | 4214 | 1162 | 1404 | 4257 | 1180 | 1574 | 3623 | 1190 | 993 |
| 2016 | QGN15113.1 | FIM1_1800 | CP015056. | 5128 | 462 | 523 | 3790 | 379 | 386 | 4517 | 538 | 669 | 824 | 242 | 604 | 2221 | 195 | 545 | 1215 | 188 | 386 |
| 2017 | QGN15114.1 | FIM1_1801 | CP015056. | 2134 | 1139 | 691 | 1069 | 1692 | 781 | 1597 | 1198 | 940 | 1598 | 2065 | 1456 | 4002 | 2544 | 1631 | 2733 | 2380 | 1057 |
| 2018 | QGN15115.1 | FIM1_1802 | CP015056. | 11889 | 1970 | 1513 | 12255 | 1578 | 1161 | 10647 | 1997 | 2148 | 988 | 1057 | 2217 | 4338 | 1393 | 1944 | 1539 | 1345 | 1484 |
| 2019 | QGN15116.1 | FIM1_1803 | CP015056. | 41057 | 1543 | 1379 | 24396 | 1063 | 941 | 32852 | 1499 | 1908 | 718 | 927 | 2365 | 5221 | 1573 | 2226 | 1627 | 1476 | 1642 |
| 2020 | QGN15117.1 | FIM1_1804 | CP015056. | 4998 | 403 | 340 | 2601 | 289 | 231 | 3051 | 380 | 444 | 548 | 272 | 430 | 1380 | 287 | 515 | 723 | 365 | 310 |
| 2021 | QGN15118.1 | FIM1_1805 | CP015056. | 1030 | 36 | 24 | 671 | 23 | 22 | 364 | 24 | 32 | 235 | 35 | 51 | 167 | 28 | 46 | 216 | 37 | 26 |
| 2022 | QGN15119.1 | FIM1_1806 | CP015056. | 7130 | 1086 | 654 | 5182 | 543 | 446 | 6370 | 787 | 896 | 924 | 642 | 970 | 3793 | 775 | 1096 | 2215 | 712 | 747 |
| 2023 | QGN15120.1 | FIM1_1807 | CP015056. | 2021 | 54 | 51 | 1277 | 79 | 39 | 1056 | 55 | 64 | 129 | 83 | 69 | 234 | 81 | 75 | 140 | 75 | 70 |
| 2024 | QGN15121.1 | FIM1_1808 | CP015056. | 10154 | 1510 | 1396 | 5211 | 897 | 915 | 5368 | 1440 | 1900 | 1131 | 968 | 1668 | 3197 | 1217 | 1951 | 1875 | 951 | 1271 |
| 2025 | QGN15122.1 | FIM1_1809 | CP015056. | 7871 | 1274 | 2001 | 10938 | 1880 | 1354 | 8351 | 2254 | 2573 | 1302 | 1113 | 3273 | 6794 | 1589 | 2778 | 2680 | 1512 | 2018 |
| 2026 | QGN15123.1 | FIM1_1810 | CP015056. | 2276 | 806 | 1003 | 3773 | 938 | 836 | 2934 | 964 | 1269 | 2465 | 758 | 1507 | 4465 | 889 | 1299 | 3063 | 794 | 1032 |
| 2027 | QGN15124.1 | FIM1_1811 | CP015056. | 1770 | 1445 | 927 | 1924 | 961 | 785 | 1892 | 940 | 1204 | 5115 | 966 | 1379 | 4830 | 930 | 1420 | 4607 | 819 | 984 |
| 2028 | QGN15125.1 | FIM1_1812 | CP015056. | 2479 | 316 | 512 | 1842 | 267 | 392 | 1857 | 425 | 620 | 977 | 266 | 633 | 2132 | 232 | 632 | 1384 | 264 | 422 |
| 2029 | QGN15126.1 | FIM1_1813 | CP015056. | 1130 | 449 | 398 | 981 | 328 | 293 | 849 | 341 | 435 | 1427 | 269 | 363 | 1311 | 275 | 471 | 1252 | 271 | 259 |
| 2030 | QGN15127.1 | FIM1_1814 | CP015056. | 922 | 645 | 615 | 1465 | 562 | 583 | 1240 | 655 | 763 | 2303 | 507 | 863 | 2881 | 410 | 869 | 2284 | 436 | 596 |
| 2031 | QGN15128.1 | FIM1_1815 | CP015056. | 2007 | 572 | 363 | 2708 | 439 | 358 | 1803 | 454 | 434 | 1338 | 283 | 521 | 1652 | 281 | 503 | 1293 | 280 | 344 |
| 2032 | QGN15129.1 | FIM1_1816 | CP015056. | 1701 | 1117 | 1420 | 2663 | 1576 | 1248 | 2105 | 1491 | 1797 | 5541 | 1608 | 2265 | 8936 | 1688 | 2094 | 6632 | 1417 | 1462 |
| 2033 | QGN15130.1 | FIM1_1817 | CP015056. | 1694 | 899 | 1070 | 2182 | 1175 | 870 | 1618 | 1009 | 1389 | 3691 | 1041 | 1719 | 6995 | 1169 | 1483 | 4557 | 1057 | 1238 |
| 2034 | QGN15131.1 | FIM1_1818 | CP015056. | 18847 | 22557 | 13090 | 19246 | 18750 | 11747 | 16714 | 18108 | 15668 | 40914 | 19647 | 19652 | 48551 | 18805 | 19590 | 43859 | 17791 | 14515 |
| 2035 | QGN15132.1 | FIM1_1819 | CP015056. | 3691 | 1814 | 1941 | 4698 | 1989 | 1736 | 3741 | 2069 | 2491 | 4637 | 1582 | 2836 | 8544 | 1615 | 2468 | 5866 | 1421 | 1891 |
| 2036 | QGN15133.1 | FIM1_1820 | CP015056. | 761 | 557 | 263 | 591 | 417 | 316 | 673 | 403 | 444 | 1707 | 277 | 347 | 1453 | 310 | 345 | 1445 | 318 | 210 |
| 2037 | QGN15134.1 | FIM1_1821 | CP015056. | 1019 | 1754 | 1135 | 926 | 1377 | 1071 | 904 | 1176 | 1600 | 5406 | 1341 | 1418 | 4652 | 1205 | 1494 | 4853 | 1334 | 975 |
| 2038 | QGN15135.1 | FIM1_1822 | CP015056. | 1021 | 388 | 327 | 968 | 281 | 289 | 806 | 269 | 434 | 1576 | 226 | 382 | 1422 | 183 | 407 | 1445 | 183 | 239 |
| 2039 | QGN15136.1 | FIM1_1823 | CP015056. | 261 | 268 | 110 | 167 | 167 | 220 | 154 | 125 | 187 | 910 | 212 | 213 | 555 | 152 | 264 | 612 | 280 | 135 |
| 2040 | QGN15137.1 | FIM1_1824 | CP015056. | 2842 | 735 | 689 | 2609 | 659 | 599 | 2063 | 705 | 926 | 1079 | 534 | 889 | 2754 | 490 | 882 | 1735 | 466 | 573 |
| 2041 | QGN15138.1 | FIM1_1825 | CP015056. | 1308 | 3965 | 2903 | 2648 | 3859 | 3018 | 1769 | 3017 | 3679 | 15942 | 4054 | 4658 | 12708 | 3436 | 5033 | 14735 | 3270 | 3092 |
| 2042 | QGN15139.1 | FIM1_1826 | CP015056. | 3249 | 2900 | 4648 | 7480 | 3913 | 3510 | 4434 | 3242 | 5979 | 10673 | 3894 | 5803 | 24860 | 5109 | 5424 | 18182 | 3375 | 4040 |
| 2043 | QGN15140.1 | FIM1_1827 | CP015056. | 6071 | 5271 | 3619 | 5714 | 3872 | 3148 | 5393 | 3714 | 4672 | 11536 | 3845 | 5196 | 13951 | 3879 | 5535 | 11211 | 3171 | 3686 |
| 2044 | QGN15141.1 | FIM1_1828 | CP015056. | 5165 | 2070 | 2332 | 10550 | 1823 | 2033 | 7491 | 1671 | 3004 | 9864 | 1875 | 3592 | 12854 | 2011 | 3384 | 11023 | 1759 | 2588 |
| 2045 | QGN15142.1 | FIM1_1829 | CP015056. | 797 | 219 | 213 | 696 | 133 | 148 | 619 | 146 | 282 | 472 | 109 | 275 | 575 | 103 | 278 | 568 | 112 | 190 |
| 2046 | QGN15143.1 | FIM1_1830 | CP015056. | 2168 | 1152 | 1147 | 2523 | 1295 | 1002 | 2240 | 1230 | 1546 | 4541 | 1266 | 1642 | 7014 | 1069 | 1610 | 5128 | 1054 | 1207 |
| 2047 | QGN15144.1 | FIM1_1831 | CP015056. | 487 | 466 | 325 | 396 | 406 | 326 | 431 | 378 | 442 | 2221 | 406 | 475 | 1822 | 359 | 494 | 2068 | 402 | 260 |
| 2048 | QGN15145.1 | FIM1_1832 | CP015056. | 517 | 221 | 281 | 859 | 203 | 238 | 583 | 217 | 370 | 1865 | 158 | 428 | 1593 | 171 | 424 | 1736 | 183 | 288 |
| 2049 | QGN15146.1 | FIM1_1833 | CP015056. | 2279 | 634 | 700 | 2203 | 623 | 567 | 2143 | 663 | 843 | 2495 | 457 | 828 | 2619 | 390 | 741 | 2501 | 407 | 520 |
| 2050 | QGN15147.1 | FIM1_1834 | CP015056. | 2233 | 418 | 624 | 3292 | 390 | 481 | 3062 | 464 | 812 | 1807 | 294 | 831 | 2685 | 315 | 681 | 1997 | 248 | 525 |
| 2051 | QGN15148.1 | FIM1_1835 | CP015056. | 1533 | 384 | 563 | 1559 | 316 | 437 | 1368 | 445 | 757 | 1296 | 251 | 659 | 1864 | 259 | 648 | 1482 | 244 | 457 |
| 2052 | QGN15149.1 | FIM1_1836 | CP015056. | 1998 | 1199 | 1354 | 1706 | 1684 | 1189 | 1712 | 1656 | 1835 | 3177 | 1602 | 1948 | 7416 | 1472 | 1834 | 4448 | 1479 | 1323 |
| 2053 | QGN15150.1 | FIM1_1837 | CP015056. | 5858 | 4125 | 2249 | 4599 | 4291 | 2171 | 5782 | 4062 | 3026 | 5996 | 4928 | 4050 | 15372 | 3846 | 3813 | 9602 | 4540 | 2892 |
| 2054 | QGN15151.1 | FIM1_1838 | CP015056. | 384 | 53 | 126 | 480 | 112 | 70 | 333 | 100 | 132 | 222 | 110 | 172 | 465 | 107 | 134 | 276 | 107 | 84 |
| 2055 | QGN15152.1 | FIM1_1839 | CP015056. | 2715 | 3912 | 2661 | 4071 | 2600 | 2630 | 3235 | 2206 | 3456 | 6406 | 1859 | 3090 | 6526 | 1875 | 3157 | 5868 | 1815 | 2178 |
| 2056 | QGN15153.1 | FIM1_1840 | CP015056. | 12119 | 3842 | 3918 | 12623 | 5333 | 3004 | 10092 | 5306 | 5026 | 8069 | 5130 | 6270 | 20148 | 5453 | 6103 | 13960 | 4906 | 4329 |
| 2057 | QGN15154.1 | FIM1_1841 | CP015056. | 1773 | 978 | 886 | 2348 | 738 | 706 | 2014 | 902 | 1119 | 2133 | 490 | 901 | 2360 | 467 | 915 | 2073 | 416 | 600 |
| 2058 | QGN15155.1 | FIM1_1842 | CP015056. | 2630 | 500 | 723 | 2091 | 467 | 578 | 2151 | 537 | 978 | 974 | 406 | 976 | 4272 | 422 | 897 | 2106 | 386 | 680 |
| 2059 | QGN15156.1 | FIM1_1843 | CP015056. | 8992 | 4787 | 4240 | 12568 | 5395 | 4005 | 11105 | 4513 | 5491 | 17143 | 6229 | 6675 | 26498 | 5596 | 6495 | 20290 | 5863 | 4796 |
| 2060 | QGN15157.1 | FIM1_1844 | CP015056. | 45 | 59 | 37 | 40 | 32 | 29 | 46 | 26 | 49 | 278 | 52 | 41 | 169 | 29 | 62 | 227 | 42 | 37 |
| 2061 | QGN15158.1 | FIM1_1845 | CP015056. | 381 | 520 | 392 | 511 | 427 | 366 | 387 | 382 | 502 | 1740 | 359 | 432 | 1695 | 292 | 456 | 1532 | 333 | 290 |
| 2062 | QGN15159.1 | FIM1_1846 | CP015056. | 2121 | 3390 | 1928 | 2322 | 3173 | 1713 | 2378 | 3143 | 2609 | 7433 | 2469 | 2902 | 11685 | 2584 | 2513 | 7742 | 2703 | 1889 |
| 2063 | QGN15160.1 | FIM1_1847 | CP015056. | 7903 | 3129 | 2325 | 7767 | 1817 | 1501 | 6578 | 2366 | 2937 | 10964 | 2338 | 3203 | 13401 | 3458 | 3701 | 11020 | 2452 | 2509 |
| 2064 | QGN15161.1 | FIM1_1848 | CP015056. | 8789 | 7050 | 4158 | 8761 | 4607 | 3126 | 6220 | 4809 | 5160 | 7844 | 5207 | 6974 | 19092 | 5887 | 7136 | 11768 | 5416 | 4728 |
| 2065 | QGN15162.1 | FIM1_1849 | CP015056. | 5961 | 2417 | 3379 | 8925 | 1782 | 2683 | 7216 | 1462 | 4308 | 3347 | 1562 | 4869 | 5660 | 1865 | 5347 | 4069 | 1574 | 3494 |
| 2066 | QGN15163.1 | FIM1_1850 | CP015056. | 1267 | 485 | 490 | 1244 | 490 | 411 | 1090 | 491 | 670 | 755 | 305 | 580 | 1825 | 314 | 663 | 1216 | 330 | 394 |
| 2067 | QGN15164.1 | FIM1_1851 | CP015056. | 3472 | 3999 | 2998 | 5134 | 3244 | 2882 | 4119 | 2984 | 4043 | 8816 | 2220 | 3690 | 8803 | 2424 | 3676 | 7935 | 2151 | 2548 |
| 2068 | QGN15165.1 | FIM1_1853 | CP015056. | 81709 | 31227 | 27391 | 78659 | 35486 | 24099 | 75445 | 34374 | 33981 | 50671 | 41634 | 41505 | 103782 | 39088 | 40961 | 73115 | 38392 | 31840 |
| 2069 | QGN15166.1 | FIM1_1854 | CP015056. | 11473 | 3578 | 2841 | 22892 | 4264 | 2395 | 14129 | 4142 | 3835 | 5522 | 4101 | 4451 | 15427 | 4497 | 4250 | 8279 | 4744 | 3133 |
| 2070 | QGN15167.1 | FIM1_1855 | CP015056. | 2027 | 415 | 503 | 2839 | 384 | 395 | 3198 | 610 | 703 | 892 | 311 | 500 | 2312 | 446 | 462 | 1333 | 387 | 386 |
| 2071 | QGN15168.1 | FIM1_1856 | CP015056. | 1548 | 1278 | 1207 | 3118 | 1154 | 910 | 2237 | 1274 | 1576 | 1778 | 775 | 1620 | 5053 | 891 | 1487 | 3083 | 765 | 1078 |
| 2072 | QGN15169.1 | FIM1_1857 | CP015056. | 7830 | 3612 | 3657 | 9340 | 4973 | 3056 | 8783 | 4545 | 4789 | 14060 | 4455 | 5557 | 27752 | 3870 | 5182 | 18954 | 3689 | 3806 |
| 2073 | QGN15170.1 | FIM1_1858 | CP015056. | 1693 | 1500 | 999 | 2867 | 1178 | 876 | 2205 | 985 | 1251 | 2817 | 671 | 1019 | 3132 | 704 | 1017 | 2799 | 690 | 689 |
| 2074 | QGN15171.1 | FIM1_1859 | CP015056. | 973 | 2180 | 1206 | 1083 | 1555 | 1190 | 1018 | 1380 | 1705 | 7005 | 1502 | 1657 | 5550 | 1392 | 1743 | 6498 | 1479 | 1107 |
| 2075 | QGN15172.1 | FIM1_1860 | CP015056. | 3411 | 517 | 858 | 3982 | 635 | 730 | 3335 | 645 | 1204 | 1750 | 603 | 1469 | 5703 | 631 | 1273 | 3333 | 616 | 992 |
| 2076 | QGN15173.1 | FIM1_1861 | CP015056. | 871 | 448 | 436 | 922 | 536 | 336 | 672 | 453 | 480 | 1903 | 496 | 544 | 1925 | 337 | 470 | 1934 | 383 | 307 |
| 2077 | QGN15174.1 | FIM1_1862 | CP015056. | 1165 | 463 | 529 | 1173 | 685 | 432 | 955 | 582 | 705 | 2048 | 567 | 778 | 3395 | 555 | 705 | 2598 | 530 | 525 |
| 2078 | QGN15175.1 | FIM1_1863 | CP015056. | 352 | 358 | 428 | 378 | 434 | 393 | 317 | 405 | 585 | 2838 | 473 | 763 | 2777 | 451 | 727 | 2933 | 392 | 510 |
| 2079 | QGN15176.1 | FIM1_1864 | CP015056. | 4437 | 1008 | 1052 | 4775 | 932 | 841 | 4837 | 867 | 1346 | 5008 | 508 | 1004 | 4081 | 539 | 999 | 5046 | 487 | 688 |
| 2080 | QGN15177.1 | FIM1_1865 | CP015056. | 1962 | 394 | 537 | 1496 | 576 | 422 | 1998 | 518 | 701 | 1253 | 524 | 775 | 2592 | 605 | 685 | 1690 | 522 | 554 |
| 2081 | QGN15178.1 | FIM1_1866 | CP015056. | 6427 | 125 | 171 | 2599 | 148 | 113 | 2126 | 142 | 200 | 443 | 133 | 222 | 673 | 134 | 189 | 569 | 124 | 133 |
| 2082 | QGN15179.1 | FIM1_1867 | CP015056. | 603 | 29 | 94 | 657 | 68 | 84 | 620 | 51 | 108 | 599 | 81 | 144 | 706 | 73 | 118 | 612 | 62 | 101 |
| 2083 | QGN15180.1 | FIM1_1868 | CP015056. | 1529 | 1067 | 1049 | 2092 | 765 | 666 | 1351 | 848 | 1238 | 7304 | 1411 | 1720 | 6146 | 1307 | 1862 | 6562 | 1419 | 1109 |
| 2084 | QGN15181.1 | FIM1_1869 | CP015056. | 1735 | 1240 | 1003 | 1493 | 1474 | 963 | 1762 | 1178 | 1329 | 8175 | 1423 | 1834 | 7405 | 1409 | 1774 | 7892 | 1450 | 1310 |
| 2085 | QGN15182.1 | FIM1_1870 | CP015056. | 1377 | 629 | 946 | 2351 | 681 | 818 | 1804 | 689 | 1174 | 5557 | 617 | 1355 | 4146 | 542 | 1179 | 5463 | 526 | 813 |
| 2086 | QGN15183.1 | FIM1_1871 | CP015056. | 5434 | 525 | 1085 | 4883 | 866 | 764 | 4207 | 889 | 1286 | 4325 | 708 | 1387 | 6696 | 794 | 1364 | 5137 | 665 | 980 |
| 2087 | QGN15184.1 | FIM1_1872 | CP015056. | 3057 | 360 | 447 | 2625 | 525 | 363 | 2539 | 461 | 572 | 2248 | 459 | 712 | 3729 | 434 | 587 | 2529 | 430 | 507 |
| 2088 | QGN15185.1 | FIM1_1873 | CP015056. | 995 | 522 | 513 | 1179 | 435 | 542 | 1199 | 406 | 791 | 5136 | 521 | 943 | 3990 | 453 | 885 | 5021 | 444 | 623 |
| 2089 | QGN15186.1 | FIM1_1874 | CP015056. | 575 | 450 | 1283 | 923 | 639 | 885 | 633 | 593 | 1726 | 8513 | 950 | 1424 | 12555 | 626 | 1433 | 11136 | 606 | 1067 |
| 2090 | QGN15187.1 | FIM1_1875 | CP015056. | 88 | 6 | 52 | 74 | 25 | 29 | 62 | 23 | 52 | 265 | 45 | 35 | 415 | 45 | 48 | 316 | 33 | 32 |
| 2091 | QGN15188.1 | FIM1_1876 | CP015056. | 2314 | 661 | 373 | 701 | 463 | 325 | 1545 | 580 | 575 | 502 | 310 | 317 | 1474 | 272 | 376 | 776 | 327 | 229 |
| 2092 | QGN15189.1 | FIM1_1877 | CP015056. | 2131 | 1115 | 339 | 897 | 897 | 382 | 1858 | 751 | 489 | 2249 | 964 | 641 | 2751 | 1043 | 558 | 2318 | 1246 | 464 |
| 2093 | QGN15190.1 | FIM1_1878 | CP015056. | 10016 | 936 | 1379 | 11910 | 1350 | 944 | 10546 | 1364 | 1822 | 1271 | 930 | 2509 | 9191 | 1417 | 2177 | 2972 | 1217 | 1553 |
| 2094 | QGN15191.1 | FIM1_1879 | CP015056. | 656 | 313 | 310 | 914 | 279 | 283 | 906 | 310 | 399 | 1852 | 282 | 395 | 1693 | 269 | 407 | 1803 | 256 | 273 |
| 2095 | QGN15192.1 | FIM1_1880 | CP015056. | 3503 | 2796 | 2069 | 5340 | 2218 | 1836 | 4367 | 1942 | 2672 | 10918 | 2208 | 3663 | 11107 | 2472 | 3492 | 11806 | 2242 | 2258 |
| 2096 | QGN15193.1 | FIM1_1881 | CP015056. | 696 | 389 | 442 | 1196 | 361 | 373 | 832 | 414 | 561 | 2921 | 416 | 784 | 3064 | 355 | 741 | 3271 | 425 | 543 |
| 2097 | QGN15194.1 | FIM1_1882 | CP015056. | 6864 | 2372 | 1990 | 9148 | 2694 | 1681 | 5680 | 2374 | 2537 | 2906 | 1826 | 2548 | 8544 | 1746 | 2566 | 4814 | 1674 | 1717 |
| 2098 | QGN15195.1 | FIM1_1883 | CP015056. | 289 | 21 | 84 | 342 | 79 | 39 | 310 | 71 | 83 | 302 | 61 | 94 | 659 | 47 | 89 | 382 | 57 | 67 |

| | | | | | | | | | | | | | | | | | | | | | |
|---|---|---|---|---|---|---|---|---|---|---|---|---|---|---|---|---|---|---|---|---|---|
| 2099 | QGN15196.1 | FIM1_1884 | CP015056. | 3360 | 3280 | 3094 | 9424 | 3110 | 2620 | 5276 | 2865 | 3860 | 10666 | 2242 | 4592 | 13362 | 2737 | 4230 | 11255 | 2270 | 2904 |
| 2100 | QGN15197.1 | FIM1_1885 | CP015056. | 1740 | 905 | 1032 | 1779 | 948 | 957 | 1602 | 798 | 1407 | 6282 | 908 | 1828 | 5261 | 861 | 1697 | 6600 | 792 | 1232 |
| 2101 | QGN15198.1 | FIM1_1886 | CP015056. | 2011 | 561 | 653 | 1576 | 498 | 560 | 1792 | 497 | 861 | 3231 | 407 | 866 | 3040 | 323 | 904 | 3361 | 342 | 570 |
| 2102 | QGN15199.1 | FIM1_1887 | CP015056. | 1213 | 3996 | 3555 | 3218 | 3818 | 3186 | 2127 | 3279 | 4226 | 21559 | 4855 | 5715 | 18241 | 4420 | 5665 | 21713 | 4287 | 3793 |
| 2103 | QGN15200.1 | FIM1_1888 | CP015056. | 926 | 1604 | 1741 | 1165 | 1676 | 1575 | 1076 | 1534 | 2347 | 6943 | 1669 | 1926 | 6942 | 1483 | 1923 | 6812 | 1494 | 1317 |
| 2104 | QGN15201.1 | FIM1_1889 | CP015056. | 1305 | 742 | 835 | 1351 | 975 | 832 | 1346 | 881 | 1195 | 4298 | 855 | 1220 | 5521 | 1025 | 1111 | 4687 | 883 | 755 |
| 2105 | QGN15202.1 | FIM1_1890 | CP015056. | 2638 | 682 | 328 | 758 | 850 | 358 | 2478 | 583 | 470 | 804 | 997 | 531 | 1659 | 1364 | 541 | 1275 | 1229 | 450 |
| 2106 | QGN15203.1 | FIM1_1891 | CP015056. | 5238 | 73 | 978 | 5583 | 334 | 789 | 5763 | 415 | 1280 | 2875 | 316 | 1158 | 4986 | 333 | 1054 | 3448 | 269 | 731 |
| 2107 | QGN15204.1 | FIM1_1892 | CP015056. | 420 | 687 | 656 | 1533 | 834 | 710 | 588 | 600 | 913 | 2980 | 501 | 1007 | 2734 | 416 | 786 | 3007 | 470 | 603 |
| 2108 | QGN15205.1 | FIM1_1893 | CP015056. | 2272 | 1170 | 751 | 2883 | 1045 | 751 | 2387 | 878 | 1081 | 6577 | 1407 | 1321 | 5323 | 1386 | 1573 | 6080 | 1391 | 970 |
| 2109 | QGN15206.1 | FIM1_1894 | CP015056. | 5375 | 476 | 1070 | 4754 | 758 | 878 | 4510 | 725 | 1389 | 3499 | 608 | 2071 | 5561 | 564 | 1833 | 4187 | 559 | 1321 |
| 2110 | QGN15207.1 | FIM1_1895 | CP015056. | 449 | 765 | 721 | 825 | 780 | 631 | 528 | 718 | 1035 | 3800 | 775 | 945 | 3646 | 751 | 924 | 3784 | 617 | 667 |
| 2111 | QGN15208.1 | FIM1_1896 | CP015056. | 1033 | 600 | 709 | 1202 | 698 | 658 | 1172 | 739 | 958 | 5372 | 738 | 1290 | 4921 | 723 | 1064 | 5327 | 716 | 781 |
| 2112 | QGN15209.1 | FIM1_1897 | CP015056. | 1918 | 1453 | 1108 | 1909 | 1321 | 973 | 1779 | 1180 | 1453 | 6812 | 1385 | 1655 | 5973 | 1247 | 1560 | 6134 | 1249 | 1199 |
| 2113 | QGN15210.1 | FIM1_1898 | CP015056. | 2972 | 1825 | 1451 | 4071 | 1609 | 1364 | 3156 | 1556 | 1944 | 5554 | 1137 | 1472 | 4904 | 997 | 1624 | 4936 | 1005 | 987 |
| 2114 | QGN15211.1 | FIM1_1899 | CP015056. | 17442 | 5359 | 4682 | 24045 | 4933 | 4539 | 19531 | 4077 | 6340 | 19223 | 5013 | 5767 | 17023 | 5583 | 5760 | 18661 | 5095 | 3877 |
| 2115 | QGN15212.1 | FIM1_1900 | CP015056. | 359 | 263 | 270 | 307 | 249 | 254 | 402 | 246 | 363 | 1794 | 272 | 351 | 1654 | 241 | 339 | 1671 | 208 | 245 |
| 2116 | QGN15213.1 | FIM1_1901 | CP015056. | 4600 | 2139 | 2576 | 7301 | 2782 | 1984 | 5361 | 2446 | 3225 | 6686 | 2502 | 4071 | 14805 | 2745 | 3718 | 9841 | 2333 | 2763 |
| 2117 | QGN15214.1 | FIM1_1902 | CP015056. | 1368 | 654 | 482 | 979 | 698 | 434 | 1214 | 669 | 639 | 2870 | 625 | 744 | 2745 | 581 | 723 | 2804 | 598 | 465 |
| 2118 | QGN15215.1 | FIM1_1903 | CP015056. | 1654 | 653 | 441 | 1928 | 668 | 426 | 1712 | 562 | 620 | 3497 | 620 | 848 | 3404 | 678 | 715 | 3272 | 544 | 526 |
| 2119 | QGN15216.1 | FIM1_1904 | CP015056. | 26963 | 2246 | 1474 | 20340 | 2452 | 1334 | 19478 | 1993 | 1804 | 13399 | 3495 | 2655 | 12132 | 3812 | 2597 | 12813 | 3396 | 1862 |
| 2120 | QGN15217.1 | FIM1_1905 | CP015056. | 892 | 888 | 638 | 1091 | 754 | 662 | 998 | 729 | 907 | 4684 | 735 | 929 | 3426 | 649 | 925 | 4247 | 675 | 639 |
| 2121 | QGN15218.1 | FIM1_1907 | CP015056. | 417 | 1101 | 658 | 577 | 744 | 659 | 437 | 597 | 978 | 4853 | 861 | 1087 | 3300 | 767 | 1075 | 4624 | 859 | 624 |
| 2122 | QGN15219.1 | FIM1_1908 | CP015056. | 1544 | 2842 | 2634 | 3324 | 3225 | 2201 | 2408 | 2600 | 2998 | 13805 | 4000 | 4286 | 11389 | 3733 | 4151 | 14622 | 3496 | 2979 |
| 2123 | QGN15220.1 | FIM1_1909 | CP015056. | 8385 | 1967 | 2579 | 9936 | 2396 | 2198 | 8515 | 1976 | 3480 | 11321 | 2212 | 3957 | 16815 | 2399 | 3601 | 12982 | 2025 | 2811 |
| 2124 | QGN15221.1 | FIM1_1910 | CP015056. | 1333 | 424 | 1313 | 1760 | 719 | 979 | 1335 | 631 | 1622 | 3822 | 649 | 2229 | 6434 | 633 | 1831 | 4878 | 604 | 1493 |
| 2125 | QGN15222.1 | FIM1_1911 | CP015056. | 9398 | 6177 | 3684 | 10933 | 5117 | 3265 | 8881 | 4626 | 4753 | 22912 | 6097 | 6476 | 20634 | 5594 | 6156 | 21681 | 5556 | 4369 |
| 2126 | QGN15223.1 | FIM1_1912 | CP015056. | 7160 | 1866 | 1743 | 7514 | 2118 | 1454 | 6783 | 1930 | 2247 | 5255 | 1658 | 2658 | 9604 | 1737 | 2290 | 6614 | 1630 | 1651 |
| 2127 | QGN15224.1 | FIM1_1913 | CP015056. | 2089 | 2335 | 960 | 1462 | 2002 | 1033 | 2110 | 1492 | 1328 | 8404 | 2268 | 2078 | 6265 | 1841 | 1770 | 7722 | 2440 | 1447 |
| 2128 | QGN15225.1 | FIM1_1914 | CP015056. | 355 | 503 | 405 | 355 | 341 | 353 | 346 | 325 | 528 | 2324 | 327 | 563 | 1512 | 335 | 544 | 2025 | 371 | 384 |
| 2129 | QGN15226.1 | FIM1_1915 | CP015056. | 1462 | 789 | 1322 | 2529 | 1162 | 1008 | 2164 | 1149 | 1590 | 2190 | 1181 | 1732 | 8149 | 1277 | 1495 | 4679 | 1145 | 1080 |
| 2130 | QGN15227.1 | FIM1_1916 | CP015056. | 1390 | 478 | 746 | 1194 | 744 | 671 | 1378 | 745 | 947 | 2242 | 537 | 897 | 4005 | 489 | 852 | 3069 | 489 | 662 |
| 2131 | QGN15228.1 | FIM1_1917 | CP015056. | 883 | 803 | 419 | 566 | 718 | 487 | 922 | 614 | 586 | 3696 | 799 | 935 | 2898 | 755 | 861 | 3471 | 885 | 597 |
| 2132 | QGN15229.1 | FIM1_1918 | CP015056. | 1073 | 779 | 525 | 846 | 568 | 507 | 855 | 520 | 749 | 3716 | 582 | 793 | 2740 | 565 | 877 | 3195 | 581 | 523 |
| 2133 | QGN15230.1 | FIM1_1919 | CP015056. | 4720 | 1083 | 816 | 3568 | 1068 | 715 | 3638 | 945 | 1102 | 3412 | 846 | 959 | 4349 | 768 | 900 | 3547 | 724 | 661 |
| 2134 | QGN15231.1 | FIM1_1920 | CP015056. | 3653 | 2049 | 719 | 1690 | 1974 | 785 | 3363 | 1682 | 905 | 7364 | 2404 | 1601 | 5934 | 2350 | 1408 | 7223 | 2826 | 963 |
| 2135 | QGN15232.1 | FIM1_1921 | CP015056. | 819 | 1619 | 2123 | 643 | 2121 | 1852 | 769 | 2395 | 2749 | 16326 | 2200 | 2967 | 14143 | 1455 | 2985 | 16063 | 1592 | 1923 |
| 2136 | QGN15233.1 | FIM1_1922 | CP015056. | 3387 | 1372 | 1521 | 3768 | 1530 | 1417 | 3242 | 1395 | 2069 | 6365 | 1294 | 2130 | 7545 | 1215 | 1905 | 6358 | 1141 | 1434 |
| 2137 | QGN15234.1 | FIM1_1923 | CP015056. | 3308 | 1944 | 1258 | 4209 | 1668 | 1089 | 3925 | 1472 | 1602 | 6914 | 2061 | 2002 | 6818 | 1815 | 1949 | 6870 | 1949 | 1422 |
| 2138 | QGN15235.1 | FIM1_1924 | CP015056. | 616 | 1108 | 686 | 936 | 837 | 711 | 788 | 696 | 913 | 3038 | 753 | 969 | 2330 | 765 | 937 | 2539 | 936 | 658 |
| 2139 | QGN15236.1 | FIM1_1925 | CP015056. | 2849 | 1224 | 1592 | 2713 | 1377 | 1214 | 2721 | 1209 | 2037 | 7362 | 1374 | 2277 | 9714 | 1336 | 2035 | 8173 | 1147 | 1575 |
| 2140 | QGN15237.1 | FIM1_1926 | CP015056. | 483 | 619 | 548 | 1221 | 492 | 486 | 810 | 422 | 772 | 2377 | 318 | 461 | 1766 | 348 | 440 | 2123 | 293 | 271 |
| 2141 | QGN15238.1 | FIM1_1927 | CP015056. | 4276 | 2205 | 1229 | 2010 | 2834 | 1330 | 4032 | 2235 | 1586 | 10002 | 4050 | 2469 | 10741 | 3209 | 2390 | 9685 | 3986 | 1684 |
| 2142 | QGN15239.1 | FIM1_1928 | CP015056. | 1997 | 982 | 1086 | 2902 | 949 | 950 | 2598 | 985 | 1531 | 7549 | 736 | 1390 | 6342 | 576 | 1246 | 7219 | 591 | 956 |
| 2143 | QGN15240.1 | FIM1_1929 | CP015056. | 635 | 96 | 348 | 1010 | 347 | 286 | 829 | 337 | 455 | 3370 | 254 | 503 | 3052 | 224 | 362 | 3053 | 226 | 317 |
| 2144 | QGN15241.1 | FIM1_1930 | CP015056. | 516 | 538 | 507 | 660 | 617 | 428 | 566 | 438 | 644 | 3112 | 540 | 690 | 2850 | 393 | 611 | 2788 | 397 | 420 |
| 2145 | QGN15242.1 | FIM1_1931 | CP015056. | 10493 | 8060 | 12165 | 13542 | 10138 | 10344 | 11501 | 9275 | 14609 | 26490 | 7615 | 15822 | 29851 | 7273 | 14914 | 27236 | 6631 | 10611 |
| 2146 | QGN15243.1 | FIM1_1932 | CP015056. | 1205 | 561 | 580 | 1217 | 437 | 507 | 994 | 435 | 749 | 1769 | 306 | 590 | 1673 | 251 | 537 | 1757 | 276 | 339 |
| 2147 | QGN15244.1 | FIM1_1933 | CP015056. | 5831 | 1905 | 3251 | 8053 | 2795 | 2622 | 6147 | 2697 | 4083 | 11876 | 3083 | 4610 | 18162 | 3243 | 4312 | 14832 | 2825 | 3092 |
| 2148 | QGN15245.1 | FIM1_1934 | CP015056. | 4020 | 283 | 245 | 3005 | 226 | 211 | 4415 | 230 | 323 | 566 | 136 | 302 | 1063 | 148 | 303 | 702 | 119 | 225 |
| 2149 | QGN15246.1 | FIM1_1935 | CP015056. | 24925 | 490 | 604 | 18045 | 668 | 504 | 23519 | 620 | 711 | 2767 | 428 | 814 | 4428 | 405 | 756 | 2948 | 345 | 565 |
| 2150 | QGN15247.1 | FIM1_1936 | CP015056. | 5958 | 3648 | 2693 | 8166 | 3878 | 2479 | 7035 | 3593 | 3511 | 13132 | 3515 | 3919 | 15062 | 3600 | 3807 | 12596 | 3581 | 2608 |
| 2151 | QGN15248.1 | FIM1_1937 | CP015056. | 3463 | 2050 | 1761 | 4585 | 1725 | 1355 | 4117 | 1538 | 2071 | 7323 | 1981 | 2399 | 6238 | 1951 | 2408 | 7379 | 1882 | 1777 |
| 2152 | QGN15249.1 | FIM1_1938 | CP015056. | 537 | 1429 | 843 | 695 | 892 | 790 | 518 | 840 | 1106 | 3875 | 975 | 1053 | 2889 | 852 | 1118 | 3224 | 984 | 741 |
| 2153 | QGN15250.1 | FIM1_1939 | CP015056. | 1438 | 251 | 149 | 999 | 141 | 126 | 1322 | 125 | 197 | 687 | 170 | 149 | 511 | 166 | 138 | 730 | 141 | 77 |
| 2154 | QGN15251.1 | FIM1_1940 | CP015056. | 4659 | 1871 | 2426 | 3518 | 1956 | 1760 | 3695 | 2683 | 3041 | 4225 | 1730 | 2711 | 12154 | 1250 | 2710 | 7095 | 1464 | 1878 |
| 2155 | QGN15252.1 | FIM1_1941 | CP015056. | 1599 | 1487 | 1088 | 3264 | 1006 | 803 | 2043 | 1077 | 1379 | 1649 | 682 | 1341 | 4551 | 1232 | 1389 | 2760 | 859 | 891 |
| 2156 | QGN15253.1 | FIM1_1942 | CP015056. | 2742 | 632 | 681 | 2556 | 455 | 545 | 2198 | 542 | 925 | 1873 | 381 | 692 | 2516 | 451 | 775 | 1985 | 306 | 511 |
| 2157 | QGN15254.1 | FIM1_1943 | CP015056. | 501 | 659 | 767 | 1153 | 807 | 575 | 690 | 804 | 929 | 3469 | 762 | 1217 | 4368 | 678 | 1072 | 3764 | 638 | 801 |
| 2158 | QGN15255.1 | FIM1_1944 | CP015056. | 540 | 471 | 378 | 786 | 405 | 293 | 769 | 455 | 489 | 2138 | 290 | 523 | 2082 | 387 | 526 | 2038 | 289 | 345 |
| 2159 | QGN15256.1 | FIM1_1945 | CP015056. | 2227 | 483 | 512 | 2217 | 607 | 459 | 1418 | 503 | 656 | 1058 | 417 | 731 | 1874 | 398 | 751 | 1629 | 348 | 465 |
| 2160 | QGN15257.1 | FIM1_1946 | CP015056. | 4836 | 1359 | 1628 | 7537 | 1805 | 1203 | 4900 | 1582 | 1908 | 3824 | 894 | 1676 | 6753 | 806 | 1441 | 5045 | 731 | 1099 |
| 2161 | QGN15258.1 | FIM1_1947 | CP015056. | 563 | 995 | 661 | 1277 | 637 | 665 | 836 | 634 | 934 | 3453 | 632 | 974 | 3054 | 538 | 967 | 3305 | 524 | 648 |
| 2162 | QGN15259.1 | FIM1_1948 | CP015056. | 10540 | 1780 | 787 | 9607 | 1800 | 871 | 10051 | 1335 | 1000 | 6391 | 1928 | 1371 | 6440 | 2134 | 1249 | 6154 | 2072 | 892 |
| 2163 | QGN15260.1 | FIM1_1949 | CP015056. | 274 | 254 | 347 | 259 | 334 | 258 | 275 | 356 | 424 | 981 | 406 | 471 | 2028 | 386 | 501 | 1390 | 354 | 331 |
| 2164 | QGN15261.1 | FIM1_1950 | CP015056. | 955 | 850 | 844 | 1537 | 674 | 743 | 1032 | 709 | 1165 | 1999 | 622 | 1242 | 3151 | 617 | 1276 | 2302 | 565 | 837 |
| 2165 | QGN15262.1 | FIM1_1951 | CP015056. | 5714 | 809 | 1482 | 5589 | 649 | 1149 | 5388 | 705 | 1836 | 3958 | 424 | 1536 | 3562 | 493 | 1792 | 3361 | 342 | 1067 |
| 2166 | QGN15263.1 | FIM1_1952 | CP015056. | 5781 | 2225 | 1195 | 6403 | 1865 | 990 | 5397 | 1686 | 1576 | 4374 | 1547 | 1578 | 6073 | 1846 | 1697 | 4634 | 1690 | 1128 |
| 2167 | QGN15264.1 | FIM1_1953 | CP015056. | 12578 | 1577 | 1505 | 12140 | 1523 | 1027 | 13745 | 1502 | 1902 | 1934 | 1146 | 1764 | 6600 | 1110 | 1822 | 3500 | 1088 | 1201 |
| 2168 | QGN15265.1 | FIM1_1954 | CP015056. | 3951 | 2018 | 580 | 1666 | 1534 | 645 | 3157 | 1429 | 893 | 586 | 1766 | 1157 | 4323 | 1844 | 1190 | 2254 | 2202 | 833 |
| 2169 | QGN15266.1 | FIM1_1955 | CP015056. | 1393 | 334 | 351 | 1987 | 344 | 270 | 1521 | 317 | 398 | 1518 | 288 | 555 | 2127 | 259 | 522 | 1529 | 265 | 337 |
| 2170 | QGN15267.1 | FIM1_1956 | CP015056. | 453 | 329 | 557 | 333 | 645 | 406 | 276 | 625 | 706 | 3207 | 520 | 616 | 3321 | 387 | 544 | 3502 | 424 | 411 |
| 2171 | QGN15268.1 | FIM1_1957 | CP015056. | 1232 | 955 | 705 | 1703 | 871 | 656 | 1323 | 615 | 984 | 5532 | 883 | 1160 | 4186 | 723 | 1082 | 5950 | 762 | 698 |
| 2172 | QGN15269.1 | FIM1_1958 | CP015056. | 1432 | 412 | 820 | 3024 | 485 | 682 | 1935 | 474 | 992 | 3044 | 311 | 1208 | 3559 | 468 | 1100 | 3012 | 321 | 738 |
| 2173 | QGN15270.1 | FIM1_1959 | CP015056. | 1462 | 752 | 498 | 1367 | 608 | 421 | 1223 | 517 | 591 | 2564 | 502 | 596 | 2184 | 432 | 618 | 2253 | 468 | 393 |
| 2174 | QGN15271.1 | FIM1_1960 | CP015056. | 2637 | 2175 | 1744 | 3726 | 2137 | 1570 | 2684 | 1890 | 2158 | 7258 | 2181 | 2506 | 8878 | 2052 | 2558 | 7249 | 1778 | 1806 |
| 2175 | QGN15272.1 | FIM1_1961 | CP015056. | 23950 | 778 | 497 | 18393 | 797 | 515 | 22127 | 905 | 694 | 6081 | 460 | 843 | 9638 | 344 | 956 | 6609 | 333 | 668 |
| 2176 | QGN15273.1 | FIM1_1962 | CP015056. | 46224 | 36981 | 19046 | 42095 | 28742 | 17637 | 37155 | 26308 | 22829 | 50410 | 28886 | 29032 | 70889 | 26755 | 28633 | 56744 | 25478 | 21901 |
| 2177 | QGN15274.1 | FIM1_1963 | CP015056. | 633 | 566 | 335 | 562 | 349 | 330 | 536 | 324 | 435 | 1297 | 371 | 487 | 1381 | 288 | 515 | 1239 | 313 | 354 |
| 2178 | QGN15275.1 | FIM1_1964 | CP015056. | 2925 | 5906 | 6622 | 10799 | 9508 | 5374 | 8075 | 8349 | 7631 | 35540 | 11533 | 11283 | 39102 | 10992 | 10577 | 37746 | 9347 | 7422 |
| 2179 | QGN15276.1 | FIM1_1965 | CP015056. | 1517 | 633 | 823 | 2309 | 806 | 671 | 1822 | 801 | 1059 | 1912 | 534 | 531 | 3543 | 553 | 506 | 2800 | 516 | 365 |
| 2180 | QGN15277.1 | FIM1_1966 | CP015056. | 2460 | 992 | 1162 | 4041 | 1323 | 905 | 2965 | 1076 | 1412 | 4680 | 825 | 1380 | 4819 | 784 | 1113 | 4502 | 756 | 811 |
| 2181 | QGN15278.1 | FIM1_1967 | CP015056. | 3976 | 1048 | 1404 | 6353 | 1210 | 1079 | 4701 | 1161 | 1758 | 5609 | 887 | 1953 | 8267 | 1002 | 1714 | 6668 | 836 | 1313 |
| 2182 | QGN15279.1 | FIM1_1968 | CP015056. | 1966 | 1487 | 971 | 1414 | 1881 | 982 | 2025 | 1599 | 1280 | 3592 | 1892 | 1576 | 7685 | 1745 | 1473 | 5638 | 1885 | 1073 |

| | | | | | | | | | | | | | | | | | | | | | |
|---|---|---|---|---|---|---|---|---|---|---|---|---|---|---|---|---|---|---|---|---|---|
| 2183 | QGN15280.1 | FIM1_1969 | CP015056. | 1 | 2 | 2 | 6 | 0 | 0 | 1 | 1 | 1 | 1 | 0 | 2 | 7 | 0 | 3 | 2 | 0 | 1 |
| 2184 | QGN15281.1 | FIM1_1970 | CP015056. | 267 | 638 | 439 | 534 | 593 | 382 | 278 | 457 | 558 | 1443 | 483 | 640 | 1733 | 358 | 552 | 1538 | 321 | 386 |
| 2185 | QGN15282.1 | FIM1_1971 | CP015056. | 251 | 231 | 305 | 528 | 298 | 256 | 388 | 312 | 423 | 1570 | 257 | 412 | 1316 | 233 | 397 | 1414 | 218 | 293 |
| 2186 | QGN15283.1 | FIM1_1972 | CP015056. | 29048 | 463 | 2220 | 27015 | 365 | 1535 | 24719 | 518 | 2714 | 2485 | 223 | 2142 | 2915 | 530 | 1983 | 2477 | 318 | 1387 |
| 2187 | QGN15284.1 | FIM1_1973 | CP015056. | 19567 | 2095 | 3383 | 17408 | 4511 | 2590 | 18516 | 5196 | 4148 | 5820 | 4115 | 4764 | 16258 | 4493 | 4404 | 10271 | 4567 | 3337 |
| 2188 | QGN15285.1 | FIM1_1974 | CP015056. | 1 | 1 | 0 | 6 | 4 | 2 | 1 | 0 | 0 | 5 | 1 | 1 | 4 | 0 | 1 | 7 | 0 | 0 |
| 2189 | QGN15286.1 | FIM1_1975 | CP015056. | 1995 | 832 | 992 | 2246 | 608 | 878 | 1896 | 592 | 1325 | 3309 | 601 | 1122 | 3056 | 593 | 1285 | 3132 | 553 | 722 |
| 2190 | QGN15287.1 | FIM1_1976 | CP015056. | 5667 | 1681 | 1748 | 6397 | 2380 | 1406 | 5624 | 2083 | 2158 | 5178 | 1641 | 2054 | 10366 | 1527 | 2004 | 7395 | 1416 | 1382 |
| 2191 | QGN15288.1 | FIM1_1977 | CP015056. | 4273 | 599 | 447 | 3835 | 572 | 433 | 4325 | 481 | 653 | 2148 | 466 | 717 | 2400 | 447 | 666 | 2144 | 451 | 471 |
| 2192 | QGN15289.1 | FIM1_1978 | CP015056. | 3831 | 2277 | 1609 | 4489 | 1870 | 1294 | 4288 | 1986 | 2107 | 1669 | 1521 | 2361 | 6710 | 1680 | 1993 | 3167 | 1770 | 1501 |
| 2193 | QGN15290.1 | FIM1_1979 | CP015056. | 1655 | 516 | 542 | 1742 | 566 | 453 | 1572 | 551 | 693 | 2470 | 395 | 704 | 2916 | 381 | 661 | 2484 | 291 | 454 |
| 2194 | QGN15291.1 | FIM1_1980 | CP015056. | 2405 | 6818 | 6860 | 5672 | 8488 | 6213 | 4620 | 6312 | 8961 | 31125 | 6918 | 6910 | 29412 | 5587 | 7266 | 30308 | 5111 | 4648 |
| 2195 | QGN15292.1 | FIM1_1981 | CP015056. | 2571 | 9078 | 6553 | 6583 | 8459 | 5745 | 6401 | 7365 | 7715 | 24333 | 9839 | 9973 | 22662 | 8691 | 10103 | 24772 | 7920 | 7476 |
| 2196 | QGN15293.1 | FIM1_1982 | CP015056. | 535 | 9 | 20 | 468 | 7 | 8 | 597 | 19 | 21 | 23 | 6 | 22 | 88 | 8 | 15 | 31 | 4 | 7 |
| 2197 | QGN15294.1 | FIM1_1983 | CP015056. | 5504 | 3726 | 2427 | 10461 | 3828 | 2171 | 7313 | 2912 | 3034 | 12791 | 3202 | 3907 | 13954 | 3666 | 3932 | 13163 | 3022 | 2620 |
| 2198 | QGN15295.1 | FIM1_1984 | CP015056. | 676 | 308 | 335 | 846 | 430 | 300 | 708 | 389 | 452 | 2175 | 429 | 522 | 1955 | 359 | 467 | 2024 | 358 | 350 |
| 2199 | QGN15296.1 | FIM1_1985 | CP015056. | 3924 | 1024 | 1528 | 4412 | 1538 | 1320 | 4536 | 1240 | 1916 | 10325 | 1829 | 2407 | 12236 | 1656 | 2284 | 10710 | 1686 | 1720 |
| 2200 | QGN15297.1 | FIM1_1986 | CP015056. | 3280 | 1228 | 1142 | 3324 | 1284 | 1083 | 2875 | 1091 | 1617 | 9253 | 1399 | 2005 | 8016 | 1389 | 1788 | 8942 | 1210 | 1307 |
| 2201 | QGN15298.1 | FIM1_1987 | CP015056. | 156 | 657 | 458 | 453 | 556 | 467 | 201 | 443 | 626 | 3227 | 491 | 556 | 2812 | 489 | 571 | 2714 | 562 | 375 |
| 2202 | QGN15299.1 | FIM1_1988 | CP015056. | 840 | 1483 | 1780 | 2294 | 1874 | 1462 | 1547 | 1499 | 2276 | 8337 | 1898 | 2247 | 9975 | 1673 | 1939 | 8624 | 1583 | 1431 |
| 2203 | QGN15300.1 | FIM1_1989 | CP015056. | 7806 | 664 | 485 | 5408 | 686 | 371 | 6410 | 562 | 686 | 1683 | 699 | 653 | 3239 | 1035 | 551 | 2145 | 818 | 436 |
| 2204 | QGN15301.1 | FIM1_1990 | CP015056. | 3327 | 1243 | 941 | 2937 | 1471 | 917 | 3677 | 1252 | 1231 | 5712 | 1590 | 1854 | 6835 | 1483 | 1552 | 5900 | 1634 | 1157 |
| 2205 | QGN15302.1 | FIM1_1991 | CP015056. | 5031 | 788 | 1471 | 4294 | 1599 | 1113 | 4148 | 1617 | 1784 | 6675 | 1174 | 1795 | 9847 | 934 | 1570 | 7809 | 929 | 1228 |
| 2206 | QGN15303.1 | FIM1_1992 | CP015056. | 6981 | 1970 | 2959 | 9863 | 3433 | 2492 | 7642 | 2928 | 3851 | 4906 | 2783 | 4030 | 15324 | 2869 | 3728 | 8680 | 2479 | 2786 |
| 2207 | QGN15304.1 | FIM1_1993 | CP015056. | 2059 | 1089 | 1367 | 3721 | 824 | 1197 | 2600 | 864 | 1915 | 3274 | 563 | 1558 | 4049 | 807 | 1671 | 3219 | 555 | 1194 |
| 2208 | QGN15305.1 | FIM1_1994 | CP015056. | 16783 | 12019 | 7061 | 19900 | 12884 | 6223 | 18085 | 12577 | 8641 | 28986 | 13518 | 11060 | 39618 | 12046 | 11029 | 33051 | 11985 | 8200 |
| 2209 | QGN15306.1 | FIM1_1995 | CP015056. | 7843 | 1743 | 1792 | 8304 | 2022 | 1427 | 7717 | 2016 | 2367 | 2567 | 1796 | 2561 | 9646 | 1929 | 2153 | 4727 | 1926 | 1754 |
| 2210 | QGN15307.1 | FIM1_1996 | CP015056. | 7862 | 4437 | 3941 | 7395 | 5348 | 3658 | 8256 | 4442 | 4856 | 24057 | 6621 | 6450 | 22413 | 6518 | 5875 | 22356 | 6617 | 4127 |
| 2211 | QGN15308.1 | FIM1_1997 | CP015056. | 1477 | 850 | 1019 | 1861 | 1127 | 880 | 1538 | 862 | 1253 | 5892 | 1120 | 1398 | 5497 | 1019 | 1272 | 5401 | 967 | 913 |
| 2212 | QGN15309.1 | FIM1_1998 | CP015056. | 965 | 967 | 933 | 1422 | 937 | 825 | 1124 | 727 | 1177 | 3717 | 855 | 1164 | 4580 | 776 | 1074 | 3874 | 807 | 865 |
| 2213 | QGN15310.1 | FIM1_1999 | CP015056. | 1101 | 199 | 166 | 1596 | 303 | 145 | 842 | 251 | 228 | 730 | 195 | 234 | 1047 | 228 | 199 | 773 | 183 | 144 |
| 2214 | QGN15311.1 | FIM1_2000 | CP015056. | 1528 | 5414 | 3391 | 3852 | 4701 | 3351 | 2990 | 3969 | 4430 | 14892 | 4733 | 5263 | 15105 | 4544 | 5350 | 15330 | 4435 | 3929 |
| 2215 | QGN15312.1 | FIM1_2001 | CP015056. | 6085 | 2157 | 946 | 2650 | 2492 | 1051 | 5321 | 2051 | 1309 | 7056 | 2578 | 2118 | 8380 | 2498 | 1886 | 7267 | 3077 | 1409 |
| 2216 | QGN15313.1 | FIM1_2002 | CP015056. | 2326 | 2566 | 1802 | 2129 | 2813 | 1591 | 2598 | 2389 | 2174 | 10626 | 2580 | 2536 | 10147 | 2397 | 2306 | 10843 | 2501 | 1587 |
| 2217 | QGN15314.1 | FIM1_2003 | CP015056. | 1444 | 1600 | 3171 | 2433 | 2672 | 2623 | 1680 | 2267 | 4158 | 10494 | 3110 | 4718 | 16552 | 3260 | 4090 | 13366 | 3071 | 3106 |
| 2218 | QGN15315.1 | FIM1_2004 | CP015056. | 1830 | 614 | 432 | 1270 | 582 | 363 | 1338 | 588 | 572 | 2014 | 353 | 393 | 1912 | 254 | 401 | 1838 | 241 | 246 |
| 2219 | QGN15316.1 | FIM1_2005 | CP015056. | 11404 | 22202 | 18475 | 24126 | 29727 | 16050 | 13946 | 26275 | 22762 | 60088 | 30890 | 29927 | 100384 | 29979 | 30805 | 77453 | 23757 | 20907 |
| 2220 | QGN15317.1 | FIM1_2006 | CP015056. | 1121 | 252 | 598 | 1841 | 373 | 448 | 1491 | 342 | 695 | 2219 | 411 | 601 | 2774 | 355 | 549 | 2360 | 356 | 483 |
| 2221 | QGN15318.1 | FIM1_2007 | CP015056. | 811 | 476 | 341 | 1030 | 473 | 305 | 931 | 404 | 483 | 2639 | 412 | 483 | 2033 | 322 | 474 | 2677 | 351 | 324 |
| 2222 | QGN15319.1 | FIM1_2008 | CP015056. | 979 | 198 | 390 | 1242 | 377 | 333 | 1043 | 333 | 478 | 2622 | 288 | 466 | 2197 | 246 | 494 | 2352 | 229 | 348 |
| 2223 | QGN15320.1 | FIM1_2009 | CP015056. | 9878 | 224 | 181 | 6894 | 166 | 146 | 6871 | 174 | 290 | 956 | 150 | 260 | 775 | 143 | 261 | 871 | 106 | 179 |
| 2224 | QGN15321.1 | FIM1_2010 | CP015056. | 751 | 19 | 28 | 556 | 16 | 26 | 450 | 17 | 25 | 124 | 29 | 27 | 135 | 16 | 34 | 121 | 19 | 29 |
| 2225 | QGN15322.1 | FIM1_2011 | CP015056. | 3844 | 1379 | 1336 | 5214 | 1765 | 1163 | 4821 | 1433 | 1715 | 9948 | 1648 | 2329 | 10453 | 1599 | 2112 | 9483 | 1514 | 1582 |
| 2226 | QGN15323.1 | FIM1_2012 | CP015056. | 12813 | 1175 | 1471 | 12513 | 1762 | 1267 | 13255 | 1337 | 1752 | 8344 | 1253 | 1999 | 8996 | 1449 | 1842 | 7603 | 1346 | 1378 |
| 2227 | QGN15324.1 | FIM1_2013 | CP015056. | 3686 | 432 | 921 | 2870 | 495 | 636 | 2072 | 576 | 1156 | 2535 | 382 | 1399 | 4937 | 363 | 1180 | 3484 | 310 | 878 |
| 2228 | QGN15325.1 | FIM1_2014 | CP015056. | 3730 | 1060 | 1317 | 4342 | 940 | 1038 | 3641 | 1208 | 1635 | 4085 | 771 | 1648 | 5242 | 697 | 1536 | 4072 | 595 | 1068 |
| 2229 | QGN15326.1 | FIM1_2015 | CP015056. | 57 | 14 | 10 | 48 | 14 | 10 | 44 | 8 | 18 | 106 | 11 | 34 | 95 | 9 | 28 | 100 | 5 | 26 |
| 2230 | QGN15327.1 | FIM1_2016 | CP015056. | 6379 | 680 | 1387 | 7857 | 820 | 1199 | 7086 | 866 | 1791 | 10778 | 614 | 2558 | 10062 | 471 | 2429 | 10118 | 490 | 1747 |
| 2231 | QGN15328.1 | FIM1_2017 | CP015056. | 4949 | 716 | 831 | 3035 | 853 | 658 | 3710 | 994 | 1044 | 2511 | 570 | 904 | 4020 | 435 | 804 | 2947 | 440 | 603 |
| 2232 | QGN15329.1 | FIM1_2018 | CP015056. | 579 | 127 | 302 | 807 | 43 | 155 | 407 | 229 | 394 | 790 | 63 | 183 | 874 | 60 | 161 | 812 | 62 | 111 |
| 2233 | QGN15330.1 | FIM1_2019 | CP015056. | 4999 | 365 | 1110 | 2687 | 745 | 846 | 3988 | 837 | 1337 | 3987 | 826 | 1250 | 4725 | 739 | 1113 | 4487 | 640 | 738 |
| 2234 | QGN15331.1 | FIM1_2020 | CP015056. | 5854 | 2029 | 2428 | 6635 | 2751 | 1810 | 7011 | 2739 | 2978 | 4173 | 1777 | 2915 | 10701 | 1510 | 2248 | 7401 | 1390 | 1733 |
| 2235 | QGN15332.1 | FIM1_2021 | CP015056. | 8118 | 1545 | 2892 | 7718 | 3342 | 2341 | 6340 | 3324 | 3571 | 10550 | 2714 | 3366 | 12710 | 2468 | 3293 | 11486 | 2266 | 2227 |
| 2236 | QGN15333.1 | FIM1_2022 | CP015056. | 1086 | 1459 | 1057 | 1533 | 992 | 994 | 1311 | 854 | 1354 | 6654 | 1082 | 1556 | 4905 | 998 | 1486 | 6482 | 996 | 972 |
| 2237 | QGN15334.1 | FIM1_2023 | CP015056. | 3027 | 804 | 1645 | 4989 | 1359 | 1418 | 3470 | 1420 | 2211 | 7455 | 1086 | 1561 | 9694 | 859 | 1582 | 8406 | 804 | 1144 |
| 2238 | QGN15335.1 | FIM1_2024 | CP015056. | 430 | 291 | 752 | 7342 | 139 | 414 | 1164 | 475 | 935 | 517 | 63 | 163 | 878 | 55 | 91 | 619 | 67 | 74 |
| 2239 | QGN15336.1 | FIM1_2025 | CP015056. | 251 | 97 | 156 | 533 | 103 | 113 | 251 | 111 | 173 | 852 | 121 | 394 | 910 | 80 | 330 | 942 | 85 | 207 |
| 2240 | QGN15337.1 | FIM1_2026 | CP015056. | 2213 | 682 | 821 | 2334 | 587 | 590 | 2181 | 634 | 1016 | 5593 | 604 | 1189 | 4861 | 551 | 1067 | 5418 | 528 | 776 |
| 2241 | QGN15338.1 | FIM1_2027 | CP015056. | 220 | 575 | 745 | 356 | 598 | 588 | 207 | 519 | 919 | 4926 | 730 | 1137 | 4277 | 601 | 1014 | 4984 | 556 | 785 |
| 2242 | QGN15339.1 | FIM1_2028 | CP015056. | 1127 | 2174 | 1116 | 976 | 1954 | 1153 | 907 | 1536 | 1587 | 6645 | 2057 | 1520 | 6090 | 1581 | 1719 | 6138 | 1753 | 1055 |
| 2243 | QGN15340.1 | FIM1_2029 | CP015056. | 3082 | 1392 | 1789 | 4055 | 1075 | 1501 | 2927 | 1106 | 2297 | 6003 | 884 | 2735 | 5594 | 973 | 2685 | 5895 | 787 | 1812 |
| 2244 | QGN15341.1 | FIM1_2030 | CP015056. | 7674 | 276 | 293 | 7692 | 244 | 219 | 6857 | 337 | 422 | 1291 | 212 | 476 | 2174 | 267 | 381 | 1521 | 215 | 297 |
| 2245 | QGN15342.1 | FIM1_2031 | CP015056. | 8813 | 404 | 667 | 5974 | 302 | 437 | 3794 | 725 | 937 | 6562 | 456 | 806 | 7587 | 337 | 705 | 7228 | 380 | 680 |
| 2246 | QGN15343.1 | FIM1_2032 | CP015056. | 104 | 5 | 7 | 134 | 15 | 5 | 46 | 8 | 9 | 28 | 14 | 20 | 112 | 13 | 12 | 67 | 19 | 9 |
| 2247 | QGN15344.1 | FIM1_2033 | CP015056. | 3651 | 804 | 982 | 4511 | 902 | 782 | 3656 | 950 | 1267 | 4434 | 759 | 1384 | 6853 | 751 | 1188 | 5395 | 658 | 888 |
| 2248 | QGN15345.1 | FIM1_2034 | CP015056. | 5333 | 1058 | 1336 | 5393 | 1169 | 1035 | 5368 | 1087 | 1676 | 6507 | 1083 | 1870 | 6868 | 1109 | 1611 | 6176 | 1041 | 1267 |
| 2249 | QGN15346.1 | FIM1_2035 | CP015056. | 646 | 213 | 202 | 442 | 178 | 212 | 590 | 184 | 284 | 982 | 123 | 233 | 689 | 96 | 228 | 1127 | 103 | 103 |
| 2250 | QGN15347.1 | FIM1_2037 | CP015056. | 136 | 314 | 319 | 352 | 209 | 284 | 171 | 282 | 438 | 1065 | 147 | 339 | 923 | 99 | 307 | 1051 | 110 | 191 |
| 2251 | QGN15348.1 | FIM1_2038 | CP015056. | 3783 | 890 | 1301 | 3679 | 953 | 1070 | 2867 | 1036 | 1695 | 3109 | 1059 | 1511 | 5729 | 681 | 1517 | 5060 | 822 | 1035 |
| 2252 | QGN15349.1 | FIM1_2039 | CP015056. | 1295 | 315 | 273 | 1006 | 429 | 258 | 1157 | 265 | 321 | 1997 | 439 | 514 | 1553 | 489 | 496 | 1908 | 492 | 326 |
| 2253 | QGN15350.1 | FIM1_2040 | CP015056. | 686 | 353 | 189 | 570 | 286 | 166 | 377 | 236 | 254 | 1550 | 357 | 327 | 1383 | 341 | 290 | 1477 | 372 | 218 |
| 2254 | QGN15351.1 | FIM1_2041 | CP015056. | 8257 | 2283 | 1213 | 8230 | 2519 | 1278 | 7287 | 1527 | 1650 | 6419 | 5102 | 1292 | 7493 | 5088 | 1271 | 5858 | 5850 | 930 |
| 2255 | QGN15352.1 | FIM1_2042 | CP015056. | 3920 | 2673 | 2399 | 5261 | 3421 | 2315 | 4172 | 2520 | 2967 | 12384 | 2544 | 3257 | 11603 | 2213 | 2881 | 12218 | 2034 | 2043 |
| 2256 | QGN15353.1 | FIM1_2043 | CP015056. | 2858 | 1930 | 851 | 1938 | 1690 | 956 | 2800 | 1350 | 1182 | 7290 | 2095 | 1751 | 6631 | 1812 | 1601 | 6683 | 2246 | 1153 |
| 2257 | QGN15354.1 | FIM1_2044 | CP015056. | 1234 | 318 | 519 | 1097 | 371 | 415 | 1010 | 458 | 615 | 3342 | 502 | 819 | 4353 | 381 | 813 | 3576 | 374 | 582 |
| 2258 | QGN15355.1 | FIM1_2045 | CP015056. | 937 | 184 | 290 | 730 | 216 | 213 | 702 | 247 | 370 | 1855 | 237 | 421 | 1634 | 175 | 399 | 1676 | 165 | 265 |
| 2259 | QGN15356.1 | FIM1_2046 | CP015056. | 324 | 197 | 153 | 779 | 146 | 132 | 343 | 160 | 202 | 1031 | 125 | 280 | 922 | 159 | 230 | 888 | 138 | 173 |
| 2260 | QGN15357.1 | FIM1_2047 | CP015056. | 2566 | 405 | 442 | 3241 | 412 | 341 | 2163 | 394 | 574 | 1256 | 376 | 592 | 2159 | 433 | 566 | 1581 | 338 | 357 |
| 2261 | QGN15358.1 | FIM1_2048 | CP015056. | 2115 | 1678 | 1309 | 2250 | 1512 | 1110 | 2523 | 1222 | 1749 | 4213 | 1225 | 1256 | 4453 | 1265 | 1133 | 4366 | 1215 | 891 |
| 2262 | QGN15359.1 | FIM1_2049 | CP015056. | 452 | 310 | 269 | 460 | 244 | 240 | 511 | 207 | 373 | 1807 | 264 | 380 | 1525 | 241 | 392 | 1894 | 220 | 277 |
| 2263 | QGN15360.1 | FIM1_2050 | CP015056. | 3653 | 3269 | 2514 | 6136 | 3195 | 2167 | 4369 | 2753 | 3158 | 11859 | 2419 | 3305 | 10593 | 2475 | 3123 | 10978 | 2237 | 2164 |
| 2264 | QGN15361.1 | FIM1_2051 | CP015056. | 2292 | 1308 | 1304 | 3961 | 1156 | 1054 | 3284 | 1196 | 1644 | 6076 | 1055 | 1889 | 6271 | 1277 | 1811 | 5844 | 1142 | 1321 |
| 2265 | QGN15362.1 | FIM1_2052 | CP015056. | 6894 | 1033 | 2709 | 4294 | 1226 | 1954 | 5718 | 1546 | 3422 | 9122 | 1321 | 2844 | 10952 | 1377 | 2731 | 9470 | 1163 | 2183 |
| 2266 | QGN15363.1 | FIM1_2053 | CP015056. | 4462 | 6322 | 5704 | 8723 | 5963 | 4850 | 5931 | 5922 | 7248 | 13402 | 4543 | 4819 | 16159 | 4539 | 4832 | 13325 | 4279 | 3258 |

| | | | | | | | | | | | | | | | | | | | | | |
|---|---|---|---|---|---|---|---|---|---|---|---|---|---|---|---|---|---|---|---|---|---|
| 2267 | QGN15364.1 | FIM1_2054 | CP015056. | 1121 | 1116 | 1565 | 1235 | 863 | 1222 | 1156 | 946 | 1826 | 3079 | 716 | 1816 | 3464 | 829 | 1914 | 3411 | 597 | 1331 |
| 2268 | QGN15365.1 | FIM1_2055 | CP015056. | 1660 | 85 | 459 | 1786 | 684 | 337 | 1976 | 388 | 491 | 2449 | 486 | 545 | 4381 | 425 | 460 | 3166 | 368 | 396 |
| 2269 | QGN15366.1 | FIM1_2056 | CP015056. | 1913 | 448 | 489 | 1605 | 625 | 435 | 2013 | 311 | 613 | 1679 | 426 | 473 | 2613 | 402 | 528 | 1914 | 307 | 350 |
| 2270 | QGN15367.1 | FIM1_2057 | CP015056. | 1803 | 352 | 403 | 1948 | 439 | 345 | 1960 | 310 | 515 | 1177 | 401 | 678 | 1674 | 371 | 565 | 1333 | 326 | 370 |
| 2271 | QGN15368.1 | FIM1_2058 | CP015056. | 2123 | 65 | 181 | 2287 | 323 | 166 | 2754 | 157 | 196 | 1255 | 221 | 317 | 1434 | 218 | 269 | 1245 | 197 | 196 |
| 2272 | QGN15369.1 | FIM1_2059 | CP015056. | 1148 | 429 | 379 | 1385 | 679 | 329 | 983 | 461 | 446 | 2301 | 680 | 684 | 2436 | 661 | 619 | 2295 | 579 | 412 |
| 2273 | QGN15370.1 | FIM1_2060 | CP015056. | 1561 | 2090 | 4062 | 9057 | 2681 | 3401 | 3841 | 2358 | 4771 | 8978 | 1301 | 3501 | 7737 | 1735 | 2576 | 8827 | 1871 | 2174 |
| 2274 | QGN15371.1 | FIM1_2061 | CP015056. | 1308 | 1681 | 1031 | 2151 | 1011 | 956 | 1752 | 1093 | 1377 | 6008 | 1290 | 1630 | 4824 | 1216 | 1594 | 5471 | 1288 | 1140 |
| 2275 | QGN15372.1 | FIM1_2062 | CP015056. | 4695 | 148 | 515 | 6024 | 548 | 409 | 5697 | 525 | 641 | 2350 | 371 | 799 | 3343 | 481 | 536 | 2349 | 329 | 467 |
| 2276 | QGN15373.1 | FIM1_2063 | CP015056. | 15104 | 1951 | 1820 | 18025 | 1624 | 1514 | 17284 | 1524 | 2443 | 6058 | 1210 | 2230 | 8280 | 1321 | 2236 | 6573 | 1088 | 1547 |
| 2277 | QGN15374.1 | FIM1_2064 | CP015056. | 4865 | 402 | 783 | 8976 | 305 | 706 | 8466 | 248 | 1374 | 298 | 122 | 339 | 735 | 149 | 286 | 401 | 91 | 212 |
| 2278 | QGN15375.1 | FIM1_2065 | CP015056. | 218 | 18 | 7 | 359 | 37 | 13 | 239 | 30 | 14 | 40 | 25 | 28 | 89 | 20 | 22 | 44 | 16 | 18 |
| 2279 | QGN15376.1 | FIM1_2066 | CP015056. | 43369 | 513 | 709 | 24562 | 652 | 445 | 27822 | 634 | 868 | 545 | 301 | 739 | 2849 | 942 | 574 | 1212 | 495 | 449 |
| 2280 | QGN15377.1 | FIM1_2067 | CP015056. | 3241 | 889 | 1465 | 4546 | 989 | 1187 | 3361 | 906 | 2008 | 3517 | 614 | 2616 | 6391 | 890 | 2406 | 3784 | 805 | 1793 |
| 2281 | QGN15378.1 | FIM1_2068 | CP015056. | 5429 | 636 | 628 | 4630 | 726 | 562 | 5308 | 585 | 770 | 3315 | 749 | 1032 | 4537 | 755 | 954 | 3715 | 589 | 705 |
| 2282 | QGN15379.1 | FIM1_2069 | CP015056. | 4076 | 425 | 959 | 5187 | 670 | 803 | 4950 | 608 | 1185 | 7371 | 530 | 1364 | 6364 | 567 | 1193 | 6519 | 467 | 892 |
| 2283 | QGN15380.1 | FIM1_2070 | CP015056. | 102 | 48 | 41 | 113 | 51 | 49 | 99 | 69 | 67 | 360 | 54 | 78 | 216 | 48 | 78 | 258 | 74 | 55 |
| 2284 | QGN15381.1 | FIM1_2071 | CP015056. | 3273 | 1316 | 1137 | 3980 | 1349 | 1020 | 3764 | 1155 | 1532 | 7361 | 1201 | 2001 | 6926 | 1189 | 1735 | 7501 | 1145 | 1310 |
| 2285 | QGN15382.1 | FIM1_2072 | CP015056. | 1660 | 1320 | 909 | 1607 | 942 | 820 | 1622 | 885 | 1211 | 5527 | 982 | 1274 | 4255 | 917 | 1279 | 5292 | 965 | 880 |
| 2286 | QGN15383.1 | FIM1_2073 | CP015056. | 2567 | 2362 | 1453 | 2700 | 2711 | 1744 | 3212 | 2109 | 1952 | 11591 | 3083 | 2783 | 10231 | 2237 | 2498 | 11182 | 2826 | 1742 |
| 2287 | QGN15384.1 | FIM1_2074 | CP015056. | 1524 | 469 | 432 | 1458 | 484 | 420 | 1413 | 405 | 558 | 3296 | 549 | 711 | 2690 | 532 | 788 | 3180 | 480 | 461 |
| 2288 | QGN15385.1 | FIM1_2075 | CP015056. | 3805 | 5035 | 5408 | 8300 | 4337 | 4441 | 6044 | 3800 | 6565 | 14928 | 3417 | 7461 | 16372 | 3435 | 6972 | 15248 | 3049 | 5004 |
| 2289 | QGN15386.1 | FIM1_2076 | CP015056. | 1299 | 873 | 868 | 1844 | 723 | 890 | 1710 | 558 | 1256 | 4536 | 827 | 1379 | 4218 | 750 | 1520 | 4245 | 730 | 893 |
| 2290 | QGN15387.1 | FIM1_2077 | CP015056. | 5418 | 1442 | 1584 | 4243 | 1603 | 1217 | 4229 | 1637 | 1921 | 2915 | 1558 | 1986 | 7220 | 1765 | 1810 | 5044 | 1591 | 1249 |
| 2291 | QGN15388.1 | FIM1_2078 | CP015056. | 963 | 1155 | 935 | 1530 | 1066 | 784 | 1253 | 1072 | 1318 | 1526 | 922 | 1061 | 3731 | 839 | 1295 | 2161 | 729 | 856 |
| 2292 | QGN15389.1 | FIM1_2079 | CP015056. | 2204 | 773 | 3666 | 1877 | 263 | 2752 | 984 | 1070 | 4997 | 1623 | 166 | 2955 | 2349 | 119 | 3212 | 2446 | 110 | 2242 |
| 2293 | QGN15390.1 | FIM1_2080 | CP015056. | 64 | 11 | 12 | 157 | 26 | 21 | 80 | 35 | 17 | 47 | 18 | 23 | 73 | 24 | 23 | 64 | 24 | 17 |
| 2294 | QGN15391.1 | FIM1_2081 | CP015056. | 14636 | 8959 | 3113 | 11018 | 8442 | 3935 | 14550 | 6904 | 4275 | 16393 | 11867 | 5840 | 23156 | 9144 | 6160 | 19075 | 11951 | 4329 |
| 2295 | QGN15392.1 | FIM1_2082 | CP015056. | 24 | 11 | 3 | 23 | 11 | 9 | 11 | 10 | 5 | 14 | 8 | 4 | 31 | 8 | 1 | 17 | 11 | 7 |
| 2296 | QGN15393.1 | FIM1_2083 | CP015056. | 3335 | 334 | 474 | 1666 | 398 | 392 | 2046 | 430 | 593 | 1021 | 315 | 482 | 2410 | 302 | 470 | 1550 | 272 | 365 |
| 2297 | QGN15394.1 | FIM1_2084 | CP015056. | 3318 | 4447 | 4255 | 5048 | 4267 | 3799 | 3927 | 3965 | 5263 | 18274 | 4252 | 6167 | 18403 | 4106 | 5851 | 18118 | 3925 | 4195 |
| 2298 | QGN15395.1 | FIM1_2085 | CP015056. | 3238 | 1578 | 1785 | 4819 | 1573 | 1675 | 3512 | 1453 | 2347 | 8743 | 1534 | 2928 | 8736 | 1575 | 2846 | 8824 | 1418 | 1903 |
| 2299 | QGN15396.1 | FIM1_2086 | CP015056. | 2138 | 1912 | 2733 | 3241 | 2341 | 2212 | 2117 | 2006 | 3374 | 8588 | 1804 | 3126 | 10804 | 1733 | 2925 | 9473 | 1646 | 1892 |
| 2300 | QGN15397.1 | FIM1_2087 | CP015056. | 1706 | 3624 | 3186 | 4242 | 4646 | 3086 | 3196 | 3406 | 4320 | 2321 | 2389 | 3725 | 10006 | 2033 | 3600 | 6271 | 1819 | 2462 |
| 2301 | QGN15398.1 | FIM1_2088 | CP015056. | 2028 | 852 | 724 | 1900 | 1215 | 731 | 2084 | 879 | 952 | 3140 | 1013 | 1054 | 4217 | 1161 | 987 | 3320 | 1078 | 663 |
| 2302 | QGN15399.1 | FIM1_2089 | CP015056. | 725 | 294 | 202 | 668 | 193 | 184 | 769 | 195 | 284 | 1469 | 169 | 286 | 1049 | 198 | 326 | 1476 | 178 | 202 |
| 2303 | QGN15400.1 | FIM1_2090 | CP015056. | 1821 | 358 | 459 | 2078 | 479 | 354 | 2005 | 399 | 608 | 3481 | 442 | 714 | 2976 | 416 | 638 | 3176 | 408 | 443 |
| 2304 | QGN15401.1 | FIM1_2091 | CP015056. | 447 | 1252 | 711 | 522 | 915 | 669 | 493 | 763 | 968 | 4818 | 1045 | 1109 | 3710 | 901 | 1054 | 4552 | 938 | 691 |
| 2305 | QGN15402.1 | FIM1_2092 | CP015056. | 643 | 497 | 299 | 1022 | 549 | 402 | 589 | 372 | 447 | 2756 | 547 | 529 | 1958 | 483 | 432 | 2487 | 482 | 358 |
| 2306 | QGN15403.1 | FIM1_2093 | CP015056. | 4913 | 2013 | 1020 | 2215 | 2546 | 1157 | 4192 | 1870 | 1477 | 8157 | 3062 | 2174 | 8466 | 3244 | 2065 | 8148 | 3433 | 1533 |
| 2307 | QGN15404.1 | FIM1_2094 | CP015056. | 5720 | 6737 | 3977 | 4325 | 6393 | 3781 | 5114 | 5388 | 4969 | 26272 | 7342 | 6679 | 22790 | 6131 | 6288 | 26091 | 6727 | 4600 |
| 2308 | QGN15405.1 | FIM1_2095 | CP015056. | 2905 | 380 | 821 | 3587 | 477 | 631 | 2519 | 522 | 1045 | 3209 | 290 | 1013 | 3812 | 269 | 880 | 3235 | 199 | 605 |
| 2309 | QGN15406.1 | FIM1_2096 | CP015056. | 2405 | 447 | 1064 | 6065 | 769 | 823 | 3053 | 555 | 1434 | 2510 | 354 | 615 | 3657 | 376 | 530 | 2815 | 295 | 386 |
| 2310 | QGN15407.1 | FIM1_2097 | CP015056. | 493 | 705 | 1065 | 761 | 720 | 998 | 489 | 654 | 1415 | 2515 | 573 | 1450 | 2740 | 568 | 1312 | 2582 | 557 | 821 |
| 2311 | QGN15408.1 | FIM1_2098 | CP015056. | 17919 | 81 | 350 | 13939 | 15 | 171 | 13904 | 87 | 422 | 214 | 16 | 430 | 234 | 20 | 275 | 196 | 9 | 221 |
| 2312 | QGN15409.1 | FIM1_2099 | CP015056. | 36 | 11 | 8 | 17 | 10 | 15 | 13 | 13 | 6 | 36 | 11 | 12 | 27 | 11 | 7 | 21 | 11 | 10 |
| 2313 | QGN15410.1 | FIM1_2100 | CP015056. | 2004 | 1504 | 934 | 1666 | 1330 | 957 | 2129 | 1272 | 1234 | 5407 | 1493 | 1445 | 5066 | 1361 | 1365 | 4839 | 1410 | 982 |
| 2314 | QGN15411.1 | FIM1_2101 | CP015056. | 709 | 599 | 456 | 639 | 556 | 427 | 779 | 474 | 659 | 2712 | 491 | 590 | 2390 | 490 | 537 | 2817 | 459 | 416 |
| 2315 | QGN15412.1 | FIM1_2102 | CP015056. | 3181 | 510 | 652 | 2352 | 641 | 609 | 3056 | 548 | 905 | 3586 | 470 | 961 | 3526 | 506 | 872 | 3551 | 528 | 632 |
| 2316 | QGN15413.1 | FIM1_2103 | CP015056. | 6380 | 2896 | 4355 | 14859 | 3941 | 3741 | 8279 | 2925 | 5230 | 7065 | 2630 | 5648 | 15765 | 4125 | 4293 | 10167 | 2999 | 3517 |
| 2317 | QGN15414.1 | FIM1_2104 | CP015056. | 556 | 435 | 323 | 302 | 473 | 377 | 294 | 274 | 441 | 3304 | 807 | 904 | 2776 | 810 | 712 | 3188 | 855 | 566 |
| 2318 | QGN15415.1 | FIM1_2105 | CP015056. | 433 | 259 | 266 | 332 | 275 | 278 | 427 | 214 | 384 | 2169 | 296 | 485 | 1611 | 239 | 431 | 2146 | 269 | 308 |
| 2319 | QGN15416.1 | FIM1_2106 | CP015056. | 1044 | 429 | 637 | 1731 | 393 | 573 | 978 | 371 | 907 | 5089 | 367 | 1088 | 3906 | 344 | 839 | 5056 | 275 | 656 |
| 2320 | QGN15417.1 | FIM1_2107 | CP015056. | 986 | 554 | 390 | 1859 | 559 | 367 | 1422 | 471 | 528 | 2676 | 508 | 578 | 2831 | 458 | 583 | 2552 | 491 | 444 |
| 2321 | QGN15418.1 | FIM1_2108 | CP015056. | 609 | 273 | 498 | 918 | 346 | 386 | 738 | 406 | 643 | 3273 | 394 | 678 | 2829 | 283 | 602 | 3304 | 331 | 478 |
| 2322 | QGN15419.1 | FIM1_2109 | CP015056. | 245 | 242 | 125 | 267 | 167 | 101 | 240 | 138 | 180 | 1014 | 150 | 222 | 687 | 149 | 171 | 923 | 167 | 125 |
| 2323 | QGN15420.1 | FIM1_2110 | CP015056. | 360 | 293 | 215 | 600 | 362 | 193 | 387 | 251 | 233 | 2331 | 225 | 428 | 1747 | 178 | 316 | 2126 | 236 | 241 |
| 2324 | QGN15421.1 | FIM1_2111 | CP015056. | 41 | 8 | 26 | 112 | 42 | 24 | 56 | 23 | 33 | 174 | 18 | 41 | 170 | 21 | 42 | 160 | 16 | 22 |
| 2325 | QGN15422.1 | FIM1_2112 | CP015056. | 2992 | 676 | 861 | 4876 | 601 | 698 | 3628 | 599 | 1135 | 4149 | 361 | 961 | 3659 | 351 | 853 | 4179 | 279 | 634 |
| 2326 | QGN15423.1 | FIM1_2113 | CP015056. | 1778 | 737 | 1118 | 2297 | 875 | 860 | 1988 | 760 | 1466 | 3049 | 731 | 1476 | 5454 | 678 | 1375 | 3790 | 669 | 1118 |
| 2327 | QGN15424.1 | FIM1_2115 | CP015056. | 1690 | 886 | 749 | 1688 | 741 | 672 | 1395 | 690 | 1006 | 4753 | 885 | 1328 | 4066 | 674 | 1121 | 4580 | 711 | 730 |
| 2328 | QGN15425.1 | FIM1_2116 | CP015056. | 1772 | 644 | 746 | 2249 | 748 | 652 | 1750 | 682 | 987 | 4808 | 669 | 1256 | 5260 | 619 | 1071 | 4841 | 581 | 811 |
| 2329 | QGN15426.1 | FIM1_2117 | CP015056. | 3663 | 1395 | 1411 | 3971 | 1736 | 1212 | 3666 | 1604 | 1850 | 8562 | 1577 | 2068 | 8523 | 1443 | 1889 | 8074 | 1318 | 1351 |
| 2330 | QGN15427.1 | FIM1_2118 | CP015056. | 4313 | 1084 | 995 | 4419 | 1146 | 909 | 4211 | 977 | 1289 | 4816 | 865 | 1346 | 5291 | 804 | 1226 | 4650 | 759 | 911 |
| 2331 | QGN15428.1 | FIM1_2119 | CP015056. | 1237 | 647 | 361 | 599 | 708 | 440 | 1171 | 524 | 510 | 2405 | 784 | 668 | 2257 | 831 | 730 | 2296 | 822 | 484 |
| 2332 | QGN15429.1 | FIM1_2120 | CP015056. | 967 | 1735 | 1297 | 1250 | 1090 | 1232 | 894 | 965 | 1755 | 5084 | 1052 | 2064 | 4518 | 1262 | 1917 | 4557 | 1091 | 1303 |
| 2333 | QGN15430.1 | FIM1_2121 | CP015056. | 6067 | 2199 | 2364 | 7976 | 2780 | 2003 | 7735 | 2397 | 2932 | 9614 | 2050 | 3230 | 12306 | 2141 | 2941 | 9893 | 1809 | 2144 |
| 2334 | QGN15431.1 | FIM1_2122 | CP015056. | 1105 | 795 | 755 | 1432 | 774 | 730 | 1275 | 563 | 959 | 5873 | 817 | 1178 | 4584 | 775 | 1069 | 5413 | 821 | 805 |
| 2335 | QGN15432.1 | FIM1_2123 | CP015056. | 14711 | 3103 | 4698 | 18311 | 4179 | 4052 | 14743 | 3896 | 6141 | 14700 | 4092 | 6801 | 23658 | 4017 | 6402 | 18314 | 3630 | 4637 |
| 2336 | QGN15433.1 | FIM1_2124 | CP015056. | 8282 | 1472 | 2367 | 11646 | 2095 | 2518 | 6222 | 1669 | 3240 | 4996 | 618 | 1741 | 6255 | 853 | 1339 | 5062 | 459 | 1035 |
| 2337 | QGN15434.1 | FIM1_2125 | CP015056. | 4056 | 537 | 719 | 4279 | 851 | 593 | 4591 | 811 | 927 | 2118 | 670 | 1058 | 3612 | 582 | 984 | 2699 | 598 | 652 |
| 2338 | QGN15435.1 | FIM1_2126 | CP015056. | 2037 | 706 | 964 | 3391 | 885 | 775 | 2443 | 632 | 1162 | 3392 | 450 | 1363 | 3700 | 634 | 1120 | 3315 | 496 | 776 |
| 2339 | QGN15436.1 | FIM1_2127 | CP015056. | 2641 | 556 | 590 | 3992 | 627 | 537 | 3141 | 577 | 782 | 3120 | 536 | 754 | 3798 | 547 | 682 | 3382 | 529 | 495 |
| 2340 | QGN15437.1 | FIM1_2128 | CP015056. | 656 | 573 | 486 | 1108 | 730 | 474 | 636 | 563 | 570 | 2598 | 576 | 890 | 2453 | 403 | 710 | 2657 | 468 | 511 |
| 2341 | QGN15438.1 | FIM1_2129 | CP015056. | 1307 | 3171 | 1814 | 1429 | 2680 | 1885 | 1275 | 2364 | 2483 | 12920 | 2548 | 2717 | 10559 | 2219 | 2647 | 12189 | 2407 | 1788 |
| 2342 | QGN15439.1 | FIM1_2130 | CP015056. | 40688 | 1846 | 1317 | 30363 | 1990 | 1344 | 29359 | 1524 | 1732 | 7550 | 1330 | 1619 | 6494 | 1469 | 1527 | 6828 | 1300 | 1177 |
| 2343 | QGN15440.1 | FIM1_2131 | CP015056. | 4121 | 7743 | 6432 | 9250 | 7749 | 5545 | 6597 | 6580 | 7483 | 30410 | 8906 | 10385 | 27787 | 8432 | 10406 | 31589 | 7879 | 7406 |
| 2344 | QGN15441.1 | FIM1_2132 | CP015056. | 4582 | 1714 | 682 | 1553 | 1888 | 840 | 3876 | 1481 | 943 | 5107 | 2064 | 1429 | 5240 | 2304 | 1224 | 4881 | 2347 | 1027 |
| 2345 | QGN15442.1 | FIM1_2133 | CP015056. | 1171 | 902 | 717 | 1380 | 1034 | 590 | 1145 | 864 | 876 | 4824 | 967 | 970 | 3880 | 803 | 929 | 4521 | 855 | 571 |
| 2346 | QGN15443.1 | FIM1_2134 | CP015056. | 594 | 603 | 601 | 1081 | 535 | 589 | 824 | 490 | 796 | 4207 | 584 | 1017 | 3254 | 521 | 931 | 4081 | 547 | 672 |
| 2347 | QGN15444.1 | FIM1_2135 | CP015056. | 3682 | 1462 | 1182 | 3014 | 1493 | 1113 | 3537 | 1232 | 1534 | 9789 | 1719 | 2064 | 7876 | 1592 | 1975 | 9120 | 1624 | 1486 |
| 2348 | QGN15445.1 | FIM1_2136 | CP015056. | 1627 | 759 | 1220 | 1835 | 1089 | 1044 | 1874 | 886 | 1554 | 8491 | 1191 | 2151 | 8158 | 1109 | 1892 | 8337 | 1033 | 1411 |
| 2349 | QGN15446.1 | FIM1_2137 | CP015056. | 635 | 893 | 770 | 966 | 926 | 653 | 689 | 689 | 899 | 3589 | 618 | 780 | 2571 | 542 | 714 | 3405 | 596 | 505 |
| 2350 | QGN15447.1 | FIM1_2138 | CP015056. | 295 | 782 | 628 | 889 | 848 | 592 | 741 | 658 | 825 | 5036 | 662 | 1026 | 4138 | 664 | 937 | 4937 | 635 | 663 |

| | | | | | | | | | | | | | | | | | | | | | |
|---|---|---|---|---|---|---|---|---|---|---|---|---|---|---|---|---|---|---|---|---|---|
| 2351 | QGN15448.1 | FIM1_2139 | CP015056. | 3441 | 2630 | 2108 | 4362 | 1773 | 1759 | 3467 | 1603 | 2539 | 9154 | 1651 | 3111 | 7059 | 1816 | 2753 | 8763 | 1822 | 1968 |
| 2352 | QGN15449.1 | FIM1_2140 | CP015056. | 2266 | 815 | 605 | 2587 | 642 | 618 | 1963 | 655 | 936 | 2210 | 480 | 872 | 3041 | 525 | 842 | 2411 | 424 | 611 |
| 2353 | QGN15450.1 | FIM1_2141 | CP015056. | 474 | 1292 | 717 | 589 | 998 | 673 | 489 | 794 | 1024 | 6717 | 994 | 1227 | 5006 | 940 | 1281 | 6252 | 973 | 830 |
| 2354 | QGN15451.1 | FIM1_2142 | CP015056. | 416 | 76 | 113 | 484 | 78 | 82 | 417 | 94 | 141 | 730 | 88 | 187 | 580 | 90 | 182 | 747 | 70 | 150 |
| 2355 | QGN15452.1 | FIM1_2143 | CP015056. | 1380 | 478 | 548 | 1439 | 577 | 581 | 1190 | 446 | 684 | 3557 | 524 | 886 | 2832 | 447 | 808 | 3458 | 513 | 549 |
| 2356 | QGN15453.1 | FIM1_2144 | CP015056. | 1058 | 565 | 842 | 1898 | 738 | 748 | 1406 | 563 | 995 | 6899 | 707 | 1392 | 5167 | 678 | 1206 | 6852 | 667 | 896 |
| 2357 | QGN15454.1 | FIM1_2145 | CP015056. | 689 | 454 | 674 | 2269 | 555 | 557 | 1484 | 390 | 895 | 2070 | 240 | 628 | 2238 | 384 | 557 | 2159 | 242 | 408 |
| 2358 | QGN15455.1 | FIM1_2146 | CP015056. | 11918 | 8426 | 9236 | 13673 | 8738 | 8167 | 10315 | 7059 | 11350 | 19982 | 7960 | 12045 | 26762 | 7484 | 13786 | 23415 | 6347 | 9215 |
| 2359 | QGN15456.1 | FIM1_2147 | CP015056. | 605 | 403 | 308 | 955 | 628 | 274 | 614 | 435 | 398 | 3426 | 577 | 604 | 2642 | 451 | 578 | 3201 | 540 | 405 |
| 2360 | QGN15457.1 | FIM1_2148 | CP015056. | 372 | 300 | 380 | 867 | 320 | 291 | 455 | 293 | 454 | 3152 | 302 | 626 | 2101 | 291 | 493 | 2977 | 267 | 357 |
| 2361 | QGN15458.1 | FIM1_2149 | CP015056. | 1121 | 588 | 598 | 1667 | 654 | 517 | 1275 | 525 | 805 | 3425 | 542 | 964 | 3071 | 515 | 832 | 3464 | 469 | 568 |
| 2362 | QGN15459.1 | FIM1_2150 | CP015056. | 1146 | 667 | 999 | 2305 | 685 | 859 | 1416 | 466 | 1310 | 4253 | 530 | 1335 | 3530 | 717 | 1229 | 4042 | 599 | 804 |
| 2363 | QGN15460.1 | FIM1_2151 | CP015056. | 2560 | 433 | 346 | 1956 | 632 | 328 | 2151 | 441 | 437 | 2293 | 769 | 649 | 2359 | 776 | 626 | 2245 | 924 | 444 |
| 2364 | QGN15461.1 | FIM1_2152 | CP015056. | 812 | 116 | 95 | 798 | 128 | 68 | 643 | 121 | 100 | 773 | 159 | 186 | 768 | 167 | 146 | 719 | 171 | 115 |
| 2365 | QGN15462.1 | FIM1_2153 | CP015056. | 982 | 212 | 393 | 1105 | 174 | 277 | 887 | 207 | 451 | 1556 | 178 | 497 | 1269 | 146 | 457 | 1480 | 173 | 302 |
| 2366 | QGN15463.1 | FIM1_2154 | CP015056. | 1334 | 1145 | 391 | 595 | 921 | 496 | 1119 | 698 | 614 | 4456 | 1068 | 868 | 3070 | 985 | 759 | 4681 | 1379 | 576 |
| 2367 | QGN15464.1 | FIM1_2155 | CP015056. | 1004 | 444 | 476 | 1540 | 533 | 425 | 1359 | 437 | 643 | 2520 | 400 | 676 | 2753 | 423 | 634 | 2670 | 382 | 456 |
| 2368 | QGN15465.1 | FIM1_2156 | CP015056. | 981 | 1103 | 738 | 1251 | 813 | 738 | 1250 | 765 | 1006 | 4631 | 844 | 1342 | 3235 | 843 | 1283 | 4076 | 912 | 872 |
| 2369 | QGN15466.1 | FIM1_2157 | CP015056. | 2600 | 1973 | 3648 | 3892 | 2381 | 3113 | 3015 | 2185 | 4656 | 7723 | 1642 | 4869 | 10816 | 1636 | 4383 | 8412 | 1553 | 3191 |
| 2370 | QGN15467.1 | FIM1_2158 | CP015056. | 2706 | 1409 | 1301 | 2807 | 2124 | 1168 | 2680 | 1672 | 1596 | 6932 | 2351 | 2474 | 9451 | 2150 | 2191 | 7512 | 2236 | 1589 |
| 2371 | QGN15468.1 | FIM1_2159 | CP015056. | 147 | 34 | 31 | 450 | 71 | 33 | 157 | 22 | 37 | 430 | 58 | 57 | 339 | 47 | 54 | 405 | 38 | 36 |
| 2372 | QGN15469.1 | FIM1_2160 | CP015056. | 4409 | 1117 | 1087 | 4306 | 879 | 1013 | 3865 | 651 | 1437 | 6023 | 993 | 1645 | 4600 | 1036 | 1736 | 5558 | 859 | 1107 |
| 2373 | QGN15470.1 | FIM1_2161 | CP015056. | 3374 | 850 | 1108 | 3717 | 978 | 987 | 3197 | 880 | 1427 | 4944 | 718 | 1561 | 5580 | 656 | 1420 | 4972 | 617 | 1033 |
| 2374 | QGN15471.1 | FIM1_2162 | CP015056. | 1045 | 598 | 414 | 1202 | 613 | 377 | 1025 | 433 | 558 | 2729 | 629 | 781 | 2266 | 538 | 747 | 2425 | 508 | 475 |
| 2375 | QGN15472.1 | FIM1_2163 | CP015056. | 373 | 397 | 537 | 673 | 502 | 428 | 284 | 388 | 616 | 3723 | 549 | 974 | 3316 | 424 | 813 | 3807 | 419 | 647 |
| 2376 | QGN15473.1 | FIM1_2164 | CP015056. | 740 | 670 | 535 | 1044 | 569 | 483 | 787 | 501 | 719 | 2446 | 485 | 698 | 2476 | 411 | 731 | 2265 | 407 | 451 |
| 2377 | QGN15474.1 | FIM1_2165 | CP015056. | 2494 | 2495 | 1469 | 2193 | 2187 | 1236 | 2362 | 2053 | 1940 | 6780 | 2115 | 1994 | 8468 | 2022 | 1922 | 6752 | 1964 | 1374 |
| 2378 | QGN15475.1 | FIM1_2166 | CP015056. | 156 | 230 | 216 | 506 | 307 | 217 | 417 | 254 | 288 | 2431 | 235 | 391 | 1694 | 205 | 313 | 2396 | 213 | 237 |
| 2379 | QGN15476.1 | FIM1_2167 | CP015056. | 4300 | 3250 | 3406 | 7946 | 3353 | 2840 | 5114 | 3111 | 4346 | 11629 | 2694 | 5382 | 14489 | 2958 | 4765 | 12036 | 2544 | 3543 |
| 2380 | QGN15477.1 | FIM1_2168 | CP015056. | 5630 | 2820 | 2367 | 6020 | 3040 | 2357 | 6080 | 2455 | 3128 | 14810 | 3560 | 4329 | 14559 | 3242 | 4109 | 14314 | 3327 | 2941 |
| 2381 | QGN15478.1 | FIM1_2169 | CP015056. | 1137 | 1331 | 1590 | 1725 | 1748 | 1465 | 1459 | 1575 | 2062 | 7347 | 1784 | 2149 | 9004 | 1755 | 2122 | 7698 | 1518 | 1443 |
| 2382 | QGN15479.1 | FIM1_2170 | CP015056. | 3274 | 2158 | 2693 | 6528 | 1924 | 2263 | 4312 | 1914 | 3450 | 7577 | 1397 | 3432 | 8764 | 1674 | 3449 | 7752 | 1399 | 2292 |
| 2383 | QGN15480.1 | FIM1_2171 | CP015056. | 2039 | 674 | 899 | 1463 | 864 | 723 | 1829 | 949 | 1186 | 1368 | 568 | 967 | 2496 | 470 | 903 | 1891 | 437 | 672 |
| 2384 | QGN15481.1 | FIM1_2172 | CP015056. | 5166 | 1583 | 1114 | 6247 | 1733 | 1063 | 6332 | 1314 | 1483 | 2605 | 1152 | 1179 | 4114 | 980 | 1130 | 3107 | 1123 | 735 |
| 2385 | QGN15482.1 | FIM1_2173 | CP015056. | 613 | 791 | 575 | 932 | 461 | 566 | 885 | 426 | 811 | 2862 | 375 | 678 | 1862 | 354 | 704 | 2534 | 344 | 442 |
| 2386 | QGN15483.1 | FIM1_2174 | CP015056. | 2770 | 738 | 914 | 3864 | 743 | 655 | 2887 | 692 | 1101 | 3925 | 544 | 1128 | 3972 | 795 | 1082 | 3973 | 591 | 679 |
| 2387 | QGN15484.1 | FIM1_2175 | CP015056. | 5498 | 1900 | 1916 | 6664 | 2170 | 1692 | 5641 | 1791 | 2583 | 6017 | 2347 | 3013 | 11823 | 2277 | 2747 | 8347 | 2252 | 2187 |
| 2388 | QGN15485.1 | FIM1_2176 | CP015056. | 763 | 485 | 984 | 1189 | 389 | 874 | 747 | 468 | 1340 | 4620 | 280 | 1415 | 3691 | 223 | 1193 | 4679 | 195 | 828 |
| 2389 | QGN15486.1 | FIM1_2177 | CP015056. | 188 | 56 | 24 | 329 | 58 | 32 | 274 | 43 | 43 | 297 | 52 | 49 | 220 | 53 | 52 | 346 | 40 | 40 |
| 2390 | QGN15487.1 | FIM1_2178 | CP015056. | 2647 | 2491 | 3200 | 6799 | 2487 | 2569 | 3768 | 2579 | 4193 | 11789 | 1659 | 3956 | 13694 | 3443 | 3579 | 11808 | 1867 | 2811 |
| 2391 | QGN15488.1 | FIM1_2179 | CP015056. | 9114 | 4604 | 7074 | 16733 | 5851 | 6148 | 11765 | 5080 | 8665 | 25232 | 4909 | 9871 | 28107 | 5266 | 9533 | 25121 | 4491 | 6773 |
| 2392 | QGN15489.1 | FIM1_2180 | CP015056. | 3697 | 2347 | 1006 | 1986 | 2385 | 1039 | 3529 | 1852 | 1352 | 6991 | 2645 | 2071 | 7587 | 2439 | 1833 | 7208 | 2946 | 1400 |
| 2393 | QGN15490.1 | FIM1_2181 | CP015056. | 518 | 2533 | 1656 | 1941 | 2361 | 1377 | 1475 | 2188 | 2055 | 5915 | 2128 | 2482 | 8910 | 2135 | 2275 | 6922 | 2013 | 1594 |
| 2394 | QGN15491.1 | FIM1_2182 | CP015056. | 9822 | 542 | 1142 | 14437 | 351 | 969 | 10627 | 605 | 1396 | 2115 | 189 | 497 | 1347 | 220 | 290 | 1866 | 414 | 306 |
| 2395 | QGN15492.1 | FIM1_2183 | CP015056. | 5664 | 2133 | 1650 | 5760 | 1925 | 1430 | 5482 | 1837 | 2186 | 4513 | 1498 | 1780 | 7308 | 1380 | 1790 | 5455 | 1162 | 1263 |
| 2396 | QGN15493.1 | FIM1_2184 | CP015056. | 4450 | 2353 | 1968 | 6544 | 1646 | 1828 | 5277 | 1701 | 2595 | 3521 | 1507 | 2573 | 6314 | 1676 | 2670 | 4634 | 1516 | 1885 |
| 2397 | QGN15494.1 | FIM1_2185 | CP015056. | 12793 | 1160 | 2115 | 18970 | 1434 | 1611 | 14750 | 1374 | 2611 | 4035 | 736 | 1802 | 5865 | 1049 | 1588 | 4230 | 811 | 1129 |
| 2398 | QGN15495.1 | FIM1_2186 | CP015056. | 1237 | 1150 | 924 | 1415 | 1098 | 878 | 1359 | 946 | 1172 | 5996 | 1098 | 1396 | 4881 | 1032 | 1378 | 6003 | 1119 | 943 |
| 2399 | QGN15496.1 | FIM1_2187 | CP015056. | 2866 | 1731 | 736 | 1797 | 1328 | 805 | 2710 | 1030 | 1097 | 5618 | 1440 | 1187 | 4730 | 1296 | 1135 | 5392 | 1375 | 863 |
| 2400 | QGN15497.1 | FIM1_2188 | CP015056. | 5326 | 4668 | 3203 | 8010 | 4096 | 2671 | 6695 | 3582 | 4055 | 17584 | 3003 | 4061 | 15492 | 3264 | 3863 | 16838 | 2763 | 2692 |
| 2401 | QGN15498.1 | FIM1_2189 | CP015056. | 375 | 1165 | 554 | 526 | 776 | 553 | 487 | 730 | 750 | 4258 | 777 | 823 | 3267 | 656 | 765 | 3914 | 810 | 566 |
| 2402 | QGN15499.1 | FIM1_2190 | CP015056. | 5031 | 1326 | 1282 | 5390 | 1332 | 1100 | 4763 | 1370 | 1677 | 3619 | 1352 | 1843 | 6589 | 1160 | 1700 | 4892 | 1139 | 1270 |
| 2403 | QGN15500.1 | FIM1_2191 | CP015056. | 12268 | 4750 | 4525 | 13336 | 3900 | 3669 | 11303 | 3926 | 5827 | 23948 | 5106 | 7406 | 21703 | 5591 | 7575 | 23274 | 5062 | 5092 |
| 2404 | QGN15501.1 | FIM1_2192 | CP015056. | 1719 | 507 | 456 | 1679 | 545 | 383 | 2204 | 415 | 552 | 3773 | 490 | 853 | 2744 | 515 | 664 | 3546 | 594 | 545 |
| 2405 | QGN15502.1 | FIM1_2193 | CP015056. | 6005 | 2699 | 1339 | 2689 | 3382 | 1347 | 5196 | 2604 | 1657 | 8739 | 4186 | 2776 | 10626 | 4077 | 2551 | 9120 | 4761 | 1976 |
| 2406 | QGN15503.1 | FIM1_2194 | CP015056. | 1226 | 525 | 255 | 972 | 504 | 365 | 1060 | 339 | 420 | 1934 | 491 | 387 | 1588 | 529 | 401 | 1806 | 601 | 272 |
| 2407 | QGN15504.1 | FIM1_2195 | CP015056. | 2522 | 638 | 432 | 2053 | 543 | 411 | 2584 | 451 | 584 | 2210 | 457 | 616 | 1820 | 437 | 645 | 2004 | 439 | 363 |
| 2408 | QGN15505.1 | FIM1_2196 | CP015056. | 1018 | 70 | 137 | 1304 | 125 | 107 | 1331 | 137 | 148 | 1510 | 124 | 284 | 1237 | 153 | 202 | 1406 | 161 | 165 |
| 2409 | QGN15506.1 | FIM1_2197 | CP015056. | 5799 | 760 | 811 | 4993 | 814 | 622 | 6477 | 817 | 1019 | 4268 | 840 | 1166 | 4257 | 738 | 1051 | 4431 | 828 | 759 |
| 2410 | QGN15507.1 | FIM1_2198 | CP015056. | 2986 | 1439 | 2013 | 8057 | 2834 | 1644 | 4147 | 2019 | 2158 | 6326 | 1186 | 1485 | 7039 | 894 | 1000 | 6237 | 1090 | 874 |
| 2411 | QGN15508.1 | FIM1_2199 | CP015056. | 251 | 28 | 38 | 226 | 53 | 44 | 221 | 54 | 49 | 277 | 48 | 52 | 226 | 74 | 33 | 227 | 69 | 48 |
| 2412 | QGN15509.1 | FIM1_2200 | CP015056. | 1215 | 599 | 840 | 1963 | 993 | 841 | 1156 | 722 | 1272 | 2422 | 765 | 827 | 3106 | 951 | 807 | 2954 | 850 | 590 |
| 2413 | QGN15510.1 | FIM1_2201 | CP015056. | 1671 | 764 | 551 | 2186 | 730 | 519 | 1906 | 610 | 731 | 3655 | 669 | 857 | 3233 | 640 | 727 | 3615 | 657 | 555 |
| 2414 | QGN15511.1 | FIM1_2202 | CP015056. | 7371 | 1635 | 3378 | 8140 | 2420 | 2849 | 6968 | 2295 | 4312 | 7561 | 1819 | 5988 | 12656 | 2039 | 4820 | 8943 | 1864 | 3541 |
| 2415 | QGN15512.1 | FIM1_2203 | CP015056. | 2178 | 655 | 854 | 2255 | 489 | 601 | 2014 | 492 | 1029 | 3011 | 432 | 1008 | 2814 | 440 | 1125 | 2598 | 355 | 677 |
| 2416 | QGN15513.1 | FIM1_2204 | CP015056. | 2253 | 294 | 928 | 2423 | 427 | 623 | 1974 | 538 | 1109 | 3200 | 321 | 1121 | 3830 | 271 | 967 | 3244 | 288 | 721 |
| 2417 | QGN15514.1 | FIM1_2205 | CP015056. | 2829 | 542 | 620 | 1845 | 552 | 568 | 2103 | 498 | 778 | 3340 | 519 | 907 | 3092 | 500 | 913 | 3123 | 486 | 645 |
| 2418 | QGN15515.1 | FIM1_2206 | CP015056. | 1559 | 1408 | 420 | 936 | 1105 | 493 | 1502 | 863 | 569 | 3398 | 1184 | 773 | 2782 | 1026 | 775 | 3271 | 1289 | 532 |
| 2419 | QGN15516.1 | FIM1_2207 | CP015056. | 304 | 240 | 279 | 1301 | 323 | 273 | 697 | 266 | 408 | 407 | 158 | 372 | 1893 | 222 | 306 | 824 | 161 | 237 |
| 2420 | QGN15517.1 | FIM1_2208 | CP015056. | 2 | 0 | 0 | 1 | 0 | 1 | 2 | 1 | 0 | 0 | 1 | 0 | 0 | 1 | 0 | 0 | 0 | 0 |
| 2421 | QGN15518.1 | FIM1_2209 | CP015056. | 3493 | 1264 | 1420 | 3672 | 1787 | 1059 | 2636 | 1556 | 1801 | 1570 | 1079 | 1774 | 4831 | 746 | 1411 | 2926 | 796 | 934 |
| 2422 | QGN15519.1 | FIM1_2210 | CP015056. | 6039 | 2949 | 1279 | 3964 | 3226 | 1419 | 6202 | 2701 | 1767 | 11565 | 4270 | 2693 | 12847 | 3780 | 2497 | 11607 | 4905 | 1811 |
| 2423 | QGN15520.1 | FIM1_2211 | CP015056. | 1742 | 3666 | 3215 | 2654 | 4534 | 2805 | 2117 | 3692 | 3878 | 17210 | 4152 | 4571 | 16772 | 3742 | 4399 | 17002 | 3449 | 3272 |
| 2424 | QGN15521.1 | FIM1_2212 | CP015056. | 1296 | 741 | 1000 | 1325 | 670 | 760 | 957 | 861 | 1259 | 3190 | 486 | 1181 | 4395 | 411 | 1112 | 3428 | 369 | 759 |
| 2425 | QGN15522.1 | FIM1_2213 | CP015056. | 1133 | 532 | 496 | 1095 | 569 | 485 | 1345 | 528 | 673 | 3658 | 601 | 783 | 3450 | 580 | 753 | 3648 | 558 | 526 |
| 2426 | QGN15523.1 | FIM1_2214 | CP015056. | 911 | 290 | 456 | 825 | 318 | 386 | 888 | 321 | 575 | 2363 | 334 | 670 | 2354 | 324 | 599 | 2438 | 297 | 420 |
| 2427 | QGN15524.1 | FIM1_2215 | CP015056. | 4740 | 5227 | 2880 | 3919 | 4660 | 3076 | 4958 | 3794 | 3807 | 20008 | 6245 | 5035 | 16229 | 4404 | 5146 | 19972 | 5657 | 3654 |
| 2428 | QGN15525.1 | FIM1_2216 | CP015056. | 762 | 502 | 573 | 873 | 571 | 559 | 772 | 521 | 746 | 3895 | 550 | 917 | 3502 | 494 | 805 | 4107 | 546 | 558 |
| 2429 | QGN15526.1 | FIM1_2217 | CP015056. | 983 | 397 | 490 | 2684 | 514 | 447 | 1553 | 482 | 696 | 1277 | 369 | 734 | 3389 | 338 | 584 | 2165 | 296 | 472 |
| 2430 | QGN15527.1 | FIM1_2218 | CP015056. | 6552 | 4041 | 6135 | 9711 | 5076 | 5121 | 7601 | 4460 | 7453 | 18382 | 4953 | 10197 | 23518 | 5223 | 9041 | 20459 | 4482 | 6756 |
| 2431 | QGN15528.1 | FIM1_2219 | CP015056. | 1305 | 2452 | 1579 | 1416 | 2089 | 1458 | 1222 | 1981 | 2146 | 8011 | 2011 | 2148 | 8311 | 1931 | 2018 | 7872 | 1854 | 1425 |
| 2432 | QGN15529.1 | FIM1_2220 | CP015056. | 2748 | 795 | 925 | 2913 | 778 | 803 | 2954 | 753 | 1178 | 5159 | 838 | 1352 | 5275 | 754 | 1311 | 5016 | 703 | 979 |
| 2433 | QGN15530.1 | FIM1_2221 | CP015056. | 757 | 607 | 518 | 1093 | 509 | 495 | 769 | 476 | 725 | 2604 | 441 | 845 | 2710 | 463 | 791 | 2623 | 416 | 503 |
| 2434 | QGN15531.1 | FIM1_2222 | CP015056. | 1148 | 371 | 513 | 2085 | 320 | 388 | 1418 | 336 | 679 | 2050 | 227 | 602 | 2225 | 193 | 516 | 1909 | 204 | 426 |

| | | | | | | | | | | | | | | | | | | | | | |
|---|---|---|---|---|---|---|---|---|---|---|---|---|---|---|---|---|---|---|---|---|---|
| 2435 | QGN15532.1 | FIM1_2223 | CP015056. | 1492 | 358 | 592 | 1849 | 533 | 502 | 1677 | 567 | 766 | 5701 | 461 | 942 | 4660 | 504 | 825 | 5723 | 477 | 636 |
| 2436 | QGN15533.1 | FIM1_2224 | CP015056. | 528 | 3669 | 2229 | 799 | 3694 | 2065 | 718 | 2811 | 2900 | 14773 | 3576 | 3336 | 13639 | 3086 | 3170 | 14723 | 3162 | 2321 |
| 2437 | QGN15534.1 | FIM1_2225 | CP015056. | 3569 | 446 | 581 | 3397 | 739 | 483 | 2542 | 567 | 765 | 1409 | 533 | 784 | 4009 | 774 | 862 | 2245 | 512 | 563 |
| 2438 | QGN15535.1 | FIM1_2226 | CP015056. | 11943 | 7191 | 6189 | 16022 | 7380 | 5111 | 12548 | 6805 | 7592 | 26991 | 6972 | 9338 | 29271 | 7806 | 8477 | 27669 | 7176 | 6134 |
| 2439 | QGN15536.1 | FIM1_2227 | CP015056. | 2600 | 808 | 869 | 2306 | 1304 | 640 | 3314 | 1202 | 1026 | 3572 | 1683 | 1228 | 5071 | 2082 | 1182 | 3775 | 2005 | 849 |
| 2440 | QGN15537.1 | FIM1_2228 | CP015056. | 1419 | 594 | 620 | 1777 | 631 | 538 | 1632 | 664 | 809 | 366 | 525 | 770 | 2702 | 482 | 749 | 952 | 507 | 512 |
| 2441 | QGN15538.1 | FIM1_2229 | CP015056. | 1464 | 856 | 448 | 1424 | 744 | 385 | 1373 | 650 | 589 | 774 | 593 | 670 | 2205 | 587 | 679 | 1102 | 609 | 432 |
| 2442 | QGN15539.1 | FIM1_2230 | CP015056. | 1121 | 212 | 195 | 3243 | 121 | 135 | 1347 | 224 | 272 | 1000 | 155 | 267 | 1140 | 204 | 278 | 950 | 172 | 193 |
| 2443 | QGN15540.1 | FIM1_2231 | CP015056. | 978 | 544 | 701 | 1252 | 574 | 606 | 1153 | 557 | 923 | 2500 | 594 | 867 | 3279 | 475 | 859 | 2857 | 492 | 589 |
| 2444 | QGN15541.1 | FIM1_2232 | CP015056. | 8970 | 151 | 207 | 8814 | 282 | 206 | 4766 | 286 | 212 | 495 | 206 | 265 | 1072 | 204 | 264 | 717 | 186 | 167 |
| 2445 | QGN15542.1 | FIM1_2233 | CP015056. | 706 | 1835 | 1179 | 779 | 1515 | 1078 | 706 | 1293 | 1514 | 6576 | 1274 | 1399 | 5226 | 1260 | 1379 | 6253 | 1255 | 899 |
| 2446 | QGN15543.1 | FIM1_2234 | CP015056. | 986 | 269 | 361 | 1197 | 292 | 255 | 985 | 314 | 423 | 1468 | 220 | 469 | 2140 | 175 | 449 | 1600 | 198 | 306 |
| 2447 | QGN15544.1 | FIM1_2235 | CP015056. | 494 | 377 | 308 | 802 | 362 | 363 | 611 | 282 | 429 | 2404 | 322 | 487 | 1562 | 303 | 441 | 2194 | 334 | 299 |
| 2448 | QGN15545.1 | FIM1_2236 | CP015056. | 1672 | 749 | 816 | 1740 | 665 | 761 | 1738 | 599 | 1156 | 6471 | 661 | 1367 | 4684 | 596 | 1272 | 6454 | 615 | 946 |
| 2449 | QGN15546.1 | FIM1_2237 | CP015056. | 146 | 24 | 14 | 372 | 22 | 18 | 178 | 16 | 16 | 120 | 13 | 26 | 122 | 7 | 16 | 119 | 20 | 17 |
| 2450 | QGN15547.1 | FIM1_2238 | CP015056. | 1894 | 506 | 674 | 3120 | 512 | 570 | 2332 | 439 | 895 | 3015 | 350 | 811 | 2588 | 358 | 686 | 2876 | 289 | 503 |
| 2451 | QGN15548.1 | FIM1_2239 | CP015056. | 574 | 554 | 718 | 623 | 341 | 576 | 584 | 412 | 939 | 2751 | 320 | 930 | 1973 | 268 | 866 | 2603 | 290 | 581 |
| 2452 | QGN15549.1 | FIM1_2240 | CP015056. | 803 | 433 | 559 | 1197 | 299 | 471 | 807 | 264 | 685 | 2794 | 312 | 739 | 1999 | 284 | 702 | 2794 | 284 | 494 |
| 2453 | QGN15550.1 | FIM1_2241 | CP015056. | 469 | 1588 | 1823 | 1514 | 1894 | 1682 | 990 | 1679 | 2510 | 9434 | 1825 | 2753 | 11021 | 1780 | 2704 | 10384 | 1816 | 1893 |
| 2454 | QGN15551.1 | FIM1_2242 | CP015056. | 2 | 2 | 1 | 5 | 4 | 1 | 5 | 3 | 3 | 42 | 7 | 2 | 40 | 3 | 6 | 40 | 5 | 5 |
| 2455 | QGN15552.1 | FIM1_2243 | CP015056. | 1362 | 1035 | 714 | 1710 | 963 | 759 | 1380 | 832 | 1017 | 4039 | 833 | 929 | 3666 | 674 | 862 | 3749 | 744 | 658 |
| 2456 | QGN15553.1 | FIM1_2244 | CP015056. | 7903 | 2568 | 2493 | 7356 | 3819 | 2306 | 7872 | 3468 | 3384 | 12880 | 3650 | 3831 | 17894 | 3334 | 3743 | 14306 | 3139 | 2606 |
| 2457 | QGN15554.1 | FIM1_2245 | CP015056. | 4285 | 900 | 788 | 5877 | 643 | 578 | 5648 | 674 | 1005 | 5283 | 850 | 1232 | 4824 | 1276 | 1278 | 5093 | 981 | 779 |
| 2458 | QGN15555.1 | FIM1_2246 | CP015056. | 9316 | 9198 | 5861 | 8267 | 9085 | 5561 | 9058 | 8222 | 7561 | 26720 | 9239 | 10540 | 30497 | 8148 | 9630 | 26296 | 8369 | 7150 |
| 2459 | QGN15556.1 | FIM1_2247 | CP015056. | 1405 | 516 | 703 | 1507 | 597 | 570 | 1781 | 601 | 953 | 4512 | 623 | 1110 | 4844 | 599 | 904 | 4549 | 546 | 704 |
| 2460 | QGN15557.1 | FIM1_2248 | CP015056. | 3246 | 1757 | 884 | 1678 | 2198 | 974 | 3327 | 1645 | 1136 | 8603 | 2774 | 1853 | 7196 | 2835 | 1827 | 8706 | 3093 | 1397 |
| 2461 | QGN15558.1 | FIM1_2249 | CP015056. | 6131 | 3062 | 1513 | 3471 | 3264 | 1606 | 6153 | 2488 | 1986 | 11451 | 3842 | 2626 | 11586 | 4275 | 2674 | 11868 | 4194 | 1986 |
| 2462 | QGN15559.1 | FIM1_2250 | CP015056. | 923 | 288 | 153 | 441 | 251 | 168 | 900 | 187 | 171 | 691 | 279 | 269 | 441 | 355 | 255 | 630 | 437 | 167 |
| 2463 | QGN15560.1 | FIM1_2251 | CP015056. | 1364 | 320 | 288 | 1638 | 329 | 265 | 1427 | 308 | 366 | 1317 | 294 | 477 | 1745 | 315 | 385 | 1317 | 303 | 322 |
| 2464 | QGN15561.1 | FIM1_2252 | CP015056. | 4806 | 1974 | 2461 | 9442 | 2190 | 2051 | 6199 | 2135 | 3088 | 8103 | 1659 | 3745 | 10712 | 2157 | 3240 | 8388 | 1724 | 2431 |
| 2465 | QGN15562.1 | FIM1_2253 | CP015056. | 19285 | 10241 | 6865 | 21957 | 11463 | 6313 | 20361 | 10088 | 8805 | 35657 | 12435 | 13170 | 47969 | 11479 | 11808 | 36605 | 11833 | 8866 |
| 2466 | QGN15563.1 | FIM1_2254 | CP015056. | 2819 | 1756 | 1610 | 5330 | 1445 | 1678 | 3679 | 1033 | 2219 | 8979 | 1587 | 3004 | 7090 | 1542 | 2836 | 9377 | 1437 | 1875 |
| 2467 | QGN15564.1 | FIM1_2255 | CP015056. | 22463 | 10197 | 8674 | 23129 | 9434 | 7446 | 21050 | 9150 | 10773 | 23872 | 10017 | 12297 | 30049 | 10126 | 11305 | 25475 | 8772 | 8457 |
| 2468 | QGN15565.1 | FIM1_2256 | CP015056. | 2236 | 276 | 1773 | 3510 | 529 | 1217 | 2675 | 390 | 1849 | 2579 | 506 | 2380 | 2665 | 652 | 2296 | 2746 | 589 | 1476 |
| 2469 | QGN15566.1 | FIM1_2257 | CP015056. | 4574 | 181 | 1000 | 13698 | 200 | 641 | 8865 | 290 | 1196 | 1715 | 98 | 932 | 1717 | 254 | 732 | 1600 | 149 | 534 |
| 2470 | QGN15567.1 | FIM1_2258 | CP015056. | 2321 | 2517 | 1047 | 2152 | 2145 | 1077 | 1542 | 1558 | 1397 | 7567 | 1731 | 1754 | 6689 | 1305 | 1634 | 7178 | 1508 | 1087 |
| 2471 | QGN15568.1 | FIM1_2259 | CP015056. | 414 | 330 | 350 | 445 | 259 | 282 | 378 | 323 | 444 | 1967 | 246 | 418 | 1522 | 182 | 378 | 1937 | 188 | 255 |
| 2472 | QGN15569.1 | FIM1_2260 | CP015056. | 2848 | 962 | 1167 | 3572 | 1235 | 1042 | 3116 | 957 | 1527 | 8839 | 1019 | 1857 | 7246 | 1110 | 1685 | 8535 | 961 | 1278 |
| 2473 | QGN15570.1 | FIM1_2261 | CP015056. | 4186 | 1505 | 1766 | 4292 | 1988 | 1566 | 4044 | 1483 | 2264 | 15523 | 2198 | 3363 | 11879 | 2166 | 2897 | 14068 | 2233 | 2273 |
| 2474 | QGN15571.1 | FIM1_2262 | CP015056. | 1788 | 2703 | 3214 | 4542 | 1838 | 2274 | 3178 | 1673 | 3605 | 8711 | 1563 | 3914 | 7988 | 1811 | 3270 | 8953 | 1735 | 2194 |
| 2475 | QGN15572.1 | FIM1_2263 | CP015056. | 2636 | 2892 | 3479 | 5856 | 2590 | 3051 | 3948 | 2161 | 4299 | 10659 | 2179 | 5454 | 9808 | 2388 | 4901 | 10444 | 2265 | 3527 |
| 2476 | QGN15573.1 | FIM1_2264 | CP015056. | 1379 | 829 | 793 | 1582 | 905 | 715 | 907 | 698 | 1086 | 4169 | 953 | 1365 | 5113 | 921 | 1257 | 4246 | 850 | 911 |
| 2477 | QGN15574.1 | FIM1_2265 | CP015056. | 1173 | 744 | 572 | 1934 | 701 | 541 | 1394 | 659 | 795 | 2745 | 454 | 851 | 3086 | 539 | 799 | 2577 | 452 | 480 |
| 2478 | QGN15575.1 | FIM1_2266 | CP015056. | 215 | 1495 | 885 | 455 | 1344 | 844 | 339 | 1026 | 1182 | 2435 | 969 | 949 | 3358 | 941 | 1046 | 2536 | 970 | 617 |
| 2479 | QGN15576.1 | FIM1_2267 | CP015056. | 3209 | 1067 | 1126 | 2539 | 935 | 942 | 2518 | 961 | 1411 | 4101 | 966 | 1379 | 4511 | 918 | 1287 | 3884 | 803 | 906 |
| 2480 | QGN15577.1 | FIM1_2268 | CP015056. | 900 | 944 | 882 | 890 | 882 | 734 | 960 | 805 | 1111 | 5845 | 968 | 1164 | 5099 | 865 | 1025 | 5738 | 784 | 692 |
| 2481 | QGN15578.1 | FIM1_2270 | CP015056. | 1030 | 554 | 439 | 751 | 529 | 416 | 1002 | 510 | 633 | 3660 | 573 | 738 | 2760 | 585 | 672 | 3211 | 575 | 482 |
| 2482 | QGN15579.1 | FIM1_2271 | CP015056. | 2253 | 830 | 925 | 2197 | 1075 | 798 | 1574 | 824 | 1228 | 5848 | 1030 | 1514 | 5968 | 915 | 1401 | 5666 | 758 | 991 |
| 2483 | QGN15580.1 | FIM1_2272 | CP015056. | 767 | 384 | 774 | 1687 | 317 | 664 | 800 | 375 | 1009 | 1826 | 270 | 971 | 1974 | 249 | 967 | 1763 | 211 | 649 |
| 2484 | QGN15581.1 | FIM1_2273 | CP015056. | 204 | 63 | 153 | 609 | 125 | 125 | 332 | 92 | 194 | 484 | 67 | 208 | 464 | 79 | 211 | 493 | 56 | 134 |
| 2485 | QGN15582.1 | FIM1_2274 | CP015056. | 4127 | 899 | 2171 | 4211 | 1436 | 1803 | 3533 | 1286 | 3057 | 3089 | 1115 | 2526 | 7410 | 822 | 2142 | 5089 | 798 | 1588 |
| 2486 | QGN15583.1 | FIM1_2275 | CP015056. | 2090 | 1962 | 1721 | 4022 | 1569 | 1508 | 2484 | 1389 | 2141 | 6783 | 1108 | 1836 | 6305 | 1508 | 1734 | 6649 | 1034 | 1216 |
| 2487 | QGN15584.1 | FIM1_2276 | CP015056. | 1177 | 988 | 493 | 854 | 807 | 473 | 804 | 689 | 709 | 2113 | 693 | 561 | 2304 | 626 | 654 | 1783 | 613 | 441 |
| 2488 | QGN15585.1 | FIM1_2277 | CP015056. | 32 | 3 | 3 | 45 | 7 | 8 | 18 | 1 | 3 | 5 | 10 | 4 | 9 | 2 | 1 | 9 | 2 | 2 |
| 2489 | QGN15586.1 | FIM1_2278 | CP015056. | 7963 | 800 | 1011 | 5186 | 1008 | 980 | 5216 | 820 | 1316 | 2040 | 716 | 1607 | 5156 | 898 | 1404 | 3237 | 823 | 1026 |
| 2490 | QGN15587.1 | FIM1_2279 | CP015056. | 999 | 252 | 215 | 1341 | 187 | 163 | 1213 | 218 | 322 | 810 | 142 | 228 | 855 | 122 | 206 | 837 | 130 | 128 |
| 2491 | QGN15588.1 | FIM1_2280 | CP015056. | 1109 | 1083 | 1516 | 1881 | 1203 | 1236 | 1190 | 942 | 1860 | 5738 | 1250 | 2024 | 6081 | 1367 | 1823 | 5651 | 1262 | 1275 |
| 2492 | QGN15589.1 | FIM1_2281 | CP015056. | 1755 | 433 | 516 | 1583 | 483 | 433 | 1043 | 545 | 706 | 1764 | 382 | 808 | 3202 | 284 | 768 | 2216 | 309 | 506 |
| 2493 | QGN15590.1 | FIM1_2282 | CP015056. | 536 | 197 | 282 | 713 | 177 | 202 | 550 | 178 | 352 | 1000 | 103 | 315 | 1199 | 70 | 330 | 966 | 76 | 199 |
| 2494 | QGN15591.1 | FIM1_2283 | CP015056. | 3185 | 717 | 1212 | 2522 | 552 | 1084 | 2825 | 678 | 1607 | 3546 | 445 | 1606 | 3299 | 409 | 1675 | 3347 | 375 | 1088 |
| 2495 | QGN15592.1 | FIM1_2284 | CP015056. | 7149 | 1236 | 1663 | 7109 | 1315 | 1428 | 5629 | 1440 | 2240 | 2057 | 1077 | 2112 | 7172 | 1101 | 2156 | 4136 | 948 | 1515 |
| 2496 | QGN15593.1 | FIM1_2285 | CP015056. | 6600 | 1405 | 2058 | 8708 | 2622 | 2173 | 5847 | 1610 | 2691 | 12138 | 2342 | 4376 | 14404 | 2994 | 3332 | 13641 | 2243 | 3007 |
| 2497 | QGN15594.1 | FIM1_2286 | CP015056. | 1384 | 770 | 609 | 1842 | 636 | 585 | 1517 | 491 | 832 | 4287 | 645 | 984 | 3256 | 675 | 999 | 4209 | 642 | 666 |
| 2498 | QGN15595.1 | FIM1_2287 | CP015056. | 2547 | 574 | 472 | 2700 | 457 | 475 | 2777 | 366 | 716 | 3592 | 425 | 728 | 2790 | 387 | 731 | 3467 | 393 | 489 |
| 2499 | QGN15596.1 | FIM1_2288 | CP015056. | 3276 | 922 | 1188 | 3544 | 1261 | 997 | 3030 | 1192 | 1428 | 4267 | 1212 | 1629 | 6193 | 1151 | 1542 | 5176 | 1134 | 1106 |
| 2500 | QGN15597.1 | FIM1_2289 | CP015056. | 2153 | 2083 | 2059 | 1572 | 2617 | 1903 | 1910 | 2390 | 2492 | 10468 | 2851 | 2721 | 9432 | 2598 | 2899 | 10687 | 2521 | 1891 |
| 2501 | QGN15598.1 | FIM1_2290 | CP015056. | 10696 | 725 | 1014 | 7961 | 1196 | 967 | 8613 | 841 | 1293 | 5891 | 1183 | 1479 | 6780 | 1938 | 1463 | 6043 | 1470 | 1041 |
| 2502 | QGN15599.1 | FIM1_2291 | CP015056. | 3968 | 2419 | 1878 | 7078 | 2048 | 1911 | 5238 | 1696 | 2558 | 8430 | 2353 | 2880 | 7975 | 3024 | 3160 | 8151 | 2573 | 2069 |
| 2503 | QGN15600.1 | FIM1_2292 | CP015056. | 260 | 897 | 539 | 446 | 692 | 507 | 405 | 518 | 678 | 3843 | 679 | 784 | 2755 | 580 | 729 | 3675 | 606 | 492 |
| 2504 | QGN15601.1 | FIM1_2293 | CP015056. | 104 | 545 | 512 | 237 | 472 | 442 | 133 | 347 | 644 | 2715 | 396 | 673 | 1959 | 363 | 630 | 2588 | 377 | 403 |
| 2505 | QGN15602.1 | FIM1_2294 | CP015056. | 8237 | 7225 | 5194 | 9763 | 6838 | 4749 | 8619 | 5837 | 6530 | 26487 | 7610 | 7483 | 27774 | 6527 | 7682 | 25598 | 6496 | 5397 |
| 2506 | QGN15603.1 | FIM1_2295 | CP015056. | 187 | 452 | 556 | 463 | 545 | 513 | 279 | 452 | 713 | 3441 | 481 | 803 | 2972 | 397 | 773 | 3408 | 388 | 514 |
| 2507 | QGN15604.1 | FIM1_2297 | CP015056. | 1904 | 652 | 726 | 1918 | 788 | 696 | 1784 | 653 | 970 | 4789 | 826 | 1152 | 4396 | 659 | 1113 | 4620 | 682 | 790 |
| 2508 | QGN15605.1 | FIM1_2298 | CP015056. | 4004 | 651 | 857 | 4071 | 822 | 653 | 3798 | 747 | 1103 | 3658 | 858 | 1218 | 6760 | 634 | 1168 | 4936 | 621 | 828 |
| 2509 | QGN15606.1 | FIM1_2299 | CP015056. | 1444 | 745 | 1116 | 2441 | 950 | 830 | 1750 | 896 | 1311 | 2114 | 882 | 1797 | 4088 | 846 | 1637 | 2784 | 731 | 1079 |
| 2510 | QGN15607.1 | FIM1_2300 | CP015056. | 877 | 229 | 403 | 1008 | 288 | 331 | 706 | 259 | 459 | 2460 | 314 | 520 | 2368 | 224 | 469 | 2365 | 267 | 303 |
| 2511 | QGN15608.1 | FIM1_2301 | CP015056. | 1945 | 1834 | 1635 | 5876 | 1778 | 1568 | 3641 | 1340 | 2056 | 9750 | 1907 | 2849 | 8390 | 1993 | 2490 | 9251 | 1949 | 1825 |
| 2512 | QGN15609.1 | FIM1_2302 | CP015056. | 3692 | 1585 | 2885 | 4248 | 2085 | 2624 | 3185 | 1538 | 3629 | 9217 | 1845 | 5155 | 8627 | 1469 | 4736 | 9156 | 1404 | 3374 |
| 2513 | QGN15610.1 | FIM1_2303 | CP015056. | 937 | 92 | 125 | 1796 | 207 | 114 | 838 | 150 | 129 | 403 | 131 | 174 | 656 | 118 | 172 | 430 | 122 | 136 |
| 2514 | QGN15611.1 | FIM1_2304 | CP015056. | 668 | 167 | 359 | 875 | 110 | 229 | 629 | 207 | 390 | 447 | 82 | 443 | 604 | 91 | 376 | 494 | 83 | 278 |
| 2515 | QGN15612.1 | FIM1_2305 | CP015056. | 1722 | 446 | 460 | 1577 | 481 | 355 | 1498 | 407 | 574 | 2396 | 409 | 737 | 2601 | 400 | 710 | 2310 | 402 | 501 |
| 2516 | QGN15613.1 | FIM1_2306 | CP015056. | 632 | 130 | 266 | 950 | 149 | 169 | 654 | 132 | 309 | 1132 | 65 | 272 | 1289 | 55 | 256 | 1030 | 50 | 160 |
| 2517 | QGN15614.1 | FIM1_2307 | CP015056. | 1486 | 2623 | 1366 | 1562 | 2199 | 1324 | 1505 | 1953 | 1769 | 8573 | 1841 | 1989 | 7513 | 1792 | 1767 | 7925 | 1839 | 1308 |
| 2518 | QGN15615.1 | FIM1_2308 | CP015056. | 1218 | 1763 | 885 | 745 | 1491 | 893 | 1161 | 1210 | 1181 | 6883 | 1725 | 1505 | 5903 | 1391 | 1411 | 6985 | 1646 | 1049 |

| | | | | | | | | | | | | | | | | | | | | | |
|---|---|---|---|---|---|---|---|---|---|---|---|---|---|---|---|---|---|---|---|---|---|
| 2519 | QGN15616.1 | FIM1_2309 | CP015056. | 855 | 131 | 281 | 1553 | 212 | 223 | 754 | 161 | 340 | 497 | 91 | 271 | 651 | 96 | 238 | 568 | 71 | 144 |
| 2520 | QGN15617.1 | FIM1_2310 | CP015056. | 314 | 235 | 262 | 465 | 260 | 276 | 325 | 234 | 347 | 237 | 140 | 300 | 579 | 235 | 265 | 365 | 234 | 215 |
| 2521 | QGN15618.1 | FIM1_2311 | CP015056. | 6271 | 522 | 1067 | 5834 | 820 | 812 | 6634 | 912 | 1439 | 600 | 492 | 1312 | 2497 | 544 | 1084 | 1001 | 582 | 872 |
| 2522 | QGN15619.1 | FIM1_2312 | CP015056. | 4816 | 3057 | 3102 | 7850 | 3972 | 2830 | 6043 | 3110 | 4044 | 15451 | 3071 | 4095 | 15763 | 3031 | 3710 | 15124 | 2753 | 2622 |
| 2523 | QGN15620.1 | FIM1_2313 | CP015056. | 3308 | 888 | 554 | 2468 | 848 | 523 | 3034 | 761 | 788 | 2350 | 818 | 864 | 2753 | 1109 | 768 | 2448 | 984 | 577 |
| 2524 | QGN15621.1 | FIM1_2314 | CP015056. | 1803 | 772 | 907 | 2285 | 823 | 715 | 2243 | 747 | 1181 | 3970 | 514 | 909 | 3557 | 477 | 874 | 3659 | 450 | 576 |
| 2525 | QGN15622.1 | FIM1_2315 | CP015056. | 3715 | 430 | 543 | 3923 | 434 | 493 | 5523 | 582 | 698 | 2009 | 297 | 449 | 3701 | 421 | 454 | 2634 | 313 | 360 |
| 2526 | QGN15623.1 | FIM1_2316 | CP015056. | 1305 | 441 | 542 | 1587 | 522 | 439 | 1327 | 503 | 749 | 875 | 520 | 794 | 3192 | 577 | 764 | 1879 | 530 | 563 |
| 2527 | QGN15624.1 | FIM1_2317 | CP015056. | 7551 | 1050 | 1149 | 6172 | 1247 | 1062 | 6696 | 1198 | 1561 | 4226 | 1107 | 1405 | 7136 | 1057 | 1419 | 5587 | 880 | 990 |
| 2528 | QGN15625.1 | FIM1_2318 | CP015056. | 13325 | 15588 | 11233 | 7756 | 12310 | 8381 | 9093 | 16936 | 14477 | 33599 | 11714 | 8174 | 33603 | 9449 | 6977 | 32677 | 10772 | 6172 |
| 2529 | QGN15626.1 | FIM1_2319 | CP015056. | 202 | 38 | 131 | 266 | 113 | 80 | 184 | 74 | 150 | 366 | 76 | 146 | 616 | 86 | 116 | 446 | 57 | 96 |
| 2530 | QGN15627.1 | FIM1_2320 | CP015056. | 309 | 185 | 136 | 309 | 151 | 99 | 210 | 145 | 155 | 519 | 159 | 230 | 734 | 137 | 197 | 595 | 156 | 136 |
| 2531 | QGN15628.1 | FIM1_2321 | CP015056. | 2240 | 959 | 1046 | 2503 | 1105 | 887 | 2299 | 978 | 1350 | 5150 | 1132 | 1571 | 6149 | 986 | 1502 | 5504 | 937 | 1052 |
| 2532 | QGN15629.1 | FIM1_2322 | CP015056. | 792 | 685 | 698 | 595 | 584 | 578 | 736 | 629 | 909 | 2505 | 468 | 796 | 2940 | 379 | 868 | 2653 | 364 | 554 |
| 2533 | QGN15630.1 | FIM1_2323 | CP015056. | 2682 | 2429 | 1543 | 3136 | 2254 | 1589 | 2665 | 1856 | 2088 | 11955 | 2200 | 2252 | 9950 | 1917 | 1989 | 11671 | 2186 | 1541 |
| 2534 | QGN15631.1 | FIM1_2324 | CP015056. | 986 | 3055 | 2427 | 2067 | 3279 | 2144 | 1980 | 3527 | 3225 | 8493 | 2715 | 3291 | 12132 | 2326 | 3049 | 8696 | 2388 | 2373 |
| 2535 | QGN15632.1 | FIM1_2325 | CP015056. | 80 | 22 | 31 | 91 | 24 | 16 | 95 | 29 | 41 | 111 | 26 | 42 | 129 | 20 | 36 | 115 | 22 | 29 |
| 2536 | QGN15633.1 | FIM1_2326 | CP015056. | 700 | 283 | 278 | 1004 | 268 | 264 | 701 | 232 | 363 | 2052 | 240 | 394 | 1687 | 220 | 349 | 1869 | 219 | 252 |
| 2537 | QGN15634.1 | FIM1_2327 | CP015056. | 7763 | 1622 | 1944 | 9143 | 2253 | 1691 | 7070 | 1767 | 2511 | 8550 | 1564 | 2738 | 9720 | 1681 | 2464 | 9002 | 1505 | 1876 |
| 2538 | QGN15635.1 | FIM1_2328 | CP015056. | 4651 | 4146 | 3755 | 5938 | 5149 | 3473 | 5399 | 4534 | 4869 | 16269 | 5388 | 6513 | 23752 | 5228 | 5973 | 18497 | 5018 | 4411 |
| 2539 | QGN15636.1 | FIM1_2329 | CP015056. | 925 | 118 | 203 | 643 | 223 | 167 | 916 | 247 | 259 | 1554 | 185 | 257 | 1591 | 179 | 252 | 1468 | 180 | 159 |
| 2540 | QGN15637.1 | FIM1_2330 | CP015056. | 1401 | 314 | 259 | 660 | 380 | 233 | 810 | 342 | 370 | 1194 | 304 | 359 | 1401 | 338 | 308 | 1158 | 298 | 211 |
| 2541 | QGN15638.1 | FIM1_2331 | CP015056. | 2289 | 2081 | 1966 | 5482 | 1467 | 1703 | 3200 | 1307 | 2617 | 8085 | 1482 | 2229 | 6363 | 1791 | 2462 | 7318 | 1516 | 1485 |
| 2542 | QGN15639.1 | FIM1_2332 | CP015056. | 8829 | 4626 | 6463 | 8530 | 3971 | 4455 | 9572 | 4727 | 8246 | 18443 | 5475 | 9067 | 25721 | 8619 | 10460 | 19370 | 6534 | 6847 |
| 2543 | QGN15640.1 | FIM1_2333 | CP015056. | 594 | 129 | 285 | 556 | 233 | 185 | 505 | 241 | 330 | 1801 | 212 | 334 | 1578 | 182 | 330 | 1837 | 177 | 189 |
| 2544 | QGN15641.1 | FIM1_2334 | CP015056. | 2152 | 349 | 413 | 2476 | 348 | 301 | 1809 | 376 | 519 | 1831 | 260 | 387 | 1797 | 238 | 430 | 1860 | 191 | 265 |
| 2545 | QGN15642.1 | FIM1_2335 | CP015056. | 8475 | 6930 | 4092 | 14706 | 4900 | 3870 | 9577 | 4030 | 5362 | 7081 | 3978 | 5241 | 13839 | 3940 | 4968 | 10433 | 3686 | 3677 |
| 2546 | QGN15643.1 | FIM1_2336 | CP015056. | 8529 | 238 | 303 | 10310 | 381 | 300 | 11618 | 273 | 385 | 1139 | 343 | 652 | 2781 | 550 | 541 | 1805 | 386 | 460 |
| 2547 | QGN15644.1 | FIM1_2337 | CP015056. | 850 | 35 | 85 | 926 | 117 | 66 | 784 | 119 | 107 | 372 | 104 | 137 | 577 | 116 | 127 | 433 | 103 | 95 |
| 2548 | QGN15645.1 | FIM1_2338 | CP015056. | 1338 | 1107 | 1161 | 1759 | 1371 | 991 | 1438 | 1419 | 1627 | 4336 | 1017 | 954 | 5371 | 862 | 899 | 4681 | 850 | 612 |
| 2549 | QGN15646.1 | FIM1_2339 | CP015056. | 1170 | 240 | 372 | 1872 | 252 | 325 | 1287 | 320 | 551 | 2477 | 216 | 579 | 2959 | 197 | 497 | 2432 | 200 | 364 |
| 2550 | QGN15647.1 | FIM1_2340 | CP015056. | 2230 | 916 | 868 | 1063 | 698 | 857 | 1714 | 963 | 1285 | 3004 | 715 | 1520 | 3407 | 483 | 1610 | 3187 | 610 | 980 |
| 2551 | QGN15648.1 | FIM1_2341 | CP015056. | 3769 | 536 | 333 | 3812 | 320 | 200 | 4163 | 433 | 415 | 894 | 307 | 417 | 1516 | 333 | 457 | 1077 | 231 | 298 |
| 2552 | QGN15649.1 | FIM1_2342 | CP015056. | 1122 | 1725 | 1170 | 1714 | 1355 | 961 | 1028 | 1071 | 1461 | 6241 | 1123 | 1679 | 5012 | 1267 | 1656 | 5944 | 1017 | 1055 |
| 2553 | QGN15650.1 | FIM1_2343 | CP015056. | 460 | 42 | 38 | 343 | 33 | 22 | 364 | 31 | 40 | 198 | 11 | 25 | 128 | 4 | 23 | 207 | 12 | 22 |
| 2554 | QGN15651.1 | FIM1_2344 | CP015056. | 5254 | 158 | 391 | 5751 | 105 | 241 | 4870 | 158 | 473 | 1506 | 99 | 382 | 1659 | 155 | 369 | 1554 | 114 | 232 |
| 2555 | QGN15652.1 | FIM1_2345 | CP015056. | 2884 | 471 | 808 | 4225 | 586 | 588 | 2703 | 468 | 1030 | 1989 | 390 | 545 | 2317 | 362 | 520 | 2028 | 334 | 355 |
| 2556 | QGN15653.1 | FIM1_2346 | CP015056. | 4865 | 1854 | 2188 | 5842 | 2768 | 2027 | 5229 | 2047 | 2873 | 18212 | 2642 | 3754 | 14986 | 2313 | 3388 | 17256 | 2249 | 2582 |
| 2557 | QGN15654.1 | FIM1_2347 | CP015056. | 824 | 549 | 369 | 904 | 489 | 374 | 896 | 369 | 527 | 2331 | 391 | 469 | 1566 | 354 | 513 | 1799 | 401 | 351 |
| 2558 | QGN15655.1 | FIM1_2348 | CP015056. | 1895 | 2475 | 1618 | 2044 | 2009 | 1464 | 1851 | 1802 | 2184 | 11752 | 2160 | 2478 | 10033 | 1992 | 2245 | 11427 | 1996 | 1628 |
| 2559 | QGN15656.1 | FIM1_2349 | CP015056. | 3554 | 989 | 1439 | 4576 | 1167 | 1250 | 4045 | 1019 | 1889 | 9625 | 1112 | 2521 | 9515 | 1059 | 2173 | 9805 | 981 | 1674 |
| 2560 | QGN15657.1 | FIM1_2350 | CP015056. | 2592 | 2265 | 2266 | 4402 | 1868 | 1994 | 3406 | 1684 | 2845 | 7890 | 1617 | 3059 | 7662 | 1776 | 3118 | 7484 | 1492 | 2121 |
| 2561 | QGN15658.1 | FIM1_2351 | CP015056. | 1289 | 840 | 728 | 3055 | 608 | 693 | 1736 | 530 | 1040 | 5211 | 655 | 998 | 3975 | 808 | 971 | 4744 | 560 | 643 |
| 2562 | QGN15659.1 | FIM1_2352 | CP015056. | 3121 | 436 | 624 | 3300 | 505 | 478 | 2965 | 519 | 794 | 1286 | 532 | 816 | 4450 | 520 | 803 | 2733 | 493 | 597 |
| 2563 | QGN15660.1 | FIM1_2353 | CP015056. | 1398 | 922 | 743 | 1568 | 779 | 670 | 1396 | 664 | 1024 | 3589 | 550 | 882 | 2817 | 462 | 807 | 3552 | 453 | 610 |
| 2564 | QGN15661.1 | FIM1_2354 | CP015056. | 1386 | 140 | 486 | 1183 | 318 | 341 | 1298 | 285 | 578 | 1893 | 215 | 661 | 1941 | 201 | 586 | 2029 | 211 | 474 |
| 2565 | QGN15662.1 | FIM1_2355 | CP015056. | 780 | 222 | 176 | 527 | 156 | 145 | 711 | 188 | 229 | 533 | 115 | 126 | 485 | 73 | 145 | 467 | 105 | 97 |
| 2566 | QGN15663.1 | FIM1_2356 | CP015056. | 1364 | 196 | 355 | 1449 | 216 | 224 | 1117 | 226 | 456 | 293 | 141 | 383 | 1087 | 197 | 401 | 551 | 147 | 261 |
| 2567 | QGN15664.1 | FIM1_2357 | CP015056. | 3858 | 1481 | 832 | 2438 | 1548 | 860 | 3697 | 1246 | 1078 | 6612 | 1807 | 1471 | 6168 | 1597 | 1415 | 6458 | 1973 | 1047 |
| 2568 | QGN15665.1 | FIM1_2358 | CP015056. | 783 | 786 | 992 | 1025 | 955 | 882 | 877 | 836 | 1259 | 3841 | 630 | 1144 | 3045 | 519 | 1058 | 3732 | 555 | 717 |
| 2569 | QGN15666.1 | FIM1_2359 | CP015056. | 2321 | 3732 | 2683 | 2172 | 3221 | 2422 | 2041 | 2982 | 3445 | 10713 | 3375 | 3702 | 12412 | 3166 | 3572 | 11087 | 3132 | 2442 |
| 2570 | QGN15667.1 | FIM1_2360 | CP015056. | 5324 | 1393 | 1505 | 8922 | 1230 | 1168 | 6698 | 1188 | 1966 | 8436 | 782 | 1600 | 7264 | 1238 | 1489 | 7979 | 909 | 1058 |
| 2571 | QGN15668.1 | FIM1_2361 | CP015056. | 3594 | 2777 | 2104 | 4072 | 2273 | 1871 | 3263 | 1981 | 2681 | 9067 | 1886 | 2975 | 9371 | 2300 | 3062 | 8897 | 1875 | 2009 |
| 2572 | QGN15669.1 | FIM1_2362 | CP015056. | 2933 | 959 | 866 | 2677 | 1066 | 734 | 2818 | 911 | 1115 | 6432 | 1069 | 1397 | 5703 | 989 | 1292 | 6127 | 915 | 934 |
| 2573 | QGN15670.1 | FIM1_2363 | CP015056. | 4627 | 1328 | 1033 | 2750 | 1664 | 981 | 3722 | 1300 | 1312 | 5744 | 1467 | 1798 | 6694 | 1272 | 1602 | 5798 | 1346 | 1158 |
| 2574 | QGN15671.1 | FIM1_2364 | CP015056. | 3681 | 1466 | 1523 | 3628 | 1838 | 1316 | 3259 | 1621 | 1973 | 7442 | 1503 | 2062 | 8569 | 1325 | 1980 | 7611 | 1331 | 1465 |
| 2575 | QGN15672.1 | FIM1_2365 | CP015056. | 14579 | 18324 | 11815 | 14168 | 14167 | 11256 | 12330 | 11868 | 13949 | 44515 | 17551 | 16723 | 37601 | 14904 | 16960 | 44077 | 15244 | 12181 |
| 2576 | QGN15673.1 | FIM1_2366 | CP015056. | 719 | 346 | 319 | 1404 | 260 | 285 | 898 | 288 | 466 | 2392 | 255 | 538 | 2117 | 288 | 533 | 2442 | 258 | 369 |
| 2577 | QGN15674.1 | FIM1_2367 | CP015056. | 578 | 310 | 257 | 1240 | 379 | 262 | 1244 | 276 | 338 | 1110 | 254 | 387 | 1407 | 225 | 338 | 1149 | 233 | 275 |
| 2578 | QGN15675.1 | FIM1_2368 | CP015056. | 8638 | 690 | 1283 | 6677 | 1062 | 1109 | 7370 | 1010 | 1819 | 3003 | 1096 | 2114 | 6391 | 979 | 1994 | 3589 | 1037 | 1474 |
| 2579 | QGN15676.1 | FIM1_2369 | CP015056. | 1815 | 489 | 851 | 1827 | 760 | 728 | 1429 | 634 | 1070 | 3655 | 673 | 1246 | 4332 | 524 | 1204 | 3803 | 513 | 892 |
| 2580 | QGN15677.1 | FIM1_2370 | CP015056. | 755 | 205 | 192 | 592 | 214 | 181 | 1035 | 222 | 291 | 1993 | 252 | 364 | 1679 | 140 | 320 | 1795 | 199 | 197 |
| 2581 | QGN15678.1 | FIM1_2371 | CP015056. | 21731 | 803 | 1898 | 25295 | 2025 | 1408 | 20877 | 2304 | 2475 | 2412 | 830 | 2689 | 7901 | 1001 | 1954 | 3394 | 1039 | 1605 |
| 2582 | QGN15679.1 | FIM1_2372 | CP015056. | 5946 | 5298 | 2525 | 4644 | 4466 | 2452 | 4183 | 3966 | 3491 | 8131 | 3895 | 3252 | 12457 | 3790 | 3200 | 9591 | 3354 | 2368 |
| 2583 | QGN15680.1 | FIM1_2373 | CP015056. | 737 | 251 | 1704 | 1673 | 223 | 951 | 611 | 251 | 1582 | 753 | 78 | 1062 | 1083 | 110 | 695 | 1037 | 72 | 546 |
| 2584 | QGN15681.1 | FIM1_2374 | CP015056. | 745 | 813 | 733 | 1270 | 775 | 637 | 940 | 686 | 921 | 5634 | 735 | 905 | 4793 | 649 | 843 | 5566 | 559 | 569 |
| 2585 | QGN15682.1 | FIM1_2375 | CP015056. | 754 | 972 | 1037 | 1359 | 1429 | 1005 | 937 | 965 | 1361 | 5314 | 1111 | 1525 | 5970 | 1062 | 1386 | 5683 | 1084 | 1068 |
| 2586 | QGN15683.1 | FIM1_2376 | CP015056. | 1315 | 647 | 944 | 2449 | 858 | 887 | 1515 | 836 | 1234 | 4222 | 734 | 1523 | 4215 | 673 | 1531 | 3904 | 614 | 1032 |
| 2587 | QGN15684.1 | FIM1_2377 | CP015056. | 13704 | 4158 | 4490 | 13019 | 6460 | 3488 | 11663 | 5267 | 5273 | 18647 | 7772 | 6695 | 26281 | 7244 | 6422 | 23652 | 7036 | 4688 |
| 2588 | QGN15685.1 | FIM1_2378 | CP015056. | 804 | 450 | 517 | 1094 | 509 | 442 | 542 | 487 | 712 | 1678 | 352 | 729 | 2623 | 334 | 660 | 1979 | 312 | 464 |
| 2589 | QGN15686.1 | FIM1_2379 | CP015056. | 3321 | 867 | 915 | 3907 | 1190 | 780 | 3564 | 977 | 1105 | 7715 | 955 | 1238 | 6739 | 843 | 1125 | 7273 | 849 | 730 |
| 2590 | QGN15687.1 | FIM1_2380 | CP015056. | 1028 | 734 | 449 | 1004 | 586 | 503 | 951 | 433 | 626 | 3346 | 573 | 769 | 2380 | 482 | 780 | 3058 | 564 | 542 |
| 2591 | QGN15688.1 | FIM1_2381 | CP015056. | 2295 | 1338 | 1061 | 2484 | 1087 | 898 | 2531 | 976 | 1378 | 4958 | 1062 | 1487 | 4473 | 999 | 1459 | 4581 | 938 | 981 |
| 2592 | QGN15689.1 | FIM1_2382 | CP015056. | 636 | 380 | 284 | 1095 | 409 | 301 | 973 | 300 | 402 | 2666 | 279 | 433 | 2103 | 314 | 459 | 2407 | 311 | 306 |
| 2593 | QGN15690.1 | FIM1_2383 | CP015056. | 3677 | 352 | 442 | 3816 | 400 | 377 | 3711 | 464 | 634 | 2141 | 256 | 465 | 2486 | 193 | 479 | 2043 | 215 | 366 |
| 2594 | QGN15691.1 | FIM1_2384 | CP015056. | 6898 | 7253 | 5638 | 6653 | 7855 | 5415 | 6924 | 6386 | 6988 | 27578 | 8445 | 9285 | 25931 | 7038 | 8414 | 26852 | 7789 | 6163 |
| 2595 | QGN15692.1 | FIM1_2385 | CP015056. | 1189 | 688 | 635 | 1360 | 592 | 536 | 1081 | 566 | 810 | 3327 | 516 | 862 | 3061 | 431 | 853 | 3300 | 411 | 593 |
| 2596 | QGN15693.1 | FIM1_2386 | CP015056. | 392 | 442 | 778 | 1262 | 436 | 668 | 602 | 326 | 1017 | 2804 | 198 | 709 | 2117 | 227 | 678 | 2479 | 165 | 478 |
| 2597 | QGN15694.1 | FIM1_2387 | CP015056. | 774 | 253 | 299 | 993 | 278 | 261 | 850 | 241 | 398 | 1926 | 267 | 529 | 1722 | 227 | 453 | 1986 | 215 | 294 |
| 2598 | QGN15695.1 | FIM1_2388 | CP015056. | 3414 | 5169 | 5134 | 5549 | 5878 | 4523 | 4066 | 4790 | 6407 | 15793 | 4359 | 5949 | 14705 | 4144 | 5653 | 15326 | 3705 | 4046 |
| 2599 | QGN15696.1 | FIM1_2389 | CP015056. | 5797 | 654 | 1064 | 5149 | 1131 | 927 | 5812 | 878 | 1336 | 3622 | 1018 | 1433 | 5210 | 1233 | 1364 | 4016 | 1036 | 948 |
| 2600 | QGN15697.1 | FIM1_2390 | CP015056. | 1622 | 664 | 648 | 2008 | 691 | 508 | 1829 | 643 | 788 | 4086 | 538 | 932 | 3840 | 591 | 841 | 4020 | 509 | 627 |
| 2601 | QGN15698.1 | FIM1_2391 | CP015056. | 599 | 823 | 511 | 964 | 961 | 598 | 725 | 669 | 685 | 6022 | 825 | 1003 | 4314 | 868 | 912 | 5426 | 917 | 563 |
| 2602 | QGN15699.1 | FIM1_2392 | CP015056. | 875 | 867 | 645 | 877 | 572 | 634 | 941 | 540 | 897 | 4531 | 705 | 1068 | 3244 | 668 | 1062 | 3886 | 747 | 734 |

| | | | | | | | | | | | | | | | | | | | | | |
|---|---|---|---|---|---|---|---|---|---|---|---|---|---|---|---|---|---|---|---|---|---|
| 2603 | QGN15700.1 | FIM1_2393 | CP015056. | 148 | 301 | 334 | 286 | 224 | 253 | 204 | 155 | 415 | 1413 | 198 | 512 | 889 | 190 | 449 | 1247 | 210 | 325 |
| 2604 | QGN15701.1 | FIM1_2394 | CP015056. | 3363 | 326 | 752 | 3726 | 324 | 561 | 2702 | 415 | 1001 | 3975 | 206 | 828 | 4540 | 215 | 696 | 3926 | 157 | 524 |
| 2605 | QGN15702.1 | FIM1_2395 | CP015056. | 17679 | 9401 | 8126 | 19935 | 12872 | 7961 | 17065 | 10571 | 10123 | 35411 | 16084 | 14859 | 43496 | 14369 | 14552 | 37640 | 14863 | 10776 |
| 2606 | QGN15703.1 | FIM1_2396 | CP015056. | 4573 | 1473 | 1803 | 5397 | 1827 | 1575 | 4418 | 1482 | 2289 | 7412 | 1191 | 2315 | 8184 | 1203 | 1872 | 7324 | 1211 | 1506 |
| 2607 | QGN15704.1 | FIM1_2397 | CP015056. | 7017 | 1194 | 1807 | 7388 | 1611 | 1529 | 6414 | 1172 | 2277 | 12103 | 1285 | 2664 | 11975 | 967 | 2421 | 12128 | 1084 | 1886 |
| 2608 | QGN15705.1 | FIM1_2398 | CP015056. | 1287 | 1534 | 2121 | 2257 | 1276 | 1953 | 1707 | 1165 | 2720 | 6048 | 759 | 3055 | 4354 | 721 | 2491 | 6274 | 774 | 1970 |
| 2609 | QGN15706.1 | FIM1_2399 | CP015056. | 746 | 591 | 1131 | 745 | 452 | 922 | 700 | 393 | 1373 | 2121 | 268 | 1184 | 1516 | 244 | 1175 | 2223 | 257 | 744 |
| 2610 | QGN15707.1 | FIM1_2400 | CP015056. | 713 | 328 | 393 | 659 | 253 | 324 | 689 | 294 | 525 | 2832 | 229 | 586 | 2211 | 157 | 531 | 3005 | 188 | 397 |
| 2611 | QGN15708.1 | FIM1_2401 | CP015056. | 839 | 636 | 495 | 1162 | 392 | 448 | 883 | 403 | 655 | 2816 | 417 | 699 | 2212 | 381 | 694 | 2399 | 408 | 412 |
| 2612 | QGN15709.1 | FIM1_2402 | CP015056. | 1110 | 1122 | 911 | 1407 | 1026 | 824 | 1410 | 946 | 1173 | 7450 | 1124 | 1676 | 5383 | 926 | 1456 | 7278 | 1024 | 1029 |
| 2613 | QGN15710.1 | FIM1_2403 | CP015056. | 3902 | 904 | 1147 | 4647 | 1071 | 1049 | 4299 | 877 | 1517 | 10998 | 1139 | 1945 | 8389 | 980 | 1677 | 10390 | 1032 | 1315 |
| 2614 | QGN15711.1 | FIM1_2404 | CP015056. | 6761 | 4655 | 3342 | 8918 | 4875 | 3396 | 5499 | 3339 | 4296 | 13837 | 4133 | 4747 | 15006 | 3965 | 3472 | 13462 | 3429 | 3037 |
| 2615 | QGN15712.1 | FIM1_2405 | CP015056. | 1193 | 1360 | 1121 | 2045 | 1375 | 1124 | 1570 | 1175 | 1560 | 6579 | 1289 | 1662 | 5886 | 1108 | 1622 | 6182 | 1144 | 1136 |
| 2616 | QGN15713.1 | FIM1_2406 | CP015056. | 14839 | 2857 | 3930 | 40690 | 4923 | 3942 | 25837 | 3189 | 5284 | 18121 | 4282 | 5463 | 28584 | 9032 | 5898 | 21479 | 3911 | 4184 |
| 2617 | QGN15714.1 | FIM1_2407 | CP015056. | 2915 | 786 | 510 | 2394 | 1059 | 542 | 3388 | 769 | 696 | 2837 | 1039 | 959 | 3691 | 1297 | 862 | 3148 | 1196 | 698 |
| 2618 | QGN15715.1 | FIM1_2408 | CP015056. | 247 | 228 | 196 | 511 | 182 | 191 | 256 | 168 | 251 | 2294 | 151 | 302 | 1598 | 98 | 258 | 2235 | 137 | 194 |
| 2619 | QGN15716.1 | FIM1_2409 | CP015056. | 380 | 579 | 400 | 591 | 404 | 361 | 331 | 456 | 539 | 2293 | 328 | 549 | 2123 | 261 | 519 | 2258 | 308 | 366 |
| 2620 | QGN15717.1 | FIM1_2410 | CP015056. | 5322 | 2952 | 3366 | 6802 | 4173 | 3113 | 5505 | 3317 | 4426 | 21248 | 4363 | 6206 | 18429 | 3838 | 5450 | 20352 | 3856 | 4002 |
| 2621 | QGN15718.1 | FIM1_2411 | CP015056. | 953 | 464 | 366 | 1249 | 419 | 350 | 951 | 304 | 496 | 3369 | 403 | 556 | 2464 | 338 | 541 | 3301 | 338 | 359 |
| 2622 | QGN15719.1 | FIM1_2412 | CP015056. | 599 | 600 | 337 | 467 | 448 | 362 | 490 | 284 | 485 | 2165 | 434 | 565 | 1987 | 308 | 540 | 2208 | 395 | 404 |
| 2623 | QGN15720.1 | FIM1_2413 | CP015056. | 3164 | 559 | 1157 | 4558 | 1311 | 1057 | 3740 | 1051 | 1474 | 9199 | 1200 | 1896 | 8735 | 1090 | 1683 | 8995 | 986 | 1260 |
| 2624 | QGN15721.1 | FIM1_2414 | CP015056. | 1906 | 1196 | 869 | 2833 | 1022 | 848 | 2031 | 782 | 1125 | 5926 | 808 | 1288 | 4534 | 842 | 1253 | 5206 | 767 | 837 |
| 2625 | QGN15722.1 | FIM1_2415 | CP015056. | 1352 | 587 | 452 | 2100 | 650 | 396 | 1858 | 539 | 534 | 3805 | 695 | 766 | 2837 | 584 | 683 | 3598 | 571 | 470 |
| 2626 | QGN15723.1 | FIM1_2416 | CP015056. | 3862 | 2412 | 2367 | 4543 | 2901 | 2142 | 4000 | 2408 | 3072 | 16716 | 2969 | 3841 | 15253 | 2504 | 3905 | 15785 | 2487 | 2609 |
| 2627 | QGN15724.1 | FIM1_2417 | CP015056. | 1699 | 460 | 893 | 1907 | 656 | 821 | 1568 | 631 | 1093 | 4918 | 592 | 1107 | 4850 | 446 | 994 | 4689 | 459 | 745 |
| 2628 | QGN15725.1 | FIM1_2418 | CP015056. | 22254 | 9054 | 7204 | 20951 | 10523 | 6483 | 19813 | 8562 | 8765 | 42447 | 12354 | 13189 | 44825 | 11509 | 12482 | 42554 | 10521 | 9227 |
| 2629 | QGN15726.1 | FIM1_2419 | CP015056. | 2016 | 483 | 476 | 2783 | 618 | 356 | 1808 | 432 | 618 | 2389 | 443 | 609 | 2051 | 447 | 542 | 2290 | 391 | 399 |
| 2630 | QGN15727.1 | FIM1_2420 | CP015056. | 4301 | 710 | 906 | 3649 | 1050 | 847 | 4112 | 860 | 1227 | 5873 | 970 | 1507 | 6021 | 1117 | 1411 | 5480 | 1018 | 1070 |
| 2631 | QGN15728.1 | FIM1_2422 | CP015056. | 8857 | 2195 | 2394 | 11655 | 2257 | 2077 | 10570 | 1831 | 3130 | 16216 | 2419 | 3958 | 16844 | 2380 | 3463 | 16087 | 2276 | 2839 |
| 2632 | QGN15729.1 | FIM1_2423 | CP015056. | 735 | 176 | 337 | 1492 | 414 | 279 | 761 | 334 | 419 | 2014 | 251 | 429 | 2492 | 257 | 367 | 2027 | 233 | 287 |
| 2633 | QGN15730.1 | FIM1_2424 | CP015056. | 9307 | 825 | 873 | 13434 | 721 | 697 | 10922 | 708 | 1129 | 1709 | 554 | 947 | 2827 | 588 | 984 | 2002 | 496 | 663 |
| 2634 | QGN15731.1 | FIM1_2425 | CP015056. | 730 | 548 | 394 | 691 | 261 | 367 | 665 | 331 | 543 | 1780 | 344 | 505 | 1224 | 290 | 460 | 1533 | 353 | 316 |
| 2635 | QGN15732.1 | FIM1_2426 | CP015056. | 13966 | 1166 | 2145 | 18093 | 1500 | 1632 | 14435 | 1451 | 2734 | 6976 | 1352 | 2686 | 10600 | 1306 | 2539 | 8262 | 1164 | 1791 |
| 2636 | QGN15733.1 | FIM1_2427 | CP015056. | 245 | 157 | 160 | 332 | 140 | 149 | 201 | 112 | 167 | 1336 | 204 | 291 | 1006 | 207 | 254 | 1259 | 187 | 174 |
| 2637 | QGN15734.1 | FIM1_2428 | CP015056. | 6710 | 297 | 413 | 4116 | 104 | 238 | 4116 | 272 | 538 | 941 | 106 | 379 | 1055 | 128 | 299 | 833 | 101 | 217 |
| 2638 | QGN15735.1 | FIM1_2429 | CP015056. | 2197 | 955 | 665 | 2347 | 842 | 558 | 1960 | 750 | 849 | 3552 | 747 | 1100 | 3618 | 701 | 1071 | 3646 | 625 | 741 |
| 2639 | QGN15736.1 | FIM1_2430 | CP015056. | 3914 | 802 | 1025 | 3406 | 757 | 833 | 3319 | 806 | 1280 | 7522 | 775 | 1629 | 6540 | 714 | 1534 | 7852 | 557 | 1055 |
| 2640 | QGN15737.1 | FIM1_2431 | CP015056. | 3869 | 986 | 1490 | 3492 | 1062 | 1317 | 1962 | 944 | 1894 | 4745 | 1001 | 2684 | 5786 | 1086 | 2233 | 4562 | 865 | 1629 |
| 2641 | QGN15738.1 | FIM1_2432 | CP015056. | 2197 | 349 | 757 | 3027 | 419 | 594 | 2042 | 400 | 990 | 3707 | 391 | 1342 | 5192 | 465 | 1157 | 4381 | 331 | 877 |
| 2642 | QGN15739.1 | FIM1_2433 | CP015056. | 5469 | 14561 | 18198 | 23408 | 18203 | 15096 | 19497 | 18502 | 19824 | 79311 | 22780 | 28285 | 72892 | 29022 | 30046 | 81805 | 26015 | 22031 |
| 2643 | QGN15740.1 | FIM1_2434 | CP015056. | 3372 | 2483 | 3235 | 9621 | 2496 | 2784 | 7271 | 2605 | 4178 | 21556 | 3616 | 4555 | 16707 | 4573 | 5036 | 21622 | 3963 | 3447 |
| 2644 | QGN15741.1 | FIM1_2435 | CP015056. | 1076 | 1086 | 1124 | 1768 | 1135 | 943 | 1197 | 799 | 1482 | 6076 | 1115 | 1334 | 6224 | 1023 | 1149 | 5937 | 977 | 928 |
| 2645 | QGN15742.1 | FIM1_2436 | CP015056. | 934 | 96 | 218 | 1165 | 205 | 177 | 730 | 172 | 256 | 564 | 161 | 265 | 785 | 137 | 245 | 727 | 132 | 176 |
| 2646 | QGN15743.1 | FIM1_2437 | CP015056. | 1528 | 61 | 89 | 816 | 31 | 39 | 1185 | 83 | 127 | 416 | 50 | 103 | 471 | 33 | 105 | 447 | 39 | 53 |
| 2647 | QGN15744.1 | FIM1_2438 | CP015056. | 5470 | 689 | 796 | 5614 | 1011 | 606 | 3701 | 847 | 907 | 3642 | 747 | 1165 | 4282 | 755 | 1038 | 3686 | 667 | 714 |
| 2648 | QGN15745.1 | FIM1_2439 | CP015056. | 2989 | 82 | 240 | 2103 | 234 | 157 | 1413 | 231 | 233 | 628 | 142 | 298 | 1004 | 168 | 265 | 792 | 132 | 177 |
| 2649 | QGN15746.1 | FIM1_2440 | CP015056. | 314 | 81 | 151 | 2787 | 121 | 267 | 717 | 89 | 198 | 554 | 55 | 154 | 416 | 46 | 87 | 584 | 74 | 76 |
| 2650 | QGN15747.1 | FIM1_2441 | CP015057. | 0 | 0 | 0 | 0 | 0 | 0 | 0 | 0 | 0 | 0 | 0 | 0 | 0 | 0 | 0 | 0 | 0 | 0 |
| 2651 | QGN15748.1 | FIM1_2444 | CP015057. | 20182 | 8281 | 6519 | 62662 | 19641 | 5816 | 23538 | 7510 | 6883 | 19482 | 21377 | 20003 | 61459 | 19938 | 18502 | 35133 | 17894 | 16084 |
| 2652 | QGN15749.1 | FIM1_2445 | CP015057. | 718 | 782 | 284 | 1222 | 313 | 159 | 409 | 261 | 250 | 2380 | 883 | 661 | 2993 | 690 | 737 | 2595 | 658 | 523 |
| 2653 | QGN15750.1 | FIM1_2446 | CP015057. | 898 | 158 | 170 | 1214 | 188 | 126 | 709 | 137 | 210 | 756 | 179 | 268 | 785 | 184 | 263 | 744 | 159 | 175 |
| 2654 | QGN15751.1 | FIM1_2447 | CP015057. | 1254 | 198 | 289 | 1608 | 250 | 299 | 984 | 212 | 385 | 1270 | 197 | 519 | 1255 | 243 | 461 | 1255 | 182 | 278 |
| 2655 | QGN15752.1 | FIM1_2448 | CP015057. | 23 | 16 | 26 | 193 | 30 | 24 | 54 | 10 | 23 | 170 | 21 | 71 | 224 | 31 | 37 | 173 | 16 | 34 |
| 2656 | QGN15753.1 | FIM1_2449 | CP015057. | 49 | 14 | 18 | 180 | 36 | 22 | 63 | 14 | 23 | 169 | 16 | 35 | 165 | 28 | 32 | 164 | 15 | 19 |
| 2657 | QGN15754.1 | FIM1_2450 | CP015057. | 1767 | 104 | 179 | 2472 | 208 | 145 | 1479 | 114 | 235 | 2796 | 196 | 327 | 3129 | 280 | 301 | 2854 | 192 | 229 |
| 2658 | QGN15755.1 | FIM1_2451 | CP015057. | 198 | 19 | 149 | 1152 | 60 | 116 | 138 | 25 | 186 | 6516 | 633 | 577 | 5398 | 1547 | 514 | 6635 | 735 | 401 |
| 2659 | QGN15756.1 | FIM1_2452 | CP015057. | 58 | 22 | 30 | 196 | 30 | 26 | 92 | 33 | 31 | 225 | 27 | 46 | 203 | 27 | 32 | 206 | 24 | 16 |
| 2660 | QGN15757.1 | FIM1_2453 | CP015057. | 359 | 105 | 122 | 601 | 69 | 76 | 454 | 64 | 162 | 1021 | 90 | 187 | 892 | 96 | 174 | 896 | 85 | 129 |
| 2661 | QGN15758.1 | FIM1_2454 | CP015057. | 899 | 94 | 206 | 1869 | 144 | 150 | 816 | 137 | 217 | 2133 | 119 | 234 | 1947 | 121 | 251 | 2045 | 114 | 196 |
| 2662 | QGN15759.1 | FIM1_2455 | CP015057. | 695 | 186 | 436 | 5706 | 285 | 347 | 2290 | 185 | 503 | 1968 | 317 | 685 | 5077 | 600 | 760 | 3039 | 268 | 588 |
| 2663 | QGN15760.1 | FIM1_2456 | CP015057. | 550 | 83 | 218 | 540 | 64 | 99 | 461 | 124 | 299 | 1315 | 101 | 307 | 1144 | 109 | 284 | 1245 | 68 | 189 |
| 2664 | QGN15761.1 | FIM1_2457 | CP015057. | 970 | 173 | 192 | 1330 | 171 | 172 | 1084 | 229 | 326 | 707 | 187 | 274 | 1395 | 160 | 279 | 916 | 155 | 212 |
| 2665 | QGN15762.1 | FIM1_2458 | CP015057. | 15692 | 962 | 1209 | 12629 | 811 | 866 | 9383 | 812 | 1391 | 5999 | 780 | 1803 | 6152 | 1136 | 1573 | 5834 | 867 | 1060 |
| 2666 | QGN15763.1 | FIM1_2459 | CP015057. | 4709 | 1860 | 783 | 2959 | 1773 | 887 | 3622 | 1518 | 1059 | 4789 | 1673 | 1288 | 5011 | 1589 | 1405 | 4551 | 1860 | 944 |
| 2667 | QGN15764.1 | FIM1_2460 | CP015057. | 8962 | 3643 | 1491 | 5179 | 3211 | 1587 | 9005 | 2516 | 1953 | 8059 | 3755 | 2673 | 8353 | 4192 | 2588 | 8434 | 4190 | 1866 |
| 2668 | QGN15765.1 | FIM1_2461 | CP015057. | 4106 | 1388 | 1638 | 5457 | 1840 | 1543 | 4724 | 1691 | 2160 | 8158 | 1737 | 2177 | 10176 | 1667 | 2256 | 8417 | 1522 | 1476 |
| 2669 | QGN15766.1 | FIM1_2462 | CP015057. | 8150 | 3113 | 2718 | 7507 | 3752 | 2222 | 7257 | 3624 | 3354 | 14807 | 4174 | 4569 | 18292 | 3738 | 4724 | 16696 | 3447 | 3064 |
| 2670 | QGN15767.1 | FIM1_2463 | CP015057. | 2740 | 490 | 568 | 2503 | 472 | 451 | 2476 | 507 | 700 | 2623 | 387 | 749 | 3113 | 435 | 758 | 2686 | 389 | 510 |
| 2671 | QGN15768.1 | FIM1_2464 | CP015057. | 2971 | 1232 | 1427 | 2305 | 1080 | 1199 | 1672 | 1063 | 1912 | 6276 | 888 | 1598 | 6674 | 905 | 1882 | 5936 | 668 | 1152 |
| 2672 | QGN15769.1 | FIM1_2465 | CP015057. | 750 | 423 | 541 | 636 | 350 | 472 | 649 | 447 | 728 | 1865 | 289 | 648 | 1985 | 231 | 705 | 1921 | 212 | 416 |
| 2673 | QGN15770.1 | FIM1_2466 | CP015057. | 2258 | 156 | 498 | 3374 | 275 | 351 | 2761 | 351 | 726 | 1036 | 218 | 769 | 3337 | 256 | 672 | 1802 | 241 | 515 |
| 2674 | QGN15771.1 | FIM1_2467 | CP015057. | 272 | 219 | 248 | 411 | 172 | 190 | 220 | 190 | 319 | 1616 | 354 | 346 | 1681 | 396 | 307 | 1647 | 397 | 250 |
| 2675 | QGN15772.1 | FIM1_2468 | CP015057. | 2198 | 1545 | 1359 | 3117 | 1171 | 1204 | 2692 | 1049 | 1889 | 9158 | 1340 | 2213 | 7165 | 1193 | 2148 | 9409 | 1195 | 1361 |
| 2676 | QGN15773.1 | FIM1_2469 | CP015057. | 2436 | 1565 | 1416 | 3416 | 1492 | 1281 | 2664 | 1334 | 1921 | 5020 | 904 | 1836 | 5145 | 884 | 1637 | 4863 | 959 | 1139 |
| 2677 | QGN15774.1 | FIM1_2470 | CP015057. | 10421 | 2954 | 4496 | 14331 | 3585 | 3600 | 12127 | 3239 | 5736 | 14938 | 3747 | 6495 | 23307 | 3459 | 6144 | 18444 | 3504 | 4613 |
| 2678 | QGN15775.1 | FIM1_2471 | CP015057. | 257 | 567 | 915 | 548 | 828 | 743 | 271 | 608 | 1157 | 4428 | 746 | 1262 | 3996 | 694 | 1128 | 4523 | 692 | 835 |
| 2679 | QGN15776.1 | FIM1_2472 | CP015057. | 4554 | 1005 | 1305 | 4920 | 984 | 1048 | 4138 | 1001 | 1732 | 7536 | 909 | 2140 | 7041 | 771 | 1793 | 7192 | 721 | 1254 |
| 2680 | QGN15777.1 | FIM1_2473 | CP015057. | 2211 | 1331 | 623 | 1636 | 1065 | 616 | 2069 | 691 | 823 | 4509 | 1154 | 1038 | 3623 | 1113 | 990 | 4321 | 1441 | 654 |
| 2681 | QGN15778.1 | FIM1_2474 | CP015057. | 8464 | 2409 | 1586 | 7951 | 2163 | 1434 | 7274 | 1858 | 2010 | 9651 | 2089 | 2348 | 9510 | 2146 | 2328 | 9219 | 1987 | 1628 |
| 2682 | QGN15779.1 | FIM1_2475 | CP015057. | 1429 | 541 | 1122 | 1716 | 677 | 826 | 1566 | 717 | 1382 | 4655 | 555 | 1480 | 5356 | 459 | 1312 | 4781 | 439 | 996 |
| 2683 | QGN15780.1 | FIM1_2476 | CP015057. | 7803 | 10095 | 6697 | 8806 | 8080 | 5438 | 6525 | 7628 | 8059 | 18746 | 8157 | 9184 | 20303 | 7488 | 8777 | 20318 | 6652 | 6866 |
| 2684 | QGN15781.1 | FIM1_2477 | CP015057. | 3442 | 1436 | 1315 | 3043 | 1318 | 1210 | 3249 | 1434 | 1869 | 4699 | 1523 | 1830 | 7011 | 1222 | 1882 | 5265 | 1180 | 1232 |
| 2685 | QGN15782.1 | FIM1_2478 | CP015057. | 967 | 820 | 653 | 830 | 753 | 654 | 870 | 620 | 912 | 4746 | 790 | 1247 | 3601 | 670 | 1114 | 4659 | 739 | 804 |
| 2686 | QGN15783.1 | FIM1_2479 | CP015057. | 1145 | 515 | 465 | 1262 | 527 | 473 | 1020 | 472 | 631 | 3539 | 513 | 867 | 2916 | 433 | 720 | 3524 | 431 | 561 |

| | | | | | | | | | | | | | | | | | | | | | |
|---|---|---|---|---|---|---|---|---|---|---|---|---|---|---|---|---|---|---|---|---|---|
| 2687 | QGN15784.1 | FIM1_2480 | CP015057. | 2838 | 1117 | 1157 | 3703 | 1274 | 962 | 3123 | 1121 | 1449 | 7416 | 1232 | 2005 | 8066 | 1196 | 1713 | 7639 | 1108 | 1388 |
| 2688 | QGN15785.1 | FIM1_2481 | CP015057. | 2176 | 493 | 722 | 2301 | 647 | 616 | 2228 | 585 | 951 | 5675 | 739 | 1274 | 5704 | 600 | 1134 | 5498 | 628 | 851 |
| 2689 | QGN15786.1 | FIM1_2482 | CP015057. | 2236 | 760 | 853 | 2235 | 841 | 741 | 2115 | 780 | 1151 | 4544 | 788 | 1376 | 4506 | 714 | 1223 | 4531 | 742 | 890 |
| 2690 | QGN15787.1 | FIM1_2483 | CP015057. | 2892 | 561 | 718 | 2950 | 619 | 608 | 3303 | 525 | 955 | 5653 | 521 | 1153 | 5304 | 525 | 970 | 5955 | 518 | 784 |
| 2691 | QGN15788.1 | FIM1_2484 | CP015057. | 686 | 218 | 180 | 1043 | 230 | 184 | 897 | 191 | 236 | 1487 | 164 | 229 | 1140 | 123 | 235 | 1412 | 125 | 154 |
| 2692 | QGN15789.1 | FIM1_2485 | CP015057. | 4273 | 812 | 1194 | 4861 | 823 | 862 | 3870 | 918 | 1479 | 5333 | 548 | 1394 | 6149 | 589 | 1219 | 5641 | 453 | 846 |
| 2693 | QGN15790.1 | FIM1_2486 | CP015057. | 7874 | 11464 | 7098 | 7046 | 8504 | 6861 | 6834 | 7106 | 8531 | 29326 | 10511 | 10972 | 23299 | 8801 | 10586 | 29933 | 9625 | 8381 |
| 2694 | QGN15791.1 | FIM1_2487 | CP015057. | 5015 | 1801 | 1415 | 4763 | 1726 | 1276 | 3980 | 1513 | 1842 | 5972 | 1420 | 2108 | 7465 | 1469 | 2066 | 6016 | 1388 | 1352 |
| 2695 | QGN15792.1 | FIM1_2488 | CP015057. | 567 | 181 | 166 | 779 | 304 | 174 | 550 | 290 | 244 | 438 | 190 | 322 | 996 | 183 | 323 | 634 | 187 | 205 |
| 2696 | QGN15793.1 | FIM1_2489 | CP015057. | 786 | 648 | 510 | 836 | 357 | 410 | 798 | 444 | 686 | 672 | 294 | 704 | 1296 | 256 | 774 | 818 | 250 | 479 |
| 2697 | QGN15794.1 | FIM1_2490 | CP015057. | 1186 | 1041 | 720 | 1952 | 810 | 692 | 1576 | 697 | 913 | 1616 | 687 | 923 | 2693 | 671 | 1032 | 1884 | 625 | 674 |
| 2698 | QGN15795.1 | FIM1_2491 | CP015057. | 9347 | 212 | 232 | 8067 | 71 | 125 | 7591 | 246 | 343 | 648 | 55 | 302 | 580 | 77 | 262 | 578 | 92 | 178 |
| 2699 | QGN15796.1 | FIM1_2492 | CP015057. | 7767 | 1683 | 1364 | 6722 | 1855 | 1239 | 6378 | 1459 | 1791 | 1926 | 1750 | 1928 | 6759 | 1619 | 1934 | 3756 | 1580 | 1365 |
| 2700 | QGN15797.1 | FIM1_2493 | CP015057. | 2416 | 1097 | 1011 | 2666 | 1095 | 921 | 2488 | 969 | 1254 | 4493 | 1200 | 1618 | 5315 | 1029 | 1726 | 4474 | 1047 | 1120 |
| 2701 | QGN15798.1 | FIM1_2494 | CP015057. | 1548 | 550 | 539 | 1749 | 483 | 480 | 1596 | 503 | 660 | 1796 | 444 | 728 | 3071 | 366 | 695 | 1976 | 362 | 530 |
| 2702 | QGN15799.1 | FIM1_2495 | CP015057. | 1978 | 3010 | 3955 | 4647 | 3731 | 3462 | 3139 | 3622 | 5278 | 12402 | 4509 | 5796 | 23241 | 4032 | 5495 | 17858 | 3667 | 4012 |
| 2703 | QGN15800.1 | FIM1_2496 | CP015057. | 1863 | 641 | 889 | 1510 | 528 | 723 | 1625 | 539 | 1199 | 5388 | 372 | 1211 | 4935 | 381 | 1083 | 5339 | 258 | 744 |
| 2704 | QGN15801.1 | FIM1_2497 | CP015057. | 1151 | 290 | 286 | 1045 | 479 | 252 | 817 | 311 | 333 | 2986 | 432 | 517 | 2395 | 307 | 473 | 3176 | 295 | 310 |
| 2705 | QGN15802.1 | FIM1_2498 | CP015057. | 231 | 186 | 58 | 318 | 98 | 66 | 196 | 73 | 95 | 529 | 105 | 115 | 376 | 74 | 121 | 449 | 90 | 78 |
| 2706 | QGN15803.1 | FIM1_2499 | CP015057. | 3195 | 586 | 673 | 3689 | 736 | 629 | 2872 | 548 | 860 | 3063 | 638 | 1089 | 3732 | 739 | 1012 | 3118 | 569 | 756 |
| 2707 | QGN15804.1 | FIM1_2500 | CP015057. | 10111 | 653 | 724 | 7361 | 768 | 518 | 5722 | 861 | 965 | 2654 | 512 | 1024 | 4672 | 548 | 953 | 3131 | 306 | 665 |
| 2708 | QGN15805.1 | FIM1_2501 | CP015057. | 35 | 0 | 4 | 29 | 3 | 7 | 16 | 3 | 5 | 3 | 4 | 8 | 23 | 4 | 6 | 16 | 0 | 5 |
| 2709 | QGN15806.1 | FIM1_2502 | CP015057. | 7205 | 2320 | 2544 | 6815 | 2011 | 2183 | 4784 | 1674 | 3374 | 3894 | 1630 | 3597 | 6540 | 2045 | 4287 | 5032 | 1515 | 2574 |
| 2710 | QGN15807.1 | FIM1_2503 | CP015057. | 1384 | 496 | 346 | 886 | 414 | 311 | 933 | 409 | 499 | 537 | 381 | 511 | 1192 | 338 | 612 | 764 | 382 | 392 |
| 2711 | QGN15808.1 | FIM1_2504 | CP015057. | 1883 | 145 | 567 | 1996 | 536 | 481 | 1492 | 511 | 698 | 2019 | 461 | 1069 | 2921 | 404 | 884 | 2377 | 404 | 682 |
| 2712 | QGN15809.1 | FIM1_2505 | CP015057. | 1040 | 370 | 463 | 844 | 442 | 435 | 507 | 336 | 603 | 1036 | 388 | 764 | 1574 | 346 | 736 | 1381 | 354 | 483 |
| 2713 | QGN15810.1 | FIM1_2506 | CP015057. | 1551 | 142 | 395 | 2336 | 168 | 291 | 1523 | 215 | 496 | 2173 | 122 | 477 | 2203 | 126 | 371 | 2103 | 92 | 285 |
| 2714 | QGN15811.1 | FIM1_2507 | CP015057. | 2221 | 709 | 627 | 2162 | 658 | 540 | 2205 | 484 | 766 | 3984 | 519 | 798 | 3016 | 540 | 767 | 3978 | 540 | 524 |
| 2715 | QGN15812.1 | FIM1_2508 | CP015057. | 12 | 11 | 9 | 57 | 21 | 4 | 18 | 17 | 13 | 87 | 5 | 18 | 60 | 11 | 13 | 81 | 11 | 8 |
| 2716 | QGN15813.1 | FIM1_2509 | CP015057. | 8 | 4 | 5 | 39 | 9 | 8 | 8 | 2 | 3 | 41 | 4 | 11 | 41 | 4 | 5 | 39 | 6 | 12 |
| 2717 | QGN15814.1 | FIM1_2510 | CP015057. | 134 | 144 | 146 | 282 | 117 | 157 | 160 | 84 | 197 | 1187 | 149 | 239 | 888 | 144 | 252 | 1095 | 159 | 160 |
| 2718 | QGN15815.1 | FIM1_2511 | CP015057. | 9291 | 3486 | 1829 | 6846 | 4542 | 1987 | 8982 | 3793 | 2453 | 10055 | 5152 | 3707 | 15146 | 4609 | 3334 | 10984 | 5453 | 2584 |
| 2719 | QGN15816.1 | FIM1_2512 | CP015057. | 1000 | 659 | 833 | 1869 | 796 | 778 | 1319 | 667 | 1134 | 4740 | 668 | 1518 | 4970 | 625 | 1300 | 4652 | 539 | 977 |
| 2720 | QGN15817.1 | FIM1_2513 | CP015057. | 2247 | 4925 | 3674 | 6120 | 4062 | 3401 | 4747 | 3390 | 4586 | 19534 | 5407 | 6292 | 16569 | 5203 | 6022 | 19579 | 4899 | 4264 |
| 2721 | QGN15818.1 | FIM1_2514 | CP015057. | 1193 | 249 | 271 | 1582 | 169 | 246 | 1546 | 178 | 412 | 1985 | 168 | 372 | 1748 | 193 | 332 | 1834 | 160 | 251 |
| 2722 | QGN15819.1 | FIM1_2515 | CP015057. | 379 | 60 | 50 | 341 | 36 | 41 | 451 | 38 | 66 | 305 | 29 | 54 | 261 | 27 | 63 | 302 | 41 | 41 |
| 2723 | QGN15820.1 | FIM1_2516 | CP015057. | 4272 | 1969 | 2377 | 5664 | 2857 | 2058 | 4773 | 2271 | 2999 | 12649 | 2986 | 4020 | 17624 | 2757 | 3410 | 14532 | 2635 | 2767 |
| 2724 | QGN15821.1 | FIM1_2517 | CP015057. | 8287 | 1095 | 1549 | 6338 | 1336 | 1263 | 6484 | 1347 | 1944 | 6867 | 805 | 1478 | 6234 | 1130 | 1484 | 6157 | 897 | 952 |
| 2725 | QGN15822.1 | FIM1_2518 | CP015057. | 2035 | 1213 | 1178 | 2762 | 1309 | 1127 | 2104 | 1190 | 1651 | 7808 | 1398 | 1872 | 7430 | 1263 | 1754 | 7751 | 1257 | 1148 |
| 2726 | QGN15823.1 | FIM1_2519 | CP015057. | 4681 | 1670 | 844 | 1684 | 2392 | 803 | 4158 | 2148 | 1100 | 3381 | 2557 | 1915 | 8635 | 2747 | 1665 | 5388 | 3133 | 1355 |
| 2727 | QGN15824.1 | FIM1_2520 | CP015057. | 1088 | 423 | 326 | 1035 | 557 | 324 | 1019 | 369 | 392 | 2264 | 458 | 524 | 1975 | 512 | 490 | 2103 | 544 | 328 |
| 2728 | QGN15825.1 | FIM1_2521 | CP015057. | 723 | 353 | 337 | 773 | 460 | 321 | 1112 | 442 | 465 | 1227 | 555 | 606 | 2371 | 565 | 557 | 1563 | 545 | 452 |
| 2729 | QGN15826.1 | FIM1_2522 | CP015057. | 926 | 367 | 336 | 629 | 284 | 277 | 810 | 276 | 384 | 1788 | 298 | 440 | 1579 | 290 | 432 | 1786 | 299 | 276 |
| 2730 | QGN15827.1 | FIM1_2523 | CP015057. | 5345 | 2441 | 2746 | 7194 | 2809 | 2374 | 6117 | 2537 | 3451 | 13022 | 2900 | 4095 | 14845 | 2781 | 3770 | 13037 | 2473 | 2853 |
| 2731 | QGN15828.1 | FIM1_2524 | CP015057. | 6862 | 696 | 735 | 5267 | 841 | 653 | 6099 | 958 | 1102 | 748 | 658 | 1296 | 2951 | 729 | 1067 | 1266 | 847 | 867 |
| 2732 | QGN15829.1 | FIM1_2525 | CP015057. | 533 | 906 | 643 | 695 | 538 | 672 | 513 | 499 | 933 | 1679 | 399 | 597 | 1605 | 393 | 652 | 1436 | 342 | 422 |
| 2733 | QGN15830.1 | FIM1_2526 | CP015057. | 1673 | 3731 | 4722 | 2316 | 3954 | 4934 | 1272 | 2985 | 6518 | 10852 | 3692 | 8017 | 11758 | 3155 | 8866 | 11052 | 2894 | 5982 |
| 2734 | QGN15831.1 | FIM1_2527 | CP015057. | 127 | 124 | 87 | 451 | 126 | 82 | 128 | 153 | 106 | 117 | 124 | 278 | 488 | 106 | 289 | 229 | 130 | 188 |
| 2735 | QGN15832.1 | FIM1_2528 | CP015057. | 30313 | 25359 | 12547 | 29168 | 20486 | 11969 | 26734 | 19198 | 15552 | 38041 | 19984 | 18808 | 46013 | 17606 | 19246 | 41804 | 18055 | 14374 |
| 2736 | QGN15833.1 | FIM1_2529 | CP015057. | 11704 | 4206 | 4549 | 45873 | 4487 | 3940 | 21667 | 4035 | 5941 | 11800 | 3099 | 6026 | 22412 | 6985 | 5826 | 15173 | 3095 | 4345 |
| 2737 | QGN15834.1 | FIM1_2530 | CP015057. | 6514 | 673 | 673 | 6027 | 735 | 579 | 5140 | 748 | 919 | 507 | 387 | 814 | 3529 | 327 | 766 | 1354 | 365 | 647 |
| 2738 | QGN15835.1 | FIM1_2531 | CP015057. | 6331 | 2707 | 1757 | 6589 | 2480 | 1399 | 6235 | 2542 | 2237 | 2501 | 1767 | 2007 | 6709 | 2014 | 2205 | 4171 | 1906 | 1559 |
| 2739 | QGN15836.1 | FIM1_2532 | CP015057. | 235 | 482 | 370 | 271 | 473 | 292 | 280 | 538 | 436 | 648 | 306 | 430 | 1471 | 215 | 471 | 950 | 259 | 331 |
| 2740 | QGN15837.1 | FIM1_2533 | CP015057. | 3499 | 2266 | 2162 | 4017 | 1637 | 1676 | 3778 | 1844 | 2852 | 2866 | 1326 | 2745 | 5433 | 1211 | 2969 | 3500 | 1137 | 1934 |
| 2741 | QGN15838.1 | FIM1_2534 | CP015057. | 5892 | 2100 | 1214 | 5039 | 1308 | 1127 | 4990 | 1537 | 1824 | 2938 | 1115 | 1719 | 5374 | 1178 | 1856 | 3598 | 1028 | 1298 |
| 2742 | QGN15839.1 | FIM1_2535 | CP015057. | 514 | 641 | 223 | 659 | 340 | 239 | 614 | 366 | 334 | 1446 | 334 | 403 | 1329 | 233 | 366 | 1227 | 307 | 256 |
| 2743 | QGN15840.1 | FIM1_2536 | CP015057. | 409 | 395 | 300 | 1155 | 434 | 250 | 487 | 514 | 459 | 390 | 394 | 420 | 1529 | 345 | 334 | 729 | 408 | 370 |
| 2744 | QGN15841.1 | FIM1_2537 | CP015057. | 1304 | 1436 | 489 | 1343 | 1478 | 536 | 1049 | 1026 | 751 | 927 | 1309 | 613 | 2681 | 941 | 728 | 1649 | 924 | 466 |
| 2745 | QGN15842.1 | FIM1_2538 | CP015057. | 1220 | 971 | 692 | 2126 | 648 | 476 | 1974 | 754 | 865 | 589 | 427 | 600 | 1310 | 398 | 730 | 801 | 416 | 424 |
| 2746 | QGN15843.1 | FIM1_2539 | CP015057. | 580 | 623 | 418 | 913 | 412 | 310 | 719 | 378 | 524 | 629 | 439 | 506 | 1410 | 415 | 604 | 1022 | 468 | 360 |
| 2747 | QGN15844.1 | FIM1_2540 | CP015057. | 5901 | 1678 | 2028 | 8951 | 1966 | 1584 | 7555 | 1866 | 2541 | 3922 | 1659 | 3052 | 10626 | 1952 | 2827 | 6390 | 1733 | 2144 |
| 2748 | QGN15845.1 | FIM1_2541 | CP015057. | 1262 | 1654 | 800 | 1330 | 1107 | 739 | 1424 | 1117 | 964 | 2260 | 1074 | 1069 | 2951 | 853 | 1229 | 2313 | 925 | 772 |
| 2749 | QGN15846.1 | FIM1_2542 | CP015057. | 218 | 430 | 272 | 307 | 287 | 267 | 216 | 243 | 387 | 1262 | 317 | 398 | 1080 | 286 | 382 | 1056 | 301 | 247 |
| 2750 | QGN15847.1 | FIM1_2543 | CP015057. | 2341 | 605 | 719 | 2490 | 644 | 566 | 2072 | 798 | 885 | 1676 | 615 | 951 | 3532 | 603 | 938 | 2044 | 595 | 715 |
| 2751 | QGN15848.1 | FIM1_2544 | CP015057. | 2251 | 465 | 529 | 2274 | 468 | 491 | 1972 | 492 | 733 | 1465 | 447 | 629 | 2210 | 506 | 630 | 1673 | 386 | 452 |
| 2752 | QGN15849.1 | FIM1_2546 | CP015057. | 691 | 1392 | 977 | 814 | 939 | 844 | 745 | 1029 | 1322 | 2082 | 562 | 1327 | 3093 | 676 | 1238 | 2245 | 577 | 835 |
| 2753 | QGN15850.1 | FIM1_2547 | CP015057. | 9133 | 851 | 1109 | 8027 | 913 | 727 | 7660 | 1057 | 1339 | 892 | 632 | 1895 | 3161 | 472 | 1554 | 1458 | 690 | 1230 |
| 2754 | QGN15851.1 | FIM1_2548 | CP015057. | 8602 | 3895 | 3187 | 12001 | 5001 | 2758 | 9413 | 4400 | 4037 | 2033 | 4201 | 3031 | 13529 | 2419 | 3014 | 6778 | 3419 | 2196 |
| 2755 | QGN15852.1 | FIM1_2549 | CP015057. | 14765 | 3292 | 4099 | 16171 | 3444 | 2948 | 12695 | 3901 | 4867 | 4608 | 2702 | 5152 | 17400 | 2977 | 4640 | 9569 | 2545 | 3410 |
| 2756 | QGN15853.1 | FIM1_2550 | CP015057. | 2371 | 1513 | 1196 | 2364 | 1411 | 1170 | 2177 | 1445 | 1650 | 5866 | 1168 | 1735 | 5025 | 725 | 1933 | 5312 | 758 | 1110 |
| 2757 | QGN15854.1 | FIM1_2551 | CP015057. | 55 | 17 | 5 | 109 | 6 | 4 | 58 | 7 | 8 | 16 | 2 | 8 | 36 | 5 | 7 | 15 | 7 | 3 |
| 2758 | QGN15855.1 | FIM1_2552 | CP015057. | 9645 | 1162 | 1513 | 18352 | 579 | 757 | 18858 | 1023 | 1587 | 2440 | 848 | 1611 | 4688 | 1093 | 1136 | 2870 | 848 | 893 |
| 2759 | QGN15856.1 | FIM1_2553 | CP015057. | 1247 | 75 | 341 | 1763 | 262 | 264 | 1387 | 265 | 412 | 1170 | 223 | 498 | 2098 | 243 | 466 | 1353 | 185 | 335 |
| 2760 | QGN15857.1 | FIM1_2554 | CP015057. | 1800 | 630 | 976 | 1245 | 845 | 714 | 1408 | 1033 | 1233 | 1787 | 610 | 1373 | 6367 | 441 | 1204 | 3382 | 439 | 897 |
| 2761 | QGN15858.1 | FIM1_2555 | CP015057. | 95497 | 4300 | 4138 | 104541 | 2295 | 2641 | 96198 | 3774 | 4840 | 8972 | 2650 | 4535 | 14242 | 11513 | 3835 | 9520 | 5235 | 2578 |
| 2762 | QGN15859.1 | FIM1_2556 | CP015057. | 5046 | 36 | 55 | 2731 | 107 | 50 | 2865 | 78 | 76 | 118 | 52 | 120 | 415 | 67 | 106 | 214 | 53 | 71 |
| 2763 | QGN15860.1 | FIM1_2557 | CP015057. | 32196 | 6013 | 6995 | 43785 | 8058 | 5260 | 27165 | 8392 | 8464 | 12532 | 7668 | 8762 | 31638 | 7713 | 6891 | 22981 | 8233 | 5704 |
| 2764 | QGN15861.1 | FIM1_2558 | CP015057. | 1700 | 1033 | 551 | 952 | 938 | 617 | 1389 | 783 | 762 | 3707 | 1012 | 995 | 2940 | 933 | 932 | 3371 | 1029 | 619 |
| 2765 | QGN15862.1 | FIM1_2559 | CP015057. | 516 | 261 | 232 | 1970 | 280 | 232 | 671 | 383 | 309 | 868 | 197 | 438 | 1565 | 143 | 318 | 1014 | 169 | 234 |
| 2766 | QGN15863.1 | FIM1_2560 | CP015057. | 7408 | 2212 | 1984 | 8198 | 1406 | 1342 | 7571 | 1666 | 2280 | 12930 | 2585 | 3306 | 11932 | 3334 | 3668 | 11587 | 2765 | 2252 |
| 2767 | QGN15864.1 | FIM1_2561 | CP015057. | 719 | 102 | 201 | 1427 | 159 | 112 | 804 | 133 | 251 | 1210 | 127 | 360 | 1224 | 148 | 332 | 1238 | 119 | 243 |
| 2768 | QGN15865.1 | FIM1_2562 | CP015057. | 6240 | 1329 | 1625 | 8855 | 1437 | 1302 | 7320 | 1536 | 2024 | 5772 | 1240 | 2617 | 9589 | 1406 | 2288 | 7079 | 1184 | 1757 |
| 2769 | QGN15866.1 | FIM1_2563 | CP015057. | 5800 | 2052 | 1526 | 6970 | 2014 | 1322 | 6142 | 2145 | 1960 | 5528 | 1842 | 2206 | 8643 | 1994 | 1949 | 6342 | 1905 | 1483 |
| 2770 | QGN15867.1 | FIM1_2564 | CP015057. | 9781 | 763 | 611 | 10918 | 774 | 655 | 12585 | 741 | 830 | 2099 | 510 | 973 | 3154 | 407 | 923 | 2356 | 438 | 679 |

| | | | | | | | | | | | | | | | | | | | | | |
|---|---|---|---|---|---|---|---|---|---|---|---|---|---|---|---|---|---|---|---|---|---|
| 2771 | QGN15868.1 | FIM1_2565 | CP015057. | 2722 | 890 | 766 | 2348 | 605 | 700 | 2360 | 579 | 1032 | 3559 | 690 | 1011 | 2866 | 637 | 1106 | 2926 | 601 | 747 |
| 2772 | QGN15869.1 | FIM1_2566 | CP015057. | 4151 | 874 | 1381 | 7215 | 1146 | 1070 | 4895 | 1240 | 1734 | 4874 | 798 | 1924 | 9509 | 913 | 1727 | 6379 | 732 | 1262 |
| 2773 | QGN15870.1 | FIM1_2567 | CP015057. | 3088 | 550 | 513 | 2504 | 653 | 474 | 3527 | 545 | 666 | 2684 | 529 | 766 | 2999 | 614 | 693 | 2700 | 548 | 530 |
| 2774 | QGN15871.1 | FIM1_2568 | CP015057. | 1445 | 1322 | 610 | 825 | 896 | 639 | 1234 | 697 | 818 | 3505 | 1131 | 1117 | 2872 | 783 | 1123 | 3150 | 1089 | 710 |
| 2775 | QGN15872.1 | FIM1_2570 | CP015057. | 1898 | 135 | 201 | 2107 | 144 | 145 | 1591 | 177 | 257 | 456 | 125 | 295 | 1352 | 131 | 295 | 639 | 121 | 221 |
| 2776 | QGN15873.1 | FIM1_2571 | CP015057. | 1413 | 133 | 140 | 886 | 301 | 179 | 1196 | 227 | 171 | 1223 | 222 | 285 | 1098 | 394 | 275 | 1078 | 300 | 216 |
| 2777 | QGN15874.1 | FIM1_2572 | CP015057. | 888 | 223 | 100 | 662 | 185 | 118 | 812 | 150 | 133 | 741 | 185 | 185 | 607 | 262 | 190 | 661 | 232 | 130 |
| 2778 | QGN15875.1 | FIM1_2573 | CP015057. | 711 | 383 | 341 | 1349 | 283 | 323 | 949 | 256 | 501 | 1685 | 195 | 447 | 1417 | 219 | 495 | 1628 | 198 | 275 |
| 2779 | QGN15876.1 | FIM1_2574 | CP015057. | 1737 | 1907 | 1134 | 1639 | 1392 | 1012 | 1514 | 1236 | 1416 | 5748 | 1252 | 1262 | 5326 | 1130 | 1348 | 5554 | 1206 | 907 |
| 2780 | QGN15877.1 | FIM1_2575 | CP015057. | 608 | 868 | 566 | 917 | 859 | 566 | 711 | 659 | 744 | 2938 | 675 | 749 | 2916 | 518 | 713 | 2818 | 545 | 533 |
| 2781 | QGN15878.1 | FIM1_2576 | CP015057. | 11417 | 10287 | 7022 | 14735 | 10161 | 6477 | 11976 | 8567 | 8895 | 33785 | 11003 | 12489 | 38420 | 10431 | 11520 | 33313 | 10456 | 8538 |
| 2782 | QGN15879.1 | FIM1_2577 | CP015057. | 26200 | 2343 | 1906 | 15775 | 2514 | 1471 | 21804 | 2700 | 2385 | 10466 | 2948 | 2925 | 13619 | 2440 | 3466 | 11356 | 2885 | 2138 |
| 2783 | QGN15880.1 | FIM1_2578 | CP015057. | 249 | 75 | 38 | 219 | 66 | 43 | 239 | 49 | 57 | 56 | 41 | 43 | 119 | 34 | 40 | 64 | 34 | 27 |
| 2784 | QGN15881.1 | FIM1_2579 | CP015057. | 1091 | 239 | 140 | 1377 | 154 | 120 | 1484 | 143 | 200 | 285 | 87 | 98 | 437 | 97 | 104 | 285 | 91 | 85 |
| 2785 | QGN15882.1 | FIM1_2580 | CP015057. | 9197 | 540 | 442 | 7307 | 370 | 368 | 6851 | 440 | 584 | 862 | 294 | 485 | 1948 | 390 | 518 | 1402 | 316 | 343 |
| 2786 | QGN15883.1 | FIM1_2581 | CP015057. | 54660 | 13106 | 11112 | 50596 | 11430 | 9990 | 52204 | 9543 | 13382 | 36798 | 12660 | 16316 | 37173 | 15418 | 16897 | 36951 | 12905 | 11760 |
| 2787 | QGN15884.1 | FIM1_2582 | CP015057. | 1029 | 820 | 675 | 1043 | 440 | 496 | 1250 | 613 | 836 | 1952 | 279 | 526 | 1966 | 228 | 533 | 1875 | 226 | 343 |
| 2788 | QGN15885.1 | FIM1_2583 | CP015057. | 62 | 610 | 534 | 134 | 361 | 437 | 70 | 361 | 731 | 2222 | 315 | 373 | 1756 | 213 | 389 | 1987 | 232 | 240 |
| 2789 | QGN15886.1 | FIM1_2584 | CP015057. | 668 | 43 | 16 | 222 | 20 | 12 | 900 | 44 | 18 | 305 | 43 | 64 | 226 | 68 | 37 | 249 | 45 | 47 |
| 2790 | QGN15887.1 | FIM1_2585 | CP015057. | 3936 | 2800 | 3232 | 6574 | 4428 | 2598 | 4761 | 3697 | 3862 | 17185 | 3592 | 4674 | 17441 | 3588 | 3981 | 17031 | 3173 | 3021 |
| 2791 | QGN15888.1 | FIM1_2586 | CP015057. | 996 | 505 | 759 | 1189 | 726 | 570 | 876 | 622 | 941 | 3898 | 708 | 1132 | 4136 | 672 | 1004 | 4005 | 669 | 709 |
| 2792 | QGN15889.1 | FIM1_2587 | CP015057. | 6601 | 2939 | 2583 | 7417 | 4016 | 2261 | 7803 | 3374 | 3232 | 14970 | 3035 | 3202 | 16726 | 2660 | 2831 | 14974 | 2643 | 2102 |
| 2793 | QGN15890.1 | FIM1_2588 | CP015057. | 8128 | 1234 | 5667 | 7487 | 1865 | 3656 | 7460 | 2331 | 6769 | 3987 | 1857 | 12130 | 9797 | 2162 | 10999 | 6138 | 1724 | 8401 |
| 2794 | QGN15891.1 | FIM1_2589 | CP015057. | 6 | 4 | 1 | 7 | 5 | 4 | 7 | 0 | 8 | 22 | 2 | 3 | 8 | 2 | 4 | 12 | 1 | 3 |
| 2795 | QGN15892.1 | FIM1_2590 | CP015057. | 14909 | 760 | 1472 | 11964 | 562 | 998 | 10800 | 845 | 1888 | 609 | 252 | 1326 | 2148 | 432 | 1073 | 980 | 268 | 842 |
| 2796 | QGN15893.1 | FIM1_2591 | CP015057. | 2666 | 527 | 467 | 1445 | 687 | 381 | 2491 | 757 | 601 | 716 | 585 | 663 | 2029 | 535 | 681 | 1118 | 562 | 407 |
| 2797 | QGN15894.1 | FIM1_2592 | CP015057. | 4996 | 1064 | 875 | 3611 | 972 | 781 | 4641 | 844 | 1173 | 5556 | 1112 | 1556 | 6285 | 993 | 1515 | 5434 | 917 | 1089 |
| 2798 | QGN15895.1 | FIM1_2593 | CP015057. | 3526 | 549 | 944 | 4781 | 622 | 796 | 4073 | 663 | 1215 | 5809 | 706 | 1451 | 6596 | 738 | 1281 | 5627 | 615 | 1061 |
| 2799 | QGN15896.1 | FIM1_2594 | CP015057. | 1944 | 1291 | 1805 | 3540 | 1686 | 1446 | 2809 | 1568 | 2107 | 5817 | 1240 | 2288 | 7119 | 1253 | 1989 | 6267 | 1132 | 1580 |
| 2800 | QGN15897.1 | FIM1_2595 | CP015057. | 1632 | 857 | 730 | 1647 | 834 | 623 | 1681 | 768 | 960 | 4958 | 973 | 1219 | 4599 | 819 | 1034 | 4678 | 818 | 789 |
| 2801 | QGN15898.1 | FIM1_2596 | CP015057. | 8218 | 1816 | 1276 | 5160 | 934 | 1037 | 6445 | 1348 | 1649 | 2375 | 1071 | 1241 | 3439 | 683 | 1432 | 2828 | 852 | 877 |
| 2802 | QGN15899.1 | FIM1_2597 | CP015057. | 1698 | 516 | 628 | 1264 | 677 | 526 | 1728 | 602 | 754 | 1417 | 534 | 836 | 3079 | 684 | 721 | 1960 | 609 | 557 |
| 2803 | QGN15900.1 | FIM1_2598 | CP015057. | 1307 | 702 | 784 | 1179 | 603 | 733 | 1041 | 659 | 1052 | 4076 | 584 | 1067 | 3500 | 426 | 1106 | 4229 | 472 | 733 |
| 2804 | QGN15901.1 | FIM1_2599 | CP015057. | 679 | 725 | 492 | 396 | 754 | 469 | 773 | 684 | 642 | 3396 | 749 | 824 | 3436 | 756 | 711 | 3427 | 733 | 526 |
| 2805 | QGN15902.1 | FIM1_2600 | CP015057. | 463 | 1358 | 759 | 554 | 1065 | 864 | 473 | 859 | 1073 | 5637 | 929 | 1060 | 4078 | 839 | 1041 | 5540 | 999 | 713 |
| 2806 | QGN15903.1 | FIM1_2601 | CP015057. | 937 | 1884 | 1254 | 1573 | 1672 | 1227 | 1079 | 1543 | 1719 | 5305 | 1380 | 1692 | 6489 | 1419 | 1457 | 5594 | 1469 | 1052 |
| 2807 | QGN15904.1 | FIM1_2602 | CP015057. | 2280 | 1088 | 1027 | 3656 | 1387 | 1027 | 2705 | 1189 | 1362 | 5612 | 940 | 1615 | 6826 | 978 | 1403 | 5829 | 1038 | 1070 |
| 2808 | QGN15905.1 | FIM1_2603 | CP015057. | 312 | 33 | 211 | 629 | 143 | 138 | 438 | 171 | 269 | 679 | 114 | 316 | 1481 | 117 | 268 | 884 | 107 | 189 |
| 2809 | QGN15906.1 | FIM1_2604 | CP015057. | 1109 | 407 | 433 | 1359 | 324 | 316 | 890 | 361 | 541 | 569 | 322 | 537 | 1671 | 279 | 507 | 984 | 279 | 376 |
| 2810 | QGN15907.1 | FIM1_2605 | CP015057. | 34 | 0 | 0 | 38 | 0 | 1 | 26 | 1 | 1 | 6 | 1 | 1 | 2 | 0 | 2 | 5 | 0 | 2 |
| 2811 | QGN15908.1 | FIM1_2606 | CP015057. | 48 | 35 | 14 | 182 | 24 | 16 | 115 | 22 | 23 | 47 | 18 | 17 | 88 | 9 | 20 | 65 | 12 | 11 |
| 2812 | QGN15909.1 | FIM1_2607 | CP015057. | 2946 | 1944 | 1367 | 1629 | 2440 | 1182 | 2572 | 2325 | 1807 | 7569 | 2778 | 2296 | 9742 | 2154 | 2280 | 7828 | 2257 | 1586 |
| 2813 | QGN15910.1 | FIM1_2608 | CP015057. | 996 | 596 | 433 | 1079 | 525 | 444 | 1064 | 472 | 651 | 2494 | 524 | 689 | 2441 | 456 | 710 | 2432 | 518 | 424 |
| 2814 | QGN15911.1 | FIM1_2609 | CP015057. | 865 | 318 | 336 | 1527 | 252 | 276 | 962 | 238 | 491 | 2589 | 208 | 499 | 1869 | 171 | 450 | 2352 | 169 | 301 |
| 2815 | QGN15912.1 | FIM1_2610 | CP015057. | 4792 | 849 | 521 | 4521 | 725 | 517 | 5723 | 642 | 718 | 3017 | 609 | 876 | 3482 | 530 | 744 | 2877 | 576 | 569 |
| 2816 | QGN15913.1 | FIM1_2611 | CP015057. | 5189 | 796 | 1022 | 2827 | 796 | 1038 | 3885 | 768 | 1357 | 5759 | 802 | 2259 | 7022 | 647 | 1970 | 6134 | 706 | 1504 |
| 2817 | QGN15914.1 | FIM1_2612 | CP015057. | 12252 | 2142 | 1641 | 10604 | 1852 | 1290 | 11498 | 2034 | 2230 | 707 | 1694 | 2369 | 7704 | 1665 | 2305 | 2705 | 1726 | 1642 |
| 2818 | QGN15915.1 | FIM1_2613 | CP015057. | 5453 | 1337 | 1898 | 7052 | 2040 | 1441 | 4530 | 1901 | 2350 | 1987 | 1523 | 2925 | 8992 | 1677 | 2511 | 3741 | 1493 | 1944 |
| 2819 | QGN15916.1 | FIM1_2614 | CP015057. | 411 | 606 | 701 | 470 | 380 | 495 | 366 | 435 | 806 | 2302 | 340 | 894 | 3411 | 283 | 809 | 2531 | 278 | 618 |
| 2820 | QGN15917.1 | FIM1_2615 | CP015057. | 57 | 21 | 49 | 65 | 35 | 34 | 80 | 45 | 75 | 90 | 39 | 69 | 248 | 43 | 58 | 143 | 49 | 53 |
| 2821 | QGN15918.1 | FIM1_2616 | CP015057. | 1296 | 352 | 373 | 1266 | 447 | 316 | 1147 | 390 | 470 | 2536 | 481 | 793 | 2725 | 447 | 704 | 2660 | 452 | 523 |
| 2822 | QGN15919.1 | FIM1_2617 | CP015057. | 530 | 749 | 830 | 645 | 651 | 712 | 598 | 717 | 1068 | 5106 | 672 | 1305 | 4919 | 770 | 1099 | 5034 | 632 | 845 |
| 2823 | QGN15920.1 | FIM1_2618 | CP015057. | 442 | 424 | 307 | 405 | 277 | 246 | 327 | 243 | 395 | 1276 | 227 | 437 | 1242 | 182 | 392 | 1160 | 209 | 229 |
| 2824 | QGN15921.1 | FIM1_2619 | CP015057. | 348 | 226 | 190 | 453 | 121 | 196 | 298 | 159 | 292 | 1381 | 130 | 297 | 1003 | 103 | 295 | 1301 | 91 | 191 |
| 2825 | QGN15922.1 | FIM1_2620 | CP015057. | 200 | 829 | 601 | 328 | 614 | 479 | 212 | 548 | 703 | 2598 | 491 | 726 | 2074 | 396 | 660 | 2590 | 387 | 419 |
| 2826 | QGN15923.1 | FIM1_2621 | CP015057. | 20665 | 4530 | 4275 | 18971 | 4844 | 3154 | 18891 | 4896 | 5428 | 11418 | 5429 | 7145 | 28809 | 5904 | 6372 | 18144 | 5441 | 4817 |
| 2827 | QGN15924.1 | FIM1_2622 | CP015057. | 937 | 360 | 491 | 1239 | 388 | 383 | 958 | 434 | 638 | 1561 | 314 | 766 | 2443 | 279 | 661 | 1732 | 280 | 532 |
| 2828 | QGN15925.1 | FIM1_2623 | CP015057. | 2266 | 1249 | 970 | 2318 | 1085 | 883 | 2328 | 912 | 1292 | 7514 | 1194 | 1665 | 5831 | 1090 | 1524 | 7554 | 1055 | 1050 |
| 2829 | QGN15926.1 | FIM1_2624 | CP015057. | 13029 | 3985 | 4338 | 18428 | 4450 | 3752 | 14893 | 4169 | 5511 | 13017 | 3991 | 8539 | 21387 | 4721 | 7679 | 15985 | 4003 | 5637 |
| 2830 | QGN15927.1 | FIM1_2625 | CP015057. | 2034 | 491 | 335 | 1508 | 280 | 338 | 1591 | 340 | 510 | 1254 | 270 | 460 | 1644 | 218 | 501 | 1272 | 230 | 325 |
| 2831 | QGN15928.1 | FIM1_2626 | CP015057. | 1602 | 801 | 1391 | 1944 | 928 | 1136 | 1611 | 877 | 1674 | 2284 | 825 | 1807 | 4264 | 895 | 1670 | 3045 | 789 | 1204 |
| 2832 | QGN15929.1 | FIM1_2627 | CP015057. | 623 | 685 | 1194 | 603 | 468 | 1145 | 478 | 456 | 1601 | 2861 | 430 | 1723 | 1925 | 362 | 1700 | 2454 | 453 | 1088 |
| 2833 | QGN15930.1 | FIM1_2628 | CP015057. | 644 | 603 | 547 | 1115 | 529 | 542 | 782 | 468 | 792 | 3084 | 453 | 805 | 2818 | 467 | 738 | 2936 | 476 | 533 |
| 2834 | QGN15931.1 | FIM1_2629 | CP015057. | 10568 | 4499 | 2296 | 6105 | 5020 | 2517 | 10714 | 4090 | 3333 | 15611 | 8025 | 4852 | 25623 | 3639 | 4692 | 18498 | 6091 | 3575 |
| 2835 | QGN15932.1 | FIM1_2630 | CP015057. | 840 | 369 | 317 | 550 | 346 | 301 | 760 | 259 | 404 | 1966 | 311 | 463 | 1758 | 431 | 439 | 1816 | 383 | 299 |
| 2836 | QGN15933.1 | FIM1_2631 | CP015057. | 553 | 70 | 100 | 558 | 126 | 100 | 505 | 112 | 143 | 674 | 97 | 195 | 816 | 141 | 149 | 670 | 109 | 124 |
| 2837 | QGN15934.1 | FIM1_2632 | CP015057. | 678 | 229 | 419 | 1066 | 261 | 371 | 730 | 295 | 528 | 2903 | 321 | 641 | 2101 | 303 | 583 | 2728 | 306 | 438 |
| 2838 | QGN15935.1 | FIM1_2633 | CP015057. | 1211 | 375 | 573 | 2061 | 510 | 503 | 1012 | 499 | 684 | 2678 | 405 | 516 | 2587 | 332 | 458 | 2761 | 360 | 334 |
| 2839 | QGN15936.1 | FIM1_2634 | CP015057. | 3792 | 2564 | 4639 | 17343 | 2252 | 4766 | 9391 | 1975 | 5860 | 9059 | 1674 | 4519 | 8190 | 1932 | 4264 | 8545 | 2075 | 3247 |
| 2840 | QGN15937.1 | FIM1_2635 | CP015057. | 33455 | 27620 | 22169 | 34037 | 22347 | 18837 | 29475 | 20473 | 24665 | 47652 | 26746 | 25216 | 54917 | 25339 | 27230 | 51683 | 23333 | 20831 |
| 2841 | QGN15938.1 | FIM1_2636 | CP015057. | 6190 | 2510 | 2256 | 6272 | 2636 | 1998 | 5682 | 2417 | 3030 | 8538 | 2460 | 3605 | 11529 | 2079 | 3549 | 9318 | 2026 | 2411 |
| 2842 | QGN15939.1 | FIM1_2637 | CP015057. | 2547 | 2895 | 4748 | 6914 | 3365 | 3784 | 3929 | 2656 | 5675 | 11876 | 2317 | 5811 | 13435 | 2462 | 5033 | 11784 | 2043 | 3842 |
| 2843 | QGN15940.1 | FIM1_2638 | CP015057. | 3120 | 650 | 890 | 2465 | 774 | 798 | 2854 | 792 | 1163 | 5761 | 956 | 1395 | 5592 | 875 | 1285 | 5660 | 881 | 943 |
| 2844 | QGN15941.1 | FIM1_2639 | CP015057. | 992 | 3184 | 3138 | 1536 | 3918 | 3056 | 928 | 3122 | 4080 | 4534 | 2027 | 3446 | 9872 | 1508 | 2505 | 7214 | 1526 | 2084 |
| 2845 | QGN15942.1 | FIM1_2640 | CP015057. | 5509 | 2295 | 2486 | 9534 | 3113 | 2571 | 5713 | 2380 | 3276 | 6166 | 1484 | 2563 | 9354 | 1331 | 2394 | 7366 | 1203 | 1753 |
| 2846 | QGN15943.1 | FIM1_2641 | CP015057. | 1118 | 758 | 635 | 1115 | 625 | 555 | 1090 | 590 | 889 | 4405 | 528 | 1034 | 3942 | 489 | 1003 | 4257 | 513 | 693 |
| 2847 | QGN15944.1 | FIM1_2642 | CP015057. | 1170 | 274 | 191 | 1393 | 202 | 160 | 1307 | 264 | 269 | 1008 | 189 | 285 | 1223 | 131 | 260 | 1153 | 134 | 188 |
| 2848 | QGN15945.1 | FIM1_2643 | CP015057. | 6043 | 1190 | 1615 | 6764 | 1272 | 1388 | 6037 | 1661 | 2122 | 6046 | 1199 | 2261 | 8076 | 975 | 2484 | 6309 | 911 | 1580 |
| 2849 | QGN15946.1 | FIM1_2644 | CP015057. | 2097 | 1876 | 1351 | 2779 | 1284 | 1243 | 2329 | 1317 | 1787 | 4115 | 1026 | 1696 | 4784 | 949 | 1885 | 4065 | 880 | 1166 |
| 2850 | QGN15947.1 | FIM1_2645 | CP015057. | 1471 | 673 | 449 | 1391 | 414 | 385 | 1459 | 558 | 618 | 2025 | 360 | 596 | 1851 | 343 | 553 | 1964 | 270 | 421 |
| 2851 | QGN15948.1 | FIM1_2646 | CP015057. | 3716 | 1983 | 1816 | 5765 | 1588 | 1603 | 4689 | 1526 | 2303 | 7522 | 1465 | 2584 | 7026 | 1551 | 2511 | 6690 | 1376 | 1804 |
| 2852 | QGN15949.1 | FIM1_2647 | CP015057. | 5935 | 1546 | 2132 | 7708 | 2053 | 1784 | 6217 | 1813 | 2675 | 9843 | 1856 | 3352 | 11910 | 1890 | 3018 | 9826 | 1637 | 2243 |
| 2853 | QGN15950.1 | FIM1_2648 | CP015057. | 472 | 211 | 169 | 748 | 181 | 161 | 539 | 156 | 237 | 1047 | 152 | 236 | 1066 | 157 | 225 | 987 | 146 | 171 |
| 2854 | QGN15951.1 | FIM1_2649 | CP015057. | 1193 | 616 | 509 | 1500 | 617 | 417 | 1260 | 578 | 631 | 2714 | 575 | 756 | 2461 | 572 | 741 | 2525 | 543 | 482 |

| | | | | | | | | | | | | | | | | | | | | | |
|---|---|---|---|---|---|---|---|---|---|---|---|---|---|---|---|---|---|---|---|---|---|
| 2855 | QGN15952.1 | FIM1_2650 | CP015057. | 10363 | 11834 | 8908 | 8920 | 9863 | 7360 | 8042 | 8611 | 9695 | 29970 | 12787 | 12337 | 27003 | 10954 | 12839 | 30843 | 10248 | 9783 |
| 2856 | QGN15953.1 | FIM1_2651 | CP015057. | 1131 | 141 | 181 | 1855 | 356 | 196 | 1126 | 223 | 223 | 544 | 190 | 221 | 824 | 181 | 250 | 637 | 179 | 169 |
| 2857 | QGN15954.1 | FIM1_2652 | CP015057. | 4490 | 5630 | 3564 | 5920 | 6700 | 3387 | 5271 | 5409 | 4620 | 18306 | 5979 | 6729 | 23749 | 5289 | 6003 | 19036 | 5448 | 4481 |
| 2858 | QGN15955.1 | FIM1_2653 | CP015057. | 1799 | 1422 | 1062 | 1766 | 1110 | 1019 | 1619 | 1108 | 1413 | 7899 | 1059 | 1570 | 6133 | 872 | 1567 | 7617 | 909 | 1096 |
| 2859 | QGN15956.1 | FIM1_2654 | CP015057. | 5139 | 1324 | 1425 | 5831 | 1419 | 1252 | 4854 | 1231 | 1731 | 4003 | 1029 | 2002 | 6773 | 1308 | 1976 | 5014 | 1094 | 1368 |
| 2860 | QGN15957.1 | FIM1_2655 | CP015057. | 2757 | 1094 | 829 | 2462 | 982 | 626 | 2486 | 979 | 1169 | 320 | 749 | 1188 | 3810 | 868 | 1173 | 975 | 838 | 785 |
| 2861 | QGN15958.1 | FIM1_2656 | CP015057. | 519 | 74 | 84 | 857 | 89 | 95 | 406 | 81 | 107 | 1055 | 62 | 258 | 1027 | 229 | 206 | 1052 | 90 | 169 |
| 2862 | QGN15959.1 | FIM1_2657 | CP015057. | 1931 | 692 | 495 | 2258 | 453 | 387 | 1704 | 510 | 665 | 3549 | 554 | 706 | 2787 | 488 | 685 | 3459 | 513 | 461 |
| 2863 | QGN15960.1 | FIM1_2658 | CP015057. | 2300 | 8 | 51 | 2948 | 2 | 17 | 1555 | 9 | 44 | 190 | 4 | 30 | 286 | 5 | 24 | 189 | 4 | 28 |
| 2864 | QGN15961.1 | FIM1_2659 | CP015057. | 2317 | 707 | 1183 | 1904 | 710 | 1040 | 1482 | 838 | 1630 | 6252 | 878 | 1279 | 6409 | 790 | 1430 | 6262 | 695 | 931 |
| 2865 | QGN15962.1 | FIM1_2660 | CP015057. | 440 | 1179 | 2418 | 848 | 1605 | 2180 | 211 | 1423 | 3546 | 3290 | 1389 | 2482 | 5475 | 879 | 1652 | 4369 | 889 | 1549 |
| 2866 | QGN15963.1 | FIM1_2661 | CP015057. | 2412 | 403 | 825 | 4719 | 759 | 582 | 2817 | 699 | 1031 | 940 | 606 | 1105 | 4872 | 588 | 1025 | 2479 | 598 | 712 |
| 2867 | QGN15964.1 | FIM1_2662 | CP015057. | 7827 | 2903 | 3172 | 8636 | 4274 | 2601 | 8299 | 3613 | 4130 | 10456 | 3462 | 4144 | 18395 | 3262 | 3937 | 14250 | 2873 | 2936 |
| 2868 | QGN15965.1 | FIM1_2663 | CP015057. | 526 | 342 | 291 | 234 | 413 | 215 | 595 | 462 | 318 | 302 | 310 | 317 | 1005 | 333 | 295 | 568 | 319 | 218 |
| 2869 | QGN15966.1 | FIM1_2664 | CP015057. | 2322 | 1356 | 2341 | 2963 | 1780 | 1838 | 2241 | 1479 | 2856 | 8354 | 1488 | 2692 | 8516 | 1204 | 2561 | 8030 | 1140 | 1821 |
| 2870 | QGN15967.1 | FIM1_2665 | CP015057. | 2746 | 178 | 242 | 4080 | 128 | 186 | 3032 | 138 | 300 | 1185 | 80 | 305 | 1117 | 102 | 288 | 1100 | 73 | 195 |
| 2871 | QGN15968.1 | FIM1_2666 | CP015057. | 4659 | 281 | 371 | 6158 | 279 | 330 | 6628 | 324 | 493 | 996 | 219 | 478 | 1961 | 289 | 434 | 1374 | 222 | 293 |
| 2872 | QGN15969.1 | FIM1_2667 | CP015057. | 3537 | 2060 | 1774 | 5030 | 2077 | 1505 | 4457 | 2068 | 2352 | 7972 | 1849 | 2770 | 10393 | 1782 | 2369 | 8850 | 1686 | 1767 |
| 2873 | QGN15970.1 | FIM1_2668 | CP015057. | 7397 | 804 | 2117 | 8606 | 1760 | 1696 | 8444 | 1882 | 2784 | 2580 | 1270 | 1909 | 9556 | 1366 | 1881 | 5557 | 1184 | 1291 |
| 2874 | QGN15971.1 | FIM1_2669 | CP015057. | 147 | 29 | 26 | 339 | 30 | 25 | 243 | 23 | 52 | 58 | 6 | 18 | 78 | 7 | 25 | 46 | 8 | 11 |
| 2875 | QGN15972.1 | FIM1_2671 | CP015057. | 306 | 38 | 40 | 272 | 35 | 40 | 145 | 24 | 54 | 84 | 17 | 60 | 136 | 16 | 37 | 123 | 17 | 35 |
| 2876 | QGN15973.1 | FIM1_2672 | CP015057. | 964 | 386 | 511 | 1499 | 488 | 451 | 1264 | 449 | 661 | 3600 | 452 | 899 | 3641 | 500 | 756 | 3647 | 433 | 570 |
| 2877 | QGN15974.1 | FIM1_2674 | CP015057. | 10840 | 1593 | 3665 | 12933 | 2709 | 3383 | 12223 | 2022 | 4695 | 10669 | 1956 | 6464 | 14054 | 2065 | 5575 | 10979 | 1843 | 4364 |
| 2878 | QGN15975.1 | FIM1_2675 | CP015057. | 122722 | 219 | 811 | 103215 | 95 | 399 | 102603 | 458 | 1063 | 1459 | 74 | 294 | 4026 | 954 | 228 | 2076 | 112 | 248 |
| 2879 | QGN15976.1 | FIM1_2676 | CP015057. | 105842 | 6377 | 9276 | 101588 | 3316 | 6153 | 99229 | 5801 | 11509 | 23016 | 7078 | 7492 | 36086 | 23962 | 6821 | 28051 | 14857 | 5796 |
| 2880 | QGN15977.1 | FIM1_2677 | CP015057. | 1893 | 3202 | 2270 | 1808 | 3251 | 2206 | 1570 | 2588 | 2935 | 11198 | 3065 | 3152 | 12374 | 2875 | 3017 | 11290 | 3035 | 2080 |
| 2881 | QGN15978.1 | FIM1_2678 | CP015057. | 695 | 582 | 377 | 895 | 429 | 312 | 600 | 359 | 476 | 1969 | 385 | 572 | 1585 | 322 | 545 | 1770 | 345 | 369 |
| 2882 | QGN15979.1 | FIM1_2679 | CP015057. | 3773 | 516 | 781 | 4851 | 640 | 651 | 3729 | 638 | 1006 | 2521 | 488 | 1037 | 5409 | 468 | 1008 | 3564 | 476 | 699 |
| 2883 | QGN15980.1 | FIM1_2680 | CP015057. | 4029 | 2954 | 1893 | 3432 | 3039 | 1930 | 3720 | 2345 | 2452 | 14279 | 3907 | 3218 | 13124 | 3236 | 3036 | 14356 | 3820 | 2127 |
| 2884 | QGN15981.1 | FIM1_2681 | CP015057. | 26677 | 2073 | 1630 | 29465 | 1415 | 1323 | 26210 | 1467 | 1985 | 5643 | 1645 | 1933 | 6799 | 1955 | 2078 | 5444 | 1606 | 1299 |
| 2885 | QGN15982.1 | FIM1_2682 | CP015057. | 7450 | 173 | 243 | 5497 | 345 | 149 | 6602 | 385 | 296 | 494 | 277 | 349 | 1417 | 271 | 387 | 587 | 274 | 205 |
| 2886 | QGN15983.1 | FIM1_2683 | CP015057. | 9142 | 2118 | 1630 | 5737 | 2097 | 1681 | 6763 | 1630 | 2247 | 2843 | 862 | 2177 | 4482 | 1018 | 2063 | 3096 | 918 | 1443 |
| 2887 | QGN15984.1 | FIM1_2684 | CP015057. | 10292 | 2936 | 1898 | 7035 | 3227 | 1778 | 11757 | 2729 | 2433 | 12474 | 4593 | 3402 | 13355 | 4219 | 3136 | 12329 | 5279 | 2378 |
| 2888 | QGN15985.1 | FIM1_2685 | CP015057. | 1445 | 867 | 862 | 2031 | 723 | 772 | 1747 | 636 | 1138 | 4235 | 558 | 979 | 3328 | 508 | 853 | 3931 | 493 | 608 |
| 2889 | QGN15986.1 | FIM1_2686 | CP015057. | 178 | 117 | 113 | 370 | 98 | 104 | 219 | 101 | 167 | 825 | 118 | 164 | 956 | 89 | 180 | 839 | 61 | 123 |
| 2890 | QGN15987.1 | FIM1_2687 | CP015057. | 827 | 199 | 471 | 2386 | 480 | 315 | 1492 | 448 | 538 | 1380 | 316 | 626 | 3660 | 382 | 566 | 2059 | 336 | 432 |
| 2891 | QGN15988.1 | FIM1_2688 | CP015057. | 4404 | 1042 | 950 | 5311 | 1173 | 856 | 5710 | 971 | 1251 | 3098 | 1075 | 1441 | 5662 | 1144 | 1385 | 3815 | 1067 | 970 |
| 2892 | QGN15989.1 | FIM1_2689 | CP015057. | 3907 | 1309 | 1160 | 4110 | 986 | 1116 | 3808 | 862 | 1616 | 5883 | 876 | 1729 | 4988 | 867 | 1747 | 5710 | 798 | 1201 |
| 2893 | QGN15990.1 | FIM1_2690 | CP015057. | 3713 | 4047 | 3915 | 6192 | 4099 | 3455 | 4897 | 3514 | 5100 | 18852 | 4524 | 6758 | 19698 | 4534 | 5919 | 19048 | 4282 | 4477 |
| 2894 | QGN15991.1 | FIM1_2691 | CP015057. | 1032 | 799 | 930 | 1318 | 951 | 734 | 1214 | 754 | 1140 | 5188 | 896 | 1612 | 5860 | 840 | 1326 | 5221 | 803 | 971 |
| 2895 | QGN15992.1 | FIM1_2692 | CP015057. | 838 | 534 | 422 | 1214 | 463 | 442 | 898 | 406 | 601 | 3907 | 446 | 733 | 3093 | 397 | 622 | 3886 | 401 | 529 |
| 2896 | QGN15993.1 | FIM1_2693 | CP015057. | 1918 | 1584 | 1455 | 2670 | 1503 | 1252 | 2016 | 1301 | 1838 | 9438 | 1579 | 2303 | 8115 | 1432 | 2061 | 9276 | 1472 | 1512 |
| 2897 | QGN15994.1 | FIM1_2694 | CP015057. | 1295 | 454 | 308 | 1764 | 347 | 229 | 1684 | 367 | 360 | 1730 | 343 | 546 | 1762 | 361 | 501 | 1747 | 331 | 330 |
| 2898 | QGN15995.1 | FIM1_2695 | CP015057. | 1182 | 2621 | 1562 | 1207 | 2432 | 1599 | 1108 | 1980 | 2043 | 9962 | 2310 | 2442 | 9002 | 2195 | 2317 | 9935 | 2217 | 1589 |
| 2899 | QGN15996.1 | FIM1_2696 | CP015057. | 5949 | 5960 | 3021 | 3813 | 5027 | 2817 | 5226 | 4221 | 3929 | 14368 | 7163 | 4741 | 17096 | 5121 | 4868 | 15372 | 6064 | 3608 |
| 2900 | QGN15997.1 | FIM1_2697 | CP015057. | 323 | 452 | 255 | 155 | 360 | 257 | 287 | 315 | 353 | 1534 | 374 | 381 | 1483 | 252 | 324 | 1544 | 300 | 245 |
| 2901 | QGN15998.1 | FIM1_2698 | CP015057. | 3310 | 1618 | 2400 | 5098 | 1988 | 1854 | 3609 | 1704 | 2966 | 9871 | 1933 | 4077 | 12684 | 1975 | 3661 | 10531 | 1721 | 2673 |
| 2902 | QGN15999.1 | FIM1_2699 | CP015057. | 3519 | 1542 | 748 | 2284 | 1581 | 856 | 3374 | 1212 | 1040 | 7426 | 2155 | 1490 | 6137 | 2004 | 1546 | 7010 | 2243 | 1098 |
| 2903 | QGN16000.1 | FIM1_2700 | CP015057. | 2091 | 257 | 285 | 3313 | 348 | 250 | 1941 | 266 | 383 | 764 | 221 | 391 | 1630 | 259 | 371 | 1066 | 217 | 268 |
| 2904 | QGN16001.1 | FIM1_2701 | CP015057. | 2400 | 1112 | 1709 | 3829 | 1717 | 1601 | 2370 | 1514 | 2242 | 4376 | 1133 | 1706 | 8252 | 776 | 1624 | 5800 | 841 | 1158 |
| 2905 | QGN16002.1 | FIM1_2702 | CP015057. | 405 | 93 | 103 | 699 | 113 | 66 | 547 | 107 | 136 | 767 | 100 | 147 | 579 | 106 | 148 | 692 | 100 | 90 |
| 2906 | QGN16003.1 | FIM1_2703 | CP015057. | 381 | 176 | 548 | 477 | 50 | 221 | 631 | 128 | 486 | 763 | 60 | 262 | 509 | 65 | 128 | 825 | 57 | 111 |
| 2907 | QGN16004.1 | FIM1_2704 | CP015057. | 4237 | 694 | 945 | 4332 | 955 | 808 | 4619 | 820 | 1197 | 3486 | 723 | 1213 | 5708 | 829 | 1061 | 4189 | 813 | 840 |
| 2908 | QGN16005.1 | FIM1_2705 | CP015057. | 6918 | 1157 | 1750 | 5808 | 2007 | 1627 | 6324 | 1302 | 2255 | 5655 | 1578 | 1982 | 5823 | 2892 | 1830 | 5615 | 2236 | 1286 |
| 2909 | QGN16006.1 | FIM1_2706 | CP015057. | 83 | 18 | 7 | 193 | 15 | 8 | 93 | 14 | 6 | 68 | 9 | 17 | 51 | 8 | 3 | 62 | 13 | 5 |
| 2910 | QGN16007.1 | FIM1_2707 | CP015057. | 2 | 2 | 0 | 7 | 1 | 1 | 3 | 0 | 0 | 3 | 2 | 0 | 3 | 2 | 1 | 3 | 1 | 1 |
| 2911 | QGN16008.1 | FIM1_2708 | CP015057. | 5647 | 3872 | 1754 | 3293 | 4031 | 2189 | 5357 | 3085 | 2444 | 15414 | 5525 | 3632 | 15017 | 4623 | 3469 | 15302 | 6096 | 2763 |
| 2912 | QGN16009.1 | FIM1_2709 | CP015057. | 1811 | 3131 | 1802 | 2387 | 2343 | 1613 | 2025 | 1767 | 2431 | 10117 | 2379 | 3143 | 9477 | 2337 | 3179 | 10046 | 2249 | 2073 |
| 2913 | QGN16010.1 | FIM1_2710 | CP015057. | 4044 | 7789 | 5128 | 9450 | 6014 | 4624 | 7807 | 5239 | 6296 | 25048 | 7047 | 8602 | 20383 | 6673 | 7903 | 25122 | 6471 | 5767 |
| 2914 | QGN16011.1 | FIM1_2711 | CP015057. | 344 | 334 | 200 | 454 | 271 | 175 | 376 | 247 | 271 | 730 | 242 | 260 | 1017 | 188 | 318 | 874 | 180 | 188 |
| 2915 | QGN16012.1 | FIM1_2712 | CP015057. | 1739 | 2360 | 1268 | 1996 | 1807 | 1297 | 2003 | 1481 | 1660 | 8802 | 2187 | 2464 | 7247 | 1597 | 2233 | 8418 | 1972 | 1688 |
| 2916 | QGN16013.1 | FIM1_2713 | CP015057. | 1862 | 1008 | 1263 | 2561 | 1225 | 1086 | 1776 | 1058 | 1638 | 8641 | 1145 | 2218 | 8016 | 938 | 1882 | 8252 | 881 | 1348 |
| 2917 | QGN16014.1 | FIM1_2714 | CP015057. | 3146 | 718 | 1160 | 3931 | 864 | 971 | 3752 | 795 | 1507 | 4421 | 673 | 1372 | 4628 | 677 | 1341 | 4225 | 615 | 988 |
| 2918 | QGN16015.1 | FIM1_2715 | CP015057. | 2036 | 1812 | 1577 | 2007 | 2244 | 1469 | 2064 | 1944 | 2030 | 8733 | 2261 | 2340 | 9454 | 1986 | 2282 | 8469 | 2004 | 1593 |
| 2919 | QGN16016.1 | FIM1_2716 | CP015057. | 5205 | 3865 | 4331 | 4613 | 3937 | 3071 | 4646 | 4141 | 5531 | 26197 | 5502 | 6558 | 26580 | 6639 | 8289 | 25924 | 5612 | 4718 |
| 2920 | QGN16017.1 | FIM1_2717 | CP015057. | 6 | 5 | 10 | 11 | 4 | 2 | 7 | 3 | 4 | 48 | 4 | 10 | 31 | 4 | 5 | 35 | 5 | 3 |
| 2921 | QGN16018.1 | FIM1_2718 | CP015057. | 12880 | 2485 | 2796 | 13743 | 2285 | 2412 | 14488 | 2266 | 3759 | 11028 | 2936 | 3580 | 16306 | 2550 | 3357 | 12604 | 2774 | 2774 |
| 2922 | QGN16019.1 | FIM1_2719 | CP015057. | 2140 | 972 | 1447 | 3540 | 1649 | 1369 | 2154 | 1301 | 1806 | 6783 | 1338 | 1648 | 6620 | 1069 | 1390 | 6985 | 1127 | 1050 |
| 2923 | QGN16020.1 | FIM1_2720 | CP015057. | 5888 | 1525 | 2655 | 6696 | 1580 | 2303 | 6542 | 1460 | 3443 | 6469 | 1453 | 3824 | 8924 | 1445 | 3510 | 7223 | 1273 | 2759 |
| 2924 | QGN16021.1 | FIM1_2721 | CP015057. | 1349 | 188 | 288 | 1312 | 429 | 275 | 1292 | 387 | 409 | 491 | 324 | 317 | 1252 | 398 | 257 | 888 | 353 | 228 |
| 2925 | QGN16022.1 | FIM1_2722 | CP015057. | 619 | 2159 | 1520 | 865 | 2032 | 1466 | 735 | 1649 | 2005 | 10813 | 1986 | 2437 | 8240 | 1791 | 2183 | 10685 | 1876 | 1585 |
| 2926 | QGN16023.1 | FIM1_2723 | CP015057. | 1757 | 520 | 680 | 666 | 486 | 533 | 1897 | 483 | 846 | 1986 | 401 | 1077 | 2723 | 312 | 986 | 2227 | 285 | 694 |
| 2927 | QGN16024.1 | FIM1_2724 | CP015057. | 2741 | 506 | 1423 | 3441 | 787 | 1169 | 3382 | 748 | 1790 | 7092 | 696 | 2626 | 7921 | 752 | 2363 | 7304 | 585 | 1675 |
| 2928 | QGN16025.1 | FIM1_2725 | CP015057. | 4 | 1 | 1 | 1 | 1 | 0 | 1 | 0 | 1 | 5 | 1 | 2 | 2 | 1 | 3 | 0 | 0 | 0 |
| 2929 | QGN16026.1 | FIM1_2726 | CP015057. | 1756 | 430 | 575 | 2786 | 399 | 537 | 1916 | 342 | 790 | 1659 | 233 | 828 | 2021 | 280 | 818 | 1716 | 217 | 539 |
| 2930 | QGN16027.1 | FIM1_2727 | CP015057. | 333 | 88 | 82 | 678 | 87 | 71 | 321 | 109 | 113 | 456 | 57 | 184 | 480 | 35 | 130 | 461 | 48 | 121 |
| 2931 | QGN16028.1 | FIM1_2728 | CP015057. | 4243 | 143 | 549 | 1977 | 224 | 471 | 2338 | 225 | 734 | 1089 | 155 | 512 | 1636 | 98 | 465 | 1348 | 110 | 320 |
| 2932 | QGN16029.1 | FIM1_2729 | CP015057. | 224 | 19 | 162 | 288 | 16 | 95 | 233 | 14 | 282 | 774 | 56 | 195 | 1876 | 119 | 197 | 1663 | 67 | 177 |
| 2933 | QGN16030.1 | FIM1_2730 | CP015057. | 1158 | 41 | 83 | 1585 | 55 | 56 | 923 | 38 | 122 | 109 | 23 | 54 | 272 | 25 | 54 | 182 | 17 | 41 |
| 2934 | QGN16031.1 | FIM1_2731 | CP015057. | 6581 | 2297 | 2395 | 7931 | 2808 | 2025 | 7426 | 2503 | 2952 | 8251 | 2604 | 3751 | 13620 | 2609 | 3489 | 9779 | 2419 | 2599 |
| 2935 | QGN16032.1 | FIM1_2732 | CP015057. | 13049 | 4897 | 5094 | 19028 | 5529 | 4196 | 16712 | 4842 | 6178 | 20199 | 5642 | 8061 | 30646 | 5710 | 7313 | 25151 | 5149 | 5577 |
| 2936 | QGN16033.1 | FIM1_2733 | CP015057. | 556 | 178 | 179 | 632 | 172 | 175 | 579 | 94 | 216 | 1427 | 163 | 226 | 1064 | 162 | 185 | 1322 | 152 | 162 |
| 2937 | QGN16034.1 | FIM1_2734 | CP015057. | 206 | 207 | 187 | 162 | 192 | 145 | 154 | 163 | 242 | 819 | 171 | 189 | 745 | 173 | 189 | 931 | 143 | 119 |
| 2938 | QGN16035.1 | FIM1_2735 | CP015057. | 649 | 462 | 402 | 742 | 461 | 324 | 624 | 336 | 506 | 2455 | 513 | 699 | 2046 | 422 | 601 | 2464 | 473 | 418 |

| | | | | | | | | | | | | | | | | | | | | | |
|---|---|---|---|---|---|---|---|---|---|---|---|---|---|---|---|---|---|---|---|---|---|
| 2939 | QGN16036.1 | FIM1_2736 | CP015057. | 646 | 474 | 375 | 706 | 388 | 413 | 578 | 402 | 536 | 2695 | 351 | 609 | 1946 | 304 | 541 | 2416 | 339 | 392 |
| 2940 | QGN16037.1 | FIM1_2737 | CP015057. | 1616 | 2131 | 1629 | 1459 | 2246 | 1332 | 1500 | 2164 | 2120 | 6554 | 1988 | 2194 | 7937 | 1786 | 1875 | 7569 | 1855 | 1369 |
| 2941 | QGN16038.1 | FIM1_2738 | CP015057. | 235 | 145 | 426 | 611 | 271 | 368 | 292 | 321 | 567 | 1437 | 243 | 573 | 2208 | 193 | 478 | 1776 | 205 | 405 |
| 2942 | QGN16039.1 | FIM1_2739 | CP015057. | 840 | 1242 | 3077 | 4464 | 1291 | 2174 | 1778 | 1692 | 3774 | 6159 | 690 | 2658 | 7021 | 869 | 2327 | 6545 | 620 | 1637 |
| 2943 | QGN16040.1 | FIM1_2740 | CP015057. | 2753 | 3700 | 4107 | 4474 | 3519 | 3580 | 3422 | 3380 | 5246 | 17193 | 4269 | 6022 | 17581 | 4141 | 5646 | 16821 | 3770 | 4073 |
| 2944 | QGN16041.1 | FIM1_2741 | CP015057. | 504 | 606 | 693 | 737 | 742 | 610 | 490 | 590 | 861 | 3872 | 698 | 1093 | 3879 | 541 | 1005 | 3967 | 521 | 658 |
| 2945 | QGN16042.1 | FIM1_2742 | CP015057. | 3174 | 2148 | 4081 | 8472 | 4023 | 3382 | 4544 | 2677 | 5057 | 20982 | 3992 | 6509 | 26258 | 6475 | 5606 | 22979 | 4165 | 4240 |
| 2946 | QGN16043.1 | FIM1_2743 | CP015057. | 1160 | 724 | 393 | 476 | 628 | 293 | 766 | 544 | 471 | 923 | 583 | 472 | 2039 | 524 | 523 | 1525 | 591 | 317 |
| 2947 | QGN16044.1 | FIM1_2744 | CP015057. | 3668 | 4267 | 5601 | 5193 | 4955 | 5162 | 4261 | 4208 | 7598 | 13551 | 3375 | 8516 | 16674 | 3289 | 8040 | 14794 | 3087 | 5707 |
| 2948 | QGN16045.1 | FIM1_2745 | CP015057. | 161 | 1403 | 864 | 623 | 1203 | 831 | 384 | 874 | 1146 | 5115 | 1132 | 1474 | 4230 | 895 | 1350 | 5027 | 1026 | 952 |
| 2949 | QGN16046.1 | FIM1_2746 | CP015057. | 1714 | 878 | 1037 | 2705 | 686 | 952 | 2083 | 651 | 1443 | 5281 | 650 | 1739 | 3942 | 643 | 1526 | 5148 | 652 | 1108 |
| 2950 | QGN16047.1 | FIM1_2747 | CP015057. | 2117 | 2864 | 1424 | 2304 | 2207 | 1582 | 2375 | 1755 | 2051 | 11210 | 2606 | 2421 | 9050 | 2035 | 2306 | 10259 | 2315 | 1669 |
| 2951 | QGN16048.1 | FIM1_2748 | CP015057. | 1676 | 278 | 277 | 1606 | 383 | 307 | 1523 | 246 | 396 | 2462 | 245 | 433 | 2047 | 210 | 390 | 2419 | 227 | 261 |
| 2952 | QGN16049.1 | FIM1_2749 | CP015057. | 3756 | 319 | 495 | 3531 | 376 | 363 | 3209 | 392 | 630 | 3329 | 287 | 702 | 3958 | 262 | 643 | 3248 | 224 | 431 |
| 2953 | QGN16050.1 | FIM1_2750 | CP015057. | 280 | 54 | 61 | 401 | 72 | 41 | 274 | 57 | 58 | 237 | 47 | 56 | 216 | 37 | 58 | 244 | 47 | 33 |
| 2954 | QGN16051.1 | FIM1_2751 | CP015057. | 646 | 504 | 442 | 1477 | 420 | 328 | 851 | 336 | 447 | 2136 | 319 | 562 | 1594 | 393 | 478 | 1939 | 393 | 329 |
| 2955 | QGN16052.1 | FIM1_2752 | CP015057. | 1070 | 816 | 898 | 1588 | 830 | 770 | 1195 | 829 | 1162 | 5659 | 698 | 1393 | 5515 | 727 | 1220 | 5405 | 614 | 941 |
| 2956 | QGN16053.1 | FIM1_2753 | CP015057. | 3218 | 951 | 683 | 3625 | 685 | 417 | 3098 | 761 | 747 | 2689 | 856 | 1044 | 3939 | 1172 | 1118 | 2719 | 985 | 719 |
| 2957 | QGN16054.1 | FIM1_2754 | CP015057. | 27706 | 733 | 1198 | 21842 | 1087 | 892 | 22353 | 971 | 1530 | 3763 | 1002 | 1706 | 7581 | 1054 | 1501 | 5091 | 973 | 1128 |
| 2958 | QGN16055.1 | FIM1_2755 | CP015057. | 1247 | 656 | 757 | 2233 | 786 | 648 | 1769 | 685 | 980 | 4424 | 545 | 985 | 3962 | 607 | 885 | 4099 | 592 | 570 |
| 2959 | QGN16056.1 | FIM1_2756 | CP015057. | 262 | 847 | 557 | 1206 | 688 | 657 | 549 | 457 | 795 | 3628 | 617 | 821 | 2778 | 552 | 858 | 3469 | 576 | 530 |
| 2960 | QGN16057.1 | FIM1_2757 | CP015057. | 7438 | 2758 | 1748 | 3254 | 3885 | 1974 | 6069 | 3102 | 2443 | 12914 | 5582 | 3693 | 15859 | 4943 | 3391 | 13805 | 6112 | 2607 |
| 2961 | QGN16058.1 | FIM1_2758 | CP015057. | 483 | 650 | 361 | 493 | 485 | 359 | 432 | 411 | 501 | 983 | 427 | 420 | 1439 | 420 | 435 | 1040 | 416 | 291 |
| 2962 | QGN16059.1 | FIM1_2759 | CP015057. | 398 | 237 | 176 | 527 | 176 | 198 | 567 | 193 | 279 | 666 | 188 | 141 | 448 | 210 | 126 | 668 | 186 | 86 |
| 2963 | QGN16060.1 | FIM1_2760 | CP015057. | 13796 | 3534 | 4016 | 16950 | 5387 | 3548 | 15185 | 4627 | 5144 | 8579 | 5294 | 6679 | 26995 | 5081 | 5902 | 16328 | 4933 | 4763 |
| 2964 | QGN16061.1 | FIM1_2761 | CP015057. | 1768 | 903 | 858 | 2229 | 926 | 765 | 2059 | 914 | 1167 | 5595 | 845 | 1293 | 5584 | 789 | 1166 | 5617 | 716 | 821 |
| 2965 | QGN16062.1 | FIM1_2762 | CP015057. | 4753 | 798 | 1588 | 4973 | 1868 | 1201 | 4579 | 1736 | 1861 | 5116 | 1672 | 2162 | 8225 | 1643 | 1978 | 6729 | 1291 | 1370 |
| 2966 | QGN16063.1 | FIM1_2763 | CP015057. | 5663 | 9 | 8 | 6770 | 4 | 4 | 6789 | 18 | 7 | 205 | 8 | 18 | 297 | 122 | 10 | 114 | 23 | 18 |
| 2967 | QGN16064.1 | FIM1_2764 | CP015057. | 1696 | 1061 | 1712 | 3181 | 824 | 1476 | 2207 | 870 | 2093 | 1789 | 323 | 1814 | 2362 | 487 | 1683 | 1768 | 405 | 1225 |
| 2968 | QGN16065.1 | FIM1_2765 | CP015057. | 818 | 1248 | 1101 | 736 | 875 | 967 | 854 | 999 | 1340 | 3111 | 809 | 1404 | 3021 | 824 | 1513 | 2850 | 760 | 1010 |
| 2969 | QGN16066.1 | FIM1_2766 | CP015057. | 6818 | 3409 | 2007 | 6761 | 2269 | 1845 | 6352 | 2216 | 2812 | 8666 | 2306 | 2648 | 8862 | 2358 | 3174 | 8722 | 2259 | 1881 |
| 2970 | QGN16067.1 | FIM1_2767 | CP015057. | 5652 | 1151 | 1962 | 4481 | 1185 | 1470 | 4849 | 1429 | 2447 | 2676 | 1019 | 2506 | 7220 | 888 | 2256 | 4112 | 885 | 1702 |
| 2971 | QGN16068.1 | FIM1_2768 | CP015057. | 1677 | 1053 | 943 | 3424 | 756 | 854 | 1987 | 639 | 1232 | 4096 | 475 | 1027 | 3777 | 754 | 1032 | 3939 | 534 | 676 |
| 2972 | QGN16069.1 | FIM1_2769 | CP015057. | 2345 | 1662 | 1882 | 2511 | 1300 | 1675 | 2142 | 1211 | 2495 | 6715 | 1215 | 2549 | 5669 | 1284 | 2523 | 6070 | 1182 | 1725 |
| 2973 | QGN16070.1 | FIM1_2770 | CP015057. | 4403 | 532 | 1056 | 4463 | 593 | 873 | 4398 | 730 | 1244 | 3248 | 476 | 1465 | 4968 | 434 | 1296 | 3896 | 390 | 901 |
| 2974 | QGN16071.1 | FIM1_2771 | CP015057. | 78 | 298 | 327 | 288 | 342 | 326 | 146 | 337 | 462 | 1210 | 293 | 501 | 1372 | 245 | 426 | 1326 | 252 | 262 |
| 2975 | QGN16072.1 | FIM1_2772 | CP015057. | 3118 | 718 | 841 | 3855 | 728 | 668 | 2860 | 746 | 1110 | 6677 | 803 | 1215 | 7207 | 677 | 1042 | 6711 | 652 | 776 |
| 2976 | QGN16073.1 | FIM1_2773 | CP015057. | 613 | 350 | 300 | 525 | 216 | 284 | 548 | 200 | 356 | 1343 | 226 | 375 | 1081 | 176 | 424 | 1218 | 202 | 243 |
| 2977 | QGN16074.1 | FIM1_2774 | CP015057. | 5052 | 332 | 383 | 3998 | 793 | 481 | 2222 | 466 | 562 | 1630 | 441 | 407 | 2291 | 551 | 389 | 1790 | 443 | 280 |
| 2978 | QGN16075.1 | FIM1_2775 | CP015057. | 3402 | 2083 | 2211 | 4095 | 2433 | 1932 | 4393 | 2178 | 2756 | 12987 | 2792 | 4010 | 13786 | 2457 | 3741 | 13168 | 2538 | 2861 |
| 2979 | QGN16076.1 | FIM1_2776 | CP015057. | 1162 | 296 | 422 | 1218 | 375 | 345 | 1185 | 341 | 559 | 3390 | 365 | 621 | 3055 | 349 | 524 | 3221 | 317 | 409 |
| 2980 | QGN16077.1 | FIM1_2777 | CP015057. | 1301 | 378 | 876 | 1545 | 522 | 613 | 1144 | 514 | 1126 | 3488 | 307 | 768 | 3503 | 327 | 717 | 3314 | 271 | 489 |
| 2981 | QGN16078.1 | FIM1_2778 | CP015057. | 397 | 338 | 461 | 440 | 352 | 328 | 285 | 346 | 576 | 2487 | 349 | 560 | 2288 | 343 | 508 | 2363 | 351 | 374 |
| 2982 | QGN16079.1 | FIM1_2779 | CP015057. | 641 | 574 | 635 | 783 | 841 | 587 | 939 | 622 | 732 | 3008 | 733 | 862 | 3082 | 592 | 755 | 2782 | 621 | 560 |
| 2983 | QGN16080.1 | FIM1_2780 | CP015057. | 1026 | 506 | 384 | 1600 | 282 | 282 | 1062 | 316 | 457 | 316 | 116 | 203 | 392 | 62 | 135 | 323 | 69 | 81 |
| 2984 | QGN16081.1 | FIM1_2781 | CP015057. | 6183 | 397 | 1078 | 9479 | 616 | 739 | 8114 | 742 | 1291 | 1366 | 281 | 813 | 5007 | 411 | 708 | 2809 | 323 | 597 |
| 2985 | QGN16082.1 | FIM1_2782 | CP015057. | 685 | 1779 | 1065 | 775 | 1322 | 949 | 699 | 1152 | 1271 | 5568 | 1240 | 1369 | 4162 | 1103 | 1310 | 5167 | 1178 | 945 |
| 2986 | QGN16083.1 | FIM1_2783 | CP015057. | 8604 | 3362 | 2060 | 4514 | 4339 | 2233 | 8038 | 3380 | 2706 | 14576 | 6080 | 4026 | 16522 | 5892 | 3751 | 14872 | 6599 | 2927 |
| 2987 | QGN16084.1 | FIM1_2784 | CP015057. | 1760 | 622 | 568 | 1702 | 537 | 491 | 1736 | 535 | 782 | 3015 | 487 | 849 | 3170 | 368 | 790 | 2967 | 409 | 619 |
| 2988 | QGN16085.1 | FIM1_2785 | CP015057. | 2727 | 2354 | 2049 | 3005 | 1691 | 1985 | 2835 | 1571 | 2690 | 10350 | 1736 | 3096 | 8285 | 1570 | 3083 | 9636 | 1465 | 2136 |
| 2989 | QGN16086.1 | FIM1_2786 | CP015057. | 1833 | 1178 | 1618 | 3247 | 1347 | 1315 | 2504 | 1313 | 2160 | 4547 | 1330 | 2805 | 9016 | 1382 | 2384 | 6479 | 1357 | 1848 |
| 2990 | QGN16087.1 | FIM1_2787 | CP015057. | 8668 | 3203 | 3173 | 11014 | 3567 | 2568 | 9162 | 3574 | 4278 | 3345 | 3230 | 4837 | 17953 | 3225 | 4630 | 7673 | 3358 | 3386 |
| 2991 | QGN16088.1 | FIM1_2788 | CP015057. | 677 | 139 | 257 | 1243 | 119 | 249 | 775 | 135 | 356 | 1153 | 116 | 351 | 1331 | 128 | 305 | 1072 | 93 | 247 |
| 2992 | QGN16089.1 | FIM1_2789 | CP015057. | 182 | 236 | 191 | 327 | 153 | 199 | 295 | 155 | 293 | 1287 | 123 | 287 | 814 | 155 | 299 | 1092 | 147 | 179 |
| 2993 | QGN16090.1 | FIM1_2790 | CP015057. | 2214 | 600 | 997 | 2655 | 732 | 744 | 2295 | 655 | 1207 | 4439 | 485 | 913 | 3932 | 432 | 832 | 4426 | 370 | 602 |
| 2994 | QGN16091.1 | FIM1_2791 | CP015057. | 1150 | 119 | 91 | 1574 | 154 | 73 | 724 | 108 | 120 | 645 | 117 | 181 | 610 | 116 | 170 | 563 | 93 | 103 |
| 2995 | QGN16092.1 | FIM1_2792 | CP015057. | 245 | 640 | 616 | 568 | 487 | 523 | 407 | 428 | 782 | 2254 | 406 | 815 | 1931 | 343 | 808 | 2240 | 340 | 569 |
| 2996 | QGN16093.1 | FIM1_2793 | CP015057. | 6773 | 908 | 983 | 6098 | 1102 | 836 | 6221 | 935 | 1325 | 5042 | 847 | 1134 | 5583 | 819 | 967 | 5373 | 784 | 706 |
| 2997 | QGN16094.1 | FIM1_2794 | CP015057. | 1507 | 1936 | 1320 | 2018 | 1244 | 927 | 1552 | 1358 | 1600 | 8932 | 1566 | 1906 | 8031 | 2007 | 2030 | 8323 | 1746 | 1324 |
| 2998 | QGN16095.1 | FIM1_2795 | CP015057. | 1671 | 326 | 360 | 1237 | 381 | 300 | 1711 | 379 | 481 | 552 | 432 | 565 | 2555 | 427 | 466 | 1313 | 396 | 363 |
| 2999 | QGN16096.1 | FIM1_2796 | CP015057. | 7281 | 105 | 150 | 7393 | 166 | 109 | 4012 | 163 | 189 | 371 | 149 | 179 | 815 | 111 | 160 | 551 | 108 | 122 |
| 3000 | QGN16097.1 | FIM1_2797 | CP015057. | 74 | 33 | 45 | 150 | 16 | 28 | 125 | 20 | 35 | 258 | 19 | 55 | 175 | 24 | 32 | 309 | 20 | 33 |
| 3001 | QGN16098.1 | FIM1_2798 | CP015057. | 1434 | 1045 | 1503 | 1103 | 906 | 1305 | 1400 | 816 | 1917 | 5033 | 944 | 2085 | 4581 | 994 | 1888 | 4987 | 810 | 1370 |
| 3002 | QGN16099.1 | FIM1_2800 | CP015057. | 5640 | 1542 | 1350 | 6840 | 1754 | 1166 | 6713 | 1465 | 1756 | 11695 | 2206 | 2182 | 11126 | 2852 | 2186 | 10708 | 2402 | 1477 |
| 3003 | QGN16100.1 | FIM1_2801 | CP015057. | 4112 | 3926 | 2777 | 6491 | 3044 | 2559 | 5025 | 2627 | 3621 | 15407 | 2829 | 3942 | 12205 | 2708 | 3665 | 15587 | 2579 | 2582 |
| 3004 | QGN16101.1 | FIM1_2802 | CP015057. | 1583 | 332 | 463 | 2542 | 472 | 366 | 1771 | 428 | 585 | 3353 | 479 | 832 | 2907 | 453 | 761 | 2993 | 442 | 547 |
| 3005 | QGN16102.1 | FIM1_2803 | CP015057. | 2071 | 535 | 643 | 2606 | 749 | 539 | 2150 | 607 | 837 | 5127 | 710 | 1182 | 5292 | 761 | 988 | 5111 | 730 | 774 |
| 3006 | QGN16103.1 | FIM1_2804 | CP015057. | 4669 | 1275 | 1567 | 4795 | 1699 | 1401 | 4829 | 1336 | 2056 | 6159 | 1372 | 1904 | 7492 | 1180 | 1820 | 7132 | 1154 | 1306 |
| 3007 | QGN16104.1 | FIM1_2805 | CP015057. | 2908 | 1835 | 1379 | 2468 | 1689 | 1339 | 2344 | 1417 | 1778 | 6240 | 1228 | 1516 | 5343 | 1032 | 1487 | 6214 | 983 | 930 |
| 3008 | QGN16105.1 | FIM1_2806 | CP015057. | 350 | 138 | 146 | 396 | 120 | 125 | 374 | 100 | 204 | 777 | 93 | 145 | 638 | 55 | 165 | 741 | 88 | 112 |
| 3009 | QGN16106.1 | FIM1_2807 | CP015057. | 1331 | 1180 | 828 | 1876 | 828 | 746 | 1584 | 709 | 1101 | 4239 | 608 | 962 | 3040 | 591 | 879 | 4022 | 625 | 609 |
| 3010 | QGN16107.1 | FIM1_2808 | CP015057. | 651 | 905 | 942 | 1417 | 1198 | 820 | 888 | 880 | 1281 | 6601 | 1215 | 1319 | 6088 | 1165 | 1209 | 6279 | 1031 | 840 |
| 3011 | QGN16108.1 | FIM1_2809 | CP015057. | 193 | 60 | 143 | 517 | 80 | 120 | 219 | 66 | 168 | 787 | 61 | 176 | 791 | 62 | 177 | 837 | 56 | 89 |
| 3012 | QGN16109.1 | FIM1_2810 | CP015057. | 879 | 152 | 86 | 424 | 136 | 83 | 464 | 89 | 116 | 815 | 193 | 322 | 1009 | 118 | 134 | 1195 | 179 | 159 |
| 3013 | QGN16110.1 | FIM1_2811 | CP015057. | 727 | 195 | 268 | 818 | 249 | 189 | 751 | 252 | 332 | 1799 | 275 | 476 | 1556 | 295 | 402 | 1659 | 278 | 265 |
| 3014 | QGN16111.1 | FIM1_2812 | CP015057. | 6688 | 2008 | 1990 | 6878 | 1978 | 1661 | 6122 | 1745 | 2573 | 9409 | 1729 | 2635 | 8653 | 1914 | 2461 | 8899 | 1658 | 1694 |
| 3015 | QGN16112.1 | FIM1_2813 | CP015057. | 4172 | 626 | 754 | 3104 | 464 | 599 | 3301 | 497 | 1039 | 3184 | 399 | 803 | 3002 | 481 | 748 | 2927 | 366 | 558 |
| 3016 | QGN16113.1 | FIM1_2814 | CP015057. | 8959 | 1427 | 1296 | 6001 | 1468 | 1034 | 6482 | 1322 | 1624 | 6647 | 1003 | 1571 | 6906 | 976 | 1531 | 6390 | 837 | 1068 |
| 3017 | QGN16114.1 | FIM1_2815 | CP015057. | 385 | 394 | 289 | 434 | 316 | 271 | 379 | 264 | 362 | 1137 | 207 | 278 | 783 | 163 | 305 | 983 | 169 | 161 |
| 3018 | QGN16115.1 | FIM1_2816 | CP015057. | 489 | 203 | 304 | 457 | 320 | 232 | 349 | 272 | 364 | 1125 | 275 | 405 | 1164 | 231 | 370 | 1148 | 197 | 218 |
| 3019 | QGN16116.1 | FIM1_2817 | CP015057. | 969 | 557 | 335 | 729 | 463 | 317 | 746 | 361 | 447 | 1361 | 387 | 477 | 1582 | 348 | 505 | 1360 | 335 | 317 |
| 3020 | QGN16117.1 | FIM1_2818 | CP015057. | 1016 | 134 | 270 | 1091 | 171 | 237 | 1014 | 168 | 340 | 1990 | 159 | 379 | 1732 | 149 | 322 | 1958 | 176 | 276 |
| 3021 | QGN16118.1 | FIM1_2819 | CP015057. | 2438 | 1606 | 1519 | 2526 | 1489 | 1278 | 2530 | 1533 | 1961 | 7782 | 1247 | 1980 | 7345 | 1161 | 1750 | 7489 | 1275 | 1272 |
| 3022 | QGN16119.1 | FIM1_2820 | CP015057. | 5152 | 1115 | 1017 | 10725 | 1025 | 889 | 4465 | 876 | 1446 | 3003 | 774 | 1160 | 6574 | 1167 | 1023 | 4630 | 532 | 839 |

| | | | | | | | | | | | | | | | | | | | | | |
|---|---|---|---|---|---|---|---|---|---|---|---|---|---|---|---|---|---|---|---|---|---|
| 3023 | QGN16120.1 | FIM1_2821 | CP015057. | 2835 | 714 | 1245 | 3000 | 811 | 957 | 3074 | 794 | 1531 | 3902 | 863 | 1592 | 6958 | 883 | 1618 | 5175 | 780 | 1042 |
| 3024 | QGN16121.1 | FIM1_2822 | CP015057. | 748 | 231 | 251 | 1306 | 197 | 258 | 881 | 171 | 352 | 2112 | 260 | 584 | 1828 | 261 | 504 | 1923 | 269 | 384 |
| 3025 | QGN16122.1 | FIM1_2823 | CP015057. | 1806 | 3266 | 1884 | 2823 | 2852 | 1647 | 2175 | 2466 | 2390 | 8682 | 2590 | 3449 | 10388 | 2763 | 2884 | 8826 | 2443 | 2111 |
| 3026 | QGN16123.1 | FIM1_2824 | CP015057. | 2461 | 594 | 593 | 2784 | 689 | 549 | 2554 | 656 | 831 | 4739 | 752 | 1026 | 4409 | 618 | 926 | 4465 | 668 | 661 |
| 3027 | QGN16124.1 | FIM1_2825 | CP015057. | 1054 | 590 | 723 | 1124 | 606 | 634 | 742 | 685 | 925 | 4585 | 582 | 1110 | 4678 | 500 | 1025 | 4341 | 499 | 737 |
| 3028 | QGN16125.1 | FIM1_2826 | CP015057. | 1857 | 456 | 904 | 1809 | 453 | 603 | 1027 | 456 | 1113 | 2115 | 305 | 835 | 3105 | 423 | 926 | 2115 | 266 | 601 |
| 3029 | QGN16126.1 | FIM1_2827 | CP015057. | 4054 | 2636 | 1454 | 2127 | 2202 | 1426 | 3410 | 1931 | 1908 | 11206 | 3030 | 2247 | 9547 | 2695 | 2337 | 10788 | 3023 | 1614 |
| 3030 | QGN16127.1 | FIM1_2828 | CP015057. | 2054 | 297 | 470 | 2589 | 377 | 456 | 2601 | 286 | 622 | 3056 | 245 | 501 | 2730 | 275 | 503 | 2893 | 249 | 331 |
| 3031 | QGN16128.1 | FIM1_2829 | CP015057. | 812 | 1176 | 1289 | 1316 | 1582 | 1290 | 1040 | 1252 | 1771 | 6027 | 1603 | 1740 | 7943 | 1508 | 1738 | 6331 | 1349 | 1223 |
| 3032 | QGN16129.1 | FIM1_2830 | CP015057. | 7414 | 1408 | 1288 | 7415 | 1855 | 932 | 8067 | 1468 | 1517 | 3877 | 1621 | 2065 | 5980 | 1298 | 1695 | 4207 | 1272 | 1214 |
| 3033 | QGN16130.1 | FIM1_2831 | CP015057. | 1704 | 179 | 179 | 3787 | 96 | 105 | 2903 | 217 | 268 | 152 | 46 | 99 | 311 | 35 | 66 | 221 | 53 | 49 |
| 3034 | QGN16131.1 | FIM1_2832 | CP015057. | 5068 | 595 | 790 | 4636 | 676 | 635 | 4986 | 609 | 973 | 3176 | 647 | 927 | 4892 | 710 | 910 | 3821 | 632 | 670 |
| 3035 | QGN16132.1 | FIM1_2833 | CP015057. | 3383 | 1444 | 734 | 1373 | 1539 | 822 | 2585 | 1222 | 987 | 5184 | 1566 | 1197 | 4949 | 1725 | 1197 | 4790 | 1741 | 910 |
| 3036 | QGN16133.1 | FIM1_2834 | CP015057. | 6830 | 396 | 714 | 5797 | 637 | 533 | 5921 | 803 | 1120 | 837 | 420 | 1311 | 4552 | 483 | 1051 | 1269 | 485 | 778 |
| 3037 | QGN16134.1 | FIM1_2835 | CP015057. | 1380 | 1205 | 658 | 1251 | 722 | 545 | 1183 | 839 | 923 | 544 | 551 | 941 | 2871 | 526 | 904 | 1086 | 586 | 628 |
| 3038 | QGN16135.1 | FIM1_2836 | CP015057. | 2443 | 498 | 548 | 1939 | 463 | 408 | 2438 | 490 | 619 | 1940 | 341 | 683 | 2574 | 374 | 598 | 1927 | 382 | 476 |
| 3039 | QGN16136.1 | FIM1_2837 | CP015057. | 1443 | 455 | 806 | 1803 | 292 | 547 | 1468 | 393 | 1054 | 308 | 293 | 1132 | 1423 | 346 | 1139 | 655 | 305 | 772 |
| 3040 | QGN16137.1 | FIM1_2838 | CP015057. | 5211 | 2232 | 1522 | 6188 | 2622 | 1294 | 4734 | 2442 | 1936 | 4742 | 1806 | 1936 | 8459 | 1782 | 1893 | 6039 | 1649 | 1226 |
| 3041 | QGN16138.1 | FIM1_2839 | CP015057. | 1650 | 337 | 423 | 2140 | 300 | 378 | 1794 | 336 | 555 | 3538 | 275 | 615 | 3132 | 270 | 615 | 3234 | 245 | 420 |
| 3042 | QGN16139.1 | FIM1_2840 | CP015057. | 228 | 47 | 64 | 324 | 69 | 71 | 235 | 54 | 92 | 507 | 51 | 97 | 369 | 71 | 101 | 391 | 81 | 61 |
| 3043 | QGN16140.1 | FIM1_2841 | CP015057. | 2296 | 901 | 556 | 2232 | 739 | 470 | 2701 | 704 | 728 | 2715 | 689 | 1013 | 3039 | 675 | 909 | 2853 | 750 | 634 |
| 3044 | QGN16141.1 | FIM1_2842 | CP015057. | 1541 | 2899 | 1907 | 1498 | 2709 | 1751 | 1455 | 2550 | 2410 | 7448 | 2695 | 2697 | 9891 | 2532 | 2477 | 8772 | 2448 | 1732 |
| 3045 | QGN16142.1 | FIM1_2843 | CP015057. | 321 | 76 | 141 | 541 | 112 | 125 | 407 | 105 | 169 | 1132 | 84 | 269 | 770 | 86 | 206 | 1015 | 104 | 169 |
| 3046 | QGN16143.1 | FIM1_2844 | CP015057. | 1423 | 448 | 385 | 1101 | 304 | 315 | 728 | 308 | 496 | 1782 | 241 | 527 | 1572 | 214 | 526 | 1796 | 228 | 357 |
| 3047 | QGN16144.1 | FIM1_2845 | CP015057. | 118028 | 27866 | 21663 | 114553 | 22277 | 18874 | 111899 | 21808 | 26170 | 60103 | 24887 | 26539 | 71048 | 32220 | 28692 | 61583 | 27068 | 20040 |
| 3048 | QGN16145.1 | FIM1_2846 | CP015057. | 3921 | 1042 | 417 | 1101 | 1041 | 420 | 2909 | 899 | 595 | 1684 | 860 | 650 | 2288 | 880 | 609 | 1866 | 903 | 519 |
| 3049 | QGN16146.1 | FIM1_2847 | CP015057. | 28224 | 695 | 986 | 13936 | 990 | 859 | 10086 | 1228 | 1302 | 7053 | 1003 | 1710 | 10205 | 257 | 1456 | 7553 | 520 | 1153 |
| 3050 | QGN16147.1 | FIM1_2848 | CP015057. | 2107 | 720 | 484 | 2107 | 452 | 390 | 2315 | 485 | 654 | 2282 | 308 | 494 | 1768 | 269 | 436 | 2142 | 310 | 291 |
| 3051 | QGN16148.1 | FIM1_2849 | CP015057. | 12549 | 15083 | 8806 | 12099 | 11508 | 8850 | 10425 | 10094 | 10503 | 27433 | 12561 | 12365 | 26746 | 11214 | 12201 | 27451 | 11577 | 9101 |
| 3052 | QGN16149.1 | FIM1_2850 | CP015057. | 1884 | 2122 | 1353 | 1425 | 1768 | 1268 | 1656 | 1546 | 1752 | 7136 | 1531 | 1622 | 6029 | 1421 | 1688 | 6363 | 1571 | 1104 |
| 3053 | QGN16150.1 | FIM1_2851 | CP015057. | 4052 | 1035 | 1637 | 5306 | 1517 | 1247 | 4331 | 1604 | 2153 | 4535 | 1267 | 2138 | 9862 | 1366 | 2013 | 7420 | 1117 | 1491 |
| 3054 | QGN16151.1 | FIM1_2852 | CP015057. | 1210 | 762 | 874 | 1481 | 571 | 721 | 1174 | 508 | 994 | 3233 | 413 | 862 | 2359 | 572 | 869 | 2794 | 494 | 519 |
| 3055 | QGN16152.1 | FIM1_2853 | CP015057. | 5597 | 3091 | 1064 | 2848 | 3093 | 1211 | 4775 | 2232 | 1413 | 11983 | 4140 | 2470 | 11575 | 3484 | 2381 | 11646 | 4418 | 1704 |
| 3056 | QGN16153.1 | FIM1_2854 | CP015057. | 1020 | 1819 | 1223 | 2103 | 1484 | 1084 | 1352 | 1219 | 1648 | 6061 | 1158 | 1332 | 4695 | 1121 | 1320 | 5954 | 1039 | 948 |
| 3057 | QGN16154.1 | FIM1_2855 | CP015057. | 197 | 169 | 101 | 215 | 74 | 70 | 156 | 69 | 116 | 349 | 89 | 130 | 367 | 59 | 143 | 339 | 60 | 79 |
| 3058 | QGN16155.1 | FIM1_2856 | CP015057. | 1440 | 161 | 439 | 2044 | 310 | 329 | 1434 | 343 | 559 | 2484 | 259 | 564 | 2669 | 266 | 535 | 2450 | 234 | 359 |
| 3059 | QGN16156.1 | FIM1_2857 | CP015057. | 813 | 754 | 510 | 587 | 578 | 416 | 643 | 540 | 620 | 2170 | 484 | 578 | 1887 | 503 | 578 | 2177 | 555 | 363 |
| 3060 | QGN16157.1 | FIM1_2858 | CP015057. | 1220 | 795 | 749 | 1226 | 715 | 599 | 796 | 663 | 986 | 2040 | 662 | 717 | 3053 | 675 | 768 | 2272 | 643 | 516 |
| 3061 | QGN16158.1 | FIM1_2859 | CP015057. | 1752 | 3335 | 2828 | 2145 | 2074 | 1943 | 1879 | 2244 | 3635 | 14672 | 3262 | 3727 | 12427 | 3746 | 3853 | 15085 | 3093 | 2552 |
| 3062 | QGN16159.1 | FIM1_2860 | CP015057. | 8884 | 11293 | 9245 | 3471 | 11941 | 7836 | 2878 | 13738 | 12694 | 19053 | 11825 | 7475 | 23719 | 10663 | 5132 | 24181 | 11146 | 5777 |
| 3063 | QGN16160.1 | FIM1_2862 | CP015057. | 133 | 178 | 166 | 173 | 137 | 130 | 150 | 136 | 184 | 875 | 161 | 193 | 506 | 122 | 213 | 813 | 123 | 128 |
| 3064 | QGN16161.1 | FIM1_2863 | CP015057. | 4410 | 399 | 638 | 3666 | 654 | 576 | 4807 | 496 | 883 | 1121 | 507 | 933 | 3011 | 552 | 822 | 1898 | 563 | 608 |
| 3065 | QGN16162.1 | FIM1_2864 | CP015057. | 1359 | 2686 | 1607 | 1231 | 2786 | 1574 | 1336 | 2626 | 1973 | 7246 | 2570 | 2621 | 8236 | 2657 | 2152 | 6999 | 2970 | 1673 |
| 3066 | QGN16163.1 | FIM1_2865 | CP015057. | 509 | 679 | 583 | 397 | 842 | 591 | 479 | 717 | 751 | 4035 | 834 | 952 | 3728 | 730 | 824 | 3852 | 831 | 656 |
| 3067 | QGN16164.1 | FIM1_2866 | CP015057. | 1657 | 393 | 424 | 2060 | 380 | 363 | 1914 | 277 | 552 | 2728 | 255 | 655 | 2189 | 300 | 536 | 2809 | 263 | 364 |
| 3068 | QGN16165.1 | FIM1_2867 | CP015057. | 868 | 883 | 851 | 1561 | 895 | 824 | 1226 | 730 | 1117 | 6309 | 795 | 1306 | 5265 | 686 | 1155 | 5609 | 656 | 849 |
| 3069 | QGN16166.1 | FIM1_2868 | CP015057. | 150 | 29 | 22 | 329 | 47 | 36 | 141 | 26 | 24 | 146 | 24 | 38 | 162 | 26 | 27 | 140 | 20 | 20 |
| 3070 | QGN16167.1 | FIM1_2869 | CP015057. | 2362 | 40 | 90 | 2208 | 40 | 99 | 2071 | 49 | 175 | 640 | 24 | 113 | 646 | 23 | 84 | 571 | 15 | 68 |
| 3071 | QGN16168.1 | FIM1_2870 | CP015057. | 1211 | 2924 | 2815 | 3136 | 3387 | 2394 | 2205 | 2937 | 3476 | 15960 | 3278 | 4652 | 16879 | 3349 | 3946 | 16421 | 3177 | 2896 |
| 3072 | QGN16169.1 | FIM1_2871 | CP015057. | 1247 | 688 | 582 | 1439 | 595 | 512 | 1396 | 548 | 705 | 5184 | 613 | 911 | 3991 | 554 | 815 | 4973 | 582 | 573 |
| 3073 | QGN16170.1 | FIM1_2872 | CP015057. | 18649 | 690 | 909 | 23075 | 961 | 787 | 27380 | 870 | 1216 | 4918 | 862 | 1113 | 7112 | 670 | 993 | 5285 | 655 | 783 |
| 3074 | QGN16171.1 | FIM1_2873 | CP015057. | 846 | 327 | 274 | 876 | 294 | 198 | 881 | 276 | 315 | 256 | 166 | 251 | 667 | 188 | 221 | 438 | 163 | 156 |
| 3075 | QGN16172.1 | FIM1_2874 | CP015057. | 12380 | 4396 | 4467 | 16381 | 5325 | 3663 | 12158 | 4746 | 5487 | 5667 | 2413 | 3352 | 13054 | 2855 | 3231 | 8199 | 2310 | 2413 |
| 3076 | QGN16173.1 | FIM1_2875 | CP015057. | 5922 | 1934 | 1196 | 5726 | 1917 | 990 | 5452 | 1565 | 1459 | 4887 | 1712 | 1710 | 6028 | 1916 | 1666 | 5273 | 1730 | 1141 |
| 3077 | QGN16174.1 | FIM1_2876 | CP015057. | 3176 | 1818 | 1275 | 3588 | 1394 | 985 | 3091 | 1533 | 1716 | 9039 | 1697 | 1718 | 7902 | 1574 | 2008 | 7913 | 1405 | 1198 |
| 3078 | QGN16175.1 | FIM1_2877 | CP015057. | 2628 | 2662 | 1255 | 1417 | 3165 | 1331 | 2221 | 2473 | 1620 | 8914 | 3378 | 2296 | 8699 | 3297 | 2164 | 8929 | 3787 | 1656 |
| 3079 | QGN16176.1 | FIM1_2878 | CP015057. | 1388 | 765 | 749 | 2109 | 888 | 623 | 1775 | 716 | 1003 | 7084 | 648 | 1180 | 5286 | 676 | 948 | 6653 | 557 | 769 |
| 3080 | QGN16177.1 | FIM1_2879 | CP015057. | 4988 | 611 | 598 | 8162 | 895 | 471 | 5889 | 640 | 742 | 3991 | 808 | 954 | 5318 | 826 | 873 | 4542 | 693 | 610 |
| 3081 | QGN16178.1 | FIM1_2880 | CP015057. | 751 | 716 | 590 | 1563 | 763 | 561 | 1247 | 617 | 808 | 4185 | 679 | 984 | 3472 | 631 | 857 | 3837 | 577 | 592 |
| 3082 | QGN16179.1 | FIM1_2881 | CP015057. | 1382 | 1096 | 615 | 975 | 976 | 628 | 1151 | 750 | 857 | 4858 | 937 | 1158 | 3783 | 845 | 1078 | 4894 | 844 | 760 |
| 3083 | QGN16180.1 | FIM1_2882 | CP015057. | 3106 | 574 | 712 | 3844 | 854 | 608 | 3279 | 644 | 843 | 5905 | 698 | 1248 | 5844 | 879 | 1002 | 6105 | 787 | 856 |
| 3084 | QGN16181.1 | FIM1_2883 | CP015057. | 2929 | 1225 | 1498 | 4303 | 1268 | 1282 | 3530 | 1162 | 1997 | 10665 | 1239 | 2221 | 9065 | 1226 | 1814 | 9781 | 1164 | 1397 |
| 3085 | QGN16182.1 | FIM1_2884 | CP015057. | 1581 | 339 | 324 | 1588 | 388 | 323 | 1483 | 325 | 497 | 2350 | 319 | 643 | 2762 | 410 | 494 | 2568 | 332 | 375 |
| 3086 | QGN16183.1 | FIM1_2885 | CP015057. | 2330 | 1177 | 555 | 1060 | 1277 | 657 | 2236 | 952 | 684 | 4000 | 1462 | 1163 | 3841 | 1611 | 1040 | 3945 | 1612 | 848 |
| 3087 | QGN16184.1 | FIM1_2886 | CP015057. | 1490 | 693 | 221 | 679 | 630 | 273 | 1472 | 418 | 331 | 3296 | 836 | 571 | 2142 | 825 | 527 | 3323 | 992 | 381 |
| 3088 | QGN16185.1 | FIM1_2887 | CP015057. | 5804 | 3550 | 1628 | 2786 | 3663 | 1726 | 4985 | 2818 | 2187 | 12374 | 4429 | 3042 | 10451 | 3924 | 2939 | 11968 | 4742 | 2173 |
| 3089 | QGN16186.1 | FIM1_2888 | CP015057. | 951 | 193 | 385 | 1088 | 180 | 275 | 692 | 189 | 424 | 2016 | 106 | 482 | 1537 | 84 | 404 | 1944 | 79 | 244 |
| 3090 | QGN16187.1 | FIM1_2889 | CP015057. | 2368 | 1269 | 1235 | 2270 | 1467 | 1095 | 2104 | 1306 | 1587 | 7478 | 1399 | 1911 | 7921 | 1077 | 1833 | 7137 | 1114 | 1273 |
| 3091 | QGN16188.1 | FIM1_2890 | CP015057. | 4122 | 3353 | 3222 | 6194 | 2843 | 2756 | 4724 | 2520 | 3888 | 13554 | 2761 | 4617 | 12621 | 2995 | 4757 | 13120 | 2435 | 3121 |
| 3092 | QGN16189.1 | FIM1_2891 | CP015057. | 6735 | 4921 | 2707 | 6292 | 4285 | 2328 | 5741 | 3946 | 3293 | 12063 | 2995 | 2891 | 13314 | 3270 | 2814 | 11743 | 2833 | 1925 |
| 3093 | QGN16190.1 | FIM1_2892 | CP015057. | 1080 | 248 | 209 | 1405 | 186 | 183 | 768 | 170 | 277 | 775 | 113 | 187 | 877 | 103 | 205 | 768 | 104 | 117 |
| 3094 | QGN16191.1 | FIM1_2893 | CP015057. | 2464 | 591 | 469 | 1900 | 548 | 400 | 2031 | 559 | 598 | 1040 | 273 | 376 | 1394 | 189 | 383 | 1083 | 229 | 216 |
| 3095 | QGN16192.1 | FIM1_2894 | CP015057. | 4006 | 462 | 482 | 4178 | 535 | 401 | 3564 | 476 | 653 | 2172 | 464 | 758 | 3440 | 419 | 643 | 2698 | 389 | 501 |
| 3096 | QGN16193.1 | FIM1_2895 | CP015057. | 3759 | 2204 | 2075 | 3536 | 1928 | 1925 | 3434 | 1660 | 2649 | 12704 | 1917 | 3178 | 10037 | 1880 | 2866 | 12154 | 1699 | 1957 |
| 3097 | QGN16194.1 | FIM1_2896 | CP015057. | 2677 | 825 | 768 | 3323 | 982 | 658 | 2941 | 896 | 1012 | 4284 | 768 | 1113 | 4120 | 813 | 994 | 4557 | 782 | 676 |
| 3098 | QGN16195.1 | FIM1_2897 | CP015057. | 11947 | 2914 | 3372 | 20762 | 3824 | 2743 | 14933 | 2859 | 4287 | 7705 | 2784 | 4317 | 17355 | 3643 | 3583 | 11553 | 3299 | 2679 |
| 3099 | QGN16196.1 | FIM1_2898 | CP015057. | 13857 | 16473 | 9289 | 14050 | 12281 | 8577 | 11606 | 10662 | 10845 | 31799 | 13238 | 14334 | 31289 | 11902 | 14019 | 32677 | 12032 | 10859 |
| 3100 | QGN16197.1 | FIM1_2899 | CP015057. | 11940 | 6463 | 2435 | 10100 | 7204 | 2693 | 13673 | 5942 | 3270 | 15529 | 9836 | 5364 | 23181 | 7571 | 5226 | 16074 | 9806 | 3864 |
| 3101 | QGN16198.1 | FIM1_2900 | CP015057. | 21320 | 2376 | 1933 | 26406 | 4110 | 1809 | 26239 | 3233 | 2477 | 10065 | 4189 | 2540 | 16988 | 5037 | 2276 | 13203 | 4668 | 1768 |
| 3102 | QGN16199.1 | FIM1_2901 | CP015057. | 3790 | 1152 | 974 | 2900 | 1067 | 756 | 2862 | 1016 | 1223 | 4015 | 1021 | 1512 | 5119 | 1056 | 1446 | 4652 | 1025 | 952 |
| 3103 | QGN16200.1 | FIM1_2903 | CP015057. | 5987 | 1584 | 1433 | 8440 | 1461 | 1211 | 6296 | 1332 | 1948 | 4931 | 977 | 1878 | 7255 | 1260 | 1856 | 5742 | 928 | 1265 |
| 3104 | QGN16201.1 | FIM1_2904 | CP015057. | 7332 | 1512 | 985 | 6247 | 1179 | 780 | 5347 | 1213 | 1355 | 1765 | 842 | 1166 | 4468 | 876 | 1135 | 2747 | 749 | 856 |
| 3105 | QGN16202.1 | FIM1_2905 | CP015057. | 1756 | 317 | 464 | 2127 | 571 | 310 | 1982 | 738 | 599 | 462 | 375 | 597 | 2584 | 453 | 626 | 1036 | 416 | 398 |
| 3106 | QGN16203.1 | FIM1_2906 | CP015057. | 519 | 88 | 168 | 595 | 110 | 109 | 466 | 98 | 139 | 857 | 96 | 125 | 654 | 61 | 118 | 792 | 66 | 128 |

| | | | | | | | | | | | | | | | | | | | | | |
|---|---|---|---|---|---|---|---|---|---|---|---|---|---|---|---|---|---|---|---|---|---|
| 3107 | QGN16204.1 | FIM1_2907 | CP015057. | 746 | 1055 | 795 | 416 | 919 | 694 | 519 | 899 | 1053 | 2391 | 682 | 821 | 2838 | 486 | 785 | 2364 | 590 | 550 |
| 3108 | QGN16205.1 | FIM1_2908 | CP015057. | 2243 | 1389 | 1628 | 2118 | 1667 | 1292 | 2213 | 1627 | 2043 | 7971 | 1554 | 2103 | 9254 | 1488 | 1998 | 8290 | 1448 | 1383 |
| 3109 | QGN16206.1 | FIM1_2909 | CP015057. | 2357 | 696 | 652 | 2153 | 895 | 581 | 2334 | 760 | 813 | 4168 | 837 | 1008 | 4113 | 817 | 932 | 4047 | 766 | 644 |
| 3110 | QGN16207.1 | FIM1_2911 | CP015057. | 194 | 277 | 401 | 799 | 208 | 334 | 471 | 227 | 579 | 2378 | 124 | 432 | 1680 | 151 | 393 | 2301 | 152 | 282 |
| 3111 | QGN16208.1 | FIM1_2912 | CP015057. | 14734 | 622 | 1426 | 9942 | 1333 | 1049 | 11772 | 1506 | 1812 | 2459 | 1359 | 1745 | 8001 | 1376 | 1622 | 4465 | 1356 | 1185 |
| 3112 | QGN16209.1 | FIM1_2913 | CP015057. | 31 | 2 | 10 | 35 | 8 | 5 | 42 | 6 | 12 | 21 | 8 | 7 | 43 | 9 | 4 | 24 | 7 | 7 |
| 3113 | QGN16210.1 | FIM1_2914 | CP015057. | 3197 | 1910 | 886 | 1232 | 1267 | 790 | 2387 | 1529 | 1176 | 963 | 1203 | 1388 | 3039 | 1261 | 1320 | 1525 | 1598 | 1000 |
| 3114 | QGN16211.1 | FIM1_2915 | CP015057. | 815 | 1406 | 2145 | 1633 | 941 | 1998 | 1069 | 845 | 2858 | 2510 | 436 | 2101 | 2359 | 479 | 2071 | 2467 | 449 | 1438 |
| 3115 | QGN16212.1 | FIM1_2916 | CP015057. | 6938 | 3352 | 4832 | 6663 | 3289 | 3871 | 5967 | 3191 | 6190 | 5966 | 2830 | 5722 | 13045 | 2122 | 5414 | 9663 | 2349 | 3817 |
| 3116 | QGN16213.1 | FIM1_2917 | CP015057. | 3084 | 1072 | 1061 | 4557 | 1044 | 844 | 3499 | 978 | 1293 | 5053 | 563 | 1107 | 4860 | 508 | 1062 | 4963 | 437 | 701 |
| 3117 | QGN16214.1 | FIM1_2918 | CP015057. | 2597 | 994 | 554 | 2807 | 809 | 467 | 2756 | 771 | 662 | 2867 | 542 | 668 | 2220 | 504 | 558 | 2977 | 474 | 375 |
| 3118 | QGN16215.1 | FIM1_2919 | CP015057. | 4448 | 1338 | 1672 | 7297 | 1649 | 1465 | 6467 | 1679 | 2372 | 10114 | 1794 | 2955 | 11747 | 1757 | 2690 | 10175 | 1592 | 1983 |
| 3119 | QGN16216.1 | FIM1_2920 | CP015057. | 627 | 303 | 209 | 372 | 218 | 175 | 508 | 205 | 275 | 1311 | 256 | 236 | 904 | 200 | 224 | 1171 | 166 | 165 |
| 3120 | QGN16217.1 | FIM1_2921 | CP015057. | 3001 | 1984 | 1131 | 1269 | 2537 | 1244 | 3000 | 2118 | 1540 | 8096 | 2627 | 2139 | 8494 | 2060 | 2076 | 7881 | 2487 | 1511 |
| 3121 | QGN16218.1 | FIM1_2922 | CP015057. | 518 | 400 | 721 | 1313 | 387 | 575 | 929 | 402 | 865 | 5375 | 355 | 1026 | 4821 | 346 | 944 | 5101 | 277 | 669 |
| 3122 | QGN16219.1 | FIM1_2923 | CP015057. | 2699 | 2612 | 2329 | 4193 | 2603 | 1702 | 4130 | 2305 | 2700 | 12137 | 3327 | 3335 | 11175 | 3268 | 3456 | 13349 | 2629 | 2538 |
| 3123 | QGN16220.1 | FIM1_2924 | CP015057. | 3268 | 4811 | 3659 | 5876 | 4722 | 3272 | 5114 | 4076 | 4419 | 18725 | 5792 | 5573 | 16456 | 5308 | 5576 | 19215 | 4989 | 4236 |
| 3124 | QGN16221.1 | FIM1_2925 | CP015057. | 65 | 33 | 25 | 326 | 46 | 41 | 98 | 37 | 34 | 161 | 29 | 34 | 160 | 26 | 30 | 152 | 35 | 25 |
| 3125 | QGN16222.1 | FIM1_2926 | CP015057. | 944 | 1948 | 1511 | 2790 | 1759 | 1230 | 1577 | 1572 | 1947 | 6612 | 1404 | 2052 | 7699 | 1677 | 2130 | 6746 | 1332 | 1405 |
| 3126 | QGN16223.1 | FIM1_2927 | CP015057. | 6599 | 4556 | 3747 | 8173 | 3865 | 3246 | 7435 | 3309 | 4988 | 12529 | 3411 | 5585 | 13508 | 3347 | 5332 | 12966 | 3334 | 3712 |
| 3127 | QGN16224.1 | FIM1_2928 | CP015057. | 7831 | 528 | 575 | 10046 | 574 | 410 | 7507 | 621 | 743 | 629 | 267 | 547 | 2022 | 457 | 435 | 1093 | 311 | 382 |
| 3128 | QGN16225.1 | FIM1_2929 | CP015057. | 4678 | 528 | 1098 | 5397 | 801 | 768 | 3688 | 863 | 1336 | 3014 | 622 | 1541 | 7367 | 605 | 1357 | 4544 | 603 | 1006 |
| 3129 | QGN16226.1 | FIM1_2930 | CP015057. | 603 | 206 | 160 | 209 | 260 | 137 | 495 | 170 | 202 | 894 | 213 | 232 | 762 | 298 | 213 | 769 | 227 | 167 |
| 3130 | QGN16227.1 | FIM1_2931 | CP015057. | 2821 | 2911 | 1471 | 2557 | 2429 | 1490 | 2833 | 1943 | 1928 | 9665 | 2657 | 2411 | 7852 | 2175 | 2313 | 8912 | 2472 | 1695 |
| 3131 | QGN16228.1 | FIM1_2932 | CP015057. | 1057 | 734 | 686 | 1475 | 502 | 568 | 1176 | 556 | 904 | 3651 | 444 | 912 | 2687 | 561 | 858 | 3472 | 465 | 580 |
| 3132 | QGN16229.1 | FIM1_2933 | CP015057. | 4292 | 1440 | 1595 | 5353 | 1633 | 1332 | 5022 | 1358 | 2004 | 6807 | 1558 | 2429 | 10533 | 1402 | 2331 | 8181 | 1304 | 1714 |
| 3133 | QGN16230.1 | FIM1_2934 | CP015057. | 6116 | 2038 | 1250 | 3758 | 2681 | 1304 | 5267 | 1999 | 1619 | 8208 | 2836 | 2213 | 9022 | 2777 | 2120 | 8045 | 2957 | 1600 |
| 3134 | QGN16231.1 | FIM1_2935 | CP015057. | 5135 | 8745 | 7856 | 12642 | 10498 | 6414 | 9471 | 9188 | 9257 | 30975 | 13509 | 13671 | 39838 | 13037 | 13865 | 33219 | 11429 | 9836 |
| 3135 | QGN16232.1 | FIM1_2936 | CP015057. | 9336 | 1442 | 2051 | 10694 | 3087 | 1918 | 9088 | 2654 | 2606 | 9286 | 3182 | 3161 | 14851 | 3596 | 2910 | 11605 | 3685 | 2147 |
| 3136 | QGN16233.1 | FIM1_2937 | CP015057. | 862 | 920 | 758 | 1276 | 1030 | 575 | 1066 | 862 | 869 | 999 | 522 | 790 | 2716 | 614 | 689 | 1680 | 501 | 503 |
| 3137 | QGN16234.1 | FIM1_2939 | CP015057. | 1068 | 301 | 494 | 2139 | 348 | 384 | 1101 | 341 | 693 | 2156 | 182 | 542 | 2227 | 162 | 467 | 2175 | 147 | 304 |
| 3138 | QGN16235.1 | FIM1_2940 | CP015057. | 17 | 6 | 4 | 12 | 5 | 2 | 9 | 4 | 5 | 33 | 9 | 11 | 28 | 9 | 11 | 38 | 6 | 4 |
| 3139 | QGN16236.1 | FIM1_2941 | CP015057. | 894 | 1345 | 993 | 1556 | 1212 | 943 | 1140 | 1090 | 1312 | 5651 | 1075 | 1334 | 4482 | 985 | 1296 | 5245 | 1008 | 852 |
| 3140 | QGN16237.1 | FIM1_2942 | CP015057. | 5157 | 4118 | 7893 | 9786 | 5405 | 6786 | 8186 | 4944 | 9614 | 13973 | 3443 | 9987 | 17821 | 3485 | 9116 | 14734 | 3254 | 6766 |
| 3141 | QGN16238.1 | FIM1_2943 | CP015057. | 1041 | 1001 | 777 | 1514 | 827 | 673 | 1416 | 829 | 1040 | 5177 | 668 | 1126 | 4314 | 635 | 1056 | 4967 | 572 | 719 |
| 3142 | QGN16239.1 | FIM1_2944 | CP015057. | 1251 | 719 | 440 | 1114 | 649 | 392 | 1330 | 633 | 618 | 4011 | 681 | 837 | 3071 | 514 | 674 | 4134 | 604 | 489 |
| 3143 | QGN16240.1 | FIM1_2945 | CP015057. | 1046 | 606 | 408 | 813 | 619 | 419 | 911 | 539 | 479 | 3118 | 614 | 669 | 2157 | 452 | 664 | 2905 | 587 | 448 |
| 3144 | QGN16241.1 | FIM1_2946 | CP015057. | 4133 | 816 | 1264 | 5375 | 1063 | 1036 | 4303 | 885 | 1576 | 5975 | 947 | 1978 | 7746 | 789 | 1708 | 6573 | 817 | 1351 |
| 3145 | QGN16242.1 | FIM1_2947 | CP015057. | 2666 | 3127 | 2084 | 3878 | 2260 | 1722 | 2954 | 2192 | 2741 | 13601 | 2478 | 3472 | 11430 | 2937 | 3440 | 13466 | 2535 | 2270 |
| 3146 | QGN16243.1 | FIM1_2948 | CP015057. | 1891 | 464 | 859 | 1869 | 636 | 634 | 1889 | 596 | 1134 | 4509 | 589 | 1335 | 5709 | 493 | 1074 | 4919 | 485 | 840 |
| 3147 | QGN16244.1 | FIM1_2949 | CP015057. | 3170 | 3071 | 2601 | 6283 | 2198 | 2303 | 3981 | 2096 | 3382 | 8947 | 1664 | 3355 | 9433 | 1753 | 3355 | 8831 | 1518 | 2216 |
| 3148 | QGN16245.1 | FIM1_2950 | CP015057. | 246 | 31 | 47 | 470 | 45 | 42 | 289 | 41 | 63 | 163 | 30 | 54 | 233 | 33 | 35 | 190 | 35 | 35 |
| 3149 | QGN16246.1 | FIM1_2951 | CP015057. | 3926 | 1261 | 1540 | 4142 | 1470 | 1178 | 3340 | 1427 | 1957 | 7806 | 2129 | 2315 | 13318 | 2733 | 2203 | 9118 | 1890 | 1549 |
| 3150 | QGN16247.1 | FIM1_2952 | CP015057. | 2763 | 527 | 639 | 3022 | 495 | 535 | 2555 | 464 | 847 | 4034 | 368 | 943 | 3529 | 456 | 839 | 3982 | 345 | 563 |
| 3151 | QGN16248.1 | FIM1_2953 | CP015057. | 1427 | 805 | 1136 | 3421 | 1151 | 945 | 2439 | 1044 | 1420 | 4185 | 885 | 1760 | 6072 | 1089 | 1554 | 4592 | 949 | 1141 |
| 3152 | QGN16249.1 | FIM1_2954 | CP015057. | 1152 | 395 | 372 | 1309 | 425 | 295 | 1180 | 346 | 464 | 2523 | 414 | 594 | 2496 | 358 | 608 | 2423 | 353 | 444 |
| 3153 | QGN16250.1 | FIM1_2955 | CP015057. | 531 | 952 | 642 | 615 | 767 | 669 | 540 | 638 | 900 | 4933 | 847 | 1179 | 3420 | 777 | 1027 | 4640 | 936 | 711 |
| 3154 | QGN16251.1 | FIM1_2956 | CP015057. | 352 | 173 | 207 | 460 | 297 | 160 | 327 | 233 | 209 | 1583 | 263 | 353 | 1389 | 196 | 338 | 1410 | 234 | 213 |
| 3155 | QGN16252.1 | FIM1_2957 | CP015057. | 6027 | 2341 | 1234 | 2952 | 2911 | 1248 | 5037 | 2426 | 1592 | 10404 | 2996 | 2689 | 11010 | 3144 | 2401 | 10896 | 3984 | 1816 |
| 3156 | QGN16253.1 | FIM1_2958 | CP015057. | 6689 | 993 | 892 | 5880 | 1129 | 783 | 6770 | 831 | 1163 | 9646 | 1206 | 1894 | 8048 | 1157 | 1708 | 9878 | 1107 | 1272 |
| 3157 | QGN16254.1 | FIM1_2959 | CP015057. | 4688 | 4085 | 1264 | 2936 | 3538 | 1600 | 4023 | 2605 | 1630 | 13126 | 4661 | 3111 | 10161 | 3264 | 3060 | 12241 | 4181 | 2274 |
| 3158 | QGN16255.1 | FIM1_2960 | CP015057. | 5795 | 2456 | 1036 | 3720 | 2439 | 1152 | 5328 | 1841 | 1461 | 9282 | 2677 | 2063 | 9052 | 2751 | 1759 | 9425 | 2959 | 1518 |
| 3159 | QGN16256.1 | FIM1_2961 | CP015057. | 6038 | 2201 | 1511 | 4068 | 3022 | 1559 | 5790 | 2477 | 1998 | 12946 | 3620 | 3240 | 13089 | 3332 | 2816 | 12783 | 3853 | 2168 |
| 3160 | QGN16257.1 | FIM1_2962 | CP015057. | 1554 | 238 | 219 | 2010 | 211 | 185 | 1652 | 185 | 260 | 761 | 134 | 227 | 1131 | 140 | 253 | 964 | 133 | 162 |
| 3161 | QGN16258.1 | FIM1_2963 | CP015057. | 645 | 261 | 276 | 661 | 274 | 237 | 506 | 268 | 382 | 1027 | 206 | 352 | 1479 | 194 | 302 | 1176 | 180 | 225 |
| 3162 | QGN16259.1 | FIM1_2964 | CP015057. | 4162 | 7659 | 4410 | 5753 | 5266 | 4079 | 4476 | 4474 | 5549 | 18175 | 5923 | 6843 | 15574 | 5614 | 6831 | 17215 | 5711 | 4691 |
| 3163 | QGN16260.1 | FIM1_2965 | CP015057. | 1236 | 1432 | 959 | 1870 | 1059 | 840 | 1677 | 962 | 1247 | 4939 | 877 | 1191 | 4250 | 857 | 1230 | 4781 | 755 | 761 |
| 3164 | QGN16261.1 | FIM1_2966 | CP015057. | 142 | 356 | 217 | 232 | 292 | 239 | 180 | 239 | 331 | 1862 | 215 | 281 | 1235 | 224 | 289 | 1771 | 238 | 176 |
| 3165 | QGN16262.1 | FIM1_2967 | CP015057. | 1786 | 1148 | 1143 | 2413 | 1272 | 970 | 2283 | 1155 | 1443 | 4108 | 891 | 1420 | 5397 | 882 | 1364 | 4032 | 794 | 923 |
| 3166 | QGN16263.1 | FIM1_2968 | CP015057. | 2246 | 1231 | 1515 | 2533 | 1166 | 1383 | 2099 | 1233 | 2016 | 4681 | 935 | 2161 | 7235 | 821 | 2108 | 5137 | 738 | 1490 |
| 3167 | QGN16264.1 | FIM1_2969 | CP015057. | 2122 | 4883 | 3467 | 4062 | 4026 | 2986 | 3296 | 3571 | 4009 | 16532 | 4920 | 5538 | 13626 | 4417 | 5489 | 17270 | 4377 | 4055 |
| 3168 | QGN16265.1 | FIM1_2970 | CP015057. | 1516 | 543 | 437 | 1975 | 549 | 436 | 1543 | 617 | 528 | 2032 | 461 | 475 | 2318 | 359 | 539 | 2144 | 366 | 321 |
| 3169 | QGN16266.1 | FIM1_2971 | CP015057. | 479 | 717 | 273 | 400 | 598 | 277 | 454 | 503 | 393 | 1884 | 567 | 449 | 1608 | 442 | 537 | 1869 | 579 | 281 |
| 3170 | QGN16267.1 | FIM1_2972 | CP015057. | 779 | 626 | 414 | 1494 | 506 | 336 | 1347 | 543 | 539 | 2204 | 315 | 557 | 2784 | 314 | 460 | 2137 | 283 | 348 |
| 3171 | QGN16268.1 | FIM1_2973 | CP015057. | 8932 | 147 | 166 | 5569 | 95 | 161 | 7122 | 144 | 266 | 2656 | 72 | 150 | 2540 | 100 | 158 | 2188 | 83 | 108 |
| 3172 | QGN16269.1 | FIM1_2974 | CP015057. | 4084 | 777 | 1065 | 3999 | 1212 | 946 | 3874 | 1049 | 1467 | 3047 | 1083 | 1947 | 8169 | 978 | 1751 | 5387 | 1025 | 1323 |
| 3173 | QGN16270.1 | FIM1_2975 | CP015057. | 3060 | 2066 | 1124 | 2367 | 2275 | 1242 | 2981 | 1669 | 1551 | 7465 | 2190 | 1985 | 7307 | 2316 | 1889 | 7279 | 2497 | 1316 |
| 3174 | QGN16271.1 | FIM1_2976 | CP015057. | 679 | 500 | 363 | 1205 | 481 | 330 | 851 | 350 | 403 | 2001 | 447 | 545 | 1973 | 438 | 587 | 1845 | 405 | 406 |
| 3175 | QGN16272.1 | FIM1_2977 | CP015057. | 1478 | 416 | 717 | 1986 | 603 | 637 | 1328 | 583 | 983 | 2436 | 393 | 971 | 3060 | 388 | 835 | 2544 | 292 | 598 |
| 3176 | QGN16273.1 | FIM1_2978 | CP015057. | 1670 | 2823 | 1691 | 2832 | 1933 | 1559 | 2006 | 1695 | 2204 | 8382 | 2005 | 2641 | 6872 | 1935 | 2677 | 8121 | 2052 | 1693 |
| 3177 | QGN16274.1 | FIM1_2979 | CP015057. | 249 | 96 | 105 | 413 | 131 | 111 | 381 | 109 | 148 | 1277 | 126 | 212 | 910 | 123 | 154 | 1120 | 138 | 132 |
| 3178 | QGN16275.1 | FIM1_2980 | CP015057. | 1096 | 290 | 562 | 1964 | 466 | 512 | 1388 | 390 | 770 | 5761 | 414 | 951 | 4680 | 374 | 763 | 5367 | 341 | 592 |
| 3179 | QGN16276.1 | FIM1_2981 | CP015057. | 6925 | 2816 | 1746 | 4047 | 3965 | 1866 | 6331 | 3026 | 2173 | 14484 | 4835 | 3410 | 13527 | 4104 | 3234 | 13241 | 5007 | 2494 |
| 3180 | QGN16277.1 | FIM1_2982 | CP015057. | 4763 | 1950 | 1988 | 5729 | 2069 | 1676 | 5710 | 1680 | 2547 | 5596 | 1544 | 2724 | 8229 | 1888 | 2424 | 6501 | 1571 | 1813 |
| 3181 | QGN16278.1 | FIM1_2983 | CP015057. | 3288 | 5465 | 3339 | 1991 | 5155 | 3290 | 2682 | 4553 | 4375 | 16215 | 5247 | 5104 | 16530 | 4547 | 4950 | 15660 | 4946 | 3458 |
| 3182 | QGN16279.1 | FIM1_2984 | CP015057. | 482 | 500 | 252 | 251 | 567 | 310 | 509 | 394 | 375 | 1722 | 593 | 511 | 1408 | 518 | 515 | 1687 | 532 | 358 |
| 3183 | QGN16280.1 | FIM1_2985 | CP015057. | 4808 | 5931 | 11478 | 19341 | 12111 | 10503 | 15720 | 9994 | 14956 | 19838 | 9476 | 15351 | 57297 | 12060 | 13872 | 35887 | 9890 | 10504 |
| 3184 | QGN16281.1 | FIM1_2986 | CP015057. | 976 | 745 | 730 | 1221 | 529 | 649 | 1178 | 532 | 942 | 2994 | 458 | 831 | 3208 | 449 | 799 | 2856 | 407 | 588 |
| 3185 | QGN16282.1 | FIM1_2987 | CP015057. | 2271 | 1715 | 1098 | 3045 | 1745 | 1074 | 2519 | 1306 | 1397 | 5544 | 1198 | 1766 | 5768 | 1123 | 1607 | 5061 | 1103 | 1172 |
| 3186 | QGN16283.1 | FIM1_2988 | CP015057. | 1684 | 361 | 689 | 2801 | 495 | 634 | 2662 | 426 | 969 | 2244 | 373 | 360 | 2691 | 350 | 371 | 2464 | 338 | 267 |
| 3187 | QGN16284.1 | FIM1_2989 | CP015057. | 578 | 1216 | 605 | 616 | 803 | 605 | 599 | 765 | 855 | 2855 | 796 | 713 | 2210 | 695 | 767 | 2638 | 755 | 503 |
| 3188 | QGN16285.1 | FIM1_2990 | CP015057. | 57419 | 9489 | 5709 | 51342 | 9152 | 5272 | 59675 | 9070 | 7389 | 19804 | 6434 | 2762 | 22098 | 6573 | 2593 | 19053 | 5877 | 1852 |
| 3189 | QGN16286.1 | FIM1_2991 | CP015057. | 364 | 83 | 260 | 995 | 118 | 276 | 535 | 119 | 350 | 669 | 79 | 1420 | 757 | 69 | 1441 | 628 | 73 | 876 |
| 3190 | QGN16287.1 | FIM1_2992 | CP015057. | 251 | 45 | 41 | 271 | 38 | 38 | 142 | 54 | 50 | 168 | 66 | 156 | 289 | 48 | 133 | 203 | 44 | 66 |

| | | | | | | | | | | | | | | | | | | | | | |
|---|---|---|---|---|---|---|---|---|---|---|---|---|---|---|---|---|---|---|---|---|---|
| 3191 | QGN16288.1 | FIM1_2993 | CP015057. | 1346 | 635 | 672 | 1565 | 569 | 623 | 1464 | 584 | 928 | 4043 | 601 | 1186 | 3925 | 526 | 1034 | 3859 | 491 | 742 |
| 3192 | QGN16289.1 | FIM1_2994 | CP015057. | 2134 | 651 | 697 | 1610 | 612 | 594 | 1517 | 635 | 863 | 3537 | 500 | 1066 | 3409 | 468 | 964 | 3357 | 447 | 633 |
| 3193 | QGN16290.1 | FIM1_2995 | CP015057. | 620 | 171 | 223 | 491 | 110 | 168 | 319 | 161 | 316 | 319 | 96 | 288 | 774 | 66 | 317 | 490 | 70 | 206 |
| 3194 | QGN16291.1 | FIM1_2996 | CP015057. | 4478 | 173 | 429 | 5266 | 107 | 293 | 4069 | 213 | 524 | 925 | 83 | 513 | 1239 | 173 | 497 | 1032 | 85 | 296 |
| 3195 | QGN16292.1 | FIM1_2997 | CP015057. | 3345 | 1048 | 1064 | 3926 | 846 | 817 | 3047 | 815 | 1255 | 4074 | 831 | 1391 | 5151 | 811 | 1326 | 4396 | 715 | 943 |
| 3196 | QGN16293.1 | FIM1_2998 | CP015057. | 3408 | 2139 | 3658 | 6278 | 2106 | 2897 | 4537 | 2352 | 4349 | 7665 | 1568 | 4645 | 8731 | 1688 | 4155 | 7701 | 1489 | 3001 |
| 3197 | QGN16294.1 | FIM1_2999 | CP015057. | 813 | 401 | 561 | 1744 | 422 | 464 | 1296 | 418 | 682 | 1660 | 357 | 1029 | 2378 | 403 | 965 | 1738 | 376 | 667 |
| 3198 | QGN16295.1 | FIM1_3000 | CP015057. | 6378 | 844 | 673 | 5719 | 758 | 678 | 5607 | 660 | 812 | 4154 | 896 | 934 | 3658 | 1441 | 995 | 3801 | 1204 | 630 |
| 3199 | QGN16296.1 | FIM1_3001 | CP015057. | 945 | 380 | 743 | 1327 | 882 | 654 | 1254 | 717 | 853 | 3490 | 716 | 1051 | 4173 | 692 | 998 | 3754 | 726 | 701 |
| 3200 | QGN16297.1 | FIM1_3002 | CP015057. | 514 | 266 | 286 | 724 | 282 | 314 | 456 | 217 | 393 | 1401 | 217 | 481 | 1231 | 199 | 476 | 1363 | 193 | 336 |
| 3201 | QGN16298.1 | FIM1_3003 | CP015057. | 1210 | 555 | 615 | 635 | 662 | 513 | 951 | 536 | 751 | 3760 | 687 | 904 | 3486 | 579 | 722 | 3655 | 665 | 594 |
| 3202 | QGN16299.1 | FIM1_3004 | CP015057. | 6271 | 3097 | 3325 | 8455 | 3791 | 3062 | 7752 | 3121 | 4526 | 22073 | 3456 | 5306 | 19016 | 3271 | 4610 | 21698 | 2933 | 3538 |
| 3203 | QGN16300.1 | FIM1_3005 | CP015057. | 1216 | 1814 | 866 | 722 | 1572 | 1072 | 1047 | 1254 | 1190 | 6874 | 2139 | 1673 | 4608 | 1529 | 1617 | 6145 | 2362 | 1154 |
| 3204 | QGN16301.1 | FIM1_3006 | CP015057. | 4906 | 3292 | 1792 | 5052 | 3812 | 1946 | 6084 | 3076 | 2415 | 14196 | 4067 | 3586 | 15484 | 3164 | 3342 | 13734 | 3882 | 2418 |
| 3205 | QGN16302.1 | FIM1_3007 | CP015057. | 971 | 2696 | 2057 | 1550 | 3279 | 1979 | 1387 | 2768 | 2696 | 14414 | 2780 | 3112 | 13149 | 2659 | 2815 | 14230 | 2590 | 2085 |
| 3206 | QGN16303.1 | FIM1_3008 | CP015057. | 2902 | 1207 | 1244 | 3307 | 1358 | 1153 | 2951 | 1151 | 1690 | 9577 | 1418 | 2103 | 8176 | 1194 | 1990 | 9154 | 1155 | 1398 |
| 3207 | QGN16304.1 | FIM1_3009 | CP015057. | 2068 | 768 | 854 | 2184 | 943 | 847 | 2319 | 760 | 1255 | 4933 | 909 | 1655 | 6721 | 880 | 1525 | 5634 | 850 | 1156 |
| 3208 | QGN16305.1 | FIM1_3010 | CP015057. | 878 | 426 | 437 | 1001 | 399 | 401 | 897 | 375 | 576 | 2915 | 371 | 552 | 2281 | 354 | 552 | 2978 | 361 | 384 |
| 3209 | QGN16306.1 | FIM1_3011 | CP015057. | 1580 | 341 | 300 | 3627 | 388 | 224 | 2065 | 356 | 338 | 1205 | 303 | 426 | 1443 | 287 | 383 | 1112 | 268 | 253 |
| 3210 | QGN16307.1 | FIM1_3012 | CP015057. | 1782 | 1846 | 1343 | 2265 | 1735 | 1261 | 2212 | 1587 | 1764 | 8977 | 1951 | 2112 | 7930 | 1985 | 2020 | 8762 | 1989 | 1407 |
| 3211 | QGN16308.1 | FIM1_3013 | CP015057. | 3543 | 1040 | 1472 | 3942 | 1298 | 1178 | 3907 | 1092 | 1804 | 5710 | 1281 | 2390 | 8641 | 1193 | 2195 | 6761 | 1120 | 1713 |
| 3212 | QGN16309.1 | FIM1_3014 | CP015057. | 9197 | 3707 | 1765 | 4066 | 4485 | 1720 | 8706 | 3927 | 2262 | 10961 | 5821 | 3835 | 17140 | 4917 | 3449 | 12960 | 6699 | 2564 |
| 3213 | QGN16310.1 | FIM1_3015 | CP015057. | 2753 | 4019 | 3118 | 3771 | 3110 | 2648 | 3060 | 2661 | 3597 | 14435 | 3406 | 4918 | 11127 | 3322 | 4737 | 13943 | 3301 | 3379 |
| 3214 | QGN16311.1 | FIM1_3016 | CP015057. | 2373 | 327 | 418 | 1720 | 394 | 345 | 1988 | 348 | 494 | 3021 | 284 | 639 | 3111 | 275 | 566 | 2941 | 224 | 427 |
| 3215 | QGN16312.1 | FIM1_3017 | CP015057. | 22422 | 22703 | 11480 | 20572 | 16515 | 10688 | 17163 | 15486 | 13874 | 35071 | 16433 | 17766 | 39980 | 15403 | 17024 | 36521 | 15678 | 13135 |
| 3216 | QGN16313.1 | FIM1_3018 | CP015057. | 291 | 505 | 296 | 247 | 460 | 341 | 275 | 414 | 433 | 2161 | 417 | 464 | 1512 | 329 | 423 | 2088 | 391 | 305 |
| 3217 | QGN16314.1 | FIM1_3019 | CP015057. | 4228 | 3225 | 3838 | 6084 | 2770 | 3170 | 4782 | 2516 | 4855 | 7345 | 2727 | 5311 | 13027 | 2502 | 4872 | 9771 | 2337 | 3676 |
| 3218 | QGN16315.1 | FIM1_3020 | CP015057. | 211 | 92 | 31 | 218 | 88 | 57 | 251 | 80 | 78 | 292 | 33 | 33 | 230 | 26 | 33 | 277 | 32 | 17 |
| 3219 | QGN16316.1 | FIM1_3021 | CP015057. | 8528 | 2469 | 2206 | 7863 | 3270 | 2148 | 7499 | 2436 | 2822 | 8341 | 2363 | 3040 | 9988 | 2699 | 2844 | 8527 | 2554 | 1908 |
| 3220 | QGN16317.1 | FIM1_3022 | CP015057. | 552 | 30 | 117 | 480 | 204 | 151 | 474 | 151 | 179 | 914 | 136 | 183 | 896 | 166 | 150 | 757 | 194 | 111 |
| 3221 | QGN16318.1 | FIM1_3023 | CP015057. | 2573 | 490 | 772 | 2503 | 356 | 477 | 2546 | 586 | 973 | 1334 | 187 | 667 | 2518 | 245 | 582 | 1812 | 131 | 387 |
| 3222 | QGN16319.1 | FIM1_3024 | CP015057. | 706 | 26 | 13 | 308 | 23 | 11 | 165 | 21 | 16 | 293 | 50 | 72 | 386 | 12 | 14 | 525 | 35 | 33 |
| 3223 | QGN16320.1 | FIM1_3025 | CP015057. | 618 | 495 | 701 | 2230 | 444 | 538 | 1357 | 564 | 887 | 3361 | 437 | 805 | 3688 | 414 | 738 | 3358 | 394 | 557 |
| 3224 | QGN16321.1 | FIM1_3026 | CP015057. | 2727 | 1028 | 1470 | 4297 | 2000 | 1137 | 3646 | 1931 | 1858 | 7805 | 1824 | 2140 | 11491 | 1541 | 1781 | 8905 | 1285 | 1447 |
| 3225 | QGN16322.1 | FIM1_3027 | CP015057. | 16 | 3 | 0 | 31 | 8 | 1 | 11 | 4 | 5 | 69 | 5 | 3 | 33 | 4 | 5 | 59 | 6 | 4 |
| 3226 | QGN16323.1 | FIM1_3028 | CP015057. | 4292 | 2412 | 2275 | 6853 | 2580 | 2027 | 5951 | 2038 | 3036 | 9174 | 2810 | 3602 | 14786 | 2338 | 3432 | 11174 | 2509 | 2645 |
| 3227 | QGN16324.1 | FIM1_3029 | CP015057. | 1097 | 750 | 1067 | 1865 | 888 | 839 | 1182 | 781 | 1364 | 3917 | 808 | 1469 | 4624 | 822 | 1360 | 3901 | 718 | 953 |
| 3228 | QGN16325.1 | FIM1_3030 | CP015057. | 4835 | 2044 | 3506 | 5351 | 2397 | 3113 | 4622 | 2129 | 4469 | 8972 | 1859 | 4320 | 9671 | 1591 | 3731 | 8929 | 1560 | 2837 |
| 3229 | QGN16326.1 | FIM1_3031 | CP015057. | 4030 | 977 | 1442 | 4393 | 1247 | 1181 | 3654 | 1232 | 1732 | 4307 | 709 | 1420 | 4823 | 714 | 1300 | 4243 | 570 | 878 |
| 3230 | QGN16327.1 | FIM1_3032 | CP015057. | 5605 | 3689 | 1792 | 4282 | 2334 | 1544 | 4027 | 2241 | 2273 | 3835 | 2076 | 2635 | 6574 | 2011 | 2481 | 4682 | 1956 | 1917 |
| 3231 | QGN16328.1 | FIM1_3033 | CP015057. | 1639 | 1076 | 919 | 2155 | 936 | 847 | 2246 | 898 | 1231 | 2427 | 635 | 1247 | 3435 | 724 | 1371 | 2707 | 670 | 929 |
| 3232 | QGN16329.1 | FIM1_3034 | CP015057. | 9773 | 1063 | 2475 | 11488 | 1073 | 1739 | 9124 | 1527 | 3144 | 1151 | 644 | 3581 | 6123 | 699 | 3092 | 2672 | 625 | 2420 |
| 3233 | QGN16330.1 | FIM1_3035 | CP015057. | 4134 | 9943 | 8942 | 9948 | 9051 | 7236 | 6694 | 8427 | 10850 | 22431 | 7509 | 11607 | 24659 | 8580 | 10666 | 23276 | 7573 | 7980 |
| 3234 | QGN16331.1 | FIM1_3036 | CP015057. | 6510 | 2320 | 2195 | 9172 | 2663 | 1729 | 7103 | 2381 | 2793 | 5485 | 1606 | 2352 | 9804 | 2152 | 2383 | 6699 | 1531 | 1610 |
| 3235 | QGN16332.1 | FIM1_3037 | CP015057. | 157 | 661 | 1190 | 924 | 715 | 1003 | 166 | 656 | 1529 | 3139 | 573 | 1011 | 3636 | 421 | 1078 | 3807 | 490 | 692 |
| 3236 | QGN16333.1 | FIM1_3038 | CP015057. | 12829 | 16008 | 10114 | 12925 | 11895 | 9514 | 10807 | 9630 | 11754 | 39271 | 15867 | 14952 | 32934 | 13395 | 15074 | 39923 | 13485 | 11310 |
| 3237 | QGN16334.1 | FIM1_3039 | CP015057. | 1527 | 2164 | 1721 | 1863 | 2255 | 1696 | 1344 | 1915 | 2241 | 9793 | 2227 | 2223 | 8636 | 2234 | 2186 | 8873 | 2393 | 1509 |
| 3238 | QGN16335.1 | FIM1_3040 | CP015057. | 4109 | 2162 | 844 | 1912 | 2295 | 1115 | 2904 | 1619 | 1263 | 8641 | 2575 | 1899 | 6052 | 2657 | 1711 | 8214 | 2822 | 1323 |
| 3239 | QGN16336.1 | FIM1_3041 | CP015057. | 4241 | 1037 | 1057 | 3732 | 1000 | 864 | 3677 | 971 | 1394 | 6394 | 968 | 1572 | 5559 | 818 | 1430 | 6279 | 820 | 1120 |
| 3240 | QGN16337.1 | FIM1_3043 | CP015057. | 413 | 140 | 101 | 1078 | 134 | 74 | 592 | 128 | 115 | 654 | 88 | 116 | 566 | 75 | 113 | 662 | 107 | 82 |
| 3241 | QGN16338.1 | FIM1_3044 | CP015057. | 46249 | 304 | 259 | 54714 | 282 | 237 | 57562 | 223 | 364 | 2396 | 281 | 351 | 2476 | 708 | 404 | 2181 | 287 | 255 |
| 3242 | QGN16339.1 | FIM1_3045 | CP015057. | 952 | 135 | 375 | 1280 | 285 | 313 | 866 | 215 | 486 | 2253 | 203 | 615 | 2671 | 274 | 557 | 2237 | 185 | 391 |
| 3243 | QGN16340.1 | FIM1_3046 | CP015057. | 3202 | 574 | 1061 | 4570 | 879 | 818 | 3434 | 767 | 1400 | 3163 | 670 | 1411 | 6590 | 749 | 1275 | 4449 | 631 | 1050 |
| 3244 | QGN16341.1 | FIM1_3047 | CP015057. | 1503 | 650 | 608 | 1443 | 579 | 455 | 1351 | 549 | 690 | 2536 | 500 | 716 | 2371 | 410 | 667 | 2591 | 455 | 467 |
| 3245 | QGN16342.1 | FIM1_3048 | CP015057. | 1145 | 2527 | 1505 | 1471 | 2317 | 1476 | 1307 | 1899 | 1987 | 11484 | 2983 | 2539 | 8627 | 2440 | 2545 | 10938 | 2792 | 1802 |
| 3246 | QGN16343.1 | FIM1_3049 | CP015057. | 864 | 434 | 668 | 1323 | 541 | 604 | 912 | 425 | 884 | 3248 | 452 | 1090 | 2831 | 364 | 941 | 3211 | 479 | 642 |
| 3247 | QGN16344.1 | FIM1_3050 | CP015057. | 487 | 771 | 579 | 746 | 681 | 564 | 627 | 576 | 805 | 2350 | 545 | 700 | 2050 | 519 | 604 | 2128 | 565 | 428 |
| 3248 | QGN16345.1 | FIM1_3051 | CP015057. | 1457 | 474 | 980 | 1577 | 611 | 793 | 1435 | 596 | 1242 | 3766 | 382 | 1421 | 3931 | 375 | 1129 | 3631 | 323 | 859 |
| 3249 | QGN16346.1 | FIM1_3052 | CP015057. | 636 | 552 | 496 | 1257 | 420 | 470 | 895 | 399 | 718 | 3310 | 457 | 846 | 2787 | 414 | 772 | 3157 | 415 | 555 |
| 3250 | QGN16347.1 | FIM1_3053 | CP015057. | 1124 | 2319 | 1940 | 2574 | 2263 | 1863 | 1843 | 2017 | 2436 | 6377 | 1304 | 2153 | 5485 | 1208 | 2143 | 5645 | 1112 | 1449 |
| 3251 | QGN16348.1 | FIM1_3054 | CP015057. | 7300 | 642 | 590 | 7435 | 660 | 555 | 8031 | 606 | 895 | 452 | 565 | 533 | 2488 | 783 | 605 | 1243 | 733 | 393 |
| 3252 | QGN16349.1 | FIM1_3055 | CP015057. | 4 | 7 | 3 | 40 | 33 | 17 | 4 | 7 | 4 | 20 | 5 | 3 | 15 | 4 | 6 | 14 | 1 | 4 |
| 3253 | QGN16350.1 | FIM1_3056 | CP015057. | 6766 | 1644 | 2150 | 7244 | 2617 | 1727 | 7335 | 2391 | 2741 | 3187 | 2750 | 3118 | 12438 | 2693 | 3178 | 6678 | 2654 | 2187 |
| 3254 | QGN16351.1 | FIM1_3057 | CP015057. | 501 | 405 | 371 | 1293 | 457 | 409 | 567 | 251 | 464 | 1691 | 290 | 531 | 1085 | 193 | 499 | 1430 | 266 | 329 |
| 3255 | QGN16352.1 | FIM1_3058 | CP015057. | 5987 | 863 | 1886 | 6895 | 1572 | 1660 | 5494 | 1361 | 2338 | 10578 | 1421 | 2989 | 12346 | 1245 | 2599 | 10867 | 1124 | 2026 |
| 3256 | QGN16353.1 | FIM1_3059 | CP015057. | 1885 | 347 | 509 | 2504 | 367 | 511 | 1879 | 329 | 777 | 3712 | 304 | 869 | 3111 | 318 | 726 | 3553 | 284 | 539 |
| 3257 | QGN16354.1 | FIM1_3060 | CP015057. | 1350 | 395 | 424 | 1758 | 457 | 457 | 1468 | 402 | 592 | 2647 | 321 | 697 | 2492 | 344 | 552 | 2520 | 381 | 426 |
| 3258 | QGN16355.1 | FIM1_3061 | CP015057. | 5158 | 1806 | 2923 | 8310 | 2465 | 2437 | 6030 | 1991 | 3708 | 10987 | 1838 | 4545 | 12394 | 1939 | 4126 | 11275 | 1706 | 3080 |
| 3259 | QGN16356.1 | FIM1_3062 | CP015057. | 1989 | 1348 | 938 | 1454 | 1184 | 1060 | 1919 | 1016 | 1300 | 4584 | 1226 | 1536 | 4359 | 1062 | 1605 | 4052 | 1067 | 1099 |
| 3260 | QGN16357.1 | FIM1_3063 | CP015057. | 2539 | 553 | 787 | 2879 | 590 | 710 | 2519 | 483 | 1134 | 3927 | 350 | 1229 | 3977 | 358 | 1012 | 3994 | 375 | 752 |
| 3261 | QGN16358.1 | FIM1_3064 | CP015057. | 9317 | 3642 | 3180 | 11259 | 4193 | 2751 | 9278 | 3913 | 4108 | 9073 | 4756 | 4563 | 14796 | 4174 | 4701 | 12223 | 4316 | 3347 |
| 3262 | QGN16359.1 | FIM1_3065 | CP015057. | 1646 | 1316 | 1537 | 1957 | 1888 | 1411 | 1612 | 1514 | 1942 | 6590 | 1596 | 2480 | 9168 | 1628 | 2039 | 7234 | 1564 | 1579 |
| 3263 | QGN16360.1 | FIM1_3066 | CP015057. | 3302 | 4921 | 3864 | 5496 | 3999 | 3559 | 4116 | 3647 | 4942 | 13026 | 2979 | 5248 | 11882 | 2930 | 4892 | 13095 | 2867 | 3382 |
| 3264 | QGN16361.1 | FIM1_3067 | CP015057. | 796 | 676 | 415 | 1068 | 486 | 349 | 721 | 491 | 569 | 2369 | 520 | 548 | 1849 | 449 | 613 | 2111 | 476 | 342 |
| 3265 | QGN16362.1 | FIM1_3068 | CP015057. | 12362 | 2840 | 3201 | 10505 | 3732 | 2436 | 10380 | 3650 | 4153 | 5274 | 3264 | 4617 | 15149 | 3047 | 4390 | 8323 | 2999 | 3218 |
| 3266 | QGN16363.1 | FIM1_3069 | CP015057. | 525 | 1312 | 716 | 486 | 893 | 730 | 481 | 792 | 925 | 4381 | 911 | 925 | 3188 | 757 | 863 | 4067 | 860 | 655 |
| 3267 | QGN16364.1 | FIM1_3070 | CP015057. | 6378 | 5744 | 4009 | 7641 | 5198 | 3664 | 6542 | 4502 | 5099 | 24558 | 6698 | 6211 | 21379 | 5527 | 6132 | 22953 | 5441 | 4300 |
| 3268 | QGN16365.1 | FIM1_3071 | CP015057. | 2986 | 2397 | 2536 | 5231 | 2156 | 2184 | 3766 | 1745 | 3090 | 12920 | 1696 | 3869 | 10356 | 1751 | 3353 | 12742 | 1603 | 2433 |
| 3269 | QGN16366.1 | FIM1_3072 | CP015057. | 1885 | 582 | 1391 | 2959 | 726 | 1174 | 1815 | 689 | 1681 | 4046 | 529 | 2295 | 4399 | 724 | 2014 | 4106 | 521 | 1423 |
| 3270 | QGN16367.1 | FIM1_3074 | CP015057. | 970 | 504 | 544 | 1604 | 444 | 501 | 1347 | 400 | 720 | 3132 | 461 | 966 | 2701 | 496 | 820 | 3166 | 469 | 614 |
| 3271 | QGN16368.1 | FIM1_3075 | CP015057. | 3530 | 873 | 528 | 2863 | 846 | 574 | 3488 | 683 | 713 | 3906 | 845 | 823 | 3104 | 878 | 742 | 3767 | 876 | 624 |
| 3272 | QGN16369.1 | FIM1_3076 | CP015057. | 2149 | 515 | 537 | 2160 | 582 | 477 | 1937 | 540 | 689 | 2816 | 630 | 890 | 3804 | 512 | 779 | 3179 | 562 | 535 |
| 3273 | QGN16370.1 | FIM1_3077 | CP015057. | 2795 | 944 | 1294 | 3143 | 1588 | 1059 | 3070 | 1435 | 1686 | 8120 | 1433 | 1876 | 9655 | 1195 | 1631 | 9052 | 1197 | 1258 |
| 3274 | QGN16371.1 | FIM1_3078 | CP015057. | 937 | 427 | 504 | 1046 | 531 | 396 | 856 | 479 | 630 | 2860 | 504 | 810 | 2523 | 489 | 702 | 3112 | 473 | 534 |

| | | | | | | | | | | | | | | | | | | | | | |
|---|---|---|---|---|---|---|---|---|---|---|---|---|---|---|---|---|---|---|---|---|---|
| 3275 | QGN16372.1 | FIM1_3079 | CP015057. | 1988 | 1389 | 1116 | 2257 | 1238 | 931 | 2283 | 861 | 1428 | 7999 | 1486 | 1845 | 5625 | 1414 | 1641 | 8328 | 1593 | 1160 |
| 3276 | QGN16373.1 | FIM1_3080 | CP015057. | 3375 | 1113 | 740 | 3904 | 1003 | 698 | 3381 | 764 | 1045 | 5285 | 887 | 1000 | 4339 | 794 | 1120 | 4856 | 818 | 736 |
| 3277 | QGN16374.1 | FIM1_3081 | CP015057. | 6577 | 988 | 432 | 5545 | 1236 | 423 | 6963 | 781 | 602 | 5170 | 814 | 1175 | 5948 | 686 | 1101 | 5768 | 747 | 728 |
| 3278 | QGN16375.1 | FIM1_3082 | CP015057. | 4444 | 960 | 1072 | 4608 | 1290 | 919 | 3881 | 1141 | 1410 | 6136 | 1021 | 1543 | 6719 | 935 | 1376 | 6063 | 817 | 1031 |
| 3279 | QGN16376.1 | FIM1_3083 | CP015057. | 7746 | 3261 | 1549 | 3789 | 3888 | 1763 | 6330 | 3102 | 2082 | 10544 | 4712 | 3029 | 11573 | 4703 | 2774 | 10545 | 5471 | 2163 |
| 3280 | QGN16377.1 | FIM1_3085 | CP015057. | 159 | 17 | 6 | 282 | 21 | 19 | 121 | 10 | 18 | 165 | 26 | 25 | 159 | 22 | 21 | 200 | 21 | 20 |
| 3281 | QGN16378.1 | FIM1_3086 | CP015057. | 8254 | 490 | 448 | 7045 | 350 | 321 | 6762 | 406 | 660 | 5167 | 504 | 565 | 3862 | 889 | 503 | 4945 | 665 | 381 |
| 3282 | QGN16379.1 | FIM1_3087 | CP015057. | 5722 | 5206 | 3265 | 5404 | 7917 | 4095 | 5307 | 5087 | 3993 | 19993 | 5972 | 4932 | 20133 | 6129 | 4419 | 19533 | 6939 | 3535 |
| 3283 | QGN16380.1 | FIM1_3088 | CP015057. | 10270 | 232 | 357 | 5520 | 102 | 237 | 6888 | 172 | 445 | 1224 | 128 | 467 | 1181 | 162 | 461 | 1162 | 125 | 304 |
| 3284 | QGN16381.1 | FIM1_3089 | CP015057. | 889 | 307 | 360 | 1979 | 365 | 291 | 863 | 296 | 374 | 1357 | 312 | 492 | 1311 | 244 | 479 | 1369 | 243 | 281 |
| 3285 | QGN16382.1 | FIM1_3090 | CP015057. | 810 | 260 | 177 | 1338 | 286 | 135 | 940 | 219 | 206 | 1113 | 224 | 300 | 1106 | 247 | 269 | 1147 | 279 | 195 |
| 3286 | QGN16383.1 | FIM1_3092 | CP015057. | 2182 | 195 | 174 | 3195 | 280 | 158 | 1845 | 209 | 228 | 1117 | 197 | 271 | 1159 | 277 | 226 | 1073 | 281 | 176 |
| 3287 | QGN16384.1 | FIM1_3097 | CP015057. | 1 | 0 | 0 | 1 | 1 | 0 | 1 | 1 | 0 | 1 | 1 | 0 | 2 | 1 | 2 | 0 | 1 | 0 |
| 3288 | QGN16385.1 | FIM1_3098 | CP015058. | 666 | 447 | 833 | 1493 | 429 | 534 | 719 | 353 | 935 | 2624 | 819 | 1860 | 3496 | 936 | 1353 | 2841 | 746 | 1342 |
| 3289 | QGN16386.1 | FIM1_3099 | CP015058. | 1755 | 1382 | 1552 | 8639 | 841 | 1133 | 5773 | 972 | 2073 | 8715 | 1104 | 2029 | 7022 | 1639 | 2012 | 8938 | 1352 | 1337 |
| 3290 | QGN16387.1 | FIM1_3100 | CP015058. | 2354 | 625 | 595 | 2614 | 481 | 467 | 2386 | 451 | 736 | 4321 | 493 | 860 | 3030 | 539 | 767 | 4376 | 472 | 586 |
| 3291 | QGN16388.1 | FIM1_3101 | CP015058. | 1671 | 693 | 596 | 2447 | 675 | 557 | 1834 | 517 | 823 | 4345 | 612 | 975 | 3254 | 528 | 891 | 4342 | 614 | 575 |
| 3292 | QGN16389.1 | FIM1_3102 | CP015058. | 9659 | 17138 | 9551 | 9688 | 12714 | 8996 | 7711 | 11196 | 11308 | 36168 | 14173 | 14543 | 28858 | 12471 | 14544 | 34252 | 12852 | 10782 |
| 3293 | QGN16390.1 | FIM1_3103 | CP015058. | 129 | 77 | 75 | 652 | 131 | 60 | 213 | 46 | 87 | 682 | 116 | 117 | 520 | 129 | 126 | 563 | 130 | 94 |
| 3294 | QGN16391.1 | FIM1_3104 | CP015058. | 4330 | 951 | 1304 | 3798 | 1186 | 1080 | 2835 | 1097 | 1664 | 6537 | 1076 | 1928 | 7307 | 977 | 1784 | 7067 | 835 | 1226 |
| 3295 | QGN16392.1 | FIM1_3105 | CP015058. | 1 | 13 | 12 | 10 | 5 | 15 | 4 | 6 | 10 | 66 | 12 | 10 | 57 | 8 | 9 | 62 | 15 | 16 |
| 3296 | QGN16393.1 | FIM1_3106 | CP015058. | 861 | 815 | 964 | 1566 | 940 | 802 | 966 | 827 | 1277 | 3924 | 829 | 1461 | 5169 | 709 | 1349 | 4377 | 696 | 1005 |
| 3297 | QGN16394.1 | FIM1_3107 | CP015058. | 146 | 1833 | 1330 | 1334 | 3672 | 1279 | 256 | 2834 | 1741 | 3127 | 436 | 1211 | 3665 | 312 | 925 | 3104 | 259 | 667 |
| 3298 | QGN16395.1 | FIM1_3108 | CP015058. | 3436 | 1989 | 1796 | 4293 | 1850 | 1297 | 3641 | 1906 | 2267 | 7921 | 1102 | 2145 | 9182 | 1521 | 1882 | 8062 | 1010 | 1396 |
| 3299 | QGN16396.1 | FIM1_3109 | CP015058. | 1910 | 3089 | 1556 | 1731 | 2818 | 1719 | 2088 | 2124 | 2012 | 10129 | 2852 | 2393 | 7731 | 2578 | 2479 | 10048 | 2939 | 1657 |
| 3300 | QGN16397.1 | FIM1_3110 | CP015058. | 37472 | 1145 | 1515 | 31945 | 442 | 717 | 29835 | 1384 | 1738 | 4181 | 266 | 739 | 4024 | 1455 | 600 | 3531 | 719 | 499 |
| 3301 | QGN16398.1 | FIM1_3111 | CP015058. | 8208 | 260 | 507 | 10173 | 201 | 309 | 8807 | 253 | 637 | 1453 | 240 | 531 | 1552 | 516 | 499 | 1368 | 347 | 329 |
| 3302 | QGN16399.1 | FIM1_3112 | CP015058. | 1406 | 385 | 567 | 738 | 296 | 393 | 986 | 371 | 675 | 2968 | 286 | 717 | 2456 | 325 | 715 | 2709 | 309 | 536 |
| 3303 | QGN16400.1 | FIM1_3113 | CP015058. | 7044 | 466 | 753 | 5206 | 502 | 467 | 3951 | 513 | 993 | 1800 | 558 | 1196 | 2779 | 562 | 1276 | 2053 | 460 | 956 |
| 3304 | QGN16401.1 | FIM1_3114 | CP015058. | 2748 | 1020 | 1372 | 3310 | 1293 | 1068 | 2475 | 1450 | 1850 | 4518 | 908 | 1668 | 5487 | 945 | 1329 | 5540 | 1088 | 1108 |
| 3305 | QGN16402.1 | FIM1_3115 | CP015058. | 239 | 157 | 223 | 386 | 149 | 173 | 201 | 198 | 301 | 1194 | 142 | 312 | 1210 | 111 | 234 | 1137 | 95 | 199 |
| 3306 | QGN16403.1 | FIM1_3116 | CP015058. | 2900 | 486 | 958 | 7053 | 912 | 819 | 4475 | 713 | 1210 | 4298 | 706 | 1842 | 6177 | 904 | 1529 | 5017 | 817 | 1027 |
| 3307 | QGN16404.1 | FIM1_3117 | CP015058. | 29 | 25 | 13 | 33 | 9 | 11 | 27 | 13 | 20 | 60 | 7 | 29 | 55 | 8 | 26 | 48 | 13 | 19 |
| 3308 | QGN16405.1 | FIM1_3118 | CP015058. | 4801 | 4404 | 4419 | 6511 | 5634 | 5059 | 4603 | 3548 | 6225 | 9299 | 1242 | 3808 | 9682 | 1968 | 3085 | 9598 | 998 | 2337 |
| 3309 | QGN16406.1 | FIM1_3119 | CP015058. | 372 | 628 | 580 | 1047 | 625 | 504 | 654 | 515 | 803 | 5034 | 547 | 975 | 4121 | 576 | 900 | 4933 | 464 | 617 |
| 3310 | QGN16407.1 | FIM1_3120 | CP015058. | 3053 | 1989 | 1666 | 4093 | 1757 | 1588 | 3553 | 1525 | 2361 | 11225 | 1746 | 2724 | 9413 | 1500 | 2574 | 11313 | 1488 | 1865 |
| 3311 | QGN16408.1 | FIM1_3121 | CP015058. | 2008 | 603 | 624 | 2631 | 646 | 566 | 1921 | 496 | 876 | 4902 | 647 | 1122 | 4588 | 541 | 1009 | 4916 | 548 | 768 |
| 3312 | QGN16409.1 | FIM1_3122 | CP015058. | 573 | 1605 | 1827 | 1280 | 1542 | 1628 | 810 | 1321 | 2493 | 6069 | 841 | 2372 | 6015 | 1110 | 2198 | 6032 | 823 | 1522 |
| 3313 | QGN16410.1 | FIM1_3123 | CP015058. | 3977 | 958 | 2040 | 4722 | 1247 | 1500 | 4426 | 1105 | 2583 | 5069 | 1071 | 2784 | 7322 | 1112 | 2499 | 6126 | 941 | 1911 |
| 3314 | QGN16411.1 | FIM1_3124 | CP015058. | 4437 | 3088 | 2225 | 5973 | 3163 | 1984 | 5335 | 3025 | 2934 | 13798 | 3165 | 3680 | 15420 | 3269 | 3212 | 14413 | 2971 | 2352 |
| 3315 | QGN16412.1 | FIM1_3125 | CP015058. | 2102 | 329 | 395 | 1927 | 396 | 317 | 1829 | 474 | 520 | 1238 | 338 | 537 | 1521 | 267 | 456 | 1301 | 299 | 318 |
| 3316 | QGN16413.1 | FIM1_3126 | CP015058. | 4099 | 1141 | 1528 | 4960 | 1395 | 1093 | 3606 | 1333 | 1879 | 2531 | 1030 | 2272 | 7107 | 1126 | 2047 | 4215 | 1032 | 1509 |
| 3317 | QGN16414.1 | FIM1_3127 | CP015058. | 3954 | 869 | 1185 | 3840 | 882 | 1011 | 3235 | 859 | 1634 | 1040 | 636 | 1835 | 2951 | 733 | 1829 | 1613 | 744 | 1257 |
| 3318 | QGN16415.1 | FIM1_3128 | CP015058. | 5696 | 6524 | 6541 | 8946 | 6859 | 5430 | 6574 | 6413 | 8119 | 19989 | 6871 | 12143 | 27516 | 7105 | 10753 | 21263 | 6722 | 8137 |
| 3319 | QGN16416.1 | FIM1_3129 | CP015058. | 3639 | 2458 | 1957 | 3680 | 2790 | 1805 | 3627 | 2299 | 2477 | 14487 | 3076 | 3796 | 12371 | 2759 | 3559 | 14220 | 2697 | 2528 |
| 3320 | QGN16417.1 | FIM1_3130 | CP015058. | 713 | 661 | 657 | 761 | 832 | 579 | 661 | 740 | 863 | 5236 | 848 | 840 | 4392 | 799 | 782 | 4985 | 859 | 585 |
| 3321 | QGN16418.1 | FIM1_3131 | CP015058. | 1633 | 731 | 675 | 1537 | 548 | 506 | 1094 | 599 | 895 | 5016 | 621 | 927 | 3841 | 698 | 1019 | 4566 | 541 | 606 |
| 3322 | QGN16419.1 | FIM1_3132 | CP015058. | 1035 | 2952 | 2041 | 1383 | 2898 | 1908 | 1171 | 2503 | 2623 | 12367 | 2562 | 2887 | 11506 | 2655 | 2609 | 11889 | 2564 | 1718 |
| 3323 | QGN16420.1 | FIM1_3133 | CP015058. | 2012 | 282 | 260 | 1919 | 483 | 220 | 1502 | 357 | 308 | 1468 | 356 | 451 | 1731 | 247 | 329 | 1653 | 343 | 251 |
| 3324 | QGN16421.1 | FIM1_3134 | CP015058. | 19201 | 2809 | 6457 | 26271 | 6765 | 5242 | 21649 | 6145 | 7987 | 22175 | 7274 | 8029 | 35648 | 8137 | 8234 | 28698 | 7291 | 5630 |
| 3325 | QGN16422.1 | FIM1_3135 | CP015058. | 937 | 729 | 688 | 1232 | 657 | 629 | 1104 | 515 | 885 | 5329 | 720 | 1189 | 4221 | 621 | 992 | 5260 | 624 | 735 |
| 3326 | QGN16423.1 | FIM1_3136 | CP015058. | 6193 | 2575 | 1627 | 4705 | 3605 | 1646 | 5978 | 2513 | 2174 | 11319 | 4184 | 3334 | 13565 | 3721 | 3032 | 11762 | 4472 | 2351 |
| 3327 | QGN16424.1 | FIM1_3137 | CP015058. | 34 | 13 | 16 | 98 | 6 | 9 | 61 | 16 | 9 | 144 | 13 | 9 | 76 | 14 | 11 | 136 | 11 | 7 |
| 3328 | QGN16425.1 | FIM1_3138 | CP015058. | 7683 | 3874 | 2112 | 4335 | 4240 | 2214 | 7955 | 3345 | 2875 | 17230 | 5630 | 3637 | 15933 | 5004 | 3453 | 16823 | 6002 | 2717 |
| 3329 | QGN16426.1 | FIM1_3139 | CP015058. | 508 | 492 | 490 | 327 | 624 | 477 | 550 | 495 | 701 | 1529 | 311 | 395 | 1209 | 376 | 336 | 1367 | 332 | 270 |
| 3330 | QGN16427.1 | FIM1_3140 | CP015058. | 471 | 242 | 310 | 759 | 256 | 225 | 500 | 222 | 380 | 2173 | 309 | 452 | 1759 | 244 | 423 | 2126 | 239 | 335 |
| 3331 | QGN16428.1 | FIM1_3141 | CP015058. | 538 | 422 | 751 | 585 | 633 | 636 | 495 | 534 | 900 | 3049 | 692 | 1038 | 3251 | 678 | 848 | 3165 | 641 | 628 |
| 3332 | QGN16429.1 | FIM1_3142 | CP015058. | 780 | 118 | 152 | 1317 | 128 | 99 | 821 | 140 | 168 | 755 | 93 | 175 | 896 | 105 | 175 | 761 | 94 | 101 |
| 3333 | QGN16430.1 | FIM1_3143 | CP015058. | 4761 | 4032 | 3075 | 6804 | 4440 | 2921 | 6015 | 3690 | 3863 | 13916 | 4338 | 4450 | 13374 | 3993 | 4434 | 13221 | 4105 | 3094 |
| 3334 | QGN16431.1 | FIM1_3144 | CP015058. | 431 | 364 | 306 | 400 | 210 | 277 | 354 | 241 | 415 | 1569 | 194 | 351 | 1173 | 129 | 376 | 1439 | 173 | 213 |
| 3335 | QGN16432.1 | FIM1_3145 | CP015058. | 1328 | 619 | 810 | 1271 | 778 | 647 | 1540 | 650 | 1048 | 5267 | 861 | 1180 | 5457 | 817 | 1082 | 5427 | 791 | 741 |
| 3336 | QGN16433.1 | FIM1_3146 | CP015058. | 1728 | 401 | 379 | 1574 | 436 | 300 | 1728 | 484 | 492 | 2855 | 381 | 567 | 2483 | 345 | 510 | 2711 | 331 | 384 |
| 3337 | QGN16434.1 | FIM1_3147 | CP015058. | 3636 | 361 | 727 | 4270 | 365 | 685 | 3529 | 456 | 948 | 4009 | 349 | 899 | 4451 | 347 | 845 | 4035 | 290 | 572 |
| 3338 | QGN16435.1 | FIM1_3148 | CP015058. | 1422 | 407 | 629 | 1744 | 362 | 636 | 1487 | 375 | 924 | 3777 | 287 | 1047 | 3041 | 258 | 905 | 3735 | 287 | 694 |
| 3339 | QGN16436.1 | FIM1_3149 | CP015058. | 1076 | 588 | 656 | 1687 | 541 | 635 | 1381 | 479 | 888 | 4791 | 468 | 1215 | 3483 | 570 | 1053 | 4856 | 547 | 745 |
| 3340 | QGN16437.1 | FIM1_3150 | CP015058. | 6011 | 2464 | 1489 | 2907 | 3331 | 1537 | 5506 | 2611 | 1889 | 13449 | 4943 | 3654 | 14959 | 4390 | 2903 | 13495 | 5396 | 2499 |
| 3341 | QGN16438.1 | FIM1_3151 | CP015058. | 405 | 677 | 584 | 693 | 687 | 478 | 400 | 865 | 806 | 2815 | 597 | 504 | 2829 | 524 | 499 | 2903 | 482 | 311 |
| 3342 | QGN16439.1 | FIM1_3152 | CP015058. | 559 | 205 | 127 | 512 | 180 | 131 | 415 | 141 | 181 | 1100 | 154 | 229 | 849 | 128 | 228 | 878 | 129 | 141 |
| 3343 | QGN16440.1 | FIM1_3153 | CP015058. | 1021 | 464 | 540 | 1517 | 431 | 462 | 1151 | 438 | 705 | 4002 | 351 | 830 | 3092 | 355 | 701 | 3751 | 300 | 495 |
| 3344 | QGN16441.1 | FIM1_3154 | CP015058. | 1081 | 1759 | 1221 | 837 | 1874 | 1193 | 1129 | 1750 | 1633 | 6204 | 1579 | 1967 | 7544 | 1438 | 1700 | 6029 | 1483 | 1219 |
| 3345 | QGN16442.1 | FIM1_3155 | CP015058. | 11484 | 965 | 1088 | 12820 | 1395 | 978 | 12199 | 1243 | 1380 | 4714 | 1347 | 2051 | 7049 | 1528 | 1838 | 5291 | 1332 | 1383 |
| 3346 | QGN16443.1 | FIM1_3156 | CP015058. | 4746 | 2824 | 2354 | 6532 | 2549 | 2219 | 5324 | 1804 | 3025 | 14451 | 2715 | 4174 | 11565 | 2886 | 3974 | 14413 | 2717 | 2810 |
| 3347 | QGN16444.1 | FIM1_3157 | CP015058. | 2922 | 313 | 1455 | 3806 | 489 | 1154 | 2836 | 455 | 1696 | 1051 | 244 | 2326 | 1766 | 420 | 2202 | 1352 | 278 | 1488 |
| 3348 | QGN16445.1 | FIM1_3158 | CP015058. | 772 | 242 | 202 | 960 | 227 | 181 | 918 | 253 | 308 | 1792 | 158 | 268 | 1279 | 152 | 247 | 1567 | 152 | 176 |
| 3349 | QGN16446.1 | FIM1_3159 | CP015058. | 413 | 346 | 357 | 637 | 319 | 298 | 334 | 304 | 436 | 1431 | 278 | 412 | 1381 | 224 | 399 | 1364 | 223 | 266 |
| 3350 | QGN16447.1 | FIM1_3160 | CP015058. | 430 | 27 | 157 | 851 | 107 | 127 | 639 | 155 | 252 | 888 | 112 | 280 | 1377 | 88 | 248 | 983 | 101 | 201 |
| 3351 | QGN16448.1 | FIM1_3161 | CP015058. | 4119 | 988 | 686 | 8112 | 921 | 422 | 11341 | 1846 | 904 | 634 | 332 | 372 | 1119 | 313 | 431 | 898 | 301 | 232 |
| 3352 | QGN16449.1 | FIM1_3162 | CP015058. | 2398 | 1157 | 1118 | 5820 | 2670 | 1740 | 2583 | 1217 | 1495 | 6870 | 1393 | 1963 | 8261 | 1896 | 1955 | 6957 | 1545 | 1344 |
| 3353 | QGN16450.1 | FIM1_3163 | CP015058. | 225 | 673 | 526 | 378 | 472 | 517 | 306 | 488 | 735 | 3085 | 518 | 831 | 2402 | 351 | 776 | 2931 | 450 | 500 |
| 3354 | QGN16451.1 | FIM1_3164 | CP015058. | 4508 | 1648 | 1402 | 5400 | 1901 | 1305 | 4764 | 1561 | 1874 | 8682 | 1737 | 2450 | 9976 | 1682 | 2163 | 9354 | 1567 | 1606 |
| 3355 | QGN16452.1 | FIM1_3165 | CP015058. | 444 | 860 | 534 | 458 | 573 | 425 | 394 | 552 | 738 | 2673 | 601 | 677 | 2364 | 525 | 733 | 2646 | 559 | 440 |
| 3356 | QGN16453.1 | FIM1_3166 | CP015058. | 937 | 550 | 825 | 1356 | 984 | 644 | 1071 | 886 | 1060 | 3967 | 750 | 1189 | 4479 | 710 | 996 | 4252 | 653 | 734 |
| 3357 | QGN16454.1 | FIM1_3167 | CP015058. | 6661 | 2200 | 1087 | 2826 | 2771 | 1178 | 5671 | 2210 | 1531 | 6591 | 2970 | 1940 | 8063 | 3216 | 1930 | 6922 | 3473 | 1437 |
| 3358 | QGN16455.1 | FIM1_3168 | CP015058. | 1246 | 1018 | 775 | 1916 | 889 | 792 | 1363 | 846 | 1048 | 2438 | 559 | 1024 | 3084 | 536 | 962 | 2482 | 557 | 638 |

| | | | | | | | | | | | | | | | | | | | | | |
|---|---|---|---|---|---|---|---|---|---|---|---|---|---|---|---|---|---|---|---|---|---|
| 3359 | QGN16456.1 | FIM1_3169 | CP015058. | 1524 | 1093 | 711 | 1296 | 772 | 708 | 1305 | 648 | 986 | 3460 | 660 | 818 | 2775 | 635 | 913 | 3301 | 613 | 534 |
| 3360 | QGN16457.1 | FIM1_3170 | CP015058. | 1206 | 43 | 36 | 610 | 40 | 22 | 1004 | 44 | 54 | 195 | 43 | 60 | 199 | 29 | 50 | 192 | 36 | 36 |
| 3361 | QGN16458.1 | FIM1_3171 | CP015058. | 933 | 1810 | 1077 | 827 | 1489 | 1039 | 907 | 1244 | 1442 | 5227 | 1283 | 1512 | 5141 | 1134 | 1409 | 5413 | 1309 | 936 |
| 3362 | QGN16459.1 | FIM1_3172 | CP015058. | 522 | 1007 | 1075 | 1689 | 841 | 905 | 1016 | 778 | 1462 | 5676 | 817 | 1724 | 5181 | 1058 | 1556 | 5241 | 865 | 1122 |
| 3363 | QGN16460.1 | FIM1_3173 | CP015058. | 7203 | 594 | 761 | 5696 | 778 | 674 | 6704 | 777 | 1033 | 3415 | 683 | 1168 | 5128 | 771 | 1026 | 4048 | 707 | 800 |
| 3364 | QGN16461.1 | FIM1_3174 | CP015058. | 485 | 313 | 360 | 614 | 450 | 236 | 491 | 422 | 365 | 428 | 297 | 432 | 1213 | 285 | 394 | 752 | 301 | 261 |
| 3365 | QGN16462.1 | FIM1_3175 | CP015058. | 1497 | 1718 | 1047 | 1615 | 1163 | 979 | 1331 | 1061 | 1387 | 2332 | 843 | 1314 | 3628 | 715 | 1263 | 2794 | 783 | 864 |
| 3366 | QGN16463.1 | FIM1_3176 | CP015058. | 1197 | 298 | 263 | 2110 | 323 | 277 | 1242 | 293 | 388 | 1063 | 175 | 281 | 1276 | 176 | 255 | 1072 | 166 | 182 |
| 3367 | QGN16464.1 | FIM1_3177 | CP015058. | 3314 | 1463 | 1300 | 3245 | 1493 | 1186 | 3332 | 1301 | 1750 | 7116 | 1370 | 1924 | 7835 | 1294 | 1730 | 7206 | 1270 | 1245 |
| 3368 | QGN16465.1 | FIM1_3178 | CP015058. | 2175 | 1587 | 1328 | 3846 | 1406 | 1119 | 2722 | 985 | 1661 | 7106 | 1244 | 2017 | 5866 | 1396 | 1809 | 6517 | 1336 | 1239 |
| 3369 | QGN16466.1 | FIM1_3179 | CP015058. | 2001 | 242 | 255 | 2216 | 193 | 194 | 1562 | 176 | 345 | 1602 | 145 | 345 | 1541 | 139 | 318 | 1482 | 152 | 236 |
| 3370 | QGN16467.1 | FIM1_3180 | CP015058. | 4500 | 593 | 966 | 6699 | 765 | 782 | 5753 | 670 | 1209 | 4215 | 615 | 1309 | 5292 | 892 | 1146 | 4222 | 545 | 855 |
| 3371 | QGN16468.1 | FIM1_3181 | CP015058. | 4266 | 1844 | 1024 | 1915 | 2479 | 1154 | 3411 | 2044 | 1534 | 6398 | 2603 | 2016 | 7479 | 2939 | 1825 | 6552 | 2895 | 1442 |
| 3372 | QGN16469.1 | FIM1_3182 | CP015058. | 1177 | 1675 | 1086 | 1407 | 1394 | 1069 | 1461 | 1229 | 1539 | 7915 | 1462 | 1573 | 6142 | 1295 | 1538 | 7476 | 1306 | 1103 |
| 3373 | QGN16470.1 | FIM1_3183 | CP015058. | 6979 | 5915 | 4686 | 12013 | 6509 | 3700 | 9318 | 6218 | 5592 | 10422 | 4256 | 5382 | 20573 | 4968 | 5099 | 15503 | 4125 | 3663 |
| 3374 | QGN16471.1 | FIM1_3184 | CP015058. | 93 | 837 | 502 | 306 | 640 | 433 | 239 | 511 | 649 | 2146 | 441 | 656 | 1887 | 485 | 658 | 1957 | 465 | 447 |
| 3375 | QGN16472.1 | FIM1_3185 | CP015058. | 1354 | 1397 | 545 | 815 | 854 | 502 | 1286 | 989 | 704 | 1859 | 713 | 741 | 2291 | 683 | 678 | 1870 | 801 | 492 |
| 3376 | QGN16473.1 | FIM1_3186 | CP015058. | 4145 | 1549 | 2567 | 6368 | 2218 | 2406 | 4479 | 1933 | 3402 | 6067 | 1775 | 4052 | 9091 | 1621 | 3867 | 7340 | 1625 | 2723 |
| 3377 | QGN16474.1 | FIM1_3187 | CP015058. | 630 | 398 | 405 | 722 | 441 | 354 | 548 | 317 | 568 | 3216 | 451 | 790 | 2385 | 349 | 664 | 3103 | 393 | 520 |
| 3378 | QGN16475.1 | FIM1_3188 | CP015058. | 732 | 306 | 435 | 1015 | 403 | 372 | 912 | 336 | 544 | 3477 | 398 | 715 | 2638 | 363 | 589 | 3441 | 338 | 453 |
| 3379 | QGN16476.1 | FIM1_3189 | CP015058. | 579 | 293 | 230 | 542 | 210 | 205 | 596 | 187 | 326 | 1875 | 235 | 412 | 1273 | 229 | 375 | 1626 | 233 | 253 |
| 3380 | QGN16477.1 | FIM1_3190 | CP015058. | 972 | 421 | 415 | 1097 | 397 | 398 | 1020 | 444 | 585 | 3720 | 347 | 570 | 2571 | 339 | 514 | 3182 | 323 | 387 |
| 3381 | QGN16478.1 | FIM1_3191 | CP015058. | 189 | 684 | 542 | 481 | 670 | 540 | 351 | 625 | 758 | 4099 | 608 | 850 | 3179 | 552 | 735 | 3963 | 572 | 522 |
| 3382 | QGN16479.1 | FIM1_3192 | CP015058. | 961 | 397 | 434 | 987 | 536 | 435 | 983 | 406 | 610 | 2235 | 434 | 718 | 2633 | 419 | 626 | 2230 | 408 | 436 |
| 3383 | QGN16480.1 | FIM1_3193 | CP015058. | 47 | 34 | 29 | 60 | 31 | 17 | 30 | 21 | 16 | 142 | 39 | 27 | 151 | 36 | 31 | 124 | 28 | 21 |
| 3384 | QGN16481.1 | FIM1_3194 | CP015058. | 5593 | 795 | 1402 | 5910 | 1506 | 1111 | 5369 | 1165 | 1800 | 3177 | 1307 | 2104 | 7855 | 1624 | 1962 | 5133 | 1377 | 1494 |
| 3385 | QGN16482.1 | FIM1_3195 | CP015058. | 2848 | 3052 | 1129 | 1811 | 2423 | 1266 | 2483 | 2121 | 1572 | 9722 | 3158 | 2138 | 8497 | 2430 | 2093 | 9177 | 3206 | 1532 |
| 3386 | QGN16483.1 | FIM1_3196 | CP015058. | 415 | 323 | 251 | 572 | 341 | 266 | 355 | 232 | 388 | 2531 | 297 | 504 | 1759 | 288 | 405 | 2564 | 271 | 321 |
| 3387 | QGN16484.1 | FIM1_3197 | CP015058. | 12298 | 191 | 334 | 10643 | 270 | 262 | 16579 | 436 | 463 | 469 | 222 | 409 | 1393 | 394 | 395 | 767 | 281 | 266 |
| 3388 | QGN16485.1 | FIM1_3198 | CP015058. | 1213 | 23 | 267 | 1591 | 175 | 183 | 1548 | 209 | 366 | 394 | 109 | 454 | 1543 | 170 | 341 | 683 | 173 | 278 |
| 3389 | QGN16486.1 | FIM1_3199 | CP015058. | 1643 | 1419 | 1364 | 3435 | 1650 | 1129 | 2894 | 1308 | 1661 | 10166 | 1488 | 2211 | 9415 | 1455 | 1930 | 9747 | 1448 | 1465 |
| 3390 | QGN16487.1 | FIM1_3200 | CP015058. | 8250 | 5620 | 4783 | 9697 | 5943 | 4151 | 8003 | 4710 | 5977 | 25616 | 7860 | 9183 | 29022 | 7189 | 8577 | 25561 | 7116 | 6169 |
| 3391 | QGN16488.1 | FIM1_3201 | CP015058. | 349 | 341 | 341 | 508 | 302 | 322 | 470 | 267 | 456 | 2472 | 276 | 401 | 1805 | 259 | 366 | 2281 | 294 | 288 |
| 3392 | QGN16489.1 | FIM1_3202 | CP015058. | 1433 | 286 | 323 | 1391 | 305 | 295 | 1547 | 334 | 409 | 1319 | 232 | 431 | 1766 | 233 | 367 | 1401 | 195 | 320 |
| 3393 | QGN16490.1 | FIM1_3203 | CP015058. | 3159 | 697 | 840 | 3429 | 742 | 669 | 2885 | 726 | 1086 | 2905 | 640 | 1160 | 5380 | 709 | 963 | 3980 | 561 | 779 |
| 3394 | QGN16491.1 | FIM1_3204 | CP015058. | 687 | 478 | 499 | 801 | 465 | 446 | 788 | 411 | 635 | 2497 | 447 | 732 | 2691 | 478 | 721 | 2412 | 462 | 486 |
| 3395 | QGN16492.1 | FIM1_3205 | CP015058. | 5635 | 3738 | 2879 | 8671 | 3089 | 2554 | 5976 | 2647 | 3721 | 12903 | 3163 | 4588 | 13208 | 3548 | 4485 | 12216 | 2934 | 3228 |
| 3396 | QGN16493.1 | FIM1_3206 | CP015058. | 2122 | 847 | 1162 | 2682 | 1232 | 969 | 2611 | 989 | 1380 | 6404 | 1277 | 1893 | 7558 | 1098 | 1608 | 6515 | 1151 | 1227 |
| 3397 | QGN16494.1 | FIM1_3207 | CP015058. | 1246 | 379 | 601 | 1981 | 451 | 537 | 1262 | 446 | 830 | 1220 | 405 | 880 | 3154 | 417 | 841 | 2183 | 382 | 598 |
| 3398 | QGN16495.1 | FIM1_3208 | CP015058. | 2363 | 1733 | 2336 | 3223 | 1939 | 1794 | 2441 | 1468 | 2709 | 10468 | 2332 | 3778 | 8981 | 2208 | 3843 | 10734 | 2044 | 2774 |
| 3399 | QGN16496.1 | FIM1_3209 | CP015058. | 4456 | 3677 | 5135 | 8467 | 4958 | 3973 | 5997 | 4408 | 6172 | 18868 | 4750 | 7732 | 25377 | 4642 | 6563 | 20855 | 4189 | 4781 |
| 3400 | QGN16497.1 | FIM1_3210 | CP015058. | 257 | 273 | 719 | 225 | 230 | 555 | 196 | 328 | 900 | 2072 | 209 | 809 | 2960 | 142 | 801 | 2216 | 169 | 578 |
| 3401 | QGN16498.1 | FIM1_3211 | CP015058. | 363 | 239 | 270 | 863 | 253 | 266 | 412 | 194 | 378 | 2163 | 217 | 533 | 1680 | 224 | 492 | 2049 | 194 | 314 |
| 3402 | QGN16499.1 | FIM1_3212 | CP015058. | 2713 | 2743 | 1797 | 3610 | 2127 | 1596 | 2571 | 1810 | 2289 | 12044 | 2110 | 3036 | 9569 | 2157 | 2829 | 11618 | 2068 | 1979 |
| 3403 | QGN16500.1 | FIM1_3213 | CP015058. | 5632 | 1684 | 2552 | 7593 | 2529 | 2204 | 6720 | 2242 | 3269 | 11677 | 2062 | 3508 | 14099 | 2067 | 3060 | 12052 | 2001 | 2264 |
| 3404 | QGN16501.1 | FIM1_3214 | CP015058. | 3096 | 3146 | 1176 | 1060 | 2593 | 1282 | 2373 | 2114 | 1616 | 9675 | 3369 | 2057 | 8037 | 3211 | 2094 | 9650 | 3655 | 1429 |
| 3405 | QGN16502.1 | FIM1_3215 | CP015058. | 4922 | 1524 | 1438 | 6088 | 1797 | 1170 | 6104 | 1520 | 1822 | 6341 | 1492 | 1910 | 8775 | 1440 | 1628 | 7030 | 1448 | 1254 |
| 3406 | QGN16503.1 | FIM1_3216 | CP015058. | 217 | 52 | 74 | 243 | 49 | 62 | 333 | 59 | 99 | 165 | 26 | 67 | 215 | 28 | 77 | 184 | 11 | 37 |
| 3407 | QGN16504.1 | FIM1_3217 | CP015058. | 4527 | 1493 | 1475 | 3057 | 1646 | 1108 | 3886 | 1841 | 1905 | 1494 | 1166 | 1809 | 5633 | 1417 | 1774 | 2952 | 1306 | 1170 |
| 3408 | QGN16505.1 | FIM1_3218 | CP015058. | 344 | 97 | 415 | 294 | 97 | 195 | 177 | 137 | 487 | 280 | 94 | 368 | 579 | 84 | 324 | 427 | 101 | 213 |
| 3409 | QGN16506.1 | FIM1_3219 | CP015058. | 1914 | 3442 | 2909 | 4143 | 2364 | 2263 | 3643 | 2484 | 3455 | 8224 | 1876 | 4048 | 9027 | 1586 | 3693 | 8138 | 1479 | 2499 |
| 3410 | QGN16507.1 | FIM1_3220 | CP015058. | 14069 | 544 | 565 | 14210 | 607 | 491 | 13347 | 595 | 632 | 1805 | 523 | 635 | 3683 | 891 | 577 | 2449 | 728 | 480 |
| 3411 | QGN16508.1 | FIM1_3221 | CP015058. | 5219 | 4299 | 4252 | 8566 | 4368 | 3792 | 6394 | 4111 | 5374 | 11730 | 4237 | 6306 | 18343 | 4357 | 5612 | 14111 | 3996 | 4291 |
| 3412 | QGN16509.1 | FIM1_3222 | CP015058. | 5660 | 2345 | 2756 | 6549 | 2082 | 2328 | 5728 | 2101 | 3484 | 8960 | 2267 | 3468 | 9208 | 2158 | 3452 | 8828 | 1783 | 2497 |
| 3413 | QGN16510.1 | FIM1_3223 | CP015058. | 6290 | 3889 | 2641 | 6214 | 3808 | 2433 | 7197 | 3141 | 3409 | 14475 | 3664 | 4281 | 16070 | 3371 | 3979 | 14135 | 3496 | 2812 |
| 3414 | QGN16511.1 | FIM1_3224 | CP015058. | 4576 | 735 | 1203 | 4557 | 1218 | 1059 | 3457 | 1003 | 1583 | 1423 | 756 | 1905 | 6442 | 722 | 1695 | 3042 | 651 | 1284 |
| 3415 | QGN16512.1 | FIM1_3225 | CP015058. | 368 | 127 | 82 | 138 | 71 | 83 | 144 | 90 | 105 | 266 | 84 | 128 | 311 | 50 | 124 | 284 | 50 | 94 |
| 3416 | QGN16513.1 | FIM1_3226 | CP015058. | 1261 | 714 | 570 | 1921 | 476 | 493 | 1509 | 515 | 750 | 1386 | 400 | 825 | 2186 | 370 | 826 | 1549 | 324 | 591 |
| 3417 | QGN16514.1 | FIM1_3227 | CP015058. | 1554 | 1743 | 1223 | 1747 | 1857 | 1242 | 1648 | 1321 | 1685 | 9357 | 1875 | 1950 | 8504 | 1541 | 1956 | 8368 | 1539 | 1292 |
| 3418 | QGN16515.1 | FIM1_3228 | CP015058. | 1468 | 1326 | 1335 | 2313 | 1100 | 1214 | 1759 | 914 | 1599 | 6096 | 938 | 1753 | 4803 | 842 | 1653 | 5941 | 819 | 1160 |
| 3419 | QGN16516.1 | FIM1_3229 | CP015058. | 2330 | 3600 | 2102 | 3158 | 2991 | 2006 | 2349 | 2783 | 2729 | 9246 | 3196 | 3628 | 9070 | 2670 | 3438 | 9296 | 2800 | 2421 |
| 3420 | QGN16517.1 | FIM1_3230 | CP015058. | 3040 | 3715 | 2620 | 3681 | 2877 | 2238 | 3433 | 2329 | 3287 | 13901 | 3594 | 4260 | 12362 | 3200 | 4271 | 14425 | 3119 | 3067 |
| 3421 | QGN16518.1 | FIM1_3231 | CP015058. | 2488 | 4878 | 2782 | 2991 | 4056 | 2430 | 2372 | 3865 | 3549 | 12777 | 4135 | 4406 | 14563 | 4065 | 4063 | 13169 | 3666 | 3060 |
| 3422 | QGN16519.1 | FIM1_3232 | CP015058. | 912 | 74 | 61 | 2196 | 137 | 79 | 1361 | 69 | 101 | 261 | 71 | 90 | 433 | 68 | 76 | 298 | 86 | 48 |
| 3423 | QGN16520.1 | FIM1_3233 | CP015058. | 3200 | 715 | 2939 | 3695 | 627 | 2376 | 3112 | 715 | 3583 | 2302 | 288 | 3005 | 2113 | 313 | 2600 | 2332 | 251 | 1759 |
| 3424 | QGN16521.1 | FIM1_3234 | CP015058. | 6273 | 2563 | 1173 | 2965 | 3307 | 1363 | 5250 | 2318 | 1588 | 9513 | 3567 | 2391 | 9014 | 3918 | 2268 | 9127 | 4113 | 1652 |
| 3425 | QGN16522.1 | FIM1_3235 | CP015058. | 1474 | 995 | 1059 | 2024 | 860 | 923 | 1559 | 892 | 1443 | 6938 | 679 | 1512 | 5552 | 564 | 1374 | 6985 | 540 | 1037 |
| 3426 | QGN16523.1 | FIM1_3236 | CP015058. | 20008 | 5352 | 3649 | 18487 | 5015 | 3482 | 20258 | 5162 | 5057 | 5320 | 5831 | 6839 | 19914 | 5532 | 6503 | 8715 | 7049 | 4845 |
| 3427 | QGN16524.1 | FIM1_3237 | CP015058. | 6366 | 4705 | 3816 | 10205 | 3810 | 3366 | 7196 | 3758 | 4769 | 8952 | 2863 | 4894 | 15419 | 3129 | 5154 | 11360 | 2647 | 3391 |
| 3428 | QGN16525.1 | FIM1_3238 | CP015058. | 561 | 524 | 419 | 633 | 364 | 350 | 515 | 363 | 514 | 1537 | 219 | 470 | 1438 | 190 | 456 | 1366 | 204 | 304 |
| 3429 | QGN16526.1 | FIM1_3239 | CP015058. | 6944 | 3881 | 2351 | 7452 | 2740 | 2086 | 6450 | 2528 | 2801 | 7253 | 1969 | 2274 | 7330 | 1848 | 2524 | 6695 | 1643 | 1635 |
| 3430 | QGN16527.1 | FIM1_3240 | CP015058. | 3619 | 2945 | 1522 | 2972 | 1951 | 1362 | 3177 | 1978 | 2049 | 8444 | 2165 | 2016 | 7899 | 1692 | 2251 | 7785 | 1701 | 1395 |
| 3431 | QGN16528.1 | FIM1_3241 | CP015058. | 1852 | 753 | 1007 | 3589 | 567 | 841 | 2451 | 506 | 1344 | 4331 | 419 | 1302 | 3536 | 631 | 1366 | 3800 | 435 | 867 |
| 3432 | QGN16529.1 | FIM1_3242 | CP015058. | 1419 | 359 | 434 | 2202 | 368 | 352 | 1473 | 401 | 554 | 2776 | 343 | 576 | 2467 | 331 | 574 | 2591 | 283 | 413 |
| 3433 | QGN16530.1 | FIM1_3243 | CP015058. | 1450 | 1370 | 1365 | 1702 | 1666 | 1119 | 1074 | 1187 | 1649 | 6986 | 1634 | 2173 | 7357 | 1273 | 1913 | 7002 | 1361 | 1343 |
| 3434 | QGN16531.1 | FIM1_3244 | CP015058. | 1814 | 773 | 2267 | 1706 | 481 | 1582 | 1461 | 1303 | 2884 | 1607 | 340 | 1674 | 1864 | 400 | 1496 | 1690 | 444 | 1129 |
| 3435 | QGN16532.1 | FIM1_3245 | CP015058. | 1556 | 712 | 914 | 2085 | 848 | 771 | 1614 | 738 | 1242 | 2301 | 665 | 1268 | 4741 | 638 | 1109 | 3036 | 567 | 865 |
| 3436 | QGN16533.1 | FIM1_3246 | CP015058. | 7108 | 492 | 651 | 5807 | 719 | 496 | 7763 | 675 | 761 | 3073 | 477 | 685 | 4144 | 589 | 674 | 2981 | 371 | 456 |
| 3437 | QGN16534.1 | FIM1_3247 | CP015058. | 10971 | 2072 | 3353 | 16220 | 3731 | 2594 | 12758 | 3524 | 4309 | 5272 | 3062 | 5553 | 22498 | 3544 | 4336 | 10686 | 2896 | 3383 |
| 3438 | QGN16535.1 | FIM1_3248 | CP015058. | 897 | 214 | 252 | 812 | 152 | 179 | 869 | 151 | 286 | 870 | 85 | 232 | 708 | 79 | 221 | 781 | 79 | 144 |
| 3439 | QGN16536.1 | FIM1_3249 | CP015058. | 2576 | 3111 | 2575 | 4772 | 2409 | 2344 | 3144 | 2072 | 3330 | 7520 | 1719 | 3462 | 8212 | 1994 | 3545 | 7874 | 1778 | 2384 |
| 3440 | QGN16537.1 | FIM1_3250 | CP015058. | 930 | 878 | 613 | 609 | 665 | 564 | 622 | 610 | 789 | 2940 | 646 | 859 | 2500 | 536 | 937 | 2502 | 493 | 620 |
| 3441 | QGN16538.1 | FIM1_3251 | CP015058. | 2472 | 357 | 415 | 2127 | 372 | 298 | 2458 | 394 | 510 | 2252 | 342 | 637 | 2514 | 404 | 592 | 2233 | 354 | 413 |
| 3442 | QGN16539.1 | FIM1_3252 | CP015058. | 116 | 15 | 12 | 410 | 19 | 19 | 182 | 19 | 17 | 95 | 23 | 34 | 136 | 18 | 34 | 114 | 13 | 35 |

| | | | | | | | | | | | | | | | | | | | | | |
|---|---|---|---|---|---|---|---|---|---|---|---|---|---|---|---|---|---|---|---|---|---|
| 3443 | QGN16540.1 | FIM1_3253 | CP015058. | 10106 | 889 | 2491 | 30178 | 1197 | 2514 | 16890 | 974 | 3512 | 3261 | 541 | 3023 | 6047 | 654 | 3124 | 3960 | 449 | 1959 |
| 3444 | QGN16541.1 | FIM1_3254 | CP015058. | 2948 | 3186 | 2987 | 4917 | 3333 | 2515 | 3134 | 2960 | 3761 | 13270 | 2850 | 3856 | 13414 | 2952 | 3519 | 12929 | 2578 | 2626 |
| 3445 | QGN16542.1 | FIM1_3255 | CP015058. | 2315 | 477 | 434 | 2556 | 449 | 350 | 2282 | 428 | 546 | 3289 | 409 | 660 | 2937 | 348 | 585 | 3239 | 391 | 407 |
| 3446 | QGN16543.1 | FIM1_3256 | CP015058. | 84 | 53 | 24 | 64 | 38 | 17 | 53 | 27 | 20 | 139 | 32 | 43 | 115 | 14 | 40 | 115 | 30 | 15 |
| 3447 | QGN16544.1 | FIM1_3258 | CP015058. | 153 | 340 | 277 | 167 | 286 | 235 | 97 | 254 | 379 | 829 | 230 | 376 | 680 | 129 | 387 | 918 | 186 | 233 |
| 3448 | QGN16545.1 | FIM1_3259 | CP015058. | 233 | 134 | 480 | 197 | 288 | 314 | 168 | 341 | 468 | 1744 | 227 | 618 | 1395 | 170 | 527 | 1690 | 201 | 327 |
| 3449 | QGN16546.1 | FIM1_3260 | CP015058. | 3358 | 803 | 750 | 3505 | 945 | 670 | 3175 | 753 | 1005 | 5066 | 838 | 1301 | 5774 | 783 | 1179 | 5038 | 794 | 853 |
| 3450 | QGN16547.1 | FIM1_3261 | CP015058. | 712 | 567 | 367 | 967 | 438 | 412 | 656 | 358 | 602 | 1279 | 315 | 484 | 1967 | 320 | 501 | 1446 | 296 | 349 |
| 3451 | QGN16548.1 | FIM1_3262 | CP015058. | 22944 | 28044 | 16775 | 23277 | 22359 | 15735 | 19805 | 19553 | 19558 | 67619 | 26635 | 25754 | 60381 | 22768 | 25740 | 70579 | 22306 | 19060 |
| 3452 | QGN16549.1 | FIM1_3263 | CP015058. | 1946 | 1173 | 1085 | 2791 | 1173 | 1009 | 2730 | 1153 | 1551 | 8480 | 1079 | 1745 | 7274 | 979 | 1716 | 8783 | 1005 | 1258 |
| 3453 | QGN16550.1 | FIM1_3264 | CP015058. | 1589 | 758 | 987 | 1972 | 1044 | 927 | 1881 | 902 | 1358 | 8758 | 1020 | 1762 | 6813 | 972 | 1443 | 8113 | 917 | 1149 |
| 3454 | QGN16551.1 | FIM1_3265 | CP015058. | 1054 | 881 | 752 | 1486 | 1074 | 741 | 1055 | 809 | 990 | 4324 | 945 | 1254 | 4870 | 882 | 1004 | 4522 | 805 | 790 |
| 3455 | QGN16552.1 | FIM1_3266 | CP015058. | 327 | 634 | 433 | 165 | 618 | 379 | 195 | 507 | 528 | 2110 | 488 | 411 | 1647 | 424 | 423 | 1981 | 513 | 305 |
| 3456 | QGN16553.1 | FIM1_3267 | CP015058. | 2031 | 3309 | 2111 | 2829 | 2986 | 2016 | 2300 | 2823 | 2802 | 12389 | 2547 | 3050 | 12108 | 2438 | 2721 | 11746 | 2542 | 2021 |
| 3457 | QGN16554.1 | FIM1_3268 | CP015058. | 727 | 781 | 563 | 692 | 530 | 523 | 620 | 467 | 735 | 3787 | 657 | 743 | 2717 | 523 | 683 | 3272 | 567 | 493 |
| 3458 | QGN16555.1 | FIM1_3269 | CP015058. | 797 | 225 | 463 | 1232 | 353 | 334 | 897 | 279 | 523 | 1877 | 320 | 887 | 2883 | 317 | 690 | 2050 | 255 | 489 |
| 3459 | QGN16556.1 | FIM1_3270 | CP015058. | 8258 | 4755 | 3434 | 7515 | 5184 | 3421 | 4965 | 3837 | 4808 | 1476 | 575 | 1883 | 5838 | 1705 | 1354 | 3854 | 749 | 1004 |
| 3460 | QGN16557.1 | FIM1_3271 | CP015058. | 8362 | 2900 | 2771 | 6912 | 2488 | 2597 | 7911 | 3049 | 3899 | 11729 | 2082 | 3115 | 10573 | 1593 | 3548 | 11397 | 1751 | 2253 |
| 3461 | QGN16558.1 | FIM1_3272 | CP015058. | 2274 | 422 | 886 | 2302 | 639 | 654 | 2027 | 662 | 1108 | 4366 | 531 | 1244 | 5402 | 469 | 1080 | 4735 | 484 | 895 |
| 3462 | QGN16559.1 | FIM1_3273 | CP015058. | 1027 | 2334 | 966 | 558 | 1772 | 1025 | 963 | 1538 | 1233 | 6039 | 1938 | 1705 | 4566 | 1351 | 1696 | 5304 | 1793 | 1177 |
| 3463 | QGN16560.1 | FIM1_3274 | CP015058. | 3950 | 1344 | 1833 | 4341 | 1500 | 1535 | 3992 | 1426 | 2294 | 7855 | 1226 | 2972 | 8945 | 1281 | 2747 | 7771 | 1089 | 1905 |
| 3464 | QGN16561.1 | FIM1_3275 | CP015058. | 6527 | 1626 | 2016 | 7689 | 2455 | 1841 | 5801 | 2144 | 2531 | 6383 | 2701 | 2726 | 8349 | 2782 | 2674 | 6815 | 2538 | 2105 |
| 3465 | QGN16562.1 | FIM1_3276 | CP015058. | 697 | 151 | 315 | 1472 | 153 | 256 | 780 | 218 | 416 | 1926 | 101 | 439 | 2498 | 132 | 323 | 1980 | 94 | 288 |
| 3466 | QGN16563.1 | FIM1_3277 | CP015058. | 2093 | 1205 | 1414 | 3007 | 1649 | 1244 | 2496 | 1243 | 1860 | 7198 | 1747 | 2401 | 8894 | 1598 | 2255 | 7632 | 1526 | 1725 |
| 3467 | QGN16564.1 | FIM1_3278 | CP015058. | 12086 | 3556 | 4254 | 11930 | 4014 | 3674 | 10627 | 3414 | 5331 | 13674 | 4867 | 5948 | 21680 | 4343 | 5722 | 15942 | 4184 | 4178 |
| 3468 | QGN16565.1 | FIM1_3279 | CP015058. | 4035 | 4041 | 2146 | 3455 | 3203 | 2008 | 3180 | 2811 | 2709 | 11566 | 2855 | 3050 | 11149 | 2684 | 2964 | 11162 | 2776 | 2065 |
| 3469 | QGN16566.1 | FIM1_3280 | CP015058. | 3506 | 1140 | 1738 | 5253 | 1459 | 1472 | 3757 | 1486 | 2229 | 3485 | 1100 | 2226 | 5193 | 1150 | 2049 | 4109 | 905 | 1476 |
| 3470 | QGN16567.1 | FIM1_3281 | CP015058. | 3159 | 3785 | 3752 | 5213 | 2684 | 3491 | 3720 | 2109 | 4761 | 9149 | 2597 | 4756 | 7584 | 2534 | 5128 | 8422 | 2515 | 3438 |
| 3471 | QGN16568.1 | FIM1_3282 | CP015058. | 9256 | 2332 | 3443 | 10988 | 3448 | 2491 | 10300 | 3817 | 4576 | 2298 | 3034 | 5304 | 17109 | 3256 | 4732 | 6680 | 3387 | 3473 |
| 3472 | QGN16569.1 | FIM1_3283 | CP015058. | 819 | 789 | 582 | 1615 | 640 | 533 | 1248 | 607 | 788 | 3562 | 450 | 639 | 2609 | 459 | 650 | 3156 | 411 | 431 |
| 3473 | QGN16570.1 | FIM1_3284 | CP015058. | 2241 | 763 | 1040 | 2509 | 922 | 813 | 1664 | 979 | 1364 | 4886 | 660 | 1498 | 4935 | 475 | 1348 | 4600 | 435 | 904 |
| 3474 | QGN16571.1 | FIM1_3286 | CP015058. | 2069 | 1166 | 844 | 2819 | 1093 | 836 | 2563 | 978 | 1195 | 6583 | 850 | 1215 | 5825 | 630 | 1133 | 5919 | 708 | 775 |
| 3475 | QGN16572.1 | FIM1_3287 | CP015058. | 423 | 316 | 250 | 732 | 373 | 232 | 614 | 283 | 294 | 2052 | 298 | 345 | 1743 | 251 | 302 | 1752 | 246 | 266 |
| 3476 | QGN16573.1 | FIM1_3288 | CP015058. | 35007 | 8890 | 17328 | 42778 | 16938 | 14221 | 40072 | 14667 | 21820 | 23926 | 17419 | 25186 | 65572 | 17466 | 23445 | 43016 | 17317 | 17383 |
| 3477 | QGN16574.1 | FIM1_3289 | CP015058. | 45 | 11 | 8 | 77 | 28 | 19 | 37 | 9 | 16 | 79 | 5 | 23 | 63 | 2 | 25 | 78 | 5 | 17 |
| 3478 | QGN16575.1 | FIM1_3290 | CP015058. | 1666 | 1490 | 1116 | 2697 | 1355 | 974 | 1627 | 1082 | 1427 | 2936 | 870 | 1341 | 4236 | 886 | 1543 | 3141 | 708 | 991 |
| 3479 | QGN16576.1 | FIM1_3291 | CP015058. | 43 | 15 | 10 | 85 | 14 | 6 | 67 | 31 | 14 | 20 | 9 | 13 | 50 | 3 | 6 | 49 | 12 | 6 |
| 3480 | QGN16577.1 | FIM1_3292 | CP015058. | 28324 | 29188 | 16857 | 24917 | 21627 | 15447 | 21910 | 19903 | 20177 | 48997 | 24769 | 22921 | 53671 | 21805 | 24427 | 48259 | 21235 | 17791 |
| 3481 | QGN16578.1 | FIM1_3293 | CP015058. | 432 | 339 | 264 | 389 | 351 | 186 | 328 | 282 | 277 | 1422 | 341 | 273 | 1085 | 270 | 298 | 1414 | 303 | 192 |
| 3482 | QGN16579.1 | FIM1_3294 | CP015058. | 669 | 298 | 174 | 359 | 310 | 178 | 490 | 260 | 254 | 844 | 279 | 231 | 876 | 199 | 253 | 855 | 263 | 158 |
| 3483 | QGN16580.1 | FIM1_3295 | CP015058. | 3953 | 423 | 623 | 5307 | 499 | 425 | 4513 | 626 | 877 | 550 | 396 | 787 | 1403 | 490 | 782 | 1016 | 442 | 557 |
| 3484 | QGN16581.1 | FIM1_3297 | CP015058. | 3121 | 1312 | 1544 | 3202 | 1332 | 1406 | 2729 | 1144 | 1923 | 5877 | 1216 | 2192 | 5221 | 1305 | 2236 | 5310 | 1294 | 1445 |
| 3485 | QGN16582.1 | FIM1_3298 | CP015058. | 4659 | 1558 | 1313 | 3173 | 2718 | 1316 | 4797 | 2044 | 1797 | 9294 | 2923 | 2701 | 11620 | 3126 | 2536 | 9611 | 3429 | 1931 |
| 3486 | QGN16583.1 | FIM1_3299 | CP015058. | 7800 | 2604 | 2496 | 10440 | 3445 | 2251 | 8427 | 2836 | 3069 | 11635 | 2973 | 3883 | 16370 | 2850 | 3480 | 12915 | 2849 | 2602 |
| 3487 | QGN16584.1 | FIM1_3300 | CP015058. | 9290 | 12883 | 7356 | 8426 | 9168 | 7364 | 7053 | 8005 | 9024 | 26949 | 10540 | 11247 | 22120 | 9339 | 10861 | 25256 | 9828 | 8133 |
| 3488 | QGN16585.1 | FIM1_3301 | CP015058. | 3738 | 1622 | 1489 | 3662 | 1205 | 1136 | 3684 | 1267 | 1855 | 7064 | 1328 | 1824 | 5998 | 1613 | 1608 | 7114 | 1561 | 1117 |
| 3489 | QGN16586.1 | FIM1_3302 | CP015058. | 628 | 510 | 259 | 202 | 461 | 293 | 450 | 427 | 344 | 2639 | 632 | 534 | 1733 | 599 | 432 | 2465 | 701 | 324 |
| 3490 | QGN16587.1 | FIM1_3303 | CP015058. | 480 | 463 | 367 | 845 | 327 | 296 | 731 | 347 | 439 | 2590 | 348 | 664 | 1772 | 336 | 560 | 2500 | 345 | 387 |
| 3491 | QGN16588.1 | FIM1_3304 | CP015058. | 12087 | 6362 | 3847 | 8001 | 7951 | 3973 | 12108 | 6746 | 5063 | 14993 | 10185 | 7164 | 27202 | 8970 | 6462 | 19894 | 10832 | 4989 |
| 3492 | QGN16589.1 | FIM1_3305 | CP015058. | 34663 | 5637 | 7705 | 43777 | 6967 | 7365 | 39613 | 4363 | 10130 | 17170 | 5009 | 7818 | 22116 | 5650 | 8249 | 19806 | 4245 | 5731 |
| 3493 | QGN16590.1 | FIM1_3306 | CP015058. | 746 | 366 | 291 | 517 | 344 | 380 | 432 | 311 | 475 | 787 | 222 | 373 | 931 | 174 | 307 | 721 | 167 | 225 |
| 3494 | QGN16591.1 | FIM1_3307 | CP015058. | 7784 | 1000 | 743 | 8853 | 773 | 607 | 5572 | 867 | 965 | 2293 | 730 | 915 | 3340 | 826 | 1002 | 2548 | 585 | 679 |
| 3495 | QGN16592.1 | FIM1_3308 | CP015058. | 21729 | 281 | 210 | 10958 | 326 | 186 | 13394 | 272 | 264 | 568 | 231 | 240 | 994 | 212 | 251 | 711 | 206 | 154 |
| 3496 | QGN16593.1 | FIM1_3309 | CP015058. | 4873 | 2064 | 2144 | 5334 | 3446 | 1976 | 5117 | 2847 | 2561 | 8768 | 3210 | 3603 | 13824 | 2740 | 3188 | 10425 | 2810 | 2402 |
| 3497 | QGN16594.1 | FIM1_3310 | CP015058. | 1692 | 1057 | 1048 | 2813 | 896 | 959 | 2410 | 725 | 1388 | 4931 | 912 | 1526 | 5657 | 834 | 1341 | 5033 | 864 | 990 |
| 3498 | QGN16595.1 | FIM1_3311 | CP015058. | 4114 | 4244 | 3407 | 5129 | 4386 | 2962 | 4010 | 3585 | 4217 | 16529 | 4524 | 5688 | 16754 | 4006 | 5339 | 16227 | 3755 | 3887 |
| 3499 | QGN16596.1 | FIM1_3312 | CP015058. | 15494 | 1321 | 1274 | 12070 | 1499 | 1062 | 11041 | 1574 | 1853 | 1231 | 895 | 1784 | 5690 | 928 | 1705 | 1972 | 1001 | 1163 |
| 3500 | QGN16597.1 | FIM1_3313 | CP015058. | 716 | 29 | 271 | 2054 | 163 | 161 | 1372 | 213 | 323 | 325 | 92 | 312 | 959 | 124 | 280 | 545 | 125 | 245 |
| 3501 | QGN16598.1 | FIM1_3314 | CP015058. | 1148 | 565 | 402 | 1118 | 421 | 371 | 1071 | 369 | 578 | 2244 | 356 | 451 | 1908 | 342 | 527 | 1916 | 270 | 306 |
| 3502 | QGN16599.1 | FIM1_3315 | CP015058. | 171 | 785 | 411 | 370 | 531 | 354 | 257 | 453 | 569 | 1710 | 442 | 450 | 1513 | 435 | 482 | 1650 | 441 | 326 |
| 3503 | QGN16600.1 | FIM1_3316 | CP015058. | 981 | 1560 | 1002 | 1354 | 1771 | 875 | 973 | 1479 | 1259 | 6253 | 1741 | 1550 | 6990 | 1401 | 1592 | 6164 | 1318 | 1045 |
| 3504 | QGN16601.1 | FIM1_3317 | CP015058. | 108 | 127 | 162 | 104 | 82 | 135 | 98 | 116 | 215 | 820 | 73 | 180 | 624 | 45 | 183 | 690 | 67 | 120 |
| 3505 | QGN16602.1 | FIM1_3318 | CP015058. | 907 | 891 | 309 | 458 | 634 | 316 | 785 | 512 | 396 | 2450 | 818 | 686 | 2003 | 760 | 666 | 2351 | 940 | 465 |
| 3506 | QGN16603.1 | FIM1_3319 | CP015058. | 565 | 1065 | 966 | 803 | 1115 | 723 | 784 | 1094 | 1149 | 2983 | 1132 | 1305 | 5201 | 1024 | 1201 | 3782 | 1042 | 904 |
| 3507 | QGN16604.1 | FIM1_3320 | CP015058. | 10308 | 1247 | 1277 | 19678 | 1188 | 950 | 17971 | 1018 | 1783 | 5734 | 2243 | 1944 | 8839 | 4407 | 2596 | 7128 | 2605 | 1476 |
| 3508 | QGN16605.1 | FIM1_3321 | CP015058. | 9998 | 1488 | 828 | 6859 | 2289 | 725 | 7895 | 1976 | 1120 | 1162 | 2145 | 1004 | 5237 | 3037 | 948 | 2755 | 2817 | 681 |
| 3509 | QGN16606.1 | FIM1_3322 | CP015058. | 2169 | 311 | 334 | 1677 | 413 | 211 | 1140 | 423 | 395 | 627 | 379 | 385 | 2107 | 297 | 375 | 1162 | 373 | 268 |
| 3510 | QGN16607.1 | FIM1_3323 | CP015058. | 2936 | 831 | 1319 | 3771 | 690 | 1039 | 3185 | 748 | 1761 | 5101 | 459 | 1263 | 4703 | 409 | 1220 | 4727 | 350 | 833 |
| 3511 | QGN16608.1 | FIM1_3324 | CP015058. | 11274 | 601 | 1107 | 11112 | 814 | 876 | 12657 | 836 | 1430 | 4016 | 497 | 1389 | 4664 | 625 | 1220 | 4070 | 569 | 917 |
| 3512 | QGN16609.1 | FIM1_3325 | CP015058. | 48 | 13 | 2 | 43 | 2 | 9 | 12 | 3 | 5 | 51 | 7 | 20 | 63 | 0 | 19 | 78 | 9 | 10 |
| 3513 | QGN16610.1 | FIM1_3326 | CP015058. | 4892 | 5243 | 5338 | 12394 | 7715 | 4590 | 8828 | 7441 | 6422 | 26318 | 8561 | 10225 | 32764 | 8811 | 8831 | 27946 | 8114 | 6772 |
| 3514 | QGN16611.1 | FIM1_3327 | CP015058. | 4037 | 752 | 720 | 2972 | 1077 | 653 | 2834 | 795 | 977 | 3137 | 1038 | 1462 | 5166 | 1128 | 1358 | 3811 | 1008 | 1005 |
| 3515 | QGN16612.1 | FIM1_3328 | CP015058. | 271 | 300 | 171 | 402 | 173 | 160 | 289 | 202 | 238 | 1181 | 136 | 174 | 792 | 84 | 188 | 1087 | 108 | 125 |
| 3516 | QGN16613.1 | FIM1_3329 | CP015058. | 882 | 252 | 171 | 853 | 259 | 116 | 614 | 224 | 216 | 1330 | 226 | 191 | 961 | 275 | 177 | 1239 | 274 | 106 |
| 3517 | QGN16614.1 | FIM1_3331 | CP015058. | 249 | 367 | 381 | 206 | 424 | 292 | 179 | 362 | 494 | 1522 | 309 | 389 | 1326 | 307 | 339 | 1536 | 295 | 217 |
| 3518 | QGN16615.1 | FIM1_3332 | CP015058. | 204 | 522 | 407 | 278 | 415 | 356 | 209 | 390 | 547 | 1658 | 409 | 508 | 1448 | 445 | 517 | 1624 | 380 | 335 |
| 3519 | QGN16616.1 | FIM1_3333 | CP015058. | 4496 | 2284 | 1535 | 3850 | 2871 | 1753 | 4377 | 2212 | 2172 | 9747 | 3647 | 2543 | 9198 | 4274 | 2524 | 9354 | 4134 | 1795 |
| 3520 | QGN16617.1 | FIM1_3334 | CP015058. | 44221 | 32018 | 17877 | 39576 | 25096 | 16455 | 34788 | 23750 | 22154 | 40890 | 25790 | 27581 | 62598 | 24187 | 26578 | 47325 | 24013 | 20905 |
| 3521 | QGN16618.1 | FIM1_3335 | CP015058. | 1908 | 1090 | 633 | 1441 | 1161 | 605 | 2217 | 905 | 799 | 3694 | 1081 | 1079 | 4159 | 977 | 1029 | 3804 | 1079 | 740 |
| 3522 | QGN16619.1 | FIM1_3336 | CP015058. | 537 | 1121 | 735 | 520 | 951 | 649 | 607 | 800 | 959 | 4463 | 841 | 1051 | 3058 | 791 | 912 | 4134 | 902 | 646 |
| 3523 | QGN16620.1 | FIM1_3337 | CP015058. | 5362 | 843 | 2235 | 6019 | 1080 | 2103 | 5424 | 919 | 2831 | 5684 | 633 | 2492 | 6235 | 775 | 2245 | 5661 | 603 | 1710 |
| 3524 | QGN16621.1 | FIM1_3338 | CP015058. | 44 | 3 | 12 | 46 | 10 | 9 | 28 | 7 | 7 | 30 | 1 | 16 | 28 | 2 | 17 | 50 | 7 | 7 |
| 3525 | QGN16622.1 | FIM1_3339 | CP015058. | 1654 | 704 | 925 | 2882 | 863 | 880 | 1804 | 762 | 1211 | 1815 | 530 | 1350 | 2913 | 514 | 1360 | 2154 | 446 | 973 |
| 3526 | QGN16623.1 | FIM1_3340 | CP015058. | 1250 | 2186 | 7634 | 3808 | 3018 | 6573 | 2249 | 2973 | 9008 | 8459 | 1616 | 10660 | 11078 | 1940 | 9073 | 9964 | 1596 | 6713 |

| | | | | | | | | | | | | | | | | | | | | | |
|---|---|---|---|---|---|---|---|---|---|---|---|---|---|---|---|---|---|---|---|---|---|
| 3527 | QGN16624.1 | FIM1_3341 | CP015058. | 620 | 686 | 803 | 929 | 687 | 637 | 636 | 603 | 1023 | 3531 | 696 | 1278 | 4422 | 583 | 1165 | 3656 | 599 | 857 |
| 3528 | QGN16625.1 | FIM1_3342 | CP015058. | 188 | 96 | 143 | 471 | 77 | 100 | 286 | 87 | 190 | 851 | 47 | 133 | 603 | 67 | 122 | 632 | 55 | 76 |
| 3529 | QGN16626.1 | FIM1_3343 | CP015058. | 12656 | 5035 | 6666 | 18130 | 5361 | 5098 | 17919 | 5178 | 9121 | 3719 | 4483 | 7203 | 18678 | 4926 | 7485 | 10469 | 4585 | 5103 |
| 3530 | QGN16627.1 | FIM1_3344 | CP015058. | 48913 | 5972 | 13155 | 58707 | 7932 | 10277 | 55986 | 9345 | 18006 | 5360 | 5891 | 19809 | 38681 | 7528 | 20150 | 20625 | 6564 | 13976 |
| 3531 | QGN16628.1 | FIM1_3345 | CP015058. | 3360 | 3788 | 6341 | 4328 | 4741 | 4589 | 2755 | 3817 | 8365 | 2193 | 3172 | 8787 | 16083 | 4321 | 8404 | 6626 | 4046 | 6106 |
| 3532 | QGN16629.1 | FIM1_3346 | CP015058. | 351 | 85 | 129 | 798 | 133 | 118 | 500 | 141 | 170 | 260 | 101 | 201 | 622 | 125 | 159 | 421 | 127 | 105 |
| 3533 | QGN16630.1 | FIM1_3347 | CP015058. | 6201 | 647 | 933 | 7463 | 1063 | 886 | 4804 | 833 | 1203 | 2412 | 499 | 1039 | 5178 | 645 | 903 | 3467 | 467 | 713 |
| 3534 | QGN16631.1 | FIM1_3348 | CP015058. | 3960 | 555 | 523 | 3137 | 485 | 423 | 2526 | 469 | 690 | 3860 | 353 | 764 | 3745 | 219 | 665 | 3551 | 182 | 455 |
| 3535 | QGN16632.1 | FIM1_3349 | CP015058. | 4178 | 780 | 1248 | 5495 | 885 | 1068 | 4170 | 892 | 1639 | 4469 | 696 | 1463 | 5431 | 748 | 1416 | 4528 | 692 | 975 |
| 3536 | QGN16633.1 | FIM1_3350 | CP015058. | 18315 | 4701 | 5024 | 41167 | 5105 | 4697 | 32871 | 4369 | 6636 | 18265 | 3870 | 4468 | 17765 | 6889 | 4380 | 17750 | 4431 | 3172 |
| 3537 | QGN16634.1 | FIM1_3351 | CP015058. | 2376 | 824 | 1364 | 2460 | 878 | 1094 | 1726 | 754 | 1726 | 3763 | 874 | 1857 | 5802 | 890 | 1923 | 4657 | 834 | 1343 |
| 3538 | QGN16635.1 | FIM1_3353 | CP015058. | 5511 | 8830 | 4974 | 5178 | 6156 | 4639 | 4565 | 5162 | 5854 | 19304 | 7368 | 7312 | 15924 | 6315 | 7365 | 19430 | 6477 | 5432 |
| 3539 | QGN16636.1 | FIM1_3355 | CP015058. | 717 | 1386 | 1168 | 2055 | 1056 | 1066 | 1292 | 943 | 1577 | 5585 | 714 | 1198 | 4507 | 847 | 1155 | 5444 | 753 | 781 |
| 3540 | QGN16637.1 | FIM1_3356 | CP015058. | 1341 | 533 | 984 | 3495 | 691 | 820 | 2274 | 469 | 1290 | 4396 | 468 | 1152 | 5307 | 661 | 1099 | 4612 | 385 | 831 |
| 3541 | QGN16638.1 | FIM1_3357 | CP015058. | 3146 | 295 | 316 | 2905 | 440 | 324 | 3339 | 322 | 419 | 1455 | 246 | 412 | 1255 | 251 | 383 | 1319 | 246 | 296 |
| 3542 | QGN16639.1 | FIM1_3358 | CP015058. | 56 | 3 | 1 | 80 | 20 | 14 | 49 | 9 | 4 | 69 | 9 | 20 | 46 | 6 | 13 | 57 | 8 | 17 |
| 3543 | QGN16640.1 | FIM1_3359 | CP015058. | 4904 | 4350 | 2885 | 5530 | 3697 | 2580 | 5455 | 3431 | 3750 | 20185 | 4164 | 4340 | 16417 | 3671 | 4073 | 19576 | 3638 | 3010 |
| 3544 | QGN16641.1 | FIM1_3360 | CP015058. | 6701 | 2300 | 2472 | 9486 | 1942 | 2418 | 5807 | 1724 | 3259 | 11256 | 1892 | 4251 | 9546 | 1815 | 4307 | 10907 | 1627 | 2934 |
| 3545 | QGN16642.1 | FIM1_3361 | CP015058. | 6082 | 3949 | 4281 | 7582 | 5630 | 3572 | 6638 | 4620 | 5066 | 19786 | 5206 | 5531 | 20566 | 5041 | 5052 | 20258 | 4613 | 3626 |
| 3546 | QGN16643.1 | FIM1_3362 | CP015058. | 9460 | 928 | 1621 | 11177 | 1635 | 1311 | 9665 | 1421 | 2123 | 5646 | 1476 | 2322 | 12191 | 1523 | 2032 | 8096 | 1333 | 1510 |
| 3547 | QGN16644.1 | FIM1_3363 | CP015058. | 2202 | 1212 | 897 | 3396 | 978 | 863 | 2667 | 918 | 1229 | 3573 | 780 | 1467 | 3923 | 932 | 1310 | 3797 | 881 | 925 |
| 3548 | QGN16645.1 | FIM1_3364 | CP015058. | 2822 | 916 | 1238 | 4024 | 1016 | 1098 | 3253 | 921 | 1681 | 6485 | 783 | 1714 | 6994 | 780 | 1478 | 6167 | 728 | 1059 |
| 3549 | QGN16646.1 | FIM1_3365 | CP015058. | 483 | 195 | 234 | 412 | 161 | 168 | 574 | 227 | 263 | 1579 | 236 | 332 | 1431 | 144 | 284 | 1547 | 167 | 216 |
| 3550 | QGN16647.1 | FIM1_3366 | CP015058. | 1492 | 30 | 255 | 1837 | 81 | 136 | 1624 | 135 | 303 | 252 | 55 | 317 | 1919 | 59 | 241 | 749 | 61 | 243 |
| 3551 | QGN16648.1 | FIM1_3367 | CP015058. | 964 | 621 | 559 | 1338 | 745 | 549 | 897 | 513 | 742 | 1549 | 433 | 574 | 2234 | 279 | 731 | 1722 | 291 | 427 |
| 3552 | QGN16649.1 | FIM1_3368 | CP015058. | 467 | 426 | 330 | 673 | 419 | 262 | 378 | 289 | 397 | 1906 | 482 | 613 | 1534 | 502 | 558 | 2305 | 421 | 471 |
| 3553 | QGN16650.1 | FIM1_3369 | CP015058. | 1079 | 835 | 920 | 2537 | 1379 | 905 | 1123 | 934 | 1181 | 3288 | 583 | 901 | 3351 | 776 | 737 | 3129 | 586 | 581 |
| 3554 | QGN16651.1 | FIM1_3370 | CP015058. | 461 | 570 | 633 | 1113 | 471 | 583 | 825 | 394 | 783 | 2720 | 443 | 1121 | 1971 | 514 | 985 | 2333 | 597 | 664 |
| 3555 | QGN16652.1 | FIM1_3371 | CP015058. | 411 | 872 | 649 | 416 | 508 | 646 | 367 | 525 | 888 | 2908 | 602 | 737 | 2265 | 481 | 843 | 2897 | 434 | 496 |
| 3556 | QGN16653.1 | FIM1_3372 | CP015058. | 1048 | 609 | 553 | 1383 | 542 | 479 | 1345 | 541 | 731 | 2592 | 481 | 694 | 2388 | 363 | 620 | 2540 | 350 | 373 |
| 3557 | QGN16654.1 | FIM1_3373 | CP015058. | 1479 | 870 | 263 | 737 | 656 | 381 | 1320 | 584 | 393 | 1834 | 756 | 481 | 1568 | 693 | 528 | 1724 | 1005 | 283 |
| 3558 | QGN16655.1 | FIM1_3374 | CP015058. | 10127 | 2328 | 2434 | 30672 | 2727 | 2020 | 18880 | 2170 | 3187 | 3624 | 2691 | 3075 | 13289 | 5467 | 3167 | 6771 | 2698 | 2309 |
| 3559 | QGN16656.1 | FIM1_3375 | CP015058. | 27892 | 6711 | 3855 | 22226 | 6367 | 3457 | 20472 | 5397 | 4892 | 13318 | 5514 | 3758 | 18496 | 5993 | 4101 | 15775 | 4745 | 2912 |
| 3560 | QGN16657.1 | FIM1_3376 | CP015058. | 16 | 16 | 14 | 28 | 19 | 9 | 20 | 27 | 14 | 49 | 17 | 18 | 87 | 4 | 16 | 66 | 11 | 5 |
| 3561 | QGN16658.1 | FIM1_3377 | CP015058. | 4138 | 1231 | 715 | 1586 | 1178 | 611 | 3914 | 1212 | 913 | 1735 | 1158 | 1132 | 3715 | 1074 | 1145 | 2349 | 1176 | 799 |
| 3562 | QGN16659.1 | FIM1_3378 | CP015058. | 59 | 18 | 8 | 71 | 18 | 11 | 71 | 17 | 18 | 76 | 20 | 20 | 88 | 10 | 20 | 67 | 11 | 16 |
| 3563 | QGN16660.1 | FIM1_3379 | CP015058. | 1466 | 394 | 391 | 1356 | 290 | 389 | 1485 | 284 | 563 | 1194 | 214 | 681 | 1428 | 222 | 691 | 1173 | 228 | 481 |
| 3564 | QGN16661.1 | FIM1_3380 | CP015058. | 958 | 100 | 262 | 952 | 222 | 200 | 931 | 211 | 321 | 625 | 152 | 478 | 911 | 122 | 434 | 708 | 145 | 307 |
| 3565 | QGN16662.1 | FIM1_3381 | CP015058. | 13123 | 17608 | 10972 | 10846 | 14341 | 9301 | 10444 | 12466 | 12582 | 39656 | 17034 | 15369 | 33852 | 14847 | 16086 | 41017 | 13866 | 11900 |
| 3566 | QGN16663.1 | FIM1_3382 | CP015058. | 2395 | 320 | 322 | 1820 | 370 | 307 | 1984 | 351 | 410 | 1183 | 212 | 324 | 1348 | 331 | 310 | 1036 | 290 | 205 |
| 3567 | QGN16664.1 | FIM1_3383 | CP015058. | 4054 | 1276 | 1707 | 6321 | 1802 | 1322 | 5315 | 1432 | 2186 | 5899 | 1213 | 2009 | 8706 | 1431 | 1796 | 6491 | 1160 | 1269 |
| 3568 | QGN16665.1 | FIM1_3384 | CP015058. | 10137 | 4617 | 2974 | 5976 | 6269 | 3090 | 11111 | 5213 | 3842 | 25843 | 10885 | 6312 | 29988 | 8875 | 5716 | 26402 | 11678 | 4553 |
| 3569 | QGN16666.1 | FIM1_3385 | CP015058. | 114 | 105 | 40 | 105 | 69 | 35 | 154 | 55 | 53 | 290 | 47 | 73 | 256 | 32 | 56 | 280 | 58 | 39 |
| 3570 | QGN16667.1 | FIM1_3386 | CP015058. | 138 | 531 | 423 | 89 | 561 | 394 | 135 | 408 | 531 | 2614 | 609 | 668 | 2033 | 650 | 564 | 2512 | 664 | 440 |
| 3571 | QGN16668.1 | FIM1_3387 | CP015058. | 4052 | 1141 | 991 | 4541 | 1039 | 807 | 4365 | 954 | 1351 | 1179 | 826 | 1319 | 5063 | 893 | 1243 | 2583 | 827 | 1010 |
| 3572 | QGN16669.1 | FIM1_3388 | CP015058. | 9304 | 1211 | 1911 | 7424 | 1678 | 1447 | 7157 | 1924 | 2449 | 2449 | 1563 | 3061 | 8648 | 1626 | 2656 | 4065 | 1871 | 1941 |
| 3573 | QGN16670.1 | FIM1_3389 | CP015058. | 683 | 808 | 1282 | 2054 | 546 | 883 | 1234 | 562 | 1598 | 4417 | 525 | 1486 | 4231 | 1079 | 1399 | 4262 | 785 | 945 |
| 3574 | QGN16671.1 | FIM1_3390 | CP015058. | 3882 | 2530 | 1967 | 5781 | 2109 | 1579 | 4781 | 2217 | 2501 | 8408 | 2143 | 3014 | 11831 | 3067 | 2964 | 8739 | 2520 | 2120 |
| 3575 | QGN16672.1 | FIM1_3391 | CP015058. | 603 | 965 | 928 | 1721 | 1221 | 769 | 1509 | 1093 | 1200 | 4689 | 1080 | 1170 | 5723 | 990 | 1026 | 4680 | 991 | 754 |
| 3576 | QGN16673.1 | FIM1_3392 | CP015058. | 1596 | 556 | 923 | 1562 | 635 | 704 | 1333 | 812 | 1159 | 3324 | 512 | 1022 | 3824 | 400 | 940 | 3488 | 382 | 706 |
| 3577 | QGN16674.1 | FIM1_3393 | CP015058. | 165 | 224 | 200 | 385 | 121 | 164 | 454 | 175 | 330 | 346 | 73 | 129 | 337 | 49 | 131 | 359 | 67 | 74 |
| 3578 | QGN16675.1 | FIM1_3394 | CP015058. | 13581 | 1334 | 1529 | 8531 | 1860 | 1625 | 8617 | 1367 | 2060 | 3569 | 933 | 2191 | 7092 | 1117 | 2016 | 4901 | 929 | 1440 |
| 3579 | QGN16676.1 | FIM1_3395 | CP015058. | 2706 | 2118 | 1211 | 2161 | 1287 | 1021 | 2412 | 1282 | 1639 | 1402 | 1358 | 1924 | 3401 | 1444 | 1835 | 1983 | 1589 | 1346 |
| 3580 | QGN16677.1 | FIM1_3396 | CP015058. | 3681 | 1233 | 665 | 1642 | 764 | 580 | 1851 | 808 | 943 | 1970 | 644 | 817 | 2517 | 503 | 918 | 1823 | 527 | 589 |
| 3581 | QGN16678.1 | FIM1_3397 | CP015058. | 5685 | 3979 | 5402 | 8295 | 6432 | 4211 | 4477 | 5012 | 6801 | 6088 | 5250 | 8297 | 29250 | 7038 | 8426 | 12726 | 3879 | 5849 |
| 3582 | QGN16679.1 | FIM1_3398 | CP015058. | 3 | 8 | 6 | 12 | 5 | 5 | 3 | 10 | 5 | 22 | 9 | 15 | 31 | 2 | 10 | 39 | 4 | 5 |
| 3583 | QGN16680.1 | FIM1_3399 | CP015058. | 2664 | 530 | 762 | 1812 | 619 | 632 | 2103 | 789 | 1002 | 1676 | 590 | 993 | 3124 | 419 | 913 | 2033 | 478 | 667 |
| 3584 | QGN16681.1 | FIM1_3400 | CP015058. | 3814 | 904 | 948 | 3476 | 822 | 842 | 2896 | 828 | 1298 | 6121 | 856 | 1216 | 5310 | 780 | 1296 | 5502 | 755 | 845 |
| 3585 | QGN16682.1 | FIM1_3401 | CP015058. | 2235 | 1009 | 853 | 4278 | 1016 | 786 | 3916 | 1081 | 1239 | 2942 | 604 | 547 | 3594 | 647 | 667 | 3069 | 659 | 404 |
| 3586 | QGN16683.1 | FIM1_3402 | CP015058. | 2494 | 3012 | 2420 | 3204 | 2820 | 2050 | 3139 | 2534 | 2928 | 9365 | 2528 | 3535 | 9707 | 2459 | 3343 | 8775 | 2545 | 2342 |
| 3587 | QGN16684.1 | FIM1_3403 | CP015058. | 7317 | 3134 | 1466 | 4105 | 3631 | 1580 | 6919 | 3282 | 2029 | 8740 | 4558 | 2859 | 12921 | 4060 | 2754 | 9051 | 4946 | 2058 |
| 3588 | QGN16685.1 | FIM1_3404 | CP015058. | 1379 | 90 | 30 | 826 | 26 | 20 | 511 | 104 | 41 | 169 | 39 | 34 | 168 | 28 | 40 | 171 | 30 | 26 |
| 3589 | QGN16686.1 | FIM1_3405 | CP015058. | 10484 | 7541 | 3989 | 8545 | 7513 | 3863 | 9918 | 6854 | 5335 | 18841 | 9973 | 6958 | 27606 | 7856 | 7245 | 21990 | 8912 | 5368 |
| 3590 | QGN16687.1 | FIM1_3407 | CP015058. | 1102 | 1417 | 946 | 1927 | 928 | 835 | 1469 | 832 | 1180 | 4329 | 933 | 1194 | 3744 | 833 | 1374 | 3808 | 831 | 882 |
| 3591 | QGN16688.1 | FIM1_3408 | CP015058. | 2390 | 923 | 671 | 2977 | 644 | 567 | 2369 | 692 | 891 | 2365 | 504 | 863 | 2959 | 505 | 832 | 2310 | 485 | 578 |
| 3592 | QGN16689.1 | FIM1_3409 | CP015058. | 5483 | 7860 | 5440 | 12302 | 6903 | 4674 | 6679 | 5721 | 6740 | 12146 | 8231 | 9409 | 22746 | 7276 | 9337 | 17359 | 7464 | 6935 |
| 3593 | QGN16690.1 | FIM1_3410 | CP015058. | 2314 | 1108 | 723 | 1487 | 1088 | 700 | 2231 | 1099 | 910 | 3543 | 1131 | 1037 | 4808 | 1035 | 1030 | 3619 | 1066 | 736 |
| 3594 | QGN16691.1 | FIM1_3411 | CP015058. | 1612 | 2311 | 882 | 970 | 1557 | 943 | 1569 | 1497 | 1289 | 4850 | 1640 | 1501 | 4797 | 1357 | 1635 | 4435 | 1558 | 1053 |
| 3595 | QGN16692.1 | FIM1_3412 | CP015058. | 1830 | 1347 | 1005 | 2259 | 1016 | 854 | 2093 | 944 | 1280 | 3869 | 882 | 1198 | 3929 | 794 | 1312 | 3583 | 774 | 859 |
| 3596 | QGN16693.1 | FIM1_3413 | CP015058. | 3098 | 2644 | 1652 | 2131 | 2521 | 1366 | 3012 | 2683 | 2201 | 3696 | 2053 | 2257 | 8807 | 1921 | 2197 | 5784 | 1837 | 1505 |
| 3597 | QGN16694.1 | FIM1_3414 | CP015058. | 3339 | 2381 | 2187 | 5976 | 1375 | 1829 | 4267 | 2143 | 2802 | 2723 | 1556 | 2728 | 4116 | 1063 | 3072 | 3515 | 1100 | 1894 |
| 3598 | QGN16695.1 | FIM1_3415 | CP015058. | 4841 | 397 | 536 | 3402 | 696 | 356 | 4817 | 964 | 696 | 349 | 458 | 645 | 1869 | 574 | 583 | 743 | 579 | 405 |
| 3599 | QGN16696.1 | FIM1_3416 | CP015058. | 530 | 353 | 328 | 634 | 286 | 190 | 546 | 382 | 373 | 364 | 214 | 393 | 1293 | 245 | 328 | 615 | 259 | 265 |
| 3600 | QGN16697.1 | FIM1_3417 | CP015058. | 11757 | 13517 | 8664 | 10662 | 10593 | 7671 | 9472 | 9206 | 10136 | 31414 | 13522 | 12792 | 30476 | 11739 | 13079 | 31249 | 11448 | 9906 |
| 3601 | QGN16698.1 | FIM1_3418 | CP015058. | 1278 | 2567 | 1448 | 1632 | 2283 | 1520 | 1238 | 1852 | 1970 | 9223 | 2028 | 2089 | 7915 | 1800 | 1988 | 9048 | 1998 | 1379 |
| 3602 | QGN16699.1 | FIM1_3419 | CP015058. | 9561 | 1669 | 1634 | 10533 | 2097 | 1374 | 10474 | 1754 | 2133 | 8714 | 1647 | 2533 | 10827 | 1793 | 2231 | 8772 | 1602 | 1635 |
| 3603 | QGN16700.1 | FIM1_3420 | CP015058. | 257 | 30 | 112 | 728 | 34 | 69 | 226 | 40 | 151 | 254 | 16 | 69 | 298 | 31 | 68 | 265 | 18 | 41 |
| 3604 | QGN16701.1 | FIM1_3421 | CP015058. | 100 | 95 | 102 | 599 | 97 | 74 | 174 | 89 | 110 | 335 | 66 | 112 | 350 | 71 | 89 | 330 | 59 | 58 |
| 3605 | QGN16702.1 | FIM1_3422 | CP015058. | 193 | 32 | 63 | 840 | 24 | 38 | 433 | 31 | 84 | 648 | 51 | 121 | 689 | 151 | 122 | 610 | 61 | 98 |
| 3606 | QGN16703.1 | FIM1_3423 | CP015058. | 5702 | 141 | 284 | 2202 | 241 | 163 | 2336 | 252 | 349 | 733 | 127 | 315 | 1518 | 100 | 294 | 934 | 98 | 210 |
| 3607 | QGN16704.1 | FIM1_3424 | CP015058. | 1837 | 447 | 464 | 2979 | 489 | 407 | 1999 | 420 | 642 | 550 | 427 | 682 | 1550 | 458 | 708 | 827 | 479 | 492 |
| 3608 | QGN16705.1 | FIM1_3425 | CP015058. | 3663 | 409 | 682 | 3677 | 601 | 525 | 2992 | 662 | 951 | 6037 | 924 | 575 | 7916 | 1142 | 688 | 6646 | 823 | 415 |
| 3609 | QGN16706.1 | FIM1_3426 | CP015058. | 12671 | 1003 | 1158 | 27645 | 1254 | 808 | 18461 | 1459 | 1682 | 2137 | 2513 | 1764 | 13131 | 5776 | 2075 | 5230 | 2675 | 1358 |
| 3610 | QGN16707.1 | FIM1_3427 | CP015058. | 4558 | 427 | 641 | 5118 | 610 | 419 | 5032 | 701 | 900 | 1385 | 437 | 867 | 3115 | 472 | 896 | 1869 | 469 | 635 |

| | | | | | | | | | | | | | | | | | | | | | |
|---|---|---|---|---|---|---|---|---|---|---|---|---|---|---|---|---|---|---|---|---|---|
| 3611 | QGN16708.1 | FIM1_3428 | CP015058. | 373 | 67 | 24 | 354 | 39 | 24 | 627 | 86 | 36 | 89 | 17 | 25 | 134 | 14 | 27 | 122 | 27 | 12 |
| 3612 | QGN16709.1 | FIM1_3429 | CP015058. | 2542 | 2628 | 1815 | 2012 | 2270 | 1627 | 2358 | 2135 | 2359 | 9303 | 2588 | 2620 | 9949 | 2307 | 2723 | 9378 | 2350 | 1768 |
| 3613 | QGN16710.1 | FIM1_3430 | CP015058. | 2263 | 1700 | 1211 | 3329 | 1542 | 1154 | 2978 | 1430 | 1547 | 4913 | 1182 | 1257 | 4654 | 1078 | 1434 | 4528 | 1053 | 884 |
| 3614 | QGN16711.1 | FIM1_3431 | CP015058. | 1381 | 114 | 88 | 1742 | 140 | 82 | 1512 | 107 | 110 | 517 | 233 | 238 | 681 | 289 | 170 | 509 | 379 | 148 |
| 3615 | QGN16712.1 | FIM1_3432 | CP015058. | 518 | 1088 | 661 | 1372 | 904 | 573 | 1011 | 808 | 857 | 2633 | 597 | 697 | 2880 | 583 | 691 | 2554 | 535 | 477 |
| 3616 | QGN16713.1 | FIM1_3433 | CP015058. | 4582 | 2609 | 2075 | 5553 | 2506 | 1927 | 5179 | 2445 | 2962 | 9275 | 2251 | 2994 | 13099 | 2032 | 2823 | 9725 | 2029 | 2091 |
| 3617 | QGN16714.1 | FIM1_3434 | CP015058. | 733 | 385 | 528 | 877 | 384 | 451 | 642 | 387 | 680 | 2822 | 364 | 822 | 2806 | 312 | 747 | 2714 | 303 | 542 |
| 3618 | QGN16715.1 | FIM1_3435 | CP015058. | 1496 | 1683 | 1856 | 3032 | 2106 | 1536 | 3007 | 1920 | 2458 | 9767 | 2390 | 2440 | 14114 | 1948 | 2233 | 11341 | 1912 | 1789 |
| 3619 | QGN16716.1 | FIM1_3437 | CP015058. | 4830 | 13521 | 14256 | 11692 | 11306 | 11479 | 8488 | 10589 | 16524 | 29250 | 12024 | 20955 | 53040 | 11626 | 17443 | 41800 | 10201 | 13356 |
| 3620 | QGN16717.1 | FIM1_3438 | CP015058. | 797 | 3309 | 1754 | 961 | 2634 | 1361 | 1087 | 3076 | 2367 | 2180 | 1964 | 2621 | 5228 | 1739 | 2884 | 3127 | 1797 | 1912 |
| 3621 | QGN16718.1 | FIM1_3439 | CP015058. | 4234 | 719 | 991 | 7234 | 678 | 1030 | 4530 | 637 | 1560 | 3546 | 791 | 1159 | 6290 | 1085 | 1487 | 4933 | 740 | 956 |
| 3622 | QGN16719.1 | FIM1_3440 | CP015058. | 3254 | 1873 | 1638 | 6621 | 1840 | 1349 | 4902 | 1828 | 1972 | 7299 | 1709 | 2528 | 9497 | 2137 | 2390 | 7809 | 1822 | 1771 |
| 3623 | QGN16720.1 | FIM1_3441 | CP015058. | 3163 | 342 | 630 | 3567 | 365 | 491 | 3302 | 480 | 750 | 4660 | 321 | 935 | 3858 | 386 | 843 | 4128 | 334 | 600 |
| 3624 | QGN16721.1 | FIM1_3442 | CP015058. | 6560 | 2871 | 2790 | 8204 | 4140 | 2798 | 5231 | 3124 | 3473 | 9865 | 3653 | 5029 | 16715 | 3251 | 4153 | 12015 | 2932 | 3310 |
| 3625 | QGN16722.1 | FIM1_3443 | CP015058. | 273 | 4 | 39 | 220 | 23 | 25 | 136 | 29 | 36 | 133 | 31 | 152 | 276 | 28 | 135 | 225 | 22 | 79 |
| 3626 | QGN16723.1 | FIM1_3444 | CP015058. | 2748 | 4341 | 2544 | 2398 | 3568 | 2393 | 2812 | 3629 | 3423 | 7117 | 3392 | 3273 | 11555 | 3163 | 3372 | 8551 | 3213 | 2308 |
| 3627 | QGN16724.1 | FIM1_3445 | CP015058. | 1037 | 3712 | 2113 | 2481 | 2709 | 1983 | 1978 | 2280 | 2656 | 8843 | 3008 | 3185 | 8114 | 2787 | 3240 | 7885 | 3027 | 2207 |
| 3628 | QGN16725.1 | FIM1_3446 | CP015058. | 1351 | 1022 | 783 | 1661 | 1170 | 690 | 1145 | 975 | 1021 | 4137 | 898 | 1366 | 4558 | 852 | 1247 | 4047 | 820 | 884 |
| 3629 | QGN16726.1 | FIM1_3447 | CP015058. | 2157 | 1508 | 1630 | 2222 | 1750 | 1444 | 2361 | 1388 | 2149 | 9244 | 1808 | 2805 | 9567 | 1604 | 2724 | 8537 | 1562 | 1836 |
| 3630 | QGN16727.1 | FIM1_3448 | CP015058. | 1285 | 411 | 430 | 1964 | 432 | 351 | 1550 | 426 | 560 | 2835 | 351 | 684 | 2995 | 375 | 581 | 2767 | 330 | 493 |
| 3631 | QGN16728.1 | FIM1_3449 | CP015058. | 988 | 737 | 750 | 1527 | 827 | 662 | 1216 | 684 | 998 | 3737 | 545 | 931 | 3240 | 498 | 757 | 3281 | 460 | 564 |
| 3632 | QGN16729.1 | FIM1_3450 | CP015058. | 6534 | 3259 | 1361 | 3390 | 3840 | 1488 | 5589 | 3327 | 1861 | 8010 | 4246 | 2943 | 11101 | 4275 | 2800 | 8570 | 4695 | 2066 |
| 3633 | QGN16730.1 | FIM1_3451 | CP015058. | 0 | 0 | 0 | 4 | 2 | 0 | 5 | 0 | 0 | 1 | 0 | 0 | 0 | 0 | 0 | 1 | 0 | 0 |
| 3634 | QGN16731.1 | FIM1_3452 | CP015058. | 3516 | 876 | 1104 | 3223 | 1114 | 937 | 4025 | 1027 | 1460 | 4888 | 973 | 1745 | 7089 | 856 | 1624 | 5723 | 784 | 1242 |
| 3635 | QGN16732.1 | FIM1_3453 | CP015058. | 13 | 2 | 0 | 34 | 0 | 0 | 17 | 6 | 0 | 12 | 2 | 1 | 18 | 1 | 2 | 17 | 2 | 0 |
| 3636 | QGN16733.1 | FIM1_3454 | CP015058. | 108191 | 5121 | 1217 | 79010 | 5196 | 889 | 85740 | 5993 | 1637 | 27225 | 2411 | 3601 | 34024 | 3043 | 3449 | 24916 | 1771 | 2544 |
| 3637 | QGN16734.1 | FIM1_3455 | CP015058. | 17241 | 55 | 30 | 11232 | 80 | 20 | 12779 | 153 | 37 | 844 | 31 | 123 | 1348 | 45 | 109 | 834 | 31 | 76 |
| 3638 | QGN16735.1 | FIM1_3456 | CP015058. | 4062 | 26 | 14 | 1426 | 22 | 7 | 2073 | 34 | 13 | 51 | 16 | 28 | 105 | 18 | 18 | 78 | 17 | 17 |
| 3639 | QGN16736.1 | FIM1_3457 | CP015058. | 4707 | 356 | 184 | 6225 | 334 | 205 | 5874 | 261 | 273 | 191 | 226 | 296 | 1068 | 201 | 314 | 465 | 275 | 167 |
| 3640 | QGN16737.1 | FIM1_3458 | CP015058. | 35539 | 1623 | 1569 | 53293 | 3648 | 1820 | 35896 | 2344 | 2248 | 17116 | 2842 | 3208 | 26115 | 2809 | 3200 | 19249 | 2823 | 2224 |
| 3641 | QGN16738.1 | FIM1_3460 | CP015058. | 10324 | 3180 | 3536 | 14221 | 3900 | 2561 | 10591 | 4115 | 4137 | 9774 | 4171 | 5103 | 17655 | 4621 | 5098 | 13755 | 3906 | 3529 |
| 3642 | QGN16739.1 | FIM1_3461 | CP015058. | 588 | 657 | 774 | 1369 | 735 | 673 | 971 | 786 | 1138 | 514 | 715 | 1140 | 3951 | 709 | 1064 | 1518 | 777 | 730 |
| 3643 | QGN16740.1 | FIM1_3462 | CP015058. | 966 | 397 | 513 | 1430 | 528 | 460 | 1103 | 447 | 668 | 2142 | 411 | 891 | 3003 | 404 | 786 | 2337 | 349 | 569 |
| 3644 | QGN16741.1 | FIM1_3463 | CP015058. | 12291 | 1098 | 1381 | 16505 | 1346 | 1002 | 13766 | 1657 | 1892 | 1131 | 914 | 2467 | 4592 | 1453 | 1843 | 2197 | 1410 | 1449 |
| 3645 | QGN16742.1 | FIM1_3464 | CP015058. | 869 | 242 | 169 | 2004 | 168 | 156 | 649 | 166 | 232 | 808 | 163 | 302 | 878 | 153 | 271 | 692 | 150 | 209 |
| 3646 | QGN16743.1 | FIM1_3465 | CP015058. | 426 | 137 | 211 | 1593 | 79 | 133 | 410 | 117 | 271 | 578 | 57 | 206 | 738 | 74 | 208 | 632 | 59 | 144 |
| 3647 | QGN16744.1 | FIM1_3466 | CP015058. | 5020 | 1905 | 1583 | 5907 | 1849 | 1507 | 4642 | 1798 | 2217 | 5953 | 1781 | 2640 | 8178 | 1690 | 2597 | 5903 | 1667 | 1803 |
| 3648 | QGN16745.1 | FIM1_3467 | CP015058. | 1963 | 396 | 429 | 3541 | 276 | 270 | 2280 | 385 | 586 | 220 | 176 | 813 | 1653 | 347 | 602 | 369 | 308 | 522 |
| 3649 | QGN16746.1 | FIM1_3468 | CP015058. | 3067 | 2812 | 1476 | 2397 | 1869 | 1329 | 2213 | 1642 | 1880 | 4587 | 2034 | 2182 | 4545 | 1755 | 2298 | 4539 | 1790 | 1594 |
| 3650 | QGN16747.1 | FIM1_3469 | CP015058. | 2957 | 1053 | 1305 | 3700 | 1228 | 1008 | 2697 | 1381 | 1696 | 1262 | 864 | 2137 | 6071 | 990 | 1821 | 2085 | 969 | 1452 |
| 3651 | QGN16748.1 | FIM1_3470 | CP015058. | 912 | 434 | 450 | 860 | 486 | 334 | 912 | 553 | 588 | 618 | 427 | 703 | 2494 | 486 | 613 | 1181 | 519 | 408 |
| 3652 | QGN16749.1 | FIM1_3471 | CP015058. | 2158 | 2032 | 1359 | 2589 | 1573 | 1348 | 2418 | 1356 | 1806 | 6134 | 1848 | 2298 | 5782 | 1516 | 2080 | 5779 | 1858 | 1521 |
| 3653 | QGN16750.1 | FIM1_3472 | CP015058. | 1091 | 213 | 398 | 1411 | 336 | 326 | 1329 | 358 | 514 | 2163 | 273 | 632 | 2547 | 337 | 551 | 2308 | 296 | 440 |
| 3654 | QGN16751.1 | FIM1_3473 | CP015058. | 5152 | 2416 | 1085 | 2367 | 2084 | 1081 | 4919 | 2056 | 1498 | 3417 | 3072 | 1918 | 8388 | 2565 | 1689 | 5124 | 3509 | 1426 |
| 3655 | QGN16752.1 | FIM1_3474 | CP015058. | 3957 | 80 | 223 | 4727 | 63 | 160 | 4999 | 141 | 250 | 312 | 52 | 264 | 1367 | 43 | 206 | 610 | 74 | 193 |
| 3656 | QGN16753.1 | FIM1_3475 | CP015058. | 9147 | 2125 | 2287 | 7833 | 2660 | 1859 | 7943 | 2526 | 2800 | 8601 | 2523 | 3366 | 11237 | 2452 | 3236 | 8806 | 2204 | 2258 |
| 3657 | QGN16754.1 | FIM1_3476 | CP015058. | 4393 | 710 | 919 | 4194 | 761 | 800 | 4770 | 766 | 1213 | 5065 | 688 | 1468 | 5544 | 631 | 1291 | 4954 | 665 | 911 |
| 3658 | QGN16755.1 | FIM1_3477 | CP015058. | 3125 | 2119 | 1199 | 3766 | 1124 | 977 | 4103 | 1290 | 1614 | 5542 | 1201 | 1813 | 5853 | 1163 | 1780 | 4949 | 1111 | 1201 |
| 3659 | QGN16756.1 | FIM1_3478 | CP015058. | 123 | 90 | 187 | 227 | 142 | 130 | 130 | 148 | 229 | 345 | 125 | 250 | 526 | 101 | 235 | 374 | 113 | 188 |
| 3660 | QGN16757.1 | FIM1_3479 | CP015058. | 9541 | 4743 | 5501 | 10524 | 4169 | 4350 | 9419 | 3904 | 6565 | 17187 | 4738 | 10128 | 19138 | 4415 | 9538 | 17128 | 3960 | 6495 |
| 3661 | QGN16758.1 | FIM1_3480 | CP015058. | 2039 | 2842 | 1059 | 666 | 2167 | 1145 | 1536 | 1787 | 1368 | 8604 | 3151 | 2295 | 7605 | 2346 | 2157 | 8387 | 3333 | 1531 |
| 3662 | QGN16759.1 | FIM1_3481 | CP015058. | 4054 | 2278 | 1943 | 4247 | 2470 | 1629 | 4556 | 2098 | 2455 | 7820 | 2231 | 2919 | 11697 | 2296 | 2761 | 9516 | 2045 | 2050 |
| 3663 | QGN16760.1 | FIM1_3482 | CP015058. | 1341 | 2596 | 3652 | 3148 | 1685 | 3210 | 2570 | 1848 | 4662 | 7663 | 1101 | 4393 | 6152 | 1240 | 4450 | 7717 | 1190 | 2951 |
| 3664 | QGN16761.1 | FIM1_3483 | CP015058. | 827 | 594 | 679 | 971 | 571 | 589 | 818 | 539 | 874 | 4658 | 550 | 1137 | 3698 | 434 | 1044 | 4250 | 428 | 698 |
| 3665 | QGN16762.1 | FIM1_3484 | CP015058. | 2120 | 880 | 1189 | 2585 | 757 | 959 | 2204 | 803 | 1470 | 5073 | 718 | 1797 | 5561 | 674 | 1713 | 5103 | 640 | 1209 |
| 3666 | QGN16763.1 | FIM1_3485 | CP015058. | 736 | 313 | 360 | 829 | 327 | 339 | 684 | 291 | 424 | 1835 | 304 | 588 | 1277 | 340 | 571 | 1424 | 381 | 382 |
| 3667 | QGN16764.1 | FIM1_3486 | CP015058. | 2127 | 2515 | 2943 | 5063 | 2350 | 2588 | 3700 | 2146 | 3550 | 11023 | 1818 | 3967 | 10163 | 1881 | 3551 | 10359 | 1789 | 2626 |
| 3668 | QGN16765.1 | FIM1_3487 | CP015058. | 714 | 825 | 555 | 887 | 590 | 556 | 859 | 555 | 804 | 3530 | 571 | 894 | 2261 | 472 | 812 | 3339 | 577 | 570 |
| 3669 | QGN16766.1 | FIM1_3488 | CP015058. | 530 | 307 | 239 | 812 | 236 | 233 | 606 | 277 | 387 | 1350 | 211 | 430 | 1579 | 210 | 403 | 1240 | 215 | 244 |
| 3670 | QGN16767.1 | FIM1_3489 | CP015058. | 1122 | 404 | 416 | 1339 | 487 | 313 | 988 | 498 | 554 | 2844 | 539 | 817 | 2794 | 557 | 728 | 2674 | 514 | 496 |
| 3671 | QGN16768.1 | FIM1_3490 | CP015058. | 1608 | 800 | 652 | 1878 | 650 | 582 | 1734 | 599 | 830 | 3585 | 725 | 973 | 3279 | 653 | 1013 | 3203 | 701 | 631 |
| 3672 | QGN16769.1 | FIM1_3491 | CP015058. | 4845 | 1073 | 1320 | 3984 | 1282 | 1106 | 4677 | 1272 | 1783 | 7780 | 1093 | 1934 | 7754 | 886 | 1691 | 7470 | 878 | 1193 |
| 3673 | QGN16770.1 | FIM1_3492 | CP015058. | 1057 | 576 | 684 | 638 | 467 | 587 | 749 | 618 | 885 | 2342 | 440 | 888 | 3071 | 279 | 866 | 2480 | 303 | 585 |
| 3674 | QGN16771.1 | FIM1_3493 | CP015058. | 143 | 88 | 64 | 221 | 83 | 59 | 161 | 84 | 94 | 58 | 59 | 90 | 170 | 50 | 57 | 93 | 57 | 58 |
| 3675 | QGN16772.1 | FIM1_3494 | CP015058. | 13617 | 5618 | 5039 | 11137 | 5020 | 4089 | 10957 | 4310 | 6058 | 18508 | 8178 | 8010 | 23213 | 7557 | 7738 | 21107 | 7110 | 5687 |
| 3676 | QGN16773.1 | FIM1_3495 | CP015058. | 1018 | 117 | 122 | 1622 | 75 | 98 | 1358 | 91 | 164 | 481 | 57 | 145 | 547 | 65 | 119 | 428 | 82 | 92 |
| 3677 | QGN16774.1 | FIM1_3496 | CP015058. | 1106 | 482 | 260 | 1017 | 414 | 303 | 1008 | 375 | 358 | 1730 | 426 | 525 | 1529 | 373 | 507 | 1419 | 508 | 344 |
| 3678 | QGN16775.1 | FIM1_3497 | CP015058. | 12366 | 998 | 1409 | 5068 | 574 | 702 | 3539 | 1218 | 1695 | 3404 | 701 | 1366 | 5583 | 633 | 1208 | 3835 | 418 | 925 |
| 3679 | QGN16776.1 | FIM1_3498 | CP015058. | 111 | 141 | 105 | 89 | 116 | 72 | 102 | 182 | 136 | 413 | 97 | 142 | 431 | 62 | 150 | 402 | 99 | 80 |
| 3680 | QGN16777.1 | FIM1_3499 | CP015058. | 3740 | 1509 | 1883 | 5131 | 1712 | 1496 | 3860 | 1617 | 2316 | 6046 | 1469 | 2794 | 8090 | 1367 | 2604 | 6569 | 1308 | 1925 |
| 3681 | QGN16778.1 | FIM1_3500 | CP015058. | 1200 | 1506 | 1310 | 3452 | 1163 | 1112 | 2523 | 1205 | 1667 | 4397 | 850 | 1740 | 4444 | 1035 | 1765 | 4282 | 953 | 1155 |
| 3682 | QGN16779.1 | FIM1_3501 | CP015058. | 11503 | 1619 | 1679 | 8987 | 1709 | 1306 | 9314 | 2123 | 2238 | 2552 | 1224 | 2320 | 10019 | 1034 | 2180 | 5731 | 980 | 1561 |
| 3683 | QGN16780.1 | FIM1_3503 | CP015058. | 649 | 52 | 283 | 1313 | 171 | 126 | 573 | 200 | 272 | 166 | 111 | 380 | 1147 | 243 | 379 | 442 | 147 | 263 |
| 3684 | QGN16781.1 | FIM1_3504 | CP015058. | 1683 | 311 | 272 | 852 | 245 | 206 | 1641 | 435 | 351 | 608 | 279 | 369 | 1317 | 471 | 314 | 874 | 450 | 288 |
| 3685 | QGN16782.1 | FIM1_3505 | CP015058. | 3085 | 3957 | 2910 | 2964 | 3846 | 2685 | 2396 | 3334 | 3604 | 6236 | 2576 | 2661 | 8863 | 2505 | 2839 | 6379 | 2157 | 1943 |
| 3686 | QGN16783.1 | FIM1_3506 | CP015058. | 518 | 632 | 436 | 1420 | 489 | 370 | 1008 | 444 | 580 | 2149 | 409 | 659 | 2105 | 366 | 666 | 2089 | 327 | 432 |
| 3687 | QGN16784.1 | FIM1_3507 | CP015058. | 2288 | 1797 | 1670 | 2211 | 1758 | 1515 | 2436 | 1656 | 2230 | 4722 | 1199 | 1501 | 4742 | 984 | 1662 | 4623 | 941 | 1061 |
| 3688 | QGN16785.1 | FIM1_3508 | CP015058. | 703 | 651 | 467 | 656 | 726 | 460 | 753 | 629 | 642 | 2969 | 695 | 893 | 3417 | 682 | 736 | 2956 | 680 | 590 |
| 3689 | QGN16786.1 | FIM1_3509 | CP015058. | 972 | 12 | 11 | 854 | 18 | 10 | 602 | 17 | 8 | 65 | 10 | 17 | 82 | 12 | 22 | 65 | 12 | 16 |
| 3690 | QGN16787.1 | FIM1_3510 | CP015058. | 144 | 3 | 0 | 141 | 1 | 2 | 185 | 2 | 2 | 4 | 0 | 2 | 17 | 1 | 0 | 5 | 2 | 1 |
| 3691 | QGN16788.1 | FIM1_3511 | CP015058. | 2793 | 6130 | 4049 | 4922 | 4035 | 3654 | 3748 | 3704 | 5193 | 11055 | 3037 | 4965 | 11236 | 2889 | 5044 | 11482 | 2853 | 3342 |
| 3692 | QGN16789.1 | FIM1_3512 | CP015058. | 6396 | 2806 | 3079 | 15033 | 2943 | 2752 | 9536 | 2755 | 4000 | 6408 | 2363 | 4450 | 13239 | 2548 | 4336 | 9225 | 2175 | 3087 |
| 3693 | QGN16790.1 | FIM1_3513 | CP015058. | 1344 | 354 | 454 | 10330 | 739 | 692 | 1801 | 357 | 596 | 1791 | 435 | 876 | 2697 | 459 | 889 | 2201 | 422 | 559 |
| 3694 | QGN16791.1 | FIM1_3514 | CP015058. | 5632 | 2813 | 2120 | 7137 | 2727 | 1627 | 6695 | 2758 | 2759 | 1860 | 1255 | 2056 | 7526 | 1277 | 1880 | 3341 | 1253 | 1384 |

| | | | | | | | | | | | | | | | | | | | | | |
|---|---|---|---|---|---|---|---|---|---|---|---|---|---|---|---|---|---|---|---|---|---|
| 3695 | QGN16792.1 | FIM1_3515 | CP015058. | 1 | 0 | 0 | 1 | 0 | 0 | 1 | 1 | 0 | 0 | 3 | 0 | 2 | 0 | 0 | 0 | 0 | 0 |
| 3696 | QGN16793.1 | FIM1_3516 | CP015058. | 10513 | 5138 | 3490 | 9006 | 6052 | 2892 | 10669 | 5960 | 4508 | 3295 | 4952 | 5659 | 17777 | 4979 | 5136 | 6856 | 5971 | 3929 |
| 3697 | QGN16794.1 | FIM1_3517 | CP015058. | 989 | 456 | 497 | 1415 | 374 | 388 | 1074 | 355 | 652 | 835 | 222 | 438 | 1090 | 211 | 493 | 816 | 195 | 267 |
| 3698 | QGN16795.1 | FIM1_3518 | CP015058. | 6179 | 3601 | 992 | 3303 | 2050 | 946 | 5051 | 1951 | 1396 | 900 | 2263 | 2011 | 5528 | 2001 | 1920 | 2316 | 2910 | 1329 |
| 3699 | QGN16796.1 | FIM1_3519 | CP015058. | 1189 | 741 | 418 | 592 | 585 | 355 | 836 | 647 | 603 | 632 | 471 | 744 | 1754 | 452 | 738 | 958 | 545 | 518 |
| 3700 | QGN16797.1 | FIM1_3520 | CP015058. | 2020 | 1011 | 758 | 2018 | 620 | 638 | 1938 | 1037 | 1163 | 381 | 434 | 962 | 1336 | 499 | 845 | 606 | 557 | 622 |
| 3701 | QGN16798.1 | FIM1_3522 | CP015058. | 6051 | 3005 | 965 | 2321 | 3077 | 1152 | 5056 | 2533 | 1366 | 3051 | 3161 | 2290 | 8316 | 3287 | 2082 | 4903 | 3847 | 1638 |
| 3702 | QGN16799.1 | FIM1_3523 | CP015058. | 11913 | 5168 | 3440 | 12685 | 5005 | 2816 | 11834 | 4655 | 4330 | 7646 | 5385 | 6147 | 21924 | 5714 | 5411 | 12287 | 5589 | 4175 |
| 3703 | QGN16800.1 | FIM1_3524 | CP015058. | 2298 | 1567 | 994 | 1956 | 1263 | 1001 | 2281 | 1171 | 1253 | 6423 | 1607 | 1531 | 5186 | 1168 | 1624 | 5944 | 1392 | 1004 |
| 3704 | QGN16801.1 | FIM1_3525 | CP015058. | 102 | 389 | 312 | 407 | 341 | 335 | 315 | 356 | 429 | 1483 | 309 | 395 | 1929 | 264 | 381 | 1549 | 252 | 280 |
| 3705 | QGN16802.1 | FIM1_3526 | CP015058. | 4940 | 3616 | 2248 | 5812 | 2564 | 2064 | 5148 | 2645 | 2994 | 7362 | 1923 | 2368 | 7382 | 1841 | 2579 | 7109 | 1855 | 1605 |
| 3706 | QGN16803.1 | FIM1_3527 | CP015058. | 2916 | 1940 | 1547 | 3341 | 1407 | 1178 | 2947 | 1550 | 2059 | 1041 | 849 | 1594 | 4057 | 856 | 1617 | 2412 | 864 | 1061 |
| 3707 | QGN16804.1 | FIM1_3528 | CP015058. | 17996 | 630 | 1137 | 20267 | 969 | 800 | 20470 | 1049 | 1542 | 1174 | 678 | 1785 | 6709 | 1086 | 1531 | 2950 | 913 | 1132 |
| 3708 | QGN16805.1 | FIM1_3529 | CP015058. | 113 | 443 | 416 | 280 | 397 | 418 | 276 | 377 | 595 | 2670 | 469 | 615 | 2292 | 368 | 621 | 2685 | 372 | 414 |
| 3709 | QGN16806.1 | FIM1_3530 | CP015058. | 318 | 327 | 210 | 264 | 265 | 244 | 313 | 251 | 306 | 2086 | 297 | 466 | 1506 | 234 | 374 | 1751 | 273 | 273 |
| 3710 | QGN16807.1 | FIM1_3531 | CP015058. | 1178 | 669 | 899 | 1621 | 783 | 789 | 1352 | 694 | 1179 | 6804 | 900 | 1432 | 6430 | 757 | 1333 | 6634 | 729 | 1020 |
| 3711 | QGN16808.1 | FIM1_3532 | CP015058. | 666 | 633 | 562 | 1112 | 726 | 510 | 816 | 617 | 771 | 4619 | 740 | 1050 | 3595 | 561 | 929 | 4194 | 544 | 636 |
| 3712 | QGN16809.1 | FIM1_3533 | CP015058. | 3761 | 203 | 263 | 3154 | 256 | 218 | 4737 | 237 | 345 | 1745 | 203 | 341 | 1782 | 161 | 313 | 1633 | 183 | 238 |
| 3713 | QGN16810.1 | FIM1_3534 | CP015058. | 6182 | 1236 | 1709 | 6327 | 1438 | 1370 | 6389 | 1265 | 2071 | 7463 | 1351 | 2934 | 8609 | 1410 | 2523 | 7583 | 1263 | 1869 |
| 3714 | QGN16811.1 | FIM1_3535 | CP015058. | 859 | 1060 | 1008 | 1215 | 1094 | 903 | 845 | 933 | 1275 | 5190 | 1041 | 1527 | 5292 | 1154 | 1431 | 4816 | 1058 | 1012 |
| 3715 | QGN16812.1 | FIM1_3536 | CP015058. | 352 | 367 | 307 | 421 | 298 | 277 | 346 | 260 | 401 | 2211 | 330 | 450 | 1724 | 240 | 435 | 2189 | 239 | 296 |
| 3716 | QGN16813.1 | FIM1_3537 | CP015058. | 6860 | 5404 | 3022 | 3119 | 5332 | 3128 | 6130 | 4473 | 3787 | 21886 | 8094 | 6147 | 21253 | 5916 | 6038 | 19998 | 7678 | 4434 |
| 3717 | QGN16814.1 | FIM1_3538 | CP015058. | 598 | 606 | 636 | 1036 | 535 | 587 | 787 | 514 | 856 | 4059 | 488 | 951 | 3488 | 437 | 929 | 3691 | 422 | 597 |
| 3718 | QGN16815.1 | FIM1_3539 | CP015058. | 814 | 477 | 391 | 881 | 329 | 363 | 845 | 300 | 538 | 2032 | 334 | 618 | 1484 | 336 | 582 | 1892 | 317 | 372 |
| 3719 | QGN16816.1 | FIM1_3540 | CP015058. | 6601 | 3262 | 2190 | 7166 | 2515 | 1866 | 6871 | 2361 | 2725 | 7505 | 2074 | 2667 | 8188 | 2184 | 2806 | 6887 | 1876 | 1988 |
| 3720 | QGN16817.1 | FIM1_3541 | CP015058. | 581 | 135 | 173 | 597 | 140 | 103 | 559 | 149 | 212 | 646 | 157 | 282 | 647 | 377 | 273 | 582 | 322 | 168 |
| 3721 | QGN16818.1 | FIM1_3542 | CP015058. | 931 | 228 | 277 | 903 | 253 | 206 | 907 | 249 | 339 | 1647 | 291 | 386 | 1699 | 268 | 347 | 1630 | 249 | 307 |
| 3722 | QGN16819.1 | FIM1_3543 | CP015058. | 10 | 0 | 24 | 61 | 8 | 14 | 29 | 12 | 19 | 28 | 4 | 23 | 48 | 14 | 29 | 30 | 5 | 11 |
| 3723 | QGN16820.1 | FIM1_3544 | CP015058. | 919 | 2495 | 1746 | 2238 | 2170 | 1367 | 2038 | 2089 | 2144 | 8245 | 1889 | 2515 | 7610 | 1682 | 2447 | 7914 | 1609 | 1669 |
| 3724 | QGN16821.1 | FIM1_3545 | CP015058. | 881 | 93 | 75 | 1028 | 55 | 26 | 827 | 95 | 79 | 263 | 30 | 56 | 435 | 79 | 66 | 310 | 44 | 39 |
| 3725 | QGN16822.1 | FIM1_3546 | CP015058. | 1794 | 671 | 786 | 2094 | 1383 | 664 | 1870 | 1074 | 938 | 6741 | 1278 | 1053 | 8371 | 1217 | 890 | 7126 | 1170 | 772 |
| 3726 | QGN16823.1 | FIM1_3547 | CP015058. | 29 | 639 | 833 | 57 | 745 | 661 | 25 | 635 | 1078 | 3072 | 799 | 750 | 2657 | 747 | 757 | 2840 | 658 | 462 |
| 3727 | QGN16824.1 | FIM1_3548 | CP015058. | 87 | 170 | 161 | 443 | 66 | 108 | 226 | 82 | 190 | 1449 | 329 | 440 | 1188 | 506 | 436 | 1374 | 362 | 289 |
| 3728 | QGN16825.1 | FIM1_3549 | CP015058. | 440 | 98 | 97 | 567 | 147 | 88 | 507 | 141 | 137 | 291 | 95 | 124 | 501 | 93 | 115 | 313 | 78 | 75 |
| 3729 | QGN16826.1 | FIM1_3550 | CP015058. | 911 | 1568 | 1732 | 2354 | 1414 | 1804 | 970 | 1147 | 2297 | 3710 | 631 | 1908 | 3502 | 509 | 1339 | 3341 | 419 | 1022 |
| 3730 | QGN16827.1 | FIM1_3551 | CP015058. | 341 | 1913 | 1084 | 970 | 1449 | 983 | 686 | 1299 | 1407 | 3132 | 1284 | 1467 | 4032 | 1254 | 1303 | 3220 | 1326 | 938 |
| 3731 | QGN16828.1 | FIM1_3552 | CP015058. | 658 | 519 | 458 | 642 | 370 | 400 | 492 | 370 | 647 | 2056 | 351 | 607 | 1541 | 300 | 661 | 1796 | 294 | 375 |
| 3732 | QGN16829.1 | FIM1_3553 | CP015058. | 645 | 295 | 309 | 598 | 301 | 228 | 470 | 426 | 376 | 347 | 245 | 434 | 1135 | 196 | 385 | 576 | 246 | 271 |
| 3733 | QGN16830.1 | FIM1_3554 | CP015058. | 262 | 550 | 448 | 487 | 397 | 426 | 337 | 348 | 616 | 2231 | 342 | 752 | 1770 | 338 | 666 | 2098 | 353 | 464 |
| 3734 | QGN16831.1 | FIM1_3555 | CP015058. | 593 | 456 | 357 | 902 | 328 | 365 | 845 | 270 | 526 | 2578 | 283 | 654 | 2158 | 204 | 596 | 2587 | 218 | 424 |
| 3735 | QGN16832.1 | FIM1_3556 | CP015058. | 4276 | 1083 | 1380 | 4849 | 1397 | 1012 | 4310 | 1352 | 1609 | 4748 | 1322 | 2042 | 7091 | 1433 | 1897 | 5504 | 1169 | 1347 |
| 3736 | QGN16833.1 | FIM1_3557 | CP015058. | 5088 | 3776 | 1890 | 3382 | 3977 | 2015 | 4575 | 3299 | 2379 | 10822 | 5174 | 3535 | 12143 | 4045 | 3546 | 9881 | 4716 | 2416 |
| 3737 | QGN16834.1 | FIM1_3558 | CP015058. | 2410 | 924 | 696 | 3281 | 639 | 621 | 2423 | 572 | 864 | 4361 | 546 | 1065 | 3325 | 576 | 935 | 4319 | 516 | 652 |
| 3738 | QGN16835.1 | FIM1_3559 | CP015058. | 591 | 196 | 196 | 673 | 136 | 150 | 525 | 149 | 223 | 454 | 155 | 244 | 787 | 139 | 191 | 545 | 100 | 162 |
| 3739 | QGN16836.1 | FIM1_3560 | CP015058. | 3172 | 1276 | 875 | 3570 | 890 | 778 | 2818 | 817 | 1154 | 3713 | 645 | 1047 | 3503 | 650 | 968 | 3778 | 615 | 711 |
| 3740 | QGN16837.1 | FIM1_3561 | CP015058. | 2730 | 228 | 558 | 4387 | 305 | 414 | 2688 | 242 | 658 | 3110 | 201 | 734 | 3326 | 239 | 645 | 3004 | 162 | 441 |
| 3741 | QGN16838.1 | FIM1_3562 | CP015058. | 3404 | 761 | 816 | 4825 | 831 | 646 | 3828 | 795 | 991 | 5831 | 1103 | 1537 | 7074 | 1354 | 1284 | 5751 | 1089 | 943 |
| 3742 | QGN16839.1 | FIM1_3563 | CP015058. | 2982 | 1105 | 1102 | 2218 | 1214 | 972 | 2598 | 1102 | 1380 | 7146 | 1430 | 1966 | 7363 | 1154 | 1712 | 7297 | 1149 | 1279 |
| 3743 | QGN16840.1 | FIM1_3565 | CP015058. | 198 | 57 | 30 | 314 | 92 | 57 | 177 | 52 | 61 | 326 | 38 | 43 | 296 | 19 | 52 | 351 | 51 | 25 |
| 3744 | QGN16841.1 | FIM1_3566 | CP015058. | 464 | 167 | 130 | 734 | 192 | 121 | 413 | 122 | 157 | 1209 | 182 | 129 | 995 | 132 | 146 | 1148 | 119 | 113 |
| 3745 | QGN16842.1 | FIM1_3567 | CP015058. | 2167 | 4717 | 3926 | 5328 | 5641 | 3060 | 4523 | 4880 | 4672 | 19059 | 6407 | 6341 | 20658 | 5909 | 6134 | 20059 | 5472 | 4325 |
| 3746 | QGN16843.1 | FIM1_3568 | CP015058. | 2775 | 2033 | 3110 | 5898 | 2604 | 2298 | 3491 | 2628 | 4022 | 5526 | 2511 | 3881 | 15448 | 2057 | 3637 | 8949 | 2175 | 2724 |
| 3747 | QGN16844.1 | FIM1_3569 | CP015058. | 3581 | 2418 | 679 | 1045 | 1614 | 665 | 2360 | 1605 | 950 | 1246 | 1818 | 1139 | 3277 | 1829 | 1171 | 1718 | 2088 | 882 |
| 3748 | QGN16845.1 | FIM1_3570 | CP015058. | 1349 | 1559 | 812 | 878 | 977 | 726 | 1077 | 1119 | 1082 | 3123 | 889 | 1139 | 3351 | 897 | 1247 | 2838 | 844 | 777 |
| 3749 | QGN16846.1 | FIM1_3571 | CP015058. | 3371 | 4053 | 2584 | 3724 | 3347 | 2432 | 3315 | 3270 | 3552 | 9504 | 3481 | 3554 | 12114 | 2734 | 3602 | 9856 | 2681 | 2330 |
| 3750 | QGN16847.1 | FIM1_3572 | CP015058. | 675 | 394 | 368 | 933 | 348 | 306 | 626 | 287 | 463 | 1886 | 299 | 559 | 1498 | 255 | 521 | 1767 | 252 | 314 |
| 3751 | QGN16848.1 | FIM1_3573 | CP015058. | 3185 | 3358 | 2561 | 3005 | 3695 | 2146 | 2834 | 3455 | 3214 | 12138 | 3571 | 3899 | 15472 | 3188 | 3568 | 11439 | 2974 | 2619 |
| 3752 | QGN16849.1 | FIM1_3574 | CP015058. | 10995 | 1036 | 1485 | 18102 | 1300 | 1020 | 12581 | 1218 | 1996 | 969 | 690 | 1698 | 7242 | 1291 | 1319 | 2416 | 852 | 1073 |
| 3753 | QGN16850.1 | FIM1_3575 | CP015058. | 2668 | 711 | 1443 | 5603 | 1810 | 1149 | 3534 | 1246 | 1711 | 4059 | 1004 | 894 | 5536 | 1174 | 715 | 5067 | 761 | 492 |
| 3754 | QGN16851.1 | FIM1_3576 | CP015058. | 2011 | 3662 | 2759 | 3693 | 2753 | 2355 | 2604 | 2709 | 3523 | 9896 | 2072 | 3766 | 10751 | 2339 | 3164 | 9346 | 1849 | 2265 |
| 3755 | QGN16852.1 | FIM1_3577 | CP015058. | 494 | 317 | 456 | 728 | 262 | 313 | 535 | 368 | 502 | 1457 | 242 | 598 | 1968 | 176 | 519 | 1552 | 200 | 420 |
| 3756 | QGN16853.1 | FIM1_3579 | CP015058. | 3791 | 1220 | 1385 | 5212 | 1542 | 1206 | 4492 | 1566 | 1908 | 2786 | 1391 | 1828 | 7620 | 1163 | 1690 | 4583 | 1115 | 1247 |
| 3757 | QGN16854.1 | FIM1_3580 | CP015058. | 6952 | 3040 | 1637 | 3371 | 4252 | 1830 | 6193 | 3427 | 2184 | 12739 | 5199 | 3546 | 13296 | 4785 | 3228 | 11840 | 5358 | 2490 |
| 3758 | QGN16855.1 | FIM1_3581 | CP015058. | 956 | 3257 | 3354 | 2229 | 3018 | 2450 | 1064 | 2366 | 3949 | 19000 | 2703 | 5408 | 18118 | 2757 | 5295 | 20130 | 2718 | 3184 |
| 3759 | QGN16856.1 | FIM1_3582 | CP015058. | 5254 | 1399 | 1126 | 7590 | 1679 | 1164 | 5271 | 1469 | 1633 | 6812 | 1427 | 1286 | 7551 | 1350 | 1282 | 6510 | 1144 | 867 |
| 3760 | QGN16857.1 | FIM1_3583 | CP015058. | 1963 | 1490 | 1645 | 1504 | 1780 | 1255 | 1433 | 2011 | 2011 | 1661 | 894 | 948 | 6436 | 736 | 824 | 3320 | 798 | 672 |
| 3761 | QGN16858.1 | FIM1_3584 | CP015058. | 101 | 794 | 488 | 138 | 551 | 384 | 96 | 588 | 668 | 1487 | 425 | 482 | 1598 | 271 | 504 | 1308 | 232 | 325 |
| 3762 | QGN16859.1 | FIM1_3585 | CP015058. | 680 | 923 | 934 | 903 | 937 | 763 | 431 | 768 | 1202 | 3400 | 782 | 1640 | 5564 | 586 | 1194 | 3963 | 483 | 966 |
| 3763 | QGN16860.1 | FIM1_3586 | CP015058. | 2484 | 1721 | 1439 | 4397 | 1888 | 1163 | 3733 | 1894 | 1885 | 2467 | 2120 | 2545 | 6379 | 2126 | 2361 | 3585 | 2250 | 1630 |
| 3764 | QGN16861.1 | FIM1_3587 | CP015058. | 111 | 30 | 28 | 148 | 57 | 49 | 154 | 43 | 50 | 132 | 38 | 68 | 188 | 54 | 61 | 138 | 77 | 28 |
| 3765 | QGN16862.1 | FIM1_3588 | CP015058. | 2633 | 605 | 691 | 1392 | 721 | 479 | 2009 | 791 | 868 | 781 | 591 | 923 | 3121 | 556 | 866 | 1593 | 589 | 609 |
| 3766 | QGN16863.1 | FIM1_3589 | CP015058. | 13810 | 1860 | 1801 | 18536 | 1725 | 1385 | 16611 | 1661 | 2315 | 3118 | 1543 | 2548 | 9109 | 2028 | 2169 | 4872 | 1566 | 1648 |
| 3767 | QGN16864.1 | FIM1_3590 | CP015058. | 5 | 0 | 0 | 11 | 0 | 1 | 6 | 0 | 1 | 3 | 3 | 0 | 1 | 0 | 0 | 2 | 0 | 1 |
| 3768 | QGN16865.1 | FIM1_3591 | CP015058. | 893 | 324 | 702 | 2112 | 598 | 633 | 828 | 407 | 893 | 1436 | 219 | 610 | 2627 | 214 | 531 | 1895 | 172 | 343 |
| 3769 | QGN16866.1 | FIM1_3592 | CP015058. | 439 | 1160 | 2391 | 544 | 1313 | 1870 | 519 | 3052 | 3314 | 2710 | 1239 | 1575 | 2491 | 2255 | 1630 | 2673 | 900 | 1156 |
| 3770 | QGN16867.1 | FIM1_3593 | CP015058. | 1 | 0 | 1 | 4 | 2 | 1 | 0 | 2 | 0 | 4 | 0 | 0 | 1 | 1 | 2 | 1 | 1 | 0 |
| 3771 | QGN16868.1 | FIM1_3594 | CP015058. | 4921 | 2381 | 1593 | 5333 | 2207 | 1473 | 5160 | 2259 | 2217 | 4654 | 2206 | 2281 | 7714 | 2088 | 2491 | 5096 | 2159 | 1568 |
| 3772 | QGN16869.1 | FIM1_3595 | CP015058. | 12076 | 2599 | 4047 | 9514 | 3214 | 3400 | 10470 | 3227 | 5178 | 6264 | 2811 | 5942 | 12874 | 2735 | 5735 | 9280 | 2456 | 4008 |
| 3773 | QGN16870.1 | FIM1_3596 | CP015058. | 87 | 42 | 16 | 79 | 16 | 23 | 64 | 14 | 33 | 88 | 19 | 17 | 70 | 17 | 39 | 44 | 23 | 16 |
| 3774 | QGN16871.1 | FIM1_3597 | CP015058. | 10008 | 10152 | 7532 | 13712 | 7541 | 6955 | 9978 | 6727 | 8889 | 20512 | 8163 | 10637 | 19783 | 7344 | 10698 | 19299 | 6944 | 7419 |
| 3775 | QGN16872.1 | FIM1_3598 | CP015058. | 4753 | 2458 | 822 | 2785 | 1853 | 832 | 4531 | 1750 | 1134 | 4517 | 2358 | 1482 | 5243 | 1976 | 1604 | 4394 | 2509 | 1062 |
| 3776 | QGN16873.1 | FIM1_3599 | CP015058. | 14419 | 4513 | 2655 | 11785 | 3211 | 2060 | 11457 | 3306 | 3413 | 14030 | 4686 | 5257 | 17727 | 4618 | 5356 | 14735 | 4111 | 3691 |
| 3777 | QGN16874.1 | FIM1_3600 | CP015058. | 71 | 316 | 222 | 163 | 346 | 277 | 98 | 296 | 360 | 1492 | 271 | 390 | 1238 | 205 | 394 | 1555 | 224 | 246 |
| 3778 | QGN16875.1 | FIM1_3601 | CP015058. | 3599 | 763 | 1317 | 3936 | 1290 | 1043 | 3394 | 1227 | 1720 | 6603 | 1246 | 2206 | 9878 | 1195 | 1883 | 7844 | 1088 | 1440 |

| | | | | | | | | | | | | | | | | | | | | |
|---|---|---|---|---|---|---|---|---|---|---|---|---|---|---|---|---|---|---|---|---|
| 3779 | QGN16876.1 | FIM1_3603 | CP015058. | 1014 | 287 | 464 | 1156 | 424 | 360 | 921 | 357 | 607 | 2364 | 400 | 697 | 2558 | 332 | 631 | 2563 | 314 | 428 |
| 3780 | QGN16877.1 | FIM1_3604 | CP015058. | 3799 | 3106 | 1486 | 3386 | 3135 | 1605 | 4061 | 2354 | 1948 | 10394 | 3479 | 3183 | 10556 | 2861 | 2836 | 10351 | 3510 | 2012 |
| 3781 | QGN16878.1 | FIM1_3605 | CP015058. | 3128 | 820 | 821 | 3458 | 692 | 761 | 3217 | 675 | 1189 | 4058 | 707 | 1356 | 4374 | 575 | 1263 | 3878 | 570 | 847 |
| 3782 | QGN16879.1 | FIM1_3606 | CP015058. | 1000 | 367 | 443 | 635 | 212 | 342 | 836 | 275 | 571 | 3009 | 275 | 607 | 2046 | 176 | 619 | 2735 | 224 | 345 |
| 3783 | QGN16880.1 | FIM1_3607 | CP015058. | 1437 | 760 | 788 | 1560 | 998 | 761 | 1388 | 902 | 996 | 6846 | 1165 | 1318 | 5753 | 1194 | 1086 | 6171 | 1083 | 816 |
| 3784 | QGN16881.1 | FIM1_3608 | CP015058. | 1160 | 152 | 265 | 1489 | 241 | 225 | 1396 | 255 | 332 | 1365 | 324 | 493 | 1946 | 289 | 415 | 1523 | 304 | 341 |
| 3785 | QGN16882.1 | FIM1_3609 | CP015058. | 6358 | 1027 | 1296 | 8816 | 1209 | 1491 | 3140 | 964 | 1832 | 2394 | 376 | 984 | 2975 | 288 | 765 | 2542 | 274 | 543 |
| 3786 | QGN16883.1 | FIM1_3610 | CP015058. | 1043 | 984 | 1211 | 1440 | 1213 | 1089 | 1176 | 1129 | 1685 | 6830 | 1026 | 1842 | 6210 | 863 | 1665 | 6648 | 844 | 1176 |
| 3787 | QGN16884.1 | FIM1_3611 | CP015058. | 1460 | 424 | 500 | 1427 | 494 | 381 | 1348 | 426 | 638 | 3096 | 538 | 931 | 3115 | 476 | 735 | 3232 | 441 | 498 |
| 3788 | QGN16885.1 | FIM1_3612 | CP015058. | 1969 | 1014 | 1086 | 2925 | 1006 | 850 | 2506 | 914 | 1426 | 6323 | 911 | 1664 | 5890 | 857 | 1423 | 5756 | 763 | 1060 |
| 3789 | QGN16886.1 | FIM1_3613 | CP015058. | 1630 | 326 | 853 | 3033 | 363 | 638 | 2470 | 315 | 1145 | 1244 | 313 | 2066 | 3081 | 308 | 1719 | 2257 | 282 | 1221 |
| 3790 | QGN16887.1 | FIM1_3614 | CP015058. | 231 | 218 | 194 | 212 | 189 | 195 | 248 | 156 | 316 | 1585 | 177 | 450 | 991 | 153 | 362 | 1499 | 188 | 268 |
| 3791 | QGN16888.1 | FIM1_3615 | CP015058. | 653 | 1189 | 586 | 724 | 1100 | 607 | 936 | 886 | 785 | 5618 | 1518 | 1149 | 4625 | 1154 | 1130 | 5225 | 1358 | 720 |
| 3792 | QGN16889.1 | FIM1_3616 | CP015058. | 3023 | 2664 | 1431 | 1907 | 2835 | 1540 | 3231 | 2316 | 1915 | 12030 | 3315 | 2743 | 10652 | 2684 | 2516 | 11823 | 3038 | 1799 |
| 3793 | QGN16890.1 | FIM1_3617 | CP015058. | 97 | 169 | 126 | 87 | 190 | 88 | 100 | 175 | 135 | 903 | 205 | 215 | 624 | 162 | 185 | 792 | 188 | 124 |
| 3794 | QGN16891.1 | FIM1_3618 | CP015058. | 3539 | 1174 | 2031 | 4041 | 1405 | 1760 | 3411 | 1149 | 2633 | 10790 | 1305 | 3409 | 9794 | 1200 | 2872 | 10672 | 1038 | 2175 |
| 3795 | QGN16892.1 | FIM1_3619 | CP015058. | 517 | 347 | 246 | 499 | 263 | 289 | 439 | 228 | 350 | 1220 | 245 | 327 | 956 | 195 | 347 | 1085 | 254 | 205 |
| 3796 | QGN16893.1 | FIM1_3620 | CP015058. | 2569 | 536 | 1041 | 3692 | 783 | 851 | 3269 | 746 | 1291 | 5639 | 680 | 1550 | 6407 | 693 | 1262 | 5674 | 563 | 981 |
| 3797 | QGN16894.1 | FIM1_3621 | CP015058. | 1858 | 1382 | 1086 | 2306 | 951 | 1001 | 1660 | 899 | 1385 | 5906 | 998 | 1589 | 4662 | 741 | 1575 | 5783 | 831 | 946 |
| 3798 | QGN16895.1 | FIM1_3622 | CP015058. | 2959 | 1535 | 1467 | 4325 | 1476 | 1274 | 3680 | 1451 | 1873 | 8857 | 1410 | 2305 | 8317 | 1354 | 2011 | 8751 | 1298 | 1470 |
| 3799 | QGN16896.1 | FIM1_3623 | CP015058. | 6129 | 72 | 220 | 4322 | 58 | 129 | 4814 | 80 | 243 | 1005 | 62 | 221 | 1007 | 70 | 193 | 969 | 39 | 138 |
| 3800 | QGN16897.1 | FIM1_3624 | CP015058. | 1591 | 142 | 533 | 2739 | 70 | 301 | 1566 | 183 | 623 | 345 | 70 | 549 | 1024 | 85 | 456 | 505 | 57 | 324 |
| 3801 | QGN16898.1 | FIM1_3625 | CP015058. | 18483 | 15730 | 8954 | 22403 | 14559 | 7953 | 18017 | 13412 | 10996 | 31437 | 15117 | 14594 | 39613 | 13702 | 13830 | 32891 | 13198 | 10548 |
| 3802 | QGN16899.1 | FIM1_3626 | CP015058. | 191 | 144 | 236 | 149 | 116 | 166 | 147 | 142 | 284 | 587 | 84 | 271 | 814 | 80 | 252 | 596 | 55 | 168 |
| 3803 | QGN16900.1 | FIM1_3627 | CP015058. | 13011 | 509 | 480 | 9273 | 420 | 308 | 8227 | 471 | 612 | 764 | 296 | 571 | 1625 | 323 | 536 | 1038 | 260 | 363 |
| 3804 | QGN16901.1 | FIM1_3628 | CP015058. | 14705 | 6080 | 5798 | 22692 | 7325 | 4223 | 22517 | 7790 | 7623 | 4013 | 6349 | 10058 | 29216 | 7737 | 9268 | 11350 | 7328 | 6518 |
| 3805 | QGN16902.1 | FIM1_3629 | CP015058. | 6764 | 3000 | 2869 | 11578 | 2677 | 2387 | 9161 | 2507 | 3862 | 4488 | 2660 | 4895 | 13858 | 2699 | 4254 | 7634 | 2627 | 3144 |
| 3806 | QGN16903.1 | FIM1_3630 | CP015058. | 391 | 133 | 152 | 800 | 113 | 99 | 483 | 94 | 159 | 447 | 77 | 163 | 445 | 57 | 140 | 502 | 71 | 79 |
| 3807 | QGN16904.1 | FIM1_3631 | CP015058. | 770 | 381 | 357 | 1123 | 338 | 368 | 889 | 319 | 455 | 2390 | 328 | 537 | 1885 | 266 | 457 | 2274 | 263 | 366 |
| 3808 | QGN16905.1 | FIM1_3632 | CP015058. | 9188 | 5304 | 5127 | 9687 | 5719 | 4613 | 9599 | 5255 | 6718 | 16491 | 6709 | 8021 | 25618 | 5746 | 7256 | 18196 | 5921 | 5448 |
| 3809 | QGN16906.1 | FIM1_3633 | CP015058. | 10481 | 1683 | 3035 | 10544 | 698 | 1524 | 10284 | 1317 | 3415 | 4494 | 598 | 2648 | 6340 | 742 | 1782 | 5134 | 479 | 1392 |
| 3810 | QGN16907.1 | FIM1_3634 | CP015058. | 9504 | 5345 | 4167 | 4791 | 1776 | 2715 | 9118 | 3836 | 5595 | 8258 | 1518 | 2136 | 8633 | 1364 | 2205 | 8745 | 1432 | 1594 |
| 3811 | QGN16908.1 | FIM1_3635 | CP015058. | 5370 | 7956 | 5604 | 2614 | 2991 | 3321 | 5320 | 6348 | 7127 | 12878 | 3220 | 3584 | 11985 | 2834 | 3192 | 12758 | 2952 | 2119 |
| 3812 | QGN16909.1 | FIM1_3636 | CP015058. | 8735 | 2583 | 3604 | 3802 | 1087 | 1925 | 7809 | 3210 | 4556 | 6874 | 1062 | 1888 | 8310 | 924 | 1573 | 7355 | 865 | 1255 |
| 3813 | QGN16910.1 | FIM1_3637 | CP015058. | 7098 | 4299 | 4389 | 2856 | 1661 | 2557 | 6456 | 4331 | 5686 | 8386 | 1443 | 2356 | 8136 | 1171 | 1993 | 7790 | 1228 | 1502 |
| 3814 | QGN16911.1 | FIM1_3638 | CP015058. | 7818 | 10968 | 8597 | 3263 | 4483 | 5403 | 7830 | 8089 | 10711 | 19500 | 5919 | 6071 | 19279 | 4676 | 6549 | 18843 | 4173 | 4105 |
| 3815 | QGN16912.1 | FIM1_3639 | CP015058. | 1347 | 617 | 490 | 556 | 175 | 242 | 1112 | 376 | 624 | 1423 | 190 | 269 | 898 | 143 | 232 | 1527 | 139 | 147 |
| 3816 | QGN16913.1 | FIM1_3640 | CP015058. | 3174 | 1498 | 1343 | 1325 | 533 | 783 | 3235 | 1253 | 1819 | 3822 | 499 | 797 | 2899 | 426 | 719 | 3656 | 436 | 520 |
| 3817 | QGN16914.1 | FIM1_3641 | CP015058. | 2809 | 772 | 1023 | 1233 | 521 | 520 | 2764 | 1142 | 1219 | 2916 | 485 | 733 | 2455 | 479 | 623 | 2713 | 452 | 433 |
| 3818 | QGN16915.1 | FIM1_3642 | CP015058. | 6888 | 1410 | 2985 | 3653 | 734 | 1467 | 8522 | 1620 | 3612 | 8861 | 839 | 2195 | 7566 | 805 | 1762 | 8806 | 779 | 1431 |
| 3819 | QGN16916.1 | FIM1_3643 | CP015058. | 2860 | 1693 | 1896 | 1331 | 493 | 962 | 3084 | 1306 | 2435 | 3929 | 481 | 883 | 3425 | 401 | 793 | 4090 | 390 | 552 |
| 3820 | QGN16917.1 | FIM1_3644 | CP015058. | 6440 | 3797 | 3772 | 3886 | 1606 | 2062 | 7982 | 4031 | 4757 | 7115 | 1158 | 1968 | 6933 | 1029 | 1539 | 6944 | 1008 | 1108 |
| 3821 | QGN16918.1 | FIM1_3645 | CP015058. | 8014 | 2968 | 4071 | 3865 | 1441 | 2205 | 7734 | 3868 | 5280 | 10886 | 1256 | 2303 | 9922 | 1094 | 1826 | 10512 | 1067 | 1422 |
| 3822 | QGN16919.1 | FIM1_3646 | CP015058. | 2742 | 1585 | 1460 | 1623 | 710 | 873 | 3148 | 1859 | 1995 | 2993 | 576 | 750 | 2604 | 489 | 668 | 3160 | 506 | 478 |
| 3823 | QGN16920.1 | FIM1_3647 | CP015058. | 26306 | 36240 | 23017 | 15217 | 15740 | 16233 | 20065 | 24100 | 25390 | 46539 | 20037 | 18921 | 38178 | 16964 | 19497 | 47222 | 17303 | 14150 |
| 3824 | QGN16921.1 | FIM1_3648 | CP015058. | 7447 | 3114 | 1665 | 1292 | 1390 | 1082 | 6036 | 1942 | 2281 | 5017 | 1856 | 1302 | 4258 | 2001 | 1130 | 4879 | 1957 | 895 |
| 3825 | QGN16922.1 | FIM1_3649 | CP015058. | 5338 | 1772 | 1848 | 2216 | 999 | 1033 | 5866 | 2056 | 2461 | 3466 | 640 | 825 | 3059 | 783 | 782 | 3377 | 613 | 539 |
| 3826 | QGN16923.1 | FIM1_3650 | CP015058. | 8607 | 2500 | 1956 | 3092 | 1033 | 1066 | 6647 | 2320 | 2406 | 3034 | 731 | 995 | 3106 | 673 | 884 | 2852 | 650 | 634 |
| 3827 | QGN16924.1 | FIM1_3651 | CP015058. | 4838 | 1457 | 1755 | 1626 | 762 | 851 | 4101 | 1502 | 2066 | 3099 | 753 | 1165 | 3268 | 667 | 961 | 3021 | 613 | 666 |
| 3828 | QGN16925.1 | FIM1_3652 | CP015058. | 1921 | 448 | 677 | 1054 | 93 | 259 | 1930 | 279 | 799 | 832 | 86 | 214 | 712 | 167 | 207 | 817 | 114 | 164 |
| 3829 | QGN16926.1 | FIM1_3653 | CP015058. | 18849 | 7909 | 5327 | 6839 | 3136 | 3133 | 16224 | 6398 | 6857 | 9930 | 3279 | 3231 | 10461 | 2784 | 2987 | 9775 | 3079 | 2033 |
| 3830 | QGN16927.1 | FIM1_3654 | CP015058. | 8843 | 10125 | 9992 | 6273 | 3999 | 5537 | 10261 | 9466 | 12306 | 20123 | 4503 | 6485 | 18768 | 4161 | 5469 | 20233 | 4086 | 4047 |
| 3831 | QGN16928.1 | FIM1_3655 | CP015058. | 8542 | 4499 | 6194 | 4759 | 2188 | 3326 | 9602 | 5233 | 7850 | 13842 | 1975 | 3580 | 11768 | 1806 | 3127 | 12886 | 1650 | 2214 |
| 3832 | QGN16929.1 | FIM1_3656 | CP015058. | 7735 | 7437 | 6972 | 3964 | 2885 | 3851 | 8278 | 6698 | 8480 | 14347 | 3175 | 4667 | 13683 | 2856 | 4024 | 14107 | 2887 | 2785 |
| 3833 | QGN16930.1 | FIM1_3657 | CP015058. | 2822 | 848 | 882 | 1228 | 309 | 473 | 2855 | 879 | 1201 | 2319 | 296 | 428 | 1967 | 286 | 347 | 2181 | 254 | 225 |
| 3834 | QGN16931.1 | FIM1_3658 | CP015058. | 5331 | 3287 | 1909 | 1481 | 1210 | 1174 | 4733 | 2618 | 2470 | 6493 | 1420 | 1343 | 4768 | 1235 | 1236 | 6268 | 1556 | 871 |
| 3835 | QGN16932.1 | FIM1_3659 | CP015058. | 6984 | 2011 | 2454 | 3191 | 897 | 1291 | 7554 | 2477 | 3160 | 2696 | 736 | 1325 | 4124 | 725 | 1108 | 3048 | 743 | 814 |
| 3836 | QGN16933.1 | FIM1_3660 | CP015058. | 4236 | 2501 | 1750 | 1759 | 885 | 890 | 3927 | 1947 | 2002 | 4155 | 883 | 998 | 3134 | 678 | 937 | 4300 | 783 | 568 |
| 3837 | QGN16934.1 | FIM1_3661 | CP015058. | 9396 | 3082 | 4466 | 3537 | 1519 | 2192 | 8356 | 3719 | 5363 | 6031 | 1228 | 2175 | 6029 | 952 | 1698 | 6092 | 932 | 1272 |
| 3838 | QGN16935.1 | FIM1_3662 | CP015058. | 4327 | 1024 | 1856 | 2257 | 462 | 975 | 4687 | 1350 | 2375 | 5582 | 419 | 905 | 4863 | 340 | 762 | 5190 | 318 | 560 |
| 3839 | QGN16936.1 | FIM1_3663 | CP015058. | 3964 | 4413 | 4475 | 1095 | 2151 | 2804 | 4289 | 5369 | 6039 | 15379 | 2493 | 2330 | 13997 | 2768 | 1987 | 14781 | 2677 | 1649 |
| 3840 | QGN16937.1 | FIM1_3664 | CP015058. | 5667 | 2423 | 1818 | 1854 | 1022 | 918 | 4594 | 1850 | 2149 | 4559 | 1048 | 1182 | 4146 | 953 | 1031 | 4912 | 881 | 721 |
| 3841 | QGN16938.1 | FIM1_3665 | CP015058. | 3544 | 1387 | 1711 | 1397 | 430 | 944 | 3356 | 1359 | 2393 | 4978 | 441 | 959 | 3627 | 317 | 792 | 4822 | 346 | 499 |
| 3842 | QGN16939.1 | FIM1_3666 | CP015058. | 2098 | 1997 | 1820 | 842 | 710 | 951 | 2097 | 1929 | 2356 | 5334 | 737 | 1027 | 4592 | 626 | 839 | 5368 | 663 | 617 |
| 3843 | QGN16940.1 | FIM1_3667 | CP015058. | 3743 | 2779 | 2835 | 2066 | 1073 | 1611 | 4452 | 2269 | 3547 | 6949 | 1003 | 1872 | 5348 | 767 | 1463 | 6594 | 830 | 1099 |
| 3844 | QGN16941.1 | FIM1_3668 | CP015058. | 1263 | 2158 | 2074 | 867 | 815 | 1093 | 1918 | 2222 | 2772 | 4739 | 712 | 1067 | 4070 | 602 | 1025 | 4555 | 607 | 719 |
| 3845 | QGN16942.1 | FIM1_3669 | CP015058. | 4098 | 1590 | 1603 | 1854 | 479 | 764 | 4637 | 1411 | 1965 | 4004 | 404 | 821 | 3289 | 343 | 761 | 3651 | 343 | 525 |
| 3846 | QGN16943.1 | FIM1_3670 | CP015058. | 1929 | 1190 | 1103 | 881 | 332 | 637 | 1856 | 850 | 1419 | 2568 | 313 | 594 | 2065 | 244 | 540 | 2422 | 244 | 356 |
| 3847 | QGN16944.1 | FIM1_3671 | CP015058. | 2476 | 925 | 837 | 735 | 266 | 494 | 2211 | 748 | 1188 | 1810 | 233 | 441 | 1527 | 180 | 382 | 1782 | 195 | 279 |
| 3848 | QGN16945.1 | FIM1_3672 | CP015058. | 11198 | 2936 | 2206 | 5511 | 964 | 1256 | 12318 | 2460 | 2762 | 3095 | 773 | 1183 | 3895 | 827 | 1116 | 3249 | 777 | 738 |
| 3849 | QGN16946.1 | FIM1_3673 | CP015058. | 10853 | 2277 | 1882 | 2900 | 961 | 1185 | 10375 | 2111 | 2609 | 6590 | 1147 | 1325 | 5537 | 1445 | 1099 | 6060 | 1358 | 863 |
| 3850 | QGN16947.1 | FIM1_3674 | CP015058. | 308 | 109 | 90 | 178 | 23 | 41 | 272 | 88 | 97 | 129 | 27 | 50 | 164 | 29 | 42 | 124 | 25 | 23 |
| 3851 | QGN16948.1 | FIM1_3675 | CP015058. | 7993 | 7187 | 5909 | 4100 | 2324 | 3260 | 7502 | 5386 | 7351 | 10457 | 2443 | 3351 | 9417 | 2044 | 3089 | 9463 | 1981 | 2204 |
| 3852 | QGN16949.1 | FIM1_3676 | CP015058. | 1030 | 289 | 154 | 544 | 104 | 87 | 1304 | 229 | 203 | 327 | 48 | 66 | 316 | 32 | 42 | 280 | 46 | 41 |
| 3853 | QGN16950.1 | FIM1_3677 | CP015058. | 2900 | 1830 | 1910 | 1339 | 663 | 1071 | 3060 | 1480 | 2692 | 5040 | 646 | 1037 | 4363 | 591 | 917 | 4871 | 600 | 654 |
| 3854 | QGN16951.1 | FIM1_3678 | CP015058. | 6070 | 5926 | 2423 | 838 | 2010 | 1510 | 5152 | 4584 | 3306 | 6533 | 1974 | 1501 | 5462 | 1737 | 1342 | 6402 | 2017 | 1009 |
| 3855 | QGN16952.1 | FIM1_3679 | CP015058. | 102756 | 81744 | 55431 | 56624 | 39483 | 36167 | 89855 | 74842 | 64443 | 69272 | 43873 | 42911 | 100085 | 40468 | 40403 | 80018 | 38517 | 32251 |
| 3856 | QGN16953.1 | FIM1_3680 | CP015058. | 576 | 196 | 116 | 274 | 79 | 73 | 664 | 166 | 155 | 689 | 101 | 88 | 534 | 70 | 77 | 706 | 83 | 52 |
| 3857 | QGN16954.1 | FIM1_3681 | CP015058. | 5474 | 1384 | 1509 | 2469 | 585 | 843 | 4164 | 1399 | 1984 | 3525 | 541 | 808 | 2904 | 494 | 705 | 3278 | 502 | 536 |
| 3858 | QGN16955.1 | FIM1_3682 | CP015058. | 25808 | 2895 | 3083 | 8167 | 1352 | 1514 | 20770 | 3108 | 3708 | 5736 | 1205 | 1522 | 5939 | 1372 | 1277 | 5776 | 1206 | 896 |
| 3859 | QGN16956.1 | FIM1_3683 | CP015058. | 2040 | 2015 | 1565 | 1268 | 715 | 1056 | 2197 | 1371 | 2133 | 4012 | 905 | 1195 | 2746 | 791 | 1089 | 3645 | 941 | 753 |
| 3860 | QGN16957.1 | FIM1_3684 | CP015058. | 2645 | 580 | 752 | 1113 | 294 | 445 | 2433 | 776 | 979 | 1753 | 287 | 420 | 1830 | 238 | 398 | 2000 | 214 | 273 |
| 3861 | QGN16958.1 | FIM1_3685 | CP015058. | 3140 | 1233 | 1254 | 1321 | 353 | 745 | 2826 | 1126 | 1794 | 3162 | 317 | 658 | 2700 | 259 | 582 | 3234 | 264 | 381 |
| 3862 | QGN16959.1 | FIM1_3686 | CP015058. | 2269 | 1353 | 979 | 518 | 400 | 543 | 2298 | 943 | 1349 | 3709 | 506 | 689 | 2692 | 521 | 566 | 3683 | 571 | 449 |

| | | | | | | | | | | | | | | | | | | | | | |
|---|---|---|---|---|---|---|---|---|---|---|---|---|---|---|---|---|---|---|---|---|---|
| 3863 | QGN16960.1 | FIM1_3687 | CP015058. | 1434 | 1132 | 1013 | 846 | 340 | 586 | 1501 | 787 | 1337 | 2252 | 242 | 601 | 1707 | 256 | 530 | 1924 | 240 | 400 |
| 3864 | QGN16961.1 | FIM1_3688 | CP015058. | 4458 | 1375 | 2194 | 1423 | 565 | 1067 | 3591 | 1744 | 2795 | 1704 | 498 | 957 | 2972 | 446 | 842 | 2347 | 408 | 575 |
| 3865 | QGN16962.1 | FIM1_3689 | CP015058. | 6865 | 1890 | 2004 | 3068 | 729 | 1148 | 6208 | 1557 | 2719 | 4857 | 634 | 1267 | 4719 | 576 | 1123 | 4909 | 527 | 826 |
| 3866 | QGN16963.1 | FIM1_3690 | CP015058. | 4327 | 469 | 1569 | 2741 | 206 | 763 | 4577 | 593 | 2052 | 3382 | 167 | 897 | 2762 | 201 | 748 | 3039 | 178 | 540 |
| 3867 | QGN16964.1 | FIM1_3691 | CP015058. | 8374 | 3998 | 4491 | 4637 | 1629 | 2478 | 8130 | 4046 | 5841 | 11448 | 1815 | 2624 | 9878 | 1542 | 2354 | 11307 | 1457 | 1632 |
| 3868 | QGN16965.1 | FIM1_3692 | CP015058. | 3860 | 4033 | 2741 | 1958 | 1481 | 1542 | 4574 | 3260 | 3585 | 6928 | 1210 | 1397 | 5610 | 1057 | 1187 | 6830 | 1079 | 858 |
| 3869 | QGN16966.1 | FIM1_3693 | CP015058. | 4163 | 2770 | 2802 | 2092 | 860 | 1549 | 4178 | 1962 | 3534 | 7108 | 797 | 1554 | 5966 | 815 | 1460 | 7572 | 671 | 1026 |
| 3870 | QGN16967.1 | FIM1_3694 | CP015058. | 2277 | 1546 | 1483 | 1109 | 625 | 773 | 2080 | 1390 | 1857 | 4099 | 712 | 1025 | 3488 | 478 | 921 | 4101 | 518 | 580 |
| 3871 | QGN16968.1 | FIM1_3695 | CP015058. | 41663 | 4107 | 4260 | 22118 | 1208 | 2435 | 39942 | 3111 | 5605 | 10455 | 1384 | 2679 | 8752 | 1387 | 2483 | 10267 | 1241 | 1731 |
| 3872 | QGN16969.1 | FIM1_3696 | CP015058. | 18221 | 5720 | 6104 | 7409 | 2318 | 3365 | 16658 | 5322 | 7778 | 9212 | 1918 | 2793 | 8776 | 1648 | 2422 | 9170 | 1590 | 1709 |
| 3873 | QGN16970.1 | FIM1_3697 | CP015058. | 28515 | 10908 | 14377 | 25500 | 6719 | 8860 | 39429 | 9542 | 17409 | 24517 | 5356 | 7793 | 22004 | 7594 | 7641 | 23465 | 5125 | 5324 |
| 3874 | QGN16971.1 | FIM1_3698 | CP015058. | 11745 | 3550 | 3504 | 5514 | 1608 | 1862 | 15130 | 3634 | 4459 | 6856 | 1602 | 1916 | 7732 | 1609 | 1774 | 7267 | 1576 | 1307 |
| 3875 | QGN16972.1 | FIM1_3699 | CP015058. | 12500 | 286 | 402 | 6987 | 63 | 183 | 18007 | 219 | 554 | 701 | 51 | 121 | 598 | 49 | 77 | 692 | 40 | 67 |
| 3876 | QGN16973.1 | FIM1_3700 | CP015058. | 2400 | 491 | 1062 | 5930 | 200 | 518 | 4665 | 552 | 1516 | 2390 | 159 | 473 | 1763 | 189 | 340 | 2395 | 157 | 288 |
| 3877 | QGN16974.1 | FIM1_3701 | CP015058. | 1733 | 2005 | 1110 | 730 | 575 | 678 | 1319 | 1133 | 1522 | 3328 | 585 | 727 | 2427 | 429 | 616 | 3257 | 453 | 438 |
| 3878 | QGN16975.1 | FIM1_3702 | CP015058. | 3063 | 482 | 1077 | 1560 | 240 | 459 | 2309 | 545 | 1256 | 2281 | 159 | 482 | 1848 | 121 | 408 | 2066 | 112 | 268 |
| 3879 | QGN16976.1 | FIM1_3703 | CP015058. | 8806 | 2306 | 3434 | 4507 | 1081 | 1770 | 10274 | 2778 | 4372 | 9363 | 1073 | 1952 | 9063 | 1020 | 1659 | 9061 | 950 | 1322 |
| 3880 | QGN16977.1 | FIM1_3704 | CP015058. | 1268 | 1353 | 2166 | 578 | 513 | 1063 | 1232 | 1415 | 2566 | 6734 | 578 | 1388 | 5194 | 438 | 1058 | 6794 | 537 | 842 |
| 3881 | QGN16978.1 | FIM1_3705 | CP015058. | 6054 | 3440 | 3437 | 2258 | 1199 | 1928 | 6276 | 2908 | 4435 | 9972 | 1326 | 2416 | 8147 | 1147 | 2091 | 9719 | 1150 | 1502 |
| 3882 | QGN16979.1 | FIM1_3706 | CP015058. | 12645 | 6739 | 6217 | 5282 | 2727 | 3525 | 11333 | 6810 | 8103 | 12313 | 2784 | 3242 | 11410 | 2267 | 3279 | 11962 | 2168 | 2109 |
| 3883 | QGN16980.1 | FIM1_3707 | CP015058. | 1768 | 901 | 1158 | 800 | 251 | 706 | 2021 | 710 | 1586 | 2320 | 247 | 688 | 1844 | 294 | 607 | 2141 | 236 | 411 |
| 3884 | QGN16981.1 | FIM1_3708 | CP015058. | 9609 | 6712 | 6724 | 7787 | 3271 | 4025 | 13462 | 6742 | 8624 | 16838 | 2862 | 4803 | 16190 | 2653 | 4249 | 17190 | 2562 | 3176 |
| 3885 | QGN16982.1 | FIM1_3709 | CP015058. | 3736 | 3689 | 2763 | 1764 | 1428 | 1703 | 3849 | 3405 | 3714 | 5674 | 1370 | 1455 | 4826 | 1000 | 1320 | 5067 | 1120 | 940 |
| 3886 | QGN16983.1 | FIM1_3710 | CP015058. | 5670 | 2618 | 4147 | 3281 | 1096 | 2173 | 7394 | 3119 | 5476 | 7191 | 873 | 2115 | 7777 | 775 | 1684 | 7720 | 833 | 1320 |
| 3887 | QGN16984.1 | FIM1_3711 | CP015058. | 15437 | 3471 | 4006 | 7144 | 1398 | 2192 | 17076 | 3822 | 5089 | 9867 | 1144 | 2086 | 8565 | 972 | 1868 | 9894 | 990 | 1206 |
| 3888 | QGN16985.1 | FIM1_3712 | CP015058. | 14172 | 3494 | 4265 | 6326 | 1678 | 2202 | 14944 | 3755 | 4752 | 11058 | 1279 | 2733 | 9567 | 1994 | 2456 | 10908 | 1924 | 1879 |
| 3889 | QGN16986.1 | FIM1_3713 | CP015058. | 8652 | 1975 | 4404 | 4309 | 1353 | 2228 | 9631 | 3301 | 5357 | 10862 | 1283 | 2453 | 9794 | 1139 | 2059 | 10455 | 1082 | 1522 |
| 3890 | QGN16987.1 | FIM1_3714 | CP015058. | 1998 | 1390 | 2015 | 3063 | 402 | 1004 | 7289 | 1167 | 2609 | 1664 | 196 | 444 | 1687 | 299 | 402 | 1403 | 183 | 270 |
| 3891 | QGN16988.1 | FIM1_3715 | CP015058. | 1167 | 109 | 603 | 540 | 228 | 280 | 1168 | 550 | 698 | 978 | 165 | 303 | 1037 | 163 | 254 | 921 | 133 | 182 |
| 3892 | QGN16989.1 | FIM1_3716 | CP015058. | 15385 | 5856 | 8124 | 8810 | 2319 | 4657 | 15930 | 6030 | 10128 | 8445 | 1942 | 5336 | 9829 | 1951 | 4667 | 8990 | 1764 | 3404 |
| 3893 | QGN16990.1 | FIM1_3717 | CP015058. | 14230 | 9383 | 12197 | 13521 | 4224 | 6751 | 17531 | 9289 | 14650 | 26916 | 4069 | 7523 | 22414 | 4481 | 5878 | 25649 | 4011 | 4728 |
| 3894 | QGN16991.1 | FIM1_3718 | CP015058. | 4149 | 2552 | 2827 | 1755 | 835 | 1357 | 4284 | 2416 | 3581 | 6953 | 866 | 1509 | 5455 | 610 | 1373 | 6538 | 623 | 836 |
| 3895 | QGN16992.1 | FIM1_3719 | CP015058. | 6084 | 1177 | 2151 | 1722 | 402 | 974 | 4698 | 1209 | 2633 | 4696 | 443 | 1310 | 4511 | 469 | 1008 | 4392 | 399 | 813 |
| 3896 | QGN16993.1 | FIM1_3720 | CP015058. | 6592 | 2455 | 2201 | 3175 | 680 | 1195 | 6180 | 1690 | 2871 | 4551 | 563 | 1165 | 4013 | 726 | 1030 | 4151 | 576 | 653 |
| 3897 | QGN16994.1 | FIM1_3721 | CP015058. | 111 | 47 | 20 | 38 | 8 | 7 | 95 | 15 | 21 | 27 | 12 | 5 | 33 | 5 | 13 | 41 | 3 | 6 |
| 3898 | QGN16995.1 | FIM1_3722 | CP015058. | 686 | 206 | 160 | 316 | 67 | 88 | 633 | 154 | 198 | 168 | 48 | 91 | 249 | 58 | 76 | 165 | 48 | 73 |
| 3899 | QGN16996.1 | FIM1_3723 | CP015058. | 2370 | 262 | 373 | 1136 | 136 | 163 | 2058 | 386 | 444 | 86 | 78 | 214 | 462 | 120 | 210 | 173 | 75 | 131 |
| 3900 | QGN16997.1 | FIM1_3725 | CP015058. | 354 | 25 | 38 | 883 | 83 | 20 | 516 | 31 | 40 | 58 | 55 | 49 | 473 | 43 | 71 | 95 | 39 | 59 |
| 3901 | QGN16998.1 | FIM1_3726 | CP015058. | 0 | 0 | 0 | 0 | 0 | 0 | 0 | 0 | 0 | 0 | 0 | 0 | 0 | 0 | 0 | 0 | 0 | 0 |
| 3902 | QGN16999.1 | FIM1_3727 | CP015058. | 1365 | 1241 | 623 | 853 | 176 | 279 | 1635 | 626 | 805 | 1070 | 311 | 300 | 931 | 207 | 286 | 769 | 230 | 223 |
| 3903 | QGN17000.1 | FIM1_3728 | CP015058. | 1 | 0 | 0 | 3 | 0 | 0 | 2 | 0 | 0 | 0 | 0 | 0 | 3 | 0 | 1 | 0 | 0 | 0 |
| 3904 | QGN17001.1 | FIM1_3730 | CP015059. | 2 | 0 | 0 | 1 | 4 | 2 | 1 | 2 | 1 | 1 | 4 | 1 | 7 | 5 | 1 | 3 | 2 | 0 |
| 3905 | QGN17002.1 | FIM1_3731 | CP015059. | 22 | 114 | 55 | 49 | 111 | 25 | 19 | 73 | 69 | 71 | 153 | 97 | 346 | 123 | 127 | 193 | 104 | 78 |
| 3906 | QGN17003.1 | FIM1_3732 | CP015059. | 249 | 457 | 132 | 854 | 214 | 181 | 341 | 165 | 211 | 706 | 214 | 245 | 682 | 156 | 207 | 539 | 226 | 171 |
| 3907 | QGN17004.1 | FIM1_3733 | CP015059. | 2085 | 547 | 1237 | 2599 | 696 | 869 | 3298 | 755 | 1584 | 2648 | 555 | 978 | 2493 | 514 | 868 | 2543 | 505 | 600 |
| 3908 | QGN17005.1 | FIM1_3734 | CP015059. | 789 | 418 | 798 | 850 | 449 | 504 | 510 | 398 | 974 | 2151 | 498 | 988 | 2314 | 393 | 917 | 2436 | 383 | 625 |
| 3909 | QGN17006.1 | FIM1_3735 | CP015059. | 2548 | 717 | 1218 | 2483 | 883 | 1035 | 2295 | 767 | 1629 | 5544 | 862 | 1992 | 5733 | 869 | 1821 | 5837 | 833 | 1335 |
| 3910 | QGN17007.1 | FIM1_3736 | CP015059. | 3568 | 678 | 668 | 2343 | 956 | 561 | 2824 | 829 | 856 | 4433 | 982 | 1066 | 4263 | 892 | 1025 | 4363 | 936 | 691 |
| 3911 | QGN17008.1 | FIM1_3737 | CP015059. | 5826 | 168 | 285 | 5047 | 139 | 204 | 6790 | 157 | 395 | 1672 | 86 | 265 | 1857 | 106 | 243 | 1645 | 63 | 180 |
| 3912 | QGN17009.1 | FIM1_3738 | CP015059. | 1709 | 247 | 245 | 2281 | 314 | 169 | 1938 | 266 | 279 | 1259 | 146 | 233 | 1199 | 157 | 222 | 1342 | 118 | 156 |
| 3913 | QGN17010.1 | FIM1_3739 | CP015059. | 2309 | 1106 | 960 | 4698 | 1025 | 786 | 3539 | 757 | 1274 | 5579 | 540 | 1220 | 4910 | 872 | 1261 | 5194 | 542 | 751 |
| 3914 | QGN17011.1 | FIM1_3740 | CP015059. | 5465 | 1946 | 1476 | 7972 | 1305 | 1150 | 5389 | 1230 | 1790 | 6473 | 1298 | 2145 | 5414 | 1657 | 2061 | 5417 | 1493 | 1308 |
| 3915 | QGN17012.1 | FIM1_3741 | CP015059. | 8899 | 1257 | 1884 | 9125 | 1349 | 1560 | 7959 | 1291 | 2384 | 9743 | 1380 | 2528 | 12251 | 1123 | 2429 | 10495 | 1049 | 1746 |
| 3916 | QGN17013.1 | FIM1_3742 | CP015059. | 2175 | 929 | 1533 | 4037 | 1118 | 1237 | 3073 | 1063 | 1990 | 8344 | 1052 | 2457 | 9048 | 1018 | 2125 | 8420 | 928 | 1729 |
| 3917 | QGN17014.1 | FIM1_3743 | CP015059. | 5825 | 1610 | 2107 | 5849 | 2176 | 1629 | 5366 | 2122 | 2581 | 10630 | 2176 | 2599 | 11834 | 2111 | 2442 | 10822 | 1867 | 1777 |
| 3918 | QGN17015.1 | FIM1_3744 | CP015059. | 2230 | 782 | 1173 | 2971 | 897 | 1037 | 2284 | 964 | 1589 | 3479 | 853 | 1818 | 5569 | 604 | 1839 | 4541 | 638 | 1238 |
| 3919 | QGN17016.1 | FIM1_3745 | CP015059. | 16267 | 6634 | 8003 | 20684 | 10007 | 6851 | 15792 | 8337 | 9421 | 26927 | 8631 | 9314 | 28420 | 8556 | 9002 | 27512 | 7957 | 6351 |
| 3920 | QGN17017.1 | FIM1_3746 | CP015059. | 3332 | 1460 | 1394 | 4925 | 1320 | 1207 | 3171 | 1052 | 1799 | 5837 | 956 | 1825 | 5527 | 1167 | 1696 | 5385 | 913 | 1141 |
| 3921 | QGN17018.1 | FIM1_3747 | CP015059. | 10674 | 8421 | 8451 | 20316 | 14868 | 7492 | 15937 | 11884 | 10470 | 50421 | 15491 | 15019 | 64453 | 14266 | 14044 | 55950 | 12773 | 9746 |
| 3922 | QGN17019.1 | FIM1_3748 | CP015059. | 8636 | 5914 | 5064 | 7101 | 4715 | 4450 | 6837 | 3714 | 6115 | 21466 | 5529 | 7102 | 17089 | 6081 | 6964 | 21794 | 5528 | 5216 |
| 3923 | QGN17020.1 | FIM1_3749 | CP015059. | 5863 | 1813 | 2932 | 6026 | 2353 | 2323 | 5486 | 2181 | 3695 | 16277 | 2767 | 4325 | 17945 | 2676 | 3697 | 16909 | 2560 | 2902 |
| 3924 | QGN17021.1 | FIM1_3750 | CP015059. | 2040 | 1152 | 471 | 1031 | 1050 | 561 | 1764 | 755 | 641 | 5085 | 1140 | 996 | 3594 | 978 | 930 | 4759 | 1279 | 641 |
| 3925 | QGN17022.1 | FIM1_3751 | CP015059. | 2110 | 870 | 1233 | 2837 | 1016 | 1132 | 2321 | 864 | 1603 | 8470 | 981 | 1747 | 7522 | 861 | 1554 | 7962 | 861 | 1212 |
| 3926 | QGN17023.1 | FIM1_3752 | CP015059. | 4688 | 1037 | 1617 | 4039 | 1160 | 1374 | 4045 | 1002 | 2143 | 6803 | 1253 | 2369 | 9342 | 1094 | 2211 | 7778 | 1080 | 1797 |
| 3927 | QGN17024.1 | FIM1_3753 | CP015059. | 10116 | 800 | 890 | 7589 | 1133 | 810 | 9625 | 835 | 1193 | 4018 | 1016 | 1186 | 5617 | 1219 | 1131 | 4763 | 1225 | 838 |
| 3928 | QGN17025.1 | FIM1_3754 | CP015059. | 3361 | 1505 | 1176 | 3831 | 1860 | 974 | 3185 | 1591 | 1615 | 1891 | 1567 | 1568 | 8018 | 1751 | 1598 | 4028 | 1699 | 1142 |
| 3929 | QGN17026.1 | FIM1_3755 | CP015059. | 87 | 196 | 117 | 175 | 215 | 157 | 101 | 137 | 196 | 660 | 163 | 151 | 503 | 164 | 172 | 610 | 153 | 101 |
| 3930 | QGN17027.1 | FIM1_3756 | CP015059. | 19317 | 22564 | 12881 | 18654 | 16910 | 12187 | 15839 | 14634 | 15310 | 47862 | 20082 | 19362 | 44595 | 17413 | 19466 | 49005 | 17311 | 14428 |
| 3931 | QGN17028.1 | FIM1_3757 | CP015059. | 825 | 465 | 456 | 1078 | 393 | 406 | 832 | 415 | 555 | 2637 | 345 | 616 | 2108 | 311 | 629 | 2478 | 358 | 393 |
| 3932 | QGN17029.1 | FIM1_3758 | CP015059. | 661 | 1431 | 1113 | 616 | 1092 | 1076 | 556 | 941 | 1372 | 4540 | 1331 | 1455 | 3266 | 1128 | 1454 | 4011 | 1411 | 893 |
| 3933 | QGN17030.1 | FIM1_3759 | CP015059. | 4264 | 9231 | 5877 | 10973 | 7112 | 4328 | 8556 | 7150 | 7355 | 10361 | 8342 | 10095 | 31319 | 8729 | 8858 | 21915 | 8059 | 6745 |
| 3934 | QGN17031.1 | FIM1_3760 | CP015059. | 332 | 752 | 425 | 476 | 584 | 414 | 297 | 482 | 587 | 1888 | 362 | 477 | 1936 | 381 | 453 | 1884 | 366 | 331 |
| 3935 | QGN17032.1 | FIM1_3761 | CP015059. | 958 | 420 | 487 | 945 | 494 | 417 | 1026 | 424 | 607 | 2784 | 430 | 632 | 2720 | 422 | 531 | 2846 | 394 | 401 |
| 3936 | QGN17033.1 | FIM1_3762 | CP015059. | 647 | 1301 | 1159 | 1598 | 1365 | 1009 | 1108 | 1111 | 1505 | 5946 | 1271 | 1356 | 6110 | 1343 | 1453 | 6097 | 1211 | 908 |
| 3937 | QGN17034.1 | FIM1_3763 | CP015059. | 3453 | 1784 | 2333 | 3608 | 2004 | 1954 | 3026 | 1869 | 3149 | 3072 | 2036 | 3173 | 9210 | 1674 | 3395 | 6322 | 1759 | 2188 |
| 3938 | QGN17035.1 | FIM1_3764 | CP015059. | 218 | 311 | 489 | 268 | 251 | 360 | 200 | 221 | 604 | 1237 | 245 | 582 | 1201 | 227 | 571 | 1202 | 198 | 376 |
| 3939 | QGN17036.1 | FIM1_3765 | CP015059. | 11476 | 4918 | 6960 | 14373 | 5816 | 5463 | 12202 | 5443 | 8412 | 12152 | 5192 | 10799 | 26993 | 5501 | 9340 | 18850 | 5046 | 7176 |
| 3940 | QGN17037.1 | FIM1_3766 | CP015059. | 285 | 74 | 54 | 379 | 79 | 54 | 281 | 52 | 47 | 281 | 59 | 82 | 314 | 39 | 58 | 360 | 35 | 45 |
| 3941 | QGN17038.1 | FIM1_3767 | CP015059. | 2158 | 1659 | 1762 | 4187 | 1372 | 1379 | 3035 | 1448 | 2230 | 8980 | 1236 | 2080 | 8110 | 1470 | 2108 | 8886 | 1259 | 1432 |
| 3942 | QGN17039.1 | FIM1_3768 | CP015059. | 902 | 23344 | 22936 | 4896 | 28703 | 19222 | 10184 | 31469 | 25903 | 70550 | 20094 | 21105 | 72449 | 19421 | 22133 | 70188 | 19632 | 15586 |
| 3943 | QGN17040.1 | FIM1_3769 | CP015059. | 6704 | 1152 | 1009 | 5875 | 1004 | 876 | 5366 | 952 | 1325 | 5530 | 1449 | 1690 | 5975 | 1397 | 1663 | 5327 | 1385 | 1182 |
| 3944 | QGN17041.1 | FIM1_3770 | CP015059. | 4978 | 4541 | 4440 | 6644 | 4141 | 3784 | 5608 | 3785 | 5380 | 16043 | 4528 | 6333 | 19205 | 4509 | 6065 | 16422 | 4113 | 4299 |
| 3945 | QGN17042.1 | FIM1_3771 | CP015059. | 681 | 1040 | 890 | 1484 | 1132 | 761 | 1301 | 903 | 1117 | 4275 | 908 | 1395 | 4532 | 844 | 1257 | 4450 | 827 | 879 |
| 3946 | QGN17043.1 | FIM1_3772 | CP015059. | 2723 | 1096 | 1582 | 5138 | 1485 | 1192 | 3558 | 1128 | 1932 | 6164 | 1103 | 2207 | 7370 | 1531 | 1925 | 6564 | 1089 | 1506 |

| | | | | | | | | | | | | | | | | | | | | | |
|---|---|---|---|---|---|---|---|---|---|---|---|---|---|---|---|---|---|---|---|---|---|
| 3947 | QGN17044.1 | FIM1_3773 | CP015059. | 12456 | 3092 | 5795 | 13381 | 6320 | 5053 | 10786 | 5989 | 7691 | 19106 | 6457 | 8532 | 34079 | 6350 | 8062 | 23757 | 6029 | 5843 |
| 3948 | QGN17045.1 | FIM1_3774 | CP015059. | 2055 | 3993 | 5228 | 4791 | 3288 | 3873 | 3521 | 3171 | 5987 | 15045 | 2431 | 6964 | 14498 | 2367 | 5614 | 15191 | 2042 | 4120 |
| 3949 | QGN17046.1 | FIM1_3775 | CP015059. | 302 | 93 | 180 | 373 | 134 | 135 | 272 | 157 | 207 | 964 | 125 | 228 | 936 | 145 | 228 | 964 | 110 | 161 |
| 3950 | QGN17047.1 | FIM1_3776 | CP015059. | 627 | 220 | 334 | 677 | 162 | 328 | 632 | 132 | 461 | 1369 | 132 | 617 | 1054 | 95 | 636 | 1369 | 127 | 394 |
| 3951 | QGN17048.1 | FIM1_3777 | CP015059. | 10366 | 3564 | 6883 | 15133 | 5452 | 4761 | 12106 | 4948 | 8424 | 2731 | 4309 | 11912 | 26694 | 5052 | 10465 | 11234 | 4564 | 7660 |
| 3952 | QGN17049.1 | FIM1_3778 | CP015059. | 2082 | 108 | 204 | 1602 | 223 | 140 | 1866 | 205 | 228 | 392 | 139 | 253 | 1181 | 174 | 214 | 669 | 130 | 163 |
| 3953 | QGN17050.1 | FIM1_3779 | CP015059. | 797 | 1115 | 1084 | 1049 | 1308 | 991 | 813 | 1132 | 1368 | 2244 | 997 | 1372 | 4531 | 817 | 1467 | 3021 | 795 | 962 |
| 3954 | QGN17051.1 | FIM1_3780 | CP015059. | 5707 | 5819 | 6559 | 8027 | 6893 | 5207 | 6180 | 5595 | 7487 | 28206 | 8021 | 11572 | 26231 | 7968 | 10883 | 28973 | 7256 | 8086 |
| 3955 | QGN17052.1 | FIM1_3782 | CP015059. | 3144 | 1323 | 2217 | 5062 | 1943 | 1813 | 3446 | 1794 | 2827 | 11493 | 1360 | 2985 | 12267 | 1408 | 2647 | 11395 | 1147 | 1941 |
| 3956 | QGN17053.1 | FIM1_3783 | CP015059. | 3333 | 576 | 787 | 4257 | 648 | 692 | 3779 | 708 | 1082 | 3676 | 529 | 1318 | 4872 | 533 | 1289 | 3796 | 483 | 873 |
| 3957 | QGN17054.1 | FIM1_3784 | CP015059. | 2576 | 2330 | 2971 | 3996 | 1783 | 2638 | 3002 | 1548 | 3573 | 11174 | 2258 | 5609 | 10018 | 2229 | 5295 | 10879 | 2159 | 3765 |
| 3958 | QGN17055.1 | FIM1_3785 | CP015059. | 233 | 259 | 331 | 516 | 206 | 265 | 255 | 217 | 476 | 1657 | 154 | 453 | 1302 | 154 | 507 | 1633 | 137 | 345 |
| 3959 | QGN17056.1 | FIM1_3786 | CP015059. | 6192 | 3124 | 3286 | 6557 | 2999 | 2752 | 5662 | 3032 | 4274 | 14923 | 2765 | 4226 | 15270 | 2637 | 4344 | 13988 | 2105 | 3030 |
| 3960 | QGN17057.1 | FIM1_3787 | CP015059. | 323 | 54 | 29 | 368 | 65 | 52 | 292 | 60 | 42 | 190 | 29 | 36 | 195 | 30 | 36 | 202 | 37 | 27 |
| 3961 | QGN17058.1 | FIM1_3788 | CP015059. | 5785 | 4251 | 4429 | 7553 | 4376 | 3774 | 5194 | 3479 | 5441 | 19546 | 4023 | 6867 | 20973 | 4985 | 6329 | 19139 | 3961 | 4572 |
| 3962 | QGN17059.1 | FIM1_3789 | CP015059. | 900 | 520 | 505 | 1425 | 385 | 405 | 1100 | 432 | 682 | 2464 | 316 | 543 | 2116 | 342 | 606 | 2404 | 325 | 375 |
| 3963 | QGN17060.1 | FIM1_3790 | CP015059. | 481 | 354 | 408 | 477 | 299 | 303 | 303 | 345 | 509 | 1386 | 255 | 378 | 1596 | 145 | 373 | 1414 | 184 | 254 |
| 3964 | QGN17061.1 | FIM1_3791 | CP015059. | 4674 | 1940 | 2124 | 5838 | 2120 | 1732 | 4347 | 1881 | 2714 | 11188 | 2009 | 3375 | 11626 | 1902 | 3032 | 10757 | 1792 | 2167 |
| 3965 | QGN17062.1 | FIM1_3792 | CP015059. | 5680 | 2159 | 1866 | 5130 | 1394 | 1294 | 2944 | 1765 | 2495 | 11116 | 1519 | 2558 | 10840 | 3009 | 3260 | 9916 | 1777 | 1917 |
| 3966 | QGN17063.1 | FIM1_3793 | CP015059. | 12833 | 4940 | 4370 | 12020 | 4336 | 3963 | 11989 | 3288 | 5363 | 17324 | 3892 | 6511 | 13758 | 4541 | 6047 | 16895 | 4315 | 4405 |
| 3967 | QGN17064.1 | FIM1_3794 | CP015059. | 7351 | 9645 | 5479 | 6302 | 7542 | 4513 | 5772 | 6454 | 6487 | 18562 | 8535 | 8288 | 18525 | 7411 | 8248 | 17765 | 7327 | 6231 |
| 3968 | QGN17065.1 | FIM1_3795 | CP015059. | 3268 | 160 | 444 | 2490 | 295 | 284 | 3022 | 368 | 460 | 2216 | 379 | 563 | 3024 | 420 | 485 | 2487 | 389 | 380 |
| 3969 | QGN17066.1 | FIM1_3796 | CP015059. | 626 | 104 | 234 | 800 | 202 | 166 | 642 | 213 | 292 | 1054 | 207 | 327 | 1277 | 173 | 275 | 1093 | 160 | 191 |
| 3970 | QGN17067.1 | FIM1_3797 | CP015059. | 3517 | 3122 | 3133 | 7152 | 3189 | 2726 | 5170 | 2748 | 3957 | 13580 | 2244 | 3357 | 12049 | 2524 | 3272 | 12730 | 2029 | 2130 |
| 3971 | QGN17068.1 | FIM1_3798 | CP015059. | 3644 | 1187 | 1477 | 3742 | 1196 | 1331 | 3371 | 1519 | 2221 | 5109 | 990 | 1625 | 5786 | 993 | 1674 | 5607 | 1064 | 1140 |
| 3972 | QGN17069.1 | FIM1_3799 | CP015059. | 11563 | 8767 | 11424 | 26219 | 8864 | 9252 | 15566 | 8102 | 13846 | 32834 | 8061 | 18759 | 41565 | 10131 | 16826 | 34800 | 7769 | 12477 |
| 3973 | QGN17070.1 | FIM1_3801 | CP015059. | 2305 | 1846 | 1356 | 6350 | 1506 | 1178 | 3972 | 1270 | 1798 | 6065 | 1480 | 1896 | 6904 | 1811 | 1938 | 5581 | 1339 | 1348 |
| 3974 | QGN17071.1 | FIM1_3802 | CP015059. | 2843 | 1306 | 1979 | 9627 | 1032 | 1455 | 5023 | 1097 | 2400 | 6216 | 769 | 2341 | 5325 | 1161 | 1809 | 6046 | 988 | 1343 |
| 3975 | QGN17072.1 | FIM1_3803 | CP015059. | 373 | 327 | 211 | 456 | 231 | 181 | 249 | 199 | 278 | 1237 | 190 | 323 | 920 | 209 | 323 | 1025 | 195 | 180 |
| 3976 | QGN17073.1 | FIM1_3804 | CP015059. | 410 | 129 | 239 | 429 | 102 | 205 | 341 | 127 | 347 | 1172 | 75 | 296 | 1142 | 65 | 306 | 1127 | 69 | 224 |
| 3977 | QGN17074.1 | FIM1_3805 | CP015059. | 2253 | 306 | 271 | 1707 | 206 | 217 | 1914 | 229 | 364 | 1077 | 186 | 305 | 1307 | 179 | 319 | 1213 | 198 | 211 |
| 3978 | QGN17075.1 | FIM1_3806 | CP015059. | 1201 | 723 | 477 | 555 | 776 | 559 | 906 | 640 | 746 | 1728 | 710 | 762 | 2171 | 618 | 761 | 1713 | 644 | 506 |
| 3979 | QGN17076.1 | FIM1_3807 | CP015059. | 14059 | 7761 | 4857 | 15647 | 7659 | 4510 | 13901 | 6257 | 6245 | 20932 | 7516 | 8713 | 27252 | 7213 | 8090 | 21899 | 7035 | 6090 |
| 3980 | QGN17077.1 | FIM1_3808 | CP015059. | 1714 | 390 | 398 | 1893 | 357 | 291 | 1785 | 343 | 513 | 1535 | 244 | 469 | 2233 | 299 | 433 | 1545 | 264 | 301 |
| 3981 | QGN17078.1 | FIM1_3809 | CP015059. | 2069 | 1756 | 808 | 915 | 1622 | 829 | 1396 | 1263 | 995 | 6217 | 1904 | 1361 | 4763 | 1651 | 1306 | 5717 | 1992 | 965 |
| 3982 | QGN17079.1 | FIM1_3810 | CP015059. | 653 | 2013 | 1197 | 976 | 1660 | 1188 | 829 | 1332 | 1686 | 6159 | 1500 | 1810 | 5395 | 1411 | 1754 | 6085 | 1469 | 1142 |
| 3983 | QGN17080.1 | FIM1_3811 | CP015059. | 18566 | 3005 | 2184 | 22176 | 3483 | 2170 | 19311 | 2918 | 2950 | 13169 | 2272 | 2054 | 11933 | 3436 | 2222 | 11981 | 2367 | 1464 |
| 3984 | QGN17081.1 | FIM1_3812 | CP015059. | 43 | 8 | 10 | 147 | 16 | 4 | 105 | 10 | 12 | 55 | 8 | 20 | 57 | 11 | 21 | 45 | 13 | 11 |
| 3985 | QGN17082.1 | FIM1_3813 | CP015059. | 2866 | 3092 | 2473 | 3399 | 3530 | 2029 | 2934 | 2959 | 2974 | 14633 | 3528 | 3376 | 13682 | 3181 | 3253 | 14609 | 2924 | 2343 |
| 3986 | QGN17083.1 | FIM1_3814 | CP015059. | 75 | 10 | 16 | 76 | 12 | 13 | 39 | 8 | 29 | 133 | 12 | 31 | 115 | 11 | 14 | 127 | 15 | 10 |
| 3987 | QGN17084.1 | FIM1_3815 | CP015059. | 10355 | 8259 | 12285 | 25715 | 8368 | 10558 | 24030 | 8496 | 16343 | 12348 | 6583 | 12322 | 37007 | 6901 | 10772 | 22691 | 5870 | 8718 |
| 3988 | QGN17085.1 | FIM1_3816 | CP015059. | 4581 | 1547 | 965 | 2782 | 1763 | 1167 | 3240 | 1243 | 1381 | 5434 | 1798 | 1997 | 5438 | 1589 | 1987 | 5346 | 2011 | 1347 |
| 3989 | QGN17086.1 | FIM1_3817 | CP015059. | 1784 | 1019 | 2504 | 2814 | 1078 | 2104 | 2175 | 1163 | 3332 | 9618 | 1863 | 5244 | 15509 | 1466 | 4592 | 12240 | 1507 | 3364 |
| 3990 | QGN17087.1 | FIM1_3818 | CP015059. | 5575 | 52 | 49 | 2102 | 42 | 34 | 3847 | 62 | 60 | 225 | 54 | 74 | 275 | 56 | 64 | 261 | 43 | 33 |
| 3991 | QGN17088.1 | FIM1_3819 | CP015059. | 5110 | 984 | 1777 | 8616 | 1448 | 1449 | 6237 | 1176 | 2365 | 3613 | 754 | 1773 | 6925 | 1064 | 1643 | 5023 | 746 | 1184 |
| 3992 | QGN17089.1 | FIM1_3820 | CP015059. | 1697 | 1068 | 633 | 832 | 1228 | 615 | 1351 | 972 | 772 | 6975 | 1703 | 997 | 5230 | 1419 | 1023 | 6364 | 1639 | 710 |
| 3993 | QGN17090.1 | FIM1_3821 | CP015059. | 1681 | 421 | 462 | 1078 | 634 | 399 | 1432 | 564 | 647 | 2777 | 675 | 900 | 3626 | 706 | 769 | 2923 | 603 | 578 |
| 3994 | QGN17091.1 | FIM1_3822 | CP015059. | 14789 | 4050 | 3086 | 19711 | 3132 | 2498 | 13707 | 3034 | 3869 | 7348 | 2881 | 3988 | 12186 | 3143 | 3894 | 8764 | 2648 | 2783 |
| 3995 | QGN17092.1 | FIM1_3823 | CP015059. | 579 | 63 | 37 | 627 | 48 | 37 | 518 | 49 | 64 | 254 | 56 | 36 | 228 | 82 | 34 | 174 | 86 | 17 |
| 3996 | QGN17093.1 | FIM1_3824 | CP015059. | 96578 | 5268 | 6640 | 110659 | 7177 | 5364 | 116683 | 6743 | 8147 | 19702 | 6340 | 8098 | 37105 | 11899 | 7191 | 26322 | 8347 | 5605 |
| 3997 | QGN17094.1 | FIM1_3825 | CP015059. | 167 | 133 | 246 | 570 | 168 | 240 | 124 | 139 | 337 | 1046 | 126 | 302 | 835 | 116 | 320 | 1031 | 107 | 185 |
| 3998 | QGN17095.1 | FIM1_3826 | CP015059. | 4681 | 4110 | 3238 | 5516 | 4991 | 2907 | 4955 | 4417 | 4145 | 17998 | 5266 | 5332 | 21160 | 5008 | 4800 | 17999 | 4879 | 3572 |
| 3999 | QGN17096.1 | FIM1_3827 | CP015059. | 427 | 568 | 354 | 562 | 385 | 346 | 424 | 442 | 491 | 1246 | 315 | 393 | 1367 | 227 | 431 | 1178 | 234 | 245 |
| 4000 | QGN17097.1 | FIM1_3828 | CP015059. | 2377 | 3243 | 1872 | 2768 | 2856 | 1754 | 2557 | 2269 | 2282 | 10353 | 2255 | 2330 | 8485 | 2128 | 2351 | 9605 | 2105 | 1583 |
| 4001 | QGN17098.1 | FIM1_3829 | CP015059. | 421 | 2184 | 1809 | 383 | 1973 | 1539 | 295 | 1989 | 2433 | 6371 | 1617 | 1857 | 7261 | 1389 | 2064 | 6889 | 1396 | 1276 |
| 4002 | QGN17099.1 | FIM1_3830 | CP015059. | 4934 | 2083 | 2446 | 7445 | 2704 | 1943 | 5923 | 2216 | 3099 | 5782 | 1468 | 2800 | 9776 | 1898 | 2725 | 7078 | 1439 | 1910 |
| 4003 | QGN17100.1 | FIM1_3831 | CP015059. | 2450 | 385 | 575 | 2632 | 299 | 447 | 1781 | 391 | 792 | 2000 | 201 | 721 | 2602 | 226 | 771 | 2064 | 155 | 474 |
| 4004 | QGN17101.1 | FIM1_3832 | CP015059. | 658 | 793 | 602 | 817 | 560 | 567 | 744 | 522 | 812 | 1855 | 489 | 795 | 1865 | 459 | 839 | 1861 | 424 | 501 |
| 4005 | QGN17102.1 | FIM1_3833 | CP015059. | 4514 | 54 | 47 | 2208 | 43 | 33 | 1502 | 37 | 57 | 204 | 32 | 79 | 220 | 27 | 56 | 200 | 29 | 48 |
| 4006 | QGN17103.1 | FIM1_3834 | CP015059. | 6021 | 3078 | 2214 | 7495 | 2398 | 1877 | 6469 | 2185 | 2805 | 8998 | 1653 | 2512 | 8715 | 1709 | 2333 | 8749 | 1431 | 1690 |
| 4007 | QGN17104.1 | FIM1_3835 | CP015059. | 682 | 248 | 240 | 917 | 250 | 167 | 710 | 193 | 263 | 1230 | 211 | 274 | 1230 | 220 | 295 | 1275 | 174 | 168 |
| 4008 | QGN17105.1 | FIM1_3836 | CP015059. | 2614 | 5043 | 2258 | 2780 | 3109 | 2123 | 2870 | 3013 | 3031 | 15233 | 3709 | 3597 | 12358 | 2980 | 3295 | 14669 | 3262 | 2345 |
| 4009 | QGN17106.1 | FIM1_3837 | CP015059. | 3353 | 1973 | 1652 | 3646 | 1964 | 1527 | 2717 | 1544 | 2212 | 4902 | 1873 | 2490 | 7069 | 1436 | 2482 | 6356 | 1371 | 1633 |
| 4010 | QGN17107.1 | FIM1_3838 | CP015059. | 3070 | 960 | 1114 | 3298 | 1064 | 968 | 3036 | 1133 | 1512 | 3248 | 1122 | 1460 | 5795 | 1054 | 1358 | 4378 | 984 | 974 |
| 4011 | QGN17108.1 | FIM1_3839 | CP015059. | 4914 | 866 | 1846 | 4177 | 1943 | 1468 | 3190 | 1700 | 2207 | 8684 | 1002 | 2308 | 11096 | 663 | 1984 | 9437 | 877 | 1567 |
| 4012 | QGN17109.1 | FIM1_3840 | CP015059. | 857 | 252 | 438 | 856 | 667 | 367 | 855 | 473 | 466 | 2597 | 773 | 800 | 2848 | 940 | 650 | 2577 | 819 | 507 |
| 4013 | QGN17110.1 | FIM1_3841 | CP015059. | 6908 | 1020 | 1297 | 7001 | 1325 | 1029 | 6292 | 1500 | 1704 | 2468 | 1022 | 1926 | 7131 | 1105 | 1742 | 4057 | 939 | 1285 |
| 4014 | QGN17111.1 | FIM1_3842 | CP015059. | 428 | 89 | 86 | 1431 | 170 | 101 | 457 | 67 | 98 | 554 | 96 | 119 | 560 | 73 | 102 | 519 | 99 | 78 |
| 4015 | QGN17112.1 | FIM1_3843 | CP015059. | 11412 | 8826 | 6198 | 11731 | 9174 | 5854 | 10951 | 7935 | 7868 | 28850 | 10259 | 9582 | 32256 | 8903 | 9507 | 29252 | 9026 | 6685 |
| 4016 | QGN17113.1 | FIM1_3844 | CP015059. | 5006 | 421 | 550 | 2531 | 721 | 561 | 4945 | 574 | 719 | 1920 | 703 | 818 | 2803 | 939 | 680 | 2083 | 770 | 556 |
| 4017 | QGN17114.1 | FIM1_3845 | CP015059. | 1588 | 519 | 422 | 1124 | 860 | 426 | 1323 | 608 | 523 | 3180 | 731 | 726 | 2702 | 967 | 666 | 2986 | 1005 | 497 |
| 4018 | QGN17115.1 | FIM1_3846 | CP015059. | 2885 | 794 | 488 | 2481 | 755 | 477 | 2261 | 599 | 644 | 2382 | 579 | 611 | 2461 | 522 | 592 | 2188 | 616 | 462 |
| 4019 | QGN17116.1 | FIM1_3847 | CP015059. | 27270 | 23321 | 17278 | 27322 | 23048 | 15109 | 24140 | 20126 | 19513 | 66350 | 27971 | 25552 | 65735 | 25049 | 25713 | 70134 | 23902 | 19480 |
| 4020 | QGN17117.1 | FIM1_3848 | CP015059. | 219 | 540 | 501 | 666 | 778 | 479 | 259 | 564 | 635 | 3196 | 481 | 665 | 3186 | 414 | 579 | 3149 | 446 | 401 |
| 4021 | QGN17118.1 | FIM1_3849 | CP015059. | 12011 | 18544 | 18096 | 26117 | 24245 | 14678 | 18922 | 21921 | 20643 | 56154 | 29011 | 29144 | 73866 | 27960 | 28052 | 62712 | 24728 | 21433 |
| 4022 | QGN17119.1 | FIM1_3850 | CP015059. | 3871 | 2384 | 1564 | 5191 | 1597 | 1228 | 3762 | 1819 | 2087 | 2794 | 1419 | 2058 | 6804 | 1864 | 2052 | 4061 | 1409 | 1464 |
| 4023 | QGN17120.1 | FIM1_3851 | CP015059. | 7782 | 499 | 534 | 5834 | 680 | 456 | 6620 | 593 | 603 | 3531 | 561 | 701 | 4188 | 424 | 628 | 3774 | 465 | 500 |
| 4024 | QGN17121.1 | FIM1_3852 | CP015059. | 1025 | 1378 | 963 | 1635 | 1142 | 818 | 1357 | 862 | 1233 | 5486 | 870 | 1045 | 3732 | 788 | 901 | 5007 | 894 | 636 |
| 4025 | QGN17122.1 | FIM1_3853 | CP015059. | 2093 | 782 | 929 | 3539 | 927 | 834 | 2448 | 849 | 1238 | 8461 | 992 | 1472 | 7271 | 823 | 1338 | 8491 | 806 | 937 |
| 4026 | QGN17123.1 | FIM1_3854 | CP015059. | 5292 | 3831 | 3878 | 10754 | 3497 | 3420 | 6963 | 3096 | 5101 | 20270 | 3152 | 6811 | 18313 | 3743 | 5969 | 20794 | 3113 | 4370 |
| 4027 | QGN17124.1 | FIM1_3855 | CP015059. | 3245 | 2488 | 2836 | 6317 | 2656 | 2553 | 4564 | 2377 | 3680 | 11776 | 1955 | 3996 | 11936 | 2091 | 3550 | 12181 | 1922 | 2496 |
| 4028 | QGN17125.1 | FIM1_3856 | CP015059. | 2038 | 1033 | 2008 | 3083 | 943 | 1702 | 2019 | 884 | 2483 | 2899 | 439 | 1787 | 2683 | 512 | 1929 | 2893 | 389 | 1289 |
| 4029 | QGN17126.1 | FIM1_3857 | CP015059. | 5134 | 5159 | 3819 | 5984 | 4328 | 3423 | 5038 | 3683 | 4670 | 18182 | 5525 | 6406 | 17274 | 5091 | 6556 | 16682 | 4631 | 4537 |
| 4030 | QGN17127.1 | FIM1_3858 | CP015059. | 2731 | 588 | 904 | 2484 | 647 | 723 | 2307 | 593 | 1145 | 3403 | 462 | 1175 | 3225 | 469 | 1041 | 3339 | 373 | 741 |

| | | | | | | | | | | | | | | | | | | | | | |
|---|---|---|---|---|---|---|---|---|---|---|---|---|---|---|---|---|---|---|---|---|---|
| 4031 | QGN17128.1 | FIM1_3859 | CP015059. | 30 | 21 | 22 | 33 | 21 | 28 | 36 | 24 | 30 | 73 | 15 | 33 | 92 | 17 | 13 | 60 | 10 | 22 |
| 4032 | QGN17129.1 | FIM1_3860 | CP015059. | 10416 | 926 | 1870 | 5514 | 1592 | 1271 | 6961 | 1775 | 2068 | 6817 | 1394 | 1771 | 10016 | 1038 | 1687 | 8346 | 904 | 1155 |
| 4033 | QGN17130.1 | FIM1_3861 | CP015059. | 1466 | 455 | 626 | 1365 | 334 | 493 | 1238 | 402 | 794 | 505 | 303 | 716 | 1691 | 278 | 601 | 993 | 274 | 414 |
| 4034 | QGN17131.1 | FIM1_3862 | CP015059. | 2905 | 2117 | 4056 | 14885 | 1680 | 3523 | 5517 | 1690 | 5011 | 11353 | 1421 | 4089 | 10374 | 2345 | 3454 | 11035 | 2183 | 2779 |
| 4035 | QGN17132.1 | FIM1_3863 | CP015059. | 7619 | 2193 | 3235 | 8415 | 4148 | 2676 | 8050 | 3691 | 4003 | 15439 | 4354 | 4914 | 21373 | 3998 | 4694 | 17726 | 3420 | 3350 |
| 4036 | QGN17133.1 | FIM1_3864 | CP015059. | 406 | 18 | 172 | 459 | 13 | 89 | 281 | 17 | 240 | 816 | 43 | 104 | 1620 | 62 | 115 | 1497 | 36 | 110 |
| 4037 | QGN17134.1 | FIM1_3865 | CP015059. | 435 | 177 | 567 | 1345 | 176 | 581 | 321 | 198 | 838 | 3403 | 84 | 577 | 3256 | 86 | 691 | 3230 | 57 | 398 |
| 4038 | QGN17135.1 | FIM1_3866 | CP015059. | 1939 | 1048 | 949 | 2205 | 1087 | 1106 | 1511 | 937 | 1393 | 4750 | 738 | 1069 | 5996 | 557 | 1035 | 4951 | 542 | 765 |
| 4039 | QGN17136.1 | FIM1_3867 | CP015059. | 1063 | 1583 | 928 | 1803 | 1086 | 826 | 1606 | 1068 | 1216 | 5397 | 1104 | 1237 | 4443 | 955 | 1218 | 5437 | 976 | 753 |
| 4040 | QGN17137.1 | FIM1_3868 | CP015059. | 3443 | 306 | 270 | 2411 | 172 | 257 | 2841 | 226 | 419 | 1071 | 118 | 276 | 1012 | 93 | 306 | 962 | 128 | 223 |
| 4041 | QGN17138.1 | FIM1_3869 | CP015059. | 5337 | 11760 | 8576 | 19372 | 11892 | 7572 | 12102 | 11431 | 10999 | 35939 | 12482 | 14495 | 43516 | 12891 | 13695 | 36909 | 11500 | 9791 |
| 4042 | QGN17139.1 | FIM1_3870 | CP015059. | 4460 | 1167 | 2020 | 5276 | 1181 | 1562 | 5176 | 1342 | 2543 | 4043 | 801 | 1714 | 3640 | 668 | 1565 | 4184 | 610 | 1036 |
| 4043 | QGN17140.1 | FIM1_3871 | CP015059. | 2218 | 1923 | 1055 | 1479 | 1827 | 1029 | 2214 | 1580 | 1447 | 8201 | 2140 | 1976 | 7888 | 1951 | 1799 | 8141 | 2155 | 1427 |
| 4044 | QGN17141.1 | FIM1_3872 | CP015059. | 12853 | 3206 | 2652 | 17246 | 4052 | 2304 | 16449 | 3688 | 3345 | 17889 | 3423 | 3280 | 17448 | 3588 | 3140 | 16942 | 3090 | 2210 |
| 4045 | QGN17142.1 | FIM1_3873 | CP015059. | 723 | 617 | 1213 | 1254 | 789 | 1053 | 680 | 742 | 1528 | 2584 | 635 | 2512 | 4666 | 517 | 2230 | 3350 | 504 | 1617 |
| 4046 | QGN17143.1 | FIM1_3874 | CP015059. | 1291 | 238 | 347 | 1665 | 244 | 285 | 1437 | 292 | 438 | 1980 | 218 | 494 | 2067 | 187 | 471 | 1895 | 150 | 334 |
| 4047 | QGN17144.1 | FIM1_3875 | CP015059. | 254 | 209 | 214 | 406 | 131 | 158 | 293 | 162 | 264 | 1310 | 139 | 323 | 1282 | 138 | 288 | 1273 | 114 | 227 |
| 4048 | QGN17145.1 | FIM1_3876 | CP015059. | 100 | 78 | 105 | 229 | 49 | 90 | 137 | 91 | 153 | 342 | 29 | 108 | 424 | 30 | 98 | 357 | 23 | 68 |
| 4049 | QGN17146.1 | FIM1_3877 | CP015059. | 1105 | 58 | 82 | 1024 | 26 | 60 | 720 | 43 | 117 | 471 | 29 | 89 | 475 | 45 | 90 | 433 | 29 | 72 |
| 4050 | QGN17147.1 | FIM1_3878 | CP015059. | 356 | 24 | 13 | 199 | 18 | 9 | 149 | 26 | 20 | 108 | 10 | 24 | 108 | 11 | 18 | 111 | 1 | 17 |
| 4051 | QGN17148.1 | FIM1_3879 | CP015059. | 118505 | 6798 | 8439 | 93832 | 8773 | 6235 | 99447 | 8139 | 10364 | 16633 | 5830 | 12707 | 44294 | 8317 | 11943 | 30409 | 5405 | 9002 |
| 4052 | QGN17149.1 | FIM1_3880 | CP015059. | 2404 | 2536 | 2278 | 6179 | 2036 | 2176 | 4168 | 1855 | 3052 | 11478 | 1544 | 3412 | 9055 | 1788 | 3153 | 11376 | 1686 | 2372 |
| 4053 | QGN17150.1 | FIM1_3881 | CP015059. | 7181 | 3667 | 5175 | 9311 | 3976 | 4744 | 8239 | 3439 | 6637 | 17995 | 3884 | 8126 | 18518 | 4000 | 8161 | 18046 | 3550 | 5721 |
| 4054 | QGN17151.1 | FIM1_3882 | CP015059. | 127 | 14 | 42 | 384 | 24 | 28 | 210 | 23 | 51 | 206 | 26 | 60 | 160 | 20 | 46 | 193 | 31 | 35 |
| 4055 | QGN17152.1 | FIM1_3883 | CP015059. | 648 | 196 | 395 | 831 | 160 | 317 | 617 | 222 | 494 | 2341 | 138 | 447 | 1908 | 117 | 418 | 2095 | 134 | 274 |
| 4056 | QGN17153.1 | FIM1_3884 | CP015059. | 2196 | 277 | 350 | 1857 | 256 | 253 | 2000 | 310 | 400 | 1354 | 164 | 397 | 1403 | 167 | 321 | 1338 | 154 | 224 |
| 4057 | QGN17154.1 | FIM1_3885 | CP015059. | 2775 | 1490 | 1795 | 2854 | 1543 | 1425 | 2334 | 1751 | 2219 | 6363 | 1445 | 2480 | 9474 | 1187 | 2328 | 7090 | 1178 | 1633 |
| 4058 | QGN17155.1 | FIM1_3886 | CP015059. | 3506 | 2726 | 3001 | 6435 | 2408 | 2604 | 4574 | 2239 | 3759 | 10508 | 2328 | 4360 | 10597 | 2653 | 4346 | 10458 | 2255 | 3015 |
| 4059 | QGN17156.1 | FIM1_3887 | CP015059. | 460 | 270 | 324 | 626 | 229 | 311 | 520 | 267 | 470 | 1181 | 159 | 463 | 1137 | 158 | 455 | 1157 | 145 | 292 |
| 4060 | QGN17157.1 | FIM1_3888 | CP015059. | 1459 | 3008 | 2131 | 2486 | 2524 | 1779 | 2142 | 2275 | 2771 | 10887 | 2715 | 3014 | 9538 | 2325 | 3451 | 10089 | 2160 | 2130 |
| 4061 | QGN17158.1 | FIM1_3889 | CP015059. | 1289 | 821 | 984 | 1332 | 620 | 690 | 1017 | 774 | 1274 | 1328 | 587 | 1538 | 3451 | 535 | 1650 | 2161 | 537 | 1016 |
| 4062 | QGN17159.1 | FIM1_3890 | CP015059. | 4877 | 2350 | 1774 | 5557 | 1707 | 1393 | 4633 | 1619 | 2241 | 4252 | 1678 | 2319 | 6315 | 2101 | 2518 | 4600 | 1747 | 1593 |
| 4063 | QGN17160.1 | FIM1_3891 | CP015059. | 1962 | 228 | 235 | 1713 | 171 | 135 | 1480 | 244 | 282 | 662 | 157 | 250 | 1062 | 197 | 302 | 783 | 115 | 161 |
| 4064 | QGN17161.1 | FIM1_3892 | CP015059. | 2554 | 665 | 966 | 5036 | 764 | 622 | 3316 | 956 | 1210 | 1746 | 750 | 1208 | 7189 | 727 | 1384 | 3802 | 406 | 918 |
| 4065 | QGN17162.1 | FIM1_3893 | CP015059. | 240 | 42 | 82 | 258 | 61 | 61 | 292 | 74 | 102 | 295 | 76 | 114 | 407 | 87 | 113 | 312 | 81 | 77 |
| 4066 | QGN17163.1 | FIM1_3894 | CP015059. | 2828 | 547 | 512 | 3076 | 364 | 404 | 2942 | 367 | 660 | 2561 | 320 | 662 | 2375 | 300 | 653 | 2409 | 324 | 446 |
| 4067 | QGN17164.1 | FIM1_3895 | CP015059. | 315 | 126 | 98 | 332 | 113 | 67 | 259 | 83 | 121 | 699 | 110 | 161 | 458 | 88 | 141 | 572 | 111 | 88 |
| 4068 | QGN17165.1 | FIM1_3896 | CP015059. | 1542 | 558 | 797 | 2610 | 660 | 652 | 1729 | 646 | 1040 | 4876 | 578 | 1250 | 5271 | 568 | 1144 | 4959 | 510 | 782 |
| 4069 | QGN17166.1 | FIM1_3897 | CP015059. | 900 | 383 | 320 | 777 | 323 | 239 | 999 | 312 | 383 | 1321 | 341 | 429 | 1207 | 342 | 432 | 1253 | 362 | 300 |
| 4070 | QGN17167.1 | FIM1_3898 | CP015059. | 6327 | 3271 | 1246 | 2120 | 4187 | 1479 | 4688 | 3536 | 1912 | 9015 | 4451 | 2999 | 11991 | 4506 | 2940 | 9397 | 5262 | 2222 |
| 4071 | QGN17168.1 | FIM1_3899 | CP015059. | 1922 | 389 | 559 | 2690 | 448 | 466 | 2159 | 486 | 752 | 3710 | 421 | 801 | 3813 | 404 | 727 | 3739 | 377 | 523 |
| 4072 | QGN17169.1 | FIM1_3900 | CP015059. | 658 | 297 | 209 | 569 | 228 | 170 | 464 | 207 | 261 | 1831 | 264 | 315 | 1452 | 226 | 308 | 1810 | 202 | 197 |
| 4073 | QGN17170.1 | FIM1_3901 | CP015059. | 3239 | 596 | 851 | 3287 | 867 | 696 | 3854 | 846 | 1134 | 3922 | 876 | 1244 | 6033 | 943 | 1056 | 5117 | 904 | 830 |
| 4074 | QGN17171.1 | FIM1_3902 | CP015059. | 8982 | 343 | 387 | 5255 | 349 | 384 | 5339 | 280 | 537 | 1641 | 273 | 458 | 2166 | 415 | 376 | 1659 | 279 | 353 |
| 4075 | QGN17172.1 | FIM1_3903 | CP015059. | 518 | 408 | 318 | 923 | 471 | 370 | 646 | 350 | 468 | 3081 | 405 | 502 | 2753 | 493 | 492 | 3268 | 377 | 321 |
| 4076 | QGN17173.1 | FIM1_3904 | CP015059. | 4743 | 6530 | 4658 | 6417 | 5209 | 4233 | 5102 | 4421 | 5599 | 22051 | 6683 | 7171 | 17610 | 5923 | 7360 | 21818 | 5887 | 5072 |
| 4077 | QGN17174.1 | FIM1_3905 | CP015059. | 7484 | 2278 | 4085 | 13782 | 4280 | 3530 | 11098 | 3317 | 5211 | 13415 | 2706 | 5851 | 22640 | 3161 | 5241 | 16461 | 2728 | 3936 |
| 4078 | QGN17175.1 | FIM1_3906 | CP015059. | 745 | 333 | 638 | 926 | 467 | 626 | 639 | 355 | 836 | 2975 | 236 | 919 | 3236 | 195 | 843 | 2832 | 209 | 593 |
| 4079 | QGN17176.1 | FIM1_3907 | CP015059. | 2336 | 1253 | 1688 | 3687 | 965 | 1472 | 2518 | 925 | 2116 | 4068 | 635 | 2187 | 3870 | 644 | 2180 | 3804 | 549 | 1410 |
| 4080 | QGN17177.1 | FIM1_3908 | CP015059. | 1661 | 822 | 1049 | 2244 | 1005 | 859 | 1854 | 913 | 1393 | 4586 | 840 | 1384 | 5004 | 777 | 1355 | 4834 | 812 | 961 |
| 4081 | QGN17178.1 | FIM1_3909 | CP015059. | 937 | 681 | 615 | 485 | 934 | 593 | 716 | 870 | 770 | 3253 | 1049 | 1002 | 3672 | 1028 | 862 | 3289 | 1109 | 610 |
| 4082 | QGN17179.1 | FIM1_3910 | CP015059. | 7655 | 4501 | 3247 | 8520 | 4324 | 3229 | 6174 | 3479 | 4245 | 16676 | 4836 | 5466 | 17914 | 4739 | 5490 | 16047 | 4340 | 3659 |
| 4083 | QGN17180.1 | FIM1_3911 | CP015059. | 400 | 124 | 106 | 371 | 74 | 94 | 365 | 67 | 184 | 950 | 79 | 222 | 750 | 93 | 202 | 968 | 70 | 86 |
| 4084 | QGN17181.1 | FIM1_3912 | CP015059. | 22807 | 364 | 903 | 21989 | 530 | 925 | 29673 | 524 | 1305 | 10728 | 538 | 1213 | 13128 | 1260 | 1129 | 9653 | 730 | 944 |
| 4085 | QGN17182.1 | FIM1_3913 | CP015059. | 2281 | 120 | 248 | 2810 | 191 | 180 | 2537 | 201 | 321 | 719 | 142 | 285 | 1138 | 240 | 262 | 727 | 230 | 204 |
| 4086 | QGN17183.1 | FIM1_3914 | CP015059. | 1030 | 299 | 234 | 714 | 317 | 226 | 621 | 241 | 332 | 1549 | 226 | 418 | 1432 | 281 | 350 | 1548 | 240 | 261 |
| 4087 | QGN17184.1 | FIM1_3915 | CP015059. | 4061 | 1961 | 1579 | 3502 | 2241 | 1316 | 3792 | 2054 | 1905 | 7642 | 2309 | 2426 | 10304 | 2171 | 2179 | 8514 | 1959 | 1631 |
| 4088 | QGN17185.1 | FIM1_3916 | CP015059. | 810 | 314 | 548 | 1325 | 563 | 458 | 814 | 366 | 682 | 3993 | 462 | 914 | 4304 | 449 | 797 | 3984 | 343 | 557 |
| 4089 | QGN17186.1 | FIM1_3917 | CP015059. | 2852 | 851 | 1009 | 2752 | 1151 | 891 | 2452 | 1090 | 1401 | 3711 | 1290 | 1591 | 7342 | 1144 | 1441 | 5651 | 1152 | 1110 |
| 4090 | QGN17187.1 | FIM1_3918 | CP015059. | 700 | 849 | 1131 | 1664 | 868 | 925 | 1505 | 935 | 1357 | 4057 | 593 | 1932 | 4224 | 497 | 1750 | 3925 | 427 | 1309 |
| 4091 | QGN17188.1 | FIM1_3919 | CP015059. | 1715 | 157 | 196 | 1394 | 96 | 156 | 1809 | 123 | 267 | 1140 | 81 | 300 | 1015 | 56 | 239 | 1007 | 54 | 191 |
| 4092 | QGN17189.1 | FIM1_3920 | CP015059. | 391 | 128 | 205 | 539 | 126 | 144 | 394 | 108 | 261 | 899 | 93 | 298 | 637 | 92 | 235 | 807 | 94 | 167 |
| 4093 | QGN17190.1 | FIM1_3921 | CP015059. | 2990 | 1266 | 1679 | 5348 | 1537 | 1406 | 4306 | 1294 | 2067 | 12119 | 1520 | 2629 | 11086 | 1607 | 2215 | 12131 | 1386 | 1802 |
| 4094 | QGN17191.1 | FIM1_3922 | CP015059. | 263 | 466 | 332 | 442 | 311 | 276 | 324 | 353 | 458 | 2538 | 332 | 579 | 2269 | 223 | 512 | 2337 | 293 | 361 |
| 4095 | QGN17192.1 | FIM1_3923 | CP015059. | 972 | 3496 | 1745 | 1653 | 2836 | 1510 | 1259 | 2314 | 2191 | 9107 | 2819 | 2611 | 8583 | 2518 | 2683 | 9282 | 2517 | 1783 |
| 4096 | QGN17193.1 | FIM1_3924 | CP015059. | 1081 | 378 | 179 | 355 | 667 | 203 | 1011 | 498 | 195 | 1124 | 645 | 352 | 1461 | 779 | 349 | 1185 | 859 | 263 |
| 4097 | QGN17194.1 | FIM1_3925 | CP015059. | 1095 | 74 | 147 | 1211 | 54 | 97 | 767 | 73 | 233 | 800 | 43 | 170 | 719 | 45 | 161 | 752 | 38 | 97 |
| 4098 | QGN17195.1 | FIM1_3926 | CP015059. | 5205 | 1021 | 990 | 7700 | 1084 | 835 | 6464 | 1087 | 1396 | 4250 | 672 | 1205 | 5494 | 906 | 1126 | 4429 | 693 | 849 |
| 4099 | QGN17196.1 | FIM1_3927 | CP015059. | 387 | 239 | 614 | 728 | 175 | 366 | 519 | 240 | 805 | 692 | 120 | 613 | 905 | 139 | 538 | 714 | 82 | 367 |
| 4100 | QGN17197.1 | FIM1_3928 | CP015059. | 2397 | 1152 | 1006 | 3978 | 1634 | 977 | 2574 | 1435 | 1500 | 5484 | 1508 | 1696 | 7974 | 1085 | 1611 | 5842 | 1236 | 1120 |
| 4101 | QGN17198.1 | FIM1_3929 | CP015059. | 459 | 875 | 941 | 948 | 1082 | 867 | 715 | 873 | 1179 | 4906 | 1002 | 1204 | 3926 | 911 | 1152 | 4759 | 954 | 805 |
| 4102 | QGN17199.1 | FIM1_3930 | CP015059. | 1537 | 532 | 643 | 1680 | 757 | 548 | 1543 | 636 | 813 | 4008 | 694 | 1052 | 3903 | 651 | 925 | 3685 | 586 | 652 |
| 4103 | QGN17200.1 | FIM1_3931 | CP015059. | 2 | 0 | 0 | 5 | 2 | 2 | 0 | 2 | 0 | 11 | 1 | 0 | 7 | 0 | 0 | 2 | 0 | 1 |
| 4104 | QGN17201.1 | FIM1_3932 | CP015059. | 2648 | 1546 | 2152 | 3964 | 1958 | 1927 | 3227 | 1610 | 2781 | 7342 | 1172 | 2225 | 6927 | 902 | 2129 | 6780 | 860 | 1458 |
| 4105 | QGN17202.1 | FIM1_3933 | CP015059. | 555 | 416 | 442 | 709 | 346 | 403 | 584 | 292 | 586 | 2078 | 259 | 510 | 1502 | 263 | 498 | 1819 | 267 | 332 |
| 4106 | QGN17203.1 | FIM1_3934 | CP015059. | 3013 | 605 | 489 | 3655 | 616 | 427 | 3726 | 493 | 569 | 4158 | 577 | 721 | 3543 | 522 | 615 | 4159 | 527 | 410 |
| 4107 | QGN17204.1 | FIM1_3935 | CP015059. | 3786 | 1128 | 991 | 4426 | 868 | 781 | 4037 | 723 | 1241 | 5844 | 844 | 1316 | 4671 | 1000 | 1365 | 5829 | 835 | 861 |
| 4108 | QGN17205.1 | FIM1_3936 | CP015059. | 1514 | 558 | 661 | 1657 | 536 | 574 | 1227 | 564 | 975 | 4144 | 363 | 753 | 4260 | 354 | 674 | 3717 | 293 | 446 |
| 4109 | QGN17206.1 | FIM1_3937 | CP015059. | 76 | 18 | 26 | 83 | 7 | 43 | 52 | 12 | 35 | 249 | 8 | 32 | 151 | 2 | 29 | 203 | 7 | 20 |
| 4110 | QGN17207.1 | FIM1_3938 | CP015059. | 102 | 90 | 49 | 159 | 63 | 43 | 67 | 55 | 57 | 210 | 55 | 57 | 196 | 20 | 46 | 217 | 34 | 34 |
| 4111 | QGN17208.1 | FIM1_3939 | CP015059. | 2016 | 326 | 375 | 3615 | 240 | 242 | 2801 | 267 | 529 | 806 | 135 | 237 | 958 | 246 | 186 | 915 | 205 | 138 |
| 4112 | QGN17209.1 | FIM1_3940 | CP015059. | 7120 | 510 | 1128 | 5723 | 1065 | 888 | 6734 | 1051 | 1489 | 1338 | 1085 | 1309 | 5907 | 1204 | 1237 | 2534 | 1266 | 983 |
| 4113 | QGN17210.1 | FIM1_3941 | CP015059. | 303 | 4 | 23 | 312 | 15 | 16 | 252 | 15 | 30 | 16 | 6 | 10 | 49 | 3 | 5 | 19 | 4 | 4 |
| 4114 | QGN17211.1 | FIM1_3942 | CP015059. | 2109 | 431 | 558 | 1759 | 407 | 482 | 1672 | 456 | 657 | 2204 | 234 | 526 | 2128 | 182 | 468 | 2041 | 205 | 372 |

| | | | | | | | | | | | | | | | | | | | | | |
|---|---|---|---|---|---|---|---|---|---|---|---|---|---|---|---|---|---|---|---|---|---|
| 4115 | QGN17212.1 | FIM1_3943 | CP015059. | 6541 | 2155 | 2499 | 8217 | 2838 | 2135 | 7509 | 2242 | 3153 | 12463 | 2931 | 3833 | 16148 | 2729 | 3455 | 13938 | 2767 | 2610 |
| 4116 | QGN17213.1 | FIM1_3944 | CP015059. | 879 | 537 | 675 | 911 | 605 | 558 | 759 | 592 | 948 | 3723 | 613 | 1098 | 4061 | 519 | 961 | 4002 | 455 | 746 |
| 4117 | QGN17214.1 | FIM1_3945 | CP015059. | 281 | 842 | 529 | 305 | 644 | 588 | 367 | 582 | 792 | 4033 | 752 | 836 | 2653 | 653 | 758 | 3537 | 776 | 569 |
| 4118 | QGN17215.1 | FIM1_3946 | CP015059. | 5901 | 985 | 792 | 1341 | 1177 | 725 | 5235 | 986 | 1034 | 4872 | 1228 | 1222 | 5144 | 1113 | 1164 | 4980 | 1118 | 823 |
| 4119 | QGN17216.1 | FIM1_3947 | CP015059. | 2308 | 600 | 1003 | 2511 | 823 | 783 | 2584 | 856 | 1290 | 4349 | 815 | 1327 | 5648 | 610 | 1269 | 4684 | 675 | 943 |
| 4120 | QGN17217.1 | FIM1_3948 | CP015059. | 3550 | 604 | 990 | 4033 | 772 | 863 | 3826 | 767 | 1380 | 5805 | 640 | 1467 | 6021 | 630 | 1386 | 5556 | 538 | 994 |
| 4121 | QGN17218.1 | FIM1_3949 | CP015059. | 3294 | 664 | 879 | 4561 | 721 | 805 | 3892 | 652 | 1106 | 5976 | 631 | 1451 | 5802 | 581 | 1213 | 5882 | 503 | 915 |
| 4122 | QGN17219.1 | FIM1_3950 | CP015059. | 1892 | 1038 | 918 | 2648 | 893 | 792 | 2144 | 784 | 1186 | 4966 | 766 | 1395 | 4800 | 706 | 1293 | 4431 | 640 | 885 |
| 4123 | QGN17220.1 | FIM1_3951 | CP015059. | 2776 | 1337 | 1226 | 3635 | 1432 | 1076 | 3136 | 1407 | 1677 | 5281 | 1132 | 1880 | 6482 | 1250 | 1788 | 5315 | 1124 | 1179 |
| 4124 | QGN17221.1 | FIM1_3952 | CP015059. | 4832 | 686 | 906 | 4720 | 896 | 729 | 4630 | 956 | 1189 | 903 | 821 | 1218 | 5671 | 845 | 1143 | 2550 | 805 | 874 |
| 4125 | QGN17222.1 | FIM1_3953 | CP015059. | 2469 | 947 | 1419 | 4192 | 1238 | 1258 | 2878 | 1065 | 1882 | 4158 | 827 | 1874 | 5302 | 751 | 1854 | 4104 | 769 | 1294 |
| 4126 | QGN17223.1 | FIM1_3954 | CP015059. | 567 | 482 | 463 | 563 | 376 | 379 | 454 | 441 | 616 | 2748 | 383 | 542 | 2577 | 516 | 627 | 2447 | 359 | 373 |
| 4127 | QGN17224.1 | FIM1_3955 | CP015059. | 1215 | 289 | 543 | 1103 | 389 | 421 | 809 | 471 | 669 | 2750 | 358 | 670 | 3624 | 226 | 613 | 2956 | 223 | 391 |
| 4128 | QGN17225.1 | FIM1_3956 | CP015059. | 2259 | 2214 | 1115 | 1344 | 2621 | 1104 | 1805 | 1843 | 1443 | 10272 | 3280 | 2764 | 8941 | 2796 | 2211 | 9783 | 3352 | 1687 |
| 4129 | QGN17226.1 | FIM1_3957 | CP015059. | 3431 | 1657 | 1476 | 4392 | 1511 | 1327 | 3501 | 1329 | 1992 | 6763 | 1275 | 1896 | 8118 | 1193 | 1931 | 6659 | 1203 | 1313 |
| 4130 | QGN17227.1 | FIM1_3958 | CP015059. | 2331 | 1185 | 896 | 7980 | 1938 | 911 | 11222 | 1413 | 1254 | 12593 | 3494 | 1814 | 14251 | 4568 | 2135 | 12767 | 3818 | 1423 |
| 4131 | QGN17228.1 | FIM1_3959 | CP015059. | 1406 | 430 | 591 | 1789 | 508 | 520 | 1394 | 568 | 769 | 3137 | 328 | 542 | 3804 | 283 | 471 | 3236 | 253 | 377 |
| 4132 | QGN17229.1 | FIM1_3960 | CP015059. | 150 | 102 | 144 | 361 | 239 | 83 | 250 | 199 | 135 | 759 | 189 | 145 | 835 | 166 | 139 | 824 | 156 | 111 |
| 4133 | QGN17230.1 | FIM1_3961 | CP015059. | 1298 | 151 | 601 | 2471 | 788 | 515 | 1221 | 625 | 782 | 2983 | 572 | 867 | 3825 | 465 | 752 | 3006 | 431 | 563 |
| 4134 | QGN17231.1 | FIM1_3962 | CP015059. | 2105 | 1765 | 1297 | 2127 | 1122 | 1044 | 2038 | 1214 | 1692 | 3506 | 608 | 964 | 3127 | 744 | 994 | 3210 | 646 | 686 |
| 4135 | QGN17232.1 | FIM1_3963 | CP015059. | 7929 | 1152 | 943 | 4446 | 1228 | 792 | 5830 | 1140 | 1265 | 885 | 701 | 1007 | 3210 | 677 | 1127 | 1572 | 675 | 758 |
| 4136 | QGN17233.1 | FIM1_3964 | CP015059. | 6036 | 694 | 1278 | 4534 | 1535 | 891 | 6248 | 1618 | 1536 | 1976 | 1308 | 1652 | 8630 | 1544 | 1598 | 5201 | 1398 | 1143 |
| 4137 | QGN17234.1 | FIM1_3965 | CP015059. | 643 | 556 | 405 | 669 | 470 | 414 | 518 | 411 | 585 | 1671 | 397 | 503 | 1657 | 343 | 585 | 1487 | 343 | 356 |
| 4138 | QGN17235.1 | FIM1_3966 | CP015059. | 58 | 77 | 244 | 305 | 99 | 107 | 98 | 120 | 253 | 1073 | 63 | 292 | 1657 | 369 | 301 | 1224 | 104 | 240 |
| 4139 | QGN17236.1 | FIM1_3967 | CP015059. | 1350 | 1580 | 931 | 1172 | 979 | 862 | 1254 | 827 | 1181 | 3914 | 930 | 1290 | 3463 | 810 | 1444 | 3711 | 909 | 929 |
| 4140 | QGN17237.1 | FIM1_3968 | CP015059. | 2235 | 619 | 801 | 2026 | 790 | 675 | 2051 | 805 | 1093 | 3934 | 772 | 1122 | 4878 | 671 | 1077 | 4093 | 721 | 818 |
| 4141 | QGN17238.1 | FIM1_3969 | CP015059. | 811 | 1135 | 1531 | 1994 | 1577 | 1285 | 1361 | 1168 | 1949 | 9294 | 1488 | 1746 | 8727 | 1399 | 1501 | 9177 | 1315 | 1164 |
| 4142 | QGN17239.1 | FIM1_3970 | CP015059. | 1310 | 535 | 455 | 921 | 699 | 363 | 1236 | 585 | 613 | 1723 | 594 | 845 | 3996 | 518 | 750 | 2627 | 582 | 551 |
| 4143 | QGN17240.1 | FIM1_3971 | CP015059. | 491 | 35 | 242 | 713 | 127 | 181 | 587 | 151 | 273 | 558 | 96 | 368 | 1419 | 104 | 276 | 774 | 97 | 234 |
| 4144 | QGN17241.1 | FIM1_3972 | CP015059. | 1170 | 272 | 352 | 1090 | 245 | 311 | 1104 | 314 | 478 | 2000 | 218 | 500 | 2397 | 175 | 443 | 2217 | 185 | 337 |
| 4145 | QGN17242.1 | FIM1_3973 | CP015059. | 5941 | 5385 | 2992 | 5443 | 5500 | 2965 | 5511 | 4613 | 3936 | 17035 | 5355 | 4952 | 15370 | 4523 | 4831 | 15978 | 4985 | 3252 |
| 4146 | QGN17243.1 | FIM1_3974 | CP015059. | 2620 | 918 | 740 | 3331 | 747 | 586 | 2787 | 589 | 995 | 4338 | 672 | 1216 | 3551 | 708 | 1129 | 4192 | 602 | 759 |
| 4147 | QGN17244.1 | FIM1_3975 | CP015059. | 2947 | 481 | 856 | 4469 | 625 | 621 | 3143 | 604 | 1017 | 2752 | 291 | 717 | 3395 | 311 | 597 | 2978 | 240 | 470 |
| 4148 | QGN17245.1 | FIM1_3976 | CP015059. | 29229 | 2977 | 4620 | 31957 | 4618 | 3686 | 29603 | 4174 | 5723 | 21486 | 4438 | 7557 | 25896 | 5803 | 7339 | 23276 | 4102 | 5096 |
| 4149 | QGN17246.1 | FIM1_3977 | CP015059. | 16409 | 323 | 273 | 8659 | 254 | 172 | 11184 | 315 | 354 | 277 | 118 | 433 | 2241 | 94 | 389 | 747 | 80 | 267 |
| 4150 | QGN17247.1 | FIM1_3978 | CP015059. | 1080 | 1131 | 532 | 844 | 1029 | 559 | 1049 | 1014 | 788 | 2754 | 835 | 972 | 3480 | 730 | 862 | 2753 | 768 | 654 |
| 4151 | QGN17248.1 | FIM1_3979 | CP015059. | 268 | 554 | 458 | 367 | 565 | 507 | 288 | 453 | 682 | 2642 | 385 | 790 | 2756 | 419 | 663 | 2741 | 407 | 484 |
| 4152 | QGN17249.1 | FIM1_3980 | CP015059. | 14789 | 13731 | 9140 | 14273 | 11708 | 8329 | 12269 | 9613 | 10695 | 34104 | 12321 | 12878 | 29484 | 11101 | 12321 | 35627 | 10962 | 9643 |
| 4153 | QGN17250.1 | FIM1_3981 | CP015059. | 2261 | 134 | 238 | 5514 | 102 | 138 | 3618 | 150 | 277 | 1033 | 62 | 224 | 899 | 85 | 194 | 1124 | 107 | 116 |
| 4154 | QGN17251.1 | FIM1_3982 | CP015059. | 4666 | 2479 | 2417 | 7770 | 2866 | 2185 | 5277 | 2254 | 2938 | 13922 | 2114 | 3498 | 12260 | 2112 | 3287 | 13378 | 1817 | 2325 |
| 4155 | QGN17252.1 | FIM1_3983 | CP015059. | 1877 | 1591 | 1113 | 2923 | 1593 | 1162 | 1835 | 1310 | 1486 | 5993 | 1140 | 1818 | 5560 | 1070 | 1847 | 5628 | 1059 | 1140 |
| 4156 | QGN17253.1 | FIM1_3984 | CP015059. | 308 | 315 | 198 | 214 | 191 | 178 | 190 | 175 | 253 | 1495 | 179 | 280 | 993 | 179 | 267 | 1292 | 176 | 141 |
| 4157 | QGN17254.1 | FIM1_3985 | CP015059. | 735 | 438 | 521 | 789 | 416 | 430 | 642 | 381 | 664 | 3111 | 389 | 819 | 2932 | 349 | 741 | 2922 | 314 | 500 |
| 4158 | QGN17255.1 | FIM1_3986 | CP015059. | 374 | 1369 | 1058 | 990 | 1504 | 863 | 715 | 1469 | 1452 | 3264 | 1064 | 1517 | 5236 | 1092 | 1530 | 3741 | 945 | 943 |
| 4159 | QGN17256.1 | FIM1_3987 | CP015059. | 1624 | 603 | 529 | 1612 | 599 | 428 | 1361 | 585 | 646 | 1469 | 485 | 640 | 2318 | 407 | 696 | 1649 | 427 | 463 |
| 4160 | QGN17257.1 | FIM1_3988 | CP015059. | 1318 | 767 | 763 | 1293 | 582 | 688 | 1240 | 598 | 1026 | 4663 | 481 | 1011 | 3717 | 365 | 1027 | 4556 | 386 | 626 |
| 4161 | QGN17258.1 | FIM1_3989 | CP015059. | 6901 | 4803 | 5809 | 9210 | 7386 | 5169 | 7664 | 5390 | 7399 | 35624 | 6706 | 9720 | 39044 | 6484 | 8384 | 36336 | 5982 | 6365 |
| 4162 | QGN17259.1 | FIM1_3990 | CP015059. | 2591 | 796 | 1536 | 2020 | 861 | 1006 | 1372 | 1196 | 2019 | 3009 | 660 | 1327 | 7791 | 560 | 1363 | 4407 | 439 | 938 |
| 4163 | QGN17260.1 | FIM1_3991 | CP015059. | 3851 | 1725 | 578 | 2129 | 1528 | 677 | 3114 | 1457 | 905 | 1585 | 931 | 904 | 2563 | 1175 | 873 | 1698 | 1191 | 717 |
| 4164 | QGN17261.1 | FIM1_3992 | CP015059. | 1516 | 628 | 468 | 1651 | 445 | 415 | 1454 | 449 | 632 | 1941 | 369 | 586 | 2002 | 352 | 624 | 1995 | 329 | 374 |
| 4165 | QGN17262.1 | FIM1_3993 | CP015059. | 7856 | 5745 | 4093 | 9086 | 6080 | 3482 | 8275 | 5435 | 5402 | 19737 | 6371 | 7022 | 24881 | 6144 | 6482 | 20952 | 5633 | 4866 |
| 4166 | QGN17263.1 | FIM1_3994 | CP015059. | 362 | 39 | 33 | 475 | 26 | 27 | 444 | 26 | 48 | 301 | 34 | 73 | 292 | 24 | 37 | 352 | 34 | 33 |
| 4167 | QGN17264.1 | FIM1_3995 | CP015059. | 224 | 173 | 105 | 812 | 234 | 110 | 276 | 144 | 136 | 395 | 107 | 110 | 573 | 89 | 117 | 475 | 70 | 78 |
| 4168 | QGN17265.1 | FIM1_3996 | CP015059. | 4990 | 1077 | 646 | 2064 | 1390 | 598 | 4025 | 1176 | 849 | 1154 | 1048 | 1035 | 3655 | 1283 | 950 | 2001 | 1230 | 732 |
| 4169 | QGN17266.1 | FIM1_3997 | CP015059. | 3062 | 1057 | 1268 | 2701 | 1188 | 1095 | 3501 | 1255 | 1652 | 4046 | 954 | 1400 | 6297 | 863 | 1331 | 5082 | 933 | 995 |
| 4170 | QGN17267.1 | FIM1_3998 | CP015059. | 1268 | 359 | 992 | 1556 | 693 | 824 | 1356 | 628 | 1331 | 5146 | 517 | 1287 | 4815 | 464 | 1043 | 5306 | 468 | 804 |
| 4171 | QGN17268.1 | FIM1_3999 | CP015059. | 149 | 346 | 209 | 171 | 217 | 179 | 97 | 162 | 309 | 614 | 211 | 279 | 565 | 110 | 217 | 563 | 113 | 169 |
| 4172 | QGN17269.1 | FIM1_4000 | CP015059. | 1121 | 798 | 623 | 1040 | 679 | 593 | 991 | 590 | 872 | 2816 | 465 | 547 | 1993 | 371 | 520 | 2481 | 382 | 325 |
| 4173 | QGN17270.1 | FIM1_4001 | CP015059. | 174 | 148 | 59 | 119 | 85 | 55 | 134 | 73 | 82 | 413 | 89 | 113 | 407 | 65 | 97 | 344 | 94 | 71 |
| 4174 | QGN17271.1 | FIM1_4002 | CP015059. | 5373 | 3592 | 2479 | 4749 | 3535 | 2175 | 4836 | 3044 | 3158 | 11746 | 3556 | 4244 | 13003 | 3388 | 4116 | 11640 | 3275 | 2912 |
| 4175 | QGN17272.1 | FIM1_4003 | CP015059. | 4966 | 3916 | 2860 | 5449 | 3110 | 2791 | 4504 | 2509 | 3536 | 11153 | 2978 | 4341 | 10388 | 2932 | 4463 | 10752 | 2859 | 2977 |
| 4176 | QGN17273.1 | FIM1_4004 | CP015059. | 999 | 1049 | 940 | 1071 | 809 | 814 | 845 | 718 | 1242 | 3223 | 733 | 1257 | 3230 | 721 | 1392 | 3025 | 641 | 890 |
| 4177 | QGN17274.1 | FIM1_4005 | CP015059. | 3487 | 1537 | 2420 | 6152 | 1511 | 1972 | 4731 | 1734 | 3224 | 4051 | 1182 | 3494 | 7605 | 1269 | 3287 | 5314 | 1083 | 2512 |
| 4178 | QGN17275.1 | FIM1_4006 | CP015059. | 2407 | 1698 | 1896 | 3828 | 1857 | 1497 | 2866 | 1692 | 2370 | 9375 | 1719 | 2722 | 9546 | 1723 | 2621 | 9685 | 1506 | 1810 |
| 4179 | QGN17276.1 | FIM1_4007 | CP015059. | 2495 | 1226 | 1302 | 2677 | 1251 | 998 | 2632 | 1143 | 1636 | 5194 | 1068 | 1619 | 5448 | 936 | 1472 | 4763 | 895 | 1042 |
| 4180 | QGN17277.1 | FIM1_4008 | CP015059. | 1272 | 169 | 253 | 1072 | 301 | 123 | 1130 | 283 | 268 | 1948 | 453 | 367 | 1735 | 466 | 305 | 1869 | 452 | 329 |
| 4181 | QGN17278.1 | FIM1_4009 | CP015059. | 8413 | 2452 | 1591 | 7217 | 1639 | 1190 | 7238 | 1700 | 1997 | 9134 | 2374 | 2399 | 7998 | 2781 | 2270 | 8661 | 2711 | 1602 |
| 4182 | QGN17279.1 | FIM1_4010 | CP015059. | 3937 | 830 | 1145 | 5589 | 1070 | 945 | 4518 | 906 | 1444 | 5139 | 849 | 1721 | 6014 | 860 | 1538 | 5515 | 812 | 1093 |
| 4183 | QGN17280.1 | FIM1_4011 | CP015059. | 3390 | 969 | 1251 | 4086 | 1114 | 1127 | 3776 | 930 | 1707 | 7445 | 1052 | 2039 | 7688 | 926 | 1714 | 7303 | 911 | 1350 |
| 4184 | QGN17281.1 | FIM1_4012 | CP015059. | 3133 | 1851 | 1239 | 3217 | 1826 | 1233 | 3025 | 1553 | 1733 | 9044 | 1682 | 1985 | 8011 | 1386 | 1747 | 8755 | 1504 | 1302 |
| 4185 | QGN17282.1 | FIM1_4013 | CP015059. | 1899 | 1080 | 1009 | 2382 | 1097 | 905 | 1969 | 985 | 1397 | 5761 | 1266 | 1656 | 6213 | 1229 | 1429 | 5463 | 1182 | 996 |
| 4186 | QGN17283.1 | FIM1_4014 | CP015059. | 627 | 204 | 323 | 523 | 193 | 243 | 790 | 220 | 390 | 1500 | 209 | 384 | 1060 | 131 | 313 | 1456 | 189 | 217 |
| 4187 | QGN17284.1 | FIM1_4015 | CP015059. | 4787 | 1074 | 2531 | 2772 | 1461 | 2117 | 3805 | 1676 | 3170 | 11312 | 1374 | 2920 | 12380 | 1114 | 2629 | 11789 | 1263 | 1920 |
| 4188 | QGN17285.1 | FIM1_4016 | CP015059. | 5162 | 1506 | 1654 | 8218 | 1522 | 1509 | 6139 | 1205 | 2046 | 8010 | 1268 | 2416 | 7396 | 1584 | 2268 | 7749 | 1441 | 1548 |
| 4189 | QGN17286.1 | FIM1_4017 | CP015059. | 341 | 272 | 190 | 294 | 306 | 225 | 378 | 213 | 278 | 1461 | 270 | 340 | 1321 | 333 | 246 | 1323 | 300 | 208 |
| 4190 | QGN17287.1 | FIM1_4018 | CP015059. | 6004 | 2780 | 1221 | 2247 | 3471 | 1334 | 5191 | 2772 | 1597 | 8950 | 4044 | 2780 | 11624 | 3632 | 2495 | 9336 | 4460 | 1923 |
| 4191 | QGN17288.1 | FIM1_4019 | CP015059. | 1713 | 393 | 452 | 1659 | 361 | 385 | 1673 | 338 | 579 | 3689 | 460 | 745 | 3222 | 405 | 654 | 3477 | 398 | 559 |
| 4192 | QGN17289.1 | FIM1_4020 | CP015059. | 3936 | 2993 | 2260 | 5488 | 3811 | 2003 | 4964 | 3128 | 2861 | 12525 | 2828 | 2798 | 12506 | 2709 | 2453 | 12744 | 2581 | 1810 |
| 4193 | QGN17290.1 | FIM1_4021 | CP015059. | 1025 | 741 | 752 | 992 | 777 | 643 | 1155 | 610 | 968 | 4651 | 681 | 1136 | 4214 | 559 | 1042 | 4220 | 515 | 732 |
| 4194 | QGN17291.1 | FIM1_4022 | CP015059. | 11 | 20 | 16 | 49 | 32 | 21 | 13 | 17 | 25 | 138 | 32 | 43 | 100 | 20 | 33 | 114 | 20 | 15 |
| 4195 | QGN17292.1 | FIM1_4023 | CP015059. | 8722 | 4704 | 4262 | 11444 | 5301 | 3708 | 9148 | 4430 | 5216 | 19971 | 4935 | 6505 | 24061 | 5103 | 5920 | 21172 | 4617 | 4333 |
| 4196 | QGN17293.1 | FIM1_4024 | CP015059. | 2582 | 1565 | 2049 | 17256 | 1555 | 1839 | 6245 | 1454 | 2452 | 9073 | 1236 | 2394 | 7710 | 1466 | 1918 | 8330 | 1525 | 1416 |
| 4197 | QGN17294.1 | FIM1_4025 | CP015059. | 339 | 151 | 161 | 393 | 92 | 159 | 230 | 106 | 249 | 1343 | 114 | 372 | 939 | 129 | 245 | 1261 | 110 | 188 |
| 4198 | QGN17295.1 | FIM1_4026 | CP015059. | 4495 | 79 | 62 | 2492 | 83 | 66 | 2121 | 73 | 94 | 589 | 118 | 249 | 1177 | 59 | 63 | 1080 | 115 | 94 |

| | | | | | | | | | | | | | | | | | | | | | |
|---|---|---|---|---|---|---|---|---|---|---|---|---|---|---|---|---|---|---|---|---|---|
| 4199 | QGN17296.1 | FIM1_4027 | CP015059. | 1424 | 203 | 596 | 8633 | 193 | 343 | 3045 | 187 | 546 | 1191 | 125 | 574 | 1247 | 178 | 270 | 1149 | 133 | 308 |
| 4200 | QGN17297.1 | FIM1_4028 | CP015059. | 1474 | 762 | 715 | 1400 | 699 | 592 | 1088 | 634 | 829 | 3374 | 875 | 1323 | 3519 | 771 | 1182 | 3453 | 767 | 845 |
| 4201 | QGN17298.1 | FIM1_4029 | CP015059. | 191 | 23 | 26 | 539 | 82 | 31 | 282 | 48 | 29 | 232 | 43 | 51 | 324 | 55 | 51 | 261 | 56 | 28 |
| 4202 | QGN17299.1 | FIM1_4030 | CP015059. | 7614 | 3071 | 1646 | 3481 | 4089 | 1846 | 7072 | 3286 | 2273 | 13175 | 5705 | 3809 | 17020 | 4662 | 3369 | 14736 | 6269 | 2710 |
| 4203 | QGN17300.1 | FIM1_4031 | CP015059. | 1867 | 3193 | 1829 | 2112 | 2477 | 1785 | 1798 | 2049 | 2517 | 12160 | 2677 | 3092 | 9717 | 2468 | 2822 | 11239 | 2590 | 2009 |
| 4204 | QGN17301.1 | FIM1_4033 | CP015059. | 1030 | 713 | 547 | 1392 | 592 | 549 | 1030 | 604 | 785 | 3457 | 435 | 668 | 2689 | 443 | 731 | 3442 | 402 | 451 |
| 4205 | QGN17302.1 | FIM1_4034 | CP015059. | 358 | 247 | 267 | 477 | 398 | 225 | 469 | 327 | 318 | 2034 | 450 | 484 | 1772 | 512 | 402 | 1987 | 535 | 289 |
| 4206 | QGN17303.1 | FIM1_4035 | CP015059. | 190 | 396 | 214 | 237 | 351 | 191 | 177 | 259 | 267 | 1404 | 365 | 368 | 1376 | 415 | 347 | 1388 | 438 | 257 |
| 4207 | QGN17304.1 | FIM1_4036 | CP015059. | 2254 | 1388 | 996 | 2857 | 1147 | 947 | 2709 | 871 | 1326 | 8499 | 1201 | 1783 | 6007 | 1154 | 1532 | 8296 | 1250 | 1130 |
| 4208 | QGN17305.1 | FIM1_4037 | CP015059. | 1527 | 565 | 834 | 1869 | 742 | 701 | 1680 | 733 | 1030 | 7563 | 774 | 1378 | 6608 | 643 | 1125 | 7186 | 653 | 887 |
| 4209 | QGN17306.1 | FIM1_4038 | CP015059. | 1179 | 423 | 423 | 1253 | 439 | 340 | 1200 | 380 | 576 | 3306 | 527 | 766 | 2618 | 497 | 630 | 3371 | 488 | 464 |
| 4210 | QGN17307.1 | FIM1_4039 | CP015059. | 4135 | 519 | 457 | 3783 | 896 | 445 | 3686 | 719 | 560 | 2145 | 947 | 690 | 3191 | 847 | 640 | 2378 | 911 | 501 |
| 4211 | QGN17308.1 | FIM1_4040 | CP015059. | 24 | 1 | 2 | 39 | 7 | 2 | 24 | 5 | 3 | 8 | 12 | 0 | 20 | 3 | 3 | 10 | 5 | 2 |
| 4212 | QGN17309.1 | FIM1_4041 | CP015059. | 3420 | 655 | 372 | 1571 | 772 | 343 | 3098 | 693 | 463 | 700 | 965 | 717 | 2577 | 1065 | 708 | 1460 | 1202 | 500 |
| 4213 | QGN17310.1 | FIM1_4042 | CP015059. | 4579 | 803 | 707 | 3814 | 1052 | 679 | 4162 | 750 | 988 | 5697 | 1064 | 1370 | 5942 | 983 | 1128 | 5536 | 965 | 898 |
| 4214 | QGN17311.1 | FIM1_4043 | CP015059. | 4032 | 712 | 904 | 2968 | 956 | 826 | 4070 | 838 | 1223 | 2714 | 893 | 1371 | 4873 | 913 | 1206 | 3421 | 910 | 939 |
| 4215 | QGN17312.1 | FIM1_4044 | CP015059. | 523 | 30 | 164 | 400 | 183 | 121 | 556 | 179 | 166 | 656 | 174 | 217 | 1172 | 155 | 180 | 785 | 159 | 162 |
| 4216 | QGN17313.1 | FIM1_4045 | CP015059. | 4811 | 2485 | 2131 | 3855 | 3060 | 1984 | 4680 | 2672 | 2669 | 13058 | 3431 | 3435 | 12953 | 3235 | 3352 | 12312 | 3370 | 2514 |
| 4217 | QGN17314.1 | FIM1_4046 | CP015059. | 998 | 1120 | 813 | 866 | 875 | 852 | 870 | 651 | 1113 | 4826 | 950 | 1208 | 3431 | 878 | 1187 | 4103 | 979 | 789 |
| 4218 | QGN17315.1 | FIM1_4047 | CP015059. | 6313 | 3255 | 3924 | 8736 | 5740 | 3415 | 7752 | 4548 | 4757 | 24525 | 6173 | 7016 | 28662 | 6043 | 5702 | 25497 | 5750 | 4346 |
| 4219 | QGN17316.1 | FIM1_4048 | CP015059. | 132 | 470 | 266 | 98 | 339 | 233 | 134 | 326 | 341 | 1119 | 273 | 275 | 1213 | 303 | 275 | 1015 | 263 | 184 |
| 4220 | QGN17317.1 | FIM1_4049 | CP015059. | 1262 | 1257 | 587 | 405 | 1326 | 617 | 1022 | 983 | 785 | 4932 | 1600 | 1346 | 3793 | 1412 | 1200 | 4334 | 1647 | 830 |
| 4221 | QGN17318.1 | FIM1_4050 | CP015059. | 5015 | 1899 | 2457 | 6159 | 2296 | 2009 | 5639 | 2346 | 3107 | 11322 | 2127 | 3234 | 14724 | 2058 | 2958 | 11947 | 1913 | 2230 |
| 4222 | QGN17319.1 | FIM1_4051 | CP015059. | 8692 | 5288 | 9706 | 14050 | 5674 | 8736 | 9832 | 5886 | 12218 | 15321 | 2578 | 10337 | 17641 | 2656 | 10375 | 15971 | 2447 | 6771 |
| 4223 | QGN17320.1 | FIM1_4052 | CP015059. | 166 | 57 | 31 | 231 | 69 | 26 | 124 | 66 | 25 | 149 | 52 | 58 | 223 | 41 | 46 | 134 | 33 | 29 |
| 4224 | QGN17321.1 | FIM1_4053 | CP015059. | 2053 | 628 | 381 | 1882 | 371 | 267 | 1556 | 528 | 590 | 257 | 187 | 551 | 2192 | 291 | 483 | 772 | 242 | 364 |
| 4225 | QGN17322.1 | FIM1_4054 | CP015059. | 9784 | 1179 | 1827 | 8183 | 1599 | 1357 | 7695 | 1706 | 2337 | 4469 | 1424 | 2058 | 8809 | 1275 | 2071 | 5640 | 1278 | 1428 |
| 4226 | QGN17323.1 | FIM1_4055 | CP015059. | 2969 | 925 | 993 | 5111 | 945 | 648 | 4012 | 912 | 1245 | 5011 | 594 | 948 | 5453 | 986 | 803 | 4560 | 550 | 560 |
| 4227 | QGN17324.1 | FIM1_4056 | CP015059. | 5631 | 818 | 744 | 6826 | 798 | 656 | 6260 | 746 | 1002 | 2913 | 593 | 1229 | 4921 | 648 | 1016 | 3256 | 502 | 840 |
| 4228 | QGN17325.1 | FIM1_4057 | CP015059. | 1869 | 891 | 722 | 2816 | 734 | 676 | 2353 | 691 | 1001 | 4466 | 671 | 1037 | 3741 | 604 | 978 | 3845 | 540 | 718 |
| 4229 | QGN17326.1 | FIM1_4058 | CP015059. | 84 | 367 | 289 | 166 | 291 | 271 | 84 | 218 | 380 | 1543 | 237 | 331 | 1426 | 282 | 302 | 1671 | 265 | 241 |
| 4230 | QGN17327.1 | FIM1_4059 | CP015059. | 23969 | 1410 | 1938 | 18152 | 2535 | 1422 | 22564 | 2590 | 2374 | 2088 | 1936 | 2504 | 8694 | 2407 | 2144 | 4027 | 2333 | 1740 |
| 4231 | QGN17328.1 | FIM1_4060 | CP015059. | 2415 | 2877 | 931 | 1399 | 1928 | 1003 | 2055 | 1559 | 1137 | 5964 | 2553 | 1587 | 5029 | 1905 | 1668 | 5042 | 2632 | 1095 |
| 4232 | QGN17329.1 | FIM1_4061 | CP015059. | 7422 | 2404 | 1819 | 5536 | 3086 | 2310 | 7075 | 3041 | 2541 | 11813 | 5337 | 3185 | 12735 | 4921 | 3272 | 11884 | 6384 | 2399 |
| 4233 | QGN17330.1 | FIM1_4062 | CP015059. | 1599 | 1163 | 483 | 1200 | 751 | 648 | 1300 | 746 | 709 | 2865 | 1475 | 674 | 2872 | 1231 | 867 | 2536 | 1776 | 546 |
| 4234 | QGN17331.1 | FIM1_4063 | CP015059. | 4015 | 930 | 1371 | 6486 | 1604 | 1227 | 6126 | 1129 | 1616 | 5988 | 1199 | 2083 | 8625 | 1235 | 1870 | 6530 | 1065 | 1380 |
| 4235 | QGN17332.1 | FIM1_4064 | CP015059. | 3087 | 4176 | 3237 | 6545 | 3723 | 3457 | 5102 | 3191 | 4066 | 15230 | 4986 | 4665 | 12502 | 4282 | 5447 | 14616 | 4204 | 3394 |
| 4236 | QGN17333.1 | FIM1_4065 | CP015059. | 4785 | 4307 | 1442 | 2449 | 3383 | 1427 | 4651 | 3300 | 1962 | 4844 | 4213 | 2453 | 8406 | 3824 | 2842 | 6042 | 4714 | 1927 |
| 4237 | QGN17334.1 | FIM1_4066 | CP015059. | 17835 | 2200 | 1608 | 19680 | 1640 | 1484 | 17476 | 1857 | 2146 | 4403 | 1434 | 1990 | 6220 | 1618 | 2127 | 4706 | 1419 | 1460 |
| 4238 | QGN17335.1 | FIM1_4067 | CP015059. | 4118 | 395 | 490 | 5009 | 547 | 462 | 4382 | 524 | 598 | 2015 | 478 | 656 | 2431 | 529 | 693 | 1928 | 491 | 455 |
| 4239 | QGN17336.1 | FIM1_4068 | CP015059. | 1348 | 375 | 196 | 1171 | 238 | 211 | 1175 | 200 | 281 | 861 | 267 | 289 | 838 | 193 | 336 | 840 | 291 | 188 |
| 4240 | QGN17337.1 | FIM1_4069 | CP015059. | 5316 | 1316 | 1287 | 6942 | 1392 | 1007 | 6259 | 1345 | 1749 | 2202 | 1268 | 2161 | 10580 | 1381 | 2024 | 5844 | 1300 | 1448 |
| 4241 | QGN17338.1 | FIM1_4070 | CP015059. | 5149 | 1203 | 1587 | 6422 | 1535 | 1130 | 5886 | 1659 | 1936 | 3498 | 1571 | 2393 | 12257 | 2728 | 2849 | 6931 | 1334 | 1782 |
| 4242 | QGN17339.1 | FIM1_4071 | CP015059. | 218 | 47 | 29 | 646 | 59 | 39 | 383 | 40 | 50 | 141 | 72 | 48 | 259 | 63 | 75 | 202 | 64 | 45 |
| 4243 | QGN17340.1 | FIM1_4072 | CP015059. | 1955 | 634 | 1028 | 2448 | 938 | 873 | 1975 | 733 | 1313 | 6932 | 931 | 1750 | 6646 | 786 | 1488 | 6724 | 812 | 1190 |
| 4244 | QGN17341.1 | FIM1_4073 | CP015059. | 1869 | 410 | 674 | 1748 | 538 | 587 | 2154 | 456 | 895 | 4236 | 533 | 1203 | 3690 | 567 | 1050 | 3995 | 503 | 732 |
| 4245 | QGN17342.1 | FIM1_4074 | CP015059. | 491 | 702 | 893 | 1852 | 359 | 549 | 1385 | 559 | 1176 | 3381 | 648 | 1078 | 4124 | 1330 | 1077 | 3215 | 1052 | 690 |
| 4246 | QGN17343.1 | FIM1_4075 | CP015059. | 988 | 453 | 634 | 1697 | 243 | 409 | 864 | 416 | 957 | 1143 | 212 | 432 | 2777 | 204 | 532 | 2263 | 158 | 313 |
| 4247 | QGN17344.1 | FIM1_4076 | CP015059. | 75 | 11 | 3 | 112 | 24 | 18 | 69 | 13 | 7 | 22 | 4 | 5 | 26 | 7 | 13 | 18 | 8 | 5 |
| 4248 | QGN17345.1 | FIM1_4077 | CP015059. | 109024 | 18015 | 15913 | 110396 | 18411 | 12077 | 109246 | 16676 | 19244 | 12382 | 20098 | 20965 | 66360 | 28758 | 20634 | 41886 | 24999 | 15628 |
| 4249 | QGN17346.1 | FIM1_4078 | CP015059. | 1553 | 33 | 91 | 951 | 74 | 72 | 1108 | 50 | 122 | 113 | 28 | 142 | 779 | 53 | 109 | 145 | 86 | 85 |
| 4250 | QGN17347.1 | FIM1_4079 | CP015059. | 133 | 2 | 1 | 88 | 1 | 3 | 187 | 5 | 5 | 12 | 6 | 4 | 11 | 6 | 1 | 10 | 9 | 5 |
| 4251 | QGN17348.1 | FIM1_4080 | CP015059. | 13522 | 476 | 1635 | 17799 | 602 | 1240 | 21108 | 1041 | 1986 | 8647 | 522 | 1764 | 16406 | 1646 | 1694 | 10598 | 897 | 1274 |
| 4252 | QGN17349.1 | FIM1_4081 | CP015059. | 119 | 16 | 12 | 130 | 23 | 12 | 81 | 12 | 18 | 34 | 10 | 13 | 61 | 19 | 14 | 38 | 14 | 8 |
| 4253 | QGN17350.1 | FIM1_4082 | CP015059. | 383 | 19 | 4 | 361 | 55 | 12 | 186 | 17 | 10 | 48 | 22 | 7 | 57 | 34 | 10 | 42 | 27 | 6 |
| 4254 | QGN17351.1 | FIM1_4083 | CP015059. | 1492 | 486 | 337 | 1672 | 487 | 358 | 1860 | 431 | 444 | 1069 | 447 | 400 | 1154 | 435 | 424 | 999 | 431 | 266 |
| 4255 | QGN17352.1 | FIM1_4084 | CP015059. | 3490 | 2779 | 1848 | 2689 | 2622 | 1824 | 2762 | 2337 | 2469 | 9638 | 2840 | 2803 | 9249 | 2484 | 2947 | 8858 | 2483 | 2022 |
| 4256 | QGN17353.1 | FIM1_4085 | CP015059. | 9162 | 5530 | 4760 | 10089 | 6829 | 4299 | 8878 | 6345 | 6268 | 18059 | 8045 | 8063 | 30841 | 7719 | 7741 | 21484 | 7716 | 5697 |
| 4257 | QGN17354.1 | FIM1_4086 | CP015059. | 12039 | 6900 | 4193 | 9608 | 8638 | 4104 | 12783 | 7250 | 5421 | 14528 | 11025 | 8019 | 30036 | 9973 | 7090 | 20240 | 11976 | 5559 |
| 4258 | QGN17355.1 | FIM1_4087 | CP015059. | 471 | 257 | 189 | 775 | 412 | 243 | 458 | 205 | 236 | 346 | 274 | 418 | 1024 | 175 | 341 | 723 | 200 | 276 |
| 4259 | QGN17356.1 | FIM1_4088 | CP015059. | 1965 | 1103 | 1161 | 3993 | 1507 | 886 | 2895 | 1353 | 1609 | 1859 | 1228 | 1586 | 7551 | 1399 | 1376 | 4126 | 1400 | 1060 |
| 4260 | QGN17357.1 | FIM1_4089 | CP015059. | 901 | 214 | 143 | 1234 | 140 | 96 | 911 | 141 | 157 | 822 | 87 | 152 | 626 | 110 | 156 | 693 | 87 | 91 |
| 4261 | QGN17358.1 | FIM1_4090 | CP015059. | 634 | 146 | 129 | 640 | 120 | 101 | 678 | 140 | 136 | 690 | 93 | 185 | 692 | 91 | 176 | 694 | 112 | 136 |
| 4262 | QGN17359.1 | FIM1_4091 | CP015059. | 3549 | 2169 | 851 | 1671 | 2044 | 972 | 3207 | 1867 | 1127 | 5773 | 2541 | 1730 | 6134 | 2589 | 1683 | 5617 | 3117 | 1248 |
| 4263 | QGN17360.1 | FIM1_4092 | CP015059. | 2021 | 1038 | 1062 | 2406 | 797 | 904 | 2122 | 710 | 1449 | 4596 | 711 | 1205 | 4052 | 756 | 1242 | 4323 | 632 | 812 |
| 4264 | QGN17361.1 | FIM1_4093 | CP015059. | 5741 | 454 | 784 | 3888 | 723 | 604 | 4173 | 828 | 1054 | 1076 | 502 | 1050 | 5215 | 503 | 993 | 2297 | 528 | 721 |
| 4265 | QGN17362.1 | FIM1_4094 | CP015059. | 29029 | 3185 | 1958 | 40152 | 4162 | 1944 | 36003 | 3364 | 2647 | 1855 | 2420 | 1205 | 7869 | 3370 | 1168 | 3521 | 3409 | 928 |
| 4266 | QGN17363.1 | FIM1_4095 | CP015059. | 3428 | 1755 | 1127 | 2908 | 1125 | 1049 | 2991 | 1033 | 1519 | 3806 | 1030 | 1201 | 3017 | 869 | 1438 | 3533 | 1056 | 901 |
| 4267 | QGN17364.1 | FIM1_4096 | CP015059. | 1124 | 1959 | 1387 | 1325 | 1198 | 1549 | 1151 | 1201 | 1846 | 4234 | 1196 | 1835 | 3194 | 1134 | 1861 | 3361 | 1289 | 1318 |
| 4268 | QGN17365.1 | FIM1_4097 | CP015059. | 2699 | 2528 | 614 | 752 | 1885 | 673 | 1761 | 1813 | 887 | 1719 | 1735 | 1151 | 3369 | 1585 | 1107 | 2176 | 1940 | 844 |
| 4269 | QGN17366.1 | FIM1_4098 | CP015059. | 1829 | 2140 | 1730 | 1173 | 1539 | 1351 | 1025 | 1978 | 2358 | 1619 | 1000 | 1900 | 3885 | 831 | 2100 | 2485 | 837 | 1236 |
| 4270 | QGN17367.1 | FIM1_4099 | CP015059. | 70 | 489 | 327 | 75 | 309 | 302 | 34 | 303 | 453 | 574 | 241 | 359 | 966 | 206 | 442 | 653 | 223 | 267 |
| 4271 | QGN17368.1 | FIM1_4100 | CP015059. | 759 | 257 | 195 | 948 | 216 | 168 | 756 | 267 | 269 | 903 | 130 | 280 | 991 | 195 | 261 | 935 | 166 | 150 |
| 4272 | QGN17369.1 | FIM1_4101 | CP015059. | 2729 | 373 | 513 | 2622 | 475 | 392 | 2530 | 497 | 634 | 936 | 326 | 759 | 2408 | 273 | 648 | 1335 | 291 | 495 |
| 4273 | QGN17370.1 | FIM1_4102 | CP015059. | 4558 | 775 | 1523 | 5359 | 1102 | 1194 | 4883 | 1020 | 1928 | 4951 | 828 | 2185 | 8146 | 863 | 1856 | 5570 | 780 | 1435 |
| 4274 | QGN17371.1 | FIM1_4103 | CP015059. | 967 | 1982 | 6635 | 2788 | 2158 | 5339 | 1576 | 2308 | 7922 | 5520 | 1524 | 10288 | 8977 | 1438 | 8916 | 6714 | 1513 | 6733 |
| 4275 | QGN17372.1 | FIM1_4104 | CP015059. | 5390 | 912 | 827 | 3357 | 862 | 687 | 5248 | 924 | 1040 | 786 | 795 | 1074 | 2360 | 936 | 1094 | 1276 | 998 | 873 |
| 4276 | QGN17373.1 | FIM1_4105 | CP015059. | 97 | 326 | 182 | 122 | 228 | 156 | 75 | 418 | 275 | 100 | 128 | 152 | 290 | 94 | 182 | 177 | 115 | 129 |
| 4277 | QGN17374.1 | FIM1_4106 | CP015059. | 14623 | 6869 | 3492 | 10886 | 8386 | 2553 | 9527 | 8353 | 5075 | 6440 | 8114 | 2800 | 21185 | 5871 | 2503 | 13093 | 6735 | 2015 |
| 4278 | QGN17375.1 | FIM1_4107 | CP015059. | 1662 | 772 | 682 | 1584 | 781 | 612 | 1500 | 791 | 916 | 4016 | 909 | 1061 | 3941 | 743 | 1014 | 3478 | 793 | 668 |
| 4279 | QGN17376.1 | FIM1_4108 | CP015059. | 2184 | 2389 | 1710 | 2391 | 2110 | 1532 | 2251 | 1869 | 2183 | 10956 | 2178 | 2627 | 9503 | 1904 | 2384 | 10168 | 1874 | 1707 |
| 4280 | QGN17377.1 | FIM1_4109 | CP015059. | 498 | 288 | 202 | 312 | 205 | 196 | 485 | 218 | 273 | 1319 | 206 | 263 | 956 | 152 | 305 | 1133 | 182 | 197 |
| 4281 | QGN17378.1 | FIM1_4110 | CP015059. | 1095 | 885 | 530 | 948 | 747 | 478 | 1039 | 868 | 765 | 1499 | 654 | 721 | 2621 | 631 | 730 | 1986 | 648 | 488 |
| 4282 | QGN17379.1 | FIM1_4111 | CP015059. | 3398 | 1376 | 1522 | 3532 | 1541 | 1294 | 3162 | 1521 | 1961 | 7279 | 1625 | 2248 | 7822 | 1511 | 2006 | 7116 | 1471 | 1457 |

| | | | | | | | | | | | | | | | | | | | | | |
|---|---|---|---|---|---|---|---|---|---|---|---|---|---|---|---|---|---|---|---|---|---|
| 4283 | QGN17380.1 | FIM1_4112 | CP015059. | 2210 | 1664 | 1447 | 4171 | 1014 | 912 | 2520 | 1482 | 1832 | 4569 | 1335 | 1806 | 7596 | 2269 | 2010 | 4992 | 1562 | 1285 |
| 4284 | QGN17381.1 | FIM1_4113 | CP015059. | 6918 | 953 | 516 | 10755 | 1170 | 481 | 5650 | 1730 | 823 | 1548 | 1465 | 1056 | 5702 | 951 | 1135 | 3227 | 1168 | 695 |
| 4285 | QGN17382.1 | FIM1_4114 | CP015059. | 4735 | 2003 | 540 | 1089 | 1703 | 465 | 3217 | 1714 | 760 | 911 | 1667 | 1092 | 2974 | 1498 | 1206 | 1536 | 2155 | 836 |
| 4286 | QGN17383.1 | FIM1_4115 | CP015059. | 1258 | 612 | 659 | 1387 | 747 | 470 | 1698 | 902 | 819 | 1902 | 526 | 938 | 3471 | 603 | 879 | 2099 | 618 | 616 |
| 4287 | QGN17384.1 | FIM1_4116 | CP015059. | 303 | 164 | 160 | 244 | 241 | 127 | 249 | 215 | 243 | 110 | 145 | 208 | 656 | 130 | 186 | 257 | 168 | 133 |
| 4288 | QGN17385.1 | FIM1_4117 | CP015059. | 493 | 159 | 119 | 593 | 150 | 90 | 429 | 122 | 168 | 51 | 99 | 127 | 345 | 81 | 121 | 119 | 98 | 94 |
| 4289 | QGN17386.1 | FIM1_4118 | CP015059. | 4723 | 1542 | 1277 | 5908 | 1159 | 1053 | 4733 | 1092 | 1682 | 1125 | 984 | 1636 | 4815 | 1322 | 1709 | 3109 | 1032 | 1165 |
| 4290 | QGN17387.1 | FIM1_4119 | CP015059. | 1940 | 1302 | 1113 | 2409 | 1484 | 864 | 1918 | 1337 | 1319 | 2185 | 1437 | 1328 | 4233 | 1325 | 1231 | 3230 | 1167 | 1078 |
| 4291 | QGN17388.1 | FIM1_4120 | CP015059. | 3127 | 2204 | 982 | 1257 | 2510 | 1165 | 2305 | 2308 | 1464 | 5025 | 2320 | 1642 | 7396 | 2268 | 1706 | 5064 | 2612 | 1148 |
| 4292 | QGN17389.1 | FIM1_4121 | CP015059. | 2296 | 527 | 424 | 1096 | 735 | 422 | 2132 | 514 | 574 | 2529 | 660 | 850 | 2652 | 786 | 727 | 2258 | 794 | 559 |
| 4293 | QGN17390.1 | FIM1_4122 | CP015059. | 660 | 441 | 485 | 1418 | 349 | 408 | 819 | 368 | 638 | 2112 | 280 | 645 | 2351 | 298 | 615 | 2103 | 265 | 421 |
| 4294 | QGN17391.1 | FIM1_4123 | CP015059. | 2450 | 704 | 662 | 2069 | 609 | 601 | 2105 | 689 | 994 | 1438 | 616 | 960 | 2850 | 459 | 983 | 1854 | 449 | 645 |
| 4295 | QGN17392.1 | FIM1_4124 | CP015059. | 52 | 55 | 43 | 151 | 29 | 42 | 71 | 38 | 46 | 181 | 25 | 69 | 135 | 36 | 52 | 145 | 34 | 27 |
| 4296 | QGN17393.1 | FIM1_4125 | CP015059. | 9104 | 6162 | 3502 | 11922 | 4624 | 3009 | 11486 | 3931 | 4731 | 8183 | 5239 | 6241 | 20197 | 5600 | 6180 | 13753 | 4722 | 4400 |
| 4297 | QGN17394.1 | FIM1_4126 | CP015059. | 1036 | 208 | 289 | 1285 | 270 | 186 | 1165 | 284 | 397 | 647 | 149 | 203 | 782 | 91 | 171 | 674 | 115 | 152 |
| 4298 | QGN17395.1 | FIM1_4127 | CP015059. | 169 | 22 | 46 | 311 | 32 | 31 | 159 | 34 | 54 | 58 | 32 | 40 | 147 | 16 | 40 | 99 | 14 | 29 |
| 4299 | QGN17396.1 | FIM1_4128 | CP015059. | 1340 | 1413 | 731 | 1051 | 1097 | 611 | 1337 | 1023 | 912 | 3189 | 952 | 954 | 3891 | 961 | 933 | 3290 | 958 | 714 |
| 4300 | QGN17397.1 | FIM1_4129 | CP015059. | 845 | 2643 | 1586 | 943 | 1920 | 1515 | 905 | 1719 | 2005 | 7947 | 1919 | 2372 | 6060 | 1749 | 2226 | 7279 | 2008 | 1536 |
| 4301 | QGN17398.1 | FIM1_4130 | CP015059. | 8261 | 1113 | 1031 | 6566 | 1042 | 884 | 7533 | 1013 | 1498 | 2656 | 893 | 1284 | 6559 | 751 | 1444 | 4058 | 810 | 1059 |
| 4302 | QGN17399.1 | FIM1_4131 | CP015059. | 19 | 8 | 6 | 70 | 14 | 7 | 14 | 8 | 11 | 13 | 3 | 13 | 18 | 3 | 4 | 17 | 6 | 6 |
| 4303 | QGN17400.1 | FIM1_4132 | CP015059. | 225 | 107 | 159 | 294 | 133 | 134 | 275 | 131 | 215 | 598 | 64 | 171 | 782 | 89 | 180 | 611 | 70 | 162 |
| 4304 | QGN17401.1 | FIM1_4133 | CP015059. | 2420 | 605 | 667 | 3062 | 425 | 570 | 2520 | 547 | 916 | 2746 | 305 | 881 | 2274 | 359 | 848 | 2468 | 268 | 581 |
| 4305 | QGN17402.1 | FIM1_4134 | CP015059. | 2272 | 1752 | 1369 | 2883 | 1343 | 1151 | 2542 | 1217 | 1732 | 6359 | 1008 | 1557 | 5092 | 978 | 1570 | 6275 | 829 | 1068 |
| 4306 | QGN17403.1 | FIM1_4135 | CP015059. | 4163 | 600 | 567 | 3552 | 668 | 444 | 4203 | 760 | 747 | 1904 | 521 | 790 | 3521 | 566 | 722 | 2351 | 501 | 536 |
| 4307 | QGN17404.1 | FIM1_4136 | CP015059. | 4041 | 2272 | 1532 | 5056 | 1603 | 1371 | 4156 | 1511 | 2038 | 5525 | 1249 | 2017 | 5551 | 1338 | 2191 | 5405 | 1218 | 1444 |
| 4308 | QGN17405.1 | FIM1_4137 | CP015059. | 7889 | 1093 | 1634 | 6769 | 1908 | 1386 | 7654 | 1789 | 2198 | 4679 | 1818 | 2674 | 11207 | 1823 | 2298 | 7147 | 1742 | 1814 |
| 4309 | QGN17406.1 | FIM1_4138 | CP015059. | 2321 | 1285 | 744 | 2383 | 876 | 665 | 2437 | 878 | 977 | 3965 | 811 | 1123 | 4211 | 833 | 1046 | 4253 | 819 | 708 |
| 4310 | QGN17407.1 | FIM1_4139 | CP015059. | 4237 | 1932 | 1208 | 2168 | 2230 | 1311 | 3940 | 1904 | 1642 | 11465 | 3275 | 2500 | 10632 | 2696 | 2235 | 10878 | 3531 | 1767 |
| 4311 | QGN17408.1 | FIM1_4140 | CP015059. | 303 | 634 | 668 | 1039 | 774 | 559 | 498 | 711 | 870 | 2907 | 773 | 1037 | 5276 | 632 | 965 | 4163 | 694 | 717 |
| 4312 | QGN17409.1 | FIM1_4141 | CP015059. | 3179 | 7526 | 5007 | 6390 | 5810 | 4576 | 4255 | 4939 | 6150 | 24314 | 7627 | 8896 | 22593 | 7206 | 8637 | 24758 | 6952 | 6244 |
| 4313 | QGN17410.1 | FIM1_4142 | CP015059. | 1088 | 592 | 505 | 1981 | 545 | 423 | 1369 | 504 | 655 | 2457 | 472 | 693 | 2943 | 440 | 662 | 2478 | 426 | 439 |
| 4314 | QGN17411.1 | FIM1_4143 | CP015059. | 1408 | 741 | 580 | 1350 | 585 | 514 | 1356 | 664 | 707 | 1974 | 507 | 735 | 2373 | 471 | 729 | 1992 | 488 | 530 |
| 4315 | QGN17412.1 | FIM1_4144 | CP015059. | 356 | 374 | 309 | 319 | 244 | 293 | 320 | 235 | 432 | 2017 | 241 | 413 | 1320 | 228 | 419 | 1603 | 215 | 261 |
| 4316 | QGN17413.1 | FIM1_4145 | CP015059. | 1363 | 411 | 376 | 1173 | 233 | 348 | 886 | 293 | 445 | 2186 | 187 | 455 | 1897 | 187 | 552 | 1848 | 163 | 306 |
| 4317 | QGN17414.1 | FIM1_4146 | CP015059. | 1678 | 1470 | 625 | 489 | 1326 | 689 | 1347 | 1195 | 844 | 3346 | 1384 | 1155 | 3878 | 1281 | 1042 | 3523 | 1705 | 795 |
| 4318 | QGN17415.1 | FIM1_4147 | CP015059. | 416 | 715 | 680 | 1219 | 529 | 587 | 809 | 563 | 902 | 1669 | 445 | 791 | 2146 | 510 | 801 | 1796 | 437 | 549 |
| 4319 | QGN17416.1 | FIM1_4148 | CP015059. | 13502 | 1193 | 1834 | 12238 | 1744 | 1335 | 11943 | 2039 | 2491 | 1920 | 1348 | 2429 | 9042 | 1464 | 2241 | 3841 | 1449 | 1679 |
| 4320 | QGN17417.1 | FIM1_4149 | CP015059. | 6689 | 8325 | 8772 | 11563 | 8087 | 7682 | 9078 | 7131 | 10266 | 31431 | 9677 | 13873 | 29650 | 9462 | 14573 | 31497 | 8313 | 10591 |
| 4321 | QGN17418.1 | FIM1_4150 | CP015059. | 3606 | 1831 | 1442 | 2742 | 1369 | 1212 | 2736 | 1292 | 1734 | 3055 | 1533 | 1874 | 5245 | 1487 | 2230 | 4120 | 1364 | 1392 |
| 4322 | QGN17419.1 | FIM1_4151 | CP015059. | 6405 | 1341 | 1820 | 10879 | 1236 | 1532 | 8381 | 977 | 2421 | 5806 | 1386 | 2613 | 9086 | 1629 | 2466 | 7080 | 1273 | 1919 |
| 4323 | QGN17420.1 | FIM1_4152 | CP015059. | 2516 | 1657 | 526 | 1030 | 1498 | 589 | 2202 | 1404 | 738 | 5000 | 2006 | 1200 | 4706 | 1968 | 1190 | 4544 | 2104 | 939 |
| 4324 | QGN17421.1 | FIM1_4153 | CP015059. | 3424 | 713 | 1050 | 3135 | 956 | 838 | 3398 | 1006 | 1389 | 3791 | 779 | 1578 | 7311 | 813 | 1323 | 4761 | 739 | 1076 |
| 4325 | QGN17422.1 | FIM1_4154 | CP015059. | 440 | 600 | 819 | 1170 | 505 | 600 | 673 | 628 | 1034 | 3507 | 394 | 777 | 3714 | 477 | 740 | 3302 | 242 | 502 |
| 4326 | QGN17423.1 | FIM1_4155 | CP015059. | 627 | 3714 | 2277 | 1726 | 2976 | 2272 | 1175 | 2552 | 2849 | 11596 | 3431 | 3604 | 8732 | 3007 | 3551 | 10714 | 3349 | 2518 |
| 4327 | QGN17424.1 | FIM1_4156 | CP015059. | 4408 | 3502 | 1315 | 2063 | 2830 | 1488 | 3376 | 2710 | 1817 | 6132 | 3343 | 2447 | 8057 | 2781 | 2305 | 6488 | 3401 | 1760 |
| 4328 | QGN17425.1 | FIM1_4157 | CP015059. | 9057 | 2072 | 1326 | 6740 | 1978 | 1139 | 8647 | 1874 | 1730 | 2952 | 1822 | 2299 | 7725 | 2072 | 2190 | 4798 | 2025 | 1586 |
| 4329 | QGN17426.1 | FIM1_4158 | CP015059. | 4993 | 3333 | 2980 | 6676 | 4471 | 2387 | 5738 | 4312 | 3684 | 4688 | 3022 | 3719 | 13321 | 2921 | 3238 | 7432 | 2720 | 2357 |
| 4330 | QGN17427.1 | FIM1_4159 | CP015059. | 2516 | 1368 | 999 | 2306 | 1011 | 799 | 2207 | 1008 | 1254 | 2417 | 684 | 1059 | 3273 | 637 | 1142 | 2427 | 592 | 708 |
| 4331 | QGN17428.1 | FIM1_4160 | CP015059. | 509 | 370 | 279 | 510 | 197 | 245 | 488 | 259 | 388 | 920 | 184 | 306 | 980 | 170 | 355 | 835 | 199 | 222 |
| 4332 | QGN17429.1 | FIM1_4161 | CP015059. | 22475 | 14417 | 9192 | 26277 | 10289 | 7810 | 21182 | 10322 | 11346 | 23728 | 10724 | 14899 | 30749 | 11451 | 14483 | 24966 | 11029 | 10532 |
| 4333 | QGN17430.1 | FIM1_4162 | CP015059. | 2305 | 1113 | 1162 | 1026 | 169 | 566 | 2196 | 737 | 1456 | 3977 | 820 | 1755 | 3317 | 908 | 1992 | 3580 | 867 | 1110 |
| 4334 | QGN17431.1 | FIM1_4163 | CP015059. | 14026 | 7300 | 5982 | 12815 | 6355 | 5492 | 10821 | 5453 | 7591 | 9027 | 4492 | 5880 | 13640 | 3780 | 6340 | 11681 | 4182 | 4608 |
| 4335 | QGN17432.1 | FIM1_4164 | CP015059. | 1373 | 45 | 60 | 335 | 20 | 32 | 416 | 37 | 64 | 173 | 43 | 86 | 205 | 40 | 87 | 178 | 36 | 55 |
| 4336 | QGN17433.1 | FIM1_4165 | CP015059. | 617 | 400 | 353 | 666 | 329 | 296 | 441 | 286 | 474 | 1063 | 288 | 348 | 947 | 193 | 380 | 1320 | 168 | 258 |
| 4337 | QGN17434.1 | FIM1_4166 | CP015059. | 16542 | 358 | 773 | 12804 | 474 | 570 | 14903 | 624 | 1060 | 894 | 400 | 994 | 2433 | 588 | 938 | 1307 | 588 | 715 |
| 4338 | QGN17435.1 | FIM1_4167 | CP015059. | 2014 | 732 | 742 | 2011 | 748 | 586 | 1600 | 1020 | 1102 | 467 | 467 | 1070 | 3104 | 465 | 877 | 1040 | 487 | 690 |
| 4339 | QGN17436.1 | FIM1_4169 | CP015059. | 783 | 365 | 433 | 1015 | 400 | 313 | 846 | 385 | 543 | 1907 | 369 | 691 | 2482 | 396 | 586 | 2056 | 343 | 402 |
| 4340 | QGN17437.1 | FIM1_4170 | CP015059. | 289 | 539 | 611 | 544 | 533 | 524 | 255 | 577 | 778 | 3108 | 375 | 1081 | 3280 | 422 | 1099 | 2842 | 294 | 733 |
| 4341 | QGN17438.1 | FIM1_4171 | CP015059. | 149 | 49 | 74 | 95 | 24 | 51 | 122 | 57 | 122 | 158 | 29 | 479 | 230 | 22 | 524 | 168 | 18 | 271 |
| 4342 | QGN17439.1 | FIM1_4172 | CP015059. | 3377 | 687 | 733 | 5512 | 603 | 582 | 3642 | 682 | 989 | 684 | 394 | 802 | 2416 | 532 | 866 | 1265 | 366 | 564 |
| 4343 | QGN17440.1 | FIM1_4173 | CP015059. | 3607 | 1024 | 837 | 3357 | 875 | 643 | 2740 | 824 | 1148 | 1944 | 589 | 870 | 2802 | 533 | 981 | 2072 | 466 | 610 |
| 4344 | QGN17441.1 | FIM1_4174 | CP015059. | 7124 | 3074 | 1702 | 4586 | 4011 | 1918 | 7541 | 3025 | 2235 | 11950 | 5867 | 4185 | 18251 | 4096 | 3790 | 13924 | 6378 | 3073 |
| 4345 | QGN17442.1 | FIM1_4175 | CP015059. | 1367 | 1140 | 1023 | 2093 | 869 | 927 | 1712 | 749 | 1480 | 7016 | 901 | 1728 | 5681 | 877 | 1698 | 6854 | 820 | 1029 |
| 4346 | QGN17443.1 | FIM1_4176 | CP015059. | 1087 | 793 | 1013 | 1538 | 823 | 810 | 953 | 891 | 1283 | 5172 | 736 | 1415 | 5056 | 679 | 1396 | 4606 | 606 | 942 |
| 4347 | QGN17444.1 | FIM1_4177 | CP015059. | 5682 | 6375 | 5815 | 4224 | 4487 | 5236 | 4849 | 4170 | 7335 | 16189 | 4988 | 5876 | 13305 | 3997 | 6381 | 16254 | 4316 | 4258 |
| 4348 | QGN17445.1 | FIM1_4178 | CP015059. | 4520 | 5201 | 5556 | 3687 | 4301 | 4543 | 3518 | 4184 | 7084 | 15223 | 4424 | 5526 | 13938 | 3544 | 5401 | 16299 | 3857 | 3727 |
| 4349 | QGN17446.1 | FIM1_4180 | CP015059. | 22 | 28 | 17 | 52 | 20 | 13 | 39 | 22 | 22 | 101 | 14 | 16 | 77 | 9 | 13 | 124 | 17 | 14 |
| 4350 | QGN17447.1 | FIM1_4181 | CP015059. | 15617 | 1030 | 1689 | 20279 | 1346 | 1457 | 19389 | 1129 | 2062 | 6324 | 1345 | 2629 | 8453 | 1648 | 2370 | 6871 | 1496 | 1850 |
| 4351 | QGN17448.1 | FIM1_4182 | CP015059. | 7800 | 6456 | 4278 | 8191 | 6308 | 4167 | 7117 | 4905 | 5531 | 27012 | 8007 | 8856 | 26861 | 7188 | 8424 | 26450 | 7135 | 5873 |
| 4352 | QGN17449.1 | FIM1_4183 | CP015059. | 1873 | 1116 | 921 | 2063 | 984 | 908 | 1714 | 883 | 1410 | 5854 | 947 | 1348 | 5339 | 873 | 1369 | 5781 | 823 | 983 |
| 4353 | QGN17450.1 | FIM1_4184 | CP015059. | 1379 | 1284 | 1229 | 1753 | 1283 | 1120 | 1385 | 1287 | 1670 | 5975 | 1250 | 1880 | 8712 | 1299 | 1624 | 7342 | 1212 | 1293 |
| 4354 | QGN17451.1 | FIM1_4185 | CP015059. | 839 | 477 | 622 | 1168 | 524 | 483 | 871 | 560 | 778 | 3255 | 453 | 695 | 2513 | 377 | 623 | 3236 | 354 | 430 |
| 4355 | QGN17452.1 | FIM1_4186 | CP015059. | 1868 | 1108 | 1848 | 2375 | 1152 | 1609 | 1649 | 1398 | 2478 | 3184 | 1395 | 3361 | 8708 | 744 | 3529 | 5373 | 985 | 2177 |
| 4356 | QGN17453.1 | FIM1_4187 | CP015059. | 1117 | 482 | 438 | 1419 | 431 | 416 | 1443 | 570 | 643 | 1281 | 335 | 448 | 1694 | 249 | 429 | 1168 | 367 | 327 |
| 4357 | QGN17454.1 | FIM1_4188 | CP015059. | 1471 | 936 | 668 | 1380 | 1117 | 528 | 1537 | 1127 | 821 | 2016 | 969 | 1076 | 4197 | 917 | 941 | 2654 | 905 | 720 |
| 4358 | QGN17455.1 | FIM1_4189 | CP015059. | 6796 | 1458 | 2060 | 6249 | 2033 | 1675 | 4937 | 1634 | 2453 | 6606 | 1893 | 3736 | 12117 | 1794 | 3195 | 9093 | 1616 | 2389 |
| 4359 | QGN17456.1 | FIM1_4190 | CP015059. | 1412 | 1099 | 1606 | 2336 | 1175 | 1335 | 1515 | 948 | 2115 | 8611 | 1076 | 2254 | 6850 | 1034 | 2036 | 7985 | 920 | 1431 |
| 4360 | QGN17457.1 | FIM1_4191 | CP015059. | 2268 | 1035 | 1212 | 4435 | 1339 | 1166 | 2944 | 1154 | 1590 | 5521 | 949 | 2087 | 6914 | 1164 | 1865 | 5623 | 1055 | 1368 |
| 4361 | QGN17458.1 | FIM1_4192 | CP015059. | 859 | 834 | 411 | 534 | 670 | 443 | 875 | 583 | 591 | 2840 | 759 | 632 | 2399 | 666 | 659 | 3076 | 803 | 433 |
| 4362 | QGN17459.1 | FIM1_4193 | CP015059. | 166 | 216 | 170 | 313 | 188 | 163 | 167 | 98 | 231 | 1261 | 135 | 274 | 1006 | 157 | 267 | 1342 | 135 | 190 |
| 4363 | QGN17460.1 | FIM1_4194 | CP015059. | 1424 | 915 | 1045 | 1633 | 840 | 793 | 1073 | 940 | 1298 | 2214 | 578 | 1089 | 4757 | 573 | 1074 | 3363 | 468 | 737 |
| 4364 | QGN17461.1 | FIM1_4195 | CP015059. | 4376 | 2067 | 1336 | 4233 | 1328 | 1140 | 4041 | 1363 | 1946 | 4811 | 1297 | 2044 | 5839 | 1384 | 1983 | 5052 | 1249 | 1252 |
| 4365 | QGN17462.1 | FIM1_4196 | CP015059. | 17035 | 1064 | 2013 | 15690 | 1531 | 1544 | 15327 | 1574 | 2663 | 5047 | 1293 | 3135 | 10759 | 1433 | 2775 | 6915 | 1253 | 2157 |
| 4366 | QGN17463.1 | FIM1_4197 | CP015059. | 838 | 996 | 1922 | 1654 | 683 | 1104 | 730 | 994 | 1979 | 3647 | 555 | 1998 | 5213 | 494 | 1450 | 3784 | 412 | 1087 |

| | | | | | | | | | | | | | | | | | | | | | |
|---|---|---|---|---|---|---|---|---|---|---|---|---|---|---|---|---|---|---|---|---|---|
| 4367 | QGN17464.1 | FIM1_4198 | CP015059. | 3965 | 841 | 1362 | 4825 | 961 | 1002 | 3589 | 1324 | 1751 | 2287 | 724 | 1393 | 6134 | 538 | 1350 | 3607 | 589 | 914 |
| 4368 | QGN17465.1 | FIM1_4199 | CP015059. | 1564 | 175 | 202 | 1286 | 179 | 149 | 1369 | 228 | 239 | 359 | 167 | 232 | 778 | 152 | 241 | 596 | 153 | 176 |
| 4369 | QGN17466.1 | FIM1_4201 | CP015059. | 1142 | 338 | 398 | 1229 | 347 | 366 | 1061 | 319 | 550 | 1603 | 225 | 541 | 1813 | 190 | 442 | 1675 | 205 | 352 |
| 4370 | QGN17467.1 | FIM1_4203 | CP015059. | 219 | 146 | 185 | 671 | 197 | 138 | 410 | 199 | 234 | 1950 | 206 | 332 | 1374 | 193 | 277 | 1973 | 162 | 187 |
| 4371 | QGN17468.1 | FIM1_4204 | CP015059. | 4068 | 578 | 799 | 3568 | 526 | 637 | 3515 | 685 | 1067 | 4817 | 457 | 1082 | 3962 | 352 | 857 | 4770 | 351 | 686 |
| 4372 | QGN17469.1 | FIM1_4205 | CP015059. | 25423 | 331 | 858 | 20579 | 806 | 663 | 22748 | 1019 | 1173 | 1173 | 739 | 1017 | 4818 | 1466 | 885 | 1848 | 1255 | 667 |
| 4373 | QGN17470.1 | FIM1_4206 | CP015059. | 1887 | 1257 | 831 | 2420 | 1093 | 609 | 2176 | 1008 | 1105 | 686 | 968 | 1047 | 2699 | 1282 | 1049 | 1287 | 1290 | 771 |
| 4374 | QGN17471.1 | FIM1_4207 | CP015059. | 1613 | 1505 | 1786 | 4008 | 1565 | 1453 | 2247 | 1416 | 2237 | 4431 | 1207 | 2330 | 7623 | 1105 | 2166 | 5352 | 1110 | 1636 |
| 4375 | QGN17472.1 | FIM1_4208 | CP015059. | 925 | 418 | 723 | 2109 | 525 | 506 | 1365 | 623 | 890 | 2848 | 394 | 1002 | 4215 | 420 | 834 | 3052 | 330 | 622 |
| 4376 | QGN17473.1 | FIM1_4209 | CP015059. | 1025 | 291 | 463 | 1765 | 335 | 380 | 1170 | 383 | 595 | 2396 | 334 | 704 | 2237 | 302 | 725 | 2191 | 250 | 469 |
| 4377 | QGN17474.1 | FIM1_4210 | CP015059. | 3943 | 1270 | 1233 | 3532 | 1431 | 1053 | 3399 | 1293 | 1686 | 2548 | 1240 | 1711 | 5950 | 1196 | 1587 | 3760 | 1236 | 1229 |
| 4378 | QGN17475.1 | FIM1_4211 | CP015059. | 780 | 923 | 392 | 445 | 673 | 415 | 620 | 678 | 585 | 1505 | 510 | 542 | 1713 | 435 | 586 | 1409 | 479 | 362 |
| 4379 | QGN17476.1 | FIM1_4212 | CP015059. | 5859 | 1884 | 1575 | 7382 | 1912 | 1374 | 5236 | 1647 | 2011 | 7583 | 2048 | 2690 | 11260 | 1993 | 2589 | 8551 | 1689 | 1868 |
| 4380 | QGN17477.1 | FIM1_4213 | CP015059. | 905 | 279 | 236 | 1729 | 246 | 246 | 947 | 203 | 330 | 728 | 190 | 366 | 1056 | 210 | 381 | 828 | 170 | 263 |
| 4381 | QGN17478.1 | FIM1_4214 | CP015059. | 2485 | 971 | 714 | 3043 | 583 | 526 | 2417 | 697 | 885 | 2105 | 633 | 988 | 2680 | 679 | 967 | 2115 | 629 | 705 |
| 4382 | QGN17479.1 | FIM1_4215 | CP015059. | 13758 | 2045 | 2809 | 24471 | 2507 | 2274 | 17053 | 2351 | 3603 | 4719 | 2635 | 3724 | 15044 | 4475 | 3932 | 8739 | 2754 | 2702 |
| 4383 | QGN17480.1 | FIM1_4216 | CP015059. | 1864 | 474 | 570 | 2222 | 356 | 338 | 1513 | 419 | 691 | 1646 | 296 | 696 | 2208 | 464 | 672 | 1791 | 331 | 465 |
| 4384 | QGN17481.1 | FIM1_4217 | CP015059. | 2553 | 496 | 696 | 3150 | 376 | 495 | 2340 | 448 | 883 | 1799 | 264 | 803 | 2541 | 466 | 969 | 1916 | 224 | 542 |
| 4385 | QGN17482.1 | FIM1_4218 | CP015059. | 110 | 42 | 27 | 105 | 21 | 11 | 88 | 26 | 39 | 71 | 22 | 43 | 104 | 17 | 48 | 72 | 18 | 32 |
| 4386 | QGN17483.1 | FIM1_4219 | CP015059. | 927 | 399 | 281 | 831 | 326 | 278 | 910 | 336 | 400 | 865 | 306 | 373 | 1131 | 262 | 447 | 852 | 313 | 263 |
| 4387 | QGN17484.1 | FIM1_4220 | CP015059. | 1111 | 743 | 486 | 777 | 597 | 407 | 1092 | 602 | 635 | 1121 | 517 | 662 | 1851 | 461 | 746 | 1399 | 468 | 453 |
| 4388 | QGN17485.1 | FIM1_4221 | CP015059. | 116 | 247 | 176 | 120 | 124 | 182 | 118 | 151 | 274 | 847 | 125 | 325 | 611 | 108 | 302 | 778 | 96 | 156 |
| 4389 | QGN17486.1 | FIM1_4222 | CP015059. | 690 | 1982 | 935 | 1774 | 1213 | 851 | 999 | 1168 | 1396 | 1175 | 1002 | 1359 | 4194 | 1115 | 1445 | 2592 | 1006 | 1002 |
| 4390 | QGN17487.1 | FIM1_4223 | CP015059. | 1399 | 2502 | 1665 | 2645 | 1821 | 1584 | 1843 | 1812 | 2279 | 4298 | 1901 | 2429 | 6632 | 1913 | 2784 | 4998 | 1723 | 1708 |
| 4391 | QGN17488.1 | FIM1_4224 | CP015059. | 1933 | 4955 | 3816 | 4477 | 3934 | 3710 | 2523 | 3184 | 4929 | 12289 | 3866 | 5184 | 9163 | 3438 | 5250 | 11197 | 3693 | 3519 |
| 4392 | QGN17489.1 | FIM1_4225 | CP015059. | 537 | 71 | 328 | 1487 | 317 | 208 | 919 | 409 | 371 | 437 | 233 | 450 | 1215 | 279 | 425 | 511 | 289 | 344 |
| 4393 | QGN17490.1 | FIM1_4226 | CP015059. | 149800 | 11467 | 15068 | 165078 | 14309 | 12922 | 153690 | 11421 | 18903 | 37885 | 12526 | 15898 | 56723 | 31762 | 14932 | 41379 | 22245 | 11094 |
| 4394 | QGN17491.1 | FIM1_4227 | CP015059. | 13069 | 1378 | 790 | 11445 | 937 | 618 | 5912 | 1510 | 1324 | 1143 | 934 | 1058 | 5270 | 372 | 1075 | 1700 | 686 | 684 |
| 4395 | QGN17492.1 | FIM1_4228 | CP015059. | 175 | 8 | 6 | 121 | 13 | 14 | 116 | 11 | 15 | 17 | 17 | 13 | 47 | 16 | 13 | 20 | 9 | 9 |
| 4396 | QGN17493.1 | FIM1_4229 | CP015059. | 18288 | 330 | 267 | 10352 | 224 | 173 | 8364 | 353 | 288 | 402 | 237 | 383 | 1226 | 217 | 389 | 677 | 240 | 264 |
| 4397 | QGN17494.1 | FIM1_4230 | CP015059. | 3478 | 2759 | 1327 | 2071 | 2280 | 1101 | 3116 | 2501 | 1679 | 1498 | 1600 | 1641 | 4168 | 1400 | 1730 | 2347 | 1686 | 1190 |
| 4398 | QGN17495.1 | FIM1_4231 | CP015059. | 8159 | 4332 | 2815 | 7234 | 5045 | 2555 | 8409 | 4534 | 3618 | 6019 | 4272 | 4125 | 14303 | 3796 | 3976 | 8455 | 4083 | 2867 |
| 4399 | QGN17496.1 | FIM1_4232 | CP015059. | 942 | 326 | 402 | 1302 | 365 | 353 | 750 | 341 | 594 | 644 | 229 | 444 | 1514 | 216 | 377 | 976 | 235 | 333 |
| 4400 | QGN17497.1 | FIM1_4233 | CP015059. | 3330 | 218 | 466 | 3015 | 281 | 358 | 2590 | 331 | 553 | 1827 | 298 | 511 | 2279 | 295 | 474 | 1865 | 323 | 407 |
| 4401 | QGN17498.1 | FIM1_4234 | CP015059. | 678 | 1074 | 674 | 712 | 828 | 708 | 577 | 718 | 963 | 5026 | 908 | 1018 | 3676 | 744 | 952 | 4825 | 859 | 643 |
| 4402 | QGN17499.1 | FIM1_4235 | CP015059. | 1476 | 512 | 626 | 2038 | 715 | 589 | 1504 | 504 | 795 | 3471 | 675 | 993 | 3235 | 646 | 1034 | 3367 | 580 | 683 |
| 4403 | QGN17500.1 | FIM1_4236 | CP015059. | 4271 | 1363 | 1758 | 5005 | 1762 | 1448 | 5353 | 1557 | 2398 | 5847 | 1323 | 2523 | 9221 | 1318 | 2239 | 7217 | 1213 | 1700 |
| 4404 | QGN17501.1 | FIM1_4237 | CP015059. | 8341 | 3586 | 4139 | 9449 | 5466 | 3516 | 8464 | 4342 | 5147 | 28867 | 6278 | 7734 | 30375 | 5976 | 6975 | 29155 | 5419 | 5209 |
| 4405 | QGN17502.1 | FIM1_4238 | CP015059. | 1129 | 2619 | 1809 | 1299 | 2208 | 1634 | 1212 | 1870 | 2235 | 11864 | 2329 | 2603 | 9574 | 2190 | 2413 | 10727 | 2280 | 1672 |
| 4406 | QGN17503.1 | FIM1_4239 | CP015059. | 2195 | 2622 | 1349 | 1951 | 2030 | 1250 | 2164 | 1948 | 1746 | 6830 | 1819 | 1774 | 8043 | 1794 | 1846 | 6868 | 1787 | 1282 |
| 4407 | QGN17504.1 | FIM1_4240 | CP015059. | 2348 | 1179 | 536 | 1241 | 1109 | 638 | 1894 | 847 | 767 | 3278 | 1178 | 1027 | 3021 | 1205 | 951 | 3146 | 1299 | 698 |
| 4408 | QGN17505.1 | FIM1_4241 | CP015059. | 4158 | 1369 | 1468 | 4335 | 1658 | 1093 | 3930 | 2088 | 1885 | 1647 | 1106 | 1712 | 6599 | 1173 | 1617 | 2810 | 1200 | 1190 |
| 4409 | QGN17506.1 | FIM1_4242 | CP015059. | 1252 | 174 | 118 | 1142 | 64 | 108 | 1001 | 117 | 197 | 703 | 65 | 146 | 738 | 51 | 174 | 606 | 52 | 108 |
| 4410 | QGN17507.1 | FIM1_4243 | CP015059. | 876 | 620 | 498 | 652 | 534 | 431 | 733 | 575 | 628 | 1715 | 385 | 631 | 1922 | 396 | 623 | 1759 | 408 | 377 |
| 4411 | QGN17508.1 | FIM1_4244 | CP015059. | 964 | 393 | 656 | 2072 | 261 | 488 | 1274 | 427 | 865 | 1145 | 211 | 735 | 1924 | 227 | 788 | 1325 | 167 | 555 |
| 4412 | QGN17509.1 | FIM1_4245 | CP015059. | 226 | 10 | 15 | 303 | 21 | 21 | 345 | 28 | 29 | 78 | 20 | 20 | 88 | 4 | 21 | 87 | 10 | 17 |
| 4413 | QGN17510.1 | FIM1_4246 | CP015059. | 1279 | 441 | 392 | 1254 | 360 | 302 | 1135 | 356 | 515 | 1606 | 264 | 550 | 2365 | 265 | 526 | 1787 | 256 | 370 |
| 4414 | QGN17511.1 | FIM1_4247 | CP015059. | 1197 | 141 | 237 | 1210 | 352 | 170 | 1120 | 334 | 273 | 1146 | 281 | 380 | 1969 | 295 | 330 | 1476 | 321 | 240 |
| 4415 | QGN17512.1 | FIM1_4248 | CP015059. | 1308 | 1125 | 1021 | 1770 | 1136 | 945 | 1521 | 991 | 1317 | 5068 | 758 | 1216 | 4352 | 680 | 1087 | 4807 | 656 | 746 |
| 4416 | QGN17513.1 | FIM1_4249 | CP015059. | 8542 | 3297 | 3015 | 13653 | 3843 | 2403 | 11582 | 3482 | 3775 | 11503 | 3132 | 4245 | 18077 | 4058 | 3683 | 13746 | 3197 | 2805 |
| 4417 | QGN17514.1 | FIM1_4250 | CP015059. | 1621 | 889 | 615 | 1919 | 828 | 568 | 1887 | 714 | 840 | 2756 | 766 | 748 | 2831 | 764 | 775 | 2729 | 693 | 514 |
| 4418 | QGN17515.1 | FIM1_4251 | CP015059. | 1830 | 2552 | 2405 | 2336 | 2017 | 2109 | 1901 | 1830 | 3198 | 10790 | 2007 | 3160 | 10567 | 2206 | 2931 | 10212 | 1939 | 2089 |
| 4419 | QGN17516.1 | FIM1_4252 | CP015059. | 1487 | 815 | 715 | 1965 | 683 | 654 | 1241 | 558 | 973 | 4248 | 700 | 1098 | 3795 | 647 | 1170 | 3656 | 600 | 783 |
| 4420 | QGN17517.1 | FIM1_4253 | CP015059. | 1351 | 817 | 1316 | 2797 | 1236 | 1269 | 1679 | 857 | 1809 | 10280 | 1401 | 2314 | 12915 | 1441 | 1905 | 11361 | 1349 | 1644 |
| 4421 | QGN17518.1 | FIM1_4254 | CP015059. | 2645 | 567 | 561 | 3666 | 614 | 464 | 2982 | 523 | 767 | 1865 | 590 | 745 | 2815 | 587 | 692 | 2198 | 581 | 545 |
| 4422 | QGN17519.1 | FIM1_4255 | CP015059. | 653 | 1602 | 1093 | 1888 | 1410 | 1121 | 1132 | 1076 | 1495 | 6114 | 1197 | 1647 | 5325 | 1023 | 1507 | 5710 | 1073 | 1105 |
| 4423 | QGN17520.1 | FIM1_4256 | CP015059. | 5191 | 5 | 5 | 3498 | 4 | 2 | 4397 | 4 | 2 | 349 | 7 | 7 | 287 | 7 | 1 | 175 | 4 | 5 |
| 4424 | QGN17521.1 | FIM1_4257 | CP015059. | 30952 | 1001 | 1491 | 22033 | 1198 | 1134 | 23234 | 1265 | 2009 | 2503 | 799 | 1855 | 8556 | 1061 | 1707 | 3848 | 844 | 1397 |
| 4425 | QGN17522.1 | FIM1_4258 | CP015059. | 133 | 23 | 16 | 106 | 34 | 22 | 78 | 16 | 26 | 129 | 27 | 35 | 112 | 17 | 26 | 133 | 22 | 33 |
| 4426 | QGN17523.1 | FIM1_4259 | CP015059. | 3194 | 1293 | 2071 | 3632 | 1570 | 1674 | 2824 | 1467 | 2663 | 2703 | 1392 | 3091 | 8438 | 1423 | 2660 | 4761 | 1296 | 2032 |
| 4427 | QGN17524.1 | FIM1_4260 | CP015059. | 829 | 573 | 534 | 1475 | 449 | 413 | 1248 | 507 | 680 | 699 | 326 | 738 | 2410 | 363 | 725 | 1185 | 356 | 573 |
| 4428 | QGN17525.1 | FIM1_4261 | CP015059. | 5058 | 368 | 302 | 4367 | 338 | 275 | 5807 | 242 | 449 | 654 | 334 | 392 | 1312 | 593 | 382 | 899 | 418 | 278 |
| 4429 | QGN17526.1 | FIM1_4262 | CP015059. | 3590 | 2179 | 1568 | 4101 | 2225 | 1389 | 4051 | 1894 | 2000 | 3258 | 2040 | 2075 | 7830 | 2147 | 2063 | 5912 | 2094 | 1512 |
| 4430 | QGN17527.1 | FIM1_4263 | CP015059. | 1087 | 462 | 521 | 1190 | 385 | 412 | 910 | 463 | 687 | 1472 | 214 | 550 | 1984 | 205 | 553 | 1511 | 212 | 385 |
| 4431 | QGN17528.1 | FIM1_4264 | CP015059. | 0 | 0 | 0 | 0 | 0 | 0 | 0 | 0 | 0 | 2 | 0 | 0 | 0 | 0 | 0 | 0 | 0 | 0 |
| 4432 | QGN17529.1 | FIM1_4265 | CP015059. | 419 | 531 | 542 | 531 | 611 | 478 | 534 | 561 | 726 | 4620 | 557 | 818 | 3868 | 575 | 635 | 4516 | 527 | 528 |
| 4433 | QGN17530.1 | FIM1_4266 | CP015059. | 11637 | 3037 | 4531 | 21096 | 5909 | 4296 | 14339 | 4518 | 6207 | 11418 | 6691 | 7322 | 31605 | 6544 | 7129 | 20284 | 5828 | 5661 |
| 4434 | QGN17531.1 | FIM1_4267 | CP015059. | 2103 | 788 | 920 | 1863 | 1193 | 874 | 1944 | 900 | 1146 | 3941 | 1011 | 1255 | 4900 | 871 | 1163 | 4157 | 788 | 953 |
| 4435 | QGN17532.1 | FIM1_4268 | CP015059. | 6379 | 1857 | 2053 | 6799 | 2746 | 1770 | 5681 | 2297 | 2718 | 4082 | 2371 | 2802 | 11563 | 2147 | 2622 | 6379 | 2064 | 1837 |
| 4436 | QGN17533.1 | FIM1_4269 | CP015059. | 717 | 324 | 239 | 569 | 292 | 141 | 607 | 325 | 280 | 557 | 288 | 272 | 933 | 270 | 280 | 678 | 231 | 208 |
| 4437 | QGN17534.1 | FIM1_4270 | CP015059. | 7889 | 1298 | 1385 | 7994 | 1400 | 1137 | 7032 | 1281 | 1771 | 5386 | 1002 | 1569 | 6799 | 1120 | 1519 | 5407 | 1027 | 1015 |
| 4438 | QGN17535.1 | FIM1_4271 | CP015059. | 1038 | 575 | 460 | 1411 | 683 | 430 | 1031 | 468 | 638 | 2214 | 558 | 799 | 2628 | 549 | 706 | 2302 | 530 | 464 |
| 4439 | QGN17536.1 | FIM1_4272 | CP015059. | 1680 | 359 | 496 | 2480 | 234 | 307 | 2287 | 312 | 570 | 2200 | 306 | 707 | 2056 | 444 | 622 | 2159 | 352 | 427 |
| 4440 | QGN17537.1 | FIM1_4273 | CP015059. | 2950 | 1473 | 654 | 1321 | 1537 | 686 | 2276 | 1158 | 894 | 3960 | 1654 | 1302 | 4003 | 1936 | 1180 | 3751 | 1925 | 913 |
| 4441 | QGN17538.1 | FIM1_4274 | CP015059. | 2920 | 98 | 119 | 3218 | 165 | 120 | 3298 | 110 | 138 | 1299 | 100 | 177 | 1347 | 113 | 148 | 1222 | 130 | 129 |
| 4442 | QGN17539.1 | FIM1_4275 | CP015059. | 469 | 5 | 6 | 168 | 8 | 13 | 175 | 11 | 16 | 69 | 7 | 25 | 75 | 2 | 12 | 72 | 6 | 11 |
| 4443 | QGN17540.1 | FIM1_4276 | CP015059. | 2886 | 1803 | 1539 | 2554 | 1609 | 1444 | 2771 | 1448 | 2064 | 7967 | 1662 | 2104 | 6729 | 1468 | 2197 | 7258 | 1382 | 1546 |
| 4444 | QGN17541.1 | FIM1_4277 | CP015059. | 319 | 126 | 120 | 614 | 112 | 116 | 369 | 85 | 166 | 935 | 73 | 251 | 746 | 108 | 217 | 886 | 66 | 123 |
| 4445 | QGN17542.1 | FIM1_4278 | CP015059. | 3159 | 1978 | 2056 | 4485 | 1940 | 1807 | 3734 | 1667 | 2515 | 7990 | 1768 | 2944 | 9389 | 1863 | 2970 | 7624 | 1749 | 1967 |
| 4446 | QGN17543.1 | FIM1_4279 | CP015059. | 2712 | 3541 | 1198 | 906 | 2729 | 1479 | 2263 | 2283 | 1638 | 8776 | 3897 | 2393 | 7617 | 2566 | 2501 | 8286 | 3638 | 1734 |
| 4447 | QGN17544.1 | FIM1_4280 | CP015059. | 5559 | 2606 | 2229 | 6754 | 1737 | 1956 | 5205 | 1512 | 2894 | 5691 | 1662 | 3117 | 6429 | 1792 | 3306 | 5468 | 1647 | 2172 |
| 4448 | QGN17545.1 | FIM1_4281 | CP015059. | 6823 | 4144 | 2964 | 8759 | 3617 | 2747 | 7117 | 3030 | 3821 | 7721 | 3553 | 4661 | 13894 | 3664 | 4514 | 9372 | 3443 | 3129 |
| 4449 | QGN17546.1 | FIM1_4282 | CP015059. | 17075 | 570 | 1031 | 20057 | 501 | 673 | 19074 | 836 | 1349 | 2098 | 328 | 552 | 2591 | 984 | 623 | 2085 | 603 | 404 |
| 4450 | QGN17547.1 | FIM1_4283 | CP015059. | 523 | 36 | 41 | 420 | 21 | 17 | 518 | 37 | 40 | 72 | 33 | 37 | 136 | 22 | 41 | 81 | 25 | 26 |

| | | | | | | | | | | | | | | | | | | | | | |
|---|---|---|---|---|---|---|---|---|---|---|---|---|---|---|---|---|---|---|---|---|---|
| 4451 | QGN17548.1 | FIM1_4284 | CP015059. | 3777 | 3480 | 4642 | 6884 | 3361 | 4574 | 4940 | 3066 | 5929 | 11014 | 2409 | 6263 | 12252 | 2538 | 6284 | 11037 | 2574 | 4364 |
| 4452 | QGN17549.1 | FIM1_4285 | CP015059. | 9935 | 1522 | 1497 | 8242 | 1293 | 1248 | 6068 | 1153 | 1852 | 4956 | 1313 | 2009 | 5473 | 1231 | 1935 | 5087 | 1257 | 1304 |
| 4453 | QGN17550.1 | FIM1_4286 | CP015059. | 3105 | 641 | 1665 | 2456 | 1767 | 1311 | 2906 | 1692 | 2294 | 2141 | 2008 | 2326 | 7020 | 2113 | 2323 | 5030 | 2092 | 1569 |
| 4454 | QGN17551.1 | FIM1_4287 | CP015059. | 15 | 6 | 4 | 37 | 14 | 10 | 18 | 3 | 7 | 38 | 9 | 10 | 42 | 6 | 10 | 29 | 8 | 5 |
| 4455 | QGN17552.1 | FIM1_4288 | CP015059. | 1586 | 998 | 831 | 1776 | 927 | 695 | 1852 | 976 | 1204 | 780 | 791 | 855 | 3192 | 894 | 980 | 1933 | 785 | 575 |
| 4456 | QGN17553.1 | FIM1_4289 | CP015059. | 800 | 1926 | 1598 | 1728 | 1566 | 1414 | 924 | 1261 | 2146 | 6699 | 1442 | 1866 | 6314 | 1444 | 1859 | 6574 | 1317 | 1318 |
| 4457 | QGN17554.1 | FIM1_4290 | CP015059. | 2243 | 568 | 830 | 2408 | 881 | 758 | 2209 | 839 | 1028 | 6381 | 738 | 1352 | 5311 | 736 | 1157 | 6052 | 667 | 868 |
| 4458 | QGN17555.1 | FIM1_4291 | CP015059. | 978 | 557 | 476 | 1003 | 569 | 468 | 917 | 475 | 637 | 2584 | 570 | 792 | 2697 | 544 | 772 | 2525 | 476 | 560 |
| 4459 | QGN17556.1 | FIM1_4292 | CP015059. | 470 | 862 | 606 | 400 | 762 | 489 | 448 | 599 | 660 | 3374 | 699 | 883 | 3166 | 719 | 744 | 3221 | 676 | 549 |
| 4460 | QGN17557.1 | FIM1_4293 | CP015059. | 1095 | 486 | 383 | 1104 | 440 | 394 | 1018 | 359 | 523 | 2157 | 329 | 624 | 2018 | 331 | 536 | 2274 | 300 | 400 |
| 4461 | QGN17558.1 | FIM1_4294 | CP015059. | 1450 | 706 | 805 | 1902 | 679 | 602 | 1275 | 651 | 972 | 3017 | 495 | 1121 | 3436 | 601 | 1066 | 3129 | 399 | 736 |
| 4462 | QGN17559.1 | FIM1_4295 | CP015059. | 1276 | 602 | 617 | 1417 | 581 | 573 | 1261 | 546 | 897 | 3885 | 493 | 1041 | 3431 | 495 | 879 | 3838 | 439 | 657 |
| 4463 | QGN17560.1 | FIM1_4296 | CP015059. | 2300 | 904 | 916 | 2461 | 993 | 791 | 2253 | 841 | 1221 | 3677 | 694 | 1112 | 3982 | 633 | 957 | 3795 | 649 | 724 |
| 4464 | QGN17561.1 | FIM1_4297 | CP015059. | 2036 | 111 | 481 | 2640 | 216 | 374 | 2442 | 183 | 546 | 3922 | 147 | 709 | 3969 | 213 | 577 | 3699 | 166 | 505 |
| 4465 | QGN17562.1 | FIM1_4298 | CP015059. | 617 | 33 | 157 | 783 | 213 | 87 | 470 | 182 | 179 | 85 | 107 | 226 | 713 | 111 | 197 | 198 | 131 | 148 |
| 4466 | QGN17563.1 | FIM1_4299 | CP015059. | 841 | 158 | 174 | 857 | 259 | 101 | 745 | 178 | 181 | 939 | 265 | 254 | 1319 | 226 | 229 | 1242 | 192 | 167 |
| 4467 | QGN17564.1 | FIM1_4300 | CP015059. | 384 | 177 | 126 | 757 | 171 | 89 | 496 | 143 | 161 | 566 | 135 | 153 | 575 | 113 | 146 | 549 | 101 | 80 |
| 4468 | QGN17565.1 | FIM1_4301 | CP015059. | 22 | 1 | 5 | 40 | 3 | 2 | 23 | 8 | 5 | 41 | 5 | 6 | 32 | 3 | 6 | 47 | 4 | 3 |
| 4469 | QGN17566.1 | FIM1_4303 | CP015059. | 124 | 43 | 27 | 255 | 62 | 31 | 172 | 35 | 34 | 332 | 63 | 67 | 252 | 55 | 35 | 305 | 62 | 40 |
| 4470 | QGN17567.1 | FIM1_4304 | CP015059. | 35 | 8 | 10 | 134 | 5 | 6 | 83 | 13 | 17 | 138 | 7 | 29 | 162 | 15 | 15 | 178 | 8 | 20 |
| 4471 | QGN17568.1 | FIM1_4306 | CP015059. | 0 | 0 | 0 | 0 | 0 | 0 | 0 | 0 | 0 | 0 | 0 | 0 | 0 | 0 | 0 | 0 | 0 | 0 |
| 4472 | QGN17569.1 | FIM1_4307 | CP015059. | 980 | 264 | 209 | 1691 | 349 | 225 | 1057 | 182 | 279 | 2207 | 294 | 383 | 1538 | 311 | 318 | 2085 | 276 | 215 |
| 4473 | QGN17570.1 | FIM1_4308 | CP015059. | 118 | 51 | 54 | 280 | 62 | 55 | 199 | 34 | 57 | 452 | 51 | 86 | 398 | 81 | 98 | 423 | 68 | 68 |
| 4474 | QGN17571.1 | FIM1_4314 | CP015059. | 371 | 241 | 240 | 507 | 449 | 217 | 538 | 315 | 231 | 1834 | 484 | 432 | 1768 | 740 | 526 | 1676 | 720 | 308 |
| 4475 | QGN17572.1 | FIM1_4315 | CP015059. | 0 | 0 | 0 | 0 | 0 | 0 | 0 | 0 | 0 | 0 | 0 | 0 | 0 | 0 | 0 | 0 | 0 | 0 |
| 4476 | QGN17573.1 | FIM1_4779 | CP015060. | 0 | 0 | 0 | 0 | 0 | 0 | 0 | 0 | 0 | 0 | 0 | 1 | 5 | 1 | 0 | 1 | 0 | 1 |
| 4477 | QGN17574.1 | FIM1_4780 | CP015060. | 45 | 32 | 20 | 27 | 29 | 18 | 96 | 35 | 24 | 156 | 41 | 37 | 217 | 101 | 44 | 143 | 106 | 24 |
| 4478 | QGN17575.1 | FIM1_4781 | CP015060. | 671 | 43 | 25 | 706 | 28 | 23 | 474 | 31 | 38 | 180 | 24 | 41 | 168 | 24 | 30 | 199 | 28 | 19 |
| 4479 | QGN17576.1 | FIM1_4782 | CP015060. | 909 | 116 | 151 | 1467 | 153 | 127 | 978 | 130 | 173 | 700 | 112 | 215 | 907 | 107 | 200 | 710 | 92 | 147 |
| 4480 | QGN17577.1 | FIM1_4783 | CP015060. | 4125 | 404 | 1189 | 7519 | 466 | 1164 | 5860 | 377 | 1651 | 4660 | 336 | 2748 | 4485 | 405 | 2488 | 4637 | 340 | 1668 |
| 4481 | QGN17578.1 | FIM1_4785 | CP015060. | 4189 | 374 | 433 | 5343 | 541 | 480 | 4058 | 426 | 549 | 1482 | 420 | 630 | 1989 | 456 | 600 | 1681 | 463 | 374 |
| 4482 | QGN17579.1 | FIM1_4786 | CP015060. | 8332 | 640 | 570 | 23176 | 1993 | 561 | 11517 | 1435 | 753 | 661 | 990 | 695 | 5296 | 1658 | 545 | 1550 | 1765 | 484 |
| 4483 | QGN17580.1 | FIM1_4787 | CP015060. | 1246 | 47 | 73 | 1343 | 178 | 74 | 995 | 147 | 94 | 294 | 119 | 122 | 491 | 102 | 99 | 343 | 102 | 66 |
| 4484 | QGN17581.1 | FIM1_4788 | CP015060. | 3992 | 109 | 140 | 9405 | 169 | 115 | 7417 | 98 | 161 | 592 | 137 | 306 | 720 | 163 | 280 | 580 | 112 | 190 |
| 4485 | QGN17582.1 | FIM1_4789 | CP015060. | 104 | 278 | 124 | 300 | 297 | 118 | 97 | 287 | 197 | 483 | 227 | 251 | 1213 | 76 | 271 | 741 | 111 | 121 |
| 4486 | QGN17583.1 | FIM1_4790 | CP015060. | 1997 | 395 | 313 | 1130 | 267 | 217 | 1277 | 274 | 431 | 1510 | 364 | 488 | 1447 | 324 | 605 | 1410 | 329 | 365 |
| 4487 | QGN17584.1 | FIM1_4791 | CP015060. | 930 | 2729 | 566 | 860 | 2372 | 756 | 990 | 1891 | 717 | 4414 | 2039 | 1165 | 5335 | 1501 | 1254 | 4228 | 1975 | 774 |
| 4488 | QGN17585.1 | FIM1_4792 | CP015060. | 608 | 245 | 296 | 1237 | 339 | 228 | 1234 | 312 | 407 | 420 | 156 | 383 | 1475 | 265 | 320 | 788 | 208 | 248 |
| 4489 | QGN17586.1 | FIM1_4793 | CP015060. | 4610 | 632 | 1150 | 6569 | 994 | 927 | 4671 | 853 | 1447 | 4764 | 926 | 1699 | 8394 | 1086 | 1657 | 6109 | 775 | 1169 |
| 4490 | QGN17587.1 | FIM1_4794 | CP015060. | 7893 | 1363 | 1875 | 10063 | 1146 | 1225 | 5926 | 1555 | 2281 | 4074 | 866 | 1889 | 8160 | 1525 | 1672 | 5381 | 820 | 1351 |
| 4491 | QGN17588.1 | FIM1_4795 | CP015060. | 12230 | 5162 | 4969 | 7685 | 5345 | 3361 | 6375 | 6468 | 6193 | 6962 | 3042 | 4050 | 22432 | 2722 | 3946 | 12740 | 2246 | 2932 |
| 4492 | QGN17589.1 | FIM1_4796 | CP015060. | 9880 | 1764 | 2121 | 8079 | 1845 | 1533 | 9789 | 3050 | 2705 | 2798 | 1041 | 1628 | 6425 | 932 | 1805 | 4026 | 914 | 1169 |
| 4493 | QGN17590.1 | FIM1_4797 | CP015060. | 6988 | 5721 | 1600 | 2917 | 4243 | 1803 | 5454 | 4120 | 2159 | 4974 | 4418 | 2632 | 8844 | 3449 | 2854 | 6785 | 4315 | 2027 |
| 4494 | QGN17591.1 | FIM1_4798 | CP015060. | 729 | 1180 | 764 | 316 | 822 | 634 | 512 | 810 | 1062 | 1030 | 620 | 874 | 1989 | 592 | 939 | 1404 | 602 | 653 |
| 4495 | QGN17592.1 | FIM1_4799 | CP015060. | 1146 | 665 | 639 | 1064 | 596 | 553 | 1086 | 687 | 870 | 925 | 542 | 1204 | 2902 | 456 | 1124 | 1678 | 472 | 816 |
| 4496 | QGN17593.1 | FIM1_4800 | CP015060. | 18 | 9 | 5 | 33 | 5 | 10 | 14 | 7 | 9 | 66 | 11 | 11 | 58 | 9 | 10 | 52 | 17 | 11 |
| 4497 | QGN17594.1 | FIM1_4801 | CP015060. | 258 | 376 | 238 | 729 | 266 | 197 | 313 | 305 | 282 | 684 | 314 | 407 | 1082 | 179 | 299 | 879 | 294 | 291 |
| 4498 | QGN17595.1 | FIM1_4802 | CP015060. | 394 | 397 | 496 | 639 | 512 | 406 | 532 | 441 | 636 | 3606 | 536 | 750 | 3902 | 474 | 657 | 3668 | 481 | 529 |
| 4499 | QGN17596.1 | FIM1_4803 | CP015060. | 2964 | 950 | 1243 | 3347 | 1209 | 870 | 2930 | 1287 | 1566 | 1730 | 1014 | 1830 | 6212 | 1063 | 1643 | 3324 | 1026 | 1234 |
| 4500 | QGN17597.1 | FIM1_4804 | CP015060. | 2200 | 4711 | 3373 | 4549 | 4200 | 2954 | 3371 | 3418 | 3909 | 14282 | 4921 | 5321 | 12391 | 4423 | 5364 | 14226 | 4375 | 3848 |
| 4501 | QGN17598.1 | FIM1_4805 | CP015060. | 3866 | 2236 | 1520 | 3873 | 2931 | 1691 | 4616 | 2045 | 2014 | 8657 | 3544 | 3621 | 10819 | 3442 | 3198 | 8512 | 4105 | 2425 |
| 4502 | QGN17599.1 | FIM1_4806 | CP015060. | 2458 | 1416 | 1049 | 3570 | 1364 | 923 | 3224 | 1300 | 1384 | 5541 | 1268 | 1661 | 5822 | 1222 | 1563 | 5839 | 1212 | 1100 |
| 4503 | QGN17600.1 | FIM1_4807 | CP015060. | 3512 | 2557 | 1397 | 3090 | 2136 | 1281 | 3543 | 2015 | 1929 | 2588 | 1922 | 2247 | 7008 | 1655 | 2317 | 3568 | 1902 | 1647 |
| 4504 | QGN17601.1 | FIM1_4808 | CP015060. | 2986 | 1668 | 1387 | 4837 | 1413 | 1167 | 3934 | 1386 | 1788 | 1755 | 1076 | 1831 | 5200 | 1154 | 1857 | 3290 | 1019 | 1345 |
| 4505 | QGN17602.1 | FIM1_4809 | CP015060. | 1691 | 719 | 440 | 1178 | 533 | 273 | 1237 | 531 | 506 | 1531 | 518 | 521 | 1521 | 448 | 646 | 1526 | 418 | 357 |
| 4506 | QGN17603.1 | FIM1_4810 | CP015060. | 12713 | 10564 | 6791 | 20470 | 10130 | 6279 | 14203 | 8385 | 8587 | 31594 | 10668 | 12659 | 36676 | 9720 | 12795 | 31920 | 8581 | 8597 |
| 4507 | QGN17604.1 | FIM1_4811 | CP015060. | 8398 | 1060 | 1314 | 6089 | 1022 | 944 | 4546 | 1074 | 1678 | 845 | 684 | 1703 | 4690 | 1108 | 1551 | 2093 | 880 | 1120 |
| 4508 | QGN17605.1 | FIM1_4812 | CP015060. | 371 | 87 | 140 | 531 | 88 | 133 | 652 | 99 | 231 | 390 | 78 | 210 | 586 | 67 | 207 | 410 | 77 | 183 |
| 4509 | QGN17606.1 | FIM1_4813 | CP015060. | 3355 | 2090 | 493 | 1295 | 1542 | 503 | 2783 | 1430 | 677 | 1560 | 1573 | 869 | 2784 | 1537 | 995 | 1920 | 1854 | 719 |
| 4510 | QGN17607.1 | FIM1_4814 | CP015060. | 1024 | 606 | 758 | 1138 | 764 | 606 | 898 | 704 | 1001 | 3893 | 871 | 1187 | 5126 | 786 | 1125 | 4274 | 772 | 798 |
| 4511 | QGN17608.1 | FIM1_4815 | CP015060. | 1527 | 388 | 500 | 1730 | 354 | 405 | 1784 | 453 | 695 | 3954 | 339 | 748 | 3656 | 339 | 713 | 3697 | 300 | 535 |
| 4512 | QGN17609.1 | FIM1_4816 | CP015060. | 2198 | 427 | 707 | 2147 | 479 | 525 | 2216 | 427 | 866 | 4248 | 570 | 1255 | 3929 | 480 | 1146 | 4147 | 482 | 898 |
| 4513 | QGN17610.1 | FIM1_4817 | CP015060. | 2469 | 1562 | 897 | 1987 | 1167 | 886 | 2329 | 1119 | 1161 | 5041 | 1288 | 1478 | 4952 | 1051 | 1520 | 4682 | 1087 | 977 |
| 4514 | QGN17611.1 | FIM1_4818 | CP015060. | 1790 | 453 | 702 | 1894 | 572 | 536 | 1857 | 537 | 853 | 4492 | 537 | 1215 | 4678 | 445 | 1000 | 4404 | 421 | 793 |
| 4515 | QGN17612.1 | FIM1_4819 | CP015060. | 4211 | 3755 | 5331 | 5680 | 2660 | 4654 | 4724 | 2701 | 6703 | 9394 | 2483 | 8256 | 10975 | 2716 | 7940 | 9599 | 2550 | 5831 |
| 4516 | QGN17613.1 | FIM1_4820 | CP015060. | 2928 | 1249 | 1360 | 4272 | 1276 | 1150 | 3677 | 1278 | 1742 | 5595 | 947 | 1627 | 5322 | 1001 | 1566 | 5487 | 861 | 1064 |
| 4517 | QGN17614.1 | FIM1_4821 | CP015060. | 411 | 624 | 855 | 2185 | 780 | 869 | 789 | 583 | 1046 | 5306 | 606 | 1362 | 3981 | 706 | 1030 | 4597 | 601 | 721 |
| 4518 | QGN17615.1 | FIM1_4822 | CP015060. | 1371 | 1137 | 1441 | 1347 | 976 | 1175 | 1122 | 903 | 1937 | 7183 | 849 | 1750 | 6335 | 697 | 1742 | 7312 | 598 | 1133 |
| 4519 | QGN17616.1 | FIM1_4823 | CP015060. | 1843 | 390 | 690 | 2070 | 492 | 576 | 1926 | 499 | 953 | 5182 | 522 | 1310 | 4228 | 554 | 1114 | 5195 | 538 | 835 |
| 4520 | QGN17617.1 | FIM1_4824 | CP015060. | 6030 | 2016 | 1300 | 4073 | 2315 | 1370 | 5875 | 2028 | 1796 | 11048 | 2675 | 2386 | 10368 | 2517 | 2210 | 10920 | 2751 | 1720 |
| 4521 | QGN17618.1 | FIM1_4825 | CP015060. | 1653 | 183 | 307 | 2900 | 256 | 338 | 2462 | 158 | 436 | 1777 | 82 | 243 | 1359 | 145 | 231 | 1527 | 97 | 136 |
| 4522 | QGN17619.1 | FIM1_4826 | CP015060. | 1195 | 733 | 587 | 1688 | 638 | 497 | 1615 | 587 | 803 | 4296 | 604 | 886 | 3800 | 547 | 787 | 4171 | 610 | 553 |
| 4523 | QGN17620.1 | FIM1_4827 | CP015060. | 1163 | 931 | 816 | 1231 | 917 | 687 | 1472 | 825 | 1083 | 4735 | 834 | 1190 | 5007 | 786 | 993 | 4732 | 725 | 785 |
| 4524 | QGN17621.1 | FIM1_4828 | CP015060. | 1873 | 749 | 613 | 1999 | 777 | 575 | 1670 | 506 | 796 | 2567 | 498 | 868 | 2510 | 529 | 929 | 2411 | 389 | 623 |
| 4525 | QGN17622.1 | FIM1_4829 | CP015060. | 5967 | 726 | 1735 | 7230 | 1445 | 1230 | 5813 | 1235 | 2235 | 2151 | 605 | 2526 | 8660 | 1312 | 2216 | 3843 | 673 | 1588 |
| 4526 | QGN17623.1 | FIM1_4830 | CP015060. | 6098 | 781 | 1544 | 7386 | 1238 | 1248 | 5801 | 1178 | 2206 | 2008 | 830 | 2637 | 8862 | 805 | 2404 | 4082 | 759 | 1841 |
| 4527 | QGN17624.1 | FIM1_4831 | CP015060. | 554 | 1321 | 1452 | 699 | 1473 | 1203 | 491 | 1364 | 1822 | 8438 | 1538 | 1924 | 7728 | 1292 | 1768 | 8681 | 1207 | 1254 |
| 4528 | QGN17625.1 | FIM1_4832 | CP015060. | 2919 | 2326 | 2797 | 3845 | 2670 | 2345 | 2970 | 2477 | 3706 | 10980 | 2099 | 4107 | 12565 | 2094 | 3718 | 11002 | 1713 | 2787 |
| 4529 | QGN17626.1 | FIM1_4833 | CP015060. | 9448 | 3984 | 2295 | 6352 | 2795 | 1826 | 7120 | 2928 | 3000 | 12199 | 3352 | 3397 | 12848 | 3347 | 4104 | 10957 | 2951 | 2467 |
| 4530 | QGN17627.1 | FIM1_4834 | CP015060. | 4016 | 249 | 647 | 6548 | 567 | 409 | 5081 | 749 | 801 | 439 | 305 | 1006 | 2287 | 606 | 840 | 971 | 548 | 636 |
| 4531 | QGN17628.1 | FIM1_4835 | CP015060. | 2545 | 3753 | 2202 | 3146 | 2708 | 2017 | 2737 | 2312 | 2687 | 9267 | 3230 | 3417 | 7729 | 2728 | 3426 | 8551 | 3004 | 2473 |
| 4532 | QGN17629.1 | FIM1_4836 | CP015060. | 173 | 415 | 524 | 410 | 286 | 324 | 314 | 402 | 756 | 1699 | 1022 | 1417 | 3727 | 901 | 1428 | 2706 | 867 | 1008 |
| 4533 | QGN17630.1 | FIM1_4837 | CP015060. | 9 | 97 | 52 | 34 | 23 | 30 | 16 | 25 | 64 | 136 | 88 | 141 | 163 | 51 | 168 | 187 | 63 | 99 |
| 4534 | QGN17631.1 | FIM1_4838 | CP015060. | 65 | 0 | 5 | 69 | 5 | 2 | 36 | 5 | 6 | 14 | 8 | 4 | 28 | 6 | 5 | 22 | 1 | 5 |

| | | | | | | | | | | | | | | | | | | | | | |
|---|---|---|---|---|---|---|---|---|---|---|---|---|---|---|---|---|---|---|---|---|---|
| 4535 | QGN17632.1 | FIM1_4839 | CP015060. | 395 | 308 | 536 | 1324 | 325 | 415 | 593 | 382 | 641 | 597 | 217 | 641 | 1795 | 202 | 608 | 1027 | 189 | 455 |
| 4536 | QGN17633.1 | FIM1_4840 | CP015060. | 6034 | 7460 | 4790 | 6883 | 6088 | 4337 | 6261 | 5840 | 6062 | 19459 | 7834 | 7197 | 21868 | 6519 | 7927 | 19356 | 6562 | 5384 |
| 4537 | QGN17634.1 | FIM1_4841 | CP015060. | 1439 | 179 | 127 | 1345 | 110 | 122 | 1205 | 150 | 188 | 416 | 77 | 133 | 486 | 80 | 145 | 378 | 62 | 77 |
| 4538 | QGN17635.1 | FIM1_4842 | CP015060. | 127137 | 42058 | 36871 | 117655 | 33506 | 33165 | 111063 | 31758 | 42837 | 62239 | 39372 | 46654 | 86713 | 42283 | 52243 | 75308 | 37977 | 37432 |
| 4539 | QGN17636.1 | FIM1_4843 | CP015060. | 71 | 0 | 8 | 314 | 10 | 7 | 98 | 4 | 7 | 3 | 8 | 5 | 11 | 6 | 9 | 16 | 8 | 14 |
| 4540 | QGN17637.1 | FIM1_4844 | CP015060. | 5737 | 158 | 377 | 22790 | 150 | 424 | 9590 | 159 | 463 | 353 | 57 | 145 | 312 | 73 | 67 | 386 | 90 | 100 |
| 4541 | QGN17638.1 | FIM1_4845 | CP015060. | 2147 | 315 | 265 | 3639 | 236 | 165 | 2558 | 253 | 344 | 269 | 250 | 428 | 1296 | 373 | 389 | 619 | 328 | 241 |
| 4542 | QGN17639.1 | FIM1_4846 | CP015060. | 2031 | 36 | 257 | 2346 | 157 | 188 | 1996 | 211 | 331 | 390 | 159 | 618 | 1244 | 125 | 480 | 669 | 146 | 421 |
| 4543 | QGN17640.1 | FIM1_4847 | CP015060. | 4922 | 94 | 256 | 3791 | 269 | 194 | 3713 | 308 | 349 | 262 | 182 | 341 | 757 | 220 | 360 | 426 | 195 | 265 |
| 4544 | QGN17641.1 | FIM1_4848 | CP015060. | 7995 | 307 | 404 | 7899 | 696 | 387 | 6495 | 508 | 584 | 1645 | 1011 | 1205 | 5862 | 1069 | 1207 | 3371 | 948 | 855 |
| 4545 | QGN17642.1 | FIM1_4849 | CP015060. | 2461 | 2886 | 1746 | 4445 | 2249 | 1190 | 4720 | 2825 | 2437 | 2054 | 2057 | 2829 | 7288 | 2961 | 2422 | 3456 | 2676 | 1773 |
| 4546 | QGN17643.1 | FIM1_4850 | CP015060. | 7513 | 282 | 295 | 5325 | 180 | 176 | 7220 | 359 | 417 | 124 | 114 | 435 | 1002 | 122 | 425 | 328 | 92 | 284 |
| 4547 | QGN17644.1 | FIM1_4851 | CP015060. | 1275 | 873 | 644 | 1022 | 558 | 615 | 1054 | 516 | 871 | 2937 | 523 | 788 | 2302 | 566 | 848 | 2594 | 525 | 527 |
| 4548 | QGN17645.1 | FIM1_4852 | CP015060. | 510 | 427 | 388 | 798 | 281 | 324 | 662 | 432 | 535 | 1026 | 300 | 468 | 1576 | 237 | 600 | 1090 | 222 | 355 |
| 4549 | QGN17646.1 | FIM1_4853 | CP015060. | 1720 | 169 | 211 | 1618 | 232 | 202 | 1853 | 236 | 256 | 550 | 162 | 302 | 1220 | 246 | 300 | 765 | 214 | 218 |
| 4550 | QGN17647.1 | FIM1_4854 | CP015060. | 10817 | 3127 | 1852 | 6910 | 3067 | 1430 | 8401 | 3366 | 2556 | 904 | 1826 | 1934 | 5857 | 1894 | 1933 | 2088 | 2147 | 1375 |
| 4551 | QGN17648.1 | FIM1_4855 | CP015060. | 2145 | 88 | 418 | 2208 | 539 | 273 | 2259 | 713 | 519 | 430 | 448 | 615 | 1740 | 469 | 648 | 854 | 518 | 439 |
| 4552 | QGN17649.1 | FIM1_4856 | CP015060. | 204 | 46 | 92 | 302 | 39 | 55 | 283 | 63 | 106 | 90 | 32 | 88 | 227 | 27 | 71 | 129 | 33 | 58 |
| 4553 | QGN17650.1 | FIM1_4857 | CP015060. | 848 | 11 | 9 | 1468 | 45 | 33 | 618 | 18 | 19 | 42 | 12 | 17 | 62 | 18 | 16 | 51 | 6 | 13 |
| 4554 | QGN17651.1 | FIM1_4858 | CP015060. | 2483 | 1277 | 616 | 1107 | 1297 | 695 | 2132 | 1042 | 845 | 3793 | 1898 | 1412 | 3852 | 1567 | 1525 | 3716 | 2042 | 956 |
| 4555 | QGN17652.1 | FIM1_4859 | CP015060. | 3730 | 846 | 851 | 4183 | 889 | 756 | 3930 | 930 | 1180 | 4717 | 916 | 1220 | 6390 | 784 | 1081 | 5014 | 704 | 810 |
| 4556 | QGN17653.1 | FIM1_4860 | CP015060. | 384 | 261 | 509 | 764 | 358 | 386 | 426 | 407 | 565 | 605 | 268 | 600 | 2021 | 249 | 525 | 1065 | 221 | 370 |
| 4557 | QGN17654.1 | FIM1_4861 | CP015060. | 737 | 826 | 528 | 896 | 690 | 514 | 844 | 680 | 737 | 1486 | 520 | 753 | 2268 | 530 | 840 | 1783 | 491 | 575 |
| 4558 | QGN17655.1 | FIM1_4862 | CP015060. | 1029 | 2005 | 930 | 828 | 1300 | 933 | 869 | 1237 | 1275 | 5300 | 1292 | 1277 | 4851 | 1241 | 1421 | 4743 | 1253 | 918 |
| 4559 | QGN17656.1 | FIM1_4863 | CP015060. | 4754 | 4985 | 1713 | 2424 | 4150 | 2085 | 3891 | 3540 | 2454 | 11553 | 5265 | 3201 | 11789 | 4197 | 3494 | 11740 | 5266 | 2289 |
| 4560 | QGN17657.1 | FIM1_4864 | CP015060. | 55 | 67 | 38 | 73 | 40 | 33 | 87 | 62 | 72 | 308 | 36 | 58 | 230 | 40 | 49 | 258 | 30 | 28 |
| 4561 | QGN17658.1 | FIM1_4865 | CP015060. | 2121 | 495 | 461 | 1676 | 528 | 399 | 1440 | 664 | 594 | 851 | 346 | 709 | 2614 | 348 | 613 | 1405 | 365 | 478 |
| 4562 | QGN17659.1 | FIM1_4866 | CP015060. | 1425 | 581 | 1582 | 3657 | 586 | 1091 | 2733 | 761 | 1976 | 1901 | 353 | 1823 | 4109 | 485 | 1531 | 2492 | 366 | 1158 |
| 4563 | QGN17660.1 | FIM1_4867 | CP015060. | 2609 | 1712 | 682 | 1601 | 1480 | 662 | 2453 | 1360 | 916 | 3388 | 1385 | 1287 | 4711 | 1093 | 1233 | 3385 | 1267 | 848 |
| 4564 | QGN17661.1 | FIM1_4868 | CP015060. | 1114 | 1417 | 873 | 1521 | 935 | 706 | 1339 | 834 | 1075 | 3213 | 904 | 1157 | 3151 | 887 | 1199 | 2858 | 913 | 752 |
| 4565 | QGN17662.1 | FIM1_4869 | CP015060. | 495 | 3193 | 1999 | 650 | 1964 | 1649 | 647 | 1789 | 2383 | 7482 | 1638 | 2702 | 5556 | 1611 | 2422 | 7135 | 1576 | 1531 |
| 4566 | QGN17663.1 | FIM1_4870 | CP015060. | 2726 | 692 | 328 | 1473 | 578 | 284 | 2347 | 576 | 387 | 361 | 333 | 385 | 1277 | 388 | 367 | 666 | 371 | 299 |
| 4567 | QGN17664.1 | FIM1_4871 | CP015060. | 2109 | 393 | 282 | 3109 | 395 | 269 | 1954 | 238 | 366 | 1076 | 137 | 201 | 936 | 175 | 236 | 923 | 136 | 160 |
| 4568 | QGN17665.1 | FIM1_4872 | CP015060. | 5513 | 288 | 194 | 3824 | 140 | 146 | 5232 | 196 | 287 | 621 | 100 | 159 | 685 | 59 | 132 | 591 | 80 | 91 |
| 4569 | QGN17666.1 | FIM1_4873 | CP015060. | 3273 | 1200 | 701 | 2580 | 1042 | 747 | 3138 | 811 | 1019 | 4259 | 1013 | 987 | 3384 | 1025 | 997 | 3831 | 1076 | 687 |
| 4570 | QGN17667.1 | FIM1_4874 | CP015060. | 1941 | 1677 | 1281 | 1294 | 2133 | 1153 | 2379 | 1942 | 1682 | 6626 | 2311 | 2090 | 9244 | 1827 | 2060 | 6619 | 2211 | 1543 |
| 4571 | QGN17668.1 | FIM1_4875 | CP015060. | 1774 | 466 | 1234 | 4076 | 826 | 972 | 1937 | 781 | 1546 | 4658 | 366 | 1679 | 5797 | 578 | 1537 | 4596 | 375 | 1113 |
| 4572 | QGN17669.1 | FIM1_4876 | CP015060. | 47 | 46 | 60 | 138 | 58 | 48 | 47 | 43 | 72 | 195 | 28 | 107 | 221 | 37 | 107 | 208 | 21 | 67 |
| 4573 | QGN17670.1 | FIM1_4877 | CP015060. | 3274 | 295 | 503 | 2920 | 512 | 379 | 2767 | 670 | 623 | 1311 | 431 | 675 | 3099 | 391 | 613 | 1738 | 398 | 424 |
| 4574 | QGN17671.1 | FIM1_4878 | CP015060. | 13910 | 17485 | 10788 | 13962 | 13203 | 10611 | 11722 | 11160 | 12832 | 45938 | 17135 | 16402 | 39826 | 14550 | 16825 | 46689 | 15197 | 12275 |
| 4575 | QGN17672.1 | FIM1_4879 | CP015060. | 3574 | 1571 | 1524 | 4001 | 1759 | 1339 | 3957 | 1684 | 1941 | 8309 | 1806 | 2475 | 9466 | 1689 | 2288 | 8252 | 1724 | 1743 |
| 4576 | QGN17673.1 | FIM1_4880 | CP015060. | 3302 | 3820 | 2429 | 3712 | 3083 | 2052 | 3350 | 2964 | 3037 | 15389 | 4191 | 4449 | 16123 | 4138 | 4569 | 14779 | 3779 | 3238 |
| 4577 | QGN17674.1 | FIM1_4881 | CP015060. | 2662 | 1036 | 1156 | 3236 | 1365 | 1076 | 2612 | 1287 | 1371 | 4012 | 1343 | 1643 | 7311 | 1358 | 1569 | 5360 | 1178 | 1160 |
| 4578 | QGN17675.1 | FIM1_4882 | CP015060. | 20366 | 30 | 24 | 11071 | 24 | 15 | 8135 | 73 | 35 | 53 | 15 | 46 | 156 | 14 | 20 | 88 | 11 | 25 |
| 4579 | QGN17676.1 | FIM1_4883 | CP015060. | 8930 | 1567 | 2119 | 9715 | 2279 | 1840 | 9592 | 2052 | 2738 | 5507 | 1867 | 3047 | 12809 | 1870 | 2768 | 7988 | 1843 | 2063 |
| 4580 | QGN17677.1 | FIM1_4884 | CP015060. | 2480 | 1104 | 1496 | 3573 | 1351 | 1195 | 3064 | 1265 | 1928 | 7564 | 1397 | 2477 | 9506 | 1311 | 2229 | 8180 | 1227 | 1608 |
| 4581 | QGN17678.1 | FIM1_4885 | CP015060. | 934 | 787 | 717 | 1422 | 666 | 621 | 1125 | 797 | 951 | 2477 | 426 | 978 | 3429 | 503 | 964 | 2431 | 430 | 639 |
| 4582 | QGN17679.1 | FIM1_4886 | CP015060. | 194 | 275 | 217 | 359 | 160 | 172 | 318 | 202 | 255 | 835 | 123 | 284 | 877 | 153 | 260 | 742 | 139 | 177 |
| 4583 | QGN17680.1 | FIM1_4887 | CP015060. | 256 | 304 | 295 | 230 | 217 | 261 | 290 | 287 | 402 | 1060 | 200 | 332 | 1289 | 198 | 323 | 1032 | 190 | 234 |
| 4584 | QGN17681.1 | FIM1_4888 | CP015060. | 762 | 764 | 654 | 1297 | 526 | 579 | 942 | 684 | 963 | 1030 | 451 | 872 | 2426 | 401 | 945 | 1453 | 382 | 541 |
| 4585 | QGN17682.1 | FIM1_4889 | CP015060. | 1085 | 381 | 551 | 1371 | 286 | 451 | 1085 | 387 | 752 | 2539 | 269 | 752 | 1885 | 457 | 791 | 2321 | 290 | 447 |
| 4586 | QGN17683.1 | FIM1_4890 | CP015060. | 622 | 755 | 431 | 462 | 420 | 296 | 529 | 441 | 478 | 1208 | 423 | 526 | 1931 | 408 | 562 | 1463 | 332 | 349 |
| 4587 | QGN17684.1 | FIM1_4891 | CP015060. | 1578 | 1256 | 692 | 1399 | 843 | 639 | 1520 | 868 | 864 | 3940 | 779 | 751 | 3035 | 763 | 897 | 3273 | 785 | 508 |
| 4588 | QGN17685.1 | FIM1_4892 | CP015060. | 8566 | 3196 | 2682 | 8764 | 3957 | 2054 | 7446 | 3941 | 3473 | 3030 | 2854 | 4351 | 12463 | 3375 | 3713 | 5898 | 3272 | 2617 |
| 4589 | QGN17686.1 | FIM1_4893 | CP015060. | 9860 | 7638 | 4641 | 12134 | 7531 | 3751 | 10750 | 7465 | 6118 | 6275 | 7202 | 7959 | 25307 | 7202 | 7823 | 10877 | 7636 | 5680 |
| 4590 | QGN17687.1 | FIM1_4894 | CP015060. | 3168 | 1116 | 1062 | 3512 | 837 | 963 | 3282 | 1139 | 1397 | 1301 | 695 | 1417 | 3538 | 680 | 1440 | 2396 | 656 | 954 |
| 4591 | QGN17688.1 | FIM1_4895 | CP015060. | 2051 | 1433 | 959 | 2611 | 1524 | 889 | 2384 | 1469 | 1334 | 1222 | 988 | 1508 | 4276 | 871 | 1450 | 2076 | 951 | 1088 |
| 4592 | QGN17689.1 | FIM1_4896 | CP015060. | 4796 | 628 | 704 | 3582 | 438 | 462 | 4175 | 632 | 870 | 683 | 373 | 916 | 2021 | 418 | 883 | 1054 | 444 | 614 |
| 4593 | QGN17690.1 | FIM1_4897 | CP015060. | 3024 | 830 | 613 | 2627 | 527 | 479 | 2925 | 798 | 843 | 564 | 412 | 865 | 1845 | 528 | 908 | 1040 | 542 | 654 |
| 4594 | QGN17691.1 | FIM1_4898 | CP015060. | 686 | 923 | 721 | 1334 | 677 | 547 | 859 | 879 | 940 | 673 | 488 | 811 | 2712 | 514 | 836 | 1213 | 405 | 586 |
| 4595 | QGN17692.1 | FIM1_4899 | CP015060. | 7252 | 8012 | 6950 | 8123 | 6431 | 6214 | 5365 | 5502 | 8462 | 17561 | 5932 | 6771 | 15804 | 4867 | 7278 | 18875 | 4969 | 4865 |
| 4596 | QGN17693.1 | FIM1_4900 | CP015060. | 4833 | 4634 | 3866 | 4160 | 2859 | 3171 | 3743 | 3261 | 4973 | 6419 | 3024 | 3810 | 8084 | 2762 | 3807 | 6863 | 2974 | 2741 |
| 4597 | QGN17694.1 | FIM1_4901 | CP015060. | 674 | 314 | 312 | 1229 | 181 | 238 | 674 | 196 | 452 | 427 | 159 | 334 | 827 | 185 | 375 | 540 | 194 | 222 |
| 4598 | QGN17695.1 | FIM1_4902 | CP015060. | 2533 | 944 | 700 | 2295 | 706 | 574 | 2003 | 706 | 888 | 1608 | 486 | 886 | 2856 | 381 | 875 | 2080 | 429 | 595 |
| 4599 | QGN17696.1 | FIM1_4903 | CP015060. | 3503 | 1125 | 1186 | 4356 | 1205 | 1005 | 2916 | 1211 | 1467 | 2069 | 575 | 983 | 3724 | 451 | 885 | 2488 | 457 | 616 |
| 4600 | QGN17697.1 | FIM1_4904 | CP015060. | 1645 | 950 | 1230 | 1292 | 884 | 1319 | 1143 | 879 | 1734 | 1239 | 658 | 1749 | 2305 | 444 | 1662 | 1478 | 529 | 1151 |
| 4601 | QGN17698.1 | FIM1_4905 | CP015060. | 981 | 191 | 309 | 955 | 196 | 213 | 767 | 275 | 472 | 527 | 147 | 400 | 1622 | 147 | 375 | 945 | 130 | 266 |
| 4602 | QGN17699.1 | FIM1_4906 | CP015060. | 3840 | 1288 | 1195 | 3705 | 1139 | 978 | 3852 | 1257 | 1512 | 2622 | 831 | 1178 | 4042 | 835 | 1216 | 3033 | 780 | 860 |
| 4603 | QGN17700.1 | FIM1_4907 | CP015060. | 5971 | 1323 | 577 | 2657 | 1611 | 634 | 5590 | 1514 | 766 | 1924 | 1547 | 1230 | 4238 | 1691 | 1229 | 2501 | 1964 | 878 |
| 4604 | QGN17701.1 | FIM1_4908 | CP015060. | 650 | 517 | 361 | 547 | 375 | 385 | 667 | 357 | 551 | 1984 | 358 | 509 | 1315 | 285 | 567 | 1538 | 385 | 332 |
| 4605 | QGN17702.1 | FIM1_4909 | CP015060. | 523 | 2815 | 1689 | 820 | 2265 | 1465 | 594 | 1985 | 2180 | 10226 | 2291 | 2567 | 9184 | 2047 | 2402 | 9756 | 2211 | 1634 |
| 4606 | QGN17703.1 | FIM1_4910 | CP015060. | 1467 | 317 | 270 | 1275 | 294 | 314 | 1737 | 307 | 409 | 2758 | 232 | 397 | 2321 | 238 | 381 | 2395 | 269 | 292 |
| 4607 | QGN17704.1 | FIM1_4911 | CP015060. | 1602 | 832 | 707 | 1265 | 858 | 652 | 1510 | 778 | 889 | 1828 | 797 | 1096 | 4124 | 871 | 1031 | 2544 | 748 | 751 |
| 4608 | QGN17705.1 | FIM1_4912 | CP015060. | 1160 | 726 | 634 | 1105 | 420 | 493 | 823 | 555 | 832 | 3548 | 531 | 974 | 3196 | 851 | 1103 | 3167 | 627 | 697 |
| 4609 | QGN17706.1 | FIM1_4913 | CP015060. | 2187 | 1314 | 516 | 1328 | 1467 | 574 | 2148 | 1091 | 646 | 3745 | 1614 | 1090 | 3985 | 1670 | 1031 | 3645 | 1916 | 747 |
| 4610 | QGN17707.1 | FIM1_4914 | CP015060. | 4502 | 1246 | 986 | 5043 | 1682 | 1015 | 5270 | 1429 | 1312 | 3885 | 1545 | 1786 | 6495 | 1778 | 1675 | 4888 | 1655 | 1253 |
| 4611 | QGN17708.1 | FIM1_4915 | CP015060. | 4431 | 946 | 1168 | 6071 | 1040 | 917 | 4994 | 1079 | 1552 | 1483 | 882 | 1804 | 6779 | 942 | 1593 | 3136 | 915 | 1205 |
| 4612 | QGN17709.1 | FIM1_4916 | CP015060. | 3057 | 978 | 446 | 1031 | 893 | 469 | 2623 | 869 | 631 | 1254 | 1068 | 922 | 1832 | 1324 | 911 | 1431 | 1429 | 709 |
| 4613 | QGN17710.1 | FIM1_4917 | CP015060. | 5376 | 782 | 800 | 4739 | 916 | 678 | 5507 | 843 | 1068 | 2766 | 768 | 1251 | 4647 | 732 | 1056 | 3117 | 766 | 776 |
| 4614 | QGN17711.1 | FIM1_4918 | CP015060. | 170 | 111 | 121 | 313 | 168 | 109 | 169 | 132 | 158 | 771 | 146 | 135 | 602 | 105 | 140 | 705 | 103 | 82 |
| 4615 | QGN17712.1 | FIM1_4919 | CP015060. | 1067 | 1234 | 668 | 962 | 1132 | 717 | 1149 | 987 | 923 | 5230 | 1423 | 1193 | 4692 | 1010 | 1138 | 4977 | 1206 | 786 |
| 4616 | QGN17713.1 | FIM1_4920 | CP015060. | 1635 | 198 | 310 | 1554 | 193 | 298 | 1213 | 194 | 437 | 2211 | 124 | 546 | 2024 | 129 | 445 | 2198 | 98 | 336 |
| 4617 | QGN17714.1 | FIM1_4921 | CP015060. | 1895 | 9151 | 5692 | 3492 | 8175 | 4995 | 3689 | 7769 | 7263 | 29494 | 11175 | 11010 | 37098 | 10029 | 10897 | 31209 | 9298 | 7469 |
| 4618 | QGN17715.1 | FIM1_4922 | CP015060. | 948 | 1238 | 580 | 681 | 754 | 528 | 903 | 753 | 723 | 3247 | 725 | 846 | 3014 | 687 | 861 | 3228 | 705 | 567 |

| | | | | | | | | | | | | | | | | | | | | | |
|---|---|---|---|---|---|---|---|---|---|---|---|---|---|---|---|---|---|---|---|---|---|
| 4619 | QGN17716.1 | FIM1_4923 | CP015060. | 793 | 730 | 559 | 1021 | 739 | 430 | 750 | 542 | 705 | 1876 | 613 | 691 | 2390 | 657 | 629 | 2000 | 591 | 412 |
| 4620 | QGN17717.1 | FIM1_4924 | CP015060. | 749 | 4084 | 3814 | 964 | 2609 | 3478 | 420 | 3094 | 4966 | 2183 | 1052 | 2277 | 2710 | 512 | 1739 | 2405 | 612 | 1325 |
| 4621 | QGN17718.1 | FIM1_4925 | CP015060. | 699 | 81 | 96 | 2815 | 200 | 101 | 716 | 134 | 106 | 159 | 97 | 122 | 443 | 77 | 117 | 254 | 111 | 63 |
| 4622 | QGN17719.1 | FIM1_4926 | CP015060. | 1516 | 689 | 871 | 2050 | 840 | 661 | 1697 | 710 | 1119 | 5899 | 755 | 1307 | 5382 | 725 | 1179 | 5760 | 634 | 860 |
| 4623 | QGN17720.1 | FIM1_4927 | CP015060. | 2022 | 906 | 555 | 890 | 1055 | 584 | 1562 | 807 | 666 | 3852 | 1093 | 967 | 3139 | 1200 | 888 | 3377 | 1242 | 675 |
| 4624 | QGN17721.1 | FIM1_4928 | CP015060. | 1496 | 720 | 399 | 726 | 820 | 441 | 1506 | 727 | 542 | 3259 | 976 | 744 | 2575 | 1027 | 686 | 3026 | 1142 | 540 |
| 4625 | QGN17722.1 | FIM1_4929 | CP015060. | 417 | 403 | 647 | 346 | 379 | 611 | 352 | 509 | 879 | 2587 | 347 | 807 | 2947 | 269 | 765 | 2664 | 292 | 537 |
| 4626 | QGN17723.1 | FIM1_4931 | CP015060. | 4849 | 3814 | 2575 | 5050 | 3634 | 2350 | 5173 | 3419 | 3338 | 3356 | 2455 | 2975 | 8983 | 2562 | 2582 | 5434 | 2434 | 2002 |
| 4627 | QGN17724.1 | FIM1_4932 | CP015060. | 15000 | 1614 | 2483 | 15439 | 3169 | 2095 | 14712 | 2687 | 3250 | 10019 | 2261 | 2757 | 15659 | 2488 | 2579 | 11902 | 2014 | 1773 |
| 4628 | QGN17725.1 | FIM1_4933 | CP015060. | 981 | 1197 | 1357 | 2205 | 1496 | 1163 | 1972 | 1264 | 1797 | 8367 | 932 | 1740 | 6898 | 935 | 1375 | 8215 | 799 | 1094 |
| 4629 | QGN17726.1 | FIM1_4935 | CP015060. | 13437 | 530 | 745 | 11017 | 645 | 571 | 12420 | 742 | 992 | 3324 | 464 | 836 | 4689 | 552 | 860 | 3572 | 433 | 600 |
| 4630 | QGN17727.1 | FIM1_4936 | CP015060. | 2123 | 459 | 930 | 4237 | 674 | 628 | 3050 | 787 | 1186 | 674 | 344 | 1246 | 2819 | 429 | 1124 | 1276 | 409 | 830 |
| 4631 | QGN17728.1 | FIM1_4937 | CP015060. | 6704 | 3858 | 5371 | 11250 | 3901 | 4176 | 8553 | 3846 | 7031 | 2995 | 3259 | 7941 | 19412 | 3791 | 7457 | 9753 | 3254 | 5430 |
| 4632 | QGN17729.1 | FIM1_4938 | CP015060. | 2429 | 902 | 533 | 1863 | 902 | 559 | 2353 | 737 | 740 | 4795 | 841 | 953 | 3630 | 754 | 853 | 4411 | 889 | 660 |
| 4633 | QGN17730.1 | FIM1_4939 | CP015060. | 700 | 557 | 762 | 1008 | 494 | 676 | 905 | 504 | 1023 | 4737 | 528 | 1174 | 3307 | 459 | 1047 | 4847 | 485 | 772 |
| 4634 | QGN17731.1 | FIM1_4940 | CP015060. | 1356 | 1336 | 630 | 832 | 1275 | 743 | 1362 | 1112 | 852 | 2912 | 1199 | 1029 | 3031 | 1207 | 976 | 2990 | 1293 | 767 |
| 4635 | QGN17732.1 | FIM1_4941 | CP015060. | 5717 | 5106 | 4891 | 8981 | 6846 | 4043 | 8062 | 5753 | 5792 | 24280 | 7947 | 7610 | 23815 | 8049 | 7688 | 24209 | 6826 | 5442 |
| 4636 | QGN17733.1 | FIM1_4942 | CP015060. | 2488 | 669 | 689 | 2075 | 804 | 506 | 2251 | 910 | 932 | 494 | 686 | 1054 | 4026 | 722 | 926 | 1386 | 765 | 661 |
| 4637 | QGN17734.1 | FIM1_4943 | CP015060. | 6553 | 1376 | 1589 | 5754 | 1223 | 1406 | 5828 | 1171 | 2087 | 2744 | 1026 | 2502 | 5753 | 833 | 2734 | 3979 | 842 | 1665 |
| 4638 | QGN17735.1 | FIM1_4944 | CP015060. | 6220 | 351 | 516 | 5134 | 467 | 398 | 5677 | 561 | 693 | 335 | 311 | 700 | 1699 | 383 | 584 | 627 | 421 | 468 |
| 4639 | QGN17736.1 | FIM1_4945 | CP015060. | 5907 | 6302 | 4889 | 11391 | 5207 | 4214 | 9676 | 4909 | 6464 | 8009 | 5397 | 6468 | 18738 | 5494 | 6575 | 14798 | 4575 | 4713 |
| 4640 | QGN17737.1 | FIM1_4946 | CP015060. | 4029 | 1559 | 1641 | 5096 | 2013 | 1365 | 4575 | 1808 | 2159 | 4490 | 1878 | 2402 | 10524 | 1739 | 2179 | 6738 | 1821 | 1670 |
| 4641 | QGN17738.1 | FIM1_4947 | CP015060. | 1818 | 1204 | 1242 | 1792 | 893 | 907 | 1274 | 1064 | 1610 | 8242 | 1180 | 1988 | 7495 | 1380 | 2071 | 7705 | 1056 | 1323 |
| 4642 | QGN17739.1 | FIM1_4948 | CP015060. | 1146 | 362 | 333 | 1066 | 207 | 275 | 1164 | 228 | 460 | 1896 | 223 | 470 | 1613 | 191 | 466 | 1675 | 210 | 308 |
| 4643 | QGN17740.1 | FIM1_4949 | CP015060. | 2431 | 807 | 1252 | 2722 | 745 | 1095 | 2171 | 771 | 1528 | 3629 | 723 | 1573 | 4346 | 736 | 1532 | 3674 | 630 | 1138 |
| 4644 | QGN17741.1 | FIM1_4950 | CP015060. | 1883 | 2585 | 2682 | 1983 | 1988 | 2432 | 1647 | 2062 | 3630 | 8533 | 2287 | 3789 | 9786 | 1934 | 4110 | 8906 | 1645 | 2710 |
| 4645 | QGN17742.1 | FIM1_4951 | CP015060. | 3790 | 1396 | 1562 | 5864 | 1543 | 1210 | 4618 | 1511 | 1917 | 3192 | 1241 | 2198 | 8015 | 1262 | 1954 | 4762 | 1155 | 1476 |
| 4646 | QGN17743.1 | FIM1_4952 | CP015060. | 5530 | 3974 | 2238 | 3019 | 5023 | 2556 | 4859 | 4045 | 2996 | 17708 | 7378 | 4886 | 18306 | 6372 | 4960 | 17027 | 7832 | 3614 |
| 4647 | QGN17744.1 | FIM1_4953 | CP015060. | 3572 | 1605 | 660 | 1466 | 1804 | 647 | 3035 | 1388 | 833 | 4842 | 2064 | 1380 | 5093 | 2338 | 1221 | 4813 | 2633 | 964 |
| 4648 | QGN17745.1 | FIM1_4954 | CP015060. | 199 | 210 | 116 | 94 | 160 | 124 | 206 | 164 | 165 | 712 | 196 | 246 | 681 | 131 | 246 | 627 | 144 | 170 |
| 4649 | QGN17746.1 | FIM1_4955 | CP015060. | 6410 | 3576 | 5281 | 8098 | 7574 | 4415 | 7009 | 5962 | 5814 | 32633 | 10134 | 9737 | 32465 | 8311 | 9215 | 33620 | 8373 | 6583 |
| 4650 | QGN17747.1 | FIM1_4956 | CP015060. | 18467 | 479 | 471 | 15904 | 657 | 358 | 19168 | 512 | 571 | 1735 | 596 | 609 | 2267 | 893 | 553 | 1833 | 774 | 418 |
| 4651 | QGN17748.1 | FIM1_4957 | CP015060. | 5590 | 232 | 180 | 4837 | 308 | 162 | 5046 | 298 | 224 | 342 | 328 | 217 | 908 | 335 | 225 | 589 | 404 | 156 |
| 4652 | QGN17749.1 | FIM1_4958 | CP015060. | 161 | 182 | 381 | 545 | 125 | 208 | 191 | 165 | 479 | 157 | 34 | 167 | 627 | 83 | 180 | 248 | 31 | 129 |
| 4653 | QGN17750.1 | FIM1_4959 | CP015060. | 555 | 3 | 0 | 185 | 1 | 1 | 137 | 3 | 2 | 254 | 50 | 111 | 329 | 9 | 5 | 365 | 37 | 22 |
| 4654 | QGN17751.1 | FIM1_4960 | CP015060. | 1643 | 2323 | 2169 | 2146 | 3080 | 1766 | 1580 | 2480 | 2516 | 15221 | 3769 | 4296 | 15821 | 3681 | 4197 | 15078 | 3126 | 2896 |
| 4655 | QGN17752.1 | FIM1_4961 | CP015060. | 4571 | 2054 | 2005 | 5713 | 1747 | 1090 | 4126 | 2461 | 2484 | 7122 | 3426 | 3357 | 16648 | 4226 | 3426 | 10114 | 3554 | 2491 |
| 4656 | QGN17753.1 | FIM1_4962 | CP015060. | 2269 | 590 | 347 | 2254 | 436 | 296 | 2108 | 333 | 437 | 2307 | 469 | 586 | 1715 | 442 | 547 | 2227 | 581 | 405 |
| 4657 | QGN17754.1 | FIM1_4963 | CP015060. | 2102 | 414 | 532 | 2130 | 474 | 388 | 1819 | 534 | 639 | 2091 | 344 | 515 | 2226 | 415 | 531 | 2003 | 381 | 339 |
| 4658 | QGN17755.1 | FIM1_4964 | CP015060. | 351 | 51 | 29 | 442 | 42 | 25 | 306 | 37 | 39 | 220 | 37 | 47 | 285 | 30 | 49 | 215 | 24 | 25 |
| 4659 | QGN17756.1 | FIM1_4965 | CP015060. | 6036 | 6445 | 3250 | 4911 | 5748 | 3332 | 5951 | 5338 | 4358 | 15607 | 5694 | 5742 | 19033 | 5035 | 5332 | 15137 | 5477 | 3812 |
| 4660 | QGN17757.1 | FIM1_4966 | CP015060. | 4881 | 3907 | 3908 | 6204 | 3141 | 3412 | 5582 | 2888 | 4865 | 10903 | 2011 | 4685 | 10174 | 2381 | 4547 | 10216 | 1885 | 3117 |
| 4661 | QGN17758.1 | FIM1_4967 | CP015060. | 5062 | 1090 | 1181 | 5164 | 996 | 904 | 5174 | 914 | 1581 | 5251 | 658 | 1502 | 5563 | 987 | 1489 | 5278 | 743 | 973 |
| 4662 | QGN17759.1 | FIM1_4968 | CP015060. | 2069 | 1675 | 817 | 751 | 1519 | 883 | 1313 | 1510 | 1146 | 2396 | 1389 | 1291 | 4157 | 1272 | 1357 | 2581 | 1491 | 953 |
| 4663 | QGN17760.1 | FIM1_4969 | CP015060. | 19755 | 9363 | 9501 | 18995 | 9623 | 8298 | 17308 | 9031 | 11608 | 31412 | 8652 | 13019 | 31047 | 8393 | 13088 | 31414 | 7473 | 9119 |
| 4664 | QGN17761.1 | FIM1_4970 | CP015060. | 13202 | 6673 | 7410 | 15491 | 9449 | 6347 | 13560 | 8417 | 9187 | 33253 | 10828 | 12000 | 42737 | 10223 | 10774 | 35488 | 10207 | 8131 |
| 4665 | QGN17762.1 | FIM1_4971 | CP015060. | 10496 | 1528 | 3198 | 14191 | 3676 | 2986 | 8636 | 2858 | 4137 | 13861 | 2208 | 4307 | 17736 | 2550 | 3868 | 15130 | 1986 | 2883 |
| 4666 | QGN17763.1 | FIM1_4972 | CP015060. | 2337 | 1009 | 1355 | 5898 | 1566 | 1320 | 2623 | 1135 | 1861 | 2392 | 838 | 1433 | 7182 | 812 | 1238 | 3997 | 707 | 943 |
| 4667 | QGN17764.1 | FIM1_4973 | CP015060. | 1318 | 618 | 1273 | 3206 | 1172 | 1213 | 2154 | 932 | 1757 | 5147 | 753 | 1690 | 6440 | 590 | 1636 | 5721 | 661 | 1136 |
| 4668 | QGN17765.1 | FIM1_4974 | CP015060. | 830 | 269 | 395 | 880 | 273 | 270 | 666 | 297 | 503 | 1827 | 256 | 532 | 1829 | 247 | 424 | 1829 | 248 | 287 |
| 4669 | QGN17766.1 | FIM1_4975 | CP015060. | 82 | 19 | 73 | 88 | 10 | 44 | 50 | 26 | 94 | 109 | 5 | 115 | 121 | 4 | 95 | 101 | 6 | 48 |
| 4670 | QGN17767.1 | FIM1_4976 | CP015060. | 5862 | 4432 | 4249 | 8727 | 5698 | 3709 | 7017 | 4975 | 5330 | 14154 | 5368 | 6910 | 23556 | 5313 | 6003 | 15963 | 5252 | 4614 |
| 4671 | QGN17768.1 | FIM1_4977 | CP015060. | 1829 | 1928 | 535 | 670 | 1569 | 612 | 1695 | 1313 | 754 | 4646 | 1513 | 831 | 3453 | 1416 | 930 | 4351 | 1829 | 596 |
| 4672 | QGN17769.1 | FIM1_4978 | CP015060. | 844 | 572 | 571 | 1011 | 457 | 546 | 755 | 416 | 809 | 2927 | 714 | 1873 | 4365 | 525 | 1415 | 3433 | 564 | 1067 |
| 4673 | QGN17770.1 | FIM1_4979 | CP015060. | 17946 | 10972 | 10971 | 27146 | 14205 | 9551 | 21801 | 12806 | 13878 | 37162 | 11284 | 15161 | 46239 | 12564 | 15118 | 39423 | 10501 | 10655 |
| 4674 | QGN17771.1 | FIM1_4980 | CP015060. | 1863 | 156 | 355 | 3381 | 226 | 336 | 1913 | 153 | 460 | 974 | 154 | 542 | 1542 | 106 | 454 | 1169 | 138 | 362 |
| 4675 | QGN17772.1 | FIM1_4981 | CP015060. | 14696 | 13242 | 10393 | 15879 | 12478 | 8395 | 14099 | 12588 | 12248 | 33610 | 13066 | 12664 | 39311 | 12583 | 13351 | 35703 | 11408 | 9524 |
| 4676 | QGN17773.1 | FIM1_4982 | CP015060. | 1974 | 564 | 641 | 2198 | 601 | 505 | 1900 | 580 | 726 | 3482 | 524 | 942 | 3393 | 501 | 852 | 3380 | 471 | 594 |
| 4677 | QGN17774.1 | FIM1_4983 | CP015060. | 1874 | 637 | 554 | 2557 | 587 | 582 | 1816 | 558 | 795 | 3937 | 450 | 870 | 3255 | 401 | 790 | 3805 | 411 | 582 |
| 4678 | QGN17775.1 | FIM1_4984 | CP015060. | 4789 | 419 | 895 | 7467 | 547 | 719 | 5351 | 482 | 1211 | 4107 | 359 | 1295 | 6037 | 475 | 1068 | 4489 | 334 | 857 |
| 4679 | QGN17776.1 | FIM1_4985 | CP015060. | 3753 | 1422 | 1701 | 5052 | 1586 | 1416 | 4018 | 1571 | 2138 | 8565 | 1323 | 2553 | 10652 | 1426 | 2234 | 8960 | 1261 | 1706 |
| 4680 | QGN17777.1 | FIM1_4986 | CP015060. | 6380 | 2400 | 2876 | 8807 | 2685 | 2406 | 6868 | 2191 | 3577 | 12618 | 2616 | 4804 | 15411 | 2691 | 4183 | 13295 | 2373 | 3106 |
| 4681 | QGN17778.1 | FIM1_4987 | CP015060. | 1536 | 2532 | 3093 | 4640 | 2311 | 2846 | 3049 | 2447 | 4118 | 6328 | 1181 | 3251 | 6655 | 1148 | 3106 | 6244 | 1016 | 2217 |
| 4682 | QGN17779.1 | FIM1_4988 | CP015060. | 290 | 126 | 142 | 414 | 109 | 101 | 283 | 81 | 153 | 890 | 78 | 183 | 589 | 62 | 145 | 767 | 68 | 109 |
| 4683 | QGN17780.1 | FIM1_4989 | CP015060. | 12643 | 2976 | 4139 | 19527 | 4766 | 3325 | 16013 | 3668 | 5296 | 14705 | 4047 | 5198 | 25590 | 5518 | 4204 | 18863 | 4507 | 3183 |
| 4684 | QGN17781.1 | FIM1_4990 | CP015060. | 2 | 3 | 0 | 11 | 0 | 0 | 3 | 0 | 2 | 5 | 0 | 0 | 4 | 0 | 2 | 7 | 1 | 2 |
| 4685 | QGN17782.1 | FIM1_4991 | CP015060. | 2014 | 2352 | 1950 | 3658 | 2479 | 1569 | 2547 | 1903 | 2610 | 4599 | 2355 | 2987 | 12801 | 2475 | 2686 | 8577 | 2309 | 2041 |
| 4686 | QGN17783.1 | FIM1_4992 | CP015060. | 437 | 816 | 877 | 1215 | 702 | 747 | 591 | 651 | 1145 | 3597 | 655 | 1319 | 3310 | 643 | 1376 | 3551 | 516 | 920 |
| 4687 | QGN17784.1 | FIM1_4993 | CP015060. | 4736 | 1088 | 1314 | 4845 | 1565 | 1111 | 4736 | 1583 | 1768 | 4063 | 1462 | 1858 | 9239 | 1312 | 1766 | 5833 | 1370 | 1285 |
| 4688 | QGN17785.1 | FIM1_4994 | CP015060. | 1 | 0 | 0 | 0 | 0 | 0 | 0 | 0 | 0 | 1 | 1 | 0 | 1 | 0 | 0 | 1 | 0 | 0 |
| 4689 | QGN17786.1 | FIM1_4995 | CP015060. | 3713 | 785 | 1128 | 7510 | 1151 | 974 | 4897 | 754 | 1371 | 3231 | 863 | 1279 | 4622 | 1250 | 1176 | 3447 | 1030 | 901 |
| 4690 | QGN17787.1 | FIM1_4996 | CP015060. | 710 | 315 | 332 | 1511 | 330 | 270 | 1004 | 290 | 380 | 2282 | 271 | 490 | 1975 | 265 | 441 | 2161 | 221 | 292 |
| 4691 | QGN17788.1 | FIM1_4997 | CP015060. | 2563 | 1313 | 1185 | 2420 | 991 | 925 | 2364 | 941 | 1488 | 7403 | 782 | 1571 | 6001 | 668 | 1563 | 7525 | 680 | 1061 |
| 4692 | QGN17789.1 | FIM1_4998 | CP015060. | 4809 | 1435 | 1633 | 8095 | 1323 | 1366 | 5470 | 1196 | 2030 | 5492 | 1123 | 2353 | 7370 | 1625 | 2126 | 5997 | 1202 | 1453 |
| 4693 | QGN17790.1 | FIM1_4999 | CP015060. | 4128 | 3215 | 1845 | 3930 | 2697 | 1816 | 4516 | 2432 | 2441 | 11303 | 3142 | 2968 | 9964 | 2607 | 2953 | 10709 | 2921 | 2014 |
| 4694 | QGN17791.1 | FIM1_5000 | CP015060. | 2502 | 303 | 262 | 2273 | 308 | 220 | 2357 | 289 | 346 | 508 | 219 | 440 | 2120 | 204 | 377 | 989 | 215 | 281 |
| 4695 | QGN17792.1 | FIM1_5001 | CP015060. | 6744 | 258 | 391 | 5905 | 402 | 345 | 7125 | 341 | 504 | 1820 | 239 | 377 | 2797 | 523 | 347 | 1904 | 307 | 259 |
| 4696 | QGN17793.1 | FIM1_5002 | CP015060. | 3342 | 735 | 1069 | 6302 | 1071 | 923 | 4570 | 839 | 1396 | 3990 | 824 | 1594 | 6899 | 715 | 1346 | 5215 | 612 | 1052 |
| 4697 | QGN17794.1 | FIM1_5003 | CP015060. | 1430 | 1005 | 1036 | 1708 | 1040 | 853 | 1197 | 1195 | 1256 | 4116 | 748 | 971 | 4479 | 558 | 1081 | 3702 | 521 | 657 |
| 4698 | QGN17795.1 | FIM1_5004 | CP015060. | 7966 | 3266 | 1405 | 4357 | 2185 | 1321 | 3761 | 2400 | 1967 | 8836 | 2690 | 2314 | 8175 | 2098 | 2435 | 8039 | 2627 | 1720 |
| 4699 | QGN17796.1 | FIM1_5005 | CP015060. | 17326 | 19137 | 12239 | 17611 | 15161 | 11214 | 15919 | 13339 | 14083 | 43466 | 17589 | 17676 | 39192 | 15344 | 17525 | 45473 | 15430 | 13489 |
| 4700 | QGN17797.1 | FIM1_5006 | CP015060. | 329 | 156 | 207 | 414 | 138 | 169 | 340 | 121 | 276 | 1510 | 117 | 309 | 1144 | 111 | 328 | 1584 | 107 | 192 |
| 4701 | QGN17798.1 | FIM1_5007 | CP015060. | 7871 | 1014 | 1171 | 6800 | 1580 | 903 | 6868 | 1553 | 1597 | 1032 | 1129 | 1867 | 6665 | 1300 | 1598 | 2092 | 1374 | 1245 |
| 4702 | QGN17799.1 | FIM1_5008 | CP015060. | 5055 | 4159 | 2844 | 7260 | 4924 | 2967 | 4216 | 3598 | 3755 | 6883 | 2413 | 3106 | 10173 | 2700 | 3374 | 7627 | 2367 | 2118 |

| | | | | | | | | | | | | | | | | | | | | | |
|---|---|---|---|---|---|---|---|---|---|---|---|---|---|---|---|---|---|---|---|---|---|
| 4703 | QGN17800.1 | FIM1_5009 | CP015060. | 777 | 273 | 116 | 377 | 141 | 66 | 255 | 135 | 116 | 127 | 63 | 129 | 227 | 26 | 84 | 143 | 31 | 53 |
| 4704 | QGN17801.1 | FIM1_5010 | CP015060. | 2844 | 902 | 742 | 2228 | 861 | 632 | 2202 | 898 | 998 | 894 | 576 | 809 | 2634 | 501 | 785 | 1664 | 493 | 544 |
| 4705 | QGN17802.1 | FIM1_5011 | CP015060. | 110 | 75 | 149 | 152 | 150 | 80 | 120 | 140 | 149 | 430 | 114 | 174 | 637 | 107 | 160 | 476 | 124 | 113 |
| 4706 | QGN17803.1 | FIM1_5012 | CP015060. | 525 | 777 | 865 | 1056 | 613 | 874 | 654 | 514 | 1066 | 2874 | 517 | 1250 | 1943 | 522 | 1197 | 2455 | 539 | 848 |
| 4707 | QGN17804.1 | FIM1_5013 | CP015060. | 1997 | 1081 | 900 | 2038 | 812 | 752 | 1566 | 785 | 1217 | 1871 | 689 | 1097 | 3248 | 674 | 1219 | 2187 | 644 | 775 |
| 4708 | QGN17805.1 | FIM1_5014 | CP015060. | 3366 | 1286 | 1811 | 4371 | 1233 | 1787 | 4153 | 1394 | 2435 | 6984 | 1040 | 3002 | 7796 | 848 | 3148 | 6849 | 829 | 2185 |
| 4709 | QGN17806.1 | FIM1_5015 | CP015060. | 1176 | 2897 | 1972 | 2974 | 2439 | 1833 | 2126 | 2225 | 2638 | 8806 | 2007 | 2569 | 8407 | 1837 | 2590 | 8375 | 1792 | 1792 |
| 4710 | QGN17807.1 | FIM1_5016 | CP015060. | 1912 | 183 | 558 | 3389 | 435 | 500 | 1911 | 299 | 718 | 2077 | 183 | 505 | 2768 | 290 | 433 | 2261 | 172 | 319 |
| 4711 | QGN17808.1 | FIM1_5017 | CP015060. | 1169 | 1438 | 1582 | 2057 | 1963 | 1348 | 1619 | 1621 | 2083 | 2918 | 1602 | 1695 | 8227 | 1619 | 1539 | 4741 | 1614 | 1151 |
| 4712 | QGN17809.1 | FIM1_5018 | CP015060. | 399 | 329 | 321 | 540 | 224 | 272 | 316 | 197 | 423 | 1691 | 193 | 325 | 1236 | 185 | 337 | 1478 | 170 | 250 |
| 4713 | QGN17810.1 | FIM1_5019 | CP015060. | 3830 | 545 | 578 | 2668 | 531 | 416 | 3242 | 708 | 749 | 653 | 509 | 777 | 2097 | 646 | 665 | 919 | 634 | 481 |
| 4714 | QGN17811.1 | FIM1_5020 | CP015060. | 5034 | 418 | 501 | 3582 | 455 | 319 | 4524 | 529 | 699 | 423 | 380 | 801 | 2172 | 484 | 647 | 809 | 498 | 484 |
| 4715 | QGN17812.1 | FIM1_5021 | CP015060. | 3219 | 389 | 438 | 2945 | 342 | 325 | 2704 | 312 | 559 | 2275 | 241 | 639 | 2326 | 252 | 534 | 2280 | 222 | 382 |
| 4716 | QGN17813.1 | FIM1_5022 | CP015060. | 9622 | 979 | 1372 | 9716 | 1296 | 1213 | 9233 | 1212 | 1847 | 5991 | 1060 | 1779 | 7295 | 1005 | 1596 | 6324 | 995 | 1152 |
| 4717 | QGN17814.1 | FIM1_5023 | CP015060. | 2829 | 1603 | 1090 | 2429 | 1651 | 983 | 2201 | 1414 | 1368 | 4541 | 1658 | 1596 | 5187 | 1297 | 1630 | 4355 | 1333 | 1067 |
| 4718 | QGN17815.1 | FIM1_5024 | CP015060. | 1745 | 2184 | 1818 | 2131 | 2756 | 1638 | 2109 | 2160 | 2409 | 14950 | 2627 | 2897 | 12968 | 2438 | 2435 | 14687 | 2579 | 1780 |
| 4719 | QGN17816.1 | FIM1_5025 | CP015060. | 3372 | 2186 | 2022 | 3909 | 1218 | 1797 | 4085 | 2010 | 3030 | 10750 | 1745 | 2428 | 10615 | 926 | 2252 | 10896 | 1371 | 1579 |
| 4720 | QGN17817.1 | FIM1_5026 | CP015060. | 14988 | 889 | 632 | 13854 | 1068 | 595 | 16037 | 902 | 896 | 5130 | 990 | 1080 | 4880 | 1378 | 975 | 4811 | 1240 | 722 |
| 4721 | QGN17818.1 | FIM1_5027 | CP015060. | 5275 | 2690 | 1999 | 4143 | 3697 | 1655 | 4587 | 2890 | 2620 | 7106 | 2713 | 1989 | 6939 | 2699 | 1773 | 7271 | 2697 | 1308 |
| 4722 | QGN17819.1 | FIM1_5028 | CP015060. | 6324 | 1638 | 1699 | 9187 | 1999 | 1497 | 8813 | 1802 | 2189 | 12121 | 2357 | 2748 | 12690 | 2609 | 2669 | 11501 | 1886 | 1840 |
| 4723 | QGN17820.1 | FIM1_5029 | CP015060. | 4046 | 1228 | 1820 | 4209 | 1354 | 1455 | 4320 | 1504 | 2292 | 9711 | 1728 | 2857 | 10721 | 1474 | 2556 | 10077 | 1458 | 1976 |
| 4724 | QGN17821.1 | FIM1_5030 | CP015060. | 2028 | 2730 | 3436 | 2586 | 2528 | 2912 | 2575 | 2418 | 4358 | 9484 | 2158 | 4204 | 9389 | 2094 | 4286 | 8758 | 1878 | 2938 |
| 4725 | QGN17822.1 | FIM1_5031 | CP015060. | 5767 | 3534 | 2662 | 5750 | 4563 | 2270 | 4916 | 3469 | 3371 | 13519 | 5103 | 3796 | 15878 | 3859 | 3582 | 14976 | 4386 | 2547 |
| 4726 | QGN17823.1 | FIM1_5032 | CP015060. | 2730 | 1508 | 815 | 1379 | 1841 | 948 | 2096 | 1545 | 1149 | 4779 | 2095 | 1308 | 5302 | 2353 | 1251 | 4425 | 2268 | 979 |
| 4727 | QGN17824.1 | FIM1_5033 | CP015060. | 2270 | 997 | 1324 | 2866 | 933 | 995 | 2751 | 1021 | 1668 | 5028 | 970 | 1721 | 5805 | 917 | 1756 | 5092 | 842 | 1218 |
| 4728 | QGN17825.1 | FIM1_5034 | CP015060. | 9089 | 2213 | 4430 | 11039 | 1809 | 3579 | 9998 | 2385 | 5543 | 7566 | 1909 | 7258 | 9123 | 2427 | 7093 | 7563 | 2011 | 5015 |
| 4729 | QGN17826.1 | FIM1_5035 | CP015060. | 2263 | 959 | 664 | 1905 | 767 | 609 | 1944 | 846 | 893 | 918 | 601 | 922 | 2940 | 588 | 906 | 1623 | 552 | 651 |
| 4730 | QGN17827.1 | FIM1_5036 | CP015060. | 1686 | 1474 | 952 | 1665 | 906 | 885 | 1462 | 837 | 1239 | 3873 | 1010 | 1061 | 3005 | 796 | 1186 | 3209 | 871 | 809 |
| 4731 | QGN17828.1 | FIM1_5037 | CP015060. | 2668 | 814 | 763 | 3755 | 788 | 727 | 2827 | 785 | 1000 | 4178 | 661 | 1240 | 4439 | 729 | 1148 | 4079 | 654 | 845 |
| 4732 | QGN17829.1 | FIM1_5038 | CP015060. | 3154 | 1682 | 2007 | 4465 | 2313 | 1742 | 3653 | 1854 | 2459 | 14659 | 2389 | 3257 | 12235 | 2118 | 2908 | 14028 | 2234 | 2088 |
| 4733 | QGN17830.1 | FIM1_5039 | CP015060. | 2389 | 635 | 901 | 2730 | 816 | 838 | 2146 | 866 | 1261 | 4025 | 731 | 1326 | 5800 | 664 | 1243 | 4614 | 560 | 886 |
| 4734 | QGN17831.1 | FIM1_5040 | CP015060. | 12817 | 15030 | 8549 | 11557 | 11595 | 8390 | 11398 | 9762 | 10368 | 35858 | 14904 | 13508 | 32251 | 12637 | 14584 | 35947 | 13081 | 10241 |
| 4735 | QGN17832.1 | FIM1_5041 | CP015060. | 622 | 288 | 265 | 1231 | 305 | 235 | 857 | 276 | 279 | 888 | 197 | 356 | 1098 | 245 | 330 | 892 | 228 | 214 |
| 4736 | QGN17833.1 | FIM1_5042 | CP015060. | 23124 | 8927 | 9099 | 28567 | 12800 | 7565 | 23781 | 12125 | 11546 | 28897 | 12831 | 16523 | 56663 | 13732 | 15842 | 39408 | 12115 | 11382 |
| 4737 | QGN17834.1 | FIM1_5043 | CP015060. | 1501 | 1054 | 575 | 1357 | 611 | 571 | 1208 | 520 | 764 | 1676 | 685 | 644 | 2205 | 613 | 852 | 1734 | 645 | 528 |
| 4738 | QGN17835.1 | FIM1_5044 | CP015060. | 314 | 212 | 182 | 282 | 167 | 138 | 334 | 244 | 251 | 185 | 104 | 195 | 485 | 100 | 192 | 234 | 120 | 110 |
| 4739 | QGN17836.1 | FIM1_5045 | CP015060. | 6790 | 1230 | 2164 | 7536 | 1517 | 1809 | 9452 | 1316 | 2671 | 4858 | 998 | 1912 | 6750 | 570 | 1998 | 6308 | 861 | 1213 |
| 4740 | QGN17837.1 | FIM1_5046 | CP015060. | 12450 | 1560 | 1952 | 13849 | 1612 | 1492 | 11452 | 1685 | 2564 | 1958 | 968 | 1690 | 9316 | 1480 | 1483 | 4878 | 1081 | 1085 |
| 4741 | QGN17838.1 | FIM1_5047 | CP015060. | 660 | 1392 | 1099 | 675 | 1433 | 891 | 544 | 1613 | 1519 | 3566 | 1385 | 1205 | 5619 | 1375 | 1082 | 4488 | 1323 | 899 |
| 4742 | QGN17839.1 | FIM1_5048 | CP015060. | 534 | 423 | 557 | 565 | 382 | 610 | 547 | 339 | 691 | 2311 | 316 | 777 | 1915 | 304 | 791 | 2284 | 291 | 531 |
| 4743 | QGN17840.1 | FIM1_5049 | CP015060. | 2304 | 1625 | 1495 | 3634 | 1535 | 1310 | 2965 | 1382 | 1878 | 4819 | 1509 | 2306 | 7385 | 1397 | 2109 | 5958 | 1352 | 1590 |
| 4744 | QGN17841.1 | FIM1_5050 | CP015060. | 6437 | 882 | 1271 | 7049 | 1159 | 983 | 6597 | 993 | 1641 | 3687 | 815 | 1650 | 7913 | 816 | 1589 | 5709 | 691 | 1180 |
| 4745 | QGN17842.1 | FIM1_5051 | CP015060. | 126 | 11 | 10 | 149 | 13 | 13 | 120 | 10 | 14 | 47 | 10 | 13 | 86 | 7 | 18 | 74 | 7 | 8 |
| 4746 | QGN17843.1 | FIM1_5052 | CP015060. | 1695 | 591 | 580 | 1617 | 694 | 621 | 1464 | 576 | 828 | 2288 | 497 | 571 | 2066 | 514 | 594 | 2035 | 457 | 400 |
| 4747 | QGN17844.1 | FIM1_5053 | CP015060. | 749 | 1187 | 846 | 1101 | 937 | 814 | 969 | 827 | 1148 | 4376 | 945 | 1194 | 3652 | 898 | 1155 | 3766 | 949 | 738 |
| 4748 | QGN17845.1 | FIM1_5054 | CP015060. | 634 | 538 | 398 | 944 | 376 | 371 | 573 | 406 | 532 | 3015 | 490 | 615 | 2533 | 379 | 721 | 3095 | 381 | 394 |
| 4749 | QGN17846.1 | FIM1_5055 | CP015060. | 1571 | 2926 | 2080 | 2355 | 2591 | 1843 | 1887 | 2277 | 2554 | 13946 | 2608 | 3034 | 12023 | 2268 | 2806 | 13687 | 2319 | 2006 |
| 4750 | QGN17847.1 | FIM1_5056 | CP015060. | 2073 | 580 | 898 | 3224 | 786 | 702 | 2380 | 681 | 1081 | 6667 | 666 | 1473 | 5793 | 673 | 1355 | 6405 | 524 | 982 |
| 4751 | QGN17848.1 | FIM1_5057 | CP015060. | 6870 | 3512 | 1907 | 2801 | 4417 | 2019 | 5678 | 3368 | 2471 | 18198 | 6218 | 3996 | 16743 | 5723 | 3692 | 17915 | 6887 | 2960 |
| 4752 | QGN17849.1 | FIM1_5058 | CP015060. | 3233 | 1735 | 3432 | 1951 | 1843 | 2586 | 1699 | 2216 | 4057 | 6753 | 1528 | 2917 | 6709 | 992 | 2780 | 7508 | 1206 | 1924 |
| 4753 | QGN17850.1 | FIM1_5059 | CP015060. | 920 | 588 | 444 | 1171 | 439 | 371 | 1002 | 466 | 619 | 2029 | 356 | 602 | 2201 | 330 | 562 | 2128 | 332 | 425 |
| 4754 | QGN17851.1 | FIM1_5060 | CP015060. | 60 | 6 | 6 | 25 | 11 | 5 | 13 | 5 | 2 | 63 | 8 | 6 | 46 | 4 | 8 | 41 | 12 | 5 |
| 4755 | QGN17852.1 | FIM1_5061 | CP015060. | 3540 | 1813 | 1643 | 2473 | 1798 | 1407 | 2356 | 1468 | 2018 | 7830 | 1480 | 1786 | 6081 | 1451 | 1668 | 7048 | 1367 | 1162 |
| 4756 | QGN17853.1 | FIM1_5062 | CP015060. | 1005 | 308 | 500 | 1607 | 296 | 373 | 1172 | 316 | 624 | 2652 | 229 | 643 | 2348 | 241 | 587 | 2508 | 199 | 436 |
| 4757 | QGN17854.1 | FIM1_5063 | CP015060. | 1360 | 1989 | 1347 | 1565 | 1308 | 1122 | 1445 | 1069 | 1629 | 6356 | 1611 | 2153 | 5413 | 1474 | 2055 | 6032 | 1449 | 1416 |
| 4758 | QGN17855.1 | FIM1_5064 | CP015060. | 543 | 354 | 407 | 687 | 503 | 350 | 705 | 440 | 515 | 662 | 422 | 584 | 1734 | 406 | 570 | 1218 | 424 | 438 |
| 4759 | QGN17856.1 | FIM1_5065 | CP015060. | 11346 | 1508 | 2316 | 9837 | 2122 | 1820 | 9894 | 1950 | 2927 | 4448 | 2374 | 2975 | 11980 | 2305 | 3103 | 8126 | 2040 | 2138 |
| 4760 | QGN17857.1 | FIM1_5066 | CP015060. | 1225 | 362 | 292 | 1261 | 322 | 303 | 1112 | 282 | 406 | 1008 | 224 | 358 | 1030 | 191 | 409 | 1042 | 220 | 227 |
| 4761 | QGN17858.1 | FIM1_5067 | CP015060. | 1977 | 3208 | 1959 | 2287 | 2894 | 2024 | 2087 | 2489 | 2691 | 12398 | 2638 | 2750 | 11695 | 2459 | 2652 | 11691 | 2579 | 1793 |
| 4762 | QGN17859.1 | FIM1_5068 | CP015060. | 397 | 1238 | 950 | 1110 | 978 | 845 | 703 | 932 | 1267 | 3699 | 640 | 996 | 3414 | 516 | 932 | 3578 | 510 | 676 |
| 4763 | QGN17860.1 | FIM1_5069 | CP015060. | 9426 | 78 | 99 | 8380 | 60 | 83 | 14270 | 104 | 140 | 351 | 53 | 138 | 979 | 137 | 177 | 430 | 66 | 132 |
| 4764 | QGN17861.1 | FIM1_5070 | CP015060. | 130 | 41 | 55 | 203 | 43 | 67 | 162 | 26 | 70 | 108 | 23 | 56 | 116 | 29 | 86 | 111 | 25 | 62 |
| 4765 | QGN17862.1 | FIM1_5071 | CP015060. | 4407 | 8077 | 6969 | 6060 | 5981 | 6273 | 5220 | 6035 | 8819 | 8648 | 4357 | 7094 | 12757 | 4659 | 7566 | 10615 | 4234 | 5226 |
| 4766 | QGN17863.1 | FIM1_5072 | CP015060. | 91 | 106 | 130 | 179 | 70 | 98 | 98 | 110 | 157 | 199 | 45 | 175 | 430 | 44 | 146 | 228 | 46 | 82 |
| 4767 | QGN17864.1 | FIM1_5073 | CP015060. | 413 | 661 | 495 | 656 | 423 | 404 | 449 | 424 | 601 | 2145 | 357 | 660 | 1901 | 285 | 691 | 2020 | 305 | 446 |
| 4768 | QGN17865.1 | FIM1_5074 | CP015060. | 834 | 696 | 425 | 1482 | 476 | 338 | 1181 | 476 | 535 | 2444 | 300 | 586 | 1939 | 389 | 603 | 2145 | 373 | 421 |
| 4769 | QGN17866.1 | FIM1_5075 | CP015060. | 3857 | 461 | 558 | 5417 | 704 | 468 | 4607 | 578 | 754 | 1121 | 513 | 882 | 2292 | 531 | 974 | 1480 | 519 | 671 |
| 4770 | QGN17867.1 | FIM1_5076 | CP015060. | 2388 | 3538 | 2720 | 2611 | 2276 | 2389 | 2161 | 2216 | 3451 | 7196 | 2748 | 3927 | 8028 | 2809 | 4240 | 7078 | 2753 | 2804 |
| 4771 | QGN17868.1 | FIM1_5077 | CP015060. | 2474 | 1832 | 1753 | 3630 | 1879 | 1402 | 3029 | 1919 | 2206 | 2560 | 1159 | 1928 | 5857 | 1187 | 2005 | 3586 | 1127 | 1360 |
| 4772 | QGN17869.1 | FIM1_5078 | CP015060. | 878 | 227 | 292 | 1619 | 279 | 275 | 1236 | 236 | 357 | 1297 | 167 | 419 | 1491 | 181 | 336 | 1190 | 158 | 259 |
| 4773 | QGN17870.1 | FIM1_5079 | CP015060. | 4938 | 5793 | 3087 | 7461 | 6256 | 3418 | 6458 | 4511 | 3983 | 17149 | 4951 | 5136 | 15882 | 4354 | 5179 | 16092 | 4476 | 3574 |
| 4774 | QGN17871.1 | FIM1_5080 | CP015060. | 1252 | 864 | 744 | 1477 | 839 | 667 | 1381 | 770 | 950 | 3418 | 535 | 956 | 3559 | 473 | 909 | 3393 | 498 | 674 |
| 4775 | QGN17872.1 | FIM1_5081 | CP015060. | 2013 | 1351 | 1027 | 3376 | 1080 | 880 | 2713 | 1048 | 1348 | 4454 | 869 | 1421 | 4316 | 1009 | 1406 | 4243 | 842 | 1008 |
| 4776 | QGN17873.1 | FIM1_5082 | CP015060. | 357 | 394 | 350 | 533 | 334 | 276 | 437 | 307 | 388 | 1986 | 441 | 589 | 1490 | 381 | 494 | 1969 | 408 | 403 |
| 4777 | QGN17874.1 | FIM1_5083 | CP015060. | 5160 | 3051 | 3094 | 6005 | 3085 | 2487 | 5337 | 2820 | 4101 | 3851 | 3061 | 4535 | 14866 | 3340 | 4647 | 7372 | 3302 | 3316 |
| 4778 | QGN17875.1 | FIM1_5084 | CP015060. | 2116 | 469 | 444 | 863 | 632 | 428 | 1525 | 499 | 541 | 821 | 536 | 638 | 1546 | 778 | 662 | 1099 | 659 | 469 |
| 4779 | QGN17876.1 | FIM1_5085 | CP015060. | 2544 | 694 | 1128 | 3570 | 916 | 855 | 2115 | 865 | 1523 | 2688 | 883 | 1426 | 7831 | 1245 | 1546 | 4407 | 756 | 1037 |
| 4780 | QGN17877.1 | FIM1_5086 | CP015060. | 4756 | 1227 | 1621 | 4950 | 1503 | 1403 | 4883 | 1443 | 2164 | 5534 | 1613 | 2404 | 10627 | 1520 | 2247 | 8218 | 1450 | 1636 |
| 4781 | QGN17878.1 | FIM1_5087 | CP015060. | 604 | 894 | 1315 | 1627 | 783 | 1266 | 895 | 596 | 1682 | 4007 | 686 | 2220 | 3197 | 709 | 2029 | 3733 | 694 | 1438 |
| 4782 | QGN17879.1 | FIM1_5088 | CP015060. | 10879 | 5885 | 6086 | 20088 | 5603 | 5084 | 12744 | 5015 | 7634 | 6818 | 5996 | 8976 | 23201 | 6463 | 8012 | 15667 | 5943 | 5837 |
| 4783 | QGN17880.1 | FIM1_5089 | CP015060. | 2376 | 2754 | 2494 | 1241 | 1684 | 1794 | 1642 | 2723 | 3266 | 2444 | 1409 | 2504 | 6924 | 1100 | 2673 | 4451 | 1262 | 1766 |
| 4784 | QGN17881.1 | FIM1_5090 | CP015060. | 753 | 111 | 278 | 971 | 214 | 168 | 762 | 300 | 330 | 253 | 183 | 377 | 918 | 184 | 347 | 438 | 154 | 258 |
| 4785 | QGN17882.1 | FIM1_5091 | CP015060. | 247 | 784 | 398 | 850 | 581 | 365 | 259 | 559 | 622 | 632 | 241 | 520 | 926 | 246 | 602 | 711 | 210 | 384 |
| 4786 | QGN17883.1 | FIM1_5092 | CP015060. | 27 | 4 | 4 | 25 | 4 | 4 | 20 | 2 | 8 | 41 | 7 | 14 | 37 | 2 | 4 | 48 | 1 | 6 |

| | | | | | | | | | | | | | | | | | | | | | |
|---|---|---|---|---|---|---|---|---|---|---|---|---|---|---|---|---|---|---|---|---|---|
| 4787 | QGN17884.1 | FIM1_5093 | CP015060. | 2078 | 1084 | 1163 | 2268 | 867 | 1029 | 2164 | 870 | 1545 | 2522 | 490 | 1135 | 3124 | 586 | 1196 | 2477 | 487 | 822 |
| 4788 | QGN17885.1 | FIM1_5094 | CP015060. | 93744 | 74124 | 58561 | 95472 | 65965 | 51529 | 79499 | 59914 | 65833 | 117024 | 75746 | 80865 | 163915 | 70192 | 79707 | 139621 | 63518 | 63244 |
| 4789 | QGN17886.1 | FIM1_5095 | CP015060. | 303 | 372 | 345 | 478 | 369 | 280 | 319 | 291 | 472 | 2045 | 326 | 520 | 1708 | 301 | 454 | 2038 | 282 | 303 |
| 4790 | QGN17887.1 | FIM1_5096 | CP015060. | 1330 | 1198 | 678 | 1123 | 1013 | 631 | 1250 | 746 | 886 | 6061 | 942 | 1180 | 4498 | 1021 | 1013 | 6346 | 1109 | 715 |
| 4791 | QGN17888.1 | FIM1_5097 | CP015060. | 10870 | 6756 | 4468 | 14105 | 6375 | 3884 | 10558 | 5775 | 5660 | 19686 | 6917 | 8625 | 27786 | 7027 | 7647 | 21146 | 6727 | 5484 |
| 4792 | QGN17889.1 | FIM1_5098 | CP015060. | 516 | 84 | 196 | 621 | 200 | 139 | 518 | 221 | 219 | 691 | 161 | 231 | 1133 | 157 | 195 | 865 | 147 | 154 |
| 4793 | QGN17890.1 | FIM1_5099 | CP015060. | 528 | 1816 | 1129 | 535 | 1420 | 1099 | 521 | 1239 | 1545 | 5211 | 1356 | 1520 | 5040 | 1313 | 1541 | 5394 | 1381 | 991 |
| 4794 | QGN17891.1 | FIM1_5100 | CP015060. | 1678 | 584 | 719 | 1964 | 578 | 633 | 1544 | 543 | 979 | 3747 | 549 | 915 | 4053 | 486 | 907 | 3655 | 518 | 614 |
| 4795 | QGN17892.1 | FIM1_5101 | CP015060. | 4454 | 741 | 756 | 3861 | 661 | 671 | 4021 | 650 | 1007 | 1848 | 505 | 1015 | 2990 | 571 | 959 | 2327 | 454 | 672 |
| 4796 | QGN17893.1 | FIM1_5102 | CP015060. | 23212 | 1255 | 1392 | 22255 | 1348 | 1214 | 22049 | 1385 | 1965 | 1496 | 782 | 1622 | 5442 | 1060 | 1393 | 2869 | 918 | 1056 |
| 4797 | QGN17894.1 | FIM1_5103 | CP015060. | 481 | 581 | 470 | 521 | 399 | 407 | 333 | 376 | 650 | 1078 | 290 | 558 | 1285 | 284 | 566 | 1023 | 313 | 366 |
| 4798 | QGN17895.1 | FIM1_5104 | CP015060. | 1086 | 525 | 514 | 1264 | 509 | 413 | 1077 | 541 | 666 | 2415 | 452 | 649 | 2179 | 429 | 677 | 2205 | 449 | 448 |
| 4799 | QGN17896.1 | FIM1_5105 | CP015060. | 3007 | 2115 | 1387 | 3522 | 1882 | 1339 | 2996 | 1561 | 1888 | 7704 | 1967 | 2208 | 7613 | 1864 | 2329 | 7386 | 1809 | 1577 |
| 4800 | QGN17897.1 | FIM1_5106 | CP015060. | 1584 | 407 | 595 | 2413 | 382 | 511 | 1555 | 385 | 885 | 2960 | 298 | 768 | 3341 | 384 | 722 | 3057 | 266 | 462 |
| 4801 | QGN17898.1 | FIM1_5107 | CP015060. | 2501 | 2994 | 2510 | 5469 | 3130 | 2038 | 4574 | 2835 | 2905 | 10914 | 3846 | 3746 | 10548 | 3700 | 3927 | 10892 | 3240 | 2969 |
| 4802 | QGN17899.1 | FIM1_5108 | CP015060. | 786 | 808 | 605 | 590 | 668 | 589 | 680 | 758 | 860 | 2264 | 570 | 623 | 3274 | 586 | 689 | 2582 | 503 | 479 |
| 4803 | QGN17900.1 | FIM1_5109 | CP015060. | 880 | 664 | 588 | 696 | 820 | 523 | 1046 | 573 | 739 | 4669 | 871 | 864 | 3959 | 758 | 791 | 4462 | 700 | 566 |
| 4804 | QGN17901.1 | FIM1_5110 | CP015060. | 7979 | 736 | 705 | 6775 | 649 | 653 | 7195 | 636 | 954 | 6662 | 698 | 1215 | 5232 | 681 | 1075 | 6457 | 680 | 760 |
| 4805 | QGN17902.1 | FIM1_5111 | CP015060. | 1961 | 907 | 1052 | 2378 | 1132 | 836 | 2225 | 1007 | 1237 | 8358 | 1178 | 1554 | 7127 | 1011 | 1417 | 7567 | 1059 | 1104 |
| 4806 | QGN17903.1 | FIM1_5112 | CP015060. | 2264 | 655 | 573 | 1831 | 531 | 530 | 1982 | 538 | 774 | 2811 | 500 | 694 | 2447 | 433 | 736 | 2709 | 432 | 463 |
| 4807 | QGN17904.1 | FIM1_5113 | CP015060. | 563 | 150 | 65 | 484 | 119 | 79 | 422 | 88 | 119 | 756 | 104 | 111 | 522 | 110 | 135 | 826 | 97 | 84 |
| 4808 | QGN17905.1 | FIM1_5114 | CP015060. | 1176 | 271 | 211 | 623 | 337 | 178 | 1073 | 337 | 305 | 1351 | 322 | 295 | 1318 | 322 | 294 | 1341 | 325 | 215 |
| 4809 | QGN17906.1 | FIM1_5115 | CP015060. | 916 | 1435 | 864 | 1369 | 1304 | 753 | 1375 | 1122 | 1102 | 5410 | 1507 | 1396 | 5345 | 1561 | 1453 | 5028 | 1498 | 994 |
| 4810 | QGN17907.1 | FIM1_5116 | CP015060. | 822 | 1337 | 1033 | 1401 | 1138 | 752 | 945 | 1125 | 1233 | 7143 | 1154 | 1632 | 5641 | 1186 | 1504 | 6772 | 1095 | 1083 |
| 4811 | QGN17908.1 | FIM1_5117 | CP015060. | 1124 | 400 | 427 | 614 | 215 | 305 | 1209 | 456 | 590 | 1588 | 211 | 374 | 1676 | 204 | 377 | 1555 | 261 | 261 |
| 4812 | QGN17909.1 | FIM1_5118 | CP015060. | 2367 | 3395 | 2058 | 2339 | 2125 | 1991 | 2020 | 2196 | 2583 | 7695 | 1903 | 2506 | 6236 | 1810 | 2637 | 7141 | 2071 | 1729 |
| 4813 | QGN17910.1 | FIM1_5119 | CP015060. | 1725 | 1448 | 1240 | 1931 | 1038 | 991 | 1629 | 1002 | 1524 | 4243 | 776 | 1268 | 3691 | 842 | 1354 | 3935 | 624 | 864 |
| 4814 | QGN17911.1 | FIM1_5120 | CP015060. | 1509 | 730 | 841 | 1932 | 794 | 625 | 1758 | 810 | 1040 | 1224 | 524 | 1230 | 3724 | 671 | 955 | 2112 | 741 | 753 |
| 4815 | QGN17912.1 | FIM1_5121 | CP015060. | 1018 | 2206 | 1944 | 917 | 1490 | 1744 | 919 | 1412 | 2434 | 6652 | 1526 | 2555 | 4967 | 1450 | 2565 | 6175 | 1425 | 1676 |
| 4816 | QGN17913.1 | FIM1_5122 | CP015060. | 15487 | 1355 | 2167 | 17451 | 1884 | 1497 | 15542 | 1937 | 2855 | 1877 | 1444 | 2839 | 13426 | 1659 | 2601 | 5081 | 1666 | 1977 |
| 4817 | QGN17914.1 | FIM1_5123 | CP015060. | 1064 | 3453 | 2695 | 6023 | 3435 | 3100 | 1603 | 3264 | 3636 | 6007 | 2733 | 4104 | 9081 | 1611 | 2873 | 7053 | 3118 | 2623 |
| 4818 | QGN17915.1 | FIM1_5124 | CP015060. | 453 | 621 | 648 | 610 | 598 | 548 | 606 | 568 | 937 | 3621 | 728 | 963 | 3715 | 729 | 931 | 3657 | 590 | 697 |
| 4819 | QGN17916.1 | FIM1_5125 | CP015060. | 2528 | 1426 | 1766 | 1121 | 2294 | 1360 | 1996 | 2940 | 2204 | 4519 | 2386 | 2303 | 9529 | 1929 | 2368 | 6912 | 1977 | 1589 |
| 4820 | QGN17917.1 | FIM1_5126 | CP015060. | 1736 | 1239 | 931 | 2285 | 1130 | 821 | 2210 | 1143 | 1192 | 4080 | 943 | 1224 | 4593 | 810 | 1093 | 3970 | 814 | 843 |
| 4821 | QGN17918.1 | FIM1_5127 | CP015060. | 7702 | 2243 | 3010 | 12783 | 2833 | 2429 | 9335 | 2909 | 3784 | 9206 | 2548 | 3998 | 14003 | 2918 | 3871 | 10898 | 2538 | 2622 |
| 4822 | QGN17919.1 | FIM1_5128 | CP015060. | 4676 | 2029 | 2120 | 5468 | 1803 | 1626 | 4522 | 2059 | 2735 | 8016 | 2176 | 3146 | 10549 | 2612 | 3183 | 8598 | 2078 | 2175 |
| 4823 | QGN17920.1 | FIM1_5129 | CP015060. | 3339 | 1126 | 691 | 3451 | 820 | 691 | 3684 | 794 | 1075 | 3957 | 816 | 978 | 3831 | 745 | 984 | 3748 | 857 | 623 |
| 4824 | QGN17921.1 | FIM1_5130 | CP015060. | 3383 | 1959 | 1964 | 5544 | 1698 | 1536 | 3981 | 1776 | 2522 | 3459 | 930 | 2313 | 5836 | 1242 | 2175 | 4048 | 951 | 1377 |
| 4825 | QGN17922.1 | FIM1_5131 | CP015060. | 1138 | 879 | 3284 | 7882 | 820 | 1968 | 4090 | 1737 | 4163 | 6967 | 911 | 2761 | 12409 | 2158 | 2568 | 8595 | 919 | 1984 |
| 4826 | QGN17923.1 | FIM1_5132 | CP015060. | 666 | 113 | 124 | 547 | 109 | 73 | 521 | 121 | 165 | 49 | 67 | 160 | 438 | 63 | 136 | 131 | 87 | 84 |
| 4827 | QGN17924.1 | FIM1_5133 | CP015060. | 3912 | 1050 | 1146 | 3696 | 784 | 905 | 3632 | 825 | 1536 | 2915 | 638 | 1571 | 5765 | 693 | 1526 | 3631 | 638 | 1180 |
| 4828 | QGN17925.1 | FIM1_5134 | CP015060. | 304 | 676 | 427 | 226 | 480 | 383 | 284 | 403 | 545 | 2023 | 416 | 559 | 1599 | 358 | 602 | 1843 | 400 | 361 |
| 4829 | QGN17926.1 | FIM1_5135 | CP015060. | 911 | 1204 | 873 | 889 | 895 | 805 | 787 | 726 | 1120 | 5905 | 1026 | 1307 | 4242 | 902 | 1359 | 5254 | 975 | 772 |
| 4830 | QGN17927.1 | FIM1_5136 | CP015060. | 2055 | 2559 | 2122 | 3747 | 1822 | 1892 | 2460 | 1745 | 2783 | 8049 | 1640 | 2838 | 7215 | 1798 | 2903 | 7515 | 1472 | 1980 |
| 4831 | QGN17928.1 | FIM1_5137 | CP015060. | 723 | 823 | 703 | 1626 | 799 | 678 | 835 | 859 | 911 | 1593 | 537 | 838 | 2873 | 490 | 790 | 1955 | 436 | 536 |
| 4832 | QGN17929.1 | FIM1_5138 | CP015060. | 2197 | 2054 | 1439 | 2844 | 1982 | 1104 | 1828 | 1777 | 1870 | 1023 | 1336 | 1704 | 4804 | 1338 | 1830 | 2244 | 1411 | 1211 |
| 4833 | QGN17930.1 | FIM1_5139 | CP015060. | 1470 | 1685 | 2329 | 4214 | 2466 | 1847 | 2396 | 2010 | 3039 | 2474 | 2107 | 4266 | 8975 | 2138 | 4069 | 4783 | 1943 | 2890 |
| 4834 | QGN17931.1 | FIM1_5140 | CP015060. | 725 | 2715 | 2467 | 4039 | 2277 | 2133 | 2262 | 2428 | 3140 | 9252 | 2123 | 3326 | 10388 | 2838 | 3112 | 9054 | 2640 | 2168 |
| 4835 | QGN17932.1 | FIM1_5141 | CP015060. | 8735 | 1008 | 1268 | 10146 | 1481 | 987 | 10042 | 1303 | 1665 | 5520 | 1254 | 1739 | 9868 | 906 | 1669 | 6903 | 963 | 1200 |
| 4836 | QGN17933.1 | FIM1_5142 | CP015060. | 2721 | 752 | 1294 | 3079 | 996 | 1021 | 2405 | 1232 | 1689 | 2280 | 818 | 1689 | 6524 | 640 | 1395 | 3770 | 701 | 1056 |
| 4837 | QGN17934.1 | FIM1_5143 | CP015060. | 4836 | 1765 | 1419 | 4398 | 1727 | 1271 | 3921 | 1493 | 1802 | 7532 | 1595 | 1806 | 7267 | 1481 | 1880 | 7564 | 1340 | 1230 |
| 4838 | QGN17935.1 | FIM1_5144 | CP015060. | 2587 | 1649 | 1474 | 2537 | 1342 | 1351 | 2223 | 1292 | 1950 | 7238 | 877 | 1967 | 6366 | 638 | 1889 | 6448 | 680 | 1306 |
| 4839 | QGN17936.1 | FIM1_5145 | CP015060. | 3037 | 1264 | 2061 | 3417 | 1367 | 1706 | 3004 | 1136 | 2545 | 6119 | 952 | 2952 | 5805 | 1363 | 2537 | 6103 | 1102 | 1843 |
| 4840 | QGN17937.1 | FIM1_5146 | CP015060. | 1561 | 437 | 241 | 1096 | 346 | 191 | 1207 | 266 | 299 | 1759 | 357 | 460 | 1252 | 538 | 405 | 1565 | 561 | 234 |
| 4841 | QGN17938.1 | FIM1_5147 | CP015060. | 22447 | 26953 | 17078 | 22830 | 21443 | 16178 | 18965 | 17874 | 19713 | 60355 | 26315 | 25299 | 56060 | 22785 | 25752 | 62136 | 22290 | 19281 |
| 4842 | QGN17939.1 | FIM1_5148 | CP015060. | 378 | 277 | 308 | 395 | 192 | 224 | 346 | 221 | 387 | 140 | 93 | 387 | 505 | 111 | 383 | 288 | 115 | 290 |
| 4843 | QGN17940.1 | FIM1_5149 | CP015060. | 7766 | 4223 | 7545 | 6793 | 2485 | 3533 | 9444 | 5160 | 7834 | 1793 | 1397 | 6516 | 8204 | 983 | 3720 | 4465 | 1017 | 3154 |
| 4844 | QGN17941.1 | FIM1_5150 | CP015060. | 13153 | 6629 | 4228 | 15163 | 5749 | 3450 | 12666 | 5533 | 5340 | 7315 | 6603 | 7771 | 25865 | 6643 | 7147 | 14154 | 6674 | 5387 |
| 4845 | QGN17942.1 | FIM1_5151 | CP015060. | 253 | 229 | 128 | 332 | 163 | 133 | 232 | 130 | 180 | 748 | 144 | 156 | 665 | 121 | 189 | 772 | 139 | 107 |
| 4846 | QGN17943.1 | FIM1_5152 | CP015060. | 444 | 115 | 326 | 411 | 169 | 250 | 219 | 150 | 404 | 1487 | 106 | 371 | 1431 | 101 | 330 | 1515 | 74 | 231 |
| 4847 | QGN17944.1 | FIM1_5153 | CP015060. | 8010 | 2931 | 3156 | 9504 | 3767 | 2689 | 8976 | 3218 | 4145 | 15234 | 4415 | 5516 | 22926 | 4068 | 4824 | 18931 | 4173 | 3745 |
| 4848 | QGN17945.1 | FIM1_5154 | CP015060. | 4 | 2 | 0 | 17 | 7 | 2 | 11 | 0 | 7 | 43 | 2 | 3 | 19 | 0 | 2 | 45 | 3 | 3 |
| 4849 | QGN17946.1 | FIM1_5155 | CP015060. | 1231 | 558 | 623 | 1295 | 526 | 550 | 1426 | 518 | 849 | 5411 | 586 | 1017 | 4253 | 458 | 825 | 5128 | 557 | 657 |
| 4850 | QGN17947.1 | FIM1_5156 | CP015060. | 796 | 1065 | 532 | 516 | 513 | 519 | 796 | 495 | 752 | 2908 | 774 | 650 | 2225 | 596 | 755 | 2940 | 601 | 445 |
| 4851 | QGN17948.1 | FIM1_5157 | CP015060. | 2378 | 1676 | 1414 | 3062 | 1599 | 1277 | 2412 | 1349 | 1866 | 7614 | 1691 | 2152 | 7397 | 1458 | 2103 | 7461 | 1433 | 1536 |
| 4852 | QGN17949.1 | FIM1_5158 | CP015060. | 3855 | 546 | 978 | 3179 | 824 | 751 | 3037 | 891 | 1298 | 2240 | 682 | 1301 | 5616 | 609 | 1205 | 3257 | 564 | 836 |
| 4853 | QGN17950.1 | FIM1_5159 | CP015060. | 999 | 903 | 828 | 1834 | 959 | 753 | 1534 | 975 | 1101 | 2369 | 449 | 711 | 2633 | 465 | 701 | 2319 | 420 | 488 |
| 4854 | QGN17951.1 | FIM1_5160 | CP015060. | 9035 | 7344 | 4061 | 4360 | 9530 | 4632 | 8298 | 7699 | 5422 | 30024 | 12618 | 8416 | 32384 | 8565 | 7912 | 28874 | 11362 | 5869 |
| 4855 | QGN17952.1 | FIM1_5161 | CP015060. | 1539 | 761 | 921 | 1840 | 1034 | 694 | 1512 | 1100 | 1301 | 1576 | 975 | 1369 | 7368 | 948 | 1071 | 3461 | 1083 | 877 |
| 4856 | QGN17953.1 | FIM1_5162 | CP015060. | 2751 | 2130 | 1534 | 3129 | 1718 | 1363 | 2868 | 1496 | 2161 | 10247 | 1867 | 2265 | 8559 | 1651 | 2405 | 10350 | 1643 | 1597 |
| 4857 | QGN17954.1 | FIM1_5163 | CP015060. | 6970 | 1481 | 3025 | 5910 | 2378 | 2486 | 6571 | 2209 | 3762 | 4417 | 1740 | 3381 | 9080 | 1545 | 3093 | 6652 | 1472 | 2358 |
| 4858 | QGN17955.1 | FIM1_5164 | CP015060. | 2365 | 188 | 168 | 2789 | 252 | 167 | 1920 | 191 | 207 | 969 | 162 | 191 | 901 | 159 | 152 | 892 | 138 | 135 |
| 4859 | QGN17956.1 | FIM1_5165 | CP015060. | 3829 | 435 | 711 | 4762 | 652 | 546 | 3613 | 620 | 834 | 3059 | 559 | 880 | 4671 | 526 | 910 | 3414 | 415 | 658 |
| 4860 | QGN17957.1 | FIM1_5166 | CP015060. | 3721 | 1203 | 2142 | 4602 | 1782 | 1602 | 3615 | 1808 | 2717 | 3407 | 1906 | 3175 | 10693 | 1951 | 2666 | 6565 | 1914 | 2040 |
| 4861 | QGN17958.1 | FIM1_5167 | CP015060. | 365 | 544 | 489 | 585 | 460 | 402 | 487 | 486 | 610 | 1484 | 417 | 618 | 2286 | 352 | 623 | 1871 | 323 | 415 |
| 4862 | QGN17959.1 | FIM1_5168 | CP015060. | 2436 | 167 | 470 | 2280 | 297 | 336 | 2722 | 298 | 561 | 2847 | 271 | 733 | 3266 | 283 | 664 | 2946 | 234 | 501 |
| 4863 | QGN17960.1 | FIM1_5169 | CP015060. | 727 | 157 | 164 | 963 | 189 | 173 | 424 | 139 | 205 | 919 | 182 | 318 | 1119 | 117 | 194 | 1137 | 150 | 175 |
| 4864 | QGN17961.1 | FIM1_5170 | CP015060. | 5368 | 1186 | 1323 | 4944 | 1298 | 1201 | 4721 | 1060 | 1711 | 5998 | 1042 | 2064 | 7869 | 1027 | 1854 | 6655 | 991 | 1350 |
| 4865 | QGN17962.1 | FIM1_5171 | CP015060. | 3599 | 327 | 493 | 4131 | 728 | 415 | 3181 | 425 | 603 | 2960 | 501 | 685 | 2769 | 517 | 680 | 2868 | 438 | 444 |
| 4866 | QGN17963.1 | FIM1_5172 | CP015060. | 4652 | 7865 | 3740 | 5491 | 5929 | 3700 | 4863 | 5004 | 4719 | 20444 | 6363 | 5897 | 17072 | 5528 | 6055 | 18701 | 5732 | 4217 |
| 4867 | QGN17964.1 | FIM1_5173 | CP015060. | 2656 | 970 | 1361 | 4233 | 1370 | 1040 | 2752 | 1065 | 1687 | 5006 | 884 | 1492 | 6877 | 886 | 1307 | 5605 | 669 | 968 |
| 4868 | QGN17965.1 | FIM1_5174 | CP015060. | 8776 | 2668 | 2979 | 8287 | 2911 | 2692 | 8006 | 2689 | 3842 | 7241 | 2112 | 3703 | 9407 | 2209 | 3800 | 7196 | 1904 | 2514 |
| 4869 | QGN17966.1 | FIM1_5175 | CP015060. | 2978 | 1917 | 1830 | 2573 | 1645 | 1290 | 2282 | 1612 | 2250 | 15536 | 2381 | 3500 | 13062 | 3400 | 4031 | 13676 | 2650 | 2447 |
| 4870 | QGN17967.1 | FIM1_5176 | CP015060. | 1155 | 668 | 317 | 600 | 649 | 246 | 1034 | 448 | 353 | 1832 | 433 | 342 | 1378 | 481 | 366 | 2008 | 405 | 246 |

| | | | | | | | | | | | | | | | | | | | | | |
|---|---|---|---|---|---|---|---|---|---|---|---|---|---|---|---|---|---|---|---|---|---|
| 4871 | QGN17968.1 | FIM1_5177 | CP015060. | 1549 | 666 | 599 | 583 | 1280 | 568 | 1232 | 1061 | 744 | 4079 | 1299 | 1113 | 4085 | 1508 | 1051 | 3937 | 1575 | 840 |
| 4872 | QGN17969.1 | FIM1_5178 | CP015060. | 2147 | 815 | 672 | 1928 | 640 | 587 | 2124 | 597 | 902 | 3169 | 642 | 1023 | 3222 | 722 | 966 | 2965 | 769 | 659 |
| 4873 | QGN17970.1 | FIM1_5179 | CP015060. | 5319 | 2943 | 1423 | 2522 | 3168 | 1599 | 4592 | 2397 | 2054 | 12568 | 4333 | 3017 | 11803 | 3806 | 2869 | 12728 | 4693 | 2076 |
| 4874 | QGN17971.1 | FIM1_5180 | CP015060. | 1569 | 382 | 434 | 2437 | 590 | 443 | 1646 | 376 | 590 | 2963 | 403 | 721 | 2656 | 400 | 639 | 2863 | 411 | 446 |
| 4875 | QGN17972.1 | FIM1_5181 | CP015060. | 3660 | 3341 | 3398 | 7187 | 3300 | 3085 | 4769 | 2933 | 4440 | 14899 | 2839 | 5706 | 15171 | 3366 | 5012 | 15264 | 2927 | 3668 |
| 4876 | QGN17973.1 | FIM1_5182 | CP015060. | 816 | 1154 | 933 | 1127 | 927 | 884 | 814 | 874 | 1263 | 3724 | 906 | 1502 | 3663 | 892 | 1381 | 3500 | 857 | 980 |
| 4877 | QGN17974.1 | FIM1_5183 | CP015060. | 4228 | 2994 | 2576 | 4088 | 3599 | 2256 | 4361 | 3096 | 3330 | 12602 | 4480 | 4710 | 17206 | 3755 | 4494 | 14810 | 4186 | 3274 |
| 4878 | QGN17975.1 | FIM1_5184 | CP015060. | 723 | 502 | 357 | 952 | 372 | 332 | 910 | 330 | 484 | 3682 | 465 | 586 | 2852 | 376 | 520 | 3870 | 403 | 378 |
| 4879 | QGN17976.1 | FIM1_5185 | CP015060. | 2686 | 2167 | 2141 | 4190 | 2425 | 1983 | 3000 | 1971 | 2746 | 13113 | 2276 | 3297 | 11394 | 2393 | 2866 | 12932 | 2086 | 2022 |
| 4880 | QGN17977.1 | FIM1_5186 | CP015060. | 1883 | 1094 | 1144 | 3092 | 927 | 931 | 2391 | 920 | 1386 | 6024 | 979 | 1697 | 5241 | 944 | 1646 | 5619 | 845 | 1085 |
| 4881 | QGN17978.1 | FIM1_5187 | CP015060. | 4483 | 875 | 1410 | 8893 | 904 | 1107 | 6671 | 733 | 1851 | 4699 | 674 | 1848 | 7236 | 948 | 1703 | 5398 | 641 | 1293 |
| 4882 | QGN17979.1 | FIM1_5188 | CP015060. | 5393 | 4513 | 4613 | 6115 | 7098 | 3975 | 5332 | 5533 | 5724 | 31875 | 7434 | 7956 | 29344 | 5553 | 7586 | 32155 | 4726 | 5421 |
| 4883 | QGN17980.1 | FIM1_5189 | CP015060. | 201 | 176 | 150 | 504 | 233 | 162 | 190 | 111 | 212 | 1143 | 155 | 284 | 787 | 128 | 253 | 1196 | 113 | 162 |
| 4884 | QGN17981.1 | FIM1_5190 | CP015060. | 1517 | 792 | 662 | 2160 | 616 | 645 | 1575 | 501 | 949 | 4029 | 578 | 1058 | 3166 | 477 | 1007 | 3986 | 499 | 655 |
| 4885 | QGN17982.1 | FIM1_5191 | CP015060. | 8284 | 880 | 825 | 6091 | 1224 | 770 | 5964 | 885 | 1010 | 1923 | 660 | 1035 | 2951 | 782 | 1020 | 2010 | 788 | 686 |
| 4886 | QGN17983.1 | FIM1_5192 | CP015060. | 32319 | 432 | 210 | 14107 | 567 | 207 | 8682 | 531 | 312 | 1557 | 472 | 394 | 2334 | 303 | 400 | 1713 | 310 | 270 |
| 4887 | QGN17984.1 | FIM1_5193 | CP015060. | 18141 | 14494 | 8566 | 18386 | 11314 | 7779 | 14671 | 10264 | 10260 | 26225 | 12920 | 13230 | 30961 | 11946 | 12888 | 26755 | 11684 | 10047 |
| 4888 | QGN17985.1 | FIM1_5194 | CP015060. | 6260 | 3948 | 2705 | 6862 | 2862 | 2263 | 6597 | 2009 | 3407 | 11859 | 2007 | 3376 | 10287 | 2482 | 2768 | 12809 | 2040 | 2080 |
| 4889 | QGN17986.1 | FIM1_5195 | CP015060. | 9795 | 2991 | 3346 | 13725 | 4452 | 3293 | 10720 | 3066 | 4441 | 15824 | 3320 | 3839 | 21929 | 3757 | 3362 | 18015 | 2941 | 2688 |
| 4890 | QGN17987.1 | FIM1_5196 | CP015060. | 562 | 155 | 237 | 591 | 169 | 191 | 411 | 170 | 333 | 972 | 123 | 272 | 1022 | 124 | 269 | 961 | 101 | 199 |
| 4891 | QGN17988.1 | FIM1_5197 | CP015060. | 589 | 297 | 203 | 719 | 312 | 223 | 592 | 207 | 302 | 945 | 272 | 295 | 1018 | 192 | 314 | 882 | 206 | 227 |
| 4892 | QGN17989.1 | FIM1_5199 | CP015060. | 3928 | 1940 | 1532 | 4647 | 2061 | 1270 | 4108 | 1627 | 1804 | 13616 | 2146 | 2768 | 11385 | 2016 | 2379 | 13695 | 1953 | 1727 |
| 4893 | QGN17990.1 | FIM1_5200 | CP015060. | 1206 | 52 | 140 | 1814 | 256 | 117 | 715 | 159 | 149 | 531 | 101 | 130 | 792 | 103 | 132 | 531 | 79 | 103 |
| 4894 | QGN17991.1 | FIM1_5201 | CP015060. | 2411 | 90 | 205 | 2347 | 183 | 147 | 1970 | 219 | 284 | 783 | 179 | 305 | 1311 | 150 | 236 | 834 | 173 | 150 |
| 4895 | QGN17992.1 | FIM1_5202 | CP015060. | 1513 | 626 | 600 | 2106 | 554 | 461 | 1601 | 623 | 777 | 3247 | 405 | 587 | 2618 | 363 | 529 | 2818 | 355 | 373 |
| 4896 | QGN17993.1 | FIM1_5203 | CP015060. | 2261 | 260 | 442 | 2172 | 471 | 342 | 1820 | 456 | 531 | 1566 | 410 | 638 | 2241 | 339 | 577 | 1793 | 358 | 395 |
| 4897 | QGN17994.1 | FIM1_5204 | CP015060. | 1906 | 271 | 293 | 1971 | 224 | 209 | 2064 | 250 | 376 | 449 | 148 | 186 | 1016 | 110 | 227 | 726 | 111 | 140 |
| 4898 | QGN17995.1 | FIM1_5205 | CP015060. | 1703 | 858 | 913 | 2252 | 901 | 618 | 1945 | 866 | 1068 | 5681 | 1066 | 1472 | 5308 | 1087 | 1303 | 5804 | 976 | 935 |
| 4899 | QGN17996.1 | FIM1_5206 | CP015060. | 2267 | 594 | 569 | 2359 | 531 | 380 | 2300 | 561 | 665 | 2432 | 504 | 844 | 2329 | 439 | 716 | 2549 | 447 | 549 |
| 4900 | QGN17997.1 | FIM1_5207 | CP015060. | 588 | 176 | 192 | 1022 | 129 | 123 | 770 | 140 | 254 | 1023 | 91 | 230 | 960 | 92 | 202 | 1027 | 60 | 125 |
| 4901 | QGN17998.1 | FIM1_5208 | CP015060. | 24 | 6 | 6 | 228 | 20 | 10 | 41 | 8 | 10 | 51 | 14 | 15 | 59 | 13 | 19 | 65 | 5 | 15 |
| 4902 | QGN17999.1 | FIM1_5209 | CP015060. | 60 | 16 | 4 | 235 | 20 | 1 | 61 | 12 | 10 | 168 | 18 | 24 | 130 | 25 | 20 | 151 | 27 | 19 |
| 4903 | QGN18000.1 | FIM1_4316 | CP015061. | 6423 | 2523 | 1885 | 11192 | 3966 | 1535 | 7684 | 2282 | 2137 | 4185 | 3921 | 3196 | 9391 | 3240 | 3687 | 6042 | 3069 | 2740 |
| 4904 | QGN18001.1 | FIM1_4317 | CP015061. | 0 | 0 | 0 | 0 | 0 | 0 | 0 | 0 | 0 | 0 | 0 | 0 | 0 | 0 | 0 | 0 | 0 | 0 |
| 4905 | QGN18002.1 | FIM1_4318 | CP015061. | 1612 | 101 | 129 | 1689 | 90 | 93 | 1440 | 94 | 158 | 1192 | 91 | 218 | 966 | 81 | 182 | 1171 | 65 | 149 |
| 4906 | QGN18003.1 | FIM1_4319 | CP015061. | 454 | 284 | 404 | 1714 | 612 | 397 | 683 | 286 | 501 | 4925 | 683 | 690 | 4172 | 962 | 813 | 4314 | 485 | 558 |
| 4907 | QGN18004.1 | FIM1_4320 | CP015061. | 7862 | 1445 | 2135 | 7720 | 2091 | 1812 | 7400 | 1545 | 2718 | 10478 | 2264 | 3459 | 13599 | 2429 | 3167 | 11781 | 2140 | 2390 |
| 4908 | QGN18005.1 | FIM1_4321 | CP015061. | 1537 | 836 | 1097 | 2767 | 914 | 999 | 2045 | 690 | 1404 | 5047 | 790 | 2269 | 4670 | 710 | 2112 | 4918 | 712 | 1402 |
| 4909 | QGN18006.1 | FIM1_4322 | CP015061. | 2662 | 719 | 661 | 3206 | 902 | 586 | 3459 | 718 | 903 | 4421 | 790 | 1126 | 3714 | 956 | 1077 | 4206 | 943 | 798 |
| 4910 | QGN18007.1 | FIM1_4323 | CP015061. | 39553 | 7983 | 7426 | 41277 | 9375 | 6032 | 39677 | 8223 | 9165 | 37450 | 11518 | 13729 | 48923 | 11867 | 12042 | 41102 | 10594 | 9096 |
| 4911 | QGN18008.1 | FIM1_4324 | CP015061. | 75 | 39 | 22 | 94 | 29 | 12 | 48 | 34 | 30 | 224 | 22 | 45 | 127 | 20 | 27 | 202 | 18 | 25 |
| 4912 | QGN18009.1 | FIM1_4325 | CP015061. | 3372 | 1313 | 1181 | 3387 | 1120 | 1000 | 2990 | 1054 | 1563 | 8597 | 1091 | 2011 | 7328 | 846 | 1848 | 8971 | 904 | 1392 |
| 4913 | QGN18010.1 | FIM1_4326 | CP015061. | 367 | 786 | 870 | 723 | 892 | 750 | 474 | 708 | 1082 | 5492 | 900 | 1458 | 4990 | 959 | 1260 | 5192 | 831 | 899 |
| 4914 | QGN18011.1 | FIM1_4327 | CP015061. | 3217 | 1566 | 1389 | 5229 | 2169 | 1209 | 4489 | 1452 | 1854 | 6494 | 1502 | 1238 | 5647 | 1504 | 1082 | 6404 | 1626 | 720 |
| 4915 | QGN18012.1 | FIM1_4328 | CP015061. | 1285 | 726 | 432 | 1066 | 743 | 443 | 1322 | 539 | 598 | 2909 | 688 | 683 | 2353 | 595 | 639 | 2565 | 658 | 489 |
| 4916 | QGN18013.1 | FIM1_4329 | CP015061. | 1912 | 400 | 432 | 1648 | 377 | 419 | 1225 | 312 | 583 | 2527 | 194 | 1135 | 2032 | 200 | 963 | 2446 | 171 | 681 |
| 4917 | QGN18014.1 | FIM1_4330 | CP015061. | 1606 | 1473 | 1544 | 3434 | 1645 | 1321 | 2388 | 1598 | 2006 | 9642 | 1498 | 1878 | 9145 | 1267 | 1684 | 9026 | 1259 | 1266 |
| 4918 | QGN18015.1 | FIM1_4331 | CP015061. | 11552 | 2340 | 2987 | 19971 | 4142 | 2404 | 14088 | 2926 | 3339 | 17450 | 3639 | 3156 | 16072 | 4606 | 2891 | 16312 | 3976 | 1978 |
| 4919 | QGN18016.1 | FIM1_4332 | CP015061. | 3563 | 966 | 1043 | 4512 | 910 | 960 | 3627 | 830 | 1427 | 6219 | 1001 | 1535 | 5511 | 941 | 1434 | 5968 | 943 | 973 |
| 4920 | QGN18017.1 | FIM1_4333 | CP015061. | 1889 | 1946 | 1928 | 2619 | 1812 | 1749 | 2330 | 1545 | 2462 | 12269 | 1932 | 3330 | 10456 | 1945 | 2888 | 11788 | 1884 | 2066 |
| 4921 | QGN18018.1 | FIM1_4334 | CP015061. | 2047 | 762 | 561 | 1943 | 511 | 538 | 1754 | 393 | 744 | 2985 | 470 | 862 | 2160 | 510 | 906 | 2604 | 545 | 567 |
| 4922 | QGN18019.1 | FIM1_4335 | CP015061. | 4125 | 1145 | 1732 | 4879 | 1301 | 1540 | 4163 | 1097 | 2271 | 10835 | 1493 | 3915 | 10740 | 1453 | 3517 | 10957 | 1276 | 2590 |
| 4923 | QGN18020.1 | FIM1_4336 | CP015061. | 1183 | 750 | 825 | 1282 | 653 | 766 | 1554 | 699 | 1185 | 3786 | 510 | 782 | 3267 | 335 | 757 | 3785 | 408 | 475 |
| 4924 | QGN18021.1 | FIM1_4337 | CP015061. | 1980 | 1066 | 886 | 2252 | 874 | 804 | 2155 | 734 | 1222 | 6533 | 1005 | 1439 | 5283 | 779 | 1391 | 6202 | 868 | 917 |
| 4925 | QGN18022.1 | FIM1_4338 | CP015061. | 3395 | 2829 | 2388 | 4167 | 3071 | 2251 | 4010 | 2751 | 3040 | 16148 | 2935 | 4407 | 16078 | 3039 | 3812 | 15906 | 2999 | 2820 |
| 4926 | QGN18023.1 | FIM1_4339 | CP015061. | 223 | 378 | 460 | 320 | 283 | 419 | 205 | 318 | 650 | 2802 | 224 | 654 | 1887 | 166 | 578 | 2563 | 190 | 427 |
| 4927 | QGN18024.1 | FIM1_4340 | CP015061. | 2845 | 7369 | 9524 | 4202 | 5866 | 8688 | 3223 | 4825 | 11556 | 21696 | 6303 | 15530 | 19161 | 5880 | 14402 | 21180 | 5666 | 10669 |
| 4928 | QGN18025.1 | FIM1_4341 | CP015061. | 382 | 61 | 269 | 508 | 246 | 218 | 381 | 172 | 301 | 1193 | 197 | 475 | 971 | 210 | 384 | 1224 | 218 | 266 |
| 4929 | QGN18026.1 | FIM1_4342 | CP015061. | 920 | 376 | 494 | 1067 | 332 | 387 | 881 | 261 | 619 | 1861 | 307 | 695 | 1374 | 324 | 655 | 1829 | 344 | 475 |
| 4930 | QGN18027.1 | FIM1_4343 | CP015061. | 21377 | 25350 | 15448 | 22290 | 20823 | 14352 | 18657 | 18341 | 18015 | 53550 | 22336 | 22358 | 48744 | 20401 | 22207 | 54666 | 20263 | 17116 |
| 4931 | QGN18028.1 | FIM1_4344 | CP015061. | 3116 | 2248 | 1045 | 2130 | 1801 | 1095 | 2898 | 1398 | 1389 | 6493 | 1947 | 1579 | 5108 | 2108 | 1643 | 6146 | 2406 | 1133 |
| 4932 | QGN18029.1 | FIM1_4345 | CP015061. | 161 | 345 | 541 | 373 | 404 | 439 | 220 | 310 | 677 | 2347 | 242 | 732 | 1969 | 297 | 658 | 2283 | 236 | 418 |
| 4933 | QGN18030.1 | FIM1_4346 | CP015061. | 661 | 1556 | 478 | 279 | 1314 | 645 | 554 | 1026 | 782 | 3729 | 1373 | 954 | 2931 | 1025 | 852 | 3720 | 1397 | 608 |
| 4934 | QGN18031.1 | FIM1_4347 | CP015061. | 3922 | 627 | 1061 | 5006 | 914 | 1124 | 4253 | 779 | 1417 | 4466 | 596 | 1854 | 5037 | 720 | 1589 | 4713 | 622 | 1193 |
| 4935 | QGN18032.1 | FIM1_4348 | CP015061. | 5337 | 1186 | 1440 | 5436 | 1417 | 1243 | 5431 | 1221 | 1875 | 7945 | 1453 | 2053 | 9283 | 1222 | 1952 | 8073 | 1213 | 1541 |
| 4936 | QGN18033.1 | FIM1_4349 | CP015061. | 7568 | 62 | 91 | 4359 | 34 | 55 | 4531 | 51 | 94 | 303 | 34 | 120 | 268 | 43 | 67 | 275 | 42 | 69 |
| 4937 | QGN18034.1 | FIM1_4350 | CP015061. | 3561 | 1902 | 1569 | 4288 | 1865 | 1310 | 3701 | 1952 | 2040 | 9461 | 1543 | 2013 | 9513 | 1528 | 1812 | 8463 | 1435 | 1348 |
| 4938 | QGN18035.1 | FIM1_4351 | CP015061. | 1732 | 3574 | 2614 | 2125 | 3717 | 2182 | 2000 | 3298 | 3356 | 9431 | 4169 | 4544 | 17637 | 3853 | 3961 | 12717 | 3691 | 3164 |
| 4939 | QGN18036.1 | FIM1_4352 | CP015061. | 1061 | 1049 | 607 | 1058 | 822 | 613 | 1123 | 687 | 803 | 5541 | 1038 | 1082 | 3502 | 858 | 1090 | 5211 | 1042 | 722 |
| 4940 | QGN18037.1 | FIM1_4353 | CP015061. | 3476 | 1021 | 1118 | 3402 | 1539 | 984 | 3834 | 1227 | 1425 | 8373 | 1856 | 2001 | 8992 | 1726 | 1625 | 8730 | 1688 | 1293 |
| 4941 | QGN18038.1 | FIM1_4354 | CP015061. | 1395 | 1314 | 1629 | 2241 | 908 | 1384 | 1564 | 880 | 2150 | 5605 | 658 | 2186 | 4202 | 687 | 1902 | 5762 | 610 | 1341 |
| 4942 | QGN18039.1 | FIM1_4355 | CP015061. | 31 | 17 | 19 | 90 | 23 | 12 | 51 | 26 | 25 | 91 | 5 | 29 | 86 | 13 | 16 | 88 | 9 | 20 |
| 4943 | QGN18040.1 | FIM1_4356 | CP015061. | 22380 | 16279 | 15259 | 29575 | 15938 | 12786 | 23796 | 13638 | 17559 | 51238 | 17710 | 24014 | 52718 | 17758 | 22872 | 52379 | 16479 | 17198 |
| 4944 | QGN18041.1 | FIM1_4357 | CP015061. | 1303 | 1771 | 1655 | 2256 | 1326 | 1589 | 1888 | 1243 | 2244 | 5728 | 784 | 2841 | 4822 | 756 | 2915 | 5815 | 761 | 1781 |
| 4945 | QGN18042.1 | FIM1_4358 | CP015061. | 3428 | 3843 | 3049 | 4937 | 2998 | 2830 | 3583 | 2298 | 3947 | 14637 | 2675 | 4842 | 11839 | 2821 | 4563 | 14861 | 2487 | 3235 |
| 4946 | QGN18043.1 | FIM1_4359 | CP015061. | 1677 | 2498 | 2143 | 3204 | 2238 | 1819 | 2327 | 1729 | 2662 | 6754 | 1299 | 2366 | 6456 | 1246 | 2239 | 6927 | 1115 | 1529 |
| 4947 | QGN18044.1 | FIM1_4360 | CP015061. | 5298 | 6442 | 5136 | 9001 | 7136 | 4451 | 7609 | 6694 | 6584 | 27390 | 6881 | 7928 | 28680 | 6611 | 7316 | 27233 | 5825 | 5405 |
| 4948 | QGN18045.1 | FIM1_4361 | CP015061. | 5069 | 1168 | 1129 | 4709 | 2072 | 1002 | 5621 | 1528 | 1351 | 6886 | 2280 | 1883 | 7798 | 2638 | 1670 | 6839 | 2683 | 1297 |
| 4949 | QGN18046.1 | FIM1_4363 | CP015061. | 2074 | 633 | 338 | 1708 | 637 | 328 | 2059 | 449 | 410 | 2897 | 635 | 559 | 2381 | 818 | 487 | 2566 | 889 | 428 |
| 4950 | QGN18047.1 | FIM1_4364 | CP015061. | 102 | 310 | 302 | 225 | 193 | 259 | 159 | 184 | 358 | 1823 | 236 | 498 | 1279 | 217 | 464 | 1813 | 218 | 327 |
| 4951 | QGN18048.1 | FIM1_4365 | CP015061. | 191 | 935 | 696 | 297 | 928 | 764 | 233 | 682 | 963 | 4693 | 790 | 1280 | 4121 | 772 | 1084 | 4473 | 835 | 770 |
| 4952 | QGN18049.1 | FIM1_4366 | CP015061. | 16082 | 5111 | 5807 | 15284 | 5191 | 4618 | 14048 | 4892 | 7297 | 21300 | 6179 | 7823 | 22356 | 5642 | 9435 | 22086 | 5487 | 5495 |
| 4953 | QGN18050.1 | FIM1_4367 | CP015061. | 1735 | 1402 | 907 | 3236 | 1199 | 853 | 2832 | 984 | 1270 | 6751 | 1156 | 1511 | 5852 | 1204 | 1559 | 6061 | 1087 | 1125 |
| 4954 | QGN18051.1 | FIM1_4368 | CP015061. | 2230 | 1257 | 935 | 2524 | 1127 | 948 | 2224 | 817 | 1236 | 5505 | 1146 | 1526 | 4613 | 977 | 1579 | 5353 | 1009 | 1062 |

| | | | | | | | | | | | | | | | | | | | | | |
|---|---|---|---|---|---|---|---|---|---|---|---|---|---|---|---|---|---|---|---|---|---|
| 4955 | QGN18052.1 | FIM1_4369 | CP015061. | 2973 | 1972 | 1059 | 1579 | 2398 | 1044 | 2763 | 1806 | 1371 | 11576 | 3061 | 2224 | 8974 | 2695 | 2003 | 11072 | 3298 | 1582 |
| 4956 | QGN18053.1 | FIM1_4370 | CP015061. | 1251 | 550 | 380 | 1163 | 539 | 383 | 1320 | 343 | 532 | 3446 | 571 | 673 | 2471 | 681 | 649 | 3468 | 784 | 422 |
| 4957 | QGN18054.1 | FIM1_4371 | CP015061. | 10453 | 1342 | 2063 | 6927 | 2179 | 1953 | 10256 | 1711 | 2659 | 8436 | 1640 | 3039 | 11057 | 1844 | 2663 | 9122 | 1653 | 1918 |
| 4958 | QGN18055.1 | FIM1_4372 | CP015061. | 2453 | 2225 | 1483 | 2426 | 1819 | 1623 | 1887 | 1365 | 1722 | 7297 | 1921 | 2250 | 4919 | 1774 | 2005 | 6047 | 2153 | 1532 |
| 4959 | QGN18056.1 | FIM1_4373 | CP015061. | 9987 | 5064 | 2719 | 6339 | 6764 | 2816 | 9913 | 5357 | 3524 | 22037 | 7580 | 5499 | 23691 | 6518 | 5260 | 21584 | 7685 | 3781 |
| 4960 | QGN18057.1 | FIM1_4374 | CP015061. | 2995 | 585 | 744 | 3910 | 561 | 753 | 3186 | 531 | 959 | 4218 | 497 | 935 | 3641 | 446 | 874 | 4051 | 502 | 646 |
| 4961 | QGN18058.1 | FIM1_4376 | CP015061. | 368 | 243 | 167 | 306 | 202 | 151 | 256 | 154 | 210 | 1267 | 187 | 259 | 934 | 154 | 237 | 1298 | 177 | 166 |
| 4962 | QGN18059.1 | FIM1_4377 | CP015061. | 5522 | 3697 | 2371 | 4251 | 4479 | 2211 | 5892 | 3777 | 3042 | 7307 | 4684 | 4090 | 17002 | 3921 | 3790 | 11245 | 4295 | 2862 |
| 4963 | QGN18060.1 | FIM1_4378 | CP015061. | 124 | 225 | 153 | 163 | 153 | 142 | 129 | 127 | 207 | 564 | 108 | 222 | 647 | 151 | 249 | 586 | 106 | 165 |
| 4964 | QGN18061.1 | FIM1_4379 | CP015061. | 1952 | 1475 | 2627 | 3177 | 1770 | 2104 | 2630 | 1427 | 3264 | 11226 | 1833 | 4729 | 12325 | 1933 | 3807 | 12019 | 1733 | 3195 |
| 4965 | QGN18062.1 | FIM1_4380 | CP015061. | 2375 | 698 | 810 | 3034 | 925 | 674 | 2407 | 678 | 1009 | 5991 | 985 | 1429 | 6237 | 865 | 1222 | 5839 | 804 | 894 |
| 4966 | QGN18063.1 | FIM1_4381 | CP015061. | 673 | 827 | 965 | 1053 | 1121 | 779 | 762 | 933 | 1114 | 5411 | 1088 | 1450 | 6515 | 1212 | 1226 | 5567 | 1111 | 861 |
| 4967 | QGN18064.1 | FIM1_4382 | CP015061. | 1012 | 532 | 581 | 1408 | 701 | 478 | 1148 | 532 | 702 | 4422 | 658 | 983 | 3448 | 577 | 787 | 4289 | 561 | 573 |
| 4968 | QGN18065.1 | FIM1_4383 | CP015061. | 3491 | 3776 | 3097 | 4847 | 4260 | 2957 | 3772 | 3530 | 4005 | 16683 | 4313 | 5053 | 17073 | 3988 | 4901 | 16428 | 3773 | 3454 |
| 4969 | QGN18066.1 | FIM1_4384 | CP015061. | 6325 | 477 | 716 | 7609 | 553 | 525 | 8699 | 615 | 854 | 2937 | 401 | 956 | 2953 | 875 | 949 | 2676 | 608 | 675 |
| 4970 | QGN18067.1 | FIM1_4385 | CP015061. | 2676 | 747 | 1032 | 3391 | 970 | 875 | 2798 | 928 | 1371 | 4094 | 914 | 1558 | 6102 | 760 | 1381 | 4707 | 790 | 1009 |
| 4971 | QGN18068.1 | FIM1_4386 | CP015061. | 7256 | 12262 | 10861 | 13978 | 11894 | 9325 | 9922 | 10755 | 13370 | 30635 | 11425 | 15293 | 34769 | 11908 | 14734 | 31475 | 10282 | 10999 |
| 4972 | QGN18069.1 | FIM1_4387 | CP015061. | 55 | 31 | 17 | 54 | 39 | 22 | 27 | 22 | 15 | 102 | 24 | 23 | 94 | 17 | 24 | 77 | 18 | 21 |
| 4973 | QGN18070.1 | FIM1_4388 | CP015061. | 10341 | 2004 | 631 | 4340 | 2099 | 744 | 5119 | 1825 | 958 | 2394 | 2640 | 854 | 4478 | 2167 | 714 | 3071 | 2419 | 650 |
| 4974 | QGN18071.1 | FIM1_4389 | CP015061. | 315 | 141 | 234 | 618 | 138 | 127 | 490 | 110 | 262 | 973 | 80 | 322 | 946 | 111 | 247 | 1006 | 93 | 177 |
| 4975 | QGN18072.1 | FIM1_4390 | CP015061. | 257 | 64 | 69 | 242 | 53 | 77 | 288 | 74 | 96 | 523 | 44 | 112 | 377 | 67 | 120 | 480 | 48 | 81 |
| 4976 | QGN18073.1 | FIM1_4391 | CP015061. | 11870 | 8338 | 4102 | 7627 | 9551 | 4178 | 11741 | 7671 | 5199 | 24259 | 11426 | 8951 | 27590 | 8576 | 8398 | 23597 | 10105 | 6528 |
| 4977 | QGN18074.1 | FIM1_4392 | CP015061. | 5516 | 2456 | 1359 | 8807 | 1969 | 1329 | 5233 | 1255 | 2018 | 3599 | 1676 | 1664 | 4385 | 1054 | 1484 | 3657 | 1034 | 1006 |
| 4978 | QGN18075.1 | FIM1_4393 | CP015061. | 89 | 115 | 72 | 177 | 83 | 66 | 178 | 95 | 85 | 116 | 49 | 94 | 212 | 52 | 106 | 130 | 59 | 43 |
| 4979 | QGN18076.1 | FIM1_4394 | CP015061. | 710 | 621 | 462 | 540 | 520 | 490 | 624 | 571 | 669 | 1344 | 273 | 453 | 1583 | 243 | 416 | 1354 | 211 | 274 |
| 4980 | QGN18077.1 | FIM1_4396 | CP015061. | 1979 | 2628 | 2699 | 3879 | 2004 | 2354 | 2540 | 1828 | 3415 | 12583 | 1824 | 3915 | 10118 | 2132 | 3900 | 12310 | 1855 | 2643 |
| 4981 | QGN18078.1 | FIM1_4397 | CP015061. | 1369 | 829 | 998 | 2258 | 893 | 837 | 1517 | 773 | 1272 | 5409 | 605 | 1130 | 5651 | 667 | 1091 | 5269 | 464 | 762 |
| 4982 | QGN18079.1 | FIM1_4398 | CP015061. | 285 | 56 | 83 | 1112 | 49 | 63 | 1034 | 65 | 119 | 932 | 37 | 64 | 771 | 26 | 53 | 812 | 39 | 46 |
| 4983 | QGN18080.1 | FIM1_4399 | CP015061. | 4890 | 6284 | 7840 | 14665 | 10929 | 6532 | 10714 | 9028 | 9099 | 41035 | 11744 | 12757 | 42881 | 11946 | 11771 | 41581 | 10510 | 8656 |
| 4984 | QGN18081.1 | FIM1_4400 | CP015061. | 1189 | 840 | 743 | 1351 | 723 | 648 | 1256 | 861 | 981 | 2248 | 706 | 968 | 3225 | 586 | 994 | 2581 | 585 | 671 |
| 4985 | QGN18082.1 | FIM1_4401 | CP015061. | 1249 | 2483 | 1413 | 1309 | 2204 | 1226 | 1444 | 1937 | 1790 | 6052 | 1709 | 2063 | 7824 | 1675 | 1994 | 6487 | 1739 | 1355 |
| 4986 | QGN18083.1 | FIM1_4402 | CP015061. | 213 | 247 | 130 | 190 | 180 | 129 | 222 | 165 | 170 | 436 | 95 | 88 | 363 | 76 | 78 | 393 | 96 | 58 |
| 4987 | QGN18084.1 | FIM1_4403 | CP015061. | 15670 | 2599 | 2752 | 24276 | 2619 | 2071 | 18720 | 2694 | 3653 | 2666 | 1920 | 3839 | 13517 | 2720 | 3469 | 5379 | 2317 | 2595 |
| 4988 | QGN18085.1 | FIM1_4404 | CP015061. | 666 | 296 | 265 | 646 | 270 | 287 | 551 | 260 | 401 | 2577 | 249 | 416 | 1739 | 221 | 393 | 2473 | 218 | 285 |
| 4989 | QGN18086.1 | FIM1_4405 | CP015061. | 256 | 476 | 470 | 569 | 405 | 394 | 350 | 438 | 602 | 3100 | 399 | 635 | 2357 | 305 | 608 | 2824 | 291 | 381 |
| 4990 | QGN18087.1 | FIM1_4406 | CP015061. | 1334 | 493 | 793 | 2105 | 665 | 644 | 1393 | 647 | 1024 | 3553 | 524 | 1151 | 4291 | 480 | 1024 | 3705 | 409 | 795 |
| 4991 | QGN18088.1 | FIM1_4407 | CP015061. | 1573 | 38 | 51 | 695 | 45 | 34 | 850 | 45 | 56 | 153 | 26 | 53 | 178 | 28 | 53 | 162 | 21 | 36 |
| 4992 | QGN18089.1 | FIM1_4408 | CP015061. | 1328 | 166 | 464 | 1350 | 195 | 327 | 1334 | 438 | 602 | 336 | 76 | 531 | 1159 | 65 | 406 | 699 | 60 | 334 |
| 4993 | QGN18090.1 | FIM1_4409 | CP015061. | 1941 | 5008 | 3287 | 2865 | 4278 | 2319 | 2132 | 7936 | 4827 | 21856 | 6138 | 3370 | 20671 | 4907 | 2587 | 20967 | 7149 | 2592 |
| 4994 | QGN18091.1 | FIM1_4410 | CP015061. | 1259 | 755 | 650 | 1380 | 513 | 534 | 1266 | 556 | 938 | 1633 | 364 | 803 | 2253 | 427 | 872 | 2000 | 314 | 553 |
| 4995 | QGN18092.1 | FIM1_4412 | CP015061. | 527 | 71 | 139 | 1933 | 133 | 89 | 726 | 126 | 169 | 497 | 76 | 137 | 814 | 102 | 123 | 594 | 61 | 96 |
| 4996 | QGN18093.1 | FIM1_4413 | CP015061. | 1916 | 947 | 948 | 1853 | 968 | 810 | 1934 | 904 | 1209 | 5445 | 902 | 1272 | 4964 | 778 | 1293 | 5083 | 762 | 849 |
| 4997 | QGN18094.1 | FIM1_4414 | CP015061. | 2089 | 1461 | 2300 | 2482 | 2265 | 1791 | 2325 | 2141 | 2878 | 8773 | 2100 | 2342 | 12177 | 1910 | 2175 | 9456 | 1835 | 1689 |
| 4998 | QGN18095.1 | FIM1_4415 | CP015061. | 10811 | 818 | 1894 | 10724 | 1550 | 1484 | 9390 | 1443 | 2575 | 2540 | 1064 | 2864 | 12109 | 1205 | 2634 | 5661 | 1033 | 1879 |
| 4999 | QGN18096.1 | FIM1_4416 | CP015061. | 4806 | 631 | 1257 | 5567 | 1108 | 1021 | 4156 | 871 | 1597 | 2859 | 814 | 1959 | 7083 | 827 | 1700 | 4386 | 747 | 1263 |
| 5000 | QGN18097.1 | FIM1_4417 | CP015061. | 1536 | 571 | 727 | 1798 | 741 | 608 | 1333 | 685 | 944 | 4949 | 906 | 1118 | 5924 | 818 | 1122 | 5269 | 721 | 800 |
| 5001 | QGN18098.1 | FIM1_4418 | CP015061. | 5383 | 4622 | 3198 | 5640 | 3905 | 2989 | 5060 | 3500 | 4251 | 15523 | 4701 | 4718 | 16871 | 4179 | 4809 | 15140 | 4373 | 3173 |
| 5002 | QGN18099.1 | FIM1_4419 | CP015061. | 962 | 944 | 795 | 1550 | 678 | 684 | 1341 | 648 | 1053 | 2596 | 540 | 1016 | 2844 | 522 | 1037 | 2438 | 457 | 737 |
| 5003 | QGN18100.1 | FIM1_4420 | CP015061. | 1605 | 540 | 759 | 2511 | 531 | 659 | 1933 | 652 | 1003 | 2756 | 421 | 1093 | 3524 | 465 | 963 | 2797 | 396 | 708 |
| 5004 | QGN18101.1 | FIM1_4421 | CP015061. | 1486 | 780 | 656 | 1535 | 702 | 578 | 1605 | 601 | 903 | 2798 | 399 | 709 | 2366 | 371 | 708 | 2515 | 366 | 481 |
| 5005 | QGN18102.1 | FIM1_4422 | CP015061. | 16 | 5 | 11 | 88 | 6 | 9 | 40 | 6 | 11 | 63 | 7 | 14 | 30 | 7 | 15 | 54 | 8 | 8 |
| 5006 | QGN18103.1 | FIM1_4423 | CP015061. | 584 | 1892 | 1204 | 1752 | 1369 | 1178 | 1153 | 1085 | 1512 | 5991 | 1185 | 1615 | 4534 | 1081 | 1636 | 5217 | 1197 | 1090 |
| 5007 | QGN18104.1 | FIM1_4424 | CP015061. | 7608 | 539 | 484 | 3771 | 309 | 441 | 3085 | 471 | 637 | 3669 | 479 | 871 | 3129 | 121 | 660 | 3895 | 306 | 606 |
| 5008 | QGN18105.1 | FIM1_4425 | CP015061. | 504 | 543 | 400 | 614 | 460 | 388 | 553 | 479 | 527 | 1960 | 437 | 567 | 2098 | 377 | 510 | 1822 | 370 | 337 |
| 5009 | QGN18106.1 | FIM1_4426 | CP015061. | 432 | 591 | 441 | 720 | 572 | 399 | 513 | 400 | 595 | 3275 | 439 | 684 | 2755 | 489 | 591 | 3068 | 444 | 416 |
| 5010 | QGN18107.1 | FIM1_4427 | CP015061. | 6635 | 3614 | 3312 | 9267 | 3229 | 2821 | 7017 | 2943 | 4254 | 23103 | 3129 | 5336 | 18215 | 3444 | 4714 | 23505 | 2873 | 3459 |
| 5011 | QGN18108.1 | FIM1_4428 | CP015061. | 6555 | 5381 | 2796 | 12973 | 4832 | 3477 | 6344 | 3557 | 4196 | 24052 | 5389 | 4640 | 26285 | 5361 | 6377 | 24210 | 2848 | 4168 |
| 5012 | QGN18109.1 | FIM1_4429 | CP015061. | 2151 | 1555 | 1253 | 3301 | 1558 | 959 | 2586 | 1538 | 1568 | 2679 | 1291 | 1678 | 6430 | 1211 | 1703 | 3864 | 1212 | 1196 |
| 5013 | QGN18110.1 | FIM1_4430 | CP015061. | 403 | 299 | 232 | 308 | 229 | 219 | 385 | 198 | 303 | 1279 | 236 | 255 | 883 | 193 | 259 | 1141 | 233 | 175 |
| 5014 | QGN18111.1 | FIM1_4431 | CP015061. | 9313 | 1098 | 949 | 9530 | 1131 | 886 | 8268 | 1194 | 1271 | 3616 | 821 | 1290 | 5084 | 828 | 1260 | 3767 | 823 | 822 |
| 5015 | QGN18112.1 | FIM1_4432 | CP015061. | 5756 | 4815 | 2866 | 8644 | 4511 | 2941 | 6164 | 3706 | 3820 | 16670 | 4564 | 5134 | 17803 | 4236 | 4934 | 16192 | 4216 | 3440 |
| 5016 | QGN18113.1 | FIM1_4433 | CP015061. | 542 | 453 | 336 | 598 | 360 | 347 | 502 | 259 | 452 | 4025 | 363 | 535 | 2641 | 350 | 454 | 3992 | 328 | 344 |
| 5017 | QGN18114.1 | FIM1_4434 | CP015061. | 4496 | 883 | 1143 | 4496 | 1040 | 995 | 4409 | 858 | 1493 | 5956 | 1088 | 2148 | 6549 | 1077 | 1930 | 5977 | 990 | 1373 |
| 5018 | QGN18115.1 | FIM1_4435 | CP015061. | 9953 | 1618 | 2499 | 10488 | 2229 | 1935 | 9889 | 2082 | 3130 | 11093 | 2123 | 4081 | 15368 | 2107 | 3509 | 12202 | 1903 | 2681 |
| 5019 | QGN18116.1 | FIM1_4436 | CP015061. | 2645 | 820 | 1141 | 3337 | 876 | 958 | 2862 | 975 | 1411 | 5582 | 836 | 1553 | 6001 | 768 | 1426 | 5374 | 719 | 1024 |
| 5020 | QGN18117.1 | FIM1_4437 | CP015061. | 1108 | 380 | 448 | 747 | 334 | 390 | 853 | 336 | 613 | 2334 | 367 | 717 | 2826 | 276 | 677 | 2299 | 306 | 522 |
| 5021 | QGN18118.1 | FIM1_4438 | CP015061. | 3022 | 640 | 831 | 2447 | 739 | 701 | 2381 | 685 | 1070 | 3223 | 645 | 1214 | 5084 | 429 | 1185 | 3703 | 488 | 843 |
| 5022 | QGN18119.1 | FIM1_4439 | CP015061. | 1418 | 1418 | 1265 | 2080 | 1379 | 1184 | 1485 | 1102 | 1582 | 5742 | 1089 | 1543 | 4797 | 935 | 1531 | 4979 | 893 | 1058 |
| 5023 | QGN18120.1 | FIM1_4440 | CP015061. | 3093 | 615 | 1633 | 4049 | 770 | 1118 | 2699 | 825 | 2018 | 2120 | 478 | 2021 | 5418 | 512 | 1885 | 3121 | 404 | 1280 |
| 5024 | QGN18121.1 | FIM1_4441 | CP015061. | 6713 | 1814 | 2202 | 8607 | 1879 | 1717 | 7570 | 1913 | 2913 | 1908 | 1531 | 3272 | 9339 | 1693 | 2969 | 4856 | 1449 | 2146 |
| 5025 | QGN18122.1 | FIM1_4443 | CP015061. | 3724 | 242 | 333 | 1190 | 294 | 230 | 2575 | 430 | 418 | 491 | 137 | 419 | 1560 | 121 | 337 | 824 | 113 | 228 |
| 5026 | QGN18123.1 | FIM1_4444 | CP015061. | 9756 | 18265 | 16885 | 22130 | 22759 | 14417 | 16082 | 20563 | 20244 | 63538 | 25898 | 28316 | 74041 | 25932 | 26779 | 65568 | 23879 | 19741 |
| 5027 | QGN18124.1 | FIM1_4445 | CP015061. | 1702 | 923 | 779 | 2480 | 1040 | 731 | 1718 | 900 | 1037 | 4306 | 832 | 1267 | 4914 | 696 | 1101 | 4393 | 724 | 802 |
| 5028 | QGN18125.1 | FIM1_4446 | CP015061. | 4976 | 2574 | 3988 | 6642 | 3049 | 3137 | 4971 | 3164 | 4685 | 10938 | 3047 | 6187 | 17596 | 3474 | 5338 | 13581 | 3036 | 4145 |
| 5029 | QGN18126.1 | FIM1_4447 | CP015061. | 3246 | 1028 | 939 | 3438 | 1018 | 833 | 3189 | 837 | 1262 | 5420 | 1000 | 1722 | 6237 | 786 | 1489 | 5641 | 847 | 1175 |
| 5030 | QGN18127.1 | FIM1_4448 | CP015061. | 625 | 71 | 181 | 987 | 168 | 118 | 537 | 164 | 209 | 230 | 118 | 178 | 726 | 110 | 161 | 371 | 116 | 128 |
| 5031 | QGN18128.1 | FIM1_4449 | CP015061. | 678 | 558 | 515 | 2000 | 487 | 495 | 1063 | 464 | 720 | 1656 | 274 | 604 | 2013 | 514 | 671 | 1733 | 343 | 430 |
| 5032 | QGN18129.1 | FIM1_4450 | CP015061. | 2618 | 2411 | 2590 | 3221 | 2234 | 2214 | 2268 | 2128 | 3317 | 4420 | 1797 | 4179 | 9212 | 1718 | 3795 | 6746 | 1511 | 2844 |
| 5033 | QGN18130.1 | FIM1_4451 | CP015061. | 7797 | 1072 | 1920 | 5340 | 931 | 1623 | 5891 | 1131 | 2531 | 4028 | 798 | 2644 | 5450 | 647 | 2725 | 4382 | 601 | 1919 |
| 5034 | QGN18131.1 | FIM1_4452 | CP015061. | 8354 | 10601 | 5858 | 7945 | 7466 | 5911 | 6740 | 6145 | 7191 | 24731 | 8883 | 8685 | 19943 | 7487 | 8813 | 24072 | 7981 | 6163 |
| 5035 | QGN18132.1 | FIM1_4454 | CP015061. | 5078 | 3001 | 1766 | 7090 | 3032 | 1818 | 5635 | 2805 | 2547 | 7873 | 2909 | 2962 | 11251 | 2793 | 3006 | 8435 | 2673 | 2070 |
| 5036 | QGN18133.1 | FIM1_4455 | CP015061. | 492 | 1709 | 844 | 441 | 1159 | 794 | 509 | 963 | 1105 | 5591 | 980 | 1166 | 4301 | 858 | 1097 | 5384 | 995 | 790 |
| 5037 | QGN18134.1 | FIM1_4456 | CP015061. | 2976 | 1296 | 1636 | 3503 | 1120 | 1265 | 2804 | 1293 | 1980 | 2067 | 556 | 1380 | 4061 | 576 | 1374 | 2955 | 462 | 884 |
| 5038 | QGN18135.1 | FIM1_4457 | CP015061. | 1308 | 1709 | 1519 | 1773 | 1735 | 1411 | 1390 | 1520 | 1990 | 3376 | 1351 | 1760 | 5441 | 1114 | 1803 | 3963 | 1141 | 1191 |

| | | | | | | | | | | | | | | | | | | | | | |
|---|---|---|---|---|---|---|---|---|---|---|---|---|---|---|---|---|---|---|---|---|---|
| 5039 | QGN18136.1 | FIM1_4458 | CP015061. | 1260 | 1033 | 1018 | 1649 | 832 | 944 | 1254 | 800 | 1302 | 6394 | 990 | 1807 | 5921 | 862 | 1663 | 6169 | 830 | 1292 |
| 5040 | QGN18137.1 | FIM1_4460 | CP015061. | 1071 | 156 | 370 | 854 | 271 | 338 | 881 | 288 | 444 | 1103 | 203 | 545 | 1398 | 279 | 502 | 1184 | 193 | 385 |
| 5041 | QGN18138.1 | FIM1_4461 | CP015061. | 517 | 164 | 179 | 331 | 155 | 174 | 392 | 134 | 243 | 474 | 125 | 274 | 605 | 121 | 267 | 563 | 126 | 167 |
| 5042 | QGN18139.1 | FIM1_4462 | CP015061. | 1158 | 694 | 621 | 1596 | 876 | 575 | 1436 | 652 | 830 | 4214 | 689 | 1055 | 3720 | 575 | 908 | 3963 | 568 | 694 |
| 5043 | QGN18140.1 | FIM1_4463 | CP015061. | 6486 | 297 | 492 | 6013 | 458 | 428 | 6481 | 430 | 742 | 3960 | 332 | 623 | 4475 | 303 | 605 | 3665 | 278 | 459 |
| 5044 | QGN18141.1 | FIM1_4464 | CP015061. | 3361 | 1067 | 664 | 2947 | 603 | 591 | 2920 | 660 | 884 | 2567 | 487 | 759 | 2599 | 388 | 841 | 2333 | 375 | 528 |
| 5045 | QGN18142.1 | FIM1_4465 | CP015061. | 1697 | 1068 | 1043 | 1641 | 964 | 939 | 1273 | 793 | 1321 | 4997 | 922 | 1406 | 3916 | 842 | 1534 | 4382 | 817 | 946 |
| 5046 | QGN18143.1 | FIM1_4466 | CP015061. | 952 | 871 | 904 | 1449 | 838 | 786 | 1080 | 873 | 1154 | 3199 | 653 | 1259 | 3888 | 845 | 1280 | 3325 | 638 | 907 |
| 5047 | QGN18144.1 | FIM1_4467 | CP015061. | 1526 | 2575 | 1632 | 2119 | 2406 | 1513 | 1634 | 2108 | 2105 | 9622 | 2099 | 2665 | 9646 | 2105 | 2308 | 9562 | 2086 | 1600 |
| 5048 | QGN18145.1 | FIM1_4468 | CP015061. | 623 | 254 | 214 | 609 | 176 | 170 | 691 | 188 | 276 | 1109 | 133 | 270 | 896 | 90 | 258 | 1088 | 114 | 169 |
| 5049 | QGN18146.1 | FIM1_4469 | CP015061. | 4784 | 2211 | 1246 | 1681 | 2582 | 1113 | 4170 | 2315 | 1523 | 5650 | 2582 | 2333 | 7976 | 2189 | 2081 | 6361 | 2708 | 1679 |
| 5050 | QGN18147.1 | FIM1_4470 | CP015061. | 7025 | 1911 | 1309 | 3733 | 1946 | 1128 | 4883 | 1924 | 1882 | 2928 | 1681 | 2387 | 7342 | 1895 | 2223 | 3880 | 1967 | 1591 |
| 5051 | QGN18148.1 | FIM1_4471 | CP015061. | 1214 | 592 | 529 | 1510 | 387 | 492 | 1179 | 388 | 712 | 1892 | 432 | 845 | 1613 | 501 | 941 | 1824 | 491 | 606 |
| 5052 | QGN18149.1 | FIM1_4472 | CP015061. | 3295 | 1348 | 1475 | 4225 | 1319 | 1201 | 3844 | 1409 | 1962 | 7487 | 1226 | 2369 | 8673 | 1120 | 2046 | 7455 | 1032 | 1550 |
| 5053 | QGN18150.1 | FIM1_4473 | CP015061. | 1574 | 556 | 610 | 1653 | 708 | 487 | 1475 | 729 | 797 | 1662 | 526 | 739 | 2896 | 388 | 645 | 2182 | 429 | 463 |
| 5054 | QGN18151.1 | FIM1_4474 | CP015061. | 6958 | 6231 | 8023 | 10069 | 9026 | 6946 | 5390 | 7966 | 10968 | 15675 | 8824 | 8657 | 27104 | 8793 | 7802 | 20736 | 8541 | 5816 |
| 5055 | QGN18152.1 | FIM1_4475 | CP015061. | 263 | 636 | 853 | 619 | 474 | 775 | 402 | 442 | 1135 | 1829 | 307 | 1160 | 1993 | 312 | 1125 | 1675 | 289 | 787 |
| 5056 | QGN18153.1 | FIM1_4476 | CP015061. | 1233 | 1310 | 995 | 893 | 1166 | 855 | 1152 | 1052 | 1265 | 3729 | 933 | 1412 | 3259 | 828 | 1412 | 3483 | 866 | 933 |
| 5057 | QGN18154.1 | FIM1_4477 | CP015061. | 2313 | 1475 | 1446 | 3877 | 1326 | 1333 | 2801 | 1373 | 1914 | 5591 | 1221 | 2402 | 6541 | 1317 | 2081 | 5485 | 1195 | 1533 |
| 5058 | QGN18155.1 | FIM1_4478 | CP015061. | 2702 | 1709 | 1397 | 4162 | 1154 | 1289 | 3083 | 1075 | 1779 | 5741 | 896 | 1772 | 4788 | 964 | 1822 | 5539 | 866 | 1156 |
| 5059 | QGN18156.1 | FIM1_4479 | CP015061. | 2072 | 1138 | 1257 | 4041 | 947 | 829 | 2325 | 1213 | 1620 | 2601 | 1468 | 2076 | 7167 | 1581 | 1901 | 3847 | 1555 | 1300 |
| 5060 | QGN18157.1 | FIM1_4480 | CP015061. | 686 | 411 | 317 | 774 | 326 | 271 | 607 | 337 | 439 | 500 | 203 | 413 | 1094 | 231 | 406 | 684 | 255 | 310 |
| 5061 | QGN18158.1 | FIM1_4481 | CP015061. | 3372 | 1124 | 1440 | 3344 | 1081 | 1260 | 2885 | 1091 | 1965 | 4456 | 1121 | 1761 | 5859 | 1009 | 1650 | 4502 | 960 | 1228 |
| 5062 | QGN18159.1 | FIM1_4482 | CP015061. | 37020 | 61 | 48 | 31256 | 30 | 31 | 33169 | 51 | 94 | 511 | 31 | 60 | 847 | 53 | 59 | 431 | 49 | 47 |
| 5063 | QGN18160.1 | FIM1_4483 | CP015061. | 44 | 6 | 8 | 67 | 4 | 4 | 30 | 4 | 5 | 37 | 6 | 8 | 33 | 4 | 8 | 37 | 2 | 2 |
| 5064 | QGN18161.1 | FIM1_4484 | CP015061. | 1384 | 11 | 6 | 247 | 6 | 8 | 828 | 11 | 3 | 19 | 5 | 22 | 59 | 5 | 9 | 23 | 8 | 6 |
| 5065 | QGN18162.1 | FIM1_4485 | CP015061. | 33 | 7 | 6 | 47 | 11 | 6 | 22 | 9 | 7 | 12 | 7 | 6 | 19 | 10 | 2 | 18 | 4 | 4 |
| 5066 | QGN18163.1 | FIM1_4486 | CP015061. | 7164 | 1022 | 812 | 4709 | 1204 | 614 | 5999 | 866 | 961 | 853 | 774 | 1224 | 4052 | 1360 | 927 | 2056 | 934 | 800 |
| 5067 | QGN18164.1 | FIM1_4487 | CP015061. | 3810 | 139 | 222 | 2237 | 139 | 117 | 3173 | 222 | 242 | 227 | 87 | 226 | 689 | 101 | 168 | 331 | 85 | 117 |
| 5068 | QGN18165.1 | FIM1_4488 | CP015061. | 50 | 15 | 22 | 29 | 28 | 16 | 42 | 22 | 32 | 43 | 18 | 42 | 102 | 30 | 47 | 46 | 30 | 29 |
| 5069 | QGN18166.1 | FIM1_4489 | CP015061. | 1622 | 1298 | 729 | 1473 | 745 | 546 | 1875 | 804 | 968 | 766 | 664 | 1076 | 3587 | 761 | 1046 | 2047 | 646 | 722 |
| 5070 | QGN18167.1 | FIM1_4490 | CP015061. | 21381 | 2254 | 3717 | 22874 | 1834 | 2975 | 27623 | 3557 | 5399 | 11036 | 3355 | 4663 | 16271 | 3968 | 5082 | 12905 | 3307 | 3422 |
| 5071 | QGN18168.1 | FIM1_4491 | CP015061. | 1928 | 496 | 539 | 2444 | 607 | 462 | 1699 | 521 | 709 | 1238 | 405 | 704 | 2889 | 619 | 651 | 1953 | 500 | 494 |
| 5072 | QGN18169.1 | FIM1_4492 | CP015061. | 979 | 857 | 490 | 1083 | 755 | 501 | 1014 | 640 | 632 | 2663 | 582 | 566 | 1992 | 519 | 602 | 2593 | 538 | 371 |
| 5073 | QGN18170.1 | FIM1_4493 | CP015061. | 20174 | 2623 | 2872 | 34645 | 3732 | 3084 | 26179 | 2687 | 4325 | 10496 | 2768 | 1817 | 17548 | 3597 | 1868 | 12966 | 3048 | 1557 |
| 5074 | QGN18171.1 | FIM1_4494 | CP015061. | 56 | 27 | 9 | 111 | 31 | 21 | 47 | 30 | 19 | 123 | 34 | 20 | 127 | 44 | 20 | 116 | 36 | 10 |
| 5075 | QGN18172.1 | FIM1_4495 | CP015061. | 5096 | 389 | 630 | 4546 | 464 | 516 | 4300 | 466 | 779 | 3202 | 488 | 757 | 3507 | 568 | 750 | 3191 | 473 | 466 |
| 5076 | QGN18173.1 | FIM1_4496 | CP015061. | 615 | 621 | 396 | 600 | 437 | 332 | 577 | 406 | 486 | 1018 | 297 | 397 | 1646 | 331 | 421 | 1367 | 318 | 277 |
| 5077 | QGN18174.1 | FIM1_4497 | CP015061. | 5801 | 2950 | 1812 | 6267 | 2068 | 1482 | 6010 | 1878 | 2195 | 11120 | 3788 | 3979 | 10504 | 4039 | 4170 | 11327 | 4168 | 2850 |
| 5078 | QGN18175.1 | FIM1_4498 | CP015061. | 15287 | 15487 | 11232 | 13488 | 13212 | 10329 | 12539 | 10865 | 12921 | 42551 | 16863 | 15733 | 37498 | 14615 | 16034 | 43995 | 14652 | 11936 |
| 5079 | QGN18176.1 | FIM1_4499 | CP015061. | 4553 | 1346 | 3068 | 6412 | 1958 | 2762 | 5267 | 1821 | 4052 | 7574 | 1445 | 4354 | 9869 | 1367 | 4000 | 8267 | 1301 | 2773 |
| 5080 | QGN18177.1 | FIM1_4500 | CP015061. | 5772 | 1903 | 1719 | 7122 | 1956 | 1597 | 5615 | 1767 | 2300 | 7666 | 1750 | 2470 | 9894 | 1589 | 2536 | 8358 | 1356 | 1682 |
| 5081 | QGN18178.1 | FIM1_4501 | CP015061. | 301 | 1126 | 1084 | 382 | 1314 | 1062 | 452 | 1072 | 1354 | 3729 | 1028 | 1833 | 5041 | 1009 | 1663 | 3981 | 1013 | 1262 |
| 5082 | QGN18179.1 | FIM1_4502 | CP015061. | 201 | 210 | 146 | 216 | 116 | 138 | 165 | 110 | 185 | 884 | 125 | 198 | 611 | 106 | 197 | 730 | 145 | 143 |
| 5083 | QGN18180.1 | FIM1_4503 | CP015061. | 720 | 413 | 537 | 1176 | 385 | 449 | 884 | 371 | 679 | 2726 | 355 | 791 | 2706 | 319 | 678 | 2417 | 287 | 482 |
| 5084 | QGN18181.1 | FIM1_4504 | CP015061. | 2723 | 409 | 712 | 2706 | 585 | 531 | 2523 | 521 | 882 | 1909 | 565 | 993 | 4703 | 521 | 920 | 2996 | 497 | 650 |
| 5085 | QGN18182.1 | FIM1_4505 | CP015061. | 5274 | 2040 | 905 | 2257 | 2295 | 1025 | 4417 | 2023 | 1213 | 4921 | 2308 | 1531 | 5993 | 2344 | 1502 | 4849 | 2696 | 1165 |
| 5086 | QGN18183.1 | FIM1_4506 | CP015061. | 706 | 1524 | 810 | 799 | 1099 | 860 | 834 | 949 | 1168 | 5488 | 1114 | 1386 | 4197 | 920 | 1215 | 5013 | 1099 | 858 |
| 5087 | QGN18184.1 | FIM1_4507 | CP015061. | 2276 | 1803 | 1828 | 4458 | 2148 | 1634 | 4285 | 2000 | 2258 | 10172 | 1563 | 2617 | 8976 | 1583 | 2376 | 9641 | 1587 | 1653 |
| 5088 | QGN18185.1 | FIM1_4508 | CP015061. | 1766 | 751 | 695 | 1877 | 781 | 703 | 2054 | 750 | 941 | 3880 | 673 | 1158 | 3788 | 645 | 1042 | 3585 | 642 | 797 |
| 5089 | QGN18186.1 | FIM1_4509 | CP015061. | 3589 | 2344 | 2590 | 4226 | 2248 | 2125 | 3806 | 2154 | 3187 | 8676 | 1769 | 2947 | 8059 | 1877 | 2873 | 7577 | 1637 | 2027 |
| 5090 | QGN18187.1 | FIM1_4510 | CP015061. | 4508 | 3516 | 2605 | 5242 | 3479 | 2291 | 5591 | 2955 | 3261 | 14840 | 4234 | 4457 | 13621 | 3726 | 4232 | 14016 | 3824 | 3048 |
| 5091 | QGN18188.1 | FIM1_4512 | CP015061. | 1120 | 557 | 827 | 1386 | 548 | 655 | 1200 | 539 | 960 | 2255 | 413 | 1201 | 3510 | 364 | 980 | 2957 | 398 | 668 |
| 5092 | QGN18189.1 | FIM1_4513 | CP015061. | 221 | 167 | 128 | 210 | 127 | 95 | 156 | 156 | 194 | 892 | 172 | 201 | 688 | 134 | 192 | 833 | 162 | 130 |
| 5093 | QGN18190.1 | FIM1_4514 | CP015061. | 538 | 224 | 256 | 463 | 166 | 199 | 495 | 200 | 334 | 1487 | 134 | 305 | 1183 | 101 | 296 | 1323 | 128 | 210 |
| 5094 | QGN18191.1 | FIM1_4515 | CP015061. | 1099 | 866 | 960 | 1694 | 803 | 851 | 1182 | 813 | 1246 | 4678 | 636 | 1211 | 4335 | 642 | 1201 | 4390 | 575 | 799 |
| 5095 | QGN18192.1 | FIM1_4516 | CP015061. | 1501 | 503 | 318 | 546 | 637 | 269 | 815 | 481 | 359 | 1477 | 809 | 442 | 1404 | 843 | 506 | 1304 | 760 | 291 |
| 5096 | QGN18193.1 | FIM1_4517 | CP015061. | 1610 | 497 | 1032 | 2626 | 679 | 804 | 2069 | 607 | 1221 | 3186 | 275 | 1187 | 4447 | 380 | 939 | 3866 | 218 | 743 |
| 5097 | QGN18194.1 | FIM1_4518 | CP015061. | 1126 | 333 | 432 | 1006 | 416 | 430 | 830 | 349 | 586 | 1665 | 460 | 806 | 3060 | 409 | 799 | 2224 | 420 | 622 |
| 5098 | QGN18195.1 | FIM1_4519 | CP015061. | 949 | 804 | 745 | 1019 | 740 | 548 | 1028 | 625 | 991 | 2050 | 720 | 974 | 3094 | 640 | 1028 | 2514 | 623 | 683 |
| 5099 | QGN18196.1 | FIM1_4520 | CP015061. | 4138 | 1258 | 1100 | 3298 | 1600 | 915 | 3778 | 1464 | 1372 | 2586 | 1602 | 1801 | 5890 | 1439 | 1828 | 3808 | 1455 | 1196 |
| 5100 | QGN18197.1 | FIM1_4521 | CP015061. | 3753 | 1053 | 1476 | 4059 | 1386 | 1286 | 3383 | 1262 | 1937 | 7720 | 1142 | 2300 | 8934 | 1092 | 2109 | 7699 | 991 | 1564 |
| 5101 | QGN18198.1 | FIM1_4522 | CP015061. | 3729 | 257 | 404 | 2811 | 438 | 294 | 1400 | 306 | 483 | 2336 | 175 | 425 | 2945 | 256 | 426 | 2215 | 129 | 293 |
| 5102 | QGN18199.1 | FIM1_4523 | CP015061. | 5213 | 2488 | 1669 | 6446 | 2295 | 1518 | 5526 | 2229 | 2186 | 7582 | 2115 | 2799 | 9949 | 2075 | 2447 | 8188 | 2024 | 1780 |
| 5103 | QGN18200.1 | FIM1_4524 | CP015061. | 2479 | 2229 | 1632 | 3239 | 1682 | 1477 | 2344 | 1583 | 2099 | 6324 | 1209 | 1812 | 4985 | 1346 | 1872 | 5495 | 1255 | 1173 |
| 5104 | QGN18201.1 | FIM1_4525 | CP015061. | 508 | 243 | 189 | 349 | 140 | 168 | 433 | 219 | 298 | 591 | 127 | 180 | 621 | 107 | 222 | 546 | 119 | 130 |
| 5105 | QGN18202.1 | FIM1_4526 | CP015061. | 9428 | 4509 | 4286 | 10057 | 4869 | 3715 | 9543 | 4619 | 5449 | 13914 | 4512 | 6125 | 16302 | 4629 | 6496 | 14264 | 4198 | 4405 |
| 5106 | QGN18203.1 | FIM1_4527 | CP015061. | 11849 | 11361 | 11272 | 28896 | 9897 | 10625 | 23755 | 8568 | 15667 | 8258 | 8782 | 14503 | 33789 | 8882 | 15483 | 15342 | 8682 | 11044 |
| 5107 | QGN18204.1 | FIM1_4528 | CP015061. | 2545 | 65 | 110 | 1146 | 90 | 92 | 1567 | 101 | 148 | 58 | 29 | 57 | 222 | 22 | 62 | 117 | 27 | 37 |
| 5108 | QGN18205.1 | FIM1_4529 | CP015061. | 4115 | 198 | 412 | 2170 | 197 | 280 | 2569 | 252 | 559 | 686 | 117 | 335 | 1195 | 125 | 323 | 914 | 109 | 218 |
| 5109 | QGN18206.1 | FIM1_4530 | CP015061. | 4095 | 1174 | 1382 | 4024 | 1496 | 1228 | 3917 | 1513 | 1823 | 5326 | 1191 | 2294 | 7630 | 1235 | 2055 | 5837 | 1179 | 1416 |
| 5110 | QGN18207.1 | FIM1_4531 | CP015061. | 1613 | 2818 | 4031 | 2632 | 2494 | 3841 | 2027 | 2300 | 5025 | 7714 | 1637 | 4912 | 6917 | 1658 | 4777 | 7461 | 1533 | 3397 |
| 5111 | QGN18208.1 | FIM1_4532 | CP015061. | 205 | 1015 | 1176 | 257 | 1121 | 978 | 150 | 1058 | 1661 | 2111 | 1078 | 1422 | 3880 | 1150 | 1401 | 2875 | 998 | 994 |
| 5112 | QGN18209.1 | FIM1_4533 | CP015061. | 8295 | 4372 | 3050 | 4013 | 6707 | 3022 | 7171 | 5727 | 4208 | 9242 | 7168 | 5817 | 21976 | 7124 | 5191 | 15731 | 7969 | 4003 |
| 5113 | QGN18210.1 | FIM1_4534 | CP015061. | 907 | 566 | 684 | 881 | 493 | 587 | 722 | 560 | 878 | 4128 | 458 | 918 | 3100 | 312 | 843 | 3958 | 311 | 546 |
| 5114 | QGN18211.1 | FIM1_4535 | CP015061. | 1361 | 2089 | 841 | 1264 | 1438 | 897 | 1369 | 1139 | 1191 | 6191 | 1467 | 1584 | 4684 | 1230 | 1555 | 5853 | 1363 | 1094 |
| 5115 | QGN18212.1 | FIM1_4536 | CP015061. | 1211 | 1289 | 1821 | 3263 | 1039 | 1283 | 1978 | 1102 | 2299 | 7795 | 877 | 2702 | 7916 | 1553 | 2281 | 7886 | 1105 | 1670 |
| 5116 | QGN18213.1 | FIM1_4537 | CP015061. | 976 | 478 | 531 | 1030 | 708 | 470 | 898 | 542 | 657 | 3181 | 662 | 961 | 3823 | 591 | 787 | 3378 | 615 | 596 |
| 5117 | QGN18214.1 | FIM1_4538 | CP015061. | 943 | 961 | 1147 | 1999 | 397 | 635 | 1051 | 1059 | 1604 | 1074 | 231 | 730 | 1411 | 312 | 580 | 1237 | 282 | 402 |
| 5118 | QGN18215.1 | FIM1_4539 | CP015061. | 762 | 738 | 691 | 866 | 768 | 616 | 693 | 735 | 938 | 4257 | 737 | 997 | 3883 | 554 | 979 | 4285 | 560 | 646 |
| 5119 | QGN18216.1 | FIM1_4540 | CP015061. | 352 | 362 | 547 | 387 | 283 | 420 | 419 | 450 | 693 | 2717 | 296 | 712 | 2211 | 276 | 614 | 2486 | 287 | 446 |
| 5120 | QGN18217.1 | FIM1_4542 | CP015061. | 2847 | 2763 | 3071 | 6281 | 2615 | 2771 | 3641 | 2474 | 4062 | 15567 | 2403 | 4886 | 15475 | 2881 | 4481 | 16079 | 2449 | 3215 |
| 5121 | QGN18218.1 | FIM1_4543 | CP015061. | 5355 | 2051 | 1062 | 2229 | 3066 | 1174 | 4823 | 2415 | 1489 | 7521 | 3594 | 2523 | 9930 | 4251 | 2303 | 8588 | 4371 | 1718 |
| 5122 | QGN18219.1 | FIM1_4544 | CP015061. | 1231 | 246 | 538 | 2346 | 371 | 412 | 1401 | 307 | 702 | 2899 | 171 | 673 | 2924 | 216 | 590 | 2774 | 153 | 456 |

| | | | | | | | | | | | | | | | | | | | | | |
|---|---|---|---|---|---|---|---|---|---|---|---|---|---|---|---|---|---|---|---|---|---|
| 5123 | QGN18220.1 | FIM1_4545 | CP015061. | 4213 | 1325 | 795 | 2681 | 1689 | 853 | 3948 | 1279 | 1075 | 3802 | 1624 | 1546 | 5409 | 1919 | 1438 | 4110 | 1892 | 1119 |
| 5124 | QGN18221.1 | FIM1_4546 | CP015061. | 7878 | 3494 | 1366 | 4163 | 3939 | 1567 | 7184 | 2875 | 1933 | 13752 | 4756 | 2987 | 13613 | 4869 | 3021 | 13280 | 5156 | 2196 |
| 5125 | QGN18222.1 | FIM1_4547 | CP015061. | 63 | 124 | 60 | 87 | 105 | 53 | 94 | 65 | 59 | 235 | 70 | 73 | 186 | 56 | 81 | 243 | 71 | 35 |
| 5126 | QGN18223.1 | FIM1_4548 | CP015061. | 1591 | 841 | 964 | 1476 | 883 | 781 | 1387 | 1051 | 1297 | 2354 | 758 | 1070 | 3745 | 598 | 1119 | 3397 | 541 | 668 |
| 5127 | QGN18224.1 | FIM1_4549 | CP015061. | 3291 | 1011 | 1985 | 6017 | 1641 | 2037 | 3712 | 1103 | 2612 | 4036 | 605 | 2229 | 6052 | 455 | 2086 | 4684 | 438 | 1462 |
| 5128 | QGN18225.1 | FIM1_4550 | CP015061. | 123 | 62 | 41 | 137 | 65 | 33 | 71 | 60 | 58 | 52 | 29 | 46 | 201 | 29 | 33 | 105 | 26 | 24 |
| 5129 | QGN18226.1 | FIM1_4551 | CP015061. | 311 | 126 | 103 | 377 | 109 | 85 | 193 | 88 | 121 | 523 | 103 | 179 | 523 | 88 | 222 | 505 | 86 | 127 |
| 5130 | QGN18227.1 | FIM1_4552 | CP015061. | 696 | 694 | 691 | 1226 | 735 | 583 | 1320 | 731 | 943 | 1982 | 535 | 886 | 3055 | 672 | 771 | 2161 | 558 | 609 |
| 5131 | QGN18228.1 | FIM1_4553 | CP015061. | 1430 | 478 | 273 | 957 | 338 | 218 | 1246 | 326 | 354 | 1984 | 351 | 401 | 1594 | 265 | 370 | 1740 | 329 | 288 |
| 5132 | QGN18229.1 | FIM1_4554 | CP015061. | 4084 | 947 | 1413 | 5321 | 1371 | 1406 | 3874 | 1284 | 1924 | 3541 | 1154 | 2336 | 7550 | 961 | 2394 | 5432 | 929 | 1682 |
| 5133 | QGN18230.1 | FIM1_4555 | CP015061. | 6977 | 1267 | 2631 | 7340 | 3905 | 2002 | 6542 | 3225 | 3145 | 15055 | 3789 | 4328 | 19412 | 3703 | 4258 | 16368 | 3275 | 3002 |
| 5134 | QGN18231.1 | FIM1_4556 | CP015061. | 1162 | 1765 | 1784 | 4362 | 1636 | 1414 | 2886 | 1422 | 2265 | 9477 | 1792 | 3160 | 9549 | 2156 | 2843 | 9938 | 1887 | 1900 |
| 5135 | QGN18232.1 | FIM1_4557 | CP015061. | 3727 | 3912 | 2990 | 5559 | 3317 | 2811 | 4078 | 2795 | 4125 | 14411 | 3536 | 4819 | 13409 | 3235 | 4917 | 13463 | 3003 | 3260 |
| 5136 | QGN18233.1 | FIM1_4558 | CP015061. | 2185 | 584 | 1670 | 3443 | 829 | 1357 | 2588 | 784 | 2099 | 4119 | 525 | 2558 | 5608 | 504 | 2080 | 4721 | 450 | 1569 |
| 5137 | QGN18234.1 | FIM1_4559 | CP015061. | 1181 | 515 | 989 | 1730 | 401 | 919 | 1221 | 466 | 1284 | 2080 | 344 | 1742 | 2426 | 376 | 1686 | 2145 | 314 | 1177 |
| 5138 | QGN18235.1 | FIM1_4560 | CP015061. | 134 | 111 | 171 | 147 | 55 | 120 | 116 | 81 | 219 | 1014 | 71 | 184 | 623 | 40 | 219 | 1056 | 43 | 115 |
| 5139 | QGN18236.1 | FIM1_4561 | CP015061. | 9575 | 1518 | 2403 | 10984 | 2682 | 1709 | 9160 | 2774 | 2970 | 4775 | 1896 | 3527 | 13201 | 2148 | 3460 | 7317 | 1809 | 2329 |
| 5140 | QGN18237.1 | FIM1_4562 | CP015061. | 8203 | 66 | 246 | 3678 | 109 | 129 | 3159 | 178 | 283 | 235 | 98 | 299 | 531 | 243 | 248 | 283 | 221 | 165 |
| 5141 | QGN18238.1 | FIM1_4563 | CP015061. | 1781 | 2158 | 1078 | 1148 | 1422 | 1241 | 1600 | 1169 | 1493 | 6994 | 2056 | 1556 | 5083 | 1586 | 1552 | 6053 | 1972 | 1047 |
| 5142 | QGN18239.1 | FIM1_4564 | CP015061. | 296 | 292 | 386 | 365 | 230 | 400 | 233 | 234 | 509 | 1195 | 160 | 356 | 692 | 109 | 394 | 963 | 126 | 228 |
| 5143 | QGN18240.1 | FIM1_4565 | CP015061. | 7521 | 1974 | 3338 | 12313 | 2635 | 2634 | 9606 | 2412 | 4410 | 2876 | 1687 | 4625 | 12458 | 1834 | 4346 | 6238 | 1495 | 3240 |
| 5144 | QGN18241.1 | FIM1_4566 | CP015061. | 2581 | 2243 | 10203 | 7428 | 3787 | 6975 | 3740 | 3316 | 13193 | 40629 | 5496 | 9821 | 46739 | 6647 | 8107 | 44726 | 4971 | 7110 |
| 5145 | QGN18242.1 | FIM1_4567 | CP015061. | 3200 | 4854 | 3628 | 6224 | 4771 | 3191 | 4934 | 4717 | 4657 | 16323 | 4718 | 5945 | 19030 | 4416 | 5306 | 17467 | 4101 | 4156 |
| 5146 | QGN18243.1 | FIM1_4568 | CP015061. | 5935 | 3312 | 3295 | 7997 | 3584 | 2732 | 6497 | 3329 | 4257 | 15886 | 3687 | 5160 | 17358 | 3889 | 4569 | 15743 | 3081 | 3266 |
| 5147 | QGN18244.1 | FIM1_4569 | CP015061. | 1309 | 2004 | 1857 | 2816 | 1834 | 1702 | 1927 | 1648 | 2330 | 9737 | 1466 | 2425 | 7508 | 1376 | 2232 | 9170 | 1433 | 1591 |
| 5148 | QGN18245.1 | FIM1_4570 | CP015061. | 3528 | 699 | 882 | 3567 | 798 | 804 | 2898 | 630 | 1119 | 6846 | 882 | 1530 | 6012 | 787 | 1415 | 6947 | 726 | 1086 |
| 5149 | QGN18246.1 | FIM1_4571 | CP015061. | 977 | 791 | 717 | 1406 | 833 | 641 | 1347 | 710 | 928 | 5780 | 784 | 1186 | 4569 | 744 | 1008 | 5436 | 683 | 791 |
| 5150 | QGN18247.1 | FIM1_4572 | CP015061. | 8917 | 3885 | 1924 | 7414 | 3305 | 1538 | 9339 | 3648 | 2644 | 2524 | 3001 | 3479 | 12451 | 3031 | 2901 | 5497 | 3527 | 2258 |
| 5151 | QGN18248.1 | FIM1_4573 | CP015061. | 3620 | 666 | 1036 | 7630 | 933 | 968 | 6052 | 668 | 1485 | 3912 | 868 | 1786 | 7067 | 743 | 1683 | 4839 | 764 | 1284 |
| 5152 | QGN18249.1 | FIM1_4574 | CP015061. | 1209 | 447 | 455 | 1650 | 375 | 355 | 1438 | 422 | 624 | 1753 | 318 | 667 | 2208 | 275 | 716 | 1886 | 296 | 445 |
| 5153 | QGN18250.1 | FIM1_4575 | CP015061. | 8925 | 7651 | 5100 | 9326 | 5615 | 4314 | 7651 | 4848 | 6312 | 22395 | 6992 | 8538 | 21025 | 6620 | 8596 | 21382 | 6002 | 5889 |
| 5154 | QGN18251.1 | FIM1_4576 | CP015061. | 837 | 1502 | 1029 | 928 | 1139 | 1079 | 881 | 1022 | 1432 | 5431 | 1099 | 1459 | 4313 | 987 | 1428 | 5000 | 1147 | 1045 |
| 5155 | QGN18252.1 | FIM1_4577 | CP015061. | 163 | 424 | 376 | 475 | 280 | 281 | 346 | 313 | 461 | 1695 | 288 | 517 | 1698 | 332 | 586 | 1568 | 280 | 368 |
| 5156 | QGN18253.1 | FIM1_4578 | CP015061. | 629 | 484 | 294 | 617 | 320 | 231 | 776 | 376 | 405 | 982 | 258 | 404 | 1310 | 352 | 406 | 1037 | 307 | 259 |
| 5157 | QGN18254.1 | FIM1_4579 | CP015061. | 401 | 99 | 233 | 440 | 259 | 176 | 441 | 286 | 261 | 706 | 182 | 297 | 1101 | 200 | 265 | 778 | 204 | 186 |
| 5158 | QGN18255.1 | FIM1_4580 | CP015061. | 2173 | 2776 | 1368 | 1573 | 2565 | 1317 | 2005 | 2236 | 1761 | 7332 | 2857 | 2294 | 8331 | 2597 | 2159 | 7292 | 2821 | 1588 |
| 5159 | QGN18256.1 | FIM1_4582 | CP015061. | 3302 | 963 | 848 | 4644 | 999 | 701 | 4080 | 953 | 1128 | 1535 | 888 | 1325 | 5815 | 1014 | 1273 | 3977 | 893 | 1029 |
| 5160 | QGN18257.1 | FIM1_4583 | CP015061. | 3003 | 686 | 837 | 3360 | 850 | 736 | 2550 | 753 | 1098 | 5046 | 880 | 1224 | 5532 | 805 | 1202 | 4917 | 727 | 853 |
| 5161 | QGN18258.1 | FIM1_4584 | CP015061. | 3856 | 1343 | 1038 | 5543 | 1303 | 985 | 3986 | 1038 | 1380 | 5337 | 934 | 1505 | 5002 | 1033 | 1414 | 5224 | 961 | 975 |
| 5162 | QGN18259.1 | FIM1_4585 | CP015061. | 6036 | 1310 | 1282 | 6960 | 1305 | 1079 | 5989 | 1193 | 1654 | 6947 | 1290 | 2150 | 8236 | 1463 | 1879 | 6916 | 1199 | 1360 |
| 5163 | QGN18260.1 | FIM1_4586 | CP015061. | 15064 | 10520 | 7258 | 14794 | 9484 | 6091 | 13106 | 9408 | 9220 | 34892 | 11220 | 13847 | 43047 | 11818 | 14270 | 36049 | 10822 | 9995 |
| 5164 | QGN18261.1 | FIM1_4587 | CP015061. | 3795 | 1914 | 762 | 1324 | 2057 | 902 | 2917 | 1809 | 1134 | 4988 | 2122 | 1395 | 5585 | 1887 | 1296 | 4904 | 2263 | 1014 |
| 5165 | QGN18262.1 | FIM1_4588 | CP015061. | 1011 | 1775 | 2360 | 1950 | 2040 | 2173 | 1579 | 1635 | 3120 | 12113 | 2081 | 4128 | 10106 | 2056 | 3683 | 11702 | 1980 | 2697 |
| 5166 | QGN18263.1 | FIM1_4589 | CP015061. | 1264 | 1822 | 938 | 848 | 1752 | 975 | 1253 | 1361 | 1200 | 8845 | 2498 | 1626 | 7691 | 2375 | 1741 | 8997 | 2583 | 1274 |
| 5167 | QGN18264.1 | FIM1_4590 | CP015061. | 605 | 296 | 331 | 845 | 403 | 362 | 695 | 318 | 408 | 2646 | 406 | 597 | 2355 | 384 | 488 | 2637 | 425 | 390 |
| 5168 | QGN18265.1 | FIM1_4591 | CP015061. | 1228 | 2208 | 1389 | 1318 | 2149 | 1527 | 1196 | 1701 | 1828 | 10214 | 2236 | 2671 | 9208 | 2024 | 2492 | 9354 | 2040 | 1737 |
| 5169 | QGN18266.1 | FIM1_4592 | CP015061. | 1282 | 1051 | 830 | 1644 | 1297 | 683 | 1504 | 1036 | 997 | 4951 | 998 | 1166 | 4464 | 906 | 1036 | 4723 | 808 | 733 |
| 5170 | QGN18267.1 | FIM1_4593 | CP015061. | 1161 | 493 | 479 | 995 | 368 | 487 | 1191 | 363 | 756 | 3501 | 327 | 765 | 2894 | 244 | 714 | 3209 | 303 | 500 |
| 5171 | QGN18268.1 | FIM1_4594 | CP015061. | 892 | 257 | 228 | 2110 | 205 | 237 | 1051 | 241 | 387 | 3466 | 402 | 404 | 3584 | 266 | 554 | 3414 | 231 | 336 |
| 5172 | QGN18269.1 | FIM1_4595 | CP015061. | 4200 | 1982 | 2706 | 4239 | 2681 | 2023 | 4796 | 2548 | 3317 | 11224 | 3034 | 2798 | 13232 | 2376 | 3005 | 11908 | 2367 | 1991 |
| 5173 | QGN18270.1 | FIM1_4596 | CP015061. | 4981 | 2003 | 1253 | 2837 | 2352 | 1343 | 4772 | 1799 | 1637 | 8851 | 3280 | 2249 | 8481 | 3361 | 2260 | 8475 | 3705 | 1622 |
| 5174 | QGN18271.1 | FIM1_4597 | CP015061. | 550 | 1000 | 1351 | 932 | 1213 | 1135 | 668 | 1032 | 1845 | 6942 | 1253 | 1554 | 6869 | 1265 | 1439 | 7002 | 1222 | 1044 |
| 5175 | QGN18272.1 | FIM1_4598 | CP015061. | 576 | 350 | 314 | 1150 | 283 | 284 | 717 | 228 | 432 | 2008 | 258 | 463 | 1599 | 212 | 426 | 2064 | 208 | 300 |
| 5176 | QGN18273.1 | FIM1_4599 | CP015061. | 256 | 278 | 175 | 565 | 309 | 139 | 477 | 255 | 213 | 936 | 197 | 215 | 792 | 150 | 218 | 881 | 156 | 150 |
| 5177 | QGN18274.1 | FIM1_4600 | CP015061. | 5399 | 2190 | 1876 | 7025 | 2786 | 1656 | 5925 | 2360 | 2328 | 6992 | 2582 | 2944 | 11007 | 2457 | 2742 | 8041 | 2332 | 1952 |
| 5178 | QGN18275.1 | FIM1_4601 | CP015061. | 994 | 751 | 584 | 1238 | 571 | 511 | 975 | 593 | 815 | 1905 | 393 | 664 | 2160 | 375 | 722 | 1774 | 350 | 479 |
| 5179 | QGN18276.1 | FIM1_4602 | CP015061. | 209 | 539 | 292 | 210 | 290 | 324 | 139 | 286 | 435 | 1555 | 342 | 458 | 944 | 278 | 453 | 1146 | 397 | 234 |
| 5180 | QGN18277.1 | FIM1_4603 | CP015061. | 4514 | 3162 | 2456 | 4690 | 3071 | 2190 | 4030 | 3067 | 3302 | 6331 | 2775 | 3377 | 10174 | 2469 | 3666 | 7101 | 2378 | 2234 |
| 5181 | QGN18278.1 | FIM1_4604 | CP015061. | 340 | 51 | 153 | 335 | 44 | 76 | 388 | 71 | 108 | 625 | 67 | 68 | 526 | 54 | 87 | 570 | 42 | 54 |
| 5182 | QGN18279.1 | FIM1_4605 | CP015061. | 1337 | 780 | 571 | 1324 | 824 | 530 | 1599 | 668 | 748 | 4388 | 752 | 1028 | 3831 | 748 | 969 | 4237 | 718 | 620 |
| 5183 | QGN18280.1 | FIM1_4606 | CP015061. | 839 | 584 | 444 | 1080 | 524 | 447 | 990 | 387 | 652 | 2200 | 379 | 604 | 1922 | 400 | 650 | 2135 | 401 | 441 |
| 5184 | QGN18281.1 | FIM1_4607 | CP015061. | 4472 | 1790 | 1473 | 4983 | 1506 | 1513 | 4258 | 1470 | 2039 | 5056 | 1234 | 1957 | 5903 | 1242 | 2115 | 5172 | 1145 | 1305 |
| 5185 | QGN18282.1 | FIM1_4608 | CP015061. | 1033 | 255 | 327 | 1150 | 225 | 249 | 539 | 255 | 413 | 1104 | 114 | 476 | 1545 | 129 | 501 | 1214 | 88 | 291 |
| 5186 | QGN18283.1 | FIM1_4609 | CP015061. | 1137 | 2697 | 1866 | 1373 | 2643 | 1557 | 1591 | 2752 | 2448 | 6489 | 3122 | 2863 | 10530 | 2310 | 2676 | 7537 | 2613 | 1971 |
| 5187 | QGN18284.1 | FIM1_4610 | CP015061. | 896 | 4 | 6 | 609 | 4 | 1 | 300 | 4 | 8 | 18 | 7 | 16 | 38 | 2 | 1 | 36 | 6 | 9 |
| 5188 | QGN18285.1 | FIM1_4611 | CP015061. | 0 | 1 | 0 | 5 | 0 | 1 | 2 | 0 | 1 | 0 | 1 | 0 | 0 | 0 | 0 | 3 | 0 | 1 |
| 5189 | QGN18286.1 | FIM1_4612 | CP015061. | 9610 | 3370 | 4387 | 12203 | 4662 | 3950 | 9647 | 4124 | 5665 | 12963 | 4336 | 6929 | 26043 | 4121 | 6191 | 16596 | 4036 | 4597 |
| 5190 | QGN18287.1 | FIM1_4613 | CP015061. | 3859 | 19115 | 15874 | 6869 | 16221 | 13507 | 14143 | 19594 | 18868 | 27400 | 9682 | 9506 | 32162 | 8899 | 10228 | 26978 | 9237 | 7148 |
| 5191 | QGN18288.1 | FIM1_4614 | CP015061. | 955 | 395 | 392 | 1069 | 367 | 305 | 937 | 458 | 517 | 1741 | 374 | 543 | 2192 | 273 | 448 | 1701 | 329 | 306 |
| 5192 | QGN18289.1 | FIM1_4615 | CP015061. | 3156 | 786 | 900 | 3005 | 769 | 828 | 2998 | 610 | 1185 | 5789 | 781 | 1305 | 5332 | 871 | 1175 | 5518 | 808 | 788 |
| 5193 | QGN18290.1 | FIM1_4616 | CP015061. | 1122 | 2882 | 1867 | 1387 | 2913 | 1725 | 1113 | 2675 | 2404 | 7894 | 2434 | 2531 | 10294 | 2306 | 2403 | 8419 | 2395 | 1772 |
| 5194 | QGN18291.1 | FIM1_4617 | CP015061. | 4245 | 1320 | 2493 | 5489 | 2759 | 2154 | 4980 | 2396 | 2964 | 8724 | 2125 | 3204 | 12224 | 2110 | 3178 | 10099 | 1659 | 2293 |
| 5195 | QGN18292.1 | FIM1_4618 | CP015061. | 2627 | 2410 | 3232 | 2771 | 2703 | 2564 | 2221 | 2805 | 3792 | 9975 | 2673 | 4521 | 16616 | 2516 | 3918 | 12238 | 2364 | 2947 |
| 5196 | QGN18293.1 | FIM1_4619 | CP015061. | 586 | 750 | 635 | 842 | 567 | 532 | 669 | 473 | 792 | 2442 | 555 | 868 | 2626 | 557 | 848 | 2525 | 496 | 592 |
| 5197 | QGN18294.1 | FIM1_4620 | CP015061. | 1710 | 598 | 446 | 1282 | 526 | 389 | 1287 | 493 | 588 | 634 | 464 | 619 | 2221 | 419 | 563 | 1157 | 399 | 442 |
| 5198 | QGN18295.1 | FIM1_4621 | CP015061. | 184 | 16 | 69 | 140 | 101 | 36 | 159 | 111 | 57 | 78 | 85 | 111 | 507 | 85 | 95 | 203 | 92 | 55 |
| 5199 | QGN18296.1 | FIM1_4622 | CP015061. | 4852 | 611 | 704 | 5744 | 616 | 601 | 5156 | 697 | 982 | 2584 | 470 | 890 | 3868 | 489 | 813 | 2796 | 383 | 610 |
| 5200 | QGN18297.1 | FIM1_4623 | CP015061. | 2233 | 611 | 855 | 1816 | 832 | 718 | 2259 | 737 | 1073 | 4681 | 927 | 1501 | 5265 | 841 | 1409 | 4889 | 882 | 1023 |
| 5201 | QGN18298.1 | FIM1_4624 | CP015061. | 1143 | 1890 | 2022 | 1621 | 1366 | 1768 | 1177 | 1392 | 2575 | 8186 | 1459 | 2759 | 6574 | 1498 | 2842 | 7968 | 1308 | 1836 |
| 5202 | QGN18299.1 | FIM1_4625 | CP015061. | 1275 | 729 | 565 | 1321 | 715 | 484 | 1155 | 754 | 779 | 1848 | 542 | 664 | 2439 | 562 | 555 | 1967 | 572 | 388 |
| 5203 | QGN18300.1 | FIM1_4626 | CP015061. | 1046 | 159 | 309 | 1367 | 214 | 229 | 751 | 203 | 406 | 1298 | 185 | 404 | 1681 | 188 | 356 | 1377 | 169 | 234 |
| 5204 | QGN18301.1 | FIM1_4627 | CP015061. | 1665 | 1254 | 1853 | 3480 | 1515 | 1614 | 1965 | 1593 | 2555 | 7252 | 1061 | 2675 | 8980 | 1435 | 2263 | 7135 | 985 | 1593 |
| 5205 | QGN18302.1 | FIM1_4628 | CP015061. | 585 | 179 | 303 | 649 | 177 | 248 | 379 | 260 | 432 | 723 | 122 | 307 | 989 | 102 | 300 | 845 | 100 | 186 |
| 5206 | QGN18303.1 | FIM1_4629 | CP015061. | 1740 | 830 | 504 | 1243 | 502 | 483 | 1361 | 593 | 719 | 1923 | 445 | 597 | 2007 | 269 | 712 | 1914 | 361 | 428 |

| | | | | | | | | | | | | | | | | | | | | | |
|---|---|---|---|---|---|---|---|---|---|---|---|---|---|---|---|---|---|---|---|---|---|
| 5207 | QGN18304.1 | FIM1_4630 | CP015061. | 4882 | 904 | 760 | 4789 | 667 | 776 | 4947 | 754 | 1064 | 2191 | 469 | 1099 | 2774 | 433 | 1113 | 2154 | 477 | 768 |
| 5208 | QGN18305.1 | FIM1_4631 | CP015061. | 2544 | 2038 | 1695 | 3991 | 1430 | 1351 | 3380 | 1468 | 2064 | 2943 | 1101 | 1993 | 4902 | 1230 | 2230 | 3610 | 1034 | 1429 |
| 5209 | QGN18306.1 | FIM1_4632 | CP015061. | 9941 | 1079 | 1238 | 7398 | 1164 | 1008 | 8857 | 1329 | 1692 | 1866 | 915 | 1855 | 5595 | 1036 | 1632 | 2940 | 1056 | 1280 |
| 5210 | QGN18307.1 | FIM1_4633 | CP015061. | 314 | 55 | 63 | 194 | 33 | 35 | 330 | 66 | 90 | 188 | 32 | 36 | 251 | 48 | 56 | 179 | 31 | 35 |
| 5211 | QGN18308.1 | FIM1_4634 | CP015061. | 10723 | 1115 | 1883 | 10907 | 1336 | 1584 | 9549 | 1330 | 2702 | 930 | 749 | 3046 | 6631 | 873 | 2816 | 2905 | 773 | 2020 |
| 5212 | QGN18309.1 | FIM1_4635 | CP015061. | 1722 | 493 | 811 | 2678 | 585 | 633 | 2037 | 454 | 1092 | 5031 | 522 | 1256 | 4858 | 547 | 1134 | 4838 | 442 | 745 |
| 5213 | QGN18310.1 | FIM1_4636 | CP015061. | 670 | 388 | 399 | 830 | 394 | 333 | 656 | 350 | 558 | 2271 | 388 | 609 | 2135 | 369 | 607 | 2201 | 285 | 413 |
| 5214 | QGN18311.1 | FIM1_4637 | CP015061. | 1379 | 660 | 780 | 2441 | 606 | 548 | 1890 | 615 | 954 | 4611 | 576 | 1071 | 3673 | 675 | 912 | 4678 | 583 | 658 |
| 5215 | QGN18312.1 | FIM1_4638 | CP015061. | 1584 | 741 | 860 | 2111 | 694 | 739 | 1597 | 762 | 1094 | 5606 | 644 | 1061 | 4897 | 570 | 1061 | 5389 | 558 | 715 |
| 5216 | QGN18313.1 | FIM1_4639 | CP015061. | 4483 | 943 | 1308 | 6650 | 1395 | 1151 | 5033 | 1248 | 1789 | 5768 | 1002 | 1835 | 7479 | 977 | 1673 | 5783 | 960 | 1214 |
| 5217 | QGN18314.1 | FIM1_4640 | CP015061. | 994 | 956 | 513 | 911 | 641 | 564 | 771 | 605 | 740 | 3052 | 621 | 638 | 2636 | 513 | 725 | 2673 | 553 | 504 |
| 5218 | QGN18315.1 | FIM1_4641 | CP015061. | 8896 | 4389 | 4354 | 7266 | 4190 | 3584 | 7874 | 4360 | 5371 | 15910 | 4504 | 4690 | 13781 | 3789 | 4969 | 15895 | 3793 | 3469 |
| 5219 | QGN18316.1 | FIM1_4642 | CP015061. | 694 | 3075 | 1961 | 1649 | 2867 | 1790 | 1188 | 2478 | 2674 | 5859 | 2110 | 2124 | 7374 | 2230 | 2295 | 6049 | 2103 | 1527 |
| 5220 | QGN18317.1 | FIM1_4643 | CP015061. | 772 | 562 | 335 | 625 | 364 | 288 | 695 | 365 | 448 | 1321 | 330 | 367 | 1178 | 344 | 437 | 1172 | 323 | 250 |
| 5221 | QGN18318.1 | FIM1_4644 | CP015061. | 1159 | 405 | 657 | 1066 | 524 | 488 | 1154 | 481 | 756 | 3729 | 493 | 1036 | 4013 | 385 | 905 | 3803 | 462 | 683 |
| 5222 | QGN18319.1 | FIM1_4645 | CP015061. | 1089 | 546 | 637 | 1025 | 843 | 578 | 1096 | 830 | 856 | 4497 | 736 | 1065 | 4661 | 688 | 932 | 4409 | 749 | 720 |
| 5223 | QGN18320.1 | FIM1_4646 | CP015061. | 824 | 1432 | 1100 | 718 | 1332 | 901 | 770 | 1313 | 1369 | 5577 | 1104 | 1546 | 5940 | 1020 | 1332 | 5444 | 1066 | 919 |
| 5224 | QGN18321.1 | FIM1_4647 | CP015061. | 388 | 1927 | 1320 | 490 | 1586 | 1231 | 367 | 1342 | 1668 | 7840 | 1487 | 1776 | 6382 | 1353 | 1714 | 7241 | 1316 | 1124 |
| 5225 | QGN18322.1 | FIM1_4648 | CP015061. | 2061 | 1437 | 1073 | 3255 | 1038 | 1001 | 2540 | 997 | 1421 | 5948 | 1034 | 1437 | 4526 | 984 | 1416 | 5416 | 940 | 941 |
| 5226 | QGN18323.1 | FIM1_4649 | CP015061. | 4240 | 1043 | 1522 | 5748 | 1359 | 1280 | 4641 | 1253 | 2040 | 7241 | 1347 | 2396 | 9003 | 1467 | 2209 | 7687 | 1345 | 1596 |
| 5227 | QGN18324.1 | FIM1_4650 | CP015061. | 2195 | 1496 | 847 | 1693 | 1165 | 885 | 1934 | 1116 | 1175 | 4589 | 989 | 1327 | 4388 | 972 | 1344 | 4110 | 971 | 919 |
| 5228 | QGN18325.1 | FIM1_4651 | CP015061. | 11652 | 2741 | 2423 | 11607 | 3306 | 1870 | 11894 | 2982 | 3073 | 2711 | 2257 | 3211 | 12013 | 2969 | 2527 | 6232 | 2820 | 1967 |
| 5229 | QGN18326.1 | FIM1_4652 | CP015061. | 376 | 952 | 925 | 622 | 1268 | 895 | 448 | 981 | 1329 | 4440 | 979 | 1063 | 4410 | 958 | 991 | 4279 | 956 | 703 |
| 5230 | QGN18327.1 | FIM1_4653 | CP015061. | 846 | 433 | 521 | 1470 | 657 | 479 | 952 | 620 | 692 | 1735 | 711 | 1042 | 4286 | 719 | 912 | 2821 | 752 | 732 |
| 5231 | QGN18328.1 | FIM1_4654 | CP015061. | 1356 | 1364 | 905 | 1743 | 783 | 755 | 1439 | 897 | 1195 | 3306 | 520 | 1012 | 2661 | 679 | 1065 | 2935 | 686 | 656 |
| 5232 | QGN18329.1 | FIM1_4655 | CP015061. | 8002 | 1524 | 1108 | 6220 | 1071 | 915 | 6375 | 1322 | 1569 | 1152 | 760 | 1165 | 4163 | 628 | 1218 | 2137 | 660 | 891 |
| 5233 | QGN18330.1 | FIM1_4656 | CP015061. | 4718 | 1547 | 1500 | 5880 | 1317 | 1068 | 5741 | 1901 | 2065 | 888 | 936 | 1921 | 4241 | 1127 | 1753 | 1603 | 1162 | 1344 |
| 5234 | QGN18331.1 | FIM1_4657 | CP015061. | 1542 | 889 | 643 | 1430 | 826 | 486 | 1425 | 805 | 881 | 1642 | 610 | 757 | 2793 | 567 | 762 | 1892 | 567 | 484 |
| 5235 | QGN18332.1 | FIM1_4658 | CP015061. | 4022 | 17797 | 12549 | 8389 | 18888 | 9842 | 5943 | 18607 | 14982 | 25165 | 18829 | 19337 | 47440 | 18299 | 18009 | 36297 | 17041 | 14164 |
| 5236 | QGN18333.1 | FIM1_4659 | CP015061. | 18 | 5 | 1 | 9 | 1 | 1 | 8 | 2 | 7 | 6 | 0 | 1 | 8 | 0 | 1 | 5 | 4 | 3 |
| 5237 | QGN18334.1 | FIM1_4660 | CP015061. | 1689 | 86 | 62 | 1131 | 60 | 46 | 1256 | 53 | 67 | 172 | 50 | 48 | 239 | 51 | 62 | 144 | 57 | 32 |
| 5238 | QGN18335.1 | FIM1_4661 | CP015061. | 736 | 335 | 100 | 473 | 363 | 123 | 457 | 247 | 168 | 264 | 282 | 145 | 695 | 282 | 106 | 475 | 258 | 87 |
| 5239 | QGN18336.1 | FIM1_4662 | CP015061. | 2518 | 2774 | 1136 | 1048 | 2435 | 1110 | 1926 | 2130 | 1509 | 8898 | 3518 | 1830 | 7172 | 2953 | 2105 | 8375 | 3449 | 1376 |
| 5240 | QGN18337.1 | FIM1_4664 | CP015061. | 83 | 30 | 49 | 63 | 25 | 29 | 66 | 30 | 46 | 148 | 23 | 43 | 88 | 15 | 49 | 126 | 16 | 21 |
| 5241 | QGN18338.1 | FIM1_4665 | CP015061. | 4306 | 1830 | 1305 | 4712 | 2469 | 1099 | 4926 | 2589 | 1691 | 2083 | 2713 | 2273 | 10012 | 3023 | 1908 | 4924 | 3256 | 1467 |
| 5242 | QGN18339.1 | FIM1_4666 | CP015061. | 614 | 253 | 617 | 1563 | 336 | 471 | 773 | 355 | 898 | 363 | 386 | 1008 | 1247 | 342 | 971 | 675 | 355 | 643 |
| 5243 | QGN18340.1 | FIM1_4667 | CP015061. | 9162 | 3970 | 1218 | 3931 | 3953 | 1384 | 7565 | 3494 | 1805 | 4653 | 4748 | 2878 | 12613 | 4522 | 2601 | 6711 | 5707 | 2015 |
| 5244 | QGN18341.1 | FIM1_4668 | CP015061. | 1798 | 432 | 260 | 2011 | 357 | 214 | 1628 | 295 | 374 | 191 | 276 | 291 | 592 | 294 | 235 | 280 | 321 | 194 |
| 5245 | QGN18342.1 | FIM1_4669 | CP015061. | 93 | 13 | 25 | 98 | 13 | 15 | 91 | 32 | 23 | 48 | 34 | 28 | 169 | 23 | 23 | 133 | 19 | 16 |
| 5246 | QGN18343.1 | FIM1_4670 | CP015061. | 2442 | 1189 | 1599 | 3780 | 1702 | 1250 | 3234 | 1670 | 2065 | 3492 | 1406 | 2306 | 9273 | 1299 | 1984 | 5362 | 1287 | 1550 |
| 5247 | QGN18344.1 | FIM1_4671 | CP015061. | 2027 | 747 | 868 | 3058 | 655 | 705 | 2676 | 760 | 1089 | 4458 | 516 | 1186 | 4622 | 557 | 1054 | 4531 | 469 | 716 |
| 5248 | QGN18345.1 | FIM1_4672 | CP015061. | 3429 | 975 | 746 | 3344 | 685 | 658 | 3372 | 773 | 1028 | 3615 | 559 | 858 | 3013 | 518 | 888 | 3227 | 521 | 584 |
| 5249 | QGN18346.1 | FIM1_4673 | CP015061. | 726 | 173 | 228 | 1155 | 175 | 166 | 1161 | 220 | 269 | 1257 | 198 | 331 | 1346 | 189 | 330 | 1199 | 166 | 244 |
| 5250 | QGN18347.1 | FIM1_4674 | CP015061. | 928 | 196 | 136 | 929 | 171 | 153 | 975 | 176 | 211 | 721 | 139 | 215 | 769 | 132 | 195 | 733 | 119 | 146 |
| 5251 | QGN18348.1 | FIM1_4675 | CP015061. | 12601 | 3556 | 2170 | 9505 | 3064 | 2067 | 9808 | 2752 | 2820 | 11044 | 3960 | 3735 | 12390 | 3437 | 3975 | 10643 | 3439 | 2623 |
| 5252 | QGN18349.1 | FIM1_4676 | CP015061. | 2319 | 1695 | 1132 | 2537 | 1626 | 940 | 2570 | 1513 | 1474 | 6666 | 1565 | 1923 | 7881 | 1530 | 1659 | 6632 | 1432 | 1215 |
| 5253 | QGN18350.1 | FIM1_4677 | CP015061. | 0 | 0 | 0 | 1 | 1 | 0 | 0 | 1 | 0 | 3 | 0 | 0 | 0 | 0 | 1 | 1 | 1 | 2 |
| 5254 | QGN18351.1 | FIM1_4678 | CP015061. | 3527 | 2176 | 949 | 1733 | 2021 | 829 | 3050 | 2113 | 1215 | 1118 | 1704 | 1589 | 5900 | 1423 | 1489 | 3029 | 1744 | 1125 |
| 5255 | QGN18352.1 | FIM1_4679 | CP015061. | 6281 | 1145 | 622 | 3613 | 438 | 343 | 3705 | 673 | 803 | 4780 | 1339 | 1267 | 5555 | 1382 | 1619 | 4975 | 1361 | 952 |
| 5256 | QGN18353.1 | FIM1_4680 | CP015061. | 1376 | 445 | 347 | 1870 | 344 | 266 | 1632 | 336 | 484 | 1336 | 383 | 443 | 1658 | 412 | 506 | 1371 | 385 | 289 |
| 5257 | QGN18354.1 | FIM1_4681 | CP015061. | 4414 | 1744 | 1559 | 9231 | 1784 | 1342 | 6415 | 1749 | 1966 | 8767 | 1880 | 2052 | 10729 | 2664 | 2014 | 8786 | 2075 | 1311 |
| 5258 | QGN18355.1 | FIM1_4682 | CP015061. | 3768 | 4629 | 4239 | 5506 | 5110 | 4078 | 4072 | 4361 | 5489 | 16102 | 4819 | 5860 | 19606 | 4467 | 5705 | 16345 | 4135 | 4026 |
| 5259 | QGN18356.1 | FIM1_4683 | CP015061. | 37 | 246 | 123 | 16 | 115 | 114 | 33 | 157 | 157 | 433 | 101 | 148 | 260 | 59 | 156 | 349 | 121 | 95 |
| 5260 | QGN18357.1 | FIM1_4684 | CP015061. | 2195 | 1378 | 827 | 1486 | 1208 | 825 | 1843 | 1105 | 1162 | 6066 | 1243 | 1082 | 5430 | 1058 | 1222 | 5897 | 1115 | 809 |
| 5261 | QGN18358.1 | FIM1_4685 | CP015061. | 2156 | 1678 | 2107 | 5800 | 1367 | 1623 | 1566 | 1163 | 3119 | 5780 | 1249 | 1458 | 6235 | 1046 | 546 | 4956 | 583 | 975 |
| 5262 | QGN18359.1 | FIM1_4686 | CP015061. | 1582 | 83 | 152 | 4127 | 128 | 131 | 1417 | 140 | 186 | 170 | 45 | 108 | 376 | 85 | 97 | 212 | 58 | 57 |
| 5263 | QGN18360.1 | FIM1_4687 | CP015061. | 8 | 13 | 4 | 9 | 8 | 8 | 3 | 8 | 12 | 13 | 2 | 3 | 16 | 2 | 5 | 14 | 3 | 2 |
| 5264 | QGN18361.1 | FIM1_4688 | CP015061. | 410 | 1211 | 623 | 443 | 979 | 458 | 656 | 1001 | 787 | 2301 | 579 | 706 | 3386 | 716 | 710 | 2429 | 682 | 539 |
| 5265 | QGN18362.1 | FIM1_4689 | CP015061. | 2705 | 732 | 745 | 2096 | 776 | 649 | 2032 | 662 | 991 | 4657 | 763 | 1242 | 4614 | 674 | 1060 | 4623 | 614 | 752 |
| 5266 | QGN18363.1 | FIM1_4690 | CP015061. | 8211 | 1251 | 1778 | 6791 | 2180 | 1592 | 6470 | 1853 | 2370 | 8809 | 2004 | 2970 | 11235 | 1962 | 2745 | 9080 | 1824 | 2024 |
| 5267 | QGN18364.1 | FIM1_4691 | CP015061. | 2708 | 1410 | 345 | 2245 | 1048 | 349 | 2557 | 982 | 510 | 477 | 912 | 649 | 1748 | 1023 | 559 | 835 | 1363 | 459 |
| 5268 | QGN18365.1 | FIM1_4692 | CP015061. | 1668 | 732 | 896 | 2495 | 690 | 1014 | 1956 | 502 | 1237 | 4190 | 1141 | 1259 | 2956 | 1279 | 1308 | 3551 | 1708 | 877 |
| 5269 | QGN18366.1 | FIM1_4693 | CP015061. | 1136 | 229 | 203 | 1058 | 133 | 140 | 1106 | 202 | 299 | 496 | 184 | 250 | 980 | 209 | 259 | 582 | 166 | 227 |
| 5270 | QGN18367.1 | FIM1_4694 | CP015061. | 3031 | 480 | 640 | 2104 | 465 | 494 | 2208 | 577 | 892 | 577 | 314 | 708 | 2637 | 352 | 684 | 1241 | 319 | 506 |
| 5271 | QGN18368.1 | FIM1_4695 | CP015061. | 1357 | 485 | 404 | 612 | 481 | 292 | 724 | 532 | 572 | 216 | 273 | 397 | 650 | 328 | 369 | 353 | 317 | 293 |
| 5272 | QGN18369.1 | FIM1_4696 | CP015061. | 2 | 1 | 3 | 14 | 8 | 5 | 7 | 7 | 5 | 4 | 2 | 5 | 11 | 2 | 2 | 6 | 6 | 4 |
| 5273 | QGN18370.1 | FIM1_4697 | CP015061. | 2830 | 1492 | 416 | 1108 | 1239 | 385 | 2344 | 1436 | 586 | 385 | 979 | 870 | 2294 | 1205 | 720 | 795 | 1539 | 615 |
| 5274 | QGN18371.1 | FIM1_4698 | CP015061. | 2740 | 956 | 993 | 2487 | 1004 | 775 | 2356 | 1048 | 1289 | 1595 | 748 | 1054 | 3431 | 678 | 1049 | 2292 | 613 | 667 |
| 5275 | QGN18372.1 | FIM1_4699 | CP015061. | 2316 | 756 | 468 | 1971 | 529 | 352 | 2128 | 689 | 656 | 284 | 278 | 623 | 1224 | 381 | 546 | 425 | 418 | 378 |
| 5276 | QGN18373.1 | FIM1_4700 | CP015061. | 16 | 21 | 27 | 19 | 27 | 21 | 14 | 21 | 32 | 111 | 28 | 23 | 101 | 31 | 19 | 66 | 31 | 13 |
| 5277 | QGN18374.1 | FIM1_4701 | CP015061. | 174 | 1133 | 1116 | 108 | 781 | 1001 | 67 | 816 | 1652 | 2673 | 537 | 808 | 2252 | 516 | 1010 | 2773 | 534 | 552 |
| 5278 | QGN18375.1 | FIM1_4702 | CP015061. | 5098 | 1790 | 1546 | 4582 | 1532 | 1356 | 4391 | 1490 | 2165 | 920 | 1408 | 2117 | 6842 | 1306 | 2167 | 3748 | 1170 | 1413 |
| 5279 | QGN18376.1 | FIM1_4703 | CP015061. | 10976 | 5260 | 6382 | 15270 | 5623 | 4933 | 11671 | 6233 | 8178 | 4005 | 4234 | 9590 | 19752 | 4633 | 8431 | 9038 | 4220 | 6361 |
| 5280 | QGN18377.1 | FIM1_4704 | CP015061. | 3290 | 1293 | 1041 | 3848 | 831 | 675 | 3174 | 1015 | 1343 | 970 | 728 | 1324 | 3990 | 946 | 1326 | 2041 | 892 | 881 |
| 5281 | QGN18378.1 | FIM1_4705 | CP015061. | 1226 | 1351 | 706 | 1219 | 758 | 608 | 1310 | 790 | 957 | 3226 | 1000 | 1252 | 4201 | 921 | 1305 | 3481 | 888 | 826 |
| 5282 | QGN18379.1 | FIM1_4706 | CP015061. | 1425 | 1506 | 890 | 1448 | 1052 | 813 | 1092 | 952 | 1145 | 2556 | 988 | 1042 | 3429 | 905 | 1269 | 2800 | 901 | 767 |
| 5283 | QGN18380.1 | FIM1_4707 | CP015061. | 1747 | 679 | 458 | 1795 | 455 | 363 | 1172 | 492 | 608 | 510 | 311 | 563 | 1495 | 308 | 608 | 853 | 282 | 384 |
| 5284 | QGN18381.1 | FIM1_4708 | CP015061. | 3389 | 1407 | 636 | 1624 | 1122 | 543 | 2762 | 1215 | 891 | 536 | 766 | 852 | 2535 | 727 | 855 | 1213 | 738 | 588 |
| 5285 | QGN18382.1 | FIM1_4709 | CP015061. | 2302 | 1483 | 1025 | 2079 | 1143 | 709 | 2466 | 1423 | 1471 | 910 | 943 | 1145 | 3535 | 852 | 1015 | 1515 | 969 | 783 |
| 5286 | QGN18383.1 | FIM1_4710 | CP015061. | 2483 | 1028 | 838 | 1516 | 1056 | 714 | 1763 | 1020 | 1084 | 1400 | 1067 | 1371 | 4445 | 841 | 1311 | 2277 | 818 | 883 |
| 5287 | QGN18384.1 | FIM1_4711 | CP015061. | 8701 | 1301 | 1515 | 9067 | 1452 | 1138 | 9727 | 1448 | 1794 | 1540 | 1039 | 2177 | 5510 | 1179 | 2093 | 2856 | 1042 | 1488 |
| 5288 | QGN18385.1 | FIM1_4712 | CP015061. | 1159 | 378 | 320 | 1112 | 218 | 232 | 1205 | 361 | 422 | 231 | 187 | 420 | 582 | 217 | 322 | 285 | 221 | 243 |
| 5289 | QGN18386.1 | FIM1_4713 | CP015061. | 4210 | 820 | 1698 | 3834 | 1480 | 1150 | 3912 | 1519 | 2061 | 5717 | 1628 | 2152 | 8815 | 1315 | 2414 | 6453 | 1200 | 1533 |
| 5290 | QGN18387.1 | FIM1_4714 | CP015061. | 3578 | 1236 | 1142 | 5621 | 1274 | 954 | 4676 | 975 | 1479 | 1098 | 898 | 1158 | 3108 | 804 | 1058 | 1919 | 843 | 766 |

| | | | | | | | | | | | | | | | | | | | | | |
|---|---|---|---|---|---|---|---|---|---|---|---|---|---|---|---|---|---|---|---|---|---|
| 5291 | QGN18388.1 | FIM1_4715 | CP015061. | 11528 | 2484 | 3050 | 10324 | 2739 | 2194 | 9914 | 2813 | 3811 | 5151 | 2933 | 4420 | 15959 | 3186 | 4081 | 9505 | 2650 | 3049 |
| 5292 | QGN18389.1 | FIM1_4716 | CP015061. | 11232 | 542 | 1565 | 9133 | 1520 | 1100 | 11816 | 1602 | 2023 | 1428 | 1475 | 2717 | 11461 | 1706 | 2570 | 4286 | 1661 | 1773 |
| 5293 | QGN18390.1 | FIM1_4717 | CP015061. | 4509 | 3828 | 2201 | 1819 | 3643 | 2392 | 3338 | 3347 | 2975 | 6741 | 4916 | 3792 | 9416 | 3888 | 4235 | 7309 | 4906 | 2776 |
| 5294 | QGN18391.1 | FIM1_4718 | CP015061. | 3872 | 1900 | 632 | 1252 | 1722 | 706 | 2943 | 1532 | 899 | 1097 | 1643 | 989 | 2786 | 1923 | 1280 | 1618 | 2106 | 808 |
| 5295 | QGN18392.1 | FIM1_4719 | CP015061. | 788 | 372 | 162 | 594 | 218 | 159 | 681 | 218 | 203 | 375 | 235 | 270 | 853 | 258 | 282 | 457 | 286 | 192 |
| 5296 | QGN18393.1 | FIM1_4720 | CP015061. | 3678 | 1768 | 749 | 2009 | 1929 | 802 | 3555 | 1796 | 931 | 3251 | 2360 | 1729 | 6407 | 2348 | 1592 | 4100 | 2893 | 1265 |
| 5297 | QGN18394.1 | FIM1_4721 | CP015061. | 962 | 659 | 894 | 1658 | 511 | 882 | 1308 | 550 | 1273 | 2304 | 330 | 1368 | 2316 | 441 | 1338 | 2276 | 383 | 880 |
| 5298 | QGN18395.1 | FIM1_4722 | CP015061. | 4184 | 1813 | 537 | 1187 | 1216 | 446 | 2977 | 1350 | 751 | 409 | 1012 | 1237 | 2154 | 1408 | 951 | 630 | 1858 | 805 |
| 5299 | QGN18396.1 | FIM1_4723 | CP015061. | 233 | 10 | 9 | 188 | 11 | 13 | 168 | 9 | 20 | 65 | 9 | 18 | 64 | 9 | 17 | 61 | 6 | 8 |
| 5300 | QGN18397.1 | FIM1_4724 | CP015061. | 996 | 485 | 331 | 1428 | 502 | 344 | 814 | 460 | 476 | 2265 | 445 | 458 | 1924 | 312 | 436 | 2291 | 323 | 295 |
| 5301 | QGN18398.1 | FIM1_4725 | CP015061. | 461 | 258 | 179 | 2 | 0 | 53 | 491 | 188 | 251 | 702 | 128 | 179 | 669 | 101 | 169 | 675 | 125 | 127 |
| 5302 | QGN18399.1 | FIM1_4726 | CP015061. | 103 | 44 | 43 | 0 | 1 | 23 | 130 | 44 | 80 | 275 | 45 | 59 | 235 | 35 | 41 | 235 | 45 | 42 |
| 5303 | QGN18400.1 | FIM1_4727 | CP015061. | 1196 | 636 | 403 | 0 | 2 | 124 | 1106 | 430 | 518 | 1288 | 257 | 371 | 1151 | 224 | 348 | 1253 | 230 | 221 |
| 5304 | QGN18401.1 | FIM1_4728 | CP015061. | 849 | 56 | 134 | 0 | 2 | 35 | 817 | 66 | 155 | 314 | 46 | 129 | 404 | 62 | 115 | 296 | 35 | 80 |
| 5305 | QGN18402.1 | FIM1_4729 | CP015061. | 1515 | 908 | 691 | 1 | 2 | 246 | 1677 | 668 | 934 | 2759 | 441 | 752 | 2582 | 382 | 717 | 2609 | 355 | 481 |
| 5306 | QGN18403.1 | FIM1_4730 | CP015061. | 151 | 582 | 249 | 278 | 475 | 307 | 213 | 416 | 369 | 2063 | 452 | 320 | 1609 | 351 | 317 | 1702 | 453 | 250 |
| 5307 | QGN18404.1 | FIM1_4731 | CP015061. | 2731 | 1861 | 607 | 1260 | 1318 | 610 | 2292 | 1309 | 841 | 3118 | 1373 | 1055 | 3367 | 1328 | 1079 | 2769 | 1443 | 729 |
| 5308 | QGN18405.1 | FIM1_4732 | CP015061. | 7230 | 3097 | 2644 | 9106 | 2166 | 1937 | 7041 | 2542 | 3390 | 1614 | 1407 | 3362 | 7270 | 1737 | 3290 | 2886 | 1578 | 2141 |
| 5309 | QGN18406.1 | FIM1_4733 | CP015061. | 5597 | 1073 | 919 | 4268 | 881 | 693 | 5072 | 1132 | 1168 | 623 | 500 | 922 | 1441 | 528 | 826 | 806 | 505 | 667 |
| 5310 | QGN18407.1 | FIM1_4734 | CP015061. | 2722 | 322 | 257 | 1787 | 300 | 173 | 2393 | 358 | 321 | 388 | 214 | 279 | 868 | 182 | 234 | 547 | 217 | 201 |
| 5311 | QGN18408.1 | FIM1_4736 | CP015061. | 969 | 248 | 129 | 958 | 193 | 92 | 588 | 220 | 154 | 127 | 100 | 93 | 429 | 91 | 85 | 201 | 81 | 77 |
| 5312 | QGN18409.1 | FIM1_4737 | CP015061. | 1695 | 190 | 379 | 2889 | 450 | 258 | 2701 | 488 | 490 | 2634 | 482 | 762 | 3706 | 494 | 688 | 2841 | 431 | 513 |
| 5313 | QGN18410.1 | FIM1_4738 | CP015061. | 242 | 33 | 61 | 502 | 62 | 62 | 204 | 51 | 70 | 223 | 44 | 92 | 342 | 28 | 74 | 213 | 31 | 59 |
| 5314 | QGN18411.1 | FIM1_4739 | CP015061. | 723 | 314 | 203 | 992 | 294 | 201 | 973 | 285 | 302 | 759 | 207 | 291 | 1303 | 305 | 271 | 845 | 242 | 198 |
| 5315 | QGN18412.1 | FIM1_4740 | CP015061. | 5947 | 378 | 861 | 6508 | 466 | 700 | 6464 | 445 | 1112 | 1664 | 272 | 1141 | 3365 | 285 | 1081 | 2188 | 262 | 759 |
| 5316 | QGN18413.1 | FIM1_4741 | CP015061. | 570 | 315 | 393 | 1534 | 252 | 299 | 1013 | 212 | 494 | 2555 | 193 | 589 | 2360 | 240 | 479 | 2322 | 195 | 357 |
| 5317 | QGN18414.1 | FIM1_4742 | CP015061. | 991 | 172 | 169 | 888 | 125 | 110 | 827 | 152 | 202 | 120 | 127 | 194 | 560 | 96 | 186 | 277 | 112 | 141 |
| 5318 | QGN18415.1 | FIM1_4743 | CP015061. | 47 | 12 | 10 | 111 | 14 | 8 | 29 | 12 | 9 | 24 | 19 | 22 | 64 | 18 | 29 | 58 | 14 | 19 |
| 5319 | QGN18416.1 | FIM1_4744 | CP015061. | 1271 | 2606 | 1837 | 3808 | 2031 | 1535 | 2592 | 1925 | 2284 | 8049 | 2016 | 2754 | 7909 | 2103 | 2986 | 7197 | 1896 | 1895 |
| 5320 | QGN18417.1 | FIM1_4745 | CP015061. | 586 | 647 | 897 | 1068 | 737 | 712 | 676 | 758 | 1133 | 3450 | 534 | 1192 | 3923 | 553 | 1174 | 3597 | 459 | 767 |
| 5321 | QGN18418.1 | FIM1_4746 | CP015061. | 2799 | 1461 | 1371 | 7065 | 2607 | 2004 | 3199 | 1385 | 1781 | 5398 | 1436 | 1570 | 5732 | 1391 | 1175 | 5141 | 1301 | 1016 |
| 5322 | QGN18419.1 | FIM1_4747 | CP015061. | 845 | 483 | 391 | 1256 | 483 | 447 | 904 | 355 | 559 | 2255 | 410 | 603 | 1992 | 373 | 513 | 2201 | 363 | 389 |
| 5323 | QGN18420.1 | FIM1_4748 | CP015061. | 3699 | 3288 | 3273 | 6034 | 2763 | 2709 | 4478 | 2666 | 4056 | 12383 | 2239 | 4446 | 12151 | 2360 | 4416 | 11415 | 2021 | 3029 |
| 5324 | QGN18421.1 | FIM1_4749 | CP015061. | 1075 | 260 | 309 | 1217 | 240 | 262 | 948 | 184 | 400 | 1423 | 164 | 442 | 1310 | 216 | 420 | 1318 | 158 | 281 |
| 5325 | QGN18422.1 | FIM1_4750 | CP015061. | 5664 | 2335 | 2630 | 6893 | 3118 | 2218 | 5953 | 2729 | 3274 | 14620 | 3192 | 3952 | 16958 | 2982 | 3621 | 15252 | 2812 | 2847 |
| 5326 | QGN18423.1 | FIM1_4751 | CP015061. | 400 | 124 | 145 | 487 | 130 | 95 | 295 | 154 | 172 | 526 | 152 | 157 | 532 | 106 | 135 | 521 | 138 | 85 |
| 5327 | QGN18424.1 | FIM1_4752 | CP015061. | 22 | 4 | 1 | 45 | 0 | 3 | 28 | 6 | 3 | 2 | 2 | 4 | 13 | 1 | 2 | 8 | 5 | 5 |
| 5328 | QGN18425.1 | FIM1_4753 | CP015061. | 1733 | 1689 | 707 | 1544 | 1477 | 805 | 1975 | 1112 | 1049 | 6987 | 1812 | 1399 | 5844 | 1880 | 1289 | 7037 | 2140 | 1045 |
| 5329 | QGN18426.1 | FIM1_4754 | CP015061. | 1377 | 1243 | 1094 | 2228 | 1096 | 902 | 1812 | 999 | 1390 | 8509 | 1328 | 1791 | 6634 | 1170 | 1621 | 8578 | 1147 | 1105 |
| 5330 | QGN18427.1 | FIM1_4755 | CP015061. | 453 | 293 | 289 | 871 | 225 | 269 | 504 | 272 | 405 | 1973 | 193 | 393 | 1433 | 182 | 345 | 1776 | 168 | 249 |
| 5331 | QGN18428.1 | FIM1_4756 | CP015061. | 2605 | 1114 | 754 | 2228 | 1358 | 724 | 2495 | 1081 | 922 | 5812 | 1210 | 1499 | 6044 | 1204 | 1311 | 5964 | 1296 | 927 |
| 5332 | QGN18429.1 | FIM1_4757 | CP015061. | 2367 | 1027 | 762 | 1284 | 1283 | 865 | 2135 | 957 | 994 | 5850 | 1238 | 1609 | 4737 | 1351 | 1517 | 5615 | 1275 | 1069 |
| 5333 | QGN18430.1 | FIM1_4758 | CP015061. | 1526 | 443 | 452 | 2181 | 397 | 372 | 1682 | 370 | 596 | 2540 | 244 | 546 | 2439 | 275 | 507 | 2340 | 186 | 345 |
| 5334 | QGN18431.1 | FIM1_4759 | CP015061. | 7551 | 4048 | 4012 | 16361 | 3750 | 3134 | 12513 | 3140 | 4942 | 12296 | 3699 | 5087 | 16052 | 4652 | 4842 | 13283 | 3931 | 3472 |
| 5335 | QGN18432.1 | FIM1_4760 | CP015061. | 9149 | 2383 | 2506 | 27367 | 6791 | 2589 | 12296 | 3806 | 3149 | 1690 | 739 | 2626 | 4120 | 715 | 2372 | 2457 | 730 | 1685 |
| 5336 | QGN18433.1 | FIM1_4761 | CP015061. | 1380 | 304 | 220 | 923 | 249 | 234 | 782 | 274 | 324 | 667 | 182 | 271 | 643 | 184 | 297 | 609 | 183 | 164 |
| 5337 | QGN18434.1 | FIM1_4762 | CP015061. | 288 | 136 | 69 | 227 | 134 | 60 | 184 | 127 | 103 | 185 | 85 | 71 | 319 | 69 | 81 | 242 | 59 | 51 |
| 5338 | QGN18435.1 | FIM1_4763 | CP015061. | 303 | 77 | 100 | 339 | 167 | 82 | 214 | 214 | 136 | 340 | 116 | 113 | 545 | 88 | 109 | 435 | 87 | 78 |
| 5339 | QGN18436.1 | FIM1_4764 | CP015061. | 303 | 280 | 244 | 894 | 413 | 188 | 256 | 232 | 273 | 735 | 236 | 278 | 836 | 156 | 251 | 768 | 175 | 189 |
| 5340 | QGN18437.1 | FIM1_4765 | CP015061. | 715 | 142 | 241 | 1591 | 126 | 194 | 708 | 98 | 335 | 1270 | 116 | 346 | 922 | 160 | 321 | 1141 | 116 | 194 |
| 5341 | QGN18438.1 | FIM1_4766 | CP015061. | 287 | 100 | 119 | 718 | 146 | 86 | 427 | 96 | 132 | 604 | 111 | 198 | 662 | 132 | 186 | 595 | 118 | 135 |
| 5342 | QGN18439.1 | FIM1_4767 | CP015061. | 3081 | 308 | 476 | 2606 | 464 | 398 | 2336 | 357 | 583 | 2659 | 409 | 730 | 2235 | 450 | 611 | 2519 | 390 | 456 |
| 5343 | QGN18440.1 | FIM1_4769 | CP015061. | 320 | 40 | 39 | 1276 | 56 | 43 | 455 | 41 | 36 | 112 | 50 | 60 | 262 | 60 | 79 | 150 | 45 | 41 |
| 5344 | QGN18441.1 | FIM1_4770 | CP015061. | 524 | 137 | 180 | 1945 | 608 | 127 | 696 | 212 | 187 | 528 | 462 | 463 | 1946 | 469 | 473 | 1162 | 407 | 379 |
| 5345 | QGN18442.1 | FIM1_4771 | CP015061. | 597 | 458 | 214 | 1244 | 500 | 255 | 456 | 388 | 296 | 2146 | 793 | 652 | 2982 | 654 | 654 | 2553 | 566 | 511 |
| 5346 | QGN18443.1 | FIM1_4772 | CP015061. | 393 | 11 | 34 | 492 | 17 | 23 | 435 | 24 | 38 | 273 | 19 | 49 | 237 | 19 | 36 | 306 | 11 | 45 |
| 5347 | QGN18444.1 | FIM1_4773 | CP015061. | 2443 | 1586 | 2595 | 4728 | 1349 | 2342 | 2990 | 1071 | 3390 | 8002 | 1158 | 4494 | 6318 | 1321 | 4155 | 8331 | 1331 | 2964 |
| 5348 | QGN18445.1 | FIM1_4776 | CP015061. | 0 | 0 | 0 | 0 | 0 | 0 | 0 | 0 | 0 | 0 | 0 | 0 | 0 | 0 | 0 | 0 | 0 | 0 |

**Batch**

| sample | batch |
|---|---|
| SC-RI-1 | b1 |
| SC-RI-2 | b1 |
| SC-RI-3 | b1 |
| KS-1-1 | b1 |
| KS-1-2 | b1 |
| KS-1-3 | b1 |
| Km-V-1 | b1 |
| Km-V-2 | b1 |
| Km-V-3 | b1 |
| SC-RIII-1 | b2 |
| SC-RIII-2 | b2 |
| SC-RIII-3 | b2 |
| KS-3-1 | b2 |
| KS-3-2 | b2 |
| KS-3-3 | b2 |
| SC-RI-1_NaCl | b3 |
| SC-RI-2_NaCl | b3 |
| SC-RI-3_NaCl | b3 |
| SC-RI-1_TM | b3 |
| SC-RI-2_TM | b3 |
| SC-RI-3_TM | b3 |
| SC-RIII-1_NaCl | b3 |
| SC-RIII-2_NaCl | b3 |
| SC-RIII-3_NaCl | b3 |
| SC-RIII-1_TM | b3 |
| SC-RIII-2_TM | b3 |
| SC-RIII-3_TM | b3 |
| KS-1-1_NaCl | b3 |
| KS-1-2_NaCl | b3 |
| KS-1-3_NaCl | b3 |
| KS-1-1_TM | b3 |
| KS-1-2_TM | b3 |
| KS-1-3_TM | b3 |
| KS-3-1_NaCl | b3 |
| KS-3-2_NaCl | b3 |
| KS-3-3_NaCl | b3 |
| KS-3-1_TM | b3 |
| KS-3-2_TM | b3 |
| KS-3-3_TM | b3 |
| Km-V-1_NaCl | b3 |
| Km-V-2_NaCl | b3 |
| Km-V-3_NaCl | b3 |
| Km-V-1_TM | b3 |
| Km-V-2_TM | b3 |
| Km-V-3_TM | b3 |
