## Supplementary material for "Intergeneric chromosomal transfer in yeast results in improved phenotypes and widespread transcriptional responses": TableS1

**Table S1: Primers used in this study.**

| Application | Primer name | Primer sequence(5'-3') | Description |
|---|---|---|---|
| chr1 circularization | P1 | AAGTGAGACAATATCGTAACGTCCG | amplification of upstream donor sequence |
| | P2 | ACGGATGCTATTTCAGAATATTTCGT | |
| | P3 | TATTCTGAAATAGCATCCGTGGCGCGCCTAGTGCAACCTTTTACTTGT | amplification of further downstream donor sequence |
| | P4 | AATATGGTGGGGCCTTTGACTTCCA | |
| | P5 | [illegible]CAAAGGCCCCACCATATTCCCGCAGCTTCGCTAGTAATCAGCGAATTCTGATTGGAAAG | amplification of *KmURA3* cassete |
| | P6 | ACATATTTCACTCTTGCTCGTGCCGGTAGAGGTGTGGTCA | |
| | P7 | CGAGCAAGAGTGAAATATGTTCTTTTTATAAGC | amplification of downstream donor sequence |
| | P8 | CAGAGGTATTGTGGATCCTTCTACAGT | |
| | gA1 F | ATCTCAAGCAAATTGCAATATTG | gRNA for deletion of chr1 left telomere |
| | gA1 R | AACCAATATTGCAATTTGCTTGA | |
| | gA2 F | ATCTTTGATGTAAGCTCTCTTCC | gRNA for deletio of chr1 right telomere and repeat sequence |
| | gA2 R | AACGGAAGAGAGCTTACATCAAA | |
| chr1 left arm ARS insertion | PL A1 | TCCATACTCTTCAGGATATCATATAATAACAACTCTAAT | amplification of upstream donor sequence |
| | PL A2 | AATGTTTTTTTCTTGAATTATTTAGTATTTGCTGTA | |
| | PL A3 | CTAAATAATTCAAGAAAAAAACATTATCGATTGAAGTTTTGTCCAACTAT | amplification of *KmARS1* |
| | PL A4 | TGACCCGGCGGGGACAAGGCAAGCTCTTTTAACGATATATCATTTTCAAT | |
| | PL A5 | ATTGAAAATGATATATCGTTAAAAGAGCTTGCCTTGTCCCCGCCGGGTCA | amplification of *hyg* $^{R}$ cassete |
| | PL A6 | CTTGAACATCTGCCCCTCTCTATCTACTGGATGGCGGCGTTAGTATCGAA | |
| | PL A7 | AGATAGAGAGGGGCAGATGTTCAAG | amplification of downstream donor sequence |
| | PL A8 | GATCAACTTCCTCTGTGGATTGTGA | |
| | gAL F | ATCAAAAAACATTAGATAGAGAG | gRNA for insertion of chr1 left arm ARS |
| | gAL R | AACCTCTCTATCTAATGTTTTTT | |
| chr1 right arm ARS&CEN insertion | PR A1 | GTTGATAGATATAAATTGGGACTTC | amplification of upstream donor sequence |
| | PR A2 | AGTTGCCTATTTAAACTTTCTCCTTTT | |
| | PR A3 | AAGGAGAAAGTTTAAATAGGCAACTATCGATTGAAGTTTTGTCCAACTAT | amplification of *KmARS1* &*KmCEN5* |
| | PR A4 | TGACCCGGCGGGGACGAGGCAAGCTGAGCTCCTTTCATTTCTGATAAAAG | |
| | PR A5 | CTTTTATCAGAAATGAAAGGAGCTCAGCTTGCCTCGTCCCCGCCGGGTCA | amplification of *kan* $^{R}$ cassete |
| | PR A6 | GGCATGACTATACCTCTTCAGTGTCACTGGATGGCGGCGTTAGTATCGAA | |
| | PR A7 | GACACTGAAGAGGTATAGTCATGCCTA | amplification of downstream donor sequence |
| | PR A8 | TCTATTTCGTCAAGTTTTTGAAGGG | |
| | gAR F | ATCATAGGCAACTGACACTGAAG | gRNA for insertion of chr1 right arm ARS&CEN |
| | gAR R | AACCTTCAGTGTCAGTTGCCTAT | |
| R1 PCR test | PA TF | GGTACCAAACTCTCATTTAGAGACCCT | verifying marker 4 in R1 (circularization site, 207968-1630 bp of R1) |
| | PA TR | GTAACCTAGGCTAATACGCGAAACA | |
| | PAL TF | GACCACTTCATCAAGTTTCGAAAGTGAA | verifying marker 5 in R1 (left arm ARS insert site, 65682-66542 bp of R1) |
| | PAL TR | AAGATTATTTCCTGTTAGACCCATGGC | |
| | PAR TF | TTAGTCGCTTCTACACATGGTTCTT | verifying marker 6 in R1 (right arm ARS&CEN insert site, 156872-157772 bp of R1) |
| | PAR TR | CAAATAATCAAGTACTATTTAATCTATGTAACATA | |
| | I T1F | AAAACAGCCACCATAAATACGCAGA | verifying marker 1 in R1 (37143-38132 bp of R1) |
| | I T1R | AATGTGGGAACAAAGACGACAAAAG | |
| | I T2F | TTTGAATGCGAACCACTCATGTTTT | verifying marker 2 in R1 (114231-115612 bp of R1) |
| | I T2R | CGATGAGAATTAATGCTGAATGGAC | |
| | I T3F | GCGTCGTGCAAGAGCTGAATTTT | verifying marker 3 in R1 (191213-192795 bp of R1) |
| | I T3R | ATGCCAAACTTAAAGAGACTACCCAT | |
| | PC A1 | TGGAATGATTCTTTTAACTCGGGTAAGAAGG | amplification of upstream donor sequence |
| | PC A2 | GGTATTCGTTCTTTCGGCGCGCCTGTTGAATGTGGTAACCCAATAGCA | |

| | | | |
|---|---|---|---|
| | PC A3 | TTACCACATTCAACAGGCGCGCCGAAAGAACGAATACCCTCTCCATTC | amplification of further downstream donor sequence |
| | PC A4 | TCCAATCAGAATTCGGAAGCATATCCATCGTGCTTAAAAT | |
| | PC A5 | CGATGGATATGCTTCCGAATTCTGATTGGAAAGACCATT | amplification of *KmURA3* cassete |
| chr3 circularization | PC A6 | GATGCTTTCATCCCTTGCCGGTAGAGGTGTGGTCA | |
| | PC A7 | ACACCTCTACCGGCAAGGGATGAAAGCATCTCTAAATATTATCATGT | amplification of downstream donor sequence |
| | PC A8 | GTTGGCGCCTCCGAACAAAC | |
| | gC1 F | ATCAATATTGCAAATTGCTTGAA | gRNA for deletion of chr3 left telomere |
| | gC1 R | AACTTCAAGCAATTTGCAATATT | |
| | gC2 F | ATCTTTCATCCCTCTGACGAAGG | gRNA for deletion of chr3 right telomere |
| | gC2 R | AACCCTTCGTCAGAGGGATGAAA | |
| | PCL A1 | GTGCAGCTATGCATGGGAGTGATTTCAC | amplification of upstream donor sequence |
| | PCL A2 | AAAACTTCAATCGATGTGGTCGTGTTGAATGAATCATTATTATAATGC | |
| | PCL A3 | ATTCAACACGACCACATCGATTGAAGTTTTGTCCAACTATCC | amplification of *KmARS1* &*KmCEN5* |
| | PCL A4 | GGGACAAGGCAAGCTGAGCTCCTTTCATTTCTGATAAAAGTAAGGCTT | |
| chr3 left arm ARS&CEN insertion | PCL A5 | AAATGAAAGGAGCTCAGCTTGCCTTGTCCCCGC | amplification of *hyg* $^{R}$ cassete |
| | PCL A6 | AAGTGTCAGACCTATACTGGATGGCGGCGTTAGT | |
| | PCL A7 | ACGCCGCCATCCAGTATAGGTCTGACACTTACCAAAACTCAT | amplification of downstream donor sequence |
| | PCL A8 | TTGTAGGGACTTAGTGATCGTGTTCAAATAATTGCCA | |
| | gCL F | ATCCGATGAACCCACTGATATTT | gRNA for insertion of chr3 left arm ARS&CEN |
| | gCL R | AACAAATATCAGTGGGTTCATCG | |
| | PCR A1 | TCTGTGGGTATTTCCGTGTATATGG | amplification of upstream donor sequence |
| | PCR A2 | AAAACTTCAATCGATCTAAGAGCCGGTAAAAAAAATCCGG | |
| | PCR A3 | TTTACCGGCTCTTAGATCGATTGAAGTTTTGTCCAACTATCC | amplification of *KmARS1* |
| | PCR A4 | GGGACGAGGCAAGCTCTTTTAACGATATATCATTTTCAATTTATTTATT | |
| chr3 right arm ARS insertion | PCR A5 | ATATATCGTTAAAAGAGCTTGCCTCGTCCCCGC | amplification of *kan* $^{R}$ cassete |
| | PCR A6 | CCACTCCCCTTATAAACTGGATGGCGGCGTTAGT | |
| | PCR A7 | ACGCCGCCATCCAGTTTATAAGGGGAGTGGCAGCGG | amplification of downstream donor sequence |
| | PCR A8 | TTTTTGAACACGGACAATGCCATTACG | |
| | gCR F | ATCCGGCTCTTAGCGGACCGTTT | gRNA for insertion of chr3 right arm ARS |
| | gCR R | AACAAACGGTCCGCTAAGAGCCG | |
| | PC TF | GAGCATGGTTAGGCAGGCGGAATGTAATAG | verifying marker 13 in R3 (circularization site, 332349-1732 bp of R3) |
| | PC TR | ATGCATCTGAGACCGAAGATGATGAATTCGA | |
| | PCL TF | AGGGCTTCATCAGCAACTGTAGGAG | verifying marker 14 in R3 (left arm ARS&CEN insert site, 108060-109506 bp of R3) |
| | PCL TR | CACCAACTCAGTCTCGCTTTTGCCT | |
| | PCR TF | CAATATCGTCTAGAGCTGGCACAGG | verifying marker 15 in R3 (right arm ARS insert site, 242412-243858 bp of R3) |
| | PCR TR | GAAGTAGCACAGAAGTGGTGCCATAGA | |
| | LTR F | ATGCCTCGTATGGCAACCAACCGTA | verifying 29 kb deletion in R3 in KS-R3 |
| | LTR R | ACGACACCATGTCTGATGCATTTTAG | |
| | III T1F | CATCATGCGCAGTCTGTGGTGGT | verifying marker 1 in R3 (21000-21645 bp of R3) |
| | III T1R | CCCAATGACATAGACAATCCGCCA | |
| | III T2F | CTAGCTACAGTCACTGCTATCAACGC | verifying marker 2 in R3 (41169-42072 bp of R3) |
| | III T2R | GATCATGAGACGATTCAATGACGTGC | |
| | III T3F | CCACATCACGTGAATCAGGAGGC | verifying marker 3 in R3 (60978-61740 bp of R3) |
| | III T3R | CCTCGCATTCTCAAGGTTGTTCTGC | |
| | III T4F | GCTCTGCTGCAGGATACTGAAAAG | verifying marker 4 in R3 (80745-81549 bp of R3) |
| R3 PCR test | III T4R | GGGCATTCACTGTGTCGTTATCTT | |
| | III T5F | TGGCCTGTGGGTGATAACTTCCAG | verifying marker 5 in R3 (126458-127310 bp of R3) |
| | III T5R | GCCAGCGATGATATAATTTGCCTGC | |

| | | | |
|---|---|---|---|
| | III T6F | GGCACTTCTGTATATACCGATGAGCAT | verifying marker 6 in R3 (147947-148614 bp of R3) |
| | III T6R | GTTGTGCACAAGGAACTAGTCGGTC | |
| | III T7F | CGACGTATTGGTGGAGACTTGAAACA | verifying marker 7 in R3 (168329-169115 bp of R3) |
| | III T7R | GAGATTAGGCCGTGGAGTGTGGAG | |
| | III T8F | CAGGTGGTGTATCGGCAGTAACAG | verifying marker 8 in R3 (193002-193806 bp of R3) |
| | III T8R | GAAGAGGCAAGAAGGCAATGGGA | |
| | III T9F | CTAGTTGGGCCTCTATGTCCCCAT | verifying marker 9 in R3 (220536-221222 bp of R3) |
| | III T9R | CTTCATCGCATTTTCAACATGTTTGCGA | |
| | III T10F | GTTGACGACGATGCTCTTACGGAC | verifying marker 10 in R3 (265609-266253 bp of R3) |
| | III T10R | CCGCAAGATGAATCCATGGTACCT | |
| | III T11F | GCTCTAGTGTGAACTCACTCACTGAT | verifying marker 11 in R3 (287295-287815 bp of R3) |
| | III T11R | CTGGGATGTGGTTGAACTCTCGC | |
| | III T12F | CATGATGTCCCCAAAGCGTGTACA | verifying marker 12 in R3 (310594-311217 bp of R3) |
| | III T12R | CCAGGAGTACCTGCGCTTATTCTC | |
| qPCR | ScFLO9 RT F | AGTTTGTCCTCCGACAGAACCCAGT | *ScFLO9* qPCR |
| | ScFLO9 RT R | GCGGCATGCCTGCCAGCAAA | |
| | KmFLO5 RT F | ATGTTCGCGCGGGCGGAACCCTATT | *KmFLO5* qPCR |
| | KmFLO5 RT R | AGCACCAGATATTACCTTTGCCACGACA | |
| | kmSPS22 RT F | TCGACTTGGGCGACCTAAAGAGCA | *KmSPS22* qPCR |
| | kmSPS22 RT R | GGGAAGCTAACAGCACTCAATGACG | |
| | scSPS22 RT F | TCAAGGTGGCTTGATGATCGCCG | *ScSPS22* qPCR |
| | scSPS22 RT R | GCGCTACCTTTCACCAGTTTCAAGC | |
| | ALPHA1 RT F | CTGCTCGCTGAAGAATGGCACGC | *ScHMLALPHA1* qPCR |
| | ALPHA1 RT R | TCCGTCACCACGTACTTCAGCATAA | |
| | ALPHA2 RT F | ACGGTTTTTGTTGGCCCTAGATAAGAA | *ScHMLALPHA2* qPCR |
| | ALPHA2 RT R | ACCTTTTAAATCCACAAATCACAGATGAGT | |
| | kmSWC4 RT F | AGCTTGCACCAGGTGTGTTCCTCA | *KmSWC4* qPCR |
| | kmSWC4 RT R | AGGGAGAGAAGGCATGGCTGGC | |
| | scSWC4 RT F | TGGTGAGAAGCTATCGCCGGGT | *ScSWC4* qPCR |
| | scSWC4 RT R | AGGGCATCACAGGGCGAGAAGG | |
| | kmTRK1 RT F | AGTCCGTCTGGCCTGAGTGCAT | *KmTRK1* qPCR |
| | kmTRK1 RT R | GGCTGGTGACCCATGCTGAAAGC | |
| | scTRK1 RT F | TCGTCCGAAACGCTCGCCGT | *ScTRK1* qPCR |
| | scTRK1 RT R | GGTCCCCTGCGGATGGCACAA | |
| | kmMPE1 RT F | TCGGCCAAACGCACTTGCCA | *KmMPE1* qPCR |
| | kmMPE1 RT R | ACCCTGGCCCATTCCGTTGCT | |
| | scMPE1 RT F | CCGTTGGAGGCATGAAGGGCA | *ScMPE1* qPCR |
| | scMPE1 RT R | TCCAACTGGCACCGCTGGCT | |

**R1.gb**

```
LOCUS       Exported             214653 bp ds-DNA     circular SYN 20-APR-2024
DEFINITION  synthetic circular DNA.
ACCESSION   .
VERSION     .
KEYWORDS    chromosome_R1
SOURCE      synthetic DNA construct
  ORGANISM  synthetic DNA construct
REFERENCE   1  (bases 1 to 214653)
  AUTHORS   Yilin Lyu
  TITLE     Direct Submission
  JOURNAL   Exported Saturday, April 20, 2024 from SnapGene 3.2.1
            http://www.snapgene.com
FEATURES             Location/Qualifiers
     source          1..214653
                     /organism="synthetic DNA construct"
                     /lab_host="Saccharomyces cerevisiae"
                     /mol_type="other DNA"
     CDS             complement(564..1367)
                     /codon_start=1
                     /label=KmURA3
                     /note="KmURA3"
                     /translation="MSTKSYSERAAAHRSPVAAKLLNLMEEKKSNLCASLDVRKTAELL
                     RLVEVLGPYICLLKTHVDILEDFSFENTIVPLKQLAEKHKFLIFEDRKFADIGNTVKLQ
                     YTSGVYRIAEWSDITNAHGVTGAGIVAGLKQGAEEVTKEPRGLLMLAELSSKGSLAHGE
                     YTRGTVEIAKSDKDFVIGFIAQNDMGGREEGYDWLIMTPGVGLDDKGDALGQQYRTVDE
                     VVAGGSDIIIVGRGLFAKGRDPVVEGERYRKAGWDAYLKRVGRSA"
     CDS             complement(3168..3530)
                     /codon_start=1
                     /label=PAU8
                     /note="PAU8"
                     /translation="MVKLTSIAAGVAAIAATASATTTLAQSDERVNLVELGVYVSDIRA
                     HLAQYYMFQAAHPTETYPVEVAEAVFNYGDFTTMLTGIAPDQVTRMITGVPWYSSRLKP
                     AISSALSKDGIYTIAN"
     CDS             3841..4068
                     /codon_start=1
                     /locus_tag="YAL067W-A"
                     /label=YAL067W-A
                     /note="YAL067W-A"
```

/translation="MPIIGVPRCLIKPFSVPVTFPFSVKKNIRILDLDPRTEAYCLSLN
SVCFKRLPRRKYFHLLNSYNIKRVLGVVYC"
CDS complement(8598..10379)
/codon_start=1
/label=SEO1
/note="SEO1"
/translation="MYSIVKEIIVDPYKRLKWGFIPVKRQVEDLPDDLNSTEIVTISNS
IQSHETAENFITTTSEKDQLHFETSSYSEHKDNVNVTRSYEYRDEADRPWWRFFDEQEY
RINEKERSHNKWYSWFKQGTSFKEKKLLIKLDVLLAFYSCIAYWVKYLDTVNINNAYVS
GMKEDLGFQGNDLVHTQVMYTVGNIIFQLPFLIYLNKLPLNYVLPSLDLCWSLLTVGAA
YVNSVPHLKAIRFFIGAFEAPSYLAYQYLFGSFYKHDEMVRRSAFYYLGQYIGILSAGG
IQSAVYSSLNGVNGLEGWRWNFIIDAIVSVVVGLIGFYSLPGDPYNCYSIFLTDDEIRL
ARKRLKENQTGKSDFETKVFDIKLWKTIFSDWKIYILTLWNIFCWNDSNVSSGAYLLWL
KSLKRYSIPKLNQLSMITPGLGMVYLMLTGIIADKLHSRWFAIIFTQVFNIIGNSILAA
WDVAEGAKWFAFMLQCFGWAMAPVLYSWQNDICRRDAQTRAITLVTMNIMAQSSTAWIS
VLVWKTEEAPRYLKGFTFTACSAFCLSIWTFVVLYFYKRDERNNAKKNGIVLYNSKHGV
EKPTSKDVETLSVSDEK"
CDS complement(12928..13314)
/codon_start=1
/label=YAL065C
/note="YAL065C"
/translation="MNSATSETTTNTGAAETTTSTGAAETKTVVTSSISRFNHAETQTA
SATDVIGHSSSVVSVSETGNTKSLITSGLSTMSQQPRSTPASSIIGSSTASLEISTYVG
IANGLLTNNGISVFISTVLLAIVW"
CDS 13409..13789
/codon_start=1
/locus_tag="YAL064W-B"
/label=YAL064W-B
/note="YAL064W-B"
/translation="MAGEAVSEHTPDSQEVTVTSVVCCLDSVVEIGHHVVYSVVTPLIV
AVLIDTMAGEAVLEHTSDSQEEIVTTVVCSVVPLVCFVVSVVCFVISVVEIGHHVVYSV
VAPLTVTVAVETIAEEMDSVHT"
CDS complement(15110..15490)
/codon_start=1
/label=TDA8
/note="TDA8"
/translation="MTGYFLPPQTSSYTFRFAKVDDSAILSVGGDVAFGCCAQEQPPIT
STNFTINGIKPWQGRLPDNIAGTVYMYAGFYCPMKIVYSNAVSWHTLPVSVELPDVTTV
SDDFAGHVYSFDDDLTAQLYYP"

```
CDS             23316..23600
                /codon_start=1
                /locus_tag="YAL064W"
                /label=YAL064W
                /note="YAL064W"
                /translation="MNPFASLEGQDNISSVFFLHMQQFESQVKDRFRFPIFRLERKTFG
                NSCYQVETLKVKCRPRHAKSCNLLTLLFKSRTQSVLVPNFGFLILNSEP"
CDS             complement(24145..24435)
                /codon_start=1
                /locus_tag="YAL063C-A"
                /label=YAL063C-A
                /note="YAL063C-A"
                /translation="MCPRTVLLIININHWFYDKNIVRIILTFRLDSGHISDICFINKNL
                ANALITADISLLKRHDIRCTKYIITYYQRYRNKEKGKFISLCKNTIISSSV"
CDS             complement(25752..29720)
                /codon_start=1
                /label=FLO9
                /note="FLO9"
                /translation="MSLAHYCLLLAIVTLLGLTNVVSATTAACLPANSRKNGMNVNFYQ
                YSLRDSSTYSNAAYMAYGYASKTKLGSVGGQTDISIDYNIPCVSSSGTFPCPQEDLYGN
                WGCKGIGACSNNPIIAYWSTDLFGFYTTPTNVTLEMTGYFLPPQTGSYTFKFATVDDSA
                ILSVGGSIAFECCAQEQPPITSTNFTINGIKPWNGSPPDNITGTVYMYAGFYYPMKIVY
                SNAVAWGTLPISVTLPDGTTVSDDFEGYVYTFDNNLSQPNCTIPDPSNYTVSTTITTTE
                PWTGTFTSTSTEMTTITGTNGVPTDETIIVVKTPTTASTIITTTEPWTGTFTSTSTEMT
                TITGTNGVPTDETIIVVKTPTTASTIITTTEPWTGTSTSTSTELTTVTGTNGLPTDETI
                IVVRTPTTASTIITTTEPWTGTSTSTSTELTTVTGTNGLPTDETIIVVRTPTTASTIIT
                TTEPWTGTSTSTSTELTTVTGTNGLPTDETIIVVRTPTTANTIITITEPWTGTSTSTST
                ELTTVTGTNGLPTDETIIVVKTPTTASTIITTTEPWTGTFTSTSTEMTTVTGTNGQPTD
                ETVIVVKTPTTANTIITTTEPWTGTFTSTSTEMTTITGTNGVPTDETIIVVKTPTTAST
                IITTTEPWTGTFTSTSTEMTTITGTNGVPTDETIIVVKTPTTASTIITTTEPWTGTFTS
                TSTEMTTITGTNGVPTDETIIVVKTPTTASTIITTTEPWTGTFTSTSTEMTTVTGTNGQ
                PTDETVIVIRTPTSEGLVTTTTEPWTGTFTSTSTEMTTVTGTNGQPTDETVIVIRTPTS
                EGLVTTTTEPWTGTFTSTSTEMTTITGTNGQPTDETVIIVKTPTTAISSSLSSSSGQIT
                SFITSSRPIITPFYPSNGTSVISSSVISSSDTSSLVISSSVTSSLVTSSPVISSSFISS
                PVISSTTTSASILSESSKSSVIPTSSSTSGSSESETGSASSASSSSSISSESPKSTYSS
                SSLPPVTSATTSQEITSSLPPVTTTKTSEQTTLVTVTSCESHVCTESISSAIVSTATVT
                VSGATTEYTTWCPISTTEITKQTTETTKQTKGTTEQTTETTKQTTVVTISSCESDVCSK
                TASPAIVSTSTATINGVTTEYTTWCPISTTESKQQTTLVTVTSCGSGVCSETTSPAIVS
                TATATVNDVVTVYSTWRPQTTNEQSVSSKMNSATSETTTNTGAAETTTSTGAAETKTVV
```

TSSISRFNHAETQTASATDVIGHSSSVVSVSETGNTKSLTSSGLSTMSQQPRSTPASSM
VGSSTASLEISTYAGSANSLLAGSGLSVFIASLLLAII"

CDS 33319..34692
/codon_start=1
/label=GDH3
/note="GDH3"
/translation="MTSEPEFQQAYDEIVSSVEDSKIFEKFPQYKKVLPIVSVPERIIQ
FRVTWENDNGEQEVAQGYRVQFNSAKGPYKGGLRFHPSVNLSILKFLGFEQIFKNALTG
LDMGGGKGGLCVDLKGKSDNEIRRICYAFMRELSRHIGKDTDVPAGDIGVGGREIGYLF
GAYRSYKNSWEGVLTGKGLNWGGSLIRPEATGFGLVYYTQAMIDYATNGKESFEGKRVT
ISGSGNVAQYAALKVIELGGIVVSLSDSKGCIISETGITSEQIHDIASAKIRFKSLEEI
VDEYSTFSESKMKYVAGARPWTHVSNVDIALPCATQNEVSGDEAKALVASGVKFVAEGA
NMGSTPEAISVFETARSTATNAKDAVWFGPPKAANLGGVAVSGLEMAQNSQKVTWTAER
VDQELKKIMINCFNDCIQAAQEYSTEKNTNTLPSLVKGANIASFVMVADAMLDQGDVF"

CDS 35200..36453
/codon_start=1
/label=BDH2
/note="BDH2"
/translation="MRALAYFGKGNIRFTNHLKEPHIVAPDELVIDIEWCGICGTDLHE
YTDGPIFFPEDGHTHEISHNPLPQAMGHEMAGTVLEVGPGVKNLKVGDKVVVEPTGTCR
DRYRWPLSPNVDKEWCAACKKGYYNICSYLGLCGAGVQSGGFAERVVMNESHCYKVPDF
VPLDVAALIQPLAVCWHAIRVCEFKAGSTALIIGAGPIGLGTILALNAAGCKDIVVSEP
AKVRRELAEKMGARVYDPTAHAAKESIDYLRSIADGGDGFDYTFDCSGLEVTLNAAIQC
LTFRGTAVNLAMWGHHKIQFSPMDITLHERKYTGSMCYTHHDFEAVIEALEEGRIDIDR
ARHMITGRVNIEDGLDGAIMKLINEKESTIKIILTPNNHGELNREADNEKKEISELSSR
KDQERLRESINEAKLRHT"

CDS 36907..38055
/codon_start=1
/label=BDH1
/note="BDH1"
/translation="MRALAYFKKGDIHFTNDIPRPEIQTDDEVIIDVSWCGICGSDLHE
YLDGPIFMPKDGECHKLSNAALPLAMGHEMSGIVSKVGPKVTKVKVGDHVVVDAASSCA
DLHCWPHSKFYNSKPCDACQRGSENLCTHAGFVGLGVISGGFAEQVVVSQHHIIPVPKE
IPLDVAALVEPLSVTWHAVKISGFKKGSSALVLGAGPIGLCTILVLKGMGASKIVVSEI
AERRIEMAKKLGVEVFNPSKHGHKSIEILRGLTKSHDGFDYSYDCSGIQVTFETSLKAL
TFKGTATNIAVWGPKPVPFQPMDVTLQEKVMTGSIGYVVEDFEEVVRAIHNGDIAMEDC
KQLITGKQRIEDGWEKGFQELMDHKESNVKILLTPNNHGEMK"

CDS complement(38248..38670)
/codon_start=1

/label=YAL059C-A
/note="YAL059C-A"
/translation="MYLAREMDLAILPSRRLVKFKAFTKRSLSMDEIELASLSSRAFLF
NFLPLLLLLAFLDIFASSNASFLAAVLIKILVKSVFSALGSSLKSFTSGSRASDCLAAL
EFFDIFLAMLCFRRYLTSIVKEKTTFCRLCSHIQYF"
CDS 38261..38899
/codon_start=1
/label=ECM1
/note="ECM1"
/translation="MWEQRRQKVVFSLTILVRYRLKQSMAKKISKNSRAARQSDALEPE
VKDLSELPRAEKTDLTNILIRTAAKNEALLEAKISKKANKSKRGKKLNKKALEDKLANS
ISSMDRDRLVKALNFTNRLDGKIAKSISRAKYIQNTRKAGWDSTNETIKKELAFLNGGL
SVQAKSASEGNAEKEDEEIPEVFDSLAEDNTVQKTPTNRFGVLPDDVEE"
CDS 39216..40724
/codon_start=1
/label=CNE1
/note="CNE1"
/translation="MKFSAYLWWLFLNLALVKGTSLLSNVTLAEDSFWEHFQAYTNTKH
LNQEWITSEAVNNEGSKIYGAQWRLSQGRLQGSAWDKGIAVRTGNAAAMIGHLLETPIN
VSETDTLVVQYEIKLDNSLTCGGAFIKLMSGFMNVEALKHYAPDTEGVELVFGPDYCAP
EINGVQFAINKVDKITHESKLRYLQEMPLSKLTDTSQSHLYTLIIDESAQSFQILIDGK
TVMVREHIEDKKKVNFEPPITPPLMIPDVSVAKPHDWDDRIRIPDPEAVKLSDRDERDP
LMIPHPDGTEPPEWNSSIPEYILDPNAQKPSWWKELEHGEWIPPMIKNPLCTAERGCGQ
QIPGLINNAKYKGPGELNEIINPNYMGEWHPPEIENPLYYEEQHPLRIENVISGVILEF
WSGSPNMLISNIYVGKNVTEAQIIGNKTWLMRDRAFRGSDGPTERKFMNSRLGNLQTTF
HNERESPNPFDRIIDRILEQPLKFVLTAAVVLLTTSVLCCVVFT"
CDS 41011..43653
/codon_start=1
/label=GPB2
/note="GPB2"
/translation="MEISSSPWNDGGYSPYERNRVAVSPFSSALEGEERIETSRSLGDH
CFEPLPYVTNYLSIFALFGKEIFGDKGNVSSRNEYLLKKYYSLKKPFVLRHNGHALKNP
DMPLQRNDILQTNFMVDKFLNRTVRSVNFNNFKIISDMQSKSGRGTKSGTNQNQSADAI
QNICLPSIPSALPYFQYYRKLLTVNTKEWDILKLHSLWVPKLRKDFKDFSLYGDKNSLK
PIDSHYDEDNTMKKNLFFERSPSRQTLDGKGCASKGYDISSGNMIIPSLFSEDKLPALT
YHCSVELNGNIYIFGGLMPCYSYEEDAPMLNDFFVDGIKNLPPPLLPQVINNPSMVNNP
HLYVASIPSCRFSKPKMGGYIPPPLLCVQGSKLTDRHIFFYGGFEIRTETRGDENGKYH
LKKRLYVNNTGYILDIMSFKFTKIDIIVQPSKYNAYPTMSSRFGHLQISIDNPNRRASV
HSSSMNEIHKMGSASMKQGSSITSGRLEKAAVLSSLPHNTVHTVIIFGGYRQTGDDRYE

AMNDLWKIEIPVIRRGKKGYCKFSETANAILLTPSEKDKSDWPEERAFSAFSVHGTSLM
DRSSLDMRLLNNLKNHFVLKPSYISQDRVVSPKPVFPMMVHGTHQDLFNSGSAAQESPK
AGASASSASAASFDPDMDDNLENYIVNPGRKSSSIPMTAIGRQRLILSQEKPVGKTVVL
HGGSNGLNVLDDMWLMDLECETWTPIETFAKADSSEDGDEKLDSVNVGLVGHRMESIGR
ICVCIGGMVQEDVDQFYSENDDEPPRKRKVDTLPLGGNFLNTIDLSTQCWEEHKITLSK
KEDDEDRQDSENEDTNSNIVVGVGGTSLQCDKSIILIGGLISRRSNVKEIYLHGTITKS
IFPSVNPSA"

CDS 43929..44471
/codon_start=1
/label=PEX22
/note="PEX22"
/translation="MPPPSRSRINKTRTLGIVGTAIAVLVTSYYIYQKVTSAKEDNGAR
PPEGDSVKENKKARKSKCIIMSKSIQGLPIKWEEYAADEVVLLVPTSHTDGSMKQAIGD
AFRKTKNEHKIIYCDSMDGLWSCVRRLGKFQCILNSRDFTSSGGSDAAVVPEDIGRFVK
FVVDSDVEDVLIDTLCN"

CDS complement(44633..46774)
/codon_start=1
/label=ACS1
/note="ACS1"
/translation="MSPSAVQSSKLEEQSSEIDKLKAKMSQSAATAQQKKEHEYEHLTS
VKIVPQRPISDRLQPAIATHYSPHLDGLQDYQRLHKESIEDPAKFFGSKATQFLNWSKP
FDKVFIPDPKTGRPSFQNNAWFLNGQLNACYNCVDRHALKTPNKKAIIFEGDEPGQGYS
ITYKELLEEVCQVAQVLTYSMGVRKGDTVAVYMPMVPEAIITLLAISRIGAIHSVVFAG
FSSNSLRDRINDGDSKVVITTDESNRGGKVIETKRIVDDALRETPGVRHVLVYRKTNNP
SVAFHAPRDLDWATEKKKYKTYYPCTPVDSEDPLFLLYTSGSTGAPKGVQHSTAGYLLG
ALLTMRYTFDTHQEDVFFTAGDIGWITGHTYVVYGPLLYGCATLVFEGTPAYPNYSRYW
DIIDEHKVTQFYVAPTALRLLKRAGDSYIENHSLKSLRCLGSVGEPIAAEVWEWYSEKI
GKNEIPIVDTYWQTESGSHLVTPLAGGVTPMKPGSASFPFFGIDAVVLDPNTGEELNTS
HAEGVLAVKAAWPSFARTIWKNHDRYLDTYLNPYPGYYFTGDGAAKDKDGYIWILGRVD
DVVNVSGHRLSTAEIEAAIIEDPIVAECAVVGFNDDLTGQAVAAFVVLKNKSSWSTATD
DELQDIKKHLVFTVRKDIGPFAAPKLIILVDDLPKTRSGKIMRRILRKILAGESDQLGD
VSTLSNPGIVRHLIDSVKL"

CDS 47651..50002
/codon_start=1
/label=FLC2
/note="FLC2"
/translation="MIFLNTFARCLLTCFVLCSGTARSSDTNDTTPASAKHLQTTSLLT
CMDNSQLTASFFDVKFYPDNNTVIFDIDATTTLNGNVTVKAELLTYGLKVLDKTFDLCS
LGQVSLCPLSAGRIDVMSTQVIESSITKQFPGIAYTIPDLDAQVRVVAYAQNDTEFETP

LACVQAILSNGKTVQTKYAAWPIAAISGVGVLTSGFVSVIGYSATAAHIASNSISLFIY
FQNLAITAMMGVSRVPPIAAAWTQNFQWSMGIINTNFMQKIFDWYVQATNGVSNVVVAN
KDVLSISVQKRAISMASSSDYNFDTILDDSNLYTTSEKDPSNYSAKILVLRGIERVAYL
ANIELSNFFLTGIVFFLFFLFVVVVSLIFFKALLEVLTRARILKETSNFFQYRKNWGSI
IKGTLFRLSIIAFPQVSLLAIWEFTQVNSPAIVVDAVVILLIITGLLVYGTIRVFIKGR
ESLRLYKNPAYLLYSDTYFLNKFGFLYVQFKADKFWWLLPLLSYAFLRSLFVAVLQNQG
KAQAMIIFVIELAYFVCLCWIRPYLDKRTNVFNIAIHLVNLINAFFFLFFSNLFKQPAV
VSSVMAVILFVLNAVFALFLLLFTIVTCTLALLHRNPDVRYQPMKDDRVSFIPKIQNDF
DGKNKNDSELFELRKAVMDTNENEEEKMFRDDTFGKNLNANTNTARLFDDETSSSSFKQ
NSSPFDASEVTEQPVQPTSAVMGTGGSFLSPQYQRASSASRTNLAPNNTSTSSLMKPES
SLYLGNSNKSYSHFNNNGSNENARNNNPYL"

CDS complement(49739..49750)
/codon_start=1
/product="Factor Xa recognition and cleavage site"
/label=Factor Xa site
/note="Factor Xa site"
/translation="IEGR"

CDS 50316..53459
/codon_start=1
/label=OAF1
/note="OAF1"
/translation="MVENSTQKAPHAGNDDNSSTKPYSEAFFLGFNNPTPGLEAEHSST
SPAPENSETHNRKRNRILFVCQACRKSKTKCDREKPECGRCVKHGLKCVYDVSKQPAPR
IPSKDAIISRLEKDMFYWKDKAMKLLTEREVNESGKRSASPINTNNASGDSPDTKKQHK
MEPIYEQSGNGDINNGTRNDIEINLYRSHPTMIMSKVMKREVKPLSENYIIIQDCFLKI
LVTSVFLDTSKNTMIPALTANANITRAQPSVANNLLKLKEMLIRQCQTEDEKNRVNEFT
DRILQNTNSNRNLKIGMLLSMLYNSVGYQYLEDHCPQGGEYSDLLRNLINECEAILPSY
EIIERYKNHFYEYVYPSLPFIELEIFEESLSQTIFPDPNNPSKVQIRMGSTHLRAKVEN
LSLLLVILKLSYMSIRFLDHSTADSSFYLSKEIIDKYPIPNDFILLSQRCLASENWCAC
ANENIISCLLYIWSFFAFSPEEGDFFLEHPTDVISSLIMMLSTSIGLHRDPSDFPQLIS
PSTSDKRTLNHRRILWLSIVTVCSFEASLKGRHSVSPISLMALFLNIKDPDSLTVYMNR
VRGDLSDINNHKLLRIHKFTFKRAQLALLLSDLDNLTMTYYGSFHLHSIEFIREKIEIF
VEENFPIVPLKSVAQDKSDLDDMNVISEMNILSSENSSSFHNRIMNKLLMLRTSMAVFL
HFETLITKDKSIFPFYKKYFMVSCMDALSLINYFNKFFNGEYRHAISSLTSFNVTKFIQ
LALSSTIFSLLGIILRIGLAIHMLSSEVQKLSGTTDPRIKELNTKVEKFSTLQRDLESA
LEGIYCSASEHLRFTYFPVFKMLALFDVIVQRMRKGELWHGIFTMIQMEQMHSRIIKTL
SITLGVKLDKKDRLLEELMACNHVANFSVEDIDELNRNIKKEIQISSGLKPPVNTIDLT
NGEPFGNAVPTFTKTWSSSLDNLEKLSSAAAVGQSLDYNSGLRQGPLAGGGSKEQTPIA
GMNNLNNSINATPIVDNSSGSQLPNGFDRGQANNTPFPGYFGGLDLFDYDFLFGNDFA"

CDS complement(53607..54347)
/codon_start=1
/label=AIM2
/note="AIM2"
/translation="MASNQPGKCCFEGVCHDGTPKGRREEIFGLDTYAAGSTSPKEKVI
VILTDVYGNKFNNVLLTADKFASAGYMVFVPDILFGDAISSDKPIDRDAWFQRHSPEVT
KKIVDGFMKLLKLEYDPKFIGVVGYCFGAKFAVQHISGDGGLANAAAIAHPSFVSIEEI
EAIDSKKPILISAAEEDHIFPANLRHLTEEKLKDNHATYQLDLFSGVAHGFAARGDISI
PAVKYAKEKVLLDQIYWFNHFSNV"
CDS complement(54553..56541)
/codon_start=1
/label=GEM1
/note="GEM1"
/translation="MTKETIRVVICGDEGVGKSSLIVSLTKAEFIPTIQDVLPPISIPR
DFSSSPTYSPKNTVLIDTSDSDLIALDHELKSADVIWLVYCDHESYDHVSLFWLPHFRS
LGLNIPVILCKNKCDSISNVNANAMVVSENSDDDIDTKVEDEEFIPILMEFKEIDTCIK
TSAKTQFDLNQAFYLCQRAITHPISPLFDAMVGELKPLAVMALKRIFLLSDLNQDSYLD
DNEILGLQKKCFNKSIDVNELNFIKDLLLDISKHDQEYINRKLYVPGKGITKDGFLVLN
KIYAERGRHETTWAILRTFHYTDSLCINDKILHPRLVVPDTSSVELSPKGYRFLVDIFL
KFDIDNDGGLNNQELHRLFKCTPGLPKLWTSTNFPFSTVVNNKGCITLQGWLAQWSMTT
FLNYSTTTAYLVYFGFQEDARLALQVTKPRKMRRRSGKLYRSNINDRKVFNCFVIGKPC
CGKSSLLEAFLGRSFSEEYSPTIKPRIAVNSLELKGGKQYYLILQELGEQEYAILENKD
KLKECDVICLTYDSSDPESFSYLVSLLDKFTHLQDLPLVFVASKADLDKQQQRCQIQPD
ELADELFVNHPLHISSRWLSSLNELFIKITEAALDPGKNTPGLPEETAAKDVDYRQTAL
IFGSTVGFVALCSFTLMKLFKSSKFSK"
CDS complement(56741..58609)
/codon_start=1
/label=SPC72
/note="SPC72"
/translation="MVRRWIPSGRHLRNNDNTGDDDDSEFTNSMDSGMSIPSLRDSMTT
RSSHNDPIKPALMNDSNKVKNLEKELTNAKIKIQVLYEYIRRIPNKDGNAPSLGNDTDF
RNSIIEGLNLEINKLKQDLKAKEVEYQDTLQFVQENLENSESIVNTINHLLSFILTHFN
EQDENAHLLDKEERETLEETLELSSDYVLEKMDTLSKFIIQFLQDFLHSKSRAESKQDK
EEFLSLAQSSPAGSQLESRDSPSSKEENTDGGYQNDEIHDSNNHIDTENVMANSTSLPI
SAVESRFEKTLDTQLEIVIENLHKEYDQFINSIRLKFEKSQKLEKIIASKLNEQSHLLD
SLELEENSSSVIEKQDHLISQLKEKIESQSVLINNLEKLKEDIIKMKQNEKVLTKELET
QTKINKLKENNWDSYINDLEKQINDLQIDKSEEFHVIQNQLDKLDLENYQLKNQLNTLD
NQKLILSQYESNFIKFNQNLLLHLDSIFNILQKILQESSIAQFDRKMKSIKSVPNALKN
LNLIQPKLESLYTFIETALESIINSYISSLISMETPEQPHQQGNELTATPNKELTLRIE

ELQRRWISERERRKLDANASEARIKALEQENESLRSKLFNLSINNP"
CDS complement(57465..57476)
/codon_start=1
/product="Factor Xa recognition and cleavage site"
/label=Factor Xa site
/note="Factor Xa site"
/translation="IEGR"
CDS complement(58781..59137)
/codon_start=1
/label=BOL3
/note="BOL3"
/translation="MKLPQTMLRSISVKHVRWPRILTGSKLWYSTQMAMTPEEKMITDK
LQQELEPEVCKVQDVSGGCGSMFAINITSKKFNGLSLIKQHQLVNRILRDDISRWHGLQ
LTTKKSTGKGPASS"
CDS 59270..59602
/codon_start=1
/label=BOL1
/note="BOL1"
/translation="MFKRAMSTDGPVARTILKRLECGFPDYKNFAFGLYNDSHKHKGHA
GVQGNVSAETHFRIEMVSKKFEGLKLPQRHRMVYSLLQDEMAQANGIHALQLSLKTPQE
YESKAK"
CDS complement(59702..60214)
/codon_start=1
/label=GCV3
/note="GCV3"
/translation="MLRTTRLWTTRMPTVSKLFLRNSSGNALNKNKLPFLYSSQGPQAV
RYTSQHEWIAVHQDKTAFVGITKYATDALGDATYVELPEVGTEIAQGESLGSIESVKSA
SEIYQPADGTVEEINTNLEENPGVVNEDPMGDGWLVKMKLGEGVNVEQVEGLMSLEQYE
KTLVHDD"
CDS complement(60447..62804)
/codon_start=1
/label=PTA1
/note="PTA1"
/translation="MSSAEMEQLLQAKTLAMHNNPTEMLPKVLETTASMYHNGNLSKLK
LPLAKFFTQLVLDVVSMDSPIANTERPFIAAQYLPLLLAMAQSTADVLVYKNIVLIMCA
SYPLVLDLVAKTSNQEMFDQLCMLKKFVLSHWRTAYPLRATVDDETDVEQWLAQIDQNI
GVKLATIKFISEVVLSQTKSPSGNEINSSTIPDNHPVLNKPALESEAKRLLDMLLNYLI
EEQYMVSSVFIGIINSLSFVIKRRPQTTIRILSGLLRFNVDAKFPLEGKSDLNYKLSKR
FVERAYKNFVQFGLKNQIITKSLSSGSGSSIYSKLTKISQTLHVIGEETKSKGILNFDP

SKGNSKKTLSRQDKLKYISLWKRQLSALLSTLGVSTKTPTPVSAPATGSSTENMLDQLK
ILQKYTLNKASHQGNTFFNNSPKPISNTYSSVYSLMNSSNSNQDVTQLPNDILIKLSTE
AILQMDSTKLITGLSIVASRYTDLMNTYINSVPSSSSSKRKSDDDDDGNDNEEVGNDGP
TANSKKIKMETEPLAEEPEEPEDDDRMQKMLQEEESAQEISGDANKSTSAIKEIAPPFE
PDSLTQDEKLKYLSKLTKKLFELSGRQDTTRAKSSSSSSILLDDDDSSSWLHVLIRLVT
RGIEAQEASDLIREELLGFFIQDFEQRVSLIIEWLNEEWFFQTSLHQDPSNYKKWSLRV
LESLGPFLENKHRRFFIRLMSELPSLQSDHLEALKPICLDPARSSLGFQTLKFLIMFRP
PVQDTVRDLLHQLKQEDEGLHKQCDSLLDRLK"

CDS 63068..64315
/codon_start=1
/label=ERV46
/note="ERV46"
/translation="MKRSTLLSLDAFAKTEEDVRVRTRAGGLITLSCILTTLFLLVNEW
GQFNSVVTRPQLVVDRDRHAKLELNMDVTFPSMPCDLVNLDIMDDSGEMQLDILDAGFT
MSRLNSEGRPVGDATELHVGGNGDGTAPVNNDPNYCGPCYGAKDQSQNENLAQEEKVCC
QDCDAVRSAYLEAGWAFFDGKNIEQCEREGYVSKINEHLNEGCRIKGSAQINRIQGNLH
FAPGKPYQNAYGHFHDTSLYDKTSNLNFNHIINHLSFGKPIQSHSKLLGNDKRHGGAVV
ATSPLDGRQVFPDRNTHFHQFSYFAKIVPTRYEYLDNVVIETAQFSATFHSRPLAGGRD
KDHPNTLHVRGGIPGMFVFFEMSPLKVINKEQHGQTWSGFILNCITSIGGVLAVGTVMD
KLFYKAQRSIWGKKSQ"

CDS 64592..67156
/codon_start=1
/label=CDC24
/note="CDC24"
/translation="MAIQTRFASGTSLSDLKPKPSATSISIPMQNVMNKPVTEQDSLFH
ICANIRKRLEVLPQLKPFLQLAYQSSEVLSERQSLLLSQKQHQELLKSNGANRDSSDLA
PTLRSSSISTATSLMSMEGISYTNSNPSATPNMEDTLLTFSMGILPITMDCDPVTQLSQ
LFQQGAPLCILFNSVKPQFKLPVIASDDLKVCKKSIYDFILGCKKHFAFNDEELFTISD
VFANSTSQLVKVLEVVETLMNSSPTIFPSKSKTQQIMNAENQHRHQPQQSSKKHNEYVK
IIKEFVATERKYVHDLEILDKYRQQLLDSNLITSEELYMLFPNLGDAIDFQRRFLISLE
INALVEPSKQRIGALFMHSKHFFKLYEPWSIGQNAAIEFLSSTLHKMRVDESQRFIINN
KLELQSFLYKPVQRLCRYPLLVKELLAESSDDNNTKELEAALDISKNIARSINENQRRT
ENHQVVKKLYGRVVNWKGYRISKFGELLYFDKVFISTTNSSSEPEREFEVYLFEKIIIL
FSEVVTKKSASSLILKKKSSTSASISASNITDNNGSPHHSYHKRHSNSSSSNNIHLSSS
SAAAIIHSSTNSSDNNSNNSSSSSLFKLSANEPKLDLRGRIMIMNLNQIIPQNNRSLNI
TWESIKEQGNFLLKFKNEETRDNWSSCLQQLIHDLKNEQFKARHHSSTSTTSSTAKSSS
MMSPTTTMNTPNHHNSRQTHDSMASFSSSHMKRVSDVLPKRRTTSSSFESEIKSISENF
KNSIPESSILFRISYNNNSNNTSSSEIFTLLVEKVWNFDDLIMAINSKISNTHNNNISP
ITKIKYQDEDGDFVVLGSDEDWNVAKEMLAENNEKFLNIRLY"

```
CDS             68267..69295
                /label=HphMX4
                /note="HphMX4"
CDS             complement(69853..71595)
                /codon_start=1
                /label=CLN3
                /note="CLN3"
                /translation="MAILKDTIIRYANARYATASGTSTATAASVSAASCPNLPLLLQKR
                RAIASAKSKNPNLVKRELQAHHSAISEYNNDQLDHYFRLSHTERPLYNLTNFNSQPQVN
                PKMRFLIFDFIMYCHTRLNLSTSTLFLTFTILDKYSSRFIIKSYNYQLLSLTALWISSK
                FWDSKNRMATLKVLQNLCCNQYSIKQFTTMEMHLFKSLDWSICQSATFDSYIDIFLFQS
                TSPLSPGVVLSAPLEAFIQQKLALLNNAAGTAINKSSSSQGPSLNINEIKLGAIMLCEL
                ASFNLELSFKYDRSLIALGAINLIKLSLNYYNSNLWENINLALEENCQDLDIKLSEISN
                TLLDIAMDQNSFPSSFKSKYLNSNKTSLAKSLLDALQNYCIQLKLEEFYRSQELETMYN
                TIFAQSFDSDSLTCVYSNATTPKSATVSSAATDYFSDHTHLRRLTKDSISPPFAFTPTS
                SSSSPSPFNSPYKTSSSMTTPDSASHHSHSGSFSSTQNSFKRSLSIPQNSSIFWPSPLT
                PTTPSLMSNRKLLQNLSVRSKRLFPVRPMATAHPCSAPTQLKKRSTSSVDCDFNDSSNL
                KKTR"
CDS             complement(72791..73600)
                /codon_start=1
                /label=CYC3
                /note="CYC3"
                /translation="MGWFWADQKTTGKDIGGAAVSSMSGCPVMHESSSSSPPSSECPVM
                QGDNDRINPLNNMPELAASKQPGQKMDLPVDRTISSIPKSPDSNEFWEYPSPQQMYNAM
                VRKGKIGGSGEVAEDAVESMVQVHNFLNEGCWQEVLEWEKPHTDESHVQPKLLKFMGKP
                GVLSPRARWMHLCGLLFPSHFSQELPFDRHDWIVLRGERKAEQQPPTFKEVRYVLDFYG
                GPDDENGMPTFHVDVRPALDSLDNAKDRMTRFLDRMISGPSSSSSAP"
gap             75003..75012
                /estimated_length="10"
CDS             75899..77401
                /codon_start=1
                /label=CDC19
                /note="CDC19"
                /translation="MSRLERLTSLNVVAGSDLRRTSIIGTIGPKTNNPETLVALRKAGL
                NIVRMNFSHGSYEYHKSVIDNARKSEELYPGRPLAIALDTKGPEIRTGTTTNDVDYPIP
                PNHEMIFTTDDKYAKACDDKIMYVDYKNITKVISAGRIIYVDDGVLSFQVLEVVDDKTL
                KVKALNAGKICSHKGVNLPGTDVDLPALSEKDKEDLRFGVKNGVHMVFASFIRTANDVL
                TIREVLGEQGKDVKIIVKIENQQGVNNFDEILKVTDGVMVARGDLGIEIPAPEVLAVQK
                KLIAKSNLAGKPVICATQMLESMTYNPRPTRAEVSDVGNAILDGADCVMLSGETAKGNY
```

PINAVTTMAETAVIAEQAIAYLPNYDDMRNCTPKPTSTTETVAASAVAAVFEQKAKAII
VLSTSGTTPRLVSKYRPNCPIILVTRCPRAARFSHLYRGVFPFVFEKEPVSDWTDDVEA
RINFGIEKAKEFGILKKGDTYVSIQGFKAGAGHSNTLQVSTV"

CDS complement(77539..77631)
/codon_start=1
/label=YAL037C-A
/note="YAL037C-A"
/translation="MSISFPKMQHLIVMTTIGDKKVNNNIILFL"

CDS 78133..78936
/codon_start=1
/label=YAL037W
/note="YAL037W"
/translation="MDMEIEDSSPIDDLKLQKLDTNVYFGPCEILTQPILLQYENIKFI
IGVNLSTEKIASFYTQYFRNSNSVVVNLCSPTTAAVATKKAAIDLYIRNNTILLQKFVG
QYLQMGKKIKTSLTQAQTDTIQSLPQFCNSNVLSGEPLVQYQAFNDLLALFKSFSHFGN
ILVISSHSYDCALLKFLISRVMTYYPLVTIQDSLQYMKATLNISISTSDEFDILNDKEL
WEFGQTQEILKRRQTSSVKRRCVNLPENSTIDNRMLMGTTKRGRF"

CDS complement(79156..80265)
/codon_start=1
/label=RBG1
/note="RBG1"
/translation="MSTTVEKIKAIEDEMARTQKNKATSFHLGQLKAKLAKLRRELLTS
ASSGSGGGAGIGFDVARTGVASVGFVGFPSVGKSTLLSKLTGTESEAAEYEFTTLVTVP
GVIRYKGAKIQMLDLPGIIDGAKDGRGRGKQVIAVARTCNLLFIILDVNKPLHHKQIIE
KELEGVGIRLNKTPPDILIKKKEKGGISITNTVPLTHLGNDEIRAVMSEYRINSAEIAF
RCDATVDDLIDVLEASSRRYMPAIYVLNKIDSLSIEELELLYRIPNAVPISSGQDWNLD
ELLQVMWDRLNLVRIYTKPKGQIPDFTDPVVLRSDRCSVKDFCNQIHKSLVDDFRNALV
YGSSVKHQPQYVGLSHILEDEDVVTILKK"

CDS 80540..83548
/codon_start=1
/label=FUN12
/note="FUN12"
/translation="MAKKSKKNQQNYWDEEFEEDAAQNEEISATPTPNPESSAGADDTS
REASASAEGAEAIEGDFMSTLKQSKKKQEKKVIEEKKDGKPILKSKKEKEKEKKEKEKQ
KKKEQAARKKAQQQAQKEKNKELNKQNVEKAAAEKAAAEKSQKSKGESDKPSASAKKPA
KKVPAGLAALRRQLELKKQLEEQEKLEREEEERLEKEEEERLANEEKMKEEAKAAKKEK
EKAKREKRKAEGKLLTRKQKEEKKLLERRRAALLSSGNVKVAGLAKKDGEENKPKKVVY
SKKKKRTTQENASEAIKSDSKKDSEVVPDDELKESEDVLIDDWENLALGDDDEEGTNEE
TQESTASHENEDQNQGEEEEEGEEEEEEEEERAHVHEVAKSTPAATPAATPTPSSASPN

KKDLRSPICCILGHVDTGKTKLLDKIRQTNVQGGEAGGITQQIGATYFPIDAIKAKTKV
MAEYEKQTFDVPGLLVIDTPGHESFSNLRSRGSSLCNIAILVIDIMHGLEQQTIESIKL
LRDRKAPFVVALNKIDRLYDWKAIPNNSFRDSFAKQSRAVQEEFQSRYSKIQLELAEQG
LNSELYFQNKNMSKYVSIVPTSAVTGEGVPDLLWLLLELTQKRMSKQLMYLSHVEATIL
EVKVVEGFGTTIDVILSNGYLREGDRIVLCGMNGPIVTNIRALLTPQPLRELRLKSEYV
HHKEVKAALGVKIAANDLEKAVSGSRLLVVGPEDDEDELMDDVMDDLTGLLDSVDTTGK
GVVVQASTLGSLEALLDFLKDMKIPVMSIGLGPVYKRDVMKASTMLEKAPEYAVMLCFD
VKVDKEAEQYAEQEGIKIFNADVIYHLFDSFTAYQEKLLEERRKDFLDYAIFPCVLQTL
QIINKRGPMIIGVDVLEGTLRVGTPICAVKTDPTTKERQTLILGKVISLEINHQPVQEV
KKGQTAAGVAVRLEDPSGQQPIWGRHVDENDTLYSLVSRRSIDTLKDKAFRDQVARSDW
LLLKKLKVVFGIE"

CDS 83831..84700
/codon_start=1
/label=MTW1
/note="MTW1"
/translation="MSAPTMRSTSILTEHLGYPPISLVDDIINAVNEIMYKCTAAMEKY
LLSKSKIGEEDYGEEIKSGVAKLESLLENSVDKNFDKLELYVLRNVLRIPEEYLDANVF
RLENQKDLVIVDENELKKSEEKLREKVNDVELAFKKNEMLLKRVTKVKRLLFTIRGFKQ
KLNELLKCKDDVQLQKILESLKPIDDTMTLLTDSLRKLYVDSESTSSTEEVEALLQRLK
TNGKQNNKDFRTRYIDIRTNNVLRKLGLLGDKEDEKQSAKPDARTQAGDIVSIDIEEPQ
LDLLDDVL"

CDS complement(84823..86064)
/codon_start=1
/label=FUN19
/note="FUN19"
/translation="MGLYSPESEKSQLNMNYIGKDDSQSIFRRLNQNLKASNNNNDSNK
NGLNMSDYSNNSPYGRSYDVRINQNSQNNGNGCFSGSIDSLVDEHIIPSPPLSPKLESK
ISHNGSPRMASSVLVGSTPKGAVENVLFVKPVWPNGLSRKRYRYATYGFLSQYKIFSNL
AQPYSKNIINRYNNLAYNARHKYSKYNDDMTPPPLPSSSSRLPSPLASPNLNRQARYNM
RKQALYNNNLGKFESDTEWIPRKRKVYSPQRRTMTTSPHRAKKFSPSASTPHTNIASIE
AIHDAPQYIPNVSWKKLPDYSPPLSTLPTDSNKSLKIEWKGSPMDLSTDPLRNELHPAE
LVLAQTLRLPCDLYLDSKRRLFLEKVYRLKKGLPFRRTDAQKACRIDVNKASRLFQAFE
KVGWLQDSNFTKYL"

CDS 86819..87340
/codon_start=1
/label=POP5
/note="POP5"
/translation="MVRLKSRYILFEIIFPPTDTNVEESVSKADILLSHHRASPADVSI
KSILQEIRRSLSLNLGDYGSAKCNSLLQLKYFSNKTSTGIIRCHREDCDLVIMALMLMS

KIGDVDGLIVNPVKVSGTIKKIEQFAMRRNSKILNIIKCSQSSHLSDNDFIINDFKKIG
RENENENEDD"
CDS complement(87448..88587)
/codon_start=1
/label=PRP45
/note="PRP45"
/translation="MFSNRLPPPKHSQGRVSTALSSDRVEPAILTDQIAKNVKLDDFIP
KRQSNFELSVPLPTKAEIQECTARTKSYIQRLVNAKLANSNNRASSRYVTETHQAPANL
LLNNSHHIEVVSKQMDPLLPRFVGKKARKVVAPTENDEVVPVLHMDGSNDRGEADPNEW
KIPAAVSNWKNPNGYTVALERRVGKALDNENNTINDGFMKLSEALENADKKARQEIRSK
MELKRLAMEQEMLAKESKLKELSQRARYHNGTPQTGAIVKPKKQTSTVARLKELAYSQG
RDVSEKIILGAAKRSEQPDLQYDSRFFTRGANASAKRHEDQVYDNPLFVQQDIESIYKT
NYEKLDEAVNVKSEGASGSHGPIQFTKAESDDKSDNYGA"
CDS complement(88862..91144)
/codon_start=1
/label=GIP4
/note="GIP4"
/translation="MVDVQKRKKLLAKAAASASIPAIKGSVPLDSYDIKIIQYKNALYK
LNELNRLLNVLVPHLKKKRDNDESYKIIPLVNFILSLCEGPIFNVSPVLAKRYHLLCRF
QLIKLSEVQQRLSTNFIDVEGWMFPEEVPLDHYKSCIYNNSLQWKILNSLSCIAQNAIK
IYNAKLRQILLERDAYKARSLPFDTSIIEDLLNPVEMTLILDLAVLINDPVRDKSTHSF
YKLQWQVMEKLNSCVHSKIFPILRTYYNQLQKFSETRPTSLSNLQKDLPHWEWTLHRIY
TFHLRVFSVLCVIISFSRQIFLPNKQHFLDIKTRLSSENVYHYDLIICELMALLSPECD
DVTALFELQENLKFWTQTARTDNNSSRTPIFHLQPGLVVELFNNHICKIIPKLRSIMGL
LSNWMDCWKYIEKNYKTFDETNDLRENLKEKLERDKALYLEVKNAKSKLKKKPSITKLP
ASSSPSPSPTSSASPSRQASLESIRTRARAHLASNSSRSPSVSPVRTTFNNKNAETKKS
VVSPEKRKLINGRRPRSSSLQSYTNKQQTSYLNSTRHPSIAPPSKLNNQRSNSLQSSTM
TLNQKIVQDTVRHLMNKSASTPNPSASSSLAPSPKVSSINNTSSGKSSSTLIANSSDTL
AIETLTLDPESNSSELSIKRVRFAGVPPMTEAENPKPTKVGWYKKPAVLHYPPIPASAM
IKPLQHKSKYNTLRQEEGFTFRKSLRDGLEWENGESGSETTMMPFGIEIKESTGHRIAS
KIRSKLR"
CDS 91399..91865
/codon_start=1
/label=SNC1
/note="SNC1"
/translation="MSSSTPFDPYALSEHDEERPQNVQSKSRTAELQAVSTESHRVPSR
KLTLY*LSTSLILMRIHSYTFFVFIFRKLMIPWE**EIT*IK*QKEVKD*RPLKIKPIT
*RSQPKALRGVPIGSEKPCGTRI*K*RCVWL**SSYCLL*SSSPLLFTLVD"
CDS complement(91968..96383)

/codon_start=1
/label=MYO4
/note="MYO4"
/translation="MSFEVGTKCWYPHKEQGWIGGEVTKNDFFEGTFHLELKLEDGETV
SIETNSFENDDDHPTLPVLRNPPILESTDDLTTLSYLNEPAVLHAIKKRYMNGQIYTYS
GIVLIAANPFDKVDHLYSREMIQNYSSKRKDELEPHLFAIAEEAYRFMVHEKANQTVVV
SGESGAGKTVSAKYIMRYFASVQESNNREGEVEMSQIESQILATNPIMEAFGNAKTTRN
DNSSRFGKYLQILFDENTTIRGSKIRTYLLEKSRLVYQPETERNYHIFYQILEGLPEPV
KQELHLSSPKDYHYTNQGGQPNIAGIDEAREYKITTDALSLVGINHETQLGIFKILAGL
LHIGNIEMKMTRNDASLSSEEQNLQIACELLGIDPFNFAKWIVKKQIVTRSEKIVTNLN
YNQALIARDSVAKFIYSTLFDWLVDNINKTLYDPELDQQDHVFSFIGILDIYGFEHFEK
NSFEQFCINYANEKLQQEFNQHVFKLEQEEYVKEEIEWSFIEFSDNQPCIDLIENKLGI
LSLLDEESRLPSGSDESWASKLYSAFNKPPSNEVFSKPRFGQTKFIVSHYAVDVEYEVE
GFIEKNRDSVSLGHLDVFKATTNPIFKQILDNRELRSDDAPEEQNTEKKIMIPARLSQK
KPTLGSMFKKSLGELMAIINSTNVHYIRCIKPNSEKKPWEFDNLMVLSQLRACGVLETI
RISCAGFPSRWTFDEFVQRYFLLTDYSLWSGILYNPDLPKEAIVNFCQSILDATISDSA
KYQIGNTKIFFKAGMLAFLEKLRTNKMNEICIIIQKKIRARYYRLQYLQTMESIKKCQS
QIRSLLVRTRVDHELKTRAAILLQTNIRALWKREYYRAAIGQIIKLQCTCKRKLILDSV
NRKFMLMAAVIIQSYIRSYGHKTDYRTLKRSSILVQSAMRMQLARRRYIVLQKEVEERN
IRASYGIGLLEEAIEFKNSFILNLEMLNDSYTRLTQLLQGDLSNIPSKQRQEYETIVNG
YNDKISKLKTLQVEIMNTLNKKNALKERKKKQSSLIQSHMQSLAAIKGNKPSRLSDEVK
SMKQELAFIENVIAQDFTTTYSANKNDKVKGLGIAGQQVKPKLVNVIRRESGNPDLLEL
LMDLNCYTLEVTEGYLKKVNVTEVNGDNVLGPIHVITTVVSSLVRNGLLIQSSKFISKV
LLTVESIVMSLPKDETMLGGIFWLSNLSRLPAFAANQKTLYEANGGDEKDKLTLIYLND
LENETLKVFDKIYSTWLVKFMKHASAHIEIFDMVLNEKLFKNSGDEKFAKLFTFLNEFD
AVLCKFQVVDSMHTKIFNDTLKYLNVMLFNDLITKCPALNWKYGYEVDRNIERLVSWFE
PRIEDVRPNLIQIIQAVKILQLKISNLNEFKLLFDFWYALNPAQIQAILLKYKPANKGE
AGVPNEILNYLANVIKRENLSLPGKMEIMLSAQFDSAKNHLRYDTSAITQNSNTEGLAT
VSKIIKLDRK"
CDS 97013..98599
/codon_start=1
/label=FRT2
/note="FRT2"
/translation="MQNAQIKSSSKGSGIDGTDRNSKDGVEKRPLEDVKQMIDAGTPDV
GHKSTVETKPNVGWQASHSNLAALHEKEQKYEMEHHHARHKLHRQVIPDYTSASTAMFS
DCMFNAAPDKVRSLSTMKSSGLSPKHPFNVVATFKGPFPQHSVESKPLDGGYSAKDHFP
SFKMLQAQQHPAHRHYKDNDKYGLKSPSRSFVKDKKRLVHRFLKSMEPSSSGQSKDSSA
LAPAFDPILPNVISKPSKRPTHHSHSSDGSSSTQTDISLQSLLYHDLESSPKKHVSPSR
PPSVASESSPAVANPIGLSPKDACNASFSQSSSSSLSSSSSSSSSTSFSQSVAVDPLEP

PGNITYSSSNLSLNSDELDYYQRHIGLQLQQTEALLKHSLKDEVLKDENDLVKNIANFD
KIVKELRDLRSRTIGWKELVEEDYLMNLKQDFDKENPESFEARLSDTINTNVAKLQDLE
KRMASCKDRLASRKEVMRKMESLLSLENSLMISKKNVTFASKYRNEALDIVFLIIIIVI
CYTFKHLVSHK"
CDS 98800..99585
/codon_start=1
/label=SAW1
/note="SAW1"
/translation="MAPSIATVKIARDMVLPLRIFVNRKQILQTNDKTSNKSNATIFEA
PLLSNNSIICLKSPNTRIYLSQQDKKNLCDEIKEDLLLIVYELASPEIISSVLSKIRVG
HSTDFQINVLPKLFAGADTDNAVTSHIQSVTRLAKFKYKLHYKHKWELDIFINSIKKIA
NLRHYLMFQTLTLNGFSLNAGPKTLLARKIEKQPQVPNLLIENGDADALDTPVEEDIKP
VIEFMYKPVINLGEIIDVHVLHRPRRHKVRTQSKQPQEE"
CDS complement(99743..103810)
/codon_start=1
/label=DRS2
/note="DRS2"
/translation="MNDDRETPPKRKPGEDDTLFDIDFLDDTTSHSGSRSKVTNSHANA
NYIPPSHVLPEETIDLDADDDNIENDVHENLFMSNNHDDQTSWNANRFDSDAYQPQSLR
AVKPPGLFARFGNGLKNAFTFKRKKGPESFEMNHYNAVTNNELDDNYLDSRNKFNIKIL
FNRYILRKNVGDAEGNGEPRVIHINDSLANSSFGYSDNHISTTKYNFATFLPKFLFQEF
SKYANLFFLCTSAIQQVPHVSPTNRYTTIGTLLVVLIVSAMKECIEDIKRANSDKELNN
STAEIFSEAHDDFVEKRWIDIRVGDIIRVKSEEPIPADTIILSSSEPEGLCYIETANLD
GETNLKIKQSRVETAKFIDVKTLKNMNGKVVSEQPNSSLYTYEGTMTLNDRQIPLSPDQ
MILRGATLRNTAWIFGLVIFTGHETKLLRNATATPIKRTAVEKIINRQIIALFTVLIVL
ILISSIGNVIMSTADAKHLSYLYLEGTNKAGLFFKDFLTFWILFSNLVPISLFVTVELI
KYYQAFMIGSDLDLYYEKTDTPTVVRTSSLVEELGQIEYIFSDKTGTLTRNIMEFKSCS
IAGHCYIDKIPEDKTATVEDGIEVGYRKFDDLKKKLNDPSDEDSPIINDFLTLLATCHT
VIPEFQSDGSIKYQAASPDEGALVQGGADLGYKFIIRKPNSVTVLLEETGEEKEYQLLN
ICEFNSTRKRMSAIFRFPDGSIKLFCKGADTVILERLDDEANQYVEATMRHLEDYASEG
LRTLCLAMRDISEGEYEEWNSIYNEAATTLDNRAEKLDEAANLIEKNLILIGATAIEDK
LQDGVPETIHTLQEAGIKIWVLTGDRQETAINIGMSCRLLSEDMNLLIINEETRDDTER
NLLEKINALNEHQLSTHDMNTLALVIDGKSLGFALEPELEDYLLTVAKLCKAVICCRVS
PLQKALVVKMVKRKSSSLLLAIGDGANDVSMIQAAHVGVGISGMEGMQAARSADIAVGQ
FKFLKKLLLVHGSWSYQRISVAILYSFYKNTALYMTQFWYVFANAFSGQSIMESWTMSF
YNLFFTVWPPFVIGVFDQFVSSRLLERYPQLYKLGQKGQFFSVYIFWGWIINGFFHSAI
VFIGTILIYRYGFALNMHGELADHWSWGVTVYTTSVIIVLGKAALVTNQWTKFTLIAIP
GSLLFWLIFFPIYASIFPHANISREYYGVVKHTYGSGVFWLTLIVLPIFALVRDFLWKY
YKRMYEPETYHVIQEMQKYNISDSRPHVQQFQNAIRKVRQVQRMKKQRGFAFSQAEEGG

QEKIVRMYDTTQKRGKYGELQDASANPFNDNNGLGSNDFESAEPFIENPFADGNQNSNR
FSSSRDDISFDI"
CDS complement(104338..105258)
/codon_start=1
/label=MAK16
/note="MAK16"
/translation="MSDEIVWQVINQSFCSHRIKAPNGQNFCRNEYNVTGLCTRQSCPL
ANSKYATVKCDNGKLYLYMKTPERAHTPAKLWERIKLSKNYTKALQQIDEHLLHWSKFF
RHKCKQRFTKLTQVMITERRLALREEERHYVGVAPKVKRREQNRERKALVAAKIEKAIE
KELMDRLKSGAYGDKPLNVDEKVWKKIMGQMEEENSQDEEEDWDEEEESDDGEVEYVAD
DGEGEYVDVDDLEKWLADSDREASSASESESDSESESDSDSDEENKNSAKRRKKGTSAK
TKRPKVEIEYEEEHEVQNAEQEVAQ"
CDS complement(105679..109986)
/codon_start=1
/label=LTE1
/note="LTE1"
/translation="MEIFSQKDYYPTPSSNVISYESDCVSKPVNSADLPALIVHLSSPL
EGVDYNASADFFLIYRNFITPQDLHDLLIYRFRWCIREITTNAAKAKRRRIGEVALVRT
FVLLRHSILNYFVQDFLPNITLRLRLIEFLNDKHIEQYPKIISSCIINLKKNWVHCSKL
VWENIELNEPDKLDFDAWLHYSLKDFTQLESLHKRGSRLSIYARQSFASPDFRNQSVLS
LYKTSDVFRLPEKLQSSNSSKNQRSPSMLLFPDNTSNVYSKHRIAKEPSVDNESEDMSD
SKQKISHLSKVTLVSTLMKGVDYPSSYAVDKIMPPTPAKKVEFILNSLYIPEDLNEQSG
TLQGTSTTSSLDNNSNSNSRSNTSSMSVLHRSAIGLLAKWMKNHNRHDSSNDKKFMSAI
KPANQKPEMDAFVKYVVSISSLNRKSSKEEEEEFLNSDSSKFDILSARTIDEVESLLHL
QNQLIEKVQTHSNNNRGPTVNVDCERREHIHDIKILQQNSFKPSNDNFSAMDNLDLYQT
VSSIAQSVISLTNTLNKQLQNNESNMQPSPSYDALQRRKVKSLTTAYYNKMHGSYSAES
MRLFDKDNSSSRTDENGPQRLLFHETDKTNSEAITNMTPRRKNHSQSQKSMTSSPLKNV
LPDLKESSPLNDSREDTESITYSYDSELSSSSPPRDTVTKKSRKVRNIVNNTDSPTLKT
KTGFLNLREFTFEDTKSLDEKKSTIDGLEKNYDNKENQESEYESTKKLDNSLDASSEAN
NYDITTRKKHSSCNHKIKQAVVRPASGRISISRVQSIAITPTKELSIVDPEQNKSNSVI
EEISEIEPLNLEYNKKSALYSDTSSTVISISTSKLFESAQNSPLKQTQNPQREFPNGTS
VSETNRIRLSIAPTIESVVSDLNSITTGSTVETFETSRDLPVPHQRIINLREEYQRGNQ
DIISNTSSLHELKTIDLSDSNNDLESPSTHAKNNKYFFSPDDGSIDVASPMKNVEELKS
KFLKNESETNSNISGSVLTMDDIDINDTSSARNTRRANSESAFTGSLNKKNLNEIANML
DDSINDDPITVALMKLEGTYEKIPEKPENTKSSDAIGIKTSKLADEVEMLNLNNLPSFQ
NSPAEKRKSLLIERRRQTIMNIPFTPDQSEKEGFTSSSPEKIDVSANVDVAVQAAQIQE
LIGQYRIHDSRLMISNNESHVPFILMYDSLSVAQQMTLIEKEILGEIDWKDLLDLKMKH
EGPQVISWLQLLVRNETLSGIDLAISRFNLTVDWIISEILLTKSSKMKRNVIQRFIHVA
DHCRTFQNFNTLMEIILALSSSVVQKFTDAWRLIEPGDLLTWEELKKIPSLDRNYSTIR

NLLNSVNPLVGCVPFIVVYLSDLSANAEKKDWILEDKVVNYNKFDTNVQIVKNFIQRVQ
WSKFYTFKVNHELLSKCVYISTLTQEEINELST"
CDS complement(110386..112665)
/codon_start=1
/label=PMT2
/note="PMT2"
/translation="MSSSSSTGYSKNNAAHIKQENTLRQRESSSISVSEELSSADERDA
EDFSKEKPAAQSSLLRLESVVMPVIFTALALFTRMYKIGINNHVVWDEAHFGKFGSYYL
RHEFYHDVHPPLGKMLVGLSGYLAGYNGSWDFPSGEIYPDYLDYVKMRLFNASFSALCV
PLAYFTAKAIGFSLPTVWLMTVLVLFENSYSTLGRFILLDSMLLFFTVASFFSFVMFHN
QRSKPFSRKWWKWLLITGISLGCTISVKMVGLFIITMVGIYTVIDLWTFLADKSMSWKT
YINHWLARIFGLIIVPFCIFLLCFKIHFDLLSHSGTGDANMPSLFQARLVGSDVGQGPR
DIALGSSVVSIKNQALGGSLLHSHIQTYPDGSNQQQVTCYGYKDANNEWFFNRERGLPS
WSENETDIEYLKPGTSYRLVHKSTGRNLHTHPVAAPVSKTQWEVSGYGDNVVGDNKDNW
VIEIMDQRGDEDPEKLHTLTTSFRIKNLEMGCYLAQTGNSLPEWGFRQQEVVCMKNPFK
RDKRTWWNIETHENERLPPRPEDFQYPKTNFLKDFIHLNLAMMATNNALVPDPDKFDYL
ASSAWQWPTLNVGLRLCGWGDDNPKYFLLGTPASTWASSVAVLAFMATVVILLIRWQRQ
YVDLRNPSNWNVFLMGGFYPLLAWGLHYMPFVIMSRVTYVHHYLPALYFALIILAYCFD
AGLQKWSRSKCGRIMRFVLYAGFMALVIGCFWYFSPISFGMEGPSSNFRYLNWFSTWDI
ADKQEA"
CDS complement(112991..114544)
/codon_start=1
/label=FUN26
/note="FUN26"
/translation="MSTSADTDTIKKPILAVPEPALADTHSEEISRSGEEHESENNEHS
DEEGDNYSEREQSVSTEPLDTLPLRKKLKNLSYITFFAIGIGLLWPWNCILSASQYFKH
DIFKDTSIWAKIFTSSMMSFSTISSMLFNIYLAKRQYKYSRRVINGLVWEIIVFTVMCF
FTILHFLLPKWFNFMFIMMLVVISSMGTAMTQNGIMAIANVFGSEYSQGVMVGQAVAGV
LPSLVLFALAFIENSSVSTTGGILLYFFTTTLVVTICVVMFSVSKISRKVNENWNVEDG
HITDVLLGSLRSNEEEIRIVGRIDQMEDEDHRRTNGTRDDNDEGEELQLKVPFEVLFAK
LKYLVLSIFTTFVVTLVFPVFASATYVTGLPLSNAQYIPLIFTLWNLGDLYGRVIADWP
MFRDQKFTPRKTFIYSLLRVAAIPLFLMFTAITSSSSGDEEHNGSVIVDLCYMLLQFLF
GVTNGHVISMSFMKVPEQLDNDDEKEAAGGFTNIFVSTGLALGSIISYVFVFIIDFIIR
"
CDS complement(114960..117473)
/codon_start=1
/label=CCR4
/note="CCR4"
/translation="MNDPSLLGYPNVGPQQQQQQQQQQHAGLLGKGTPNALQQQLHMNQ

LTGIPPPGLMNNSDVHTSSNNNSRQLLDQLANGNANMLNMNMDNNNNNNNNNNNNNNNG
GGSGVMMNASTAAVNSIGMVPTVGTPVNINVNASNPLLHPHLDDPSLLNNPIWKLQLHL
AAVSAQSLGQPNIYARQNAMKKYLATQQAQQAQQQAQQQAQQQVPGPFGPGPQAAPPAL
QPTDFQQSHIAEASKSLVDCTKQALMEMADTLTDSKTAKKQQPTGDSTPSGTATNSAVS
TPLTPKIELFANGKDEANQALLQHKKLSQYSIDEDDDIENRMVMPKDSKYDDQLWHALD
LSNLQIFNISANIFKYDFLTRLYLNGNSLTELPAEIKNLSNLRVLDLSHNRLTSLPAEL
GSCFQLKYFYFFDNMVTTLPWEFGNLCNLQFLGVEGNPLEKQFLKILTEKSVTGLIFYL
RDNRPEIPLPHERRFIEINTDGEPQREYDSLQQSTEHLATDLAKRTFTVLSYNTLCQHY
ATPKMYRYTPSWALSWDYRRNKLKEQILSYDSDLLCLQEVESKTFEEYWVPLLDKHGYT
GIFHAKARAKTMHSKDSKKVDGCCIFFKRDQFKLITKDAMDFSGAWMKHKKFQRTEDYL
NRAMNKDNVALFLKLQHIPSGDTIWAVTTHLHWDPKFNDVKTFQVGVLLDHLETLLKEE
TSHNFRQDIKKFPVLICGDFNSYINSAVYELINTGRVQIHQEGNGRDFGYMSEKNFSHN
LALKSSYNCIGELPFTNFTPSFTDVIDYIWFSTHALRVRGLLGEVDPEYVSKFIGFPND
KFPSDHIPLLARFEFMKTNTGSKKV"

CDS complement(117728..118729)
/codon_start=1
/label=ATS1
/note="ATS1"
/translation="MSCVYAFGSNGQRQLGLGHDEDMDTPQRSVPGDDGAIVRKIACGG
NHSVMLTNDGNLVGCGDNRRGELDSAQALRQVHDWRPVEVPAPVVDVACGWDTTVIVDA
DGRVWQRGGGCYEFTQQHVPLNSNDERIAVYGCFQNFVVVQGTRVYGWGSNTKCQLQEP
KSRSLKEPVLVYDTGSVAVDYVAMGKDFMVIVDEGGRIVHASGRLPTGFELKQQQKRHN
LVVLCMWTSIHLWNARLNTVESFGRGTHSQLFPQERLDFPIVGVATGSEHGILTTANQE
GKSHCYNVYCWGWGEHGNCGPQKGSQPGLQLVGQYSGKPRVFGGCATTWIVL"

CDS 118364..118933
/codon_start=1
/label=YAL019W-A
/note="YAL019W-A"
/translation="MLLSELVATASSLPYTAISIHNNCRVPAARHIHHGCRYFHGPPVM
HLPQCLRTIQFSPSVISTSYQIPVICQHHAVVPTARYLPDYCSIISWHRPLWGIHILIV
PQSQLPLPIRPKRIHTTHRYKPVIAFNDHIPSLALWICLHYQGSNGCVTPVAAKFFIIF
HFVGLKEIMSPSRNATRNLNQYWRVL"

CDS 119033..122428
/codon_start=1
/label=FUN30
/note="FUN30"
/translation="MSGSHSNDEDDVVQVPETSSPTKVASSSPLKPTSPTVPDASVASL
RSRFTFKPSDPSEGAHTSKPLPSGSPEVALVNLAREFPDFSQTLVQAVFKSNSFNLQSA
RERLTRLRQQRQNWTWNKNASPKKSETPPPVKKSLPLANTGRLSSIHGNINNKSSKITV

AKQKTSIFDRYSNVINQKQYTFELPTNLNIDSEALSKLPVNYNKKRRLVRADQHPIGKS
YESSATQLGSAREKLLANRKYGRHANDNDEEEEESMMTDDDDASGDDYTESTPQINLDE
QVLQFINDSDIVDLSDLSDTTMHKAQLIASHRPYSSLNAFVNTNFNDKDTEENASNKRK
RRAAASANESERLLDKITQSIRGYNAIESVIKKCSSYGDLVTSQMKKWGVQVEGDNSEL
DLMNLGEDDDDDNDDGNNDNNNSNNNNTAGADATSKEKEDTKAVVEGFDETSAEPTPAP
APAPVERETKRIRNTTKPKVVEDEDDDVDLEAIDDELPQSEHEDDDYEEEDEDYNDEEE
DVEYDDGDDDDDDDDEFVATRKNTHVISTTSRNGRKPIVKFFKGKPRLLSPEISLKDYQ
QTGINWLNLLYQNKMSCILADDMGLGKTCQVISFFAYLKQINEPGPHLVVVPSSTLENW
LREFQKFAPALKIEPYYGSLQEREELRDILERNAGKYDVIVTTYNLAAGNKYDVSFLKN
RNFNVVVYDEGHMLKNSTSERFAKLMKIRANFRLLLTGTPLQNNLKELMSLLEFIMPNL
FISKKESFDAIFKQRAKTTDDNKNHNPLLAQEAITRAKTMMKPFILRRRKDQVLKHLPP
KHTHIQYCELNAIQKKIYDKEIQIVLEHKRMIKDGELPKDAKEKSKLQSSSSKNLIMAL
RKASLHPLLFRNIYNDKIITKMSDAILDEPAYAENGNKEYIKEDMSYMTDFELHKLCCN
FPNTLSKYQLHNDEWMQSGKIDALKKLLKTIIVDKQEKVLIFSLFTQVLDILEMVLSTL
DYKFLRLDGSTQVNDRQLLIDKFYEDKDIPIFILSTKAGGFGINLVCANNVIIFDQSFN
PHDDRQAADRAHRVGQTKEVNITTLITKDSIEEKIHQLAKNKLALDSYISEDKKSQDVL
ESKVSDMLEDIIYDENSKPKGTKE"

CDS 120402..120419
/codon_start=1
/product="HQHQHQ metal affinity tag"
/label=HQ tag
/note="HQ tag"
/note="The HQ tag binds to nickel and other metal ions much
like the 6xHis tag, but may allow elution at lower
concentrations of imidazole (Godat et al., 2008)."
/translation="HQHQHQ"

CDS complement(122678..123655)
/codon_start=1
/label=LDS1
/note="LDS1"
/translation="MSFTGSLALAGIGGLVYKFGGGQSYEKLPYVNIPFNQYLDKVYKK
HFSKVMSRTRYVLMNFFKDAFTGGAFMYPFKGFLEFNTNKSSYSTTMLGILSSYLIMFA
LVSFVYWATITPMYTAFLIVLGPIGLFIAIFHSFLQANVFTLLFMRLSHFNNHLVEVCL
EKNGLEENLSEVKPIKYYAPINSIYFWAYYFPFKLVKYMLGLSVLFVLLVISFFPLIGP
ILFHILISPFITQIYFTKVLRLQNFDNIQRRENIYLHAGQYASFGFLAGLIESVPILAG
FAISTNTIGSVLFNLDHPMVPENLVETQAEIEAAPQDINQQPNQ"

CDS 124339..128409
/codon_start=1
/label=PSK1

/note="PSK1"
/translation="MPYIGASNLSEHSFVNLKEKHAITHKGTSSSVASLQTPPSPDQEN
HIDNELENYDTSLSDVSTPNKKEGDEFEQSLRDTFASFRKTKPPPPLDFEQPRLPSTAS
SSVDSTVSSPLTDEDIKELEFLPNESTHSYSYNPLSPNSLAVRLRILKRSLEIIIQNPS
MLLEPTPDDLPPLKEFAGRRSSLPRTSASANHLMNRNKSQIWNTTSATLNAFVNNTSSS
SAASSALSNKKPGTPVFPNLDPTHSQTFHRANSLAYLPSILPEQDPLLKHNNSLFRGDY
GNNISPERPSFRQPFKDQTSNLRNSSLLNERAYQEDETFLPHHGPSMDLLNEQRANLKS
LLNLLNETLEKNTSERASDLHMISLFNLNKLMLGDPKKNNSERDKRTEKLKKILLDSLA
EPFFEHYNFIGDNPIADTDELKEEIDEFTGSGDTTAITDIRPQQDYGRILRTFTSTKNS
APQAIFTCSQEDPWQFRAANDLACLVFGISQNAIRALTLMDLIHTDSRNFVLHKLLSTE
GQEMVFTGEIIGIVQPETLSSSKVVWASFWAKRKNGLLVCVFEKVPCDYVDVLLNLDDF
GAENIVDKCELLSDGPTLSSSSTLSLPKMASSPTGSKLEYSLERKILEKSYTKPTSTEN
RNGDENQLDGDSHSEPSLSSSPVRSKKSVKFANDIKDVKSISQSLAKLMDDVRNGVVFD
PDDDLLPMPIKVCNHINETRYFTLNHLSYNIPCAVSSTVLEDELKLKIHSLPYQAGLFI
VDSHTLDIVSSNKSILKNMFGYHFAELVGKSITEIIPSFPKFLQFINDKYPALDITLHK
NKGLVLTEHFFRKIQAEIMGDRKSFYTSVGIDGLHRDGCEIKIDFQLRVMNSKVILLWV
THSRDVVFEEYNTNPSQLKMLKESELSLMSSASSSASSSKKSSSRISTGTLKDMSNLST
YEDLAHRTNKLKYEIGDDSRAHSQSTLSEQEQVPLENDKDSGEMMLADPEMKHKLELAR
IYSRDKSQFVKEGNFKVDENLIISKISLSPSTESLADSKSSGKGLSPLEEEKLIDENAT
ENGLAGSPKDEDGIIMTNKRGNQPVSTFLRTPEKNIGAQKHVKKFSDFVSLQKMGEGAY
GKVNLCIHKKNRYIVVIKMIFKERILVDTWVRDRKLGTIPSEIQIMATLNKKPHENILR
LLDFFEDDDYYYIETPVHGETGCIDLFDLIEFKTNMTEFEAKLIFKQVVAGIKHLHDQG
IVHRDIKDENVIVDSKGFVKIIDFGSAAYVKSGPFDVFVGTIDYAAPEVLGGNPYEGQP
QDIWAIGILLYTVVFKENPFYNIDEILEGDLKFNNAEEVSEDCIELIKSILNRCVPKRP
TIDDINNDKWLVI"

CDS 128993..130900
/codon_start=1
/label=TPD3
/note="TPD3"
/translation="MSGARSTTAGAVPSAATTSTTSTTSNSKDSDSNESLYPLALLMDE
LKHDDIANRVEAMKKLDTIALALGPERTRNELIPFLTEVAQDDEDEVFAVLAEQLGKFV
PYIGGPQYATILLPVLEILASAEETLVREKAVDSLNNVAQELSQEQLFSDFVPLIEHLA
TADWFSSKVSACGLFKSVIVRIKDDSLRKNILALYLQLAQDDTPMVKRAVGKNLPILID
LLTQNLGLSTDEDWDYISNIFQKIINDNQDSVKFLAVDCLISILKFFNAKGDESHTQDL
LNSAVKLIGDEAWRVRYMAADRFSDLASQFSSNQAYIDELVQPFLNLCEDNEGDVREAV
AKQVSGFAKFLNDPSIILNKILPAVQNLSMDESETVRSALASKITNIVLLLNKDQVINN
FLPILLNMLRDEFPDVRLNIIASLKVVNDVIGIELLSDSLLPAITELAKDVNWRVRMAI
IEYIPILAEQLGMQFFDQQLSDLCLSWLWDTVYSIREAAVNNLKRLTEIFGSDWCRDEI
ISRLLKFDLQLLENFVSRFTILSALTTLVPVVSLDVVTEQLLPFISHLADDGVPNIRFN

VAKSYAVIVKVLIKDEAKYDALIKNTILPSLQTLCQDEDVDVKYFAKKSLAECQELLKN
"
CDS complement(131019..132218)
/codon_start=1
/label=NTG1
/note="NTG1"
/translation="MQKISKYSSMAILRKRPLVKTETGPESELLPEKRTKIKQEEVVPQ
PVDIDWVKSLPNKQYFEWIVVRNGNVPNRWATPLDPSILVTPASTKVPYKFQETYARMR
VLRSKILAPVDIIGGSSIPVTVASKCGISKEQISPRDYRLQVLLGVMLSSQTKDEVTAM
AMLNIMRYCIDELHSEEGMTLEAVLQINETKLDELIHSVGFHTRKAKYILSTCKILQDQ
FSSDVPATINELLGLPGVGPKMAYLTLQKAWGKIEGICVDVHVDRLTKLWKWVDAQKCK
TPDQTRTQLQNWLPKGLWTEINGLLVGFGQIITKSRNLGDMLQFLPPDDPRSSLDWDLQ
SQLYKEIQQNIMSYPKWVKYLEGKRELNVEAEINVKHEEKTVEETMVKLENDISVKVED
"
CDS complement(132368..133135)
/codon_start=1
/label=SYN8
/note="SYN8"
/translation="MDVLKLGYELDQLSDLVEERTRLVSVLKLAPTSNDNVTLKRQLGS
ILELLQKCAPNDELISRYNTILDKIPDTAVDKELYRFQQQVARNTDEVSKESLKKVRFK
NDDELTVMYKDDDEQDEESPLPSTHTPYKDEPLQSQLQSQSQPQPPQPMVSNQELFINQ
QQQLLEQDSHLGALSQSIGRTHDISLDLNNEIVSQNDSLLVDLENLIDNNGRNLNRASR
SMHGFNNSRFKDNGNCVIILVLIVVLLLLLLVL"
CDS 133386..134603
/codon_start=1
/label=DEP1
/note="DEP1"
/translation="MSQQTPQESEQTTAKEQDLDQESVLSNIDFNTDLNHNLNLSEYCI
SSDAGTEKMDSDEEKSLANLPELKYAPKLSSLVKQETLTESLKRPHEDEKEAIDEAKKM
KVPGENEDESKEEEKSQELEEAIDSKEKSTDARDEQGDEGDNEEENNEEDNENENEHTA
PPALVMPSPIEMEEQRMTALKEITDIEYKFAQLRQKLYDNQLVRLQTELQMCLEGSHPE
LQVYYSKIAAIRDYKLHRAYQRQKYELSCINTETIATRTFIHQDFHKKVTDLRARLLNR
TTQTWYDINKERRDMDIVIPDVNYHVPIKLDNKTLSCITGYASAAQLCYPGEPVAEDLA
CESIEYRYRANPVDKLEVIVDRMRLNNEISDLEGLRKYFHSFPGAPELNPLRDSEINDD
FHQWAQ"
CDS 134915..136099
/codon_start=1
/label=CYS3
/note="CYS3"

/translation="MTLQESDKFATKAIHAGEHVDVHGSVIEPISLSTTFKQSSPANPI
GTYEYSRSQNPNRENLERAVAALENAQYGLAFSSGSATTATILQSLPQGSHAVSIGDVY
GGTHRYFTKVANAHGVETSFTNDLLNDLPQLIKENTKLVWIETPTNPTLKVTDIQKVAD
LIKKHAAGQDVILVVDNTFLSPYISNPLNFGADIVVHSATKYINGHSDVVLGVLATNNK
PLYERLQFLQNAIGAIPSPFDAWLTHRGLKTLHLRVRQAALSANKIAEFLAADKENVVA
VNYPGLKTHPNYDVVLKQHRDALGGGMISFRIKGGAEAASKFASSTRLFTLAESLGGIE
SLLEVPAVMTHGGIPKEAREASGVFDDLVRISVGIEDTDDLLEDIKQALKQATN"

CDS 136315..138192
/codon_start=1
/label=SWC3
/note="SWC3"
/translation="MPAVLRTRSKESSIEQKPASRTRTRSRRGKRGRDDDDDDDDEESD
DAYDEVGNDYDEYASRAKLATNRPFEIVAGLPASVELPNYNSSLTHPQSIKNSGVLYDS
LVSSRRTWVQGEMFELYWRRPKKIVSESTPAATESPTSGTIPLIRDKMQKMCDCVMSGG
PHTFKVRLFILKNDKIEQKWQDEQELKKKEKELKRKNDAEAKRLRMEERKRQQMQKKIA
KEQKLQLQKENKAKQKLEQEALKLKRKEEMKKLKEQNKNKQGSPSSSMHDPRMIMNLNL
MAQEDPKLNTLMETVAKGLANNSQLEEFKKFIEIAKKRSLEENPVNKRPSVTTTRPAPP
SKAKDVAEDHRLNSITLVKSSKTAATEPEPKKADDENAEKQQSKEAKTTAESTQVDVKK
EEEDVKEKGVKSEDTQKKEDNQVVPKRKRRKNAIKEDKDMQLTAFQQKYVQGAEIILEY
LEFTHSRYYLPKKSVVEFLEDTDEIIISWIVIHNSKEIEKFKTKKIKAKLKADQKLNKE
DAKPGSDVEKEVSFNPLFEADCPTPLYTPMTMKLSGIHKRFNQIIRNSVSPMEEVVKEM
EKILQIGTRLSGYNLWYQLDGYDDEALSESLRFELNEWEHAMRSRRHKR"

CDS complement(138300..139781)
/codon_start=1
/label=MDM10
/note="MDM10"
/translation="MLPYMDQVLRAFYQSTHWSTQNSYEDITATSRTLLDFRIPSAIHL
QISNKSTPNTFNSLDFSTRSRINGSLSYLYSDAQQLEKFMRNSTDIPLQDATETYRQLQ
PNLNFSVSSANTLSSDNTTVDNDKKLLHDSKFVKKSLYYGRMYYPSSDLEAMIIKRLSP
QTQFMLKGVSSFKESLNVLTCYFQRDSHRNLQEWIFSTSDLLCGYRVLHNFLTTPSKFN
TSLYNNSSLSLGAEFWLGLVSLSPGCSTTLRYYTHSTNTGRPLTLTLSWNPLFGHISST
YSAKTGTNSTFCAKYDFNLYSIESNLSFGCEFWQKKHHLLETNKNNNDKLEPISDELVD
INPNSRATKLLHENVPDLNSAVNDIPSTLDIPVHKQKLLNDLTYAFSSSLRKIDEERST
IEKFDNKINSSIFTSVWKLSTSLRDKTLKLLWEGKWRGFLISAGTELVFTRGFQESLSD
DEKNDNAISISATDTENGNIPVFPAKFGIQFQYST"

CDS 139970..140749
/codon_start=1
/label=SPO7
/note="SPO7"

/translation="MEPESIGDVGNHAQDDSASIVSGPRRRSTSKTSSAKNIRNSSNIS
PASMIFRNLLILEDDLRRQAHEQKILKWQFTLFLASMAGVGAFTFYELYFTSDYVKGLH
RVILQFTLSFISITVVLFHISGQYRRTIVIPRRFFTSTNKGIRQFNVKLVKVQSTWDEK
YTDSVRFVSRTIAYCNIYCLKKFLWLKDDNAIVKFWKSVTIQSQPRIGAVDVKLVLNPR
AFSAEIREGWEIYRDEFWAREGARRRKQAHELRPKSE"
CDS 141030..141626
/codon_start=1
/label=FUN14
/note="FUN14"
/translation="MTLAFNMQRLVFRNLNVGKRMFKNVPLWRFNVANKLGKPLTRSVG
LGGAGIVAGGFYLMNRQPSKLIFNDSLGAAVKQQGPLEPTVGNSTAITEERRNKISSHK
QMFLGSLFGVVLGVTVAKISILFMYVGITSMLLCEWLRYKGWIRINLKNIKSVIVLKDV
DLKKLLIDGLLGTEYMGFKVFFTLSFVLASLNANK"
CDS complement(141814..142461)
/codon_start=1
/label=ERP2
/note="ERP2"
/translation="MIKSTIALPSFFIVLILALVNSVAASSSYAPVAISLPAFSKECLY
YDMVTEDDSLAVGYQVLTGGNFEIDFDITAPDGSVITSEKQKKYSDFLLKSFGVGKYTF
CFSNNYGTALKKVEITLEKEKTLTDEHEADVNNDDIIANNAVEEIDRNLNKITKTLNYL
RAREWRNMSTVNSTESRLTWLSILIIIIIAVISIAQVLLIQFLFTGRQKNYV"
CDS complement(143619..145547)
/codon_start=1
/label=SSA1
/note="SSA1"
/translation="MSKAVGIDLGTTYSCVAHFANDRVDIIANDQGNRTTPSFVAFTDT
ERLIGDAAKNQAAMNPSNTVFDAKRLIGRNFNDPEVQADMKHFPFKLIDVDGKPQIQVE
FKGETKNFTPEQISSMVLGKMKETAESYLGAKVNDAVVTVPAYFNDSQRQATKDAGTIA
GLNVLRIINEPTAAAIAYGLDKKGKEEHVLIFDLGGGTFDVSLLSIEDGIFEVKATAGD
THLGGEDFDNRLVNHFIQEFKRKNKKDLSTNQRALRRLRTACERAKRTLSSSAQTSVEI
DSLFEGIDFYTSITRARFEELCADLFRSTLDPVEKVLRDAKLDKSQVDEIVLVGGSTRI
PKVQKLVTDYFNGKEPNRSINPDEAVAYGAAVQAAILTGDESSKTQDLLLLDVAPLSLG
IETAGGVMTKLIPRNSTIPTKKSEIFSTYADNQPGVLIQVFEGERAKTKDNNLLGKFEL
SGIPPAPRGVPQIEVTFDVDSNGILNVSAVEKGTGKSNKITITNDKGRLSKEDIEKMVA
EAEKFKEEDEKESQRIASKNQLESIAYSLKNTISEAGDKLEQADKDTVTKKAEETISWL
DSNTTASKEEFDDKLKELQDIANPIMSKLYQAGGAPGGAAGGAPGGFPGGAPPAPEAEG
PTVEEVD"
CDS 146290..147276
/codon_start=1

/label=EFB1
/note="EFB1"
/translation="MASTDFSKIETLKQLNASLADKSYIEGYVPI*FTL*IVVFLSFFF
FLWLHVKGS*LIMITFFRLCE**I*FFGPCL*WEVRPPI*VTKEMWLTITVSDDSTEQS
IIISYLLFFFPFLFKKRKTESKDCITRKKFSLLTSKMFCFSF*NSTAVSQADVTVFKAF
QSAYPEFSRWFNHIASKADEFDSFPAASAAAAEEEEDDDVDLFGSDDEEADAEAEKLKA
ERIAAYNAKKAAKPAKPAAKSIVTLDVKPWDDETNLEEMVANVKAIEMEGLTWGAHQFI
PIGFGIKKLQINCVVEDDKVSLDDLQQSIEEDEDHVQSTDIAAMQKL"

CDS 147823..151647
/codon_start=1
/label=VPS8
/note="VPS8"
/translation="MEQNGLDHDSRSSIDTTINDTQKTFLEFRSYTQLSEKLASSSSYT
APPLNEDGPKGVASAVSQGSESVVSWTTLTHVYSILGAYGGPTCLYPTATYFLMGTSKG
CVLIFNYNEHLQTILVPTLSEDPSIHSIRSPVKSIVICSDGTHVAASYETGNICIWNLN
VGYRVKPTSEPTNGMTPTPALPAVLHIDDHVNKEITGLDFFGARHTALIVSDRTGKVSL
YNGYRRGFWQLVYNSKKILDVNSSKEKLIRSKLSPLISREKISTNLLSVLTTTHFALIL
LSPHVSLMFQETVEPSVQNSLVVNSSISWTQNCSRVAYSVNNKISVISISSSDFNVQSA
SHSPEFAESILSIQWIDQLLLGVLTISHQFLVLHPQHDFKILLRLDFLIHDLMIPPNKY
FVISRRSFYLLTNYSFKIGKFVSWSDITLRHILKGDYLGALEFIESLLQPYCPLANLLK
LDNNTEERTKQLMEPFYNLSLAALRFLIKKDNADYNRVYQLLMVVVRVLQQSSKKLDSI
PSLDVFLEQGLEFFELKDNAVYFEVVANIVAQGSVTSISPVLFRSIIDYYAKEENLKVI
EDLIIMLNPTTLDVDLAVKLCQKYNLFDLLIYIWNKIFDDYQTPVVDLIYRISNQSEKC
VIFNGPQVPPETTIFDYVTYILTGRQYPQNLSISPSDKCSKIQRELSAFIFSGFSIKWP
SNSNHKLYICENPEEEPAFPYFHLLLKSNPSRFLAMLNEVFEASLFNDDNDMVASVGEA
ELVSRQYVIDLLLDAMKDTGNSDNIRVLVAIFIATSISKYPQFIKVSNQALDCVVNTIC
SSRVQGIYEISQIALESLLPYYHSRTTENFILELKEKNFNKVLFHIYKSENKYASALSL
ILETKDIEKEYNTDIVSITDYILKKCPPGSLECGKVTEVIETNFDLLLSRIGIEKCVTI
FSDFDYNLHQEILEVKNEETQQKYLDKLFSTPNINNKVDKRLRNLHIELNCKYKSKREM
ILWLNGTVLSNAESLQILDLLNQDSNFEAAAIIHERLESFNLAVRDLLSFIEQCLNEGK
TNISTLLESLRRAFDDCNSAGTEKKSCWILLITFLITLYGKYPSHDERKDLCNKLLQEA
FLGLVRSKSSSQKDSGGEFWEIMSSVLEHQDVILMKVQDLKQLLLNVFNTYKLERSLSE
LIQKIIEDSSQDLVQQYRKFLSEGWSIHTDDCEICGKKIWGAGLDPLLFLAWENVQRHQ
DMISVDLKTPLVIFKCHHGFHQTCLENLAQKPDEYSCLICQTESNPKIV"

CDS complement(151710..155282)
/codon_start=1
/label=TFC3
/note="TFC3"
/translation="MVLTIYPDELVQIVSDKIASNKGSMFMSHSPFRLRLGDKRTIL*K

IIY*RHIEEITLNQLWDISGKYFDLSDKKVKQFVLSCVILKKDIEVYCDGAITTKNVTD
IIGDANHSYSVGITEDSLWTLLTGYTKKESTIGNSAFELLLEVAKSGEKGINTMDLAQV
TGQDPRSVTGRIKKINHLLTSSQLIYKGHVVKQLKLKKFSHDGVDSNPYINIRDHLATI
VEVVKRSKNGIRQIIDLKRELKFDKEKRLSKAFIAAIAWLDEKEYLKKVLVVSPKNPAI
KIRCVKYVKDIPDSKGSPSFEYDSNSADEDSVSDSKAAFEDEDLVEGLDNFNATDLLQN
QGLVMEEKEDAVKNEVLLNRFYPLQNQTYDIADKSGLKGISTMDVVNRITGKEFQRAFT
KSSEYYLESVDKQKENTGGYRLFRIYDFEGKKKFFRLFTAQNFQKLTNAEDEISVPKGF
DELGKSRTDLKTLNEDNFVALNNTVRFTTDSDGQDIFFWHGELKIPPNSKKTPNKNKRK
RQVKNSTNASVAGNISNPKRIKLEQHVSTAQEPKSAEDSPSSNGGTVVKGKVVNFGGFS
ARSLRSLQRQRAILKVMNTIGGVAYLREQFYESVSKYMGSTTTLDKKTVRGDVDLMVES
EKLGARTEPVSGRKIIFLPTVGEDAIQRYILKEKDSKKATFTDVIHDTEIYFFDQTEKN
RFHRGKKSVERIRKFQNRQKNAKIKASDDAISKKSTSVNVSDGKIKRRDKKVSAGRTTV
VVENTKEDKTVYHAGTKDGVQALIRAVVVTKSIKNEIMWDKITKLFPNNSLDNLKKKWT
ARRVRMGHSGWRAYVDKWKKMLVLAIKSEKISLRDVEELDLIKLLDIWTSFDEKEIKRP
LFLYKNYEENRKKFTLVRDDTLTHSGNDLAMSSMIQREISSLKKTYTRKISASTKDLSK
SQSDDYIRTVIRSILIESPSTTRNEIEALKNVGNESIDNVIMDMAKEKQIYLHGSKLEC
TDTLPDILENRGNYKDFGVAFQYRCKVNELLEAGNAIVINQEPSDISSWVLIDLISGEL
LNMDVIPMVRNVRPLTYTSRRFEIRTLTPPLIIYANSQTKLNTARKSAVKVPLGKPFSR
LWVNGSGSIRPNIWKQVVTMVVNEIIFHPGITLSRLQSRCREVLSLHEISEICKWLLER
QVLITTDFDGYWVNHNWYSIYEST"

misc_feature 155581..155698
/label=CEN1
/note="CEN1"

gap 156560..156569
/estimated_length="10"

CDS 156721..158340
/codon_start=1
/label=NUP60
/note="NUP60"
/translation="MHRKSLRRASATVPSAPYRKQIISNAHNKPNLFSKIKTFFTQKDS
ARVSPRNNVANKQPRNESFNRRISSMPGGYFHSEISPDSTVNRSVVVSAVGEARNDIEN
KEEEYDETHETNISNAKLANFFSKKGNEPLSEIEIEGVMSLLQKSSKSMITSEGEQKSA
EGNNIDQSLILKESGSTPISISNAPTFNPKYDTSNASMNTTLGSIGSRKYSFNYSSLPS
PYKTTVYRYSAAKKIPDTYTANTSAQSIASAKSVRSGVSKSAPSKKISNTAAALVSLLD
ENDSKKNNAASELANPYSSYVSQIRKHKRVSPNAAPRQEISEEETTVKPLFQNVPEQGE
EPMKQLNATKISPSAPSKDSFTKYKPARSSSLRSNVVVAETSPEKKDGGDKPPSSAFNF
SFNTSRNVEPTENAYKSENAPSASSKEFNFTNLQAKPLVGKPKTELTKGDSTPVQPDLS
VTPQKSSSKGFVFNSVQKKSRSNLSQENDNEGKHISASIDNDFSEEKAEEFDFNVPVVS
KQLGNGLVDENKVEAFKSLYTF"

```
CDS             complement(158529..159188)
                /codon_start=1
                /label=ERP1
                /note="ERP1"
                /translation="MLLTSLLQVFACCLVLPAQVTAFYYYTSGAERKCFHKELSKGTLF
                QATYKAQIYDDQLQNYRDAGAQDFGVLIDIEETFDDNHLVVHQKGSASGDLTFLASDSG
                EHKICIQPEAGGWLIKAKTKIDVEFQVGSDEKLDSKGKATIDILHAKVNVLNSKIGEIR
                REQKLMRDREATFRDASEAVNSRAMWWIVIQLIVLAVTCGWQMKHLGKFFVKQKIL"
CDS             159469..160749
                /codon_start=1
                /label=SWD1
                /note="SWD1"
                /translation="MNILLQDPFAVLKEHPEKLTHTIENPLRTECLQFSPCGDYLALGC
                ANGALVIYDMDTFRPICVPGNMLGAHVRPITSIAWSPDGRLLLTSSRDWSIKLWDLSKP
                SKPLKEIRFDSPIWGCQWLDAKRRLCVATIFEESDAYVIDFSNDPVASLLSKSDEKQLS
                STPDHGYVLVCTVHTKHPNIIIVGTSKGWLDFYKFHSLYQTECIHSLKITSSNIKHLIV
                SQNGERLAINCSDRTIRQYEISIDDENSAVELTLEHKYQDVINKLQWNCILFSNNTAEY
                LVASTHGSSAHELYIWETTSGTLVRVLEGAEEELIDINWDFYSMSIVSNGFESGNVYVW
                SVVIPPKWSALAPDFEEVEENVDYLEKEDEFDEVDEAEQQQGLEQEEEIAIDLRTREQY
                DVRGNNLLVERFTIPTDYTRIIKMQSS"
CDS             162412..163221
                /codon_start=1
                /label=Kan R
                /note="Kan R"
                /translation="MGKEKTHVSRPRLNSNMDADLYGYKWARDNVGQSGATIYRLYGKP
                DAPELFLKHGKGSVANDVTDEMVRLNWLTEFMPLPTIKHFIRTPDDAWLLTTAIPGKTA
                FQVLEEYPDSGENIVDALAVFLRRLHSIPVCNCPFNSDRVFRLAQAQSRMNNGLVDASD
                FDDERNGWPVEQVWKEMHKLLPFSPDSVVTHGDFSLDNLIFDEGKLIGCIDVGRVGIAD
                RYQDLAILWNCLGEFSPSLQKRLFQKYGIDNPDMNKLQFHLMLDEFF"
CDS             complement(163888..165753)
                /codon_start=1
                /label=RFA1
                /note="RFA1"
                /translation="MSSVQLSRGDFHSIFTNKQRYDNPTGGVYQVYNTRKSDGANSNRK
                NLIMISDGIYHMKALLRNQAASKFQSMELQRGDIIRVIIAEPAIVRERKKYVLLVDDFE
                LVQSRADMVNQTSTFLDNYFSEHPNETLKDEDITDSGNVANQTNASNAGVPDMLHSNSN
                LNANERKFANENPNSQKTRPIFAIEQLSPYQNVWTIKARVSYKGEIKTWHNQRGDGKLF
                NVNFLDTSGEIRATAFNDFATKFNEILQEGKVYYVSKAKLQPAKPQFTNLTHPYELNLD
                RDTVIEECFDESNVPKTHFNFIKLDAIQNQEVNSNVDVLGIIQTINPHFELTSRAGKKF
```

DRRDITIVDDSGFSISVGLWNQQALDFNLPEGSVAAIKGVRVTDFGGKSLSMGFSSTLI
PNPEIPEAYALKGWYDSKGRNANFITLKQEPGMGGQSAASLTKFIAQRITIARAQAENL
GRSEKGDFFSVKAAISFLKVDNFAYPACSNENCNKKVLEQPDGTWRCEKCDTNNARPNW
RYILTISIIDETNQLWLTLFDDQAKQLLGVDANTLMSLKEEDPNEFTKITQSIQMNEYD
FRIRAREDTYNDQSRIRYTVANLHSLNYRAEADYLADELSKALLA"

CDS 166100..166927
/codon_start=1
/label=SEN34
/note="SEN34"
/translation="MPPLVFDIDHIKLLRKWGICGVLSGTLPTAAQQNVFLSVPLRLML
EDVLWLHLNNLADVKLIRQEGDEIMEGITLERGAKLSKIVNDRLNKSFEYQRKFKKDEH
IAKLKKIGRINDKTTAEELQRLDKSSNNDQLIESSLFIDIANTSMILRDIRSDSDSLSR
DDISDLLFKQYRQAGKMQTYFLYKALRDQGYVLSPGGRFGGKFIAYPGDPLRFHSHLTI
QDAIDYHNEPIDLISMISGARLGTTVKKLWVIGGVAEETKETHFFSIEWAGFG"

CDS complement(167731..171321)
/codon_start=1
/label=YAR009C
/note="YAR009C"
/translation="METFTGYLKSTCFHQISPYPPSIMSIQVKVHANILILSFIECLRM
PMHRQIRYSLKNNTITYFNESDVDWSSAIDYQCPDCLIGKSTKHRHIKGSRLKYQNSYE
PFQYLHTDIFGPVHNLPKSAPSYFISFTDETTKFRWVYPLHDRREDSILDVFTTILAFI
KNQFQASVLVIQMDRGSEYTNRTLHKFLEKNGITPCYTTTADSRAHGVAERLNRTLLDD
CRTQLQCSGLPNHLWFSAIEFSTIVRNSLASPKSKKSARQHAGLAGLDISTLLPFGQPV
IVNDHNPNSKIHPRGIPGYALHPSRNSYGYIIYLPSLKKTVDTTNYVILQGKESRLDQF
NYDALTFDEDLNRLTASYHSFIASNEIQESNDLNIESDHDFQSDIELHPEQPRNVLSKA
VSPTDSTPPSTHTEDSKRVSKTNIRAPREVDPNISESNILPSKKRSSTPQISNIESTGS
GGMHKLNVPLLAPMSQSNTHESSHASKSKDFRHSDSYSENETNHTNVPISSTGGTNNKT
VPQISDQETEKRIIHRSPSIDASPPENNSSHNIVPIKTPTTVSEQNTEESIIADLPLPD
LPPESPTEFPDPFKELPPINSHQTNSSLGGIGDSNAYTTINSKKRSLEDNETEIKVSRD
TWNTKNMRSLEPPRSKKRIHLIAAVKAVKSIKPIRTTLRYDEAITYNKDIKEKEKYIEA
YHKEVNQLLKMNTWDTDKYYDRKEIDPKRVINSMFIFNKKRDGTHKARFVARGDIQHPD
TYDTGMQSNTVHHYALMTSLSLALDNNYYITQLDISSAYLYADIKEELYIRPPPHLGMN
DKLIRLKKSLYGLKQSGANWYETIKSYLIKQCGMEEVRGWSCVFKNSQVTICLFVDDMI
LFSKDLNANKKIITTLKKQYDTKIINLGESDNEIQYDILGLEIKYQRGKYMKLGMEKSL
TEKLPKLNVPLNPKGKKLRAPGQPGLYIDQDELEIDEDEYKEKVHEMQKLIGLASYVGY
KFRFDLLYYINTLAQHILFPSRQVLDMTYELIQFMWDTRDKQLIWHKNKPTEPDNKLVA
ISDASYGNQPYYKSQIGNIYLLNGKVIGGKSTKASLTCTSTTEAEIHAISESVPLLNNL
SYLIQELNKKPIIKGLLTDSRSTISIIKSTNEEKFRNRFFGTKAMRLRDEVSGNNLYVY
YIETKKNIADVMTKPLPIKTFKLLTNKWIH"

CDS complement(171678..173000)
/codon_start=1
/label=YAR010C
/note="YAR010C"
/translation="MESQQLSQHSPISHGSACASVTSKEVHTNQDPLDVSASKTEECEK
ASTKANSQQTTTPASSAVPENPHHASPQTAQSHSPQNGPYPQQCMMTQNQANPSGWSFY
GHPSMIPYTPYQMSPMYFPPGPQSQFPQYPSSVGTPLSTPSPESGNTFTDSSSADSDMT
STKKYVRPPPMLTSPNDFPNWVKTYIKFLQNSNLGGIIPTVNGKPVRQITDDELTFLYN
TFQIFAPSQFLPTWVKDILSVDYTDIMKILSKSIEKMQSDTQEANDIVTLANLQYNGST
PADAFETKVTNIIDRLNNNGIHINNKVACQLIMRGLSGEYKFLRYTRHRHLNMTVAELF
LDIHAIYEEQQGSRNSKPNYRRNPSDEKNDSRSYTNTTKPKVIARNPQKTNNSKSKTAR
AHNVSTSNNSPSTDNDSISKSTTEPIQLNNKHDLHLRPETY"
CDS complement(173876..176005)
/codon_start=1
/label=BUD14
/note="BUD14"
/translation="MSNKEEHVDETSASGVKEVSSIAARHDNGYAPSLITSTSGMDSFQ
SHALLNDPTLIEDYSDIINNRPTSGSKLTLGNEDSESMGGSVVVTPTSNKSSPFNSKLN
ILSNAAEKGHDVLRNRDDDKELEEENVEKHMHSNSKRDQRHYKENSSELPDSYDYSDSE
FEDNLERRLQEIETDSVDSADKDEVHFSVNNTMNPDVDDFSDGLKYAISEDEDEEENYS
DDDDFDRKFQDSGFQGEKDDLEEENDDYQPLSPPRELDPDKLYALYAFNGHDSSHCQLG
QDEPCILLNDQDAYWWLVKRITDGKIGFAPAEILETFPERLARLNCWKNENMSSQSVAS
SDSKDDSISSGNKNQSDAESIIPTPALNGYGKGNKSVSFNDVVGYADRFIDDAIEDTSL
DSNDDGGEGNGQSYDDDVDNDKETKVTHRDEYTEAKLNFGKFQDDDTSDVVSDVSFSTS
LNTPLNVKKVRRQDNKNESEPKTSSSKDREDDYNANRYVGQEKSEPVDSDYDTDLKKVF
EAPRMPFANGMAKSDSQNSLSTIGEFSPSSSEWTNESPSTPIVEESSSIPSSRAIKDIS
QYIHAKSKIEETTNVENTEGQIQASLGSSGGMANQTDAEQPKEELEKHHSTPEEEKQST
LSLHSSSEEDFYMDEQRAVSSASINSSLSGSRALSNTNMSDPASKPNSLVQHLYAPVFD
RMDVLMKQLDEIIRK"
CDS 175706..175717
/codon_start=1
/product="Factor Xa recognition and cleavage site"
/label=Factor Xa site
/note="Factor Xa site"
/translation="IEGR"
CDS 176509..177429
/codon_start=1
/label=ADE1
/note="ADE1"

/translation="MSITKTELDGILPLVARGKVRDIYEVDAGTLLFVATDRISAYDVI
MENSIPEKGILLTKLSEFWFKFLSNDVRNHLVDIAPGKTIFDYLPAKLSEPKYKTQLED
RSLLVHKHKLIPLEVIVRGYITGSAWKEYVKTGTVHGLKQPQGLKESQEFPEPIFTPST
KAEQGEHDENISPAQAAELVGEDLSRRVAELAVKLYSKCKDYAKEKGIIIADTKFEFGI
DEKTNEIILVDEVLTPDSSRFWNGASYKVGESQDSYDKQFLRDWLTANKLNGVNGVKMP
QDIVDRTRAKYIEAYETLTGSKWSH"

CDS complement(177530..178837)

/codon_start=1
/label=KIN3
/note="KIN3"
/translation="MHRRQFFQEYRSPQQQQGHPPRSEYQVLEEIGRGSFGSVRKVIHI
PTKKLLVRKDIKYGHMNSKERQQLIAECSILSQLKHENIVEFYNWDFDEQKEVLYLYME
YCSRGDLSQMIKHYKQEHKYIPEKIVWGILAQLLTALYKCHYGVELPTLTTIYDRMKPP
VKGKNIVIHRDLKPGNIFLSYDDSDYNINEQVDGHEEVNSNYYRDHRVNSGKRGSPMDY
SQVVVKLGDFGLAKSLETSIQFATTYVGTPYYMSPEVLMDQPYSPLSDIWSLGCVIFEM
CSLHPPFQAKNYLELQTKIKNGKCDTVPEYYSRGLNAIIHSMIDVNLRTRPSTFELLQD
IQIRTARKSLQLERFERKLLDYENELTNIEKILEKQAIEYERELSQLKEQFTQAVEERA
REVISGKKVGKVPESINGYYGKKFAKPAYHWQTRYR"

CDS complement(179345..182269)

/codon_start=1
/label=CDC15
/note="CDC15"
/translation="MNSMADTDRVNLTPIQRASEKSVQYHLKQVIGRGSYGVVYKAINK
HTDQVVAIKEVVYENDEELNDIMAEISLLKNLNHNNIVKYHGFIRKSYELYILLEYCAN
GSLRRLISRSSTGLSENESKTYVTQTLLGLKYLHGEGVIHRDIKAANILLSADNTVKLA
DFGVSTIVNSSALTLAGTLNWMAPEILGNRGASTLSDIWSLGATVVEMLTKNPPYHNLT
DANIYYAVENDTYYPPSSFSEPLKDFLSKCFVKNMYKRPTADQLLKHVWINSTENVKVD
KLNKFKEDFTDADYHWDADFQEEKLNISPSKFSLAAAPAAWAENNQELDLMPPTESQLL
SQLKSSSKPLTDLHVLFSVCSLENIADTIIECLSRTTVDKRLITAFGSIFVYDTQHNHS
RLRLKFIAMGGIPLIIKFEHLAKEFVIDYPQTLIECGIMYPPNFASLKTPKYILELVYR
FYDLTSTAFWCRWCFKHLDISLLLNNIHERRAQSILLKLSSYAPWSFEKILPSLIDSKL
KKKILISPQITYVVFKSINYMITTNDDKIHKSAIPSSSSLPLSSSPTRNSPVNSVQSPS
RSPVHSLMATRPSSPMRHKSISNFPHLTISSKSRLLIELPEGFFTWLTSFFVDMAQIKD
LSVLKYFTKLCYLTVHINSTFLNDLLDNDAFFAFIRNIDTIIPFIDDAKTAAFIWKQIT
AICVEMSLDMDQMSASLFSTAMNFIRKKNNTSISGLEIILNCLHFTLRNVNDDVAPTVG
SSESHSVFLIKVNNDAAIELPIDQLVDLFYALNDDDVNLSKLISIFTKICSLPGFENLT
INIIFHPNFYEKIVSFFDTYFNSLLIQIDLLKFIKLIFSKSLLKLYDYTGQPDPIKQTE
PNRRNKATVFKLRAILVQITEFLNNNWNKDVPKRNSNQVGGDSVLICQLCEDIRSLSKK
GSLQKVSSVTAAIGSSPTKDERSNLRSSKDKSDGFSVPITTFQT"

CDS complement(183990..184157)
/codon_start=1
/label=PAU7
/note="PAU7"
/translation="MVKLTSIAAGVAAIAAGASAAATTTLSQSDERVNLVELGVYVSDI
RAHLAEYYSF"
CDS complement(186415..186954)
/codon_start=1
/locus_tag="YAR023C"
/label=YAR023C
/note="YAR023C"
/translation="MINFLLFVLTILATLTNIWFSGVLSPAMVIRICLGGSMVVLQIWS
FSRPISNETFRTKLLLEVITHRPSIAGKEWKTITYNMNQYLFKAGLWKTPYHFFCEHQC
YEFFKDLIKGKYPDVQWDTANTQPFISVPENQAATQNSDVEPTVKWCLFKAAEIQAHAV
REYWQSQYPDVGIPAI"
CDS complement(187070..187081)
/codon_start=1
/product="Factor Xa recognition and cleavage site"
/label=Factor Xa site
/note="Factor Xa site"
/translation="IEGR"
CDS 190900..191607
/codon_start=1
/label=UIP3
/note="UIP3"
/translation="MQTPSENTDVKLDTLDEPSAHLIEENVALPEDTFNSYWSYILNEI
ARCKPLMIMFLIPVCLVLLITFFHDIKGILVFLVISLILSIIILLIGITAFVSETLNKG
FIIKLLVEVITRKPAVGGKEWRIIAYNMNQYLFDHGIWHTPYYFFCEHRCHKFFKSLIK
QTRSNAHLSSPTNGAENTQSNTPAKEVSNEMVKPYIFSSDPVLEAYLIKAAEIHKEAEF
EYWRKQYPEVDLP"
CDS 192022..192726
/codon_start=1
/label=YAR028W
/note="YAR028W"
/translation="MQTPSENTDVKMDTLDEPSAHLIEENVALPEDTFSSHLSYVLYEI
AHCKPIMFMIIIIVSLISLIVLFHDNDGCTVILVMSLIVASMALMVVAAFTFGKAITEQ
EFMIKLLVEVIARKPAGKEWGTVAYNMNQYLFMKRLWYTPYYFYSGKKCHEFFTTLIKE
VNSGSHSDSSSNSAEDTQSPVSAGKTSNGLNNFYSIRSDPILMAYVLKATQIEKEAQSE
YWRKQYPDADLP"

```
CDS             193451..193675
                /codon_start=1
                /label=YAR029W
                /note="YAR029W"
                /translation="MNKYLFDHKIWSTPYYFYCEEDCHRLFLSFIEGRTFEKPTSNAEE
                NVQETEAGESFTLNPGEDFQNCFPRQRIL"
CDS             193541..193552
                /codon_start=1
                /product="Factor Xa recognition and cleavage site"
                /label=Factor Xa site
                /note="Factor Xa site"
                /translation="IEGR"
CDS             193966..194862
                /codon_start=1
                /label=PRM9
                /note="PRM9"
                /translation="MSPQYHFYFVSFRNLVLNEKCLRSKKQVMKSFNWYKTDRYFDPHN
                ILQHHSRAIEKTRYKLGMQTSSESTDAKSDFLDEPSAYLIEKNVALPKDIFGSYLSYWI
                YEVTRHKAAVILLVLIVTSILLLVFFYNTEFCVAFEILLFSFCFPGTCMVVIAFSEPIG
                DREFKVKLLMEIITRKPAVKGKEWRTITYKMNQYLFDHGLWDTPYYFYRDEDCHRYFLS
                LIKGRTFKKQKESSASNVKDAQSNDETAGTPNEAAESSSFSAGPNFIKLLTKAAEIEQQ
                FQKEYWRQEYPGVDEFF"
CDS             195237..195941
                /codon_start=1
                /label=MST28
                /note="MST28"
                /translation="MQTPPESTDVKLDTLNEPSAHLIEKNVALPKDIFRSYLSYWIYEI
                ARYTPVMILSLVIGVLVLLIIFFNDNEACVFNSAIFAFTSLVGLLIILSDGNPKLVSRR
                NFRTELLVDVITRKPAVEGKEWRIITYNMNQYLFNHGQWHTPYYFYSDEDCYRYFLRLV
                EGVTPKKQTATSIGNSPVTAKPEDAIESASPSSRLNYQNFLLKAAEIERQAQENYWRRR
                HPNIDALLKKTE"
CDS             197122..197139
                /codon_start=1
                /product="HQHQHQ metal affinity tag"
                /label=HQ tag
                /note="HQ tag"
                /note="The HQ tag binds to nickel and other metal ions much
                like the 6xHis tag, but may allow elution at lower
                concentrations of imidazole (Godat et al., 2008)."
```

/translation="HQHQHQ"
CDS 197323..199386
/codon_start=1
/label=YAT1
/note="YAT1"
/translation="MPNLKRLPIPPLQDTLNRYLARVEPLQDERQNRRTRRTVLSAENL
DALNTLHERLLEYDARLAESNPESSYIEQFWYDAYLLYDATVVLNVNPYFQLQDDPTIK
DTPETAAQGPYGAHTVQVRRAARLTTSILKFIRQIRHGTLRTDTVRGKTPLSMDQYERL
FGSSRIPPGPGEPSCHLQTDATSHHVVAMYRGQFYWFDVLDTRNEPIFATPEQLEWNLY
SIIMDAESAGSGSAPFGVFTTESRRVWSNIRDYLFHADDCTNWRNLKLIDSALFVVCLD
DVAFAADQQDELTRSMLCGTSTINLDPHQHQPPLNVQTGTCLNRWYDKLQLIVTKNGKA
GINFEHTGVDGHTVLRLATDIYTDSILSFARGVTKNVVDIFSDDDGKPSSSSLASAAHS
ANLITIPRKLEWRTDNFLQSSLHFAETRISDLISQYEFVNLDFSNYGASHIKTVFKCSP
DAFVQQVFQVAYFALYGRFETVYEPAMTKAFQNGRTEAIRSVTGQSKLFVKSLLDQDAS
DATKIQLLHDACTAHSQITRECSQGLGQDRHLYALYCLWNQWYKDKLELPPIFRDKSWT
TMQNNVLSTSNCGNPCLKSFGFGPVTANGFGIGYIIRDHSVSVVVSSRHRQTARFASLM
EKSLLEIDRIFKRQQARAAKPAARTTASANTKSEDMKYLLSGYDYFDVSVSG"
CDS complement(199467..199547)
/codon_start=1
/locus_tag="YAR035C-A"
/label=YAR035C-A
/note="YAR035C-A"
/translation="MRLNYSRCYYSSQRRRQSLPKRFPLI"
CDS 199749..203315
/codon_start=1
/label=SWH1
/note="SWH1"
/translation="MEQPDLSSVAISKPLLKLKLLDALRQGSFPNLQDLLKKQFQPLDD
PNVQQVLHLMLHYAVQVAPMAVIKEIVHHWVSTTNTTFLNIHLDLNERDSNGNTPLHIA
AYQSRGDIVAFLLDQPTINDCVLNNSHLQAIEMCKNLNIAQMMQVKRSTYVAETAQEFR
TAFNNRDFGHLESILSSPRNAELLDINGMDPETGDTVLHEFVKKRDVIMCRWLLEHGAD
PFKRDRKGKLPIELVRKVNENDTATNTKIAIDIELKKLLERATREQSVIDVTNNNLHEA
PTYKGYLKKWTNFAQGYKLRWFILSSDGKLSYYIDQADTKNACRGSLNMSSCSLHLDSS
EKLKFEIIGGNNGVIRWHLKGNHPIETNRWVWAIQGAIRYAKDREILLHNGPYSPSLAL
SHGLSSKVSNKENLHATSKRLTKSPHLSKSTLTQNDHDNDDDSTNNNNNKSNNDYDDNN
NNNNNDDDDYDDDDESRPLIEPLPLISSRSQSLSEITPGPHSRKSTVSSTRAADIPSDD
EGYSEDDSDDDGNSSYTMENGGENDGDEDLNAIYGPYIQKLHMLQRSISIELASLNELL
QDKQQHDEYWNTVNTSIETVSEFFDKLNRLTSQREKRMIAQMTKQRDVNNVWIQSVKDL
EMELVDKDEKLVALDKERKNLKKMLQKKLNNQPQVETEANEESDDANSMIKGSQESTNT

LEEIVKFIEATKESDEDSDADEFFDAEEAASDKKANDSEDLTTNKETPANAKPQEEAPE
DESLIVISSPQVEKKNQLLKEGSFVGYEDPVRTKLALDEDNRPKIGLWSVLKSMVGQDL
TKLTLPVSFNEPTSLLQRVSEDIEYSHILDQAATFEDSSLRMLYVAAFTASMYASTTNR
VSKPFNPLLGETFEYARTDGQYRFFTEQVSHHPPISATWTESPKWDFYGECNVDSSFNG
RTFAVQHLGLWYITIRPDHNISVPEETYSWKKPNNTVIGILMGKPQVDNSGDVKVTNHT
TGDYCMLHYKAHGWTSAGAYEVRGEVFNKDGKKLWVLGGHWNDSIYGKKVTARGGELTL
DRIKTANSATGGPKLDGSKFLIWKANERPSVPFNLTSFALTLNALPPHLIPYLAPTDSR
LRPDQRAMENGEYDKAAAEKHRVEVKQRAAKKEREQKGEEYRPKWFVQEEHPVTKSLYW
KFNGEYWNKRKNHDFKDCADIF"

ORIGIN

1 cagaggtatt gtggatcctt ctacagtact tctgaataca cctaaaaggt tgttggatgc
61 taaatttagc aaaagtcttt tttagctcac tattaggctt gttaaagtct gaaattgttg
121 aaaggcactc aaaagataaa ttgacaatta gcattaacgg cacagttgaa agagtcaccc
181 acttgaaatt agctcggtta tcaaatataa ttatctccgg taaagagctc tgcagcaggg
241 ttaatctatt cctatactta cgctgtagga acattttatt attaggatcc gactactgcc
301 tacatattta ttcggaaggc ttgatgtcga aaatttttga gcttataaaa agaacatatt
361 tcactcttgc tcgtgccggt agaggtgtgg tcaataagag cgacctcata ctatacctga
421 gaaagcaacc tgacctacag gaaagagtta ctcaagaata agaattttcg ttttaaaacc
481 taagagtcac tttaaaattt gtatacactt atttttttta taacttattt aataataaaa
541 atcataaatc ataagaaatt cgcttaagcg gatctgccta ctctcttcaa gtaagcgtcc
601 catcccgcct ttctgtatct ctcaccttcc actacaggat ctcttccctt tgcgaaaaga
661 cctctaccaa caataatgat gtctgatcca ccggcaacaa cttcatccac agttctgtat
721 tgttgtccca aagcatcacc tttgtcatca agaccaacac ctggcgtcat gatcaaccaa
781 tcgtagccct cttctcttcc acccatatcg ttttgagcaa taaatccaat aacaaagtcc
841 ttatcactct tggcaatttc cacggtccca cgagtgtatt caccgtgcgc tagagacccc
901 ttggacgata actcggcaag cattaacaac cctctaggtt cttttgtaac ttcctcggca
961 ccttgcttca aaccagcaac aatgcccgca ccagtcacac cgtgtgcatt ggtgatatca
1021 gaccattcgg cgatacggta tacaccagac gtgtattgta atttaacagt gttcccaatg
1081 tcggcaaact tcctgtcttc aaatatcaaa aacttgtgtt tctctgctaa ttgcttcaac
1141 ggcacaatgg tattctcaaa gctgaaatcc tccaagatat ctacatgtgt cttcaataga
1201 cagatatatg gacccaaaac ctcaactaat cttaacaact ctgctgtttt acgaacatca
1261 agagaagcac ataagtttga cttcttctct tccatcaagt ttaaaagctt ggcagcaact
1321 ggacttctat gagcagctgc tctttccgag taactcttag tcgacatcct cctttgatta
1381 gtttatgggg ctagctgatg atacttcctc ttgctgttcc agtttacaaa tgatttaaac
1441 gaacttttac acttttacgg tagaatttct tgttggtgat gagctcagat tcaatatgac
1501 tgcttaatca gtgaaaatta atggaaacag ccagacacac agcctttgaa aaatgttaaa
1561 aacgatcaca tatatgtagt cacgtggtta gtgcatagag tcagtcgatg gtggttagtt
1621 ccctagccag agccattccc ctcttgtttc gaaatgcgat gactgaatgg ggaagaagct
1681 cttagtattt ctgtagtttt ttttgccgat ttctattgtt gcctgccccg gactttcttt

1741 taccttttgt agctacaact attaattata atgcataata gatctagaag cgtattgtga
1801 tgagcatcct tggaaataag aaaggaagtc aaaggtacag taaaaaaaga gccgtgacag
1861 gcacctaatt tcagttttga ttgaggtata atgagctcag aagaccaaga tgctctaaaa
1921 gtaaagcaga atggtctttc caatcagaat tcgctgatta ctagcgaagc tgcgggaata
1981 tggtggggcc tttgacttcc agaatgcttt cacggaatta ttttagatgt acatttagct
2041 ccatttcctg tgcctccgat aggaaggcat catggtacta ccgtgacgga gaatacgtag
2101 gctgactttt tcgtcagttt gttgtccgtt tacaaaattg gtgaatgaat tctagccttt
2161 gctcattaat tgccctcaca agaatttgga agtgcgtaga acaagtaaaa ggttgcacta
2221 ggcgcgccac ggatgctatt tcagaatatt tcgtacttac acaggccata cattagaata
2281 atatgtcaca tcactgtcgt aacactcttt attcaccgag caataatacg gtagtggctc
2341 aaactcatgc gggtgctatg atacaattat atcttatttc cattcccata tgctaaccgc
2401 aatatcctaa aagcataact gatgcatctt taatcttgta tgtgacacta ctcatacgaa
2461 gggactatat ctagtcaaga cgatactgtg ataggtacgt tatttaatag gatctataac
2521 gaaatgtcaa ataattttac ggtaatataa cttatcagcg gcgtatacta aaacggacgt
2581 tacgatattg tctcacttca tcttaccacc ctctatctta ttgctgatag aacactaacc
2641 cctcagcttt atttctagtt acagttacac aaaaaactat gccaacccag aaatcttgat
2701 attttacgtg tcaaaaaatg agggtctcta aatgagagtt tggtaccatg acttgtaact
2761 cgcactgccc tgatctgcaa tcttgttctt agaagtgacg catattctat acggcccgac
2821 gcgacgcgcc aaaaaatgaa aaacgaagca gcgactcatt tttatttaag gacaaaggtt
2881 gcgaagccgc acatttccaa tttcattgtt gtttattgga catacactgt tagctttatt
2941 accgtccacg ttttttctac aatagtgtag aagtttcttt cttatgttca tcgtattcat
3001 aaaatgcttc acgaacaccg tcattgatca aataggtcta taatattaat atacatttat
3061 ataatctacg gtatttatat catcaaaaaa aagtagtttt tttattttat tttgttcgtt
3121 aattttcaat ttctatggaa acccgttcgt aaaattggcg tttgtctcta gtttgcgata
3181 gtgtagatac cgtccttgga tagagcactg gagatggctg gctttaatct gctggagtac
3241 catggaacac cggtgatcat tctggtcact tggtctggag caataccggt caacatggtg
3301 gtgaagtcac cgtagttgaa aacggcttca gcaacttcga ctgggtaggt ttcagttggg
3361 tgggcggctt ggaacatgta gtattgggct aagtgagctc tgatatcaga gacgtagaca
3421 cccaattcca ccaagttgac tctttcgtca gattgagcta gagtggtggt tgcagaagca
3481 gtagcagcga tggcagcgac accagcggcg attgaagtta atttgaccat tgtatttgtt
3541 ttgtttgtta gtgctgatat aagcttaaca ggaaaggaaa gaataaagac atattctcaa
3601 aggcatatag ttgaagcagc tctatttata cccattccct catgggttgt tgctatttaa
3661 acgatcgctg actggcacca gttcctcatc aaatattctc tatatctcat ctttcacaca
3721 atctcattat ctctatggag atgctcttgt ttctgaacga atcataaatc tttcataggt
3781 ttcgtatgtg gagtactgtt ttatggcgct tatgtgtatt cgtatgcgca gaatgtggga
3841 atgccaatta taggggtgcc gaggtgcctt ataaaaccct tttctgtgcc tgtgacattt
3901 cctttttcgg tcaaaaagaa tatccgaatt ttagatttgg accctcgtac agaagcttat
3961 tgtctaagcc tgaattcagt ctgctttaaa cggcttccgc ggaggaaata tttccatctc
4021 ttgaattcgt acaacattaa acgtgtgttg ggagtcgtat actgttaggg tctgtaaact

4081 tgtgaactct cggcaaatgc cttggtgcaa ttacgtaatt ttagccgctg agaagcggat
4141 ggtaatgaga caagttgata tcaaacagat acatatttaa aagagggtac cgctaattta
4201 gcagggcagt attattgtag tttgatatgt acggctaact gaacctaagt agggatatga
4261 gagtaagaac gttcggctac tcttctttct aagtgggatt tttcttaatc cttggattct
4321 taaaaggtta ttaaagttcc gcacaaagaa cgcttggaaa tcgcattcat caaagaacaa
4381 ctcttcgttt tccaaacaat cttcccgaaa aagtagccgt tcatttccct tccgatttca
4441 ttcctagact gccaaatttt tcttgctcat ttataatgat tgataagaat tgtatttgtg
4501 tcccattctc gtagataaaa ttcttggatg ttaaaaaatt attattttct tcataaagaa
4561 gctttcaaga tataagatac gaaatagggg ttgataattg catgacagta gctttagatc
4621 aaaaaggaaa gcatggaggg aaacagtaaa cagtgaaaat tctcttgaga accaaagtaa
4681 accttcattg aagagcttcc ttaaaaaatt tagaatctcc catgtcaacg ggtttccata
4741 cctccccagc atcatacatc ttttttcaaa gaaacttcaa atgcctcttt tatgcaaggg
4801 gcaaaatcct gaaatgactt aaacttagca gtttcgtctt ttttcaaaga gaatggttga
4861 agaagaattg ttttggacgc ttattgacaa tctgttgcat tgataaagta cctactatcc
4921 cagactatat ttgtatacaa gtacaaaatt aggtttgttg aaacaacttt ccgatcattg
4981 gtgcccgtat ctgatgtttt tttagtaatt tctttgtaaa tacagggagt tgtttcgaaa
5041 gcttatgaga aaaatacatg aatgacaggt aaaaatattg gctcgaaaaa gaggacaaaa
5101 agagaaatca taaatgagta aacccacttg ctggacatta tccagtaaag gcttggtagt
5161 aaccataata ttacccaggt acgaaacgct aagaacttga aagactcata aaacttccag
5221 gttaagctat ttttgaaaat attctgaggt aaaagccatt aaggtccaga taaccaaggg
5281 acaataaacc tatgcttttc ttgtcttcaa tttcagtatc tttccatttt gataatgagc
5341 tagtgatccg gaaagctact ttatgatgtt tcaaggcctg aagtttgaat atttatgtag
5401 ttcaacatca aatgtgtcta ttttgtgatg aggcaaccgt cgacaacctt attatcgaaa
5461 aagaacaaca agttcacatg cttgttactc tctataacta gagagtactt tttttggaag
5521 caagtaagaa taagtcaatt tctacttacc tcattaggga aaaatttaat agcagttgtt
5581 ataacgacaa atacaggccc taaaaaaattc actgtattca atggtctacg aatcgtcaat
5641 cgcttgcggt tatggcacga agaacaatgc aatagctctt acaagccact acatgacaag
5701 caactcataa tttaagtgga tagcttgtga taaattgaat tttctctgtt tagtacttgc
5761 cgaatagtta cttgttagtt gcagatgctt tttgatgaca aagttatcaa tctcaatatt
5821 aaacttttta ggctttcagg tttaatcttt ctttgagggt gtattaattt tcatacaaat
5881 atttgattca ttattcgttt tactgttaca ttagacctgc tcattacatg gagtaactta
5941 agttttctca aacgcttgat agcatgattt gatgtagtaa aaaaaaaggc agagtttcca
6001 aaaaaaattg ttaatcgaca aagttaatat tatggtggta gtatctcaaa tatctggata
6061 accagatcgt acatctctga taaacaatct ttgccactgc tttatccttt taaattgtat
6121 tgagtgcttc agtcattgca aaattttacg agatttaaaa tttgtgaacc cgaccttacc
6181 gagaaatgat gagctaattt ttataggtcg acccttctgt cgcttactgg gttgattatc
6241 ttgtgctttc ttagtatcta tcacaaagga gacaaaatcg ttgataaaaa gtgcatcaac
6301 attcccagcc agaaaatgca catcataaag acatgttatt caagagccac gaccgtcttc
6361 aatttatctt ttataaaaaa cccttgttct actgacagga tggaatagat attaaatata

6421 cattttgcat tttttttttt ttctgtattg aagatttgta tatgaaagat gtttatacat
6481 caaatgcttt gaataaagcc atcttaattt caatttcatg ccctccttca ccgttttctg
6541 ttggtctaga ggtagcttgt tgtggtcact aatgagaact taaatagttt tcaactgctg
6601 gtgataaatc aataatttat gttcttaacc taacatttga tgacctttga tgcgttggtt
6661 atgttgaaga caaattgcct ctaatcagtt ccattaagaa atcttcttaa ctcctccaaa
6721 tattctgccc atacgatacc tatttgttta ctttgtcatt ttgccataag attggtatcc
6781 acttcttgtc tgtaaaataa ttagaaagta gcacaatttt tacagtaatg tagcacgcgt
6841 aactcctaaa ctttgtcata atggttgaaa tgaatgtatg atataaaaac tcggaccctg
6901 ttttacttct tttatagaac cttatttttg acgcagggag gcgacattta tccaaattaa
6961 gttttgacat ggcgcatcag ggaataaaaa aaactttatt atgtggccga atcaacatta
7021 atcaaatgca ctaatattgt aacgttctta caaagggcag acaacttgag aactttcatg
7081 cgtgcaacag tattaatatt ttactgtctt gatatcgtta tcctcatcgt aacgtgaatt
7141 tttttgtctc atacgttaag gtaaattttg atgacccccg ttgtccttgt ttgccttact
7201 gtataaagca cccttttatt gtttagaata ctagaatgat aactgcattc ggactatgaa
7261 agaaaaaatg gtagtagcaa aggataggca tcgccgtatt tactactttg taaaccagtg
7321 gatttttgct caacatataa aaaactaaag accttttttt catcaatata cttctgagac
7381 gtgcagatgt gatattcggg tttgagcttg tagtcaacga agcgggttca tgggcaaatt
7441 ttcttttttt tcccttttttt tttgtctaga ttatttcgaa tatgagttaa tcatacgttg
7501 attagtactg ttggtctctc attgaaattt tacgtgacac catcatttta cttccacata
7561 agttctaatg ttacgtagtt caattttagt cgacctagct tcatatttat tttagaagca
7621 attcgtaatt atcattttgc tttcgaagaa aattaagact tcatttacta ttctcgtgat
7681 attttagtag gcgcttcttt tgtatcgaac cattttattg caatggccct taagttaccg
7741 ttattcatac caatttgacg ttaattttaa atgcgttctg aagtttctta aataacccgg
7801 attggttagg ttcagccatg cctggcgcgt acattgaggc attagaagat ccgcagataa
7861 ataataagct tagtaaatcc taaagataac aactaaaatt atatttccat cagctcaata
7921 ccgcagtact ttgaaacctg atttatatat tgcagaactt aattaaaagt acattgtagt
7981 tcaaaaaata aatatcaaac ttttggaccc tctcttattg cctcccaatt aattaaaaca
8041 tcttttcttc caatctacag gtttgaaaag gtaataagta atataaactt gagaaccaaa
8101 aaaaaaaaaa aaaaaaaata ctgatcctta caggttttaa ggttgcaaag ggaacattta
8161 ttgaaaggag ctaacaatag tgggtatgag taaagatata tagatcgata ttttgaattc
8221 taaatgatga actagggaag taatttaggt gaaacattgc aaccaatcat tttacacttt
8281 tggttgcacg taatgtacct ttttatgata tttttttttt atagtagtag tgtgaaaatt
8341 tcttcaggac ttgcaaaaag aatctaactg atcttcggat gagcctttat cgattatttt
8401 tttcctaaat ataatacttt acaagcgaat gttttgttag gagaaagata taaaaattat
8461 gcggcatagg catattatcc aataaaaagg aaatttatat ataaacttca tttacgtcat
8521 aagaaaatgt taagttctct taacgaaaac tgtgcgaatt ttgtgttaaa gctggatgat
8581 gagaaattat tctcgtatta tttttcatca gatactgata aggtttcaac gtcttttgac
8641 gttggctttt ccacaccatg tttagagtta taaagcacaa taccgttctt cttggcattg
8701 ttcctttcat cacgtttata gaagtagagt acaacaaaag tccaaatgga gagacaaaaa

8761 gcagaacatg cagtgaaagt aaaccccttt aaatacctgg gagcttcttc tgttttccaa
8821 accaaaacac ttatccatgc ggtagatgat tgagccataa tattcattgt aactaaagta
8881 atagctctag tttgagcatc tcggcgacaa atatcgtttt gccaagagta taaaacagga
8941 gccatagccc aaccaaaaca ttgcagcata aatgcaaacc atttggctcc ttctgcgacg
9001 tcccaagcgg ctaatatgga gttaccaatg atattgaaaa cctgagtaaa aataatcgca
9061 aaccaacgag agtgtaattt atctgcaata ataccagtaa gcatcaaata aaccatacct
9121 aaacccggag taatcatgga taactgattg agcttaggaa tagagtatct tttcaaagat
9181 ttcaaccata gtaggtatgc cccagatgaa acattactgt cattccaaca gaaaatattc
9241 cataaagtta aaatgtatat tttccaatca ctgaaaattg ttttccacag tttaatatcg
9301 aatacttttg tttcaaaatc acttttacct gtttggtttt cttttaatct tttcctcgcc
9361 aacctaattt catcatcagt taagaaaata gaataacagt tgtatgggtc acctggcagg
9421 gagtaaaatc caataaggcc cactacgaca gacacaatag cgtcaataat aaagttccat
9481 ctccatccct ctaaaccatt tacaccattt aacgatgaat atacggctga ctggatccca
9541 ccagcggata gaataccgat atactggccc aaatagtaaa aagcagaacg acgcaccatt
9601 tcatcatgtt tgtaaaagga accaaacaaa tattggtatg ccaaataact tggcgcttca
9661 aaagccccaa tgaaaaacct aattgctttc aagtgtggta cagaattgac atatgcagca
9721 ccaacggtta aaagcgacca acataagtcg aggcttggta aaacatagtt taatgggagc
9781 ttgttcaggt aaatcaaaaa tggcaattga aatataatat taccaactgt gtacattact
9841 tgagtatgca ccaaatcatt accttgaaag cctaaatctt ccttcattcc cgaaacgtaa
9901 gcgttgttta tattaaccgt atccagatat ttcacccaat aagcaataca agaataaaag
9961 gctaaaagga catccaattt aattaataat tttttttctt tgaaagaggt accctgtttg
10021 aaccaactat accatttatt gtgagatctt tccttttcat tgatccgata ctcttgttca
10081 tcgaaaaatc tccaccatgg cctatcggct tcatctctat attcataact tctagtaacg
10141 ttcacattgt ctttatgttc actatagcta ctagtctcaa aatgtagttg atctttttca
10201 cttgtagtcg tgatgaaatt ttcagctgtt tcatgactct ggatactgtt ggagatagtg
10261 acaatttctg ttgaatttaa gtcatctggc aggtcttcca cctgccgctt tactggaata
10321 aaaccccatt ttagtctttt gtaaggatct acaataatct ctttaacaat tgaatacatg
10381 tatgttattt atatatgcag tagttctctt tgtaattttt tttaacaaat agagagtaag
10441 atatgtagcg aatgtccatt catcataaca ggtaaactag atgctctttt atatagtctg
10501 gttgtataaa taatttatat cctcatctaa acgcatgtct ccgcttggtt tcttatctat
10561 ttgtggaggc acccgtagta ctgtgctttc gtattttttt atttatttat tatttgtagc
10621 agttttttc cagtgacaca atctttacca ttacacagtt tttactattt ctaatgatta
10681 cattggacca tcggaaaact gcgctaactt tggataacgc cacagaacgt gatctgacta
10741 ttgtaaaggt gtcattcggt aaggtaaacc ttcaaggcgt ccacaccttt aaccatataa
10801 cagcgtaaac tcttgttaaa atacaggggt aagacattgg tggatattca acaagatccg
10861 atacctccaa ttccgatttc tacaaattgt gctctacata ttactgcgat gcaattatcc
10921 acacataaaa tcgatgcctt tagaaaaaga cataaagcag acggcattgt agatatttga
10981 ccaattagaa ttgaatgaga tgaatatctc accaagctat tcgatattat agataaagtt
11041 tgtatcttag ttatgcattg tgagttggtt gctctactcg cggttaccag tctcttctaa

11101 aaaatctaag gccaatggta tccatgacat ttctgcactt tttgtaattg gtttagttat
11161 gaaaggaacg tcaaaccaaa tggtttttca gataagaaat tgacagtatc tgagaatttg
11221 ctatcaaagc tcagaggatt tacatatttt aacgtaatta aaacattttt atgttcgata
11281 tattagcaaa tagcgtatta atatacagct gttgcgctca tggtaaaatt tagcgatata
11341 ctttgcatct tggctgcaaa gaagaatgaa tcggatatac tatttttgat cataatgacg
11401 gacatcatga tataataacg ttatacggat aactttattt caaaagcacc atcatgttat
11461 ctcttgtaaa aagaagtatt cttcattcaa taccaattac tcgtcacatt cttccaatcc
11521 aattaatatt ggttaaaatg aaccatgtgc aaatcagaaa cataaaatta tatcacttta
11581 tttcatatgg tttcatgctt acaaagctta ctgtctttct ctttaactta tttttctaca
11641 ggctacgaat tctttgcagg cttactttac tcatattatc attacctgta caaatatata
11701 ttaaagaaat ccaaacaaaa atgcttgaaa agcatacagc ttccgataca tcatgtatat
11761 agaaaatcac agtacaaaaa ttttgaattt atgtataacc gtttcgcctg atatatgtaa
11821 gagctcttga ttgtcggaat agttcgggaa ttctgggtgg aactagtagc tggagatgcg
11881 ttctaaagga tctaaaatca gactcacccc aaaaaccaaa attttgatat tcaactttag
11941 tattagccag tcttaaaatg attttttgcc aagaaagccg aagtttaacg agcgttttaa
12001 aatatgcaca agtccattaa ttaaatttga tcacagtgat taccaatttt gttgaaagca
12061 gtaatttgtg acgtccgttt ttggcacaag taaaaaatat ttgttttgaa gtctagttac
12121 aagaagctac tgaaaacaca gggctggata tagatgttta taagctcctc ccatggtcta
12181 agagtttctc caaacgccta atgtatgcaa tttgttaaat aaaaatgaag aatagatgca
12241 taaacaagcg tcattaaagt tcttatcatt attccatgta gaaatgacct gcaagtacga
12301 gctggttatg cgaaactgag cttaacaaca tagttttctt ttccattggc ttggatataa
12361 atttttcgct gagaaacttc tctgcatttt ttaagcatta agcgtacata acataaagtc
12421 actgagatat tagaggttat aaaaaaagac tacgtgggtg tttacctaag attttcacaa
12481 cttgtttatt aattggataa atagtatttt tcttggcatt tacaaacttt atgtttttag
12541 gtgtatttgc agttccatta agggaaataa gtcataagct tttttggaaa acagattttt
12601 tgccccgctt aacctgaaat tgatattaaa agaatgtgtg aatggccatt aactaacaag
12661 agaattaata atgttaaaac acagatacct cgaaacaaac tctatgtaaa cacttatttt
12721 attgtggtaa tattttttga taacaacaca tctgaaacaa aataatgcaa agccgaatag
12781 ttaggctaaa aatgtactct tagacattta aaaaggttta tgaatcctat ggtatttaat
12841 atattaaaga acgaagtaaa tgggaaaaaa tgtgtaaaca ctataagcgt gatgatagaa
12901 ttattaatat aagatgatgc cgtgcgttta ccatacgatt gccagcaata cggtggaaat
12961 aaaaacactt atgccattat tggtcaacag accattggca ataccaacgt aggttgagat
13021 ttctaaagag gcagtactag atcctattat gctacttgct ggtgtgctac gaggctgttg
13081 cgacatagta cttaacccgg aagttattag actcttggtg ttgccagttt cggatacaga
13141 aacaacacta ctgctgtgac caatcacatc ggtcgcggaa gccgtctgtg tttcagcatg
13201 attgaatctt gaaattgaag aggtgactac tgttttcgtc tcagcagctc cagtactggt
13261 agttgtctca gcagctccag tattggttgt tgtctcactg gtagcactgt tcattttaga
13321 gctgacagac tcttcattcg tagtctgtgg cctccatgtt ggatagaccg taacaacatc
13381 attcacagta gccgtggccg tcgaaacaat ggcaggtgaa gcagtttcgg aacacacacc

13441 agattcgcag gaagtaacag taactagcgt agtttgttgc ctcgattctg tggtggaaat
13501 aggacaccat gtcgtgtatt ctgtggtaac gccgttaata gtagcagtgc ttatagatac
13561 aatggctgga gaagcagtct tagagcatac gtcagattca caagaagaaa ttgtaactac
13621 tgtggtttgc tctgttgtcc ccttggtttg ctttgttgtc tccgtagttt gctttgttat
13681 ctctgtggta gaaatagggc accatgtggt atactctgtt gtggcaccgc taacagtaac
13741 ggtggccgtg gaaacaattg cagaggagat ggattcagtg cacacatgag attcgcagga
13801 tgtcacggta accaaagtgg tttgttcgct cgtttttgta gtggtagcag gtggtaatga
13861 agaagcagtt tcctggcttg ttgttgtact gataacaggt ggtaatgatg aagtaatttc
13921 ctgacttgct gtcgcactgg taacaggtgg taatgaagaa gtaatttcct gacttgttgt
13981 tgtactggta acaggtggta atgatgaagt aatttcctga cttgttgttg tactggtaac
14041 aggtggtaat gaagaagtaa tttcctgact tgttgttgca ctggtaacag gtggtaatga
14101 tgaagtaatt tcctgacttg ttgttgtact ggtaacaggt ggtaatgatg aagtaatttc
14161 ctgacttgtt gttgtactgg taacaggtgg taatgaagaa gtaatttcct gacttgttgt
14221 tgcactggta acaggtggta atgatgaagt aatttcctga cttgttgttg tactggtaac
14281 aggtggtaat gaagaagtaa tttcctgact tgttgttgta ctggtaacag gtggtaatga
14341 tgaagtaatt tcctgacttg ctgtcgcact ggtaacaggt ggtaatgaag aagtaatttc
14401 ctgacttgtt gttgcactgg taacaggtgg taatgatgaa gtaatttcct gacttgttgt
14461 tgcactggta acaggtggta atgatgaagt aatttcctga cttgttgttg tactggtaac
14521 aggtggtaat gatgaagcag tttcctggct tgttgttgca gtggtaatag gtggtaatga
14581 tgaagacgaa tatgtagact ttggtgattc agaagagata gaagaggaag aagagacaga
14641 actagctgaa ctagtttcgc tctcagaaga accagaggtg gaactactgg ttggaatgac
14701 ggatgattta aatgattcag agaatataga agtggaggtt gttgtagaag aaatgacagg
14761 ggaagaaatg aatgaagaag aaatgactgg agaagaagtg actagagaag aagtgactga
14821 ggaagaaatg actagagaag aagtgtctga ggaagaaatt actgaggagg aaatcacaga
14881 agttccattg ctaggataga atggggtaat aattggatgc taagacgtga tagagctggt
14941 tatttgttct gaagaagatg atgacaaact ggatgaggtg gcagtggttg gacttttagc
15001 aacaataacg gtttcgtcag ttggttgacc attagtgcca gtgacagtag tcatctcagt
15061 agaagtcgaa gtggtagttg cgatggtgct agtagtatgt tttgaaggat cagggatagt
15121 acaattgggc tgttaggtca tcgtcaaaag agtaaacgtg ccctgcaaaa tcatcgctaa
15181 cagtagtaac atcaggcaat tccacgctaa ctggcagtgt atgccaagaa acagcgtttg
15241 agtagacaat cttcattgga cagtagaatc cagcatacat atacacagtc cctgcaatgt
15301 tatcgggcaa tcttccttgc catggcttga taccattaat cgtaaagttt gtggatgtaa
15361 ttggaggttg ctcttgtgca cagcatccga atgcaacgtc gccaccgact gatagaattg
15421 cagagtcatc gaccttagca aacctgaacg tgtaagaact tgtttgtggt ggtaaaaagt
15481 aacctgtcat ttctagagta gcgttttttt gggatagtat agaaaccaag taaatcagta
15541 ctccaatagg tctttcctgg ctattttgaa cattcactat gacctctgca tccccagttt
15601 ccataagcat ctgattgagc acatttaaaa gttcctgaag agataacaca agaaagatcg
15661 tagtcaattg agaaatccgt ctgtccgcca acggaaccta attccaattc gaggcatatc
15721 cagaagtcat atattgcggg taggaatacg ttgttgaatt cagtaatgag tacaaaagtt

15781 gacattcatc ccatttttcc tcgggtccgt tggcaagcat gcttgtgtac ttcctgaaac
15841 aacattggac gtggccaggt aggtgaagag tgccaaaaat aaatagtgat gaggcatttc
15901 ttggttcgtc aatcaggtat gggtatatta aacaataact ttaagccatg ctgttaatga
15961 tgaaaattct gatgctggtc tcttcatatg tcattgtttt gggaactgtc ttttttcgat
16021 cagatgtatg cgcaacgatt tgtttagcgt agtgatgttc ccgcgtagta cttaggaaat
16081 tttggtatcg aaacgtccct acaacacttt ttgtcgctga tatctgcaca tgtgagtaac
16141 gcagattcca ttgcattatt accttcccct tcccagaaac gatttattac gactcaacaa
16201 gttcaagaaa aaaattaaaa aggttacaac cgcattttgc gtttaggtgc agcgagactt
16261 accttgatgc agtcctaaag ggtcctgcag cggttgtaac gaatcctatt tgcttttgtg
16321 cattcctaaa cgggctgaag taaaagtctt gaggaactcc aggagaagat acgataagct
16381 ccgaagtgcc aaattttatt atcaaaaaaa aatatttgta cgagcggaaa aggattatac
16441 tcttcctctt ttgcttcatc acaagaggat tattttgacg tcgtctgttg tactacttct
16501 ttatcggtgt gcgcaggaaa agaaagaaaa aggaaggaaa aatagatgtc aaaaatcgat
16561 gtcaaaaaat ctcgtgagac ctccggaatt ttgacgctgc aagtcaatct acgggaaaga
16621 agaaattttt taaacctaat gcaaaataag cttttcttgg aaaataagat tttcggcaat
16681 aaaaggtaaa tgcagccaaa aatcaaaata cttcagaaga agtcgtagcg aggactgcta
16741 aggggaagcg gatttgaaga tcctttccag aacaagaagg agccgaaagc tgtcaggaac
16801 tgttcctgat tttttaggaa aacaattaat aggtatctcg tctagagtag tatctcgagc
16861 ttccagaagt tgcagataat caaaatcatt gttttatccc tttttttaga ttacagctta
16921 gaagagtaga gagcaagttt actgaaacgg ttccttgttt acaataatat tcctaacaaa
16981 ctttacgaat taggatgcag catgattttt tatattgctt cacttcctaa agtatgaatt
17041 tttatccgta gtcgcaaaca aaacagctac tggaaatctg cagcttgtta aaaaccggta
17101 gtttccgaat actcctcgtc cttgagttgt ataccgttaa acttcctagg gtgtcatgtg
17161 tctggcccaa ttggcccaca aaatctggtc ctattgacgg ttttcttttg attttcagca
17221 tcttcctcta agaaggacag aaaattatgt aatatatggg agaaacggcc tcccaactgc
17281 taagtgtccc cggcagcacg agtaagcaaa attcaggcaa actattgcat taagaagccg
17341 tacataattc agcgtgatat gatgaaattt tgttaattgc aaattttagt acgatttggt
17401 tgttagtgtg tgtttatgca agtaattatt gaaccctaag tagttactgt cttcttttgc
17461 tgtaattcgt ggattcacgg ccctccagca acatggattg aaaggttctt taaagttttc
17521 aatccgtaaa gttctgaaat gtattttaat catgtcaata attttactgg tgagtagcat
17581 ttatgaccaa aagcgtactt aaattagcag caaaaaaatt tttaataacg aaactataag
17641 gaaaatacga ggtactgatt atgagagtcc ccgtttctca tttttgagac atgatctgaa
17701 caaggctgaa aacagcaatc tttttcgata acttttgcaa aaatttcaaa cattgttgtt
17761 tgaatgcagc caatttttat agggtacaga gcttaatgct ttacatgtgc tttattttcg
17821 gtactttcct taaagtgtct acattatctc tcaggacttg aatgtcttcg gctgaattac
17881 tataaaatct tgagttttct ctgaagttta atcctaagac aatagtggtg agtgatgtag
17941 ttcacgtgtg tgccactggt aataatagag ataactatct cagttaagtt tgaaaaggta
18001 aaaaatagtt taagtagtca ttttttgcga cggtcattct tctctgatgc acgttcttta
18061 gactacctat aaacaccatt cttacggaat tataatggaa ataaaacatc agtacgtgtt

18121 gctgtcggtg atagaggggt aacagaacct taattgaaaa attagcacag tgcataattt
18181 attaacatga ttgttttctg tggaaaataa gaaattcagc accagtaaaa gacgagaaat
18241 ataggcacat aaatgcgctc ttactcgtat gttccaggat gaaaatgttt agggcatcaa
18301 gtattgccga aagggcaata tgctttaaca ccagaaaatc cactgtatac tcgttacggg
18361 taaacaaagc aaaacgcagt gcgtgataat gtttctaaaa tctctgcaca ctgttgaaat
18421 gcggctctga tactttagcc cttagtacct gacggtgcct aaaatgagga taagtatatg
18481 agtttacaac gaatataatg ctttttacct catcatataa agaataatat taatttaaaa
18541 gtaaaaatta gacctaggtg gtaatgagga atgtgtccta atgaatttat gtagttgatg
18601 cttttgcagc gttttttttt ttttgtatta atcaaagtgc actacgtgac aaaaataaaa
18661 agtggatgaa aacaaaaaaa tcacagattt tcataaaaat aacatggtat tcttaatcgt
18721 atatcgtaat ggattgtcaa aaataaagag aatatagtcg gttcaaacca ctttgtacgt
18781 ctacagagta tagtgtagaa attatagtta ggattgtgac atcccttttt ctttaacata
18841 ataagtttgc ttcaaattaa ttgatcaatt aggataataa ctctcaacac ctttattatt
18901 tatgtttaga gagtttatgg tcagatggtg attggtattg atacttatac tttttttctc
18961 attagtataa tacgctatat tccatgtttg aatagatctt attttttggc tgtctagata
19021 ataaacatac cgtttttaac acacacacac caatgagtag acgttattgc gactgtatgc
19081 cctttaagat attctgttca aagagataag gtatttattc gttttcttgc aatttcttct
19141 catctatcat tatcatacca gattgaatct tattttttaa tttttagtct atttgatgac
19201 ggctactgaa tctttacatt acttgagctg agaagtaaaa ttgcattatt catattaaac
19261 aaaaaaatca atttcccata aagacaatag ctcaacttca tcacgtcctt ggtttttacc
19321 gagtacatat atttgatgaa ttctgatttg aaaatatcaa tttgttcctc tagaatattt
19381 ccactcgcac attcaaaagc aggtattgga acgcacaaga agcgatcgtc accttccatt
19441 ggggttagta gcaggatata tatttccatt ctatattcca ggttctcttt cacaatttta
19501 ttgaatgaaa tacctgagct accaaatcag tcttaataac accatctttt acaccggtac
19561 tacaacaaac catactgagt ctgaaattgg acttctgcaa tgaaaaattt aagccctgat
19621 ctacgcataa tgcccgctat aaaccttatt ttttatatgg gggtctggcg cttcgggaaa
19681 agagaggaaa acttgtaact caatatatct cgatacaaca ttacgttttg taaatttatc
19741 acaaaagcca aatgatgata tctctcttgc aagttatcga acattgattg gtaatttgtt
19801 tgaaaattgt taatttattg aatatttctt ttgcaaaaga aatagtctca gcgaaagctg
19861 gttacaaaat ttacatcatg agtttacggg atttgtaaat acgctttttg cataaaaata
19921 ctttgccgtt tcccaccctt gcatattcac ttactccccc cttcatatac tctatgtaat
19981 gatgattaag ctttggccgc taagtctctc aattagtgtt gattttggtt ttattcatat
20041 gattcttctt tagtgaagta ttgatcaatt acgtgagtca gctttttgaa aaccccattt
20101 ggaaggaatt aggaaattat tttgcttact acgaccacta atttaccgcc atttctgggc
20161 ctttttattg actattttga ccatgtgctc gactagaaga acggcatcat aatctgctgg
20221 tagagttagt ctataatgat tgttgaaaat aaaggcataa gagatattcc acctaaaatt
20281 caagttattg actttattat caggatctta gtatcctttt ttggtaagtc atattcaatg
20341 aactaggtct cgcaaacttt ttgttcgaaa agcggtagtg catagttatg ctaactctgg
20401 atatatggca taaaccgtac aacactagcc cattttttg gaagtagtga gggcagctag

20461 actgtatgat gaatattcgc ctgcatactg agttttttgg tccttttttt tatgtggctg
20521 gccttacgat atgatgcaca attcataatt tggaagaagg gcagaacaat tgcatctgtg
20581 cttggctctc aagaacggtg tttggtgcat caaaagtttt cgactgctta tttggtcgga
20641 aatataaaaa ctcgatcctc ttatctaagc agtatacatt cttcttttg aaatgaatgt
20701 actccgtaat atcttcttat ttggcatttt catccttaac ttttgcatgg ctctgaacta
20761 gtcagatagt tgccctttc agcaaacctc ttattattga aagcatggtg tacatccgtt
20821 atactattat attataagaa attgggatgc caattttttt gcttttgttt tgcctgtttt
20881 ccttcttttc gcaaaagtaa ttgcagattt aatagcagga tattataccg ttggtaaaac
20941 ttaaggattt tatgaacaat agcttcaagt acagcattca tagaaccaac tactaaggat
21001 gaaactagta tgtttttgtc aaaatatttt cttgaccttg ctgtaacatc aagatctgtt
21061 tctctaagat attaaagttg agtaaaaaca aagctgatat gagaaaaata cgtaattgct
21121 ccacataata cgtgggtcag acataaaggt agaatacttg atacagaaga gattattcgg
21181 tactcttgat ggcgtgcttg aactggtgcc tcttaacaac cggtaatata gtcagatgag
21241 tcactacgag tgtgtgtagt agcaagtgtt ttacctacgt ggcagtaaga gtagctctat
21301 ggttgtgtaa tagtggtgct tattcctaat gctctgaagt ctgaagcggt acagttggtc
21361 tggtctatat catggtcaaa ggagcaaaca tatcttctga agtgaccgca aatagtacta
21421 tgatgtggtt ggcaatataa cttaaaagga aataaccaca aggaattgca cccatgtaca
21481 cagtttttcc cggaaattgg gaaaccagta cgaacatata tcaattactt ttacattatc
21541 ttgattggaa aactggcaca attgactgtg acgttagtag tatgttgttc ttctctacat
21601 cagttaaata tacgtgtcaa catcaactaa tcaactttcc cccgtttacc acattgaagc
21661 tgggtgtgga agatttattt gaagaaacta aaacgtaccc tgtcatttcc tgagtccccc
21721 ttttcaactt agtgtgaaag ccgaacaatt ataatcctcg gtagacaaca gatttattgt
21781 actaaagtta ctcttcctgt tatcttcctt gattttactg ttatagcaat gacccaccgc
21841 aatcaggaga gccgccgtat ggaatagcat accaagtcat aaaatcgtca acctattaac
21901 ggggttcagg ttctttttca gcgtagtagc cctttaacaa gcgctgacaa agttgacact
21961 cagagaaaat tcaggattta ttgtaatcca gctactcatc cttagatccg cttgcaggca
22021 tggttttttt caccttgaga ggctattttg ggtaagccag gaaggctgaa aaatcccaaa
22081 aggacacagt aataagaaat tgttgttgtt gtatgatgca tttagaactc aaaagacgag
22141 tttctgaaaa tgcttacaat actccatagg taacatgatt tttttattaa aaaagtatac
22201 tgttcctttg ggtaaaaatt atgcaaccct tgagtgtccg atgaagataa gactacgaaa
22261 caatttgcgg taaatttttt ctgctattga catttacaca tgctccaatc cattaccctt
22321 tccattctcg taataaaacc tcgaactgtt atttcatatt tacatctaga cgggtatcgg
22381 cctcaacaac tccaaacaaa agtaaataga aaagagccag acctatcgca ccgggtagag
22441 ccagaaaata ttttaaacta tagttgacgt attctacggc tgttgtttag gacaatactt
22501 tttccttcac aggcttcgaa ttacgcacat gcagaactcc tgtagaaacc acgaagaaaa
22561 gtttaattaa ctttcaaatg ccagaactaa agattgatga aacagttata cgaatttaag
22621 gggaacttgc tttttccttt tgcttcatca taagcgcaat attcgcagcg gtggcgtcgc
22681 gtcttaccat ttatcggcac atttttccga aaaggaagga aaatggcggt gtcaaacggt
22741 ctggtgtgac atctagaagt tcggcattgc aagttaatct acgggaaaaa gaaaattttc

22801 ttataaccct aatgtataat aaggttttcg gagaaagcaa ctcacgggga acaattggtt
22861 agaagagacc gtagaagcgc cccactaacc tggagttgat tatcgaggat ctttctcaga
22921 aaagcagcta ataaatgagc acagtactca aaaattactc atagtattat aattacttac
22981 ttagtcaata tggctgcgct tttaaaatga gtatctttta ttttttgaca aggaagaaaa
23041 agatgcaagc aaagagaaat ttactaagta caagctcatt tccaaaaatt cagtttgcct
23101 gtaacaatgg gtagcgatct tcaacatatg ggattatgcc ttgaatgaga tttttgaacg
23161 tagaggaaaa aatcgcagac aggctttatt gtgctgtttg acatagtata ctcagccgtt
23221 gtggtgttaa tgaatactat cgatgtgtat acaaacgtac ttcaaataag caatgcgaat
23281 atactgcaac ttttcggcct ttgcaacgat ttgtaatgaa tccttttgca tcgctagaag
23341 gacaagataa tatttcttct gttttttttt tacatatgca acaatttgaa agtcaggtca
23401 aagacagatt ccggttcccc atattcagat tggagagaaa aacttttggc aactcatgtt
23461 accaagtcga gacgcttaaa gttaagtgtc ggccaagaca cgcaaaatct tgtaatcttt
23521 taacgctgct cttcaaatca cggacgcaat cagtacttgt acctaatttt ggttttctaa
23581 tattgaatag cgaaccatag gtcgaaaatt taaggccaca taaatccaga gcccgcaact
23641 tggataggta tttacttgat ttttagtttg ctttcaatag tgtcgtgaaa ttataaagta
23701 cgccgcatat atatcttgat tagttaaaaa tagcagtgtt caatgatgat ttgatagggt
23761 tcataactgg taccagcgta gtacaattac gattatccat gaacataaaa gtggtttaag
23821 tactatatat cagtgaagct tcaaagtaag caaacgagat accaagatct tgtaggacca
23881 cgatatataa gaagcctagt tccgttgtag catcacatag aagagagcga caatgcctca
23941 gaaattctac agcaacagaa agcacagtga gtcattaact gttggaataa gactcaactg
24001 cgagctatcg actagtactt atattactag tatattatat acggtgtgaa aagacgacat
24061 gaagattgag gaaatggtcc caagatctaa tggacactga aatgcaaaag ttgataatgt
24121 aacatgaaaa ttaaggacag catattaaac agaagaggaa ataatggtat ttttacatag
24181 agatatgaat ttcccttttt ctttgttcct atatctctga taataagtta taatatactt
24241 tgtgcatctt atatcatgcc ttttcaacaa tgaaatatca gcagttatca aagcattcgc
24301 caaattttta ttaataaaac aaatgtcaga aatatggccg gaatctaatc tgaaagttaa
24361 aataattctt acgatattct tatcataaaa ccaatgatta atgtttatga tgagcaaaac
24421 tgtgcgaggg cacatacata cagtgggacc tcatgagatc cagaaaaaaa gcaaaggaaa
24481 gttcgatcga aacattagct cttagatatt acagacgtga ggtatctctt ccttgaagag
24541 aatctggccc tgatcaaaaa ctttattttg atactgttcg aaaagagaaa taaggacaac
24601 gttctaattt atacaaatat aaccagtagg ctttatcgta aaatcctcga acttgcttct
24661 catcacagaa gtataaaaag taaagaggtc aagcaagaaa cgaataacca ccaaattttt
24721 ttcaattgaa aagcgaaacg taatggaaat tcacctgctt tgaatcgttg ttgaattctt
24781 atcatcagtc attattttcc tcttttaaag cttaagtcaa ccgcatctaa gtttaacact
24841 tcgttcccgt tttgaagttt atcaaaattc caaaatacag tagttaggtg ctcgtgacaa
24901 acctttttcc aaagtgtcat tagacttcat ggaactacta cacttctggg gaacgaaaag
24961 ctaccatttg ctcatcattg gataatttct tttttttttt ttgatgcatc caaacttgga
25021 cccctttga atgtcgagta tgaagtatca attttagggc aagtagtttt acaataattt
25081 tggtgcactt ttacgtctct gctgatattt ttaatactag tgcaaaagaa ttcgaaataa

25141 atataaaaga acaaggagga ttaacatact ttaaataact aattatcacc attatttcaa
25201 gtttcaagta ctgcgtatgg catgcacagc agatttctac tcaaaaacat cctaagcgaa
25261 ccacactaga tcttacgtta gtactgctga aaatggcaat aattgttaaa gcaaaaattg
25321 gcgatggttc attaattgct aaacaaaata gggcattttt cctcgttaat gataagtact
25381 catacttctg tttctataag tgttagttgt aacttatgaa atatcgattt aaattcgaag
25441 tgttgtttgc aggatataaa tcaaaaaaaa aaaaaaaaaa aaaaaaaatt aaataaataa
25501 ataaaataaa tacaatgata tcgataacgg tgaaattctt ttcatggatt tttgttgccc
25561 aagaaaataa caataacgtt ttctttatga tacatatatc tacttttca aaaaaggaag
25621 cgcaagaatt atcatttagt tcaatttgaa tatttgaaag tttggaggag aaacagttaa
25681 aaaataattc atgtcagcgt atatttagca aagaaaagat acacagatac gtaaaaagaa
25741 cgcgaatttt attaaataat tgccagcaat aaggacgcaa tgaagacact taaaccacta
25801 ccggccagta agctgttggc actgccagca tacgttgaaa tttctaaaga agctgtacta
25861 gatcctacca tgctacttgc tggtgtgcta cgaggctgtt gcgacatagt actcaacccg
25921 gaacttgtta gactcttggt gttgccagtt tcggatacag aaacaacact actgctgtga
25981 ccaatcacat cggtcgcgga agccgtctgt gtttcagcat gattgaatct tgaaattgaa
26041 gaggtgacta ctgttttcgt ctcagcagct ccagtactgg tagttgtctc agcagctcca
26101 gtattggttg ttgtctcact ggtagcactg ttcattttag agctgacaga ctgttcattc
26161 gtagtctgtg gcctccatgt agaatagacc gtaacaacat cattcacagt agccgtggcc
26221 gtcgaaacaa tggcaggtga agtagtttcg gaacacacac cagatccgca ggaagtaaca
26281 gtaactagcg tagtttgttg cttcgattct gtggtggaaa taggacacca tgttgtgtat
26341 tctgtggtaa cgccattaat agtagcagtg cttgtagata caatggctgg agaagcagtc
26401 ttagagcata cgtcagattc acaagaagaa attgtaacta ctgtggtttg ttttgttgtt
26461 tctgtggttt gctctgttgt ccccttggtt tgctttgttg tctccgtagt ttgctttgtt
26521 atctctgtgg tagaaatagg gcaccatgtg gtatactctg ttgtggcacc gctaacagta
26581 acggtggccg tggaaacaat cgcagaggag atagattcag tgcacacatg agattcgcag
26641 gatgtcacgg taaccaaagt ggtttgttcg ctcgtttttg tagtggtaac aggtggtaat
26701 gaagaagtaa tttcctgact tgttgttgca ctggtaacag gtggtaatga tgaagacgaa
26761 tatgtagact ttggtgattc agaagagata gaagaggaag aagaggcaga actagctgaa
26821 cccgtttcgc tctcagaaga accagaggtg gaactactgg ttggaatgac ggatgattta
26881 gatgattcag agagtataga agcggaggtt gttgtagaag aaatgacagg ggaagaaatg
26941 aatgaagaag aaatgactgg agaagaagtg actagagaag aagtgactga ggaagaaatg
27001 actagagaag aagtgtctga ggaagaaatt actgaggagg aaatcacaga agttccattg
27061 ctaggataga atggggtaat aattggacgc gaagacgtga taaagctggt gatttgtcct
27121 gaagaagatg acaaactgga tgagatggca gtagttggag ttttgacaat aatgacagtt
27181 tcatcagttg gttgaccgtt ggttccagtg atggtggtca tctcagtaga tgtagaggta
27241 aaagtaccag tccatggttc agtggtggtt gtaaccaaac cttcactagt tggggttctg
27301 ataacaatca cagtttcgtc agttggttga ccgttggtac cggtgacggt ggtcatctca
27361 gtagatgtag aggtaaaagt accagtccat ggttcagtgg tggttgtaac caaaccttca
27421 ctagttgggg ttctgataac aatcacagtt tcgtcagttg gttgaccgtt ggtaccggtg

27481 acagtagtca tttctgtgga tgtagatgtg aaagtaccgg tccatggttc ggtcgtagtt
27541 atgatggtgc tagcagttgt tggtgttttg acaacaatga tggtttcgtc agttggtaca
27601 ccgttggtgc cagtgatagt agtcatctca gtagacgtag aggtgaaagt gccagtccat
27661 ggctcagttg tagttatgat ggtgctagca gttgttggtg ttttgacaac aatgatggtt
27721 tcgtcagttg gtacaccgtt ggtgccagtg atagtagtca tttctgtgga tgtagatgtg
27781 aaagtaccgg tccatggttc ggtcgtagtt atgatggtgc tagcagttgt tggtgttttg
27841 acaacaatga tggtttcgtc agttggtaca ccgttggtgc cagtgatagt agtcatctca
27901 gtagacgtag aggtgaaagt gccagtccat ggctcagttg tagttatgat ggtgttagca
27961 gttgtaggtg ttttaacaac aatgacagtt tcatcagttg gttgaccgtt ggtaccagtg
28021 acagtagtca tttctgtgga tgtagatgtg aaagtaccgg tccatggttc ggtcgtagtt
28081 atgatggtgc tagcagttgt tggtgttttg acaacaatga tggtttcatc agttggcaag
28141 ccattggtac cggtgactgt ggtcaattcg gtagaagtag aagtggaagt gccagtccat
28201 ggctcagtta tagttatgat ggtgttagca gttgttggtg ttctgacaac aatgatggtt
28261 tcatcagttg gcaagccatt ggtaccggtg actgtggtca attcggtaga agtagaagtg
28321 gaagtgccag tccatggctc agttgtagtt atgatggtgc tagcagttgt tggtgttctg
28381 acaacaatga tggtttcatc agttggcaag ccattggtac cggtgactgt ggtcaattcg
28441 gtagaagtag aagtggaagt gccagtccat ggctcagttg tagttatgat ggtgctagca
28501 gttgttggtg ttctgacaac aatgatggtt tcgtcagttg gcaagccatt agtaccggtg
28561 actgtggtca attcggtaga agtagaagtg gaagtgccag tccatggctc agttgtagtt
28621 atgatggtgc tagcagttgt tggtgttttg acaacaatga tggtttcgtc agttggtaca
28681 ccgttggtgc cagtgatagt agtcatttct gtggatgtag atgtgaaagt accggtccat
28741 ggttcggtcg tagttatgat ggtgctagca gttgttggtg ttttgacaac aatgatggtt
28801 tcgtcagttg gtacaccgtt ggtgccagtg atagtagtca tctcagtaga cgtagaggtg
28861 aaagtaccgg tccatggctc ggttgtagtt atggtagtac tgacagtata atttgaaggg
28921 tctggaatgg tacagtttgg ctggcttaga ttgttgtcaa aagtatatac gtacccttca
28981 aagtcatcac taacggtagt gccatctggt agtgtcacac taattggaag tgtaccccag
29041 gcaacggcat ttgagtaaac aatcttcatt ggataataga aaccagcata catgtagaca
29101 gtccctgtaa tattatcagg gggacttcca ttccatggct tgataccatt gatggtgaag
29161 ttagtcgacg tgatgggagg ttgttcttgt gcacaacatt cgaacgcaat gctaccaccg
29221 actgatagaa ttgcagagtc gtcaactgta gcaaacttga atgtgtaaga acccgtctgt
29281 ggtggtaaaa aataacctgt catttctagg gttacgtttg ttggggtagt atagaaacca
29341 aataaatcag tactccagta tgcaattatt ggattattag aacaagcacc aattcctttg
29401 catccccaat taccatataa atcttcttga ggacaaggaa atgtgcctga tgaactaaca
29461 caaggaatat tataatcaat cgagatatca gtttgtcctc cgacagaacc cagtttagtt
29521 tttgaggcat atccataagc catatatgct gcattcgaat atgtggagga atctctcaat
29581 gaatactggt aaaagtttac attcatacca ttcttccttg agtttgctgg caggcatgcc
29641 gctgtagtcg cagagacaac attagttaat cccagcaatg tgacgatggc tagtagtaaa
29701 caataatgtg ccagagacat ttttggggct tttattgtac aattgttctt tttaaattgc
29761 aatttaaaga gcgtacctgt aaataagaag gaagaacgtt atgttattaa tggactttta

29821 gtgtcatcga attttatgta atatataaga aggtagaata atttggcagg ataatgtgtt
29881 agcaaaggag gaaatcgaat acctttaaaa gagaaaaaat tttttagctg cttaaatttc
29941 tgtgttatac cacccgatag attttgagtt atgctttcta attgatctga ctgcgaacgt
30001 tttctttatg ccatctgaat tgtcaggaac aaagaagaaa aagaaaagtt tttaaaaaat
30061 ctgtggtcgt gtgtgatgta cctttccttt acatgcatta atgcgctctg aaatgtggta
30121 cgatatcctt acagagaata tattttctgt atatcgtgca atgttgaata acctatgaag
30181 gaaagtaccc atcgctcaag gtaagcattc caggagggtc gccagaaact taaactagtt
30241 ttagcgacag atccgaaaat tgatagagac attgaaaaaa tcactactcc gtccttttta
30301 gtgctttctc aatgcataat tttggtgcac gactaaaaaa ttctagaaca ctatagttgc
30361 attttttggg ccggaagaag aaaaacgcat gtaactttaa tgtcaaataa agttttcacc
30421 tagtaagcgc gatacaaaaa aaacacagaa atagccatag gaaagtgaat tttgtcagcc
30481 gactaaaatt aaggttagct tacaaagcag caaaaaattt gacatcgcac ggtattccct
30541 gaaaaaggag caggcaggtg ctgtatattt ttttcggttc ctgcctctta catggcgtcg
30601 gtgtatctta aatactaaag tgagctgact accctttga gtgccctatg tgacctctga
30661 tctcgaaagt aaacaagaga tacctaattt cacagccact ttttgttgcg gacactgacg
30721 ggatgtgttg tgaatatttt aaaccttaaa agtatttatt ggttagttat acttaattct
30781 tatacgtcct ttaaaaccag tgtgcagtaa gtctgtcaca caaaaatatg attgatgcat
30841 tttcaaaaaa taatggcttt gaggtaattc gttccacttt tgagacttat tcacatgtcc
30901 cagctgggtg caatcaaaat ataccggcat ttcagcaagg taactatata cgcttttttt
30961 ctcatggttt ttgaaatcat acctgccaag cttgctgcac ttggaaggca actccgtttt
31021 tcagagatta gatccttttt tttttgaaga aaaggcagcc aagttacgtc atagagaaaa
31081 ctccctagag cgctgcaaac acttgttagt ttgcagctta cttcacgtgt ggctggaaaa
31141 ttgaaaagtt acaatacggt aaaaatactt tgaagacact acactataac tgcttgacca
31201 aatttttact cgtgaaaaat ttgttccagc cggtctgtta ctgatattaa gttaaggaat
31261 agtaaaccgt atatttttca tccacgtttt tgtagtttat tttcggttta ctagctatgc
31321 attgattgtc tatcagagca tatcaaggtg gttatggagg tgctgatatt taatcaataa
31381 aattatagaa attttgagaa aacaataatg attgtttcta caatgaaaat cagttcatgg
31441 aatagttgcc ttcgcactac tttttcacca gattaatggt gcatcagagc tttccctttt
31501 tgtctttttc ttgtttcctt tttcgattgg tttatcacgt aagtttctta tgacataaga
31561 agaggcaaga aacgaaaagg gaaatattgc ctactttttt cttttcgaac gtaaataatg
31621 acgggatttt ggtgtcattt tttttcgtct ggattttctc agagaagtac gtttcgtcgc
31681 acaagctaaa atcattgaca gggcatgagt tacgtcaatc tctggtcaat ccatgtccaa
31741 aaaaatttct tgatttgttc aaagtttttg cataggcaca ttaattggtt tagcgaagta
31801 tacattctga agaacatttt tgggtgtatt ttccacatag aaaattcgat tttttttttt
31861 caatgcacca cttgtcgagt atacgttaac tttaatgtat tgaagatgca aaaatgaaaa
31921 gcctacttgg catatggaat tacaataaat ggtcaattcc tgtttaagct agcagcggca
31981 ctgtcggtgt aaatttgggt ttaaactatt gtttacttac tttgagaaat ttatgtatcg
32041 cctatatttt tcaataagtg cgatatcttt agctttccac agaacccccc ctcaggctat
32101 ggacgtgcgg ttatatgatc ttctaataaa aatctcttag actacggttc atggaatact

32161 tcttgacttc ttggcgaagg aaaggcgcat aaatgttgtg atcgaatgac cagtaatacg
32221 tcgtttgttc tcttgctgaa cagcgaggaa gagatattca tccaaatgca tgaacgcaga
32281 atcctgtaaa agtcgtaaaa atgttaatcg caatgttgta aaaaccgtca aggcatttat
32341 cgttcaaaat ggcgatcatt gttactacta aacacttact gttaaattaa aaagctgatg
32401 ttgcgtaatc cattgaccaa ttcataggca atgtatttca atgacgcaca agattcataa
32461 caaatttttt atttacgatg tacctgtaca ttgtgcagaa ggtcttcaga gtgagtttaa
32521 gctaggctgt aaatatttta atgttaagat gaaatttaag tgagctggta atatcaagtg
32581 aggcataatt tgttatatgt agctgaactt caactttaaa tagaaaaatt acaactaaca
32641 agcaccggat tgtttcagaa ttcaaagtgt agaagctatt attcttgcaa aataaaacgc
32701 tttcaaagtt ttcttctata aacatacttg tggcagcttg tttttttgt tttattttta
32761 agttttgttg ggtctctgag aactttcaaa aaaagaaaaa gtaaagtatg ataaaacgga
32821 gcacttgcca aagtaattaa cgcccattaa aaagaaggca taggaggcat atacatatat
32881 atatatatat atatatatgg ctgctgacag atattctgca cttaaaaact aaaaatatta
32941 taccaacttt tctttttctt cccgttcagt ttgcttgatt ggcccagctc tttgaagaaa
33001 ggaaaaaatg cggagaggga gccaatgaga ttttaaaggg tatattactt atcttatcga
33061 taagcagtat tgatattaaa gggacagttt tatcgttggt taatatggaa aaagtgatga
33121 ccatgatgcc tttcttaaaa agagtatttc tttttatttc actttcacat aaacagttaa
33181 tgacttctga ctttgagccg ttcgaactca gttatataaa ggtacataca taggccacac
33241 acacacacac acacacatat atatatatat atagggaagt agcaacagtc accgaaaaga
33301 aaaggtaaaa agtaaaaaat gacaagcgaa ccagagtttc agcaggctta cgatgagatc
33361 gtttcttctg tggaggattc caaaattttt gaaaaattcc cacagtataa aaaagtgtta
33421 cctattgttt ctgtcccgga gaggatcatt caattcaggg tcacgtggga aaatgataat
33481 ggcgagcaag aagtggctca aggatacagg gtgcagttca attcagccaa gggcccttac
33541 aagggtggcc tacgcttcca cccatcagtg aacctgtcta tcctaaaatt tttgggtttt
33601 gaacagatct tcaagaatgc gctcactggg ctagatatgg gcggtggtaa gggtggcctg
33661 tgtgtggact tgaaaggcaa gtctgacaac gagatcagaa ggatttgtta tgcgttcatg
33721 agagaactga gcaggcatat tggtaaggac acagacgtgc ccgcaggaga tattggtgtc
33781 ggtggccgtg aaattggcta cctattcggc gcttacagat catacaagaa ctcctgggaa
33841 ggtgtgttga ctggtaaggg tttaaactgg ggtggctcac ttatcaggcc ggaggccacc
33901 gggttcggct tagtttacta tacgcaagca atgatcgatt atgcaacaaa cggcaaggag
33961 tcgtttgagg gcaaacgtgt gacaatctcc ggaagtggca atgttgcgca atatgcagct
34021 ttgaaagtga tcgagctggg tggtattgtg gtgtctttat ccgattcgaa ggggtgcatc
34081 atctctgaga cgggcattac ttctgagcaa attcacgata tcgcttccgc caagatccgt
34141 ttcaagtcgt tagaggaaat cgttgatgaa tactctactt tcagcgaaag taagatgaag
34201 tacgttgcag gagcacgccc atggacgcat gtgagcaacg tcgacattgc cttgccctgt
34261 gccacccaaa acgaggtcag tggtgacgaa gccaaggccc tagtggcatc tggcgttaag
34321 ttcgttgccg aaggtgctaa catgggttct acacccgagg ctatttctgt tttcgaaaca
34381 gcgcgtagca ctgcaaccaa tgcaaaggat gcagtttggt ttgggccacc aaaggcagct
34441 aacctgggcg gcgtggcagt atccggtctg gaaatggctc agaattctca aaaagtaact

34501 tggactgccg agcgggtcga tcaagaacta aagaagataa tgatcaactg cttcaacgac
34561 tgcatacagg ccgcacaaga gtactctacg gaaaaaaata caaacacctt gccatcattg
34621 gtcaaggggg ccaacattgc cagcttcgtc atggtggctg acgcaatgct tgaccaggga
34681 gacgtttttt agccgtaagc gctattttct ttttgttcgt aactatctgt gtatgtagta
34741 gtgtaatcta cttttaattt actatgcaaa tagggttcag cattacggaa gaaactgaaa
34801 tcccttccgc ggaagtttct tagtagtggc cgtgcggggt gaggagatta catgtcggta
34861 attagatgat taacctaggc aatttgaagg gggatagtgg cattggttag ctcagatatg
34921 ataaggagaa ctaagcaagg gggttaacca ccacggctgt agcacaagac cggcagatgc
34981 gattattagc aacacattag ttaatgcttt tgataaaatg tatataaagg ctgtcgtaat
35041 gtgcagtagt aaggacctga ctgtgtttgt ggttctcttc attcttgaac cttgtcattg
35101 gtaaaagacc atcgtcaaga tatttgaaag ttaatagaca gttaacaata ataacaacag
35161 caataagaat aacaataaat tcattgaaca tatttcagaa tgagagcctt agcgtatttc
35221 ggtaaaggta acatcagatt caccaaccat ttaaaggagc cacatattgt ggcgcccgat
35281 gagcttgtga ttgatatcga atggtgtggt atttgcggta cggacctgca tgagtacaca
35341 gatggtccta tctttttccc agaagatgga cacacacatg agattagtca taacccattg
35401 ccacaggcga tgggccacga aatggctggt accgttttgg aggtgggccc tggtgtgaaa
35461 aacttgaaag tgggagacaa ggtagttgtc gagcccacag gtacatgcag agaccggtat
35521 cgttggcccc tgtcgccaaa cgttgacaag gaatggtgcg ctgcttgcaa aaagggctac
35581 tataacattt gttcatattt ggggctttgt ggtgcgggtg tgcagagcgg tggatttgca
35641 gaacgtgttg tgatgaacga atctcactgc tacaaagtac cggacttcgt gcccttagac
35701 gttgcagctt tgattcaacc gttggctgtg tgctggcatg caattagagt ctgcgagttc
35761 aaagcaggct ctacggcttt gatcattggt gctggcccca tcggactggg cacgatactg
35821 gcgttgaacg ctgcaggttg caaggacatc gtcgtttcag agcctgccaa ggtaagaaga
35881 gaactggctg aaaaaatggg tgccagggtt tacgacccaa ctgcgcacgc tgccaaggag
35941 agcattgatt atctgaggtc gattgctgat ggtggagacg gcttcgatta cacatttgat
36001 tgctccgggt tggaagtcac attgaatgct gctattcagt gtctcacttt cagaggcacc
36061 gcagtgaact tggccatgtg gggccatcac aagatacagt tttctccgat ggacatcaca
36121 ttgcatgaaa gaaagtacac agggtccatg tgctacacac accacgattt tgaggcagta
36181 atagaagctt tggaagaagg caggattgac attgatagag caagacatat gataacgggc
36241 agagtcaaca ttgaggacgg ccttgatggc gccatcatga agctgataaa cgagaaggag
36301 tctacaatca agattattct gactccaaac aatcacggag agttgaacag ggaagccgat
36361 aatgagaaga aagaaatttc cgagctgagc agtcggaaag atcaagaaag actacgagaa
36421 tcaataaacg aggctaaact gcgtcacaca tgattgtgat tgagtactca cgttctcgtg
36481 ttaatcccgc ggtcttcttg ttttactaac ttttctttct ctcatagcat tctcttgaca
36541 gtgttttata tacatcatat gtacatttat cgagccaatc gagggcagca gtttaacatc
36601 aagccggatt tgctcacgct actttgaccc cttttcgttt cgacggagag aagaaaccgg
36661 tgttttccta tccttgccta ttctttcctc cttacggggt cctagcctgt ttctcttgat
36721 atgataatag gtggaaacgt agaaaaaaaa atcgacatat aaaagtgggg cagatacttc
36781 gtgtgacaat ggccaattca agccctttgg gcagatgttg cccttcttct ttcttaaaaa

36841 gtcttagtac gattgaccaa gtcagaaaaa aaaaaaaaaa ggaactaaaa aaagttttaa
36901 ttaattatga gagctttggc atatttcaag aagggtgata ttcacttcac taatgatatc
36961 cctaggccag aaatccaaac cgacgatgag gttattatcg acgtctcttg gtgtgggatt
37021 tgtggctcgg atcttcacga gtacttggat ggtccaatct tcatgcctaa agatggagag
37081 tgccataaat tatccaacgc tgctttacct ctggcaatgg gccatgagat gtcaggaatt
37141 gtttccaagg ttggtcctaa agtgacaaag gtgaaggttg gcgaccacgt ggtcgttgat
37201 gctgccagca gttgtgcgga cctgcattgc tggccacact ccaaatttta caattccaaa
37261 ccatgtgatg cttgtcagag gggcagtgaa aatctatgta cccacgccgg ttttgtagga
37321 ctaggtgtga tcagtggtgg ctttgctgaa caagtcgtag tctctcaaca tcacattatc
37381 ccggttccaa aggaaattcc tctagatgtg gctgctttag ttgagcctct ttctgtcacc
37441 tggcatgctg ttaagatttc tggtttcaaa aaaggcagtt cagccttggt tcttggtgca
37501 ggtcccattg ggttgtgtac cattttggta cttaagggaa tgggggctag taaaattgta
37561 gtgtctgaaa ttgcagagag aagaatagaa atggccaaga aactgggcgt tgaggtgttc
37621 aatccctcca agcacggtca taaatctata gagatactac gtggtttgac caagagccat
37681 gatgggtttg attacagtta tgattgttct ggtattcaag ttactttcga aacctctttg
37741 aaggcattaa cattcaaggg gacagccacc aacattgcag tttggggtcc aaaacctgtc
37801 ccattccaac caatggatgt gactctccaa gagaaagtta tgactggttc gatcggctat
37861 gttgtcgaag acttcgaaga agttgttcgt gccatccaca acggagacat cgccatggaa
37921 gattgtaagc aactaatcac tggtaagcaa aggattgagg acggttggga aaagggattc
37981 caagagttga tggatcacaa ggaatccaac gttaagattc tattgacgcc taacaatcac
38041 ggtgaaatga agtaatgaca aaataatatt tggggcccct cgcggctcat ttgtagtatc
38101 taagattatg tattttcttt tataatattt gttgttatga aacagacaga agtaagtttc
38161 tgcgactata ttattttttt ttttcttctt tttttttcct ttattcaact tggcgatgag
38221 ctgaaaattt ttttggttaa ggacccttta gaagtattga atgtgggaac aaagacgaca
38281 aaaggtagtt ttttccttga ctatactggt aagatatcgt ctaaaacaaa gcatggccaa
38341 gaaaatatca aagaattcaa gagctgctag acaatcggat gctcttgaac cagaggtaaa
38401 ggatttaagt gaactaccta gagctgaaaa aaccgatttg actaatattt tgattagaac
38461 agcagccaag aatgaggcat tgctggaagc aaagatatct aagaaagcca ataaaagtaa
38521 gaggggcaag aagttaaata aaaaggctct ggaagacaaa ctggccaact ctatttcatc
38581 catggacagg gatcgtttag tgaaggcctt gaattttacc aatcgtctgg acggtaaaat
38641 tgccaagtcc atttctcgtg ccaagtacat tcaaaataca agaaaggctg gctgggatag
38701 caccaatgag actataaaaa aagagctggc tttttgaac ggagggttgt ctgtgcaggc
38761 aaaaagtgct agtgaaggta atgctgaaaa ggaagatgag gagatcccag aagtttttga
38821 ctctttagca gaggataaca cagtgcagaa gactcctaca aatagattcg gtgtcctgcc
38881 agacgatgtt gaagaataga aaattttcat atgaaaggtc ctaggaatac acgattcttg
38941 tacgcattct tcttttttct atcttctttc attctttgta cattagataa catggtttta
39001 gcttagtttt attttatttt ttatatatct ggatgtatac tattattgaa aaacttcatt
39061 aatagttaca actttttcaa tatcaagttg attaagaaaa agaaaattat tatgggttag
39121 ctgaaaaccg tgtgatgcat gtcgtttaag gattgtgtaa aaaagtgaac ggcaacgcat

39181 ttctaatata gataacggcc acacaaagta gtactatgaa attttctgcg tatttatggt
39241 ggctgttttt gaatctagcg ttggtgaaag gcacttcatt gctatccaac gttacattag
39301 cggaagattc tttctgggag cattttcagg cttacactaa tacaaagcat ttaaaccaag
39361 agtggatcac aagtgaagcc gtcaacaatg aaggctctaa aatatatggt gcacaatggc
39421 gactatcaca gggtcgattg caaggatccg catgggataa aggaatcgca gttcgaacag
39481 gcaatgccgc agctatgata ggacatctct tggagacacc tattaatgtt tcagaaacgg
39541 ataccttggt tgtccagtac gaaattaagt tggacaattc tttgacgtgc ggcggtgcgt
39601 ttattaagtt aatgtctggt ttcatgaatg ttgaagcatt aaaacactat gcacccgata
39661 cagagggtgt cgagttagtt tttggtccgg attattgtgc tcctgaaata aatggcgtgc
39721 aatttgccat caataaggtt gacaagatca cacatgaatc taaactaaga tatttgcaag
39781 agatgcccct gtcaaaatta actgatacct cgcaatctca tctgtatacg ctcataatag
39841 atgaatcagc gcagtctttt caaattctta tagacggtaa gacggttatg gtaagagaac
39901 atatcgaaga caagaaaaag gtcaattttg agccacccat tacaccgcct ttaatgattc
39961 ctgatgtttc agtagcgaaa ccgcatgatt gggatgatcg catccgaatc ccagatcctg
40021 aggcggtgaa gctcagtgat cgggatgaac gagacccatt gatgattcca catccagatg
40081 gcactgaacc accagaatgg aacagctcca tccccgaata cattcttgac ccaaatgctc
40141 aaaagccctc gtggtggaag gaacttgagc acggggaatg gataccgccc atgattaaaa
40201 atcctctttg cactgcagaa cgtggttgtg gccagcagat accagggctg ataaataatg
40261 ccaagtacaa aggtccaggc gaactcaatg aaatcataaa tcccaattac atgggggaat
40321 ggcatccacc ggaaattgaa aacccgctat actacgaaga gcagcaccca ttgcgcatcg
40381 aaaacgttat cagtggtgtg atcctcgagt tttggagtgg atctccaaac atgttgataa
40441 gcaacattta tgttggtaaa aatgtaacag aggcgcaaat tattgggaat aagacttggc
40501 tgatgagaga ccgcgcgttt agaggctccg atggccccac agaacgcaaa ttcatgaata
40561 gcagactagg aaatctacaa acaactttcc ataacgaaag agaatcccct aatccatttg
40621 accgcattat agatcgcata ttagagcaac ctctgaaatt tgtgcttact gcggccgtcg
40681 tgctcttgac gacgtcggtt ctttgttgtg tagtatttac atagtggaca agtgttagtt
40741 tataacatgg tctcaataat tgcaccacaa cggcttctct tttatagatg gttaacatta
40801 tagtatcaat attatcatca tgattaaatg atgatgtata atacttaccc gatgttaaat
40861 cttatttttt catgcagtaa gtaatcatgc aacaagaaaa acccgtaatt aagcgaacat
40921 agaacaacta gcatccccga taagacggaa tagaatagta aagattgtga ttcattggca
40981 ggtccattgt cgcattacta aatcataggc atggaaattt ccagttcacc atggaacgac
41041 ggtggataca gcccctatga gagaaacaga gtcgctgtat caccattttc atcagcgttg
41101 gaaggcgaag aacgaataga aacctctcga tctttgggtg atcattgctt tgaacctttg
41161 ccatacgtga cgaattatct ttctattttc gcgctttttg gtaaagagat atttggtgac
41221 aagggaaatg tgagctcaag aaatgaatat ttgctaaaaa aatactactc tttgaaaaag
41281 ccatttgtat tgcgacataa tgggcatgcg ttgaagaatc ccgacatgcc actccagagg
41341 aatgacatat tgcaaaccaa tttcatggtt gacaaatttc tgaatcgtac tgtgcggtca
41401 gtgaatttta ataatttcaa gataatatca gatatgcaaa gtaaaagcgg tcgaggaaca
41461 aagtcaggca caaatcagaa tcaaagtgcc gacgctattc aaaatatttg tctaccatct

41521 ataccgtcgg cgttgcctta tttccagtat tataggaagc tattgacagt taataccaaa
41581 gaatgggata ttttaaaact gcacagttta tgggtaccaa agctaaggaa ggattttaaa
41641 gatttttcgt tgtatggtga taaaaactct ttaaagccga tcgatagtca ctatgatgag
41701 gataatacca tgaagaaaaa tttatttttt gaaagatctc caagtcgaca gactctagat
41761 ggtaaagggt gtgcctctaa ggggtatgac atttcttccg gtaatatgat tatcccatcc
41821 ctattttctg aagataagct gccggcttta acttatcatt gttccgtaga attaaatgga
41881 aacatttaca tatttggggg attgatgcca tgctacagct atgaggagga tgcgccgatg
41941 ctgaacgatt tttttgtaga cggaataaag aacttacctc cgcctttact acctcaagtg
42001 attaataatc catcaatggt caataatcct catctttatg tcgcttctat accatcatgc
42061 cggtttagca aacctaaaat ggggggttat ataccgcctc cattgctatg tgttcaagga
42121 tccaaattaa cggaccgaca tattttcttt tatggcggat ttgaaatcag gacagaaacc
42181 cgtggtgatg aaaatgggaa gtatcatctc aagaaaagat tatatgtgaa taacactggt
42241 tacatactcg atattatgtc gttcaagttc actaaaatag atatcatagt acaaccttcc
42301 aaatataatg catatccgac aatgtcatcg aggtttggtc acttacaaat ttctattgat
42361 aatccaaata ggagagctag cgttcattct tcaagcatga acgaaattca taaaatgggg
42421 agtgcttcca tgaaacaagg tagcagcatc acttccgggc ggcttgaaaa agcagcagta
42481 ctttcatcat tacctcataa tactgtgcac acggttataa tatttggtgg ttacagacaa
42541 accggtgatg atcgttacga agcaatgaat gatttgtgga agatagagat acccgtgata
42601 cgtcgcggta aaaaaggtta ttgtaagttt tcagagacag ctaacgcgat actactgacg
42661 ccaagcgaaa aggacaaatc ggattggccc gaagaaagag ccttttctgc cttttctgtt
42721 catgggactt cgttaatgga taggagttct cttgacatga gactattgaa caacttaaaa
42781 aaccattttg ttttaaaacc gtcatatata tcacaggatc gcgttgttag tcctaaaccg
42841 gttttcccca tgatggttca tggcacgcat caagatcttt tcaatagtgg ctctgcggca
42901 caagaatcgc ccaaagctgg tgcctcggcc agcagcgcaa gtgctgcgag ctttgatccc
42961 gatatggacg ataatttgga aaattatata gtcaatccag ggagaaaatc gtcatctatt
43021 ccaatgactg cgatagggag acagagatta attttaagcc aagagaagcc agtaggtaaa
43081 actgttgtat tgcatggtgg gtctaacggt ctcaacgttc ttgatgatat gtggttgatg
43141 gacttagagt gtgagacatg gactccaata gagacatttg caaaggcaga ttcgagcgaa
43201 gacggtgatg aaaaattgga tagtgtgaac gtaggtctcg ttggccacag aatggaaagt
43261 attggacgaa tatgtgtatg tataggtggt atggtacaag aggatgttga ccaattttac
43321 tcggagaatg atgatgagcc tcctcgaaaa cgcaaggtcg atacattacc gttgggtggt
43381 aattttttga acacaattga tttaagcacg cagtgttggg aagaacataa aattactctg
43441 tccaagaagg aagacgatga ggacagacaa gatagcgaaa atgaagatac aaattcaaat
43501 atagtagttg gtgtcggtgg cacttctttg caatgtgaca aaagtattat tttgattggc
43561 ggattgatat ctagacggag caatgtaaaa gaaatatatt tacatggtac cataacgaaa
43621 agtatttttc ctagcgtaaa tcctagtgca taaaaaggca gttttcaatg ctttcacttt
43681 gtaaactttg tttagtagta gaatataata tattcagttt tgttttatag tcacataaca
43741 ctttgtcttt caaagaataa tctccttcgc aataccagcg aaatattttg gcaaaaaatt
43801 aacaattagg ttcatagtcc cctaattcaa ttaatcgaaa aaaaaaaaat aaaatataag

43861 ggaagattgt gctgatgaaa tagacaatga aacaataatg aagaataaag aagaagaaga
43921 tataaaacat gccaccacca tcaagaagta gaataaacaa aacaagaaca ttaggaatag
43981 tgggtacagc tatagcagtg ttggtcacgt cctactatat atatcaaaag gtgacaagtg
44041 caaaggaaga taatggggca cgacctccag agggtgattc agtaaaagag aacaaaaagg
44101 caaggaagag caaatgtatt ataatgagca agtcgataca aggactgccc ataaagtggg
44161 aggagtacgc cgctgatgaa gtggttttgc tggtacctac gagccacact gatggatcaa
44221 tgaaacaagc cattggggat gcctttcgca agacgaaaaa cgaacacaaa atcatatatt
44281 gcgatagcat ggatggatta tggtcatgtg taagacggct aggtaaattt cagtgcatat
44341 tgaactccag ggacttcaca agtagtggtg gtagcgatgc agcagtcgtt cctgaagata
44401 taggcaggtt tgtcaaattt gttgttgata gcgatgtaga ggatgtgctg attgacactt
44461 tatgcaatta atgtagaaaa gagtttcttg taacagtatg taaagaataa ataattataa
44521 gtataaataa aaagagaagg tgaaataata ataagtaagc agctcggtta taagagaaca
44581 aaaacacacg aaaaaaaaaa agtcgtcaat ataaaaagga aagaaatcat cattacaact
44641 tgaccgaatc aattagatgt ctaacaatgc cagggtttga caatgtagaa acgtcgccta
44701 gttggtcact ttctcctgct aggatttttc ttaaaatacg tctcataatt ttgccggatc
44761 ttgtcttggg caagtcatcc actaaaatga tcaattttgg tgcggcaaat ggcccgatgt
44821 cttttctaac agtaaagacc aaatgcttct tgatatcttg taattcatca tctgttgcgg
44881 tggaccaact agatttgttt ttcaacacca caaatgcagc aactgcttga ccagtcaagt
44941 catcgttgaa tccgacaaca gcacactcgg ccacaattgg atcttcgata atagcagcct
45001 caatttcagc ggtagacaga cggtgaccag agacgttcac cacatcgtct acacgaccca
45061 aaatccagat ataaccatcc ttatcctttg cagcaccatc accagtgaaa tagtagccag
45121 ggtaagggtt caaataagtg tctagatacc tatcatgatt tttccaaata gttcttgcaa
45181 atgatggcca tgcagctttg acggcaagga caccctctgc gtggctggtg ttaagttctt
45241 caccagtgtt agggtcaaga acaactgcat caataccgaa gaaggggaat gaggcagaac
45301 ccggtttcat tggtgtaaca ccaccagcca gcggggtgac cagatgcgaa ccagattctg
45361 tttgccagta ggtgtctaca atggggattt catttttacc tattttttca gagtaccact
45421 cccaaacttc agcagcaatt ggctcaccga ccgaacccaa gcaacgcaaa gattttaagg
45481 aatgattttc gatgtaggaa tcaccagctc ttttcaacaa acgcaaagca gttggcgcaa
45541 cataaaattg ggtgactttg tgttcatcaa taatatccca ataacgggag taatttgggt
45601 acgcaggagt cccttcaaag accaaagtgg cacaaccata tagtaaggga ccataaacca
45661 cataagtgtg gcctgtaatc cagccaatgt ctccagctgt gaagaaaacg tcttcttggt
45721 gagtgtcaaa agtgtagcgc atggtcaaca aagctcccag caagtaacct gcggtagaat
45781 gttgaacacc cttgggggca ccagtagaac cagacgtata caacaagaat aatggatcct
45841 cagaatcaac gggtgtgcat ggatagtagg tcttgtattt cttcttttct gttgcccaat
45901 ccaaatctct gggggcatgg aaagcaacag atggattgtt ggtctttcta taaaccaaga
45961 cgtgtctcac gcctggggtc tctcttagcg cgtcatcaac aattctttta gtctcaatga
46021 ctttaccacc tctgttggat tcatctgtag tgatgacaac tttagagtcc ccatcgttga
46081 tacgatctct caaggagttg gaagaaaacc cggcaaagac tacggagtga atggcaccga
46141 tacgggaaat ggccaacaag gttatgattg cttctgggac cataggcatg tacacggcaa

46201 cagtatcgcc cttgcgaacg cccatagagt aagtcagcac ttgtgccact tgacaaactt
46261 cttcaagtag ttccttgtag gtaatggaat agccttggcc aggctcgtca ccttcgaaaa
46321 taatggcttt cttgttagga gtcttcaagg catgtctgtc aacacagttg taacaggcgt
46381 ttaattggcc gttgaggaac catgcattgt tctggaagga gggcctgccc gttttagggt
46441 ctgggatgaa caccttatcg aatggcttag accagtttaa aaattgggta gctttagaac
46501 cgaagaactt agcagggtct tcaatagact ccttgtgcaa gcgctgatag tcctgcaacc
46561 cgtccaagtg tggagaatag tgggtagcaa ttgcgggctg cagtctatct gagatgggcc
46621 gttgtggcac gatcttgacc gaagtcaaat gttcatactc atgttccttc ttctgctgcg
46681 cagtggcggc agactgggac atttttgctt tcaacttgtc aatttcactt gactgttctt
46741 ctagttttga tgattgtacg gcagagggcg acatagcaca gtgggcaatg tctttctagt
46801 agttttgata tgtttggttt tgcttataga tagaaaatat aagaacaaga tataacgtac
46861 taccagataa cctaagggag aaatatgctt agaatagccg cccagtttat atacaaaatg
46921 aagggagaac tatttgccac cgaggaactg taccccaact gcaataccca ttgaataatg
46981 gcatcggagg ctcggcggca attcgtaccc caaccttttt tttttacttt tctttggatc
47041 ttagagataa cagaaaaaaa ggatgacccc aatcatttgc cacggcatgt caacaggtga
47101 gtgccttttg aggggggggg gtcatctcga catccggcga aatggagcag tcacacgtga
47161 acatttttag gggatggaga gtgctacgcc gttcgtccga gatgattatc atatttacac
47221 agccgtacat acacgtgcca tttatcttga tatcattctg gacgtatgtg cacatgtgat
47281 ttgcttttgt tttttaaga atgtcgggta ataaacagat tgtttttctg ggaggataat
47341 cttttctttt ttcctgttgg tattctaaaa ttaaccttgc tgtttctttt tttttttt
47401 ttcgcgcgac tactcagcca tcttgcattt ttaaagaaaa agataatcat taatgccttc
47461 acgggaatac gtatagaaca ttattaaaag tatatgaatg gcatatatat atagaacacc
47521 acccttggaa aacatttata ccccttaaac taaaacaatt tgctgcgcta taccgtgttt
47581 cagtgtatta taatacattc atttctgttt cattacgatt atattgacgt gataaaaaga
47641 ttatatagcc atgatcttcc taaacacctt cgcaaggtgc cttttaacgt gtttcgtact
47701 gtgcagcggt acagcacgtt cctctgacac aaacgacact actccggcgt ctgcaaagca
47761 tttgcagacc acttctttat tgacgtgtat ggacaattcg caattaacgg catcattctt
47821 tgatgtgaaa ttttaccccg ataataatac tgttatcttt gatattgacg ctacgacgac
47881 gcttaatggg aacgtcactg tgaaggctga gctgcttact tacggactga aagtcctgga
47941 taagactttt gatttatgtt ccttgggcca agtatcgctt tgccccctaa gtgctgggcg
48001 tattgatgtc atgtccacac aggtgatcga atcatccatt accaagcaat ttcccggcat
48061 tgcttacacc attccagatt tggacgcaca agtacgtgtg gtggcatacg ctcagaatga
48121 cacggaattc gaaactccgc tggcttgtgt ccaggctatc ttgagtaacg ggaagacagt
48181 gcaaacaaag tatgcggcct ggcccattgc cgctatctca ggtgtcggtg tacttacctc
48241 agggtttgtg tctgtgatcg gttactcagc cactgctgct cacattgcgt ccaactccat
48301 ctcattgttc atatacttcc aaaatctagc tatcactgca atgatgggtg tctcaagggt
48361 tccacccatt gctgccgcgt ggacgcagaa tttccaatgg tccatgggta tcatcaatac
48421 aaacttcatg caaaagattt ttgattggta cgtacaggcc actaatggtg tctcaaatgt
48481 tgtggtagct aacaaggacg tcttgtccat tagtgtgcaa aaacgtgcta tctctatggc

48541 atcgtctagt gattacaatt ttgacaccat tttagacgat tcgaatctgt acaccacttc
48601 tgagaaggat ccaagcaatt actcagccaa gattctcgtg ttaagaggta tagaaagagt
48661 tgcttatttg gctaatattg agctatctaa tttcttttg accggtattg tgttttttct
48721 attcttccta tttgtagttg tcgtctcttt gattttcttt aaggcgctat tggaagttct
48781 tacaagagca agaatattga aagagacttc caatttcttc caatatagga agaactgggg
48841 gagtattatc aaaggcaccc ttttcagatt atctatcatc gccttccctc aagtttctct
48901 tctggcgatt tgggaattta ctcaggtcaa ctctccagcg attgttgttg atgcggtagt
48961 aatattactg atcatcacgg gacttctggt ttatggaact ataagggttt tcatcaaggg
49021 aagagagtct ctcagattat acaagaatcc tgcgtaccta ctttacagtg atacctactt
49081 cttgaacaag tttgggttct tatacgttca attcaaagca gataagtttt ggtggctttt
49141 acccttatta agttatgcgt tcttaagatc cctgtttgtt gccgttttac aaaaccaagg
49201 taaggctcaa gcaatgatca tctttgtcat tgaactagct tacttcgttt gtctctgttg
49261 gataagacca tatttggaca agagaactaa tgttttcaat attgctattc atttggtgaa
49321 tttgatcaat gcatttttct ttttgttttt cagtaatttg ttcaagcaac cagcagtggt
49381 ttcgtcagtg atggcggtta ttctgttcgt tttgaacgcg gtgtttgctc tattcctatt
49441 attgttcact attgtcacct gtacactggc attactacac agaaacccag atgtccgtta
49501 ccaaccaatg aaagatgacc gtgtgtcatt cattcctaag attcaaaatg atttcgatgg
49561 caaaaacaaa aatgattctg aactgtttga attgagaaaa gctgttatgg acaccaatga
49621 aaatgaggaa gaaaaaatgt tccgtgacga cactttcggc aagaacctga atgcaaacac
49681 aaatacagca agactctttg atgatgagac tagttcatcc tcttttaagc aaaattcctc
49741 tcccttcgat gcctcggaag taacggagca acctgtgcaa ccaacctccg ctgtcatggg
49801 tacgggtggc agcttcttgt ctccacagta ccaacgtgcg tcatctgctt ctcgtactaa
49861 tctagcgccg aataatacaa gcacctccag tttaatgaag cctgaatcaa gtctctacct
49921 ggggaattcc aataaatcat attcgcattt taacaacaac ggcagcaacg aaaacgcccg
49981 caacaacaac ccatatttgt aatccaatat atactcacat gtaacaactt attatataaa
50041 tatttaaggg caaggatatc ctacattata tttcatagaa aaccgctcaa aaaggtgtat
50101 tatctccatt acatcccaac accacacata tttcagcgat aaaaacctta aatgtgaaat
50161 tcgctttggc tctgcttcct taaatgtacg caattgccgc ttttttctga catctttttt
50221 gacgtgtaga gaaggaaaca gatcctccag aagggattta ctgttggcta ttttgtgtta
50281 gaagcaggtt aataatagat taggttgcgt aagtcatggt cgaaaatagt acgcagaagg
50341 ccccacatgc cggaaatgat gataatagct ctaccaagcc atattcggag gcgtttttct
50401 tagggttcaa taacccaacg cctggattag aagctgagca ctcaagcaca tcgcctgccc
50461 ccgagaactc cgaaacacat aataggaaaa gaaatagaat attgtttgtc tgccaggctt
50521 gtaggaagtc aaaaacaaag tgtgatagag aaaaacctga atgtggtcga tgcgtcaagc
50581 atgggttaaa atgtgtttat gacgtatcaa aacagccagc accacgaatt ccgagtaaag
50641 acgccattat atcaaggttg gaaaaagata tgttttattg gaaagataaa gctatgaagc
50701 tactaacaga gagagaggtg aatgaatcag gcaagagatc agcaagtccg atcaatacaa
50761 acaatgctag cggggacagt cctgatacca agaagcagca taaaatggaa cctatatatg
50821 aacaaagtgg taacggggat ataaacaatg gtaccagaaa tgatattgaa atcaacttgt

50881 atagaagtca tccaaccatg atcatgagta aagtcatgaa aagagaagtt aagccgttat
50941 ctgaaaatta tattataatt caggactgtt ttctaaaaat cctggtcact tcagtgttcc
51001 ttgacacttc aaagaacacg atgataccgg cattgacggc aaacgcgaat attacaagag
51061 cccagcctag cgtagcaaat aacctttga aattgaagga aatgctaatc agacagtgtc
51121 aaaccgaaga tgaaaaaaat cgtgtaaacg aattcactga tagaatacta caaaatacaa
51181 attcaaatag aaacttgaaa atcggtatgc tattatcaat gctttacaat tctgtcggtt
51241 accaatatct ggaggatcat tgccctcaag gtggcgaata ttcggattta ttgagaaatt
51301 tgatcaatga atgtgaagct attttgccat cttacgaaat tattgaacgc tacaagaacc
51361 acttttatga gtacgtttat ccaagtctac ctttcatcga attagaaatt tttgaagaat
51421 cattaagtca aacaattttt ccggacccaa acaacccctc caaggtgcaa atacgtatgg
51481 gtagcacaca tttgagagct aaggtggaaa acttgagtct tctattggtt atcttgaaac
51541 tctcatacat gtcaataagg tttttagatc atagtacagc agactcgagt ttttatcttt
51601 caaaggaaat aattgataaa tacccaatac cgaacgattt tattttattg agtcaaagat
51661 gtctagcatc ggaaaattgg tgtgcatgcg ctaatgaaaa catcatatca tgtttactat
51721 atatctggtc tttttttgct ttttctcctg aagagggtga tttctttctc gagcacccca
51781 ccgatgttat cagtagtttg ataatgatgc tttccacctc gattggtctc cacagagatc
51841 cttcagattt ccctcaattg atttccccgt ccacctcaga taaaagaacc ttgaatcaca
51901 gaagaatact ctggttgagt atcgttaccg tttgttcgtt tgaagcaagt ctcaaaggta
51961 gacattctgt ctcaccgata tctttaatgg ccttattcct aaatattaag gatcctgatt
52021 ctctgacggt atatatgaac cgagttaggg gcgatctaag cgatatcaat aatcacaaac
52081 ttttgagaat tcataaattt acattcaaga gagcccagct tgcgttactc ctgtcggact
52141 tagataactt gacgatgaca tactatggta gtttccattt gcattcaatt gaattcataa
52201 gagaaaagat tgagattttt gtggaggaaa actttcccat agtaccatta aaaagtgtcg
52261 cacaggataa gtcagacctt gatgacatga atgtgatttc agaaatgaat atattatctt
52321 cagaaaattc ttcttcattt cacaatcgaa taatgaataa actattgatg ttgagaactt
52381 caatggccgt attcttgcat tttgaaacac ttatcactaa ggataaaagt atcttcccat
52441 tctacaagaa atactttatg gttagctgta tggatgcgtt gtcactaata aattatttca
52501 ataagttttt caacggagaa tatcgacacg caatatcttc tttaaccagt tttaatgtta
52561 caaaatttat tcagttagca ctatccagca caatcttcag cctattaggg attatactaa
52621 gaataggttt agccatccat atgttatctt ctgaagtaca aaagttatcg ggaacgacag
52681 atccaagaat aaaggagtta aataccaaag tcgaaaaatt tagtaccctg caaagagatc
52741 tcgagtctgc tttagaaggt atatattgct ctgcttcgga acatttaaga ttcacatact
52801 tccccgtttt taagatgttg gctttattcg atgtcattgt acaaaggatg agaaagggtg
52861 aattatggca cggcatattt acgatgattc aaatggaaca aatgcattct aggataatca
52921 agacattaag cattacctta ggagtcaaac tggacaaaaa ggataggcta ttagaggaat
52981 tgatggcatg caatcacgtt gcgaatttta gcgttgaaga tatagatgag ctgaaccgta
53041 atatcaaaaa agagattcaa atttcttcag gattgaagcc gcctgtaaac acaattgact
53101 taaccaacgg cgaaccattc ggaaatgctg ttcctacctt cacaaagaca tggagttcat
53161 ccttagataa tttagaaaaa ctatcatcgg ccgctgcagt tggtcagagc ttggactaca

53221 acagtggttt acgtcagggt cctttggcgg gtggtggttc aaaagagcaa acgccaatag
53281 ccgggatgaa taacttgaac aattcaatca atgctacacc aattgtcgat aactcatctg
53341 gatcacaact tcctaatggt ttcgatagag gccaagcgaa taatactcct tttccaggtt
53401 attttggagg tttggattta tttgattatg actttttgtt tggcaatgac tttgcttaaa
53461 aattttcttt ccaaactcct acctattcat ttcatcaatt aattaatatt atatagccac
53521 gaatttatga aactgaccga taatataaag tgctcaatat atatatatat gtatataacg
53581 gttaacgtaa gaagagctct tccctcttaa acattcgaaa aatgattgaa ccagtatatt
53641 tggtcgagca agactttctc cttcgcatat tttacggcag gtatggatat atcgcctctt
53701 gctgcaaacc cgtgagccac accactgaag aggtctaact ggtaagtagc gtgattatcc
53761 tttaattttt cctccgttaa gtgtcttaag tttgccggaa agatgtgatc ctcttccgct
53821 gctgaaatca atattggttt cttgctatca attgcttcaa tttcctcgat gctgacgaaa
53881 gatggatgtg caatggctgc agcattggca agaccccgt cgccactaat gtgttggacg
53941 gcaaactttg caccaaaaca gtaacccaca acgccaataa actttgggtc atattcaagt
54001 tttaacaact tcatgaatcc atcaacaatt ttcttggtga cttcaggaga atgtctttga
54061 aaccaggcat cacgatcaat tggtttgtcc gatgagatag catcgccgaa taaaatatcg
54121 ggaacaaaga ccatgtaccc agcactagca aatttgtcgg ccgttaataa aacattgttg
54181 aatttattgc catacacatc tgtcaagata actataactt tttccttggg agatgtagag
54241 cctgctgcat aagtatctaa accgaagatt tcttcacgac gacccttggg tgttccatcg
54301 tgacaaactc cttcaaagca acacttgcca ggttgattag atgccatttg attgaaataa
54361 ttttgttctg ctgtagttag acgtagtgga aaactttaag tctactgagt cttgagacct
54421 tatcaccctt tgaaggtttc ttgcaaatga gcgtggtttg gcattttta tcggaaagaa
54481 aaaaagggct ccgccttagg ccagatatca tagaaatgca acacttccct aatatagaaa
54541 tttgggcatt aattattttg agaattttga tgatttgaat aatttcatta acgtaaagga
54601 acatagtgct acgaatccaa cagtggaccc aaaaatgaga gccgtttgtc tgtagtcgac
54661 atcttttgct gctgtttctt ctggcaatcc cggagtgttt ttgccaggat caagagcagc
54721 ttctgtgatt ttaataaaca attcattaag ggaacttagc catctggatg atatgtgcag
54781 tgggtggttc acaaatagct cgtctgccag ttcatctggt tggatttgac acctttgttg
54841 ctgcttatcc aaatctgcct tagaagctac aaataccaac ggtagatctt gtaaatgtgt
54901 gaatttgtct agaagcgaaa ctaagtagga gaatgattct gggtcgctgg aatcgtatgt
54961 tagacagatt acgtcacatt cttttaactt atccttattc tctagtatgg cgtattcctg
55021 ttctccaagt tcttgcaaaa tcaaatagta ctgtttccca cctttgagtt ctaaactatt
55081 gactgcaatt cttggtttga ttgtcggaga atactcctcc gagaaagatc tgcccaagaa
55141 ggcctctagc aaagagcttt tgccgcaaca tggctttcca atgacaaagc aattgaacac
55201 ttttctgtca ttgatattgg atctgtaaag tttcccggaa cggcgtctca ttttccttgg
55261 cttggttact tgtagggcta gtcttgcatc ttcctgaaag ccaaaataca ccaagtaagc
55321 ggtagttgtg ctatagttca agaaagtcgt catactccat tgtgctagcc agccttgtaa
55381 ggtgatgcaa cccttgttgt ttacgacagt ggagaagggg aaattcgttg aggtccatag
55441 tttaggcagc cctggtgtgc acttaaatag acgatgtaat tcttgattat tcaaaccacc
55501 atcattgtcg atatcaaact tcaaaaaaat atctacaaga aatctgtagc ccttggggct

55561 caattccaca ctggaagtgt cagggacaac caacctcgga tggagaattt tgtcattaat
55621 acacaaggaa tctgtgtaat ggaaagttct taggatagcc catgtagttt cgtgtctccc
55681 cctttcagcg tatattttgt tcagtacaag gaaaccatct ttggtgatgc cttttcccgg
55741 tacgtatagc ttgcggttaa tgtactcttg atcgtgcttg gaaatatcca aaagcaaatc
55801 tttaataaaa ttcagttcgt ttacatcgat actcttattg aagcactttt tttgtaagcc
55861 caagatttcg ttgtcatcta aatatgagtc ctggtttaaa tcgcttaaaa gaaaaattct
55921 ttttaaagcc atgacagcca atggctttag ttcacctacc atggcatcaa ataaaggtga
55981 tattgggtgt gttatagccc tttggcaaag ataaaacgct tggttaagat caaactgtgt
56041 cttggcactt gtcttaatgc aagtgtcgat ttctttaaac tccattaata ttgggataaa
56101 ttcttcatcc tccactttgg tatcgatatc atcatcactg ttctctgaca cgaccattgc
56161 attggcatta acattcgata tggaatcaca tttatttttg cagagaatga caggaatatt
56221 caaccccagg gatctgaaat gaggcaacca aaagagagaa acatggtcat acgattcgtg
56281 atcgcaatac acaagccaaa ttacgtcggc ggacttcaac tcatggtcta aagctatgag
56341 gtccgaatct gaagtgtcta taagtactgt attcttagga gaatatgtag gtgatgatga
56401 gaaatctctt gggatactga tgggtggcag cacgtcctgt atggtcggta tgaattcagc
56461 ttttgttaat gatacaatca gactggattt accaacccct tcatcaccgc aaataactac
56521 ccgaatcgtt tctttagtca ttgtgttgtt caacacatta gtatttagaa gtccgctatt
56581 tttgttttca atttttattt gtttaatata caaatttcat tcttgttttg agggtaaacc
56641 actataccaa atgttgaaga tctaaaggta tcgatcaaat atgttgctag agagtgactg
56701 agtgttacat taaatatatt tatatataaa cgtatgatat ttagggattg ttgattgata
56761 ggttgaaaag tttcgatctc aatgactcat tttcctgttc taaagccttg attcgagctt
56821 ctgaagcgtt agcatcgagc ttccgtcttt ctctttcgga tatccatctt ctttgtaact
56881 cctctattcg taaggtcagt tctttatttg gagttgccgt taattcattg ccctgttgat
56941 gtggctgctc tggggtttcc atggaaatca gtgaagagat atatgagttt attatggact
57001 ctaatgcagt ctctataaaa gtgtagagcg attctagttt gggctgaatc aaattcaagt
57061 ttttcaacgc atttggtaca gattttatgg atttcatttt cctgtcaaat tgggcaatag
57121 aactttcttg taagattttc tgcagaatat tgaaaatact gtcaaggtgt agtaaaaggt
57181 tctgattaaa tttgataaaa ttgctttcat attgggatag tattaacttt tggttgtcca
57241 aagtatttaa ctgatttttt aattggtaat tctccaagtc taatttgtct aactgatttt
57301 gtattacgtg gaattcctct gatttatcga tttgaaggtc attgatttgt ttttccaagt
57361 cattgatgta gctgtcccaa ttattttcct ttagtttatt gattttggtc tgagtttcca
57421 gttctttggt taaaactttt tcattttgtt tcattttaat gatgtcttcc ttcaattttt
57481 ccaaattgtt tatcaagaca gattgcgatt cgatcttttc cttcagttgg gaaatcaaat
57541 gatcttgttt ttctataacg gaactagaat tttcttccaa ttctaaggaa tctagtagat
57601 gagactgctc atttagtttg gacgctatta ttttttctaa tttttgcgat ttctcgaatt
57661 tcaatctaat ggaatttata aattgatcat attctttgtg caaattttct atgacaatct
57721 ctaattgagt gtctaaggtt ttctcaaaac gagactcgac ggctgaaatt ggtaaggaag
57781 tactgtttgc cattacattt tctgtatcaa tgtggttgtt actgtcatga atttcgtcat
57841 tctggtatcc accatcagta ttctcctctt tacttgatgg tgagtcccta ctttctaact

57901 gtgatcctgc tggtgaggac tgggccagtg aaagaaattc ttccttatcc tgctttgatt
57961 ccgctcgact tttggaatgc aaaaagtcct gcaagaattg gatgatgaac ttggacaaag
58021 tatccatttt ttcaagaaca taatccgagc tcagctctaa agtttcctcc agggtctccc
58081 tctcttcttt atcaagaaga tgggcatttt catcttgctc attaaaatgc gtcaatataa
58141 aggacaatag gtgattgatc gtattcacga tactttccga attttctaaa ttctcttgaa
58201 cgaattgtag cgtatcttgg tactcgactt ccttcgcctt taaatcctgt ttcaatttgt
58261 ttatttcaag atttagaccc tcgataatcg aatttctaaa atcagtgtca ttgcccagcg
58321 atggtgcatt gccgtcttta ttagggattc tgcgaatgta ttcatagagt acttgaatct
58381 tgattttggc attagtcaac tccttctcca aatttttgac tttgttggaa tcgttcatca
58441 gagcaggctt gatgggatcg ttatgagatg acctcgtggt catggagtcc ctgagtgatg
58501 gtatggacat cccagaatcc atcgagtttg tgaactcgct gtcgtcatca tcaccagtgt
58561 tgtcattatt gcgaagatgc ctgccactag gaatccatcg acgtaccatg gctataactt
58621 tccttatgtt gtttgcttag ttttttgata ttagtgttgc ttatgtgaaa tttcgcgatt
58681 tcaattaaaa taataaatac atatataaag aatatacaca gagggaagca aaagtaaact
58741 aaaagtgata cttacacgag cttttttggt tccaaactgt tcatgatgat gccggaccct
58801 tcccagttga cttcttagtg gtcaattgta ggccatgcca tctggaaata tcgtccctca
58861 aaattctgtt caccagctgg tgctgcttga tgagactcag tccgttgaac ttcttgcttg
58921 ttatgttgat agcaaacatg gatccgcagc caccggaaac gtcttgcact ttacacactt
58981 caggttccag ttcctgttgt agtttatcgg tgatcatctt ctcctccgga gtcattgcca
59041 tctgcgttga gtaccaaagc tttgagcccg tcagaatcct tggccaccgg acatgcttca
59101 cagatataga acgtagcatg gtctgtggga gcttcatttc tatgttttac cttctctttt
59161 cgcttttatg gttctcagtg accaaataaa gaaacttata tatgttccgg aatgacgaat
59221 caaaaagaga atagcatcgt tagcagcaaa cgaaagtgga aagagaataa tgttcaagag
59281 agcaatgagc acagatggtc ccgtggcacg taccatcctg aagagactgg aatgcggctt
59341 tccagattac aagaactttg cgtttggcct ctacaacgat tctcacaagc ataagggcca
59401 tgctggtgta cagggaaatg tctctgctga gacacatttc cggattgaga tggtcagtaa
59461 aaagttcgaa ggcctgaaac ttccacaacg ccatcgtatg gtttattccc tcttgcaaga
59521 cgagatggct caggcgaacg gtatccatgc tttacaattg tcactaaaga ccccacagga
59581 gtatgaatcc aaagcgaaat agaatgcata agcataagtg tacacgttga gtttattgtt
59641 ttatttcccc tacatatata tacatatata tgaaattact ttacgtacgt ataagctttg
59701 ttcagtcatc atgaaccagt gtcttttcgt actgttctaa ggacattaga ccctcgacct
59761 gttccacatt aacgccctca ccaagcttca ttttgactag ccagccgtca cccataggat
59821 cttcgttcac cacacctgga ttttcctcaa gattagtgtt aatttcctct acggtaccat
59881 cggcaggctg gtagatctcg gaggctgact tgacggactc aatggaccct agcgactcac
59941 cttgggcaat ctcagtgccc acttctggca actcaacata ggtagcgtcc cctaaggcat
60001 cagtggcgta ttttgtaatt ccgacaaagg cagtcttgtc ctgatgcaca gctatccact
60061 catgttggga agtgtacctc acggcttgag gtccttggga tgagtacaaa aatggtagtt
60121 tattcttgtt tagggcattg ccggagctgt ttctcaaaaa caatttgctc acagtgggca
60181 tgcgggtggt ccatagtcta gtagtgcgta acattgtcga tgtggtatgc ttcatgtgga

60241 gattcccttt cccattagat acttgtttgt tggtctgtat atatagaaga aagagttagc
60301 gaaagtgact ccgccgctga atgactcctt acggaagtgt caaaattgcg aggtccctat
60361 agcacagaat gatagataaa acattgattt gcaagttgaa ggaagaccct acacatgcgt
60421 atatatgatg tatgtaatgg ttgtgatcat tttagcctgt caagcagtga atcgcactgc
60481 ttgtgtaagc cttcatcttc ttgctttagc tgatgcagca ggtcgcgaac agtgtcctgc
60541 actgggggtc taaacataat gagaaacttt agcgtttgga aaccaaggga acttcttgcc
60601 ggatccaggc agataggctt cagtgcctca agatgatcgc tttgaagact gggcagttcg
60661 ctcataagtc tgatgaagaa tcgtctgtgt ttgttttcaa ggaatggacc cagagactcg
60721 agaactctta aggaccattt tttgtagtta gagggatctt gatgcagcga ggtttggaag
60781 aaccattctt cattgagcca ttcaatgatc agactgacac gttgctcgaa atcctggatg
60841 aagaagccaa gcagttcttc acgaatcagg tcactggcct cttgtgcttc gattcctctc
60901 gtaaccaatc tgattaagac gtgtaaccat gacgaggagt cgtcatcgtc cagtaatatg
60961 gaggaggaag acgaagattt ggcccgggta gtatcctggc gaccggataa ttcaaacagt
61021 ttcttggtca gctttgagag gtactttagt ttttcatcct gcgtcaatga gtcaggttca
61081 aagggggggtg cgatctcctt aatggcagaa gttgatttgt tggcatctcc tgagatttct
61141 tgggcgcttt cctcttcttg aagcatcttc tgcattcggt catcgtcttc gggctcctct
61201 ggttcctccg ctagtggttc tgtttccatt ttgattttct tgctattagc cgttgggcca
61261 tcgtttccaa cttcttcatt gtcgttgccg tcgtcatcat cgtcggattt cctctttgat
61321 gatgacgagg acggtacaga attgatgtac gtattcatta aatccgtgta cctcgaagca
61381 acgatagaca atccggtgat cagtttcgtg ctgtccattt gcaagatggc ctctgtggac
61441 agcttgataa gtatgtcatt gggtagctgg gtcacatcct ggttggagtt cgaactgttc
61501 atcaatgagt acacagatga gtaggtgttg ctgattggtt tgggtgagtt gttgaaaaaa
61561 gtattgccct ggtgtgaagc cttgttgagg gtgtattttt gcaatatctt cagttgatca
61621 agcatgtttt cggttgaaga gcccgttgca ggtgcggaca caggcgtggg ggtctttgtg
61681 gacaccccta gagtagacaa taacgcggat aattgccttt tccatagtga gatgtatttt
61741 agtttgtcct gcctggacaa cgttttcttg ctattgccct tggaagggtc gaagttcaaa
61801 attcccttgc tcttggtctc ttcgccaata acgtgtaaag tttgagaaat cttggtcagc
61861 ttggagtaga tcgatgaccc tgatccggat gagagggatt ttgtaatgat ttgatttttt
61921 agcccaaatt gcacaaagtt cttgtacgcc ctttcaacaa atctcttgga tagtttgtag
61981 ttcaagtcag acttgccctc taggggaaac ttggcgtcga cgttgaaacg caacagcccg
62041 gaaagaattc ttattgttgt ctgcggcctt cttttgatga cgaaggataa agaattgatg
62101 ataccaatga aaacggacga gaccatgtac tgttcctcaa ttaggtagtt tagcaacata
62161 tcaagaagcc tcttagcctc gctctccaaa gccggtttgt tcaacacagg gtggttatcc
62221 gggatggtag atgaattaat ctcgttgccg ctgggtgatt tagtttgcga cagcacgacc
62281 tcagatatga acttgatggt cgctaatttc acgccgatat tttggtcaat ctgcgccagc
62341 cattgttcga catccgtttc atcgtcaacg gtggcacgca aaggatatgc agttctccag
62401 tgcgagagca cgaacttctt cagcatacac aactgatcaa acatttcctg gtttgatgtc
62461 ttagcaacca gatccaacac cagcgggtat gaagcgcaca taataagcac gatattcttg
62521 tacactagta cgtccgcggt ggattgcgcc atagcaagaa gtagtggcag atattgagca

62581 gcaataaacg gtctctcagt attcgcaatt ggagagtcca tcgacaccac gtctagaact
62641 aactgtgtaa aaaacttggc caaaggcaac ttcagcttgc tgagattacc gttgtggtac
62701 atggatgccg tagtttcgag caccttgggc agcatctccg ttggattgtt gtgcatggcc
62761 agtgtcttgg cctgtaacaa ttgttccatc tctgcagatg acattgcgct gcttagtggt
62821 agttatatgc ttcttgccac gatttaacca tttgttcagt caagtactaa cggttaaaag
62881 gtatcgaaat atggcaactt ttcactttta gatcaagtca ctatatacga cttgaacatc
62941 agaacggcga ttttccatca agatggagtg gaaaccacgc cattataaag gaaagctagt
63001 tttatgtctc gtatacatgc ggagtaggac agtgatataa cacacatagc tagacacaat
63061 agacatcatg aaaaggtcca cgttgctgtc gctggacgca ttcgctaaga ccgaagagga
63121 cgtacgagtc cgcaccaggg ccggcgggct gatcacttta tcgtgcatct tgaccacgtt
63181 atttctgctg gtgaacgagt ggggacagtt caattctgtg gtaacaaggc cacaattggt
63241 ggtggaccgt gaccgacacg caaagctgga gcttaatatg gatgtgacat ttccatcgat
63301 gccatgtgac ctggtgaatc tcgatattat ggacgactct ggagagatgc aactagacat
63361 tcttgacgca gggttcacga tgtctaggtt gaatagcgag ggtcgccccg tgggagatgc
63421 tactgagttg catgtgggtg ggaacggcga cggaaccgcg ccggttaata acgatcctaa
63481 ctattgtggg ccatgttacg gtgccaaaga tcagtcgcag aatgagaatc tagcacagga
63541 agagaaggtt tgctgccaag actgtgatgc agtgagatca gcatacttgg aggcaggctg
63601 ggcttttttc gacgggaaga atatcgagca gtgtgaaaga gagggctatg tcagcaagat
63661 taacgagcac ttgaatgaag gctgcaggat caaaggttct gcacaaatta acagaattca
63721 ggggaatctt cactttgccc ctggaaaacc ctaccagaat gcatatggac attttcatga
63781 tacttctttg tacgacaaga cttcgaattt gaacttcaac cacatcatca atcatttgag
63841 ctttgggaag ccgatccagt cccacagtaa gttgttagga aacgataagc gccacggcgg
63901 cgccgtagtt gccacttctc ccttggacgg acgccaggtg ttcccggaca ggaacacaca
63961 ctttcaccag ttctcgtatt ttgccaagat tgtccccacc agatatgagt acttggataa
64021 tgttgtcatt gagaccgcgc agttcagcgc cacttttcat tcccgacctc ttgccggtgg
64081 aagggacaag gatcatccaa acacacttca cgttaggggt ggtatccctg gtatgttcgt
64141 ctttttcgaa atgtctccat tgaaagtcat caataaggaa cagcacgggc agacttggtc
64201 gggcttcatc ttgaattgta tcaccagcat tggtggtgtc ctagctgtgg gcactgtcat
64261 ggacaagcta ttctacaaag cacagagatc gatctggggc aagaagagcc agtagaggaa
64321 gagactgtca tagggaagag ccctttctac atactactac ataatatata tatatagtat
64381 agaaattggt atatcactac ttgtacaaat atcatattgt acgataatcg cgaagaacga
64441 cgcactggtg ggaagaagtg gaaaacagaa gctttaaggt agaaacagaa caagaatgtg
64501 gctatggtag gatagcagaa gagtaccatt gctgttatca tttgttgcct agccctatca
64561 agacctgtct gctaatccaa cccgagagat catggcgatc caaacccgtt ttgcctcggg
64621 cacatcttta tccgatttga aaccaaaacc aagtgcaact tccatctcca tacccatgca
64681 aaatgtcatg aacaagcctg tcacggaaca ggactcactg ttccatatat gcgcaaacat
64741 ccggaaaaga ctggaggtgt tacctcaact caaacctttt ttacaattgg cctaccaatc
64801 gagcgaggtt ttgagtgaaa ggcaatctct tttgctatcc caaaagcagc atcaggaact
64861 gctcaagtcc aatggcgcta accgggacag tagcgacttg gcaccaactt taaggtctag

64921 ctctatctcc acagctacca gtctcatgtc gatggaaggt atatcataca cgaattcgaa
64981 tccctcggcc accccaaata tggaggacac tttactgact tttagtatgg gtattttgcc
65041 cattaccatg gattgcgacc ctgtgacaca actatcacag ctgtttcaac aaggtgcgcc
65101 cctctgtata cttttcaact ctgtgaagcc gcaatttaaa ttaccggtaa tagcatctga
65161 cgatttgaaa gtctgtaaaa aatccattta tgactttata ttgggctgca agaaacactt
65221 tgcatttaac gatgaggagc ttttcactat atccgacgtt tttgccaact ctacttccca
65281 gctggtcaaa gtgctagaag tagtagaaac gctaatgaat tccagcccta ctattttccc
65341 ctctaagagt aagacacagc aaatcatgaa cgcagaaaac caacaccgac atcagcctca
65401 gcagtcttcg aagaagcata acgagtatgt taaaattatc aaggaattcg ttgcaacgga
65461 aagaaaatat gttcacgatt tggaaatttt ggataaatat agacagcagt tattagacag
65521 caatctaata acgtctgaag agttgtacat gttgttccct aatttgggtg atgctataga
65581 ttttcaaaga agatttctaa tatccttgga aataaatgct ttagtagaac cttccaagca
65641 aagaatcggg gctcttttca tgcattccaa acattttttt aagttgtatg agccttggtc
65701 tattggccaa aatgcagcca tcgaatttct ctcttcaact ttgcacaaga tgagggttga
65761 tgaatcgcag cggttcataa ttaacaataa actggaattg caatccttcc tttataaacc
65821 cgtgcaaagg ctttgtagat atcccctgtt ggtcaaagaa ttgcttgctg aatcgagtga
65881 cgataataat acgaaagaac ttgaagctgc tttagatatt tctaaaaata ttgcgagaag
65941 tatcaacgaa aatcaaagaa gaacagaaaa tcatcaagtg gtgaagaaac tttatggtag
66001 agtggtcaac tggaagggtt atagaatttc caagttcggt gagttattat atttcgataa
66061 agtgttcatt tcaacaacaa atagctcctc ggaacctgaa agagaatttg aggtttatct
66121 ttttgaaaaa atcatcatcc ttttttcaga ggtagtgact aagaaatctg catcatcact
66181 aatccttaag aagaaatcct caacctcagc atcaatctcc gcctcgaaca taacggacaa
66241 caatggcagc cctcaccaca gttaccataa gaggcatagc aatagtagta gcagtaataa
66301 tatccattta tcttcgtctt cagcagcggc gataatacat tccagtacca atagtagtga
66361 caacaattcc aacaattcat catcatcctc attattcaag ctgtccgcta acgaacctaa
66421 gctggatcta agaggtcgaa ttatgataat gaatctgaat caaatcatac cgcaaaacaa
66481 ccggtcatta aatataacat gggaatccat aaaagagcaa ggtaatttcc ttttgaaatt
66541 caaaaatgag gaaacaagag ataattggtc atcgtgttta caacagttga ttcatgatct
66601 gaaaaatgag cagtttaagg caagacatca ctcttcaaca tcgacgactt catcgacagc
66661 caaatcatct tcaatgatgt cacccaccac aactatgaat acaccgaatc atcacaacag
66721 ccgccagaca cacgatagta tggcttcttt ctcaagttct catatgaaaa gggtttcgga
66781 tgtcctgcct aaacggagga ccacttcatc aagtttcgaa agtgaaatta aatccatttc
66841 agaaaatttc aagaactcta ttccagaatc ttccatactc ttcaggatat catataataa
66901 caactctaat aatacctcta gtagcgagat cttcacactt ttggtagaaa aagtttggaa
66961 ttttgacgac ttgataatgg cgatcaattc taaaatttcg aatacacata ataacaacat
67021 ttcaccaatc accaagatca aatatcagga cgaagatggg gattttgttg tgttaggtag
67081 cgatgaagat tggaatgttg ctaaagaaat gttggcggaa aacaatgaga aattcttgaa
67141 cattcgtctg tattgataaa taaaactagt atacagcaaa tactaaataa ttcaagaaaa
67201 aaacattatc gattgaagtt ttgtccaact atccactatg gatatgcgtt ttgttgatta

67261 accttaaata acacgtattt cgcattttcc aaaagccttt tttcataact acaaactaac
67321 tatttttgtt tattttacat cagtaaatta tgcgcagaaa atatgtaagg ctatatactc
67381 aatatagtgg aagagcctct ggctacttaa tctgggttca tatattctgt cagtggtata
67441 gtaaatttaa atagtgaatt tgggcgcatg tagagagatc ttttgattaa taacccagta
67501 tttgattctt gaatgttatt tgtctcttat taagtatttc agtttgattt atatattttg
67561 aaagtaattg ttgctttacc ttccaaacaa taaaaaaata taaaaaaatg taaaaaatat
67621 aataaatatt aaataaaata ctacttgttt gaaaatcaga gaaaatccac caaaatatca
67681 atcatttcaa ggatttccga accaagttcg agatagtcct ttaaggtcag taaaattcag
67741 ttgcacgtat aacagatatt tttcattttg ttccaattta aaagtccccc atttttaaaa
67801 ttattgaaaa taaaaattaa aaaattaaaa ggaatctctc tatgtcactt taaaataaat
67861 aaattgaaaa tgatatatcg ttaaaagagc ttgccttgtc cccgccgggt cacccggcca
67921 gcgacatgga ggcccagaat accctccttg acagtcttga cgtgcgcagc tcaggggcat
67981 gatgtgactg tcgcccgtac atttagccca tacatcccca tgtataatca tttgcatcca
68041 tacattttga tggccgcacg gcgcgaagca aaaattacgg ctcctcgctg cagacctgcg
68101 agcagggaaa cgctcccctc acagacgcgt tgaattgtcc ccacgccgcg cccctgtaga
68161 gaaatataaa aggttaggat ttgccactga ggttcttctt tcatatactt ccttttaaaa
68221 tcttgctagg atacagttct cacatcacat ccgaacataa acaaccatgg gtaaaaagcc
68281 tgaactcacc gcgacgtctg tcgagaagtt tctgatcgaa aagttcgaca gcgtctccga
68341 cctgatgcag ctctcggagg gcgaagaatc tcgtgctttc agcttcgatg taggagggcg
68401 tggatatgtc ctgcgggtaa atagctgcgc cgatggtttc tacaaagatc gttatgttta
68461 tcggcacttt gcatcggccg cgctcccgat tccggaagtg cttgacattg gggaattcag
68521 cgagagcctg acctattgca tctcccgccg tgcacagggt gtcacgttgc aagacctgcc
68581 tgaaaccgaa ctgcccgctg ttctgcagcc ggtcgcggag gccatggatg cgatcgctgc
68641 ggccgatctt agccagacga gcgggttcgg cccattcgga ccgcaaggaa tcggtcaata
68701 cactacatgg cgtgatttca tatgcgcgat tgctgatccc catgtgtatc actggcaaac
68761 tgtgatggac gacaccgtca gtgcgtccgt cgcgcaggct ctcgatgagc tgatgctttg
68821 ggccgaggac tgccccgaag tccggcacct cgtgcacgcg gatttcggct ccaacaatgt
68881 cctgacggac aatggccgca taacagcggt cattgactgg agcgaggcga tgttcgggga
68941 ttcccaatac gaggtcgcca acatcttctt ctggaggccg tggttggctt gtatggagca
69001 gcagacgcgc tacttcgagc ggaggcatcc ggagcttgca ggatcgccgc ggctccgggc
69061 gtatatgctc cgcattggtc ttgaccaact ctatcagagc ttggttgacg gcaatttcga
69121 tgatgcagct tgggcgcagg gtcgatgcga cgcaatcgtc cgatccggag ccgggactgt
69181 cgggcgtaca caaatcgccc gcagaagcgc ggccgtctgg accgatggct gtgtagaagt
69241 actcgccgat agtggaaacc gacgccccag cactcgtccg agggcaaagg aataatcagt
69301 actgacaata aaaagattct tgttttcaag aacttgtcat ttgtatagtt tttttatatt
69361 gtagttgttc tattttaatc aaatgttagc gtgatttata ttttttttcg cctcgacatc
69421 atctgcccag atgcgaagtt aagtgcgcag aaagtaatat catgcgtcaa tcgtatgtga
69481 atgctggtcg ctatactgct gtcgattcga tactaacgcc gccatccagt agatagagag
69541 gggcagatgt tcaagctata cccattatat tgatccacac ttagtattaa gatacgtctg

69601 tgaaggatga aaaaaaatgt ataatgtgac tagaggaagt aaggagaaaa aacgatagta
69661 atcgtatttt aggttgtgcg tttttataat tttttttttt ttgtaattct atgcaaatgt
69721 aatataagta tatttaaaga aataatgagt cctgtgaaaa caaaaagaaa aaaagatcat
69781 taatgtatgt taacgtattt gctttgcaaa ttttaattta tttgttgtta aatgcatttt
69841 ttttttgtcg tttcagcgag ttttcttgag gttgctacta tcattaaaat cacaatccac
69901 agaggaagtt gatctctttt tcagttgggt gggggcagag catgggtgag cagtggccat
69961 gggtctaaca ggaaataatc tttttgaacg cacagataaa ttttgtaata attttctatt
70021 tgacattaga gatggggtgg tgggagttag tgggcttggc caaaagatgc ttgaattttg
70081 tgggatgctc agtgaccttt taaaagaatt ttgggtagaa gagaacgaac ctgaatgtga
70141 atggtgtgat gcagagtctg gggtcgtcat tgaacttgaa gtcttgtaag gggaattgaa
70201 tggagatgga gaggatgaag atgaggttgg agtgaaggca aatggtggag aaatgctatc
70261 tttggtcaac cttcttaaat gagtgtgatc cgagaaatag tctgtggccg cagatgaaac
70321 cgtagcgctc tttggagtag tagcatttga gtaaacacaa gtcaatgaat cgctgtcaaa
70381 ggactgagca aagatagtat tgtacatggt ttccaattct tgtgaacggt agaattcttc
70441 cagtttcaat tgaatacaat agttttgtaa tgcgtctaag agagattttg ctaaagatgt
70501 cttattgcta ttcaaatatt ttgatttgaa actggagggg aaagaatttt ggtccattgc
70561 tatatccaat aaagtattag agatttctga caatttaata tctaggtctt ggcagttttc
70621 ctccaaagcc agattgatat tttcccaaag gtttgaatta tagtagttca aagataattt
70681 gatgaggtta attgcaccca gcgcaattag tgaacgatca tatttaaatg ataattcgag
70741 attgaaggaa gctaactcgc acaacataat ggcacccaat ttgatctcgt tgatgttcaa
70801 agaggggcct tgagaagagg acgatttatt aatagcagta ccagcagcgt tatttaataa
70861 ggccagtttc tgttgaatga aagcttccaa aggggcagaa aggacaacgc caggcgataa
70921 cggggacgta gattggaaca agaagatgtc gatgtaggag tcgaatgttg ccgactgaca
70981 gatggaccaa tcgagtgatt tgaaaagatg catttccata gtcgtgaatt gctttataga
71041 atattgattg caacacaagt tttgcaagac tttcaaagtg gccattctat tcttggagtc
71101 ccaaaatttg gacgaaatcc aaagcgcggt caaggacaag agctggtagt tgtaactctt
71161 gataatgaac cgcgaggaat acttgtccaa gatagtgaaa gtaaggaaca aagtcgaggt
71221 cgatagattg agtcttgtgt gacagtacat gatgaagtca aagatcaaga aacgcatctt
71281 cggattaact tgtggctgag agttgaagtt agtcaggttg tacagcggcc tttctgtgtg
71341 ggaaagacgg aaatagtggt ccaattgatc attattgtat tcgctgatcg ctgagtgatg
71401 tgcttgcaat tctcttttaa cgagattagg gtttttagac tttgcactag caatggcccg
71461 cctcttttgc aagagcaagg gcaaattagg acatgaggca gcgctgacag aggcggcagt
71521 ggcggtggaa gtgccactag cggtagcata ccttgcatta gcgtatctaa ttatggtatc
71581 cttcaatatg gccatcgtac agaaagcgta tcaaatcagt gtcttgaacg agagtaaaag
71641 ggagatgcaa tggaatgtaa gaaatgctat gggtcagaaa agaaatgcag aggagttaaa
71701 ccgaatgagg aaatgcaatg gatacgttaa attggagatg tgagattgcg actgggactc
71761 ggatggatgc ttgcttttgg atgctagtac agagaaaaaa gagggagaaa agaaaagaga
71821 aatagaaaaa ggttggttta agtcggcaga ggagactact cgggcaattc gtttcctaat
71881 gggtatagtc ctctttcctc agaaatccat ttgactggca gactcagtag tagaagaaaa

71941 aatcaagaaa gaaattgtgt gaaagtatac tacacaagaa aatgtttaag aagaagatta
72001 aaagctcgag gaaagtacag atatacaaat tataaatagg taggaggaag aaaaaaaaaa
72061 gatgagggca aaaacccaag gccaaatatg gaaatgtggc agagggacac actaatccga
72121 tagtaaattt atgtaacttg atcattacag tgagagcagg cttggtaatt tctttttctt
72181 gcctgataat ttttcacttt ttcttttttt ttttcttgaa agttcacaat ttagggtttg
72241 agcacagcgt ttggttgcga cacttcccca gaaggaaaac cggcctcttt ggctggggag
72301 ggaagaaggg ggaggagagc tcagaaagcc ctatcgcgat atgagggacc gatctgtatc
72361 agcatacttt cggtatcatc acggtcgcag gatagagata gtgaatgagt tagctgtttt
72421 acaggccaag cgttcaaacg agacgaatgg cgacatttgc ccgtcaagaa aacccgccga
72481 gtttttttcct aaacgggtaa aacagccatg caccgcagca cgtcgacgag gtgatatttc
72541 caatttggga aatttcccaa atcagtaatg tagcctctac gggtgtctct gtcagccccg
72601 tggtcgccag cacagaatgt atcgtacccc tgaaggtagt tttttaccgc cgtggcacac
72661 gataaaggtg caccttgtga taataaggtg gaaaaatata tatgaaaaag tgaaattgat
72721 tgtggctgca ctaggacatc attatttctt acttggctat ttacacgtac ttacgctggc
72781 tgtatatcat ttaaggggcg gaggacgaag aggacggacc cgagatcatc cggtccaaga
72841 aacgggtcat ccggtcctta gcattgtcta gactatctag ggcaggacgg acatccacgt
72901 ggaaagtagg cattccgttt tcgtcgtcgg gccctccgta gaaatccaag acgtatctaa
72961 cttccttgaa ggttggaggt tgttgttccg ctttgcgctc gcctcggagt acaatccagt
73021 cgtgcctgtc gaatggtagt tcttggctaa aatgggacgg aaacagtagg ccgcacaggt
73081 gcatccagcg agcacgaggg ctcaatacgc ccggtttccc catgaatttc agcaacttag
73141 gctgcacgtg gctttcatct gtgtgcggtt tttcccattc gagcacttcc tgccagcacc
73201 cttcatttag aaagttgtgg acctgcacca tggactccac tgcatcttcg gcgacttcgc
73261 cgctaccgcc aatcttgccc tttctaacca tagcattgta catctgttgt ggagaaggat
73321 actcccagaa ctcgttactg tctggactct tggggatgct ggagatggtc cgatcaacgg
73381 gcaagtccat cttttggcca ggctgtttgg atgctgccaa ctccggcata ttgttcagcg
73441 ggtttattct atcgttatct ccctgcataa cggggcactc agaggatggt ggcgacgacg
73501 acgacgactc gtgcatgact gggcaccctg acatggatga tactgctgcc ccaccaatat
73561 ctttgcccgt agttttttga tctgcccaaa accaacccat tttttgtagt ttctgttgta
73621 tttgctctgc tttattcgtg ctactcgtat gtgtatgcta tcttattagc tgcctatttc
73681 atttctctta gtattcgcat ttaaagctga gaaatttttt gataatcatt tcccgatgaa
73741 aagaaaaaag ggaaaaagtc gataaaaaga ggtaagcgaa aagaaaaaga aaaaatagaa
73801 aatttgggtg gggggggcgg aagatcccac gccgcgcaag agatatttca atattactac
73861 tacatagtat atgcggcgct accatacgta caactttttt ttcttttttt ttttttttgc
73921 cttctaaatt tgtaattcgg tcacactttt gtcgcagtgt tgcaaacgtc ttgaaagaat
73981 tgtaggtgtt gtaaaccaca acttgctccc ttgaaagcgt tgctgattat ttcctttgaa
74041 cctgttgcat tgttgtattg ttgtattgct gctgatgttt taggcactta gtattagata
74101 ttcctaagcc tccctcacca taaattacct ttattacttg catgactatt attagcagag
74161 catgtagtat gggactcaag accgatatga tacacaccaa agacgtaggc accggcgatt
74221 aaatcaaagg ctccgatagc cgaaaagtga gaagaaaaaa aaaggaaaaa aaggaattgt

74281 cctaatgagc ggtgtggccg acttgccata atatcagtta gggctactat caatgtttta
74341 tctacgttgg agtaagatcg tttatcactt ccatatttgg accaaatgaa aagttcaatc
74401 ggccaagtat ttcatggatg gaatgacgtt tggtaaggaa gtgctttttc tttttccaca
74461 tattttccct ttctctcggg gaaattttgt ttctaaacat aaaaaataaa gcaacagcaa
74521 aaaagagggt ctgtccagcg aataagaaga aaacctcctt ttcggctttt gaagataggt
74581 tgcagttgtc tgcgggcaca aaatgggcaa tttttttaat actttttacg tatgagacaa
74641 gatttttttc gccaattata tcgcatgaag aataaccaga gtttttctcc gaacgttaag
74701 ggagttgaag taaaaataaa gaaaggacca aatgagaatg ggtatgcttg gtcttagtct
74761 tcgaatcaaa ttctgcttcc ctgttcatgg caacgtcacc tcaattattt ggaaaggggg
74821 ggttttccga ctttatttga gatgacttga gatgtgtgtc aatgctagta ttttggagat
74881 taatctcagt acaaaacaat attaaaaaga ggtgaattat ttttcccccc ttattttttt
74941 tttgttagaa ttgatccaaa tgtaaataaa caatcacaag gaaaaaaaaa aaaaaaaaaa
75001 aannnnnnnn nnaaaaaaaa aaaaaaaaaa aaaaaaaaaa atagccgcca tgaccccgga
75061 tcgtcggttg tgatacggtc agggtagcgc cctggtcaaa cttcagaact aaaaaaataa
75121 taaggaagaa aaaaatagct aatttttccg gcagaaagat tttcgctacc cgaaagtttt
75181 tccggcaagc taaatggaaa aaggaaagat tattgaaaga gaaagaaaga aaaaaaaaaa
75241 atgtacaccc agacatcggg cttccacaat ttcggctcta ttgttttcca tctctcgcaa
75301 cggcgggatt cctctatggc gtgtgatgtc tgtatctgtt acttaatcca gaaactggca
75361 cttgacccaa ctctgccacg tgggtcgttt tgccatcgac agattgggag attttcatag
75421 tagaattcag catgatagct acgtaaatgt gttccgcacc gtcacaaagt gttttctact
75481 gttctttctt ctttcgttca ttcagttgag ttgagtgagt gctttgttca atggatctta
75541 gctaaaatgc atattttttc tcttggtaaa tgaatgcttg tgatgtcttc caagtgattt
75601 cctttccttc ccatatgatg ctaggtacct ttagtgtctt cctaaaaaaa aaaaaaggct
75661 cgccatcaaa acgatattcg ttggcttttt tttctgaatt ataaatactc tttggtaact
75721 tttcatttcc aagaacctct tttttccagt tatatcatgg tcccctttca aagttattct
75781 ctactctttt tcatattcat tctttttcat cctttggttt tttattctta acttgtttat
75841 tattctctct tgtttctatt tacaagacac caatcaaaac aaataaaaca tcatcacaat
75901 gtctagatta gaaagattga cctcattaaa cgttgttgct ggttctgact tgagaagaac
75961 ctccatcatt ggtaccatcg gtccaaagac caacaaccca gaaaccttgg ttgctttgag
76021 aaaggctggt ttgaacattg tccgtatgaa cttctctcac ggttcttacg aataccacaa
76081 gtctgtcatt gacaacgcca gaaagtccga agaattgtac ccaggtagac cattggccat
76141 tgctttggac accaagggtc cagaaatcag aactggtacc accaccaacg atgttgacta
76201 cccaatccca ccaaaccacg aaatgatctt caccaccgat gacaagtacg ctaaggcttg
76261 tgacgacaag atcatgtacg ttgactacaa gaacatcacc aaggtcatct ccgctggtag
76321 aatcatctac gttgatgatg gtgttttgtc tttccaagtt ttggaagtcg ttgacgacaa
76381 gactttgaag gtcaaggctt tgaacgccgg taagatctgt tcccacaagg gtgtcaactt
76441 accaggtacc gatgtcgatt tgccagcttt gtctgaaaag gacaaggaag atttgagatt
76501 cggtgtcaag aacggtgtcc acatggtctt cgcttctttc atcagaaccg ccaacgatgt
76561 tttgaccatc agagaagtct tgggtgaaca aggtaaggac gtcaagatca ttgtcaagat

76621 tgaaaaccaa caaggtgtta acaacttcga cgaaatcttg aaggtcactg acggtgttat
76681 ggttgccaga ggtgacttgg gtattgaaat cccagcccca gaagtcttgg ctgtccaaaa
76741 gaaattgatt gctaagtcta acttggctgg taagccagtt atctgtgcta cccaaatgtt
76801 ggaatccatg acttacaacc caagaccaac cagagctgaa gtttccgatg tcggtaacgc
76861 tatcttggat ggtgctgact gtgttatgtt gtctggtgaa accgccaagg gtaactaccc
76921 aatcaacgcc gttaccacta tggctgaaac cgctgtcatt gctgaacaag ctatcgctta
76981 cttgccaaac tacgatgaca tgagaaactg tactccaaag ccaacctcca ccaccgaaac
77041 cgtcgctgcc tccgctgtcg ctgctgtttt cgaacaaaag gccaaggcta tcattgtctt
77101 gtccacttcc ggtaccaccc caagattggt ttccaagtac agaccaaact gtccaatcat
77161 cttggttacc agatgcccaa gagctgctag attctctcac ttgtacagag gtgtcttccc
77221 attcgttttc gaaaaggaac ctgtctctga ctggactgat gatgttgaag cccgtatcaa
77281 cttcggtatt gaaaaggcta aggaattcgg tatcttgaag aagggtgaca cttacgtttc
77341 catccaaggt ttcaaggccg gtgctggtca ctccaacact ttgcaagtct ctaccgttta
77401 aaaaaagaat catgattgaa tgaagatatt atttttttga attatatttt ttaaatttta
77461 tataaagaca tggtttttct tttcaactca aataaagatt tataagttac ttaaataaca
77521 tacattttat aaggtattct ataaaaagag tattatgtta ttgttaacct ttttgtctcc
77581 aattgtcgtc ataacgatga ggtgttgcat ttttggaaac gagattgaca tagagtcaaa
77641 atttgctaaa tttgatccct cccatcgcaa gataatcttc cctcaaggtt atcatgatta
77701 tcaggatggc gaaaggatac gctaaaaatt caataaaaaa ttcaatataa ttttcgtttc
77761 ccaagaacta acttggaagg ttatacatgg gtacataaat gcagatgcca gtgaactatg
77821 ttcagcttct ggccttcgtt tggtggttta atctattttt tataaaaaat gacgcgggca
77881 gattcaatta gtgtcctaaa tttattcgcg tttcaagatt tcaaaggatt gatcctctta
77941 tcagaaacga taagtgctac tccgtcctat tcttctagcc atctagtacg tattcttttc
78001 ataacataat cccttattta cagaatgtgt ttcgaagaaa aattaattag atgggaagaa
78061 aactgaagtg gcttatataa tcagtgacat agtgccaata attacgcaaa aagcaaagga
78121 aataacactg ctatggatat ggaaatcgaa gattcaagcc ccatagatga cctgaagtta
78181 caaaaactgg ataccaatgt ttattttgga ccctgtgaga tattgacaca acctattctt
78241 ttgcaatatg aaaatattaa gttcatcatt ggtgtcaatc taagtactga aaagatagcg
78301 tcgttttata cccagtattt caggaactct aattcggtag tcgtgaatct ttgctcacca
78361 actacagcag cagtagcaac aaagaaggcc gcaattgatt tgtatatacg aaacaataca
78421 atactactac agaaattcgt tggacagtac ttgcagatgg gcaaaaagat aaaaacatct
78481 ttaacacagg cacaaaccga tacaatccaa tcactgcccc agttttgtaa ttcgaatgtc
78541 ctcagtggtg agcccttggt acagtaccag gcattcaacg atctgttggc actctttaag
78601 tcatttagtc attttggaaa tatcttggtt atatcatcac attcctatga ttgcgcactt
78661 ctcaaatttc ttatttccag ggtgatgacc tactatccac tagtgaccat ccaggattct
78721 ttgcaatata tgaaagcaac cctgaacata tccatcagta catccgatga gttcgatatt
78781 ctgaatgata aagaactgtg ggagtttggc caaacccagg aaattctaaa acgtaggcag
78841 acgagctcag tcaagaggag atgtgtcaat ttaccagaaa actctacgat cgataacaga
78901 atgcttatgg gtaccacaaa gcgaggtcgc ttttgaagag ccctcggtag cataacattt

78961 ttaattatta cgactgtttt ttttattcat tatgtagaga taattaaatg ttatagatgc
79021 tctatactca aacggtggaa gaaaaacagc gaaaaaaaat aaccgatacc cccttttcga
79081 atacaaatgc ttgtatattc aattatgaat tatttttttt ttttttcatt tcttatatta
79141 ttttttgttc gagaatcact ttttcaagat ggtaacaaca tcttcgtctt ccaaaatgtg
79201 actcaacccc acgtattgag gttgatgttt gacactgcta ccgtaaacca gagcatttct
79261 aaagtcgtcc actaaagatt tatgaatttg gttacaaaaa tccttgacac tgcaacggtc
79321 tgatcttagc accacagggt cggtaaaatc tggtatttgg ccctttggtt tagtgtaaat
79381 acggactaga tttagtctat cccacatgac ttgcaacagc tcgtccaagt tccaatcttg
79441 accagacgaa ataggcacgg cattaggaat tcggtaaagt aattccaatt cctctattga
79501 cagagaatca atcttgttta acacatagat ggcaggcatg tatcttcttg acgaagcttc
79561 caaaacatca atcaaatcat ccacagtggc atcacacctg aaggcaatct cagcgctatt
79621 tattctgtac tcgctcataa cggctctgat ttcgtcattc cccagatggg tcaatgggac
79681 tgtgtttgtg atggaaatac cacctttctc tttttttttg atcaagatat ctggcggagt
79741 tttattcaga cgaatcccca caccttccag ttccttctca atgatttgct tatgatgcaa
79801 gggtttgttc acatctagga tgataaataa caggttacag gttcttgcca cggcaataac
79861 ttgcttacct ctacctctac catccttagc accatcgata ataccaggta aatccaacat
79921 ttggatcttg gcacctttat aacgaatgac accggggacg gtaaccaggg tggtaaactc
79981 gtactcagct gcttcagact cagtaccagt caacttggac agtaatgtag atttccccac
80041 cgacgggaac ccgacaaacc ccacactggc cacaccagtt ctagccacat caaaaccaat
80101 accagcacca ccaccgctgc cggatgaagc actggtcaac aattctcttc tcagtttggc
80161 cagcttggcc ttcagttgac ccaaatggaa agatgtggcc ttgttctttt gggtacgggc
80221 catttcatct tcgatagctt tgatttttc aactgtagta gacattttgc tcaatcaaca
80281 actctacgct tgcacctact gcatctagct tcaaacactt cctatcattg cgccctcatc
80341 acaccgtaat atcccatctt aaaagtggaa aactcttata gctcatcgat gaaaaaaacg
80401 ggccctcgtc gcttgtgatg tgaaaaaatt tttcaagctt taagcccatt gaaagcaaga
80461 gatcttgcac tagaataagt ggcaaaggtg aactttgagg ggataagaag ggcaatcagg
80521 aacatcagat aagtgaaaga tggcgaaaaa gagtaaaaag aaccaacaga actactggga
80581 tgaggaattc gaagaagacg ccgcccagaa cgaagaaatc agtgccacgc caactccaaa
80641 tccagaaagc agcgcaggtg cagatgacac ttccagagaa gcaagtgcaa gtgctgaagg
80701 tgctgaggcc attgaaggcg acttcatgtc tactttgaag caatcgaaga agaagcaaga
80761 aaagaaggtt attgaagaga agaaggatgg taagcctata ctaaagtcca agaaggaaaa
80821 ggagaaggaa aaaaaggaaa aggagaagca gaagaagaaa gaacaagctg ccaggaagaa
80881 ggcccaacag caagctcaaa aggagaagaa caaggagttg aacaagcaaa atgttgaaaa
80941 agctgctgct gagaaggctg ctgctgagaa atcccaaaaa tctaaaggtg aaagtgataa
81001 accaagtgct agtgctaaga agccagccaa gaaagtacct gccggtttgg ctgctttgag
81061 acgtcaatta gaattgaaga aacaacttga agaacaagaa aagttggaaa gagaggaaga
81121 agaaagattg gagaaagaag aggaggaaag attggccaac gaagaaaaaa tgaaggaaga
81181 agctaaagca gctaaaaagg aaaaggagaa ggcaaagcgt gaaaaacgaa aggctgaagg
81241 taagctattg accagaaagc aaaaagaaga aaagaaatta ttggaaagaa gacgtgccgc

81301 tttattgtct tccggtaacg tcaaagttgc cggtctggcc aagaaggatg gagaagaaaa
81361 caaaccaaag aaggttgttt acagcaagaa gaagaagaga acaacccagg aaaacgcctc
81421 cgaagccatt aaatctgact ctaagaaaga ctcggaagtt gtacctgatg acgaactcaa
81481 agaatccgaa gatgttttga ttgatgattg ggaaaatttg gctcttggtg atgatgacga
81541 ggagggaacc aacgaagaaa cgcaagaatc caccgcaagc catgaaaatg aagaccaaaa
81601 tcaaggcgaa gaagaagaag aaggagaaga agaagaagaa gaagaagaag aaagagcaca
81661 tgtgcatgaa gttgccaaaa gcacaccagc agctacacca gcagctactc caactccatc
81721 cagcgcttct ccaaacaaaa aagatcttcg ttccccaatt tgttgtattt tgggtcatgt
81781 cgataccggt aagactaaat tgttagacaa aatcagacaa accaacgttc aaggtggtga
81841 agctggtggc atcacccaac agattggtgc cacttatttc cccatcgacg ctattaaggc
81901 aaaaactaaa gttatggctg aatatgaaaa acaaactttt gatgtcccag gtcttttggt
81961 tattgatacc ccaggtcacg aatccttctc taacttacgt tcaagaggtt cttcattgtg
82021 taacatcgca attttggtta ttgacattat gcatggtttg gaacaacaga ctattgaatc
82081 tatcaaactg ttaagagata gaaaggctcc atttgtcgtt gccctaaaca aaattgatag
82141 attatatgac tggaaagcca ttccaaacaa ttcattcaga gactcctttg caaagcaatc
82201 aagagctgtt caagaggaat ttcaatctag gtattctaag attcaattgg aattagctga
82261 acaaggtttg aattcggaat tgtatttcca aaacaaaaat atgtctaagt atgtctccat
82321 tgtcccaaca tctgccgtca ccggtgaggg tgttccagat ttattgtggt tgctattaga
82381 attgacccaa aagaggatgt ccaaacaatt gatgtacttg tctcacgtgg aagcaaccat
82441 tttggaagtg aaagtcgtag aaggttttgg taccacaatt gatgttatct tgtccaacgg
82501 ttacttgaga gaaggtgacc gtattgtact gtgtggtatg aatggtccaa ttgtaacgaa
82561 tatcagagca ttactaacac cacaaccatt acgtgaacta cgtttgaaat ctgaatatgt
82621 ccaccacaaa gaagtcaagg ctgctttagg tgtcaagatt gccgctaatg atttagaaaa
82681 agccgtttct ggttctaggc tgctagttgt cggtcctgaa gatgacgaag atgaattgat
82741 ggacgacgtt atggatgatt tgactggttt gttggactcc gttgacacaa ctggtaaagg
82801 tgttgtggtc caagcatcca ccttgggttc tttggaagct ttgttggatt tcttgaaaga
82861 catgaaaatc cctgtgatgt ctatcgggtt aggtccagtg tacaagcgtg atgttatgaa
82921 agcctccact atgttggaaa aggctccaga gtatgccgtg atgttatgtt ttgatgttaa
82981 agtggataag gaagctgaac aatacgctga acaagaagga attaagatct ttaatgcaga
83041 cgtcatctat catttatttg attcatttac agcataccaa gaaaagttat tggaagaacg
83101 tcgtaaagat ttcctagatt acgctatttt cccatgtgtc ttacaaacct tacaaattat
83161 taacaaacgt ggtccaatga ttattggtgt agacgttctg gaaggtactc tacgtgtggg
83221 aactcctatt tgcgctgtga aaaccgaccc tactacaaag gaaagacaaa ctttgatatt
83281 aggtaaagtc atctctttag aaatcaacca tcaacctgtc caagaagtaa agaagggcca
83341 aaccgctgct ggtgttgccg tccgtctaga agatccctcc ggtcaacaac ctatctgggg
83401 tcgtcatgtt gacgagaatg atacattata ctccttggtt tcaagaagat ctattgacac
83461 tttgaaggat aaagctttta gggaccaagt tgctagatcc gattggctgc tattgaagaa
83521 gctgaaggtc gttttcggca tcgaatgagc atggcatacg ctgacttgtc aacccaatca
83581 cattctacaa aatttaatga attaaatagg taattgtata taaaaatgtg aacctttgtg

83641 tattagtttc aattctatct tacttttcat tgccatttta cttctttcac cttgctgtct
83701 ttcaaccttg gaaattttta tagtacgcgt aaacaaaaaa ggtaaataag aggcattgaa
83761 tataagttgg catttattag gaagttgagt aataacacgt tgaaactggg ttaagacgat
83821 caaaacaacc atgtctgctc ccactatgag atccacctca atattgacag agcatttggg
83881 atatccgccc atctcgcttg ttgatgatat cattaatgct gtaaatgaaa ttatgtacaa
83941 gtgcactgct gccatggaaa aatatctgct atccaagagc aaaatcggcg aggaagatta
84001 tggagaagag atcaaaagtg gagttgctaa gttggaatca cttttggaaa actccgtgga
84061 taagaatttt gacaaactag aactatatgt tttgaggaac gtccttcgaa tccctgaaga
84121 gtatttggac gccaatgttt ttagattgga gaaccaaaag gatctggtca ttgtagatga
84181 gaatgagttg aagaaaagtg aggagaaact tcgagagaaa gtgaacgacg tggagttagc
84241 gttcaaaaag aatgaaatgc tattgaaaag agttacaaaa gtgaaaagac tgttgtttac
84301 gataagagga ttcaaacaaa agctaaacga gttactgaaa tgcaaagacg atgtacaatt
84361 gcagaaaatt ttggagtcgt taaaacctat agatgacaca atgactctac tgactgattc
84421 attacgtaaa ctatatgttg atagtgaaag taccagttca acagaggagg tagaggcact
84481 actgcagaga ttgaagacca acgggaagca aaataataag gatttcagaa cacgatatat
84541 cgatataagg acgaataatg tcctacgaaa attggggcta ctaggtgata aagaggacga
84601 aaaacagtct gccaagccgg atgcgaggac gcaagcaggg gatatagtta gtatagatat
84661 tgaagagcct caattggatt tacttgatga tgtgttataa tataaagtgg gaaaaagtat
84721 gtgctatgat atgatgtatg tattcacgaa tgtattatgt agaaaaatgc taaaaaattg
84781 gataaaagaa aaccatgttt aaaatgcata ccaccatgtg tattataagt acttcgtaaa
84841 attcgaatcc tgtagccagc caaccttctc gaaagcttgg aatagtcttg atgctttatt
84901 aacgtcgatc ctacaggctt tttgggcgtc ggtccttcta aacggcaacc ctttctttag
84961 tctataaact ttttccaaaa ataaccttct cttagaatcc agatacaaat cacaaggtaa
85021 ccttagagtt tgagcgagaa ctagttcagc agggtgtagc tcgttcctca gcgggtctgt
85081 ggacaggtcc attggagacc ccttccactc gatcttgagt gacttgttac tgtctgtagg
85141 aagcgtagat aagggcggag agtaatctgg taattttttc catgagacgt ttggtatata
85201 ttgaggagca tcatgaatcg cctcaatgga tgcaatgtta gtgtgaggag tggatgcaga
85261 gggtgagaac ttcttggcgc gatgtggact ggtagtcatc gttcttcttt gtggtgagta
85321 taccttgcgt ttccgtggta tccattcagt gtcggattca aactttccta gattgttatt
85381 gtagagagcc tgtttcctca tattatatct tgcttgtcta ttcaaattcg gagatgctaa
85441 cggggaaggt aatctagaag aagaggaagg cagaggagga ggagtcatat catcattgta
85501 tttggaatat ttatgtctag cattataggc cagattgttg taccggttga taatgttctt
85561 agaatatggt tgggccaaat tgctgaaaat tttgtattga gacagaaacc cgtaggtggc
85621 gtagcggtac ctttttcttg ataacccatt gggccataca ggcttcacga acagcacatt
85681 ctctacagcg cctttaggcg tagatcccac tagcactgaa gaggccatgc ggggtgagcc
85741 attgtggcta attttcgact ccagcttggg cgacaaaggt ggcgatggta ttatatgttc
85801 atcaaccaag gagtcaatgc tgccagaaaa gcatccattg ccattatttt gtgagttctg
85861 gttaattctt acgtcgtacg agcgcccata gggtgaatta ttgctataat cactcatgtt
85921 taaaccgttt ttattactat cgttattgtt gttacttgct ttcaaattct gattcagacg

85981 tctgaaaatg gactgcgaat catccttacc aatgtaattc atatttaatt gagacttttc
86041 agattcagga gaatataatc ccatgtttcc cctataaatg ttacgatgcg cgaatatcct
86101 gtttatcgat gcccagtgta tgagccataa cggggatgtt atgaaccatg tggctacttt
86161 tataagcggt tctcttagga agaagggggt tcttgataga cccctgcacc tcatctagcg
86221 gaggtgcacg gatgtaccaa cagttttagt gaacattatt cactaaagaa gcattgggca
86281 tactcagagc caatggcaag ctcgtttacc agttcaaata tgtcgtttca ttatctgtat
86341 gactgtcgta actttgaatc gatctaatgt gttgaccctg tctcaggctc acccatggcg
86401 gcgcctgcac ctgtgggtga aggaagaaag acgatgtttg tgagggaact gaattgggtt
86461 gaagttcata tcctaaacaa acacttcacc agccatggat gcatgccttg tcttttcgca
86521 gttggtggca tgaaaatata tatcacccac caaaccctct tactcttttc ttaccaagta
86581 actccagtaa gtgctcgttt ttttcttctt ccattcaaac ctgcttaaaa acctcgacaa
86641 acgagccccc aacgtactac cacagcaacc actctggttt ttctatcttg ttgtctttaa
86701 ttgcctcctg actttgtttt gttttgttct tgcttagcgc tcttgaaaaa tattttactt
86761 ttcactatca gattaatgtg atagcaatag ttagtgcaac aaagaaacaa gtcgataaat
86821 ggtacgttta aaaagtagat atatcctttt tgaaattata ttcccaccta cagacaccaa
86881 cgttgaggaa tctgtgtcga aagcagacat cttgctttcg catcacagag catcgcctgc
86941 ggatgtgtcc ataaagtcga tactccaaga gatacgacgc tcgctgtcgt tgaatctggg
87001 cgactatggg tctgcaaaat gtaactctct cttgcagttg aaatactttt caaataagac
87061 gtctacgggg ataatccgat gccatcgaga ggattgcgac cttgttatca tggcattgat
87121 gttgatgtcg aaaattggcg acgtcgatgg actgatcgtg aaccccgtca aggtaagtgg
87181 gaccatcaag aaaatagagc agtttgctat gagaaggaat tctaaaattc tgaacataat
87241 caagtgtagt caatcatcac acctcagcga taatgacttt attatcaatg atttcaagaa
87301 aattggaagg gaaaacgaaa acgaaaacga ggacgattag aatatattaa tatatagatg
87361 tacacgtata tgcagtagtt ttattttttt atctataata caactcaagc acaagaatgc
87421 tttgttttcc tagtgctcat cctgggccta ggcgccatag ttatccgatt tatcatcgga
87481 ttcagcttta gtaaactgaa tggggccgtg agaaccactg gcaccttcac tcttaacatt
87541 gaccgcttcg tccagctttt cgtagttggt cttgtatatg ctttcaatat cttgttggac
87601 gaacagtggg ttgtcataaa cctggtcttc atgccttttg gcggaggcat ttgcccctct
87661 tgtgaaaaat cttgaatcgt actgcagatc cggttgttct gaacgctttg ctgcgcccag
87721 aattatcttt tcggatacgt ctcttccttg agagtacgcc agctctttta gtctggccac
87781 tgtgctcgtt tgctttttgg gcttaactat tgctcccgtc tgcggagtcc cgttgtggta
87841 tctggctcgt tggctcaatt ctttcaattt agattcttta gcaagcattt cctgttccat
87901 agcaagccgc ttcaattcca ttttggacct gatctcttgt cttgccttct tgtcagcgtt
87961 ttctaacgct tcggagagct tcataaaccc atcgttgatg gtattatttt cgttgtcaag
88021 agctttacct acacgtcttt ccaaggccac ggtataacca tttggatttt tccagtttga
88081 cacagctgca ggtatcttcc actcatttgg atcagcttct cccctatcat tgctgccatc
88141 catatggaga acaggcacga cttcgtcgtt ttctgtgggt gctacaacct ttctcgcctt
88201 cttcccaacg aaccttggca acaatggatc catttgcttg gacactacct caatatggtg
88261 gctgttgttc aataatagat tcgcgggtgc ctgatgtgtt tcggtgacgt accttgatga

88321 tgccctgtta tttgagttgg ctagtttcgc attcacaagc cgctgaatgt atgacttggt
88381 tcttgctgta cattcttgga tttctgcttt cgttggcaaa ggaaccgata gttcgaaatt
88441 agactgtctc tttggaataa aatcatcgag cttaacgttt ttagcgattt ggtcagtcaa
88501 tattgccggc tcaacgcgat ctgagctcaa agccgtcgaa actcgtcctt gagaatgttt
88561 tggaggtggt agtctgttac taaacatatt tggttgaggt tgctgatacg tcctctcggc
88621 tcagagctgc catgtgtcaa taacccccata atttgaactt ctttcatcaa ctttttagct
88681 ggggaaaaat caaattcgtg aagaattaca ataatacgtt aaggtaaaag attaaatatt
88741 aaaaaatagt atgagtactt ttgaatcatc agacaagaac aatgaaggat atgaatagta
88801 ttagatatgt attctttttt ttttccaggg acataaagag ttgtttttat aaggtgcgga
88861 gttatctcaa tttgcttctg attttagaag ctattctatg cccggtcgac tctttgattt
88921 cgatcccaaa cggcatcatg gtagtttcgg agccagattc gccattttcc cactctaatc
88981 catctcgaag actttttcga aaggtaaacc cttcttcttg tctcagcgtg ttatatttgg
89041 acttatgctg tagcggcttg atcattgccg aagcaggtat tggcggataa tgtagcacag
89101 ccggcttctt gtaccaaccc acctttgttg gtttaggatt ttctgcctct gtcattggcg
89161 gtacaccagc aaatctcacc cttttgatgg acaattcact ggagtttgac tccggatcta
89221 gagttaatgt ctctatagcc aaagtatcac tactattggc aatcagggta cttgatgatt
89281 ttcctgagga ggtgttgttt atactggata cttttggtga aggcgctaac gatgaagatg
89341 cagaaggatt gggtgtagat gcacttttgt tcatcaaatg ccgcactgta tcttggacga
89401 ttttttgatt taacgtcatg gttgaggact gcaaagaatt ggacctttga ttgttgagct
89461 ttgaaggtgg agctattgac ggatgtcttg tagaattcaa atatgaggtt tgttgtttgt
89521 ttgtatagga ttgcaaagat gaggatctag gccttctgcc gttgatcagt tttcttttct
89581 ccggagacac cacgcttttc ttcgtttctg cgttttttatt gttgaaagtg gtccttacag
89641 gcgatactga gggagaccta gaagaatttg aggctaaatg cgcccttgcc cttgttctta
89701 tgctctctaa agaggcctgt cttgatgggc ttgctgaaga tgtgggtgag ggggatggag
89761 aagaagaggc tggcagcttt gttattgacg gcttcttctt tagcttactt ttcgcgtttt
89821 taacctctaa atacagagcc ttatctcttt ctagcttttc tttcaaattc tcccgcaagt
89881 cattggtttc atcgaaggtt ttataatttt tctcaatata tttccagcaa tccatccaat
89941 tggataataa tcccataatg gacctcaatt tcggtataat cttgcagata tgattattaa
90001 acaattccac aactaaacca ggttgaaggt ggaaaatggg agttcttgag gaattgttat
90061 cggtacgcgc cgtttgagtc caaaatttta aattctcctg cagttcgaat aaagctgtga
90121 catcatcaca ttctggagag agcaaagcca tcagctcaca aattatcaaa tcgtaatggt
90181 acacgttttc gctggataat cttgtcttga tgtccaagaa atgttgttta ttcggtagga
90241 aaatttgcct agagaaactg ataataacac acaagacaga aaaaaacccgc aagtggaaag
90301 tgtatatcct gtgtagagtc cattcccaat gtggtagatc cttctgtaaa tttgacagtg
90361 aagtaggtct tgtttccgaa aatttctgca attggttgta atatgtcctt agaatgggga
90421 agattttcga atgaacacag gagttcagtt tctccatcac ttgccattgc aatttgtaaa
90481 aggaatgggt ggacttatcc cttactggat cattaatcaa aacggctaaa tctagaatca
90541 atgtcatttc aactggattt aacaagtctt caattattga agtgtcaaat ggaagtgacc
90601 gagctttgta cgcatccctt tccaataaaa tctgtctcaa cttggcatta taaattttta

90661 tagcattttg agcaatgcaa gacagcgaat ttaaaatctt ccattgtaga gaattattgt
90721 atatgcacga tttatagtga tctaggggta cctcctcggg aaacatccag ccctctacat
90781 caataaaatt agttgataat ctctgctgca cttcgcttag ctttatcagc tggaatcgac
90841 agagcaaatg gtacctctta gccaatacgg ggcttacatt aaatatagga ccctcacaaa
90901 gtgataagat gaaattaacg aggggaataa ttttgtagga ctcgtcattg tctcttttct
90961 tcttcaaatg cggtaccaga acgtttagta gcctattcag ttcattcagc ttgtaaagag
91021 cgttcttata ttgaatgatt tttatatcat acgagtccaa cggaacagaa cctttgattg
91081 ccggaataga agcgctggca gcagctttgg ctagcaactt ttttctcttt tgcacatcta
91141 ccatgttgta atataaaact agtctaaatc ctcaaccttc agcttccaat cactcctttt
91201 atcattgttc attcgctaac aaacttcaaa catatgcctt tattatgcgt cttcctgctg
91261 tttcaaccat tgattgcagg gtaacagaca tttttaaggg tctttcccac agcatctata
91321 agaaagatcg tcaaaagtat tagttaaaca ttgaaaattt gcgccaaaga catagcaagc
91381 gcaacgtatt cattgtccat gtcgtcatct actccctttg acccttatgc tctatccgag
91441 cacgatgaag aacgacccca gaatgtacag tctaagtcaa ggactgcgga actacaagct
91501 gtaagtacag aaagccacag agtaccatct aggaaattaa cattatacta actttctaca
91561 tcgttgatac ttatgcgtat acattcatat acgttcttcg tgtttatttt taggaaattg
91621 atgataccgt gggaataatg agagataaca taaataaagt agcagaaaga ggtgaaagat
91681 taacgtccat tgaagataaa gccgataacc tagcggtctc agcccaaggc tttaagaggg
91741 gtgccaatag ggtcagaaaa gccatgtggt acaaggatct aaaaatgaag atgtgtctgg
91801 ctttagtaat catcatattg cttgttgtaa tcatcgtccc cattgctgtt cactttagtc
91861 gatagaagtt cactcgcaat gctatatata tagggagctt ccatatgtag tgtaggatat
91921 atgtatatat acatatatac atatatgggc gtatatttac tttgttctta ttttctgtct
91981 aattttataa ttttactgac agtagctaag ccctctgtat tgctgttctg tgttattgca
92041 ctagtgtcat aacgcagatg gtttttagca gagtcaaatt gggcagaaag cataatttcc
92101 atcttccctg gcaaagacag attttctctt ttgatcacgt tagctaaata attcaagatt
92161 tcgttcggta ctccagcttc gcccttgtta gcaggcttgt atttcaacag aattgcttgt
92221 atttgggctg gattcaaggc ataccagaaa tcgaatagta acttgaactc gttcaagttg
92281 cttatcttca actgtaatat cttgacagcc tgaattattt gaatcaaatt gggacgaacg
92341 tcttcaattc ttggttcaaa ccagctcacc aaccgctcga tatttctatc cacttcgtac
92401 ccatacttcc aatttaacgc gggacatttg gttattaggt cattaaatag cattacgttc
92461 agatacttca gggtgtcgtt aaaaattttt gtatgcattg agtcaacaac ttgaaattta
92521 cacaagacag catcaaattc atttaagaat gtgaataatt tagcgaactt ctcatcacca
92581 gaatttttaa atagcttctc gtttaaaacc atatcgaaga tctcgatgtg ggctgaagca
92641 tgcttcatga atttcactaa ccaagtggaa taaatcttgt cgaatacttt taaagtttca
92701 ttctccaaat cattcaaata tatcagtgtt aacttatctt tttcatcacc accattcgct
92761 tcatataatg ttttttgatt tgcagcaaat gcgggaagtc tggaaagatt acttaaccaa
92821 aagataccac caagcattgt ttcgtccttt ggtagactca taacaatgct ttctacagtc
92881 aataggactt tcgagatgaa tttgctactt tgaatcaata acccatttct cactaagcta
92941 cttaccactg tagtaatgac gtgtatagga cccagaacat tgtctccatt cacctcagta

93001 acatttactt ttttcaagta accttctgtg acttctaatg tgtaacaatt caggtccatt
93061 agtaattcta gaagatcagg atttcctgat tctctacgaa ttacattgac tagcttcggt
93121 tttacttgtt gtcctgcgat tcccaaacct ttcactttat cattcttatt agcagaatat
93181 gtggtggtga aatcttgtgc tattacgttt tcaataaacg ctagttcttg cttcattgat
93241 ttgacttcat cacttaatct tgaaggtttg ttacctttta tagcagccaa actttgcata
93301 tgtgactgta ttaaagaaga ttgcttcttt tttctctcct tcaaggcatt cttcttgttt
93361 aaagtattca tgatttcaac ttgtaacgtc ttcagcttgg aaatcttatc attataacca
93421 tttactatcg tttcgtactc ttgcctttgc ttagatggaa tatttgacaa atctccctgc
93481 aacaactgtg ttaatcttgt atatgagtca tttaacatct ctagattcaa aatgaaagag
93541 ttcttaaatt cgatagcttc ctccagtaat ccaatcccgt aactagccct aatattccgt
93601 tcttcaactt ctttttgcag aactatatac cttcgtctag ctaactgcat tctcatggcg
93661 ctttgtacaa gaatggagct ccttttcaat gttctgtaat ctgttttgtg accatatgat
93721 ctgatgtagc tttggataat tactgccgcc atgagcatga atttcctgtt aacactatct
93781 aaaatcaatt ttcttttaca ggtacattgt agttttatta tttgcccaat ggcagctcta
93841 tagtattccc gtttccataa tgctctaatg ttcgtttgta gtaatatggc agctcttgtc
93901 ttgagctcat gatcgaccct tgtacgaacc aacaaacttc taatttggct ttggcacttc
93961 tttattgatt ccatggtctg caaatactga agcctatagt atcttgccct tatctttttc
94021 tgtattatga tgcatatttc attcatttta ttagtcctta gcttttccaa aaatgcaagc
94081 attcctgctt taaagaaaat tttggtatta ccaatttggt attttgctga atcagagata
94141 gtagcatcta gaatagattg acagaagttg acgattgcct cttttgggag gtctggatta
94201 taaagaattc cactccataa gctgtagtcc gtcaggagga aatatctttg aacgaattcg
94261 tcaaaagtcc atcttgatgg aaagcctgca catgaaatcc tgattgtttc cagcacacca
94321 caagctctta attgcgataa gaccatcaaa ttatcgaact cccatggctt tttttcagaa
94381 tttggtttta tgcaacgaat ataatgaaca ttagtagaat ttatgatggc catcaattcc
94441 ccgagcgatt ttttgaacat agatcccagg gttggtttct tttgacttaa tcttgctggg
94501 atcattatct ttttttcagt attttgctct tcaggagcat cgtcactcct aagttctctg
94561 ttgtctaaaa tttgtttgaa aattggattt gtcgttgctt tgaaaacatc cagatgacct
94621 agggaaacgc tgtctctatt cttttcaata aacccttcaa cttcatattc tacatcaaca
94681 gcataatggc ttactataaa cttcgtttgt ccgaatcttg gcttcgaaaa aacttcattt
94741 gagggcggtt tattaaaggc agaatatagt tttgaggccc atgattcatc tgagcctgat
94801 ggtaatctac tttcttcatc taatagagaa agtattccaa gtttattctc aatcaagtct
94861 atgcaaggtt ggttgtcgct gaactcaatg aaagaccact caatttcctc ctttacatat
94921 tcttcttgct ccaatttgaa aacgtgctgg ttgaactctt gttgtaattt ttcattcgca
94981 tagtttatac agaattgttc gaatgagttt ttctcaaaat gctcaaaccc gtagatatct
95041 aaaataccaa taaaggaaaa gacatgatct tgttgatcca gctcaggatc gtataatgtc
95101 ttgttaatgt tgtctactag ccaatcaaaa agcgtggaat aaataaattt cgccacagag
95161 tccctagcaa ttagcgcttg attgtaattc aaattagtga caattttttc agaccttgtg
95221 acgatttgtt tcttgacaat ccattttgca aaattgaagg gatcaatacc caataattca
95281 catgcgattt gcaggttttg ctcttccgat gatagtgatg cgtcgtttct agtcattttc

95341 atctcgatat tacctatatg tagcaagcct gcaagaattt tgaaaattcc aagctgagtc
95401 tcgtggttta tacctactaa tgacagggca tcagtggtta tcttgtattc tcgagcttca
95461 tctataccag ctatgttcgg ctgtccacct tggttagtat aatggtagtc cttgggagat
95521 gacaaatgca gttcttgttt taccggttct ggcaatcctt ccaaaatttg gtaaaaaata
95581 tggtaatttc tttctgtctc tggctgataa actaacctgg atttttccaa gagataggtt
95641 ctaattttgg accccctgat ggtagtattc tcatcaaata aaatttgcag atatttgccg
95701 aaacgagagg aattatcatt tctggtagtt ttggcgttac caaaggcttc catgatcggg
95761 ttagtagcta aaatttggct ctcgatttga gacatttcca cttccccttc acgattgttg
95821 ctttcttgga ctgaggcaaa gtatctcata atgtatttag cagagacggt cttaccagca
95881 ccggattcac cactgactac aacagtctgg ttagcctttt catggaccat gaacctgtac
95941 gcctcctctg ctatggcaaa aaggtgcggc tccagctcat ctttacgctt actagaataa
96001 ttctgtatca tctcgcgaga gtacaaatgg tcgactttat caaagggatt agcggcaata
96061 aggacaatac cagagtaagt atatatctgt ccattcatgt atcttttttt tatggcatgc
96121 agcactgccg gttcgttcag atatgatagg gtggttaagt catcagtaga ctctaaaata
96181 ggtgggtttc gtagtacagg tagcgtggga tgatcatcat cattttcgaa gctatttgtt
96241 tcaatggata cagtttctcc atcctccaat ttcaactcta agtggaacgt tccttcaaag
96301 aagtcattct tggtgacttc gccgcctatc cagccttgtt ctttgtgagg gtaccaacac
96361 ttagttccta cttcaaatga catgcgggag aactggttat aggatttttt ttgttttttt
96421 gtgttttaga attagattga attataaaaa gaagaacaaa agggtatcgt attgaaaata
96481 aattgtctcg ccaaactggt aacaatgttt tcagctagaa caataagaaa agaagagaag
96541 gtaaaaaaaa aggtgataac tccgtaggaa ttgaggaatt gagtatgcag aatcagaata
96601 aaggctgact ttcaaaaaaa ggttgtatta caattgcagg ttttcgataa aagagaccct
96661 attctcatct actactgcta aacttcgaga tattttcgaa tttttcagtc ttttcttttt
96721 ttttttcgc attagttcag aaccctaaag aatggtaaac attctatgga taacccggag
96781 agtgagtttc ttaaagacct agttttattt taagggtttt aactcaatct tgatgttttc
96841 attgtgtacc ctaaagaaag tttaagaata gccctaactg ttaccttttg aaataaaata
96901 aggggaaggt caaaaagcta ttgttctatt gttatgaaac attgtctcaa agagtaagaa
96961 taacacaaat tgatggcagt tttttacgta gtccagtagt tgtccaggta caatgcaaaa
97021 tgctcaaata aagagctctt ctaaaggcag cggaatagat ggtacagatc gcaatagcaa
97081 agatggtgta gaaaagagac ccctggaaga tgtaaagcaa atgattgacg ctggaacacc
97141 agatgttggc cacaaatcta ctgttgaaac taagccaaac gttggatggc aagcctctca
97201 cagtaatttg gctgcattac acgaaaaaga gcagaaatat gaaatggagc accatcatgc
97261 tcgtcataaa ctgcatcgtc aagttattcc ggattacacg tctgcctcga ccgcaatgtt
97321 cagcgattgt atgttcaacg cagcaccaga taaagtacga agtctcagta cgatgaagtc
97381 ttctggactc tcgccaaaac acccatttaa cgtagtcgcc acctttaaag gaccattccc
97441 gcagcatagt gtagaatcaa agcctctcga tggtggatac tctgccaaag accattttcc
97501 ctcatttaag atgttgcaag cccagcagca cccagcccat cgccattaca aagacaacga
97561 caagtacggt cttaaatcac cttcccggtc cttcgtgaag gacaagaaaa ggttggttca
97621 ccggtttttg aaatccatgg agccttcttc gtctgggcaa tctaaggatt cgtctgcact

97681 ggcgccggct ttcgatccaa tattgcccaa tgttatatct aagccttcca agcgacccac
97741 acatcattcg cattcatcag acgggagttc tagcacgcag acagatatat cgttacagag
97801 cttgctttac catgatcttg aaagctcacc aaagaaacat gtttcgccct caagaccgcc
97861 ctctgtagct tccgaatcct ctcctgccgt tgctaatccc attgggcttt cgccaaaaga
97921 cgcctgcaat gcatcgtttt cgcagtcgtc ctcatcttcg ttgtcttctt cttcatcgtc
97981 ttcatcatcg acgtcattct cacagtcagt ggctgttgat cctcttgaac ctcctggaaa
98041 tatcacatat agtagttcga atctttcgct aaattcagat gaattagact actatcagcg
98101 tcatatcgga ttgcagttac agcagacaga agctttacta aagcacagtt tgaaagatga
98161 ggttctgaaa gatgaaaatg accttgttaa aaacattgca aattttgaca agatcgttaa
98221 agagctaagg gacttaagat ccaggaccat tggatggaaa gagcttgttg aagaggatta
98281 tttaatgaat ttgaaacagg attttgacaa ggaaaacccc gaatcatttg aggcacgttt
98341 gagtgataca ataaatacaa acgtggcaaa attacaagat ttagagaaaa gaatggcttc
98401 ttgcaaagac aggttggcct ctaggaagga agtaatgagg aaaatggaaa gtttattgtc
98461 tttggagaat tccttaatga tatccaaaaa aaatgtaaca ttcgcatcta aataccgcaa
98521 cgaggccctt gatattgtct ttttaattat catcatcgtc atatgctata ccttcaagca
98581 tctagtatcg cataaataaa aaatagtatt tgtatatcaa aaaatgatcc tgtgattttt
98641 tcatatgtaa cgtataaatg taaaaatgtg cttcttctgg tatttttaat caagtggaaa
98701 gatgagtgga aaaaagggca atgaaataga aaaggacagg cctgaaaggg aagaatacaa
98761 gaagattgag tatattggac ttcacagtaa ccgtgaaaaa tggcaccaag tatagcaacg
98821 gtaaagatag ccagggacat ggttttgcca ttacgtatat ttgtcaatag aaagcagatc
98881 cttcaaacca atgataagac tagcaataag tcgaatgcca ctatatttga agcaccatta
98941 ttatcaaata actccataat ctgcttaaaa tcaccaaata caagaatata tttatcgcaa
99001 caagataaga agaatctttg tgacgagatc aaggaggacc tgttattgat tgtttacgaa
99061 ctagcgtccc cggaaatcat cagttccgta ctcagcaaaa taagagttgg tcattctact
99121 gatttccaaa tcaacgttct gcccaaactt tttgcaggtg ccgatacgga taatgcggta
99181 acttctcaca tccagtctgt gacaaggctg gctaaattca aatacaagtt gcactacaaa
99241 cataagtggg agctcgacat attcatcaac agcattaaga agatcgccaa tttaaggcac
99301 tatttgatgt ttcaaacatt aacattaaac ggtttctcat taaatgcagg acccaaaacg
99361 ttattagcta ggaaaataga aaaacagccc caggtaccta atttgttaat agaaaatggg
99421 gacgctgatg ccctggatac accggtggaa gaggatataa aacctgtaat agaatttatg
99481 tacaagcctg ttattaattt aggtgaaatt attgatgtac atgtgttgca taggcctaga
99541 agacataagg tacgtaccca gtcgaagcaa ccccaggagg aatgaaaaac cgataacaaa
99601 gtgatggctt aatattataa cttctatata acggatatat tttatggtaa atgtacatat
99661 ttcagtaatg gtaataatga cttttctttt tatcttattt ttatttttgt attttttgtc
99721 ttctgctctt tgtttctgtg cctcatatat caaatgaaat atcatctctc gaagaactga
99781 atctatttga attttgatta ccatcagcaa atggattttc aatgaacggt tctgcacttt
99841 caaagtcgtt acttcccagt ccattattgt cattaaaggg gttcgccgaa gcatcttgta
99901 attcaccgta cttacctctc ttttgagtag tatcatacat tctgacaatt ttttcttgcc
99961 caccctcttc agcctgtgaa aaggcaaatc ctctttgttt tttcattctt tgcacttgcc

100021 tcaccttcct gatggcattt tgaaattgct gaacgtgcgg cctagagtcg ctgatattgt
100081 atttctgcat ttcttgaata acatgatacg tttctggttc atacattctt ttatagtact
100141 tccatagaaa atctcttacc agtgcaaaaa ttggtaaaac gatcaaagtt aaccaaaata
100201 caccggatcc atacgtgtgt ttaaccacac cataatactc tcttgagatg ttagcatgag
100261 gaaatataga agcataaatt gggaagaata ttaaccaaaa taagagggaa ccaggaatgg
100321 caattaaggt gaatttcgtc cattgattag ttaccaaagc agcctttccc aaaacaataa
100381 tgacacttgt agtgtagaca gttacacccc aagaccaatg atcagctagt tcgccgtgca
100441 tatttaaggc aaacccatat ctgtaaatca atatggtgcc aataaatact attgcagaat
100501 ggaaaaagcc attaataatc catccccaga agatgtaaac agagaaaaat tgacccttct
100561 gtcctaattt gtacaattgt gggtaccgct caagtaatct actactaaca aattgatcaa
100621 atacaccaat gacaaaaggg ggccaaacag tgaagaataa gttgtagaaa ctcattgtcc
100681 atgattccat aatggattga cctgaaaagg cattggcaaa aacgtaccaa aactgcgtca
100741 tgtataatgc tgtattcttg taaaaagagt acaaaattgc gacagaaatt ctttgataag
100801 accaggagcc atggacaagt aatagttttt ttaaaaattt aaattggcca acagctatat
100861 cagctgaacg agccgcttgc ataccttcca taccactaat accgacacca acatgagctg
100921 cctgtatcat actaacatcg ttggcaccat cgccaatggc tagcagtagt gaagacgact
100981 ttctttttac cattttaaca accaaggctt tctgtagtgg agatacacgg caacatataa
101041 ccgctttaca aagcttcgcc acagttagca aataatcttc taattctggt tccaaggcaa
101101 agcctaatga cttcccatca atgacgagcg ctaaggtatt catatcatgt gttgacaatt
101161 gatgctcgtt tagagcgtta atcttctcta acaaatttct ctcagtatca tctctggttt
101221 cctcgttgat gatcaacaaa ttcatgtctt cactcaataa acggcaactc ataccaatgt
101281 taatagcagt ttcctgtctg tcaccagtta aaacccaaat tttaataccc gcttcttgta
101341 atgtgtggat agtttctgga acaccgtcct gtaacttatc ttcaatagca gttgcaccaa
101401 ttaatattag atttttctcg attagatttg cggcttcgtc tagcttctcg gctctgttat
101461 ctaatgttgt ggcagcctca ttataaatgc tattccattc ttcatattct ccttcagaga
101521 tatctctcat cgccaaacac aacgtccgca aaccctcaga tgcataatct tctaaatgtc
101581 tcattgtagc ttctacatat tgattagctt catcatccaa tctttccaga atgacagtat
101641 cagcaccttt acagaataac tttatcgaac catccggaaa tctaaatata gcgctcattc
101701 tcttcctggt ggaattaaat tcacaaatgt taagtagttg atactctttt tcctcgccag
101761 tttcctccaa taaaacagtt acagagtttg gtttacggat gataaactta taccctaaat
101821 ctgcaccacc ttgaacgagg gcaccttcat ctggagaggc tgcttgatat ttaatagatc
101881 catcgctttg aaattctgga atgacagtat gacaggtagc cagtaatgtt aagaagtcat
101941 tgataatagg tgaatcctca tcagagggat catttaactt tttcttcaaa tcgtcaaatt
102001 ttctataacc aacttcaatc ccatcttcaa cagtggccgt tttatcttca ggtattttat
102061 caatatagca atggcctgca atagagcagg atttaaattc cataatattt cttgttaaag
102121 ttcctgtctt gtcactgaat atatattcta tttgaccaag ttcttcaacc aaagaggatg
102181 tacgaaccac agttggggta tcagtttttt cgtagtacaa atctagatct gaacctatca
102241 taaaagcctg ataatatttg attaattcaa cggtgacaaa tagagaaata ggaactagat
102301 tcgaaaatag aatccaaaat gttaaaaagt ctttgaagaa taagccagcc ttgttggtac

102361 cctccaggta aaggtacgat aaatgtttgg catctgcagt agacataata acattaccaa
102421 ttgaagaaat taaaattagc acgattaaaa ctgtgaacaa tgcaataatc tgtctgttga
102481 taattttctc aaccgcggtt cttttaattg gggttgcagt agcattacgc aataacttag
102541 tttcatgacc agtgaagata actaaaccaa atatccatgc agtatttctt aaagttgcac
102601 ctcttaaaat catttggtca ggagataacg gtatttgacg atcatttaaa gtcattgtac
102661 cttcataagt atacaagcta gagttcggct gttcggaaac aacttttccg ttcatgttct
102721 tcaaagtttt aacgtctata aatttggcag tttctactct cgactgtttg attttcaaat
102781 ttgtttcacc atccaagttg gcagtttcaa tgtagcaaag accctccggt tccgaagatg
102841 acaaaattat ggtatcagca ggaataggtt cctctgattt tactctaatt atgtcaccta
102901 cacgaatatc aatccatcgt ttctcaacaa agtcatcatg tgcttctgaa aatatttctg
102961 ctgtcgaatt atttaattct ttatcagaat tagctctctt gatatcttcg atacattcct
103021 tcatggcaga aacaatcaaa accactaata aagtaccaat tgtggtgtat ctattagttg
103081 gcgagacgtg aggcacctgt tgaatggcag atgtacataa aaaaaacaga ttagcgtatt
103141 tggaaaattc ttggaacaaa aatttaggca gaaaggtcgc aaaattatac ttggtggtag
103201 atatatggtt gtcactataa ccgaaggagg aattagcgag agaatcgttg atgtggatga
103261 ctcttggttc accattgccc tctgcgtcgc caacgttctt tcttaaaatg taccggttga
103321 aaagaatttt aatgttgaac ttattcctcg agtctagata attatcgtct aattcgttat
103381 tagtcaccgc attgtaatga ttcatttcga aactttcggg ccctttcttc cgtttgaacg
103441 taaaggcatt tttgagacca ttaccaaacc tagcaaataa accgggaggc ttgactgctc
103501 gtagggattg tggttgatat gcatctgagt cgaatctatt ggcattccaa gacgtttggt
103561 cgtcatggtt attgctcata aatagatttt catggacatc attttcaatg ttatcatcat
103621 cagcgtctaa gtcgatagtc tcttcgggaa gtacatgact tggtggtata taattcgcgt
103681 tcgcatgtga gttggtgacc tttgaacggg atccagaatg agaagtcgtg tcgtctaaga
103741 agtcaatatc gaaaagcgtg tcgtcctccc caggtttcct ctttgggggg gtttctctgt
103801 cgtcattcat ggtaaaatca gggaatgaaa gaacctacca gtaaattatt gcttggcgct
103861 aacctctatt tggcgttact ggcttcttgt tgcactattc cccttagaat tgccaacaat
103921 tgttgattat atgtttcttc aacttagtgg cattaaacag atttgggttt tctggcaaaa
103981 aaagccaatc acgtgatcag gaagatttca gtgatttacg cttcaaataa cctttacctc
104041 aataatcccg taatagtaag cttctaatag taagcttcta atttgcaatg ttattagccc
104101 aacagctact aaaggtatta tttttttttt ttttgataag aaatttaagt gttacagaat
104161 gggccatctt acaaaaataa tagtctttat gtatttttat atatgtaaaa gaattgaaat
104221 attttataac tggttgttat tatggtacag tgcgctgccc aatccacgtg gaaaaatcct
104281 gttcattcaa taatagaaac tgaaatcatg taatgttgcg tagtatagtg cgtgagctta
104341 ttgtgccact tcttgctcag cattttgaac ttcgtgttcc tcctcgtatt cgatttcgac
104401 tttaggacgt ttagttttgg cactagttcc cttcttacgt ctcttggctg aattcttatt
104461 ttcttcatca ctatcgctgt cgctctcgct ttcactgtcg ctttcgcttt cactggcgct
104521 ggaagcttct ctgtcagagt cagctaacca cttttctaag tcatccacgt caacgtattc
104581 accttcgcca tcatctgcaa cgtattcaac ttccccatca tcactttcct cttcttcatc
104641 ccagtcttct tcctcgtcct gagaattctc ctcttccatt tgacccatta tcttcttcca

104701 gaccttttcg tcaacattta atggtttatc accgtaagca ccactcttta atctgtccat
104761 caattctttt tcgatggctt tttcaatctt ggcagctacc aatgcctttc tctccctgtt
104821 ttgttctctt cttttgacct ttggtgctac accaacgtag tgtctctcct cctcccttaa
104881 tgctaaacgt ctctctgtta tcatgacctg cgtcaatttt gtaaatctct gtttacactt
104941 atgacggaaa aacttgctcc aatgtagtag gtgttcgtcg atctgttgta aagccttcgt
105001 gtaatttttg gatagtttga ttctttccca taacttggca ggagtgtgcg ctctttcagg
105061 cgtcttcata tacaagtaca gtttcccatt gtcacacttc actgttgcat acttggagtt
105121 ggcaagtggg catgattgcc ttgtacacaa cccagtgacg ttatactcat ttctgcaaaa
105181 attttgacca ttaggtgcct taattctatg agagcagaaa ctttgattaa tcacttgcca
105241 aacaatttcg tcggacatat cctcgttgta ttctaaccgt tgtagttatg tactgaagag
105301 aacactgtca aaagaaagaa ctaagcaatg caatatctgc ctctatacca atcacttttt
105361 catttttttt caaaagctca tcggaaaatt tttcaaaaaa aaaaaaaaaa aaaaaaaaag
105421 gtttattacc ctactgcatt ttgataatct gaacataatg agctaatgaa agcaattctc
105481 atttaaaaac aagtattctc tcttattgaa gtatgcatta tctatcatta taaattcttt
105541 tattctgttc gagtccatgt ttttaaaaaa aaaaaaacat gtatgtatgc tccatctata
105601 tatgctccat ctgtatattt tatatgcaaa gttttttaca agaggaattt gggaactgga
105661 ggaaagtggc acaatacctc atgtggatag ttcattaatc tcttcttgtg ttaatgtgct
105721 aatataaaca cacttactca gcaattcgtg gttaactttg aaagtgtaaa acttggacca
105781 ttgaaccctt tgaatgaaat tcttgactat ttgaacgttc gtatcgaatt tattgtaatt
105841 cactacttta tcttctaaga tccagtcttt cttctctgcg tttgctgata ggtccgaaag
105901 atatacgacg atgaaaggga cgcaacccac cagtggattc acagagttaa gtagatttct
105961 tattgtagaa taattcctgt ctagcgaggg aatcttcttt agctcttccc aagtcagtaa
106021 gtcgcctggt tcaatcagac gccatgcatc agtgaatttt tgaacaactg aagaacttaa
106081 ggctagaatg atttccatca atgtgttaaa gttctgaaat gtcctgcagt ggtctgcaac
106141 gtgtataaac ctttgaatga catttctttt cattttgctg ctcttggtaa gaagtatttc
106201 tgatataatc caatccacag ttaagttaaa tctggatata gctaaatcaa tgccagataa
106261 ggtctcgttc cttactagta gttgcaacca acttataact tgtgggcctt catgcttcat
106321 ctttaaatct aataaatctt tccagtctat ttctcctaaa atttccttct ctatcaaggt
106381 catttgttgc gctacagaca gggaatcgta cattagtata aatggaacgt gactttcatt
106441 attagaaatc attaatctcg aatcgtgaat cctgtattga ccgattaatt cctggatttg
106501 cgccgcttgc acagcaacat caacattagc tgacacgtct attttttcag gagaggagga
106561 agtaaagcct tccttctcgg attgatctgg agtaaagggg atattcatta tagtttgtct
106621 ccttctttct attaataacg attttctttt ttctgcgggg ctattttgga atgacggcaa
106681 gttgttaagg tttagcattt ccacttcatc tgctaacttg ctcgttttaa ttccaatagc
106741 gtcactactc ttggtgttct ctggtttttc gggaattttc tcgtatgtac cttccagttt
106801 cattaatgcc acggttattg gatcgtcatt aattgaatca tcgagcatat tagcaatttc
106861 atttaaattt ttcttattca aggagcctgt aaaagccgac tcagagtttg ctctccgtgt
106921 gttccgggca ctagatgtgt cattaatgtc gatatcgtcc attgtcaata cacttcctga
106981 tatgttacta ttagtttcac tctcgttctt cagaaattta cttttcagct cttcaacgtt

107041 tttcattgga gaagcaacgt caatgctacc gtcatctggt gaaaaaaaat acttgttgtt
107101 tttcgcatgg gtacttggac tttctaggtc attattggaa tcgcttaaat caattgtttt
107161 taactcatgc aatgaagaag tatttgagat tatatcttga ttaccccttt gatattcttc
107221 tctcaaattt attatccttt ggtgtggcac tggtaaatct ctggacgttt cgaaagtttc
107281 aactgtgcta ccagtggtaa tcgaattcaa atcacttaca acagattcta ttgtgggtgc
107341 tatggataat ctgattctat tggtttcact cacgctggtc ccgttgggaa attctctctg
107401 tggattttga gtttgtttta gaggactatt ctgagcagat tcaaaaagct tgctcgttga
107461 gatactgata acggtggacg atgtatcaga gtacaacgcg cttttttat tatattccag
107521 gttgagaggt tctatttcac ttatttcctc tataactgag tttgacttgt tttgctcagg
107581 atctactatg gataactctt tggttggtgt tatggcaatg ctttgaactc ttgaaatgct
107641 gatccttcca ctagccggtc ttacaactgc ttgtttgatc ttgtgattgc aagatgaatg
107701 ttttttctt gtagttatat cataattatt ggcttctgat gatgcatcga gtgagttgtc
107761 cagctttttt gtgctctcat attctgattc ctgattttct ttgttatcat aattcttttc
107821 aaggccatct atagtactct ttttctcatc taaagacttg gtatcttcaa atgtaaactc
107881 tcttagattt aaaaatccag tttttgtttt tagagttgga ctatctgtat tattcacgat
107941 gtttctgact tttcttgact ttttcgttac agtatctctg ggtggagagc tggatgacaa
108001 ctcggaatcg tatgagtatg ttattgattc tgtgtcttcc ctactatcgt ttaagggaga
108061 cgattcttta agatcaggta gaacattttt cagaggcgaa gaagtcatgc tttttggga
108121 ttgggaatga tttttcctcc ttggtgtcat atttgtaatc gcttctgagt tggttttatc
108181 agtttcatgg aataacaatc tttgaggtcc attctcgtct gtacgtgaag aactattatc
108241 tttatcaaag agcctcatac tttctgctga gtagctaccg tgcattttat tgtaatatgc
108301 tgtcgtgaga ctttttactt tccttcgttg cagtgcatca tacgagggtg atggctgcat
108361 attcgattca ttgttctgca gttgcttgtt taaagtatta gttaaagaga tgactgactg
108421 agcaattgaa cttacggttt gatacagatc caaattgtcc atagcgctaa aattatcatt
108481 actcggttta aagctatttt gttgtaaaat ctttatatca tgtatgtgtt cccgacgttc
108541 acaatcaaca ttaactgtag gacctcggtt gttattcgaa tgtgtttgaa ctttttctat
108601 cagctggttt tgtaaatgca gtagggactc cacttcatct atggttcttg cacttaggat
108661 gtcaaattta gaagaatcag agttaagaaa ctcttcctct tcttccttgg aagattttct
108721 attgagagat gatattgata caacgtattt gacaaacgca tccatttcag gtttctggtt
108781 tgctggttta atagcactca taaatttttt atcattactg ctatcatggc gattatggtt
108841 tttcatccac ttagctaaaa gaccaatagc gctgcgatgt aacacagaca ttgaagatgt
108901 attagatcta ctattactgt tgctattgtt atctagagat gatgtcgtcg atgtaccttg
108961 aagcgttccg ctttgttcat ttaaatcttc aggaatatat aaggaattca gaataaattc
109021 aactttttta gccggtgttg gaggcattat tttatcgact gcatatgatg acgggtagtc
109081 tacgcctttc attagtgtgg atactagagt gacttttgac aagtggctga tcttttgttt
109141 actatctgac atgtcctcgc tttcattatc aactgaaggt tctttcgcta ttcggtgctt
109201 cgagtaaaca tttgatgtat tatcgggaaa taaaagcata gaagggcttc tctggttttt
109261 ggaagaattt gaagactgta gcttctccgg cagtctaaat acatcagaag ttttatataa
109321 tgataaaacg ctttggttac gaaaatcagg actggcaaaa ctttgcctgg cgtaaatgct

109381 cagtcggctt cctcttttgt gtaggctttc tagttgcgtg aaatccttta acgaataatg
109441 cagccatgca tcaaagtcta atttatcagg ttcatttagt tcaatattct cccaaaccaa
109501 cttagaacaa tgcacccagt ttttttcag attgattata cagctggaaa ttatttttgg
109561 gtactgttca atgtgtttgt cgtttagaaa ctcgatcaac cgaagtctta atgttatgtt
109621 cggtaaaaaa tcttgcacaa aataattcag tatggaatgc ctcaacaaca caaacgtcct
109681 cacgagggca acttcaccaa ttcgccttct cttagctttt gcagcatttg ttgtgatctc
109741 cctaatacac catcgaaatc tatagatgag taagtcgtgt aagtcttgag gcgttataaa
109801 attcctataa ataaggaaaa aatctgcaga tgcgttgtag tctacacctt ccaaagggga
109861 agaaaggtgt acaattaacg cgggcaggtc tgcactattc actggtttgg atacacaatc
109921 gctttcatag ctaataacat ttgaggatgg agtcgggtag taatctttct ggctaaatat
109981 ttccatggtt cagggtagcg aatctttgaa tggctagagg ctatgtaaag caaacaaaaa
110041 ggttcgcgta aatcaacgag cgaataacac gagtacggtt ggggtgggct aaagggttca
110101 agaaattatc agtttctgtt tacagttttt ttttcattgt tgataaaaag atcaagaatt
110161 tcattattcg cgaaaatcaa aatgcagaaa ggaaaaaata gaagggtaat aaaacagcat
110221 cggatcgcaa agatgagtaa ggagaacagc ctggttaaca attaaagagt gtttatcgaa
110281 attcattata tagtggttta tatagaccac ttcttctgct ggttgatata gaaattttat
110341 ttaattcttg ttttttactt atgtacttac tacgatgatc ttaattcatg cttcttgctt
110401 gtcggcaatg tcccaagtgg aaaaccagtt taagtagcgg aagttactac ttggtccctc
110461 cataccaaat gatattgggg agaagtacca gaagcaacca attacaagtg ccatgaatcc
110521 ggcgtatagg acgaaccgca tgatacggcc gcacttagat ctggaccatt tttgcaaacc
110581 ggcgtcgaaa cagtacgcca aaatgatcag tgcaaaatac aaggcaggca agtaatgatg
110641 aacgtaggtg actctagaca tgataacgaa tggcatgtag tgtaggcccc aagctagtag
110701 tgggtagaac ccgcccatta agaaaacgtt ccagttagat ggatttctta ggtccacata
110761 ttgtctttgc catctgatca gtaagataac gaccgtggcc atgaatgcga ggacggcaac
110821 actagaagcc cacgtggaag ctggggtacc caataggaag tattttggat tatcatcacc
110881 ccagccacat agtctcaaac ccacattcaa agttggccat tgccatgctg aggaagctaa
110941 gtaatcaaat ttgtctggat ctggcaccaa agcgttatta gtggccatca tggctagatt
111001 taaatgaatg aagtctttta agaagttggt ctttgggtat tgaaaatctt cgggtcttgg
111061 tggcaacctt tcattttcgt gggtctcgat gttccaccag gtcctcttgt ccctcttgaa
111121 tgggtttttc atgcagacaa cctcttgttg tctgaaaccc cattcgggca aactgttacc
111181 ggtttgagcc aagtaacagc ccatctccaa gttcttgata cggaaagagg tggtcaatgt
111241 gtgcaacttc tcagggtctt catctcctct ttggtccatg atctcaataa cccaattgtc
111301 tttgttgtca ccaacaacat tgtcaccgta accagaaacc tcccattgtg tctttgacac
111361 tggtgcagca actgggtggg tgtgcaagtt tctgcccgtg cttttgtgta ccaatctata
111421 ggaggtacct ggcttcaaat actcgatgtc agtttcgttt tctgaccatg atggtaagcc
111481 tctttctctg ttgaaaaacc attcgttgtt agcatctttg taaccataac aggttacttg
111541 ttgttggttg gacccatctg gataagtttg tatatgtgag tgcaatagag atcctccaag
111601 agcttggttt ttgatggaaa caacggagga acctagagca atgtcacggg ggccttgtcc
111661 gacgtcagaa cccactaatc ttgcttggaa aagagatggc atgttagcat cacctgtacc

111721 agaatgcgat aataggtcaa aatgtatttt gaagcacaat aggaaaatgc agaaggggac
111781 gataataaga ccaaatattc ttgccaacca gtggttaata taggttttcc atgacatgga
111841 tttatctgcc aaaaaggtcc ataagtcaat cacagtatag ataccgacca tagtgatgat
111901 aaatagaccc accattttga cggaaatagt gcaacccaaa gaaataccag tgatcaacag
111961 ccatttccac cactttctag agaacggctt ggacctctgg ttgtggaaca taacaaaact
112021 aaagaacgat gcgacagtga agaaaagtag catggagtcc aaaagaatga acctgcccaa
112081 agtactatac gagttttcaa acaaaaccaa cacggtcatc agccaaactg ttggtaaaga
112141 aaatccaata gctttggcag tgaagtaggc caatggcaca cagagcgcgg aaaatgacgc
112201 gttgaacagt ctcattttaa cataatccaa atagtctggg taaatttccc cagaagggaa
112261 gtcccaagaa ccgttgtagc ctgccaaata accagacaac ccgaccagca tttttcctag
112321 gggaggatgg acatcgtggt aaaattcgtg tctcaagtaa taagaaccaa atttaccaaa
112381 gtgcgcctca tcccaaacaa catggttgtt gatgccgatt ttgtacatcc tggtaaacaa
112441 cgccaatgca gtaaagatca ccggcattac aacggattcc aggcgtaaca gtgagctttg
112501 tgcagcgggc ttttccttcg agaaatcttc cgcgtctctc tcatcagcgc tcgaaagttc
112561 ctcactgacg ctgatggaag acgattctct ttgtctcagt gtattctctt gcttaatgtg
112621 ggcggcattg tttttgctgt acccggtaga cgaagacgag gacatgattg ctggaccacg
112681 gttcgaaaca gaatgacagt agcgatgtgg actaagccgg ttctccttgg taatgttgtt
112741 agtctcgaga aatgaccttt tttttaccct caaaaagaat gcaacactat taataaacag
112801 tacacgaaac ggatctttcc gtaagttctt tcgttctttt ttttcgtatt cttttcttat
112861 gttttttttt ttcagggcga cgtgtccaat aatatgtatg tttgtcgcta tgtacgagat
112921 attattgcta agtgacagta aatcgtatca cgactgtaaa tgatggcgtt tcggtatgta
112981 cagtatctat ctacctgata ataaagtcaa ttatgaagac gaatacgtag cttatgatgc
113041 tgcccagagc tagtcctgta gaaacgaaaa tattagtgaa tccaccggcc gcttccttct
113101 cgtcgtcgtt gtccagttgc tctggcacct tcatgaaact catagagatg acgtgcccgt
113161 tggttacgcc gaaaaggaac tgcagtagca tgtaacacaa gtcaacgatt acggagccat
113221 tgtgctcttc atcgccgctt gaagaggagg tgatggctgt gaacatcaag aacagtggta
113281 ttgcggccac ccgcaacaat gagtagatga aggttttgcg tggcgtaaat ttctggtcac
113341 ggaacatggg ccagtcggca atgactcttc cgtaaaggtc gcctaggttc cacagcgtga
113401 atatgagagg tatgtactgt gcgttgctta aaggaagccc tgtcacgtag gtggcagacg
113461 caaatacagg aaaaacaaga gttacgacaa acgtggtgaa tatggaaaga accaggtact
113521 ttagtttggc aaataagacc tcgaaaggca cttttagttg gagttcctca ccttcgtcgt
113581 tgtcgtcgcg agtgccgttg gtgcggcggt ggtcttcatc ctccatttgg tcaatgcggc
113641 caacaatacg gatttcctcc tcattggagc gcagagaccc taacaacaca tcagtgatat
113701 gtccgtcttc cacattccaa ttctcgttca ctttccgact gattttgctc acgctgaaca
113761 tgaccacaca aatggtgacc acgagtgttg tggtaaaaaa gtatagaaga atcccgcccg
113821 tagtagacac agaagagttc tcgatgaaag ctagggcaaa aagcactagg gagggcagga
113881 caccagcaac ggcttgcccc accatgacac cttgactgta ctcggaaccg aagacgttgg
113941 ctatggccat gataccattc tgtgtcatgg ctgtccccat ggaactgatc actacaagca
114001 tcattataaa catgaaattg aaccatttag gtaaaaggaa atgcaaaatt gtaaagaagc

114061 acatgacagt aaagacaata atctcccaca caagcccgtt tatgaccctt ctcgagtatt
114121 tgtactgtct tttggccaag tagatgttga acagcattga cgatatggta gaaaaggaca
114181 tcatagagct tgtgaagatc tttgcccaaa tggaggtgtc tttgaaaata tcgtgcttaa
114241 aatattgcga ggcactgagg atacagttcc acggccataa aagacctatt cctatggcga
114301 aaaatgtaat atatgaaaga tttttcaact tttttcttaa cggcaatgta tccagtggtt
114361 cggttgacac agattgctct ctttcagaat aattatcgcc ttcttcatct gagtgctcgt
114421 tgttctctga ttcatgttct tctccagagc gtgatatctc ctctgaatgc gtatcggcca
114481 gtgcaggctc tggcaccgca aggattggct tcttgatggt atcagtgtcc gcactagtac
114541 tcatttttgc aatctggtat gctttcttgc gtttgatgat aagctgtcgc cgtaacgtaa
114601 atcattcatc ctttcctatg attttttaag tcaatttttt tttctttttc gggaaattaa
114661 ttctgttttt ggcattaata acaacaagaa gaattaagga ttcgttatag aatacaagat
114721 tcttggtttg gttaaataat tttggctctc ttaaatatta taaaaagata cttaagtaag
114781 aaagaaacag gacgaaaaag atacgcaaaa gagagagaga gagagagaga aaaaaaagat
114841 gagaaaaaaa agtataacgt gaaggcttta tgggaagtgc ggtgagattg ggatcgtttc
114901 aattttataa tgaggtagtg tacagagagg agggagggag tgggatgaaa gtgtgcggtt
114961 tatactttct tactgcctgt gtttgtcttc ataaattcaa atcttgctaa caatggtata
115021 tggtcactgg ggaatttgtc gttggggaac ccgataaact tactcacgta ttcagggtcc
115081 acttcaccca atagcccacg cacccttaga gcatgtgtag aaaaccatat atagtcgata
115141 acatctgtga atgacggtgt gaaattggtg aatggtagtt ctccgataca attataactt
115201 gatttgagag ccaagttatg tgagaaattt ttctccgaca tgtaaccgaa atctctacca
115261 tttccctctt gatgtatttg gacacggcct gtatttatca attcgtatac ggcggagttg
115321 atgtatgaat tgaagtcacc acaaatgagc acaggaaatt tcttaatgtc ctgtctaaaa
115381 ttgtgcgatg tctcctcctt tagcagcgtt tccagatgat ctaacaggac acctacttgg
115441 aatgtcttga catcattaaa ttttggatcc cagtgcaaat gcgtggtgac cgcccatatg
115501 gtgtcgccac taggaatgtg ttgtagcttt aagaacagtg caacgttgtc tttgttcatt
115561 gcacggttta aataatcttc agttctttgg aacttcttgt gtttcatcca agcaccgctg
115621 aaatccatgg cgtctttggt gatcaacttg aattggtccc ttttgaaaaa aatgcaacac
115681 ccgtccactt tcttggagtc cttggaatgc atggtcttgg ctcttgcctt tgcatggaag
115741 atgcctgtat aaccgtgctt gtccaatagg ggcacccaat actcttcaaa agtcttagac
115801 tccacttctt gtaaacacaa cagatcactg tcgtacgaga gaatctgctc ctttaattta
115861 ttgcgcctgt aatcccaact taacgcccac gacggtgtgt aacggtacat ttttggggtg
115921 gcatagtgtt gacataaggt gttgtaggat aagacggtga acgtcctctt ggctaaatcg
115981 gtggccagat gctcagtgga ttgctgcaaa gaatcgtact ccctctgtgg ttccccatcg
116041 gtattgattt cgatgaacct acgttcatgc ggtaagggaa tctctggtct gttgtccctt
116101 aggtagaaaa tcaatcccgt gacagatttt tctgtaagga tctttaaaaa ctgtttttct
116161 aaggggtttc cttctacacc aagaaactga aggttacaca ggttcccaaa ctcccatggt
116221 aatgtggtga ccatgttatc aaaaaagtag aagtatttca attggaaaca tgagcctagt
116281 tccgctggta gagatgttaa cctattatgc gacaggtcca aaacgcgtag gttgcttagg
116341 ttcttgatct ccgctggcag ttccgtgagg ctattgccat tcaaatatag tctcgttaga

116401 aaatcgtact tgaagatgtt ggcgctgata ttgaagattt gcaagttgga caaatctagc
116461 gcgtgccata attggtcgtc gtatttcgag tccttgggca tgaccattct gttttcaatg
116521 tcgtcatctt cgtcgatgct gtactgagac agtttcttgt gttgtaagag cgcctggttg
116581 gcttcgtcct tgccattagc aaacagctct atcttgggag tcaatggtgt agaaactgca
116641 ctgttagttg ccgtgcctga gggagtgcta tctcccgtag gttgttgttt ctttgctgtc
116701 ttgctgtcgg tgagagtgtc ggccatttcc atcaaggctt gctttgtgca gtctaccagt
116761 gatttggagg cttctgcaat gtgagattgc tggaaatcgg tgggctgcaa agctggtggt
116821 gcagcctgag gtccggggcc aaatgggccc gggacctgct gctgtgcctg ttgctgcgct
116881 tgttgctgag cttgctgagc ctgttgtgta gccaaatact tcttcatagc gttttgtcta
116941 gcataaatgt tgggttgccc cagagattgt gcggacactg cagccagatg caattgaagc
117001 ttccagatcg gattgtttag taaagaaggg tcatccaagt gcgggtgtaa taaagggtta
117061 ctggcgttca cattgatgtt cacaggcgta ccaacggtag gaaccatccc tatagaattt
117121 actgcggctg tggaggcatt catcatcacc ccgcttccac cgccgttgtt gttgttgttg
117181 ttgttgttgt tattgttgtt attgttgtta tccatgttca tattgagcat gttcgcgttc
117241 ccattagcca attggtcgag caactggcgc gagttgttgt tgctggaagt gtgtacatcg
117301 ctgttgttca tgagtcccgg tggaggtatc ccggtgagct ggttcatgtg tagctgctgc
117361 tgtagagcgt ttggtgtacc ctttcctagt agcccggcat gctgctgttg ctgctgttgc
117421 tgttgctgct gcgggcccac attagggtag cctagtaaag aagggtcgtt cattagcggt
117481 agtttgcagg gataacgtca gtcggagttc ccttgctggt gtcccttatg ctgtgccctt
117541 gtgcttgaga accttctcta aggtggaaac tttttcaaat tttttcgttg ttggcccgtt
117601 tttcgcaagt acgggcgata acaaaaggcc taaacggcta ctttaccttt gagatctacc
117661 acgatgagaa ttaatgctga atggacatct atgtatatga tagtgggtat atagttactt
117721 atcagtgcta gagcacgatc cacgtggtgg cacatccgcc aaacacgcga ggttttccag
117781 agtattggcc cacgagctgc agtccaggct gggagccctt ttgcgggccg cagttgccat
117841 gctctcccca gccccagcag tatacattgt aacagtgaga cttgccttct tgattagcag
117901 tagttagaat accgtgctca ctcccggttg caacaccgac aatagggaag tctaggcgct
117961 cttgcgggaa gagttgggaa tgtgtgcccc taccaaacga ctctaccgta ttgaggcgcg
118021 cattccacag gtggatcgag gtccacatgc acagtaccac tagattgtgt cttttttgct
118081 gttgtttgag ctcgaaccca gtgggcaggc gaccggatgc gtgcactatg cggccgccct
118141 cgtccactat gaccatgaag tccttgccca tggccacgta gtctacggcc acagacccgg
118201 tatcgtacac caatacgggc tctttcagtg atcgggattt gggctcttgc aattgacact
118261 ttgtgttgct gccccagccg tatactcggg tgccttgcac caccacaaag ttctggaaac
118321 atccgtatac tgcgatgcgc tcatcgttgg aattcaatgg cacatgttgc tgagtgaact
118381 cgtagcaacc gcctcctctc tgccatacac ggccatcagc atccacaata actgtcgtgt
118441 cccagccgca cgccacatcc accacgggtg ccggtacttc cacgggcctc cagtcatgca
118501 cctgccgcag tgcttgcgca ctatccagtt ctccccgtct gttatctcca catcctacca
118561 gattcccgtc atttgtcagc atcacgctgt ggttcccacc gcacgctatc ttcctgacta
118621 ttgctccatc atctcctggc acagacctct gtggggtatc catatcctca tcgtgcccca
118681 gtcccagttg cctttgccca ttagacccaa acgcatacac acaactcatc gatacaagcc

118741 tgttatagcc tttaatgatc acattccatc acttgcgctt tggatctgcc tgcattatca
118801 aggctcaaac ggctgcgtta cccccgtcgc cgcgaaattt ttcataattt ttcactttgt
118861 aggattaaaa gagatcatga gcccatctcg caatgcaaca cgtaacttaa atcagtactg
118921 gcgtgtgcta tagtgctcta tattcgagtt tgttgctact ggtggacacc cgactatcta
118981 cagtaaggaa cgtaaacaag aaaaagagag aaaatacgct atagttgaaa acatgagtgg
119041 ttcgcattca aatgatgagg atgacgtagt gcaagtgccc gagacgtcct ctcccaccaa
119101 ggtagcatcg tcgtctccct taaagcctac ttcgccaaca gttccggatg caagtgtggc
119161 gtctttgaga agcaggttta ctttcaagcc ttcagatccc agcgaaggag ctcatacttc
119221 gaagccgctc ccatctggga gtcctgaggt agcactggtt aaccttgcga gagagttccc
119281 cgatttctct caaactctgg tgcaggctgt tttcaaatct aactctttta acttacagtc
119341 tgccagggaa cgtcttacaa gattgaggca gcaaagacaa aattggacat ggaacaagaa
119401 cgcatctccc aagaagtcag aaaccccgcc acctgtcaag aagtcattac cactggcaaa
119461 cacaggccgt ttatcatcta tccatggcaa tatcaacaac aaatcctcca agattaccgt
119521 ggccaaacag aaaacgtcca tttttgaccg ttactcaaac gtcatcaacc agaaacaata
119581 cacttttgag ctgccaacta acttgaatat agactcggag gcactgagca agttgcccgt
119641 gaactacaac aaaaagagaa ggctggtaag ggcagatcag catccaattg gcaagtctta
119701 tgagtcatcc gctacacaat taggttctgc aagagagaaa ctactggcga accgcaaata
119761 cggtcgtcat gcaaacgaca acgatgaaga ggaggaagag agtatgatga cggacgatga
119821 cgatgcaagt ggcgacgact acacagaatc cacgccgcag ataaatctgg atgaacaagt
119881 tttacagttt attaatgact ctgatattgt cgatctctcg gacctctcag ataccacgat
119941 gcataaggct caactcatag cctcacatag gccatattct tctttaaatg cctttgtaaa
120001 cacaaatttc aatgataagg acactgagga gaacgcatcg aacaagagaa aaagacgtgc
120061 ggctgcatcc gccaatgaga gtgagaggct gctcgataaa atcacccaaa gtataagagg
120121 ttacaatgca attgagtctg tgatcaagaa atgttcttcc tacggtgatt tggtcacttc
120181 gcaaatgaag aaatggggtg tgcaagtgga aggcgataac tctgagttgg acctgatgaa
120241 ccttggggaa gacgatgacg acgacaatga tgatggcaat aacgataata ataatagcaa
120301 caataacaat accgctggcg cagacgccac tagcaaggaa aaagaagata caaaggccgt
120361 agtggaaggt tttgatgaaa ctagcgcaga acctactcca gcaccagcac cagcaccagt
120421 ggaaagagaa acaaaacgaa ttagaaacac aactaagcca aaagtggtcg aagatgaaga
120481 tgacgatgta gatttggagg caatcgatga cgaattgccg cagtctgagc atgaagatga
120541 tgactatgaa gaggaggacg aagactataa cgatgaggaa gaagatgtgg aatatgatga
120601 tggtgacgat gatgacgatg atgacgatga atttgtcgct accagaaaaa acacacacgt
120661 gatctccacc acgagcagaa atggccgtaa acctattgtc aagttcttca agggcaaacc
120721 cagactgtta agcccggaaa tttcactaaa agactaccaa caaacgggta taaactggtt
120781 gaatctgcta taccaaaaca agatgtcatg tatccttgca gacgacatgg gtctaggtaa
120841 aacatgtcaa gtcatttcat ttttcgcata tttgaaacaa ataaacgaac cgggtcctca
120901 cttggttgtt gtgccatcat cgacgctaga aaattggtta agggagttcc agaaattcgc
120961 acctgctttg aagattgaac cctactatgg ctctttacaa gaaagggaag aattgcgtga
121021 tatcctggaa aggaacgctg ggaaatatga tgttatcgtg accacgtata acttggctgc

121081 aggtaataaa tacgacgttt cgtttttgaa aaatagaaac ttcaatgttg tggtttatga
121141 tgaaggtcat atgttgaaaa attccacttc agagagattt gccaaactga tgaaaattcg
121201 tgccaatttc cgccttttat taactggtac gccattacaa aataacttga aggaactaat
121261 gtcgctgttg gaatttatca tgccaaatct tttcatttcc aaaaaggaat catttgacgc
121321 aatcttcaaa caacgtgcca agaccacaga cgataacaaa aatcacaacc cgctattagc
121381 gcaagaagcc attacaagag ctaaaacgat gatgaagcca tttattttga gaagacgtaa
121441 ggatcaagtg ttgaaacatt tgccaccaaa gcacacgcat attcagtatt gtgaattgaa
121501 cgcaatacaa aaaaaaatat atgataagga aatacaaatc gtgttagaac ataagagaat
121561 gattaaagat ggcgaattgc caaaagatgc aaaagaaaag tctaaattac aatcttcaag
121621 ttccaaaaat ttaataatgg cattgcgaaa ggcctctctg catccacttt tgttcagaaa
121681 tatctataat gataaaatca tcactaaaat gagtgatgcc atattggatg aacctgctta
121741 tgctgaaaac ggtaacaaag agtatattaa ggaagatatg agctatatga cggattttga
121801 gttgcacaaa ctatgctgca atttcccgaa cacgttatcc aaataccaac ttcataatga
121861 cgagtggatg caatctggga agatagacgc tttgaaaaaa ttgctgaaaa caatcattgt
121921 tgacaaacag gaaaaggtgc tgatattttc cttattcact caagtcctgg atattctaga
121981 gatggttttg tccaccttag attataaatt tttaagatta gatggttcca cgcaagtgaa
122041 tgatagacaa ctactaatag ataagtttta tgaagataag gatattccca ttttcatctt
122101 atcaacaaag gcaggtggat tcggtattaa tttggtgtgc gcaaataatg ttattatatt
122161 cgatcaaagt tttaacccac atgatgacag acaagctgct gatagggcac atcgtgtggg
122221 acaaacaaag gaagttaata taaccacttt aattactaag gattccatag aggaaaagat
122281 tcatcaactg gccaaaaata aactagcttt agattcgtat atcagtgaag ataaaaaatc
122341 tcaagatgtg ttggaaagta aagttagtga tatgttggag gatataattt atgatgaaaa
122401 ctcgaaaccg aagggaacca aagaataaat aaataaaaat atagtaaaaa agtaaataga
122461 taagcagaaa ataaaccaaa aaaataacat atttattttc ttctgcgtga acgcaccatg
122521 tggaaaaaga gagttgatag aacaaaggac aaagaaattt agaacggtaa aaatattttc
122581 tagagtataa tatacatata tgtgtatata tacataggtt agtatgtata gctaggtaat
122641 tttaatctgg ggagagaaat ggtgaacttt tttcaattta ttgattcggt tgttgattga
122701 tatcttgtgg tgctgcttca atttcagctt gagtttctac taaattttca gggaccattg
122761 gatgatcaag gttaaataaa acgctaccaa tagtatttgt agaaatggca aagcccgcta
122821 aaatgggcac ggactcaatc aaaccggcca agaaaccaaa tgaggcatat tggccagcgt
122881 ggagataaat attttcacgt ctttgtatgt tgtcgaaatt ttgtaaacgt aaaactttcg
122941 tgaagtaaat ttgagtgatg aaaggagaaa ttaaaatatg aaaaaggatt ggtccaataa
123001 gtgggaaaaa agatataacc aacagcacaa aaagaaccga caggcctaac atgtacttga
123061 ccaacttgaa tgggaaatag taagcccaga aatatatcga gttaataggt gcatagtact
123121 tgattggttt gacttcactc aaattctcct ctaaaccatt cttctctaag cacacctcca
123181 ctaaatgatt attaaaatga ctgagtctca taaacaataa agtgaaaaca tttgcttgca
123241 aaaaggaatg aaaaatggca atgaaaagcc caatagggcc cagcacaatc aaaaaagctg
123301 tgtacattgg cgtaattgta gcccaataca caaagcttac cagggcaaac attatcaagt
123361 aactcgataa tatacctaac atagtcgttg agtaactgct cttattggtg ttgaactcta

123421 aaaacccttt gaacgggtac atgaatgcac ctcctgtgaa tgcatctttg aaaaagttca
123481 ttagcacgta tctggtacgg ctcataactt tgctgaagtg cttcttgtaa actttatcca
123541 agtactgatt gaaggggatg ttgacatatg gcagtttttc atacgattgg cctccgccaa
123601 acttgtacac caaaccacca atacctgcca gcgctaatga acctgtaaaa ctcatttcct
123661 gttgttgtta ttgtttgtat aatgtacctt tgttccctac actataacaa cacttattcg
123721 tggcaagaac gaaaataaaa aatgatttga aacttccttt tccacatcta tatctatata
123781 tatatgtata tgtataatac ttattactat gatattattc catgtgtcac aaaaccagaa
123841 agatcccgtc gtattattga gtcacgtgct ttattccatg gcgggtaata aatattaaaa
123901 caagggtttt ttaagtaaat gattacatgc taaggaagtg gtgaataaga tttggcaagg
123961 ggcaggtcgc taaccacaac atagcattcg ttgaaaagga aatcaacgtt acaaagtgca
124021 gttttttgta ttattttcct attatcctct tcttttcctt tgtttcaggc gtcgttgcac
124081 ctttttgcta tacaaaagct cataaaagat tagaaacccc ggatatcgtc acaacagcgg
124141 tatattatag ttattttgca tctttttggt aagtcagaaa ttagaaacag tagcttacca
124201 attgagtgaa agtttccgtt cgtcatctcc ctcttttgtt ttttttcacc tttgtttttc
124261 tgtttcttct atttttgttt tgttttgttt tgttttgttc tctccctaat ttgcatatag
124321 gtaaacatca aagaagtaat gccctacatc ggtgcttcca acctctcaga acattcattt
124381 gttaatttga aggaaaaaca tgcgattaca cataaaggta cgagcagttc tgtagcatct
124441 ttgcagacac caccgagccc cgatcaagag aaccatattg acaatgaatt agaaaattac
124501 gatacgtctt taagtgacgt ttcaaccccg aataaaaagg aaggtgatga gttcgagcaa
124561 agtttaagag atacatttgc gagctttcgg aagactaaac ccccacctcc tttagatttt
124621 gaacaaccaa gacttccttc gacagcttct tcatccgttg attcaaccgt atcatcgccc
124681 ttaacggatg aagacataaa ggagttagag tttcttccga atgaatcaac tcattcttat
124741 tcgtacaatc cactttcgcc aaattccctg gcagtcagat tgaggatttt gaagagatca
124801 ttggaaatca taatacaaaa cccaagtatg ctactggagc ctactccaga tgatttgcct
124861 cctttgaaag agtttgcggg ccgtaggagc agtttaccaa ggacatcggc ttctgcaaac
124921 catttaatga acagaaataa gagccagatt tggaacacta cttccgctac tttaaatgca
124981 tttgtaaata atacctcttc ctcctcagca gcatcttctg ctttatctaa caaaaaaccg
125041 ggcaccccag ttttccctaa tttggatcca acacattctc aaacattcca tagagccaac
125101 tcgttggctt atttaccttc catcttacct gagcaagatc cgctgctcaa acataataat
125161 tctttatttc gtggcgacta tggaaacaac ataagtcctg aaaggccaag ttttagacaa
125221 cccttcaagg atcaaactag caatctccgc aatagcagtt tactcaatga gagggcatat
125281 caggaagatg aaactttttt accgcaccat ggaccctcta tggatctatt gaatgaacaa
125341 agagcgaact tgaaaagtct tctgaattta ttaaacgaaa cactggagaa aaatacttcc
125401 gagagagctt cggatcttca tatgatatcg ttgttcaatt tgaataaact aatgcttgga
125461 gatcccaaga aaaataattc agaacgcgat aaaagaactg aaaagcttaa aaagattttg
125521 ctggatagtc ttgcagaacc attctttgag cactataatt ttattggaga taatccgatc
125581 gcagatacag atgaactaaa ggaggaaatt gatgaattta caggttctgg agatacgaca
125641 gcgataacag atatacggcc ccaacaggac tatggccgca tattgaggac attcacttct
125701 accaaaaatt ccgccccaca ggcaattttt acatgtagtc aggaagaccc ttggcaattc

125761 agagctgcga atgatctagc gtgcttagta ttcggtatct cacagaatgc cattcgcgct
125821 ttaaccttga tggatttaat tcacaccgat agcagaaatt ttgttttaca caaattactt
125881 tctacggagg gtcaagaaat ggttttcaca ggcgaaatca ttggtatagt tcaaccagaa
125941 acactcagct catccaaagt agtatgggca tcgttttggg caaaaaggaa aaacggctta
126001 ttagtttgtg ttttcgaaaa ggttccttgc gattatgttg atgtactttt gaacctggat
126061 gattttgggg ccgagaatat tgtagacaaa tgtgagttat tatcagatgg acccacattg
126121 tcttcctctt ctacattatc gctacctaag atggcttctt caccaactgg tagtaaatta
126181 gagtattcgt tggagaggaa aatcctggaa aagagttata ctaagcctac ttcaacagag
126241 aatcgcaacg gcgatgaaaa ccaacttgat ggagatagtc attctgaacc atcgctgtca
126301 tcatcgccag taaggtcgaa gaaaagtgta aagttcgcaa atgatattaa agacgtcaag
126361 agtataagcc aatcgttagc caaattaatg gatgatgtga ggaatggggt tgtatttgat
126421 cccgatgacg accttttgcc tatgcccatc aaagtttgca accacattaa tgaaacaaga
126481 tattttactc taaatcatct atcttataat atcccatgcg cggtttcctc cactgtgttg
126541 gaggatgagc tgaaattaaa gattcacagt ttaccttacc aggcgggttt gtttattgtg
126601 gattcgcata ctttagatat tgtaagttcc aataaatcta ttttaaaaaa catgtttggt
126661 tatcattttg ctgagctggt gggaaaatcc attactgaaa taattccttc tttcccaaaa
126721 ttcctccaat ttataaatga caaatatcct gcgttggata tcacactcca taaaaataaa
126781 ggtttggtat taacagaaca ttttttagg aaaattcagg cagagattat gggtgatcgt
126841 aaaagctttt atacgtcggt gggtattgat ggccttcata gggatggctg tgaaatcaaa
126901 attgatttcc agctgcgtgt catgaattct aaagtgattt tgctttgggt tacacattcg
126961 agagacgtgg tatttgaaga atataataca aatccatctc aattgaagat gctgaaggag
127021 agtgaattaa gtttaatgag cagtgcaagt agttctgcca gctcttccaa aaaatcttcg
127081 tctaggatat ccaccgggac attaaaggac atgagtaatc tgtcaacata tgaggatttg
127141 gcccaccgaa cgaataagct taagtatgaa atcggagatg attctagagc acattctcaa
127201 tctactttgt ccgagcagga acaagttccc ctggaaaacg ataaggacag tggcgagatg
127261 atgcttgcag accccgaaat gaagcacaag ttagaattgg ccagaattta ctcaagagat
127321 aaatctcaat ttgtgaaaga aggaaatttt aaagttgacg aaaatttgat tattagcaaa
127381 atttcacttt ccccaagcac tgaatcctta gcagattcta aaagttctgg gaaagggctt
127441 tctccacttg aagaggaaaa gctaattgac gaaaacgcta cagaaaacgg attagcggga
127501 tcacctaaag acgaagacgg aatcataatg actaacaagc gaggaaacca acctgttagt
127561 actttcctac gcacccccga aaagaacatc ggtgctcaaa agcatgttaa gaagttttcg
127621 gacttcgtaa gtctgcaaaa aatgggtgaa ggtgcatatg gtaaggtcaa cctatgtatt
127681 cataagaaga ataggtatat tgtggtgatt aagatgattt ttaaagaaag aatccttgta
127741 gatacatggg ttagagatag gaaattaggc actatacctt ctgagatcca aattatggcc
127801 acgttgaaca aaaaaccaca tgagaatatt ttacggttac tggatttttt tgaagacgac
127861 gattactatt atatcgaaac tcccgtacat ggtgaaacag gatgtataga tcttttcgat
127921 ctaattgaat ttaaaaccaa catgaccgaa tttgaagcaa aattgatatt caagcaggtt
127981 gtagcgggaa taaaacatct acacgaccag ggtattgttc acagagatat caaggatgag
128041 aatgttatcg tagattctaa aggctttgtt aagattattg attttggatc tgctgcgtat

128101 gtcaaaagcg gaccatttga tgtttttgtt gggacaatag attatgctgc ccctgaagtc
128161 ttaggaggaa acccttatga gggccaacca caggatattt gggctattgg tattctattg
128221 tatacggtgg tcttcaaaga aaaccccttc tacaatatag atgaaatatt agaaggcgac
128281 ctgaaattca ataatgcaga ggaagttagt gaagattgca ttgagttaat caagagtatt
128341 ttgaaccgtt gcgtaccgaa gagacccacc attgacgaca taaataacga caaatggttg
128401 gttatttgaa ggattaatga tcattctagg acattagtaa ttggaaaaga aaatgcaaaa
128461 aaatatccag tagaaacaat gcttggtatg tcgttcttct tactttcttc agtaatgagt
128521 tggtagtttt ctctatttaa atatgaataa atcaatatgt actttctttc tttaatcaaa
128581 atgttaatat gataaaaata cagcatccaa agcagttaat caagacttaa atataaaaaa
128641 tttacatatt tagaaaacaa agatagggta aatattgaaa ttacatagta aaccatacct
128701 taaaagcaga aatactgata ccacatgaac tattgatcaa tactactgct attctcttcc
128761 taatgcaaag agcttattac cattattaaa tcactacaga cgataatacc cggaatgccc
128821 tttttgcagg gaaagcgaaa aaggtgaaag agttaacagg agaaagtgtt agggaactaa
128881 gttacgaagg aaggcagata gcaacaatac taacgtggaa cttatttcga cgactataac
128941 aactggtatc tgattatcga agaataattg agtggtcaga aaggagcgaa atatgtctgg
129001 agcaagatca acaacggcag gtgccgtgcc ctcggcagca acaacatcaa caacatcaac
129061 aacgtcaaac tctaaggact cagactctaa cgagtcatta tatcccttgg ctctgcttat
129121 ggatgagtta aaacatgatg atattgctaa tagggtagaa gccatgaaaa aactagatac
129181 catcgcgttg gcactcggtc ccgaaagaac aagaaacgag ttgattccct ttttaacgga
129241 agttgcacaa gatgatgaag atgaggtgtt tgccgtttta gccgaacagt taggaaaatt
129301 tgtcccctac attggcggtc ctcaatacgc cacaatccta ttaccagttt tggaaatttt
129361 ggcatctgca gaagaaactt tggttagaga aaaggccgta gattctctga ataacgtggc
129421 ccaagaactt tctcaagaac aattatttag tgacttcgtc cctttaattg aacatttagc
129481 tactgcagat tggttttctt caaaagtttc tgcttgtggt cttttcaagt ctgttattgt
129541 tagaatcaaa gatgattcat tgagaaagaa tatcctggct ttatacttac aactcgctca
129601 agatgatact ccaatggtga aaagggccgt cggtaaaaac ctgcccatct tgatcgatct
129661 gttgactcaa aatttgggat tatctacaga cgaagattgg gattacattt ctaacatttt
129721 ccagaaaatc attaacgata atcaagattc tgtcaagttt ctggcagttg attgtttaat
129781 ttccatcttg aaatttttta acgctaaagg tgatgagtct cacactcaag atttattgaa
129841 ctctgctgtc aaattaattg gtgacgaagc gtggagggta cgttacatgg ctgccgatag
129901 attttcagat ttagcctcgc aattcagttc caaccaagca tatatcgatg aattagtaca
129961 accatttttg aacctttgtg aggacaacga gggagatgtt agggaagctg tggctaaaca
130021 agtttctggg tttgccaagt tcctaaatga tccttcaatt atattgaata agatcttacc
130081 tgctgtgcag aatttgagta tggacgaaag tgaaacagtg agatctgctt tggcttctaa
130141 gattacaaat attgtattac tgttgaataa agatcaagtc attaacaatt ttcttccgat
130201 tttactgaat atgctaagag atgagttccc tgacgttcgt ttaaatatca ttgccagctt
130261 gaaggttgtc aatgacgtaa taggaattga gctgctatca gactctttgt tacctgccat
130321 aacagaatta gccaaggacg tgaattggag agttagaatg gctataattg agtacatacc
130381 tatcttggca gaacaattag gtatgcaatt ttttgaccaa cagttaagcg atttatgtct

130441 ttcatggttg tgggatactg tttattctat cagagaagcc gcagtgaata atttaaaaag
130501 gttaacggaa atatttggct ctgattggtg tcgtgatgaa attatttcaa gactgctcaa
130561 atttgatcta caattactgg aaaattttgt ctcgaggttc acaatactct ctgctctaac
130621 cactttggtg cccgtggtat cgttagatgt cgttactgag caactattac cattcatttc
130681 tcacttggct gatgacggtg ttccaaatat taggtttaat gtggccaaat cctacgctgt
130741 gatagtgaag gttttaatta aggacgaggc caaatatgat gcattaatta agaacacaat
130801 tttaccctca ttgcaaacgc tgtgtcaaga cgaagatgtt gatgtaaaat acttcgctaa
130861 gaaaagtttg gcagaatgtc aagaactttt aaaaaattga tactagttca aatatataca
130921 tacatacaca tatgtacact tgaatgaata aaaagtatac atattaacaa ctaggcctgc
130981 tttcttttct tttctttttt tttttttttt taccacactt agtcctctac tttaacagaa
131041 atatcatttt ccagtttgac catagtttct tcaactgttt tctcctcgtg tttaacattg
131101 atttccgcct ccacgttcag ttcacgtttt ccttccaggt atttcaccca ctttggataa
131161 ctcatgatat tttgctggat ttccttgtac aattgagact gcaaatccca atctaaggag
131221 cttctgggat catccggtgg caagaattgt aacatgtcac caaggttcct cgatttagtg
131281 attatttgtc caaatccaac cagcaaccca tttatttctg tccaaagccc cttaggcagc
131341 caattttgta attgagttct tgtctgatct ggagtcttgc atttctgcgc atctacccat
131401 ttccaaagct tggtcaatct atccacgtga acgtcaacgc agataccttc aatcttgccc
131461 catgcctttt gtaatgttag ataagccatt ttgggcccta caccgggcag gcccaatagc
131521 tcattaatcg tagctggaac atcactcgaa aactgatctt gaaggatttt gcaggttgac
131581 aaaatatact tggccttcct tgtgtggaac ccaactgaat gaatcaattc gtctaatttg
131641 gtctcattga tttgtaaaac ggcctctaat gtcatacctt cttcgctgtg cagttcgtct
131701 atacaatacc gcattatgtt aagcattgcc attgcggtaa cttcatcttt tgtttgcgat
131761 gatagcatca ccccaagaag gacctgtaat ctgtagtccc tcggtgatat ctgctctttc
131821 gagataccac acttagaggc aaccgttaca ggaatggatg atccaccaat tatatcgacg
131881 ggggctagaa tcttagatct cagtactcgc attctagcgt atgtttcttg aaacttgtaa
131941 gggactttcg tcgaggccgg agtgacaagg atcgaggggt ccaatggtgt ggcccacctg
132001 ttgggcacat tgccgtttct aaccacaatc cattcgaagt actgcttatt tggcagcgat
132061 ttaacccagt cgatatccac gggttgaggg acaacctctt cttgtttgat tttggtcctt
132121 ttctccggta ggagttctga ttctggccca gtttcagtct ttaccagcgg tcttttcctc
132181 agaattgcca tagatgagta tttactgatc ttttgcatat tttttttttt tttgggctat
132241 aaagtatata tagatacaaa tatatgatga atcattaaag aggaggttat tactaagtga
132301 aagaaaaaga aaaaaaaaaa gatcaaaacc aaacttcgta ttcgagccta aaaaacagaa
132361 tataatgtta taatactaat agaagcaaca ggagcaccac tatcagcaca agaataatca
132421 cgcagttgcc gttatccttg aacctagaat tgttaaagcc atgcatggac cgagaggccc
132481 tgttcaaatt cctaccatta ttatcaatca ggttttctaa atcaaccagc agcgagtcgt
132541 tctgagaaac gatctcatta ttaaggtcca gcgaaatgtc atgagtcctc ccaatagact
132601 gagacagtgc gcccaagtgg gaatcttgct ctaacaactg ctgttgctgg ttaataaata
132661 gttcttggtt ggaaaccatg ggttgtggtg gctgtggctg tgactgtgac tgtaactgcg
132721 attgcaatgg ttcatccttg tatggagtat gcgtcgaagg cagcggcgac tcctcgtctt

132781 gttcatcgtc atccttatac atgaccgtaa gctcgtcatc attcttgaac cgcacttttt
132841 tcaaagactc tttggacacc tcatcagtat ttcttgccac ctgctgttgg aacctataca
132901 actccttgtc aaccgccgta tcaggaattt tgtccagtat ggtattgtat cgggatatta
132961 gttcatcatt gggcgcacac ttctgcaata actccaatat tgagcccaat tgccgtttca
133021 aagtaacgtt atcgtttgag gtcggggcga gcttcaacac cgacactagc cttgttcttt
133081 cttcgaccag atcagagagc tggtccagtt cataacccag cttcaacaca tccataccgt
133141 gtgtgcttac gcatctattt gctgtcgtga gatctgtctc tatgctttat tcgttttcca
133201 ttgtaaagtc tcagcattta tttcttgttc tttacttgtt tttgcccttt cgggtgatca
133261 aagtcgtgct gggaaatttt attcttataa aatgattttt agaaataata aactcataac
133321 agtgcaacgg caaagtacaa gggaaggaag cacagaagca agaggaggcg catcgatcgt
133381 ggcagatgag tcagcaaaca ccacaggaaa gtgaacagac cacagcgaaa gaacaggacc
133441 ttgatcaaga gagcgtgttg agcaacattg acttcaatac ggatttgaat cacaatttga
133501 atttatcgga atactgtata tccagtgacg caggaacaga gaagatggat agcgacgagg
133561 agaagtcgtt ggccaatctg ccggagttga aatacgctcc caagctatcc agcctggtga
133621 agcaagagac gctcaccgag agcttgaaaa gaccacacga agatgagaaa gaggcgatag
133681 atgaggccaa gaagatgaaa gtgccgggag agaacgagga cgaaagcaag gaagaggaaa
133741 agagtcaaga actggaagag gcaattgaca gcaaggagaa gagcaccgac gccagggacg
133801 agcaagggga cgaaggtgat aatgaggagg aaaacaacga ggaggataat gaaaacgaaa
133861 acgagcatac agcaccgcct gcgctggtga tgccctcccc catcgaaatg gaggaacaga
133921 ggatgactgc gctgaaggaa atcaccgaca tcgagtacaa gttcgcgcaa ttgcgccaaa
133981 aactatatga caatcaattg gtgcggttgc aaacggagct gcagatgtgt ctggaagggt
134041 cacacccgga attgcaggtc tactactcga agattgccgc gatccgtgac tacaagctac
134101 accgagcgta ccagcgacag aagtacgagc tttcatgcat caacacagaa acaatcgcta
134161 ccaggacatt cattcaccag gacttccaca agaaggtcac cgacctgcga gccaggctgc
134221 tgaacagaac cacgcagacc tggtacgata tcaacaagga gcgccgcgat atggatatag
134281 tcatcccaga tgtcaattac cacgtcccca tcaaacttga taacaagacg ctgagctgta
134341 tcacgggcta cgccagcgca gcacagctgt gctatcccgg cgagcccgtg gcagaggacc
134401 tcgcttgcga aagcatcgag taccgctaca gagccaaccc ggtggacaaa ctcgaagtca
134461 ttgtggaccg aatgaggctc aataacgaga ttagcgacct cgaaggcctg cgcaaatatt
134521 tccactcctt cccgggtgct cctgagttga acccgcttag agactccgaa atcaacgacg
134581 acttccacca gtgggcccag tgaccgccac actggacccc ataccacttc tttttgttat
134641 tcttaaatat gttgtaacgc tatgtaattc cacccttcat tactaataat tagccattca
134701 cgtgatctca gccagttgtg gcgccacact ttttttcca taaaaatcct cgaggaaaag
134761 aaaagaaaaa aatatttcag ttatttaaag cataagatgc caggtagatg gaacttgtgc
134821 cgtgccagat tgaattttga aagtacaatt gaggcctata cacatagaca tttgcacctt
134881 atacatatac acacaagaca aaaccaaaaa aaatatgact ctacaagaat ctgataaatt
134941 tgctaccaag gccattcatg ccggtgaaca tgtggacgtt cacggttccg tgatcgaacc
135001 catttctttg tccaccactt tcaaacaatc ttctccagct aaccctatcg gtacttacga
135061 atactccaga tctcaaaatc ctaacagaga gaacttggaa agagcagttg ccgctttaga

135121 gaacgctcaa tacgggttgg ctttctcctc tggttctgcc accaccgcca caatcttgca
135181 atcgcttcct cagggctccc atgcggtctc tatcggtgat gtgtacggtg gtacccacag
135241 atacttcacc aaagtcgcca acgctcacgg tgtggaaacc tccttcacta acgatttgtt
135301 gaacgatcta cctcaattga taaaggaaaa caccaaattg gtctggatcg aaaccccaac
135361 caacccaact ttgaaggtca ccgacatcca aaaggtggca gaccttatca agaagcacgc
135421 tgccggccaa gacgtgatct tggttgtcga caacaccttc ttgtccccat atatctccaa
135481 tccattgaac ttcggtgcag acatcgttgt ccactccgct acaaagtaca tcaacggtca
135541 ctcagacgtt gtgctcggtg tcctggccac taataacaag ccattgtacg agcgtctgca
135601 gttcttacaa aacgccattg gtgctatccc atctcctttc gatgcttggt tgacccacag
135661 aggtttgaag actttgcatc tacgtgtcag acaagctgcc ctcagcgcca acaaaatcgc
135721 tgaattcttg gcagcagaca aggaaaacgt tgtcgcagtc aactacccag gtttgaagac
135781 acaccctaac tacgacgtag tgttaaagca acaccgtgat gcccttggtg gtggtatgat
135841 ctccttcaga atcaagggtg gtgctgaagc tgcttccaag ttcgcctcct ccacaagact
135901 gttcacattg gccgaatccc ttggtggtat cgaatctcta ttggaagtgc ccgctgtgat
135961 gacccacggt ggtatcccaa aggaggccag agaggcctct ggtgtttttg acgacttggt
136021 tagaatctct gtcggtattg aagacactga cgatcttttg gaagacatca agcaagcctt
136081 gaaacaagcc accaactaat cgccagtgcc acgtctctgc cttcgaccgg acctttttaa
136141 gtacgataaa tatcctttta taaatatata gtctaaaata tccattaata ctgtgctcaa
136201 tcaatcgtgt tagatgattt agttttttcc aaatcgttat tatagtgcag aagtagtata
136261 cataaaggca tatgcatgcg atttggaagt aacgctcgcc gtagacaagt aagaatgcct
136321 gctgtcttga gaaccaggtc caaagaatcc tctatagagc agaagcctgc ttccagaact
136381 agaacgagat caagaagggg caagcgtggt cgtgacgatg atgatgatga cgacgatgag
136441 gaaagcgatg atgcatacga tgaagtaggt aatgactatg acgagtatgc ttcaagagcg
136501 aagctggcca ccaataggcc cttcgaaata gtcgcgggac tgcctgctag tgtggagctg
136561 cccaactata actcttcgct tactcatccg caatcaatta aaaattctgg ggtgctttac
136621 gactctctgg tcagttccag aagaacctgg gttcagggtg agatgtttga actgtattgg
136681 cgaagaccta agaaaattgt tagtgaatct accccagcag cgacggagag tccaacatct
136741 ggaacgattc ctttgattcg agataagatg cagaaaatgt gcgattgtgt aatgagtgga
136801 ggtcctcaca cgttcaaagt tagacttttc atactgaaga atgacaaaat cgaacagaaa
136861 tggcaagatg agcaagagtt gaagaaaaag gaaaaggaac tgaaacgaaa gaacgatgca
136921 gaggccaaaa gattgaggat ggaggaaagg aaaaggcagc agatgcaaaa gaaaatagcc
136981 aaggaacaaa aacttcaatt gcagaaggaa aataaagcca agcagaagtt ggaacaggag
137041 gcgctgaagc taaaaagaaa ggaagaaatg aaaaaactaa aggaacaaaa taaaaataaa
137101 cagggttcac cttcttcctc catgcatgac ccaagaatga taatgaattt gaatttgatg
137161 gcacaagaag atccaaaact aaacacttta atggaaaccg tcgcaaaggg tcttgccaat
137221 aatagtcaac tggaggaatt taaaaagttc attgaaattg ccaaaaaaag gtcactagag
137281 gagaacccag ttaataagcg tccatctgtc acaacaacgc gacctgcacc tccctctaaa
137341 gctaaagacg tagccgaaga tcaccggtta aactcgataa ccttggtgaa aagttccaaa
137401 accgctgcca cggaacctga accaaaaaaa gctgatgacg agaatgcaga gaagcaacag

137461 tccaaagagg caaagacaac tgccgaatcg actcaagtag atgtcaagaa agaagaagaa
137521 gatgtgaagg aaaagggtgt gaaatcagag gatacacaaa agaaagaaga taatcaagtg
137581 gtaccgaaaa ggaaaagaag aaagaacgca ataaaggaag ataaagatat gcaattgacc
137641 gcgttccaac agaaatacgt tcaaggtgcg gagatcatcc tggagtattt agaattcacc
137701 cattcgaggt attacctgcc taagaaatca gtagtagaat ttttggagga tacggatgag
137761 attataatat cttggattgt tatacacaat tctaaagaaa ttgagaagtt caaaaccaag
137821 aaaataaaag ctaaactgaa agccgaccaa aaactaaaca aggaggatgc caagccaggc
137881 tctgatgtgg agaaggaagt cagctttaat cctcttttg aagccgattg ccctacccct
137941 ctctacaccc caatgacaat gaagttatcg gggattcaca aaagatttaa ccaaatcatc
138001 cgaaatagcg tttctccaat ggaagaagtt gttaaagaaa tggaaaaaat tctgcaaatt
138061 ggtactagat tgtctggcta taatctgtgg taccaattgg atggatacga tgatgaagct
138121 ttgagcgaaa gtttgcggtt cgaactaaat gagtgggagc acgccatgag aagcagaaga
138181 cacaaaagat aaggtgttcg gttactttat tctgctttaa cgccattatg attatacaca
138241 ttgtattact tatttttaa cctgtatatt aaaacctta ttttatttca cattactcat
138301 catgtggagt actggaattg tatgccaaac tttgccggga aaactggtat attgccgttt
138361 tctgtatcag ttgctgatat agatattgca ttatcattct tttcatcatc ggataaactt
138421 tcttgaaagc ctctagtgaa taccagctct gtcccggcag atattaaaaa tcccctccat
138481 ttgccttccc ataatagttt taaagtcttg tcacgtaatg acgtgcttaa tttccaaaca
138541 ctggtaaaaa tggaactatt tattttgtta tcaaattttt cgatggtaga tctttcttca
138601 tcgatttttc ttaatgacga cgagaatgca taagttaaat catttaatag cttttgtttg
138661 tgaacaggta tatctagtgt agaaggaata tcgttaacag ctgaattcag atccggtaca
138721 ttttcgtgca gtagtttagt cgctctgctg tttggattta tatcaaccaa ttcgtcggag
138781 attggttcta atttatcatt attgttttta ttggtttcaa gcaaatgatg cttttttgc
138841 caaaattcgc acccaaatga aagatttgat tcaatcgaat aaagattaaa atcatacttc
138901 gcgcaaaaag tagaatttgt ccctgtcttg gccgaatatg tggaggatat atggccgaat
138961 aatggattcc aagataatgt caaagttagt ggtcgtcctg tgtttgtaga atgtgtgtaa
139021 tatcttaaag ttgtcgaaca accggggctt aaacttacta accctaacca aaattcagca
139081 ccaagcgaca acgaagaatt attgtacagt gaggtgttaa acttggaagg cgtggtaagg
139141 aaattgtgta atactctata accacataat agatcactgg tggaaaatat ccactcctgt
139201 aaattgcggt gagaatctct ttgaaaatag cacgttaaaa cgtttaagct ttctttgaaa
139261 ctactgacac ccttaagcat aaattgggtt tgtggactta gtcgttttat tatcattgct
139321 tctaaatcag agctggggta gtacattcta ccataataaa gggatttttt aacaaatttc
139381 gagtcatgta gtaatttctt gtcattgtcg actgtggtgt tgtcactact caacgtattc
139441 gcactactaa cactgaaatt gaggtttggt tgcaattgtc tgtatgtttc ggtggcatct
139501 tgtaatggga tatcagtaga gttgcgcatg aatttctcca attgctgtgc atcggagtat
139561 aaataactca gagaaccatt tatcctggac ctcgtagaaa aatctaaaga attgaatgta
139621 ttgggagtag atttgttgga aatttgcagg tgtattgctg agggaattcg gaaatctaat
139681 aatgttctcg atgtggccgt tatatcctcg tagctatttt gcgtactcca atgggtgctc
139741 tgataaaatg cccttagtac ttggtccata tagggtagca tcaagatcgg tcttctctgt

139801 tcgtgtcttt ttcctaacgt atatttgctt tgtttcttca ctcaacaata aagtcaaagt
139861 aaaattaaat actaattatt cttaaaaggg aagatgcgaa atttagcgaa aatctattga
139921 ttatacacac aaaggaagaa aggtagtgga aagctaaata aaggaggtca tggagccaga
139981 gagcataggc gatgtgggga accatgccca ggatgatagt gccagtatag tgtccgggcc
140041 tcgcaggcgt tctactagca agacatccag tgcgaagaat atacggaact ccagtaatat
140101 ctctccagca tcgatgattt tcaggaattt gttgatactg gaggatgatt taagacgcca
140161 agctcacgaa caaaagatac tgaagtggca attcactttg ttcttagcgt ctatggccgg
140221 tgtaggcgca tttaccttct acgaacttta tttcacttca gattatgtca agggcctcca
140281 tagggttatt ttgcaattca ctctttcttt catttccatt actgtagttc tttttcatat
140341 cagtggacaa tatagaagaa ctatcgtcat tccaagaaga ttttttacct ctactaataa
140401 agggattagg cagtttaatg tgaagctagt taaagtacag tctacgtggg acgagaaata
140461 cacagattca gtaagatttg tgagtcgaac aattgcttat tgtaatattt attgtttgaa
140521 aaaatttctg tggcttaaag acgataatgc cattgtgaaa ttttggaaaa gtgtcacgat
140581 acaatcccaa ccgaggatcg gagctgtgga tgtgaaatta gtcctcaacc ccagagcatt
140641 tagtgcagag attagagaag gatgggagat ttatagagac gagttttggg ccagggaagg
140701 tgctagaaga cgcaaacaag cgcacgaact ccgacctaaa tcagaatgaa agagttggag
140761 ggcttcttcc ttcgaataag aggtcatatt tacctatgta aaattgtaac catctatgtt
140821 cacacataaa ttatatttta tacattatta gaagtgaagc tgttgtgtcg tgaaaatttt
140881 acaaatccgt catttcatat ttaagttttc caacaagtgc tagaaaacct aggggttgtt
140941 gaaattggtt aaacaaggca tcttattata catacaacag cataacgcta gaggggcaag
141001 aaggaagaac ttaaaataat aggtgtaaaa tgactttggc ttttaatatg caacggttgg
141061 tgtttcgtaa tttgaatgtt gggaagcgca tgttcaagaa cgtcccctta tggaggttta
141121 atgtcgccaa taaattagga aagcccttaa ctcgctctgt agggttaggc ggtgctggca
141181 tagttgctgg tggcttttac ttgatgaatc gccagccttc taagttgata ttcaatgatt
141241 ctttaggggc agctgtcaaa caacagggtc ccttggaacc aactgtgggc aacagtacgg
141301 caattaccga ggaaaggagg aacaaaataa gtagtcacaa gcagatgttt ttgggatcat
141361 tattcggtgt tgttttagga gttacggtgg ctaagatatc aattttgttt atgtatgtcg
141421 gtattacaag catgcttctt tgtgaatggt tacggtacaa gggatggatt cgcattaatt
141481 tgaaaaatat caaatctgta attgttttga aagatgtaga cttgaagaaa ctgcttattg
141541 atgggttatt gggtacagaa tacatgggtt ttaaagtatt ctttacattg agtttcgtat
141601 tagcaagttt aaatgctaac aaatgagcaa gacaaatgac cagatataaa cgagggttat
141661 attctttcgt tttatacttt tttatttttg gtatttcatt tatcctatac agtaaatata
141721 catagggcta aggaagaaaa aaaaatcacg tcgaatataa acctaattgt gttctatatt
141781 gcggacatat atttttcgta gattgaaaag ttcttaaacg taattttttt gacgaccagt
141841 gaagaggaat tgaataagta gaacttgggc aatacttata acggcaatga taatgataat
141901 caatatagat aaccaagtca accttgattc ggtggaattg acggtagaca tgtttctcca
141961 ttctctggct ctcaaatagt tcaaagtctt cgtgattttg ttcaagtttc tatctatttc
142021 ctccacggcg ttattggcaa taatgtcatc gttattgaca tcagcttcat gctcgtcagt
142081 caaagttttt tccttttcta gtgtaatttc tacctttttc aacgctgtac catagttatt

142141 ggaaaaacaa aaggtatact tccccactcc aaacgatttt aataaaaagt ctgagtactt
142201 cttttgtttc tcactagtaa tcacagatcc atcaggagca gtaatatcaa aatcaatctc
142261 aaaattacca ccggttagaa cttggtaacc cacagccagg gaatcatcct cagtaaccat
142321 atcgtagtac aggcattctt tgctgaatgc tggtaaactg atagcaactg gtgcataact
142381 ggaagatgca gccactgaat tgaccaacgc caaaattaaa acaatgaaga aagagggtag
142441 agcaattgta gatttgatca tggttttcct ttacaatgag tgtactccac tgtttatctt
142501 attctagttt gggtcacctg atgcagaatt ggcggtgtcg atccctaagg ttaatgaatc
142561 gaaatcactg gagattttgt tagtttttca ttttaactag ttcaagtttt ggcagaatgt
142621 actttttcgt gtttcgggac acgtcgctga aaaggattga aatataacca aacgccatca
142681 tgtaaggtgc agtgaacatt agcgctaaga ggattgagga ttgctttatg aaagtttatg
142741 aacattttct tgaattaaat aaataagtct gaaaacactt gaattcgaga tgctgtagca
142801 atttgtaata ttatttaaag ttcacagagt ggcttccttg gcggatctga aaaataggta
142861 gtgatttaag gctatataat attagataca caaaacatac cagaaatttt cctcgaagac
142921 cagcagaagg gaaccaatat cacgacatta aaatattgtt atctcttttt attcagtgct
142981 tgtatttcag cttccattga aaacgatgac tgtcttctca atcttcatgt cgagccctca
143041 cactgtacat gataatatac tagtaccatg aaaactagtc gatagatgat aattgatttt
143101 tatttgaaat agaatcttta atgatcacag tggatctttt aaaaacttgg tgagatagtg
143161 ctagtgatcc gtagtctaaa tgagttacgt acgaaaggga gccaaaagca atctgaccaa
143221 tttgtatata tatacatcta tccgaaaagg aagacatcaa ttagtacggg cgtgtggtct
143281 agtggtatga ttctcgcttt gggcgacttc ctgattaaac aggaagacaa agcatgcgag
143341 aggccctggg ttcaattccc agctcgcccc gaataatttt tttttgccta tctataaaat
143401 taaagtagca gtacttcaac cattagtgtt agcgataatc aagaagtgaa actctttctc
143461 tatttttttt ttaattgaaa aatttccttt ctctatagcg tatagaatat atgttacatg
143521 tatatatata tataaagtaa aaacgttcgg aaaattcctc attataccca gatcattaaa
143581 agacattttc gttattatca attgccgcac caattggctt aatcaacttc ttcaacggtt
143641 ggaccttcag cctctggagc tggaggagca ccacctggga aaccgcctgg agcaccacct
143701 gcagcgccac ctggagcacc accagcttgg tacaacttag acatgattgg gttggcaatg
143761 tcttgcaact ccttcaactt gtcatcgaat tcttccttgc tggcagtggt gttgctgtct
143821 aaccaagaaa tagtctcttc agccttcttg gtgacggtgt ccttgtcagc ttgttccaat
143881 ttgtcaccag cttcagaaat ggtgttcttc aaagagtaag caatggattc caattggttc
143941 ttggaagcaa ttctttgaga ttccttttca tcttcttcct tgaatttttc ggcttcagca
144001 accatctttt cgatatcttc cttggacaat ctacccttgt cgttggtaat agtgatcttg
144061 ttagacttac cagtaccctt ttcgacggcg gaaacattca aaataccgtt agagtcgaca
144121 tcgaaagtga cttcaatttg tgggacacct cttggagctg gtggaatacc actcaattcg
144181 aacttaccca acaagttgtt gtccttagtc ttggctcttt caccttcaaa gacttgaatc
144241 aagacacctg gttggttatc agcataagtg gaaaagatct cggacttctt tgttggaatg
144301 gtagagtttc ttggaatcaa cttggtcatg acaccaccag cagtttcaat acccaaggat
144361 aatggagcga catccaacaa caatagatct tgagtcttgg aagattcgtc accagtcaaa
144421 atagcagctt gaacagcagc accgtaagca acagcttcat ctgggttgat agatctgttt

144481 ggttccttac cgttgaagta gtcagtgacc aatttttgga cctttggaat tctggtagaa
144541 ccaccgacca agacaatttc atcgacttga gatttgtcca atttagcatc tctcaagacc
144601 ttttcaactg ggtccaaagt agatctgaac aagtcagcac acaattcttc gaatctggct
144661 ctggtgatgg aagtgtagaa atcgatacct tcgaacaaag agtcaatttc aacggaagtt
144721 tgagcggagg aagacaaagt tctcttggct ctttcacaag cggttcttaa tcttctcaaa
144781 gctctttggt tggtagacaa gtccttcttg ttctttctct tgaattcttg gatgaagtgg
144841 ttgaccaatc tgttgtcaaa atcttcacca cccaaatggg tgtcaccagc ggtggcctta
144901 acttcaaaga taccgtcttc aatggacaac aaagagacat cgaaagtacc accacccaag
144961 tcgaaaatca agacgtgttc ttccttaccc ttcttgtcca aaccgtaagc aatggcagcg
145021 gcggtaggtt cgttaataat acgcaagaca ttcaaaccag caatggtacc agcatccttg
145081 gtagcttgtc tttgagaatc gttgaagtaa gctgggacag tgacgacagc gtcattgacc
145141 ttggcaccca agtaagattc ggcagtttcc ttcatcttac ccaagaccat ggaggagatt
145201 tgttctgggg taaagttctt ggtttcaccc ttaaattcaa cttgaatttg tggcttaccg
145261 tcaacatcga tcaacttgaa tgggaagtgc ttcatgtcag cctgcacttc tgggtcgttg
145321 aagtttctac cgatcaaacg cttagcgtcg aaaacggtat tcgaaggatt catagcagct
145381 tgattcttag cagcatcacc aatcaatctt tcagtgtcag tgaaagcgac aaaagatgga
145441 gtggttctgt taccttgatc gttggcaata atgtccacac gatcattagc aaagtgagca
145501 acacacgagt atgttgtacc taaatcaata ccgacagctt ttgacatatt atctgttatt
145561 tacttgaatt tttgtttctt gtaatacttg attacttttc ttttgatgtg cttatcttac
145621 aaatagagaa aataaaacaa cttaagtaag aattgggaaa cgaaactaca actcaatccc
145681 ttctcgaaga tacatcaatc caccccttat ataaccttga agtcctcgaa acgatcagct
145741 aatctaaatg gcccccccttc tttttgggtt ctttctctcc cttttgccgc cgatggaacg
145801 ttctggaaaa agaagaataa tttaattact ttctcaacta aaatctggag aaaaaacgca
145861 aatgacagct tctaaacgtt ccgtgtgctt tctttctaga atgttctgga aagtttacaa
145921 caatccacaa gaacgaaaat gccgttgaca atgatgaaac catcatccac acaccgcgca
145981 cacgtgcttt atttcttttt ctgaattttt tttttccgcc attttcaacc aaggaaattt
146041 tttttcttag ggctcagaac ctgcaggtga agaagcgctt tagaaatcaa agcacaacgt
146101 aacaatttgt cgacaaccga gcctttgaag aaaaaatttt tcacattgtc gcctctaaat
146161 aaatagttta aggttatcta cccactatat ttagttggtt cttttttttt tccttctact
146221 ctttatcttt ttacctcatg ctttctacct ttcagcactg aagagtccaa ccgaatatat
146281 acacacataa tggcatccac cgatttctcc aagattgaaa ctttgaaaca attaaacgct
146341 tctttggctg acaagtcata cattgaaggg tatgttccga tttagtttac tttatagatc
146401 gttgtttttc tttctttttt ttttttccta tggttacatg taaagggaag ttaactaata
146461 atgattactt tttttcgctt atgtgaatga tgaatttaat tctttggtcc gtgtttatga
146521 tgggaagtaa gacccccgat atgagtgaca aaagagatgt ggttgactat cacagtatct
146581 gacgatagca cagagcagag tatcattatt agttatctgt tatttttttt tcctttttttg
146641 ttcaaaaaaa gaaagacaga gtctaaagat tgcattacaa gaaaaaagtt ctcattacta
146701 acaagcaaaa tgttttgttt ctccttttaa aatagtactg ctgtttctca agctgacgtc
146761 actgtcttca aggctttcca atctgcttac ccagaattct ccagatggtt caaccacatc

146821 gcttccaagg ccgatgaatt cgactctttc ccagctgcct ctgctgccgc tgccgaagaa
146881 gaagaagatg acgatgtcga tttattcggt tccgacgatg aagaagctga cgctgaagct
146941 gaaaagttga aggctgaaag aattgccgca tacaacgcta agaaggctgc taagccagct
147001 aagccagctg ctaagtccat tgtcactcta gatgtcaagc catgggatga tgaaaccaat
147061 ttggaagaaa tggttgctaa cgtcaaggcc atcgaaatgg aaggtttgac ctggggtgct
147121 caccaattta tcccaattgg tttcggtatc aagaagttgc aaattaactg tgttgtcgaa
147181 gatgacaagg tttccttgga tgacttgcaa caaagcattg aagaagacga agaccacgtc
147241 caatctaccg atattgctgc tatgcaaaaa ttataaaagg cttttttata aactttttat
147301 aattaacatt aaagcaaaaa caacattgta aagattaaca aataaatgaa aaaaacaacg
147361 aaataactta ggttttaggc taaaaaaaac agaaggaatt ttgaacgata aacttttcga
147421 ctgcacacga aacattatta ctaatttgtg taaccactat ataaggaatc gtgtttatta
147481 attgaattta ttccgggaat attcaagtta tgtatatctc ttttcatatt cttaaataca
147541 catactcata atatcttgtc gaaaatacgc ggtgtaggga gttatggtgg ataacttttt
147601 cacgattaga agaaaaggaa aatttcatta ttcgtagctt aacatggcaa aaacgagaaa
147661 gacatataat caaaacgtga gtttcctgtg gaaaaaaaaa aaagggaacc tctggttacg
147721 atgatatacc tgcgtgaaaa aggacagtta ttaccaatac atacaaaggc ttaataagtg
147781 taaaatatat atctgccgag accattactc attacaccta gaatggagca aaatggcctt
147841 gaccacgaca gcagatctag catcgatacg actattaatg acactcaaaa gactttccta
147901 gaatttagat cgtataccca attaagtgaa aaactggcat ctagttcttc atatacggca
147961 cctcccctga acgaagatgg tcctaaaggg gtagcttctg cagtgtcaca aggctccgaa
148021 tccgtagtct catggacaac tttaacacac gtatattcca tcctgggtgc ttatggaggg
148081 cccacgtgct tgtatccgac agccacgtat tttttgatgg gcacttctaa aggatgcgta
148141 ctcattttta attataatga acatttgcag acaatcctag tgccgacctt atctgaggac
148201 ccttctattc actcaataag aagtccagtg aaatcaattg tcatatgttc cgatggtact
148261 catgtagctg cctcatacga gaccggaaat atatgcattt ggaacttgaa cgtagggtat
148321 agagtgaaac ccacttctga accaacaaat ggtatgaccc caacgcctgc cttaccggca
148381 gtcttacaca tcgatgacca tgtgaacaag gaaatcacag ggttagactt ttttggtgct
148441 cggcatacag ccctgattgt tagtgatagg acaggtaaag tatcactcta taacggttac
148501 agaagaggct tttggcagtt ggtgtataat tcaaaaaaaa ttttagatgt gaactcttcc
148561 aaggagaaat taataaggtc aaagttgtct ccactaatat cacgggagaa aatttccact
148621 aatttgttga gtgtactcac aactacacat tttgccctta ttttattatc gccacacgtt
148681 tctttgatgt ttcaagaaac tgttgaaccc tcagtacaaa attctctagt cgtgaatagc
148741 tctatttcat ggactcaaaa ctgttccagg gttgcttatt ccgtaaataa taaaatttct
148801 gttatttcca tatcttcatc agacttcaat gttcagtccg ctagccattc tcctgaattt
148861 gcagaatcta tattatccat tcaatggatt gaccagctcc tacttggtgt tttaaccata
148921 tcgcaccaat ttttggtatt gcaccccaa catgacttca agatcctgtt aagattggat
148981 tttctgattc acgatttgat gatcccacct aataaatatt ttgtaataag tagaagaagt
149041 ttctacctgt taacaaacta ctcatttaaa attggcaaat ttgtgtcttg gtcagatatt
149101 actttaagac atattttgaa aggcgactac ttgggtgcat tggagttcat agaatcactt

149161 ttgcaacctt actgtccact ggcaaacttg ttgaagctag ataataatac ggaagagagg
149221 actaagcaac ttatggaacc attttacaat ctgtccttgg ctgccctaag gtttcttata
149281 aaaaaagata atgccgacta caatagggtt taccaattat taatggtagt tgttcgtgtt
149341 ttgcagcaat cttccaaaaa actagactca attccttctc tagacgtctt tttggaacag
149401 ggtctggagt tctttgaatt gaaggacaac gcggtatatt ttgaagttgt agcaaatatt
149461 gttgcccaag gatcagttac gtcaatttcc ccagttcttt tcaggtccat aattgattac
149521 tatgctaagg aggagaattt aaaagtaatt gaagacttaa tcatcatgtt aaatcctact
149581 acgcttgatg ttgatcttgc cgtcaaacta tgccaaaagt ataatttgtt cgatttatta
149641 atatatattt ggaacaagat ctttgatgat tatcaaaccc cagtggtgga cttgatatac
149701 aggatttcta accaaagtga aaaatgtgtg atcttcaatg gtcctcaagt acctcctgaa
149761 acgactatat ttgattacgt aacgtatatc cttactggca ggcaatatcc acaaaacttg
149821 tctatatcac caagtgataa atgctccaaa atacaaaggg aactttcagc atttattttt
149881 agtggcttct ccataaaatg gccgtcgaac agcaatcata aactttacat atgcgaaaat
149941 ccagaagaag agccagcatt tccttacttt cacctttttat tgaaatcgaa tccgagtagg
150001 ttcttagcaa tgctcaatga agtgtttgaa gcgtccttgt ttaacgatga caatgacatg
150061 gttgcatcag ttggagaagc agaattggta agtaggcaat atgttattga tctactattg
150121 gatgctatga aagatacggg aaattcagac aacatcaggg tacttgttgc aattttcatt
150181 gcaactagta tatcaaaata tcctcaattt attaaagtgt ctaaccaagc cctcgactgc
150241 gttgttaata ccatatgctc ctctagggtt caaggtatat atgaaatttc tcaaatagct
150301 ctggagtcgc ttttacccta ttatcattca agaacaacag aaaattttat actggaacta
150361 aaagaaaaaa atttcaataa agttcttttc catatctata aaagtgaaaa taagtacgcc
150421 agtgcgcttt cacttatttt agaaactaag gacatcgaaa aagaatataa cacggacatt
150481 gtatccataa ccgactacat actcaaaaaa tgcccacctg gaagtttaga atgtggcaaa
150541 gttactgaag ttatcgagac gaactttgat cttcttctct caaggatcgg tatcgaaaaa
150601 tgcgtcacaa tttttctga ctttgactat aatcttcatc aagaaatcct ggaagtaaag
150661 aatgaggaga ctcagcaaaa gtatttggat aagctttttt ctacgccaaa tatcaacaat
150721 aaggtcgata agcgtttaag aaatttacac atcgaattga actgtaaata caagagcaaa
150781 agggaaatga ttctttggct taatggtaca gttctcagca acgctgagag cttacaaatt
150841 ctggacttat tgaatcaaga ctctaatttt gaagctgcag ctataattca cgaacgcttg
150901 gaaagtttta acctagcagt cagggattta ttaagtttta ttgaacaatg tctaaatgaa
150961 gggaaaacaa atatatctac tttattggaa tctttgagga gggcctttga tgattgtaat
151021 tctgctggta ccgagaaaaa atcgtgttgg atattattga ttacattcct gatcactcta
151081 tatgggaaat atccttcaca cgatgaaagg aaagatttat gtaataaact acttcaagaa
151141 gcatttttgg gattggttag gtccaagagt tcctctcaga aggattcagg tggggaattc
151201 tgggaaataa tgtcttctgt tcttgagcac caagacgtta ttttaatgaa agttcaggat
151261 ttaaagcaac tgctactgaa tgtttttaat acttataaat tggaaagatc tctttctgag
151321 ttgattcaaa agattataga ggattcttcg caagatcttg ttcaacagta tagaaaattt
151381 ctgagtgaag ggtggtctat acacaccgac gactgcgaaa tctgcgggaa aaaaatatgg
151441 ggagctggtc tggacccatt actttttcta gcttgggaaa atgtacagcg ccaccaagat

151501 atgattagtg tagatctcaa aactcccctt gtcatattca aatgtcacca tggctttcac
151561 cagacttgcc tcgaaaactt ggcccagaaa cccgatgaat attcttgttt aatttgccag
151621 acggaatcta acccaaaaat agtataacat ttctaaatat ttaatacaac tttggttaca
151681 taaaagtaaa atttatacac ctcatttcat tatgtagatt catatataga ataccaatta
151741 tgattgaccc aatagccatc aaaatcagta gttattaata cttgtctttc taggagccat
151801 ttgcatattt ctgatatttc atgaagcgaa agtacttcac gacacctaga ttgcaatcta
151861 ctcaatgtta tccctggatg aaatattatt tcgttaacga ccatagtaac tacctgcttc
151921 catatgtttg gcctaatgga accagatcca ttcacccata aacgagaaaa tggtttgccc
151981 agtggaactt tgacagcaga cttccttgct gtattcaatt ttgtctgaga attggcatat
152041 ataatcagag ggggagttaa tgttcgtatt tcaaatctcc ttgaagtata cgttaaaggt
152101 cgaacatttc tcaccattgg aattacatcc atattcaata gctctcccga aatcaaatca
152161 attaaaaccc aagaggatat atcggacggc tcttgattga taacaatagc gtttccggcc
152221 tccaataatt cattaacctt acatctatac tgaaaagcta caccaaaatc tttataattt
152281 cctctatttt ccaaaatgtc tggtaaagta tcagtacatt caagttttga gccatggaga
152341 taaatttgct tttccttagc catatccatg atgacgttat ctattgattc gtttccaacg
152401 ttcttcaacg cctctatttc atttctagtg gtcgaaggac tttctattaa tatggaccgg
152461 atcactgtgc gaatataatc gtcgctttga ctcttcgata agtccttagt agaagcggaa
152521 atctttctag tgtaagtttt ttttaaagaa gagatctctc tttgaatcat agaagacatg
152581 gccagatcgt tgccagaatg tgtaagtgtg tcatcacgta ccagagtaaa tttttttcta
152641 ttctcttcgt agttcttata aagaaagagc ggccttttta tttccttttc atcaaaagag
152701 gtccaaatat caagcaattt gataagatct agttcttcaa catccctcag tgaaatcttt
152761 tcacttttaa tggctagaac gagcattttt ttccacttat cgacatatgc cctccaacca
152821 ctatgaccca ttcttactcg ccgtgccgtc catttctttt ttaggttatc taaagaatta
152881 ttaggaaata attttgttat tttgtcccac attatttcat ttttaatact tttagtaact
152941 acaacagctc tgattaaagc ctgaacacca tctttagtgc ctgcatgata gacagttttg
153001 tcttctttag tattttccac gaccaccgtt gtcctaccag cagacacttt tttgtctctc
153061 cttttgatct ttccatctga tacgttgacc gacgtactct tcttagagat agcgtcatct
153121 gaagctttga tcttagcatt cttttggcgg ttctgaaact tacgaattct ttcaactgat
153181 ttctttcctc tgtgaaacct attttttttcc gtttggtcaa agaagtatat ttccgtatca
153241 tgtataacat cagtaaaggt tgctttttta ctatcttttt ctttcaggat gtacctttgg
153301 atagcgtcct ctccaacagt gggcaaaaaa ataatttttc ttcctgatac aggctctgtt
153361 ctggctccta atttttcgct ttctaccatc aaatcaacat caccacggac agtcttttta
153421 tctaatgtcg ttgtggagcc catatattta gaaacgcttt cgtaaaattg ttctctcagg
153481 tatgctaccc caccaatcgt attcataact ttcaaaatgg ctctctgtct ctgtagtgaa
153541 cgcaaagagc gggcagaaaa gccgccgaag ttaaccacct tgcctttgac aacagtgcct
153601 ccgtttgaac ttggactatc ttcagcagat ttcggctcct gtgcagtgct gacatgctgc
153661 tctagcttaa tccttttggg attcgaaatg tttcctgcaa cagaagcatt agtactgttt
153721 ttaacctgcc tcttccgttt gtttttattc ggagtttttt ttgagtttgg gggaattttt
153781 aattcaccgt gccagaagaa tatatcctgt ccatcgctgt ccgttgtaaa tctaacagtg

153841 ttgttgagtg cgacgaaatt atcctcgttg agagttttca aatcggtacg agatttgcct
153901 agctcatcaa acccttttgg aacggatatt tcgtcttccg catttgttaa cttttgaaag
153961 ttctgagctg tgaacagcct aaaaaacttc ttctttccct caaaatcgta tatgcgaaaa
154021 agcctatacc cccctgtatt ttctttttgc ttatccacac tttctaaata atattcgctt
154081 gatttggtaa aagctcgctg aaattctttt ccggtaattc gatttacaac atccatagtt
154141 gaaattcctt taaggccaga cttatctgca atgtcataag tctgattttg aagtggataa
154201 aatcgattaa gaagaacttc attctttaca gcatcctctt tctcttccat aacaaggcct
154261 tgattttgta ataaatcagt cgcattgaaa ttatctaaac cttcgactaa gtcttcatct
154321 tcgaaagctg ccttgctatc tgatacagaa tcttcatccg cgctattgct atcatactca
154381 aatgaaggcg agcctttaga gtctggaata tctttcacgt attttacaca tctgatttta
154441 atggcaggat tcttgggtga tactacaagc actttcttta agtactcctt ttcatctaac
154501 catgcaatag ctgcaataaa agctttagaa agtcttttct ctttgtcaaa tttcaattca
154561 cgctttaaat caattatctg gcgaatacca ttttttgatc gttttaccac ctcaactatt
154621 gttgctaaat gatccctaat attaatatag ggattactat ccaccccgtc atggctgaat
154681 ttttttagct tcaattgctt cacgacgtgt cccttataaa tcagttgtga acttgttaac
154741 aggtggttta ttttcttgat acgtccagtc acacttctag gatcttgccc agttacctgc
154801 gccaaatcca tagtattgat cccttttttct cctgatttgg caacttcgag aagtagttca
154861 aatgcagaat ttccaatagt tgactccttt tttgtgtatc ccgttaataa tgtccatagg
154921 ctgtcctcag taatcccaac cgagtatgaa tgattagcgt cgcctataat atcagtcaca
154981 tttttagttg ttatagcacc atcacaatac acctcaatgt cctttttcaa tatcacgcat
155041 gaaagcacga actgtttaac ttttttatca gacaaatcaa aatatttacc agatatatcc
155101 cacagctgat tcaaagtgat ttcttcaatg tgtcgttagt aaataatctt tcacaatata
155161 gtacgtttat cacctaaacg gagccgaaaa ggagaatgag acatgaacat acttccctta
155221 tttgaagcaa ttttatcaga cactatttgt acgagttcgt caggataaat cgtcagtacc
155281 atttttcttg tggctagttg gcttcaacca aacgtcctct tctctcttat ggcaagaaga
155341 aagttatatg tgtgactggt tgtttatttc actttcgcga ctgaaagcgc cgccctttat
155401 gatgcaaaaa accaagcggt atttgaaaat gcagatttgc agaaataaaa gaaataaaaa
155461 taaaaatctt ataattttac cagcttcaat aactcgattt gcataagtgt gccttagtat
155521 gcttattata ttgactttga cattgaactt caaaaccttt tatgtaatga tttaagtctt
155581 gtcacatgac ataataataa ataattttaa aaatataaaa tatttttaat agtttttaaa
155641 tattttacag tttatttttt aaatttattt atatgttttt gttttccgaa gcagtcaaag
155701 tattttaatt ttcggagctt tcatttcaag cgcctttttt ttacacataa tacgatcaag
155761 ataaattatt atagtggtac agaactctta agcactaggc ggtggacctt attagctcaa
155821 tcttgagtca gtccatgtat cgttttataa tacttttttta agcactttct ttaataaata
155881 ttccattgaa gtactgttac tgaaatgaga tgaactgttc agaatgtaga aatggcgcca
155941 gaaatcaatc attgtttagc aaaaacacct ttcgtctgct gcctcgggtg ttttttcaaa
156001 ttatttcagc aggtaaatta agatagttat tcgagtgatt gccaaatatc atgttctact
156061 tcgaagactt atagctaatt aattttttca taatgaaggt gtcgttaatt gttctgatta
156121 gtaacatgaa actcaaaaat catcaaaaaa agaaaagcta aatgtatact tttttgtcta

156181 cattagttac cttttattac atgagaaagt tatttttctt cttttttttt ttttttttga
156241 aactttttcc tctcggaaaa taaaagatat atttacaagt gaaagcttat tgtaatgtgt
156301 acttttaaac atcaaataac agacctttac atcaaataag caccgcaaga tatcctaaaa
156361 tcgacatcca atgcatcgta aatcattgag gagggctagc gctactgtgc cttccgctcc
156421 ctatcgaaag cagattatta gcaatgcaca caataaacca aaccttttct ctaaaattaa
156481 aactttcttt acccaaaaag attcagccag agtgagtcca aggaataatg ttgctaataa
156541 acaaccacgc aatgagtctn nnnnnnnnnt tttttttttt ttttttttga aactttttcc
156601 tctcggaaaa taaaagatat atttacaagt gaaagcttat tgtaatgtgt acttttaaac
156661 atcaaataac agacctttac atcaaataag caccgcaaga tatcctaaaa tcgacatcca
156721 atgcatcgta aatcattgag gagggctagc gctactgtgc cttccgctcc ctatcgaaag
156781 cagattatta gcaatgcaca caataaacca aaccttttct ctaaaattaa aactttcttt
156841 acccaaaaag attcagccag agtgagtcca aggaataatg ttgctaataa acaaccacgc
156901 aatgagtctt ttaacagaag aatctcaagt atgcctggag gttatttcca ttctgagata
156961 tccccagatt ctactgtaaa ccgttccgta gttgtttctg cagtgggtga agccagaaac
157021 gacattgaga ataaagaaga ggagtatgat gaaacacatg aaactaacat ctccaatgca
157081 aagcttgcaa actttttag taaaaaaggt aatgagcctt tatcagaaat tgaaatagag
157141 ggtgtgatgt cattgttaca aaaatcaagc aaatccatga taacttcgga aggagaacaa
157201 aaatcagccg aaggtaataa tatcgaccag tcgcttatct tgaaggagtc aggaagtaca
157261 ccaatcagca tatctaatgc gccgaccttc aacccaaaat atgatacttc aaatgcgtca
157321 atgaatacga ctttgggaag cattggttca agaaaataca gtttcaatta ttctagcctg
157381 ccctcaccat acaaaacaac cgtttataga tatagtgcag cgaaaaagat ccccgataca
157441 tacacagcca acacatctgc tcaaagtata gcatctgcta aatcggtaag aagtggtgtt
157501 tcaaagtcag ctcctagtaa gaaaataagt aatacagctg cggcattggt ctcactatta
157561 gatgaaaatg acagtaagaa gaataatgca gcttcagaac ttgctaatcc atactcctca
157621 tatgtaagcc aaatacgcaa acataagaga gtttctccaa atgctgcacc aaggcaagag
157681 atcagtgaag aagaaactac tgttaagcca ttatttcaaa acgttcctga acaaggcgaa
157741 gaaccaatga aacaactgaa cgccaccaaa atttcaccat ctgcgccaag caaagattct
157801 tttactaaat acaaacctgc aaggtcctca tccttacgct caaatgtcgt cgtagctgaa
157861 acctcacctg aaaagaagga tggtggagat aaacctccat cctctgcttt taacttctcg
157921 tttaatactt caagaaacgt tgaacctact gagaatgctt ataagagcga gaacgcacca
157981 tctgcatcat caaaggaatt caattttacc aacctacagg cgaagccgtt agttggaaag
158041 ccaaaaaccg aacttacaaa gggcgattct actcccgtcc aaccagatct ttcggttact
158101 cctcaaaaaa gttcatcgaa aggctttgtt tttaatagtg ttcaaaagaa atcacggtcc
158161 aatctttcac aagaaaacga taatgaaggt aaacatatca gcgcctcaat tgataacgac
158221 ttttcagagg aaaaggcgga agagtttgat ttcaatgttc ccgtggtgtc taagcagcta
158281 ggaaatggct tggttgatga aaataaagtt gaggctttca agtccctata taccttttga
158341 taatgaaaat tttagccgtg acataattac cgtatagccc aactcaatac gtaagtttgt
158401 gtaaaatacc attccaagat gatattattt agtcattttt tttccacttt ctcaaaaaga
158461 aggaatacct ttagcggctc ttataaacta taaatttcta gaagatacat aaaaggtttt

158521 tagtctgatc ataaaatttt ttgcttaaca aaaaatttgc ccaggtgttt catttgccag
158581 ccacaagtaa cagcgagaac aattaattga atgacaatcc accacatagc acgagaatta
158641 acagcttcag aggcgtctct aaaggtagct tcacgatctc tcatcaattt ttgctctctt
158701 ctaatttcgc cgatcttgga gtttaggacg ttaaccttgg catgtagaat gtcaatagtg
158761 gctttaccct tagaatctaa cttttcatca gagcccactt ggaattcaac gtcaatcttc
158821 gttttagcct taatcaacca gccaccagct tcgggctgaa tacagatttt atgttcaccc
158881 gaatcagacg caaggaaagt taaatcacca cttgctgaac ctttctgatg aacaaccagg
158941 tggttatcat caaaagtttc ctcaatatca atcaagacac caaaatcttg cgcaccagcg
159001 tctctgtaat tttgtaattg gtcatcgtaa atttgtgcct tgtaagttgc ttggaacaaa
159061 gtacctttag acaattcctt gtggaagcac ttacgttcag caccagaagt ataataatag
159121 aacgcagtaa cttgagctgg taaaactaga cagcaggcaa aaacctgtaa aagagaggtt
159181 aaaagcatga ttagtagaga gattggttac ctttaaatac ttttccaaac tacagaggga
159241 agatagagta agttttgtat gtacacattt ctgctgatgt gttttttttt ttcaacttat
159301 tacgcgattc gttttttttt tacggtaaca gaatacagaa taaattcacg tacaaaaata
159361 gagaatatat aaaataatag gttgacgatt atattggatc ttcccctggg gttcaagagt
159421 cgagaccgag tccttttagt ttgtgtatat cagctggttc ttttcgttat gaacatcctt
159481 ttacaggatc cattcgctgt tcttaaggaa catcctgaga agctcacaca tacgattgag
159541 aaccctttac gcactgaatg tctccagttc agtccttgcg gtgattacct ggctcttggg
159601 tgtgccaatg gagcacttgt tatttacgat atggatacgt tcaggcctat ttgtgtccca
159661 ggaaatatgt tgggagcaca tgttcgaccc attacatcta tcgcatggtc tccagatggt
159721 agattgttgc ttacaagctc tagagactgg tcaataaaac tgtgggatct ttcaaagcca
159781 agtaagcctt tgaaagaaat acgattcgat tctccaattt ggggttgcca atggctggat
159841 gctaaaaggc ggctttgtgt agctacgata tttgaggaaa gtgacgcata tgttattgac
159901 ttcagcaatg atccggtcgc aagccttctc agtaaatcag acgaaaaaca attgagttcg
159961 acacctgatc atggatatgt tcttgtttgt acagtacata ccaaacatcc aaatattatt
160021 attgttggaa cttcaaaagg ttggctagac ttctataaat tccattctct atatcaaaca
160081 gaatgtattc attcccttaa aatcacgagt tctaatatca aacatttaat tgtctcgcaa
160141 aatggtgaaa gattagctat taactgctcc gatagaacaa taagacaata cgaaataagt
160201 attgatgatg aaaactctgc ggttgagttg accttagagc ataagtacca ggatgtgatt
160261 aataaattac agtggaactg tatcctcttt agtaataata ctgccgaata cttagtcgct
160321 tctacacatg gttcttctgc acatgaacta tacatctggg aaacgactag tggaacgttg
160381 gtgagagtcc tggaaggggc tgaagaggag ttgatagata taaattggga cttctatagt
160441 atgagtatag tgagtaatgg ttttgaatct gggaacgtgt atgtgtggtc tgttgttatt
160501 ccgccaaagt ggagtgcttt ggcgccagat tttgaagaag tagaagagaa tgtcgactat
160561 ttggagaagg aagatgaatt tgatgaggtc gatgaggcag aacagcagca aggactagaa
160621 caagaggaag aaatagctat cgatcttcgg acgagagagc aatatgatgt tagaggtaat
160681 aacttgcttg tagaacggtt cacaatccct acagattata cgaggataat taagatgcag
160741 tcatcatagg tttctcttca aaaggagaaa gtttaaatag gcaactatcg attgaagttt
160801 tgtccaacta tccactatgg atatgcgttt tgttgattaa ccttaaataa cacgtatttc

160861 gcattttcca aaagcctttt ttcataacta caaactaact atttttgttt attttacatc
160921 agtaaattat gcgcagaaaa tatgtaaggc tatatactca atatagtgga agagcctctg
160981 gctacttaat ctgggttcat atattctgtc agtggtatag taaatttaaa tagtgaattt
161041 gggcgcatgt agagagatct tttgattaat aacccagtat ttgattcttg aatgttattt
161101 gtctcttatt aagtatttca gtttgattta tatattttga aagtaattgt tgctttacct
161161 tccaaacaat aaaaaaatat aaaaaaatgt aaaaaatata ataaatatta aataaaatac
161221 tacttgtttg aaaatcagag aaaatccacc aaaatatcaa tcatttcaag gatttccgaa
161281 ccaagttcga gatagtcctt taaggtcagt aaaattcagt tgcacgtata acagatattt
161341 ttcattttgt tccaatttaa aagtccccca tttttaaaat tattgaaaat aaaaattaaa
161401 aaattaaaag gaatctctct atgtcacttt aaaataaata aattgaaaat gatatatcgt
161461 taaaaggatc ccgcaacacc acctatttct aagaggagag ttctaaaaaa atcatagtac
161521 cacacaggta aactaaaaca gctaaattca acataatacc tgtggatata attacataca
161581 aaaatataat taaaaaaata cattaaaaat aatttatttt tgtaaaaccc ataaaatata
161641 ttttactttc ggaacaactt tttaacttat aattttgttt taaataaaaa cgttgtattt
161701 aaaaataata aaatattaag taaaaattta aagcatattt atttaaaata taaaatacta
161761 ctaaaattac tctaaacttc aaaataaaaa aataataaat tatatagttt taaaattaat
161821 aatttgtata cacgtgacca gaccataata gttttctttt cttgaaactg ccatgaattt
161881 aatagatttt ttttacacat atttacagta agttttgttt ttacgttgaa ttaatattgt
161941 atacgtactt agaaggataa cttccaacta tgaatatgta ggttaaaagg taaatagaga
162001 agccttactt ttatcagaaa tgaaaggagc tcagcttgcc tcgtccccgc cgggtcaccc
162061 ggccagcgac atggaggccc agaataccct ccttgacagt cttgacgtgc gcagctcagg
162121 ggcatgatgt gactgtcgcc cgtacattta gcccatacat ccccatgtat aatcatttgc
162181 atccatacat tttgatggcc gcacggcgcg aagcaaaaat tacggctcct cgctgcagac
162241 ctgcgagcag ggaaacgctc ccctcacaga cgcgttgaat tgtccccacg ccgcgcccct
162301 gtagagaaat ataaaaggtt aggatttgcc actgaggttc ttctttcata tacttccttt
162361 taaaatcttg ctaggataca gttctcacat cacatccgaa cataaacaac catgggtaag
162421 gaaaagactc acgtttcgag gccgcgatta aattccaaca tggatgctga tttatatggg
162481 tataaatggg ctcgcgataa tgtcgggcaa tcaggtgcga caatctatcg attgtatggg
162541 aagcccgatg cgccagagtt gtttctgaaa catggcaaag gtagcgttgc caatgatgtt
162601 acagatgaga tggtcagact aaactggctg acggaattta tgcctcttcc gaccatcaag
162661 cattttatcc gtactcctga tgatgcatgg ttactcacca ctgcgatccc cggcaaaaca
162721 gcattccagg tattagaaga atatcctgat tcaggtgaaa atattgttga tgcgctggca
162781 gtgttcctgc gccggttgca ttcgattcct gtttgtaatt gtccttttaa cagcgatcgc
162841 gtatttcgtc tcgctcaggc gcaatcacga atgaataacg gtttggttga tgcgagtgat
162901 tttgatgacg agcgtaatgg ctggcctgtt gaacaagtct ggaaagaaat gcataagctt
162961 ttgccattct caccggattc agtcgtcact catggtgatt tctcacttga taaccttatt
163021 tttgacgagg ggaaattaat aggttgtatt gatgttggac gagtcggaat cgcagaccga
163081 taccaggatc ttgccatcct atggaactgc ctcggtgagt tttctccttc attacagaaa
163141 cggctttttc aaaaatatgg tattgataat cctgatatga ataaattgca gtttcatttg

163201 atgctcgatg agtttttcta atcagtactg acaataaaaa gattcttgtt ttcaagaact
163261 tgtcatttgt atagtttttt tatattgtag ttgttctatt ttaatcaaat gttagcgtga
163321 tttatatttt ttttcgcctc gacatcatct gcccagatgc gaagttaagt gcgcagaaag
163381 taatatcatg cgtcaatcgt atgtgaatgc tggtcgctat actgctgtcg attcgatact
163441 aacgccgcca tccagtgaca ctgaagaggt atagtcatgc ctaccgcgat ttctttgaca
163501 cagaattgaa aaattttgca ttttttggta atttcctaat aatacgaagt gcaataatct
163561 cactttgata ggagcacgtc atgatggtaa tttcatacta ctgaacgtaa atgttgaagg
163621 tgaatttgta aagctttgtt atttgaagct gtacacctaa agggcggtaa attgttgggt
163681 gttagtacca tgccattaat tagaatttgt tagcattttt attcatttgt gcattatggg
163741 ttcaaattca tatgactgaa cgtgtagttt catatccagt catcaagaga tgctgaaccg
163801 cccttcaaaa acttgacgaa atagaagttt ttttttaca tttctcatat gttacataga
163861 ttaaatagta cttgattatt tgatacatta agctaacaaa gccttggata actcatcggc
163921 aagatagtcg gcttcagccc tgtaattcaa gctgtgtagg ttagcaacgg tatatctaat
163981 tctgctttga tcattgtatg tatcctcacg cgctctaatc ctaaagtcat attcgttcat
164041 ttggatactt tgagtaattt ttgtgaattc gttggggtct tcttccttca aagacattaa
164101 tgtattagca tcaacaccca ataattgttt agcttggtcg tcaaataaag tgagccatag
164161 ttgattggtt tcgtcaataa ttgatattgt caagatgtat ctccaatttg gccttgcatt
164221 attggtgtcg cacttctcac atctccaagt accatcaggc tgttccagaa ctttcttatt
164281 acaattctca ttagaacagg caggatatgc aaaattatca acttttaaga aacttatagc
164341 agctttaaca ctaaaaaagt cacctttctc gcttcttcct agattttcag cttgagctct
164401 agcaatagta atacgctgag caatgaattt tgttaagcta gcagccgatt gaccacccat
164461 accgggttct tgctttaaag tgatgaagtt tgcgttgcgg cccttggaat cataccaacc
164521 ctttaaggca tatgcctcag gaatttctgg attcggaatc agggtactag aaaatcccat
164581 agacaaagat ttgccaccaa aatccgtcac acgaacacct ttaatggcag caacagaacc
164641 ttcaggaagg ttgaaatcaa gggcttgctg attccatagg ccaacagaga tagaaaaccc
164701 agagtcgtca acaattgtga tgtcacgacg atcgaatttc ttcccagccc ttgaagttag
164761 ctcaaaatgt gggtttatag tttggataat accgaggacg tctacgttgg aatttacttc
164821 ctggttctga atagcatcta gtttgatgaa attgaaatgg gttttcggaa cattactttc
164881 atcgaaacat tcttctataa cagtgtctct atccaaattc agttcataag ggtgtgttag
164941 attagtaaat tggggcttag ctggttggag ttttgccttt gatacatagt atactttgcc
165001 ttcttgtaaa atttcgttaa attttgtagc aaaatcatta aacgccgtgg ctcggatttc
165061 tccagaggta tccaagaagt tgacattgaa tagtttacca tcacctcttt gattgtgcca
165121 cgttttaatt tctcccttgt aggaaactct tgctttgata gtccaaacgt tttggtatgg
165181 agacagttgt tcgatggcaa aaattggtct ggttttttgc gaattagggt tttcattggc
165241 gaattttctc tcatttgcat tcaagtttga gtttgaatgc agcatatcag ggacaccagc
165301 attgctggcg tttgtttgat tggcaacatt accactgtca gttatatctt cgtcttttaa
165361 ggtttcattt ggatgctctg agaaatagtt atccaaaaaa gtactagttt ggttgaccat
165421 atcagcacgc gactggacca actcaaagtc atctactaaa agaacgtatt tctttctttc
165481 cctgacaata gcaggttctg caattatcac gcgaatgata tcaccccttt gtagttccat

165541 tgactggaac ttggatgcag cttggtttct caacagagcc ttcatatggt aaataccatc
165601 ggaaatcatg atcaaattct ttctgttgct gttagcccca tcagatttcc tggtgttata
165661 aacttgataa acgccaccgg tgggattatc gtacctttgc ttattggtga agatgctatg
165721 aaaatcgccc ctcgaaagtt gaacactgct catctcttgt aagtataatc tggtcttctt
165781 gctggtttcg cctttaccgt aataagaaga gtgaatagtt tttgttttac gtgtagaact
165841 taaagtgata acatttgttc aagtaaacct ttatgttagt tcacgcgtct tttgtcgcct
165901 cgtctaattt ttacgcgtga catttttcca agcagagata ttttattgag cagcgaagaa
165961 gagttagaga ataagaaagt gatgcgataa gaaatccacc caattagcat agatcctttc
166021 gtatatggct gaagaaggtg gtacgcgcat agctattaac atatatgcaa aaagaacggc
166081 aaaaggcgag gaggttttta tgccaccgct agtatttgac atagatcaca tcaaacttct
166141 aaggaaatgg ggtatttgtg gtgtgttatc tggaactttg cctactgcag cacagcaaaa
166201 tgtatttttg tcggtacctt tgaggcttat gttagaagat gtgctgtggc tgcatttgaa
166261 caatcttgcc gatgtgaaat taataagaca agagggagat gagattatgg agggaataac
166321 attagagcgg ggcgccaaac tatctaaaat tgtcaacgat cgtttgaaca agtcatttga
166381 atatcagaga aagttcaaaa aggatgaaca cattgcaaaa ttaaagaaaa tcggtagaat
166441 caatgataaa accacagctg aagaattgca acggcttgat aaatctagca ataatgacca
166501 gctaattgaa tcttctttgt tcattgacat tgctaatacc tctatgattt taagagacat
166561 acggagtgat tcagacagct tatcccgcga tgatatcagt gatttgttat ttaagcagta
166621 cagacaggca ggaaaaatgc agacctattt cttatacaag gcattgagag atcaagggta
166681 cgttttgtcc ccaggtggac gttttggtgg gaagtttata gcataccctg gtgatcctct
166741 tcgtttccat tcacatctga cgatacaaga tgcgattgat tatcataatg agccgattga
166801 cctaatatcc atgataagtg gtgcaagact aggaacgact gtgaaaaaac tttgggtcat
166861 aggcggtgtt gcggaagaga caaaggaaac tcatttcttc tcaatagaat gggctggatt
166921 tggttaagct gggaatcagt catgtataat tattttctca gaatttatgt atttataagg
166981 tttttcagaa gcatacatat gtgtaataca atttttaaat ttgcattgat attttgatgc
167041 attcagcggg aaagtagttg tttatcacta gacatataat tatgtttatt tatatttagt
167101 gggagcaaaa cagtttattg aatgtttacc agaaccgaaa aaaaagctct tctaaactgt
167161 tgacatccag ttcatttact tccacgtgta gatgtgaagg aacaaatatt ttagcatcgt
167221 tcatacaagt aattatgcta tattatcgat cctcggattt cagcttccgt tatatcggat
167281 gattgttact cgacctttat gtcgtctttt tacatcatat atgataatat gctagcagtt
167341 ttaatacaaa ttgatcgaag atagttggtt ctgagaaatg ggtgaatgtt gagataattg
167401 ttgggattcc attgttgata aaggctataa tattaggtat acagaatata ctagaagttc
167461 tcctcgagga tataggaatc ctcaaaatgg aatctatatt tctacatact aatattacga
167521 ttattcctca ttccgtttta tatgtttata ttcattgatc ctattacatt atcaatcctt
167581 gcgtttcagc ttccactaat ttagatgact atttctcatc atttgcgtca tcttctaaca
167641 ccgtatatga taatatacta gtaatgtaaa tactagttag tagatgatag ttgatttcta
167701 ttccaacata ccacccataa tgtaatagat ctaatgaatc catttgttag ttaatagttt
167761 aaatgttttt atcggaagag gttttgtcat cacatcagca atgttcttct tggtctcgat
167821 gtagtatacg tataaattat tacctgatac ttcatctcta agtctcattg cctttgtgcc

167881 aaaaaatctg tttctaaatt tctcttcatt tgtagactta attatactga tcgttgatct
167941 actatcagta agtaagcctt taataattgg tttcttgtta agttcttgta tcaggtaact
168001 tagattattt aataatggga cagattcact tatcgcgtgt atttctgctt ccgtagttga
168061 agtacatgtt aatgaagcct tggtggactt tcctccaatt acctttccat taagtaaata
168121 tatgttgcca atttgtgatt tataatacgg ttggttgcca tacgaagcat cacttattgc
168181 gactagttta ttatctggct cggtaggttt gtttttgtgc catatcagtt gtttatctct
168241 agtgtcccac atgaattgta tcaactcata tgtcatgtct aaaacttgcc tagaggggaa
168301 tagtatatgt tgagcaagtg tgttgatgta gtatagtaag tcaaatctaa atttatatcc
168361 aacatatgaa gctagaccaa tcaacttttg catttcatgt accttctctt tgtattcatc
168421 ttcatctatt tctagttcat cctggtctat ataaagacct ggttgacctg gagctctaag
168481 tttctttcct tttgggttca aaggtacgtt tagtttgggt aatttttctg tcaaggattt
168541 ttccatacct aatttcatgt atttacctct ttgatatttg atttctaagc caagtatgtc
168601 gtactgaatt tcgttatcac tttcacccag atttattatc tttgtatcgt attgtttctt
168661 gagtgttgtt atgattttct tatttgcatt taagtctttg ctgaacaata tcatatcatc
168721 aacgaataag caaattgtta cttgactatt cttaaatacg catgaccatc cacgaacttc
168781 ttccatacca cactgtttta tcaggtatga tttgatagtt tcgtaccagt tcgctccact
168841 ttgtttcaat ccataaagtg atttcttcaa acgtatcaac ttatcattca ttcctaaatg
168901 tggtggaggt cttatgtata attcttcttt gatgtctgca tacaaatatg ccgaagatat
168961 gtctaattgt gtaatatagt agttattgtc taatgcaagt gacagggatg tcatcaacgc
169021 ataatgatgt acagtgttgg attgcatacc tgtatcgtac gtatcaggat gctgaatgtc
169081 acctcttgca acaaatctag ctttatgagt tccgtcacgt ttcttgttga agataaacat
169141 tgagtttatt actcttttag ggtctatttc ttttctgtca taatatttgt cagtgtccca
169201 agtattcatt ttcaatagtt ggttgacttc tttgtggtat gcctcgatat atttttcttt
169261 ttctttaata tctttattat aggtgattgc ctcatcgtat cgtaaggttg tccgtattgg
169321 tttgattgat tttactgctt ttacagctgc aatcaggtga attcgtttct tcgatctcgg
169381 aggttctaaa ctacgcatat tcttagtatt ccatgtgtct cgtgatacct taatttcagt
169441 ttcattatct tctaatgatc ttttcttact gttgatagta gtataggcat tagagtcacc
169501 aataccaccc aaactggaat tagtttgatg agaatttatc ggtgggagtt ctttaaatgg
169561 gtcagggaat tcggtaggag attctggagg tagatcaggg agtgggagat cagcgatgat
169621 agattcctcg gtattctgtt cagaaacagt agttggcgtt ttgataggaa caatattgtg
169681 cgatgaatta ttttccggtg gagaagcatc gattgaaggt gaacggtgta taatcctttt
169741 ctcagtctct tggtcactta tctgcggaac agttttgttg ttggtaccac ccgtactgga
169801 tattggtacg tttgtatgat tagtctcatt ttcactgtac gagtctgagt gtctgaaatc
169861 tttagattta ctggcgtgcg acgactcatg tgtgttagat tgggacatgg gagcaagtaa
169921 aggaacattt aatttatgca taccacccga accggtactc tcgatattgg aaatttgggg
169981 ggtgctagat ctcttctttg atggaagaat attagattca gatatgttgg ggtcaacttc
170041 tctgggtgcg cgaatattgg ttttagaaac acgtttcgaa tcttcagtat gagttgacgg
170101 aggtgtggaa tcggttggac tcacagcttt tgaaaggaca tttctcggtt gctcaggatg
170161 tagttcaatg tcggattgga agtcatggtc agattctatg ttaagatcat tggattcttg

170221 gatctcattt gacgcaatga acgaatgata tgaagcagtt aaacggttta agtcttcatc
170281 gaaagtgagt gcgtcgtaat tgaattgatc taatctggat tccttgccct gaagaataac
170341 atagttagtt gtatctactg tcttctttaa ggatggaaga tagatgatat atccataaga
170401 gtttcgagac ggatgtagag cgtagcctgg gatgccacga ggatgtattt tggagttagg
170461 gttgtgatca ttgacgataa caggttgacc gaaaggtaac aaagtactga tatcaagtcc
170521 tgccaagcca gcatgttgtc ttgcagattt tttgctttta ggtgaagcta gtgaatttct
170581 cacaatagta gaaaattcga ttgcagagaa ccataaatgg ttcggtaaac cactacattg
170641 cagttgagta cggcagtcat ctaataaggt acggtttagc cgttcagcga ctccatgtgc
170701 tcgggaatcc gctgtggttg tatagcatgg agttatacca ttttttcaa ggaatttatg
170761 gagagttctg ttagtatact cagaaccacg gtccatttgt ataaccaaga cactggcctg
170821 aaactggttt ttaataaaag ctagtatcgt agtaaaaaca tcgaggatag agtcctcgcg
170881 acggtcgtgt aatggataaa cccaacggaa ttttgttgtc tcatcagtaa atgagatgaa
170941 ataggatggt gcactttttg gtaggttgtg aactggacca aatatgtcag tatgtaggta
171001 ttgaaagggt tcgtatgaat tttggtattt tagtcgtgaa cctttgatat gtctgtgttt
171061 ggtgcttttg ccgattaaac aatcaggaca ttgatagtca atagcactag accagtcgac
171121 atctgattcg ttaaaatacg tgatggtgtt atttttaagt gagtatcgaa tttgtctgtg
171181 cattggcatg cgcaagcatt cgatgaatga aaggataagg atatttgcgt gtactttcac
171241 ttgtatggac attattgatg gtgggtacgg agatatttga tggaagcaag tactttttag
171301 atacccagta aaagtctcca tattttacga taggtgcaag tacagtgccg tcagaccgtt
171361 ctaagacgtt tttggtaaag catgctgtga tatctactgc agccaattca ttcaaactga
171421 gtaagtcata ggctatgtta ggagtgtgca atacctttat tgatgttttg gtgttgtcct
171481 ggaagtgaaa ttgtaggtca ccaatagcgt taattggtat atttcttttt tgagcatcaa
171541 ctacgtttat gtcaggatta gatgatgctg agtgtatgtg atgagcagat cttataaggg
171601 ttcgtgatgc tcctgaatcg agaaggaggt gtccagggag ttcatcatca gaatgattag
171661 tatgatttac tgtagattca gtaagtttct ggcctaagat gaaggtcgtg cttattgttc
171721 aattgaatcg gttcagtagt tgatttactg atggaatcgt tgtccgtgct gggagagtta
171781 ttagatgtgg atacattgtg agccctggct gttttcgatt tcgaattatt tgtttttttga
171841 ggattccgag ctataacttt gggtttggtt gtattcgtat agctgcgaga atcattcttc
171901 tcatcactcg gatttctcct gtaattaggt ttactgtttc tcgatccctg ttgttcttca
171961 taaatagcat ggatatctaa gaacagttca gcgactgtca tatttagatg tcgatgacgt
172021 gtgtagcgta aaaatttata ttcgccagat agacctctca taattaattg gcatgcgacc
172081 ttgttattga tatgaatgcc attattgttc agtctgtcga taatgtttgt gacttttgtt
172141 tcaaatgcat ctgcaggtgt actgccatta tattgcaaat ttgccagggt cacaatgtcg
172201 tttgcctctt gggtatcaga ttgcattttt tcaatacttt tggaaagaat tttcatgata
172261 tccgtataat caacggatag gatgtctttg acccaggtag gtaggaattg agagggagca
172321 aatatttgaa aagtgttata caagaaggtg agttcatcat cagtgatctg acgtacgggt
172381 tttccgttta ctgtcggaat aataccaccg agattcgagt tttgtaaaaa tttgatgtat
172441 gttttaaccc aatttggaaa gtcattaggt gaggttaaca ttggtggtgg tctgacatat
172501 tttttagtgg atgtcatatc agagtccgct gaggatgaat cagtaaatgt attacctgac

172561 tcaggtgatg gagtgctcag aggcgttcca actgatgatg gatactgcgg aaactgtgat
172621 tgtggcccag gtggaaagta cataggcgac atttgataag gtgtatacgg aatcatagat
172681 gggtgtccgt aaaatgacca accagatgga ttggcttggt tttgggtcat catgcactgc
172741 tgtgggtacg gcccattctg tggtgaatgt gactgagcag tttgaggaga ggcatgatgg
172801 gggttctctg gaacagctga tgaagcaggt gttgttgtct gttgagagtt agccttagtg
172861 gaagccttct cacattcttc tgttttggaa gctgaaacgt ctaacggatc ttgatttgtg
172921 tggacttcct tagaagtaac cgaagcacag gcgctaccat gagaaatggg tgaatgttga
172981 gataattgtt gggattccat tgttgataaa ggctataata ttaggtatac agaatatact
173041 agaagttctc ctcgaggata taggaatcct caaaatggaa tctatatttc tacatactaa
173101 tattacgatt attcctcatt ccgttttata tgtttatatt cattgatcct attacattat
173161 caatccttgc gtttcagctt ccactaattt agatgactat ttctcatcat ttgcgtcatc
173221 ttctaacacc gtatatgata atatactagt aatgtaaata ctagttagta gatgatagtt
173281 gatttctatt ccaacaggtt cttattttca acaatgtaat tgatggcctt aaatctctac
173341 tacatcataa agcttctaag cacttaccat tccttcataa gtctagtatt gtaatgagtt
173401 gggcacatgg cgcagttggt agcgcgcttc ccttgcaagg aagaggtcat cggttcgatt
173461 ccggttgcgt ccaaattttt ttgttaatcc aacacaattg aattcgtgaa tagctgactg
173521 tcatcagtaa tgttcgtgga aagtacctat ccatactgtt gtatcacgac taagtagttg
173581 tcgactacta cctcctcaac cccagttata tccctatgac acattggagg atgctgaata
173641 atgacagaat tttattcctc cttttcatta tcataatctg aagcaaagtt aaaaaaataga
173701 agaagtaaga taaactttgt agatacgata tatagttgtt tgttttagct atcatatatg
173761 ctgaactgtt actaccttat tttttccgaa atgtttctaa aacaaataaa tattcatgaa
173821 tatgatgcaa gttcgttgga tgagaaaaag accaggcttt attgtaagga caatatcatt
173881 tacgaataat ttcatccaat tgtttcatca acacatccat tctgtcaaaa actggagcat
173941 agagatgttg aaccaaagaa ttgggcttcg aagccggatc ggacatattg gtattagaca
174001 atgcccttga tccggaaagt gaactattta tgcttgctga tgataccgcc ctttgttcat
174061 ccatataaaa atcttcttca gatgatgagt gtaacgataa ggttgattgc ttttcttctt
174121 cgggggtact gtgatgtttt tctagctcct ctttaggttg ttctgcgtca gtttgatttg
174181 ccattcctcc actagacccc aagctggctt gaatttgccc ctctgtgttt tccacgtttg
174241 ttgtctcttc aattttcgat tttgcatgaa tatattgtga aatgtccttt attgctctag
174301 atgatggaat actgctgctt tcttcaacta ttggagtgga tggtgattcg tttgtccatt
174361 ctgacgacga cggtgaaaac tcgccaattg ttgaaaggga attttgagag tctgattttg
174421 ccataccatt cgcaaatggc atacgaggcg cttcaaatac ttttttcaga tcggtatcgt
174481 aatcactgtc aaccggttca gacttttctt gccccacata ccgatttgca ttataatcat
174541 cttctctgtc tttactcgat gatgtcttag gctcactttc gtttttgttg tcttgtctcc
174601 gaactttttt cacatttagt ggtgtattca aagatgtact aaaggagacg tcgctcacta
174661 catcgctcgt atcatcatcc tggaatttgc caaaatttaa cttcgcttca gtgtattcgt
174721 cccggtgcgt tacttttgtt tccttgtcgt tatcaacatc atcgtcgtac gactgtccat
174781 tgccttcacc gccatcatcg ttactatcta acgaggtatc ttcaattgca tcatctataa
174841 acctgtctgc ataaccgaca acatcattga aactaacaga tttgttcccc ttaccatatc

174901 cattcagtgc aggtgtcgga attatactct ctgcatcact ttgattcttg ttaccagagc
174961 tgatggaatc atctttcgaa tcggaggaag caaccgattg agaagacatg ttttcatttt
175021 tccagcaatt caatcgagct agtctttctg gaaaggtttc tagaatttcc gctggcgcaa
175081 acccgatttt accatcagtg atcctcttaa ccagccacca ataggcatcc tggtcattca
175141 aaagtataca aggttcgtct tgccctaatt gacaatgtga agaatcatgg ccattgaacg
175201 catataaagc atatagttta tcagggtcca gttctcttgg cggcgataag ggttggtaat
175261 cgtcgttttc ctcctccaaa tcgtctttct ccccttgaaa ccctgaatct tgaaattttc
175321 tgtcaaagtc atcgtcatca gagtagtttt cctcttcatc ctcgtcctcc gagatggcgt
175381 atttcaagcc atcactaaaa tcgtcaacgt cagggttcat tgtattattc actgaaaaat
175441 gcacctcatc cttatcggcg ctatccacgg aatcagtttc gatctcttgt agccttcttt
175501 ccaaattgtc ctcaaattca gaatccgaat aatcatatga gtcgggcaat tccgaagaat
175561 tttctttgta atgtcgctga tcccttttac tgttgctatg catatgtttt tccacgttct
175621 cctcctctaa ctctttgtca tcatctctat ttcgcagaac atcatggccc ttttctgccg
175681 cattactcag tatattaagt ttcgaattga agggcgaact cttattcgaa gtcggagtca
175741 ccacaacact tccgcccata ctctccgaat cctcgtttcc taaagtaagt ttacttccac
175801 ttgtaggcct attattaatg atatctgaat aatcctctat tagggttgga tcattcagta
175861 gcgcgtgcga ttgaaaggag tccatgcccg acgtcgacgt gattagcgaa ggcgcgtaac
175921 cattgtcatg tctagcagct atagaactaa cctccttgac accacttgcg gaagtctcat
175981 caacatgctc ttccttatta ctcattctct taccaagcag agaatgttat ctaaaaacta
176041 cgtgtatttc acctctttct cgacttgaac acgtccaact ccttaagtac taccacagcc
176101 aggaaagaat ggatccagtt ctacacgata gcaaagcaga aaacacaacc agcgtacccc
176161 tgtagaagct tctttgttta cagcacttga tccatgtagc catactcgaa atttcaactc
176221 atctgaaact tttcctgaag gttgaaaaag aatgccataa gggtcacccg aagcttattc
176281 acgagtcagt ctgactcttg cgagagatga ggatgtaata atactaatct cgaagatgcc
176341 atctaataca tatagacata catatatata tatatacatt ctatatattc ttacccagat
176401 tctttgaggt aagacggttg ggttttatct tttgcagttg gtactattaa gaacaatcga
176461 atcataagca ttgcttacaa agaatacaca tacgaaatat taacgataat gtcaattacg
176521 aagactgaac tggacggtat attgccattg gtggccagag gtaaagttag agacatatat
176581 gaggtagacg ctggtacgtt gctgtttgtt gctacggatc gtatctctgc atatgacgtt
176641 attatggaaa acagcattcc tgaaaagggg atcctattga ccaaactgtc agagttctgg
176701 ttcaagttcc tgtccaacga tgttcgtaat catttggtcg acatcgcccc aggtaagact
176761 attttcgatt atctacctgc aaaattgagc gaaccaaagt acaaaacgca actagaagac
176821 cgctctctat tggttcacaa acataaacta attccattgg aagtaattgt cagaggctac
176881 atcaccggat ctgcttggaa agagtacgta aaaacaggta ctgtgcatgg tttgaaacaa
176941 cctcaaggac ttaaagaatc tcaagagttc ccagaaccaa tcttcacccc atcgaccaag
177001 gctgaacaag gtgaacatga cgaaaacatc tctcctgccc aggccgctga gctggtgggt
177061 gaagatttgt cacgtagagt ggcagaactg gctgtaaaac tgtactccaa gtgcaaagat
177121 tatgctaagg agaagggcat catcatcgca gacactaaat tcgaattcgg tattgacgaa
177181 aagaccaatg aaattattct agtggacgag gtgctaacgc cagactcctc tagattctgg

177241 aacggtgcct cttataaggt aggagaatcc caagattctt acgataagca atttttaaga
177301 gactggctta ctgctaataa gttgaacggt gttaacggcg tcaaaatgcc ccaagacatt
177361 gtcgacagga caagggccaa atatatagag gcttatgaaa cattgacagg gtctaaatgg
177421 tctcactaac gtgatttaca tatactacaa gtcgccagtg taactcctca ctgaatatga
177481 ttcatacata cccgtatgta ttaatgtata aatgttctca gagcaaattt tatcgatatc
177541 ttgtttgcca gtggtatgca ggtttggcaa attttttacc ataatatccg tttatagatt
177601 ctggaacctt accaactttc ttaccgctaa ttacttccct ggctcgctcc tccactgcct
177661 gggtaaattg ttccttcaac tgactcagtt ctctttcgta ttcaatagct tgcttctcga
177721 ggattttttc aatgtttgtc agctcatttt catagtccag taacttcctt tcaaatctct
177781 ctaattgcaa cgactttctt gcagttcgta tctgaatatc ttgcagtaat tcaaaagtgg
177841 aaggcctggt tcttaagttc acatctatca ttgaatgtat tatggcatta agccctctag
177901 agtaatactc agggacggtg tcacatttcc cgtttttaat cttagtttgt agctcgagat
177961 aattttttgc ctgaaatggg gggtgcaacg aacacatctc aaaaataaca caacctagtg
178021 accagatgtc ggatagtggg gagtatggtt ggtccatcaa cacttcaggc gacatgtaat
178081 atggtgtacc gacgtatgtt gtggcaaatt gaatactagt ttccagagat ttggctaacc
178141 caaaatcacc taactttacc acaacttgac tatagtccat agggctcccc cttttccctg
178201 aattcactct atggtctctg taataattac tattcacttc ctcgtgaccg tctacttgtt
178261 cattaatatt gtaatcgcta tcatcatagc ttaagaatat atttcctggt ttcagatcac
178321 gatggataac gatgtttttg ccttttaccg gtggtttcat ccggtcatat attgtggtca
178381 aagttggcaa ttcaacacca taatgacatt tatagagcgc agtcaataat tgggccagga
178441 taccccacac aattttttct ggtatatatt tatgctcctg tttgtagtgc ttaatcatct
178501 gggataaatc acccctggaa caatattcca tataaaggta taacacttct ttttgttcat
178561 cgaagtccca gttataaaat tctacaatat tttcatgctt caactgcgat agaatgctac
178621 attcagcgat cagctgttgt ctctctttgc tattcatatg gccatatttg atatcctttc
178681 taaccaaaag tttcttggta ggtatatgga tgacttttcg tacagaccca aatgaacctc
178741 tcccaatttc ttcgagaact tggtattctg accttggtgg gtgtccctgc tgctgctgag
178801 gactacggta ttcttggaaa aactgtcgtc tatgcatact cacacagaga attgattcaa
178861 ttatcaaata gcactctcat tgaaattagt attgtgaatc ttgctctttt catgttatat
178921 gatttgatat tcttttgaaa agtcgctttt atttacgttt aacctaatta ggaaacgtaa
178981 tgaaaaaatt cagaaacctt aaaaaaaaaa cttggctgta acctatcgga agactgtgcc
179041 actgcaatca tgtcagatat cgtatttcag atttattgat ctatagctag aaacatttaa
179101 caaaatgcac tttgagtcgt tcatacattt aatccccaat tgaaaaaaaa aagaaaagaa
179161 aaaaagcata tatatgtata tgctttttta tcattactgg cctctttaaa ttcaaaaact
179221 tttctgatct cttttccaaa cagatgcgtt ttcagtattg gaaggttcac aattctatat
179281 atagtgttaa tgtaatgctg tattatttct ctatatatgt atgtatgcac atgcaattcc
179341 tacattatgt ttgaaatgtt gtaatgggga cggaaaagcc gtcactttta tctttggagg
179401 atcgcaaatt actacgctca tcttttgttg gagaactacc aattgctgca gtgacgcttg
179461 aaactttttg caggcttcct ttttttgata atgaacggat atcctcacat agctgacaga
179521 tcaaaactga gtcccccca acttgatttg aattcctttt tgggacatcc ttgttccagt

179581 tgttgtttaa aaactccgtt atttgtacta aaatggcacg aagtttgaaa acagtggcct
179641 tattacgacg atttggttcg gtttgcttta taggatccgg ctgtcctgtg tagtcatata
179701 gcttcaataa cgattttgag aatattagct ttatgaattt caatagatcg atttgaataa
179761 gtaaactgtt gaaataggta tcaaagaaag aaacaatctt ttcataaaag ttcgggtgaa
179821 atatgatatt aattgtaagg ttttcaaaac cgggcaatga gcatatcttc gtgaaaatgc
179881 taattagttt actgaggtta acgtcatcgt cattcaatgc ataaaacagg tcaactaatt
179941 gatcaatcgg taattcaata gcagcgtcat tgttgacctt tatgaggaaa acactatgag
180001 actctgatga gcccactgta ggagccacat catcatttac attgcgtaac gtaaaatgca
180061 agcagttcag tatgatctcc agtccactta tggaggtgtt attctttttt ctgatgaaat
180121 tcatagctgt agaaaataaa gaagcactca tttggtccat atccaaactc atttcaacgc
180181 atatagcagt gatttgtttc caaatgaaag ctgctgtctt tgcgtcgtcg ataaagggaa
180241 tgatggtatc aatattccgg ataaaagcaa aaaaagcatc gttgtcaagc agatcattta
180301 aaaaagtgct gtttatatgt actgtaaggt aacaaagctt ggtgaagtac tttaaaacag
180361 ataaatcttt gatttgggcc atgtcaacaa aaaaagatgt tagccaggta aagaaaccct
180421 ccggtaattc aattagtagt cttgattttg aagatatggt cagatgggga aagtttgaaa
180481 tgctcttgtg tcgcattgga gaagagggac gcgttgccat caaagaatga acaggggacc
180541 ttgatggaga ctgtaccgaa ttcactggtg agttcctcgt gggtgaggag gataacggta
180601 gagaggaaga agaaggaata gcggacttgt gtattttatc gtcattcgtg gttatcatat
180661 agtttattga tttgaagact acgtaagtaa tttgaggact gattaaaatt ttcttttta
180721 gcttagagtc aattaaagag ggcaaaattt tctcaaaaga ccatggtgca tatgacgata
180781 gctttagtag tatggattgg gctcttcttt catggatgtt attcagaagg agtgatatat
180841 cgaggtgttt gaaacaccag cgacaccaga aggctgtgga tgttaaatcg tagaacctat
180901 agacgagttc taaaatatac tttggggttt tcagcgatgc aaaattcgga ggatacatta
180961 ttccacattc aattaaagtc tgagggtagt cgatgacgaa ctctttggct aaatgttcga
181021 atttaatgat cagtggaatt cctcccatag cgatgaattt cagtcgcaac ctagagtggt
181081 tatgctgggt atcgtaaaca aaaatggagc caaatgcagt tattaatcgt ttatcaacag
181141 ttgtgcgcga cagacactcg ataattgtat cagcgatgtt ctcgagggag caaacactga
181201 aaagcacatg caagtccgtc aaaggcttag atgaactctt caattggctc agcaattgac
181261 tttcagtggg gggcattaaa tctagttctt gattgttttc tgcccaggca gcgggagccg
181321 ctgcaagact gaatttagag ggtgatatat ttagtttctc ttcttgaaaa tcggcatccc
181381 aatgataatc agcgtcggta aagtcctcct tgaacttgtt gagcttgtcg accttcacat
181441 tttcggtaga gttgatccac acatgcttga gtaactggtc ggctgtcggc ctcttgtaca
181501 tgtttttcac aaagcattta gataagaaat cctttagtgg ctcagagaaa gagctaggtg
181561 ggtagtaggt atcattttca acagcgtagt agatattggc gtctgtcaaa ttgtggtagg
181621 gtggattctt tgtgagcatt tcaactacag tggcacctag agaccaaatg tcgctgagcg
181681 tagaagctcc cctgttgccc aggatctctg gagccatcca attgagtgtg cccgctagcg
181741 ttaaggcgct ggagttcaca atagtggaaa cgccaaaatc agcaagtttg acagtgttat
181801 cagcactcag caggatgtta gccgccttga tgtccctgtg gatgactcct tcaccgtgta
181861 aatatttcag ccccaatagt gtctgtgtca cataggtttt cgattcattt tcacttaatc

181921 cggtagagct ccttgaaatg agcctcctca aagaaccatt agcgcagtat tcgaggagga
182041 aattttttaa caagctaatt tctgccataa tgtcattaag ttcctcatca ttttcgtaca
182101 cgacctcctt tattgccacg acttggtcag tatgtttatt aatggctttg taaactaccc
182161 cgtaagaacc cctcccaatg acctgcttca agtggtattg cacggatttc tcagatgccc
182221 tctggatggg agtcaagttg actctatcgg tatcggccat actgttcatg gtatagtctt
182281 accaggaaaa tgggtagtgc ttatgtgtgt tttgtccttc ctcgagcctc caagtagaag
182341 atataccttt tgtgaggcag atctcccgta tacaaaaata acagcaagaa aagcggaaag
182401 accatcgcaa ggtggaaagg attataatgg cacagcaaag tccgcacaga gcactacagt
182461 atagcataga gtgctaatga gttgataggc ccaattttga ttatgcctct tttccataca
182521 cgacgccaga ggacattatt acattacagt agttcgccgc tagatgacaa acgacatcct
182581 taccgatatg agatgtgcaa agctacataa tggcaacaag cgttatgaac agccttgtct
182641 ttacgaccac agaaaagccg tattagagct cttcagctgc aaaattttct tctaatatga
182701 tgcaaagcca tcaaaaatca tgcatagtta tgaaatacct gatgaaacgc ttcgagttcg
182761 tgctcaagaa attactgaaa ggttaccgag aagaaaaata tctatgagac acgataaggc
182821 cccttctgaa tccattgtcc tgggcttgtt cattctattt accacttaaa attgatcctt
182881 tcaaaggaat ttttttctat ttccaatagt atatttgtac aaaaactaca aaaatggata
182941 aaaaataaca gtaatttgtg actactgtaa atatcactga tttggatttt gtaatgagta
183001 ctgctcatgc ccatgccgat gcaagtggat cataaatttt actaaacgat attcgataat
183061 gcgccaagcc tttataagga actcaaaata acccatatgg acagtttcag aaggccaaat
183121 aacgatcaag gacattcact catgtttttc aaaggcgaag agtgtaaaat tttcttctat
183181 atagttcgaa tattttatct tataaatttc agtcgtcatt ttccacattc gaactcaaat
183241 aatgataaag aacgctgcag taatggctta aaaaaacata ctttataacc cattatctct
183301 tacgtgtaat ttaaaattgt ttatagtact atttggttat gcttgtatgc ctctattatt
183361 tacttgatct ttttatgttt tcttatgatt gaattattta tattctaaat tcctcacgaa
183421 tttatactga agatttcctt ccaggcgaga ataataaaca catatttatg atgataacaa
183481 gacgaacgtg tattaagctc ccagtacgag ggaagcagta aaaattatcc caagatccat
183541 ttaaaatgga taactccacg agctacaaca aaatactaag ggaataggcc gttatttccg
183601 taaaggatgg tttaataata agaaatttat aatattaata atacatatat acaaaaattt
183661 atatttatat acatgcgcct aactattcat actattaatt tcatattatt aagctttttt
183721 tttttcattt atcatttttt ttcgtaacct ctcatacctg tacaggtttc attcgtaaag
183781 cagggactct agtttgcgat agtgtagata ccgtctacgg atagagcact agagatagct
183841 ggctttaatc tgctggagta ccatggaaca ccagtgataa ctctggtaac ttggtcggcg
183901 ggaataccag tcaacatggt ggtgaaatca ccgtagttga aaacagcttc tgcaatttca
183961 actggataag tttcagttgg gtgagcagct tagaaagagt agtattcagc caaatgagct
184021 ctgatatcgg aaacataaac acctaattca accaaattaa ctctttcgtc agattgagat
184081 aatgtagtgg ttgctgcggc ggaggcacca gcagcaatgg cggcgacacc ggcagcgatt
184141 gaagttaatt tgaccattgt atttgttgtt ttttgggtta ttgcttagtg atgatatagg
184201 cttaactgga aggaaaagaa cagagaaatg tctcaaacaa agctgatcaa gccgctgtat

184261 ttatatgaaa ctttgaacaa ctacatctgc acacatgggc tcttactggt cgcccatctc
184321 acactcatgc cttccacatt ccacttagcg actaagtcat tattactatg gggacgggtt
184381 gttcttgaac gatgctatac ttcgtatagg aagccgtttt tttatgcccc atcctttcat
184441 atgttccata gcacaagaat gttctctaca ggaaaagtgc ctatagggct gcagctgcag
184501 ttttggccaa gaaatagaac caaagccaaa tttattttgg gccctcgttc aagggccatc
184561 tcacccttgg cactaaacgg ttagtaggag ggaaatcgga cttttcccaa attagaaaca
184621 atgaaaaatt aagtgtgagc tcttagagtc gcatctgcag gaatatgcac acaaaaaggg
184681 gagctgtacg taaataatca gaccacacaa actattgcca accatttgat actcacgcta
184741 gatatgatgg gggttcttgt ttggacaaca caagtctcag agccagcgta gatatgcttg
184801 tacataaatg acgactgggg catcaattga atcgggttac attgtgcgag ctattacatg
184861 aagagaatat gcctttaggg taatttccaa atgtaggaag tctcgctaag tagggcgccc
184921 aaatctgtat agcgatgttg ttgaggccat atagtaaaat gacgtgccaa ttaccgagct
184981 tttgatggag gtaaaatcta agattaatct tgcgccttga aaccactaga aatgaaagga
185041 attggtgaaa aaataatcgc gcaatagatg acatggaacg acagaagtct tgtattgtgc
185101 acgaatccgc aatattcaaa gccgaagttc atatacgaat gcgaactatt tcttagggta
185161 gctctctgta tgggccgcca taaattagta ccaaaagata ggtttttgaa aaggctacaa
185221 tgtgcttttt tccttcttgc tttcgagtcc ggtgaacaga atattacgac gtccttgtat
185281 taagagccag acctcctgtt agcgtcacta taagagtaag tctgaaatac gcaacaacta
185341 cagtgcaatg aaaaagtgct caactcaatg acaataaaca atttaaccat ggcaggttaa
185401 aatattactg cgatcagtaa aaatggggat atcacctttt gacacataac atagcaataa
185461 agtaacagat cattagtgat cggacaacct gaaccaacga tataatgtcg aagccaccac
185521 taccttttaag attagtagcg ctgcaggggg agacaatgag agaaatttcc cgccacatga
185581 actgagtcag gagttttttt tttcttgctg gagaatcatt taatttcatg gttaaactcc
185641 tctataagca tcccattctc ccatgcctga aaacactttt gtccattcga tcctcatgca
185701 gccctcgtta atatgctaaa atggctcatt aaattgtaga ttgtatcgtt cgagaaacgt
185761 caggcatgat agatgttgca atcacaggac attgattatt taatcctgct ctcaacatgt
185821 tcaataagtt gaagagttgc tgatctcccc gtatatctta tgaaccaaag catggtgggt
185881 gaatgttatg gttatccttg ttgaaaaatg attgatagac tggattgagc ggaaaaacat
185941 gggtcaatat gctgatcttg acatttttca aaatccacgg gggatcaaat caacttctta
186001 tagcgtatga cctcttttac attgtttaat gatgttaaga ttgcgatatt atagtcagtt
186061 aagttactca aacgcacaga tttaatagaa aactgcgtct tcgttgccta gtcgatcata
186121 ataaattcgc agattatttc gaatttgatc tccttcgaaa tcaagtttat tctcttcaca
186181 acaaaaaatg cttttaactt gaacaaaact cgtaaactat ttccccactg ttgcttcggg
186241 acgacccagt tattcaatat cttgcaatgc taattttttt tgggagagca gttgcaaata
186301 ttgcaaacac atctaaagcg tacccacaat ttatgacttc ctggagccca gaacagccca
186361 aaaaaaaaaa gatgcgttct ttttatacca atatattaga tacgtaaact ctactcatat
186421 tgcaggtatg cccacatctg gatattgact ttgccaatat tcccgcacag catgggcttg
186481 aatttcggct gctttaaaga ggcaccactt tacggttggt tcaacatcag aattttgagt
186541 tgcagcctga ttttctggaa cactgatgaa cggctgtgta ttcgctgtat cccactgtac

186601 atcaggatat tttcccttta tgagatcctt gaaaaattca tagcactggt gttcacaaaa
186661 aaagtggtat ggtgttttcc ataagccagc cttgaacaaa tattgattca tgttatacgt
186721 tatggttttc cattctttcc cagctatcga tggtctatga gttattacct ctagtagaag
186781 ttttgtacgg aatgtttcat tacttattgg tctactgaat gaccatatct gaaggactac
186841 catagagcca cctaaacata tccggatcac catggcaggg gagagaacac cagaaaacca
186901 aatgtttgtt aaagtcgcca aaatcgtcaa gacaaaaagg aggaaattga tcattatata
186961 cttggcgcgt acaatttcgt aaagcaaata actctggtat gatgcaaact catcctctgg
187021 aaggacgata tcagctgaga ttaaaggact ttcagggttg tcaggagatc cttccctcaa
187081 tgggattgct ttaggatcgt ccgtgacatg agtgtttttt ttaaataaga ttgcatgttt
187141 aacaaacgat ttaacttgct tttgcttaca agtcaagtaa accttatcct gatagcttag
187201 gaaaaataga cttgaatgtg tcgaacattt caaacctcaa ttggtatttt cctttttttc
187261 aactgtacgt acatagcttt tcgctttctt tagcgccccc agatgaaagt atatatcgta
187321 acaaggatgg gaacatgaaa ggtactgaaa aaacatctgt atttattaaa agtaaatcaa
187381 aagcagactg ggaagttctg tcgtagggat ttttttttta atgttatgtg tgtaggatta
187441 ttctatttcc ttgaatttct cgatcgagat ttttcgtacc tgtgtatttt tggatataag
187501 agtgtttctg atctattgag tgagcaggtc tccagcggaa tatagagtag attgaatatg
187561 gaagaggact acattaaggc ttattgttag ttagttactg ttaggacgct tcggcgagct
187621 gatgtctgac ttctcgttgt atcaaagagc tcccaatacg ccagcgcatt taaactatga
187681 tcacggaatg ctggattagt agtatagcaa aagtaacact tgtccaccgc agactccatc
187741 acttagtcaa caccttgggt gttttaccgc tgataatggc cgtaaaatcg ccagatatat
187801 atcatcattg ttcttcgcga ataatacgta acacagtctc ttttcgaaat ttagatgagg
187861 accataggca tgacttattt actgagatgt ccctgcgtta aaacttttac tggccgattg
187921 ctaactttat atttgttaat aaaactattc acgcctgtgt cctaattgtt ggataatacc
187981 taaacaataa cgatgttgta tagctaagag gacgaccaga caaaaagtta taaactttac
188041 cctcgttgaa aatggggcag ccacctatga atcacttccc attacaatgc cgaatagcaa
188101 aatatgtagt agaacacgta cacgcatgat aattacttcc atgctgtact tattttttgg
188161 gtgtctcttc agaaagaatg ctttatataa ccatgtgttt gaattagcga tcagctaata
188221 acaagtcagt gtccaaatag ttaaaacatt gtgacccaaa tatcacaaat aagtggttgt
188281 ttggccgagc ggtctaaggc gcctgattca agaaatatct tgaccgcagt gaactgtggg
188341 aatactcagg tatcgtaaga tgcaagagtt cgaatctctt agcaaccatt atttttttctt
188401 tttcctccta tacttcataa tctacgtagg aatgaaagta ccaacattat accaatgagg
188461 gtgtgtttcg tggatgcata tactctgaag ataaaaacaa actcaagtcc gcttcctacg
188521 gtttgagtat ttcttaccac tacataataa agaatattac gttaactgta aaatcaagta
188581 gacttggaaa atacaacgag aacactttcc tgattctgca tcagcgtttt cttatcacca
188641 gctgtacttc tacattagct aactctcctt tctataaagg gcgtctttca cttcacttgt
188701 gccatgttac aaagctccaa acgcacttct aactgagtac aatgcacgat cccactgaca
188761 gacaaaacag cttcacagaa tttgatcatg ccatcgtaaa aaccacgtag taaggaataa
188821 aaaatcccga agtcgatcat actatgtaga gatgtacatg aatagtctag gaatctggtt
188881 ttccagcatg ttgctttggt ctgctccaag cgctatggaa gcgctcgcca tgagatatgc

188941 tgtttcaagg caaaatagca aagctctttg taaagaaata caattcagag aagaagctac
189001 agcattttgt ttctggatga tccctgcagg ttcatactac taagtaaatc ttgaacagtt
189061 caaatttcaa caattcagaa accgctcttt ttatatactc taccaaacga gatgaaacag
189121 catttttta ctcttataag gtaccaatat tttgacgtat gctttcttta atgttcgcga
189181 tcgggctggg ccattaaact taccttagat attatttgga acagcaccgc aagtgctgat
189241 gtcccagaaa tgggcgccgg ttcaattagg tcgtgaagtc agacatatgg agactctcgg
189301 actgaaagca ctaagggatg atagctggca tgccaattcc attttaaatt tacacatcaa
189361 gttacagggt ttgggaaaat cacgttcaaa gcctgaaaat ttgaggttgt tcacggaaat
189421 catttggtta tgtctgtcgg cctgctattt agagacattt tttattgcaa caacctactc
189481 tatgcactta cacggaatcg cagaataacg cgcgcacaac acaattggga aacgatagga
189541 ttttgaatag tgtattgctt tgtaccgatt taaataattc tttctcgtgt tgaatccgag
189601 ttgaagatga gtatgctttg aagaggtgaa atatcatcag taaaaaaaat aacgacaact
189661 gcaggactcg aacctgcgcg ggcaaagccc aaaagatttc taatctttcg ccttaaccac
189721 tcggccaagt tgccaaaatt gtatgttatt tgttgtatct caaaatgaga tatgtcagta
189781 tgacaatacg tcaccctgaa cgttcataaa acacatatga aacaacctta taacaaaacg
189841 aacaacatga gacaaaaccc gaccttccct agctgaacta cccaaagtat aaatgcctga
189901 acaattagtt tagatccgag attccgcgct tccaccactt agtatgattc atattttata
189961 taatatataa gataagtaac attccgtgaa ttaatctgat aaactgtttt gacaactggt
190021 tacttcccta agactgttta tattaggatt gtcaagacac tccggtatta ctcgagcccg
190081 taatacaaca ttattttcag tgataaaata tgtaaaccaa ttataagaaa aaggattgcg
190141 ttgcatcaca actgtaaacc attaattaaa aagagcaatt gctatttaga tttgttgctg
190201 agaattggct aaaaaatctg ataattgtag gacttctatt attgctaggg gcaatgtgtt
190261 ggaatgcaat tctgttggaa taaaaatcca ctatcgtcta tcaactaata gttatattat
190321 caatatatta tcatatacgg tgtagagatg atggcataag gtatgaaaag ctgtcatcga
190381 agttagagga agctgaagtg taaggattga taatgcaata ggataatgaa acatataaaa
190441 cggaatgagg aataatcgta agattggtat atagaaatat agactccatt atggggattc
190501 ctagaccctc gaggagaact tcaagtatat tctgtatacc taatattata gcctttatca
190561 acaatggaat cccaacaatt atctcaaaat tcccccaatt ctcaacatcc gactgccatg
190621 caatgtgctt ttctggatct cactcatgat cataatggcc ctgtaaaagg ctcgcactat
190681 tattattata tcttcactat atattatttc ggaggctgta cctatcagtg aaaaaacgcc
190741 tctaaaaatg aaaaaaaaaa agaatatgaa aggggttctg aattgctaaa atatttcgtc
190801 aaagctcaat tagtatcatg atcaagtcgt aattcgaatc agcataacaa cctccaaaac
190861 catataataa ccttacacaa gacaagatat caattcaaca tgcaaacccc ttcagaaaat
190921 accgacgtca agttggatac tctcgacgaa cccagtgcac atttaatcga ggaaaatgtg
190981 gctcttccag aggatacatt caattcgtac tggagttata tacttaatga aatcgctcgt
191041 tgtaaaccgc taatgattat gttcctaata cctgtgtgtt tggttttatt gattacgttt
191101 tttcatgata tcaaaggtat ccttgtgttt ttagtgattt ctcttatcct ctctattatc
191161 attttattga tcggtataac tgccttcgtg tctgagacct tgaataaggg tttcataatt
191221 aagcttttag tagaagtcat tacacgtaaa ccagcagtag gggggaagga atggagaata

191281 atcgcatata atatgaacca gtatctgttt gaccatggga tatggcatac tccgtattac
191341 ttttttgtg aacataggtg ccataaattt ttcaaaagcc ttatcaaaca gacaaggtcg
191401 aatgcacatt tgagttcacc aacgaacggt gcagagaata cgcagtcaaa cacaccagca
191461 aaagaggttt caaatgagat ggtaaaacct tatatcttta gttctgatcc agttttagaa
191521 gcttaccta ttaaagctgc ggaaattcac aaagaagctg aatttgagta ttggagaaag
191581 caatacccag aggttgattt gccttagggc cgaatttttg gtatttatct agtatattct
191641 aatataaaat gtacgagcat cattaacttc aagaacatta cgaagcccgc aattaagtgt
191701 cagtccatct gggtgtaaaa gttatgtacg ctcgaaacaa attttatgta gtttacttta
191761 gatgcaaatg ctattatata ttttgcttta tgatcctcgg cttgatgctc gccaacgtga
191821 gatagctggt catcacaata gatcagccgg gacgcttttc gatcacatcg aatcccttcg
191881 ggacgttgca acaatacgtg aaaaatgcct caaaaataat aaatacaatg gtgaacaacg
191941 ttaaaaaagc ataaaacagc tggctatttt gatcaggata acatctataa gtgccatatt
192001 aaggcaagat atcaattgac catgcaaaca ccttcagaaa ataccgacgt aaagatggat
192061 actctcgacg aacccagtgc acatttaatc gaagagaatg tagctcttcc cgaagacaca
192121 ttcagttcac atctgagtta tgtactttat gaaattgctc attgtaaacc gatcatgttt
192181 atgatcatca taatcgtgag tttgatctca ttgattgtgc tttttcatga taacgacggg
192241 tgcactgtga tcttagtgat gtcccttata gtagcctcca tggctttaat ggtggttgca
192301 gcattcacat tcgggaaagc gatcactgaa caggagttca tgataaagct tttagtggag
192361 gtgatcgcac gcaagcctgc ggggaaggaa tggggtactg tcgcatataa tatgaaccaa
192421 tatctattca tgaagagact atggtatacc ccgtactatt tctatagcgg caagaagtgc
192481 catgagttct tcaccactct tatcaaggaa gtgaattctg gttcgcactc ggattcctca
192541 tcgaatagtg ccgaggatac acaatcacct gtctcagcag ggaagacttc aaatggtcta
192601 aacaactttt atagtattag atcagaccct attttgatgg catatgtttt gaaggcaaca
192661 caaatagaaa aggaggctca aagtgaatac tggagaaagc aatatcctga cgctgattta
192721 ccttgaagcg gaagcatttt attcaccaag tatacttact tttctttaaa acgagaacaa
192781 gaatcgaatt caagaacatc tcgaagccag aattgagcat catatattcg agctgtacaa
192841 acatcatggc ctacaactat cgtatttgta agttttttta gaggttttca tatttgttta
192901 ataagggttc tgtcagtttt tgtcacattc tattgttgcg cttcgcataa tgcagccaag
192961 aaaatccaaa caatagaaaa agaaaaaaag gatctcaaaa agggtttggt gttgtagtta
193021 taagaataac tagtgaataa aaaagctgtt gtttggtccg tattacattc gtcaaaaatt
193081 tagtactcaa atcgtgtatg caatcgcaac cacaaaataa aaatattaga ctggatgtgt
193141 tgagtggaga tggtgccaat ttagttgagg gaaatgtggt ccttcccaaa gacatgttca
193201 attcgtactt aagttattca ctttacgtgt tacgaggggg ctcattgtaa gccgataatg
193261 attatgttcc tggcatctgt aattttgatt tcactgacta atttccgagt ataccatctc
193321 atgtcccttc tatcctcttt tttcatctcc gggacagacc gacaataaag catctaatat
193381 taggctttcg ttagaggtaa gcacacgcca gcgttcggtg aaaggggaat gaaacattat
193441 cacgtacaag atgaataaat atctatttga ccataaaata tggagtactc cttactactt
193501 ttattgcgaa gaagattgcc accgtctttt tctaagtttt attgagggaa gaactttcga
193561 gaagccaaca agcaacgctg aggaaaatgt acaggagact gaagctggcg aatctttcac

193621 attaaatccc ggagaagatt ttcaaaattg ctttccaaga cagcggatat tgtagaacaa
193681 tctcaagtga agtattggca agatattggt gcaattattt gaaaggaagg agaaatattc
193741 tgacagtacc ttgctagcaa agggatttac caatccactg acgctaaaat ggggtagtaa
193801 attagataaa ttgcattcta acgtgacttt atatagtggg aaacagatat gtagcacaca
193861 aaacggcatg attatgctta attgattcct attttttaac gtaaatactc tcccagaacg
193921 atcagaaaac ttaacccgca accatctttg ctgtgctaac aacttatgtc gcctcaatac
193981 catttttatt ttgtatcatt ccggaactta gtattgaatg aaaaatgcct ccgaagtaaa
194041 aagcaggtga tgaaaagttt caattggtat aagacagatc gctattttga tccgcataac
194101 atccttcaac accatagcag agctatagag aagacaagat ataaactggg catgcaaaca
194161 tcttcagaaa gtaccgacgc caagtcggat tttctcgacg aacccagtgc atatttaatt
194221 gagaaaaatg tggctcttcc caaggacata ttcggttcgt acttaagtta ttggatatat
194281 gaagttactc gtcataaagc ggcagtaatt ttgctcgtac ttattgtgac ttcaatttta
194341 ttattagtgt ttttttataa tacggaattt tgcgttgcct ttgagatact attgttttcc
194401 ttttgctttc caggaacatg catggttgta attgcattta gtgaaccgat cggtgatcgg
194461 gaatttaaag ttaagcttct gatggaaatt atcacacgta aaccggcggt aaaggggaaa
194521 gaatggagga caattacata caagatgaac cagtatttat ttgatcatgg gctatgggat
194581 actccctact acttttaccg tgatgaagat tgccaccgtt attttctaag tcttattaag
194641 ggaagaactt tcaagaagca aaaggaatcg tcagccagca atgttaaaga cgcacaatca
194701 aatgacgaaa ccgctggcac accaaacgaa gccgctgagt cttctagttt tagtgccgga
194761 ccgaacttta taaagctcct caccaaggca gccgaaatcg aacaacaatt tcaaaaggaa
194821 tattggcgac aagagtatcc tggtgtcgat gagttttttt agacagaaga cgggagacac
194881 tagcacacaa ctttaccagg caaggtattt gacgctagca tgtgtccaat tcagtgtcat
194941 ttatgatttt ttgtagtagg atataaatat atacagcgct ccaaatagtg cggttgcccc
195001 aaaaacacca cggaacctca tctgttctcg tactttgttg tgacaaagta gctcactgcc
195061 ttattatcac attttcatta tgcaacgctt cggaaaatac gatgttgaaa atgcctctag
195121 agatgaaaaa caatcgtaaa agggtcctgc gtaattgaaa catttgatca gtatgcagtg
195181 gcacagaaac aaccaggaat actatagtca taggcaatac aaggtatata ttggctatgc
195241 agacccctcc agaaagtacc gacgtcaagt tagatacact taacgaacct agtgcacatt
195301 taattgagaa aaatgtggct cttcctaagg acatattccg ttcgtacttg agttattgga
195361 tctatgaaat cgctcgctat acaccagtca tgattttgtc cctggtaata ggggttttgg
195421 ttttattaat tatatttttt aatgacaacg aagcttgtgt tttcaattct gcaatatttg
195481 cttttacttc tcttgtaggt ttgttaataa tattaagtga tggtaatcca aagctagtca
195541 gtcgtcgaaa ttttaggacc gagcttttag tggatgtcat cacacgtaaa ccggcggtag
195601 aagggaaaga atggaggatc atcacataca acatgaacca atatttgttt aatcatgggc
195661 aatggcatac tccgtattac ttttacagcg atgaggattg ctaccgttat tttctacgcc
195721 ttgttgaggg agtaaccccc aagaagcaaa cagccacgtc aattggcaat tctccggtca
195781 ccgctaagcc tgaagatgcc atcgagtcag cttctcctag ttccagactg aattatcaaa
195841 actttttgct caaggcagcg gagatcgaac gacaagctca ggaaaattac tggcgaaggc
195901 ggcatcccaa tatcgatgcg cttcttaaaa agacggaata gcttagagac actaccatac

195961 gtaaagcgaa cataaactag agtatgatat ataatcagca ctaactggcc ggaaaacggc
196021 cgaaggaagc ctcgaaaagt cgattcgtgt tggacccatt tgctgaacaa agtggttcat
196081 tgcctaccta ttatggtagt agtcgtgata atcgtgtggt tggttttgtc aacggtgcat
196141 ttgcattttc atgacaataa accttgcgtt ttcgttctcg ggatattact ttccctccac
196201 ttctttcgcc tcaatagctc ctataagcat tctcagggcg tatgtcggtg atcgagattt
196261 ccaagcaagc ttttagtgga aatcatcgcg cgcaagccag cggtaaaggg aaaagaacgg
196321 aggacgatta catacaagat gaacgaataa ataaattaat aataaataat aataaaaagt
196381 acagtagcat taaatattat taagtttaat gattaaaaat tggttaattg tcaagaaaat
196441 ctaaggtatt aataaataaa taatactatg acaacttgca gcgaaagcat cagccccaat
196501 gaaaattaat cagaattgaa tctgagcgta tttatttgat aacggtttac gtaactgttg
196561 gaataaaaat caactatcat ctactaacta gtgtttacgt tactagtata ttatcatata
196621 cggtgttaga agatgacgca aatgatgaga aatagtcatc gttttcaacg gaagctgaaa
196681 tacaaggatt gataatgtaa taggatcaat gaatatcaac atataaaacg atgataataa
196741 tatttataga attgtgtaga attgcagatt cccttttatg gattcctaaa tcctcgagaa
196801 gaacttctag tatatctacg tacctaatat tattgcctta ttaaaaatgg aatcccaaca
196861 attatctcaa aattccccca attctcatca gtaacacccc accccgtatt acttttaccg
196921 tgatgaagat tggcatcgtt actttctaaa cgtaggacgt gcggaatgac aaaaccatca
196981 gcagtgtcac gatctctcca gtcacaatgg caatcatgag tgcatagtcc aaagtaaagg
197041 ggcaaggaaa agcatgattg aaaggactcc ccatctggac tctatatgtc atcagcggct
197101 aaaaaaaagc atatagcaca acatcagcat cagcatcagc actagagtca tcggcccggc
197161 ggtccgcggt catccccgcg gactttccgt ccgcccggcg ggctgtatca gcgtcaactg
197221 gaacgcgcat atatatacaa gacacacata acatagaagc acacccacga caataaccac
197281 acgacaataa ccacacccgc ccacccctcc tttccgtata caatgccaaa cttaaagaga
197341 ctacccatcc cgccactgca ggacacgctc aaccgctacc tggcacgcgt ggaacccctg
197401 caggacgagc gccaaaaccg ccgtacgcgc cgcactgtgc tctccgcaga aaacctggac
197461 gcattgaaca cgctgcacga gcggctgcta gaatacgacg cacggctcgc ggaaagcaac
197521 ccagagtcct catacatcga gcagttctgg tatgacgcgt acttgctata tgatgcaact
197581 gtcgttctca acgtcaaccc gtacttccaa ctgcaggacg acccaaccat caaagacaca
197641 ccagagacgg cggcacaggg cccctatggc gcacacacgg tgcaggttcg tcgtgccgca
197701 cgactcacca cctctattct caagttcatc cgccagattc gccacggcac actccgcaca
197761 gacactgtgc gcggcaaaac gccgctgtcg atggaccagt atgagcggct attcggctcc
197821 agtagaatcc ctccgggtcc cggcgagccc tcttgccact tgcaaacaga cgccacgtcg
197881 catcacgtgg tggcgatgta tcgtggccag ttctactggt tcgacgtgct ggacacacgc
197941 aacgagccca tcttcgccac cccagaacaa ctggagtgga acctctactc gatcatcatg
198001 gacgcggaat ccgccggaag cggatccgcg ccctttggcg tgttcaccac agagtcgcgc
198061 cgggtgtggt ccaacatcag ggactatctg ttccatgcgg acgactgcac caactggcgc
198121 aatctcaagc tgatcgactc cgcgctgttc gtggtctgtc tcgacgacgt ggcgtttgcc
198181 gccgatcagc aggacgagct cacgcgttcg atgctgtgcg ggacttctac catcaatctc
198241 gacccgcacc aacaccagcc gccattgaac gtgcagacag gcacctgtct caaccgctgg

198301 tacgacaagt tacaactgat cgtgaccaag aacggtaagg cgggcatcaa cttcgaacac
198361 accggtgtgg acggccacac tgtgctgcgg ctcgccacag acatctacac agactcgatc
198421 ctgagcttcg cacgcggtgt caccaagaac gtcgtcgaca tctttagcga cgacgatgga
198481 aaaccatcgt cgtcgtcgtt ggcctcggcg gctcactccg ccaacttgat caccatccct
198541 cgtaaactgg aatggcgcac tgacaatttc ctgcaatcgt cgctgcactt tgccgagacg
198601 cgcatctcgg acttgatctc gcaatacgag tttgttaatc ttgacttctc caactacggc
198661 gcgtcccata tcaagacagt gttcaagtgc tcgccagacg ccttcgtgca gcaggtgttc
198721 caggtcgcat acttcgcgtt gtacggtcgc ttcgagaccg tgtacgagcc tgccatgacc
198781 aaggcgttcc aaaacggccg cacagaggcc atccgctccg tcacgggcca atcgaagctc
198841 tttgtcaagt cactactgga ccaggatgcc tcggacgcca ccaaaattca gctcttgcac
198901 gacgcctgta cggcgcactc gcaaatcaca agggaatgct cccaggggct cggccaggac
198961 cgtcacttgt atgcgctcta ctgcctctgg aaccaatggt acaaggacaa gttggagctc
199021 ccacccatct tccgcgacaa gtcctggact accatgcaga acaacgtctt gagcacctcc
199081 aactgcggta acccctgcct caagagcttc gggttcgggc ctgtcaccgc caacggcttc
199141 ggcatcggct acatcatcag agaccactcc gtctctgtgg tggtgtcctc aaggcatcgc
199201 cagactgctc ggtttgcgtc gctcatggaa aagtcgctgc tggagatcga ccgcatcttc
199261 aaacggcagc aagctcgcgc agcaaaaccc gctgccagga ccactgctag cgccaacacc
199321 aaatcagaag acatgaaata cctgttgtcc ggctacgatt acttcgacgt gagcgtgtcc
199381 ggttgagttt atgctgagtt tttgcgcatc aatattattt ttactactac tactactact
199441 actactacta catactatta aatatactaa ataagaggaa aacgctttgg aagtgactgg
199501 cgccgccgct ggctactata atagcagcga ctgtaattta atctcatccc gtcgtttgga
199561 ttacctcttt tactcgccga gcgaacgtgc accaaaaagg gaaaggaaaa aaagaaaaaa
199621 aaaggaaaaa ggaaactcaa aacttggata aatagaagca ctcaaactaa attaaactgc
199681 caaaaaaaaa aaaaataaaa agggaaaagt ttaaacatca aagtacacct ttcacccctc
199741 cacacaccat ggaacaacct gatctatcgt ctgtggccat cagtaagccg ctgctgaagt
199801 tgaaacttct cgacgccctt cgccagggaa gtttccccaa cctacaagat ctcctaaaga
199861 aacaattcca gccgctagac gacccaaacg tccaacaagt gctccatctc atgctccact
199921 atgccgtgca agtcgccccc atggctgtca taaaggaaat cgtccatcat tgggtctcaa
199981 ctacaaacac cacttttcta aacatccatc ttgatctaaa cgaacgggac tccaacggca
200041 acaccccatt gcacatcgcc gcctaccagt cccgcggtga tatcgtagcc ttcctcctgg
200101 accaaccaac catcaacgac tgcgtgctca acaactccca cttgcaggcc atcgaaatgt
200161 gcaagaacct aaacatcgcg cagatgatgc aggtgaaacg ctccacatac gttgcagaga
200221 ccgcccagga attcagaaca gcttttaaca acagggactt cggccaccta gaatctatcc
200281 tctccagccc tcgaaacgca gaactgctcg acatcaacgg tatggacccg gagactggcg
200341 ataccgttct gcacgaattc gtcaagaaaa gagacgtcat catgtgccgt tggttgcttg
200401 aacacggtgc tgaccccttc aagagagacc gcaagggcaa actgcccatc gagctcgtta
200461 ggaaagtcaa tgaaaacgac accgccacca acaccaagat cgccatcgac atcgaactga
200521 aaaaactatt ggaaagggcc accagggagc aaagtgtcat cgacgtcaca aacaacaact
200581 tgcacgaggc ccccacttac aaaggctacc tgaaaaaatg gaccaacttc gctcaaggct

200641 acaaattgcg ttggttcatc cttagtagcg atgggaaact atcctactac atcgatcagg
200701 ccgacactaa gaatgcctgc aggggctccc taaacatgtc ttcgtgctct ctgcatttgg
200761 attcgtctga aaagttgaaa ttcgaaatta tcggcggtaa caacggtgtt atcaggtggc
200821 atttaaaggg gaaccacccc atcgagacaa atagatgggt ttgggccatc cagggcgcca
200881 taagatacgc aaaggacaga gaaattttgc tgcacaatgg cccctattcg ccttctctgg
200941 ccttaagcca tggcttgtca tccaaagtgt ccaataaaga aaacttgcat gcaacttcaa
201001 aacggttgac caagagcccg catctgtcca aatccacact gacacaaaac gatcacgata
201061 atgacgatga cagcactaac aacaacaaca acaaaagtaa taatgattat gacgataata
201121 ataataataa taataatgac gatgatgatt atgatgatga tgatgaaagt agaccccctca
201181 tagaaccatt accgttgatt tcatccagaa gccaaagctt aagcgaaatc actcccggtc
201241 cacattctag gaagtctaca gtctcgtcta caagggcagc cgatatacca tcagatgatg
201301 agggttactc tgaggacgat tctgatgacg acggtaactc ctcttacaca atggaaaacg
201361 gcggtgaaaa cgatggcgac gaagatctaa atgccattta tggtccctat attcaaaaac
201421 tacacatgct acaaagatcc atttccatcg agttggcatc tttgaacgaa ttgctgcaag
201481 ataaacaaca acacgatgag tactggaaca ccgtcaacac ttctattgaa accgtcagcg
201541 aatttttcga caaattaaat cggttgacct ctcaaaggga aaaaagaatg attgcccaaa
201601 tgaccaagca acgggatgtt aacaatgttt ggattcaatc ggtaaaagat ctggaaatgg
201661 aactggttga taaagacgaa aaattggttg ccttggataa agaacggaaa aatctgaaaa
201721 aaatgcttca aaaaaaattg aacaatcaac cacaggttga aactgaggct aatgaagaat
201781 ccgatgatgc aaattcaatg ataaaaggat cccaagaatc aacaaatacc cttgaggaaa
201841 tcgtaaaatt tatcgaagca acaaaggaaa gtgatgagga ttctgacgcc gacgaatttt
201901 tcgacgcaga agaagctgct tccgacaaaa aagccaatga ttcggaagac ttaaccacaa
201961 acaaggagac tccagctaat gcgaaaccac aagaagaagc tcctgaagac gagagcctta
202021 ttgtgatcag ttctccacag gtggaaaaga agaaccaact attaaaagag ggatcattcg
202081 tcggatatga agacccagtg agaaccaaac tggctttaga cgaagataat cgtcccaaga
202141 ttggtctctg gtctgtttta aagtctatgg tcggtcaaga cttaaccaaa ctaactctac
202201 cggtatcgtt caatgagcca acatccttac tacagagagt atctgaagat attgagtatt
202261 ctcatattct tgaccaagct gccacttttg aagactcctc tttaagaatg ctatatgtag
202321 ctgcctttac tgcatcaatg tacgcatcta ccactaacag agtgtctaaa ccattcaacc
202381 ccttactcgg tgaaactttt gaatatgcca gaactgatgg tcagtaccga ttcttcaccg
202441 aacaagtctc tcaccaccca cctatctctg ctacttggac agaatcgccc aaatgggatt
202501 tttacggtga atgtaatgtt gattcgtcat tcaatgggcg cacgttcgcc gtgcaacatt
202561 taggattatg gtacattact atccggcctg atcataatat tagtgttccc gaggaaactt
202621 attcctggaa aaaaccaaat aacactgtta tcggtatttt aatggggaaa ccacaagtag
202681 acaacagtgg ggacgtcaaa gtcacaaacc atactacagg cgactattgt atgctgcatt
202741 acaaagccca tggctggacc tcagccggtg catatgaagt cagaggtgaa gtattcaaca
202801 aggacggtaa aaaattatgg gttcttggtg ggcattggaa tgattccatt tacgggaaaa
202861 aagtaactgc tagaggcgga gaactgacat tagacagaat aaaaacggca aattctgcca
202921 cgggaggacc aaaactagat gggtctaagt ttctgatatg gaaagcaaat gaaaggcctt

202981 cagtgccatt taatttaacg tcgtttgcat tgactttgaa tgctttgcca ccccacttga
203041 taccatattt agcacccaca gatagtcgtt taaggcccga tcaaagggct atggaaaatg
203101 gtgaatacga taaagctgcc gcggaaaagc atcgtgttga agtaaaacaa agggcagcaa
203161 aaaaagaaag ggaacaaaaa ggagaagaat acagacctaa gtggtttgtc caggaggagc
203221 accccgttac caaaagtcta tactggaaat ttaatggaga gtattggaac aaaagaaaaa
203281 atcatgactt taaagattgt gctgatattt tctaagctgt gcaatgtagt cacaataaca
203341 ctcgttcatt tgtatccatt gcgaatgccg gtacatcgga aaacaggata ggacctattt
203401 aattatatag tatgaagtat tcataacttc ttgaggcatc aatacatcat attccatgag
203461 ctgcgtggca ttcatactca ttgatttaaa ggttttttat tttcatggaa aagattaacc
203521 gggctgaacg aaatatatta aagatgctaa aacttatgct ttcattgact ttcaatagtg
203581 tccactaacc aaaaaaaaaa actactctaa caagggatcc ccatggattc aaagccgata
203641 ccaaacagat attactctgc atagaattca aaatattatc catataaaga tgggaaagaa
203701 ttccaaaagg aaaattctgc tctagaaggt cacaaaacta gtaagaagtt gacccccctg
203761 ccattaaaaa acgtttttaa cagctctagc aatattctaa tttcgaaagt gctctcaaaa
203821 gaatttattc atttgcgaaa aaaagaatat ctcaaaattt tctcgatcac gtacaacatc
203881 gtagtattta aaggattatt aagtcaacaa ataatttcca caagaaaggt acctctagtt
203941 ttggtgatga agcaagacaa taactggcaa gggctctcac taaatattaa ccccttttcaa
204001 ataaaaaaag gattatggct tgcagcgctc ccaacaatta aactctgtta ctaccaaaaa
204061 acattgagcc caagaatgga ataaaatttt cactacacct cggacatgga tttgtacatg
204121 tcctattatc ctgtaatttt gacatatact gatgtggacc ttttgtttcg tataaatcgc
204181 tatttatttc cccagataac taaagaaaat ccttcaaccc agcgttctat attactatat
204241 tctcaaccca ccattttcct gctgtgcgat aattctcatt caatactacc atttcaaacc
204301 tagaaaaggg tgtctttatt aaaactgtaa gaaaatttta tcatggactc tgatgaaaag
204361 tcttctgaag agtaactttt tgatttttca aaaatgaaag aaactgctgt ctccatttat
204421 ggcctcatat ctgaaggaaa ttcgtctgct attagtcata ccactatctt ctttttgaaa
204481 ctgcatttca aggttgatta ggaagcatct aacaatcatt cgatagggaa aacagaaagg
204541 ccgttacagt tttttaataa tatcaatatc aagagaaaga actagggaca ttgcaaaatc
204601 aaaacctctg atacttggtt ttacgtaaaa attaccaaca cttaaacgtt aaaatttcaa
204661 actatctact gaaatataaa ttttcagtcg aagagatgaa acgattattt tcttgaataa
204721 ttcgttgtca caatttaaca gaaaaattac tgttttttta agtagttcac aaagagactc
204781 aaactgagat catggagacg tcggcaagca attgattgtt agttgttatc aaaactcatt
204841 agcttcggta aaactacata gattgaaaat cagtagtaac atacaacgtc gtttgcatgg
204901 ggttacaaac cgaccaacct gatgctttca aaaattcttt acttactggt tcacagtggc
204961 caagaagcat ataaatctgt actttgcagt aaaaaacaga tgatattgct cattcccagc
205021 gattgccaaa tacgatttat aacagattga agttatcttt ttcgaaagct ctatcatatg
205081 gctttgagta ataaggtgag attttttgg tgaagaaaat aaatacggtg ccttgatgca
205141 aaaattttag ccgcaaatct catcttcagt cccacatgat tcacctgatt acgactcact
205201 atttcatctg ttccataaat ctcggctttt tatatacaga acattagaca acgggaagag
205261 aaaaacgtca gtataaccca cttttgttcg taaaaaaagt ttattcactt ctactccgta

205321 ccaatcaatg actttaatgg tgaaacaatg atgaggagag taagacgatg taattttcat
205381 aaagcaatat catcccatgt tagtttactg gcggctacga gtaagaatat atactttagg
205441 agtgtaacca taccagtaat tgtgttccaa acaagttttt gaagcgcgtg tgctgaaagt
205501 aaaaaaaact taccttaata acgcaatttc tggcgaggtt aaaatattac atataaaaag
205561 ccttccaagt aattttgccc actaaaaagt aacacaaagg tccactggtt ctcagcttct
205621 tgttctttca aaaggatctt caaatccgct tccacttaag cagctttttg ctacgactta
205681 ttctgagata cccagtttcg cgcgggcact tctacccgcc gaaaaagttt atatgtcgga
205741 aagcttattt tgaattaagg tttggaataa atttcttttc tgcagatttt catgcagcgt
205801 cgagctgcta aacgtcacaa ctatttgacg tacattctgc aaaagtatgt cgacttcgca
205861 cggaccatta tgcttgcttt actacttatt ttgctttcga cttgttgagt ccgaaaagct
205921 gttcctgggt accaatagaa aatcagctga aatcttctca acgaaaaagt aaaataaact
205981 tggcagcatg cgttaatatc agcagaagaa aattttcact gaaaacagcc tttcactatt
206041 attttgcttt ttcaactgtt taaaatttta tcaaacaccg actaagtgcg taaaaattca
206101 ccaaaaaaaa aaaagcatga ctatcataat tgcaactagg gttctcttta aagctcttct
206161 tttcatctta gccatataac cacagaatgt tatgcaaaat tgataaacct tcctcaatat
206221 gctttattga tagttgtagc ctctgtaagg ttgctcaagc tgaccttgga aaacgctaaa
206281 atgtctttgc taaccggttt aagaaatgag ggattgaaat caatctttat ctgtatttgt
206341 tgaaaggttt ttcatgaaac taatctagaa tatatggtct ataaacgccc taaaagagaa
206401 gaaaatgctc aacttttata ttatctcgta acttgcagtc agtagcgtta acacatcaga
206461 ctggtcattg ccaaggaaat tttacgtcgt gtggtgaatt tcgaccaata ttatattgct
206521 ggatgatagc gctattatga tgcatttggg ataatgcgtt ttttcaatga tggctccaac
206581 taggctaatt aaatggttca agacttttcg aaaaatgcgc actttgggaa tcatacatca
206641 tggaactgct aaatttttaa atgacttgat agaatttcta aaaatattaa cgcttgaata
206701 aggttcttgt aaatagactt tgacactttt caggaggatc tttcaggaat aatcatcaca
206761 tttacccaat tatgaatcag ctcgtccaaa caacaaaaga tatccactat caccgaatct
206821 aatggattat catctaagca gaatgtcaag actctagcta ttgtcagctc tttcaaacgc
206881 gcaagctgat atcgtgatct acgcattatc ttcccttaca tccatcacat ccttatattc
206941 agtattgaag ctttttttacc ggctatttac acaactagtc taaagggtct ggaaatcaca
207001 acacttcatt cttcttttta attagttccg tcccttccca gtcatgtata actaccgtta
207061 aaagacatat atcaactata gctcagagtt agttaacaaa caatgaaaaa attataatga
207121 actattctta ctcgattaca taacagtcat aaaatatgga taggcaatat ccgccaatta
207181 taactacaaa taatcaaaag ttatataaag ataaatgaat aggattgaga gaatatttta
207241 atggtttact aattcaacaa acacgtcttc atatcacatc ctaaatggat aatatataaa
207301 gaaacattta aacgattatt gacaacctgg ttcttgatta ctagtacaaa atcaaaatca
207361 aacagctcca ctctagaatg aggcagctat gaaaaaaaaa cagcaggact tcggaaattg
207421 aaaaaaataa ctgtctctac atcgtaaatg ataggaagtt attcatccga tgatgttatt
207481 gttcttcagt gacagcttgt ggatacttta atagaagtgg aagctactaa ggaagcataa
207541 tttcaacaat ttttgtaaga aaatagcata tttgtgcata ggccccaatc ttgtcaaagg
207601 ataaaatcta ttcgattgga ctgagatttc aatgtaatat aaacaaggat aacctaccaa

207661 aaacggcaac ttttagatga tgcaccaatc gtgaaccttt atcatacagc atggtgcagg
207721 tgatgcttac aatgtggtat ttattacagc ggttttcagc aaattttttg ttgaagtgta
207781 taccctctat catattaagg gaaattttta gcattacctt taaattttca ctgtgagaaa
207841 ctgaacttct tttttttttg ttaggaaaga tcaatttcag gataatttag catacgattt
207901 aatgcaaaga catatgtatg ttaaaaaaaa acaacatgat gcaattattt cagctcaatc
207961 ctaagcgata gactccataa cgtatatttt tatattcttg acgatcctcg aagacctctc
208021 caacggaaca tattgattct tctaattttt acattatttt ttcaagtatc tgtcattttt
208081 ttcttcttga ccctttcagc ttttttgtcg tgctgtagat tttttttatt cttattaaaa
208141 tttaatagca tagttactgc caaaaaaaaa aatccttcaa cagggtcaaa aacataaatc
208201 caaggtggct tcatcatttt ttagaatcca atatttatta tcttatttgc cttcatgcac
208261 accctgcgtt ctacctgtac atcgttgaca accaatactt aaaagtatcg gagaagaatt
208321 ccgcaatatg ctagagtttc ggccactatg agcttatttt cacttatgat aaaatgcgaa
208381 acacttgctg ctttaatgtt actttgcagt agtaatatct ttgcctgcat tacctttctg
208441 caatacaacc atgtggactg cgaatgatct ttcatgctga aatctagata gaattaaatg
208501 gttccctctg tagctgcttc ttcccttcga taacttatta tgaagtaggc atcctgtgtt
208561 ttttacaaac tgctgttcat tccaacaggt acctatgata tcattcaaat tcgcctttcg
208621 ccactagtat attgcaatag cagtatacca acctgttggt gaatcaattt atctaatcat
208681 ttccgtttta aacccacaat tcattttttt tttttttttt ttcaagtatt tcaaatatgt
208741 ctactgagga cttttatctg caactgatac aaaaccacta tacggcaact tcttaactcg
208801 tagagctttg gacgagcttt ttcttttgta atgccgttac ctatgcaaag aaagaccagc
208861 tggccattgg ttatatcatt attttttcat tgaaaccaca ctgaagagaa gtttgaagat
208921 tatccaacaa ctttaaacac ctaaaaccca aaaggcttca agttacatcc atgaatcaag
208981 aactggaaac gcagcaaaat gaatgtttta taacctttat ttcatgcgaa gatcagtaaa
209041 attcaccata atgccaaaat gtccaactct aactgagagt ttaattaggt ttacaaaaca
209101 ttctattgta tgtagattcc cagacaaaaa aaagatatgc agcaaatatt cgttatgtct
209161 aatcggttat ctaagctacc gaatgcaatt acatttcctt aggtggttca gtacaggtct
209221 taggttgagt ttacaaaaga attctatgac cagattcttg tggctcgtct aatgaggatc
209281 attgaaactt atcctttaaa aggtcaagtt gaaagttgac gaagttcaag gtaaggactt
209341 gttttgacca acttccacaa tacagacttc accacttgca aattaagatt attagtccga
209401 tatggtaaac tttgctcgaa gctaatgttt cctccaaggc gtccaatgat ttttttctga
209461 ggatacgcat tacacattac cggcgtcatt tgcagtgaca gaggatcaac ttgattaaat
209521 aaccttacgc taaaggccaa attaatcact aaagcagatg agtaaataat caagttgttt
209581 gaccagcggt gaatatcctc tagataacat tgatcatcta atcggtgcta ttcacttgtt
209641 ctcaagaaac actttgttgc taaaattcac ttacaaactt aaccaatggt cccgctgcga
209701 acgccggcta attatgagcc agaattatcc atgccttttc ctgatcatcg gagattcgta
209761 tacggaatgc taaacaaccg gaagaaaagt cattacattc catttttgtg atttaaaaag
209821 tgtaggaacc cccacctccc acgggcgaca tggatggtga aaattgccac aggatagaac
209881 atcccaacac aagaagccat ttaaaaattc ctcatcctag atatcaagtt tgcttacggc
209941 tacagatcag aatacgcata tgaaagatgg gcccaattga taatttatac tatggtctag

210001 tcacaaagtt ctctgtccct attagtataa aaacaaaaca tgcttgtgat tgattttgtc
210061 cctccgccac tctactggat aagaaaaccc ctgaaatacg atcatgttgc ctgcgatttc
210121 tgaaaaggtt gaaatattac acaataaaga ccctccactc aaatttgctt gttcggaatc
210181 aggggtagct ccgctggacc tcggctcctt cttctttcag aaggatcttc aaatgcgcct
210241 ccacttaagc agccttctgc cacgacttct tctgagacac ccggtttcgc gcgggcactt
210301 ctgcccgcca aaaaagttta tatgtcggaa agcttatttt gaattaaagt ttgaaataaa
210361 tttcttttct gcagattttc atgcagcgtc gaactcctga attctcacga gattatttga
210421 cgttgctttt cttctccctt tttccacgca ctacgccaat gcaagtgatc cgcttttgaa
210481 gaagcaaaaa atacttcttt cagctcgtac cggcattttt attatttgtc aaaactggcg
210541 tttggaaacc agctcttcac tttttcttcg tgctttcctc gtcgccatct cgagaattct
210601 ggttgagctc gttttaaaaa tgcgtaaaaa gaaagaagac gtctgctaca accctctcaa
210661 gtctcttaaa ggatagttcg aaacttgtct gattatcctt gaataacgaa ctttgatttt
210721 caatgtttta tttccggtca ttgagatatg atagcctgtt cctggatatt acgagggaat
210781 tttcatatat tgtattgtat tttttttttt tcataacgaa ttctagaaaa ggctttgttg
210841 gcatgtgagg atggcagcgg ttggaaactt tgccgagaac aatttctcac aagttaacct
210901 gagattttca cactttttac ttagtggttg ttggcactta agtagaagct aatagtagca
210961 atgatatcga aataaaatgc tgtatcacgg gcgattattc catggcgaaa tgaggtccgt
211021 aatttcttcg tcagagggag gaaagtgact ttgggaatac tcaaatggaa tagatgtgac
211081 aagttctaga atgacaatgt caacccaagg ggagctaagt ttctaacatg gaaagcaaag
211141 aaaggccctc agtaccgttc aatctaaaat catcttcatt aacttccaaa tgtcttatcg
211201 ccctttcaca tccgtatcta acacctatag atgatcttat cacgtttgaa gaattatgga
211261 gaatcgtgaa tatgggtaaa aaaacaatta tttcgtatat tgcgcctggt attttctgag
211321 cttcaaatat gcttacaaga actcgaatac agcatatata aacatattat agataaagca
211381 tatagttctg aacccaattg taatcgtagt gttttagatt ggaagagaaa tattactatt
211441 cttaactcgc aaagatttat atttttgcat cctctgaaaa gaaaaagtta gactacttta
211501 tctaatactt gtttttcctc ggttgaaata gaaagcttct caaggcgtgt ttctttaatt
211561 ccttgtagca tgataatgaa tattttcata ttaatgttta cttttcccat ttatgcattt
211621 tcgcaacaaa tatttaatta attgtaaaaa tatcatacaa aaaagaggaa ctcggatcta
211681 agccaattta gtataaacgt caagaagtaa gaagaactgg cagtttcaaa ttcaattttt
211741 taatgtggta cgtacatctc tagtccatat cgtattcagc aatttataaa aaagtctgtc
211801 acgataaatt ttcgtttgtg tgagaaattg gctacttagg aagagtttgc tacataatcg
211861 aaatttttat taagagtaac acatgtatat attcttttag aatcctctga ttattaagtt
211921 gtttgacaga ccggcgagtc taatgagagc ttattacttg ataaaaaggt tgcataacca
211981 taataaatcc acagtataat gctatatggt gcagagcgaa ggattgcgca aaaagctctt
212041 cctttcaaaa atcatgaaaa acgaaccttg tagagtaatt ttgtaagaag atttaaactc
212101 atgtctttat cactataagt cgatatcata ctcatattat aatgtcgaac caggatatcc
212161 gtacgacctc atggggagat aaaacatcca aatagttcct ttattgaata ttgtagctat
212221 gaagtgctag taagtggggt atcgacgtga tgacacccat gctaaagctc cactaaagac
212281 aaatacgaac cacgtatgcc atttataaag agctaaaaag cttccgaaca acctgctccg

212341 gaaacctgaa gaagtgtatc atatatattt attctaattt gctctcaaaa attttcatta
212401 aacaacagga gaaacgattg agacatacag caagctatag tgaaatctga atattacttt
212461 tctctttcag tgttacagta ctatattaaa tcccaaacaa agtgtacttg tttaatttgc
212521 tgcttatcct acgccaattt ttcaaatgaa tctggaacaa ttgataagat acctttatta
212581 cagcaatagc aagcgcgtga caggcttcat cgttttagat aacaatctgt atggtcaact
212641 caacgattct taggggctac aattaaactt gaaataacta tgattcctca tgtaatgttt
212701 aagctctcaa gatatttgtc ctcacttccc gtttggctct cttaacaata gaaggtatta
212761 aaagaattcg acagtatggt tgagtactag tgtctccttg tagttttttt cccttgtgat
212821 tttgggtttg ctcataatac gatgtagcgc atattcaatg actaaagttg ttattgcttc
212881 ctctgacgac aagtccgctc ccacttatag tgtcctgagt gattatttat ttttatgtga
212941 acaccaaaag gaataaagta ggtccgtgtc catttactgc ttctttcaca cttttagcca
213001 atcttgacgt ggatatcttg ttttgcctaa cccattctct gtgttgttag tataatccac
213061 catatcactt accatatcaa ccattatgtt cagttaactc acctttaaag tactgtgcac
213121 aacaccctat ttccatctga tcggtcgctt tatctggaat atctgttttc aatctacgaa
213181 gacataagtt tgaaaaatat aaccagagtt catcgagacc ctgcccgcca caaccgccat
213241 taaaatattg aaatatgagc caaaagtttc agcctggtat tctgaatgaa attgtaatcc
213301 gactatagag aacgatttat ttggcttttg ggtagtatga ttatccttgt ctggagttag
213361 atatagaaga atttaagaag ggccacgaaa tttatcctat ggtatgggaa acgagaaaaa
213421 ctccagtgct gctaattaga actaaaatat gaaaagaaaa ggaagaggcg aggatgagtg
213481 tatctaaccc acgtagcttg aaagttactc gctacatgct aaaagtttac ttaaattaca
213541 aatcttcact ttcatgaacg tcctctacta gctattatga aggcgcagag agtagtctcg
213601 ttaaccaatt ttatgcaagt cgttgaaaaa acggcgactc tttagtagac tggtcaagcg
213661 gcatcggaaa cagttctcag aacagaaaag aagagattat tgagttaagg tccgttaaag
213721 ggtcatgata agcccaataa gataccaagt agacatgtta caccgtgagt agtaaacggg
213781 gttatatttt attatgttgc tagtctcatt ttatggccct tgtagtttgg aggaaaaatg
213841 atcggataac caaaacgaaa ttccttaagt gatttgtgtc cctaaaagaa aatcttcagg
213901 tataaccgca aacgccaaat gccacaagcc caaaaatttc cgtgtctttt cccgatgcta
213961 tccttgatta aagaactttt gattgtcaaa gtgaaaatac ctatatggac tcttttgaag
214021 aattgaaaga aataggcaaa tcacttgtgt acattacact ctctactaaa cgatttcgta
214081 acctaggcta atacgcgaaa caagcgcttt ttttatttag tgaagcatca tattacctaa
214141 gcgccacttt gcttgacgtt agtaacaaaa atgatgaaat tgtaatagaa gaaactttta
214201 tccgcaaact attgaggtac catgctagtc cgtctcaagc tgtttgatgc aagagaacgt
214261 tcaactcaat ttccggctag tacgtcactt tttctacggt ggcttcgcag acgaatgttt
214321 tccgacacat gatacttatc accgaaaaac cttatttcac ggaaaaaacc ttatttacat
214381 taaagtttga aaaatttctt cttttccgca atatggtggg gcctttgact tccagaatgc
214441 tttcacggaa ttattttaga tgtacattta gctccatttc ctgtgcctcc gataggaagg
214501 catcatggta ctaccgtgac ggagaatacg taggctgact ttttcgtcag tttgttgtcc
214561 gtttacaaaa ttggtgaatg aattctagcc tttgctcatt aattgccctc acaagaattt
214621 ggaagtgcgt agaacaagta aaaggttgca cta

//

**R3.gb**

```
LOCUS       Exported              340705 bp ds-DNA     circular SYN 15-APR-2024
DEFINITION  synthetic circular DNA
ACCESSION   .
VERSION     .
KEYWORDS    W303-1A-R3
SOURCE      synthetic DNA construct
  ORGANISM  synthetic DNA construct
REFERENCE   1  (bases 1 to 340705)
  AUTHORS   Yilin Lyu
  TITLE     Direct Submission
  JOURNAL   Exported Apr 15, 2024 from SnapGene 4.2.4
            http://www.snapgene.com
FEATURES             Location/Qualifiers
     source          1..340705
                     /organism="synthetic DNA construct"
                     /lab_host="Saccharomyces cerevisiae"
                     /mol_type="other DNA"
     CDS             1..315
                     /codon_start=1
                     /locus_tag="YCR107W"
                     /label=AAD3
                     /translation="VGASEQTDAEIKISEALAKIAEEHGTESVTAIAIAYVRSKAKNFF
                     PSVEGGKIEDLKENIKALSIDLTPDNIKYLESIVPFDIGFPNNFIVLNSLTQKYGTNNV
                     "
     CDS             complement(991..1794)
                     /codon_start=1
                     /label=KmURA3
                     /translation="MSTKSYSERAAAHRSPVAAKLLNLMEEKKSNLCASLDVRKTAELL
                     RLVEVLGPYICLLKTHVDILEDFSFENTIVPLKQLAEKHKFLIFEDRKFADIGNTVKLQ
                     YTSGVYRIAEWSDITNAHGVTGAGIVAGLKQGAEEVTKEPRGLLMLAELSSKGSLAHGE
                     YTRGTVEIAKSDKDFVIGFIAQNDMGGREEGYDWLIMTPGVGLDDKGDALGQQYRTVDE
                     VVAGGSDIIIVGRGLFAKGRDPVVEGERYRKAGWDAYLKRVGRSA"
     misc_feature    3189..6332
                     /label=YCLWTy5-1
     CDS             complement(8489..10336)
                     /codon_start=1
                     /locus_tag="YCL073C"
                     /label=GEX1
```

/translation="MSSSVVGASSNKKSGIRQSCEIIERERHSNDDTYSMTSTFFKLKE
NEIMSAQFDSLKYKILLISTAFVCGFGISLDYTLRSTYTGYATNSYSEHSLLSTVQVIN
AVVSVGSQVVYSRLSDHFGRLRLFLVATIFYIMGTIIQSQATRLTMYAAGSVFYNCGYV
GTNLLLTLILSDFSSLKWRMFYQYASYWPYIIIPWISGNIITAANPQKNWSWNIAMWAF
IYPLSALPIIFLILYMKYKSSKTAEWRSLKEQARKERTGGLFENLVFLFWKLDIVGILL
ITVSLGCILVPLTLANETSQKWHNSKIIATLVSGGCLFFIFLYWEAKFAKSPLLPFKLL
SDRGIWAPLGVTFFNFFTFFISCDYLYPVLLVSMKESSTSAARIVNLPDFVAATASPFY
SLLVAKTRKLKLSVIGGCAAWMVCMGLFYKYRGGSGSHEGVIAASVIMGLSGLLCSNSV
IVILQAMTTHSRMAVITGIQYTFSKLGAAIGASVSGAIWTQTMPNQLYKNLGNDTLAEI
AYASPYTFISDYPWGSPERDAVVESYRYVQRIIMTVGLACTVPFFAFTMFMRDPELIDK
ATHEEFTEDGLVVLPDEENIFSQIKALFRHNRSNKKLGC"
CDS 11716..13092
/codon_start=1
/locus_tag="YCL069W"
/label=VBA3
/translation="MNMLIVGRVVASVGGSGLQTLCFVIGCTMVGERSRPLVISILSCA
FAVAAIVGPIIGGAFTTHVTWRWCFYINLPIGGLAIIMFLLTYKAENKGILQQIKDAIG
TISSFTFSKFRHQVNFKRLMNGIIFKFDFFGFALCSAGLVLFLLGLTFGGNKYSWNSGQ
VIAYLVLGVLLFIFSLVYDFFLFDKFNPEPDNISYRPLLLRRLVAKPAIIIINMVTFLL
CTGYNGQMIYSVQFFQLIFASSAWKAGLHLIPIVITNVIAAIASGVITKKLGLVKPLLI
FGGVLGVIGAGLMTLMTNTSTKSTQIGVLLLPGFSLGFALQASLMSAQLQITKDRPEAA
MDFIEVTAFNTFMKSLGTTLGGVLSTTVFSASFHNKVSRAHLEPYEGKTVDDMILYRLQ
NYDGSHSTIGNILSDSIKNVFWMDLGFYALGFLFCSFSSNKKLIIPKKDETPEDNLEDK
"
CDS complement(13513..14295)
/codon_start=1
/locus_tag="YCL068C"
/label=YCL068C
/translation="MFVLIDNVLAYLLEQDDLFVTARFAIQGQIVSRRVNKIHISNITD
VLLQQFISHTLPYNDNIVPKKILDSMRTAVRQLLEATACVSRECPLVKRSQDIKRARKR
LLSDWYRLGADANMDAVLLVVNSAWRFLAVWRPFVNSIQHATQELYQNIAHYLLHGNVN
IQRVTALLQLVMGQDDLLFSMDDVLQEVFRIQLYLNKMLPHNSHKWQKPSPFDSANLLL
NFRDWTTDNALLQELLLSYPTINKNKHKNHSVPRLIQV"
CDS complement(14396..15028)
/codon_start=1
/locus_tag="YCL067C"
/label=HMLALPHA2
/translation="MNKIPIKDLLNPQITDEFKSSILDINKKLFSICCNLPKLPESVTT
EEEVELRDILGFLSRANKNRKISDEEKKLLQTTSQLTTTITVLLKEMRSIENDRSNYQL

TQKNKSADGLVFNVVTQDMINKSTKPYRGHRFTKENVRILESWFAKNIENPYLDTKGLE
NLMKNTSLSRIQIKNWVSNRRRKEKTITIAPELADLLSGEPLAKKKE"
CDS 15292..15819
/codon_start=1
/locus_tag="YCL066W"
/label=HMLALPHA1
/translation="MFTSKPAFKIKNKASKSYRNTAVSKKLKEKRLAEHVRPSCFNIIR
PLKKDIQIPVPSSRFLNKIQIHRIASGSQNTQFRQFNKTSIKSSKKYLNSFMAFRAYYS
QFGSGVKQNVLSSLLAEEWHADKMQHGIWDYFAQQYNFINPGFGFVEWLTNNYAEVRGD
GYWEDVFVHLAL"
CDS complement(17808..18890)
/codon_start=1
/locus_tag="YCL064C"
/label=CHA1
/translation="MSIVYNKTPLLRQFFPGKASAQFFLKYECLQPSGSFKSRGIGNLI
MKSAIRIQKDGKRSPQVFASSGGNAGFAAATACQRLSLPCTVVVPTATKKRMVDKIRNT
GAQVIVSGAYWKEADTFLKTNVMNKIDSQVIEPIYVHPFDNPDIWEGHSSMIDEIVQDL
KSQHISVNKVKGIVCSVGGGGLYNGIIQGLERYGLADRIPIVGVETNGCHVFNTSLKIG
QPVQFKKITSIATSLGTAVISNQTFEYARKYNTRSVVIEDKDVIETCLKYTHQFNMVIE
PACGAALHLGYNTKILENALGSKLAADDIVIIIACGGSSNTIKDLEEALDSMRKKDTPV
IEVADNFIFPEKNIVNLKSA"
CDS 19300..20571
/codon_start=1
/locus_tag="YCL062W"
/label=VAC17
/translation="MATQALEDITERLLIRSQEAILQLDLWIQRQQRSSICQTTDQESL
DKLSQQYNQYMSQLNSLYVRSESVRDKLSKEQQRRLITEDNEHQRIEDLVREFQDITLR
LNELATVPNEAPNDSPQSQSTRSSLGSFQPRPLKIIERQRLCMVTPSKPPKKSVGFNPI
NEVDCPSKTNSLPCSPKKQPARNRTLRAAKSHDTGLNKSKKPSSSDTYESFFKNRQRLS
LTFFDEMDDEDFDSDQDTIILPNISTPPHVGVTAKGAEFEPLRRYNSHESILSNKPAPS
KSLNLGSFSASFFRPSNPTFGTSISNVQVNCHPTVAATMAPSRNGPRISSSKALLSSFI
ARSDTHTVKENNTNLKHASFMDKFNSSLSTISESFQSKRGRKNKGMNEERISNHNVAQE
QKNNMDISVSIEELQDALNTELLF"
CDS complement(20826..24116)
/codon_start=1
/locus_tag="YCL061C"
/label=MRC1
/translation="MDDALHALSSLTAKKRTTTYKKVAVPILDENDNTNGNGPNDIDNP
PELTGNGFLFANATLNRVKNRLEGKKAPEQNHNNGKDRSENSLPTQLISNLYDGGEELE

KSEVKDNSYSEKNVSSSFTQTQRIPVSIQQDKVFNVPIHSVNDGKPTQLIKEDGLVNET
SQALKTPLTTGRPGATQRIDSSGATSQTQPIKSIEPQSQIITTSSNHSNALSPKIPIIP
TELIGTSPLFQSIQNRGPDTQMDVPPQTAHDEDKTQAIGIPQATHQEQKTQIDTVAQTL
QDEVPHTLKIREIQSELASEDSKREKARNVEYKKPQKPIPTKKFFSKESFLADFDDSSS
NEDDDIKLENAHPKPVQNDDELHENKSVELNLTDETRINEKRVPLLSSYANNLKREIDS
SKCITLDLDSDSDEYGDDDMDSIKLSKDESVLPISQLSKATILNLKARLSKQNQKLSQR
PNKSKDPKVDHNVLLNTLRKASRKQILDHQKEVIETKGLKLEDMAKEKEIVENLLEQEI
LRNKRIRQKEKRREKLEENDFQLNAHDSGSDSGSESSGFALSGNEIADYESSGSENDNR
RESDSEKEDDEIILKQKKSHHVKHIINESDSDTEVEAKPKEKADESLPKRIAINLGHYG
DNIGEDTDKFQETNVLDTQNIEEVMAERNTIENEVKDDVYVNEEADEAIRRQLIDKEKL
QLKQKEKEHEAKIKELKKRGVTNFFEMEAEESEDEWHGIGGADGEGSDDYDSDLEKMID
DYSKNNFNPHEIREMLAAENKEMDIKMINKILYDIKNGGFRNKRAKNSLELELSDDDED
DVLQQYRLKRRELMRKRRLEIGDDAKLVKNPKSSAFFESMVEDIIEYKNPFGAEEEYNL
DITSTATDLDTQDNSINVGDNTGNNEQKPVDQKNKKVIISEDFVQKSLSFLKSNNYEDF
ETDKELSRIQHGNDEAIEDLYTLKQNSSIKSFTNSQTDSTTSKTVNTIIDLEKRPEDED
EVENGDTSLVGVFKHPSIIKSFASRTDINDKFKEGNKTVKILKSYKTVGSSKASITYMG
KTRKLIAPKRKTEGSHRYHHDHHNKKMKMKTKTKSNKLFESGQDSFDN"

CDS complement(24439..25389)
/codon_start=1
/locus_tag="YCL059C"
/label=KRR1
/translation="MVSTHNRDKPWDTDDIDKWKIEEFKEEDNASGQPFAEESSFMTLF
PKYRESYLKTIWNDVTRALDKHNIACVLDLVEGSMTVKTTRKTYDPAIILKARDLIKLL
ARSVPFPQAVKILQDDMACDVIKIGNFVTNKERFVKRRQRLVGPNGNTLKALELLTKCY
ILVQGNTVSAMGPFKGLKEVRRVVEDCMKNIHPIYHIKELMIKRELAKRPELANEDWSR
FLPMFKKRNVARKKPKKIRNVEKKVYTPFPPAQLPRKVDLEIESGEYFLSKREKQMKKL
NEQKEKQMEREIERQEERAKDFIAPEEEAYKPNQN"

CDS complement(25533..25991)
/codon_start=1
/locus_tag="YCL058C"
/label=FYV5
/translation="MQYHSALYVYIYVTFTTIPYKEKPDIISICFSMLSFVFDFSVRIC
SRTLESFSWSLISSSAFKVVSAFSLAGSCVLASRSSVGIIVSLLLFNFSTCNFVLFLSA
VLIDLFFCTFLPTPTFLPTPFFFMLHLPIFSLLNALELLYLIIAGLHI"

CDS 25594..25935
/codon_start=1
/locus_tag="YCL058W-A"
/label=ADF1
/translation="MGKCSMKKKGVGKNVGVGKKVQKKRSISTAERKRTKLQVEKLNKS

SETMIPTLLREASTQEPAKLKAETTLKAEELIKDQEKDSKVREQIRTEKSKTNDSMLKQ
IEMISGFSL"
CDS complement(26042..26335)
/codon_start=1
/locus_tag="YCL057C-A"
/label=MIC10
/translation="MSEQAQTQQPAKSTPSKDSNKNGSSVSTILDTKWDIVLSNMLVKT
AMGFGVGVFTSVLFFKRRAFPVWLGIGFGVGRGYAEGDAIFRSSAGLRSSKV"
CDS 26778..28916
/codon_start=1
/locus_tag="YCL057W"
/label=PRD1
/translation="MRLLLCKNWFASPVISPLLYTRSLYSMANTTSFPIAPQAPPNWSF
TPSDISGKTNEIINNSNNFYDSMSKVESPSVSNFVEPFMKFENELGPIINQLTFLQHVS
SDKEIRDASVNSSMKLDELNIDLSLRHDIFLQFARVWQDVQSKADSVERETFKYVEKSY
KDYIHSGLELDEGNRLKIKEIKKKISVNSINFSKNLGEQKEYITFTKEQLEGVPDSILT
QFETIKSDKDSNETLYKVTFKYPDIFPVMKLASSAQTRKQAFLADQNKVPENEAILLDT
LKLRDELASLLGYDTYANYNLYDKMAEDSTTVMNFLNDLKDKLIPLGRKELQVLQDMKA
EDVKKLNQGADPNYYIWDHRYYDNKYLLENFNVDLEKISEYFPLEATITGMLEIYETLF
NLKFIETKDSQNKSVWHDDVKQIAVWNMDDPKSPNFVGWIYFDLHPRDGKYGHAANFGL
SSSFMIDDTTRSYPVTALVCNFSKSTKDKPSLLKHNEIVTFFHELGHGIHDLVGQNKES
RFNGPGSVPWDFVEAPSQMLEFWTWNKNELINLSSHYKTGEKIPESLINSLIKTKHVNG
ALFTLRQLHFGLFDMKVHTCKDLQNLSICDTWNQLRQDISLISNGGTLSKGYDSFGHIM
SDSYSAGYYGYLWAEVFATDMYHTKFAKDPLNAKNGIQYRDIVLARGGLYDINDNLKEF
LGREPSKDAFLKELGLQN"
CDS 27212..27223
/codon_start=1
/product="Factor Xa recognition and cleavage site"
/label=Factor Xa site
/translation="IEGR"
CDS complement(28935..29369)
/codon_start=1
/locus_tag="YCL056C"
/label=PEX34
/translation="MVSKKNTAEISAKDIWENIWSGVSSLLDFFAVLENLGVVNDKLYV
SGLLRKVWLCYSCISVIKCVWKLIKLCKVKFKIDQRLDGEGNGLVKDKLINFKKKYNEH
IRHITAALLQDLSYLMVLIYPGTRLFKRLSNIITLCRIIV"
CDS 29940..30947
/codon_start=1

/locus_tag="YCL055W"
/label=KAR4
/translation="MAFQDPTYDQNKSRHINNSHLQGPNQETIEMKSKHVSFKPSRDFH
TNDYSNNYIHGKSLPQQHVTNIENRVDGYPKLQKLFQAKAKQINQFATTPFGCKIGIDS
IVPTLNHWIQNENLTFDVVMIGCLTENQFIYPILTQLPLDRLISKPGFLFIWANSQKIN
ELTKLLNNEIWAKKFRRSEELVFVPIDKKSPFYPGLDQDDETLMEKMQWHCWMCITGTV
RRSTDGHLIHCNVDTDLSIETKDTTNGAVPSHLYRIAENFSTATRRLHIIPARTGYETP
VKVRPGWVIVSPDVMLDNFSPKRYKEEIANLGSNIPLKNEIELLRPRSPVQKAQ"
CDS 32921..33007
/codon_start=1
/locus_tag="YCL054W-A"
/label=RDT1
/translation="MIRQKIFVFIVKSRRNSICPAIRRKEDY"
CDS 33460..35985
/codon_start=1
/locus_tag="YCL054W"
/label=SPB1
/translation="MGKTQKKNSKGRLDRYYYLAKEKGYRARSSFKIIQINEKYGHFLE
KSKVVIDLCAAPGSWCQVASKLCPVNSLIIGVDIVPMKPMPNVITFQSDITTEDCRSKL
RGYMKTWKADTVLHDGAPNVGLGWVQDAFTQSQLTLQALKLAVENLVVNGTFVTKIFRS
KDYNKLIWVFQQLFEKVEATKPPASRNVSAEIFVVCKGFKAPKRLDPRLLDPKEVFEEL
PDGQQNMESKIYNPEKKVRKRQGYEEGDNLLYHETSILDFVRTEDPISMLGEMNKFTID
ENDHEWKILKKLKQTTDEFRSCIEDLKVLGKKDFKMILRWRKIAREILGIEVKDDAKTE
IEVVPLTEEEQIEKDLQGLQEKQRLNVKRERRRKNEMKQKELQRMQMNMITPTDIGIEA
ASLGKESLFNLKTAEKTGILNDLAKGKKRMIFTDDELAKDNDIYIDENIMIKDKDSAAD
ADDLESELNAMYSDYKTRRSERDAKFRAKQARGGDNEEEWTGFNEGSLEKKEEEGKDYI
EDNDDEGVEGDSDDDEAITNLISKLKGQEGDHKLSSKARMIFNDPIFNNVEPDLPVNTV
NDGIMSSESVGDISKLNKKRKHEEMHQKQDEADSSDESSSDDSDFEIVANDNASEEFDS
DYDSEEEKNQTKKEKHSRDIDIATVEAMTLAHQLALGQKNKHDLVDEGFNRYTFRDTEN
LPDWFLEDEKEHSKINKPITKEAAMAIKEKIKAMNARPIKKVAEAKARKRMRAVARLEK
IKKKAGLINDDSDKTEKDKAEEISRLMRKVTKKPKTKPKVTLVVASGRNKGLAGRPKGV
KGKYKMVDGVMKNEQRALRRIAKKHHKKK"
CDS complement(36154..37404)
/codon_start=1
/locus_tag="YCL052C"
/label=PBN1
/translation="MVTRHRVTVLYNAPEDIGNHMRQNDTHLTVRGGSGVVLQQRWLLE
RTGSLDKSFTRITWRPRADLARSLSVIENELSAGFSVYSNSSDVPERFITNPVYNSFHS
EKFDIEQYLPPEVDLNLSWNPEDFTYDISVEPTQIQIVEYRLLKQGEEFTIARVKDEKL

EVGVFFVDASDESDVDIGGIRCNWRMDDGKMERCQKTSLLYKQGHIAYNHSTTTTSLYL
NEPIGLHPKIMIDLTDFEERPKCMYLMHLQLPLELFIDKFQSSPLLLFGEDDLELPEYS
LRDKAWGSESIFELKAGTMNEVTLHTRYIEPSNNKGDKLEVSFDPEVILACDTGDNKVS
RNPFYKKGLGYESLFTDDTTFRHLNSTTLLVPIPRPDTKDYSKIKNGTLLCLLISIIYI
FSKVFGNNKKKRSVKRE"

CDS 37876..39627
/codon_start=1
/locus_tag="YCL051W"
/label=LRE1
/translation="MPNTHTQHVQISEPNPVNTLSTPSKRGHRHRRSLAISGDFDFLKQ
PAAIVNLPPPQAAENCPSTAPTAVSSTLSPIRYNRFPCKTNEDAGTLDLPEPRFYPLSP
KNNLQTPSPRFFISEEPSFSSPVKGVPDAIINLDDALKTRPRSFKSHRRSESAPPDLEV
MVDKGNCAAGSNSMIKEEEDSLIEPESKNEYYEQKLPTALLSPLRPSLCVSEQAIDVDD
SALNGSPTHHNHGMQNANARNSNTFNSLKIKGQKQRYYHYTKQLPLTVGCDSQSPKEQR
SAASMTINQAMTPSSLAYTPSKLASTPATPVSFYDSNADINLESDNFPLKDNPRYAKDG
YPKKCGNSQLNRVLDSDKRQDFSGESRRRRSGSPISHMQHRNLIDNMKGRRNSNTINSI
FNYKSQHYEMPYDDMMKNENINAQSMPFSVNGVNNENSIGGVITRADDAPLQHSVVKSC
TPDGKEEMNRLKSNDSNEYSKSEGQIRTNSQLSKDILMGEPGDMVDLSSFVNAQRKASN
ETGDLVFSLSQDDDALKTFHASNSAATSNESWCISDDALGKQAQDSEVRRKRKSKLGLF
RHIFSRK"

CDS complement(39847..40812)
/codon_start=1
/locus_tag="YCL050C"
/label=APA1
/translation="MSIPADIASLISDKYKSAFDNGNLKFIQTETTKTKDPKTSMPYLI
SHMPSLIEKPERGQTPEGEDPLGKPEEELTVIPEFGGADNKAYKLLLNKFPVIPEHTLL
VTNEYQHQTDALTPTDLLTAYKLLCALDNEESDKRHMVFYNSGPASGSSLDHKHLQILQ
MPEKFVTFQDRLCNGKEHFLPTFNTEPLQDAKVSFAHFVLPMPESEETVDEDLLAMCYI
SILQRALTFFQDWLNENPELKKSYNLMLTKEWICVVPRSKAFSDEMKIGFNSTGYCGMI
LTKNDEVFSKITEKPELINDILLECGFPNTSGQKPNEYNY"

CDS complement(41797..42735)
/codon_start=1
/locus_tag="YCL049C"
/label=YCL049C
/translation="MFSKYLVTASSLFVALTSAASTVDLDALLLLPGVESHDGVDTVFS
TKDFYQVSFVKSIAPAIVNSSVIFHDVSRGVAMGNVKSRASIFNPEETYYDWEQYQVVN
NGDWRTEWAPASDCIWREEKDNSDETPDRFPISVPYNWTSQYSIVDYDTDANEDNLDFR
FIKSLLDKKNWLKKINQTVSQSSIMVAPMIRPYNVVQLWYSKDMVWANVQRQYCSGVYP
GGTQCSAWSRYYHVDAPTCDEPVASYMTKMLENEVQCPNERNATTLEPLRLNKQGDSDF

SLTFEEEEEEETGSKSLWSTLKKIFSKRSIS"
CDS 43499..43738
/codon_start=1
/locus_tag="YCL048W-A"
/label=YCL048W-A
/translation="MQIKNIVAVLATVTAINAQVGIEPNATTPNATQPNATQPNTTLPT
ASVTTTVSIGEAVVNTMAAGAFGAAIAAGVAFLF"
CDS 44174..45565
/codon_start=1
/label=SPS22
/translation="MNRITRKSCLFAIIFASLFVTHALGAAIDPPRRPHNVKPFHNGNL
ELQRRANEPFFEIDVKSLNTNSPISELCKKDLHVIESSHDLFHLQNQCEFILGSLKVTN
YDSNILDLNSLRAIGGDLIIQDSPELIRIQAGNLNKIEGLFQLQGLTSLVSVEIPTLKF
CQSLEWKVVPILNYVSMDSQNIEIIKDIVISDTSLANIENFNKVQEIDTFNINNNRFLE
TIHSNVKTIRGQFSVHANAKELELEMPHLREVENITIRDTSLVYLPQLTKVKSSLEFIE
NYFYELNLNNLQKIGGTLGIINNVNLIKVNLENLTDIQGGLMIADNESLEDITFLPNLK
QIGGAIFFEGSFKDIMFDSLKLVKGSAFIKSSSNVLDCNKWTNPSNGRSIIRGGKFTCI
SGKKENTLNVKQDGTIIEKGYKDLTQEGEDSKKRVISKYANSANPSMQLDPLLFGTCLV
AMLLF"
CDS complement(45670..46446)
/codon_start=1
/locus_tag="YCL047C"
/label=POF1
/translation="MKKTFEQFRKSNLLFQVLKGPQHLECQKLFVLDSSFNPPHLAHFQ
LLSQTIKNFKLKDTRSHVLLLLAVNNADKLPKPASFPTRLEMMCLFADYLQEKLPQSVV
SVGLTVFSKFIDKDKILHEQFVKGCSADIGYLVGFDTIARIFDEKYYHPLKISDVMESF
MSGSQLYCLARGDCHLSAESQLRYASDILEGKFEPVIPREWGARIHVMQNDYPALRNVS
SSEIRNKLKNGQVESLKDELPLCIYDYLINNKTIFD"
CDS complement(46632..48914)
/codon_start=1
/locus_tag="YCL045C"
/label=EMC1
/translation="MKITCTDLVYVFILLFLNTSCVQAVFSDDAFITDWQLANLGPWEK
VIPDSRDRNRVLILSNPTETSCLVSSFNVSSGQILFRNVLPFTIDEIQLDSNDHNAMVC
VNSSSNHWQKYDLHDWFLLEEGVDNAPSTTILPQSSYLNDQVSIKNNELHILDEQSKLA
EWKLELPQGFNKVEYFHREDPLALVLNVNDTQYMGFSANGTELIPVWQRDEWLTNVVDY
AVLDVFDSRDVELNKDMKAELDSNSLWNAYWLRLTTNWNRLINLLKENQFSPGRVFTKL
LALDAKDTTVSDLKFGFAKILIVLTHDGFIGGLDMVNKGQLIWKLDLEIDQGVKMFWTD
KNHDELVVFSHDGHYLTIEVTKDQPIIKSRSPLSERKTVDSVIRLNEHDHQYLIKFEDK

DHLLFKLNPGKNTDVPIVANNHSSSHIFVTEHDTNGIYGYIIENDTVKQTWKKAVNSKE
KMVAYSKRETTNLNTLGITLGDKSVLYKYLYPNLAAYLIANEEHHTITFNLIDTITGEI
LITQEHKDSPDFRFPMDIVFGEYWVVYSYFSSEPVPEQKLVVVELYESLTPDERLSNSS
DNFSYDPLTGHINKPQFQTKQFIFPEIIKTMSISKTTDDITTKAIVMELENGQITYIPK
LLLNARGKPAEEMAKDKKKEFMATPYTPVIPINDNFIITHFRNLLPGSDSQLISIPTNL
ESTSIICDLGLDVFCTRITPSGQFDLMSPTFEKGKLLITIFVLLVITYFIRPSVSNKKL
KSQWLIK"
CDS complement(49120..50373)
/codon_start=1
/locus_tag="YCL044C"
/label=MGR1
/translation="MAVFTPPSGNSNSTDHTHTQDDHDKDDNDIKKFYIRPSLGLKLWG
PLVPAPDNLPGLYTLITIQSAVGFFALWRLRRLYKLPPPRRIATGTHSDLSFGELPSEM
IVNGKTKIKKDIADFPTLNRFSTTHGDIVLAPPPIIPRQSRFVSVRKLLWGLFGSLLLS
QSLLELTRLNFLKYDPWCDEMKSVRDKKFFNNIVKYYHEGIDPTKIKVKDAMNGTPLST
NIPEVKQSVALARAQVEAQNPIIKWFGPLEYKPMSFNEYLNRMEFHLDMFEFFQNKRNI
RENSIELINSISHNPQSSSTGLEGLSESKKLHLQNVEKRLHFLASSGDSISAPVKKRSS
TTLSRGVILPHDTKGPQDIDLDTIRSLYDPWMTLALETSLSIKFIPTTMPSHTKTPTST
DQPLPGPTPKALTNEKTH"
CDS complement(50662..52230)
/codon_start=1
/locus_tag="YCL043C"
/label=PDI1
/translation="MKFSAGAVLSWSSLLLASSVFAQQEAVAPEDSAVVKLATDSFNEY
IQSHDLVLAEFFAPWCGHCKNMAPEYVKAAETLVEKNITLAQIDCTENQDLCMEHNIPG
FPSLKIFKNSDVNNSIDYEGPRTAEAIVQFMIKQSQPAVAVVADLPAYLANETFVTPVI
VQSGKIDADFNATFYSMANKHFNDYDFVSAENADDDFKLSIYLPSAMDEPVVYNGKKAD
IADADVFEKWLQVEALPYFGEIDGSVFAQYVESGLPLGYLFYNDEEELEEYKPLFTELA
KKNRGLMNFVSIDARKFGRHAGNLNMKEQFPLFAIHDMTEDLKYGLPQLSEEAFDELSD
KIVLESKAIESLVKDFLKGDASPIVKSQEIFENQDSSVFQLVGKNHDEIVNDPKKDVLV
LYYAPWCGHCKRLAPTYQELADTYANATSDVLIAKLDHTENDVRGVVIEGYPTIVLYPG
GKKSESVVYQGSRSLDSLFDFIKENGHFDVDGKALYEEAQEKAAEEADADAELADEEDA
IHDEL"
CDS 52593..52952
/codon_start=1
/locus_tag="YCL042W"
/label=YCL042W
/translation="MEKKKILDAALAPRIISGVPTDGQPLSGGPLSWAWCHTTLKRWAL
MKTRPYKYPLSTEPPTPPSVPNSASVNHNTTTNTTLSYTRCHSTTYTKPLRERSSRPWT

RSATISRLPPRSWTN"
CDS 52847..54349
/codon_start=1
/locus_tag="YCL040W"
/label=GLK1
/translation="MSFDDLHKATERAVIQAVDQICDDFEVTPEKLDELTAYFIEQMEK
GLAPPKEGHTLASDKGLPMIPAFVTGSPNGTERGVLLAADLGGTNFRICSVNLHGDHTF
SMEQMKSKIPDDLLDDENVTSDDLFGFLARRTLAFMKKYHPDELAKGKDAKPMKLGFTF
SYPVDQTSLNSGTLIRWTKGFRIADTVGKDVVQLYQEQLSAQGMPMIKVVALTNDTVGT
YLSHCYTSDNTDSMTSGEISEPVIGCIFGTGTNGCYMEEINKITKLPQELRDKLIKEGK
THMIINVEWGSFDNELKHLPTTKYDVVIDQKLSTNPGFHLFEKRVSGMFLGEVLRNILV
DLHSQGLLLQQYRSKEQLPRHLTTPFQLSSEVLSHIEIDDSTGLRETELSLLQSLRLPT
TPTERVQIQKLVRAISRRSAYLAAVPLAAILIKTNALNKRYHGEVEIGCDGSVVEYYPG
FRSMLRHALALSPLGAEGERKVHLKIAKDGSGVGAALCALVA"
CDS 54654..56891
/codon_start=1
/locus_tag="YCL039W"
/label=GID7
/translation="MSHTNKIAYVLNNDTEETASPSSVGCFDKKQLTKLLIHTLKELGY
DSAANQLLLESGGYQNESNHIQTFFKLIKTGQFHLINWQIVCSLPLAHSSPLRSEWLQR
LLIPTPTPATTSLFDHMLLQLQYLQQLMSSVNSSTCSDAEIATLRNYVEIMILVNRQIF
LEFFHPVTNSASHKGPHTALPVLYLRKILKNFIEIWDSLLVSNDQFLNEENIFNPETTL
RELSTYLTNPKLTAQLNLERDHLIDAISKYIDPNELVPKGRLLHLLKQAIKYQQSQDIF
NIIDPDDDASFSSPPHRINLLQDNFSHDLTVTFQEWKTIQDTTDEIWFLTFSPNGKYLA
SATSESSRGYFITVYDVEQDFKIYKTCVSLSQSVLYLMFSPDSRYLVACPFSEDVTIYD
MNATSLPDASATDSFLLYPSTRLSPMDSFKLDTTTYPDDTESSASSSSRPANANSNQSR
VWCCDAFHTAERAGWMVVGSPDREAIVHSLTTKESLFSLKGRTCIALGHDENISGRKSI
DPAKVLYKPTSSNGNWQYVEDDETFPRVHDVKISYDDKYVLLMTHQGVIDVYDFSGFPS
KEELSKQTVDPKNFLIPRIARLDVGKNMTCISLPLNTTHQGFHRQQISESQHLVLVSLQ
DNELQMWDYKENILIQKYFGQKQQHFIIRSCFAYGNKLVMSGSEDGKIYIWDRIRGNLV
SVLSGHSTVMSNSTKPMGKNCNVVASNPADKEMFASGGDDGKIKIWKISRN"
CDS complement(56950..58536)
/codon_start=1
/locus_tag="YCL038C"
/label=ATG22
/translation="MSYGTINDMNESVTNYRIKKAQNNIKGWYAYSFSSEPFVVSAVST
YIPLLLQQFASINGVKVHDHSIPCLSETGSDSDKCVLGLFNNRIFVDTSSFALYVFSLS
VLFQTIIVISVSGIVDLWGSVKFKGRILVWFGIVGALSTVAISKLNDTQIYSLAGLYIV
ANGCFGVINVVGNSLLPIFVKDSLKCQSQGAYEPDKVDSLTTVISGRGASLGYSSALIV

QIVSMFLVASKKGSKQDVQVAVLFVGIWWFVWQLPMIWLIDDVTIPIRVDDSTLASARS
PYPGEQDALGQLNWKNYLSYGWVSLFESFKHARLLKDVMIFLIAWFIISDSITTINSTA
VLFSKAELHMSTLNLIMISVLTVVNAMLGAFMIPQFLATKFRWTSSQTLMYIIIWASFI
PFYGILGFFFNAFGLKHKFEMFLLAIWYGLSLGGLSAVSRSVFSLIVPPGKESTFFSMF
SITDKGSSILGPFLVGLLTDKTHNIRYSFYFFFLLLMLSLPVLNCLDVKRGRREAEELS
QVLPESERRLD"

CDS complement(59382..60686)
/codon_start=1
/locus_tag="YCL037C"
/label=SRO9
/translation="MSAETAAANTATAPVPEVQEQESSKSKQVNLTPAPLPTSSPWKLA
PTEIPVSTISIEDLDATRKKKNRTPTPKSSTATKWVPIKASITVSGTKRSGSKNGASNG
NSNKSKNNKTAASSTSSSNANRKKKHHQHNAKKQQQMKKDGFESAVGEEDSKDATSQEN
GQSTQQQQPPHHRNHHHSHHHNSNGPQRRKFHNSNNAGMPQNQGFPPQFKPYQGRNARN
NNNNRSKYHNHFHHNQQHPQQPMVKLQQQFYPVQPVLMAINNIARQIEYYFSEENLTVD
NYLRSKLSKDGFAPLSLISKFYRVVNMSFGGDTNLILAALREIVANEAATVNVAEGTLA
AKEGDNVTGEAKEPSPLDKYFVRSKSWSNWLPETFETEINIEKELVGDALDQFMISLPP
VPQQEEESSTELASQEQETKEDSAPVAAGESESSL"

CDS 61034..62734
/codon_start=1
/locus_tag="YCL036W"
/label=GFD2
/translation="MQVQKMVRDNSNNGSDKSVHWERRNNNGAGPRYRSRSGNTGALAT
KLSNGTLSVRGLVKDRTGSGKIAGCVEAFLDARTQLNTPWDRAKCNWLDQIGYYVQLRK
TAFSKELDQLRKPMIDAYVAEMRQKFDASYGQSRAQLEAKLAQVDSEWHMVHGDVHAKL
EKLVEERRFLKRLSDTIVPPRSKRSQRLSPLTKEDRANCICPQPKGMSDTAWFEAIQKK
MLGMNGTIKLLETEQKLLADEKNSVRKTFWPMVEAHSRSNEFAYLEKCIRLMASQRAIC
FCLDIEAFETNQNVITEIGISIYDPRENMVPSMVPITKNYHLIIEESLELRNQKWVCDY
KDCYLLGESYVLSLKECVHFIQSLINYYLVPVTEEDKTWSRAFVGHHVSGDLKWLETIG
VKFPGRGYEGHLDHTLLLAETPGDLDVFILDTEQFYRKSYGEKGSSLGKILRLFEIPHA
FLHNAGNDAYYTLHLFMKFCDVNFRKISGMDDVLKVMGQVKVWGERDVREPKVVPMSYA
ISIEEAVKNRTYRKGVKSSRKERVCQTEFGGLTYFGTAKDAFTSTLPTH"

CDS complement(62849..63181)
/codon_start=1
/locus_tag="YCL035C"
/label=GRX1
/translation="MVSQETIKHVKDLIAENEIFVASKTYCPYCHAALNTLFEKLKVPR
SKVLVLQLNDMKEGADIQAALYEINGQRTVPNIYINGKHIGGNDDLQELRETGELEELL
EPILAN"

CDS 63666..64730
/codon_start=1
/locus_tag="YCL034W"
/label=LSB5
/translation="MGFLSDHPHTAITETIFRIVSSRDYTLEVELAPLIQLIKADHNDY
NYTVNQEEAARALRKKIKYGNRLQQSRTLDLLDLFISQGVKFTVMYNDDKLLQRLRGMA
TNSENSGSGEKYEPRIIKKCAAYAISWLNYITQNNLENARAYSGLYQLGQTVKQRYSKS
SRSRRSGGGSGGRSNFMDDSADDTLYQSNSLTSADRLYRIPQINMNKEAPRIRLIISDA
LASAVSLQNSLIGLPKGKFSTDDEEATSKFIQARAIRRKVLRYLQLVTEGEFLGSLIHA
NDELVAALTAYDDRSAQDDSSDESDHGSYDDGIYDENEQDNSRYIDSESSEEESLSSYQ
PSTISNPFGDHNKI"
CDS complement(64784..65290)
/codon_start=1
/locus_tag="YCL033C"
/label=MXR2
/translation="MNKWSRLYVITVRRTFPGRRNIVLTQYWNKSKKMSDESNDVKWND
ALTPLQLMVLRDKATERPNTGAYLHTNESGVYHCANCDRPLYSSKAKFDARCGWPAFYE
EVSPGAITYHRDNSLMPARVEICCARCGGHLGHVFEGEGWKQLLNLPKDTRHCVNSASL
NLKKD"
CDS 65449..66489
/codon_start=1
/locus_tag="YCL032W"
/label=STE50
/translation="MEDGKQAINEGSNDASPDLDVNGTILMNNEDFSQWSVDDVITWCI
STLEVEETDPLCQRLRENDIVGDLLPELCLQDCQDLCDGDLNKAIKFKILINKMRDSKL
EWKDDKTQEDMITVLKNLYTTTSAKLQEFQSQYTRLRMDVLDVMKTSSSSSPINTHGVS
TTVPSSNNTIIPSSDGVSLSQTDYFDTVHNRQSPSRRESPVTVFRQPSLSHSKSLHKDS
KNKVPQISTNQSHPSAVSTANTPGPSPNEALKQLRASKEDSCERILKNAMKRHNLADQD
WRQYVLVICYGDQERLLELNEKPVIIFKNLKQQGLHPAIMLRRRGDFEEVAMMNGSDNV
TPGGRL"
CDS complement(66683..67576)
/codon_start=1
/locus_tag="YCL031C"
/label=RRP7
/translation="MGIEDISAMKNGFIVVPFKLPDHKALPKSQEASLHFMFAKRHQSS
NSNESDCLFLVNLPLLSNIEHMKKFVGQLCGKYDTVSHVEELLYNDEFGLHEVDLSALT
SDLMSSTDVNEKRYTPRNTALLKFVDAASINNCWNALKKYSNLHAKHPNELFEWTYTTP
SFTTFVNFYKPLDIDYLKEDIHTHMAIFEQREAQAQEDVQSSIVDEDGFTLVVGKNTKS
LNSIRKKILNKNPLSKHENKAKPISNIDKKAKKDFYRFQVRERKKQEINQLLSKFKEDQ

ERIKVMKAKRKFNPYT"
CDS complement(67942..70341)
/codon_start=1
/locus_tag="YCL030C"
/label=HIS4
/translation="MVLPILPLIDDLASWNSKKEYVSLVGQVLLDGSSLSNEEILQFSK
EEEVPLVALSLPSGKFSDDEIIAFLNNGVSSLFIASQDAKTAEHLVEQLNVPKERVVVE
ENGVFSNQFMVKQKFSQDKIVSIKKLSKDMLTKEVLGEVRTDRPDGLYTTLVVDQYERC
LGLVYSSKKSIAKAIDLGRGVYYSRSRNEIWIKGETSGNGQKLLQISTDCDSDALKFIV
EQENVGFCHLETMSCFGEFKHGLVGLESLLKQRLQDAPEESYTRRLFNDSALLDAKIKE
EAEELTEAKGKKELSWEAADLFYFALAKLVANDVSLKDVENNLNMKHLKVTRRKGDAKP
KFVGQPKAEEEKLTGPIHLDVVKASDKVGVQKALSRPIQKTSEIMHLVNPIIENVRDKG
NSALLEYTEKFDGVKLSNPVLNAPFPEEYFEGLTEEMKEALDLSIENVRKFHAAQLPTE
TLEVETQPGVLCSRFPRPIEKVGLYIPGGTAILPSTALMLGVPAQVAQCKEIVFASPPR
KSDGKVSPEVVYVAEKVGASKIVLAGGAQAVAAMAYGTETIPKVDKILGPGNQFVTAAK
MYVQNDTQALCSIDMPAGPSEVLVIADEDADVDFVASDLLSQAEHGIDSQVILVGVNLS
EKKIQEIQDAVHNQALQLPRVDIVRKCIAHSTIVLCDGYEEALEMSNQYAPEHLILQIA
NANDYVKLVDNAGSVFVGAYTPESCGDYSSGTNHTLPTYGYARQYSGANTATFQKFITA
QNITPEGLENIGRAVMCVAKKEGLDGHRNAVKIRMSKLGLIPKDFQ"
CDS complement(69383..69394)
/codon_start=1
/product="Factor Xa recognition and cleavage site"
/label=Factor Xa site
/translation="IEGR"
CDS complement(70607..71929)
/codon_start=1
/locus_tag="YCL029C"
/label=BIK1
/translation="MDRYQRKIGCFIQIPNLGRGQLKYVGPVDTKAGMFAGVDLLANIG
KNDGSFMGKKYFQTEYPQSGLFIQLQKVASLIEKASISQTSRRTTMEPLSIPKNRSIVR
LTNQFSPMDDPKSPTPMRSFRITSRHSGNQQSMDQEASDHHQQQEFGYDNREDRMEVDS
ILSSDRKANHNTTSDWKPDNGHMNDLNSSEVTIELREAQLTIEKLQRKQLHYKRLLDDQ
RMVLEEVQPTFDRYEATIQEREKEIDHLKQQLELERRQQAKQKQFFDAENEQLLAVVSQ
LHEEIKENEERNLSHNQPTGANEDVELLKKQLEQLRNIEDQFELHKTKWAKEREQLKMH
NDSLSKEYQNLSKELFLTKPQDSSSEEVASLTKKLEEANEKIKQLEQAQAQTAVESLPI
FDPPAPVDTTAGRQQWCEHCDTMGHNTAECPHHNPDNQQFF"
CDS 72158..73375
/codon_start=1
/label=RNQ1

/translation="MDTDKLISEAESHFSQGNHAEAVAKLTSAAQSNPNDEQMSTIESL
IQKIAGYVMDNRSGGSDASQDRAAGGGSSFMNTLMADSKGSSQTQLGKLALLATVMTHS
SNKGSSNRGFDVGTVMSMLSGSGGGSQSMGASGLAALASQFFKSGNNSQGQGQGQGQGQ
GQGQGQGQGSFTALASLASSFMNSNNNNQQGQNQSSGGSSFGALASMASSFMHSNNNQN
SNNSQQGYNQSYQNGNQNSQGYNNQQYQGGNGGYQQQQGQSGGAFSSLASMAQSYLGGG
QTQSNQQQYNQQGQNNQQQYQQQGQNYQHQQQGQQQQQGHSSSFSALASMASSYLGNNS
NSNSSYGGQQQANEYGRPQQNGQQQSNEYGRPQYGGNQNSNGQHESFNFSGNFSQQNNN
GNQNRY"
CDS 73811..75349
/codon_start=1
/locus_tag="YCL027W"
/label=FUS1
/translation="MVATIMQTTTTVLTTVAAMSTTLASNYISSQASSSTSVTTVTTIA
TSIRSTPSNLLFSNVAAQPKSSSASTIGLSIGLPIGIFCFGLLILLCYFYLKRNSVSIS
NPPMSATIPREEEYCRRTNWFSRLFWQSKCEDQNSYSNRDIEKYNDTQWTSGDNMSSKI
QYKISKPIIPQHILTPKKTVKNPYAWSGKNISLDPKVNEMEEEKVVDAFLYTKPPNIVH
IESSMPSYNDLPSQKTVSSKKTALKTSEKWSYESPLSRWFLRGSTYFKDYGLSKTSLKT
PTGAPQLKQMKMLSRISKGYFNESDIMPDERSPILEYNNTPLDANDSVNNLGNTTPDSQ
ITSYRNNNIDLITARPHSVIYGTTAQQTLETNFNDHHDCNKSTEKHELIIPTPSKPLKK
RKKRRQSKMYQHLQHLSRSKPLPLTPNSKYNGEASVQLGKTYTVIQDYEPRLTDEIRIS
LGEKVKILATHTDGWCLVEKCNTQKGSIHVSVDDKRYLNEDRGIVPGDCLQEYD"
CDS complement(75413..75994)
/codon_start=1
/locus_tag="YCL026C-B"
/label=HBN1
/translation="MSAVATYLKTLTARRTIYALKPELPGEITINDIQSVVQTIIKETP
TAFNSQPNRAVILTGETHKKVWDEVTKAIESPAGQKRPASARDEAFGSVIFFTDDKVTE
KLKADFPAYAAAFPSFADHTSGAAQINSWVALEAMGLGGHLQHYNGYIKAALPSKIPES
WTVQAQLVFGTPAAPPGEKTYIKNDVEIFN"
CDS complement(76712..77293)
/codon_start=1
/locus_tag="YCL026C-A"
/label=FRM2
/translation="MSPTGNYLNAITNRRTIYNLKPELPQGVGLDDVKRTVHVILKNTP
TAFNSQVNRAVIIVGDTHKRIWDAVASAMPTAEAKKRPESCRDEAYGSVIFFTDEGPTE
KLQRDFPALAAAFPTCAAHTTGAVQIQSWTALELLGLGANLQHYNDYVKSALPQDVPIA
WTVQSQLVFGVPTALPEEKTFINNVINVYH"
CDS complement(78026..79927)
/codon_start=1

/locus_tag="YCL025C"
/label=AGP1
/translation="MSSSKSLYELKDLKNSSTEIHATGQDNEIEYFETGSNDRPSSQPH
LGYEQHNTSAVRRFFDSFKRADQGPQDEVEATQMNDLTSAISPSSRQAQELEKNESSDN
IGANTGHKSDSLKKTIQPRHVLMIALGTGIGTGLLVGNGTALVHAGPAGLLIGYAIMGS
ILYCIIQACGEMALVYSNLTGGYNAYPSFLVDDGFGFAVAWVYCLQWLCVCPLELVTAS
MTIKYWTTSVNPDVFVIIFYVLVITINIFGARGYAEAEFFFNCCKILMMTGFFILGIII
DVGGAGNDGFIGGKYWHDPGAFNGKHAIDRFKGVAATLVTAAFAFGGSEFIAITTAEQS
NPRKAIPGAAKQMIYRILFLFLATIILLGFLVPYNSDQLLGSTGGGTKASPYVIAVASH
GVRVVPHFINAVILLSVLSMANSSFYSSARLFLTLSEQGYAPKVFSYIDRAGRPLIAMG
VSALFAVIAFCAASPKEEQVFTWLLAISGLSQLFTWTAICLSHLRFRRAMKVQGRSLGE
LGFKSQTGVWGSAYACIMMILILIAQFWVAIAPIGEGKLDAQAFFENYLAMPILIALYV
GYKVWHKDWKLFIRADKIDLDSHRQIFDEELIKQEDEEYRERLRNGPYWKRVVAFWC"
CDS 81170..84283
/codon_start=1
/locus_tag="YCL024W"
/label=KCC4
/translation="MTVANTETHSAAKPSSTIGPWKLGETLGFGSTGKVQLAQHERTGH
RTAVKVISKSIFNNNGNHSNDDSVLPYNIEREIVIMKLLSHPNVLSLYDVWETNNNLYL
ILEYAEKGELFNLLVDHGPLPEREAINCFRQIIIGISYCHALGIVHRDLKPENLLLDSF
YNIKIADFGMAALQTDADLLETSCGSPHYAAPEIVSGLPYEGFASDVWSCGVILFALLT
GRLPFDEENGNVRDLLLKVQKGQFEMPNDTEISRDAQDLIGKILVVDPRQRIKIRDILS
HPLLKKYQTIKDSKSIKDLPRENTYLYPLADSNNHTSASIDDSILQNLVVLWHGRHADD
IVSKLKENGTNKEKILYALLYRFKLDSVRGSNKKNRNKIKKTKKNKRSSTLSSSSSLLL
NNRSIQSTPRRRTSKRHSREFSSSRKRSSFLLSSNPTDSSPIPLRSSKRITHINVASAN
TQATPSGVPNPHKRNSKKRSSKRLSYMPNTKRSSLTSKSLSNFTNLIDDDDWEYIEKDA
KRTSSNFATLIDEIFEPEKFELAKREKAELQRKVQEAKRQSVNAQKINEDEFGSEVSDG
MKELKKINDKVSSPLINYEFSQQELLQDIDTLLTNRYQLSSYTRPISRLDPGLTPVTET
LPNNLKEKTALLQDTEKKIIETIRRSKFLGSLLNVRGGLSPGKSELAPIEESPIVSTTP
LIYNDRMEPRRISDVEVPHFTRKSKHFTTANNRRSVLSLYAKDSIKDLNEFLIKEDPDL
PPQGSTDNESRSEDPEIAESITDSRNIQYDEDDSKDGDNVNNDNILSDFPQGVGISQEY
DMKDKNPNQSPISKSAEPTLVVKLPSLSSFQGKNASGLGLYQREPSKVTLPSLTSNNSS
VGENIEDGAEKGTESEKIAASLSDDDLKEDNDKKDNDTVNAPTTVKKPPNSVLLKKFSK
GKILELEIHAKIPEKRLYEGLHKLLEGWKQYGLKNLVFNITNMIITGKLVNDSILFLRS
TLFEIMVLPNGDGRSLIKFNKKTGSTKTLTKLATEIQIILQKEGVLDK"
gap 85045..85144
/estimated_length=100
CDS 86068..86445
/codon_start=1

/locus_tag="YCL021W-A"
/label=YCL021W-A
/translation="MVLTDAEELRSPVITSDMSFFDLESNHSSDSVHLLCEKYTHKLPI
ESESQTTFRLAPTKQRLYRQSTLYVPLSLKQRVFLFTERVKSIWAGLPRCKPNKYFKVA
FALAVLTPLAIWIFYIDFRVH"
CDS join(87550..88839,88841..92863)
/codon_start=1
/locus_tag="YCL019W"
/label=YCL019W
/translation="MESQQLHQNPRSLHGSAYASVTSKEVPSNQDPLAVSASNLPEFDR
DSTKVNSQQETTPGTSAVPENHHHVSPQPASVPPPQNGQYQQHGMMTPNKAMASNWAHY
QQPSMMTCSHYQTSPAYYQPDPHYPLPQYIPPLSTSSPDPIDSQNQHSEVPQAETKVRN
NVLPPHTLTSEENFSTWVKFYIRFLKNSNLGDIIPNDQGEIKRQMTYEEHAYIYNTFQA
FAPFHLLPTWVKQILEINYADILTVLCKSVSKMQTNNQELKDWIALANLEYDGSTSADT
FEITVSTIIQRLKENNINVSDRLACQLILKGLSGDFKYLRNQYRTKTNMKLSQLFAEIQ
LIYDENKIMNLNKPSQYKQHSEYKNVSRTSPNTTNTKVTTRNYQRTNSSKPRAAKAHNI
ATSSKFSRVNNDHINESTVSSQYLSDDNELSLGQQQKESKPTHTIDSNDELPDHLLIDS
GASQTLVRSAHYLHHATPNSEINIVDAQKQDIPINAIGNLHFNFQNGTKTSIKALHTPN
IAYDLLSLSELANQNITACFTRNTLERSDGTVLAPIVKHGDFYWLSKKYLIPSHISKLT
INNVNKSKSVNKYPYPLIHRMLGHANFRSIQKSLKKNAVTYLKESDIEWSNASTYQCPD
CLIGKSTKHRHVKGSRLKYQESYEPFQYLHTDIFGPVHHLPKSAPSYFISFTDEKTRFQ
WVYPLHDRREESILNVFTSILAFIKNQFNARVLVIQMDRGSEYTNKTLHKFFTNRGITA
CYTTTADSRAHGVAERLNRTLLNDCRTLLHCSGLPNHLWFSAVEFSTIIRNSLVSPKND
KSARQHAGLAGLDITTILPFGQPVIVNNHNPDSKIHPRGIPGYALHPSRNSYGYIIYLP
SLKKTVDTTNYVILQDKQSKLDQFNYDTLTFDDDLNRLTAHNQSFIEQNETEQSYDQNT
ESDHDYQSEIEINSDPLVNDFSSQSINPLQLDKEPVQKVRAPKEVDADISEYNILPSPV
RSRTPHIINKESTEMGGTVESDTTSPRHSSTFTARNQKRPGSPNDMIDLTSQDRVNYGL
ENIKTTRLGGTEEPYIQRNSDTNIKYRTTNSTPSIDDRSSNSESTTPIISIETKAVCDN
TPSIDTDPPEYRSSDHATPNIMPDKSSKNVTADSILDDLPLPDLTHQSPTDTSDVSKDI
PHIHSRQTNSSLGGMDDSNVLTTTKSKKRSLEDNETEIEVSRDTWNNKNMRSLEPPRSK
KRINLIAAIKGVKSIKPVRTTLRYDEAITYNKDNKEKDRYVEAYHKEISQLLKMNTWDT
NKYYDRNDIDPKKVINSMFIFNKKRDGTHKARFVARGDIQHPDTYDSDMQSNTVHHYAL
MTSLSIALDNDYYITQLDISSAYLYADIKEELYIRPPPHLGLNDKLLRLRKSLYGLKQS
GANWYETIKSYLINCCDMQEVRGWSCVFKNSQVTICLFVDDMILFSKDLNANKKIITTL
KKQYDTKIINLGESDNEIQYDILGLEIKYQRSKYMKLGMEKSLTEKLPKLNVPLNPKGK
KLRAPGQPGHYIDQDELEIDEDEYKEKVHEMQKLIGLASYVGYKFRFDLLYYINTLAQH
ILFPSRQVLDMTYELIQFMWDTRDKQLIWHKNKPTKPDNKLVAISDASYGNQPYYKSQI
GNIFLLNGKVIGGKSTKASLTCTSTTEAEIHAVSEAIPLLNNLSHLVQELNKKPIIKGL
LTDSRSTISIIKSTNEEKFRNRFFGTKAMRLRDEVSGNNLYVYYIETKKNIADVMTKPL

PIKTFKLLTNKWIH"
CDS 87550..88866
/codon_start=1
/locus_tag="YCL020W"
/label=YCL020W
/translation="MESQQLHQNPRSLHGSAYASVTSKEVPSNQDPLAVSASNLPEFDR
DSTKVNSQQETTPGTSAVPENHHHVSPQPASVPPPQNGQYQQHGMMTPNKAMASNWAHY
QQPSMMTCSHYQTSPAYYQPDPHYPLPQYIPPLSTSSPDPIDSQNQHSEVPQAETKVRN
NVLPPHTLTSEENFSTWVKFYIRFLKNSNLGDIIPNDQGEIKRQMTYEEHAYIYNTFQA
FAPFHLLPTWVKQILEINYADILTVLCKSVSKMQTNNQELKDWIALANLEYDGSTSADT
FEITVSTIIQRLKENNINVSDRLACQLILKGLSGDFKYLRNQYRTKTNMKLSQLFAEIQ
LIYDENKIMNLNKPSQYKQHSEYKNVSRTSPNTTNTKVTTRNYQRTNSSKPRAAKAHNI
ATSSKFSRVNNDHINESTVSSQYLSDDNELSLRPATERI"
promoter 93364..93771
/gene="S. cerevisiae LEU2"
/label=LEU2 promoter
CDS 93772..94868
/codon_start=1
/locus_tag="YCL018W"
/label=LEU2
/translation="MSAPKKIVVLPGDHVGQEITAEAIKVLKAISDVRSNVKFDFENHL
IGGAAIDATGVPLPDEALEASKKADAVLLGAVGGPKWGYR*C*T*TRFTKNP*RTSIVR
QLKTM*LCIRLSFRLISNQATIC*RY*LRCCQRISGRYLLW*EKGRRW*WCRLG**TIH
RSRSAKNHKNGRFHGPTT*ATIAYLVLG*S*CFGLFKIMEKNCGGNHQERIPYIEGSTS
ID*FCRHDPS*EPNPPKWYYNHQQHVW*YHLR*SLRYPRFLGVCCHLRPWPLCQTRTPH
LVCTNHATVLLQICQRIRSTLSPLSCLLQ*C*NCH*TCLKKVRPLKMQLKRFWMQVSEL
VI*VVPTVPPKSVMLSPKKLRKSLL"
CDS complement(95227..96720)
/codon_start=1
/locus_tag="YCL017C"
/label=NFS1
/translation="MLKSTATRSITRLSQVYNVPAATYRACLVSRRFYSPPAAGVKLDD
NFSLETHTDIQAAAKAQASARASASGTTPDAVVASGSTAMSHAYQENTGFGTRPIYLDM
QATTPTDPRVLDTMLKFYTGLYGNPHSNTHSYGWETNTAVENARAHVAKMINADPKEII
FTSGATESNNMVLKGVPRFYKKTKKHIITTRTEHKCVLEAARAMMKEGFEVTFLNVDDQ
GLIDLKELEDAIRPDTCLVSVMAVNNEIGVIQPIKEIGAICRKNKIYFHTDAAQAYGKI
HIDVNEMNIDLLSISSHKIYGPKGIGAIYVRRRPRVRLEPLLSGGGQERGLRSGTLAPP
LVAGFGEAARLMKKEFDNDQAHIKRLSDKLVKGLLSAEHTTLNGSPDHRYPGCVNVSFA
YVEGESLLMALRDIALSSGSACTSASLEPSYVLHALGKDDALAHSSIRFGIGRFSTEEE

VDYVVKAVSDRVKFLRELSPLWEMVQEGIDLNSIKWSGH"
CDS complement(97071..98213)
/codon_start=1
/locus_tag="YCL016C"
/label=DCC1
/translation="MSINLHSAPEYDPSYKLIQLTPELLDIIQDPVQNHQLRFKSLDKD
KSEVVLCSHDKTWVLKQRKHSNTVLLMREFVPEQPITFDETLLFGLSKPYMDVVGFAKT
ESEFETRETHGELNLNSVPIYNGELDFSDKIMKRSSTKVIGTLEELLENSPCSALEGIS
KWHKIGGSVKDGVLCILSQDFLFKALHVLLMSAMAESLDLQHLNVEDTHHAVGKDIEDE
FNPYTREIIETVLNKFAVQEQEAENNTWRLRIPFIAQWYGIQALRKYVSGISMPIDEFL
IKWKSLFPPFFPCDIDIDMLRGYHFKPTDKTVQYIAKSTLPMDPKERFKVLFRLQSQWD
LEDIKPLIEELNSRGMKIDSFIMKYARRKRLGKKTVVTSR"
CDS 98731..103641
/codon_start=1
/locus_tag="YCL014W"
/label=BUD3
/translation="MEKDLSSLYSEKKDKENDETLFNIKLSKSVVETTPLNGHSLFDDD
KSLSDWTDNVFTQSVFYHGSDDLIWGKFFVCVYKSPNSNKLNAIIFDKLGTSCFESVDI
SSNSQYYPAIENLSPSDQESNVKKCIAVILLQRYPLLSPSDLSQILSNKSENCDYDPPY
AGDLASSCQLITAVPPEDLGKRFFTSGLLQNRFVSSTLLDVIYENNESTIELNNRLVFH
LGEQLEQLFNPVTEYSPEQTEYGYKAPEDELPTESDDDLVKAICNELLQLQTNFTFNLV
EFLQKFLIALRVRVLNEEINGLSTTKLNRLFPPTIDEVTRINCIFLDSLKTAIPYGSLE
VLKACSITIPYFYKAYTRHEAATKNFSKDIKLFIRHFSNVIPEREVYTEMKIESIIKGP
QEKLLKLKLIIERLWKSKKWRPKNQEMAKKCYNNIIDVIDSFGKLDSPLHSYSTRVFTP
SGKILTELAKCWPVELQYKWLKRRVVGVYDVVDLNDENKRNLLVIFSDYVVFINILEAE
SYYTSDGSNRPLISDILMNSLINEVPLPSKIPKLKVERHCYIDEVLVSILDKSTLRFDR
LKGKDSFSMVCKLSSAFISSSSVADLITKARILEKDTAFHLFKASRSHFTLYSTAHELC
AYDSEKIKSKFALFLNIPPSKEILEVNNLHLAFFARFCSNDGRDNIVILDVLTKHDDKH
IEVTSDNIVFTIINQLAIEIPICFSSLNSSMAKDLLCVNENLIKNLEHQLEEVKHPSTD
EHRAVNSKLSGASDFDATHEKKRSYGTITTFRSYTSDLKDSPSGDNSNVTKETKEILPV
KPTKKSSKKPREIQKKTKTNASKAEHIEKKKPNKGKGFFGVLKNVFGSKSKSKPSPVQR
VPKKISQRHPKSPVKKPMTSEKKSSPKRAVVSSPKIKKKSTSFSTKESQTAKSSLRAVE
FKSDDLIGKPPDVGNGAHPQENTRISSVVRDTKYVSYNPSQPVTENTSNEKNVEPKADQ
STKQDNISNFADVEVSASSYPEKLDAETDDQIIGKATNSSSVHGNKELPDLAEVTTANR
VSTTSAGDQRIDTQSEFLRAADVENLSDDDEHRQNESRVFNDDLFGDFIPKHYRNKQEN
INSSSNLFPEGKVPQEKGVSNENTNISLKTNEDASTLTQKLSPQASKVLTENSNELKDT
NNEGKDAKDIKLGDDYSDKETAKEITKPKNFVEGITERKEIFPTIPRLAPPASKINFQR
SPSYIELFQGMRVVLDKHDAHYNWKRLASQVSLSEGLKVNTEEDAAIINKSQDDAKAER
MTQISEVIEYEMQQPIPTYLPKAHLDDSGIEKSDDKFFEIEEELKEELKGSKTGNEDVG

NNNPSNSIPKIEKPPAFKVIRTSPVRIIGRTFEDTRKYENGSPSDISFTYDTHNNDEPD
KRLMELKFPSQDEIPDDRFYTPAEEPTAEFPVEELPNTPRSINVTTSNNKSTDDKLSSG
NIDQKPTELLDDLEFSSFNIAFGNTSMSTDNMKISSDLSSNKTVLGNAQKVQESPSGPL
IYVLPQSSTKHEKEGFLRKKQKDEPIWVSPSKIDFADLSRRTKALTPERNTVPLKNNDS
RKYKYTGEGSIGNMTNMLLTKDASYAYLKDFVALSDDEDEDGKQNCAVGGPEKLKFY"
CDS complement(join(103767..104083,104151..104238))
/codon_start=1
/locus_tag="YCL012C"
/label=YCL012C
/translation="MKSLFYLKLLLWVVLLSLCLLMAHRKTKVADKFRALRSRIQLRFN
RHIRLNDSFADDLENGLHSRNFDIISENSNDVRGGLDDVSKNEIKQIMENDNVDFDKAR
LLYMERKFGQNGIAPDGTPIDPKAFTFDSR"
CDS complement(104525..105808)
/codon_start=1
/locus_tag="YCL011C"
/label=GBP2
/translation="MERELGMYGNDRSRSRSPVRRRLSDDRDRYDDYNDSSSNNGNGSR
RQRRDRGSRFNDRYDQSYGGSRYHDDRNWPPRRGGRGRGGSRSFRGGRGGGRGRTLGPI
VERDLERQFDATKRNFENSIFVRNLTFDCTPEDLKELFGTVGEVVEADIITSKGHHRGM
GTVEFTKNESVQDAISKFDGALFMDRKLMVRQDNPPPEAAKEFSKKATREEIDNGFEVF
IINLPYSMNWQSLKDMFKECGHVLRADVELDFNGFSRGFGSVIYPTEDEMIRAIDTFNG
MEVEGRVLEVREGRFNKRKNNDRYNQRREDLEDTRGTEPGLAQDAAVHIDETAAKFTEG
VNPGGDRNCFIYCSNLPFSTARSDLFDLFGPIGKINNAELKPQENGQPTGVAVVEYENL
VDADFCIQKLNNYNYGGCSLQISYARRD"
CDS complement(106021..106800)
/codon_start=1
/locus_tag="YCL010C"
/label=SGF29
/translation="MDGYWDVVVSSLQDIYNANEVIPFDDELQTKKLNFLNMSKDQLQL
HLNTFQEHMENVNRVHRILDNVRSNLSLMLNQSREEKSEENTEDAEEGEGTRMALSQGK
KAVGKVGRSYWTSEYNPNAPILVGSEVAYKPRRGSADGEWIQCEVLKVVADGTRFEVRD
PEPDELGNSGKVYKCNRKELLLIPPGFPTKNYPPGTKVLARYPETTTFYPAIVIGTKRD
GTCRLRFDGEEEVDKETEVTRRLVLPSPTALANLARK"
CDS complement(107069..107998)
/codon_start=1
/locus_tag="YCL009C"
/label=ILV6
/translation="MLRSLLQSGHRRVVASSCATMVRCSSSSTSALAYKQMHRHATRPP
LPTLDTPSWNANSAVSSIIYETPAPSRQPRKQHVLNCLVQNEPGVLSRVSGTLAARGFN

IDSLVVCNTEVKDLSRMTIVLQGQDGVVEQARRQIEDLVPVYAVLDYTNSEIIKRELVM
ARISLLGTEYFEDLLLHHHTSTNAGAADSQELVAEIREKQFHPANLPASEVLRLKHEHL
NDITNLTNNFGGRVVDISETSCIVELSAKPTRISAFLKLVEPFGVLECARSGMMALPRT
PLKTSTEEAADEDEKISEIVDISQLPPG"

CDS complement(108146..109303)
/codon_start=1
/locus_tag="YCL008C"
/label=STP22
/translation="MSANGKISVPEAVVNWLFKVIQPIYNDGRTTFHDSLALLDNFHSL
RPRTRVFTHSDGTPQLLLSIYGTISTGEDGSSPHSIPVIMWVPSMYPVKPPFISINLEN
FDMNTISSSLPIQEYIDSNGWIALPILHCWDPAAMNLIMVVQELMSLLHEPPQDQAPSL
PPKPNTQLQQEQNTPPLPPKPKSPHLKPPLPPPPPPQPASNALDLMDMDNTDISPTNHH
EMLQNLQTVVNELYREDVDYVADKILTRQTVMQESIARFHEIIAIDKNHLRAVEQAIEQ
TMHSLNAQIDVLTANRAKVQQFSSTSHVDDEDVNSIAVAKTDGLNQLYNLVAQDYALTD
TIECLSRMLHRGTIPLDTFVKQGRELARQQFLVRWHIQRITSPLS"

CDS join(109473..109483,109561..109641,109738..109867)
/codon_start=1
/label=VMA9
/translation="MSSFYTVVGVFIVVSAMSVLFWIMAPKNNQAVWRSTVILTLAMMF
LMWAITFLCQLHPLVAPRRSDLRPEFAE"

CDS complement(111452..111463)
/label=ARS core sequence

CDS 112701..113729
/label=HphMX4

misc_feature 114037..114370
/label=ARS307

CDS 114415..115980
/codon_start=1
/locus_tag="YCL004W"
/label=PGS1
/translation="MTTRLLQLTRPHYRLLSLPLQKPFNIKRQMSAANPSPFGNYLNTI
TKSLQQNLQTCFHFQAKEIDIIESPSQFYDLLKTKILNSQNRIFIASLYLGKSETELVD
CISQALTKNPKLKVSFLLDGLRGTRELPSACSATLLSSLVAKYGSERVDCRLYKTPAYH
GWKKVLVPKRFNEGLGLQHMKIYGFDNEVILSGANLSNDYFTNRQDRYYLFKSRNFSNY
YFKLHQLISSFSYQIIKPMVDGSINIIWPDSNPTVEPTKNKRLFLREASQLLDGFLKSS
KQSLPITAVGQFSTLVYPISQFTPLFPKYNDKSTEKRTILSLLSTITSNAISWTFTAGY
FNILPDIKAKLLATPVAEANVITASPFANGFYQSKGVSSNLPGAYLYLSKKFLQDVCRY
RQDHAITLREWQRGVVNKPNGWSYHAKGIWLSARDKNDANNWKPFITVIGSSNYTRRAY
SLDLESNALIITRDEELRKKMKAELDNLLQYTKPVTLEDFQSDPERHVGTGVKIATSIL

GKKL"
CDS complement(join(116118..116867,116944..116985))
/codon_start=1
/locus_tag="YCL002C"
/label=YCL002C
/translation="MLVIVLQGLAGFLSIIAILCQKRYNKLHRSIYGLSYDLFLLDFVG
NGLYLYCALHYCYSSLVREQLSQRFPLFYPLNDARSIPISSFLILKDFCVSCCCMMVLR
QLYYYRSTKHIYQGISITSIIIISVFLVLGIFTYGCSISNLPLKNSGKFGVFYLEHINY
LWVMANLLKCFKYVPQMSINWMGCSTVGLSSKFALISFLAESIDLLGRLVIPTNALFYE
IPFNSTPFWVKLIQFVTLLVILCQVQYVYVGRKPRLPKGKL"
CDS 117226..117792
/codon_start=1
/locus_tag="YCL001W"
/label=RER1
/translation="MDYDSSDTMNGGSSNPLITKMNTMKLLYQHYLDKVTPHAKERWAV
LGGLLCLFMVRITMAEGWYVICYGLGLFLLNQFLAFLTPKFDMSLQQDEENNELEAGEK
SEEFRPFIRRLPEFKFWYNSIRATVISLLLSLFSIFDIPVFWPILLMYFILLFFLTMRR
QIQHMIKYRYIPLDIGKKKYSHSSN"
CDS 118390..118851
/codon_start=1
/locus_tag="YCL001W-A"
/label=YCL001W-A
/translation="MTFLQFINNNRQEGQGYISEKLFKTKKNEMIRKTVTNLVAVRLKN
LSHEFDVIENYLRYIASTSEHLFTAIKRHFNKCARKLLKEAIDSKSNSETATVVLQEGF
SGICLLKASSIILKLKLKFPKKKDRTDISKLCDKKERMTQWLEISILMN"
CDS 119080..119334
/codon_start=1
/locus_tag="YCL001W-B"
/label=YCL001W-B
/translation="MDDFLEHLSKDDNKAWYGAEETERAAKLDAIETLLITDSVLKRND
VKKREKYLDLIENSGNNNGKIFVLSTSKITVSNLTNQQI"
misc_feature 119695..119811
/label=CEN3
CDS 120995..121309
/codon_start=1
/locus_tag="YCR001W"
/label=YCR001W
/translation="MLYLLKKLKHIDSYGESNLSMFYLKFYCPWKLCPLLPILYGRNKS
VQNAWSCHKLHRFIDFPTPSANMNFFFFIRVCYGTVSLFISTFLKKISFFIYVNFTVYF

"
CDS complement(122690..123658)
/codon_start=1
/locus_tag="YCR002C"
/label=CDC10
/translation="MDPLSSVQPASYVGFDTITNQIEHRLLKKGFQFNIMVVGQSGLGK
STLINTLFASHLIDSATGDDISALPVTKTTEMKISTHTLVEDRVRLNINVIDTPGFGDF
IDNSKAWEPIVKYIKEQHSQYLRKELTAQRERFITDTRVHAILYFLQPNGKELSRLDVE
ALKRLTEIANVIPVIGKSDTLTLDERTEFRELIQNEFEKYNFKIYPYDSEELTDEELEL
NRSVRSIIPFAVVGSENEIEINGETFRGRKTRWSAINVEDINQCDFVYLREFLIRTHLQ
DLIETTSYIHYEGFRARQLIALKENANSRSSAHMSSNAIQR"
CDS 123930..124481
/codon_start=1
/locus_tag="YCR003W"
/label=MRPL32
/translation="MNSLIFGKQLAFHKIVPTTAIGWLVPLGNPSLQIPGQKQLGSIHR
WLREKLQQDHKDTEDKDFFSNNGILLAVPKKKVSHQKKRQKLYGPGKKQLKMIHHLNKC
PSCGHYKRANTLCMYCVGQISHIWKTHTAKEEIKPRQEEELSELDQRVLYPGRRDTKYT
KDLKDKDNYLERRVRTLKKD"
CDS complement(124885..125628)
/codon_start=1
/locus_tag="YCR004C"
/label=YCP4
/translation="MVKIAIITYSTYGHIDVLAQAVKKGVEAAGGKADIYRVEETLPDE
VLTKMNAPQKPEDIPVATEKTLLEYDAFLFGVPTRFGNLPAQWSAFWDKTGGLWAKGSL
NGKAAGIFVSTSSYGGGQESTVKACLSYLAHHGIIFLPLGYKNSFAELASIEEVHGGSP
WGAGTLAGPDGSRTASPLELRIAEIQGKTFYETAKKLFPAKEAKPSTEKKTTTSDAAKR
QTKPAAATTAEKKEDKGLLSCCTVM"
CDS complement(126256..127638)
/codon_start=1
/locus_tag="YCR005C"
/label=CIT2
/translation="MTVPYLNSNRNVASYLQSNSSQEKTLKERFSEIYPIHAQDVRQFV
KEHGKTKISDVLLEQVYGGMRGIPGSVWEGSVLDPEDGIRFRGRTIADIQKDLPKAKGS
SQPLPEALFWLLLTGEVPTQAQVENLSADLMSRSELPSHVVQLLDNLPKDLHPMAQFSI
AVTALESESKFAKAYAQGISKQDYWSYTFEDSLDLLGKLPVIAAKIYRNVFKDGKMGEV
DPNADYAKNLVNLIGSKDEDFVDLMRLYLTIHSDHEGGNVSAHTSHLVGSALSSPYLSL
ASGLNGLAGPLHGRANQEVLEWLFALKEEVNDDYSKDTIEKYLWDTLNSGRVIPGYGHA
VLRKTDPRYMAQRKFAMDHFPDYELFKLVSSIYEVAPGVLTEHGKTKNPWPNVDAHSGV

LLQYYGLKESSFYTVLFGVSRAFGILAQLITDRAIGASIERPKSYSTEKYKELVKNIES
KL"
CDS complement(127840..128313)
/codon_start=1
/locus_tag="YCR006C"
/label=YCR006C
/translation="MFIRVCNRLIYTPTNVLLITVEDGRISVLLWFRYAIPAELCYTRL
ARILRGKHCADFPQSCCHNISRFGVLNKSVLGSFNQWLGGISKERKLIARTFDAFIRWS
STRGEEVTTYFFLQKKSVTFSVARRLSGRQQWEAQRKNNNNGKRNYLLSVTFV"
CDS complement(131321..132040)
/codon_start=1
/locus_tag="YCR007C"
/label=YCR007C
/translation="MQPHLDNNSNNDDVKLDTLGEQNVLSSAENITLPEDTFKSYMTYL
LYEMAHYKPMIFSFLALSVSILIVVIFHNVKACDVVFGFSIFVTSILFLSTLIPFNVYI
SDEGFRIKLLLEVITHRPAVKGKEWRAITDNMNQYLLDNGLWSTRYYFYSSERCYKFFR
FLVKEKPPGVNVNSSVKDATSTQIDAPANEASNEVIKCFSFSSDPIFEAYFVKAVEVEK
QAQQEYWRKQYPDADIP"
CDS 133780..135591
/codon_start=1
/locus_tag="YCR008W"
/label=SAT4
/translation="MTGMNDNNAAIPQQTPRKHALSSKVMQLFRSGSRSSRQGKASSNI
QPPSNINTNVPSASKSAKFGLHTPTTATPRVVSNPSNTAGVSKPGMYMPEYYQSASPSH
SSSSASLNNHIDINTSKSSSAASLTSSVSALSLSPTSAINISSKSLSPKFSHHSNSNTA
ITPAPTPTASNINNVNKITNTSAPICGRFLVHKDGTHEHHLKNAKRQEKLSTMIKNMVG
ASKLRGEAKSAVPDIIMDPKTTLKSNKNPPTLFAGFMKQVVDMDDKYPEGAPTSGALNC
PERDIYRSDQKDSKNNTHNITTTKKDRQCFAEKYGRCQEVLGKGAFGVVRICQKKNVSS
QDGNKSEKLYAVKEFKRRTSESAEKYSKRLTSEFCISSSLHHTNIVTTLDLFQDAKGEY
CEVMEYCAGGDLFTLVVAAGKLEYMEADCFFKQLIRGVVYMHEMGVCHRDLKPENLLLT
HDGVLKITDFGNSECFKMAWEKNIHLSGGVCGSSPYIAPEEYIKEEFDPRPVDIWACGV
IYMAMRTGRQLWSSAEKDDPFYMNYLKGRKEKGGYEPIESLKRARCRNVIYSMLDPVPY
RRINGKQILNSEWGREIKCCHNGRALK"
CDS complement(136055..136852)
/codon_start=1
/locus_tag="YCR009C"
/label=RVS161
/translation="MSWEGFKKAINRAGHSVIIKNVDKTIDKEYDMEERRYKVLQRAGE
ALQKEAKGFLDSLRAVTASQTTIAEVISNLYDDSKYVAGGGYNVGNYYLQCVQDFDSET

VKQLDGPLRETVLDPITKFSTYFKEIEEAIKKRDHKKQDFDAAKAKVRRLVDKPAKDAS
KLPRAEKELSLAKDIFENLNNQLKTELPQLVSLRVPYFDPSFEALIKIQLRFCTDGYTR
LAQIQQYLDQQSRDDYANGLLDTKIEELLGQMTSLDICALGIK"
CDS complement(137583..138434)
/codon_start=1
/locus_tag="YCR010C"
/label=ADY2
/translation="MSDKEQTSGNTDLENAPAGYYSSHDNDVNGVAEDERPSHDSLGKI
YTGGDNNEYIYIGRQKFLKSDLYQAFGGTLNPGLAPAPVHKFANPAPLGLSAFALTTFV
LSMFNARAQGITVPNVVVGCAMFYGGLVQLIAGIWEIALENTFGGTALCSYGGFWLSFA
AIYIPWFGILEAYEDNESDLNNALGFYLLGWAIFTFGLTVCTMKSTVMFFLLFFLLALT
FLLLSIGHFANRLGVTRAGGVLGVVVAFIAWYNAYAGVATKQNSYVLARPFPLPSTERV
IF"
CDS complement(139034..142183)
/codon_start=1
/locus_tag="YCR011C"
/label=ADP1
/translation="MGSHRRYLYYSILSFLLLSCSVVLAKQDKTPFFEGTSSKNSRLTA
QDKGNDTCPPCFNCMLPIFECKQFSECNSYTGRCECIEGFAGDDCSLPLCGGLSPDESG
NKDRPIRAQNDTCHCDNGWGGINCDVCQEDFVCDAFMPDPSIKGTCYKNGMIVDKVFSG
CNVTNEKILQILNGKIPQITFACDKPNQECNFQFWIDQLESFYCGLSDCAFEYDLEQNT
SHYKCNDVQCKCVPDTVLCGAKGSIDISDFLTETIKGPGDFSCDLETRQCKFSEPSMND
LILTVFGDPYITLKCESGECVHYSEIPGYKSPSKDPTVSWQGKLVLALTAVMVLALFTF
ATFYISKSPLFRNGLGSSKSPIRLPDEDAVNNFLQNEDDTLATLSFENITYSVPSINSD
GVEETVLNEISGIVKPGQILAIMGGSGAGKTTLLDILAMKRKTGHVSGSIKVNGISMDR
KSFSKIIGFVDQDDFLLPTLTVFETVLNSALLRLPKALSFEAKKARVYKVLEELRIIDI
KDRIIGNEFDRGISGGEKRRVSIACELVTSPLVLFLDEPTSGLDASNANNVIECLVRLS
SDYNRTLVLSIHQPRSNIFYLFDKLVLLSKGEMVYSGNAKKVSEFLRNEGYICPDNYNI
ADYLIDITFEAGPQGKRRRIRNISDLEAGTDTNDIDNTIHQTTFTSSDGTTQREWAHLA
AHRDEIRSLLRDEEDVEGTDGRRGATEIDLNTKLLHDKYKDSVYYAELSQEIEEVLSEG
DEESNVLNGDLPTGQQSAGFLQQLSILNSRSFKNMYRNPKLLLGNYLLTILLSLFLGTL
YYNVSNDISGFQNRMGLFFFILTYFGFVTFTGLSSFALERIIFIKERSNNYYSPLAYYI
SKIMSEVVPLRVVPPILLSLIVYPMTGLNMKDNAFFKCIGILILFNLGISLEILTIGII
FEDLNNSIILSVLVLLGSLLFSGLFINTKNITNVAFKYLKNFSVFYYAYESLLINEVKT
LMLKERKYGLNIEVPGATILSTFGFVVQNLVFDIKILALFNVVFLIMGYLALKWIVVEQ
K"
CDS 143056..144306
/codon_start=1
/locus_tag="YCR012W"

/label=PGK1
/translation="MSLSSKLSVQDLDLKDKRVFIRVDFNVPLDGKKITSNQRIVAALP
TIKYVLEHHPRYVVLASHLGRPNGERNEKYSLAPVAKELQSLLGKDVTFLNDCVGPEVE
AAVKASAPGSVILLENLRYHIEEEGSRKVDGQKVKASKEDVQKFRHELSSLADVYINDA
FGTAHRAHSSMVGFDLPQRAAGFLLEKELKYFGKALENPTRPFLAILGGAKVADKIQLI
DNLLDKVDSIIIGGGMAFTFKKVLENTEIGDSIFDKAGAEIVPKLMEKAKAKGVEVVLP
VDFIIADAFSADANTKTVTDKEGIPAGWQGLDNGPESRKLFAATVAKAKTIVWNGPPGV
FEFEKFAAGTKALLDEVVKSSAAGNTVIIGGGDTATVAKKYGVTDKISHVSTGGGASLE
LLEGKELPGVAFLSEKK"

terminator 144307..144577
/gene="S. cerevisae PGK1"
/label=PGK1 terminator
/note="transcription terminator for 3-phosphoglycerate
kinase gene"

CDS complement(144495..146243)
/codon_start=1
/locus_tag="YCR014C"
/label=POL4
/translation="MSLKGKFFAFLPNPNTSSNKFFKSILEKKGATIVSSIQNCLQSSR
KEVVILIEDSFVDSDMHLTQKDIFQREAGLNDVDEFLGKIEQSGIQCVKTSCITKWVQN
DKFAFQKDDLIKFQPSIIVISDNADDGQSSTDKESEISTDVESERNDDSNNKDMIQASK
PLKRLLQGDKGRASLVTDKTKYKNNELIIGALKRLTKKYEIEGEKFRARSYRLAKQSME
NCDFNVRSGEEAHTKLRNIGPSIAKKIQVILDTGVLPGLNDSVGLEDKLKYFKNCYGIG
SEIAKRWNLLNFESFCVAAKKDPEEFVSDWTILFGWSYYDDWLCKMSRNECFTHLKKVQ
KALRGIDPECQVELQGSYNRGYSKCGDIDLLFFKPFCNDTTELAKIMETLCIKLYKDGY
IHCFLQLTPNLEKLFLKRIVERFRTAKIVGYGERKRWYSSEIIKKFFMGVKLSPRELEE
LKEMKNDEGTLLIEEEEEEETKLKPIDQYMSLNAKDGNYCRRLDFFCCKWDELGAGRIH
YTGSKEYNRWIRILAAQKGFKLTQHGLFRNNILLESFNERRIFELLNLKYAEPEHRNIE
WEKKTA"

CDS complement(146527..147480)
/codon_start=1
/locus_tag="YCR015C"
/label=CTO1
/translation="MKTIIISDFDETITRVDTICTIAKLPYLLNPRLKPEWGHFTKTYM
DGYHKYKYNGTRSLPLLSSGVPTIISQSNFNKLFADELKYQNHNRVVELNSVNEITKQQ
IFKSISLDQMKTFARDQNHEDCLLRDGFKTFCSSVVKNFESDFYVLSINWSKEFIHEVI
GDRRLKNSHIFCNDLKKVSDKCSQSYNGEFDCRLLTGSDKVKILGEILDKIDSGCNKEG
NSCSYWYIGDSETDLLSILHPSTNGVLLINPQENPSKFIKITEKIIGIPKDKISSFEAD
NGPAWLQFCEKEGGKGAYLVKSWDSLKDLIMQVTKM"

CDS 148944..149816
/codon_start=1
/locus_tag="YCR016W"
/label=YCR016W
/translation="MSENHVPAWKRIALKRQTISSGDESKEKGQSNLIDDDPLNITTHL
STGNLTKKEKKRIINGESKSSTKKGKRVSKPGTKKKEKLSKDEKNSKKNKILKDQLRYL
IEFFRTKSESKFPTGILELESVKENYGDSLIKDEPSESGVVEVWKFSKQKQNWLIKHFF
NLDEIPSVYNDLLLLYFRDLQGKSKEELISKCKGKLKQWNDYVEDQETKIKALIAEDKA
SEPINGEEKEEGEKDGNAEQGKQKEVQDEQEEVQMPNKELVQRSLKLLEIWKNDDSEQI
ELKNFFVDV"
CDS complement(150083..152944)
/codon_start=1
/locus_tag="YCR017C"
/label=CWH43
/translation="MLIINGKIIPIAHTICAFSAFFAALVTGYSLHFHKIVTNAHYTYP
DEWFPSVSATIGDRYPERSIFQILIALTAFPRFLLLLGHYYLNQSKVCFLVGVLRTVSC
GGWVYITSTDDHDIHDIFMITYIVLTLPWDIMITRYSSPLTSKNKGLTATIFFGTLFPM
IYWYIQHSVQQRAGAYSIYAYFEWSLILLDIAFDAFAYADFKKIDIVLAFNEKPGNTSF
FQIRDSSPINYGEEKSSELQKSGEKKVEKEKPVARSATGSYFRFDSFFYLLTNIFNGFL
FWSNVTSLLCSIWHFPLWYMGISGYEAAILGYLGPIFLYLPFVSEAFTQYGVLLGGIIA
IGAYIVQMPELRLISVAVGTSITVATFVQNLRYITNAETSFSFALTWLLGLVASVILKM
GFYTNNPTWVILDERNGGYNKTALVLTVLFGMLSPYVNSINFEGKRNAQAKSASLIGKL
FLAVGFGSLLFGIHQLLTDSSTTIYWAWEGYNESHGPLPWPWGALTCTVMLFASLSSVK
FMGKPLVPCLLLLISTAVLSARSITQWPKYIFGGLLYAIAMLWLVPSYFSALGQVQNIW
VYVLSFSVYIIFVLAHVWVVAYAFVPMGWVLREKIETVLAFSSTFIIIGALTCKNLNIQ
LVTMGKKFFIYVFFFAVALLSLTARFVYDIRPTGIPQPYHPDSQLITAGIWTIHFGLDN
DMWASEDRMINLIKDMELDVVGLLETDTQRITMGNRDLTSKLAHDLNMYADFGPGPNKH
TWGCVLLSKFPIVNSTHHLLPSPVGELAPAIHATLQTYNDTLVDVFVFHSGQEEDEEDR
RLQSNYMAKLMGNTTRPAILLSYLVVDPGEGNYNTYVSETSGMHDIDPSDDDRWCEYIL
YRGLRRTGYARVARGTITDTELQVGKFQVLSEQALVEHSDSMYEYGHMSEPEYEDMKFP
DKFLGEGERGHFYHVFDEPRYYL"
CDS complement(153548..153928)
/codon_start=1
/label=SRD1
/translation="NKITPSKNRVTSACNSERRTTSQEANNLEGYHSCAQGTSRSASIT
KKYSKKTTSRPKREKRQTILPNGEIKECSKCKDTWTIQWRSGPDQNRELCSPCGLAYGK
RLKKENEKKRQAADKRIDRNNP"
misc_feature 153837..154833
/label=W-IIIR-WTy1-2

gap 153990..154028
/estimated_length=39
misc_feature 154835..156887
/label=W-IIIR-WTy1-3
misc_feature 154835..156887
/label=W-IIIR-WTy1-3
misc_feature 162984..192029
/label=lost region
LTR 163256..163774
/label=Ty1 LTR
CDS 164760..165851
/codon_start=1
/locus_tag="YCR019W"
/label=MAK32
/translation="MMNEEDSTETKSLVITNGMFIIDDIERSKYNIHYKNVPGGGGTFA
ILGACIISSGNVTSKGLKWIVDRGSDFPKEVIREIDSWGTDVRFRDDFSRLTTKGLNYY
EGSDDLRKFKFLTPKKQINVDDWISTFGQKIIDEMHAFHLLCSGSRCLDIINDLLRVKS
SKGTKPIVIWEPFPDLCDFDHQNDIKSVMQRNDVTVILSPNAEESSRLFGLSSKEPTSL
EECLALAHRFDDFMDENNMCILRCGALGSISVSEKFKNGRTYDHFPAYHFKTQSKVLDP
TGGGNSFLGGFAVSYALTKSLDIASICGNIAAGAIIEQFGIPRYDPIAKTWNGITFLDR
LKFYLSQSGLQYNINDLYKSLTR"
CDS complement(165935..166582)
/codon_start=1
/locus_tag="YCR020C"
/label=PET18
/translation="MSCTTDKLIQKYDALVRKTTEHKFAKELCAGTLKDRSLYIYLSQD
LQFFETSLRLICKTTSLAPTTHALITLAKKIGFFSNDENSYFHDCLELLAPSLTKEERD
NFDNKAIPGVDAYINFLDELRKDASITWPSLVTSLWVAEELYWRWARDTPRAPGLHWKY
QKWIDLHDGEHFQTWCEFLKAEVDKFPVEEVESIFVKVSQFEFEFFESCYNA"
CDS complement(166753..167019)
/codon_start=1
/locus_tag="YCR020C-A"
/label=MAK31
/translation="MDILKLSDFIGNTLIVSLTEDRILVGSLVAVDAQMNLLLDHVEER
MGSSSRMMGLVSVPRRSVKTIMIDKPVLQELTANKVELMANIV"
CDS 167243..167479
/codon_start=1
/locus_tag="YCR020W-B"
/label=HTL1

/translation="MSQNNTISSMNPERAYNNVTLKNLTAFQLLSQRENICELLNLVES
TERHNSIINPERQRMSLEEMKKMLDALKNERKK"
CDS complement(168032..169030)
/codon_start=1
/locus_tag="YCR021C"
/label=HSP30
/translation="MNDTLSSFLNRNEALGLNPPHGLDMHITKRGSDWLWAVFAVFGFI
LLCYVVMFFIAENKGSRLTRYALAPAFLITFFEFFAFFTYASDLGWTGVQAEFNHVKVS
KSITGEVPGIRQIFYSKYIAWFLSWPCLLFLIELAASTTGENDDISALDMVHSLLIQIV
GTLFWVVSLLVGSLIKSTYKWGYYTIGAVAMLVTQGVICQRQFFNLKTRGFNALMLCTC
MVIVWLYFICWGLSDGGNRIQPDGEAIFYGVLDLCVFAIYPCYLLIAVSRDGKLPRLSL
TGGFSHHHATDDVEDAAPETKEAVPESPRASGETAIHEPEPEAEQAVEDTA"
CDS complement(169344..169688)
/codon_start=1
/locus_tag="YCR022C"
/label=YCR022C
/translation="MENRCSNPCYIQSDPSDLLSLRLPARFCWPFSRSSRIFQGSSRRR
ILSEGKNASKACFVRIFGMAKVSKAISRSRSRLLFRSLPPHPACLFEYQISSSSLFHHP
FLLWQGQVFF"
CDS complement(170461..172296)
/codon_start=1
/locus_tag="YCR023C"
/label=YCR023C
/translation="MARQKLTFKEQMDGFPWVQLVVVSLVRFSEPIAFSSLFPYVYFMV
RDFNIAPNDAQVSKYSGYLSSSFALCQVISAYHWGRFSEKHGRKITLTCGLIGTSVSLL
ILGFSRNFYQALVARSLMGLLNGNVGVIRTIIGEIATERKHQALAFSTMPLLFQFGAVV
GPMIGGFLVFRDGTMNEVPLWFPHFAKRIIRSYPYALPNVVVCMFLMFGLTNATLFLEE
THPAFKDRRDYGLEVGDFIKKNIFGIQPKRRPWQKRIQDDSENIHHRNENVNSIRGQDS
EEDENSPLVNTTNDDDTESIQSIDPILTRRQSVGLIRTYSLHEPTDAVHANIDTAPDGC
KESSIFHHVFHTKVFYPISVNFIMALHLIVYNEFLPVFLAYDLAVDPENPKKLASKFPW
KISGGIGYEPEQTGTLLSTTGIFGCFVVIFIFPIVDRNFDCLTIFRTLVKLYPIMYVMV
PYVVFLQNERIPSWYTVVYLYIITGIKTFCGALTSPQIMLLIHNSSPLSCRSVINGATI
SISASARFIGPLVWGYIMSWSQQNDVAWVSWWSLSLFCMVALYQSYKIAPIDDNENELH
GQGSEDAYNSQSQSSDLRMAHRSSLSSLSNQRCTT"
CDS complement(172667..174145)
/codon_start=1
/locus_tag="YCR024C"
/label=SLM5
/translation="MFHAFTFLKGGRFYSSLTVKSLYEQVHHTSHDPISINGWIKSIRL

LKRIAFLDLQDGTSVNPLRIVIPLTNTDEVQFLKILKTGQTLSISNATWQSTPNRKQPF
ELQIKNPVKSIKLVGPVSENYPLQKKYQTLRYLRSLPTLKYRTAYLSAILRLRSFVEFQ
FMLYFQKNHFTKVSPPILTSNDCEGAGELFQVSTNTSPTASSYFGKPTYLTVSTQLHLE
ILALSLSRCWTLSPCFRAEKSDTPRHLSEFWMLEVEMCFVNSVNELTSFVETTIKHIIK
ACIDNQQELLPKQFISSQENNASSELSINQETQQIKTRWEDLINEKWHNITYTNAIEIL
KKRHNEVSHFKYEPKWGQPLQTEHEKFLAGEYFKSPVFVTDYPRLCKPFYMKQNSTPDD
TVGCFDLLVPGMGEIIGGSLREDDYDKLCREMKARGMNRSGELDWYVSLRKEGSAPHGG
FGLGFERFISYLYGNHNIKDAIPFYRTSAESIDF"

misc_feature    complement(174154..174424)
/label=W-IIIR-CTy1-4

misc_feature    complement(174154..174424)
/label=W-IIIR-CTy1-4

CDS    complement(174522..174786)
/codon_start=1
/locus_tag="YCR024C-B"
/label=YCR024C-B
/translation="MCVCAIPFFEFFLPFIPHYAFLLFVSSVRFTVNERCYYLVCVLKL
NCAFFYGYDI*IKTSVCFVFRQIP*NPNCSLFSFYHYYFFPLL"

CDS    complement(174866..174988)
/codon_start=1
/locus_tag="YCR024C-A"
/label=PMP1
/translation="MTLPGGVILVFILVGLACIAIIATIIYRKWQARQRGLQRF"

CDS    complement(175369..175779)
/codon_start=1
/locus_tag="YCR025C"
/label=YCR025C
/translation="MSSFGHRAFFSCCPLLEGETLHSIKLRKEATYQTQSGWPVRWPIR
YPRALPLESHVHPYHMIVKLPHPRTQPRLFLWLSLHWTGIPFLGKPGIGRNRVKSKQFS
HFQQSKIKTTMRKKTTNRKQKAERCQYRLVTA"

CDS    complement(176034..178262)
/codon_start=1
/locus_tag="YCR026C"
/label=NPP1
/translation="MELQNDLESLDNELNDFSEDPFRDDFITDEDAVRSGWRSAWTRMK
YWFYKNRLKWTNNPIVIGDAKDSRDGSNFRRGIPLYELDANGQPIDTELVDENELSFGT
GFHSKVPFKIIFRTLFGSLVFAIFLILMINIAKPHHSTRVLSHFGSPEFDPYVKYFNGT
HEFFPLTIVISLDGFHPSLISKRNTPFLHDLYELKYDGGMNITSTPFMVPSFPTETFPN
HWTLVTGQYPIHHGIVSNVFWDPDLNEEFHPGVLDPRIWNNNDTEPIWQTVQSAFDGDI

PFKAATHMWPGSDVNYTKYNEEKLQPEHKNPIARERTPFYFDEFNAKEPLSQKLSKIIE
YVDMSTLNERPQLILGYVPNVDAFGHKHGYPSESEYYYEDFTETLGEVDTFLKQLVESL
QERNLTSFTNLVIVSDHGMSDIVVPSNVIIWEDLLDEKLRKDYVSHAYLEGPMMAISLK
DSGNINEVYHNLKTSIDEDKYTVYVNGNFPKEWNFNDGKNHHMASIWIVPEPGYAVMKK
EQLKKVAKGDHKDKNEDNVFTIGSHGYDNNAIDMRSVFIGMGPYFPQGYIEPFQNTEIY
NLLCDICGVAEKDRNSNDGTGMLMNQLREPQSSEEVEIEDDFDYLVSKFGEFSTYNIIW
GGYPEETEQDNVDNDNDDNDDGNTDEIAAMPSSSLTIKLEMTTSIPSATETLLGETSPS
SRSSSSSSIQASATASTVGDWLQDIINDAKDLIDDIIDSIDDLVDSDT"

CDS complement(179293..179922)
/codon_start=1
/locus_tag="YCR027C"
/label=RHB1
/translation="MEYATMSSSNSTHNFQRKIALIGARNVGKTTLTVRFVESRFVESY
YPTIENEFTRIIPYKSHDCTLEILDTAGQDEVSLLNIKSLTGVRGIILCYSIINRASFD
LIPILWDKLVDQLGKDNLPVILVGTKADLGRSTKGVKRCVTKAEGEKLASTIGSQDKRN
QAAFIECSAELDYNVEETFMLLLKQMERVEGTLGLDAENNNKCSIM"

LTR 192668..192983
/label=Ty1 LTR

CDS complement(193981..195519)
/codon_start=1
/locus_tag="YCR028C"
/label=FEN2
/translation="MMKESKSITQHEVERESVSSKRAIKKRLLLFKIDLFVLSFVCLQY
WINYVDRVGFTNAYISGMKEDLKMVGNDLTVSNTVFMIGYIVGMVPNNLMLLCVPPRIW
LSFCTFAWGLLTLGMYKVTSFKHICAIRFFQALFESCTFSGTHFVLGSWYKEDELPIRS
AIFTGSGLVGSMFSGFMQTSIFTHLNGRNGLAGWRWLFIIDFCITLPIAIYGFIFFPGL
PDQTSAVSKFSMTRYIFNEQELHYARRRLPARDESTRLDWSTIPRVLKRWHWWMFSLVW
VLGGENLGFASNSTFALWLQNQKYTLAQRNNYPSGIFAVGIVSTLCSAVYMSKIPRARH
WHVSVFISLVMVIVAVLIRADPLNPKVVFSAQYLGGVAYAGQAVFFSWANIICHADLQE
RAIVLASMNMFSGAVNAWWSILFFASDMVPKFERGCYALLATAISSGIVSVVIRSLQIK
ENLSKKQVPYIDANDMPGEDDDDDNQDNENDGDDESMEVELHNEEMAEISNPFR"

CDS complement(join(196045..196210,196294..196535))
/codon_start=1
/locus_tag="YCR028C-A"
/label=RIM1
/translation="MFLRTQARFFHATTKKMDFSKMSIVGRIGSEFTEHTSANNNRYLK
YSIASQPRRDGQTNWYNITVFNEPQINFLTEYVRKGALVYVEADAANYVFERDDGSKGT
TLSLVQKDINLLKNGKKLEDAEGQENAASSE"

CDS complement(196921..199533)

/codon_start=1
/locus_tag="YCR029C-A"
/label=SYP1
/translation="MTEQRTKYADSILTTKSPYEATETIRIRLSQVKLLNKDFYLLFKE
LANLKRNYAQQLRKIIAENEDITKILNAQMIESNVLTPQEMSAFRFNSLGELRNVWDTV
IEELKSDLKSSTEYYNTLDQQVVRELKESVENNTSWRESKDLHSKLSKNAASIEHYSKN
NENSSHLEEARRQWDQQSPYLFELFETIDYNRLDTLKNCMLRFQTSFSDYLLNTTKECE
TVMTKFLAFEPQSEIDRFAKDASQYNFQLSSSSKEVVPNNASPASATGARPVSVSNGAA
NTEREKKSPQKDKRKSAFGNIGHRLASASSSLTHNDLMNNEFSDSTNNSSLKSKKSSHT
LRSKVGSIFGRNKTKNKRQQQSSSNSHIQASITETPNNSSTRVSSTATSSIYQKQRRPT
YSSSKSNNWTPGEASDTPPLPPHATPKNVDAPVTADTPPAQTFTPSEVPPSTPQQSSPP
TAKEPDSSNLPKTVPISISQPPLQPQSKTKPLPVEPASPSISLPTATVDNQPSGQVDSR
PLHIRAPALPPSRKQNFIHNRDSQLYDSLPNHGSGATPTSSSLSSIPQERPVSTLSSQI
TGELRELNPQATGSSTSLVGQSLFQHSSLDTSQFGLNASIAEVLNASFKDGMLQNSQLI
GEIALNYLPNSVMNSPLPIGINLRINNGAKFEKVILNQAFIERVAPEEFKVNPSFIDSR
TLGAIKYSIKEPIAPIVIHPVWRFESHQASVVLTVKMSPSLPDEISQIVIEDLVVFVNI
DGANATSALSKPQGSFSKEKKRITWRFKEPVVLTRNGEGQRLIARFITDGLAHESAKGV
ITKFTISETDNVALPHSGAGSGITLTCQELDENNPFGGEWLDVNTKRTLTTGNYHGLA"
CDS complement(join(200595..201001,201309..201315))
/codon_start=1
/locus_tag="YCR031C"
/label=RPS14A
/translation="MSNVVQARDNSQVFGVARIYASFNDTFVHVTDLSGKETIARVTGG
MKVKADRDESSPYAAMLAAQDVAAKCKEVGITAVHVKIRATGGTRTKTPGPGGQAALRA
LARSGLRIGRIEDVTPVPSDSTRKKGGRRGRRL"
CDS 202615..209118
/codon_start=1
/locus_tag="YCR032W"
/label=BPH1
/translation="MNSIINAASKVLRLQDDVKKATIILGDILILQPINHEVEPDVENL
VQHELTKIIQGYPIQDNMIINSKKGTVEDDLCELNNYTCFALSKSFDLCHDSRNFNIAQ
PKRWIQLLETLTDSVSFAVIVQIILTLSNISLINKQTLGKLKKLRIRIFEILSNKNDSW
KSTLLQKNLIEWYIFMLSVDCTPLELQNLYLHKELKFCNGILNSLTLQVSDPRSQNYLQ
FENTYKLFQIQKSSRINNSFLFYIEFNSVTSNRIMTIERHIYLEIKEGQFCISNDNYII
GLFENFEFEAGTLYFIGVLIDHNNRITLYVDGSMINQLTLFENSICQLSTCELGSMICS
IKVYRFYLWDGLLTEFAINILQAIGTNYQYTFSKKKEGPEVLSLCQDFLIAKAHLMARP
TTEISSTKYIDEIELLEMENIIIDVNPNDILQDFTESSNFTVKFEESTNSKNIPEVGKC
YFYRSSNLVSKFVSIDSIRLAFLNMTESGSIDDLFHHVSHLMNLLRNIDILNWFKKDFG
FPLFAYTLKQKITQDLSQPLNIQFFNLFLEFCGWDFNDISKSIILDTDAYENIVLNLDL

WYMNEDQSSLASGGLEIIRFLFFQISSLMEASIYSKFNSNKFNDMNILEKLCLSYQAVT
KRENQNSKFNELSSDLISVFVTLLKSNTDKRHLQWFLHLSYYFIKRKDVRSTEIILQAV
DQLFSFYLDQGSDENAKILSEIIPLKLILMIMDQIVENNESNPITCLNILFKVVLTNKP
LFKQFYKNDGLKLILTMLCKVGKSYREEIISLLLTYSIGNYTTANEIFSGAEDMIGGIS
NDKITAKEIIYLAVNFIEWHVINSNASDSSSVLDLNNHILRFVEDLKSLSAVPINESVF
DPKKSYVMVSLLDLSIALNESEDISKFKSSSKVISELIKGNIMCALTKYAAYDFEVYMS
TFFCHSTEYKLVYPKTVMNNSSYLELSFIVTLLPEILNDLVDSNNNLNLMMLKHPYTMS
NLLYFLRKFRPDTSQIVMPKDFYFSSYTCLLHCVIQIDKSSFYHFKNVSKSQLLQDFKI
CIMNLIYSNTLKQIIWEKEEYEMFSESLMAHQEVLFAHGACDNETVGLLLIFFANRLRD
CGYNKAVFNCMKVIIKNKERKLKEVACFFDPANKSEVLEGLSNILSCNNSETMNLITEQ
YPFFFNNTQQVRFINIVTNILFKNNNFSPISVRQIKNQVYEWKNARSEYVTQNNKKCLI
LFRKDNTSLDFKIKKSISRYTYNLKTDREENAVFYRNNLNLLIFHLKHTLEIQSNPNSS
CKWSSDFAEDFDGMKRRLLPAWEPKYEPLINEEDANQDTITGGNRQRRESGSILSYEFI
EHMETLESEPVGDLNENRKILRLLKDNDSIATIWNCSLIIGLEIKEGILIHGSNYLYFV
SDYYFSLEDKKILKLSEVSQESRDMTVSLINGPDVKRVSTFLKHEVFVWKLLDITFVTK
RPFLLRDVAIELLFKERVSAFFSFYNKRVRDDVLRVLNKIPKHLPADPIFSSVLQEIND
RGNSIVARNGIGKASIASKFTSVFSANNSLIDGFEISKKWVRGEISNFYYLLSINILAG
RSFNDLTQYPVFPWVIADYESNVLDLENPKTYRDLSKPMGAQSEKRKLQFIERYEALAS
LENADSAPFHYGTHYSSAMIVSSYLIRLKPFVESFLLLQGGSFGPADRLFSSLERAWSS
ASSENTTDVRELTPEFFFLPEFLINVNSYDFGTDQSGKKVDDVVLPPWANGDPKVFIQK
NREALESPYVSAHLHEWIDLIFGYKQKGDIAVKSVNVFNRLSYPGAVNLDNIDDENERR
AITGIIHNFGQTPLQIFQEPHPEKIACNVQQLTTEVWRKVPMKPIFEKTIFNLNEKNRS
VDYVIHDPSYFDSLYWRGFAFPNLFFRTEESLVSLRIVHKNWLKIGLDIFKKTHMAQIT
SFAYWKLGEFITGDKNGLIKVWKYRKDKHSVSGNLENKKTMFGHLCELKEMRCYHDYNT
LLTLDISGLVYVWDMINFELVRQITNDAQKVAISQHAGSIMVLTKNNAISIFNLNGQIY
TSKKFEPAKIVSSIDFFDFTKLDAGYRKHIYWKEMEILLVGFEDGTIEIYELFLNFHNE
WAIKLLKQLCTEKGKAITSIKGQGKTYLSQKRRKDTAEPHEIEVIAGTLDGRLAIWY"

CDS 209584..213264
/codon_start=1
/locus_tag="YCR033W"
/label=SNT1
/translation="MGYPPPTRRLGDKKRYHYSNNPNRRHPSAVYSKNSFPKSSNNGFV
SSPTADNSTNPSVTPSTASVPLPTAAPGSTFGIEAPRPSRYDPSSVSRPSSSSYSSTRK
IGSRYNPDVERSSSTTSSTPESMNTSTITHTNTDIGNSRYSRKTMSRYNPQSTSSTNVT
HFPSALSNAPPFYVANGSSRRPRSMDDYSPDVTNKLETNNVSSVNNNSPHSYYSRSNKW
RSIGTPSRPPFDNHVGNMTTTSNTNSIHQREPFWKANSTTILKSTHSQSSPSLHTKKFH
DANKLDKPEASVKVETPSKDETKAISYHDNNFPPRKSVSKPNAPLEPDNIKVGEEDALG
KKEVHKSGREIAKEHPTPVKMKEHDELEARAKKVSKINIDGKQDEIWTTAKTVASAVEV
SKESQKELTRSVERKESPEIRDYERAYDPKALKTDVTKLTVDNDNKSYEEPLEKVEGCI

FPLPKAETRLWELKNQKRNKIISEQKYLLKKAIRNFSEYPFYAQNKLIHQQATGLILTK
IISKIKKEEHLKKINLKHDYFDLQKKYEKECEILTKLSENLRKEEIENKRKEHELMEQK
RREEGIETEKEKSLRHPSSSSSSRRRNRADFVDDAEMENVLLQIDPNYKHYQAAATIPP
LILDPIRKHSYKFCDVNNLVTDKKLWASRILKDASDNFTDHEHSLFLEGYLIHPKKFGK
ISHYMGGLRSPEECVLHYYRTKKTVNYKQLLIDKNKKRKMSAAAKRRKRKERSNDEEVE
VDESKEESTNTIEKEEKSENNAEENVQPVLVQGSEVKGDPLGTPEKVENMIEQRGEEFA
GELENAERVNDLKRAHDEVGEESNKSSVIETNNGVQIMDPKGAVQNGYYPEETKELDFS
LENALQRKKHKSAPEHKTSYWSVRESQLFPELLKEFGSQWSLISEKLGTKSTTMVRNYY
QRNAARNGWKLLVDETDLKRDGTSSESVQQSQILIQPERPNINAYSNIPPQQRPALGYF
VGQPTHGHNTSISSIDGSIRPFGPDFHRDTFSKISAPLTTLPPPRLPSIQFPRSEMAEP
TVTDLRNRPLDHIDTLADAASSVTNNQNFSNERNAIDIGRKSTTISNLLNNSDRSMKSS
FQSASRHEAQLEDTPSMNNIVVQEIKPNITTPRSSSISALLNPVNGNGQSNPDGRPLLP
FQHAISQGTPTFPLPAPRTSPISRAPPKFNFSNDPLAALAAVASAPDAMSSFLSKKENN
N"

CDS 213687..214730
/codon_start=1
/locus_tag="YCR034W"
/label=ELO2
/translation="MNSLVTQYAAPLFERYPQLHDYLPTLERPFFNISLWEHFDDVVTR
VTNGRFVPSEFQFIAGELPLSTLPPVLYAITAYYVIIFGGRFLLSKSKPFKLNGLFQLH
NLVLTSLSLTLLLLMVEQLVPIIVQHGLYFAICNIGAWTQPLVTLYYMNYIVKFIEFID
TFFLVLKHKKLTFLHTYHHGATALLCYTQLMGTTSISWVPISLNLGVHVVMYWYYFLAA
RGIRVWWKEWVTRFQIIQFVLDIGFIYFAVYQKAVHLYFPILPHCGDCVGSTTATFAGC
AIISSYLVLFISFYINVYKRKGTKTSRVVKRAHGGVAAKVNEYVNVDLKNVPTPSPSPK
PQHRRKR"

CDS complement(214929..216113)
/codon_start=1
/locus_tag="YCR035C"
/label=RRP43
/translation="MAESTTLETIEIHPITFPPEVLARISPELSLQRHLSLGIRPCLRK
YEEFRDVAIENNTLSRYADAGNIDTKNNILGSNVLKSGKTIVITSITGGIIEETSAAIK
DLDDFGEEELFEVTKEEDIIANYASVYPVVEVERGRVGACTDEEMTISQKLHDSILHSR
ILPKKALKVKAGVRSANEDGTFSVLYPDELEDDTLNETNLKMKRKWSYVLYAKIVVLSR
TGPVFDLCWNSLMYALQSVKLPRAFIDERASDLRMTIRTRGRSATIRETYEIICDQTKS
VPLMINAKNIAFASNYGIVELDPECQLQNSDNSEEEEVDIDMDKLNTVLIADLDTEAEE
TSIHSTISILAAPSGNYKQLTLVGGGAKITPEMIKRSLLLSRVRADDLSTRFNI"

CDS 216392..217393
/codon_start=1
/locus_tag="YCR036W"

/label=RBK1
/translation="MGITVIGSLNYDLDTFTDRLPNAGETFRANHFETHAGGKGLNQAA
AIGKLKNPSSRYSVRMIGNVGNDTFGKQLKDTLSDCGVDITHVGTYEGINTGTATILIE
EKAGGQNRILIVEGANSKTIYDPKQLCEIFPEGKEEEEYVVFQHEIPDPLSIIKWIHAN
RPNFQIVYNPSPFKAMPKKDWELVDLLVVNEIEGLQIVESVFDNELVEEIREKIKDDFL
GEYRKICELLYEKLMNRKKRGIVVMTLGSRGVLFCSHESPEVQFLPAIQNVSVVDTTGA
GDTFLGGLVTQLYQGETLSTAIKFSTLASSLTIQRKGAAESMPLYKDVQKDA"
CDS complement(217509..220280)
/codon_start=1
/locus_tag="YCR037C"
/label=PHO87
/translation="MRFSHFLKYNAVPEWQNHYLDYNELKNLIYTLQTDELKQETPTGD
LNDDADSQTPGPIADIESNIAAGEPSSSKRRFTHKLKRKLFGSKTPSGSKRGDSDEKAI
DGNNINEETIELDELSPQGKTTSFNKNFIRKKFFESRSSSVSSEGKTLFSSYDTFVTNL
SDEKLKVDDFYKRMEAKFYERFDHLINDLEKEGIVTRLNETFNPEIQALPPLREIISGT
SETHSSNNPFEIHSSNIDSELRNRFDYSEEEMDEDDDVDVFADTTDNTALLNYSQFNIK
SQKKSLLKQTIINLYIDLCQLKSFIELNRMGFSKITKKSDKVLHMNTRQELIESEEFFK
DTYIFQHETLSSLNSKIAQLIEFYAVLMGQPGNVDSCKQELKSYLHDHIVWERSNTWKD
MLGLSSQNNDIITIEDEAEKLMQEKLQIEYFKYPLPKPINLKFTKIENLAVPKLFFGKR
AMKIGFIIIVTGVLLGVKTFNDPVEHRCMALVECCAFLWASEAIPLHITGLLVPLLTVL
FRVLKDDDGKVMGAAAASTEILGTMWSSTIMILLAGFTLGEALSQYNVAKVLASWLLAL
AGTKPRNVLLMAMSVVFFLSMWISNVASPVLTYSLLTPLLDPLDYTSPFAKALVMGVAL
SADIGGMASPISSPQNIISMQYLKPYGIGWGQFFAVALPTGILSMLCSWALMILTFKIG
KTKLEKFKPIRTRFTIKQYFIIIVTIATILLWCVESQIESAFGSSGEIAVIPIVLFFGT
GLLSTKDFNTFPWSIVVLAMGGIALGKAVSSSGLLVTIARALQKKIQNDGVFAILCIFG
ILMLVVGTFVSHTVSAIIIIPLVQEVGDKLSDPKAAPILVFGCALLASCGMGLASSGFP
NVTAISMTDKKGNRWLTVGAFISRGVPASLLAFVCVITLGYGISSSVLKGST"
CDS complement(220716..222644)
/codon_start=1
/locus_tag="YCR038C"
/label=BUD5
/translation="MSPKNKYVYICVEYIYIYFAKIHKQSTLSSDTTKMFVLIDNVLAY
LLEQDDLFVTARFAIQGQIVSRRVNKIHISNITDVLLQQFISHTLPYNDNIVPKKILDS
MRTAVRQLLEATACVSRECPLVKRSQDIKRARKRLLSDWYRLGADANMDAVLLVVNSAW
RFLAVWRPFVNSIQHATQELYQNIAHYLLHGNVNIQRVTALLQLVMGQDDLLFSMDDVL
QEVFRIQLYLNKMLPHNSHKWQKPSPFDSANLLLNFRDWTTDNALLQELLLSYPTINKN
KHKNHSVPRLIQIWVESYWQDSETTLKDILNFWYSHLAEYYEYQELFADIVQLFINKKR
TRQLKIHYIGLTDKEIEENKPPLDYENLFLQYEIDKTNANDELCGATDLSDLLFQWKQG
ELLEVEAFALNVSPWSLAKTLTLLESSLYLDIETIEFTRHFKHNDTTIDSVFTLSNQLS

SYVLETTLQQTHTISYWLQVALSCLYLRNLNSLASIITSLQNHSIERLSLPIDVKSDHL
FQRLKVVVHPNNNYNVYRRTIKHIFHSQLPCVPFTSLLIRDITFIRDGNDTFTKDGNNV
NMQKFNQITKIVAFAQYLQQKQYEDIHCSNTTARSLLGAMIKVHTLYNDNKDRAYQVSI
AKVPRLT"

CDS complement(222641..223000)
/codon_start=1
/label=putative MATa2
/translation="MRSIENDRSNYQLTQKNKSADGLVFNVVTQDMINKSTKPYRGHRF
TKENVRILESWFAKNIENPYLDTKGLENLMKNTSLSRIQIKNWVSNRRRKEKTITIAPE
LADLLSGEPLAKKKE"

CDS complement(222988..223068)
/codon_start=1
/label=putative MATa2
/translation="MRRRSCCKQHLNSLLPLLYYSKKCAA"

CDS join(223297..223401,223456..223695,223754..223783)
/codon_start=1
/locus_tag="YCR097W"
/label=MATa1
/translation="MDGICSMAENINRTLFNILGTEIDEINLNTNNLYNFIMESNLTKV
EQHTLHKNISNNRLEIYHHIKKEKSPKGKSSISPQARAFLEQVFRRKQSLNSKEKEEVA
KKCGITPLQVRFINKRMRSK"

CDS 223901..224233
/codon_start=1
/locus_tag="YCR041W"
/label=YCR041W
/translation="MLKYVVTDIGKMCLYIWPYRVWSWRRLFIFRVLNVVSIAILFETP
HRLALVPNVCLYTHMAIPLSTCLFCLCLCICIKYDITQTQANNQFLASFFVLILTINDL
DVTFVI"

CDS complement(224164..228387)
/codon_start=1
/locus_tag="YCR042C"
/label=TAF2
/translation="MMSFSKNATPRAIVSESSTLHEMKFRNFRVAHEKISLDIDLATHC
ITGSATIIIIPLIQNLEYVTFDCKEMTIKDVLVENRRCDQFIHDDPLQTNLNGLTSQNV
LYSDNSIEQSHFLRSKFASLNEYPETDSKSQLTIKIPSSIKISLEDANALSNYTPITPS
IKTTPGFQESVFTPITLQIEYEIRNPKSGIKFDTVYADKPWLWNVYTSNGEICSSASYW
VPCVDLLDEKSTWELEFSVPRLVKNIGTSKLIGQNGEESEKEKEDTPEHDEEEEGKPAR
VIKDEDKDSNLKNDEEGKNSKSKDAQDNDEEEEEGESDEEEEEGEEERRNIEESNNPSL
RDVIVCCSEYSNIKELPHPIDLTKKKCIFQIINPVAPHHIGWAIGAFNSWSLPLISPPS

VDAEDEVEEDKLRENVVDNVNDTMDDDIGSDIIPIQIFTLPTQETDELTVINSTVVCQK
IIDFYSKEFGSYPFTCYSMVFLPTAPSKHMDFAALGICNTRLLYPLEVIDKAFSTTNEL
AWALANQWSCVNITPLDMNDYWCCLGIAGYMVFQVTKKLMGNNTYKYQLKRNSEAIVEQ
DFEKPPIGSTFTGSSRPISWSSKDLSFIQLKAPMILHILDRRMTKTERSFGMSRVLPKI
FLQAMSGDLPNNSLTSSHFQHVCERVNKSKLENFFNEWVYGSGVPILRVTQRFNRKRMV
IELGIRQVQDEELGHEKVVGEEGFFKSALDHLEHPDLNRTECFTGSMTIRIHEHDGTPY
EHIVEIKDTFTKIDIQYNTKYRRLRKRGGGANDENGVENNNEEKPIVVDVNCLGNVYMS
PEECSRFSLTEFNRTSESNELLKQNEAFEWIRIDSDLEWICQMHINQPDYMFSSQLRQD
GDIEAQLEAIRYYEDVVVNGGVKSLVYSSILFRTAIDERYFFGIRLAACEALSKYVYDP
DFTGGVKHLIQIFQILFCLEDSNIPKSNNFENPKLYFLQCNIPKYLAKVKNENGKCPKL
VKQFLLDILVYNENGENKYSDDAYVRSLIENVVKVALNEYKDKAYMEKVKTQLLRYENL
VNWLSSYESLIKTTIMYAKYKLHKVGAYDFTELTGMIMHTLTLGINNGDISRESFQNEF
LMVLKIMLLEGGLKNKDALVLFTEILCFHEDSYIRDKSVDVLSECVNLVVMDGSLDTIS
DDIKSSVQSVHNEVKNIKSEDDIELFLSGHYVDDMKIKIEKIGRQNISGLIQICRDMFK
GYSPLKILLWDVLNLPVLSLYQRKQIHDLVRVMYTLINSFVVRLETPRERRLVAKMNSN
EEGKLDIVIKRESILKVHIKKEVTSTVEAPKKANKIKISLKGDKPVRKVEKQIVKPKVT
SKQRKVKSHVNRMGSLPLRFVKIQQQPRVMVHLSSVPYSQFVQITKVTSRSFMVKIRTK
NDAKN"
CDS complement(229250..229633)
/codon_start=1
/locus_tag="YCR043C"
/label=YCR043C
/translation="MIPAPLDASLLREHAFQGTNDLSTVLSPSTFTDEGGYKPVLKYGL
GYFNYGLVIDDEVYDYSVCDIIRGHVYDHFWCYFCCFMILFTIWLISLNWCPSSKKSKF
DWSKKKDDFKMEGGDLEYQHVKI"
CDS complement(229867..230940)
/codon_start=1
/locus_tag="YCR044C"
/label=PER1
/translation="MRLAVVVTLLVHCFLVTCSPGDNLDEFIDCTYACEYNRRCPNSQI
NYIDPETNMFHDIEFFDTPPLYSKLLFWDCISDCDYQCQHIITRWRIDEEEEIYQFHGK
WPFLRVLGTQEFFSTIFSIGNFIPHYKGFVKFSRIIREEGDRRRKNSRSILIWNYLYVT
VAGMLAWTASSVFHCRDLIITEKLDYFFAGLTVLTGFHAIFARMTSMFLYPKIAQAFTA
SVAAIFALHILRLYVDWSYTYNMRFNIFFGVLQYILLIMLSCQNYHALQKQKLMGEFKK
TAYSSFKRQIFKLCVIPILLVIVTTMAMSLELFDFFSYEWQIDAHALWHLCTIWPSWVL
YDFFLEDYAYWGNRQLY"
CDS complement(231125..232600)
/codon_start=1
/locus_tag="YCR045C"

/label=RRT12
/translation="MKPQCILISLLVNLAYAEEYLVRFKNPTAFQQFTSNSNRSWRQFI
DNKIEKKFSIGSFRGVTMNLSKNLVNKLKKSPLVADIVPNFRFEAFEGDSVNSAESSYT
FNATAKYSYEDVEEEQNITYQPDAPRHLARISRHYQLPFDVGDKDRYKSWFNYYYEHDY
QGQDVNAYIMDTGIFADHPEFEDRVIQGIDLTKEGFGDQNGHGTHVAGLVGSKTYGAAK
RVNLVEVKVLGKDGSGEASNVLSGLEFIVEHCTKVSRPQGKKCVANLSLGSFRSPIINM
AVEGAIEEGIVFVAAAGNFNLDAYWASPASAENVITVGAFDDHIDTIAKFSNWGPCVNI
FAPGVEIESLSHLNYNDTLILSGTSMSTPIVTGVAAILLSKGIEPEMIAQEIEYLSTRN
VFHRRTLFFKPSTPNQILYNGVDKLDDPYDDETFPRLNIEAIAKELEEYNATLQTPMSE
NLQSGSKLWGWNNDVTLPLGEIRLKRRDFMKNL"
CDS complement(232904..233413)
/codon_start=1
/locus_tag="YCR046C"
/label=IMG1
/translation="MWSRNVRLLGSWTRSYMVPATKRKTIPVYPPVQRIASSQIMKQVA
LSEIESLDPGAVKRKLISKKNKDRLKAGDVVRIVYDSSKCSYDTFVGYILSIDRKQLVQ
DASLLLRNQIAKTAVEIRVPLFSPLIERIDLLTPHVSSRQRNKHYYIRGTRLDVGDLEA
GLRRKK"
CDS complement(233708..234535)
/codon_start=1
/locus_tag="YCR047C"
/label=BUD23
/translation="MSRPEELAPPEIFYNDSEAHKYTGSTRVQHIQAKMTLRALELLNL
QPCSFILDIGCGSGLSGEILTQEGDHVWCGLDISPSMLATGLSRELEGDLMLQDMGTGI
PFRAGSFDAAISISAIQWLCNADTSYNDPKQRLMRFFNTLYAALKKGGKFVAQFYPKND
DQVDDILQSAKVAGFSGGLVVDDPESKKNKKYYLVLSSGAPPQGEEQVNLDGVTMDEEN
VNLKKQLRQRLKGGKDKESAKSFILRKKELMKRRGRKVAKDSKFTGRKRRHRF"
CDS 234919..236751
/codon_start=1
/locus_tag="YCR048W"
/label=ARE1
/translation="MTETKDLLQDEEFLKIRRLNSAEANKRHSVTYDNVILPQESMEVS
PRSSTTSLVEPVESTEGVESTEAERVAGKQEQEEEYPVDAHMQKYLSHLKSKSRSRFHR
KDASKYVSFFGDVSFDPRPTLLDSAINVPFQTTFKGPVLEKQLKNLQLTKTKTKATVKT
TVKTTEKTDKADAPPGEKLESNFSGIYVFAWMFLGWIAIRCCTDYYASYGSAWNKLEIV
QYMTTDLFTIAMLDLAMFLCTFFVVFVHWLVKKRIINWKWTGFVAVSIFELAFIPVTFP
IYVYYFDFNWVTRIFLFLHSVVFVMKSHSFAFYNGYLWDIKQELEYSSKQLQKYKESLS
PETREILQKSCDFCLFELNYQTKDNDFPNNISCSNFFMFCLFPVLVYQINYPRTSRIRW
RYVLEKVCAIIGTIFLMMVTAQFFMHPVAMRCIQFHNTPTFGGWIPATQEWFHLLFDMI

PGFTVLYMLTFYMIWDALLNCVAELTRFADRYFYGDWWNCVSFEEFSRIWNVPVHKFLL
RHVYHSSMGALHLSKSQATLFTFFLSAVFHEMAMFAIFRRVRGYLFMFQLSQFVWTALS
NTKFLRARPQLSNVVFSFGVCSGPSIIMTLYLTL"
CDS complement(236454..236762)
/codon_start=1
/locus_tag="YCR050C"
/label=YCR050C
/translation="MVAVHKVRYNVIMILGPEQTPNEKTTLDNCGLARRNLVLLKAVHT
NCDSWNMNRYPLTLLKMANMAISWNTALKKKVNNVAWLLLKCNAPMELWYTCLSKNL"
CDS 237061..237729
/codon_start=1
/locus_tag="YCR051W"
/label=YCR051W
/translation="MNANIWVAASDGNLDRVEHILRESKGAMTPQSKDINGYTPMHAAA
AYGHLDLLKKMCNEYNGDINVLDNDGDTPLHHVEDVATARLIVEELGGDFTIRNVEGQT
PYDSFVENGEDGELIEYMRIKSGVADVHGVDGVQGEGVIDSKLLEEFKDNVRYTLENDP
EEGADEATLQRRRQLEQIITGDNAEEELERYIRAMVREQMLGQGSMAGSGDEPDSKRRK
"
CDS 237984..239435
/codon_start=1
/locus_tag="YCR052W"
/label=RSC6
/translation="MVTQTNPVPVTYPTDAYIPTYLPDDKVSNLADLKKLIEMDSRLDL
YLTRRRLDTSINLPTNTKTKDHPPNKEMLRIYVYNTTESSPRSDSGTPADSGKTTWTLR
IEGKLLHESANGKHPFSEFLEGVAVDFKRLKPLGMGKKRKRDSSLSLPLNLQQPEYNDQ
DSTMGDNDNGEDEDSAEAESREEIVDALEWNYDENNVVEFDGIDIKRQGKDNLRCSITI
QLRGVDGGKVQYSPNLATLIGMQTGSVNDAVYSIYKYILINNLFVTEQTEAQDGSNDAE
DSSNENNNKNGAGDDDGVEGSTPKDKPELGEVKLDSLLQKVLDTNAAHLPLMNVVQTVN
KLVSPLPPIILDYTIDLSKDTTYGATTLDVDVSHILHQPQPQPNLQKEEETDAEDTAKL
REITKLALQLNSSAQKYQFFHELSLHPRETLTHYLWSSKQNELVLQGDQYFNEDAARTS
DIYSNNNNDRSLMGNISLLYSQGRL"
CDS 239687..241231
/codon_start=1
/locus_tag="YCR053W"
/label=THR4
/translation="MPNASQVYRSTRSSSPKTISFEEAIIQGLATDGGLFIPPTIPQVD
QATLFNDWSKLSFQDLAFAIMRLYIAQEEIPDADLKDLIKRSYSTFRSDEVTPLVQNVT
GDKENLHILELFHGPTYAFKDVALQFVGNLFEYFLQRTNANLPEGEKKQITVVGATSGD
TGSAAIYGLRGKKDVSVFILYPTGRISPIQEEQMTTVPDENVQTLSVTGTFDNCQDIVK

AIFGDKEFNSKHNVGAVNSINWARILAQMTYYFYSFFQATNGKDSKKVKFVVPSGNFGD
ILAGYFAKKMGLPIEKLAIATNENDILDRFLKSGLYERSDKVAATLSPAMDILISSNFE
RLLWYLAREYLANGDDLKAGEIVNNWFQELKTNGKFQVDKSIIEGASKDFTSERVSNEE
TSETIKKIYESSVNPKHYILDPHTAVGVCATERLIAKDNDKSIQYISLSTAHPAKFADA
VNNALSGFSNYSFEKDVLPEELKKLSTLKKKLKFIERADVELVKNAIEEELAKMKL"

CDS complement(241366..243057)
/codon_start=1
/locus_tag="YCR054C"
/label=CTR86
/translation="MPMNNFLDEFNLFDSIITMMKNDPCCVEDYEPIVENLNRIFQRTF
NDEEHRKSMANSQLFWERLRDTLEAMLLPASLNENSSIPYTRTVRGLILMMRNLAAENQ
EIPQKLLLQNLVIRGFLHATSEYVVDTPLIKHLYIACLTCLFNIQQNYSTVDMTTFPAL
LQFLQYPYGIKLEDGEEEEHFWLPYLFLFKTYLNNDEFSNEFFRDNDTPQKDYYCVRDR
IFFDIVTAKFIQDQENSFLIEKGRNYLDDSKLEITSIDLSVLECISKSLTTASFGKYLN
GLEERQPGKFTTLLQILQLVVTSKEDWNTYELTAIMSWCYPILQRLACKDIPAFFNKSC
NDYAPSVAIQLHSTLLSCLDIISDLCKFNHVRKFLISYDSVKILVSLLDTFQKNLLRIN
FLKGNGDTVNEIKITDHEGNKIEDRLLIFNRVNTNESFIRADNFPHCKLVIIEILASLV
YAHPEIQDQIRELGGLALILSNCVIDDNDPFIKERSIVCLKFLLKNNAKNQEYVKKMEA
QDVVQDDALSKAGFEISVEKGGKVRLVSKEEDPGNENSEIISIDED"

CDS complement(243447..246218)
/codon_start=1
/locus_tag="YCR057C"
/label=PWP2
/translation="MKSDFKFSNLLGTVYRQGNITFSDDGKQLLSPVGNRVSVFDLINN
KSFTFEYEHRKNIAAIDLNKQGTLLISIDEDGRAILVNFKARNVLHHFNFKEKCSAVKF
SPDGRLFALASGRFLQIWKTPDVNKDRQFAPFVRHRVHAGHFQDITSLTWSQDSRFILT
TSKDLSAKIWSVDSEEKNLAATTFNGHRDYVMGAFFSHDQEKIYTVSKDGAVFVWEFTK
RPSDDDDNESEDDDKQEEVDISKYSWRITKKHFFYANQAKVKCVTFHPATRLLAVGFTS
GEFRLYDLPDFTLIQQLSMGQNPVNTVSVNQTGEWLAFGSSKLGQLLVYEWQSESYILK
QQGHFDSTNSLAYSPDGSRVVTASEDGKIKVWDITSGFCLATFEEHTSSVTAVQFAKRG
QVMFSSSLDGTVRAWDLIRYRNFRTFTGTERIQFNCLAVDPSGEVVCAGSLDNFDIHVW
SVQTGQLLDALSGHEGPVSCLSFSQENSVLASASWDKTIRIWSIFGRSQQVEPIEVYSD
VLALSMRPDGKEVAVSTLKGQISIFNIEDAKQVGNIDCRKDIISGRFNQDRFTAKNSER
SKFFTTIHYSFDGMAIVAGGNNNSICLYDVPNEVLLKRFIVSRNMALNGTLEFLNSKKM
TEAGSLDLIDDAGENSDLEDRIDNSLPGSQRGGDLSTRKMRPEVRVTSVQFSPTANAFA
AASTEGLLIYSTNDTILFDPFDLDVDVTPHSTVEALREKQFLNALVMAFRLNEEYLINK
VYEAIPIKEIPLVASNIPAIYLPRILKFIGDFAIESQHIEFNLIWIKALLSASGGYINE
HKYLFSTAMRSIQRFIVRVAKEVVNTTTDNKYTYRFLVSTDGSMEDGAADDDEVLLKDD
ADEDNEENEENDVVMESDDEEGWIGFNGKDNKLPLSNENDSSDEEENEKELP"

CDS complement(246444..247220)
/codon_start=1
/locus_tag="YCR059C"
/label=YIH1
/translation="MDDDHEQLVEELEAVEAIYPDLLSKKQEDGSIIVVKVPQHEYMTL
QISFPTHYPSEEAPNVIEVGVCTSLAKRDLYDTKYLQHLFQEVMDSVFHRGSVCLFDFL
TELDGVLYVEPEEETEPVQQSDIPTDPFEGWTASDPITDRGSTFMAFAAHVTSEEQAFA
MLDLLKTDSKMRKANHVMSAWRIKQDGSAATYQDSDDDGETAAGSRMLHLITIMDVWNV
IVVVARWFGGAHIGPDRFKHINSTAREAVVRAGFDS"
CDS 247389..247724
/codon_start=1
/locus_tag="YCR060W"
/label=TAH1
/translation="MSQFEKQKEQGNSLFKQGLYREAVHCYDQLITAQPQNPVGYSNKA
MALIKLGEYTQAIQMCQQGLRYTSTAEHVAIRSKLQYRLELAQGAVGSVQIPVVEVDEL
PEGYDRS"
misc_feature 247806..248051
/label=ARS315
CDS 249460..250269
/codon_start=1
/label=Kan R
/translation="MGKEKTHVSRPRLNSNMDADLYGYKWARDNVGQSGATIYRLYGKP
DAPELFLKHGKGSVANDVTDEMVRLNWLTEFMPLPTIKHFIRTPDDAWLLTTAIPGKTA
FQVLEEYPDSGENIVDALAVFLRRLHSIPVCNCPFNSDRVFRLAQAQSRMNNGLVDASD
FDDERNGWPVEQVWKEMHKLLPFSPDSVVTHGDFSLDNLIFDEGKLIGCIDVGRVGIAD
RYQDLAILWNCLGEFSPSLQKRLFQKYGIDNPDMNKLQFHLMLDEFF"
CDS 250644..252539
/codon_start=1
/locus_tag="YCR061W"
/label=TVS1
/translation="MVRFVSILSLFGCAATLVTAHDDMDMDMDMDMDMDMNIDTTTSQS
IDVSSTASIVPVPHEPKHLHGLPILQSPSLTPAERLYWENYNTTTYFTTQAGNRSALRY
HIITLLLVAFVLYPVSLALSAARSRWYLPLLFVNLCICISSVMALSVFKNTFPEEDWYA
HNIYGTTSVLLLVFMLVHFFAAVLSVPVSLASKKEYRPVDTIPLNDLESTPVMVNSARG
SPSPSSNRDTLFSLSSDTTTATATNNNKRRRAEGEDEGDNTSNHDTLRDEDYDNDDDEI
ASIEAPPLLPQDIPVFRILFTNTKYQMLAAHLSCVANVVFHMLTYPLFMYIFVDLIIGF
AVGNLLGKGIRIFNLLAHWIKGGVFFTLGVVSLARYCGFAAKYGWAWNNISFTSQLTQT
RSSNLLFRFAPAGTFTMEFVESFLIFFYGSTNIFLEHLAGNGGAWTAKDLQHVSIAFMF
IGTGLCGLLTEYKLNHWRFEHARKRPQTDVVAATPGYSPNPFPAFTIFWTGILMSQHAQ

SSQFSTTIHTQWGYLLSYGSFFRLLTFLILFLVPNTNSAASKPFTELITSFCLLCGGLV
FMESTDQSIEAMEYRGFTPMFTFNLSVGFVSLLMAWEMILFIWKDWLIKTRKTSL"
CDS 253399..253872
/codon_start=1
/locus_tag="YCR063W"
/label=BUD31
/translation="MPRIKTRRSKPAPDGFEKIKPTLTDFEIQLRDAQKDKSSKLAAKS
NEQLWEIMQLHHQRSRYIYTLYYKRKAISKDLYDWLIKEKYADKLLIAKWRKTGYEKLC
CLRCIQKNETNNGSTCICRVPRAQLEEEARKKGTQVSFHQCVHCGCRGCASTD"
CDS 254391..256085
/codon_start=1
/locus_tag="YCR065W"
/label=HCM1
/translation="MMNEDISIIDGHNSFLTEKSTVLLTQAKRTLEDEKEMITPPSSTV
RKTMKEVNKRPSHPLSPDHSSPIAPSKAKRQRSDTCARSNGNLTLEEILQSLERRRING
ELAKKPPYSYATLICLAILQSQEGKLTLSQIYHWIHVHFPYYKQKDASWQNSIRHNLSL
NDAFIKTEKSCDGKGHFWEVRPGAETKFFKGENRGYEFVKDSLQDIGKYFEIDSTLDEL
EQVESGEGNDDLPDEEEREEAGKFPSIEIQLNSSPILRVSQLHHIPQLKTDNSVLNPHE
NLESMRNMIENDVNNIDSLEPPYVMKKYHTSLGLPSLVNAKDHFQAGVKNNNITQANRF
NTLPITSAKSPQNFRKYFTSFNSNFEDLSPLRSNVGAGSLLDPLPYSPLKLYDQKNLAL
MSKPQSQQSYSNSQLPPPPSSHGSDLLKTPKMRHSDGLEKTPSRLISTPKDGNSILRKW
QTPSHLFEDLYCSPLFRAIETPIRYITTPGGTLETQISPRKSSAPDVLTSATNSKFASS
GLFGVDVYSVWKRATEKISDGNNTTDSNQKHHPYHNHPSNDSGNEKN"
CDS 256581..258044
/codon_start=1
/locus_tag="YCR066W"
/label=RAD18
/translation="MDHQITTASDFTTTSIPSLYQLDTLLRCHICKDFLKVPVLTPCGH
TFCSLCIRTHLNNQPNCPLCLFEFRESLLRSEFLVSEIIQSYTSLRSSLLDALRIPKPT
PVPENEEVPGPENSSWIELISESESDSVNAADDDLQIVATSERKLAKRSMTDILPLSSK
PSKRNFAMFRSERIKKKSKPNEQMAQCPICQQFYPLKALEKTHLDECLTLQSLGKKPKI
STTFPTESNPHNKSSSRFKVRTPEVDKSSCGETSHVDKYLNSMMSAEHQRLPKINFTSM
TQSQIKQKLSSLGLSTNGTRQNMIKRYNHYEMLWNSNFCDSLEPVDEAELKRQLLSWDV
SHNKTPQNSSNKGGISKLMIMKSNGKSSSYRKLLENFKNDKFNRKGWMVMFRKDFARLI
REAKMKIKTGSSDSSGSVGHSNDGDGVEKVQSDQGTEDQQMEKDQDTVINEDRVAGERN
LPNEDSTDADLSRELMDLNEYSKDPPGNN"
CDS complement(258206..261403)
/codon_start=1
/locus_tag="YCR067C"

/label=SED4
/translation="MSGNSANYDVGYPIYGAKFINEGTLLVAGGGGQFNSSFPNKITAL
KVNFQKKKHIRRFREITLDSIDDAPTSLDCNNNLILVGCNELFNDSSMENVNHHLRKFV
FEQEHLKFVASIDFNRTTDPSVFTKFVYINQRATVAAIASSEVPTVIRIIDPRNLTENY
EIETGREVNDLHFAPNGILLSYITSNSLEVASVRDGNFVARKTDFDKNLVLSNIRFLND
NTLLVAASLSNSDGVSLLKLGVSSKGVKILKTASFMFDLNGITSMDVSPNKKFVALSSN
DNLVAIVSVEKLKLVQLVPRVHESTITKVTFSPDSRYLASTSMGNTINVLKLSGTSSSI
LRNIWKFFLNFVLLVVLAGAIQLGYKHNVHGFIYKHAHDIYKSKFKENTTIDQGSSSYF
TINDDYRGITESADIISATDVASDIETEFSSFDTSTMRTTTEDEQKFVWISSSADSQFT
SADIPTSASSSSSSSSSFYEESVTNEPIVSSPTSEITKPLASPTEPNIVEKPSLPLNS
ESIDLLSSSSNSITEYPEPTPDLEEKLSSLIVEQSESEITTDRESVSKLLSTESPSLSH
MPSSSSSSLSLSSSLTTSPTTALSTSTATAVTTTQTNPTNDAANTSFLDNSKPASTREI
YKTKIITEVITKIEYRNIPASDSNAEAEQYVTTSSSMLLTPTDTMVSSPVSEIDPIASE
LERMVETPTHSISIASEFDSVASNLIPNEEILSTSASQDSISSHPSTFSDSSITSGFQS
IEVSTVTSSVLASESIPSISDSTFSKFHSISEPVSSAIVETATSSFSKTETKTSRVIAF
STEDSERSSALIDNSEYTSVLADNLEPTSVLADNSEPTSVLADSSEPTSVFTDAVQSPK
TSVGQSSLSESTNIEGTSMASMIFSSSGASIGALSDIGKGTLSVESASSTVAQPMPGVT
TTAPSFVSSPHKISASSIDASGFVQKEIMIEVQSSKDSSEAFGVRHKISENVNTPVSRM
LTTEMQASGTVDVTEDVSLSSEVISALNVEITSLPNPVAPPQTIAAPLNNNSNTNIVND
DNAVAGTVNYAGLHDEL"
CDS 262295..263857
/codon_start=1
/locus_tag="YCR068W"
/label=ATG15
/translation="MLHKSPSRKRFASPLHLGCILTLTVLCLIAYYFALPDYLSVGKSS
SRGAMDQKSDGTFRLKSIYRHGVGANHRLHQRLEVTPEVISAAGMLYQETTTQGQDFED
QEPLWTTNAEYATTNPFDFEFELRRMPLLMKRMKERDPEFIESYIYGETYMTEEEEHAM
WIDDDIVAPNITDRGTVVSLALMSSNAYVRIPQTGDWRNVTEPWNETEPEDFGWDGDGI
RGHVFYNEVENIVVLSIKGTSAQGLPGSGEDETTGNDKINDNLLFSCCCARVSYLWTTV
CDCYVKSYICDESCLEKELRRKDRFYSAVVDIYKGVLKEYPDAAIWVTGHSLGGALASL
LGRTFGLPAVAFESPGELLPSKRLHLPFPPGLPSYMEGIWHFGHNADPIFMGTCNGASS
SCSLVGYAMETACHTGRVCVYDVVNDKGWSVNMFNHRIHKVIDEVLLGYEQAAKCVEPE
PCVDCYNWKFIPSRDWESSSRLITKTKSHAAPTTTTRTTATTTSSSTCVGRNWLGFCTK
YEL"
CDS 264136..265092
/codon_start=1
/locus_tag="YCR069W"
/label=CPR4
/translation="MWLKSLLLCLYSLVLCQVHAAPSSGKQITSKDVDLQKKYEPSPPA

THRGIITIEYFDPVSKSMKEADLTFELYGTVVPKTVNNFAMLAHGVKAVIEGKDPNDIH
TYSYRKTKINKVYPNKYIQGGVVAPDVGPFTVYGPKFDDENFYLKHDRPERLAMAYFGP
DSNTSEFIITTKADGNEELDGKSVVFGQITSGLDQLMDAIQYTETDEYGKPQHELRFLY
FVLEILKISNILDLHAAYTEKVEKFRNGDVSVGSTLENIFRNDKAYTPLTTSTGTTAYD
LNHPISRALMCLTVLGLCFIAYKGMHEKPHTVSLRHK"

CDS complement(265184..265624)
/codon_start=1
/locus_tag="YCR071C"
/label=IMG2
/translation="MISSCVTRCFGRGKCLPGPATASIYQTIRCISTNSNKAAEAPIFP
KLEDVKMHELIGNNNFGKKTYYVERSRTGNLPVYSAYKNGGNKIITEIRKIEGDVIQLR
NDLQEQLPFIPKKSWSVVMQSKKIIIKGNAVEAVKRVLTKKF"

CDS complement(265886..267433)
/codon_start=1
/locus_tag="YCR072C"
/label=RSA4
/translation="MSTLIPPPSKKQKKEAQLPREVAIIPKDLPNVSIKFQALDTGDNV
GGALRVPGAISEKQLEELLNQLNGTSDDPVPYTFSCTIQGKKASDPVKTIDITDNLYSS
LIKPGYNSTEDQITLLYTPRAVFKVKPVTRSSSAIAGHGSTILCSAFAPHTSSRMVTGA
GDNTARIWDCDTQTPMHTLKGHYNWVLCVSWSPDGEVIATGSMDNTIRLWDPKSGQCLG
DALRGHSKWITSLSWEPIHLVKPGSKPRLASSSKDGTIKIWDTVSRVCQYTMSGHTNSV
SCVKWGGQGLLYSGSHDRTVRVWDINSQGRCINILKSHAHWVNHLSLSTDYALRIGAFD
HTGKKPSTPEEAQKKALENYEKICKKNGNSEEMMVTASDDYTMFLWNPLKSTKPIARMT
GHQKLVNHVAFSPDGRYIVSASFDNSIKLWDGRDGKFISTFRGHVASVYQVAWSSDCRL
LVSCSKDTTLKVWDVRTRKLSVDLPGHKDEVYTVDWSVDGKRVCSGGKDKMVRLWTH"

CDS complement(267669..271664)
/codon_start=1
/locus_tag="YCR073C"
/label=SSK22
/translation="MMMDILNTQQQKAAEGGRVLAPHTISSKLVKRLSSHSSHKLSRSD
LKALGGSETISDGPSQLTFKDRYVFNESLYLKKLKKTALDDYYTRGIKLTNRYEEDDGD
DEIIRLSNGDRIDEDLHSGVKFFSTTPYCRKMRSDSDELAWNEIATERFKWQSMLARVL
KGDIVKGEKTRIANQVKKPGLNKELSDEIWLELKAWLNGRTMQEMEQSLTYLRDSSDSV
FEEIMKFQIPQGKILSLDALEAILQDLMNRYHSVVSYWPNLKKMYKDKPITNTAEFTAR
IDVMNSWLNFKTNLTLRRQELDDWINRFSPISSSDNCQEDFDGVPQWNCKMKILAEQLM
KEKNIESIFQKKIFYPLSPWMFKLKLHFIVYRETLTKMNIKYPYERLRSLLAFPVYLIK
EVILTRLSYARKLKNPTMMMIDQMIDDFNAFIRLSVQLKYTLTKYCSNLPFDVDFDPTF
ENTVIEAIRYLFFLLNLKLIDSSKQNFKAPDLLLKYWDHLKNTGHYINGAETVIPNEFL
KLTLRLVHKLQFYLLKQQNFPPTFANASEAEKWLSSIFENLGAMKRKLNRFSNILVKAF

QNSAVYQINHNAQLVKKLKDAHYFLVYSGNTFESSGVYMFAAPELLGCDNDTILRILRN
KSIGCDLVPKLDIGNNLNVYDITTKETDLNILVSKGEDSKGIPYYRVVANSSSDLDRHA
HQSKKKNFSTDPFDQHLDEKNNEVFELEVALSSLGALVVLYPGEPVVWDGPVYKLPGNN
LFASNEMDLGKIGNPNTLILLNQGSNYALTYQIDKFNQTVGDSVSFIEKRCSLNSIESS
LQKINKAYYKLTYTVLNNYKGILGSFMKQCPGNELLNSIFMFGRDFGRSFLKYNAFSSK
RKYVIIFLMVKLGMNWLKFLVEECDPTDQRTFRWCVLAMDFAMQMTSGYNILALNVKQF
QELKERVSVCMSLLISHFDVMGARATEAENGMQQARLNIDTEENIDEEATLEINSRLRL
EAIKTLEKTMKRNPRQMGKVLDATDQGNKYLLSLASSLSNVSMRWQKRSFIGGGTFGQV
YSAINLENGEILAVKEIKIHDTTTMKKIFPLIKEEMTVLEMLNHPNIVQYYGVEVHRDK
VNIFMEYCEGGSLASLLDHGRIEDEMVTQVYTFELLEGLAYLHQSGVVHRDIKPENILL
DFNGIIKYVDFGTARTVVGSRTRTVRNAAVQDFGVETKSLNEMMGTPMYMAPETISGSA
VKGKLGADDVWALGCVVLEMATGRRPWSNLDNEWAIMYHVAAGRIPQLPNRDEMTAAGR
AFLERCLVQDPTMRATAVELLIDPWMIQIREIAFGNSEKDQVPILSS"

CDS 272044..272991
/codon_start=1
/locus_tag="YCR073W-A"
/label=SOL2
/translation="MTTTVPKIFAFHEFSDVAEAVADHVVHAQDGALAPKNERKHSVPN
ISMNALDMTREASCKSTASAAEGKSGSSGSGSGSSKPKKEKRFKIALSGGSLIEVLHEG
LLKRDDVRWGDWDIYFADERLVPFSSNESNYGCAKRKILDLIDTAKYGTPKVYHIDESL
IDDPQECADNYEKVLIRGFAGRDSVKLPMFDLFLLGCAPDGHIASLFPNFQDNLREKLA
WVVPVENAPSGPSTRISLTIPVICHSHRVTFVVEGATKAPIIKTIMERPEKGLPSSIVN
EGAAGRVSWFVDDDALTDVLVTKKKYKFHQGLSI"

CDS complement(273114..273896)
/codon_start=1
/locus_tag="YCR075C"
/label=ERS1
/translation="MVSLDDILGIVYVTSWSISMYPPIITNWRHKSASAISMDFVMLNT
AGYSYLVISIFLQLYCWKMTGDESDLGRPKLTQFDFWYCLHGCLMNVVLLTQVVAGARI
WRFPGKGHRKMNPWYLRILLASLAIFSLLTVQFMYSNYWYDWHNSRTLAYCNNLFLLKI
SMSLIKYIPQVTHNSTRKSMDCFPIQGVFLDVTGGIASLLQLIWQLSNDQGFSLDTFVT
NFGKVGLSMVTLIFNFIFIMQWFVYRSRGHDLASEYPL"

CDS 274056..274283
/codon_start=1
/locus_tag="YCR075W-A"
/label=EGO2
/translation="MEAEKQSDIKGTIAFDTHGNVIESTGVGSQRIEDIGDLSKVTLDA
EGFAQVQGDSLLVHLYKRNDITLAVYTSAQ"

CDS complement(274374..275126)

/codon_start=1
/locus_tag="YCR076C"
/label=FUB1
/translation="MIENKVELVAELVLESIGKTEVVSRHTEGTKSCQVSFRIKDSPSE
KGSTSFLSELVVIQTLDDNDKYTVVIRHGTSITMACVVGYSDFKLPTELKWPLERESLP
VEPDLKPIMTQLKRQTAGSADMPKFDDEYQAQARQNQGTAPLNPYPGLTVTEPSFANPA
GGYADGDLYPVGTSHPDWSGGLPNPLGNPSSQGGMIFDPNRRPAPRREDMPPGWMPGSK
YDEPFGPGSGGFGGSGSGGFGGSGSGFI"
CDS complement(275319..277709)
/codon_start=1
/locus_tag="YCR077C"
/label=PAT1
/translation="MSFFGLENSGNARDGPLDFEESYKGYGEHELEENDYLNDETFGDN
VQVGTDFDFGNPHSSGSSGNAIGGNGVGATARSYVAATAEGISGPRTDGTAAAGPLDLK
PMESLWSTAPPPAMAPSPQSTMAPAPAPQQMAPLQPILSMQDLERQQRQMQQQFMNFHA
MGHPQGLPQGPPQQQFPMQPASGQPGPSQFAPPPPPPGVNVNMNQMPMGPVQVPVQASP
SPIGMSNTPSPGPVVGATKMPLQSGRRSKRDLSPEEQRRLQIRHAKVEKILKYSGLMTP
RDKDFITRYQLSQIVTEDPYNEDFYFQVYKIIQRGGITSESNKGLIARAYLEHSGHRLG
GRYKRTDIALQRMQSQVEKAVTVAKERPSKLKDQQAAAGNSSQDNKQANTVLGKISSTL
NSKNPRRQLQIPRQQPSDPDALKDVTDSLTNVDLASSGSSSTGSSAAAVASKQRRRSSY
AFNNGNGATNLNKSGGKKFILELIETVYEEILDLEANLRNGQQTDSTAMWEALHIDDSS
YDVNPFISMLSFDKGIKIMPRIFNFLDKQQKLKILQKIFNELSHLQIIILSSYKTTPKP
TLTQLKKVDLFQMIILKIIVSFLSNNSNFIEIMGLLLQLIRNNNVSFLTTSKIGLNLIT
ILISRAALIKQDSSRSNILSSPEISTWNEIYDKLFTSLESKIQLIFPPREYNDHIMRLQ
NDKFMDEAYIWQFLASLALSGKLNHQRIIIDEVRDEIFATINEAETLQKKEKELSVLPQ
RSQELDTELKSIIYNKEKLYQDLNLFLNVMGLVYRDGEISELK"
CDS 277926..279254
/codon_start=1
/locus_tag="YCR079W"
/label=PTC6
/translation="MRLGNAYAYCKPSQNVGLKLDLLRGLPGYVGHATSRINRLENQDN
YSIKMMRSWPNAYGSALNCSVFDGHGEKGAQLSQLLADKLCSSLDFPEPSWDKQDLKKL
VQEYARRFPEGNYWKHKLSTFEKFYNKFIKNCNSKQELLLMKEGDSAILGQNGGRMIFD
KMGNIIDKIALLTELDRLRLFYGFARFDLDQCCGLGTAAGSTASSIFLYPYDDPNAPID
EGKDDDSWIISHSGLLKLIVTQVGDSKIILCDQDGIAHALTTTHHINSSRERHRLSIDP
SRLDPDAFGETRFLNNFANTRSFGDVAGKPYGISSEPDIFSFLVGNTLHLPRSERSKLP
FNGDECFLALVTDGITNKLADQEVVDLITSTVNSWGLKKATPQFVAEETIKFIQAIATK
HSDNATCVVVRLSNWGNWPNVDRTGPQRETKLMNAQSNETKLN"
CDS 279452..283735

/codon_start=1
/locus_tag="YCR081W"
/label=SRB8
/translation="MNNGSGRYLLTPPDDLHPYVPSSKPQEQVYPDFKPWEHTAAEDQI
LANFVAKGFYHTPMVNFESISARSSVHESLVTQSNILSQQFDKIIKIREDHINKIPSNS
TTTLHGPGFQLPNRITLTDHRKETWLHELSSSHTSLVKIGKFIPHGLKRRQVIEQCYLK
FIPLKRAIWLIKCCYFIEWKSNHKKKRSNAAGADDAISMHLLKDWTDTFVYILEKLIFD
MTNHYNDSQQLRTWKRQISYFLKLLGNCYSLRLINKEIFHHWLVEFINKMENFEFLPLS
LHILMIFWNDICQIDTNAPVAATITSSQKEPFFLVTKITDMLLHKYYIVSSSKSMINDE
NYIINDIKKNNKIKLNILKILSSLILKIFQEQSLEVFIFPTSNWEIYKPLLFEIVSNAD
TNQNSDMKKKLELISYRNESLKNNSSIRNVIMSASNANDFQLTIVTCKQFPKLSCIQLN
CIDTQFTKLLDDNPTEFDWPTYVDQNPLTMHKIIQLILWSIHPSRQFDHYESNQLVAKL
LLLRINSTDEDLHEFQIEDAIWSLVFQLAKNFSAQKRVVSYMMPSLYRLLNILITYGII
KVPTYIRKLISSGLLYLQDSNDKFVHVQLLINLKISPLMKSQYNMVLRNVMEYDVKFYE
IFNFDQLVEITEQIKMRILSNDITNLQLSKTPLSIKIMVAEWYLSHLCSGILSSVNRTV
LLKIFKIFCIDLEVFHHFFKWIEFIVYHQLLSDIESLEALMDILLCYQKLFSQFINDHI
LFTKTFIFIYKKVLKEKDVPAYNVTSFMPFWKFFMKNFPFVLKVDNDLRIELQSVYNDE
KLKTEKLKNDKSEVLKVYSMINNSNQAVGQTWNFPEVFQVNIRFLLHNSEIIDTNTSKQ
FQKARNNVMLLIATNLKEYNKFMSIFLKRKDFTNKNLIQLISLKLLTFEVTQNVLGLEY
IIRLLPINLENNDGSYGLFLKYHKEQFIKSNFEKILLTCYELEKKYHGNECEINYYEIL
LKILITYGSSPKLLATSTKIIMLLLNDSVENSSNILEDILYYSTCPSETDLNDIPLGSG
QPDNDTVVTNDDKSDDDDHTVDEIDHVEYYVMMDFANLWVFQAFTCFCIKKIMENNEPA
MAMEDLKNFIFQIIEITNSNDLCSQIFDQLKDMQTIEMITQIVEKDFCTSCLQNNNQKI
DDNYIVVVIEIITSLSMRFQRETSGMIVISTENYHLLIKIIRQLSELNEGNLSKREIQI
DAVLKIFSFHQDSIFQRIIADLSADKPTSPFIDSICKLFDKISFNLRLKLFLYEILSSL
KSFAIYSSTIDAPAFHTSGKVELPKKLLNLPPFQVSSFVKETKLHSGDYGEEEDADQEE
SFSLNLGIGIVEIAHENEQKWLIYDKKDHKYVCTFSMEPYHFISNYNTKYTDDMATGSN
DTTAFNDSCVNLSLFDARFERKNPH"
CDS 283964..284350
/codon_start=1
/locus_tag="YCR082W"
/label=AHC2
/translation="MITPKGTHDAVAKFQKTDLHQDLDYIVLQQRRTQLETLINERESF
VKNLCSLFHKIQNTKNYQEFVDVLAENRDLLREIFTVENGFQKQKWISNDDIPQIDWDK
FALDINAYIAENDQLLALYEDGLL"
CDS 284658..285041
/codon_start=1
/locus_tag="YCR083W"
/label=TRX3

/translation="MLFYKPVMRMAVRPLKSIRFQSSYTSITKLTNLTEFRNLIKQNDK
LVIDFYATWCGPCKMMQPHLTKLIQAYPDVRFVKCDVDESPDIAKECEVTAMPTFVLGK
DGQLIGKIIGANPTALEKGIKDL"
CDS complement(285392..287533)
/codon_start=1
/locus_tag="YCR084C"
/label=TUP1
/translation="MTASVSNTQNKLNELLDAIRQEFLQVSQEANTYRLQNQKDYDFKM
NQQLAEMQQIRNTVYELELTHRKMKDAYEAEIKHLKLGLEQRDHQIASLTVQQQQQQQQ
QQQVQQHLQQQQQQLAAASASVPVAQQPPATTSATATPAANTTTGSPSAFPVQASRPNL
VGSQLPTTTLPVVSSNAQQQLPQQQLQQQQLQQQQPPPQVSVAPLSNTAINGSPTSKET
TTLPSVKAPESTLKETEPENNNTSKINDTGSATTATTTTATETEIKPKEEDATPASLHQ
DHYLVPYNQRANHSKPIPPFLLDLDSQSVPDALKKQTNDYYILYNPALPREIDVELHKS
LDHTSVVCCVKFSNDGEYLATGCNKTTQVYRVSDGSLVARLSDDSAANNHRNSITENNT
TTSTDNNTMTTTTTTTITTTAMTSAAELAKDVENLNTSSSPSSDLYIRSVCFSPDGKFL
ATGAEDRLIRIWDIENRKIVMILQGHEQDIYSLDYFPSGDKLVSGSGDRTVRIWDLRTG
QCSLTLSIEDGVTTVAVSPGDGKYIAAGSLDRAVRVWDSETGFLVERLDSENESGTGHK
DSVYSVVFTRDGQSVVSGSLDRSVKLWNLQNANNKSDSKTPNSGTCEVTYIGHKDFVLS
VATTQNDEYILSGSKDRGVLFWDKKSGNPLLMLQGHRNSVISVAVANGSSLGPEYNVFA
TGSGDCKARIWKYKKIAPN"
CDS 287997..288350
/codon_start=1
/locus_tag="YCR085W"
/label=YCR085W
/translation="MLFLYTYVYVFLCTNNDVYNETSVMLSKTSAHCFIAEEVTTDNGL
ICGLAMLGKTKYQFYELFTVYSIQSLTQLASRVKKGGLIMARLILFTLCALPVLFHFIL
FMLQYLVFVYIEK"
CDS 288475..289047
/codon_start=1
/locus_tag="YCR086W"
/label=CSM1
/translation="MDPLTVYKNSVKQQIDSADLLVANLVNENFVLSEKLDTKATEIKQ
LQKQIDSLNAQVKELKTQTSQQAENSEVIKDLYEYLCNVRVHKSYEDDSGLWFDISQGT
HSGGSSDDYSIMDYKLGFVKGQAQVTEVIYAPVLKQRSTEELYSLQSKLPEYLFETLSF
PLSSLNQFYNKIAKSLNKKREKKDETE"
CDS complement(289089..289535)
/codon_start=1
/locus_tag="YCR087C-A"
/label=YCR087C-A

/translation="MVTFNCEVCNDTVPKKNTEKHYYRCPNAYYTCIDCSKTFEDGVSY
KNHTSCISEDEKYQKALYKGNKKQKQKQQQKQQQQPVATPAKKVEKPVIKKAEKVEKTS
NGIELHKGKSLYKILKTMKDKGAKKTFLKSLVVDSEGQIRYAKE"

CDS 290136..291914
/codon_start=1
/locus_tag="YCR088W"
/label=ABP1
/translation="MALEPIDYTTHSREIDAEYLKIVRGSDPDTTWLIISPNAKKEYEP
ESTGSSFHDFLQSFDETKVQYGLARVSPPGSDVEKIIIIGWCPDSAPLKTRASFAANFA
AVANNLFKGYHVQVTARDEDDLDENELLMKISNAAGARYSIQTSSKQQGKASTPPVKKS
FTPSKSPAPVSKKEPVKTPSPAPAAKISSRVNDNNDDDDWNEPELKERDFDQAPLKPNQ
SSYKPIGKIDLQKVIAEEKAKEDPRLVQKPTAAGSKIDPSSDIANLKNESKLKRDSEFN
SFLGTTKPPSMTESSLKNDDDKVIKGFRNEKSPAQLWAERKAKQNSGNAETKAEAPKPE
VPEDEPEGEPDVKDLKSKFEGLAASEKEEEEMENKFAPPPKKSEPTIISPKSFSKPQEP
VKTEEAQQPKTDYKKIGNPLPGMHIEADNEEEPEENDDDWDDDEEEAAQPPLPSRNVAS
GAPVQKEEPEQEEIAPSLPSRNSIPAPKQEEAPEQAPEEEIEEEAEEAAPQLPSRSSAA
PPPPPRRATPEKKPKENPWATAEYDYDAAEDNELTFVENDKIINIEFVDDDWWLGELEK
DGSKGLFPSNYVSLGN"

CDS 292502..297346
/codon_start=1
/locus_tag="YCR089W"
/label=FIG2
/translation="MNSFASLGLIYSVVNLLTRVEAQIVFYQNSSTSLPVPTLVSTSIA
DFHESSSTGEVQYSSSYSYVQPSIDSFTSSSFLTSFEAPTETSSSYAVSSSLITSDTFS
SYSDIFDEETSSLISTSAASSEKASSTLSSTAQPHRTSHSSSSFELPVTAPSSSSLPSS
TSLTFTSVNPSQSWTSFNSEKSSALSSTIDFTSSEISGSTSPKSLESFDTTGTITSSYS
PSPSSKNSNQTSLLSPLEPLSSSSGDLILSSTIQATTNDQTSKTIPTLVDATSSLPPTL
RSSSMAPTSGSDSISHNFTSPPSKTSGNYDVLTSNSIDPSLFTTTSEYSSTQLSSLNRA
SKSETVNFTASIASTPFGTDSATSLIDPISSVGSTASSFVGISTANFSTQGNSNYVPES
TASGSSQYQDWSSSSLPLSQTTWVVINTTNTQGSVTSTTSPAYVSTATKTVDGVITEYV
TWCPLTQTKSQAIGVSSSISSVPQASSFSGSSILSSNSSTLAASNNVPESTASGSSQYQ
DWSSSSLPLSQTTWVVINTTNTQGSVTSTTSPAYVSTATKTVDGVITEYVTWCPLTQTK
SQAIGISSSTISATQTSKPSSILTLGISTLQLSDATFKGTETINTHLMTESTSITEPTY
FSGTSDSFYLCTSEVNLASSLSSYPNFSSSEGSTATITNSTVTFGSTSKYPSTSVSNPT
EASQHVSSSVNSLTDFTSNSTETIAVISNIHKTSSNKDYSLTTTQLKTSGMQTLVLSTV
TTTVNGAATEYTTWCPASSIAYTTSISYKTLVLTTEVCSHSECTPTVITSVTATSSTIP
LLSTSSSTVLSSTVSEGAKNPAASEVTINTQVSATSEATSTSTQVSATSATATASESST
TSQVSTASETISTLGTQNFTTTGSLLFPALSTEMINTTVVSRKTLIISTEVCSHSKCVP
TVITEVVTSKGTPSNGHSSQTLQTEAVEVTLSSHQTVTMSTEVCSNSICTPTVITSVQM

RSTPFPYLTSSTSSSSLASTKKSSLEASSEMSTFSVSTQSLPLAFTSSEKRSTTSVSQW
SNTVLTNTIMSSSSNVISTNEKPSSTTSPYNFSSGYSLPSSSTPSQYSLSTATTTINGI
KTVYTTWCPLAEKSTVAASSQSSRSVDRFVSSSKPSSSLSQTSIQYTLSTATTTISGLK
TVYTTWCPLTSKSTLGATTQTSSTAKVRITSASSATSTSISLSTSTESESSSGYLSKGV
CSGTECTQDVPTQSSSPASTLAYSPSVSTSSSSSFSTTTASTLTSTHTSVPLLPSSSSI
SASSPSSTSLLSTSLPSAAFTSSTLPTATAVSSSTFIASSLPLSSKSSLSLSPVSSSIL
MSQFSSSSSSSSSSSSSSLASLPSLSISPTVDTVSVLQPTTSIATLTCTDSQCQQEVST
ICNGSNCDDVTSTATTPPSTVTDTMTCTGSECQKTTSSSCDGYSCKVSETYKSSATISA
CSGEGCQASATSELNSQYVTMTSVITPSAITTTSVEVHSTESTISITTVKPVTYTSSDT
NGELITITSSSQTVIPSVTTIITRTKVAITSAPKPTTTTYVEQRLSSSGIATSFVAAAS
STWITTPIVSTYAGSASKFLCSKFFMIMVMVINFI"

CDS 296262..296279
/codon_start=1
/product="6xHis affinity tag"
/label=6xHis
/translation="HHHHHH"

CDS 296544..296573
/codon_start=1
/product="10xHis affinity tag"
/label=10xHis
/translation="HHHHHHHHHH"

CDS 296547..296576
/codon_start=1
/product="10xHis affinity tag"
/label=10xHis
/translation="HHHHHHHHHH"

CDS 296550..296579
/codon_start=1
/product="10xHis affinity tag"
/label=10xHis
/translation="HHHHHHHHHH"

CDS 296553..296582
/codon_start=1
/product="10xHis affinity tag"
/label=10xHis
/translation="HHHHHHHHHH"

CDS 296556..296585
/codon_start=1
/product="10xHis affinity tag"

/label=10xHis
/translation="HHHHHHHHHH"
CDS complement(297398..297946)
/codon_start=1
/locus_tag="YCR090C"
/label=YCR090C
/translation="MPLFLVLKATLSENVTKVSIENTNESRAEFAFDLQCTSCRELHDS
KVIINTFEEYAMPASKGTASFLMKCKFCSKELSVNLCAFEDEYLTDQSDDKWAKIKDVR
KKHGLSKVKEDSFIPLSLDCRGCELIKFYPDTITFEVSLSSGKVMSCQLEDNEWYDYDD
NLGEEVTMTDFSSSIIKGK"
CDS 299486..301648
/codon_start=1
/locus_tag="YCR091W"
/label=KIN82
/translation="MTQQEYRSPSQRLSKGRSMSLPKIFARNLRSLQNNAPPGKNIDVN
CLNVNSCSLSASPSSQINMACNGNKQDLPIPFPLHVECNDSWSSSKLNKFKSMFNHNRS
KSNGTTDASTSEKGTHKREPRSTIHTELLQSSIIGEPNVHSTTSSTLIPNEAICSTPNE
ISGSSSPDAELFTFDMPTDPSSFHTPSSPSYIAKDSRNLSNGSLNDINENEELQNFHRK
ISENGSASPLANLSLSNSPIDSPRKNSETRKDQIPMNITPRLRRAASEPFNTAKDGLMR
EDYIALKQPPSLGDIVEPRRSRRLRTKSFGNKFQDITVEPQSFEKIRLLGQGDVGKVYL
VRERDTNQIFALKVLNKHEMIKRKKIKRVLTEQEILATSDHPFIVTLYHSFQTKDYLYL
CMEYCMGGEFFRALQTRKSKCIAEEDAKFYASEVVAALEYLHLLGFIYRDLKPENILLH
QSGHVMLSDFDLSIQATGSKKPTMKDSTYLDTKICSDGFRTNSFVGTEEYLAPEVIRGN
GHTAAVDWWTLGILIYEMLFGCTPFKGDNSNETFSNILTKDVKFPHDKEVSKNCKDLIK
KLLNKNEAKRLGSKSGAADIKRHPFFKKVQWSFLRNQDPPLIPALNDNGCELPFILSCN
KHPKRNSVSEQETKMFCEKVANDDEIDEADPFHDFNSMSLTKKDHNILTYSENYTYGKI
LYKATCTRPRHNSSHRSFFKDIIPEL"
CDS complement(301846..304902)
/codon_start=1
/locus_tag="YCR092C"
/label=MSH3
/translation="MAGQPTISRFFKKAVKSELTHKQEQEVAVGNGAGSESICLDTDEE
DNLSSVASTTVTNDSFPLKGSVSSKNSKNSEKTSGTSTTFNDIDFAKKLDRIMKRRSDE
NVEAEDDEEEGEEDFVKKKARKSPTAKLTPLDKQVKDLKMHHRDKVLVIRVGYKYKCFA
EDAVTVSRILHIKLVPGKLTIDESSPQDCNHRQFAYCSFPDVRLNVHLERLVHHNLKVA
VVEQAETSAIKKHDPGASKSSVFERKISNVFTKATFGVNSTFVLRGKRILGDTNSIWAL
SRDVHQGKVAKYSLISVNLNNGEVVYDEFEEPNLADEKLQIRIKYLQPIEVLVNTDDLP
LHVAKFFKDISCPLIHKQEYDLEDHVVQAIKVMNEKIQLSPSLIRLVSKLYSHMVEYNN
EQVMLIPSIYSPFASKIHMLLDPNSLQSLDIFTHDGGKGSLFWLLDHTRTSFGLRMLRE

WILKPLVDVHQIEERLDAIECITSEINNSIFFESLNQMLNHTPDLLRTLNRIMYGTTSR
KEVYFYLKQITSFVDHFKMHQSYLSEHFKSSDGRIGKQSPLLFRLFSELNELLSTTRLP
HFLTMINVSAVMEKNSDKQVMDFFNLNNYDCSEGIIKIQRESESVRSQLKEELAEIRKY
LKRPYLNFRDEVDYLIEVKNSQIKDLPDDWIKVNNTKMVSRFTTPRTQKLTQKLEYYKD
LLIRESELQYKEFLNKITAEYTELRKITLNLAQYDCILSLAATSCNVNYVRPTFVNGQQ
AIIAKNARNPIIESLDVHYVPNDIMMSPENGKINIITGPNMGGKSSYIRQVALLTIMAQ
IGSFVPAEEIRLSIFENVLTRIGAHDDIINGDSTFKVEMLDILHILKNCNKRSLLLLDE
VGRGTGTHDGIAISYALIKYFSELSDCPLILFTTHFPMLGEIKSPLIRNYHMDYVEEQK
TGEDWMSVIFLYKLKKGLTYNSYGMNVAKLARLDKDIINRAFSISEELRKESINEDALK
LFSSLKRILKSDNITATDKLAKLLSLDIH"

gap 305154..305163
/estimated_length=10

CDS 305719..312045
/codon_start=1
/locus_tag="YCR093W"
/label=CDC39
/translation="MLSATYRDLNTASNLETSKEKQAAQIVIAQISLLFTTLNNDNFES
VEREIRHILDRSSVDIYIKVWERLLTLSSRDILQAGKFLLQENLLHRLLLEFAKDLPKK
STDLIELLKERTFNNQEFQKQTGITLSLFIDLFDKSANKDIIESLDRSSQINDFKTIKM
NHTNYLRNFFLQTTPETLESNLRDLLHSLEGESLNDLLALLLSEILSPGSQNLQNDPTR
SWLTPPMVLDATNRGNVIARSISSLQANQINWNRVFNLMSTKYFLSAPLMPTTASLSCL
FAALHDGPVIDEFFSCDWKVIFKLDLAIQLHKWSVQNGCFDLLNAEGTRKVSETIPNTK
QSLLYLLSIASLNLELFLQREELSDGPMLAYFQECFFEDFNYAPEYLILALVKEMKRFV
LLIENRTVIDEILITLLIQVHNKSPSSFKDVISTITDDSKIVDAAKIIINSDDAPIANF
LKSLLDTGRLDTVINKLPFNEAFKILPCARQIGWEGFDTFLKTKVSPSNVDVVLESLEV
QTKMTDTNTPFRSLKTFDLFAFHSLIEVLNKCPLDVLQLQRFESLEFSLLIAFPRLINF
GFGHDEAILANGDIAGINNDIEKEMQNYLQKMYSGELAIKDVIELLRRLRDSDLPRDQE
VFTCITHAVIAESTFFQDYPLDALATTSVLFGSMILFQLLRGFVLDVAFRIIMRFAKEP
PESKMFKFAVQAIYAFRIRLAEYPQYCKDLLRDVPALKSQAQVYQSIVEAATLANAPKE
RSRPVQEMIPLKFFAVDEVSCQINQEGAPKDVVEKVLFVLNNVTLANLNNKVDELKKSL
TPNYFSWFSTYLVTQRAKTEPNYHDLYSKVIVAMGSGLLHQFMVNVTLRQLFVLLSTKD
EQAIDKKHLKNLASWLGCITLALNKPIKHKNIAFREMLIEAYKENRLEIVVPFVTKILQ
RASESKIFKPPNPWTVGILKLLIELNEKANWKLSLTFEVEVLLKSFNLTTKSLKPSNFI
NTPEVIETLSGALGSITLEQQQTEQQRQIILMQQHQQQMLIYQQRQQQQQQRQQQQQHH
ISANTIADQQAAFGGEGSISHDNPFNNLLGSTIFVTHPDLKRVFQMALAKSVREILLEV
VEKSSGIAVVTTTKIILKDFATEVDESKLKTAAIIMVRHLAQSLARATSIEPLKEGIRS
TMQSLAPNLMSLSSSPAEELDTAINENIGIALVLIEKASMDKSTQDLADQLMQAIAIRR
YHKERRADQPFITQNTNPYSLSLPEPLGLKNTGVTPQQFRVYEEFGKNIPNLDVIPFAG
LPAHAPPMTQNVGLTQPQQQQAQMPTQILTSEQIRAQQQQQQLQKSRLNQPSQSAQPPG

VNVPNPQGGIAAVQSDLEQNQRVLVHLMDILVSQIKENATKNNLAELGDQNQIKTIIFQ
ILTFIAKSAQKDQLALKVSQAVVNSLFATSESPLCREVLSLLLEKLCSLSLVARKDVVW
WLVYALDSRKFNVPVIRSLLEVNLIDATELDNVLVTAMKNKMENSTEFAMKLIQNTVLS
DDPILMRMDFIKTLEHLASSEDENVKKFIKEFEDTKIMPVRKGTKTTRTEKLYLVFTEW
VKLLQRVENNDVITTVFIKQLVEKGVISDTDNLLTFVKSSLELSVSSFKESDPTDEVFI
AIDALGSLIIKLLILQGFKDDTRRDYINAIFSVIVLVFAKDHSQEGTTFNERPYFRLFS
NILYEWATIRTHNFVRISDSSTRQELIEFDSVFYNTFSGYLHALQPFAFPGFSFAWVTL
LSHRMLLPIMLRLPNKIGWEKLMLLIIDLFKFLDQYTSKHAVSDAVSVVYKGTLRVILG
ISNDMPSFLIENHYELMNNLPPTYFQLKNVILSAIPKNMTVPNPYDVDLNMEDIPACKE
LPEVFFDPVIDLHSLKKPVDNYLRIPSNSLLRTILSAIYKDTYDIKKGVGYDFLSVDSK
LIRAIVLHVGIEAGIEYKRTSSNAVFNTKSSYYTLLFNLIQNGSIEMKYQIILSIVEQL
RYPNIHTYWFSFVLMNMFKSDEWNDQKLEVQEIILRNFLKRIIVNKPHTWGVSVFFTQL
INNNDINLLDLPFVQSVPEIKLILQQLVKYSKKYTTSEQDDQSATINRRQTPLQSNA"

CDS 312364..313539
/codon_start=1
/locus_tag="YCR094W"
/label=CDC50
/translation="MVSLFKRGKAPPLTKEGPTSKKPPNTAFRQQRLKAWQPILSPQSV
LPLLIFVACIFTPIGIGLIVSATKVQDLTIDYSHCDTKASTTAFEDIPKKYIKYHFKSK
VENKPQWRLTENENGEQSCELQFEIPNDIKKSIFIYYKITNFYQNHRRYVQSFDTKQIL
GEPIKKDDLDTSCSPIRSREDKIIYPCGLIANSMFNDTFSQVLSGIDDTEDYNLTNKHI
SWSIDRHRFKTTKYNASDIVPPPNWMKKYPDGYTDENLPDIHTWEEFQVWMRTAAFPKF
YKLTLKNESASLPKGKYQMNIELNYPISLFGGTKSFVLTTNGAIGGRNMSLGVLYLIVA
GLCALFGIIFLVKLIFQPRAMGDHTYLNFDDEENEDYEDVHAENTTLREIL"

CDS complement(313772..314860)
/codon_start=1
/locus_tag="YCR095C"
/label=OCA4
/translation="MLVPPANFGIAEEGIYRCSKVETLNLSFLETLNLKTAIFIGGQEP
SKFFKDFFTRSSIKWIVLRMSDFSAAAVPVKSSSVSNANLYSNNNSTLSLQEEKKKSTA
NGSQNSTTGDPVIQEELAYHLTDNDDLMLIKSTCLKRTFKTLLNVDNYNVLLVDKTALV
IGILRKIQKWNIASIINEYRLFSGKNRNYFAETFLEIINIEIEQEKDNKTIVDNKAKKL
PLENNRTHSIEYKANSGKLIRVNEDDLCREPEVPQRLLTLINQIETKVKNNKVLQVSGV
LGDDLKKTSSDLGIFGHRYRLAFNKKENGDYGYYKARGKDNVKIRIPCDSELPDWFRFQ
RDLWEKENVPEEHHFYREHIFT"

CDS 315241..315399
/codon_start=1
/locus_tag="YCR095W-A"
/label=YCR095W-A

/translation="MTVLIKLGLRILHVYKGFFRKVILKYFFFSSEHTKVNKKSSMHAF
LCKIYKR"
CDS complement(318778..319137)
/codon_start=1
/locus_tag="YCR096C"
/product="YCR096C"
/label=HMRA2
/translation="MRSIENDRSNYQLTQKNKSADGLVFNVVTQDMINKSTKPYRGHRF
TKENVRILESWFAKNIENPYLDTKGLENLMKNTSLSRIQIKNWVSNRRRKEKTITIAPE
LADLLSGEPLAKKKE"
CDS join(319434..319538,319593..319838,319891..319920)
/codon_start=1
/locus_tag="YCR097W"
/label=HMRA1
/translation="MDGICSMAENINRTLFNILGTEIDEINLNTNNLYNFIMESNLTKV
EQHTLHKNISNNRLEIYHHIKKEKSPKGKSSISPQARAFLEQVFRRKQSLNSKEKEEVA
KKCGITPLQVRVWFINKRMRSK"
CDS complement(323090..324646)
/codon_start=1
/locus_tag="YCR098C"
/label=GIT1
/translation="MEDKDITSVNEKEVNENTNSRIIKYDAERRATRTETSKKDKWKNI
VTIIASGFALISDGYVNGSMSMLNKVFVMEYGKKNYSSKVSTRVSNAALVGIIFGQFFM
GIAADYYSRKSCILVATAILVIGSALCAASHGTTVPGMFWMLTVMRGLVGIGVGAEYPT
STLSANESANEYTTTKRGGILVMVTNLPLAFGGPFATIIFLIVYKICSGTKHLEAIWRT
VFAIGCFWPLSVFYFRWKTATTEVYEKGRIKRNIPYFLALKFYWKRLLGTCGTWFMYDF
VTFPNGIFSSTIISSVIKDQNDLVKVAEWNLLLGVLAVLGVPIGAYLSDRIGRKYTLMF
GFSGYIIFGLIIGCAYDQLKKITPLFIIFYAFMNMLGNAGPGDMLGVISSEASATAVRG
VFYGLSAVTGKIGSVVGVECFQPIRDNLGARWTFIIAAICGLIGIIITYFFVPHSLESD
LMKQDVEFHNYLVSNGWTGKMGFDETDEESMVRTIEVEENGTNCSKKNAEIISVRQVDQ
S"
CDS complement(326873..327340)
/codon_start=1
/locus_tag="YCR099C"
/label=YCR099C
/translation="MRSIFLLHFDYKTCEEEDFEDWNLADGKCLNGAKYMYKRRKQDAR
CLVKRTFKDMILHEIPCDSCTESDYECSSEFVRDAKGDCIPDYDQIALSDICDKANGET
VSLEPLQLIKGDKCKKPMEIEAMNIPCEKILRESSNGKKIATIENKFDFEI"
CDS complement(327312..328262)

/codon_start=1
/locus_tag="YCR100C"
/label=EMA35
/translation="MSSTDIWISNDASTFQKAQLPTQLRHVKVIKIREDSIGRIILLIS
TEITNEENADPDLSEIFISDSQGLKFSPVEWTPNHQFGNFRLTFPDFLKGTIFGSFHPS
IDYSNHQVNYTENIAGGETKISVDNGLTWSNLKVVDEENADSFGCDITRPERCSLQGYF
YNLKLSNPSAGIILMTGSVGDDNEFDRKDRKTFISRDGGLTWRVAHNSSGLYATGDLGN
IIVYIPSPSYKDGDVQSKLYFSLDQGRTWNQYELADALFYIHPLELINTTPDGSGSKFI
LSGHLITTASQEGNNTNISYIARSVLYAIDFSAAF"
CDS complement(328523..329071)
/codon_start=1
/locus_tag="YCR101C"
/label=YCR101C
/translation="MILLHAIYTLWVIILLPLLNAEKFVPKVTEAPIETSFNLVSFDDS
NTSIRLDGWGVVWISFDAGENWETVKEIEERIFRFTVDPFHGQERGFAFICESPKFYIT
DDRGESWRALTIPSSEEYLDGDCFITTHPRNKELLIANCYSYMIDADVLYDPSEIYLSN
DGNPFLKLNLPWKRKKTTI"
CDS complement(330401..331507)
/codon_start=1
/locus_tag="YCR102C"
/label=YCR102C
/translation="MKAVVIEDGKAVVKEGVPIPELEEGFVLIKTLAVAGNPTDWAHID
YKVGPQGSILGCDAAGQIVKLGPAVDPKDFSIGDYIYGFIHGSSVRFPSNGAFAEYSAI
STVVAYKSPNELKFLGEDVLPAGPVRSLEGAATIPVSLTTAGLVLTYNLGLNLKWEPST
PQRNGPILLWGGATAVGQSLIQLANKLNGFTKIIVVASRKHEKLLKEYGADELFDYHDI
DVVEQIKHKYNNISYLVDCVANQNTLQQVYKCAADKQDATVVELTNLTEENVKKENRRQ
NVTIDRTRLYSIGGHEVPFGGITFPADPEARRAATEFVKFINPKISDGQIHHIPARVYK
NGLYDVPRILEDIKIGKNSGEKLVAVLN"
CDS 333839..334213
/codon_start=1
/locus_tag="YCR104W"
/label=PAU3
/translation="MVKLTSIAAGVAAIAAGVAAAPATTTLSPSDERVNLVELGVYVSD
IRAHLAQYYLFQAAHPTETYPVEIAEAVFNYGDFTTMLTGIPAEQVTRVITGVPWYSTR
LRPAISSALSKDGIYTAIPK"
CDS 335109..336194
/codon_start=1
/locus_tag="YCR105W"
/label=ADH7

/translation="MLYPEKFQGIGISNAKDWKHPKLVSFDPKPFGDHDVDVEIEACGI
CGSDFHIAVGNWGPVPENQILGHEIIGRVVKVGSKCHTGVKIGDRVGVGAQALACFECE
RCKSDNEQYCTNDHVLTMWTPYKDGYISQGGFASHVRLHEHFAIQIPENIPSPLAAPLL
CGGITVFSPLLRNGCGPGKRVGIVGIGGIGHMGILLAKAMGAEVYAFSRGHSKREDSMK
LGADHYIAMLEDKGWTEQYSNALDLLVVCSSSLSKVNFDSIVKIMKIGGSIVSIAAPEV
NEKLVLKPLGLMGVSISSSAIGSRKEIEQLLKLVSEKNVKIWVEKLPISEEGVSHAFTR
MESGDVKYRFTLVDYDKKFHK"
CDS 336997..339495
/codon_start=1
/locus_tag="YCR106W"
/label=RDS1
/translation="MDSITVKKPRLRLVCLQCKKIKRKCDKLRPACSRCQQNSLQCEYE
ERTDLSANVAANDSDGFNSSHKLNFEQQPVLERTGLRYSLQVPEGVVNATLSIWNAEDM
LVIVGLVTFLDYPFAAHSLAQHDQYIRALCASLYGMALVDFSNYANGIPCEDTSRSILG
PLSFIEKAIFRRIEHSKQFRVQPAALGLLYNAFSMEEENFSTLLPSLIAEVEDVLMQKK
DCEILLRCFYQNIYPFYPFMDISLFESDLTSLLLQDDNNRWKISTEVKNVRKKIETLSL
LTIVMAMALMHSKLDANLLSMVKENASESARKLSLLCHKLLCLLDVFRYPNENTFTCLL
YFYVSEHLDPESPDCVLSPTNLLTLHHLLNLSMTLGLQYEPSKYKRFKDPEVIRQRRIL
WLGVQSLLFQISLAEGDAGKSNSEYMEAYLTDFEEYIEASSEYEKSSASESNVQMNDIV
WNKYKFHVILSKLMSDCTSVIQHPQLFHILENIKRSEDFMAENFPTSSIYQPLHEKEPN
AIKVGKSTVLDVMDIQKTEIFLTNIVGSMCFLNIFDVLSLHFEKKCVMHWEEYEKNYHF
LTLKSFNAYLKLAGLISDYLENKFQGNILESRGYIIDKQICFMLVRIWMFQCRILLRFS
YKQESQKKLASSSISTNDNEKEDEMIVILERLIKHIRNQMAHLVDLAKGKLQDSYFGAY
QTVPMFRYVVYLIDVGGLVSVTNGFWDKISSDGEIPPKVQQAVRLKWGLDCNNSRRIKQ
KLISSQSLQSFNQVLLCQMEDAVLSSSFAIKANTAMSQNTAEEFFNISEEEALNQLLEN
NNFDAFWDLLGENLSDMPSL"
CDS 339929..340705
/codon_start=1
/locus_tag="YCR107W"
/label=AAD3
/translation="MIGSASDSPSKLGRLRFLSETAAIKVSPLILGEVSYDGARSDFLK
SMNKNRAFELLDTFYEAGGNFIDAANNCQNEQSEEWIGEWIQSRRLRDQIVIATKFIKS
DKKYKAGESNTANYCGNHKRSLHVSVRDSLRKLQTDWIDILYVHWWDYMSSIEEFMDSL
HILVQQGKVLYLGVSDTPAWVVSAANYYATSYGKTPFSIYQGKWNVLNRDFERDIIPMA
RHFGMALAPWDVMGGGRFQSKKAMEERRKNGEGIRSF"
ORIGIN
1 gttggcgcct ccgaacaaac agatgcagaa atcaagatta gtgaagcatt ggccaagatt
61 gctgaggaac atggcactga gtctgttact gctattgcta ttgcctatgt tcgctctaag
121 gcgaaaaatt tttttccgtc ggttgaagga ggaaaaattg aggatctcaa agagaacatt

181 aaggctctca gtatcgatct aacgccagac aatataaaat acttagaaag tatagttcct
241 tttgacatcg gatttcctaa taattttatc gtgttaaatt ccttgactca aaaatatggt
301 acgaataatg tttagataat ttttcagtaa tcaactacgc aagtaaagca gtaaatacgt
361 tactgctggt attaatgtca tgtattgagg caatgatgct atgctcttaa cgacatgtag
421 cttacaaaac gttcctatgg ttttactgca tggtatttac aaattagata gcagttccat
481 ccgcccttga catattatta ttcagacaag gtgtatatga gcataaatat gtatatatgc
541 acatgaggtt tgtattaacc ttgcaactgt accaaaatac aatcttcttt gctgctattt
601 aacggcgtat gtgcagttca taaatgcgtg ttctgtatag tacatccgcg cacattcttc
661 ctagcggaag acatattaac gtagcccgta cgccctggtt aagacttggt atagttccaa
721 taattggaat attactgcat agtggtccaa tagcaggatt tagcataaac atgataatat
781 ttagagatgc tttcatccct tgccggtaga ggtgtggtca ataagagcga cctcatacta
841 tacctgagaa agcaacctga cctacaggaa agagttactc aagaataaga attttcgttt
901 taaaacctaa gagtcacttt aaaatttgta tacacttatt ttttttataa cttatttaat
961 aataaaaatc ataaatcata agaaattcgc ttaagcggat ctgcctactc tcttcaagta
1021 agcgtcccat cccgcctttc tgtatctctc accttccact acaggatctc ttccctttgc
1081 gaaaagacct ctaccaacaa taatgatgtc tgatccaccg gcaacaactt catccacagt
1141 tctgtattgt tgtcccaaag catcaccttt gtcatcaaga ccaacacctg gcgtcatgat
1201 caaccaatcg tagccctctt ctcttccacc catatcgttt tgagcaataa atccaataac
1261 aaagtcctta tcactcttgg caatttccac ggtcccacga gtgtattcac cgtgcgctag
1321 agaccccttg gacgataact cggcaagcat taacaaccct ctaggttctt ttgtaacttc
1381 ctcggcacct tgcttcaaac cagcaacaat gcccgcacca gtcacaccgt gtgcattggt
1441 gatatcagac cattcggcga tacggtatac accagacgtg tattgtaatt taacagtgtt
1501 cccaatgtcg gcaaacttcc tgtcttcaaa tatcaaaaac ttgtgtttct ctgctaattg
1561 cttcaacggc acaatggtat tctcaaagct gaaatcctcc aagatatcta catgtgtctt
1621 caatagacag atatatggac ccaaaacctc aactaatctt aacaactctg ctgttttacg
1681 aacatcaaga gaagcacata agtttgactt cttctcttcc atcaagttta aaagcttggc
1741 agcaactgga cttctatgag cagctgctct ttccgagtaa ctcttagtcg acatcctcct
1801 ttgattagtt tatggggcta gctgatgata cttcctcttg ctgttccagt ttacaaatga
1861 tttaaacgaa cttttacact tttacggtag aatttcttgt tggtgatgag ctcagattca
1921 atatgactgc ttaatcagtg aaaattaatg gaaacagcca gacacacagc ctttgaaaaa
1981 tgttaaaaac gatcacatat atgtagtcac gtggttagtg catagagtca gtcgatggtg
2041 gttagttccc tagccagagc cattcccctc ttgtttcgaa atgcgatgac tgaatgggga
2101 agaagctctt agtatttctg tagttttttt tgccgatttc tattgttgcc tgccccggac
2161 tttcttttac cttttgtagc tacaactatt aattataatg cataatagat ctagaagcgt
2221 attgtgatga gcatccttgg aaataagaaa ggaagtcaaa ggtacagtaa aaaaagagcc
2281 gtgacaggca cctaatttca gttttgattg aggtataatg agctcagaag accaagatgc
2341 tctaaaagta aagcagaatg gtctttccaa tcagaattcg gaagcatatc catcgtgctt
2401 aaaatgattg ggtccgcgtc cgactcacct agcaagttag gacgcctccg atttctttct
2461 gaaactgccg ctattaaagt atccccgtta atcctaggag aagtctcata cgatggagca

2521 cgttcggatt ttctcaaatc aatgaacaag aatcgagctt ttgaattgct tgatactttt
2581 tacgaggcag gtggaaattt cattgatgcc gcaaacaact gccaaaacga gcaatcagaa
2641 gaatggattg gtgaatggat acagtccaga aggttacgtg atcaaattgt cattgcaacc
2701 aagtttataa aaagcgataa aaagtataaa gcaggtgaaa gtaacactgc caactactgt
2761 ggtaatcaca agcgtagttt acatgtgagt gtgagggatt ctctccgcaa attgcaaact
2821 gattggattg atatacttta cgttcactgg tgggattata tgagttcaat cgaagaattt
2881 atggatagtt tgcatattct ggtccagcag ggcaaggtcc tctatttggg tgtatctgat
2941 acacctgctt gggttgtttc tgcggcaaat tactacgcta catcttatgg taaaactccc
3001 tttagtatct accaaggtaa atggaacgtg ttgaacagag attttgagcg tgatattatt
3061 ccaatggcta ggcatttcgg tatggccctc gccccatggg atgtcatggg aggtggaaga
3121 tttcagagta aaaaagcaat ggaggaacgg aggaagaatg gagagggtat tcgttctttc
3181 ggcgcgcctg ttgaatgtgg taacccaata gcatgatatg agtaatgctt tagtattgtt
3241 tcagagttgt ttcagtaatg ttttagacaa ggagaacata tagtagcaaa cctctaatcc
3301 ggtagtactt aagaaactac agtttctatg tacgaaagca gtaactatgt aattattaca
3361 tttacatgac atataggaag gtccaataaa cttactacat tatgacctat aagctagatc
3421 gtaattcatt acgtcaacag gttatgagcc ctcagagcaa tgcttctgag aacataatca
3481 atctatctag ccccaacaat tataaacagt ggctgtacgg tatcgagacc gctgctgaat
3541 atgctaacga atatatgaac gaattcgttc ataccggaga tatccaatca atgaaaaggg
3601 attacaatct cagcgcgaat gatgaaagct ttgtcaaaac cgtatttaac agtttcctgg
3661 taaagctcta caagaaaact atcgtgggtg aagctgcatg tgaaatgaac tggatatgtg
3721 atgattcgct tggaagggtc tctgcttatg atattttctc gcacttcgaa gaaaactata
3781 atgaagtcac tattggatcc aggcttactc ttatagagga cctaccaaat atatcctcca
3841 agcctgtaga tgaaattgct tcctttttga aaactctatt cacgatgctt gaagacaata
3901 gcgaagaaca ggacaaaaaa aaaagacgcg acaccaatat cgcgttgtta ttaatgacct
3961 tcttacccga gttaaaagaa tcattccacg agaaattcgg tgactctaag gctcttcagc
4021 tatcacaagt cattagattc tgtaaattaa atgcgtcatc gaattcatca tcttcggtct
4081 cagatgcatt ggttgcacaa gacagaagaa actatcaaaa gaaaggaaat aagggatgta
4141 tataatttac ggagctgatc atcgcataag caactgttct ctgcttaaac gaagaatacc
4201 agaagcacga atctttaaat tatatcctaa tgacaaaacg agtagatctt catctgctag
4261 tgtcgcgatt cctgactatg aaacgcaagg ccaaacagca ggacagataa caccgaagtc
4321 ctggctctgt atgttatctt cgactgttcc agctaccaaa tcctcagaat ggatttttga
4381 cacaggatgt acttcccaca tgtgccacga ccgttccatt ttttcatcat ttactagatc
4441 ctctaggaaa gactttgtca gaggagttgg cggttccata cccatcatgg gctccggaac
4501 tgtaaacatt ggcactgttc aattacatga cgtatcttac gttcctgatt taccagttaa
4561 cctgatatcc gtttgaaaac tatgcactaa atcaaactct tctgttacgt tcacaaaaga
4621 gggtgtcact gtgcaatcac ctgatgatgt ggtttctaca gctgggtatt cacaataaaa
4681 gactctggga agtacaaagc tcgccttgtc gcccaaggac atactcaaaa ggctggtatt
4741 gactatcagg aaacctttgc accagtcatt cgatatgact ctgttagatt atttctggcc
4801 cttgctagct gcctcaaact aatagtatat cagatggacg ttgacactgc gtttctaaac

4861 tcaaccatgg atgagccgat ctacgtaaaa caaccacccg ggtttgttaa cgagaggaat
4921 cccgactatg tatgggaact atacggcggt atgtatggac tcaaacaagc cccattacta
4981 tggaacgaac atatcaacaa tactcttaaa aagattggtt tctgtcgaca tgaaggcgaa
5041 catggcttat attttcgttc cacatctgat ggtcccatct acattgccgt atacgtagac
5101 gacttacttg ttgctgctcc ctctcctaaa atatatgaca gggttaagca agaattaacg
5161 aaattatact caatgaagga tctcggtaaa gttgacaaat tcctcggtct taacattcat
5221 caatcgtcaa acggagacat cactctctcc cttcaagact atattgctaa agctgcatct
5281 gaaagcgaaa taaacacatt taaacttaca cagactccgc tctgcaactc aaagcctctt
5341 ttcgaaacaa cttccccgca tctaaaagac atcactcctt atcagagcat agttggtcag
5401 cttctctttt gtgcaaatac tggtcgtccg gacatatcgt atccagtctc attactctcc
5461 aggttccttc gagaacctcg cgcaatccat ttggagtctg ctcggcgggt tctacggtac
5521 ctatatacca ccagaagtat gtgtctcaag tatcgttctg gatctcagtt ggcattaact
5581 gtatattgtg atgcatctca tggagctatt cacgatctac cacactctac tggggggtac
5641 gtgactctac ttgctggtgc tcccgttacg tggtcatcga agaagctcaa gggtgtgatt
5701 cctgtaccat ctactgaggc agaatacatt actgcaagtg aaactgtcat ggagatataa
5761 tggattcaaa acttgtttga acacttaggc cagccactta tctcatcaac atcatatgta
5821 gataataaac ctgctataaa actgtctaaa catcctgtat ttcacacgag agcaaaacac
5881 attgctttga gataccacaa gctaagaaat gcagtggcag caggcataat taccatagaa
5941 catgttatca caaagaaaca agttgctgac atatttacaa aaatccttcc agctgaatca
6001 tttaaaacac atagggctgt catgataagg gaaccagaaa ctacaaaata accatactca
6061 tgcgtattca gttatggggg gatgttaaat gtggtaacct aatagcatga tatgagtaat
6121 gctttagtat tgtttcagag ttgtttcagt aatgttttag acaaagaaaa catataatag
6181 taaacctgta atcaggtagt acttaagaaa ctatactttc tgtgtacaaa acactaacta
6241 tgtaattctt acatttacat aacatgtaga aaggtccaat aaacttacta tattatgaca
6301 tataagttag atcgtaattc actacgtcaa catatcccac gataaaaaaa ataccttgac
6361 tcaagtgccg aaacaggaaa taatgaaaga tttcatacaa ggttgtttct aaggtgctat
6421 atgtccctac ggcctcgttt agccccatct cgcatgcaat accgtaacat aggtatatac
6481 acacagagaa gtagtaattt tacgtgtctt atacttattt tatatagaaa tgtccgacac
6541 ttgcgctaca ctaaggcata taagatagat cttaattcct tgtattaaca tataaacaga
6601 caacataaca tagccagttt atcttactta tcttttcctg gaagagatgg cattatttgt
6661 gtgtatcaaa gaggcctttt tcaattttca ttttaatatt tttttaatat tcttatattg
6721 aacacatttt tctcccctag atttatgttt cctattgctc tccatatcac attatatcta
6781 tgaatgaata tgcattgctc ttctgatact atggtatgag aggccttggt aaagggcatc
6841 gataaccgaa agaatcatcc gcagagaagt agaatcttag agcatatcgt ctcgttgaga
6901 ccgtatagat tcgcaaaaat aacagaaact tgcgttaaca ttgaatttca aactcagttt
6961 tccataatta actgttaaaa aaggctacta taagggtgtc aattactgtt ctgtttcgca
7021 ggattaattc cgttatgtgc tcttgtaatt gagttagcgc ttcttatatc tcttcagtaa
7081 tgcagcagta ataaaacaca ttggcttcgc tttcgctttt tttccaaatc attatcgaag
7141 attctttgct tatgatatac tggtgctcga ggatgatgag aggacacaac tttttgaggc

7201 ttatcgttat cgtctgagtc acaacaacat acattaactg tccacgattc agggatgtgc
7261 tatgagttca attttcgctg tagtgctgcc attggcgctg ctactggtgt ctggaaaaac
7321 tccacagctt tcaatggaga tgtgactgca cgagtgagac attatctgta ttagcttcac
7381 aaccatactc tgttagtaag gagccatagt cgacatctgt ccagctcaat cgggtatgat
7441 atttgatatc cccaaaatac atggctgtac catgtaaaat gagcggtata tcgaagatca
7501 gtatatttca gcctatacag gatcctacta tttcaacaaa aatgatttca tcaaaattaa
7561 atcacgctgc aacattggtg aagaatctca aaaggttagt aacatcaata tatcgcggag
7621 tagctatcat tgtggcagtt aacttaaagg atatacagtt aattcaacaa gattttcaag
7681 tccatttttt ttgggcacgc tgaacacgct caatgtagta cttgttttgg aaatcactcg
7741 caagtttaat gggacgaatt atatcagtaa tgtgtaaatt ccgacacaat gttcggttag
7801 tagatttact gctggattat tgggttcacc cgtagactgt tgccgactcc aagaaaacta
7861 tgatctcaat acaggcataa agaattgttg tatctgtgcc catgtgatct gtaccttttt
7921 tgcggtacca aaatcgcatg ttgcactcac actacggaac acatcaactt gtaacgacaa
7981 caatacaaaa taccaggaga catatcactt tttttcgcct agaatgattt acaatggcat
8041 acacttagaa aagtgctatg acatgacgcc cttactagaa gttacatgat tctccttttt
8101 tttctttctt tctttctttt tcggtacaac ataattaatg cattcataca gagcgcattt
8161 ccagcatcat ttattggata gtaactcctc ccgcactttc aatatcaatg acgctgactc
8221 ttttactgac gctgagtctt taccgacgct gagtctttac cgacgctgag tctttaccga
8281 cgctgagtct ttaccgacgc ttgtgaaaat aaaaagtacg acacagctat atgcgtcgca
8341 agttcagatg tgggtctatc gagcatgttt atcctatcct gttatgtcat ttttatcgga
8401 agtaataaaa gtgtagaaat cacttgtttt tgcctttttt gtgattttgc tttcattttt
8461 ttttgccctt ttttcaccag cttggggatc aacatcctaa tttcttgtta cttcgattat
8521 gtctaaaaag tgccttgatt tgagagaaaa tattttcctc atctggcaag acaaccaaac
8581 catcttcagt gaattcttcg tgtgtcgcct tgtctattag ttccgggtct ctcatgaaca
8641 ttgtaaacgc aaagaacggt accgtacatg ccaagccaac cgtcattatt attcgttgaa
8701 catatctgta agattcaacc acagcatctc tttcgggtga gccccaagga taatcactaa
8761 tgaatgtata aggtgatgca tatgctattt ctgccaatgt atcgtttcca aggttcttgt
8821 agagttggtt aggcatggtt tgtgtccata tggcaccaga aacggaggca ccgatagcag
8881 cgcctagctt tgaaaaggta tactggatgc cggttattac agccatccta ctatgcgtag
8941 tcatggcttg cagtatgacg atcactgaat tgctgcatag gagaccactc aaacccatga
9001 taacagatgc agcgataaca ccttcatgag acccagatcc tcctctgtat ttgtaaaaaa
9061 ggcccatgca caccatccat gcagcacaac ctccgattac agaaagtttc agtttccttg
9121 tctttgccac caacaaactg tagaatggag atgcagtagc agcaacaaag tcaggaaggt
9181 ttactatccg agcagccgaa gtggacgatt ctttcatcga taccagcaaa acaggataca
9241 gatagtcaca cgaaatgaaa aaggtgaaaa aattgaaaaa agtaacacca aggggtgccc
9301 aaattccacg atcacttagt aatttgaacg gtagaagagg agatttggca aatttggcct
9361 cccaatataa aaaaatgaaa aataaacagc cacctgaaac taaagtggca attatttttg
9421 aattgtgcca cttctgtgat gtctcattag ccaacgtcaa agggacaagg atacacccta
9481 gcgacacagt tattaataat atgccaacaa tatcgagttt ccagaataga aacaccaaat

9541 tctcaaataa tccgcccgtt ctttcctttc tagcctgttc tttgagagat ctccactcag
9601 cagtctttga agatttgtac ttcatataaa gaataagaaa tataattggc aaggcagaga
9661 gtgggtaaat aaaagcccac attgcaatat tccaggacca gtttttctga ggatttgctg
9721 ctgtgataat attacctgaa atccatggta ttatgatata tggccaatat gaggcgtact
9781 ggtaaaacat tctccacttc aaggaggaga aatcagaaag tattaatgtc aggagcagat
9841 ttgttccgac gtatccacag ttatagaaaa ccgatcctgc tgcatacatt gtgagacggg
9901 tcgcctgtga ttgaatgatg gttcccatta tataaaaaat agttgcaact aaaaaaagcc
9961 ttagtcttcc gaagtggtca gagagtctgg agtagacaac ttgggatccg acacttacaa
10021 cagcattgat aacttggaca gttgaaagta aggagtgttc tgaatatgag ttcgtcgcat
10081 agcccgtata ggtcgatcta agtgtgtagt ctaaactaat tccaaaccca catacaaacg
10141 cggtacttat cagtagaatt ttatatttca aggaatcaaa ctgtgcagac ataatttcgt
10201 tttctttgag cttaaaaaag gtcgatgtca tagagtacgt gtcatcgttt gagtgacgtt
10261 ctcgctcaat aatctcacag gactgcctaa ttccactttt tttattactt gaggcaccaa
10321 caacactaga actcatccga taatttattt acttgattat tcctttttt ttttttctc
10381 tttccagtcc ataaaactat tttacttttc ttatttatgt aaatagtatt taagattcta
10441 cggtcgctac atctgctaga aatggaatta ctttaataca ttccatgttc tcatatatac
10501 ttttactgac acctttcgtc actatttttt gtttaacttt ttttttggaa gtactaaaat
10561 tatatgctcg taattattga gtatcaaaaa aaaaaaaaaa aaaaattgct gtgacacccc
10621 ttcaatgtgg tgtctataca cctggtgact ttaactttaa ctttgacctg tcgtcgtaca
10681 atcaacttag tttccgaaaa tccacgacga accagttata ccgaagactt ttgtacgaca
10741 agaatactgt accgtcgaaa ctatctcttt tcaaactgat gagtgaagct tgtgtttatg
10801 tctgttgaaa caaggaaaaa aatgcggtgc ctgtcgacag gatatgcgca caagtgcaaa
10861 atcatcaaaa aattgagtac ctagatccgg tggtgtagga cgagcaaagt tccttcataa
10921 acaaaccgta agggttactg atacacaatt tcctttttgt aaagagtatt tgagcaaatt
10981 agtcagcgat tgaaaacaac accctttttt ctatatatag taggtgaaag ttaacatgcg
11041 agtaaaagta gtgccacatt tcctttgcag caggtgcaaa ttggtacata tttaacagat
11101 agcgccgcac ctcatatgca gcgcgtacat gatacgagac gacaagatat gcaaaagata
11161 atagtgtcat cacaccttta tgagaagcga attttttttt ttttttggtt taaatatata
11221 tatatacata tatatagata tagatataca catgtataga tgtattctag ttatgcttat
11281 acctagaact gatcagacca gatcatacct ggtatagagc ttaatcgata gaaattcaaa
11341 aatatggagg aaactaagta ctcttcgcag caggagatag aagaagcatg tggttcagac
11401 gcttcattga atgctagagg tagcaatgat tctccaatgg gactttcctt gtacctctgc
11461 ctggcttcgt gaactcttgt actattcata actgcactgg atattttgat agtgggaact
11521 attattgacg tggtcgcaga acagttcgga aactactcca aaacaggttg gctcgttaca
11581 ggctacagtt taccaaatgc tattctgagt ctcatttggg gaagattcgc atctatcata
11641 ggtttccagc atagtctcat tttagcaata cttatttttg aagccggatc cctaattgct
11701 gcccttgcct cttcaatgaa tatgctcatt gtcggtagag ttgttgctag tgttggggga
11761 agcggacttc aaacgctttg ctttgttatt ggttgtacga tggttggtga aaggtcacgt
11821 ccattggtga tttccatcct aagttgtgca tttgctgtag ctgctatcgt tggtcctata

11881 atcggaggtg cctttacaac ccatgttacc tggaggtggt gcttctatat caatcttcct
11941 atcggtggtc ttgccattat tatgttttta ctcacatata aggccgagaa taagggtata
12001 cttcaacaaa ttaaagatgc tataggaaca atctcgagct ttacttttag taagttcaga
12061 caccaagtta attttaaaag acttatgaat ggcataatct tcaagtttga cttctttggt
12121 tttgccctct gctctgcagg gctggtcctt ttcctactgg ggctaacctt tggtggtaat
12181 aaatatagtt ggaactctgg ccaagtcatc gcatatttgg ttttgggtgt cttacttttt
12241 attttttcat tggtgtacga tttcttctta ttcgataaat tcaacccgga acctgataat
12301 atatcctaca ggcctctcct tctaagaaga ttggtagcaa aaccagccat aataataata
12361 aacatggtaa catttctatt atgtaccggt tacaatgggc aaatgatata ctctgtccag
12421 tttttccaac ttatatttgc gtcgagtgca tggaaagccg gtcttcactt gataccaatc
12481 gttattacca acgttattgc ggccattgca agtggtgtga ttaccaaaaa gctcggttta
12541 gttaaaccac tcttaatatt tggaggcgtt cttggggtaa ttggagcagg gcttatgaca
12601 cttatgacaa atacgtccac gaagtcaact caaattggtg ttttgctatt accggggttt
12661 tcccttggat ttgctctaca agcatcgctc atgagtgcac agcttcaaat taccaaagat
12721 cgtccagaag ctgctatgga ctttattgaa gtaacagctt tcaatacatt catgaagtca
12781 ttaggtacaa ctcttggtgg tgtgctttca accactgttt tttccgcctc ctttcacaac
12841 aaagtatcac gagctcatct agagccttac gaaggaaaaa cggttgatga catgattttg
12901 tatcgtcttc aaaactacga cggttctcat tcgactattg gaaacatttt aagcgactcc
12961 attaagaacg tattttggat ggatctaggg ttttatgcct taggattttt gttttgtagt
13021 ttttcatcca ataagaaatt aatcatacca aaaaaggacg agacaccaga agataattta
13081 gaagacaagt agcgcagtta ttttcttatt ttcattttat ttttttcgcc ttttatacag
13141 acttcaacac aatcagaatc aaataggtgt atcgcaatgg aatgtaattt cttaagtatt
13201 ctatatgtac ttaaaaccta ttaatatatg gatcaacaca gtatcttatg aatgggtttt
13261 tgattttttt atgttttttt aaaacattaa agttttcggc acggacttat ttggaattca
13321 aattattaat gaaagaacaa ttaactaatt aatgtactta gtatttggcc attattatcg
13381 atttcggggg ccaaatctaa ccaaattcaa cctacatttt ttcaaattga ttcaaacacc
13441 tttcacaata agatttttat atctagcgca catagaatga aatgtaaaca aagatttcag
13501 aaaaatcgtc attcaaactt gtattagacg agggacggag tgatttttgt gtttgttttt
13561 attaattgtg ggataggata gtagcaactc ttggaggaga gcattgtcag ttgtccagtc
13621 tctgaagtta agtagtaagt ttgcggagtc aaagggggat ggcttttgcc atttgtgaga
13681 gttgtgcggc agcatcttat tcaaatagag ctgtattctg aagacctctt gtagaacatc
13741 atccatacta aaaagtaaat cgtcctgtcc cattacgagc tgtagtagtg ctgtgaccct
13801 ctgtatattt acgttgccat gaagaaggta atgggcgata ttttgataca attcctgagt
13861 tgcatgttgg attgagttta cgaagggtcg ccagacggcc agaaacctcc aggcggagtt
13921 aacaactagt aatacggcat ccatgtttgc atcagcgccg agcctatacc agtcactgag
13981 tagacgtttt cttgctcttt ttatgtcctg acttcttttg acgagggggc attctctaga
14041 gacacaggca gttgcttcca gcaactgccg tacggccgtt ctcatgctgt cgaggatttt
14101 ttttgggacg atattgtcat tatagggcag tgtgtgactt atgaattgtt gtagaaggac
14161 gtctgtgatg ttggagatat gtattttgtt aactcttctt gagacaattt ggccctggat

14221 agcgaagcgt gcggttacaa ataggtcgtc ttgttcaaga aggtaggcga ggacattatc
14281 tatcagtaca aacatcttag tagtgtctga ggagagggtt gattgtttat gtatttttgc
14341 gaaatatata tatatatata ttctacacag atatatacat atttgttttt cgggctcatt
14401 ctttcttctt tgccagaggc tcaccgctca agaggtccgc taattctgga gcgattgtta
14461 ttgttttttc ttttcttctt ctattcgaaa cccagttttt gatttgaatg cgagataaac
14521 tggtattctt cattagattc tctaggccct tggtatctag atatgggttc tcgatgttct
14581 ttgcaaacca actttctagt attcggacat tttcttttgt aaaccggtgt cctctgtaag
14641 gtttagtact tttgtttatc atatcttgag ttaccacatt aaataccaac ccatccgccg
14701 atttattttt ctgtgtaagt tgataattac ttctatcgtt ttctatgctg cgcatttctt
14761 tgagtaatac agtaatggta gtagtgagtt gagatgttgt ttgcaacaac ttcttctcct
14821 catcactaat cttacggttt ttgttggccc tagataagaa tcctaatata tcccttaatt
14881 caacttcttc ttctgttgtt acactctctg gtaacttagg taaattacag caaatagaaa
14941 agagcttttt atttatgtct agtatgctgg atttaaactc atctgtgatt tgtggattta
15001 aaaggtcttt tatgggtatt ttattcattt tttcttgctt atcttccttt ttttcttgcc
15061 cacttctaag ctgatttcaa tctctccttt atatatattt ttaagttcca acattttatg
15121 tttcaaaaca ttaatgatgt ctgggttttg tttgggatgc aatttattgc ttcccaatgt
15181 agaaaagtac atcatatgaa acaacttaaa ctcttaacta cttcttttaa ccttcacttt
15241 ttatgaaatg tatcaaccat atataataac ttaatagacg acattcacaa tatgtttact
15301 tcgaagcctg ctttcaaaat taagaacaaa gcatccaaat catacagaaa cacagcggtt
15361 tcaaaaaagc tgaaagaaaa acgtctagct gagcatgtga ggccaagctg cttcaatatt
15421 attcgaccac tcaagaaaga tatccagatt cctgttcctt cctctcgatt tttaaataaa
15481 atccaaattc acaggatagc gtctggaagt caaaatactc agtttcgaca gttcaataag
15541 acatctataa aatcttcaaa gaaatattta aactcattta tggcttttag agcatattac
15601 tcacagtttg gctccggtgt aaaacaaaat gtcttgtctt ctctgctcgc tgaagaatgg
15661 cacgcggaca aaatgcagca cggaatatgg gactacttcg cgcaacagta taattttata
15721 aaccctggtt ttggttttgt agagtggttg acgaataatt atgctgaagt acgtggtgac
15781 ggatattggg aagatgtgtt tgtacatttg gccttataga gtgtggtcgt ggcggaggtt
15841 gtttatcttt cgagtactga atgttgtcag tatagctatc ctatttgaaa ctccccatcg
15901 tcttgctctt gttcccaatg tttgtttata cactcatatg gctataccct tatctacttg
15961 cctcttttgt ttatgtctat gtatttgtat aaaatatgat attactcaga ctcaagcaaa
16021 caatcaaaga aatctttcac tgctcttttc tgtgttccat ttagttttta gtacgattgc
16081 attgtctata tactgtattt accaaatctt aattttagtc aaacattaat gtttactttc
16141 gtatttaagg atgtgtatac taagccttgg gaaaatgcgt gttccttgtg ccatccgaaa
16201 gctacttcat aaaaaagata ctgttcattt tcaagcccgc atagttggat caaatattaa
16261 ctatatatca acatggtcca aagatcgtaa gtttggtcta atgttgtttt agcaaaaaaa
16321 aatactatac ctttcattaa cgtttattga atttcttgaa aaaaaaaccc taaattatta
16381 taaaattata gcaggatagt tcttagtgga agatattaag caaactattt tctgaattgc
16441 tatgcgaaat ttctctaatg ccagctgagt aactaactct catggtacat aaaaatacgc
16501 atcttaaaaa atttgatatt tcatgattta tttatagaaa taacgatgct tattgtgctt

16561 tgttgggtgt ttgacttcta tgttaactta cttcaacatg aaagcccgac gtttgcaatg
16621 atatcctctt tttcacccga ttattttcat ttattagtaa gtggggttct gggtgaaaaa
16681 atcaaaacaa aaatatgaat ttgtaatgat ttttatattt tcgccggagg tgctggaaat
16741 ggcaaacgaa aatactatga cataaaagct gggcacacta tgtacgttct ttttaatttt
16801 ttatcaacat gagaaaaatt catgaacact gtaccttata ttatctaata aatgaagtaa
16861 gctttgcatc tccgtaacag ctgcatatat ataattaagc gggagctttc tttcgatctc
16921 tttcattcca tatgtttatt atgtactgat gaaaagcatt ttactaagtt gaggtctctg
16981 taagtatatt gtgaccatta tgctaaaata ctggggtctt catttgtgtc aaattctacg
17041 taatagttcc cctttttctt acatttttat actaaagagt tacggtattt atacaaacga
17101 ggggctttac tgtaatatat gattaagatt caagctgaaa cattctaagt tagaggtata
17161 aaatgattat tgatgaaaga gctgcaaatg tcggcaaaat ataaatacct tagatgttac
17221 cagctgggaa gattattacg gaacacattc ttataaatct ataggaatac ataggattgt
17281 atttctattg acaaagtaca acctatattt tatgttatgt tatgttatgt ttgctatgac
17341 tatagttgct tttcttcact tagtaaggat taactaactc tcgagctagt aaatttgacg
17401 tgagaagcat aattatatta ctaatagaac gatatctggt gtgttttatt acagactgca
17461 ctgagtgtta gaggtagaaa ttataagtta aaaagcagta gtttatgctt tatgctcgag
17521 tatcaagtga atttgaacag gctagtgctt cattggtact tctttcatgg ataattttga
17581 gcaaatttct gcagcatgtc cccctttata caaattctgt gcattgccgg cctagaaata
17641 tgtcaacgtt ttggatatgt tgatgcttac ttcgagaaat cttacactaa tacttctgga
17701 aaaaatcaat actagcaaaa tagtgatata tgagtaaaat gtatgtagta catgtatgaa
17761 aattatcaag ggcaaattga tgcttcaacg aaaaagttat tggattttca agcacttttt
17821 aaattcacaa tatttttttc tggaaatatg aaattgtcag cgacttctat tacaggagtg
17881 tctttttttc tcatgctatc caacgcttct tccaagtcct ttatagtatt agaggagccg
17941 ccacaagcaa taattatcac aatgtcatcc gcagctaatt ttgagcccag tgcattttct
18001 aggatcttag tgttgtaacc caaatgcaat gcggcgccac atgccggttc aatcaccata
18061 ttgaattgat gtgtatattt aagacaggtt tcaataacat ctttgtcctc tattacaacg
18121 gatctggtgt tgtatttgcg agcgtattcg aaagtttgat tagagatcac ggccgttcct
18181 agagaagtag caatacttgt tatcttcttg aattgaactg gttggcctat tttcaaagaa
18241 gtattgaaaa catgacatcc attcgtttcc acccccacaa tagggatcct atcagctaaa
18301 ccataccttt ccaaaccttg aataatacca ttgtataaac cacctccacc aacgctgcat
18361 actatgcctt taaccttatt cacggaaata tgttgcgatt tcaaatcttg tactatttca
18421 tctatcatag atgaatgtcc ttcccaaata tccggattat cgaagggatg aacataaatg
18481 ggctcaatga cctgagagtc tattttattc atgacatttg tttttaaaaa agtatctgct
18541 tctttccagt aggcaccact cacgataacc tgggcaccgg tgttcctgat tttatctacc
18601 attctcttct ttgtcgctgt aggaaccacg actgtacatg gtagagacag tctttgacat
18661 gctgttgcag cagcaaaacc ggcattaccg ccagaactag cgaaaacctg aggagatctt
18721 ttaccgtcct tttgaattcg aatggcactt ttcatgatga gattaccgat tcctctactt
18781 ttgaaggagc cacttggttg aaggcattca tatttcaaga aaaattgtgc agaagccttt
18841 ccggggaaga attgacgtaa taatggtgtt ttattgtaga ctatcgacat ctcgctggtt

18901 aattttcctg tctcttgtct atccagcact taaaaaagag aaaactgtat aaacattttc
18961 cttttatttg gcgatattta tatacctgtg ttcctcctca tattgtccct tttatcttat
19021 ctcatcgcaa acaatttgat aaagctttgg cactatttaa tgacactcag tggaattaca
19081 tttccgctgg ggcttaacag gagccgccca tgcggaatca tgtctccgcg cagaggactc
19141 attagtttcc gttcgcggtg attagtactc gcgaaaagag aaaaagaaaa atagtaggaa
19201 catttcgcaa ttgatcaaaa tgtaaacagt gcagtaaaga atgggaccat aagatagata
19261 agaaacagct cgcataagga aacaaggaca catcgattaa tggcaaccca agccctagag
19321 gatatcacgg agaggctttt aataaggtcg caagaggcta tcttgcaatt agatctgtgg
19381 atacaacgtc agcagagatc atcaatatgc cagacaacag atcaagagtc attggataag
19441 ttatcccaac agtacaacca gtatatgtct caactgaact ctttgtatgt tagatcggaa
19501 tctgttcgag acaagttgag caaggaacaa caacgcagat tgatcacaga ggataatgag
19561 catcaacgca tagaagactt ggttcgtgaa ttccaggata tcactttgag gttgaacgag
19621 ctggccaccg tcccaaatga agcgcctaat gattctccac aatcgcaatc caccagaagt
19681 agtttagggt catttcaacc tcgaccattg aaaataattg agaggcaacg tctgtgtatg
19741 gtaactccat cgaaaccacc aaaaaaatcg gtaggcttta accccatcaa tgaagtcgat
19801 tgtccttcga aaactaactc tttaccgtgc tcacctaaaa aacaacctgc aagaaatcgc
19861 actttacgtg cagccaaatc acatgatact ggcttgaaca aaagtaagaa accgtcttcc
19921 tcggatacgt atgagtcttt cttcaaaaat agacaaagac tttcgttgac cttctttgat
19981 gaaatggatg atgaagattt tgattctgat caagatacta tcattctacc aaacataagt
20041 accccctccac atgttggtgt gaccgcaaag ggtgctgaat tcgaaccttt aaggagatat
20101 aactctcacg aaagtatact atctaacaaa ccagcacctt ctaagtcgct caatctggga
20161 agtttctccg cctcattctt caggccatct aatccgacgt ttggaacttc gatatcaaat
20221 gtccaagtga actgtcatcc aactgttgca gcgacaatgg ctcctagtcg taatggtcct
20281 cgtatttcaa gttctaaggc gttgttatca tcattcattg cacgatcaga tacacatacg
20341 gtaaaagaga acaacacaaa tcttaagcat gcatctttta tggataagtt taattcatcg
20401 ttaagcacaa tatcagaatc ttttcaaagt aagaggggga gaaagaataa gggcatgaat
20461 gaagaacgaa tatcaaatca taatgtagca caggaacaaa aaaataatat ggatataagc
20521 gtctctatag aagaattgca agatgctttg aatacagaac tgctgtttta attaatgcct
20581 cctttaccta acctactctt cttttgctcc aaatgtttat tctttttttt tttattcaga
20641 atatcttcaa gttttttttgt atctttttgt attattatta aacttcttta taatatttgt
20701 cttataaaat cctatcataa catgactatg gcttggccta gactcgggtg ccatcttttt
20761 taatgcgact acttcaagac agcttctgga gttcaatcaa cttcttcgga aaagataaaa
20821 aaccactaat tatcaaagct atcttgtccg ctttcaaaaa gtttattcga ttttgttttc
20881 gttttcattt tcattttttt attatgatgg tcatggtggt agcggtggct cccttcagtt
20941 ttcctctttg gtgctattaa ttttctcgtt ttccccatat atgtgataga agcttttgaa
21001 ctccccactg tcttgtaaga ctttagaatc ttgactgtct tgttaccttc tttaaactta
21061 tcatttatgt ctgttctgga cgcaaaggat ttgattattg atggatgttt gaaaacgcca
21121 actagtgatg tatctccatt ttccacctca tcttcatcct ctgggcgctt ttctaagtca
21181 attattgtgt ttactgtttt tgaagtggtc gaatcagttt gagaatttgt gaatgacttt

21241 atcgaactat tctgttttag ggtatataaa tcttcaattg cttcatcatt gccatgttgt
21301 atccttgata gctccttatc cgtttcaaaa tcctcataat tattactttt taggaaggat
21361 aagcttttct gcacaaagtc ttcagatatg ataacctttt tgttcttttg atctacaggt
21421 ttctgttcat tatttcccgt attatcacct acattaatgc tattgtcctg agtatctagg
21481 tcagtagccg tacttgtaat gtctaggtta tattcctctt cagccccgaa ggggttttta
21541 tattcgataa tatcttccac catactttca aaaaatgcgc tcgactttgg gtttttaaca
21601 agctttgcat catcgccaat ttctaatctc ctttttctca tcaattctct ccttttgaga
21661 cgatattgtt gtagtacgtc gtcttcatca tcatcactta gttctagctc taagctattt
21721 tttgctcttt tattacgaaa tcctccattt ttaatgtcat aaagaatttt gtttatcatt
21781 ttgatatcca tttccttatt ttctgcggca agcatttctc tgatttcatg aggattgaaa
21841 ttgttcttgg aataatcatc gatcattttt tctaggtcag aatcatagtc gtcagatcct
21901 tctccatctg caccaccgat accatgccat tcatcctcgg attcttcggc ttccatttca
21961 aagaagtttg tgacacctct tttcttcaac tcttttatct tagcctcgtg ctccttttct
22021 ttctgtttta gttgcaattt ctctttgtca atcagctgac ggcgaattgc ttcatcagct
22081 tcttcattga cgtatacatc atctttgact tcattttcga ttgtatttct ttcggccatc
22141 acctcttcaa tgttttgtgt atctagaaca ttggtttctt gaaatttgtc ggtatcctct
22201 ccgatattat caccgtaatg gcccagatta atggcgatcc ttttcggtag actttcatca
22261 gctttctctt tgggtttggc ttcgacttcg gtatcagagt ctgattcgtt aatgatatgt
22321 ttcacgtggt gggatttttt ttgctttaga ataatttcat catcttcctt ttcactatcg
22381 gattctctac gattgtcatt ttcactacca gatgattcat aatcggcaat ttcattacca
22441 ctcaatgcaa atccagaaga ctctgaccct gagtcagaac cggaatcatg ggcgttcaat
22501 tgaaagtcat tttcttcaag tttttccctt ctcttctctt tttgtctgat tcttttattt
22561 ctaagaattt cttgttctag caagttctct actatttcct tttctttggc catatcctcc
22621 agcttcagac ctttcgtttc aatcacttct ttctgatgat ccagaatttg tttcctgcta
22681 gctttcctta aggtatttaa tagtacatta tgatcaactt ttgggtcttt acttttattc
22741 ggcctttgag acaacttttg attttgttta gagagtcttg ccttcaggtt taatatagtc
22801 gcctttgata actgagagat tggcaaaacg ctttcatctt tggataattt aatgctatcc
22861 atatcatcat ctccatattc gtcactgtcg ctgtcgaggt ccagtgttat gcatttcgag
22921 gagtcaattt cccgtttcaa attattagca taggagctca atagtggaac ccttttctca
22981 tttattcttg tctcatcagt tagatttaat tcaacacttt tgttctcatg cagttcatcg
23041 tcattctgta ctggttttgg atgtgcattt tccagcttta tatcatcgtc ctcatttgag
23101 gacgaatcgt caaaatcagc aagaaaagac tctttagaaa aaaatttttt ggttggtatt
23161 ggcttttgcg gttttttgta ttcgacattt ctagcttttt cccttttaga atcttctgaa
23221 gccaattcgc tttgtatttc ccgaattttt aatgtgtgtg gtacttcatc ttgaagtgtt
23281 tgtgcgactg tatctatttg ggttttctgt tcctgatgcg tagcctgtgg aataccaatt
23341 gcttgtgtct tatcctcatc atgcgcagtc tgtggtggta catccatttg ggtgtcagga
23401 ccacgatttt gaatggactg aaataaagga ctagttccaa taagttccgt tggaattata
23461 ggaattttcg gcgacagagc attagaatga ttgctggaag tcgtaattat ttgggattgt
23521 ggttcaattg attttatagg ctgagtttgt gaggttgcac cgctgctatc tatacgctgt

23581 gtagcccctg gccgtcctgt cgtgagaggt gttttcagcg cttgagaagt ttcgttgacg
23641 agaccatctt ctttgattaa ttgggtgggt ttcccgtcat taacggaatg gataggtacg
23701 ttaaacacct tgtcttgttg gatgctgaca ggtattcttt gcgtctgcgt gaatgaagaa
23761 gagacatttt tctcactgta actattatct ttaacttcgg atttctccag ctcttcacca
23821 ccatcgtaaa gattagagat taattgagtt ggtaacgaat tctcactcct atcttttcca
23881 ttattgtgat tctgttcggg tgctttcttg ccttctaatc tgttcttaac acgatttaat
23941 gtggcattgg caaataaaaa cccgttccca gtcaactctg gcggattgtc tatgtcattg
24001 ggcccatttc cattagtatt atcgttctca tctagtatgg gaacagcaac ttttttgtag
24061 gtggtagttc tcttctttgc agtcaacgag gacaaagcat gcaaggcatc atccatcact
24121 aaaatatttg gtgataagtt caaaagcgaa taacgaactt ccttagttgt ttgtctgttc
24181 ttggtttttt tccaacataa tgtgtttgaa atgctattga agcaaacgcg ttttacgcat
24241 tctatctctt catgacgcgt cgaagaagga taaggaaaat ttcaagtatt gatgtatatc
24301 tctctgcata tgagagccag tagagttggt gtcctagtaa gaatttcttt cactttacaa
24361 acatatctaa gtaaacgaaa tgtgtgtgtg tttctatata tatatagaca tatatgaagg
24421 attccgtagc ggtgtaaact aattttggtt tggcttgtat gcttcttctt ccggagctat
24481 gaaatctttt gctctctctt cctgcctttc gatttctctt tccatttgct tttccttttg
24541 ctcatttaat ttcttcattt gcttttctct cttgcttaag aaatactcac cactttcaat
24601 ttccaaatca acctttctag gcaattgggc aggaggaaat ggagtataga ccttcttttc
24661 gacgtttctg atcttcttgg gtttctttct ggccacattc ctcttcttaa acatgggcaa
24721 gaatcttgac caatcttcat tggctaactc tggccttttt gccaattctc tttttatcat
24781 taattccttg atatgataga tagggtgaat atttttcata caatcttcta ctactcgacg
24841 gacctccttc aagcccttga agggacccat ggcacttact gtgttacctt gtactagaat
24901 gtaacattta gttagaagtt ccaaagcctt taaagtatta ccgttagggc ctacaagacg
24961 ttgtcttctc ttgacaaacc tttctttgtt agtaacgaaa ttaccaattt taataacgtc
25021 gcatgccatg tcatcttgta ggatcttaac ggcttgcggg aaaggaacgg atctcgccaa
25081 taatttgatc aaatctctgg ctttcaaaat gatagcggga tcgtatgttt ttctagttgt
25141 ttttactgtc atagaacctt cgactaaatc tagaacacac gctatgttgt gtttgtctag
25201 agcccttgtt acatcattcc aaatcgtctt caagtaactt tctctgtatt taggaaacaa
25261 agtcataaaa ctggactctt cagcaaaagg ttgaccggat gcgttatcct cttccttaaa
25321 ctcctctatc ttccatttat caatatcatc cgtatcccaa ggtttatctc tgttatgtgt
25381 agacaccatc gtttgccaat ttggatattt gtgtgaccct tttgtttgct gtctacttta
25441 caatagttaa cctcatcatc tcttttttttt gaaaattttc atatctcatc gctaaaagaa
25501 ttagaaatat aaggaaaaaa aaattttcgt tttcagatgt gcaagcctgc tataataagg
25561 tacaataact caagggcatt tagcaaggaa aaaatgggca agtgtagcat gaaaaagaaa
25621 ggtgtgggca agaatgttgg tgttggcaag aaagtacaaa aaaagaggtc gatcagcacc
25681 gctgaaagga agagaacaaa gttacaagtg gaaaagttaa acaaaagtag tgaaacaatg
25741 ataccgacgc tgctgcggga ggcaagtaca caagagccag ctaaactgaa agctgagact
25801 actttgaaag ccgaggagct gatcaaggac caggaaaagg actccaaggt acgagagcaa
25861 attcggacag aaaaatcaaa aacaaacgac agcatgctga agcagatcga aatgatatcc

25921 ggcttttcct tatagggaat agtggtgaaa gttacgtaaa tatatacata tagagcggaa
25981 tggtactgca tatttttatt attttttttt ttgaatatat ataaagcatc gtcgcttaag
26041 actaaacctt cgaggatctg aggccagcgc tggatctaaa gatggcgtcg ccctcggcgt
26101 agcctcttcc aacaccaaat ccaatgccta gccatacagg aaatgcacgg cgcttgaaga
26161 ataatactga ggtgaacaca cccacgccaa aacccatggc agtcttgacc agcatgttgg
26221 acaggacaat atcccactta gtgtcgagga tcgtggacac ggagctgcca ttcttgttag
26281 aatccttgga gggagtgctt ttcgctggtt gttgtgtttg tgcttgttcg gacatggtat
26341 attttgtctt tttccgtgtt tattccctct cgtagcaaag ttaatctgcc aattgacagt
26401 agtttaatat atggtattat cgttgtttga agccctttaa aaggaaaaaa cgggttaggg
26461 ccacccggcg cgaagtaata gctgctgatt ggtcagaata gtaaaacggc ttgaattgat
26521 agtatatcgt actgtatata ccttatattt gggctaataa tcatctccca aatgctccag
26581 tatacatgtc acctagctat gtatgaggtt gactacctta tatgcagcta ctacacaata
26641 actgtagtat cgcgctgcac gagcgtgacg gcggttaaat ctggattttt tgccaccttt
26701 tttatacgaa aataagaact aggtaaatag gaaggaatgg ttggatttga gtttaaaaat
26761 agagggaata cttggagatg cgattgttgc tgtgcaagaa ttggtttgcg tcacctgtaa
26821 tctcaccact actgtatacc cgctccttat attcaatggc taacactact agtttcccta
26881 ttgctcccca ggccccgcct aattggtcgt tcactcccag cgatattagt gggaaaacca
26941 acgaaatcat caacaacagc aacaatttct atgattctat gagtaaggta gagagccctt
27001 ccgtgagtaa ttttgtggag cctttcatga agtttgaaaa tgaattgggc ccaataatta
27061 accaattaac tttcttacag catgtgtcgt ctgataaaga aattagggac gcatctgtga
27121 actcctcaat gaaactggat gagttgaaca tcgatctatc tctgcgtcac gacatctttt
27181 tgcaattcgc ccgcgtctgg caggatgttc aatcgaaggc agattctgtg gaaagagaaa
27241 ctttcaaata cgttgagaag tcttacaagg actacattca ttctggtttg gaacttgacg
27301 agggaaaccg attgaaaatc aaagagatca aaaagaagat ctccgttaac tctattaatt
27361 tttcgaagaa tctgggagaa caaaaggaat acatcacttt caccaaagaa caattggaag
27421 gtgtgccgga ttctattttg acgcagttcg agacaataaa atctgacaaa gatagcaatg
27481 aaaccttgta taaagtcacc ttcaaatatc cggacatttt tcccgtgatg aaattggcat
27541 cctcagctca gactagaaag caggcctttt tggccgacca aaataaggtc cctgaaaatg
27601 aagctatact gttggataca ttgaagctgc gtgacgaatt ggcctcgtta ttgggctatg
27661 acacgtatgc gaactacaac ctgtatgata aaatggctga agatagcact acggtaatga
27721 actttttgaa tgatttgaag gacaagctaa ttccgctggg cagaaaggaa ctacaggtct
27781 tgcaagatat gaaagccgaa gatgttaaga aacttaacca gggtgcagat ccaaactact
27841 acatttggga ccaccgttac tacgataaca aatatttgtt agaaaacttc aatgtggacc
27901 tagaaaagat ttctgaatat tttccactag aggctacgat tactggtatg ctggaaatat
27961 acgaaacatt gtttaatttg aagtttatcg agacgaaaga ttctcaaaac aaatctgttt
28021 ggcatgacga cgtcaaacaa atcgccgttt ggaatatgga tgatccaaag tctccaaact
28081 ttgttggttg gatttatttc gatttacatc ctcgtgatgg taaatatggc cacgctgcca
28141 attttggttt atcgtcatca ttcatgattg atgacaccac aagatcgtat ccggttactg
28201 cgttggtttg caatttctcc aaatctacga aggataaacc ttctctactg aagcataacg

28261 aaatagtgac ctttttccat gaattgggcc atggtatcca tgacctggtg ggacaaaaca
28321 aggaatcgag gtttaatggc cccggatctg ttccatggga ttttgtggag gcaccttccc
28381 aaatgttaga attttggact tggaataaga atgaattaat caacctctca tcacattaca
28441 aaacgggcga aaaaattcca gaatctttga tcaattcatt gatcaaaact aaacacgtaa
28501 atggtgcttt attcactcta agacaattac attttgggtt atttgatatg aaagtacata
28561 cttgtaaaga cttgcaaaac ctgtcaattt gcgatacctg gaaccaattg agacaggata
28621 tttctttgat ttctaatggt ggtacgttat ccaagggtta tgattcattt ggccatataa
28681 tgtcagactc ttactctgcc ggttattacg gttatctatg ggcggaagtc tttgcaactg
28741 atatgtatca caccaaattc gctaaggatc cgttaaatgc caagaatggg atacaatacc
28801 gtgatattgt gttggctcgt ggtggccttt atgatattaa tgataatctg aaagaatttt
28861 tgggtaggga accttctaag gatgctttct tgaaggagct gggcttacag aactaaattc
28921 gttcaagcat aaaattatac aattattcta caaagtgtta ttatgttaga taatctttta
28981 aataacctcg tccctggata gattaaaacc attaaatagc tcaaatcttg gagtaaagca
29041 gcagtaatat gtctgatatg ttcgttatat tttttcttga agtttatcaa cttatccttt
29101 actagaccat ttccctcccc atccaatcgt tgatcgattt taaatttcac cttgcataat
29161 tttattaatt tccatacaca ttttatcact gaaatgcacg agtaacacag ccaaactttc
29221 cgtaatagcc cacttacata taacttatcg ttgaccacgc ctaggttttc caatactgcg
29281 aagaaatcaa gcagcgaact tacaccactc catatattct cccatatatc ttttgcactg
29341 atttcagccg tatttttctt cgaaaccatt caactttctt ttcttcttcc tttactttta
29401 ttcatctcct ctttacttct tacctcagag tcgaacattg aaatcatttc ccgcgaattt
29461 ttcgattata attacccttt ctacttaaca agaagcaata agagcaaagg ggcgaaatgc
29521 ttgtcccata ctttggcttt tggatgtgac attaggagcc ttatctcctc ttaatgcctt
29581 aagtcaaaaa atagtttttg ggatatatct tgtgatcaaa tgctttgcaa aagggtaaaa
29641 cagcgtttac aataggatag ccaaaagtaa aaaaaaaaaa aaaaaaaaaa aagaaggaag
29701 ttagtatcga gctcaaacaa ttttagtttt gctggacata ttttcagttt cattactgta
29761 tatttgcagc ttggtttttg tcgctctgta gtcttgtttc acacttcgtc atttccgagt
29821 ttcatcttgc gacataactt gtttcaactt ttcaagtgcg actttttgtt aaatattatg
29881 acatagaaga aagtagaaca gccattataa aacaaaagag aagtgagaat actattaaga
29941 tggcattcca agatccaact tacgaccaga ataaaagcag acacatcaac aacagtcact
30001 tgcaagggcc aaaccaggaa acaatagaaa tgaaatctaa acacgtatca ttcaaaccct
30061 ctagagactt ccatacaaac gattactcga ataactacat tcatgggaag tcgctaccgc
30121 aacagcatgt tactaatatt gagaataggg ttgatggcta tccaaaactt cagaaattat
30181 ttcaggcgaa agctaaacaa ataaatcaat ttgccactac gccatttggg tgtaaaatcg
30241 gaatagattc cattgttcca acgttgaatc actggataca gaacgaaaat ttgactttcg
30301 acgtggtgat gattggctgc ttaacagaaa atcagtttat ttacccaatt ttaacccaat
30361 tgccattgga tagattgatc tccaaaccag gtttcctgtt catctgggcc aattctcaaa
30421 aaatcaatga acttactaaa cttttgaata atgaaatatg ggctaaaaag tttagaagaa
30481 gtgaagaatt ggtttttgtt cctattgaca agaaatcacc gttttatcca ggtttagatc
30541 aggacgatga aacgttgatg gaaaaaatgc aatggcactg ttggatgtgt atcacaggta

30601 cagtaaggag gtctacagat ggacatctta ttcattgtaa cgtagacact gacttgagta
30661 tcgaaacgaa ggacaccact aatggtgctg taccatccca tttgtatcgt attgcagaaa
30721 acttctctac cgcgactaga cgattacata ttattcctgc aaggactggt tacgagacac
30781 ccgtcaaagt aagacctggc tgggttatag tgagcccaga tgttatgttg gataacttct
30841 cacccaagag atataaagaa gagatagcta atttaggttc gaatatccca ttaaaaaatg
30901 agattgagct gttaagacca agaagtccag tacaaaaagc acaataaagc atcaacacat
30961 aaatccttgc ttagctcaat taaatatact agtaaataag tatataaaca ataattttgc
31021 attttatttt actggaaact cttctcaaaa ccaaattgcg caaggattga ttcagtacaa
31081 ttatgcaaac tcgaaaagta aataaacaaa aagatacagt tgagtgaaag gtaaataaac
31141 attattctac ttgattttcc tctgtttttt ccatttttct tttttgtctt ttcgtgttct
31201 agcgtgttca ctttttcagc catgtaatta cttaaatagg aagtttacac gatgttctcg
31261 cataaaaagc tggaagtgca gaacaaagag gaagaaaata ggaacagaga atttgtttgt
31321 ttatttgttt gtttgcttgt ttgtttgttt gttttgattt gaagtcgagg tacttattcg
31381 aaataattat tttgcccagg gttttggata gttttttgtt ttccaattag atttttagaa
31441 atattttatc atagtttttt gtttatttat ttgtttattt atttgttgat ttgtttattt
31501 atttgttggt ttgtttattt aagtttgtaa atgtttaata aacaaaacaa tatttgcaag
31561 ccattgaaaa ctttgtaaat ttgtttgaga atttacgaaa caatgaaaat taaggactct
31621 ttgatggtga acgaattact actttgcagc aggtaattct ctaaaactca gaaacaaggc
31681 atgtattttt cactaggaaa aagaaggaaa atgggactac agtattaggg aaacaggatt
31741 aaaattcgta ggtaataata acccatactt tcattgtagc agtactttcg ttaataaaat
31801 ggatattgtt gtacttcggt agatgttata ttctaaagag ttaaattatc caattccaaa
31861 ttctagggac ggtttccaaa gatttgtatt gtccagtgac ataattttca ctaaaacatt
31921 gatctgttta ataggtttgc tataaatgca caataatttt tcacccatga tacccaattt
31981 caagagaagc aattgctaca tataattatt taggctttac tatctactac tcattgactg
32041 tgccctttta cacaattata acaaatatgt caaagcagat gccatgaact ttgtatctga
32101 atttttgatt tccttttaat tctaattgca gacgacgtaa atatagttct gaatttcaaa
32161 gtcactgtta attaattgtt ctaattgttt ggttttttta atataaatca ctagtgctta
32221 agttctgttg acgcacacag tacctatctt tgattccttc gtgcaaacag tattccggca
32281 cgttaattga taccaaaaag atttcaaaat cagatgcttc aagactaatt gtatacttta
32341 tgtgacttct ggagtttaat ttgttgtatt gatgagataa tattgtctat tatattgcca
32401 ataacaacca gcgccaggtg tactaagtcg agatgacaga aaatttataa taagtgtaaa
32461 tacaataaat tttgtaaata aataactact tcaatagaaa tctcaaagtc gaagaaacac
32521 agaacacaga atgaaactgg actaaatgat tttcaataaa ggaaaagcac gaaaaatttt
32581 ttgaactttt ttgggacagc gtggatcgtg taattactgt attgggaaat ttacacgatt
32641 ctcgacaaat aaagaaaaag gcgcgtttaa agacttacaa gcacaccttt gaattatttt
32701 tgttctctat gattatcaaa tgtgctagtg tttgcaagat ggtgcttttt tttctttaga
32761 ttgccacgtt tctgctcctc tttgtggttt tctttttttt tttcaatctt acttataaaa
32821 tgaatatttc aattgatgaa tagctatata ttggatacaa aaattagcat ttaatcgaaa
32881 actgcagcat gtatttaatc gaaaactaca gcatgtagct atgatacgac agaagatttt

32941 tgtttttata gttaagtcaa gaagaaattc tatttgtcca gcaatccggc gcaaagaaga
33001 ctactaaagg gtattgaaaa atttctagaa ccagaaataa tcgaaagcat tacttttttt
33061 ggctctataa aggagttctg gacagcttga acttccgaag tctgtctgtt tagctcgtta
33121 agataaatct caaaagagta gcaagcaggt gggagggata atttggatta aagtgtatat
33181 attgcgtata tatagccacg taaacctcac actattgagt ttaaacattt ttcaccttac
33241 gttattattc ttttgtctaa ttttaatctc ctccagtata ctaaaaccgc aaaagcgggt
33301 tacccagctt gtttcttttt gtctgaaaaa ttttttgcga tgcccattat gaaaaattgg
33361 caataatagt atagtagtta gtttaagttt agattctttc aatactcatt ctgcttcagt
33421 ttgtagttag atttaactca atagaggtga ttggcaaaaa tgggtaagac acaaaagaag
33481 aatagtaagg gacgtttaga tagatactat tatctagcaa aggagaaagg ttatcgtgct
33541 cgttcatcct ttaagattat tcagattaat gaaaagtatg gccacttctt agaaaaatcg
33601 aaggttgtta ttgatctgtg tgctgctcct ggttcatggt gtcaagttgc atccaaactc
33661 tgtcctgtca actccttaat tattggtgtt gatattgttc caatgaagcc gatgcccaat
33721 gttataactt ttcaaagtga tattacaaca gaagattgta gatcaaaatt gaggggttat
33781 atgaagactt ggaaagctga tacagtgttg catgatggtg ctcctaatgt tggtttgggt
33841 tgggttcagg atgctttcac ccaatctcaa ttaaccttac aagctttgaa gttggccgtc
33901 gaaaatttgg tggtgaacgg tacttttgtt actaaaattt tcagatccaa ggattataat
33961 aaattgattt gggttttcca acaattgttt gagaaagttg aagccacaaa accacccgca
34021 tcaagaaatg tttctgcaga aatttttgta gtgtgtaaag gtttcaaggc accaaagagg
34081 ctggacccaa gattactgga tccaaaagaa gtttttgaag aattgccaga tgggcaacag
34141 aatatggagt ccaagattta taatcccgaa aaaaaagtta gaaaaagaca aggttatgag
34201 gaaggtgata atttactgta tcatgaaacc tcaattttgg attttgtcag gacggaagac
34261 ccaataagca tgcttggaga aatgaataag tttacaattg atgagaatga ccatgaatgg
34321 aagatcttga agaaattgaa acaaaccaca gacgagttcc gttcctgtat tgaagattta
34381 aaagtcttgg gtaaaaagga ttttaaaatg atcctaagat ggagaaagat tgcaagggaa
34441 atcttgggca tcgaagttaa ggacgacgct aaaacagaaa ttgaagtagt accgttaaca
34501 gaagaggagc aaattgaaaa agatttgcaa ggtttacagg agaagcagcg tctaaatgtc
34561 aagcgcgaaa gaagaaggaa gaacgaaatg aagcaaaagg aactacagag aatgcaaatg
34621 aacatgataa cccccactga tattggtatt gaagccgcaa gtttgggtaa agaatcgttg
34681 tttaatttga aaactgcaga aaagactggt atcttgaacg acttggcgaa gggtaagaaa
34741 agaatgattt ttacagacga tgaattagcc aaagataatg atatttacat tgatgaaaac
34801 atcatgatca aagataagga ttctgccgct gatgcggacg atttagaaag cgagttgaat
34861 gccatgtata gtgattataa aactagaagg tcggaaaggg atgccaagtt tagagctaag
34921 caagcacgtg gtggcgataa tgaggaagaa tggaccggtt ttaatgaggg aagtctagaa
34981 aagaaagaag aggaaggtaa agattatatc gaagacaatg acgatgaagg tgtcgaagga
35041 gactccgatg atgatgaagc catcaccaat ttgattagca aattgaaggg acaagaaggt
35101 gatcacaaat tgagtagtaa ggcacgtatg attttcaatg atccgatatt taataacgtt
35161 gaacctgatt tgccagtaaa taccgtcaat gatggcataa tgagttccga gtctgttggt
35221 gatatttcta aattaaataa gaaaagaaaa catgaagaaa tgcaccagaa acaagatgaa

35281 gcagattctt cggatgagag ttcaagtgat gactccgatt tcgaaattgt ggccaacgat
35341 aatgcatcag aagaattcga ttctgattat gattcagagg aagaaaaaaa tcaaacaaag
35401 aaagaaaagc attccagaga cattgatatt gctactgttg aagccatgac tttggcacat
35461 cagttagcat tgggtcagaa aaacaagcat gatcttgttg atgaaggttt caatagatac
35521 accttccgtg acaccgaaaa tttgccagat tggtttttag aagatgaaaa ggaacattca
35581 aagataaata agccgattac taaggaggca gcgatggcaa ttaaagaaaa aataaaggca
35641 atgaacgccc gtcctattaa gaaggttgct gaagctaagg caagaaagag gatgcgtgcc
35701 gtagctcggt tagaaaagat caagaagaag gcgggcttga ttaacgatga ttcagacaag
35761 acagagaagg acaaggctga agaaatttct agattgatgc gtaaggttac caaaaaacca
35821 aagaccaagc caaaggttac tttggttgtt gcctcaggta ggaacaaagg tttggcaggt
35881 agaccaaagg gcgttaaggg taagtataag atggttgatg gtgtcatgaa gaatgaacaa
35941 agagccctaa gacgtatcgc aaagaagcat cacaagaaaa agtagtaaca aagagcatta
36001 tttttccatt cccttgtata atattgaaaa taaaggaaaa gaaagcaata gatatattat
36061 gtatacatag aattaatcaa attatcatca aaaaacttat aggaaaccta tcgttttgaa
36121 ttcagaaaat acgagtgcac agtagaacaa tttttattcc cgttttactg atcttttctt
36181 cttgttgtta ccaaatacct tggagaaaat gtatatgatg gagatgagta agcatagtaa
36241 cgtaccattt ttgatcttgg aataatcctt tgtgtcaggc cttggaattg gtactagaag
36301 agttgtcgag ttcaaatggc ggaatgtagt atcgtctgta aagagagatt catatcctag
36361 acctttttta taaaatggat tacgggaaac tttattgtca cctgtgtcgc aggctaatat
36421 aacttctgga tcaaatgaaa cttctaattt atccccttta ttattagaag gctcaatata
36481 tctagtatgc aatgtcactt cattcattgt gccggctttc aattcaaaga tactttcaga
36541 accccatgcc ttatctcgaa gagagtattc tggtaattct aagtcgtctt ctccaaaaag
36601 tagtaaggga gaggattgga atttatcgat aaataattct aacggcaatt gcaggtgcat
36661 tagatacatg catttagggc gttcttcgaa atctgtgaga tcaatcatga tttttggatg
36721 caaaccgata ggttcattca gatatagtga tgtcgtagtc gtcgagtgat tgtatgcgat
36781 atggccctgt ttatacaata aggatgtttt ctgacatctt tccattttac catcgtccat
36841 cctccaatta caacgtattc caccaatatc gacatcactt tcatcacttg catccacaaa
36901 gaatacacct acttcgagtt tctcatcttt cactcttgca attgtaaatt cttcaccctg
36961 tttcaacaga cgatattcaa caatttggat ttgtgtgggc tccactgata tatcatatgt
37021 aaaatcttct ggattccatg acagattcaa atctacttcg ggaggcaagt actgctctat
37081 gtcaaacttc tcactgtgaa atgaattgta gactgggtta gtaataaacc tttccggcac
37141 atccgaagaa tttgagtaaa ctgaaaagcc agcactcagt tcattttcta taacgcttaa
37201 acttctagcc aagtccgctc tgggcctcca agtgattctc gtaaaggatt tatccaagct
37261 tccagtcctc tctaatagcc acctttgttg taaaaccaca ccagaacctc cacgaacagt
37321 caaatgagtg tcattttggc gcatatgatt accgatatcc tcaggggcat tgtagagtac
37381 agtcactcta tgtcttgtca ccattactca gctagaaacg cacacttctt gcgtttattc
37441 tcctcccgat ttgattatga ttagcgtttc cattatttga tctttttat cagggggatc
37501 ttcaaagaaa ggaacttttt ttaggaaaaa aaaaataatc atcttttatt acatgtgtaa
37561 tacttcaagc aatcataaac aagagttaag aaggcaaagt tctgctgtta gtgtgtcctc

37621 caatactctt gcacgacaaa gttttagtta tccccttcta cgtgtattgt tcctccgtgg
37681 tcgattgtgt tcaaagattg ggagctcttt tcgtgttttg cactaagagc atactgttaa
37741 tccgcaaagg ccactaaaac ttacctgcat ttgtctactt aaagatacat tccaactatt
37801 tacattcatt tttatttcta aaagagcatt ctgaaccctc ggcaattatt tcatactcct
37861 tctcagaata ggaaaatgcc caatacgcat actcaacatg tgcaaatatc agagccaaat
37921 cctgtaaata ctttgtctac accatccaaa agaggtcacc gccatcgcag atcgctagca
37981 atatcaggag attttgattt tttgaaacag cctgcagcaa ttgtgaattt accacctcca
38041 caggcggctg aaaattgtcc ttcaactgcc ccaactgctg tatcaagtac attatcgcca
38101 atacgctaca atagatttcc ttgcaaaacc aatgaagacg ctggaacgtt agatttgcct
38161 gaaccaagat tttatccgtt atcaccaaag aacaatctgc aaacaccaag tccacgattt
38221 ttcattagtg aagagccaag tttttcatcg ccagttaaag gcgtcccaga tgccattatt
38281 aaccttgacg atgcgttgaa gacaaggcct aggtcattta aatcacatag aagatctgaa
38341 tccgctcctc ctgatttgga ggttatggta gataagggca attgtgcagc cggttctaac
38401 tctatgatta aagaagaaga ggactcctta attgaaccag aatcgaaaaa tgaatattat
38461 gagcaaaagc ttccaacagc actattatcc ccactgcggc cttccctttg tgtatctgaa
38521 caggccattg atgtagatga ttcagctctc aatgggtcac cgacccatca caaccatggg
38581 atgcaaaacg ccaatgcacg gaattccaac acattcaatt cgttgaagat caaaggccaa
38641 aagcaaagat attatcatta tacgaagcag ctacctttga ccgtaggctg tgactcgcaa
38701 tctccaaaag aacaaaggtc ggctgcttca atgacaatca atcaggcaat gacaccttct
38761 tccctggcct atacccccttc taaactagca tctactcccg caacaccagt atccttttat
38821 gacagcaatg cggacattaa cttagaaagt gataattttc cactaaaaga taaccctaga
38881 tatgccaagg atggttatcc taaaaagtgc ggcaattcac agcttaatcg tgtgctggat
38941 agcgataaaa gacaggattt tagtggagaa tcgagaagaa gaagatcggg cagtcctatc
39001 tcccacatgc aacaccgcaa cctgattgat aatatgaaag gtagacgaaa cagtaacacg
39061 ataaactcaa tcttcaacta caagagtcaa cattatgaaa tgccatatga tgatatgatg
39121 aaaaatgaaa acattaatgc acagtccatg cccttttcag tcaacggtgt caacaatgaa
39181 aatagtatcg gaggggttat tacgagagcg gacgatgcac cccttcaaca ctctgtggtc
39241 aaatcctgta cgcctgatgg caaggaagaa atgaataggc ttaaaagtaa tgacagtaat
39301 gaatattcca agtctgaagg gcagatcaga accaattcgc aactaagtaa ggacattctc
39361 atgggtgaac caggtgatat ggttgatctg tcctcttttg tcaacgcgca gagaaaagcc
39421 tcaaatgaaa ctggtgactt agtctttagt ttatcccagg atgatgacgc actgaaaacg
39481 ttccatgcga gcaacagcgc agcaacaagc aatgaaagct ggtgtattag cgatgatgcg
39541 ttaggaaagc aggcgcagga cagtgaagtt aggaggaaaa gaaaatccaa attaggactc
39601 tttagacata ttttttcaag gaagtaatac aacaatatta ataataagta ataaaaaggt
39661 gaatcgtaac cttaacggca ttgattgcat gtagttttaa taataaagtg tagaataata
39721 aatataaaat ttcaaagtca tcagaaataa aaattgaata gacatataca tatagaaatt
39781 acttctatca tttaaataca tatatataca tacatgtata cgtgtatact tgtacttgtg
39841 aaggactcaa tagttgtatt cgtttggttt ttgaccagaa gtgtttggga aaccacattc
39901 caataagata tcgttaatca attcaggttt ttcagtaatc ttggagaaaa cttcatcatt

39961 tttggttaag atcataccac aataacctgt ggagttgaaa cctatcttca tttcatcaga
40021 aaaggccttc gagcgtggaa cgacacagat ccattccttg gttaacataa gattgtagga
40081 tttctttagt tctggatttt cgttcaacca gtcctggaaa aaggtcaaag ctctttgcaa
40141 tatggagatg taacacatag ctaataggtc ttcatcaaca gtttcttcag actccggcat
40201 tggcaagaca aaatgagcga acgagacttt agcatcttgc aaaggttcag tattgaaagt
40261 tggtaggaaa tgttctttac cattacatag tctatcttgg aaagtgacga actttttcagg
40321 catttgcaga atttgcaaat gtttgtggtc caatgaagaa ccactggctg gaccagaatt
40381 gtaaaagacc atgtgtctct tgtcggattc ttcattgtcc aaggcacaca gcaacttata
40441 agcagtcaat aaatcggttg gggtcaaggc atcagtttga tgttggtatt cgttagttac
40501 caataaagtg tgttcaggga ttacagggaa tttgtttaat agcaatttat acgctttgtt
40561 atcggcacca ccaaattctg ggataaccgt taattcttcc tcaggtttgc ctagtggatc
40621 ctctccttct ggagtttggc cacgctctgg cttttcgatc agacttggca tgtgactaat
40681 caagtatggc atgctggttt ttgggtcctt tgttttcgtt gtttcagtct ggataaattt
40741 taagttacca ttatcgaagg cacttttgta cttgtcacta attaaagatg caatgtcagc
40801 ggggatactc atttttattt taatttttac ttttctgttt gttctaaaat ctatctaaac
40861 tggctttcaa gatcaatcta ttgtctttta aggtaaactt taaattggaa ataatagtaa
40921 tgttagttcc ttcattttaa ccttgtattg tatttccttt gcgtgatgaa aaaaaaactg
40981 aaaaagagaa aaataagaaa atcttctaga acgttccgaa acaggacact tagcacacaa
41041 atacagaata ggaaagtaaa aggcaatata tgaatgcagt gcttgtaact ggtgcttgta
41101 tccaagaata gcttcttgct gtaggttatg ggaatatcgt gtaagctggg gtgacttttg
41161 agctattcgc gacgcccgac gccgtaataa ctactttcga cagaccactt atgacagtat
41221 ttcaggccgc tcttataaaa tgacatgtta acaaacagtt ctgattattc gccttttgac
41281 aggacgataa tgtaaatagt tgtggtagta tcattcaggt atgtaactgt ttactttgta
41341 tcgcttgaaa aaaataagca tttcagagcc ttctttggag ctcaagtgga ttgaggccac
41401 agcaagaccg gccagtttga atgctcaact cttcaaaaga aattcctcaa atatgtccag
41461 tttcatgtac tgtccggtgt gatttattat tttttattta ctttgtagtt cttaaagcta
41521 agattttttt ctttgataaa ttcttgtttt catatcctaa aattaaaggg aaaataaaca
41581 atacataaca aaacatataa aaaccaacac aataaaaaaa aggatcaaat actcattaaa
41641 gtaacttaca cgggggctaa aaacggagtt tgatgaatat tcacaagata aaaatcatat
41701 gtatgtttct gatatatcga tatacaatca aacactttca agaatttgtt tgtagacttt
41761 ttgctagaga cctcatcaaa gtgctaccaa ctaagatcaa cttatacttc ttttagagaa
41821 aattttttc aatgtactcc aaagagattt agatcctgtc tcttcctctt cctcttcctc
41881 gaaagtcaaa gaaaaatcag agtctccctg cttattcagg cggagaggct ctagggtagt
41941 tgcgtttctc tcattgggac actgaacctc attttccaac attttggtca tgtaagaggc
42001 gacaggctca tcgcaggtgg gtgcatcaac atggtagtac ctggaccaag cgctacattg
42061 agtccctcct ggataaacac cgctacaata ttgtctttgg acgtttgccc aaaccatatc
42121 ttttgaatac caaagctgga ccacattgta tggcctaatc attggtgcta ccataatact
42181 ggattgggaa acagtctggt taattttttt caaccaattt ttcttatcta gcaatgattt
42241 aataaacctg aaatctaaat tgtcttcgtt agcgtctgtg tcataatcta caattgagta

42301 ctgtgacgtc caattatatg gcaccgagat ggggaatctg tccggtgttt cgtcgctgtt
42361 atccttctcc tccctccaaa tgcagtcaga ggcaggtgcc cattcggttc gccagtctcc
42421 gttatttact acttggtact gttcccaatc gtaatacgtt tcctctgggt tgaagatact
42481 tgctctgctc ttgacattgc ccatagccac accacgagaa acatcgtgga agattacgga
42541 gctgtttacg atagcaggag caatggattt gacgaatgac acttgataaa agtctttggt
42601 cgaaaataca gtatcaacgc cgtcgtggga ctcgacccct ggtagaagaa gcagagcatc
42661 tagatcaacg gtagatgctg cagaggtcaa agccacaaag agggaagatg cagttacgag
42721 gtatttacta aacattggtt cgaagtgatt acagattttt ctctcttgaa ccctggaatt
42781 gcaatagctt ggttcctatt gggaccggac cctcttgcgt aaagtatata tttagaagca
42841 gctatttaaa tctgaatgca agacgaaaaa aagctcgttt ataggagaaa aataccaaaa
42901 aaaggaaaac catacacatt cgggtatccc agctttgctt gcaaaatcaa tcgatgagca
42961 agacaaacag ggccagctga tgcatatgtt ttgtgttgct ttcctacgat cagctaatgc
43021 gagtgttata aatattcccc tagttttctt ctcctccttt atgtaacaat aacttgtgga
43081 cttccccaac ataagactaa aagcaaaggg ataacaaggg atttatctta ttccatccct
43141 ttgcctagcg cactgttacg cagctgatct tatcaacaag tgtggttctt gcctctaaga
43201 tttacattgc cttttcccaa agggaaggtg ggacaaaact cattttccct aaggggaaac
43261 gtaagatacc ctttagggaa gataagggaa gttagtagat tgacgacgta aatctgtgat
43321 gagatgaaaa aaaaagatta gcatctatat aagaaagtac ataacctcca tttaagttga
43381 aaaaaatttg tatgtaatgt cttttcttcc tcttttcttt atttcattct caccgctaaa
43441 tcgataacgg ctaaaacaat taaagctccc ttttcaaacc ttatacttcc gttaaataat
43501 gcaaatcaaa aacattgtcg ctgttctagc tacagtcact gctatcaacg cccaagttgg
43561 cattgaacct aacgccacta ctccaaatgc tacgcagcca aatgctacgc agccaaatac
43621 tactcttcct accgcttcgg tgactacgac agtttcgatt ggcgaagccg tagttaatac
43681 catggcggcg ggtgcctttg gagcagcaat cgctgccggt gttgccttct tattctagga
43741 acatttttat ttcctttatt tttttctttc tttctaattc cataaccgtt gttcaaaatt
43801 ggggagcttt ttaactttcg ttccattaag ctttttcttt gttctttacg tttgattcta
43861 tttcaaccat aatttatttt ttcgttattt aataattact aatatttaat actagatgta
43921 tatacgttgt gtgtatgcgc tactatagta tagcttaact gttttattat acccttttttt
43981 ctctctttac tgtacagaag atgcgtcact aattcaaaaa aaaaaaaaaa aaaaaaaatat
44041 aaaaaagaaa gattttggag aagctatatc cttttgtgac atacaataga gaaacttgtt
44101 ctgttggtaa gatatctcat ctgaaataca acaaggaaaa ccaaaagcca ttttttcatt
44161 acttgatcga cttatgaacc gtattactag gaaaagttgt ttattcgcga ttatatttgc
44221 atcattattt gtgacacatg cattgggtgc cgctattgat ccgccaaggc gaccacataa
44281 tgtgaagcct tttcataacg gtaatctcga acttcaaaga agagcaaatg aaccgttttt
44341 tgaaatagat gtcaagagtc tgaacacaaa ctcaccgata tcagagttgt gtaaaaaaga
44401 tttgcacgtc attgaatcgt ctcatgatct ttttcattta caaaaccaat gtgaattcat
44461 cttggggtca ttaaaagtca caaactatga ttctaacatt ttggatttga acagcttgag
44521 ggccattggt ggtgacctga ttattcagga ttcacctgaa ctgatcagaa tccaagccgg
44581 gaacttgaat aaaatcgaag ggctcttcca attacaggga ctaacctctt tggtttctgt

44641 tgaaattcca actttgaaat tttgtcagtc actggagtgg aaagttgttc ccatcttgaa
44701 ctacgtctcc atggattctc agaatattga gattataaag gatattgtca tatcggatac
44761 ttcattagca aacatcgaga atttcaacaa ggttcaggaa attgatactt tcaatatcaa
44821 taataacaga tttttagaaa ctattcattc gaacgttaaa accattaggg gacaattcag
44881 tgtacatgcg aacgctaagg agctagaact tgaaatgcca cacttgagag aagtggaaaa
44941 cataacgatt agggacacat cattggtcta ccttccacaa ttaacaaaag tgaaaagctc
45001 tttagagttc atcgaaaatt acttttacga attgaacctg aacaatttgc agaagattgg
45061 tggaacatta ggaattatca acaatgtaaa tttaataaaa gttaatttgg agaacttaac
45121 agacattcaa ggtggcttga tgatcgccga taacgaatcc ctcgaggata ttactttcct
45181 gccaaacttg aagcagattg gaggtgctat tttctttgaa ggttcgttca aagatatcat
45241 gttcgatagc ttgaaactgg tgaaaggtag cgcttttatt aagagttcat caaacgtgtt
45301 ggattgcaat aaatggacaa acccatcaaa tggaagatca atcatcaggg gtgggaaatt
45361 cacttgtatt tctggtaaga aggaaaatac gctgaatgtt aaacaggatg gtacaatcat
45421 agaaaaaggg tacaaagatt taacgcaaga aggtgaagac tccaagaaaa gagtgatttc
45481 aaaatacgcg aactcagcaa atccaagcat gcaattggac ccccttcttt ttggtacatg
45541 ccttgttgct atgttattgt tttaagatta ctagtctagt ataacagaaa gtttttttg
45601 ttcctaatat tatctttatc tatatacata tatctccatg aaaggcattt gaatttctat
45661 ctttctcttt caatcaaata ttgtcttatt attgatcaaa taatcgtata tgcacaatgg
45721 caactcgtct ttcaaactct ccacttgccc attcttcagt ttgttcctaa tctcggatga
45781 tgaaacattt cttaatgctg gataatcatt ttgcataaca tgaatcctag cgccccattc
45841 tcttggtatt accggttcga attttccctc aaggatgtca ctggcgtatc ttagttgcga
45901 ttcagcactg agatggcaat cgcctctcgc caagcaatat aattgagatc ccgacatgaa
45961 gctctccatt acatcactga tttttaaagg atgataatat ttttcatcaa agatcctagc
46021 aattgtatca aaaccaacta agtagcctat atctgcactg catcctttaa caaattgctc
46081 atgtaatatt ttgtccttgt cgatgaattt cgagaaaaca gtcaacccga cagatactac
46141 agattggggg agcttctcct gaaggtagtc agcgaataag cacatcattt ccagacgagt
46201 tggaaaagat gccggcttag gcaacttatc tgcattattc accgctaaca gtaataaaac
46261 atgcgaacgg gtgtccttca atttgaagtt tttaatggtc tgcgatagta gttgaaaatg
46321 ggccagatgt ggtggattga atgaagaatc aaggacaaat aacttctgac attctagatg
46381 ctggggtcct ttgagaacct gaaatagtaa attgcttttt cgaaactgct cgaacgtctt
46441 cttcatatta gtctgctagt gcacgtactg ctatcaaccg tcattgatta agtgtccatt
46501 ttccaaagaa gtaggagatt atataaccac gagtatgacg tcttccaggg aaaaaaaaaa
46561 aaaaaaaaaa gaaagccaaa ataaggagcc ttgaaatgtg tatctgatat atataatgtg
46621 tatgtaaata tctatttaat tagccattgg gatttcaact tcttgtttga aacagaagga
46681 cggataaaat acgtgatcac caacaagacg aatatagtaa taagcaattt accccttttcg
46741 aaagtaggac tcattaaatc aaattggccc gaaggtgtga tccttgtaca aaatacatca
46801 aggcctagat cacatataat gcttgtggat tccagattgg ttgggatgga gatcaactgc
46861 gaatcggatc ctggcaatag atttctgaaa tgagtgataa tgaaattatc attaattggg
46921 ataactggcg tgtatggggt agccataaac tcttttttct tatccttggc catttcttct

46981 gctggtttac ctcttgcatt caataaaagc tttggtatgt aggtgatttg tccattttct
47041 aattccataa cgattgcctt tgtggtaata tcatccgttg tcttggaaat ggacattgtt
47101 ttgataatct cgggaaaaat gaattgttta gtttgaaatt gaggtttgtt aatgtgtcca
47161 gtcaatggat cataagaaaa attgtcgctt gagttagaca aacgctcatc tggggttagt
47221 gactcatata attccaccac tactaacttt tgttctggaa caggttcaga actgaaatag
47281 gaataaacga cccaatattc accgaaaaca atatccattg gaaacctaaa atccggagaa
47341 tccttgtgct cttgggtaat gaggatttct cctgtaatgg tatcaattaa gttaaaagtg
47401 attgtatgat gttcttcatt agcgatcaga taagccgcta ggttggggta caaatattta
47461 taaagaaccg atttgtcacc tagtgtaata ccaagagtgt ttaggtttgt tgtttccctc
47521 ttgctatatg ccaccatttt ctctttcgaa tttacggctt ttttccaagt ttgttttacc
47581 gtatcgtttt cgattatgta gccataaatg ccattcgtgt catgctctgt gacgaatatg
47641 tgggaactag aatggttgtt ggcaactatt ggtacatccg tattcttgcc gggattcaat
47701 ttgaacagta aatgatcctt atcctcaaac ttaatcagat actggtgatc atgttcattc
47761 agcctaataa cggaatcaac agttttcctt tcagataggg gggatcttga tttgataatc
47821 ggttgatctt tagtaacttc aattgtcaaa taatgcccat catgcgaaaa aacaacaagt
47881 tcgtcatggt ttttatccgt ccagaacatt ttgacgccct gatcaatttc taaatcgagt
47941 ttccagataa gttggccctt attgaccata tcaaggccgc cgataaagcc atcatgcgtc
48001 aaaacaatta agattttggc gaatccgaac ttcaaatctg ataccgtggt atccttagcg
48061 tctagagcta ggagtttagt gaagacacgt cctggtgaga attggttttc tttcaataag
48121 ttgataaggc gattccaatt agttgtcaat ctaagccagt aagcattcca aagcgaattt
48181 gaatcaagtt ccgctttcat atctttgttc aactccacat ccctagaatc gaagacgtcc
48241 aatacagcat agtctaccac gttagtcaac cattcatctc tttgccaaac ggggatcaat
48301 tctgtgccat tggcagagaa tcccatatat tgggtatcat taacgttcaa cactaacgcc
48361 aggggatctt cacgatgaaa atattccact ttattgaacc cttgaggtaa ctccaatttc
48421 cattctgcca gttttgactg ctcatcgaga atatgtagtt cattgttctt aatagatact
48481 tgatcgttta aatatgagga ttgaggtaaa atggtcgtag aaggggcatt atctacgcct
48541 tcctctagta aaaaccaatc gtgtaaatca tatttctgcc aatggttgct tgaagagttc
48601 acacaaacca ttgcgttatg gtcattacta tccagttgaa tctcatcaat ggtaaagggt
48661 aaaacgtttc tgaaaagaat ctgtccggaa gaaacgttaa acgaagaaac taagcaggaa
48721 gtttcggtag ggttcgataa gatgagaacc ctgttgcggt ctcgagaatc agggatgact
48781 ttctcccaag gacctaagtt agccagttgc caatcagtga taaatgcatc atctgaaaaa
48841 acggcttgga cacaactcgt gtttaggaag agtaaaatga agacgtacac caagtctgta
48901 cacgttatct tcattgctat gggggaaggg gaggatgaaa gtgttgatat gaatgtaggt
48961 attagttatt aatggagtgt atatatatat atgttattat atatttgcat atataatatg
49021 aaatcccagc catattttct ctggtagccg tctgaaaaat cacggtgtac gaagaaggat
49081 ttaatatacg cacggtacaa ctaagcaatc cgcaaagacc taatgtgtct tttcattagt
49141 gagagccttg gggtaggcc ctggtaacgg ctggtccgtg ctagtgggtg tcttggtatg
49201 ggagggcatg gtagttggta tgaatttgat gcttagcgaa gtttctaagg ccaaagtcat
49261 ccatggatca taaagcgatc ttattgtatc gagatcaata tcttgcgggc ctttcgtgtc

49321 atggggcaaa atgacacctc gggagagtgt ggtgctggat ctcttcttta ctggtgcgga
49381 aatggaatct cccgaagatg ctaagaaatg cagtcttttt tccacatttt gtagatggag
49441 ttttttggac tcggaaagac cttcaaggcc agtagaagaa gactgcggat tgtgggatat
49501 ggaattgatg agttcaatgg aattttctct aatgtttctt ttattttgaa aaaactcgaa
49561 catgtccaag tgaaattcca tgcgattgag gtactcgttg aaagacatgg gcttgtattc
49621 caagggtccg aaccatttaa taatgggatt ctgcgcctca acttgcgctc tagcgagagc
49681 gacgctttgt ttgacctcag ggatatttgt cgagagagga gtaccgttca tagcatcctt
49741 gacttttatt ttggtggggt ctatgccctc gtgataatat ttgacaatat tgttgaaaaa
49801 cttcttgtca cgtacggatt tcatttcgtc gcaccagggg tcgtatttaa gaaagttcag
49861 gcgagtaagc tccaacagtg actgagaaag tagcaaagag ccaaacaacc cccataagag
49921 ctttctgacg ctgacgaatc gagattggcg aggtatgatg ggaggagggg cgagcacaat
49981 gtcaccatgg gtggtggaga agcggttcaa agttggaaag tcagcaatat cctttttgat
50041 tttagtcttg ccattgacaa tcatttcact gggtagttcg ccaaaggata aatccgagtg
50101 agtgccagtg gcaatgcggc gcggtggcgg tagtttgtag agccttctca gtctccaaag
50161 ggcaaagaaa cccactgcag attggatagt gattagagtg tatagtcccg gtaggttatc
50221 aggagcgggt acgagcggac cccacagttt taagcctaaa cttggcctta tgtagaattt
50281 cttgatatca ttatcatctt tgtcgtggtc atcttgtgtg tgagtatggt cggtggaatt
50341 gctattacct gatggaggag taaatacagc catcttttat tacggtaatt ggaaaaggag
50401 agggaatgga ggaggatgag ataagtagtt tcctaagcag taaagcaagt gtccacattt
50461 aagtaaaaga cgatgccatg cacttattat tactacgtgg tcaattttct ctttccctaa
50521 ggcgttcgtg cagtgtgacg aatatagcga acaactattg tgtttgaatt ttaacgttta
50581 tctttttatg atttttttaa aaaaacttcc tagaaaattt cttatatatc tctatttaat
50641 gaaaaaccaa agtgatcaga attacaattc atcgtgaatg gcatcttctt cgtcagccaa
50701 ttcagcgtca gcatcggctt cctcagcagc tttttcctgg gcttcttcgt acaaggcctt
50761 accgtcgacg tcgaagtgac cgttttcctt gatgaagtcg aataaagagt ccaaggatct
50821 tgaaccttgg tacacaacag attcggactt cttaccacct gggtataaga cgattgttgg
50881 gtaaccttca attacgacgc ctctgacatc gttttcagtg tggtctagtt tagcaatcaa
50941 aacgtcggat gtggcgttgg cgtaggtatc agctagttct tggtaagttg gggccaatct
51001 cttacagtga ccacaccatg ggcatagta caaaacaaga acgtccttct ttgggtcgtt
51061 gacgatttcg tcatggttct taccgaccaa ttggaagaca gaggaatctt ggttctcgaa
51121 gatctcttgg gacttcacga ttggggaggc atcacctttc aagaagtcct taaccaaaga
51181 ttcaatagcc ttagactcca acacgatctt gtcgctcaat tcgtcaaacg cctcttcaga
51241 gagttgaggc aaaccgtact tcaagtcttc agtcatgtcg tggatggcaa atagagggaa
51301 ttgttccttc atgttcaagt tgccggcgtg tctgccgaat tttctggcat cgatgctaac
51361 aaagttcatt agacctctgt tctttttggc caactcggta aagagaggct tgtattcttc
51421 caattcttcc tcgtcattgt agaataagta acccaaaggc aaaccgcttt cgacgtattg
51481 ggcgaaaacg gaaccgtcga tttcaccaaa gtagggcaag gcttccactt gcaaccattt
51541 ttcaaaaaca tcagcgtcag cgatatcggc tttcttaccg ttgtatacta caggctcgtc
51601 catggcggag ggcaagtaaa tagaaagctt gaaatcatcg tctgcgtttt cagcggagac

51661 aaagtcgtag tcgttgaagt gtttgttggc catggagtaa aaggtggcgt tgaagtcggc
51721 gtcaatctta ccggattgga cgataactgg agtgacaaaa gtctcgttag caaggtaagc
51781 tggtagatca gcaacaacgg cgacagccgg ttggctttgc ttgatcatga attggacaat
51841 ggcctcggca gttctaggtc cctcgtaatc gatcgagttg ttaacatcgc tgtttttgaa
51901 aatcttcaag cttgggaacc ctggaatgtt gtgttccata cacagatcct ggttttcagt
51961 acagtcgatc tgggccaagg taatgttttt ctcaactaaa gtctcggcgg ctttaacgta
52021 ttcaggagcc atgttcttac agtggccaca ccatggagca aaaaactccg caagcaccaa
52081 gtcgtgcgac tgaatgtact cattgaagga gtcggtggcc aacttaacga cagcggagtc
52141 ttcaggggcc acagcctctt gttgggcgaa aacagaggag gcgagcagca gggaggacca
52201 tgacaggacg gcaccagcag aaaacttcat aacgggatag atgtatgttt ttctttatgt
52261 agagcttggc agctttaaaa ttaccaatta gacatgctgc ttgcaacaag aaaatgcacg
52321 cgtaacaaaa tatatatata tatatatata tatatgtatg tccatacggg tttttcgttt
52381 tgttgtggtt gtaacagcac aacaaaatgc tacacggtgg caaattggaa aaagagacta
52441 gtgagagaaa agggaaaaag aggcgccgcc cgacagggta acatattatc acgtgcagcc
52501 caggataatt ttcaggacac gtgtttcgaa aggtttgtcg ctccgaaaaa atcaaaaaaa
52561 acaaaaaacg ggaaataaca ataacgacaa aaatggaaaa aaaaaaaatt ttagacgcgg
52621 cgcttgcacc ccgcattata agtggtgtgc cgacggacgg tcaacccctt tcaggggggcc
52681 ccctatcttg ggcgtggtgt cacaccacgc tcaaacggtg ggcactaatg aaaacgcgcc
52741 catataaata tccgctatca acagaacccc caacccccccc atcagtgccc aactcagctt
52801 ccgtaaacca caacaccacc actaatacaa ctctatcata cacaagatgt cattcgacga
52861 cttacacaaa gccactgaga gagcggtcat ccaggccgtg gaccagatct gcgacgattt
52921 cgaggttacc cccgagaagc tggacgaatt aactgcttac ttcatcgaac aaatggaaaa
52981 aggtctagct ccaccaaagg aaggccacac attggcctcg gacaaaggtc ttcctatgat
53041 tccggcgttc gtcaccgggt cacccaacgg gacggagcgc ggtgttttac tagccgccga
53101 cctgggtggt accaatttcc gtatatgttc tgttaacttg catggagatc atactttctc
53161 catggagcaa atgaagtcca agattcccga tgatttgcta gacgatgaga acgtcacatc
53221 tgacgacctg tttgggtttc tagcacgtcg tacactggcc tttatgaaga agtatcaccc
53281 ggacgagttg gccaagggta aagacgccaa gcccatgaaa ctggggttca ctttctcata
53341 ccctgtagac cagacctctc taaactccgg gacattgatc cgttggacca agggtttccg
53401 catcgcggac accgtcggaa aggatgtcgt gcaattgtac caggagcaat taagcgctca
53461 gggtatgcct atgatcaagg ttgttgcatt aaccaacgac accgtcggaa cgtacctatc
53521 gcattgctac acgtccgata acacggactc aatgacgtcc ggagaaatct cggagccggt
53581 catcggatgt attttcggta ccggtaccaa tgggtgctat atggaggaga tcaacaagat
53641 cacgaagttg ccacaggagt tgcgtgacaa gttgataaag gagggtaaga cacacatgat
53701 catcaatgtc gaatgggggt ccttcgataa tgagctcaag cacttgccta ctactaagta
53761 tgacgtcgta attgaccaga aactgtcaac gaacccggga tttcacttgt ttgaaaaacg
53821 tgtctcaggg atgttcttgg gtgaggtgtt gcgtaacatt ttagtggact tgcactcgca
53881 aggcttgctt ttgcaacagt acaggtccaa ggaacaactt cctcgccact tgactacacc
53941 tttccagttg tcatccgaag tgctgtcgca tattgaaatt gacgactcga caggtctacg

54001 tgaaacagag ttgtcattat tacagagtct cagactgccc accactccaa cagagcgtgt
54061 tcaaattcaa aaattggtgc gcgcgatttc taggagatct gcgtatttag ccgccgtgcc
54121 gcttgccgcg atattgatca agacaaatgc tttgaacaag agatatcatg gtgaagtcga
54181 gatcggttgt gatggttccg ttgtggaata ctaccccggt ttcagatcta tgctgagaca
54241 cgccttagcc ttgtcaccct tgggtgccga gggtgagagg aaggtgcact tgaagattgc
54301 caaggatggt tccggagtgg gtgccgcctt gtgtgcgctt gtagcatgat ctttttacat
54361 ttttttggtt tgtgtacgta tcccaccgta cttaccatct tctctccttt atatatatat
54421 atatatatgt atattttcag tgtatataca tacattctta tacaataccg tataagaacg
54481 tatgtatgta tgtatgattt tctcacatta atgaacaata tcatcggaaa cgggcctttg
54541 gtttttaagg gaaaatgaag aacaataaat gccttgctag gtcttccttt atcatttttg
54601 gaagtcaaat atgaaaaaca cttccaaggg ggcgtactac ttcaactaat aaaatgtcac
54661 acactaataa gatcgcatac gtgttgaaca atgacacgga ggaaacagcc tcgccctctt
54721 ccgttggttg ttttgacaag aaacagctca ctaaattact gatacatact ctaaaggagc
54781 tgggctacga ctccgccgct aaccagttac tgctagagag cggcggatat caaaatgagt
54841 ctaaccacat ccagactttc ttcaaactca tcaagaccgg ccaattccat cttattaatt
54901 ggcagattgt ttgctcgctg ccccttgccc atagctctcc gctaaggtcc gaatggcttc
54961 aaaggctgct cattcccacg ccgacgcccg ccacgacttc acttttcgac cacatgctcc
55021 tgcaactgca atacctgcaa cagctgatga gctctgtaaa ttcatctact tgttcggacg
55081 cagagatcgc cacgctcaga aattacgtag agatcatgat tctagttaat agacaaatat
55141 tcctcgaatt cttccatcca gtcacaaatt ccgcctctca caagggccct cacactgcac
55201 tgcccgtcct gtatttgcgc aaaatcctca aaaacttcat cgagatatgg gattccctgc
55261 tggtgtctaa cgatcagttc ctaaatgaag aaaacatctt caatccggaa acgaccttga
55321 gagaactgtc gacgtacctg accaacccaa aactaaccgc gcagttgaac ctcgagcgag
55381 accatctgat tgacgccatc tccaaatata tcgatccgaa cgaactcgtt cccaagggtc
55441 gtctcttgca tctcttgaaa caggccatca agtatcaaca atcacaagac atttttaata
55501 tcatcgatcc ggatgatgac gcctctttct cctctccacc ccaccggatc aaccttttgc
55561 aggataactt ctcacacgat ctgactgtga cctttcaaga atggaagact atccaagaca
55621 ccacagacga aatttggttt ttgacatttt cgcccaatgg caagtatttg gcttctgcga
55681 cttccgagtc ttcaagaggc tacttcatta ctgtttatga tgtggaacaa gacttcaaaa
55741 tttataagac ttgcgtgagc ttgtcacaat ccgtattgta cctcatgttt tctcctgata
55801 gccggtatct ggttgcttgc cctttcagcg aagatgttac catttacgac atgaatgcca
55861 cctccctgcc cgatgcgtct gccacagact cgttcctact atacccttct accaggctct
55921 cgcccatgga ctcgtttaaa ctggacacca ccacgtaccc agatgatacc gaatcatcag
55981 catcatcttc ctcgaggccg gcaaacgcga attctaatca atcaagagta tggtgctgcg
56041 atgccttcca cactgcggaa cgtgcgggct ggatggtggt tggatcgccc gacagagaag
56101 ctattgtaca ctcactcacc acgaaagagt cactttttag cttaaaaggc aggacttgta
56161 tcgcgttggg ccacgatgaa aacatctctg ggagaaaatc aatcgatcct gcaaaagtcc
56221 tttacaaacc aacaagtagc aatggtaact ggcaatatgt ggaggacgac gaaacctttc
56281 cacgtgtgca cgacgtgaaa atcagctatg acgataaata cgtgctttta atgacccacc

56341 agggcgttat agatgtctac gatttcagtg ggttcccctc caaagaggag ctatctaagc
56401 aaacagtgga tcccaaaaac tttctgattc caaggatcgc aagattggac gtgggcaaaa
56461 acatgacttg tatctcatta ccgctaaata ccacgcatca aggcttccat aggcaacaaa
56521 tatccgagtc tcaacatttg gtattagtta gccttcaaga taacgaatta caaatgtggg
56581 actataaaga aaatattctc atacaaaagt actttggaca gaagcaacag cactttatta
56641 tcagatcttg cttcgcctac ggtaataagc ttgtcatgag cggatcagaa gatggtaaaa
56701 tttacatttg ggatagaatt agaggtaacc tagtatctgt tttatccggt cattccacgg
56761 taatgagtaa ttccacaaaa cccatgggga aaaattgtaa cgtagttgca tcaaaccctg
56821 cagataaaga aatgtttgct tctggcggtg atgacggtaa gataaaaatc tggaaaattt
56881 caagaaatta atactccacc ttatatatat acgtaagcaa agttttatgt aacaaaaaaa
56941 aaaaaagaac taatccaacc ttctttcact ttcaggtaaa acttgactga gttcttcagc
57001 ctctcttcta cctctcttga catccaaaca gtttagcaca ggcaatgata gcatcaaaag
57061 caaaaagaag aaataaaacg aatagcgaat attatgcgtt ttatcggtaa gcagtccaac
57121 aaggaagggt cccaggatgg acgacccctt atctgtgata ctgaacatac taaaaaacgt
57181 ggattctttt cctggaggta caatcaaact gaaaactgat cttgaaaccg cggacaggcc
57241 acctagtgat aatccatacc aaatggccaa taagaacatt tcaaacttat gctttaaacc
57301 gaacgcattg aagaaaaatc caagaatacc ataaaatggt atgaaacttg cccaaatgat
57361 aatgtacatc aaagtttgac tagaagtcca ccgaaacttt gtggcaagaa attgtggaat
57421 cataaaggca cccagcattg catttacaac ggtcaaaaca cttatcatga ttaaattgag
57481 ggtactcatg tgcagttctg ccttggagaa caaaaccgct gtagaattta tagttgtaat
57541 ggaatcacta ataataaacc acgcaataag aaaaatcatc acatctttca atagtctggc
57601 atgtttaaac gattcgaaaa gcgaaaccca accatatgaa aggtaattct tccagtttag
57661 ttgacccaag gcgtcttgct caccgggata cggactgcgg gcggatgcta atgtagaatc
57721 gtcaactctt atcggtattg tcacatcgtc aatcaaccag atcatgggca gttgccacac
57781 aaaccaccaa atcccaacga aaagaacagc cacttgaaca tcctgcttac tgcccttttt
57841 agatgcgact aagaacatag atacaatctg aacaatgagg gcacttgaat aacctaaaga
57901 tgcacctcta ccgctaataa cagtagttaa cgagtctacc ttatcaggtt cataagctcc
57961 ttgactttga catttcaaag aatccttgac aaaaatgggc agaagagaat tcccaacaac
58021 attgataacg ccaaaacaac cattggccac tatataaagc ccagccagag aataaatctg
58081 ggtatcattc aattttgaaa tcgcaacagt cgacaatgcg cccacaatac caaaccaaac
58141 cagaattctg cctttgaatt taacgctccc ccagagatct actatccctg aaacggaaat
58201 gactattata gtttggaata aaacgctaag ggaaaagaca tataatgcaa aacttgaagt
58261 atctacgaag atccgattgt tgaacaaacc aagaacacac ttatctgaat cactacccgt
58321 ttctgacagg cagggtatgg agtgatcgtg aacttttaca ccatttatac tcgcaaattg
58381 ctgcagtagt aagggaatat acgttgaaac cgcagaaacg acaaatggtt cgctagaaaa
58441 tgagtaagcg taccatccct tgatattgtt ttgggctttt tttattcgat agttcgttac
58501 cgattcattc atatcattta tagttccata gctcatatct aatgtagaga atatgttatg
58561 agcaaactat tgcttataag ggcaaaaacc gtggtaatat aggtacagtt tgaaatcttt
58621 ccttttgttt acactacgca tctttcaata gctgaggtat tttcgtttag gaaaataccg

58681 ttttaaggaa ttaattttag caatttccta aaattgttga gcggcgctat caatgccgca
58741 caatacaaca attaataatc taagacacgt tgcattatac atgacagaag ctaaagcatg
58801 gtcttagatg acaataaaaa ggaatttgac agcacaaaag agaaaaaaaa aactatacat
58861 aaaatttttt acatatgaat tgcttagagt aaaataggca tttttcatat cttcttccca
58921 attattaaat tagcaaaaaa ttgaataaaa aaagcaggtg gtttaagaat tgcgcaggga
58981 gaagaaagaa tatcatatct catgatggta gccacttaga cagagagaaa acgaagtgag
59041 aagaggctag tatttttata tcttaaaaaa aaaaaaaaac aaaaaaaaaa aaaacataaa
59101 tatttgtatc ttattgctta tgatgataat gtacaatgaa ttcaaaacat gaaaattgtt
59161 caaagttttt atttttattt tctctttcgt attaaactga tggaacaaat gaaaaaaatg
59221 gataaaaaaa caaaaaaaaa aaaaaacaaa aaaaaaacag ctttgctgga ggatgattaa
59281 cttttaattt ttttaattat taggtttttc aattttttat ggttttttgt gtaaaatgca
59341 atgaacgaat ccattttttta ttttgcaagt gtgagaggcc tttataagga agactcggat
59401 tcaccggcag caaccggcgc agagtcttct ttggtttctt gttcttgaga agcgagttca
59461 gtggatgatt cctcttcttg ttgaggaaca ggtggtaggg atatcatgaa ttggtccaat
59521 gcatcgccga ccagttcttt ttcaatatta atttcagttt caaaagtttc tggtaaccag
59581 tttgaccagc tcttggaacg aacgaagtac ttatccaatg gagatggttc tttggcttca
59641 ccggtaacgt tatcaccttc cttggcggcc aaagtacctt ctgcaacatt gacggtagcg
59701 gcttcgttag cgacaatttc tctcaatgcg gctaaaatca ggttagtgtc acctccgaag
59761 gacatgttaa caactctgta aaacttagag attaaagaca atggagcaaa accatccttg
59821 gagagtttgg accttaagta attgtcgacg gtcaagtttt cttcgctgaa atagtattca
59881 atttgtctag caatgttgtt gatggccatt aacactggtt ggactggata aaactgttgc
59941 tgtaatttga ccataggttg ttgaggatgt tgttggttgt gatggaagtg gttgtggtat
60001 ttagagcggt tgttgttgtt attacgagcg ttgcgtcctt ggtaaggttt aaactgtggt
60061 gggaagcctt ggttctgagg cataccggcg ttattactgt tgtggaactt tctcctttga
60121 ggaccattgc tgttatggtg atgactgtgg tgatgattac ggtggtgagg tggttgttgc
60181 tgttgtgtag attgaccatt ttcttgagag gtagcgtctt ttgaatcttc ctcacctact
60241 gccgattcaa agccatcttt cttcatttgt tgttgcttct tagcattatg ttggtgatgc
60301 ttctttttcc tgttagcatt actcgacgat gtcgacgatg ctgcagtttt gttgtttttg
60361 ctcttgttgc tgttgccatt acttgcacca ttcttggaac cggatctttt ggtgccagag
60421 acggtaatgg aggccttgat gggaacccac ttggtagcag tcgatgattt cggagtgggt
60481 gttctgttct tcttctttct tgtggcatcc aagtcttcta ttgagatagt agaaacaggg
60541 atctcagtag gagcaagttt ccatggggaa gatgtgggca atggtgccgg cgtcaagttg
60601 acttgcttgc tcttggagct ctcttgttct tgcacttctg ggactggggc agtagcagtg
60661 tttgcggcgg cggtttcagc agacatcgtg ggcagagaag tgaaaagaag atgaaacacg
60721 ttagcaggtc cagagtaatc ctgatgttct attaccgatc taggatccca aaagatcttc
60781 ataaaaccga aaaagatgtt aagaaatgtt tacacttgta cttttctttc tttgttcttg
60841 acctaccaga agtgaaaaaa atttttcatt gcgtgggtac atccgtgcac catttctgaa
60901 aaatttgcga tgagcatccc tcaaaatccg tattatccgc tagtgttttg aggttcaaca
60961 agacaacata taagaaggga agaaggaaag gaggcaatga gaacttgggt cttgctcgag

61021 agtccagatc cacatgcagg ttcaaaaaat ggtgagagat aacagtaata acggtagcga
61081 taaaagcgtc cattgggaga ggaggaataa taacggcgca ggccccccgtt atcgttccag
61141 aagcggtaat accggtgctt tggcaacaaa actaagtaat gggacgctct ctgtcagagg
61201 attagtgaag gaccgaacag gaagcggcaa gatcgcgggc tgtgtggagg cgtttctgga
61261 tgccaggacc caattgaata cgccctggga ccgtgctaag tgcaattggc tggaccagat
61321 aggttactat gtacagttga gaaagaccgc gttttctaag gaattggacc aactaaggaa
61381 gcccatgatc gatgcatatg tggcggagat gaggcagaag tttgatgcct cctatggaca
61441 atccagggcg caattggaag ccaaactggc gcaggtggac agtgaatggc atatggtaca
61501 tggtgatgtg catgcaaaac tggaaaaact cgtggaagaa cgccggtttt tgaaaagatt
61561 aagcgacacg atcgtaccac ccaggtccaa aagatcacag cggctgtctc cattgaccaa
61621 agaggaccga gccaactgta tctgtccgca gcccaaagga atgagcgaca ccgcttggtt
61681 cgaagccatt cagaagaaaa tgttaggaat gaatggtacc atcaagctcc tagagacaga
61741 acagaaacta ctggctgacg agaaaaacag cgtgaggaag acgttctggc ccatggtgga
61801 agcacattca cgctcgaatg aatttgctta tctggagaaa tgcatcaggc tgatggcctc
61861 tcagagagca atatgctttt gtcttgatat agaggctttc gaaacaaacc agaacgtaat
61921 caccgaaatt gggatttcaa tttatgaccc cagggaaaat atggtgccgt caatggttcc
61981 aattacaaag aattaccacc taattatcga ggagtccctg gaacttagaa accaaaaatg
62041 ggtctgtgac tacaaggatt gctacttatt gggagaaagc tatgttttga gcttgaaaga
62101 gtgcgtgcat ttcattcaat cactaataaa ctattacttg gtcccggtga ccgaagaaga
62161 caagacatgg tcaagggcat ttgttggtca tcacgtgagc ggggatctta agtggctgga
62221 gactattggt gtcaaattcc ctggcagagg gtatgaaggc catctggacc atacgctgct
62281 tttggctgaa actcccggtg atctagacgt gttcatcttg gacactgagc agttttacag
62341 gaaatcgtat ggcgaaaagg gcagcagtct gggcaagatt ctgcggttgt tcgagatacc
62401 gcatgcgttt ctacacaatg ccggtaacga tgcctactat accctgcatt tgttcatgaa
62461 gttttgcgat gttaatttca ggaaaataag cggcatggac gatgttctta aagtaatggg
62521 ccaagtaaaa gtttggggag aacgagacgt acgagagcct aaagtggtgc ccatgtcgta
62581 tgccatctcc atcgaggagg cagtcaaaaa tcggacgtac cgcaagggcg tcaagagcag
62641 taggaaggaa agagtctgcc aaacggaatt cggtgggtta acgtatttcg gaactgctaa
62701 agacgccttc acaagcactc ttccgacaca ctaaatatat tcatcgattt ttttcctgta
62761 aatacaaaca catgcttgta tacatataca tatacatata tataagctca ttataaacct
62821 gtgtgcatgg aaaaaacttt gtctgccctt aatttgcaag aataggttct aacaattcct
62881 ccaattcacc agtctccctc aattcctgca agtcgtcgtt gcctccaata tgtttaccat
62941 taatatagat gtttggcacg gttctttggc cattaatctc atataacgca gcctgaatgt
63001 ctgcgccttc cttcatgtca ttcaattgca aaaccagaac tttggacctg ggaaccttta
63061 acttttcaaa aagcgtgttt agggctgcat ggcagtatgg acagtacgtt ttggatgcga
63121 cgaagatctc gttttctgca ataaggtcct tgacgtgctt gatagtttct tgagatacca
63181 tttttttcc ttctgaggtt ttgtctattt gtataattat gcgaatgctc gtataagtac
63241 tgcacgaagt tctttatttt atgctaaagt cccttttat attatataaa cttttcattt
63301 attttatata tttactcctt tcaagacaaa attcccacat cacgtgaatc aggaggcgca

63361 aggggggtag taaggcagta gtagataaag aaaacaagca tcgcaatctg gaaacgccgt
63421 tgcggctgct cgttatctgt agacagctca cttgtgcatg aagttcatgt tttatagatg
63481 atggagccat acttcaaggt gataactatg aaactgtttt gaccttgacg tgtaaagtga
63541 tcaaggtaat agttaaactt tatatatgtt gtttatcaat gtggtatcct ttctaaggat
63601 gaagacaaca cgattgtgac agtgtgagag agtgataaag cattggaggc aggcaaaaca
63661 agaacatggg gtttctttcg gatcatccgc atacagctat caccgagacg atctttcgaa
63721 ttgtctcttc aagggactat acgctagaag tggagttagc tcccttgatt cagcttatca
63781 aagcagatca caacgattat aactataccg tgaaccaaga agaagctgcg cgagcactta
63841 gaaagaaaat aaagtatggg aaccggctgc agcagtccag gactctggac ctcttggatc
63901 tgttcatctc acaaggcgtg aagtttacag tcatgtacaa tgatgacaag ctgctgcaaa
63961 ggctgagagg gatggctacg aactctgaaa acagcgggtc cggtgaaaag tatgagccta
64021 ggattattaa gaagtgcgcg gcgtatgcta tctcgtggct aaactatatt acgcagaaca
64081 accttgagaa tgcgagggcg tattcaggtc tttatcaatt gggtcaaaca gtcaaacaga
64141 ggtactcgaa gagcagccgt tcccgccgta gcggcggagg ctctggcgga cggtctaact
64201 tcatggatga tagcgccgac gacactttgt accagtccaa ctcgctgact agtgccgata
64261 gattgtatag gattccacag atcaacatga acaaagaggc acccaggatt cggctcataa
64321 tcagcgatgc tctcgcgtct gctgtgtcgt tacagaactc gctgatcggg ctgcccaagg
64381 gcaagttctc cacagacgat gaagaggcca cgtccaagtt catccaggca agggcgatca
64441 gaaggaaagt tcttagatat ctgcaattgg tcacggaggg tgagtttctg ggtagcttaa
64501 ttcatgccaa tgatgaactg gtggctgcgt tgactgcata cgacgaccga agcgctcagg
64561 acgatagcag cgatgaaagt gatcacggct cgtacgatga cggtatttat gatgaaaacg
64621 agcaggataa cagcaggtat attgatagcg agtcctcaga ggaagagagc ttgtcttctt
64681 atcagccgtc cacgatctca aaccctttcg gtgatcataa caaaatttga agaagtatgt
64741 acgtatgcat acacacatat atatatatat atatatagta catttaatcc ttcttgaggt
64801 ttaaagacgc actgttcaca cagtgtctgg tgtccttggg caagtttagc aactgtttcc
64861 agccttcacc ttcaaacaca tgtcccaagt gtccaccaca ccttgcacaa catatctcca
64921 ccctcgcagg cattaaagaa ttgtcacgat gatatgtgat ggctccaggg gatacctctt
64981 cgtagaatgc gggccatcca caacgagcgt cgaacttggc cttgctcgaa tacaacggtc
65041 tgtcgcagtt ggcacaatgg tagacaccgg actcgttggt gtgtaaatac gcaccggtgt
65101 tgggcctttc agtggcctta tctctcagca ccatcagctg caatggtgtc agggcatcgt
65161 tccacttcac gtcattcgat tcgtcactca ttttcttgct cttattccag tactgcgtca
65221 gtacaatgtt tcttctccct ggaaaagtcc tgcgtacagt tataacgtac agcctgctcc
65281 acttattcat tttttgatat tgcttccttt ctccctttat ttaaactatc atcggcgaca
65341 tatttcccaa taggcaaagc tgaaaaatac tatcgctaat tataacaaag aagctaggtc
65401 gaaggactcg aaaagacaga ggtcgtacta gcagagatag caaatcagat ggaggacggt
65461 aaacaggcca tcaatgaggg atcaaacgat gcttcgccgg atctggacgt gaatggcaca
65521 atattgatga ataatgaaga cttttcccag tggtcggttg atgatgtgat aacttggtgt
65581 atatccacgc tggaggtgga agaaaccgat ccattatgtc agagactgcg agaaaatgat
65641 attgtaggag atcttttgcc ggaattgtgc ttgcaagatt gccaggactt gtgtgacggt

65701 gatttgaata aggccataaa attcaagata ctgatcaata agatgagaga cagcaagttg
65761 gagtggaagg acgacaagac tcaagaggac atgataacgg tactgaaaaa cttgtacact
65821 actacatctg cgaaattgca agaatttcaa tcgcagtaca caaggctgag gatggatgtc
65881 ttggacgtaa tgaagaccag ctcaagctct tctccgatta acacacatgg agtgtccact
65941 acggtacctt cttcaaacaa cacaattata cccagtagtg acggtgtgtc tctttcacaa
66001 acagactatt tcgacacagt tcataaccga caatcaccgt caaggagaga atccccggta
66061 acggtattta ggcaacccag tctttcccac tcaaaatctt tgcacaagga tagcaaaaac
66121 aaagtacccc aaatatctac aaaccaatct cacccatctg ccgtttcaac agcgaacaca
66181 ccggggccat cacctaacga ggcgttaaaa cagttgcgtg catctaaaga agactcctgc
66241 gaacggatct tgaaaaacgc aatgaaaaga cataacttag cagatcagga ttggagacaa
66301 tatgtcttgg tcatttgcta tggggatcaa gagaggctgt tagaattgaa cgaaaagcct
66361 gtgatcatat tcaagaactt aaagcaacag ggtttgcacc ccgccattat gttaagaaga
66421 agaggtgatt tcgaagaagt agcaatgatg aacggaagtg acaatgtcac ccccggtgga
66481 agactctaat gtgcagttgt catgcacatc atcatactaa acttacacga ataggataac
66541 atgtatgcta gcagaatata tatgaaagaa aaaaattatt gatgccttta aacttatact
66601 attatactat attatgttat attatattat tagttttata gatatattga gatatgttga
66661 atatgatgac gaggatggtg gtttaagtgt atggattgaa ttttctctta gctttcatta
66721 ccttgattct ttcttgatct tccttaaatt tacttaacag ttgattgatc tcctgttttt
66781 tacgttctct gacctgaaat ctatagaaat ctttctttgc ctttttatct atatttgaaa
66841 ttggcttggc cttattttca tgtttggata atggattttt gtttaatatt ttctttctta
66901 tggaattcaa tgacttggtg ttctttccta caactaatgt gaatccatcc tcatccacta
66961 tagaactttg aacatcctct tgtgcttgag cttcacgctg ttcaaaaatt gccatatgtg
67021 tatgaatatc ttctttcaaa taatcaatat ccagtggttt gtagaagtta acaaaagttg
67081 tgaatgatgg agtcgtatat gtccattcaa atagttcatt tggatgtttg gcatgcaaat
67141 tcgagtattt tttcaaagca ttccagcagt tatttatact tgcagcatca acaaatttta
67201 atagcgccgt gtttcttggt gtgtatctct tctcgttgac gtcagtggag gacatcagat
67261 cggaggtcaa tgccgataaa tctacttcat gtaatccaaa ttcatcgtta tatagtagtt
67321 cctctacatg cgatactgta tcgtatttcc cacagagctg cccgacaaat ttcttcatgt
67381 gctctatgtt agataataat ggaaggttga ccaaaaacaa acagtcagac tcgttggaat
67441 ttgaactctg gtgtctttta gcaaacatga aatgcaacga agcttcctgg cttttgggta
67501 gtgccttgtg atccggtaat ttgaacggca ccactataaa cccgttcttc atggcgctaa
67561 tgtcttcaat acccatcccg taacactcta tctgaggttg ttatctgtta cctagttcgt
67621 tcgccctaaa tgcctcttgc tttaccatcc tctcaatgtt ctcttctttt tcctgcttat
67681 ttcttaggtg accaaaaagt gaaaaatttt ccaatttaac ttacgtcgtt cgaagtgatg
67741 acaataagga tattcattta ttaatcgcta tttgataccc actcttgcta ctacctctct
67801 tttaatccaa aattacaatt tttacgttac cattcattac atagtgtatc tctattcatt
67861 caagactgta ttttgtttga tatatatatg tatatataca tattattttc gttagtgttc
67921 ggtttccaag ttagaaataa tctactggaa atcctttggg atcaacccaa gcttactcat
67981 tctgattttc acagcgtttc tgtgaccgtc tagaccctcc ttcttggcaa cgcacataac

68041 agctctaccg atgttttcta aaccttcagg ggtaatgttt tgggcagtga taaacttttg
68101 gaaggttgca gtgttggcac cactgtactg cctagcgtaa ccataggttg gtaatgtatg
68161 gttagtacca cttgaatagt caccgcacga ttctggagtg taagcaccca caaatacgga
68221 ccctgcattg tcaaccaatt taacataatc gttagcattg gcgatttgta gaatcaaatg
68281 ttctggtgca tattggttgg acatttcaag ggcttcttcg taaccgtcac aaagaacgat
68341 cgtactgtga gcaatacatt tacgaacaat atccacacgt ggcagttgta aagcttgatt
68401 gtggacagca tcttgaatct cttgaatttt cttttcgctc aagttaacac caacaaggat
68461 aacttgggag tcaataccgt gttcagcttg cgatagcaaa tcacttgcaa caaaatccac
68521 atcggcatct tcatcggcaa taaccaaaac ttcacttggg ccagctggca tatcaatgga
68581 acatagagct tgagtgtcat tttgaacata cattttggcg gcagtcacaa attgattacc
68641 tggacccaag atcttatcca ctttaggaat agtttctgtc ccgtaagcca tagcagcaac
68701 ggcttgggca ccaccagcta gaacaatctt ggaagcgcca actttttctg cgacataaac
68761 aacttcgggt gaaactttac catcagattt tcttggtgga gatgcaaaca caatctcctt
68821 acattgggca acttgtgctg gaacaccaag cattaatgca gtacttggta aaatggcagt
68881 gccaccaggg atatacaaac caactttttc aataggacga gggaatctgg aacacaagac
68941 accaggttgg gtttcaactt caagagtctc tgttggcaat tgagcagcat ggaatttgcg
69001 gacgttttca attgaaaggt ccaaagcttc cttcatttcc tcggttaaac cttcaaagta
69061 ttcttctggg aatggagcat taagaacagg attggataat tttacaccat caaacttttc
69121 tgtgtactcc aaaagggcag agttaccttt gtctctaaca ttttcgatga tcggattgac
69181 taaatgcata atttcagaag tcttttggat tggtctgctc aaagccttct gcacaccaac
69241 tttgtcggaa gccttcacca cgtccaagtg aattggaccg gtcagttttt cttcttcagc
69301 ctttggttgt ccaacaaact ttggcttagc atcacctttc cgtcttgtaa ccttcagatg
69361 cttcatattc agattattct cgacgtcctt caatgaaaca tcgttggcca ctaatttggc
69421 cagtgcaaag tagaacaaat cggcagcctc ccaagaaagc tccttcttac cctttgcctc
69481 agtcagttct tcagcttctt ccttgatctt ggcatctaac aatgcagagt cgttgaatag
69541 tcttctagta taagattcct ctggagcgtc ctgtagcctt tgttttagta aagattctag
69601 ccccaccaaa ccatgcttga attcaccaaa gcaagacatg gtctccaagt ggcaaaatcc
69661 aacgttttct tgttcaacga taaactttaa ggcatccgaa tcacagtcag tagagatttg
69721 taaaagcttt tggccattgc cagaagtttc acccttgatc cagatttcat tcctagaacg
69781 agaataataa acgccacgac ccaaatcgat ggcctttgct atagatttct tcgaagaata
69841 caccaaccct agacaacgct catattggtc gacaactagg gtggtatata aaccgtcagg
69901 acggtctgta cgtacttcac caagcacttc tttggtcaac atatccttgc ttaatttctt
69961 tatggacaca attttatctt gcgagaattt ttgttttacc atgaattgat tggagaaaac
70021 accgttctct tccacaacaa cacgctcctt tggtacattc aattgttcaa ccaagtgttc
70081 ggctgtttta gcatcttggc tagcaatgaa cagagaagaa actccgttgt tcaagaaggc
70141 aatgatttca tcatcgctga atttaccact tggcaaggac aaagccacca atggaacttc
70201 ttcctctttg gagaactgga gaatctcttc attactcagg ctcgagccat ccaaaagtac
70261 ctgaccaaca agtgaaacgt attccttctt actattccat gaggccagat catcaattaa
70321 cggtagaatc ggcaaaacca ttattcagaa aaaaaatttt gtaaactatt gtattactat

70381 tacacagcgc agttgtgcta tgatattttt atgtatgtac aacacacatc ggaggtgaat
70441 ataacgttcc atatctatta tatacacagt atactactgt tcatagtcat atccttttct
70501 taccttctat atcgaatgac tgataaaaaa acgtgagtca ctgtgcatgg gtttagcaat
70561 taattaacta atttaccgga gtcactatta gagtcagttc gactgcctag aagaactgct
70621 ggttgtcagg attgtgatgg gggcattctg ctgtattatg acccatcgta tcgcaatgct
70681 cacaccactg ttgtcttcct gccgtggtat cgactggtgc aggggggtcg aaaattggca
70741 acgattccac ggctgtttgt gcttgagcct gttccaactg tttgattttt tcattagcct
70801 cttcaagttt tttcgttaag gatgccacct cttccgatga ggaatcttgt ggttttgtca
70861 aaaatagttc cttgctcaaa ttttggtatt ctttactgag cgaatcgtta tgcattttca
70921 attgttcgcg ttctttagcc cactttgtct tgtgtaactc aaattggtct tctatgttgc
70981 gtaattgttc cagctgtttt ttcaggagtt cgacatcttc gttggcacca gtgggttgat
71041 tatgagaaag atttctctct tcgttttctt tgatctcttc gtgtagttgg cttacgacag
71101 caagtagctg ttcattctca gcgtcaaaaa actgcttttg tttggcttgc tgtctgcgtt
71161 cgagctccaa ttgttgcttg agatggtcta tctctttctc tctttcttgt attgtggctt
71221 catacctatc aaaagtcggt tgcacttctt cgaggaccat tctttggtca tcgagtagcc
71281 ttttgtagtg tagttgtttc ctttgtagct tttcgatggt caattgggct tctcgtaatt
71341 caattgtaac ttcgctgcta ttgaggtcat tcatgtggcc attgtccggt ttccaatcgc
71401 tggtggtgtt gtgattagcc tttctgtctg atgacaggat agagtcgacc tccattctgt
71461 cttctctgtt atcgtaacca aattcttgct gttgatggtg atccgatgcc tcctggtcca
71521 tcgactgttg attaccgctg tgccgactgg tgatccggaa acttctcatg ggtgtggggg
71581 atttaggatc atccatggga gagaactggt tagtgagcct cacaatagat ctgtttttgg
71641 gtattgatag cggttccatc gtcgttcttc tcgaggtttg cgatatcgat gccttctcga
71701 tcaatgatgc gactttttgc aactgaataa atagtccact ttgaggatac tccgtttgaa
71761 aatacttctt ccccatgaat gatccatcgt tcttaccaat gttggcaagt aagtctacac
71821 cagcaaacat tccagctttc gtgtccactg gacccacgta tttcagttgt ccgcgcccca
71881 aatttgggat ttgtatgaaa catcctatct ttctttgata tctatccatg gtatttcaaa
71941 cgcatacaca gccagccaca gctcaacgcc ttttaccttg tcctttgatg cctgtctcgt
72001 ccaaacgttt ttggtgtctt ggccaattgc ccttctgaaa aatctcactg tccgcaactc
72061 attaaaagat acccaagcaa gctacacgat aaagaaagga gaaagttcat taaaagaacg
72121 tacatatagc gatacaaacg tatagcaaag atctgaaatg gatacggata agttaatctc
72181 agaggctgag tctcattttt ctcaaggaaa ccatgcagaa gctgttgcga agttgacatc
72241 cgcagctcag tcgaacccca atgacgagca aatgtcaact attgaatcat taattcaaaa
72301 aatcgcagga tacgtcatgg acaaccgtag tggtggtagt gacgcctcgc aagatcgtgc
72361 tgctggtggt ggttcatctt ttatgaacac tttaatggca gactctaagg gttcttccca
72421 aacgcaacta ggaaaactag ctttgttagc cacagtgatg acacactcat caaataaagg
72481 ttcttctaac agagggtttg acgtagggac tgtcatgtca atgctaagtg gttctggcgg
72541 cgggagccaa agtatgggtg cttccggcct ggctgccttg gcttctcaat tctttaagtc
72601 aggtaacaat tcccaaggtc agggacaagg tcaaggtcaa ggtcaaggtc aaggacaagg
72661 tcaaggtcaa ggttctttta ctgctttggc gtctttggct tcatctttca tgaattccaa

72721 caacaataat cagcaaggtc aaaatcaaag ctccggtggt tcctcctttg gagcactggc
72781 ttctatggca agctctttta tgcattccaa taataatcag aactccaaca atagtcaaca
72841 gggctataac caatcctatc aaaacggtaa ccaaaatagt caaggttaca ataatcaaca
72901 gtaccaaggt ggcaacggtg gttaccaaca acaacaggga caatctggtg gtgctttttc
72961 ctcattggcc tccatggctc aatcttactt aggtggtgga caaactcaat ccaaccaaca
73021 gcaatacaat caacaaggcc aaaacaacca gcagcaatac cagcaacaag gccaaaacta
73081 tcagcatcaa caacagggtc agcagcagca acaaggccac tccagttcat tctcagcttt
73141 ggcttccatg gcaagttcct acctgggcaa taactccaat tcaaattcga gttatggggg
73201 ccagcaacag gctaatgagt atggtagacc gcaacagaat ggtcaacagc aatccaatga
73261 gtacggaaga ccgcaatacg gcggaaacca gaactccaat ggacagcacg aatccttcaa
73321 tttttctggc aacttttctc aacagaacaa taacggcaac cagaaccgct actgaacgat
73381 gattcagttc gccttctatc ctttgtttac gtatttgttt atatatataa ctttattttt
73441 ttttattaat tgggctgcaa gacaattttg ttgtcagtga tgcctcaatc cttcttttgc
73501 ttccatattt accatgtgga ccctttcaaa acagagttgt atctctgcag gatgcccttt
73561 ttgacgtatt gaatggcata attgcactgt cacttttcgc gctgtctcat tttggtgcga
73621 tgatgaaaca aacatgaaac gtctgtaatt tgaaacaaat aacgtaattc tcgggattgg
73681 ttttatttaa atgacaatgt aagagtggct ttgtaaggta tgtgttgctc ttaaaatatt
73741 tggatacgac atcctttatc ttttttcctt taagagcagg atataagcca tcaagtttct
73801 gaaaatcaaa atggtagcaa caataatgca gacgacaaca actgtgctga cgacagtcgc
73861 cgcaatgtct actaccttag catcaaatta catatcttcg caagctagtt cctcgacgag
73921 tgtaacaaca gtaacgacaa tagcgacatc aatacgctct acaccgtcta atctactctt
73981 ttctaatgtg gcggctcagc caaaatcatc ttcagcaagc acaattgggc tttcaatcgg
74041 acttcccatc ggaatattct gtttcggatt acttatcctt ttgtgttatt tctaccttaa
74101 aaggaattcg gtgtccattt caaatccacc catgtcagct acgattccaa gggaagagga
74161 atattgtcgc cgcactaatt ggttctcacg gttattttgg cagagtaagt gtgaggatca
74221 gaattcatat tctaatcgtg atattgagaa gtataacgac acccagtgga cctcgggtga
74281 taacatgtct tcaaaaatac agtacaaaat ttccaaaccc ataataccgc agcatatact
74341 gacacctaag aaaacggtga agaacccata tgcttggtct ggtaaaaaca tttcgttaga
74401 ccccaaagtg aacgaaatgg aggaagagaa agttgtggat gcattcctgt atactaaacc
74461 accgaatatt gtccatattg aatccagcat gccctcgtat aatgatttac cttctcaaaa
74521 aacggtgtcc tcaaagaaaa ctgcgttaaa aacgagtgag aaatggagtt acgaatctcc
74581 actatctcga tggttcttga ggggttctac atactttaag gattatggct tatcaaagac
74641 ctctttaaag actccaactg gggctccaca actgaagcaa atgaaaatgc tctcccggat
74701 aagtaagggt tacttcaatg agtcagatat aatgcctgac gaacgatcgc ccatcttgga
74761 gtataataac acgcctctgg atgcaaatga cagtgtgaat aacttgggta ataccacgcc
74821 agattcacaa atcacatctt atcgcaacaa taacatcgat ctaatcacgg caagacccca
74881 ttcagtgata tacggtacta ctgcacaaca aactttggaa accaacttca atgatcatca
74941 tgactgcaat aaaagcactg agaaacacga gttgataata cccaccccat caaaaccact
75001 aaagaaaagg aaaaaaagaa gacaaagtaa aatgtatcag catttacaac atttgtcacg

75061 ttctaaacca ttgccgctta ctccaaactc caaatataat ggggaggcta gcgtccaatt
75121 agggaagaca tatacagtta ttcaggatta cgagcctaga ttgacagacg aaataagaat
75181 ctcgctgggt gaaaaagtta aaattctggc cactcatacc gatggatggt gtctggtaga
75241 aaagtgtaat acacaaaagg gttctattca cgtcagtgtt gacgataaaa gatacctcaa
75301 tgaagataga ggcattgtgc ctggtgactg tctccaagaa tacgactgat gaaaataata
75361 ttgacgttcg catttaatct atacctataa ttctgtactt atatactgtt ccttaattga
75421 agatttcaac atcgtttttg atgtaggtct tttcacctgg aggtgcggct ggggtaccga
75481 agactaattg agcttgtacg gtccaagact cagggatttt gcttggcaaa gcagctttta
75541 tgtaaccatt gtagtgttgt aggtgaccac ccaggcccat tgcctccaag gcaacccacg
75601 agttgatttg agcggcacca gaggtatggt ccgcgaaact agggaatgca gctgcgtacg
75661 ctgggaagtc agcctttagc ttttcagtta ccttgtcgtc ggtgaagaag attacagaac
75721 caaaggcctc atcccttgct gaagcaggcc tcttttgacc ggcagggctt tctatagcct
75781 tagtcacttc gtcccaaact tttttgtgag tttcaccagt caagataaca gcgcgatttg
75841 gctgggagtt gaaagcggtg ggtgtttctt taatgatggt ttggacgacg gattggatgt
75901 cgttgatagt aatttcacca ggtaactccg gtttcaaagc gtaaatagta cgacgagcag
75961 ttaaagtttt caaataagtt gcaacagcag acatgatatt ggattgtttt tttgtatata
76021 tgttgatata ggatacttca gtctacgaaa aaagtacaaa ttatgtagtc agttccttca
76081 gtatggtgtc cttatatact gtagtttgga caaggtgcaa atgccaagac cctagcccga
76141 aaagctcgag gcaccccagg atctttttct ttacgtaatt ttcacgtaaa acgccacagt
76201 ccgatttttc tcgaataatc attagtaaaa gcggtatact ggattattgt acgataacaa
76261 ggtagagctt tattactaag ctaagacgtt cttacatcaa tagtgctgtt cgttattgat
76321 gttaggagaa ggagcgggtc tggtgaatag tgtaagcagt gtttctgaac tttttcttcg
76381 tctaagtcct tgtaatgtaa ggtaagaatg caagcatctt gtttgtaacg cgattgtacg
76441 ttgacgttag taagtcacaa acccaagctt aacttcttcg tgaggaagga aagtgttgtc
76501 tcctactttt ttcaaatttt cgaattgtat ttatatttat ttagtacttc ttgagtttac
76561 atatccttcg taaaaatgca acttttgtcg aaaaacactt ccaaaaaaaa ataataatga
76621 atttatgaag catactaacg agcgagcaca tcgctgaggt atcattactt catgagataa
76681 attaagatct cctcatatgc gaatttcctg ttcagtgata aacgttgatt acgttattga
76741 taaaagtctt ttcttctggc aaggcagttg gaacaccaaa gaccaattga gattgtacag
76801 tccacgcaat aggaacatct tgaggcaaag cagatttgac gtagtcatta tagtgttgca
76861 aattagcccc caatcccaat agttcgaggg cagtccaaga ctgaatttgc acagcaccgg
76921 tcgtatgagc ggcgcatgtt gggaaagcgg ctgccaaggc tggaaaatct ctttgcagtt
76981 tttcagttgg tccttcatca gtgaagaaaa tgactgaacc gtaagcctca tctctgcaag
77041 actctggtct cttcttggct tcagcagttg gcattgcgct cgcaacagca tcccatatcc
77101 ttttgtgtgt atcaccaacg ataatgacag cgcgattcac ttgtgagtta aaagctgttg
77161 gcgtattctt gagaataacg tgtacagttc tctttacatc atccaaaccg acaccttgtg
77221 gtaattcggg cttcaaattg tagatggtac gacggtttgt aatagcgttt aagtagtttc
77281 cagttgggga catttctttg gcttggaggt ctggtgttct tgattttgat ggtgtatata
77341 gctttaaaaa accaaaaatg atcaaccttt atatctctct tcgatcacaa catgggtgga

77401 gacaaacaca aagtagtgat gacccactca ggtgttaaag tgtgtcatat ctgcaaatga
77461 cactgtattt ttttttcctt tgccatttat tgtcgcagta aggaaaagcg cagagcaaag
77521 ctgacaaatt gaaaaattat tcggaagagc attgacaaat attgattaat aatcaagtaa
77581 cactcgcaga cattgttatt tcagaagtaa aaatatcgca gaacaagttg gttaaactga
77641 atgcttcagt cattctaatg ctaattacac cttaacaaaa ttaaaaattc ttaacgtaga
77701 tttttaaaaa aggcaagaag atcgttcctg atgcacgacg aaaacggttg caccccctact
77761 aaggcaaaat tgcatatgcg gatgaaatca aacctcttct gttgcattaa gaacaaaagg
77821 aaatcatttt tctctcactt catattattt cacttatatt ttcctctcca cttccatcac
77881 gcaatcgtta gtgcttttta ttttttttct tcgctcataa aggactagaa ttaaaatgaa
77941 aatcgtctat cattacacgt atgctattac tacaaaacag aagtacaaaa atgaataaat
78001 ataaaagaag taaatgcttt tttttttaac accagaaggc aacgaccctt ttccaataag
78061 gtccgttcct caaacgttcc ctatattctt cgtcttcttg cttgattaat tcttcatcaa
78121 agatttgtct atgagaatct aggtcgatct tgtctgccct gatgaacagt ttccaatcct
78181 tgtgccagac cttgtagccg acataaagtg caatcaagat tggcatagcc aagtagtttt
78241 cgaaaaaggc ttgtgcatcc agcttacctt caccaatggg ggcgatagcg acccaaaatt
78301 gggcaataag aattaaaatc atcataatgc aagcgtaggc agatccccaa acaccagttt
78361 gagatttgaa acccaattct ccaagagatc tcccttggac tttcatggct cttctaaatc
78421 taagatggga taaacaaatg gcagtccatg tgaaaagctg agacaaacca gaaatggcca
78481 ataaccaagt gaaaacttgt tcttccttgg gagatgcagc acagaaggca ataacggcaa
78541 acaatgcaga aacacccatg gcaatcaatg gtctaccggc tctgtcgatg taggagaaaa
78601 ccttaggagc gtaaccttgc tcggatagag ttaaaaataa acgagcactg gagtagaagg
78661 aggagttagc catggacagc acggaaagta gaataacggc gttaatgaag tgtgggacga
78721 cacggacacc gtgggatgca acagcaatga catatggcga ggctttagta ccaccaccgg
78781 tagaacccaa taattgatcg gaattgtatg gcaccaagaa acccagtaga ataatggtag
78841 ccaagaatag gaataagatt ctgtagatca tttgtttggc cgcacctgga atggcctttc
78901 ttggattaga ttgttctgca gtggtgatgg caataaactc tgaaccacca aaggcaaaag
78961 cagcagtcac taatgttgca gcaacacctt taaatctgtc aatggcatgt ttaccattga
79021 aagcgcccgg atcgtgccag tatttaccac caataaaacc atcattacca gcgccaccaa
79081 catcgatgat aataccaaga atgaagaacc cagtcatcat caaaattttg caacagttga
79141 agaagaactc agcttctgca taaccacgag caccgaaaat attaatagta atcaccaaaa
79201 catagaaaat aatgacgaac acatccgggt tcacagatgt cgtccaatat ttgatagtca
79261 tggatgcggt caccaattcc agaggacaca cacacagcca ttgcaaacaa taaacccaag
79321 cgactgcaaa cccaaaacca tcatccacaa ggaaactggg gtatgcattg tagccaccag
79381 tcaagttact atacactagc gccatttcac cacatgcttg aataatacag tacaagatag
79441 aacccataat agcgtaacca ataagtagtc cagctggacc cgcatgaacc aacgcggtac
79501 cgttaccgac caataaccca gtaccgatac ccgtacccaa cgcaatcatc agaacatgtc
79561 taggctgaat ggttttcttc agcgagtccg acttatgacc tgtattagcg cctatgttgt
79621 ccgaactttc atttttttct agttcttgag cctgtctaga agaaggtgag atagccgacg
79681 taagatcgtt catttgtgtt gcttctactt catcctgtgg accctgatcc gctcttttaa

79741 aggagtcgaa aaacctacgc acggcagaag tgttatgctg ttcgtaacct aaatgaggtt
79801 gtgaggatgg acggtcattg gagcctgttt cgaaatattc aatttcatta tcctgccccg
79861 tggcatgtat ttctgtggag ctatttttca agtctttcag ttcgtataga gacttcgacg
79921 acgacattgt gcgaagctat ctttgtctat attagcgtgc ttcttctata aagattacat
79981 aataaacttg ttctacattg ttgtgggcag accatcaaag acaaagcact tatatatgta
80041 tgtaaaacaa atgagtatta cgaggaaaat aacaaaaaga aactaatcac gaaggatatg
80101 agtcttgtta gcggacgagg ggaagatagc gcaagacaca cgtatagccg acctatccta
80161 tgcatcttag ccgtcttatc aggattgatt tgcggttcag tgaaagagat aaccactgcc
80221 aagctggaag aaatggctag aaggcactaa aaaaagagcc tcgcagcgca acttaagaac
80281 tgtggactac taaggagcag caatccgctg gacgtggggg agttgaaagt tggccggata
80341 ctgcaacctg cttatctatg actcgatgag accgttattt agtgcgatgg tgccactggg
80401 agtgccatca acggcaaagt tgcagctttt tgcagcagat cagtgccgct tagacggcac
80461 cctcaagcgg cactgaagaa cggcgcgctt ttttttttct tctattatgc cggtcaagga
80521 aaaaaagaaa ccaaaaaaaa atgccgtggg aggcacggat gcaagatttc tccaaagtcc
80581 ctttttagcc gccgaggaaa tgaggaaata aggggttagc cgccgagtga agccgccgag
80641 gtataaaagg gagaatgcac accttcgtaa gaagattcta tttgaaagag tgctaggtaa
80701 agtcatcaaa aggcttattg atataataat tgctttgcaa aatggaaaac aaatgacttt
80761 gaaaatggga gaaaacaaaa attatatagg taatgttgag ttcttgattt tttttttgtt
80821 tctgtccttg ccacagctct tttcatggaa cattgggctc taggataatt tttttatgtt
80881 ttctgttatg tttgccatgg taggttgagg taaaggcgct ctttcagcaa tgacgattta
80941 tgctaatatg ttttccttaa aaggaaaaca tctcgttctt tttttcgcgt gctgtgcaag
81001 aaaacgcgta acgcagtctt agtaaaaggg ctctcttgaa aactacatga aaactaaaaa
81061 agaagatgta aatttggata caacagcgta gaatagcata atagagtgtg caataaacgt
81121 gtataggctt gataatagtg ctgacgtaaa ttaccagaac tactgcagta tgactgtggc
81181 gaataccgag acccattctg ctgctaaacc atcgagtact ataggcccat ggaagcttgg
81241 tgaaacactg ggctttggta gcaccggcaa ggtccaactt gcgcaacatg agcgcacggg
81301 ccatagaaca gccgtcaaag taatctcaaa gtccattttc aacaacaacg gaaaccacag
81361 taacgacgat tcagtgctgc catacaacat cgagcgtgag attgttatta tgaaactttt
81421 gagccaccca aacgttctga gtctttacga cgtttgggaa accaacaata acctgtacct
81481 tatcctggaa tacgccgaaa agggagaact gttcaacttg cttgtggacc acggtccctt
81541 gcccgagcgt gaggccatca attgcttcag acagataatt ataggcattt catactgcca
81601 tgcgctagga atagtacatc gggatttaaa gccagaaaat cttttgctcg atagtttcta
81661 taatatcaaa attgcggatt ttggtatggc agctttacag acagacgccg acctgctaga
81721 aacttcttgt gggtctcctc attatgccgc tcctgagatc gtatcaggtc tgccctatga
81781 aggatttgcc agcgacgttt ggtcttgcgg tgtaatcctc tttgcccttt taacgggtag
81841 actacccttc gatgaagaaa acggtaatgt tagggatttg ttattgaaag tccaaaaggg
81901 ccagtttgaa atgcccaatg acaccgaaat ttccagagat gcacaagatt taatcggcaa
81961 gattctcgtt gtagatccca ggcaaagaat aaaaatcaga gacatcctta gtcatccact
82021 gctaaaaaaa tatcaaacca ttaaggattc caaaagtatc aaagatttac ctcgtgaaaa

82081 tacctatcta tatccactgg ctgactcgaa caaccatact agcgcctcta tcgatgattc
82141 aatcctacaa aacttggtgg tactttggca tggtagacac gccgatgaca tcgtatctaa
82201 actgaaagag aatggaacca acaaagaaaa aattctatat gcgctactat accgtttcaa
82261 actggactct gtaaggggat cgaataagaa gaaccgcaat aagataaaga aaaccaaaaa
82321 aaataaaaga tctagcacac tttcgtcatc gtcatcatta ttattgaaca accgaagcat
82381 acaatctacg ccacgcagac gtacatccaa gaggcattcc agggaattct cttctagcag
82441 gaagagatcg tcctttttac tgtcctcgaa tcctaccgat agttctccaa tacctttgag
82501 aagtagtaaa agaattacac atattaacgt agcgtctgca aatacccaag caacgccaag
82561 tggtgtcccg aatcctcaca agaggaactc aaaaaaacgg tcaagcaaaa gattatctta
82621 tatgcctaat acaaaaagaa gctcactaac atcaaaatca ttatcaaact ttactaacct
82681 aatagatgat gatgattggg agtacattga aaaggatgca aagagaacaa gttccaactt
82741 cgctacactg attgatgaaa tatttgagcc tgagaaattt gaattggcga aaagagaaaa
82801 ggctgaactt caaagaaaag ttcaggaagc aaaaaggcaa tcagtgaatg cacagaagat
82861 taatgaggac gagtttggat ccgaagtttc tgatggaatg aaagagctga aaaaaataaa
82921 tgacaaagtg tcgtccccgt tgataaatta cgaattttcg caacaagaac tattgcaaga
82981 tatagacacc ttactaacga atcgttatca actttcgtca tatactaggc ctatttcgag
83041 actggatccc ggattaacgc ctgttactga gacacttcct aacaacctaa aagaaaaaac
83101 agctctgctg caggatactg aaaagaaaat aatagaaacg atacgcagat ccaaattttt
83161 aggatcgcta ctaaatgtta gagggggact atcgccaggg aaaagtgaac tggcacctat
83221 cgaagagtct cccatagttt caaccacacc actaatatat aatgatcgaa tggaacctcg
83281 taggatatcc gatgtggaag tcccacattt cacgaggaaa tcaaaacact ttactactgc
83341 taataatcgg cgctcagtct tatctttgta tgcgaaggat tcaatcaaag acttaaacga
83401 atttttaata aaggaagatc ctgatttgcc tccacaagga agcactgata acgaaagtag
83461 gagcgaagat cccgaaatag ccgagagtat cactgattca aggaatatac aatatgatga
83521 ggatgatagt aaggatggtg ataatgtgaa taatgataat atattgagcg actttcctca
83581 aggcgttggc atatcacagg aatacgacat gaaggataaa aatccaaacc aatctccaat
83641 atcaaaaagc gcagagccca cactggtagt gaaacttccg tctttgagtt ctttccaagg
83701 aaaaaacgcc agtgggttgg gcctatacca aagagagcct tctaaggtaa ccttaccgag
83761 ccttacaagt aataacagca gcgtcggaga aaacatagag gatggggcgg aaaaagggac
83821 tgagagtgag aaaatcgctg cttccctgtc agacgatgac ttgaaggaag ataatgacaa
83881 gaaagataac gacacagtga atgcccctac tacagtaaaa aagccaccta acagcgtact
83941 cttgaaaaaa ttctcaaaag gtaaaatctt agaacttgaa atacatgcca aaatacctga
84001 aaagagacta tacgagggcc tacataaact actagaaggc tggaagcaat acgggttgaa
84061 aaacctggtg tttaacatca ctaatatgat cataacagga aaattggtga atgacagtat
84121 tttattttta cgatctacgc tttttgaaat aatggtttta ccaaatggag acggtagaag
84181 tttaattaaa ttcaataaga aaacaggatc tactaaaacg ctgaccaagc ttgctacaga
84241 aatccaaatt attttacaaa aagaaggtgt tttggacaaa taaaaaaaag ggagggataa
84301 tcgatcccca aatggacaat acgatgtaac accgttatat aattaataaa tgtatatatg
84361 taactatttt ttgaaaaaaa taaaatggta atttattaaa tatttggacc ggtcaacaat

84421 ctcagatata gttgaaaaaa tctttaattg taatagcaga aaaaaaaaac tccgatacgg
84481 ggagtcgaac cccggtctcc acggtgaaag cgtgatgtga tagccgttac actatatcgg
84541 ataacaaatt attaaaatta tttaataaaa agatgaacta atggtgtata aaaacatcat
84601 aaaaactcaa ctcaactgtt ccaataaaat ctgcaattta aattaattat tctgaaaaaa
84661 cacgtaaatt gttggaatgg aaccctaaaa tatttcctct ttagtactgt tggaatagaa
84721 atcaactatc atctactaac tagtatttac attactagta tattatcata tacggtgtta
84781 gaagatgacg caaatgatga gaaatagtca tctaaattag tggaagctga aacgcaagga
84841 ttgataatgt aataggatca atgaatataa acatataaaa cggaatgagg aataatcgta
84901 atattagtat gtagaaatat agattccatt ttgaggattc ctatatcctc gaggagaact
84961 tctagtatat tctgtatacc taatattata gcctttatca acaatggaat cccaacaatt
85021 atctaattac ccacacattt ctcannnnnn nnnnnnnnnn nnnnnnnnnn nnnnnnnnnn
85081 nnnnnnnnnn nnnnnnnnnn nnnnnnnnnn nnnnnnnnnn nnnnnnnnnn nnnnnnnnnn
85141 nnnngtatgt tggaatagaa atcaactatc atctactaac tagtatttac attactagta
85201 tattatcata tacggtgtta gaagatgacg caaatgatga gaaatagtca tctaaattag
85261 tggaagctga aacgcaagga ttgataatgt aataggatca atgaatataa acatataaaa
85321 cggaatgagg aataatcgta atattagtat gtagaaatat agattccatt ttgaggattc
85381 ctatatcctc gaggagaact tctagtatat tctgtatacc taatattata gcctttatca
85441 acaatggaat cccaacaatt atctcaacat tcacccattt ctcaagtact attcatgtta
85501 cttgagaaat gggtgaattt tgagatagtt gttgggattc cattgttgat aaaggctaat
85561 aatattaggt atacagaata tgctagaggt tctcctcgag catataggaa tcctcaaaat
85621 ggaattagca tttctacata ataacctcaa ttccctttttg tggattcaca tattctcgat
85681 cggaactcct agcatagtgc agacaacact cacatagctc tcccatttat caagttggaa
85741 ttcactccaa gaaattttct tgcttctctc tttcccacag aaatgaaaag gaacagtttg
85801 gcggctattg ttttagccct tactttttgg accctttgaa aaaatgccaa gaatgcgtga
85861 cggtttccct ccatgctcga gaaccacaaa aaccctatgt ggtattcgat tatgcgactc
85921 acggcaaata tttgacttgt caaagttatt cgggatttcc ttgaatccct gagaagtgaa
85981 atggaaagaa tttaagggca taaatacctt tcaagcaccc agaagaacag taaaataaag
86041 caaggtacgt gaaattaata tttttaaatg gttctaaccg atgccgaaga actgcgcagt
86101 ccggttataa cgtctgacat gtcctttttt gatttggaat ccaaccactc aagtgactct
86161 gttcatttac tttgcgaaaa atatacccac aaattgccca tcgaaagtga atcgcaaacc
86221 accttcagac tggcaccgac aaagcaaaga ttatacagac agagtacttt atacgtaccg
86281 ttaagtctca agcaaagggt tttcttattt actgaacggg taaagagtat ctgggccggc
86341 ttgccaagat gcaaaccgaa taagtatttc aaagttgcat ttgccttagc cgtcctgaca
86401 ccattggcta tttggatatt ttatattgac tttcgtgtac attgatcaca tcgactgttc
86461 tattggcaaa tgaaccacgg gcattgacta tttttcaggt tactactata tattattgtt
86521 ggaataaaaa tcaactatca tctactaact agtatttacg ttactagtat attatcatat
86581 acggtgttag aagatgacgc aaatgatgag aaatagtcat ctaaattagt ggaagctgaa
86641 acacaaggat tgataatgta ataggatcaa tgaatattaa catataaaat gatgataata
86701 atatttatag aattgtgtag aattgcagat tcccttttat ggtgttggaa taaaaatcaa

86761 ctatcatcta ctaactagta tttacgttac tagtatatta tcatatacgg tgttagaaga
86821 tgacgcaaat gatgagaaat agtcatctaa attagtggaa gctgaaacac aaggattgat
86881 aatgtaatag gatcaatgaa tattaacata taaaatgatg ataataatat ttatagaatt
86941 gtatagaatt gcagattccc ttttatggat tcctaaatcc tcgaggagaa cttctagtat
87001 atctacatac ctaatattat tgccttataa aaaatggaat cccaacaatt acatcaaaat
87061 ccacgttctt ttcatatgga ttcctaaatc ctcgaggaga atttataata tagtctgtat
87121 acataatatt atagcttcta acgacaatgg aattctaaca attatcaaat tgtccgccag
87181 ttcctcaaaa agatccatgt ataatcttca ttattacagc cctcttgacc tctaatcatg
87241 aatgttctca aataaatctg ttggaataaa aatcaactat catctactaa ctagtattta
87301 cgttactagt atattatcat atacggtgtt agaagatgac gcaaatgatg agaaatagtc
87361 atctaaatta gtggaagctg aaacgcaagg attgataatg taataggatc aatgaatatt
87421 aacatataaa atgatgataa taatatttat agaattgtgt agaattgcag attccctttt
87481 atggattcct aaatcctcga ggagaacttc tagtatatct acatacctaa tattattgcc
87541 ttataaaaaa tggaatccca acaattacat caaaatccac gttctcttca tggtagcgcc
87601 tatgcttcgg ttacttctaa ggaagtccca tcaaatcaag atccgttagc cgtttcagct
87661 tccaatttac cggaatttga tagagattcc actaaggtta attctcaaca agagacaaca
87721 cctgggacat cagctgttcc agagaaccat catcatgtct ctcctcaacc tgcttcagta
87781 ccacctccac agaatggaca gtaccaacag cacggcatga tgaccccaaa caaagctatg
87841 gcctctaact gggcacatta ccaacaaccg tctatgatga cgtgttcaca ttatcaaacg
87901 tcacctgcgt attatcaacc ggacccacac tatccgttgc cacagtatat cccaccactg
87961 agtacttcct cacctgatcc aatcgattca cagaatcaac actctgaagt acctcaagct
88021 gagacaaagg tgagaaataa cgtcttacca ccacacactt taacatcaga agaaaacttt
88081 tctacatggg ttaaatttta catcagattt ttgaagaact ctaatctcgg tgacatcatt
88141 ccaaatgacc agggtgaaat caaaagacaa atgacttatg aagaacatgc gtatatatac
88201 aataccttcc aagcatttgc cccatttcat ttattgccaa catgggtaaa acaaatttta
88261 gaaattaatt atgctgacat ccttacagtc ctttgtaaaa gtgtgtccaa aatgcaaact
88321 aacaatcaag aattaaagga ttggatagct cttgccaacc ttgagtacga cggaagtaca
88381 tctgctgata catttgaaat tacagtcagt acgatcattc agaggctaaa agaaaacaat
88441 atcaatgtta gcgacagatt ggcctgtcaa ctaatactta aaggtctatc cggtgacttc
88501 aaatacctac gtaatcaata tcgtaccaaa acgaacatga aactttccca attattcgct
88561 gaaattcagt taatatatga cgaaaataaa atcatgaatc taaataaacc gtcccaatac
88621 aaacaacaca gcgaatacaa aaatgtttct cgcacatctc caaacacgac taacacaaag
88681 gttacaactc gtaattatca gagaacaaat agttcaaaac caagagcagc aaaagctcac
88741 aatattgcta catctagtaa attctcaagg gtgaacaatg atcacattaa tgaatcaacc
88801 gtttcatcac aatacttaag cgatgacaac gaacttagtc ttaggccagc aacagaaaga
88861 atctaagcca acacacacaa tagactcgaa tgacgaacta cctgatcacc ttcttattga
88921 ttcaggagct tcgcaaacgc ttgtcagatc agcccattat ttacaccatg caacacccaa
88981 ttctgaaata aacatagtcg atgctcaaaa acaagacatt cctataaatg ccattggtaa
89041 tcttcacttc aactttcaga acggcaccaa aacatcaata aaagcactac acacaccaaa

89101 catagcctat gatctattaa gtttgagtga gctggctaat caaaatatta ctgcctgctt
89161 taccagaaac actttagaaa gatcggatgg tacagtccta gctcccatag tcaaacatgg
89221 agacttttac tggttatcta aaaaatacct aattccttcg cacatttcaa agctaacaat
89281 aaacaacgtc aacaaaagca aaagcgtaaa taaatatcca tatccgttaa tacatcgaat
89341 gcttggacat gctaacttcc gaagtattca gaagtctctt aagaagaatg cagttacata
89401 tttgaaggaa tcggatattg aatggtctaa cgctagcaca tatcaatgtc ctgactgtct
89461 aatcggcaaa agcacgaaac atagacatgt caaaggatca cgactaaagt accaagaatc
89521 atatgagcct tttcagtact tgcataccga tatatttggt cctgtacatc acttaccgaa
89581 aagtgcacct tcttacttta tatcgtttac agatgagaaa accagattcc aatgggtgta
89641 cccattacac gaccgtcgtg aagaatctat cctcaatgtt tttacatcga tattagcatt
89701 tattaagaac caattcaatg ctcgcgttct agttatccag atggatcgtg gctccgagta
89761 cactaacaaa actcttcata agttctttac gaacagaggt attactgcat gctatacaac
89821 cacggcagat tctagagcac acggtgtcgc tgaacgatta aatcgtactt tattaaacga
89881 ttgtcgcaca ctgcttcatt gcagtggtct accaaatcat ctatggttct cagcagtcga
89941 attttctact ataatcagaa attcattagt ctcaccaaaa aacgataaat ccgcaagaca
90001 acatgcaggt ttagctggac tggacattac tactatacta cctttcggtc aaccggttat
90061 agttaacaac cataatcccg actcgaaaat acatcctcgt ggcattccag gttacgcctt
90121 acatccgtca cgaaactctt atggctatat tatctatctt ccatcattaa aaaagacagt
90181 agatactacc aattacgtta tattacaaga caagcaatcc aaattggacc aattcaatta
90241 tgatacactc acctttgatg atgatctcaa tcgtttaaca gcccataacc aatcttttat
90301 tgaacaaaat gaaacggagc agtcatatga tcaaaataca gaatctgatc atgactatca
90361 atcggagatt gaaataaact ctgatcctct agtgaacgac ttctcgtccc aatcaataaa
90421 ccctttacaa ttagacaagg aaccagtcca aaaagtacgt gcaccaaaag aagttgatgc
90481 cgacatatct gaatacaata ttcttccatc tcctgtacga tctcgtacac cccatatcat
90541 taataaagag agtaccgaaa tgggtggtac cgttgaatca gatactactt cacctagaca
90601 ctcgtctacc ttcactgcac gaaaccaaaa gcgacctggt agtcccaatg atatgattga
90661 tttgacctca caggatagag ttaattatgg acttgaaaac atcaaaacta cacgtttggg
90721 tggtacggag gaaccatata ttcaacgaaa tagtgataca aatatcaaat acaggactac
90781 aaatagtacg ccctcaatag atgaccgttc gtccaacagt gaatccacta ctcccatcat
90841 ctccatagaa acaaaggctg tatgtgataa tacaccctcc attgatacgg atccgccaga
90901 atatcgatct tctgaccatg cgactcctaa tataatgcct gacaaatcct caaaaaatgt
90961 tacggctgat tctattcttg acgacctccc acttcctgac ttaacccatc aatctcctac
91021 ggacacttct gatgtttcaa aagatattcc acacatacac tctcgtcaga ctaattccag
91081 tttgggtggt atggatgatt ctaatgttct gactactacc aaaagtaaga aaagatcatt
91141 agaagataat gaaactgaaa ttgaggtatc ccgagacaca tggaataata agaatatgag
91201 aagtctggaa ccaccaagat cgaagaaacg cataaattta attgcagcaa taaaaggagt
91261 gaaatcgatc aaaccagttc gaacgacctt aagatatgat gaagcaatta cttataataa
91321 agacaacaaa gaaaaagaca gatatgttga agcttatcat aaagaaatta gccaactatt
91381 gaaaatgaac acttgggata caaacaaata ttatgataga aatgacatag atcctaaaaa

91441 agtaataaac tcaatgttta tatttaacaa gaaacgtgat ggtacacaca aagctagatt
91501 tgttgcaaga ggcgacattc aacaccccga tacatatgat tctgatatgc aatccaatac
91561 cgtacatcac tatgcactga tgacgtcact gtcaatcgca ttagacaacg actattatat
91621 cacacagctg gacatatcct ctgcttactt atatgctgat atcaaagaag aattatacat
91681 aagacctcca ccacatttag gtttgaatga taaattacta cgtttgagaa aatcactcta
91741 tggtttgaaa caaagtggtg caaactggta tgaaaccatt aaatcatatt taataaattg
91801 ttgcgacatg caagaagttc gcggatggtc atgcgtattt aagaatagtc aagtaacaat
91861 ttgcttattc gttgatgata tgatattatt cagcaaagac ttaaatgcaa ataagaaaat
91921 cataacaaca ctcaagaaac aatacgatac aaagataata aatctgggtg aaagtgataa
91981 cgaaattcag tacgacatac ttggattaga gatcaaatat caaagaagca agtacatgaa
92041 attaggtatg gaaaaatcct tgacagaaaa attacccaaa ctaaacgtac ctttgaaccc
92101 aaaaggaaag aaacttagag ctccaggtca accaggtcat tatatagacc aggatgaact
92161 agaaatagat gaagatgaat acaaagagaa agtacatgaa atgcaaaagt tgattggtct
92221 agcttcatat gttggatata aatttagatt tgacttacta tactacatca acacacttgc
92281 tcaacatata ctattcccct ctaggcaagt tttagacatg acatatgagt taatacaatt
92341 catgtgggac actagagata aacaattaat atggcacaaa aacaaaccta ccaagccaga
92401 taataaacta gtcgcaataa gcgatgcttc atatggtaac caaccatatt acaagtcaca
92461 aattggtaac attttcctac tcaacggaaa agtgattgga ggaaagtcga caaaggcttc
92521 gttaacatgc acttcaacta cagaagcaga aatacacgca gtcagtgaag ctattccgct
92581 attgaataac ctcagtcacc ttgtgcaaga acttaacaag aaaccaatta ttaaaggctt
92641 acttactgat agtagatcaa cgatcagtat aattaagtct acaaatgaag agaaatttag
92701 aaacagattt tttggcacaa aggcaatgag acttagagat gaagtatcag gtaataattt
92761 atacgtatac tacatcgaga ccaagaagaa cattgctgat gtgatgacaa aacctcttcc
92821 gataaaaaca tttaaactat taactaacaa atggattcat tagatctatt acattatggg
92881 tggtatgttg gaataaaaat caactatcat ctactaacta gtatttacgt tactagtata
92941 ttatcatata cggtgttaga agatgacgca aatgatgaga aatagtcatc taaattagtg
93001 gaagctgaaa cgcaaggatt gataatgtaa taggatcaat gaatattaac atataaaatg
93061 atgataataa tatttataga attgtgtaga attgcagatt cccttttatg gattcctaaa
93121 tcctcgagga gaacttctag tatatctaca tacctaatat tattgcctta ttaaaaatgg
93181 aatcccaaca attacatcaa aatccacatt ctcttcaaaa tcaattgtcc tgtacttcct
93241 tgttcatgtg tgttcaaaaa cgttatattt ataggataat tatactctat ttctcaacaa
93301 gtaattggtt gtttggccga gcggtctaag gcgcctgatt caagaaatat cttgaccgca
93361 gttaactgtg ggaatactca ggtatcgtaa gatgcaagag ttcgaatctc ttagcaacca
93421 ttattttttt cctcaacata acgagaacac acaggggcgc tatcgcacag aatcaaattc
93481 gatgactgga aattttttgt taatttcaga ggtcgcctga cgcatatacc tttttcaact
93541 gaaaaattgg gagaaaaagg aaaggtgaga gcgccggaac cggcttttca tatagaatag
93601 agaagcgttc atgactaaat gcttgcatca caatacttga agttgacaat attatttaag
93661 gacctattgt tttttccaat aggtggttag caatcgtctt actttctaac ttttcttacc
93721 ttttacattt cagcaatata tatatatata tttcaaggat ataccattct aatgtctgcc

93781 cctaagaaga tcgtcgtttt gccaggtgac cacgttggtc aagaaatcac agccgaagcc
93841 attaaggttc ttaaagctat ttctgatgtt cgttccaatg tcaagttcga tttcgaaaat
93901 catttaattg gtggtgctgc tatcgatgct acaggtgttc cacttccaga tgaggcgctg
93961 gaagcctcca agaaggctga tgccgttttg ttaggtgctg tgggtggtcc taaatggggg
94021 taccggtagt gttagacctg aacaaggttt actaaaaatc cgtaaagaac ttcaattgta
94081 cgccaactta agaccatgta actttgcatc cgactctctt ttagacttat ctccaatcaa
94141 gccacaattt gctaaaggta ctgacttcgt tgttgtcaga gaattagtgg gaggtattta
94201 ctttggtaag agaaaggaag acgatggtga tggtgtcgct tgggatagtg aacaatacac
94261 cgttccagaa gtgcaaagaa tcacaagaat ggccgctttc atggccctac aacatgagcc
94321 accattgcct atttggtcct tggataaagc taatgttttg gcctcttcaa gattatggag
94381 aaaaactgtg gaggaaacca tcaagaacga attccctaca ttgaaggttc aacatcaatt
94441 gattgattct gccgccatga tcctagttaa gaacccaacc cacctaaatg gtattataat
94501 caccagcaac atgtttggtg atatcatctc cgatgaagcc tccgttatcc caggttcctt
94561 ggggtttgtt gccatctgcg tccttggcct ctttgccaga caagaacacc gcatttggtt
94621 tgtacgaacc atgccacggt tctgctccag atttgccaaa gaataaggtc aaccctatcg
94681 ccactatctt gtctgctgca atgatgttga aattgtcatt gaacttgcct gaagaaggta
94741 aggccattga agatgcagtt aaaaaggttt tggatgcagg tatcagaact ggtgatttag
94801 gtggttccaa cagtaccacc gaagtcggtg atgctgtcgc cgaagaagtt aagaaaatcc
94861 ttgcttaaaa agattctctt tttttatgat atttgtacat aaactttata aatgaaattc
94921 ataatagaaa cgacacgaaa ttacaaaatg gaatatgttc atagggtaga cgaaactata
94981 tacgcaatct acatacattt atcaagaagg agaaaaagga ggatgtaaag gaatacaggt
95041 aagcaaattg atactaatgg ctcaacgtga taaggaaaaa gaattgcact ttaacattaa
95101 tattgacaag gaggagggca ccacacaaaa agttaggtgt aacagaaaat catgaaacta
95161 tgattcctaa tttatatatt ggaggatttt ctctaaaaaa aaaaaaatac aacaaataaa
95221 aaacactcaa tgacctgacc atttgatgga gtttaagtca ataccttctt gaaccatttc
95281 ccataatggt gaaagttccc tcaagaattt tactctgtca gaaacggcct taacgacgta
95341 gtcgacctcc tcttcagtac taaatctacc aataccaaat ctgatggaag aatgggctaa
95401 tgcatcatcc ttacccagcg catgtaaaac ataagaaggt tctagggaag cagatgtaca
95461 ggctgaaccc gaggataatg cgatatccct tagtgccatc aataaagatt ctccttccac
95521 gtaggcgaaa gaaacgttaa cacaccctgg ataacgatga tctggagatc cgttcaacgt
95581 ggtatgttca gcggataata gacctttgac taatttatcg gatagtcttt tgatgtgagc
95641 ttggtcgttg tcaaattctt tcttcatcaa tctcgcagct tcaccaaatc ccgctaccaa
95701 tggggggggcc aaagtaccag atctcaatcc tctctcttgg ccaccaccgg atagtaaagg
95761 ttctaatcta actcttggtc tccttcttac atagatggca cctattccct ttggaccgta
95821 aatcttgtga gaagaaattg atagtaaatc aatgttcatt tcattgacat caatgtgaat
95881 cttaccatag gcttgtgcgg cgtcagtatg aaagtagatc ttattctttc tacaaattgc
95941 accaatttct ttaataggtt gaatgacacc gatttcatta ttgacagcca tcacagagac
96001 gagacaggta tctggtctaa tggcatcttc caattccttc aaatcgataa gaccttgatc
96061 gtccacattt aggaaagtga cttcaaatcc ctccttcatc atggcccgtg cggcttccaa

96121 gacacacttg tgttccgttc tagtggtgat gatgtgtttc ttagtcttct tataaaatct
96181 tgggacaccc ttaagaacca tattattaga ttcggtcgct cccgaagtga atattatttc
96241 cttggggtcg gcattgatca tctttgctac gtgagctcta gcattttcca cagcagtatt
96301 tgtttcccaa ccgtaagagt gagtgttgga atgaggatta ccataaagtc ccgtataaaa
96361 cttcaacatc gtatccaaaa ccctagggtc tgttggtgta gtggcttgca tgtcaagata
96421 tatgggacga gtaccaaaac ctgtgttttc ttgataagca tggctcattg cagtgctacc
96481 agaagctact acagcatctg gggtggtacc ggatgcactc gcacgggcac tagcctgtgc
96541 ctttgcagca gcctgaatat cggtatgcgt ttccagagag aagttgtcgt ctaacttcac
96601 gcctgctgca ggaggggaat agaatctcct gcttaccaaa caagccctat atgtggccgc
96661 tggaacgttg taaacttgag ataatcttgt tatcgatctt gtagcagttg atttcaacat
96721 cttccagttg tagcagttcc ttcgtgtatg cgatatattt taatgaaaat gatgtatctg
96781 aaatagaaaa tataacacac taatgaactc tgtcggtctt tactctgagt gagttactta
96841 gatagtcatc atatatccct gggcgatggc cttactattt gtcagcgacc gtatctaaaa
96901 ttggtaggtc actatcaact attgcgggta atagggtgca taatagtatt gtaatatata
96961 taaatatata aagctactat gatagatatt tgttgctatt cacgtcatcc actattactt
97021 tagtggttcc gctatatatt cgcattgccc atcgaaggca gttttgtgtg ctacctgctc
97081 gtgaccacgg tctttttgcc cagtctttta cggcgggcat acttcatgat gaaactgtct
97141 attttcatac ctcttgaatt tagttcttca attagaggct tgatatcctc caagtcccac
97201 tgtgactgta gcctaaacag gactttaaac cgttctttgg ggtccattgg tagtgtgctt
97261 ttcgctatat actggacagt cttatcggta ggcttgaaat gatagcctcg cagcatgtca
97321 atgtcaatgt cacatgggaa gaaaggtggg aaaagggact tccacttgat gaggaactca
97381 tcaattggca tgcttattcc agaaacatat ttccttagcg cttgaatccc gtaccactga
97441 gctataaacg gtattctcaa gcgccacgta ttgttttcag cctcttgctc ttgaacagca
97501 aatttattca gcactgtttc aatgatttct cttgtgtatg gattgaactc gtcctcaatg
97561 tccttcccca cagcgtgatg tgtatcctca acattcagat gctgtagatc gagtgattct
97621 gccattgcgc tcatcagtag tacatgcagt gctttgaaaa ggaagtcttg tgaaagaata
97681 cacaacacac cgtctttcac agatccacca atcttatgcc attttgatat accttctagc
97741 gcagaacatg gtgagttctc aagtagttct tccagggtcc cgataacctt tgtagatgac
97801 ctcttcatga ttttgtcgga gaaatccagt tctccgttgt atattggtac tgaattcaag
97861 ttcaattcgc catgtgtctc tctggtctca aattctgatt cagtcttggc gaatcccacg
97921 acgtccatgt acggcttgga cagtccaaac aagagcgttt cgtcgaaagt aataggttgt
97981 tcaggaacaa attctctcat tagtagaact gtgtttgaat gtttgcgctg cttcagcacc
98041 caagtcttgt cgtgcgaaca cagtacaact tcagacttgt ctttgtccaa tgacttaaac
98101 cttaactggt gattttgaac cggatcctgt attatatcca gtaactctgg tgtcaactgg
98161 atcagcttat aagatggatc atactcgggt gcggaatgta ggttgatgga catggtacag
98221 gattattagt gagcaaatct ctggtggatt tttaccgttc atctcatagt gctttttttag
98281 tttatttttc actttttct ggttatacta tacgctgcct actgaagcaa taagcaagaa
98341 acacaacagt ataaagggac aaagataata aaccgcctac tactaaataa tatcgctgtg
98401 gtatgctact cctagagatt tcgatcacca tcgtggtgct ctttgtcata cgcatagaat

98461 tgacaaaaaa gaaccctaag ctataaatta ggtcatgttg agcgattgcc agttgccaag
98521 acctagggtt ttgttcgaac aattctaaaa aggtaaataa aaacaatggt aaacaaaggc
98581 tttgatcttc aatgaggaag atttctctca ctgcggatag taaaaacaac actagtttgt
98641 ctgtactcgc attctactgc tgctacctgc tactcattac gtgttcgggc tgttatctgg
98701 ttgctaaaag agtatattta cacctcacca atggagaaag acctgtcgtc tctttactct
98761 gaaaagaaag acaaagagaa cgatgaaacc ttatttaaca tcaaactatc caaatctgtt
98821 gtcgagacca caccgctaaa tggtcattca ttgtttgatg atgataaatc actttcagac
98881 tggacggata atgtgttcac tcaatcagta ttctatcacg ggtcagatga cttgatatgg
98941 gggaagttct ttgtctgcgt gtacaagtcc cccaacagca ataagttgaa cgctataata
99001 ttcgacaaat taggaacatc atgcttcgaa tccgtcgata tatcttccaa ctcgcaatac
99061 tatccggcca ttgagaattt gagtccaagt gatcaggaaa gcaatgttaa gaaatgcatt
99121 gctgtcattc tgttacagcg ctatccatta ctttcaccat cagacttatc acaaatattg
99181 tccaataaat cggaaaattg cgactatgac cccccttatg ctggagattt ggctagtagt
99241 tgccagttga taacagcagt tcctccagaa gatctgggga agcgcttctt tacatcagga
99301 cttctgcaaa atagatttgt cagctctacc ctgttagatg ttatttatga aaacaatgaa
99361 tccaccatcg aactaaataa taggttggta ttccatctgg gtgaacaact tgaacaactt
99421 tttaacccag tcacagaata ctcaccggaa cagacagaat atggttataa ggcgccagag
99481 gacgaattac ccacagaatc ggatgatgat cttgtcaagg ccatttgcaa cgagttatta
99541 caactacaaa caaattttac tttcaatttg gtagaatttt tgcaaaaatt cctgatcgcc
99601 ttgagagtca gagtactcaa tgaagaaatt aatgggttat ccacaaccaa attaaatcga
99661 ctcttcccac ctacaataga tgaagtcaca agaatcaatt gtatttttct agactcgcta
99721 aagacagcaa tcccttacgg ttccctcgaa gtactgaagg catgcagcat tactattcct
99781 tatttctaca aagcatatac aagacacgag gcggccacaa agaacttcag caaagatatt
99841 aaattgttta ttaggcattt cagcaatgta attccagaaa gagaggtcta cacggaaatg
99901 aaaatcgaga gtataattaa gggacctcag gaaaaactac tgaagctaaa gttaattata
99961 gagagattgt ggaagtcgaa aaaatggaga ccgaaaaatc aagaaatggc aaaaaaatgc
100021 tacaacaata tcattgatgt cattgattcg tttggaaaat tagattcccc acttcattct
100081 tatagtacca gagtatttac tccatcggga aaaatcctta cagaattagc caaatgctgg
100141 cccgtagaac tgcaatacaa atggctgaag agaagggtag tcggtgtgta tgatgtagtg
100201 gatttgaatg atgaaaataa gagaaattta ttagtcatat tcagtgatta tgtggttttc
100261 atcaatatac tggaggcaga aagttactac acttcagatg gatcaaacag gcccttaatc
100321 tcagatattt taatgaactc attgatcaac gaagttccgt tgccctccaa gatccctaag
100381 ttgaaagtgg agcgtcattg ctatatagat gaggttctag tttctatatt agacaaaagc
100441 actctacgtt ttgatcgatt gaagggaaaa gattctttct caatggtatg taaattatcc
100501 tctgcattta tctcttcttc gtcagttgct gacttgatta cgaaggctag aattttggaa
100561 aaagacactg catttcattt atttaaagct agtagaagcc attttacatt atattctact
100621 gctcacgagc tttgcgctta tgattccgaa aaaataaaat caaaatttgc cttattcctg
100681 aacataccac catccaagga gatattggag gtcaacaacc ttcatttggc tttttttgca
100741 agattttgca gtaacgatgg tagagataac atcgtaatct tagacgtctt aaccaaacat

100801 gacgataaac atatagaagt tacatccgat aacattgttt tcaccataat taatcaattg
100861 gccattgaaa taccgatatg cttttcttcc ttaaactcat cgatggccaa agatttactc
100921 tgtgtaaatg agaatttgat aaaaaactta gaacatcaat tggaagaggt caagcaccct
100981 tcaacagacg aacatagggc tgttaatagc aaactttccg gtgcatccga tttcgatgct
101041 actcacgaga agaaaagatc atacggtacc ataacaacat ttagaagcta tacaagcgac
101101 ttgaaggaca gtccatcagg cgataatagt aatgtcacca aggaaactaa ggaaatttta
101161 ccagtgaaac ctacgaaaaa gtcttcaaaa aaaccaagag aaattcaaaa gaagaccaag
101221 acaaacgcct ctaaagcaga gcacatagaa aagaagaagc ctaacaaagg caaagggttt
101281 tttggcgtgt taaaaaatgt ttttggaagt aaaagcaaga gcaagccttc accagttcaa
101341 agagtgccta aaaaaatatc gcagaggcat cctaagtctc cagtgaagaa gccaatgacc
101401 tcagaaaaga aatcctcccc taaaagggca gtcgtttcat ctcccaaaat taaaaagaaa
101461 agtacttctt tttccacaaa agaatcacaa actgctaaat cttctcttcg agcagttgaa
101521 ttcaaatctg atgacttgat cggaaaacca cctgatgttg gaaatggcgc acatcctcaa
101581 gaaaatacca gaatatcttc agtagtaagg gatacaaaat atgtctccta caatccctct
101641 cagcctgtga cagaaaatac cagtaacgaa aaaaatgtcg aaccaaaagc ggatcaatcc
101701 acaaagcagg ataacatttc caattttgca gatgtagagg tatctgcgtc ttcttatcct
101761 gaaaaacttg atgcagaaac agatgatcaa ataattggga aggcgacgaa ttcgtcatca
101821 gttcatggaa ataaagagct gccagacctt gctgaggtga ctacagcaaa tagggtttct
101881 acaacatcgg ctggggacca acgtattgat acccaaagcg aatttttacg tgcagctgat
101941 gttgaaaact taagtgatga cgatgaacac agacagaatg aaagtagagt ttttaacgat
102001 gacctctttg gtgattttat tcctaagcat taccgtaata aacaggagaa cattaacagc
102061 tcgagtaatt tgtttccaga gggaaaggtg ccccaagaaa agggcgtatc aaatgaaaac
102121 actaacatat ctctcaaaac taatgaagat gcatctacat tgacgcagaa actctctcca
102181 caagcgagta aagtgctgac agaaaattct aatgaattaa aagataccaa caatgaaggg
102241 aaggacgcaa aggacataaa attaggagat gattacagtg ataaagaaac agcgaaagaa
102301 ataactaaac caaaaaattt tgttgaagga ataactgaac ggaaagaaat attccccact
102361 attcctaggt tagcgccgcc agcttcaaaa attaactttc aaaggtcacc atcctatatt
102421 gagctctttc aaggaatgag ggtggtttta gataagcatg atgcccatta taactggaaa
102481 cgcttggcta gtcaagtctc cttaagtgag ggactaaaag tcaatactga ggaagatgcg
102541 gcaattataa ataaaagtca ggatgatgcc aaggcggaaa gaatgactca aatttctgaa
102601 gtgattgagt atgaaatgca gcaacctatc ccaacttatt tgcctaaggc gcatctagat
102661 gactcgggta ttgaaaaaag tgatgacaaa ttcttcgaaa ttgaagaaga acttaaggaa
102721 gaattgaagg gcagcaaaac cggtaatgaa gatgtcggta ataataatcc atccaattct
102781 attccaaaaa tcgagaagcc cccagcattc aaagttatta gaacatcgcc tgtgagaatt
102841 atcggaagga cttttgaaga cactagaaaa tatgaaaatg gctctccatc tgatatttcg
102901 ttcacttatg atactcacaa caatgatgaa cctgacaaaa ggctgatgga attaaaattt
102961 ccatcccaag atgaaattcc ggatgacaga ttctatactc cagcagagga acccactgct
103021 gaatttccgg tggaagaact tccaaatact ccgcgaagta ttaacgttac aacttcaaat
103081 aacaagagca cagacgataa gttgagtagc ggtaatattg atcaaaaacc taccgaactg

103141 ttagatgatt tagaattcag ttcatttaat atagcatttg gaaatacctc catgagtact
103201 gacaatatga aaatatcatc cgacttaagt tcgaataaaa ccgtgttagg aaatgctcag
103261 aaagttcaag agtctcctag tggaccatta atctatgttt tgcctcagag tagcacaaag
103321 catgagaaag aggggttcct tcgaaagaaa caaaaagacg agcccatttg ggtttcccct
103381 agcaaaattg actttgctga tctaagtagg agaactaaag cattgacgcc agagcgtaat
103441 actgttcctt tgaaaaacaa cgacagtaga aaatacaaat atactggaga gggatctatc
103501 ggtaatatga caaatatgct gttaactaaa gatgcttcgt acgcgtactt aaaagatttt
103561 gttgcgttga gtgacgatga agatgaagat gggaaacaga actgcgctgt tggtggccca
103621 gagaaactga aattttattg agtgttttat tgattttttt tttcttttta atttaatgca
103681 atgtatacat ttacataaaa tatggacaaa taagaaaaag ggggaaaaac ctcccactta
103741 agtaaaactg aatatattag atacctttac ctactatcaa aggtgaatgc ttttggatca
103801 ataggggttc catcgggggc aataccattt tggccgaact ttctttccat atataatagt
103861 ctagccttat cgaaatctac attatcgttt tccataattt gtttaatctc attcttggag
103921 acatcatcca aaccacctct tacgtcgttg ctattctccg agataatatc gaaattcctg
103981 gaatggagac cgttttctag gtcatcagca aaggaatcat ttaaacgtat atgtctatta
104041 aaacgtaatt gtattcttga acgcagtgca cgaaatttgt cggctattac agctgttagt
104101 atttagatac ggtttgcact ttgcattttc aaatgcttaa atatacatac caactttggt
104161 tttccggtgt gccataagaa ggcataaaga caatagcaca acccagagga gcaatttgag
104221 atagaacaaa ctcttcatac tcgtatttgc aatttttgga aattaattgt gtagcttttc
104281 ttctcttaat tggtggcagt ttttatatat tcagtttaga ataccctttt tttttttttt
104341 tttttcttcg cttttcgcat ctcatcgctg gaagtggtgc tcttttacag ggattaaata
104401 aggttattct ttttggtcaa aatgataaat ataactatta aagttattta taacccgccc
104461 gcttccttat tatttatacg ttatcataaa gtacacaggt catggttcgg ttggtgctta
104521 ggaattaatc acgtctagca taagagatct gtaaactaca accaccataa ttataattat
104581 ttaatttttg aatacaaaaa tctgcatcta ctaaattttc atattctaca acagcaacac
104641 cagttggttg accattttcc tgtggtttca attccgcgtt attgattttg ccgataggcc
104701 caaacaagtc gaataaatcg cttcttgctg ttgagaatgg taaattacta caataaatga
104761 aacagtttct atcccctcct ggattgacac cttcagtaaa ttttgctgca gtttcatcaa
104821 tgtggacagc ggcatcctgc gcaagacctg gttcagtacc tctggtatct tcaaggtcct
104881 cacgcctttg attataacga tcattgttct ttctcttgtt gaaacgccct tctctaactt
104941 ccaaaactct accttctact tccatgccgt tgaatgtatc gatagctcta atcatttcat
105001 cctcagtagg ataaatgaca gaaccgaatc ctcttgaaaa tccgttgaaa tccaattcta
105061 catcggcacg caagacatga ccacattctt taaacatatc ttttaaggat tgccaattca
105121 tagagtacgg taaattgatg atgaacactt caaacccatt atctatttct tccctagtag
105181 ctttcttaga aaattccttg gcagcttcag gaggaggatt atcctgtctt accattagtt
105241 tccggtccat aaagagggca ccatcaaact tcgatatggc atcttggaca gattcgtttt
105301 tggtaaattc cacagtcccc ataccacggt gatggccctt tgatgtgata atgtcagcct
105361 ccacaacttc gcccactgta ccaaacaatt ccttaaggtc ttcaggggta caatcaaaag
105421 tcaagtttct cacgaagata ctattttcaa aatttctctt ggtcgcgtca aattgccttt

105481 ctaagtctct ttcaacaatt ggacctaaag tacgacccct accgccaccg cgtccccctc
105541 tgaatgatct gcttcctcct ctgccacggc ctcctcggcg agggggccag ttcctatcat
105601 cgtggtagcg gctgccacca taactctgat cgtaccgatc attgaaacgg gagcctcggt
105661 cgcgtctctg acgacgactg ccattaccat tattactgct actatcgtta taatcatcgt
105721 acctgtctct gtcgtcgctc aaacgacgac gtacaggtga tcttgatcta ctcctatcat
105781 ttccatacat ccctagctct ctctccatct tggcgaaaaa atccagctga tgtttgtttc
105841 cttttcgccc atagcttatt tctaatggga aatcctcaat ataattgatt tcatccaagc
105901 tttcattttt tttcaatttt tttcaccttc aagtgaatac atcacttccg acgataccaa
105961 aataatgaca aaatgccatg agaagatctt atgatatgta gtaaatgtta accaccattg
106021 ctatttcctt gctaggtttg ccaaagccgt tggcgaaggc agcactagac ggcgagtcac
106081 ttccgtctct ttgtccactt cctcttcacc atcaaatcga agcctgcaag tgccgtccct
106141 cttggttccg atgacaattg caggataaaa agtggtcgtt tcaggatatc ttgccagaac
106201 cttagtacct ggcgggtagt tcttagtggg aaagcctggc gggatcaaca gtagttcttt
106261 ccgattacat ttatacacct tgcctgagtt gcccaattca tcaggttcag gatctctcac
106321 ttcaaatcgt gtaccatctg ctacgacttt gagtacttca cactgtatcc attcgccgtc
106381 ggcactgccc cttctgggct tataggcaac ctctgagccc accaaaatag gtgcatttgg
106441 gttgtattcg ctggtccagt acgatctgcc aactttcccc acagctttct tgccctgcga
106501 caatgccatg cgtgtccctt ctccttcttc tgcatcttca gtattttcct cactcttttc
106561 ctctcgtgac tgattcaaca tcaacgacaa attggaacgt acattgtcaa ggattcggtg
106621 cactctgttg acgttttcca tgtgttcttg gaaagtgttg agatgcaatt gcagctgatc
106681 ctttgacata ttaaggaaat ttagtttttt tgtttgcagc tcatcatcaa agggaatgac
106741 ctcgttggca ttgtatatgt cttgcaagga cgacactaca acatcccaat atccgtccat
106801 aataagaaag gtgaccgtgt gttttgctgt gaaaaactcc ccccacttag agaagccacc
106861 aaggtattgt gtctttaacc tttttcgtat ctggcaaaat cgaagaggaa agcagaaaac
106921 caaaagaaaa aaatagttta gtaacgcgtc acctcgtaca aacgtacaat catagcaact
106981 tatgagtaaa taggctcatg gttatagata cgttatatag atgtatagag gagagtcccg
107041 agggcgatcg caaggccgag agactaacct aaccaggtgg tagttgggaa atgtcgacga
107101 tttcgctgat cttttcgtct tcgtcggcag cttcctcggt gcttgtcttc aaaggagttc
107161 ttggcaatgc catcataccg cttcttgcac actctaggac accgaatggc tcgaccaact
107221 tcaagaaggc agagatacgt gtgggttttg cagacaactc cacaatacag cttgtttcgc
107281 tgatgtcgac gacacgacct ccaaagttgt tggtcaagtt ggtgatatcg ttcaaatgct
107341 cgtgcttcaa ccttaatacc tcactggcgg gcaagttggc agggtggaat tgcttttctc
107401 tgatttcggc gaccaattct tgggagtcag cggcgccagc attggtggaa gtgtggtggt
107461 gcaatagtag gtcttcgaag tattcagtac ccaatagaga gattctggcc atcactagct
107521 ctcttttgat gatctcagaa ttggtatagt ctaggacggc gtagacgggg accaagtctt
107581 cgatttgtct gcgtgcttgt tcgactacgc catcttgccc ttgcaaaaca atggtcattc
107641 tacttaggtc tttgacctcg gtgttgcaca cgaccaacga atcgatgtta aagcctctgg
107701 cagctaacgt acccgagact ctggacaaga caccgggttc gttttgcacc aaacagttca
107761 agacatgctg ttttcttggt tgacgagaag gcgctggtgt ttcgtaaatg atggatgaaa

107821 cggcactgtt ggcattccag gaaggagtgt ctagtgtggg caagggaggt cttgttgcgt
107881 gtctgtgcat ctgcttgtac gccaacgcgg aggtcgacga ggaactgcaa cgcaccatgg
107941 tagcacatga agaagcaacc accctgcggt ggccgctttg caataacgat ctcagcatta
108001 gaaagactgg gttagtgagc tcagatgttc taaagattct gtatatacga actatgtact
108061 tccatatata ccattccccg ttcttgttag gtaaaggtta aaaaaatatt ttttatggca
108121 cttcggcgat gcgaaagaaa gtgagtcacg ataacggtga ggtgattcgt tggatgtgcc
108181 atcgcacaag gaattgctgg cgtgctaatt cacgaccttg cttgacaaaa gtatcgaggg
108241 gaatggtgcc acggtgcagc atcctagata agcactcaat ggtgtccgtg agagcgtagt
108301 cctgggcaac gaggttgtat agttgattga gtccatctgt tttggccact gctatggagt
108361 tcacgtcctc gtcatccaca tgcgaagttg agctaaactg ttggactttg gccctatttg
108421 cggtcaggac gtctatctgt gcgtttagcg agtgcatggt ctgctcaata gcctgctcaa
108481 cggctcgtaa atggttcttg tcgatggcga ttatttcatg gaatctggca attgattctt
108541 gcatgacggt ctgtctcgtc agtatcttgt ccgctacata gtccacatcc tctctgtata
108601 actcattgac tacggtttgt agattttgta gcatttcatg atggttggtt ggggagatat
108661 ccgtgttgtc catgtccata aggtcgagtg cgttagatgc cggttgaggt ggtggaggtg
108721 gaggcaatgg cggttttagg tgtggagact tgggctttgg gggcagtggg ggagtattct
108781 gctcctgctg gagctgcgta ttgggcttag ggggtagact gggcgcctga tcttgtggcg
108841 gctcgtgtaa gagactcatt aattcttgta cgaccattat aaggttcatc gcggcaggat
108901 cccagcagtg gagtatgggc agcgcaatcc acccattgga gtcgatgtat tcttgaatgg
108961 gcaaggaact actgatggtg ttcatgtcga aattttccaa gtttatactt ataaagggag
109021 gttttaccgg gtacatactg ggtacccaca tgatcacagg aatactgtga ggagatgaac
109081 catcctcacc agtactaatg gtaccatata tggacaacag tagttgtgga gtgccgtcag
109141 agtgcgtaaa taccettgtt cttggcctta aactgtgaaa attgtccagc agcgctaaag
109201 aatcgtggaa agttgttctt ccgtcgttat aaatgggttg aatgactttg aacagccaat
109261 tgaccaccgc ctcgggaaca gagatcttgc cgtttgcaga catcgttgct cttcactctc
109321 tctcttttct tggccgttaa gataccaact gcccacaaaa ttttgccgta actttttatt
109381 ctcgctctgt gaaaaaagac tactacattt tttgaccttt tatctatact ttaagaaaca
109441 aaggtccaac gaataataaa gaacaaagag taatgagtag tttgtatgtg tcataatctc
109501 ttctctccgc ttccctttc ttccttcttc ttctactaac tctctgcgcg tccaatttag
109561 ctatactgtt gttggtgtat tcatagtggt tagtgccatg tccgtgctgt tctggataat
109621 ggctccaaag aacaaccaag cgtatgtgca acattttttt tacgatgtgt ctccatatta
109681 ctagtttccc tttcttttt actaacaaga gtttgtttgt tttgttttgt cgtacagtgt
109741 ttggagaagt acagtgattt tgacattggc catgatgttt ttgatgtggg ccattacgtt
109801 cctttgccag ctccatccct tggtggctcc acgccgttca gatttgagac ctgaatttgc
109861 agaataaagc gtcgagtata tcaagccaag tgacgtttcc tgagtataag aaaaataagc
109921 aaaataagta tacataaaaa taaatacgaa tagaaatata gtggcatcta aacaacccag
109981 agattttttg aaaaacacaa tgagtatata ctgatagaca accacacggt ctttgaaacg
110041 agacgccgtc tacggttgta tcaaaatgtg taaacgggcc atttcaaagc ttgatcttct
110101 ccataattag aacccatgtc ccgtggaagc gttttaaaag acatgaggca gaaacaggag

110161 tgtgatatcc ttgcatgcgc aagttggtaa tacactgcac aaaacacgcg tgtgcacata
110221 cacttatgtg gttcaccgtg ccgctgctgt gtttatctgt tgctcgacat gtgctgctaa
110281 tgattatgtt ggaatagaca aacaatttaa agtaatttct tcttttttta tagtgagaaa
110341 tctcaaaaga aaaaaaaaaa aataaaaaaa actaaagagc tgccgggtga gcaacgacta
110401 aaagaagaaa ccatttaggg cttcatcagc aactgtagga ggagaaagca ggtatataac
110461 tagccgcaat atgtttgtgg tggattggag cgtgcagcta tgcatgggag tgatttcacc
110521 tctcttccgt gccctcgtgc aactgccttt atcaattttc gtatggaatg ggtttcagct
110581 tgtggcattg cctatcaaca tcccattgag gttgttcctg gggacgtcgc tgagccgttt
110641 agttgcacaa acaagcacat tagacttcta tgtggttttg acgttgtttc aatattttgc
110701 tgtgctttgt gcttttggca gcatcatagg actcatcttt ggatttatat tgggtgtgtt
110761 ccactcaatc tgcggggtac ccagtgtata cataagtcta gaatggaaac ggtggtttgc
110821 tccgatacgt acggtccttg aacgtgcttc cactagtatt gtcaacatta tgcgaggaca
110881 aactattgcg ccaataccca tgcctaagcc caatcccacg catatatcaa agcctaacat
110941 gaaaaaattc catgatgagc ctggagctga tgatatgact ataacgcatg atgtgaactg
111001 ctacatcacc ccttgccaaa cgcctactaa cgaaaaaatt cagcattata ataatgattc
111061 attcaacacg accacatcga ttgaagtttt gtccaactat ccactatgga tatgcgtttt
111121 gttgattaac cttaaataac acgtatttcg cattttccaa aagccttttt tcataactac
111181 aaactaacta tttttgttta ttttacatca gtaaattatg cgcagaaaat atgtaaggct
111241 atatactcaa tatagtggaa gagcctctgg ctacttaatc tgggttcata tattctgtca
111301 gtggtatagt aaatttaaat agtgaatttg ggcgcatgta gagagatctt ttgattaata
111361 acccagtatt tgattcttga atgttatttg tctcttatta agtatttcag tttgatttat
111421 atattttgaa agtaattgtt gctttacctt ccaaacaata aaaaaatata aaaaaatgta
111481 aaaaatataa taaatattaa ataaaatact acttgtttga aaatcagaga aaatccacca
111541 aaatatcaat catttcaagg atttccgaac caagttcgag atagtccttt aaggtcagta
111601 aaattcagtt gcacgtataa cagatatttt tcattttgtt ccaatttaaa agtcccccat
111661 ttttaaaatt attgaaaata aaaattaaaa aattaaaagg aatctctcta tgtcacttta
111721 aaataaataa attgaaaatg atatatcgtt aaaaggatcc cgcaacacca cctatttcta
111781 agaggagagt tctaaaaaaa tcatagtacc acacaggtaa actaaaacag ctaaattcaa
111841 cataatacct gtggatataa ttacatacaa aaatataatt aaaaaaatac attaaaaata
111901 atttattttt gtaaaaccca taaaatatat tttactttcg gaacaacttt ttaacttata
111961 attttgtttt aaataaaaac gttgtattta aaaataataa aatattaagt aaaaatttaa
112021 agcatattta tttaaaatat aaaatactac taaaattact ctaaacttca aaataaaaaa
112081 ataataaatt atatagtttt aaaattaata atttgtatac acgtgaccag accataatag
112141 ttttcttttc ttgaaactgc catgaattta atagattttt tttacacata tttacagtaa
112201 gttttgtttt tacgttgaat taatattgta tacgtactta gaaggataac ttccaactat
112261 gaatatgtag gttaaaaggt aaatagagaa gccttacttt tatcagaaat gaaaggagct
112321 cagcttgcct tgtccccgcc gggtcacccg gccagcgaca tggaggccca gaataccctc
112381 cttgacagtc ttgacgtgcg cagctcaggg gcatgatgtg actgtcgccc gtacatttag
112441 cccatacatc cccatgtata atcatttgca tccatacatt ttgatggccg cacggcgcga

112501 agcaaaaatt acggctcctc gctgcagacc tgcgagcagg gaaacgctcc cctcacagac
112561 gcgttgaatt gtccccacgc cgcgcccctg tagagaaata taaaaggtta ggatttgcca
112621 ctgaggttct tctttcatat acttcctttt aaaatcttgc taggatacag ttctcacatc
112681 acatccgaac ataaacaacc atgggtaaaa agcctgaact caccgcgacg tctgtcgaga
112741 agtttctgat cgaaaagttc gacagcgtct ccgacctgat gcagctctcg gagggcgaag
112801 aatctcgtgc tttcagcttc gatgtaggag ggcgtggata tgtcctgcgg gtaaatagct
112861 gcgccgatgg tttctacaaa gatcgttatg tttatcggca ctttgcatcg gccgcgctcc
112921 cgattccgga agtgcttgac attggggaat tcagcgagag cctgacctat tgcatctccc
112981 gccgtgcaca gggtgtcacg ttgcaagacc tgcctgaaac cgaactgccc gctgttctgc
113041 agccggtcgc ggaggccatg gatgcgatcg ctgcggccga tcttagccag acgagcgggt
113101 tcggcccatt cggaccgcaa ggaatcggtc aatacactac atggcgtgat ttcatatgcg
113161 cgattgctga tccccatgtg tatcactggc aaactgtgat ggacgacacc gtcagtgcgt
113221 ccgtcgcgca ggctctcgat gagctgatgc tttgggccga ggactgcccc gaagtccggc
113281 acctcgtgca cgcggatttc ggctccaaca atgtcctgac ggacaatggc cgcataacag
113341 cggtcattga ctggagcgag gcgatgttcg gggattccca atacgaggtc gccaacatct
113401 tcttctggag gccgtggttg gcttgtatgg agcagcagac gcgctacttc gagcggaggc
113461 atccggagct tgcaggatcg ccgcggctcc gggcgtatat gctccgcatt ggtcttgacc
113521 aactctatca gagcttggtt gacggcaatt tcgatgatgc agcttgggcg cagggtcgat
113581 gcgacgcaat cgtccgatcc ggagccggga ctgtcgggcg tacacaaatc gcccgcagaa
113641 gcgcggccgt ctggaccgat ggctgtgtag aagtactcgc cgatagtgga aaccgacgcc
113701 ccagcactcg tccgagggca aaggaataat cagtactgac aataaaaaga ttcttgtttt
113761 caagaacttg tcatttgtat agttttttta tattgtagtt gttctatttt aatcaaatgt
113821 tagcgtgatt tatatttttt ttcgcctcga catcatctgc ccagatgcga agttaagtgc
113881 gcagaaagta atatcatgcg tcaatcgtat gtgaatgctg gtcgctatac tgctgtcgat
113941 tcgatactaa cgccgccatc cagtataggt ctgacactta ccaaaactca ttcgtcacca
114001 atgaaacttt gatgtctctt tctaatagag ctaagcttcg aagaaatgcc agtgatgcgg
114061 acatcgttaa tataaagatt ttacgaagga attctaggtg atattgcaat tacttcttct
114121 catgcactaa caagtgaatg atagaaatat gttgagttcc taactgcctg attttaaata
114181 agtttcatat tataatcttt tagcatatat atatatatat tgatcctctc tcttctttat
114241 tttctgccag taacccatgt gtgaagaaga aaacataaat aaaaaagcag tagcacatgg
114301 acacattcac gcccgaacac tcctaaaaag cagcccacac aagaaagtag atataatgta
114361 ggacacccag cttgtccata attgctaata gcatactcag gataacatat attaatgacg
114421 actcgtttgc tccaactcac tcgtcctcat tacagattat tatccctacc tctccagaaa
114481 cccttcaata taaaaaggca gatgtccgct gcgaaccctt ctccatttgg caattatttg
114541 aacacgatca ctaagtccct acaacagaat ttacaaacat gctttcattt ccaagcaaaa
114601 gaaatcgata taatcgaatc tccatctcag ttttacgatc tcttgaagac aaaaatactt
114661 aattcacaaa atagaatatt cattgcgtct ctgtatttag gcaaaagcga gactgagttg
114721 gtggactgca tatcccaggc attgaccaag aaccccaagt tgaaagtttc ttttctactt
114781 gatggccttc gaggaacaag agaattgcct tccgcctgtt ccgccacttt attatcgtct

114841 ttagtagcca aatatgggtc agagagagtg gattgccgat tgtacaagac gcctgcttat
114901 catggttgga aaaaagtctt ggttcccaag agatttaatg aaggtttagg cttacaacat
114961 atgaaaatat atgggtttga taacgaggtc attctttcgg gagccaacct ttcgaacgac
115021 tatttcacca acagacaaga tagatactat ctctttaaat ctcgaaactt ctccaactat
115081 tattttaaat tacatcaact cataagttcc ttcagttatc agattataaa gccaatggtg
115141 gatggtagca tcaacatcat ttggccagat tcgaatccta ctgttgaacc gacgaaaaat
115201 aaaaggctgt ttttaaggga agcatctcaa ttactagatg gctttttaaa gagttctaaa
115261 caaagcctcc cgattactgc cgtgggtcaa ttctccacat tagtttaccc aatttctcaa
115321 ttcactccac tttttcccaa atataatgac aaatcgaccg aaaaaagaac aatattgtca
115381 ttgctttcca ctataacaag caatgccatt tcttggacgt tcactgcagg atacttcaat
115441 attttgccag acatcaaagc aaaactgctg gcaacgccgg ttgctgaggc aaatgtaata
115501 acagcttccc cctttgcaaa cggcttttac caatcaaagg gcgtctcatc aaatttacct
115561 ggtgcttact tgtacctgtc aaaaaaattt ctacaagatg tatgtaggta cagacaagat
115621 catgctatta cattaagaga atggcaaaga ggcgtagtaa ataagccgaa tggttggtca
115681 tatcacgcaa aaggtatttg gctttccgct cgtgataaaa atgatgctaa caattggaaa
115741 ccctttatca cggttatagg atcttcaaac tatacgagaa gggcgtattc attagatttg
115801 gaatcgaatg ctctcattat tacaagagat gaagagctaa gaaaaaaaat gaaagcagag
115861 ttagataatt tattacaata tacaaaacct gtaactctag aagactttca atcagaccca
115921 gaaagacatg ttggcactgg tgtaaagata gctacctcca ttttgggtaa aaaactttag
115981 attctattct atcatagtat gtcgcctgta aatatggaaa aataaaagaa aaaatagtaa
116041 taataaatgc attatctaaa actccttaaa tttgcaaaca tcgcaaagcc attaaggtag
116101 agaatgagct gaccttgtta tagttttcct tttggcaacc gtggtttccg tcctacgtaa
116161 acatactgaa cttgacaaag tatgaccaat aaggtgacaa attgtatcaa ttttacccaa
116221 aagggcgttg aattaaatgg tatctcgtaa aacaatgcat tggtcggtat taccagtcta
116281 cctaataaat cgatagattc agccaaaaat gatatcaatg caaactttga tgataaacct
116341 actgttgagc atcccatcca gtttatactc atttgaggca catacttgaa acatttaagc
116401 agatttgcca ttacccacaa ataatttatg tgttccaaat agaaaacgcc aaatttacct
116461 gaatttttca acggtaaatt agagatggaa cagccatatg taaatatgcc taatacaagg
116521 aacacactta taattattat ggaagtaatg gaaatgccct gataaatatg tttagttgac
116581 ctataatagt ataactgcct taaaaccatc atgcagcaac aggaaacgca aaaatctttt
116641 agaatgagga aactggatat aggtatacta cgggcatcat tcaacggata aaaaagtggg
116701 aatctttgag acagttgttc cctcaccaga gatgaataac aataatgcaa cgcgcaatac
116761 aagtacagac cattaccaac aaaatctagc agaaataaat catatgaaag cccgtaaata
116821 gatcgatgaa gtttattata gcgcttttga catagaatgg cgatgatcta tgtaagaaat
116881 ggaagttttc tcccttagtt agtcaaacct ttcatgtata ctttctcctt caattatgca
116941 tactgataaa aaacctgcca agccctgcag aacaataaca agcatgtgaa tgattcaata
117001 taattgcttc gcagggctcc tcaacctctc ctctggaaaa ccttacctca atgaggttaa
117061 aaattgtggg tttttcgttc gcagaggaaa aaaaaacttg tcaaaacacg cctctgcagt
117121 agaacacgaa ttatacataa gaaaaataac gatacaagaa aggcgatttc gatttgggac

117181 taagtgttac acgaaaggca tagagtatca aaaaaaggaa gcaagatgga ttacgatagc
117241 tctgatacaa tgaacggtgg ttcaagtaac cccttaatca ctaagatgaa tacaatgaaa
117301 ttattatatc aacactattt ggataaagtc actcctcacg ctaaggagag gtgggctgta
117361 ttgggtggtt tgttatgttt gtttatggtt cgtattacaa tggccgaagg ctggtatgtg
117421 atttgttatg gtctaggtct atttttattg aatcaatttt tagccttttt gaccccaaaa
117481 ttcgatatgt ccttacagca agatgaagaa aacaacgaat tggaagctgg agaaaaatca
117541 gaagaattcc gtccattcat cagaagatta ccagagttca aattctggta taacagcatt
117601 agagccactg tcatttccct cttgttgtcg ctattttcaa tcttcgatat tccagtattt
117661 tggcccatct tattgatgta tttcatatta ttgttttttt taactatgag aaggcagatt
117721 caacatatga taaaatatag atatataccc ttagatatcg gtaagaagaa atattctcat
117781 tcttctaact gatataccct gactccacat acctattcaa cgaacacatt cgtatcttaa
117841 ttatatgtaa acaaaattat atgatagtta cagaataatt gtgattattt tctccttcta
117901 ctcatcttct cttatattgg aagaatacat gaagttattg attaagcaaa attgaaggga
117961 agaggctcat tttcttccat ttataaatgg agagcttgag atgcctccaa ttgataaatt
118021 tttgtaaaga ctttcaagaa acatgtctaa aaagaattga cagcttcgaa cctttctgca
118081 taattactaa taccaaatta gtgagaaatg gggtatgtcg aggaataaag gatgcaaaag
118141 agagagtaat gtcatgcaag ttgtgagttt gaaatgttca cacgtttata atgtccatac
118201 tccgtggcgt gcatgtgttt tagtgtatta gttttccaag ttattaaaga acttagtaaa
118261 tttatcatcg aacgcatttt gattaagtgg cacaatagac ttcaatggaa aatgaaaatc
118321 atatctgtag caaaaacgac agcgaagata acggttacac aattatcacg ttgattcgtc
118381 aacttaaaga tgaccttttt acaatttatc aataataata gacaagaagg acaaggttat
118441 atttcagaaa aattattcaa aactaagaag aatgagatga taagaaaaac agttaccaat
118501 ttagtagctg taagattaaa aaacttatcc cacgaatttg atgtaataga gaattatctg
118561 cgctatatag ctagcaccag tgaacatcta tttactgcta ttaagcgcca ctttaacaaa
118621 tgtgccagaa aacttttgaa agaagcaatc gactccaaat caaactcaga aactgctacg
118681 gtggttctgc aagaaggatt ttctggcatt tgcttattga aagcgtcttc aattatatta
118741 aaactaaaat tgaagtttcc aaaaaagaaa gatagaactg atattagcaa attgtgtgac
118801 aagaaagaac ggatgacaca gtggttagaa atttcaattt tgatgaactg agaataataa
118861 tattatgctc ccctggattt tatgcgaaga cactgctgaa aaaatttctg atatatgcag
118921 agaaggagca aataagccag ttttaaggaa ccgagataag ctgttttcgc ccattgttct
118981 agttctacaa aatatctaca gggaatcgat gaggttgtaa gaaatcctga atactctatg
119041 atagtgcata acactaaaaa gttgaaagaa tcccgtatca tggacgattt ccttgaacat
119101 ttgagcaaag acgataacaa agcatggtat ggcgcggaag aaaccgagag agctgcaaaa
119161 ttagatgcaa tagaaacact acttattaca gatagtgtac taaaaaggaa cgacgtgaaa
119221 aaacgtgaaa aatacctaga cctaatagag aatagtggaa acaacaatgg caaaatattc
119281 gtactcagta cttcaaaaat cacagtgagc aacttgacaa accaacagat ataggctgtg
119341 tcttaaaata taccatccaa taccttgatg aacttttcaa agatgactaa aataagtgaa
119401 atttcaacat taacttcgaa tttttttctt tttatctaac acttgtcaaa cagaatataa
119461 ggattacttg tcttctttgc tacatattgc taccacttct attacacaat agtttcaata

119521 gcttgcagcg tagctaaact ctaaaattta tctaaatcac tcatataaac cgaacccttc
119581 cccttccgct tatagtacag tacctataca tttcataaac atggcatggc gatcagcgcc
119641 aaacaatatg gaaaatccac agaaagctat tcattgaaaa aatagtacaa ataagtcaca
119701 tgatgatatt tgattttatt atatttttaa aaaaagtaaa aaataaaaag tagtttattt
119761 ttaaaaaata aaatttaaaa tattagtgta tttgatttcc gaaagttaaa aaagaaatag
119821 taagaaatat atatttcatt gaatggatat atgaaacgtt tactggtgga agttttgctc
119881 atatattatt attcaataga agtaataaag aaaaagttgg taaagcaact taacagtaaa
119941 aaggtaatga ttgaaaaagt ttttgaacat ctaagctata tgttgatggg tttacaattt
120001 taccattagt actcatgcct atacttttct gttcgtcctt aatgtccgcg atttagagca
120061 atcattgaaa gtactagata cattttagcc agagaggact cgttgacgta gaattaaaat
120121 tcaaatgaat ttccgcccca ttcatatacc ccaaataaca aacatattaa aacttcataa
120181 ttattcaaaa tgtggagtag tatagaagag cagtaccttc aaaattgatt tcttcagttt
120241 cccacccggg atccacttgt catgcggtga gaatcgtata ttgcgtataa tccgtgtttc
120301 atcacccaca ttatagtaca aacctactgg tgtaaccatt atcatattca tgacttgtag
120361 ttaaaaatct ttcacgataa attgaggaaa aaatcacgtt aaatagtttc atatcattct
120421 ttagttaaag tctgtgcata aaaagaggat cgggtgttgt actcgttcta tcaaaatatt
120481 gttattgttg cattatcaca cgaattacaa tcaatttttt tatctacagg ccagagtacc
120541 tgaaataatg gaattagatt caaaaaggat agcttaatcg gtctttatac taattatttt
120601 acgcccagaa cgaaacataa cgtgcattaa atctcactgt cacaacattg tatataatgg
120661 tagcatcgta ctaactattg gcattcattc aatattatac aattatctta ttaatacttt
120721 ctcaagtagt cttgaagacc agcatgtagg aaggtgatga tatgctccgt aaataattgg
120781 aaatattgag atataataac gatgacaata ctaacaaaag taataggtga gctccgccaa
120841 ttgattgttt tgttttgaat atatattgat gcttaacaat ttagtcgtca aatagcacct
120901 gcaaaaccta cctcttacag gctttttttc aatttactcc ccatgtggga gttctacgta
120961 aaccacttag aatgcagttt ctcttctaaa taacatgtta tatcttctga agaaattgaa
121021 acatattgat tcttatggtg agagcaattt atcaatgttt tatcttaagt tctactgccc
121081 ttggaaatta tgtcctttgc ttcccatttt atatggaaga aacaaatcag tacaaaatgc
121141 ttggtcatgt cataagcttc atcgatttat tgacttccca actccttcgg ctaatatgaa
121201 tttttttttc tttataagag tttgctatgg aactgtctcg ctgtttattt caacgtttct
121261 aaagaaaata tcctttttta tttatgtgaa cttcactgta tacttttagt acctttcggt
121321 ggatttcttc tattttactc gatcactttt cttgactgtg cactgtcggt acaatcctta
121381 ttgtattaac ttgcgaagaa tatttagtcg ttcagtaggg ttcttttgga agaaatgcta
121441 gcttaacatg agtgtgatct cttctagaat taactctttt gtttatccat gtaatctgtt
121501 accaacacat aaaatgcatt gatagaccaa atatcaaaag cgtgcaggca tctcttatct
121561 aacagtaagc aaaattttgc ctgcgcttta cttcaatttt ccattctttt tctgagtctt
121621 cgtcagtgca tattttttta ctgacagcac cattaatcaa tcagctcacc tgcgcatttc
121681 caataatcgt gctcatgaag atcataaaga aacgtccaaa catgaaagtg ctcctttaga
121741 atttctaaat ccaaacgata tatgtacccc tcgttgattt gtcctaaaat gagtatagct
121801 tgattttcgt tatatatttt ttcatagtat tcagaagagg agagaggatg tgatttatca

121861 atatctgcat taagtaatga actttgccct ctactgagga tattcccact agaaaactat
121921 aaaaaatggg agaaagtcaa tcgctgttca ggattcaaca tagttgttga caacgtatga
121981 taagaaggcc agtccagtcc tcctaagtta ggttaaggga ggtagggaaa aactgaagtg
122041 taatgataat cgaggatgaa agtgtgcggc cagaaaaatc ttttcttttt atattaatct
122101 gtaataaata caagacctat atattaaagt aaaacaacat tgaatcagat cgcatactgg
122161 tgtagacgtg aagtagacat atgataatcg agcagggtgc tcaatttggt gtgggacacg
122221 tttgggaagg ttaaaagaag aggtatatcc ggattacatt tatttttggc gtattttaat
122281 aatttactaa aagcgcgaat agcttcccat tagcaacgct cagggactct ttctaatgtt
122341 tatagtaata ctttaataag atgagtacca attgcttctc tatgaggcac aatccctaac
122401 caaactttta tttagaatta ccggtaaact ccgttttctt catagattaa atatttggct
122461 aagacaacaa aatgttcttt cattttacca aacgagaagg tgatagctgt acgccagaaa
122521 agaaggtaaa aactctcgcc aatccagcaa tacctgtacc tctagttttc gatcgcatta
122581 tgtcattatg taaaaccaag gagagaagca agttacggtg ttttcatata gaataatttg
122641 agaattctta ataacataag atatataatc accaccattc ttatgagatt caacgttgaa
122701 tggcgttgct agacatatga gctgaggaac gactattcgc attttctttc aaggcaatta
122761 attgtcttgc tctgaaccct tcataatgaa tgtaggaagt tgtttcgatt aagtcttgga
122821 gatgagttcg aatcaaaaat tcccttaaat atacaaaatc acactggttg atatcctcaa
122881 cattaatagc gctccaacga gtttttcttc ccctgaaggt ttcaccgttt atttcaatct
122941 cattctcaga accaaccact gcaaacggaa tgatagatct aacacttctg tttagttcta
123001 attcctcgtc agttagttct tccgaatcat aaggataaat cttgaaattg tatttttcga
123061 attcattttg aataagctcc ctaaactccg ttctttcatc taaagtcaat gtatccgact
123121 tgccaataac tggtataaca tttgctattt ctgtcaatct tttcaaggct tcaacgtcaa
123181 ggcggctcaa ctcctttcca tttggttgca ggaaataaag aattgcatga actcttgtat
123241 cagtaataaa cctttcacgt tgggctgtca attctttacg taagtattga gaatgttgtt
123301 ccttaatgta cttcacaata ggctcccaag ctttagaatt gtcaataaag tcaccaaatc
123361 caggtgtatc tataacatta atattcaagc gaacgcggtc ctccacaaga gtatgagtag
123421 aaattttcat ttcagttgtt tttgtaacag gcagggcaga aatatcatca ccagtagcag
123481 aatcaatcaa atgtgaggca aataacgtat ttattagagt acttttaccc aatccggatt
123541 ggccaacaac cattatatta aattgaaaac ctttcttcaa cagacgatgt tcgatctgat
123601 tcgtgatggt atcaaaacca acataagaag caggctgtac tgagctgaga ggatccatta
123661 atatatttat agagtgcttg tagtaaccgt ggggcttggc cttgtatttt cttgtctttg
123721 cctgaaaaaa gttaagtatt actgatacat gtttacatat attctattca gcgattctga
123781 tctatttgtg gtctctcgag gaaatctcat cgtatcatac gattatttcc gcaagcattc
123841 agaaagcaaa tcttaatata aggcacaatg ttcccaaata ataccttaac ttggagacaa
123901 aaacaccagg gaatatcagc aaacaaacta tgaattcttt gatttttggt aaacaattag
123961 catttcacaa aattgtgcct accactgcaa ttgggtggtt ggtaccgcta ggaaatcctt
124021 cactgcagat tccaggccaa aaacaactgg gatctatcca ccgttggttg agagaaaagc
124081 tacaacaaga tcataaggac actgaagata aagatttttt ctctaataat ggtattctac
124141 tagcagttcc taaaaaaaaa gtatcacacc aaaaaaaaag gcaaaaactt tacggtccag

124201 gtaagaagca attgaagatg attcaccatt tgaataagtg cccatcatgc ggccattata
124261 agagagccaa tacactgtgt atgtattgtg ttggacaaat aagtcatata tggaaaacgc
124321 ataccgctaa agaagaaatt aagccgagac aagaggagga actttccgaa ctagaccaaa
124381 gagtcctata tcctggtaga agagatacca aatataccaa ggatttgaaa gataaagata
124441 actatttgga acgtcgcgtt cggactttaa aaaaggacta ggctcatctt tcatctagtg
124501 gtttcctttt taaataaact taagcctgta tatagaaact taatttcaaa ctgtatattt
124561 attgtttaag acagcatcaa tggtagaata caaggagcgg aaacagaaag aaaagtcgaa
124621 atagaatttt tcgcttttgg cttaacaaaa tgaggtcttt gaaatcaaga gttttgcttg
124681 acaaaaaaca actatatttc aaaatattag gaaatgatat tattagtaca aggacttaca
124741 gaaatatact tttaatataa aaaggaaaaa ataaaataaa aagagttaat tctttattaa
124801 ttttccaacg tcctacattg atatatctgc tgagatagaa ttaatgttag taagaactaa
124861 atattagacg cgaactgttg tttcttacat gacagtacag caggataata atcctttgtc
124921 ctccttcttt tctgcagttg tagctgctgc aggtttagtt tgtctcttag ccgcatcaga
124981 agtagtggtc ttcttttcag tggagggctt ggcttctttt gcagggaaaa gttttttggc
125041 ggtttcgtag aatgttttac cttgaatttc agcaattctc aattcaagtg gagacgcagt
125101 tcttgagccg tcaggtcctg caagggtacc agcaccccat ggagagccac cgtgtacctc
125161 ttctatactg gctaactcag caaatgaatt cttataaccc agtggtaaaa agataattcc
125221 gtgatgagct aaataagaca aacaggcttt aacggtactt tcttgaccac ctccgtaact
125281 ggaagtacta acgaatatcc ccgcagcttt gccgttcaaa gagcccttgg cccataatcc
125341 accggtttta tcccaaaagg cggaccattg agccggcaaa ttaccaaacc tagttggaac
125401 accgaacaaa aaggcgtcat attcgagcaa cgttttctca gtggcaacag gaatatcttc
125461 aggtttctga ggagcgttca tcttggtgag gacttcatca ggtaaagttt cctcgaccct
125521 gtatatatca gctttaccac cagctgcctc cacacctttc ttaacagctt gggctaaaac
125581 gtctatgtgc ccgtaggtag agtaagtaat tatcgcaatc tttaccattc ttatagtgct
125641 tcaattgagt ctgtattgtt ttgtttactc tctttttttt tattgatact aacctgaccc
125701 ctcttttcct ttttcctctt cttgttttct ttctgataaa gtttacaaac ttcaaataaa
125761 aatgaacgta aaacaagtaa aaatgtagga tgttatagta tcaactagct gccggaatag
125821 tgcaaattgt atgaatcgaa ataagtacgc tagccaaggc agtaaggcct aatgccctta
125881 ggcaaattta ttatttatat taagttccaa gaaggggaag cgccttcttt ccccgcagta
125941 cagtatttta cccggaggtt cttaataagt tattactaaa agggtcataa cggatttatt
126001 aataatggga ttataaaaat aatgcgtgtg tacctagtta tcgcgtgata gcttccgcaa
126061 ttttccaacc attagtaaat tttatgtaac taaaacgagg aaggaaatag taacgttcat
126121 tttcttaatt ataaatatca ttatcatagt gtatggttaa aagaatatat tattgtatta
126181 aataagtgct aaatactaaa tggtcatgag gaaagaaaaa tatgcagagg ggtgtaaaag
126241 taggatgtaa tccaactata gtttgctttc aatgtttttg accaattcct tgtatttctc
126301 agtagaatag gactttggcc tttcaatgga agcaccgatg gccctatcag tgatcaattg
126361 agcaagaata ccaaatgccc ttgaaacgcc aaataaaacg gtatagaaag aagattcttt
126421 tagtccataa tattgtaata agacaccaga gtgagcatct acatttggcc atggattttt
126481 ggttttacca tgttcagtca atacgccagg tgctacctcg tatattgatg aaactaactt

126541 gaataattca taatctggaa aatggtccat ggcaaactta cgctgagcca tataacgagg
126601 atcagttttc cttagcacag catgaccata accgggaatg actcttcctg agtttagagt
126661 atcccataaa tatttttcga tcgtatcttt agagtagtca tcatttacct cttctttaag
126721 tgcaaataac cattctagta cttcttgatt agcacgccca tgaagtgggc cagccaaccc
126781 gttcaaacct gatgcaaggg acagataagg tgatgatagt gctgagccca caagatggga
126841 tgtatgtgca gatacattac caccttcgtg atccgaatga atggttaaat aaagtctcat
126901 caagtccacg aaatcttcat ccttagaacc aatcaagttg accagatttt tagcataatc
126961 ggcatttggg tccacttcac ccattttgcc atctttgaat acattacgat aaattttagc
127021 tgcaataact ggcaatttac ccagcaagtc tagtgaatct tcaaaagtat aactccaata
127081 atcttgcttg gaaattcctt gagcataagc cttagcaaac tttgactcgc tttccaaggc
127141 agttacagca atagagaatt gagccattgg gtgtaagtcc tttggtaaat tatccaaaag
127201 ttgaacgaca tgactaggta gttccgatct tgacattaga tcagctgata agttttcaac
127261 ttgcgcttga gttggaacct cgccagttag caataaccaa aagagagctt ctggtagtgg
127321 ttgtgagctt ccttttgcct tgggcaggtc cttttgaatg tcggcgatcg tacgacctct
127381 gaaacgaata ccgtcttctg ggtccaaaac ggaaccttcc catacgctcc ctggaatacc
127441 tctcatacca ccatatacct gttctaatag aacatcgcta attttagttt tgccatgctc
127501 tttaacgaat tgccttacat cttgagcatg gatggggtag atttcgctaa atctctcttt
127561 tagagtcttt tcttggcttg aatttgattg taaatatgat gcaacatttc tgtttgaatt
127621 tagataagga actgtcattt ttcttgttac tagtattatt aaaacaaaaa gttttgagaa
127681 cctgttatga tatgtgttga tattgttccc tgaaacttaa taatgaggaa cgaacaccat
127741 atctaaagaa ccaagtttta atagtcttat atacgctcaa cgtcgagaat aatgggccct
127801 tttgagagca tgactatctt tagattaacg tcaccaattt catacgaatg taacgctcaa
127861 aaggtaattg cgcttaccat tgttattgtt tttccgctgt gcctcccatt gctgacgccc
127921 tgatagacgt ctagctacgg aaaaggtcac acttttttc tgcaagaaaa aataggtcgt
127981 gacctcttcg cccctagtcg atgaccaccg gataaatgcg tcgaatgtgc gcgctattag
128041 tttgcgctct tttgagatac cgccaagcca ttgattgaat gacccaagga cggatttgtt
128101 caacactcca aagcggctta tgttgtggca acaactctgg ggaaagtcag cgcaatgttt
128161 tcccctaagg attcgcgcga gccgtgtgta acaaagttcc gccggtatag catatctaaa
128221 ccacagaagg acgcttatac ggccgtcttc cacagtaatt agaaggacat tagtcggagt
128281 gtaaataaga cggttgcata cgcgtatgaa catgggccat tcattttacg gaaatggcaa
128341 cacattttct tgtttttaat gtaaatagta cggagttcct ttttggtgcg ctgaatgtat
128401 gccgtagttt ggtttcttgc caggcgtgag aagagaggtc acacatgctg ttggggttca
128461 tgattttcag ccttcgctct gctaagtaaa aggtccattg atccgaagcg cttcctccaa
128521 tatcgcagtg ataaacggtc atgtaagcct acagtcttta aactgtagat gatttaatgg
128581 attatctcat acattgaata tcacacttca gcagttcaag ttttcattaa cattggtctt
128641 ctccagctta ctctgtaaag tccgctgcag ttttctagaa atcataatct gtacctccat
128701 tcagccatct gaggaacctc cgaaataacg agtaaatgtt caaacatgcc atgcgccttg
128761 gtttattgta cgagttttag catattgcaa aaaaaaactg ctcctgagta cacttgtttt
128821 acgtcggaaa gataaatgcc gtaccatttc tgccagtagc gacaccacac attgtaaaaa

128881 caatgtgggg cgagccggga ctcgaacccg ggacctcccg caccctaagc gagaatcata
128941 cctctagacc acacgccctc ctttgaaaaa tcccggcaga acagcgcctg aagtctggga
129001 tacagcccta ttttccctgc catcgtcatt gaccttttc ttaacgacca gattaacagc
129061 cagttattga aagtttgcga acgtgaagta aaactgcaaa atggtcgttt taagtcaaaa
129121 gtaaccctgc gtcacacatg agaaacatta tcgctaagtt gttgttacta cttcttgtaa
129181 ttaacttact gtcgcattcc aaatggactg cgaaacagac acgaaacaat tacgaaagtg
129241 acgacagagt aaatattgtc attaaagaga atgatgacct ccgggtaaac tgtgagaata
129301 agtctagacc attcgttaaa cttcatcttt aattgagaag catagtaaaa aataattaac
129361 aagttaacac aactatcttg acccatgcta ccaaggacat aagtaatatc ctccaccact
129421 tttactcaaa gtatatctca ttctgagaaa tgggtgaatg ttgagataat tgttgggatt
129481 ccattgttga taaaggctat aatattaggt atacagaata tactagaagt tctcctcgag
129541 ggtctaggaa tccccataat ggaatctata tttctatata ccaatattac gattattcct
129601 cattccgttt tatatgtttc attatcctat tacattatca atccttacac ttcagcttcc
129661 tctaacttcg atgacagctt ctcataactt atgtcatcat cttaacaccg tatatgataa
129721 tatattgata atataactat tagttgatag acgatagtgg atttttattc caacacattc
129781 gagctagtat aaaattgtga ggaattatcg ggcatcttgt ttattaggta cgcaaattgt
129841 aataattgcc aagttgtgct ttttagttga gaagtattcc gtcctcatga aatggctaat
129901 ataagtatgg atttcataaa tattatccaa agtaattata cacacttgtt ttttgctatg
129961 tcttgggttt aagagaaaat gctacataga ccgtaagaac gtcttcaaga atgcaaggat
130021 gctgcaagcg aagttgttcc cgttatgaag cagaaagggc acatcaacat ggttcaaaga
130081 tttaaatttc ggcacttgtt cttgttcact atatgtttta agattcttat cgtgcaagta
130141 ttcaattata tcctctaagt gaccgaacaa tcaacacatc aagatttttg tagttttgaa
130201 gttccacaac ctacaggtta gtgctatttg gtctaattta tacatgggaa ctatgtgaaa
130261 cagtaatgtt agcacaatgg acctctcaaa gcctaagaaa attttgctgg ggacaaaaag
130321 tggttcttga tggtaaagaa gtagtttagc ttcataataa tttctttgta aaatttttct
130381 cgtagcacac taagctttca gaaatcaaat aaggcagagg aagtggtttt gtcgtattca
130441 atgactaatt gtaagactcg agtataactt atgcagtttc caagaagtaa gcaaacagtt
130501 tctctcaacc tttggtgaca cactatgcgc atgcaagtcg taatgcttct ctatttatgt
130561 gtgggtttat taaaatagtt agaattatct aggtttgtgg gaatttggcc atgaatattg
130621 aatagtaaca agaacaaagc gtgaaaatgt tgaatataat gaaagctgaa gacatcaaat
130681 catgttctcc aaatctttct gtctcgtata ttttctttga tatttgtttt atgcggaaat
130741 tgcagtggaa acaatcgcga aacattcgag aaagaaacta ggtagctaca aaagtatgga
130801 aaaaagggta ataagcattc cttttttttt agccgcaaag cctgcttaac catacatgaa
130861 gcactttaac atatagatgc tgctcttcac tgttacatta tatacagaaa aaagtgctcg
130921 caataagaaa caaaccttca gaaaaagaag cttatcttga gcacaagtca ttcacagcaa
130981 aatttctttt ttcagttgct acatgttaat tatggagtcc tgagagtaag taactagcca
131041 ttagaattaa tttccttccc ttaaaaatga tagggaatgc acgacaatac gttaatgtgc
131101 acccaacttg caccacagca tattctatcc tttagtttgt acaatattat tagctgtgag
131161 aaagcccctt cgtgtggcac aaatttatta atctctgtca aaatacttct acttgtttat

131221 actgtataaa cgattaagtt ccattatgaa ttgtgaattt agaatcggtt tggtacatcg
131281 ctcagtataa gtgctaaata aaattatcaa gatttatatt tcatggtata tcggcgtcag
131341 gatattgctt tctccaatat tcctgttgtg cttgtttctc aacttctact gctttaacaa
131401 agtatgcttc gaatattggg tcagaactga aactaaagca ttttattacc tcatttgaag
131461 cctcatttgc tggtgcatct atctgcgtac ttgtggcgtc ctttaccgat gaatttacat
131521 tcacacctgg gggtttttct ttcacaagaa atctgaagaa tttgtagcat ctttcactac
131581 tataaaagta atagcgagta ctccataaac cattatcaag taaatattga ttcatattgt
131641 ctgtgattgc tctccattct tttccctta ccgctggcct gtgggtgata acttccagca
131701 aaagcttaat tctgaaaccc tcatccgaga tatacacatt aaacggaatc aacgtagaca
131761 aaaacaaaat agaagtgacg aaaattgaaa aaccaaaaac gacatcacaa gctttaacat
131821 tatgaaagat cacaactatt aaaattgaaa ctgaaagtgc caagaaggaa aatatcatcg
131881 gtttgtaatg agccatctcg tacagcaagt aggtcatata tgatttaaag gtgtcttcag
131941 gtaaagtgat attttctgcg gatgataaca cattttgttc ccctaatgta tccaatttga
132001 catcgtcatt attactgttg ttgtctaaat gaggctgcat gcagatacgg agttttgttt
132061 cttgtcttga tatggtatag acaaccgact ggtgtatata aacagaacac agacttacat
132121 tttcaggttc gaagttttat tgaagttcat aaactctttt taaattattt tgtaaacttg
132181 agctattcaa gcagtattct tatttaacct aagcatagcc ccggatacaa ggcgtataat
132241 acccaactaa cttaaaatga gagccactta ctgaacacct tggaaggtca ttgtttggtt
132301 gaaaggaata ttattaacta agccttagaa caaggcgtca cctaaatgat aatggctttt
132361 cttctgggcg cggtcggggc ttcgactcat ttgatactag gttaattacc cttttacgca
132421 tacctaaata tttgtaactt tagatctgtc cactaaaagc gtttcgatga cttagttgtg
132481 ctactaatag taaactcttt gtgcaggcaa attatatcat cgctggctag actttttgtc
132541 aaaagttctg tttatacagt gggcagctag agtgcttaat aacaaagata ttgtccttta
132601 gacttacaca agagcagaag caaagataag cactgtttgt tctgcttcca tcgtttttgc
132661 tgaaaatttt gcatgcatat ccaaccaata tttaacacct ggattatctc ggttcaaaga
132721 ctacaataaa cgccccaata attgtgatgc ttttttcaat ccagcccgca caaatgaata
132781 ccgctgctat tggatgccat cttcagatgt acgtccaaac tgtggttggt atagttttta
132841 gaacctcgtg gtgaggagtg ttccattctt tcaaaggcaa tgagaaaaaa aaacagttgc
132901 gatattagtc aactgttttg catggtttct ttgacgcagt tgtaaaatca tactcgaagg
132961 gtagttggaa aattttcaac atatattagt aatacttttt ctgtaaaagt gacacaagat
133021 aaaatgactc catggccaag ttggttaagg cgtgcgactg ttaatcgcaa gatcgtgagt
133081 tcaaccctca ctggggtcgt taatttttct tttctttttt tttttttatc tccgacggag
133141 aaaaaatttt ttctcaaccc ccggcatggc gaatggaaaa atcacctggc acgtcatttc
133201 atcaaatcaa catcggcctc cccgtgctaa tatgttaaca ggccgagctc ttgccatctc
133261 acgcattttc cgctatacaa atagtatata taatattaca gcttttaaag agggaagagg
133321 aaacggagaa gaaaatatac tactttatag agatattcgg caattttttt tagcaaaagt
133381 ttattaaacc ctaatataga cctttacttg gagtttcgat tttaagaatt ttttatcatt
133441 ttttttttt gacgatttcc tgtcggtttc gttataacct gttgtgttgt tgttgttgtt
133501 gttattgctg ggttgttgtt attcacttta acattatcga ataaaatttg gactttcaaa

133561 gtcctgttca ataagttgta ctaagccttt aagcgttttt aagttacaaa taaaaactgg
133621 aaagcttaag gtagctttct aatattaggc tctcaacctt tttttactct ccatttaccg
133681 ctagatatta actagtagta agtattgtga ctactattcc ctgtaatcaa aaaaaaagta
133741 atcagatttt attttatttc gacattaccc ctcaaatata tgactggtat gaatgataat
133801 aatgccgcta ttcctcagca aactccaagg aaacatgcgc tatcttctaa agttatgcaa
133861 ctttttagaa gcggttcaag atcatctagg cagggaaagg cctcatcgaa tatccagcca
133921 ccttctaata taaacacaaa cgttccatcg gcgtctaaat cagccaaatt tggtttacat
133981 accccaacca ctgctactcc tagggtagtt tctaatcctt ctaatactgc aggtgtgagt
134041 aaaccgggca tgtatatgcc cgaatattac cagtcggcat caccatcgca ctctagttca
134101 tccgcatcat taaacaacca tattgatatt aacacctcta agtcatcatc agctgcttct
134161 ttaacttcgt cagtatcagc tttatcctta tcacccacat cagccataaa tattagctcc
134221 aaaagtttga gcccaaagtt ctctcatcat agtaacagca atactgctat tacacccgcg
134281 cctactccca ctgcttcaaa tattaataat gtaaataaga taaccaatac aagtgcacct
134341 atttgtggga ggtttcttgt gcataaagat ggtacccatg aacatcactt aaaaaatgct
134401 aagagacaag aaaagctaag cacaatgatt aaaaacatgg ttggtgcgag caaattacgt
134461 ggtgaggcaa aatctgctgt ccctgatata ataatggatc caaagacgac tttaaaatcc
134521 aacaagaatc ctcctactct ttttgcaggc ttcatgaagc aggtcgtgga tatggatgat
134581 aaatatccag aaggcgctcc cacaagtggc gctttaaatt gtcctgaaag ggatatatac
134641 aggtcagatc aaaaagattc caaaaataat acgcataata tcactactac taaaaaagat
134701 aggcaatgtt ttgccgaaaa gtatggtcgc tgtcaagaag tccttggtaa aggtgctttt
134761 ggtgtagtaa gaatatgtca aaagaaaaat gtttcttctc aagatggtaa taaaagtgaa
134821 aagctttatg cagtgaaaga gttcaagcgt agaacatccg aatcagcaga aaagtattct
134881 aagaggttga cttctgaatt ttgcatttct tcttcattac accatacaaa tattgttact
134941 acactagatc ttttccaaga tgccaaaggc gagtactgtg aagtaatgga atattgtgca
135001 ggtggcgatc tattcacttt ggtcgttgcc gccggaaaat tagaatatat ggaagcagat
135061 tgtttcttca agcagcttat tagaggtgtt gtttatatgc atgaaatggg tgtttgtcat
135121 agagatttga agcctgagaa cttactgctt acgcacgatg gtgtgctaaa aattacagac
135181 tttggtaaca gcgaatgttt caagatggca tgggaaaaaa atattcacct tagtggaggc
135241 gtttgcggtt catcgccgta catcgcccca gaggaatata tcaaagaaga gtttgatcca
135301 agacccgtag atatatgggc atgtggtgtc atttatatgg caatgagaac tggtagacaa
135361 ttgtggagtt ctgctgaaaa agacgatcca ttttatatga attatttaaa aggacgtaag
135421 gaaaagggag gctatgagcc aatcgaaagt ttaaaaagag ccaggtgtag gaatgttata
135481 tattcgatgt tagatcccgt tccgtacaga agaattaacg ggaaacaaat tttgaacagt
135541 gaatggggaa gggagataaa atgctgccat aatgggcgcg cattgaaata aacgagtact
135601 tcactttcaa atatcacgat attttggaac aattcggtat tttttactta atctagtaca
135661 ctaaggaatg ctttgttatc cggcattcgt atcttattcc tcgcttctat tgttctactt
135721 ttatatcccg tttggctgat tacggatcac gttcaaattg gtaaatccca ttaattaaaa
135781 aagaattgta accttattta aaaaaaaaaa tagtacataa caataaaaaa aaaaaagata
135841 ataattttga atttattgct agacattctt acgtttattg tgttcatgtt aacaaaaaca

135901 catgcccgcg gttattatcg ctttttaatt gctttcttaa aaagagtcgg ttggtcagat
135961 agagcagata gtaaattata gtgctattta ggtctgttaa tttaaatgaa agaaacataa
136021 atgaccgtaa aaaactaaag gcaaaagcat taatttattt tatcccgagc gcacaaatat
136081 ctaggcttgt catttgtcct aatagttctt cgattttagt gtctaataac ccattggcat
136141 agtcgtctct tgattgttgg tccaaatatt gttgaatctg cgctaaacga gtgtaaccat
136201 cagtacagaa ccttagctga atcttgatta aagcttcaaa acttgggtca aagtaaggta
136261 ctcttaatga aactaactgt ggtagttcag ttttcaattg gttattaaga ttttcgaaaa
136321 tatctttagc taagctcaat tctttttcag cccttggcag tttagaggca tctttagcag
136381 gtttgtccac taatctacga acttttgcct tcgcagcatc gaagtcttgt ttcttatggt
136441 ctctcttttt tatggcctcc tcaatttctt tgaaatacgt cgaaaacttt gttattggat
136501 ctagtacggt ttctcttaag ggcccgtcta attgcttaac agtttcgcta tcaaaatctt
136561 gaacacattg caaataatag ttaccaacgt tgtaaccacc accagcaaca tattttgaat
136621 cgtcatagag gttagagatg acctcggcaa tggtagtctg tgatgctgtc acagctctca
136681 atgagtccaa gaaacctttg gcttcctttt gtaatgcctc acctgctctt tgaagaactt
136741 tataacgacg ttcttccatg tcatactctt tatcaatggt cttgtcgaca ttcttaatta
136801 tcacactgtg accagctctg ttgatagctt tcttaaaacc ttcccaactc atcttgtatc
136861 agctttcttc tggcttctta taataggatt tgtatagttt tctatatcac agcgtttcct
136921 tttcttctct cacgtctgcc tgttattaaa tttgttttcc ctttctttta attcggattc
136981 tcataattgt tcaaaatccg cgcatttgga tcgtccagcc cctgcagcgc ccagttcagc
137041 caacaaaaat tcatagggac gtaaaggaaa cgtttcatgt ctgttacaat gaaaaagaca
137101 agtcgaatca cattatgaat ggcaaacgat agacctttcg ttgcctttca attacatgat
137161 attcgtggca acatcttctc cgctaaccta aaagttttat cagaaacata tatcttattt
137221 tttctgaagt aaaatcaaag tgcagggaag acaagtaaag gagtcagcaa aaaagtgaag
137281 taaacttata tgttaactgt ttataaccgt accttgctta catcatctat attttcgtaa
137341 tcatttttgt acattgttta tatcttgttt tgaagtttta gctttgaaaa acatttaatt
137401 gcgaagtaac ataatgttat taatagtgta cgactgaaac agccattcta tttaaacata
137461 taaaagtacg attagttggt gtggttatat aagtcataaa ataattatca tattaaaaaa
137521 ttgaaaaata aaaaactact ctttttttatt tcaatagttc tcgttattag taggtcgtgc
137581 tcttaaaaga ttaccctttc agtagatggt aatgggaatg gacgagccag tacatatgaa
137641 ttctgctttg tagcaacacc tgcatatgcg ttgtaccaag caatgaaagc aacaacaact
137701 cccaggacac caccagctct tgtgacacca agtctattag caaagtgacc aatagacaac
137761 agtaggaaag ttaatgctag taagaagaac aacaaaaaga acataacagt ggatttcatg
137821 gtacaaacgg ttaaaccaaa cgtaaagatg gcccacccca acaaataaaa tcctaaagca
137881 ttattcaaat cagattcatt gtcttcgtaa gcttccaaga taccaaacca aggaatgtaa
137941 attgcagcga aactcaacca aaacccaccg taagaacata atgcggtacc accaaaagta
138001 ttttccaaag ctatctccca aataccagca atcaattgca ccaaaccacc ataaaacata
138061 gcacaaccga cgacaacatt aggaacagtg atcccttgcg ctctcgcatt gaacatggac
138121 agcacaaatg tcgtcaacgc gaaggctgaa agacctaagg gcgcaggatt agcaaatttg
138181 tgcactggag caggagctaa ccctggattc aaggtaccac caaaggcttg gtataagtcg

138241 ctcttcaaaa acttttgacg cccaatatag atatattcat tgttatcacc tccagtgtaa
138301 atcttgccca acgaatcatg agatggacgt tcatcttctg caacgccatt aacgtcgtta
138361 tcatgggaac tatagtatcc tgctggtgca ttctccaaat ctgtgtttcc gctcgtttgt
138421 tccttgtcag acatattatt tgtttgtata tgagttgttt tgtggttgtt tagttatatc
138481 tgtgttagct gtcgactatt tgcctttttc ttataaacag atgctagtag tttctgttgt
138541 attgttattg ctattggacg atatataaaa tgaacctata gcaagatctc agggaaatac
138601 gatccaacca aggtgcttta tatacagctt ggtgtctctg acaggttgga gtacagagga
138661 gaaggcgcca tagccgccca acccgcgata acgaaaaaaa gctgtaagaa aatgcagtct
138721 tccccaacgt ttgttacccc aacaataccg aaatagccgc ccaaccgcaa atctggcaaa
138781 aaatcttttc gtatctcgct gcagatgacc ttccaacccg cccctttttg tagtcacgcc
138841 tttaagggcc cttgtttggc ttgctcacat ggggctacac gtaaaaaccg tagcttggag
138901 aaccactcag ccgctcgcct attccacgga gctccggatc aacgggcagt ggaccctctc
138961 gaaaataagg aaagttgcgg ggccaaaaat tattatttct tatgtaaaag aagagactat
139021 atacgacgcc tatctacttt tgttccacaa ctatccactt aagggctaga taccccatta
139081 ttaaaaacac cacattaaac agagccagga tcttgatgtc aaatacaagg ttttggacaa
139141 caaatccaaa tgtgctcaag atagtagcgc ctggaacttc aatatttaag ccgtactttc
139201 tctctttcag catcaatgtt ttgacctcat tgatcaataa agattcgtag gcgtagtaaa
139261 acacagagaa gtttttcagg tacttgaagg ccacgtttgt aatattctta gtattgataa
139321 atagtccgct aaacagtagt gagcccaaaa gcaccagcac gcttaatatt atggagttat
139381 tcaagtcttc aaaaattatg ccgatggtta ggatttccaa cgatatccca aggttaaaca
139441 gtataaggat tccaatacat ttaaaaaaag cattgtcttt catgtttaaa ccagtcattg
139501 ggtaaacaat caatgacaag agtataggtg gtacaacacg tagcgggacc acttcgctca
139561 ttatcttact aatgtagtat gcaagtggcg agtaatagtt attggatctt tcttttatga
139621 aaatgatcct ttccagagcg aacgagctga gacctgtgaa tgtaacaaaa ccgaagtacg
139681 ttagtataaa gaagaacagc cccattctgt tctgaaaacc gctgatatca ttggagacgt
139741 tgtaatatag tgttcccaag aataaactca ataggatcgt cagtaaataa ttacccaata
139801 atagtttagg gtttctgtac atgtttttaa aacttcttga attcaatatc gataactgtt
139861 gcagaaaacc agcagattgt tgacctgtgg gtaaatctcc attcaaaacg ttactttcct
139921 catcaccttc gcttaaaact tcctcgatct cctgtgaaag ctctgcataa tagacgctat
139981 ctttatattt atcgtgtagt agtttggtat ttaagtcaat ctcagtagca cctcgccttc
140041 catctgttcc ctctacatct tcttcatctc tgagtaaaga tctgatctca tctctatgag
140101 ctgcaagatg agcccactct ctctgtgttg taccatcact gctagtaaat gttgtttggt
140161 gtatcgtatt atcaatatcg ttagtatccg taccagcttc taaatcggaa atgtttctga
140221 ttcttctcct tttcccctga ggaccggctt caaaagtaat atcaatcaaa taatcagcaa
140281 tattatagtt gtccggacag atatatccct catttctcaa aaattctgac actttttttgg
140341 catttccgga atagaccatc tcacctttac ttaacaggac caatttatcg aataaataaa
140401 atatatttga tcttggctga tgaatagata gcaccaatgt cctgttatag tcgctggata
140461 accttaccaa acattcaata acattattgg cattactagc atctaaacca gatgtaggtt
140521 catccaaaaa taaaaccaat ggagatgtca ctaattcaca tgcaatggaa actcggcgtt

140581 tttcacctcc actaatacca cgatcaaatt cattaccaat aatacgatct ttgatatcaa
140641 taattcttag ttcttccaac accttataaa ctcttgcctt cttggcctcg aatgacaatg
140701 cttttggcaa tcttaacagc gcactattta atacggtttc aaaaacagtc aaagtgggca
140761 gcaaaaagtc atcttgatcg acgaacccga ttattttcga gaaagattta cggtccatac
140821 taataccgtt aacttttatg gaacccgaaa cgtgacctgt tttccgtttc attgctagga
140881 tatctaataa agtagtttta cccgcaccag atccacccat gatagctaat atttggccgg
140941 gcttcacgat accacttatt tcattcagca cagtttcttc aacaccatct gaatttatcg
141001 aggggacact ataagtgata ttttcaaaac ttaatgtcgc cagtgtgtca tcttcatttt
141061 gtaagaaatt attcaccgca tcttcatctg gcaaacgaat gggagacttt gaggaaccca
141121 atccatttct gaataacgga gatttagaaa tgtaaaaggt agcaaatgta aaaagtgcca
141181 ggaccatcac agcagtcaat gccaacacca atttcccttg ccatgacact gttggatctt
141241 ttgaaggaga tttgtaacct ggaatctcac tataatgaac acattcaccg gattcacact
141301 tcaaagtaat ataagggtca ccaaacacgg tcaatatcaa atcattcata gaaggctcac
141361 tgaatttaca ttgccttgtt tctaaatcac agctgaaatc tcctggccct tttattgtct
141421 ctgtcaggaa atccgagata tctatagacc ccttagcacc acacaacaca gtgtcgggaa
141481 cgcatttgca ttgaacgtca ttacacttat aatgggaggt attctgttcc aagtcgtatt
141541 caaaggcaca atcacttaag ccacaataga agctttctaa ctgatctatc caaaactgaa
141601 aattacattc ttgattaggt ttatcacagg caaatgtaat ttgtggtatt ttgccgttca
141661 aaatctgtag aattttctca ttggtcacat tacaacctga aaatacttta tctacaatca
141721 taccattctt ataacatgtc cccttaatac taggatcagg catgaacgca tcacagacaa
141781 aatcttcttg acaaacgtca caattgatcc ctccccatcc gttatcacaa tgacaggtgt
141841 cattttgtgc tcttatggga cgatccttat taccgctttc atccggtgat agaccgccac
141901 agaggggcag agagcaatca tcacctgcaa acccttctat acactcacat ctaccagtgt
141961 acgaattgca ttcagaaaac tgtttgcatt caaaaatagg tagcatacaa ttaaaacatg
142021 gcgggcatgt atcattgccc ttatcttgtg cagttagacg cgaatttttc gaagaagtac
142081 cttcaaagaa tggggtctta tcttgttttg caagtaccac tgagcaggat aataatagaa
142141 atgataatat actatagtag agataacgtc gatgacttcc catactgtaa ttgcttttag
142201 ttgtgtattt ttagtgtgca agtttctgta aatcgattaa ttttttttc tttcctcttt
142261 ttattaacct taatttttat tttagattcc tgacttcaac tcaagacgca cagatattat
142321 aacatctgca taataggcat ttgcaagaat tactcgtgag taaggaaaga gtgaggaact
142381 atcgcatacc tgcatttaaa gatgccgatt tgggcgcgaa tcctttattt tggcttcacc
142441 ctcatactat tatcagggcc agaaaaagga agtgtttccc tccttcttga attgatgtta
142501 ccctcataaa gcacgtggcc tcttatcgag aaagaaatta ccgtcgctcg tgatttgttt
142561 gcaaaaagaa caaaactgaa aaaacccaga cacgctcgac ttcctgtctt cctattgatt
142621 gcagcttcca atttcgtcac acaacaaggt cctagcgacg gctcacaggt tttgtaacaa
142681 gcaatcgaag gttctggaat ggcgggaaag ggtttagtac cacatgctat gatgcccact
142741 gtgatctcca gagcaaagtt cgttcgatcg tactgttact ctctctcttt caaacagaat
142801 tgtccgaatc gtgtgacaac aacagcctgt tctcacacac tcttttcttc taaccaaggg
142861 ggtggtttag tttagtagaa cctcgtgaaa cttacattta catatatata aacttgcata

142921 aattggtcaa tgcaagaaat acatatttgg tcttttctaa ttcgtagttt ttcaagttct
142981 tagatgcttt ctttttctct tttttacaga tcatcaagga agtaattatc tacttttttac
143041 aacaaatata aaacaatgtc tttatcttca aagttgtctg tccaagattt ggacttgaag
143101 gacaagcgtg tcttcatcag agttgacttc aacgtcccat tggacggtaa gaagatcact
143161 tctaaccaaa gaattgttgc tgctttgcca accatcaagt acgttttgga acaccaccca
143221 agatacgttg tcttggcttc tcacttgggt agaccaaacg gtgaaagaaa cgaaaaatac
143281 tctttggctc cagttgctaa ggaattgcaa tcattgttgg gtaaggatgt caccttcttg
143341 aacgactgtg tcggtccaga agttgaagcc gctgtcaagg cttctgcccc aggttccgtt
143401 attttgttgg aaaacttgcg ttaccacatc gaagaagaag gttccagaaa ggtcgatggt
143461 caaaaggtca aggcttccaa ggaagatgtt caaaagttca gacacgaatt gagctctttg
143521 gctgatgttt acatcaacga tgccttcggt accgctcaca gagctcactc ttctatggtc
143581 ggtttcgact tgccacaacg tgctgccggt ttcttgttgg aaaaggaatt gaagtacttc
143641 ggtaaggctt tggagaaccc aaccagacca ttcttggcca tcttaggtgg tgccaaggtt
143701 gctgacaaga ttcaattgat tgacaacttg ttggacaagg tcgactctat catcattggt
143761 ggtggtatgg ctttcacctt caagaaggtt ttggaaaaca ctgaaatcgg tgactccatc
143821 ttcgacaagg ctggtgctga aatcgttcca aagttgatgg aaaaggccaa ggccaagggt
143881 gtcgaagtcg tcttgccagt cgacttcatc attgctgatg ctttctctgc tgatgccaac
143941 accaagactg tcactgacaa ggaaggtatt ccagctggct ggcaagggtt ggacaatggt
144001 ccagaatcta gaaagttgtt tgctgctact gttgcaaagg ctaagaccat tgtctggaac
144061 ggtccaccag gtgttttcga attcgaaaag ttcgctgctg gtactaaggc tttgttagac
144121 gaagttgtca agagctctgc tgctggtaac accgtcatca ttggtggtgg tgacactgcc
144181 actgtcgcta agaagtacgg tgtcactgac aagatctccc atgtctctac tggtggtggt
144241 gcttctttgg aattattgga aggtaaggaa ttgccaggtg ttgctttctt atccgaaaag
144301 aaataaattg aattgaattg aaatcgatag atcaattttt ttcttttctc tttccccatc
144361 ctttacgcta aaataatagt ttattttatt ttttgaatat tttttattta tatacgtata
144421 tatagactat tatttatctt ttaatgatta ttaagatttt tattaaaaaa aaattcgctc
144481 ctcttttaat gcctttatgc agtttttttt tcccattcga tatttctatg ttcgggttca
144541 gcgtatttta agtttaataa ctcgaaaatt ctgcgttcgt taaagctttc gagaaggata
144601 ttatttcgaa ataaaccgtg ttgtgtaagc ttgaagcctt tttgcgctgc caatattctt
144661 atccatctat tgtactcttt agatccagta tagtgtattc ttcctgctcc aagctcatcc
144721 cacttgcaac aaaaaaagtc taatcttctg caataatttc catccttggc attcagagac
144781 atatattggt caatcggttt taattttgtt tcttcttctt cttcttcttc aattaacaat
144841 gtgccttcat catttttcat ttctttcagt tcttctaatt ctcttggaga caatttgact
144901 cccatgaaaa atttcttgat tatctcagaa gaataccacc tctttctttc tccataccca
144961 acaatcttcg ctgtacgaaa tctctccact attcttttta agaatagctt ttccaagttt
145021 ggcgttagct gtaaaaaaca atggatatag ccatccttgt acaacttaat acaaagcgtt
145081 tccatgattt ttgccaactc ggtcgtgtca ttacaaaacg gcttgaaaaa taaaagatca
145141 atgtcaccac acttggaata gcccctatta taacttccct gtaattcgac ttggcattca
145201 ggatcaatgc cacgcagcgc tttttgaacc ttctttaaat gtgtgaaaca ttcattccga

145261 gacatcttgc ataaccaatc gtcgtaatat gaccaaccaa ataaaattgt ccaatctgat
145321 acaaactcct ctgggtcctt cttagctgca acacaaaagc tttcaaaatt tagaagattc
145381 cagcgtttag caatttccga cccaatgccg taacaatttt tgaagtattt caacttgtct
145441 tctaatccca ctgaatcatt taaacctggt aaaactcccg tatctaatat aacttgtatt
145501 tttttggcaa tactaggccc gatattcctt aatttagtat gtgcttcttc accggaacga
145561 acattgaaat cgcaattttc catcgactgt ttagccagtc tataacttct tgcacgaaat
145621 ttctcacctt cgatctcata tttttttgtt aaccttttca acgctccgat aatcaattca
145681 ttgtttttgt acttcgtttt gtcagtaaca agggaagctc ttcctttatc cccctgtaag
145741 agtcgcttaa gaggttttga agcttgtatc atatctttat tgttgctgtc atcattcctt
145801 tcactttcta cgtcagttga aatctcactc tctttatcag tagaactttg tccgtcatca
145861 gcgttatctg atataacgat aatggatggt tgaaatttaa tcaaatcatc tttttgaaac
145921 gcaaatttat cattctggac ccactttgtg atgcaactgg ttttcacaca ttgaatgcct
145981 gactgttcaa tcttaccaag aaattcatcg acatcattta agcctgcttc cctttggaaa
146041 atatctttct gagtcaaatg catatcagaa tcaacaaagg agtcctcaat caaaatgacc
146101 acttccttac ggctagattg aagacaattt tgaatacttg acacaattgt ggcgcccttt
146161 ttctccaata tactcttaaa gaacttattg gaagatgtgt taggattagg taaaaaggcg
146221 aaaaatttac cctttagaga catgtgcgat ttgtctaaca ggtcgcatgt ttatccttta
146281 cttattatca ctgtgacata taagttcttt gcaagccctt aaaattatta gccccatgta
146341 tttgaacatc tcaaagtaat tttagaccga gaccgattac ggagatagaa gaatagtgca
146401 agtgaaatac tattttatct ttacttaaaa aagaaaaaac aggaggaaaa ttagcggtat
146461 tccattatta ttagcactgc aacgtcctgg ttatataaat ttgaatatac aggatttcat
146521 ccgtttctac atttttgtta cctgcatgat caaatctttt aaagagtccc aggatttcac
146581 aaggtaggca cctttaccgc cttccttttc acaaaactgt aaccacgctg gaccattatc
146641 agcttcaaaa ctcgagattt tatcctttgg gataccaata atcttttcgg taatttttat
146701 gaatttggaa ggattttctt gcgggtttat tagcagtaca ccattagtag atggatgtaa
146761 tatggacaac aagtcagtct cactatcccc tatataccaa tagctgcaag aatttccttc
146821 tttattgcaa cctgagtcga tcttatctaa tatttcaccc agtatcttga ccttatcaga
146881 gcctgtcaat aaccgacaat caaattcacc attgtagctt tgagaacact tgtcactaac
146941 tttttttaga tcgttacaaa aaatatgact atttttaagt cttctgtcac caatgacttc
147001 atgaataaac tcttttgacc aatttataga taaaacgtaa aaatcacttt caaaattttt
147061 gacaactgaa gagcaaaacg ttttgaaacc atctcttagt aaacaatctt cgtgattctg
147121 atctctggcg aacgttttca tttgatccaa agaaatcgac ttaaaaatct gttgttttgt
147181 aatttcattt acactattta actcaacaac cctgttatga ttttgatatt ttaattcgtc
147241 cgcaaacaac ttattgaagt tagactgcga aattatcgta ggtacgcctg aagataacaa
147301 cggcagtgat ctcgttccat tgtatttgta tttatggtat ccgtccatat atgtcttggt
147361 aaaatgaccc cactcaggtt tcaaccgcgg gttcagtaga tacggtaatt tagcaatggt
147421 gcagattgtg tcaactcttg tgattgtttc atcaaaatct gaaataataa tggttttcat
147481 atcctccact gttcgcttct ggttactgca attgttaaat gcattggctc gattgtcaac
147541 attgctttgt gcgaagaaag gtcgctttta aagaaaacga taagaactaa cctctaacta

147601 cataataata agaaaaaagt atacagataa acaagctcag tagtaataca taaaataaaa
147661 agttttgcaa atcgattgtc cacacacttc tatatcaatt ccaactaaga aaacgaagga
147721 accgactcaa accggtgtaa aagctaggct ttcaatctct gctcctccaa acaaaactag
147781 tgcttgatat cacatccaat taaattacgt aaataatttt agagtgacac catcgtacaa
147841 agagggcatt aagcttattt ttttaattct tatgaaaaaa aaaatgtgaa agagagtaga
147901 aaattttcat aacgaatctc ttctattctt atgtaccgtt ccgccaaatt tttcatgaaa
147961 aaatttggga aaaagaaaaa aaaaaaaaat agtatataaa gtaaaaaaga tgcgcaagcc
148021 cggaatcgaa ccgggggccc aacgatggca acgttggatt ttaccactaa accacttgcg
148081 ctagtaatct tgatacatga ttcgagaaaa gcaacaatat tatgtatacg gaatatacta
148141 ggagttctct tcaaggatag aggaatcctc aaaatggaat ccatatttct acatactaat
148201 attacgatta tcccttcttc tattttatat gtttcattat cctattacat tatcaatcct
148261 tgcacttcag cttcctctaa cttcgatgac agcttctcat aacttatgtc atcatcttaa
148321 caccgtatat gataatatat tgataatata actattagtt gatagacgat agtggatttc
148381 tattccaaca agaacatcct tgaaaggtcg actattaaca taagaataag agagagtgtg
148441 cggccgatgc tgttgctata gctcggtcgt agtttaacac cacggcatta ttcagcttta
148501 aaacaattga attataaacc tgcttttctt actacgaaaa gtgtttccct ttaataaatc
148561 tattgtacat actttacagt tgatagtacc ttcaccataa aaatatgcaa acgcttttgt
148621 tattgggaac aggaatacca accgttctca tggaaaatag caaagtgtaa aagaatttat
148681 tatcatcgag tgataaagaa tatcgccaga tctttacgga tatatatcgc taaaaatatt
148741 tgtcgccaat aaaagatatg tctaacgctc cctttgcagg tagtttttta ctggcatcgc
148801 ctttttactg caaattattg aaaattttcg agcgagaaaa aggaacgaaa agagatgagc
148861 taaattcaaa tatcaaagta ctgtcataag tgctttcttt atcgacaaat tacaagcaga
148921 agaaacattc acttcacaat taaatgtccg aaaatcacgt tcctgcctgg aaaagaattg
148981 ctttgaagag acagactata agcagtggtg acgaaagtaa agaaaaaggc caatccaatc
149041 taatagatga tgatccacta aatatcacta cgcatttatc cactggtaat ttgacaaaga
149101 aagagaaaaa aagaatcatt aatggtgaga gcaaatcctc taccaaaaaa gggaagcgtg
149161 tgtcgaaacc agggacgaag aagaaggaaa aattgtcaaa ggatgaaaag aattcgaaaa
149221 aaaataaaat tcttaaagat caattacgtt acctgataga atttttcagg acaaagtctg
149281 agagcaagtt tcccaccgga atcctagaat tggaaagtgt aaaagaaaat tacggcgact
149341 ccctgatcaa agacgagcca tcagagtctg gtgttgttga agtttggaaa ttttccaagc
149401 aaaagcaaaa ttggcttatt aagcattttt tcaacttgga cgaaatcccc tcagtatata
149461 atgatctttt gcttttatat ttcagggact tacaaggtaa atcaaaagaa gaactaatat
149521 caaaatgcaa aggaaaactg aagcaatgga acgactatgt ggaagatcaa gaaacaaaaa
149581 taaaagcgtt aattgcagag gataaggcta gcgaaccaat taatggtgaa gaaaaagaag
149641 aaggtgaaaa agacggtaat gccgaacaag ggaagcaaaa agaggtacaa gacgagcaag
149701 aggaagtaca aatgccgaat aaagagcttg ttcaaagaag tctgaaatta ctagaaattt
149761 ggaagaatga tgactcggag cagatagaac ttaagaattt cttcgttgat gtttaacata
149821 gagttcaaca acgttgttca gttgtagtat atattttact gtgaaatacg ctcaaaaagt
149881 agcgggataa atcgtttcag gaagtataat ttcttacaaa actataacaa gatcgtctaa

149941 gtaacataca taaagaaatc acataaattt ttaaaattcg aggtcaaaaa aatcaggtgg
150001 atacataact taccttaagt tatcaagatt ttcagtacac acaatgtgat tacactgatt
150061 tataaaacca ccttacggcc tcttataagt aataacgtgg ctcatcaaaa acatggtaga
150121 agtgacccct ctcaccttcg cctaaaaact tatctggaaa tttcatgtcc tcatattccg
150181 gttcactcat atgaccgtat tcatacatag aatccgagtg ctctactaac gcttgctcac
150241 tcaaaacttg gaacttacca acttgtagct ccgtatcggt tatcgttcct cttgcaactc
150301 tagcatatcc tgttcttctc aagcccctat acaagatata ctcacaccat ctatcatcgt
150361 cagaaggatc aatgtcgtgc attccggatg tttcactaac gtacgtattg tagttgcctt
150421 cacctggatc aacaactaag taacttaata aaatagctgg gcgagtcgta ttgcccatga
150481 gcttagccat gtagttactt tgcagtcttc tatcctcttc atcctcttct tgtccactat
150541 ggaatacaaa gacgtcaacg agagtgtcat tgtacgtttg aagtgtggca tgaatggctg
150601 gcgcaagttc cccaactgga gagggcaata aatgatgcgt agaatttacg atagggaatt
150661 tagaaagaag aacacagccc caggtatgtt tatttggacc tggtccgaaa tctgcataca
150721 tattcaaatc atgagctagt ttgctagtta gatccctgtt ccccatggta attctttgtg
150781 tatctgtttc tagtagacct accacatcta gttccatatc tttaataagg ttgatcattc
150841 tgtcttcaga tgcccacata tcattatcga gaccaaagtg gatagtccaa ataccagctg
150901 taatcaactg agaatctgga tgataaggct gaggaattcc tgtaggtcta atatcataca
150961 cgaacctagc tgttagtgat agcagggcca cggcaaagaa gaaaacataa atgaagaatt
151021 ttttgcccat agtcaccagt tgaatgttaa ggtttttgca tgttaaagca ccaataatga
151081 taaatgtgga agaaaaggca agaaccgtct caatcttctc cctcagtacc cagcccattg
151141 gaacaaatgc gtatgcaacg acccaaacat gggcaaggac aaagataata taaacggaga
151201 atgataggac ataaacccat atgttttgaa cttggcctaa tgcagaaaaa tacgaaggaa
151261 ctaaccaaag catagcgata gcgtacaata aaccaccgaa aatatattta ggccattgtg
151321 taatgcttct agctgaaagt acagcagtgg atatgagaag caacaaacat ggaactaatg
151381 gcttgcccat aaacttcaca gaactcaaag aagcaaataa catgaccgta caagttaagg
151441 cgccccaagg ccatggcaag ggaccgtgtg attcattgta accttcccat gcccaataaa
151501 tagtagtaga agaatccgtc aataactgat gaattccgaa taacaacgag ccaaaaccaa
151561 cagccaaaaa taatttgcca atcaacgaag cagattttgc ttgagcattc cttttccctt
151621 cgaaattaat tgaattaaca taaggcgaca gcatgccgaa taaaacagta agcacgagag
151681 ctgtcttatt atacccacca ttacgttcat ctaaaatgac ccaagttgga ttgttggtat
151741 agaaccccat tttcaagatc acagatgcaa caagacctaa cagccaagtt agagcaaaag
151801 agaaactagt ctccgcattt gtgatatatc ttagattttg tacaaacgtt gcaacggtaa
151861 tggaagttcc cacagctaca gaaatcaacc ttaattctgg catctgaaca atataggcac
151921 caatggcaat aatacctcct aaaagtacac catattgcgt gaaggcttca gaaacgaacg
151981 gcagatataa gaaaatgggt cccaaataac ccaatattgc agcttcataa cctgagattc
152041 ccatatacca tagcgggaaa tgccaaatac tacataacag ggacgtaacg ttcgaccaga
152101 aaagaaaacc gttaaaaata tttgttagta agtaaaaaaa agagtcaaac ctgaaatatg
152161 aaccagttgc gcttctagca acgggttttt ccttttcaac cttcttttca ccacttttct
152221 gcaattctga actttttct tctccataat ttatgggact agagtctcta atttggaaaa

152281 aactggtatt accgggtttc tcattaaaag cgagaacaat atctatcttt ttgaaatcag
152341 cgtaagcaaa tgcatcaaat gcaatatcta aaagaatcag agaccattcg aaataagcat
152401 atatagaata tgccccagct ctctgttgga cggagtgctg aatgtaccag taaatcatcg
152461 ggaacaatgt tccaaaaaaa attgtagcag tcaacccttt gttcttcgaa gttaaaggac
152521 tagaatagcg ggtaatcatt atatcccatg gtaacgttaa aacaatgtat gtgatcataa
152581 atatatcatg aatatcgtgg tcatctgtac ttgtaatgta tacccaacca ccgcaagaga
152641 ctgtccggag tacaccgaca aggaagcata ccttagattg gttcaagtag tagtgaccta
152701 gtagcagtaa aaatcttgga aaagcagtta gagctattaa gatttggaaa atagaacgtt
152761 ccggatagcg gtccccgata gtggctgata cactaggaaa ccactcatct ggatacgtat
152821 aatgtgcatt ggttacaatt ttatgaaaat gtaatgaata accagtgacc aaagctgcaa
152881 agaaggcgga gaatgcacaa atagtatgag ctatagggat gatcttccca ttgatgatca
152941 gcataactgg cggtggatac cttctgtatt cctggttact tattcctcga gaaatcggac
153001 attaaaaggg tgcaaatact gtggagaaat tataagttac caagtcaaaa agcgaatgag
153061 aatctattgt tgcgattttt ttggttacta cttcttaccc gaactaactt tacataataa
153121 aaaagcatga agcgcatatt cagggactca aaaatcccaa actggcactt ctgtatatac
153181 cgatgagcat actttgattg taaaaaacaa cattaaaaag aactgcaaac ctgaaccaag
153241 aatacttaat ttcaaaaccg agaaatgagg tacaacttgt agaagttcac ttaaacttat
153301 tgagcataca atctcaacaa aactaactgc aaagtttgtt gctatagtca tatagtaata
153361 ctagcaggaa gccattaatc aaaaagaaac tccatataaa acgtaataaa tattatgaat
153421 aatagtacaa tataagaaca aagacgaaga aaatacatta taactgaagt gccaatatta
153481 gaaaaaaaaa aggtggaata tcctagtgga aaaatatcat ctatgcggtg aaagaattaa
153541 atagatacta tggattgttt cgatctatcc ttttatctgc tgcttgcctt tttttttcat
153601 tctccttctt cagtcttttt ccataggcga gtccacaggg actacaaagt tccctgtttt
153661 ggtcgggtcc actacgccat tgaattgtcc aagtgtcttt acatttagag cattccttta
153721 tctcaccatt tgggaggatt gtttgtctct tttctctttt aggacgacta gtagtctttt
153781 tgctgtattt cttcgtaata ctggcagacc gactagttcc ttgtgcacaa ctatggtagc
153841 cttcaagatt gtttgcttct tggctagtcg tacgtctctc ggaattgcat gcactagtta
153901 cacgattctt tgagggtgta attttgttgg aataaaaatc aactatcatc tactaactag
153961 tatttacgtt actagtatat tatcatatan nnnnnnnnnn nnnnnnnnnn nnnnnnnnnn
154021 nnnnnnnntc tgacggcact gtacttgcac ctatcgtaaa atatggagac ttttactggg
154081 tatctaaaaa gtacttgctt ccatcaaata tctccgtacc caccatcaat aatgtccata
154141 caagtgaaag tacacgcaaa tatccttatc ctttcattca tcgaatgctt gcgcatgcca
154201 atgcacagac aattcgatac tcacttaaaa ataacaccat cacgtattta acgaatcaga
154261 tgtcgactgg tctagtgcta ttgactatca atgtcctgat tgtttaatcg gcaaaagcac
154321 caaacacaga catatcaaag gttcacgact aaaataccaa aattcatacg aacccttca
154381 atacctacat actgacatat ttggtccagt tcacaaccta ccaaaagtgc accatcctat
154441 ttcatctcat ttactgatga gacaacaaaa ttccgttggg ttatccatta cacgaccgtc
154501 gcgaggactc tatcctcgat gtttttacta cgatactagc ttttattaag aaccagtttc
154561 aggccagtgt cttggttata caaatggacc gtggttctga gtatactaac agaactctcc

154621 ataaattcct tgaaaaaaat ggtataactc catgctatac aaccacagcg gattcccgag
154681 cacatggagt cgctgaacgg ctcaaccgta ccttattaga tgactgccgt actcaactgc
154741 aatgtagtgg tttaccgaac catttatggt tctctgcaat cgaattttct actattgtga
154801 gaaattcact agcttcacct aaaagcaaaa aatctgcaag acaacatgct ggcttggcag
154861 gacttgatat cagtactttg ttacctttcg gtcaacctgt tatcgtcaat gatcacaacc
154921 ctaactccaa aatacatcct cgtggcatcc caggttacgc tctacatccg tctcgaaact
154981 cttatggata tatcatctat cttccatcct taaagaagac agtagataca actaactatg
155041 ttattcttca gggcaaggaa tccagattag atcaattcaa ttacgacgca ctcactttcg
155101 atgaagactt aaaccgttta actgcttcat atcaatcgtt cattgcgtca aatgagatcc
155161 aacaatccga tgatcttaac atagaatctg accatgactt ccaatctgac atcgaactac
155221 atcctgagca accgagaaat gtcctttcaa aagctgtgag tccaaccgat tccacacctc
155281 cgtcaactca tactgaagat tcgaaacgtg tttctaaaac caatattcgc gcacccagag
155341 aagttgaccc caacatatct gaatctaata ttcttccatc aaagaagaga tctagcaccc
155401 cccaaatttc caatatcgag agtaccggtt cgggtggtat gcataaatta aatgttcctt
155461 tactttgctc ccatgtccca atctaacaca catgagtcgt cgcacgccag taaatctaaa
155521 gatttcagac actcagactc gtacagtgaa aatgagacta atcatacaaa cgtaccaata
155581 tccagtacgg gtggtaccaa caacaaaact gttccgcaga taagtgacca agagactgag
155641 aaaaggatta tacaccgttc accttcaatc gatgcttctc caccggaaaa taattcatcg
155701 cacaatattg ttcctatcaa aacgccaact actgtttctg aacagaatac cgaggaatct
155761 atcatcgctg atctcccact ccctgatcta cctccagaat ctcctaccga attccctgac
155821 ccatttaaag aactcccacc gataaattct cgtcaaacta attccagttt gggtggtatt
155881 ggtgactcta atgcctatac tactatcaac agtaagaaaa gatcattaga agataatgaa
155941 actgaaatta aggtatcacg agacacatgg aatactaaga atatgcgtag tttagaacct
156001 ccgagatcga agaaacgaaa ttcacctgat tgcagctgta aaagcagtaa aatcaatcaa
156061 accaatacgg acaaccttac gatacgatga ggcaatcacc tataataaag atattaaaga
156121 aaaagaaaaa tatatcgagg cataccacaa agaagtcaat caactgttga agatgaaaac
156181 ttgggacact gacgaatatt atgacagaaa agaaatagac cctaaaagag taataaactc
156241 aatgtttatc ttcaacaaga aacgtgacgg tactcataaa gctagatttg ttgcaagagg
156301 tgatattcag catcctgaca cttacgactc aggcatgcaa tccaataccg tacatcacta
156361 tgcattaatg acatccctgt cacttgcatt agacaataac tactatatta cacaattaga
156421 catatcttcg gcatatttgt atgcagacat caaagaagaa ttatacataa gacctccacc
156481 acatttagga atgaatgata agttgatacg tttgaagaaa tcactttatg gattgaaaca
156541 aagtggagcg aactggtacg aaactatcaa atcatacctg atacaacaat gtggtatgga
156601 agaagttcgt ggatggtcat gcgtatttaa aaacagtcaa gtgacaattt gtttattcgt
156661 agatgatatg gtattgttta gcaaaaatct aaattcaaac aaaagaatta tagagaagct
156721 taagatgcaa tacgacacca agattataaa tctaggcgaa agtgatgagg aaattcaata
156781 tgacatactt ggcttagaaa tcaaatatca aagaggtaaa tacatgaaat taggtatgga
156841 aaactcatta actgagaaaa tacccaaatt aaacgtacct ttgaatccaa aaggaagaaa
156901 acttagcgct ccaggtcaac caggtcttta tatagaccag gatgaactag aaatagatga

156961 agatgaatac aaagagaagg tacatgaaat gcaaaagttg attggtctag cttcatatgt
157021 tggatataaa tttagatttg acttactata ctacatcaac acacttgctc aacatatact
157081 attcccctct aggcaagttt tagacatgac atatgagtta atacaattca tgtgggacac
157141 tagagataaa caattaatat ggcacaaaaa caaacctacc aagccagata ataaactagt
157201 cgcaataagc gatgcttcat atggtaacca accatattac aagtcacaaa ttggtaacat
157261 tttcctactc aacggaaaag tgattggagg aaagtcgaca aaggcttcgt taacatgcac
157321 ttcaactaca gaagcagaaa tacacgcagt cagtgaagct ataccgctat tgaataacct
157381 cagtcacctt gtgcaagaac ttaacaagaa accaattatt aaaggcttac ttactgatag
157441 tagatcaacg atcagtataa ttaagtctac aaatgaagag aaatttagaa acagattttt
157501 tggcacaaag gcaatgagac ttagagatga agtatcaggt aataatttat acgtatacta
157561 catcgagacc aagaagaaca ttgctgatgt gatgacaaaa cctcttccga taaaaacatt
157621 taaactatta acaaacaaat ggattcatta gatctattac attatgggtg gtatgttgga
157681 ataaaaatca actatcatct actaactagt atttacgtta ctagtatatt atcatatacg
157741 gtgttagaag atgacgcaaa tgatgagaaa tagtcatcta aattagtgga agctgaaacg
157801 caaggattga taatgtaata ggatcaatga atattaacat ataaaacgat gataataata
157861 tttatagaat tgtgtagaat tgcagattcc cttttatgga ttcctaaatc ctcgaggaga
157921 acttctagta tatctacata cctaatatta tagccttaat cacaatggaa tcccaacaat
157981 tacatcaaaa tccacattct ctacatggta gcgcctgtgc ttcggttact tctaaggaag
158041 tccacacaaa tcaagatccg ttagacgttt cagcttccaa aattcaagaa tatgataagg
158101 cttccactaa ggctaactct caacagacaa caacacctgc ttcatcagct gttccagaga
158161 acccccatca tgcctctcct caacctgctt cagtaccacc tccacagaat gggccgtacc
158221 cacagcagtg catgatgacc caaaaccaag ccaatccatc tggttggtca ttttacggac
158281 acccatctat gattccgtat acaccttatc aaatgtcgcc tatgtacttt ccacctgggc
158341 cacaatcaca gtttccgcag tatccatcat cagttggaac gcctctgagc actccatcac
158401 ctgagtcagg taatacattt actgattcat cctcagcgga ctctgatatg acatccacta
158461 aaaaatatgt cagaccacca ccaatgttaa cctcacctaa tgactttcca aattgggtta
158521 aaacatacat caaattttta caaaactcga atctcggtgg tattattccg acagtaaacg
158581 gaaaacccgt acgtcagatc actgatgatg aactcacctt cttgtataac acttttcaaa
158641 tatttgctcc ctctcaattc ctacctacct gggtcaaaga catcctatcc gttgattata
158701 cggatatcat gaaaattctt tccaaaagta ttgaaaaaat gcaatctgat acccaagagg
158761 caaacgacat tgtgaccctg gcaaatttgc aatataatgg cagtacacct gcagatgcat
158821 ttgaaacaaa agtcacaaac attatcgaca gactgaacaa taatggcatt catatcaata
158881 acaaggtcgc atgccaatta attatgagag gtctatctgg cgaatataaa tttttacgct
158941 acacacgtca tcgacatcta aatatgacag tcgctgaact gttcttagat atccatgcta
159001 tttatgaaga acaacaggga tcgagaaaca gcaaacctaa ttacaggaga aatccgagtg
159061 atgagaagaa tgattctcgc agctatacga atacaaccaa acccaaagtt atagctcgga
159121 atcctcaaaa aacaaataat tcgaaatcga aaacagccag ggctcacaat gtatccacat
159181 ctaataactc tcccagcacg gacaacgatt ccatcagtaa atcaactact gaaccgattc
159241 aattgaacaa taagcacgac cttcatctta ggccagaaac ttactgaatc tacagtaaat

159301 catactaatc attctgatga tgaactccct ggacacctcc ttctcgattc aggagcatca
159361 cgaaccctta taagatctgc tcatcacata cactcagcat catctaatcc tgacataaac
159421 gtagttgatg ctcaaaaaag aaatatacca attaacgcta ttggtgacct acaatttcac
159481 ttccaggaca acaccaaaac atcaataaag gtattgcaca ctcctaacat agcctatgac
159541 ttactcagtt tgaatgaatt ggctgcagta gatatcacag catgctttac caaaaacgtc
159601 ttagaacggt ctgacggcac tgtacttgca cctatcgtaa aatatggaga cttttactgg
159661 gtatctaaaa agtacttgct tccatcaaat atctccgtac ccaccatcaa taatgtccat
159721 acaagtgaaa gtacacgcaa atatccttat cctttcattc atcgaatgct tgcgcatgcc
159781 aatgcacaga caattcgata ctcacttaaa aataacacca tcacgtattt taacgaatca
159841 gatgtcgact ggtctagtgc tattgactat caatgtcctg attgtttaat cggcaaaagc
159901 accaaacaca gacatatcaa aggttcacga ctaaaatacc aaaattcata cgaacccttt
159961 caatacctac atactgacat atttggtcca gttcacaacc taccaaaaag tgcaccatcc
160021 tatttcatct catttactga tgagacaaca aaattccgtt gggtttatcc attacacgac
160081 cgtcgcgagg actctatcct cgatgttttt actacgatac tagcttttat taagaaccag
160141 tttcaggcca gtgtcttggt tatacaaatg gaccgtggtt ctgagtatac taacagaact
160201 ctccataaat tccttgaaaa aaatggtata actccatgct atacaaccac agcggattcc
160261 cgagcacatg gagtcgctga acggctcaac cgtaccttat tagatgactg ccgtactcaa
160321 ctgcaatgta gtggtttacc gaaccattta tggttctctg caatcgaatt ttctactatt
160381 gtgagaaatt cactagcttc acctaaaagc aaaaaatctg caagacaaca tgctggcttg
160441 gcaggacttg atatcagtac tttgttacct ttcggtcaac ctgttatcgt caatgatcac
160501 aaccctaact ccaaaataca tcctcgtggc atcccaggct acgctctaca tccgtctcga
160561 aactcttatg gatatatcat ctatcttcca tccttaaaga agacagtaga tacaactaac
160621 tatgttattc ttcagggcaa ggaatccaga ttagatcaat tcaattacga cgcactcact
160681 ttcgatgaag acttaaaccg tttaactgct tcatatcaat cgttcattgc gtcaaatgag
160741 atccaacaat ccgatgatct taacatagaa tctgaccatg acttccaatc tgacatcgaa
160801 ctacatcctg agcaaccgag aaatgtcctt tcaaaagctg tgagtccaac cgattccaca
160861 cctccgtcaa ctcatactga agattcgaaa cgtgtttcta aaaccaatat tcgcgcaccc
160921 agagaagttg accccaacat atctgaatct aatattcttc catcaaagaa gagatctagc
160981 accccccaaa tttccaatat cgagagtacc ggttcgggtg gtatgcataa attaaatgtt
161041 cctttacttg ctcccatgtc ccaatctaac acacatgagt cgtcgcacgc cagtaaatct
161101 aaagatttca gacactcaga ctcgtacagt gaaaatgaga ctaatcatac aaacgtacca
161161 atatccagta cgggtggtac caacaacaaa actgttccgc agataagtga ccaagagact
161221 gagaaaagga ttatacaccg ttcaccttca atcgatgctt ctccaccgga aaataattca
161281 tcgcacaata ttgttcctat caaaacgcca actactgttt ctgaacagaa taccgaggaa
161341 tctatcatcg ctgatctccc actccctgat ctacctccag aatctcctac cgaattccct
161401 gacccattta aagaactccc accgataaat tctcatcaaa ctaattccag tttgggtggt
161461 attggtgact ctaatgccta tactactatc aacagtaaga aaagatcatt agaagataat
161521 gaaactgaaa ttaaggtatc acgagacaca tggaatacta agaatatgcg tagtttagaa
161581 cctccgagat cgaagaaacg aattcacctg attgcagctg taaaagcagt aaaatcaatc

161641 aaaccaatac gaacaacctt acgatacgat gaggcaatca cctataataa agatattaaa
161701 gaaaaagaaa aatatatcga agcataccac aaagaagtca accaactgtt gaagatgaaa
161761 acttgggaca ctgacagata ttatgacaga aaagaaatag accccaaaag agtaataaat
161821 tcaatgttta tctttaacag gaaacgtgac ggtactcata aagctagatt tgttgcaaga
161881 ggtgatattc agcatcctga cacttacgac tcaggcatgc aatccaatac cgtacatcac
161941 tatgcattaa tgacatccct gtcacttgca ttagacaata actactatat tacacaatta
162001 gacatatctt cggcatattt gtatgcagac atcaaagaag aattatacat aagacctcca
162061 ccacatttag gaatgaatga taagttgata cgtttgaaga aatcacttta tggattgaaa
162121 caaagtggag cgaactggta cgaaactatc aaatcatacc tgataaaaca gtgtggtatg
162181 gaagaagttc gtggatggtc atgcgtattt aagaatagtc aagtaacaat ttgcttattc
162241 gttgatgata tgatattgtt cagcaaggac ttaaattcga ataaaagaat tatagcaaaa
162301 ctcaagatgc aatacgatac caagattata aatctaggtg aaagcgatga tgaaattcaa
162361 tacgacatcc ttggcttaga aatcaaatat caaagaggta aatacatgaa attaggtatg
162421 gaaaactcat taactgagaa aatacccaaa ttaaacgtac ctttgaatcc aaacggaaga
162481 aaacttggtg ccccaggcca accaggtctc tatataaacc agcaagaact agagctagaa
162541 gaagatgatt acaaaatgaa ggtacatgaa atgcaaaagc tgataggtct agcatcatat
162601 gttggatata aatttagatt tgacctatta tactacatca acacacttgc acaacatata
162661 ctatttccgt ccaagcaagt gttagatatg acatatgaat tgatacagtt catatggaat
162721 acgagagata agcaattaat atggcacaaa agcaaacctg ttaagccaac aaataaatta
162781 gttgttataa gcgatgcctc gtatggcaac caaccgtatt ataaatcaca aattggcaac
162841 atatatttac ttaatggaaa ggtaattgga ggaaagtcca ccaaggcttc attaacatgt
162901 acttcaacta cggaagcaga aatacacgca ataagtgaat ctgtcccatt attaaataat
162961 ctaagttacc tgatacaaga actcgacaag aaaccaatta ctaaaggatt attaaccgac
163021 agtaaatcta caatcagtat aattatatcc aataatgaag agaaatttag gaacagattt
163081 tttggtacta aagcaatgag attgagagat gaagtatcag gaaatcatct gcacgtatgc
163141 tatatcgaaa ccaaaaagaa tattgcagac gtaatgacca aacctcttcc gataaaaaca
163201 ttcaaactat taacaaacaa atggattcat tagatctatt acattatggg tggtatgttg
163261 gaataaaaat ccactatcgt ctatcaacta atagttatat tatcaatata ttatcatata
163321 cggtgttaag atgatgacat aagttatgag aagctgtcat cgatgttaga ggaagctgaa
163381 acgcaaggat tgataatgta ataggatcaa tgaatataaa catataaaac ggaatgagga
163441 ataatcgtaa tattagtatg tagaaatata gattccattt tgaggattcc tatatcctcg
163501 aggagaactt ctagtatatt ctgtatacct aatattatag cctttatcaa caatggaatc
163561 ccaacaatta tctaattacc cacctatatc tcataatgaa acatataaaa cggaatgagg
163621 aataatcgta atattagtat gtagaaatat agattccatt ttgaggattc ctatatcctc
163681 gaggagaact tctagtatat tctgtataca taatattact gcctttatca ataatggaat
163741 cccaacaact atctgaaaat tcacgtgatt ctcatgtgag tacatcagaa aacaaccacc
163801 taacaaccgt gtttgaatgt tgatctgaaa acaaaaatta atctctcgta cttattatgg
163861 atattgtatt ttctcaaaat tccaaatgat tttcttgtta gatggtattc acggtcatcg
163921 gaatctcagt tcaatatcac caatttgctt ataaggttta tgcaagataa catgtttggt

163981 cttgttccca agtataactt aagaaaaaaa aattacaatc caagaaaggt aatacagaat
164041 tcatagcaat aatggggtcc attttcaata gataatgtac ctggaagaaa agggctgact
164101 aggaaacaaa aagaaaaaca aaactatgat actccatcaa acatagggaa atttactcga
164161 atatgaaaag tacaaatctg tcaaaagcac atcgtatatt gtcttggttc cgcttttatt
164221 cagaagaatt actgaaatga agaggaatga cgtaataaac aaacaacgat tctgatatca
164281 gaaagacatc ttgaatattc ctatgtagta attaactatc ccagtactta caaacataac
164341 ttgacgatgc acgctacgat ttttgtgtgt ttaaactttt gttggttggg actcaagaag
164401 atcttctctg agataattgt aattaaatgt actttaggag tgttcatgta taatttgcct
164461 ctgcaaaaac ttaataatgt attttaaaac ttgcatgttg ctctgtgaat ctgaatgtat
164521 gataatgcta cgattactct agggaacttc ctttttacat ttaaaatttc cgagatcctc
164581 agctgacaaa taaatgtcat atcaagacaa taaacggtag tgctaaagta attgcatatt
164641 aaacaggcgc gactaaagct ttcatccctt ctattattgc ataacctggc tatacatgaa
164701 aacacttctc acatttgatt taagaagttt actgttctaa gaaaataaag aacgtgataa
164761 tgatgaatga agaggattct acagaaacga aaagcctagt cataactaat ggcatgttta
164821 tcatagacga catcgagcgt agtaaatata atattcacta taagaatgtc ccaggaggcg
164881 gagggacttt tgccattttg ggtgcatgca taatatcttc cggcaatgtc acatccaaag
164941 gtttgaagtg gatagtggac agaggctctg actttccaaa ggaagttata agggaaatag
165001 actcatgggg tactgatgtg aggtttcgag atgactttag cagattaact accaaagggt
165061 tgaattatta cgagggaagt gatgatttga gaaagttcaa gtttttgacg ccgaagaagc
165121 agattaacgt cgatgactgg atttccacat ttgggcagaa gataattgat gaaatgcatg
165181 cgtttcattt gctatgttct gggtctagat gcttagacat aataaacgat ctgctacggg
165241 tgaaaagttc aaagggcaca aaaccaatcg tgatttggga gccattccca gatctttgcg
165301 actttgatca tcaaaatgac attaaaagtg taatgcagag gaacgatgtt acggtaatat
165361 tatctccaaa tgccgaagaa tcaagtcgct tatttggttt aagtagcaag gaaccgacta
165421 gtttggaaga atgtctagca ttagcgcatc gtttcgatga tttcatggat gaaaacaata
165481 tgtgtattct acgatgcggt gccctcggaa gcatatcggt aagtgagaag tttaagaacg
165541 gacgaaccta tgaccatttc cccgcctacc atttcaaaac tcagtctaaa gtactagatc
165601 ctactggcgg gggaaactcg ttccttggcg gctttgcagt ttcttatgcc ctaacgaaaa
165661 gcttagatat tgctagtata tgtgggaaca tcgctgcagg cgcaataatt gaacaattcg
165721 gaataccgag gtacgatcca attgctaaaa cctggaacgg aatcacattc ttggatagac
165781 tgaaatttta cctttcacag tccggtcttc aatataatat aaacgatctt tacaaaagtc
165841 taacacgatg atcatatcag cttgttcatt gctcttattt ttctgcaaaa atatacatat
165901 atatatataa gtaagtgtta tatactactt gagattaggc gttgtaacaa gattcaaaaa
165961 attcgaactc gaactgtgaa accttcacaa atatgctttc cacttcttcg acgggaaact
166021 tgtcaacttc agcctttaga aattcacacc aagtttgaaa atgctcacca tcatgtaaat
166081 caatccattt ttgatatttc caatgcaacc ctggggctct aggagtatca cgagcccatc
166141 tccaatagag ttcctcagca acccataagc tggttactaa ggatggccat gtaattgagg
166201 cgtcctttct cagctcatct aagaaattaa tatacgcatc aacgccgggg atcgctttat
166261 tgtcaaaatt atctctttct tccttggtga gggatggtgc caataattct aagcagtcat

166321 gaaagtatga gttttcatca ttagaaaaaa atccaatctt tttggctaag gttattaaag
166381 cgtgagtagt tggtgctaaa gaagtcgtct tacatatcaa ccttaagcta gtttcaaaaa
166441 attgcagatc ttgtgataaa tagatgtaca aactacggtc cttcaatgtt ccggcacata
166501 gttccttagc gaatttatgt tctgtggttt tcctaacaag ggcgtcgtac ttttgtatta
166561 acttatcagt ggtacagctc attgttcttc cttgcagata ttgttgatat tgttattatt
166621 tgcgctaaat tttagttaat acagaaaaaa atatctaagt tcagtccagt tcatattcgt
166681 ttatatatct ttcgaagcca gaccttttt gccaagcaat tgttactatg gaaactatta
166741 gaaccatatg cgctaaacaa tattagccat caattcaact ttattcgcag taagctcctg
166801 cagaacaggc ttatcaatca ttatggtctt aacggaacgc ctagggacgc tgactaggcc
166861 catcattcta ctactggagc ccatacgttc ctcaacatga tctaatagca aattcatttg
166921 ggcatctaca gcaaccaagc ttccaactaa aatacgatct tctgtaaggg aaactattaa
166981 agtatttcca ataaaatctg acagtttcaa gatgtccatg acgctttggt gccccttact
167041 aagtttgttg aagactatct tcaccctatc aatgatatca tcgcatttaa aaaatagtca
167101 gaactcatgc acttcgcgag atattttttg aaaaaaaaaa aattaatccg ccacataaga
167161 tgcacaaaat gagcctataa cattgaattt tatgatagaa cgaaaggatt gtgttattta
167221 gaacagagta tcagtatatc gtatgtcaca gaacaacaca atcagctcaa tgaatcctga
167281 aagagcttac aacaatgtaa cgctgaaaaa tttaacagca tttcagttat tatctcaaag
167341 agaaaacata tgcgagttat tgaacttggt agaaagtacg gaaagacaca atagtattat
167401 caatcctgaa aggcaaagga tgagtttgga agaaatgaaa aaaatgctcg atgctttgaa
167461 aaatgaaagg aaaaagtagg aaaacgcaac attcacaatt gtaagctaca aaatattgaa
167521 tgtgaatcct gcttcattac aaatgaagaa tttcaaagcg tatctagttc accaaaagtc
167581 acaatatctt attattattt ggaagctttt atatgcatct aatggtcata aatcccatct
167641 cactttataa aacaagttac cgcgaatgta tccagtgata cagaatagag tgtaacaaat
167701 gagaataaaa ataccatatt atcgtttaat tttggtaggg tacgctaaaa taaagtggtg
167761 tttgcgtctc catataacta aaaagaattg aaatggccta ctatcgatta gactattcag
167821 gcatggaagc agtaagaaca tttcttgatt ttggaaaaat gagattagat tattaattat
167881 tgtgtttgtg tatataattt attttgcata tttcataaag aaattaaaat tagattatta
167941 aatattaagt ttcattagtg acattaaaag aagaaaactg atgttttgaa atgtgttaag
168001 caaagaatga ttaagacaat ctcaagctgc tctaagcagt atcttcgaca gcttgctctg
168061 cttcaggttc gggttcgtgg attgcagtct ctccagatgc tcttgggctc tctggaacag
168121 cttcttttgt ttcaggagcc gcatcttcca catcgtccgt agcatggtga tgagagaatc
168181 ctcctgtcaa agatagcctt ggcaatttgc catcacggct gactgcaatt agcaagtaac
168241 atggataaat ggcaaataca cataaatcca aaacaccata aaagatagcc tcaccgtctg
168301 gttgaatacg gttaccacca tcacttagac cccaacagat aaagtacaac caaacgatta
168361 ccatgcaggt acacagcata agtgcattga accctctagt tttcaaattg aagaattgac
168421 gttggcatat cacaccttgg gtaaccaaca tagcgacagc accaatggtg taataacccc
168481 acttgtaggt ggacttgatc aatgaaccaa ctaatagcga aacaacccag aataaggtac
168541 ccacgatttg aattaacagc gaatgtacca tatccaaggc ggaaatgtcg tcattctcac
168601 cagtagtact agcggctaac tcgattaaaa ataaaaggca tggccaggac aagaaccagg

168661 caatatattt cgagtaaaag atttgtctaa taccgggaac ttcacctgtg atagacttgc
168721 taaccttgac gtggttaaat tcagcttgaa caccagtcca acctaaatca gaagcataag
168781 tgaagaaagc aaaaaattca aagaaagtga tcaaaaatgc aggagctaag gcatatctag
168841 tcaatctgga gcccttgttc tccgcaatga agaacatcac aacatagcat agcaatataa
168901 agccaaagac tgcaaacact gcccataacc aatccgaacc tctcttggta atgtgcatat
168961 ccaggccatg tggtggatta agccctaaag cctcgttacg atttaaaaag cttgatagcg
169021 tatcgttcat ttgaaatttg ttgtttttag taatcaaaag atattaaagt ctcaaacttg
169081 ttgttttgtt tttgtaatta taaataataa taattgagtt aatatttgtt gttgtgttgt
169141 attgcagaat gtaaaaacaa gaaacaagat tagaaagtga ataggtgtta cgctaaataa
169201 ttgaagactc tcacagactt atatatgatt tttttagtgt gatacaactt gtaatgaggt
169261 aagggattga tattatactt tctaactgat gatgaatatg aaaactttgg ccaattaggg
169321 agaggaaagt tttcgatgag atgctagaaa aaaacttgtc cttgccatag cagaaaggga
169381 tgatgaaata acgaagatga gctaatttgg tattcgaata aacatgcagg atgtggcgga
169441 agggatcgga ataagagtcg ggatcgggat cgggatatgg ctttgcttac tttagccata
169501 ccaaatattc ttacaaagca cgctttcgat gcgtttttgc cttctgagag aatcctccta
169561 cggctcgacc cttgaaaaat tctagaggat ctggaaaatg gccaacaaaa tcgcgcaggg
169621 agacgcaagg aaagtaagtc agatggatcc gactgtatgt aacaagggtt gctgcagcgg
169681 ttttccatcc ctttcggcac cctgcacaag tagttattct tacttatgca aaacggatat
169741 gagctacata gcgttcattt tttaggtttt cagaagctat tacaacgcag taaaaaggat
169801 acacaacttt cttacgggga agaagaatac ggctcaatgc acccacttca acccccggta
169861 tcacgcggca ctaaagttgt ttgattgtaa tagtaatact gctgttgagg ttttcttctt
169921 cagggctgct cacaacgtga tatatttgaa ttgacaggac gtattcagcg cagttgagga
169981 aaattaacat atgacatgcg caggaaataa gtgctcaagt tttctacttt tgaagaaagg
170041 actttggttt acttgcctac tatctccatg tcaagtactc ggttgtatga tgttaaggag
170101 cgcagggaac actaatcgat aaaaaaccct tattggctat tactaaatgt aacggccgtg
170161 ctctagtgaa atttgtatag aaaaatgggt tctctcgctc aaaatgggcg ccattacttg
170221 tctggtcaaa ttgagttcct ggatataatt acataaacgg attgaaattt acataaggag
170281 tataataatt ccggaaacta tcttgagtgc aatgccatca cgcggccatg ccttgacata
170341 acatgctctc atcaaaaagg tgcaaatttt cgccagttat ggttgatatc taaaatctac
170401 tggagttcat tttttttttt attagatatc ctttttgcaa gccttctgaa aaaaaatttc
170461 tcatgtggta cagcgttggt tacttaagct gcttaaacta gatcgatgag ccattcttaa
170521 atcagaagac tgtgactgcg aattgtaggc atcttcacta ccctgtccat gaagctcatt
170581 ttcgttatca tcaattggtg ctatcttata actttgataa agagctacca tacaaaaaag
170641 acttaacgac caccaactga cccaggcgac gtcattttgc tgggaccaag acataatata
170701 gccccatact aagggaccta tgaaacgagc agaggcagaa atactaatgg tggcgccatt
170761 gatgactgat ctacaactca agggactcga attatgaatt aataacataa tttgtggtga
170821 cgttaaagcg ccacaaaatg tttttatccc tgtgattatg tacaagtaga cgacagtata
170881 ccagctagga atccgttcat tctgtagaaa aacaacgtaa ggaaccataa cgtacataat
170941 agggtacagc ttgactaaag ttctgaaaat tgttaaacaa tcgaaatttc gatcaactat

171001 gggaaaaatg aaaataacca caaaacaacc aaagatacct gttgtcgaca aaagagtacc
171061 ggtttgttct ggttcataac ctataccgcc agatattttc cacggaaatt ttgaagccag
171121 cttctttgga ttttctggat ctacggctaa atcataagct aaaaaaacag gcaaaaattc
171181 gttgtataca atcaaatgta aagccataat aaaattcacc gatatagggt aaaatacttt
171241 tgtatgaaaa acgtgatgaa atatactact ttctttacaa ccgtctggag ctgtatctat
171301 attggcatgc acagcgtctg ttggttcatg cagagaatat gtcctaatca ggcctacaga
171361 ctgtcttctt gttaaaatag gatcaatcga ttgtatgctt tcagtatcat cgtcattggt
171421 agtattcact aggggactat tttcatcctc ttcactatct tgtcctcgaa tgctgttcac
171481 attctcatta cggtggtgaa tgttttccga atcatcctga atgcgctttt gccagggtct
171541 tcttttcggc tgtataccaa atatattctt cttaataaaa tcaccgacct ctaaaccgta
171601 atctcttcta tctttaaaag caggatgtgt ttcttccaaa aacaatgttg cattagttaa
171661 accaaacatc aaaaacatgc acactaccac gtttggcaag gcgtacggat atgacctaat
171721 tattcttttt gcaaaatgtg gaaaccatag tggcacttca ttcattgttc catctctaaa
171781 tacaagaaac ccaccgatca taggcccaac aacggcacca aattgaaata ataaaggcat
171841 agtactgaaa gctaaagcct gatgttttct ttcagttgct atttcaccaa taatggttct
171901 aataacgccg acgttaccat ttagcaatcc cattaaactt cttgccacca aagcctgata
171961 gaaattgcgt gaaaatccca gtattaacaa tgatacagat gttcctataa gcccgcaagt
172021 caatgttatt tttctgccat gcttttcaga gaatctaccc cagtggtacg cagatatgac
172081 ttggcataac gcaaatgatg aagataaata acctgaatat ttggacactt gagcatcatt
172141 gggagcaata ttaaaatctc taaccatgaa ataaacataa ggaaatagtg acgaaaacgc
172201 aattggttcg ctgaacctaa ctaaggacac aacaacaagt tggacccagg ggaaaccatc
172261 catttgttct ttgaaagtaa gcttttgacg cgccatatct tttatattcg tgtgttcata
172321 tgcttaccac ttcctagttc ctttgctagc attcctttag gtgatctaaa ttcaagctct
172381 gtggataagg accattatag tattttttac taatgataga tactttttgt tatattttgt
172441 tttggcattt tcgtatccga gatttccaat gagcggaatg tgtcattctt tgtcaattaa
172501 cataggtcag gccgaatgcc aaatggagtt tagtaccttc atattgaaga aaaatgctac
172561 attattatat ttacaaggat ggtcttaaat aaagcttact tttaaggggg gaagaatggt
172621 gaacagtgag taaatacata gaaggagata agaaacaaat ggaatatcaa aaatcgatgg
172681 attctgcaga tgttctataa aagggtatgg catcctttat attatggttg ccatataagt
172741 atgagataaa tctctcaaac cctagaccaa agcctccgtg tggtgcactt ccttcttttc
172801 tcagagaaac ataccagtcc aattctccag atctattcat cccgcgtgct ttcatttctc
172861 tacataactt gtcatagtca tcttccctta aactcccacc aattatttca cccattccag
172921 gaaccagtag atcaaagcat ccaacagtat cgtcaggagt ggaattttgt ttcatgtaga
172981 atggtttaca aagacgtgga tagtcggtaa cgaaaactgg ggacttaaaa tactctccgg
173041 ctaaaaattt ttcatgttca gtttgcaaag gctgtcccca tttaggttca tacttaaagt
173101 gtgaaacttc attgtggcgt ttcttgagaa tttctattgc attggtatac gttatattgt
173161 gccatttttc atttattaaa tcttcccatc gtgttttaat ttgttgtgtc tcttgattta
173221 ttgatagctc tgacgatgca ttattttctt gtgaagagat aaattgcttc ggcaagagtt
173281 cttgttggtt atctatacaa gctttaatta tgtgttttat tgtagtctca acaaacgatg

173341 ttagctcgtt gacgctatta acaaagcaca tttccacttc aagcatccaa aactccgaaa
173401 ggtgtcttgg agtatcactc ttttcggctc taaagcaagg agataacgtc caacaccttg
173461 acagtgataa cgctaaaatt tccaagtgca attgagtgga cacagtcaaa taagtcggct
173521 tcccaaagta cgaggatgca gttggcgacg tattggtgga gacttgaaac aactcgccgg
173581 caccttcaca atcgtttgaa gttaatattg gtggtgaaac tttggtgaag tggtttttct
173641 ggaaatatag catgaactgg aattctacaa aggatcttaa ccgtaaaatt gcacttaagt
173701 aagcggttct gtattttagt gtaggtaagg accttaaata acgtaaggtt tgatatttct
173761 tttgtaatgg atagttttct gaaacgggac ccacaagttt aattgacttg acaggatttt
173821 tgatttgcaa ttcaaaaggt tgttttctat taggggtgct ttgccaggta gcattagata
173881 tagataaagt ttgaccagtt ttcagaattt ttaggaactg tacttcatca gtatttgtga
173941 gtggaataac tattcttaat gggttcacag aagtcccatc ttgtaaatcc aaaaacgcta
174001 tacgttttaa tagtcttatg gatttgatcc atccattaat tgaaatggga tcatggctag
174061 tatggtgtac ctgctcgtac aatgatttaa ctgttagtga agagtaaaat ctaccacctt
174121 taaggaaggt gaaagcatga aacattagct gtttgtattc gcttgttctc tttctttctc
174181 tcttagtaag cgataagtcg agctctactt aaattgtggt acgttttctt tttttcctat
174241 aatataaaaa agataaaaaa tcaggacgag aaaaagtaga tcacaactat aagttgtgca
174301 tactcatact ccacactcca cggcctaatc tctgtaagga ttgtatttta ttttttattt
174361 atttattatt tagattttct aaaaataaaa aaaaatcata cactattttg attacactac
174421 tatgtaattt atctttgaaa acaaccatga aaaactcaat tttacacaca catgagttat
174481 catgaggagt catcagccag acttaacaat cacacacatt gttaagagtg gaaaaaaata
174541 atagtgataa aaggaaaaaa gcgaacaatt tggattttac gggatctgtc taaatacgaa
174601 acacacactc gttttaattc aaatatcata accataaaaa aaagcacaat tcaacttgag
174661 cacgcacact aagtagtaac aacgttcatt tacagtaaag cgaacggacg aaacaaataa
174721 aagaaaggca tagtgaggga taaaaggtag aaagaattca aaaaagggta tcgcacacac
174781 acacatagta gtctaattca tgttctagaa tggatataaa caaagatgaa ggggacagga
174841 aataaaatgg ctcgctccgc cctttttaga atctttgcaa tcctctttgt ctagcttgcc
174901 attttctgta gataatggta gcaataatgg caatacaagc caaaccgacc aaaatgaaaa
174961 ctaaaataac accacctggc aaagtcattt atgatagatt aaaagtgctg aaatgggaag
175021 aggagaaaca aacaaaggat atgtaggaac aaaagtttgc ttgagtgact agtttgaagt
175081 tgagaaaaaa aaaagagatg aaaaagttaa cttattttgc tcttccttca aagtgtgtag
175141 cgtcttcctt taaatactat tgaaagggaa aaagaataaa tggaaaaagg gtagtaatga
175201 aaaactctaa aggaagataa gacgcaataa gagaaaaaaa gggaagttaa gattacaaga
175261 gaaacaataa attgctttca aagctttata tactggcaag cacggaaggc taaatttgaa
175321 aaaaaaaaat gggaaattat tatcggaaaa aattgaagct atcttctatt acgcagtgac
175381 aagtctatat tgacatcttt ctgctttctg ctttctgttt gttgtttttt ttctcatagt
175441 tgttttgatt ttactctgtt ggaaatgcga aaattgtttt gattttaccc gattgcgacc
175501 tattcccggc tttcccaaaa aaggaattcc tgtccaatgc aacgacagcc atagaaacag
175561 ccgcggttgg gtgcgtgggt gcggtaattt aacgatcata tggtatgggt gcacgtggga
175621 ttctagggga agtgcgcgcg gataccgaat cggccatcga accggccacc cggattgagt

175681 ttggtatgtt gcctctttgc gcaattttat tgaatgcaaa gtttctcctt ctagtaaagg
175741 acaacagctg aaaaatgctc gatgtccaaa tgacgacata aacctcctcg tttaggaaag
175801 agttaacacg cctagtttca gcttgtttct ttttattgtg taattagttt tcttacttag
175861 tacatcttat tgaccccact gtcatcggat gtgagagagt ggttcacttc tgaattaagt
175921 ttagagatat actaaatcta aataaaagtc tctttctgcg tgcatatgtt ttcttgcgcg
175981 ggactcacaa ggatgcgtct tttatttcat aacgcggttg tcccacctta agcttaggta
176041 tcagaatcga ctaaatcgtc gatgctgtca attatgtcgt caatgagatc ttttgcgtcg
176101 ttgattatgt cttgaagcca atcccccact gtgctagcag tagcgctagc ttgtatgctg
176161 ctgctgctgc tgcttcttga tgatggtgat gtttcgccca gtagagtctc agttgctgat
176221 ggtattgaag ttgtcatttc tagttttatc gttaatgacg aagatggcat agcggctatt
176281 tcatcagtgt ttccatcgtc gttgtcatca ttatcattgt caacattgtc ttgttctgtc
176341 tcttcggggt acccgcccca aattatatta taagtgctga attcaccaaa cttactgacc
176401 aaataatcaa agtcatcttc aatctctact tcttcgctgc tctggggttc gcggagttgg
176461 ttcataagca tcccagtccc atcattggaa tttctgtcct tttctgccac accgcaaata
176521 tcacacaaaa ggttgtaaat ttcggtattt tggaacggct caatgtatcc ctgtggaaaa
176581 tatggcccca taccaataaa tacagatctc atatcgatcg cgttattgtc gtatccatgt
176641 gatccaatcg tgaacacatt gtcttcgttt ttgtccttat gatcaccttt tgccaccttc
176701 ttcaattgtt ctttcttcat cactgcatac ccaggctcgg gcacaatcca gattgacgcc
176761 atgtgatgat tttttccatc attaaagttc cattctttgg ggaaatttcc attaacgtaa
176821 accgtatact tgtcttcatc tatagaagtc tttaaattgt ggtaaacctc attgatgttt
176881 ccggaatctt tcaacgatat agccatcatc ggaccctcta gatatgcgtg cgatacataa
176941 tccttcctca atttttcgtc cagtaagtct tcccatataa taacatttga gggaactacg
177001 atatcgctca taccatgatc gctaacaatg accaaattag taaagctggt taaatttctt
177061 tcttgcagcg attccactag ttgcttcaga aatgtatcta cttcccccag tgtttcagtg
177121 aagtcttcat aatagtattc cgactctgac ggatatccat gcttatgtcc aaaggcatct
177181 acgttcggta cataaccgag aattagctgt ggtctttcgt tcagtgtact catatccaca
177241 tattcaataa tcttggataa tttttgcgaa agtggttctt tagcattgaa ttcgtcgaag
177301 taaaatggag ttctctctct agcaatagga tttttatgtt caggttgtag tttctcttca
177361 ttatacttgg tataattcac atcgctacct ggccacatat gggtagcagc tttgaatggt
177421 atatcaccgt caaatgcaga ctgaacagtt tgccatattg gttctgtatc attattgttc
177481 catattcgag ggtccaatac acctggatgg aattcctcat taagatcagg atcccaaaat
177541 acgttagaga ctataccgtg gtgtattggg tattgtccag taaccaacgt ccaatggttg
177601 ggaaaggtct ccgtagggaa gctgggtacc ataaaaggtg tggacgtgat attcatacct
177661 ccatcatatt tcaattcata taagtcatgt aaaaacggtg tgttcctctt agatatgagt
177721 gaaggatgga aaccgtctag tgaaattact atcgttaagg ggaaaaattc atgcgtaccg
177781 ttaaaatact tcacgtaagg gtcaaattca ggactgccaa agtgcgatag cactctcgtg
177841 gagtgatggg gttttgctat gttaatcatc agaattaaaa aaatggcaaa caccagcgag
177901 ccaaacaatg tgcgaaatat tattttaaaa ggcactttgg aatgaaatcc cgttccaaaa
177961 gaaagttcat tctcatcaac aagttcagta tcaatgggtt gaccattcgc gtctaattca

178021 tatagcggta taccccttct aaagttagaa ccatccctac tatctttcgc gtcgccaatc
178081 actatgggat tgtttgtcca cttcagtcta ttcttataaa accaatattt catcctggtc
178141 cacgcagatc gccaccccga tcttacagca tcttcatccg ttatgaaatc atcacgaaat
178201 ggatcttcac taaaatcatt cagctcgtta tcgagcgact ctaaatcatt ctgaagttcc
178261 atctatagag agtgttatac tgtacctctt ttcagatata tgtaattacc tcgcctcact
178321 tattttgctt atttgtggtt tgttaggcag tttgcatatc gtaccaatgt ttttttataa
178381 cttcatgtct tttctttttt ttcattttta gcatcgtacg cagatatctt gtctcgttat
178441 gatattttgt gagataggta tccaaaatta ataagagaaa atgtcgtatt gtgttcacaa
178501 ttcacctgat tgtttgacta gtccaacccc tggtataatt gtcacttaat atattcttag
178561 cttccaggtg atgttttatt ccattattta tgttatgtat atataatcgt aactcttaga
178621 gcaaaaagta aaaaaaaagg tggtacttac cgagggagaa tatatacaaa tttattcgta
178681 aatgtaaata attaattgaa gcagagctaa aagaaaaata caagaaaatt tatttacact
178741 cgctgtttgg ttctgatctt ttggtattca caattttagt gctgtccaaa aatattgcga
178801 tgtgctgata aagtacatag tgtgatccga cttcgtccca gtcaaagaaa ggatgaaata
178861 ttagttagta catttactgg ttgctactgg atgtcctatc tgaatcttcc atcttaggta
178921 caaaaagatt aagaaagtca attgtcctta gttttaaaag acagttactt catgtcttta
178981 acatcaacgg caaattactt tgaaaaaatg cactaaaagt taacacatgt aggagcattc
179041 taaacatgaa atttttaatg gtacttaggt ttgctaacgc atcttgaaaa ctagcaaaaa
179101 ccccataatt aaaatattct tgcttgttaa acgaggcgtc caagccattg aaaggtttta
179161 tcaatattca acctcttact ataaatgcag cacaccgtaa caggtatgag aaagatgtgg
179221 tattattata caaattgtac tgaccgcttt cctactaata aaaccaatga tatcctaggg
179281 acgctcgagt tttcacatta tagaacattt attattattt tcggcatcaa gccccagagt
179341 tccttcgaca cgttccattt gtttcaaaag gagcataaaa gtttcttcaa cattataatc
179401 taactcggca ctgcattcta taaatgctgc ctggttcctc ttatcttgac tgccaattgt
179461 cgaagctagt ttctctcctt cagctttcgt gacacacctt tttacacctt ttgtactcct
179521 tcccaaatca gctttggtac ccacaagtat taccgggagg ttatccttac ccagctgatc
179581 taccagcttg tcccagagaa tgggaataag atcaaagcta gcacgattta ttatactata
179641 gcacagtatt atgcctcgta cgcccgtcaa cgatttaatg tttaatagag aaacttcatc
179701 ttggcctgca gtatctagaa tttccagagt acagtcatga cttttataag gaattatcct
179761 ggtaaattca ttttcaatag tgggataata ggattcaacg aaccgcgatt ctacgaagcg
179821 aaccgttaat gtggttttgc cgacatttct agctcctata agagcaatct ttctctgaaa
179881 gttatgtgtg gagttcgaag aagacatagt ggcgtattcc attatgtcac aggttatatc
179941 ttttctcgcc gtgccccaat aaaactgttt cggtaaagag acaattcgaa ccagcccaaa
180001 agaatgaata tgaaaatatc aatatgcgtc aatgaaaatg gaggcaagtt caataatttc
180061 actggaaata taactagtaa taattcgtgt aactgctgta gaaattgtta gatattaaac
180121 ctggcagaag cgtcttgtta atacttatag agaaaccacc agtagcgttc ttctgtatct
180181 gaagatagtg aaaaatcaaa aaaataaatc tgtattcaaa aaaaggtcct acccggattc
180241 gaaccggggt tgtccggatc aaaaccgaaa gtgataacca ctacactata ggaccaggac
180301 ttcttgtaat attttgtatc tcaaaatgag gtatgtcagt atgacaatac gtcatcctaa

180361 acgttcataa aacacatatg aaacaacctt ataacaaaac gaacaacatg agataaaacc
180421 cggccttccc tagctgaact acccaaaagt ataaatgcct gaacaattag tttagatccg
180481 agattccgcg cttccaccac ttagtatgat tcatatttta tatgagaaat gggtgaatgt
180541 tgagataatt gttgggattc cattgttgat aaaggctata atattaggta tacagaatat
180601 actagaagtt ctcctcgagg atttaggaat ccataaaagg gaatctgcaa ttctacacaa
180661 ttctataaat attattatca tcgttttata tgttaatatt cattgatcct attacattat
180721 caatccttgc gtttcagctt ccactaattt agatgactat ttctcatcat ttgcgtcatc
180781 ttctaacacc gtatatgata atatactagt aacgtaaata ctagttagta gatgatagtt
180841 gatttttatt ccaacatacc acccataatg taatagatct aatgaatcca tttgtttgtt
180901 aatagtttaa atgtttttat cggaagaggt tttgtcatca catcagcaat gttcttcttg
180961 gtctcgatgt agtatacgta taaattatta cctgatactt catctctaag tctcattgcc
181021 tttgtgccaa aaaatctgtt tctaaatttc tcttcatttg tagacttaat tatactgatc
181081 gttgatctac tatcagtaag taagccttta ataattggtt tcttgttaag ttcttgcaca
181141 aggtgactga ggttattcaa tagcggtata gcttcactga ctgcgtgtat ttctgcttct
181201 gtagttgaag tgcatgttaa cgaagccttt gtcgactttc ctccaatcac ttttccgttg
181261 agtaggaaaa tgttaccaat ttgtgacttg taatatggtt ggttaccata tgaagcatcg
181321 cttattgcga ctagtttatt atctggcttg gtaggtttgt ttttgtgcca tattaattgt
181381 ttatctctag tgtcccacat gaattgtatt aactcatatg tcatgtctaa aacttgccta
181441 gaggggaata gtatatgttg agcaagtgtg ttgatgtagt atagtaagtc aaatctaaat
181501 ttatatccaa catatgaagc tagaccaatc aacttttgca tttcatgtac cttctctttg
181561 tattcatctt catctatttc tagttcatcc tggtctatat aaagacctgg ttgacctgga
181621 gcgctaagtt ttcttccttt tggattcaaa ggtacgttta atttgggtat tttctcagtt
181681 aatgagtttt ccatacctaa tttcatgtat ttacctcttt gatatttgat ttctaagcca
181741 agtatgtcat attgaatttc ctcatcactt tcgcctagat ttataatctt ggtgtcgtat
181801 tgcatcttaa gcttctctat aattcttttg tttgaattta gatttttgct aaacaatacc
181861 atatcatcta cgaataaaca aattgtcact tgactgtttt taaatacgca tgaccatcca
181921 cgaacttctt ccataccaca ttgttgtatc aggtatgatt tgatagtttc gtaccagttc
181981 gctccacttt gtttcaatcc ataaagtgat ttcttcaaac gtatcaactt atcattcatt
182041 cctaaatgtg gtggaggtct tatgtataat tcttctttga tgtctgcata caaatatgcc
182101 gaagatatgt ctaattgtgt aatatagtag ttattgtcta atgcaagtga cagggatgtc
182161 attaatgcat agtgatgtac ggtattggat tgcatgcctg agtcgtaagt gtcaggatgc
182221 tgaatatcac ctcttgcaac aaatctagct ttatgagtac cgtcacgttt cttgttgaag
182281 ataaacattg agtttattac tcttttaggg tctatttctt ttctgtcata atattcgtca
182341 gtgtcccaag ttttcatctt caacagttga ttgacttctt tgtggtatgc ctcgatatat
182401 ttttcttttt ctttaatatc tttattatag gtgattgcct catcgtatcg taaggttgtc
182461 cgtattggtt tgattgattt tactgctttt acagctgcaa tcaggtgaat tcgtttcttc
182521 gatctcggag gttctaaact acgcatattc ttagtattcc atgtgtctcg tgatacctta
182581 atttcagttt cattatcttc taatgatctt ttcttactgt tgatagtagt ataggcatta
182641 gagtcaccaa taccacccaa actggaatta gtttgacgag aatttatcgg tgggagttct

182701 ttaaatgggt cagggaattc ggtaggagat tctggaggta gatcagggag tgggagatca
182761 gcgatgatag attcctcggt attctgttca gaaacagtag ttggcgtttt gataggaaca
182821 atattgtgcg atgaattatt ttccggtgga gaagcatcga ttgaaggtga acggtgtata
182881 atccttttct cagtctcttg gtcacttatc tgcggaacag ttttgttgtt ggtaccaccc
182941 gtactggata ttggtacgtt tgtatgatta gtctcatttt cactgtacga gtctgagtgt
183001 ctgaaatctt tagatttact ggcgtgcgac gactcatgtg tgttagattg ggacatggga
183061 gcaagtaaag gaacatttaa tttatgcata ccacccgaac cggtactctc gatattggaa
183121 atttgggggg tgctagatct cttctttgat ggaagaatat tagattcaga tatgttgggg
183181 tcaacttctc tgggtgcacg aagattgatt tcagatatag gttttgaatc ttcagtatga
183241 gttgacggag gtgtggaatc ggttggactc acagcttttg aaaggacatt tctcggttgc
183301 tcaggatgta gttcaatgtc agattggaag tcatggtcag attctatgtt aagatcattg
183361 gattcttgga tctcatttga cgcaatgaac gaatgatatg aagcagttaa acggtttaag
183421 tcttcatcga aagtgagtgc gtcgtaattg aattgatcta atctggattc cttgccctga
183481 agaataacat agttagttgt atctactgtc ttctttaagg atggaagata gatgatatat
183541 ccataagagt ttcgagacgg atgtagagcg taacctggga tgccacgagg atgtattttg
183601 gagttagggt tgtgatcatt gacgataaca ggttgaccga aaggtaacaa agtactgata
183661 tcaagtcctg ccaagccagc atgttggctt gcagattttt tgcttttagg tgaagctagt
183721 gaatttctca caatagtaga aaattcgatt gcagagaacc ataaatggtt cggtaaacca
183781 ctacattgca gttgagtacg gcagtcatct aataaggtac ggttgagccg ttcagcgact
183841 ccatgtgctc gggaatccgc tgtggttgta tagcatggag ttataccatt tttttcaagg
183901 aatttatgga gagttctgtt agtatactca gaaccacggt ccatttgtat aaccaagaca
183961 ctggcctgaa attggttttt aatgaaagct agtatcgtgg taaatacatc gaggatagag
184021 tcctcgcgac ggtcgtgcaa tggataaacc caacggaatt ttgttgtctc atcagtaaat
184081 gagatgaaat aggatggtgc actttttggt aggttgtgaa ctggaccaaa tatgtcagta
184141 tgtaggtatt gaaagggttc gtatgaattt tggtatttta gtcgtgaacc tttgatatgt
184201 ctgtgtttgg tgcttttgcc gattaaacaa tcaggacatt gatagtcaat agcactagac
184261 cagtcgacat ctgattcgtt aaaatacgtg atggtgttat ttttaagtga gtatcgaatt
184321 gtctgtgcat tggcatgcgc aagcattcga tgaatgaaag gataaggata tttgcgtgta
184381 ctttcacttg tatggacatt attgatggtg ggtacggaga tatttgatgg aagcaagtac
184441 cttttagata cccagtaaaa gtctccatat tgtacgatag gtgcaagtac agtgccgtca
184501 gatcgttcta agacgttttt ggtaaagcat gctgtgatat ctactgcagc caattcattc
184561 aaactgagta agtcataggc tatgttagga gtgtgcaata cctttattga tgttttggtg
184621 ttgtcctgga agtgaaattg taggtcacca atagcgttaa ttggtatatt tcttttttga
184681 gcatcaacta cgtttatgtc aggattagat gatgctgagt gtatgtgatg agcagatctt
184741 ataagggttc gtgatgctcc tgaatcgaga aggaggtgtc cagggagttc atcatcagaa
184801 tgattagtgt gatttaccgt agattcagta agttcctggc ctaaggtgaa ggtcgtgctt
184861 attgttcaat tgaatcggtt cagtagttga tttactgatg gaatcgttgt ccgtgctggg
184921 agagttatta gatgtggata cattgtgagc cctggctgtt ttcgatttcg aattatttgt
184981 tttttgagga ttccgagcta taactttggg tttggttgta ttcgtatagc tgcgagaatc

185041 attcttctca tcactcagat ttctcctgta attaggtttg ctgtttctcg atccctgttg
185101 ttcttcataa atagcatgga tatctaagaa cagttcagcg actgtcatat ttagatgtcg
185161 atgacgtgtg tagcgtaaaa atttatattc gccagataga cctctcataa ttaattggca
185221 tgcgaccttg ttattgatat gaatgccatt attgttcagt ctgtcgataa tgtttgtgac
185281 ttttgtttca aatgcatctg caggtgtact gccattatat tgcaaatttg ccagggtcac
185341 aatgtcgttt gcctcttggg tatcagattg cattttttca atacttttgg aaagaatttt
185401 catgatatcc gtataatcaa cggataggat gtctttgacc caggtaggta ggaattgaga
185461 gggagcaaat atttgaaaag tgttatacaa gaaggtgagt tcatcatcag tgatctgacg
185521 tacgggtttt ccgtttactg tcggaataat accaccgaga ttcgagtttt gtaaaaattt
185581 gatgtatgtt ttaacccaat ttggaaagtc attaggtgag gttaacattg gtggtggtct
185641 gacatatttt ttagtggatg tcatatcaga gtccgctgag gatgaatcag taaatgtatt
185701 acctgactca ggtgatggag tgctcagagg cgttccaact gatgatggat actgcggaaa
185761 ctgtgattgt ggcccaggtg gaaagtacat aggcgacatt tgataaggtg tatacggaat
185821 catagatggg tgtccgtaaa atgaccaacc agatggattt gcttggtttt gggtcatcat
185881 gcactgctgt gggtacggcc cattctgtgg aggtggtact gaagcaggtt gaggagaggc
185941 atgatggggg ttctctggaa cagctgatga agcaggtgtt gttgtctgtt gagagttagc
186001 cttagtggaa gccttatcat attcttgaat tttggaagct gaaacgtcta acggatcttg
186061 atttgtgtgg acttccttag aagtaaccga agcacaggcg ctaccatgag aaatttgtgg
186121 gtaattagat aattgttggg attccattgt tgataaaggc tataatatta ggtatacaga
186181 atatactaga agttctcctc gaggatttag gaatccataa aagggaatct gcaattctac
186241 acaattctat aaatattatt atcatcgttt tatatgttaa tattcattga tcctattaca
186301 ttatcaatcc ttgcgtttca gcttccacta atttagatga ctatttctca tcatttgcgt
186361 catcttctaa caccgtatat gataatatac tagtaacgta aatactagtt agtagatgat
186421 agttgatttt tattccaaca ttatataata tataagataa gtaacattcc gtgaattaat
186481 ctgataaact gttttgacaa ctggttactt ccctaagact gtttatatta ggattgtcaa
186541 gacactccgg tattactcga gcccgtaata caacataata tctacattct agataccgac
186601 atatcaatat aaaaatactg aatattgaga catcttttag attacgtttt ggttgccgca
186661 aaccaaaaat gcaatttaaa tgggtgaata atttgaatgg ttagaaatct atttgttgga
186721 ataaaaatcc actatcgtct atcaactaat agttatatta tcaatatatt atcatatacg
186781 gtgttaagat gatgacataa gttatgagaa gctgtcatcg atgttagagg aagctgaaac
186841 gcaaggattg ataatgtaat aggatcaatg aatataaaca tataaaacgg aatgaggaat
186901 aatcgtaata ttagtatgta gaaatataga ttccattttg aggattccta tatcctcgag
186961 gagaacttct agtatattct gtatacctaa tattatagcc tttatcaaca atggaatccc
187021 aacaattatc taattaccca cacatttctc atggtagcgc ctgtgcttcg gttacttcta
187081 aggaagtcca cacaaatcaa gatccgttag acgtttcagc ttccaaaaca gaagaatgtg
187141 agaaggcttc cactaaggct aactctcaac agacaacaac acctgcttca tcagctgttc
187201 cagagaaccc ccatcatgcc tctcctcaaa ctgctcagtc acattcacca cagaatgggc
187261 cgtacccaca gcagtgcatg atgacccaaa accaagccaa tccatctggt tggtcatttt
187321 acggacaccc atctatgatt ccgtatacac cttatcaaat gtcgcctatg tactttccac

187381 ctgggccaca atcacagttt ccgcagtatc catcatcagt tggaacgcct ctgagcactc
187441 catcacctga gtcaggtaat acatttactg attcatcctc agcagactct gatatgacat
187501 ccactaaaaa atatgtcaga ccaccaccaa tgttaacctc acctaatgac tttccaaatt
187561 gggttaaaac atacatcaaa tttttacaaa actcgaatct cggtggtatt attccgacag
187621 taaacggaaa acccgtacgt cagatcactg atgatgaact caccttcttg tataacactt
187681 ttcaaatatt tgctccctct caattcctac ctacctgggt caaagacatc ctatccgttg
187741 attatacgga tatcatgaaa attctttcca aaagtattga aaaaatgcaa tctgataccc
187801 aagaggcaaa cgacattgtg accctggcaa atttgcaata taatggcagt acacctgcag
187861 atgcatttga aacaaaagtc acaaacatta tcgacagact gaacaataat ggcattcata
187921 tcaataacaa ggtcgcatgc caattaatta tgagaggtct atctggcgaa tataaatttt
187981 tacgctacac acgtcatcga catctaaata tgacagtcgc tgaactgttc ttagatatcc
188041 atgctattta tgaagaacaa cagggatcga gaaacagcaa acctaattac aggagaaatc
188101 cgagtgatga gaagaatgat tctcgcagct atacgaatac aaccaaaccc aaagttatag
188161 ctcggaatcc tcaaaaaaca aataattcga aatcgaaaac agccagggct cacaatgtat
188221 ccacatctaa taactctccc agcacggaca acgattccat cagtaaatca actactgaac
188281 cgattcaatt gaacaataag cacgaccttc atcttaggcc agaaacttac tgaatctaca
188341 gtaaatcata ctaatcattc tgatgatgaa ctccctggac acctccttct cgattcagga
188401 gcatcacgaa cccttataag atctgctcat cacatacact cagcatcatc taatcctgac
188461 ataaacgtag ttgatgctca aaaaagaaat ataccaatta acgctattgg tgacctacaa
188521 tttcacttcc aggacaacac caaaacatca ataaaggtat tgcacactcc taacatagcc
188581 tatgacttac tcagtttgaa tgaattggct gcagtagata tcacagcatg ctttaccaaa
188641 aacgtcttag aacggtctga cggcactgta cttgcaccta tcgtaaaata tggagacttt
188701 tactgggtat ctaaaaagta cttgcttcca tcaaatatct ccgtacccac catcaataat
188761 gtccatacaa gtgaaagtac acgcaaatat ccttatcctt tcattcatcg aatgcttgcg
188821 catgccaatg cacagacaat tcgatactca cttaaaaata acaccatcac gtattttaac
188881 gaatcagatg tcgactggtc tagtgctatt gactatcaat gtcctgattg tttaatcggc
188941 aaaagcacca aacacagaca tatcaaaggt tcacgactaa aataccaaaa ttcatacgaa
189001 ccctttcaat acctacatac tgacatattt ggtccagttc acaacctacc aaaaagtgca
189061 ccatcctatt tcatctcatt tactgatgag acaacaaaat tccgttgggt ttatccatta
189121 cacgaccgtc gcgaggactc tatcctcgat gtttttacta cgatactagc ttttattaaa
189181 aaccagtttc aggccagtgt cttggttata caaatggacc gtggttctga gtatactaac
189241 agaactctcc ataaattcct tgaaaaaaat ggtataactc catgctatac aaccacagcg
189301 gattcccgag cacatggagt cgctgaacgg ctaaaccgta ccttattaga tgactgccgt
189361 actcaactgc aatgtagtgg tttaccgaac catttatggt tctctgcaat cgaattttct
189421 actattgtga gaaattcact agcttcacct aaaagcaaaa aatctgcaag acaacatgct
189481 ggcttggcag gacttgatat cagtactttg ttacctttcg gtcaacctgt tatcgtcaat
189541 gatcacaacc ctaactccaa aatacatcct cgtggcatcc caggttacgc tctacatccg
189601 tctcgaaact cttatggata tatcatctat cttccatcct taaagaagac agtagataca
189661 actaactatg ttattcttca gggcaaggaa tccagattag atcaattcaa ttacgacgca

189721 ctcactttcg atgaagactt aaaccgttta actgcttcat atcaatcgtt cattgcgtca
189781 aatgagatcc aagaatccaa tgatcttaac atagaatctg accatgactt ccaatccgac
189841 attgaactac atcctgagca accgagaaat gtcctttcaa aagctgtgag tccaaccgat
189901 tccacacctc cgtcaactca tactgaagat tcgaaacgtg tttctaaaac caatattcgc
189961 gcacccagag aagttgaccc caacatatct gaatctaata ttcttccatc aaagaagaga
190021 tctagcaccc cccaaatttc caatatcgag agtaccggtt cgggtggtat gcataaatta
190081 aatgttcctt tacttgctcc catgtcccaa tctaacacac atgagtcgtc gcacgccagt
190141 aaatctaaag atttcagaca ctcagactcg tacagtgaaa atgagactaa tcatacaaac
190201 gtaccaatat ccagtacggg tggtaccaac aacaaaactg ttccgcagat aagtgaccaa
190261 gagactgaga aaaggattat acaccgttca ccttcaatcg atgcttctcc accggaaaat
190321 aattcatcgc acaatattgt tcctatcaaa acgccaacta ctgtttctga acagaatacc
190381 gaggaatcta tcatcgctga tctcccactc cctgatctac ctccagaatc tcctaccgaa
190441 ttccctgacc catttaaaga actcccaccg ataaattctc atcaaactaa ttccagtttg
190501 ggtggtattg gtgactctaa tgcctatact actatcaaca gtaagaaaag atcattagaa
190561 gataatgaaa ctgaaattaa ggtatcacga gacacatgga atactaagaa tatgcgtagt
190621 ttagaacctc cgagatcgaa gaaacgaatt cacctgattg cagctgtaaa agcagtaaaa
190681 tcaatcaaac caatacggac aaccttaaga tacgatgagg caatcaccta taataaagat
190741 attaaagaaa aggaaaaata tatcgaagca taccacaaag aagtcaacca actattgaaa
190801 atgaatactt gggacactga caaatattat gacagaaaag aaatagaccc taaaagagta
190861 ataaattcaa tgtttatctt caacaggaaa cgtgacggta ctcataaagc tagatttgtt
190921 gcaagaggtg atattcagca tcctgacact tacgactcag gcatgcaatc caataccgta
190981 catcactatg cattgatgac atccctatca ctttcattag acaataacta ctatattaca
191041 caattagaca tatcttcggc atatttgtat gcagacatca aagaagaatt atacataaga
191101 cctccaccac atttaggaat gaatgataag ttgatacgtt tgaagaaatc actttatgga
191161 ttgaaacaaa gtggagcgaa ctggtacgaa actatcaaat catacctgat aaaacagtgt
191221 ggtatggaag aagttcgtgg atggtcatgc gtatttaaga atagtcaagt aacaatatgt
191281 ttatttgttg acgacatgat tttatttagc aaagacttaa atgcaaataa gaaaatcata
191341 acaacactca agaaacaata cgatacaaag ataataaatc tgggtgaaag tgataacgaa
191401 attcagtacg acatacttgg cttagaaatc aaatatcaaa gaggtaaata catgaaatta
191461 ggtatggaaa aatctttgac agaaaaatta cccaaattaa acgtaccttt gaatccaaaa
191521 ggaagaaaac ttagcgctcc aggtcaacca ggtctttata tagaccagga tgaactagaa
191581 atagatgaag atgaatacaa agagaaggta catgaaatgc aaaagttgat tggtctagct
191641 tcatatgttg gatataaatt tagatttgac ttactatact acatcaacac acttgctcaa
191701 catatactat tcccctctag gcaagtttta gacatgacat atgagttaat acaattcatg
191761 tgggacacta gagataaaca actgatatgg cacaaaaaca aacctaccga gccagataat
191821 aaactagtcg caataagtga tgcttcatat ggtaaccaac catattacaa gtcacaaatt
191881 ggcaacatat atttacttaa tggaaaggta attggaggaa agtccaccaa ggcttcatta
191941 acatgtactt caactacgga agcagaaata cacgcgataa gtgaatctgt cccattatta
192001 aataacctca gtcaccttgt gcaagaactt aacaagaaac caattactaa aggattacta

192061 accgacagta aatctacaat cagtataatt atatccaata atgaagagaa atttagaaac
192121 agattttttg gtactaaagc aatgagacta agagatgaag tatcaggaaa tcatctgcac
192181 gtatgctata tcgaaaccaa aaagaatatt gcagacgtaa tgaccaaacc tcttccgata
192241 aaaacattca aactattaac aaacaaatgg attcattaga tctattacat tatgggtggt
192301 atgttggaat aaaaatccac tatcgtctat caactaatag ttatattatc aatatattat
192361 catatacggt gttaagatga tgacataagt tatgagaagc tgtcatcgat gttagaggaa
192421 gctgaaacgc aaggattgat aatgtaatag gatcaatgaa tataaacata taaaacggaa
192481 tgaggaataa tcgtaatatt agtatgtaga aatatagatt ccattttgag gattcctata
192541 tcctcgagga gaacttctag tatattctgt atacctaata ttatagcctt tatcaacaat
192601 ggaatcccaa caattatcta attacccaca catttctcac tattctcggt aaaggttata
192661 atattagtgt tggaataaaa ctctaaaata ccatccattc agaagtattc atgctactag
192721 cattttatca catgaggtgt aagaagatga cataaagatt gagatacagt catcaaacct
192781 aatggaagcg aaatgcaggg attgataatg taataagata atgaaacata agaaacagaa
192841 ggtaggagta ttattatgta gaaataccga ttccatttta aggattccta tgtcctcgag
192901 tagaacgtct agtatatacg aaatattata ccctttaaaa aacagtggac ccaacgataa
192961 tcacaacatt caccgatttc tcacctgaac agatcacgat catccttcag aacatcttgc
193021 tgtagccata ttaaattcct ttgtctctgg actctttctc gcgcgactga tctcccaatg
193081 aagaactgcg tcaggtcatc tcgtcctcaa tttctaacca aaattggatg attttagttc
193141 tcactgacgc tgcttgaacg ttaacaccga aagagcatgc aataatcaca gcacggtaaa
193201 ccaacttgtg tcatatttaa gagaaatatt aaaacgctaa gaagatagct cccataatag
193261 aagtacaagt tatatttatc gttgtccaac tgcagaagtt aaataaaggt aaataaatcg
193321 agctaagaga aaaaaaaagt ataggcaaaa ttccgaccgg agctagcttt atcttgctcc
193381 tccacctaag tcacctgctc aggtcactca agttcacaat cttctgagag gatagataaa
193441 ggaaaaattg gaacagatct tttcatggcc tggcaattgt ctgtggattg gtaaagctct
193501 gtcagtgaaa aaaaaagttc tcgctccttc ttggtctctt ctattgcctg agattctaat
193561 gctttcctta cattgtttca taaaataacc aatttttcgt gtcacaataa tcttgaaatt
193621 tgaagtttct aaaatgcatc agacatggtg tcgtaattag ttgacttact ttaacttcta
193681 ccaagaagta gaaaatagca gagtgcgtga acgcatagca gagaacaaaa tacttactaa
193741 accttttcga ataagttagc aagaagcttt taaaaacata gacttagaaa ttgtcctgca
193801 cttggcaagg aagaaaagaa agattagtta aacagaaatc gaaaaatgca tgatgaaaaa
193861 ctaagtagac ctctttagaa gtgtcttagc ggctattgtt catgatactt ttgaaaagag
193921 ccctgagatt ttccattacc atcaacagga ccaatgaata gctaaaagaa tcgaatgtat
193981 ctatcggaaa ggatttgaaa tttcggccat ttcctcatta tgaagttcaa cttccatact
194041 ctcgtcgtcg ccatcatttt cattatcctg gttgtcgtca tcgtcatctt ctccgggcat
194101 gtcattagca tctatataag gaacctgttt cttagacaaa ttctctttta tctgtagtga
194161 gcgtatgacg accgagacaa ttccgcttga tattgccgta gccaagaggg cgtagcaacc
194221 tctctcaaac ttgggcacca tatctgaagc aaagaataat atagaccacc atgcgttaac
194281 ggcccctgaa aacatattca tggaagcaag aacgatagca cgttcttgaa gatctgcatg
194341 acaaataatg tttgcccacg aaaaaaaaac cgcttgtcca gcgtatgcta cgcctccaag

194401 atactgtgca gaaaagacga cttttggatt tagtgggtct gcacgtatta gtaccgcaac
194461 aataaccatt accaatgata tgaaaacaga aacatgccaa tgcctagctc ttgggatctt
194521 actcatatat acagcagaac aaagcgtaga aactatacct acggcaaata tccccgaagg
194581 ataattattt ctttgcgcca acgtatattt ttggttttgt aaccataatg caaatgtaga
194641 attagatgcg aaacccaagt tctcacctcc cagaacccaa acaagagaga acatccacca
194701 gtgccacctt tttaagactc taggaatagt cgaccagtct aaccgggtgc tttcgtccct
194761 agcggggagc cttctcctag cataatgtag ctcttgttca ttaaaaatgt atctcgtcat
194821 agaaaatttg ctaacagcac ttgtttgatc aggaaggccg gggaagaaaa taaacccata
194881 aattgcaatg ggtaatgtga tacaaaaatc aataatgaat aaccatctcc aacccgccaa
194941 gccattccgc ccattcaaat gagtaaagat acttgtttgc ataaatccac tgaacataga
195001 tcccaccaaa ccgctacctg taaaaatagc acttctaatg ggcaattcgt cttctttata
195061 ccacgaaccc aaaacaaaat gtgttcctga aaatgtgcaa ctctcaaata aggcttgaaa
195121 gaatctaatt gcgcaaatat gtttgaacga tgtaactttg tacattccca aggtcaataa
195181 accccaagca aacgtacaaa aacttagcca tatcctgggc ggaacacaca ataacattaa
195241 attattgggg accataccta caatgtaacc aatcatgaaa actgtgttag acacggtcaa
195301 atcgtttccg accatcttaa gatcttcctt cataccggat atatatgcat tggtgaaacc
195361 gacacggtcg acataattaa tccagtattg caagcaaaca aatgatagca caaacaagtc
195421 tattttaaac agaagtaatc tctttttaat ggcacgcttg gaagaaacag attctctctc
195481 aacctcatgt tgagtgatag atttcgattc cttcatcatg acttaggccc aggctctctc
195541 caatcaaaaa ttctgaaaac agcttctcaa tcttgaataa atcggaatac ttttatatca
195601 gaattacagg tccgcctcat ttctcttaag agccgatgtc ttcctttaca atatacgcat
195661 gtagaagatt tgatcgatgc tgagttcttg taattcgtca aaaatgcctt tcaagatcga
195721 ctactcaaaa tctacaataa tcggactgca tggatcaatg gaggtgtagg caattttgta
195781 acaatgaagg atatgttaaa ttcgctaaag cttcctcctc ttttgctaaa tttcatcatc
195841 aaaaagcatt ttgcataaca cacgttaacc aatatattat agtatatata cattctattt
195901 gagaataagc atagttttac ataactagtt cgttattcca aatgtatttt tcctatttgc
195961 cataaatatt ttaagtaaaa gaaagcatgc tctagataaa aaatatcgag gaagagtcga
196021 aataagcaag cgtaaatatt acaattattc tgaagaagca gcattttctt ggccctcagc
196081 atcttctaat ttcttcccat tcttcaataa attaatgtcc ttttgaacta agctcaaagt
196141 agtaccctta gaaccgtcgt ctctctcgaa gacatagtta gcagcatctg cttcaacata
196201 taccaaagcg ctgaataaat tatggaaagg aaatgttagt aaattataac gcgtaattgt
196261 aaagtaagta aaattgaaga ataaataaca taccctttc taacatattc tgtcaaaaaa
196321 ttgatttgag gttcattgaa aacggtgata ttataccaat tggtttggcc atctcttctt
196381 ggttgcgaag cgatactata tttcaaataa cgattgttat tagcagaagt atgttcagtg
196441 aattcagagc caatgcggcc gacgatggac attttcgaga agtccatctt cttggtagta
196501 gcatggaaga aacgagcttg agtacgtaaa aacattacaa tggattagtg gatttttgtt
196561 tgttttgtat ttgtcaagga cacacaatta gcaaatgact tgttatcaaa ttcgtggtat
196621 ttgtctttct caactactgt gttagctttt tttttttcgt ctgggcgcaa acgcctttta
196681 agcactcctc cacaaataat aagtattaga tgaaaagtcc taaaagtccg aaaaatggaa

196741 tagaaaagaa cgaaaataca gaatctttac agaacattat agtacataag tgtagttcta
196801 cgtaagtctt aaacaaaggt ttcaaaagga aaatatgatt tagcggcgcc acctccataa
196861 attatggatt agcaaacaaa agaacaatta atattgaaaa atagaagcat agcagaaata
196921 tcaagcgaga ccatggtagt taccagtggt caaagttctt tttgtgttta catcgagcca
196981 ctcaccacca aatggattgt tttcatccag ctcctgacaa gtcaaggtga ttccactgcc
197041 ggctccgcta tggggtagag ccacgttatc cgtctcactg atagtgaatt tggtaatcac
197101 gcctttagcg gattcatgtg ccaggccgtc tgtaataaat ctggcgatta atctttgacc
197161 ttcaccattt cttgtcaaaa caactggttc tttaaatctc cacgttattc tcttcttttc
197221 tttactgaag gagccctgag gttttgataa agcgcttgtt gcattagcac catcgatatt
197281 cacaaagaca acgaggtctt ctattacgat ttgtgatatt tcatcgggta agcttggtga
197341 catcttgaca gtcaatacaa cacttgcctg gtgagattca aatctccaga ccggatgaat
197401 aacaataggc gcgatcggct cctttatgga atatttaata gctcctaagg ttctagaatc
197461 gataaatgaa gggttaactt tgaattcttc tggagcaaca cgctcaataa aggcttggtt
197521 cagaatgacc ttttcgaact tagcgccatt attgattctc aaattaatac cgataggtaa
197581 aggcgaattc atgacagaat tgggtaaata gttgagggca atttccccaa tcagttggga
197641 gttctgcaac ataccgtctt tgaaagaagc attgagcact tctgcgatac tagcattcaa
197701 cccaaattga gatgtatcta aagaggagtg ctgaaacaac gattgaccaa ctaaagatgt
197761 tgaagagccc gtggcttgag gatttagttc tcttagctcg ccagtaatct gagaggagag
197821 tgtggaaact ggacgttcct ggggaataga ggacaaggaa gaagaagtag gggtagcgcc
197881 tgagccgtga ttaggcaaag agtcatagag ttgagaatcc ctgttatgaa tgaaattttg
197941 tttccttgag ggtggtaaag caggtgcgcg gatatgcaag ggccttgaat caacttgacc
198001 tgaaggttgg ttatcaaccg tagcagtagg aagtgaaata ctgggagagg caggctcaac
198061 tggtaatggt tttgtttttg attgtggctg aagtggaggt tgagaaattg atatcggtac
198121 tgtcttgggt aagttacttg agtctggttc tttcgcagtg ggtggagaac tttgttgtgg
198181 cgttgaagga ggcacttcag aaggagtaaa agtttgcgca ggtggtgtat cggcagtaac
198241 aggtgcatct acgtttttag gagtagcatg gggaggaaga ggaggagtgt cactagcttc
198301 acccggggtc caattgtttg attttgatga tgaataggtg ggtcgacgtt gcttctgata
198361 aatagaagaa gttgcagttg aagacactct ggtggaagaa ttattcggag tctcagtgat
198421 ggatgcctga atatgagagt ttgatgatga ttgttgttgt ctcttgttct tggtcttatt
198481 tctaccgaaa attgagccca ctttagatct tagggtgtgg gacgactttt ttgatttcaa
198541 ggaagaatta ttagtagagt ctgaaaattc gttattcatg agatcattat gagtgagact
198601 ggaagatgca gaggcaagac gatgtccaat gtttccaaat gcactcttac gcttatcctt
198661 ttgtggggat ttcttttctc tttccgtatt agcggcacca ttggatacag agaccgggcg
198721 ggcaccagtg gcagaagctg gcgaagcatt attaggaact acttctttgg aagaagagga
198781 tagttggaag ttgtattgac tggcatcttt ggcaaaacga tcaatttcac tttgcggttc
198841 gaatgccagg aatttcgtca tcaccgtttc gcattctttg gtagtattta gtaaataatc
198901 gctaaaactg gtttgaaatc taagcataca gttttttaaa gtgtccaaac ggttatagtc
198961 aatagtctcg aacagctcaa acaaataggg gctttgctgg tcccattgcc ttcttgcctc
199021 ttctaagtga ctagagtttt cgttattctt actatagtgc tcaattgaag ctgcattttt

199081 actaagcttg gaatgtaagt ccttgctttc tctccagcta gtattgttct ccacggattc
199141 cttcaattca cgaacgactt gctggtccaa agtgttgtag tattctgtac ttgacttcaa
199201 atctgatttt agttcttcaa ttacagtatc ccatacattt cttagctctc ccagcgagtt
199261 gaatctgaat gcactcattt cctgcggagt caaaacatta ctttcaatca tttgagcgtt
199321 taagatcttt gttatatctt cattttcagc tattattttt ctcaattgct gagcatagtt
199381 cctcttcaga tttgccaatt ccttaaataa caggtaaaag tctttattca acaatttgac
199441 ctgcgataac ctgattctaa tagtttcggt agcctcgtaa ggactcttag tagtcaatat
199501 gctatctgca tatttggttc tttgttccgt catgagaggt gattattcgc aattttcaaa
199561 taagcttgct tcaatacgac tacagcaaaa gtaattcctt tattaaggca ataactacga
199621 gaccttttct ttgtatttcg tgaaaacttg aatgcaatat agagtttttc aatctgcaag
199681 tacttaaggg gcccttttag ctacacataa tatagaaatg aaagatcacg ggctacgaaa
199741 agccattcat aatcatatac aaatatcacc tgtaatatat aagaatatat gagctattga
199801 ttttcttcta ccgcctgatt tagttgtttt cactcttcaa aactaaattc agtattagcg
199861 acagtggagg tgctagtact ctgtaaatga aagtaaacgc tggtattcat gtagagcacc
199921 aacctgactg tgaatgcgta agctgaaaca tgatagccgc ctgtaaaata ttgttatcaa
199981 taagtagacc ggcttacacg aataggattc gctttattct ttactcccca ctcaacccta
200041 agcaatgttc aaaatagtgt gagattattt tatgcgcgcc tccttcaaaa aaaaatttac
200101 atatataaca taggtggaaa aacttccctc cgatgtctaa taagttaatc aatagaatga
200161 aatggacacg aaaaggcgta gacatggaag atgacaatga cgagaacgtc attttttttt
200221 ccacgtattt taaaaagggg tagcctgctt caccagatgt tcttgtcaac aaaattctaa
200281 aatgatgatt tttttaaaca caatttatgc tagatagtat ctgaaagcat tcaaacttta
200341 tgattacagt gttttcgaca gttttaaatc tctgaaccga ttttgtaact cttggtaatt
200401 gtgtccttac atttccgtaa tgacgttaat caaatcgcaa tcttgaatct tcttaactaa
200461 aaggttataa gggctttttt ttattactaa attgacgtat gagctgattt gattatgggc
200521 gctatataat tatatgtaca cattttttaa ctgtaaaaaa taatacggca atacaagtac
200581 aatacatgca taactcataa tcttctacct cttctaccac ccttctttct ggtggagtca
200641 gatggaactg gggtaacatc ttcgatacgg ccaattctca aaccagatct ggccaaagct
200701 ctcaaagcag cttgaccacc tggacctgga gtcttggttc tagtaccacc ggtagctctg
200761 atcttaacgt gaacggcagt gataccgact tccttacact tagcggcaac atcttgggca
200821 gccaacatag cagcgtatgg agaagattca tctctgtcag ccttaacctt cataccacca
200881 gtaactctgg cgatggtttc cttaccagat aaatcggtaa catgaacgaa agtatcgttg
200941 aaagaagcgt aaattctagc aacaccaaaa acttgggaat tgtcacgagc ttgaacaacg
201001 tctaaatgga aatagtaaat cgatatcgga cgatatcaaa cgtaatgttg ttagtatcca
201061 aaagtttttt tttttaattc ttacatagta tcttttgcttt tccatcgaaa aaaaagttga
201121 caaaatatac tgccattgac acaataggct caactaatcg ctatgaatta ctttggaggc
201181 aatgaattcg ttgaattgtc tttatgtttc tgactttcaa tgtatcttgc acgtcgtcag
201241 aaacattcaa caacaatcaa tggaaaatag agatgcttta aaaatgttca ctatgtgatt
201301 aaacatacta gacatggcgg gttcttgtgg tttcttactc tccttgggtt taaaatagaa
201361 tcttcattaa aaagtgcttt ctgtaatata tagtgattaa taaccaattc atttcaactg

201421 tctacctgag gaagccctgc accatatgga atgagccagg aggcgggatg gtctgccgca
201481 agcagacaaa tttgccaaaa tgcagtagag aatggcaatg ctcgtgctga aggtgaaaaa
201541 aagaacgcaa aaaaaaagag tagtaaaatg tacagatgtt tattaagtcc gtacaccgca
201601 cgtaagtact ccaaagcagt ctctattaca ttttaaatag ggagaaagaa gatacataac
201661 aagtatgaac aagaacgata atcagtgagt aacaagaata gagatgtcag gcccgtgtga
201721 agtacttgaa agtaataagt gagtatatag aatataccag aacttgtaag ctttcgaata
201781 acaaaaagag ccatttgtac ttattctcga actcggttgt agagaggacg ttgccttgaa
201841 agtcacaaat cccctacaat aataccggtc cttggaatgt aaattatttt gttaattttc
201901 ttataagttc acagttgaaa aaacaacggt cacagtgaat caataaaaga aggatctcac
201961 tttataaaga aaagaacagg aaaaaaaaat gtatatctgg gtatattacc cggcccttga
202021 taatattttt caataatttt gccaggtaac aatgtgattc atgcatggct taaagaaaat
202081 acattgctga agcaaaagct caccaaaaat ttgcgttgca aagctttgtt tgcggtatgt
202141 taataaagtt accatgaagc ttttttactg ctctcagacc ataaagacag gaagcaactt
202201 gtcatatcta tttcgacttc agcggttatt ttcttcagcc cattgataag gcgatatatg
202261 cctagaagaa aggaagagtg cagacttgat aataaagcaa gaccaagagc aactaaatag
202321 actaaaatat agagccttcc aaactgttag ttattattac ttaatactcg tacgggtgat
202381 gaaggtagtg actttgtaat ccgggtaatg gggaaaataa gaagccttca aagctatagc
202441 atttgtccct actgcatcgg cccctttcca gtacaataga atctcgttga tgtgattact
202501 gattcaactg acccaaataa atcatttgaa gaactcataa aaggactata gcagcaagaa
202561 ctgcaagcca attggttctt tctcctttct gccaaaaaaa aacacaatac ggatatgaat
202621 tcaattatta atgctgcttc gaaagtctta agactccaag acgatgtgaa gaaggctact
202681 ataatattag gagatatact gatattacaa ccaattaatc acgaagttga accagatgta
202741 gaaaacttgg tacagcatga actaaccaag ataatacaag gttatcccat acaggataat
202801 atgattatta atagcaaaaa aggcacagtt gaagatgact tatgcgaact caataactat
202861 acctgttttg cactttcgaa aagctttgat ttatgccatg atagcagaaa tttcaacata
202921 gcgcagccga aacgatggat acaattatta gagacattaa ctgactcagt tagtttcgca
202981 gttattgttc aaattattct cactttatct aacatttcgc taataaataa acaaaccttg
203041 gggaagttaa aaaaactgag gattcgaatt ttcgaaatac tatcaaataa aaacgatagt
203101 tggaaatcta cattactaca gaaaaatctt atagaatggt acattttat gctttccgtg
203161 gattgcacac ctttagaatt gcaaaactta tatctccata aggagttgaa attctgtaac
203221 ggtatcttga attcattaac actccaagtt tctgatcctc gctcacaaaa ttacctgcaa
203281 tttgagaaca cgtataagct ttttcaaata caaaagtcat ctagaattaa caactcgttc
203341 cttttttaca tagaattcaa ttccgttacc tcaaatagga taatgaccat agaaagacac
203401 atttatttgg aaattaagga aggccagttt tgtatttcaa atgataacta cataatcggt
203461 ttatttgaaa acttcgaatt cgaagcgggc actttgtact ttattggagt tttaattgat
203521 cacaataatc gaataactct ttatgttgat ggaagtatga tcaatcagct cacgttattt
203581 gaaaactcta tatgccaatt aagcacttgt gaactgggat ccatgatttg ttcaattaaa
203641 gtatatagat tttatttgtg ggatggatta ttaacagaat ttgcgataaa tatacttcaa
203701 gctatcggca ccaattacca atatacattt agcaagaaaa aagaagggcc tgaagtttta

203761 tcgctctgcc aagacttttt gatcgctaag gctcatttaa tggccaggcc tacaacagaa
203821 atatcttcca caaaatacat cgatgagatt gaacttcttg aaatggaaaa tatcattatt
203881 gatgttaacc caaatgatat tcttcaagat ttcaccgaat cgtctaattt tacggtaaaa
203941 tttgaggaaa gcacgaactc gaaaaatatt ccggaagtgg gtaagtgcta tttctatagg
204001 agttcaaact tggtttcaaa atttgtgtcc attgattcta tacggcttgc gtttttaaac
204061 atgacagaat ccggtagtat agacgatctg tttcatcatg tatcacatct gatgaatctt
204121 ttacgaaata ttgatattct taattggttt aaaaaagact ttggctttcc tttatttgct
204181 tatactttaa aacaaaaaat aacacaagat ttatctcagc ctctgaatat ccaatttttc
204241 aatttattct tagaattttg cgggtgggat ttcaacgata tttccaaatc cataattcta
204301 gatactgatg cctacgaaaa catagtcctt aacttggatt tatggtatat gaatgaggat
204361 caaagttctc tggcgtcagg cggattagaa attatcagat ttcttttctt ccaaatttca
204421 agtttgatgg aagcctctat ttattctaag ttcaattcca ataaattcaa tgatatgaat
204481 atcctagaaa aactatgttt aagctatcag gctgtcacaa aaagagaaaa tcagaacagt
204541 aaatttaatg agctatcaag tgatttaatt tctgtatttg ttactttatt gaaaagcaat
204601 actgataaac gacatctgca gtggttttta catctctcat attactttat taagagaaaa
204661 gatgtacgtt ctacagagat tatacttcaa gcggtagatc aacttttttc gttttactta
204721 gatcaaggta gtgacgaaaa tgcgaagata ctttcagaga ttataccact taagctaata
204781 ctgatgatta tggatcaaat agtggaaaat aatgaatcaa accctattac gtgcttgaat
204841 atcttattta aggtagttct gaccaataaa ccgcttttca aacaatttta caaaaatgat
204901 ggtttgaaac tcatattgac tatgctttgt aaggtaggga aaagctatcg agaggagatt
204961 atttctttgc ttctcacata ttctattggc aattatacca cagctaacga aatattttca
205021 ggtgctgaag acatgattgg aggaatttca aacgacaaga taactgcaaa agaaattatt
205081 tatttggctg tcaacttcat tgagtggcat gtgattaatt ctaatgccag tgattcttct
205141 tctgtattgg acctgaacaa ccatatatta agattcgtcg aagatctgaa atcgctgagc
205201 gctgttccga ttaatgaatc tgtatttgat cctaaaaaaa gttatgtgat ggtttcatta
205261 ttagatctct cgatagcttt gaatgaatcg gaggacatct caaagttcaa gagctcttca
205321 aaagtgattt cagagctcat taaaggtaat ataatgtgtg ctcttacgaa atatgccgct
205381 tatgatttcg aagtctatat gagcacattt ttttgtcaca gtacagaata caaactggtt
205441 tatccaaaaa ctgtaatgaa caattccagt tacttagagc tatcatttat agtgacactc
205501 ctacccgaaa tacttaatga cctggtagat agcaataaca atttgaacct gatgatgttg
205561 aagcatccat acacgatgtc aaatctcctt tattttcttc gcaaatttcg acctgatacg
205621 tcacagatag ttatgcctaa agatttttat ttctcaagtt atacatgtct cttgcattgt
205681 gttattcaga ttgataaatc atcattttac catttcaaaa acgtttctaa gtcgcaactg
205741 ttacaggact tcaaaatctg cataatgaac ttaatatatt ccaatactct aaagcagata
205801 atctgggaga aagaagaata cgagatgttt tctgagtcac tgatggcgca tcaggaagtt
205861 ttatttgcac atggagcatg tgataatgag accgttggct tattgttaat attttttgcc
205921 aacagattac gtgattgtgg atacaacaaa gcagtcttca attgtatgaa agtgatcatt
205981 aagaacaagg aaaggaaact aaaggaggtg gcgtgttttt ttgacccagc gaataaaagt
206041 gaagtactcg aaggtttaag taatatcctc tcatgcaata actctgaaac aatgaacctc

206101 ataactgaac aatacccatt ttttttcaac aatacacaac aggtacggtt cataaacatt
206161 gtcaccaata tcttgtttaa gaacaacaat ttttctccaa taagcgttag acagatcaaa
206221 aaccaagttt acgaatggaa aaatgcaaga tcagaatacg tcacccaaaa caataaaaag
206281 tgccttattt tatttagaaa agacaacaca tccttagatt ttaaaatcaa aaagtccata
206341 tcaagataca cttacaacct caaaacggat agagaagaaa atgcagtttt ctatcgaaat
206401 aatttaaatc ttttgatttt tcatctgaaa catacactgg agatacaatc aaatccaaat
206461 tcgtcctgca agtggtcatc ggactttgca gaagattttg atgggatgaa acggaggctt
206521 ttgcctgctt gggaaccaaa atatgaacca ctcattaacg aggaagatgc taatcaagat
206581 actataacag gtggtaacag acaaaggaga gaaagtggaa gcattttatc ctacgaattt
206641 atcgaacata tggagactct tgagtcggag ccagttggag atttgaatga gaatagaaaa
206701 attcttagac ttttgaagga taacgattcc attgcaacta tttggaattg cagtttgatt
206761 attggattag aaattaagga ggggatttta attcatggca gtaattacct ttactttgta
206821 agtgattact attttagttt agaggataaa aagattctaa aattatcaga agtatcgcaa
206881 gaatcacggg atatgacggt tagcttaatt aacggccctg atgttaaaag ggtatcaact
206941 ttcctaaagc acgaagtctt tgtttggaaa cttctcgata tcactttcgt taccaaacga
207001 ccctttctac ttcgggatgt cgccatcgaa ttattgttca aagagagagt tagcgctttt
207061 tttagttttt acaacaaaag agtgagagat gacgttttac gggtactgaa taagatcccg
207121 aagcaccttc cagcagatcc aattttttca agcgttttac aagaaataaa cgaccgagga
207181 aatagtatag tggcaagaaa tggaatagga aaggcaagca ttgcttctaa attcactagc
207241 gtcttctcag cgaacaacag cctaatagat ggatttgaga tcagcaaaaa atgggttagg
207301 ggagagattt ctaattttta ttacctgttg agtatcaaca tcctagcggg aaggtcattc
207361 aacgatttga cccaatatcc agtgtttccg tgggttattg cagattacga aagtaacgta
207421 ctcgatttag agaatcctaa aacttaccgg gacctatcga aacctatggg cgctcaaagt
207481 gagaaaagga aattacagtt tatagagcgt tatgaagctt tggcttccct ggaaaatgct
207541 gattccgcac catttcatta tggcacgcat tattcctcag ctatgatagt atcttcatat
207601 ctgataaggc tgaagccctt tgtcgaatcc tttttgttat tgcaaggcgg aagttttggc
207661 cctgcagatc gtttatttag ttcgcttgaa agggcctgga gctctgcttc ttctgaaaat
207721 acaacggatg tcagggaatt gacacctgaa ttttttttc tacctgaatt tttgatcaac
207781 gttaatagtt atgactttgg tacagaccaa agcggtaaaa aagttgacga cgtcgtactt
207841 ccaccctggg caaatggtga cccaaaggtt ttcattcaaa agaatagaga agctttagaa
207901 agtccttatg tatcagcaca tttacatgaa tggattgatt tgatatttgg ttacaaacaa
207961 aagggggaca ttgctgtgaa atctgttaac gtattcaaca gattgagtta cccaggcgct
208021 gtaaatctag ataacattga cgatgaaaat gagcgcagag ctatcacagg cattattcac
208081 aactttggtc aaacgccttt acaaatattt caggaacctc atccggaaaa aatagcctgc
208141 aatgttcaac agctaacaac agaggtatgg cgtaaggttc caatgaagcc aatatttgag
208201 aagacaatct ttaatttgaa tgaaaagaac aggtccgtcg attatgttat acacgatcct
208261 agttacttcg attcattata ctggaggggc ttcgctttcc caaacttgtt tttcagaacg
208321 gaagaatcgt tagtgtcatt gagaattgtg cataaaaatt ggttaaaaat tggactagat
208381 atttttaaaa agacgcatat ggctcagatt acatcgtttg cgtactggaa gttgggcgaa

208441 ttcataactg gcgataaaaa tgggctgata aaagtttgga aatatcgtaa agataagcat
208501 tcggtttcag gtaaccttga gaacaaaaaa acaatgtttg ggcacctatg cgagctaaag
208561 gagatgcgct gttatcacga ctacaatacg cttttaacct tagacatcag cggcttagta
208621 tatgtctggg acatgattaa tttcgaacta gtaaggcaaa taacaaatga tgcgcaaaag
208681 gtcgcaatat ctcaacatgc agggagcatt atggtattga ctaagaataa cgccatttcg
208741 atcttcaatc taaatggaca aatatataca tcaaagaaat tcgaaccagc taaaattgta
208801 agctcaattg atttttttga cttcactaag ttagacgcag gttacagaaa gcatatctat
208861 tggaaagaga tggaaatact actagtgggc tttgaagatg gaactataga aatttacgag
208921 ctctttttga attttcataa tgaatgggcg ataaagctac tgaagcagct ctgtaccgaa
208981 aaagggaaag ccataactag cattaaggga caggggaaga catacctgtc ccagaaaaga
209041 cgcaaggata cagcagagcc tcatgagata gaagtgattg cgggaacatt agatggcaga
209101 ttagctattt ggtactaggc atcacatcgt aacgcctttc tttaaatgat tcaatttttg
209161 tagtttatat ctttactttt gaaactgatt tctcatccca tctagtattg taattgcgta
209221 cgtatccaat atcattacca acgccgggta ttttttcta gtatttcttc tccatttcgc
209281 ctatggaaaa cggcaaaagg gtaaaagaaa aaaacaaacg attaattctt ctttgaatta
209341 tgtaaaaatc aaaacgcaac cgcagattta atagagacca gaaattcgga ttactattga
209401 ctttgtgcac caccttcaaa tttactcatt gtttaagaca ggcagtggga aagaagccgt
209461 catattgctc gaatccttaa caagcaaaat atacaaccac taaattattc cgaaagggcc
209521 tgcttaataa tttgcctact aacttgtgca tagaacagca aacagaaaca aagcgtaaga
209581 aacatggggt atccgccacc tacacgaagg cttggagata agaaaaggta ccattattcc
209641 aataatccta accgaaggca tccttccgct gtttattcca agaatagctt tccaaaatca
209701 agcaataatg gatttgtatc ttctcctact gccgataatt caacaaatcc gtctgtaact
209761 cccagtactg catctgtacc tcttcctaca gcggcacctg gaagcacgtt tggtatcgaa
209821 gcacccaggc catctcgata tgatccgagc tcagtcagta ggccttcgtc atcatcttat
209881 tcgtcaacaa gaaaaattgg aagccgttat aacccagatg tggaaagatc ctcttcaacc
209941 actagttcaa ctccggaaag tatgaatacg agcaccataa cacacaccaa tacggatatc
210001 ggaaactcac gctattctcg aaaaaccatg agcagatata atcctcaatc tactagttct
210061 acaaacgtta cccactttcc ctcggcatta tcaaacgctc caccgtttta tgttgccaac
210121 gggagttctc ggagacctcg atcaatggat gattatagtc ctgatgtaac gaacaagctc
210181 gaaacaaata atgtttcatc tgttaataat aacagccctc attcttatta ctctaggagc
210241 aacaaatgga gatccattgg aacgccttcc agaccaccat ttgataatca tgtcggcaat
210301 atgacgacca ccagcaatac taactcgatc catcaaaggg aaccttttg gaaagcaaat
210361 agtactacta ttttaaaatc aactcattca cagtcatcgc cttcccttca tactaaaaaa
210421 tttcacgatg cgaataaatt ggacaaacca gaggcttcag ttaaagttga aacacccagt
210481 aaagatgaga caaaagccat atcgtaccat gataacaatt ttccaccaag aaaatcagtt
210541 tctaaaccta atgcaccttt agaacccgat aatatcaagg ttggcgaaga agatgcattg
210601 gggaaaaaag aagtacataa aagtgggcgt gagatagcaa aggaacatcc tactcctgta
210661 aaaatgaaag agcatgatga actagaagct cgcgctaaaa aagtaagtaa aatcaatatt
210721 gatggaaagc aggacgaaat ttggacgacg gcaaaaacag tggccagtgc agtcgaagtt

210781 tccaaagaaa gtcaaaagga actaacacgc tctgttgaaa ggaaggaaag cccagaaatt
210841 agagattatg aaagagcata cgatccgaaa gccctgaaaa cagacgtaac aaagttgaca
210901 gtagacaatg ataataaaag ttacgaagaa cctcttgaaa aagtggaggg gtgtattttc
210961 ccattaccaa aagcagaaac gagattatgg gaattgaaaa accagaaaag aaacaaaata
211021 ataagtgaac aaaagtactt actgaaaaag gcaattagga atttctcaga gtatcctttt
211081 tacgcacaga acaaacttat acatcagcag gctaccggac ttatcttgac gaaaattata
211141 tcaaagataa aaaaggaaga acatttgaaa aaaataaatt taaaacatga ttatttcgat
211201 ctccagaaga agtatgaaaa agaatgcgaa attttgacta aactgagtga aaatttaagg
211261 aaggaagaaa tcgaaaataa acgtaaagag cacgaactaa tggagcagaa aagacgtgaa
211321 gaaggtatcg aaacagaaaa ggaaaaaagc ttacggcatc catcctcgtc ttcctcatct
211381 cgtcgcagaa atagggctga cttcgttgat gatgcggaaa tggaaaatgt attgctacaa
211441 atcgacccaa attataaaca ttatcaggct gctgcaacaa ttcctccgct aattttagat
211501 ccaatccgca aacactctta caaattctgt gatgtaaata acttggttac agacaaaaag
211561 ctttgggcgt ctagaatatt gaaagacgcc tctgacaact ttactgacca tgagcactct
211621 ttatttttgg agggttattt aattcatcct aaaaaatttg gtaaaatttc tcactacatg
211681 ggcggcttaa gaagtcctga agagtgtgtc ctacattatt atagaacaaa gaaaactgtg
211741 aattataaac aacttcttat cgataagaac aagaaaagaa aaatgtcagc cgctgccaag
211801 cgccgcaaga ggaaggaaag aagtaatgac gaggaagtcg aagttgatga gagtaaagaa
211861 gagtcaacga acacgataga gaaggaagaa aaaagtgaga acaatgccga ggaaaatgtt
211921 cagccggttc tagttcaagg ttctgaagtg aaaggtgatc cattaggtac accggaaaaa
211981 gttgaaaata tgattgaaca gagaggcgaa gagtttgcag gtgaattgga aaatgctgag
212041 agggtaaacg acttaaaaag ggcgcatgat gaagttggag aagagagcaa taagtccagt
212101 gtaatagaaa ccaacaatgg ggtacaaata atggatccaa aaggagctgt tcagaatggt
212161 tattatccag aggagaccaa agaacttgac ttcagtttag agaatgcgtt acagagaaag
212221 aaacacaaat ctgcaccaga gcataaaaca agttattgga gtgttcgtga atctcaactc
212281 tttccagaat tgttgaagga gtttggctct caatggtctc tcatatcaga aaaactgggt
212341 accaaatcta ctacaatggt aaggaattac taccaaagaa atgcagctcg caatggatgg
212401 aaattactgg ttgatgaaac cgacttaaag cgagatggga ctagttcaga atctgtacaa
212461 caatctcaaa ttttgataca accagaacga ccaaacatca atgcctatag taatattcct
212521 cctcaacaaa gaccggcttt gggttatttt gttggacaac caactcatgg gcataataca
212581 tctatttcat ctatcgatgg ctctataaga ccatttgggc ctgattttca tcgtgatacc
212641 ttttctaaaa ttagtgctcc tttaaccact ttaccaccac caagactacc atctattcag
212701 tttcctcgtt cagaaatggc agaacctaca gtgacagatt tgcgtaacag gcccttagac
212761 catattgaca cgttggctga tgcagcttcg tcagtaacaa ataatcaaaa cttcagtaat
212821 gaaaggaatg caattgacat tggccgtaaa tcgacgacaa tcagcaatct attgaataat
212881 tcggatcgaa gcatgaaatc ttctttccaa agcgcttcaa gacacgaagc acagctcgaa
212941 gacactccca gcatgaacaa tattgtagta caagaaataa aaccgaatat tactacgcca
213001 agatcgagtt ctatttctgc attactaaat cctgtaaatg ggaatgggca atcaaaccca
213061 gatggaaggc cgttgctgcc atttcagcat gctatttctc aaggcactcc tactttccct

213121 ttaccggccc ctcgcactag tccaataagt cgtgcgcctc caaagttcaa tttttcgaat
213181 gatccgttgg cagctttggc tgcggttgcc tccgcgccag atgcaatgag cagtttttta
213241 tctaaaaagg aaaataataa ttgaacaaac ggctgagacg ggcaatacat atgctctact
213301 tcttttccat ccaatggttg gtgaaactct cgagcataca ttaccttacg tgtgttagtg
213361 tactatatta tatatatata tatgtatata tataaaggga ggagttttta attataattg
213421 taatttcgta ttttttctgc attatacagt tttttccgat tttaaacgac tttatttaag
213481 tgtcgtgtaa atatgtcaca ttttattttt gtacgtattc acatgtcctg gcgtgcggcc
213541 attgctgaaa atcgcaaaac ccacagagaa ataaacatcg cgaaaaagtc aatgaaaaat
213601 tggaaaatat ttttcatttc actattatcc acaagcaatt ttgtacaaag tgaaaaggtt
213661 gaactaatta tcttcgtcta gaagccatga attcactcgt tactcaatat gctgctccgt
213721 tgttcgagcg ttatccccaa cttcatgact atttaccaac tttggagcga ccattttta
213781 atatttcgtt gtgggaacat ttcgatgatg tcgtcactcg tgtaactaac ggtagatttg
213841 ttccaagcga attccaattc attgcaggtg aattaccatt aagcactttg ccccctgtgc
213901 tatacgccat cactgcctat tacgttatta tttttggtgg caggtttttg ttaagtaagt
213961 cgaaaccatt taaattaaat ggccttttcc aattgcataa tttggtttta acttcacttt
214021 cattgacgct tttattgctt atggttgaac aattagtgcc aattattgtt cagcacgggt
214081 tatacttcgc tatctgtaat attggtgctt ggactcaacc gctcgttaca ttatattaca
214141 tgaattacat tgtcaagttt attgaattta tagacacctt tttcttggtg ctaaaacata
214201 aaaaattgac atttttgcat acttatcacc atggcgctac tgccttatta tgttacaccc
214261 aattgatggg caccacatct atttcttggg tccctatttc attgaacctt ggtgttcacg
214321 tggttatgta ttggtactat ttcttggctg ccagaggcat cagggtctgg tggaaggaat
214381 gggttaccag atttcaaatt atccaatttg ttttggatat cggtttcata tattttgctg
214441 tctaccaaaa agcagttcac ttgtatttcc caattttgcc acattgtggt gactgtgtgg
214501 gttcaacaac tgccaccttt gcaggttgtg ccattatttc ttcatatttg gtactattta
214561 tttcatttta cattaacgtt tataaacgta aaggcaccaa aaccagtaga gtggtaaagc
214621 gtgcccacgg cggtgttgcc gcaaaggtta atgagtatgt taacgttgac ttgaaaaacg
214681 ttcctactcc atctccatca ccaaaacctc aacacagaag aaaaaggtaa gtgtaaaatc
214741 tttgaaagaa tttaagtatt caactttcgt atattcgttt tttcttagtg gatctattgt
214801 tactattatc actattatta tattgtaaaa gaccggatgg ttttgttata tattacatac
214861 acatgttatc gttgaaaaaa gttttccgtt tcctttcgac agtcatcaga taattttatc
214921 cgagtctttt atatgttaaa tcttgttgac aaatcgtccg ctctaaccct agataacaac
214981 aatgatcttt ttatcatttc tggcgttatt ttagcgcctc ctcccactaa tgttaattgc
215041 ttatagttac ctgaaggagc agcgaggatg gagatcgtac tgtgtatact tgtttcttcc
215101 gcttcagtat ctaggtctgc tattagcaca gtgttcaatt tatccatatc aatgtccact
215161 tcctcttctt cagagttatc agagttttgt aattggcatt cggggtctaa ctccactatc
215221 ccataatttg aagcaaatgc aatattcttt gcgtttatca ttaacggtac tgacttagtt
215281 tggtcgcaaa taatttcata cgtttctctt atggtggcac ttcttcctct tgttcttata
215341 gtcattctta aatcggacgc acgctcatct ataaatgctc taggtaactt tacgctctgt
215401 aaagcgtaca tcaaagaatt ccaacacaaa tcgaagactg gaccagtgcg actcaagact

215461 acaattttcg catacaaaac gtatgaccat tttcttttca ttttcaggtt tgtttcattc
215521 agcgtatcat cttctaattc atcgggatac aagacggaaa acgtaccgtc ttcattcgca
215581 ctacgaacac cagcttttac cttcaaggct tttttaggca atatcctaga gtgcaatatg
215641 gaatcgtgta gtttttgtga tatagtcatt tcttcatcgg tgcaagcgcc cactctacct
215701 ctttcaactt ccaccaccgg atagacagaa gcataatttg caattatgtc ctcctctttg
215761 gtcacttcga acaattcttc ctcaccgaaa tcatccaaat ctttaatcgc ggcgctagtt
215821 tcttctatta ttccacccgt aatagaagtt atgactatgg ttttcccact tttcaaaaca
215881 tttgagccta aaatgttgtt tttagtgtct atattaccag catccgcata acgcgataaa
215941 gtattatttt ctatagcaac atctctaaat tcttcatatt ttcttagaca tggcctgatt
216001 cctagggata agtgtctttg taaagacaat tctggagata ttctcgcaag aacttcaggt
216061 ggaaaagtaa ttgggtgtat ctcaatagtt tctaacgtgg tactttcagc catttccaat
216121 attagatgcc tgtgaaacta ctgtcaaact tatttgtgta acaatactca tcgtctaatt
216181 tgtccattat tgttatatac tctgtcataa aaaaagaaag aaaaatctta tactttcgtc
216241 gactgtgatt tgaaaatctc tccagatcaa acgggaataa aagggtttat atccgaaggt
216301 ggccagaaca atcatgaagc aaataggata aaaatagcca ataagatgtt tgttgtcttt
216361 catcccaaag ttccaacagc atatttcaaa gatgggtatt acagtaatag gttctctaaa
216421 ctatgatttg gacacattta cggatagatt acctaacgct ggagaaactt tcagggctaa
216481 ccacttcgaa acacatgctg gtggtaaggg attgaaccaa gctgcggcca ttggtaaatt
216541 aaaaaacccc agcagcagat atagtgttcg aatgattggt aatgttggaa atgatacatt
216601 tggtaaacaa ttgaaggaca ctttatccga ttgcggagtc gatatcactc acgtcggtac
216661 ttacgaaggc attaatacgg gtaccgctac catactaatt gaagagaaag ctggtggcca
216721 aaataggata ttgattgtag aaggtgctaa cagcaagact atttatgacc cgaaacagtt
216781 gtgtgaaatt tttccagagg gcaaggagga agaagagtat gttgtttttc aacacgaaat
216841 tcctgatcct ctttccatta ttaaatggat acatgcgaac aggccgaatt ttcagatcgt
216901 atataacccc tcacctttca aggccatgcc taagaaagat tgggagttgg tagacctttt
216961 ggtcgttaat gaaattgagg gtcttcaaat cgtggaaagt gtatttgata atgaacttgt
217021 tgaagaaata agggagaaga taaaggacga ctttttagga gaatatcgta aaatttgtga
217081 gcttttgtat gaaaaactca tgaatcgaaa gaaaagagga attgtggtta tgactttggg
217141 ttcgagaggg gtgcttttct gttcgcacga aagccctgaa gtacaattcc ttccggctat
217201 tcaaaatgtt tcggttgttg atactacagg agctggagat actttcctgg gcggtttggt
217261 tactcaattg tatcaaggag agaccttgtc tacggctata aagttctcta cattagctag
217321 ttcattgacc attcaaagaa aaggtgctgc tgaaagcatg ccactgtata aagatgttca
217381 gaaagatgca taaaatctac tgcaattttt tacttttagt ttgttaaatt ttagttttcg
217441 tcatatatat aaagtttaaa aaggtacaaa gaatgcagta ataagaaatg taattacatg
217501 agggcagttt aagtgctacc ttttaagacg gaagaactaa taccataacc gagagtaatt
217561 acgcagacaa acgctaacaa cgaagcagga acacctctgg agataaaagc gcctacagtt
217621 agccatctat taccctttttt atcggtcata gaaatagcag taacgtttgg aaatccagat
217681 gaagccaacc ccataccgca tgaggctaac aaggcgcaac cgaacacaag aattggagct
217741 gcctttggat cggataattt gtcaccaact tcttgcacca agggaataat gatgattgct

217801 gacacagtat gtgagacaaa agtgcccaca actaacatta aaataccgaa aatacataag
217861 atagcaaaaa caccatcgtt ctgaattttc ttttgtaatg ctcttgcaat agttaccaac
217921 aagcctgaag atgaaactgc cttaccaagg gctataccac ccatagcaag aacaacaatt
217981 gaccaaggga atgtgttgaa atcctttgtt gatagtagac ctgtaccaaa aaacaggact
218041 atcggtatta ctgcaatttc accggacgat ccaaaagcac tttctatttg tgactctaca
218101 caccatagaa gaatagtagc aatagttaca atgatgataa aatattgctt tatagtaaat
218161 ctggtcctta ttggtttaaa tttttccagt ttagttttgc ctattttaaa ggtgagtatc
218221 atcaaggccc aggagcacag catcgataga atacctgtag gcagagcgac agcaaaaaat
218281 tgcccccagc cgattccata aggttttaag tactgcatgg agatgatatt ctgtggcgaa
218341 gaaataggtg aagccatacc accaatatct gccgaaagtg caacacccat gactaatgcc
218401 ttagcaaacg gtgaagtgta gtccagcgga tctagtaagg gtgttaatag agaatatgtc
218461 aatactgggg aggcaacgtt ggaaatccac atcgaaagaa agaatacaac actcattgcc
218521 attaaaagga catttcttgg cttggtacct gcaagggcca ataaccacga tgccaaaact
218581 ttcgcaacgt tatattgcga caaggcttca cccaatgtga aacctgctaa taaaatcata
218641 attgttgacg accacattgt acctaagatt tctgtagagg cagctgctgc tcccattacc
218701 ttaccgtcat cgtcttttag taccctaaaa aggacagtta gaaggggaac caataaacct
218761 gtgatgtgta atggaatggc ttcactagcc cataagaaag cacagcattc taccaatgcc
218821 atacaccggt gttcgacagg gtcattgaaa gttttaacac ccaacaaaac acctgtgaca
218881 atgataatga agcctatttt cattgctctt ttcccaaaaa atagcttagg aactgccaaa
218941 ttttcaattt tagtaaactt caaattaatt ggcttaggca atggatactt gaaatattca
219001 atttgaagct tttcttgcat aagtttctca gcttcatctt caatagttat tatatcgtta
219061 ttttgcgaag agaggcccaa catgtctttc catgtgttgc ttctttccca aacaatgtgg
219121 tcgtgcaggt acgactttaa ctcttgcttg catgaatcta cgttcccagg ctgacccatg
219181 agaacagcat aaaattcaat aagttgtgca attttactgt ttaaactgct taaagtttca
219241 tgctggaaga tgtaggtgtc tttgaaaaat tcttcacttt ctattaattc ttgcctagtg
219301 ttcatgtgca atactttatc agacttctta gtaattttac tgaaacccat tctgttcaat
219361 tcgataaaag atttcaactg gcaaaggtct atgtaaagat ttattattgt ctgttttaat
219421 aatgactttt tctgagattt aatgttaaat tgcgaataat tcaagagggc ggtattgtcg
219481 gtagtgtcag caaacacgtc aacgtcatca tcttcatcca tttcttcttc gctgtaatca
219541 aacctatttc tcaattcact gtcgatgttt gaagagtgta tttcaaatgg gttattagat
219601 gaatgtgtct ctgatgtacc agaaataatt tctcttaaag gaggcaatgc ttgaatttca
219661 ggattgaaag tttcattcaa tcttgttaca atgccttcct tctccaaatc attaatcaag
219721 tggtcaaatc tttcatagaa cttagcttcc attcttttgt agaaatcatc tactttcaat
219781 ttctcgtcgc tcaggttagt tacgaatgta tcataagaac tgaagagcgt ctttccctcg
219841 ctactcacag atgagctgcg tgattcaaag aatttcttac gtataaaatt cttattgaaa
219901 gaggtggttt tcccttgagg agataactcg tctaactcaa ttgtttcctc gttaatattg
219961 ttcccatcta tggccttttc gtcggagtct ccccttttgc ttcctgaagg tgttttagaa
220021 ccaaagagct tacgcttgag tttatgtgta aatcttcttt tcgatgaaga tggttctcct
220081 gcagctatgt tgctttctat atcagcgatt ggacctggag tctgagagtc agcgtcatcg

220141 tttaagtcac cggttggcgt ttcttgtttc aattcatctg tctgtaatgt gtagatcaaa
220201 tttttcaatt cgttataatc taggtaatga ttctgccatt cagggacagc gttgtatttg
220261 agaaagtgtg agaatctcat ttcgaggcac aataatagcc gcctttaccg tagttttgct
220321 gcacctttat ctgagagctg actgcttttt tggtgtgaaa cactgttttc tggtaataat
220381 ttttcaatgc atcggattac ttttcccacg tgcgaaatca tcaattaatt agattgaaaa
220441 aagggtaagg gaaaataaga aagaggcggt agcctaaaga tacggtaatt gaaacgtttc
220501 ctatgcacaa tcttaaacct ttttaggtaa ttgattaagt tgactgtaat atctgtaaaa
220561 gattacatct aatttacgtc tggtttctta tcaaaaaata ttatctactg tagttatgca
220621 agtattctta attttcagta ctaaacatga atattaatta tttgtgagcg tacaagaagc
220681 aaaaggaagt catctttctt tgaacagttc tgtttttagg taagccttgg aaccttagct
220741 atactgactt gatacgccct gtctttgttg tcgttatata aagtgtgcac ctttatcata
220801 gcccctaata agcttcttgc agtagtattt gaacagtgta tatcttcata ttgcttttgt
220861 tgtaaatatt gcgcaaaagc gactatcttt gtgatttggt tgaacttttg catattcacg
220921 ttattaccat ctttagtgaa tgtatcgttt ccgtctctta tgaaggtaat gtccctgata
220981 agcagtgatg taaaaggtac acaaggaagc tgactgtgga aaatatgttt aattgttctt
221041 ctataaacgt tgtagttgtt gtttggatgt acgacgactt ttaggcgctg aaaaaggtgg
221101 tctgatttaa catctatcgg gagagatagt ctttctattg aatgattttg caatgatgta
221161 atgattgaag caagtgagtt taagtttcgt aagtatagac aagacagtgc aacttgtaac
221221 cagtaggaaa tggtgtgcgt ttgctgcaaa gttgtctcaa gaacgtagga cgataactgg
221281 ttggaaagcg taaacacgga gtcaattgtt gtatcgttgt gtttgaaatg tcttgtgaat
221341 tctattgttt caatatccaa gtaaagagaa gattctaaga gagtcaatgt ctttgcaagt
221401 gaccatggag atacgtttag agcgaaggct tcgacttcta gaagttcacc ctgtttccat
221461 tggaaaagta aatcactgag gtcagttgca ccgcacaatt catcatttgc gttcgttttg
221521 tctatctcgt attggagaaa taagttttcg tagtccaggg gcggtttatt ttcttcgatt
221581 tccttatcag ttagaccaat gtaatgaatc ttcaattgcc tcgttctttt tttgtttata
221641 aacagctgaa ctatgtctgc aaacagttct tggtattcat aatattcagc caagtgactg
221701 taccaaaaat tgaggatatc ttttaatgtt gtctcactat cttgccaata agactctacc
221761 cagatttgta ttagacgagg gacggagtga tttttgtgtt tgtttttatt aattgtggga
221821 taggatagta gcaactcttg gaggagagca ttgtcagttg tccagtctct gaagttaagt
221881 agtaagtttg cggagtcaaa gggggatggc ttttgccatt tgtgagagtt gtgcggcagc
221941 atcttattca aatagagctg tattctgaag acctcttgta gaacatcatc catactaaaa
222001 agtaaatcgt cctgtcccat tacgagctgt agtagtgctg tgaccctctg tatatttacg
222061 ttgccatgaa gaaggtaatg ggcgatattt tgatacaatt cctgagttgc atgttggatt
222121 gagtttacga agggtcgcca gacggccaga aacctccagg cggagttaac aactagtaat
222181 acggcatcca tgtttgcatc agcgccgagc ctataccagt cactgagtag acgttttctt
222241 gctcttttta tgtcctgact tcttttgacg agggggcatt ctctagagac acaggcagtt
222301 gcttccagca actgccgtac ggccgttctc atgctgtcga ggattttttt tgggacgata
222361 ttgtcattat agggcagtgt gtgacttatg aattgttgta gaaggacgtc tgtgatgttg
222421 gagatatgta ttttgttaac tcttcttgag acaatttggc cctggatagc gaagcgtgcg

222481 gttacaaata ggtcgtcttg ttcaagaagg taggcgagga cattatctat cagtacaaac
222541 atcttagtag tgtctgagga gagggttgat tgtttatgta tttttgcgaa atatatatat
222601 atatattcta cacagatata tacatatttg tttttcgggc tcattctttc ttctttgcca
222661 gaggctcacc gctcaagagg tccgctaatt ctggagcgat tgttattgtt ttttcttttc
222721 ttcttctatt cgaaacccag tttttgattt gaatgcgaga taaactggta ttcttcatta
222781 gattctctag gcccttggta tctagatatg ggttctcgat gttctttgca aaccaacttt
222841 ctagtattcg gacattttct tttgtaaacc ggtgtcctct gtaaggttta gtacttttgt
222901 ttatcatatc ttgagttacc acattaaata ccaacccatc cgccgattta tttttctgtg
222961 taagttgata attacttcta tcgttttcta tgctgcgcat ttctttgagt aatacagtaa
223021 tggtagtagt gagttgagat gttgtttgca acaacttctt ctcctcatca ctaatcttac
223081 ggtttttgtt ggccctagat aagaatccta atatatccct taattcaact tcttcttctg
223141 ttgttacact ctctggtaac ttaggtaaat tacagcaaat agaaaagagc tttttattct
223201 tgatttttgt tctttcgggg aaactgtata aaacttccaa aaaggaaaag taaaacaata
223261 catctcctta tatcaaagaa aatcaagaag gacaacatgg atggtatttg tagtatggcg
223321 gaaaacataa acagaactct gtttaacatt ctaggtactg agattgatga aatcaatctc
223381 aatactaata atctttataa tgtatgtttt catttcaagg atagcctttg aatcaattta
223441 ctaacaatac ttcagtttat aatggaaagt aatttgacta aagtagagca acatacatta
223501 cacaaaaata tttctaacaa taggttagaa atataccacc acattaaaaa agagaagagc
223561 ccaaagggaa aatcatcaat atcaccccaa gcacgggcat ttttagaaca ggtttttaga
223621 agaaagcaaa gccttaattc caaggaaaaa gaagaagttg caaagaaatg tggcattact
223681 ccacttcaag taagagtttg ggtatgtaat atgagaatca aacttaaata tatcctatac
223741 taacaatttg tagttcataa ataaacgtat gagatctaaa taaattcgtt ttcaatgatt
223801 aaaatagcat agtcgggttt ttcttttagt ttcagctttc cgcaacagta taattttata
223861 aaccctggtt ttggttttgt agagtggttg acgaataatt atgctgaagt acgtggtgac
223921 ggatattggg aagatgtgtt tgtacatttg gccttataga gtgtggtcgt ggcggaggtt
223981 gtttatcttt cgagtactga atgttgtcag tatagctatc ctatttgaaa ctccccatcg
224041 tcttgctctt gttcccaatg tttgtttata cactcatatg gctataccct tatctacttg
224101 cctcttttgt ttatgtctat gtatttgtat aaaatatgat attactcaga ctcaagcaaa
224161 caatcaattc ttagcatcat tctttgttct tatcttaacc ataaacgatc ttgatgtgac
224221 ttttgtaatt tgaacgaatt ggctatacgg gacggatgac aaatgcacca ttactctagg
224281 ttgttgttgg atcttaacaa accgtaaagg taaactgccc atgcggttca catgactttt
224341 gactttcctt tgtttgctag ttaccttcgg cttcacaatt tgtttttcca cttttctaac
224401 aggtttatca cctttcaaac ttatctttat cttattcgcc ttcttgggtg cctccacagt
224461 agaggttact tcctttttaa tatgtacttt taggatactt tcacgcttta taacaatatc
224521 aagtttacct tcttcattac tattcatctt cgccacaagt cttctctccc ttggtgtttc
224581 caatctaact acaaaactgt tgattagggt gtacatcacc ctaacaagat catgtatttg
224641 cttcctctgg tacaagctaa gaacaggtaa attcaaaaca tcccagagta atatcttcaa
224701 agggctatac cctttaaaca tatctcggca tatttgtatt aacccactaa tattttgacg
224761 gccaatcttt tctattttta ttttcatatc atcgacgtaa tgaccactta aaaacaactc

224821 aatatcatcc tcacttttta tatttttaac ttcattgtgc acagattgga cggaggactt
224881 aatatcgtca cttatggtat ccaaactacc atccataaca actagattta cacattcaga
224941 aagcacatca acacttttat ccctaatata agaatcctca tggaagcaaa gtatttcagt
225001 aaacaaaaca agggcatcct tgttttttaa accaccttct aaaagcatga ttttcaaaac
225061 cattaaaaac tcattctgaa agctttccct ggaaatatct ccgttattta tacctaatgt
225121 taatgtatgc attatcattc ctgtcaattc cgtaaagtca taagcaccca ctttatgcaa
225181 tttgtactta gcatacatga tagtagtctt aatcaaagac tcgtatgatg aaagccaatt
225241 caccaaattt tcgtacctca ataactgagt cttaactttt tccatatatg ctttatcttt
225301 atactcattt aaagcaactt taacaacatt ttcaatcaag ctgcggacgt acgcatcatc
225361 actgtattta ttttcaccat tctcattata aacaagaata tccagtaaaa attgcttcac
225421 caattttgga catttaccat tttcattttt cactttagcc aaatatttgg gaatattaca
225481 ctgtaagaaa tacaacttag gattctcaaa gttattactc tttggaatat tagagtcttc
225541 taggcaaaac aaaatctgaa aaatctgaat taaatgctta acaccgccag taaaatccgg
225601 atcatatacg tatttactaa gcgcttcgca cgccgcgagt cttatgccaa aaaagtaacg
225661 ctcgtcgatc gccgttctaa acaaaatact tgaataaaca agtgatttca caccaccatt
225721 aacaacgacg tcctcataat atcgtatggc ttctagttgg gcctctatgt ccccatcttg
225781 tctcaactga gaagaaaaca tgtaatccgg ctgattaatg tgcatttggc aaatccattc
225841 cagatcagag tctatgcgta tccactcaaa tgcttcgttt tgcttaagca attcattact
225901 ctcagacgta cgattaaatt ccgtcaaact gaatcgggaa cactcttcgg gcgacatgta
225961 tacatttcct agacaattca catccacaac aataggcttc tcctcattat tgttttcaac
226021 accattttca tcatttgcac caccacctct tttccttaat cttctgtact ttgtattgta
226081 ctgaatatct atttttgtga atgtatcttt gatttccaca atatgctcat acggagtacc
226141 atcgtgttca tggatcctta tagtcatcga gcccgtgaag cattcggttc ggttcaaatc
226201 tggatgttct aagtggtcta gtgcactttt gaaaaatcct tcctccccta ccactttttc
226261 gtggccaagt tcttcatctt gaacttgcct tatacccagt tctataacca tcctcttcct
226321 attaaatctt tgggtgacac gtaatatggg taccccagac ccatataccc attcgttgaa
226381 aaaattctct aatttacttt tattaactct ttcgcaaaca tgttgaaaat gcgatgaagt
226441 caacgaatta ttcggtaaat caccagacat agcttgaagg aaaattttag gtaatactcg
226501 agacataccg aaagatcgtt ctgttttagt catccttctg tcaagtatgt gtagtatcat
226561 cggtgccttc aattgtataa aggacaaatc tttagaagac caagatattg gcctagaact
226621 gccggtaaaa gtgctcccaa taggcggttt ctcgaagtct tgttccacaa tcgcctcact
226681 attacgcttt aattgatatt tatacgtgtt attacccatt aattttttgg ttacctgaaa
226741 caccatataa ccagcaatac caagacagca ccagtagtcg ttcatatcta aaggagttat
226801 attcacacaa gaccattggt tagcaagtgc ccatgctaac tcattcgtag tactgaatgc
226861 tttatcaata acttctagag ggtacaataa tctggtatta caaatgccta atgctgcaaa
226921 atccatatgc ttagaaggtg cggtaggtaa aaacaccata gagtaacaag tgaaaggata
226981 agacccaaat tctttcgagt agaaatctat aattttttgg cagacaactg tcgaatttat
227041 aactgttaac tcatctgttt cctgcgtcgg aagtgtgaaa atttgaatgg gtataatatc
227101 cgaaccaatg tcgtcatcca tagtatcgtt aacattgtcc acaacattct ctctcaactt

227161 gtcttcctct acttcgtcct cggcatcaac acttggaggt gatatcaaag gtaaagacca
227221 tgaattaaag gcgcctatag cccaaccaat gtgatgtgga gccacaggat taattatctg
227281 aaatatgcat tttttttcg tcaaatcaat cgggtgcgga agttctttaa tatttgaata
227341 ttctgaacaa cacacaatca catccctcaa actcggattg ttgctttcct caatattccg
227401 cctttcctct tccccttcct cttcctcttc gtcactttcg ccttcctctt cttcttcatc
227461 attatcttgt gcatctttgc ttttactatt tttgccttct tcgtcatttt tcaagttaga
227521 atccttatct tcgtctttga taactcttgc cggcttcccc tcttcctctt catcgtgctc
227581 aggcgtatcc tccttctctt tttcactctc ttctccattt tgtccgatta atttcgaagt
227641 acctatattt ttaaccaatc tcggtacgct gaattctaac tcccatgtag atttttcatc
227701 aagcaaatcg acacatggga cccaatatga tgcagaactg caaatctcac catttgaagt
227761 gtaaacgttc cataaccagg gcttgtcagc atacacagta tcgaatttaa tacccgactt
227821 tgggtttctg atttcatatt caatttgtaa tgtaattgga gtgaaaacag attcttgaaa
227881 cccaggggta gtcttaattg aaggagtaat cggagtgtaa ttacttaatg cattggcgtc
227941 ctccaaagat attttgatgg aagatggtat ttttatagtt aactgggatt tagagtccgt
228001 ttctgggtat tcattcaagc tagcaaactt agatctcaaa aaatgtgact gttcaatgga
228061 attgtcgctg tataatacat tttgtgaagt caatccattc aaatttgttt gaagtgggtc
228121 gtcatgaata aattgatcac atcgacgatt ttcgaccaga acatctttaa tagtcatttc
228181 cttgcaatca aaagttacat attctaggtt ttggatcaac ggaatgatta ttatagtagc
228241 gctaccggta atgcagtgag tagctagatc tatatccaac gagatttttt catgggcaac
228301 tctaaaattt ctaaacttca tctcatgcaa agtgctagat tcactaacaa tggctctagg
228361 agtggcgttt ttggaaaagg acatcatgaa gtccacctat ttatcctgtt gtcacctctt
228421 cactgtccaa aacttcatca aaaaatactt ttacatccgt tccattttac tatagttttc
228481 atcttacgca atttaccaat tgccccattc ggcaaaagta tttaagtgtt acccggttac
228541 ttaaggtaat accaaaaata ttcactatat tgaaagttct atgtatttta ttgaattaaa
228601 taagactgat tttgtagcac gatatccgca agaatgattc acaaccattt tagatgtctg
228661 cagctttttt tttttgattt ttttactaat tttagtgcga ttttagaata actcttggat
228721 taaataaata aaaaaaatac aattatcact cagattttga agcaatctga taaaaattgc
228781 agcatattat tataacaatt ctagttcact acaaacagca cattatcaag aaacttatgt
228841 ttgatggaca gtttcttcaa ataattgaat gcagtaaaaa ataaaaaaaa taaccatgga
228901 atatgtttct aagaattcgc aatccttgtt tatataacta tagctttttt tatgttataa
228961 gaaaacattt attcctttaa ccaaaaggaa aaaggtaaca accccactta cccctctgtc
229021 aatagttaat cattttatta ttttcaactt taaacatcca aaaaaaataa aataaattaa
229081 attaaaatca aatagaaaat gcattattac tataataata tctctatgta gacatataca
229141 tattatttca ttgctttttg taataaacga ggaaaatggt gggataaaaa aatgcattgt
229201 tattccttcg aaaccagcag taataagtcg tcctgagaac gatgtagatt tagatcttta
229261 catgttggta ctctaaatca cctccttcca tcttaaagtc gtccttcttc ttgctccaat
229321 caaacttgct tttcttgctg ctgggacacc aatttaaact tattaaccag atagtaaaca
229381 aaatcatgaa acaacagaaa taacaccaaa aatgatcgta aacgtgaccc cttatgatat
229441 cacacactga ataatcatac acttcatcgt ctataaccag tccataattg aaataaccga

229501 gtccgtattt caatacaggt ttatatccac cttcatcggt aaacgtactt ggactcaaaa
229561 ctgtgcttag atcattagtt ccttggaatg catgttctct caataaggaa gcatcaagtg
229621 gagctgggat catcctttgc caaagtacgt taagagcgct ttcactgcct actgttgccc
229681 caaaaatttc ttggccttat cttggcacaa atattctttt gcatttagtt ctatacatac
229741 agaaagagcg aaaaaaaaag tgaaaaataa acagaatgtg ataacggtaa aagaaaatca
229801 aaaataaaca aaaaaaaata catatataac tattattcat gatagtaaac aggtgaatta
229861 agatatttag tacaattgtc tattacccca ataggcataa tcctcgagga aaaagtcgta
229921 caatacccac gagggccata ttgtgcacag atgccatagc gcatgggcat caatctgcca
229981 ttcgtaacta aaaaagtcaa acagttccaa tgacatggcc attgtggtta caatcacaag
230041 aagaattgga atgacacaca gtttgaagat ttgccgcttg aagctggagt acgcggtttt
230101 cttgaattcg cccattagct tttgcttttg cagagcatgg tagttttggc atgataacat
230161 tatcaataaa atatactgta aaacaccgaa aaaaatgttg aatctcatgt tgtatgtgta
230221 cgaccagtca acatagagtc tcaagatgtg cagggcaaag attgccgcaa ctgacgcagt
230281 gaacgcttgt gctatcttgg ggtacaggaa catggaagtc attcttgcaa atattgcatg
230341 aaaccctgtt aggacagtta aacctgcgaa aaagtaatct agcttctccg ttatgatcaa
230401 atcacgacag tgaaagaccg agcttgctgt ccaagccaac attcccgcca cagtaacgta
230461 aaggtagttc caaatcagta tacttctgct gttttttctc ctcctatctc cctcttcgcg
230521 tatgattcta gaaaacttta caaatccctt ataatgtgga ataaagttac ctatactgaa
230581 tatcgtcgag aagaattcct gagtccccaa tactctcaaa aatggccatt tcccatggaa
230641 ttggtatatt tcttcctctt cgtcaattct ccagcgcgta atgatgtgct gacattggta
230701 atcacaatct gagatacaat cccagaacaa taacttagag tacaaaggcg gggtatcgaa
230761 aaactcaata tcatgaaaca tgttggtttc agggtcaatg tagtttattt gggaattcgg
230821 acatcttctg ttatactcgc acgcatacgt acagtctata aactcatcta aattatctcc
230881 tggagagcat gtcactagga aacagtgaac aagtagggtc acaaccacag ctaacctcat
230941 atttcttgcc acacttctgt ttcttttctc ccgaagggta tggtttcaca ccttaggttg
231001 ctcttttatt cgctttatga aaaatgaaaa atgaaaagta caagaatacg caagagaaga
231061 gcgaaaaaac taatgttgac atgacattat gcaataattt atttatgaga gaaagagaaa
231121 aacactacaa atttttcata aaatcacgcc tcttcaatcg aatctcacca agaggtagtg
231181 tgacatcgtt attccaaccc cacagttttg aaccagattg aagattctca gacataggag
231241 tttgtaaagt ggcattgtac tcctccagtt ccttagcaat tgcctctatg ttcaatcgag
231301 ggaacgtttc gtcgtcatat ggatcgtcca gtttatcgac gccgttgtaa agaatctggt
231361 ttggcgtaga aggcttgaaa aacaacgttc ttctatggaa aacattacgc gtggacaaat
231421 actcaatctc ctgtgctatc atttcaggct caattccctt cgagagtagg atcgctgcaa
231481 ctccggtgac aatgggcgtc gacatagatg tacctgacaa aattaaagtg tcgttgtagt
231541 tcagatgaga tagcgactca atttccacgc ctggggcaaa gatgtttaca cagggccccc
231601 aattgctgaa cttggcaatc gtgtcaatgt ggtcatcaaa ggcccctacg gtgataacgt
231661 tttctgcaga cgcaggtgag gcccagtagg cgtctaaatt gaagttcccc gccgcggcaa
231721 caaatacaat accttcttca atggccccct ccactgccat gttgattatg gggctcctga
231781 aactccctag acttagattg gccacgcatt ttttaccctg tgggcgactg acctttgtgc

231841 aatgttccac gatgaactcc agaccactaa gaacgttact ggcctcgcca gacccgtctt
231901 tgcccaagac tttgacctcc acaagattga ccctctttgc cgctccatac gttttggaac
231961 ctacgagtcc cgccacgtgc gttccgtggc cattctggtc gccaaaccct tctttggtca
232021 agtcaatccc ctggatgact ctgtcttcga attccggatg gtccgcgaag atacccgtat
232081 ccatgatata ggcgttgacg tcttgacctt gatagtcgtg ttcatagtag taattgaacc
232141 agcttttgta gcggtccttg tccccaacgt cgaatgggag ttggtagtgg cgggaaatcc
232201 gggccaagtg acggggtgcg tctggttgat acgttatatt ttgctcttcc tcgacgtctt
232261 cgtacgagta tttggcggta gcgttaaacg tataactcga ctcggcgcta tttacactgt
232321 cgccttcaaa agcttcgaac ctgaagttgg gcacaatatc agccaccagt gggcttttct
232381 tcagcttgtt cactaagttc ttggacaggt tcatggtcac gccgcggaag gatccgatgg
232441 agaatttctt ctcaattttg ttgtcgatga actgtctcca tgacctgttg gaattcgaag
232501 tgaattgttg gaatgctgtg ggatttttga acctcaccaa atactcctct gcgtatgcga
232561 ggttgaccag caaagagatg agtatgcact gtggcttcat gaccagtgtt tttttgctgt
232621 ttttcttggc ggatgttaga ggatgtctag cacttgagta tacctctata tgctgtgtaa
232681 aatagaaaat aaaaaagcca atgagaacaa tattttgtgt cgtttttttt attttcgctt
232741 caattctcta tacacgtacg tcatcactac gggaaaccaa aaaaaaaaaa aagaaaacag
232801 tacaaaaagg tttactttac ccatagttag tatacgtaga ttttatacac gcactatttt
232861 tcttatatac aggagatggg tggccacaga acccgcgcct agcctatttc tttcttctta
232921 gacctgcctc gaggtcgccg acatccaacc ttgtacctct gatgtagtag tgtttgtttc
232981 tttgtctgct cgagacgtgg ggggttagca agtcgattct ctcgatcagc ggcgaaaaca
233041 atggcactct aatctcgacg gccgtcttgg ctatctggtt ccgcaacagc aacgaggcgt
233101 cttgcaccag ttgtttgcgg tctatggaaa ggatgtagcc aacaaaggtg tcgtacgagc
233161 acttggacga gtcgtacaca atccggacca cgtcgcctgc cttcaagcgg tccttgttct
233221 ttttactgat gagcttcctc ttaacggccc cgggatccag agactctatt tctgagaggg
233281 ccacctgctt cataatctgc gacgaagcta tgcgctgcac aggtgggtac acggggatgg
233341 tttttctctt ggtggcgggg accatgtagg accttgtcca tgatccaagc aatctgacgt
233401 tcctgctcca catcgctatg gttctctctc cctttcagca gctgctcaga gggaagactc
233461 gattcttctt cttcttcttt actgggagtt ccaaagcacc aaaattgaaa accaagaaaa
233521 aaaggaaaat acatcactac ataaaagtgc gcggagccat tgaccactca catacagata
233581 taagtgtata tatacaagta tacatataca agtatacatg tagaaggaga cagagacaga
233641 gggaaatgta taagtgtgta tttttatagt atacaaaaga atatgcgtgt gaggatgtgt
233701 gtttcttcta gaacctgtgt cttcttttcc tcccggtgaa cttggagtcc ttcgcaactt
233761 tcctcccacg tcttttcatg agctccttct ttcttagaat gaaactcttg gcagactcct
233821 tgtctttgcc tcccttcaag cgctggcgca gttgtttctt caagttgacg ttctcctcgt
233881 ccatggtcac accgtccaaa ttcacctgct cctccccctg cggtggggcc ccactgctca
233941 acacaaggta gtacttctta ttcttttag actctgggtc gtccaccaca agcccgccac
234001 tgaaccctgc caccttggca gactgcagta tgtcgtccac ctggtcgtcg tttttcgggt
234061 agaactgggc cacaaatttc cctcccttct tcagtgcagc atacaatgtg ttgaaaaacc
234121 tcatcaaccg ctgtttagga tcgttgtatg aagtgtccgc attgcacagc cattggatcg

234181 cactgatact aatagccgcg tcaaacgagc ccgcccggaa cggtatcccg gtgcccatat
234241 cctgcaacat caagtcgccc tccagctctc tactaagacc ggtcgcaagc atgctgggcg
234301 atatatccaa accacaccac acatggtctc cctcctgcgt caaaatctcc ccagacagtc
234361 cggacccgca cccgatatcc agaatgaaac tgcacggctg cagattcaaa agctccaacg
234421 ccctcagcgt catcttcgcc tggatatgct gcactctggt cgaacccgtg tacttgtgtg
234481 cttcgctatc attatagaaa atctccggtg gtgccaactc ctcaggacgt gacattattt
234541 cttctctgat atatttcctg tgtttccgta ccgcaccttt ttagcactac ttttttacta
234601 tgctcttctt cttctgcttc ttctgctttt ttcctcttta tcacactatg tatgtgctgc
234661 tcatctcttc tttttatcga taaaattgaa aaatgtgaga tggtgtagag tgaaaaaaaa
234721 aaaaaaatct ggcttggcca tcaaataccc ggccgtggtt ggactcgttt agcgaacaat
234781 agcacccagc agaccctggc aacatgcgga tgatataaga aggacgagcg tggtggagga
234841 aaggggcgcc attggcacac tcacgcaggt ggttgttcag cacggcttgc agcaagagcg
234901 ccaaaacaga ttgcaagaat gacggagact aaggatttgt tgcaagacga agagtttctt
234961 aagatccgca gactcaattc cgcagaagcc aacaaacggc attcggtcac gtacgataac
235021 gtgatcctgc cacaggagtc catggaggtt tcgccacggt cgtctaccac gtcgctggtg
235081 gagccagtgg agtcgactga aggagtggag tcgactgagg cggaacgtgt ggcagggaag
235141 caggagcagg aggaggagta ccctgtggac gcccacatgc aaaagtacct ttcacacctg
235201 aagagcaagt ctcggtcgag gttccaccga aaggatgcta gcaagtatgt gtcgtttttt
235261 ggggacgtga gttttgatcc tcgccccacg ctcctggaca gcgccatcaa cgtgcccttc
235321 cagacgactt tcaaaggtcc ggtgctggag aaacagctca aaaatttaca gttgacaaag
235381 accaagacca aggccacggt gaagactacg gtgaagacta cggagaaaac ggacaaggca
235441 gatgcccccc caggagaaaa actggagtcg aacttttcag ggatctacgt gttcgcatgg
235501 atgttcttgg gctggatagc catcaggtgc tgcacagatt actatgcgtc gtacggcagt
235561 gcatggaata agctggaaat cgtgcagtac atgacaacgg acttgttcac gatcgcaatg
235621 ttggacttgg caatgttcct gtgcactttc ttcgtggttt tcgtgcactg gctggtgaaa
235681 aagcggatca tcaactggaa gtggactggg ttcgttgcag tgagcatctt cgagttggct
235741 ttcatccccg tgacgttccc catttacgtc tactactttg atttcaactg ggtcacgaga
235801 atcttcctgt tcctgcactc cgtggtgttt gttatgaaga gccactcgtt tgccttttac
235861 aacgggtatc tttgggacat aaagcaggaa ctcgagtact cttccaaaca gttgcaaaaa
235921 tacaaggaat ctttgtcccc agagacccgc gagattctgc aaaaaagttg cgacttttgc
235981 cttttcgaat tgaactacca gaccaaggat aacgacttcc ccaacaacat cagttgcagc
236041 aatttcttca tgttctgttt gttccccgtc ctcgtgtacc agatcaacta cccaagaacg
236101 tcgcgcatca gatggaggta tgtgttggag aaggtgtgcg ccatcattgg caccatcttc
236161 ctcatgatgg tcacggcaca gttcttcatg cacccggtgg ccatgcgctg tatccagttc
236221 cacaacacgc ccaccttcgg cggctggatc cccgccacgc aagagtggtt ccacctgctc
236281 ttcgacatga ttccgggctt cactgttctg tacatgctca cgttttacat gatatgggac
236341 gctttattga attgcgtggc ggagttgacc aggtttgcgg acagatattt ctacggcgac
236401 tggtggaatt gcgtttcgtt tgaagagttt agcagaatct ggaacgtccc cgttcacaaa
236461 tttttactaa gacacgtgta ccacagctcc atgggcgcat tgcatttgag caagagccaa

236521 gctacattat ttactttttt cttgagtgcc gtgttccacg aaatggccat gttcgccatt
236581 ttcagaaggg ttagaggata tctgttcatg ttccaactgt cgcagtttgt gtggactgct
236641 ttgagcaaca ccaagtttct acgggcaaga ccgcagttgt ccaacgttgt cttttcgttt
236701 ggtgtctgtt cagggcccag tatcattatg acgttgtacc tgaccttatg aactgccacc
236761 ataccacgtg tgtccctcgc aagcccttga tagatataca atagggaatg ggcgtccgct
236821 ccaccgtggt caaagacagg ggcaaagagc tcctaggtct atatatatat ctatatacat
236881 atttatatat attattagaa ctttacaata tagtatatac cattcattgt ttaagtttcg
236941 ggtaatactt tttttttcct tgtcataacc ccaaaaattt tcgatgcctt tgatataatt
237001 gagaacaaga agagtttgca ggtgacaaaa atcgatgatt ataggtgttg tgacgacaaa
237061 atgaacgcta atatatgggt ggctgcttca gatggtaatt tggaccgagt ggaacatatc
237121 ctccgcgaga gtaaaggcgc catgaccccg caatccaagg acattaacgg ctacactcca
237181 atgcatgctg ccgccgcata cggccacctg gatttgctga agaaaatgtg caatgagtac
237241 aatggagaca ttaatgtgtt ggacaacgac ggcgataccc cgttgcacca tgtggaggat
237301 gtggccactg ccaggttgat cgtggaagag ctgggtggag acttcactat caggaatgtg
237361 gagggccaaa cgccatacga ctcgttcgtc gagaacggtg aagatggtga gctaatcgag
237421 tacatgagga ttaagtccgg cgtggccgat gttcacggag tggacggcgt gcagggtgag
237481 ggtgtcatcg acagcaaatt gctggaagag ttcaaggaca acgtgagata caccttggaa
237541 aatgaccctg aggaaggagc cgatgaggcc actctgcaac gcaggaggca gttggaacag
237601 atcattacgg gagacaacgc tgaggaggag ttggaaaggt acatccgtgc tatggtcaga
237661 gagcagatgc tgggccaggg ctccatggcg ggttccgggg acgaaccaga ttccaagaga
237721 agaaaataac gacccagcac aaaggctctt acagcttgct aaaagaaatt gaacgcgacg
237781 ctacatgaac tacttctttc tcttacatag tcttttccct tatgtatctt ttctgtacat
237841 taatagacgt tcttacaagg taaaatttca ccgcgttttt aaatagaatg aaaaaaacgt
237901 tgtagagtga aagaaaagca acaaatatac agttcacaag gcagcttcgt atagtaatac
237961 agcacgaaaa acagctcata gaaatggtaa cacagaccaa tccggtccct gttacatatc
238021 caacggatgc ttatatcccc acgtatctgc ccgatgataa ggtctccaat ctggcagatt
238081 tgaaaaaatt gatagaaatg gattccagac tagatttgta tctgacaaga aggaggctgg
238141 atacgtccat caatttacct acaaacacca agaccaagga ccatcccccc aataaagaga
238201 tgctgaggat ttacgtctac aacactacgg aaagcagccc tcgcagcgat tctggcaccc
238261 cagcggactc aggcaagact acatggacac tgagaataga aggtaagctt ctgcacgagt
238321 ccgcaaacgg aaagcaccca tttagtgagt ttttggaagg tgtcgcggtc gactttaaaa
238381 gactgaaacc gctgggcatg ggcaagaaga ggaaacgcga ttcgtcattg agccttcctt
238441 tgaatctgca acaacccgaa tacaatgatc aagatagcac catgggcgat aacgacaacg
238501 gcgaggatga ggacagtgca gaggcagaat ccagggagga aattgtagac gcactggaat
238561 ggaactacga tgaaaacaac gttgtggagt ttgatggtat cgacatcaag aggcaaggca
238621 aggataattt gcgatgcagt ataaccatcc agttgagggg tgtcgacggt ggaaaagtac
238681 agtactcgcc caacttagct accttgatag gtatgcaaac gggctccgtt aatgacgcgg
238741 tttattcgat ctacaagtac attttgatca acaatctgtt tgttacggaa caaacagagg
238801 ctcaagatgg ttccaacgat gccgaagaca gcagtaacga gaataacaat aaaaacggtg

238861 ctggtgacga tgatggcgtc gagggaagta ctccaaagga taagcccgaa ttgggtgaag
238921 tgaagctaga ttcactctta caaaaggtat tggatacaaa cgccgcgcac ctccccttga
238981 tgaatgttgt gcaaaccgtg aacaaactgg tatcacccct accgcccatc atcctagatt
239041 atacaattga tctttccaaa gataccacct atggtgctac caccttggat gtagatgtgt
239101 cgcacattct ccaccagcct caaccccagc caaatttaca aaaagaggaa gaaacagatg
239161 ctgaagacac agcaaaacta cgtgaaatca caaagcttgc cttgcagttg aactctagtg
239221 ctcaaaaata ccagtttttc cacgaactgt ctttgcatcc aagagaaacg ctgactcact
239281 acttatggtc ttccaagcaa aacgagcttg tgctgcaggg cgaccaatac ttcaatgaag
239341 atgctgcaag aacgagtgac atatacagta acaacaacaa tgacaggtca ctaatgggca
239401 atatctcact actgtactcc caaggaagac tataacatca ctgtatgatc tcattctttc
239461 tgcaattatt gcacgcctcc tcgtgagtag tgaccgtgcg aacaaaagag tcattacaac
239521 gaggaaatag aagaaaatga aaaattttcg acaaaatgta tagtcatttc tatcaataac
239581 gtatacacgt ttttttaata ctatctatat tgtgtatctt atatttttcc atttggtagt
239641 atagcagcaa atatttgttc ttagacggcc accatccctc acaccaatgc ctaacgcttc
239701 ccaagtttac agatctacca gatccagctc tccaaagaca atctcttttg aagaggctat
239761 cattcaaggt ctggccactg acggtggtct tttcattcca ccaactattc cacaagtgga
239821 ccaagccact cttttcaatg attggtcaaa gctctccttc caagacttag cctttgctat
239881 catgagacta tacattgccc aagaagagat tccagatgct gatctaaagg acttgatcaa
239941 gagatcttat tctactttcc gttctgatga agtcaccccc ttggtgcaaa acgtcactgg
240001 tgacaaggag aatttgcaca ttttagaatt attccacggt cctacctacg ctttcaaaga
240061 cgttgcttta caatttgtcg gtaatctttt tgaatacttc ttacaaagaa ccaacgccaa
240121 tttacctgaa ggcgagaaaa agcaaatcac tgtggtcggt gctacttccg gtgacactgg
240181 ttctgcagcc atctacggtt taagaggcaa aaaggacgtt tccgttttca tcttatatcc
240241 aaccggtaga atttccccaa ttcaagaaga acaaatgacc accgttccag atgaaaacgt
240301 ccagactttg tctgttaccg gtactttcga caactgtcaa gatatcgtca aagctatttt
240361 cggtgacaaa gaattcaact ctaaacacaa cgtcggtgct gttaactcca tcaactgggc
240421 aagaatcttg gcccaaatga cctattactt ttattcattc ttccaagcca ccaacggtaa
240481 ggactccaag aaggtcaagt tcgttgtgcc aagtgggaac ttcggtgata tattggccgg
240541 ttattttgcc aagaaaatgg gtttgcctat tgaaaaactg gccatcgcta ccaatgaaaa
240601 cgacattttg gacagatttt tgaaatctgg tctatacgaa agatcagaca aggttgctgc
240661 tactttatcc ccagcaatgg atatcttaat ctcttctaac tttgaaagac tactatggta
240721 cctagctcgt gaatacctag ctaatggtga tgatttgaaa gccggtgaaa tcgtcaacaa
240781 ttggttccag gaattgaaga ccaacggtaa gttccaagtt gacaaatcca tcattgaagg
240841 cgcatcaaag gactttacat cagaaagagt ttccaatgaa gaaacatctg aaacaatcaa
240901 gaagatctac gaatcatctg taaatccaaa acattacatc ttagatcctc acacagctgt
240961 cggtgtttgc gccacagaaa gattgattgc aaaagataat gacaagtcca tccaatacat
241021 ttctctatct accgctcacc cagctaaatt tgccgatgct gtaaacaatg cattgtctgg
241081 attttccaat tattcatttg aaaaggatgt tttgcctgag gaattgaaga aactatccac
241141 attaaagaag aaattaaaat tcatcgaaag agctgacgtt gaattggtca aaaacgctat

241201 tgaagaagaa cttgctaaaa tgaaattata agatcttcct ttttaatgaa tatattttat
241261 cttatataac atatagaatt ctataattaa tagagagtat gaagtacgat tatccttaat
241321 gttaatcaaa gtgaatcatg attgcgagag taaggaacca taaaattaat cttcatctat
241381 gctaataatc tcagaattct cgttcccagg gtcttcttct ttagatacta atctaacttt
241441 cccgcccttt tcaactgata tttcaaaccc agctttgctc aatgcatcgt cttgaacaac
241501 gtcttgagct tccatttttt tgacatattc ctgattcttg gcattattct ttaacaaaaa
241561 cttcaagcaa acaatagatc tttccttgat aaacggatca ttatcatcga tgacacaatt
241621 ggaaagaatt aatgcaagac cacctaattc tcttatttga tcttggattt caggatgtgc
241681 atacactaac gatgccaata tttcgattat tactaattta caatggggaa aattatcagc
241741 cctaataaag gattcgttgg tattaacacg gttgaaaatt aataaccggt cctcgatttt
241801 gttaccttca tgatctgtga ttttaatttc attcaccgta tcaccgtttc ctttcaaaaa
241861 attaatcctc aacaaattct tttggaaagt atccaagaga gataccaata ttttcacaga
241921 gtcatacgaa attaagaatt ttctaacatg attgaatttg cacaagtcag aaattatgtc
241981 caggcaagaa agtaaagtgg agtgtaattg gatggcaact gaaggagcat aatcgttaca
242041 acttttattg aaaaaggcag gaatatcctt gcatgcaaga cgttgcagaa tggggtagca
242101 ccatgacata attgcagtca actcataggt attccaatct tctttactcg ttacaaccaa
242161 ttgcaatatc tgcaacaaag tggtgaattt tcctggctgt ctttcttcta acccattgag
242221 gtatttacca aaagaagcag ttgtaagact tttgctaata cattctaaga cagataggtc
242281 aatagaagtt atttccaatt ttgaatcatc cagatagttt ctgcccttct caattaaaaa
242341 ggaattctct tgatcctgga tgaatttggc tgtcactata tcgaaaaata ttctatccct
242401 aacacaataa tagtctttct ggggtgtatc attatccctg aaaaattcgt tggaaaattc
242461 atcattgttg agatacgtct tgaaaagaaa taaatatggt agccagaaat gctcttcttc
242521 ttcaccgtct tccaatttga tcccataagg gtattgaaga aattgtaaaa gagctggaaa
242581 agtagtcata tccactgtag agtagttctg ctgtatattg aaaaggcacg ttaaacatgc
242641 gatgtataga tgtttgatta gcggagtgtc aacgacatac tcactagttg catgcagaaa
242701 accacgaatt acgaggtttt gtagtaaaag cttttggggt atttcctggt tttcagcggc
242761 aaggtttctc atcattaaga taaggcccct cactgttctt gtatacggta ttgagctatt
242821 ctcatttaac gacgctggca acagcattgc ttccaaggtg tctcttaatc gttcccaaaa
242881 aagctgggag ttagccattg atttcctatg ttcttcatca ttaaacgtcc tttgaaatat
242941 acggttcagg ttttcgacga ttggctcata atcctcgacg caacatgggt cgttcttcat
243001 catggtaatg attgaatcaa ataaattgaa ttcatctaga aaattgttca taggcattgt
243061 tatttagtta tcttcttttt tgatgaatca ataattttat aaggcttagt gtaattgttc
243121 tcccgatgat agtatgaaac ggtagtcgat aaatttgtgt caaatttttt cctcatccca
243181 tcaattcttc attttttccc ttcctatttt ttttttact gaagcgatga tgccttatat
243241 ataaggaaaa tagcctcgtc agtcaaagga cgactcccgt aatggtagcc tcagtttata
243301 ttaggcgctc tcatcatgat gtatatcgat attggcgatg atattttttt ttgttgatat
243361 taaatacaaa tgtaatgctt tatatagttc tatacttggg gaatacatga tgtgaatgtg
243421 cataaataga ggacagtgaa tatttttcaa ggaagctctt tctcattttc ttcttcatca
243481 ctggaatcat tttcattaga caagggtaat ttgttatcct tcccattgaa accaatccat

243541 ccttcctcgt cgtcagattc catgactaca tcgttctctt cgttctcttc gttatcttca
243601 tctgcgtcat cttttaatag aacctcgtca tcatcagccg cgccatcttc catggaccca
243661 tcagttgata ccaaaaatct ataggtgtat ttgttatcag tagtggtatt gactacttcc
243721 ttagccactc taacaataaa tctttgtatc gacctcatag ccgtcgagaa gagatatttg
243781 tgttcattta tgtaaccacc gctcgcagat aatagagctt tgatccaaat taggttaaac
243841 tcaatgtgtt gggattcaat ggcaaaatca ccgatgaact tcagaatcct cggtaaatat
243901 attgcaggaa tattacttgc aaccaagggg atttccttaa taggtatggc ttcatagact
243961 ttattgatca aatattcttc atttaacctg aacgccatta ctaatgcatt taaaaactgc
244021 ttttctcgta gcgcctctac agtagaatgg ggggtgacgt ccacatccag atcaaaggga
244081 tcaaataata tcgtgtcatt ggtggaatat atcaataaac cttccgttga agcagcggca
244141 aatgcattcg ccgttgggga gaattgcacc gaagtaactc taacctctgg tctcattttt
244201 cttgtggaca ggtcgccacc tctttgagac cctggtaaag aattatcaat acgatcctcc
244261 aaatctgaat tttcgcctgc atcgtcaatc aaatctaatg aacctgcttc agtcattttc
244321 ttactgttta aaaattcgag agtaccattc aaagccatgt ttctggacac aatgaatctt
244381 tttaacaaga cttcatttgg aacatcatat agacaaatgg agttattatt accaccagcc
244441 acaatagcca taccatcaaa actgtaatgt attgtagtaa aaaatttgga tcgttcagaa
244501 tttttggcag tgaacctatc ttgattaaac ctaccagata ttatatcctt tctacagtca
244561 atgttgccca cctgcttggc atcttctatg ttgaaaatgg atatttgacc ctttaaggta
244621 gatactgcaa cttctttacc atctggtctc attgataagg ctaaaacatc ggaataaact
244681 tctataggtt ctacttgttg gcttctacca aatatggacc agattctaat tgttttatcc
244741 catgatgcag aagctaagac actgttctct tgactaaatg aaagacacga aacagggcct
244801 tcatgtccgg acaaagcatc taataattga ccagtttgca cggaccaaac atgaatgtca
244861 aaattgtcca gggacccggc acaaaccact tcacctgatg gatccaccgc taaacaattg
244921 aattggattc tttcagtacc agtgaatgtt ctaaaattac gatacctgat taagtcccac
244981 gctctcaccg taccatctaa cgatgatgag aacatgacct gaccccttt cgcaaactgt
245041 acagcagtaa ctgaagaggt gtgttcttca aaagtggcca aacaaaaccc tgatgtaatg
245101 tcccaaactt tgattttccc atcttcggat gctgtcacta cacgtgaacc atccggagag
245161 tatgcaagac tatttgtgga atcgaaatgg ccctgctgct tcaagatata cgattccgat
245221 tgccattcgt aaactagtaa ttggcccagt ttgctggaac caaacgccag ccattcgccg
245281 gtttggttga cgctaacggt gttgactggg ttttgcccca tagaaagctg ttgaatcaaa
245341 gtgaaatcag gcaaatcgta aagacggaat tccccactag taaatccgac agctaaaagc
245401 cttgttgctg gatggaaggt gacacacttt actttggctt ggtttgcgta aaaaaaatgt
245461 ttctttgtga ttctccagct gtatttcgaa atatctactt cttcttgctt gtcgtcgtct
245521 tcactttcat tgtcgtcgtc atcggatggc ctcttggtaa attcccagac aaagacagca
245581 ccgtctttgc ttacagtgta gattttttcc tgatcatgac tgaagaacgc acccataacg
245641 tagtctctgt gcccattaaa tgttgtcgcc gcaaggttct tttcctctga atctacggac
245701 catatttttg cgcttaagtc tttggaagtc gtaaggataa atctggaatc ttgtgaccac
245761 gtcaaagacg ttatgtcttg aaagtgtccc gcatgcaccc tatggcggac gaagggagca
245821 aactgtctgt ctttattaac atctggagtc ttccaaatct gtaaaaacct gcctgaggct

245881 aatgcaaaga gtctcccatc agggctgaac ttcacagcgg agcatttttc tttgaagttg
245941 aaatggtgaa gcacgttacg ggctttgaaa ttgacaagga tggcgcgacc gtcctcgtca
246001 atagaaatca gcaatgtgcc ttgtttgttc agatcaatgg cagcaatatt tttgcgatgc
246061 tcgtattcaa acgtgaacga tttgttgttg attaagtcaa acacgctgac cctattcccc
246121 accggtgaga gtagttgctt gccatcatcg gaaaaggtga tgttaccttg cctgtagacc
246181 gtacctaaaa ggttagagaa cttgaaatcg gatttcattg ggatgcggtt caatctgcgg
246241 tctgctctcc tgtctacctg ttgaaaacga tactattatt actattattg atattatact
246301 agttattgta catacctaga actatactcg aaactttttt ttttttcatc tttcagctca
246361 tcgcaaaatt tttccaaaaa atttcaaaaa gaactcccgt cgcatgtgat caaggttaca
246421 ggtgcttgac ataatcataa tgattacgag tcgaagccgg ccctgacaac agcttctctt
246481 gccgtagaat tgatgtgttt aaaccggtcg ggacctatgt gggcaccgcc gaaccaacgg
246541 gccaccacaa cgatgacgtt ccacacatcc atgatggtga tgaggtgcag cattctggag
246601 ccggcggccg tttcaccgtc atcatcggaa tcttgatatg ttgccgcaga gccatcctgc
246661 ttgattcgcc atgcactcat gacatggttt gccttacgca tcttggagtc ggtcttcagt
246721 aggtctagca tggcaaacgc ttgttcctcg gaggtaacat gtgctgcaaa ggccatgaaa
246781 gtcgagcctc tatcagtaat ggggtccgac gcggtccagc cctcgaaggg gtctgtggga
246841 atgtcactct gctggaccgg ttctgtctcc tcctctggtt caacgtacaa gacaccgtcg
246901 agttctgtga ggaagtcaaa tagacagaca gatccgcggt ggaaaacaga gtccatcact
246961 tcctggaaca aatgctgaag gtacttggta tcgtagagat cgcgcttagc caaagaagtg
247021 cagacaccaa cttcgatgac attaggagcc tcctcggagg ggtagtgtgt cgggaaggag
247081 atctgcagtg tcatgtattc atgctgcggc actttcacaa cgatgatgct tccgtcttcc
247141 tgcttcttgg agagaagatc cggatagatg gcctcgacgg cctccagttc ttcgaccaac
247201 tgttcgtgat cgtcatccat tgagcttttc tttcctctct cttttttttt tcttgttaca
247261 tattcctata tatatgtata tatatctata tatatatata tcccagcgta tatgactgca
247321 caagacgcaa ttttcttgaa cggtttttttc ttttcagcac tgcgaaagag agagcaacgg
247381 taaagagcat gagccaattt gaaaagcaga aggaacaggg caattctttg ttcaaacagg
247441 gcctgtatcg cgaggctgtg cactgttatg accaactaat tactgctcaa ccgcagaacc
247501 cggtcgggta cagcaacaaa gccatggcgc tgatcaaact gggtgaatat acacaggcta
247561 ttcaaatgtg ccagcaagga ctgcggtaca cctcaacggc agagcatgta gctatcagat
247621 ccaaattgca atatcgtcta gagctggcac agggagcggt aggttcagta cagatccctg
247681 ttgtagaggt tgatgaacta ccggagggat acgaccggtc ctgagaaaag agaaagggaa
247741 cacttctaat ttatctttaa aattatatat atatatatat atatatgtgt gtttgtatac
247801 tctgtgggta tttccgtgta tatggttaat aatagtagta tcttgtcagt tttttttatg
247861 tttttcttcg cgcgtcaact ttctaccaag agaaaaacaa tataaggtct ccttactcta
247921 taggagaata aaacaaacaa aaataaaaag cacatcgtag cgccaagaaa atactgcaaa
247981 taccaaatac cacaataata ctacaattat ctatacacaa gtgttttgcc gcttaaaact
248041 tcgattttca tagtacgaaa ctatacaccc ttggtttttc tcttttctaa atacatatct
248101 accttgtaaa gaatttcccg cacatctttg cgggcataca gttcatgtat tggcaactaa
248161 cggaactaag gcaacatatc ttgcatattg caatgttcac tatatagatg aaaacttata

248221 tctaggtttc acgacacgag aataactaag aagcacgatc catgatatag aaaaatcagt
248281 tacgacgaag cacggcaaat tagccgccga agaccgatat ttttgccacc ggtcacagtt
248341 ttcttttcca cggagctctt cgcggttttt ttgttccgga ttttttttac cggctcttag
248401 atcgattgaa gttttgtcca actatccact atggatatgc gttttgttga ttaaccttaa
248461 ataacacgta tttcgcattt tccaaaagcc ttttttcata actacaaact aactattttt
248521 gtttatttta catcagtaaa ttatgcgcag aaaatatgta aggctatata ctcaatatag
248581 tggaagagcc tctggctact taatctgggt tcatatattc tgtcagtggt atagtaaatt
248641 taaatagtga atttgggcgc atgtagagag atcttttgat taataaccca gtatttgatt
248701 cttgaatgtt atttgtctct tattaagtat ttcagtttga tttatatatt ttgaaagtaa
248761 ttgttgcttt accttccaaa caataaaaaa atataaaaaa atgtaaaaaa tataataaat
248821 attaaataaa atactacttg tttgaaaatc agagaaaatc caccaaaata tcaatcattt
248881 caaggatttc cgaaccaagt tcgagatagt cctttaaggt cagtaaaatt cagttgcacg
248941 tataacagat atttttcatt ttgttccaat ttaaaagtcc cccattttta aaattattga
249001 aaataaaaat taaaaaatta aaaggaatct ctctatgtca ctttaaaata aataaattga
249061 aaatgatata tcgttaaaag agcttgcctc gtccccgccg ggtcacccgg ccagcgacat
249121 ggaggcccag aataccctcc ttgacagtct tgacgtgcgc agctcagggg catgatgtga
249181 ctgtcgcccg tacatttagc ccatacatcc ccatgtataa tcatttgcat ccatacattt
249241 tgatggccgc acggcgcgaa gcaaaaatta cggctcctcg ctgcagacct gcgagcaggg
249301 aaacgctccc ctcacagacg cgttgaattg tccccacgcc gcgcccctgt agagaaatat
249361 aaaaggttag gatttgccac tgaggttctt ctttcatata cttcctttta aaatcttgct
249421 aggatacagt tctcacatca catccgaaca taaacaacca tgggtaagga aaagactcac
249481 gtttcgaggc cgcgattaaa ttccaacatg gatgctgatt tatatgggta taaatgggct
249541 cgcgataatg tcgggcaatc aggtgcgaca atctatcgat tgtatgggaa gcccgatgcg
249601 ccagagttgt ttctgaaaca tggcaaaggt agcgttgcca atgatgttac agatgagatg
249661 gtcagactaa actggctgac ggaatttatg cctcttccga ccatcaagca ttttatccgt
249721 actcctgatg atgcatggtt actcaccact gcgatccccg gcaaaacagc attccaggta
249781 ttagaagaat atcctgattc aggtgaaaat attgttgatg cgctggcagt gttcctgcgc
249841 cggttgcatt cgattcctgt ttgtaattgt ccttttaaca gcgatcgcgt atttcgtctc
249901 gctcaggcgc aatcacgaat gaataacggt ttggttgatg cgagtgattt tgatgacgag
249961 cgtaatggct ggcctgttga acaagtctgg aaagaaatgc ataagctttt gccattctca
250021 ccggattcag tcgtcactca tggtgatttc tcacttgata accttatttt tgacgagggg
250081 aaattaatag gttgtattga tgttggacga gtcggaatcg cagaccgata ccaggatctt
250141 gccatcctat ggaactgcct cggtgagttt tctccttcat tacagaaacg gctttttcaa
250201 aaatatggta ttgataatcc tgatatgaat aaattgcagt ttcatttgat gctcgatgag
250261 tttttctaat cagtactgac aataaaaaga ttcttgtttt caagaacttg tcatttgtat
250321 agttttttta tattgtagtt gttctatttt aatcaaatgt tagcgtgatt tatatttttt
250381 ttcgcctcga catcatctgc ccagatgcga agttaagtgc gcagaaagta atatcatgcg
250441 tcaatcgtat gtgaatgctg gtcgctatac tgctgtcgat tcgatactaa cgccgccatc
250501 cagtttataa ggggagtggc agcggcggta gacactgcgc tctataagaa tacttgcaag

250561 ggtcttgtct attgtataat tcgctagtat ttgttttgca ttgtactctt aataccccaa
250621 ccaaaaacaa aatagtgaga gtaatggtgc gttttgtttc aattttaagt ttattcggct
250681 gcgcggcgac gcttgtcacg gcccatgatg acatggacat ggacatggat atggacatgg
250741 atatggacat gaatatcgat acgacaacgt ctcaatccat agatgtctca tccacggctt
250801 caatcgtccc cgtgccacat gaaccaaaac atttgcatgg ccttcctata ctgcaatcgc
250861 cctcgcttac ccctgcggag agattgtact gggaaaacta caacaccaca acctacttta
250921 ctacacaggc tgggaatagg tctgcccttc gctaccacat tattacgctg ctcttggttg
250981 catttgtgct ctaccctgtg tccctggcgc taagcgccgc ccgttctagg tggtacttac
251041 ccctgctgtt tgttaatcta tgcatttgta tttcgtccgt aatggcattg tccgtgttca
251101 aaaatacttt cccggaagaa gactggtatg cgcataatat ctatggcacc acttctgtgc
251161 tacttctcgt ttttatgctt gttcacttct tcgctgcggt gctttctgtc cccgtctcat
251221 tagcatcgaa aaaggagtac cgtccggttg acaccatccc tctgaatgat cttgaatcta
251281 cgcccgtcat ggtgaatagt gcacgtggct ctccaagtcc ttcttccaac agagacacgt
251341 tgttctcgct ctcttcagac accacgaccg ccacggccac caataataat aaacggagac
251401 gcgctgaagg cgaagacgag ggtgataaca cctccaacca cgacactttg cgcgacgaag
251461 actacgataa tgatgacgac gaaattgctt ccattgaagc gccacctctg cttcctcaag
251521 acatacccgt tttccgaatc ttgtttacca acacgaagta ccagatgctt gccgcgcacc
251581 tctcgtgcgt cgccaacgtg gtctttcaca tgcttaccta cccgctattc atgtacatct
251641 ttgtagacct aatcatcggc ttcgctgtag gtaacttgct cggcaagggc atccgcatct
251701 ttaatctctt ggcccactgg attaagggcg gcgtattttt tactctgggc gttgtctctt
251761 tagcaagata ctgcggtttc gcagctaagt acggctgggc atggaacaac atcagcttca
251821 cctctcaact cacacaaacg cgttcctcca atcttctttt ccggtttgct cctgcgggga
251881 ctttcaccat ggaattcgtt gaatccttcc tcattttctt ttacgggtcc accaacatct
251941 tcttggagca cctggcagga aacggcggcg catggactgc caaggattta cagcatgtgt
252001 cgatcgcgtt tatgttcata ggaaccggat tgtgtgggct actcacggag tacaagctca
252061 accattggcg attcgagcat gcccgcaaac ggccacagac cgatgtagtt gctgccacac
252121 cggggtactc tccaaacccg ttccccgctt tcaccatatt ttggactggg attctgatgt
252181 cccagcacgc acagtcctcg caattttcta ctaccattca cacgcaatgg ggatacttgt
252241 tgtcctatgg gtccttcttc cgtctgctaa catttttgat tctgtttttg gtgcccaaca
252301 ccaacagtgc cgcatccaag cctttcacgg agttgatcac ctcgttctgt ctcctctgtg
252361 gtggtctggt atttatggag tccacggatc agtccattga agccatggaa tatagggggt
252421 ttacccccat gttcactttc aacctcagtg ttggattcgt ttctttgttg atggcttggg
252481 aaatgatttt atttatttgg aaagactggc tcatcaaaac caggaaaacc agtctttaac
252541 cgccataatg gcttttcttt tttctttttt ttttcacccc cccctcatct ttgagtgaga
252601 agcttatatg ctttactttt cctcacctac tttttaattc ttaattatta gtttatccag
252661 cattcagtca attctgaatg cgttatcgct tcctcaaaat ttataaaaat gaataagcaa
252721 gatctcattg gaattgagtt caatgacgat tattgagggc ccatacctga taaaagtgaa
252781 tatattatgt ataatgactc tactggaggt tctctctgag aatgtaaaat aatccacaaa
252841 agacatgaaa attagttact ctacataatg tataataagt tcttccttcc ttttttcttt

252901 ttttagtttt tctactcacc atcctaccta cgccaagctg ctctactgtc ttggatctgt
252961 gaaaaaacgc ctagaaacat gtataaaacc taccgcttta aaccataatt tcaaatcgaa
253021 atggcactat ggccgagtgg ttaaggcgag agactcgaat ggaataaaaa gttcggctat
253081 ctcttgggct ctgcccgcgc tggttcaaat cctgctggtg tcgtttaatt ttttttaaat
253141 aacatcgttg attaaaacaa tcctattcac tgcagtcaca gacttagcac atttacaacc
253201 agtacaagat caatcacacg cgttgtggta ggaactttga cgatgatact gcgttggctc
253261 ttgaacgcat agtacatttg tcgttcacaa cggaaagtga aatactgaaa aaaaaagaaa
253321 aattttctta gggactcatc atcaatgtaa acacctctac agccgtcaag actagcaaag
253381 ctcacctcac aaaacgacat gccgcgcata aagaccagaa gatccaagcc tgcacctgac
253441 gggttcgaaa aaatcaagcc aaccctcaca gatttcgaaa tccaactcag agatgcccaa
253501 aaggacaagt cgtctaagct cgcagcaaag tccaatgagc agctctggga gataatgcaa
253561 ctccaccacc agcgctctag atacatatat actctgtact acaagagaaa ggccatctcc
253621 aaagaccttt acgattggtt gataaaggaa aagtatgctg ataaattgct aattgccaaa
253681 tggcgcaaaa ccgggtatga aaaactgtgc tgtctgcgct gcattcaaaa gaacgaaact
253741 aacaacggta gcacttgcat ctgcagggtg cctcgtgcac agttagagga agaagcacgc
253801 aaaaagggca cacaggtgtc cttccatcag tgcgtccact gcggctgccg tggatgtgca
253861 agcacagact aacgactata tgcacgttgc agtattgttc tggccatgta attatctctc
253921 tcccaacgtt ttagaagagg ttttcgtctc cttcgtcaaa gacgaccact ttcccatttt
253981 ggtcattgga acgcgaaaaa acaaaacaga cgcgagaacg cgacgcgaaa aaatatattt
254041 ttccgttttt ttttttttc ggaacaaaga taacgcgtta aaacctgtgg atgtaataag
254101 gatctgaaac actgactaaa acgcgtcctc atttaaaatc aaaaaattac ttcttctcct
254161 cccttacggt gccctttgat ccttctcaaa ctttaaatat atgttgcatg attaattgaa
254221 aatcatatag agtatatctt aacagaggtt ccaacgcttt aaatcgtatt ctattctttt
254281 cccccctttt attatagtta catctactat ttgagcttct tttattgacc gaaactacct
254341 cttatttcga agcaaaaaaa aaaaaaatca acaaaaattc caacttttca atgatgaatg
254401 aagacatatc catcattgat ggccataata gttttttaac ggaaaaaagc accgtgctat
254461 taacccaagc caagagaaca ctagaagacg aaaaggaaat gattactccc ccgagctcaa
254521 ctgtgagaaa aacaatgaag gaagtaaata agaggccgtc gcatcccctc tcaccggatc
254581 actcgtcccc aattgctcca tctaaggcca agcgccaaag atcggacaca tgcgctcggt
254641 ccaatggtaa cctaaccttg gaagaaattc ttcaatcttt ggaaagaaga agaataaatg
254701 gtgaactcgc caagaaacct ccatattcgt atgcaacttt gatttgcttg gccattttgc
254761 aatctcagga gggaaagcta acgctatccc agatatatca ttggatccac gttcacttcc
254821 cttattacaa gcagaaagat gctagttggc aaaattcaat aagacataac ttgtctttaa
254881 atgatgcgtt catcaagact gaaaagtcct gcgatggtaa gggtcatttc tgggaggtca
254941 gaccgggtgc cgaaacaaaa tttttcaaag gtgaaaatcg tggttatgaa tttgtaaagg
255001 actccttaca agacattggg aagtattttg aaatagattc tacacttgat gaattagaac
255061 aagttgagag tggagaaggc aatgatgatc ttcctgacga ggaagaaaga gaggaagcag
255121 ggaaattccc ttccattgaa attcaattga actcctcccc tatactgaga gtttcccagt
255181 tacatcacat accgcaattg aaaacagaca acagtgtact gaaccctcac gaaaacctag

255241 aatcgatgcg gaacatgata gaaaacgatg tcaacaatat agattccttg gaacctcctt
255301 atgtcatgaa gaaatatcat acttctttag gcttaccgtc gctggtgaat gccaaagatc
255361 atttccaggc gggtgtgaaa aacaataata tcacccaggc aaatagattt aatacactcc
255421 ctataactag cgcaaagtct cctcagaatt tcagaaaata tttcacctca ttcaattcaa
255481 attttgaaga tttatctcca cttcgaagta atgtaggggc tggttctcta ctcgacccac
255541 ttccgtattc cccattgaag ctgtacgatc agaaaaatct tgcgctcatg tcgaaaccac
255601 aatctcagca atcatattcc aattctcaac ttccacctcc accttcctct catggttcgg
255661 acttacttaa aacacccaag atgaggcatt ccgatggctt agagaaaacc ccatcgcggt
255721 tgataagcac acctaaggac ggtaactcga ttttgaggaa atggcagact ccttcacacc
255781 tttttgaaga tttgtactgt tctccgctat ttagagctat agagactcca atcaggtata
255841 tcacgacgcc ggggggcact ttggaaaccc aaatttcacc aagaaagtcc tctgcacccg
255901 atgtcctcac aagcgcaacg aattccaaat ttgcttcaag cgggctgttt ggcgtggatg
255961 tttattctgt ttggaagcgc gcaactgaaa agatttctga tggtaacaat actacggata
256021 gcaatcaaaa acatcatcct tatcataatc acccttccaa cgatagcggt aatgaaaaga
256081 attgactttt tttttttttt tttttcattt tcttttcctc accgaaaact aattattctt
256141 aatctgatgg attcatcgca aacggttgga tgagtcctaa actgggctgg agttatcata
256201 tcacatagca ttttcttttt aatcttattg tattgtcggg tcaactaact tcttttcccg
256261 ctcttctatg gtataatcaa ttgcagaaga ggcaggaaaa aattgtaatc ataacaaaaa
256321 tgatacaaat ttttttcta agttttatat tttacatatc tttatttttg atccttacgg
256381 atttttaata cagtatttag ctgaagtttc cattcacttc acttaccttt ttgatctctt
256441 ttatttttac ttgccccgtt gccttgcatt tttcaataga aagtgaaaaa ctaagtacca
256501 cttgagcaat gccacattag aagaaaaacc atccgcaagt gagcatcaca gctactaaga
256561 aaaggccatt tttactactc atggaccacc aaataaccac tgcaagcgac ttcacgacta
256621 cttcaatacc gagcctgtac caattggata cacttttgag atgtcacatt tgtaaagatt
256681 ttctaaaagt ccccgtctta acaccttgtg gccatacatt ttgttccctt tgtattagaa
256741 cacatttgaa taaccaacca aattgtcctc tctgcctttt cgagttcaga gagtccttgc
256801 tgagaagtga gttcctggtc agtgaaataa ttcaaagtta tacatcccta cgatcttcct
256861 tactagatgc actaaggata ccgaagccta cccctgtccc tgagaatgag gaagtaccag
256921 gtcctgaaaa ttcttcatgg atagaactca tatcagagtc tgaaagtgac agtgtaaatg
256981 ccgctgatga tgacttgcaa attgttgcaa caagtgaaag aaaacttgcc aaaagatcca
257041 tgactgatat attaccactg agttccaaac catccaaaag gaattttgca atgttcagaa
257101 gtgaacgtat caagaaaaaa tcaaagccaa atgaacaaat ggcccagtgc cccatatgtc
257161 aacaatttta tcctcttaaa gcccttgaaa aaacacattt ggatgaatgc ctaactttac
257221 aatcactagg caaaaaacca aaaatttcta ccactttccc tacagagtca aatccacata
257281 acaaaagttc atccagattc aaggtacgaa ctccagaagt cgacaaaagc tcatgtggtg
257341 agacctcaca tgtggataag tatttaaact caatgatgag tgcagaacac caaagattgc
257401 cgaagatcaa ttttacgtct atgactcaat cccaaataaa acaaaaactg tcatcgttgg
257461 gactgtcaac taatggtact aggcaaaaca tgattaaaag atacaatcac tacgaaatgc
257521 tttggaattc taatttttgt gattctctag aacctgttga tgaagctgaa ctaaaaagac

257581 agttgttaag ctgggatgtt tcacacaata aaacccccca aaatagtagc aacaagggtg
257641 gaatttctaa attaatgata atgaagagta atgggaaatc ttcttcatat aggaaattac
257701 ttgaaaattt caaaaacgat aaatttaata ggaaaggatg gatggttatg tttcggaagg
257761 attttgctag gcttatcagg gaagcaaaaa tgaaaataaa aacaggttca tcggacagtt
257821 caggttcagt gggacattct aatgatggag atggtgttga aaaagttcaa agtgaccagg
257881 gaaccgagga tcagcaaatg gagaaggatc aggacactgt tatcaacgaa gatagagttg
257941 ctggtgaaag aaatttgcct aacgaagatt caactgatgc tgacttatca agagaattaa
258001 tggacttgaa tgaatatagt aaagacccac ccggtaacaa ttaaggtgtg tatatgtaat
258061 caggcctgtt tgttagcttg tgcacatttg ttaattaata atttatttaa aaatgggcaa
258121 aatataaatg cataaaagat gacatagaga tatgtgcaca aagctatgta atcatattta
258181 acattaatat atattatttc gttattcaca attcgtcatg aaggccagcg tagtttacgg
258241 ttcctgctac tgcattatca tcgttgacaa tgtttgtatt cgaattatta ttcaaaggtg
258301 ctgcaattgt ttgcggaggg gccactggat taggcaaaga agtgatctcc acgttaagtg
258361 ctgaaataac ttcactagat agagagacgt cttcggtaac atctaccgta ccagatgcct
258421 gcatttccgt cgtaagcatt cgggaaacag gagtatttac attttcgctg attttgtggc
258481 gtaccccaaa cgcttcagat gaatcttttg atgactgtac ttctatcatt atttctttct
258541 gtacaaatcc tgatgcatca attgaactag cagatatttt gtgagggcta cttacaaaag
258601 atggtgctgt tgtggttaca cctggcatcg gctgagctac tgtgcttgac gcgctctcca
258661 cggataaagt tcccttgcct atatcagaca gagcgccaat tgaagcacca ctactactaa
258721 agatcataga tgccatagaa gtcccttcaa tatttgtgga ttctgatagg gaagattgac
258781 caacactcgt cttaggtgat tgtacagcat cagtgaaaac actagtgggt tccgaactgt
258841 cagccaaaac actagtgggt tccgaattgt cagccaaaac actagtgggt tccaaattgt
258901 cagccaacac actggtgtat tccgaattat caatcaaggc actggagcgt tctgaatcct
258961 cggtagagaa agcaattact ctcgaagtct tcgtttctgt ttttgagaat gagctagttg
259021 ccgtctccac aatagctgat gaaacaggtt cagagatgga atgaaattta gaaaaagtgg
259081 aatctgagat tgaagggatt gattcagaag caagaacaga agaagttaca gtcgatactt
259141 ctattgactg gaagccagag gttatagatg aatcagaaaa tgtactcgga tgcgaggaaa
259201 tagagtcctg ggatgctgat gttgacaaaa tttcttcatt tgggatcaaa ttagacgcaa
259261 ctgagtcaaa ttcggatgct atagaaatag aatgcgtagg tgtttcaacc atacgctcta
259321 attctgatgc tatgggatca atttcggaga cgggcgaaga aaccatagtg tctgtgggcg
259381 tcaacaacat tgaagaagat gttgtaacat actgctcggc ttctgcattg gaatctgatg
259441 caggtatatt cctatactca atttttgtaa tgacttcagt gattattttt gtcttataaa
259501 tctctctcgt ggatgccggt tttgagttgt ctagaaaaga tgtgttggct gcatcattgg
259561 tggggtttgt ttgtgtggtg gtaaccgcag tcgcggtact tgtggataat gctgtagtag
259621 ggctagttgt taacgatgag gaaagagata atgaagatga agatgaagac ggcatgtgtg
259681 atagcgaagg tgattcagta cttaagagtt ttgaaacgct ttctctatct gttgttattt
259741 cgctttctga ttgttctact attaacgagg acaatttctc ctccaagtca ggagtcggct
259801 caggatattc tgtgatagaa tttgaagatg aggataacaa atctatcgac tctgaattca
259861 gtggaagcga tggtttttcg acaatattcg gttctgtagg agaggctagc ggcttcgtaa

259921 tttcagaggt aggtgaagac acaattggtt cattagtgac actctcttca tagaaagaag
259981 aagaagaaga agaagaagaa gaagaagctg atgttgggat atctgcagaa gtgaattgtg
260041 aatcagctga tgacgatatc caaacaaatt tttgctcatc ttctgtggtg gttctcatag
260101 tgctagtgtc aaatgaggaa aattcagtct ctatatcgct agcaacatct gtagcgctga
260161 tgatatctgc actttcagta atgcctctgt aatcgtcatt aatagtgaaa taagaagaag
260221 atccttggtc tatggtagtg ttttccttga atttcgactt atatatatca tgagcatgtt
260281 tgtaaataaa gccgtgcaca ttatgtttgt agcctagttg aatagcgcca gccaaaacca
260341 ctaacaaaac aaaattcaag aaaaatttcc aaatatttcg taaaattgat gacgatgtac
260401 cggaaagctt cagaacgttg atcgtgttcc ccatagaagt gcttgccaaa tatctagaat
260461 ctggagaaaa agtaactttt gttatagtag attcatgtac tcttggaacc aactgaacaa
260521 gctttagctt ttcaacgctg acgatggcaa ccaagttatc gttagatgat aatgcaacga
260581 acttcttatt tggggagaca tccatcgatg ttattccatt caaatcgaac atgaaggatg
260641 ctgtttttaa aattttaaca cctttggaac tcacacctag ttttaacaag gaaactccat
260701 cagagttcga taatgatgct gccactaata gcgtattgtc attgagaaac ctgatattag
260761 aaagaacaag atttttatca aaatccgttt tccttgctac aaaatttccg tccctaacag
260821 acgctacttc caaagaatta gaagtgatat atgacaaaag aataccgtta ggggcgaagt
260881 gcaaatcatt aacttcccta cccgtttcaa tttcgtaatt ttccgtcaaa ttacgcgggt
260941 caatgattct tatcacagtg ggcacttcag aggacgcaat tgcagctact gtggcccttt
261001 gattaatgta aacaaactta gtgaaaacgg atgggtctgt agttctgttg aaatcaatac
261061 tggccacgaa ttttagatgt tcttgttcaa acacaaactt cctcaagtgg tggttaacgt
261121 tctccatgga ggaatcatta aacagttcat tgcaaccaac tagaatcagg ttattattac
261181 aatccaatga agtgggcgca tcgtcaatcg aatctaatgt gatctcacga aaccttctaa
261241 tgtgtttctt tttctggaag ttcaccttta aagctgtaat tttgtttgga aacgaggaat
261301 tgaactggcc tccaccacca gccaccagta gagtgccttc atttataaat ttagcaccat
261361 atatcggata tccgacgtca tagtttgcag agttgccact cattattatg ctagtttttc
261421 taagtttaaa aggagagaaa aaaaaggaag agaggacgta tcgtagggaa ttgtgatgct
261481 tgttgacaaa cagcctccgt cgatcgaagc agctataagc gtaggttact tgaagcgttc
261541 acagttctca tttacactgt acaattctcg agatataaat gaaaattact acaaccaaga
261601 aaaaaggaca ggagggctaa aaggataata aaatagggca gttacaaatg tggtcaaaaa
261661 tactagggct caaaattgct aagtaagggc ttgacagatg taacataata cgaaataaac
261721 tggatatcgt ttaataatta tcaaggtgat ttaaatcgta aatacatagg ctgggccata
261781 tacactaaca tgtgtcgtga ccaatgtgca gcagatagac ttgctcatta aatatattcc
261841 aggtaggatt ctctaagggt ttttttttt ctctatagct gtcacttttt tttttttctc
261901 ttatcaatca cagatataat aagcataaca tcggaaagaa aataatgata gggcaaaagg
261961 gaaagcattt ccagttgaat acagtatctc tactatattg ttgttcgctg aagacagccg
262021 tttccggaaa aaagtgattg tgtgatcgca gttacccgca cctttgttca acagtacata
262081 ataagaatct gtagatatag atgaactttc tctattacat aagactagcc taaaagcctt
262141 gtttctatca tttaaaggct tttgagacac tcaaatcagt gtcaaccttt tggataagtc
262201 aaagaaagta cgccgctgct cagaaaattg ttatcgtgat tgagaactga tctaggcatt

262261 acaattaaag gaaacaaggg aaatattcta ttgaatgttg cataaaagcc cttcaagaaa
262321 gagatttgct tctcctttgc atctaggatg cattctaacg cttacagtgc tctgccttat
262381 tgcttactac tttgctttac cggattattt atcggtaggg aagagctcat ccaggggtgc
262441 catggatcaa aaatctgatg gaacctttag acttaagtct atctacaggc atggcgtcgg
262501 agcaaaccat cggctacacc agaggctgga ggtcactcca gaagttattt ctgccgctgg
262561 aatgctgtat caagaaacca caactcaagg acaggatttc gaagaccaag aacccctatg
262621 gactactaat gccgaatatg ccaccactaa cccattcgat tttgagtttg agctacgaag
262681 gatgccgcta ttgatgaagc ggatgaagga aagagatcct gaattcatag agtcgtacat
262741 ttacggagaa acatacatga cggaggagga agaacacgcc atgtggatag atgatgacat
262801 tgtggcaccc aatatcacag atagaggtac agtagtgtcg ttagcattaa tgtcgtctaa
262861 tgcttacgtg agaataccgc aaacgggaga ctggcgtaac gtcacggagc cgtggaatga
262921 aacagagccg gaagattttg gttgggatgg tgatggtata cgtggtcacg tcttttacaa
262981 tgaagtggag aatatcgtgg tactctcgat aaagggaacc agcgctcagg gtttgccagg
263041 gtctggcgaa gatgagacaa ctggaaacga caagatcaac gataatttgt tattttcgtg
263101 ttgttgtgca agagtgagct acctgtggac taccgtgtgc gattgctatg tgaagtcgta
263161 tatatgcgat gagtcttgcc tggaaaagga actgagacgt aaggatagat tttactctgc
263221 agtcgttgac atatacaaag gcgtcctcaa agaataccccc gatgcggcca tatgggtcac
263281 aggccactca ctgggaggcg cattggccag tttactgggc cgcacttttg gattgcctgc
263341 ggtcgcattc gagtcccctg gagagctact accttcaaaa agacttcact taccattccc
263401 accgggactt ccctcataca tggagggtat ctggcacttc ggtcacaacg cagacccaat
263461 cttcatgggt acatgcaacg gagctagttc aagttgctca ctggtgggct acgctatgga
263521 aaccgcgtgc cacactggta gagtctgtgt ctacgatgtg gtcaacgaca agggctggag
263581 tgtaaacatg ttcaaccaca gaatccacaa agtcattgac gaagttctcc ttggatacga
263641 gcaggctgcc aagtgcgttg aaccagagcc ctgcgtagat tgctacaact ggaagtttat
263701 tccaagcaga gactgggaat cctcatcgag gctcattacc aagactaaaa gccacgctgc
263761 accaaccacg acgactcgca ccactgctac tactacctct tcatctacct gtgtaggccg
263821 caattggctt ggcttctgca ccaaatacga gttgtaagac ccatacatct attatgaccc
263881 tagtgttgtt ttagggccta tgcgcccttt ttttcttcca ttgtctgata aataatgtat
263941 cattttgaca acttaaaggt tcagctaaac gacacgagaa gatatattaa tactgcttta
264001 aagtgaaagg cgtcagtcag gtgcaattaa taggtcacat attctttgct tcaattgctc
264061 tctagcaaac tcacacactc gtagattgga agaccgaaga aaatagatag actttcgaat
264121 aagttcagca acaacatgtg gttgaaatcc ttgctgctct gcctgtactc cttagtactc
264181 tgccaagtcc acgctgcacc ttcatcaggg aagcagatta cctccaagga tgttgatctt
264241 cagaaaaaat atgagcccag tccccccgcc acacatcgtg gaataatcac tatcgaatac
264301 tttgatcccg tttcgaagtc gatgaaagag gcggatctga cttttgagtt gtacggtact
264361 gtcgtgccca aaactgtgaa caactttgct atgctggccc atggtgttaa ggcagttatc
264421 gaagggaaag atcccaatga tatacatact tactcgtacc gtaagaccaa aatcaacaag
264481 gtttacccta acaagtatat ccagggtggt gtggttgccc cagatgtggg tcctttcacc
264541 gtctatgggc ccaaatttga tgacgaaaac ttttacttaa aacatgacag gcctgaaaga

264601 ctcgcaatgg cctattttgg acctgattct aacacctcgg aattcatcat caccactaaa
264661 gccgatggaa atgaggaatt ggatggcaaa agtgtcgtgt ttggtcaaat aacttctggt
264721 ctagatcaac taatggatgc tattcaatac acagaaacag acgaatatgg aaagcctcag
264781 catgaattac ggttcctgta tttcgttcta gaaatcttaa aaattagtaa catcttagat
264841 ttgcacgctg cgtacacaga aaaagtcgag aagtttagaa atggcgatgt gtctgttggc
264901 tccactttgg aaaacatctt ccgtaacgat aaagcctaca cacctttaac cacctccact
264961 ggaaccaccg cctatgattt aaaccaccca atttccagag ccttgatgtg tttaactgtt
265021 cttggccttt gtttcattgc ctacaagggc atgcacgaaa agcctcatac ggtttcatta
265081 agacacaagt aagcaacttc ttttcgatgg cttgatgacc acatggctat gattttaatt
265141 aatatatata catgacaatt atttttttat acattagcga tacctaaaat ttcttggtta
265201 ataccctttt gacagcctct actgcattgc ccttgataat aattttttttc gattgcatca
265261 ccacagacca tgattttttg ggtatgaaag gcagttgctc ctgcaagtca tttcttagtt
265321 gaattacatc tccttcaatc tttctgatct ccgtgataat cttgttacct ccatttttat
265381 aagcggaata caccggtaga tttccggtcc tgcttctctc cacgtagtag gtctttttac
265441 caaaattgtt gtttcctatg agctcatgca tcttcacgtc ttccagcttt ggaaatattg
265501 gcgcctcagc agctttattt gaattagtgg atatacatct tatcgtttgg tatatcgagg
265561 cagtggcagg ccctggaagg catttacccc taccaaaaca tctagtaaca cacgaactga
265621 tcatagtttt gctcacacgc taccttggag taagttgctt taggtaaacg ggcacttctt
265681 tcaaaaggtt ttccttcttg ctggagctat actgcttata gtaccatatt ttaaattcat
265741 accaagccat tcgtacgcga tgaccattat agatcgacct tgcgacacca tcatatttaa
265801 tattcaaaaa gaaaaaaatg cagttctata tcacagttat ataatattta tatgtgcatg
265861 tatgttactt gtctttgcac ataactcaat gcgtccacaa tcttaccatc ttgtctttcc
265921 caccactaca cactctttta ccgtcgacac tccagtcgac ggtataaact tcgtctttat
265981 gaccagggag gtcaacagaa agttttctag ttcttacatc ccacactttc aacgtggtat
266041 ctttggaaca tgacaccagt agtcggcagt ccgatgacca cgcaacctgg tatacgctgg
266101 ctacatgccc tctaaatgtg gagataaact taccatctct accgtcccaa agtttgatag
266161 agttatcaaa agacgctgag acaatatacc taccatcagg gctgaacgcc acatgattga
266221 ctaatttttg gtgaccggtc attcttgcta taggcttggt agattttagt gggttccata
266281 aaaacatggt ataatcatcg cttgcagtaa ccatcatttc ttctgaattt ccattctttt
266341 tacagatttt ttcataattt tccaatgcct ttttctgggc ttcttctggt gtagaaggct
266401 tcttacctgt atgatcgaat gcaccaatgc gcaatgcgta atctgtagat aaagataagt
266461 gattaaccca gtgcgcatgc gacttcaaaa tgttgataca tctgccctgc gaattgatgt
266521 cccatacacg tacggttcta tcgtgagagc cactatacaa tagaccttgg ccgccccatt
266581 tgacacaaga cactgaattt gtgtgaccac tcatcgtata ctggcaaact ctgctcacag
266641 tgtcccaaat cttaatagta ccatccttag aagatgaagc taatcttggt ttggagcccg
266701 gcttcacaag atgtataggt tcccaactta aagaagtgat ccacttggaa tgacctctga
266761 gagcatcacc tagacactga ccgctttttg ggtcccataa tcttatggta ttgtccatgg
266821 atcccgtagc aattacttct ccatcggggg accaggaaac gcagagaacc caattgtagt
266881 gaccctttag agtatgcatt ggcgtttggg tgtcacagtc ccaaatcctt gcagtattat

266941 cacctgcacc ggttaccatc ctagaactcg tatgtggtgc gaaggcagaa cacaaaattg
267001 tggaaccgtg acctgcaatg gctgatgaac ttctagttac cggcttgact ttgaaaactg
267061 ctcttggcgt atacagtagc gtgatctgat cttctgtact gttatagcct ggttttatta
267121 atgaagaata taggttatct gttatatcaa tcgtcttcac agggtcactg gccttcttac
267181 cttgaattgt acagctgaag gtatatggca ctggatcgtc tgaagtaccg ttcaattgat
267241 ttaaaagttc ttctaactgt ttctcggaga tagcaccggg aactctcagg gcgccaccta
267301 cattgtcacc agtatctaaa gcttggaact tgattgaaac attgggtaaa tctttcggaa
267361 taatagctac ttctctggga agttgagcct ctttcttttg tttcttagaa ggtggaggaa
267421 tcagggtgga catctgtata ttgtgttagt gtgtgtttat cgaggttaca aatataaaaa
267481 tcttgtgata tatgtctttt aaagatactc tttttgaagc tcatcgcatt tttttttttt
267541 ttttcatttt aaaatttttc accggtttct cgggtaacag ctctaaatga aatcttgcct
267601 tcatcaccat tattgaatta cattacatta tatatcgtag tatatcatat ttttagacgt
267661 tgaccactct atgagcttag gataggtact tgatctttct ctgagttgcc aaatgctatt
267721 tcacggattt gtatcatcca agggtctatc agtagttcca cagcagtagc cctcatagtg
267781 gggtcttgaa ccaaacacct ttccaagaag gctcttcccg ctgcagtcat ttcgtctcta
267841 ttgggtagtt gcggtattcg acctgcagca acgtggtaca tgatggccca ttcattatcc
267901 aagttagacc aaggtcgtct acctgtggcc atttctagca caacacatcc taatgcccat
267961 acatcgtccg ctccaagttt tcccttaact gccgagcctg aaatagtctc tggagccata
268021 tacatcggtg tccccatcat ttcattgagg gactttgttt ctactccaaa atcttgaacg
268081 gctgcgttcc gcacagttct agtcctagat cctacaacgg tacgtgccgt accaaaatcc
268141 acatatttta tgattccatt gaaatccagc aagatattct ccggtttaat gtcgcgatgc
268201 accacgccag attggtgcaa atatgccaaa ccttctaata gttcgaatgt gtacacttgt
268261 gttaccattt catcttcaat tcttccatga tccaataacg aggctaaaga accaccctca
268321 cagtattcca tgaagatgtt aactttatcg cgatgtactt cgacaccata gtactggaca
268381 atattaggat ggtttaacat ttccaatacg gtcatctctt ctttaatcag gggaaaaatc
268441 ttcttcattg ttgtggtatc gtgtatcttt atttccttaa cagctaagat ttcaccgttt
268501 tccagattaa ttgcagagta tacctgtcca aatgttccac cgccaatgaa gcttcttttt
268561 tgccacctca ttgatacatt cgataatgag gatgctagcg atagtaggta tttgtttccc
268621 tgatctgtag catccaatac cttacccatt tgcctgggat tcctcttcat agtcttttcc
268681 aacgtcttta tagcttccag tctcaacctg ctgtttattt ctagggtggc ctcttcatca
268741 atattctctt cagtatcaat attcaatctt gcctgttgca tgccattttc agcttcagtg
268801 gctcgtgcac ccataacgtc gaaatgtgaa attaataatg acatacatac tgataccctc
268861 tccttcagtt cttgaaattg ctttacattc agcgccagga tattataacc actagtcatc
268921 tgcatcgcaa aatccattgc aagaacgcac catcggaaag ttcgctgatc ggtaggatca
268981 cactcttcaa caaggaattt caaccagttc attcctaatt taaccatcag aaagatgata
269041 acgtacttcc tctttgagct aaaggcgtta tatttaagga aacttcttcc aaaatccctt
269101 ccaaacatga atatcgaatt taacaactca tttcccggac attgcttcat aaagctacct
269161 agaattcctt tgtagttgtt caatactgta taagtaagtt tgtaatatgc cttattgatt
269221 ttttgtaggg aggattcaat tgaattgagt gaacaacgtt tctctatgaa tgaaacagaa

269281 tcacctaccg tttgattaaa cttgtcgatt tgataagtca gtgcataatt agaaccttga
269341 ttgagtaaaa tcaacgtatt tgggttacca attttcccta aatccatttc gttggatgca
269401 aaaaggttgt tacctggaag cttatatact ggtccatccc aaactactgg ctctccagga
269461 tatagtacaa ctagtgcacc caatgagctc aaagcaactt ccaattcaaa aacttcattg
269521 ttcttttcat caaggtgctg atcaaaaggg tctgttgaaa aattcttctt tttggactga
269581 tgagcatgcc tgtccaaatc acttgacgaa tttgctacta ctcggtagta aggaattcct
269641 ttggaatcct cccctttcga tactagaatg ttcaaatctg tttcttttgt tgttatatca
269701 tacacattca aattatttcc aatgtcaagc tttgggacca aatcacagcc aatggattta
269761 tttcgcaaaa ttcttaagat ggtatcattg tcacaaccta ataattcagg agcagcaaac
269821 atatatacac cactagactc aaaagtgtta ccggagtata ccaaaaaata gtgagcatct
269881 tttaactttt taacaagttg tgcattatga ttaatctgat aaacagcaga attttggaac
269941 gccttgacta gaatattgct gaacctgttc agctttcttt tcatggcacc caaattttcg
270001 aaaatggaac ttagccattt ttctgcttct gaagcgttag caaatgttgg tgggaagttt
270061 tgttgtttca aaagatagaa ttgcaattta tgtacgagtc tcaaagttaa cttgagaaat
270121 tcatttggaa tcacggtttc tgcaccgtta atatagtgac cggtgttttt taggtgatcc
270181 cagtatttca agagtagatc gggtgctttg aaattttgtt tactggaatc aatcaacttt
270241 aaattcaaca gaaaaaataa ataacgaatg gcttctatta cagtattttc gaacgtcggg
270301 tcaaaatcca catcgaacgg caaattggag caatattttg tcagtgtgta cttcaattgc
270361 acagaaagtc gaataaaagc gttaaaatca tcgatcattt gatcgatcat catcattgtt
270421 ggatttttaa gctttcgtgc atatgacaat ctagtcaaaa taacttcttt gattaaatag
270481 acggggaacg ccagtagtga tcttaacctt tcataaggat attttatgtt catctttgtc
270541 aaagtttctc tgtagactat aaaatgtagt ttcagtttga acatccaagg tgatagcgga
270601 tagaaaattt ttttttggaa tatagactcg atgttctttt ccttcatcaa ttgttctgca
270661 agaatcttca ttttgcagtt ccattggggc acaccatcaa aatcctcttg gcaattatcc
270721 gaactactta tcggtgagaa acggtttatc cagtcgtcca actcttgcct cctcaacgtt
270781 aagttcgttt taaagttcag ccaagaattc attacgtcta ttctagcggt aaattctgca
270841 gtattggtga ttggtttatc cttatacatt tttttcaagt taggccaata agagacaacg
270901 ctgtgatatc tgttcatgag gtcttgtaag atggcctcca gtgcatccag gctcaatatc
270961 ttgccctgtg gaatttgaaa cttcattatc tcttcaaaaa cggaatctga actatctctt
271021 aaatatgtaa gcgactgttc catctcttgc atggtcctcc cattcagcca tgccttcaat
271081 tcgagccata tctcatctga gagctcctta tttaaccctg gtttcttgac ttggttagca
271141 atcctcgtct tttcaccttt aacaatatct cccttcagca ctctggccag cattgactgc
271201 catttgaacc gttcggtcgc aatttcattc caagctagtt catcactgtc tgacctcatt
271261 ttcctgcaat aaggtgtagt ggagaaaaac ttgacacctg agtgcaggtc ttcatcaatt
271321 ctgtcgccat tagacaaccg aataatttcg tcatcaccgt cgtcttcctc gtagcggtta
271381 gtgagtttta tgcccctcgt gtagtagtca tctaaagcgg tctttttag ctttttcaag
271441 tatagcgatt cattgaaaac gtatcggtcc ttaaaagtca gctgactggg gccgtcgctt
271501 attgtttccg agccacccaa tgctttcaaa tcagatcttg atagtttatg gctggaatga
271561 cttgataatc tcttcacgag cttacttgag atggtatgag gagccagaac tctcccgcct

271621 tcagccgctt tttgttgctg tgtattcagt atatccatca tcattttcac ctacaaggaa
271681 ctacctttta tagccaccct aagtaaaaca acattagctt agcatcctca attcttatcg
271741 tatgttgttg ctgctatttt tatcctattg ttcctggcat cgctttttta cataaggtac
271801 caaggcaaga gaaaagaccc gcgaaatttt caattcgaga catagggtta atacgaaata
271861 tgttaaggtc tagtttccaa aaaatgaaga aaatgtgatt agacatcctg gggaaattag
271921 gtttaaatag ggcgggcgct acaggggttt tcctaacaaa tttcaatgat aatagtggca
271981 tcatcatcgt catatccagt gtaggtatgg actagaacag aaagcaattt gcagcaagac
272041 aatatgacta cgacggtacc caagatattc gcgtttcacg agttttcaga cgtggcagag
272101 gccgtagctg accatgtagt ccacgcgcaa gacggtgcat tggctccaaa gaacgagagg
272161 aaacactctg ttcccaacat cagcatgaat gcactggata tgacgagaga ggcctcttgc
272221 aaaagcacag catctgccgc ggaagggaaa agtggtagca gtggtagtgg cagtggtagc
272281 agtaagccca aaaaggagaa acggttcaag attgctctct ccggtgggtc attgatcgaa
272341 gtgctacacg aaggtctgct aaaacgagac gatgtacggt ggggagactg ggacatttac
272401 tttgcagacg agagacttgt acccttcagc tcgaatgaaa gcaattatgg atgcgccaaa
272461 aggaagattt tggacctgat agacacggcg aagtatggaa ctccgaaggt gtaccacatt
272521 gacgagtcat tgattgacga cccgcaagaa tgcgctgata actatgaaaa ggtgctaatc
272581 cgcgggtttg ccggtagaga ttccgtcaaa cttccgatgt tcgacttgtt cctgcttggt
272641 tgtgcccccg atggtcatat cgcatcactc ttccctaact tccaggacaa tctacgtgag
272701 aaacttgcat gggtggtgcc cgtggagaac gctcctagtg ggccctcgac cagaatttcg
272761 ctgactatac ctgtaatctg ccattctcac agggttactt tcgttgtcga aggtgcaacc
272821 aaggcgccca tcatcaagac cattatggaa aggcctgaaa agggcctacc tagcagtatt
272881 gtcaacgaag gtgctgctgg tcgtgtatca tggtttgttg acgacgatgc tcttacggac
272941 gtcctcgtca ccaaaaaaaa gtataaattc caccaaggtt tgtctattta aaccatgcat
273001 gtgtgtacat acatacatac acacatacat acatacatac atatatttat gtatatttat
273061 gtatatatat atatatatat atatgcgtaa ttatgcagat ttctccttgt gtgctacagc
273121 gggtactctg acgccagatc atggcctcga gatcgatata caaaccactg catgataaag
273181 atgaagttga atattaaagt taccattgac agtcccactt ttccaaaatt tgtcacgaac
273241 gtatccagac tgaaaccttg atcgttagac aactgccaaa tcaattggag cagcgaggcg
273301 ataccgccag tgacatctag aaacacaccc tgaatgggga aacaatccat agattttctt
273361 gtcgagttat gcgtcacttg tgggatgtac ttgattagtg acatcgatat tttgagtaaa
273421 aacaaattgt tgcaatacgc cagagttctt gagttatgcc aatcgtacca gtagttggag
273481 tacataaatt gtacggttag cagtgaaaaa atggccagtg atgcgagtaa aatccttagg
273541 taccatggat tcatcttgcg gtgaccttta cctggaaatc gccagattct cgctccagct
273601 accacctggg tcaataagac aacattcatc aagcacccat gcaggcaata ccagaaatca
273661 aattgcgtca acttgggcct gcccaagtca gactcatcac ccgtcatttt ccagcagtac
273721 aattgcaaaa atatggatat gaccaggtaa gagtaacctg ccgtatttaa catgacaaaa
273781 tccatcgata tcgcgctcgc tgacttatgg cgccaattgg tgattatcgg tggatacatc
273841 gatatcgacc atgacgtaac atacacgata cctagtatat cgtctaacga caccattttg
273901 cttcgcgttt gcttttgttt gcttttagtt tttcttccct tgtttacata tggaaactca

273961 ccttaaaatt tcaagattat caaaaaatat aaaggattcc gctattaaag ggcaatacca
274021 caacctcaaa ccaaatcacc taaaccaaat caaagatgga agcggaaaaa caatctgata
274081 ttaagggaac cattgctttc gatactcacg gtaacgttat agagtctacc ggcgtgggca
274141 gtcaaaggat cgaagatatc ggtgatttat cgaaagttac gctggatgca gaaggttttg
274201 ctcaagttca aggagattcg ctactcgtac acctgtacaa acgcaatgat atcactttag
274261 ctgtatatac cagcgctcaa taaattcttt ccttcccttt cgacatagct atgtctgtat
274321 actatctaaa aactaaatat cacaataaac aaagtactaa aataaacacc catttaaata
274381 aagccagatc ctgaccctcc aaatccgccg gatcctgaac ctccaaatcc gccagatcca
274441 ggaccaaatg gttcgtcgta cttggatcca ggcatccatc ctggcggcat atcttcccgt
274501 cttggggcag gtcttctatt tggatcgaaa atcataccgc cttggctact tggattcccc
274561 agaggattgg gaagccctcc agaccaatcc ggatggcttg tgccaacagg atacagatca
274621 ccatccgcat accctcctgc tgggttagcg aagctcggtt cagttaccgt aagtccggga
274681 tatgggttca acggtgctgt tccctggttt tgtcttgctt gtgcctggta ctcatcgtca
274741 aattttggca tgtccgcact acctgcagtt tgtcttttaa gttgtgtcat tataggtttc
274801 aggtctggct ccacaggtaa gctttctctt tctagtggcc atttcaattc tgtgggcaat
274861 ttaaaatcgc tgtaacccac cacacatgcc atggtgatcg acgtaccatg acgaataaca
274921 acagtatact tgtcattgtc atccagagtc tgaatcacca ctagctcaga gagaaacgat
274981 gttgaaccct tctcactggg agaatcttta atgcgaaaac tcacttggca tgatttggtt
275041 ccctctgtat ggcgtgacac cacctcagtt ttaccgatgg actccaacac tagttctgct
275101 accagttcaa ccttattctc aatcattgtt taactcttgc gttactatgg tggtattatt
275161 gatgctaata cgggtttttt atgaaaaaaa aaggataaca aacttatata aacttatata
275221 aaaacatata caataaatga tctacaaagg gtaggaaata aaaaataagg gagaaaaaaa
275281 aatacatgcg taagtacatt aaaattacag gaaaaatctt actttagttc tgatatttca
275341 ccatcgcgat acaccaaccc cataacgttt aggaacaaat tcaaatcttg gtatagtttc
275401 tctttattat aaataataga ttttaactct gtgtctaatt cttgagacct ctgaggtaat
275461 acactcaatt ctttctcttt cttttgtaag gtctccgcct cgttaatagt ggcaaagatt
275521 tcatcacgta cttcatcaat aatgattctc tggtggttta gctttccact tagtgctaaa
275581 ctagctagga actgccaaat gtatgcttca tccataaact tgtcattttg taaacgcatg
275641 atgtggtcgt tatattccct tggagggaaa atcagctgaa tcttactttc caatgaagtg
275701 aataatttat cataaatctc attccatgtg gagatttcag gagaggaaag aatattagat
275761 cttgatgaat cttgcttgat taatgcggca cgagaaatca aaatggtgat caaatttaga
275821 ccaattttgg aggtggtcaa gaacgaaacg ttgttgtttc tgattaactg tagcaacaga
275881 cccataattt cgataaaatt ggagttatta gacaaaaacg agacaatgat ctttaatatg
275941 atcatttgga acagatcgac tttcttcaat tgtgtcaaag ttggttttgg tgtagtcttg
276001 taggaactca atatgatgat ttgcaagtgt gataattcat tgaagatttt ttgcaggatt
276061 ttcaattttt gctgcttatc caagaaatta aaaattctag gcataatctt gatacctttta
276121 tcaaatgata gcatcgaaat gaaagggttt acgtcatatg aactgtcgtc gatgtgaagg
276181 gcctcccaca ttgcagtgct gtcagtttgc tggccattcc tcaagttagc ttccaagtct
276241 aaaatctctt cataaactgt ttcaattaac tcaagaatga attttttgcc cccagatttg

276301 ttcaaatttg tggcaccatt accgttgttg aacgcgtatg aagatcttct tctttgctta
276361 gaagcaacag cagcggcaga agagcccgta gaggaggacc ctgaagaggc caagtccacg
276421 ttggtcagag agtcagtgac gtcttttagc gcatcggggt cagaaggctg ttgtctgggg
276481 atctgcagtt gtcttcttgg attcttgctg ttcaaagtgg aagagatttt gcccagaacc
276541 gtgtttgctt gcttattatc ctggctagag ttaccagcag ccgcttgttg atccttcaac
276601 ttagaaggtc tttccttagc cacagtgaca gccttttcta cttgactttg cattctctgt
276661 agggcaatat cggttctctt atagcgacca ccgagtctgt gtccagaatg ttccaaatac
276721 gccctagcaa tcaaaccttt gttggattcg gacgtgatac cgcctctttg gataatcttg
276781 tagacctgga agtagaaatc ctcattgtaa gggtcctcag tgacaatttg agacaactga
276841 tatctggtga tgaagtcctt atcacgagga gtcattaaac ctgagtattt caagattttc
276901 tccactttgg catgacgaat ctgcaaacgt ctttgctctt caggcgacaa atctctcttc
276961 gatctgcgtc cactttgcag aggcatttta gttgcgccaa ccacagggcc tggagaagga
277021 gtgttggaca taccgatggg tgaaggcgaa gcttgaactg gaacttgtac aggacccatt
277081 ggcatttgat tcatattcac attaacgcca ggaggtggag gtggaggcgc aaattgtgag
277141 ggacctggtt gacccgacgc aggctgcatt ggaaattgct gctgaggcgg accctgtggg
277201 agaccctgtg gatgacccat ggcgtggaaa ttcataaact gttgctgcat ttgacgttgt
277261 tgtctttcca agtcttgcat cgacaagatt ggctgtaggg gggccatttg ctgcggagca
277321 ggagccggag ccattgtact ttggggtgaa ggcgccattg ctggaggtgg tgcagtagac
277381 cacaaagatt ccattggctt caggtctaga ggtcctgctg ctgccgttcc atcggtccta
277441 gggccgctaa ttccttctgc agtagctgca acatatgatc tagccgtggc accgacgcca
277501 ttaccaccaa ttgcgttgcc gctgctgccg ctgctgtgag gatttccaaa atcaaagtcg
277561 gtaccaacct gaacattatc accaaatgtt tcgtcgttca aatagtcgtt ctcctcaagt
277621 tcgtgctcgc catagccctt gtaactctct tcaaagtcca gaggaccatc ccgcgcatta
277681 ccgctatttt ctaacccaaa gaaggacatt gctagtgctt cttgctgctc ttacttccgg
277741 ttaaaacctt tgcttccttc aaggtgtaca tgtttctttt cattcacctt gtttctttct
277801 ttaaaaagag gaagaggaaa aaaaaaattt cgtgtaatgc gagaagaaca aatagggaag
277861 tgataacaat agtatcaata acaggctgca atcggggcaa ttaagcatca gaagagggga
277921 atttgatgcg gctggggaat gcttatgcgt attgcaagcc gtcgcaaaat gttgggctga
277981 agctggactt gttacggggt cttccagggt acgtgggtca cgccacgagt aggatcaacc
278041 ggttagagaa ccaggacaat tactccatta aaatgatgcg gtcgtggcct aatgcgtatg
278101 gaagtgccct gaactgctct gtattcgatg gccatgggga gaaaggtgcg caactgtcac
278161 agttgttagc cgataaattg tgcagtagcc tggatttccc tgaaccctcc tgggacaagc
278221 aggacttgaa gaagttggta caggaatatg caagaaggtt tcccgaggga aattactgga
278281 aacataagct gtccacgttt gaaaaattct acaacaagtt tataaagaac tgcaactcga
278341 aacaagaact gctgctgatg aaggagggcg atagtgcgat ccttggacaa aatggtggca
278401 ggatgatttt tgataaaatg gggaatatta ttgacaaaat tgcactgctc actgagctag
278461 accgtctgcg attgttctac gggtttgcca ggttcgatct cgaccaatgc tgcggcctgg
278521 gcaccgcagc ggggtccaca gcgtcgtcca tatttctata tccgtatgat gatccgaatg
278581 cacccattga tgaggggaaa gacgatgact cgtggatcat atcgcactct ggattgctca

278641 aactcattgt cacgcaggtg ggcgactcta agatcatcct ctgcgatcaa gacgggattg
278701 cacacgcatt aacaacaact caccacatca attccagtag ggaacgacac cgtttgagca
278761 tagacccttc tcgattggac ccagacgcct ttggagagac aaggtttttg aataatttcg
278821 ccaacactag atcgtttggt gatgtcgctg ggaaaccgta tggtatatct agtgagccgg
278881 atattttctc gtttcttgtc ggaaatacct tgcatttgcc gcgatctgaa aggtcgaagc
278941 tgcccttcaa tggtgatgaa tgcttcctgg cgcttgtcac cgacggcatc accaataaac
279001 ttgcagacca agaggttgtt gacctcatca cctctacggt gaactcctgg ggactgaaaa
279061 aggcgactcc tcaatttgtt gcagaggaga caatcaaatt tattcaggca atagccacta
279121 aacactcaga caacgccact tgcgtggtgg ttaggctttc caattggggc aactggccca
279181 atgttgacag aactggcccc caaagagaaa ccaaactaat gaatgcacaa tctaacgaaa
279241 caaaattaaa ctaaacaccc ccctgggtgg aaacaagtcc tagataatca tcttttgtat
279301 ttttttcttt ccttttatta gtatgtatac atatatatta ctttttatat tctttatagc
279361 gcattttaaa cacctttctg aaaaaatgct gagatttttg atgctaatcg aaagattaaa
279421 aaaaggtcta tagaggtaag ccaataacgc gatgaataac ggttctggtc gatacttgct
279481 gactccccca gatgatcttc acccctatgt gccaagctcg aaacctcagg aacaagtata
279541 ccctgatttc aagccttggg agcacactgc agcagaagat caaatcctag caaactttgt
279601 ggctaagggc ttttaccata caccaatggt aaatttcgag tccatatctg cgagatcatc
279661 tgttcatgaa tcattagtca ctcaatccaa cattctttcc cagcaattcg acaaaattat
279721 caagattaga gaagaccaca ttaataagat cccctcaaat tccacgacga cattacacgg
279781 gcctggtttt cagttgccta atagaataac ccttactgat catagaaagg aaacgtggtt
279841 gcatgaattg agttcgtctc acacttcgct ggtcaaaatt ggcaagttta tacctcacgg
279901 cttgaaaaga aggcaagtca tcgagcagtg ctatttaaaa tttataccat tgaaaagggc
279961 gatttggttg ataaagtgct gctattttat cgaatggaaa tcgaaccaca aaaagaagag
280021 gtcaaatgct gctggggcag atgatgccat ttccatgcac ctgctaaagg actggacgga
280081 tacctttgta tacatcctgg aaaagctcat ctttgatatg acaaatcact ataacgattc
280141 tcaacaactg cgtacgtgga agaggcagat ttcttatttt ttaaaacttt tggggaattg
280201 ctactcacta agattgatca ataaggaaat ctttcatcat tggcttgtag agtttataaa
280261 taagatggaa aacttcgaat ttttgccatt atctttacat attttgatga ttttttggaa
280321 cgacatctgc caaattgata caaatgctcc tgttgcggct acaataacat caagtcaaaa
280381 agagcccttc tttctggtaa caaaaatcac tgatatgcta ttgcacaaat attatattgt
280441 ttccagcagc aaatcaatga taaatgacga gaactacatc atcaatgata taaagaaaaa
280501 caacaagata aagttgaata ttctcaaaat attatccagt ttaattttga aaatttttca
280561 agaacaatct ttagaggtgt ttatatttcc cacatctaac tgggaaattt acaagccctt
280621 actttttgaa atagtctcaa acgccgacac taatcaaaat tctgatatga agaaaaaatt
280681 agagttaatt agttacagaa acgagtcatt gaagaataat tcttctatac gaaacgtaat
280741 aatgtctgcc agcaacgcaa atgactttca attaactatc gtcacctgta aacaatttcc
280801 aaaactatca tgcattcaat taaattgtat agatactcag ttcaccaagc tactggacga
280861 taaccctaca gaattcgatt ggcccactta cgttgaccaa aatcccctta caatgcataa
280921 aattattcaa ttaattctct ggtccataca tccatcaagg caatttgatc actatgaatc

280981 taatcaactg gtagcgaaat tattactatt gcgaataaat tcaacagatg aggatttgca
281041 cgaattccag atagaagatg ccatttggtc attggttttc caattagcca aaaatttttc
281101 ggcccaaaag agggtggtat catatatgat gccttctttg tatcgcctgc ttaatatact
281161 aattacttat ggcatcatta aggtccctac gtatatcaga aagctaatca gttccggcct
281221 actttatctc caagattcca atgataagtt tgtgcatgtc cagctgttaa ttaacttgaa
281281 aatttcaccg ttgatgaaaa gtcaatacaa tatggtattg aggaacgtta tggaatatga
281341 cgttaaattt tatgaaattt ttaatttcga ccaactcgtg gaaatcacag aacaaatcaa
281401 aatgcgaata ctctccaatg atataactaa tttgcaactg tcgaaaactc ctctgagcat
281461 taaaatcatg gttgcagaat ggtacttatc acatttatgt tccggtattt tatctagtgt
281521 taaccgcaca gtgttgctaa aaatattcaa gattttttgt atcgatctgg aggttttcca
281581 ccactttttt aagtggatcg agtttattgt ctaccatcaa ttgctaagtg atatagaatc
281641 tctggaggca ttgatggaca tcttgctatg ctaccaaaaa ttgttctcac aattcattaa
281701 tgaccatatt ctttttacga agacgttcat attcatttac aagaaagttt tgaaagaaaa
281761 agacgtgcct gcttataatg tgacttcatt tatgccattc tggaaatttt ttatgaaaaa
281821 cttccctttt gttttaaagg tggataacga tttaaggatt gagttacaat ctgtttacaa
281881 tgatgagaaa ttgaaaactg agaagctgaa gaatgataaa tcagaagtct tgaaggtgta
281941 ttccatgatc aataattcaa accaagctgt tggacagact tggaattttc ccgaggtgtt
282001 tcaagtaaac atcaggtttc tactacacaa ctccgagatc attgatacaa atacaagcaa
282061 acagttccag aaagcacgaa acaatgtcat gcttttgatt gccactaact tgaaggagta
282121 caataaattt atgtccattt tcttgaaaag gaaagacttt actaacaaaa atttaattca
282181 attgatctct ctaaaacttc taacttttga agtgacgcag aatgtgttgg ggctcgagta
282241 tattattcga ttattaccaa taaacttgga aaataatgac ggctcatatg gtctgttttt
282301 gaagtatcat aaagaacaat tcataaagtc aaattttgag aaaattttac ttacatgtta
282361 tgaattagaa aaaaaatatc atggcaacga atgtgaaata aattattatg agatcctatt
282421 gaaaatttta ataacttatg ggtcatctcc caaattactt gcaacatcta caaaaatcat
282481 tatgttgtta ttgaatgata gcgtggaaaa ctcatctaat attttggagg atattttgta
282541 ctactcaact tgtccgtcgg aaaccgatct taacgatatt ccattgggta gtggacaacc
282601 agacaatgac actgttgtaa ccaacgatga taaaagtgac gatgatgatc acacagtcga
282661 cgaaattgat catgtagaat attacgttat gatggacttt gccaatcttt gggttttcca
282721 agcgtttacc tgtttctgca tcaaaaaaat catggagaat aatgagccag caatggcaat
282781 ggaagacttg aagaacttca tattccaaat tatcgaaata actaattcta atgatttatg
282841 ttcacaaata tttgaccaac tgaaggatat gcagaccatt gagatgataa cccaaatagt
282901 ggagaaagat ttctgcactt cttgtttgca aaacaacaac caaaagatag atgataatta
282961 catcgttgtg gtgatcgaga ttataacgtc attatcgatg aggtttcaaa gagaaacttc
283021 tggtatgata gttatttcca cggagaacta tcatttacta ataaagatca taagacaatt
283081 aagtgaactg aacgaaggaa atttatctaa gagagaaatc caaatagatg ccgtcttgaa
283141 aatttttagc tttcatcagg attccatttt ccaacgcatc atcgctgatt tatcagctga
283201 taaacccaca agtccattca ttgatagcat atgcaagctg tttgataaaa tatcatttaa
283261 tttaagattg aagctgttct tgtacgaaat tttgtcttca ttgaaatcat tcgccatcta

283321 ttcatccaca attgatgccc cagcattcca cacaagcggt aaggtcgaac taccgaagaa
283381 attgctgaac ttaccaccat tccaagtgtc ctctttcgtt aaggaaacaa aacttcatag
283441 tggcgactac ggggaagaag aagatgcaga ccaagaagaa tcgtttagtt taaatttagg
283501 aatcggcata gttgaaatag cgcacgaaaa cgaacagaaa tggctcattt atgacaagaa
283561 agatcataaa tatgtctgca cattttccat ggagccgtac cacttcatct ccaactataa
283621 taccaagtac acagatgaca tggctacagg cagtaatgat acgactgcgt ttaacgattc
283681 ctgtgtaaac ctgagtcttt ttgatgctcg gtttgagagg aaaaatccac attgatctca
283741 gaatatatcc aaatggataa attataaatt taccaataac agtaattatg tgtcagtttt
283801 aatacccaac caattgattc ctctaaatga taccgtacca aatgacatgt ttgccaccga
283861 gtataaaact ggggtttatg agcgatggtt aaggaggatt aagacaaact gctcatccgc
283921 ttacaagatc agtactagcg tgtaccagtc tatttaaact ggaatgatca ccccaaaggg
283981 aacgcacgat gctgtagcca agtttcaaaa aactgacctg catcaagatc tcgattacat
284041 cgtactgcaa caacgaagaa cacagctaga gacgcttatt aacgaaagag aatcttttgt
284101 caagaatctg tgttctcttt tccacaaaat tcaaaatacc aagaattacc aggaatttgt
284161 tgatgtattg gcggagaata gggatttact gcgagaaatt ttcactgtag agaacggatt
284221 tcaaaagcaa aaatggatta gcaacgacga cattccccag atagactggg acaagtttgc
284281 cctagatatc aacgcttata tagcagagaa cgatcaattg ttggctttgt atgaagatgg
284341 cttattatga tcaaatagcg gctatggaca tttttaaaca tatatttata tatatataca
284401 tatatgcgta tattgcataa atcacaaaga aaaacaacgc cctagatata gtgacccaaa
284461 atattatgtt taagttactg gttggggttc atgtacattt ttcactatct tttaattcca
284521 aatgtggatt gtgcttctga gattttgttg cctagtattc tgtgatggga aaaatcgccc
284581 cgatgagata taaaatgagc tagcccccta accaaagaaa gcaggattgg ttagtacata
284641 gagaaaccaa agctgttatg ttgttctata agcctgtgat gaggatggcg gtgagaccgc
284701 taaaaagcat aagattccag tcctcataca ccagtattac taaattgacg aacctaacag
284761 aatttaggaa tttgatcaag caaaatgata aactagtcat cgacttttat gctacttggt
284821 gtggcccctg taagatgatg caaccacact taacgaaatt aattcaggct tatccagatg
284881 taagatttgt caagtgcgac gtggacgaat caccagatat tgccaaagag tgtgaagtga
284941 cggctatgcc cacctttgtt cttggcaagg atggccaact catcggcaag atcattggag
285001 ctaaccctac tgctttagag aagggaatca aagatctata aattttatat tcataagcac
285061 atgtagtttt tttttttttt agttttaaat gtaagaaaaa agatagtata tatatatata
285121 tatatacata tacatataaa aggaaagtaa gcttatacaa tacacaaata aattcatatc
285181 aaagaataat gaaccgcaat attcagaaac acaggaaaag gaggggaagg gatgaatggt
285241 gaggaaagta actgtttgtt ttaaagcgta cctggatcat aacataacaa ataagtagat
285301 aaaaaagaat agagagaaag atataatgac aatgaattga agaatagttt agttagttac
285361 atttgtaaag tgttcctttt gtgttctgtt cttaatttgg cgctattttt ttatacttcc
285421 aaatccttgc tttacaatca ccgctaccag tagcaaaaac gttatattct ggacccagcg
285481 aagacccgtt tgccacagcc acagatataa ctgaattcct atgaccttgc aacatcaata
285541 acggattgcc ggatttctta tcccaaaaca ggacaccacg atctttggaa ccggacaaga
285601 tgtactcatc attttgtgtg gtggccacgg acaatacaaa gtctttatgc ccgatatacg

285661 taacttcaca agtgccggaa tttggagttt tcgaatcgct cttgttgttt gcattctgca
285721 aattccagag cttaacagat ctatctaatg agccggatac aacgctttgt ccatctctcg
285781 tgaagacaac gctataaaca gagtccttgt ggcctgtacc ggattcgttt tccgaatcta
285841 gtctttccac caagaatccg gtctcggaat cccaaactct cacagcacga tctagagaac
285901 cagcagcgat gtatttacca tcacctggtg atacagcgac ggtggtaaca ccatcttcaa
285961 tggataaagt caatgaacac tggcctgtac gtaagtccca aatacgaacg gtacggtcac
286021 cagaaccgga gactaatttg tcacctgagg gaaagtagtc caatgaataa atatcttgtt
286081 cgtggccttg aagaatcata acaatctttc tattttcaat atcccaaatt ctaatcagtc
286141 tgtcttcagc acctgttgcc aaaaatttcc catctggaga aaaacacact gaacggatat
286201 acaagtcgga tgatggggaa gacgaagtgt tcaggttttc cacatctttt gccaattctg
286261 ctgccgaagt catcgctgtg gtagtaattg tggtggtagt agtggttgtc attgtattgt
286321 tatccgtgga cgtggtggtg ttattttcag tgatcgaatt tcgatggtta ttggcagcag
286381 aatcgtcaga taaacgggcc accagagaac cgtctgaaac gcgatacact tgagtagttt
286441 tgttgcagcc tgtggctaag tattcaccat cgttactgaa cttcacgcaa caaacaactg
286501 aagtatgatc caaagatttg tgtaactcaa cgtcaatttc tcttggtagt gccgggttgt
286561 ataaaatata ataatcattt gtttgcttct tcagagcatc gggaacagac tgggaatcta
286621 gatccaaaag gaaaggtggg ataggtttag agtggtttgc tctttgatta taagggacta
286681 agtagtgatc ctggtgcaaa ctagccgggg tggcgtcttc ctccttaggt ttgatttcag
286741 tttcagttgc ggtggtagtg gtggccgtgg tggcggatcc ggtgtcattt atcttcgagg
286801 tattattatt ttccggttca gtttctttca acgtagattc aggtgccttg acagagggta
286861 aagtagtggt ctctttagaa gtaggagatc cgttgatggc tgtgttactc aatggtgcca
286921 cggaaacctg gggaggtggt tgctgttgtt gaagttgctg ctgttgcagt tgctgttgtg
286981 gtagttgttg ttgggcgttt gaggacacca caggcaaagt ggtggtaggc aactgtgagc
287041 caaccagatt aggacggcta gcttgtactg ggaaggccga tggcgaacca gtagttgtgt
287101 ttgctgctgg agtggcggtg gccgaagtag tagccggtgg ttgttgcgca actggaacag
287161 atgcagatgc agcggctagc tgctgttgtt gctgttgtaa atgctgctgg acctgttgct
287221 gctgttgctg ttgctgctgt tgctggacgg tcaaagatgc aatttgatgg tctctttgct
287281 ccagccctag tttcaagtgc ttgatctctg cttcgtacgc gtccttcatt ttcctgtgag
287341 ttagctccag ttcgtagacg gtgtttctta tctgctgcat ctcagccagc tgctggttca
287401 ttttgaaatc gtaatccttt tggttttgaa gacggtaggt atttgcctct tgtgagactt
287461 ggagaaactc ctgtctgatg gcatcgagaa gctcattcag cttattctgc gtattcgaaa
287521 cgctggcagt catattggtt tggatggaaa gctgatttct ttcttcccct gcttatcaaa
287581 aaagacaaaa aaacaataga aaaagagtag tggtgtgcga aaggtaactt ggcgtgaaag
287641 aaaattaaac aattgcagaa ggacagatag aggggcacgt gttcagccag ttgttcttgc
287701 ttcgctgatg tttatcaatt gggggtagat gggggaattt ctctctattt gtttacttat
287761 gtacttgtta ttttgttttg ttgtttgttg ctttgacggg agagcttttt aattttttcg
287821 cagtttttca gtgtttttttg agaagtacga aaaattaagg tccgtcggta gcgataatgt
287881 aagagggttc tgtctctaaa aatgttaggg ggaaagttcc aaaaaattat taactcctag
287941 attgctcagc caaaaagaag ggtaaattag ggctttcctg gtttatttat ttgtttatgt

288001 tgttcttata tacatatgtt tatgtctttt tatgtacaaa taatgatgtt tacaatgaaa
288061 catctgttat gctatcaaaa acgtcagcgc attgttttat agctgaagag gttacaacgg
288121 ataacggtct gatttgcggc ctggcaatgc ttggtaaaac taaatatcaa ttttacgaat
288181 tatttactgt atatagtatt cagtccctca ctcaactggc gtcaagagtg aagaagggcg
288241 gcctcattat ggctcgcctt attctgttca ccctctgcgc tcttcccgta ttatttcatt
288301 ttattttgtt tatgcttcaa tatcttgtat ttgtttacat tgaaaaatga ttggaaaaag
288361 gttacattga aatcaaaaca gtgaagaagc agacagacaa gaaggtcatt agatagataa
288421 acaggaacaa taaaaaaaaa agaaagcgaa acaagacaat ccaacgtgcg caaaatggat
288481 ccattgactg tatacaaaaa ctcagtgaaa caacagatcg attccgcaga cttactagtg
288541 gcaaatttgg taaatgagaa tttcgtactg tcggagaagc tggatacaaa agcaaccgag
288601 atcaaacagc tgcagaagca gatagactcg ctaaacgcac aagtcaagga gctgaagact
288661 cagacttccc aacaggcaga aaattcagaa gtcataaagg acctttacga gtacctctgt
288721 aacgtacgtg tacataaaag ctacgaggat gattctgggt tgtggttcga catctcgcag
288781 ggcacccact cagggggatc ttccgatgat tattcgataa tggactataa actcggattt
288841 gtcaagggcc aggcccaagt cacagaagtc atctatgcgc ccgttcttaa acagcgatcc
288901 accgaggaac tatactcgct acagtcgaaa ctaccggaat acctcttcga aacgttgagt
288961 ttccccctct cgtcgctaaa ccaattctat aacaaaatcg ctaaaagcct gaataagaaa
289021 agagagaaaa aagatgaaac cgagtaagct gctacataat gtctatatat ctacacataa
289081 aattccgatt attcctttgc atacctgatt tgcccctcag aatccacaac cagacttttc
289141 aagaaggtct tttttgcccc tttatccttc atggttttca aaattttgta caacgacttg
289201 cccttgtgaa gctcgattcc gttcgaggtc ttttccactt tctctgcctt cttgatcaca
289261 ggcttctcca ctttctttgc aggagttgcc actggctgct gctgctgctt ctgctgctgc
289321 ttctgcttct gcttcttgtt gcccttgtat aacgctttct ggtacttctc gtcctcgctg
289381 atgcaagacg tgtgattctt gtaactcacg ccatcttcaa acgtcttgga gcaatctatg
289441 catgtatagt acgcgttagg acatctataa taatgctttt cggtattctt cttgggcaca
289501 gtatcattgc acacctcaca gttgaacgta accatcctgt aataacaaat atttcttgac
289561 tgagaccgtt tgctgttgta tacagaatac tcttagagct catcgcaagt taaaaatttt
289621 caattttttt tcactttttc ccgtcaaggc aaaaagcaac caaaagagaa tgaacctcta
289681 tttttgattt atttattatg aaatgctgcc agtccactca tccgcatcaa tgtagtagtc
289741 acaaagctaa tatttagacg ttactttgat atctctgtcc atgagagcac ttattttagg
289801 aagttaaatg agacagtcaa tagttcacaa tatcccgtca gcaatggagg gaaaaggcat
289861 tccttttcca taggatttta atcgttttca agcatcatac gccctcgagg aactcttgtt
289921 ttcgctttac tatgcaacca ttgatgtatt tctgcataaa tgtgcgtcac gtggccttgt
289981 gtctcttatt tccacttgtt ttttcacaat gcggaaaacc tcgattaaag tagaaaaaaa
290041 ggatataata ggagtatacc atattggata gttcaatcta taaacaaaca atcgcataac
290101 cgcacgtata tatacgcaca cacctatcaa tcacaatggc tttggaacct attgattaca
290161 ctactcactc gagagagatc gacgcagagt acctgaagat tgtcagaggc tccgatcctg
290221 acaccacctg gttgattatt tcacccaatg cgaaaaaaga atacgaacct gagtctaccg
290281 gttcctcctt tcacgatttc ttgcaatcgt ttgatgaaac caaggtccag tacggactgg

290341 cacgtgtgtc cccaccaggg tcagacgttg agaagattat tatcattggt tggtgtcctg
290401 attctgcgcc attgaagaca agggcctctt tcgccgccaa ttttgctgca gttgctaata
290461 atctgttcaa gggttaccac gttcaagtta ccgccagaga cgaggacgat cttgacgaaa
290521 atgaactgtt gatgaaaatc agtaacgcgg ccggtgcccg ttattctatt cagacttcct
290581 ccaagcaaca ggggaaggct tccactcctc ccgtgaagaa atccttcaca ccttccaaga
290641 gccctgctcc agtttctaag aaggaaccag tcaagactcc ttccccagca cctgctgcta
290701 agatttcttc ccgtgttaac gacaacaatg acgacgacga ttggaatgag cctgaattaa
290761 aggaacgcga cttcgatcag gctcccctga aaccaaatca atcatcttac aaaccaattg
290821 gcaaaatcga cttgcaaaaa gtgattgctg aagaaaaggc taaggaggac ccacgtcttg
290881 ttcaaaagcc aaccgctgct ggttccaaga ttgatcctag ttctgatatc gctaatttaa
290941 agaacgaatc aaaattaaag agggactccg agtttaactc ctttttgggc accactaaac
291001 ccccctccat gacggaatct tcattaaaga atgatgatga taaagtcatt aagggtttta
291061 gaaacgagaa atcacctgct caattatggg ccgaaagaaa ggcaaagcaa aacagtggca
291121 acgccgaaac taaggctgag gcaccaaaac ctgaagttcc agaagatgag cctgaaggtg
291181 aacctgacgt caaagatttg aaatcaaaat ttgaaggatt ggccgcttca gaaaaagagg
291241 aggaagaaat ggaaaacaaa tttgctcctc ctccaaagaa atcagaacca actattatct
291301 caccaaaatc cttctccaag ccacaagaac ctgtgaaaac tgaagaagct cagcagccta
291361 agactgatta caagaagatc ggcaacccat tacccggtat gcacattgaa gcggataatg
291421 aggaagaacc agaagagaat gatgatgact gggatgatga tgaagaggag gctgctcaac
291481 ctcctttgcc ttcgaggaat gttgcgtcag gagcaccagt gcaaaaagaa gagcctgaac
291541 aagaagagat cgccccaagc ttaccttcta gaaactcgat cccagctcca aaacaagaag
291601 aagcacctga acaagcacct gaagaagaaa ttgaagaaga agctgaggaa gccgctccac
291661 agctgccatc aagaagctct gcagctcctc ctccgcctcc aagacgagca actccagaga
291721 aaaagccaaa ggaaaatcct tgggccacag cagaatatga ttacgatgct gcagaagata
291781 acgaactgac ctttgtggaa aatgacaaga ttatcaatat tgaatttgtc gacgatgact
291841 ggtggctagg ggaactagag aaagacggct caaaaggtct cttccccagc aattatgtgt
291901 ctttgggcaa ctagaatcac acgtcagcgt catgctatta tattattctt acgtaaaaaa
291961 atacttacag agacgttaaa gcttttgtat agatactttc acgagccttc cctttcagta
292021 aaatgaacaa attattttgc cttgtccttc tttgtttcaa gcgtttatat gactcctcta
292081 cagtattatt tttcaacaga cggtgtgccc gtacggcgtt gtcaaaacct cgaacataaa
292141 acaacactga aacaaactta ttgcactcct gcatcgcata cagatctgat acttactaca
292201 accataatgc tgttttagaa tcgcatcagg tgtttaccag aagtgtatgg tacatggtgc
292261 tttccttatg ctcactattt cggatttact tctagtctat gtatagattt gttagagacc
292321 agttttaagc tacaatggaa cgttaagatt aagaaatgca tgaagagtct cccagtagat
292381 atataaagct gaattaaatt cagtacaatg gaagtttcca cgtctttctg tattttcata
292441 cccttaatta taggaaaagc acactacaaa aatacgctat catctaccga atataactgc
292501 aatgaactca tttgcgtcat taggtctgat atattcagta gtaaaccttt taactagagt
292561 agaggctcaa attgtgttct accagaatag tagtacttca ctgcctgtcc ctactttagt
292621 atccacctca atagcagatt ttcacgagtc ctcatcaact ggcgaagtgc agtattcatc

292681 ctcttattcg tatgtgcagc cctcaataga ctccttcact tcatctagct tcttaacaag
292741 ttttgaagct cctaccgaaa cttcttccag ctatgcagtt tcttcctcat tgataacttc
292801 tgatactttt tcttcatact ctgatatctt cgatgaagaa acaagttcat taatatcaac
292861 ctcagctgcc tcatcggaga aagcctcgtc caccctttct tcaactgcac aacctcatag
292921 gacatctcac tcttcctctt cattcgagct accagtcact gctccatcat cctctagttt
292981 accgtcctca acttcattga catttacgtc agttaatcca tctcaaagtt ggacttcatt
293041 taactcagaa aaatctagcg ctctttcctc aaccatagat tttacttctt ctgagatttc
293101 aggttcaaca tctccaaaga gcctggaaag tttcgatacc accggtacta taacttcatc
293161 ttattctcct tctccttctt caaaaaattc taaccagacc tcactactca gcccattgga
293221 gcctctgtcc agttcttcag gagatttaat attgagttca actattcaag ctactaccaa
293281 tgaccaaact tcaaaaacta ttccaactct tgttgacgcc acatcgtcat taccaccaac
293341 attgaggtca tccagtatgg caccaacaag tggttctgat tcaatctcac acaactttac
293401 gagccccccc tctaaaacaa gtggtaacta cgatgttttg acttcaaact caatagatcc
293461 ttctctattt acaactacga gtgaatattc atctacacaa ttgtcgagtt taaatcgggc
293521 ctcaaaaagt gaaacagtta atttcactgc ttctattgct tccacaccat ttggtacaga
293581 ttcggctact tctctaatag accccattag ttcagtgggt tctacagcat ctagctttgt
293641 gggaatttca accgccaatt ttagtacaca agggaactcg aactatgttc ctgaatcaac
293701 tgcaagtgga agttcacaat accaggactg gtcaagctct tctcttccgc tgtcacaaac
293761 cacttgggtt gtcatcaaca caactaatac acaagggtct gtaacgtcaa ccacatcccc
293821 ggcttatgtt tctacggcca ccaaaacggt tgacggggtg atcaccgaat atgttacatg
293881 gtgtcctcta acacaaacca aatcacaagc aattggggtc agttcgtcca tttctagcgt
293941 tccacaagcc tcctcattta gcggtagttc tattttgagc tccaattcca gcactcttgc
294001 tgcctcgaac aacgttcctg aatcaactgc aagcggaagt tcacaatacc aggactggtc
294061 aagctcttct cttccgctgt cacaaaccac ttgggttgtc atcaacacaa ctaatacaca
294121 agggtctgta acgtcaacca catccccggc ttatgtttct acggccacca aaacggttga
294181 cggggtgatc accgaatatg ttacatggtg tcctctaaca caaaccaaat cacaagcaat
294241 tgggatcagt tcatccacga ttagcgccac acaaacctct aaaccatctt caatattaac
294301 attggggata tcgaccttac aattgtctga tgccacattt aaggggactg aaactataaa
294361 cacccatctc atgaccgaaa gtacttcaat cacagagccc acctatttta gtggcacatc
294421 ggatagtttt tatttgtgca ccagtgaagt taatcttgcg tcttccttat cttcttatcc
294481 aaatttttca tcttcagaag gctctacggc gaccattact aactctaccg ttacatttgg
294541 atcgaccagc aagtatccat ctactagtgt atctaaccca acagaagcca gccaacatgt
294601 gagctctagt gtgaactcac tcactgattt tacttcaaat tcaaccgaaa ccatcgcagt
294661 tatatctaat attcacaaaa cttcgtcaaa taaagactat tcattgacga ctacgcaatt
294721 aaagaccagc ggaatgcaaa cgcttgtgct ttctactgtc acaacaacgg tgaacggtgc
294781 tgctacggaa tacacaacgt ggtgcccggc atcaagtatt gcttatacga catccatatc
294841 atataaaaca ttagttttga ccactgaagt ctgctctcat tctgagtgta ctccaacggt
294901 tattaccagt gttactgcaa caagctctac aatccccctt ttatcaacct ctagctctac
294961 ggtattatct tctacagtat ccgaaggtgc aaaaaatccc gctgcttctg aagtaactat

295021 taatacccaa gtttctgcta cttccgaagc tactagtact agcactcaag tgtctgctac
295081 ttctgcgacg gccactgcta gcgagagttc aaccacatcc caggtttcta ctgcttccga
295141 aactattagc actctcggta ctcaaaactt taccactact ggaagcttac ttttcccggc
295201 tttgtctact gaaatgataa atactactgt ggtttcccga aaaaccctaa ttattagtac
295261 agaggtatgt tcccattcca aatgtgtccc aacagtcatt accgaggttg ttacttcgaa
295321 aggcacgcct tctaatggac attcttctca aactctacaa acggaggcag tagaggtgac
295381 attgtcatcc catcaaaccg taactatgag taccgaagta tgttctaatt cgatttgcac
295441 accgactgtt attacatctg tgcaaatgag aagtactcct tttccatact taacttcttc
295501 aacgtcaagt tcctctttag cctccaccaa aaaaagttcc ttagaagcct cctcagaaat
295561 gtccaccttt tctgtcagta cgcaaagttt gcctttggca ttcacaagtt cagaaaaacg
295621 ctccaccaca tctgtctctc aatggtcaaa taccgtttta actaatacaa taatgtcctc
295681 ttcttctaat gtcatatcaa caaatgaaaa gcccagtagt actacctctc catacaactt
295741 ctcttcgggg tactctttac cttcttcttc tacaccttcc caatattcac tatctacagc
295801 tactacaaca atcaacggaa tcaaaactgt gtacacaact tggtgtccat tggcagaaaa
295861 atctactgta gctgcttctt ctcaatcttc ccgcagtgtt gacaggtttg tttcgtcgtc
295921 aaaaccatcc tcatctttat ctcagacctc tattcaatat acattatcta ctgctaccac
295981 caccataagt ggtttgaaga ctgtatacac gacttggtgt ccattaacaa gtaaatcgac
296041 tttaggtgct actactcaaa cttcctcgac agccaaagtt agaattactt ccgcttcatc
296101 tgcaacatct acttctattt ctttgagcac ttcaacagaa tcagaatctt catctggata
296161 tttgtcgaaa ggagtatgct caggtactga atgtacgcaa gatgtgccaa cacaatcatc
296221 ctcacctgct tcaacgttag catattcccc ctctgtttct acatcatcat catcatcatt
296281 ctcaacaaca actgcatcaa cactaacgtc aacacacacc tctgtcccgt tattaccatc
296341 atctagctct atatcagcat cttcgccatc atcaacttcg ttgttatcca cttctttacc
296401 atctgccgct tttacgtcat caacacttcc aacagcaaca gcagtatctt cctccacttt
296461 catagcgtct tctctaccat tgtcctctaa atcatcattg tcgttatcgc cagtctcgtc
296521 gtctattttg atgtctcagt tttcatcatc atcatcatca tcatcatcat catcatcatc
296581 atcattggca tcgttgccat ctctttctat atcaccaact gttgacactg tttctgttct
296641 acaaccaact acttccatcg caacactaac ttgcacagac tcacaatgcc aacaggaggt
296701 atccactatc tgtaatggat ccaactgtga cgatgtgact tcaactgcca ctactcctcc
296761 atctacggtt actgatacta tgacatgtac tggatctgag tgccagaaaa ccacatctag
296821 cagctgtgat ggttactcgt gtaaagtatc cgaaacgtat aaatcaagcg ctacaatatc
296881 tgcatgtagt ggagaaggat gccaagcttc cgctacaagt gagctaaatt ctcaatacgt
296941 cacgatgacg tctgtcatta ccccaagtgc cataacaaca acatcagtgg aagtgcattc
297001 aactgaatcc actatatcaa ttactacagt gaagccagtt acatatacat ccagtgatac
297061 taatggagaa ctgataacca taacaagttc cagccaaact gtaattccat cagtaacgac
297121 gataataacg agaacaaaag tggccataac ttcagcacca aagccaacaa ctacgaccta
297181 tgttgagcaa cgactttcct ccagtgggat tgctacttct tttgttgctg ctgcatcctc
297241 aacttggatt actacaccca ttgtcagtac gtatgctggt tcggcgtcaa aatttctctg
297301 tagtaagttc tttatgataa tggtaatggt gatcaacttc atttaatttc ccgtatatat

297361 tatatgtacg tatatatgct ataaacagac actcttttta cttaccttta ataatgcttg
297421 atgagaaatc agtcattgtt acttcttccc caaggttgtc atcataatca taccattcat
297481 tatcctctag ttggcatgac attactttcc ccgaacttag agaaacttca aacgttatgg
297541 tgtcgggata aaatttgatt aattcacaac ctctgcaatc aaggcttaaa ggaatgaaag
297601 aatcttcttt gaccttcgat aaaccatgct tcttccgcac atcttttatc ttggcccatt
297661 tgtcgtcact ctgatcagtt aaatattcat cctcaaaagc gcacaggtta actgacaatt
297721 cttttgaaca aaatttacac ttcatcaaga aagatgcggt accttttgat gccggcatag
297781 catattcttc aaatgtattg atgatgactt tagaatcgtg taattctcta caactggtac
297841 attgtagatc gaaggcaaat tctgcccttg attcgtttgt attctcaatt gaaaccttgg
297901 tcacgttttc tgataatgtt gctttcagaa ccaaaaataa cggcattatc gtacacaacc
297961 tggttttctt tattggttaa gttgtcttta ttcaaaattt ttcattactt cttataaaca
298021 gcttcttgac gcttgtaatt tcataagttt ttgatacaag caatggaaac aaaattaaga
298081 aaatgtaggc aacgaagaac taaatttgaa aaaaaaaaaa aaaaaattta ctgccaagat
298141 catctaaatt atcttatagt agcaaagctt gagcataaaa ggatcaatct tttagttttt
298201 tatggcaaag tacatttgca gtaagtaatt agtgatgaga taaacaaaga ataataatcc
298261 aattcgaaat caaaaattca aacaacaaaa aacataattt cacatatata tctaattcat
298321 tcacgaaaaa cacgtcagtt ccttctgtct gttgtaaata ggatgcatcc gcagtgaatc
298381 atggaaatat ttcaaaactt tcagcccaat cattttattt ggtcaatgaa gccataaacg
298441 ctccgtaatt ctaagattaa aaaaaattca taaaaataag gaccaatata aatatatata
298501 tttaatgtgc tagtgacaat tctagtggta atagtgcaaa ttatactaca attggccatg
298561 gtgaagccta aacacaccat tacgtcttca cagtgttatc cattgtattt tcaaaggtat
298621 gctactacta atgcaccagt gtataccatg atacggtgaa aagggcatgt aatattgcgt
298681 taatttgtcg ataacctttc tttacgttta aatttttttt ttataactgg cgacactaaa
298741 cttctcgata aaatacaaca taaaaagcag tacctagcta agcgaacgac agataataaa
298801 tagcgaaaca tctaggttaa tagccaatta gcaatacatt taaggattga tagtgtggta
298861 tttacaaggt aatagcccgc taggctataa ggaaactact ctaaattcac cttccatata
298921 tgtgactgcc aagaaaatat tactagttat caagcttttt ttgtgtgcat tcaacttcaa
298981 cgaatccttc gtcatcaatg aaacaaaaga acaagccaga actgaaaaaa aaaaatcaaa
299041 accactaaga atgtcacatc cttccaaaac tcctgctggc tcttcaactc ttcaaagaat
299101 atcaagataa atatcattat ctccagtgtg gccgacccat gcatgagtgc gaggagcatt
299161 aagttggatc ctgggaaaac atgtcaagga cattcaataa agtacttttc atactacaac
299221 agacgctaga agaactagaa tgtaaccgtg aactgcctct caaacacata attatgatag
299281 cgccgtcttc ggcatccttt ttcggaaata tactttcctt tttccctctt tcagctatat
299341 tgttcaagga gtaacataag acataataac cccagttttt ttagagaaag ttcttcttct
299401 ttaattatag tatgtttgaa acgtacaaga attcattgag tgccctaagg attaccgcaa
299461 acaaggttca acagtgttgg tacacatgac tcagcaagaa taccgttccc cctcacaacg
299521 cttatccaag gggaggagca tgtcgctacc caaaatattt gctcgtaatt tgagatctct
299581 gcaaaacaat gcacctcctg gcaaaaacat cgatgtcaat tgtttgaacg tcaattcttg
299641 ttcgttgtcc gcaagcccaa gctcacaaat taatatggct tgtaatggaa acaagcaaga

299701 tcttcccata ccgtttcccc tgcatgtaga atgcaacgat agctggtcaa gctccaaact
299761 taacaagttc aaatcaatgt ttaatcataa cagatcaaag agcaatggta ctacagatgc
299821 gtcaacttca gaaaaaggta cgcataagcg tgaaccccgg tcgacgatac atacagagct
299881 gttacaaagt tccattatcg gtgagccaaa tgtccatagt actacaagta gcacacttat
299941 acccaatgag gcgatatgct ccacacctaa tgagatctca ggtagctctt ctccggacgc
300001 ggagttattt acctttgaca tgcccacaga cccgtcatcc ttccacactc ctagctcccc
300061 aagttatata gcaaaggaca gtagaaacct gagtaatgga tctttgaatg atattaacga
300121 aaatgaagag ctccaaaatt tccatagaaa aatcagcgaa aatggcagtg cctccccccct
300181 ggctaacttg tcattatcca attcaccaat tgattcccca aggaaaaata gcgaaaccag
300241 aaaggatcaa atacctatga acataacacc acgtttaagg agggccgctt ccgaaccgtt
300301 caatacggca aaggatgggt taatgcggga agattacatt gccttgaaac aacctccaag
300361 cttgggagat attgtagaac cgaggagatc tcgtcgttta agaaccaagt cattcggtaa
300421 caagttccaa gacattactg tcgaacctca atccttcgaa aaaattagac tacttggcca
300481 aggtgacgta ggtaaagtgt atttagtgag ggaacgcgat accaaccaga tattcgccct
300541 gaaagttttg aataaacatg agatgatcaa gaggaagaaa attaaacgag tactcactga
300601 acaggaaatt ctcgcgacaa gtgatcatcc atttattgtg acactgtatc attcctttca
300661 aaccaaagac tatttgtatc tctgtatgga atactgcatg ggaggggaat tctttagagc
300721 cttacaaaca agaaaaagta aatgcattgc agaagaagat gcgaagtttt acgccagtga
300781 agtagtagca gctttggaat atttacacct actgggcttc atatacagag atttgaaacc
300841 cgaaaacata ttactgcatc aatctggtca tgtcatgctt tctgactttg atttatccat
300901 ccaagcaacg ggatcaaaaa aacccaccat gaaagactct acgtatttag atacaaaaat
300961 ttgttcagat ggattcagaa ctaattcctt tgttggtact gaagagtatt tagctccaga
301021 agtaatcaga gggaatggcc acactgcagc agtagactgg tggactttag gaatattgat
301081 ttacgagatg ctatttggct gtactccatt taaaggagat aattcaaatg aaacattctc
301141 taacatttta accaaggacg tcaaatttcc acatgataag gaagtttcga agaattgtaa
301201 agacctgata aagaaactac taaacaaaaa cgaggcaaaa aggcttggtt ccaaatcagg
301261 agctgcagac ataaagagac atcccttctt caaaaaagtt cagtggtcgt tcttaagaaa
301321 ccaagacccc cctctaatac ctgcattaaa tgataacggc tgcgaacttc cttttatatt
301381 gtcttgcaat aaacacccga aaaggaactc agtgagtgaa caggaaacca aaatgttctg
301441 tgagaaagtt gcaaacgatg atgaaattga tgaggctgat ccattccatg attttaattc
301501 tatgagttta acgaagaaag atcacaatat cttaacctac tctgaaaatt atacctacgg
301561 aaaaattcta tacaaagcaa cttgtacaag gccaaggcat aacagctcac atagaagttt
301621 ctttaaagac atcatacctg aactataaca tgtttacaga aagataaacc catggccaaa
301681 tttctgtatt gtttctctca atttatctta tgataaaaat gtaaaaaatt atttaaaagt
301741 aaacaaataa acaaaggaac aattcaaaaa cgagaaagtg aatgaaaaat ataatatcat
301801 ttattgtctg ataatgctgc atttagaaca tacgtaccat ccgcatcagt ggatatccaa
301861 tgatagtaat ttcgcgagtt tatccgttgc tgttatatta tcacttttta atattctttt
301921 caaagagctg aataatttca acgcgtcttc gttaatggat tccttccgca attcttctga
301981 aatactgaat gcccgattta taatatcttt gtccaggcgt gccaatttcg ccacattcat

302041 cccataacta ttataagtca atcccttttt taacttatat agaaaaatta cactcatcca
302101 gtcctcgcca gttttttgtt cttccacgta atccatatga taattcctta ttaacggaga
302161 tttgatttct cccagcatgg gaaaatgggt agtaaataat atcaaggggc agtcacttaa
302221 ctcagaaaaa tactttatta aagcataaga aattgctata ccatcgtgcg tgccagtacc
302281 tcttcccact tcgtctaata atagtaaaga ccgtttattg caatttttca agatgtgtag
302341 gatatcaagc atttccactt taaaagtaga atcaccgttt ataatatcat cgtgcgcacc
302401 gattcgagtg agtacgtttt caaatatgct taatctgatc tcttctgcgg ggacaaatga
302461 gccgatctgt gccattatag taagcagtgc cacttgtcta atataagatg atttcccacc
302521 catattcggc cccgttataa tattgatttt accgttttct ggggacatca tgatatcatt
302581 tggtacataa tgaacatcca gcgactcgat aattggattt cttgcatttt ttgcgattat
302641 ggcttgttga ccattcacaa aagttggtct aacataattt acgttgcatg atgtggctgc
302701 taacgacaaa atacagtcat actgcgccaa attgagtgta attttacgga gctctgtata
302761 ttctgccgta attttgttca agaattcttt atactgtagt tcagattccc gaattaataa
302821 gtccttgtaa tattctagct tttgagtcag tttctgggtt ctgggagtgg taaatctact
302881 gaccatcttc gtattgttaa cttttatcca atcatctggc aagtccttaa tttgcgagtt
302941 tttcacttcg attaagtaat caacttcatc tctaaaattt agatatggac gtttgagata
303001 ttttcgtatt tctgccaatt cttcctttaa ctgtgaccgt actgattcgc tttccctttg
303061 aatttttatt ataccctctg aacaatcata gttatttaaa ttaaaaaaat ccattacttg
303121 cttatctgaa tttttttcca ttaccgcaga aacgttgatc atggtcaaaa aatgaggcaa
303181 ccgagtggta gaaagtagtt cattcaattc actaaatagt ctaaaaagta aaggagattg
303241 tttgcctatc cttccatctg atgacttgaa atgttctgac aggtaagatt gatgcatctt
303301 gaagtgatca acgaaagaag ttatttgctt taaatagaaa tagacttctt ttctagaagt
303361 tgtaccatac attatgcgat ttaaagttct taataagtca ggggtatgat tcaacatttg
303421 attcaacgat tcaaaaaata tactgttgtt gatttcggat gtaatgcact caatggcatc
303481 aagccgctct tcaatttggt gtacatcaac caaaggtttg agaatccatt ctctcaacat
303541 tcttaatcca aacgatgtcc ttgtatggtc caataaccaa aacaaagaac ctttaccacc
303601 atcatgggta aaaatgtcca aactttgcag ggagttagga tcaagtaaca tatgtatttt
303661 tgatgcgaag ggcgaataga tagaaggaat caacatcacc tgctcattat tgtactcaac
303721 catatgcgaa tataacttag aaactaagcg tatgagagac ggcgagagtt gaattttctc
303781 attcattact tttattgcct gaactacatg atcttccaaa tcatactcct gcttgtgtat
303841 taaaggacat gaaatatctt tgaaaaattt cgctacatgt aatggaagat catctgtatt
303901 taccagtact tctatgggct gtaaatattt gattcgtatc tgtagtttct catcagcaag
303961 attaggctct tcaaattcat catacacgac ttccccgtta tttaaattga cagaaattaa
304021 ggaatattta gccacctttc cctgatgtac gtcacgggac aaagcccata tactgtttgt
304081 atcaccgaga atacgtttcc ccctaaggac aaaggtggaa ttaacaccaa atgtagcttt
304141 ggtaaagaca tttgaaatct ttctttcaaa aacgcttgat ttgctggcac ctggatcatg
304201 cttcttaata gcgcttgttt ctgcttgctc taccacggca acctttaaat tatgatgcac
304261 aagtctctct aggtgaacgt ttaatctgac atccgggaaa gaacagtacg caaactgcct
304321 atgattgcaa tcttgaggac tagactcatc gatagtcaat tttccaggca caagtttgat

304381 gtgaagtatt ctgctaaccg ttactgcatc ctctgcaaaa catttgtact tgtagcctac
304441 tctaataaca agcactttat ctctatgatg cattttcagg tccttcacct gtttgtccaa
304501 gggagtaagt ttcgctgtag gggactttct ggcttttttt tttacgaaat cttcctcacc
304561 ctcttcctca tcatcttcag cctcaacatt ttcatcactt cgtcttttca taatcctatc
304621 caatttctta gcaaagtcaa tatcattaaa tgttgtcgaa gtaccactag tcttttctga
304681 atttttcgaa ttcttggaag aaacactgcc tttgagtgga aagctatcat tagttactgt
304741 tgtgcttgca acagaagata aattgtcctc ttcatcagtg tcaaggcaga tggattcgct
304801 accagcgcca tttccaaccg caacttcttg ttcttgctta tgcgtcagct ctgattttac
304861 cgccttcttg aaaaaacctgc ttattgtggg ttgtcccgcc attattgtta atagattcaa
304921 caaaataaat ctatttgcac tgcttttggc tctcaaaagt accagtttag gttcattacc
304981 tatcaccatt gttccagctt atttatcatt gaaaattcac gtgatcgcaa tattccgtca
305041 ggagtgataa atacacgaaa atttttcttt ttttttcggt ggcaaccaaa gtcaaaccat
305101 cgaattcgaa taagataaac aggggcaaag gtgagtcggg gaaacactat tggnnnnnnn
305161 nnntcattaa atgttgtcga agtaccacta gtcttttctg aatttttcga attcttggaa
305221 gaaacactgc ctttgagtgg aaagctatca ttagttactg ttgtgcttgc aacagaagat
305281 aaattgtcct cttcatcagt gtcaaggcag atggattcgc taccagcgcc atttccaacc
305341 gcaacttctt gttcttgctt atgcgtcagc tctgatttta ccgccttctt gaaaaacctg
305401 cttattgtgg gttgtcccgc cattattgtt aatagattca acaaaataaa tctatttgca
305461 ctgcttttgg ctctcaaaag taccagttta ggttcattac ctatcaccat tgttccagct
305521 tatttatcat tgaaaattca cgtgatcgca atattccgtc aggagtgata aatacacgaa
305581 aatttttctt tttttcggtg gcaaccaaag tcaaaccatc gaattcgaat aagataaaca
305641 ggggcaaagg tgagtcgggg aaacactatt ggtaaaagat aagcaattga gaaacgtcac
305701 tcagggacaa ttgattacat gctatcggcc acataccgtg atttgaacac agcatctaat
305761 ttagaaacat caaaggaaaa acaggccgct caaatcgtca ttgcacaaat tagtttatta
305821 ttcacgactc ttaacaacga caattttgaa tccgtggaaa gagaaattag acatatttta
305881 gacaggtcgt ccgtagatat ttacataaaa gtttgggaac gattattaac cttaagttct
305941 cgggatattt tacaagcggg aaaattttta cttcaagaaa atctactaca cagactacta
306001 ttagaatttg cgaaggattt accgaagaaa agcacagacc ttattgagct gttgaaagaa
306061 cgaaccttca ataaccagga gtttcaaaaa caaacaggaa ttacattatc acttttcatt
306121 gatctatttg ataaatctgc aaacaaggac attatagagt cacttgaccg ctcctctcag
306181 attaacgatt tcaagacaat taagatgaat catacaaatt atttaaggaa tttttttctt
306241 caaaccacac cagaaacact agagtccaat ctacgcgact tattgcattc cttggaaggt
306301 gaaagtctaa atgacttatt agctctttta ctgtccgaaa tactttcacc tgggtctcag
306361 aatttacaaa atgatcccac acggagttgg ttgacacctc cgatggtttt agacgcaacg
306421 aaccgtggga acgttatagc aagatctata agttctctgc aagccaacca gataaattgg
306481 aatcgtgtgt ttaatttaat gtcaacaaag tatttcttga gcgcaccatt gatgcctact
306541 acagcatctt tgagttgctt atttgcagca ttgcacgatg gtccagttat tgatgaattt
306601 ttcagttgcg actggaaagt tattttcaaa ctagatttgg ccattcaact tcataagtgg
306661 tcggtacaga atggttgctt tgacttatta aatgcagaag gtaccaggaa agtttctgaa

306721 accatcccaa acacaaagca atctttactc tacttattat ccattgcatc attgaattta
306781 gaattgttcc tacaaaggga ggaattgtct gatggtccta tgctagctta ttttcaagag
306841 tgcttctttg aagatttcaa ctacgcccct gaatatctta ttttagcatt agtcaaagaa
306901 atgaagcggt tcgttttatt gatagaaaac aggacagtca tagacgaaat acttattacc
306961 ttattgattc aagtgcataa taaatcaccg tcatcgttca aggacgttat ttctacaata
307021 accgatgatt ctaaaatcgt agatgcagca aaaatcataa tcaactcgga tgacgcacct
307081 attgccaact ttttaaaatc gttgttagat acgggaagat tagatacggt cattaataaa
307141 cttcctttca atgaagcttt taaaattttg ccatgcgcaa gacaaattgg ttgggagggg
307201 ttcgatactt tcttaaaaac aaaagtttct ccatctaatg tcgatgtagt gctggaatca
307261 ctagaggttc aaacgaaaat gactgataca aacactccat ttaggtcatt aaagacattt
307321 gacttattcg cttttcattc attaattgaa gtactgaaca aatgcccact agatgttctc
307381 caattacaaa ggtttgaatc cttggaattt tccttattaa ttgcatttcc tagattgatc
307441 aattttggtt ttggacacga tgaagctatt ttagccaatg gtgacatcgc agggattaat
307501 aatgatattg aaaaggagat gcagaactat ttacagaaaa tgtatagtgg tgagttagcc
307561 attaaagatg taatcgaact tctgagaagg ttaagagata gcgacttgcc aagggaccag
307621 gaagtcttca catgtattac ccatgccgtt atagcagaat cgacattctt ccaagattat
307681 ccattggatg cattggctac tacatctgtt ctttttggat ccatgattct ctttcaactg
307741 ttacgtggat tcgtattaga cgtcgcattt aggataatca tgaggtttgc caaggagcct
307801 ccagagtcca agatgtttaa gtttgctgta caagctattt atgcatttag gatacgtttg
307861 gccgaatatc cacagtattg taaggacctc ttgagagatg ttccggcttt gaagtctcag
307921 gctcaagttt accaatctat cgtcgaagct gctaccctag caaatgctcc aaaggaaagg
307981 tcaagacccg tccaggaaat gatcccatta aaattttttg ctgtagatga agtttcatgt
308041 cagatcaatc aagaaggtgc tcctaaagat gtcgtagaaa aagttctttt tgttctcaac
308101 aacgttactc tggctaactt gaataataag gttgatgaat tgaaaaaaag tttgacacca
308161 aattattttt cttggttttc cacatattta gttacgcaaa gggctaaaac agaacctaac
308221 tatcatgatc tttatagcaa ggttatagtt gctatggggt cagggttgct acatcagttc
308281 atggtcaacg ttactttgag acaattattt gtcctactat ctacaaaaga cgagcaagcc
308341 atcgataaaa agcacctaaa gaatttggct tcatggttag gatgtatcac attagctttg
308401 aataaaccaa ttaaacacaa gaatatcgca ttcagggaaa tgttaatcga agcttataag
308461 gaaaatagac ttgaaatagt tgtgcctttt gtaacaaaga ttttacaaag ggcttctgaa
308521 tcaaaaattt tcaagcctcc aaatccctgg actgttggca tattaaagct gttgattgag
308581 ttgaacgaaa aagcaaactg gaaattaagt ttgactttcg aagttgaggt tttattaaaa
308641 tcttttaatt tgaccaccaa atctctcaag ccctcgaatt tcatcaatac tccggaagtt
308701 atagaaactt tatccggtgc tttgggatca atcactctgg agcaacaaca aacagagcaa
308761 caaaggcaaa ttatactaat gcaacaacac cagcaacaga tgctaatata tcaacagaga
308821 caacaacaac aacaacaaag gcaacaacaa caacaacatc atattagtgc aaatacaatc
308881 gcagaccaac aagcggcatt tggcggcgag ggttcaattt cacacgacaa tccttttaac
308941 aacttacttg gttctactat ttttgtaacc caccctgact tgaagagggt atttcaaatg
309001 gctttagcca agtcagttcg cgaaattttg ttggaagtag tcgaaaagtc atcaggaatt

309061 gctgttgtta cgacgacaaa aataatactt aaagactttg ccactgaagt tgatgagtct
309121 aagttgaaga cggctgcaat cattatggta aggcatttgg cacaaagttt agctcgagct
309181 acttcaattg aaccattgaa agaaggcata cgttctacta tgcaatcact agcaccgaat
309241 ttaatgtctc tttcttcttc acctgcagag gagcttgaca cggcaataaa tgaaaatatt
309301 ggcattgctc tagttttgat tgagaaagca tctatggaca agtctactca agatttagca
309361 gaccaattga tgcaagcgat tgctattcgt cgttatcaca aggaaagaag ggcagaccaa
309421 ccatttatta cgcaaaatac caatccatat tcactgtctt taccagaacc tcttggtttg
309481 aaaaacactg gtgttactcc tcaacaattc agggtatacg aagaatttgg taagaatatt
309541 ccaaacttgg atgttattcc gtttgcagga ttgcccgctc acgctccacc gatgactcaa
309601 aatgtgggtt taactcagcc tcagcaacaa caagcgcaaa tgcctaccca aatcctaacc
309661 tccgaacaaa taagagctca acaacaacag cagcaattac agaaaagccg tttgaatcag
309721 ccatcccagt cggctcaacc tccaggagtg aatgtcccaa atcctcaagg tgggattgct
309781 gcagttcaat cagatttgga acagaatcaa cgtgttctcg ttcacctcat ggacatttta
309841 gtttctcaaa ttaaagaaaa tgctacgaag aataacttag ctgaattagg cgatcaaaac
309901 caaattaaaa ccatcatttt tcaaattttg acattcattg caaaaagcgc acaaaaggat
309961 caattagctt taaaggtatc ccaagctgtc gttaatagcc tttttgccac tagtgagagt
310021 cctctctgca gagaagtttt gtccctactt ttggaaaagt tatgttcttt atccctcgtt
310081 gctagaaaag acgttgtctg gtggttagtt tatgccttgg acagtaggaa attcaatgtt
310141 cccgttatca gatcccttct agaagttaat ttaattgatg ctacagaatt agataacgtt
310201 ttagttactg caatgaaaaa taaaatggag aactcaactg aatttgctat gaaattaatt
310261 cagaatactg tcttgtctga tgatccaatt ttgatgagaa tggacttcat taaaacctta
310321 gaacacttgg cctcttcgga agatgaaaat gtaaagaaat tcatcaaaga gttcgaagat
310381 actaagataa tgccagtgag gaaaggtacc aaaaccacaa gaacagaaaa gctttactta
310441 gtatttacgg aatgggtaaa attacttcaa agagttgaga ataacgacgt aatcacaact
310501 gtttttatca agcaattagt cgaaaagggt gttatcagcg atactgataa tttacttaca
310561 tttgtcaaaa gttctcttga gctatcagtt tcttcattca aagaaagtga cccgactgat
310621 gaggttttca tcgctattga tgctctagga tcgctaatta taaaattgtt gattttacag
310681 ggtttcaaag atgatacaag aagagattac ataaatgcaa tattttctgt gatcgtttta
310741 gtgtttgcta aggatcatag ccaagagggt accacattca atgaacgacc atatttcaga
310801 ctattttcta acatcttata cgaatgggct accatcagga cgcacaattt tgttagaata
310861 tctgattcca gcactaggca ggagctgatc gaatttgatt ctgtatttta caacactttc
310921 tcaggatatt tgcacgctct gcaaccattt gccttccctg gattctcatt tgcatgggtg
310981 acactattat cacacagaat gttattacca attatgctaa gattacccaa taaaataggt
311041 tgggaaaagt taatgctttt gattatcgat ttgtttaaat ttttggacca atacacaagt
311101 aaacatgcag tctctgacgc tgtttcggtt gtttataagg gaacactgcg tgttatttta
311161 ggcatttcga atgatatgcc atcctttttg attgaaaatc actatgaatt aatgaacaat
311221 ctacctccaa catatttcca actaaagaat gttattttat ctgctattcc taagaatatg
311281 accgttccca acccatatga cgtggatctt aatatggagg atattccagc atgtaaagaa
311341 ctacctgaag tcttctttga tcctgtaatt gatttacact cattgaaaaa gccagttgac

311401 aactacctac gtattccctc aaattcatta ttaagaacaa tactaagcgc tatttacaag
311461 gatacctatg acataaaaaa gggcgtaggc tacgactttt tatctgttga tagtaaatta
311521 attcgcgcta ttgtattaca tgtgggcatt gaagctggaa tagagtataa gagaacttct
311581 tcaaatgcgg tatttaatac gaagtcttct tattatactt tattgttcaa tctgattcaa
311641 aatggtagca tcgaaatgaa atatcaaatt attctgtcta ttgtggaaca attgcggtat
311701 ccaaacatcc acacctattg gttcagcttt gtgttaatga atatgttcaa aagtgacgaa
311761 tggaatgatc aaaaacttga agtccaagaa attattttaa gaaacttttt aaaaagaatt
311821 attgttaaca aaccacatac ctggggtgtt tcagttttct ttactcagtt gataaacaat
311881 aacgatatta atcttttaga cctgcccttt gtacaaagtg ttcccgaaat taaactaatt
311941 ttacaacaat tagtaaaata ttccaaaaaa tacacaacca gtgaacaaga tgaccaatcc
312001 gccaccatca atagaaggca aacccctcta caatccaacg cataaaaaaa ctgcatttat
312061 ttatatacga aaatcatgat ttcagaaaaa aaaatataac ttgtataata ctatatttca
312121 gagaatacta ttttgaataa gaatagaagc aatttcgatt aacttttctt actgtatatg
312181 ggtcgctttc acttttgaac tactgaaaat tagtgttacc cgtattattg ttttggagaa
312241 taataataat aattagcatc aatttatatc aaacaagagc tgcaattacc ttatgtccat
312301 gacgcactgc gaacaaagaa taattgagta atcgaacttg aagttctata gaacagcttg
312361 tatatggttt cattgttcaa aagaggtaag gctccaccgc tcacgaaaga aggcccccact
312421 tctaaaaagc ctcctaacac agcgtttaga caacaaaggc ttaaggcatg gcaaccaata
312481 ctgtctcctc aaagtgtgct tccgttgtta atattcgttg catgtatatt tactcctatt
312541 ggtattggac tcattgtaag cgctactaag gtacaagatc taacaattga ttatagtcat
312601 tgtgatacaa aagcatctac aactgctttt gaagatatac caaagaagta cattaaatat
312661 cactttaaaa gtaaagttga aaataaacca caatggaggc taaccgaaaa tgaaaatggc
312721 gaacaatcat gcgaactgca gttcgaaatc ccaaacgata tcaagaaatc catttttata
312781 tattataaaa taaccaattt ttatcaaaat catcgcagat atgtccaatc gtttgacaca
312841 aagcaaatat taggggagcc tatcaaaaaa gatgatctgg atacaagctg tagtccaata
312901 agaagtaggg aagacaaaat aatatatccc tgtgggttga tcgctaattc catgtttaat
312961 gatacatttt ctcaggtgtt gagtggtata gatgacacag aagactataa tttaactaac
313021 aagcatatat catggagtat tgatcgtcac agatttaaaa ccaccaagta taatgctagc
313081 gatattgttc caccgccaaa ctggatgaag aagtatcccg atgggtatac agatgaaaat
313141 cttcctgata tccatacttg ggaagagttc caggtatgga tgaggactgc agcctttccc
313201 aagttttaca agttgacgtt gaaaaatgaa tctgcttctt taccgaaggg taaatatcaa
313261 atgaacattg agttgaatta tccgatttca ctctttggtg gcacaaaatc atttgtactg
313321 actacaaatg gagctattgg tggtagaaat atgtcactag gcgtactgta cctcatcgtt
313381 gcagggcttt gcgccttatt tggcatcatt tttttggtta aattaatctt ccaaccaaga
313441 gcgatgggtg atcacactta tttgaatttt gatgatgaag aaaacgagga ttatgaggat
313501 gtacacgcag agaatacaac attgagggaa attttataga ggccaaaaaa caaactttag
313561 tgcctgtagg tatttgtgtt tataagtaaa caagttttgt tgacttcctg aaacgtaaac
313621 aaaagaaaaa ttgaagtatg cctttttacg cttggaaaat agtgcgacaa agaactccca
313681 gggtttgcta ttcccattta gatgatgaaa tgatagaaaa gtggaaaacg tttatcatga

313741 aatcatctat acatatacgt aaaaatgtaa actatgtaaa aatatgttcc ctataaaagt
313801 gatgctcttc cggaacgttt tctttttccc acaaatctcg ttgaaacctg aaccaatctg
313861 gtaattcaga atcacagggt atccttattt tgacgttatc ttttcccctt gctttataat
313921 agccataatc tccattttct tttttattaa atgctaatct atacctatgg ccaaaaattc
313981 ctaggtcaga agaggttttc ttcaaatcgt ctcctagaac tccacttact tgtaatactt
314041 tattattttt cactttggtt tcgatttgat ttattaatgt caaaagccgc tgaggcactt
314101 ctggctctct gcaaaggtca tcttcattta ctctaataag tttgccagaa ttggccttat
314161 actctatcga atgcgtccta ttattctcaa ggggcaattt ttttgctttg ttgtccacaa
314221 ttgtcttatt atccttctcc tgttctatct caatgttaat tatttctaga aatgtttccg
314281 caaaataatt tctattctta ccagaaaata atctgtactc atttatgatt gatgcaatat
314341 tccatttctg aatttttcga agtataccaa tgaccaacgc ggtcttatct accaataaaa
314401 cattataatt atccacattc aataatgttt tgaaagttct ttttaaacac gtgctcttaa
314461 tcagcattaa gtcatcattg tcagtcaagt ggtaagccaa ttcctcctga attacaggat
314521 cccctgtagt agaattttgt gacccatttg cagtggattt tttcttttct tcctgtaatg
314581 atagagtact gttattattc gaatataaat ttgcgttaga tacagaagag ctttttaccg
314641 gaactgccgc agccgagaaa tcagacatcc ttaaaacaat ccattttatg gatgatcttg
314701 tgaaaaaatc tttgaaaaac ttggatggtt cttgacctcc aataaatata gctgtcttta
314761 aattgagagt ttccagaaag gataaattta atgtctccac tttagaacat ctatatattc
314821 cttcttcagc aatcccaaag ttagcaggtg gaaccagcat tatgaattca ggttatggat
314881 cgcaccagtg ctgtaagtgt ctctcttttt attctacgat ctccatataa caaccacatc
314941 tcatttcttt accttactgt ccatttaatg ctagaaaact tccgaatatt gtattattat
315001 tacccggata tgtcaaggcg gtatattaag atgttccttt caaaagtttc attataccgc
315061 atacggtaag gacagccata agtagtttta ctgataaaat atggggctaa agaactgtat
315121 atttatggat agcactaaaa gtaaagattt tccccgtgtc ctgacttgga gaaacaaaat
315181 ttcattgtca taaaaaactt ttaaaatatg atattcaaag cttgagcatt gggcttctgg
315241 atgacagttc taatcaagtt gggtctaagg atattgcatg tatacaaagg attctttcgg
315301 aaagtaatcc tgaaatattt tttttttcg tctgaacata cgaaagtaaa taaaaaaagt
315361 tcaatgcatg ccttcctatg caagatttat aaaaggtgaa caggtcagga gcagccacat
315421 ctcaataaag gcagttaaag acggtgatgt aggtgcagaa agagaaagtg aacaaaggat
315481 gatttgagat gtgtaatgat tcgtttgagc agcaacaact gctaagatat tcatactaat
315541 aggccacttc acacattatg ccttggagaa tatcctgcat tgctcctttt atagtatagt
315601 ctattcagca ctaacaagca aagcgaacat aatggtaaaa ctaacttcaa ctgttgctgg
315661 tgtcgcctgc atcgctggcc ctcccctcca ctacggccat tttagcacaa cttgaatgag
315721 tcaacttggt tgagttgcat gttaacgtct ctgatatcag agctcacttg ctcaataata
315781 catgttacaa actgcccact aaattgaaca ctctaacatt atattgacct gaattatttg
315841 aaacgaagtc aacagaatga tcactgattt tccaatgtac tccgctatgt tgaagctagc
315901 catcttaagc gatcaatgac ggtgctcaat cattcttcga aatgggctaa aaatgacagt
315961 tgaaatatat tctttacgtt cattttttt catactgcaa acccattcta gaaccaatgg
316021 taaaagtgac gggagttgct ttcttttact atttcaatga tggctatagc ctgcatacac

316081 acaaaagaat gtaagtgatt gcgtgaaagc ttaaagtctt aaaactgtat ttttttcaac
316141 tcctcccata tattaaactg gtcctcacag ttcgcaggat aactaacaat taaaaaaatt
316201 gttcatatgg ttgtatttct ttccgtctca cattattaaa attcaacgaa gaagtttcta
316261 caataacaga caagggccta cgattactat gtactggaag gtgagtcttg aattcatcct
316321 cagagattgt tctaaagacc aacagaaatc catatagatt accgtagtta ctctgatgga
316381 atttcagagt aagacccttg tcaacttcat cactttaagt tgagatgtag aaagacactt
316441 ttgtaaggtc tccaacaagt atatttgata aatagactgt tgagctacaa aaactcaatg
316501 tgaagtaaaa gacctcataa atgatcgatt atctgtttgg caggtactaa taaaactgca
316561 ttggtgtctt ttttcactta cgttaaaaaa aactttttgc aatgatttaa gcgtgcgtga
316621 agataacact acaatccatt ttaaagcaac atccacattg agtgtataca ccacaaaggt
316681 tttttcaggg cgtttttctc gccactttat gttgaccaaa attattaatg gaacttacaa
316741 cgtttccaaa agttagttaa atacatacgt ctatttacta agcaagaaat atatcatgac
316801 aagcccaaat attatattgt tatgtttaca aaaaaaaaat ggctatatac atcaagtctg
316861 gaggcttttt ataacaagca agtggggtaa cttagacata agattgactt ctttgaattc
316921 aacaaaaata catacttttg atgatttcaa tggtagaagc agtagtaact atgccgatga
316981 aataaaatcc tgatatcatc tatatagtag tactcacgtt actagtatat catctataag
317041 atgtaggaaa atgtcataag aaatgagaaa cagtcattaa atttaatgga aacgaaatgc
317101 aaggattggt gatgagataa gataatgaaa catagaaaac gaaaggagaa ataacagtag
317161 tattattatt attattatta tgtagaaata ttgactctct cttgtagatt catatatctt
317221 caaggggaac ttcttgtaca ttttgtatac ctatattata gcgtttaaca acaatggaat
317281 cccaacaatt accaaaattt tcacacattc ctcatataaa atctaagcaa tgactagaga
317341 actatcgcta aattatatag agcaaggaaa ccgaataatt acatgtgtca gataagaaaa
317401 aaataaagaa aacaacgcgt catgaaaaag agttactgaa ccttcagatc ctacttattg
317461 taatgcttcg cgacatccaa tccatttaat aatcaattta aaactagagt tggtagagtt
317521 ccttgttgaa cgtgataacc caaaagcata atacgagtaa tgtttcagta ttgctattat
317581 atgtttacac aaggaaaaca tataataaca aacctctaat ccggtagtac ttaagaaact
317641 atagtttcta tgtacaaaaa ggtaactatg taattcttac atttacataa catatagaag
317701 ggtccaataa acttactaaa cttactacct tgttgtatat aggctagatc gtaatccact
317761 acgtcaacat aaaaaaaaaa cttaagaagt ttgaatttta tgtacaaaca gattgttaaa
317821 atataatata agattatgga aacgaacttg gtctaaaaaa aatttaaagt tttataaaat
317881 cctcgaacta tcgctgttat acatgatgtc cccaaagcgt gtacaaataa ttttgtagta
317941 ttgtatcggt aatatcatac aagaagaaaa tagttgcaaa tctctaaccc actagtactt
318001 aaaaaaactg tagtttcagt gcaaaaaagt tttaacatta cgtatcttgt acccttttta
318061 ttgcatatag aaaggtcaaa taatccttca catcatgaaa tataagctaa atcgcatttc
318121 ttttcgtcca catttgcaaa caaaactttt caataataat tttataaata gtatcaatat
318181 atatatatat atatttattt gtttactttt tctatcagtg ttttcaattt tttattaaac
318241 aatgtttgat tttttaaatc gcaatttaat acctaaatat aaaaaatgtt attatattgc
318301 aaaaacccat caaccttgaa aaaaaagtag aaacgtttta tttaattcta tcaatacatc
318361 ataaaatacg aacgatcccc gtccaagtta tgagcttaat cttccataaa aatatttgaa

318421 agcaatagat catgtactaa actaaaatca gggaaattaa gactcctttt gaagtaatac
318481 ctattactta ctaatacgtt tgagaataag cgcaggtact cctggttttt gttaaaatta
318541 caaatttata cttagcatta cgaagattct cgattccgaa aaacaaaaat tttatcgtca
318601 tatacaaatc tagaaattac cagagctatc catcttgttc aagaaggtag gcgaggacat
318661 tatctatcag tacaaacatc ttagtagtgt ctgaggagag ggttgattgt ttatgtattt
318721 ttgcgaaata tatatatata tattctacac agatatatac atatttgttt ttcgggctca
318781 ttctttcttc tttgccagag gctcaccgct caagaggtcc gctaattctg gagcgattgt
318841 tattgttttt tcttttcttc ttctattcga aacccagttt ttgatttgaa tgcgagataa
318901 actggtattc ttcattagat tctctaggcc cttggtatct agatatgggt tctcgatgtt
318961 ctttgcaaac caactttcta gtattcggac attttctttt gtaaaccggt gtcctctgta
319021 aggtttagta cttttgttta tcatatcttg agttaccaca ttaaatacca acccatccgc
319081 cgatttattt ttctgtgtaa gttgataatt acttctatcg ttttctatgc tgcgcatttc
319141 tttgagtaat acagtaatgg tagtagtgag ttgagatgtt gtttgcaaca acttcttctc
319201 ctcatcacta atcttacggt ttttgttggc cctagataag aatcctaata tatcccttaa
319261 ttcaacttct tcttctgttg ttacactctc tggtaactta ggtaaattac agcaaataga
319321 aaagagcttt ttattcttga tttttgttct ttcggggaaa ctgtataaaa cttccaaaaa
319381 ggaaaagtaa aacaatacat ctccttatat caaagaaaat caagaaggac aacatggatg
319441 gtatttgtag tatggcggaa aacataaaca gaactctgtt taacattcta ggtactgaga
319501 ttgatgaaat caatctcaat actaataatc tttataatgt atgttttcat ttcaaggata
319561 gcctttgaat caatttacta acaatacttc agtttataat ggaaagtaat ttgactaaag
319621 tagagcaaca tacattacac aaaaatattt ctaacaatag gttagaaata taccaccaca
319681 ttaaaaaaga gaagagccca aagggaaaat catcaatatc accccaagca cgggcatttt
319741 tagaacaggt ttttagaaga aagcaaagcc ttaattccaa ggaaaaagaa gaagttgcaa
319801 agaaatgtgg cattactcca cttcaagtaa gagtttgggt atgtaatatg agaatcaaac
319861 ttaaatatat cctatactaa caatttgtag ttcataaata aacgtatgag atctaaataa
319921 attcgttttc aatgattaaa atagcatagt cgggtttttc ttttagtttc agctttccgc
319981 aacagtataa ttttataaac cctggttttg gttttgtaga gtggttgacg aataattatg
320041 ctgaagtacg tggtgacgga tattgggaag atgtgtttgt acatttggcc ttatagagtg
320101 tggtcgtggc ggaggttgtt tatctttcga gtactgaatg ttgtcagtat agctatccta
320161 tttgaaactc cccatcgtct tgctcttgtt cccaatgttt gtttatatac tcatatttct
320221 atgtgtttat acaattgcta ttgtttatat aatgtagtga cattttctct taatcttata
320281 ctaatttcta tgacatttat ataagaagag acttatgatc aacataattt tgcaaacttt
320341 gagagaaata tgtctttcta ctgcgataaa gttattattt agattacatg tcaccaacat
320401 tttcgtatat ggcgatataa tttatcatgt tttggtatga taatttaatt tttaaaaaaa
320461 caaatttaat tgacctcatt aattaatatt tattaatacc tttaaatgtt gaggtaaata
320521 gctattttct ctcttctttt cctttagttg gaatttgcac aagaaaatgt ttttccacac
320581 actttagcgt tttttcctaa atgttggaat aaaaaacaac tatcatctat caactagtag
320641 tcacactacc aatgtgttat cattatactg tgttaaacaa tgacataagg tatgaaaatt
320701 tgtcaacgaa gttagagaaa gctggatgca aggattgata atgtggtagg aaaatgaaac

320761 atataacgga atgaggaata atcgtaatat cagtatatag aaatatagat tcccttttga
320821 ggattcctat atcctcgagg agaacttcta gtatattcgg tatacattgt agagaatgtg
320881 gattttgatg taattgttgg gattccattg tgattaaggc tataatatta ggtatgtaga
320941 tatactagaa gttctcctcg aggatttagg aatccataaa agggaatctg caattctata
321001 caattctata aatattatta tcatcatttt atatgttaat attcattgat cctattacat
321061 tatcaatcct tgcgtttcag cttccactaa tttagatgac tatttctcat catttgcgtc
321121 atcttctaac accgtatatg ataatatact agtaacgtaa atactagtta gtagatgata
321181 gttgattttt attccaacat acataatatt atagctttta tcaacaatga aattccaaca
321241 attatctcaa catccaccta ttcctcacct atattctcga ggagaacttc tactatattc
321301 tatataccta atattattac ttttatctac aatgcaaccc cacaataata taaaaattca
321361 ccaattccgc atctgcagat tactttccta aatttgcata tagaattgtc aagcgcaaat
321421 ccgacgtcga ttccgcggcg gatgggtcat tctaggtcat tctaccaatt ttatttgaga
321481 ccaggtttat tcaaccggta acatagaaat attcatacaa ttaagcttct atggccaagt
321541 tggtaaggcg ccacactagt aatgtggaga tcatcggttc aaatccgatt ggaagcattt
321601 tttatcacgt tattcggtga cacccaggtt gccgccgcgt tcgcgtccat cgtcatctga
321661 aaaataatga atattaatgg accttgtgcc ccataaaggt tccatgttcc ataagtcttc
321721 aataatactt ttgtatatta aatttaccgt tattcggaga tctcttacgg cttatgattt
321781 tctttacatt ccaggccgcc ttttgatgac attcagtgcg tcacgacgcg aatagggctt
321841 tgaaagcctt aatgtggagt aaataaatat atgaatgcaa taagtaaaca taagacgagt
321901 tcttctatat ccggtgtacc taataatata gattttgcca acaatggaat ctcattaccc
321961 atcccaagat aaaatattga tcaaggcatg ggccaccaac ataggaatac gcaaaatagg
322021 attgatattt ttacatatta tattatcttt cttcttctgt ctcaaatata gctcttctcc
322081 tatttaaaaa attatcaatt cttgaatctc aacttccatt aaatttgatg attgtttccc
322141 aatatcgatg tcatcttttt acactgtaaa ttataattta ctaggaacat gaatactgat
322201 caatggatga tagttgattc ttattgcaat aaaaagtact ttctaagacg ctaggacttc
322261 taaacacaga tattatgaaa atgccaaaaa atacagaata ttttccttct tttcactata
322321 acaagtagta agaaattaag caatatggta atttgagctt atcagataaa agatcctggt
322381 agtcttgcaa aggtgtgagt acttttgatg aaactaagtt tcagcgcagc tatttcaatt
322441 ttggaacctt taaagaaagt ttaatcgaaa cattgaacga tccaaagcac aactttgtag
322501 cgggaagtca caaacaaata taaaaagtgt taataaactg ccatcctatg ttagaattgt
322561 gaaacttttt ggactaataa aaggttttca acatggtgtt ccaaagcact tatttctagg
322621 ttagctatgg taacgagttt taacgtaatc aaaacaatat acaattcaag atctgtatca
322681 tttacactaa cacaccccat tttaatagag ctgagtactt cctacattta ttttgcatat
322741 gatatcgcca ctgcatcatt tctgtagtcg taaaatttcc ttctcattaa gacaccgaaa
322801 atagctaagg attataacaa aacaaacaaa tgctcaaaca aaacttttag taaatctttt
322861 gaaatgtaaa agttacttag aataactaaa aaaaaaatat tacgctatac catatatttc
322921 cattcccaat tttatcgttt tataataata ttcagtttca agaacaatga atatatttca
322981 aacattacat gcactcttgg gagtaagctg catcataata aattaaaatt aacgctttag
323041 aaaaagaaat tttgaggtta aaggatggag gaagaaatcg ataccaattt caactttgat

323101 cgacctgtct gactgaaatt atttctgcgt ttttcttact acaattagta ccattctctt
323161 caacttcaat agttctaacc attgattctt catctgtctc atcaaatccc atcttaccag
323221 tccagccatt ggataccaaa tagttgtgaa attcaacgtc ttgcttcatt aaatcgcttt
323281 caagagaatg tggaacaaag aaatatgtaa taatgatacc aataagacca caaattgcag
323341 caataataaa agtccatctt gcacccaaat tatccctaat gggttggaaa cattcgacgc
323401 ctactacaga accgatttta ccagtcacag cagataaacc atagaaaaca cctctaacag
323461 cggttgctga cgcttcacta ctaataacac caagcatatc acctggtcca gcattaccta
323521 acatattcat gaatgcgtag aagataataa acaagggggt gattttttc aattggtcgt
323581 acgcacatcc aatgattaga ccaaagatga tgtacccaga gaaaccaaac atcaacgtat
323641 atttacgacc aatacgatcg gacagataag caccaattgg tacacccagt acagctaaaa
323701 ctcccaacag taagttccac tctgccactt ttactaaatc attttggtcc ttgataacgg
323761 aactgataat tgttgaactg aaaataccat ttgggaaggt aacaaaatca tacataaacc
323821 atgtaccaca tgtaccaagt aaccttttcc aataaaattt caatgctagg aaatatggta
323881 tatttctctt gattctacct ttttcataga cttctgtagt agcagtcttc catctaaaat
323941 agaacacact caatggccaa aagcacccta ttgcaaaaac agtcctccag atcgcctcta
324001 aatgttttgt tcctgaacag attttgtaga cgattaaaaa gatgatcgta gcaaatggac
324061 caccgaaggc tagtggcaaa tttgtcacca taaccaggat accacctctt ttggtagtgg
324121 tatattcatt agcagactca ttagcactta atgtactagt aggatattct gcaccaacac
324181 caatacctac caaacctctc ataactgtta acatccaaaa catgccaggt acagtagtac
324241 cgtgagaggc agcacacaga gcactaccaa taaccaagat agcagtggcc acaaggatac
324301 aagattttct actataataa tcagcagcga tacccataaa gaattggcca aaaataatac
324361 caactaaggc tgcgttggaa actctagtcg acacttttga gctatagttt ttcttaccgt
324421 actccataac aaaaaccttg ttcagcatgc tcattgaacc atttacgtaa ccatcactta
324481 ttagagcaaa accggacgca atgattgtaa ctatgttttt ccatttatct ttctttgagg
324541 tttcagtacg ggttgcacgc ctctcggcat catattttat tattctagaa ttagtgttct
324601 cgttcacttc cttctcattt accgatgtga tatctttatc ttccattttt tatcctattc
324661 tattttttg attataagca aaaagatcaa ctcacgtctg cggctggttg actagtcaca
324721 agaaacagat aataaatact gtaacgaaca tctgctacaa ctacatgcat cggggtttat
324781 atattcacga gtttcctact tcattcagac acattctgga tgtcttccag cagatgtccc
324841 tcatcaaaga attaagatag ccagattgaa aaactaggtt aaccagagaa ctaagttaac
324901 taatgttgga aaataatcgt acgatacagt ggacggaatg cccatatgcc tatataatca
324961 atccatgcta aggtgggggt ccagtaaggc aattcatgaa tatgaagata aatgtggcac
325021 caaacggcaa cgcggaggac cgtaggatcc aatcatataa cccctattat tctaacatca
325081 ttgctgttgg gtacaacatt ctacgccgct attactacgt gctgcttctt ttgattcttt
325141 gctttcttta ccgtcatgca cgtgcgccgc gaaaagtatt aatcatgcgg acactgtggc
325201 atttatggga tactgtcaaa aagaatatgc tcaatactgt catattcata tgcacgttct
325261 ctttttaatt tatcttggca aaacgtgggg ctggaacttt tggaacgcac tgcaacatct
325321 gtggctgtgc agtcaagaaa agtaagtact catcgtcatg ctctaatgtg tatatggtat
325381 ttgtggtgca gacaccatta gaagtatagt tcacgaaatg atgacacgat agcatatgct

325441 tttcgacggc tgataacctt gtattctctt atttttgtat tcttttcgta agcggcattt
325501 gtttgcagaa ccttatcgca gatatcgata aacatttgtt gacaattctt tttcgctccc
325561 cttctttttg tttatcttcg ttatattcaa tagtctacat acactaaccg ttgaattgtt
325621 gcaaacaaat actaataaaa atctatatat agtaaattga agatgtactg atcagtctat
325681 cattgcttag tgcacccaga ctcgaatctt aaataccact ttacacatct actaaatttt
325741 gccctcacaa aatgaagaca ggattcaaaa ccgattaata gtagcagaaa ctaaaaaagt
325801 acgaatatta gtaaaattca tgttcttgaa tcgagctact atctttgtcg ggagggtaaa
325861 cgattataac tcaaaatgac tggaactggt gattattaat ttttacgttt cctgtgccaa
325921 taagcggaag ataagaggat agaagaaaag aaaggcggca cttggcgaac tacaatggcg
325981 attatattca tggcgattat attcatacaa aggtaatgga ggcctcggat aatggacaat
326041 attgagaaaa tccttatgct tacttctctt aataaaaaat agacacagcc atttattatg
326101 cgtaaaaaag attacccact tgtcttcgat gcgtgctgct gccaatcaac cttttgagcg
326161 gaacttcgag ctcgcaatgc gtctggaatg ttgctagaga cagtcttggt tatctgtgac
326221 atgtgtttcg ttcaggcgtg tgagcatctt cttgttcgat ttcaaaatta ccgccttgac
326281 tcgtgaaact ggataattcg ttggcgtttt catataagtc gtctgatggc gaaaactttt
326341 cctttactta gcatacagca aatatcccca tttgacggat ttttgaaaaa tgagcccgct
326401 aacccagaat gaactgcatt accaagcatt tatgtaaacg ttccgccacc atctttggta
326461 aggtatacta ttatgttctg gatttaaggt tgattcacaa tttttcatca ccaaaatctg
326521 gtggcatgcc tagttgtctg gtttcaggca atttagccat catagaaaag catcctctgt
326581 cttgagttga gaaaatgtta ctcatagagc caaacaaata aaccctggaa ttaaaatata
326641 gtttaaacac aacttcatat tacactgtaa atacatatca caataataaa ttaacgaaat
326701 agtaagacct tatgtttcta tatgacatca gtcccaaata gtgtcaattt tttgtcgtca
326761 gtgtcgaaca tttttactgt ttattcatca tatcgtgagg tatgtaggca tcttctattg
326821 acttaagcat gataaaagat tcacgcaaaa ctgtatcgaa atattgataa aattaaattt
326881 caaagtcaaa cttgttttca atggttgcta ttttcttacc attcgagctc tccctcagga
326941 ttttttcaca tggaatgttc atggcctcaa tttccatggg ctttttgcat ttatctcctt
327001 taattaattg taatggctct aatgatacag tttcgccatt agctttgtca catatatcgg
327061 aaagggcaat ctgatcataa tctggtatac agtcaccctt tgcatctcta acaaattcag
327121 aagagcattc gtagtcggat tcagtacaac tgtcacaagg tatttcatgc aaaatcatat
327181 ctttgaatgt cctttttacc aaacaacggg cgtcctgttt cctcctttta tacatgtatt
327241 tggcgccatt aaggcatttc ccatctgcca aattccagtc ttcgaaatct tcctcttcac
327301 acgttttata gtcaaaatgc agcagaaaaa tcgatcgcat acaggacact tcttgcaata
327361 tatgagatgt tggtgttgtt tccttcttga ctagccgtag taatgagatg tccgcttaaa
327421 ataaattttg agcctgatcc atctggcgtt gtattaatca actctaatgg atggatataa
327481 aataaagcgt cagcaagctc atattgattc catgttctac cttggtccaa ggaaaaataa
327541 agtttggatt gtacatcacc atctttatat gaaggagacg ggatatatac aataatattt
327601 cccagatcac cagtagcata taatccagaa gaattatggg ccaccctcca tgttagacca
327661 ccgtctctag aaatgaaagt ttttcggtcc ttccgatcga attcattgtc atcgccaaca
327721 gaacctgtca ttaatatgat cccagcagaa ggatttgaaa gttttagatt gtaaaagtaa

327781 ccctgaagtg aacatctctc aggcctagtg atatcacagc cgaacgaatc ggcattttct
327841 tcatcaacaa ctttcaaatt tgaccatgtg aggccgttat caacggatat tttggtttct
327901 cctccggcta tattttcagt atagtttact tggtgattag aatagtcaat ggaaggatga
327961 aacgatccaa atattgtccc tttcaagaaa tcaggaaaag tgagcctaaa atttccaaac
328021 tgatggtttg gtgtccattc aacaggtgag aatttcaacc cttgcgaatc tgatatgaaa
328081 atctctgaga gatctggatc agcattttcc tcatttgtga tttctgtcga tataagaagg
328141 atgatccttc cgatagaatc ttcacgaatt ttaatcactt tgacgtgccg taattgagta
328201 ggcagctgtg ccttttgaaa agtagatgca tcattggata tccagatgtc cgtagatgac
328261 atctcgttat gcccgtcatc ttgcgttaaa acgaccacat atgacttcaa tatcttgtat
328321 tgagtaacaa tcttatcttt aaattcatca aaatctttaa acgttttgcc tccatcacta
328381 ctgaaagcca actgtgtgta ggtagcagtt aagcgatatt cttcatcgta ttcacggtta
328441 gagaacaaac agagtatcga agtatcatta ccagtaaagt cagaatcctt gctagactta
328501 atgaaatgac agctttaagt tgttatatcg tcgtcttttt tcttttccaa ggaaggttta
328561 attttaaaaa aggattccca tcattgctca agtaaatttc acttgggtca tataaaacgt
328621 ctgcgtctat catatagcta tagcaattcg caataagaag ttctttgttt ctaggatgag
328681 tagttataaa acagtcgcca tctaaatatt cttctgatga gggtatagtt aaagccctcc
328741 atgactcccc acggtcgtcg gtaatgtaga atttgggtga ttcacatata aaagcgaaac
328801 ctctttcctg tccatggaaa ggatcaacag taaatctgaa aatgcgctct tcaatttctt
328861 tgaccgtttc ccaattttct ccagcgtcga aacttatcca tacaaccccc caaccatcta
328921 atctgataga agtgttcgaa tcatcaaaac tcactagatt aaatgatgtt tctataggag
328981 cctccgttac ttttgggaca aatttctctg cattgagtaa cggaagtagt ataattaccc
329041 aaagagtata tatggcatga agtaatatca taattgagga tctcttacct tttcttagtt
329101 caaagttttt tttcaaagct cggaacttta tgctacaatt caatgactaa aaaaaaggta
329161 gcggataatc ttttggttgc aaattggcgg catttaaata tattttgggt tgcaagtatt
329221 ctaaaaatat tctcataata ttctaaaaat atccagtcta ggtaattgca attattttga
329281 gtccggtgac tttgagttga ccatgttcac ttcctgtaca gccatcaagg taacgtgttt
329341 accatctaat ggagaaaacc aagagataag actagaaacc tagttaggat acttggattt
329401 gctaatgtcg ccagtagtac gataatacca taaagatacg ctgttgaacc tgatttgtat
329461 atttaaagaa tgggaatctt taagggatca tagtctagaa tggtattctg cagaatagat
329521 tttaaatggt gaaaactttt cctcactatg tccaattgat gctttttagg atgacgttag
329581 aattttattc tggtgaaggc acttttaata tgaagattcg ttgaagtatc aataaccttt
329641 tattggcata cccttaccta actggtctct ctcacctcgt gatgttacga acgccacgat
329701 tattctcaaa ctaggtaaag attttttctt caaaattagt gggaaagact cgacagaccc
329761 gcttaccagg tattctttct aaacaatagg gtttaccagt acccctgaaa atgataatgc
329821 ccatgtgcgg ctgatcgcat taactacgta tgcagttgaa tcaaagccat accacagaaa
329881 atggtacaag agcccactgg atagaatagt gctggctatc aaaccaatat tttctttcca
329941 gctcttgggg gacctctact gcgttatatc gtagtttcca ttcagcctgc ttatcgcctt
330001 aggaaaaggg aatgtaagag gcagaaaagt gcggcgcaat cgttcggaca gcgagccgct
330061 attataggta agaattctta aagagcttcg ccggatatta ttacagtgta ttaaatttcg

330121 ccaagtcttt gccaatggaa atgggtgcat aattttatgt gcttggtaca tatgagttca
330181 atagatatgt agtagttgga tagtagatag taaggatata taatcatgga agggtgacgc
330241 acaacaaatc acataatgag ttatgatttt cactgttctc atctgtttcc cctacaaggc
330301 gtgatcaata acaaacaatt tcattaaaaa aaaaaaacca agtcactcta acatacgtaa
330361 taactgtgta tacaaaagaa aaataatatc tgttcgtcat ctagtttaat acggcaacta
330421 gtttttcacc agagttctta ccgattttaa tgtcttccag gatacgagga acatcgtaaa
330481 gcccgttctt atagaccctt gctggaatat ggtgaatttg cccatcacta atctttggat
330541 tgatgaactt gacgaattcg gtggcagctc tcctggcttc tgggtcagca gggaaagtaa
330601 tgccaccaaa tggtacttca tggccgccta ttgaatacag tcttgttctg tcaatagtga
330661 cattttgcct cctattctcc tttttgacgt tttcttctgt caaattagtt aattcgacaa
330721 cggtagcatc ctgtttatcg gccgcacatt tgtacacttg ttgaagcgta ttttgattcg
330781 cgacacagtc gactaaatac gagatattgt tgtacttgtg tttaatttgt tctaccacgt
330841 caatatcatg gtaatcaaat agttcatcag caccatattc tttcaacagt ttttcgtgtt
330901 tccgagaagc cacaacaatg atcttggtga agccattcaa tttattggct aattggatga
330961 gcgactgacc tactgcagtt gcaccgcccc ataataagat ggggccgttt ctttgtgggg
331021 ttgatggctc ccacttcagg ttcaagccca agttataggt caacaccaag ccagctgtgg
331081 tcagtgacac tgggatagtg gctgcccctt ccaaagacct gacagggccg gcaggtagaa
331141 catcttcacc caaaaatttg agttcattgg gtgatttgta ggcaaccaca gttgaaatag
331201 cagaatattc agcaaaagca ccattggaag gaaaccttac ggaagatccg tgaatgaacc
331261 cataaatata atcaccaata gaaaagtctt taggatcgac ggctgggccc aatttgacaa
331321 tttggccggc agcgtcacat cccagaatag atccttgagg cccgaccttg tagtcaatgt
331381 gtgcccaatc agtcgggtta ccagcaacag cgagtgtctt aatcaatacg aatccttctt
331441 ccaattcagg aatgggaacg ccctctttga caaccgcttt accgtcttca atgacgacag
331501 ccttcatggt ttctggaatt gcaacttgcc tgtttgtagg ttatatgtaa ttgtcgttga
331561 taatctgtat tcagcttgcc attttataga atatggttag aaggatctta agctagataa
331621 agaaaaagaa aaaaatagaa aaaagcctgc ccgctttata agaacttttc tcaatatctg
331681 ttgaggaaca cttagtaata ttgtctatca tggttgcttt attcttggtg atattactct
331741 ctgggattcg atgtgttctc atcatctgaa gggtttccta tgtgttgaaa gcctatccca
331801 ctagacggag ccgcgcgcca gccgggcgga tccgagacat cttagtcagc tgagacaaga
331861 aaggcacagc tctcggacag gaactgctgt catatgaaaa cctaaaaatg tttcctggaa
331921 attccatgac tgcgcaaagt ttgatacgtt tattgttctt ctctactacc gcactttcta
331981 tctcggaaac ctttcaactg aatgaactat tgttaattgt atggcgtgtg gccttgctat
332041 ttcagcaaat ttagagtaaa gtaaactccg tcggaccagt atagtaacga aacaagccat
332101 tgattctgca gcatattgcg accaaaacct ggaaattggt ggctatgagt tttaatttat
332161 aattatatga agacaaagct tggaatgctg agtacttttt cttcaatgct gctcactgcc
332221 cagaagcgtt cagaacaact ctactggagc cattctgctt gaaggaggat aaaaatgcca
332281 agatcaagtc ttttaaagac tctatccctt ttcatcggaa aagcactaca agtttattct
332341 acagaagtgg acaaactgtg gaacttgatc catactgagt aagtatggaa gactgttaac
332401 ttggaggatt tttctctgct ttctaaagaa ggttcactgc tttacactca cctaggttct

332461 catgagtatt gctagctagt gcgagtttga tctatacttc acagactttt aacaccccgt
332521 tctttaagtt ttccctgatt atcaagcgga tttccaattc ttattcccca ctatcgcacc
332581 tttcttatct gaattgtatt cttgagtggc ctaattgtaa tcttttattt tgttttctaa
332641 ttattgaagg cggtttctcg aaacatttca cgatttccaa aattcaagaa ttgaaattat
332701 gaacatagaa gacaaaatgc cttttttatc aactattatg aattctctag aaagtggttg
332761 caaaggaaat aaatatgtat ttgctgagag gaaaagtttg gaagaacaaa aagtatttct
332821 ttaatggagc tgattatcaa gcattcttta cggggagctt tcgaaatttt ttatattcaa
332881 ggacatttat gttgtgtggg ttattaaacg tgggactttg gaaatacatc caagaagcac
332941 aattgagctc taacgatgaa ccgttaccat aaagatagag acatttcttt tgacttagaa
333001 gctgcaagga gtatcaatta ttaatacacc tttcctagag aacattttta aaataaaata
333061 aaataaagta gcgctctcta ctacccttac actttttcc ttcatatctg gctagaaaag
333121 agggattaca tctcggaatc aaagttttaa tactgctact acgaactggt tacaattcta
333181 tttttatctt acacgatact ttgaacatca acttctctca ataccatccg ccttcagcgg
333241 ctacaattac gtaattgcac caaggcattc gccgagagtt tacatatttg cagaccctaa
333301 cagtatacga ctcccaacac acgtttaatg ttgtacgaat tcaagagatg gaaatatttc
333361 ttccgcggaa gccgtttaaa gcagactgaa tttaggctta gacaataagc ttctgaacga
333421 gggtccaaat ctaaaattcg gatattcttt ttgaccgaaa aaggaaatgt cacaggcgcg
333481 cgcaaaaggg tttttcaag gcacatctcg acaccccctac aattggcatt ctcacattct
333541 gcgcatacga atacacatga gtgccataaa acaatactcc acatacgaaa cctatgaaag
333601 atgtatgatt cgttcagaaa caagagcatc tccatagaga taatgagatt gtgtgaaaga
333661 tgagatatgg agaatatgtg atgagaaact ggtgccagtc agcgatcgtt taaatagcaa
333721 caacccatga gggaatgggt ataaatagag ctgcttctgt cccatgtctt aaagatatgt
333781 cttttttaat ttctttcctg ctaagcttat atcagcacta acaaacaaaa caaatacaat
333841 ggtcaaatta acttcaatcg ctgctggtgt tgccgccatc gctgccggtg ttgccgctgc
333901 cccagccact accactctat ctccatctga cgaaagggtc aacttggtcg aattgggtgt
333961 ttacgtctcc gatatcagag ctcatttggc tcaatactac ttgtttcaag cagctcatcc
334021 aactgagacc tacccagttg agattgctga agctgttttc aactatggtg acttcaccac
334081 tatgttgact ggtattccag ctgaacaagt caccagagtc atcactggtg tcccatggta
334141 ctccactaga ttgagaccag ccatctccag tgctctatct aaggacggta tctacactgc
334201 tattccaaaa tagagacatg cctttactaa tgaacttcca tagacattga aaaataacga
334261 aagaataaaa aaaaaatctt tttttttttt tgatataaat acagttgact atataaacat
334321 atatcaatac tataagccta tatttgatca agacgacatt atttaggtgc atcaacaacg
334381 gaaaagaaac tttcttctag gctatccttc tggggaagca gttgaaagtg ctatccagta
334441 ttattgcata tattcttaat gagtagtgct gaagtttgat acgtttggct ctgttgcttt
334501 tatttctccc acgaaaataa attgatttct ttatttttca gtgtgttggc ccggagttgt
334561 ttacacacat gtctcttttt ggattaatgc tgtagatcag ggactatgcg agcgacaagt
334621 cagagtaacc atattgaatt ttcgtctcct gttatacgaa tatttgcatt tctgtgccta
334681 cttatgtgtg tatatgtttt tattagtcag aaaatggagt aagtgttaaa gtaaggatag
334741 tgttaaggag gtaataacta aagatttgta ggataaggcg gtgttatttg ttgcaatttt

334801 ctcaggggtg gagtggaaat ggtttgattc aattatcagt taagcaagga agagttgaag
334861 ggcgaacatt tgcaataatc ccaatgtcat ttaatgcggg tcggctgaga cgctcaatat
334921 tcggccgaaa agtatcattc ggccgcataa taaaatggaa tagaagataa agcagatgtc
334981 actaattgag caccaaaact ctgagtagaa attatataaa gaagcacgaa catgacctgc
335041 tttagatgtt gcttgatatt attgaaataa aaaaaaaaca tagaaccact gaaaaataca
335101 aaaaaaaaat gctttaccca gaaaaatttc agggcatcgg tatttccaac gcaaaggatt
335161 ggaagcatcc taaattagtg agttttgacc caaaaccctt tggcgatcat gacgttgatg
335221 ttgaaattga agcctgtggt atctgcggat ctgattttca tatagccgtt ggtaattggg
335281 gtccagtccc agaaaatcaa atccttggac atgaaataat tggccgcgtg gtgaaggttg
335341 gatccaagtg ccacactggg gtaaaaatcg gtgaccgtgt tggtgttggt gcccaagcct
335401 tggcgtgttt tgagtgtgaa cgttgcaaaa gtgacaacga gcaatactgt accaatgacc
335461 acgttttgac tatgtggact ccttacaagg acggctacat ttcacaagga ggctttgcct
335521 cccacgtgag gcttcatgaa cactttgcta ttcaaatacc agaaaatatt ccaagtccgc
335581 tagccgctcc attattgtgt ggtggtatta cagttttctc tccactacta agaaatggct
335641 gtggtccagg taagagggta ggtattgttg gcatcggtgg tattgggcat atggggattc
335701 tgttggctaa agctatggga gccgaggttt atgcgttttc gcgaggccac tccaagcggg
335761 aggattctat gaaactcggt gctgatcact atattgctat gttggaggat aaaggctgga
335821 cagaacaata ctctaacgct ttggaccttc ttgtcgtttg ctcatcatct ttgtcgaaag
335881 ttaattttga cagtatcgtt aagattatga agattggagg ctccatcgtt tcaattgctg
335941 ctcctgaagt taatgaaaag cttgttttaa aaccgttggg cctaatggga gtatcaatct
336001 caagcagtgc tatcggatct aggaaggaaa tcgaacaact attgaaatta gtttccgaaa
336061 agaatgtcaa aatatgggtg gaaaaacttc cgatcagcga agaaggcgtc agccatgcct
336121 ttacaaggat ggaaagcgga gacgtcaaat acagatttac tttggtcgat tatgataaga
336181 aattccataa atagtctata tacgtaatat ttttcagaat ctcttaaaat gcatgtcaaa
336241 tctcggaatt tatcatgacc cattacataa caagagaaaa gtggtttgca ttgtatgtat
336301 accttaaacc aattcttcgg tagtcacatg acctatatat agggacattc atttcttcaa
336361 taggatattt cttaaactta agggaaattt tgatgaaaat ttagaatctc tcgacaacag
336421 tgcagagcag taagttttat tcaatacgaa atgtgtgcag ggttttaccc attcatgtcg
336481 agtacgtcta catttgttct gcgatttaac ttaatactta gcctactcat atgtatcgaa
336541 gccaaactat taaatctgaa aactgaatac ccattaagta ggtacattaa tgcaattttc
336601 gtccgaaagg tagagtgtta atcatgcttt tacttcgaaa tctagagtcc gagcatgtct
336661 gatgaggtcc tcttaaagca cttcaatttt aaaagagaag agcgcaaatt tttacaaaga
336721 atgcctatat taatatactg tgctatctaa gaggatggtt caatggtttg aaatgagttg
336781 aaaaataaag atttacatag tagcgtgtat tcttcacatt tttgcattcg gtagattttc
336841 tactcaagac ccgtgcagat cactattaca ctcattatta caagtcattg agaacgaaat
336901 tttgacatca ttactaataa tgttactcat taatacttta ttaatgtaag cggtagataa
336961 acaaatgaac atgaaattgg ctgctgaaat gctctgatgg attcgattac agtaaaaaaa
337021 cctcggttaa gattggtttg cctgcaatgc aaaaagatca aacggaaatg tgataaactg
337081 cggcctgctt gctcgcgatg ccaacaaaat tcattacagt gtgaatatga agagagaaca

337141 gatttatctg ccaatgttgc agcaaacgac tctgatggat tcaattcctc tcataagctc
337201 aatttcgaac agcaacctgt acttgaaagg actgggctta gatattcctt acaagtgcct
337261 gaaggtgtcg ttaatgctac gctgtcgata tggaacgccg aagatatgct agttatagta
337321 ggattagtta catttctgga ttatcctttt gctgcgcata gtctggcgca acatgaccag
337381 tatatcaggg cactttgtgc ttcgttgtac ggcatggcgc ttgttgactt tagcaattat
337441 gctaatggta ttccttgtga agacacatca agaagtatac taggaccatt gtcattcata
337501 gaaaaggcca tttttagacg gatagaacat agtaagcaat ttcgagttca gcctgccgcc
337561 ttagggttat tatacaatgc attttcaatg gaagaagaaa acttctcgac tcttctaccg
337621 tcactcatcg ctgaagtgga agacgtgttg atgcaaaaaa aagactgtga aatacttttg
337681 aggtgtttct atcaaaatat ttatcccttc tatcctttta tggacatttc actctttgag
337741 agcgatctca ctagtttgct tttacaagac gacaataatc gttggaaaat tagtactgaa
337801 gttaaaaatg tgcgcaaaaa aatagaaact ttgtcattac ttacaatagt aatggccatg
337861 gccttgatgc attcaaaatt ggatgcaaat cttctttcaa tggtaaaaga aaatgcctcc
337921 gaaagtgcca ggaaactttc tcttttatgt cataaactat tatgcctcct ggatgtattt
337981 cgctatccaa atgagaacac ttttacttgc cttttatatt tctacgtttc agagcattta
338041 gatcccgaga gtcccgattg tgtactgagc cccactaact tgcttactct gcaccatctt
338101 ttaaatttgt ccatgacctt aggtcttcaa tatgagcctt cgaagtacaa acgtttcaaa
338161 gatccagaag tgataaggca gagacggata ttatggttag gagttcagtc attacttttt
338221 caaatttctc ttgctgaagg tgatgctggt aaatcaaata gtgaatatat ggaggcatat
338281 ttaacagact tcgaagaata tattgaagct tcctcagagt atgaaaaaag ttctgcgagt
338341 gaatcgaacg tgcaaatgaa tgatattgtt tggaataagt acaaatttca cgtcattttg
338401 agtaaactaa tgtctgattg cacttcagtt atacaacatc cgcagctttt ccacatttta
338461 gaaaatatta aaagatctga agattttatg gctgagaact ttcctacaag ttcgatttac
338521 caaccccttc atgaaaagga accaaatgcg atcaaagttg gcaaaagtac ggttctcgat
338581 gtcatggata ttcaaaaaac tgaaatattt cttacaaata ttgtgggaag tatgtgtttt
338641 ttaaacattt ttgatgtcct atcgttacat tttgaaaaaa aatgtgttat gcactgggaa
338701 gaatatgaaa agaactatca tttccttact ttgaaaagtt tcaatgcata cttaaagcta
338761 gcagggttga tatctgatta tctcgagaat aagtttcaag ggaacatttt agagagtcgc
338821 ggttatatca tagataaaca aatatgtttt atgcttgtaa ggatctggat gttccaatgt
338881 cgtattttgt taaggttttc atacaagcaa gaaagtcaga aaaaattggc ctcttccagt
338941 atatccacta acgataatga aaaagaagat gaaatgattg tcattttaga aagacttatt
339001 aaacacattc gtaaccaaat ggcacattta gtggatctag caaagggaaa acttcaagat
339061 agttactttg gtgcttacca aactgttccc atgtttagat acgttgtgta tttgatcgat
339121 gttggcggct tagtatctgt gacaaatggg ttttgggata agatttccag tgatggtgaa
339181 ataccgccaa aagtacaaca agccgtgaga ttgaaatggg gattggactg caataattcg
339241 agaagaatca aacaaaagtt aataagcagc cagagtttgc agagtttcaa tcaagttctg
339301 ttgtgccaga tggaggatgc agttctctcc agttccttcg caataaaagc caataccgct
339361 atgtcccaaa acacggctga agaatttttc aatatcagcg aagaagaggc tttaaatcaa
339421 ctattggaaa acaacaattt tgatgccttc tgggatttat taggtgaaaa tctgagcgat

339481 atgccttctt tgtgagaata cggatatttg gaaaggctaa aaaaaaaaaa acaagccaaa
339541 aacaagaaac actgttatcc tccacgtttt ttccactgtt ttaagactcg acaagacgga
339601 aagattcctc acatttcgca gtgtttcttt taacctgctc gtcaatttga aaaaatgagc
339661 atggttaggc aggcggaatg taataggtgc ggtgcgtata aataagtatc tcggccgttc
339721 ctctattgcg ctcggccgag atgtcctaca tgcataaaat agattaaaaa aagggatcac
339781 gatactcaaa tatgcatata ataacaacaa cacactttgc ggtaaataga aggataatat
339841 cgtaatcttt tgctggtttc gatgatgact tgaaaaaatt caattactct ttcgatcatc
339901 tagaagaagc atatccatcg tgcttaaaat gattgggtcc gcgtccgact cacctagcaa
339961 gttaggacgc ctccgatttc tttctgaaac tgccgctatt aaagtatccc cgttaatcct
340021 aggagaagtc tcatacgatg gagcacgttc ggattttctc aaatcaatga acaagaatcg
340081 agcttttgaa ttgcttgata ctttttacga ggcaggtgga aatttcattg atgccgcaaa
340141 caactgccaa aacgagcaat cagaagaatg gattggtgaa tggatacagt ccagaaggtt
340201 acgtgatcaa attgtcattg caaccaagtt tataaaaagc gataaaaagt ataaagcagg
340261 tgaaagtaac actgccaact actgtggtaa tcacaagcgt agtttacatg tgagtgtgag
340321 ggattctctc cgcaaattgc aaactgattg gattgatata ctttacgttc actggtggga
340381 ttatatgagt tcaatcgaag aatttatgga tagtttgcat attctggtcc agcagggcaa
340441 ggtcctctat ttgggtgtat ctgatacacc tgcttgggtt gtttctgcgg caaattacta
340501 cgctacatct tatggtaaaa ctccctttag tatctaccaa ggtaaatgga acgtgttgaa
340561 cagagatttt gagcgtgata ttattccaat ggctaggcat ttcggtatgg ccctcgcccc
340621 atgggatgtc atgggaggtg gaagatttca gagtaaaaaa gcaatggagg aacggaggaa
340681 gaatggagag ggtattcgtt ctttc
//
