## Supplementary material for "Intergeneric chromosomal transfer in yeast results in improved phenotypes and widespread transcriptional responses": TableS2

**Table S2: SNPs and INDELs on Km chromosomes of KS-R1 & KS-R3.**

**KS-R1**

| chromosome | position | reference | alteration | type | FIM1_Gene | Description |
|---|---|---|---|---|---|---|
| contig1 | 7457 | G | C,A | exonic | FIM1_3 | hypothetical protein |
| contig1 | 332172 | CT | C,CAT | intergenic | | |
| contig1 | 382798 | CTTTTTTTTTTTTT | C,CT | intergenic | | |
| contig1 | 577448 | A | ATTTTTTTTTTTTTTTTTTTTTTTTT | intergenic | | |
| contig1 | 642124 | ATTTTTTTTTTTTTTT | A,AT | intergenic | | |
| contig1 | 1073978 | GTTTTTTTTTTTTTTTTTTTTTAAGCACAATAAAAGGAATCTTTTTT | G,GT | intergenic | | |
| contig1 | 1083707 | CTTTTTTTTTTTTTTTTT | C,CT | intergenic | | |
| contig1 | 1105436 | C | CT,CTT | intergenic | | |
| contig1 | 1250108 | T | TAAAAAAAAAAAAAAAAAAAAAA | intergenic | | |
| contig1 | 1341130 | CA | C | intergenic | | |
| contig1 | 1341379 | CAA | C,CA | intergenic | | |
| contig1 | 1405194 | GTTTTTTTTTTTTTTTTTTTTTTTTTTTTTTTTTTTTTTTTTTTTTTTTTTTTTTTTTTTTTTTTTTTTTTTTTTTTTTTTT | G,GT | intergenic | | |
| contig1 | 1410607 | CAA | C,CA | intergenic | | |
| contig1 | 1461905 | CAA | C,CA | intergenic | | |
| contig1 | 1591046 | GCTTCTT | G,GCTT | intergenic | | |
| contig1 | 1602175 | GTT | G,GT | intergenic | | |
| contig2 | 86383 | TAAA | T,TA | intergenic | | |
| contig2 | 413453 | C | T | exonic | FIM1_1044 | DBF4, regulatory subunit for Cdc7p protein kinase |
| contig2 | 942132 | GAAAAAAAAAAAAAAAAAA | G,GA | intergenic | | |
| contig2 | 1224699 | GTTTTTTTTTTTTTTTTTT | G | intergenic | | |
| contig2 | 1224717 | T | TAAAAAAAAAAAAAAAAAA,TAAAAAAAAAAAAAAAAAAA | intergenic | | |
| contig2 | 1226529 | CAA | C,CA | intergenic | | |
| contig2 | 1267050 | A | AGGGCTTG | intergenic | | |
| contig2 | 1267052 | A | ACGGTTGTTG | intergenic | | |
| contig2 | 1314570 | A | C | exonic | FIM1_1502 | hypothetical protein |

| | | | | | | |
|---|---|---|---|---|---|---|
| contig2 | 1622960 | GA | G,GTA | intergenic | | |
| contig2 | 1671429 | CAAA | C,CA | intergenic | | |
| contig2 | 1679077 | TAAAAAA | T | intergenic | | |
| contig2 | 1693829 | AG | A | intergenic | | |
| contig3 | 549040 | T | TAAAAAAAAAAAAAAAAAAAAAAAAAAA | intergenic | | |
| contig3 | 855910 | CTTTTTTTTTTTTTTTTTTTTTTTT | C | intergenic | | |
| contig3 | 1106742 | T | A | exonic | FIM1_2208 | hypothetical protein |
| contig3 | 1274847 | CAAAAAAAAAAAAAAAA | C | intergenic | | |
| contig3 | 1274864 | A | ATTTTTTTTTTTTTTTTT,ATTTTTTTTTTTTTTTTTT | intergenic | | |
| contig3 | 1436059 | GTTTTTTTTT | G,GT | intergenic | | |
| contig4 | 236003 | GTTTT | G | intergenic | | |
| contig4 | 364985 | GAA | G,GA | intergenic | | |
| contig4 | 467471 | CTTTTT | C,CT | intergenic | | |
| contig4 | 574179 | ATTTTTTTTTTTTTTTTTTTTTTTTTTGACGTAAAAATCGAGTAGTAAGTAAGCCAAATTTTTTTTTT | A,ATT | intergenic | | |
| contig4 | 652557 | A | ACATGTTCCTACACTACTTTGCCAGAG | exonic | FIM1_2725 | hypothetical protein |
| contig4 | 970716 | G | GTTTTTTTTTT,GTTTTTTTTTTT | intergenic | | |
| contig4 | 1067695 | GAAAA | G,GAA | intergenic | | |
| contig4 | 1093979 | GAAAAA | G,GAA | intergenic | | |
| contig5 | 81301 | GAAA | G,GA | intergenic | | |
| contig5 | 193212 | ATTTTTTTTTTTTTTTTTTTTTTTGGGACCTAGACTAGACATCTTTTAGTAGCCAGATAGCCAGCTAGCTTTTGTGGGATT | A,AT | intergenic | | |
| contig5 | 236370 | ATT | A,AT | intergenic | | |
| contig5 | 236643 | GTTTTTTTTTTTTTTTTTTTTTTTTT | G,GT | intergenic | | |
| contig5 | 375159 | CAAAAAAAAAAAAAAAAAAAAAACAAAAAAA | C,CA | intergenic | | |
| contig5 | 565928 | GTTTTTTTTTTTTTTTTTTT | G,GT | intergenic | | |

| | | | | | | |
|---|---|---|---|---|---|---|
| contig5 | 916266 | CTTTTTTTTTTTTTTTTTTTTTTTTTTTTTTTCCTTCCTTCGATGAAAATAAAAAAAAAAAAAAAAAAAAAATTTTTTTTTTTTTT | C,CT | intergenic | | |
| contig5 | 973407 | T | TAAAAAAAAAAAAAAAAAAAAAAAA,TAAAAAAAAAAAAAAAAAAAAAAAA | intergenic | | |
| contig5 | 1065616 | T | TGCAGCCATAGTAGACAGAGATA | exonic | FIM1_3590 | hypothetical protein |
| contig5 | 1070324 | T | TAAAAAAAAAAAAAAAAAAAA,TAAAAAAAAAAAAAAAAAAAAA | exonic | FIM1_3593 | hypothetical protein |
| contig5 | 1073886 | A | T | exonic | FIM1_3596 | hypothetical protein |
| contig5 | 1308056 | C | T | intergenic | | |
| contig5 | 1318492 | GTTTTTTTTTTTTTTTTTTTTTTTTTTTTTTTTATTT | G,GT | intergenic | | |
| contig6 | 69168 | ATTTTTTTTTTTTTTT | A,ATT | intergenic | | |
| contig6 | 126957 | GAAAAAAAAAAAAAAAAAAAAAAATTAAAAAAATTAAAAAAAAAAAAATA | G | intergenic | | |
| contig6 | 252427 | TAAATAAAAAAAA | T | intergenic | | |
| contig6 | 414552 | A | ATTTTTTTTTTTTTTTTTTTTTTTTTTTTTTT | intergenic | | |
| contig6 | 567820 | TAAAAAAAAAAAAAAAAAAAAAAAAAAAA | T,TA | intergenic | | |
| contig6 | 632215 | GT | G,GAT | intergenic | | |
| contig6 | 689584 | A | ATTTTTTTTTTTTTTTTT,ATTTTTTTTTTTTTTTTTT | intergenic | | |
| contig6 | 705087 | A | G | intergenic | | |
| contig6 | 705940 | TAAAAAAAAAAAAAAAAAAAAAAAAAAAC | TC,T | intergenic | | |
| contig6 | 707571 | CGGTTAAGGGATAATACCAAAACAATTCCTAAAACTCCAAGTAAGAATCATGTAGCTGGATCTATGTGCTCTTCAACTTCATCCTCTGTTA | C,CTT | intergenic | | |
| contig6 | 1113708 | ATTTTTTTTTTTTTTTTTTTGTTTTTTTTTTTTTTTT | G | intergenic | | |

| | | | | | | |
|---|---|---|---|---|---|---|
| contig6 | 1173150 | G | A | | | |
| contig6 | 1173158 | T | C | | | |
| contig6 | 1173162 | A | T | | | |
| contig6 | 1173164 | C | G | | | |
| contig6 | 1173172 | G | T | | | |
| contig6 | 1173182 | A | C | | | |
| contig6 | 1173185 | C | T | | | |
| contig6 | 1173188 | C | T | exonic | FIM1_4298 | flocculation protein FLO9 |
| contig6 | 1173197 | A | T | | | |
| contig6 | 1173201 | G | A | | | |
| contig6 | 1173202 | T | A | | | |
| contig6 | 1173206 | C | G | | | |
| contig6 | 1173209 | T | C | | | |
| contig6 | 1173212 | T | C | | | |
| contig6 | 1173215 | T | C | | | |
| contig6 | 1192035 | T | TGAATGCGGTGACGCTCCTAA | intergenic | | |
| contig7 | 1227 | T | TTC,TC | intergenic | | |
| contig7 | 259279 | A | ATTTTTTTTTTTTTTTTT,ATTTTTTTTTTTTTTTTTT | intergenic | | |
| contig7 | 453069 | ATTTTTTTTTTTTTTT | A | intergenic | | |
| contig7 | 453086 | T | A | intergenic | | |
| contig7 | 453088 | T | TAAAAAAAAAAAAAAAAAAA | intergenic | | |
| contig7 | 487037 | CAA | C,CA | intergenic | | |
| contig7 | 505277 | CTT | C,CT | intergenic | | |
| contig7 | 615461 | T | A | | | |
| contig7 | 615465 | T | C | | | |
| contig7 | 615466 | T | A | exonic | FIM1_4611 | hypothetical protein |
| contig7 | 615471 | T | C | | | |
| contig7 | 615474 | T | G | | | |
| contig7 | 615475 | T | G | | | |
| contig7 | 778183 | CTT | C | intergenic | | |
| contig7 | 957564 | CA | C | intergenic | | |
| contig8 | 109596 | T | TAGATAACCCGGACATATCACGTGACGACAAA | intergenic | | |
| contig8 | 109904 | T | C | intergenic | | |
| contig8 | 212748 | CTTT | C,CTT | intergenic | | |
| contig8 | 381045 | CTTTTTTTTTTTTT | C,CTT | intergenic | | |

| | | | | | | |
|---|---|---|---|---|---|---|
| contig8 | 468936 | A | AGCCGTG | exonic | FIM1_4990 | hypothetical protein |
| contig8 | 468939 | A | ACGGTAGTACCCT | | | |
| contig8 | 468942 | A | AGGGGAGG | | | |
| contig8 | 468945 | A | ATGCAACAACAAC | | | |
| contig8 | 653029 | T | G | intergenic | | |
| contig8 | 653032 | T | TAAA | intergenic | | |
| contig8 | 653837 | G | A | intergenic | | |
| contig8 | 653841 | G | A | intergenic | | |
| contig8 | 653844 | G | A | intergenic | | |
| contig8 | 680260 | ATTTTTTTTTTTTTTTT | A,ATT | intergenic | | |
| contig8 | 700246 | CT | C,CTT | intergenic | | |
| contig8 | 848873 | GAA | G,GA | intergenic | | |

**KS-R3**

| chromosome | position | reference | alteration | Type | FIM1_Gene |
|---|---|---|---|---|---|
| contig1 | 382798 | CTTTTTTTTTTTTT | C,CT | intergenic | |
| contig1 | 1073978 | GTTTTTTTTTTTTTTTTTTTTTAAGCACAATAAAAGGAATCTTTTT | G,GT | intergenic | |
| contig1 | 1083707 | CTTTTTTTTTTTTTTTTTT | C,CTT | intergenic | |
| contig1 | 1105436 | C | CT,CTT | intergenic | |
| contig1 | 1405194 | GTTTTTTTTTTTTTTTTTTTTTTTTTTTTTTTTTTTTTTTTTTTTTTTTTTTTTTTTTTTTTTTTTTTTTTTTTTTTTTTT | G,GT | intergenic | |
| contig2 | 286979 | T | TAAAAAAAAAAAAAAAAAA,TAAAAAAAAAAAAAAAAAAA | intergenic | |
| contig2 | 319249 | T | TTA | intergenic | |
| contig2 | 586809 | A | ACTT | intergenic | |
| contig2 | 586813 | A | T | intergenic | |
| contig2 | 942132 | GAAAAAAAAAAAAAAAAAA | G,GA | intergenic | |
| contig2 | 1224699 | GTTTTTTTTTTTTTTTTTT | G | intergenic | |
| contig2 | 1224717 | T | TAAAAAAAAAAAAAAAAAA,TAAAAAAAAAAAAAAAAAAA | intergenic | |
| contig2 | 1267050 | A | AGGGCTTG | intergenic | |
| contig2 | 1267052 | A | ACGGTTGTTG | intergenic | |
| contig2 | 1495529 | A | ATTTTTTTTTTTTTTTTT,ATTTTTTTTTTTTTTTT | intergenic | |
| contig2 | 1622960 | GA | G,GTA | intergenic | |
| contig2 | 1679077 | TAAAAAA | T | intergenic | |
| contig2 | 1693829 | AG | A | intergenic | |
| contig3 | 549040 | T | TAAAAAAAAAAAAAAAAAAAAAAAAA,TAAAAAAAAAAAAAAAAAAAAAAAAAAA | intergenic | |
| contig3 | 855910 | CTTTTTTTTTTTTTTTTTTTTTTTTT | C | intergenic | |

| | | | | | | |
|---|---|---|---|---|---|---|
| contig4 | 574179 | ATTTTTTTTTTTTTTTTTTTTTTTTTTGACGTAAAAATCGAGTAGTAAGTAAGCCAAATTTTTTTTT | A,ATT | intergenic | | |
| contig4 | 962204 | CTTTTTTTTTTTTTTTTTTTTTTTTTCCAAAAATCTATGTGGGCGCGACTGTCAACATTATGATTTTTTTTTTAAT | C | intergenic | | |
| contig4 | 970716 | G | GTTTTTTTTTT | intergenic | | |
| contig4 | 1234285 | TG | T | intergenic | | |
| contig5 | 178689 | G | GT,GTT | intergenic | | |
| contig5 | 236643 | GTTTTTTTTTTTTTTTTTTTTTTTTTT | G,GTT | intergenic | | |
| contig5 | 486371 | A | G | exonic | QDR3 | uncharacterized protein |
| contig5 | 890442 | TA | T | intergenic | | |
| contig5 | 1073886 | A | T | exonic | FIM1_3596 | hypothetical protein |
| contig5 | 1318492 | GTTTTTTTTTTTTTTTTTTTTTTTTTTTTTTTTATTT | G,GT | intergenic | | |
| contig6 | 69168 | ATTTTTTTTTTTTTT | A,AT | intergenic | | |
| contig6 | 252427 | TAAATAAAAAAAA | T | intergenic | | |
| contig6 | 264753 | A | ATAGCGATGAGCCGTCGT | intergenic | | |
| contig6 | 567820 | TAAAAAAAAAAAAAAAAAAAAAAAAAA | T | intergenic | | |
| contig6 | 567866 | A | T | intergenic | | |
| contig6 | 567871 | A | T | intergenic | | |
| contig6 | 567875 | A | T | intergenic | | |
| contig6 | 632215 | GT | G,GAT | intergenic | | |
| contig6 | 689584 | A | ATTTTTTTTTTTTTTTTT,ATTTTTTTTTTTTTTTT | intergenic | | |
| contig6 | 705940 | TAAAAAAAAAAAAAAAAAAAAAAAAAAAC | TC,T | intergenic | | |
| contig6 | 1113708 | GTTTTTTTTTTTTTTTT | G | intergenic | | |
| contig6 | 1208163 | G | T | intergenic | | |
| contig7 | 1227 | T | TC,C | intergenic | | |
| contig7 | 218179 | T | TAAAAAAAAAAAAAAAAAAAA,TAAAAAAAAAAAAAAAAAAAAA | intergenic | | |

| | | | | | | |
|---|---|---|---|---|---|---|
| contig7 | 259262 | GAAAAAAAAAAAAAAAAAC | GC,G | intergenic | | |
| contig7 | 350134 | ATTTTTTTTTTTTTTTTTTTTTTTT | A | intergenic | | |
| contig7 | 487037 | CAAA | C,CAA | intergenic | | |
| contig7 | 933511 | C | A | exonic | FIM1_4767 | hypothetical protein |
| contig8 | 8824 | T | A | intergenic | | |
| contig8 | 623290 | C | CAA,CAAA | intergenic | | |
| contig8 | 640583 | A | T | exonic | FIM1_5077 | ALG1, beta-1,4-mannosyltransferase |
