## Supplementary material for "Intergeneric chromosomal transfer in yeast results in improved phenotypes and widespread transcriptional responses": TableS3

Table S3. Quantification of spot assays.

| control | | column | fold | count_1 | count_2 | count_3 | count_1/fold | count_2/fold | count_3/fold | average_YPD |
|---|---|---|---|---|---|---|---|---|---|---|
| YPD_1 | Km-V | 5 | 5 | 13308 | 10216 | 12517 | 2661.6 | 2043.2 | 2503.4 | 2231.866667 |
| | KS-R1 | 5 | 5 | 9622 | 7253 | 11432 | 1924.4 | 1450.6 | 2286.4 | 1962.866667 |
| | KS-R3 | 5 | 5 | 6824 | 6327 | 7106 | 1364.8 | 1265.4 | 1421.2 | 1335.1 |
| | Sc-V | 4 | 25 | 4933 | 5177 | 4994 | 197.32 | 207.08 | 199.76 | 226.72 |
| | Sc-R1 | 4 | 25 | 8799 | 8849 | 9923 | 351.96 | 353.96 | 396.92 | 354.86 |
| | Sc-R3 | 4 | 25 | 7768 | 7829 | 5531 | 310.72 | 313.16 | 221.24 | 296.5866667 |
| YPD_2 | Km-V | 5 | 5 | 9060 | 9220 | 12635 | 1812 | 1844 | 2527 | |
| | KS-R1 | 5 | 5 | 10842 | 8912 | 10825 | 2168.4 | 1782.4 | 2165 | |
| | KS-R3 | 5 | 5 | 5312 | 7301 | 7183 | 1062.4 | 1460.2 | 1436.6 | |
| | Sc-V | 4 | 25 | 6149 | 7044 | 5711 | 245.96 | 281.76 | 228.44 | |
| | Sc-R1 | 4 | 25 | 8077 | 8094 | 9487 | 323.08 | 323.76 | 379.48 | |
| | Sc-R3 | 4 | 25 | 7107 | 9487 | 6766 | 284.28 | 379.48 | 270.64 | |

| treatment | strain | column | fold | count_1 | count_2 | count_3 | count_1/fold | count_2/fold | count_3/fold | average_YPD | norm_count_1 | norm_count_2 | norm_count_3 | log2(norm_count_1) | log2(norm_count_2) | log2(norm_count_3) | strain | average | t.test | Ftest | p value | direction |
|---|---|---|---|---|---|---|---|---|---|---|---|---|---|---|---|---|---|---|---|---|---|---|
| 0.2 µg/mL TM | Km-V | 5 | 5 | 2258 | 3098 | 2669 | 451.6 | 619.6 | 533.8 | 2231.866667 | 0.202341836 | 0.27761515 | 0.239171994 | -2.305133451 | -1.848841793 | -2.063879632 | Km-V | -2.072618292 | Km-V vs Sc-V | 0.094815794 | 0.017237 | |
| | KS-R1 | 5 | 5 | 2535 | 2302 | 2527 | 507 | 460.4 | 505.4 | 1962.866667 | 0.25829569 | 0.234554903 | 0.257480556 | -1.952904525 | -2.092002438 | -1.957464608 | KS-R1 | -2.000790524 | KS-R1 vs Km-V | 0.214040088 | 0.633693 | |
| | KS-R3 | 5 | 5 | 695 | 448 | 359 | 139 | 89.6 | 71.8 | 1335.1 | 0.104112052 | 0.067111078 | 0.053778743 | -3.263791017 | -3.897305262 | -4.216820151 | KS-R3 | -3.79263881 | KS-R3 vs Km-V | 0.362627452 | 0.005133 | KS<Km |
| | Sc-V | 3 | 125 | 1933 | 1984 | 573.4 | 15.464 | 15.872 | 4.5872 | 226.72 | 0.068207481 | 0.070007057 | 0.020232886 | -3.873926216 | -3.836355827 | -5.627154043 | Sc-V | -4.445812029 | | | | |
| | Sc-R1 | 4 | 25 | 721 | 952 | 94.4 | 28.84 | 38.08 | 3.776 | 354.86 | 0.081271487 | 0.107309925 | 0.010640816 | -3.621106893 | -3.220144578 | -6.554247387 | Sc-R1 | -4.465166286 | | | | |
| | Sc-R3 | 3 | 125 | 1177 | 1595 | 291.6 | 9.416 | 12.76 | 2.3328 | 296.5866667 | 0.031747887 | 0.043022838 | 0.007865492 | -4.977195611 | -4.538753507 | -6.990247307 | Sc-R3 | -5.502065475 | | | | |
| 0.3 µg/mL TM | Km-V | 5 | 5 | 1728 | 1834 | 1665 | 345.6 | 366.8 | 333 | 2231.866667 | 0.15484796 | 0.164346735 | 0.149202461 | -2.691075719 | -2.605185298 | -2.74465676 | Km-V | -2.680305926 | Km-V vs Sc-V | 0.217907584 | 1.47E-05 | |
| | KS-R1 | 5 | 5 | 615 | 2403 | 2636 | 123 | 480.6 | 527.2 | 1962.866667 | 0.062663451 | 0.244845974 | 0.268586761 | -3.996231956 | -2.030053623 | -1.896539901 | KS-R1 | -2.640941827 | KS-R1 vs Km-V | 0.007137721 | 0.959065 | |
| | KS-R3 | 4 | 25 | 592 | 385 | 268 | 23.68 | 15.4 | 10.72 | 1335.1 | 0.017736499 | 0.011534717 | 0.008029361 | -5.817134914 | -6.437873644 | -6.960499089 | KS-R3 | -6.405169215 | KS-R3 vs Km-V | 0.029768313 | 0.007097 | KS<Km |
| | Sc-V | 2 | 625 | 2620 | 2216 | 2923 | 4.192 | 3.5456 | 4.6768 | 226.72 | 0.018489767 | 0.015638673 | 0.020628088 | -5.757129136 | -5.998738067 | -5.599246119 | Sc-V | -5.785037774 | | | | |
| | Sc-R1 | 2 | 625 | 2451 | 1952 | 2540 | 3.9216 | 3.1232 | 4.064 | 354.86 | 0.011051119 | 0.008801217 | 0.011452404 | -6.499663763 | -6.828081194 | -6.44820575 | Sc-R1 | -6.591983569 | | | | |
| | Sc-R3 | 2 | 625 | 1173 | 176 | 194 | 1.8768 | 0.2816 | 0.3104 | 296.5866667 | 0.006327999 | 0.00094947 | 0.001046574 | -7.304035013 | -10.04059069 | -9.900109469 | Sc-R3 | -9.081578391 | | | | |
| 0.4 µg/mL TM | Km-V | 5 | 5 | 1966 | 2709 | 805 | 393.2 | 541.8 | 161 | 2231.866667 | 0.176175399 | 0.242756437 | 0.072136926 | -2.504915615 | -2.042418543 | -3.793118249 | Km-V | -2.780150802 | Km-V vs Sc-V | 0.09995248 | 0.005998 | |
| | KS-R1 | 5 | 5 | 3597 | 3395 | 1779 | 719.4 | 679 | 355.8 | 1962.866667 | 0.366504772 | 0.34592263 | 0.181265496 | -1.448096112 | -1.531478697 | -2.463823761 | KS-R1 | -1.81446619 | KS-R1 vs Km-V | 0.557363507 | 0.192446 | |
| | KS-R3 | 4 | 25 | 1641 | 591 | 602 | 65.64 | 23.64 | 24.08 | 1335.1 | 0.049164857 | 0.017706539 | 0.018036102 | -4.346228756 | -5.819573959 | -5.792968603 | KS-R3 | -5.319590439 | KS-R3 vs Km-V | 0.926782417 | 0.023766 | KS<Km |
| | Sc-V | 2 | 625 | 2406 | 3067 | 3109 | 3.8496 | 4.9072 | 4.9744 | 226.72 | 0.016979534 | 0.021644319 | 0.02194072 | -5.880059306 | -5.529867781 | -5.510245331 | Sc-V | -5.640057473 | | | | |
| | Sc-R1 | 2 | 625 | 2414 | 1995 | 2439 | 3.8624 | 3.192 | 3.9024 | 354.86 | 0.010884292 | 0.008995097 | 0.010997013 | -6.521608571 | -6.7966455 | -6.506744489 | Sc-R1 | -6.608332853 | | | | |
| | Sc-R3 | 2 | 625 | 1726 | 1191 | 55.8 | 2.7616 | 1.9056 | 0.08928 | 296.5866667 | 0.009311275 | 0.006425103 | 0.000301025 | -6.746805562 | -7.282064613 | -11.69782909 | Sc-R3 | -8.575566423 | | | | |
| 0.6 M NaCl | Km-V | 5 | 5 | 2868 | 1491 | 2424 | 573.6 | 298.2 | 484.8 | 2231.866667 | 0.2570046 | 0.133610132 | 0.217217277 | -1.960133913 | -2.903898679 | -2.202789238 | Km-V | -2.355607277 | Km-V vs Sc-V | 0.177122276 | 0.00027 | |
| | KS-R1 | 5 | 5 | 1935 | 1450 | 2308 | 387 | 290 | 461.6 | 1962.866667 | 0.197160615 | 0.147743097 | 0.235166253 | -2.342556705 | -2.758837372 | -2.088247048 | KS-R1 | -2.396547042 | KS-R1 vs Km-V | 0.646054184 | 0.910978 | |
| | KS-R3 | 5 | 5 | 2183 | 1292 | 2090 | 436.6 | 258.4 | 418 | 1335.1 | 0.327016703 | 0.193543555 | 0.313085162 | -1.612563769 | -2.36926983 | -1.675372957 | KS-R3 | -1.885735519 | KS-R3 vs Km-V | 0.846714678 | 0.275842 | |
| | Sc-V | 2 | 625 | 2455 | 2050 | 2470 | 3.928 | 3.28 | 3.952 | 226.72 | 0.017325335 | 0.014467184 | 0.017431193 | -5.850972923 | -6.111072038 | -5.842184906 | Sc-V | -5.934743289 | | | | |
| | Sc-R1 | 2 | 625 | 2299 | 1072 | 2481 | 3.6784 | 1.7152 | 3.9696 | 354.86 | 0.010365778 | 0.004833455 | 0.011186383 | -6.592027781 | -7.692729341 | -6.482112512 | Sc-R1 | -6.922289878 | | | | |
| | Sc-R3 | 2 | 625 | 2649 | 2942 | 2644 | 4.2384 | 4.7072 | 4.2304 | 296.5866667 | 0.014290595 | 0.015871246 | 0.014263622 | -6.128790182 | -5.97744078 | -6.131515849 | Sc-R3 | -6.079248937 | | | | |
| 0.8 M NaCl | Km-V | 5 | 5 | 359 | 187 | 289.8 | 71.8 | 37.4 | 57.96 | 2231.866667 | 0.032170381 | 0.016757273 | 0.025969293 | -4.958123188 | -5.899068762 | -5.267049437 | Km-V | -5.374747129 | Km-V vs Sc-V | 0.080345879 | 0.000823 | |
| | KS-R1 | 5 | 5 | 301 | 387 | 389 | 60.2 | 77.4 | 77.8 | 1962.866667 | 0.030669429 | 0.039432123 | 0.039635907 | -5.02705488 | -4.6644848 | -4.657048212 | KS-R1 | -4.782862631 | KS-R1 vs Km-V | 0.325614849 | 0.122154 | |
| | KS-R3 | 5 | 5 | 1740 | 1569 | 1760 | 348 | 313.8 | 352 | 1335.1 | 0.260654633 | 0.235038574 | 0.263650663 | -1.939788594 | -2.089030547 | -1.923300471 | KS-R3 | -1.984039871 | KS-R3 vs Km-V | 0.069932727 | 0.000274 | KS>Km>Sc |
| | Sc-V | 1 | 3125 | 2897 | 3098 | 2704 | 0.92704 | 0.99136 | 0.86528 | 226.72 | 0.00408892 | 0.004372618 | 0.003816514 | -7.934064358 | -7.837286899 | -8.033528891 | Sc-V | -7.934960049 | KS-R3 vs Sc-V | 0.928016267 | 1.71E-07 | |
| | Sc-R1 | 1 | 3125 | 1109 | 546 | 450 | 0.35488 | 0.17472 | 0.144 | 354.86 | 0.001000056 | 0.000492363 | 0.000405794 | -9.965702976 | -10.98798949 | -11.26696544 | Sc-R1 | -10.7402193 | | | | |
| | Sc-R3 | 2 | 625 | 612 | 154 | 405 | 0.9792 | 0.2464 | 0.648 | 296.5866667 | 0.003301564 | 0.000830786 | 0.002184859 | -8.242634468 | -10.23323577 | -8.838244213 | Sc-R3 | -9.104704817 | | | | |
| 1.0 M NaCl | Km-V | 5 | 5 | 184 | 272 | 397 | 36.8 | 54.4 | 79.4 | 2231.866667 | 0.01648844 | 0.024374216 | 0.035575602 | -5.922401266 | -5.35850038 | -4.812968024 | Km-V | -5.364623223 | Km-V vs Sc-V | 0.287254256 | 0.001394 | |
| | KS-R1 | 5 | 5 | 217 | 225 | 655 | 43.4 | 45 | 131 | 1962.866667 | 0.022110519 | 0.022925653 | 0.066739123 | -5.499123324 | -5.446893365 | -3.90532346 | KS-R1 | -4.950446716 | KS-R1 vs Km-V | 0.545813672 | 0.536363 | |
| | KS-R3 | 5 | 5 | 2339 | 1532 | 1984 | 467.8 | 306.4 | 396.8 | 1335.1 | 0.350385739 | 0.229495918 | 0.297206202 | -1.512984038 | -2.123459603 | -1.750463874 | KS-R3 | -1.795635838 | KS-R3 vs Km-V | 0.470632583 | 0.000621 | KS>Km>Sc |
| | Sc-V | 1 | 3125 | 2159 | 2836 | 2836 | 0.69088 | 0.90752 | 0.90752 | 226.72 | 0.003047283 | 0.004002823 | 0.004002823 | -8.3582608 | -7.96476651 | -7.96476651 | Sc-V | -8.095931273 | KS-R3 vs Sc-V | 0.705508888 | 8.98E-06 | |
| | Sc-R1 | 1 | 3125 | 249 | 672 | 1425 | 0.07968 | 0.21504 | 0.456 | 354.86 | 0.000224539 | 0.000605985 | 0.001285014 | -12.12074469 | -10.6884292 | -9.604000423 | Sc-R1 | -10.80439144 | | | | |
| | Sc-R3 | 1 | 3125 | 2116 | 1791 | 2871 | 0.67712 | 0.57312 | 0.91872 | 296.5866667 | 0.002283043 | 0.001932386 | 0.003097644 | -8.774826494 | -9.015400784 | -8.334612791 | Sc-R3 | -8.708280023 | | | | |
| 2% ethanol | Km-V | 5 | 5 | 6206 | 7200 | 7745 | 1241.2 | 1440 | 1549 | 2231.866667 | 0.556126411 | 0.645199833 | 0.694037876 | -0.84651524 | -0.63218203 | -0.526913698 | Km-V | -0.66853699 | Km-V vs Sc-V | 0.099185717 | 0.025065 | |
| | KS-R1 | 5 | 5 | 7221 | 8194 | 6888 | 1444.2 | 1638.8 | 1377.6 | 1962.866667 | 0.735760622 | 0.834901335 | 0.701830656 | -0.442691629 | -0.260322379 | -0.510805129 | KS-R1 | -0.404606379 | KS-R1 vs Km-V | 0.774732141 | 0.092978 | |
| | KS-R3 | 5 | 5 | 1807.6 | 3964 | 2199.8 | 361.52 | 792.8 | 439.96 | 1335.1 | 0.270781215 | 0.593813198 | 0.329533368 | -1.884800438 | -0.751918937 | -1.601503536 | KS-R3 | -1.41274097 | KS-R3 vs Km-V | 0.141818297 | 0.102787 | |
| | Sc-V | 4 | 25 | 1192 | 2173 | 815.4 | 47.68 | 86.92 | 32.616 | 226.72 | 0.210303458 | 0.383380381 | 0.143860268 | -2.249455522 | -1.383151584 | -2.797259896 | Sc-V | -2.143289001 | | | | |
| | Sc-R1 | 4 | 25 | 2872 | 3409 | 1776.8 | 114.88 | 136.36 | 71.072 | 354.86 | 0.323733303 | 0.384264217 | 0.200281801 | -1.627122308 | -1.379829458 | -2.319896759 | Sc-R1 | -1.775616175 | | | | |
| | Sc-R3 | 4 | 25 | 1421 | 2931 | 838 | 56.84 | 117.24 | 33.52 | 296.5866667 | 0.191647186 | 0.395297608 | 0.113019241 | -2.383475282 | -1.338988868 | -3.145359687 | Sc-R3 | -2.289274613 | | | | |
| 3% glycerol | Km-V | 5 | 5 | 9568 | 5884 | 10138 | 1913.6 | 1176.8 | 2027.6 | 2231.866667 | 0.857398889 | 0.527271641 | 0.908477209 | -0.221961547 | -0.92338169 | -0.138477773 | Km-V | -0.427940337 | Km-V vs Sc-V | 0.885127577 | 0.034088 | |
| | KS-R1 | 5 | 5 | 9744 | 5927 | 8092 | 1948.8 | 1185.4 | 1618.4 | 1962.866667 | 0.992833611 | 0.603912645 | 0.824508372 | -0.010376139 | -0.727588214 | -0.278393952 | KS-R1 | -0.338786102 | KS-R1 vs Km-V | 0.828152645 | 0.797515 | |
| | KS-R3 | 5 | 5 | 4905 | 3034 | 2556 | 981 | 606.8 | 511.2 | 1335.1 | 0.734776421 | 0.45449779 | 0.382892667 | -0.444622763 | -1.137654814 | -1.384988064 | KS-R3 | -0.989088547 | KS-R3 vs Km-V | 0.877707174 | 0.209583 | |
| | Sc-V | 4 | 25 | 2017 | 2655 | 1559 | 80.68 | 106.2 | 62.36 | 226.72 | 0.355857445 | 0.468419195 | 0.275052929 | -1.490628674 | -1.094127897 | -1.86221883 | Sc-V | -1.482325134 | | | | |
| | Sc-R1 | 4 | 25 | 3681 | 3776 | 3516 | 147.24 | 151.04 | 140.64 | 354.86 | 0.414924195 | 0.425632644 | 0.396325311 | -1.269080307 | -1.232319292 | -1.335242987 | Sc-R1 | -1.278880862 | | | | |
| | Sc-R3 | 4 | 25 | 2798 | 3324 | 506 | 111.92 | 132.96 | 20.24 | 296.5866667 | 0.377360187 | 0.448300665 | 0.068243122 | -1.405985874 | -1.157461454 | -3.873172546 | Sc-R3 | -2.145539958 | | | | |
| Thr | Km-V | 5 | 5 | 12787 | 11211 | 11472 | 2557.4 | 2242.2 | 2294.4 | 2231.866667 | 1.145856981 | 1.004629906 | 1.0280184 | 0.196426986 | 0.006664127 | 0.039866087 | Km-V | 0.080985734 | Km-V vs Sc-V | 0.216161553 | 0.168813 | |
| | KS-R1 | 5 | 5 | 9794 | 8876 | 10166 | 1958.8 | 1775.2 | 2033.2 | 1962.866667 | 0.9979282 | 0.904391536 | 1.035831946 | -0.002992076 | -0.144980604 | 0.050789959 | KS-R1 | -0.03239424 | KS-R1 vs Km-V | 0.998014154 | 0.242104 | |
| | KS-R3 | 5 | 5 | 4386 | 7476 | 4552 | 877.2 | 1495.2 | 910.4 | 1335.1 | 0.657029436 | 1.119916111 | 0.681896487 | -0.605970088 | 0.163390669 | -0.552375342 | KS-R3 | -0.331651587 | KS-R3 vs Km-V | 0.105453733 | 0.18068 | |
| | Sc-V | 4 | 25 | 6085 | 4101 | 4642 | 243.4 | 164.04 | 185.68 | 226.72 | 1.073570924 | 0.723535639 | 0.818983769 | 0.102417505 | -0.466864015 | -0.288093236 | Sc-V | -0.217513248 | | | | |
| | Sc-R1 | 4 | 25 | 6235 | 5203 | 6173 | 249.4 | 208.12 | 246.92 | 354.86 | 0.702812377 | 0.586484811 | 0.695823705 | -0.508788497 | -0.76983435 | -0.523206266 | Sc-R1 | -0.600609704 | | | | |
| | Sc-R3 | 4 | 25 | 5264 | 4920 | 4225 | 210.56 | 196.8 | 169 | 296.5866667 | 0.709944255 | 0.663549721 | 0.56981658 | -0.494222347 | -0.591723521 | -0.811430495 | Sc-R3 | -0.632458788 | | | | |
| 42°C | Km-V | 5 | 5 | 11677 | 7566 | 8491 | 2335.4 | 1513.2 | 1698.2 | 2231.866667 | 1.046388673 | 0.677997491 | 0.760887747 | 0.065418829 | -0.560648161 | -0.394244464 | Km-V | -0.296491265 | Km-V vs Sc-V | / | / | |
| | KS-R1 | 4 | 25 | 9725 | 5696 | 7386 | 389 | 227.84 | 295.44 | 1962.866667 | 0.198179533 | 0.116075128 | 0.150514554 | -2.335120117 | -3.10686922 | -2.732025104 | KS-R1 | -2.72467148 | KS-R1 vs Km-V | 0.827691454 | 0.001128 | KS<Km |
| | KS-R3 | 5 | 5 | 6280 | 4182 | 4424 | 1256 | 836.4 | 884.8 | 1335.1 | 0.940753502 | 0.626469927 | 0.662721893 | -0.088111341 | -0.674682838 | -0.593524514 | KS-R3 | -0.452106231 | KS-R3 vs Km-V | 0.979915584 | 0.584717 | |
| | Sc-V | 0 | | | | | | | | 226.72 | | | | | | | Sc-V | | | | | |
| | Sc-R1 | 0 | | | | | | | | 354.86 | | | | | | | Sc-R1 | | | | | |
| | Sc-R3 | 0 | | | | | | | | 296.5866667 | | | | | | | Sc-R3 | | | | | |
| 37°C | Km-V | 5 | 5 | 13176 | 17794 | 13742 | 2635.2 | 3558.8 | 2748.4 | 2231.866667 | 1.180715694 | 1.594539698 | 1.23143557 | 0.239661618 | 0.673140015 | 0.300341146 | Km-V | 0.404380927 | Km-V vs Sc-V | 0.592654897 | 0.000441 | |
| | KS-R1 | 5 | 5 | 9679 | 8497 | 9501 | 1935.8 | 1699.4 | 1900.2 | 1962.866667 | 0.986210644 | 0.865774547 | 0.968073906 | -0.020032271 | -0.207936707 | -0.046810904 | KS-R1 | -0.091593294 | KS-R1 vs Km-V | 0.315814567 | 0.028344 | KS<Km |
| | KS-R3 | 5 | 5 | 11434 | 11982 | 9836 | 2286.8 | 2396.4 | 1967.2 | 1335.1 | 1.7128305 | 1.794921729 | 1.473447682 | 0.77638239 | 0.843920934 | 0.559195835 | KS-R3 | 0.72649972 | KS-R3 vs Km-V | 0.573198321 | 0.115105 | |
| | Sc-V | 3 | 125 | 6424 | 7351 | 4523 | 51.392 | 58.808 | 36.184 | 226.72 | 0.226676076 | 0.259386027 | 0.159597742 | -2.14129596 | -1.946827332 | -2.647487857 | Sc-V | -2.245203716 | | | | |
| | Sc-R1 | 3 | 125 | 9612 | 9453 | 7811 | 76.896 | 75.624 | 62.488 | 354.86 | 0.216693907 | 0.213109395 | 0.17609198 | -2.206269504 | -2.230333897 | -2.505598891 | Sc-R1 | -2.314067431 | | | | |
| | Sc-R3 | 3 | 125 | 6378 | 2025.2 | 2418 | 51.024 | 16.2016 | 19.344 | 296.5866667 | 0.172037403 | 0.054626866 | 0.065222082 | -2.539205834 | -4.194245542 | -3.938495687 | Sc-R3 | -3.557315687 | | | | |
| 20°C | Km-V | 5 | 5 | 4836 | 3939 | 4681 | 967.2 | 787.8 | 936.2 | 2231.866667 | 0.433359221 | 0.352978075 | 0.419469502 | -1.206364692 | -1.50234952 | -1.253362172 | Km-V | -1.320692128 | Km-V vs Sc-V | 0.019737152 | 0.043035 | |
| | KS-R1 | 5 | 5 | 6434 | 5552 | 3306 | 1286.8 | 1110.4 | 661.2 | 1962.866667 | 0.655571783 | 0.565703223 | 0.336854261 | -0.609174336 | -0.821882704 | -1.569803547 | KS-R1 | -1.000286862 | KS-R1 vs Km-V | 0.180807545 | 0.353373 | |
| | KS-R3 | 5 | 5 | 255 | 1762 | 814 | 51 | 352.4 | 162.8 | 1335.1 | 0.038199386 | 0.263950266 | 0.121938432 | -4.710306748 | -1.921661976 | -3.0357752 | KS-R3 | -3.222581308 | KS-R3 vs Km-V | 0.025357418 | 0.141797 | |
| | Sc-V | 3 | 125 | 170 | 1051 | 1246.8 | 1.36 | 8.408 | 9.9744 | 226.72 | 0.005998589 | 0.037085392 | 0.043994354 | -7.381161202 | -4.753005184 | -4.506537793 | Sc-V | -5.546901393 | | | | |
| | Sc-R1 | 3 | 125 | 5893 | 6841 | 2531.6 | 47.144 | 54.728 | 20.2528 | 354.86 | 0.132852392 | 0.154224201 | 0.057072648 | -2.912103886 | -2.696898922 | -4.131056679 | Sc-R1 | -3.246686496 | | | | |
| | Sc-R3 | 3 | 125 | 695 | 2208 | 1160 | 5.56 | 17.664 | 9.28 | 296.5866667 | 0.018746628 | 0.059557634 | 0.031289336 | -5.737225048 | -4.069569759 | -4.998185126 | Sc-R3 | -4.934993311 | | | | |
| Sorbitol | Km-V | 5 | 5 | 4110 | 5740 | 4887 | 822 | 1148 | 977.4 | 2231.866667 | 0.368301571 | 0.514367644 | 0.437929386 | -1.441040543 | -0.9591282 | -1.191229832 | Km-V | -1.197132859 | Km-V vs Sc-V | 0.280950129 | 0.424965 | |
| | KS-R1 | 5 | 5 | 4694 | 4319 | 4527 | 938.8 | 863.8 | 905.4 | 1962.866667 | 0.478280067 | 0.440070645 | 0.461264137 | -1.06407243 | -1.184192955 | -1.116334965 | KS-R1 | -1.12153345 | KS-R1 vs Km-V | 0.117559448 | 0.626007 | |
| | KS-R3 | 5 | 5 | 3760 | 5350 | 3001 | 752 | 1070 | 600.2 | 1335.1 | 0.563253689 | 0.801438095 | 0.44955434 | -0.828143238 | -0.319337008 | -1.153432581 | KS-R3 | -0.766970942 | KS-R3 vs Km-V | 0.494725011 | 0.198982 | |
| | Sc-V | 4 | 25 | 1259 | 2846 | 2125 | 50.36 | 113.84 | 85 | 226.72 | 0.222124206 | 0.502117149 | 0.374911785 | -2.170561475 | -0.993904096 | -1.415376917 | Sc-V | -1.526614163 | | | | |
| | Sc-R1 | 4 | 25 | 4654 | 4669 | 4801 | 186.16 | 186.76 | 192.04 | 354.86 | 0.524601251 | 0.526292059 | 0.541171166 | -0.930706846 | -0.926064469 | -0.885843121 | Sc-R1 | -0.914204812 | | | | |
| | Sc-R3 | 4 | 25 | 3591 | 4336 | 1837 | 143.64 | 173.44 | 73.48 | 296.5866667 | 0.484310376 | 0.584786909 | 0.247752203 | -1.045996183 | -0.77401708 | -2.01303021 | Sc-R3 | -1.277681158 | | | | |
| Ser | Km-V | 6 | 1 | 8163 | 9448 | 7385 | 8163 | 9448 | 7385 | 2231.866667 | 3.657476552 | 4.233227791 | 3.30888942 | 1.870848615 | 2.081758123 | 1.726347079 | Km-V | 1.892984606 | Km-V vs Sc-V | 0.438900144 | 0.000289 | |
| | KS-R1 | 6 | 1 | 10358 | 5556 | 8911 | 10358 | 5556 | 8911 | 1962.866667 | 5.276975852 | 2.830553952 | 4.539788744 | 2.399711382 | 1.501084422 | 2.182625164 | KS-R1 | 2.027806989 | KS-R1 vs Km-V | 0.253741151 | 0.66588 | |
| | KS-R3 | 6 | 1 | 4542 | 6341 | 3372 | 4542 | 6341 | 3372 | 1335.1 | 3.40199236 | 4.74945697 | 2.525653509 | 1.766379901 | 2.247762572 | 1.336656731 | KS-R3 | 1.783599735 | KS-R3 vs Km-V | 0.266558804 | 0.718479 | |
| | Sc-V | 4 | 25 | 3766 | 4092 | 2636 | 150.64 | 163.68 | 105.44 | 226.72 | 0.664431898 | 0.721947777 | 0.465067043 | -0.589806758 | -0.470033613 | -1.104489388 | Sc-V | -0.721443253 | | | | |
| | Sc-R1 | 4 | 25 | 4353 | 1807 | 4111 | 174.12 | 72.28 | 164.44 | 354.86 | 0.490672378 | 0.203685961 | 0.463394015 | -1.027168037 | -2.295581551 | -1.109688685 | Sc-R1 | -1.477479424 | | | | |
| | Sc-R3 | 4 | 25 | 4313 | 5071 | 3542 | 172.52 | 202.84 | 141.68 | 296.5866667 | 0.581684949 | 0.683914764 | 0.477701852 | -0.78169012 | -0.548111562 | -1.065817624 | Sc-R3 | -0.798539769 | | | | |
| 1/4N | Km-V | 5 | 5 | 7587 | 5839 | 7566 | 1517.4 | 1167.8 | 1513.2 | 2231.866667 | 0.679879324 | 0.523239142 | 0.677997491 | -0.556649399 | -0.934457626 | -0.560648161 | Km-V | -0.683918395 | Km-V vs Sc-V | 0.268366071 | 0.000169 | |

Average growth relative to YPD 30°C, subtracted by Km-V

| | Km-V | KS-R1 | KS-R3 | Sc-V | Sc-R1 | Sc-R3 |
|---|---|---|---|---|---|---|
| 2% ethanol | 0 | 0.263930611 | -0.74 | -1.475 | -1.11 | -1.62073762 |
| 3% glycerol | 0 | 0.089154235 | -0.56 | -1.054 | -0.85 | -1.71759962 |
| 0.02% Glu | 0 | 0.333595763 | -0.76 | -0.933 | -0.46 | -1.27905444 |
| 1% Glu | 0 | 0.283500843 | -0.25 | -0.572 | -0.78 | -0.51642161 |
| 3% Glu | 0 | 0.376174163 | -0.45 | 0.153 | 0.26 | 0.06963234 |
| 5% Glu | 0 | 0.106672442 | -0.47 | -0.195 | -0.22 | -0.5615968 |
| Thr | 0 | -0.113379974 | -0.41 | -0.298 | -0.68 | -0.71344452 |
| Ser | 0 | 0.134822383 | -0.11 | -2.614 | -3.37 | -2.69152438 |
| 1/4N | 0 | -0.009429224 | -2.06 | -4.649 | -3.65 | -6.45988863 |
| 1/2AA1/2N | 0 | 0.16987529 | -1.63 | -2.492 | -2.27 | -2.82481039 |
| 20°C | 0 | 0.320405266 | -1.9 | -4.226 | -1.93 | -3.61430118 |
| 37°C | 0 | -0.495974221 | 0.322 | -2.65 | -2.72 | -3.96169661 |
| 42°C | 0 | -2.428180215 | -0.16 | NA | NA | NA |
| DTT | 0 | 0.501347256 | -0.23 | -0.402 | -0.72 | -1.00639391 |
| 5-FU | 0 | -0.278174621 | -0.64 | 0.7873 | 0.73 | -0.4827989 |
| Sorbitol | 0 | 0.075599409 | 0.43 | -0.329 | 0.28 | -0.0805483 |
| 0.012% H2O2 | 0 | -0.428540282 | -0.19 | -1.917 | -1.28 | -1.28735028 |
| 0.016% H2O2 | 0 | -0.580664345 | -1.52 | -4.626 | -5.72 | -4.44843473 |
| 0.024% H2O2 | 0 | -0.36135861 | -1.84 | NA | NA | NA |
| 55 mM AcOH | 0 | 0.040163016 | 0.498 | 0.9092 | 0.1 | 0.57367737 |
| 57.5 mM AcOH | 0 | -0.071452478 | 0.715 | 1.2008 | 0.5 | 1.02548512 |
| 60 mM AcOH | 0 | -0.125797082 | 1.694 | 4.6063 | 3.08 | 4.57039281 |
| 0.2 µg/mL TM | 0 | 0.071827768 | -1.72 | -2.373 | -2.39 | -3.42944718 |
| 0.3 µg/mL TM | 0 | 0.039364099 | -3.72 | -3.105 | -3.91 | -6.40127247 |
| 0.4 µg/mL TM | 0 | 0.965684612 | -2.54 | -2.86 | -3.83 | -5.79541562 |
| 0.6 M NaCl | 0 | -0.040939765 | 0.47 | -3.579 | -4.57 | -3.72364166 |
| 0.8 M NaCl | 0 | 0.591884498 | 3.391 | -2.56 | -5.37 | -3.72995769 |
| 1.0 M NaCl | 0 | 0.414176507 | 3.569 | -2.731 | -5.44 | -3.3436568 |

| | | | | | | | | | | | | | | | | | | | | | | |
|---|---|---|---|---|---|---|---|---|---|---|---|---|---|---|---|---|---|---|---|---|---|---|
| | KS-R1 | 5 | 5 | 7956 | 4693 | 5988 | 1591.2 | 938.6 | 1197.6 | 1962.866667 | 0.810651089 | 0.478178175 | 0.610128044 | -0.302846996 | -1.064379811 | -0.71281605 | KS-R1 | -0.693347619 | KS-R1 vs Km-V | 0.489542326 | 0.972079 | |
| | KS-R3 | 4 | 25 | 5388 | 7680 | 3009 | 215.52 | 307.2 | 120.36 | 1335.1 | 0.16142611 | 0.230095124 | 0.090150551 | -2.631054144 | -2.119697684 | -3.471519888 | KS-R3 | -2.740757239 | KS-R3 vs Km-V | 0.183566446 | 0.007629 | KS<Km |
| | Sc-V | 3 | 125 | 469 | 739 | 1002 | 3.752 | 5.912 | 8.016 | 226.72 | 0.016549047 | 0.026076217 | 0.035356387 | -5.917108025 | -5.261121584 | -4.821885345 | Sc-V | -5.333371651 | | | | |
| | Sc-R1 | 3 | 125 | 1600 | 2804 | 2389 | 12.8 | 22.432 | 19.112 | 354.86 | 0.036070563 | 0.063213662 | 0.053857859 | -4.793034247 | -3.983619803 | -4.214699198 | Sc-R1 | -4.330451116 | | | | |
| | Sc-R3 | 3 | 125 | 238 | 682 | 111 | 1.904 | 5.456 | 0.888 | 296.5866667 | 0.006419709 | 0.018395972 | 0.002994066 | -7.283276453 | -5.764466287 | -8.38367835 | Sc-R3 | -7.14380703 | | | | |
| 5% Glu | Km-V | 5 | 5 | 11562 | 11820 | 10438 | 2312.4 | 2364 | 2087.6 | 2231.866667 | 1.036083398 | 1.059203059 | 0.935360535 | 0.051140135 | 0.082979193 | -0.096405535 | Km-V | 0.012571265 | Km-V vs Sc-V | 0.17499332 | 0.355385 | |
| | KS-R1 | 5 | 5 | 10433 | 12177 | 9535 | 2086.6 | 2435.4 | 1907 | 1962.866667 | 1.063037055 | 1.240736338 | 0.971538226 | 0.088191886 | 0.311196569 | -0.041657333 | KS-R1 | 0.119243707 | KS-R1 vs Km-V | 0.446739051 | 0.413192 | |
| | KS-R3 | 5 | 5 | 3577 | 7652 | 4210 | 715.4 | 1530.4 | 842 | 1335.1 | 0.535840012 | 1.146281177 | 0.63066437 | -0.900125781 | 0.196960974 | -0.665055667 | KS-R3 | -0.456073491 | KS-R3 vs Km-V | 0.053442108 | 0.237908 | |
| | Sc-V | 4 | 25 | 5941 | 5335 | 3931 | 237.64 | 213.4 | 157.24 | 226.72 | 1.048165138 | 0.941249118 | 0.693542696 | 0.06786603 | -0.087351487 | -0.527943394 | Sc-V | -0.182476284 | | | | |
| | Sc-R1 | 4 | 25 | 6946 | 7944 | 8192 | 277.84 | 317.76 | 327.68 | 354.86 | 0.782956659 | 0.895451727 | 0.923406414 | -0.352995646 | -0.159312434 | -0.114962342 | Sc-R1 | -0.209090141 | | | | |
| | Sc-R3 | 4 | 25 | 6327 | 7456 | 2759 | 253.08 | 298.24 | 110.36 | 296.5866667 | 0.853308757 | 1.005574537 | 0.372100342 | -0.228860241 | 0.008020024 | -1.42623638 | Sc-R3 | -0.549025532 | | | | |
| 3% Glu | Km-V | 5 | 5 | 10807 | 7001 | 10864 | 2161.4 | 1400.2 | 2172.8 | 2231.866667 | 0.968427027 | 0.627367226 | 0.973534859 | -0.046284752 | -0.67261793 | -0.038695457 | Km-V | -0.252532713 | Km-V vs Sc-V | 0.835365625 | 0.607965 | |
| | KS-R1 | 5 | 5 | 11009 | 9781 | 11353 | 2201.8 | 1956.2 | 2270.6 | 1962.866667 | 1.121726726 | 0.996603607 | 1.156777502 | 0.165721251 | -0.004908299 | 0.210111399 | KS-R1 | 0.12364145 | KS-R1 vs Km-V | 0.177432034 | 0.162524 | |
| | KS-R3 | 5 | 5 | 3922 | 5001 | 3492 | 784.4 | 1000.2 | 698.4 | 1335.1 | 0.587521534 | 0.749157366 | 0.523106883 | -0.767286364 | -0.416659295 | -0.934822341 | KS-R3 | -0.706256 | KS-R3 vs Km-V | 0.691270379 | 0.155504 | |
| | Sc-V | 4 | 25 | 5868 | 6097 | 4138 | 234.72 | 243.88 | 165.52 | 226.72 | 1.035285815 | 1.075688073 | 0.730063514 | 0.050029113 | 0.105259788 | -0.453906113 | Sc-V | -0.099539071 | | | | |
| | Sc-R1 | 4 | 25 | 8139 | 9242 | 9391 | 325.56 | 369.68 | 375.64 | 354.86 | 0.917432227 | 1.041762949 | 1.058558305 | -0.124326509 | 0.059027032 | 0.082100734 | Sc-R1 | 0.005600419 | | | | |
| | Sc-R3 | 4 | 25 | 6893 | 7935 | 5095 | 275.72 | 317.4 | 203.8 | 296.5866667 | 0.929643949 | 1.070176227 | 0.687151591 | -0.105249821 | 0.097848387 | -0.54129969 | Sc-R3 | -0.182900375 | | | | |
| 0.02% Glu | Km-V | 5 | 5 | 8135 | 7425 | 8552 | 1627 | 1485 | 1710.4 | 2231.866667 | 0.7289862 | 0.665362327 | 0.766354024 | -0.456036591 | -0.587787911 | -0.383917084 | Km-V | -0.475913862 | Km-V vs Sc-V | 0.538045602 | 0.001258 | |
| | KS-R1 | 5 | 5 | 9288 | 8728 | 8674 | 1857.6 | 1745.6 | 1734.8 | 1962.866667 | 0.946370954 | 0.889311551 | 0.883809394 | -0.0795223 | -0.16923917 | -0.178192829 | KS-R1 | -0.142318099 | KS-R1 vs Km-V | 0.435798777 | 0.007801 | KS>Km>Sc |
| | KS-R3 | 5 | 5 | 2679 | 5566 | 1534 | 535.8 | 1113.2 | 306.8 | 1335.1 | 0.401318253 | 0.833795221 | 0.229795521 | -1.317181319 | -0.262234991 | -2.121577417 | KS-R3 | -1.233664576 | KS-R3 vs Km-V | 0.024283283 | 0.293979 | |
| | Sc-V | 4 | 25 | 2061 | 2435 | 1938 | 82.44 | 97.4 | 77.52 | 226.72 | 0.363620325 | 0.429604799 | 0.341919548 | -1.459495253 | -1.218917986 | -1.548271188 | Sc-V | -1.408894809 | KS-R1 vs Sc-V | 0.186001159 | 0.000254 | |
| | Sc-R1 | 4 | 25 | 4597 | 5196 | 4169 | 183.88 | 207.84 | 166.76 | 354.86 | 0.518176182 | 0.585695767 | 0.469931804 | -0.948485391 | -0.771776626 | -1.089476685 | Sc-R1 | -0.936579567 | | | | |
| | Sc-R3 | 4 | 25 | 2684 | 3737 | 1057 | 107.36 | 149.48 | 42.28 | 296.5866667 | 0.361985254 | 0.504001079 | 0.142555296 | -1.465997165 | -0.988501273 | -2.81040646 | Sc-R3 | -1.754968299 | | | | |
| 1% Glu | Km-V | 6 | 1 | 4499 | 1784 | 6300 | 4499 | 1784 | 6300 | 2231.866667 | 2.015801422 | 0.799330904 | 2.822749268 | 1.011353525 | -0.323135227 | 1.497100987 | Km-V | 0.728439762 | Km-V vs Sc-V | 0.433489104 | 0.404706 | |
| | KS-R1 | 6 | 1 | 5523 | 2578 | 4356 | 5523 | 2578 | 4356 | 1962.866667 | 2.813741806 | 1.313385185 | 2.219203206 | 1.49248995 | 0.393290087 | 1.150041777 | KS-R1 | 1.011940605 | KS-R1 vs Km-V | 0.525212305 | 0.67776 | |
| | KS-R3 | 6 | 1 | 1699 | 2366 | 1587 | 1699 | 2366 | 1587 | 1335.1 | 1.272563853 | 1.772151899 | 1.188675006 | 0.347738048 | 0.825502269 | 0.249354323 | KS-R3 | 0.474198213 | KS-R3 vs Km-V | 0.193182642 | 0.679953 | |
| | Sc-V | 4 | 25 | 8454 | 6893 | 4324 | 338.16 | 275.72 | 172.96 | 226.72 | 1.491531404 | 1.216125618 | 0.762879323 | 0.576794354 | 0.282292257 | -0.390473235 | Sc-V | 0.156204459 | | | | |
| | Sc-R1 | 4 | 25 | 8437 | 8920 | 8403 | 337.48 | 356.8 | 336.12 | 354.86 | 0.951022939 | 1.005466945 | 0.947190441 | -0.072447956 | 0.007865653 | -0.078273573 | Sc-R1 | -0.047618625 | | | | |
| | Sc-R3 | 4 | 25 | 10182 | 9457 | 6579 | 407.28 | 378.28 | 263.16 | 296.5866667 | 1.37322424 | 1.275445064 | 0.88729545 | 0.457567229 | 0.35100076 | -0.172513524 | Sc-R3 | 0.212018155 | | | | |
| 5-FU | Km-V | 5 | 5 | 8220 | 6521 | 7101 | 1644 | 1304.2 | 1420.2 | 2231.866667 | 0.736603142 | 0.584353904 | 0.636328335 | -0.441040543 | -0.775085717 | -0.65215673 | Km-V | -0.622760997 | Km-V vs Sc-V | 0.708607862 | 0.008622 | |
| | KS-R1 | 5 | 5 | 6064 | 6993 | 3424 | 1212.8 | 1398.6 | 684.8 | 1962.866667 | 0.61787182 | 0.712529294 | 0.348877492 | -0.694620518 | -0.488978767 | -1.51920757 | KS-R1 | -0.900935618 | KS-R1 vs Km-V | 0.175221502 | 0.446138 | |
| | KS-R3 | 5 | 5 | 1678.2 | 6215 | 2047.2 | 335.64 | 1243 | 409.44 | 1335.1 | 0.251396899 | 0.931016403 | 0.306673657 | -1.99196124 | -0.103121509 | -1.705223847 | KS-R3 | -1.266768865 | KS-R3 vs Km-V | 0.053622273 | 0.340496 | |
| | Sc-V | 4 | 25 | 7135 | 6772 | 5306 | 285.4 | 270.88 | 212.24 | 226.72 | 1.258821454 | 1.194777699 | 0.936132675 | 0.332073671 | 0.256742215 | -0.095215083 | Sc-V | 0.164533601 | | | | |
| | Sc-R1 | 4 | 25 | 9126 | 9300 | 10318 | 365.04 | 372 | 412.72 | 354.86 | 1.02868737 | 1.048300738 | 1.163050217 | 0.040804597 | 0.068052659 | 0.217913389 | Sc-R1 | 0.108923548 | | | | |
| | Sc-R3 | 4 | 25 | 4319 | 5408 | 1751.6 | 172.76 | 216.32 | 70.064 | 296.5866667 | 0.582494156 | 0.729365222 | 0.23623449 | -0.77968452 | -0.455286685 | -2.081708482 | Sc-R3 | -1.105559895 | | | | |
| DTT | Km-V | 5 | 5 | 4248 | 4347 | 5921 | 849.6 | 869.4 | 1184.2 | 2231.866667 | 0.380667901 | 0.389539399 | 0.530587251 | -1.393395171 | -1.360158841 | -0.914338083 | Km-V | -1.222630698 | Km-V vs Sc-V | 0.113497465 | 0.064911 | |
| | KS-R1 | 5 | 5 | 5394 | 7298 | 5359 | 1078.8 | 1459.6 | 1071.8 | 1962.866667 | 0.54960432 | 0.74360629 | 0.546038108 | -0.86353475 | -0.427389121 | -0.872926456 | KS-R1 | -0.721283442 | KS-R1 vs Km-V | 0.950448313 | 0.078383 | |
| | KS-R3 | 5 | 5 | 2149 | 4376 | 1527 | 429.8 | 875.2 | 305.4 | 1335.1 | 0.321923451 | 0.655531421 | 0.22874691 | -1.635210417 | -0.609263162 | -2.128175838 | KS-R3 | -1.457549805 | KS-R3 vs Km-V | 0.212972609 | 0.645682 | |
| | Sc-V | 4 | 25 | 1842 | 1922 | 1755 | 73.68 | 76.88 | 70.2 | 226.72 | 0.324982357 | 0.339096683 | 0.309633028 | -1.621566697 | -1.560231422 | -1.691368728 | Sc-V | -1.624388949 | | | | |
| | Sc-R1 | 4 | 25 | 2332.6 | 2239.8 | 2341.8 | 93.304 | 89.592 | 93.672 | 354.86 | 0.26293186 | 0.252471397 | 0.263968889 | -1.927239126 | -1.985808143 | -1.921560189 | Sc-R1 | -1.944869152 | | | | |
| | Sc-R3 | 4 | 25 | 1981.8 | 1978.4 | 1009 | 79.272 | 79.136 | 40.36 | 296.5866667 | 0.267281065 | 0.266822514 | 0.13608164 | -1.903570461 | -1.906047692 | -2.877455662 | Sc-R3 | -2.229024605 | | | | |
| 1/2AA1/2N | Km-V | 5 | 5 | 7547 | 7147 | 7592 | 1509.4 | 1429.4 | 1518.4 | 2231.866667 | 0.67629488 | 0.640450445 | 0.680327379 | -0.564275663 | -0.642841149 | -0.555698945 | Km-V | -0.587605252 | Km-V vs Sc-V | 0.04753482 | 0.004266 | |
| | KS-R1 | 5 | 5 | 6892 | 6927 | 8307 | 1378.4 | 1385.4 | 1661.4 | 1962.866667 | 0.702238223 | 0.705804436 | 0.846415107 | -0.50996757 | -0.502659598 | -0.240562717 | KS-R1 | -0.417729962 | KS-R1 vs Km-V | 0.178386616 | 0.141286 | |
| | KS-R3 | 4 | 25 | 6890 | 11183 | 4844 | 275.6 | 447.32 | 193.76 | 1335.1 | 0.206426485 | 0.335046064 | 0.145127706 | -2.276300012 | -1.577568636 | -2.78460513 | KS-R3 | -2.212824592 | KS-R3 vs Km-V | 0.012483093 | 0.042585 | KS<Km |
| | Sc-V | 3 | 125 | 2740 | 4191 | 3278 | 21.92 | 33.528 | 26.224 | 226.72 | 0.096683133 | 0.147882851 | 0.115666902 | -3.37059196 | -2.757473332 | -3.111951999 | Sc-V | -3.080005763 | | | | |
| | Sc-R1 | 3 | 125 | 5523 | 6656 | 6270 | 44.184 | 53.248 | 50.16 | 354.86 | 0.124511075 | 0.150053542 | 0.141351519 | -3.005654025 | -2.736450719 | -2.822640709 | Sc-R1 | -2.854915151 | | | | |
| | Sc-R3 | 3 | 125 | 2699 | 4744 | 3297 | 21.592 | 37.952 | 26.376 | 296.5866667 | 0.072801654 | 0.127962597 | 0.088931847 | -3.779884955 | -2.966205921 | -3.491156044 | Sc-R3 | -3.41241564 | | | | |
| 0.012% H2O2 | Km-V | 5 | 5 | 3085 | 1674 | | 617 | 334.8 | 0 | 2231.866667 | 0.276450206 | 0.150008961 | | -1.854908448 | -2.736879409 | | Km-V | -2.295893928 | Km-V vs Sc-V | 0.576295809 | 0.197631 | |
| | KS-R1 | 5 | 5 | 2434 | 906 | | 486.8 | 181.2 | 0 | 1962.866667 | 0.248004619 | 0.092313963 | | -2.011561104 | -3.437307316 | | KS-R1 | -2.72443421 | KS-R1 vs Km-V | 0.705356925 | 0.660034 | |
| | KS-R3 | 5 | 5 | 2115 | 674 | | 423 | 134.8 | 0 | 1335.1 | 0.3168302 | 0.10096622 | | -1.658218236 | -3.308055403 | | KS-R3 | -2.48313682 | KS-R3 vs Km-V | 0.6250688 | 0.859851 | |
| | Sc-V | 4 | 25 | 573 | 163 | | 22.92 | 6.52 | 0 | 226.72 | 0.10109386 | 0.028757939 | | -3.306232714 | -5.119895889 | | Sc-V | -4.213064301 | | | | |
| | Sc-R1 | 4 | 25 | 1079 | 516 | | 43.16 | 20.64 | 0 | 354.86 | 0.12162543 | 0.058163783 | | -3.039483192 | -4.103735086 | | Sc-R1 | -3.571609139 | | | | |
| | Sc-R3 | 4 | 25 | 567.8 | 674 | | 22.712 | 26.96 | 0 | 296.5866667 | 0.076577954 | 0.090900917 | | -3.706927082 | -3.45956134 | | Sc-R3 | -3.583244211 | | | | |
| 0.016% H2O2 | Km-V | 5 | 5 | 2287 | 1959 | | 457.4 | 391.8 | 0 | 2231.866667 | 0.204940558 | 0.175548121 | | -2.286722571 | -2.510061539 | | Km-V | -2.398392055 | Km-V vs Sc-V | 0.169593357 | 0.031492 | |
| | KS-R1 | 5 | 5 | 1735 | 893 | | 347 | 178.6 | 0 | 1962.866667 | 0.176782282 | 0.090989369 | | -2.499954609 | -3.458158191 | | KS-R1 | -2.9790564 | KS-R1 vs Km-V | 0.291562351 | 0.359228 | |
| | KS-R3 | 5 | 5 | 628 | 311 | | 125.6 | 62.2 | 0 | 1335.1 | 0.09407535 | 0.046588271 | | -3.410039436 | -4.423889414 | | KS-R3 | -3.916964425 | KS-R3 vs Km-V | 0.276070136 | 0.099677 | |
| | Sc-V | 2 | 625 | 611 | 1940 | | 0.9776 | 3.104 | 0 | 226.72 | 0.004311927 | 0.013690896 | | -7.857451663 | -6.190639296 | | Sc-V | -7.024045479 | | | | |
| | Sc-R1 | 2 | 625 | 672 | 943 | | 1.0752 | 1.5088 | 0 | 354.86 | 0.003029927 | 0.004251818 | | -8.366501109 | -7.877704571 | | Sc-R1 | -8.12210284 | | | | |
| | Sc-R3 | 2 | 625 | 1540 | 1684 | | 2.464 | 2.6944 | 0 | 296.5866667 | 0.008307858 | 0.009084697 | | -6.911307675 | -6.782345888 | | Sc-R3 | -6.846826782 | | | | |
| 0.024% H2O2 | Km-V | 4 | 25 | 1738 | 2206 | | 69.52 | 88.24 | 0 | 2231.866667 | 0.031148814 | 0.039536412 | | -5.00467895 | -4.660674241 | | Km-V | -4.832676595 | Km-V vs Sc-V | / | / | |
| | KS-R1 | 4 | 25 | 1194 | 1505 | | 47.76 | 60.2 | 0 | 1962.866667 | 0.02433176 | 0.030669429 | | -5.36101553 | -5.02705488 | | KS-R1 | -5.194035205 | KS-R1 vs Km-V | 0.981138333 | 0.270711 | |
| | KS-R3 | 3 | 125 | 1275 | 2103 | | 10.2 | 16.824 | 0 | 1335.1 | 0.007639877 | 0.012601303 | | -7.032234842 | -6.31028324 | | KS-R3 | -6.671259041 | KS-R3 vs Km-V | 0.566165809 | 0.044187 | KS<Km |
| | Sc-V | | | | | | | | | 226.72 | | | | | | | Sc-V | | | | | |
| | Sc-R1 | | | | | | | | | 354.86 | | | | | | | Sc-R1 | | | | | |
| | Sc-R3 | | | | | | | | | 296.5866667 | | | | | | | Sc-R3 | | | | | |
| 55 mM AcOH | Km-V | 4 | 25 | 1619 | 1584 | 1598 | 64.76 | 63.36 | 63.92 | 2231.866667 | 0.02901607 | 0.028388793 | 0.028639704 | -5.107004046 | -5.138534696 | -5.125839624 | Km-V | -5.123792789 | Km-V vs Sc-V | 0.00672245 | 0.028448 | |
| | KS-R1 | 4 | 25 | 1581 | 1386 | 1383 | 63.24 | 55.44 | 55.32 | 1962.866667 | 0.032218184 | 0.028244404 | 0.028183269 | -4.955980999 | -5.145891109 | -5.14901721 | KS-R1 | -5.083629773 | KS-R1 vs Km-V | 0.040351408 | 0.594738 | |
| | KS-R3 | 4 | 25 | 1454 | 1338 | 1270 | 58.16 | 53.52 | 50.8 | 1335.1 | 0.04356228 | 0.040086885 | 0.038049584 | -4.520776725 | -4.640725879 | -4.715975498 | KS-R3 | -4.625826034 | KS-R3 vs Km-V | 0.050621859 | 0.000983 | KS=Sc>Km |
| | Sc-V | 3 | 125 | 1891 | 1425 | 1320 | 15.128 | 11.4 | 10.56 | 226.72 | 0.066725476 | 0.050282287 | 0.046577276 | -3.90561849 | -4.313805934 | -4.424229924 | Sc-V | -4.214551449 | KS-R3 vs Sc-V | 0.229889054 | 0.070206 | |
| | Sc-R1 | 3 | 125 | 1770 | 1073 | 1331 | 14.16 | 8.584 | 10.648 | 354.86 | 0.03990306 | 0.024189821 | 0.0300062 | -4.647356792 | -5.369456076 | -5.058595581 | Sc-R1 | -5.025136149 | | | | |
| | Sc-R3 | 3 | 125 | 1774 | 1560 | 1432 | 14.192 | 12.48 | 11.456 | 296.5866667 | 0.047851106 | 0.042078763 | 0.038626146 | -4.385303922 | -4.570763902 | -4.694278439 | Sc-R3 | -4.550115421 | | | | |
| 57.5 mM AcOH | Km-V | 4 | 25 | 988 | 934 | 988 | 39.52 | 37.36 | 39.52 | 2231.866667 | 0.017707151 | 0.016739351 | 0.017707151 | -5.819524085 | -5.900612577 | -5.819524085 | Km-V | -5.846553582 | Km-V vs Sc-V | 0.111693983 | 0.000465 | |
| | KS-R1 | 4 | 25 | 612 | 1024 | 853 | 24.48 | 40.96 | 34.12 | 1962.866667 | 0.012471555 | 0.020867439 | 0.01738274 | -6.325214809 | -5.582602651 | -5.84620072 | KS-R1 | -5.91800606 | KS-R1 vs Km-V | 0.030456847 | 0.774407 | |
| | KS-R3 | 4 | 25 | 968 | 882 | 1010 | 38.72 | 35.28 | 40.4 | 1335.1 | 0.029001573 | 0.026424987 | 0.030259906 | -5.107725042 | -5.241953434 | -5.046448702 | KS-R3 | -5.132042393 | KS-R3 vs Km-V | 0.359579359 | 0.000361 | Sc>KS>Km |
| | Sc-V | 3 | 125 | 1067 | 1031 | 1319 | 8.536 | 8.248 | 10.552 | 226.72 | 0.037649965 | 0.036379675 | 0.04654199 | -4.731207677 | -4.78072352 | -4.425323289 | Sc-V | -4.645751495 | | | | |
| | Sc-R1 | 3 | 125 | 1272 | 921 | 1102 | 10.176 | 7.368 | 8.816 | 354.86 | 0.028676098 | 0.020763118 | 0.0248436 | -5.124007481 | -5.589833091 | -5.330981928 | Sc-R1 | -5.348274167 | | | | |
| | Sc-R3 | 3 | 125 | 1419 | 1029 | 1545 | 11.352 | 8.232 | 12.36 | 296.5866667 | 0.03827549 | 0.027755799 | 0.041674159 | -4.707435342 | -5.171066949 | -4.584703093 | Sc-R3 | -4.821068461 | | | | |
| 60 mM AcOH | Km-V | 3 | 125 | 530 | 279 | | 4.24 | 2.232 | 0 | 2231.866667 | 0.001899755 | 0.00100006 | 0 | -9.039970862 | -9.9656981 | | Km-V | -9.502834481 | Km-V vs Sc-V | 0.333736845 | 0.010651 | |
| | KS-R1 | 3 | 125 | 739 | 130 | | 5.912 | 1.04 | 0 | 1962.866667 | 0.003011921 | 0.000529837 | 0 | -8.375100192 | -10.88216293 | | KS-R1 | -9.628631563 | KS-R1 vs Km-V | 0.450367842 | 0.933579 | |
| | KS-R3 | 3 | 125 | 1325 | 418 | | 10.6 | 3.344 | 0 | 1335.1 | 0.00793948 | 0.002504681 | 0 | -6.97673973 | -8.641157242 | | KS-R3 | -7.808948486 | KS-R3 vs Km-V | 0.646271981 | 0.217242 | |
| | Sc-V | 3 | 125 | 1037 | 873 | | 8.296 | 6.984 | 0 | 226.72 | 0.03659139 | 0.030804517 | 0 | -4.772351959 | -5.020714294 | | Sc-V | -4.896533127 | | | | |
| | Sc-R1 | 3 | 125 | 777 | 346 | | 6.216 | 2.768 | 0 | 354.86 | 0.017516767 | 0.007800259 | 0 | -5.835119648 | -7.002262209 | | Sc-R1 | -6.418690929 | | | | |
| | Sc-R3 | 3 | 125 | 1340 | 1100 | | 10.72 | 8.8 | 0 | 296.5866667 | 0.036144578 | 0.029670922 | 0 | -4.790076931 | -5.074806408 | | Sc-R3 | -4.932441669 | | | | |

p<0.05
