## Supplementary material for "Intergeneric chromosomal transfer in yeast results in improved phenotypes and widespread transcriptional responses": TableS4

Table S4. Differential expression of Sc and Km genes upon chromosomal transfers.

SC_vs_KS1

| w303_id | W303_annotation | chr | mean_SC_count | mean_KS_count | baseMean | log2FoldChange | lfcSE | pvalue | padj | DEG | treatment | S288C_id | gene | Ambiguous gene model |
|---|---|---|---|---|---|---|---|---|---|---|---|---|---|---|
| W303.asm02.HP0_G0000040 | YAL067C | chrI | 646.1884976 | 1401.967721 | 1098.537085 | 1.092526342 | 0.24597763 | 6.9573601542 | 1.0063324508 | KS>SC | YPD | YAL067C | SEO1 | n |
| W303.asm02.HP0_G0000050 | YAL065C | chrI | 44.99133383 | 544.13142 | 136.961162 | 3.427384063 | 0.40120631 | 1.0086396416 | 2.7233270324 | KS>SC | YPD | YAL065C | | n |
| W303.asm02.HP0_G0000060 | YAL064W-B | chrI | 49.31043001 | 845.1541944 | 195.7591587 | 3.991339684 | 0.30749452 | 4.9560342764 | 3.3453231365 | KS>SC | YPD | YAL064W-B | | n |
| W303.asm02.HP0_G0000070 | YAL064C-A | chrI | 132.2287944 | 1494.365292 | 417.3870198 | 3.435338158 | 0.24175882 | 5.5427114932 | 4.4895963094 | KS>SC | YPD | YAL064C-A | TDA8 | n |
| W303.asm02.HP0_G0000080 | YAL064W | chrI | 27.21449065 | 259.6531411 | 146.485206 | 3.179637884 | 0.27372467 | 3.5858756013 | 1.3831234462 | KS>SC | YPD | YAL064W | | n |
| W303.asm02.HP0_G0000090 | YAL063C-A | chrI | 54.56128855 | 433.9524795 | 236.6762418 | 2.942715391 | 0.22777164 | 5.0189246046 | 2.9038063783 | KS>SC | YPD | YAL063C-A | | n |
| W303.asm02.HP0_G0000100 | YAL063C | chrI | 1722.016798 | 15385.90726 | 5924.23424 | 3.13949327 | 0.13802085 | 1.5116150524 | 6.1220409623 | KS>SC | YPD | YAL063C | FLO9 | n |
| W303.asm02.HP0_G0000110 | YAL062W | chrI | 760.9216576 | 5658.519119 | 2406.447531 | 2.869006123 | 0.16025449 | 1.6857617799 | 4.5515568059 | KS>SC | YPD | YAL062W | GDH3 | n |
| W303.asm02.HP0_G0000120 | YAL061W | chrI | 2921.14749 | 1130.717137 | 1060.83464 | -1.352098954 | 0.1794212 | 3.3096813568 | 7.2455186460 | KS<SC | YPD | YAL061W | BDH2 | n |
| W303.asm02.HP0_G0000130 | YAL060W | chrI | 8789.91532 | 7211.120029 | 4241.749612 | -0.270745398 | 0.15901518 | 0.060546915 | 0.069074649 | ns | YPD | YAL060W | BDH1 | n |
| W303.asm02.HP0_G0000140 | YAL059W | chrI | 618.543906 | 633.267829 | 1876.370328 | 0.027043585 | 0.19888225 | 0.881132746 | 0.898099418 | ns | YPD | YAL059W | ECM1 | n |
| W303.asm02.HP0_G0000150 | YAL058W | chrI | 760.6859137 | 2840.208678 | 1307.663517 | 1.890914758 | 0.11241052 | 7.6632601926 | 1.0345401260 | KS>SC | YPD | YAL058W | CNE1 | n |
| W303.asm02.HP0_G0000160 | YAL056W | chrI | 2134.453445 | 2219.025581 | 2028.760874 | 0.043516809 | 0.10680497 | 0.634048543 | 0.658435025 | ns | YPD | YAL056W | GPB2 | n |
| W303.asm02.HP0_G0000170 | YAL055W | chrI | 729.7471292 | 346.67575 | 734.5986017 | -1.063628747 | 0.16469491 | 8.3634621993 | 1.4726966046 | KS<SC | YPD | YAL055W | PEX22 | n |
| W303.asm02.HP0_G0000180 | YAL054C | chrI | 783.3129532 | 743.2156152 | 732.8340172 | -0.059417448 | 0.11397969 | 0.542358707 | 0.585747403 | ns | YPD | YAL054C | ACS1 | n |
| W303.asm02.HP0_G0000190 | YAL053W | chrI | 5192.11692 | 7774.117919 | 5068.030492 | 0.578920826 | 0.11923816 | 1.1180521458 | 1.6770782188 | ns | YPD | YAL053W | FLC2 | n |
| W303.asm02.HP0_G0000200 | YAL051W | chrI | 2655.621531 | 799.7123022 | 1219.365232 | -1.700678965 | 0.21513187 | 1.4587571275 | 3.5805856766 | KS<SC | YPD | YAL051W | OAF1 | n |
| W303.asm02.HP0_G0000210 | YAL049C | chrI | 2056.938888 | 834.0657519 | 1631.072868 | -1.294572196 | 0.12701228 | 1.4978191904 | 4.8529341769 | KS<SC | YPD | YAL049C | AIM2 | n |
| W303.asm02.HP0_G0000220 | YAL048C | chrI | 967.3908679 | 1524.761507 | 810.8036844 | 0.647888004 | 0.17869343 | 0.000255493 | 0.000350762 | ns | YPD | YAL048C | GEM1 | n |
| W303.asm02.HP0_G0000230 | YAL047C | chrI | 675.2431705 | 852.3931997 | 828.619211 | 0.329563792 | 0.13380926 | 0.01003325 | 0.01212975 | ns | YPD | YAL047C | SPC72 | n |
| W303.asm02.HP0_G0000240 | YAL046C | chrI | 399.5082559 | 190.9124572 | 681.1992836 | -1.046713674 | 0.18592388 | 1.4359204144 | 2.3736643586 | KS<SC | YPD | YAL046C | BOL3 | n |
| W303.asm02.HP0_G0000250 | YAL044W-A | chrI | 807.2948419 | 358.7915403 | 1054.73425 | -1.160447985 | 0.15578786 | 7.0946455963 | 1.5122797192 | KS<SC | YPD | YAL044W-A | BOL1 | n |
| W303.asm02.HP0_G0000260 | YAL044C | chrI | 815.7671796 | 118.2897206 | 1121.410942 | -2.746893758 | 0.18584671 | 3.5753026887 | 3.2177724198 | KS<SC | YPD | YAL044C | GCV3 | n |
| W303.asm02.HP0_G0000270 | YAL043C | chrI | 1301.282415 | 1914.86694 | 1358.908101 | 0.547114361 | 0.19289069 | 0.003724213 | 0.004788274 | ns | YPD | YAL043C | PTA1 | n |
| W303.asm02.HP0_G0000280 | YAL042W | chrI | 5114.206376 | 4572.192158 | 4269.30249 | -0.139411335 | 0.15948518 | 0.316978116 | 0.351715444 | ns | YPD | YAL042W | ERV46 | n |
| W303.asm02.HP0_G0000290 | YAL041W | chrI | 1597.527084 | 1663.664667 | 1652.152003 | 0.044452895 | 0.09747204 | 0.588940403 | 0.627686482 | ns | YPD | YAL041W | CDC24 | n |
| W303.asm02.HP0_G0000300 | YAL040C | chrI | 613.1664848 | 2140.628797 | 1508.764537 | 1.765956053 | 0.2401826 | 1.0137051540 | 2.0290213773 | KS>SC | YPD | YAL040C | CLN3 | n |
| W303.asm02.HP0_G0000310 | YAL039C | chrI | 7708.644603 | 1206.058578 | 2707.865039 | -2.629944391 | 0.22846665 | 2.5269639347 | 9.3038217596 | KS<SC | YPD | YAL039C | CYC3 | n |
| W303.asm02.HP0_G0000320 | YAL038W | chrI | 63841.77937 | 20499.49123 | 31732.09897 | -1.627049229 | 0.13718089 | 1.0725039826 | 4.3436411297 | KS<SC | YPD | YAL038W | CDC19 | n |
| W303.asm02.HP0_G0000330 | YAL037W | chrI | 104.5850803 | 166.3580723 | 208.9327674 | 0.651422675 | 0.26728483 | 0.011399542 | 0.013578866 | ns | YPD | YAL037W | | n |
| W303.asm02.HP0_G0000340 | YAL036C | chrI | 288.4721144 | 1108.51633 | 2307.264795 | 1.918187184 | 0.17942958 | 5.1291553381 | 1.7310899266 | KS>SC | YPD | YAL036C | RBG1 | n |
| W303.asm02.HP0_G0000350 | YAL035W | chrI | 10284.86682 | 4782.44939 | 6843.984459 | -1.101092238 | 0.09000659 | 1.5811560141 | 7.1152020638 | KS<SC | YPD | YAL035W | FUN12 | n |
| W303.asm02.HP0_G0000360 | YAL034W-A | chrI | 457.3600612 | 68.0682289 | 488.2756898 | -2.721339908 | 0.15986279 | 1.0236870013 | 1.6583729421 | KS<SC | YPD | YAL034W-A | MTW1 | n |
| W303.asm02.HP0_G0000370 | YAL034C | chrI | 2073.699827 | 412.7681603 | 842.6948489 | -2.30570285 | 0.16241246 | 2.9632807106 | 2.1820521596 | KS<SC | YPD | YAL034C | FUN19 | n |
| W303.asm02.HP0_G0000380 | YAL033W | chrI | 465.2148579 | 327.9019126 | 763.0905409 | -0.499894515 | 0.14631192 | 0.000565041 | 0.000762806 | ns | YPD | YAL033W | POP5 | n |
| W303.asm02.HP0_G0000390 | YAL032C | chrI | 450.9918738 | 801.6752738 | 598.5062741 | 0.822920859 | 0.14807545 | 2.3727964898 | 3.7685591309 | ns | YPD | YAL032C | PRP45 | n |
| W303.asm02.HP0_G0000400 | YAL031C | chrI | 1164.214228 | 1281.018936 | 923.1956263 | 0.116150034 | 0.15295966 | 0.384057221 | 0.420386958 | ns | YPD | YAL031C | GIP4 | n |
| W303.asm02.HP0_G0000410 | YAL030W | chrI | 636.1953396 | 167.3881183 | 527.2552146 | -1.89498032 | 0.19588865 | 1.8168026012 | 5.6600388731 | KS<SC | YPD | YAL030W | SNC1 | n |
| W303.asm02.HP0_G0000420 | YAL029C | chrI | 3500.641257 | 3453.86607 | 3037.468219 | -0.015235722 | 0.12206952 | 0.887011771 | 0.898099418 | ns | YPD | YAL029C | MYO4 | n |
| W303.asm02.HP0_G0000430 | YAL028W | chrI | 683.7974808 | 536.323384 | 534.3758909 | -0.338171099 | 0.17000776 | 0.032739103 | 0.037883819 | ns | YPD | YAL028W | FRT2 | n |
| W303.asm02.HP0_G0000440 | YAL027W | chrI | 1049.021973 | 355.1318799 | 817.4231696 | -1.55503053 | 0.1233208 | 1.1075077450 | 5.2769486674 | KS<SC | YPD | YAL027W | SAW1 | n |
| W303.asm02.HP0_G0000450 | YAL026C | chrI | 2238.605094 | 972.0076749 | 1676.835418 | -1.194164184 | 0.13124021 | 6.7307159441 | 1.8799585912 | KS<SC | YPD | YAL026C | DRS2 | n |
| W303.asm02.HP0_G0000460 | YAL025C | chrI | 1189.728187 | 2537.356372 | 4676.537724 | 1.080602589 | 0.16608334 | 6.0298307319 | 1.0853695317 | KS>SC | YPD | YAL025C | MAK16 | n |
| W303.asm02.HP0_G0000470 | YAL024C | chrI | 505.3728723 | 1052.909689 | 849.8290653 | 1.044104707 | 0.1922953 | 4.4985296037 | 7.0073249596 | KS>SC | YPD | YAL024C | LTE1 | n |
| W303.asm02.HP0_G0000480 | YAL023C | chrI | 3982.465814 | 2733.876595 | 6301.491672 | -0.539444091 | 0.11989537 | 6.3817861622 | 9.3986305298 | ns | YPD | YAL023C | PMT2 | n |
| W303.asm02.HP0_G0000490 | YAL022C | chrI | 950.4082586 | 2074.064 | 1953.875514 | 1.115505668 | 0.14748269 | 3.0164660249 | 6.7870485561 | KS>SC | YPD | YAL022C | FUN26 | n |
| W303.asm02.HP0_G0000500 | YAL021C | chrI | 7167.433054 | 9698.539315 | 4492.707965 | 0.428681719 | 0.16573692 | 0.0074981 | 0.009202214 | ns | YPD | YAL021C | CCR4 | n |
| W303.asm02.HP0_G0000510 | YAL020C | chrI | 700.3761758 | 379.6819576 | 663.7932739 | -0.876840162 | 0.15557528 | 1.4795227924 | 2.3968269237 | ns | YPD | YAL020C | ATS1 | n |
| W303.asm02.HP0_G0000520 | YAL019W | chrI | 965.5002586 | 2696.791641 | 1540.105575 | 1.459889825 | 0.19785959 | 1.0270355120 | 2.0290213773 | KS>SC | YPD | YAL019W | FUN30 | n |
| W303.asm02.HP0_G0000530 | YAL018C | chrI | 254.8534682 | 1497.241721 | 1484.090881 | 2.535504585 | 0.14691449 | 2.2471348587 | 4.5504480888 | KS>SC | YPD | YAL018C | LDS1 | n |
| W303.asm02.HP0_G0000540 | YAL017W | chrI | 5425.193297 | 1275.054215 | 1926.275039 | -2.065753852 | 0.17371731 | 5.2000178354 | 2.2168497088 | KS<SC | YPD | YAL017W | PSK1 | n |
| W303.asm02.HP0_G0000550 | YAL016W | chrI | 5524.886188 | 1309.302962 | 3509.625116 | -2.063513004 | 0.12946276 | 1.3180296481 | 1.3345050187 | KS<SC | YPD | YAL016W | TPD3 | n |
| W303.asm02.HP0_G0000560 | YAL015C | chrI | 1958.664 | 1972.632653 | 1842.53876 | 0.007970827 | 0.15324531 | 0.953679845 | 0.953679845 | ns | YPD | YAL015C | NTG1 | n |
| W303.asm02.HP0_G0000570 | YAL014C | chrI | 1247.600781 | 996.5963213 | 1512.310717 | -0.320540365 | 0.1212316 | 0.006155785 | 0.007790916 | ns | YPD | YAL014C | SYN8 | n |
| W303.asm02.HP0_G0000580 | YAL013W | chrI | 2664.124965 | 2278.642293 | 1856.402874 | -0.205737324 | 0.15725896 | 0.137400006 | 0.154575007 | ns | YPD | YAL013W | DEP1 | n |
| W303.asm02.HP0_G0000590 | YAL012W | chrI | 10272.40909 | 8126.876455 | 7210.026308 | -0.328818026 | 0.15097931 | 0.020620285 | 0.024206422 | ns | YPD | YAL012W | CYS3 | n |
| W303.asm02.HP0_G0000600 | YAL011W | chrI | 269.3568168 | 670.4752903 | 531.0723606 | 1.300330414 | 0.16760873 | 6.0474490898 | 1.3995525036 | KS>SC | YPD | YAL011W | SWC3 | n |
| W303.asm02.HP0_G0000610 | YAL010C | chrI | 1251.712342 | 872.3769597 | 1116.447715 | -0.513557928 | 0.16403912 | 0.00149298 | 0.001950506 | ns | YPD | YAL010C | MDM10 | n |
| W303.asm02.HP0_G0000620 | YAL009W | chrI | 547.4568274 | 391.531871 | 735.512986 | -0.474412538 | 0.18069815 | 0.006794331 | 0.008466781 | ns | YPD | YAL009W | SPO7 | n |
| W303.asm02.HP0_G0000630 | YAL008W | chrI | 2236.241241 | 1054.212956 | 1555.162474 | -1.075127521 | 0.14925822 | 4.6147253984 | 8.8998275542 | KS<SC | YPD | YAL008W | FUN14 | n |
| W303.asm02.HP0_G0000640 | YAL007C | chrI | 1685.467062 | 914.1588415 | 3003.066846 | -0.877192053 | 0.13250602 | 3.0515463029 | 5.6176193304 | ns | YPD | YAL007C | ERP2 | n |
| W303.asm02.HP0_G0000650 | YAL005C/YLL024C | chrI | 28749.44004 | 1669.695398 | 17010.88815 | -4.069334207 | 0.17570253 | 2.7779265071 | 2.2501204708 | KS<SC | YPD | YAL005C | SSA1 | y |
| W303.asm02.HP0_G0000660 | YAL003W | chrI | 7558.466792 | 2122.292595 | 13415.7194 | -1.807200981 | 0.1949616 | 9.3667500609 | 2.7096669819 | KS<SC | YPD | YAL003W | EFB1 | n |
| W303.asm02.HP0_G0000670 | YAL002W | chrI | 1588.526395 | 2484.334454 | 2106.952717 | 0.642466409 | 0.09961774 | 1.0275750367 | 1.7709271909 | ns | YPD | YAL002W | VPS8 | n |
| W303.asm02.HP0_G0000680 | YAL001C | chrI | 2759.167151 | 2007.796184 | 1647.5821 | -0.455858785 | 0.11627874 | 8.2727793946 | 0.000117561 | ns | YPD | YAL001C | TFC3 | n |

| | | | | | | | | | | | | | | |
|---|---|---|---|---|---|---|---|---|---|---|---|---|---|---|
| W303.asm02.HP0_G0000700 | YAR002W | chrI | 2361.476763 | 1309.750714 | 2219.195198 | -0.844340879 | 0.14060006 | 1.642981425 | 2.772531154 | ns | YPD | YAR002W | NUP60 | n |
| W303.asm02.HP0_G0000710 | YAR002C-A | chrI | 2953.411269 | 1333.520644 | 4700.018699 | -1.137830043 | 0.14381329 | 1.930087755 | 4.598150241 | KS<SC | YPD | YAR002C-A | ERP1 | n |
| W303.asm02.HP0_G0000720 | YAR003W | chrI | 1839.607768 | 974.7788958 | 2590.541736 | -0.911834577 | 0.11383953 | 9.618340774 | 2.434642508 | ns | YPD | YAR003W | SWD1 | n |
| W303.asm02.HP0_G0000730 | YAR007C | chrI | 2786.706799 | 723.53921 | 2357.015013 | -1.922709567 | 0.17008884 | 5.651102363 | 1.990170832 | KS<SC | YPD | YAR007C | RFA1 | n |
| W303.asm02.HP0_G0000740 | YAR008W | chrI | 285.9493502 | 179.7235197 | 909.8659521 | -0.653557061 | 0.19631291 | 0.000751581 | 0.000998001 | ns | YPD | YAR008W | SEN34 | n |
| W303.asm02.HP0_G0000750 | YAR014C | chrI | 1559.4092 | 1168.306552 | 1596.564728 | -0.414621767 | 0.10853721 | 0.0001247 | 0.000174151 | ns | YPD | YAR014C | BUD14 | n |
| W303.asm02.HP0_G0000760 | YAR015W | chrI | 2352.033607 | 326.7750907 | 1256.944613 | -2.799790791 | 0.21661378 | 5.635817565 | 3.043341485 | KS<SC | YPD | YAR015W | ADE1 | n |
| W303.asm02.HP0_G0000770 | YAR018C | chrI | 1435.061314 | 423.8511447 | 1043.818519 | -1.750863396 | 0.10884759 | 1.638127576 | 1.895547624 | KS<SC | YPD | YAR018C | KIN3 | n |
| W303.asm02.HP0_G0000780 | YAR019C | chrI | 1162.229327 | 2288.020403 | 2087.379567 | 0.969557751 | 0.145371 | 2.105238640 | 3.965682089 | ns | YPD | YAR019C | CDC15 | n |
| W303.asm02.HP0_G0000790 | W303.asm02.HP0_G0 | chrI | 81.77419327 | 815.4504084 | 341.2808402 | 3.253095454 | 0.25057751 | 1.427884207 | 8.896816984 | KS>SC | YPD | YFL020C | PAU5 | y |
| W303.asm02.HP0_G0000800 | YAR023C | chrI | 226.0523151 | 1371.089668 | 527.8578251 | 2.565747992 | 0.19916143 | 1.309357898 | 6.628624362 | KS>SC | YPD | YAR023C | DFP1 | n |
| W303.asm02.HP0_G0000810 | YAR027W | chrI | 2374.734698 | 1105.08951 | 1918.000045 | -1.095025626 | 0.13679677 | 9.290203694 | 2.427440320 | KS<SC | YPD | YAR027W | UIP3 | n |
| W303.asm02.HP0_G0000820 | YAR028W | chrI | 877.8402581 | 929.9308401 | 679.5324071 | 0.067090916 | 0.15373427 | 0.622065496 | 0.654380587 | ns | YPD | YAR028W | KTD1 | n |
| W303.asm02.HP0_G0000830 | W303.asm02.HP0_G0 | chrI | 1420.281864 | 3633.618779 | 1972.532087 | 1.344948992 | 0.14022059 | 5.904105574 | 1.771231672 | KS>SC | YPD | YAR031W | PRM9 | y |
| W303.asm02.HP0_G0000840 | YAR035W | chrI | 377.1768531 | 1168.713179 | 546.8210326 | 1.602627316 | 0.2170248 | 8.954036216 | 1.859684444 | KS>SC | YPD | YAR035W | YAT1 | n |
| W303.asm02.HP0_G0000850 | YAR042W | chrI | 3377.459068 | 6318.009409 | 2622.801709 | 0.892007915 | 0.18071024 | 6.742948069 | 1.030526025 | ns | YPD | YAR042W | SWH1 | n |
| W303.asm02.HP0_G0000040 | YAL067C | chrI | 471.2711194 | 1512.29761 | 1098.537085 | 1.637878354 | 0.25056379 | 3.160190632 | 7.756831551 | KS>SC | TM | YAL067C | SEO1 | n |
| W303.asm02.HP0_G0000050 | YAL065C | chrI | 24.4795143 | 96.46866127 | 136.961162 | 1.822078262 | 0.45087546 | 2.047414739 | 3.685346530 | KS>SC | TM | YAL065C | | n |
| W303.asm02.HP0_G0000060 | YAL064W-B | chrI | 29.62750363 | 109.0209728 | 195.7591587 | 1.7847649 | 0.36204076 | 3.551336464 | 6.849006039 | KS>SC | TM | YAL064W-B | | n |
| W303.asm02.HP0_G0000070 | YAL064C-A | chrI | 80.61836269 | 369.0244362 | 417.3870198 | 2.133763696 | 0.26799757 | 5.348211867 | 1.666173697 | KS>SC | TM | YAL064C-A | TDA8 | n |
| W303.asm02.HP0_G0000080 | YAL064W | chrI | 15.88177805 | 208.7310256 | 146.485206 | 3.546066481 | 0.37200688 | 8.234056123 | 4.446390306 | KS>SC | TM | YAL064W | | n |
| W303.asm02.HP0_G0000090 | YAL063C-A | chrI | 70.49295036 | 378.1843193 | 236.6762418 | 2.360416734 | 0.25043439 | 1.036559036 | 4.664515663 | KS>SC | TM | YAL063C-A | | n |
| W303.asm02.HP0_G0000100 | YAL063C | chrI | 1479.009027 | 7518.692446 | 5924.23424 | 2.327002015 | 0.14035046 | 2.300569238 | 3.105768471 | KS>SC | TM | YAL063C | FLO9 | n |
| W303.asm02.HP0_G0000110 | YAL062W | chrI | 504.4178307 | 2860.539211 | 2406.447531 | 2.474751161 | 0.16669346 | 1.475483943 | 1.327935549 | KS>SC | TM | YAL062W | GDH3 | n |
| W303.asm02.HP0_G0000120 | YAL061W | chrI | 470.0814333 | 707.6298489 | 1060.83464 | 0.55493825 | 0.19595944 | 0.001826681 | 0.00279172 | ns | TM | YAL061W | BDH2 | n |
| W303.asm02.HP0_G0000130 | YAL060W | chrI | 2347.303983 | 1875.826057 | 4241.749612 | -0.292181102 | 0.15306052 | 0.036675418 | 0.048700145 | ns | TM | YAL060W | BDH1 | n |
| W303.asm02.HP0_G0000140 | YAL059W | chrI | 3369.657466 | 1483.453915 | 1876.370328 | -1.157922338 | 0.21667532 | 5.266522064 | 1.122600755 | KS<SC | TM | YAL059W | ECM1 | n |
| W303.asm02.HP0_G0000150 | YAL058W | chrI | 675.2771567 | 1219.780765 | 1307.663517 | 0.847112942 | 0.12100019 | 1.339681137 | 3.391067880 | ns | TM | YAL058W | CNE1 | n |
| W303.asm02.HP0_G0000160 | YAL056W | chrI | 2113.023626 | 1793.685046 | 2028.760874 | -0.216782759 | 0.11891748 | 0.052450825 | 0.068524465 | ns | TM | YAL056W | GPB2 | n |
| W303.asm02.HP0_G0000170 | YAL055W | chrI | 1078.360716 | 625.4353749 | 734.5986017 | -0.766269696 | 0.17179472 | 3.351828494 | 6.313909489 | ns | TM | YAL055W | PEX22 | n |
| W303.asm02.HP0_G0000180 | YAL054C | chrI | 940.0322422 | 561.4531559 | 732.8340172 | -0.733342638 | 0.13247513 | 1.301498123 | 3.012038513 | ns | TM | YAL054C | ACS1 | n |
| W303.asm02.HP0_G0000190 | YAL053W | chrI | 3144.656613 | 7053.769607 | 5068.030492 | 1.157368524 | 0.12093042 | 6.999770991 | 3.335185001 | KS>SC | TM | YAL053W | FLC2 | n |
| W303.asm02.HP0_G0000200 | YAL051W | chrI | 1106.372349 | 967.4858789 | 1219.365232 | -0.158053895 | 0.2027308 | 0.378599005 | 0.431922809 | ns | TM | YAL051W | OAF1 | n |
| W303.asm02.HP0_G0000210 | YAL049C | chrI | 2889.844655 | 495.1242036 | 1631.072868 | -2.526682589 | 0.132669 | 1.308022615 | 3.531661061 | KS<SC | TM | YAL049C | AIM2 | n |
| W303.asm02.HP0_G0000220 | YAL048C | chrI | 458.8653408 | 755.6081909 | 810.8036844 | 0.692383065 | 0.19350872 | 0.000129894 | 0.00022386 | ns | TM | YAL048C | GEM1 | n |
| W303.asm02.HP0_G0000230 | YAL047C | chrI | 683.3768833 | 852.6100917 | 828.619211 | 0.292235829 | 0.13594398 | 0.020255099 | 0.027344383 | ns | TM | YAL047C | SPC72 | n |
| W303.asm02.HP0_G0000240 | YAL046C | chrI | 1353.424551 | 501.5429555 | 681.1992836 | -1.412146823 | 0.18624447 | 1.999437232 | 5.398480527 | KS<SC | TM | YAL046C | BOL3 | n |
| W303.asm02.HP0_G0000250 | YAL044W-A | chrI | 1868.171286 | 668.3769917 | 1054.73425 | -1.466692244 | 0.15767088 | 7.859497131 | 3.183096338 | KS<SC | TM | YAL044W-A | BOL1 | n |
| W303.asm02.HP0_G0000260 | YAL044C | chrI | 2826.116256 | 268.6903747 | 1121.410942 | -3.350789676 | 0.187816 | 2.147424249 | 4.348534104 | KS<SC | TM | YAL044C | GCV3 | n |
| W303.asm02.HP0_G0000270 | YAL043C | chrI | 1229.382228 | 1462.954467 | 1358.908101 | 0.212863705 | 0.18487367 | 0.197706969 | 0.23209079 | ns | TM | YAL043C | PTA1 | n |
| W303.asm02.HP0_G0000280 | YAL042W | chrI | 5047.448314 | 2663.309723 | 4269.30249 | -0.908095765 | 0.16381399 | 1.531931621 | 3.446846148 | ns | TM | YAL042W | ERV46 | n |
| W303.asm02.HP0_G0000290 | YAL041W | chrI | 1607.978592 | 2174.551414 | 1652.152003 | 0.41801296 | 0.11304353 | 9.440210387 | 0.00016623 | ns | TM | YAL041W | CDC24 | n |
| W303.asm02.HP0_G0000300 | YAL040C | chrI | 1667.186768 | 1828.776255 | 1508.764537 | 0.105130129 | 0.21804356 | 0.583560217 | 0.621952336 | ns | TM | YAL040C | CLN3 | n |
| W303.asm02.HP0_G0000310 | YAL039C | chrI | 2738.352442 | 761.2219871 | 2707.865039 | -1.80592521 | 0.2304107 | 2.004640861 | 5.799139634 | KS<SC | TM | YAL039C | CYC3 | n |
| W303.asm02.HP0_G0000320 | YAL038W | chrI | 48023.11875 | 12140.03979 | 31732.09897 | -1.967904075 | 0.13744184 | 6.065851901 | 4.913340040 | KS<SC | TM | YAL038W | CDC19 | n |
| W303.asm02.HP0_G0000330 | YAL037W | chrI | 196.5662541 | 223.8051068 | 208.9327674 | 0.145064353 | 0.24552369 | 0.494327142 | 0.548499979 | ns | TM | YAL037W | | n |
| W303.asm02.HP0_G0000340 | YAL036C | chrI | 2936.696181 | 2980.355873 | 2307.264795 | 0.017848281 | 0.16244652 | 0.90444314 | 0.915748679 | ns | TM | YAL036C | RBG1 | n |
| W303.asm02.HP0_G0000350 | YAL035W | chrI | 5694.066937 | 6658.458015 | 6843.984459 | 0.213591774 | 0.09019394 | 0.013725513 | 0.018843501 | ns | TM | YAL035W | FUN12 | n |
| W303.asm02.HP0_G0000360 | YAL034W-A | chrI | 836.4371054 | 372.8360102 | 488.2756898 | -1.15102425 | 0.15725585 | 1.584993685 | 4.141435114 | KS<SC | TM | YAL034W-A | MTW1 | n |
| W303.asm02.HP0_G0000370 | YAL034C | chrI | 523.2669376 | 803.6163821 | 842.6948489 | 0.59073153 | 0.17603328 | 0.000294775 | 0.000487281 | ns | TM | YAL034C | FUN19 | n |
| W303.asm02.HP0_G0000380 | YAL033W | chrI | 1308.970686 | 539.3758281 | 763.0905409 | -1.266656262 | 0.14813567 | 7.767974590 | 2.735678007 | KS<SC | TM | YAL033W | POP5 | n |
| W303.asm02.HP0_G0000390 | YAL032C | chrI | 497.6145786 | 693.7684839 | 598.5062741 | 0.448908794 | 0.1598827 | 0.002201268 | 0.003301902 | ns | TM | YAL032C | PRP45 | n |
| W303.asm02.HP0_G0000400 | YAL031C | chrI | 818.8463311 | 832.3998659 | 923.1956263 | 0.020515338 | 0.15132057 | 0.883195787 | 0.905555174 | ns | TM | YAL031C | GIP4 | n |
| W303.asm02.HP0_G0000410 | YAL030W | chrI | 932.4174431 | 271.2999825 | 527.2552146 | -1.749324817 | 0.20077165 | 1.356572550 | 5.232494122 | KS<SC | TM | YAL030W | SNC1 | n |
| W303.asm02.HP0_G0000420 | YAL029C | chrI | 2602.738175 | 2757.555821 | 3037.468219 | 0.073681382 | 0.13215406 | 0.551123298 | 0.595213162 | ns | TM | YAL029C | MYO4 | n |
| W303.asm02.HP0_G0000430 | YAL028W | chrI | 175.9798916 | 713.4363203 | 534.3758909 | 1.990971917 | 0.18381183 | 8.764388374 | 5.915962152 | KS>SC | TM | YAL028W | FRT2 | n |
| W303.asm02.HP0_G0000440 | YAL027W | chrI | 1458.005481 | 278.9399634 | 817.4231696 | -2.365776829 | 0.13292368 | 1.636122192 | 2.650517952 | KS<SC | TM | YAL027W | SAW1 | n |
| W303.asm02.HP0_G0000450 | YAL026C | chrI | 2012.913305 | 1534.708699 | 1676.835418 | -0.365289823 | 0.13531023 | 0.003621787 | 0.005333905 | ns | TM | YAL026C | DRS2 | n |
| W303.asm02.HP0_G0000460 | YAL025C | chrI | 5877.924154 | 5010.6326 | 4676.537724 | -0.20053805 | 0.15932951 | 0.167207615 | 0.199173776 | ns | TM | YAL025C | MAK16 | n |
| W303.asm02.HP0_G0000470 | YAL024C | chrI | 627.1226234 | 1178.648287 | 849.8290653 | 0.887858379 | 0.1999869 | 4.121735093 | 7.587739604 | ns | TM | YAL024C | LTE1 | n |
| W303.asm02.HP0_G0000480 | YAL023C | chrI | 9487.064908 | 9552.360226 | 6301.491672 | 0.008879185 | 0.1141859 | 0.934701003 | 0.934701003 | ns | TM | YAL023C | PMT2 | n |
| W303.asm02.HP0_G0000490 | YAL022C | chrI | 1551.50767 | 2288.778138 | 1953.875514 | 0.536942164 | 0.15405631 | 0.00018551 | 0.000313049 | ns | TM | YAL022C | FUN26 | n |
| W303.asm02.HP0_G0000500 | YAL021C | chrI | 2629.157552 | 2870.133402 | 4492.707965 | 0.108604794 | 0.15416484 | 0.443151116 | 0.498545006 | ns | TM | YAL021C | CCR4 | n |
| W303.asm02.HP0_G0000510 | YAL020C | chrI | 841.9275344 | 417.6500311 | 663.7932739 | -0.997143707 | 0.16283713 | 5.285857536 | 1.259277824 | ns | TM | YAL020C | ATS1 | n |
| W303.asm02.HP0_G0000520 | YAL019W | chrI | 1652.110828 | 1587.872007 | 1540.105575 | -0.046716979 | 0.18196187 | 0.775952975 | 0.80579732 | ns | TM | YAL019W | FUN30 | n |
| W303.asm02.HP0_G0000530 | YAL018C | chrI | 113.4561361 | 3653.640953 | 1484.090881 | 4.960201424 | 0.1761452 | 1.778298439 | 1.440421736 | KS>SC | TM | YAL018C | LDS1 | n |
| W303.asm02.HP0_G0000540 | YAL017W | chrI | 1270.503821 | 1455.479126 | 1926.275039 | 0.167216362 | 0.16766632 | 0.271469564 | 0.314129067 | ns | TM | YAL017W | PSK1 | n |
| W303.asm02.HP0_G0000550 | YAL016W | chrI | 5468.99044 | 1301.820018 | 3509.625116 | -2.05501276 | 0.13153264 | 1.634918622 | 1.891834406 | KS<SC | TM | YAL016W | TPD3 | n |

| | | | | | | | | | | | | | | |
|---|---|---|---|---|---|---|---|---|---|---|---|---|---|---|
| W303.asm02.HP0_G0000560 | YAL015C | chrI | 1681.73886 | 1122.473832 | 1842.53876 | -0.552565018 | 0.17958608 | 0.000807017 | 0.001281732 | ns | TM | YAL015C | NTG1 | n |
| W303.asm02.HP0_G0000570 | YAL014C | chrI | 1805.385767 | 1547.234235 | 1512.310717 | -0.202957488 | 0.11800166 | 0.067606895 | 0.085564977 | ns | TM | YAL014C | SYN8 | n |
| W303.asm02.HP0_G0000580 | YAL013W | chrI | 1043.812525 | 1821.066261 | 1856.402874 | 0.788540209 | 0.15840724 | 2.7932113402 | 5.656252964 | ns | TM | YAL013W | DEP1 | n |
| W303.asm02.HP0_G0000590 | YAL012W | chrI | 6024.242868 | 7050.421049 | 7210.026308 | 0.202029591 | 0.1415779 | 0.122419384 | 0.147999553 | ns | TM | YAL012W | CYS3 | n |
| W303.asm02.HP0_G0000600 | YAL011W | chrI | 501.4114315 | 626.104021 | 531.0723606 | 0.283627012 | 0.17013841 | 0.064555812 | 0.08300033 | ns | TM | YAL011W | SWC3 | n |
| W303.asm02.HP0_G0000610 | YAL010C | chrI | 862.822549 | 1283.833731 | 1116.447715 | 0.54446084 | 0.17309293 | 0.000638965 | 0.001035124 | ns | TM | YAL010C | MDM10 | n |
| W303.asm02.HP0_G0000620 | YAL009W | chrI | 950.3327372 | 497.211599 | 735.512986 | -0.916901971 | 0.18379985 | 3.0170205407 | 5.960455214 | ns | TM | YAL009W | SPO7 | n |
| W303.asm02.HP0_G0000630 | YAL008W | chrI | 1889.315584 | 1062.440034 | 1555.162474 | -0.816628316 | 0.1545759 | 5.8316791945 | 1.211194909 | ns | TM | YAL008W | FUN14 | n |
| W303.asm02.HP0_G0000640 | YAL007C | chrI | 6483.561286 | 2664.527796 | 3003.066846 | -1.272177777 | 0.13237375 | 4.6003438751 | 2.328924086 | KS<SC | TM | YAL007C | ERP2 | n |
| W303.asm02.HP0_G0000650 | YAL005C/YLL024C | chrI | 25422.42971 | 8080.824419 | 17010.88815 | -1.631442899 | 0.1750248 | 5.8120954458 | 2.477788058 | KS<SC | TM | YAL005C | SSA1 | y |
| W303.asm02.HP0_G0000660 | YAL003W | chrI | 27897.98419 | 5211.197733 | 13415.7194 | -2.383738931 | 0.19514637 | 5.9123937241 | 4.353671742 | KS<SC | TM | YAL003W | EFB1 | n |
| W303.asm02.HP0_G0000670 | YAL002W | chrI | 981.7556535 | 2970.625858 | 2106.952717 | 1.588650843 | 0.10534878 | 1.1226659624 | 1.136699286 | KS>SC | TM | YAL002W | VPS8 | n |
| W303.asm02.HP0_G0000680 | YAL001C | chrI | 1190.17365 | 1244.174368 | 1647.5821 | 0.057495456 | 0.11624077 | 0.60149353 | 0.632739948 | ns | TM | YAL001C | TFC3 | n |
| W303.asm02.HP0_G0000700 | YAR002W | chrI | 1951.271281 | 2076.171494 | 2219.195198 | 0.0791319 | 0.1348576 | 0.529519488 | 0.57960917 | ns | TM | YAR002W | NUP60 | n |
| W303.asm02.HP0_G0000710 | YAR002C-A | chrI | 9121.505591 | 3982.258502 | 4700.018699 | -1.184298849 | 0.14389849 | 1.2086156246 | 4.079077733 | KS<SC | TM | YAR002C-A | ERP1 | n |
| W303.asm02.HP0_G0000720 | YAR003W | chrI | 4489.510037 | 2257.184499 | 2590.541736 | -0.985271667 | 0.11466003 | 5.3317756837 | 1.963062865 | ns | TM | YAR003W | SWD1 | n |
| W303.asm02.HP0_G0000730 | YAR007C | chrI | 3114.527557 | 2226.913713 | 2357.015013 | -0.448238669 | 0.17539222 | 0.004823192 | 0.006854009 | ns | TM | YAR007C | RFA1 | n |
| W303.asm02.HP0_G0000740 | YAR008W | chrI | 1856.898286 | 610.5882376 | 909.8659521 | -1.57923338 | 0.19330677 | 1.6334529639 | 5.292387603 | KS<SC | TM | YAR008W | SEN34 | n |
| W303.asm02.HP0_G0000750 | YAR014C | chrI | 1873.583459 | 1447.667876 | 1596.564728 | -0.352508122 | 0.11257005 | 0.000909454 | 0.001416649 | ns | TM | YAR014C | BUD14 | n |
| W303.asm02.HP0_G0000760 | YAR015W | chrI | 1793.19412 | 367.7702655 | 1256.944613 | -2.240454912 | 0.22037079 | 7.6956221005 | 4.452467072 | KS<SC | TM | YAR015W | ADE1 | n |
| W303.asm02.HP0_G0000770 | YAR018C | chrI | 934.2356705 | 811.3674994 | 1043.818519 | -0.186070558 | 0.11230478 | 0.079835609 | 0.099487451 | ns | TM | YAR018C | KIN3 | n |
| W303.asm02.HP0_G0000780 | YAR019C | chrI | 454.8669457 | 4014.552236 | 2087.379567 | 3.112834029 | 0.15390062 | 4.7974912291 | 1.942983947 | KS>SC | TM | YAR019C | CDC15 | n |
| W303.asm02.HP0_G0000790 | W303.asm02.HP0_G0 | chrI | 55.46532045 | 463.3365528 | 341.2808402 | 2.97670219 | 0.28456186 | 1.4451636587 | 9.004481258 | KS>SC | TM | YFL020C | PAU5 | y |
| W303.asm02.HP0_G0000800 | YAR023C | chrI | 265.3174896 | 403.8677989 | 527.8578251 | 0.559732165 | 0.21852198 | 0.004244749 | 0.006139727 | ns | TM | YAR023C | DFP1 | n |
| W303.asm02.HP0_G0000810 | YAR027W | chrI | 2757.988495 | 1303.564748 | 1918.000045 | -1.070919294 | 0.13932103 | 9.6107620825 | 2.684385271 | KS<SC | TM | YAR027W | UIP3 | n |
| W303.asm02.HP0_G0000820 | YAR028W | chrI | 793.0058335 | 294.8863075 | 679.5324071 | -1.410389561 | 0.1755639 | 5.5977279593 | 1.679318387 | KS<SC | TM | YAR028W | KTD1 | n |
| W303.asm02.HP0_G0000830 | W303.asm02.HP0_G0 | chrI | 1115.78397 | 1926.960327 | 1972.532087 | 0.775727643 | 0.14645363 | 5.1546873579 | 1.122600755 | ns | TM | YAR031W | PRM9 | y |
| W303.asm02.HP0_G0000840 | YAR035W | chrI | 387.4309188 | 501.7885402 | 546.8210326 | 0.317301462 | 0.22114288 | 0.099647807 | 0.122295035 | ns | TM | YAR035W | YAT1 | n |
| W303.asm02.HP0_G0000850 | YAR042W | chrI | 1351.138875 | 1927.301468 | 2622.801709 | 0.472817037 | 0.18969582 | 0.005637073 | 0.007872464 | ns | TM | YAR042W | SWH1 | n |
| W303.asm02.HP0_G0000040 | YAL067C | chrI | 466.8384348 | 2092.659125 | 1098.537085 | 2.1204143 | 0.25318775 | 1.5443792609 | 5.003788805 | KS>SC | NaCl | YAL067C | SEO1 | n |
| W303.asm02.HP0_G0000050 | YAL065C | chrI | 26.21707467 | 85.47896771 | 136.961162 | 1.55186071 | 0.46910119 | 0.000290743 | 0.000436114 | KS>SC | NaCl | YAL065C | | n |
| W303.asm02.HP0_G0000060 | YAL064W-B | chrI | 26.30230414 | 115.1395469 | 195.7591587 | 2.028635602 | 0.3845196 | 3.3668759121 | 6.060376641 | KS>SC | NaCl | YAL064W-B | | n |
| W303.asm02.HP0_G0000070 | YAL064C-A | chrI | 65.53189521 | 362.5533382 | 417.3870198 | 2.407035173 | 0.28063283 | 2.7395587186 | 9.246010675 | KS>SC | NaCl | YAL064C-A | TDA8 | n |
| W303.asm02.HP0_G0000080 | YAL064W | chrI | 25.65079793 | 341.7700014 | 146.485206 | 3.616364184 | 0.33765681 | 1.3260356207 | 6.318169722 | KS>SC | NaCl | YAL064W | | n |
| W303.asm02.HP0_G0000090 | YAL063C-A | chrI | 64.46934561 | 418.3970674 | 236.6762418 | 2.643445226 | 0.25923395 | 5.6527281408 | 2.543727663 | KS>SC | NaCl | YAL063C-A | | n |
| W303.asm02.HP0_G0000100 | YAL063C | chrI | 1280.920688 | 8158.859216 | 5924.23424 | 2.654676377 | 0.14134103 | 2.9998865202 | 6.074770203 | KS>SC | NaCl | YAL063C | FLO9 | n |
| W303.asm02.HP0_G0000110 | YAL062W | chrI | 469.1617559 | 4185.125609 | 2406.447531 | 3.13046723 | 0.16811722 | 4.6354415394 | 6.257846078 | KS>SC | NaCl | YAL062W | GDH3 | n |
| W303.asm02.HP0_G0000120 | YAL061W | chrI | 589.3545883 | 546.0773412 | 1060.83464 | -0.106646955 | 0.18656047 | 0.559731256 | 0.573901667 | ns | NaCl | YAL061W | BDH2 | n |
| W303.asm02.HP0_G0000130 | YAL060W | chrI | 2959.796592 | 2266.535689 | 4241.749612 | -0.375103511 | 0.15299599 | 0.012860687 | 0.016535169 | ns | NaCl | YAL060W | BDH1 | n |
| W303.asm02.HP0_G0000140 | YAL059W | chrI | 3115.185563 | 2038.113288 | 1876.370328 | -0.586002596 | 0.21408938 | 0.004866842 | 0.006570237 | ns | NaCl | YAL059W | ECM1 | n |
| W303.asm02.HP0_G0000150 | YAL058W | chrI | 748.4850525 | 1601.543536 | 1307.663517 | 1.085767356 | 0.1203615 | 7.8692999760 | 2.771362165 | KS>SC | NaCl | YAL058W | CNE1 | n |
| W303.asm02.HP0_G0000160 | YAL056W | chrI | 2182.093285 | 1730.284259 | 2028.760874 | -0.328706135 | 0.12130513 | 0.006235477 | 0.008279895 | ns | NaCl | YAL056W | GPB2 | n |
| W303.asm02.HP0_G0000170 | YAL055W | chrI | 1280.734322 | 346.6383177 | 734.5986017 | -1.86682606 | 0.17228342 | 6.4867415965 | 3.283912933 | KS<SC | NaCl | YAL055W | PEX22 | n |
| W303.asm02.HP0_G0000180 | YAL054C | chrI | 874.9145114 | 494.0756253 | 732.8340172 | -0.813008094 | 0.1330115 | 6.0395933505 | 1.322181247 | ns | NaCl | YAL054C | ACS1 | n |
| W303.asm02.HP0_G0000190 | YAL053W | chrI | 2937.525221 | 4305.99667 | 5068.030492 | 0.543484968 | 0.1212494 | 5.9857849586 | 1.010101211 | ns | NaCl | YAL053W | FLC2 | n |
| W303.asm02.HP0_G0000200 | YAL051W | chrI | 818.1301831 | 968.8691481 | 1219.365232 | 0.231722451 | 0.21637955 | 0.271358438 | 0.29702748 | ns | NaCl | YAL051W | OAF1 | n |
| W303.asm02.HP0_G0000210 | YAL049C | chrI | 2965.707092 | 544.7566189 | 1631.072868 | -2.432986036 | 0.1330323 | 3.0966589780 | 3.135367215 | KS<SC | NaCl | YAL049C | AIM2 | n |
| W303.asm02.HP0_G0000220 | YAL048C | chrI | 419.5442362 | 738.6519633 | 810.8036844 | 0.792069705 | 0.18885856 | 1.7541196071 | 2.841673763 | ns | NaCl | YAL048C | GEM1 | n |
| W303.asm02.HP0_G0000230 | YAL047C | chrI | 876.9645862 | 1031.127335 | 828.619211 | 0.229560988 | 0.13472715 | 0.084897515 | 0.096854912 | ns | NaCl | YAL047C | SPC72 | n |
| W303.asm02.HP0_G0000240 | YAL046C | chrI | 1129.620785 | 512.1866967 | 681.1992836 | -1.114976734 | 0.18871546 | 1.4643178491 | 3.121309099 | KS<SC | NaCl | YAL046C | BOL3 | n |
| W303.asm02.HP0_G0000250 | YAL044W-A | chrI | 1863.647843 | 762.1229979 | 1054.73425 | -1.272604826 | 0.15891377 | 4.0014399492 | 1.200431984 | KS<SC | NaCl | YAL044W-A | BOL1 | n |
| W303.asm02.HP0_G0000260 | YAL044C | chrI | 2474.708675 | 224.8934463 | 1121.410942 | -3.425925961 | 0.19048133 | 4.2267853578 | 3.804106822 | KS<SC | NaCl | YAL044C | GCV3 | n |
| W303.asm02.HP0_G0000270 | YAL043C | chrI | 874.2467684 | 1370.715787 | 1358.908101 | 0.625352217 | 0.19424878 | 0.000975637 | 0.00141119 | ns | NaCl | YAL043C | PTA1 | n |
| W303.asm02.HP0_G0000280 | YAL042W | chrI | 4676.925075 | 3541.733293 | 4269.30249 | -0.390050254 | 0.16104812 | 0.013851304 | 0.017260856 | ns | NaCl | YAL042W | ERV46 | n |
| W303.asm02.HP0_G0000290 | YAL041W | chrI | 1343.681447 | 1525.508815 | 1652.152003 | 0.180287756 | 0.11275154 | 0.107033934 | 0.120413176 | ns | NaCl | YAL041W | CDC24 | n |
| W303.asm02.HP0_G0000300 | YAL040C | chrI | 777.3567635 | 2025.472152 | 1508.764537 | 1.337997627 | 0.24605495 | 1.8212454443 | 3.352747295 | KS>SC | NaCl | YAL040C | CLN3 | n |
| W303.asm02.HP0_G0000310 | YAL039C | chrI | 2749.989394 | 1082.923232 | 2707.865039 | -1.305897588 | 0.23276818 | 6.9953308645 | 1.382004390 | KS<SC | NaCl | YAL039C | CYC3 | n |
| W303.asm02.HP0_G0000320 | YAL038W | chrI | 37929.02882 | 7959.135846 | 31732.09897 | -2.238789563 | 0.13785398 | 8.0365336428 | 6.509592250 | KS<SC | NaCl | YAL038W | CDC19 | n |
| W303.asm02.HP0_G0000330 | YAL037W | chrI | 282.0852838 | 280.1968069 | 208.9327674 | -0.008638975 | 0.26096761 | 0.972597801 | 0.972597801 | ns | NaCl | YAL037W | | n |
| W303.asm02.HP0_G0000340 | YAL036C | chrI | 2762.785737 | 3766.762532 | 2307.264795 | 0.432617785 | 0.17514746 | 0.01182975 | 0.015454996 | ns | NaCl | YAL036C | RBG1 | n |
| W303.asm02.HP0_G0000350 | YAL035W | chrI | 5561.955622 | 8082.109969 | 6843.984459 | 0.534183749 | 0.09133912 | 4.0426938348 | 8.186455015 | ns | NaCl | YAL035W | FUN12 | n |
| W303.asm02.HP0_G0000360 | YAL034W-A | chrI | 884.2200895 | 310.7326439 | 488.2756898 | -1.491001062 | 0.16064383 | 4.8204190337 | 1.859304484 | KS<SC | NaCl | YAL034W-A | MTW1 | n |
| W303.asm02.HP0_G0000370 | YAL034C | chrI | 532.42593 | 710.3918565 | 842.6948489 | 0.403614456 | 0.17002611 | 0.015673733 | 0.019235945 | ns | NaCl | YAL034C | FUN19 | n |
| W303.asm02.HP0_G0000380 | YAL033W | chrI | 1167.666256 | 769.4137055 | 763.0905409 | -0.589335895 | 0.14748836 | 5.0413376565 | 7.852852888 | ns | NaCl | YAL033W | POP5 | n |
| W303.asm02.HP0_G0000390 | YAL032C | chrI | 428.2230326 | 718.7644016 | 598.5062741 | 0.729827563 | 0.15836364 | 2.7666777575 | 4.768104220 | ns | NaCl | YAL032C | PRP45 | n |
| W303.asm02.HP0_G0000400 | YAL031C | chrI | 688.3096169 | 754.3847794 | 923.1956263 | 0.128539226 | 0.16363029 | 0.424851987 | 0.452802775 | ns | NaCl | YAL031C | GIP4 | n |
| W303.asm02.HP0_G0000410 | YAL030W | chrI | 957.8796084 | 198.3507958 | 527.2552146 | -2.242819854 | 0.20554394 | 2.9915661463 | 1.615445719 | KS<SC | NaCl | YAL030W | SNC1 | n |
| W303.asm02.HP0_G0000420 | YAL029C | chrI | 2685.838447 | 3224.169542 | 3037.468219 | 0.257670669 | 0.13904973 | 0.060767335 | 0.071335568 | ns | NaCl | YAL029C | MYO4 | n |

| | | | | | | | | | | | | | | |
|---|---|---|---|---|---|---|---|---|---|---|---|---|---|---|
| W303.asm02.HP0_G0000430 | YAL028W | chrI | 133.3763888 | 963.3418803 | 534.3758909 | 2.820746353 | 0.19289758 | 5.2504731557 | 3.8662575056 | KS>SC | NaCl | YAL028W | FRT2 | n |
| W303.asm02.HP0_G0000440 | YAL027W | chrI | 1497.145666 | 266.2940531 | 817.4231696 | -2.477137987 | 0.13512997 | 1.3962552064 | 1.6156667388 | KS<SC | NaCl | YAL027W | SAW1 | n |
| W303.asm02.HP0_G0000450 | YAL026C | chrI | 1679.849834 | 1622.927901 | 1676.835418 | -0.048486676 | 0.1337557 | 0.714048708 | 0.722974317 | ns | NaCl | YAL026C | DRS2 | n |
| W303.asm02.HP0_G0000460 | YAL025C | chrI | 6979.693275 | 6463.891759 | 4676.537724 | -0.107450101 | 0.16400487 | 0.505665226 | 0.525113888 | ns | NaCl | YAL025C | MAK16 | n |
| W303.asm02.HP0_G0000470 | YAL024C | chrI | 483.3380353 | 1251.582885 | 849.8290653 | 1.343240822 | 0.20061704 | 7.0430856844 | 1.6779115895 | KS>SC | NaCl | YAL024C | LTE1 | n |
| W303.asm02.HP0_G0000480 | YAL023C | chrI | 5185.123458 | 6868.059032 | 6301.491672 | 0.39907051 | 0.12022928 | 0.000807013 | 0.00118851 | ns | NaCl | YAL023C | PMT2 | n |
| W303.asm02.HP0_G0000490 | YAL022C | chrI | 1528.869739 | 3329.62528 | 1953.875514 | 1.106040568 | 0.1501985 | 7.4833220384 | 2.0901692590 | KS>SC | NaCl | YAL022C | FUN26 | n |
| W303.asm02.HP0_G0000500 | YAL021C | chrI | 1775.320438 | 2815.664031 | 4492.707965 | 0.648149026 | 0.16490787 | 6.2957932482 | 9.6218727001 | ns | NaCl | YAL021C | CCR4 | n |
| W303.asm02.HP0_G0000510 | YAL020C | chrI | 764.264694 | 878.8592505 | 663.7932739 | 0.195282089 | 0.15908422 | 0.212693605 | 0.236002493 | ns | NaCl | YAL020C | ATS1 | n |
| W303.asm02.HP0_G0000520 | YAL019W | chrI | 1026.158254 | 1312.200463 | 1540.105575 | 0.339862505 | 0.19906932 | 0.080515947 | 0.093168452 | ns | NaCl | YAL019W | FUN30 | n |
| W303.asm02.HP0_G0000530 | YAL018C | chrI | 87.27548741 | 3298.077524 | 1484.090881 | 5.182831994 | 0.19031324 | 4.9710913495 | 4.0265839931 | KS>SC | NaCl | YAL018C | LDS1 | n |
| W303.asm02.HP0_G0000540 | YAL017W | chrI | 878.8664014 | 1252.553372 | 1926.275039 | 0.495187644 | 0.17797845 | 0.004541064 | 0.006234342 | ns | NaCl | YAL017W | PSK1 | n |
| W303.asm02.HP0_G0000550 | YAL016W | chrI | 6329.990985 | 1122.760106 | 3509.625116 | -2.480716498 | 0.13227057 | 5.3165912182 | 8.6128777735 | KS<SC | NaCl | YAL016W | TPD3 | n |
| W303.asm02.HP0_G0000560 | YAL015C | chrI | 2575.44193 | 1744.281285 | 1842.53876 | -0.545536261 | 0.17115457 | 0.001164375 | 0.00162611 | ns | NaCl | YAL015C | NTG1 | n |
| W303.asm02.HP0_G0000570 | YAL014C | chrI | 2081.122286 | 1395.92491 | 1512.310717 | -0.56759171 | 0.12091989 | 2.1302008498 | 3.7510058442 | ns | NaCl | YAL014C | SYN8 | n |
| W303.asm02.HP0_G0000580 | YAL013W | chrI | 1375.068468 | 1955.702734 | 1856.402874 | 0.49547438 | 0.15437443 | 0.001118601 | 0.001589591 | ns | NaCl | YAL013W | DEP1 | n |
| W303.asm02.HP0_G0000590 | YAL012W | chrI | 4557.216226 | 7228.992163 | 7210.026308 | 0.651800286 | 0.1462706 | 6.1611135691 | 1.0184697940 | ns | NaCl | YAL012W | CYS3 | n |
| W303.asm02.HP0_G0000600 | YAL011W | chrI | 476.2854094 | 642.8011948 | 531.0723606 | 0.418471513 | 0.17293529 | 0.013709631 | 0.017260856 | ns | NaCl | YAL011W | SWC3 | n |
| W303.asm02.HP0_G0000610 | YAL010C | chrI | 1143.961104 | 1283.979605 | 1116.447715 | 0.162072609 | 0.16504427 | 0.31832147 | 0.343787187 | ns | NaCl | YAL010C | MDM10 | n |
| W303.asm02.HP0_G0000620 | YAL009W | chrI | 1360.760585 | 665.7842966 | 735.512986 | -1.007292192 | 0.18078231 | 1.2258186286 | 2.3091002074 | KS<SC | NaCl | YAL009W | SPO7 | n |
| W303.asm02.HP0_G0000630 | YAL008W | chrI | 2181.916582 | 906.8484444 | 1555.162474 | -1.24863794 | 0.15396093 | 1.7788841052 | 5.5419081739 | KS<SC | NaCl | YAL008W | FUN14 | n |
| W303.asm02.HP0_G0000640 | YAL007C | chrI | 4434.352267 | 1836.333821 | 3003.066846 | -1.258977027 | 0.13412661 | 2.1283640143 | 8.6198742579 | KS<SC | NaCl | YAL007C | ERP2 | n |
| W303.asm02.HP0_G0000650 | YAL005C/YLL024C | chrI | 28748.81954 | 9394.1198 | 17010.88815 | -1.592200178 | 0.17634334 | 4.8159166485 | 1.7731329478 | KS<SC | NaCl | YAL005C | SSA1 | y |
| W303.asm02.HP0_G0000660 | YAL003W | chrI | 30035.57096 | 7668.804106 | 13415.7194 | -1.943650252 | 0.19602109 | 9.9653644473 | 4.2483922117 | KS<SC | NaCl | YAL003W | EFB1 | n |
| W303.asm02.HP0_G0000670 | YAL002W | chrI | 908.9734307 | 3707.500511 | 2106.952717 | 2.019937749 | 0.10674451 | 2.2158469897 | 5.9827868722 | KS>SC | NaCl | YAL002W | VPS8 | n |
| W303.asm02.HP0_G0000680 | YAL001C | chrI | 1455.680177 | 1228.501068 | 1647.5821 | -0.23955313 | 0.12132746 | 0.046305659 | 0.055158211 | ns | NaCl | YAL001C | TFC3 | n |
| W303.asm02.HP0_G0000700 | YAR002W | chrI | 2708.132366 | 2908.368572 | 2219.195198 | 0.100551658 | 0.14109199 | 0.470774035 | 0.495229829 | ns | NaCl | YAR002W | NUP60 | n |
| W303.asm02.HP0_G0000710 | YAR002C-A | chrI | 7064.673432 | 3744.742758 | 4700.018699 | -0.900565896 | 0.14439407 | 2.4635792715 | 5.7014263142 | ns | NaCl | YAR002C-A | ERP1 | n |
| W303.asm02.HP0_G0000720 | YAR003W | chrI | 4399.05792 | 1583.111295 | 2590.541736 | -1.465168794 | 0.11591911 | 3.6711446507 | 2.2874055131 | KS<SC | NaCl | YAR003W | SWD1 | n |
| W303.asm02.HP0_G0000730 | YAR007C | chrI | 3592.047499 | 1698.355304 | 2357.015013 | -1.058534427 | 0.17197612 | 3.3914744870 | 7.6308175957 | KS<SC | NaCl | YAR007C | RFA1 | n |
| W303.asm02.HP0_G0000740 | YAR008W | chrI | 1828.058209 | 697.9781098 | 909.8659521 | -1.362263064 | 0.19533477 | 9.8930278953 | 2.5849524500 | KS<SC | NaCl | YAR008W | SEN34 | n |
| W303.asm02.HP0_G0000750 | YAR014C | chrI | 2229.576025 | 1300.845255 | 1596.564728 | -0.768716136 | 0.11200321 | 4.3360543647 | 1.0643042531 | ns | NaCl | YAR014C | BUD14 | n |
| W303.asm02.HP0_G0000760 | YAR015W | chrI | 2416.975906 | 284.9186865 | 1256.944613 | -3.041623304 | 0.22232282 | 3.0254282577 | 2.0421640739 | KS<SC | NaCl | YAR015W | ADE1 | n |
| W303.asm02.HP0_G0000770 | YAR018C | chrI | 1209.2635 | 1449.131984 | 1043.818519 | 0.256819373 | 0.11240889 | 0.0212817 | 0.025728623 | ns | NaCl | YAR018C | KIN3 | n |
| W303.asm02.HP0_G0000780 | YAR019C | chrI | 312.4394551 | 4292.169034 | 2087.379567 | 3.750582479 | 0.15952666 | 3.7569575536 | 1.5215678092 | KS>SC | NaCl | YAR019C | CDC15 | n |
| W303.asm02.HP0_G0000790 | W303.asm02.HP0_G0 | chrI | 52.68530567 | 578.9732608 | 341.2808402 | 3.376102665 | 0.2910789 | 7.5171464862 | 4.3492061813 | KS>SC | NaCl | YFL020C | PAU5 | y |
| W303.asm02.HP0_G0000800 | YAR023C | chrI | 270.7256693 | 630.09401 | 527.8578251 | 1.18524569 | 0.21140309 | 8.1006589265 | 1.5622699358 | KS>SC | NaCl | YAR023C | DFP1 | n |
| W303.asm02.HP0_G0000810 | YAR027W | chrI | 2641.160226 | 1325.462591 | 1918.000045 | -0.980573808 | 0.13995762 | 1.2095997943 | 3.0617994794 | ns | NaCl | YAR027W | UIP3 | n |
| W303.asm02.HP0_G0000820 | YAR028W | chrI | 850.3941242 | 331.137079 | 679.5324071 | -1.337907392 | 0.17719783 | 1.4114768411 | 4.0832008618 | KS<SC | NaCl | YAR028W | KTD1 | n |
| W303.asm02.HP0_G0000830 | W303.asm02.HP0_G0 | chrI | 1200.776224 | 2537.771361 | 1972.532087 | 1.063436024 | 0.14475611 | 9.0017128400 | 2.4304624668 | KS>SC | NaCl | YAR031W | PRM9 | y |
| W303.asm02.HP0_G0000840 | YAR035W | chrI | 284.447348 | 561.3693566 | 546.8210326 | 0.943053737 | 0.2290732 | 2.0687871266 | 3.2857207306 | ns | NaCl | YAR035W | YAT1 | n |
| W303.asm02.HP0_G0000850 | YAR042W | chrI | 871.0308568 | 1891.870574 | 2622.801709 | 1.092566412 | 0.18700189 | 2.2396741769 | 4.6516309828 | KS>SC | NaCl | YAR042W | SWH1 | n |

SC_vs_KS3

| w303_id | W303_annotation | chr | mean_SC_count | mean_KS_count | baseMean | log2FoldChange | lfcSE | pvalue | padj | DEG | treatment | S288C_id | gene | Ambiguous gene model |
|---|---|---|---|---|---|---|---|---|---|---|---|---|---|---|
| W303.asm02.HP0_G0004980 | YCL064C | chrIII | 1891.584368 | 1380.001821 | 1211.469492 | -0.431425243 | 0.22404095 | 0.047837223 | 0.059579086 | ns | YPD | YCL064C | CHA1 | n |
| W303.asm02.HP0_G0004990 | YCL063W | chrIII | 3207.233821 | 1228.884491 | 1887.140178 | -1.376308979 | 0.07360681 | 1.5019050791 | 3.4293499306 | KS<SC | YPD | YCL063W | VAC17 | n |
| W303.asm02.HP0_G0005000 | YCL061C | chrIII | 3468.676227 | 4284.193677 | 3053.729675 | 0.301357854 | 0.09445356 | 0.001340428 | 0.002186174 | ns | YPD | YCL061C | MRC1 | n |
| W303.asm02.HP0_G0005010 | YCL059C | chrIII | 7439.453364 | 6538.059948 | 6586.286877 | -0.183532971 | 0.11516103 | 0.108318361 | 0.12792772 | ns | YPD | YCL059C | KRR1 | n |
| W303.asm02.HP0_G0005020 | YCL058W-A | chrIII | 1511.391075 | 461.7867759 | 1078.478048 | -1.654418324 | 0.19301764 | 1.8837499897 | 8.3249596322 | KS<SC | YPD | YCL058W-A | ADF1 | n |
| W303.asm02.HP0_G0005030 | YCL057C-A | chrIII | 2688.531571 | 398.4603712 | 2041.01948 | -2.716788437 | 0.15327114 | 2.7744946785 | 5.4300824423 | KS<SC | YPD | YCL057C-A | MIC10 | n |
| W303.asm02.HP0_G0005040 | YCL057W | chrIII | 3497.284443 | 1112.635328 | 3546.262488 | -1.552924179 | 0.26288944 | 7.9866991169 | 2.2330158755 | KS<SC | YPD | YCL057W | PRD1 | n |
| W303.asm02.HP0_G0005050 | YCL056C | chrIII | 1270.865847 | 1030.206509 | 1042.157888 | -0.293844418 | 0.16599781 | 0.072278292 | 0.088728343 | ns | YPD | YCL056C | PEX34 | n |
| W303.asm02.HP0_G0005060 | YCL055W | chrIII | 2667.033106 | 1889.818764 | 2117.2546 | -0.484560569 | 0.15260171 | 0.001294532 | 0.002172223 | ns | YPD | YCL055W | KAR4 | n |
| W303.asm02.HP0_G0005070 | YCL054W | chrIII | 13124.33178 | 6747.54423 | 8401.514345 | -0.950729429 | 0.09391335 | 2.4857022274 | 1.4189216881 | ns | YPD | YCL054W | SPB1 | n |
| W303.asm02.HP0_G0005080 | YCL052C | chrIII | 5366.612362 | 1912.647669 | 3501.202677 | -1.471671762 | 0.10750618 | 3.0006067680 | 2.7405541814 | KS<SC | YPD | YCL052C | PBN1 | n |
| W303.asm02.HP0_G0005090 | YCL051W | chrIII | 2354.867088 | 1145.088729 | 1712.553656 | -1.027700312 | 0.10730396 | 5.1944307234 | 2.8465480364 | KS<SC | YPD | YCL051W | LRE1 | n |
| W303.asm02.HP0_G0005100 | YCL050C | chrIII | 11509.13943 | 8204.027875 | 10811.35482 | -0.482267271 | 0.10564259 | 4.3306320394 | 8.2492646469 | ns | YPD | YCL050C | APA1 | n |
| W303.asm02.HP0_G0005110 | YCL049C | chrIII | 3427.849392 | 2648.244046 | 3002.426132 | -0.362343097 | 0.15735928 | 0.019561722 | 0.026534216 | ns | YPD | YCL049C | | n |
| W303.asm02.HP0_G0005120 | YCL048W-A | chrIII | 90.27780996 | 757.9672036 | 424.7356513 | 2.493367873 | 0.64754626 | 1.1498598305 | 2.4235507197 | KS>SC | YPD | YCL048W-A | | n |
| W303.asm02.HP0_G0005130 | YCL048W | chrIII | 198.6942518 | 7768.307601 | 3835.883642 | 5.272659512 | 0.23209366 | 7.2360393799 | 3.3044579835 | KS>SC | YPD | YCL048W | SPS22 | n |
| W303.asm02.HP0_G0005140 | YCL047C | chrIII | 1910.953363 | 524.858982 | 1285.045077 | -1.844320987 | 0.1099657 | 4.1118795594 | 6.2591944405 | KS<SC | YPD | YCL047C | POF1 | n |
| W303.asm02.HP0_G0005150 | YCL045C | chrIII | 7480.391473 | 4066.57248 | 6014.488977 | -0.866459451 | 0.11723546 | 9.2536636630 | 3.1693798045 | ns | YPD | YCL045C | EMC1 | n |
| W303.asm02.HP0_G0005160 | YCL044C | chrIII | 816.9830754 | 1009.098825 | 1679.464284 | 0.275047523 | 0.31206796 | 0.353483289 | 0.381316618 | ns | YPD | YCL044C | MGR1 | n |
| W303.asm02.HP0_G0005170 | YCL043C | chrIII | 12219.76034 | 7561.244162 | 15515.30826 | -0.676706599 | 0.14668721 | 3.0072854355 | 5.9709870242 | ns | YPD | YCL043C | PDI1 | n |
| W303.asm02.HP0_G0005180 | YCL040W | chrIII | 3559.43431 | 1056.743571 | 6276.778436 | -1.688961076 | 0.202557 | 1.2887859369 | 5.5176147926 | KS<SC | YPD | YCL040W | GLK1 | n |
| W303.asm02.HP0_G0005190 | YCL039W | chrIII | 1247.455413 | 1324.348238 | 1508.467595 | 0.084875727 | 0.1203714 | 0.477129811 | 0.505376044 | ns | YPD | YCL039W | GID7 | n |
| W303.asm02.HP0_G0005200 | YCL038C | chrIII | 1707.423537 | 1817.790516 | 1815.784429 | 0.088601931 | 0.13107871 | 0.494920379 | 0.513667363 | ns | YPD | YCL038C | ATG22 | n |
| W303.asm02.HP0_G0005210 | YCL037C | chrIII | 383.8907431 | 583.6946461 | 527.5743471 | 0.566296117 | 0.24837661 | 0.018401176 | 0.025724093 | ns | YPD | YCL037C | SRO9 | n |
| W303.asm02.HP0_G0005220 | YCL036W | chrIII | 3300.921667 | 4466.638884 | 4059.236207 | 0.416094961 | 0.21068318 | 0.043039639 | 0.054596579 | ns | YPD | YCL036W | GFD2 | n |
| W303.asm02.HP0_G0005230 | YCL035C | chrIII | 4070.123388 | 2056.054673 | 3711.433848 | -0.955919855 | 0.17158614 | 1.4624178888 | 3.6427500140 | ns | YPD | YCL035C | GRX1 | n |
| W303.asm02.HP0_G0005240 | YCL034W | chrIII | 2711.772307 | 1567.939325 | 2898.909704 | -0.779763385 | 0.11058713 | 1.2266949183 | 3.9083070654 | ns | YPD | YCL034W | LSB5 | n |
| W303.asm02.HP0_G0005250 | YCL033C | chrIII | 2185.391445 | 558.8292854 | 1683.466772 | -1.875515893 | 0.23887875 | 4.4264801118 | 1.7326507866 | KS<SC | YPD | YCL033C | MXR2 | n |
| W303.asm02.HP0_G0005260 | YCL032W | chrIII | 2525.651411 | 1485.150781 | 1936.000195 | -0.761252055 | 0.07472863 | 1.5817666568 | 9.4218274778 | ns | YPD | YCL032W | STE50 | n |
| W303.asm02.HP0_G0005270 | YCL031C | chrIII | 7362.093747 | 6391.659955 | 6114.993332 | -0.200804201 | 0.11645696 | 0.082272728 | 0.099746581 | ns | YPD | YCL031C | RRP7 | n |
| W303.asm02.HP0_G0005280 | YCL030C | chrIII | 4579.873203 | 2309.679613 | 5097.396452 | -0.966675247 | 0.14381 | 1.0205804342 | 3.1071004332 | ns | YPD | YCL030C | HIS4 | n |
| W303.asm02.HP0_G0005290 | YCL029C | chrIII | 2772.466758 | 1306.448459 | 2021.981501 | -1.074722839 | 0.09658241 | 4.4967020924 | 3.2423588771 | KS<SC | YPD | YCL029C | BIK1 | n |
| W303.asm02.HP0_G0005300 | YCL028W | chrIII | 8392.188895 | 2267.769511 | 5753.045622 | -1.879344217 | 0.06847364 | 6.4085693004 | 8.7797399416 | KS<SC | YPD | YCL028W | RNQ1 | n |
| W303.asm02.HP0_G0005310 | YCL027W | chrIII | 529.0990067 | 403.2613749 | 483.300876 | -0.369109152 | 0.23793648 | 0.109748275 | 0.128282371 | ns | YPD | YCL027W | FUS1 | n |
| W303.asm02.HP0_G0005320 | YCL026C-B | chrIII | 648.7915482 | 1307.44374 | 1007.388914 | 0.969602656 | 0.20355851 | 1.0828896020 | 2.3180605543 | ns | YPD | YCL026C-B | HBN1 | n |
| W303.asm02.HP0_G0005330 | YCL026C-A | chrIII | 201.2099991 | 7992.565613 | 4678.985084 | 5.294740982 | 0.23558269 | 1.7838273434 | 6.1096086513 | KS>SC | YPD | YCL026C-A | FRM2 | n |
| W303.asm02.HP0_G0005340 | YCL025C | chrIII | 7031.986016 | 3498.875403 | 5731.653313 | -0.975829472 | 0.17568129 | 1.5711823633 | 3.8437854246 | ns | YPD | YCL025C | AGP1 | n |
| W303.asm02.HP0_G0005350 | YCL024W | chrIII | 2892.989377 | 1558.170337 | 1965.314321 | -0.882955782 | 0.10060489 | 1.0457716254 | 4.9403694027 | ns | YPD | YCL024W | KCC4 | n |
| W303.asm02.HP0_G0005360 | YCL021W-A | chrIII | 261.9458267 | 491.8467912 | 383.3625747 | 0.888669895 | 0.14658455 | 8.3304208415 | 2.2825353105 | ns | YPD | YCL021W-A | | n |
| W303.asm02.HP0_G0005370 | YCL018W | chrIII | 1734.472912 | 1133.576911 | 1447.963445 | -0.600999785 | 0.13842107 | 1.1353955358 | 2.1308108000 | ns | YPD | YCL018W | LEU2 | n |
| W303.asm02.HP0_G0005380 | YCL017C | chrIII | 3337.034821 | 2240.767563 | 2929.661665 | -0.566768259 | 0.11173481 | 3.2309795278 | 7.3774032552 | ns | YPD | YCL017C | NFS1 | n |
| W303.asm02.HP0_G0005390 | YCL016C | chrIII | 1018.706238 | 1222.989281 | 950.4497386 | 0.260505215 | 0.10317665 | 0.011072287 | 0.016137269 | ns | YPD | YCL016C | DCC1 | n |
| W303.asm02.HP0_G0005400 | YCL014W | chrIII | 1563.234755 | 3862.288476 | 2400.236527 | 1.266408294 | 0.17430014 | 1.4062317568 | 4.6988719678 | KS>SC | YPD | YCL014W | BUD3 | n |
| W303.asm02.HP0_G0005410 | YCL012C | chrIII | 751.5134763 | 767.8171501 | 655.9406413 | 0.02892287 | 0.25682511 | 0.907146306 | 0.907146306 | ns | YPD | YCL012C | | n |
| W303.asm02.HP0_G0005420 | YCL011C | chrIII | 3979.549235 | 4071.261363 | 4931.803851 | 0.032051249 | 0.14419508 | 0.822083375 | 0.834262388 | ns | YPD | YCL011C | GBP2 | n |
| W303.asm02.HP0_G0005430 | YCL010C | chrIII | 535.2840997 | 663.7944366 | 870.6100674 | 0.29186826 | 0.23905887 | 0.208046254 | 0.235556502 | ns | YPD | YCL010C | SGF29 | n |
| W303.asm02.HP0_G0005440 | YCL009C | chrIII | 2175.095464 | 3095.298557 | 4104.665729 | 0.437925927 | 0.3822612 | 0.216076989 | 0.242643832 | ns | YPD | YCL009C | ILV6 | n |
| W303.asm02.HP0_G0005450 | YCL008C | chrIII | 179.3214277 | 395.6839651 | 549.3495235 | 1.065689952 | 0.26390394 | 2.7059537261 | 5.0096710876 | KS>SC | YPD | YCL008C | STP22 | n |
| W303.asm02.HP0_G0005460 | W303.asm02.HP0_G0 | chrIII | 1089.460815 | 500.5766475 | 782.3178261 | -1.097346995 | 0.14843016 | 6.9245038931 | 2.4324539316 | KS<SC | YPD | YCL005W-A | VMA9 | y |
| W303.asm02.HP0_G0005470 | YCL005W | chrIII | 839.7819582 | 1715.732643 | 1352.864291 | 1.02171327 | 0.09196293 | 5.8568415657 | 4.0119364725 | KS>SC | YPD | YCL005W | LDB16 | n |
| W303.asm02.HP0_G0005480 | YCL004W | chrIII | 745.5783334 | 1596.299791 | 1125.326985 | 1.089177225 | 0.08968865 | 2.9401215092 | 2.5174790422 | KS>SC | YPD | YCL004W | PGS1 | n |
| W303.asm02.HP0_G0005490 | YCL002C | chrIII | 1695.131095 | 1544.08109 | 1274.186365 | -0.131076271 | 0.15470809 | 0.390582656 | 0.418045499 | ns | YPD | YCL002C | | n |
| W303.asm02.HP0_G0005500 | YCL001W | chrIII | 2597.132363 | 1474.15295 | 1826.473456 | -0.800773185 | 0.1374828 | 3.8918734704 | 1.0253589720 | ns | YPD | YCL001W | RER1 | n |
| W303.asm02.HP0_G0005510 | YCL001W-A | chrIII | 80.55560742 | 374.5275331 | 172.3737645 | 2.143658468 | 0.20932255 | 5.5911917636 | 3.6475870077 | KS>SC | YPD | YCL001W-A | | n |
| W303.asm02.HP0_G0005520 | YCL001W-B | chrIII | 90.13392278 | 1147.528118 | 612.1595912 | 3.652759626 | 0.17288397 | 1.0045922642 | 2.7525828040 | KS>SC | YPD | YCL001W-B | | n |
| W303.asm02.HP0_G0005530 | YCR002C | chrIII | 3406.446436 | 2849.798701 | 3389.825775 | -0.254973272 | 0.09024782 | 0.0045292 | 0.006971914 | ns | YPD | YCR002C | CDC10 | n |
| W303.asm02.HP0_G0005540 | YCR003W | chrIII | 2111.194375 | 1864.0999 | 1711.638185 | -0.173435639 | 0.17859067 | 0.323029354 | 0.35123033 | ns | YPD | YCR003W | MRPL32 | n |
| W303.asm02.HP0_G0005550 | YCR004C | chrIII | 3737.282882 | 1707.320615 | 2809.503227 | -1.096310262 | 0.17436796 | 1.5666512017 | 4.6658959705 | KS<SC | YPD | YCR004C | YCP4 | n |
| W303.asm02.HP0_G0005560 | YCR005C | chrIII | 2793.374778 | 970.0860815 | 3416.448465 | -1.48069933 | 0.17870391 | 3.0725591185 | 1.2755775734 | KS<SC | YPD | YCR005C | CIT2 | n |
| W303.asm02.HP0_G0005570 | W303.asm02.HP0_G0 | chrIII | 85.38702634 | 230.3378202 | 126.1214297 | 1.396002759 | 0.16097211 | 1.2868711832 | 5.8767117366 | KS>SC | YPD | YCR006C | | y |
| W303.asm02.HP0_G0005580 | YCR007C | chrIII | 307.6000019 | 557.1285756 | 451.0619636 | 0.832194186 | 0.17058224 | 7.0496738606 | 1.5330243157 | ns | YPD | YCR007C | DFP3 | n |
| W303.asm02.HP0_G0005590 | YCR008W | chrIII | 3579.43148 | 5932.824609 | 4030.58458 | 0.718164738 | 0.11664817 | 5.4365300506 | 1.5516762852 | ns | YPD | YCR008W | SAT4 | n |
| W303.asm02.HP0_G0005600 | YCR009C | chrIII | 4790.328124 | 2497.74116 | 3351.187025 | -0.922473762 | 0.13189438 | 1.5962632935 | 4.9701834366 | ns | YPD | YCR009C | RVS161 | n |
| W303.asm02.HP0_G0005610 | YCR010C | chrIII | 135.2415438 | 587.9794923 | 376.2484175 | 2.062493667 | 0.18478706 | 3.6080853371 | 2.7461538399 | KS>SC | YPD | YCR010C | ADY2 | n |
| W303.asm02.HP0_G0005620 | YCR011C | chrIII | 2490.224722 | 4380.586905 | 4008.211727 | 0.803617005 | 0.11219363 | 5.3408103743 | 1.7421214792 | ns | YPD | YCR011C | ADP1 | n |
| W303.asm02.HP0_G0005630 | YCR012W | chrIII | 33855.14442 | 9396.766953 | 31329.98713 | -1.828857052 | 0.11021242 | 8.4530606019 | 1.0527902749 | KS<SC | YPD | YCR012W | PGK1 | n |

| | | | | | | | | | | | | | | |
|---|---|---|---|---|---|---|---|---|---|---|---|---|---|---|
| W303.asm02.HP0_G0005640 | YCR014C | chrIII | 1049.485496 | 2391.892116 | 1377.405532 | 1.164292276 | 0.14294158 | 1.66018282904 | 6.68956022292 | KS>SC | YPD | YCR014C | POL4 | n |
| W303.asm02.HP0_G0005650 | YCR015C | chrIII | 983.7153399 | 1224.677573 | 873.6436017 | 0.308024701 | 0.15392361 | 0.042586053 | 0.054526068 | ns | YPD | YCR015C | CTO1 | n |
| W303.asm02.HP0_G0005660 | YCR016W | chrIII | 3686.984619 | 4683.497967 | 3476.357041 | 0.336952905 | 0.14792251 | 0.021111783 | 0.028356022 | ns | YPD | YCR016W | RBP95 | n |
| W303.asm02.HP0_G0005670 | YCR017C | chrIII | 3401.750859 | 2802.31772 | 3263.383474 | -0.271951056 | 0.16085684 | 0.08633364 | 0.103751831 | ns | YPD | YCR017C | CWH43 | n |
| W303.asm02.HP0_G0005680 | YCR018C | chrIII | 1176.406547 | 1030.842177 | 1067.342272 | -0.185231594 | 0.16039648 | 0.241444476 | 0.268925961 | ns | YPD | YCR018C | SRD1 | n |
| W303.asm02.HP0_G0005800 | YCR028C | chrIII | 1705.427788 | 2196.279125 | 1627.578649 | 0.35980789 | 0.10835714 | 0.000828342 | 0.001436491 | ns | YPD | YCR028C | FEN2 | n |
| W303.asm02.HP0_G0005810 | YCR028C-A | chrIII | 3293.694205 | 1214.170286 | 2422.230694 | -1.405370358 | 0.15871759 | 2.50972676024 | 1.22797345055 | KS<SC | YPD | YCR028C-A | RIM1 | n |
| W303.asm02.HP0_G0005820 | YCR030C | chrIII | 1276.371526 | 2518.295333 | 2071.875465 | 0.892320886 | 0.30789527 | 0.002283005 | 0.00363688 | ns | YPD | YCR030C | SYP1 | n |
| W303.asm02.HP0_G0005830 | YCR031C/YJL191W | chrIII | 10107.47746 | 6725.403915 | 10562.69539 | -0.579767812 | 0.11075165 | 1.34626760051 | 3.12607900458 | ns | YPD | YCR031C | RPS14A | y |
| W303.asm02.HP0_G0005840 | YCR032W | chrIII | 2212.830974 | 3173.699201 | 2250.104759 | 0.510521534 | 0.13131163 | 8.62458841215 | 0.00015547 | ns | YPD | YCR032W | BPH1 | n |
| W303.asm02.HP0_G0005850 | YCR033W | chrIII | 1954.038036 | 3553.832445 | 2530.895856 | 0.834626827 | 0.17846034 | 1.91708843680 | 3.97941084621 | ns | YPD | YCR033W | SNT1 | n |
| W303.asm02.HP0_G0005860 | YCR034W | chrIII | 6905.427524 | 4239.581821 | 5411.274469 | -0.692671001 | 0.12104644 | 7.86029507719 | 2.03181212372 | ns | YPD | YCR034W | ELO2 | n |
| W303.asm02.HP0_G0005870 | YCR035C | chrIII | 3730.982892 | 3405.040448 | 3548.457755 | -0.130280539 | 0.1017907 | 0.197904202 | 0.22594063 | ns | YPD | YCR035C | RRP43 | n |
| W303.asm02.HP0_G0005880 | YCR036W | chrIII | 2938.470054 | 3748.604585 | 3412.094514 | 0.342935704 | 0.1481502 | 0.019108457 | 0.026234863 | ns | YPD | YCR036W | RBK1 | n |
| W303.asm02.HP0_G0005890 | YCR037C | chrIII | 2971.987699 | 1858.845396 | 1909.606422 | -0.666655016 | 0.11916684 | 1.69287958057 | 4.06885092172 | ns | YPD | YCR037C | PHO87 | n |
| W303.asm02.HP0_G0005900 | YCR038C | chrIII | 322.2524251 | 503.6292836 | 368.4914114 | 0.634278782 | 0.11078543 | 8.08974986637 | 2.05239950311 | ns | YPD | YCR038C | BUD5 | n |
| W303.asm02.HP0_G0005920 | YCR042C | chrIII | 2539.57308 | 2969.500913 | 2384.956457 | 0.222732144 | 0.10305326 | 0.029659777 | 0.039071052 | ns | YPD | YCR042C | TAF2 | n |
| W303.asm02.HP0_G0005930 | YCR043C | chrIII | 2799.016325 | 697.0480663 | 1395.218446 | -1.944044476 | 0.19283113 | 5.74216169980 | 3.57580069491 | KS<SC | YPD | YCR043C | | n |
| W303.asm02.HP0_G0005940 | YCR044C | chrIII | 1826.555885 | 456.4165094 | 1021.454551 | -1.979107075 | 0.11191095 | 3.74974423847 | 6.42143700830 | KS<SC | YPD | YCR044C | PER1 | n |
| W303.asm02.HP0_G0005950 | YCR045C | chrIII | 319.9986195 | 1763.883403 | 1329.097377 | 2.382647108 | 0.21414508 | 1.28732619767 | 1.03743345618 | KS>SC | YPD | YCR045C | RRT12 | n |
| W303.asm02.HP0_G0005960 | YCR046C | chrIII | 1979.77157 | 731.4420674 | 1569.660751 | -1.403106543 | 0.15644742 | 9.00096543240 | 4.56715653422 | KS<SC | YPD | YCR046C | IMG1 | n |
| W303.asm02.HP0_G0005970 | YCR047C | chrIII | 1633.058537 | 1476.534112 | 2026.885355 | -0.13923852 | 0.20131159 | 0.47955391 | 0.505376044 | ns | YPD | YCR047C | BUD23 | n |
| W303.asm02.HP0_G0005980 | YCR048W | chrIII | 1251.750677 | 2499.358497 | 2380.807295 | 0.913474599 | 0.29809285 | 0.001300163 | 0.002172223 | ns | YPD | YCR048W | ARE1 | n |
| W303.asm02.HP0_G0005990 | YCR051W | chrIII | 3363.424002 | 2785.251184 | 3541.59545 | -0.269389931 | 0.09341666 | 0.003750077 | 0.005905293 | ns | YPD | YCR051W | | n |
| W303.asm02.HP0_G0006000 | YCR052W | chrIII | 2211.890338 | 2465.93589 | 2937.406491 | 0.148828259 | 0.21916024 | 0.485876616 | 0.508130506 | ns | YPD | YCR052W | RSC6 | n |
| W303.asm02.HP0_G0006010 | YCR053W | chrIII | 9823.61546 | 9531.925275 | 10002.50807 | -0.04318444 | 0.07153387 | 0.544824895 | 0.561210606 | ns | YPD | YCR053W | THR4 | n |
| W303.asm02.HP0_G0006020 | YCR054C | chrIII | 1863.386136 | 2272.430675 | 1473.674431 | 0.280034516 | 0.14121525 | 0.044926561 | 0.056467329 | ns | YPD | YCR054C | CTR86 | n |
| W303.asm02.HP0_G0006030 | YCR057C | chrIII | 3819.418325 | 5396.603631 | 4820.360301 | 0.462695105 | 0.2677575 | 0.072537039 | 0.088728343 | ns | YPD | YCR057C | PWP2 | n |
| W303.asm02.HP0_G0006040 | YCR059C | chrIII | 2549.852428 | 2786.394228 | 2933.921788 | 0.125775465 | 0.12627778 | 0.314884651 | 0.345113577 | ns | YPD | YCR059C | YIH1 | n |
| W303.asm02.HP0_G0006050 | YCR060W | chrIII | 1337.92505 | 998.5901159 | 1398.70489 | -0.413724623 | 0.13303339 | 0.001681881 | 0.002710796 | ns | YPD | YCR060W | TAH1 | n |
| W303.asm02.HP0_G0006060 | YCR061W | chrIII | 419.4896794 | 385.0906728 | 707.4412929 | -0.113988409 | 0.26935086 | 0.660075808 | 0.674853624 | ns | YPD | YCR061W | TVS1 | n |
| W303.asm02.HP0_G0006070 | YCR063W | chrIII | 258.5957888 | 493.4714422 | 390.7163254 | 0.921315416 | 0.10178301 | 8.35449932970 | 4.40217849295 | ns | YPD | YCR063W | BUD31 | n |
| W303.asm02.HP0_G0006080 | YCR065W | chrIII | 826.7433616 | 1330.51583 | 1190.631669 | 0.65814396 | 0.19881798 | 0.000715267 | 0.001256303 | ns | YPD | YCR065W | HCM1 | n |
| W303.asm02.HP0_G0006090 | YCR066W | chrIII | 728.7201745 | 1185.421019 | 777.1183729 | 0.695887593 | 0.0875785 | 1.43228487632 | 5.45063966825 | ns | YPD | YCR066W | RAD18 | n |
| W303.asm02.HP0_G0006100 | YCR067C | chrIII | 2780.451816 | 3359.033924 | 2928.065972 | 0.264874743 | 0.1615882 | 0.096296872 | 0.114718883 | ns | YPD | YCR067C | SED4 | n |
| W303.asm02.HP0_G0006110 | YCR068W | chrIII | 319.512554 | 1503.688276 | 1068.08497 | 2.01557105 | 0.36677523 | 2.68377314817 | 7.20935139804 | KS>SC | YPD | YCR068W | ATG15 | n |
| W303.asm02.HP0_G0006120 | YCR069W | chrIII | 1631.926787 | 1031.5996 | 1523.61182 | -0.6260681 | 0.22994984 | 0.005071239 | 0.007719552 | ns | YPD | YCR069W | CPR4 | n |
| W303.asm02.HP0_G0006130 | YCR071C | chrIII | 1697.896998 | 927.9053535 | 1242.703506 | -0.848517933 | 0.16070689 | 8.38628890063 | 1.98089927480 | ns | YPD | YCR071C | IMG2 | n |
| W303.asm02.HP0_G0006140 | YCR072C | chrIII | 1960.982008 | 1997.42013 | 1902.591114 | 0.025704429 | 0.16184789 | 0.871991898 | 0.878403603 | ns | YPD | YCR072C | RSA4 | n |
| W303.asm02.HP0_G0006150 | YCR073C | chrIII | 1867.194776 | 4942.131614 | 3202.971991 | 1.359207729 | 0.18437898 | 5.49047801437 | 1.97946181044 | KS>SC | YPD | YCR073C | SSK22 | n |
| W303.asm02.HP0_G0006160 | YCR073W-A | chrIII | 672.0987988 | 958.7974866 | 1202.246529 | 0.488280796 | 0.2136977 | 0.019149535 | 0.026234863 | ns | YPD | YCR073W-A | SOL2 | n |
| W303.asm02.HP0_G0006170 | YCR075C | chrIII | 568.7618073 | 353.4773585 | 427.6849427 | -0.665856557 | 0.16785271 | 5.56807787085 | 0.00010171 | ns | YPD | YCR075C | ERS1 | n |
| W303.asm02.HP0_G0006180 | YCR075W-A | chrIII | 254.5276017 | 168.7189212 | 192.5681641 | -0.558027866 | 0.24287257 | 0.017677857 | 0.024967695 | ns | YPD | YCR075W-A | EGO2 | n |
| W303.asm02.HP0_G0006190 | YCR076C | chrIII | 676.4300415 | 850.2153977 | 771.2515063 | 0.323862843 | 0.13081024 | 0.012422372 | 0.01772776 | ns | YPD | YCR076C | FUB1 | n |
| W303.asm02.HP0_G0006200 | YCR077C | chrIII | 1170.606902 | 1534.374767 | 2278.318609 | 0.367761049 | 0.23753957 | 0.110491385 | 0.128282371 | ns | YPD | YCR077C | PAT1 | n |
| W303.asm02.HP0_G0006210 | YCR079W | chrIII | 531.5054472 | 1826.412978 | 1459.1903 | 1.75643778 | 0.1196953 | 1.28044007585 | 1.25300207423 | KS>SC | YPD | YCR079W | PTC6 | n |
| W303.asm02.HP0_G0006220 | YCR081W | chrIII | 1746.664173 | 4568.328614 | 2437.626666 | 1.377164551 | 0.08354424 | 1.41214709152 | 1.61220126282 | KS>SC | YPD | YCR081W | SRB8 | n |
| W303.asm02.HP0_G0006230 | YCR082W | chrIII | 2286.052037 | 1375.51945 | 1710.527445 | -0.717142489 | 0.14249958 | 3.54971021963 | 7.97230000140 | ns | YPD | YCR082W | AHC2 | n |
| W303.asm02.HP0_G0006240 | YCR083W | chrIII | 1180.526868 | 645.2411282 | 839.7655168 | -0.840823253 | 0.18663471 | 4.33537996041 | 8.24926464690 | ns | YPD | YCR083W | TRX3 | n |
| W303.asm02.HP0_G0006250 | YCR084C | chrIII | 1238.100775 | 3161.464543 | 3483.041071 | 1.173456566 | 0.38909009 | 0.001068223 | 0.001829332 | KS>SC | YPD | YCR084C | TUP1 | n |
| W303.asm02.HP0_G0006260 | YCR086W | chrIII | 529.3858552 | 650.6937872 | 587.5539029 | 0.291654367 | 0.13709405 | 0.031553355 | 0.041169616 | ns | YPD | YCR086W | CSM1 | n |
| W303.asm02.HP0_G0006270 | YCR087C-A | chrIII | 2145.334854 | 1934.916126 | 2059.819745 | -0.1460516 | 0.13358458 | 0.269445869 | 0.297694226 | ns | YPD | YCR087C-A | | n |
| W303.asm02.HP0_G0006280 | YCR088W | chrIII | 964.5910826 | 2196.171772 | 2590.006483 | 0.988381138 | 0.43605532 | 0.012163943 | 0.017541687 | ns | YPD | YCR088W | ABP1 | n |
| W303.asm02.HP0_G0006290 | YCR089W | chrIII | 396.1650769 | 1118.358389 | 788.1735371 | 1.387044379 | 0.28703784 | 4.18749688869 | 9.25301731847 | KS>SC | YPD | YCR089W | FIG2 | n |
| W303.asm02.HP0_G0006300 | YCR090C | chrIII | 1675.918765 | 2252.996149 | 1873.774891 | 0.416420959 | 0.15180841 | 0.005457009 | 0.008215498 | ns | YPD | YCR090C | | n |
| W303.asm02.HP0_G0006310 | YCR091W | chrIII | 198.2301641 | 1085.685686 | 551.4549167 | 2.406079421 | 0.15982275 | 3.31406730738 | 3.49251708540 | KS>SC | YPD | YCR091W | KIN82 | n |
| W303.asm02.HP0_G0006320 | YCR092C | chrIII | 952.3499219 | 1570.551419 | 1151.072338 | 0.703377839 | 0.15253631 | 2.97124789534 | 5.97098702422 | ns | YPD | YCR092C | MSH3 | n |
| W303.asm02.HP0_G0006330 | YCR093W | chrIII | 4409.686105 | 5938.605692 | 4487.939788 | 0.419200567 | 0.14777578 | 0.004081804 | 0.006354626 | ns | YPD | YCR093W | CDC39 | n |
| W303.asm02.HP0_G0006340 | YCR094W | chrIII | 1921.520431 | 2472.635724 | 1837.138468 | 0.359732418 | 0.10012076 | 0.000301525 | 0.000536479 | ns | YPD | YCR094W | CDC50 | n |
| W303.asm02.HP0_G0006350 | YCR095C | chrIII | 1559.972645 | 1088.304321 | 1025.215307 | -0.499292059 | 0.1926251 | 0.008154647 | 0.01201276 | ns | YPD | YCR095C | OCA4 | n |
| W303.asm02.HP0_G0006360 | YCR095W-A | chrIII | 99.48177086 | 159.1178534 | 117.3444562 | 0.661995702 | 0.14405082 | 3.31266094397 | 6.48335070464 | ns | YPD | YCR095W-A | | n |
| W303.asm02.HP0_G0006380 | YCR098C | chrIII | 1800.491566 | 2533.116589 | 2477.889803 | 0.43826947 | 0.33625451 | 0.166485277 | 0.191667924 | ns | YPD | YCR098C | GIT1 | n |
| W303.asm02.HP0_G0006390 | W303.asm02.HP0_G00 | chrIII | 497.908802 | 674.770168 | 543.2631845 | 0.419558106 | 0.20417955 | 0.035536779 | 0.045929611 | ns | YPD | YNR065C | | y |
| W303.asm02.HP0_G0006400 | YCR100C | chrIII | 1313.946046 | 1896.211497 | 1452.925958 | 0.499405361 | 0.23508705 | 0.028600256 | 0.038041118 | ns | YPD | YCR100C | EMA35 | n |
| W303.asm02.HP0_G0006410 | YCR101C | chrIII | 841.4404356 | 1295.083827 | 842.7599021 | 0.589045519 | 0.22889532 | 0.00809518 | 0.01201276 | ns | YPD | YCR101C | | n |
| W303.asm02.HP0_G0006420 | YCR102C | chrIII | 846.4111934 | 3286.605261 | 2418.09028 | 1.862972345 | 0.24227929 | 1.64724176978 | 6.09924655298 | KS>SC | YPD | YCR102C | | n |
| W303.asm02.HP0_G0006430 | YCR104W/YIR041W/Y | chrIII | 39.55013524 | 308.4814256 | 196.0184986 | 2.91316744 | 0.17849461 | 1.94624444059 | 2.66635488360 | KS>SC | YPD | YMR325W | PAU19 | y |
| W303.asm02.HP0_G0006440 | YCR105W | chrIII | 877.1424231 | 1386.335466 | 1036.798106 | 0.652314185 | 0.10404951 | 2.80064846132 | 8.16359232343 | ns | YPD | YCR105W | ADH7 | n |

| | | | | | | | | | | | | | | |
|---|---|---|---|---|---|---|---|---|---|---|---|---|---|---|
| W303.asm02.HP0_G0006450 | YCR106W | chrIII | 2326.280465 | 1834.9041 | 1741.030874 | -0.338246019 | 0.10611266 | 0.001334711 | 0.002186174 | ns | YPD | YCR106W | RDS1 | n |
| W303.asm02.HP0_G0006460 | YCR107W | chrIII | 2230.781828 | 1525.072855 | 2595.446155 | -0.540633681 | 0.11586218 | 2.5686461392 | 5.2523062848 | ns | YPD | YCR107W | AAD3 | n |
| W303.asm02.HP0_G0004930 | YCL073C/YKR106W | chrIII | 59.73540741 | 6322.131391 | 2987.979787 | 6.682444337 | 0.31279583 | 3.8955113904 | 1.3342126512 | KS>SC | TM | YCL073C | GEX1 | y |
| W303.asm02.HP0_G0004980 | YCL064C | chrIII | 794.0181977 | 992.4409624 | 1211.469492 | 0.293519306 | 0.23365838 | 0.169750906 | 0.200481674 | ns | TM | YCL064C | CHA1 | n |
| W303.asm02.HP0_G0004990 | YCL063W | chrIII | 2586.893519 | 887.3927944 | 1887.140178 | -1.538210745 | 0.08080372 | 3.1372030508 | 6.1399545424 | KS<SC | TM | YCL063W | VAC17 | n |
| W303.asm02.HP0_G0005000 | YCL061C | chrIII | 2732.695889 | 2700.822875 | 3053.729675 | -0.01340153 | 0.08603945 | 0.858472555 | 0.890990455 | ns | TM | YCL061C | MRC1 | n |
| W303.asm02.HP0_G0005010 | YCL059C | chrIII | 5176.165946 | 6273.903934 | 6586.286877 | 0.269868022 | 0.12049888 | 0.018036295 | 0.02353307 | ns | TM | YCL059C | KRR1 | n |
| W303.asm02.HP0_G0005020 | YCL058W-A | chrIII | 1565.496866 | 555.5310963 | 1078.478048 | -1.45263594 | 0.19666137 | 6.5884702376 | 1.8804592136 | KS<SC | TM | YCL058W-A | ADF1 | n |
| W303.asm02.HP0_G0005030 | YCL057C-A | chrIII | 3995.688544 | 342.7912816 | 2041.01948 | -3.513406283 | 0.16085088 | 1.2770268825 | 5.8317560969 | KS<SC | TM | YCL057C-A | MIC10 | n |
| W303.asm02.HP0_G0005040 | YCL057W | chrIII | 7028.762077 | 1752.618689 | 3546.262488 | -1.913997229 | 0.26477918 | 1.0491937019 | 2.9334599422 | KS<SC | TM | YCL057W | PRD1 | n |
| W303.asm02.HP0_G0005050 | YCL056C | chrIII | 1083.366484 | 721.0218142 | 1042.157888 | -0.573105667 | 0.1721956 | 0.000730276 | 0.001136908 | ns | TM | YCL056C | PEX34 | n |
| W303.asm02.HP0_G0005060 | YCL055W | chrIII | 2526.787825 | 1406.60288 | 2117.2546 | -0.830269474 | 0.15568131 | 7.3831738257 | 1.6055473240 | ns | TM | YCL055W | KAR4 | n |
| W303.asm02.HP0_G0005070 | YCL054W | chrIII | 8894.629355 | 5563.610769 | 8401.514345 | -0.672859886 | 0.09511786 | 1.2576165280 | 3.3783032224 | ns | TM | YCL054W | SPB1 | n |
| W303.asm02.HP0_G0005080 | YCL052C | chrIII | 5458.041619 | 1543.081657 | 3501.202677 | -1.805846435 | 0.11027157 | 6.8977976448 | 9.4499827734 | KS<SC | TM | YCL052C | PBN1 | n |
| W303.asm02.HP0_G0005090 | YCL051W | chrIII | 2547.242988 | 1121.60713 | 1712.553656 | -1.171848674 | 0.11127426 | 3.6014058587 | 1.9735704105 | KS<SC | TM | YCL051W | LRE1 | n |
| W303.asm02.HP0_G0005100 | YCL050C | chrIII | 12257.99193 | 9997.957597 | 10811.35482 | -0.289427041 | 0.10855165 | 0.005946519 | 0.00831299 | ns | TM | YCL050C | APA1 | n |
| W303.asm02.HP0_G0005110 | YCL049C | chrIII | 2850.908363 | 2951.130274 | 3002.426132 | 0.042217135 | 0.14798117 | 0.752903335 | 0.799595015 | ns | TM | YCL049C | | n |
| W303.asm02.HP0_G0005120 | YCL048W-A | chrIII | 129.4191502 | 638.412592 | 424.7356513 | 1.868971338 | 0.58746631 | 0.000283082 | 0.000467255 | KS>SC | TM | YCL048W-A | | n |
| W303.asm02.HP0_G0005130 | YCL048W | chrIII | 153.2120005 | 7456.453689 | 3835.883642 | 5.579023395 | 0.24206923 | 2.5960784588 | 1.7783137443 | KS>SC | TM | YCL048W | SPS22 | n |
| W303.asm02.HP0_G0005140 | YCL047C | chrIII | 2195.419232 | 754.3826715 | 1285.045077 | -1.526779505 | 0.11434319 | 4.4136933209 | 3.5569175586 | KS<SC | TM | YCL047C | POF1 | n |
| W303.asm02.HP0_G0005150 | YCL045C | chrIII | 7887.900966 | 2959.747411 | 6014.488977 | -1.399401681 | 0.11885295 | 2.4160313879 | 1.5761728578 | KS<SC | TM | YCL045C | EMC1 | n |
| W303.asm02.HP0_G0005160 | YCL044C | chrIII | 3272.572703 | 2355.247578 | 1679.464284 | -0.426745802 | 0.32188191 | 0.148130587 | 0.178016583 | ns | TM | YCL044C | MGR1 | n |
| W303.asm02.HP0_G0005170 | YCL043C | chrIII | 30967.18743 | 15496.42519 | 15515.30826 | -0.983077538 | 0.14724673 | 1.6828884646 | 4.2695503641 | ns | TM | YCL043C | PDI1 | n |
| W303.asm02.HP0_G0005180 | YCL040W | chrIII | 13560.57547 | 1871.437925 | 6276.778436 | -2.794666464 | 0.20634587 | 6.7560798821 | 6.1705529590 | KS<SC | TM | YCL040W | GLK1 | n |
| W303.asm02.HP0_G0005190 | YCL039W | chrIII | 1777.338368 | 1485.459792 | 1508.467595 | -0.249962743 | 0.12880889 | 0.036647306 | 0.046061293 | ns | TM | YCL039W | GID7 | n |
| W303.asm02.HP0_G0005200 | YCL038C | chrIII | 1846.884533 | 2025.76896 | 1815.784429 | 0.117035706 | 0.13285501 | 0.318175875 | 0.360248718 | ns | TM | YCL038C | ATG22 | n |
| W303.asm02.HP0_G0005210 | YCL037C | chrIII | 393.6841511 | 833.2466178 | 527.5743471 | 1.032638439 | 0.25504783 | 3.3798486139 | 6.3430035631 | KS>SC | TM | YCL037C | SRO9 | n |
| W303.asm02.HP0_G0005220 | YCL036W | chrIII | 2098.925714 | 5504.529248 | 4059.236207 | 1.34657436 | 0.21351035 | 1.4072628892 | 3.2803257792 | KS>SC | TM | YCL036W | GFD2 | n |
| W303.asm02.HP0_G0005230 | YCL035C | chrIII | 6113.069232 | 1977.865301 | 3711.433848 | -1.593682518 | 0.17349327 | 1.4761136679 | 6.3196116409 | KS<SC | TM | YCL035C | GRX1 | n |
| W303.asm02.HP0_G0005240 | YCL034W | chrIII | 4388.682776 | 2129.822715 | 2898.909704 | -1.032588638 | 0.1124435 | 2.7505674216 | 1.1419022326 | KS<SC | TM | YCL034W | LSB5 | n |
| W303.asm02.HP0_G0005250 | YCL033C | chrIII | 2938.771479 | 673.6575734 | 1683.466772 | -2.045420566 | 0.24120696 | 3.6470752065 | 1.3504035224 | KS<SC | TM | YCL033C | MXR2 | n |
| W303.asm02.HP0_G0005260 | YCL032W | chrIII | 2458.712688 | 1290.7474 | 1936.000195 | -0.925556722 | 0.08003502 | 4.4226895698 | 2.7541294139 | ns | TM | YCL032W | STE50 | n |
| W303.asm02.HP0_G0005270 | YCL031C | chrIII | 5945.107322 | 4594.43074 | 6114.993332 | -0.36734902 | 0.11892379 | 0.001705445 | 0.002512322 | ns | TM | YCL031C | RRP7 | n |
| W303.asm02.HP0_G0005280 | YCL030C | chrIII | 10104.17787 | 3612.134844 | 5097.396452 | -1.461948618 | 0.14477504 | 2.4050699091 | 1.2203502872 | KS<SC | TM | YCL030C | HIS4 | n |
| W303.asm02.HP0_G0005290 | YCL029C | chrIII | 2877.648916 | 1669.266224 | 2021.981501 | -0.781273548 | 0.09959885 | 3.4219772801 | 1.0418019719 | ns | TM | YCL029C | BIK1 | n |
| W303.asm02.HP0_G0005300 | YCL028W | chrIII | 9411.002022 | 2735.218495 | 5753.045622 | -1.775841254 | 0.0705504 | 2.0041064720 | 2.7456258666 | KS<SC | TM | YCL028W | RNQ1 | n |
| W303.asm02.HP0_G0005310 | YCL027W | chrIII | 543.4907711 | 537.8340884 | 483.300876 | -0.014373434 | 0.23056485 | 0.946148349 | 0.967330775 | ns | TM | YCL027W | FUS1 | n |
| W303.asm02.HP0_G0005320 | YCL026C-B | chrIII | 780.610552 | 1286.992644 | 1007.388914 | 0.701021397 | 0.20779662 | 0.000588587 | 0.000948663 | ns | TM | YCL026C-B | HBN1 | n |
| W303.asm02.HP0_G0005330 | YCL026C-A | chrIII | 363.4542235 | 11420.06381 | 4678.985084 | 4.943860579 | 0.238425 | 9.5475257556 | 2.6160220570 | KS>SC | TM | YCL026C-A | FRM2 | n |
| W303.asm02.HP0_G0005340 | YCL025C | chrIII | 9572.93282 | 3161.143724 | 5731.653313 | -1.56352409 | 0.17716359 | 4.1198983712 | 1.5678502135 | KS<SC | TM | YCL025C | AGP1 | n |
| W303.asm02.HP0_G0005350 | YCL024W | chrIII | 2746.586291 | 968.9234037 | 1965.314321 | -1.492555119 | 0.10574702 | 1.2442310118 | 1.2175689187 | KS<SC | TM | YCL024W | KCC4 | n |
| W303.asm02.HP0_G0005360 | YCL021W-A | chrIII | 370.5978202 | 490.1932254 | 383.3625747 | 0.394948154 | 0.15894828 | 0.010004422 | 0.013437312 | ns | TM | YCL021W-A | | n |
| W303.asm02.HP0_G0005370 | YCL018W | chrIII | 2087.796808 | 1106.478643 | 1447.963445 | -0.901007132 | 0.14195612 | 1.6003857946 | 3.6542142311 | ns | TM | YCL018W | LEU2 | n |
| W303.asm02.HP0_G0005380 | YCL017C | chrIII | 3680.200312 | 2186.397936 | 2929.661665 | -0.744472608 | 0.11405647 | 5.3872475737 | 1.3179516385 | ns | TM | YCL017C | NFS1 | n |
| W303.asm02.HP0_G0005390 | YCL016C | chrIII | 829.9158866 | 907.6483235 | 950.4497386 | 0.110098022 | 0.11187774 | 0.256778952 | 0.298124716 | ns | TM | YCL016C | DCC1 | n |
| W303.asm02.HP0_G0005400 | YCL014W | chrIII | 1633.626573 | 2446.235451 | 2400.236527 | 0.569004224 | 0.17718736 | 0.001092978 | 0.001663756 | ns | TM | YCL014W | BUD3 | n |
| W303.asm02.HP0_G0005410 | YCL012C | chrIII | 438.1617116 | 788.0276121 | 655.9406413 | 0.807520122 | 0.26568006 | 0.001724951 | 0.002514024 | ns | TM | YCL012C | | n |
| W303.asm02.HP0_G0005420 | YCL011C | chrIII | 4928.95881 | 4715.033732 | 4931.803851 | -0.053259999 | 0.13568346 | 0.662130297 | 0.719935323 | ns | TM | YCL011C | GBP2 | n |
| W303.asm02.HP0_G0005430 | YCL010C | chrIII | 992.2169738 | 983.4132606 | 870.6100674 | -0.011512234 | 0.22922157 | 0.95660292 | 0.970774815 | ns | TM | YCL010C | SGF29 | n |
| W303.asm02.HP0_G0005440 | YCL009C | chrIII | 4617.18268 | 4644.823678 | 4104.665729 | 0.007155367 | 0.37554887 | 0.983338255 | 0.983528582 | ns | TM | YCL009C | ILV6 | n |
| W303.asm02.HP0_G0005450 | YCL008C | chrIII | 471.2455331 | 828.2430157 | 549.3495235 | 0.771282748 | 0.26829247 | 0.002953535 | 0.00421494 | ns | TM | YCL008C | STP22 | n |
| W303.asm02.HP0_G0005460 | W303.asm02.HP0_G0 | chrIII | 1126.009569 | 413.6863114 | 782.3178261 | -1.420245149 | 0.15612826 | 4.2005807056 | 1.6925869313 | KS<SC | TM | YCL005W-A | VMA9 | y |
| W303.asm02.HP0_G0005470 | YCL005W | chrIII | 764.4638669 | 2141.431591 | 1352.864291 | 1.474594065 | 0.09822375 | 2.4843887077 | 2.6181634843 | KS>SC | TM | YCL005W | LDB16 | n |
| W303.asm02.HP0_G0005480 | YCL004W | chrIII | 629.9654093 | 1498.833416 | 1125.326985 | 1.243506009 | 0.09810706 | 4.3714230466 | 3.1520260915 | KS>SC | TM | YCL004W | PGS1 | n |
| W303.asm02.HP0_G0005490 | YCL002C | chrIII | 956.0613775 | 1045.996621 | 1274.186365 | 0.113354429 | 0.15599307 | 0.41751944 | 0.468853797 | ns | TM | YCL002C | | n |
| W303.asm02.HP0_G0005500 | YCL001W | chrIII | 2271.774029 | 987.7217839 | 1826.473456 | -1.183684657 | 0.14159949 | 3.6473558924 | 1.2187506274 | KS<SC | TM | YCL001W | RER1 | n |
| W303.asm02.HP0_G0005510 | YCL001W-A | chrIII | 55.91301164 | 202.0053451 | 172.3737645 | 1.772046543 | 0.24415904 | 1.0944070701 | 2.9986753720 | KS>SC | TM | YCL001W-A | | n |
| W303.asm02.HP0_G0005520 | YCL001W-B | chrIII | 69.22515836 | 938.3120811 | 612.1595912 | 3.704423942 | 0.20386098 | 1.3717733579 | 2.3491618755 | KS>SC | TM | YCL001W-B | | n |
| W303.asm02.HP0_G0005530 | YCR002C | chrIII | 3797.599014 | 2681.797571 | 3389.825775 | -0.49827483 | 0.09267015 | 6.8578583898 | 1.5153654829 | ns | TM | YCR002C | CDC10 | n |
| W303.asm02.HP0_G0005540 | YCR003W | chrIII | 1847.710584 | 1108.690222 | 1711.638185 | -0.718203434 | 0.18258598 | 6.7697494752 | 0.000122034 | ns | TM | YCR003W | MRPL32 | n |
| W303.asm02.HP0_G0005550 | YCR004C | chrIII | 4307.589669 | 1347.263231 | 2809.503227 | -1.640555469 | 0.17688774 | 6.0299230277 | 2.7536648493 | KS<SC | TM | YCR004C | YCP4 | n |
| W303.asm02.HP0_G0005560 | YCR005C | chrIII | 7645.507948 | 746.9132263 | 3416.448465 | -3.313576047 | 0.18572934 | 1.0104363070 | 1.5381086006 | KS<SC | TM | YCR005C | CIT2 | n |
| W303.asm02.HP0_G0005570 | W303.asm02.HP0_G0 | chrIII | 68.54920082 | 142.384222 | 126.1214297 | 1.027634376 | 0.2028608 | 2.7049435282 | 5.5310039309 | KS>SC | TM | YCR006C | | y |
| W303.asm02.HP0_G0005580 | YCR007C | chrIII | 243.0513532 | 459.8521694 | 451.0619636 | 0.899818121 | 0.1834763 | 6.8495732986 | 1.3599877419 | ns | TM | YCR007C | DFP3 | n |
| W303.asm02.HP0_G0005590 | YCR008W | chrIII | 3118.840158 | 3897.971131 | 4030.58458 | 0.316323269 | 0.12103786 | 0.006947445 | 0.009614141 | ns | TM | YCR008W | SAT4 | n |
| W303.asm02.HP0_G0005600 | YCR009C | chrIII | 4446.945735 | 1994.34009 | 3351.187025 | -1.141206625 | 0.13405982 | 1.0214308991 | 3.4984008294 | KS<SC | TM | YCR009C | RVS161 | n |
| W303.asm02.HP0_G0005610 | YCR010C | chrIII | 99.82918514 | 505.29405 | 376.2484175 | 2.290605514 | 0.20708279 | 1.5499941722 | 8.8478833999 | KS>SC | TM | YCR010C | ADY2 | n |
| W303.asm02.HP0_G0005620 | YCR011C | chrIII | 3824.551048 | 5781.116306 | 4008.211727 | 0.590495264 | 0.11337082 | 1.6571558163 | 3.4398537400 | ns | TM | YCR011C | ADP1 | n |

| | | | | | | | | | | | | | | |
|---|---|---|---|---|---|---|---|---|---|---|---|---|---|---|
| W303.asm02.HP0_G0005630 | YCR012W | chrIII | 65412.71647 | 14724.67882 | 31329.98713 | -2.133403359 | 0.11046795 | 4.7264888362 | 1.0792149509 | KS<SC | TM | YCR012W | PGK1 | n |
| W303.asm02.HP0_G0005640 | YCR014C | chrIII | 736.9820261 | 1789.112286 | 1377.405532 | 1.259115019 | 0.14797625 | 9.4437353939 | 3.3174147409 | KS>SC | TM | YCR014C | POL4 | n |
| W303.asm02.HP0_G0005650 | YCR015C | chrIII | 696.2880282 | 822.3446303 | 873.6436017 | 0.223322733 | 0.16558321 | 0.136097857 | 0.165003596 | ns | TM | YCR015C | CTO1 | n |
| W303.asm02.HP0_G0005660 | YCR016W | chrIII | 2050.838723 | 3093.006458 | 3476.357041 | 0.582489009 | 0.15067255 | 9.5339817035 | 0.000163269 | ns | TM | YCR016W | RBP95 | n |
| W303.asm02.HP0_G0005670 | YCR017C | chrIII | 3981.610279 | 2165.22292 | 3263.383474 | -0.861945488 | 0.16308083 | 9.4008312446 | 2.0123654383 | ns | TM | YCR017C | CWH43 | n |
| W303.asm02.HP0_G0005680 | YCR018C | chrIII | 1307.545467 | 619.2442427 | 1067.342272 | -1.055848725 | 0.16631517 | 1.4126950435 | 3.2803257792 | KS<SC | TM | YCR018C | SRD1 | n |
| W303.asm02.HP0_G0005800 | YCR028C | chrIII | 1213.352337 | 1573.091079 | 1627.578649 | 0.369707683 | 0.1141276 | 0.001040519 | 0.001601697 | ns | TM | YCR028C | FEN2 | n |
| W303.asm02.HP0_G0005810 | YCR028C-A | chrIII | 3886.372054 | 1419.383901 | 2422.230694 | -1.426189086 | 0.16069231 | 3.1270024678 | 1.2239981088 | KS<SC | TM | YCR028C-A | RIM1 | n |
| W303.asm02.HP0_G0005820 | YCR030C | chrIII | 2278.023127 | 2288.30131 | 2071.875465 | 0.00565018 | 0.29688517 | 0.983528582 | 0.983528582 | ns | TM | YCR030C | SYP1 | n |
| W303.asm02.HP0_G0005830 | YCR031C/YJL191W | chrIII | 13299.38215 | 7849.410538 | 10562.69539 | -0.753555439 | 0.11150598 | 1.1189506852 | 2.8923819599 | ns | TM | YCR031C | RPS14A | y |
| W303.asm02.HP0_G0005840 | YCR032W | chrIII | 1631.453176 | 3406.751737 | 2250.104759 | 1.048983166 | 0.13409165 | 3.3613812295 | 1.0418019719 | KS>SC | TM | YCR032W | BPH1 | n |
| W303.asm02.HP0_G0005850 | YCR033W | chrIII | 1971.2986 | 3034.502847 | 2530.895856 | 0.607058088 | 0.18113954 | 0.000664557 | 0.001046487 | ns | TM | YCR033W | SNT1 | n |
| W303.asm02.HP0_G0005860 | YCR034W | chrIII | 5746.023683 | 3390.754253 | 5411.274469 | -0.751800581 | 0.1226538 | 7.0648817821 | 1.5867029576 | ns | TM | YCR034W | ELO2 | n |
| W303.asm02.HP0_G0005870 | YCR035C | chrIII | 3431.542107 | 3997.769497 | 3548.457755 | 0.210699826 | 0.10870695 | 0.03536689 | 0.044863555 | ns | TM | YCR035C | RRP43 | n |
| W303.asm02.HP0_G0005880 | YCR036W | chrIII | 2301.839828 | 4821.535053 | 3412.094514 | 1.050072641 | 0.15019282 | 1.7728897611 | 4.6708826400 | KS>SC | TM | YCR036W | RBK1 | n |
| W303.asm02.HP0_G0005890 | YCR037C | chrIII | 2288.892579 | 1152.5131 | 1909.606422 | -0.978643635 | 0.1234247 | 1.5179752423 | 4.8363397254 | ns | TM | YCR037C | PHO87 | n |
| W303.asm02.HP0_G0005900 | YCR038C | chrIII | 309.8929684 | 392.2175221 | 368.4914114 | 0.331920752 | 0.13023064 | 0.008326674 | 0.011407543 | ns | TM | YCR038C | BUD5 | n |
| W303.asm02.HP0_G0005920 | YCR042C | chrIII | 2353.119475 | 2395.260359 | 2384.956457 | 0.020204556 | 0.09482642 | 0.808590857 | 0.845625552 | ns | TM | YCR042C | TAF2 | n |
| W303.asm02.HP0_G0005930 | YCR043C | chrIII | 1890.486994 | 333.6651168 | 1395.218446 | -2.441949358 | 0.20075194 | 2.2016902984 | 1.5081578544 | KS<SC | TM | YCR043C | | n |
| W303.asm02.HP0_G0005940 | YCR044C | chrIII | 1362.833895 | 364.0721763 | 1021.454551 | -1.87837728 | 0.12239094 | 7.9301345722 | 9.8766221490 | KS<SC | TM | YCR044C | PER1 | n |
| W303.asm02.HP0_G0005950 | YCR045C | chrIII | 526.6409424 | 2298.254089 | 1329.097377 | 2.062857571 | 0.21439885 | 9.9666356198 | 4.8765324282 | KS>SC | TM | YCR045C | RRT12 | n |
| W303.asm02.HP0_G0005960 | YCR046C | chrIII | 2582.204624 | 987.4977049 | 1569.660751 | -1.360859464 | 0.15906543 | 5.6811069798 | 2.0481885690 | KS<SC | TM | YCR046C | IMG1 | n |
| W303.asm02.HP0_G0005970 | YCR047C | chrIII | 1996.272178 | 1908.70054 | 2026.885355 | -0.055562966 | 0.19226236 | 0.752594258 | 0.799595015 | ns | TM | YCR047C | BUD23 | n |
| W303.asm02.HP0_G0005980 | YCR048W | chrIII | 1616.381411 | 3158.648989 | 2380.807295 | 0.906088044 | 0.30288265 | 0.001889203 | 0.002724429 | ns | TM | YCR048W | ARE1 | n |
| W303.asm02.HP0_G0005990 | YCR051W | chrIII | 3369.314736 | 3334.391003 | 3541.59545 | -0.011086183 | 0.084086 | 0.879846574 | 0.906308125 | ns | TM | YCR051W | | n |
| W303.asm02.HP0_G0006000 | YCR052W | chrIII | 3020.9416 | 3312.612766 | 2937.406491 | 0.115014407 | 0.21352365 | 0.555836057 | 0.614109192 | ns | TM | YCR052W | RSC6 | n |
| W303.asm02.HP0_G0006010 | YCR053W | chrIII | 9944.662419 | 10098.60835 | 10002.50807 | 0.015695091 | 0.0626072 | 0.765781505 | 0.807015894 | ns | TM | YCR053W | THR4 | n |
| W303.asm02.HP0_G0006020 | YCR054C | chrIII | 1040.33212 | 1252.686272 | 1473.674431 | 0.250860596 | 0.1522808 | 0.072152607 | 0.089862793 | ns | TM | YCR054C | CTR86 | n |
| W303.asm02.HP0_G0006030 | YCR057C | chrIII | 3485.821897 | 5926.461458 | 4820.360301 | 0.724492675 | 0.27379322 | 0.005943283 | 0.00831299 | ns | TM | YCR057C | PWP2 | n |
| W303.asm02.HP0_G0006040 | YCR059C | chrIII | 2207.42815 | 3027.868581 | 2933.921788 | 0.450549303 | 0.12906887 | 0.000424612 | 0.000692521 | ns | TM | YCR059C | YIH1 | n |
| W303.asm02.HP0_G0006050 | YCR060W | chrIII | 1477.00347 | 1414.166857 | 1398.70489 | -0.050746537 | 0.1267411 | 0.654158372 | 0.716957576 | ns | TM | YCR060W | TAH1 | n |
| W303.asm02.HP0_G0006060 | YCR061W | chrIII | 781.9384501 | 724.3268772 | 707.4412929 | -0.095072756 | 0.26302506 | 0.694535374 | 0.749223199 | ns | TM | YCR061W | TVS1 | n |
| W303.asm02.HP0_G0006070 | YCR063W | chrIII | 272.1883472 | 377.8462117 | 390.7163254 | 0.469291647 | 0.1208113 | 9.2272979298 | 0.000160018 | ns | TM | YCR063W | BUD31 | n |
| W303.asm02.HP0_G0006080 | YCR065W | chrIII | 1026.989692 | 1436.301509 | 1190.631669 | 0.465745883 | 0.20658448 | 0.018257534 | 0.023597001 | ns | TM | YCR065W | HCM1 | n |
| W303.asm02.HP0_G0006090 | YCR066W | chrIII | 565.6104313 | 897.8544491 | 777.1183729 | 0.656570949 | 0.0988475 | 2.6029259826 | 6.4836519931 | ns | TM | YCR066W | RAD18 | n |
| W303.asm02.HP0_G0006100 | YCR067C | chrIII | 2827.053433 | 3324.05984 | 2928.065972 | 0.214409971 | 0.16791479 | 0.157338526 | 0.18743807 | ns | TM | YCR067C | SED4 | n |
| W303.asm02.HP0_G0006110 | YCR068W | chrIII | 678.9986962 | 1657.723692 | 1068.08497 | 1.177346077 | 0.36165099 | 0.000629114 | 0.001002193 | KS>SC | TM | YCR068W | ATG15 | n |
| W303.asm02.HP0_G0006120 | YCR069W | chrIII | 2167.631732 | 1449.629839 | 1523.61182 | -0.554322624 | 0.23687248 | 0.014490065 | 0.019087874 | ns | TM | YCR069W | CPR4 | n |
| W303.asm02.HP0_G0006130 | YCR071C | chrIII | 1881.359891 | 614.2855984 | 1242.703506 | -1.582905376 | 0.16602543 | 5.5358253808 | 2.6152002661 | KS<SC | TM | YCR071C | IMG2 | n |
| W303.asm02.HP0_G0006140 | YCR072C | chrIII | 1272.853426 | 2058.491999 | 1902.591114 | 0.677734996 | 0.1654319 | 3.4743367266 | 6.4322179939 | ns | TM | YCR072C | RSA4 | n |
| W303.asm02.HP0_G0006150 | YCR073C | chrIII | 1166.741042 | 6661.170974 | 3202.971991 | 2.461521853 | 0.18819905 | 1.7731299765 | 1.3495489265 | KS>SC | TM | YCR073C | SSK22 | n |
| W303.asm02.HP0_G0006160 | YCR073W-A | chrIII | 1083.279852 | 2236.802018 | 1202.246529 | 1.011550199 | 0.21656374 | 2.0164456418 | 3.8908880695 | KS>SC | TM | YCR073W-A | SOL2 | n |
| W303.asm02.HP0_G0006170 | YCR075C | chrIII | 565.0057816 | 299.9791498 | 427.6849427 | -0.885634929 | 0.1783177 | 5.0254473958 | 1.0124798429 | ns | TM | YCR075C | ERS1 | n |
| W303.asm02.HP0_G0006180 | YCR075W-A | chrIII | 243.6097861 | 151.4774649 | 192.5681641 | -0.640881148 | 0.26188135 | 0.010660254 | 0.014179173 | ns | TM | YCR075W-A | EGO2 | n |
| W303.asm02.HP0_G0006190 | YCR076C | chrIII | 732.1492753 | 786.2748023 | 771.2515063 | 0.086729759 | 0.13223974 | 0.45985225 | 0.512193156 | ns | TM | YCR076C | FUB1 | n |
| W303.asm02.HP0_G0006200 | YCR077C | chrIII | 3009.740662 | 2510.663117 | 2278.318609 | -0.23414516 | 0.24031091 | 0.285385468 | 0.325815076 | ns | TM | YCR077C | PAT1 | n |
| W303.asm02.HP0_G0006210 | YCR079W | chrIII | 835.1604876 | 2210.400606 | 1459.1903 | 1.389065529 | 0.1230804 | 7.1291200956 | 4.2464758830 | KS>SC | TM | YCR079W | PTC6 | n |
| W303.asm02.HP0_G0006220 | YCR081W | chrIII | 1184.531814 | 3064.644951 | 2437.626666 | 1.361705902 | 0.08869708 | 1.5996919449 | 1.8263149704 | KS>SC | TM | YCR081W | SRB8 | n |
| W303.asm02.HP0_G0006230 | YCR082W | chrIII | 2154.530233 | 1441.089526 | 1710.527445 | -0.569755769 | 0.14558866 | 7.8938666502 | 0.000138649 | ns | TM | YCR082W | AHC2 | n |
| W303.asm02.HP0_G0006240 | YCR083W | chrIII | 1080.346405 | 451.9250724 | 839.7655168 | -1.22246967 | 0.19323866 | 1.4089522700 | 3.2803257792 | KS<SC | TM | YCR083W | TRX3 | n |
| W303.asm02.HP0_G0006250 | YCR084C | chrIII | 3830.951126 | 5477.946272 | 3483.041071 | 0.445825161 | 0.3945701 | 0.212128921 | 0.248390274 | ns | TM | YCR084C | TUP1 | n |
| W303.asm02.HP0_G0006260 | YCR086W | chrIII | 563.8482967 | 470.4506023 | 587.5539029 | -0.245060293 | 0.15264572 | 0.078924987 | 0.096878838 | ns | TM | YCR086W | CSM1 | n |
| W303.asm02.HP0_G0006270 | YCR087C-A | chrIII | 1359.477546 | 1949.004875 | 2059.819745 | 0.511427602 | 0.13751353 | 0.000175871 | 0.000293833 | ns | TM | YCR087C-A | | n |
| W303.asm02.HP0_G0006280 | YCR088W | chrIII | 2653.93402 | 3770.397115 | 2590.006483 | 0.423705902 | 0.44208305 | 0.284742544 | 0.325815076 | ns | TM | YCR088W | ABP1 | n |
| W303.asm02.HP0_G0006290 | YCR089W | chrIII | 523.4606261 | 1211.618055 | 788.1735371 | 1.141089565 | 0.2902882 | 5.0457302095 | 9.2168671828 | KS>SC | TM | YCR089W | FIG2 | n |
| W303.asm02.HP0_G0006300 | YCR090C | chrIII | 1464.229305 | 2062.397116 | 1873.774891 | 0.485117211 | 0.15592658 | 0.001561469 | 0.002325232 | ns | TM | YCR090C | | n |
| W303.asm02.HP0_G0006310 | YCR091W | chrIII | 238.4694632 | 817.615165 | 551.4549167 | 1.74549174 | 0.16860487 | 1.1630374695 | 6.1283128201 | KS>SC | TM | YCR091W | KIN82 | n |
| W303.asm02.HP0_G0006320 | YCR092C | chrIII | 1020.060042 | 1354.528331 | 1151.072338 | 0.398449673 | 0.15962958 | 0.009707915 | 0.013168162 | ns | TM | YCR092C | MSH3 | n |
| W303.asm02.HP0_G0006330 | YCR093W | chrIII | 4222.073642 | 5390.578998 | 4487.939788 | 0.34210069 | 0.15345988 | 0.019165921 | 0.024539543 | ns | TM | YCR093W | CDC39 | n |
| W303.asm02.HP0_G0006340 | YCR094W | chrIII | 1506.214334 | 1978.664163 | 1837.138468 | 0.391257631 | 0.10457933 | 0.000167163 | 0.000282732 | ns | TM | YCR094W | CDC50 | n |
| W303.asm02.HP0_G0006350 | YCR095C | chrIII | 1067.845165 | 836.8103643 | 1025.215307 | -0.330619906 | 0.20364386 | 0.079200218 | 0.096878838 | ns | TM | YCR095C | OCA4 | n |
| W303.asm02.HP0_G0006360 | YCR095W-A | chrIII | 76.57338353 | 151.849323 | 117.3444562 | 0.975721171 | 0.18534348 | 9.7334127285 | 2.0515039135 | ns | TM | YCR095W-A | | n |
| W303.asm02.HP0_G0006380 | YCR098C | chrIII | 3765.439691 | 1689.995764 | 2477.889803 | -1.064973194 | 0.34412149 | 0.001197319 | 0.001802557 | KS<SC | TM | YCR098C | GIT1 | n |
| W303.asm02.HP0_G0006390 | W303.asm02.HP0_G0( | chrIII | 489.2193902 | 880.3584836 | 543.2631845 | 0.82219951 | 0.21033294 | 7.0488395398 | 0.000125414 | ns | TM | YNR065C | | y |
| W303.asm02.HP0_G0006400 | YCR100C | chrIII | 804.368301 | 2957.479806 | 1452.925958 | 1.808918964 | 0.24033674 | 1.3591921012 | 4.0480286494 | KS>SC | TM | YCR100C | EMA35 | n |
| W303.asm02.HP0_G0006410 | YCR101C | chrIII | 423.5789992 | 1513.84701 | 842.7599021 | 1.772764699 | 0.23621272 | 1.7058347288 | 4.9723267629 | KS>SC | TM | YCR101C | | n |
| W303.asm02.HP0_G0006420 | YCR102C | chrIII | 1454.732219 | 2972.503703 | 2418.09028 | 0.990380044 | 0.24197204 | 2.8936950642 | 5.5060586638 | ns | TM | YCR102C | | n |
| W303.asm02.HP0_G0006430 | YCR104W/YIR041W/Y | chrIII | 74.55587382 | 264.5757899 | 196.0184986 | 1.802752508 | 0.19525074 | 6.8780176969 | 3.0396400789 | KS>SC | TM | YMR325W | PAU19 | y |

| | | | | | | | | | | | | | | |
|---|---|---|---|---|---|---|---|---|---|---|---|---|---|---|
| W303.asm02.HP0_G0006440 | YCR105W | chrIII | 767.2364569 | 1117.236365 | 1036.798106 | 0.536502645 | 0.11135105 | 1.2908943494( | 2.5264646552( | ns | TM | YCR105W | ADH7 | n |
| W303.asm02.HP0_G0006450 | YCR106W | chrIII | 2235.636399 | 1202.015337 | 1741.030874 | -0.886939715 | 0.11058763 | 7.74541121198 | 2.52647937152 | ns | TM | YCR106W | RDS1 | n |
| W303.asm02.HP0_G0006460 | YCR107W | chrIII | 4149.924619 | 1320.926181 | 2595.446155 | -1.634057103 | 0.11884498 | 1.69835503725 | 1.45421650065 | KS<SC | TM | YCR107W | AAD3 | n |
| W303.asm02.HP0_G0004930 | YCL073C/YKR106W | chrIII | 33.15413562 | 5711.027746 | 2987.979787 | 7.329954921 | 0.34029006 | 3.25231122878 | 8.91133276687 | KS>SC | NaCl | YCL073C | GEX1 | y |
| W303.asm02.HP0_G0004980 | YCL064C | chrIII | 938.8599975 | 1271.911606 | 1211.469492 | 0.416244778 | 0.22881387 | 0.062664217 | 0.076651766 | ns | NaCl | YCL064C | CHA1 | n |
| W303.asm02.HP0_G0004990 | YCL063W | chrIII | 2005.637196 | 1406.799247 | 1887.140178 | -0.508711925 | 0.08087843 | 2.79096247761 | 6.48070948191 | ns | NaCl | YCL063W | VAC17 | n |
| W303.asm02.HP0_G0005000 | YCL061C | chrIII | 1580.622199 | 3555.367187 | 3053.729675 | 1.162557406 | 0.09939575 | 6.59772758644 | 4.51944339671 | KS>SC | NaCl | YCL061C | MRC1 | n |
| W303.asm02.HP0_G0005010 | YCL059C | chrIII | 6441.010101 | 7649.127971 | 6586.286877 | 0.244619955 | 0.11641466 | 0.03446702 | 0.04413067 | ns | NaCl | YCL059C | KRR1 | n |
| W303.asm02.HP0_G0005020 | YCL058W-A | chrIII | 1559.518111 | 817.1443654 | 1078.478048 | -0.903395822 | 0.19616037 | 2.75610299152 | 5.17241246354 | ns | NaCl | YCL058W-A | ADF1 | n |
| W303.asm02.HP0_G0005030 | YCL057C-A | chrIII | 4439.299385 | 381.3457272 | 2041.01948 | -3.514732318 | 0.16185152 | 1.51141803092 | 6.90214234121 | KS<SC | NaCl | YCL057C-A | MIC10 | n |
| W303.asm02.HP0_G0005040 | YCL057W | chrIII | 6923.368715 | 962.9056787 | 3546.262488 | -2.751106869 | 0.27249847 | 8.33411806043 | 4.39143913184 | KS<SC | NaCl | YCL057W | PRD1 | n |
| W303.asm02.HP0_G0005050 | YCL056C | chrIII | 1140.151563 | 1007.335111 | 1042.157888 | -0.175337868 | 0.17131668 | 0.299730265 | 0.3311536 | ns | NaCl | YCL056C | PEX34 | n |
| W303.asm02.HP0_G0005060 | YCL055W | chrIII | 2742.370756 | 1470.91427 | 2117.2546 | -0.880908275 | 0.15624059 | 1.1729203104( | 2.5506362305( | ns | NaCl | YCL055W | KAR4 | n |
| W303.asm02.HP0_G0005070 | YCL054W | chrIII | 8228.568137 | 7850.401799 | 8401.514345 | -0.067129927 | 0.09513847 | 0.478557051 | 0.512975268 | ns | NaCl | YCL054W | SPB1 | n |
| W303.asm02.HP0_G0005080 | YCL052C | chrIII | 5459.115651 | 1267.717105 | 3501.202677 | -2.088849035 | 0.1122966 | 2.55277646327 | 4.3716296933( | KS<SC | NaCl | YCL052C | PBN1 | n |
| W303.asm02.HP0_G0005090 | YCL051W | chrIII | 2244.906317 | 861.6096846 | 1712.553656 | -1.367636401 | 0.11451741 | 2.6469367094( | 1.90858067994 | KS<SC | NaCl | YCL051W | LRE1 | n |
| W303.asm02.HP0_G0005100 | YCL050C | chrIII | 13554.2371 | 9344.774977 | 10811.35482 | -0.530931268 | 0.10649416 | 5.37171805997 | 1.06655851335 | ns | NaCl | YCL050C | APA1 | n |
| W303.asm02.HP0_G0005110 | YCL049C | chrIII | 3589.78311 | 2546.641604 | 3002.426132 | -0.483693259 | 0.15963338 | 0.002173246 | 0.003134049 | ns | NaCl | YCL049C | | n |
| W303.asm02.HP0_G0005120 | YCL048W-A | chrIII | 190.7522344 | 741.5849177 | 424.7356513 | 1.567518873 | 0.58751758 | 0.001986749 | 0.002895582 | KS>SC | NaCl | YCL048W-A | | n |
| W303.asm02.HP0_G0005130 | YCL048W | chrIII | 181.2415336 | 7257.392774 | 3835.883642 | 5.289954306 | 0.24282027 | 1.49657642959 | 5.1257742713( | KS>SC | NaCl | YCL048W | SPS22 | n |
| W303.asm02.HP0_G0005140 | YCL047C | chrIII | 1836.337807 | 488.3184031 | 1285.045077 | -1.892434796 | 0.12075431 | 3.17848140143 | 3.62876626663 | KS<SC | NaCl | YCL047C | POF1 | n |
| W303.asm02.HP0_G0005150 | YCL045C | chrIII | 9546.814684 | 4145.506846 | 6014.488977 | -1.189233035 | 0.11877746 | 6.50691160084 | 3.18373889041 | KS<SC | NaCl | YCL045C | EMC1 | n |
| W303.asm02.HP0_G0005160 | YCL044C | chrIII | 1444.729908 | 1178.153614 | 1679.464284 | -0.270359205 | 0.31646017 | 0.372565842 | 0.405091431 | ns | NaCl | YCL044C | MGR1 | n |
| W303.asm02.HP0_G0005170 | YCL043C | chrIII | 16171.25112 | 10675.98134 | 15515.30826 | -0.587488003 | 0.14735691 | 5.6460422900( | 9.54947893494 | ns | NaCl | YCL043C | PDI1 | n |
| W303.asm02.HP0_G0005180 | YCL040W | chrIII | 15815.97866 | 1796.500687 | 6276.778436 | -3.086436988 | 0.20765017 | 2.22945305289 | 2.3495005249( | KS<SC | NaCl | YCL040W | GLK1 | n |
| W303.asm02.HP0_G0005190 | YCL039W | chrIII | 1869.136316 | 1347.067444 | 1508.467595 | -0.466000892 | 0.12488768 | 0.000170851 | 0.00027865 | ns | NaCl | YCL039W | GID7 | n |
| W303.asm02.HP0_G0005200 | YCL038C | chrIII | 1988.292546 | 1508.546484 | 1815.784429 | -0.390673454 | 0.13523649 | 0.003576437 | 0.005051257 | ns | NaCl | YCL038C | ATG22 | n |
| W303.asm02.HP0_G0005210 | YCL037C | chrIII | 165.3515716 | 805.5783531 | 527.5743471 | 2.189609095 | 0.26861789 | 2.72523976161 | 1.0371051315( | KS>SC | NaCl | YCL037C | SRO9 | n |
| W303.asm02.HP0_G0005220 | YCL036W | chrIII | 2457.544256 | 6526.85747 | 4059.236207 | 1.361457363 | 0.21404459 | 7.87077688754 | 1.89174812911 | KS>SC | NaCl | YCL036W | GFD2 | n |
| W303.asm02.HP0_G0005230 | YCL035C | chrIII | 6021.873388 | 2029.617107 | 3711.433848 | -1.533754826 | 0.17431326 | 4.06353330183 | 1.59058303529 | KS<SC | NaCl | YCL035C | GRX1 | n |
| W303.asm02.HP0_G0005240 | YCL034W | chrIII | 4946.20551 | 1649.035589 | 2898.909704 | -1.5687624 | 0.11400017 | 1.15325645032 | 1.0533075579( | KS<SC | NaCl | YCL034W | LSB5 | n |
| W303.asm02.HP0_G0005250 | YCL033C | chrIII | 3261.071912 | 483.0789347 | 1683.466772 | -2.674879975 | 0.24770347 | 6.26911792182 | 3.73421371865 | KS<SC | NaCl | YCL033C | MXR2 | n |
| W303.asm02.HP0_G0005260 | YCL032W | chrIII | 2846.584862 | 1009.154031 | 1936.000195 | -1.487573561 | 0.08314243 | 4.01973734681 | 6.11893351682 | KS<SC | NaCl | YCL032W | STE50 | n |
| W303.asm02.HP0_G0005270 | YCL031C | chrIII | 5596.943649 | 6799.724582 | 6114.993332 | 0.277048354 | 0.11789518 | 0.018020763 | 0.024204359 | ns | NaCl | YCL031C | RRP7 | n |
| W303.asm02.HP0_G0005280 | YCL030C | chrIII | 7920.800726 | 2057.712461 | 5097.396452 | -1.916111684 | 0.14665581 | 6.87317079285 | 5.88515249138 | KS<SC | NaCl | YCL030C | HIS4 | n |
| W303.asm02.HP0_G0005290 | YCL029C | chrIII | 2075.18733 | 1430.87132 | 2021.981501 | -0.531345626 | 0.10204978 | 1.67032815764 | 3.41544712832 | ns | NaCl | YCL029C | BIK1 | n |
| W303.asm02.HP0_G0005300 | YCL028W | chrIII | 9227.938775 | 2484.156037 | 5753.045622 | -1.886140717 | 0.07181525 | 6.09868064623 | 8.35519248534 | KS<SC | NaCl | YCL028W | RNQ1 | n |
| W303.asm02.HP0_G0005310 | YCL027W | chrIII | 682.7832852 | 203.3367298 | 483.300876 | -1.673640196 | 0.25453914 | 1.14201523005 | 3.06776640231 | KS<SC | NaCl | YCL027W | FUS1 | n |
| W303.asm02.HP0_G0005320 | YCL026C-B | chrIII | 1087.054726 | 933.4402743 | 1007.388914 | -0.211248717 | 0.20748376 | 0.299433859 | 0.3311536 | ns | NaCl | YCL026C-B | HBN1 | n |
| W303.asm02.HP0_G0005330 | YCL026C-A | chrIII | 400.7608909 | 7695.855974 | 4678.985084 | 4.220398443 | 0.24134228 | 3.44932552422 | 4.72557596819 | KS>SC | NaCl | YCL026C-A | FRM2 | n |
| W303.asm02.HP0_G0005340 | YCL025C | chrIII | 7635.121487 | 3489.860428 | 5731.653313 | -1.100631579 | 0.17732248 | 2.95693680112 | 6.75167236255 | KS<SC | NaCl | YCL025C | AGP1 | n |
| W303.asm02.HP0_G0005350 | YCL024W | chrIII | 2307.600196 | 1317.616319 | 1965.314321 | -0.799215617 | 0.10618667 | 3.78946853182 | 1.17990270195 | ns | NaCl | YCL024W | KCC4 | n |
| W303.asm02.HP0_G0005360 | YCL021W-A | chrIII | 275.7437486 | 409.8480363 | 383.3625747 | 0.556276425 | 0.1636362 | 0.000578769 | 0.000921993 | ns | NaCl | YCL021W-A | | n |
| W303.asm02.HP0_G0005370 | YCL018W | chrIII | 1616.583255 | 1008.872141 | 1447.963445 | -0.670025846 | 0.1439055 | 2.5852516354( | 4.91915936181 | ns | NaCl | YCL018W | LEU2 | n |
| W303.asm02.HP0_G0005380 | YCL017C | chrIII | 4055.831372 | 2077.737983 | 2929.661665 | -0.95320278 | 0.11483089 | 6.5443005901( | 2.29889533552 | ns | NaCl | YCL017C | NFS1 | n |
| W303.asm02.HP0_G0005390 | YCL016C | chrIII | 896.4733121 | 826.9653908 | 950.4497386 | -0.118574064 | 0.11345799 | 0.29309199 | 0.329127891 | ns | NaCl | YCL016C | DCC1 | n |
| W303.asm02.HP0_G0005400 | YCL014W | chrIII | 1190.373718 | 3705.66019 | 2400.236527 | 1.602877686 | 0.17808726 | 5.84994832721 | 2.4286149116( | KS>SC | NaCl | YCL014W | BUD3 | n |
| W303.asm02.HP0_G0005410 | YCL012C | chrIII | 508.6515372 | 681.4723606 | 655.9406413 | 0.398667164 | 0.26437077 | 0.120061314 | 0.141796552 | ns | NaCl | YCL012C | | n |
| W303.asm02.HP0_G0005420 | YCL011C | chrIII | 6016.394343 | 5879.625624 | 4931.803851 | -0.032554575 | 0.14536101 | 0.821112856 | 0.858721078 | ns | NaCl | YCL011C | GBP2 | n |
| W303.asm02.HP0_G0005430 | YCL010C | chrIII | 1031.400772 | 1017.550862 | 870.6100674 | -0.018489179 | 0.24278469 | 0.937726167 | 0.944621213 | ns | NaCl | YCL010C | SGF29 | n |
| W303.asm02.HP0_G0005440 | YCL009C | chrIII | 3985.987587 | 6109.606409 | 4104.665729 | 0.544551946 | 0.38791337 | 0.134472342 | 0.157459067 | ns | NaCl | YCL009C | ILV6 | n |
| W303.asm02.HP0_G0005450 | YCL008C | chrIII | 581.8493072 | 839.7538919 | 549.3495235 | 0.500038036 | 0.26592161 | 0.052456561 | 0.065332263 | ns | NaCl | YCL008C | STP22 | n |
| W303.asm02.HP0_G0005460 | W303.asm02.HP0_G0( | chrIII | 1054.970115 | 509.2034984 | 782.3178261 | -1.029436116 | 0.15721796 | 3.4476267472( | 8.74675674753 | KS<SC | NaCl | YCL005W-A | VMA9 | y |
| W303.asm02.HP0_G0005470 | YCL005W | chrIII | 1150.147101 | 1505.628587 | 1352.864291 | 0.386452412 | 0.09867227 | 8.33027605135 | 0.000139177 | ns | NaCl | YCL005W | LDB16 | n |
| W303.asm02.HP0_G0005480 | YCL004W | chrIII | 776.5030285 | 1504.781934 | 1125.326985 | 0.944562302 | 0.09875806 | 7.1760512532( | 3.0722469428( | ns | NaCl | YCL004W | PGS1 | n |
| W303.asm02.HP0_G0005490 | YCL002C | chrIII | 1087.780605 | 1316.0674 | 1274.186365 | 0.268392442 | 0.16011014 | 0.089919525 | 0.107121521 | ns | NaCl | YCL002C | | n |
| W303.asm02.HP0_G0005500 | YCL001W | chrIII | 2448.24807 | 1179.81154 | 1826.473456 | -1.035622684 | 0.14204757 | 1.80450842432 | 5.15036779443 | KS<SC | NaCl | YCL001W | RER1 | n |
| W303.asm02.HP0_G0005510 | YCL001W-A | chrIII | 105.9339465 | 215.3071431 | 172.3737645 | 0.975665318 | 0.2328022 | 1.73585861285 | 3.0488798713( | ns | NaCl | YCL001W-A | | n |
| W303.asm02.HP0_G0005520 | YCL001W-B | chrIII | 143.5715085 | 1284.186758 | 612.1595912 | 3.113500801 | 0.18678378 | 7.38825842634 | 9.20174004009 | KS>SC | NaCl | YCL001W-B | | n |
| W303.asm02.HP0_G0005530 | YCR002C | chrIII | 4603.186384 | 3000.126542 | 3389.825775 | -0.612512023 | 0.09291643 | 3.5974515565( | 8.96092478619 | ns | NaCl | YCR002C | CDC10 | n |
| W303.asm02.HP0_G0005540 | YCR003W | chrIII | 1980.808168 | 1357.325864 | 1711.638185 | -0.529333721 | 0.18242935 | 0.003223626 | 0.004600383 | ns | NaCl | YCR003W | MRPL32 | n |
| W303.asm02.HP0_G0005550 | YCR004C | chrIII | 4262.424021 | 1495.138942 | 2809.503227 | -1.475433356 | 0.17762583 | 3.20496202087 | 1.18670215367 | KS<SC | NaCl | YCR004C | YCP4 | n |
| W303.asm02.HP0_G0005560 | YCR005C | chrIII | 7859.694165 | 483.1145904 | 3416.448465 | -3.997263763 | 0.18841289 | 8.48905330055 | 1.93833383695 | KS<SC | NaCl | YCR005C | CIT2 | n |
| W303.asm02.HP0_G0005570 | W303.asm02.HP0_G0( | chrIII | 95.22338038 | 134.8469286 | 126.1214297 | 0.467502594 | 0.20297598 | 0.018978013 | 0.0252426 | ns | NaCl | YCR006C | | y |
| W303.asm02.HP0_G0005580 | YCR007C | chrIII | 570.727844 | 568.0118377 | 451.0619636 | -0.009380583 | 0.1780607 | 0.957388336 | 0.957388336 | ns | NaCl | YCR007C | DFP3 | n |
| W303.asm02.HP0_G0005590 | YCR008W | chrIII | 2441.655002 | 5212.785099 | 4030.58458 | 1.080141538 | 0.11931117 | 7.68376110161 | 3.0961037380( | KS>SC | NaCl | YCR008W | SAT4 | n |
| W303.asm02.HP0_G0005600 | YCR009C | chrIII | 4574.875507 | 1802.891537 | 3351.187025 | -1.324125435 | 0.13508819 | 4.45936939478 | 2.03644535695 | KS<SC | NaCl | YCR009C | RVS161 | n |
| W303.asm02.HP0_G0005610 | YCR010C | chrIII | 149.5762347 | 779.569999 | 376.2484175 | 2.324878975 | 0.20246029 | 7.93898701382 | 5.17924390901 | KS>SC | NaCl | YCR010C | ADY2 | n |

| | | | | | | | | | | | | | | |
|---|---|---|---|---|---|---|---|---|---|---|---|---|---|---|
| W303.asm02.HP0_G0005620 | YCR011C | chrIII | 3273.29894 | 4299.492442 | 4008.211727 | 0.388878435 | 0.11422272 | 0.000613789 | 0.000955558 | ns | NaCl | YCR011C | ADP1 | n |
| W303.asm02.HP0_G0005630 | YCR012W | chrIII | 52533.48658 | 12057.12955 | 31329.98713 | -2.106469917 | 0.11084074 | 1.2093567812 | 2.3668839862 | KS<SC | NaCl | YCR012W | PGK1 | n |
| W303.asm02.HP0_G0005640 | YCR014C | chrIII | 729.1886587 | 1567.77261 | 1377.405532 | 1.080465263 | 0.14968906 | 2.9403690539 | 8.2210318448 | KS>SC | NaCl | YCR014C | POL4 | n |
| W303.asm02.HP0_G0005650 | YCR015C | chrIII | 749.947748 | 764.8882909 | 873.6436017 | 0.028515881 | 0.16218429 | 0.858790863 | 0.89132082 | ns | NaCl | YCR015C | CTO1 | n |
| W303.asm02.HP0_G0005660 | YCR016W | chrIII | 2213.544713 | 5130.269768 | 3476.357041 | 1.189788022 | 0.15066384 | 1.3968923415 | 4.5565297806 | KS>SC | NaCl | YCR016W | RBP95 | n |
| W303.asm02.HP0_G0005670 | YCR017C | chrIII | 4582.46544 | 2646.933629 | 3263.383474 | -0.773721841 | 0.16303872 | 1.5493287265 | 2.9895497962 | ns | NaCl | YCR017C | CWH43 | n |
| W303.asm02.HP0_G0005680 | YCR018C | chrIII | 1352.777496 | 917.2377006 | 1067.342272 | -0.548224424 | 0.1658163 | 0.00081399 | 0.001252996 | ns | NaCl | YCR018C | SRD1 | n |
| W303.asm02.HP0_G0005800 | YCR028C | chrIII | 1348.391906 | 1728.929662 | 1627.578649 | 0.35367142 | 0.11406624 | 0.001811488 | 0.002668536 | ns | NaCl | YCR028C | FEN2 | n |
| W303.asm02.HP0_G0005810 | YCR028C-A | chrIII | 3681.639237 | 1038.124481 | 2422.230694 | -1.792997577 | 0.16310217 | 7.3039092592 | 4.1693148688 | KS<SC | NaCl | YCR028C-A | RIM1 | n |
| W303.asm02.HP0_G0005820 | YCR030C | chrIII | 2009.140917 | 2061.120577 | 2071.875465 | 0.034117519 | 0.30964697 | 0.908618744 | 0.928960955 | ns | NaCl | YCR030C | SYP1 | n |
| W303.asm02.HP0_G0005830 | YCR031C/YJL191W | chrIII | 14104.56611 | 11289.93214 | 10562.69539 | -0.317333339 | 0.11138387 | 0.004162125 | 0.00581848 | ns | NaCl | YCR031C | RPS14A | y |
| W303.asm02.HP0_G0005840 | YCR032W | chrIII | 1006.095674 | 2069.79779 | 2250.104759 | 1.024977374 | 0.13724056 | 4.7969884795 | 1.4604164926 | KS>SC | NaCl | YCR032W | BPH1 | n |
| W303.asm02.HP0_G0005850 | YCR033W | chrIII | 1410.657897 | 3261.045311 | 2530.895856 | 1.177609308 | 0.18188728 | 4.7569109476 | 1.1637442854 | KS>SC | NaCl | YCR033W | SNT1 | n |
| W303.asm02.HP0_G0005860 | YCR034W | chrIII | 6951.309029 | 5234.550508 | 5411.274469 | -0.403673715 | 0.12233573 | 0.000891145 | 0.00135652 | ns | NaCl | YCR034W | ELO2 | n |
| W303.asm02.HP0_G0005870 | YCR035C | chrIII | 2436.741578 | 4288.670006 | 3548.457755 | 0.806927986 | 0.10483955 | 1.0079369415 | 3.2113339765 | ns | NaCl | YCR035C | RRP43 | n |
| W303.asm02.HP0_G0005880 | YCR036W | chrIII | 2736.590683 | 3925.526883 | 3412.094514 | 0.510777634 | 0.15032579 | 0.000596294 | 0.000938991 | ns | NaCl | YCR036W | RBK1 | n |
| W303.asm02.HP0_G0005890 | YCR037C | chrIII | 2046.686446 | 1138.713313 | 1909.606422 | -0.834553614 | 0.12495843 | 1.7077589935 | 4.4992881175 | ns | NaCl | YCR037C | PHO87 | n |
| W303.asm02.HP0_G0005900 | YCR038C | chrIII | 256.2072575 | 426.7490116 | 368.4914114 | 0.718855144 | 0.13430473 | 6.7299846137 | 1.4184736801 | ns | NaCl | YCR038C | BUD5 | n |
| W303.asm02.HP0_G0005920 | YCR042C | chrIII | 1716.637287 | 2335.647629 | 2384.956457 | 0.440086469 | 0.10779644 | 4.0421507502 | 6.9221831598 | ns | NaCl | YCR042C | TAF2 | n |
| W303.asm02.HP0_G0005930 | YCR043C | chrIII | 2016.61867 | 634.4755033 | 1395.218446 | -1.621823227 | 0.19781303 | 6.1671224756 | 2.2234099451 | KS<SC | NaCl | YCR043C | | n |
| W303.asm02.HP0_G0005940 | YCR044C | chrIII | 1470.380552 | 648.4682898 | 1021.454551 | -1.168742514 | 0.11992355 | 9.5546711776 | 4.2225482301 | KS<SC | NaCl | YCR044C | PER1 | n |
| W303.asm02.HP0_G0005950 | YCR045C | chrIII | 520.6853984 | 2545.121807 | 1329.097377 | 2.226652866 | 0.21707771 | 7.2346573283 | 3.9645922159 | KS>SC | NaCl | YCR045C | RRT12 | n |
| W303.asm02.HP0_G0005960 | YCR046C | chrIII | 2192.962741 | 944.0857981 | 1569.660751 | -1.190276737 | 0.16063672 | 6.1982726656 | 1.8460072939 | KS<SC | NaCl | YCR046C | IMG1 | n |
| W303.asm02.HP0_G0005970 | YCR047C | chrIII | 2550.988955 | 2595.75781 | 2026.885355 | 0.023846525 | 0.20337862 | 0.904952837 | 0.928960955 | ns | NaCl | YCR047C | BUD23 | n |
| W303.asm02.HP0_G0005980 | YCR048W | chrIII | 1904.318955 | 3854.385239 | 2380.807295 | 0.945926644 | 0.3013209 | 0.001079713 | 0.001625502 | ns | NaCl | YCR048W | ARE1 | n |
| W303.asm02.HP0_G0005990 | YCR051W | chrIII | 4296.19793 | 4100.993848 | 3541.59545 | -0.067265153 | 0.09548902 | 0.479276163 | 0.512975268 | ns | NaCl | YCR051W | | n |
| W303.asm02.HP0_G0006000 | YCR052W | chrIII | 3025.852769 | 3587.205585 | 2937.406491 | 0.23552189 | 0.22120404 | 0.276671487 | 0.315866614 | ns | NaCl | YCR052W | RSC6 | n |
| W303.asm02.HP0_G0006010 | YCR053W | chrIII | 9685.052021 | 10931.18489 | 10002.50807 | 0.173959566 | 0.07276384 | 0.016534546 | 0.022428048 | ns | NaCl | YCR053W | THR4 | n |
| W303.asm02.HP0_G0006020 | YCR054C | chrIII | 850.6948098 | 1562.516573 | 1473.674431 | 0.858418438 | 0.1479275 | 4.5314334198 | 1.0013006105 | ns | NaCl | YCR054C | CTR86 | n |
| W303.asm02.HP0_G0006030 | YCR057C | chrIII | 3583.422912 | 6710.433583 | 4820.360301 | 0.852632728 | 0.27094534 | 0.001146895 | 0.001707876 | ns | NaCl | YCR057C | PWP2 | n |
| W303.asm02.HP0_G0006040 | YCR059C | chrIII | 2812.519836 | 4219.467503 | 2933.921788 | 0.576489079 | 0.12841124 | 6.0609401365 | 1.1220929712 | ns | NaCl | YCR059C | YIH1 | n |
| W303.asm02.HP0_G0006050 | YCR060W | chrIII | 1873.195565 | 1291.34828 | 1398.70489 | -0.527455326 | 0.1371391 | 0.000104511 | 0.000172506 | ns | NaCl | YCR060W | TAH1 | n |
| W303.asm02.HP0_G0006060 | YCR061W | chrIII | 772.7435059 | 1161.058572 | 707.4412929 | 0.55177958 | 0.2735297 | 0.037134127 | 0.046673169 | ns | NaCl | YCR061W | TVS1 | n |
| W303.asm02.HP0_G0006070 | YCR063W | chrIII | 244.7932866 | 697.4028762 | 390.7163254 | 1.49372095 | 0.12282699 | 1.5245746795 | 1.1603707282 | KS>SC | NaCl | YCR063W | BUD31 | n |
| W303.asm02.HP0_G0006080 | YCR065W | chrIII | 1011.594742 | 1511.644882 | 1190.631669 | 0.558426543 | 0.20267196 | 0.005007326 | 0.00692933 | ns | NaCl | YCR065W | HCM1 | n |
| W303.asm02.HP0_G0006090 | YCR066W | chrIII | 477.4541993 | 807.6499647 | 777.1183729 | 0.748102781 | 0.10449474 | 6.1402075754 | 1.6824168756 | ns | NaCl | YCR066W | RAD18 | n |
| W303.asm02.HP0_G0006100 | YCR067C | chrIII | 2550.842273 | 2726.954543 | 2928.065972 | 0.094473308 | 0.16407648 | 0.560035272 | 0.594766142 | ns | NaCl | YCR067C | SED4 | n |
| W303.asm02.HP0_G0006110 | YCR068W | chrIII | 847.4843569 | 1401.102244 | 1068.08497 | 0.65299362 | 0.35863783 | 0.054309256 | 0.067030342 | ns | NaCl | YCR068W | ATG15 | n |
| W303.asm02.HP0_G0006120 | YCR069W | chrIII | 1931.577387 | 929.3055762 | 1523.61182 | -1.009517171 | 0.2346227 | 1.0194764882 | 1.8377405116 | KS<SC | NaCl | YCR069W | CPR4 | n |
| W303.asm02.HP0_G0006130 | YCR071C | chrIII | 1534.768691 | 800.0045069 | 1242.703506 | -0.917670547 | 0.16647369 | 2.3378696068 | 5.0045021272 | ns | NaCl | YCR071C | IMG2 | n |
| W303.asm02.HP0_G0006140 | YCR072C | chrIII | 1225.584766 | 2900.214353 | 1902.591114 | 1.215336755 | 0.1660356 | 1.1799110386 | 3.4393151550 | KS>SC | NaCl | YCR072C | RSA4 | n |
| W303.asm02.HP0_G0006150 | YCR073C | chrIII | 961.4292362 | 3619.164306 | 3202.971991 | 1.869846195 | 0.1892522 | 7.7786442628 | 3.6747388414 | KS>SC | NaCl | YCR073C | SSK22 | n |
| W303.asm02.HP0_G0006160 | YCR073W-A | chrIII | 1047.975304 | 1214.525714 | 1202.246529 | 0.204354737 | 0.21738691 | 0.337178976 | 0.369548157 | ns | NaCl | YCR073W-A | SOL2 | n |
| W303.asm02.HP0_G0006170 | YCR075C | chrIII | 577.7741372 | 201.1114221 | 427.6849427 | -1.484579282 | 0.18749615 | 7.7591285463 | 2.5926844167 | KS<SC | NaCl | YCR075C | ERS1 | n |
| W303.asm02.HP0_G0006180 | YCR075W-A | chrIII | 205.1655085 | 131.9097023 | 192.5681641 | -0.598065652 | 0.26578394 | 0.020281871 | 0.026463013 | ns | NaCl | YCR075W-A | EGO2 | n |
| W303.asm02.HP0_G0006190 | YCR076C | chrIII | 877.8778833 | 704.5616375 | 771.2515063 | -0.313110781 | 0.13923951 | 0.023280947 | 0.030089526 | ns | NaCl | YCR076C | FUB1 | n |
| W303.asm02.HP0_G0006200 | YCR077C | chrIII | 2331.0404 | 3113.485805 | 2278.318609 | 0.397714952 | 0.23967759 | 0.088815143 | 0.106733987 | ns | NaCl | YCR077C | PAT1 | n |
| W303.asm02.HP0_G0006210 | YCR079W | chrIII | 900.7812042 | 2450.881077 | 1459.1903 | 1.427995357 | 0.1240065 | 3.7581217340 | 2.3402848980 | KS>SC | NaCl | YCR079W | PTC6 | n |
| W303.asm02.HP0_G0006220 | YCR081W | chrIII | 782.2967732 | 3279.293673 | 2437.626666 | 2.057465991 | 0.0928923 | 8.9804636375 | 6.1516175917 | KS>SC | NaCl | YCR081W | SRB8 | n |
| W303.asm02.HP0_G0006230 | YCR082W | chrIII | 1880.576794 | 1125.396629 | 1710.527445 | -0.726396837 | 0.14736924 | 6.3747562868 | 1.2476308732 | ns | NaCl | YCR082W | AHC2 | n |
| W303.asm02.HP0_G0006240 | YCR083W | chrIII | 1197.981758 | 482.5718699 | 839.7655168 | -1.273047563 | 0.19482772 | 2.8285389618 | 7.3115063731 | KS<SC | NaCl | YCR083W | TRX3 | n |
| W303.asm02.HP0_G0006250 | YCR084C | chrIII | 2324.611679 | 4865.17203 | 3483.041071 | 0.945044376 | 0.3923718 | 0.010015981 | 0.013721893 | ns | NaCl | YCR084C | TUP1 | n |
| W303.asm02.HP0_G0006260 | YCR086W | chrIII | 668.5820755 | 642.3628007 | 587.5539029 | -0.057571885 | 0.14634778 | 0.691197807 | 0.72841615 | ns | NaCl | YCR086W | CSM1 | n |
| W303.asm02.HP0_G0006270 | YCR087C-A | chrIII | 1751.18312 | 3219.001949 | 2059.819745 | 0.862525264 | 0.13674209 | 1.9610077319 | 4.6320355048 | ns | NaCl | YCR087C-A | | n |
| W303.asm02.HP0_G0006280 | YCR088W | chrIII | 2315.600089 | 3639.34402 | 2590.006483 | 0.557503309 | 0.43921799 | 0.168430953 | 0.195551191 | ns | NaCl | YCR088W | ABP1 | n |
| W303.asm02.HP0_G0006290 | YCR089W | chrIII | 330.5302771 | 1148.908799 | 788.1735371 | 1.693874147 | 0.29647965 | 2.5234790845 | 5.6674858127 | KS>SC | NaCl | YCR089W | FIG2 | n |
| W303.asm02.HP0_G0006300 | YCR090C | chrIII | 1454.84796 | 2332.260054 | 1873.774891 | 0.666601275 | 0.15544044 | 1.4458552925 | 2.5724957801 | ns | NaCl | YCR090C | | n |
| W303.asm02.HP0_G0006310 | YCR091W | chrIII | 214.2360016 | 754.4930198 | 551.4549167 | 1.785745264 | 0.17437124 | 2.3558800232 | 1.1953909747 | KS>SC | NaCl | YCR091W | KIN82 | n |
| W303.asm02.HP0_G0006320 | YCR092C | chrIII | 760.0836347 | 1248.860679 | 1151.072338 | 0.701107978 | 0.1590893 | 8.2318854085 | 1.5036910679 | ns | NaCl | YCR092C | MSH3 | n |
| W303.asm02.HP0_G0006330 | YCR093W | chrIII | 2045.534415 | 4921.159876 | 4487.939788 | 1.244663037 | 0.15079253 | 6.9583550904 | 2.3832366184 | KS>SC | NaCl | YCR093W | CDC39 | n |
| W303.asm02.HP0_G0006340 | YCR094W | chrIII | 1364.347749 | 1779.448407 | 1837.138468 | 0.379860658 | 0.10627681 | 0.000326448 | 0.000526158 | ns | NaCl | YCR094W | CDC50 | n |
| W303.asm02.HP0_G0006350 | YCR095C | chrIII | 928.5480111 | 669.8113359 | 1025.215307 | -0.456513342 | 0.19957137 | 0.019884935 | 0.026194578 | ns | NaCl | YCR095C | OCA4 | n |
| W303.asm02.HP0_G0006360 | YCR095W-A | chrIII | 77.37279219 | 139.6716143 | 117.3444562 | 0.829635768 | 0.19731115 | 1.8637802410 | 3.2321252281 | ns | NaCl | YCR095W-A | | n |
| W303.asm02.HP0_G0006380 | YCR098C | chrIII | 2877.254042 | 2201.041164 | 2477.889803 | -0.350272368 | 0.34025993 | 0.279472249 | 0.316427258 | ns | NaCl | YCR098C | GIT1 | n |
| W303.asm02.HP0_G0006390 | W303.asm02.HP0_G0 | chrIII | 356.3787917 | 360.9434712 | 543.2631845 | 0.021367084 | 0.2172582 | 0.920024472 | 0.933654464 | ns | NaCl | YNR065C | | y |
| W303.asm02.HP0_G0006400 | YCR100C | chrIII | 719.1045461 | 1026.445551 | 1452.925958 | 0.491495393 | 0.24093419 | 0.036372344 | 0.046138992 | ns | NaCl | YCR100C | EMA35 | n |
| W303.asm02.HP0_G0006410 | YCR101C | chrIII | 438.7533058 | 543.8558352 | 842.7599021 | 0.298364109 | 0.23818073 | 0.199103647 | 0.229220165 | ns | NaCl | YCR101C | | n |
| W303.asm02.HP0_G0006420 | YCR102C | chrIII | 1762.565289 | 4185.724016 | 2418.09028 | 1.192855028 | 0.24228279 | 4.1860989178 | 8.4337581138 | KS>SC | NaCl | YCR102C | | n |

| | | | | | | | | | | | | | | |
|---|---|---|---|---|---|---|---|---|---|---|---|---|---|---|
| W303.asm02.HP0_G0006430 | YCR104W/YIR041W/Y | chrIII | 67.28483803 | 421.6629289 | 196.0184986 | 2.609741312 | 0.20657551 | 2.8787650529! | 2.3199459544 | KS>SC | NaCl | YMR325W | PAU19 | y |
| W303.asm02.HP0_G0006440 | YCR105W | chrIII | 820.9707044 | 1251.867219 | 1036.798106 | 0.598402338 | 0.11250149 | 8.737214693 | 1.8136339591( | ns | NaCl | YCR105W | ADH7 | n |
| W303.asm02.HP0_G0006450 | YCR106W | chrIII | 1523.043331 | 1324.30561 | 1741.030874 | -0.199382159 | 0.11254451 | 0.074737852 | 0.090611378 | ns | NaCl | YCR106W | RDS1 | n |
| W303.asm02.HP0_G0006460 | YCR107W | chrIII | 4855.254826 | 1490.716619 | 2595.446155 | -1.685236174 | 0.11951578 | 7.9734928453 | 7.8026322843 | KS<SC | NaCl | YCR107W | AAD3 | n |

Km_vs_KS

| gene | mean_KS_counts | mean_km_counts | baseMean | log2FoldChange | lfcSE | pvalue | padj | DEG | treatment | strain |
|---|---|---|---|---|---|---|---|---|---|---|
| FIM1_3 | 2063.260472 | 4134.440481 | 3037.529789 | 0.970061597 | 0.136787345 | 2.66991854813 | 1.92784394440 | ns | YPD | KS.1 |
| FIM1_5 | 1149.474673 | 763.0394869 | 633.8041188 | -0.479972722 | 0.287783609 | 0.055617485 | 0.086198439 | ns | YPD | KS.1 |
| FIM1_6 | 532.1724261 | 350.1681196 | 405.4975614 | -0.540647162 | 0.208252116 | 0.004903013 | 0.009554859 | ns | YPD | KS.1 |
| FIM1_7 | 166.5375086 | 64.94534338 | 60.8195808 | -1.261462146 | 0.23758131 | 1.25848905740 | 5.69698185676 | Km<KS.1 | YPD | KS.1 |
| FIM1_8 | 540.6127825 | 563.0297939 | 375.3172182 | 0.047864742 | 0.234181924 | 0.821700046 | 0.861348025 | ns | YPD | KS.1 |
| FIM1_9 | 3913.942673 | 3312.941117 | 2901.971283 | -0.176709652 | 0.31741557 | 0.512897223 | 0.588619073 | ns | YPD | KS.1 |
| FIM1_10 | 13845.03734 | 4944.338791 | 2943.370395 | -1.300894418 | 0.324695992 | 6.22978223367 | 1.95171074386 | Km<KS.1 | YPD | KS.1 |
| FIM1_11 | 805.0526724 | 350.1619611 | 340.6940978 | -1.145858969 | 0.176207021 | 1.14110927537 | 7.09627330625 | Km<KS.1 | YPD | KS.1 |
| FIM1_12 | 9606.178975 | 1523.123224 | 1729.337393 | -2.52108004 | 0.280207359 | 1.03967048125 | 1.53028421427 | Km<KS.1 | YPD | KS.1 |
| FIM1_13 | 12274.6475 | 21659.26469 | 35774.55391 | 0.567972343 | 0.413113628 | 0.081396763 | 0.120305673 | ns | YPD | KS.1 |
| FIM1_14 | 18.52721746 | 12.99318278 | 16.92066841 | -0.370067894 | 0.37404816 | 0.22268707 | 0.289563035 | ns | YPD | KS.1 |
| FIM1_15 | 3176.756493 | 7427.593607 | 14849.47076 | 0.767687859 | 0.519744841 | 0.041956832 | 0.066328325 | ns | YPD | KS.1 |
| FIM1_16 | 4322.928405 | 6312.094667 | 13864.7932 | 0.45021284 | 0.27139305 | 0.05988768 | 0.092213311 | ns | YPD | KS.1 |
| FIM1_17 | 3455.578564 | 11058.2523 | 21161.65642 | 1.209430154 | 0.515289661 | 0.002062439 | 0.004333039 | Km>KS.1 | YPD | KS.1 |
| FIM1_18 | 1489.208762 | 1292.096227 | 2232.947735 | -0.165084463 | 0.256013408 | 0.469816743 | 0.547635123 | ns | YPD | KS.1 |
| FIM1_19 | 1172.91905 | 1085.087411 | 965.1753006 | -0.10780367 | 0.093991916 | 0.24062264 | 0.308848718 | ns | YPD | KS.1 |
| FIM1_20 | 2460.97914 | 2515.191789 | 3333.389345 | 0.029842686 | 0.091819796 | 0.739904441 | 0.793495278 | ns | YPD | KS.1 |
| FIM1_21 | 606.8020785 | 403.9863633 | 585.7584931 | -0.557245996 | 0.136499351 | 2.23347232567 | 6.46018884898 | ns | YPD | KS.1 |
| FIM1_22 | 1494.97523 | 1539.353995 | 1520.407378 | 0.041304768 | 0.072784376 | 0.564764626 | 0.637410167 | ns | YPD | KS.1 |
| FIM1_23 | 303.1081367 | 220.2223719 | 230.7427362 | -0.422944113 | 0.174420813 | 0.009947332 | 0.018187084 | ns | YPD | KS.1 |
| FIM1_24 | 517.4208745 | 670.9427168 | 1199.112786 | 0.304185816 | 0.265263879 | 0.19509773 | 0.258967772 | ns | YPD | KS.1 |
| FIM1_25 | 401.7564951 | 240.3485949 | 454.9892691 | -0.647029597 | 0.247271199 | 0.003695817 | 0.00739015 | ns | YPD | KS.1 |
| FIM1_26 | 358.04022 | 178.6962462 | 284.9825584 | -0.91655496 | 0.224193604 | 9.93156073315 | 3.01277528338 | ns | YPD | KS.1 |
| FIM1_27 | 471.4449456 | 553.6774636 | 657.296323 | 0.20976057 | 0.160918952 | 0.168543715 | 0.228102552 | ns | YPD | KS.1 |
| FIM1_28 | 6029.437664 | 11598.73299 | 11506.37819 | 0.881838518 | 0.192179265 | 1.09356995873 | 3.84216846377 | ns | YPD | KS.1 |
| FIM1_29 | 4271.181802 | 3778.58307 | 2774.21492 | -0.156003726 | 0.184845616 | 0.366530098 | 0.442164704 | ns | YPD | KS.1 |
| FIM1_30 | 1941.233347 | 1306.459053 | 2049.9388 | -0.546225947 | 0.128793714 | 1.14013450192 | 3.43976297578 | ns | YPD | KS.1 |
| FIM1_31 | 4488.773119 | 4982.144893 | 4258.824352 | 0.145147824 | 0.088619985 | 0.094390664 | 0.13714765 | ns | YPD | KS.1 |
| FIM1_32 | 2024.093003 | 2129.344564 | 2794.011073 | 0.067717446 | 0.129468673 | 0.587131893 | 0.659363695 | ns | YPD | KS.1 |
| FIM1_33 | 477.0656006 | 632.0316917 | 732.2157693 | 0.362396589 | 0.188632119 | 0.039239175 | 0.062368976 | ns | YPD | KS.1 |
| FIM1_34 | 1035.911722 | 1557.310613 | 1643.975651 | 0.578595508 | 0.078642551 | 8.74731558490 | 6.51465494534 | ns | YPD | KS.1 |
| FIM1_35 | 136.6160636 | 41.13337872 | 43.59138275 | -1.573164242 | 0.296265751 | 6.57463853433 | 3.10036272116 | Km<KS.1 | YPD | KS.1 |
| FIM1_36 | 183.3461564 | 3078.585804 | 6124.361802 | 3.737526587 | 0.497694008 | 6.36317836430 | 6.43431145577 | Km>KS.1 | YPD | KS.1 |
| FIM1_37 | 285.5179616 | 560.4785223 | 700.1965119 | 0.929865406 | 0.153769902 | 3.23624357565 | 1.75003389009 | ns | YPD | KS.1 |
| FIM1_38 | 405.4592773 | 552.2381282 | 1220.934355 | 0.40376337 | 0.18071693 | 0.01708803 | 0.029641894 | ns | YPD | KS.1 |
| FIM1_39 | 1148.877104 | 1202.720579 | 1622.369305 | 0.055522173 | 0.221689379 | 0.784201899 | 0.830379381 | ns | YPD | KS.1 |
| FIM1_40 | 1302.125406 | 719.2125459 | 1421.516646 | -0.827120399 | 0.129940631 | 5.26301329181 | 3.06957691990 | ns | YPD | KS.1 |
| FIM1_41 | 969.5257971 | 952.4846551 | 1004.945188 | -0.023997199 | 0.131590005 | 0.849748543 | 0.88367454 | ns | YPD | KS.1 |
| FIM1_42 | 335.1705468 | 650.8567564 | 1088.647994 | 0.909075656 | 0.167540566 | 1.32515796294 | 5.98245087625 | ns | YPD | KS.1 |
| FIM1_43 | 899.6377664 | 2339.777019 | 2347.041188 | 1.365403845 | 0.080317228 | 7.63309679414 | 2.37341603442 | Km>KS.1 | YPD | KS.1 |
| FIM1_44 | 107.8144527 | 117.9407358 | 223.7839843 | 0.106646183 | 0.207112884 | 0.576818806 | 0.649394333 | ns | YPD | KS.1 |
| FIM1_45 | 623.2069111 | 612.2322902 | 909.3249672 | -0.024549838 | 0.13680921 | 0.851761114 | 0.88558235 | ns | YPD | KS.1 |
| FIM1_46 | 5421.128929 | 4137.760307 | 3895.846385 | -0.367066504 | 0.136901139 | 0.005228974 | 0.010098658 | ns | YPD | KS.1 |

| | | | | | | | | | | |
|---|---|---|---|---|---|---|---|---|---|---|
| FIM1_47 | 3350.114928 | 2309.26507 | 1453.094101 | -0.519042995 | 0.108278929 | 8.86961989759 | 3.16317985595 | ns | YPD | KS.1 |
| FIM1_48 | 1947.705994 | 1614.614981 | 2574.536415 | -0.265025472 | 0.075418435 | 0.000361519 | 0.000865106 | ns | YPD | KS.1 |
| FIM1_49 | 5574.020894 | 3476.932013 | 9518.429149 | -0.558750957 | 0.290929527 | 0.027433403 | 0.045297438 | ns | YPD | KS.1 |
| FIM1_50 | 1506.482747 | 1725.64094 | 4848.123799 | 0.166653816 | 0.216149824 | 0.400315129 | 0.477022219 | ns | YPD | KS.1 |
| FIM1_51 | 4036.535494 | 2076.197936 | 3374.321267 | -0.917208957 | 0.156774866 | 1.10498621907 | 5.69669061131 | ns | YPD | KS.1 |
| FIM1_52 | 2469.236806 | 2534.745798 | 3261.822567 | 0.034637428 | 0.135788989 | 0.790605746 | 0.835620052 | ns | YPD | KS.1 |
| FIM1_53 | 15761.20152 | 21185.86383 | 32534.50092 | 0.408807693 | 0.116980118 | 0.000316951 | 0.000767313 | ns | YPD | KS.1 |
| FIM1_54 | 7097.419807 | 5434.387673 | 3260.43686 | -0.332825368 | 0.218583184 | 0.095754542 | 0.138966991 | ns | YPD | KS.1 |
| FIM1_55 | 802.7904983 | 327.4623658 | 884.141548 | -1.077955462 | 0.355312199 | 0.000383649 | 0.000914544 | Km<KS.1 | YPD | KS.1 |
| FIM1_56 | 768.7301318 | 27.04428853 | 110.1015889 | -4.656232167 | 0.348080972 | 3.67079283748 | 2.17407075791 | Km<KS.1 | YPD | KS.1 |
| FIM1_57 | 390.3454888 | 336.4064384 | 411.2940193 | -0.171763592 | 0.264590493 | 0.46402145 | 0.541648689 | ns | YPD | KS.1 |
| FIM1_58 | 2618.753278 | 2028.808259 | 3663.533176 | -0.327889807 | 0.191448988 | 0.06554143 | 0.099776198 | ns | YPD | KS.1 |
| FIM1_59 | 8510.909878 | 8563.48328 | 4886.402825 | 0.006956719 | 0.268675914 | 0.976656899 | 0.983775678 | ns | YPD | KS.1 |
| FIM1_60 | 889.3570127 | 1908.538004 | 2097.26308 | 1.074405745 | 0.122905292 | 3.78670654626 | 4.59484513846 | Km>KS.1 | YPD | KS.1 |
| FIM1_61 | 2098.858613 | 2073.944035 | 1514.364944 | -0.016649926 | 0.091498936 | 0.852574801 | 0.885872939 | ns | YPD | KS.1 |
| FIM1_62 | 678.3690306 | 743.5706172 | 1248.976291 | 0.119747489 | 0.160878758 | 0.432198576 | 0.50988568 | ns | YPD | KS.1 |
| FIM1_63 | 81.02393205 | 51.5932734 | 61.0388668 | -0.524129075 | 0.307163125 | 0.047095766 | 0.073818978 | ns | YPD | KS.1 |
| FIM1_64 | 1187.012315 | 1375.720175 | 2027.713438 | 0.203667297 | 0.097300833 | 0.031971726 | 0.051844634 | ns | YPD | KS.1 |
| FIM1_65 | 471.573438 | 337.6699113 | 253.6940693 | -0.461819111 | 0.12350817 | 0.000112179 | 0.000290369 | ns | YPD | KS.1 |
| FIM1_66 | 2842.528475 | 2934.197612 | 4634.515481 | 0.041525396 | 0.154537575 | 0.777675323 | 0.825630715 | ns | YPD | KS.1 |
| FIM1_67 | 709.706274 | 582.0396414 | 516.1440968 | -0.255541331 | 0.181301182 | 0.131972286 | 0.184067353 | ns | YPD | KS.1 |
| FIM1_68 | 6377.431843 | 10465.51643 | 15105.22591 | 0.664808886 | 0.177641286 | 7.24598239376 | 0.000193498 | ns | YPD | KS.1 |
| FIM1_69 | 6424.422755 | 9133.372778 | 14277.90827 | 0.482520308 | 0.132951123 | 0.000165953 | 0.000420804 | ns | YPD | KS.1 |
| FIM1_70 | 1431.796492 | 791.5354286 | 1316.552668 | -0.793137075 | 0.193899348 | 1.28460848160 | 3.83839471229 | ns | YPD | KS.1 |
| FIM1_71 | 748.7618266 | 604.3997611 | 738.570201 | -0.287049846 | 0.148714278 | 0.042833565 | 0.067628368 | ns | YPD | KS.1 |
| FIM1_72 | 438.9935728 | 467.3562419 | 603.4281888 | 0.081457297 | 0.158529195 | 0.588328226 | 0.660409053 | ns | YPD | KS.1 |
| FIM1_73 | 5701.336768 | 11192.96914 | 13298.10691 | 0.955849503 | 0.098652051 | 6.84827224915 | 1.11340373985 | ns | YPD | KS.1 |
| FIM1_74 | 296.6010098 | 680.3822034 | 775.8522912 | 1.178445129 | 0.10011996 | 7.50826557554 | 2.50695444552 | Km>KS.1 | YPD | KS.1 |
| FIM1_75 | 46316.33021 | 20240.17744 | 22825.45264 | -1.156194832 | 0.146498415 | 4.22811164846 | 3.91744040963 | Km<KS.1 | YPD | KS.1 |
| FIM1_76 | 640.6233909 | 580.358411 | 347.4029841 | -0.129282319 | 0.157859122 | 0.388176564 | 0.464919454 | ns | YPD | KS.1 |
| FIM1_77 | 1294.995225 | 1022.312154 | 859.8821874 | -0.327992903 | 0.105397913 | 0.001405064 | 0.003024748 | ns | YPD | KS.1 |
| FIM1_78 | 3334.184388 | 3728.374536 | 3222.916234 | 0.150832803 | 0.128211204 | 0.221768049 | 0.28851884 | ns | YPD | KS.1 |
| FIM1_79 | 13828.65966 | 6988.052883 | 3178.933275 | -0.900540855 | 0.224125773 | 1.38827649159 | 4.12585158045 | ns | YPD | KS.1 |
| FIM1_80 | 323.1431676 | 211.9830141 | 115.301577 | -0.553168286 | 0.193743812 | 0.002180441 | 0.004550206 | ns | YPD | KS.1 |
| FIM1_81 | 2491.000081 | 2309.586261 | 1455.405965 | -0.099094698 | 0.155438503 | 0.50246798 | 0.57878634 | ns | YPD | KS.1 |
| FIM1_82 | 1551.08305 | 1954.378665 | 2001.056325 | 0.322717566 | 0.098920219 | 0.000841462 | 0.00188741 | ns | YPD | KS.1 |
| FIM1_83 | 345.6723769 | 327.0989009 | 385.1962856 | -0.071186192 | 0.154079125 | 0.627332154 | 0.695330547 | ns | YPD | KS.1 |
| FIM1_84 | 918.8772712 | 905.1894011 | 813.4542519 | -0.019094727 | 0.143879472 | 0.889702389 | 0.916082721 | ns | YPD | KS.1 |
| FIM1_85 | 600.6806804 | 655.6841529 | 812.4313926 | 0.119319983 | 0.103032952 | 0.234345919 | 0.302216453 | ns | YPD | KS.1 |
| FIM1_86 | 887.6945378 | 416.6530377 | 890.901293 | -1.065822891 | 0.120020191 | 1.11604878130 | 1.44856980361 | Km<KS.1 | YPD | KS.1 |
| FIM1_87 | 786.9337735 | 378.0873804 | 533.8379842 | -0.986838305 | 0.202546426 | 2.17208329267 | 8.42253654018 | ns | YPD | KS.1 |
| FIM1_88 | 1553.639015 | 868.393944 | 812.0837853 | -0.78585458 | 0.180632309 | 4.10398316610 | 1.31385561463 | ns | YPD | KS.1 |
| FIM1_89 | 1896.637598 | 1281.570918 | 613.585864 | -0.427247373 | 0.331425651 | 0.125583634 | 0.176491124 | ns | YPD | KS.1 |
| FIM1_90 | 1340.738817 | 3375.724576 | 2907.03043 | 1.320196366 | 0.081085805 | 1.36252922658 | 2.42092246509 | Km>KS.1 | YPD | KS.1 |
| FIM1_91 | 968.1616052 | 550.4821056 | 2110.39745 | -0.699873335 | 0.270231752 | 0.003562612 | 0.007138136 | ns | YPD | KS.1 |

| | | | | | | | | | | |
|---|---|---|---|---|---|---|---|---|---|---|
| FIM1_92 | 288.5658923 | 329.8429015 | 320.0306234 | 0.179129908 | 0.143234276 | 0.190657976 | 0.254311037 | ns | YPD | KS.1 |
| FIM1_93 | 258.0748703 | 244.1331045 | 258.6469668 | -0.07388295 | 0.110313427 | 0.490516477 | 0.567912375 | ns | YPD | KS.1 |
| FIM1_94 | 457.1450531 | 372.8673838 | 357.0574842 | -0.273118625 | 0.147266288 | 0.051811445 | 0.080752487 | ns | YPD | KS.1 |
| FIM1_95 | 3187.51497 | 6631.442945 | 5406.626742 | 1.040838454 | 0.095260346 | 1.4993799928 | 3.805824216 | Km>KS.1 | YPD | KS.1 |
| FIM1_96 | 3112.818784 | 916.5668963 | 692.0664714 | -1.627278659 | 0.274453096 | 1.6594353702 | 9.224235717 | Km<KS.1 | YPD | KS.1 |
| FIM1_97 | 6724.650123 | 2432.687887 | 3895.884372 | -1.287489401 | 0.320187604 | 6.0336561472 | 1.893844752 | Km<KS.1 | YPD | KS.1 |
| FIM1_98 | 176.60813 | 121.21455 | 194.9117998 | -0.462446647 | 0.244623856 | 0.035459186 | 0.057035063 | ns | YPD | KS.1 |
| FIM1_99 | 238.9205426 | 462.4020717 | 463.1981537 | 0.897032483 | 0.18402747 | 2.5837778684 | 9.956851197 | ns | YPD | KS.1 |
| FIM1_100 | 2927.235846 | 4066.46233 | 3763.677311 | 0.449213019 | 0.135245792 | 0.000556484 | 0.001293695 | ns | YPD | KS.1 |
| FIM1_101 | 5606.194843 | 2099.312944 | 1500.413589 | -1.353829893 | 0.187429654 | 4.9933107859 | 3.786847737 | Km<KS.1 | YPD | KS.1 |
| FIM1_102 | 321.8385876 | 203.8843009 | 221.6806448 | -0.612771041 | 0.175275496 | 0.000210702 | 0.000528106 | ns | YPD | KS.1 |
| FIM1_103 | 1021.659981 | 1469.509133 | 1767.344721 | 0.504017686 | 0.11688028 | 9.0386599341 | 2.758732096 | ns | YPD | KS.1 |
| FIM1_104 | 547.8770044 | 705.5163397 | 640.3642589 | 0.357867438 | 0.072821824 | 6.4715597995 | 2.360411290 | ns | YPD | KS.1 |
| FIM1_105 | 3621.818511 | 2902.468069 | 3825.164874 | -0.286104513 | 0.182333497 | 0.093330844 | 0.135885557 | ns | YPD | KS.1 |
| FIM1_107 | 569.7837211 | 806.8111545 | 944.1502495 | 0.471145812 | 0.146237973 | 0.000759811 | 0.001719773 | ns | YPD | KS.1 |
| FIM1_108 | 643.933358 | 1136.380394 | 2134.243348 | 0.772863693 | 0.16819427 | 1.3412877751 | 4.633962973 | ns | YPD | KS.1 |
| FIM1_109 | 1269.391971 | 7670.872049 | 3009.743906 | 2.539967784 | 0.176357147 | 1.7920082129 | 1.592007296 | Km>KS.1 | YPD | KS.1 |
| FIM1_110 | 1048.463951 | 1367.227637 | 762.545038 | 0.361487228 | 0.136450069 | 0.005801026 | 0.011091509 | ns | YPD | KS.1 |
| FIM1_111 | 3941.435451 | 5141.25739 | 5944.23296 | 0.37089906 | 0.10001715 | 0.000148253 | 0.000378041 | ns | YPD | KS.1 |
| FIM1_112 | 473.4507552 | 561.4800064 | 919.1620486 | 0.221425834 | 0.174840878 | 0.177590961 | 0.238608402 | ns | YPD | KS.1 |
| FIM1_113 | 1497.895693 | 1887.810799 | 1147.542679 | 0.316183481 | 0.124653604 | 0.008601275 | 0.015907562 | ns | YPD | KS.1 |
| FIM1_114 | 829.1733003 | 841.0831378 | 985.3718464 | 0.019668049 | 0.128313062 | 0.873681958 | 0.90308908 | ns | YPD | KS.1 |
| FIM1_115 | 2211.438933 | 1759.383423 | 1055.82097 | -0.274519353 | 0.242659746 | 0.208520947 | 0.274255054 | ns | YPD | KS.1 |
| FIM1_116 | 580.1850401 | 411.4142746 | 432.800489 | -0.461311115 | 0.16231974 | 0.00272085 | 0.005570464 | ns | YPD | KS.1 |
| FIM1_117 | 4290.028125 | 8173.885227 | 11163.9285 | 0.860993109 | 0.203245155 | 5.8436291875 | 1.835357020 | ns | YPD | KS.1 |
| FIM1_118 | 340.5681398 | 154.7915164 | 186.7466403 | -1.046807784 | 0.229314488 | 8.6970343891 | 3.103855529 | Km<KS.1 | YPD | KS.1 |
| FIM1_119 | 2733.880823 | 4691.96105 | 7368.886416 | 0.712528054 | 0.204473696 | 0.000176749 | 0.000446813 | ns | YPD | KS.1 |
| FIM1_120 | 1154.468423 | 945.9292213 | 1362.100667 | -0.27445101 | 0.112347932 | 0.011823514 | 0.02129688 | ns | YPD | KS.1 |
| FIM1_121 | 3534.636448 | 1914.838044 | 3572.133845 | -0.756020855 | 0.28321075 | 0.002489441 | 0.005143259 | ns | YPD | KS.1 |
| FIM1_122 | 2561.942261 | 2899.752655 | 3375.700208 | 0.171696654 | 0.096539023 | 0.068446332 | 0.103344614 | ns | YPD | KS.1 |
| FIM1_123 | 2466.882946 | 2256.05394 | 4792.686076 | -0.121281685 | 0.120562679 | 0.297957378 | 0.370214275 | ns | YPD | KS.1 |
| FIM1_124 | 307.4897198 | 1069.808266 | 724.0275505 | 1.769138133 | 0.126151237 | 4.3909510772 | 3.260445016 | Km>KS.1 | YPD | KS.1 |
| FIM1_125 | 2658.530481 | 425.0830414 | 653.6319812 | -2.59139147 | 0.174286458 | 1.5813576559 | 1.639011320 | Km<KS.1 | YPD | KS.1 |
| FIM1_126 | 621.9974935 | 1258.114467 | 1309.752804 | 0.999542567 | 0.096432534 | 6.6990555181 | 1.406236337 | ns | YPD | KS.1 |
| FIM1_127 | 973.6347809 | 483.2104785 | 779.7362617 | -0.941528097 | 0.203010309 | 7.6106909668 | 2.737757596 | ns | YPD | KS.1 |
| FIM1_128 | 5852.805603 | 11511.76441 | 15811.42649 | 0.951659967 | 0.118082738 | 1.5942939062 | 1.547020551 | ns | YPD | KS.1 |
| FIM1_129 | 1515.851118 | 809.6710211 | 1036.64876 | -0.830542608 | 0.21137698 | 2.3474633216 | 6.770220304 | ns | YPD | KS.1 |
| FIM1_130 | 12181.03414 | 7714.854907 | 9887.24238 | -0.63552545 | 0.120909674 | 6.1439635134 | 2.557842550 | ns | YPD | KS.1 |
| FIM1_131 | 1587.792988 | 3409.52995 | 2248.066324 | 1.054543019 | 0.165507131 | 3.2180425079 | 1.915043238 | Km>KS.1 | YPD | KS.1 |
| FIM1_132 | 8948.254204 | 14058.47575 | 21182.2828 | 0.638481952 | 0.08999345 | 5.2741352301 | 3.695608840 | ns | YPD | KS.1 |
| FIM1_133 | 3825.8222 | 5414.202402 | 4961.324517 | 0.470452481 | 0.148262752 | 0.000900573 | 0.002009125 | ns | YPD | KS.1 |
| FIM1_134 | 3454.751697 | 4379.839363 | 4336.865324 | 0.326077171 | 0.118710818 | 0.004559643 | 0.008951943 | ns | YPD | KS.1 |
| FIM1_135 | 796.0765207 | 1320.464963 | 1364.517315 | 0.714946029 | 0.093833891 | 8.6775851009 | 7.008276928 | ns | YPD | KS.1 |
| FIM1_136 | 1391.116394 | 330.4870471 | 244.4891628 | -1.920962868 | 0.288866066 | 7.9922935068 | 5.537835681 | Km<KS.1 | YPD | KS.1 |
| FIM1_137 | 2976.773895 | 1368.526563 | 852.1868397 | -1.033895997 | 0.225568088 | 8.1720508818 | 2.931215078 | Km<KS.1 | YPD | KS.1 |

| | | | | | | | | | | |
|---|---|---|---|---|---|---|---|---|---|---|
| FIM1_138 | 4914.540043 | 6154.98843 | 11142.29983 | 0.29886606 | 0.157038627 | 0.044876169 | 0.070629213 | ns | YPD | KS.1 |
| FIM1_140 | 8070.105418 | 11064.6161 | 17517.12602 | 0.430513078 | 0.136376521 | 0.001024682 | 0.002266693 | ns | YPD | KS.1 |
| FIM1_141 | 200.5349474 | 829.0550853 | 718.8621058 | 2.002279422 | 0.150163861 | 3.1070742541 | 1.7767464844 | Km>KS.1 | YPD | KS.1 |
| FIM1_142 | 5867.559189 | 9988.056968 | 12027.67333 | 0.7594892 | 0.066568803 | 1.1250562945 | 3.2541599216 | ns | YPD | KS.1 |
| FIM1_143 | 797.3213273 | 320.3346053 | 247.9532581 | -1.186953837 | 0.271292743 | 1.5811505471 | 5.4100577524 | Km<KS.1 | YPD | KS.1 |
| FIM1_144 | 946.040132 | 456.465688 | 911.1906525 | -0.822025352 | 0.37477362 | 0.00794714 | 0.014780194 | ns | YPD | KS.1 |
| FIM1_145 | 14164.29621 | 14942.18624 | 13754.70225 | 0.067427438 | 0.187599988 | 0.700423396 | 0.760333056 | ns | YPD | KS.1 |
| FIM1_146 | 2594.431723 | 2616.238942 | 2501.115526 | 0.011608094 | 0.089940451 | 0.895200549 | 0.920169986 | ns | YPD | KS.1 |
| FIM1_147 | 1945.964752 | 2055.110362 | 2010.46277 | 0.076403931 | 0.082829512 | 0.347490032 | 0.422577098 | ns | YPD | KS.1 |
| FIM1_148 | 766.563299 | 899.6969456 | 787.1053614 | 0.225259038 | 0.087736368 | 0.008795512 | 0.016224572 | ns | YPD | KS.1 |
| FIM1_149 | 240.5385193 | 410.6816991 | 567.6685807 | 0.74849474 | 0.120082334 | 1.4649693441 | 8.1890140304 | ns | YPD | KS.1 |
| FIM1_150 | 949.8474017 | 966.3632629 | 1020.816971 | 0.023761442 | 0.087636861 | 0.782161229 | 0.829337621 | ns | YPD | KS.1 |
| FIM1_151 | 1168.266187 | 960.2194645 | 929.2554157 | -0.268002477 | 0.122218973 | 0.023179505 | 0.038906221 | ns | YPD | KS.1 |
| FIM1_153 | 216.2655241 | 242.6887445 | 573.2743016 | 0.148203872 | 0.183493851 | 0.387920956 | 0.464813766 | ns | YPD | KS.1 |
| FIM1_154 | 3683.512669 | 2986.613048 | 1999.653377 | -0.243594121 | 0.263244574 | 0.296698472 | 0.369018724 | ns | YPD | KS.1 |
| FIM1_155 | 3526.46766 | 4271.237087 | 3525.482726 | 0.240074934 | 0.205180905 | 0.2045816 | 0.26997174 | ns | YPD | KS.1 |
| FIM1_156 | 3038.845262 | 1407.025705 | 1443.315126 | -1.073702943 | 0.144670054 | 1.9159465183 | 1.5153949012 | Km<KS.1 | YPD | KS.1 |
| FIM1_157 | 7073.829622 | 3184.230465 | 2207.756938 | -0.96921 | 0.329400351 | 0.000661158 | 0.001516144 | ns | YPD | KS.1 |
| FIM1_158 | 3128.029844 | 5896.697703 | 6922.7706 | 0.891243449 | 0.115561836 | 2.8940514147 | 2.4611804766 | ns | YPD | KS.1 |
| FIM1_159 | 4421.798098 | 6451.798472 | 6425.530611 | 0.543111299 | 0.033820302 | 2.3902665797 | 3.7161175731 | ns | YPD | KS.1 |
| FIM1_160 | 3782.194857 | 5575.669114 | 5656.953157 | 0.550907772 | 0.073973254 | 4.7226467868 | 3.5870485136 | ns | YPD | KS.1 |
| FIM1_161 | 1148.525459 | 985.6707946 | 678.5940738 | -0.195308616 | 0.185974916 | 0.260617848 | 0.330505683 | ns | YPD | KS.1 |
| FIM1_162 | 1289.937499 | 1201.513074 | 1262.050445 | -0.093052356 | 0.156461481 | 0.531388402 | 0.607039563 | ns | YPD | KS.1 |
| FIM1_163 | 1845.943862 | 313.7656568 | 516.5720716 | -2.450057818 | 0.24489957 | 7.9636798189 | 1.6508044624 | Km<KS.1 | YPD | KS.1 |
| FIM1_164 | 1583.054349 | 1117.337087 | 893.1192894 | -0.489384731 | 0.091768128 | 5.5523545475 | 2.3232097455 | ns | YPD | KS.1 |
| FIM1_165 | 178.6153544 | 335.785821 | 587.4606674 | 0.86923913 | 0.151913812 | 2.6225477385 | 1.2930797818 | ns | YPD | KS.1 |
| FIM1_166 | 586.4843429 | 685.1402145 | 737.7645358 | 0.197144794 | 0.189427447 | 0.26394753 | 0.333962096 | ns | YPD | KS.1 |
| FIM1_167 | 1520.474685 | 467.1788089 | 515.6252124 | -1.643207434 | 0.178351585 | 1.7300303842 | 2.5166377665 | Km<KS.1 | YPD | KS.1 |
| FIM1_168 | 850.1223023 | 996.4003255 | 1298.756978 | 0.217025813 | 0.120422525 | 0.062134997 | 0.09523155 | ns | YPD | KS.1 |
| FIM1_169 | 6498.460709 | 2462.413638 | 2192.324479 | -1.369702526 | 0.128012722 | 9.7488399345 | 2.2986008850 | Km<KS.1 | YPD | KS.1 |
| FIM1_170 | 9198.08707 | 10035.32422 | 11208.52652 | 0.123843085 | 0.052096023 | 0.01649259 | 0.028769508 | ns | YPD | KS.1 |
| FIM1_171 | 469.9847572 | 1031.433784 | 1811.685558 | 1.096393019 | 0.14579374 | 8.6946798512 | 7.0107021491 | Km>KS.1 | YPD | KS.1 |
| FIM1_172 | 621.3345919 | 588.8644224 | 709.2605182 | -0.071292042 | 0.138242952 | 0.590834495 | 0.662624349 | ns | YPD | KS.1 |
| FIM1_173 | 830.3176644 | 596.9474906 | 894.9360852 | -0.455388257 | 0.122947857 | 0.00013055 | 0.000334614 | ns | YPD | KS.1 |
| FIM1_174 | 77.14449461 | 284.9532205 | 327.707724 | 1.576820173 | 0.413592456 | 7.2966968705 | 2.2617487184 | Km>KS.1 | YPD | KS.1 |
| FIM1_175 | 17.48477695 | 70.87775178 | 88.16067366 | 1.808130942 | 0.322472916 | 7.2159743181 | 3.7868641595 | Km>KS.1 | YPD | KS.1 |
| FIM1_176 | 1303.134992 | 2044.077363 | 2120.157969 | 0.619661434 | 0.137382543 | 2.8210926205 | 9.2639840181 | ns | YPD | KS.1 |
| FIM1_177 | 709.9663111 | 4471.753032 | 8182.819457 | 2.545811917 | 0.249766339 | 8.5481430993 | 1.9070408932 | Km>KS.1 | YPD | KS.1 |
| FIM1_178 | 641.4243668 | 693.5558728 | 898.2572507 | 0.109237774 | 0.087720858 | 0.203722888 | 0.269052659 | ns | YPD | KS.1 |
| FIM1_179 | 2787.846376 | 2395.718447 | 2568.32626 | -0.207371289 | 0.114994551 | 0.062612808 | 0.095845759 | ns | YPD | KS.1 |
| FIM1_180 | 2722.94977 | 1915.829243 | 1767.514357 | -0.491778866 | 0.101619129 | 7.4547580674 | 2.6836050206 | ns | YPD | KS.1 |
| FIM1_181 | 7846.670533 | 9127.008735 | 11963.46107 | 0.213236662 | 0.072738013 | 0.00292872 | 0.005972336 | ns | YPD | KS.1 |
| FIM1_182 | 1576.053086 | 5644.035703 | 5639.517076 | 1.72587764 | 0.249930411 | 2.2318920782 | 1.6281030922 | Km>KS.1 | YPD | KS.1 |
| FIM1_183 | 71.16258623 | 57.57469897 | 86.97005316 | -0.250005044 | 0.267267582 | 0.290036405 | 0.361818234 | ns | YPD | KS.1 |
| FIM1_184 | 3490.934236 | 3933.523593 | 4253.522987 | 0.160287162 | 0.134077317 | 0.213070465 | 0.279468907 | ns | YPD | KS.1 |

| | | | | | | | | | | |
|---|---|---|---|---|---|---|---|---|---|---|
| FIM1_185 | 7318.020967 | 752.193175 | 1297.052696 | -3.218404191 | 0.205582501 | 3.09451905033 | 4.66522190164 | Km<KS.1 | YPD | KS.1 |
| FIM1_186 | 876.6751078 | 428.4114147 | 342.0256655 | -0.918370267 | 0.261509468 | 0.000101487 | 0.0002649 | ns | YPD | KS.1 |
| FIM1_187 | 6295.473773 | 5157.793995 | 7421.616443 | -0.229742785 | 0.267570234 | 0.331437043 | 0.405832953 | ns | YPD | KS.1 |
| FIM1_189 | 1050.915202 | 1235.122028 | 1349.127392 | 0.221820539 | 0.108736226 | 0.035710553 | 0.057346676 | ns | YPD | KS.1 |
| FIM1_190 | 25.68603687 | 108.3574684 | 166.9618692 | 1.949484056 | 0.238841219 | 8.45370535824 | 9.18279130071 | Km>KS.1 | YPD | KS.1 |
| FIM1_191 | 1235.595112 | 907.6995334 | 673.3985639 | -0.42626719 | 0.117983812 | 0.000196226 | 0.000493292 | ns | YPD | KS.1 |
| FIM1_192 | 7680.054292 | 6725.929863 | 3739.861374 | -0.142009509 | 0.308319631 | 0.591104696 | 0.662777972 | ns | YPD | KS.1 |
| FIM1_193 | 4532.277517 | 402.7995884 | 567.2888931 | -3.303446552 | 0.361971779 | 7.27058119809 | 1.17438770975 | Km<KS.1 | YPD | KS.1 |
| FIM1_194 | 2702.479291 | 3266.036909 | 2532.895346 | 0.2385894 | 0.200790201 | 0.198865022 | 0.263336036 | ns | YPD | KS.1 |
| FIM1_195 | 681.1482093 | 721.5550002 | 1244.231713 | 0.074681393 | 0.164154622 | 0.63077723 | 0.698134977 | ns | YPD | KS.1 |
| FIM1_196 | 964.6181486 | 2336.447299 | 2924.485626 | 1.249802793 | 0.120712497 | 4.82095257113 | 1.02060591665 | Km>KS.1 | YPD | KS.1 |
| FIM1_197 | 2571.373664 | 2437.435346 | 2057.649556 | -0.065762375 | 0.209395233 | 0.733327322 | 0.787970503 | ns | YPD | KS.1 |
| FIM1_198 | 8684.306035 | 5380.113326 | 4654.542393 | -0.655400742 | 0.148937744 | 4.35579174060 | 1.39267762914 | ns | YPD | KS.1 |
| FIM1_199 | 1661.383685 | 1246.209152 | 1061.302746 | -0.377741825 | 0.175528966 | 0.021985517 | 0.037002012 | ns | YPD | KS.1 |
| FIM1_200 | 1315.839434 | 1855.767762 | 2473.384623 | 0.463857239 | 0.15358079 | 0.001526133 | 0.003279703 | ns | YPD | KS.1 |
| FIM1_201 | 1548.32637 | 1925.67517 | 2060.566516 | 0.299876164 | 0.115561353 | 0.007429131 | 0.013905163 | ns | YPD | KS.1 |
| FIM1_203 | 908.5336204 | 993.9797332 | 1224.968132 | 0.122444771 | 0.115941576 | 0.275503007 | 0.34608905 | ns | YPD | KS.1 |
| FIM1_204 | 184.8901206 | 148.0775875 | 149.289268 | -0.277612616 | 0.213163293 | 0.155728685 | 0.212609826 | ns | YPD | KS.1 |
| FIM1_205 | 1374.066594 | 1700.654105 | 1549.415675 | 0.246334609 | 0.267708374 | 0.297831899 | 0.370214275 | ns | YPD | KS.1 |
| FIM1_206 | 2064.687039 | 2949.20658 | 2888.196518 | 0.497401094 | 0.106563274 | 1.72534888673 | 5.85512326843 | ns | YPD | KS.1 |
| FIM1_207 | 325.1774993 | 472.4258233 | 798.6830117 | 0.502377735 | 0.1576635 | 0.00080883 | 0.001819137 | ns | YPD | KS.1 |
| FIM1_208 | 2292.968926 | 2395.545282 | 3150.474754 | 0.058682358 | 0.123645898 | 0.623179781 | 0.691881145 | ns | YPD | KS.1 |
| FIM1_209 | 6931.260193 | 5922.792826 | 4312.461468 | -0.213617197 | 0.12506565 | 0.076436926 | 0.113820325 | ns | YPD | KS.1 |
| FIM1_210 | 992.4099628 | 2332.706379 | 2926.221732 | 1.19370603 | 0.147005738 | 6.19218912881 | 6.28696753384 | Km>KS.1 | YPD | KS.1 |
| FIM1_211 | 1493.897331 | 1528.967784 | 1421.061327 | 0.031111806 | 0.119442138 | 0.787853292 | 0.833064851 | ns | YPD | KS.1 |
| FIM1_212 | 1004.248338 | 1695.303485 | 1343.651911 | 0.718293147 | 0.149251574 | 5.19592184847 | 1.90772776355 | ns | YPD | KS.1 |
| FIM1_213 | 599.1542452 | 1193.698622 | 1407.992985 | 0.980646405 | 0.092282868 | 4.35114966195 | 9.88446099006 | ns | YPD | KS.1 |
| FIM1_214 | 276.6009771 | 254.0867219 | 339.0121516 | -0.108210105 | 0.183472389 | 0.528613222 | 0.604479593 | ns | YPD | KS.1 |
| FIM1_215 | 782.5798007 | 897.0879065 | 763.6384068 | 0.191796278 | 0.080549315 | 0.01535165 | 0.026968383 | ns | YPD | KS.1 |
| FIM1_216 | 2399.539955 | 5279.063854 | 3833.499387 | 1.073702691 | 0.191427956 | 3.37318869241 | 1.63997836303 | Km>KS.1 | YPD | KS.1 |
| FIM1_217 | 1502.580583 | 2244.690658 | 2626.542711 | 0.551536152 | 0.134311084 | 2.03801486465 | 5.92584684490 | ns | YPD | KS.1 |
| FIM1_218 | 8332.946484 | 2061.112334 | 1854.385565 | -1.863958537 | 0.288193365 | 3.13114867836 | 2.04428670273 | Km<KS.1 | YPD | KS.1 |
| FIM1_219 | 1304.135969 | 1406.208435 | 1022.823757 | 0.104768876 | 0.090599424 | 0.237395855 | 0.305258304 | ns | YPD | KS.1 |
| FIM1_220 | 846.9149891 | 306.3119344 | 417.3276213 | -1.42593605 | 0.148663673 | 7.44767376207 | 1.19909957819 | Km<KS.1 | YPD | KS.1 |
| FIM1_221 | 669.1390216 | 815.3508122 | 1188.193571 | 0.263608243 | 0.148189027 | 0.061996844 | 0.095137045 | ns | YPD | KS.1 |
| FIM1_222 | 2699.27424 | 2127.812973 | 1793.918807 | -0.337327679 | 0.065916821 | 2.31658045038 | 8.95492442943 | ns | YPD | KS.1 |
| FIM1_224 | 528.8735741 | 1236.656803 | 1990.069359 | 1.190827087 | 0.139735956 | 2.09615556183 | 2.38635558812 | Km>KS.1 | YPD | KS.1 |
| FIM1_225 | 730.7244065 | 1357.779636 | 1133.049414 | 0.87368379 | 0.109261171 | 3.10237407657 | 2.92316496798 | ns | YPD | KS.1 |
| FIM1_226 | 656.6156846 | 413.6915672 | 393.6260039 | -0.587124162 | 0.230778382 | 0.00517924 | 0.01002206 | ns | YPD | KS.1 |
| FIM1_227 | 2166.43038 | 3364.075943 | 1808.330015 | 0.593486793 | 0.164945414 | 0.000149022 | 0.000379807 | ns | YPD | KS.1 |
| FIM1_228 | 61.3831032 | 57.22313552 | 100.5731306 | -0.094073904 | 0.14876747 | 0.507416134 | 0.583540284 | ns | YPD | KS.1 |
| FIM1_229 | 3208.891578 | 8019.635491 | 7860.361819 | 1.287939858 | 0.135438505 | 2.14212879760 | 3.35128640505 | Km>KS.1 | YPD | KS.1 |
| FIM1_230 | 1913.633164 | 1620.661268 | 1941.403259 | -0.230407016 | 0.09943518 | 0.017514561 | 0.030318351 | ns | YPD | KS.1 |
| FIM1_231 | 142.6498407 | 87.14600227 | 186.9610113 | -0.579948177 | 0.297151889 | 0.024440877 | 0.040789455 | ns | YPD | KS.1 |
| FIM1_233 | 1325.90309 | 5031.358985 | 4732.029427 | 1.87710797 | 0.156343083 | 1.00349261703 | 3.69805612574 | Km>KS.1 | YPD | KS.1 |

| | | | | | | | | | | |
|---|---|---|---|---|---|---|---|---|---|---|
| FIM1_234 | 1902.895313 | 1931.363895 | 1866.763399 | 0.021320076 | 0.061430064 | 0.72572255 | 0.781656135 | ns | YPD | KS.1 |
| FIM1_235 | 896.5374032 | 782.6480396 | 816.0938578 | -0.182792702 | 0.132613099 | 0.151316741 | 0.207326021 | ns | YPD | KS.1 |
| FIM1_236 | 8008.958932 | 3312.316013 | 2878.122841 | -1.240512343 | 0.135668496 | 7.3609753448 | 1.0315733053 | Km<KS.1 | YPD | KS.1 |
| FIM1_237 | 479.9195532 | 527.0346874 | 509.869959 | 0.105562184 | 0.272067621 | 0.66008522 | 0.723969129 | ns | YPD | KS.1 |
| FIM1_238 | 317.234311 | 439.0326101 | 439.7022536 | 0.439746061 | 0.140863233 | 0.001135705 | 0.002490141 | ns | YPD | KS.1 |
| FIM1_239 | 95.21084986 | 70.25345332 | 93.79866522 | -0.384479728 | 0.2118535 | 0.04812942 | 0.075344198 | ns | YPD | KS.1 |
| FIM1_240 | 682.0071792 | 859.986867 | 582.5653609 | 0.297421435 | 0.190607045 | 0.093529918 | 0.136135559 | ns | YPD | KS.1 |
| FIM1_241 | 4412.078769 | 3664.702981 | 3621.35305 | -0.262778058 | 0.067063753 | 7.3200525264 | 0.000195162 | ns | YPD | KS.1 |
| FIM1_242 | 33.17690948 | 42.8293546 | 58.14148934 | 0.305880677 | 0.223703676 | 0.133375852 | 0.18576284 | ns | YPD | KS.1 |
| FIM1_243 | 783.6926402 | 1532.99178 | 1207.709963 | 0.954282697 | 0.082592066 | 1.4248393625 | 4.4303598930 | ns | YPD | KS.1 |
| FIM1_245 | 2471.460068 | 2531.351929 | 4952.962543 | 0.028671415 | 0.228965874 | 0.890582928 | 0.916749444 | ns | YPD | KS.1 |
| FIM1_246 | 3933.201017 | 2771.677275 | 2323.651007 | -0.491914725 | 0.093638135 | 8.5350491097 | 3.4919300428 | ns | YPD | KS.1 |
| FIM1_247 | 2113.96917 | 2720.437793 | 1343.159356 | 0.316323353 | 0.21206435 | 0.104174165 | 0.149787997 | ns | YPD | KS.1 |
| FIM1_248 | 1834.840423 | 1006.789638 | 1115.187354 | -0.843134101 | 0.11579885 | 8.5802009430 | 6.3997750662 | ns | YPD | KS.1 |
| FIM1_249 | 643.9271894 | 380.2364369 | 995.5845502 | -0.592464635 | 0.336580717 | 0.036497477 | 0.058515936 | ns | YPD | KS.1 |
| FIM1_250 | 650.6217864 | 584.5973562 | 804.4931944 | -0.144027988 | 0.125286755 | 0.233158653 | 0.300976725 | ns | YPD | KS.1 |
| FIM1_251 | 190.6539967 | 398.3019075 | 522.5479514 | 1.031667293 | 0.134062291 | 2.5247178401 | 2.1618711281 | Km>KS.1 | YPD | KS.1 |
| FIM1_252 | 1639.402543 | 1999.713871 | 2061.09628 | 0.279254666 | 0.082352359 | 0.000561072 | 0.001303144 | ns | YPD | KS.1 |
| FIM1_253 | 4944.757054 | 4304.487521 | 4712.613198 | -0.191711024 | 0.101552159 | 0.052632944 | 0.081955835 | ns | YPD | KS.1 |
| FIM1_254 | 795.1756395 | 466.3968109 | 630.1023059 | -0.734601759 | 0.147818614 | 2.2523149915 | 8.7200522046 | ns | YPD | KS.1 |
| FIM1_255 | 3156.078246 | 4037.21134 | 3916.679253 | 0.328314117 | 0.154606855 | 0.025456252 | 0.042341977 | ns | YPD | KS.1 |
| FIM1_256 | 1876.378592 | 2614.13579 | 2945.980669 | 0.436597319 | 0.178285848 | 0.009087455 | 0.016732084 | ns | YPD | KS.1 |
| FIM1_257 | 1348.274775 | 1001.633065 | 989.1991687 | -0.411933552 | 0.114612989 | 0.000216259 | 0.00054092 | ns | YPD | KS.1 |
| FIM1_258 | 1981.806864 | 1492.973125 | 1326.343152 | -0.392310571 | 0.111725334 | 0.000306 | 0.000743698 | ns | YPD | KS.1 |
| FIM1_259 | 1417.185444 | 2303.730336 | 3182.099749 | 0.66650837 | 0.145550579 | 1.8326563200 | 6.1729622154 | ns | YPD | KS.1 |
| FIM1_260 | 450.0650025 | 477.4724951 | 531.1506979 | 0.077313326 | 0.153936611 | 0.597667241 | 0.668779695 | ns | YPD | KS.1 |
| FIM1_261 | 953.2870946 | 1672.923945 | 1417.32722 | 0.736219157 | 0.216820358 | 0.000233397 | 0.000580866 | ns | YPD | KS.1 |
| FIM1_262 | 916.9766317 | 677.8119183 | 1256.727153 | -0.386972735 | 0.200612421 | 0.037050644 | 0.05928818 | ns | YPD | KS.1 |
| FIM1_263 | 14350.43186 | 3718.377598 | 4356.538085 | -1.921921085 | 0.117204609 | 5.0572161535 | 1.0938978419 | Km<KS.1 | YPD | KS.1 |
| FIM1_264 | 1410.121351 | 1121.167717 | 998.4399574 | -0.318785332 | 0.100960316 | 0.001218327 | 0.002652593 | ns | YPD | KS.1 |
| FIM1_265 | 1171.603082 | 1885.957517 | 1462.899096 | 0.675413232 | 0.08308133 | 1.5952191816 | 1.5470205514 | ns | YPD | KS.1 |
| FIM1_266 | 1208.847205 | 1475.555528 | 972.3963985 | 0.268516648 | 0.138212761 | 0.042596616 | 0.067275609 | ns | YPD | KS.1 |
| FIM1_267 | 1510.825392 | 4575.271523 | 5139.402025 | 1.581453917 | 0.094042209 | 1.1069481420 | 2.8984563193 | Km>KS.1 | YPD | KS.1 |
| FIM1_268 | 7490.569994 | 5353.290899 | 5576.256847 | -0.455794074 | 0.145312335 | 0.001048627 | 0.002314516 | ns | YPD | KS.1 |
| FIM1_269 | 1362.693082 | 1228.726998 | 1362.757812 | -0.140951543 | 0.118399338 | 0.218396151 | 0.285134182 | ns | YPD | KS.1 |
| FIM1_270 | 53.02448969 | 82.98523194 | 135.2654486 | 0.573537861 | 0.21285238 | 0.003426732 | 0.006895643 | ns | YPD | KS.1 |
| FIM1_271 | 2845.699438 | 3540.609291 | 2806.474049 | 0.295266191 | 0.137440469 | 0.02505103 | 0.041709797 | ns | YPD | KS.1 |
| FIM1_272 | 714.2506732 | 772.9771612 | 690.4990834 | 0.105581858 | 0.138895493 | 0.427738512 | 0.505342935 | ns | YPD | KS.1 |
| FIM1_273 | 634.0982114 | 549.6631315 | 446.7911323 | -0.179575687 | 0.201152022 | 0.334619163 | 0.409427037 | ns | YPD | KS.1 |
| FIM1_274 | 981.6742962 | 997.6833822 | 1722.830084 | 0.020045705 | 0.212362203 | 0.91830459 | 0.938682008 | ns | YPD | KS.1 |
| FIM1_275 | 979.6986038 | 1074.990891 | 1602.25596 | 0.119549776 | 0.168552517 | 0.452325358 | 0.529985552 | ns | YPD | KS.1 |
| FIM1_276 | 446.3479246 | 1311.858175 | 906.7754852 | 1.521869762 | 0.134312965 | 6.5107374897 | 1.9166224267 | Km>KS.1 | YPD | KS.1 |
| FIM1_277 | 427.321698 | 380.141939 | 311.8922662 | -0.159541163 | 0.107163031 | 0.125542769 | 0.176483548 | ns | YPD | KS.1 |
| FIM1_278 | 406.2265532 | 502.993005 | 637.3066224 | 0.28579914 | 0.152816987 | 0.049212832 | 0.076943381 | ns | YPD | KS.1 |
| FIM1_279 | 2397.730751 | 2500.994628 | 3531.57243 | 0.05750219 | 0.112695377 | 0.599088556 | 0.669918087 | ns | YPD | KS.1 |

| | | | | | | | | | | |
|---|---|---|---|---|---|---|---|---|---|---|
| FIM1_280 | 14.75461542 | 17.79411007 | 26.86602388 | 0.191676112 | 0.343308028 | 0.502387638 | 0.57878634 | ns | YPD | KS.1 |
| FIM1_281 | 1905.136062 | 3846.308396 | 3055.708706 | 0.960537615 | 0.17573299 | 9.5382771056 | 4.3937896852 | ns | YPD | KS.1 |
| FIM1_282 | 921.4317083 | 643.2931873 | 392.1436837 | -0.390522709 | 0.327633138 | 0.157772323 | 0.215045838 | ns | YPD | KS.1 |
| FIM1_283 | 11546.58107 | 956.6347052 | 1789.244243 | -3.426639325 | 0.343475319 | 2.0017964532 | 4.3488809410 | Km<KS.1 | YPD | KS.1 |
| FIM1_284 | 1604.8485 | 881.7129912 | 1082.482432 | -0.826121867 | 0.149565587 | 9.0796750916 | 4.1980839759 | ns | YPD | KS.1 |
| FIM1_285 | 682.4779099 | 2117.409155 | 1445.590392 | 1.44525662 | 0.32560914 | 6.8551965430 | 2.4948502415 | Km>KS.1 | YPD | KS.1 |
| FIM1_286 | 5184.021527 | 4945.472245 | 3409.995974 | -0.060337639 | 0.173609106 | 0.712103118 | 0.770489998 | ns | YPD | KS.1 |
| FIM1_287 | 661.7134678 | 729.5544809 | 685.4018079 | 0.133151888 | 0.108184094 | 0.205232801 | 0.270615739 | ns | YPD | KS.1 |
| FIM1_288 | 106.253344 | 103.7096613 | 132.9791323 | -0.03555813 | 0.180700248 | 0.833863454 | 0.871710587 | ns | YPD | KS.1 |
| FIM1_290 | 285.8985245 | 424.5192834 | 401.1793757 | 0.53079075 | 0.163888146 | 0.000636866 | 0.001468896 | ns | YPD | KS.1 |
| FIM1_291 | 2657.547068 | 1852.090593 | 1665.079634 | -0.484327202 | 0.161582828 | 0.001582924 | 0.00339561 | ns | YPD | KS.1 |
| FIM1_292 | 5299.579073 | 1531.860867 | 1255.979238 | -1.626724213 | 0.301393203 | 3.6118584106 | 1.7496587724 | Km<KS.1 | YPD | KS.1 |
| FIM1_293 | 11699.41185 | 4198.744448 | 3668.649036 | -1.445730537 | 0.132768757 | 1.0701117253 | 2.7301568376 | Km<KS.1 | YPD | KS.1 |
| FIM1_294 | 18076.19821 | 718.4367029 | 2209.438311 | -4.428709555 | 0.406949226 | 4.3695072751 | 1.2076832607 | Km<KS.1 | YPD | KS.1 |
| FIM1_295 | 284.1402875 | 42.61914981 | 54.48768277 | -2.534451545 | 0.34363637 | 7.3989700846 | 6.6927047583 | Km<KS.1 | YPD | KS.1 |
| FIM1_296 | 386.8559322 | 78.18672128 | 99.09863344 | -2.09895524 | 0.339387752 | 1.0281099046 | 6.4256869041 | Km<KS.1 | YPD | KS.1 |
| FIM1_297 | 104.91069 | 45.49441513 | 31.93400973 | -1.004646724 | 0.34328612 | 0.000641643 | 0.001477857 | Km<KS.1 | YPD | KS.1 |
| FIM1_298 | 133.86227 | 71.99454444 | 43.81990865 | -0.808138906 | 0.23186687 | 0.000142767 | 0.000364801 | ns | YPD | KS.1 |
| FIM1_299 | 1049.552759 | 779.8605833 | 625.4742574 | -0.355212356 | 0.253899591 | 0.116881271 | 0.165948722 | ns | YPD | KS.1 |
| FIM1_300 | 1200.102538 | 845.2799995 | 1750.706662 | -0.375443448 | 0.33493606 | 0.181371872 | 0.243148764 | ns | YPD | KS.1 |
| FIM1_301 | 278.4318437 | 221.5919101 | 194.6034244 | -0.27425026 | 0.243425332 | 0.210161975 | 0.276157322 | ns | YPD | KS.1 |
| FIM1_302 | 1284.34748 | 727.9838759 | 737.3037931 | -0.773333965 | 0.16682965 | 1.1024438768 | 3.8706127645 | ns | YPD | KS.1 |
| FIM1_303 | 60248.71537 | 26552.72154 | 31660.2633 | -1.143033457 | 0.148466408 | 2.0015486169 | 1.7378192616 | Km<KS.1 | YPD | KS.1 |
| FIM1_304 | 2432.465592 | 1977.201623 | 807.8463057 | -0.260765694 | 0.202948907 | 0.164114553 | 0.222592667 | ns | YPD | KS.1 |
| FIM1_305 | 4384.960467 | 1063.967734 | 843.7830533 | -1.821405138 | 0.350795055 | 6.8822456613 | 3.2392783505 | Km<KS.1 | YPD | KS.1 |
| FIM1_306 | 1605.082556 | 298.2550638 | 334.8470566 | -2.041324449 | 0.470227319 | 2.4230396829 | 9.3591788997 | Km<KS.1 | YPD | KS.1 |
| FIM1_307 | 252.9490829 | 898.6537694 | 1116.511687 | 1.79805488 | 0.129400423 | 2.4859081077 | 1.7667704051 | Km>KS.1 | YPD | KS.1 |
| FIM1_308 | 1669.087968 | 1680.509381 | 2450.767558 | 0.009242284 | 0.146333033 | 0.947315418 | 0.962404371 | ns | YPD | KS.1 |
| FIM1_309 | 2965.56166 | 3916.551929 | 4752.360542 | 0.392049262 | 0.084348248 | 2.3071830728 | 7.6828887464 | ns | YPD | KS.1 |
| FIM1_310 | 4085.069577 | 3449.718469 | 4156.181608 | -0.222976771 | 0.156798627 | 0.133903571 | 0.186298402 | ns | YPD | KS.1 |
| FIM1_311 | 9907.964763 | 3081.680622 | 3860.09946 | -1.663624503 | 0.105357107 | 1.8163419995 | 2.5100837355 | Km<KS.1 | YPD | KS.1 |
| FIM1_312 | 2511.19033 | 3495.566578 | 3656.8811 | 0.428284569 | 0.193946109 | 0.017510438 | 0.030318351 | ns | YPD | KS.1 |
| FIM1_313 | 960.1330168 | 2725.043185 | 2693.26069 | 1.439689419 | 0.188872526 | 2.0731987024 | 1.7968926036 | Km>KS.1 | YPD | KS.1 |
| FIM1_314 | 1007.198264 | 2086.836153 | 2126.457402 | 0.971209857 | 0.216420824 | 1.4647121261 | 5.0393795488 | ns | YPD | KS.1 |
| FIM1_315 | 1019.925659 | 831.113858 | 659.2698316 | -0.286836628 | 0.08763424 | 0.000849072 | 0.00190105 | ns | YPD | KS.1 |
| FIM1_316 | 700.1452699 | 466.2663407 | 1101.901215 | -0.435776756 | 0.343867334 | 0.128282716 | 0.179599287 | ns | YPD | KS.1 |
| FIM1_317 | 104.894295 | 60.21342172 | 101.9596112 | -0.684669139 | 0.273171488 | 0.004680905 | 0.009161095 | ns | YPD | KS.1 |
| FIM1_318 | 4573.968612 | 7714.668643 | 7843.167884 | 0.748516894 | 0.057434245 | 2.4166229292 | 1.1903662448 | ns | YPD | KS.1 |
| FIM1_319 | 433.3600522 | 311.5530188 | 301.5180789 | -0.444326774 | 0.15060785 | 0.001988698 | 0.004194053 | ns | YPD | KS.1 |
| FIM1_320 | 4258.225679 | 2050.468058 | 1810.685389 | -0.957536155 | 0.239522823 | 1.3396577139 | 3.9932876733 | ns | YPD | KS.1 |
| FIM1_321 | 54.37109368 | 38.12532614 | 107.0083954 | -0.375510905 | 0.348908627 | 0.194226162 | 0.25794852 | ns | YPD | KS.1 |
| FIM1_322 | 8627.210989 | 11782.9295 | 14723.54858 | 0.434578254 | 0.105415867 | 2.3956675905 | 6.9012427694 | ns | YPD | KS.1 |
| FIM1_323 | 424.7876989 | 535.5314942 | 571.029106 | 0.285506959 | 0.223024626 | 0.160180595 | 0.217850865 | ns | YPD | KS.1 |
| FIM1_324 | 4713.384977 | 3179.098381 | 5869.511829 | -0.492670997 | 0.234396077 | 0.020313691 | 0.034574277 | ns | YPD | KS.1 |
| FIM1_325 | 30.11190622 | 15.64920357 | 17.19711774 | -0.730458468 | 0.367728258 | 0.016224876 | 0.028352217 | ns | YPD | KS.1 |

| | | | | | | | | | | |
|---|---|---|---|---|---|---|---|---|---|---|
| FIM1_326 | 38517.48323 | 5271.617922 | 4974.569664 | -2.557772452 | 0.435106819 | 1.79665687805 | 1.09404748694 | Km<KS.1 | YPD | KS.1 |
| FIM1_327 | 1530.426308 | 1553.428477 | 1256.923338 | 0.020915786 | 0.109976742 | 0.844869904 | 0.87947331 | ns | YPD | KS.1 |
| FIM1_328 | 597.583981 | 228.6456441 | 197.3835281 | -1.200076267 | 0.327949359 | 3.13193761570 | 8.87178444638 | Km<KS.1 | YPD | KS.1 |
| FIM1_329 | 979.5583435 | 1280.839033 | 674.5158635 | 0.324987814 | 0.240718404 | 0.13381965 | 0.186276653 | ns | YPD | KS.1 |
| FIM1_330 | 9954.969208 | 9076.359625 | 10680.29304 | -0.121026696 | 0.156160948 | 0.41461895 | 0.492063281 | ns | YPD | KS.1 |
| FIM1_331 | 1643.45806 | 423.7255024 | 526.2211276 | -1.824663644 | 0.267223578 | 3.01351576888 | 2.15715697125 | Km<KS.1 | YPD | KS.1 |
| FIM1_332 | 1511.009459 | 1320.845016 | 1159.272756 | -0.175036693 | 0.167007342 | 0.266825604 | 0.336542656 | ns | YPD | KS.1 |
| FIM1_333 | 1977.126343 | 1559.37928 | 1067.606057 | -0.316536085 | 0.153371706 | 0.029975823 | 0.048991366 | ns | YPD | KS.1 |
| FIM1_334 | 1215.280997 | 1813.893686 | 1901.909592 | 0.542723657 | 0.154227918 | 0.000224196 | 0.000559365 | ns | YPD | KS.1 |
| FIM1_335 | 680.8408566 | 621.1998216 | 1214.735243 | -0.100262653 | 0.291449422 | 0.692269826 | 0.752960731 | ns | YPD | KS.1 |
| FIM1_336 | 1744.461649 | 1060.512416 | 665.6980096 | -0.655608108 | 0.19977236 | 0.000419996 | 0.000992155 | ns | YPD | KS.1 |
| FIM1_337 | 3485.049206 | 3408.936383 | 3669.238901 | -0.029555118 | 0.138667453 | 0.824187765 | 0.863268033 | ns | YPD | KS.1 |
| FIM1_338 | 3631.910195 | 2235.344358 | 2051.336962 | -0.572643395 | 0.29583641 | 0.025750657 | 0.04278875 | ns | YPD | KS.1 |
| FIM1_339 | 6054.504515 | 3619.768365 | 3976.175577 | -0.697135824 | 0.167278016 | 1.13474384109 | 3.42557682611 | ns | YPD | KS.1 |
| FIM1_340 | 197.4798011 | 402.1846817 | 277.3983359 | 0.836452237 | 0.340250847 | 0.003812384 | 0.007601848 | ns | YPD | KS.1 |
| FIM1_341 | 3312.559528 | 4400.158339 | 4523.907596 | 0.391989087 | 0.118238771 | 0.000628487 | 0.001450915 | ns | YPD | KS.1 |
| FIM1_342 | 921.2064296 | 1015.968614 | 1112.983384 | 0.129104567 | 0.14354305 | 0.346813973 | 0.422064461 | ns | YPD | KS.1 |
| FIM1_343 | 2750.061184 | 3323.749721 | 2331.160854 | 0.228682977 | 0.233692079 | 0.279810748 | 0.350998101 | ns | YPD | KS.1 |
| FIM1_344 | 1976.532339 | 1434.742497 | 1776.822187 | -0.44170776 | 0.122376352 | 0.00019348 | 0.000487372 | ns | YPD | KS.1 |
| FIM1_345 | 6600.66594 | 4168.398533 | 3734.663347 | -0.583162054 | 0.232297766 | 0.005748375 | 0.011012002 | ns | YPD | KS.1 |
| FIM1_346 | 8232.320794 | 7254.895282 | 6352.248537 | -0.16611013 | 0.156100341 | 0.262406722 | 0.332435305 | ns | YPD | KS.1 |
| FIM1_347 | 2794.816722 | 1372.6274 | 1962.428707 | -1.000308 | 0.120173142 | 1.61122951123 | 1.71645970414 | Km<KS.1 | YPD | KS.1 |
| FIM1_348 | 1148.170239 | 941.8284892 | 2060.195225 | -0.218319655 | 0.296200571 | 0.394254174 | 0.471380561 | ns | YPD | KS.1 |
| FIM1_349 | 2632.285849 | 3071.891352 | 2501.27156 | 0.211026946 | 0.116966341 | 0.062258585 | 0.09539158 | ns | YPD | KS.1 |
| FIM1_350 | 514.0411539 | 671.5719642 | 551.4005951 | 0.370193865 | 0.10692901 | 0.000381129 | 0.000908973 | ns | YPD | KS.1 |
| FIM1_351 | 1902.738012 | 2255.271181 | 1800.110609 | 0.240561602 | 0.069213575 | 0.00042918 | 0.00101145 | ns | YPD | KS.1 |
| FIM1_352 | 10432.58905 | 9572.926302 | 5972.361674 | -0.093907686 | 0.292873399 | 0.711913337 | 0.770452219 | ns | YPD | KS.1 |
| FIM1_354 | 1069.215643 | 810.7991623 | 493.2727055 | -0.311917843 | 0.292048925 | 0.218273842 | 0.285091195 | ns | YPD | KS.1 |
| FIM1_355 | 1349.508554 | 2169.416754 | 2709.148435 | 0.659320234 | 0.124890691 | 5.13684802398 | 2.16025519182 | ns | YPD | KS.1 |
| FIM1_356 | 10076.0117 | 2379.496765 | 2103.957352 | -1.978824594 | 0.236640831 | 1.46180954984 | 1.68585374278 | Km<KS.1 | YPD | KS.1 |
| FIM1_357 | 4364.228928 | 7107.99115 | 3991.281274 | 0.578223287 | 0.292798549 | 0.02324011 | 0.03898164 | ns | YPD | KS.1 |
| FIM1_358 | 3921.940553 | 3509.198448 | 6074.080717 | -0.124082841 | 0.280512628 | 0.613773625 | 0.68311494 | ns | YPD | KS.1 |
| FIM1_359 | 7863.788155 | 8600.225231 | 7789.665494 | 0.127700799 | 0.049742002 | 0.009658498 | 0.017717931 | ns | YPD | KS.1 |
| FIM1_360 | 1274.25119 | 1426.531545 | 1375.551515 | 0.155779135 | 0.095822992 | 0.095914579 | 0.139148648 | ns | YPD | KS.1 |
| FIM1_361 | 640.0283546 | 651.2820704 | 560.8953301 | 0.022530819 | 0.172427928 | 0.889751097 | 0.916082721 | ns | YPD | KS.1 |
| FIM1_362 | 2375.975092 | 3771.859156 | 2519.03007 | 0.609291447 | 0.193895482 | 0.000754687 | 0.001711289 | ns | YPD | KS.1 |
| FIM1_363 | 1416.171302 | 1212.891605 | 1212.245779 | -0.206022136 | 0.145975279 | 0.139110817 | 0.192725233 | ns | YPD | KS.1 |
| FIM1_364 | 2605.52802 | 2709.633455 | 2836.126551 | 0.055145973 | 0.065683934 | 0.395352163 | 0.472459474 | ns | YPD | KS.1 |
| FIM1_365 | 1255.208687 | 1151.950493 | 1520.484683 | -0.109682824 | 0.176772364 | 0.509008705 | 0.585032844 | ns | YPD | KS.1 |
| FIM1_366 | 2478.891083 | 2204.263561 | 1946.325552 | -0.15341929 | 0.15968821 | 0.31077099 | 0.383929892 | ns | YPD | KS.1 |
| FIM1_367 | 16907.06148 | 3497.157907 | 4689.156278 | -2.096139357 | 0.313614272 | 3.91960570461 | 2.76989181540 | Km<KS.1 | YPD | KS.1 |
| FIM1_368 | 1156.216742 | 541.6402373 | 489.1536025 | -0.925736582 | 0.317568853 | 0.000796104 | 0.001791918 | ns | YPD | KS.1 |
| FIM1_369 | 46.32807459 | 29.21451799 | 35.23365655 | -0.467911327 | 0.383998547 | 0.130644226 | 0.182622935 | ns | YPD | KS.1 |
| FIM1_370 | 445.5447946 | 724.0524585 | 767.2498702 | 0.675201755 | 0.127433938 | 4.45981922029 | 1.88306435582 | ns | YPD | KS.1 |
| FIM1_371 | 167.2730593 | 203.5554891 | 98.26239274 | 0.229143114 | 0.258578183 | 0.319326477 | 0.392743936 | ns | YPD | KS.1 |

| | | | | | | | | | | |
|---|---|---|---|---|---|---|---|---|---|---|
| FIM1_372 | 19628.71866 | 15924.52562 | 15260.98212 | -0.25890377 | 0.216697866 | 0.191845761 | 0.255674434 | ns | YPD | KS.1 |
| FIM1_373 | 2194.374976 | 2066.450788 | 1422.934025 | -0.076046819 | 0.184354644 | 0.659379648 | 0.723354741 | ns | YPD | KS.1 |
| FIM1_374 | 3004.538064 | 1574.49469 | 1849.472422 | -0.902109382 | 0.131929201 | 1.85456174592 | 1.24681684945 | ns | YPD | KS.1 |
| FIM1_375 | 3102.825407 | 2415.545298 | 1675.434603 | -0.309998231 | 0.221838165 | 0.125512507 | 0.176483548 | ns | YPD | KS.1 |
| FIM1_376 | 894.7528354 | 656.3232787 | 754.4049239 | -0.362125334 | 0.273153977 | 0.131896203 | 0.184067353 | ns | YPD | KS.1 |
| FIM1_377 | 615.4265433 | 605.2706712 | 809.5753069 | -0.020795716 | 0.195994432 | 0.909143227 | 0.931806253 | ns | YPD | KS.1 |
| FIM1_378 | 79789.73581 | 36888.20518 | 46593.75577 | -1.077248023 | 0.142661289 | 7.10588340929 | 5.79537212479 | Km<KS.1 | YPD | KS.1 |
| FIM1_379 | 887.5170642 | 1009.930324 | 566.4716419 | 0.137692605 | 0.310570807 | 0.604414415 | 0.674532373 | ns | YPD | KS.1 |
| FIM1_380 | 2893.132342 | 4571.087385 | 2248.476043 | 0.555765565 | 0.268597204 | 0.01942865 | 0.033272817 | ns | YPD | KS.1 |
| FIM1_381 | 2100.589177 | 3780.805812 | 2891.447937 | 0.715310904 | 0.29019592 | 0.004916912 | 0.009570281 | ns | YPD | KS.1 |
| FIM1_383 | 29120.34426 | 5882.485825 | 5227.277974 | -2.086826988 | 0.35140161 | 4.87149530605 | 2.85125754677 | Km<KS.1 | YPD | KS.1 |
| FIM1_384 | 28903.50492 | 38327.54768 | 47056.6348 | 0.355000621 | 0.21328775 | 0.069605192 | 0.104839791 | ns | YPD | KS.1 |
| FIM1_385 | 1202.31793 | 1296.342784 | 715.9322575 | 0.096193393 | 0.174807789 | 0.558599691 | 0.631742092 | ns | YPD | KS.1 |
| FIM1_386 | 3415.536062 | 4555.373066 | 2795.587424 | 0.336475002 | 0.26960298 | 0.157154009 | 0.214320503 | ns | YPD | KS.1 |
| FIM1_387 | 498.4897564 | 725.697823 | 897.1630477 | 0.505918355 | 0.155542673 | 0.00063762 | 0.001469954 | ns | YPD | KS.1 |
| FIM1_388 | 820.6487028 | 904.0694341 | 624.7718793 | 0.127766369 | 0.142734751 | 0.349315434 | 0.424381999 | ns | YPD | KS.1 |
| FIM1_389 | 2350.198491 | 4013.631579 | 3232.834531 | 0.728369542 | 0.163344747 | 2.82006500168 | 9.26398401810 | ns | YPD | KS.1 |
| FIM1_390 | 335.9565749 | 337.9209936 | 386.9148257 | 0.008551223 | 0.206927291 | 0.964308293 | 0.973110295 | ns | YPD | KS.1 |
| FIM1_391 | 1018.308317 | 1496.483542 | 1297.739817 | 0.530683055 | 0.126505584 | 1.44570436176 | 4.28890922878 | ns | YPD | KS.1 |
| FIM1_392 | 2204.419544 | 2877.81742 | 4038.265816 | 0.362549456 | 0.13405982 | 0.004913032 | 0.009566472 | ns | YPD | KS.1 |
| FIM1_393 | 2124.918467 | 2757.420728 | 1931.746616 | 0.366252604 | 0.088745587 | 2.63319922448 | 7.53749490323 | ns | YPD | KS.1 |
| FIM1_394 | 232.2639964 | 311.0247596 | 215.3585894 | 0.359402177 | 0.232467166 | 0.087803337 | 0.128704066 | ns | YPD | KS.1 |
| FIM1_395 | 1347.868565 | 1881.597774 | 1365.180795 | 0.450979144 | 0.14908079 | 0.001538408 | 0.003304655 | ns | YPD | KS.1 |
| FIM1_396 | 801.4268899 | 1925.973514 | 2343.481647 | 1.232969249 | 0.132686822 | 1.86622965533 | 2.69898038816 | Km>KS.1 | YPD | KS.1 |
| FIM1_397 | 54.53001053 | 51.6575611 | 44.24618426 | -0.067981334 | 0.271050645 | 0.776415313 | 0.824651191 | ns | YPD | KS.1 |
| FIM1_398 | 1797.43832 | 2119.873361 | 2180.258315 | 0.223799098 | 0.126996271 | 0.067311683 | 0.10197187 | ns | YPD | KS.1 |
| FIM1_399 | 1132.708813 | 1548.607406 | 2774.821104 | 0.423575934 | 0.143430005 | 0.002043756 | 0.004297416 | ns | YPD | KS.1 |
| FIM1_401 | 845.7274615 | 1250.229733 | 1422.118783 | 0.537772753 | 0.132386363 | 2.53878183676 | 7.29240163852 | ns | YPD | KS.1 |
| FIM1_402 | 1270.212734 | 576.3867009 | 416.4323931 | -1.071634309 | 0.198322937 | 1.08652661719 | 4.98199992675 | Km<KS.1 | YPD | KS.1 |
| FIM1_403 | 2878.362283 | 1249.746177 | 967.7019184 | -1.149848586 | 0.17458462 | 6.48941927653 | 4.13378500649 | Km<KS.1 | YPD | KS.1 |
| FIM1_404 | 939.6638504 | 688.6345345 | 546.760275 | -0.389832073 | 0.219915439 | 0.052317409 | 0.081490016 | ns | YPD | KS.1 |
| FIM1_405 | 4567.990918 | 8536.484498 | 7626.635991 | 0.888586416 | 0.087221781 | 5.20664130510 | 1.01183751925 | ns | YPD | KS.1 |
| FIM1_406 | 7143.226416 | 10498.62089 | 9920.880166 | 0.545017945 | 0.081844591 | 1.39233914397 | 8.57288024910 | ns | YPD | KS.1 |
| FIM1_407 | 2842.722365 | 3839.502461 | 3459.852175 | 0.393545015 | 0.179990796 | 0.01964205 | 0.033603576 | ns | YPD | KS.1 |
| FIM1_408 | 2746.294725 | 1941.650538 | 1724.87704 | -0.490355726 | 0.081775465 | 1.15288220780 | 5.93746271619 | ns | YPD | KS.1 |
| FIM1_409 | 4452.165594 | 5433.726064 | 4668.864928 | 0.272435888 | 0.122041624 | 0.020830668 | 0.035273171 | ns | YPD | KS.1 |
| FIM1_410 | 430.6832037 | 175.0529496 | 198.6579872 | -1.221959985 | 0.207749222 | 5.06271212751 | 2.69668017498 | Km<KS.1 | YPD | KS.1 |
| FIM1_411 | 6861.265643 | 3435.535195 | 2144.197744 | -0.825554991 | 0.325133452 | 0.003113952 | 0.006310351 | ns | YPD | KS.1 |
| FIM1_412 | 1296.049458 | 855.9695325 | 728.237725 | -0.576516749 | 0.119920117 | 7.31256315632 | 2.63814370578 | ns | YPD | KS.1 |
| FIM1_413 | 5370.210224 | 4855.35031 | 4930.833769 | -0.140106258 | 0.092310995 | 0.120614384 | 0.170713104 | ns | YPD | KS.1 |
| FIM1_414 | 3767.824975 | 818.2695689 | 811.4516496 | -2.046476937 | 0.29355831 | 6.11857709445 | 4.61910789756 | Km<KS.1 | YPD | KS.1 |
| FIM1_415 | 463.5868415 | 559.4598869 | 421.903523 | 0.246289616 | 0.161549578 | 0.107151451 | 0.153369169 | ns | YPD | KS.1 |
| FIM1_416 | 2504.32819 | 1490.420017 | 1370.58872 | -0.724888306 | 0.11910916 | 3.91876715464 | 2.10538516137 | ns | YPD | KS.1 |
| FIM1_417 | 12808.55074 | 5575.960397 | 6188.655595 | -1.158234017 | 0.15319061 | 5.68600715310 | 4.69898431672 | Km<KS.1 | YPD | KS.1 |
| FIM1_418 | 1721.445377 | 464.1823986 | 901.0270708 | -1.826219238 | 0.18457778 | 1.56342684759 | 2.92407840856 | Km<KS.1 | YPD | KS.1 |

| | | | | | | | | | | |
|---|---|---|---|---|---|---|---|---|---|---|
| FIM1_419 | 1070.25791 | 1149.281182 | 844.6077708 | 0.094562628 | 0.140587993 | 0.482540874 | 0.560766375 | ns | YPD | KS.1 |
| FIM1_420 | 755.8358485 | 656.2663979 | 749.4564911 | -0.190020805 | 0.129569915 | 0.127384753 | 0.178568371 | ns | YPD | KS.1 |
| FIM1_421 | 1769.718149 | 1200.307083 | 1164.997138 | -0.543450701 | 0.103711103 | 8.2301035272 | 3.3727154075 | ns | YPD | KS.1 |
| FIM1_422 | 879.6274856 | 817.4280055 | 623.0930477 | -0.099238542 | 0.120962399 | 0.396016916 | 0.472910535 | ns | YPD | KS.1 |
| FIM1_423 | 370.3805739 | 138.5757118 | 92.95811991 | -1.066312221 | 0.45170586 | 0.002870454 | 0.005857468 | Km<KS.1 | YPD | KS.1 |
| FIM1_425 | 1474.305387 | 1698.034242 | 3568.612381 | 0.160080411 | 0.273471209 | 0.506337653 | 0.58256934 | ns | YPD | KS.1 |
| FIM1_426 | 4040.208785 | 5099.531329 | 4407.258022 | 0.319639182 | 0.120321541 | 0.006035719 | 0.011513689 | ns | YPD | KS.1 |
| FIM1_427 | 245.2543964 | 263.5827905 | 348.6366388 | 0.093204574 | 0.147352381 | 0.507607571 | 0.583625529 | ns | YPD | KS.1 |
| FIM1_428 | 98.92921532 | 355.5801363 | 345.022864 | 1.762763765 | 0.20320098 | 1.7157745103 | 2.1555004477 | Km>KS.1 | YPD | KS.1 |
| FIM1_429 | 349.7347862 | 1166.260226 | 1382.713337 | 1.709370014 | 0.117605144 | 3.3138261837 | 2.8923307481 | Km>KS.1 | YPD | KS.1 |
| FIM1_430 | 825.1109464 | 1087.300141 | 1029.537073 | 0.385229513 | 0.098901737 | 6.8237843712 | 0.000183219 | ns | YPD | KS.1 |
| FIM1_431 | 756.2914502 | 763.9370075 | 613.3133998 | 0.012660574 | 0.192756069 | 0.943792502 | 0.959609176 | ns | YPD | KS.1 |
| FIM1_432 | 168.9792167 | 123.5098894 | 229.5730587 | -0.379278681 | 0.246634076 | 0.086415033 | 0.126968337 | ns | YPD | KS.1 |
| FIM1_433 | 563.5304385 | 1397.942759 | 1934.95982 | 1.278769694 | 0.133523866 | 1.1340204275 | 1.7853644389 | Km>KS.1 | YPD | KS.1 |
| FIM1_434 | 917.5076923 | 1215.243536 | 1025.508772 | 0.383531581 | 0.13253456 | 0.002644755 | 0.005423602 | ns | YPD | KS.1 |
| FIM1_435 | 1032.563177 | 805.1187244 | 1007.262228 | -0.346283882 | 0.104901187 | 0.000709653 | 0.001616539 | ns | YPD | KS.1 |
| FIM1_436 | 470.2946789 | 247.598339 | 391.0584459 | -0.842600644 | 0.22293031 | 4.2189246979 | 0.000117652 | ns | YPD | KS.1 |
| FIM1_437 | 598.3519373 | 478.4770635 | 545.3349553 | -0.298566494 | 0.156701505 | 0.044670554 | 0.070350113 | ns | YPD | KS.1 |
| FIM1_438 | 64.96542338 | 51.8589638 | 26.92615975 | -0.223122162 | 0.364311823 | 0.453596491 | 0.53109968 | ns | YPD | KS.1 |
| FIM1_439 | 1586.088442 | 1473.412468 | 1895.423913 | -0.084906576 | 0.256264492 | 0.710535999 | 0.769128937 | ns | YPD | KS.1 |
| FIM1_440 | 1451.106134 | 587.0914628 | 823.75812 | -1.277468684 | 0.123149294 | 3.7258038727 | 7.9248613176 | Km<KS.1 | YPD | KS.1 |
| FIM1_441 | 2264.46646 | 1155.012958 | 966.3812468 | -0.949365171 | 0.110828612 | 2.2272394548 | 2.5125887274 | ns | YPD | KS.1 |
| FIM1_442 | 2040.870452 | 1749.630755 | 1211.952098 | -0.197136201 | 0.182372098 | 0.247893201 | 0.316547401 | ns | YPD | KS.1 |
| FIM1_443 | 329.8417378 | 342.6881445 | 176.328668 | 0.042330871 | 0.283067391 | 0.86432501 | 0.895300863 | ns | YPD | KS.1 |
| FIM1_444 | 1312.682876 | 1069.733842 | 1310.064377 | -0.274866273 | 0.143835634 | 0.04548203 | 0.071447142 | ns | YPD | KS.1 |
| FIM1_445 | 2690.250113 | 2239.426914 | 1874.845332 | -0.248965318 | 0.128146547 | 0.043607784 | 0.06882891 | ns | YPD | KS.1 |
| FIM1_446 | 195.3605023 | 177.7369418 | 323.4614719 | -0.119212408 | 0.172655374 | 0.463392993 | 0.541042042 | ns | YPD | KS.1 |
| FIM1_447 | 1099.277064 | 1331.487758 | 917.951041 | 0.270305468 | 0.080659693 | 0.000655252 | 0.001505718 | ns | YPD | KS.1 |
| FIM1_448 | 2468.47793 | 3321.664014 | 2575.742673 | 0.405745348 | 0.13132918 | 0.001343862 | 0.002905568 | ns | YPD | KS.1 |
| FIM1_449 | 3405.33794 | 5636.359182 | 4280.172542 | 0.662835731 | 0.202577364 | 0.000428 | 0.001009146 | ns | YPD | KS.1 |
| FIM1_450 | 34.12820246 | 48.22547615 | 132.0429841 | 0.4106155 | 0.258842612 | 0.074460201 | 0.111332664 | ns | YPD | KS.1 |
| FIM1_451 | 3013.160637 | 2068.029029 | 2000.966981 | -0.467645524 | 0.237222707 | 0.02913959 | 0.047750151 | ns | YPD | KS.1 |
| FIM1_452 | 651.8383757 | 1290.437745 | 780.8864883 | 0.936459787 | 0.169559105 | 7.2460556493 | 3.4008610241 | ns | YPD | KS.1 |
| FIM1_453 | 578.9338194 | 1647.894051 | 2775.856282 | 1.453618525 | 0.173873102 | 5.0924486499 | 5.5927002281 | Km>KS.1 | YPD | KS.1 |
| FIM1_454 | 330.0293472 | 660.0282577 | 268.9446485 | 0.769356219 | 0.377994828 | 0.013221654 | 0.02353407 | ns | YPD | KS.1 |
| FIM1_455 | 636.0939206 | 1197.221176 | 1245.188269 | 0.819394262 | 0.237032669 | 0.000154885 | 0.000393943 | ns | YPD | KS.1 |
| FIM1_456 | 2524.516406 | 1284.413455 | 1442.054858 | -0.887349885 | 0.228885599 | 2.5776906903 | 7.3998910469 | ns | YPD | KS.1 |
| FIM1_457 | 8019.999646 | 9799.66153 | 14320.3747 | 0.270447076 | 0.137130257 | 0.039670415 | 0.06301415 | ns | YPD | KS.1 |
| FIM1_458 | 10408.66933 | 13319.45607 | 20648.47836 | 0.338123735 | 0.12306426 | 0.004471932 | 0.008797098 | ns | YPD | KS.1 |
| FIM1_459 | 498.8533614 | 1236.3796 | 1542.097326 | 1.235586534 | 0.204818094 | 1.9653860004 | 1.0804193759 | Km>KS.1 | YPD | KS.1 |
| FIM1_460 | 557.9898088 | 264.1437347 | 413.5934671 | -1.021236287 | 0.18090934 | 3.0310815104 | 1.4842156018 | Km<KS.1 | YPD | KS.1 |
| FIM1_461 | 1175.124434 | 1002.533257 | 842.2776394 | -0.214365261 | 0.13110802 | 0.088958558 | 0.130205597 | ns | YPD | KS.1 |
| FIM1_462 | 668.6709535 | 687.1992629 | 709.7308589 | 0.037766388 | 0.07840214 | 0.62437719 | 0.692901298 | ns | YPD | KS.1 |
| FIM1_463 | 515.9763418 | 1011.792139 | 2168.265916 | 0.865433499 | 0.253976952 | 0.000167939 | 0.000425188 | ns | YPD | KS.1 |
| FIM1_464 | 3784.153747 | 7114.072049 | 5385.754652 | 0.863063982 | 0.167977318 | 7.0449602078 | 2.9158633139 | ns | YPD | KS.1 |

| | | | | | | | | | | |
|---|---|---|---|---|---|---|---|---|---|---|
| FIM1_465 | 12.19768512 | 17.20838134 | 27.27999917 | 0.332580369 | 0.360069177 | 0.260480814 | 0.330416127 | ns | YPD | KS.1 |
| FIM1_466 | 3167.1164 | 7713.477005 | 7328.141002 | 1.25880772 | 0.118773609 | 3.5625213177 | 8.1300658512 | Km>KS.1 | YPD | KS.1 |
| FIM1_467 | 1652.045872 | 1581.3691 | 2370.570354 | -0.053229731 | 0.215107985 | 0.787615596 | 0.832990559 | ns | YPD | KS.1 |
| FIM1_468 | 255.1197979 | 54.80100785 | 101.8914025 | -2.128694692 | 0.224431205 | 3.7116281540 | 6.1551166888 | Km<KS.1 | YPD | KS.1 |
| FIM1_469 | 1022.646028 | 518.4983104 | 815.2713542 | -0.890797648 | 0.230950503 | 2.7744346989 | 7.9235434139 | ns | YPD | KS.1 |
| FIM1_470 | 3378.352211 | 2110.73632 | 1474.85134 | -0.602597306 | 0.224778911 | 0.003359734 | 0.00677328 | ns | YPD | KS.1 |
| FIM1_471 | 1457.565052 | 1873.345164 | 924.8653515 | 0.278403967 | 0.298697677 | 0.280229405 | 0.351434659 | ns | YPD | KS.1 |
| FIM1_472 | 793.5124645 | 390.3574807 | 470.3324537 | -0.898292189 | 0.274857949 | 0.00025798 | 0.000636188 | ns | YPD | KS.1 |
| FIM1_473 | 2011.221198 | 2395.254308 | 1561.354367 | 0.221138557 | 0.19487655 | 0.22158371 | 0.288354422 | ns | YPD | KS.1 |
| FIM1_474 | 477.3568242 | 1120.448596 | 1664.142884 | 1.199066738 | 0.132786413 | 2.2576769858 | 3.0521584251 | Km>KS.1 | YPD | KS.1 |
| FIM1_475 | 1265.492143 | 943.8108878 | 707.7649955 | -0.360405049 | 0.232976425 | 0.087504667 | 0.128341897 | ns | YPD | KS.1 |
| FIM1_476 | 3479.633226 | 6344.609919 | 9708.260431 | 0.818136101 | 0.170139683 | 4.2585884257 | 1.5905763827 | ns | YPD | KS.1 |
| FIM1_477 | 8642.708385 | 7137.082783 | 8762.515758 | -0.265120917 | 0.104763658 | 0.009341852 | 0.017168716 | ns | YPD | KS.1 |
| FIM1_478 | 3299.204192 | 3623.736782 | 3891.826898 | 0.10423933 | 0.282684661 | 0.673492451 | 0.735592743 | ns | YPD | KS.1 |
| FIM1_479 | 1626.569457 | 2036.631911 | 1402.401075 | 0.274330839 | 0.229778569 | 0.188297155 | 0.251416625 | ns | YPD | KS.1 |
| FIM1_480 | 608.1961795 | 718.2917962 | 1130.206243 | 0.224902387 | 0.125588332 | 0.063174901 | 0.096676448 | ns | YPD | KS.1 |
| FIM1_481 | 3032.502104 | 1915.546252 | 1343.131751 | -0.640925486 | 0.115709852 | 1.2511011916 | 5.6790405368 | ns | YPD | KS.1 |
| FIM1_482 | 2063.049191 | 1696.940982 | 1188.457089 | -0.260028078 | 0.150326284 | 0.069233886 | 0.104362163 | ns | YPD | KS.1 |
| FIM1_483 | 1404.665151 | 2122.085855 | 2054.337398 | 0.58204194 | 0.090681292 | 6.4133909086 | 3.6928958067 | ns | YPD | KS.1 |
| FIM1_485 | 240.3738542 | 121.4393688 | 116.1817359 | -0.918624247 | 0.199625077 | 9.5017672120 | 3.3693009180 | ns | YPD | KS.1 |
| FIM1_486 | 1350.980012 | 1774.178468 | 1003.7575 | 0.35707646 | 0.174812455 | 0.029722297 | 0.048624935 | ns | YPD | KS.1 |
| FIM1_487 | 260.4601742 | 569.6346768 | 668.4113705 | 1.042866532 | 0.220837513 | 4.0942598553 | 1.5338059322 | Km>KS.1 | YPD | KS.1 |
| FIM1_488 | 3049.681764 | 4036.835686 | 5026.573853 | 0.362873742 | 0.187794478 | 0.03821793 | 0.060901411 | ns | YPD | KS.1 |
| FIM1_489 | 3658.60063 | 6505.977957 | 3569.253078 | 0.719893401 | 0.264436187 | 0.002300378 | 0.004778446 | ns | YPD | KS.1 |
| FIM1_490 | 2343.386208 | 2944.945915 | 1831.003238 | 0.31196377 | 0.126340714 | 0.010462544 | 0.019017595 | ns | YPD | KS.1 |
| FIM1_491 | 6885.564499 | 8992.756889 | 12395.74351 | 0.37414164 | 0.0935111 | 4.4624317945 | 0.000124095 | ns | YPD | KS.1 |
| FIM1_492 | 2413.74619 | 2405.770328 | 2334.251937 | -0.00503227 | 0.092733874 | 0.955785517 | 0.968044167 | ns | YPD | KS.1 |
| FIM1_493 | 716.636411 | 879.5618442 | 1321.065216 | 0.276218141 | 0.138501149 | 0.037377895 | 0.059703302 | ns | YPD | KS.1 |
| FIM1_494 | 408.0084435 | 1371.591933 | 1974.82989 | 1.720190465 | 0.121624126 | 9.1047287856 | 7.3058105981 | Km>KS.1 | YPD | KS.1 |
| FIM1_495 | 761.2425978 | 2846.315447 | 3267.866095 | 1.881999737 | 0.101625269 | 4.0476650498 | 1.8306485111 | Km>KS.1 | YPD | KS.1 |
| FIM1_496 | 3028.910791 | 5305.514006 | 5435.532296 | 0.800858927 | 0.064961541 | 1.7365656849 | 6.8566780022 | ns | YPD | KS.1 |
| FIM1_497 | 1603.686066 | 1423.854427 | 1598.935476 | -0.145090074 | 0.218663216 | 0.468584678 | 0.546334373 | ns | YPD | KS.1 |
| FIM1_498 | 448.4464121 | 487.9050938 | 630.0746192 | 0.109825064 | 0.156259658 | 0.459454931 | 0.536822048 | ns | YPD | KS.1 |
| FIM1_499 | 1981.33343 | 2603.555047 | 3312.6156 | 0.376743981 | 0.115165127 | 0.000752827 | 0.001707849 | ns | YPD | KS.1 |
| FIM1_500 | 333.8024773 | 300.6012982 | 352.4480144 | -0.137855391 | 0.154040253 | 0.346605111 | 0.422016747 | ns | YPD | KS.1 |
| FIM1_501 | 514.7297469 | 570.2093059 | 642.575356 | 0.131683827 | 0.17612207 | 0.426194962 | 0.503998083 | ns | YPD | KS.1 |
| FIM1_502 | 1737.717034 | 2136.772478 | 2460.250386 | 0.278667433 | 0.137476201 | 0.034467684 | 0.055692344 | ns | YPD | KS.1 |
| FIM1_503 | 1665.036612 | 1597.356451 | 959.0798002 | -0.053189533 | 0.17027687 | 0.740610824 | 0.794081648 | ns | YPD | KS.1 |
| FIM1_504 | 3669.529532 | 1239.889902 | 1291.733053 | -1.547701321 | 0.094843805 | 4.7178337352 | 8.6930454936 | Km<KS.1 | YPD | KS.1 |
| FIM1_505 | 1307.27535 | 803.1815069 | 1053.730224 | -0.644072318 | 0.194356636 | 0.000384081 | 0.000914697 | ns | YPD | KS.1 |
| FIM1_506 | 1606.148052 | 2042.847874 | 3207.087225 | 0.284238959 | 0.255464822 | 0.211974792 | 0.27825187 | ns | YPD | KS.1 |
| FIM1_507 | 1212.110296 | 1233.845834 | 1545.803788 | 0.023750893 | 0.133631489 | 0.853379493 | 0.886523904 | ns | YPD | KS.1 |
| FIM1_508 | 2393.049691 | 1384.296449 | 1038.605976 | -0.75633095 | 0.141695535 | 2.9977661650 | 1.3059445421 | ns | YPD | KS.1 |
| FIM1_509 | 2856.02144 | 1567.363103 | 964.2370488 | -0.782319859 | 0.226150736 | 0.000166588 | 0.000422116 | ns | YPD | KS.1 |
| FIM1_510 | 550.920056 | 794.3587464 | 854.0455006 | 0.49944606 | 0.137809393 | 0.000163754 | 0.000415864 | ns | YPD | KS.1 |

| | | | | | | | | | | |
|---|---|---|---|---|---|---|---|---|---|---|
| FIM1_511 | 3518.262907 | 1279.290511 | 1599.389843 | -1.378107865 | 0.213117854 | 9.33311616714 | 5.87750037108 | Km<KS.1 | YPD | KS.1 |
| FIM1_512 | 923.8513911 | 200.5046651 | 324.3843601 | -2.115387002 | 0.218927599 | 6.91159326696 | 1.22367176167 | Km<KS.1 | YPD | KS.1 |
| FIM1_513 | 1317.221844 | 2426.556847 | 3189.871274 | 0.852711461 | 0.130132074 | 1.44485958514 | 8.88526135487 | ns | YPD | KS.1 |
| FIM1_514 | 2418.452474 | 2518.257084 | 1894.917338 | 0.052128126 | 0.170850932 | 0.74635566 | 0.7988639 | ns | YPD | KS.1 |
| FIM1_515 | 1643.663654 | 1733.020178 | 1777.12397 | 0.073873536 | 0.072294569 | 0.299788841 | 0.372025314 | ns | YPD | KS.1 |
| FIM1_516 | 1393.354854 | 1385.524621 | 1352.961096 | -0.007833126 | 0.106836137 | 0.939950929 | 0.956485145 | ns | YPD | KS.1 |
| FIM1_517 | 896.4901725 | 851.2094237 | 1368.583645 | -0.058997357 | 0.262876322 | 0.80059971 | 0.843852449 | ns | YPD | KS.1 |
| FIM1_518 | 13959.4733 | 21284.76236 | 24314.06549 | 0.598136957 | 0.080386474 | 4.45711315291 | 3.39054096871 | ns | YPD | KS.1 |
| FIM1_519 | 3329.444654 | 784.6238849 | 587.3415799 | -1.807056407 | 0.394293843 | 1.49835053524 | 5.92551185440 | Km<KS.1 | YPD | KS.1 |
| FIM1_522 | 1338.835377 | 1930.420779 | 2603.129516 | 0.446136023 | 0.251191451 | 0.047089997 | 0.073818978 | ns | YPD | KS.1 |
| FIM1_523 | 129.0143418 | 312.9783509 | 636.3143509 | 1.20911906 | 0.199813741 | 1.84219797545 | 1.01494295989 | Km>KS.1 | YPD | KS.1 |
| FIM1_524 | 1158.981204 | 2449.020384 | 1738.364328 | 1.052023366 | 0.122638093 | 1.65565095689 | 1.88918887857 | Km>KS.1 | YPD | KS.1 |
| FIM1_525 | 1950.226996 | 1629.417485 | 1775.306378 | -0.253288215 | 0.073657918 | 0.000485346 | 0.001136816 | ns | YPD | KS.1 |
| FIM1_526 | 311.9669938 | 158.8503919 | 136.6789371 | -0.7839259 | 0.342721315 | 0.006815743 | 0.012848929 | ns | YPD | KS.1 |
| FIM1_527 | 51636.41622 | 24421.6185 | 14481.84218 | -0.871249525 | 0.35577623 | 0.003612291 | 0.007231849 | ns | YPD | KS.1 |
| FIM1_528 | 7965.439116 | 6085.953545 | 3607.913838 | -0.299155594 | 0.300021391 | 0.247498895 | 0.316287439 | ns | YPD | KS.1 |
| FIM1_529 | 40.88557307 | 27.15763008 | 26.20148965 | -0.503891262 | 0.240640116 | 0.020269944 | 0.034511625 | ns | YPD | KS.1 |
| FIM1_530 | 576.3632859 | 2522.572471 | 3718.552015 | 2.07407963 | 0.17128987 | 1.65442110921 | 6.58459601469 | Km>KS.1 | YPD | KS.1 |
| FIM1_531 | 4012.350766 | 10247.51995 | 10265.85102 | 1.199657832 | 0.296966838 | 6.72758546157 | 2.09578820734 | Km>KS.1 | YPD | KS.1 |
| FIM1_532 | 5684.35635 | 3154.766366 | 4169.976391 | -0.630048539 | 0.379929933 | 0.041605154 | 0.065835128 | ns | YPD | KS.1 |
| FIM1_533 | 1709.411048 | 1638.007497 | 1149.200053 | -0.058189273 | 0.105695439 | 0.5716871 | 0.644200073 | ns | YPD | KS.1 |
| FIM1_534 | 1066.177525 | 1217.108106 | 1090.027217 | 0.178244499 | 0.128603789 | 0.149857455 | 0.205553029 | ns | YPD | KS.1 |
| FIM1_535 | 1678.057038 | 2280.867974 | 1542.118228 | 0.328597882 | 0.328469397 | 0.235277829 | 0.3031252 | ns | YPD | KS.1 |
| FIM1_536 | 3537.32496 | 1279.668114 | 1058.344059 | -1.312036056 | 0.296786103 | 9.89483189700 | 3.49126160904 | Km<KS.1 | YPD | KS.1 |
| FIM1_537 | 1558.058101 | 2091.459743 | 995.1810894 | 0.338594831 | 0.281501138 | 0.168970599 | 0.228555935 | ns | YPD | KS.1 |
| FIM1_538 | 409.3473561 | 1076.119191 | 1018.384598 | 1.344988554 | 0.16206211 | 1.05253481522 | 1.13341140817 | Km>KS.1 | YPD | KS.1 |
| FIM1_539 | 358.4551801 | 453.210965 | 440.214433 | 0.320878358 | 0.126899035 | 0.008739677 | 0.016127557 | ns | YPD | KS.1 |
| FIM1_540 | 2714.153076 | 2969.704164 | 2288.724654 | 0.124588184 | 0.096449962 | 0.185817296 | 0.248372124 | ns | YPD | KS.1 |
| FIM1_541 | 314.4632558 | 466.8378831 | 552.7711887 | 0.562102413 | 0.07868005 | 4.38955738644 | 3.09320793166 | ns | YPD | KS.1 |
| FIM1_542 | 387.3241386 | 622.3218338 | 827.8097266 | 0.657172385 | 0.126155286 | 7.54886344891 | 3.11664694260 | ns | YPD | KS.1 |
| FIM1_543 | 913.3581704 | 879.648123 | 548.75533 | -0.046249884 | 0.207458504 | 0.809153908 | 0.850705979 | ns | YPD | KS.1 |
| FIM1_544 | 1204.170766 | 2354.934731 | 1915.488907 | 0.930214738 | 0.147423168 | 6.11585708967 | 3.52565342075 | ns | YPD | KS.1 |
| FIM1_545 | 348.587707 | 1050.270542 | 1877.281489 | 1.529639705 | 0.179792513 | 1.23629922119 | 1.43705341715 | Km>KS.1 | YPD | KS.1 |
| FIM1_546 | 9322.171143 | 4121.29287 | 4994.01369 | -1.158954814 | 0.101138687 | 2.97070274381 | 9.06702217819 | Km<KS.1 | YPD | KS.1 |
| FIM1_547 | 3198.66386 | 2971.796379 | 2150.754405 | -0.095136417 | 0.165797895 | 0.543919074 | 0.617525649 | ns | YPD | KS.1 |
| FIM1_549 | 1568.898245 | 1679.540372 | 1034.980657 | 0.087899033 | 0.169365242 | 0.582361086 | 0.654892949 | ns | YPD | KS.1 |
| FIM1_550 | 9310.503743 | 9630.006046 | 11820.81753 | 0.04536396 | 0.124249797 | 0.705399102 | 0.764733174 | ns | YPD | KS.1 |
| FIM1_551 | 117.6341125 | 374.7406252 | 566.140344 | 1.600437268 | 0.190331209 | 2.48852019339 | 2.80099275161 | Km>KS.1 | YPD | KS.1 |
| FIM1_552 | 895.7498258 | 1061.722136 | 1160.952802 | 0.23331125 | 0.111523819 | 0.031065708 | 0.050540189 | ns | YPD | KS.1 |
| FIM1_553 | 1883.219253 | 1885.440282 | 1591.648014 | 0.002151891 | 0.092671938 | 0.981073015 | 0.986083092 | ns | YPD | KS.1 |
| FIM1_554 | 1270.158771 | 1534.249084 | 1887.284174 | 0.261870691 | 0.102463446 | 0.008727819 | 0.016111651 | ns | YPD | KS.1 |
| FIM1_555 | 369.6286376 | 413.8026653 | 575.5339994 | 0.151400744 | 0.143532735 | 0.269752701 | 0.339837854 | ns | YPD | KS.1 |
| FIM1_556 | 1055.917867 | 728.9525519 | 1202.338386 | -0.511050331 | 0.128937115 | 4.07442221818 | 0.000114038 | ns | YPD | KS.1 |
| FIM1_557 | 1212.575841 | 938.0608079 | 994.4919767 | -0.357322481 | 0.101962974 | 0.000331812 | 0.000801343 | ns | YPD | KS.1 |
| FIM1_558 | 269.7188465 | 857.1748225 | 1298.567273 | 1.619256171 | 0.161326788 | 6.02844652730 | 1.16246207261 | Km>KS.1 | YPD | KS.1 |

| | | | | | | | | | | |
|---|---|---|---|---|---|---|---|---|---|---|
| FIM1_559 | 567.7860779 | 668.5182727 | 786.9371466 | 0.223448665 | 0.117281121 | 0.048948592 | 0.076578378 | ns | YPD | KS.1 |
| FIM1_560 | 810.5486631 | 1483.677333 | 2200.518531 | 0.842481277 | 0.130884282 | 3.1888457823 | 1.8999410499 | ns | YPD | KS.1 |
| FIM1_561 | 2059.446632 | 1211.674294 | 945.6953352 | -0.73139312 | 0.142046382 | 8.8360245532 | 3.6061708082 | ns | YPD | KS.1 |
| FIM1_562 | 4450.454185 | 2162.769505 | 1953.332671 | -1.023411007 | 0.09908674 | 9.4184549302 | 1.9282639209 | Km<KS.1 | YPD | KS.1 |
| FIM1_563 | 588.0535647 | 132.9936166 | 220.2215981 | -2.021697126 | 0.258272635 | 1.0451115619 | 1.0295901031 | Km<KS.1 | YPD | KS.1 |
| FIM1_564 | 445.1455662 | 1049.221585 | 1556.202246 | 1.171235772 | 0.192796347 | 1.7081967780 | 9.4741125648 | Km>KS.1 | YPD | KS.1 |
| FIM1_565 | 693.2251378 | 1167.653235 | 1520.594745 | 0.688681414 | 0.201519422 | 0.000239136 | 0.000592481 | ns | YPD | KS.1 |
| FIM1_566 | 257.570442 | 495.4362678 | 826.6143196 | 0.902306663 | 0.156741152 | 1.9949746478 | 1.0005039186 | ns | YPD | KS.1 |
| FIM1_567 | 956.0479342 | 1165.118793 | 1648.299936 | 0.270984595 | 0.117111014 | 0.01685179 | 0.029283149 | ns | YPD | KS.1 |
| FIM1_568 | 2266.717584 | 2789.139475 | 3256.95093 | 0.208280545 | 0.352512113 | 0.474288113 | 0.551949325 | ns | YPD | KS.1 |
| FIM1_569 | 1151.600868 | 1693.993532 | 2490.001199 | 0.447353212 | 0.290908919 | 0.076754185 | 0.114190212 | ns | YPD | KS.1 |
| FIM1_570 | 776.2135807 | 555.9736421 | 1243.278647 | -0.435035006 | 0.189215545 | 0.013664418 | 0.024211669 | ns | YPD | KS.1 |
| FIM1_571 | 3853.94039 | 2963.375808 | 2026.460833 | -0.290260374 | 0.303283952 | 0.265795779 | 0.335617767 | ns | YPD | KS.1 |
| FIM1_572 | 508.3506451 | 402.9712468 | 510.8466022 | -0.23074641 | 0.362519436 | 0.436799149 | 0.513608075 | ns | YPD | KS.1 |
| FIM1_573 | 6096.836673 | 4057.420743 | 4434.991654 | -0.559236362 | 0.136841714 | 2.1829000206 | 6.3212617013 | ns | YPD | KS.1 |
| FIM1_574 | 1813.943251 | 1820.936804 | 3334.420775 | 0.004235197 | 0.300831104 | 0.986984934 | 0.990568902 | ns | YPD | KS.1 |
| FIM1_575 | 2877.142291 | 1791.647022 | 2703.430966 | -0.661247159 | 0.117256973 | 6.7017790435 | 3.1573248808 | ns | YPD | KS.1 |
| FIM1_576 | 3789.543306 | 5052.20863 | 5823.24603 | 0.403268527 | 0.093051793 | 9.8974375953 | 3.0060898679 | ns | YPD | KS.1 |
| FIM1_577 | 435.042078 | 483.0034956 | 905.1260683 | 0.139483374 | 0.135114273 | 0.282131838 | 0.3536422 | ns | YPD | KS.1 |
| FIM1_578 | 2987.576117 | 3853.593222 | 4422.84059 | 0.347398184 | 0.129059928 | 0.005221407 | 0.010087961 | ns | YPD | KS.1 |
| FIM1_579 | 561.179332 | 1223.968622 | 850.9124711 | 1.095022088 | 0.12827554 | 2.2000839808 | 2.4875949556 | Km>KS.1 | YPD | KS.1 |
| FIM1_580 | 302.8794658 | 232.3991229 | 260.9150765 | -0.362924651 | 0.128496131 | 0.003399841 | 0.006845087 | ns | YPD | KS.1 |
| FIM1_581 | 931.2050661 | 874.4993565 | 835.7897162 | -0.081491125 | 0.157807637 | 0.586610396 | 0.658926782 | ns | YPD | KS.1 |
| FIM1_582 | 1425.279521 | 1297.542894 | 3226.125734 | -0.10037099 | 0.306248476 | 0.702799569 | 0.762746042 | ns | YPD | KS.1 |
| FIM1_583 | 169.6699858 | 177.4926783 | 256.7458051 | 0.054924541 | 0.199200971 | 0.76632038 | 0.816718914 | ns | YPD | KS.1 |
| FIM1_584 | 655.6220296 | 764.8691452 | 916.6185616 | 0.209673015 | 0.121828819 | 0.074659822 | 0.111541326 | ns | YPD | KS.1 |
| FIM1_585 | 2313.835606 | 10106.0461 | 10000.39067 | 2.054431108 | 0.196780368 | 3.0650576108 | 7.0924007507 | Km>KS.1 | YPD | KS.1 |
| FIM1_586 | 3316.065212 | 1772.244212 | 2018.87692 | -0.832338868 | 0.207599926 | 1.6691862984 | 4.9021262305 | ns | YPD | KS.1 |
| FIM1_587 | 434.0198758 | 453.6108029 | 743.9692453 | 0.054341188 | 0.192776459 | 0.762157798 | 0.813327981 | ns | YPD | KS.1 |
| FIM1_588 | 175.5864175 | 390.698574 | 685.0246758 | 1.101641198 | 0.172575521 | 2.7229290891 | 1.6360594466 | Km>KS.1 | YPD | KS.1 |
| FIM1_589 | 1217.597549 | 1084.693258 | 965.9544655 | -0.159732302 | 0.105080061 | 0.118103658 | 0.16758862 | ns | YPD | KS.1 |
| FIM1_590 | 1579.733014 | 1063.353644 | 896.1242943 | -0.555954988 | 0.097147209 | 5.2082075957 | 2.4890329287 | ns | YPD | KS.1 |
| FIM1_591 | 1290.934931 | 691.4247278 | 799.1978164 | -0.873415802 | 0.124757578 | 6.2064981123 | 4.3185074278 | ns | YPD | KS.1 |
| FIM1_592 | 5426.895908 | 6500.02924 | 5793.999017 | 0.242605291 | 0.138915392 | 0.068198851 | 0.10303349 | ns | YPD | KS.1 |
| FIM1_593 | 751.8962043 | 1004.429783 | 1160.875021 | 0.383967621 | 0.164645969 | 0.01365899 | 0.024211669 | ns | YPD | KS.1 |
| FIM1_594 | 772.2778212 | 1412.106949 | 2091.221904 | 0.811839552 | 0.187751969 | 4.3771905965 | 1.3986206305 | ns | YPD | KS.1 |
| FIM1_595 | 1325.62698 | 1530.697557 | 2258.61927 | 0.189249805 | 0.156470281 | 0.202536497 | 0.267841327 | ns | YPD | KS.1 |
| FIM1_596 | 854.4537799 | 2672.844908 | 3810.382358 | 1.592860039 | 0.16662699 | 7.2288295804 | 1.1714471388 | Km>KS.1 | YPD | KS.1 |
| FIM1_597 | 3860.5246 | 1798.903003 | 1847.660826 | -1.07778147 | 0.115500084 | 1.7088841607 | 2.4931667740 | Km<KS.1 | YPD | KS.1 |
| FIM1_598 | 407.1275301 | 659.6650494 | 942.4573859 | 0.660613541 | 0.149607666 | 4.0142985041 | 1.2876295975 | ns | YPD | KS.1 |
| FIM1_599 | 598.1082753 | 604.9789492 | 612.5166413 | 0.015610386 | 0.106070477 | 0.879845858 | 0.908139656 | ns | YPD | KS.1 |
| FIM1_600 | 1427.255385 | 1711.330427 | 1731.684023 | 0.234009409 | 0.178436475 | 0.161779567 | 0.219785185 | ns | YPD | KS.1 |
| FIM1_601 | 1811.413892 | 1386.563866 | 2786.680377 | -0.298529515 | 0.296583425 | 0.244306906 | 0.312817155 | ns | YPD | KS.1 |
| FIM1_602 | 2825.995219 | 2210.140202 | 2163.783996 | -0.346253681 | 0.081808895 | 1.7061293671 | 5.0076658417 | ns | YPD | KS.1 |
| FIM1_603 | 9297.93773 | 8640.828718 | 6877.477411 | -0.098812911 | 0.127085775 | 0.419885132 | 0.497600889 | ns | YPD | KS.1 |

| | | | | | | | | | | |
|---|---|---|---|---|---|---|---|---|---|---|
| FIM1_604 | 997.0520883 | 1624.668989 | 2260.077787 | 0.676736377 | 0.127997419 | 4.72939594773 | 1.99734676061 | ns | YPD | KS.1 |
| FIM1_605 | 94.61761893 | 387.7841655 | 511.9090931 | 1.963781777 | 0.189664762 | 9.82941540991 | 2.17339296285 | Km>KS.1 | YPD | KS.1 |
| FIM1_606 | 1593.046609 | 1142.201429 | 1345.168045 | -0.448331147 | 0.153528486 | 0.002174805 | 0.004542257 | ns | YPD | KS.1 |
| FIM1_607 | 582.2646326 | 321.3565338 | 715.6332677 | -0.734985382 | 0.276996749 | 0.002736036 | 0.005596948 | ns | YPD | KS.1 |
| FIM1_608 | 643.2180263 | 559.0351761 | 591.9919784 | -0.18857875 | 0.13256862 | 0.138654896 | 0.192200643 | ns | YPD | KS.1 |
| FIM1_609 | 12985.98531 | 11358.21959 | 7419.754934 | -0.165062004 | 0.212039761 | 0.397039005 | 0.473799244 | ns | YPD | KS.1 |
| FIM1_610 | 482.2612816 | 485.1908743 | 549.4681379 | 0.008590687 | 0.1135496 | 0.937865564 | 0.954949075 | ns | YPD | KS.1 |
| FIM1_611 | 308.0495369 | 372.90861 | 193.9310292 | 0.246186991 | 0.176722636 | 0.137592761 | 0.190976611 | ns | YPD | KS.1 |
| FIM1_612 | 3414.297581 | 2326.048614 | 1807.422097 | -0.543722628 | 0.078067206 | 1.66393020931 | 1.12626568589 | ns | YPD | KS.1 |
| FIM1_613 | 1372.617042 | 1553.443816 | 1297.621499 | 0.171445644 | 0.092295608 | 0.057458085 | 0.088802105 | ns | YPD | KS.1 |
| FIM1_615 | 1729.257454 | 1373.319724 | 988.5036954 | -0.316193072 | 0.121177514 | 0.006968743 | 0.013102606 | ns | YPD | KS.1 |
| FIM1_616 | 1592.216081 | 1819.454059 | 2442.610828 | 0.182642807 | 0.113447705 | 0.096675089 | 0.140067131 | ns | YPD | KS.1 |
| FIM1_617 | 333.8162822 | 213.2042887 | 457.5838486 | -0.503723512 | 0.324642074 | 0.067092648 | 0.101671009 | ns | YPD | KS.1 |
| FIM1_618 | 3049.65408 | 3105.056446 | 1419.095574 | 0.021929277 | 0.216880424 | 0.912257213 | 0.934420348 | ns | YPD | KS.1 |
| FIM1_619 | 31881.52293 | 12605.87122 | 13755.79097 | -1.289009943 | 0.166314337 | 1.02054923312 | 9.16467948519 | Km<KS.1 | YPD | KS.1 |
| FIM1_620 | 387.3313328 | 1003.379712 | 905.0081186 | 1.218730797 | 0.296817945 | 4.87142737188 | 1.54759586048 | Km>KS.1 | YPD | KS.1 |
| FIM1_621 | 301.784054 | 659.1934452 | 628.3279031 | 0.987531423 | 0.287493239 | 0.00011613 | 0.000299818 | ns | YPD | KS.1 |
| FIM1_622 | 199.2021619 | 174.3306503 | 308.1685685 | -0.179098152 | 0.125997318 | 0.140194176 | 0.193848256 | ns | YPD | KS.1 |
| FIM1_623 | 1705.293306 | 2240.739487 | 3212.491914 | 0.376951615 | 0.115582923 | 0.000779688 | 0.001760756 | ns | YPD | KS.1 |
| FIM1_625 | 11068.6351 | 1817.134974 | 1665.134305 | -2.436240996 | 0.313125857 | 4.35581743350 | 4.46808076941 | Km<KS.1 | YPD | KS.1 |
| FIM1_626 | 8118.214199 | 2240.286552 | 3460.275943 | -1.755091549 | 0.236032519 | 4.35913093130 | 3.65473775130 | Km<KS.1 | YPD | KS.1 |
| FIM1_627 | 2864.695516 | 1372.809286 | 1097.322672 | -1.029520427 | 0.133641215 | 2.37752260426 | 2.04286268674 | Km<KS.1 | YPD | KS.1 |
| FIM1_628 | 4159.769699 | 2888.22884 | 2869.052745 | -0.480553624 | 0.18228952 | 0.004905148 | 0.009554859 | ns | YPD | KS.1 |
| FIM1_629 | 3125.127553 | 2755.323417 | 2629.388905 | -0.160133711 | 0.185783845 | 0.356125537 | 0.431181442 | ns | YPD | KS.1 |
| FIM1_630 | 1068.059334 | 889.7205497 | 786.9538102 | -0.246500847 | 0.136558304 | 0.059744338 | 0.092021078 | ns | YPD | KS.1 |
| FIM1_631 | 3477.157236 | 3436.424459 | 2014.126863 | -0.013530994 | 0.256401483 | 0.952873134 | 0.96666881 | ns | YPD | KS.1 |
| FIM1_632 | 556.7858917 | 430.5800197 | 338.091691 | -0.330992478 | 0.189263637 | 0.060397401 | 0.092969391 | ns | YPD | KS.1 |
| FIM1_633 | 1271.644316 | 4263.56279 | 5665.138552 | 1.7225747 | 0.107700923 | 6.10758796860 | 9.80169359478 | Km>KS.1 | YPD | KS.1 |
| FIM1_634 | 707.3466302 | 472.3863444 | 504.229274 | -0.563872755 | 0.108565832 | 1.01045172306 | 4.07704567904 | ns | YPD | KS.1 |
| FIM1_635 | 1732.717497 | 2752.416668 | 3749.836985 | 0.647645129 | 0.11027695 | 1.72897705222 | 8.72379395011 | ns | YPD | KS.1 |
| FIM1_636 | 656.8229426 | 682.6030582 | 505.1129773 | 0.054031583 | 0.090065102 | 0.540213923 | 0.614440847 | ns | YPD | KS.1 |
| FIM1_637 | 4192.769969 | 5518.0346 | 8546.879484 | 0.377143598 | 0.123782488 | 0.00162442 | 0.003472921 | ns | YPD | KS.1 |
| FIM1_638 | 1544.92002 | 1448.305831 | 1509.720033 | -0.088354553 | 0.105609643 | 0.389938839 | 0.466872009 | ns | YPD | KS.1 |
| FIM1_639 | 178.2619192 | 67.60167514 | 79.66084673 | -1.287595102 | 0.256994284 | 5.86987872740 | 2.44578280308 | Km<KS.1 | YPD | KS.1 |
| FIM1_640 | 685.839947 | 1269.330705 | 1173.532448 | 0.876496697 | 0.081594932 | 1.52420328253 | 3.57684496727 | ns | YPD | KS.1 |
| FIM1_641 | 516.3256964 | 569.3843878 | 514.1290684 | 0.127980013 | 0.146840228 | 0.361169845 | 0.436121354 | ns | YPD | KS.1 |
| FIM1_642 | 4175.265275 | 3328.234952 | 3393.308776 | -0.280315665 | 0.22015718 | 0.163253149 | 0.221602738 | ns | YPD | KS.1 |
| FIM1_644 | 1033.127006 | 914.3304583 | 1767.328646 | -0.150860548 | 0.208858286 | 0.432974486 | 0.510507293 | ns | YPD | KS.1 |
| FIM1_645 | 1809.59857 | 2344.17339 | 1058.851261 | 0.328809212 | 0.201408385 | 0.077338106 | 0.114955804 | ns | YPD | KS.1 |
| FIM1_646 | 3124.624927 | 3736.928237 | 2889.920807 | 0.241107754 | 0.136574833 | 0.065572964 | 0.099793666 | ns | YPD | KS.1 |
| FIM1_647 | 1127.211601 | 1415.447047 | 1610.557098 | 0.290880831 | 0.193863372 | 0.106126999 | 0.152156144 | ns | YPD | KS.1 |
| FIM1_648 | 244.6081534 | 173.7376759 | 180.6262444 | -0.470995658 | 0.126566308 | 0.000118332 | 0.000304868 | ns | YPD | KS.1 |
| FIM1_650 | 1096.822803 | 787.0994267 | 655.9854887 | -0.455201261 | 0.13042642 | 0.000296888 | 0.000723318 | ns | YPD | KS.1 |
| FIM1_651 | 4488.506386 | 1671.575348 | 1317.559511 | -1.308163506 | 0.257237404 | 3.80687876263 | 1.63128525788 | Km<KS.1 | YPD | KS.1 |
| FIM1_652 | 2241.241581 | 2237.551839 | 2526.977574 | -0.002326127 | 0.096650195 | 0.980352649 | 0.98601122 | ns | YPD | KS.1 |

| | | | | | | | | | | |
|---|---|---|---|---|---|---|---|---|---|---|
| FIM1_653 | 3042.12189 | 1623.959655 | 3738.84694 | -0.748619717 | 0.313838156 | 0.005683865 | 0.010905216 | ns | YPD | KS.1 |
| FIM1_654 | 609.5088493 | 556.691956 | 618.8363942 | -0.124145882 | 0.117329431 | 0.274291136 | 0.344943984 | ns | YPD | KS.1 |
| FIM1_655 | 497.14348 | 206.6802834 | 491.6011962 | -1.215201922 | 0.166946536 | 4.2936165343( | 3.2711703305( | Km<KS.1 | YPD | KS.1 |
| FIM1_656 | 984.1317855 | 1439.505998 | 1796.04403 | 0.524085735 | 0.127432864 | 2.1009037327 | 6.0944583501 | ns | YPD | KS.1 |
| FIM1_657 | 697.0714959 | 1001.033068 | 840.9049317 | 0.483960989 | 0.164811937 | 0.001931074 | 0.004086386 | ns | YPD | KS.1 |
| FIM1_658 | 1022.366172 | 879.6906496 | 678.8625973 | -0.194535933 | 0.171457742 | 0.228095212 | 0.295360146 | ns | YPD | KS.1 |
| FIM1_659 | 1904.162892 | 2054.350476 | 2457.652158 | 0.10324868 | 0.11486625 | 0.353676738 | 0.428947287 | ns | YPD | KS.1 |
| FIM1_660 | 1723.330266 | 784.3999922 | 935.7586285 | -1.120137167 | 0.09442819 | 2.7877231193 | 9.6311961937( | Km<KS.1 | YPD | KS.1 |
| FIM1_661 | 1624.36916 | 1152.322672 | 869.1390861 | -0.446315644 | 0.192070861 | 0.012547596 | 0.022479038 | ns | YPD | KS.1 |
| FIM1_662 | 5944.530794 | 8539.970432 | 13329.56743 | 0.482073328 | 0.171064043 | 0.002819444 | 0.005755738 | ns | YPD | KS.1 |
| FIM1_663 | 1243.52997 | 1411.288538 | 728.0410642 | 0.134064409 | 0.314132888 | 0.616977537 | 0.686148358 | ns | YPD | KS.1 |
| FIM1_664 | 2296.290592 | 2128.933683 | 3029.072366 | -0.101370362 | 0.137915465 | 0.44341564 | 0.520526855 | ns | YPD | KS.1 |
| FIM1_665 | 293.6529198 | 547.8535871 | 624.5284246 | 0.879768903 | 0.107041555 | 4.8987382996 | 4.9941030820 | ns | YPD | KS.1 |
| FIM1_666 | 285.9147729 | 306.0593382 | 221.6060011 | 0.079387811 | 0.239712401 | 0.713895049 | 0.771757468 | ns | YPD | KS.1 |
| FIM1_667 | 185.0084826 | 513.9088629 | 255.920656 | 1.399065295 | 0.200873394 | 2.9511579080 | 2.1155634859 | Km>KS.1 | YPD | KS.1 |
| FIM1_668 | 3149.785786 | 4330.984739 | 3917.240784 | 0.436063431 | 0.131975156 | 0.000606277 | 0.001401593 | ns | YPD | KS.1 |
| FIM1_669 | 808.6022345 | 2036.874055 | 1242.978932 | 1.213325312 | 0.261771423 | 4.4336853516 | 1.6497819464 | Km>KS.1 | YPD | KS.1 |
| FIM1_670 | 7330.625082 | 3519.788689 | 2971.716088 | -0.806234985 | 0.392954375 | 0.011667943 | 0.021039513 | ns | YPD | KS.1 |
| FIM1_671 | 355.323041 | 630.2024262 | 622.0532644 | 0.796977941 | 0.130449936 | 2.8875422770 | 1.5648717678 | ns | YPD | KS.1 |
| FIM1_672 | 766.0861734 | 978.3549659 | 662.1761727 | 0.327351058 | 0.151738108 | 0.023427436 | 0.039269372 | ns | YPD | KS.1 |
| FIM1_673 | 1384.243179 | 433.6978126 | 349.8445058 | -1.501307163 | 0.31118974 | 9.6196476116 | 3.8972106570 | Km<KS.1 | YPD | KS.1 |
| FIM1_674 | 6995.230746 | 6986.790361 | 5215.570752 | -0.001561483 | 0.13622699 | 0.990479334 | 0.992874206 | ns | YPD | KS.1 |
| FIM1_675 | 1483.734394 | 1685.956118 | 1473.001452 | 0.174587336 | 0.118559505 | 0.127783282 | 0.179026142 | ns | YPD | KS.1 |
| FIM1_676 | 3618.604882 | 2984.742775 | 3264.916752 | -0.233176111 | 0.231647541 | 0.266901299 | 0.336542656 | ns | YPD | KS.1 |
| FIM1_677 | 6558.756161 | 2208.55517 | 1919.82398 | -1.522182108 | 0.161403209 | 2.8805438439 | 4.4643942752( | Km<KS.1 | YPD | KS.1 |
| FIM1_678 | 3300.436234 | 4387.501771 | 2766.754028 | 0.380916015 | 0.155564252 | 0.009996578 | 0.018243938 | ns | YPD | KS.1 |
| FIM1_679 | 1784.169784 | 3525.929665 | 2876.776366 | 0.870616476 | 0.260444434 | 0.000210108 | 0.000527124 | ns | YPD | KS.1 |
| FIM1_680 | 10514.44817 | 4698.550563 | 5497.70073 | -1.027910227 | 0.280983462 | 4.5676172296( | 0.000126525 | Km<KS.1 | YPD | KS.1 |
| FIM1_681 | 4042.709407 | 2765.355065 | 2011.579786 | -0.419459157 | 0.321259945 | 0.123802587 | 0.174530424 | ns | YPD | KS.1 |
| FIM1_682 | 3284.211312 | 2320.311088 | 1325.94449 | -0.400468691 | 0.288145001 | 0.110267931 | 0.157593496 | ns | YPD | KS.1 |
| FIM1_683 | 5395.533273 | 4414.514702 | 3487.012945 | -0.27346622 | 0.125589713 | 0.023939081 | 0.040072991 | ns | YPD | KS.1 |
| FIM1_684 | 3267.596128 | 2550.615572 | 1704.721099 | -0.345445053 | 0.09890823 | 0.000353017 | 0.000848025 | ns | YPD | KS.1 |
| FIM1_685 | 810.9135257 | 731.8623151 | 571.6491387 | -0.136860623 | 0.143057558 | 0.317045322 | 0.390517573 | ns | YPD | KS.1 |
| FIM1_686 | 4123.287157 | 3376.663178 | 3809.941275 | -0.282715415 | 0.071922832 | 6.7903775948( | 0.000182705 | ns | YPD | KS.1 |
| FIM1_687 | 1202.370115 | 973.8592355 | 1055.214549 | -0.287866578 | 0.120267652 | 0.013313724 | 0.023652716 | ns | YPD | KS.1 |
| FIM1_688 | 813.8756561 | 859.6056367 | 1036.037415 | 0.075117187 | 0.103336492 | 0.455553936 | 0.533015247 | ns | YPD | KS.1 |
| FIM1_689 | 123.1402828 | 80.33626059 | 143.7030848 | -0.518384895 | 0.26501265 | 0.0274023 | 0.045261103 | ns | YPD | KS.1 |
| FIM1_690 | 1422.17977 | 1429.095224 | 1524.061999 | 0.006912428 | 0.11156824 | 0.949142292 | 0.963866687 | ns | YPD | KS.1 |
| FIM1_691 | 865.9151005 | 1249.614259 | 1611.399486 | 0.503891835 | 0.133112395 | 8.5903558000 | 0.000226722 | ns | YPD | KS.1 |
| FIM1_692 | 737.3817979 | 1104.616955 | 1227.666889 | 0.552312573 | 0.140987982 | 4.5346363608 | 0.000125822 | ns | YPD | KS.1 |
| FIM1_693 | 1231.528562 | 1374.417494 | 1255.218501 | 0.151531819 | 0.100575865 | 0.122103101 | 0.172476698 | ns | YPD | KS.1 |
| FIM1_694 | 387.0273586 | 590.0253628 | 543.4684132 | 0.581088893 | 0.130020933 | 3.7289325363 | 1.2062054205 | ns | YPD | KS.1 |
| FIM1_695 | 1274.442171 | 1170.515207 | 1000.654697 | -0.117885204 | 0.082980396 | 0.147742602 | 0.203044046 | ns | YPD | KS.1 |
| FIM1_696 | 1074.634059 | 1468.974212 | 1557.699312 | 0.431963033 | 0.117114716 | 0.000144836 | 0.000369516 | ns | YPD | KS.1 |
| FIM1_697 | 1341.932652 | 999.5629268 | 898.8205216 | -0.407608307 | 0.115256547 | 0.000271316 | 0.000666565 | ns | YPD | KS.1 |

| | | | | | | | | | | |
|---|---|---|---|---|---|---|---|---|---|---|
| FIM1_698 | 130.810937 | 797.2446253 | 812.8636411 | 2.565727543 | 0.152874183 | 1.00112922148 | 2.92977522168 | Km>KS.1 | YPD | KS.1 |
| FIM1_699 | 3055.931694 | 3816.865538 | 2938.740619 | 0.272778839 | 0.225479119 | 0.183748752 | 0.245937595 | ns | YPD | KS.1 |
| FIM1_700 | 4069.815933 | 3676.266259 | 6849.827476 | -0.142056862 | 0.083475404 | 0.082818171 | 0.122188731 | ns | YPD | KS.1 |
| FIM1_701 | 1084.595246 | 921.9876778 | 1751.349444 | -0.200795234 | 0.211787609 | 0.302162372 | 0.37459701 | ns | YPD | KS.1 |
| FIM1_702 | 885.9074205 | 917.0463693 | 1186.573425 | 0.046159077 | 0.119928583 | 0.690806323 | 0.752026577 | ns | YPD | KS.1 |
| FIM1_703 | 1706.176728 | 2786.995868 | 2874.929683 | 0.670952601 | 0.152322691 | 4.12458683014 | 1.31960253890 | ns | YPD | KS.1 |
| FIM1_704 | 8230.499712 | 10192.44576 | 6127.943018 | 0.256700517 | 0.240460197 | 0.236098423 | 0.304025137 | ns | YPD | KS.1 |
| FIM1_705 | 1088.85274 | 1575.249306 | 1247.382072 | 0.439992744 | 0.268981152 | 0.063973895 | 0.097611467 | ns | YPD | KS.1 |
| FIM1_706 | 959.4752546 | 1802.744053 | 2196.0927 | 0.884202923 | 0.120526082 | 5.23317575796 | 3.96271680302 | ns | YPD | KS.1 |
| FIM1_707 | 508.2382446 | 1067.255055 | 994.1467093 | 1.053507515 | 0.093971927 | 6.11326369316 | 1.64397226343 | Km>KS.1 | YPD | KS.1 |
| FIM1_708 | 683.2010182 | 587.6301092 | 661.6962084 | -0.20363612 | 0.126704819 | 0.095229933 | 0.138294201 | ns | YPD | KS.1 |
| FIM1_709 | 1055.014547 | 1215.087688 | 1294.26645 | 0.193922371 | 0.109650039 | 0.068484842 | 0.103371387 | ns | YPD | KS.1 |
| FIM1_710 | 721.70121 | 629.9078864 | 649.6720697 | -0.183120387 | 0.133040215 | 0.151815096 | 0.20795157 | ns | YPD | KS.1 |
| FIM1_711 | 197.9533507 | 387.8043734 | 444.1594199 | 0.909291713 | 0.186628208 | 2.54725152379 | 9.82370258206 | ns | YPD | KS.1 |
| FIM1_712 | 11.51933884 | 31.09884007 | 21.06410889 | 1.08122379 | 0.444056837 | 0.002273401 | 0.00473031 | Km>KS.1 | YPD | KS.1 |
| FIM1_713 | 1007.055293 | 865.7091334 | 1449.725859 | -0.16918845 | 0.281925666 | 0.492881934 | 0.570252935 | ns | YPD | KS.1 |
| FIM1_714 | 2269.780428 | 814.131209 | 955.3241447 | -1.432385454 | 0.159406829 | 2.18252814216 | 2.95860422542 | Km<KS.1 | YPD | KS.1 |
| FIM1_715 | 769.4288395 | 1008.68623 | 1366.728225 | 0.333857557 | 0.22799298 | 0.106948906 | 0.153290351 | ns | YPD | KS.1 |
| FIM1_718 | 3967.414421 | 3526.389915 | 4056.178042 | -0.152974319 | 0.167021756 | 0.331993998 | 0.406314917 | ns | YPD | KS.1 |
| FIM1_719 | 3401.087874 | 4247.496299 | 7126.365091 | 0.299172708 | 0.142051106 | 0.027681474 | 0.045646448 | ns | YPD | KS.1 |
| FIM1_720 | 997.5921218 | 650.7440082 | 588.1164666 | -0.585164421 | 0.141018579 | 1.57159911045 | 4.63742916637 | ns | YPD | KS.1 |
| FIM1_721 | 2076.802705 | 1787.103362 | 2180.455325 | -0.202288181 | 0.133925537 | 0.11560993 | 0.164378223 | ns | YPD | KS.1 |
| FIM1_722 | 3439.779543 | 4227.877245 | 4671.506458 | 0.289151028 | 0.088670806 | 0.000883797 | 0.001972584 | ns | YPD | KS.1 |
| FIM1_723 | 1524.226981 | 1136.311525 | 877.597854 | -0.404157965 | 0.123185014 | 0.000695951 | 0.001589695 | ns | YPD | KS.1 |
| FIM1_724 | 243.4455045 | 254.569499 | 388.5643602 | 0.055930319 | 0.197116148 | 0.760026019 | 0.811227085 | ns | YPD | KS.1 |
| FIM1_725 | 3468.114842 | 3201.214316 | 5829.750812 | -0.099096761 | 0.204433855 | 0.599999045 | 0.670701579 | ns | YPD | KS.1 |
| FIM1_726 | 997.2315658 | 817.3818561 | 898.0738828 | -0.271245511 | 0.124724896 | 0.02417037 | 0.040392203 | ns | YPD | KS.1 |
| FIM1_727 | 854.7593898 | 1155.955931 | 1229.378744 | 0.409589817 | 0.140164175 | 0.00231335 | 0.004799381 | ns | YPD | KS.1 |
| FIM1_728 | 1543.191376 | 2019.86649 | 2356.280893 | 0.353173571 | 0.173939408 | 0.030790435 | 0.050125136 | ns | YPD | KS.1 |
| FIM1_729 | 533.9137608 | 1974.080618 | 2429.537675 | 1.854823818 | 0.127384896 | 1.56947059383 | 1.41965749169 | Km>KS.1 | YPD | KS.1 |
| FIM1_730 | 1525.288363 | 1688.623192 | 1867.201485 | 0.140360625 | 0.098712968 | 0.144877388 | 0.199713219 | ns | YPD | KS.1 |
| FIM1_731 | 548.5578954 | 677.2480734 | 460.353051 | 0.292947294 | 0.101808926 | 0.0031764 | 0.006431661 | ns | YPD | KS.1 |
| FIM1_732 | 4087.760266 | 5536.505528 | 8995.403339 | 0.412291589 | 0.13981708 | 0.00211385 | 0.004431692 | ns | YPD | KS.1 |
| FIM1_733 | 2261.01505 | 1564.576957 | 2077.4532 | -0.456434789 | 0.238307257 | 0.033874496 | 0.054787261 | ns | YPD | KS.1 |
| FIM1_734 | 1633.502574 | 1856.048539 | 2175.091979 | 0.181227109 | 0.060681654 | 0.002530286 | 0.005214838 | ns | YPD | KS.1 |
| FIM1_735 | 900.3596035 | 871.8382932 | 672.6776497 | -0.044845486 | 0.09416728 | 0.626307269 | 0.694423594 | ns | YPD | KS.1 |
| FIM1_736 | 2578.946642 | 3027.483096 | 2714.657046 | 0.218740636 | 0.120214867 | 0.059684178 | 0.091956886 | ns | YPD | KS.1 |
| FIM1_737 | 2373.346267 | 2340.229432 | 2322.058371 | -0.019695904 | 0.084910892 | 0.813190612 | 0.853687127 | ns | YPD | KS.1 |
| FIM1_738 | 979.8001813 | 924.1816326 | 1834.221427 | -0.07612998 | 0.165198486 | 0.626157143 | 0.6944119 | ns | YPD | KS.1 |
| FIM1_739 | 545.82408 | 312.781361 | 482.0774618 | -0.733149986 | 0.210336162 | 0.000168305 | 0.0004259 | ns | YPD | KS.1 |
| FIM1_740 | 757.1773499 | 676.7944626 | 827.4917073 | -0.150882595 | 0.133071703 | 0.237882863 | 0.30580549 | ns | YPD | KS.1 |
| FIM1_741 | 1352.737313 | 1347.900354 | 1896.080384 | -0.004628438 | 0.102317741 | 0.962998147 | 0.972577301 | ns | YPD | KS.1 |
| FIM1_742 | 955.9228752 | 2397.294305 | 1995.452123 | 1.295305974 | 0.129109748 | 1.20625491259 | 2.28545189255 | Km>KS.1 | YPD | KS.1 |
| FIM1_743 | 2060.216428 | 1601.849739 | 2245.619871 | -0.352286231 | 0.094645571 | 0.000144385 | 0.000368557 | ns | YPD | KS.1 |
| FIM1_744 | 354.1768712 | 383.3994683 | 472.0112613 | 0.101552268 | 0.162647335 | 0.509697715 | 0.585521117 | ns | YPD | KS.1 |

| | | | | | | | | | | |
|---|---|---|---|---|---|---|---|---|---|---|
| FIM1_745 | 1332.030095 | 735.5158522 | 570.5548511 | -0.838653207 | 0.102360905 | 6.60165334975 | 6.64842619737 | ns | YPD | KS.1 |
| FIM1_746 | 8064.666347 | 1724.322989 | 1985.127197 | -2.179620999 | 0.156225646 | 3.71810359269 | 2.80266140255 | Km<KS.1 | YPD | KS.1 |
| FIM1_747 | 811.6063868 | 967.0979763 | 1178.845108 | 0.23126306 | 0.154654532 | 0.115402622 | 0.164177307 | ns | YPD | KS.1 |
| FIM1_748 | 573.7166527 | 990.1426358 | 794.7275218 | 0.688089374 | 0.251780299 | 0.002389196 | 0.00494848 | ns | YPD | KS.1 |
| FIM1_749 | 1913.184876 | 1056.660035 | 1109.475622 | -0.822602989 | 0.141375083 | 1.62814589135 | 8.24010763938 | ns | YPD | KS.1 |
| FIM1_750 | 1747.032747 | 1246.905772 | 1422.150315 | -0.46980811 | 0.107334999 | 7.20089329935 | 2.23483744006 | ns | YPD | KS.1 |
| FIM1_751 | 2553.579797 | 4049.084217 | 3565.154167 | 0.654490297 | 0.07848829 | 2.91207935901 | 3.03723161657 | ns | YPD | KS.1 |
| FIM1_752 | 9661.985274 | 5034.62963 | 5517.589437 | -0.924030885 | 0.096403946 | 2.00556729147 | 3.14753857258 | ns | YPD | KS.1 |
| FIM1_753 | 1499.22255 | 3599.144296 | 2091.90095 | 1.220737748 | 0.153884974 | 2.71123048480 | 2.56921364988 | Km>KS.1 | YPD | KS.1 |
| FIM1_754 | 3406.657463 | 3659.614983 | 2116.637094 | 0.091175066 | 0.180685392 | 0.590484536 | 0.662381187 | ns | YPD | KS.1 |
| FIM1_755 | 1646.281731 | 2338.18832 | 2519.886863 | 0.499130573 | 0.066325044 | 2.92018640084 | 2.25239183631 | ns | YPD | KS.1 |
| FIM1_756 | 971.3465195 | 496.7091204 | 822.4926349 | -0.942977528 | 0.118173671 | 3.10212459524 | 2.92316496798 | ns | YPD | KS.1 |
| FIM1_757 | 745.31473 | 1699.83486 | 2005.582828 | 1.160802255 | 0.125936501 | 4.29138299537 | 6.06523590966 | Km>KS.1 | YPD | KS.1 |
| FIM1_758 | 477.709751 | 448.8682845 | 378.9891539 | -0.086233126 | 0.107817165 | 0.410714892 | 0.488128664 | ns | YPD | KS.1 |
| FIM1_759 | 546.6027017 | 544.5051435 | 738.316749 | -0.005810657 | 0.111267157 | 0.957129933 | 0.968592242 | ns | YPD | KS.1 |
| FIM1_760 | 1913.885745 | 2987.418108 | 2381.124377 | 0.603732438 | 0.158307962 | 6.21846466164 | 0.000168852 | ns | YPD | KS.1 |
| FIM1_761 | 2476.251334 | 2792.724321 | 2800.818831 | 0.155309626 | 0.169684342 | 0.331594759 | 0.405926163 | ns | YPD | KS.1 |
| FIM1_762 | 230.7511755 | 464.2605561 | 1122.594086 | 0.961747115 | 0.163608473 | 8.54372475978 | 4.45078855287 | ns | YPD | KS.1 |
| FIM1_763 | 1135.479021 | 763.4667976 | 775.0109219 | -0.548997534 | 0.124307095 | 5.10804858170 | 1.61863328891 | ns | YPD | KS.1 |
| FIM1_764 | 2182.785929 | 2205.101439 | 2454.664853 | 0.013807536 | 0.098749097 | 0.886136308 | 0.913495261 | ns | YPD | KS.1 |
| FIM1_765 | 349.6485292 | 343.8878013 | 354.7709332 | -0.018978124 | 0.238737156 | 0.929944778 | 0.948631387 | ns | YPD | KS.1 |
| FIM1_767 | 4887.66785 | 1682.343081 | 1201.766306 | -1.360903401 | 0.317546916 | 1.64188849371 | 5.59479127138 | Km<KS.1 | YPD | KS.1 |
| FIM1_768 | 528.8776525 | 376.0937411 | 370.6882992 | -0.462121508 | 0.148146711 | 0.001099169 | 0.002413224 | ns | YPD | KS.1 |
| FIM1_769 | 1762.666461 | 2760.663782 | 2602.665713 | 0.638512052 | 0.071440934 | 1.59154669293 | 2.01421676258 | ns | YPD | KS.1 |
| FIM1_770 | 3258.426081 | 2713.405737 | 2444.516323 | -0.224882076 | 0.219319468 | 0.262018882 | 0.332028512 | ns | YPD | KS.1 |
| FIM1_772 | 52.42552198 | 44.88320393 | 53.38291118 | -0.168861458 | 0.321185668 | 0.535366434 | 0.610322642 | ns | YPD | KS.1 |
| FIM1_773 | 1129.10771 | 1432.021372 | 2254.322242 | 0.301345392 | 0.201042324 | 0.104925563 | 0.15069419 | ns | YPD | KS.1 |
| FIM1_774 | 715.6477117 | 928.8811059 | 784.9324631 | 0.355685393 | 0.128595762 | 0.004118854 | 0.008154118 | ns | YPD | KS.1 |
| FIM1_776 | 985.1988162 | 2616.725958 | 2078.601108 | 1.37276769 | 0.139578011 | 7.56527357144 | 1.33465375950 | Km>KS.1 | YPD | KS.1 |
| FIM1_777 | 1612.988253 | 2804.409265 | 2381.458233 | 0.777655369 | 0.10945534 | 3.58996195116 | 2.54779753309 | ns | YPD | KS.1 |
| FIM1_778 | 808.8660092 | 899.3434543 | 665.8322517 | 0.139819834 | 0.150141959 | 0.328342688 | 0.402341102 | ns | YPD | KS.1 |
| FIM1_779 | 4065.428798 | 11760.82789 | 14674.89046 | 1.514050302 | 0.098596055 | 2.25274232516 | 2.43638979732 | Km>KS.1 | YPD | KS.1 |
| FIM1_780 | 1068.038922 | 559.8136579 | 453.4928227 | -0.829474605 | 0.249502525 | 0.000245751 | 0.00060766 | ns | YPD | KS.1 |
| FIM1_781 | 1575.338295 | 438.0751596 | 472.8245672 | -1.732226312 | 0.249222279 | 1.60007084919 | 1.17236413471 | Km<KS.1 | YPD | KS.1 |
| FIM1_782 | 2948.799865 | 3004.398038 | 1671.123472 | 0.022523987 | 0.226388128 | 0.913115114 | 0.934914117 | ns | YPD | KS.1 |
| FIM1_783 | 1413.611065 | 1129.824629 | 935.7181848 | -0.293943606 | 0.168253494 | 0.063880108 | 0.097545592 | ns | YPD | KS.1 |
| FIM1_784 | 1359.281653 | 702.7249425 | 488.4529459 | -0.86785282 | 0.224554386 | 2.82538311523 | 8.06441824342 | ns | YPD | KS.1 |
| FIM1_785 | 1826.17125 | 3874.266427 | 3305.208004 | 1.044886547 | 0.151443585 | 9.13487201120 | 6.30318834898 | Km>KS.1 | YPD | KS.1 |
| FIM1_786 | 1983.85946 | 2031.814571 | 2337.434031 | 0.032257133 | 0.113166742 | 0.769005152 | 0.81882759 | ns | YPD | KS.1 |
| FIM1_787 | 9672.339071 | 6118.064541 | 4819.528209 | -0.610987043 | 0.180208474 | 0.000312734 | 0.000758582 | ns | YPD | KS.1 |
| FIM1_788 | 366.0897021 | 220.7762667 | 200.3153645 | -0.624049566 | 0.266491095 | 0.008397956 | 0.015548876 | ns | YPD | KS.1 |
| FIM1_789 | 449.5580048 | 763.0486144 | 1008.009036 | 0.694392831 | 0.206142909 | 0.000282464 | 0.000692246 | ns | YPD | KS.1 |
| FIM1_790 | 395.1020096 | 602.8709685 | 368.2901065 | 0.540757299 | 0.21804231 | 0.006829785 | 0.01286565 | ns | YPD | KS.1 |
| FIM1_791 | 5050.651209 | 5856.602856 | 3356.780222 | 0.186421421 | 0.195848026 | 0.30511599 | 0.377787966 | ns | YPD | KS.1 |
| FIM1_792 | 694.8919265 | 1147.438558 | 1088.768169 | 0.70060144 | 0.1146887 | 3.58689847855 | 1.93544684715 | ns | YPD | KS.1 |

| | | | | | | | | | | |
|---|---|---|---|---|---|---|---|---|---|---|
| FIM1_793 | 6327.019052 | 4373.230174 | 3888.038793 | -0.511182849 | 0.121311162 | 1.3844458643( | 4.1169265839: | ns | YPD | KS.1 |
| FIM1_794 | 2204.929909 | 1025.875553 | 1027.485245 | -1.084852225 | 0.102863471 | 8.4734220603( | 1.8988862500: | Km<KS.1 | YPD | KS.1 |
| FIM1_795 | 1664.325954 | 2025.766754 | 2824.487688 | 0.262586562 | 0.147000147 | 0.061074733 | 0.0938378 | ns | YPD | KS.1 |
| FIM1_796 | 1861.199509 | 1360.537492 | 1281.943094 | -0.439406351 | 0.093740057 | 1.7527902321: | 5.9401440087( | ns | YPD | KS.1 |
| FIM1_797 | 111.9941454 | 208.58357 | 336.0054251 | 0.818607081 | 0.218073187 | 4.9321207017 | 0.000136167 | ns | YPD | KS.1 |
| FIM1_798 | 392.6803633 | 543.8688433 | 648.7373743 | 0.426034003 | 0.180843229 | 0.011937854 | 0.021479502 | ns | YPD | KS.1 |
| FIM1_799 | 4443.819183 | 1842.537038 | 1734.643283 | -1.123366049 | 0.292044867 | 1.7613621738! | 5.1606459451 | Km<KS.1 | YPD | KS.1 |
| FIM1_800 | 2438.04435 | 1809.427258 | 2792.052274 | -0.399076064 | 0.157900347 | 0.007776359 | 0.014478812 | ns | YPD | KS.1 |
| FIM1_801 | 4243.617328 | 3832.363273 | 2056.706458 | -0.117532165 | 0.25845289 | 0.609801448 | 0.679910848 | ns | YPD | KS.1 |
| FIM1_802 | 901.4950753 | 1017.128044 | 852.4081707 | 0.153805711 | 0.185191193 | 0.374131762 | 0.449808002 | ns | YPD | KS.1 |
| FIM1_803 | 1039.337319 | 1326.590399 | 1145.42991 | 0.341707527 | 0.089538835 | 0.000100573 | 0.000262789 | ns | YPD | KS.1 |
| FIM1_804 | 4278.889175 | 5938.975093 | 5144.829841 | 0.419753758 | 0.204576304 | 0.026224485 | 0.043474446 | ns | YPD | KS.1 |
| FIM1_805 | 2301.189385 | 1122.388797 | 1049.39095 | -1.003698977 | 0.136098091 | 3.1082899467( | 2.3900683902: | Km<KS.1 | YPD | KS.1 |
| FIM1_806 | 4471.551153 | 2436.050054 | 1397.046938 | -0.767821163 | 0.259431126 | 0.000981915 | 0.002180816 | ns | YPD | KS.1 |
| FIM1_807 | 3576.182852 | 203.0797604 | 604.1760257 | -4.095735328 | 0.173992245 | 6.5213377160 | 1.6221827568( | Km<KS.1 | YPD | KS.1 |
| FIM1_808 | 1658.117299 | 1302.748248 | 1277.977105 | -0.32243992 | 0.150830145 | 0.024778199 | 0.041296998 | ns | YPD | KS.1 |
| FIM1_809 | 11484.593 | 9834.221527 | 20737.53121 | -0.211048593 | 0.123747445 | 0.077060567 | 0.114577501 | ns | YPD | KS.1 |
| FIM1_810 | 1209.927688 | 1493.235508 | 1437.83775 | 0.29348835 | 0.098467656 | 0.002277786 | 0.004735473 | ns | YPD | KS.1 |
| FIM1_811 | 2028.86258 | 2912.437654 | 2368.765038 | 0.479256641 | 0.17424657 | 0.003491388 | 0.007012376 | ns | YPD | KS.1 |
| FIM1_812 | 967.7436946 | 1014.968988 | 2025.255696 | 0.062796308 | 0.139105853 | 0.637802266 | 0.704499616 | ns | YPD | KS.1 |
| FIM1_813 | 1240.592252 | 833.1840805 | 733.4134806 | -0.55564863 | 0.106865406 | 9.9778970136 | 4.0324969653: | ns | YPD | KS.1 |
| FIM1_814 | 172.4206314 | 60.70142868 | 59.6130047 | -1.351238346 | 0.296917885 | 4.9781243777! | 1.8358909399: | Km<KS.1 | YPD | KS.1 |
| FIM1_815 | 137.0730814 | 174.4707543 | 102.8411246 | 0.280482015 | 0.262425634 | 0.228218761 | 0.295443231 | ns | YPD | KS.1 |
| FIM1_816 | 623.9731087 | 311.8712775 | 391.056969 | -0.911565687 | 0.231796416 | 1.9419846980! | 5.6631734308: | ns | YPD | KS.1 |
| FIM1_817 | 2749.57651 | 1028.801533 | 1487.348653 | -1.360712515 | 0.178177946 | 2.1653013586 | 1.8702038644 | Km<KS.1 | YPD | KS.1 |
| FIM1_818 | 106.885704 | 82.87438035 | 327.2956797 | -0.325886952 | 0.197406071 | 0.074804238 | 0.111723531 | ns | YPD | KS.1 |
| FIM1_819 | 1710.374276 | 1336.877759 | 1264.18978 | -0.335146911 | 0.13158964 | 0.00812762 | 0.015098921 | ns | YPD | KS.1 |
| FIM1_820 | 15714.47479 | 6554.953797 | 6602.112733 | -1.220538282 | 0.151272739 | 9.0121378665 | 8.9670771772( | Km<KS.1 | YPD | KS.1 |
| FIM1_821 | 3014.305574 | 5768.29672 | 8074.419783 | 0.887315633 | 0.169740142 | 4.1394819423 | 1.7646891742: | ns | YPD | KS.1 |
| FIM1_822 | 838.9378878 | 785.3564833 | 963.4226905 | -0.084825264 | 0.170823564 | 0.598484007 | 0.669392521 | ns | YPD | KS.1 |
| FIM1_823 | 669.5410082 | 691.0244501 | 1053.613319 | 0.043209376 | 0.103743773 | 0.669037024 | 0.73128338 | ns | YPD | KS.1 |
| FIM1_824 | 3218.727432 | 2009.583519 | 4683.444448 | -0.524920702 | 0.332073826 | 0.060737382 | 0.093377156 | ns | YPD | KS.1 |
| FIM1_825 | 2556.28562 | 2111.729565 | 3341.939333 | -0.262214511 | 0.117078145 | 0.020696179 | 0.035105179 | ns | YPD | KS.1 |
| FIM1_826 | 140.736111 | 77.76838125 | 94.20694577 | -0.700086349 | 0.316281457 | 0.009963327 | 0.018203288 | ns | YPD | KS.1 |
| FIM1_828 | 1580.917125 | 699.66782 | 1423.450385 | -1.136979255 | 0.147365831 | 1.7796537423! | 1.5505739699: | Km<KS.1 | YPD | KS.1 |
| FIM1_829 | 232.429421 | 223.382491 | 492.427216 | -0.054481575 | 0.151118347 | 0.705282609 | 0.764733174 | ns | YPD | KS.1 |
| FIM1_830 | 2547.980771 | 2515.999249 | 2874.1411 | -0.013880059 | 0.293562274 | 0.956562322 | 0.968541301 | ns | YPD | KS.1 |
| FIM1_831 | 1324.209887 | 2395.549874 | 4234.203468 | 0.819024085 | 0.145306903 | 4.8067625510! | 2.2993888164! | ns | YPD | KS.1 |
| FIM1_832 | 690.8928424 | 522.9122983 | 754.9707349 | -0.363646805 | 0.179603038 | 0.030712184 | 0.050014113 | ns | YPD | KS.1 |
| FIM1_833 | 2952.400852 | 2379.361761 | 3921.404092 | -0.264672561 | 0.224504344 | 0.195498131 | 0.259430036 | ns | YPD | KS.1 |
| FIM1_834 | 524.2866229 | 1545.711368 | 1221.750856 | 1.531670834 | 0.11783727 | 8.5693402730: | 4.3502518222 | Km>KS.1 | YPD | KS.1 |
| FIM1_835 | 1036.274175 | 1150.619179 | 953.2250199 | 0.141092034 | 0.134470063 | 0.274462819 | 0.345072663 | ns | YPD | KS.1 |
| FIM1_836 | 1778.807 | 1339.215318 | 1089.792055 | -0.339136011 | 0.252989459 | 0.133237494 | 0.185674098 | ns | YPD | KS.1 |
| FIM1_840 | 553.9521843 | 450.7140428 | 485.8406089 | -0.27261464 | 0.157305541 | 0.067684563 | 0.102443171 | ns | YPD | KS.1 |
| FIM1_841 | 394.7635668 | 142.6329527 | 116.3551879 | -1.287235322 | 0.321618709 | 6.5283202688( | 2.0388194185! | Km<KS.1 | YPD | KS.1 |

| | | | | | | | | | | |
|---|---|---|---|---|---|---|---|---|---|---|
| FIM1_842 | 318.7234483 | 264.4475286 | 255.2486176 | -0.227600112 | 0.229659881 | 0.275037946 | 0.345621062 | ns | YPD | KS.1 |
| FIM1_843 | 915.9221636 | 193.0691429 | 1017.35679 | -1.891202552 | 0.447851874 | 6.09072865849 | 2.22804228500 | Km<KS.1 | YPD | KS.1 |
| FIM1_844 | 281.2270704 | 126.3986806 | 112.3709184 | -1.07435868 | 0.212047443 | 6.68678144189 | 2.77454025633 | Km<KS.1 | YPD | KS.1 |
| FIM1_846 | 296.1567031 | 63.09457018 | 84.39783745 | -2.095927323 | 0.271052864 | 1.79351459942 | 1.73029487680 | Km<KS.1 | YPD | KS.1 |
| FIM1_847 | 54.89932342 | 20.40088299 | 15.97427327 | -1.307305127 | 0.26294565 | 6.87511063583 | 2.84793300693 | Km<KS.1 | YPD | KS.1 |
| FIM1_850 | 1850.621401 | 641.1723534 | 698.1875307 | -1.358304426 | 0.31164483 | 1.19128653621 | 4.14451085150 | Km<KS.1 | YPD | KS.1 |
| FIM1_851 | 577.9299487 | 358.178215 | 305.0748288 | -0.592383759 | 0.257526148 | 0.010024207 | 0.018274251 | ns | YPD | KS.1 |
| FIM1_852 | 45.96431837 | 37.96604856 | 63.46054071 | -0.239317819 | 0.209049567 | 0.213417533 | 0.279850349 | ns | YPD | KS.1 |
| FIM1_853 | 212.0886829 | 67.69094032 | 101.2122781 | -1.562308928 | 0.221554203 | 1.13962052826 | 8.39942537500 | Km<KS.1 | YPD | KS.1 |
| FIM1_854 | 3605.634154 | 943.6269962 | 1559.299267 | -1.79613695 | 0.274936505 | 2.41008989077 | 1.59869296087 | Km<KS.1 | YPD | KS.1 |
| FIM1_855 | 279.9403421 | 217.3061559 | 276.8693972 | -0.325536735 | 0.196945706 | 0.074521759 | 0.111368504 | ns | YPD | KS.1 |
| FIM1_856 | 354.4350721 | 305.4317199 | 299.5086226 | -0.194932595 | 0.168486939 | 0.219818547 | 0.286431972 | ns | YPD | KS.1 |
| FIM1_857 | 378.0057913 | 202.2318897 | 194.7328761 | -0.868015323 | 0.136692863 | 5.33941464032 | 3.10685237843 | ns | YPD | KS.1 |
| FIM1_858 | 286.4540407 | 175.5648778 | 643.7882858 | -0.592302261 | 0.277973217 | 0.015484672 | 0.027182867 | ns | YPD | KS.1 |
| FIM1_859 | 740.4182668 | 338.7579533 | 460.6221006 | -1.109835559 | 0.097871919 | 1.27861625780 | 3.67694559686 | Km<KS.1 | YPD | KS.1 |
| FIM1_860 | 769.0910487 | 458.0544038 | 530.8493055 | -0.731383058 | 0.098645668 | 4.01928681865 | 3.07157479613 | ns | YPD | KS.1 |
| FIM1_861 | 528.4457155 | 713.4690368 | 689.1671746 | 0.404169901 | 0.151077376 | 0.005012278 | 0.009736854 | ns | YPD | KS.1 |
| FIM1_862 | 296.770629 | 166.2138263 | 158.4303416 | -0.779600072 | 0.190080995 | 1.26206742924 | 3.78012369685 | ns | YPD | KS.1 |
| FIM1_863 | 54.99820391 | 30.55986818 | 32.18866186 | -0.64067764 | 0.372853133 | 0.03592124 | 0.057647796 | ns | YPD | KS.1 |
| FIM1_864 | 631.1926619 | 703.0853926 | 574.4896196 | 0.125140809 | 0.253689396 | 0.581124317 | 0.653797711 | ns | YPD | KS.1 |
| FIM1_865 | 1249.16019 | 708.9528812 | 947.753543 | -0.7432149 | 0.214642859 | 0.000179408 | 0.000453074 | ns | YPD | KS.1 |
| FIM1_866 | 640.9969622 | 594.4000097 | 514.4551748 | -0.097346101 | 0.171702273 | 0.547438773 | 0.620954832 | ns | YPD | KS.1 |
| FIM1_867 | 3010.697218 | 1701.292162 | 1299.862932 | -0.756151583 | 0.203496742 | 6.58426895579 | 0.000177929 | ns | YPD | KS.1 |
| FIM1_868 | 938.5802599 | 998.2356928 | 901.8214115 | 0.079985594 | 0.155206505 | 0.587883167 | 0.660058396 | ns | YPD | KS.1 |
| FIM1_869 | 325.4714997 | 493.4099774 | 674.8613506 | 0.575285081 | 0.120992588 | 9.53855196849 | 3.37993561561 | ns | YPD | KS.1 |
| FIM1_870 | 1808.529907 | 906.0640356 | 1337.424702 | -0.919663079 | 0.214010657 | 3.92020051345 | 1.26151342525 | ns | YPD | KS.1 |
| FIM1_871 | 544.3290587 | 638.0534859 | 1085.450442 | 0.21996675 | 0.095183542 | 0.018007059 | 0.031084358 | ns | YPD | KS.1 |
| FIM1_872 | 2323.695497 | 2142.866545 | 2799.267011 | -0.111576927 | 0.101516692 | 0.259367742 | 0.329172071 | ns | YPD | KS.1 |
| FIM1_873 | 2345.073184 | 890.1929357 | 998.9041111 | -1.355831899 | 0.150404194 | 1.94511793169 | 2.65122238635 | Km<KS.1 | YPD | KS.1 |
| FIM1_874 | 17417.22103 | 8173.122793 | 4884.448618 | -1.005063792 | 0.225210911 | 1.53388780197 | 5.26282194125 | Km<KS.1 | YPD | KS.1 |
| FIM1_875 | 692.9448377 | 531.6407371 | 368.4667161 | -0.335605177 | 0.204628369 | 0.075390551 | 0.112430453 | ns | YPD | KS.1 |
| FIM1_876 | 4761.087159 | 2945.360384 | 2803.00508 | -0.586545193 | 0.268363965 | 0.013654523 | 0.024211669 | ns | YPD | KS.1 |
| FIM1_877 | 557.4391819 | 816.3062594 | 1254.645693 | 0.519140225 | 0.145645498 | 0.000198041 | 0.000497602 | ns | YPD | KS.1 |
| FIM1_878 | 2686.313558 | 3089.224025 | 3660.528863 | 0.191027401 | 0.114371677 | 0.084686367 | 0.124612445 | ns | YPD | KS.1 |
| FIM1_879 | 67.28441665 | 12.97120831 | 27.8276178 | -2.148553692 | 0.354254154 | 1.88228088359 | 1.14199358486 | Km<KS.1 | YPD | KS.1 |
| FIM1_880 | 1860.272543 | 1647.985045 | 1216.289069 | -0.166885837 | 0.101422206 | 0.091198407 | 0.133170554 | ns | YPD | KS.1 |
| FIM1_881 | 1847.127066 | 1503.877356 | 1551.362047 | -0.287274369 | 0.095214126 | 0.00203599 | 0.004286521 | ns | YPD | KS.1 |
| FIM1_882 | 743.7769766 | 482.6151194 | 510.0564056 | -0.595602034 | 0.135012908 | 4.72654969237 | 1.50445199741 | ns | YPD | KS.1 |
| FIM1_883 | 1536.725108 | 1879.324311 | 1562.433082 | 0.255749921 | 0.193328251 | 0.154446353 | 0.21103285 | ns | YPD | KS.1 |
| FIM1_884 | 288.7474588 | 360.6001647 | 330.5532116 | 0.265870246 | 0.241402629 | 0.221247769 | 0.288067953 | ns | YPD | KS.1 |
| FIM1_885 | 2475.302909 | 1782.880755 | 2050.872708 | -0.463236784 | 0.083464294 | 1.72615943272 | 7.68813176166 | ns | YPD | KS.1 |
| FIM1_886 | 2946.659967 | 5253.639298 | 4045.572158 | 0.751609915 | 0.226359919 | 0.000296228 | 0.000722065 | ns | YPD | KS.1 |
| FIM1_887 | 326.1522549 | 394.5195159 | 602.4879608 | 0.259915702 | 0.13283109 | 0.041668767 | 0.065914822 | ns | YPD | KS.1 |
| FIM1_888 | 1309.535621 | 1389.41957 | 1495.16734 | 0.078535191 | 0.136140637 | 0.547947104 | 0.621106595 | ns | YPD | KS.1 |
| FIM1_889 | 787.3802988 | 994.4368795 | 968.179931 | 0.32429066 | 0.099665589 | 0.000865515 | 0.001936122 | ns | YPD | KS.1 |

| | | | | | | | | | | |
|---|---|---|---|---|---|---|---|---|---|---|
| FIM1_890 | 181.8010008 | 214.4262103 | 335.6733714 | 0.21473072 | 0.155387896 | 0.145553899 | 0.200534657 | ns | YPD | KS.1 |
| FIM1_891 | 7412.883154 | 4744.470283 | 4113.37712 | -0.608030402 | 0.151993968 | 2.8420371234 | 8.1035287981 | ns | YPD | KS.1 |
| FIM1_892 | 282.5071408 | 128.2124082 | 170.0596042 | -0.977239669 | 0.310693265 | 0.000331604 | 0.000801229 | ns | YPD | KS.1 |
| FIM1_893 | 88.63615494 | 47.37823328 | 28.89626081 | -0.740004604 | 0.321348697 | 0.007227689 | 0.01355362 | ns | YPD | KS.1 |
| FIM1_894 | 11617.00078 | 8435.155304 | 3726.986747 | -0.376184761 | 0.270542446 | 0.114754712 | 0.163380166 | ns | YPD | KS.1 |
| FIM1_895 | 710.4527824 | 983.5472564 | 707.5695422 | 0.446964173 | 0.126569015 | 0.000258094 | 0.000636188 | ns | YPD | KS.1 |
| FIM1_896 | 156.6142162 | 132.9751846 | 140.4085589 | -0.216827251 | 0.153201275 | 0.136515724 | 0.189711097 | ns | YPD | KS.1 |
| FIM1_897 | 419.7078365 | 592.9227875 | 566.0714161 | 0.437108697 | 0.218527893 | 0.028777341 | 0.047218757 | ns | YPD | KS.1 |
| FIM1_898 | 3256.788943 | 6665.20276 | 5129.727209 | 0.969599266 | 0.192937845 | 1.0231441692 | 4.1249126757 | ns | YPD | KS.1 |
| FIM1_899 | 697.9980074 | 402.7483869 | 688.9489661 | -0.735514988 | 0.186991777 | 2.8423432668 | 8.1035287981 | ns | YPD | KS.1 |
| FIM1_900 | 1514.905273 | 2127.459216 | 1909.136672 | 0.447287869 | 0.178033164 | 0.007460715 | 0.013953781 | ns | YPD | KS.1 |
| FIM1_901 | 5164.943006 | 3949.804611 | 3456.279029 | -0.369762924 | 0.117834051 | 0.001209069 | 0.002634743 | ns | YPD | KS.1 |
| FIM1_902 | 380.035623 | 502.7182781 | 486.7253884 | 0.384944317 | 0.119460082 | 0.000882399 | 0.001970348 | ns | YPD | KS.1 |
| FIM1_903 | 707.7261801 | 832.8169207 | 1651.287398 | 0.176562978 | 0.303330403 | 0.498799033 | 0.575730761 | ns | YPD | KS.1 |
| FIM1_904 | 1215.632573 | 807.3259349 | 1256.333312 | -0.479524594 | 0.287091915 | 0.055382742 | 0.085888136 | ns | YPD | KS.1 |
| FIM1_905 | 2385.110189 | 1108.311813 | 879.7972504 | -1.072447661 | 0.13713849 | 8.7557875700 | 7.8770421629 | Km<KS.1 | YPD | KS.1 |
| FIM1_906 | 5068.820321 | 7237.935669 | 12090.97668 | 0.485617624 | 0.141443879 | 0.000346057 | 0.000832511 | ns | YPD | KS.1 |
| FIM1_907 | 9466.609364 | 12699.92252 | 16591.70845 | 0.40481905 | 0.121219369 | 0.000563853 | 0.001308382 | ns | YPD | KS.1 |
| FIM1_909 | 236.0369472 | 689.6047378 | 1077.98299 | 1.50265295 | 0.156164802 | 4.7505617712 | 7.8258426529 | Km>KS.1 | YPD | KS.1 |
| FIM1_910 | 590.4396581 | 550.999271 | 267.2848716 | -0.086266347 | 0.198039662 | 0.638840588 | 0.705333317 | ns | YPD | KS.1 |
| FIM1_912 | 529.6331261 | 108.0970424 | 111.3884135 | -2.172187609 | 0.254913219 | 2.1347342511 | 2.6550757249 | Km<KS.1 | YPD | KS.1 |
| FIM1_913 | 3279.394653 | 3060.55724 | 2001.170019 | -0.082168025 | 0.233544527 | 0.69812521 | 0.758334699 | ns | YPD | KS.1 |
| FIM1_914 | 1293.17424 | 647.9447586 | 592.0757581 | -0.970859742 | 0.120482508 | 1.5520253460 | 1.5110227194 | ns | YPD | KS.1 |
| FIM1_915 | 3199.883905 | 4815.065138 | 4187.559889 | 0.56920576 | 0.114463664 | 3.2045324725 | 1.2188493158 | ns | YPD | KS.1 |
| FIM1_916 | 1640.395882 | 1973.285955 | 1234.286868 | 0.244363765 | 0.154822003 | 0.096530411 | 0.139970502 | ns | YPD | KS.1 |
| FIM1_917 | 18153.3621 | 11065.20897 | 10820.5135 | -0.584568497 | 0.297013242 | 0.023296015 | 0.039062243 | ns | YPD | KS.1 |
| FIM1_918 | 132.5362672 | 325.2720823 | 235.7526735 | 1.185684258 | 0.251090595 | 3.0725277738 | 1.1722258953 | Km>KS.1 | YPD | KS.1 |
| FIM1_919 | 690.0110932 | 549.647938 | 300.1039835 | -0.266687072 | 0.258431809 | 0.24607183 | 0.314787183 | ns | YPD | KS.1 |
| FIM1_920 | 2573.550732 | 2455.106834 | 1881.743709 | -0.060164983 | 0.176210362 | 0.716548191 | 0.77412101 | ns | YPD | KS.1 |
| FIM1_921 | 3356.26246 | 1977.801112 | 2693.04147 | -0.728301372 | 0.14516685 | 1.7858871732 | 7.0069311410 | ns | YPD | KS.1 |
| FIM1_922 | 700.0485637 | 450.6736612 | 488.257024 | -0.586020237 | 0.180169808 | 0.000540874 | 0.001259761 | ns | YPD | KS.1 |
| FIM1_923 | 664.3129198 | 446.6180194 | 409.4636735 | -0.519278689 | 0.193539199 | 0.003959341 | 0.007863362 | ns | YPD | KS.1 |
| FIM1_924 | 1741.647689 | 893.7355064 | 926.2492243 | -0.930434485 | 0.138001352 | 3.4039058751 | 2.2107613223 | ns | YPD | KS.1 |
| FIM1_925 | 426.6250217 | 252.5947142 | 227.6234873 | -0.699808409 | 0.189377353 | 8.0863294806 | 0.000213987 | ns | YPD | KS.1 |
| FIM1_926 | 24599.85307 | 7071.783955 | 4723.754127 | -1.575474411 | 0.353425341 | 4.6889043471 | 1.7382488172 | Km<KS.1 | YPD | KS.1 |
| FIM1_927 | 3739.825691 | 5424.824215 | 6918.70648 | 0.525872221 | 0.083902386 | 1.9384496747 | 1.0667906119 | ns | YPD | KS.1 |
| FIM1_928 | 390.6615394 | 622.8359058 | 996.4788868 | 0.631230847 | 0.159761287 | 3.3240529359 | 9.3801266910 | ns | YPD | KS.1 |
| FIM1_929 | 2228.295199 | 2218.12446 | 1444.466614 | -0.005500324 | 0.220608001 | 0.978270676 | 0.984603806 | ns | YPD | KS.1 |
| FIM1_930 | 2317.763704 | 3124.705999 | 1751.919322 | 0.390380997 | 0.181751348 | 0.021761654 | 0.036712183 | ns | YPD | KS.1 |
| FIM1_931 | 753.7326712 | 1190.588016 | 1672.699267 | 0.635739183 | 0.124332087 | 1.3244604112 | 5.2840341188 | ns | YPD | KS.1 |
| FIM1_932 | 418.8915764 | 636.8274132 | 497.2585964 | 0.586989729 | 0.108805346 | 3.1904343696 | 1.3826141976 | ns | YPD | KS.1 |
| FIM1_933 | 2802.343306 | 2767.47892 | 2148.831949 | -0.017152803 | 0.078446076 | 0.824121716 | 0.863268033 | ns | YPD | KS.1 |
| FIM1_934 | 1172.407393 | 1210.862613 | 1323.273902 | 0.043661904 | 0.121181457 | 0.70948194 | 0.768268427 | ns | YPD | KS.1 |
| FIM1_935 | 1525.271213 | 2113.166175 | 1205.089316 | 0.349579077 | 0.330239081 | 0.208544095 | 0.274255054 | ns | YPD | KS.1 |
| FIM1_936 | 8870.849193 | 11655.14473 | 8369.136041 | 0.330487712 | 0.242072172 | 0.129143664 | 0.180626856 | ns | YPD | KS.1 |

| | | | | | | | | | | |
|---|---|---|---|---|---|---|---|---|---|---|
| FIM1_937 | 1859.178465 | 1290.41521 | 2593.518284 | -0.506590053 | 0.116569268 | 7.7196736853 | 2.3795152778 | ns | YPD | KS.1 |
| FIM1_938 | 3957.141464 | 3905.71978 | 3427.445661 | -0.016308301 | 0.194834809 | 0.928317311 | 0.947554087 | ns | YPD | KS.1 |
| FIM1_939 | 912.7688903 | 1564.435415 | 1869.916936 | 0.765904347 | 0.080642354 | 6.4224784687 | 9.6531209613 | ns | YPD | KS.1 |
| FIM1_940 | 1862.326076 | 3392.647767 | 2972.43512 | 0.811237265 | 0.179958478 | 1.8686769020 | 6.2900321972 | ns | YPD | KS.1 |
| FIM1_941 | 408.6716504 | 521.0899965 | 506.1071318 | 0.298518702 | 0.227546939 | 0.148820424 | 0.204412371 | ns | YPD | KS.1 |
| FIM1_942 | 5801.633993 | 1598.851913 | 1787.929814 | -1.771070101 | 0.218615875 | 2.2070070298 | 2.3115494680 | Km<KS.1 | YPD | KS.1 |
| FIM1_943 | 4171.648025 | 5579.01326 | 8646.69721 | 0.392446559 | 0.146388356 | 0.005026344 | 0.009760367 | ns | YPD | KS.1 |
| FIM1_944 | 548.4326372 | 644.6816254 | 1283.016317 | 0.190444195 | 0.250304619 | 0.395536424 | 0.472459474 | ns | YPD | KS.1 |
| FIM1_945 | 3730.545753 | 2543.879344 | 2338.598098 | -0.536564709 | 0.101672419 | 6.8167407515 | 2.8261071032 | ns | YPD | KS.1 |
| FIM1_946 | 48.02796559 | 63.15846769 | 45.99085676 | 0.355018082 | 0.181415965 | 0.036536439 | 0.05855953 | ns | YPD | KS.1 |
| FIM1_947 | 3690.375016 | 5319.610655 | 5270.519313 | 0.502652228 | 0.130902766 | 6.9087887726 | 0.00018529 | ns | YPD | KS.1 |
| FIM1_948 | 6575.985396 | 8319.42054 | 5590.866524 | 0.327080495 | 0.103130765 | 0.001148002 | 0.002515997 | ns | YPD | KS.1 |
| FIM1_949 | 1698.680743 | 3223.461806 | 1384.197803 | 0.82728071 | 0.242888788 | 0.000183689 | 0.00046365 | ns | YPD | KS.1 |
| FIM1_950 | 618.0653723 | 596.9506062 | 465.2174882 | -0.041672664 | 0.225397344 | 0.83941766 | 0.875838945 | ns | YPD | KS.1 |
| FIM1_951 | 43343.95295 | 20137.99036 | 20186.18416 | -1.067492023 | 0.148011087 | 9.2310984280 | 6.8544350267 | Km<KS.1 | YPD | KS.1 |
| FIM1_952 | 153.2091403 | 37.75023387 | 41.24879713 | -1.799046935 | 0.349264548 | 9.0530484632 | 4.1896666143 | Km<KS.1 | YPD | KS.1 |
| FIM1_953 | 5894.360902 | 4970.552135 | 3895.405561 | -0.229871789 | 0.134965679 | 0.075879648 | 0.113092045 | ns | YPD | KS.1 |
| FIM1_954 | 836.0073673 | 1848.673165 | 1865.356409 | 1.05149601 | 0.233551873 | 1.1591386819 | 4.0439796231 | Km>KS.1 | YPD | KS.1 |
| FIM1_955 | 7204.74159 | 3154.749716 | 1946.284371 | -1.033100458 | 0.305372032 | 0.000127005 | 0.000326032 | Km<KS.1 | YPD | KS.1 |
| FIM1_957 | 7286.646479 | 11872.21532 | 11679.93381 | 0.574506151 | 0.298249987 | 0.026214278 | 0.043472012 | ns | YPD | KS.1 |
| FIM1_958 | 6409.550217 | 3292.321982 | 4172.932193 | -0.905369564 | 0.181480036 | 1.4125032112 | 5.6038305230 | ns | YPD | KS.1 |
| FIM1_959 | 3220.332345 | 958.9531241 | 1630.320037 | -1.63295986 | 0.251035077 | 4.2520423035 | 2.7401438420 | Km<KS.1 | YPD | KS.1 |
| FIM1_960 | 915.2910959 | 1897.685089 | 2383.03718 | 1.024000518 | 0.126758535 | 1.1904334822 | 1.1681275294 | Km>KS.1 | YPD | KS.1 |
| FIM1_961 | 2550.901503 | 4239.775969 | 4809.67305 | 0.717541056 | 0.094138748 | 8.4358734375 | 6.8241415206 | ns | YPD | KS.1 |
| FIM1_962 | 1557.016223 | 2205.31262 | 2784.4084 | 0.48861736 | 0.099314722 | 4.9880659717 | 1.8381946821 | ns | YPD | KS.1 |
| FIM1_963 | 484.5862208 | 909.4632279 | 705.9682752 | 0.863015506 | 0.160007631 | 1.7477202654 | 7.7771988556 | ns | YPD | KS.1 |
| FIM1_964 | 107.2628873 | 177.0044773 | 229.22586 | 0.685190584 | 0.14581838 | 9.8359174625 | 3.4729375000 | ns | YPD | KS.1 |
| FIM1_965 | 1356.179205 | 1157.35601 | 1032.537501 | -0.220468905 | 0.091602173 | 0.013895884 | 0.024602144 | ns | YPD | KS.1 |
| FIM1_966 | 2023.181763 | 3094.687157 | 3950.198171 | 0.571353741 | 0.166172349 | 0.000285392 | 0.000698046 | ns | YPD | KS.1 |
| FIM1_967 | 994.9921323 | 891.7605454 | 1015.043269 | -0.148065083 | 0.116910733 | 0.19066938 | 0.254311037 | ns | YPD | KS.1 |
| FIM1_968 | 1754.329362 | 651.9151929 | 752.0076583 | -1.382924995 | 0.157080703 | 1.2349549186 | 1.5916841243 | Km<KS.1 | YPD | KS.1 |
| FIM1_969 | 1255.670163 | 525.0090973 | 440.0148115 | -1.208111372 | 0.167278735 | 6.6180474931 | 4.9810569256 | Km<KS.1 | YPD | KS.1 |
| FIM1_970 | 1003.581004 | 1051.899142 | 965.1903519 | 0.065154208 | 0.110673818 | 0.544461357 | 0.618000286 | ns | YPD | KS.1 |
| FIM1_971 | 252.1885899 | 139.3715225 | 131.9005992 | -0.79727096 | 0.185680712 | 5.1819529770 | 1.6389202835 | ns | YPD | KS.1 |
| FIM1_972 | 72.70998388 | 105.5037826 | 157.1682748 | 0.47449499 | 0.213001987 | 0.015318131 | 0.026919004 | ns | YPD | KS.1 |
| FIM1_973 | 2031.957869 | 1880.452797 | 1188.232828 | -0.107345208 | 0.086979939 | 0.20793531 | 0.273671473 | ns | YPD | KS.1 |
| FIM1_974 | 1169.818134 | 1308.371521 | 1062.195313 | 0.145919666 | 0.166342031 | 0.353034848 | 0.428273195 | ns | YPD | KS.1 |
| FIM1_975 | 2263.173228 | 3152.727793 | 2546.823218 | 0.430581779 | 0.191639265 | 0.015791579 | 0.027663065 | ns | YPD | KS.1 |
| FIM1_976 | 454.8051969 | 171.0619351 | 101.3528942 | -1.249089383 | 0.304630793 | 4.6800694700 | 1.4915660226 | Km<KS.1 | YPD | KS.1 |
| FIM1_977 | 3778.670676 | 2891.251139 | 3415.972487 | -0.294320143 | 0.306584023 | 0.26308481 | 0.333039931 | ns | YPD | KS.1 |
| FIM1_978 | 4783.797888 | 2996.806321 | 4313.41991 | -0.472453634 | 0.388483616 | 0.129951781 | 0.181706046 | ns | YPD | KS.1 |
| FIM1_979 | 1312.552012 | 1341.073082 | 2201.409289 | 0.023715766 | 0.286300619 | 0.924407931 | 0.944276281 | ns | YPD | KS.1 |
| FIM1_980 | 2671.472227 | 3987.696016 | 2473.290594 | 0.500526613 | 0.236343903 | 0.019238514 | 0.032981256 | ns | YPD | KS.1 |
| FIM1_981 | 4629.547065 | 2200.991723 | 1418.350743 | -0.872334186 | 0.34844963 | 0.003095062 | 0.006274627 | ns | YPD | KS.1 |
| FIM1_982 | 2926.294678 | 1125.401252 | 1544.672182 | -1.31436318 | 0.189837202 | 4.6554265742 | 3.2666780263 | Km<KS.1 | YPD | KS.1 |

| | | | | | | | | | | |
|---|---|---|---|---|---|---|---|---|---|---|
| FIM1_983 | 10724.38041 | 9654.306181 | 12436.09678 | -0.14471184 | 0.104371505 | 0.154183882 | 0.210790001 | ns | YPD | KS.1 |
| FIM1_984 | 575.2264558 | 50.37409379 | 91.15760918 | -2.911546441 | 0.642317885 | 1.11425689542 | 5.09506255029 | Km<KS.1 | YPD | KS.1 |
| FIM1_988 | 3152.479306 | 2385.531886 | 2634.034784 | -0.380772904 | 0.131034837 | 0.002556276 | 0.005261676 | ns | YPD | KS.1 |
| FIM1_989 | 255.7558663 | 125.7557573 | 71.54174204 | -0.946849917 | 0.211926255 | 1.69053082438 | 5.75659880308 | ns | YPD | KS.1 |
| FIM1_991 | 1684.596943 | 1604.520262 | 1301.236003 | -0.059050736 | 0.220105314 | 0.769484817 | 0.818967801 | ns | YPD | KS.1 |
| FIM1_992 | 2583.07786 | 3207.502298 | 2183.789848 | 0.271279082 | 0.20810667 | 0.156696927 | 0.213873034 | ns | YPD | KS.1 |
| FIM1_993 | 375.9467834 | 829.0908377 | 1362.119294 | 1.025320317 | 0.26175827 | 1.62518360023 | 4.78419432614 | Km>KS.1 | YPD | KS.1 |
| FIM1_994 | 1538.73244 | 1559.801645 | 2129.332173 | 0.017361395 | 0.184487331 | 0.919930099 | 0.94015042 | ns | YPD | KS.1 |
| FIM1_995 | 1883.471591 | 1176.836212 | 2185.273744 | -0.575991099 | 0.264359047 | 0.014199314 | 0.025094702 | ns | YPD | KS.1 |
| FIM1_996 | 1975.203887 | 1221.088857 | 2898.353279 | -0.514013812 | 0.359407001 | 0.082687427 | 0.122032023 | ns | YPD | KS.1 |
| FIM1_997 | 991.9482304 | 229.1652431 | 234.4595379 | -1.987449996 | 0.261398628 | 6.68216940584 | 6.08860673883 | Km<KS.1 | YPD | KS.1 |
| FIM1_998 | 1308.554433 | 1399.099743 | 1120.239119 | 0.090599939 | 0.119814053 | 0.434323634 | 0.511662742 | ns | YPD | KS.1 |
| FIM1_999 | 447.046322 | 838.5065848 | 1425.907567 | 0.874562923 | 0.138572303 | 6.78023432257 | 3.89511151903 | ns | YPD | KS.1 |
| FIM1_1000 | 2890.599182 | 2616.858682 | 961.7356256 | -0.109144671 | 0.290644113 | 0.665898036 | 0.72889829 | ns | YPD | KS.1 |
| FIM1_1001 | 116.272 | 303.4379076 | 323.4595592 | 1.358067526 | 0.120217672 | 1.32672805084 | 3.79337474306 | Km>KS.1 | YPD | KS.1 |
| FIM1_1002 | 214.8428686 | 377.3161554 | 328.5117821 | 0.780948634 | 0.144669006 | 2.03011003722 | 9.00160199213 | ns | YPD | KS.1 |
| FIM1_1003 | 3410.334455 | 2315.946315 | 3583.500302 | -0.469999926 | 0.257099639 | 0.040213057 | 0.06381498 | ns | YPD | KS.1 |
| FIM1_1004 | 3720.87164 | 3331.969342 | 3041.230843 | -0.136174853 | 0.209700772 | 0.480702534 | 0.558891121 | ns | YPD | KS.1 |
| FIM1_1005 | 1415.488952 | 1520.351021 | 1178.634649 | 0.094005092 | 0.153244243 | 0.519250014 | 0.595223231 | ns | YPD | KS.1 |
| FIM1_1006 | 2643.440715 | 5078.773316 | 3500.720483 | 0.89070524 | 0.174336661 | 7.75117119554 | 3.18958450767 | ns | YPD | KS.1 |
| FIM1_1007 | 636.650673 | 634.5610088 | 680.5163647 | -0.006402426 | 0.093402757 | 0.944152585 | 0.95977914 | ns | YPD | KS.1 |
| FIM1_1008 | 582.0786607 | 433.6454459 | 350.3893167 | -0.389956332 | 0.16687063 | 0.013353727 | 0.023709776 | ns | YPD | KS.1 |
| FIM1_1009 | 913.2264068 | 726.1198926 | 736.7478592 | -0.311554317 | 0.131326968 | 0.013665601 | 0.024211669 | ns | YPD | KS.1 |
| FIM1_1010 | 4660.363014 | 5897.020049 | 2615.082822 | 0.273222849 | 0.266945769 | 0.247012146 | 0.315746512 | ns | YPD | KS.1 |
| FIM1_1011 | 5685.824818 | 10431.27292 | 10635.097 | 0.794147068 | 0.223150018 | 0.000111536 | 0.000288855 | ns | YPD | KS.1 |
| FIM1_1012 | 6677.572103 | 3391.317456 | 2599.211892 | -0.768421855 | 0.360623508 | 0.010515554 | 0.019099994 | ns | YPD | KS.1 |
| FIM1_1013 | 1033.733002 | 2985.462321 | 2768.382107 | 1.42994291 | 0.236515162 | 1.23658320090 | 6.95141403901 | Km>KS.1 | YPD | KS.1 |
| FIM1_1014 | 888.0705075 | 2612.463114 | 2115.903474 | 1.302700425 | 0.381232512 | 6.14542596360 | 0.000167251 | Km>KS.1 | YPD | KS.1 |
| FIM1_1015 | 1912.49714 | 1531.883422 | 2155.16098 | -0.254891986 | 0.272357973 | 0.287972639 | 0.359785002 | ns | YPD | KS.1 |
| FIM1_1016 | 128.0842188 | 50.02486185 | 196.0072282 | -1.219104885 | 0.275627942 | 1.18852224713 | 4.13778738943 | Km<KS.1 | YPD | KS.1 |
| FIM1_1017 | 958.4749784 | 2584.332823 | 4189.521793 | 1.396033917 | 0.1386171 | 6.74304335506 | 1.29523709233 | Km>KS.1 | YPD | KS.1 |
| FIM1_1018 | 398.379329 | 926.366121 | 1304.181682 | 1.198606465 | 0.097328398 | 9.77917283931 | 3.98781843242 | Km>KS.1 | YPD | KS.1 |
| FIM1_1019 | 1333.224282 | 1203.561611 | 854.6122633 | -0.13292701 | 0.162621812 | 0.387866345 | 0.464813766 | ns | YPD | KS.1 |
| FIM1_1020 | 68.15778871 | 209.7832121 | 134.9193748 | 1.413524431 | 0.341527681 | 2.78038639253 | 9.14842744899 | Km>KS.1 | YPD | KS.1 |
| FIM1_1021 | 2764.99598 | 1428.700347 | 651.045516 | -0.775387109 | 0.331909492 | 0.006109041 | 0.011640168 | ns | YPD | KS.1 |
| FIM1_1022 | 8101.314737 | 7345.069818 | 7715.193246 | -0.134315699 | 0.108133318 | 0.201093798 | 0.266074906 | ns | YPD | KS.1 |
| FIM1_1024 | 3322.969609 | 3542.417339 | 1785.986354 | 0.076102984 | 0.234102396 | 0.719980079 | 0.776479708 | ns | YPD | KS.1 |
| FIM1_1025 | 5675.693887 | 3528.462071 | 5296.423224 | -0.647553763 | 0.155822572 | 1.33411975468 | 3.97916413641 | ns | YPD | KS.1 |
| FIM1_1026 | 7134.07672 | 4674.859021 | 6893.761453 | -0.572217186 | 0.15782004 | 0.00014013 | 0.000358799 | ns | YPD | KS.1 |
| FIM1_1027 | 2000.877203 | 1088.81614 | 1025.936303 | -0.814554522 | 0.195462407 | 8.76800071502 | 2.67941053791 | ns | YPD | KS.1 |
| FIM1_1028 | 760.0576602 | 288.8496327 | 216.8554548 | -1.31972124 | 0.20465091 | 1.17560148168 | 7.26975344086 | Km<KS.1 | YPD | KS.1 |
| FIM1_1029 | 4515.07646 | 9988.486035 | 12682.87503 | 1.042259568 | 0.245700923 | 3.88538981381 | 1.25193097951 | Km>KS.1 | YPD | KS.1 |
| FIM1_1030 | 2778.562479 | 2548.795885 | 4336.793453 | -0.090557513 | 0.317232873 | 0.73738485 | 0.791475648 | ns | YPD | KS.1 |
| FIM1_1031 | 105.4141566 | 60.85962026 | 81.79274124 | -0.692107477 | 0.251359844 | 0.002224341 | 0.004635986 | ns | YPD | KS.1 |
| FIM1_1032 | 1208.097296 | 636.5253293 | 743.9069697 | -0.907660374 | 0.099139513 | 1.21563528747 | 1.67528685739 | ns | YPD | KS.1 |

| | | | | | | | | | | |
|---|---|---|---|---|---|---|---|---|---|---|
| FIM1_1033 | 57.5463765 | 109.8824705 | 215.5946391 | 0.83736523 | 0.232680675 | 8.70111599729 | 0.00022928 | ns | YPD | KS.1 |
| FIM1_1034 | 752.0651786 | 1023.764195 | 648.1549468 | 0.431106758 | 0.096595491 | 5.19993454458 | 1.64356253871 | ns | YPD | KS.1 |
| FIM1_1035 | 180.7598486 | 215.349436 | 164.1932233 | 0.226867144 | 0.162380571 | 0.139652747 | 0.193260755 | ns | YPD | KS.1 |
| FIM1_1036 | 85.27980231 | 350.8314588 | 581.8946502 | 1.995449547 | 0.145993834 | 3.38471074032 | 2.21564946488 | Km>KS.1 | YPD | KS.1 |
| FIM1_1037 | 1018.477787 | 1489.436661 | 1268.132034 | 0.53992732 | 0.068436543 | 1.53846576176 | 1.35227335066 | ns | YPD | KS.1 |
| FIM1_1038 | 1473.148837 | 3044.668109 | 3371.647619 | 1.035132162 | 0.080318399 | 9.15645159961 | 4.25732212224 | Km>KS.1 | YPD | KS.1 |
| FIM1_1039 | 4743.214099 | 3869.05344 | 3203.341312 | -0.280586544 | 0.113453196 | 0.010783845 | 0.019558743 | ns | YPD | KS.1 |
| FIM1_1040 | 2695.446839 | 2571.151445 | 1696.149232 | -0.058516774 | 0.202501169 | 0.754954718 | 0.806679493 | ns | YPD | KS.1 |
| FIM1_1041 | 2751.676484 | 2889.872056 | 2064.977982 | 0.055286116 | 0.269369526 | 0.816467243 | 0.856223553 | ns | YPD | KS.1 |
| FIM1_1042 | 88.85555179 | 88.83745533 | 77.10730827 | -0.00207877 | 0.179516045 | 0.990159603 | 0.992753734 | ns | YPD | KS.1 |
| FIM1_1043 | 203.1955644 | 272.0861906 | 188.1524893 | 0.352420297 | 0.244772833 | 0.108857461 | 0.155756649 | ns | YPD | KS.1 |
| FIM1_1044 | 2743.119132 | 3369.705538 | 2014.807761 | 0.251809809 | 0.225155996 | 0.219293854 | 0.285923648 | ns | YPD | KS.1 |
| FIM1_1045 | 653.7279478 | 399.1480708 | 478.7153123 | -0.680083948 | 0.140271934 | 4.72804600408 | 1.75145412288 | ns | YPD | KS.1 |
| FIM1_1046 | 765.7686489 | 1088.74338 | 813.5084914 | 0.483906174 | 0.12697758 | 8.06297332506 | 0.000213482 | ns | YPD | KS.1 |
| FIM1_1047 | 1731.083175 | 1651.971452 | 1493.593789 | -0.06575141 | 0.070233626 | 0.342497255 | 0.417501575 | ns | YPD | KS.1 |
| FIM1_1048 | 1074.74718 | 1843.440578 | 1727.147317 | 0.757247586 | 0.109329303 | 1.34769673744 | 9.19724453881 | ns | YPD | KS.1 |
| FIM1_1049 | 1589.875377 | 1198.383798 | 2601.580577 | -0.309577973 | 0.311010987 | 0.244226224 | 0.312817155 | ns | YPD | KS.1 |
| FIM1_1050 | 171.2567964 | 90.25628926 | 107.7630703 | -0.787163618 | 0.290604041 | 0.002064478 | 0.00433549 | ns | YPD | KS.1 |
| FIM1_1051 | 615.1948052 | 494.7428289 | 591.553105 | -0.286497019 | 0.166807069 | 0.068714559 | 0.103686664 | ns | YPD | KS.1 |
| FIM1_1052 | 504.1826152 | 726.3790918 | 1052.997585 | 0.465369998 | 0.211157961 | 0.016559427 | 0.028865855 | ns | YPD | KS.1 |
| FIM1_1053 | 38.58198193 | 102.5559335 | 89.13311434 | 1.325300942 | 0.217199209 | 1.07571813509 | 6.06768449218 | Km>KS.1 | YPD | KS.1 |
| FIM1_1054 | 1055.267 | 2182.133317 | 2238.829363 | 1.030272281 | 0.100024174 | 1.24780362504 | 2.53380532024 | Km>KS.1 | YPD | KS.1 |
| FIM1_1055 | 220.0064231 | 142.3468066 | 204.8296976 | -0.581817309 | 0.17429193 | 0.000402312 | 0.000954916 | ns | YPD | KS.1 |
| FIM1_1056 | 543.3349938 | 395.8612665 | 564.8640782 | -0.426256898 | 0.151060539 | 0.003087736 | 0.006266717 | ns | YPD | KS.1 |
| FIM1_1057 | 1595.701346 | 2201.288262 | 1415.285574 | 0.449257655 | 0.103304201 | 8.51604897006 | 2.60561768918 | ns | YPD | KS.1 |
| FIM1_1058 | 1218.147543 | 1876.892885 | 2535.405671 | 0.604103757 | 0.108340953 | 1.10034677609 | 5.03608575074 | ns | YPD | KS.1 |
| FIM1_1060 | 3079.366574 | 3242.198586 | 4530.229662 | 0.064216907 | 0.198709193 | 0.727650315 | 0.783337521 | ns | YPD | KS.1 |
| FIM1_1061 | 1191.477644 | 732.803636 | 795.8899212 | -0.668424839 | 0.141857376 | 9.57890086153 | 3.39181720897 | ns | YPD | KS.1 |
| FIM1_1062 | 1325.493835 | 837.8946141 | 854.6353268 | -0.643596416 | 0.102712285 | 1.50289184238 | 8.39156780681 | ns | YPD | KS.1 |
| FIM1_1063 | 1581.828695 | 811.1304052 | 789.7390678 | -0.928585419 | 0.142676653 | 1.66696967975 | 1.01881746398 | ns | YPD | KS.1 |
| FIM1_1064 | 3304.978482 | 1336.705794 | 1260.751839 | -1.259181464 | 0.161280182 | 6.87839218799 | 6.25594170662 | Km<KS.1 | YPD | KS.1 |
| FIM1_1065 | 1057.563365 | 1372.666742 | 1242.392912 | 0.364988608 | 0.098441669 | 0.000150015 | 0.000381948 | ns | YPD | KS.1 |
| FIM1_1066 | 269.0337379 | 349.696466 | 442.4206693 | 0.359978248 | 0.122874684 | 0.002447342 | 0.005058382 | ns | YPD | KS.1 |
| FIM1_1067 | 339.3570656 | 482.1486164 | 409.8270158 | 0.480214272 | 0.136197346 | 0.000247558 | 0.000611823 | ns | YPD | KS.1 |
| FIM1_1068 | 341.6788244 | 285.1337639 | 234.1648906 | -0.216992169 | 0.239187106 | 0.3146276 | 0.387935495 | ns | YPD | KS.1 |
| FIM1_1069 | 727.4632257 | 758.2749988 | 417.8969508 | 0.045910843 | 0.281605498 | 0.852385179 | 0.885872939 | ns | YPD | KS.1 |
| FIM1_1070 | 1871.520714 | 1589.352919 | 1189.605498 | -0.22566649 | 0.105074345 | 0.027283031 | 0.045118111 | ns | YPD | KS.1 |
| FIM1_1071 | 1052.473379 | 1630.986317 | 1522.70044 | 0.623960887 | 0.067853502 | 1.53933274511 | 2.10969157216 | ns | YPD | KS.1 |
| FIM1_1072 | 3378.329263 | 1094.441452 | 1457.758092 | -1.389146477 | 0.36673836 | 1.24632823947 | 3.73523071771 | Km<KS.1 | YPD | KS.1 |
| FIM1_1074 | 713.6074092 | 1023.20979 | 1221.662544 | 0.496776228 | 0.125260258 | 4.15117866765 | 0.000115893 | ns | YPD | KS.1 |
| FIM1_1075 | 3831.679368 | 3158.365573 | 3065.067165 | -0.261624544 | 0.132581944 | 0.039998137 | 0.063514437 | ns | YPD | KS.1 |
| FIM1_1076 | 254.3634021 | 792.5810753 | 428.2304838 | 1.525894631 | 0.250638627 | 7.80674469177 | 4.44886080659 | Km>KS.1 | YPD | KS.1 |
| FIM1_1077 | 143.5787064 | 120.4085252 | 125.6029179 | -0.191853757 | 0.300125275 | 0.458694125 | 0.536079328 | ns | YPD | KS.1 |
| FIM1_1078 | 282.932149 | 316.2533383 | 294.9642808 | 0.145723629 | 0.164103173 | 0.347739598 | 0.422777248 | ns | YPD | KS.1 |
| FIM1_1079 | 398.6927772 | 748.072623 | 387.3355343 | 0.84100438 | 0.199598221 | 6.75104914747 | 2.10178157125 | ns | YPD | KS.1 |

| | | | | | | | | | | |
|---|---|---|---|---|---|---|---|---|---|---|
| FIM1_1080 | 783.1968584 | 1250.686361 | 1084.702492 | 0.58465957 | 0.247389743 | 0.008608541 | 0.015915083 | ns | YPD | KS.1 |
| FIM1_1081 | 1199.633151 | 3275.778724 | 1316.689336 | 1.28751754 | 0.303629548 | 2.3462871413 | 7.7922420082 | Km>KS.1 | YPD | KS.1 |
| FIM1_1082 | 1092.237097 | 1699.983112 | 1101.869711 | 0.564988519 | 0.222928713 | 0.005574765 | 0.010708284 | ns | YPD | KS.1 |
| FIM1_1083 | 1930.520029 | 1934.195446 | 1554.803504 | 0.001985126 | 0.101223488 | 0.983960859 | 0.988218307 | ns | YPD | KS.1 |
| FIM1_1084 | 1158.015898 | 928.8065372 | 682.8854909 | -0.288138276 | 0.173583313 | 0.077378902 | 0.114982089 | ns | YPD | KS.1 |
| FIM1_1085 | 677.6826542 | 440.8535392 | 580.9697454 | -0.601061109 | 0.108727895 | 1.4573285090 | 6.5553429767 | ns | YPD | KS.1 |
| FIM1_1086 | 320.0196282 | 489.3865975 | 635.0051541 | 0.576120973 | 0.151979649 | 7.2386387486 | 0.000193406 | ns | YPD | KS.1 |
| FIM1_1087 | 860.7246619 | 977.8072034 | 755.4464923 | 0.171149719 | 0.132525378 | 0.178870927 | 0.240054724 | ns | YPD | KS.1 |
| FIM1_1088 | 2145.696571 | 2263.206009 | 2300.985618 | 0.073931346 | 0.083498774 | 0.367070843 | 0.442709683 | ns | YPD | KS.1 |
| FIM1_1089 | 39.28641156 | 80.66082282 | 127.3354343 | 0.951655663 | 0.230998189 | 8.0416030439 | 2.4695663669 | ns | YPD | KS.1 |
| FIM1_1090 | 5878.995636 | 4528.270824 | 4998.7929 | -0.360374748 | 0.116029046 | 0.001367844 | 0.002952289 | ns | YPD | KS.1 |
| FIM1_1091 | 1900.791524 | 1467.500915 | 2368.850307 | -0.315452644 | 0.234559599 | 0.136987259 | 0.190206982 | ns | YPD | KS.1 |
| FIM1_1092 | 1130.114339 | 1120.436338 | 980.3221085 | -0.011591333 | 0.086138844 | 0.890913621 | 0.916900138 | ns | YPD | KS.1 |
| FIM1_1093 | 1988.015015 | 1005.095782 | 1159.130995 | -0.94576177 | 0.148056634 | 3.5701337165 | 2.1169744028 | ns | YPD | KS.1 |
| FIM1_1094 | 1640.707842 | 1799.66018 | 1676.028457 | 0.131330303 | 0.057906939 | 0.021929336 | 0.036944953 | ns | YPD | KS.1 |
| FIM1_1095 | 1341.844982 | 1663.736335 | 1197.186712 | 0.29859414 | 0.101410583 | 0.002544808 | 0.005240241 | ns | YPD | KS.1 |
| FIM1_1096 | 629.6847013 | 434.2669812 | 436.598392 | -0.495873036 | 0.171202522 | 0.002145455 | 0.004490382 | ns | YPD | KS.1 |
| FIM1_1097 | 16555.37416 | 11363.74178 | 8368.502534 | -0.496448476 | 0.181808226 | 0.003588213 | 0.007186537 | ns | YPD | KS.1 |
| FIM1_1098 | 1548.642065 | 2386.16359 | 3522.134248 | 0.575645762 | 0.17809277 | 0.000593736 | 0.001373878 | ns | YPD | KS.1 |
| FIM1_1099 | 1511.152518 | 1346.3165 | 822.1322244 | -0.152415602 | 0.151470225 | 0.290488508 | 0.362200583 | ns | YPD | KS.1 |
| FIM1_1100 | 5756.541838 | 7126.88063 | 7850.830426 | 0.299036001 | 0.090458856 | 0.000744503 | 0.001689737 | ns | YPD | KS.1 |
| FIM1_1101 | 207.2988247 | 92.82807377 | 78.11312331 | -0.762310448 | 0.488772496 | 0.036790102 | 0.058928125 | ns | YPD | KS.1 |
| FIM1_1102 | 573.9143403 | 944.7951678 | 1220.438134 | 0.689424204 | 0.134745809 | 1.1538324104 | 4.6181144344 | ns | YPD | KS.1 |
| FIM1_1103 | 446.3067463 | 704.4172775 | 973.7573959 | 0.570028586 | 0.246356197 | 0.010129176 | 0.018452086 | ns | YPD | KS.1 |
| FIM1_1104 | 703.2364065 | 1682.550782 | 1469.220352 | 1.196468292 | 0.186678206 | 1.9251124164 | 1.1651379892 | Km>KS.1 | YPD | KS.1 |
| FIM1_1105 | 3448.445692 | 3515.919674 | 4789.31649 | 0.027209779 | 0.072659219 | 0.704018519 | 0.763735746 | ns | YPD | KS.1 |
| FIM1_1106 | 4002.596385 | 7002.703749 | 6673.423361 | 0.762898893 | 0.163415939 | 9.6086620707 | 3.3999355477 | ns | YPD | KS.1 |
| FIM1_1107 | 652.6574487 | 780.7439686 | 984.9406452 | 0.235320159 | 0.157084034 | 0.114273306 | 0.162756856 | ns | YPD | KS.1 |
| FIM1_1108 | 552.8546932 | 992.444113 | 1788.027479 | 0.70465753 | 0.298731474 | 0.006734039 | 0.012709349 | ns | YPD | KS.1 |
| FIM1_1109 | 7738.728182 | 6625.078043 | 4698.985873 | -0.205354985 | 0.153728495 | 0.159844207 | 0.21751229 | ns | YPD | KS.1 |
| FIM1_1110 | 1039.660099 | 1022.482278 | 1348.385957 | -0.022251861 | 0.149007608 | 0.875676001 | 0.904774269 | ns | YPD | KS.1 |
| FIM1_1111 | 797.6597938 | 881.8983633 | 1127.910016 | 0.136927258 | 0.11787724 | 0.229892642 | 0.297300727 | ns | YPD | KS.1 |
| FIM1_1112 | 9572.41148 | 4507.781395 | 3978.287789 | -1.063699868 | 0.112907276 | 7.4957283687 | 1.1198573163 | Km<KS.1 | YPD | KS.1 |
| FIM1_1113 | 1454.170735 | 2560.667163 | 1419.349751 | 0.699436995 | 0.27310561 | 0.003876644 | 0.007708356 | ns | YPD | KS.1 |
| FIM1_1114 | 16306.47914 | 2080.040175 | 2368.019001 | -2.69312322 | 0.414998678 | 2.5662815070 | 1.8611152328 | Km<KS.1 | YPD | KS.1 |
| FIM1_1115 | 1414.261217 | 521.7735899 | 361.7384784 | -1.325463835 | 0.252796144 | 1.5886740612 | 7.1076020277 | Km<KS.1 | YPD | KS.1 |
| FIM1_1116 | 3002.677087 | 7290.842636 | 6624.83979 | 1.235067961 | 0.158623342 | 8.5104024125 | 7.6840747735 | Km>KS.1 | YPD | KS.1 |
| FIM1_1117 | 2358.476635 | 1830.825402 | 1887.62211 | -0.336131279 | 0.159916389 | 0.026550282 | 0.043955957 | ns | YPD | KS.1 |
| FIM1_1118 | 62.42087429 | 14.53058011 | 20.87135697 | -1.971344577 | 0.262446386 | 1.4122047169 | 1.2434899941 | Km<KS.1 | YPD | KS.1 |
| FIM1_1119 | 3667.931025 | 3651.171942 | 2937.395474 | -0.00620442 | 0.136213437 | 0.962180055 | 0.97214577 | ns | YPD | KS.1 |
| FIM1_1120 | 137.3303394 | 38.9355994 | 43.48615303 | -1.737744373 | 0.20793592 | 2.8277403151 | 3.1756225887 | Km<KS.1 | YPD | KS.1 |
| FIM1_1121 | 16329.82759 | 11939.38587 | 12996.19495 | -0.423361977 | 0.147156188 | 0.002609557 | 0.005362474 | ns | YPD | KS.1 |
| FIM1_1122 | 4398.777123 | 6550.717348 | 5340.92505 | 0.564510644 | 0.078387051 | 2.8681579840 | 2.0590311646 | ns | YPD | KS.1 |
| FIM1_1123 | 733.7364429 | 408.9848689 | 448.2279881 | -0.809098534 | 0.141274182 | 2.8867316278 | 1.4177186425 | ns | YPD | KS.1 |
| FIM1_1124 | 4203.218871 | 4900.703274 | 3235.534849 | 0.2064477 | 0.135246047 | 0.111568811 | 0.159315395 | ns | YPD | KS.1 |

| | | | | | | | | | | |
|---|---|---|---|---|---|---|---|---|---|---|
| FIM1_1125 | 11237.59309 | 15545.08444 | 21740.66123 | 0.443754286 | 0.134076098 | 0.000586134 | 0.001357382 | ns | YPD | KS.1 |
| FIM1_1126 | 1462.189174 | 762.4454135 | 764.9335658 | -0.797955491 | 0.29576527 | 0.002081617 | 0.004367797 | ns | YPD | KS.1 |
| FIM1_1127 | 719.3650804 | 525.390444 | 411.6442297 | -0.382883458 | 0.242632914 | 0.079342364 | 0.117478649 | ns | YPD | KS.1 |
| FIM1_1128 | 304.9548575 | 121.8638339 | 192.2800568 | -1.175514065 | 0.290822778 | 7.0104870088 | 2.1784617657 | Km<KS.1 | YPD | KS.1 |
| FIM1_1129 | 2319.125799 | 2958.090062 | 3127.035772 | 0.335316081 | 0.115619236 | 0.002789526 | 0.005701681 | ns | YPD | KS.1 |
| FIM1_1130 | 506.7959081 | 453.1567594 | 445.2423528 | -0.145989028 | 0.15644251 | 0.32572951 | 0.399828353 | ns | YPD | KS.1 |
| FIM1_1131 | 746.7544747 | 1314.605564 | 1327.843283 | 0.795600994 | 0.111455705 | 2.7079814481 | 1.9524938702 | ns | YPD | KS.1 |
| FIM1_1132 | 4418.300592 | 1304.712159 | 990.3518956 | -1.675201633 | 0.214252312 | 2.7033281587 | 2.5666140438 | Km<KS.1 | YPD | KS.1 |
| FIM1_1133 | 1284.939884 | 1213.204699 | 653.9017944 | -0.068142025 | 0.234987678 | 0.749007032 | 0.801529358 | ns | YPD | KS.1 |
| FIM1_1134 | 372.8753681 | 417.2348294 | 307.6470545 | 0.13263683 | 0.246653605 | 0.549488835 | 0.62271229 | ns | YPD | KS.1 |
| FIM1_1135 | 5166.79308 | 11991.18845 | 14001.82884 | 1.108442751 | 0.248097737 | 1.2121130293 | 4.2110770397 | Km>KS.1 | YPD | KS.1 |
| FIM1_1136 | 1244.74232 | 976.9438545 | 1073.687292 | -0.327471269 | 0.137742805 | 0.013182294 | 0.023480814 | ns | YPD | KS.1 |
| FIM1_1137 | 3928.747207 | 4420.996614 | 3821.963831 | 0.137625674 | 0.251446856 | 0.540997858 | 0.61519185 | ns | YPD | KS.1 |
| FIM1_1138 | 769.3039369 | 452.9505055 | 592.9896694 | -0.738346113 | 0.125086748 | 1.1769988468 | 6.0491418005 | ns | YPD | KS.1 |
| FIM1_1139 | 4826.553243 | 8767.599594 | 9084.169046 | 0.845572241 | 0.094525026 | 9.4111781140 | 1.2256704481 | ns | YPD | KS.1 |
| FIM1_1140 | 581.1560483 | 1668.263694 | 3299.452156 | 1.238653584 | 0.403167804 | 0.000233165 | 0.000580578 | Km>KS.1 | YPD | KS.1 |
| FIM1_1141 | 477.1754034 | 1865.994982 | 3765.705268 | 1.761356675 | 0.338208093 | 7.3492805945 | 3.4413700033 | Km>KS.1 | YPD | KS.1 |
| FIM1_1142 | 956.0491701 | 1027.811456 | 869.6513995 | 0.099392597 | 0.102277959 | 0.318564707 | 0.392000845 | ns | YPD | KS.1 |
| FIM1_1143 | 1408.252793 | 1645.109099 | 1700.876473 | 0.216969306 | 0.092093088 | 0.01601496 | 0.028006613 | ns | YPD | KS.1 |
| FIM1_1144 | 198.817067 | 712.5919412 | 867.3689057 | 1.737888409 | 0.235074234 | 6.2718778115 | 5.1659920716 | Km>KS.1 | YPD | KS.1 |
| FIM1_1145 | 9478.124531 | 8978.766635 | 9504.782481 | -0.074884931 | 0.093033181 | 0.410561227 | 0.488062629 | ns | YPD | KS.1 |
| FIM1_1146 | 465.8541554 | 573.4610468 | 629.5973332 | 0.280833158 | 0.140550295 | 0.036829113 | 0.058971625 | ns | YPD | KS.1 |
| FIM1_1147 | 3088.226747 | 3002.079957 | 3609.849336 | -0.038786344 | 0.103342385 | 0.700143582 | 0.760195181 | ns | YPD | KS.1 |
| FIM1_1148 | 922.5922466 | 522.4111578 | 672.4674812 | -0.780084604 | 0.156452069 | 1.8710261816 | 7.3172614812 | ns | YPD | KS.1 |
| FIM1_1149 | 217.7539198 | 225.9973659 | 286.7529395 | 0.046794439 | 0.165353337 | 0.764871131 | 0.815523762 | ns | YPD | KS.1 |
| FIM1_1150 | 510.0793792 | 657.1574469 | 929.2402497 | 0.337290736 | 0.157509399 | 0.024031729 | 0.040214548 | ns | YPD | KS.1 |
| FIM1_1151 | 7352.809732 | 5379.80644 | 6165.423123 | -0.421970822 | 0.148526882 | 0.002929146 | 0.005972336 | ns | YPD | KS.1 |
| FIM1_1152 | 1269.619067 | 1357.949767 | 803.2191854 | 0.081278017 | 0.222587074 | 0.689461894 | 0.750727275 | ns | YPD | KS.1 |
| FIM1_1153 | 94.37436835 | 104.558508 | 75.4474694 | 0.126625324 | 0.196431521 | 0.487558685 | 0.565135026 | ns | YPD | KS.1 |
| FIM1_1154 | 1063.752974 | 1143.955998 | 1354.118306 | 0.099921024 | 0.111709262 | 0.356701675 | 0.431773926 | ns | YPD | KS.1 |
| FIM1_1155 | 139.5369184 | 109.8564841 | 96.14274037 | -0.253469087 | 0.329968988 | 0.361483787 | 0.436394525 | ns | YPD | KS.1 |
| FIM1_1156 | 6538.476845 | 7445.386039 | 9766.387008 | 0.176106823 | 0.12381755 | 0.14035856 | 0.193986337 | ns | YPD | KS.1 |
| FIM1_1157 | 2775.60905 | 3622.432732 | 2947.350756 | 0.372202636 | 0.096821738 | 8.5806799744 | 0.000226586 | ns | YPD | KS.1 |
| FIM1_1158 | 1990.667526 | 1537.93467 | 1034.973562 | -0.301438677 | 0.265508639 | 0.199501704 | 0.264038568 | ns | YPD | KS.1 |
| FIM1_1159 | 1263.782172 | 1833.611092 | 1818.015142 | 0.526339144 | 0.082170231 | 7.9132320210 | 4.5043855039 | ns | YPD | KS.1 |
| FIM1_1160 | 16350.86398 | 9418.882019 | 13109.57996 | -0.783931479 | 0.082774863 | 7.9987465677 | 1.1914300651 | ns | YPD | KS.1 |
| FIM1_1161 | 419.9822388 | 107.61246 | 118.611153 | -1.729167773 | 0.36189899 | 7.1460899136 | 2.9539989756 | Km<KS.1 | YPD | KS.1 |
| FIM1_1162 | 12120.40363 | 9207.67677 | 6707.310668 | -0.306719255 | 0.298367406 | 0.233753008 | 0.301587452 | ns | YPD | KS.1 |
| FIM1_1163 | 1594.106273 | 1230.017641 | 1155.211628 | -0.362652317 | 0.092712401 | 6.6024123793 | 0.000178226 | ns | YPD | KS.1 |
| FIM1_1164 | 36.27004559 | 30.59391991 | 38.21550992 | -0.197484426 | 0.26768307 | 0.404116084 | 0.480975482 | ns | YPD | KS.1 |
| FIM1_1165 | 410.3127744 | 758.3995947 | 916.3283181 | 0.835565494 | 0.171630824 | 3.0346437346 | 1.1586609807 | ns | YPD | KS.1 |
| FIM1_1166 | 16634.12374 | 5019.702943 | 6055.591906 | -1.685363713 | 0.151499527 | 4.7060218701 | 1.2863988353 | Km<KS.1 | YPD | KS.1 |
| FIM1_1167 | 1317.990156 | 1315.760956 | 888.8186405 | -0.00261864 | 0.138184234 | 0.98424557 | 0.988218307 | ns | YPD | KS.1 |
| FIM1_1168 | 1524.64308 | 1834.032572 | 1564.529 | 0.253824501 | 0.114605514 | 0.022289732 | 0.03749126 | ns | YPD | KS.1 |
| FIM1_1169 | 273.5164415 | 576.3258722 | 944.2420511 | 1.026381523 | 0.165479218 | 1.0090931436 | 5.7113064729 | Km>KS.1 | YPD | KS.1 |

| | | | | | | | | | | |
|---|---|---|---|---|---|---|---|---|---|---|
| FIM1_1170 | 336.0529523 | 946.6741192 | 1068.3569 | 1.451867694 | 0.148322795 | 1.0292312401 | 1.7966404981 | Km>KS.1 | YPD | KS.1 |
| FIM1_1171 | 84.27567812 | 60.04447654 | 54.07909024 | -0.4370813 | 0.198932625 | 0.017736642 | 0.030670766 | ns | YPD | KS.1 |
| FIM1_1172 | 2836.453937 | 2371.345571 | 2917.458834 | -0.252764518 | 0.075509627 | 0.000677755 | 0.00155098 | ns | YPD | KS.1 |
| FIM1_1173 | 177.1439134 | 281.675665 | 611.6997115 | 0.607964946 | 0.199326839 | 0.001033533 | 0.00228424 | ns | YPD | KS.1 |
| FIM1_1174 | 1486.516797 | 721.9135957 | 552.607912 | -0.943053433 | 0.242617535 | 2.192677917 | 6.345882826 | ns | YPD | KS.1 |
| FIM1_1175 | 276.8485402 | 135.716609 | 154.079058 | -0.924607637 | 0.250880375 | 5.134151021 | 0.000141431 | ns | YPD | KS.1 |
| FIM1_1177 | 1950.340345 | 1623.249924 | 2264.533237 | -0.257731006 | 0.086549666 | 0.002400326 | 0.004967397 | ns | YPD | KS.1 |
| FIM1_1178 | 8394.717816 | 9503.614447 | 14048.49393 | 0.164510414 | 0.147465842 | 0.242129687 | 0.310454942 | ns | YPD | KS.1 |
| FIM1_1179 | 1011.584743 | 888.6889052 | 940.2419267 | -0.179309446 | 0.100659329 | 0.067545994 | 0.102264552 | ns | YPD | KS.1 |
| FIM1_1180 | 2037.272196 | 774.67973 | 1300.328035 | -1.352833952 | 0.151023438 | 3.279671620 | 4.362664789 | Km<KS.1 | YPD | KS.1 |
| FIM1_1181 | 1968.295 | 2097.935806 | 2233.461848 | 0.081292793 | 0.179482607 | 0.629355989 | 0.696986438 | ns | YPD | KS.1 |
| FIM1_1182 | 248.9319624 | 139.8836894 | 106.0153218 | -0.771560257 | 0.192147695 | 1.857889731 | 5.433863266 | ns | YPD | KS.1 |
| FIM1_1183 | 3452.112871 | 700.5234003 | 1349.595068 | -2.240373898 | 0.180362315 | 2.045885604 | 8.774380069 | Km<KS.1 | YPD | KS.1 |
| FIM1_1184 | 369.4958163 | 448.4638348 | 592.4635588 | 0.256775234 | 0.156477953 | 0.083745409 | 0.123373827 | ns | YPD | KS.1 |
| FIM1_1186 | 1096.306208 | 819.7537695 | 717.6328512 | -0.410551887 | 0.077783238 | 8.705528356 | 3.555829521 | ns | YPD | KS.1 |
| FIM1_1187 | 4239.879987 | 6424.696732 | 6179.320756 | 0.593694032 | 0.059344036 | 6.435702962 | 1.143486508 | ns | YPD | KS.1 |
| FIM1_1188 | 8626.831691 | 1021.493407 | 1370.245573 | -2.99494821 | 0.229285405 | 6.537430162 | 3.613746117 | Km<KS.1 | YPD | KS.1 |
| FIM1_1189 | 8338.344457 | 1679.572113 | 1684.525144 | -2.177332981 | 0.272201531 | 1.676345877 | 1.778213377 | Km<KS.1 | YPD | KS.1 |
| FIM1_1190 | 6018.355732 | 543.8240486 | 919.1149451 | -3.36099721 | 0.272067413 | 4.841345609 | 2.169882378 | Km<KS.1 | YPD | KS.1 |
| FIM1_1191 | 311.6904958 | 32.33762597 | 87.82668476 | -2.822350387 | 0.542486819 | 4.478540813 | 2.400941869 | Km<KS.1 | YPD | KS.1 |
| FIM1_1192 | 399.6980647 | 704.4126549 | 1024.594112 | 0.760905407 | 0.186729245 | 1.474628284 | 4.359046770 | ns | YPD | KS.1 |
| FIM1_1193 | 3976.730392 | 4738.712618 | 4984.269302 | 0.248661948 | 0.06644106 | 0.000151942 | 0.000386656 | ns | YPD | KS.1 |
| FIM1_1194 | 785.6723254 | 2282.260658 | 3125.589537 | 1.511600094 | 0.118977906 | 3.968989406 | 1.795065663 | Km>KS.1 | YPD | KS.1 |
| FIM1_1195 | 119.7719615 | 178.8976419 | 301.4078431 | 0.534374752 | 0.171015741 | 0.00093729 | 0.002086362 | ns | YPD | KS.1 |
| FIM1_1196 | 8209.19037 | 11123.96933 | 16415.82363 | 0.417545723 | 0.12566781 | 0.000586335 | 0.001357382 | ns | YPD | KS.1 |
| FIM1_1197 | 2545.599263 | 3166.673232 | 4171.618958 | 0.298609614 | 0.121987935 | 0.011305624 | 0.020445466 | ns | YPD | KS.1 |
| FIM1_1198 | 2909.075306 | 1797.106754 | 1483.304579 | -0.632419215 | 0.201005694 | 0.000708589 | 0.001616336 | ns | YPD | KS.1 |
| FIM1_1199 | 1259.134856 | 1785.087391 | 1383.38834 | 0.486110866 | 0.107309255 | 3.418402054 | 1.108640822 | ns | YPD | KS.1 |
| FIM1_1200 | 5384.901851 | 6297.055383 | 4476.768721 | 0.193882534 | 0.210031017 | 0.315661833 | 0.388982412 | ns | YPD | KS.1 |
| FIM1_1201 | 205.7816638 | 189.881163 | 205.2568071 | -0.103909332 | 0.159866868 | 0.493123063 | 0.570266676 | ns | YPD | KS.1 |
| FIM1_1202 | 466.0156284 | 373.2065154 | 329.0836406 | -0.272431159 | 0.224404324 | 0.182527927 | 0.244566775 | ns | YPD | KS.1 |
| FIM1_1203 | 1451.751786 | 997.0835861 | 1165.203984 | -0.444014734 | 0.275786914 | 0.066920004 | 0.101440287 | ns | YPD | KS.1 |
| FIM1_1204 | 8552.893133 | 5619.88014 | 4190.507741 | -0.56624245 | 0.162444192 | 0.000241819 | 0.00059883 | ns | YPD | KS.1 |
| FIM1_1205 | 1030.164296 | 847.5156794 | 847.5103943 | -0.265693117 | 0.127956479 | 0.031079618 | 0.050546289 | ns | YPD | KS.1 |
| FIM1_1206 | 857.8863759 | 760.959774 | 779.5023305 | -0.163990515 | 0.114525391 | 0.139371616 | 0.192925372 | ns | YPD | KS.1 |
| FIM1_1207 | 2426.269433 | 2392.39458 | 1871.703589 | -0.019007971 | 0.108143368 | 0.856607161 | 0.889134284 | ns | YPD | KS.1 |
| FIM1_1208 | 817.8639128 | 1532.599608 | 1630.311595 | 0.882613997 | 0.116390289 | 8.036748602 | 6.511860634 | ns | YPD | KS.1 |
| FIM1_1209 | 583.6064171 | 1044.870339 | 1031.406901 | 0.825874241 | 0.0946317 | 6.916068351 | 8.211799533 | ns | YPD | KS.1 |
| FIM1_1210 | 146.7444188 | 267.9653341 | 392.2183859 | 0.793390078 | 0.217035865 | 7.688264227 | 0.000203995 | ns | YPD | KS.1 |
| FIM1_1211 | 1198.758743 | 1354.579879 | 1425.858515 | 0.16176349 | 0.147402818 | 0.249908497 | 0.318875295 | ns | YPD | KS.1 |
| FIM1_1212 | 9258.650124 | 7656.428397 | 9933.849973 | -0.262399642 | 0.109035253 | 0.013261552 | 0.023579779 | ns | YPD | KS.1 |
| FIM1_1213 | 4768.240958 | 5972.353906 | 8591.439274 | 0.304601833 | 0.137282132 | 0.020687055 | 0.035101671 | ns | YPD | KS.1 |
| FIM1_1214 | 8616.573293 | 12635.3918 | 18090.46764 | 0.538107315 | 0.096404852 | 1.232086314 | 5.597835079 | ns | YPD | KS.1 |
| FIM1_1215 | 1099.495392 | 1062.696412 | 930.5813569 | -0.046203188 | 0.093200216 | 0.612368085 | 0.682161044 | ns | YPD | KS.1 |
| FIM1_1216 | 1207.745585 | 1396.422909 | 932.4318863 | 0.191642459 | 0.154856828 | 0.192631896 | 0.256241626 | ns | YPD | KS.1 |

| | | | | | | | | | | |
|---|---|---|---|---|---|---|---|---|---|---|
| FIM1_1217 | 2929.126472 | 1901.317831 | 3883.21751 | -0.50344506 | 0.295343142 | 0.049237803 | 0.076958238 | ns | YPD | KS.1 |
| FIM1_1218 | 523.5158622 | 285.2306018 | 430.1686544 | -0.834088441 | 0.156285374 | 2.54404320748 | 1.11610361174 | ns | YPD | KS.1 |
| FIM1_1219 | 1779.075862 | 1404.16608 | 1348.642288 | -0.328146315 | 0.105208878 | 0.001371597 | 0.002959104 | ns | YPD | KS.1 |
| FIM1_1220 | 1048.826364 | 623.0038862 | 623.5037668 | -0.684053841 | 0.206547743 | 0.000354515 | 0.000851214 | ns | YPD | KS.1 |
| FIM1_1221 | 1419.133487 | 764.3920827 | 911.9478004 | -0.878339348 | 0.088305033 | 6.16434851890 | 1.10713479716 | ns | YPD | KS.1 |
| FIM1_1222 | 5783.760169 | 6059.001404 | 5569.113365 | 0.065739012 | 0.056645729 | 0.241292965 | 0.309549382 | ns | YPD | KS.1 |
| FIM1_1223 | 537.4482235 | 593.9933444 | 358.2853896 | 0.130022462 | 0.157349725 | 0.384051995 | 0.46073274 | ns | YPD | KS.1 |
| FIM1_1224 | 292.2858325 | 218.0796891 | 283.1675351 | -0.396291855 | 0.143075807 | 0.003815903 | 0.007605815 | ns | YPD | KS.1 |
| FIM1_1225 | 4242.249082 | 3137.534234 | 2384.065711 | -0.405525774 | 0.152315886 | 0.00519567 | 0.010046039 | ns | YPD | KS.1 |
| FIM1_1227 | 7972.812526 | 7097.683232 | 4298.800342 | -0.146583307 | 0.192160355 | 0.412449842 | 0.489956533 | ns | YPD | KS.1 |
| FIM1_1228 | 1286.382394 | 1488.105213 | 1288.517643 | 0.191847075 | 0.15229755 | 0.185380414 | 0.247921387 | ns | YPD | KS.1 |
| FIM1_1230 | 1332.40502 | 1569.665654 | 2123.311934 | 0.21702114 | 0.153819579 | 0.137618322 | 0.190976611 | ns | YPD | KS.1 |
| FIM1_1231 | 3912.22209 | 3233.50911 | 3724.980557 | -0.249722823 | 0.165251543 | 0.109538645 | 0.156596195 | ns | YPD | KS.1 |
| FIM1_1232 | 2134.28127 | 1924.099179 | 3261.024632 | -0.136145637 | 0.153318758 | 0.350548163 | 0.425661769 | ns | YPD | KS.1 |
| FIM1_1233 | 477.4339415 | 657.4507141 | 1200.732231 | 0.406961944 | 0.210196939 | 0.035245471 | 0.05674635 | ns | YPD | KS.1 |
| FIM1_1234 | 4294.029302 | 7197.889622 | 8850.688557 | 0.728126398 | 0.100640318 | 1.54437763195 | 1.13322694970 | ns | YPD | KS.1 |
| FIM1_1235 | 3958.627203 | 1880.940377 | 2019.227918 | -1.044529064 | 0.1281222 | 6.21740765916 | 6.29971549986 | Km<KS.1 | YPD | KS.1 |
| FIM1_1236 | 523.1225344 | 325.2497613 | 389.0121877 | -0.637965597 | 0.173805652 | 0.000102117 | 0.000266404 | ns | YPD | KS.1 |
| FIM1_1239 | 1656.823992 | 1971.499154 | 2726.193429 | 0.220617159 | 0.191487354 | 0.215534572 | 0.282180131 | ns | YPD | KS.1 |
| FIM1_1240 | 392.9336494 | 247.3971783 | 277.3851919 | -0.626951192 | 0.162678371 | 5.01716796381 | 0.000138285 | ns | YPD | KS.1 |
| FIM1_1242 | 990.5160909 | 775.9131653 | 774.0457167 | -0.330461297 | 0.13617233 | 0.011468585 | 0.020710058 | ns | YPD | KS.1 |
| FIM1_1243 | 2992.210321 | 3440.680707 | 3450.124753 | 0.197972885 | 0.066313355 | 0.002497624 | 0.005158024 | ns | YPD | KS.1 |
| FIM1_1244 | 751.625763 | 656.7455025 | 884.1417876 | -0.184474489 | 0.115680947 | 0.09953854 | 0.143829288 | ns | YPD | KS.1 |
| FIM1_1245 | 1582.976615 | 1479.981128 | 1256.399244 | -0.089724299 | 0.133285577 | 0.483743375 | 0.561770143 | ns | YPD | KS.1 |
| FIM1_1246 | 343.7685353 | 558.3004965 | 571.5752836 | 0.679676838 | 0.103588674 | 1.98698320660 | 1.19821108519 | ns | YPD | KS.1 |
| FIM1_1247 | 1839.164502 | 673.4948328 | 887.4453676 | -1.414674452 | 0.136077034 | 2.26407334132 | 4.89728907526 | Km<KS.1 | YPD | KS.1 |
| FIM1_1248 | 854.8345766 | 1145.711231 | 1548.077761 | 0.395645341 | 0.146088487 | 0.004605579 | 0.00903734 | ns | YPD | KS.1 |
| FIM1_1249 | 5039.028405 | 6120.392181 | 4249.76047 | 0.220367007 | 0.277951867 | 0.366178986 | 0.441848279 | ns | YPD | KS.1 |
| FIM1_1250 | 8392.552302 | 4480.674867 | 2503.527131 | -0.855351679 | 0.17263764 | 1.87233645539 | 7.31726148120 | ns | YPD | KS.1 |
| FIM1_1251 | 5621.834482 | 4745.145714 | 3902.077065 | -0.233276572 | 0.111197943 | 0.030625067 | 0.049904916 | ns | YPD | KS.1 |
| FIM1_1252 | 917.5305527 | 1355.068399 | 1817.874988 | 0.528848261 | 0.152209131 | 0.000272547 | 0.000669261 | ns | YPD | KS.1 |
| FIM1_1253 | 263.6404548 | 435.9617121 | 403.9005518 | 0.706461228 | 0.10192959 | 1.46054014256 | 9.92648525853 | ns | YPD | KS.1 |
| FIM1_1254 | 597.6261048 | 825.837172 | 1261.360215 | 0.419345386 | 0.192211566 | 0.019030469 | 0.032714784 | ns | YPD | KS.1 |
| FIM1_1255 | 1433.915795 | 1290.056774 | 2162.44669 | -0.117970363 | 0.279708922 | 0.630594365 | 0.698087888 | ns | YPD | KS.1 |
| FIM1_1256 | 1468.511678 | 1512.463762 | 1269.138431 | 0.039891636 | 0.118531088 | 0.7281604 | 0.783337521 | ns | YPD | KS.1 |
| FIM1_1257 | 2912.434394 | 2246.546368 | 1885.744787 | -0.352535887 | 0.135925082 | 0.006922479 | 0.013020542 | ns | YPD | KS.1 |
| FIM1_1258 | 668.038371 | 698.5771339 | 1152.889444 | 0.061464059 | 0.125615321 | 0.612127683 | 0.682081039 | ns | YPD | KS.1 |
| FIM1_1259 | 525.0367714 | 337.8353178 | 347.9151025 | -0.575307395 | 0.200467 | 0.001988594 | 0.004194053 | ns | YPD | KS.1 |
| FIM1_1261 | 890.5371182 | 1736.280064 | 2347.620967 | 0.903132119 | 0.18791353 | 3.60008724277 | 1.35787975988 | ns | YPD | KS.1 |
| FIM1_1262 | 1016.62996 | 1075.770197 | 2501.382779 | 0.070070279 | 0.198752139 | 0.703986882 | 0.763735746 | ns | YPD | KS.1 |
| FIM1_1263 | 2132.45033 | 1872.148613 | 2066.89919 | -0.170202505 | 0.162532813 | 0.268419617 | 0.338244072 | ns | YPD | KS.1 |
| FIM1_1264 | 1017.12145 | 2052.606074 | 2953.260636 | 0.945646212 | 0.198041413 | 3.84917938476 | 1.44744273916 | ns | YPD | KS.1 |
| FIM1_1265 | 6778.130313 | 6233.538021 | 5046.128415 | -0.108365817 | 0.167193549 | 0.49261241 | 0.570073678 | ns | YPD | KS.1 |
| FIM1_1266 | 427.6889206 | 563.6415863 | 805.668196 | 0.358923018 | 0.185318615 | 0.03805658 | 0.06067937 | ns | YPD | KS.1 |
| FIM1_1267 | 2813.178523 | 2265.453488 | 4172.094383 | -0.244181719 | 0.283840855 | 0.324570126 | 0.39870034 | ns | YPD | KS.1 |

| | | | | | | | | | | |
|---|---|---|---|---|---|---|---|---|---|---|
| FIM1_1268 | 755.7747953 | 641.3556191 | 779.0367007 | -0.220470407 | 0.143180269 | 0.107072546 | 0.153335036 | ns | YPD | KS.1 |
| FIM1_1269 | 3089.798671 | 1907.793732 | 3666.06524 | -0.559788068 | 0.30674759 | 0.033950135 | 0.054873919 | ns | YPD | KS.1 |
| FIM1_1270 | 1701.570529 | 1650.587107 | 2826.111165 | -0.040649588 | 0.125444891 | 0.73709951 | 0.791475648 | ns | YPD | KS.1 |
| FIM1_1271 | 1989.827937 | 3886.390628 | 3309.465527 | 0.939723861 | 0.122014739 | 2.8590869821 | 2.4356092014 | ns | YPD | KS.1 |
| FIM1_1272 | 10043.64829 | 6677.542958 | 7662.356285 | -0.572426315 | 0.102584104 | 1.1535767510 | 5.2555351064 | ns | YPD | KS.1 |
| FIM1_1273 | 606.3030252 | 697.2375908 | 549.5565453 | 0.181302018 | 0.173804252 | 0.267247964 | 0.336852957 | ns | YPD | KS.1 |
| FIM1_1274 | 123.7215558 | 130.8949539 | 79.67623041 | 0.066866993 | 0.240603227 | 0.758217175 | 0.809643796 | ns | YPD | KS.1 |
| FIM1_1275 | 1217.127145 | 3038.496142 | 2704.232452 | 1.288244445 | 0.1322377 | 2.2298037495 | 3.7732223312 | Km>KS.1 | YPD | KS.1 |
| FIM1_1276 | 4018.213621 | 3043.997715 | 3207.325652 | -0.397494 | 0.045418694 | 1.4228377364 | 1.6461901717 | ns | YPD | KS.1 |
| FIM1_1277 | 4198.95889 | 3634.235718 | 2437.463365 | -0.187500531 | 0.169436784 | 0.240337515 | 0.308562358 | ns | YPD | KS.1 |
| FIM1_1279 | 1254.618957 | 965.5138168 | 1120.380077 | -0.359604075 | 0.12400034 | 0.002692261 | 0.005516475 | ns | YPD | KS.1 |
| FIM1_1280 | 3262.971619 | 3239.839338 | 7070.403574 | -0.009828668 | 0.113196554 | 0.928723285 | 0.947774019 | ns | YPD | KS.1 |
| FIM1_1281 | 667.3001897 | 1793.825188 | 2186.258833 | 1.402563405 | 0.1148462 | 2.3734446068 | 9.2975487552 | Km>KS.1 | YPD | KS.1 |
| FIM1_1282 | 1299.275382 | 1100.088734 | 725.1315443 | -0.189432492 | 0.27231782 | 0.429940365 | 0.507582657 | ns | YPD | KS.1 |
| FIM1_1283 | 9420.784041 | 2651.79006 | 3946.865104 | -1.672795198 | 0.293679898 | 5.9750018025 | 3.1555874700 | Km<KS.1 | YPD | KS.1 |
| FIM1_1284 | 2453.689737 | 2134.965906 | 937.7489032 | -0.170635607 | 0.215657108 | 0.388196063 | 0.464919454 | ns | YPD | KS.1 |
| FIM1_1285 | 6349.947297 | 4900.266828 | 4577.267726 | -0.360865425 | 0.102322902 | 0.00030348 | 0.000737933 | ns | YPD | KS.1 |
| FIM1_1286 | 172.7209923 | 303.4331563 | 438.3162663 | 0.759465225 | 0.178845161 | 6.9638342040 | 2.1653171978 | ns | YPD | KS.1 |
| FIM1_1287 | 699.9611044 | 1646.936294 | 1377.703421 | 1.138684997 | 0.233966586 | 1.6441685485 | 6.4713121272 | Km>KS.1 | YPD | KS.1 |
| FIM1_1288 | 327.0166777 | 793.2211736 | 608.8673932 | 1.243871874 | 0.141146461 | 1.4824718391 | 1.8934375797 | Km>KS.1 | YPD | KS.1 |
| FIM1_1289 | 428.7210391 | 1681.904445 | 2209.892379 | 1.944289351 | 0.119164283 | 1.8238644931 | 3.6294903413 | Km>KS.1 | YPD | KS.1 |
| FIM1_1290 | 4767.474258 | 4700.945698 | 4253.796382 | -0.018945961 | 0.130824667 | 0.88046367 | 0.908588832 | ns | YPD | KS.1 |
| FIM1_1291 | 173.7372642 | 602.8824066 | 1057.93371 | 1.724144482 | 0.189131456 | 3.5407487477 | 5.0329214343 | Km>KS.1 | YPD | KS.1 |
| FIM1_1292 | 158.8103901 | 160.0333791 | 242.678051 | 0.011245421 | 0.172552755 | 0.944878836 | 0.960125042 | ns | YPD | KS.1 |
| FIM1_1293 | 2009.81691 | 1352.812503 | 2514.956573 | -0.499655575 | 0.226755369 | 0.015535048 | 0.027239938 | ns | YPD | KS.1 |
| FIM1_1294 | 712.4384363 | 671.2371137 | 1168.931978 | -0.066496101 | 0.275599026 | 0.78397445 | 0.830379381 | ns | YPD | KS.1 |
| FIM1_1295 | 5810.611492 | 3929.596162 | 3331.000546 | -0.538094866 | 0.132362612 | 2.5028871721 | 7.2017719384 | ns | YPD | KS.1 |
| FIM1_1296 | 2493.583753 | 1753.079465 | 921.9107713 | -0.372444785 | 0.343030035 | 0.192477393 | 0.256237169 | ns | YPD | KS.1 |
| FIM1_1297 | 7859.162326 | 1314.506039 | 1960.959462 | -2.416135367 | 0.306035655 | 1.8579945430 | 1.9625313910 | Km<KS.1 | YPD | KS.1 |
| FIM1_1298 | 3010.965105 | 3110.58737 | 2757.110115 | 0.039769379 | 0.210592893 | 0.8375837 | 0.874864352 | ns | YPD | KS.1 |
| FIM1_1299 | 845.7900399 | 911.0939714 | 985.6162665 | 0.102535467 | 0.098244512 | 0.284843899 | 0.356413077 | ns | YPD | KS.1 |
| FIM1_1300 | 1747.404639 | 1334.122127 | 1455.409244 | -0.361149573 | 0.15320148 | 0.013235325 | 0.023549979 | ns | YPD | KS.1 |
| FIM1_1301 | 375.8849715 | 319.5256051 | 342.8589182 | -0.21910324 | 0.141205788 | 0.104746154 | 0.150523431 | ns | YPD | KS.1 |
| FIM1_1302 | 1090.908424 | 664.742357 | 736.6684656 | -0.692176925 | 0.114481514 | 5.4023914347 | 2.8683988674 | ns | YPD | KS.1 |
| FIM1_1303 | 1540.257251 | 1100.448583 | 2040.5456 | -0.428043913 | 0.210545116 | 0.027043828 | 0.04474328 | ns | YPD | KS.1 |
| FIM1_1304 | 2707.376665 | 2187.647447 | 1576.531809 | -0.24818095 | 0.261953784 | 0.28575771 | 0.357466585 | ns | YPD | KS.1 |
| FIM1_1305 | 6956.195995 | 5321.763948 | 4427.895953 | -0.358071169 | 0.154573027 | 0.014832346 | 0.026129929 | ns | YPD | KS.1 |
| FIM1_1306 | 204.0503447 | 85.49931091 | 117.1812301 | -1.174333834 | 0.21694623 | 8.4341001468 | 3.9104984371 | Km<KS.1 | YPD | KS.1 |
| FIM1_1307 | 494.6938596 | 1146.455965 | 1081.113627 | 1.179058228 | 0.133239578 | 1.2033739276 | 1.5550091661 | Km>KS.1 | YPD | KS.1 |
| FIM1_1308 | 1162.558729 | 916.4519792 | 883.5704345 | -0.326505056 | 0.118295719 | 0.004376095 | 0.008629042 | ns | YPD | KS.1 |
| FIM1_1309 | 924.14123 | 1230.276865 | 1078.806146 | 0.380738961 | 0.160369557 | 0.012263214 | 0.022027243 | ns | YPD | KS.1 |
| FIM1_1310 | 1164.439869 | 2067.574006 | 2115.586315 | 0.80752629 | 0.107856498 | 1.9611868001 | 1.5487149731 | ns | YPD | KS.1 |
| FIM1_1311 | 7888.343881 | 2573.370462 | 2063.53469 | -1.496100227 | 0.258275387 | 4.9966743183 | 2.6643574205 | Km<KS.1 | YPD | KS.1 |
| FIM1_1312 | 2504.780872 | 1635.035268 | 1879.694692 | -0.540741563 | 0.228054043 | 0.00922096 | 0.016959067 | ns | YPD | KS.1 |
| FIM1_1313 | 9236.71431 | 23926.43424 | 32699.65459 | 1.330436702 | 0.153495224 | 4.5573305399 | 5.5030872418 | Km>KS.1 | YPD | KS.1 |

| | | | | | | | | | | |
|---|---|---|---|---|---|---|---|---|---|---|
| FIM1_1314 | 155.3921954 | 886.6269292 | 895.8018927 | 2.418684209 | 0.226088813 | 6.1736072263 | 1.5831802036 | Km>KS.1 | YPD | KS.1 |
| FIM1_1315 | 1953.052871 | 2336.143938 | 3495.315727 | 0.232588958 | 0.171852129 | 0.150301778 | 0.206048869 | ns | YPD | KS.1 |
| FIM1_1316 | 250.609003 | 769.4184352 | 918.0455375 | 1.530573279 | 0.217527959 | 1.3604666728 | 9.9975209709 | Km>KS.1 | YPD | KS.1 |
| FIM1_1317 | 179.5180846 | 544.4980068 | 631.2855543 | 1.532937235 | 0.194638635 | 2.3387628504 | 2.2375663810 | Km>KS.1 | YPD | KS.1 |
| FIM1_1318 | 810.6696845 | 940.6607147 | 1423.226802 | 0.200114322 | 0.136613811 | 0.126529503 | 0.177569612 | ns | YPD | KS.1 |
| FIM1_1319 | 4267.668003 | 6828.920029 | 3635.495891 | 0.599050533 | 0.229731628 | 0.004203406 | 0.008314883 | ns | YPD | KS.1 |
| FIM1_1320 | 7203.743441 | 14453.63088 | 13061.52587 | 0.929145825 | 0.21145154 | 2.4702358190 | 8.1820394139 | ns | YPD | KS.1 |
| FIM1_1321 | 914.8671122 | 762.3547574 | 788.6044429 | -0.250606855 | 0.120323706 | 0.03115882 | 0.05064199 | ns | YPD | KS.1 |
| FIM1_1322 | 3574.379436 | 7007.529018 | 7929.863146 | 0.881138403 | 0.232367079 | 3.6910825334 | 0.000103531 | ns | YPD | KS.1 |
| FIM1_1323 | 3611.095463 | 794.3875707 | 817.2490221 | -2.123518881 | 0.180355483 | 7.9826362598 | 2.7771759015 | Km<KS.1 | YPD | KS.1 |
| FIM1_1324 | 600.9510411 | 508.5758566 | 1164.683031 | -0.194008635 | 0.257601206 | 0.397872374 | 0.474679871 | ns | YPD | KS.1 |
| FIM1_1325 | 4376.537536 | 2131.455358 | 1963.284815 | -0.995045151 | 0.157086638 | 4.5944271142 | 2.7018055429 | ns | YPD | KS.1 |
| FIM1_1326 | 1941.590755 | 1033.569224 | 1051.740856 | -0.891442838 | 0.100332054 | 1.4843028263 | 1.8934375797 | ns | YPD | KS.1 |
| FIM1_1327 | 5132.573105 | 2499.824871 | 2187.925136 | -1.017067541 | 0.107433756 | 5.2609403245 | 7.9796274740 | Km<KS.1 | YPD | KS.1 |
| FIM1_1328 | 2190.758084 | 2155.105142 | 1573.607562 | -0.022650438 | 0.070047774 | 0.74310435 | 0.796068937 | ns | YPD | KS.1 |
| FIM1_1329 | 9263.147198 | 2420.946964 | 1818.801545 | -1.828726912 | 0.241209253 | 1.1984501694 | 1.0704290112 | Km<KS.1 | YPD | KS.1 |
| FIM1_1331 | 152.4691206 | 235.6799649 | 153.9760707 | 0.515116913 | 0.284952118 | 0.038571044 | 0.061385459 | ns | YPD | KS.1 |
| FIM1_1332 | 462.5265886 | 478.1535624 | 565.8234931 | 0.045819247 | 0.129230427 | 0.712902769 | 0.770852266 | ns | YPD | KS.1 |
| FIM1_1333 | 4100.30254 | 4842.143422 | 7441.272709 | 0.204136038 | 0.218592354 | 0.307341482 | 0.380259605 | ns | YPD | KS.1 |
| FIM1_1334 | 2250.204293 | 2665.429511 | 3113.798748 | 0.213000909 | 0.199664049 | 0.249163788 | 0.31800663 | ns | YPD | KS.1 |
| FIM1_1335 | 992.5717008 | 842.8971052 | 988.3723608 | -0.211658762 | 0.170659356 | 0.187800946 | 0.250821398 | ns | YPD | KS.1 |
| FIM1_1336 | 3397.12384 | 2909.251341 | 1587.422816 | -0.178983452 | 0.261463693 | 0.441034878 | 0.518098824 | ns | YPD | KS.1 |
| FIM1_1337 | 2809.659896 | 2512.804267 | 3774.680074 | -0.123987748 | 0.28356549 | 0.617041103 | 0.686148358 | ns | YPD | KS.1 |
| FIM1_1338 | 2473.629415 | 858.1478999 | 857.7782986 | -1.451970484 | 0.20364582 | 8.1363333577 | 6.0869561586 | Km<KS.1 | YPD | KS.1 |
| FIM1_1339 | 5074.434889 | 6907.846454 | 5129.626516 | 0.401948993 | 0.185308653 | 0.020239407 | 0.034483237 | ns | YPD | KS.1 |
| FIM1_1340 | 1351.298855 | 2726.351042 | 2228.020992 | 0.985254026 | 0.123872636 | 3.5286026139 | 3.2997740610 | ns | YPD | KS.1 |
| FIM1_1341 | 285.9133879 | 664.164609 | 703.1858564 | 1.115181134 | 0.238519317 | 4.4487206254 | 1.6541393954 | Km>KS.1 | YPD | KS.1 |
| FIM1_1342 | 1214.532811 | 2575.766137 | 2388.158002 | 1.041226773 | 0.156799913 | 5.5139462788 | 3.5304868387 | Km>KS.1 | YPD | KS.1 |
| FIM1_1344 | 1182.496918 | 547.7786261 | 431.1494487 | -1.085638368 | 0.114595305 | 4.3558904576 | 6.6678630852 | Km<KS.1 | YPD | KS.1 |
| FIM1_1345 | 3058.314868 | 1479.883686 | 1529.858814 | -1.018761983 | 0.128153714 | 3.4299264104 | 3.2135374561 | Km<KS.1 | YPD | KS.1 |
| FIM1_1346 | 4503.950392 | 5590.814132 | 8008.702261 | 0.296508291 | 0.118957328 | 0.010004736 | 0.01824544 | ns | YPD | KS.1 |
| FIM1_1347 | 8122.663547 | 10100.32285 | 16702.93723 | 0.294608266 | 0.136944774 | 0.024886667 | 0.041463888 | ns | YPD | KS.1 |
| FIM1_1348 | 6494.870576 | 9616.969181 | 10458.08256 | 0.554701392 | 0.085701526 | 4.7773768295 | 2.8027652979 | ns | YPD | KS.1 |
| FIM1_1349 | 897.5830751 | 1072.167116 | 1253.590956 | 0.248136698 | 0.09266479 | 0.006200754 | 0.011783327 | ns | YPD | KS.1 |
| FIM1_1350 | 5323.428792 | 5760.463458 | 8388.563378 | 0.102562307 | 0.162661636 | 0.505525285 | 0.581782868 | ns | YPD | KS.1 |
| FIM1_1352 | 1585.493282 | 1460.389024 | 2375.419536 | -0.114308323 | 0.096679676 | 0.225896107 | 0.292970055 | ns | YPD | KS.1 |
| FIM1_1353 | 5093.737644 | 7334.052266 | 8281.703894 | 0.511732081 | 0.096538268 | 6.3216591750 | 2.6296199327 | ns | YPD | KS.1 |
| FIM1_1354 | 745.8899902 | 856.9420444 | 1056.633872 | 0.182530448 | 0.15220649 | 0.207433202 | 0.273082874 | ns | YPD | KS.1 |
| FIM1_1355 | 2179.574225 | 2137.621401 | 1650.334581 | -0.026603408 | 0.062614746 | 0.667457896 | 0.729963296 | ns | YPD | KS.1 |
| FIM1_1356 | 654.6390107 | 651.4366472 | 718.3923749 | -0.007718781 | 0.108031235 | 0.941450321 | 0.957814999 | ns | YPD | KS.1 |
| FIM1_1357 | 805.6516307 | 992.6013116 | 1025.871029 | 0.276478668 | 0.154271521 | 0.059257893 | 0.091356683 | ns | YPD | KS.1 |
| FIM1_1358 | 291.1289816 | 1239.521947 | 1315.886861 | 2.054336297 | 0.130517258 | 1.4051639695 | 1.9973402138 | Km>KS.1 | YPD | KS.1 |
| FIM1_1359 | 489.7086282 | 686.9597701 | 495.7212686 | 0.460079013 | 0.141845327 | 0.00071893 | 0.001636175 | ns | YPD | KS.1 |
| FIM1_1360 | 3385.711166 | 5355.187198 | 2675.62504 | 0.578080203 | 0.237607278 | 0.007215871 | 0.01353656 | ns | YPD | KS.1 |
| FIM1_1361 | 695.1545629 | 181.7308694 | 279.5368154 | -1.884109985 | 0.166818753 | 4.2175818999 | 1.2640042139 | Km<KS.1 | YPD | KS.1 |

| | | | | | | | | | | |
|---|---|---|---|---|---|---|---|---|---|---|
| FIM1_1362 | 914.4494773 | 320.3009724 | 484.6343487 | -1.468960491 | 0.155793917 | 3.2728864244 | 5.0410557156 | Km<KS.1 | YPD | KS.1 |
| FIM1_1363 | 2694.353652 | 1881.524005 | 3829.713585 | -0.420019827 | 0.279973642 | 0.086705492 | 0.127300915 | ns | YPD | KS.1 |
| FIM1_1364 | 181.90846 | 103.7442573 | 299.4874911 | -0.713047573 | 0.246509899 | 0.001375479 | 0.002966193 | ns | YPD | KS.1 |
| FIM1_1365 | 2720.967606 | 2922.653636 | 2515.753909 | 0.093278046 | 0.156993911 | 0.531673 | 0.607225247 | ns | YPD | KS.1 |
| FIM1_1366 | 709.8215717 | 1050.396742 | 1390.123103 | 0.533354762 | 0.147930033 | 0.00016438 | 0.000417241 | ns | YPD | KS.1 |
| FIM1_1367 | 1407.286562 | 1253.485868 | 982.0969972 | -0.162011047 | 0.089971772 | 0.065772643 | 0.099944685 | ns | YPD | KS.1 |
| FIM1_1368 | 6120.498305 | 5875.837229 | 5774.293505 | -0.056456378 | 0.093921343 | 0.538808369 | 0.612982309 | ns | YPD | KS.1 |
| FIM1_1369 | 11128.25251 | 3108.424554 | 3750.884313 | -1.766914075 | 0.198322568 | 2.1094743222 | 2.8673865445 | Km<KS.1 | YPD | KS.1 |
| FIM1_1370 | 86.25441049 | 35.33106556 | 34.13613517 | -1.130036261 | 0.307431299 | 3.4227672376 | 9.6422802984 | Km<KS.1 | YPD | KS.1 |
| FIM1_1371 | 1133.343031 | 1313.561191 | 1244.944897 | 0.203223137 | 0.104721489 | 0.046077597 | 0.072337029 | ns | YPD | KS.1 |
| FIM1_1372 | 123.1478579 | 125.0838763 | 292.5230999 | 0.022043478 | 0.177268554 | 0.894764396 | 0.920101875 | ns | YPD | KS.1 |
| FIM1_1373 | 3984.061986 | 3060.782682 | 2413.523477 | -0.365079381 | 0.111713498 | 0.000775732 | 0.001752619 | ns | YPD | KS.1 |
| FIM1_1374 | 1418.954552 | 1681.672351 | 2208.060344 | 0.234978587 | 0.102350021 | 0.018439933 | 0.03178748 | ns | YPD | KS.1 |
| FIM1_1375 | 3508.211489 | 2595.274454 | 1570.463931 | -0.341839184 | 0.29176482 | 0.176924906 | 0.237892272 | ns | YPD | KS.1 |
| FIM1_1376 | 184.9593182 | 75.9635786 | 96.17648643 | -1.228331038 | 0.174922405 | 2.7190968642 | 1.9576710418 | Km<KS.1 | YPD | KS.1 |
| FIM1_1377 | 5417.90297 | 4955.407212 | 3826.541247 | -0.122407708 | 0.106534584 | 0.237341876 | 0.305258304 | ns | YPD | KS.1 |
| FIM1_1378 | 320.5084957 | 199.2265679 | 321.4348399 | -0.638391008 | 0.171420488 | 8.2589702391 | 0.000218207 | ns | YPD | KS.1 |
| FIM1_1379 | 755.1782825 | 450.8992744 | 272.0880493 | -0.669163438 | 0.218492598 | 0.000868512 | 0.001941954 | ns | YPD | KS.1 |
| FIM1_1380 | 7688.005021 | 3376.037633 | 2315.659662 | -1.066507666 | 0.26566043 | 9.9071834373 | 3.0072140085 | Km<KS.1 | YPD | KS.1 |
| FIM1_1382 | 3765.96351 | 4493.554897 | 4035.366892 | 0.251065802 | 0.065295068 | 0.000100249 | 0.00026208 | ns | YPD | KS.1 |
| FIM1_1383 | 2443.547821 | 2616.232453 | 2587.870545 | 0.092476407 | 0.118053249 | 0.418342223 | 0.496126951 | ns | YPD | KS.1 |
| FIM1_1384 | 1226.269864 | 846.9307618 | 1200.455473 | -0.499145141 | 0.156883769 | 0.000827497 | 0.001857761 | ns | YPD | KS.1 |
| FIM1_1385 | 10206.39809 | 5447.905737 | 5001.549149 | -0.850726996 | 0.181147489 | 6.9294387747 | 2.5181853838 | ns | YPD | KS.1 |
| FIM1_1386 | 362.4371853 | 770.6605851 | 571.1931895 | 1.055331912 | 0.136344749 | 1.7011069621 | 1.4847380941 | Km>KS.1 | YPD | KS.1 |
| FIM1_1387 | 2272.834738 | 2829.414219 | 2965.730192 | 0.313950081 | 0.045990578 | 6.7163494945 | 4.2674123544 | ns | YPD | KS.1 |
| FIM1_1388 | 565.8068912 | 403.292316 | 383.3514424 | -0.475301398 | 0.098981715 | 9.2996057452 | 3.3046813273 | ns | YPD | KS.1 |
| FIM1_1389 | 492.5504746 | 371.088103 | 608.2360905 | -0.38967609 | 0.11897113 | 0.000726894 | 0.001652031 | ns | YPD | KS.1 |
| FIM1_1390 | 4298.379612 | 3748.910583 | 2724.645374 | -0.172277321 | 0.194766605 | 0.341004677 | 0.416012327 | ns | YPD | KS.1 |
| FIM1_1391 | 5090.740051 | 4825.296269 | 3696.141313 | -0.056281975 | 0.315148051 | 0.834141963 | 0.871818543 | ns | YPD | KS.1 |
| FIM1_1392 | 1933.404577 | 1079.94891 | 1101.82239 | -0.786673182 | 0.180176904 | 3.8151764532 | 1.2310609529 | ns | YPD | KS.1 |
| FIM1_1393 | 791.002718 | 931.5701392 | 1062.710524 | 0.206475521 | 0.194452794 | 0.252914274 | 0.322379839 | ns | YPD | KS.1 |
| FIM1_1394 | 2494.614215 | 340.4695958 | 587.753791 | -2.747186595 | 0.275738712 | 4.9715271419 | 1.0480232004 | Km<KS.1 | YPD | KS.1 |
| FIM1_1395 | 822.2400694 | 1176.459625 | 753.8510585 | 0.434946478 | 0.252794359 | 0.054120229 | 0.084087489 | ns | YPD | KS.1 |
| FIM1_1396 | 874.5001903 | 1786.528104 | 1338.63058 | 0.976530042 | 0.17659752 | 6.4324099623 | 3.0361707364 | ns | YPD | KS.1 |
| FIM1_1397 | 2711.993871 | 3882.900024 | 4469.632152 | 0.488140459 | 0.143885927 | 0.000399866 | 0.000949561 | ns | YPD | KS.1 |
| FIM1_1399 | 5796.812783 | 2462.868637 | 2805.973654 | -1.153193387 | 0.216490525 | 1.4165635298 | 6.3835177184 | Km<KS.1 | YPD | KS.1 |
| FIM1_1400 | 2282.536629 | 3534.900547 | 1902.072314 | 0.556646419 | 0.225694688 | 0.006871677 | 0.012929877 | ns | YPD | KS.1 |
| FIM1_1401 | 8623.119694 | 5396.174526 | 3542.249151 | -0.564552376 | 0.277492881 | 0.020750127 | 0.035172703 | ns | YPD | KS.1 |
| FIM1_1402 | 1139.14963 | 1144.134824 | 872.8792913 | 0.006486478 | 0.09370556 | 0.943596615 | 0.959606125 | ns | YPD | KS.1 |
| FIM1_1403 | 1468.340329 | 717.3107775 | 747.890367 | -0.99117899 | 0.156237869 | 4.3472994310 | 2.5625372831 | ns | YPD | KS.1 |
| FIM1_1404 | 297.4450689 | 332.3171588 | 894.6827909 | 0.12064189 | 0.297544062 | 0.639223 | 0.705486336 | ns | YPD | KS.1 |
| FIM1_1405 | 341.2084137 | 643.1197755 | 849.0081097 | 0.832516108 | 0.220704468 | 4.4463070703 | 0.000123716 | ns | YPD | KS.1 |
| FIM1_1406 | 9769.97682 | 6962.713459 | 6026.309982 | -0.418827934 | 0.235527443 | 0.04919839 | 0.076943381 | ns | YPD | KS.1 |
| FIM1_1407 | 1546.40232 | 1731.920645 | 1369.341169 | 0.15549177 | 0.109799612 | 0.144621791 | 0.199416134 | ns | YPD | KS.1 |
| FIM1_1408 | 21781.62784 | 6241.622622 | 7942.031075 | -1.771931714 | 0.127885099 | 4.6159563666 | 3.1894976283 | Km<KS.1 | YPD | KS.1 |

| | | | | | | | | | | |
|---|---|---|---|---|---|---|---|---|---|---|
| FIM1_1409 | 2711.295993 | 1295.216413 | 2107.260258 | -0.865190268 | 0.348866022 | 0.003360091 | 0.00677328 | ns | YPD | KS.1 |
| FIM1_1410 | 2361.874756 | 3541.416619 | 3987.427979 | 0.547462192 | 0.15706426 | 0.000251654 | 0.000621639 | ns | YPD | KS.1 |
| FIM1_1411 | 2126.098431 | 1259.428123 | 1205.529616 | -0.616969549 | 0.303745908 | 0.018744623 | 0.032279162 | ns | YPD | KS.1 |
| FIM1_1412 | 9823.182854 | 7384.404686 | 4924.602873 | -0.359895772 | 0.211558784 | 0.06397 | 0.097611467 | ns | YPD | KS.1 |
| FIM1_1413 | 894.5950678 | 1204.342792 | 623.2557519 | 0.340772364 | 0.28415223 | 0.169385221 | 0.228977752 | ns | YPD | KS.1 |
| FIM1_1414 | 399.8901652 | 410.1761602 | 305.0673911 | 0.03284348 | 0.178880018 | 0.845001491 | 0.87947331 | ns | YPD | KS.1 |
| FIM1_1415 | 1475.616505 | 1832.39616 | 1607.211688 | 0.303972031 | 0.087328796 | 0.000390527 | 0.000928715 | ns | YPD | KS.1 |
| FIM1_1416 | 927.6927443 | 862.5141319 | 1016.078387 | -0.100629409 | 0.102650965 | 0.314261315 | 0.387760428 | ns | YPD | KS.1 |
| FIM1_1418 | 384.9994397 | 939.1457553 | 1019.213384 | 1.266544566 | 0.096824193 | 4.8540422185 | 2.4895731997 | Km>KS.1 | YPD | KS.1 |
| FIM1_1420 | 1517.7567 | 1358.970752 | 1437.044234 | -0.150762361 | 0.114136381 | 0.172821174 | 0.23306732 | ns | YPD | KS.1 |
| FIM1_1421 | 830.6181003 | 756.1436511 | 553.3328384 | -0.112656464 | 0.229869362 | 0.589822343 | 0.661812547 | ns | YPD | KS.1 |
| FIM1_1422 | 599.7383786 | 492.9650528 | 486.6103039 | -0.250113299 | 0.191642537 | 0.160501545 | 0.218168084 | ns | YPD | KS.1 |
| FIM1_1423 | 561.753896 | 1157.055526 | 793.3963286 | 1.029702229 | 0.085164924 | 2.0840448711 | 7.5679731635 | Km>KS.1 | YPD | KS.1 |
| FIM1_1424 | 2829.955398 | 1502.0617 | 1449.15332 | -0.901942144 | 0.080047944 | 4.2504788730 | 1.1813481784 | ns | YPD | KS.1 |
| FIM1_1425 | 3346.716234 | 1576.460968 | 1331.415899 | -1.039593658 | 0.163470791 | 3.5820898483 | 2.1215353565 | Km<KS.1 | YPD | KS.1 |
| FIM1_1426 | 2727.736234 | 6542.187659 | 3731.823932 | 1.035672247 | 0.364962727 | 0.000790116 | 0.001781881 | Km>KS.1 | YPD | KS.1 |
| FIM1_1427 | 37.39896422 | 76.97271293 | 38.6833852 | 0.894833526 | 0.292053066 | 0.000524601 | 0.001224726 | ns | YPD | KS.1 |
| FIM1_1428 | 1972.836494 | 2646.29214 | 2119.551811 | 0.392995664 | 0.157530914 | 0.008616315 | 0.01592354 | ns | YPD | KS.1 |
| FIM1_1429 | 2778.379977 | 3228.336794 | 2233.881656 | 0.195927815 | 0.165573149 | 0.210212619 | 0.276157322 | ns | YPD | KS.1 |
| FIM1_1430 | 1308.97292 | 1807.370667 | 1121.095545 | 0.42481189 | 0.176811424 | 0.010537852 | 0.019133509 | ns | YPD | KS.1 |
| FIM1_1431 | 5854.282121 | 3041.925174 | 2212.056116 | -0.884162817 | 0.189284028 | 7.3019189014 | 2.6362152782 | ns | YPD | KS.1 |
| FIM1_1432 | 3481.428666 | 10338.75058 | 13282.27271 | 1.539131102 | 0.128734368 | 4.0996320659 | 1.4465013849 | Km>KS.1 | YPD | KS.1 |
| FIM1_1433 | 2659.768504 | 1715.748684 | 1941.906229 | -0.616562541 | 0.098061818 | 1.3920838064 | 7.7991181725 | ns | YPD | KS.1 |
| FIM1_1434 | 3205.39242 | 2956.436626 | 1918.794881 | -0.095376057 | 0.240704334 | 0.660648431 | 0.724427142 | ns | YPD | KS.1 |
| FIM1_1435 | 9186.610468 | 6249.95298 | 5628.031343 | -0.514151198 | 0.170183062 | 0.001380999 | 0.002976806 | ns | YPD | KS.1 |
| FIM1_1436 | 1159.80358 | 468.4672376 | 560.7691604 | -1.262896678 | 0.156754832 | 9.1913742904 | 9.1271630927 | Km<KS.1 | YPD | KS.1 |
| FIM1_1437 | 182.2346593 | 321.0676007 | 339.2170924 | 0.794298844 | 0.105378663 | 1.3703034228 | 1.0925095398 | ns | YPD | KS.1 |
| FIM1_1438 | 45343.64485 | 55519.2895 | 60172.18324 | 0.283004395 | 0.092531015 | 0.001784415 | 0.003790549 | ns | YPD | KS.1 |
| FIM1_1439 | 3224.808339 | 2112.539123 | 1220.359772 | -0.58271916 | 0.133047663 | 5.6258376541 | 1.7703062826 | ns | YPD | KS.1 |
| FIM1_1440 | 724.3851032 | 311.664688 | 401.8067455 | -1.190745322 | 0.117647426 | 5.9188248333 | 1.1457647294 | Km<KS.1 | YPD | KS.1 |
| FIM1_1441 | 1473.418407 | 399.8183899 | 322.5394479 | -1.704741691 | 0.312652303 | 2.2200919061 | 1.1011921468 | Km<KS.1 | YPD | KS.1 |
| FIM1_1442 | 4353.042475 | 470.3928111 | 578.1003268 | -3.049136623 | 0.324100027 | 6.3085872554 | 1.1249183367 | Km<KS.1 | YPD | KS.1 |
| FIM1_1443 | 496.847823 | 453.470997 | 494.6692785 | -0.121163523 | 0.148478743 | 0.392224131 | 0.469404631 | ns | YPD | KS.1 |
| FIM1_1444 | 5334.504085 | 6613.78194 | 12306.94573 | 0.288417841 | 0.144949641 | 0.037195108 | 0.059486573 | ns | YPD | KS.1 |
| FIM1_1445 | 759.1912397 | 591.5716432 | 1243.352178 | -0.286911196 | 0.274648929 | 0.234691043 | 0.302483922 | ns | YPD | KS.1 |
| FIM1_1446 | 502.8101728 | 563.9829512 | 640.2136342 | 0.149331093 | 0.155612479 | 0.312391829 | 0.385740717 | ns | YPD | KS.1 |
| FIM1_1447 | 253.3174037 | 485.0223442 | 788.7167144 | 0.845907792 | 0.233148228 | 7.6214476714 | 0.000202438 | ns | YPD | KS.1 |
| FIM1_1448 | 203.1192812 | 392.8845089 | 525.5591259 | 0.877569716 | 0.207010923 | 5.5639501320 | 1.7530495191 | ns | YPD | KS.1 |
| FIM1_1449 | 1346.560601 | 1547.458318 | 2542.263048 | 0.189425291 | 0.116880831 | 0.093965642 | 0.136689787 | ns | YPD | KS.1 |
| FIM1_1450 | 1372.909195 | 1389.782904 | 920.6746159 | 0.016365338 | 0.139589695 | 0.902700172 | 0.926057771 | ns | YPD | KS.1 |
| FIM1_1451 | 949.3927395 | 2947.160846 | 8214.938459 | 1.509883592 | 0.263026749 | 6.6052536950 | 3.4736931430 | Km>KS.1 | YPD | KS.1 |
| FIM1_1453 | 5456.636641 | 4354.90309 | 3848.212792 | -0.263637413 | 0.261009592 | 0.255357202 | 0.324413197 | ns | YPD | KS.1 |
| FIM1_1454 | 2597.346055 | 1230.329771 | 1125.145479 | -1.042604809 | 0.141363337 | 2.8783305805 | 2.2235550680 | Km<KS.1 | YPD | KS.1 |
| FIM1_1455 | 906.9825503 | 583.6127408 | 1192.667858 | -0.572741768 | 0.207408328 | 0.002802119 | 0.005725068 | ns | YPD | KS.1 |
| FIM1_1456 | 721.3658094 | 532.5604867 | 560.0456439 | -0.407046062 | 0.156636306 | 0.006249916 | 0.011872215 | ns | YPD | KS.1 |

| | | | | | | | | | | |
|---|---|---|---|---|---|---|---|---|---|---|
| FIM1_1457 | 3848.014124 | 3777.455625 | 2947.262511 | -0.02418705 | 0.151539833 | 0.867027481 | 0.897702751 | ns | YPD | KS.1 |
| FIM1_1458 | 1098.224712 | 1184.951096 | 1399.980782 | 0.101629018 | 0.130908631 | 0.419759989 | 0.497571109 | ns | YPD | KS.1 |
| FIM1_1460 | 452.78464 | 1037.595707 | 1103.395768 | 1.177652096 | 0.092570968 | 6.04936215611 | 2.66332537404 | Km>KS.1 | YPD | KS.1 |
| FIM1_1461 | 22.50383462 | 332.5079706 | 986.9451075 | 3.716313042 | 0.328446403 | 1.55632394117 | 5.09388921535 | Km>KS.1 | YPD | KS.1 |
| FIM1_1462 | 554.3580579 | 1209.203293 | 1477.699355 | 1.103176912 | 0.111353846 | 6.05657108141 | 1.09171888152 | Km>KS.1 | YPD | KS.1 |
| FIM1_1463 | 183.6613401 | 202.2934062 | 239.1789131 | 0.121197795 | 0.180102729 | 0.472949491 | 0.550906981 | ns | YPD | KS.1 |
| FIM1_1464 | 54.05778116 | 32.90270711 | 24.69833755 | -0.458550593 | 0.441152151 | 0.174373948 | 0.234716014 | ns | YPD | KS.1 |
| FIM1_1465 | 2818.390501 | 1489.299716 | 1340.776731 | -0.898427662 | 0.111185201 | 1.48538304506 | 1.44897659787 | ns | YPD | KS.1 |
| FIM1_1466 | 1136.968664 | 1060.253206 | 1195.574456 | -0.095005005 | 0.110537151 | 0.376091288 | 0.451836309 | ns | YPD | KS.1 |
| FIM1_1467 | 2662.14915 | 3338.780189 | 2349.248935 | 0.302243235 | 0.151458861 | 0.036020405 | 0.057788299 | ns | YPD | KS.1 |
| FIM1_1468 | 2976.993895 | 3212.167366 | 3250.144257 | 0.104507125 | 0.100164817 | 0.28461389 | 0.356214868 | ns | YPD | KS.1 |
| FIM1_1469 | 169.8488597 | 89.07415526 | 198.5580107 | -0.855895691 | 0.210217376 | 1.21610477401 | 3.65209062938 | ns | YPD | KS.1 |
| FIM1_1470 | 1694.627751 | 2466.067832 | 1860.483098 | 0.531951316 | 0.077906004 | 4.47300780188 | 2.87881161893 | ns | YPD | KS.1 |
| FIM1_1471 | 2454.918515 | 2221.174352 | 4424.68561 | -0.135711353 | 0.123791048 | 0.255907319 | 0.325029082 | ns | YPD | KS.1 |
| FIM1_1472 | 612.2704495 | 426.005138 | 1066.927055 | -0.488236395 | 0.159655954 | 0.001285511 | 0.002786717 | ns | YPD | KS.1 |
| FIM1_1473 | 2747.970903 | 1066.394367 | 1273.656244 | -1.333400641 | 0.1330019 | 1.20818863867 | 2.28545189255 | Km<KS.1 | YPD | KS.1 |
| FIM1_1474 | 1336.03936 | 2763.558682 | 2614.068586 | 1.004065325 | 0.15941236 | 5.69323566703 | 3.29347063296 | Km>KS.1 | YPD | KS.1 |
| FIM1_1475 | 1777.161892 | 1570.098147 | 1427.26863 | -0.171431619 | 0.09714671 | 0.070558159 | 0.105954374 | ns | YPD | KS.1 |
| FIM1_1476 | 8671.762567 | 391.3180762 | 1083.108515 | -4.377887565 | 0.258149045 | 7.45903704607 | 2.47391395361 | Km<KS.1 | YPD | KS.1 |
| FIM1_1477 | 2635.802264 | 2472.412327 | 1843.773248 | -0.074821201 | 0.246514606 | 0.735666849 | 0.790143043 | ns | YPD | KS.1 |
| FIM1_1478 | 1469.987527 | 2425.28037 | 1710.692423 | 0.62543762 | 0.252444957 | 0.005769872 | 0.011044677 | ns | YPD | KS.1 |
| FIM1_1479 | 632.5299396 | 693.716909 | 809.3494458 | 0.128883477 | 0.093138821 | 0.156912732 | 0.214105235 | ns | YPD | KS.1 |
| FIM1_1480 | 381.3532913 | 480.2560847 | 905.3788425 | 0.284531025 | 0.22205878 | 0.160066358 | 0.217755027 | ns | YPD | KS.1 |
| FIM1_1481 | 410.1289127 | 255.383519 | 507.7294437 | -0.630875098 | 0.18528972 | 0.000284245 | 0.000695583 | ns | YPD | KS.1 |
| FIM1_1482 | 11004.01083 | 9005.39149 | 6288.323436 | -0.270054497 | 0.139044975 | 0.042571476 | 0.067257255 | ns | YPD | KS.1 |
| FIM1_1483 | 222.8010055 | 396.4160975 | 398.8084871 | 0.753107266 | 0.217179459 | 0.000172221 | 0.000435586 | ns | YPD | KS.1 |
| FIM1_1484 | 475.0733888 | 833.7293311 | 742.8195055 | 0.77796793 | 0.138808608 | 6.31916509337 | 2.98838843531 | ns | YPD | KS.1 |
| FIM1_1485 | 1815.126729 | 2736.351041 | 3134.913706 | 0.581968785 | 0.081258943 | 3.68557250487 | 2.60821098317 | ns | YPD | KS.1 |
| FIM1_1486 | 939.2114158 | 1108.06897 | 2021.714419 | 0.208111806 | 0.196742024 | 0.254055686 | 0.323007166 | ns | YPD | KS.1 |
| FIM1_1487 | 1189.505075 | 791.0203591 | 1944.335533 | -0.484821544 | 0.27679418 | 0.046200305 | 0.072483922 | ns | YPD | KS.1 |
| FIM1_1488 | 3370.922443 | 5608.627759 | 5118.618575 | 0.724807131 | 0.076430332 | 8.06329901688 | 1.19746007788 | ns | YPD | KS.1 |
| FIM1_1489 | 120.6116122 | 335.6791845 | 496.6185416 | 1.425609568 | 0.163359956 | 2.25393977242 | 2.79634672513 | Km>KS.1 | YPD | KS.1 |
| FIM1_1490 | 209.8525239 | 479.622507 | 542.9241269 | 1.141012637 | 0.166859776 | 1.17370595298 | 8.06517557472 | Km>KS.1 | YPD | KS.1 |
| FIM1_1491 | 858.1843167 | 974.002563 | 1008.296104 | 0.177794747 | 0.085267459 | 0.03344652 | 0.054130266 | ns | YPD | KS.1 |
| FIM1_1492 | 1234.413319 | 1251.442699 | 1549.742596 | 0.018393478 | 0.122673614 | 0.876730189 | 0.905675392 | ns | YPD | KS.1 |
| FIM1_1493 | 664.5159584 | 662.3934597 | 639.7551531 | -0.004153542 | 0.153440189 | 0.977324581 | 0.984049745 | ns | YPD | KS.1 |
| FIM1_1494 | 212.1480236 | 159.4121936 | 226.628268 | -0.377379143 | 0.16552449 | 0.015848276 | 0.027752613 | ns | YPD | KS.1 |
| FIM1_1495 | 102.4108282 | 82.42058248 | 72.26210851 | -0.25174621 | 0.265329036 | 0.283858256 | 0.355537469 | ns | YPD | KS.1 |
| FIM1_1496 | 1158.64818 | 982.636974 | 888.9900235 | -0.225732933 | 0.110265964 | 0.034948174 | 0.056304134 | ns | YPD | KS.1 |
| FIM1_1497 | 468.673658 | 720.3091092 | 541.8235652 | 0.600030061 | 0.115582078 | 9.45049079756 | 3.84119213381 | ns | YPD | KS.1 |
| FIM1_1498 | 366.7291653 | 455.0543389 | 589.1756182 | 0.288547484 | 0.151383095 | 0.045192293 | 0.071081776 | ns | YPD | KS.1 |
| FIM1_1499 | 656.190453 | 2087.691165 | 1990.863723 | 1.589876787 | 0.20791884 | 1.26143438584 | 1.12466596229 | Km>KS.1 | YPD | KS.1 |
| FIM1_1500 | 1875.162519 | 1798.108378 | 1542.250736 | -0.056062194 | 0.132960118 | 0.661100145 | 0.72476272 | ns | YPD | KS.1 |
| FIM1_1501 | 635.7304907 | 1106.003316 | 1400.018085 | 0.766105225 | 0.142753661 | 2.49990083425 | 1.09867549915 | ns | YPD | KS.1 |
| FIM1_1503 | 941.5935748 | 2410.901237 | 2153.953742 | 1.339024919 | 0.094901318 | 3.29834429908 | 2.52450198276 | Km>KS.1 | YPD | KS.1 |

| | | | | | | | | | | |
|---|---|---|---|---|---|---|---|---|---|---|
| FIM1_1504 | 888.9758156 | 1048.932459 | 912.8558115 | 0.231313124 | 0.086842195 | 0.006591411 | 0.012463804 | ns | YPD | KS.1 |
| FIM1_1505 | 741.1551065 | 638.796214 | 530.282422 | -0.199833324 | 0.12885541 | 0.107050685 | 0.153335036 | ns | YPD | KS.1 |
| FIM1_1506 | 218.5050855 | 148.0755924 | 147.5683477 | -0.478895513 | 0.248745243 | 0.031756661 | 0.051529481 | ns | YPD | KS.1 |
| FIM1_1507 | 2853.926935 | 3856.32395 | 4989.294237 | 0.410186208 | 0.135554656 | 0.001648462 | 0.003519103 | ns | YPD | KS.1 |
| FIM1_1508 | 1446.004769 | 1846.235557 | 1713.87884 | 0.33518339 | 0.119863642 | 0.0038656 | 0.007689468 | ns | YPD | KS.1 |
| FIM1_1509 | 695.5539214 | 1134.462683 | 1673.109911 | 0.673090384 | 0.142082915 | 8.36501646715 | 2.99610920979 | ns | YPD | KS.1 |
| FIM1_1510 | 605.3547351 | 842.9893131 | 1336.410245 | 0.456873195 | 0.122955394 | 0.000124239 | 0.00031926 | ns | YPD | KS.1 |
| FIM1_1511 | 208.6548538 | 182.9299123 | 223.070927 | -0.171857621 | 0.172258785 | 0.289130707 | 0.36086936 | ns | YPD | KS.1 |
| FIM1_1512 | 400.3280495 | 528.951126 | 639.4952849 | 0.377614536 | 0.141223972 | 0.005257153 | 0.010145205 | ns | YPD | KS.1 |
| FIM1_1513 | 3419.207272 | 3917.098439 | 4753.957716 | 0.189739643 | 0.086617763 | 0.025405673 | 0.042271982 | ns | YPD | KS.1 |
| FIM1_1514 | 12244.35023 | 9486.275861 | 11778.31794 | -0.34990003 | 0.124175049 | 0.003539766 | 0.007103752 | ns | YPD | KS.1 |
| FIM1_1515 | 992.7027756 | 350.690417 | 365.2917633 | -1.452565645 | 0.162058653 | 2.57311646145 | 3.45047288294 | Km<KS.1 | YPD | KS.1 |
| FIM1_1516 | 874.6509742 | 109.6042496 | 209.8896237 | -2.689406052 | 0.436805652 | 2.25337468942 | 1.50074151002 | Km<KS.1 | YPD | KS.1 |
| FIM1_1518 | 859.2472512 | 965.4838585 | 1559.237096 | 0.156000712 | 0.138276093 | 0.239034064 | 0.307185907 | ns | YPD | KS.1 |
| FIM1_1519 | 635.2051885 | 414.1546911 | 582.3236775 | -0.586690566 | 0.140774095 | 1.44726069982 | 4.29089510229 | ns | YPD | KS.1 |
| FIM1_1520 | 3424.950041 | 3983.556015 | 4137.183329 | 0.200040835 | 0.150409504 | 0.162421976 | 0.220597688 | ns | YPD | KS.1 |
| FIM1_1521 | 3128.866424 | 2406.607496 | 4002.233212 | -0.281998303 | 0.319948632 | 0.299032602 | 0.37117944 | ns | YPD | KS.1 |
| FIM1_1523 | 355.4681934 | 397.8919837 | 369.144513 | 0.150206236 | 0.128228265 | 0.223754443 | 0.29064709 | ns | YPD | KS.1 |
| FIM1_1524 | 497.5748377 | 740.363509 | 1103.512142 | 0.544284909 | 0.137637858 | 3.95221258668 | 0.000110711 | ns | YPD | KS.1 |
| FIM1_1525 | 1376.08681 | 1654.310364 | 1670.902005 | 0.258683806 | 0.078028267 | 0.000755776 | 0.001712197 | ns | YPD | KS.1 |
| FIM1_1526 | 6477.109272 | 3579.911074 | 7536.909485 | -0.827905202 | 0.127000728 | 1.90717241355 | 1.15568608495 | ns | YPD | KS.1 |
| FIM1_1527 | 611.8278558 | 506.4861095 | 531.9941973 | -0.261865303 | 0.102649633 | 0.008846273 | 0.01631216 | ns | YPD | KS.1 |
| FIM1_1528 | 640.1627943 | 614.9800307 | 668.6270175 | -0.04283427 | 0.303035314 | 0.869663194 | 0.899869882 | ns | YPD | KS.1 |
| FIM1_1529 | 4264.048731 | 1451.77252 | 1461.92079 | -1.519953671 | 0.135489326 | 2.34854149714 | 6.60112652446 | Km<KS.1 | YPD | KS.1 |
| FIM1_1530 | 5467.916401 | 3740.314422 | 2471.08405 | -0.516431441 | 0.146785828 | 0.000237192 | 0.00058972 | ns | YPD | KS.1 |
| FIM1_1531 | 353.256182 | 964.9428233 | 741.0917151 | 1.372189596 | 0.206830122 | 3.07294870508 | 2.00892507329 | Km>KS.1 | YPD | KS.1 |
| FIM1_1532 | 139.4667838 | 708.7488111 | 768.8674388 | 2.219973323 | 0.259877252 | 1.53622854672 | 1.95466419947 | Km>KS.1 | YPD | KS.1 |
| FIM1_1533 | 305.7110844 | 594.2313169 | 731.9239523 | 0.930103518 | 0.122780751 | 7.77945797947 | 6.32398749148 | ns | YPD | KS.1 |
| FIM1_1534 | 298.1034786 | 408.4843089 | 392.2872536 | 0.43636413 | 0.114923577 | 9.32750899249 | 0.000245255 | ns | YPD | KS.1 |
| FIM1_1535 | 507.185114 | 676.8728971 | 781.0826287 | 0.406902103 | 0.082406474 | 5.30655128946 | 1.94690948857 | ns | YPD | KS.1 |
| FIM1_1536 | 934.5110503 | 753.6977818 | 720.7602547 | -0.293651172 | 0.122203593 | 0.012878552 | 0.023013936 | ns | YPD | KS.1 |
| FIM1_1537 | 1079.158027 | 2110.167921 | 1969.124807 | 0.944108746 | 0.114140033 | 2.78412362879 | 2.90987711203 | ns | YPD | KS.1 |
| FIM1_1538 | 2061.680399 | 2690.593189 | 1902.472634 | 0.366147397 | 0.121210983 | 0.001801726 | 0.003824057 | ns | YPD | KS.1 |
| FIM1_1540 | 905.0723436 | 892.6778727 | 957.5712073 | -0.017548147 | 0.150647937 | 0.902789988 | 0.926057771 | ns | YPD | KS.1 |
| FIM1_1541 | 5250.089701 | 3597.753393 | 2516.562676 | -0.49030981 | 0.199145095 | 0.007930766 | 0.014755259 | ns | YPD | KS.1 |
| FIM1_1542 | 3617.500647 | 2184.86508 | 4261.464166 | -0.611465717 | 0.27822767 | 0.012567383 | 0.022506383 | ns | YPD | KS.1 |
| FIM1_1543 | 2252.944454 | 1610.169753 | 3473.59313 | -0.369540599 | 0.316402869 | 0.170065143 | 0.229786552 | ns | YPD | KS.1 |
| FIM1_1544 | 21.62236047 | 58.19420727 | 191.583174 | 1.316442775 | 0.235194074 | 2.25125616200 | 1.11553778944 | Km>KS.1 | YPD | KS.1 |
| FIM1_1545 | 303.03609 | 918.1986947 | 1128.202135 | 1.558751764 | 0.146096941 | 9.27264474660 | 2.19673369592 | Km>KS.1 | YPD | KS.1 |
| FIM1_1546 | 169.4432996 | 171.7626837 | 298.4181561 | 0.016322759 | 0.152782937 | 0.910709392 | 0.933027024 | ns | YPD | KS.1 |
| FIM1_1547 | 858.5634429 | 643.8471452 | 968.3615341 | -0.390329127 | 0.142392395 | 0.004205074 | 0.008314883 | ns | YPD | KS.1 |
| FIM1_1548 | 803.0623538 | 917.4906204 | 1539.925275 | 0.167579478 | 0.19609034 | 0.357236993 | 0.432316721 | ns | YPD | KS.1 |
| FIM1_1549 | 781.6152702 | 1686.349073 | 1949.095125 | 1.074517705 | 0.137964849 | 1.12278474644 | 1.00465002042 | Km>KS.1 | YPD | KS.1 |
| FIM1_1550 | 720.195543 | 927.4711505 | 1478.150675 | 0.326509265 | 0.188998619 | 0.06362725 | 0.097218082 | ns | YPD | KS.1 |
| FIM1_1551 | 3574.141375 | 9944.528745 | 21078.24352 | 1.361433557 | 0.254246677 | 8.04551620119 | 3.74779429784 | Km>KS.1 | YPD | KS.1 |

| | | | | | | | | | | |
|---|---|---|---|---|---|---|---|---|---|---|
| FIM1_1552 | 334.4716223 | 216.3870526 | 398.9322626 | -0.563343529 | 0.210928654 | 0.003757343 | 0.007501117 | ns | YPD | KS.1 |
| FIM1_1553 | 1809.064221 | 1568.955451 | 3566.150704 | -0.152761437 | 0.307513773 | 0.562529539 | 0.635175773 | ns | YPD | KS.1 |
| FIM1_1554 | 880.7548747 | 975.9711036 | 1029.671585 | 0.136578392 | 0.142845273 | 0.317378141 | 0.390830755 | ns | YPD | KS.1 |
| FIM1_1556 | 134.8016369 | 267.2115936 | 412.6890585 | 0.915050479 | 0.204254285 | 1.7073090065( | 5.8018185159 | ns | YPD | KS.1 |
| FIM1_1557 | 124.5641014 | 183.499429 | 370.8202114 | 0.507806449 | 0.1824886 | 0.002993335 | 0.006090732 | ns | YPD | KS.1 |
| FIM1_1558 | 1793.321375 | 2734.625733 | 2132.708685 | 0.586700745 | 0.118104033 | 3.1665237473 | 1.2062370324( | ns | YPD | KS.1 |
| FIM1_1559 | 3292.33311 | 5793.022983 | 5562.228255 | 0.755349918 | 0.191629627 | 2.6276707856 | 7.5303353447 | ns | YPD | KS.1 |
| FIM1_1560 | 1127.979299 | 1325.039519 | 1399.809743 | 0.222346329 | 0.110375852 | 0.037948007 | 0.060529444 | ns | YPD | KS.1 |
| FIM1_1561 | 3195.493878 | 1942.094078 | 1903.390235 | -0.679880637 | 0.154262236 | 3.9970001028 | 1.2845656015 | ns | YPD | KS.1 |
| FIM1_1562 | 447.3691525 | 356.7386818 | 554.8368495 | -0.284325742 | 0.210552074 | 0.141690591 | 0.195591201 | ns | YPD | KS.1 |
| FIM1_1563 | 630.4697774 | 1433.727225 | 1104.718313 | 1.165959416 | 0.099192295 | 9.2588348087 | 3.0708468782 | Km>KS.1 | YPD | KS.1 |
| FIM1_1564 | 2316.6838 | 311.939237 | 328.3333144 | -2.725352037 | 0.318393673 | 2.8542272875 | 3.8171453643( | Km<KS.1 | YPD | KS.1 |
| FIM1_1566 | 5843.352258 | 3407.042507 | 2607.517748 | -0.720226851 | 0.189811109 | 5.2027934759 | 0.000143242 | ns | YPD | KS.1 |
| FIM1_1567 | 2230.845824 | 2254.748416 | 1116.362786 | 0.012776249 | 0.223294212 | 0.950071304 | 0.964416392 | ns | YPD | KS.1 |
| FIM1_1568 | 5325.004627 | 7477.416599 | 7098.304888 | 0.478826428 | 0.086438359 | 1.7780194417 | 7.9049568567 | ns | YPD | KS.1 |
| FIM1_1569 | 392.1018763 | 272.8936004 | 191.7269476 | -0.449771052 | 0.237921753 | 0.036294423 | 0.05820914 | ns | YPD | KS.1 |
| FIM1_1570 | 1411.573206 | 1266.059628 | 1038.993896 | -0.147726215 | 0.114687849 | 0.183647063 | 0.245867636 | ns | YPD | KS.1 |
| FIM1_1571 | 239.885189 | 291.1758269 | 365.0476731 | 0.258383966 | 0.150552887 | 0.071401536 | 0.107188486 | ns | YPD | KS.1 |
| FIM1_1572 | 1102.755019 | 505.7716884 | 549.9838849 | -1.08952123 | 0.139832308 | 1.0642681941 | 9.5400617401 | Km<KS.1 | YPD | KS.1 |
| FIM1_1573 | 941.9410712 | 994.6152067 | 1173.957086 | 0.076170046 | 0.083763542 | 0.354207 | 0.42928255 | ns | YPD | KS.1 |
| FIM1_1574 | 3181.366602 | 3511.7532 | 3203.394133 | 0.139130822 | 0.070539375 | 0.045379247 | 0.071308198 | ns | YPD | KS.1 |
| FIM1_1575 | 780.979278 | 1068.445777 | 851.4544938 | 0.434649864 | 0.111637919 | 6.3161935423( | 0.000171058 | ns | YPD | KS.1 |
| FIM1_1576 | 41.59817927 | 34.06165671 | 38.68437699 | -0.250963548 | 0.229872159 | 0.229005107 | 0.296229955 | ns | YPD | KS.1 |
| FIM1_1577 | 690.2057039 | 549.3040594 | 674.6265987 | -0.306514128 | 0.14403519 | 0.025955877 | 0.043086582 | ns | YPD | KS.1 |
| FIM1_1578 | 6352.226283 | 8810.643279 | 9826.523273 | 0.459802049 | 0.092607163 | 4.1867563355( | 1.5660987044 | ns | YPD | KS.1 |
| FIM1_1579 | 906.6891078 | 958.6861295 | 1877.861067 | 0.062952602 | 0.272306304 | 0.793311077 | 0.837589688 | ns | YPD | KS.1 |
| FIM1_1580 | 2298.482978 | 1025.898584 | 1777.236402 | -1.089970497 | 0.206321527 | 2.0383934776 | 9.0302827706 | Km<KS.1 | YPD | KS.1 |
| FIM1_1581 | 6025.791743 | 3970.818872 | 4862.540858 | -0.589935286 | 0.085146465 | 1.9388534030 | 1.2982228372( | ns | YPD | KS.1 |
| FIM1_1582 | 299.4889793 | 72.20545629 | 111.7929124 | -1.894644437 | 0.292052624 | 2.5421416027 | 1.6773414421( | Km<KS.1 | YPD | KS.1 |
| FIM1_1583 | 1540.010829 | 1058.914718 | 887.9140318 | -0.517458635 | 0.122539961 | 1.3135009996 | 3.9223694318( | ns | YPD | KS.1 |
| FIM1_1584 | 3248.821537 | 6245.201282 | 5326.866002 | 0.911452915 | 0.135052819 | 3.3758748953 | 2.1954219090 | ns | YPD | KS.1 |
| FIM1_1585 | 2054.35841 | 720.9696507 | 919.5679531 | -1.473419314 | 0.142414566 | 3.4078503264 | 7.3394179107 | Km<KS.1 | YPD | KS.1 |
| FIM1_1586 | 9348.835946 | 1829.106973 | 2013.747848 | -2.238424482 | 0.252024165 | 7.2313904895 | 1.0162759233 | Km<KS.1 | YPD | KS.1 |
| FIM1_1587 | 7700.605821 | 7625.743181 | 7941.355191 | -0.013244916 | 0.110871241 | 0.902146585 | 0.925779551 | ns | YPD | KS.1 |
| FIM1_1588 | 4682.293794 | 6623.121634 | 6573.148533 | 0.45583212 | 0.18196255 | 0.007498608 | 0.014019382 | ns | YPD | KS.1 |
| FIM1_1589 | 67.34439001 | 118.0468307 | 174.21158 | 0.778267994 | 0.147805185 | 4.2476636307 | 1.8077097145( | ns | YPD | KS.1 |
| FIM1_1590 | 332.5752815 | 463.0373491 | 907.6081619 | 0.422203762 | 0.210596675 | 0.02922618 | 0.047860515 | ns | YPD | KS.1 |
| FIM1_1591 | 79.04288588 | 107.301179 | 193.798865 | 0.369970744 | 0.244288107 | 0.091815857 | 0.133993513 | ns | YPD | KS.1 |
| FIM1_1592 | 110.7608419 | 140.6844556 | 207.5354789 | 0.307759795 | 0.183806797 | 0.073111886 | 0.109557721 | ns | YPD | KS.1 |
| FIM1_1593 | 7007.067013 | 7387.03182 | 3711.678997 | 0.056367819 | 0.305341189 | 0.829993951 | 0.868943583 | ns | YPD | KS.1 |
| FIM1_1594 | 2746.867759 | 1914.379375 | 2701.158854 | -0.432697123 | 0.262904507 | 0.063497432 | 0.097050606 | ns | YPD | KS.1 |
| FIM1_1595 | 443.2184268 | 957.787747 | 1468.553781 | 1.062183343 | 0.168645162 | 5.1007199106 | 2.9819132262 | Km>KS.1 | YPD | KS.1 |
| FIM1_1596 | 1984.940415 | 2472.448252 | 1618.955995 | 0.275411244 | 0.207778183 | 0.149911142 | 0.205569992 | ns | YPD | KS.1 |
| FIM1_1597 | 1065.981764 | 932.6195337 | 1675.69388 | -0.147212613 | 0.290206844 | 0.55973426 | 0.632737547 | ns | YPD | KS.1 |
| FIM1_1598 | 2295.708928 | 1618.614292 | 3069.22103 | -0.387749063 | 0.313336403 | 0.147063367 | 0.202278201 | ns | YPD | KS.1 |

| | | | | | | | | | | |
|---|---|---|---|---|---|---|---|---|---|---|
| FIM1_1599 | 2860.604701 | 2549.724106 | 2665.984911 | -0.160113539 | 0.088841034 | 0.065657223 | 0.099830282 | ns | YPD | KS.1 |
| FIM1_1601 | 10928.31289 | 15833.46208 | 22763.71051 | 0.508272605 | 0.135530974 | 9.8057161099 | 0.00025662 | ns | YPD | KS.1 |
| FIM1_1602 | 1586.913976 | 1325.613543 | 2256.884439 | -0.250622172 | 0.09610275 | 0.007603122 | 0.014193446 | ns | YPD | KS.1 |
| FIM1_1603 | 3101.867375 | 4018.539821 | 4657.555361 | 0.358332641 | 0.1123302 | 0.001029299 | 0.002275895 | ns | YPD | KS.1 |
| FIM1_1604 | 4706.895964 | 2769.218728 | 2197.326057 | -0.730954723 | 0.144274802 | 1.3704314114 | 5.4456040512 | ns | YPD | KS.1 |
| FIM1_1605 | 1931.180099 | 1468.385305 | 2906.898455 | -0.29855284 | 0.312929783 | 0.263640693 | 0.333658725 | ns | YPD | KS.1 |
| FIM1_1606 | 4892.328351 | 7546.060704 | 7623.839318 | 0.59571705 | 0.137533215 | 6.8212889753 | 2.1223209913 | ns | YPD | KS.1 |
| FIM1_1607 | 841.6718313 | 1329.165092 | 1263.574432 | 0.635505927 | 0.119053139 | 3.9228429905 | 1.6795304542 | ns | YPD | KS.1 |
| FIM1_1608 | 1256.158669 | 2516.448007 | 2705.623237 | 0.965878407 | 0.145740615 | 6.9616271363 | 4.4176141586 | ns | YPD | KS.1 |
| FIM1_1609 | 567.0480858 | 1126.364432 | 1307.238998 | 0.939384607 | 0.173239953 | 1.2708237103 | 5.7447072873 | ns | YPD | KS.1 |
| FIM1_1610 | 1058.192283 | 1669.320841 | 2102.081972 | 0.585986518 | 0.218443203 | 0.003461149 | 0.006957258 | ns | YPD | KS.1 |
| FIM1_1611 | 1340.771524 | 1655.45261 | 1829.231039 | 0.276936521 | 0.16747755 | 0.079563001 | 0.117770286 | ns | YPD | KS.1 |
| FIM1_1612 | 902.0628025 | 1068.241376 | 1563.798557 | 0.209966656 | 0.210796607 | 0.278611167 | 0.349581476 | ns | YPD | KS.1 |
| FIM1_1613 | 1725.687368 | 1749.029766 | 1328.538532 | 0.018894481 | 0.095798339 | 0.840107451 | 0.876032882 | ns | YPD | KS.1 |
| FIM1_1614 | 548.7695052 | 394.9026401 | 538.7302173 | -0.448871783 | 0.135974093 | 0.000598696 | 0.001384711 | ns | YPD | KS.1 |
| FIM1_1615 | 3715.025881 | 2948.694161 | 3138.809537 | -0.32137541 | 0.102267605 | 0.001278116 | 0.002774271 | ns | YPD | KS.1 |
| FIM1_1616 | 1097.617454 | 859.6609689 | 848.8679576 | -0.332360625 | 0.131392259 | 0.008575841 | 0.015866421 | ns | YPD | KS.1 |
| FIM1_1617 | 3878.64044 | 2741.61733 | 3551.417207 | -0.486348937 | 0.097733813 | 3.7505097242 | 1.4135443847 | ns | YPD | KS.1 |
| FIM1_1618 | 391.1183128 | 144.9190918 | 330.5806568 | -1.328181659 | 0.241890296 | 4.0305643915 | 1.9430288612 | Km<KS.1 | YPD | KS.1 |
| FIM1_1619 | 1387.144218 | 809.9624672 | 1644.712945 | -0.751944161 | 0.119307556 | 9.3047021118 | 5.2783230337 | ns | YPD | KS.1 |
| FIM1_1621 | 3488.486372 | 3951.261507 | 3449.992768 | 0.172983601 | 0.095417098 | 0.0633889 | 0.09694429 | ns | YPD | KS.1 |
| FIM1_1622 | 2608.510701 | 690.4604855 | 889.5684964 | -1.872560941 | 0.153495723 | 9.6406863139 | 3.9638359017 | Km<KS.1 | YPD | KS.1 |
| FIM1_1623 | 461.8507663 | 398.1940643 | 369.2683476 | -0.194706579 | 0.163297592 | 0.207328666 | 0.2730175 | ns | YPD | KS.1 |
| FIM1_1624 | 315.9125135 | 270.4646039 | 674.7249446 | -0.204187198 | 0.161627998 | 0.181910533 | 0.2438052 | ns | YPD | KS.1 |
| FIM1_1625 | 2526.627052 | 1141.055319 | 1770.134835 | -1.108586712 | 0.146856839 | 6.8289063451 | 5.5877975439 | Km<KS.1 | YPD | KS.1 |
| FIM1_1627 | 409.9853483 | 1407.057345 | 2079.215773 | 1.725498754 | 0.1704784 | 2.0051453975 | 3.9902393411 | Km>KS.1 | YPD | KS.1 |
| FIM1_1628 | 1479.451423 | 363.8348729 | 650.5022275 | -1.703667061 | 0.423882077 | 2.3262631123 | 7.7360688395 | Km<KS.1 | YPD | KS.1 |
| FIM1_1629 | 242.2344792 | 305.1744473 | 370.7286386 | 0.313558607 | 0.136504096 | 0.01666663 | 0.029032382 | ns | YPD | KS.1 |
| FIM1_1630 | 5678.914998 | 3876.523237 | 2888.50703 | -0.531450096 | 0.113509154 | 1.4991764545 | 5.1508307054 | ns | YPD | KS.1 |
| FIM1_1631 | 857.6581902 | 723.2291901 | 737.8346959 | -0.217637078 | 0.187058918 | 0.212266982 | 0.278488459 | ns | YPD | KS.1 |
| FIM1_1632 | 3182.635155 | 2263.416985 | 1564.129185 | -0.384268414 | 0.301757479 | 0.139216375 | 0.19281778 | ns | YPD | KS.1 |
| FIM1_1633 | 294.4428865 | 234.1204127 | 1191.610824 | -0.251299913 | 0.30303657 | 0.335110761 | 0.40982695 | ns | YPD | KS.1 |
| FIM1_1634 | 735.5161032 | 2495.068736 | 2116.876201 | 1.718912713 | 0.149718586 | 7.4925551192 | 2.3592064378 | Km>KS.1 | YPD | KS.1 |
| FIM1_1635 | 6626.973925 | 6520.352976 | 6104.133034 | -0.022848374 | 0.077471852 | 0.764427029 | 0.815399758 | ns | YPD | KS.1 |
| FIM1_1636 | 442.8873246 | 731.2189392 | 829.9480188 | 0.6911338 | 0.139762777 | 2.8167747920 | 1.0804513947 | ns | YPD | KS.1 |
| FIM1_1637 | 216.473977 | 200.8677147 | 238.0021329 | -0.096113733 | 0.169458209 | 0.547775548 | 0.621053635 | ns | YPD | KS.1 |
| FIM1_1638 | 358.0112096 | 42.30168279 | 70.51401508 | -2.947582348 | 0.285297711 | 7.1752181858 | 1.7161880035 | Km<KS.1 | YPD | KS.1 |
| FIM1_1640 | 2306.871409 | 2497.143978 | 2570.685489 | 0.108223164 | 0.113716258 | 0.326311073 | 0.400407279 | ns | YPD | KS.1 |
| FIM1_1641 | 1729.313107 | 4169.049667 | 4617.052727 | 1.254809965 | 0.089306595 | 8.9025355303 | 6.5132520975 | Km>KS.1 | YPD | KS.1 |
| FIM1_1642 | 2297.940365 | 2399.889323 | 1800.68227 | 0.060001465 | 0.094627611 | 0.516639111 | 0.59250336 | ns | YPD | KS.1 |
| FIM1_1643 | 1997.607101 | 1696.874336 | 1577.718136 | -0.222900578 | 0.120185328 | 0.054942387 | 0.085258383 | ns | YPD | KS.1 |
| FIM1_1644 | 749.9172625 | 624.2344026 | 640.2083275 | -0.25230618 | 0.112189164 | 0.020442834 | 0.034758407 | ns | YPD | KS.1 |
| FIM1_1645 | 854.2806775 | 636.389102 | 638.4829664 | -0.402551571 | 0.132776691 | 0.001638093 | 0.003499147 | ns | YPD | KS.1 |
| FIM1_1646 | 8789.31019 | 4914.794031 | 3165.894573 | -0.791898491 | 0.167857155 | 7.0830947669 | 2.5683962438 | ns | YPD | KS.1 |
| FIM1_1647 | 43.77076368 | 69.63323903 | 78.19150405 | 0.56821678 | 0.25562956 | 0.01289653 | 0.023037788 | ns | YPD | KS.1 |

| | | | | | | | | | | |
|---|---|---|---|---|---|---|---|---|---|---|
| FIM1_1648 | 2222.560688 | 1895.499364 | 3748.977745 | -0.207731686 | 0.165503661 | 0.183814091 | 0.245958877 | ns | YPD | KS.1 |
| FIM1_1649 | 1353.845783 | 1750.128889 | 2915.235222 | 0.331482734 | 0.187793754 | 0.058268771 | 0.089915365 | ns | YPD | KS.1 |
| FIM1_1650 | 1420.952996 | 1104.512891 | 1231.102463 | -0.302927757 | 0.244504235 | 0.167796458 | 0.227276717 | ns | YPD | KS.1 |
| FIM1_1651 | 677.2032222 | 376.0997088 | 475.7090115 | -0.796411372 | 0.177054394 | 2.02127779905 | 6.78532864393 | ns | YPD | KS.1 |
| FIM1_1652 | 826.4046724 | 2497.963421 | 2996.097767 | 1.559016824 | 0.1400791 | 5.88039825391 | 1.59863285864 | Km>KS.1 | YPD | KS.1 |
| FIM1_1653 | 977.6343669 | 787.9952417 | 589.1797007 | -0.292540262 | 0.132099015 | 0.021253249 | 0.035878831 | ns | YPD | KS.1 |
| FIM1_1654 | 343.6251877 | 648.0599735 | 472.4391557 | 0.837697887 | 0.217858247 | 3.26954706825 | 9.24729770584 | ns | YPD | KS.1 |
| FIM1_1655 | 1674.313077 | 1440.639315 | 1447.568048 | -0.206691131 | 0.107394414 | 0.047650126 | 0.074640861 | ns | YPD | KS.1 |
| FIM1_1656 | 1959.403399 | 2006.674759 | 1353.127567 | 0.031978352 | 0.125633323 | 0.792013078 | 0.836643222 | ns | YPD | KS.1 |
| FIM1_1657 | 16872.37603 | 25564.26808 | 30073.46668 | 0.583270537 | 0.101510414 | 4.27544725865 | 2.05510629099 | ns | YPD | KS.1 |
| FIM1_1658 | 748.2066176 | 705.0095419 | 1053.529141 | -0.078380604 | 0.13825592 | 0.55446593 | 0.627494996 | ns | YPD | KS.1 |
| FIM1_1659 | 3664.743334 | 4008.089814 | 2925.916133 | 0.123166234 | 0.103848937 | 0.223078894 | 0.289973049 | ns | YPD | KS.1 |
| FIM1_1660 | 1470.956863 | 1928.795025 | 2284.236219 | 0.377289714 | 0.104148009 | 0.000205139 | 0.000515178 | ns | YPD | KS.1 |
| FIM1_1661 | 510.63252 | 1158.263526 | 1659.451143 | 1.143766327 | 0.144806457 | 4.10924551424 | 3.82121428660 | Km>KS.1 | YPD | KS.1 |
| FIM1_1664 | 2209.227653 | 1859.005852 | 2194.986819 | -0.243170632 | 0.081405426 | 0.002367226 | 0.004905018 | ns | YPD | KS.1 |
| FIM1_1665 | 425.152673 | 346.976199 | 323.128888 | -0.254976191 | 0.205212342 | 0.177875085 | 0.238911595 | ns | YPD | KS.1 |
| FIM1_1666 | 878.6031302 | 830.4262484 | 919.0415446 | -0.079398581 | 0.078556258 | 0.304012239 | 0.376515033 | ns | YPD | KS.1 |
| FIM1_1667 | 2439.202616 | 609.114021 | 1963.193442 | -1.844981455 | 0.293168572 | 1.02166893085 | 6.40151502643 | Km<KS.1 | YPD | KS.1 |
| FIM1_1669 | 2657.760816 | 1956.689693 | 2080.923473 | -0.421537606 | 0.12205009 | 0.000361056 | 0.00086483 | ns | YPD | KS.1 |
| FIM1_1670 | 960.6256841 | 1065.677224 | 1932.306517 | 0.129615363 | 0.19872312 | 0.481804347 | 0.560041268 | ns | YPD | KS.1 |
| FIM1_1671 | 3572.833141 | 2783.538236 | 5657.092212 | -0.287216004 | 0.274818078 | 0.234422773 | 0.302216453 | ns | YPD | KS.1 |
| FIM1_1672 | 4599.641598 | 4177.034778 | 4201.129198 | -0.127190693 | 0.150342228 | 0.374578766 | 0.450236618 | ns | YPD | KS.1 |
| FIM1_1673 | 3826.590893 | 4229.662872 | 3566.447103 | 0.139823142 | 0.084788925 | 0.092636761 | 0.134993523 | ns | YPD | KS.1 |
| FIM1_1674 | 2168.577404 | 1455.129964 | 1673.238799 | -0.550132193 | 0.129808975 | 1.14636697293 | 3.45493764954 | ns | YPD | KS.1 |
| FIM1_1675 | 17.92376044 | 11.29405136 | 18.27940201 | -0.511494236 | 0.350109756 | 0.078526678 | 0.116513636 | ns | YPD | KS.1 |
| FIM1_1676 | 515.7984383 | 558.6510084 | 574.8902043 | 0.107693968 | 0.116025585 | 0.33792089 | 0.413012258 | ns | YPD | KS.1 |
| FIM1_1677 | 1232.287023 | 821.9229503 | 639.4036967 | -0.422336087 | 0.359815162 | 0.153343147 | 0.209755885 | ns | YPD | KS.1 |
| FIM1_1678 | 479.868421 | 246.3323357 | 212.1497853 | -0.867458218 | 0.239608917 | 7.49610276095 | 0.000199428 | ns | YPD | KS.1 |
| FIM1_1679 | 873.3083911 | 833.0292717 | 646.7479553 | -0.057200658 | 0.219870482 | 0.776316032 | 0.824651191 | ns | YPD | KS.1 |
| FIM1_1680 | 448.6483999 | 403.4910245 | 512.5961361 | -0.13729503 | 0.162670598 | 0.372566501 | 0.44825111 | ns | YPD | KS.1 |
| FIM1_1681 | 92.08570461 | 42.33250084 | 59.03730574 | -0.923925514 | 0.34543673 | 0.001676421 | 0.003571818 | ns | YPD | KS.1 |
| FIM1_1682 | 1088.33309 | 791.7438963 | 870.1066741 | -0.415734296 | 0.183832694 | 0.015600753 | 0.027338411 | ns | YPD | KS.1 |
| FIM1_1683 | 3535.690248 | 3700.175293 | 4383.462853 | 0.057879547 | 0.177556734 | 0.728671412 | 0.783644677 | ns | YPD | KS.1 |
| FIM1_1684 | 659.0738923 | 692.6406758 | 1764.440744 | 0.068058923 | 0.110083865 | 0.524544185 | 0.600461878 | ns | YPD | KS.1 |
| FIM1_1685 | 3825.583972 | 1305.090765 | 2269.185769 | -1.533309261 | 0.097709792 | 1.13973680192 | 1.49215541831 | Km<KS.1 | YPD | KS.1 |
| FIM1_1686 | 901.7347233 | 853.0955558 | 1051.0058 | -0.074502981 | 0.133058489 | 0.560356322 | 0.63315301 | ns | YPD | KS.1 |
| FIM1_1687 | 2173.424988 | 1606.893799 | 2583.761881 | -0.420558758 | 0.107383423 | 5.88787384869 | 0.000160681 | ns | YPD | KS.1 |
| FIM1_1688 | 665.5295436 | 162.4758121 | 317.1652885 | -1.910320194 | 0.258926108 | 4.56711719669 | 3.81872404261 | Km<KS.1 | YPD | KS.1 |
| FIM1_1689 | 453.6456497 | 288.7384616 | 271.523357 | -0.594054549 | 0.197013447 | 0.001194713 | 0.002605743 | ns | YPD | KS.1 |
| FIM1_1690 | 2796.4822 | 4081.174266 | 3733.747297 | 0.534854415 | 0.082384276 | 4.38044881145 | 2.57902163751 | ns | YPD | KS.1 |
| FIM1_1691 | 13543.47484 | 4037.802793 | 2389.642711 | -1.405193537 | 0.439731578 | 0.000104157 | 0.000271442 | Km<KS.1 | YPD | KS.1 |
| FIM1_1692 | 4935.443651 | 1610.252141 | 1205.142974 | -1.548959072 | 0.190973156 | 3.36203133682 | 3.47736089411 | Km<KS.1 | YPD | KS.1 |
| FIM1_1693 | 2660.967104 | 584.4841552 | 768.7516542 | -2.011119079 | 0.311213234 | 2.19637226509 | 1.46473887652 | Km<KS.1 | YPD | KS.1 |
| FIM1_1694 | 116.8076877 | 64.63699426 | 53.63782133 | -0.727543786 | 0.279931581 | 0.003264113 | 0.006593164 | ns | YPD | KS.1 |
| FIM1_1695 | 559.0715574 | 299.5049742 | 186.2138148 | -0.712661404 | 0.34514961 | 0.014022631 | 0.024817712 | ns | YPD | KS.1 |

| | | | | | | | | | | |
|---|---|---|---|---|---|---|---|---|---|---|
| FIM1_1697 | 464.4943031 | 188.0107905 | 205.3618799 | -1.179415592 | 0.269672467 | 1.6181649624 | 5.5177317943 | Km<KS.1 | YPD | KS.1 |
| FIM1_1698 | 2405.568281 | 1377.648445 | 2099.503148 | -0.694744929 | 0.264420564 | 0.003228315 | 0.006526155 | ns | YPD | KS.1 |
| FIM1_1701 | 776.139333 | 400.6872957 | 548.8457104 | -0.878747511 | 0.213084612 | 9.2253776673 | 2.8139947207 | ns | YPD | KS.1 |
| FIM1_1703 | 95.9473524 | 125.3485578 | 84.05414166 | 0.30675588 | 0.275036918 | 0.204352074 | 0.269811987 | ns | YPD | KS.1 |
| FIM1_1705 | 31.92377754 | 20.79020019 | 22.09320857 | -0.492205861 | 0.307191387 | 0.062079538 | 0.095175871 | ns | YPD | KS.1 |
| FIM1_1707 | 515.6158594 | 495.1402842 | 374.05025 | -0.05158855 | 0.168835679 | 0.746271792 | 0.7988639 | ns | YPD | KS.1 |
| FIM1_1708 | 14208.28365 | 3252.120524 | 2951.506154 | -1.734430673 | 0.470878541 | 8.0931260485 | 2.4838557736 | Km<KS.1 | YPD | KS.1 |
| FIM1_1709 | 3810.62847 | 1140.08746 | 1248.502807 | -1.602631657 | 0.276495439 | 3.8871005807 | 2.0906297718 | Km<KS.1 | YPD | KS.1 |
| FIM1_1710 | 1770.377655 | 206.154865 | 440.0704167 | -2.941139027 | 0.320266195 | 6.2600341890 | 1.0211039373 | Km<KS.1 | YPD | KS.1 |
| FIM1_1711 | 6610.532842 | 1720.395631 | 1667.795688 | -1.674811839 | 0.386714579 | 6.5739195844 | 2.3942349877 | Km<KS.1 | YPD | KS.1 |
| FIM1_1712 | 1954.497447 | 1224.070625 | 2816.95815 | -0.541422983 | 0.306825731 | 0.040193653 | 0.063804539 | ns | YPD | KS.1 |
| FIM1_1713 | 321.3188212 | 57.05571338 | 91.60946069 | -2.399690117 | 0.22804596 | 4.2994133330 | 1.0383291908 | Km<KS.1 | YPD | KS.1 |
| FIM1_1714 | 1527.106901 | 243.8436923 | 331.4930736 | -2.597514962 | 0.16562589 | 5.6843252826 | 8.3175053768 | Km<KS.1 | YPD | KS.1 |
| FIM1_1715 | 268.6796052 | 255.8395071 | 205.5618188 | -0.061564077 | 0.209305786 | 0.749681538 | 0.801906182 | ns | YPD | KS.1 |
| FIM1_1716 | 4131.011896 | 3640.648242 | 4759.1703 | -0.149010895 | 0.244674648 | 0.498377658 | 0.57540702 | ns | YPD | KS.1 |
| FIM1_1717 | 228.0127211 | 228.037934 | 180.8783533 | -0.003106388 | 0.126006132 | 0.979622964 | 0.985566076 | ns | YPD | KS.1 |
| FIM1_1718 | 204.8118944 | 160.4335291 | 178.6442181 | -0.333846471 | 0.13145758 | 0.008315631 | 0.015425155 | ns | YPD | KS.1 |
| FIM1_1719 | 420.9602096 | 264.2333167 | 232.4189604 | -0.634605277 | 0.154961891 | 1.7864508612 | 5.2310730046 | ns | YPD | KS.1 |
| FIM1_1721 | 15.48242325 | 26.07087615 | 37.01723315 | 0.53950342 | 0.37082891 | 0.074976388 | 0.11191345 | ns | YPD | KS.1 |
| FIM1_1724 | 27169.12056 | 12983.15563 | 10659.98046 | -0.964026241 | 0.244960013 | 1.7153658477 | 5.0288421286 | ns | YPD | KS.1 |
| FIM1_1725 | 3993.395015 | 12562.74025 | 18386.44368 | 1.562246421 | 0.223725171 | 1.8615406835 | 1.3599361087 | Km>KS.1 | YPD | KS.1 |
| FIM1_1727 | 1944.44352 | 2718.937076 | 1983.948707 | 0.426119194 | 0.211577102 | 0.02839745 | 0.046703244 | ns | YPD | KS.1 |
| FIM1_1728 | 5708.628792 | 6519.733609 | 6048.725339 | 0.178108226 | 0.136544121 | 0.173698099 | 0.233996221 | ns | YPD | KS.1 |
| FIM1_1729 | 2344.743989 | 3550.945846 | 3532.289846 | 0.577139487 | 0.117455484 | 4.2673673942 | 1.5926596238 | ns | YPD | KS.1 |
| FIM1_1730 | 3339.444495 | 5127.891883 | 4780.754088 | 0.61020981 | 0.070191846 | 1.5061611288 | 1.7265326304 | ns | YPD | KS.1 |
| FIM1_1731 | 2751.745171 | 4321.811962 | 4615.337124 | 0.641405203 | 0.074812244 | 4.0022751112 | 4.4247374840 | ns | YPD | KS.1 |
| FIM1_1732 | 1156.181792 | 1010.881276 | 1104.078117 | -0.177582535 | 0.151789664 | 0.218873011 | 0.285499012 | ns | YPD | KS.1 |
| FIM1_1733 | 8060.934086 | 11364.20506 | 8127.148046 | 0.400886803 | 0.278968301 | 0.10101974 | 0.145800175 | ns | YPD | KS.1 |
| FIM1_1734 | 337.7555981 | 593.8582671 | 542.6370371 | 0.730790224 | 0.226087405 | 0.000424013 | 0.001000695 | ns | YPD | KS.1 |
| FIM1_1735 | 1972.432015 | 1613.577916 | 2353.005583 | -0.265839016 | 0.158851401 | 0.077413677 | 0.114999415 | ns | YPD | KS.1 |
| FIM1_1736 | 773.7065887 | 618.5689975 | 664.8273505 | -0.299383283 | 0.147590756 | 0.033429362 | 0.054120102 | ns | YPD | KS.1 |
| FIM1_1737 | 585.8690399 | 630.916914 | 484.8333214 | 0.095584894 | 0.168197762 | 0.547275047 | 0.620910687 | ns | YPD | KS.1 |
| FIM1_1738 | 2036.918087 | 1184.916996 | 1392.724953 | -0.747090003 | 0.145125555 | 8.5837385611 | 3.5089646131 | ns | YPD | KS.1 |
| FIM1_1739 | 1966.089728 | 1935.017976 | 2986.10881 | -0.02094825 | 0.143328796 | 0.878645562 | 0.907088955 | ns | YPD | KS.1 |
| FIM1_1740 | 1006.474409 | 876.2011216 | 944.6310137 | -0.190298638 | 0.11347305 | 0.083549414 | 0.123121545 | ns | YPD | KS.1 |
| FIM1_1741 | 8751.886151 | 2999.077468 | 4603.886683 | -1.489496204 | 0.174275903 | 9.5861950389 | 1.1221487133 | Km<KS.1 | YPD | KS.1 |
| FIM1_1742 | 1361.719818 | 1004.653308 | 1223.137733 | -0.407936035 | 0.155428198 | 0.005785232 | 0.011069818 | ns | YPD | KS.1 |
| FIM1_1743 | 1512.935444 | 4116.138061 | 3528.491098 | 1.334920618 | 0.248091043 | 7.3531182182 | 3.4413700033 | Km>KS.1 | YPD | KS.1 |
| FIM1_1744 | 4376.318265 | 4605.66367 | 2700.92026 | 0.058317746 | 0.262635838 | 0.802707267 | 0.845416071 | ns | YPD | KS.1 |
| FIM1_1745 | 964.1664874 | 1480.350237 | 1396.955375 | 0.602273364 | 0.098209954 | 3.8615970677 | 2.0791607588 | ns | YPD | KS.1 |
| FIM1_1746 | 366.7680512 | 397.6036058 | 512.8983152 | 0.102869919 | 0.172373526 | 0.526377342 | 0.602144694 | ns | YPD | KS.1 |
| FIM1_1747 | 4118.584245 | 4181.701847 | 3201.610679 | 0.019680257 | 0.166145948 | 0.900347084 | 0.924314227 | ns | YPD | KS.1 |
| FIM1_1748 | 1616.432565 | 256.2501418 | 243.67275 | -2.545756735 | 0.2533021 | 3.6840659952 | 7.9000984166 | Km<KS.1 | YPD | KS.1 |
| FIM1_1749 | 6509.643462 | 4819.364251 | 4469.165477 | -0.40395084 | 0.152961804 | 0.005552262 | 0.010669179 | ns | YPD | KS.1 |
| FIM1_1750 | 2450.468508 | 4937.019857 | 4549.600814 | 0.887828117 | 0.271991782 | 0.000267808 | 0.000658274 | ns | YPD | KS.1 |

| | | | | | | | | | | |
|---|---|---|---|---|---|---|---|---|---|---|
| FIM1_1751 | 878.5449255 | 799.8779835 | 1262.995325 | -0.097784934 | 0.321126827 | 0.719695988 | 0.776341618 | ns | YPD | KS.1 |
| FIM1_1752 | 2511.612719 | 1170.097607 | 1127.180991 | -0.945810087 | 0.305348863 | 0.00041834 | 0.000988714 | ns | YPD | KS.1 |
| FIM1_1753 | 741.9740527 | 835.3731019 | 787.7175786 | 0.162048597 | 0.10886522 | 0.12532724 | 0.176280187 | ns | YPD | KS.1 |
| FIM1_1754 | 1720.873121 | 940.4424089 | 560.3280302 | -0.77329649 | 0.246731792 | 0.000542209 | 0.001261689 | ns | YPD | KS.1 |
| FIM1_1755 | 4095.817154 | 2538.370213 | 2207.912755 | -0.555845892 | 0.305654201 | 0.034709053 | 0.056009581 | ns | YPD | KS.1 |
| FIM1_1756 | 1148.07093 | 2322.235008 | 1284.85607 | 0.925592378 | 0.231803944 | 1.4628056551( | 4.3318203179( | ns | YPD | KS.1 |
| FIM1_1757 | 3668.166591 | 1443.947209 | 1650.215053 | -1.232195464 | 0.253743908 | 1.4396548086 | 5.7069981459 | Km<KS.1 | YPD | KS.1 |
| FIM1_1758 | 146.6041903 | 194.6942824 | 332.026574 | 0.348643251 | 0.2382211 | 0.104714816 | 0.150521876 | ns | YPD | KS.1 |
| FIM1_1759 | 555.7855313 | 565.7971153 | 363.4795938 | 0.021663658 | 0.24252973 | 0.921100395 | 0.941153104 | ns | YPD | KS.1 |
| FIM1_1760 | 5314.439162 | 6578.802586 | 3933.361456 | 0.25796707 | 0.235511979 | 0.225679408 | 0.292765333 | ns | YPD | KS.1 |
| FIM1_1761 | 5495.971424 | 4687.113185 | 2837.073276 | -0.184638394 | 0.258879119 | 0.422973007 | 0.500425853 | ns | YPD | KS.1 |
| FIM1_1762 | 2940.549483 | 3052.129923 | 5193.355965 | 0.042799462 | 0.258041562 | 0.852457166 | 0.885872939 | ns | YPD | KS.1 |
| FIM1_1763 | 31.17538003 | 53.76502269 | 106.1058246 | 0.642355469 | 0.311680636 | 0.016525462 | 0.028816745 | ns | YPD | KS.1 |
| FIM1_1764 | 1233.467182 | 3269.32261 | 3825.694864 | 1.385757483 | 0.104734044 | 5.3040469848( | 2.8373799730 | Km>KS.1 | YPD | KS.1 |
| FIM1_1765 | 5504.10148 | 2300.814953 | 2218.508472 | -1.23312588 | 0.117320537 | 9.4948155102 | 2.1087815697 | Km<KS.1 | YPD | KS.1 |
| FIM1_1766 | 4490.514212 | 2686.507365 | 1836.890909 | -0.619405068 | 0.284502439 | 0.013020011 | 0.023240513 | ns | YPD | KS.1 |
| FIM1_1767 | 1745.01214 | 1672.376967 | 1753.236508 | -0.05710394 | 0.131397917 | 0.651612602 | 0.716572213 | ns | YPD | KS.1 |
| FIM1_1769 | 18.83148655 | 22.58498385 | 21.2872472 | 0.16015796 | 0.414110488 | 0.619737814 | 0.688828334 | ns | YPD | KS.1 |
| FIM1_1770 | 1542.111619 | 2006.354951 | 2963.922448 | 0.36599644 | 0.105089689 | 0.000355219 | 0.000852303 | ns | YPD | KS.1 |
| FIM1_1771 | 2983.495527 | 1216.403614 | 1673.299133 | -1.256824814 | 0.143882383 | 2.8834622360( | 3.5246252147 | Km<KS.1 | YPD | KS.1 |
| FIM1_1772 | 570.911467 | 991.1836915 | 1718.984209 | 0.751218461 | 0.165319569 | 1.7956863583 | 6.0648605790 | ns | YPD | KS.1 |
| FIM1_1774 | 945.6081269 | 655.7611122 | 925.4380031 | -0.503695949 | 0.129040331 | 5.3124210653 | 0.000146018 | ns | YPD | KS.1 |
| FIM1_1775 | 4500.625809 | 6710.102414 | 6632.013889 | 0.545912398 | 0.142139134 | 6.3024144847 | 0.000170869 | ns | YPD | KS.1 |
| FIM1_1776 | 2369.547322 | 3889.664825 | 2784.014127 | 0.673311328 | 0.16170773 | 1.2163847583 | 3.6520906293 | ns | YPD | KS.1 |
| FIM1_1777 | 3327.217205 | 752.4394542 | 3183.634497 | -2.087621514 | 0.175015166 | 1.4120882577 | 5.1655434430 | Km<KS.1 | YPD | KS.1 |
| FIM1_1778 | 3574.0282 | 4053.8783 | 2756.463574 | 0.166448207 | 0.151230571 | 0.247583064 | 0.31631375 | ns | YPD | KS.1 |
| FIM1_1779 | 7555.227794 | 5734.121704 | 5361.145866 | -0.336437664 | 0.236889907 | 0.11572306 | 0.164492064 | ns | YPD | KS.1 |
| FIM1_1780 | 1067.931939 | 813.8130279 | 982.3092643 | -0.373128462 | 0.123409042 | 0.001766024 | 0.003756294 | ns | YPD | KS.1 |
| FIM1_1781 | 1231.027494 | 719.122218 | 622.2993874 | -0.740988648 | 0.144410779 | 9.5145419819 | 3.8609173213 | ns | YPD | KS.1 |
| FIM1_1782 | 1970.814425 | 1226.088987 | 1110.647824 | -0.674181371 | 0.077429462 | 1.1518927494 | 1.3420764469 | ns | YPD | KS.1 |
| FIM1_1783 | 733.2934624 | 379.5070437 | 438.0035736 | -0.909003139 | 0.156191083 | 1.3507310950( | 6.9205841379 | ns | YPD | KS.1 |
| FIM1_1784 | 3098.277981 | 2269.320736 | 2015.541673 | -0.425734634 | 0.132394196 | 0.000843271 | 0.001890614 | ns | YPD | KS.1 |
| FIM1_1785 | 572.1429516 | 824.6042798 | 875.1746038 | 0.493940216 | 0.152247561 | 0.000672077 | 0.001538766 | ns | YPD | KS.1 |
| FIM1_1786 | 58.00489302 | 46.21951437 | 53.71768876 | -0.279762372 | 0.216454443 | 0.157995868 | 0.215291549 | ns | YPD | KS.1 |
| FIM1_1787 | 1509.768499 | 1212.130796 | 2637.023263 | -0.248313959 | 0.282332907 | 0.314409241 | 0.387846758 | ns | YPD | KS.1 |
| FIM1_1788 | 399.9716359 | 473.8146029 | 541.1282185 | 0.227555325 | 0.143674625 | 0.097322875 | 0.140955255 | ns | YPD | KS.1 |
| FIM1_1789 | 1831.738052 | 1979.009944 | 1310.35748 | 0.090500411 | 0.246747529 | 0.683216523 | 0.74539522 | ns | YPD | KS.1 |
| FIM1_1790 | 1399.684117 | 833.9300008 | 819.0189478 | -0.697224358 | 0.177002884 | 3.0216239006 | 8.5949564926 | ns | YPD | KS.1 |
| FIM1_1791 | 414.345708 | 693.4742435 | 360.2382304 | 0.688885605 | 0.184308127 | 7.0023695547 | 0.000187597 | ns | YPD | KS.1 |
| FIM1_1792 | 3728.111719 | 7448.915534 | 6424.741287 | 0.893492113 | 0.251197543 | 9.0328005836 | 0.000237642 | ns | YPD | KS.1 |
| FIM1_1793 | 1660.862734 | 1697.354854 | 1819.898147 | 0.029546155 | 0.122366327 | 0.802762918 | 0.845416071 | ns | YPD | KS.1 |
| FIM1_1794 | 928.4312525 | 982.8000543 | 681.1152267 | 0.05856571 | 0.328645376 | 0.832629417 | 0.871067052 | ns | YPD | KS.1 |
| FIM1_1795 | 1025.886947 | 1115.267025 | 820.1649257 | 0.118499128 | 0.059394563 | 0.043719834 | 0.068962009 | ns | YPD | KS.1 |
| FIM1_1796 | 46.48841722 | 52.74439317 | 82.68238282 | 0.147761143 | 0.234302514 | 0.486377662 | 0.564170871 | ns | YPD | KS.1 |
| FIM1_1797 | 1164.645459 | 1686.933782 | 1607.787515 | 0.459892074 | 0.23772364 | 0.032201325 | 0.052199932 | ns | YPD | KS.1 |

| | | | | | | | | | | |
|---|---|---|---|---|---|---|---|---|---|---|
| FIM1_1798 | 5138.422663 | 3335.637511 | 1934.363292 | -0.515816225 | 0.277718249 | 0.034508202 | 0.055739709 | ns | YPD | KS.1 |
| FIM1_1799 | 1627.860947 | 1455.680175 | 1747.339532 | -0.152178656 | 0.121303447 | 0.193998683 | 0.25778404 | ns | YPD | KS.1 |
| FIM1_1800 | 2870.831773 | 941.5765772 | 846.4389477 | -1.506563231 | 0.237224448 | 1.5330729662 | 9.4044858286 | Km<KS.1 | YPD | KS.1 |
| FIM1_1801 | 1035.064379 | 699.5804225 | 2016.171127 | -0.444046178 | 0.306682726 | 0.091659382 | 0.133804408 | ns | YPD | KS.1 |
| FIM1_1802 | 7360.002182 | 5799.554942 | 3150.560927 | -0.278171226 | 0.263476251 | 0.233593003 | 0.301459194 | ns | YPD | KS.1 |
| FIM1_1803 | 21115.95011 | 4672.882288 | 4401.067168 | -1.891755464 | 0.399042514 | 5.6806218850 | 2.3728878151 | Km<KS.1 | YPD | KS.1 |
| FIM1_1804 | 2291.590895 | 934.7777886 | 737.2140736 | -1.177442262 | 0.258487782 | 6.9853956713 | 2.5366674061 | Km<KS.1 | YPD | KS.1 |
| FIM1_1805 | 440.0582784 | 39.86712325 | 87.50925761 | -3.312603607 | 0.324785879 | 2.0335623464 | 4.7497524288 | Km<KS.1 | YPD | KS.1 |
| FIM1_1806 | 3994.448599 | 1722.73539 | 1470.141121 | -1.065857767 | 0.293873329 | 4.7507551660 | 0.000131379 | Km<KS.1 | YPD | KS.1 |
| FIM1_1807 | 930.8713914 | 259.0537477 | 206.0439287 | -1.747525512 | 0.230032004 | 1.2977078661 | 1.1528743989 | Km<KS.1 | YPD | KS.1 |
| FIM1_1808 | 4459.29025 | 4693.936212 | 2362.410885 | 0.059469302 | 0.252348722 | 0.792405136 | 0.836810773 | ns | YPD | KS.1 |
| FIM1_1809 | 5700.745955 | 7051.350756 | 3503.93053 | 0.236632777 | 0.29194768 | 0.350368242 | 0.42555713 | ns | YPD | KS.1 |
| FIM1_1810 | 1879.539774 | 2095.921824 | 1536.574471 | 0.144715621 | 0.142758643 | 0.289084102 | 0.36086936 | ns | YPD | KS.1 |
| FIM1_1811 | 1182.203455 | 1364.734412 | 1497.327843 | 0.196990108 | 0.111825443 | 0.069317882 | 0.104438662 | ns | YPD | KS.1 |
| FIM1_1812 | 1318.023464 | 805.0272564 | 669.3733824 | -0.64908129 | 0.19948321 | 0.000470327 | 0.001103195 | ns | YPD | KS.1 |
| FIM1_1813 | 628.4868717 | 563.6855529 | 514.8656018 | -0.144793295 | 0.142454417 | 0.287868849 | 0.359745673 | ns | YPD | KS.1 |
| FIM1_1814 | 760.5099445 | 975.0853066 | 879.8759774 | 0.343599404 | 0.10871959 | 0.001164867 | 0.002547346 | ns | YPD | KS.1 |
| FIM1_1815 | 1366.414412 | 855.4132308 | 661.3282019 | -0.651867389 | 0.1226237 | 4.2684190797 | 1.8149901642 | ns | YPD | KS.1 |
| FIM1_1816 | 1355.332635 | 2743.509338 | 2257.114841 | 0.985927491 | 0.133598238 | 3.0965792373 | 2.3847494900 | ns | YPD | KS.1 |
| FIM1_1817 | 1154.524273 | 2317.236194 | 1715.826269 | 0.969962485 | 0.142079018 | 1.7518165338 | 1.1809332325 | ns | YPD | KS.1 |
| FIM1_1818 | 11596.60935 | 19281.94959 | 24018.39903 | 0.722278691 | 0.081479291 | 2.5398398523 | 3.1276493230 | ns | YPD | KS.1 |
| FIM1_1819 | 2552.06933 | 3331.780665 | 2817.807409 | 0.366286501 | 0.122302648 | 0.001960963 | 0.004142587 | ns | YPD | KS.1 |
| FIM1_1820 | 432.1039286 | 410.643781 | 505.4351108 | -0.064934287 | 0.175757016 | 0.694478502 | 0.755032899 | ns | YPD | KS.1 |
| FIM1_1821 | 605.0687411 | 1303.815824 | 1756.485633 | 1.073086345 | 0.140684787 | 3.9672118787 | 3.3339322798 | Km>KS.1 | YPD | KS.1 |
| FIM1_1822 | 592.0710303 | 561.9860292 | 470.8690079 | -0.070156897 | 0.126947218 | 0.566589724 | 0.639180018 | ns | YPD | KS.1 |
| FIM1_1823 | 124.4846794 | 157.0200615 | 249.6394333 | 0.244382527 | 0.32712652 | 0.375958303 | 0.451785642 | ns | YPD | KS.1 |
| FIM1_1824 | 1592.020985 | 1393.311303 | 1074.180864 | -0.171140875 | 0.178226636 | 0.305980719 | 0.378764388 | ns | YPD | KS.1 |
| FIM1_1825 | 1190.433987 | 2965.611746 | 4575.038093 | 1.2634945 | 0.171928885 | 2.3319517692 | 1.8212653142 | Km>KS.1 | YPD | KS.1 |
| FIM1_1826 | 3138.997489 | 9434.473663 | 6200.367689 | 1.489102645 | 0.233557166 | 1.3518456874 | 8.3338690147 | Km>KS.1 | YPD | KS.1 |
| FIM1_1827 | 3644.283453 | 4445.20732 | 5286.396663 | 0.274414436 | 0.109353602 | 0.009811535 | 0.01797216 | ns | YPD | KS.1 |
| FIM1_1828 | 4829.713229 | 5033.75597 | 3775.938771 | 0.055329447 | 0.129498402 | 0.657397912 | 0.722136148 | ns | YPD | KS.1 |
| FIM1_1829 | 448.4953236 | 267.659283 | 281.7080514 | -0.705442762 | 0.154445974 | 1.7765539795 | 6.0083997610 | ns | YPD | KS.1 |
| FIM1_1830 | 1462.976889 | 2009.551063 | 1821.326502 | 0.442049451 | 0.107300541 | 2.3920651962 | 6.8948576775 | ns | YPD | KS.1 |
| FIM1_1831 | 280.0085994 | 356.4340799 | 557.696713 | 0.328106442 | 0.141027079 | 0.015100052 | 0.026554528 | ns | YPD | KS.1 |
| FIM1_1832 | 408.9974494 | 434.8990835 | 410.2668753 | 0.080481495 | 0.135266683 | 0.53561066 | 0.610397842 | ns | YPD | KS.1 |
| FIM1_1833 | 1405.182877 | 736.3294282 | 903.3919552 | -0.921347883 | 0.076234971 | 2.7021324713 | 9.7413833659 | ns | YPD | KS.1 |
| FIM1_1834 | 1805.633063 | 1387.767614 | 892.8793349 | -0.347319566 | 0.167200671 | 0.027738188 | 0.045724812 | ns | YPD | KS.1 |
| FIM1_1835 | 943.8847824 | 828.4709417 | 680.737761 | -0.178214794 | 0.112971498 | 0.103652053 | 0.149123472 | ns | YPD | KS.1 |
| FIM1_1836 | 1152.052618 | 2532.675947 | 2170.345231 | 1.074532303 | 0.189251337 | 2.2571679201 | 1.1173542689 | Km>KS.1 | YPD | KS.1 |
| FIM1_1837 | 3467.764282 | 4881.067068 | 5368.555356 | 0.432295282 | 0.217414018 | 0.029852278 | 0.048821527 | ns | YPD | KS.1 |
| FIM1_1838 | 251.4906565 | 155.0549751 | 170.4799956 | -0.619611856 | 0.228394629 | 0.002943057 | 0.0059972 | ns | YPD | KS.1 |
| FIM1_1839 | 2101.419118 | 2305.570801 | 3181.795781 | 0.124132082 | 0.131722972 | 0.327058404 | 0.401063732 | ns | YPD | KS.1 |
| FIM1_1840 | 7360.492975 | 7971.918927 | 7274.870358 | 0.102199871 | 0.176098715 | 0.53697975 | 0.61132134 | ns | YPD | KS.1 |
| FIM1_1841 | 1291.055901 | 968.3307827 | 1025.017145 | -0.405623407 | 0.087233652 | 2.2351079831 | 7.4578552757 | ns | YPD | KS.1 |
| FIM1_1842 | 1464.273037 | 1650.8862 | 1034.982679 | 0.141113958 | 0.246177056 | 0.523563751 | 0.599477483 | ns | YPD | KS.1 |

| | | | | | | | | | | |
|---|---|---|---|---|---|---|---|---|---|---|
| FIM1_1843 | 6869.939892 | 9919.013227 | 8375.507969 | 0.513466536 | 0.104871843 | 5.3508410621 | 1.9602676203 | ns | YPD | KS.1 |
| FIM1_1844 | 27.88314134 | 58.53770431 | 62.51638684 | 0.947020228 | 0.27336016 | 0.000113826 | 0.000294174 | ns | YPD | KS.1 |
| FIM1_1845 | 268.6208581 | 515.5970646 | 543.7782126 | 0.912829352 | 0.122727181 | 2.3003162028 | 1.7993825643 | ns | YPD | KS.1 |
| FIM1_1846 | 1444.121814 | 3731.25973 | 3597.826933 | 1.338214506 | 0.130968513 | 1.6752780636 | 3.3742948852 | Km>KS.1 | YPD | KS.1 |
| FIM1_1847 | 4709.997501 | 5201.030763 | 4023.427754 | 0.127613015 | 0.173484655 | 0.434424809 | 0.511662742 | ns | YPD | KS.1 |
| FIM1_1848 | 5021.250041 | 7193.854954 | 7128.816569 | 0.468242307 | 0.192846008 | 0.009085565 | 0.016732084 | ns | YPD | KS.1 |
| FIM1_1849 | 4636.166627 | 1949.101291 | 3645.990782 | -1.168351222 | 0.216384676 | 9.2042626629 | 4.2477928337 | Km<KS.1 | YPD | KS.1 |
| FIM1_1850 | 762.6466723 | 816.8549521 | 680.5337264 | 0.087908465 | 0.17858091 | 0.60005884 | 0.670701579 | ns | YPD | KS.1 |
| FIM1_1851 | 2669.55319 | 2566.209182 | 3843.198155 | -0.055040444 | 0.092642294 | 0.543739667 | 0.617462872 | ns | YPD | KS.1 |
| FIM1_1853 | 50012.94979 | 48156.95198 | 51825.76471 | -0.050214842 | 0.139574351 | 0.70749086 | 0.766666745 | ns | YPD | KS.1 |
| FIM1_1854 | 10072.95616 | 8070.888285 | 6421.667845 | -0.277960708 | 0.20778823 | 0.146187413 | 0.201184614 | ns | YPD | KS.1 |
| FIM1_1855 | 1702.247219 | 1369.864013 | 833.8984346 | -0.264393672 | 0.230760186 | 0.206473784 | 0.272035772 | ns | YPD | KS.1 |
| FIM1_1856 | 1437.293113 | 2419.392199 | 1627.702602 | 0.677783979 | 0.216143435 | 0.000665469 | 0.00152497 | ns | YPD | KS.1 |
| FIM1_1857 | 5479.569339 | 9056.700607 | 6646.68315 | 0.690802896 | 0.145209673 | 7.2826487653 | 2.6311675822 | ns | YPD | KS.1 |
| FIM1_1858 | 1414.649429 | 1049.57408 | 1277.778027 | -0.409827721 | 0.127550057 | 0.000874612 | 0.001953837 | ns | YPD | KS.1 |
| FIM1_1859 | 649.9002538 | 1196.612586 | 1973.780227 | 0.834181223 | 0.166400132 | 1.4465102051 | 5.7296084955 | ns | YPD | KS.1 |
| FIM1_1860 | 2262.606772 | 2519.084851 | 1463.585236 | 0.132302823 | 0.210411713 | 0.494556672 | 0.571526003 | ns | YPD | KS.1 |
| FIM1_1861 | 520.1372166 | 542.6587762 | 621.8867677 | 0.056692283 | 0.133740683 | 0.659285479 | 0.723354741 | ns | YPD | KS.1 |
| FIM1_1862 | 696.4748308 | 1174.104699 | 889.5884481 | 0.723818364 | 0.134466733 | 2.5075346374 | 1.1010577953 | ns | YPD | KS.1 |
| FIM1_1863 | 221.2092063 | 441.3627456 | 657.76852 | 0.981471553 | 0.095042514 | 1.0357000485 | 2.1117244841 | ns | YPD | KS.1 |
| FIM1_1864 | 2974.97012 | 1190.832974 | 1432.97111 | -1.295445902 | 0.11686626 | 1.6291097468 | 4.3341288720 | Km<KS.1 | YPD | KS.1 |
| FIM1_1865 | 1166.425131 | 944.3595588 | 897.4371506 | -0.270921295 | 0.186424034 | 0.119248333 | 0.168971932 | ns | YPD | KS.1 |
| FIM1_1866 | 2409.13731 | 338.6901188 | 472.7962783 | -2.69598534 | 0.283255778 | 4.6434225568 | 8.4619147327 | Km<KS.1 | YPD | KS.1 |
| FIM1_1867 | 397.6541633 | 375.6443846 | 199.9969661 | -0.072639859 | 0.169033 | 0.64896503 | 0.714292262 | ns | YPD | KS.1 |
| FIM1_1868 | 1041.581749 | 1761.061857 | 1780.688788 | 0.726094125 | 0.137507841 | 4.3932469578 | 1.8632910157 | ns | YPD | KS.1 |
| FIM1_1869 | 1063.063844 | 1497.064664 | 2045.223674 | 0.472134635 | 0.125921762 | 0.000105605 | 0.000274782 | ns | YPD | KS.1 |
| FIM1_1870 | 1156.595085 | 1171.190368 | 1245.736782 | 0.016448913 | 0.129331832 | 0.895003738 | 0.9201578 | ns | YPD | KS.1 |
| FIM1_1871 | 3081.35542 | 2509.757105 | 1710.552191 | -0.267376812 | 0.172264216 | 0.098904303 | 0.143037474 | ns | YPD | KS.1 |
| FIM1_1872 | 1747.860808 | 1219.399192 | 908.4532218 | -0.486632757 | 0.151874488 | 0.000784222 | 0.001769391 | ns | YPD | KS.1 |
| FIM1_1873 | 712.9998984 | 1032.814294 | 969.815206 | 0.512922435 | 0.122103962 | 1.4619700991 | 4.3318203179 | ns | YPD | KS.1 |
| FIM1_1874 | 445.2860033 | 2759.865991 | 1626.170728 | 2.579179997 | 0.17112397 | 7.6742496315 | 8.1232748759 | Km>KS.1 | YPD | KS.1 |
| FIM1_1875 | 47.58042489 | 85.77271418 | 58.7017236 | 0.764254432 | 0.234594523 | 0.000360075 | 0.000862897 | ns | YPD | KS.1 |
| FIM1_1876 | 993.4300598 | 816.3589675 | 644.4371434 | -0.205699824 | 0.326646293 | 0.455706576 | 0.533068473 | ns | YPD | KS.1 |
| FIM1_1877 | 1058.656897 | 620.7889568 | 1186.719207 | -0.59594683 | 0.342676247 | 0.037896556 | 0.060466763 | ns | YPD | KS.1 |
| FIM1_1878 | 6850.914735 | 5336.723783 | 2956.918088 | -0.264398047 | 0.327213986 | 0.338199697 | 0.413198304 | ns | YPD | KS.1 |
| FIM1_1879 | 521.6723943 | 459.5680724 | 476.7881425 | -0.175250405 | 0.10661047 | 0.090809102 | 0.132796967 | ns | YPD | KS.1 |
| FIM1_1880 | 2770.498315 | 3072.632868 | 3492.685695 | 0.13959548 | 0.126997794 | 0.253934204 | 0.323007166 | ns | YPD | KS.1 |
| FIM1_1881 | 568.5533003 | 946.3490638 | 750.7455602 | 0.711695684 | 0.115027221 | 2.1315422330 | 1.1691755908 | ns | YPD | KS.1 |
| FIM1_1882 | 4544.657558 | 4152.761118 | 3247.16289 | -0.113504436 | 0.192001877 | 0.525549097 | 0.601473834 | ns | YPD | KS.1 |
| FIM1_1883 | 198.6112874 | 252.166824 | 133.49796 | 0.279279976 | 0.262843202 | 0.230822811 | 0.298348528 | ns | YPD | KS.1 |
| FIM1_1884 | 3721.954917 | 5030.201206 | 4422.520693 | 0.392925175 | 0.183057528 | 0.02174789 | 0.036701408 | ns | YPD | KS.1 |
| FIM1_1885 | 1083.975223 | 1206.195178 | 1568.879703 | 0.147678562 | 0.100594918 | 0.131966671 | 0.184067353 | ns | YPD | KS.1 |
| FIM1_1886 | 1147.574077 | 829.5793171 | 895.1978618 | -0.447368894 | 0.119343038 | 0.000110995 | 0.000287754 | ns | YPD | KS.1 |
| FIM1_1887 | 1357.10468 | 4579.717879 | 5765.02779 | 1.694953248 | 0.178436884 | 1.0318519019 | 1.6400840933 | Km>KS.1 | YPD | KS.1 |
| FIM1_1888 | 667.8169262 | 2037.99511 | 2288.547933 | 1.600779636 | 0.064267861 | 3.2988515064 | 1.6411786244 | Km>KS.1 | YPD | KS.1 |

| | | | | | | | | | | |
|---|---|---|---|---|---|---|---|---|---|---|
| FIM1_1889 | 847.9136835 | 1606.43879 | 1442.223128 | 0.897201835 | 0.120339357 | 2.07402646263 | 1.63522688615 | ns | YPD | KS.1 |
| FIM1_1890 | 1281.930821 | 431.0212285 | 1155.395381 | -1.258506698 | 0.423996235 | 0.000310959 | 0.000754645 | Km<KS.1 | YPD | KS.1 |
| FIM1_1891 | 3513.249884 | 2625.681647 | 1454.40925 | -0.331069855 | 0.28865399 | 0.187265696 | 0.250173695 | ns | YPD | KS.1 |
| FIM1_1892 | 517.7045409 | 683.8802543 | 894.0127034 | 0.339613341 | 0.23437762 | 0.109134826 | 0.156063742 | ns | YPD | KS.1 |
| FIM1_1893 | 1587.714823 | 1437.03948 | 1753.076194 | -0.135655066 | 0.12087399 | 0.24542918 | 0.31412662 | ns | YPD | KS.1 |
| FIM1_1894 | 3109.843976 | 1897.364958 | 1675.682039 | -0.675519825 | 0.152017119 | 3.40822613467 | 1.10606164514 | ns | YPD | KS.1 |
| FIM1_1895 | 375.1248681 | 1084.420754 | 1091.70638 | 1.501878924 | 0.119860716 | 3.72343283553 | 1.58325456040 | Km>KS.1 | YPD | KS.1 |
| FIM1_1896 | 720.0242039 | 934.9065197 | 1174.592518 | 0.365806857 | 0.096851461 | 0.000113716 | 0.000294043 | ns | YPD | KS.1 |
| FIM1_1897 | 1187.762657 | 1379.503471 | 1873.564261 | 0.207310682 | 0.101074994 | 0.035302207 | 0.056819308 | ns | YPD | KS.1 |
| FIM1_1898 | 2141.802701 | 1684.427793 | 1951.388759 | -0.336737892 | 0.093540832 | 0.000237918 | 0.000590346 | ns | YPD | KS.1 |
| FIM1_1899 | 12820.67254 | 7800.064091 | 8322.433107 | -0.685122036 | 0.140146459 | 3.81104479439 | 1.43418667565 | ns | YPD | KS.1 |
| FIM1_1900 | 227.807471 | 369.5966741 | 403.9192299 | 0.676854435 | 0.124398033 | 2.01178915711 | 8.92832386853 | ns | YPD | KS.1 |
| FIM1_1901 | 3611.542516 | 5538.58459 | 4115.123878 | 0.573864583 | 0.169015624 | 0.000332484 | 0.000802575 | ns | YPD | KS.1 |
| FIM1_1902 | 761.6733338 | 678.6059509 | 880.7159253 | -0.151776762 | 0.150845832 | 0.290680133 | 0.3623487 | ns | YPD | KS.1 |
| FIM1_1903 | 1117.408716 | 963.7409899 | 936.9536444 | -0.206092147 | 0.090453739 | 0.01991027 | 0.034003979 | ns | YPD | KS.1 |
| FIM1_1904 | 14236.20601 | 4014.131295 | 5400.172419 | -1.789280581 | 0.139818726 | 6.45831429327 | 3.06001081990 | Km<KS.1 | YPD | KS.1 |
| FIM1_1905 | 628.853292 | 898.8344421 | 1071.12881 | 0.490124395 | 0.132021966 | 0.00011811 | 0.000304454 | ns | YPD | KS.1 |
| FIM1_1907 | 300.3520719 | 815.2519857 | 1143.469546 | 1.377393971 | 0.185875107 | 1.18492112080 | 9.46225132584 | Km>KS.1 | YPD | KS.1 |
| FIM1_1908 | 1513.476205 | 2680.020201 | 4344.102142 | 0.78443275 | 0.153945366 | 1.04469934934 | 4.20499940369 | ns | YPD | KS.1 |
| FIM1_1909 | 5659.822861 | 5832.832482 | 4253.410281 | 0.041496752 | 0.097755311 | 0.663833225 | 0.727438391 | ns | YPD | KS.1 |
| FIM1_1910 | 930.3091069 | 1328.416401 | 1422.179399 | 0.481322422 | 0.152346523 | 0.000924677 | 0.002059208 | ns | YPD | KS.1 |
| FIM1_1911 | 6159.088181 | 6544.661007 | 7619.23478 | 0.084949106 | 0.077354457 | 0.264250234 | 0.334080153 | ns | YPD | KS.1 |
| FIM1_1912 | 4539.540343 | 4125.83294 | 3167.10355 | -0.129850129 | 0.117642916 | 0.254016724 | 0.323007166 | ns | YPD | KS.1 |
| FIM1_1913 | 1213.005605 | 1129.015483 | 2600.344044 | -0.084697114 | 0.23971106 | 0.695672033 | 0.756000079 | ns | YPD | KS.1 |
| FIM1_1914 | 223.8326806 | 280.9622141 | 540.0841743 | 0.297833965 | 0.168302451 | 0.060516752 | 0.093095498 | ns | YPD | KS.1 |
| FIM1_1915 | 1288.87643 | 3302.620642 | 2001.753845 | 1.235938179 | 0.263568181 | 3.25827787381 | 1.23739942154 | Km>KS.1 | YPD | KS.1 |
| FIM1_1916 | 843.8875525 | 1191.377335 | 1040.49571 | 0.442091145 | 0.206114011 | 0.020055323 | 0.034204742 | ns | YPD | KS.1 |
| FIM1_1917 | 509.3595918 | 459.6713368 | 992.6185635 | -0.121886306 | 0.236198953 | 0.568585652 | 0.641286243 | ns | YPD | KS.1 |
| FIM1_1918 | 591.0836932 | 672.9644055 | 883.3691243 | 0.170185788 | 0.159018896 | 0.258926481 | 0.328695903 | ns | YPD | KS.1 |
| FIM1_1919 | 2543.639629 | 1655.092134 | 1482.241881 | -0.597332448 | 0.118604869 | 2.16451643220 | 8.39974200485 | ns | YPD | KS.1 |
| FIM1_1920 | 1883.082696 | 933.5068338 | 2593.445431 | -0.837861896 | 0.326562303 | 0.002806117 | 0.005730884 | ns | YPD | KS.1 |
| FIM1_1921 | 476.1853111 | 1162.133752 | 2806.918951 | 1.239764695 | 0.165064045 | 7.17280188797 | 5.84037469602 | Km>KS.1 | YPD | KS.1 |
| FIM1_1922 | 2195.71924 | 2088.411738 | 2248.488215 | -0.07055892 | 0.060597033 | 0.239158358 | 0.307207031 | ns | YPD | KS.1 |
| FIM1_1923 | 2412.552667 | 2190.680732 | 2652.797944 | -0.134529859 | 0.089886349 | 0.126229103 | 0.177248034 | ns | YPD | KS.1 |
| FIM1_1924 | 491.0117946 | 660.4551784 | 1067.849095 | 0.392340946 | 0.165798708 | 0.012288374 | 0.02205435 | ns | YPD | KS.1 |
| FIM1_1925 | 1757.984981 | 1932.178981 | 2161.135795 | 0.131722889 | 0.090721632 | 0.137834785 | 0.191223663 | ns | YPD | KS.1 |
| FIM1_1926 | 520.6832275 | 496.405714 | 614.4443236 | -0.061935651 | 0.179206006 | 0.712763924 | 0.770852266 | ns | YPD | KS.1 |
| FIM1_1927 | 2231.285902 | 2134.329405 | 3871.751739 | -0.049562372 | 0.276543585 | 0.838539199 | 0.875419197 | ns | YPD | KS.1 |
| FIM1_1928 | 1575.908934 | 1783.403619 | 1626.853045 | 0.168957845 | 0.11410072 | 0.126432437 | 0.177483458 | ns | YPD | KS.1 |
| FIM1_1929 | 518.4851624 | 795.9643204 | 577.4460461 | 0.526940391 | 0.25415866 | 0.020380033 | 0.034675331 | ns | YPD | KS.1 |
| FIM1_1930 | 366.8369402 | 619.6749994 | 719.1019424 | 0.718110147 | 0.151473145 | 7.43394539747 | 2.67805056860 | ns | YPD | KS.1 |
| FIM1_1931 | 7480.981501 | 6965.073773 | 12554.5319 | -0.100325473 | 0.075025575 | 0.174303867 | 0.234685179 | ns | YPD | KS.1 |
| FIM1_1932 | 722.4664365 | 534.9642798 | 615.3969019 | -0.413988292 | 0.120225195 | 0.000380107 | 0.000907404 | ns | YPD | KS.1 |
| FIM1_1933 | 4204.922468 | 5926.330004 | 5025.61618 | 0.4729286 | 0.125143322 | 9.36439065916 | 0.000245846 | ns | YPD | KS.1 |
| FIM1_1934 | 2448.875138 | 720.098898 | 592.1345006 | -1.70593956 | 0.179312705 | 8.71962269465 | 1.39485925742 | Km<KS.1 | YPD | KS.1 |

| | | | | | | | | | | |
|---|---|---|---|---|---|---|---|---|---|---|
| FIM1_1935 | 14224.82745 | 2836.268204 | 2584.393175 | -2.276104893 | 0.164427485 | 1.2455376135 | 8.9805067061 | Km<KS.1 | YPD | KS.1 |
| FIM1_1936 | 4450.049985 | 6202.622277 | 5334.781665 | 0.473394215 | 0.060095685 | 1.9607233448 | 1.7053494126 | ns | YPD | KS.1 |
| FIM1_1937 | 2561.694296 | 1846.541234 | 2724.981043 | -0.45730381 | 0.103726656 | 6.3737852859 | 1.9930598238 | ns | YPD | KS.1 |
| FIM1_1938 | 367.7374293 | 768.8573708 | 1245.841524 | 1.010098915 | 0.175358636 | 1.5756420625 | 8.0069655375 | Km>KS.1 | YPD | KS.1 |
| FIM1_1939 | 804.9404304 | 207.2916126 | 307.9868761 | -1.851855941 | 0.238417535 | 2.6581922446 | 2.5285863130 | Km<KS.1 | YPD | KS.1 |
| FIM1_1940 | 2532.850634 | 3271.130658 | 3000.403003 | 0.31745111 | 0.220910565 | 0.115368081 | 0.164175116 | ns | YPD | KS.1 |
| FIM1_1941 | 1434.196023 | 2474.946948 | 1548.488475 | 0.683933652 | 0.257065458 | 0.002997039 | 0.006095775 | ns | YPD | KS.1 |
| FIM1_1942 | 1589.016812 | 1173.911448 | 919.5417893 | -0.401901194 | 0.165694436 | 0.010292187 | 0.018742177 | ns | YPD | KS.1 |
| FIM1_1943 | 485.3322849 | 1034.213847 | 1103.740139 | 1.04523848 | 0.159932663 | 1.1098275849 | 6.9103782669 | Km>KS.1 | YPD | KS.1 |
| FIM1_1944 | 440.9740919 | 616.7872795 | 568.8789386 | 0.46516716 | 0.1148732 | 3.1012256896 | 8.7962359212 | ns | YPD | KS.1 |
| FIM1_1945 | 1237.059669 | 679.4837314 | 776.3111489 | -0.822307224 | 0.158102351 | 5.4763385600 | 2.2933320148 | ns | YPD | KS.1 |
| FIM1_1946 | 3608.671454 | 3150.521775 | 2199.170287 | -0.179425539 | 0.15219201 | 0.215239817 | 0.281942625 | ns | YPD | KS.1 |
| FIM1_1947 | 555.1094238 | 828.1368288 | 995.8216166 | 0.532530999 | 0.172556765 | 0.001071936 | 0.002361772 | ns | YPD | KS.1 |
| FIM1_1948 | 6417.823218 | 2353.997189 | 2939.991438 | -1.423614021 | 0.111863589 | 3.5995530943 | 1.6429152884 | Km<KS.1 | YPD | KS.1 |
| FIM1_1949 | 171.6254456 | 281.6964339 | 448.1257848 | 0.667707997 | 0.175776385 | 5.7450526689 | 0.000156956 | ns | YPD | KS.1 |
| FIM1_1950 | 736.0903692 | 1113.124814 | 1066.766058 | 0.572648845 | 0.121967661 | 1.2870665121 | 4.4590222129 | ns | YPD | KS.1 |
| FIM1_1951 | 3539.004255 | 1110.661946 | 1505.63966 | -1.633420487 | 0.143344896 | 2.4550902076 | 7.5863812317 | Km<KS.1 | YPD | KS.1 |
| FIM1_1952 | 3712.302068 | 2554.367027 | 2539.892516 | -0.523997515 | 0.101168623 | 1.1899269938 | 4.7549291520 | ns | YPD | KS.1 |
| FIM1_1953 | 8172.662309 | 4599.213718 | 3001.487266 | -0.748341882 | 0.224395266 | 0.0002832 | 0.000693365 | ns | YPD | KS.1 |
| FIM1_1954 | 1900.19204 | 1302.833978 | 2132.792289 | -0.369179982 | 0.389614303 | 0.236367356 | 0.304250349 | ns | YPD | KS.1 |
| FIM1_1955 | 1028.190918 | 869.9790709 | 615.9159347 | -0.230737564 | 0.10659853 | 0.02600486 | 0.043153496 | ns | YPD | KS.1 |
| FIM1_1956 | 226.2559098 | 707.8495229 | 737.4129827 | 1.564031367 | 0.213160956 | 1.4036974722 | 1.1173431878 | Km>KS.1 | YPD | KS.1 |
| FIM1_1957 | 894.0471119 | 1203.758025 | 1281.058434 | 0.403640569 | 0.13996518 | 0.002640824 | 0.005420786 | ns | YPD | KS.1 |
| FIM1_1958 | 1326.906082 | 1449.010903 | 1017.167306 | 0.114955592 | 0.155101897 | 0.435587432 | 0.512559298 | ns | YPD | KS.1 |
| FIM1_1959 | 859.2508729 | 916.4667474 | 825.9505333 | 0.087579714 | 0.116654837 | 0.438272778 | 0.515219062 | ns | YPD | KS.1 |
| FIM1_1960 | 1896.725983 | 2914.819552 | 2838.54034 | 0.609524392 | 0.07634896 | 6.1445974460 | 5.6193699069 | ns | YPD | KS.1 |
| FIM1_1961 | 13767.11522 | 6427.112087 | 3122.551629 | -1.032389047 | 0.196619011 | 2.7202676385 | 1.1902666228 | Km<KS.1 | YPD | KS.1 |
| FIM1_1962 | 26618.85673 | 30581.75794 | 36346.13897 | 0.189058665 | 0.119977207 | 0.102944211 | 0.148233705 | ns | YPD | KS.1 |
| FIM1_1963 | 367.7166498 | 601.4461422 | 534.5366959 | 0.666913465 | 0.164787736 | 2.0460839611 | 5.9458339409 | ns | YPD | KS.1 |
| FIM1_1964 | 4502.585357 | 11349.10109 | 12314.88059 | 1.229369519 | 0.244004489 | 5.7015549175 | 2.3796338687 | Km>KS.1 | YPD | KS.1 |
| FIM1_1965 | 1183.927336 | 1879.484244 | 1120.310801 | 0.636089018 | 0.138283267 | 1.7759918481 | 6.0083997610 | ns | YPD | KS.1 |
| FIM1_1966 | 1979.122646 | 1559.79849 | 1545.494868 | -0.32800774 | 0.117014635 | 0.003824447 | 0.007619793 | ns | YPD | KS.1 |
| FIM1_1967 | 3144.525035 | 3256.944009 | 2172.736869 | 0.047282718 | 0.122230224 | 0.689019887 | 0.750410232 | ns | YPD | KS.1 |
| FIM1_1968 | 1157.507251 | 2365.801213 | 2275.329364 | 0.942633121 | 0.229673495 | 8.7675083360 | 2.6794105379 | ns | YPD | KS.1 |
| FIM1_1970 | 223.5069124 | 357.7665304 | 570.7668803 | 0.627348323 | 0.1747542 | 0.000142941 | 0.000365057 | ns | YPD | KS.1 |
| FIM1_1971 | 242.905432 | 373.4202946 | 411.3584587 | 0.575834394 | 0.165222506 | 0.000237472 | 0.000590122 | ns | YPD | KS.1 |
| FIM1_1972 | 17136.34734 | 3951.836387 | 3532.771367 | -1.932653639 | 0.318566611 | 3.3927764627 | 2.0166144446 | Km<KS.1 | YPD | KS.1 |
| FIM1_1973 | 11800.98484 | 8024.693175 | 7261.775774 | -0.472560336 | 0.250003915 | 0.034816551 | 0.056143894 | ns | YPD | KS.1 |
| FIM1_1975 | 1295.435909 | 1135.161968 | 1218.097813 | -0.182557321 | 0.103670068 | 0.070395914 | 0.105806547 | ns | YPD | KS.1 |
| FIM1_1976 | 3735.369852 | 4257.277149 | 2935.738703 | 0.174556189 | 0.141602438 | 0.197459449 | 0.26182323 | ns | YPD | KS.1 |
| FIM1_1977 | 2645.291481 | 1091.507188 | 1034.614862 | -1.261591682 | 0.090718403 | 6.6448938444 | 4.5285406679 | Km<KS.1 | YPD | KS.1 |
| FIM1_1978 | 2663.451614 | 3048.553565 | 2647.71356 | 0.16226029 | 0.23154514 | 0.440001794 | 0.517129442 | ns | YPD | KS.1 |
| FIM1_1979 | 1051.195706 | 1047.631883 | 814.5126033 | -0.004350434 | 0.078682262 | 0.955181 | 0.967825963 | ns | YPD | KS.1 |
| FIM1_1980 | 2643.151903 | 7695.794701 | 8603.960782 | 1.476294086 | 0.189454241 | 5.1005827944 | 4.6818080078 | Km>KS.1 | YPD | KS.1 |
| FIM1_1981 | 3246.247281 | 5718.821346 | 10239.13225 | 0.748179848 | 0.206028564 | 9.3320147660 | 0.000245255 | ns | YPD | KS.1 |

| | | | | | | | | | | |
|---|---|---|---|---|---|---|---|---|---|---|
| FIM1_1982 | 341.0480091 | 129.1408558 | 64.93626001 | -1.109746657 | 0.410742583 | 0.00100129 | 0.002219882 | Km<KS.1 | YPD | KS.1 |
| FIM1_1983 | 4849.698268 | 4928.23449 | 4897.795103 | 0.021548827 | 0.116433573 | 0.848573437 | 0.882821592 | ns | YPD | KS.1 |
| FIM1_1984 | 469.650305 | 462.3504503 | 569.3680288 | -0.020929908 | 0.109145416 | 0.843653237 | 0.878989498 | ns | YPD | KS.1 |
| FIM1_1985 | 2723.99115 | 3081.793071 | 2826.378344 | 0.168399747 | 0.116855983 | 0.136442607 | 0.189662467 | ns | YPD | KS.1 |
| FIM1_1986 | 2005.892766 | 2061.842963 | 2153.313448 | 0.038585584 | 0.083677802 | 0.638629677 | 0.705256968 | ns | YPD | KS.1 |
| FIM1_1987 | 166.1243227 | 833.4011343 | 767.2735327 | 2.240557469 | 0.210891543 | 2.4391423991 | 6.0072937801 | Km>KS.1 | YPD | KS.1 |
| FIM1_1988 | 968.6299238 | 2977.400178 | 2531.306992 | 1.560198164 | 0.179176988 | 2.0319848097 | 2.5336151449 | Km>KS.1 | YPD | KS.1 |
| FIM1_1989 | 4198.449182 | 1574.338798 | 1456.817408 | -1.342761986 | 0.200797655 | 2.2738113893 | 1.5123277622 | Km<KS.1 | YPD | KS.1 |
| FIM1_1990 | 2118.184291 | 2184.90305 | 2173.972986 | 0.042729932 | 0.109435983 | 0.687918602 | 0.749616258 | ns | YPD | KS.1 |
| FIM1_1991 | 2864.811903 | 2785.828858 | 2246.902047 | -0.035239978 | 0.184917235 | 0.838642793 | 0.875419197 | ns | YPD | KS.1 |
| FIM1_1992 | 5138.84671 | 5693.173685 | 4555.279446 | 0.125747005 | 0.212544195 | 0.519876778 | 0.595804416 | ns | YPD | KS.1 |
| FIM1_1993 | 1747.507194 | 2038.470072 | 1575.670143 | 0.202244957 | 0.158461813 | 0.178204122 | 0.239288936 | ns | YPD | KS.1 |
| FIM1_1994 | 11559.50489 | 14808.59777 | 15695.74925 | 0.349636564 | 0.079137977 | 7.3172237898 | 2.2665173570 | ns | YPD | KS.1 |
| FIM1_1995 | 5049.616988 | 5070.106995 | 3370.377933 | 0.004917611 | 0.219150859 | 0.98046181 | 0.98601122 | ns | YPD | KS.1 |
| FIM1_1996 | 4997.678313 | 5235.928093 | 7937.596766 | 0.061567452 | 0.144762853 | 0.656498031 | 0.721306913 | ns | YPD | KS.1 |
| FIM1_1997 | 1026.599267 | 1275.255539 | 1520.024576 | 0.302932848 | 0.095874844 | 0.00123524 | 0.002687065 | ns | YPD | KS.1 |
| FIM1_1998 | 736.4516545 | 1533.930545 | 1350.747153 | 1.043528129 | 0.088224481 | 4.8076499125 | 1.6382231722 | Km>KS.1 | YPD | KS.1 |
| FIM1_1999 | 738.5062124 | 849.4267994 | 411.8301405 | 0.175792274 | 0.196453326 | 0.334898313 | 0.40966784 | ns | YPD | KS.1 |
| FIM1_2000 | 1738.993557 | 4806.023105 | 5751.658379 | 1.408861083 | 0.177864769 | 2.0907481703 | 2.0080062060 | Km>KS.1 | YPD | KS.1 |
| FIM1_2001 | 3043.389715 | 2079.865286 | 3238.179242 | -0.452496723 | 0.272106422 | 0.059142004 | 0.091206284 | ns | YPD | KS.1 |
| FIM1_2002 | 1501.401332 | 2782.682575 | 3293.617611 | 0.867353749 | 0.11623704 | 2.0982447558 | 1.6517037437 | ns | YPD | KS.1 |
| FIM1_2003 | 1160.169206 | 4652.727357 | 4467.052273 | 1.957092443 | 0.156353573 | 1.4657676106 | 6.3410381415 | Km>KS.1 | YPD | KS.1 |
| FIM1_2004 | 948.5246705 | 520.0034151 | 610.1034241 | -0.818172527 | 0.170494608 | 4.4724060824 | 1.6617042763 | ns | YPD | KS.1 |
| FIM1_2005 | 10254.19801 | 31570.60103 | 32739.38233 | 1.563357067 | 0.178991929 | 1.5951787024 | 2.0142167625 | Km>KS.1 | YPD | KS.1 |
| FIM1_2006 | 932.4841974 | 1533.28484 | 869.3747403 | 0.686711568 | 0.136487997 | 1.8186779945 | 7.1299629808 | ns | YPD | KS.1 |
| FIM1_2007 | 584.2434162 | 557.692284 | 634.0629093 | -0.062540178 | 0.124476746 | 0.602804349 | 0.67316535 | ns | YPD | KS.1 |
| FIM1_2008 | 687.2604155 | 641.2980045 | 575.9856178 | -0.091202411 | 0.148960417 | 0.520977337 | 0.596928201 | ns | YPD | KS.1 |
| FIM1_2009 | 5050.3022 | 401.5404997 | 814.4961066 | -3.526422748 | 0.299848146 | 7.1935437091 | 2.6908180415 | Km<KS.1 | YPD | KS.1 |
| FIM1_2010 | 374.191829 | 81.48547142 | 78.59222337 | -2.089726312 | 0.244618275 | 2.2667237927 | 2.8052116589 | Km<KS.1 | YPD | KS.1 |
| FIM1_2011 | 2922.547989 | 3312.016743 | 2751.075001 | 0.175266655 | 0.082631008 | 0.030680097 | 0.04997822 | ns | YPD | KS.1 |
| FIM1_2012 | 8189.503002 | 3419.358242 | 3168.947475 | -1.233912211 | 0.119781363 | 8.5500573258 | 1.7638591541 | Km<KS.1 | YPD | KS.1 |
| FIM1_2013 | 1833.670321 | 1406.970151 | 1113.463587 | -0.333594126 | 0.209448834 | 0.083254228 | 0.122759272 | ns | YPD | KS.1 |
| FIM1_2014 | 2470.533923 | 2030.799517 | 1707.956662 | -0.271898273 | 0.102888603 | 0.00669296 | 0.012641411 | ns | YPD | KS.1 |
| FIM1_2015 | 31.67527642 | 9.122806003 | 21.92099029 | -1.647695063 | 0.291250081 | 7.9204382518 | 4.1304172225 | Km<KS.1 | YPD | KS.1 |
| FIM1_2016 | 4496.480046 | 2818.813133 | 2361.274511 | -0.658170872 | 0.097299065 | 5.2252626516 | 3.3542815086 | ns | YPD | KS.1 |
| FIM1_2017 | 2508.550786 | 1454.577763 | 1226.622931 | -0.725150361 | 0.194425006 | 6.6714929659 | 0.000179993 | ns | YPD | KS.1 |
| FIM1_2018 | 374.309302 | 127.3277135 | 205.6604993 | -1.3395855 | 0.351010228 | 1.2438786556 | 3.7301364145 | Km<KS.1 | YPD | KS.1 |
| FIM1_2019 | 2513.989773 | 1374.664437 | 1473.133084 | -0.783223237 | 0.231931019 | 0.000225138 | 0.000561433 | ns | YPD | KS.1 |
| FIM1_2020 | 4127.497868 | 4248.759298 | 3206.016417 | 0.036317506 | 0.192470401 | 0.839504459 | 0.875838945 | ns | YPD | KS.1 |
| FIM1_2021 | 4696.641346 | 3834.654355 | 4196.697368 | -0.260150694 | 0.18634625 | 0.134522008 | 0.187097285 | ns | YPD | KS.1 |
| FIM1_2022 | 825.9838633 | 1130.994958 | 1584.664656 | 0.422835845 | 0.151685693 | 0.003453382 | 0.006944453 | ns | YPD | KS.1 |
| FIM1_2023 | 2399.316433 | 3521.800694 | 2191.806579 | 0.513913237 | 0.165999725 | 0.001076828 | 0.00237045 | ns | YPD | KS.1 |
| FIM1_2024 | 1761.604973 | 848.0968 | 535.9085137 | -0.638930359 | 0.508612873 | 0.083387233 | 0.122918958 | ns | YPD | KS.1 |
| FIM1_2025 | 213.794976 | 171.6767325 | 215.5552265 | -0.282242491 | 0.194536016 | 0.118037199 | 0.167542102 | ns | YPD | KS.1 |
| FIM1_2026 | 1423.849373 | 1190.389436 | 1235.45582 | -0.249167479 | 0.093166182 | 0.006255677 | 0.011878623 | ns | YPD | KS.1 |

| | | | | | | | | | | |
|---|---|---|---|---|---|---|---|---|---|---|
| FIM1_2027 | 163.193664 | 670.2337077 | 972.3894514 | 2.001460086 | 0.128669101 | 3.0183118326 | 3.6624637481 | Km>KS.1 | YPD | KS.1 |
| FIM1_2028 | 639.5836655 | 1634.079369 | 2123.234624 | 1.292050384 | 0.185483612 | 3.6029811694 | 2.5533947746 | Km>KS.1 | YPD | KS.1 |
| FIM1_2029 | 2113.173896 | 1772.928283 | 2108.789834 | -0.245138918 | 0.093820829 | 0.00754014 | 0.014091734 | ns | YPD | KS.1 |
| FIM1_2030 | 4705.402503 | 1415.877072 | 1023.682246 | -1.661765689 | 0.195569857 | 1.0199503534 | 1.1911392038 | Km<KS.1 | YPD | KS.1 |
| FIM1_2031 | 3959.09797 | 3545.869193 | 1632.790141 | -0.125222595 | 0.268626706 | 0.598107704 | 0.669122066 | ns | YPD | KS.1 |
| FIM1_2032 | 59.18119181 | 68.26911354 | 27.57214771 | 0.140511657 | 0.356931325 | 0.632216285 | 0.69905468 | ns | YPD | KS.1 |
| FIM1_2033 | 2488.527587 | 3019.317229 | 1731.269197 | 0.268701774 | 0.100212548 | 0.005989256 | 0.011433826 | ns | YPD | KS.1 |
| FIM1_2034 | 3411.411398 | 2220.415314 | 2139.809632 | -0.605879454 | 0.091145013 | 1.3141094455 | 8.1112834882 | ns | YPD | KS.1 |
| FIM1_2035 | 359.4511499 | 152.5748756 | 253.7108773 | -1.156945047 | 0.210832235 | 5.7429609014 | 2.7236635352 | Km<KS.1 | YPD | KS.1 |
| FIM1_2037 | 135.6572282 | 233.9907684 | 314.2585225 | 0.694005017 | 0.238546876 | 0.001359501 | 0.00293683 | ns | YPD | KS.1 |
| FIM1_2038 | 2185.266877 | 2930.164517 | 1938.2021 | 0.385661297 | 0.175069301 | 0.019083527 | 0.032783339 | ns | YPD | KS.1 |
| FIM1_2039 | 737.9496858 | 347.0063051 | 620.846494 | -1.027682109 | 0.188021517 | 8.3529153921 | 3.8764695966 | Km<KS.1 | YPD | KS.1 |
| FIM1_2040 | 346.2573771 | 277.112252 | 447.3371595 | -0.292277864 | 0.168931468 | 0.066380246 | 0.100714157 | ns | YPD | KS.1 |
| FIM1_2041 | 5034.015205 | 3659.794729 | 5070.218881 | -0.430214494 | 0.149937518 | 0.002641601 | 0.005420786 | ns | YPD | KS.1 |
| FIM1_2042 | 2806.190994 | 3092.341865 | 3704.298972 | 0.134383319 | 0.095002183 | 0.147588794 | 0.202888713 | ns | YPD | KS.1 |
| FIM1_2043 | 1628.559729 | 1552.227375 | 2435.732405 | -0.060111365 | 0.192344625 | 0.73727412 | 0.791475648 | ns | YPD | KS.1 |
| FIM1_2044 | 709.4539269 | 905.6583137 | 810.1224928 | 0.329912969 | 0.143229158 | 0.016015842 | 0.028006613 | ns | YPD | KS.1 |
| FIM1_2045 | 504.6852188 | 311.6018064 | 395.9114986 | -0.654342596 | 0.158846829 | 1.5405068944 | 4.5483808901 | ns | YPD | KS.1 |
| FIM1_2046 | 297.612095 | 230.9793673 | 265.7848338 | -0.314654372 | 0.222824943 | 0.121335692 | 0.171538808 | ns | YPD | KS.1 |
| FIM1_2047 | 1673.312264 | 968.3478019 | 771.6282506 | -0.739409342 | 0.174532058 | 7.6108878336 | 2.3474375060 | ns | YPD | KS.1 |
| FIM1_2048 | 1460.777739 | 1512.689172 | 1837.727007 | 0.04839759 | 0.097974974 | 0.61296731 | 0.682517176 | ns | YPD | KS.1 |
| FIM1_2049 | 301.9936602 | 293.5053015 | 419.7850415 | -0.036692517 | 0.121795626 | 0.755364077 | 0.806943586 | ns | YPD | KS.1 |
| FIM1_2050 | 2957.446196 | 2664.963482 | 3690.857324 | -0.144543366 | 0.097396308 | 0.128300677 | 0.179599287 | ns | YPD | KS.1 |
| FIM1_2051 | 1996.048297 | 2038.082413 | 2042.719606 | 0.028044585 | 0.112038881 | 0.796589158 | 0.840159225 | ns | YPD | KS.1 |
| FIM1_2052 | 3627.852668 | 2173.925966 | 3036.137461 | -0.686300748 | 0.181336808 | 5.8334427124 | 0.000159283 | ns | YPD | KS.1 |
| FIM1_2053 | 3978.65666 | 6276.762863 | 6991.329792 | 0.636725404 | 0.113326662 | 7.9943249000 | 3.7274382734 | ns | YPD | KS.1 |
| FIM1_2054 | 742.6129845 | 1075.623176 | 1495.778769 | 0.513825042 | 0.118739162 | 8.2737840320 | 2.5377358544 | ns | YPD | KS.1 |
| FIM1_2055 | 1149.440631 | 1308.914436 | 893.0585356 | 0.135698417 | 0.322976201 | 0.61989208 | 0.6888459 | ns | YPD | KS.1 |
| FIM1_2056 | 1179.63798 | 1236.829843 | 844.1085238 | 0.059340104 | 0.194000199 | 0.742412549 | 0.795499124 | ns | YPD | KS.1 |
| FIM1_2057 | 1209.148435 | 749.5467653 | 693.8576477 | -0.666835778 | 0.118447586 | 6.9934778313 | 3.2854156950 | ns | YPD | KS.1 |
| FIM1_2058 | 1520.380528 | 652.5446744 | 517.6529092 | -1.084451997 | 0.281597793 | 1.8789139728 | 5.4856789994 | Km<KS.1 | YPD | KS.1 |
| FIM1_2059 | 739.5026373 | 676.1460187 | 774.0261571 | -0.119297713 | 0.139275343 | 0.371334887 | 0.447310184 | ns | YPD | KS.1 |
| FIM1_2060 | 2932.462617 | 2875.027642 | 3723.71952 | -0.021529008 | 0.295530876 | 0.933013881 | 0.95102216 | ns | YPD | KS.1 |
| FIM1_2061 | 1091.287256 | 1671.509689 | 1823.673689 | 0.580221215 | 0.1502808 | 5.3955401768 | 0.000148139 | ns | YPD | KS.1 |
| FIM1_2062 | 3461.467371 | 1859.083669 | 1165.197352 | -0.762461033 | 0.289521481 | 0.002729774 | 0.005586436 | ns | YPD | KS.1 |
| FIM1_2063 | 10646.61312 | 5335.446639 | 3667.742342 | -0.964288088 | 0.136572069 | 3.3790256781 | 2.4084029726 | ns | YPD | KS.1 |
| FIM1_2064 | 4674.155574 | 2369.872766 | 1152.213554 | -0.861606859 | 0.267740701 | 0.00033368 | 0.000805072 | ns | YPD | KS.1 |
| FIM1_2065 | 170.3441382 | 99.62845989 | 56.90405186 | -0.682811967 | 0.242382537 | 0.00186487 | 0.003953016 | ns | YPD | KS.1 |
| FIM1_2066 | 20563.06202 | 3421.722074 | 3256.199129 | -2.256727976 | 0.438069949 | 2.5948458300 | 1.2819620659 | Km<KS.1 | YPD | KS.1 |
| FIM1_2067 | 2338.40895 | 2693.88062 | 2073.776345 | 0.185568712 | 0.159079629 | 0.21842138 | 0.285134182 | ns | YPD | KS.1 |
| FIM1_2068 | 3273.321201 | 1693.866943 | 1436.187791 | -0.928303312 | 0.111705932 | 2.0713248736 | 2.1743182967 | ns | YPD | KS.1 |
| FIM1_2069 | 2997.787204 | 2074.158653 | 1555.198042 | -0.506766039 | 0.129847813 | 5.2935197137 | 0.000145579 | ns | YPD | KS.1 |
| FIM1_2070 | 66.3334094 | 57.146728 | 80.28421507 | -0.17922655 | 0.244690412 | 0.41530552 | 0.492760544 | ns | YPD | KS.1 |
| FIM1_2071 | 2325.389304 | 2109.555745 | 2107.247229 | -0.137125178 | 0.06920406 | 0.044493175 | 0.070092952 | ns | YPD | KS.1 |
| FIM1_2072 | 1037.375661 | 1109.306659 | 1485.603666 | 0.089298755 | 0.138004275 | 0.499894776 | 0.576489688 | ns | YPD | KS.1 |

| | | | | | | | | | | |
|---|---|---|---|---|---|---|---|---|---|---|
| FIM1_2073 | 1799.921104 | 2091.413116 | 3368.971596 | 0.18694986 | 0.20507686 | 0.323279463 | 0.397212973 | ns | YPD | KS.1 |
| FIM1_2074 | 932.3502216 | 734.729263 | 831.1518814 | -0.330219482 | 0.105259368 | 0.001285531 | 0.002786717 | ns | YPD | KS.1 |
| FIM1_2075 | 3774.685225 | 4687.64919 | 6104.843797 | 0.291530497 | 0.141081726 | 0.030822091 | 0.050160257 | ns | YPD | KS.1 |
| FIM1_2076 | 1021.10849 | 1350.655426 | 1345.166444 | 0.386131914 | 0.116128707 | 0.00061263 | 0.001415622 | ns | YPD | KS.1 |
| FIM1_2077 | 2959.721674 | 3052.423044 | 2543.597471 | 0.03943908 | 0.177477526 | 0.813095218 | 0.853687127 | ns | YPD | KS.1 |
| FIM1_2078 | 785.051005 | 1866.942196 | 1355.849322 | 1.185583663 | 0.190456901 | 6.4652579644( | 3.7184576153( | Km>KS.1 | YPD | KS.1 |
| FIM1_2079 | 1071.360244 | 1022.348278 | 1965.955201 | -0.050541634 | 0.298237757 | 0.844632655 | 0.87947331 | ns | YPD | KS.1 |
| FIM1_2080 | 62.07532024 | 61.4525946 | 36.46740299 | -0.014150339 | 0.287615543 | 0.955014003 | 0.967825963 | ns | YPD | KS.1 |
| FIM1_2081 | 8596.185303 | 5759.993932 | 11195.6198 | -0.499919813 | 0.237009185 | 0.01967023 | 0.033628658 | ns | YPD | KS.1 |
| FIM1_2083 | 1517.98016 | 1028.864875 | 754.0544526 | -0.486757562 | 0.232674706 | 0.021048046 | 0.035617017 | ns | YPD | KS.1 |
| FIM1_2084 | 2576.311131 | 4975.454286 | 6222.755823 | 0.940176469 | 0.069549682 | 2.5183815795 | 1.4568544602( | ns | YPD | KS.1 |
| FIM1_2085 | 2423.289649 | 2801.941878 | 2721.631889 | 0.203907116 | 0.076158241 | 0.006511219 | 0.012321534 | ns | YPD | KS.1 |
| FIM1_2086 | 1567.084793 | 3236.793428 | 3151.420719 | 1.029047209 | 0.096354436 | 2.2470300041 | 5.2238197527( | Km>KS.1 | YPD | KS.1 |
| FIM1_2087 | 1899.061738 | 3782.4712 | 3960.918562 | 0.858953793 | 0.286372764 | 0.000703776 | 0.001606093 | ns | YPD | KS.1 |
| FIM1_2088 | 1277.686323 | 1610.23268 | 1507.92372 | 0.311990819 | 0.142213386 | 0.021830146 | 0.036815246 | ns | YPD | KS.1 |
| FIM1_2089 | 459.8720268 | 287.7296404 | 349.9156491 | -0.63403545 | 0.164411353 | 4.8902773053 | 0.000135087 | ns | YPD | KS.1 |
| FIM1_2090 | 1248.026193 | 881.3095788 | 845.9294823 | -0.487295004 | 0.099042171 | 4.9965908249 | 1.8399733052 | ns | YPD | KS.1 |
| FIM1_2091 | 308.8227084 | 662.7789672 | 1214.776412 | 1.065126715 | 0.144127762 | 2.4698031509 | 1.9228905596 | Km>KS.1 | YPD | KS.1 |
| FIM1_2092 | 469.9077733 | 295.7909632 | 677.1625973 | -0.555056307 | 0.280512273 | 0.024149914 | 0.040385151 | ns | YPD | KS.1 |
| FIM1_2093 | 2448.834273 | 1731.932358 | 3388.232517 | -0.392421482 | 0.299431509 | 0.128759986 | 0.180191542 | ns | YPD | KS.1 |
| FIM1_2094 | 3237.545092 | 5270.819073 | 8107.293509 | 0.66686482 | 0.150928055 | 3.9079492804 | 1.2583849624 | ns | YPD | KS.1 |
| FIM1_2095 | 1894.018342 | 1522.39222 | 1015.029635 | -0.285037765 | 0.172653372 | 0.079160813 | 0.117275301 | ns | YPD | KS.1 |
| FIM1_2096 | 2374.657513 | 2603.837443 | 1200.383166 | 0.110459744 | 0.228921686 | 0.595821833 | 0.667315088 | ns | YPD | KS.1 |
| FIM1_2097 | 364.1604119 | 620.9209646 | 1071.318606 | 0.751702311 | 0.090973763 | 4.4318247857 | 4.5366930676 | ns | YPD | KS.1 |
| FIM1_2098 | 9752.552442 | 607.2470381 | 1283.458349 | -3.740195943 | 0.444185718 | 4.3836014231( | 5.7541997572 | Km<KS.1 | YPD | KS.1 |
| FIM1_2099 | 14.20286996 | 8.56454576 | 15.19172552 | -0.504879884 | 0.400394517 | 0.113021741 | 0.161297522 | ns | YPD | KS.1 |
| FIM1_2100 | 1237.248999 | 1207.973061 | 1852.733756 | -0.030899083 | 0.163111556 | 0.841523659 | 0.876954378 | ns | YPD | KS.1 |
| FIM1_2101 | 452.9057712 | 686.2874063 | 761.0685894 | 0.570116781 | 0.14103875 | 2.5813074782 | 7.4017318181 | ns | YPD | KS.1 |
| FIM1_2102 | 1836.777775 | 1093.040698 | 1111.61358 | -0.72215272 | 0.125610507 | 3.0657411645 | 1.4997111400( | ns | YPD | KS.1 |
| FIM1_2103 | 6102.820727 | 8470.600758 | 5729.274012 | 0.400927914 | 0.242448495 | 0.066040248 | 0.100290059 | ns | YPD | KS.1 |
| FIM1_2104 | 247.3895928 | 424.8775449 | 766.9407679 | 0.704079035 | 0.221014116 | 0.000529147 | 0.001233602 | ns | YPD | KS.1 |
| FIM1_2105 | 254.715085 | 321.2169779 | 459.8558034 | 0.307774302 | 0.163897303 | 0.046960493 | 0.073653359 | ns | YPD | KS.1 |
| FIM1_2106 | 781.4722728 | 1096.328107 | 901.2688859 | 0.444721004 | 0.180676585 | 0.008648808 | 0.015977654 | ns | YPD | KS.1 |
| FIM1_2107 | 890.4733078 | 963.796738 | 817.3913675 | 0.105960782 | 0.122921561 | 0.372036759 | 0.44793874 | ns | YPD | KS.1 |
| FIM1_2108 | 475.0092863 | 598.6009667 | 655.523916 | 0.309517918 | 0.148188252 | 0.028489512 | 0.046839167 | ns | YPD | KS.1 |
| FIM1_2109 | 158.9655566 | 155.9570422 | 233.2353216 | -0.025552177 | 0.18389001 | 0.882043706 | 0.910030576 | ns | YPD | KS.1 |
| FIM1_2110 | 280.9906122 | 405.841495 | 423.3050302 | 0.485897893 | 0.17569662 | 0.003291951 | 0.006643999 | ns | YPD | KS.1 |
| FIM1_2111 | 43.0023895 | 55.85733177 | 44.30100201 | 0.29607898 | 0.273220839 | 0.218135613 | 0.284985471 | ns | YPD | KS.1 |
| FIM1_2112 | 2404.440924 | 1027.123407 | 1126.871592 | -1.193779154 | 0.13630764 | 2.6416440509 | 3.2369899392 | Km<KS.1 | YPD | KS.1 |
| FIM1_2113 | 1276.681025 | 1677.591888 | 1409.47737 | 0.375041108 | 0.123374855 | 0.001668019 | 0.003556963 | ns | YPD | KS.1 |
| FIM1_2115 | 1009.839772 | 954.0497537 | 1230.235367 | -0.077436402 | 0.110759832 | 0.471584828 | 0.549446023 | ns | YPD | KS.1 |
| FIM1_2116 | 1213.879194 | 1301.339922 | 1207.977676 | 0.097425399 | 0.069904979 | 0.157544113 | 0.214793632 | ns | YPD | KS.1 |
| FIM1_2117 | 2389.747138 | 2570.369715 | 2470.402507 | 0.102710055 | 0.070593522 | 0.139967116 | 0.193641936 | ns | YPD | KS.1 |
| FIM1_2118 | 2741.371474 | 1740.897354 | 1670.173672 | -0.649235758 | 0.056616338 | 7.3773239675 | 2.1589521610 | ns | YPD | KS.1 |
| FIM1_2119 | 649.8585592 | 363.5399146 | 891.3692608 | -0.688599929 | 0.309639979 | 0.009950803 | 0.018187084 | ns | YPD | KS.1 |

| | | | | | | | | | | |
|---|---|---|---|---|---|---|---|---|---|---|
| FIM1_2120 | 653.3967489 | 1105.549415 | 1712.28105 | 0.724702169 | 0.142805237 | 1.3297937694 | 5.3010608999 | ns | YPD | KS.1 |
| FIM1_2121 | 4591.699428 | 3889.629655 | 3712.965937 | -0.230447284 | 0.098282008 | 0.01626815 | 0.028417853 | ns | YPD | KS.1 |
| FIM1_2122 | 802.9594051 | 1106.232358 | 1269.252022 | 0.444118341 | 0.114659727 | 6.7417621965 | 0.000181692 | ns | YPD | KS.1 |
| FIM1_2123 | 10055.52194 | 10275.02971 | 7668.136712 | 0.029438068 | 0.110407892 | 0.783791677 | 0.830379381 | ns | YPD | KS.1 |
| FIM1_2124 | 5461.684303 | 3426.273639 | 2574.161137 | -0.607986208 | 0.206563922 | 0.001467429 | 0.003154909 | ns | YPD | KS.1 |
| FIM1_2125 | 2740.177337 | 1572.232593 | 1307.919858 | -0.761420877 | 0.155224252 | 2.9542495230 | 1.1305685674 | ns | YPD | KS.1 |
| FIM1_2126 | 1644.751414 | 935.1696635 | 1140.724034 | -0.772044689 | 0.1603474 | 4.5633384435 | 1.6930023887 | ns | YPD | KS.1 |
| FIM1_2127 | 2048.999511 | 1806.343166 | 1172.311572 | -0.175118418 | 0.096048884 | 0.061835636 | 0.094918941 | ns | YPD | KS.1 |
| FIM1_2128 | 499.6542544 | 585.8950236 | 763.6469738 | 0.207918645 | 0.161068667 | 0.172683037 | 0.233048747 | ns | YPD | KS.1 |
| FIM1_2129 | 847.7536247 | 2081.566442 | 3207.282133 | 1.26851598 | 0.123019335 | 7.2220586238 | 1.5096530106 | Km>KS.1 | YPD | KS.1 |
| FIM1_2130 | 21411.94667 | 4160.797623 | 4591.296402 | -2.266727489 | 0.230600489 | 8.3067532527 | 1.5833753805 | Km<KS.1 | YPD | KS.1 |
| FIM1_2131 | 4149.42278 | 8165.558161 | 10641.61632 | 0.938553594 | 0.148374174 | 5.4319465293 | 3.1496426554 | ns | YPD | KS.1 |
| FIM1_2132 | 2178.03819 | 1001.85951 | 2400.954067 | -0.889471285 | 0.372852158 | 0.004114087 | 0.008147923 | ns | YPD | KS.1 |
| FIM1_2133 | 779.2662971 | 934.9046092 | 1195.454349 | 0.25263472 | 0.101056321 | 0.010344863 | 0.018824321 | ns | YPD | KS.1 |
| FIM1_2134 | 521.8096745 | 685.4286715 | 923.6418429 | 0.368227446 | 0.139792893 | 0.006002744 | 0.011455179 | ns | YPD | KS.1 |
| FIM1_2135 | 2181.97281 | 2034.108586 | 2432.676187 | -0.094994649 | 0.117164665 | 0.402398822 | 0.47927559 | ns | YPD | KS.1 |
| FIM1_2136 | 1129.047664 | 1581.495992 | 1894.50463 | 0.473125959 | 0.096860767 | 6.1578414713 | 2.2509376429 | ns | YPD | KS.1 |
| FIM1_2137 | 479.3090819 | 505.6713123 | 929.025873 | 0.069696377 | 0.140335977 | 0.604268695 | 0.674532373 | ns | YPD | KS.1 |
| FIM1_2138 | 399.5743611 | 1066.193075 | 1077.161676 | 1.334994992 | 0.211498 | 2.7790786274 | 1.6671200487 | Km>KS.1 | YPD | KS.1 |
| FIM1_2139 | 2371.271228 | 1446.090128 | 2822.038722 | -0.680016978 | 0.144330645 | 9.3534064260 | 3.3191561717 | ns | YPD | KS.1 |
| FIM1_2140 | 1436.475833 | 1394.787085 | 1049.093401 | -0.040433741 | 0.103371777 | 0.688140191 | 0.749616258 | ns | YPD | KS.1 |
| FIM1_2141 | 326.8719074 | 1016.346194 | 1365.113948 | 1.592499071 | 0.153282974 | 1.6913898792 | 3.6906423901 | Km>KS.1 | YPD | KS.1 |
| FIM1_2142 | 277.883247 | 185.4069424 | 184.7391978 | -0.549186068 | 0.148719747 | 0.000113226 | 0.000292928 | ns | YPD | KS.1 |
| FIM1_2143 | 847.4128023 | 735.2323788 | 887.3095082 | -0.192018626 | 0.128817513 | 0.121387182 | 0.17156285 | ns | YPD | KS.1 |
| FIM1_2144 | 910.6203076 | 1381.978496 | 1324.008097 | 0.56895939 | 0.146006178 | 4.7678480023 | 0.000131778 | ns | YPD | KS.1 |
| FIM1_2145 | 916.0870547 | 949.0784367 | 768.7166833 | 0.042105867 | 0.226560732 | 0.838468757 | 0.875419197 | ns | YPD | KS.1 |
| FIM1_2146 | 7565.875841 | 8826.024176 | 11469.12418 | 0.214028163 | 0.096589223 | 0.023232644 | 0.03898164 | ns | YPD | KS.1 |
| FIM1_2147 | 453.9972482 | 619.8347528 | 730.851883 | 0.411733507 | 0.169239698 | 0.009920828 | 0.018152136 | ns | YPD | KS.1 |
| FIM1_2148 | 349.8706725 | 476.297486 | 554.9139861 | 0.39743355 | 0.192479913 | 0.026397764 | 0.043718001 | ns | YPD | KS.1 |
| FIM1_2149 | 851.6690014 | 919.2872294 | 947.8479098 | 0.104078795 | 0.103879828 | 0.303451533 | 0.375914188 | ns | YPD | KS.1 |
| FIM1_2150 | 1010.71738 | 1214.581825 | 1219.572483 | 0.235018731 | 0.185262033 | 0.174213386 | 0.234626854 | ns | YPD | KS.1 |
| FIM1_2151 | 1422.633046 | 806.3114691 | 1008.198779 | -0.78311721 | 0.145634784 | 2.2703200313 | 9.9954355362 | ns | YPD | KS.1 |
| FIM1_2152 | 476.7144263 | 293.3803532 | 256.6004841 | -0.674244245 | 0.12449912 | 2.3335795093 | 1.0264861237 | ns | YPD | KS.1 |
| FIM1_2153 | 627.3918962 | 445.8308125 | 445.3975149 | -0.471003646 | 0.123448679 | 8.1198329805 | 0.000214645 | ns | YPD | KS.1 |
| FIM1_2154 | 659.4637224 | 464.6904448 | 1304.304847 | -0.365545665 | 0.349599767 | 0.206912266 | 0.272541309 | ns | YPD | KS.1 |
| FIM1_2155 | 819.4457528 | 949.120791 | 789.1994816 | 0.20296232 | 0.100008253 | 0.037357184 | 0.059701892 | ns | YPD | KS.1 |
| FIM1_2156 | 734.8722602 | 723.4288026 | 1257.674473 | -0.020872504 | 0.157649604 | 0.889188659 | 0.915882728 | ns | YPD | KS.1 |
| FIM1_2157 | 1992.936491 | 3273.144941 | 3731.140676 | 0.701298159 | 0.091587096 | 6.6645004661 | 5.4712689470 | ns | YPD | KS.1 |
| FIM1_2158 | 1734.913703 | 2905.782947 | 2815.198628 | 0.721789358 | 0.116193156 | 1.7798115043 | 9.8384024823 | ns | YPD | KS.1 |
| FIM1_2159 | 153.8692949 | 99.73171128 | 86.34968685 | -0.502616883 | 0.302057428 | 0.053665815 | 0.083433572 | ns | YPD | KS.1 |
| FIM1_2160 | 2665.295352 | 1460.310014 | 1771.477552 | -0.833375828 | 0.143201952 | 1.5819134562 | 8.0306320866 | ns | YPD | KS.1 |
| FIM1_2161 | 2173.131135 | 1450.837704 | 1565.997904 | -0.577255133 | 0.056209748 | 4.4166107905 | 8.6167210520 | ns | YPD | KS.1 |
| FIM1_2162 | 690.4541747 | 501.1949478 | 767.1367039 | -0.443693958 | 0.115395285 | 7.5769894913 | 0.000201365 | ns | YPD | KS.1 |
| FIM1_2163 | 275.3175572 | 693.4320239 | 761.4008034 | 1.262105235 | 0.196646527 | 1.6040863534 | 9.8159035775 | Km>KS.1 | YPD | KS.1 |
| FIM1_2164 | 539.3774216 | 732.0624401 | 769.3607597 | 0.424461128 | 0.103090401 | 2.4943740286 | 7.1814298567 | ns | YPD | KS.1 |

| | | | | | | | | | | |
|---|---|---|---|---|---|---|---|---|---|---|
| FIM1_2165 | 1499.621773 | 2573.364036 | 2668.911091 | 0.756996732 | 0.11550774 | 1.7598862381 | 1.0729698571 | ns | YPD | KS.1 |
| FIM1_2166 | 223.7096196 | 340.7652447 | 398.1753113 | 0.52775049 | 0.234490598 | 0.01302403 | 0.023240513 | ns | YPD | KS.1 |
| FIM1_2167 | 3613.775419 | 5046.419192 | 4787.92655 | 0.461195713 | 0.120530115 | 7.9112168295 | 0.000209687 | ns | YPD | KS.1 |
| FIM1_2168 | 3754.770257 | 3681.284095 | 4647.44812 | -0.027507767 | 0.074991114 | 0.709586618 | 0.768268427 | ns | YPD | KS.1 |
| FIM1_2169 | 906.7841523 | 2713.807807 | 2427.300803 | 1.55945253 | 0.106279169 | 6.1064678544 | 5.6258662177 | Km>KS.1 | YPD | KS.1 |
| FIM1_2170 | 2935.067145 | 3495.143847 | 3230.549092 | 0.234604586 | 0.138082849 | 0.076147242 | 0.113456881 | ns | YPD | KS.1 |
| FIM1_2171 | 1140.290168 | 1035.608383 | 1110.462969 | -0.123410429 | 0.174847916 | 0.452931395 | 0.530570683 | ns | YPD | KS.1 |
| FIM1_2172 | 3749.445613 | 2699.537456 | 2057.017079 | -0.456194867 | 0.112980666 | 3.3140830588 | 9.3626139794 | ns | YPD | KS.1 |
| FIM1_2173 | 510.8607491 | 585.3042579 | 774.6059075 | 0.173169606 | 0.1842361 | 0.31463713 | 0.387935495 | ns | YPD | KS.1 |
| FIM1_2174 | 1997.648848 | 1324.125225 | 1263.218158 | -0.548670916 | 0.173800176 | 0.000815519 | 0.001833351 | ns | YPD | KS.1 |
| FIM1_2175 | 3752.402367 | 4325.569016 | 3557.19596 | 0.192975541 | 0.124149479 | 0.107028896 | 0.153335036 | ns | YPD | KS.1 |
| FIM1_2176 | 563.6011443 | 739.1760371 | 925.9356295 | 0.359310572 | 0.163203702 | 0.019966553 | 0.034088402 | ns | YPD | KS.1 |
| FIM1_2177 | 165.5287265 | 71.94327174 | 79.54727981 | -1.12066516 | 0.215668224 | 3.0668984914 | 1.3313979053 | Km<KS.1 | YPD | KS.1 |
| FIM1_2178 | 2726.863727 | 3929.208992 | 4144.390667 | 0.462983044 | 0.218453287 | 0.020572132 | 0.034918579 | ns | YPD | KS.1 |
| FIM1_2179 | 7841.813911 | 9255.632956 | 9088.135023 | 0.225830462 | 0.122206416 | 0.055597679 | 0.086194595 | ns | YPD | KS.1 |
| FIM1_2180 | 1986.227797 | 1692.152274 | 2978.44097 | -0.179067753 | 0.283535968 | 0.46992047 | 0.547635123 | ns | YPD | KS.1 |
| FIM1_2181 | 812.733905 | 2917.884436 | 2827.95635 | 1.735313645 | 0.242929321 | 3.9900923084 | 3.0539552668 | Km>KS.1 | YPD | KS.1 |
| FIM1_2182 | 7310.260361 | 972.2855701 | 1562.092515 | -2.678803545 | 0.376488352 | 3.3809856419 | 2.8654861275 | Km<KS.1 | YPD | KS.1 |
| FIM1_2183 | 3581.23527 | 3353.952393 | 2582.084018 | -0.089419839 | 0.109092364 | 0.398922132 | 0.475590128 | ns | YPD | KS.1 |
| FIM1_2184 | 3414.083635 | 3646.902923 | 2910.723947 | 0.08793038 | 0.135980398 | 0.500604981 | 0.57717492 | ns | YPD | KS.1 |
| FIM1_2185 | 9752.782954 | 4100.641326 | 3145.59594 | -1.192589681 | 0.180209937 | 4.8414751697 | 3.1119300994 | Km<KS.1 | YPD | KS.1 |
| FIM1_2186 | 847.7808392 | 1156.913757 | 1562.596842 | 0.429027793 | 0.119778446 | 0.000220032 | 0.000549529 | ns | YPD | KS.1 |
| FIM1_2187 | 1583.532758 | 1212.360696 | 1825.116442 | -0.327240178 | 0.232047836 | 0.119530451 | 0.16932346 | ns | YPD | KS.1 |
| FIM1_2188 | 4203.523273 | 4147.243775 | 4842.867628 | -0.018694024 | 0.095025473 | 0.840558924 | 0.876233752 | ns | YPD | KS.1 |
| FIM1_2189 | 292.1021436 | 559.6851834 | 1001.661405 | 0.880647382 | 0.183614672 | 3.9664979220 | 1.4870630868 | ns | YPD | KS.1 |
| FIM1_2190 | 3210.212297 | 2839.818599 | 2212.447532 | -0.167652469 | 0.113366729 | 0.127051142 | 0.178150911 | ns | YPD | KS.1 |
| FIM1_2191 | 7796.647214 | 6821.289327 | 7904.87394 | -0.187435844 | 0.081107002 | 0.018614836 | 0.032077869 | ns | YPD | KS.1 |
| FIM1_2192 | 1192.19429 | 588.6639424 | 907.4031976 | -0.965031665 | 0.175302499 | 7.5773049226 | 3.5429597734 | ns | YPD | KS.1 |
| FIM1_2193 | 3005.472186 | 2211.907242 | 4372.129069 | -0.348223163 | 0.291529079 | 0.168711098 | 0.228266988 | ns | YPD | KS.1 |
| FIM1_2194 | 694.6338463 | 420.5543684 | 674.7807856 | -0.63233041 | 0.245242803 | 0.004264727 | 0.00842812 | ns | YPD | KS.1 |
| FIM1_2195 | 1527.760305 | 675.5370561 | 827.4549321 | -1.144311572 | 0.13559211 | 4.6403733074 | 5.1074905320 | Km<KS.1 | YPD | KS.1 |
| FIM1_2196 | 771.1365487 | 404.6845713 | 346.3742841 | -0.8755495 | 0.180268411 | 2.9540437899 | 1.1305685674 | ns | YPD | KS.1 |
| FIM1_2197 | 3682.700522 | 1386.253763 | 1611.03317 | -1.384871719 | 0.115396669 | 3.2401126335 | 1.1513971679 | Km<KS.1 | YPD | KS.1 |
| FIM1_2198 | 3127.706279 | 2117.271641 | 2316.489743 | -0.487621567 | 0.234871026 | 0.021845669 | 0.036828941 | ns | YPD | KS.1 |
| FIM1_2199 | 148.2715541 | 122.7851426 | 102.5282884 | -0.225797721 | 0.239512809 | 0.295893636 | 0.368201811 | ns | YPD | KS.1 |
| FIM1_2200 | 903.4891944 | 1177.542048 | 1233.819232 | 0.352860771 | 0.155925017 | 0.017211254 | 0.029845237 | ns | YPD | KS.1 |
| FIM1_2201 | 1213.339916 | 1063.233779 | 1062.087388 | -0.184257812 | 0.088449099 | 0.033399266 | 0.054088981 | ns | YPD | KS.1 |
| FIM1_2202 | 4747.719057 | 4608.141235 | 4287.758579 | -0.039751905 | 0.134722638 | 0.75884529 | 0.810140627 | ns | YPD | KS.1 |
| FIM1_2203 | 1364.166455 | 977.5533131 | 970.0864016 | -0.458727059 | 0.124589401 | 0.000141463 | 0.000362026 | ns | YPD | KS.1 |
| FIM1_2204 | 1404.485228 | 1171.171399 | 982.1773121 | -0.240262842 | 0.15569661 | 0.10403329 | 0.149628684 | ns | YPD | KS.1 |
| FIM1_2205 | 1451.206058 | 801.6412094 | 972.1536766 | -0.819243402 | 0.146598753 | 6.3508054392 | 3.0004992459 | ns | YPD | KS.1 |
| FIM1_2206 | 859.5312181 | 557.6867163 | 1280.748421 | -0.493429375 | 0.309931029 | 0.063209699 | 0.096699955 | ns | YPD | KS.1 |
| FIM1_2207 | 471.0350138 | 1024.234086 | 462.4421567 | 0.917168366 | 0.348970091 | 0.001952154 | 0.004127482 | ns | YPD | KS.1 |
| FIM1_2209 | 2068.085016 | 1738.573773 | 1758.944291 | -0.21058419 | 0.227922789 | 0.309511697 | 0.382849501 | ns | YPD | KS.1 |
| FIM1_2210 | 3478.695308 | 2645.00623 | 4726.000485 | -0.330709837 | 0.244422107 | 0.131973517 | 0.184067353 | ns | YPD | KS.1 |

| | | | | | | | | | | |
|---|---|---|---|---|---|---|---|---|---|---|
| FIM1_2211 | 1365.447489 | 3146.720704 | 4800.855103 | 1.184735156 | 0.102385948 | 7.714049768 | 2.413672804 | Km>KS.1 | YPD | KS.1 |
| FIM1_2212 | 755.5184025 | 850.2946387 | 1039.059064 | 0.160056743 | 0.127549047 | 0.19257704 | 0.256237169 | ns | YPD | KS.1 |
| FIM1_2213 | 759.9146128 | 721.7301044 | 898.7242659 | -0.069521289 | 0.113789136 | 0.528782953 | 0.604479593 | ns | YPD | KS.1 |
| FIM1_2214 | 557.9093095 | 566.1966284 | 624.8375672 | 0.021153594 | 0.107774166 | 0.840111131 | 0.876032882 | ns | YPD | KS.1 |
| FIM1_2215 | 2905.246735 | 3624.591017 | 6323.786572 | 0.282515593 | 0.192787117 | 0.114776619 | 0.163380166 | ns | YPD | KS.1 |
| FIM1_2216 | 508.1973199 | 826.1942817 | 919.6030042 | 0.677804766 | 0.119650189 | 5.560753881 | 2.647344551 | ns | YPD | KS.1 |
| FIM1_2217 | 1076.938588 | 1662.548465 | 851.7880038 | 0.542057259 | 0.242066966 | 0.013034583 | 0.02324267 | ns | YPD | KS.1 |
| FIM1_2218 | 5004.310732 | 6839.465716 | 8006.289705 | 0.438228329 | 0.094448843 | 2.214568743 | 7.394281542 | ns | YPD | KS.1 |
| FIM1_2219 | 833.1755783 | 2135.712946 | 2623.452048 | 1.340694303 | 0.096623428 | 8.868824562 | 5.962486783 | Km>KS.1 | YPD | KS.1 |
| FIM1_2220 | 1824.838002 | 1681.398327 | 1521.415233 | -0.114928856 | 0.06026307 | 0.053817388 | 0.083643083 | ns | YPD | KS.1 |
| FIM1_2221 | 548.6520008 | 777.5611118 | 801.815568 | 0.48897947 | 0.090495839 | 3.758327677 | 1.613259723 | ns | YPD | KS.1 |
| FIM1_2222 | 969.3693927 | 612.8394274 | 622.0733073 | -0.617949215 | 0.168109335 | 0.000104472 | 0.00027212 | ns | YPD | KS.1 |
| FIM1_2223 | 1058.1739 | 1252.796147 | 1068.373156 | 0.228157413 | 0.13257039 | 0.073197501 | 0.109652986 | ns | YPD | KS.1 |
| FIM1_2224 | 429.5805296 | 2395.685682 | 3951.563321 | 2.438721654 | 0.148063888 | 2.842871039 | 6.734896866 | Km>KS.1 | YPD | KS.1 |
| FIM1_2225 | 2011.727531 | 2473.754105 | 1245.852409 | 0.254029643 | 0.222388905 | 0.21042703 | 0.276366017 | ns | YPD | KS.1 |
| FIM1_2226 | 8512.583244 | 8461.806113 | 10280.44126 | -0.008625305 | 0.061102972 | 0.886550805 | 0.913553948 | ns | YPD | KS.1 |
| FIM1_2227 | 1754.080061 | 2512.974185 | 2047.478626 | 0.46898732 | 0.191534135 | 0.00857256 | 0.015866253 | ns | YPD | KS.1 |
| FIM1_2228 | 1018.077872 | 1433.300119 | 941.2334906 | 0.382768627 | 0.306838385 | 0.145864609 | 0.200851489 | ns | YPD | KS.1 |
| FIM1_2229 | 903.5668547 | 1082.649273 | 888.5419249 | 0.226825561 | 0.202626925 | 0.22565765 | 0.292765333 | ns | YPD | KS.1 |
| FIM1_2230 | 1170.037043 | 644.5179089 | 429.8878394 | -0.752480704 | 0.259849816 | 0.001247726 | 0.002713041 | ns | YPD | KS.1 |
| FIM1_2231 | 713.0997291 | 967.7516431 | 934.2259976 | 0.429421199 | 0.09226916 | 2.101091969 | 7.029544418 | ns | YPD | KS.1 |
| FIM1_2232 | 4752.937588 | 831.6928855 | 872.8070102 | -2.39659471 | 0.257653065 | 9.129958345 | 1.455818678 | Km<KS.1 | YPD | KS.1 |
| FIM1_2233 | 463.0866285 | 1032.600481 | 1744.585935 | 1.118500873 | 0.147306914 | 4.770023824 | 3.955144754 | Km>KS.1 | YPD | KS.1 |
| FIM1_2234 | 667.4629091 | 803.6460135 | 525.7881748 | 0.257526743 | 0.101756401 | 0.009422361 | 0.017310284 | ns | YPD | KS.1 |
| FIM1_2235 | 399.0342979 | 399.3761833 | 542.9644863 | -8.52E-05 | 0.188041649 | 0.999612977 | 0.999612977 | ns | YPD | KS.1 |
| FIM1_2236 | 1091.065652 | 776.7477127 | 1267.879751 | -0.465150863 | 0.132737895 | 0.000276338 | 0.000677565 | ns | YPD | KS.1 |
| FIM1_2237 | 143.2793711 | 36.52695401 | 40.526282 | -1.822043584 | 0.288118896 | 8.884609591 | 5.609255420 | Km<KS.1 | YPD | KS.1 |
| FIM1_2238 | 1536.242051 | 780.2462976 | 875.0395528 | -0.942061126 | 0.14323629 | 1.025066662 | 6.414725339 | ns | YPD | KS.1 |
| FIM1_2239 | 376.7115119 | 441.3631692 | 707.4400088 | 0.210027329 | 0.151650152 | 0.145468498 | 0.200472515 | ns | YPD | KS.1 |
| FIM1_2240 | 587.2907017 | 577.3828753 | 670.9627868 | -0.023130186 | 0.153291843 | 0.87411681 | 0.903350878 | ns | YPD | KS.1 |
| FIM1_2241 | 613.2981874 | 2822.775087 | 2657.322158 | 2.126041611 | 0.201501648 | 7.641447490 | 1.818956983 | Km>KS.1 | YPD | KS.1 |
| FIM1_2243 | 937.1006425 | 978.6510456 | 1190.464514 | 0.057464008 | 0.13305169 | 0.653378923 | 0.718038468 | ns | YPD | KS.1 |
| FIM1_2244 | 4915.114956 | 5347.256781 | 5009.643501 | 0.114006129 | 0.123374392 | 0.338461223 | 0.413358531 | ns | YPD | KS.1 |
| FIM1_2245 | 3330.402864 | 1727.022919 | 1697.103408 | -0.893427103 | 0.178637788 | 1.356972253 | 5.396432423 | ns | YPD | KS.1 |
| FIM1_2246 | 5667.694918 | 9732.843799 | 11412.85161 | 0.765696152 | 0.092358017 | 3.406813109 | 3.516368302 | ns | YPD | KS.1 |
| FIM1_2247 | 995.8023634 | 1059.808686 | 1103.334595 | 0.086355674 | 0.105157286 | 0.398828482 | 0.475590128 | ns | YPD | KS.1 |
| FIM1_2248 | 1781.463052 | 965.9373466 | 2883.141521 | -0.726009733 | 0.315219177 | 0.00744989 | 0.013938774 | ns | YPD | KS.1 |
| FIM1_2249 | 3393.956413 | 2693.252774 | 4456.848732 | -0.26845257 | 0.266289143 | 0.254434638 | 0.323406316 | ns | YPD | KS.1 |
| FIM1_2250 | 489.5171354 | 81.90408684 | 393.1173166 | -2.409267515 | 0.312466268 | 8.217211444 | 8.208961232 | Km<KS.1 | YPD | KS.1 |
| FIM1_2251 | 934.0432143 | 685.3052791 | 559.9824934 | -0.436346353 | 0.087078935 | 3.451449918 | 1.304784448 | ns | YPD | KS.1 |
| FIM1_2252 | 4252.780012 | 3999.490661 | 3579.467591 | -0.081729917 | 0.140446064 | 0.543586898 | 0.617430323 | ns | YPD | KS.1 |
| FIM1_2253 | 13015.93757 | 17300.53608 | 15903.47299 | 0.403383938 | 0.07294836 | 2.165938393 | 9.561263094 | ns | YPD | KS.1 |
| FIM1_2254 | 2463.938491 | 1793.013884 | 2607.848379 | -0.42491052 | 0.161062639 | 0.005398619 | 0.010394012 | ns | YPD | KS.1 |
| FIM1_2255 | 14105.96506 | 11290.98577 | 13851.80021 | -0.316039811 | 0.065661765 | 1.143540352 | 4.001364352 | ns | YPD | KS.1 |
| FIM1_2256 | 1763.215787 | 554.017992 | 1342.415325 | -1.597437952 | 0.198960974 | 5.931590345 | 6.034695699 | Km<KS.1 | YPD | KS.1 |

| | | | | | | | | | | |
|---|---|---|---|---|---|---|---|---|---|---|
| FIM1_2257 | 5602.914799 | 1714.294711 | 1340.549922 | -1.426782319 | 0.399622542 | 2.6313117935 | 7.5364284241 | Km<KS.1 | YPD | KS.1 |
| FIM1_2258 | 1272.910624 | 1812.955485 | 2130.826195 | 0.472329517 | 0.165893406 | 0.00262229 | 0.005386414 | ns | YPD | KS.1 |
| FIM1_2259 | 261.3865364 | 313.9779932 | 421.2064794 | 0.245837441 | 0.142074179 | 0.070328486 | 0.105806547 | ns | YPD | KS.1 |
| FIM1_2260 | 2009.372701 | 1641.875379 | 1913.190993 | -0.279404151 | 0.106559982 | 0.00704635 | 0.013238516 | ns | YPD | KS.1 |
| FIM1_2261 | 2651.809749 | 2556.655084 | 3325.419316 | -0.049602527 | 0.112055339 | 0.648447958 | 0.713881078 | ns | YPD | KS.1 |
| FIM1_2262 | 1971.12327 | 1958.266953 | 2968.684737 | -0.008356084 | 0.196497781 | 0.963509137 | 0.97267456 | ns | YPD | KS.1 |
| FIM1_2263 | 2582.861962 | 2886.183122 | 4218.55243 | 0.146625723 | 0.147968612 | 0.298789461 | 0.370971074 | ns | YPD | KS.1 |
| FIM1_2264 | 812.5255505 | 1452.637875 | 1340.425027 | 0.813565506 | 0.119329915 | 2.5582439476 | 1.6857302834 | ns | YPD | KS.1 |
| FIM1_2265 | 940.7273324 | 1307.893709 | 992.9188256 | 0.45984395 | 0.101056723 | 3.2669014693 | 1.0615829399 | ns | YPD | KS.1 |
| FIM1_2266 | 210.0498316 | 1309.28599 | 1334.817771 | 2.591581497 | 0.159935032 | 1.3666278137 | 2.8329055722 | Km>KS.1 | YPD | KS.1 |
| FIM1_2267 | 1760.880902 | 1783.401147 | 1619.757706 | 0.017437998 | 0.133597538 | 0.892046033 | 0.917875701 | ns | YPD | KS.1 |
| FIM1_2268 | 583.720934 | 1073.72644 | 1336.216251 | 0.861726103 | 0.104777809 | 4.8561080472 | 4.9608085287 | ns | YPD | KS.1 |
| FIM1_2270 | 595.4526234 | 540.7232333 | 840.3921184 | -0.120930441 | 0.193014198 | 0.500772893 | 0.577234741 | ns | YPD | KS.1 |
| FIM1_2271 | 1273.300453 | 1363.680437 | 1494.176021 | 0.092564626 | 0.12637926 | 0.447534344 | 0.524866421 | ns | YPD | KS.1 |
| FIM1_2272 | 671.7399792 | 706.8994512 | 737.2469833 | 0.064918782 | 0.163384304 | 0.674810467 | 0.736870517 | ns | YPD | KS.1 |
| FIM1_2273 | 235.6080627 | 140.0670524 | 169.1592428 | -0.664218841 | 0.238283595 | 0.002128643 | 0.004460824 | ns | YPD | KS.1 |
| FIM1_2274 | 2510.847191 | 2620.563266 | 2250.51838 | 0.051132745 | 0.230477745 | 0.807228182 | 0.848860749 | ns | YPD | KS.1 |
| FIM1_2275 | 1786.795674 | 1785.014876 | 2090.895444 | -0.001491584 | 0.173097962 | 0.99270394 | 0.994302819 | ns | YPD | KS.1 |
| FIM1_2276 | 604.7843734 | 990.4707255 | 871.5900199 | 0.677716498 | 0.14726273 | 1.6014263194 | 5.4719065516 | ns | YPD | KS.1 |
| FIM1_2278 | 3929.018844 | 2271.973648 | 1870.660216 | -0.718691392 | 0.211412929 | 0.000239123 | 0.000592481 | ns | YPD | KS.1 |
| FIM1_2279 | 748.0723892 | 383.1977654 | 338.5058139 | -0.938257523 | 0.122838493 | 4.7088839696 | 3.9240699747 | ns | YPD | KS.1 |
| FIM1_2280 | 871.4012299 | 2148.8486 | 1950.211584 | 1.278706454 | 0.109698021 | 2.3954987024 | 7.7387052238 | Km>KS.1 | YPD | KS.1 |
| FIM1_2281 | 927.0198116 | 963.8042962 | 749.2942018 | 0.049324979 | 0.184707874 | 0.775389654 | 0.82400309 | ns | YPD | KS.1 |
| FIM1_2282 | 377.9922478 | 406.9323901 | 305.8777883 | 0.098737425 | 0.124389886 | 0.410814522 | 0.488130463 | ns | YPD | KS.1 |
| FIM1_2283 | 1819.610649 | 882.054195 | 1312.087633 | -1.01402249 | 0.132521051 | 3.6678361645 | 3.0927940539 | Km<KS.1 | YPD | KS.1 |
| FIM1_2284 | 4206.023247 | 3655.488054 | 2494.53559 | -0.172533297 | 0.214210729 | 0.380167317 | 0.456402606 | ns | YPD | KS.1 |
| FIM1_2285 | 4437.544655 | 4405.904222 | 4091.590869 | -0.009241893 | 0.17038652 | 0.954167716 | 0.967561833 | ns | YPD | KS.1 |
| FIM1_2286 | 997.4540252 | 795.2160923 | 1032.886163 | -0.308852989 | 0.129487603 | 0.013246653 | 0.023561708 | ns | YPD | KS.1 |
| FIM1_2287 | 1699.950098 | 1011.229386 | 927.4687669 | -0.717635294 | 0.138690845 | 7.9528907041 | 3.2650977350 | ns | YPD | KS.1 |
| FIM1_2288 | 2081.251747 | 1639.115046 | 1886.171791 | -0.328046423 | 0.119720402 | 0.004640905 | 0.00910071 | ns | YPD | KS.1 |
| FIM1_2289 | 1204.600053 | 1815.433306 | 3305.0036 | 0.55818881 | 0.151103182 | 0.000110529 | 0.000286695 | ns | YPD | KS.1 |
| FIM1_2290 | 5821.229013 | 3092.273072 | 2575.129183 | -0.854503929 | 0.186211566 | 1.1576418051 | 4.0415915654 | ns | YPD | KS.1 |
| FIM1_2291 | 3399.782676 | 2735.580385 | 3516.64279 | -0.288891473 | 0.155696462 | 0.050667378 | 0.079043652 | ns | YPD | KS.1 |
| FIM1_2292 | 232.9156586 | 621.9164592 | 872.7159229 | 1.357128368 | 0.18017554 | 4.8806190264 | 4.0401130875 | Km>KS.1 | YPD | KS.1 |
| FIM1_2293 | 98.05118533 | 396.1996571 | 613.4994378 | 1.919842683 | 0.21946827 | 6.0637702548 | 7.9179152278 | Km>KS.1 | YPD | KS.1 |
| FIM1_2294 | 5616.195498 | 8558.028981 | 9396.152219 | 0.597923518 | 0.077317647 | 4.6550806246 | 3.8857426355 | ns | YPD | KS.1 |
| FIM1_2295 | 192.1272538 | 611.2397091 | 746.0783644 | 1.612679021 | 0.169025228 | 8.2695984565 | 1.3271371716 | Km>KS.1 | YPD | KS.1 |
| FIM1_2297 | 1187.443492 | 1054.79095 | 1216.926502 | -0.163748092 | 0.097837988 | 0.086246735 | 0.126758495 | ns | YPD | KS.1 |
| FIM1_2298 | 2514.508545 | 3145.639927 | 1665.205528 | 0.292839427 | 0.172033621 | 0.070383941 | 0.105806547 | ns | YPD | KS.1 |
| FIM1_2299 | 1177.069489 | 1603.072231 | 1448.848662 | 0.413895463 | 0.155834758 | 0.00522045 | 0.010087961 | ns | YPD | KS.1 |
| FIM1_2300 | 545.4580714 | 582.9627263 | 538.300696 | 0.087730593 | 0.14889635 | 0.536822654 | 0.611282377 | ns | YPD | KS.1 |
| FIM1_2301 | 2364.115739 | 2020.24528 | 2796.687825 | -0.199213549 | 0.192977508 | 0.266807963 | 0.336542656 | ns | YPD | KS.1 |
| FIM1_2302 | 2343.864337 | 1873.959265 | 3328.172383 | -0.309538848 | 0.108916538 | 0.003475475 | 0.006983235 | ns | YPD | KS.1 |
| FIM1_2303 | 739.1895872 | 501.0379407 | 302.1341551 | -0.491205313 | 0.225752245 | 0.016937653 | 0.029416353 | ns | YPD | KS.1 |
| FIM1_2304 | 456.0811152 | 381.0185339 | 326.5847066 | -0.235465554 | 0.165494754 | 0.13185007 | 0.184067353 | ns | YPD | KS.1 |

| | | | | | | | | | | |
|---|---|---|---|---|---|---|---|---|---|---|
| FIM1_2305 | 1018.324702 | 885.4391037 | 762.5268359 | -0.19440518 | 0.086918235 | 0.022465449 | 0.037771412 | ns | YPD | KS.1 |
| FIM1_2306 | 467.8813342 | 404.237058 | 275.4655839 | -0.196303415 | 0.140128519 | 0.143258907 | 0.197591645 | ns | YPD | KS.1 |
| FIM1_2307 | 963.9746744 | 2178.09329 | 2506.191909 | 1.151443763 | 0.117245425 | 1.3107382715 | 2.2485941037 | Km>KS.1 | YPD | KS.1 |
| FIM1_2308 | 671.4256157 | 1158.54507 | 1907.014888 | 0.712813679 | 0.216658053 | 0.000359747 | 0.000862525 | ns | YPD | KS.1 |
| FIM1_2309 | 655.682907 | 333.8406264 | 295.1554919 | -0.85731974 | 0.265679715 | 0.000326634 | 0.000790371 | ns | YPD | KS.1 |
| FIM1_2310 | 231.1326477 | 226.1537634 | 334.031678 | -0.027995514 | 0.207833473 | 0.883960614 | 0.911819211 | ns | YPD | KS.1 |
| FIM1_2311 | 3984.631584 | 2456.183863 | 1672.056037 | -0.584471633 | 0.277723743 | 0.016797093 | 0.029208507 | ns | YPD | KS.1 |
| FIM1_2312 | 3915.148989 | 4491.175741 | 4768.238598 | 0.189485147 | 0.102691789 | 0.058080513 | 0.089680495 | ns | YPD | KS.1 |
| FIM1_2313 | 1882.74375 | 1064.61535 | 1339.623606 | -0.776808049 | 0.165661884 | 8.4238848503 | 3.0150235345 | ns | YPD | KS.1 |
| FIM1_2314 | 1335.962748 | 932.0980735 | 1073.429431 | -0.505900593 | 0.096233481 | 8.1290863196 | 3.3340646694 | ns | YPD | KS.1 |
| FIM1_2315 | 2799.807593 | 1855.690033 | 1049.537091 | -0.52862845 | 0.21259287 | 0.006870795 | 0.012929877 | ns | YPD | KS.1 |
| FIM1_2316 | 889.0512426 | 1383.603195 | 886.4830386 | 0.559707604 | 0.231883281 | 0.007899997 | 0.01470351 | ns | YPD | KS.1 |
| FIM1_2317 | 4350.624335 | 3883.326995 | 2337.312787 | -0.153177846 | 0.128098775 | 0.214278106 | 0.280830763 | ns | YPD | KS.1 |
| FIM1_2318 | 6477.714603 | 5974.779484 | 13391.12303 | -0.102102813 | 0.187616332 | 0.56003327 | 0.63293174 | ns | YPD | KS.1 |
| FIM1_2319 | 136.8240303 | 328.5863503 | 164.3750103 | 1.169351049 | 0.232371748 | 6.6290804255 | 2.7551942453 | Km>KS.1 | YPD | KS.1 |
| FIM1_2320 | 174.8187815 | 128.7083598 | 214.5370624 | -0.415962521 | 0.137603003 | 0.00164885 | 0.003519103 | ns | YPD | KS.1 |
| FIM1_2321 | 1488.258794 | 1549.469557 | 1675.237695 | 0.057456236 | 0.064698215 | 0.36876503 | 0.444537442 | ns | YPD | KS.1 |
| FIM1_2322 | 453.6625876 | 623.9742978 | 811.9243081 | 0.441544685 | 0.123136589 | 0.000211908 | 0.000530569 | ns | YPD | KS.1 |
| FIM1_2323 | 1789.393099 | 2666.752752 | 3024.517997 | 0.553738664 | 0.119506483 | 1.8093842780 | 6.1028384970 | ns | YPD | KS.1 |
| FIM1_2324 | 1052.882091 | 5368.893634 | 3890.555805 | 2.294581882 | 0.173183116 | 3.8238426567 | 2.1617746837 | Km>KS.1 | YPD | KS.1 |
| FIM1_2325 | 56.29051903 | 41.49787689 | 40.96022622 | -0.395952759 | 0.181775182 | 0.019975226 | 0.034091509 | ns | YPD | KS.1 |
| FIM1_2326 | 503.8376344 | 416.3435132 | 442.7879753 | -0.26245269 | 0.120571195 | 0.024308666 | 0.040596043 | ns | YPD | KS.1 |
| FIM1_2327 | 5050.710878 | 3618.246521 | 3239.293127 | -0.468691688 | 0.093417043 | 3.1414333006 | 1.1975962199 | ns | YPD | KS.1 |
| FIM1_2328 | 3369.665713 | 8741.063122 | 7241.459513 | 1.360775433 | 0.087647428 | 2.2083874329 | 2.5550529021 | Km>KS.1 | YPD | KS.1 |
| FIM1_2329 | 532.2587055 | 424.8012942 | 371.0631883 | -0.285402432 | 0.198748724 | 0.120876539 | 0.170992857 | ns | YPD | KS.1 |
| FIM1_2330 | 619.1681652 | 338.6359558 | 471.7337333 | -0.79822598 | 0.208057232 | 3.6917492545 | 0.000103531 | ns | YPD | KS.1 |
| FIM1_2331 | 2268.942767 | 2420.94891 | 2684.208701 | 0.081558928 | 0.187946229 | 0.642153495 | 0.708048236 | ns | YPD | KS.1 |
| FIM1_2332 | 5721.622576 | 6231.433883 | 8886.548113 | 0.110333154 | 0.168684731 | 0.488294459 | 0.565603012 | ns | YPD | KS.1 |
| FIM1_2333 | 351.0009231 | 391.0014776 | 387.8348301 | 0.141442398 | 0.163297098 | 0.35995759 | 0.435035655 | ns | YPD | KS.1 |
| FIM1_2334 | 1355.759085 | 780.4330161 | 624.1776974 | -0.753582877 | 0.161431842 | 9.8237560623 | 3.4711069893 | ns | YPD | KS.1 |
| FIM1_2335 | 6828.894705 | 6919.258288 | 6406.854433 | 0.016868011 | 0.167473347 | 0.915142291 | 0.936411538 | ns | YPD | KS.1 |
| FIM1_2336 | 6445.050345 | 3996.855288 | 1689.305228 | -0.607395461 | 0.233273073 | 0.00417643 | 0.008264813 | ns | YPD | KS.1 |
| FIM1_2337 | 540.7715221 | 301.4604749 | 216.0073424 | -0.753658887 | 0.23525936 | 0.000445829 | 0.001048699 | ns | YPD | KS.1 |
| FIM1_2338 | 953.847798 | 1703.951781 | 1538.303557 | 0.819585026 | 0.099968205 | 6.5673033201 | 6.6272482794 | ns | YPD | KS.1 |
| FIM1_2339 | 904.6649777 | 1069.939271 | 640.8000692 | 0.227375373 | 0.127298307 | 0.063646089 | 0.097218082 | ns | YPD | KS.1 |
| FIM1_2340 | 1081.155077 | 840.5715929 | 1242.990799 | -0.318151413 | 0.203994935 | 0.090985411 | 0.132937568 | ns | YPD | KS.1 |
| FIM1_2341 | 2492.117404 | 688.7953218 | 716.9177179 | -1.787351615 | 0.190712729 | 2.7808412504 | 4.3233391315 | Km<KS.1 | YPD | KS.1 |
| FIM1_2342 | 806.7023289 | 1335.180858 | 1706.408292 | 0.690941419 | 0.147603237 | 1.0598254802 | 3.7288767779 | ns | YPD | KS.1 |
| FIM1_2343 | 249.0632434 | 52.33251942 | 62.07038615 | -2.077362864 | 0.307371496 | 2.4806090865 | 1.8042441821 | Km<KS.1 | YPD | KS.1 |
| FIM1_2344 | 3353.090514 | 1092.016615 | 771.224789 | -1.515559899 | 0.238477707 | 1.4626748953 | 8.9837130917 | Km<KS.1 | YPD | KS.1 |
| FIM1_2345 | 2052.173045 | 1193.547804 | 917.6130562 | -0.743703397 | 0.152211963 | 3.3900743276 | 1.2835327077 | ns | YPD | KS.1 |
| FIM1_2346 | 3361.987208 | 3651.735312 | 4009.822228 | 0.112418463 | 0.116446337 | 0.31879034 | 0.392181489 | ns | YPD | KS.1 |
| FIM1_2347 | 555.2912551 | 468.259116 | 620.2654514 | -0.221904191 | 0.171607516 | 0.169420524 | 0.228977752 | ns | YPD | KS.1 |
| FIM1_2348 | 1224.317809 | 2445.321111 | 2895.085466 | 0.978776365 | 0.105137356 | 2.5175645047 | 3.6094764873 | ns | YPD | KS.1 |
| FIM1_2349 | 2564.602083 | 2754.593198 | 2302.086355 | 0.099698025 | 0.085823735 | 0.236131282 | 0.304025137 | ns | YPD | KS.1 |

| | | | | | | | | | | |
|---|---|---|---|---|---|---|---|---|---|---|
| FIM1_2350 | 2174.87207 | 2410.000121 | 2886.681087 | 0.139402859 | 0.117456628 | 0.220002631 | 0.286596777 | ns | YPD | KS.1 |
| FIM1_2351 | 1257.060201 | 1172.789725 | 1167.699938 | -0.087487875 | 0.192321314 | 0.625243738 | 0.693708206 | ns | YPD | KS.1 |
| FIM1_2352 | 1985.327278 | 1815.536286 | 1125.829973 | -0.109319159 | 0.21446543 | 0.57888118 | 0.651568749 | ns | YPD | KS.1 |
| FIM1_2353 | 921.4367484 | 711.8570379 | 972.6931714 | -0.354541141 | 0.121906666 | 0.002642293 | 0.005420786 | ns | YPD | KS.1 |
| FIM1_2354 | 823.3179846 | 624.7732241 | 595.0128552 | -0.360226003 | 0.178534214 | 0.031400704 | 0.051001796 | ns | YPD | KS.1 |
| FIM1_2355 | 432.4252172 | 136.3877058 | 214.1355932 | -1.592826062 | 0.196314162 | 2.9933496837 | 3.1154633214 | Km<KS.1 | YPD | KS.1 |
| FIM1_2356 | 829.7160546 | 690.9203675 | 428.712158 | -0.213918712 | 0.255593543 | 0.348053417 | 0.423055399 | ns | YPD | KS.1 |
| FIM1_2357 | 2146.039933 | 1694.080906 | 2252.726222 | -0.299733179 | 0.200543915 | 0.106039052 | 0.152089691 | ns | YPD | KS.1 |
| FIM1_2358 | 565.1420146 | 564.9285889 | 1063.905746 | -0.000807009 | 0.11131581 | 0.994045712 | 0.995246009 | ns | YPD | KS.1 |
| FIM1_2359 | 1386.276658 | 3327.864729 | 4212.938516 | 1.246095884 | 0.097968651 | 5.5034584598 | 2.4446165926 | Km>KS.1 | YPD | KS.1 |
| FIM1_2360 | 4378.820703 | 2750.644253 | 2317.011115 | -0.6165105 | 0.18809662 | 0.000463894 | 0.001088619 | ns | YPD | KS.1 |
| FIM1_2361 | 2305.18448 | 2901.001442 | 3183.044409 | 0.322143087 | 0.090823074 | 0.000297216 | 0.000723764 | ns | YPD | KS.1 |
| FIM1_2362 | 1791.231108 | 1275.567328 | 1599.294081 | -0.476449065 | 0.094462151 | 2.6925492814 | 1.0343963455 | ns | YPD | KS.1 |
| FIM1_2363 | 2384.146893 | 2341.439361 | 2113.157998 | -0.023421976 | 0.150125281 | 0.87008709 | 0.899934152 | ns | YPD | KS.1 |
| FIM1_2364 | 2238.606839 | 3082.401174 | 2512.202957 | 0.452852288 | 0.07953254 | 7.6570361590 | 3.5768783935 | ns | YPD | KS.1 |
| FIM1_2365 | 8700.85646 | 12228.795 | 19621.02589 | 0.466879141 | 0.131659181 | 0.000235214 | 0.000585094 | ns | YPD | KS.1 |
| FIM1_2366 | 628.1380542 | 659.1478018 | 568.7596446 | 0.062614401 | 0.145292491 | 0.652139979 | 0.716993679 | ns | YPD | KS.1 |
| FIM1_2367 | 640.9277705 | 516.5916221 | 488.8553682 | -0.27221971 | 0.202911856 | 0.146278321 | 0.20125405 | ns | YPD | KS.1 |
| FIM1_2368 | 4839.65841 | 3314.814249 | 2404.573716 | -0.493329056 | 0.194268951 | 0.006338906 | 0.012018314 | ns | YPD | KS.1 |
| FIM1_2369 | 1072.063676 | 1062.501297 | 1138.461136 | -0.011644168 | 0.131130102 | 0.926523146 | 0.945916818 | ns | YPD | KS.1 |
| FIM1_2370 | 510.4042946 | 229.6881528 | 394.0827039 | -0.989109187 | 0.311112695 | 0.000287556 | 0.000702302 | ns | YPD | KS.1 |
| FIM1_2371 | 14319.41667 | 8909.388075 | 4375.944249 | -0.558786962 | 0.295289668 | 0.029256874 | 0.047895015 | ns | YPD | KS.1 |
| FIM1_2372 | 3145.20142 | 5666.769087 | 4975.428215 | 0.818947046 | 0.134478591 | 3.1164817979 | 1.6871052170 | ns | YPD | KS.1 |
| FIM1_2373 | 620.8301386 | 475.6246671 | 648.7126715 | -0.296322337 | 0.299590617 | 0.25148633 | 0.320724043 | ns | YPD | KS.1 |
| FIM1_2374 | 617.4707587 | 1335.636088 | 1107.153194 | 1.087302555 | 0.117094179 | 2.5838292502 | 3.6938363562 | Km>KS.1 | YPD | KS.1 |
| FIM1_2375 | 635.9201812 | 2199.768971 | 1708.348738 | 1.75710071 | 0.129797221 | 3.8490022197 | 2.2527983580 | Km>KS.1 | YPD | KS.1 |
| FIM1_2376 | 1098.050772 | 1416.78261 | 1376.278378 | 0.344829619 | 0.1368828 | 0.008672802 | 0.016016033 | ns | YPD | KS.1 |
| FIM1_2377 | 8137.338225 | 9618.226492 | 9042.424857 | 0.228195848 | 0.120997991 | 0.050890264 | 0.079366478 | ns | YPD | KS.1 |
| FIM1_2378 | 509.5757822 | 813.1722172 | 689.8052412 | 0.632500857 | 0.162340106 | 4.1662210157 | 0.000116248 | ns | YPD | KS.1 |
| FIM1_2379 | 2278.257253 | 2274.98578 | 1764.115152 | -0.001896451 | 0.123562429 | 0.987327022 | 0.99071237 | ns | YPD | KS.1 |
| FIM1_2380 | 632.3809419 | 521.2611709 | 815.8202954 | -0.252076198 | 0.171526619 | 0.118343327 | 0.167880825 | ns | YPD | KS.1 |
| FIM1_2381 | 1547.886472 | 1378.568852 | 1661.261604 | -0.161501097 | 0.084540352 | 0.051419977 | 0.080167466 | ns | YPD | KS.1 |
| FIM1_2382 | 566.6351027 | 734.4695349 | 593.4255858 | 0.349643925 | 0.142716012 | 0.010457951 | 0.019016194 | ns | YPD | KS.1 |
| FIM1_2383 | 2373.011424 | 1471.324785 | 893.42857 | -0.666094804 | 0.118907779 | 8.2403800120 | 3.8313916411 | ns | YPD | KS.1 |
| FIM1_2384 | 4346.345174 | 6183.158821 | 9984.682313 | 0.490933295 | 0.110398729 | 5.0012099668 | 1.5867627659 | ns | YPD | KS.1 |
| FIM1_2385 | 765.4220589 | 885.6472433 | 919.9138713 | 0.204295176 | 0.083180319 | 0.012333793 | 0.022127883 | ns | YPD | KS.1 |
| FIM1_2386 | 462.4611966 | 758.4979647 | 686.0131183 | 0.622603342 | 0.243839093 | 0.004680523 | 0.009161095 | ns | YPD | KS.1 |
| FIM1_2387 | 551.05085 | 465.5686306 | 480.6728324 | -0.233623562 | 0.099770298 | 0.016348574 | 0.0285383 | ns | YPD | KS.1 |
| FIM1_2388 | 2724.565844 | 3535.72872 | 6074.465123 | 0.364824843 | 0.092396868 | 5.6455966439 | 0.000154493 | ns | YPD | KS.1 |
| FIM1_2389 | 3566.238592 | 2423.26542 | 1980.459844 | -0.518868869 | 0.163002762 | 0.000791473 | 0.001783323 | ns | YPD | KS.1 |
| FIM1_2390 | 1151.260508 | 1307.948193 | 1057.863545 | 0.178620479 | 0.086361312 | 0.034826142 | 0.056143894 | ns | YPD | KS.1 |
| FIM1_2391 | 478.741047 | 811.133023 | 1136.747479 | 0.710601203 | 0.174770502 | 1.7132837217 | 5.0256995963 | ns | YPD | KS.1 |
| FIM1_2392 | 571.4396968 | 675.5481401 | 1043.287542 | 0.215031084 | 0.180521079 | 0.203319536 | 0.268619893 | ns | YPD | KS.1 |
| FIM1_2393 | 132.9248593 | 146.2372063 | 366.9729904 | 0.113259482 | 0.214937459 | 0.565992259 | 0.638650825 | ns | YPD | KS.1 |
| FIM1_2394 | 2063.739911 | 1951.986097 | 1032.077971 | -0.071015285 | 0.176696488 | 0.668987763 | 0.73128338 | ns | YPD | KS.1 |

| | | | | | | | | | | |
|---|---|---|---|---|---|---|---|---|---|---|
| FIM1_2395 | 11543.58859 | 13392.50718 | 17822.67553 | 0.205333803 | 0.103035649 | 0.040671155 | 0.06443917 | ns | YPD | KS.1 |
| FIM1_2396 | 3032.96253 | 2796.400448 | 2557.179755 | -0.112588795 | 0.090341984 | 0.202911496 | 0.268194658 | ns | YPD | KS.1 |
| FIM1_2397 | 4402.239591 | 4045.413605 | 3026.433657 | -0.115795681 | 0.106959236 | 0.265489978 | 0.335316741 | ns | YPD | KS.1 |
| FIM1_2398 | 1097.043587 | 859.7117958 | 2011.486547 | -0.328119707 | 0.146456787 | 0.018934643 | 0.032572561 | ns | YPD | KS.1 |
| FIM1_2399 | 464.22646 | 348.5260201 | 828.3418785 | -0.384694788 | 0.151080763 | 0.007544747 | 0.01409505 | ns | YPD | KS.1 |
| FIM1_2400 | 437.8895859 | 385.8088224 | 510.3101681 | -0.166118267 | 0.150951468 | 0.247717292 | 0.316403987 | ns | YPD | KS.1 |
| FIM1_2401 | 605.3384546 | 635.069909 | 714.9424682 | 0.063198365 | 0.137646045 | 0.632310765 | 0.69905468 | ns | YPD | KS.1 |
| FIM1_2402 | 828.9142489 | 1155.438707 | 1613.534404 | 0.455615078 | 0.130180048 | 0.000285988 | 0.000699161 | ns | YPD | KS.1 |
| FIM1_2403 | 2712.14622 | 2187.005066 | 2146.319139 | -0.296884397 | 0.111680555 | 0.006190629 | 0.011768582 | ns | YPD | KS.1 |
| FIM1_2404 | 4437.639252 | 5696.643875 | 5503.319899 | 0.341049737 | 0.128120363 | 0.005760805 | 0.011031564 | ns | YPD | KS.1 |
| FIM1_2405 | 1005.160048 | 1558.113226 | 1827.205237 | 0.610666237 | 0.114041285 | 3.78075969175 | 1.62148960918 | ns | YPD | KS.1 |
| FIM1_2406 | 16815.69222 | 11430.39609 | 8854.810501 | -0.454454571 | 0.280080542 | 0.06400173 | 0.097611467 | ns | YPD | KS.1 |
| FIM1_2407 | 1857.967431 | 864.7607744 | 1434.130017 | -1.039853743 | 0.191534417 | 1.00232868518 | 4.60441847532 | Km<KS.1 | YPD | KS.1 |
| FIM1_2408 | 209.7703976 | 324.8735696 | 317.70967 | 0.569340058 | 0.199983737 | 0.002164215 | 0.004523936 | ns | YPD | KS.1 |
| FIM1_2409 | 271.4633265 | 442.9728781 | 558.8150676 | 0.666359872 | 0.153317248 | 5.45308051462 | 1.71920630926 | ns | YPD | KS.1 |
| FIM1_2410 | 3709.51465 | 4250.320053 | 5763.511942 | 0.186951334 | 0.108870011 | 0.07700683 | 0.114531832 | ns | YPD | KS.1 |
| FIM1_2411 | 662.5713288 | 730.1008627 | 686.7980707 | 0.129859545 | 0.133188422 | 0.31021673 | 0.383340345 | ns | YPD | KS.1 |
| FIM1_2412 | 331.7451161 | 467.3388178 | 600.7828512 | 0.448916028 | 0.18648859 | 0.009969258 | 0.018207437 | ns | YPD | KS.1 |
| FIM1_2413 | 2406.605545 | 2543.17612 | 2181.1896 | 0.071001112 | 0.170136252 | 0.658266856 | 0.722771487 | ns | YPD | KS.1 |
| FIM1_2414 | 1417.771276 | 1573.105065 | 1461.678195 | 0.140221179 | 0.123717954 | 0.240162046 | 0.30841667 | ns | YPD | KS.1 |
| FIM1_2415 | 1114.644878 | 866.8578234 | 948.249549 | -0.344567289 | 0.125417943 | 0.004432438 | 0.008729761 | ns | YPD | KS.1 |
| FIM1_2416 | 2617.567802 | 3467.578805 | 4098.833739 | 0.395381086 | 0.08865462 | 5.61488223748 | 1.76797716021 | ns | YPD | KS.1 |
| FIM1_2417 | 1091.877484 | 1194.987279 | 1140.405553 | 0.123359815 | 0.111273479 | 0.253277839 | 0.322450333 | ns | YPD | KS.1 |
| FIM1_2418 | 13369.37318 | 13623.52768 | 15167.9165 | 0.026466666 | 0.074145564 | 0.717129888 | 0.77424505 | ns | YPD | KS.1 |
| FIM1_2419 | 1383.932012 | 880.2350971 | 811.3745993 | -0.605246757 | 0.17595661 | 0.000264126 | 0.000649543 | ns | YPD | KS.1 |
| FIM1_2420 | 2569.119903 | 1460.648335 | 1741.522806 | -0.783184974 | 0.136423642 | 2.81551025297 | 1.38547611360 | ns | YPD | KS.1 |
| FIM1_2422 | 6546.910013 | 5950.610363 | 4611.739681 | -0.134184677 | 0.074149736 | 0.066021683 | 0.100290059 | ns | YPD | KS.1 |
| FIM1_2423 | 618.5558336 | 892.8858717 | 573.2406725 | 0.469898443 | 0.208818763 | 0.014573203 | 0.025700704 | ns | YPD | KS.1 |
| FIM1_2424 | 7066.780374 | 1575.870564 | 1761.714842 | -2.106977697 | 0.17571449 | 6.25077759590 | 2.35588019239 | Km<KS.1 | YPD | KS.1 |
| FIM1_2425 | 442.5835098 | 398.5623686 | 549.0309316 | -0.13137403 | 0.197793804 | 0.47404788 | 0.55179883 | ns | YPD | KS.1 |
| FIM1_2426 | 9778.7357 | 3949.554506 | 3547.211572 | -1.277835625 | 0.128721743 | 3.60790403967 | 6.62336627210 | Km<KS.1 | YPD | KS.1 |
| FIM1_2427 | 162.8842499 | 207.1802868 | 261.1951881 | 0.312924524 | 0.16943724 | 0.049995479 | 0.078044402 | ns | YPD | KS.1 |
| FIM1_2428 | 3200.7267 | 512.5468744 | 668.4622509 | -2.509563478 | 0.276683922 | 5.51891256826 | 8.34546809334 | Km<KS.1 | YPD | KS.1 |
| FIM1_2429 | 1374.243794 | 1279.685206 | 1186.307702 | -0.099673352 | 0.077697624 | 0.192075479 | 0.255804896 | ns | YPD | KS.1 |
| FIM1_2430 | 2261.489114 | 1812.31136 | 1665.928015 | -0.305593799 | 0.110728915 | 0.004506644 | 0.008858377 | ns | YPD | KS.1 |
| FIM1_2431 | 1970.042106 | 1736.466721 | 1976.13754 | -0.166770581 | 0.150892423 | 0.24569173 | 0.314381779 | ns | YPD | KS.1 |
| FIM1_2432 | 1522.790311 | 1789.071421 | 1147.157163 | 0.214791085 | 0.145920899 | 0.122918461 | 0.173480665 | ns | YPD | KS.1 |
| FIM1_2433 | 10001.72374 | 13550.12878 | 28152.16964 | 0.35788708 | 0.265990445 | 0.128222042 | 0.179590276 | ns | YPD | KS.1 |
| FIM1_2434 | 4200.356787 | 3800.433729 | 5279.855016 | -0.121086726 | 0.223773751 | 0.553272706 | 0.626429611 | ns | YPD | KS.1 |
| FIM1_2435 | 843.831676 | 1923.89812 | 1608.339509 | 1.164536236 | 0.113715788 | 1.81697432348 | 3.64493841102 | Km>KS.1 | YPD | KS.1 |
| FIM1_2436 | 593.6696204 | 374.8631433 | 317.0982153 | -0.605267369 | 0.19477034 | 0.000856838 | 0.001917575 | ns | YPD | KS.1 |
| FIM1_2437 | 759.9516697 | 227.1933772 | 185.0976495 | -1.592082807 | 0.287802557 | 1.84003529070 | 9.25599147750 | Km<KS.1 | YPD | KS.1 |
| FIM1_2438 | 3118.717676 | 1973.632216 | 1531.349559 | -0.631303614 | 0.134569507 | 1.15187767220 | 4.02429172697 | ns | YPD | KS.1 |
| FIM1_2439 | 1385.225039 | 579.8500141 | 443.2228972 | -1.123088424 | 0.278017765 | 7.90910837030 | 2.43338368226 | Km<KS.1 | YPD | KS.1 |
| FIM1_2440 | 762.9570952 | 119.8740314 | 226.0552197 | -2.276174106 | 0.481798133 | 2.13193969168 | 9.41953815818 | Km<KS.1 | YPD | KS.1 |

| | | | | | | | | | | |
|---|---|---|---|---|---|---|---|---|---|---|
| FIM1_2444 | 21729.38658 | 13142.5068 | 20677.52651 | -0.567676559 | 0.329655923 | 0.041697829 | 0.065939828 | ns | YPD | KS.1 |
| FIM1_2445 | 485.6411456 | 995.9653859 | 805.4567149 | 0.889698333 | 0.296820581 | 0.000661313 | 0.001516144 | ns | YPD | KS.1 |
| FIM1_2446 | 590.3816705 | 236.8382462 | 299.0757199 | -1.28394715 | 0.13853411 | 2.13356982075 | 3.06777741567 | Km<KS.1 | YPD | KS.1 |
| FIM1_2447 | 806.3423648 | 415.4832542 | 473.4177694 | -0.927298334 | 0.131160145 | 3.39500344526 | 2.41539231965 | ns | YPD | KS.1 |
| FIM1_2448 | 54.07111339 | 39.01325458 | 42.2117417 | -0.313840688 | 0.3951866 | 0.318040661 | 0.391549688 | ns | YPD | KS.1 |
| FIM1_2449 | 59.39048936 | 63.33022078 | 43.40792016 | 0.061317393 | 0.351104016 | 0.832722593 | 0.871067052 | ns | YPD | KS.1 |
| FIM1_2450 | 1196.086035 | 1570.359218 | 632.294383 | 0.343445853 | 0.209385007 | 0.074475398 | 0.111332664 | ns | YPD | KS.1 |
| FIM1_2451 | 295.9256071 | 2083.40517 | 931.4556968 | 2.225774847 | 0.594795435 | 1.72236065067 | 5.84897217550 | Km>KS.1 | YPD | KS.1 |
| FIM1_2452 | 70.84891529 | 78.83334775 | 52.95915738 | 0.117103713 | 0.26024333 | 0.613099348 | 0.682517176 | ns | YPD | KS.1 |
| FIM1_2453 | 295.6639371 | 292.2138066 | 209.9167577 | -0.016016138 | 0.162764634 | 0.917301028 | 0.938063253 | ns | YPD | KS.1 |
| FIM1_2454 | 739.8405193 | 615.1117398 | 378.7276646 | -0.228515999 | 0.215329189 | 0.246893399 | 0.315675832 | ns | YPD | KS.1 |
| FIM1_2455 | 1754.189354 | 3171.671293 | 1087.383669 | 0.634370301 | 0.379903145 | 0.040257069 | 0.063844093 | ns | YPD | KS.1 |
| FIM1_2456 | 328.3889533 | 268.9123343 | 256.7192958 | -0.24337417 | 0.225930024 | 0.236530836 | 0.304382025 | ns | YPD | KS.1 |
| FIM1_2457 | 711.132606 | 743.2135515 | 402.8146455 | 0.056791796 | 0.169215491 | 0.722226847 | 0.778565236 | ns | YPD | KS.1 |
| FIM1_2458 | 8008.538137 | 2393.461461 | 2445.15704 | -1.674022586 | 0.192299687 | 1.61681873956 | 2.03637296944 | Km<KS.1 | YPD | KS.1 |
| FIM1_2459 | 2420.739682 | 1164.852363 | 2120.245717 | -0.951358838 | 0.248493562 | 2.73629925239 | 7.81912049434 | ns | YPD | KS.1 |
| FIM1_2460 | 4983.709376 | 1727.165403 | 4269.400428 | -1.362435167 | 0.307139687 | 8.30013633222 | 2.97501284242 | Km<KS.1 | YPD | KS.1 |
| FIM1_2461 | 3006.755525 | 3991.549882 | 2918.362486 | 0.398269276 | 0.088832229 | 5.00428297452 | 1.58676276598 | ns | YPD | KS.1 |
| FIM1_2462 | 4864.594294 | 5286.312222 | 5382.854023 | 0.115776505 | 0.089370316 | 0.185790162 | 0.248372124 | ns | YPD | KS.1 |
| FIM1_2463 | 1639.411228 | 1284.50557 | 944.635553 | -0.342303194 | 0.086927941 | 6.10179551010 | 0.000166246 | ns | YPD | KS.1 |
| FIM1_2464 | 1476.811008 | 2998.909477 | 1939.7883 | 0.973560438 | 0.168292469 | 1.46168755064 | 7.45071266848 | ns | YPD | KS.1 |
| FIM1_2465 | 433.0046927 | 395.2268051 | 596.242639 | -0.122484483 | 0.119419451 | 0.288833375 | 0.360679227 | ns | YPD | KS.1 |
| FIM1_2466 | 1760.923743 | 1614.572755 | 842.4749687 | -0.100527275 | 0.253974711 | 0.657978443 | 0.722614294 | ns | YPD | KS.1 |
| FIM1_2467 | 188.2855508 | 512.4577904 | 403.5880091 | 1.40338884 | 0.142485444 | 6.19238273373 | 1.10816921224 | Km>KS.1 | YPD | KS.1 |
| FIM1_2468 | 1682.962988 | 2194.575996 | 2249.658896 | 0.362932719 | 0.127852146 | 0.00325134 | 0.006570031 | ns | YPD | KS.1 |
| FIM1_2469 | 1787.675967 | 1907.351307 | 1894.960468 | 0.09023032 | 0.069767257 | 0.189708304 | 0.253164917 | ns | YPD | KS.1 |
| FIM1_2470 | 7753.68609 | 8206.543071 | 6846.182636 | 0.078277687 | 0.096996829 | 0.408610607 | 0.485859887 | ns | YPD | KS.1 |
| FIM1_2471 | 222.4121792 | 875.3948614 | 1109.614518 | 1.918678465 | 0.169706689 | 3.37106018474 | 1.02262344019 | Km>KS.1 | YPD | KS.1 |
| FIM1_2472 | 2875.661632 | 2450.692688 | 2006.773113 | -0.220747117 | 0.104598007 | 0.0300965 | 0.049156299 | ns | YPD | KS.1 |
| FIM1_2473 | 1264.813207 | 948.117284 | 1554.022244 | -0.368784526 | 0.198989661 | 0.045321435 | 0.071239855 | ns | YPD | KS.1 |
| FIM1_2474 | 5024.091848 | 3679.023118 | 3438.768156 | -0.437885898 | 0.090842323 | 9.10997577703 | 3.24192628689 | ns | YPD | KS.1 |
| FIM1_2475 | 994.126524 | 1465.423113 | 1268.99965 | 0.542574006 | 0.107120405 | 2.09911913293 | 8.15231669503 | ns | YPD | KS.1 |
| FIM1_2476 | 4875.373919 | 6437.371946 | 9869.721191 | 0.387564615 | 0.101707998 | 9.59948276154 | 0.000251487 | ns | YPD | KS.1 |
| FIM1_2477 | 2070.421534 | 2074.166437 | 2092.768011 | 0.002704662 | 0.118038995 | 0.9811279 | 0.986083092 | ns | YPD | KS.1 |
| FIM1_2478 | 567.4962395 | 711.9159432 | 1118.601481 | 0.307423597 | 0.136976293 | 0.019292495 | 0.033062406 | ns | YPD | KS.1 |
| FIM1_2479 | 723.3361532 | 725.3629696 | 825.2483291 | 0.003561027 | 0.108445883 | 0.973068034 | 0.981070462 | ns | YPD | KS.1 |
| FIM1_2480 | 2033.812821 | 2310.593547 | 2053.053403 | 0.18019824 | 0.068970419 | 0.008073571 | 0.015004115 | ns | YPD | KS.1 |
| FIM1_2481 | 1419.519681 | 1448.663061 | 1282.433671 | 0.028612508 | 0.08915784 | 0.743314333 | 0.796122456 | ns | YPD | KS.1 |
| FIM1_2482 | 1395.522547 | 1433.281154 | 1389.641121 | 0.038687319 | 0.071545479 | 0.583420782 | 0.655640025 | ns | YPD | KS.1 |
| FIM1_2483 | 1940.887068 | 1536.485816 | 1291.08699 | -0.326227293 | 0.094064862 | 0.000397424 | 0.000944666 | ns | YPD | KS.1 |
| FIM1_2484 | 551.2008724 | 422.8008684 | 351.8027649 | -0.362354114 | 0.131983498 | 0.004340155 | 0.008564963 | ns | YPD | KS.1 |
| FIM1_2485 | 2742.396922 | 1912.02465 | 1569.017362 | -0.497758804 | 0.124395354 | 3.56817748339 | 0.000100349 | ns | YPD | KS.1 |
| FIM1_2486 | 4621.254368 | 6599.899506 | 11891.10262 | 0.480746796 | 0.154639037 | 0.001099654 | 0.002413224 | ns | YPD | KS.1 |
| FIM1_2487 | 2914.38347 | 2725.062907 | 2468.994696 | -0.09332666 | 0.083208505 | 0.253168685 | 0.322450333 | ns | YPD | KS.1 |
| FIM1_2488 | 397.6258504 | 398.7325274 | 344.4512517 | 0.002850234 | 0.185407723 | 0.986891909 | 0.990568902 | ns | YPD | KS.1 |

| | | | | | | | | | | |
|---|---|---|---|---|---|---|---|---|---|---|
| FIM1_2489 | 512.1912627 | 460.1756527 | 585.8551589 | -0.138658349 | 0.169701097 | 0.386192706 | 0.463033209 | ns | YPD | KS.1 |
| FIM1_2490 | 987.0503799 | 1478.945807 | 1138.043493 | 0.547569232 | 0.154157041 | 0.000195845 | 0.000492832 | ns | YPD | KS.1 |
| FIM1_2491 | 5314.266747 | 541.6916768 | 811.3610947 | -3.046077649 | 0.406713353 | 9.3733190188 | 9.2892952428 | Km<KS.1 | YPD | KS.1 |
| FIM1_2492 | 4434.931045 | 2793.470032 | 2656.474913 | -0.592717908 | 0.222645443 | 0.003626808 | 0.007257993 | ns | YPD | KS.1 |
| FIM1_2493 | 1600.566347 | 1783.42271 | 1708.561039 | 0.152329234 | 0.069547253 | 0.026320069 | 0.043603844 | ns | YPD | KS.1 |
| FIM1_2494 | 1033.774673 | 1259.119111 | 863.5211927 | 0.268171335 | 0.128510232 | 0.030210807 | 0.049326801 | ns | YPD | KS.1 |
| FIM1_2495 | 2025.396179 | 3972.929857 | 5237.240303 | 0.915537319 | 0.181997242 | 1.1133165021 | 4.4703386589 | ns | YPD | KS.1 |
| FIM1_2496 | 1065.006593 | 1225.349579 | 1081.57116 | 0.188402119 | 0.138159898 | 0.154613372 | 0.211203055 | ns | YPD | KS.1 |
| FIM1_2497 | 638.4667252 | 512.7443283 | 585.9993221 | -0.285772123 | 0.173160556 | 0.079181455 | 0.117275301 | ns | YPD | KS.1 |
| FIM1_2498 | 155.9545672 | 88.11535909 | 134.5474941 | -0.724520135 | 0.249980634 | 0.001316833 | 0.002850847 | ns | YPD | KS.1 |
| FIM1_2499 | 2056.216546 | 1433.519789 | 1237.132266 | -0.504551445 | 0.103267889 | 5.7432094662 | 2.1024626265 | ns | YPD | KS.1 |
| FIM1_2500 | 4940.094991 | 2216.505025 | 1509.180097 | -1.071906956 | 0.221122949 | 2.0712245084 | 8.0565613210 | Km<KS.1 | YPD | KS.1 |
| FIM1_2502 | 3976.687695 | 2404.774136 | 3124.580927 | -0.684869559 | 0.159395845 | 6.5567188776 | 2.0464037902 | ns | YPD | KS.1 |
| FIM1_2503 | 685.7354445 | 456.6932976 | 574.3758891 | -0.518147182 | 0.219334346 | 0.009875619 | 0.018082887 | ns | YPD | KS.1 |
| FIM1_2504 | 1133.609848 | 917.9335211 | 853.0960671 | -0.251506532 | 0.245368561 | 0.253696125 | 0.322880077 | ns | YPD | KS.1 |
| FIM1_2505 | 506.9046422 | 398.4867203 | 596.8034414 | -0.312571219 | 0.177820769 | 0.060837531 | 0.09350223 | ns | YPD | KS.1 |
| FIM1_2506 | 1131.119398 | 907.9278073 | 537.1545035 | -0.282160051 | 0.187923445 | 0.107158036 | 0.153369169 | ns | YPD | KS.1 |
| FIM1_2507 | 1397.892999 | 774.8998331 | 1007.272581 | -0.82224159 | 0.12911527 | 5.2175803049 | 3.0466504715 | ns | YPD | KS.1 |
| FIM1_2508 | 17.58911921 | 23.28822557 | 19.23299098 | 0.263439259 | 0.371854421 | 0.382894399 | 0.459454808 | ns | YPD | KS.1 |
| FIM1_2510 | 119.3548248 | 173.2660648 | 247.8209449 | 0.476251208 | 0.204707301 | 0.011785597 | 0.021236271 | ns | YPD | KS.1 |
| FIM1_2511 | 5372.591098 | 4175.765696 | 5768.840307 | -0.323370368 | 0.192245966 | 0.070364346 | 0.105806547 | ns | YPD | KS.1 |
| FIM1_2512 | 872.9299514 | 1149.364195 | 1212.02316 | 0.37606671 | 0.127165139 | 0.002182896 | 0.00455342 | ns | YPD | KS.1 |
| FIM1_2513 | 2721.54802 | 4406.729543 | 6384.272418 | 0.637841001 | 0.192125639 | 0.000380963 | 0.000908973 | ns | YPD | KS.1 |
| FIM1_2514 | 911.0042854 | 622.5227449 | 481.8060497 | -0.519959045 | 0.140882563 | 0.000121222 | 0.000311991 | ns | YPD | KS.1 |
| FIM1_2515 | 249.582072 | 108.9222316 | 94.90197683 | -1.133713104 | 0.187888164 | 2.3660443040 | 1.2935242211 | Km<KS.1 | YPD | KS.1 |
| FIM1_2516 | 3094.535422 | 4054.01997 | 4106.495846 | 0.374713776 | 0.10896695 | 0.000412494 | 0.000976752 | ns | YPD | KS.1 |
| FIM1_2517 | 4501.270328 | 1679.211879 | 2074.916218 | -1.369673281 | 0.170755039 | 1.0019855673 | 9.8906313443 | Km<KS.1 | YPD | KS.1 |
| FIM1_2518 | 1449.137424 | 1945.05972 | 2038.925988 | 0.41475899 | 0.083068083 | 3.9376643379 | 1.4784815155 | ns | YPD | KS.1 |
| FIM1_2519 | 2287.320462 | 1920.705933 | 2929.214204 | -0.175956609 | 0.347342722 | 0.541397027 | 0.6154537 | ns | YPD | KS.1 |
| FIM1_2520 | 666.7341298 | 574.8231546 | 689.008901 | -0.193177099 | 0.166771979 | 0.21971564 | 0.286372887 | ns | YPD | KS.1 |
| FIM1_2521 | 554.8666631 | 543.0336265 | 697.9728368 | -0.027109079 | 0.177653026 | 0.871020529 | 0.900525173 | ns | YPD | KS.1 |
| FIM1_2522 | 506.5294283 | 379.8582951 | 490.9830642 | -0.384941337 | 0.151274362 | 0.007576967 | 0.014149929 | ns | YPD | KS.1 |
| FIM1_2523 | 3924.266703 | 5057.433326 | 4516.161231 | 0.361224869 | 0.059684674 | 1.0322948919 | 5.3552315822 | ns | YPD | KS.1 |
| FIM1_2524 | 3891.090776 | 3128.173135 | 1763.71944 | -0.256435317 | 0.256261225 | 0.261454854 | 0.331426633 | ns | YPD | KS.1 |
| FIM1_2525 | 365.7415514 | 494.1577453 | 706.1662017 | 0.402912958 | 0.153777239 | 0.005911791 | 0.011290273 | ns | YPD | KS.1 |
| FIM1_2526 | 1099.589102 | 2420.362649 | 5219.96843 | 1.099668307 | 0.146627882 | 1.0103392926 | 8.1202552196 | Km>KS.1 | YPD | KS.1 |
| FIM1_2527 | 143.2039211 | 137.0957021 | 199.4878147 | -0.047987488 | 0.311824108 | 0.857202377 | 0.889566505 | ns | YPD | KS.1 |
| FIM1_2528 | 18276.08689 | 18939.67743 | 23970.31229 | 0.049009795 | 0.101084548 | 0.619118452 | 0.688293698 | ns | YPD | KS.1 |
| FIM1_2529 | 16191.32163 | 11355.49136 | 8482.783516 | -0.397137422 | 0.30870044 | 0.133146882 | 0.185599815 | ns | YPD | KS.1 |
| FIM1_2530 | 3748.342529 | 2522.253084 | 1352.405008 | -0.445480532 | 0.312752566 | 0.09548744 | 0.138619788 | ns | YPD | KS.1 |
| FIM1_2531 | 4055.477556 | 3824.185566 | 3138.28937 | -0.075103444 | 0.17612194 | 0.650190947 | 0.715166916 | ns | YPD | KS.1 |
| FIM1_2532 | 166.2671091 | 327.0192798 | 477.4910332 | 0.835034196 | 0.292585483 | 0.00118522 | 0.002586171 | ns | YPD | KS.1 |
| FIM1_2533 | 2386.326229 | 2638.045542 | 2574.912595 | 0.134249014 | 0.13523138 | 0.301005842 | 0.373347027 | ns | YPD | KS.1 |
| FIM1_2534 | 3386.035742 | 2292.920152 | 2106.468145 | -0.523575796 | 0.16332498 | 0.000725137 | 0.001649546 | ns | YPD | KS.1 |
| FIM1_2535 | 376.7313518 | 422.9675878 | 461.4695118 | 0.149140212 | 0.170859986 | 0.354212034 | 0.42928255 | ns | YPD | KS.1 |

| | | | | | | | | | | |
|---|---|---|---|---|---|---|---|---|---|---|
| FIM1_2536 | 420.4957851 | 341.0857109 | 460.6005252 | -0.233089015 | 0.292908206 | 0.358864368 | 0.433969429 | ns | YPD | KS.1 |
| FIM1_2537 | 780.964169 | 1098.812901 | 1077.293257 | 0.408166276 | 0.262000654 | 0.079125056 | 0.117261589 | ns | YPD | KS.1 |
| FIM1_2538 | 1115.449637 | 730.9231442 | 815.3711556 | -0.550626837 | 0.20142452 | 0.003191689 | 0.006459989 | ns | YPD | KS.1 |
| FIM1_2539 | 463.3654911 | 712.0932631 | 615.8128797 | 0.56235511 | 0.196090747 | 0.002055718 | 0.004320743 | ns | YPD | KS.1 |
| FIM1_2540 | 4702.08959 | 4688.071214 | 3341.158667 | -0.003794582 | 0.183880364 | 0.982420291 | 0.986983228 | ns | YPD | KS.1 |
| FIM1_2541 | 851.3443983 | 1134.72884 | 1315.902532 | 0.387585428 | 0.146761251 | 0.005703905 | 0.010935233 | ns | YPD | KS.1 |
| FIM1_2542 | 155.3127309 | 263.5184139 | 401.8720866 | 0.702155386 | 0.188983997 | 7.42938796812 | 0.000197865 | ns | YPD | KS.1 |
| FIM1_2543 | 1458.623477 | 1787.045306 | 1152.744984 | 0.268515175 | 0.158692292 | 0.074214708 | 0.111009672 | ns | YPD | KS.1 |
| FIM1_2544 | 1374.991622 | 983.2761565 | 859.9406074 | -0.466743581 | 0.108377296 | 9.97944749104 | 3.02361457173 | ns | YPD | KS.1 |
| FIM1_2546 | 474.9971271 | 1170.510522 | 1183.701345 | 1.267771104 | 0.135007127 | 6.93201320502 | 1.03875800286 | Km>KS.1 | YPD | KS.1 |
| FIM1_2547 | 5273.666947 | 3524.934976 | 2021.126718 | -0.485258934 | 0.266026312 | 0.039418672 | 0.062634269 | ns | YPD | KS.1 |
| FIM1_2548 | 6302.109834 | 5952.165753 | 5162.908892 | -0.061788338 | 0.2978419 | 0.810460533 | 0.851719719 | ns | YPD | KS.1 |
| FIM1_2549 | 9207.28866 | 8824.9497 | 5865.03692 | -0.051101371 | 0.223737705 | 0.802535503 | 0.845416071 | ns | YPD | KS.1 |
| FIM1_2550 | 1464.218442 | 1348.389234 | 1720.231843 | -0.105005585 | 0.180995979 | 0.535969843 | 0.610590788 | ns | YPD | KS.1 |
| FIM1_2552 | 9828.753899 | 3154.429973 | 2587.608678 | -1.449693031 | 0.327131872 | 7.01328636694 | 2.54493797779 | Km<KS.1 | YPD | KS.1 |
| FIM1_2553 | 922.9826137 | 804.7585684 | 530.5381407 | -0.162120013 | 0.242987321 | 0.458711497 | 0.536079328 | ns | YPD | KS.1 |
| FIM1_2554 | 952.3513831 | 1898.701548 | 1249.548733 | 0.865568188 | 0.280890131 | 0.000527963 | 0.001231418 | ns | YPD | KS.1 |
| FIM1_2555 | 62633.80745 | 13413.33644 | 14232.97528 | -1.938883119 | 0.399477962 | 2.88579462038 | 1.25937089793 | Km<KS.1 | YPD | KS.1 |
| FIM1_2556 | 2286.504203 | 751.5237908 | 413.0204317 | -1.340461889 | 0.388394961 | 4.97907341404 | 0.000137311 | Km<KS.1 | YPD | KS.1 |
| FIM1_2557 | 21599.82885 | 12407.8437 | 12207.00935 | -0.724216407 | 0.217406871 | 0.000302776 | 0.000736583 | ns | YPD | KS.1 |
| FIM1_2558 | 869.4696483 | 572.1073135 | 1227.149074 | -0.51699134 | 0.248831004 | 0.020561777 | 0.034918579 | ns | YPD | KS.1 |
| FIM1_2559 | 641.380828 | 891.2021394 | 458.2771054 | 0.372429158 | 0.296292432 | 0.146070659 | 0.201079559 | ns | YPD | KS.1 |
| FIM1_2560 | 4899.51795 | 3222.7356 | 3738.299824 | -0.573931187 | 0.1412687 | 2.34595654524 | 6.77022030449 | ns | YPD | KS.1 |
| FIM1_2561 | 611.9214307 | 483.224636 | 346.8324722 | -0.293497086 | 0.217497188 | 0.140171731 | 0.193848256 | ns | YPD | KS.1 |
| FIM1_2562 | 4708.21972 | 4195.820985 | 2830.694731 | -0.155535932 | 0.127795785 | 0.206286229 | 0.271860659 | ns | YPD | KS.1 |
| FIM1_2563 | 3988.945065 | 3463.387301 | 3001.103454 | -0.197051377 | 0.090135022 | 0.025503949 | 0.042407135 | ns | YPD | KS.1 |
| FIM1_2564 | 7055.229504 | 1778.133978 | 1739.693656 | -1.945865441 | 0.149637986 | 2.90697218140 | 1.52233543184 | Km<KS.1 | YPD | KS.1 |
| FIM1_2565 | 1580.14999 | 857.4023112 | 1138.975453 | -0.848055897 | 0.141567133 | 5.42998805484 | 2.87997767301 | ns | YPD | KS.1 |
| FIM1_2566 | 3391.901305 | 4485.024875 | 2318.819173 | 0.365630313 | 0.177363848 | 0.028022127 | 0.046162278 | ns | YPD | KS.1 |
| FIM1_2567 | 1948.451502 | 1153.997438 | 1056.417571 | -0.728249569 | 0.127493372 | 3.76521705535 | 1.81863639324 | ns | YPD | KS.1 |
| FIM1_2568 | 753.7924114 | 632.3998332 | 1217.730461 | -0.211697663 | 0.232546762 | 0.315172481 | 0.38849928 | ns | YPD | KS.1 |
| FIM1_2570 | 1180.018675 | 404.9700087 | 388.1636513 | -1.434332131 | 0.246827002 | 5.09053827353 | 2.70570811013 | Km<KS.1 | YPD | KS.1 |
| FIM1_2571 | 749.9753126 | 221.0216486 | 421.0415144 | -1.617856677 | 0.28254966 | 5.68458388062 | 3.00859625596 | Km<KS.1 | YPD | KS.1 |
| FIM1_2572 | 504.7688568 | 132.5385619 | 287.6389418 | -1.850159481 | 0.206711008 | 1.18229095539 | 1.63386041752 | Km<KS.1 | YPD | KS.1 |
| FIM1_2573 | 626.9762499 | 453.7815019 | 484.8169296 | -0.434553544 | 0.157570898 | 0.003706315 | 0.007408162 | ns | YPD | KS.1 |
| FIM1_2574 | 1037.108565 | 1584.968729 | 1794.752805 | 0.587203485 | 0.126458743 | 1.60365786100 | 5.47577066472 | ns | YPD | KS.1 |
| FIM1_2575 | 468.6664218 | 884.8692886 | 933.2852499 | 0.888520759 | 0.124994538 | 2.77508132388 | 1.99509098068 | ns | YPD | KS.1 |
| FIM1_2576 | 8021.810192 | 14503.09516 | 14153.82679 | 0.848982506 | 0.053835322 | 1.21157293524 | 1.54553214175 | ns | YPD | KS.1 |
| FIM1_2577 | 13698.96962 | 5940.535255 | 5459.753975 | -1.15738979 | 0.164811162 | 3.09390073688 | 2.21151669051 | Km<KS.1 | YPD | KS.1 |
| FIM1_2578 | 150.4293175 | 77.47163407 | 73.6616089 | -0.889722928 | 0.200967611 | 2.30419028655 | 7.67806207341 | ns | YPD | KS.1 |
| FIM1_2579 | 835.129668 | 335.7273819 | 269.9166729 | -1.267698854 | 0.162511447 | 7.14201015639 | 6.48385046132 | Km<KS.1 | YPD | KS.1 |
| FIM1_2580 | 4972.245332 | 2032.660642 | 1231.593787 | -1.194660884 | 0.234145832 | 4.37116502723 | 1.85550733877 | Km<KS.1 | YPD | KS.1 |
| FIM1_2581 | 33448.79842 | 16104.9351 | 20957.63236 | -1.021076122 | 0.138150391 | 2.66297716281 | 2.06359990420 | Km<KS.1 | YPD | KS.1 |
| FIM1_2582 | 705.7209017 | 523.0790068 | 655.2949581 | -0.394127828 | 0.174011983 | 0.016035841 | 0.028031732 | ns | YPD | KS.1 |
| FIM1_2583 | 55.01559255 | 176.7867537 | 481.033384 | 1.592723476 | 0.211825897 | 3.34950336525 | 2.84364833484 | Km>KS.1 | YPD | KS.1 |

| | | | | | | | | | | |
|---|---|---|---|---|---|---|---|---|---|---|
| FIM1_2584 | 390.1987165 | 84.45564847 | 103.7048874 | -1.825881581 | 0.465037603 | 2.4823980156 | 8.2168530458 | Km<KS.1 | YPD | KS.1 |
| FIM1_2585 | 3191.138511 | 3661.138913 | 4837.266344 | 0.185985995 | 0.125885985 | 0.125248782 | 0.176219653 | ns | YPD | KS.1 |
| FIM1_2586 | 644.2484956 | 1104.097057 | 1096.603075 | 0.755121053 | 0.113768993 | 1.0044070831 | 6.3012928611 | ns | YPD | KS.1 |
| FIM1_2587 | 4619.245531 | 4590.370821 | 4513.054149 | -0.008551908 | 0.093877041 | 0.925812584 | 0.945385387 | ns | YPD | KS.1 |
| FIM1_2588 | 4900.37016 | 3492.848577 | 5707.663931 | -0.428531919 | 0.216039581 | 0.030391119 | 0.049588657 | ns | YPD | KS.1 |
| FIM1_2590 | 8016.023937 | 2402.272537 | 1955.598832 | -1.580618675 | 0.296193318 | 5.5977834809 | 2.6598827906 | Km<KS.1 | YPD | KS.1 |
| FIM1_2591 | 1422.965219 | 858.5974754 | 888.9165065 | -0.627315934 | 0.258567777 | 0.006629978 | 0.012527209 | ns | YPD | KS.1 |
| FIM1_2592 | 2834.036025 | 2011.450435 | 1822.145642 | -0.473942276 | 0.119836234 | 4.5466045328 | 0.000126083 | ns | YPD | KS.1 |
| FIM1_2593 | 2603.815358 | 2121.053069 | 1588.488274 | -0.28742964 | 0.088424238 | 0.000918656 | 0.002046715 | ns | YPD | KS.1 |
| FIM1_2594 | 1732.802561 | 1979.033541 | 2182.106981 | 0.180417832 | 0.120254674 | 0.120472313 | 0.170560546 | ns | YPD | KS.1 |
| FIM1_2595 | 1051.716045 | 1209.723766 | 1316.272804 | 0.194401163 | 0.097530501 | 0.041082914 | 0.065070836 | ns | YPD | KS.1 |
| FIM1_2596 | 4251.056873 | 2627.062021 | 2024.028705 | -0.626891905 | 0.209698497 | 0.001212677 | 0.002641449 | ns | YPD | KS.1 |
| FIM1_2597 | 1003.326723 | 946.6037037 | 977.7203132 | -0.072672098 | 0.192111536 | 0.684708992 | 0.746696019 | ns | YPD | KS.1 |
| FIM1_2598 | 748.3968969 | 693.8025034 | 1013.907709 | -0.101276604 | 0.125616082 | 0.403228162 | 0.480033526 | ns | YPD | KS.1 |
| FIM1_2599 | 398.1988962 | 725.9863503 | 964.505576 | 0.780866697 | 0.2308372 | 0.000221582 | 0.000553121 | ns | YPD | KS.1 |
| FIM1_2600 | 314.1699945 | 953.589577 | 1325.314155 | 1.537439758 | 0.187024373 | 1.3890579433 | 1.4925622609 | Km>KS.1 | YPD | KS.1 |
| FIM1_2601 | 749.27497 | 2080.661096 | 2074.916074 | 1.444283792 | 0.122790773 | 5.0312808942 | 1.7027634319 | Km>KS.1 | YPD | KS.1 |
| FIM1_2602 | 1807.712615 | 2571.545478 | 1905.859368 | 0.491792146 | 0.10584895 | 1.9370623794 | 6.5158115872 | ns | YPD | KS.1 |
| FIM1_2603 | 286.9711579 | 460.1569583 | 286.3863057 | 0.544515052 | 0.309161541 | 0.04022943 | 0.063820604 | ns | YPD | KS.1 |
| FIM1_2604 | 705.3387695 | 508.1800675 | 543.7081307 | -0.418490522 | 0.207052697 | 0.028289926 | 0.046557189 | ns | YPD | KS.1 |
| FIM1_2606 | 70.96342064 | 22.79164506 | 32.27504231 | -1.430784004 | 0.342703127 | 2.2900093793 | 7.6359226960 | Km<KS.1 | YPD | KS.1 |
| FIM1_2607 | 1538.958197 | 2523.229502 | 3002.133013 | 0.646714987 | 0.207834078 | 0.000772984 | 0.001747997 | ns | YPD | KS.1 |
| FIM1_2608 | 664.3948538 | 849.0282475 | 800.2735726 | 0.344232837 | 0.090611372 | 0.000107466 | 0.000279334 | ns | YPD | KS.1 |
| FIM1_2609 | 698.5916896 | 501.1806145 | 502.6738705 | -0.448990509 | 0.152096537 | 0.001960558 | 0.004142587 | ns | YPD | KS.1 |
| FIM1_2610 | 3200.410917 | 1202.271733 | 1233.092503 | -1.384473878 | 0.12336608 | 2.9306131390 | 8.1908990823 | Km<KS.1 | YPD | KS.1 |
| FIM1_2611 | 2560.934602 | 1928.823911 | 1871.482649 | -0.358316981 | 0.20912013 | 0.062486204 | 0.095681399 | ns | YPD | KS.1 |
| FIM1_2612 | 7311.357863 | 5229.048096 | 3394.826906 | -0.352973709 | 0.342558773 | 0.216239835 | 0.282805778 | ns | YPD | KS.1 |
| FIM1_2613 | 3572.929084 | 5982.208864 | 3064.290298 | 0.622724569 | 0.283193821 | 0.012232916 | 0.021986545 | ns | YPD | KS.1 |
| FIM1_2614 | 262.878387 | 660.7688404 | 709.2168211 | 1.291736908 | 0.143940967 | 3.1721509254 | 4.2309519716 | Km>KS.1 | YPD | KS.1 |
| FIM1_2615 | 42.84068411 | 49.95289793 | 64.47101098 | 0.177094816 | 0.267970397 | 0.454854799 | 0.532447677 | ns | YPD | KS.1 |
| FIM1_2616 | 785.8948587 | 802.2906971 | 737.3979886 | 0.029825478 | 0.089203212 | 0.732964722 | 0.787751025 | ns | YPD | KS.1 |
| FIM1_2617 | 374.1254036 | 932.0545903 | 1133.242136 | 1.302323229 | 0.089864263 | 1.4470009791 | 1.2411775640 | Km>KS.1 | YPD | KS.1 |
| FIM1_2618 | 248.802086 | 362.2103269 | 394.8937208 | 0.506929232 | 0.156603354 | 0.000672108 | 0.001538766 | ns | YPD | KS.1 |
| FIM1_2619 | 230.5391417 | 234.7672492 | 276.3989874 | 0.021838396 | 0.174908064 | 0.894479498 | 0.919999071 | ns | YPD | KS.1 |
| FIM1_2620 | 154.4235131 | 335.1964376 | 670.0593794 | 1.072359192 | 0.156443515 | 1.1890300982 | 8.1479679598 | Km>KS.1 | YPD | KS.1 |
| FIM1_2621 | 12429.97916 | 10714.86335 | 8627.981714 | -0.192717009 | 0.169943713 | 0.228739824 | 0.295963751 | ns | YPD | KS.1 |
| FIM1_2622 | 658.594268 | 702.4698861 | 652.659282 | 0.086589278 | 0.131663214 | 0.494225064 | 0.571408249 | ns | YPD | KS.1 |
| FIM1_2623 | 1464.946991 | 1516.788771 | 1779.418525 | 0.04766855 | 0.113875613 | 0.666128432 | 0.72899009 | ns | YPD | KS.1 |
| FIM1_2624 | 9733.122415 | 7802.443531 | 7410.842857 | -0.302357232 | 0.124006646 | 0.01155368 | 0.020848589 | ns | YPD | KS.1 |
| FIM1_2625 | 1095.567327 | 628.3714453 | 576.4427825 | -0.756115273 | 0.166432872 | 1.7846415544 | 6.0316519928 | ns | YPD | KS.1 |
| FIM1_2626 | 1086.632322 | 1825.784896 | 1584.582197 | 0.722054284 | 0.126842267 | 4.2819298534 | 2.0562356197 | ns | YPD | KS.1 |
| FIM1_2627 | 360.6113688 | 434.3682024 | 1047.969986 | 0.243303116 | 0.167518852 | 0.123570424 | 0.174252511 | ns | YPD | KS.1 |
| FIM1_2628 | 530.3486703 | 680.5244928 | 833.5519857 | 0.339616503 | 0.124611913 | 0.004772746 | 0.009333496 | ns | YPD | KS.1 |
| FIM1_2629 | 5897.311985 | 10759.97948 | 7639.601927 | 0.750909004 | 0.270185856 | 0.001805132 | 0.003829652 | ns | YPD | KS.1 |
| FIM1_2630 | 461.0722008 | 293.5507671 | 544.7853279 | -0.600301374 | 0.180507262 | 0.000404655 | 0.000959561 | ns | YPD | KS.1 |

| | | | | | | | | | | |
|---|---|---|---|---|---|---|---|---|---|---|
| FIM1_2631 | 342.176092 | 288.71221 | 217.1279088 | -0.225398297 | 0.145031037 | 0.103518495 | 0.148974403 | ns | YPD | KS.1 |
| FIM1_2632 | 517.1645131 | 536.4749783 | 622.236406 | 0.047976887 | 0.141747174 | 0.723805184 | 0.779928696 | ns | YPD | KS.1 |
| FIM1_2633 | 889.7982623 | 1393.963095 | 808.6597244 | 0.60707939 | 0.160767652 | 7.1833546661 | 0.000192032 | ns | YPD | KS.1 |
| FIM1_2634 | 6242.503223 | 4618.295178 | 5047.092429 | -0.355755041 | 0.264728294 | 0.129036977 | 0.180528392 | ns | YPD | KS.1 |
| FIM1_2635 | 20518.18547 | 19690.41363 | 31529.08503 | -0.056201021 | 0.109314145 | 0.596843958 | 0.668309406 | ns | YPD | KS.1 |
| FIM1_2636 | 3841.570849 | 4872.453765 | 3879.739344 | 0.336471147 | 0.074985384 | 5.40392750878 | 1.70695487975 | ns | YPD | KS.1 |
| FIM1_2637 | 2761.819422 | 3346.345359 | 4556.375614 | 0.249437133 | 0.172880954 | 0.124911379 | 0.175794656 | ns | YPD | KS.1 |
| FIM1_2638 | 1800.433588 | 1488.798507 | 1535.1718 | -0.259784291 | 0.119190384 | 0.024212524 | 0.040449062 | ns | YPD | KS.1 |
| FIM1_2639 | 721.3816222 | 3508.474355 | 3287.046383 | 2.204501034 | 0.204567234 | 5.2788772127 | 1.3607468462 | Km>KS.1 | YPD | KS.1 |
| FIM1_2640 | 4321.666593 | 3388.820626 | 3174.526742 | -0.32601019 | 0.149284257 | 0.021954383 | 0.036962965 | ns | YPD | KS.1 |
| FIM1_2641 | 704.4147984 | 774.6064705 | 974.5261912 | 0.130652651 | 0.105932519 | 0.204726438 | 0.27009123 | ns | YPD | KS.1 |
| FIM1_2642 | 817.0951295 | 519.647076 | 395.3580463 | -0.630450652 | 0.117686589 | 3.5712503946 | 1.5369351827 | ns | YPD | KS.1 |
| FIM1_2643 | 3980.928972 | 2858.272242 | 2442.68225 | -0.449466998 | 0.144748395 | 0.001180507 | 0.002578148 | ns | YPD | KS.1 |
| FIM1_2644 | 1515.712269 | 1511.133051 | 1769.190409 | -0.004595258 | 0.10354707 | 0.963680986 | 0.97267456 | ns | YPD | KS.1 |
| FIM1_2645 | 917.6799513 | 642.3659034 | 693.2951935 | -0.489717252 | 0.131141179 | 0.00010841 | 0.000281493 | ns | YPD | KS.1 |
| FIM1_2646 | 2970.608098 | 2601.676685 | 2678.289446 | -0.183596255 | 0.101086322 | 0.062302853 | 0.095430016 | ns | YPD | KS.1 |
| FIM1_2647 | 4177.411779 | 3827.780606 | 3353.209645 | -0.121422843 | 0.090468689 | 0.170232639 | 0.229950416 | ns | YPD | KS.1 |
| FIM1_2648 | 367.9225244 | 420.7258676 | 308.8158223 | 0.182951672 | 0.105007606 | 0.073264364 | 0.109687093 | ns | YPD | KS.1 |
| FIM1_2649 | 832.4870697 | 801.6437042 | 874.0597844 | -0.053559374 | 0.078170775 | 0.486110621 | 0.563992617 | ns | YPD | KS.1 |
| FIM1_2650 | 5805.21718 | 7332.141887 | 13759.61847 | 0.317098179 | 0.13365646 | 0.01355641 | 0.024052475 | ns | YPD | KS.1 |
| FIM1_2651 | 857.3035149 | 442.5065039 | 381.7676173 | -0.878310512 | 0.213253489 | 9.4499924080 | 2.8806067287 | ns | YPD | KS.1 |
| FIM1_2652 | 3301.953711 | 7616.469913 | 7485.896065 | 1.191790309 | 0.086956271 | 1.2296373451 | 7.5524022124 | Km>KS.1 | YPD | KS.1 |
| FIM1_2653 | 1098.462491 | 1312.874557 | 1690.258184 | 0.24492407 | 0.115216604 | 0.028205127 | 0.046432994 | ns | YPD | KS.1 |
| FIM1_2654 | 3339.107706 | 2526.622772 | 2235.239421 | -0.38305134 | 0.12368647 | 0.001360621 | 0.002937972 | ns | YPD | KS.1 |
| FIM1_2655 | 1637.553138 | 2021.384875 | 1444.349023 | 0.212221207 | 0.350831326 | 0.46449056 | 0.542069091 | ns | YPD | KS.1 |
| FIM1_2656 | 370.2126547 | 771.7113912 | 282.8426353 | 0.898792903 | 0.311067167 | 0.000918142 | 0.002046715 | ns | YPD | KS.1 |
| FIM1_2657 | 1241.222579 | 871.0615656 | 919.771808 | -0.486378091 | 0.131831141 | 0.000130289 | 0.000334118 | ns | YPD | KS.1 |
| FIM1_2659 | 1211.231006 | 1875.335947 | 1561.785365 | 0.602703625 | 0.135268322 | 3.7896424228 | 1.2242513671 | ns | YPD | KS.1 |
| FIM1_2660 | 307.9401421 | 732.7814153 | 1832.770411 | 1.10979902 | 0.285059983 | 1.4986999608 | 4.4275726277 | Km>KS.1 | YPD | KS.1 |
| FIM1_2661 | 2066.021066 | 2979.998306 | 1387.871653 | 0.40854445 | 0.312851421 | 0.126203143 | 0.177248034 | ns | YPD | KS.1 |
| FIM1_2662 | 5237.505743 | 7445.998347 | 5391.751476 | 0.481094916 | 0.13702396 | 0.000261351 | 0.000643675 | ns | YPD | KS.1 |
| FIM1_2663 | 293.6636454 | 184.3471795 | 406.4431412 | -0.529643851 | 0.31683935 | 0.050066666 | 0.078131011 | ns | YPD | KS.1 |
| FIM1_2664 | 1582.257878 | 2151.08802 | 2489.737755 | 0.431294418 | 0.091979891 | 1.7658532757 | 5.9803403995 | ns | YPD | KS.1 |
| FIM1_2665 | 2065.552886 | 608.1418787 | 497.1695574 | -1.679237695 | 0.214512081 | 2.4870645899 | 2.3748841334 | Km<KS.1 | YPD | KS.1 |
| FIM1_2666 | 3683.454087 | 2160.443263 | 1035.415032 | -0.711190597 | 0.190979338 | 7.0411705980 | 0.000188535 | ns | YPD | KS.1 |
| FIM1_2667 | 2738.374518 | 3968.024559 | 3083.454991 | 0.526094388 | 0.075608297 | 1.8140680025 | 1.2212433440 | ns | YPD | KS.1 |
| FIM1_2668 | 5166.992508 | 4856.371168 | 3118.802518 | -0.071485674 | 0.256028544 | 0.754527621 | 0.80639633 | ns | YPD | KS.1 |
| FIM1_2669 | 151.4355123 | 59.49190756 | 47.98920621 | -1.20641427 | 0.2855598 | 2.9818012061 | 9.7530973048 | Km<KS.1 | YPD | KS.1 |
| FIM1_2671 | 152.7746461 | 17.86693252 | 54.40823865 | -2.938539588 | 0.315195109 | 1.6947738407 | 2.8874999513 | Km<KS.1 | YPD | KS.1 |
| FIM1_2672 | 781.7385572 | 961.1537726 | 872.2161792 | 0.285766361 | 0.104271282 | 0.004903716 | 0.009554859 | ns | YPD | KS.1 |
| FIM1_2674 | 7600.474978 | 4752.933119 | 5072.522202 | -0.641133373 | 0.151326007 | 9.4553382675 | 2.8806067287 | ns | YPD | KS.1 |
| FIM1_2675 | 69899.94851 | 20644.87172 | 10561.10044 | -1.039984437 | 0.624869679 | 0.014114301 | 0.024953321 | Km<KS.1 | YPD | KS.1 |
| FIM1_2676 | 65055.69876 | 30706.5406 | 23990.20896 | -0.923203869 | 0.309780599 | 0.000648964 | 0.001492648 | ns | YPD | KS.1 |
| FIM1_2677 | 1116.883516 | 3295.122148 | 3814.504978 | 1.536067114 | 0.115360798 | 1.2752177877 | 6.9716576855 | Km>KS.1 | YPD | KS.1 |
| FIM1_2678 | 459.6198366 | 462.5861645 | 579.2869035 | 0.007040429 | 0.145322042 | 0.959596136 | 0.970126148 | ns | YPD | KS.1 |

| | | | | | | | | | | |
|---|---|---|---|---|---|---|---|---|---|---|
| FIM1_2679 | 2597.226365 | 2536.163738 | 1383.071188 | -0.031150831 | 0.153512344 | 0.831252464 | 0.870004778 | ns | YPD | KS.1 |
| FIM1_2680 | 2380.326404 | 3067.226968 | 4303.406281 | 0.342019115 | 0.143617708 | 0.012765204 | 0.022819579 | ns | YPD | KS.1 |
| FIM1_2681 | 17401.42904 | 4566.144465 | 4554.22403 | -1.865354459 | 0.18615858 | 3.97223717154 | 7.78026768835 | Km<KS.1 | YPD | KS.1 |
| FIM1_2682 | 4177.718747 | 1399.502688 | 945.3060971 | -1.414134658 | 0.303706619 | 2.63941413746 | 1.01555184330 | Km<KS.1 | YPD | KS.1 |
| FIM1_2683 | 4639.164055 | 1532.754539 | 2297.684861 | -1.537402091 | 0.18118838 | 1.47909750141 | 1.69942495832 | Km<KS.1 | YPD | KS.1 |
| FIM1_2684 | 6242.794806 | 2862.46827 | 5456.353642 | -1.019332062 | 0.249055332 | 7.83438562535 | 2.41337885362 | Km<KS.1 | YPD | KS.1 |
| FIM1_2685 | 1097.922737 | 777.399845 | 1047.779783 | -0.476397616 | 0.124442126 | 7.62694466079 | 0.000202476 | ns | YPD | KS.1 |
| FIM1_2686 | 159.0819207 | 527.3641056 | 211.4828374 | 1.651905291 | 0.200716562 | 1.00247322069 | 1.08184474467 | Km>KS.1 | YPD | KS.1 |
| FIM1_2687 | 971.3004181 | 1568.585451 | 777.3624947 | 0.568572219 | 0.290484672 | 0.024696847 | 0.041175205 | ns | YPD | KS.1 |
| FIM1_2688 | 3262.077149 | 2616.874999 | 1954.134537 | -0.299901946 | 0.129690857 | 0.016301886 | 0.02846679 | ns | YPD | KS.1 |
| FIM1_2689 | 2502.404121 | 1582.206453 | 1808.791337 | -0.641822684 | 0.10992559 | 2.15310754160 | 1.06903293607 | ns | YPD | KS.1 |
| FIM1_2690 | 3097.565153 | 5027.12779 | 6279.00711 | 0.685666269 | 0.086368494 | 7.39309703662 | 6.69270475838 | ns | YPD | KS.1 |
| FIM1_2691 | 751.2882383 | 1465.455358 | 1404.064091 | 0.953010864 | 0.078296984 | 8.89770152620 | 3.30343769349 | ns | YPD | KS.1 |
| FIM1_2692 | 618.3893231 | 647.572893 | 751.1524816 | 0.061381281 | 0.125447557 | 0.612159164 | 0.682081039 | ns | YPD | KS.1 |
| FIM1_2693 | 1385.880988 | 1976.514022 | 2325.402404 | 0.49872589 | 0.092508056 | 3.94250286156 | 1.68504740002 | ns | YPD | KS.1 |
| FIM1_2694 | 999.1446748 | 540.0110929 | 600.8518616 | -0.870690282 | 0.097938132 | 1.47452553841 | 1.89066096743 | ns | YPD | KS.1 |
| FIM1_2695 | 740.402759 | 1846.603631 | 2827.478016 | 1.291558088 | 0.121856272 | 3.33105170057 | 7.67221398627 | Km>KS.1 | YPD | KS.1 |
| FIM1_2696 | 3215.267554 | 4213.161577 | 6745.90564 | 0.343017934 | 0.20449762 | 0.069015881 | 0.104109767 | ns | YPD | KS.1 |
| FIM1_2697 | 165.298787 | 210.8437238 | 428.8738886 | 0.290343804 | 0.252546555 | 0.198068278 | 0.262490592 | ns | YPD | KS.1 |
| FIM1_2698 | 2514.315049 | 3199.597419 | 3296.22013 | 0.335185008 | 0.102242401 | 0.000781666 | 0.001764424 | ns | YPD | KS.1 |
| FIM1_2699 | 1969.437749 | 1086.718795 | 2342.557362 | -0.760137132 | 0.246307249 | 0.000657025 | 0.001508399 | ns | YPD | KS.1 |
| FIM1_2700 | 1531.720571 | 683.9826632 | 571.3223321 | -1.10056205 | 0.189665389 | 1.02747827835 | 5.33580838705 | Km<KS.1 | YPD | KS.1 |
| FIM1_2701 | 1794.348451 | 2486.125523 | 2051.847764 | 0.422052282 | 0.193826255 | 0.019144031 | 0.032864581 | ns | YPD | KS.1 |
| FIM1_2702 | 345.2227882 | 194.0049568 | 193.6273669 | -0.783807272 | 0.16994668 | 1.20648748816 | 4.19446209197 | ns | YPD | KS.1 |
| FIM1_2703 | 315.6494345 | 63.37269138 | 206.2115144 | -1.987629951 | 0.42975682 | 7.65243292649 | 3.15417181518 | Km<KS.1 | YPD | KS.1 |
| FIM1_2704 | 2797.151653 | 1671.532783 | 1576.395522 | -0.712878793 | 0.13481306 | 4.34207698597 | 1.84473381769 | ns | YPD | KS.1 |
| FIM1_2705 | 4056.916894 | 2005.427744 | 3068.511263 | -0.943865928 | 0.206915683 | 1.09244563363 | 3.84093076137 | ns | YPD | KS.1 |
| FIM1_2706 | 76.10757887 | 17.92416865 | 23.14872675 | -1.883086129 | 0.338519761 | 7.68379377535 | 4.01965026628 | Km<KS.1 | YPD | KS.1 |
| FIM1_2708 | 3076.359355 | 3252.455189 | 5778.449195 | 0.063785989 | 0.260535694 | 0.783271238 | 0.830337611 | ns | YPD | KS.1 |
| FIM1_2709 | 1309.492615 | 2461.612167 | 3182.780214 | 0.88575131 | 0.11940034 | 2.81390669419 | 2.17716731005 | ns | YPD | KS.1 |
| FIM1_2710 | 4436.109751 | 5764.260193 | 8876.301857 | 0.343280588 | 0.173652167 | 0.035501003 | 0.057083869 | ns | YPD | KS.1 |
| FIM1_2711 | 246.9530614 | 419.6450356 | 333.9744302 | 0.740947645 | 0.116720021 | 7.09598812230 | 4.06243278578 | ns | YPD | KS.1 |
| FIM1_2712 | 1213.42704 | 2535.741292 | 2749.901888 | 1.019909034 | 0.158059499 | 2.01880441666 | 1.21592638897 | Km>KS.1 | YPD | KS.1 |
| FIM1_2713 | 1299.698993 | 1874.028059 | 1923.715799 | 0.510808843 | 0.105502582 | 7.08926977096 | 2.56876308161 | ns | YPD | KS.1 |
| FIM1_2714 | 2284.823446 | 1541.073233 | 1521.290893 | -0.559437657 | 0.071974933 | 3.72962988717 | 3.13957845832 | ns | YPD | KS.1 |
| FIM1_2715 | 1295.635468 | 2575.228159 | 2877.232405 | 0.975463386 | 0.095110417 | 2.17978676093 | 4.32049367953 | ns | YPD | KS.1 |
| FIM1_2716 | 3074.442487 | 5293.860432 | 7226.098918 | 0.749372856 | 0.145180004 | 7.93309949604 | 3.26174958618 | ns | YPD | KS.1 |
| FIM1_2718 | 8711.32349 | 7320.551515 | 5181.679644 | -0.241919998 | 0.096949724 | 0.010610314 | 0.019251026 | ns | YPD | KS.1 |
| FIM1_2719 | 1633.058006 | 2008.493668 | 2146.318507 | 0.273306932 | 0.159509151 | 0.070509716 | 0.105945586 | ns | YPD | KS.1 |
| FIM1_2720 | 4043.903742 | 3663.446289 | 3393.652226 | -0.138638995 | 0.074105608 | 0.057377681 | 0.088705395 | ns | YPD | KS.1 |
| FIM1_2721 | 838.4564171 | 799.8382893 | 627.1013239 | -0.055692309 | 0.236192402 | 0.79459214 | 0.838764247 | ns | YPD | KS.1 |
| FIM1_2722 | 466.4217792 | 1490.333121 | 2572.884247 | 1.637455203 | 0.142023454 | 5.14745241053 | 1.63111947403 | Km>KS.1 | YPD | KS.1 |
| FIM1_2723 | 938.8668942 | 527.0096459 | 810.7871717 | -0.73720238 | 0.244402847 | 0.000872522 | 0.001950044 | ns | YPD | KS.1 |
| FIM1_2724 | 2018.575068 | 2143.839137 | 1911.980482 | 0.081076376 | 0.127677893 | 0.510027269 | 0.585595583 | ns | YPD | KS.1 |
| FIM1_2726 | 1349.815823 | 1077.527112 | 789.6774471 | -0.302379448 | 0.146315275 | 0.030361351 | 0.049556339 | ns | YPD | KS.1 |

| | | | | | | | | | | |
|---|---|---|---|---|---|---|---|---|---|---|
| FIM1_2727 | 275.4144734 | 102.0090504 | 134.5850001 | -1.34551199 | 0.221606588 | 1.2475609241 | 7.0052094781 | Km<KS.1 | YPD | KS.1 |
| FIM1_2728 | 1845.719499 | 500.3948589 | 592.1077316 | -1.758891523 | 0.260093847 | 5.5756588496 | 3.8904491973 | Km<KS.1 | YPD | KS.1 |
| FIM1_2729 | 156.7440492 | 1179.984883 | 331.3030649 | 2.698975844 | 0.361138712 | 2.2087820377 | 2.1172814331 | Km>KS.1 | YPD | KS.1 |
| FIM1_2730 | 766.9757078 | 146.0477801 | 150.8965523 | -2.255589869 | 0.275548609 | 2.8410383605 | 3.1833706855 | Km<KS.1 | YPD | KS.1 |
| FIM1_2731 | 4630.684122 | 5380.49334 | 4238.867516 | 0.207732629 | 0.101506527 | 0.0356788 | 0.057314185 | ns | YPD | KS.1 |
| FIM1_2732 | 10252.05958 | 9857.232712 | 8924.198354 | -0.054276663 | 0.094839259 | 0.558304402 | 0.631551705 | ns | YPD | KS.1 |
| FIM1_2733 | 373.2990107 | 324.0351655 | 314.6935133 | -0.186727128 | 0.153759316 | 0.201357786 | 0.26635336 | ns | YPD | KS.1 |
| FIM1_2734 | 111.1715028 | 210.5009785 | 239.54723 | 0.890948646 | 0.143631654 | 1.3134217763 | 7.3667117670 | ns | YPD | KS.1 |
| FIM1_2735 | 425.1824748 | 542.990162 | 651.349206 | 0.334677755 | 0.128710422 | 0.006971823 | 0.013103446 | ns | YPD | KS.1 |
| FIM1_2736 | 407.5006651 | 312.7147327 | 595.3209154 | -0.356359 | 0.148858461 | 0.012075151 | 0.021718683 | ns | YPD | KS.1 |
| FIM1_2737 | 972.3016268 | 1877.714391 | 2475.593407 | 0.92506994 | 0.119676687 | 2.3607637105 | 2.0319722249 | ns | YPD | KS.1 |
| FIM1_2738 | 234.1751066 | 550.0364457 | 482.4117522 | 1.11666835 | 0.25650995 | 2.0187286797 | 6.7813471856 | Km>KS.1 | YPD | KS.1 |
| FIM1_2739 | 1435.934839 | 2294.813826 | 2229.439946 | 0.559558617 | 0.283912229 | 0.02437322 | 0.040690191 | ns | YPD | KS.1 |
| FIM1_2740 | 2228.368059 | 4798.214126 | 5832.091995 | 1.094298665 | 0.079583829 | 7.9039021966 | 4.9721758111 | Km>KS.1 | YPD | KS.1 |
| FIM1_2741 | 362.2429731 | 673.3672114 | 950.3022383 | 0.870542032 | 0.112231004 | 2.1263878160 | 1.8397877191 | ns | YPD | KS.1 |
| FIM1_2742 | 3336.742997 | 7868.020522 | 6492.584788 | 1.127625948 | 0.252106846 | 1.1384348401 | 3.9885305139 | Km>KS.1 | YPD | KS.1 |
| FIM1_2743 | 520.0559688 | 692.1590163 | 767.070899 | 0.330507691 | 0.277729907 | 0.174761431 | 0.235173957 | ns | YPD | KS.1 |
| FIM1_2744 | 2756.070533 | 5230.309384 | 6478.222438 | 0.913076025 | 0.077563722 | 1.1901576644 | 3.9212148217 | ns | YPD | KS.1 |
| FIM1_2745 | 240.0513845 | 807.8911906 | 1426.642156 | 1.624300324 | 0.261863051 | 3.0679503915 | 1.8322993034 | Km>KS.1 | YPD | KS.1 |
| FIM1_2746 | 1361.48695 | 1011.268492 | 1398.377701 | -0.407942951 | 0.128950787 | 0.001041148 | 0.002299028 | ns | YPD | KS.1 |
| FIM1_2747 | 1439.109233 | 2282.291268 | 2989.429139 | 0.625751564 | 0.159199603 | 3.6646898946 | 0.000102899 | ns | YPD | KS.1 |
| FIM1_2748 | 1019.047672 | 554.2540459 | 558.3447268 | -0.847651765 | 0.134222457 | 6.9826230707 | 4.0067531461 | ns | YPD | KS.1 |
| FIM1_2749 | 2225.855295 | 1360.944765 | 918.7957471 | -0.680364487 | 0.133624676 | 1.3434302427 | 5.3468523659 | ns | YPD | KS.1 |
| FIM1_2750 | 200.015117 | 98.10158334 | 94.23558957 | -0.970887438 | 0.178801651 | 1.1434464858 | 5.2141578984 | ns | YPD | KS.1 |
| FIM1_2751 | 615.4405249 | 417.0678018 | 582.8865073 | -0.508119255 | 0.195926689 | 0.005271791 | 0.010169507 | ns | YPD | KS.1 |
| FIM1_2752 | 807.4881892 | 1472.651273 | 1326.891207 | 0.852609168 | 0.086923629 | 2.5710575881 | 4.3212876693 | ns | YPD | KS.1 |
| FIM1_2753 | 2098.700702 | 1797.984473 | 1404.235879 | -0.204053779 | 0.154731372 | 0.165027665 | 0.223709164 | ns | YPD | KS.1 |
| FIM1_2754 | 15322.48406 | 4702.432363 | 3605.0053 | -1.607274374 | 0.230456096 | 1.7951450495 | 1.3133597972 | Km<KS.1 | YPD | KS.1 |
| FIM1_2755 | 1097.0603 | 950.8827383 | 1085.069687 | -0.195808403 | 0.118242027 | 0.086823799 | 0.127418407 | ns | YPD | KS.1 |
| FIM1_2756 | 410.8784837 | 734.8121962 | 909.1344341 | 0.708506892 | 0.286397668 | 0.004868194 | 0.009505206 | ns | YPD | KS.1 |
| FIM1_2757 | 3627.417334 | 3180.784229 | 5736.513145 | -0.14655293 | 0.282269586 | 0.552983762 | 0.626244985 | ns | YPD | KS.1 |
| FIM1_2758 | 297.9519878 | 619.9025389 | 579.5831668 | 1.014310183 | 0.156515226 | 1.6966146186 | 1.0356635248 | Km>KS.1 | YPD | KS.1 |
| FIM1_2759 | 315.0193761 | 70.20159051 | 236.8115605 | -2.101592904 | 0.189456309 | 2.2103136800 | 6.2479037263 | Km<KS.1 | YPD | KS.1 |
| FIM1_2760 | 9685.735541 | 10559.85064 | 8012.268097 | 0.107166867 | 0.203416315 | 0.568922592 | 0.641520828 | ns | YPD | KS.1 |
| FIM1_2761 | 1276.964071 | 1856.938881 | 1486.793316 | 0.53379329 | 0.063356484 | 1.8437853965 | 1.9516664569 | ns | YPD | KS.1 |
| FIM1_2762 | 3027.328925 | 2957.799143 | 2559.5786 | -0.029139244 | 0.19066159 | 0.869845009 | 0.899870851 | ns | YPD | KS.1 |
| FIM1_2763 | 4061.788841 | 1351.86538 | 643.8802253 | -0.953043151 | 0.592039627 | 0.019997097 | 0.034117132 | ns | YPD | KS.1 |
| FIM1_2764 | 1476.044775 | 1054.799786 | 1403.205605 | -0.444583512 | 0.174415162 | 0.006754881 | 0.01273902 | ns | YPD | KS.1 |
| FIM1_2765 | 512.47189 | 625.4003634 | 1273.675358 | 0.271829204 | 0.126630118 | 0.025925186 | 0.043050001 | ns | YPD | KS.1 |
| FIM1_2766 | 4223.561636 | 3176.240986 | 3595.941157 | -0.392957313 | 0.118972415 | 0.000655658 | 0.001505955 | ns | YPD | KS.1 |
| FIM1_2767 | 3194.055945 | 2647.305843 | 2284.431879 | -0.236807576 | 0.198745104 | 0.198207392 | 0.262605001 | ns | YPD | KS.1 |
| FIM1_2768 | 1470.140881 | 1316.588219 | 1224.0717 | -0.140178929 | 0.187068632 | 0.422194206 | 0.49986106 | ns | YPD | KS.1 |
| FIM1_2769 | 1478.917954 | 1486.49305 | 2209.538832 | 0.006972691 | 0.092759055 | 0.938781305 | 0.9556859 | ns | YPD | KS.1 |
| FIM1_2770 | 2811.078098 | 1843.915533 | 1412.254494 | -0.567240327 | 0.164997894 | 0.000288359 | 0.000703919 | ns | YPD | KS.1 |
| FIM1_2771 | 104.855968 | 415.7389682 | 445.9064314 | 1.892132925 | 0.218464088 | 1.4005649584 | 1.8004678728 | Km>KS.1 | YPD | KS.1 |

| | | | | | | | | | | |
|---|---|---|---|---|---|---|---|---|---|---|
| FIM1_2772 | 2068.157194 | 2826.755913 | 1609.891903 | 0.439113478 | 0.091852086 | 1.1078689695( | 3.8869168713 | ns | YPD | KS.1 |
| FIM1_2773 | 358.7550974 | 381.4388939 | 405.4531913 | 0.079943656 | 0.162938162 | 0.60443524 | 0.674532373 | ns | YPD | KS.1 |
| FIM1_2774 | 2389.96064 | 1058.616863 | 928.7320496 | -1.055086505 | 0.264724574 | 1.1465531777( | 3.4549376495 | Km<KS.1 | YPD | KS.1 |
| FIM1_2775 | 2514.501634 | 3824.771424 | 4011.411568 | 0.594215305 | 0.082806789 | 3.2343200234 | 2.3085713223 | ns | YPD | KS.1 |
| FIM1_2776 | 754.9233367 | 883.3289303 | 718.1820608 | 0.214514877 | 0.120616997 | 0.065626739 | 0.099830282 | ns | YPD | KS.1 |
| FIM1_2777 | 839.9225632 | 850.9417793 | 817.4033035 | 0.017394411 | 0.140312739 | 0.897108512 | 0.921559952 | ns | YPD | KS.1 |
| FIM1_2778 | 236.1956094 | 586.94647 | 579.844977 | 1.287406257 | 0.11704133 | 4.2734211632 | 1.1073057441 | Km>KS.1 | YPD | KS.1 |
| FIM1_2779 | 500.3968246 | 745.7775267 | 941.1714081 | 0.545537076 | 0.142803158 | 6.8554500361 | 0.000183958 | ns | YPD | KS.1 |
| FIM1_2780 | 770.7169678 | 262.5094509 | 312.6164471 | -1.428616574 | 0.264805778 | 5.6177980490 | 2.6668459250 | Km<KS.1 | YPD | KS.1 |
| FIM1_2781 | 4989.490859 | 3092.199585 | 1642.397071 | -0.580895589 | 0.272856798 | 0.015929499 | 0.02788503 | ns | YPD | KS.1 |
| FIM1_2782 | 456.0655552 | 844.1720184 | 1563.850147 | 0.847686912 | 0.154807732 | 1.1376867821 | 5.1926529735 | ns | YPD | KS.1 |
| FIM1_2783 | 4563.38389 | 3384.945068 | 6384.891685 | -0.3412065 | 0.286690868 | 0.171825191 | 0.232038634 | ns | YPD | KS.1 |
| FIM1_2784 | 1103.096549 | 960.4646065 | 907.651831 | -0.192651785 | 0.084822956 | 0.020566644 | 0.034918579 | ns | YPD | KS.1 |
| FIM1_2785 | 1811.664186 | 2032.066634 | 2810.77649 | 0.156851458 | 0.114919688 | 0.158798079 | 0.216325423 | ns | YPD | KS.1 |
| FIM1_2786 | 1584.675534 | 2919.847331 | 2411.717556 | 0.841393604 | 0.154087141 | 1.2565638012( | 5.6934470956 | ns | YPD | KS.1 |
| FIM1_2787 | 6072.354459 | 9227.872557 | 5756.818924 | 0.500529059 | 0.274016866 | 0.038066395 | 0.06067937 | ns | YPD | KS.1 |
| FIM1_2788 | 560.7979637 | 475.3884162 | 344.3151537 | -0.213872388 | 0.176186538 | 0.195945832 | 0.259954803 | ns | YPD | KS.1 |
| FIM1_2789 | 168.4064591 | 175.4929505 | 274.6503493 | 0.049556378 | 0.192021773 | 0.781819168 | 0.829328435 | ns | YPD | KS.1 |
| FIM1_2790 | 1510.722463 | 997.7937145 | 1088.731515 | -0.573129045 | 0.127188217 | 3.1892104648( | 1.0376927444 | ns | YPD | KS.1 |
| FIM1_2791 | 719.388203 | 254.2014631 | 239.3295169 | -1.439194792 | 0.183914028 | 4.2284733667 | 3.9174404096 | Km<KS.1 | YPD | KS.1 |
| FIM1_2792 | 253.407452 | 469.287129 | 699.8149305 | 0.838778255 | 0.168564799 | 1.7367211395 | 6.8301878810( | ns | YPD | KS.1 |
| FIM1_2793 | 4057.416514 | 2071.872676 | 1821.019608 | -0.957062803 | 0.083467133 | 3.9267718712 | 1.1839812157 | ns | YPD | KS.1 |
| FIM1_2794 | 1066.55157 | 2033.902836 | 2224.607093 | 0.899627136 | 0.135027957 | 6.2479241962 | 3.9850542149( | ns | YPD | KS.1 |
| FIM1_2795 | 988.3475032 | 780.1097421 | 653.7490727 | -0.276802426 | 0.261387641 | 0.232889127 | 0.300706827 | ns | YPD | KS.1 |
| FIM1_2796 | 3935.275995 | 583.5456092 | 654.7569723 | -2.546951541 | 0.349548258 | 1.3884847692 | 1.2247715828( | Km<KS.1 | YPD | KS.1 |
| FIM1_2797 | 72.85461825 | 59.19037129 | 53.41088707 | -0.236730692 | 0.282633177 | 0.337964402 | 0.413012258 | ns | YPD | KS.1 |
| FIM1_2798 | 841.1285617 | 900.5567378 | 1657.686551 | 0.093975257 | 0.119898453 | 0.417689934 | 0.495471488 | ns | YPD | KS.1 |
| FIM1_2800 | 4053.380343 | 3162.268869 | 3329.2877 | -0.3467972 | 0.097165114 | 0.000263812 | 0.000649093 | ns | YPD | KS.1 |
| FIM1_2801 | 3272.680216 | 3497.92508 | 4352.614611 | 0.089689867 | 0.123212924 | 0.451002413 | 0.528559955 | ns | YPD | KS.1 |
| FIM1_2802 | 1232.289128 | 879.0176428 | 855.5880216 | -0.463821599 | 0.130842634 | 0.000237687 | 0.000590346 | ns | YPD | KS.1 |
| FIM1_2803 | 1437.370526 | 1545.928364 | 1290.444468 | 0.100835072 | 0.095725006 | 0.280875033 | 0.352155567 | ns | YPD | KS.1 |
| FIM1_2804 | 3029.16215 | 2192.545642 | 2299.026539 | -0.455459735 | 0.085750877 | 6.6817229776 | 2.7745402563 | ns | YPD | KS.1 |
| FIM1_2805 | 1641.45253 | 1227.561911 | 1868.510163 | -0.399825819 | 0.121714889 | 0.00069106 | 0.001579511 | ns | YPD | KS.1 |
| FIM1_2806 | 236.7267137 | 253.4416072 | 204.5735688 | 0.090484736 | 0.14550116 | 0.515181702 | 0.590968174 | ns | YPD | KS.1 |
| FIM1_2807 | 1006.783328 | 924.9949067 | 1116.620185 | -0.11297803 | 0.141753166 | 0.404982209 | 0.481775822 | ns | YPD | KS.1 |
| FIM1_2808 | 613.1657853 | 1563.486065 | 1536.198656 | 1.30793268 | 0.150387096 | 3.6675855987 | 4.4611829715 | Km>KS.1 | YPD | KS.1 |
| FIM1_2809 | 190.6724401 | 256.073446 | 172.0515214 | 0.358520208 | 0.238211198 | 0.095235363 | 0.138294201 | ns | YPD | KS.1 |
| FIM1_2810 | 380.6724185 | 67.03426035 | 237.534163 | -2.322446038 | 0.322027447 | 4.7496612801 | 3.9464311434 | Km<KS.1 | YPD | KS.1 |
| FIM1_2811 | 485.1677805 | 451.3256211 | 460.8452583 | -0.097794532 | 0.127690995 | 0.426768278 | 0.504556127 | ns | YPD | KS.1 |
| FIM1_2812 | 4166.341881 | 2407.650359 | 2959.470854 | -0.775248852 | 0.09584858 | 1.7946375003( | 1.7302948768( | ns | YPD | KS.1 |
| FIM1_2813 | 2257.507542 | 1095.392857 | 1060.003627 | -1.0078766 | 0.141989471 | 2.3702351369 | 1.7264889906 | Km<KS.1 | YPD | KS.1 |
| FIM1_2814 | 4587.067614 | 2303.887475 | 2193.081339 | -0.957554184 | 0.143673511 | 5.4868387619 | 3.5176575825( | ns | YPD | KS.1 |
| FIM1_2815 | 252.9806803 | 139.7657102 | 337.3905196 | -0.814269808 | 0.159798998 | 9.7899962611 | 3.9629968591 | ns | YPD | KS.1 |
| FIM1_2816 | 274.1704267 | 234.5042083 | 373.392304 | -0.207476497 | 0.152424752 | 0.15234416 | 0.208618826 | ns | YPD | KS.1 |
| FIM1_2817 | 521.3382262 | 324.5771366 | 534.8311899 | -0.645846304 | 0.154437291 | 1.1933621041( | 3.5894658209 | ns | YPD | KS.1 |

| | | | | | | | | | | |
|---|---|---|---|---|---|---|---|---|---|---|
| FIM1_2818 | 660.2161539 | 526.6145987 | 428.9503715 | -0.31180011 | 0.112471844 | 0.004302291 | 0.008493611 | ns | YPD | KS.1 |
| FIM1_2819 | 1587.848736 | 1391.471205 | 2042.335681 | -0.184780586 | 0.08057904 | 0.019563975 | 0.033482814 | ns | YPD | KS.1 |
| FIM1_2820 | 4196.208305 | 9963.216539 | 2772.900424 | 1.126892905 | 0.264357594 | 2.9718535475 | 9.7269548676 | Km>KS.1 | YPD | KS.1 |
| FIM1_2821 | 1887.420117 | 2495.543577 | 1743.736036 | 0.383377658 | 0.125766444 | 0.001600294 | 0.003427232 | ns | YPD | KS.1 |
| FIM1_2822 | 612.105935 | 1012.058664 | 639.5826493 | 0.689350755 | 0.146852104 | 9.9745379725 | 3.5168906033 | ns | YPD | KS.1 |
| FIM1_2823 | 1424.997782 | 2976.06935 | 3390.920902 | 1.046672522 | 0.091144133 | 2.7302975631 | 8.3847101090 | Km>KS.1 | YPD | KS.1 |
| FIM1_2824 | 1647.813969 | 1574.137944 | 1247.898865 | -0.063218262 | 0.082764934 | 0.436676559 | 0.513585314 | ns | YPD | KS.1 |
| FIM1_2825 | 615.384808 | 1089.146091 | 1050.128606 | 0.801131898 | 0.115248587 | 1.0265457741 | 7.0637140058 | ns | YPD | KS.1 |
| FIM1_2826 | 989.8057081 | 1292.42783 | 888.972011 | 0.348304518 | 0.178812717 | 0.03770092 | 0.060173909 | ns | YPD | KS.1 |
| FIM1_2827 | 2067.361741 | 2046.329918 | 3463.262847 | -0.012132947 | 0.227689276 | 0.953359928 | 0.966965472 | ns | YPD | KS.1 |
| FIM1_2828 | 1529.279317 | 982.9437768 | 768.7387174 | -0.619647397 | 0.104928697 | 1.5129381951 | 7.7040609222 | ns | YPD | KS.1 |
| FIM1_2829 | 663.2996555 | 2844.919644 | 2250.230989 | 2.065896479 | 0.133561278 | 9.6844996076 | 1.1471520368 | Km>KS.1 | YPD | KS.1 |
| FIM1_2830 | 4859.41108 | 2718.244342 | 2405.026122 | -0.801542094 | 0.147595974 | 1.6186361003 | 7.2286486527 | ns | YPD | KS.1 |
| FIM1_2831 | 1746.949347 | 309.6940551 | 339.8605135 | -2.162884815 | 0.437210275 | 9.7509420540 | 4.4834507133 | Km<KS.1 | YPD | KS.1 |
| FIM1_2832 | 3122.714426 | 2590.679347 | 1539.108163 | -0.256436927 | 0.11408314 | 0.020403364 | 0.034703158 | ns | YPD | KS.1 |
| FIM1_2833 | 1590.816328 | 1159.501855 | 1969.477907 | -0.357519709 | 0.295988177 | 0.162550924 | 0.220712567 | ns | YPD | KS.1 |
| FIM1_2834 | 3946.584689 | 4501.11582 | 1728.387067 | 0.141056932 | 0.306980126 | 0.592389902 | 0.663919748 | ns | YPD | KS.1 |
| FIM1_2835 | 809.7836244 | 1743.85764 | 1076.06063 | 0.951710714 | 0.304197967 | 0.000371645 | 0.00088763 | ns | YPD | KS.1 |
| FIM1_2836 | 1456.084712 | 1185.100267 | 864.2023526 | -0.279610604 | 0.127383922 | 0.022717686 | 0.038169702 | ns | YPD | KS.1 |
| FIM1_2837 | 992.4632894 | 1107.688338 | 858.4962492 | 0.12658614 | 0.259172471 | 0.583360386 | 0.655640025 | ns | YPD | KS.1 |
| FIM1_2838 | 3397.478043 | 3021.007675 | 2749.047553 | -0.160221615 | 0.116078112 | 0.153927882 | 0.21049786 | ns | YPD | KS.1 |
| FIM1_2839 | 1175.208939 | 940.7665874 | 734.8702793 | -0.310877074 | 0.094080207 | 0.00073673 | 0.001673621 | ns | YPD | KS.1 |
| FIM1_2840 | 164.9928268 | 124.818969 | 112.8839267 | -0.368977451 | 0.169236684 | 0.020764518 | 0.035173129 | ns | YPD | KS.1 |
| FIM1_2841 | 1537.045049 | 995.3422026 | 1116.009165 | -0.603452022 | 0.121748391 | 3.2309620813 | 1.2279630522 | ns | YPD | KS.1 |
| FIM1_2842 | 953.0664618 | 2131.422428 | 3122.825629 | 1.139947414 | 0.109641486 | 3.7274724589 | 7.9248613176 | Km>KS.1 | YPD | KS.1 |
| FIM1_2843 | 265.2938848 | 211.8583278 | 223.4923183 | -0.296311278 | 0.168435195 | 0.06202756 | 0.095154828 | ns | YPD | KS.1 |
| FIM1_2844 | 690.7754903 | 407.0823526 | 525.3170241 | -0.709879678 | 0.181161702 | 3.2011874638 | 9.0591055930 | ns | YPD | KS.1 |
| FIM1_2845 | 73062.74804 | 37649.92367 | 43296.34962 | -0.929436142 | 0.124929057 | 2.1960181096 | 1.7232161033 | ns | YPD | KS.1 |
| FIM1_2846 | 1730.549753 | 945.0436896 | 1216.290288 | -0.739911136 | 0.288940966 | 0.003547939 | 0.007114468 | ns | YPD | KS.1 |
| FIM1_2847 | 11228.50937 | 6113.877448 | 3194.810321 | -0.641462831 | 0.391929424 | 0.042365462 | 0.066953041 | ns | YPD | KS.1 |
| FIM1_2848 | 1385.8731 | 560.5771776 | 706.3170098 | -1.267196175 | 0.146232876 | 5.2184649717 | 6.2709814575 | Km<KS.1 | YPD | KS.1 |
| FIM1_2849 | 7429.564885 | 9846.525125 | 14759.95724 | 0.38749714 | 0.122042861 | 0.00103519 | 0.002286887 | ns | YPD | KS.1 |
| FIM1_2850 | 1060.209013 | 1398.207471 | 2126.038905 | 0.370852372 | 0.154049911 | 0.011381476 | 0.020575162 | ns | YPD | KS.1 |
| FIM1_2851 | 2879.377232 | 3544.770823 | 2482.05434 | 0.269660998 | 0.177274863 | 0.104817311 | 0.15058219 | ns | YPD | KS.1 |
| FIM1_2852 | 813.7813909 | 484.9322069 | 869.8070042 | -0.709216705 | 0.152304978 | 1.1467751955 | 4.0092808137 | ns | YPD | KS.1 |
| FIM1_2853 | 2851.952646 | 2108.373626 | 4109.292691 | -0.34931018 | 0.278926458 | 0.152891112 | 0.209195072 | ns | YPD | KS.1 |
| FIM1_2854 | 929.5892118 | 1464.440504 | 1743.33323 | 0.614599848 | 0.161198941 | 6.0975691606 | 0.000166221 | ns | YPD | KS.1 |
| FIM1_2855 | 119.7641229 | 99.8084287 | 121.4704813 | -0.225854471 | 0.210144146 | 0.242497532 | 0.310774143 | ns | YPD | KS.1 |
| FIM1_2856 | 1030.594571 | 877.328255 | 634.690912 | -0.20823339 | 0.174705234 | 0.204569866 | 0.26997174 | ns | YPD | KS.1 |
| FIM1_2857 | 436.4306947 | 478.8064052 | 751.6658429 | 0.121317751 | 0.162989808 | 0.431673645 | 0.509387188 | ns | YPD | KS.1 |
| FIM1_2858 | 684.1039863 | 1273.548286 | 1021.325247 | 0.868699599 | 0.127613471 | 2.4549258145 | 1.6219463382 | ns | YPD | KS.1 |
| FIM1_2859 | 1217.735339 | 2890.920803 | 3942.214549 | 1.195298798 | 0.171268517 | 3.9312816305 | 2.7742022854 | Km>KS.1 | YPD | KS.1 |
| FIM1_2860 | 3293.45793 | 7631.486614 | 11384.25993 | 1.05130133 | 0.307714144 | 0.000108054 | 0.000280715 | Km>KS.1 | YPD | KS.1 |
| FIM1_2862 | 96.00919515 | 126.0408024 | 207.0002085 | 0.343449031 | 0.202408531 | 0.06621888 | 0.100499978 | ns | YPD | KS.1 |
| FIM1_2863 | 2749.412294 | 1667.075174 | 1263.715346 | -0.646172578 | 0.221154339 | 0.001449237 | 0.003117144 | ns | YPD | KS.1 |

| | | | | | | | | | | |
|---|---|---|---|---|---|---|---|---|---|---|
| FIM1_2864 | 834.8095082 | 1775.647712 | 3071.00938 | 1.053629799 | 0.142485797 | 2.4394986033 | 1.9022735974 | Km>KS.1 | YPD | KS.1 |
| FIM1_2865 | 295.6774913 | 595.336561 | 1040.611865 | 0.956961432 | 0.177859335 | 1.5505218746 | 6.9494111049 | ns | YPD | KS.1 |
| FIM1_2866 | 1187.761571 | 771.7482988 | 682.8797352 | -0.588058043 | 0.1481664 | 3.3890202456 | 9.5526208057 | ns | YPD | KS.1 |
| FIM1_2867 | 763.7387346 | 1250.659792 | 1333.668172 | 0.676490492 | 0.145788389 | 1.3355769448 | 4.6179205016 | ns | YPD | KS.1 |
| FIM1_2868 | 127.8013422 | 73.60332361 | 53.04285322 | -0.690957944 | 0.261700311 | 0.003134238 | 0.006348874 | ns | YPD | KS.1 |
| FIM1_2869 | 1408.981582 | 257.3540757 | 263.2725796 | -2.387636328 | 0.187711936 | 2.8672823101 | 1.3985028914 | Km<KS.1 | YPD | KS.1 |
| FIM1_2870 | 1358.03325 | 4417.880089 | 4619.041052 | 1.648805817 | 0.167995695 | 5.2597562598 | 9.5501048879 | Km>KS.1 | YPD | KS.1 |
| FIM1_2871 | 862.755665 | 1040.295939 | 1028.851863 | 0.257816497 | 0.111681418 | 0.017370034 | 0.030089108 | ns | YPD | KS.1 |
| FIM1_2872 | 14628.33808 | 5464.443596 | 3341.988258 | -1.380611598 | 0.148074329 | 1.0538625194 | 1.5465976501 | Km<KS.1 | YPD | KS.1 |
| FIM1_2873 | 551.5646705 | 565.9690608 | 379.4106986 | 0.032973551 | 0.178418128 | 0.844038397 | 0.879206663 | ns | YPD | KS.1 |
| FIM1_2874 | 8594.055523 | 5931.068415 | 5304.524092 | -0.496107109 | 0.166074404 | 0.001599509 | 0.003427027 | ns | YPD | KS.1 |
| FIM1_2875 | 3626.071144 | 2106.791339 | 2510.602835 | -0.769168217 | 0.090540312 | 5.9088074759 | 6.4749597340 | ns | YPD | KS.1 |
| FIM1_2876 | 2080.627501 | 2151.569375 | 2282.743622 | 0.045657579 | 0.107304275 | 0.661856835 | 0.72543242 | ns | YPD | KS.1 |
| FIM1_2877 | 1349.791952 | 1456.079363 | 3588.738224 | 0.086933666 | 0.261188512 | 0.708270996 | 0.767344992 | ns | YPD | KS.1 |
| FIM1_2878 | 1106.240679 | 1235.202201 | 1245.67938 | 0.148807075 | 0.127165628 | 0.22446937 | 0.291423569 | ns | YPD | KS.1 |
| FIM1_2879 | 3979.877867 | 2039.895092 | 1558.259213 | -0.919525224 | 0.161722118 | 2.9244597313 | 1.4348310812 | ns | YPD | KS.1 |
| FIM1_2880 | 742.4532661 | 1188.510293 | 1078.416442 | 0.643256739 | 0.14794898 | 5.6999922532 | 1.7913746974 | ns | YPD | KS.1 |
| FIM1_2881 | 750.2218903 | 768.0033755 | 1233.208042 | 0.0301524 | 0.178974133 | 0.857628083 | 0.889780873 | ns | YPD | KS.1 |
| FIM1_2882 | 2155.317169 | 1882.202807 | 1469.445907 | -0.182411447 | 0.1317721 | 0.149737562 | 0.205450585 | ns | YPD | KS.1 |
| FIM1_2883 | 2258.77458 | 2628.472657 | 2329.138013 | 0.210830045 | 0.093617959 | 0.021228891 | 0.035849875 | ns | YPD | KS.1 |
| FIM1_2884 | 985.4748596 | 716.2517752 | 690.2018904 | -0.443325353 | 0.109210891 | 3.0958446092 | 8.7859822766 | ns | YPD | KS.1 |
| FIM1_2885 | 1217.6143 | 618.5816322 | 1656.05392 | -0.790090122 | 0.340093497 | 0.006115781 | 0.011644091 | ns | YPD | KS.1 |
| FIM1_2886 | 788.1091975 | 269.2787534 | 931.8734118 | -1.31090563 | 0.368958225 | 3.6650659005 | 0.000102899 | Km<KS.1 | YPD | KS.1 |
| FIM1_2887 | 2932.636181 | 2006.114292 | 4640.995791 | -0.450000797 | 0.274108071 | 0.062065294 | 0.095175871 | ns | YPD | KS.1 |
| FIM1_2888 | 574.3645524 | 344.5212327 | 389.7551072 | -0.690247383 | 0.171244072 | 2.0833716607 | 6.0471260282 | ns | YPD | KS.1 |
| FIM1_2889 | 1429.475362 | 2049.133447 | 2031.263372 | 0.505576862 | 0.098321647 | 1.5104902102 | 5.9687758505 | ns | YPD | KS.1 |
| FIM1_2890 | 3151.840405 | 3503.954083 | 4347.024313 | 0.144968565 | 0.109498877 | 0.172713715 | 0.233048747 | ns | YPD | KS.1 |
| FIM1_2891 | 3980.770125 | 4081.946305 | 4319.216258 | 0.035068055 | 0.087987277 | 0.684353192 | 0.746471635 | ns | YPD | KS.1 |
| FIM1_2892 | 680.8784389 | 393.2054909 | 304.0822981 | -0.757400733 | 0.146608014 | 7.6090422482 | 3.1388876604 | ns | YPD | KS.1 |
| FIM1_2893 | 1363.980949 | 665.9637145 | 642.7355356 | -0.996532929 | 0.14753209 | 2.7475070985 | 1.8056602133 | ns | YPD | KS.1 |
| FIM1_2894 | 2484.138828 | 1431.360031 | 1040.38808 | -0.766264123 | 0.131142861 | 1.5879988610 | 8.0533071701 | ns | YPD | KS.1 |
| FIM1_2895 | 2276.874665 | 2578.047771 | 3125.276002 | 0.17053678 | 0.111182965 | 0.113860442 | 0.162308223 | ns | YPD | KS.1 |
| FIM1_2896 | 1884.75055 | 1637.468953 | 1441.378662 | -0.195874956 | 0.089741601 | 0.02576372 | 0.042796163 | ns | YPD | KS.1 |
| FIM1_2897 | 9946.967177 | 7787.172924 | 5776.34956 | -0.316842085 | 0.183739653 | 0.06495698 | 0.098977327 | ns | YPD | KS.1 |
| FIM1_2898 | 8357.125827 | 11210.70432 | 16191.39412 | 0.405335041 | 0.119107648 | 0.000447405 | 0.001051413 | ns | YPD | KS.1 |
| FIM1_2899 | 7621.916468 | 6868.957979 | 9340.909001 | -0.129561966 | 0.201136376 | 0.486613015 | 0.564312296 | ns | YPD | KS.1 |
| FIM1_2900 | 15617.31156 | 6225.260431 | 6646.927741 | -1.279750765 | 0.162178162 | 3.4000216425 | 3.1915297493 | Km<KS.1 | YPD | KS.1 |
| FIM1_2901 | 2036.187178 | 1424.006224 | 1677.89953 | -0.496194277 | 0.11458323 | 8.4561975804 | 2.5888974131 | ns | YPD | KS.1 |
| FIM1_2903 | 4346.962366 | 3439.615461 | 2443.122967 | -0.318281754 | 0.132324646 | 0.01237725 | 0.022197845 | ns | YPD | KS.1 |
| FIM1_2904 | 4019.740218 | 2220.616775 | 1819.270302 | -0.79552285 | 0.191785949 | 9.9499991224 | 3.0165292890 | ns | YPD | KS.1 |
| FIM1_2905 | 1237.78572 | 1481.884257 | 846.2092377 | 0.196690718 | 0.299886123 | 0.447152689 | 0.524542473 | ns | YPD | KS.1 |
| FIM1_2906 | 333.0827086 | 168.5893108 | 192.2999731 | -0.90344948 | 0.217886262 | 7.9322205167 | 2.4374797449 | ns | YPD | KS.1 |
| FIM1_2907 | 361.3361664 | 709.7667758 | 990.5456322 | 0.903523276 | 0.206474481 | 2.8349862547 | 9.3034674255 | ns | YPD | KS.1 |
| FIM1_2908 | 1396.143003 | 2632.52336 | 2481.130868 | 0.89358119 | 0.112093994 | 3.6400285439 | 3.3912250948 | ns | YPD | KS.1 |
| FIM1_2909 | 1455.025842 | 1196.690777 | 1242.605634 | -0.26927399 | 0.107672919 | 0.010115982 | 0.018434803 | ns | YPD | KS.1 |

| | | | | | | | | | | |
|---|---|---|---|---|---|---|---|---|---|---|
| FIM1_2911 | 300.3480686 | 382.9387476 | 424.6130829 | 0.284091326 | 0.26035754 | 0.219313495 | 0.285923648 | ns | YPD | KS.1 |
| FIM1_2912 | 7801.875184 | 4348.36349 | 2982.181995 | -0.718864149 | 0.280824274 | 0.003727302 | 0.00744413 | ns | YPD | KS.1 |
| FIM1_2913 | 22.90079411 | 22.65472999 | 14.35777334 | -0.004148709 | 0.335355929 | 0.988227724 | 0.991216316 | ns | YPD | KS.1 |
| FIM1_2914 | 1478.51092 | 1907.572796 | 1880.422927 | 0.278130428 | 0.309633574 | 0.293882135 | 0.365973373 | ns | YPD | KS.1 |
| FIM1_2915 | 731.1972264 | 683.6704919 | 1489.707832 | -0.087895999 | 0.165441872 | 0.574282171 | 0.64668488 | ns | YPD | KS.1 |
| FIM1_2916 | 4147.195591 | 4144.333202 | 5132.835234 | -0.000860768 | 0.153043812 | 0.995289008 | 0.996226101 | ns | YPD | KS.1 |
| FIM1_2917 | 2335.547668 | 1705.205369 | 1446.996132 | -0.440099134 | 0.099636908 | 6.3341711279 | 1.9819183246 | ns | YPD | KS.1 |
| FIM1_2918 | 1727.113153 | 789.9485408 | 985.6773065 | -1.102396042 | 0.120682781 | 1.0276013326 | 1.4280214050 | Km<KS.1 | YPD | KS.1 |
| FIM1_2919 | 3819.392529 | 3619.205673 | 3139.017352 | -0.073667957 | 0.108674274 | 0.485572019 | 0.563499136 | ns | YPD | KS.1 |
| FIM1_2920 | 323.7465512 | 219.9556209 | 333.924461 | -0.478238325 | 0.24226019 | 0.028385375 | 0.046698822 | ns | YPD | KS.1 |
| FIM1_2921 | 1576.140179 | 1859.642615 | 2861.91268 | 0.189640402 | 0.267788233 | 0.423227464 | 0.500607854 | ns | YPD | KS.1 |
| FIM1_2922 | 572.3263156 | 1303.283675 | 940.8518398 | 1.133016296 | 0.173986674 | 1.0997304941 | 6.8560892333 | Km>KS.1 | YPD | KS.1 |
| FIM1_2923 | 2317.692436 | 3104.752331 | 3964.760454 | 0.401692514 | 0.124501918 | 0.000846475 | 0.001896942 | ns | YPD | KS.1 |
| FIM1_2924 | 2984.579217 | 4496.777926 | 6416.49765 | 0.56496386 | 0.130948058 | 7.8782469584 | 2.4253885283 | ns | YPD | KS.1 |
| FIM1_2925 | 98.69584211 | 65.6693922 | 55.21287527 | -0.44698987 | 0.334228695 | 0.111343232 | 0.159038926 | ns | YPD | KS.1 |
| FIM1_2926 | 1093.796962 | 2290.052219 | 2247.740019 | 1.002030967 | 0.192028407 | 3.4469770584 | 1.4886033737 | Km>KS.1 | YPD | KS.1 |
| FIM1_2927 | 4683.162232 | 4648.931658 | 5522.650867 | -0.010434836 | 0.069246197 | 0.878641855 | 0.907088955 | ns | YPD | KS.1 |
| FIM1_2928 | 5335.262218 | 2522.932654 | 1364.35551 | -0.93607024 | 0.293849563 | 0.000316664 | 0.000766993 | ns | YPD | KS.1 |
| FIM1_2929 | 2896.317058 | 3068.433559 | 1704.738101 | 0.070849239 | 0.210529638 | 0.714843595 | 0.772447195 | ns | YPD | KS.1 |
| FIM1_2930 | 284.2094737 | 141.4817839 | 314.4429203 | -0.81856884 | 0.338814371 | 0.004463137 | 0.008783269 | ns | YPD | KS.1 |
| FIM1_2931 | 1746.329831 | 1984.48821 | 3084.728023 | 0.169681284 | 0.147374125 | 0.227354371 | 0.294482577 | ns | YPD | KS.1 |
| FIM1_2932 | 778.7039457 | 811.4412432 | 921.3139034 | 0.054427644 | 0.137760239 | 0.680546168 | 0.742970635 | ns | YPD | KS.1 |
| FIM1_2933 | 3093.974362 | 3648.486265 | 2741.375958 | 0.229929973 | 0.092515093 | 0.011059512 | 0.020014941 | ns | YPD | KS.1 |
| FIM1_2934 | 3251.216671 | 2982.862846 | 3436.724809 | -0.110264924 | 0.175614456 | 0.50424044 | 0.580693562 | ns | YPD | KS.1 |
| FIM1_2935 | 5659.196419 | 12593.52936 | 14252.74012 | 1.105795696 | 0.165628262 | 3.8257383079 | 2.4718244262 | Km>KS.1 | YPD | KS.1 |
| FIM1_2936 | 6144.713626 | 5310.220742 | 4841.660457 | -0.187467644 | 0.179054421 | 0.264031265 | 0.333983102 | ns | YPD | KS.1 |
| FIM1_2937 | 672.5573749 | 898.1439469 | 929.7319652 | 0.372839452 | 0.19246772 | 0.037198538 | 0.059486573 | ns | YPD | KS.1 |
| FIM1_2939 | 892.1891884 | 739.5035953 | 579.7654092 | -0.245835359 | 0.168104175 | 0.120880984 | 0.170992857 | ns | YPD | KS.1 |
| FIM1_2941 | 749.7422557 | 1123.989429 | 1537.092147 | 0.554826268 | 0.136249968 | 2.3465988870 | 6.7702203044 | ns | YPD | KS.1 |
| FIM1_2942 | 4833.78842 | 5541.00267 | 7572.433439 | 0.18440879 | 0.128767145 | 0.136724178 | 0.18994772 | ns | YPD | KS.1 |
| FIM1_2943 | 835.275611 | 1266.692486 | 1210.279347 | 0.582366922 | 0.106669836 | 2.2427903803 | 9.8917394875 | ns | YPD | KS.1 |
| FIM1_2944 | 786.7732719 | 691.1271863 | 950.1685742 | -0.171886879 | 0.144039527 | 0.211617583 | 0.277856288 | ns | YPD | KS.1 |
| FIM1_2945 | 590.9754747 | 489.5896888 | 787.5940644 | -0.240565206 | 0.187281925 | 0.168340864 | 0.227890013 | ns | YPD | KS.1 |
| FIM1_2946 | 2904.671253 | 2513.761094 | 1978.107633 | -0.200650035 | 0.096848468 | 0.033794695 | 0.05467597 | ns | YPD | KS.1 |
| FIM1_2947 | 1991.683654 | 2780.874493 | 3679.15989 | 0.45981426 | 0.124366831 | 0.000132923 | 0.000340521 | ns | YPD | KS.1 |
| FIM1_2948 | 1198.163114 | 1272.288508 | 1174.689626 | 0.081763939 | 0.120447636 | 0.482695871 | 0.560815497 | ns | YPD | KS.1 |
| FIM1_2949 | 2792.020733 | 3341.686985 | 3327.628688 | 0.241067979 | 0.139963286 | 0.071933922 | 0.107955133 | ns | YPD | KS.1 |
| FIM1_2950 | 208.9177323 | 168.5598633 | 91.11159101 | -0.274079271 | 0.196693997 | 0.132797289 | 0.185164382 | ns | YPD | KS.1 |
| FIM1_2951 | 2410.092919 | 2739.178796 | 2696.25655 | 0.172642745 | 0.130158988 | 0.167869696 | 0.227314027 | ns | YPD | KS.1 |
| FIM1_2952 | 1761.555974 | 953.8452654 | 991.1695776 | -0.859164018 | 0.12193494 | 4.6213617066 | 3.2473551540 | ns | YPD | KS.1 |
| FIM1_2953 | 1512.624005 | 2066.658268 | 1696.349832 | 0.408015742 | 0.18208688 | 0.016680994 | 0.029043363 | ns | YPD | KS.1 |
| FIM1_2954 | 769.05025 | 713.5334915 | 669.8722657 | -0.104333455 | 0.079227229 | 0.180287491 | 0.241825362 | ns | YPD | KS.1 |
| FIM1_2955 | 355.860008 | 687.5371798 | 1135.911101 | 0.902811583 | 0.166089804 | 1.2713412509 | 5.7447072873 | ns | YPD | KS.1 |
| FIM1_2956 | 239.1467 | 301.3644956 | 356.0336572 | 0.29961463 | 0.177604328 | 0.072014495 | 0.10804346 | ns | YPD | KS.1 |
| FIM1_2957 | 3026.005737 | 2391.523773 | 3906.709487 | -0.278925468 | 0.252225136 | 0.21584217 | 0.282434192 | ns | YPD | KS.1 |

| | | | | | | | | | | |
|---|---|---|---|---|---|---|---|---|---|---|
| FIM1_2958 | 4117.198465 | 2267.640498 | 2248.188588 | -0.833254007 | 0.125567613 | 8.57834400565 | 5.42277781806 | ns | YPD | KS.1 |
| FIM1_2959 | 2499.623571 | 2917.55089 | 4410.205535 | 0.190867993 | 0.213088178 | 0.329314314 | 0.403432335 | ns | YPD | KS.1 |
| FIM1_2960 | 3185.212189 | 2107.977079 | 3342.8738 | -0.513288036 | 0.242372313 | 0.018768465 | 0.032309036 | ns | YPD | KS.1 |
| FIM1_2961 | 3408.006298 | 2814.081689 | 4279.333789 | -0.243468768 | 0.192529445 | 0.173677525 | 0.233996221 | ns | YPD | KS.1 |
| FIM1_2962 | 1097.521308 | 491.3121436 | 401.0509813 | -1.134148505 | 0.118059213 | 1.10736680641 | 1.74893646409 | Km<KS.1 | YPD | KS.1 |
| FIM1_2963 | 382.833436 | 371.0533819 | 393.8680425 | -0.042779103 | 0.101990478 | 0.667063524 | 0.729692399 | ns | YPD | KS.1 |
| FIM1_2964 | 3021.726301 | 4957.952302 | 7234.558437 | 0.684003354 | 0.136407635 | 1.99842751459 | 7.78557312852 | ns | YPD | KS.1 |
| FIM1_2965 | 1004.70671 | 1152.543046 | 1402.398664 | 0.186828985 | 0.119311081 | 0.105229153 | 0.151086591 | ns | YPD | KS.1 |
| FIM1_2966 | 115.9423662 | 354.3630468 | 368.3976288 | 1.549151907 | 0.17967673 | 4.41171554210 | 5.34021528514 | Km>KS.1 | YPD | KS.1 |
| FIM1_2967 | 1365.497914 | 1617.004996 | 1546.006294 | 0.237043462 | 0.083829607 | 0.003970338 | 0.007882056 | ns | YPD | KS.1 |
| FIM1_2968 | 1451.467552 | 2170.180913 | 1890.686952 | 0.565723189 | 0.096942255 | 2.60363142623 | 1.28502642316 | ns | YPD | KS.1 |
| FIM1_2969 | 1979.88963 | 3806.138917 | 5685.816099 | 0.904067379 | 0.150074877 | 3.93165772661 | 2.11003205931 | ns | YPD | KS.1 |
| FIM1_2970 | 1058.342334 | 1025.689751 | 760.6778271 | -0.043333548 | 0.102731184 | 0.665198855 | 0.728453512 | ns | YPD | KS.1 |
| FIM1_2971 | 284.0280018 | 267.9397196 | 637.3109641 | -0.069827017 | 0.230407813 | 0.73887671 | 0.792563957 | ns | YPD | KS.1 |
| FIM1_2972 | 757.4445729 | 1235.24484 | 731.7172261 | 0.670125368 | 0.147070439 | 2.02659600455 | 6.79859414879 | ns | YPD | KS.1 |
| FIM1_2973 | 4638.541503 | 1363.824684 | 901.5278038 | -1.69346238 | 0.198078369 | 6.03596157854 | 7.20117718303 | Km<KS.1 | YPD | KS.1 |
| FIM1_2974 | 2535.335551 | 2679.604191 | 1991.706768 | 0.06983966 | 0.188451847 | 0.69142576 | 0.752371644 | ns | YPD | KS.1 |
| FIM1_2975 | 1796.018454 | 1834.594747 | 2788.527927 | 0.027479263 | 0.169954422 | 0.863993925 | 0.895300863 | ns | YPD | KS.1 |
| FIM1_2976 | 570.6332057 | 571.2737722 | 629.7327363 | 0.000142398 | 0.121595998 | 0.999033857 | 0.999234708 | ns | YPD | KS.1 |
| FIM1_2977 | 1004.728435 | 708.9235851 | 839.6745868 | -0.471207448 | 0.151076117 | 0.001081847 | 0.002379394 | ns | YPD | KS.1 |
| FIM1_2978 | 1359.08527 | 2141.487127 | 2827.370299 | 0.620771005 | 0.148947163 | 1.34265158928 | 3.99742169759 | ns | YPD | KS.1 |
| FIM1_2979 | 218.841777 | 239.4857312 | 219.4068092 | 0.116029294 | 0.16357503 | 0.453529301 | 0.53109968 | ns | YPD | KS.1 |
| FIM1_2980 | 927.8752922 | 1076.905858 | 926.1346837 | 0.197469822 | 0.147235683 | 0.159598345 | 0.217237145 | ns | YPD | KS.1 |
| FIM1_2981 | 3721.944146 | 2981.985018 | 5131.577162 | -0.267849704 | 0.236567581 | 0.210071624 | 0.27611792 | ns | YPD | KS.1 |
| FIM1_2982 | 3422.959011 | 4102.152072 | 3129.856232 | 0.249737574 | 0.109326369 | 0.018691559 | 0.032198929 | ns | YPD | KS.1 |
| FIM1_2983 | 1709.414842 | 3442.764285 | 6118.843466 | 0.953167969 | 0.183120553 | 4.07982820885 | 1.74075002907 | ns | YPD | KS.1 |
| FIM1_2984 | 268.1684493 | 171.0669161 | 607.7647513 | -0.535054668 | 0.282297534 | 0.030485995 | 0.049724169 | ns | YPD | KS.1 |
| FIM1_2985 | 8243.44626 | 13404.06524 | 15548.43375 | 0.56216087 | 0.310302016 | 0.03478448 | 0.056113096 | ns | YPD | KS.1 |
| FIM1_2986 | 712.1863515 | 1053.265752 | 930.6546089 | 0.551713591 | 0.090977999 | 6.63424080933 | 3.48525322348 | ns | YPD | KS.1 |
| FIM1_2987 | 1647.564316 | 2105.487875 | 1963.239033 | 0.343909867 | 0.088061532 | 6.96502042046 | 0.000186697 | ns | YPD | KS.1 |
| FIM1_2988 | 1500.26486 | 1460.075028 | 931.4687696 | -0.036455393 | 0.132333531 | 0.774655183 | 0.823484944 | ns | YPD | KS.1 |
| FIM1_2989 | 379.5584288 | 545.6537928 | 992.0718388 | 0.475632941 | 0.186526664 | 0.006361436 | 0.012056437 | ns | YPD | KS.1 |
| FIM1_2990 | 35851.52492 | 12619.61083 | 12583.20332 | -1.494119721 | 0.07995943 | 4.58674858661 | 2.28190742184 | Km<KS.1 | YPD | KS.1 |
| FIM1_2991 | 390.2069514 | 150.9930551 | 416.6918034 | -1.285391883 | 0.222059343 | 7.83120069493 | 4.08816615501 | Km<KS.1 | YPD | KS.1 |
| FIM1_2992 | 139.5709248 | 66.09077297 | 85.80505748 | -0.97804771 | 0.244440159 | 1.26373974373 | 3.78285512942 | ns | YPD | KS.1 |
| FIM1_2993 | 924.1067902 | 1062.438674 | 1039.204649 | 0.196143531 | 0.081116393 | 0.013816055 | 0.024469518 | ns | YPD | KS.1 |
| FIM1_2994 | 1121.367646 | 1021.38283 | 988.7851955 | -0.125729986 | 0.122948969 | 0.28948834 | 0.361225105 | ns | YPD | KS.1 |
| FIM1_2995 | 303.5363114 | 401.149368 | 250.7650778 | 0.335820367 | 0.248010941 | 0.130711169 | 0.182665187 | ns | YPD | KS.1 |
| FIM1_2996 | 2909.87273 | 890.8464776 | 702.7419252 | -1.474721482 | 0.362432543 | 3.26216051698 | 1.06073520078 | Km<KS.1 | YPD | KS.1 |
| FIM1_2997 | 2173.818205 | 2255.21245 | 1624.884602 | 0.050311309 | 0.109040477 | 0.635058687 | 0.701780757 | ns | YPD | KS.1 |
| FIM1_2998 | 2966.554171 | 3054.538093 | 3567.38224 | 0.039277251 | 0.119257687 | 0.733673663 | 0.788172419 | ns | YPD | KS.1 |
| FIM1_2999 | 801.8827159 | 981.6495522 | 807.5264055 | 0.269412463 | 0.148452863 | 0.056881181 | 0.087992498 | ns | YPD | KS.1 |
| FIM1_3000 | 3761.16827 | 1561.155787 | 1759.153242 | -1.227134059 | 0.152072227 | 8.83428874927 | 8.80773277107 | Km<KS.1 | YPD | KS.1 |
| FIM1_3001 | 742.2551352 | 1018.636818 | 1124.216239 | 0.410897314 | 0.189899736 | 0.02020336 | 0.034433613 | ns | YPD | KS.1 |
| FIM1_3002 | 354.5305273 | 346.2079727 | 406.515444 | -0.031952822 | 0.132146602 | 0.801615452 | 0.844744095 | ns | YPD | KS.1 |

| | | | | | | | | | | |
|---|---|---|---|---|---|---|---|---|---|---|
| FIM1_3003 | 602.4174206 | 602.7205202 | 974.3824504 | 0.000946565 | 0.242970523 | 0.99655225 | 0.997153549 | ns | YPD | KS.1 |
| FIM1_3004 | 4733.209848 | 5671.55839 | 5703.476767 | 0.252573426 | 0.091670931 | 0.004881484 | 0.009520628 | ns | YPD | KS.1 |
| FIM1_3005 | 641.5387464 | 695.1292618 | 2134.188148 | 0.08932212 | 0.28110982 | 0.716899966 | 0.774208869 | ns | YPD | KS.1 |
| FIM1_3006 | 3407.195164 | 4597.330498 | 4840.496114 | 0.405212395 | 0.145276776 | 0.003541167 | 0.007103752 | ns | YPD | KS.1 |
| FIM1_3007 | 820.09939 | 2907.60024 | 3697.877283 | 1.796831261 | 0.12238076 | 3.06015664744 | 2.87250553226 | Km>KS.1 | YPD | KS.1 |
| FIM1_3008 | 1934.409823 | 2075.055468 | 2183.93518 | 0.098063714 | 0.08275034 | 0.227358307 | 0.294482577 | ns | YPD | KS.1 |
| FIM1_3009 | 1392.774144 | 1724.492229 | 1567.872644 | 0.299923878 | 0.089749416 | 0.000653552 | 0.001502505 | ns | YPD | KS.1 |
| FIM1_3010 | 586.2617213 | 504.6271864 | 653.7230729 | -0.205381196 | 0.115990848 | 0.067390428 | 0.102060085 | ns | YPD | KS.1 |
| FIM1_3011 | 1504.447048 | 777.6857583 | 622.8877379 | -0.906731709 | 0.163916668 | 7.32307720327 | 3.43377088466 | ns | YPD | KS.1 |
| FIM1_3012 | 1320.60215 | 1857.351866 | 2505.524487 | 0.477260868 | 0.101036478 | 1.36570911230 | 4.71179114682 | ns | YPD | KS.1 |
| FIM1_3013 | 2410.124577 | 2656.71592 | 2299.118373 | 0.135189105 | 0.094517789 | 0.143199864 | 0.197564981 | ns | YPD | KS.1 |
| FIM1_3014 | 4757.186795 | 3500.558737 | 6063.64154 | -0.357149059 | 0.274917894 | 0.139310442 | 0.192894364 | ns | YPD | KS.1 |
| FIM1_3015 | 2013.948253 | 2672.6441 | 4634.54022 | 0.389571808 | 0.120369426 | 0.000833707 | 0.001870857 | ns | YPD | KS.1 |
| FIM1_3016 | 1299.546893 | 1096.868545 | 759.6981876 | -0.22598134 | 0.144319782 | 0.101070225 | 0.145830734 | ns | YPD | KS.1 |
| FIM1_3017 | 12752.69224 | 16773.70496 | 21330.23418 | 0.377916562 | 0.118350998 | 0.000987579 | 0.002190461 | ns | YPD | KS.1 |
| FIM1_3018 | 173.1399353 | 272.7704215 | 535.4983651 | 0.605536686 | 0.183542191 | 0.000439911 | 0.001035758 | ns | YPD | KS.1 |
| FIM1_3019 | 3167.260944 | 4422.759083 | 4633.715476 | 0.462065951 | 0.1180114 | 5.48175414871 | 0.000150423 | ns | YPD | KS.1 |
| FIM1_3020 | 144.3362538 | 163.1582375 | 96.90294568 | 0.154351324 | 0.19523828 | 0.394725078 | 0.471830193 | ns | YPD | KS.1 |
| FIM1_3021 | 5070.957726 | 2585.566906 | 3834.209133 | -0.950721555 | 0.1091535 | 6.30167811505 | 7.50020301971 | ns | YPD | KS.1 |
| FIM1_3022 | 320.1725974 | 168.6261715 | 240.3611277 | -0.752896653 | 0.328246668 | 0.007191872 | 0.013506818 | ns | YPD | KS.1 |
| FIM1_3023 | 1617.26665 | 1001.67759 | 752.7178282 | -0.607748113 | 0.235104502 | 0.004407909 | 0.008684891 | ns | YPD | KS.1 |
| FIM1_3024 | 253.9177719 | 49.61633659 | 74.4569741 | -1.939223723 | 0.486451592 | 1.44125041706 | 4.96209053627 | Km<KS.1 | YPD | KS.1 |
| FIM1_3025 | 864.77734 | 1357.103319 | 931.7120277 | 0.57895774 | 0.217986637 | 0.003802179 | 0.007584539 | ns | YPD | KS.1 |
| FIM1_3026 | 2237.713559 | 2782.569034 | 2497.866158 | 0.282360517 | 0.179218278 | 0.092604843 | 0.13498655 | ns | YPD | KS.1 |
| FIM1_3028 | 3585.85615 | 5889.303416 | 4185.317805 | 0.687916899 | 0.130475374 | 4.99866332409 | 2.10570279740 | ns | YPD | KS.1 |
| FIM1_3029 | 863.8783162 | 1674.891364 | 1373.966389 | 0.927344641 | 0.124682937 | 2.24015131785 | 1.75507918209 | ns | YPD | KS.1 |
| FIM1_3030 | 3127.754099 | 2679.745078 | 3644.827954 | -0.217856994 | 0.07318794 | 0.002517849 | 0.005193325 | ns | YPD | KS.1 |
| FIM1_3031 | 2550.559867 | 1354.26194 | 1581.656361 | -0.88483628 | 0.128271978 | 1.25768221579 | 8.60655986738 | ns | YPD | KS.1 |
| FIM1_3032 | 2964.87276 | 2268.491513 | 2951.006234 | -0.35438336 | 0.164868102 | 0.022952201 | 0.038550709 | ns | YPD | KS.1 |
| FIM1_3033 | 1274.828295 | 1146.671623 | 1273.167036 | -0.143534271 | 0.125881453 | 0.236857381 | 0.304723421 | ns | YPD | KS.1 |
| FIM1_3034 | 6403.201835 | 4328.381408 | 2931.69041 | -0.454168296 | 0.291450139 | 0.072808991 | 0.109169599 | ns | YPD | KS.1 |
| FIM1_3035 | 4307.637222 | 8369.893539 | 11142.20316 | 0.910575168 | 0.167200962 | 1.18078515000 | 5.37457101673 | ns | YPD | KS.1 |
| FIM1_3036 | 4782.0918 | 4679.08057 | 3404.866561 | -0.028971043 | 0.138716373 | 0.827663447 | 0.866686097 | ns | YPD | KS.1 |
| FIM1_3037 | 249.0905384 | 684.0733955 | 1021.631874 | 1.237844347 | 0.3536558 | 5.25746356656 | 0.000144667 | Km>KS.1 | YPD | KS.1 |
| FIM1_3038 | 7734.877731 | 10401.42504 | 17148.25544 | 0.401685626 | 0.141679392 | 0.003088641 | 0.006266717 | ns | YPD | KS.1 |
| FIM1_3039 | 995.6320734 | 1910.784122 | 2869.090948 | 0.908797768 | 0.134808543 | 3.57306538087 | 2.31157350713 | ns | YPD | KS.1 |
| FIM1_3040 | 1927.375914 | 979.6639662 | 2794.51789 | -0.80016614 | 0.329703056 | 0.004516615 | 0.008874471 | ns | YPD | KS.1 |
| FIM1_3041 | 2476.099265 | 1441.011893 | 1740.792045 | -0.763335234 | 0.100483815 | 9.29457266685 | 7.48228139443 | ns | YPD | KS.1 |
| FIM1_3043 | 429.6581745 | 224.1898038 | 207.6025605 | -0.868552923 | 0.206610033 | 6.64001572484 | 2.07110208345 | ns | YPD | KS.1 |
| FIM1_3044 | 33526.26881 | 4993.571127 | 4677.982151 | -2.478492099 | 0.399302849 | 2.92669337274 | 1.91835303417 | Km<KS.1 | YPD | KS.1 |
| FIM1_3045 | 649.6413873 | 793.0541032 | 549.2223036 | 0.253635244 | 0.193998201 | 0.159159043 | 0.216698478 | ns | YPD | KS.1 |
| FIM1_3046 | 2350.279844 | 2706.05644 | 1642.342519 | 0.18372887 | 0.165193044 | 0.239080167 | 0.307185907 | ns | YPD | KS.1 |
| FIM1_3047 | 911.1452932 | 658.9816888 | 825.3510037 | -0.452686875 | 0.099712026 | 3.47938487108 | 1.12694920140 | ns | YPD | KS.1 |
| FIM1_3048 | 826.4546554 | 1526.221606 | 3099.753428 | 0.831479479 | 0.178458868 | 8.65674262634 | 3.09169379512 | ns | YPD | KS.1 |
| FIM1_3049 | 648.2280395 | 726.2930981 | 883.089918 | 0.153385889 | 0.123900735 | 0.199412945 | 0.263991325 | ns | YPD | KS.1 |

| | | | | | | | | | | |
|---|---|---|---|---|---|---|---|---|---|---|
| FIM1_3050 | 390.2229046 | 589.0043313 | 792.229839 | 0.572650672 | 0.112628131 | 1.7754781950 | 6.9770963824 | ns | YPD | KS.1 |
| FIM1_3051 | 944.9764529 | 1039.023846 | 1014.141862 | 0.127429166 | 0.135259022 | 0.326362114 | 0.400407279 | ns | YPD | KS.1 |
| FIM1_3052 | 580.6159337 | 867.8103404 | 796.9924135 | 0.548011615 | 0.142485397 | 6.1380777628 | 0.000167143 | ns | YPD | KS.1 |
| FIM1_3053 | 1151.142379 | 1289.683432 | 2143.005932 | 0.149946957 | 0.146665152 | 0.284012488 | 0.35564111 | ns | YPD | KS.1 |
| FIM1_3054 | 4829.744084 | 1685.205207 | 1456.614948 | -1.352879222 | 0.306985671 | 9.6696101141 | 3.4190696743 | Km<KS.1 | YPD | KS.1 |
| FIM1_3056 | 4520.370187 | 5109.273576 | 4038.32144 | 0.146532479 | 0.233604096 | 0.488925753 | 0.566202426 | ns | YPD | KS.1 |
| FIM1_3057 | 485.2123693 | 259.0504267 | 483.5889983 | -0.797019309 | 0.259953037 | 0.000647782 | 0.001490617 | ns | YPD | KS.1 |
| FIM1_3058 | 3874.316811 | 3420.563094 | 2914.185425 | -0.167278534 | 0.134084524 | 0.193776184 | 0.257557178 | ns | YPD | KS.1 |
| FIM1_3059 | 1316.529543 | 911.1744781 | 849.149499 | -0.514262753 | 0.106112286 | 6.8727502258 | 2.4994102612 | ns | YPD | KS.1 |
| FIM1_3060 | 962.9358253 | 613.6142867 | 717.4541639 | -0.625625056 | 0.125566309 | 2.6900535686 | 1.0342362058 | ns | YPD | KS.1 |
| FIM1_3061 | 4077.422448 | 3846.958313 | 3894.558609 | -0.079948934 | 0.104412311 | 0.431637901 | 0.509387188 | ns | YPD | KS.1 |
| FIM1_3062 | 1147.086342 | 1021.422021 | 1637.020043 | -0.147425165 | 0.184825933 | 0.393482791 | 0.470571366 | ns | YPD | KS.1 |
| FIM1_3063 | 1675.889282 | 1185.306625 | 1105.968508 | -0.487676211 | 0.087744375 | 1.5717277069 | 7.0381146195 | ns | YPD | KS.1 |
| FIM1_3064 | 6290.619756 | 5945.429525 | 6172.113327 | -0.077564604 | 0.103181321 | 0.440332315 | 0.51739567 | ns | YPD | KS.1 |
| FIM1_3065 | 1099.230199 | 2283.312715 | 2380.210882 | 1.034624406 | 0.105840821 | 2.5380017918 | 4.2801894624 | Km>KS.1 | YPD | KS.1 |
| FIM1_3066 | 2700.209616 | 3471.092703 | 4964.727518 | 0.348429633 | 0.105991517 | 0.00074277 | 0.001686573 | ns | YPD | KS.1 |
| FIM1_3067 | 542.0764179 | 565.037994 | 690.525503 | 0.054540869 | 0.130719243 | 0.664762528 | 0.728135971 | ns | YPD | KS.1 |
| FIM1_3068 | 7071.666337 | 6596.578975 | 5347.153281 | -0.086982083 | 0.195334341 | 0.63195042 | 0.699038102 | ns | YPD | KS.1 |
| FIM1_3069 | 316.7966061 | 712.842664 | 1141.648876 | 1.109431884 | 0.186861585 | 4.4896861820 | 2.4043260232 | Km>KS.1 | YPD | KS.1 |
| FIM1_3070 | 4335.383681 | 5794.133853 | 7452.029009 | 0.40287774 | 0.108729047 | 0.00014248 | 0.000364442 | ns | YPD | KS.1 |
| FIM1_3071 | 2501.536992 | 2657.51774 | 3315.501244 | 0.08081571 | 0.131232903 | 0.522211695 | 0.598204739 | ns | YPD | KS.1 |
| FIM1_3072 | 1389.834171 | 1655.159903 | 1546.247319 | 0.22518718 | 0.178900335 | 0.179221742 | 0.240460671 | ns | YPD | KS.1 |
| FIM1_3074 | 821.4621778 | 833.9002187 | 871.4180666 | 0.019824268 | 0.111745103 | 0.855066495 | 0.887905618 | ns | YPD | KS.1 |
| FIM1_3075 | 2108.088408 | 863.6901188 | 1321.472421 | -1.236900977 | 0.167848643 | 2.1063712319 | 1.6554813395 | Km<KS.1 | YPD | KS.1 |
| FIM1_3076 | 1322.513701 | 1250.260488 | 1017.158479 | -0.076934802 | 0.093474791 | 0.400011541 | 0.476774656 | ns | YPD | KS.1 |
| FIM1_3077 | 1905.011476 | 2403.932409 | 2215.349619 | 0.31717059 | 0.129418997 | 0.01093718 | 0.019815175 | ns | YPD | KS.1 |
| FIM1_3078 | 599.153461 | 680.2487666 | 790.6237241 | 0.175905515 | 0.101900951 | 0.076329238 | 0.113693999 | ns | YPD | KS.1 |
| FIM1_3079 | 1380.91879 | 1167.087161 | 2045.88673 | -0.222734607 | 0.153398891 | 0.126593576 | 0.177595049 | ns | YPD | KS.1 |
| FIM1_3080 | 2249.642974 | 1473.272898 | 1517.297272 | -0.592018349 | 0.109001254 | 2.5728550515 | 1.1277492406 | ns | YPD | KS.1 |
| FIM1_3081 | 4070.090452 | 2403.841128 | 1776.031229 | -0.720676511 | 0.154349828 | 1.0515480426 | 3.7023719123 | ns | YPD | KS.1 |
| FIM1_3082 | 2734.647144 | 2194.492827 | 1874.539366 | -0.307828954 | 0.091048769 | 0.000561436 | 0.001303379 | ns | YPD | KS.1 |
| FIM1_3083 | 3856.593666 | 2181.300354 | 5074.020734 | -0.690589862 | 0.290503078 | 0.006609039 | 0.012492389 | ns | YPD | KS.1 |
| FIM1_3085 | 116.4097665 | 35.9622264 | 43.4414597 | -1.553818291 | 0.280993106 | 2.0257229689 | 1.0118445552 | Km<KS.1 | YPD | KS.1 |
| FIM1_3086 | 4690.608799 | 1418.893893 | 1403.542008 | -1.649874751 | 0.201727746 | 1.5432546857 | 1.6475734037 | Km<KS.1 | YPD | KS.1 |
| FIM1_3087 | 3487.505474 | 4146.937442 | 7192.444108 | 0.217282406 | 0.201894667 | 0.24433808 | 0.312817155 | ns | YPD | KS.1 |
| FIM1_3088 | 4878.294494 | 432.2878809 | 879.435673 | -3.398973369 | 0.25971915 | 4.1814471873 | 2.3373819951 | Km<KS.1 | YPD | KS.1 |
| FIM1_3089 | 769.0286736 | 577.9311752 | 526.3408947 | -0.369119235 | 0.192262519 | 0.038942278 | 0.061936647 | ns | YPD | KS.1 |
| FIM1_3090 | 645.1248893 | 363.7765568 | 396.6058563 | -0.796261871 | 0.135369908 | 1.1747644082 | 6.0439016866 | ns | YPD | KS.1 |
| FIM1_3092 | 1508.434448 | 499.2549956 | 500.4249974 | -1.53449158 | 0.181730289 | 2.1209342885 | 2.4090520743 | Km<KS.1 | YPD | KS.1 |
| FIM1_3098 | 594.0314324 | 1088.553291 | 1200.264754 | 0.805080533 | 0.202877084 | 2.1068515493 | 6.1081506164 | ns | YPD | KS.1 |
| FIM1_3099 | 3319.845257 | 2031.554101 | 2350.021231 | -0.577732673 | 0.299168651 | 0.02574942 | 0.04278875 | ns | YPD | KS.1 |
| FIM1_3100 | 1554.269594 | 771.3493943 | 982.5762384 | -0.970010614 | 0.153008572 | 4.6629177117 | 2.7388448189 | ns | YPD | KS.1 |
| FIM1_3101 | 1247.643979 | 874.5382025 | 1052.947167 | -0.482455728 | 0.146474236 | 0.000577016 | 0.001337055 | ns | YPD | KS.1 |
| FIM1_3102 | 5721.886306 | 8754.849601 | 16236.58602 | 0.582100673 | 0.143353942 | 2.3260062893 | 6.7200239775 | ns | YPD | KS.1 |
| FIM1_3103 | 200.8984391 | 127.3115004 | 162.3837864 | -0.517070512 | 0.321183172 | 0.058220469 | 0.089868704 | ns | YPD | KS.1 |

| | | | | | | | | | | |
|---|---|---|---|---|---|---|---|---|---|---|
| FIM1_3104 | 2323.947895 | 2222.829016 | 1971.434572 | -0.05979027 | 0.122337994 | 0.613077113 | 0.682517176 | ns | YPD | KS.1 |
| FIM1_3105 | 3.022398144 | 8.561974445 | 14.7225111 | 1.013824144 | 0.504933761 | 0.007713454 | 0.014383221 | Km>KS.1 | YPD | KS.1 |
| FIM1_3106 | 706.0585602 | 1612.627881 | 1352.376741 | 1.164636859 | 0.118513357 | 1.2029842851 | 2.0853124803 | Km>KS.1 | YPD | KS.1 |
| FIM1_3107 | 345.087024 | 458.1981111 | 1220.887682 | 0.279847094 | 0.36947361 | 0.351958466 | 0.427175742 | ns | YPD | KS.1 |
| FIM1_3108 | 2395.092966 | 2301.650955 | 2240.122344 | -0.052631823 | 0.140732299 | 0.696328184 | 0.756547874 | ns | YPD | KS.1 |
| FIM1_3109 | 1219.615859 | 1726.870687 | 3375.384685 | 0.446993313 | 0.203762607 | 0.017576891 | 0.030415664 | ns | YPD | KS.1 |
| FIM1_3110 | 21097.81227 | 5208.819603 | 4126.047925 | -1.596886241 | 0.487649489 | 4.9522343638 | 0.000136647 | Km<KS.1 | YPD | KS.1 |
| FIM1_3111 | 5729.659288 | 1989.620525 | 1294.786463 | -1.350791691 | 0.315340565 | 1.6935289091 | 5.7628634220 | Km<KS.1 | YPD | KS.1 |
| FIM1_3112 | 673.8963822 | 523.2923483 | 703.4952814 | -0.324036276 | 0.191822198 | 0.069246131 | 0.104362163 | ns | YPD | KS.1 |
| FIM1_3113 | 3448.789636 | 3470.7287 | 1621.420472 | 0.008158952 | 0.174481627 | 0.960391822 | 0.970536119 | ns | YPD | KS.1 |
| FIM1_3114 | 1795.918562 | 1312.017004 | 1719.038682 | -0.416108455 | 0.169302647 | 0.009189293 | 0.016907075 | ns | YPD | KS.1 |
| FIM1_3115 | 171.9036836 | 277.2310175 | 260.7413305 | 0.641108654 | 0.16997475 | 6.7868152071 | 0.000182705 | ns | YPD | KS.1 |
| FIM1_3116 | 2987.632993 | 2276.183516 | 1724.004441 | -0.343115178 | 0.209727083 | 0.075165435 | 0.11216198 | ns | YPD | KS.1 |
| FIM1_3117 | 18.77524307 | 10.84940698 | 19.37440892 | -0.667404398 | 0.30308754 | 0.011023298 | 0.019963926 | ns | YPD | KS.1 |
| FIM1_3118 | 3338.647052 | 5205.469536 | 4526.624002 | 0.587937037 | 0.187118737 | 0.000790578 | 0.001782115 | ns | YPD | KS.1 |
| FIM1_3119 | 428.1064047 | 961.7934929 | 946.5580616 | 1.099346055 | 0.196266188 | 3.3537686582 | 1.6325830797 | Km>KS.1 | YPD | KS.1 |
| FIM1_3120 | 2251.35753 | 2848.575912 | 2859.777166 | 0.328352321 | 0.097160874 | 0.000548993 | 0.001276876 | ns | YPD | KS.1 |
| FIM1_3121 | 1377.75638 | 1405.850056 | 1170.818666 | 0.027549156 | 0.103794749 | 0.785326184 | 0.830745857 | ns | YPD | KS.1 |
| FIM1_3122 | 552.2246342 | 1759.101051 | 1970.926694 | 1.619647234 | 0.164630458 | 4.4426487214 | 8.1258005108 | Km>KS.1 | YPD | KS.1 |
| FIM1_3123 | 2771.265729 | 2769.11973 | 2384.973826 | -0.000924655 | 0.107293585 | 0.992933897 | 0.994332958 | ns | YPD | KS.1 |
| FIM1_3124 | 3314.272208 | 4861.669815 | 4368.791076 | 0.547535034 | 0.054890085 | 9.6233692932 | 1.6917407149 | ns | YPD | KS.1 |
| FIM1_3125 | 1243.520213 | 803.9106907 | 648.7975289 | -0.601671334 | 0.131317749 | 2.0909345955 | 7.0002689186 | ns | YPD | KS.1 |
| FIM1_3126 | 2664.376149 | 3339.633735 | 2225.153781 | 0.288309084 | 0.193668403 | 0.108939371 | 0.155829031 | ns | YPD | KS.1 |
| FIM1_3127 | 2335.356463 | 2012.494997 | 1641.556946 | -0.188273603 | 0.191969658 | 0.291791937 | 0.363643509 | ns | YPD | KS.1 |
| FIM1_3128 | 4439.764292 | 8138.579139 | 9997.73968 | 0.859969274 | 0.089735585 | 2.3079927532 | 3.5994557828 | ns | YPD | KS.1 |
| FIM1_3129 | 2319.929497 | 3380.125402 | 3927.025455 | 0.530700451 | 0.090583238 | 2.4482820243 | 1.2107557724 | ns | YPD | KS.1 |
| FIM1_3130 | 451.326046 | 724.6499644 | 1143.616367 | 0.628773378 | 0.187521102 | 0.000344757 | 0.000829785 | ns | YPD | KS.1 |
| FIM1_3131 | 901.980905 | 977.158752 | 1039.568712 | 0.10637641 | 0.14206531 | 0.433965362 | 0.511363731 | ns | YPD | KS.1 |
| FIM1_3132 | 755.002779 | 2484.38832 | 3367.068562 | 1.700913175 | 0.09324722 | 1.0967067492 | 4.1970123672 | Km>KS.1 | YPD | KS.1 |
| FIM1_3133 | 1149.99612 | 729.3705869 | 594.2050343 | -0.611121559 | 0.171711484 | 0.00016664 | 0.000422116 | ns | YPD | KS.1 |
| FIM1_3134 | 14107.98924 | 12869.25698 | 11415.25866 | -0.115319506 | 0.194361662 | 0.523438915 | 0.599472514 | ns | YPD | KS.1 |
| FIM1_3135 | 689.2961845 | 899.9590429 | 1117.0395 | 0.368846642 | 0.112910571 | 0.000774734 | 0.00175116 | ns | YPD | KS.1 |
| FIM1_3136 | 3606.377428 | 2909.112954 | 4630.98272 | -0.277523595 | 0.18169495 | 0.102499147 | 0.147678325 | ns | YPD | KS.1 |
| FIM1_3137 | 39.83985779 | 26.21453289 | 24.44135902 | -0.427880203 | 0.379738768 | 0.163295606 | 0.221602738 | ns | YPD | KS.1 |
| FIM1_3138 | 4303.940171 | 3626.321176 | 6167.117837 | -0.200824813 | 0.252569744 | 0.373771852 | 0.449483917 | ns | YPD | KS.1 |
| FIM1_3139 | 297.5980159 | 430.677284 | 667.88201 | 0.483544832 | 0.193453325 | 0.007232638 | 0.013557789 | ns | YPD | KS.1 |
| FIM1_3140 | 361.2495505 | 447.68779 | 459.0135077 | 0.288638527 | 0.13796968 | 0.029058213 | 0.047648191 | ns | YPD | KS.1 |
| FIM1_3141 | 341.7956338 | 598.1919985 | 972.2596684 | 0.772405643 | 0.145571818 | 3.4464289299 | 1.4886033737 | ns | YPD | KS.1 |
| FIM1_3142 | 608.2355925 | 425.8556594 | 264.7518978 | -0.458550083 | 0.203927335 | 0.014947273 | 0.026304451 | ns | YPD | KS.1 |
| FIM1_3143 | 3695.836119 | 4086.218302 | 5562.129101 | 0.139763292 | 0.086042712 | 0.097476042 | 0.141136003 | ns | YPD | KS.1 |
| FIM1_3144 | 251.2977973 | 226.4231547 | 358.3171778 | -0.133140512 | 0.176903671 | 0.422936446 | 0.500425853 | ns | YPD | KS.1 |
| FIM1_3145 | 880.3023814 | 1257.161642 | 1300.969336 | 0.492762889 | 0.123247856 | 3.6541286003 | 0.000102708 | ns | YPD | KS.1 |
| FIM1_3146 | 1069.579485 | 642.1225027 | 695.8146255 | -0.713564528 | 0.113248678 | 1.0244898749 | 5.7918603725 | ns | YPD | KS.1 |
| FIM1_3147 | 2411.015247 | 1592.324177 | 1134.359183 | -0.56960574 | 0.137104182 | 1.5901118783 | 4.6892748042 | ns | YPD | KS.1 |
| FIM1_3148 | 980.5899564 | 766.595052 | 864.5807955 | -0.339870461 | 0.11327176 | 0.002003229 | 0.004221121 | ns | YPD | KS.1 |

| | | | | | | | | | | |
|---|---|---|---|---|---|---|---|---|---|---|
| FIM1_3149 | 868.6675637 | 950.2194395 | 1048.570728 | 0.119814309 | 0.134648439 | 0.354111855 | 0.42928255 | ns | YPD | KS.1 |
| FIM1_3150 | 3116.714471 | 2551.878347 | 5060.087817 | -0.233461701 | 0.258450197 | 0.310074803 | 0.383307985 | ns | YPD | KS.1 |
| FIM1_3151 | 311.8056057 | 639.9279673 | 787.651183 | 0.991644051 | 0.159773816 | 1.05156303499 | 5.93816810339 | ns | YPD | KS.1 |
| FIM1_3152 | 314.9382622 | 217.2856209 | 255.1933084 | -0.503634196 | 0.148333205 | 0.000384035 | 0.000914697 | ns | YPD | KS.1 |
| FIM1_3153 | 773.1941504 | 703.1697308 | 769.2644039 | -0.129908294 | 0.114637235 | 0.242186056 | 0.310454942 | ns | YPD | KS.1 |
| FIM1_3154 | 651.2934262 | 1587.059278 | 2146.446937 | 1.213815541 | 0.201712027 | 2.24817716159 | 1.23043799548 | Km>KS.1 | YPD | KS.1 |
| FIM1_3155 | 7718.158622 | 3390.755549 | 2866.78671 | -1.15217605 | 0.139035253 | 1.66646657210 | 1.77151093936 | Km<KS.1 | YPD | KS.1 |
| FIM1_3156 | 3488.578973 | 2850.350645 | 3973.487865 | -0.275387661 | 0.126517155 | 0.023928584 | 0.040068901 | ns | YPD | KS.1 |
| FIM1_3157 | 2009.459704 | 904.9543648 | 1292.04226 | -1.028461331 | 0.268230954 | 2.26683506238 | 6.55287881195 | Km<KS.1 | YPD | KS.1 |
| FIM1_3158 | 559.1916521 | 413.9380244 | 368.4361109 | -0.409342409 | 0.139625942 | 0.002244824 | 0.004672804 | ns | YPD | KS.1 |
| FIM1_3159 | 288.3649611 | 280.4797405 | 404.0752316 | -0.039393927 | 0.141123934 | 0.770797538 | 0.820085062 | ns | YPD | KS.1 |
| FIM1_3160 | 400.2125094 | 504.984144 | 291.6823713 | 0.260645342 | 0.289446149 | 0.300339013 | 0.372615109 | ns | YPD | KS.1 |
| FIM1_3161 | 4971.766173 | 1830.251807 | 1401.267782 | -1.289597968 | 0.294560437 | 1.26095567322 | 4.37465444512 | Km<KS.1 | YPD | KS.1 |
| FIM1_3162 | 2222.888413 | 1973.300982 | 2181.720415 | -0.136193486 | 0.26565659 | 0.562923596 | 0.63547649 | ns | YPD | KS.1 |
| FIM1_3163 | 190.2699888 | 369.4597483 | 715.8975727 | 0.885886674 | 0.203648879 | 3.31380936140 | 1.07612281808 | ns | YPD | KS.1 |
| FIM1_3164 | 3094.88331 | 3444.938496 | 2822.476065 | 0.152722416 | 0.052066857 | 0.003090977 | 0.0062689 | ns | YPD | KS.1 |
| FIM1_3165 | 274.1323224 | 561.1967905 | 793.2031306 | 0.990824419 | 0.157300153 | 5.82585473597 | 3.36236975771 | ns | YPD | KS.1 |
| FIM1_3166 | 705.8588038 | 1126.691609 | 1168.677337 | 0.642539108 | 0.141665037 | 2.38027733858 | 7.89458650632 | ns | YPD | KS.1 |
| FIM1_3167 | 3283.373129 | 1924.50378 | 3498.114776 | -0.626932074 | 0.308669361 | 0.018355577 | 0.03167499 | ns | YPD | KS.1 |
| FIM1_3168 | 946.8635504 | 1115.072661 | 1110.963659 | 0.223818829 | 0.109574125 | 0.03540752 | 0.056970378 | ns | YPD | KS.1 |
| FIM1_3169 | 877.5439258 | 905.7683889 | 1077.659817 | 0.04154436 | 0.153948382 | 0.776814156 | 0.824898703 | ns | YPD | KS.1 |
| FIM1_3170 | 608.3524981 | 82.96808913 | 121.4952025 | -2.772400966 | 0.24597451 | 3.64828580128 | 1.17098205557 | Km<KS.1 | YPD | KS.1 |
| FIM1_3171 | 567.4049233 | 1096.187187 | 1716.990805 | 0.912063444 | 0.149872992 | 2.63949507213 | 1.43513529878 | ns | YPD | KS.1 |
| FIM1_3172 | 664.5059881 | 1404.628083 | 1513.119302 | 1.002785271 | 0.210929948 | 3.79712163250 | 1.43002877529 | Km>KS.1 | YPD | KS.1 |
| FIM1_3173 | 4182.834067 | 2277.336449 | 1674.368048 | -0.841680394 | 0.144156508 | 1.38632270573 | 7.07380047284 | ns | YPD | KS.1 |
| FIM1_3174 | 334.5811095 | 660.1321639 | 492.5185698 | 0.899153251 | 0.220003416 | 1.03631038760 | 3.13794533071 | ns | YPD | KS.1 |
| FIM1_3175 | 938.4288431 | 1402.893918 | 1395.309944 | 0.553212274 | 0.133177134 | 1.65098811062 | 4.85473138080 | ns | YPD | KS.1 |
| FIM1_3176 | 946.544046 | 628.4445097 | 464.5748983 | -0.566909165 | 0.128185257 | 4.77825384806 | 1.51993688581 | ns | YPD | KS.1 |
| FIM1_3177 | 2098.670428 | 2406.00131 | 2248.049981 | 0.193226538 | 0.074043718 | 0.008042893 | 0.014952689 | ns | YPD | KS.1 |
| FIM1_3178 | 1824.337172 | 1575.61504 | 2050.784747 | -0.194601285 | 0.149900612 | 0.17282108 | 0.23306732 | ns | YPD | KS.1 |
| FIM1_3179 | 1217.713815 | 763.7765666 | 479.2921944 | -0.589041046 | 0.237120829 | 0.006115355 | 0.011644091 | ns | YPD | KS.1 |
| FIM1_3180 | 3559.549254 | 1934.800653 | 1588.096631 | -0.845396709 | 0.141765776 | 6.45190637225 | 3.39663853988 | ns | YPD | KS.1 |
| FIM1_3181 | 2074.497129 | 1676.170309 | 3024.698306 | -0.246628318 | 0.266321206 | 0.295236078 | 0.367475479 | ns | YPD | KS.1 |
| FIM1_3182 | 855.1568488 | 1183.245884 | 1881.758192 | 0.446337278 | 0.127754692 | 0.000297728 | 0.000724656 | ns | YPD | KS.1 |
| FIM1_3183 | 5919.028336 | 7234.746771 | 6908.476124 | 0.267506004 | 0.150127518 | 0.061296926 | 0.09415011 | ns | YPD | KS.1 |
| FIM1_3184 | 132.0934597 | 494.5617052 | 690.2581567 | 1.799995046 | 0.234213385 | 5.75744476123 | 5.27500694053 | Km>KS.1 | YPD | KS.1 |
| FIM1_3185 | 742.84712 | 775.6363511 | 1068.012224 | 0.05317367 | 0.210943497 | 0.784312103 | 0.830379381 | ns | YPD | KS.1 |
| FIM1_3186 | 3136.635817 | 4133.589959 | 3548.73503 | 0.378179749 | 0.126444515 | 0.001950345 | 0.00412541 | ns | YPD | KS.1 |
| FIM1_3187 | 400.3939583 | 402.7310889 | 663.5778738 | 0.007860004 | 0.148380223 | 0.955758058 | 0.968044167 | ns | YPD | KS.1 |
| FIM1_3188 | 559.4495551 | 625.1260745 | 690.8064976 | 0.149736073 | 0.128461381 | 0.226001889 | 0.293030857 | ns | YPD | KS.1 |
| FIM1_3189 | 364.9178432 | 362.0240401 | 411.7370225 | -0.010194099 | 0.174493004 | 0.950526226 | 0.96468135 | ns | YPD | KS.1 |
| FIM1_3190 | 652.8023894 | 584.9953154 | 680.2043265 | -0.146630307 | 0.13884196 | 0.270302933 | 0.340444833 | ns | YPD | KS.1 |
| FIM1_3191 | 211.8355565 | 649.1753886 | 883.2861558 | 1.546345314 | 0.190817751 | 3.58349372098 | 3.69107272503 | Km>KS.1 | YPD | KS.1 |
| FIM1_3192 | 621.0924521 | 932.9138253 | 729.3971384 | 0.566894312 | 0.115309527 | 4.30673453284 | 1.60494414239 | ns | YPD | KS.1 |
| FIM1_3193 | 28.66013572 | 46.14824947 | 43.75386283 | 0.571351489 | 0.270171902 | 0.016827252 | 0.029250725 | ns | YPD | KS.1 |

| | | | | | | | | | | |
|---|---|---|---|---|---|---|---|---|---|---|
| FIM1_3194 | 3569.283159 | 2850.560208 | 2395.124519 | -0.28483436 | 0.199925029 | 0.123552091 | 0.174252511 | ns | YPD | KS.1 |
| FIM1_3195 | 1532.353002 | 1660.693211 | 3200.949999 | 0.096527831 | 0.229040781 | 0.64318942 | 0.708805868 | ns | YPD | KS.1 |
| FIM1_3196 | 280.944626 | 397.3912684 | 477.2816158 | 0.461818905 | 0.164265181 | 0.002990279 | 0.006089495 | ns | YPD | KS.1 |
| FIM1_3197 | 8444.414569 | 1848.625611 | 1536.617768 | -2.01305407 | 0.314034208 | 3.0703339588 | 2.0089250732 | Km<KS.1 | YPD | KS.1 |
| FIM1_3198 | 917.6632084 | 697.8069092 | 460.0031329 | -0.270444468 | 0.369495637 | 0.368358421 | 0.444154907 | ns | YPD | KS.1 |
| FIM1_3199 | 1663.695141 | 2343.607029 | 2359.023598 | 0.464809522 | 0.145532684 | 0.000847274 | 0.001897878 | ns | YPD | KS.1 |
| FIM1_3200 | 5471.23759 | 8008.855178 | 9302.489641 | 0.544265292 | 0.057635745 | 1.7961017572 | 2.6051330152 | ns | YPD | KS.1 |
| FIM1_3201 | 279.0753719 | 381.4859941 | 501.8279529 | 0.415392647 | 0.16326773 | 0.007209144 | 0.013529042 | ns | YPD | KS.1 |
| FIM1_3202 | 928.460243 | 612.2220341 | 534.8867205 | -0.576140229 | 0.124399585 | 1.7419071399 | 5.9072856314 | ns | YPD | KS.1 |
| FIM1_3203 | 2001.068681 | 2645.095639 | 1449.933299 | 0.38067539 | 0.133035826 | 0.002943751 | 0.0059972 | ns | YPD | KS.1 |
| FIM1_3204 | 481.0410787 | 767.3213553 | 744.7715961 | 0.66737348 | 0.067547472 | 1.8871141709 | 3.2042296929 | ns | YPD | KS.1 |
| FIM1_3205 | 4242.01211 | 4367.550759 | 4777.253445 | 0.039746164 | 0.102908701 | 0.69191794 | 0.752742565 | ns | YPD | KS.1 |
| FIM1_3206 | 1564.601745 | 1489.304019 | 1815.968829 | -0.067807592 | 0.103033854 | 0.49941209 | 0.576066577 | ns | YPD | KS.1 |
| FIM1_3207 | 937.1737555 | 1441.430063 | 891.1447567 | 0.563312221 | 0.197476666 | 0.002152985 | 0.004504248 | ns | YPD | KS.1 |
| FIM1_3208 | 1685.280509 | 2355.260972 | 3247.202603 | 0.467974458 | 0.101310351 | 2.3130128082 | 7.6971496460 | ns | YPD | KS.1 |
| FIM1_3209 | 3942.647932 | 6921.32362 | 7110.124517 | 0.783012507 | 0.130633555 | 6.1040467754 | 3.2203216020 | ns | YPD | KS.1 |
| FIM1_3210 | 143.9399065 | 192.0120844 | 590.3318852 | 0.382365405 | 0.169413809 | 0.016690447 | 0.029043363 | ns | YPD | KS.1 |
| FIM1_3211 | 337.6431538 | 368.8894625 | 437.1034821 | 0.110958767 | 0.177443207 | 0.505537153 | 0.581782868 | ns | YPD | KS.1 |
| FIM1_3212 | 1866.723393 | 2577.230211 | 3159.226961 | 0.446072297 | 0.117337798 | 8.9954063578 | 0.000236784 | ns | YPD | KS.1 |
| FIM1_3213 | 4197.682872 | 3951.450861 | 3801.11775 | -0.083223937 | 0.100569224 | 0.39610416 | 0.472910535 | ns | YPD | KS.1 |
| FIM1_3214 | 1419.731938 | 1421.938475 | 3474.386174 | 0.001634651 | 0.342006489 | 0.995425116 | 0.996226101 | ns | YPD | KS.1 |
| FIM1_3215 | 3614.072687 | 3702.042375 | 2625.623474 | 0.033549504 | 0.08445531 | 0.68572535 | 0.74747669 | ns | YPD | KS.1 |
| FIM1_3216 | 168.4817716 | 67.12606017 | 80.9259995 | -1.217663004 | 0.246223808 | 9.4268074922 | 3.8346988776 | Km<KS.1 | YPD | KS.1 |
| FIM1_3217 | 2456.62173 | 2592.32793 | 2185.918002 | 0.063545127 | 0.239619559 | 0.769121875 | 0.81882759 | ns | YPD | KS.1 |
| FIM1_3218 | 172.556982 | 150.8878672 | 239.287081 | -0.150577919 | 0.275052576 | 0.53381819 | 0.608976266 | ns | YPD | KS.1 |
| FIM1_3219 | 2024.886064 | 2711.569234 | 3158.916771 | 0.385443602 | 0.170345414 | 0.016361323 | 0.028550537 | ns | YPD | KS.1 |
| FIM1_3220 | 8817.594394 | 2902.151014 | 2141.848212 | -1.511891189 | 0.22453122 | 1.1803679868 | 8.0997665306 | Km<KS.1 | YPD | KS.1 |
| FIM1_3221 | 4220.165891 | 6452.761383 | 6394.062365 | 0.59434933 | 0.106889457 | 1.2298178699 | 5.5926361088 | ns | YPD | KS.1 |
| FIM1_3222 | 3785.806121 | 3115.77866 | 3571.471653 | -0.277061991 | 0.061117736 | 4.6889096220 | 1.4934267202 | ns | YPD | KS.1 |
| FIM1_3223 | 4185.426331 | 5126.299513 | 5186.784433 | 0.284916042 | 0.085439499 | 0.000683188 | 0.001562695 | ns | YPD | KS.1 |
| FIM1_3224 | 2661.439361 | 3340.225043 | 1877.403515 | 0.265397284 | 0.261615726 | 0.253084596 | 0.32243172 | ns | YPD | KS.1 |
| FIM1_3225 | 140.754978 | 55.71425925 | 117.7906859 | -1.176716462 | 0.302860838 | 1.3421724533 | 3.9974216975 | Km<KS.1 | YPD | KS.1 |
| FIM1_3226 | 983.2383304 | 1033.086576 | 829.6867632 | 0.065640069 | 0.133423771 | 0.608880884 | 0.679188878 | ns | YPD | KS.1 |
| FIM1_3227 | 1046.151908 | 2196.903546 | 2360.123084 | 1.039024937 | 0.133585579 | 1.3011976512 | 1.1539141381 | Km>KS.1 | YPD | KS.1 |
| FIM1_3228 | 1159.861652 | 1321.85698 | 1687.076866 | 0.176748077 | 0.123661708 | 0.138449801 | 0.192023351 | ns | YPD | KS.1 |
| FIM1_3229 | 1645.196439 | 2537.401377 | 3769.560011 | 0.609793509 | 0.095405558 | 7.1860359952 | 4.1092562156 | ns | YPD | KS.1 |
| FIM1_3230 | 2143.001812 | 3538.026591 | 4397.562342 | 0.702921376 | 0.110745589 | 7.7701647515 | 4.4330928485 | ns | YPD | KS.1 |
| FIM1_3231 | 1653.671206 | 3967.013963 | 4952.197279 | 1.251724514 | 0.074429421 | 2.0952574882 | 5.2119530020 | Km>KS.1 | YPD | KS.1 |
| FIM1_3232 | 925.1617055 | 488.1606671 | 257.3779081 | -0.834229527 | 0.231816174 | 8.7647702905 | 0.000230835 | ns | YPD | KS.1 |
| FIM1_3233 | 2111.169485 | 650.8129879 | 1744.519258 | -1.669162709 | 0.122062374 | 7.2400862709 | 4.6178755381 | Km<KS.1 | YPD | KS.1 |
| FIM1_3234 | 3130.506032 | 1817.089486 | 4010.201847 | -0.633994594 | 0.315523241 | 0.019040169 | 0.032720152 | ns | YPD | KS.1 |
| FIM1_3235 | 1061.82795 | 1303.7564 | 1430.647497 | 0.282702128 | 0.111346478 | 0.0089279 | 0.016456577 | ns | YPD | KS.1 |
| FIM1_3236 | 12489.53944 | 12666.72306 | 9277.944695 | 0.0168134 | 0.230526196 | 0.936072391 | 0.953513544 | ns | YPD | KS.1 |
| FIM1_3237 | 4968.479507 | 5453.036227 | 5182.316218 | 0.124922093 | 0.130733519 | 0.320560228 | 0.394163899 | ns | YPD | KS.1 |
| FIM1_3238 | 360.5665455 | 408.2298458 | 470.235462 | 0.167542548 | 0.121233203 | 0.152457968 | 0.208717223 | ns | YPD | KS.1 |

| | | | | | | | | | | |
|---|---|---|---|---|---|---|---|---|---|---|
| FIM1_3239 | 4406.032033 | 2887.869656 | 3239.1185 | -0.590420469 | 0.110284632 | 3.98207779643 | 1.70050103324 | ns | YPD | KS.1 |
| FIM1_3240 | 2080.742409 | 2535.058851 | 2728.842769 | 0.266456323 | 0.137812749 | 0.043638189 | 0.068855055 | ns | YPD | KS.1 |
| FIM1_3241 | 1642.968024 | 1261.157579 | 1207.718789 | -0.34134974 | 0.189284617 | 0.05282651 | 0.082231505 | ns | YPD | KS.1 |
| FIM1_3242 | 1064.771305 | 995.4804481 | 739.1983989 | -0.09058205 | 0.128833017 | 0.465444044 | 0.543054436 | ns | YPD | KS.1 |
| FIM1_3243 | 888.1655286 | 2062.852971 | 2177.611347 | 1.184169095 | 0.132063166 | 4.13060894876 | 5.45086989923 | Km>KS.1 | YPD | KS.1 |
| FIM1_3244 | 1055.632908 | 504.7802725 | 1295.076085 | -0.97550166 | 0.228663355 | 4.01002593850 | 1.28708897058 | ns | YPD | KS.1 |
| FIM1_3245 | 1104.028529 | 1588.472496 | 1235.652559 | 0.495188372 | 0.143874219 | 0.000329133 | 0.000795645 | ns | YPD | KS.1 |
| FIM1_3246 | 4415.248444 | 1693.257015 | 1352.307553 | -1.326186868 | 0.177040294 | 7.10120257094 | 5.79537212479 | Km<KS.1 | YPD | KS.1 |
| FIM1_3247 | 8378.373217 | 9148.716118 | 5932.375847 | 0.101389346 | 0.257555734 | 0.658895914 | 0.723302553 | ns | YPD | KS.1 |
| FIM1_3248 | 548.098725 | 300.0957179 | 270.241428 | -0.834843654 | 0.141952086 | 1.08830838033 | 5.62657542200 | ns | YPD | KS.1 |
| FIM1_3249 | 2184.805132 | 2742.692175 | 3260.562503 | 0.307405973 | 0.136759249 | 0.019121921 | 0.032837955 | ns | YPD | KS.1 |
| FIM1_3250 | 462.3237465 | 548.8591235 | 864.3547259 | 0.225340035 | 0.164638133 | 0.147391243 | 0.202673143 | ns | YPD | KS.1 |
| FIM1_3251 | 1503.080821 | 912.0560109 | 786.1267761 | -0.699730896 | 0.10925738 | 5.38272197270 | 3.12839273530 | ns | YPD | KS.1 |
| FIM1_3252 | 144.7145346 | 42.16223308 | 46.84830583 | -1.591277873 | 0.324912487 | 5.46397033835 | 2.29008023869 | Km<KS.1 | YPD | KS.1 |
| FIM1_3253 | 11770.72362 | 4129.633606 | 3578.117444 | -1.358985548 | 0.293524948 | 3.36598858856 | 1.27538409962 | Km<KS.1 | YPD | KS.1 |
| FIM1_3254 | 2294.000679 | 4836.870429 | 4440.083237 | 1.059136078 | 0.095759383 | 3.27902829958 | 8.63130465102 | Km>KS.1 | YPD | KS.1 |
| FIM1_3255 | 1511.547873 | 926.7846754 | 844.2191512 | -0.693859388 | 0.082915991 | 2.07161180437 | 2.17431829678 | ns | YPD | KS.1 |
| FIM1_3256 | 42.78675365 | 24.97383804 | 40.43753559 | -0.641898103 | 0.295341898 | 0.012526615 | 0.022449535 | ns | YPD | KS.1 |
| FIM1_3258 | 87.71633258 | 58.73618216 | 304.6439041 | -0.518218861 | 0.19972279 | 0.005148985 | 0.009979043 | ns | YPD | KS.1 |
| FIM1_3259 | 127.0281088 | 134.1676432 | 420.9082459 | 0.06758937 | 0.225215271 | 0.742198013 | 0.795440568 | ns | YPD | KS.1 |
| FIM1_3260 | 2124.008321 | 1958.740336 | 1532.399655 | -0.113232815 | 0.074939768 | 0.124747593 | 0.175663537 | ns | YPD | KS.1 |
| FIM1_3261 | 489.5524057 | 607.0961479 | 583.1292843 | 0.285625414 | 0.152150734 | 0.048445138 | 0.075814584 | ns | YPD | KS.1 |
| FIM1_3262 | 13969.14617 | 19807.54908 | 29873.94409 | 0.483762197 | 0.118323858 | 2.52994308209 | 7.27541435457 | ns | YPD | KS.1 |
| FIM1_3263 | 1572.116928 | 2047.945252 | 1959.690367 | 0.367670259 | 0.105191799 | 0.000338091 | 0.000814923 | ns | YPD | KS.1 |
| FIM1_3264 | 1148.350745 | 1257.755491 | 1677.565821 | 0.123465963 | 0.119989096 | 0.287152433 | 0.358940541 | ns | YPD | KS.1 |
| FIM1_3265 | 753.5834391 | 1398.843927 | 1308.290788 | 0.875218158 | 0.096266923 | 2.33520823171 | 3.13990836562 | ns | YPD | KS.1 |
| FIM1_3266 | 147.9237918 | 419.7084902 | 627.6223463 | 1.413292557 | 0.228948885 | 5.78338021440 | 3.34173246999 | Km>KS.1 | YPD | KS.1 |
| FIM1_3267 | 1504.122224 | 2893.316894 | 3631.771014 | 0.920830821 | 0.113926514 | 1.39623138095 | 1.36737226776 | ns | YPD | KS.1 |
| FIM1_3268 | 432.2245495 | 588.3674318 | 861.409376 | 0.406478922 | 0.173907189 | 0.012974577 | 0.023168888 | ns | YPD | KS.1 |
| FIM1_3269 | 612.4167225 | 599.8694959 | 594.8101771 | -0.026931277 | 0.163723775 | 0.862123315 | 0.893927364 | ns | YPD | KS.1 |
| FIM1_3270 | 4387.31239 | 6534.041553 | 3849.519547 | 0.438083427 | 0.32715808 | 0.112878031 | 0.161138652 | ns | YPD | KS.1 |
| FIM1_3271 | 4941.528442 | 2604.711203 | 3839.313539 | -0.894599192 | 0.129974635 | 1.37297924520 | 9.35694759574 | ns | YPD | KS.1 |
| FIM1_3272 | 1397.605092 | 1173.320322 | 1169.173293 | -0.235083014 | 0.137019601 | 0.073463634 | 0.10995234 | ns | YPD | KS.1 |
| FIM1_3273 | 549.165982 | 875.1800488 | 2005.529265 | 0.565608955 | 0.271411669 | 0.018367019 | 0.031683745 | ns | YPD | KS.1 |
| FIM1_3274 | 2596.796597 | 2758.194582 | 2580.125895 | 0.086219032 | 0.049434394 | 0.078805019 | 0.116891762 | ns | YPD | KS.1 |
| FIM1_3275 | 4215.421578 | 2698.153959 | 3593.207739 | -0.620572558 | 0.12105107 | 1.27685664750 | 5.09820370892 | ns | YPD | KS.1 |
| FIM1_3276 | 610.3515421 | 1023.474609 | 476.724671 | 0.67749006 | 0.20718026 | 0.000417392 | 0.000986941 | ns | YPD | KS.1 |
| FIM1_3277 | 1595.241013 | 2033.354187 | 2368.234626 | 0.336547824 | 0.106109593 | 0.001132037 | 0.002483193 | ns | YPD | KS.1 |
| FIM1_3278 | 7337.465134 | 7366.075372 | 6815.733062 | 0.005457228 | 0.097584975 | 0.954351186 | 0.967561833 | ns | YPD | KS.1 |
| FIM1_3279 | 2267.56409 | 3175.119537 | 3833.314864 | 0.46104659 | 0.133533288 | 0.000337572 | 0.000814068 | ns | YPD | KS.1 |
| FIM1_3280 | 2620.579675 | 1969.286142 | 2105.880803 | -0.392442139 | 0.125040246 | 0.001163909 | 0.002547346 | ns | YPD | KS.1 |
| FIM1_3281 | 2526.841313 | 2353.857325 | 4292.32629 | -0.094037385 | 0.145613308 | 0.499110401 | 0.575852097 | ns | YPD | KS.1 |
| FIM1_3282 | 6449.212134 | 7395.82396 | 5569.056616 | 0.147420435 | 0.305332222 | 0.574235907 | 0.64668488 | ns | YPD | KS.1 |
| FIM1_3283 | 767.9289807 | 565.5866405 | 807.0213661 | -0.409873069 | 0.158061471 | 0.006321067 | 0.011989062 | ns | YPD | KS.1 |
| FIM1_3284 | 1350.258036 | 1622.129686 | 1342.972478 | 0.243238376 | 0.153745218 | 0.095935651 | 0.139148648 | ns | YPD | KS.1 |

| | | | | | | | | | | |
|---|---|---|---|---|---|---|---|---|---|---|
| FIM1_3286 | 1568.463709 | 2242.191088 | 1644.190823 | 0.496475011 | 0.113898975 | 7.4227073509 | 2.2950881958 | ns | YPD | KS.1 |
| FIM1_3287 | 370.3070484 | 409.4558032 | 460.2756596 | 0.128165937 | 0.174718873 | 0.435374218 | 0.512559298 | ns | YPD | KS.1 |
| FIM1_3288 | 24870.84637 | 19619.91487 | 24758.36259 | -0.298220918 | 0.207609468 | 0.118684143 | 0.168268342 | ns | YPD | KS.1 |
| FIM1_3290 | 1249.081823 | 1923.232424 | 1587.834695 | 0.595154766 | 0.13116024 | 2.6187875603 | 8.6338423543 | ns | YPD | KS.1 |
| FIM1_3291 | 40.68247709 | 23.46581816 | 18.39346275 | -0.573738671 | 0.390575957 | 0.067913281 | 0.102695432 | ns | YPD | KS.1 |
| FIM1_3292 | 15954.40159 | 20513.46849 | 28547.75295 | 0.344817928 | 0.122983833 | 0.003725425 | 0.00744337 | ns | YPD | KS.1 |
| FIM1_3293 | 243.7038077 | 256.2320828 | 402.3163413 | 0.062809738 | 0.183272177 | 0.714425524 | 0.77216315 | ns | YPD | KS.1 |
| FIM1_3294 | 326.7333266 | 241.2777107 | 335.1507035 | -0.392441496 | 0.185527126 | 0.023520619 | 0.039412286 | ns | YPD | KS.1 |
| FIM1_3295 | 2897.280839 | 3168.044988 | 1289.998448 | 0.111686753 | 0.197380458 | 0.542135501 | 0.615922383 | ns | YPD | KS.1 |
| FIM1_3297 | 1914.712928 | 1631.323309 | 2133.083476 | -0.223295638 | 0.090469455 | 0.011687872 | 0.021067812 | ns | YPD | KS.1 |
| FIM1_3298 | 2708.496529 | 2401.021361 | 3671.679717 | -0.142800186 | 0.240002673 | 0.509727027 | 0.585521117 | ns | YPD | KS.1 |
| FIM1_3299 | 5605.873035 | 6849.597047 | 5009.030904 | 0.279983738 | 0.092742473 | 0.002042355 | 0.004296802 | ns | YPD | KS.1 |
| FIM1_3300 | 5252.299481 | 6389.369519 | 12154.06664 | 0.260783398 | 0.151324231 | 0.070123892 | 0.105557145 | ns | YPD | KS.1 |
| FIM1_3301 | 2351.220713 | 1505.493941 | 2184.731024 | -0.61779482 | 0.126221201 | 4.2997196645 | 1.6035311342 | ns | YPD | KS.1 |
| FIM1_3302 | 278.6035864 | 173.7271428 | 695.5945618 | -0.511356712 | 0.349699857 | 0.078355947 | 0.116294999 | ns | YPD | KS.1 |
| FIM1_3303 | 430.4840937 | 413.6920567 | 583.473781 | -0.051930112 | 0.163800959 | 0.737780935 | 0.791729972 | ns | YPD | KS.1 |
| FIM1_3304 | 6908.457234 | 6659.733309 | 10610.29114 | -0.043704601 | 0.231307753 | 0.83529421 | 0.872839465 | ns | YPD | KS.1 |
| FIM1_3305 | 24884.62626 | 11806.86105 | 11293.93362 | -1.045930433 | 0.129881056 | 1.4072293320 | 1.3754353490 | Km<KS.1 | YPD | KS.1 |
| FIM1_3306 | 361.6306308 | 304.840488 | 385.9812727 | -0.215276854 | 0.198462478 | 0.241633718 | 0.309906611 | ns | YPD | KS.1 |
| FIM1_3307 | 4670.953978 | 1312.180239 | 1497.973183 | -1.786433477 | 0.155120132 | 4.2154875998 | 1.3443622313 | Km<KS.1 | YPD | KS.1 |
| FIM1_3308 | 9924.05566 | 798.570965 | 1459.473884 | -3.565183339 | 0.222912937 | 1.9633261972 | 3.3681199418 | Km<KS.1 | YPD | KS.1 |
| FIM1_3309 | 3241.673791 | 4559.915557 | 4207.45595 | 0.462464801 | 0.147193699 | 0.001017421 | 0.002252635 | ns | YPD | KS.1 |
| FIM1_3310 | 1449.05357 | 1843.945793 | 1586.122228 | 0.333418308 | 0.109039717 | 0.001672809 | 0.003565648 | ns | YPD | KS.1 |
| FIM1_3311 | 2788.864952 | 4842.798346 | 5799.004745 | 0.788697564 | 0.063239858 | 2.9753025409 | 1.2335108450 | ns | YPD | KS.1 |
| FIM1_3312 | 8220.66541 | 6357.04407 | 3030.972899 | -0.297110647 | 0.272877907 | 0.216073052 | 0.28266196 | ns | YPD | KS.1 |
| FIM1_3313 | 856.2920415 | 1254.439054 | 449.383105 | 0.389501593 | 0.367890269 | 0.194438292 | 0.25816133 | ns | YPD | KS.1 |
| FIM1_3314 | 707.5585469 | 640.9944418 | 611.8314272 | -0.132893082 | 0.128416484 | 0.282510628 | 0.354027802 | ns | YPD | KS.1 |
| FIM1_3315 | 165.8503735 | 577.7632627 | 625.3048471 | 1.736262086 | 0.180332118 | 2.6854004301 | 4.4982717642 | Km>KS.1 | YPD | KS.1 |
| FIM1_3316 | 693.8472582 | 1650.534957 | 1921.22516 | 1.232242071 | 0.095656742 | 6.9497463382 | 3.2617913238 | Km>KS.1 | YPD | KS.1 |
| FIM1_3317 | 65.73370266 | 93.27567858 | 156.9880462 | 0.464374924 | 0.179098581 | 0.005740325 | 0.011000816 | ns | YPD | KS.1 |
| FIM1_3318 | 463.9456127 | 330.9613435 | 902.019129 | -0.386475722 | 0.291474024 | 0.126619023 | 0.177595049 | ns | YPD | KS.1 |
| FIM1_3319 | 453.1452927 | 1225.904224 | 1424.631369 | 1.393717353 | 0.15146015 | 3.2199933835 | 4.5901051814 | Km>KS.1 | YPD | KS.1 |
| FIM1_3320 | 10037.54667 | 3329.138443 | 3754.133926 | -1.43137969 | 0.300855116 | 1.5547169100 | 6.1386639900 | Km<KS.1 | YPD | KS.1 |
| FIM1_3321 | 5295.294931 | 2067.201004 | 2821.384153 | -1.163771045 | 0.334702022 | 6.7517307322 | 0.000181862 | Km<KS.1 | YPD | KS.1 |
| FIM1_3322 | 1059.343262 | 818.853801 | 576.8316103 | -0.304390896 | 0.257050767 | 0.183501396 | 0.245738746 | ns | YPD | KS.1 |
| FIM1_3323 | 2082.487398 | 1119.192527 | 1336.77278 | -0.864507784 | 0.135195318 | 4.0243327641 | 2.3777975655 | ns | YPD | KS.1 |
| FIM1_3324 | 7443.612492 | 2393.024206 | 2165.389234 | -1.586314655 | 0.165450059 | 5.5672986175 | 9.1109574415 | Km<KS.1 | YPD | KS.1 |
| FIM1_3326 | 5415.872569 | 12308.12706 | 10761.74779 | 1.126909525 | 0.180948647 | 7.0938617454 | 4.0624327857 | Km>KS.1 | YPD | KS.1 |
| FIM1_3327 | 2099.341542 | 1543.937001 | 1547.231501 | -0.412221752 | 0.154336801 | 0.005008358 | 0.00973304 | ns | YPD | KS.1 |
| FIM1_3328 | 201.4780327 | 278.4922462 | 250.7530417 | 0.420190611 | 0.188426216 | 0.016796299 | 0.029208507 | ns | YPD | KS.1 |
| FIM1_3329 | 496.630546 | 192.6317845 | 333.4106703 | -1.319820698 | 0.160265989 | 1.8879637769 | 1.9899618199 | Km<KS.1 | YPD | KS.1 |
| FIM1_3331 | 134.7640384 | 267.5864972 | 427.7347597 | 0.961192626 | 0.132880839 | 9.6462878370 | 7.1520539477 | ns | YPD | KS.1 |
| FIM1_3332 | 145.0643251 | 450.0653738 | 563.365356 | 1.602636827 | 0.121662098 | 7.3798034958 | 3.9058002544 | Km>KS.1 | YPD | KS.1 |
| FIM1_3333 | 2709.67166 | 1866.218872 | 3974.130456 | -0.484087758 | 0.197764622 | 0.008350392 | 0.015472329 | ns | YPD | KS.1 |
| FIM1_3334 | 25165.83211 | 30451.82234 | 33685.66399 | 0.259796233 | 0.125126316 | 0.031315116 | 0.050879393 | ns | YPD | KS.1 |

| | | | | | | | | | | |
|---|---|---|---|---|---|---|---|---|---|---|
| FIM1_3335 | 1191.80612 | 1113.380906 | 1422.542747 | -0.085553505 | 0.189266418 | 0.62806727 | 0.69575477 | ns | YPD | KS.1 |
| FIM1_3336 | 353.6752332 | 700.3880214 | 1146.475696 | 0.922618027 | 0.193432508 | 4.1447498705 | 1.5515523405 | ns | YPD | KS.1 |
| FIM1_3337 | 3550.397606 | 2337.360853 | 2308.848679 | -0.592863843 | 0.080264394 | 6.8029769030 | 5.1125090774 | ns | YPD | KS.1 |
| FIM1_3339 | 1320.753392 | 1041.20726 | 1165.07265 | -0.323407654 | 0.13442316 | 0.012268842 | 0.022027243 | ns | YPD | KS.1 |
| FIM1_3340 | 1505.462806 | 2840.429499 | 5574.560663 | 0.840348864 | 0.212995017 | 2.1476306273 | 6.2227503616 | ns | YPD | KS.1 |
| FIM1_3341 | 457.2035899 | 1332.80459 | 1121.771156 | 1.523113171 | 0.099096844 | 1.7848971800 | 1.9733029934 | Km>KS.1 | YPD | KS.1 |
| FIM1_3342 | 195.4241619 | 173.0887913 | 158.9391856 | -0.149148854 | 0.221366439 | 0.460940206 | 0.538430975 | ns | YPD | KS.1 |
| FIM1_3343 | 10256.31603 | 10088.28767 | 8696.620167 | -0.019079931 | 0.253901525 | 0.933053098 | 0.95102216 | ns | YPD | KS.1 |
| FIM1_3344 | 34545.28365 | 22120.68807 | 18575.06962 | -0.502972357 | 0.319406605 | 0.064214347 | 0.097905723 | ns | YPD | KS.1 |
| FIM1_3345 | 2190.272707 | 4605.67736 | 6481.108677 | 0.882335087 | 0.338954898 | 0.002275359 | 0.004732405 | ns | YPD | KS.1 |
| FIM1_3346 | 341.7842986 | 253.1786099 | 215.6425215 | -0.369274977 | 0.235195699 | 0.082418417 | 0.121698375 | ns | YPD | KS.1 |
| FIM1_3347 | 3879.979069 | 3106.18536 | 1658.200111 | -0.285763041 | 0.187570586 | 0.102175185 | 0.147254213 | ns | YPD | KS.1 |
| FIM1_3348 | 2046.12065 | 1635.5189 | 957.1650824 | -0.283944659 | 0.198709154 | 0.12272265 | 0.173253458 | ns | YPD | KS.1 |
| FIM1_3349 | 2908.623983 | 2549.287247 | 1789.716452 | -0.180471029 | 0.113594674 | 0.101126283 | 0.145869312 | ns | YPD | KS.1 |
| FIM1_3350 | 19231.89913 | 10145.1955 | 9211.614553 | -0.853377149 | 0.2040835 | 7.5717483853 | 2.3368144055 | ns | YPD | KS.1 |
| FIM1_3351 | 1384.781021 | 2054.831344 | 1694.361127 | 0.548966811 | 0.115922914 | 1.1099277256 | 3.8913956555 | ns | YPD | KS.1 |
| FIM1_3353 | 3233.862912 | 5080.21682 | 8404.574634 | 0.617684904 | 0.147525243 | 1.2419242929 | 3.7265219286 | ns | YPD | KS.1 |
| FIM1_3355 | 839.1492892 | 1316.079006 | 1440.681107 | 0.58805339 | 0.200753001 | 0.00160181 | 0.00342753 | ns | YPD | KS.1 |
| FIM1_3356 | 1471.29857 | 2157.276744 | 1282.228097 | 0.491241813 | 0.209364272 | 0.010863205 | 0.019693549 | ns | YPD | KS.1 |
| FIM1_3357 | 1997.199169 | 365.663932 | 631.5619316 | -2.401018049 | 0.161299191 | 2.3118584984 | 2.3472440877 | Km<KS.1 | YPD | KS.1 |
| FIM1_3358 | 38.69157474 | 4.315878622 | 16.18493289 | -2.6682234 | 0.566893839 | 8.1785090507 | 3.8061817144 | Km<KS.1 | YPD | KS.1 |
| FIM1_3359 | 3360.453346 | 5172.332727 | 5452.627362 | 0.607520662 | 0.095675245 | 9.4888455989 | 5.3766522613 | ns | YPD | KS.1 |
| FIM1_3360 | 4600.81256 | 3079.476534 | 3710.643881 | -0.558640354 | 0.116322185 | 7.7950757123 | 2.8020593691 | ns | YPD | KS.1 |
| FIM1_3361 | 4278.644896 | 6193.547135 | 6897.756809 | 0.521759915 | 0.088300374 | 1.8419423527 | 9.2562254593 | ns | YPD | KS.1 |
| FIM1_3362 | 6391.938971 | 5243.689001 | 3212.704601 | -0.255374531 | 0.181131163 | 0.131900796 | 0.184067353 | ns | YPD | KS.1 |
| FIM1_3363 | 1731.937269 | 1339.642741 | 1497.899917 | -0.355309721 | 0.11348017 | 0.001265783 | 0.002749374 | ns | YPD | KS.1 |
| FIM1_3364 | 2120.379498 | 2508.153454 | 1841.479941 | 0.234615964 | 0.089798713 | 0.0076329 | 0.01424369 | ns | YPD | KS.1 |
| FIM1_3365 | 313.5692222 | 273.1121293 | 341.3008944 | -0.17560591 | 0.180271594 | 0.29845253 | 0.370736913 | ns | YPD | KS.1 |
| FIM1_3366 | 1044.237046 | 1086.070007 | 426.0664148 | 0.038311451 | 0.357956611 | 0.896377484 | 0.92099917 | ns | YPD | KS.1 |
| FIM1_3367 | 670.2758819 | 679.2930117 | 713.2190129 | 0.017220864 | 0.167497274 | 0.91338522 | 0.934998245 | ns | YPD | KS.1 |
| FIM1_3368 | 317.0626315 | 346.2689071 | 621.4381627 | 0.112673962 | 0.169124819 | 0.480139426 | 0.558366911 | ns | YPD | KS.1 |
| FIM1_3369 | 975.897749 | 948.1638634 | 1156.707981 | -0.036608277 | 0.195830487 | 0.84065602 | 0.876233752 | ns | YPD | KS.1 |
| FIM1_3370 | 498.3001736 | 494.9472928 | 841.2917541 | -0.009585888 | 0.189683432 | 0.956818489 | 0.968541301 | ns | YPD | KS.1 |
| FIM1_3371 | 252.7308164 | 426.6253814 | 821.7171968 | 0.704667145 | 0.17786895 | 2.7030080329 | 7.7328723195 | ns | YPD | KS.1 |
| FIM1_3372 | 796.1142051 | 821.6743622 | 785.7021713 | 0.042961229 | 0.107843635 | 0.68213839 | 0.744532555 | ns | YPD | KS.1 |
| FIM1_3373 | 763.4107924 | 346.0853652 | 864.5272455 | -0.948829965 | 0.339080161 | 0.001089852 | 0.002393824 | ns | YPD | KS.1 |
| FIM1_3374 | 12307.06831 | 7308.220198 | 5453.42696 | -0.600924297 | 0.318901408 | 0.027288522 | 0.045118111 | ns | YPD | KS.1 |
| FIM1_3375 | 15025.11791 | 7538.107997 | 8288.966713 | -0.965196286 | 0.131037254 | 3.5778284218 | 2.7468667281 | ns | YPD | KS.1 |
| FIM1_3376 | 13.35902539 | 27.36280704 | 19.77240213 | 0.809717981 | 0.3653902 | 0.007722041 | 0.014393839 | ns | YPD | KS.1 |
| FIM1_3377 | 2092.989777 | 1247.800947 | 1614.212457 | -0.630419795 | 0.276300043 | 0.009715908 | 0.017816675 | ns | YPD | KS.1 |
| FIM1_3378 | 42.46636379 | 20.43283175 | 24.81555604 | -0.943513647 | 0.260512295 | 6.3162388823 | 0.000171058 | ns | YPD | KS.1 |
| FIM1_3379 | 915.5443205 | 533.5606987 | 602.1434 | -0.74713844 | 0.138118641 | 2.0534631942 | 9.0808705702 | ns | YPD | KS.1 |
| FIM1_3380 | 602.2856746 | 402.278301 | 384.328297 | -0.523734055 | 0.202056407 | 0.00513273 | 0.009955296 | ns | YPD | KS.1 |
| FIM1_3381 | 7322.104133 | 10168.42414 | 18287.95914 | 0.449229851 | 0.134244052 | 0.000508834 | 0.00119015 | ns | YPD | KS.1 |
| FIM1_3382 | 1322.808495 | 426.9922601 | 540.4338346 | -1.585110532 | 0.15809595 | 7.2470798602 | 1.3867008578 | Km<KS.1 | YPD | KS.1 |

| | | | | | | | | | | |
|---|---|---|---|---|---|---|---|---|---|---|
| FIM1_3383 | 3290.668697 | 2901.872513 | 2471.836453 | -0.170786296 | 0.122846464 | 0.149716038 | 0.205450585 | ns | YPD | KS.1 |
| FIM1_3384 | 5862.879783 | 4964.427175 | 10102.61494 | -0.189572268 | 0.271398173 | 0.428353921 | 0.505829754 | ns | YPD | KS.1 |
| FIM1_3385 | 79.56485182 | 80.35184585 | 83.71763438 | 0.012058404 | 0.248164771 | 0.9568604 | 0.968541301 | ns | YPD | KS.1 |
| FIM1_3386 | 77.7133014 | 190.2531321 | 715.7074906 | 1.184908622 | 0.252055163 | 3.41248733723 | 1.29103608386 | Km>KS.1 | YPD | KS.1 |
| FIM1_3387 | 2739.973234 | 2956.576377 | 1783.26767 | 0.090791001 | 0.232476345 | 0.667016509 | 0.729692399 | ns | YPD | KS.1 |
| FIM1_3388 | 5085.904488 | 3367.838819 | 3075.192619 | -0.504009837 | 0.255935264 | 0.027326172 | 0.045165351 | ns | YPD | KS.1 |
| FIM1_3389 | 818.5902224 | 985.0391982 | 1286.693949 | 0.207083166 | 0.284824471 | 0.404890886 | 0.481775822 | ns | YPD | KS.1 |
| FIM1_3390 | 3031.157456 | 4742.660828 | 3612.230617 | 0.61819058 | 0.132187852 | 1.27426625888 | 4.41775236093 | ns | YPD | KS.1 |
| FIM1_3391 | 797.0518513 | 1844.853423 | 1527.696093 | 1.131849394 | 0.212237618 | 1.42758004424 | 6.42734001822 | Km>KS.1 | YPD | KS.1 |
| FIM1_3392 | 950.928621 | 906.1499074 | 997.793671 | -0.063799419 | 0.137811873 | 0.62945931 | 0.696986438 | ns | YPD | KS.1 |
| FIM1_3393 | 210.6223583 | 91.50899234 | 168.986438 | -0.974658838 | 0.367209797 | 0.00158348 | 0.00339561 | ns | YPD | KS.1 |
| FIM1_3394 | 6579.521902 | 3246.982797 | 2718.688271 | -0.952468425 | 0.197581176 | 3.02141627470 | 1.15449661802 | ns | YPD | KS.1 |
| FIM1_3395 | 1552.069131 | 2965.18344 | 2191.632266 | 0.858555306 | 0.213054185 | 1.44572879933 | 4.28890922878 | ns | YPD | KS.1 |
| FIM1_3396 | 1548.284943 | 868.4111349 | 1060.624303 | -0.740880741 | 0.241416575 | 0.000726539 | 0.00165198 | ns | YPD | KS.1 |
| FIM1_3397 | 3852.038751 | 10464.96539 | 7705.802668 | 1.268266485 | 0.31527603 | 6.24430880745 | 1.95503060522 | Km>KS.1 | YPD | KS.1 |
| FIM1_3398 | 3.636081251 | 5.086057503 | 8.902902147 | 0.257866937 | 0.445945093 | 0.444404177 | 0.521564233 | ns | YPD | KS.1 |
| FIM1_3399 | 1407.81084 | 1297.751842 | 1046.559827 | -0.102894552 | 0.185007211 | 0.552050628 | 0.625330572 | ns | YPD | KS.1 |
| FIM1_3400 | 2159.622411 | 1865.319513 | 1585.761323 | -0.201996804 | 0.105800607 | 0.049605594 | 0.077508741 | ns | YPD | KS.1 |
| FIM1_3401 | 2182.823104 | 2150.591639 | 1416.926159 | -0.019904911 | 0.143994911 | 0.885138969 | 0.912656242 | ns | YPD | KS.1 |
| FIM1_3402 | 1864.261306 | 2808.404678 | 3669.48146 | 0.580438547 | 0.081383178 | 4.59383392072 | 3.23257761748 | ns | YPD | KS.1 |
| FIM1_3403 | 3950.185363 | 3774.675235 | 4961.124225 | -0.052792335 | 0.249755167 | 0.813729043 | 0.853970968 | ns | YPD | KS.1 |
| FIM1_3404 | 580.4679455 | 35.88215942 | 107.336423 | -3.792082555 | 0.405628113 | 1.17838466748 | 2.04981248977 | Km<KS.1 | YPD | KS.1 |
| FIM1_3405 | 6172.167387 | 8003.755084 | 10136.20419 | 0.346763358 | 0.155916282 | 0.01921602 | 0.03295405 | ns | YPD | KS.1 |
| FIM1_3407 | 939.8983948 | 1184.115536 | 1399.676544 | 0.30670919 | 0.155118793 | 0.037382068 | 0.059703302 | ns | YPD | KS.1 |
| FIM1_3408 | 1628.313704 | 1549.293788 | 1106.331544 | -0.068204719 | 0.104192917 | 0.501498927 | 0.577937725 | ns | YPD | KS.1 |
| FIM1_3409 | 5059.263835 | 7437.833248 | 9335.100371 | 0.508222216 | 0.183145894 | 0.003062762 | 0.006221822 | ns | YPD | KS.1 |
| FIM1_3410 | 1294.89975 | 1436.146289 | 1501.152017 | 0.126502581 | 0.217562228 | 0.525740363 | 0.601554348 | ns | YPD | KS.1 |
| FIM1_3411 | 892.6686286 | 1189.287587 | 1892.610748 | 0.354282246 | 0.22947578 | 0.088889699 | 0.130143099 | ns | YPD | KS.1 |
| FIM1_3412 | 1304.111767 | 1281.59493 | 1428.368822 | -0.024489385 | 0.089478682 | 0.780013949 | 0.827589976 | ns | YPD | KS.1 |
| FIM1_3413 | 1766.049777 | 2443.06009 | 2766.208354 | 0.421919488 | 0.19030431 | 0.017311474 | 0.030008565 | ns | YPD | KS.1 |
| FIM1_3414 | 2833.814912 | 1380.957057 | 2553.054635 | -0.976417969 | 0.188788386 | 4.65034171656 | 1.96730017346 | ns | YPD | KS.1 |
| FIM1_3415 | 2797.728459 | 1932.842586 | 1188.797963 | -0.408014928 | 0.320247063 | 0.133421618 | 0.185774573 | ns | YPD | KS.1 |
| FIM1_3416 | 360.6000238 | 835.9216255 | 460.4770577 | 1.098852514 | 0.257630823 | 3.11508865314 | 1.01623383930 | Km>KS.1 | YPD | KS.1 |
| FIM1_3417 | 6766.623208 | 9274.663728 | 14646.90247 | 0.435029393 | 0.121166146 | 0.000210712 | 0.000528106 | ns | YPD | KS.1 |
| FIM1_3418 | 872.0913665 | 1442.003187 | 2580.809561 | 0.698203907 | 0.127866916 | 1.71927226193 | 7.66431855120 | ns | YPD | KS.1 |
| FIM1_3419 | 6468.556059 | 3925.433217 | 3343.998621 | -0.707915068 | 0.086071055 | 6.71807390566 | 6.75200357185 | ns | YPD | KS.1 |
| FIM1_3420 | 247.0919253 | 162.0455521 | 103.0118483 | -0.46176492 | 0.33490158 | 0.100599819 | 0.145236244 | ns | YPD | KS.1 |
| FIM1_3421 | 175.6496423 | 128.1945594 | 127.9908184 | -0.350176555 | 0.31000173 | 0.186713467 | 0.249502954 | ns | YPD | KS.1 |
| FIM1_3422 | 299.6540655 | 189.627954 | 142.3187291 | -0.491074566 | 0.352934136 | 0.092889344 | 0.135321957 | ns | YPD | KS.1 |
| FIM1_3423 | 2216.174997 | 1110.905618 | 604.0889833 | -0.843323969 | 0.305402063 | 0.001550843 | 0.003329927 | ns | YPD | KS.1 |
| FIM1_3424 | 1423.279275 | 1317.822214 | 844.8276442 | -0.096092773 | 0.198945818 | 0.602544098 | 0.67316535 | ns | YPD | KS.1 |
| FIM1_3425 | 2185.366805 | 1074.239555 | 1240.169645 | -0.933034228 | 0.233126452 | 1.38439888292 | 4.11692658393 | ns | YPD | KS.1 |
| FIM1_3426 | 12205.36442 | 8884.936535 | 4545.849937 | -0.311093843 | 0.379630022 | 0.309682593 | 0.382965673 | ns | YPD | KS.1 |
| FIM1_3427 | 3110.798234 | 3149.910135 | 1391.919756 | 0.015616897 | 0.197600345 | 0.9321934 | 0.950730251 | ns | YPD | KS.1 |
| FIM1_3428 | 289.3607649 | 134.6849527 | 84.78845287 | -0.982151152 | 0.266785555 | 4.60563125715 | 0.000127436 | ns | YPD | KS.1 |

| | | | | | | | | | | |
|---|---|---|---|---|---|---|---|---|---|---|
| FIM1_3429 | 1474.775978 | 2453.051331 | 3127.203162 | 0.711753642 | 0.117445488 | 4.7354995812 | 2.5332376792 | ns | YPD | KS.1 |
| FIM1_3430 | 1801.037258 | 1347.054148 | 1770.705665 | -0.405653278 | 0.103228607 | 5.7176997736 | 0.000156294 | ns | YPD | KS.1 |
| FIM1_3431 | 976.4844884 | 327.3204839 | 368.9373469 | -1.518145809 | 0.178533377 | 1.3146633097 | 1.5245804116 | Km<KS.1 | YPD | KS.1 |
| FIM1_3432 | 601.6516143 | 830.5050699 | 922.3058643 | 0.426450145 | 0.169416702 | 0.007638891 | 0.014249525 | ns | YPD | KS.1 |
| FIM1_3433 | 3231.966483 | 4998.451283 | 3764.933484 | 0.618715721 | 0.07968426 | 3.4676349427 | 2.9339258231 | ns | YPD | KS.1 |
| FIM1_3434 | 473.9177845 | 878.6493623 | 717.5488816 | 0.875433377 | 0.092769128 | 9.1778758159 | 1.3589265530 | ns | YPD | KS.1 |
| FIM1_3435 | 1578.111801 | 4084.992185 | 3127.965965 | 1.316745811 | 0.175354738 | 6.2945028964 | 5.1760581668 | Km>KS.1 | YPD | KS.1 |
| FIM1_3437 | 5192.548067 | 16326.05492 | 16824.47536 | 1.586589582 | 0.189450916 | 3.4241207418 | 3.8109621231 | Km>KS.1 | YPD | KS.1 |
| FIM1_3438 | 602.0928596 | 2461.852939 | 2603.795388 | 1.950202969 | 0.209420016 | 3.2004065493 | 4.9447275102 | Km>KS.1 | YPD | KS.1 |
| FIM1_3439 | 3334.076686 | 3497.926985 | 1906.752553 | 0.056150368 | 0.2449964 | 0.799018604 | 0.842364389 | ns | YPD | KS.1 |
| FIM1_3440 | 3077.929565 | 3657.383504 | 3000.803043 | 0.230709508 | 0.142795287 | 0.090924561 | 0.132926739 | ns | YPD | KS.1 |
| FIM1_3441 | 2119.988762 | 973.7157878 | 987.7667858 | -1.082652569 | 0.150372959 | 9.8347672135 | 7.2809474535 | Km<KS.1 | YPD | KS.1 |
| FIM1_3442 | 4196.594298 | 6347.895541 | 5052.080606 | 0.561691233 | 0.153342462 | 0.000123958 | 0.000318704 | ns | YPD | KS.1 |
| FIM1_3444 | 1694.799116 | 3173.32888 | 4131.371059 | 0.868685397 | 0.146339995 | 7.2623715638 | 3.8071968946 | ns | YPD | KS.1 |
| FIM1_3445 | 1143.378763 | 2533.456083 | 3482.366193 | 1.082453572 | 0.193312366 | 3.5013945272 | 1.6994573437 | Km>KS.1 | YPD | KS.1 |
| FIM1_3446 | 873.6422803 | 1528.802876 | 1375.250604 | 0.796825488 | 0.072869125 | 2.1698294356 | 5.4245735890 | ns | YPD | KS.1 |
| FIM1_3447 | 1429.277963 | 2092.691855 | 2612.233532 | 0.529915182 | 0.116812719 | 3.0308768152 | 9.9071039130 | ns | YPD | KS.1 |
| FIM1_3448 | 1005.88326 | 1080.433833 | 773.1970386 | 0.098603368 | 0.090230706 | 0.264291428 | 0.334080153 | ns | YPD | KS.1 |
| FIM1_3449 | 781.9689822 | 704.7511196 | 936.3167794 | -0.142463693 | 0.112591651 | 0.191936273 | 0.255726556 | ns | YPD | KS.1 |
| FIM1_3450 | 3345.128205 | 2530.137591 | 4650.317217 | -0.324155808 | 0.27289052 | 0.177104923 | 0.238069978 | ns | YPD | KS.1 |
| FIM1_3452 | 2291.924758 | 1912.196888 | 1780.430393 | -0.246975787 | 0.121313174 | 0.035068876 | 0.056480304 | ns | YPD | KS.1 |
| FIM1_3454 | 58287.98727 | 21451.3118 | 12413.85407 | -1.338784558 | 0.241337634 | 2.8650561869 | 1.4084638863 | Km<KS.1 | YPD | KS.1 |
| FIM1_3455 | 8833.645182 | 1238.167037 | 1212.410285 | -2.694810562 | 0.2895467 | 3.4938829014 | 5.8134004798 | Km<KS.1 | YPD | KS.1 |
| FIM1_3456 | 1641.118286 | 142.1955944 | 220.5758679 | -3.357525573 | 0.34555939 | 2.6187532992 | 5.1699593904 | Km<KS.1 | YPD | KS.1 |
| FIM1_3457 | 3541.601177 | 1784.799789 | 898.4169684 | -0.825002581 | 0.316814525 | 0.002585671 | 0.005315584 | ns | YPD | KS.1 |
| FIM1_3458 | 26092.43601 | 20775.73094 | 8765.855031 | -0.287814794 | 0.202753963 | 0.124252398 | 0.175015764 | ns | YPD | KS.1 |
| FIM1_3460 | 7373.642013 | 6948.146174 | 6233.295187 | -0.07936444 | 0.135442113 | 0.541783939 | 0.615663567 | ns | YPD | KS.1 |
| FIM1_3461 | 608.0183319 | 1596.774253 | 1201.563314 | 1.170660382 | 0.358651329 | 0.000142696 | 0.000364801 | Km>KS.1 | YPD | KS.1 |
| FIM1_3462 | 733.6027937 | 983.9921135 | 799.4370233 | 0.408007388 | 0.107218047 | 9.4753090996 | 0.000248365 | ns | YPD | KS.1 |
| FIM1_3463 | 8950.814271 | 7253.78295 | 3452.321499 | -0.241255357 | 0.27158577 | 0.31350335 | 0.386921152 | ns | YPD | KS.1 |
| FIM1_3464 | 722.36491 | 269.0280004 | 332.051609 | -1.32444671 | 0.239493074 | 3.2497831302 | 1.5835133274 | Km<KS.1 | YPD | KS.1 |
| FIM1_3465 | 491.501597 | 278.0373602 | 228.7605552 | -0.677663716 | 0.306166529 | 0.010470503 | 0.019025111 | ns | YPD | KS.1 |
| FIM1_3466 | 3280.438167 | 3390.964495 | 2895.335907 | 0.046191177 | 0.075631126 | 0.535075245 | 0.610130493 | ns | YPD | KS.1 |
| FIM1_3467 | 1621.150901 | 1723.87251 | 819.1413989 | 0.060817026 | 0.349947099 | 0.833696443 | 0.871710587 | ns | YPD | KS.1 |
| FIM1_3468 | 1634.780505 | 1785.166521 | 2493.683692 | 0.115449545 | 0.154888904 | 0.433033322 | 0.510507293 | ns | YPD | KS.1 |
| FIM1_3469 | 1966.461359 | 3878.809435 | 2023.438997 | 0.840395557 | 0.291952074 | 0.001083641 | 0.002382286 | ns | YPD | KS.1 |
| FIM1_3470 | 570.1211085 | 1179.131157 | 741.3694133 | 0.935205082 | 0.26012072 | 7.1224752610 | 0.000190507 | ns | YPD | KS.1 |
| FIM1_3471 | 1512.471226 | 1560.568419 | 2447.582828 | 0.041845138 | 0.126244734 | 0.731215424 | 0.786040782 | ns | YPD | KS.1 |
| FIM1_3472 | 807.635857 | 798.832884 | 647.6004953 | -0.014795313 | 0.130363114 | 0.906206427 | 0.929179096 | ns | YPD | KS.1 |
| FIM1_3473 | 2691.266678 | 2384.58539 | 3333.603423 | -0.129985236 | 0.305038404 | 0.620120195 | 0.688945505 | ns | YPD | KS.1 |
| FIM1_3474 | 2893.502317 | 1912.929182 | 705.3237644 | -0.465131729 | 0.316209222 | 0.084390376 | 0.124213645 | ns | YPD | KS.1 |
| FIM1_3475 | 5301.682947 | 3795.106832 | 3795.78794 | -0.467393719 | 0.101950607 | 2.7357113353 | 9.0073884140 | ns | YPD | KS.1 |
| FIM1_3476 | 2838.83965 | 1876.105724 | 1593.52545 | -0.581692266 | 0.098723195 | 1.7866611604 | 9.0057135491 | ns | YPD | KS.1 |
| FIM1_3477 | 2325.471213 | 1856.346203 | 1971.95525 | -0.301312511 | 0.1499081 | 0.034849575 | 0.05616347 | ns | YPD | KS.1 |
| FIM1_3478 | 99.74037473 | 197.9015474 | 204.791761 | 0.903434277 | 0.220205862 | 9.5955160249 | 2.9215233919 | ns | YPD | KS.1 |

| | | | | | | | | | | |
|---|---|---|---|---|---|---|---|---|---|---|
| FIM1_3479 | 6230.772327 | 5793.73393 | 7511.9524 | -0.103069483 | 0.057011673 | 0.06784515 | 0.102623783 | ns | YPD | KS.1 |
| FIM1_3480 | 922.9689246 | 1177.5348 | 3057.765141 | 0.259147308 | 0.323844078 | 0.344229036 | 0.419328956 | ns | YPD | KS.1 |
| FIM1_3481 | 2725.902472 | 3956.412611 | 3386.072532 | 0.525704785 | 0.088775246 | 1.68924342819 | 8.54063623503 | ns | YPD | KS.1 |
| FIM1_3482 | 1469.650152 | 1676.56601 | 3183.553039 | 0.168205281 | 0.181461741 | 0.322178226 | 0.395957676 | ns | YPD | KS.1 |
| FIM1_3483 | 551.70755 | 807.3864864 | 933.8487455 | 0.533116981 | 0.102589238 | 1.06413620580 | 4.27631472043 | ns | YPD | KS.1 |
| FIM1_3484 | 1456.197079 | 1602.15932 | 1506.444146 | 0.13369501 | 0.083513696 | 0.102746344 | 0.147991622 | ns | YPD | KS.1 |
| FIM1_3485 | 474.5950857 | 271.1450466 | 508.4485735 | -0.764022621 | 0.166549459 | 1.41917074144 | 4.88945598248 | ns | YPD | KS.1 |
| FIM1_3486 | 2261.861939 | 2774.832523 | 3483.811659 | 0.272118803 | 0.150674025 | 0.057799498 | 0.089302019 | ns | YPD | KS.1 |
| FIM1_3487 | 519.1998288 | 599.8552819 | 881.4953144 | 0.183338606 | 0.186742218 | 0.292811302 | 0.364822497 | ns | YPD | KS.1 |
| FIM1_3488 | 407.9424379 | 538.9553553 | 422.2680436 | 0.38459077 | 0.110460606 | 0.000346288 | 0.000832666 | ns | YPD | KS.1 |
| FIM1_3489 | 725.9291702 | 661.4068922 | 767.3091327 | -0.129339437 | 0.101893245 | 0.192562525 | 0.256237169 | ns | YPD | KS.1 |
| FIM1_3490 | 1102.306582 | 970.0301308 | 1105.079606 | -0.176457547 | 0.107966539 | 0.092494424 | 0.134865111 | ns | YPD | KS.1 |
| FIM1_3491 | 2879.631929 | 2092.579839 | 2077.042796 | -0.437065842 | 0.130921666 | 0.000535362 | 0.001247505 | ns | YPD | KS.1 |
| FIM1_3492 | 524.291503 | 859.3811856 | 782.9234932 | 0.656597301 | 0.191343945 | 0.000243603 | 0.000602948 | ns | YPD | KS.1 |
| FIM1_3493 | 109.8848188 | 199.2199309 | 111.3256679 | 0.731126722 | 0.281921756 | 0.003291951 | 0.006643999 | ns | YPD | KS.1 |
| FIM1_3494 | 7601.525112 | 9530.867488 | 9553.046138 | 0.313675298 | 0.106592199 | 0.002507752 | 0.005174644 | ns | YPD | KS.1 |
| FIM1_3495 | 838.1133044 | 450.5319273 | 257.5342315 | -0.850360968 | 0.163411754 | 5.08462003801 | 2.14010022750 | ns | YPD | KS.1 |
| FIM1_3496 | 664.9069194 | 491.2103485 | 619.1826244 | -0.397626499 | 0.178127548 | 0.017353114 | 0.030070269 | ns | YPD | KS.1 |
| FIM1_3497 | 4527.212124 | 2880.875035 | 1782.456726 | -0.506623082 | 0.324809796 | 0.06565697 | 0.099830282 | ns | YPD | KS.1 |
| FIM1_3498 | 64.40574446 | 110.077283 | 150.7250461 | 0.678355302 | 0.249305368 | 0.002530376 | 0.005214838 | ns | YPD | KS.1 |
| FIM1_3499 | 2672.322964 | 3159.807664 | 2687.064173 | 0.234749822 | 0.084522546 | 0.004653106 | 0.009121042 | ns | YPD | KS.1 |
| FIM1_3500 | 1486.027486 | 1716.445195 | 1777.103672 | 0.18235276 | 0.190189554 | 0.303275545 | 0.375826944 | ns | YPD | KS.1 |
| FIM1_3501 | 6353.322978 | 4069.199091 | 2906.741389 | -0.550755782 | 0.252629222 | 0.0149027 | 0.026235291 | ns | YPD | KS.1 |
| FIM1_3503 | 523.6104814 | 753.2036394 | 375.0021449 | 0.350029968 | 0.395450816 | 0.265963477 | 0.335744302 | ns | YPD | KS.1 |
| FIM1_3504 | 901.7650232 | 312.4294834 | 551.5915906 | -1.347404974 | 0.321198419 | 2.52030610609 | 8.33124443710 | Km<KS.1 | YPD | KS.1 |
| FIM1_3505 | 1787.844536 | 2780.775566 | 3680.734522 | 0.620034193 | 0.103912213 | 1.03841075562 | 5.38134740546 | ns | YPD | KS.1 |
| FIM1_3506 | 610.2083057 | 731.1198102 | 702.3294603 | 0.231484453 | 0.181277045 | 0.172393957 | 0.232743538 | ns | YPD | KS.1 |
| FIM1_3507 | 1473.051263 | 1277.552016 | 1893.918144 | -0.196771697 | 0.10084294 | 0.0452399 | 0.071134166 | ns | YPD | KS.1 |
| FIM1_3508 | 449.1150619 | 769.071512 | 926.6437892 | 0.750210418 | 0.127764897 | 1.38241246440 | 7.06109035974 | ns | YPD | KS.1 |
| FIM1_3509 | 514.4030588 | 34.46312932 | 75.09163503 | -3.816667663 | 0.236924311 | 4.61466754403 | 8.69304549363 | Km<KS.1 | YPD | KS.1 |
| FIM1_3511 | 2394.914781 | 3904.213173 | 4938.838614 | 0.674341511 | 0.137284593 | 3.46517458508 | 1.30897825062 | ns | YPD | KS.1 |
| FIM1_3512 | 6415.720508 | 5928.165814 | 4645.559879 | -0.100275604 | 0.184790914 | 0.561820477 | 0.634663232 | ns | YPD | KS.1 |
| FIM1_3513 | 2680.354112 | 829.8550808 | 986.2109913 | -1.37105695 | 0.426954795 | 0.000107301 | 0.00027905 | Km<KS.1 | YPD | KS.1 |
| FIM1_3514 | 4104.651496 | 4142.69326 | 2916.016212 | 0.011046463 | 0.228256159 | 0.957621609 | 0.968720517 | ns | YPD | KS.1 |
| FIM1_3516 | 6432.023223 | 8409.863555 | 7410.048664 | 0.301953822 | 0.291543787 | 0.232722872 | 0.300570168 | ns | YPD | KS.1 |
| FIM1_3517 | 729.5237037 | 489.4105542 | 512.655349 | -0.556766015 | 0.115670275 | 7.41241261688 | 2.67222846152 | ns | YPD | KS.1 |
| FIM1_3518 | 3130.604455 | 2651.625398 | 3047.35086 | -0.172459168 | 0.329106672 | 0.533994063 | 0.609037245 | ns | YPD | KS.1 |
| FIM1_3519 | 564.2039159 | 796.1890494 | 757.2062454 | 0.42121083 | 0.246032444 | 0.056461953 | 0.087398324 | ns | YPD | KS.1 |
| FIM1_3520 | 1266.497641 | 2148.594045 | 1141.632281 | 0.664235366 | 0.252208367 | 0.003387326 | 0.006822651 | ns | YPD | KS.1 |
| FIM1_3522 | 2914.425552 | 2367.385377 | 3616.222836 | -0.218450147 | 0.326452306 | 0.428039758 | 0.505578774 | ns | YPD | KS.1 |
| FIM1_3523 | 7708.481934 | 9043.961976 | 7519.610076 | 0.205367762 | 0.180002681 | 0.223221018 | 0.29002992 | ns | YPD | KS.1 |
| FIM1_3524 | 1392.320518 | 1381.660243 | 1888.769054 | -0.009889491 | 0.16501396 | 0.949514403 | 0.964047787 | ns | YPD | KS.1 |
| FIM1_3525 | 170.1865218 | 567.1651989 | 478.5321531 | 1.612475808 | 0.259137334 | 2.78132591040 | 1.66712004870 | Km>KS.1 | YPD | KS.1 |
| FIM1_3526 | 3355.294256 | 2535.324368 | 3162.86158 | -0.39296869 | 0.093904723 | 1.96074378010 | 5.71452859180 | ns | YPD | KS.1 |
| FIM1_3527 | 1943.203733 | 1748.945663 | 1778.7921 | -0.127197079 | 0.225245123 | 0.535677785 | 0.610397842 | ns | YPD | KS.1 |

| | | | | | | | | | | |
|---|---|---|---|---|---|---|---|---|---|---|
| FIM1_3528 | 12425.89594 | 4814.936384 | 3188.141061 | -1.160189846 | 0.346950884 | 0.00011027 | 0.000286172 | Km<KS.1 | YPD | KS.1 |
| FIM1_3529 | 139.7152015 | 500.7956876 | 616.9667557 | 1.757376596 | 0.21113583 | 3.5873917726 | 3.9748940019 | Km>KS.1 | YPD | KS.1 |
| FIM1_3530 | 190.8086558 | 278.1519727 | 406.0561677 | 0.494448214 | 0.189870581 | 0.005247328 | 0.010130174 | ns | YPD | KS.1 |
| FIM1_3531 | 872.5411376 | 1408.263164 | 1385.990734 | 0.677759931 | 0.085556345 | 8.6370118844 | 7.7842634284 | ns | YPD | KS.1 |
| FIM1_3532 | 542.1767471 | 712.9991503 | 977.3404401 | 0.368938858 | 0.144016399 | 0.007400363 | 0.013856532 | ns | YPD | KS.1 |
| FIM1_3533 | 2489.984253 | 637.4728982 | 652.0218671 | -1.923188638 | 0.148947002 | 1.0273710697 | 5.1627990625 | Km<KS.1 | YPD | KS.1 |
| FIM1_3534 | 4005.51372 | 2722.910499 | 2703.360744 | -0.545418826 | 0.08395803 | 4.1630811896 | 2.4568598954 | ns | YPD | KS.1 |
| FIM1_3535 | 611.6649438 | 1305.03484 | 1527.632879 | 1.079737548 | 0.078186836 | 3.5592257213 | 2.2996296056 | Km>KS.1 | YPD | KS.1 |
| FIM1_3536 | 235.8297203 | 334.815763 | 460.5971985 | 0.478290574 | 0.13699803 | 0.000283031 | 0.000693293 | ns | YPD | KS.1 |
| FIM1_3537 | 3485.067106 | 4385.343327 | 8084.007475 | 0.267935535 | 0.263376335 | 0.251137617 | 0.320361448 | ns | YPD | KS.1 |
| FIM1_3538 | 505.9476804 | 925.3428985 | 903.5957502 | 0.844593045 | 0.121435355 | 9.1252922579 | 6.3031883849 | ns | YPD | KS.1 |
| FIM1_3539 | 537.4355275 | 408.310892 | 579.1258978 | -0.372981358 | 0.138912275 | 0.005122747 | 0.009939808 | ns | YPD | KS.1 |
| FIM1_3540 | 4366.491581 | 2849.307342 | 3361.474688 | -0.601297702 | 0.095127694 | 1.1609382509 | 6.5335608580 | ns | YPD | KS.1 |
| FIM1_3541 | 367.8049032 | 134.2867542 | 306.9328249 | -1.347503902 | 0.242696632 | 2.7386898672 | 1.3490081276 | Km<KS.1 | YPD | KS.1 |
| FIM1_3542 | 581.5463942 | 483.1469664 | 450.4154753 | -0.253046534 | 0.112655235 | 0.020570594 | 0.034918579 | ns | YPD | KS.1 |
| FIM1_3544 | 1083.393122 | 2491.479737 | 2705.31606 | 1.146905291 | 0.175437218 | 9.0498584657 | 5.7063429489 | Km>KS.1 | YPD | KS.1 |
| FIM1_3545 | 576.7546446 | 257.2890087 | 154.6222435 | -1.015475019 | 0.296061171 | 0.000111269 | 0.000288314 | Km<KS.1 | YPD | KS.1 |
| FIM1_3546 | 1215.4022 | 1815.564973 | 1688.710571 | 0.540870029 | 0.161070888 | 0.000407991 | 0.00096701 | ns | YPD | KS.1 |
| FIM1_3547 | 22.94384286 | 422.7587211 | 926.7482602 | 4.133211694 | 0.189662371 | 1.1218097110 | 1.8603344374 | Km>KS.1 | YPD | KS.1 |
| FIM1_3548 | 154.1940595 | 731.6832195 | 437.2404573 | 2.046945918 | 0.330836689 | 1.1709450006 | 7.2546094376 | Km>KS.1 | YPD | KS.1 |
| FIM1_3549 | 318.9625779 | 165.4598147 | 171.1134703 | -0.897983623 | 0.171288711 | 3.7320997878 | 1.6033848397 | ns | YPD | KS.1 |
| FIM1_3550 | 869.4490846 | 839.2635281 | 1459.170544 | -0.043860259 | 0.212535318 | 0.822532697 | 0.862039218 | ns | YPD | KS.1 |
| FIM1_3551 | 413.384729 | 1999.101203 | 1689.147242 | 2.193877917 | 0.2062707 | 2.4947544868 | 6.1139919074 | Km>KS.1 | YPD | KS.1 |
| FIM1_3552 | 379.0327437 | 452.7679962 | 580.9714058 | 0.236714535 | 0.148337718 | 0.094030018 | 0.136743449 | ns | YPD | KS.1 |
| FIM1_3553 | 362.836827 | 579.6629075 | 438.2540512 | 0.589668847 | 0.241223973 | 0.00682228 | 0.012856381 | ns | YPD | KS.1 |
| FIM1_3554 | 226.3107176 | 374.2415133 | 602.7025135 | 0.679971076 | 0.162957497 | 1.1523598209 | 3.4682335807 | ns | YPD | KS.1 |
| FIM1_3555 | 491.8295043 | 560.8009773 | 575.0273797 | 0.173406834 | 0.149042777 | 0.222095673 | 0.288869535 | ns | YPD | KS.1 |
| FIM1_3556 | 2837.343313 | 2526.879979 | 2232.991527 | -0.159417727 | 0.105701144 | 0.12091515 | 0.170992857 | ns | YPD | KS.1 |
| FIM1_3557 | 2796.249187 | 2469.972793 | 4774.948837 | -0.152781007 | 0.211675284 | 0.43248564 | 0.510103381 | ns | YPD | KS.1 |
| FIM1_3558 | 1703.021076 | 1243.507576 | 1198.38517 | -0.425504815 | 0.146593368 | 0.002396206 | 0.004960935 | ns | YPD | KS.1 |
| FIM1_3559 | 377.1789121 | 228.1922651 | 243.8344568 | -0.684637957 | 0.160325013 | 7.3873550046 | 2.2855778077 | ns | YPD | KS.1 |
| FIM1_3560 | 2016.352148 | 1367.025413 | 1344.499968 | -0.54209198 | 0.110974342 | 5.3342701616 | 1.9556369973 | ns | YPD | KS.1 |
| FIM1_3561 | 2045.392718 | 1582.280256 | 872.0920139 | -0.321498514 | 0.213431995 | 0.100506297 | 0.145143346 | ns | YPD | KS.1 |
| FIM1_3562 | 2531.085332 | 2111.045596 | 1735.131374 | -0.250368442 | 0.108800668 | 0.017866365 | 0.030873624 | ns | YPD | KS.1 |
| FIM1_3563 | 1665.828528 | 1603.01559 | 1925.341098 | -0.051395661 | 0.11467753 | 0.643991174 | 0.709445548 | ns | YPD | KS.1 |
| FIM1_3565 | 143.6043354 | 278.7705799 | 110.8741976 | 0.778590753 | 0.331793172 | 0.005897553 | 0.011267406 | ns | YPD | KS.1 |
| FIM1_3566 | 335.7819691 | 239.0891592 | 242.8917698 | -0.430894487 | 0.216785652 | 0.029975677 | 0.048991366 | ns | YPD | KS.1 |
| FIM1_3567 | 2502.363237 | 6427.868636 | 7172.635201 | 1.308264491 | 0.17109777 | 2.2225953018 | 1.9163625002 | Km>KS.1 | YPD | KS.1 |
| FIM1_3568 | 2521.83757 | 6564.357065 | 4147.484486 | 1.28820151 | 0.227863804 | 1.7212428366 | 8.6935869159 | Km>KS.1 | YPD | KS.1 |
| FIM1_3569 | 1522.932129 | 1074.664552 | 2003.273747 | -0.351944667 | 0.368264926 | 0.241076551 | 0.30935152 | ns | YPD | KS.1 |
| FIM1_3570 | 707.9452272 | 837.7619046 | 1301.032634 | 0.20720437 | 0.21743228 | 0.297901399 | 0.370214275 | ns | YPD | KS.1 |
| FIM1_3571 | 2199.701225 | 3636.215162 | 4121.434755 | 0.70732552 | 0.103132103 | 2.4282144636 | 1.6085708331 | ns | YPD | KS.1 |
| FIM1_3572 | 468.1298664 | 461.5158614 | 519.58149 | -0.020808691 | 0.126209676 | 0.864347747 | 0.895300863 | ns | YPD | KS.1 |
| FIM1_3573 | 1914.313688 | 3658.191195 | 4375.909811 | 0.915959622 | 0.103235595 | 1.5823951360 | 2.0082693371 | ns | YPD | KS.1 |
| FIM1_3574 | 8706.225016 | 7748.389748 | 3146.18648 | -0.119220027 | 0.334205147 | 0.670740977 | 0.73290937 | ns | YPD | KS.1 |

| | | | | | | | | | | |
|---|---|---|---|---|---|---|---|---|---|---|
| FIM1_3575 | 2450.608125 | 2288.802736 | 1718.833637 | -0.082394125 | 0.224902825 | 0.688036192 | 0.749616258 | ns | YPD | KS.1 |
| FIM1_3576 | 1732.228617 | 3309.974141 | 3522.498511 | 0.905684956 | 0.127374621 | 2.6444811557 | 1.9122519984 | ns | YPD | KS.1 |
| FIM1_3577 | 368.14621 | 403.9442473 | 487.2915457 | 0.123003724 | 0.140415365 | 0.360288524 | 0.435268433 | ns | YPD | KS.1 |
| FIM1_3579 | 2838.047185 | 3211.242204 | 2293.963276 | 0.15740095 | 0.183884049 | 0.360008401 | 0.435035655 | ns | YPD | KS.1 |
| FIM1_3580 | 3567.938351 | 2253.762247 | 5248.873732 | -0.535434949 | 0.299887908 | 0.038889311 | 0.061872184 | ns | YPD | KS.1 |
| FIM1_3581 | 876.1906947 | 2818.300267 | 4370.134313 | 1.582729724 | 0.237347119 | 1.5845445012 | 1.0754582392 | Km>KS.1 | YPD | KS.1 |
| FIM1_3582 | 3794.162774 | 2740.30653 | 2298.735857 | -0.457177786 | 0.094392305 | 7.8234039188 | 2.8102118769 | ns | YPD | KS.1 |
| FIM1_3583 | 1044.17276 | 3401.701884 | 1973.623488 | 1.579115954 | 0.262881004 | 1.1488659812 | 6.4729425328 | Km>KS.1 | YPD | KS.1 |
| FIM1_3584 | 70.24989529 | 468.0187992 | 589.4158153 | 2.670482773 | 0.190310672 | 2.3899501538 | 1.9491806583 | Km>KS.1 | YPD | KS.1 |
| FIM1_3585 | 420.6925208 | 1272.797181 | 1187.196258 | 1.520543374 | 0.20418442 | 6.7507684702 | 5.5329609785 | Km>KS.1 | YPD | KS.1 |
| FIM1_3586 | 2221.428641 | 4491.845319 | 2909.673499 | 0.949307975 | 0.197689968 | 3.3387556729 | 1.2660296854 | ns | YPD | KS.1 |
| FIM1_3587 | 87.14629084 | 178.6173008 | 85.38572007 | 0.874264812 | 0.312184542 | 0.001282236 | 0.002781999 | ns | YPD | KS.1 |
| FIM1_3588 | 1299.58316 | 1138.135003 | 1024.528335 | -0.152656095 | 0.260803185 | 0.510364181 | 0.585847208 | ns | YPD | KS.1 |
| FIM1_3589 | 10306.3597 | 6138.588115 | 3873.387259 | -0.680249239 | 0.206733708 | 0.000385861 | 0.000918497 | ns | YPD | KS.1 |
| FIM1_3591 | 787.7785723 | 683.470083 | 629.3839924 | -0.168345992 | 0.243407097 | 0.442259674 | 0.519414985 | ns | YPD | KS.1 |
| FIM1_3592 | 316.9562735 | 173.32326 | 1972.23227 | -0.70097277 | 0.329080901 | 0.012267908 | 0.022027243 | ns | YPD | KS.1 |
| FIM1_3594 | 3261.700983 | 3603.325246 | 3172.010947 | 0.135723708 | 0.115401201 | 0.224640558 | 0.291569731 | ns | YPD | KS.1 |
| FIM1_3595 | 6836.928831 | 5193.332312 | 5378.255139 | -0.366268353 | 0.159132789 | 0.015232673 | 0.026778285 | ns | YPD | KS.1 |
| FIM1_3596 | 48.75471678 | 55.84026335 | 37.01878554 | 0.13752885 | 0.334871773 | 0.624310114 | 0.692901298 | ns | YPD | KS.1 |
| FIM1_3597 | 7070.399297 | 6556.494562 | 10687.94615 | -0.103174042 | 0.111300715 | 0.339557701 | 0.414450334 | ns | YPD | KS.1 |
| FIM1_3598 | 2596.735311 | 1836.220378 | 2566.044627 | -0.436259596 | 0.221727692 | 0.031128094 | 0.050608584 | ns | YPD | KS.1 |
| FIM1_3599 | 8015.808775 | 6398.743818 | 6212.493068 | -0.310150115 | 0.115703738 | 0.005692935 | 0.010918408 | ns | YPD | KS.1 |
| FIM1_3600 | 68.75030102 | 288.6025188 | 386.4645009 | 1.981930296 | 0.208033547 | 3.7939836040 | 6.2707868539 | Km>KS.1 | YPD | KS.1 |
| FIM1_3601 | 2308.951284 | 2995.091624 | 2242.782252 | 0.352874549 | 0.138114291 | 0.00773648 | 0.014415352 | ns | YPD | KS.1 |
| FIM1_3603 | 651.8330833 | 390.1774191 | 629.563039 | -0.712776 | 0.126142686 | 5.5887667519 | 2.6581371501 | ns | YPD | KS.1 |
| FIM1_3604 | 2394.682786 | 2398.526135 | 3903.096472 | 0.002116005 | 0.16781276 | 0.989355049 | 0.992147021 | ns | YPD | KS.1 |
| FIM1_3605 | 2072.544688 | 1362.956002 | 1340.176429 | -0.597205096 | 0.06682404 | 1.7628205705 | 2.2090761557 | ns | YPD | KS.1 |
| FIM1_3606 | 530.008864 | 564.2908842 | 573.6389977 | 0.078668819 | 0.201166752 | 0.672944534 | 0.735155699 | ns | YPD | KS.1 |
| FIM1_3607 | 926.9045402 | 1083.932824 | 1506.810348 | 0.211134153 | 0.132919178 | 0.098145624 | 0.142022827 | ns | YPD | KS.1 |
| FIM1_3608 | 852.8141801 | 635.3697989 | 542.6673427 | -0.399313758 | 0.140281104 | 0.002992578 | 0.006090732 | ns | YPD | KS.1 |
| FIM1_3609 | 3809.760264 | 1616.65375 | 1486.044669 | -1.113828095 | 0.267109674 | 4.6005455150 | 1.4681022410 | Km<KS.1 | YPD | KS.1 |
| FIM1_3610 | 768.8400864 | 1313.885468 | 1668.484706 | 0.748034442 | 0.120583467 | 1.7728280397 | 9.8142397112 | ns | YPD | KS.1 |
| FIM1_3611 | 897.7570898 | 961.8462321 | 868.238066 | 0.096614107 | 0.093005308 | 0.288118997 | 0.359877481 | ns | YPD | KS.1 |
| FIM1_3612 | 1553.84236 | 1749.651248 | 1620.530005 | 0.164124425 | 0.098225362 | 0.086718151 | 0.127300915 | ns | YPD | KS.1 |
| FIM1_3613 | 1490.533758 | 1046.015457 | 1052.78488 | -0.45369435 | 0.207656592 | 0.017759692 | 0.030699954 | ns | YPD | KS.1 |
| FIM1_3614 | 147.0179785 | 102.9001274 | 312.8522437 | -0.454035303 | 0.214835855 | 0.021223711 | 0.035849875 | ns | YPD | KS.1 |
| FIM1_3615 | 491.1066312 | 878.9390075 | 1455.642772 | 0.780217034 | 0.190745768 | 1.3221637950 | 3.9458697543 | ns | YPD | KS.1 |
| FIM1_3616 | 1754.269 | 2381.052958 | 3554.016383 | 0.370827764 | 0.245079027 | 0.091935808 | 0.134129221 | ns | YPD | KS.1 |
| FIM1_3617 | 60.43832863 | 129.1860778 | 224.0788788 | 0.998656362 | 0.240067704 | 6.1138981767 | 1.9178211493 | ns | YPD | KS.1 |
| FIM1_3618 | 2319.364533 | 2894.438325 | 2714.576116 | 0.308593972 | 0.099876878 | 0.001555018 | 0.003337453 | ns | YPD | KS.1 |
| FIM1_3619 | 308.2768887 | 242.1696037 | 359.8464444 | -0.310911261 | 0.189545907 | 0.078108897 | 0.115962926 | ns | YPD | KS.1 |
| FIM1_3620 | 2003.328119 | 1951.440971 | 1503.26998 | -0.034909424 | 0.13653196 | 0.790048067 | 0.835208061 | ns | YPD | KS.1 |
| FIM1_3621 | 1224.383959 | 1285.15434 | 1545.844025 | 0.064075664 | 0.139566537 | 0.632014751 | 0.699038102 | ns | YPD | KS.1 |
| FIM1_3622 | 2302.660103 | 2640.837343 | 2433.684599 | 0.191544873 | 0.085482824 | 0.022291286 | 0.03749126 | ns | YPD | KS.1 |
| FIM1_3623 | 3262.984979 | 606.2648817 | 572.7992675 | -2.143552395 | 0.402079088 | 1.3681616142 | 6.9954820460 | Km<KS.1 | YPD | KS.1 |

| | | | | | | | | | | |
|---|---|---|---|---|---|---|---|---|---|---|
| FIM1_3624 | 1226.910738 | 679.4778869 | 462.092434 | -0.677794217 | 0.335823326 | 0.016940271 | 0.029416353 | ns | YPD | KS.1 |
| FIM1_3625 | 12406.94013 | 18030.15591 | 18717.45703 | 0.532668619 | 0.064402557 | 6.8317657696 | 6.8524263516 | ns | YPD | KS.1 |
| FIM1_3626 | 103.7702144 | 181.9095927 | 201.9656237 | 0.776887759 | 0.149164393 | 5.8128335385 | 2.4240441621 | ns | YPD | KS.1 |
| FIM1_3627 | 6508.346507 | 1127.531166 | 1302.001326 | -2.448263203 | 0.212483695 | 5.3722663018 | 1.8058800575 | Km<KS.1 | YPD | KS.1 |
| FIM1_3628 | 12601.57908 | 12066.07359 | 11043.21588 | -0.046635797 | 0.3017668 | 0.857766647 | 0.889780873 | ns | YPD | KS.1 |
| FIM1_3629 | 5752.765655 | 6281.305749 | 4731.292686 | 0.109524745 | 0.199490203 | 0.553709354 | 0.626781351 | ns | YPD | KS.1 |
| FIM1_3630 | 347.4210957 | 361.141515 | 197.6624048 | 0.047132241 | 0.206937196 | 0.805167015 | 0.847051364 | ns | YPD | KS.1 |
| FIM1_3631 | 583.6703517 | 442.6026196 | 557.9884718 | -0.380594071 | 0.125263523 | 0.00166163 | 0.003544857 | ns | YPD | KS.1 |
| FIM1_3632 | 6030.305884 | 7895.143072 | 8567.738859 | 0.374640786 | 0.106156033 | 0.000294405 | 0.000717974 | ns | YPD | KS.1 |
| FIM1_3633 | 6635.600843 | 3155.339597 | 2506.69561 | -0.914362493 | 0.308185717 | 0.000691175 | 0.001579511 | ns | YPD | KS.1 |
| FIM1_3634 | 5055.758477 | 2737.414911 | 3704.855705 | -0.769469855 | 0.268057945 | 0.001302195 | 0.002821612 | ns | YPD | KS.1 |
| FIM1_3635 | 2875.666423 | 2742.093597 | 4861.765801 | -0.055327825 | 0.248258004 | 0.804008838 | 0.846190812 | ns | YPD | KS.1 |
| FIM1_3636 | 4405.85187 | 2414.088469 | 2761.731935 | -0.733487501 | 0.291105466 | 0.004048034 | 0.008023493 | ns | YPD | KS.1 |
| FIM1_3637 | 3559.585922 | 1764.212736 | 3180.97478 | -0.870705828 | 0.293263181 | 0.000757118 | 0.001714458 | ns | YPD | KS.1 |
| FIM1_3638 | 4101.019963 | 3960.77148 | 7351.407994 | -0.039230473 | 0.269590982 | 0.869282292 | 0.899662868 | ns | YPD | KS.1 |
| FIM1_3639 | 653.3598196 | 220.5507925 | 401.9788217 | -1.354831211 | 0.346712353 | 8.3244205479 | 2.5501226740 | Km<KS.1 | YPD | KS.1 |
| FIM1_3640 | 1677.328395 | 616.7373235 | 1134.694664 | -1.276740089 | 0.308579992 | 3.7619353818 | 1.2160902225 | Km<KS.1 | YPD | KS.1 |
| FIM1_3641 | 1474.304632 | 610.3795995 | 956.6364595 | -1.138321111 | 0.278277205 | 6.1627857489 | 1.9319381916 | Km<KS.1 | YPD | KS.1 |
| FIM1_3642 | 4118.899025 | 1820.306678 | 2394.135994 | -1.046114834 | 0.278160401 | 2.9326647727 | 8.3514637919 | Km<KS.1 | YPD | KS.1 |
| FIM1_3643 | 1574.811134 | 831.7128039 | 1267.188752 | -0.766071019 | 0.31075947 | 0.004388238 | 0.008649558 | ns | YPD | KS.1 |
| FIM1_3644 | 3944.482491 | 1854.206401 | 2834.249379 | -0.966967369 | 0.26911445 | 6.7020646422 | 0.00018072 | ns | YPD | KS.1 |
| FIM1_3645 | 4239.544841 | 2204.395796 | 3118.831835 | -0.817218896 | 0.278306236 | 0.000948449 | 0.002109313 | ns | YPD | KS.1 |
| FIM1_3646 | 1618.41417 | 771.7714732 | 1263.162388 | -0.929934894 | 0.287776902 | 0.000273701 | 0.000671763 | ns | YPD | KS.1 |
| FIM1_3647 | 13233.95527 | 12671.71661 | 25119.05881 | -0.053115071 | 0.211816474 | 0.785293358 | 0.830745857 | ns | YPD | KS.1 |
| FIM1_3648 | 3245.525616 | 882.9926365 | 2447.029702 | -1.508124182 | 0.457149257 | 5.7019115421 | 0.000155948 | Km<KS.1 | YPD | KS.1 |
| FIM1_3649 | 2911.912474 | 772.4164332 | 1619.634937 | -1.750284182 | 0.300866187 | 2.4367251144 | 1.3307033418 | Km<KS.1 | YPD | KS.1 |
| FIM1_3650 | 3985.745375 | 1076.414484 | 1797.310892 | -1.728649216 | 0.296946314 | 2.5067309216 | 1.3644405180 | Km<KS.1 | YPD | KS.1 |
| FIM1_3651 | 2298.937391 | 812.7551437 | 1422.069128 | -1.341637963 | 0.299667477 | 7.1716996220 | 2.5946730423 | Km<KS.1 | YPD | KS.1 |
| FIM1_3652 | 1057.434571 | 199.7724442 | 400.6113962 | -2.096333695 | 0.418067524 | 8.3448772077 | 3.8763551922 | Km<KS.1 | YPD | KS.1 |
| FIM1_3653 | 9107.579025 | 3708.585151 | 5782.597739 | -1.150057054 | 0.290967921 | 1.0770798485 | 3.2594113421 | Km<KS.1 | YPD | KS.1 |
| FIM1_3654 | 5442.692044 | 4836.67161 | 7828.351318 | -0.142304229 | 0.227395724 | 0.491777509 | 0.569239904 | ns | YPD | KS.1 |
| FIM1_3655 | 4939.77923 | 2916.156935 | 4492.996544 | -0.657867881 | 0.257865946 | 0.004366204 | 0.008612952 | ns | YPD | KS.1 |
| FIM1_3656 | 4315.289896 | 3419.562094 | 5621.544174 | -0.275321997 | 0.253073089 | 0.223118961 | 0.289973049 | ns | YPD | KS.1 |
| FIM1_3657 | 1496.23833 | 681.7947354 | 773.3438998 | -0.966307636 | 0.315700045 | 0.00045168 | 0.001060958 | ns | YPD | KS.1 |
| FIM1_3658 | 2520.548854 | 753.55278 | 2222.721408 | -1.505886763 | 0.364521642 | 2.3343878197 | 7.7579020731 | Km<KS.1 | YPD | KS.1 |
| FIM1_3659 | 3839.297096 | 1618.887491 | 1990.138433 | -1.091485253 | 0.300194853 | 4.3411929434 | 0.000120926 | Km<KS.1 | YPD | KS.1 |
| FIM1_3660 | 2151.067435 | 786.0866104 | 1551.995642 | -1.282419176 | 0.311503209 | 4.0610758145 | 1.3009563539 | Km<KS.1 | YPD | KS.1 |
| FIM1_3661 | 4623.502222 | 1287.363207 | 2798.114036 | -1.675490964 | 0.30585544 | 2.0613672145 | 1.0265567459 | Km<KS.1 | YPD | KS.1 |
| FIM1_3662 | 2433.927551 | 1052.221449 | 1358.714328 | -1.058893964 | 0.297414063 | 6.2212151427 | 0.000168852 | Km<KS.1 | YPD | KS.1 |
| FIM1_3663 | 2041.954473 | 1537.309686 | 4056.103488 | -0.296747991 | 0.338946516 | 0.29470888 | 0.36691108 | ns | YPD | KS.1 |
| FIM1_3664 | 2637.116234 | 751.0678229 | 1737.776234 | -1.631358265 | 0.31665603 | 1.3457985525 | 6.0701249311 | Km<KS.1 | YPD | KS.1 |
| FIM1_3665 | 1800.813419 | 661.1791927 | 1234.791245 | -1.236290621 | 0.346825424 | 4.1487661942 | 0.000115891 | Km<KS.1 | YPD | KS.1 |
| FIM1_3666 | 1093.144421 | 969.4796228 | 1461.323656 | -0.132468006 | 0.287442078 | 0.597038407 | 0.668376705 | ns | YPD | KS.1 |
| FIM1_3667 | 2214.50911 | 1266.782094 | 2133.783649 | -0.688021189 | 0.274718454 | 0.004671936 | 0.009150741 | ns | YPD | KS.1 |
| FIM1_3668 | 871.1099648 | 919.3288334 | 1477.657812 | 0.061319208 | 0.266989649 | 0.795435104 | 0.83929791 | ns | YPD | KS.1 |

| | | | | | | | | | | |
|---|---|---|---|---|---|---|---|---|---|---|
| FIM1_3669 | 2294.149464 | 961.1653179 | 1245.2853 | -1.070143666 | 0.329215352 | 0.00018746 | 0.000472688 | Km<KS.1 | YPD | KS.1 |
| FIM1_3670 | 1009.776023 | 493.4855971 | 785.9254473 | -0.871992231 | 0.312308172 | 0.001322663 | 0.002862222 | ns | YPD | KS.1 |
| FIM1_3671 | 1182.524948 | 354.0360032 | 649.025862 | -1.502750772 | 0.365145246 | 2.51321620652 | 8.31333153421 | Km<KS.1 | YPD | KS.1 |
| FIM1_3672 | 6276.840077 | 1631.246173 | 2288.424868 | -1.785548414 | 0.295315442 | 5.60137340392 | 3.24410159307 | Km<KS.1 | YPD | KS.1 |
| FIM1_3673 | 5271.612296 | 1182.36581 | 2448.216564 | -1.920529225 | 0.362523771 | 3.02955527975 | 1.48421560180 | Km<KS.1 | YPD | KS.1 |
| FIM1_3674 | 163.0529396 | 43.74946248 | 71.71387351 | -1.674083365 | 0.351635728 | 8.87310938110 | 3.61833763696 | Km<KS.1 | YPD | KS.1 |
| FIM1_3675 | 4229.021975 | 2707.51418 | 4505.666315 | -0.539063139 | 0.270575027 | 0.024158469 | 0.040385881 | ns | YPD | KS.1 |
| FIM1_3676 | 621.9987249 | 119.0214875 | 187.0670644 | -2.202352942 | 0.319622596 | 6.88027354444 | 5.16279952995 | Km<KS.1 | YPD | KS.1 |
| FIM1_3677 | 1580.063384 | 1127.119734 | 1500.927836 | -0.397096543 | 0.272774203 | 0.098175698 | 0.142025035 | ns | YPD | KS.1 |
| FIM1_3678 | 2655.054 | 1018.915051 | 2981.63063 | -1.041515587 | 0.444840913 | 0.003209765 | 0.006491293 | Km<KS.1 | YPD | KS.1 |
| FIM1_3679 | 53672.74546 | 42499.28293 | 61159.05026 | -0.290527514 | 0.215435204 | 0.140964662 | 0.194643129 | ns | YPD | KS.1 |
| FIM1_3680 | 327.7181209 | 114.2657536 | 178.89611 | -1.339753313 | 0.319663133 | 2.60487108255 | 8.59365625708 | Km<KS.1 | YPD | KS.1 |
| FIM1_3681 | 2617.455381 | 784.075002 | 1295.64304 | -1.59971492 | 0.27761954 | 4.78182769208 | 2.55253142888 | Km<KS.1 | YPD | KS.1 |
| FIM1_3682 | 11923.36618 | 1774.805013 | 3493.016974 | -2.570909953 | 0.323339757 | 7.37137460449 | 8.02463646769 | Km<KS.1 | YPD | KS.1 |
| FIM1_3683 | 1183.851121 | 612.7008429 | 1390.773785 | -0.832834205 | 0.267672733 | 0.000513388 | 0.001200237 | ns | YPD | KS.1 |
| FIM1_3684 | 1341.779762 | 493.1420353 | 675.516524 | -1.29185564 | 0.294175628 | 1.18069671297 | 4.11342167160 | Km<KS.1 | YPD | KS.1 |
| FIM1_3685 | 1579.151778 | 624.8836601 | 967.9350028 | -1.149640594 | 0.330017219 | 6.79954866262 | 0.000182853 | Km<KS.1 | YPD | KS.1 |
| FIM1_3686 | 1113.467306 | 396.8795522 | 1007.071036 | -1.174966662 | 0.424921144 | 0.000709624 | 0.001616539 | Km<KS.1 | YPD | KS.1 |
| FIM1_3687 | 813.9576903 | 374.4225762 | 709.5175891 | -0.982200271 | 0.285950992 | 0.000117545 | 0.000303156 | ns | YPD | KS.1 |
| FIM1_3688 | 2062.674305 | 928.5162007 | 1342.19789 | -0.95070075 | 0.345988784 | 0.001266093 | 0.002749374 | ns | YPD | KS.1 |
| FIM1_3689 | 3493.519372 | 1495.156946 | 1805.398193 | -1.093122869 | 0.276526134 | 1.21234184118 | 3.64435085190 | Km<KS.1 | YPD | KS.1 |
| FIM1_3690 | 2502.703055 | 752.6388762 | 1028.481125 | -1.581178974 | 0.290638091 | 3.15083942919 | 1.53982575248 | Km<KS.1 | YPD | KS.1 |
| FIM1_3691 | 4555.531893 | 2739.383235 | 3698.699931 | -0.631529542 | 0.259344714 | 0.006409727 | 0.012143332 | ns | YPD | KS.1 |
| FIM1_3692 | 2246.831738 | 1271.253092 | 2354.507026 | -0.697393101 | 0.281671403 | 0.004960094 | 0.009643011 | ns | YPD | KS.1 |
| FIM1_3693 | 2253.693932 | 1603.911111 | 2060.419245 | -0.398817385 | 0.275101913 | 0.099007373 | 0.143144923 | ns | YPD | KS.1 |
| FIM1_3694 | 1180.876683 | 762.2332406 | 1198.114767 | -0.526477087 | 0.272722773 | 0.028617815 | 0.047019032 | ns | YPD | KS.1 |
| FIM1_3695 | 22370.92064 | 4399.9337 | 5682.653673 | -2.179563712 | 0.304481308 | 1.08387286513 | 8.68320048959 | Km<KS.1 | YPD | KS.1 |
| FIM1_3696 | 9171.216738 | 2449.928957 | 4593.752869 | -1.749768811 | 0.291697311 | 8.18688234049 | 4.65482737888 | Km<KS.1 | YPD | KS.1 |
| FIM1_3697 | 19954.50288 | 7208.416722 | 11978.24542 | -1.384106449 | 0.217650588 | 1.85960256689 | 1.13099300370 | Km<KS.1 | YPD | KS.1 |
| FIM1_3698 | 7017.485635 | 2388.708128 | 3455.448409 | -1.405433359 | 0.289987591 | 1.05832383927 | 4.25639539239 | Km<KS.1 | YPD | KS.1 |
| FIM1_3699 | 8101.330429 | 467.0407607 | 1113.884538 | -3.849471445 | 0.447113089 | 9.57670574371 | 1.33456893767 | Km<KS.1 | YPD | KS.1 |
| FIM1_3700 | 2701.437528 | 898.5907215 | 866.2872135 | -1.378618064 | 0.3446895 | 5.40916842344 | 1.70752619966 | Km<KS.1 | YPD | KS.1 |
| FIM1_3701 | 818.8970388 | 424.4685114 | 989.2838633 | -0.782828832 | 0.320030522 | 0.00445545 | 0.008771612 | ns | YPD | KS.1 |
| FIM1_3702 | 1494.263826 | 478.4702279 | 650.5353628 | -1.468577896 | 0.312687466 | 1.93219382161 | 7.54526237246 | Km<KS.1 | YPD | KS.1 |
| FIM1_3703 | 5098.193723 | 2383.867579 | 2849.464825 | -0.963666694 | 0.281181224 | 0.000125911 | 0.000323391 | ns | YPD | KS.1 |
| FIM1_3704 | 666.1786688 | 601.0102044 | 1314.613492 | -0.113756761 | 0.284058312 | 0.646861662 | 0.712450027 | ns | YPD | KS.1 |
| FIM1_3705 | 3170.78439 | 1785.791301 | 2843.130502 | -0.681439315 | 0.307586626 | 0.01032968 | 0.01880357 | ns | YPD | KS.1 |
| FIM1_3706 | 6341.679814 | 3101.317516 | 5158.641043 | -0.894793872 | 0.287532575 | 0.000446387 | 0.001049515 | ns | YPD | KS.1 |
| FIM1_3707 | 994.2801968 | 460.1058301 | 787.9552835 | -0.936946386 | 0.323198736 | 0.000817021 | 0.001835642 | ns | YPD | KS.1 |
| FIM1_3708 | 6607.994105 | 4420.060237 | 5903.164291 | -0.513771433 | 0.216947992 | 0.009805726 | 0.017968135 | ns | YPD | KS.1 |
| FIM1_3709 | 2022.604597 | 1071.939455 | 2293.754854 | -0.773101263 | 0.298146124 | 0.003001066 | 0.006101473 | ns | YPD | KS.1 |
| FIM1_3710 | 3526.326161 | 2449.143421 | 2787.408431 | -0.430129158 | 0.274682805 | 0.074931446 | 0.111879936 | ns | YPD | KS.1 |
| FIM1_3711 | 8586.789155 | 2842.820469 | 3620.269437 | -1.434615732 | 0.300176736 | 1.38899020814 | 5.51494516001 | Km<KS.1 | YPD | KS.1 |
| FIM1_3712 | 7677.665896 | 2286.177688 | 3898.949549 | -1.580758037 | 0.304785968 | 1.25615905758 | 5.69344709564 | Km<KS.1 | YPD | KS.1 |
| FIM1_3713 | 4884.458831 | 2634.44636 | 3243.832749 | -0.767558272 | 0.276778685 | 0.001772262 | 0.003766341 | ns | YPD | KS.1 |

| | | | | | | | | | | |
|---|---|---|---|---|---|---|---|---|---|---|
| FIM1_3714 | 2636.678524 | 565.3559854 | 1059.112359 | -1.94357381 | 0.394744225 | 2.0083495892( | 8.9210171484! | Km<KS.1 | YPD | KS.1 |
| FIM1_3715 | 622.1507731 | 301.2284024 | 396.9883538 | -0.835971595 | 0.358175292 | 0.00534863 | 0.010305745 | ns | YPD | KS.1 |
| FIM1_3716 | 8643.415238 | 3444.358806 | 5631.398114 | -1.208109183 | 0.261524738 | 4.8289764941: | 1.7848557249 | Km<KS.1 | YPD | KS.1 |
| FIM1_3717 | 9639.833194 | 5747.714533 | 8897.841964 | -0.659127794 | 0.236930594 | 0.002189611 | 0.004565514 | ns | YPD | KS.1 |
| FIM1_3718 | 2209.117441 | 797.4084115 | 1894.578112 | -1.277181603 | 0.332231237 | 1.2763886193( | 3.8184205538: | Km<KS.1 | YPD | KS.1 |
| FIM1_3719 | 2728.261865 | 963.506978 | 1434.633716 | -1.2655349 | 0.367916194 | 6.1629877406( | 0.000167546 | Km<KS.1 | YPD | KS.1 |
| FIM1_3720 | 3446.598637 | 1171.820849 | 1754.689686 | -1.375702321 | 0.319953704 | 1.4941749208 | 5.1371943546! | Km<KS.1 | YPD | KS.1 |
| FIM1_3722 | 353.6836744 | 86.82257182 | 144.6627501 | -1.822342767 | 0.335174023 | 1.8173273103: | 9.1510155555! | Km<KS.1 | YPD | KS.1 |
| FIM1_3723 | 1202.119641 | 447.0287413 | 374.7185674 | -1.139648395 | 0.408024039 | 0.000710023 | 0.001616643 | Km<KS.1 | YPD | KS.1 |
| FIM1_3725 | 362.288505 | 116.4095385 | 110.7036884 | -1.228820958 | 0.48274299 | 0.001164589 | 0.002547346 | Km<KS.1 | YPD | KS.1 |
| FIM1_3727 | 829.2157323 | 254.1102066 | 551.350287 | -1.418541888 | 0.404221949 | 3.3601587428: | 9.4766381777: | Km<KS.1 | YPD | KS.1 |
| FIM1_3731 | 18.51879776 | 10.04147328 | 113.1955211 | -0.645303303 | 0.390360951 | 0.04064019 | 0.064410622 | ns | YPD | KS.1 |
| FIM1_3732 | 294.72645 | 151.865823 | 285.7986033 | -0.790677157 | 0.322416168 | 0.004302025 | 0.008493611 | ns | YPD | KS.1 |
| FIM1_3733 | 1691.147639 | 982.2930022 | 1175.971791 | -0.745705956 | 0.151565655 | 2.8365618132: | 1.0872030062: | ns | YPD | KS.1 |
| FIM1_3734 | 452.3392111 | 533.6043884 | 827.4544815 | 0.221749874 | 0.139403643 | 0.096597125 | 0.140026427 | ns | YPD | KS.1 |
| FIM1_3735 | 1552.719449 | 1747.544814 | 1708.862647 | 0.167453954 | 0.072112497 | 0.018417066 | 0.031759065 | ns | YPD | KS.1 |
| FIM1_3736 | 1871.049899 | 1290.543403 | 1412.888503 | -0.499252054 | 0.159499751 | 0.000987056 | 0.002190278 | ns | YPD | KS.1 |
| FIM1_3737 | 3767.83283 | 1390.017254 | 793.9904804 | -1.317623735 | 0.261636161 | 4.8309878010: | 2.0367935856( | Km<KS.1 | YPD | KS.1 |
| FIM1_3738 | 1247.136985 | 648.988982 | 451.205716 | -0.909923342 | 0.136596244 | 6.1672958177: | 3.9386773675! | ns | YPD | KS.1 |
| FIM1_3739 | 2197.328178 | 1484.506045 | 1405.266701 | -0.498783467 | 0.219774245 | 0.013121552 | 0.023380988 | ns | YPD | KS.1 |
| FIM1_3740 | 3940.791778 | 1629.430597 | 2403.374038 | -1.230518028 | 0.156549259 | 4.7634647586! | 4.3967044850! | Km<KS.1 | YPD | KS.1 |
| FIM1_3741 | 5497.534394 | 4672.998636 | 3060.206979 | -0.222018549 | 0.11825188 | 0.052226327 | 0.081373622 | ns | YPD | KS.1 |
| FIM1_3742 | 1938.047459 | 2659.794453 | 2198.695666 | 0.436751818 | 0.120260001 | 0.000179101 | 0.000452528 | ns | YPD | KS.1 |
| FIM1_3743 | 3609.022054 | 3809.705795 | 3417.466024 | 0.075242917 | 0.089926972 | 0.392886432 | 0.469971146 | ns | YPD | KS.1 |
| FIM1_3744 | 1572.568244 | 1839.692119 | 1593.700646 | 0.213178873 | 0.123770438 | 0.074176351 | 0.110985668 | ns | YPD | KS.1 |
| FIM1_3745 | 11090.82863 | 9519.782953 | 12179.15248 | -0.211573923 | 0.101155766 | 0.031837794 | 0.051644286 | ns | YPD | KS.1 |
| FIM1_3746 | 2389.905973 | 2048.548592 | 1963.194845 | -0.210099194 | 0.122329952 | 0.0751959 | 0.112173794 | ns | YPD | KS.1 |
| FIM1_3747 | 9795.790736 | 16530.91034 | 16972.88969 | 0.713104977 | 0.160502702 | 3.1457586071! | 1.0255667805: | ns | YPD | KS.1 |
| FIM1_3748 | 4803.620738 | 4306.945336 | 7906.555294 | -0.145207324 | 0.14227014 | 0.285910396 | 0.357567677 | ns | YPD | KS.1 |
| FIM1_3749 | 3677.876972 | 4002.067592 | 4416.571733 | 0.118507683 | 0.078543581 | 0.12479803 | 0.175684833 | ns | YPD | KS.1 |
| FIM1_3750 | 1043.31131 | 590.4111302 | 1375.601999 | -0.671866281 | 0.311047063 | 0.012140319 | 0.021828004 | ns | YPD | KS.1 |
| FIM1_3751 | 1528.01119 | 1876.744573 | 1850.963617 | 0.288590965 | 0.084944638 | 0.000541825 | 0.001261384 | ns | YPD | KS.1 |
| FIM1_3752 | 2716.209238 | 2816.580249 | 2341.916908 | 0.050249388 | 0.100873157 | 0.60955838 | 0.679792185 | ns | YPD | KS.1 |
| FIM1_3753 | 5841.666342 | 3320.995717 | 2476.245096 | -0.77258626 | 0.159417317 | 3.8896123672: | 1.4615424113! | ns | YPD | KS.1 |
| FIM1_3754 | 2189.463392 | 2923.03459 | 2359.349226 | 0.348427208 | 0.247418046 | 0.116148856 | 0.165003015 | ns | YPD | KS.1 |
| FIM1_3755 | 75.30624454 | 131.6229343 | 213.6750868 | 0.720343134 | 0.222268564 | 0.000421085 | 0.000994256 | ns | YPD | KS.1 |
| FIM1_3756 | 11396.4646 | 14947.93472 | 22678.61819 | 0.371984964 | 0.125413212 | 0.002135391 | 0.004471199 | ns | YPD | KS.1 |
| FIM1_3757 | 574.873367 | 645.499598 | 660.2333306 | 0.157242682 | 0.11746448 | 0.166482768 | 0.225620205 | ns | YPD | KS.1 |
| FIM1_3758 | 388.7624812 | 834.4470356 | 1542.398213 | 1.038430599 | 0.191819173 | 1.0960514023( | 5.0210457889: | Km>KS.1 | YPD | KS.1 |
| FIM1_3759 | 4942.311679 | 8570.395387 | 9864.711821 | 0.698493328 | 0.246610448 | 0.001717855 | 0.003655402 | ns | YPD | KS.1 |
| FIM1_3760 | 231.141574 | 686.5638983 | 619.9495454 | 1.540098424 | 0.120401333 | 1.2328239728: | 5.6789808007( | Km>KS.1 | YPD | KS.1 |
| FIM1_3761 | 621.793512 | 851.7284161 | 735.6103109 | 0.445157628 | 0.085820173 | 1.3384946722! | 5.3314739748( | ns | YPD | KS.1 |
| FIM1_3762 | 695.3044075 | 2387.774066 | 1837.459971 | 1.732179624 | 0.157429178 | 1.6386208528( | 4.3362440122: | Km>KS.1 | YPD | KS.1 |
| FIM1_3763 | 2132.415717 | 3225.343801 | 3123.577917 | 0.536357784 | 0.205027506 | 0.004666254 | 0.00914321 | ns | YPD | KS.1 |
| FIM1_3764 | 144.3135401 | 344.0089063 | 462.6643805 | 1.229199259 | 0.113781595 | 4.1217683248! | 1.0002828007! | Km>KS.1 | YPD | KS.1 |

| | | | | | | | | | | |
|---|---|---|---|---|---|---|---|---|---|---|
| FIM1_3765 | 8015.169295 | 9899.227138 | 9241.215501 | 0.281551074 | 0.150885794 | 0.049958469 | 0.078011105 | ns | YPD | KS.1 |
| FIM1_3766 | 198.4509652 | 114.495327 | 98.51668961 | -0.747390128 | 0.170246368 | 3.7259280263 | 1.2060176923 | ns | YPD | KS.1 |
| FIM1_3767 | 1954.685887 | 1961.563658 | 2270.164745 | 0.004400472 | 0.136273873 | 0.973182458 | 0.981070462 | ns | YPD | KS.1 |
| FIM1_3768 | 3377.844453 | 11717.93451 | 26488.78546 | 1.508340114 | 0.40185672 | 1.0825275289 | 3.2719164377 | Km>KS.1 | YPD | KS.1 |
| FIM1_3769 | 3811.441212 | 2307.05451 | 2223.519758 | -0.711673495 | 0.0858523 | 3.8712773361 | 3.9792571791 | ns | YPD | KS.1 |
| FIM1_3770 | 3624.524996 | 6309.958164 | 6474.110678 | 0.793699285 | 0.057555607 | 7.9954585907 | 4.9721758111 | ns | YPD | KS.1 |
| FIM1_3771 | 723.4168568 | 1423.531125 | 1381.914983 | 0.93196862 | 0.159942927 | 1.2360012780 | 6.3392849056 | ns | YPD | KS.1 |
| FIM1_3772 | 2378.957264 | 2456.646687 | 2235.125208 | 0.041622296 | 0.160748762 | 0.784880268 | 0.830627384 | ns | YPD | KS.1 |
| FIM1_3773 | 7734.0349 | 9023.906613 | 9253.453081 | 0.198498274 | 0.178379546 | 0.235310457 | 0.3031252 | ns | YPD | KS.1 |
| FIM1_3774 | 2154.182095 | 3641.466025 | 4738.193851 | 0.714149178 | 0.162733684 | 4.0403988994 | 1.2951665286 | ns | YPD | KS.1 |
| FIM1_3775 | 199.141925 | 239.0302173 | 237.9796987 | 0.24742208 | 0.128072247 | 0.044808172 | 0.070544511 | ns | YPD | KS.1 |
| FIM1_3776 | 409.518232 | 374.89781 | 418.9431956 | -0.11781256 | 0.137734341 | 0.372370782 | 0.448232432 | ns | YPD | KS.1 |
| FIM1_3777 | 7890.693574 | 10685.40353 | 9059.563685 | 0.320852071 | 0.334812201 | 0.253073019 | 0.32243172 | ns | YPD | KS.1 |
| FIM1_3778 | 1184.754955 | 532.5812199 | 395.1968135 | -1.035355748 | 0.263024774 | 1.4686118175 | 4.3438429204 | Km<KS.1 | YPD | KS.1 |
| FIM1_3779 | 558.8465932 | 959.5119986 | 1354.296373 | 0.744694747 | 0.14497375 | 9.1575059869 | 3.7312524393 | ns | YPD | KS.1 |
| FIM1_3780 | 4179.323153 | 6991.888398 | 10131.22759 | 0.728749107 | 0.089103957 | 9.4165216634 | 9.3135577089 | ns | YPD | KS.1 |
| FIM1_3782 | 2434.342124 | 3926.379089 | 3009.537331 | 0.662840683 | 0.128409137 | 9.6114118631 | 3.8970475973 | ns | YPD | KS.1 |
| FIM1_3783 | 2395.337158 | 2098.652696 | 1425.301828 | -0.1821494 | 0.104251089 | 0.072543425 | 0.108804203 | ns | YPD | KS.1 |
| FIM1_3784 | 2004.658778 | 2254.750979 | 3653.060382 | 0.159926803 | 0.116750433 | 0.156953124 | 0.214105235 | ns | YPD | KS.1 |
| FIM1_3785 | 207.3663519 | 296.1674048 | 374.948203 | 0.459280881 | 0.196782704 | 0.011924286 | 0.021462852 | ns | YPD | KS.1 |
| FIM1_3786 | 3892.345275 | 3985.514249 | 4454.690071 | 0.033308733 | 0.070419776 | 0.631578362 | 0.698866181 | ns | YPD | KS.1 |
| FIM1_3787 | 207.2885474 | 92.6028658 | 84.05890081 | -1.111609669 | 0.17005809 | 9.7183029108 | 6.1123333731 | Km<KS.1 | YPD | KS.1 |
| FIM1_3788 | 3889.423695 | 5279.560919 | 6370.812734 | 0.424989838 | 0.108204273 | 5.5760353177 | 0.000152795 | ns | YPD | KS.1 |
| FIM1_3789 | 717.1604892 | 767.5700219 | 690.6040231 | 0.091339049 | 0.12120304 | 0.435598133 | 0.512559298 | ns | YPD | KS.1 |
| FIM1_3790 | 266.1166208 | 443.1001039 | 421.9346332 | 0.694199124 | 0.161550659 | 6.4093232343 | 2.0029135107 | ns | YPD | KS.1 |
| FIM1_3791 | 3124.728042 | 3683.910303 | 3342.107276 | 0.232370433 | 0.073129355 | 0.001262993 | 0.002745038 | ns | YPD | KS.1 |
| FIM1_3792 | 2906.809834 | 3838.76008 | 3126.315086 | 0.345590677 | 0.222719998 | 0.088721215 | 0.129934662 | ns | YPD | KS.1 |
| FIM1_3793 | 7821.471108 | 3854.887666 | 6434.595813 | -0.991231423 | 0.130015188 | 4.7515824219 | 3.9464311434 | ns | YPD | KS.1 |
| FIM1_3794 | 4127.850746 | 6052.720826 | 9539.717368 | 0.529160005 | 0.124584082 | 1.1488474237 | 3.4597554074 | ns | YPD | KS.1 |
| FIM1_3795 | 1875.349276 | 1382.79074 | 881.351855 | -0.386770472 | 0.208378583 | 0.043744708 | 0.068979373 | ns | YPD | KS.1 |
| FIM1_3796 | 435.102409 | 273.4735543 | 333.1994851 | -0.628477582 | 0.162126124 | 4.5584691423 | 0.000126342 | ns | YPD | KS.1 |
| FIM1_3797 | 3297.767193 | 4417.952448 | 4135.845504 | 0.398661257 | 0.133925777 | 0.00197553 | 0.004171588 | ns | YPD | KS.1 |
| FIM1_3798 | 2276.765558 | 2336.816109 | 2050.789803 | 0.03476551 | 0.127782948 | 0.777835409 | 0.825630715 | ns | YPD | KS.1 |
| FIM1_3799 | 11047.30646 | 12358.64804 | 14279.63957 | 0.145404705 | 0.167404574 | 0.357523746 | 0.432558521 | ns | YPD | KS.1 |
| FIM1_3801 | 2608.625001 | 2932.877274 | 2398.246948 | 0.14825588 | 0.188096703 | 0.398205079 | 0.474962903 | ns | YPD | KS.1 |
| FIM1_3802 | 3589.275304 | 1782.7854 | 2210.137681 | -0.860981718 | 0.300526979 | 0.001080289 | 0.002377019 | ns | YPD | KS.1 |
| FIM1_3803 | 225.9751138 | 233.8957646 | 313.849329 | 0.041811083 | 0.182600085 | 0.80696339 | 0.848761705 | ns | YPD | KS.1 |
| FIM1_3804 | 249.2952005 | 251.1482225 | 257.7622274 | 0.010736883 | 0.137825846 | 0.93531347 | 0.952935595 | ns | YPD | KS.1 |
| FIM1_3805 | 1253.856155 | 482.3543085 | 471.9418112 | -1.334986609 | 0.15327324 | 3.1156589293 | 3.7991184249 | Km<KS.1 | YPD | KS.1 |
| FIM1_3806 | 575.160238 | 376.1324844 | 864.3943599 | -0.514959339 | 0.264315341 | 0.028092181 | 0.046262363 | ns | YPD | KS.1 |
| FIM1_3807 | 9212.825823 | 10799.12392 | 10336.56045 | 0.225243498 | 0.064705304 | 0.00042757 | 0.001008612 | ns | YPD | KS.1 |
| FIM1_3808 | 1140.188508 | 718.9886142 | 598.9609995 | -0.647454191 | 0.104293538 | 2.1595921022 | 1.1832566860 | ns | YPD | KS.1 |
| FIM1_3809 | 946.7819004 | 803.0291366 | 2060.132763 | -0.178624275 | 0.302648624 | 0.493056374 | 0.570266676 | ns | YPD | KS.1 |
| FIM1_3810 | 516.0006389 | 1290.268622 | 1956.994185 | 1.281008393 | 0.149924438 | 1.4638971193 | 1.6858537427 | Km>KS.1 | YPD | KS.1 |
| FIM1_3811 | 12666.02459 | 5744.705454 | 5184.452121 | -1.096129205 | 0.159294258 | 9.4431027429 | 6.5068471116 | Km<KS.1 | YPD | KS.1 |

| | | | | | | | | | | |
|---|---|---|---|---|---|---|---|---|---|---|
| FIM1_3812 | 60.92608694 | 30.4474989 | 22.63589428 | -0.839160774 | 0.318755499 | 0.002303283 | 0.004782485 | ns | YPD | KS.1 |
| FIM1_3813 | 1940.255094 | 3357.497847 | 4199.466434 | 0.779275336 | 0.082073315 | 6.40331921565 | 9.65312096134 | ns | YPD | KS.1 |
| FIM1_3814 | 39.99621774 | 24.33145633 | 26.24877177 | -0.582187466 | 0.314286344 | 0.030494158 | 0.049724169 | ns | YPD | KS.1 |
| FIM1_3815 | 12537.00277 | 15281.58937 | 13481.12748 | 0.237066072 | 0.240422538 | 0.273907734 | 0.34454892 | ns | YPD | KS.1 |
| FIM1_3816 | 2274.163495 | 1299.938609 | 2349.079688 | -0.734118878 | 0.212716577 | 0.000190982 | 0.000481325 | ns | YPD | KS.1 |
| FIM1_3817 | 1418.328633 | 2439.54288 | 3144.887224 | 0.754030518 | 0.129819266 | 2.00834213311 | 1.00518129901 | ns | YPD | KS.1 |
| FIM1_3818 | 2500.103953 | 140.2857909 | 352.0551801 | -4.052807859 | 0.274291901 | 5.97426286786 | 5.94439155352 | Km<KS.1 | YPD | KS.1 |
| FIM1_3819 | 4170.822896 | 3413.562461 | 2295.003418 | -0.258504013 | 0.181318987 | 0.127579807 | 0.178791419 | ns | YPD | KS.1 |
| FIM1_3820 | 837.4172312 | 938.2888525 | 1699.424937 | 0.138181318 | 0.223956009 | 0.49888951 | 0.575730761 | ns | YPD | KS.1 |
| FIM1_3821 | 898.8301147 | 692.304093 | 934.0114174 | -0.341019944 | 0.175274314 | 0.038316199 | 0.061018915 | ns | YPD | KS.1 |
| FIM1_3822 | 10114.42336 | 6224.009231 | 5347.315536 | -0.675096264 | 0.125168207 | 2.63602473555 | 1.15442104396 | ns | YPD | KS.1 |
| FIM1_3823 | 364.1160111 | 171.0598332 | 134.5366798 | -1.048577884 | 0.156167065 | 3.28258633043 | 2.14034954048 | Km<KS.1 | YPD | KS.1 |
| FIM1_3824 | 68547.44342 | 31896.84338 | 20601.80896 | -1.012692131 | 0.231007774 | 2.17610958846 | 7.27074896057 | Km<KS.1 | YPD | KS.1 |
| FIM1_3825 | 174.1560795 | 127.6555875 | 253.0589104 | -0.348017353 | 0.304122021 | 0.183040574 | 0.245187629 | ns | YPD | KS.1 |
| FIM1_3826 | 3197.932848 | 6027.360188 | 6236.632612 | 0.906499512 | 0.065967455 | 1.25183239336 | 7.59495872804 | ns | YPD | KS.1 |
| FIM1_3827 | 296.8649876 | 398.4352785 | 477.8659757 | 0.39692775 | 0.142715301 | 0.00368009 | 0.007361661 | ns | YPD | KS.1 |
| FIM1_3828 | 1626.533919 | 2096.930225 | 3076.068311 | 0.353269443 | 0.103813207 | 0.00048624 | 0.001138374 | ns | YPD | KS.1 |
| FIM1_3829 | 232.8380316 | 1313.700322 | 2219.783123 | 2.470998612 | 0.119434073 | 1.68368877590 | 1.19662166573 | Km>KS.1 | YPD | KS.1 |
| FIM1_3830 | 3837.560377 | 4285.74868 | 3249.216534 | 0.147234281 | 0.140281889 | 0.272938031 | 0.343452303 | ns | YPD | KS.1 |
| FIM1_3831 | 1446.494835 | 1032.171096 | 740.3250707 | -0.447776678 | 0.170592795 | 0.005390369 | 0.010382146 | ns | YPD | KS.1 |
| FIM1_3832 | 467.9416252 | 549.7376528 | 785.8791234 | 0.21564902 | 0.136724227 | 0.099900588 | 0.144310519 | ns | YPD | KS.1 |
| FIM1_3833 | 1767.440286 | 105.894542 | 255.9778003 | -3.908777182 | 0.334694579 | 3.06470285345 | 1.09689904287 | Km<KS.1 | YPD | KS.1 |
| FIM1_3834 | 4211.147409 | 3161.284367 | 3235.127828 | -0.4016072 | 0.095608096 | 1.80285071232 | 5.27598958459 | ns | YPD | KS.1 |
| FIM1_3835 | 485.0676845 | 563.4331189 | 381.0551814 | 0.203790875 | 0.117610514 | 0.07325624 | 0.109687093 | ns | YPD | KS.1 |
| FIM1_3836 | 1750.769963 | 3710.282185 | 4396.370425 | 1.041566646 | 0.15529431 | 3.49971925580 | 2.26707074188 | Km>KS.1 | YPD | KS.1 |
| FIM1_3837 | 2049.427629 | 1664.743015 | 2397.157586 | -0.287125455 | 0.11011224 | 0.007280138 | 0.013641689 | ns | YPD | KS.1 |
| FIM1_3838 | 1989.072382 | 2557.971104 | 1857.047638 | 0.347034105 | 0.116155565 | 0.002067375 | 0.004339744 | ns | YPD | KS.1 |
| FIM1_3839 | 2605.798082 | 2969.794151 | 2382.466326 | 0.157453788 | 0.229641847 | 0.4505409 | 0.528143491 | ns | YPD | KS.1 |
| FIM1_3840 | 544.488885 | 632.5154996 | 898.5308475 | 0.190910604 | 0.187777765 | 0.275649719 | 0.34608905 | ns | YPD | KS.1 |
| FIM1_3841 | 4276.682396 | 3575.822865 | 2372.251608 | -0.225754079 | 0.198045252 | 0.218584222 | 0.285271906 | ns | YPD | KS.1 |
| FIM1_3842 | 471.1657315 | 242.2388884 | 197.4601246 | -0.810447009 | 0.304149445 | 0.002235764 | 0.004655893 | ns | YPD | KS.1 |
| FIM1_3843 | 7218.992287 | 10361.36401 | 11982.34472 | 0.513061575 | 0.073731083 | 1.86136580348 | 1.24970241192 | ns | YPD | KS.1 |
| FIM1_3844 | 2695.666316 | 965.02132 | 1232.990209 | -1.355676766 | 0.266897133 | 3.56766167244 | 1.53672007102 | Km<KS.1 | YPD | KS.1 |
| FIM1_3845 | 862.8484702 | 357.2207269 | 1007.305287 | -1.156898516 | 0.257924498 | 1.01108000139 | 3.56241006157 | Km<KS.1 | YPD | KS.1 |
| FIM1_3846 | 1620.564579 | 770.5844492 | 975.6377112 | -1.046379731 | 0.120342839 | 6.02219096280 | 7.20117718303 | Km<KS.1 | YPD | KS.1 |
| FIM1_3847 | 16668.85298 | 23997.02213 | 31400.47508 | 0.514226174 | 0.087255157 | 2.05728664334 | 1.02555120748 | ns | YPD | KS.1 |
| FIM1_3848 | 233.8972474 | 712.7893201 | 750.7664885 | 1.484488363 | 0.259269936 | 7.59905761031 | 3.97950648540 | Km>KS.1 | YPD | KS.1 |
| FIM1_3849 | 11864.96148 | 25561.06261 | 30737.89021 | 1.067253719 | 0.150966959 | 2.61060191249 | 1.89050138496 | Km>KS.1 | YPD | KS.1 |
| FIM1_3850 | 2691.754959 | 3223.25939 | 2505.267773 | 0.229837452 | 0.188572218 | 0.190866388 | 0.254437374 | ns | YPD | KS.1 |
| FIM1_3851 | 4320.936537 | 1561.823328 | 1348.817359 | -1.432140156 | 0.139515349 | 8.57997819683 | 1.76385915410 | Km<KS.1 | YPD | KS.1 |
| FIM1_3852 | 841.9618204 | 719.5418763 | 1300.114399 | -0.207572299 | 0.157248249 | 0.164050169 | 0.222566018 | ns | YPD | KS.1 |
| FIM1_3853 | 1686.840172 | 1912.090711 | 1715.996936 | 0.170182334 | 0.11954178 | 0.140776592 | 0.194437408 | ns | YPD | KS.1 |
| FIM1_3854 | 4781.510854 | 4966.228222 | 5777.754768 | 0.049818808 | 0.147522801 | 0.723639366 | 0.779918944 | ns | YPD | KS.1 |
| FIM1_3855 | 2943.35719 | 3353.506506 | 3691.321226 | 0.176981589 | 0.122323189 | 0.133910168 | 0.186298402 | ns | YPD | KS.1 |
| FIM1_3856 | 1492.780867 | 820.061481 | 1475.589319 | -0.835700495 | 0.130151893 | 3.59093418216 | 2.12424465591 | ns | YPD | KS.1 |

| | | | | | | | | | | |
|---|---|---|---|---|---|---|---|---|---|---|
| FIM1_3857 | 3407.744507 | 5498.170844 | 6700.010341 | 0.678535178 | 0.082959199 | 1.0502531184 | 1.0326105265 | ns | YPD | KS.1 |
| FIM1_3858 | 1596.648009 | 1028.589732 | 1106.416192 | -0.612944888 | 0.114232542 | 3.5391969833 | 1.5257803286 | ns | YPD | KS.1 |
| FIM1_3859 | 20.97527357 | 19.09640543 | 26.75156027 | -0.106844922 | 0.258769037 | 0.643047909 | 0.708805868 | ns | YPD | KS.1 |
| FIM1_3860 | 4925.957795 | 2880.629047 | 2591.085474 | -0.690829781 | 0.229870216 | 0.001002094 | 0.002220676 | ns | YPD | KS.1 |
| FIM1_3861 | 863.0577029 | 890.5125089 | 683.3851481 | 0.038381888 | 0.21393196 | 0.845266162 | 0.879564768 | ns | YPD | KS.1 |
| FIM1_3862 | 4721.074429 | 4510.944892 | 4239.368737 | -0.049721435 | 0.292200403 | 0.844742844 | 0.87947331 | ns | YPD | KS.1 |
| FIM1_3863 | 5093.70634 | 6621.678429 | 5834.400475 | 0.351795634 | 0.150929031 | 0.014414107 | 0.025429144 | ns | YPD | KS.1 |
| FIM1_3864 | 240.9867557 | 748.5822553 | 239.0951616 | 1.452953116 | 0.32014722 | 4.2458283840 | 1.5870019692 | Km>KS.1 | YPD | KS.1 |
| FIM1_3865 | 426.3765808 | 1194.605023 | 647.6237333 | 1.307778515 | 0.318116909 | 3.9444138524 | 1.2684847392 | Km>KS.1 | YPD | KS.1 |
| FIM1_3866 | 1190.48243 | 2317.403379 | 1486.262765 | 0.922244497 | 0.150064586 | 1.7734676382 | 9.8142397112 | ns | YPD | KS.1 |
| FIM1_3867 | 937.4033302 | 1427.295223 | 1536.73574 | 0.5715944 | 0.150717155 | 7.2582976790 | 0.000193723 | ns | YPD | KS.1 |
| FIM1_3868 | 1859.76048 | 479.4262092 | 508.917387 | -1.886658863 | 0.191942484 | 2.5501537320 | 4.7263615918 | Km<KS.1 | YPD | KS.1 |
| FIM1_3869 | 7574.092518 | 14015.36714 | 15450.81622 | 0.812318381 | 0.214209965 | 4.2863987490 | 0.000119467 | ns | YPD | KS.1 |
| FIM1_3870 | 3150.727033 | 1106.301391 | 1812.154242 | -1.467948126 | 0.150578227 | 1.4768745788 | 2.5248972610 | Km<KS.1 | YPD | KS.1 |
| FIM1_3871 | 1268.07047 | 1654.158436 | 2563.177529 | 0.330068628 | 0.221969008 | 0.103007572 | 0.148282023 | ns | YPD | KS.1 |
| FIM1_3872 | 9808.197652 | 5082.101657 | 5529.200178 | -0.935304702 | 0.086512359 | 6.4194637112 | 1.5428421238 | ns | YPD | KS.1 |
| FIM1_3873 | 552.3898771 | 1585.278219 | 1453.231919 | 1.467876743 | 0.168683364 | 2.5808944844 | 3.1703580395 | Km>KS.1 | YPD | KS.1 |
| FIM1_3874 | 925.0255133 | 539.6473476 | 510.1569601 | -0.751765347 | 0.122683717 | 2.8412390962 | 1.5414574158 | ns | YPD | KS.1 |
| FIM1_3875 | 199.3056488 | 331.63441 | 283.7631909 | 0.703649757 | 0.131323034 | 3.0094630384 | 1.3098931422 | ns | YPD | KS.1 |
| FIM1_3876 | 96.49487206 | 114.0598016 | 99.06440972 | 0.198575292 | 0.230108915 | 0.341890073 | 0.416888018 | ns | YPD | KS.1 |
| FIM1_3877 | 602.8125701 | 364.0877995 | 174.706943 | -0.675447025 | 0.180456947 | 7.0623431707 | 0.000189 | ns | YPD | KS.1 |
| FIM1_3879 | 66458.70108 | 33159.89998 | 20269.80526 | -0.91279754 | 0.232009963 | 1.9232603303 | 5.6118593217 | ns | YPD | KS.1 |
| FIM1_3880 | 2641.052226 | 2485.491001 | 3181.816457 | -0.077358305 | 0.181740262 | 0.649729207 | 0.714817073 | ns | YPD | KS.1 |
| FIM1_3881 | 5208.829824 | 6152.57247 | 6875.92404 | 0.236694942 | 0.059896919 | 6.5559830884 | 0.000177261 | ns | YPD | KS.1 |
| FIM1_3882 | 148.3394076 | 53.65078809 | 60.88332332 | -1.308806839 | 0.300430632 | 1.3357161008 | 4.6179205016 | Km<KS.1 | YPD | KS.1 |
| FIM1_3883 | 440.5078824 | 428.9844447 | 422.477108 | -0.035362333 | 0.142949636 | 0.796096857 | 0.839818037 | ns | YPD | KS.1 |
| FIM1_3884 | 1288.727367 | 536.7896303 | 508.2338864 | -1.194378158 | 0.197115214 | 1.8022589544 | 9.9514298537 | Km<KS.1 | YPD | KS.1 |
| FIM1_3885 | 1683.490473 | 2771.917247 | 2432.3808 | 0.699746437 | 0.109263904 | 5.3910256848 | 3.1295627517 | ns | YPD | KS.1 |
| FIM1_3886 | 3026.827909 | 2972.311745 | 3862.400914 | -0.024750274 | 0.123809336 | 0.836071823 | 0.873468568 | ns | YPD | KS.1 |
| FIM1_3887 | 337.635042 | 316.6394724 | 388.6433825 | -0.088492851 | 0.101705574 | 0.372014127 | 0.44793874 | ns | YPD | KS.1 |
| FIM1_3888 | 1275.118544 | 2056.096231 | 3161.601912 | 0.659864003 | 0.133895086 | 3.2942573052 | 1.2501090841 | ns | YPD | KS.1 |
| FIM1_3889 | 768.4595522 | 1019.494926 | 1106.781719 | 0.364380951 | 0.192232414 | 0.0414842 | 0.065664619 | ns | YPD | KS.1 |
| FIM1_3890 | 3179.150333 | 2282.214522 | 2590.63797 | -0.449278006 | 0.146195405 | 0.001315211 | 0.002848573 | ns | YPD | KS.1 |
| FIM1_3891 | 1094.388639 | 563.0426952 | 393.9541856 | -0.869160731 | 0.231747155 | 4.4759734479 | 0.000124333 | ns | YPD | KS.1 |
| FIM1_3892 | 2268.299823 | 4234.111266 | 1681.82906 | 0.756336813 | 0.300260213 | 0.003855072 | 0.007674663 | ns | YPD | KS.1 |
| FIM1_3893 | 167.5282597 | 171.7222416 | 134.945167 | 0.034676665 | 0.155264163 | 0.814397444 | 0.854413177 | ns | YPD | KS.1 |
| FIM1_3894 | 1871.465654 | 991.0279605 | 844.3097769 | -0.890597606 | 0.123299505 | 1.1963147995 | 8.8042398343 | ns | YPD | KS.1 |
| FIM1_3895 | 191.3371065 | 155.1676553 | 166.2505611 | -0.271215536 | 0.184556117 | 0.115566092 | 0.164362867 | ns | YPD | KS.1 |
| FIM1_3896 | 1226.916497 | 1574.858813 | 1224.149328 | 0.343779076 | 0.115637049 | 0.002179724 | 0.004550206 | ns | YPD | KS.1 |
| FIM1_3897 | 570.5870741 | 375.938269 | 509.3109975 | -0.571631473 | 0.137983867 | 1.6670912225 | 4.8988652288 | ns | YPD | KS.1 |
| FIM1_3898 | 2857.055001 | 2336.731104 | 4862.051882 | -0.214215358 | 0.317372937 | 0.427517788 | 0.505202137 | ns | YPD | KS.1 |
| FIM1_3899 | 1422.006379 | 1321.247739 | 974.2576244 | -0.101767987 | 0.092668609 | 0.261477293 | 0.331426633 | ns | YPD | KS.1 |
| FIM1_3900 | 358.8722423 | 298.5571499 | 381.9129481 | -0.245974251 | 0.141956654 | 0.069957607 | 0.105338709 | ns | YPD | KS.1 |
| FIM1_3901 | 2204.463031 | 1688.095803 | 1554.936371 | -0.363059535 | 0.134133607 | 0.00487509 | 0.009514936 | ns | YPD | KS.1 |
| FIM1_3902 | 4198.135411 | 1003.683672 | 1006.228109 | -1.990398471 | 0.199014562 | 3.4294999323 | 6.7437795111 | Km<KS.1 | YPD | KS.1 |

| | | | | | | | | | | |
|---|---|---|---|---|---|---|---|---|---|---|
| FIM1_3903 | 435.336923 | 435.3992745 | 596.3936974 | -0.00137846 | 0.151156489 | 0.992368972 | 0.994302819 | ns | YPD | KS.1 |
| FIM1_3904 | 3417.099988 | 5267.552358 | 7748.498376 | 0.601987976 | 0.118835672 | 1.83611471813 | 7.19265411238 | ns | YPD | KS.1 |
| FIM1_3905 | 6762.766348 | 11260.64327 | 6693.299249 | 0.682440673 | 0.18286319 | 7.26609979173 | 0.000193828 | ns | YPD | KS.1 |
| FIM1_3906 | 485.361716 | 595.3914986 | 747.9196432 | 0.269750923 | 0.157253658 | 0.070532564 | 0.105947918 | ns | YPD | KS.1 |
| FIM1_3907 | 1785.196988 | 1600.030725 | 1759.155748 | -0.150992879 | 0.106785969 | 0.145762557 | 0.200766534 | ns | YPD | KS.1 |
| FIM1_3908 | 1210.81529 | 1757.852973 | 1524.870191 | 0.52827633 | 0.075778294 | 1.64381537279 | 1.11416641412 | ns | YPD | KS.1 |
| FIM1_3909 | 460.7223572 | 487.6226762 | 1174.845268 | 0.066373515 | 0.250816198 | 0.767819933 | 0.817791515 | ns | YPD | KS.1 |
| FIM1_3910 | 4710.183773 | 5835.458202 | 6264.664992 | 0.300620964 | 0.086689225 | 0.000411646 | 0.000975208 | ns | YPD | KS.1 |
| FIM1_3911 | 241.1819616 | 176.4604776 | 182.5380654 | -0.39640113 | 0.210298383 | 0.040339261 | 0.063954054 | ns | YPD | KS.1 |
| FIM1_3912 | 15857.04623 | 10346.81547 | 4257.191824 | -0.544909211 | 0.221680684 | 0.007201841 | 0.013520437 | ns | YPD | KS.1 |
| FIM1_3913 | 1608.645295 | 1281.208663 | 592.3381723 | -0.297228752 | 0.173304188 | 0.068091439 | 0.102902463 | ns | YPD | KS.1 |
| FIM1_3914 | 504.7229106 | 627.7305144 | 465.4991353 | 0.290947305 | 0.154949639 | 0.048059939 | 0.075259111 | ns | YPD | KS.1 |
| FIM1_3915 | 2416.755947 | 3033.050498 | 2899.055429 | 0.316114606 | 0.102701745 | 0.001601448 | 0.00342753 | ns | YPD | KS.1 |
| FIM1_3916 | 614.973155 | 1056.249039 | 852.7608378 | 0.73768014 | 0.160090312 | 1.36263638637 | 4.70445247897 | ns | YPD | KS.1 |
| FIM1_3917 | 1707.102839 | 2371.560637 | 1846.392987 | 0.444394719 | 0.149434595 | 0.001841765 | 0.003905705 | ns | YPD | KS.1 |
| FIM1_3918 | 807.0175387 | 530.0350155 | 1266.487463 | -0.533300255 | 0.226997287 | 0.009924384 | 0.018152136 | ns | YPD | KS.1 |
| FIM1_3919 | 1049.881579 | 290.9923543 | 331.3315058 | -1.799276953 | 0.165019399 | 4.17656962226 | 1.08787611888 | Km<KS.1 | YPD | KS.1 |
| FIM1_3920 | 277.7788621 | 205.6178812 | 231.2274604 | -0.404406542 | 0.152708312 | 0.005428632 | 0.010447755 | ns | YPD | KS.1 |
| FIM1_3921 | 2643.422923 | 3293.901989 | 2798.983204 | 0.300509277 | 0.124592184 | 0.012452468 | 0.022324695 | ns | YPD | KS.1 |
| FIM1_3922 | 215.042711 | 338.9057336 | 500.5854168 | 0.613158457 | 0.162938079 | 7.46858973294 | 0.000198803 | ns | YPD | KS.1 |
| FIM1_3923 | 812.0032629 | 1899.020174 | 3006.232563 | 1.195836037 | 0.12648702 | 4.30265361204 | 6.60669806170 | Km>KS.1 | YPD | KS.1 |
| FIM1_3924 | 532.8577692 | 205.2521576 | 705.9763253 | -1.098428732 | 0.402003904 | 0.00093961 | 0.002090591 | Km<KS.1 | YPD | KS.1 |
| FIM1_3925 | 646.7833874 | 241.3260374 | 212.0267322 | -1.324511474 | 0.234397248 | 1.62755781685 | 8.24010763938 | Km<KS.1 | YPD | KS.1 |
| FIM1_3926 | 4066.128882 | 2601.879369 | 1880.232738 | -0.614747013 | 0.136780299 | 3.07647534517 | 1.00429559332 | ns | YPD | KS.1 |
| FIM1_3927 | 340.6016296 | 157.1293987 | 370.1716768 | -1.053703128 | 0.19090603 | 5.85537043416 | 2.77433027714 | Km<KS.1 | YPD | KS.1 |
| FIM1_3928 | 1867.056365 | 3207.272805 | 2189.577524 | 0.741667088 | 0.153422003 | 4.42698863475 | 1.64852308816 | ns | YPD | KS.1 |
| FIM1_3929 | 442.0297368 | 881.9248522 | 1378.074944 | 0.934740789 | 0.188745184 | 1.60426174630 | 6.32926422509 | ns | YPD | KS.1 |
| FIM1_3930 | 1006.424156 | 981.1035864 | 1070.974216 | -0.034780853 | 0.09434978 | 0.706236379 | 0.765474071 | ns | YPD | KS.1 |
| FIM1_3932 | 2064.072521 | 1388.82552 | 2228.120119 | -0.553534037 | 0.108705582 | 1.77841820522 | 6.98313383663 | ns | YPD | KS.1 |
| FIM1_3933 | 388.9545277 | 399.5288086 | 529.5501752 | 0.034411177 | 0.1304053 | 0.784098609 | 0.830379381 | ns | YPD | KS.1 |
| FIM1_3934 | 2196.257016 | 1258.814097 | 1094.394944 | -0.782374964 | 0.109414572 | 2.54717540283 | 1.84995585826 | ns | YPD | KS.1 |
| FIM1_3935 | 2586.157064 | 1451.770778 | 1625.570104 | -0.795532156 | 0.149461875 | 2.99061694277 | 1.30397189222 | ns | YPD | KS.1 |
| FIM1_3936 | 927.4832536 | 1065.605878 | 891.8872015 | 0.193400032 | 0.091393922 | 0.030522684 | 0.049754374 | ns | YPD | KS.1 |
| FIM1_3937 | 44.41809369 | 34.92549966 | 34.38314948 | -0.248749394 | 0.345576814 | 0.386248808 | 0.463033209 | ns | YPD | KS.1 |
| FIM1_3938 | 68.14087482 | 106.71288 | 80.08171728 | 0.538332761 | 0.270886517 | 0.024480197 | 0.040841375 | ns | YPD | KS.1 |
| FIM1_3939 | 1761.61687 | 897.0415487 | 588.5738225 | -0.864639493 | 0.25672943 | 0.000194414 | 0.000489478 | ns | YPD | KS.1 |
| FIM1_3940 | 4175.960153 | 4669.973277 | 2416.895088 | 0.124856627 | 0.280378655 | 0.611428446 | 0.681572153 | ns | YPD | KS.1 |
| FIM1_3942 | 1178.37916 | 680.9649738 | 686.7445039 | -0.76947699 | 0.111968502 | 1.92064205338 | 1.28776202366 | ns | YPD | KS.1 |
| FIM1_3943 | 4694.824779 | 5428.06726 | 4597.163711 | 0.203213865 | 0.084561478 | 0.014290665 | 0.025229262 | ns | YPD | KS.1 |
| FIM1_3944 | 538.9180278 | 675.9556709 | 950.884262 | 0.31644681 | 0.09993056 | 0.001184461 | 0.002585648 | ns | YPD | KS.1 |
| FIM1_3945 | 202.222121 | 453.7329693 | 915.1070056 | 1.071067728 | 0.234970503 | 8.54269700911 | 3.05315500146 | Km>KS.1 | YPD | KS.1 |
| FIM1_3946 | 2731.880871 | 834.5127586 | 1684.428242 | -1.514877417 | 0.331519121 | 3.20067551857 | 1.21831374941 | Km<KS.1 | YPD | KS.1 |
| FIM1_3947 | 1567.622928 | 1562.487082 | 1402.901195 | -0.004157129 | 0.127300088 | 0.972993758 | 0.981070462 | ns | YPD | KS.1 |
| FIM1_3948 | 2411.388399 | 2159.206299 | 1624.501317 | -0.153519518 | 0.088495919 | 0.076463317 | 0.113825554 | ns | YPD | KS.1 |
| FIM1_3949 | 2469.742861 | 1895.955865 | 1524.536322 | -0.370517999 | 0.092586904 | 4.47246530460 | 0.000124305 | ns | YPD | KS.1 |

| | | | | | | | | | | |
|---|---|---|---|---|---|---|---|---|---|---|
| FIM1_3950 | 1403.912589 | 1434.768224 | 1387.801633 | 0.02926363 | 0.089469035 | 0.738534488 | 0.792367711 | ns | YPD | KS.1 |
| FIM1_3951 | 2009.685898 | 2344.804463 | 2008.560255 | 0.21803221 | 0.067476326 | 0.001063996 | 0.002346356 | ns | YPD | KS.1 |
| FIM1_3952 | 3007.695175 | 2955.774996 | 1720.543302 | -0.019329019 | 0.279465278 | 0.937188181 | 0.954454699 | ns | YPD | KS.1 |
| FIM1_3953 | 1991.037685 | 1820.059155 | 1806.123057 | -0.122761791 | 0.116643264 | 0.27700997 | 0.347660091 | ns | YPD | KS.1 |
| FIM1_3954 | 335.0863931 | 621.2133134 | 635.473512 | 0.855189749 | 0.146507971 | 1.3598641705 | 6.9602101321 | ns | YPD | KS.1 |
| FIM1_3955 | 662.2204907 | 1066.689069 | 717.4408983 | 0.632208449 | 0.189767845 | 0.000369941 | 0.00088441 | ns | YPD | KS.1 |
| FIM1_3956 | 1161.589486 | 1372.157644 | 3147.383048 | 0.202404045 | 0.227113045 | 0.327269713 | 0.401223958 | ns | YPD | KS.1 |
| FIM1_3957 | 2382.247378 | 3575.422935 | 2442.72559 | 0.579023775 | 0.063712121 | 4.6283642659 | 6.0595032165 | ns | YPD | KS.1 |
| FIM1_3958 | 4518.711583 | 2756.441739 | 3999.83523 | -0.58589755 | 0.294092007 | 0.021954887 | 0.036962965 | ns | YPD | KS.1 |
| FIM1_3959 | 965.222766 | 950.1558987 | 783.6072442 | -0.021505936 | 0.109264954 | 0.839573051 | 0.875838945 | ns | YPD | KS.1 |
| FIM1_3960 | 157.8693869 | 190.0586328 | 230.82045 | 0.219387694 | 0.241083055 | 0.312491142 | 0.385767601 | ns | YPD | KS.1 |
| FIM1_3961 | 1033.925535 | 1087.826993 | 941.5538678 | 0.055710536 | 0.287625973 | 0.82422576 | 0.863268033 | ns | YPD | KS.1 |
| FIM1_3962 | 1328.46433 | 920.8446269 | 1344.310423 | -0.504508075 | 0.12937663 | 5.3892608904 | 0.000148048 | ns | YPD | KS.1 |
| FIM1_3963 | 3914.083062 | 1966.868911 | 1670.37721 | -0.878380526 | 0.262701007 | 0.000205915 | 0.000516866 | ns | YPD | KS.1 |
| FIM1_3964 | 3597.548217 | 3833.823906 | 2477.98014 | 0.070536276 | 0.282480415 | 0.775473863 | 0.82400309 | ns | YPD | KS.1 |
| FIM1_3965 | 386.5539884 | 490.7714369 | 578.3806888 | 0.323125432 | 0.134960324 | 0.012671254 | 0.022676076 | ns | YPD | KS.1 |
| FIM1_3966 | 93.0919732 | 1356.522259 | 404.0608576 | 3.526452797 | 0.49720371 | 1.2948928072 | 1.1524314339 | Km>KS.1 | YPD | KS.1 |
| FIM1_3967 | 803.4627663 | 1082.465959 | 1412.758355 | 0.397052685 | 0.162612407 | 0.009923481 | 0.018152136 | ns | YPD | KS.1 |
| FIM1_3968 | 1341.126099 | 1504.271089 | 1315.968476 | 0.156464335 | 0.120351875 | 0.178481768 | 0.239597085 | ns | YPD | KS.1 |
| FIM1_3969 | 863.6961291 | 2919.943995 | 2221.040879 | 1.701227837 | 0.172273788 | 2.5555603380 | 4.7263615918 | Km>KS.1 | YPD | KS.1 |
| FIM1_3970 | 742.3400814 | 1169.409646 | 906.8312738 | 0.586222239 | 0.216401698 | 0.003195723 | 0.006465523 | ns | YPD | KS.1 |
| FIM1_3971 | 376.0062269 | 532.7802049 | 324.094424 | 0.405678333 | 0.281617198 | 0.099535474 | 0.143829288 | ns | YPD | KS.1 |
| FIM1_3972 | 714.367566 | 804.7642915 | 550.1518357 | 0.166222646 | 0.098910138 | 0.084826574 | 0.124773636 | ns | YPD | KS.1 |
| FIM1_3973 | 3588.957562 | 4141.534735 | 6210.686297 | 0.19543018 | 0.118572046 | 0.088271384 | 0.129313938 | ns | YPD | KS.1 |
| FIM1_3974 | 1839.682209 | 1067.355804 | 1268.56398 | -0.753218909 | 0.13928447 | 2.0445749679 | 9.0496089551 | ns | YPD | KS.1 |
| FIM1_3975 | 2209.743278 | 1373.056537 | 1007.307858 | -0.655202469 | 0.139945705 | 1.1437065547 | 4.0013643529 | ns | YPD | KS.1 |
| FIM1_3976 | 19197.77669 | 9736.523534 | 8974.615497 | -0.931097274 | 0.168295314 | 6.9343930571 | 3.2607377560 | ns | YPD | KS.1 |
| FIM1_3977 | 7802.713337 | 1870.322247 | 1321.053844 | -1.765975445 | 0.406314303 | 4.9003018922 | 1.8098739356 | Km<KS.1 | YPD | KS.1 |
| FIM1_3978 | 634.8533602 | 982.3603841 | 1124.950898 | 0.589245492 | 0.164343261 | 0.000157857 | 0.000401296 | ns | YPD | KS.1 |
| FIM1_3979 | 193.8653343 | 591.7233512 | 703.252064 | 1.584220405 | 0.112271152 | 1.9813880430 | 1.5646675419 | Km>KS.1 | YPD | KS.1 |
| FIM1_3980 | 8754.792646 | 9454.024748 | 15468.15665 | 0.104088414 | 0.120426591 | 0.37130108 | 0.447310184 | ns | YPD | KS.1 |
| FIM1_3981 | 2360.544827 | 710.1758445 | 506.7927928 | -1.546711008 | 0.322557352 | 1.0055586926 | 4.0605961817 | Km<KS.1 | YPD | KS.1 |
| FIM1_3982 | 3697.968983 | 3231.006793 | 3579.231815 | -0.184971175 | 0.113848752 | 0.093593155 | 0.136187758 | ns | YPD | KS.1 |
| FIM1_3983 | 1385.515792 | 1587.2539 | 1834.252194 | 0.183783917 | 0.123914081 | 0.124135193 | 0.174900195 | ns | YPD | KS.1 |
| FIM1_3984 | 151.9659913 | 221.2744819 | 288.2195062 | 0.482336906 | 0.211171209 | 0.0130333 | 0.02324267 | ns | YPD | KS.1 |
| FIM1_3985 | 457.472522 | 732.7709166 | 708.1430085 | 0.667325947 | 0.088437554 | 1.7056734944 | 1.3512301966 | ns | YPD | KS.1 |
| FIM1_3986 | 431.0016004 | 1827.683372 | 1607.990702 | 2.005381098 | 0.204241539 | 2.0474882565 | 3.8150764330 | Km>KS.1 | YPD | KS.1 |
| FIM1_3987 | 972.9624558 | 799.3259172 | 787.8348508 | -0.270335263 | 0.115093224 | 0.01536237 | 0.026977689 | ns | YPD | KS.1 |
| FIM1_3988 | 816.4324277 | 992.2181416 | 1013.687429 | 0.266764407 | 0.122900183 | 0.024561972 | 0.040964066 | ns | YPD | KS.1 |
| FIM1_3989 | 5000.342483 | 11449.98533 | 10098.21849 | 1.175988929 | 0.102935052 | 4.2889890973 | 1.2777078298 | Km>KS.1 | YPD | KS.1 |
| FIM1_3990 | 1270.898613 | 2491.784107 | 1567.976856 | 0.856590362 | 0.263991046 | 0.00030728 | 0.000746444 | ns | YPD | KS.1 |
| FIM1_3991 | 1957.181377 | 970.2636016 | 1565.42951 | -0.892797351 | 0.268250657 | 0.000210863 | 0.000528219 | ns | YPD | KS.1 |
| FIM1_3992 | 976.6051938 | 667.3353372 | 725.4346143 | -0.53681237 | 0.089313713 | 9.5734833322 | 4.9768108232 | ns | YPD | KS.1 |
| FIM1_3993 | 5325.143667 | 8044.104023 | 8151.152732 | 0.591254811 | 0.048149478 | 5.0971045920 | 1.9506227188 | ns | YPD | KS.1 |
| FIM1_3994 | 269.9530217 | 72.65204414 | 83.69121574 | -1.812406512 | 0.208087633 | 1.1180920695 | 1.4485698036 | Km<KS.1 | YPD | KS.1 |

| | | | | | | | | | | |
|---|---|---|---|---|---|---|---|---|---|---|
| FIM1_3995 | 266.7810961 | 221.350227 | 185.7070297 | -0.219050783 | 0.259540978 | 0.342616726 | 0.417501575 | ns | YPD | KS.1 |
| FIM1_3996 | 2400.484342 | 1253.573478 | 1615.367025 | -0.7677969 | 0.325023521 | 0.005794555 | 0.011083395 | ns | YPD | KS.1 |
| FIM1_3997 | 1974.725572 | 2271.804584 | 1901.564487 | 0.190291545 | 0.126524289 | 0.118483348 | 0.168031543 | ns | YPD | KS.1 |
| FIM1_3998 | 881.1546643 | 1166.166782 | 1173.428708 | 0.373826055 | 0.159038384 | 0.013214234 | 0.023529283 | ns | YPD | KS.1 |
| FIM1_3999 | 87.58584397 | 91.42665482 | 239.0508598 | 0.049585877 | 0.227232322 | 0.810737471 | 0.851830817 | ns | YPD | KS.1 |
| FIM1_4000 | 668.9602322 | 742.7708318 | 866.1570562 | 0.137730149 | 0.15311761 | 0.344376468 | 0.419405858 | ns | YPD | KS.1 |
| FIM1_4001 | 91.3390612 | 100.0475792 | 118.8632034 | 0.112652582 | 0.217251684 | 0.571633073 | 0.644200073 | ns | YPD | KS.1 |
| FIM1_4002 | 3179.989163 | 4064.909119 | 4722.895656 | 0.346365012 | 0.081537694 | 1.59247794088 | 4.69347023452 | ns | YPD | KS.1 |
| FIM1_4003 | 3150.091599 | 2990.954976 | 4266.578467 | -0.071039086 | 0.106859448 | 0.494375218 | 0.57144905 | ns | YPD | KS.1 |
| FIM1_4004 | 615.4641146 | 867.8862871 | 1181.977227 | 0.481173937 | 0.099863091 | 8.46654974036 | 3.02811538161 | ns | YPD | KS.1 |
| FIM1_4005 | 3002.689535 | 3234.641548 | 2922.143101 | 0.097188432 | 0.15693193 | 0.514437538 | 0.590250634 | ns | YPD | KS.1 |
| FIM1_4006 | 1904.596608 | 2126.287126 | 2646.572787 | 0.15315341 | 0.085697472 | 0.068284199 | 0.103131114 | ns | YPD | KS.1 |
| FIM1_4007 | 1652.028745 | 1574.093806 | 1713.896372 | -0.067554516 | 0.078123285 | 0.379322561 | 0.45560834 | ns | YPD | KS.1 |
| FIM1_4008 | 739.5283525 | 331.819405 | 533.3020394 | -1.066210534 | 0.227311703 | 4.56344965496 | 1.69300238877 | Km<KS.1 | YPD | KS.1 |
| FIM1_4009 | 4863.820046 | 2117.33358 | 3342.353643 | -1.156877103 | 0.155774274 | 1.58427146045 | 1.25705749852 | Km<KS.1 | YPD | KS.1 |
| FIM1_4010 | 2948.923724 | 2031.576119 | 1854.404496 | -0.526654582 | 0.084522401 | 2.44859789103 | 1.33572088902 | ns | YPD | KS.1 |
| FIM1_4011 | 2374.652125 | 1841.282987 | 1968.013867 | -0.361739923 | 0.062045169 | 3.99608830795 | 1.92827733579 | ns | YPD | KS.1 |
| FIM1_4012 | 1985.257481 | 2112.171385 | 2395.085632 | 0.084701116 | 0.109791534 | 0.427118666 | 0.504850407 | ns | YPD | KS.1 |
| FIM1_4013 | 1316.006385 | 1832.017624 | 1792.394935 | 0.468708588 | 0.072875457 | 7.52433564007 | 4.29776920888 | ns | YPD | KS.1 |
| FIM1_4014 | 414.6326017 | 203.2428182 | 366.1548634 | -0.96556621 | 0.190010006 | 7.68291439458 | 3.16411416498 | ns | YPD | KS.1 |
| FIM1_4015 | 2442.525092 | 2082.952924 | 2846.782905 | -0.204899573 | 0.177578345 | 0.218658447 | 0.285293935 | ns | YPD | KS.1 |
| FIM1_4016 | 4084.583065 | 2950.281267 | 2731.732684 | -0.442602553 | 0.140890849 | 0.001056752 | 0.002331415 | ns | YPD | KS.1 |
| FIM1_4017 | 216.0010086 | 267.333346 | 389.0880982 | 0.27329407 | 0.19193692 | 0.125637335 | 0.17651673 | ns | YPD | KS.1 |
| FIM1_4018 | 2919.293797 | 2224.689898 | 4265.881815 | -0.294183151 | 0.316623011 | 0.274903352 | 0.345539206 | ns | YPD | KS.1 |
| FIM1_4019 | 1070.456891 | 781.9138016 | 802.5732301 | -0.435136536 | 0.111873997 | 6.41199073411 | 0.000173556 | ns | YPD | KS.1 |
| FIM1_4020 | 3027.202192 | 2997.166295 | 3839.847189 | -0.014106924 | 0.083629186 | 0.863621328 | 0.895107522 | ns | YPD | KS.1 |
| FIM1_4021 | 674.1796521 | 852.4702141 | 1069.033636 | 0.318695959 | 0.135109632 | 0.014035792 | 0.024832171 | ns | YPD | KS.1 |
| FIM1_4022 | 14.7311946 | 17.30497026 | 31.43776702 | 0.148130063 | 0.33061956 | 0.594468197 | 0.666098936 | ns | YPD | KS.1 |
| FIM1_4023 | 6163.962767 | 6903.339784 | 7102.690264 | 0.160625564 | 0.057268284 | 0.004606773 | 0.00903734 | ns | YPD | KS.1 |
| FIM1_4024 | 5263.507905 | 2299.811931 | 2905.728058 | -0.991669276 | 0.346811269 | 0.000817276 | 0.001835642 | ns | YPD | KS.1 |
| FIM1_4025 | 202.0733017 | 171.6202757 | 257.0080608 | -0.210123436 | 0.185078305 | 0.224108713 | 0.291031284 | ns | YPD | KS.1 |
| FIM1_4026 | 1953.561255 | 172.8398535 | 370.4821554 | -3.29023654 | 0.380939628 | 5.96241085821 | 8.40311445314 | Km<KS.1 | YPD | KS.1 |
| FIM1_4027 | 2645.575692 | 256.1562082 | 621.9376762 | -3.125817629 | 0.40530273 | 1.49143932372 | 1.59567970656 | Km<KS.1 | YPD | KS.1 |
| FIM1_4028 | 838.6501716 | 997.5352089 | 1191.808455 | 0.243089924 | 0.090663356 | 0.00617238 | 0.011738375 | ns | YPD | KS.1 |
| FIM1_4029 | 208.2748286 | 191.9897176 | 104.9791132 | -0.092710756 | 0.271924901 | 0.699218911 | 0.759356927 | ns | YPD | KS.1 |
| FIM1_4030 | 3930.678057 | 3268.998802 | 5795.456378 | -0.211492901 | 0.268648431 | 0.372914541 | 0.448561373 | ns | YPD | KS.1 |
| FIM1_4031 | 1219.450631 | 2232.038759 | 3294.908715 | 0.843442776 | 0.129425286 | 1.87192320528 | 1.13709620833 | ns | YPD | KS.1 |
| FIM1_4033 | 724.6592626 | 596.2284313 | 824.2126454 | -0.264607398 | 0.132204797 | 0.037282661 | 0.05960194 | ns | YPD | KS.1 |
| FIM1_4034 | 274.9258615 | 336.0984165 | 554.5582895 | 0.264214802 | 0.160697055 | 0.082436889 | 0.121698375 | ns | YPD | KS.1 |
| FIM1_4035 | 127.0248341 | 217.1020202 | 436.1044789 | 0.711928642 | 0.192143798 | 7.57146874472 | 0.000201326 | ns | YPD | KS.1 |
| FIM1_4036 | 1649.35673 | 1600.988442 | 1895.044124 | -0.039618679 | 0.139588055 | 0.767233071 | 0.817516498 | ns | YPD | KS.1 |
| FIM1_4037 | 1070.533196 | 1373.977545 | 1354.565124 | 0.345613692 | 0.110708282 | 0.001324859 | 0.002865728 | ns | YPD | KS.1 |
| FIM1_4038 | 768.7533866 | 713.2322834 | 775.7880711 | -0.101346232 | 0.114090437 | 0.359533888 | 0.434673412 | ns | YPD | KS.1 |
| FIM1_4039 | 2464.057697 | 814.8430103 | 1193.073608 | -1.554736286 | 0.149172036 | 1.29392228223 | 2.84834661686 | Km<KS.1 | YPD | KS.1 |
| FIM1_4041 | 1749.738533 | 605.4144176 | 1151.064925 | -1.338589536 | 0.330985482 | 4.89920791294 | 1.55542816636 | Km<KS.1 | YPD | KS.1 |

| | | | | | | | | | | |
|---|---|---|---|---|---|---|---|---|---|---|
| FIM1_4042 | 2674.211757 | 1625.682146 | 1666.45966 | -0.697123952 | 0.111415708 | 1.41055602121 | 7.89371901633 | ns | YPD | KS.1 |
| FIM1_4043 | 2368.742386 | 1736.913666 | 1573.467723 | -0.415209516 | 0.157725305 | 0.005582873 | 0.01071972 | ns | YPD | KS.1 |
| FIM1_4044 | 316.2987707 | 355.8779593 | 280.2021216 | 0.127093538 | 0.304422748 | 0.627404789 | 0.695330547 | ns | YPD | KS.1 |
| FIM1_4045 | 2847.888081 | 3482.957594 | 4361.899764 | 0.264782244 | 0.163799129 | 0.087186944 | 0.127913609 | ns | YPD | KS.1 |
| FIM1_4046 | 581.3703629 | 840.7868838 | 1270.611571 | 0.488574894 | 0.17727974 | 0.003372689 | 0.006795921 | ns | YPD | KS.1 |
| FIM1_4047 | 4796.59037 | 9102.994122 | 7888.412633 | 0.894169469 | 0.13245808 | 3.45331157846 | 2.23992504600 | ns | YPD | KS.1 |
| FIM1_4048 | 77.87402465 | 333.9561147 | 405.4900098 | 2.01610924 | 0.217795163 | 4.48591944345 | 6.82490802176 | Km>KS.1 | YPD | KS.1 |
| FIM1_4049 | 585.5247772 | 510.9399597 | 1563.26928 | -0.142289414 | 0.322744899 | 0.602796929 | 0.67316535 | ns | YPD | KS.1 |
| FIM1_4050 | 3546.528038 | 5160.043408 | 3856.464842 | 0.531184393 | 0.079906385 | 1.55358224625 | 9.51856117627 | ns | YPD | KS.1 |
| FIM1_4051 | 6807.66373 | 5301.8253 | 8292.48385 | -0.345343369 | 0.11464561 | 0.001911809 | 0.004047341 | ns | YPD | KS.1 |
| FIM1_4052 | 108.8493608 | 116.0727762 | 75.04869013 | 0.07764427 | 0.191961236 | 0.664179758 | 0.72765785 | ns | YPD | KS.1 |
| FIM1_4053 | 1163.954113 | 1890.879279 | 749.4921098 | 0.524733445 | 0.353345526 | 0.073076454 | 0.109537619 | ns | YPD | KS.1 |
| FIM1_4054 | 5457.516168 | 4047.19441 | 2907.120408 | -0.395875236 | 0.168200896 | 0.012644598 | 0.022636515 | ns | YPD | KS.1 |
| FIM1_4055 | 2528.675764 | 2204.459139 | 1475.51348 | -0.170263473 | 0.207064732 | 0.372470768 | 0.448244332 | ns | YPD | KS.1 |
| FIM1_4056 | 3949.241382 | 2514.996326 | 1623.584067 | -0.62110636 | 0.137663712 | 2.79507134557 | 9.19066751107 | ns | YPD | KS.1 |
| FIM1_4057 | 1476.873536 | 1201.692764 | 1189.612241 | -0.286067761 | 0.106082528 | 0.005598668 | 0.0107459 | ns | YPD | KS.1 |
| FIM1_4058 | 69.17322675 | 348.3913282 | 385.20337 | 2.264860938 | 0.177348694 | 2.24355870530 | 1.11617045588 | Km>KS.1 | YPD | KS.1 |
| FIM1_4059 | 13821.03279 | 6058.97723 | 4790.05668 | -1.039777325 | 0.296981089 | 8.10413735451 | 0.000214344 | Km<KS.1 | YPD | KS.1 |
| FIM1_4060 | 1262.059815 | 1265.626602 | 2445.498945 | 0.00334814 | 0.244194717 | 0.987855912 | 0.991043186 | ns | YPD | KS.1 |
| FIM1_4061 | 4282.736732 | 2433.334649 | 5649.129174 | -0.692462475 | 0.280874909 | 0.005164008 | 0.010000367 | ns | YPD | KS.1 |
| FIM1_4062 | 874.8614931 | 535.254817 | 1445.784866 | -0.562493168 | 0.318337328 | 0.038302942 | 0.061017335 | ns | YPD | KS.1 |
| FIM1_4063 | 3491.594051 | 2890.616044 | 2330.080871 | -0.25128889 | 0.15102567 | 0.080394654 | 0.11893054 | ns | YPD | KS.1 |
| FIM1_4064 | 3069.728234 | 2997.082094 | 5527.055317 | -0.03057955 | 0.182289016 | 0.85796187 | 0.889797853 | ns | YPD | KS.1 |
| FIM1_4065 | 2565.755508 | 2672.966079 | 4374.277099 | 0.045431403 | 0.281468738 | 0.853854593 | 0.886832275 | ns | YPD | KS.1 |
| FIM1_4066 | 11619.88248 | 4024.627488 | 3727.910755 | -1.504155794 | 0.116673559 | 3.61825623589 | 1.74765289063 | Km<KS.1 | YPD | KS.1 |
| FIM1_4067 | 2848.376384 | 1229.62728 | 1068.184694 | -1.192195889 | 0.10443452 | 4.61239900171 | 1.36587410913 | Km<KS.1 | YPD | KS.1 |
| FIM1_4068 | 785.4817262 | 293.3768372 | 406.8654862 | -1.362668455 | 0.179947739 | 3.54365295882 | 2.99315339051 | Km<KS.1 | YPD | KS.1 |
| FIM1_4069 | 3900.541367 | 4486.114749 | 2644.230048 | 0.164790923 | 0.246167106 | 0.456196358 | 0.533515957 | ns | YPD | KS.1 |
| FIM1_4070 | 3681.330975 | 3703.547932 | 2799.421974 | 0.006988902 | 0.248859601 | 0.975046325 | 0.982551239 | ns | YPD | KS.1 |
| FIM1_4071 | 257.0499664 | 127.6209435 | 102.4320302 | -0.90222943 | 0.256585481 | 0.000103371 | 0.000269535 | ns | YPD | KS.1 |
| FIM1_4072 | 1342.572058 | 1765.637168 | 1604.04108 | 0.380184638 | 0.108312807 | 0.0003136 | 0.000760311 | ns | YPD | KS.1 |
| FIM1_4073 | 1228.157223 | 779.9762181 | 1025.178863 | -0.631122246 | 0.120670262 | 7.14897440546 | 2.95399897568 | ns | YPD | KS.1 |
| FIM1_4074 | 769.8288638 | 1894.551666 | 1295.36021 | 1.119164 | 0.32364569 | 7.99243992956 | 0.000211727 | Km>KS.1 | YPD | KS.1 |
| FIM1_4075 | 737.3449359 | 1860.059437 | 728.1756904 | 1.208349609 | 0.269214191 | 8.97287288534 | 3.19771078829 | Km>KS.1 | YPD | KS.1 |
| FIM1_4076 | 53.49234505 | 21.9323961 | 18.61852671 | -1.058637716 | 0.366841054 | 0.000644933 | 0.001484748 | Km<KS.1 | YPD | KS.1 |
| FIM1_4077 | 69660.02309 | 39256.27338 | 36444.03178 | -0.69305894 | 0.29347995 | 0.006858369 | 0.012914605 | ns | YPD | KS.1 |
| FIM1_4078 | 774.583581 | 591.1830993 | 253.8581896 | -0.261476496 | 0.379521376 | 0.392836713 | 0.469971146 | ns | YPD | KS.1 |
| FIM1_4080 | 11087.2367 | 9574.640049 | 3950.898547 | -0.164188916 | 0.281411764 | 0.505161347 | 0.581619464 | ns | YPD | KS.1 |
| FIM1_4081 | 69.46454422 | 27.66445919 | 26.3944966 | -1.201136688 | 0.270791324 | 1.16351276179 | 4.05639522771 | Km<KS.1 | YPD | KS.1 |
| FIM1_4082 | 196.1548312 | 67.2344887 | 53.98004836 | -1.321831371 | 0.356750021 | 2.00373182887 | 5.83298177216 | Km<KS.1 | YPD | KS.1 |
| FIM1_4083 | 1064.359963 | 359.4183946 | 621.9285562 | -1.52509644 | 0.149987857 | 1.93507951297 | 3.86627332412 | Km<KS.1 | YPD | KS.1 |
| FIM1_4084 | 1906.34823 | 2643.568804 | 3451.86931 | 0.443502204 | 0.145065216 | 0.001401468 | 0.003018314 | ns | YPD | KS.1 |
| FIM1_4085 | 5943.38963 | 10488.75853 | 9779.713209 | 0.79692669 | 0.115308568 | 1.37572197136 | 9.36281369021 | ns | YPD | KS.1 |
| FIM1_4086 | 7354.349231 | 6848.24278 | 11459.64232 | -0.087037533 | 0.214887317 | 0.659186577 | 0.723354741 | ns | YPD | KS.1 |
| FIM1_4087 | 355.1026193 | 311.1483816 | 342.0848434 | -0.150178892 | 0.272936084 | 0.532394303 | 0.607909492 | ns | YPD | KS.1 |

| | | | | | | | | | | |
|---|---|---|---|---|---|---|---|---|---|---|
| FIM1_4088 | 1843.363686 | 3918.137765 | 2166.353819 | 0.964830943 | 0.27019654 | 7.3327300703( | 0.000195395 | ns | YPD | KS.1 |
| FIM1_4089 | 639.2324703 | 250.4051942 | 243.3551311 | -1.31074366 | 0.152304276 | 8.1010642383: | 9.5278474197( | Km<KS.1 | YPD | KS.1 |
| FIM1_4090 | 414.0837282 | 260.6770159 | 225.2102565 | -0.645442548 | 0.112231597 | 3.6048883979: | 1.7479843839( | ns | YPD | KS.1 |
| FIM1_4091 | 1821.835415 | 1130.043887 | 2847.784754 | -0.536139516 | 0.328550211 | 0.053603283 | 0.083362405 | ns | YPD | KS.1 |
| FIM1_4092 | 1381.59799 | 1094.556866 | 1310.075233 | -0.324259676 | 0.100648467 | 0.000968521 | 0.00215299 | ns | YPD | KS.1 |
| FIM1_4093 | 2951.823647 | 2870.326753 | 1490.982194 | -0.031033349 | 0.280019657 | 0.899471092 | 0.923605507 | ns | YPD | KS.1 |
| FIM1_4094 | 22130.90062 | 6189.265972 | 6504.902137 | -1.703037509 | 0.272612657 | 1.9364832374: | 1.1691752556( | Km<KS.1 | YPD | KS.1 |
| FIM1_4095 | 1984.73054 | 1165.34038 | 1717.111691 | -0.723284966 | 0.166426167 | 4.8113068627( | 1.5294729483( | ns | YPD | KS.1 |
| FIM1_4096 | 759.4677775 | 1041.960782 | 1803.497877 | 0.421601666 | 0.162424324 | 0.006148864 | 0.011702602 | ns | YPD | KS.1 |
| FIM1_4097 | 1137.104235 | 1204.460753 | 1921.574245 | 0.059135911 | 0.329563036 | 0.83135733 | 0.870004778 | ns | YPD | KS.1 |
| FIM1_4098 | 860.9768334 | 1252.393209 | 1806.243203 | 0.472550636 | 0.224993475 | 0.021184305 | 0.035798885 | ns | YPD | KS.1 |
| FIM1_4099 | 37.58375242 | 259.5703868 | 365.0170713 | 2.697030217 | 0.226963642 | 3.4744992188( | 1.3504401260( | Km>KS.1 | YPD | KS.1 |
| FIM1_4100 | 518.4103353 | 521.7710121 | 354.4049164 | 0.007657967 | 0.1084358 | 0.9421174 | 0.958297703 | ns | YPD | KS.1 |
| FIM1_4101 | 1671.639063 | 1042.843475 | 806.4622174 | -0.621123645 | 0.19681543 | 0.000702539 | 0.001604006 | ns | YPD | KS.1 |
| FIM1_4102 | 3124.36178 | 2679.089771 | 2062.510701 | -0.207977839 | 0.129094044 | 0.094058569 | 0.136744997 | ns | YPD | KS.1 |
| FIM1_4103 | 1098.359415 | 2484.655072 | 4905.46933 | 1.10288782 | 0.206871124 | 1.5245207207( | 6.8452081099: | Km>KS.1 | YPD | KS.1 |
| FIM1_4104 | 3006.998213 | 1118.996374 | 1512.017908 | -1.310362054 | 0.2559839 | 3.1781498531( | 1.3784913269( | Km<KS.1 | YPD | KS.1 |
| FIM1_4105 | 61.67042728 | 142.5103763 | 221.2919914 | 1.085872933 | 0.266402026 | 7.3211927492( | 2.2665173570: | Km>KS.1 | YPD | KS.1 |
| FIM1_4106 | 7465.416692 | 8173.866031 | 8180.424686 | 0.107518011 | 0.237785705 | 0.617050805 | 0.686148358 | ns | YPD | KS.1 |
| FIM1_4107 | 1006.631277 | 1037.312115 | 1193.939476 | 0.041380249 | 0.103746192 | 0.682276165 | 0.744532555 | ns | YPD | KS.1 |
| FIM1_4108 | 1443.643266 | 2052.362389 | 2811.311319 | 0.497431343 | 0.083041062 | 1.1800755284: | 6.0586953087! | ns | YPD | KS.1 |
| FIM1_4109 | 278.2052649 | 257.3276437 | 326.6748471 | -0.096468603 | 0.198722856 | 0.600765776 | 0.671340911 | ns | YPD | KS.1 |
| FIM1_4110 | 655.9937088 | 858.7666248 | 929.8709923 | 0.365400268 | 0.140714009 | 0.006725282 | 0.012697639 | ns | YPD | KS.1 |
| FIM1_4111 | 2135.322651 | 2535.706391 | 2452.532085 | 0.244783922 | 0.059262969 | 3.0311826574( | 8.6122979558( | ns | YPD | KS.1 |
| FIM1_4112 | 1850.255364 | 3566.876589 | 2371.194331 | 0.855856143 | 0.234367283 | 6.8036584577! | 0.000182864 | ns | YPD | KS.1 |
| FIM1_4113 | 4858.717562 | 1482.614249 | 1778.14431 | -1.53924835 | 0.311055606 | 4.7466714541: | 2.0029423651( | Km<KS.1 | YPD | KS.1 |
| FIM1_4114 | 1978.677718 | 1352.451956 | 2048.426331 | -0.389193189 | 0.366255808 | 0.19343057 | 0.257166511 | ns | YPD | KS.1 |
| FIM1_4115 | 921.5297279 | 1701.491941 | 1095.168122 | 0.823035173 | 0.193099777 | 5.6468471722 | 1.7757942276( | ns | YPD | KS.1 |
| FIM1_4116 | 169.5524595 | 292.3228448 | 254.7718731 | 0.617888724 | 0.334737519 | 0.028684676 | 0.047097777 | ns | YPD | KS.1 |
| FIM1_4117 | 318.7378242 | 163.6916362 | 180.9317383 | -0.769466827 | 0.346337194 | 0.008347786 | 0.015472329 | ns | YPD | KS.1 |
| FIM1_4118 | 3233.896276 | 2956.138272 | 2095.015767 | -0.105111783 | 0.247049543 | 0.635820911 | 0.702467029 | ns | YPD | KS.1 |
| FIM1_4119 | 1319.204634 | 1877.622829 | 1782.112594 | 0.480936612 | 0.141214512 | 0.000386427 | 0.000919405 | ns | YPD | KS.1 |
| FIM1_4120 | 1449.589827 | 1927.062015 | 2779.77814 | 0.300171377 | 0.334472548 | 0.284572257 | 0.356214868 | ns | YPD | KS.1 |
| FIM1_4121 | 1194.06197 | 578.5177201 | 988.6484663 | -0.910601951 | 0.284332111 | 0.000315881 | 0.00076547 | ns | YPD | KS.1 |
| FIM1_4122 | 600.2342805 | 834.9203901 | 644.6217503 | 0.445855627 | 0.145793377 | 0.001391449 | 0.002998033 | ns | YPD | KS.1 |
| FIM1_4123 | 1409.523936 | 1407.662585 | 1041.902857 | -0.001570295 | 0.152862764 | 0.991396499 | 0.993593389 | ns | YPD | KS.1 |
| FIM1_4124 | 56.25274122 | 80.01080639 | 61.31069651 | 0.413810611 | 0.263732751 | 0.076642381 | 0.114057986 | ns | YPD | KS.1 |
| FIM1_4125 | 6854.7012 | 10249.22469 | 7126.221652 | 0.529460111 | 0.187814916 | 0.002561781 | 0.005270827 | ns | YPD | KS.1 |
| FIM1_4126 | 735.0961349 | 409.3486729 | 369.1405331 | -0.806880867 | 0.148922142 | 1.7164663733: | 7.6586728316: | ns | YPD | KS.1 |
| FIM1_4127 | 132.5639972 | 68.71418263 | 56.92117253 | -0.818030537 | 0.28466989 | 0.001157229 | 0.002533985 | ns | YPD | KS.1 |
| FIM1_4128 | 796.2046343 | 1078.007936 | 1315.209878 | 0.411966572 | 0.141639437 | 0.002406953 | 0.00497697 | ns | YPD | KS.1 |
| FIM1_4129 | 569.4707042 | 1284.262807 | 2444.613147 | 1.126865942 | 0.162085853 | 5.3999651710! | 3.7784566422: | Km>KS.1 | YPD | KS.1 |
| FIM1_4130 | 4769.449486 | 3182.171424 | 2108.407292 | -0.530600421 | 0.192004057 | 0.003034642 | 0.006167215 | ns | YPD | KS.1 |
| FIM1_4132 | 167.3392031 | 242.0459046 | 198.7253864 | 0.508180553 | 0.128280946 | 4.1343285178: | 0.000115552 | ns | YPD | KS.1 |
| FIM1_4133 | 1684.513626 | 830.1654133 | 883.5089162 | -0.983037903 | 0.147684663 | 5.5323632395: | 3.5377258504( | ns | YPD | KS.1 |

| | | | | | | | | | | |
|---|---|---|---|---|---|---|---|---|---|---|
| FIM1_4134 | 1621.736043 | 1340.939306 | 1743.048971 | -0.261140055 | 0.115272582 | 0.019379574 | 0.0332002 | ns | YPD | KS.1 |
| FIM1_4135 | 2539.475875 | 1697.605675 | 1187.428661 | -0.547830807 | 0.148678149 | 0.000117214 | 0.000302458 | ns | YPD | KS.1 |
| FIM1_4136 | 2790.499865 | 2159.488576 | 2361.138264 | -0.356704135 | 0.104710129 | 0.00047733 | 0.001118565 | ns | YPD | KS.1 |
| FIM1_4137 | 4751.427481 | 3926.802748 | 3197.217042 | -0.241796785 | 0.19447801 | 0.180516712 | 0.242067558 | ns | YPD | KS.1 |
| FIM1_4138 | 1513.703726 | 1462.547984 | 1404.198415 | -0.046992407 | 0.0987011 | 0.625779487 | 0.694147814 | ns | YPD | KS.1 |
| FIM1_4139 | 2232.666331 | 1941.343752 | 3528.830738 | -0.16550153 | 0.242390302 | 0.448447178 | 0.525813036 | ns | YPD | KS.1 |
| FIM1_4140 | 376.8321007 | 1158.077847 | 1085.210523 | 1.510050396 | 0.244738722 | 4.83054740687 | 2.83062112475 | Km>KS.1 | YPD | KS.1 |
| FIM1_4141 | 2874.725057 | 6268.034419 | 8667.610626 | 1.087245575 | 0.145033836 | 1.05946611333 | 8.50136115136 | Km>KS.1 | YPD | KS.1 |
| FIM1_4142 | 925.1718155 | 1107.423568 | 850.6829663 | 0.246885252 | 0.108265162 | 0.018949297 | 0.032586503 | ns | YPD | KS.1 |
| FIM1_4143 | 873.0509116 | 768.7100939 | 875.8348652 | -0.174485402 | 0.11129105 | 0.106050078 | 0.152089691 | ns | YPD | KS.1 |
| FIM1_4144 | 211.4391251 | 246.0611788 | 395.2838946 | 0.193299847 | 0.18484722 | 0.262974098 | 0.333039931 | ns | YPD | KS.1 |
| FIM1_4145 | 725.8016487 | 512.5559358 | 511.0689433 | -0.460412827 | 0.175093029 | 0.0051948 | 0.010046039 | ns | YPD | KS.1 |
| FIM1_4146 | 766.4217287 | 759.9051001 | 1621.817164 | -0.008211041 | 0.357432304 | 0.977709399 | 0.984238013 | ns | YPD | KS.1 |
| FIM1_4147 | 505.0312496 | 1018.094677 | 825.2580876 | 0.947070337 | 0.192491465 | 1.84627302274 | 7.22675711105 | ns | YPD | KS.1 |
| FIM1_4148 | 8003.487306 | 5779.240923 | 3486.418128 | -0.381073229 | 0.274332931 | 0.114115155 | 0.16262472 | ns | YPD | KS.1 |
| FIM1_4149 | 5714.822741 | 7688.257644 | 12080.15282 | 0.411776715 | 0.110038323 | 0.000121323 | 0.00031209 | ns | YPD | KS.1 |
| FIM1_4150 | 1936.87068 | 2251.59496 | 2153.571342 | 0.199905853 | 0.149709546 | 0.160901065 | 0.218651406 | ns | YPD | KS.1 |
| FIM1_4151 | 5366.73311 | 4872.712501 | 3026.452796 | -0.129601371 | 0.132875173 | 0.310113495 | 0.383307985 | ns | YPD | KS.1 |
| FIM1_4152 | 1246.168592 | 676.9771188 | 1951.519265 | -0.697099737 | 0.342218313 | 0.015522118 | 0.027229385 | ns | YPD | KS.1 |
| FIM1_4153 | 2116.74532 | 2486.894129 | 1687.778482 | 0.211775864 | 0.160802438 | 0.164206128 | 0.22265617 | ns | YPD | KS.1 |
| FIM1_4154 | 471.1637379 | 792.3653786 | 796.973279 | 0.678310879 | 0.211781485 | 0.000527457 | 0.001230816 | ns | YPD | KS.1 |
| FIM1_4155 | 730.0920363 | 2193.143148 | 3767.699817 | 1.494834666 | 0.224629774 | 2.09430481804 | 1.40042560077 | Km>KS.1 | YPD | KS.1 |
| FIM1_4156 | 2127.122228 | 2111.597573 | 3625.250613 | -0.008227226 | 0.266348856 | 0.972202577 | 0.980678796 | ns | YPD | KS.1 |
| FIM1_4157 | 5226.153683 | 2820.263003 | 3012.997044 | -0.824213617 | 0.199125311 | 9.71286954022 | 2.95183420663 | ns | YPD | KS.1 |
| FIM1_4158 | 3662.571439 | 5188.664035 | 4488.575145 | 0.451776069 | 0.195171343 | 0.012722419 | 0.022759451 | ns | YPD | KS.1 |
| FIM1_4159 | 1492.235896 | 1241.169636 | 1271.519633 | -0.253714802 | 0.11009328 | 0.017638268 | 0.03051126 | ns | YPD | KS.1 |
| FIM1_4160 | 319.3469549 | 314.9498007 | 355.4111926 | -0.018759029 | 0.149584526 | 0.895440728 | 0.920226734 | ns | YPD | KS.1 |
| FIM1_4161 | 14741.99457 | 12341.78428 | 15530.23963 | -0.247288599 | 0.097176888 | 0.009163049 | 0.016865027 | ns | YPD | KS.1 |
| FIM1_4162 | 1196.6921 | 1161.312813 | 1415.571465 | -0.030571253 | 0.330852423 | 0.912583242 | 0.934561883 | ns | YPD | KS.1 |
| FIM1_4163 | 7985.684459 | 4687.908302 | 7457.914414 | -0.741398289 | 0.127776416 | 2.14057726668 | 1.06387331686 | ns | YPD | KS.1 |
| FIM1_4164 | 464.0051261 | 59.36251134 | 109.5779923 | -2.714675853 | 0.393523442 | 1.46067494191 | 1.16083990990 | Km<KS.1 | YPD | KS.1 |
| FIM1_4165 | 363.2374515 | 220.1237948 | 424.6697933 | -0.67642864 | 0.171396916 | 3.06620920845 | 8.70684407082 | ns | YPD | KS.1 |
| FIM1_4166 | 9444.643608 | 4057.511165 | 2248.026681 | -1.10156706 | 0.261140674 | 3.81567033055 | 1.23106095295 | Km<KS.1 | YPD | KS.1 |
| FIM1_4167 | 1189.617923 | 1724.239205 | 1085.647549 | 0.413501137 | 0.314509103 | 0.123178193 | 0.173797933 | ns | YPD | KS.1 |
| FIM1_4169 | 556.4179441 | 930.4484729 | 672.9160416 | 0.72745785 | 0.091526064 | 6.24453638592 | 5.70028780183 | ns | YPD | KS.1 |
| FIM1_4170 | 225.319734 | 559.3332399 | 738.4157999 | 1.25773261 | 0.167839261 | 7.90595906165 | 6.41633708510 | Km>KS.1 | YPD | KS.1 |
| FIM1_4171 | 78.48245721 | 24.02103982 | 117.2484734 | -1.51834345 | 0.330050354 | 2.74579231013 | 1.05403678571 | Km<KS.1 | YPD | KS.1 |
| FIM1_4172 | 2616.087609 | 1550.700812 | 1125.987572 | -0.664686254 | 0.240765968 | 0.002309382 | 0.004793148 | ns | YPD | KS.1 |
| FIM1_4173 | 2056.042573 | 867.2559454 | 1128.570396 | -1.21938407 | 0.118972452 | 1.50673866406 | 3.04716457467 | Km<KS.1 | YPD | KS.1 |
| FIM1_4174 | 4135.704153 | 4203.43632 | 5769.751469 | 0.018909013 | 0.249477901 | 0.932691419 | 0.95102216 | ns | YPD | KS.1 |
| FIM1_4175 | 1084.624276 | 1646.335083 | 1627.710598 | 0.576985391 | 0.126354166 | 2.37584391251 | 7.88513906923 | ns | YPD | KS.1 |
| FIM1_4176 | 748.5903362 | 1418.270151 | 1316.289622 | 0.902269984 | 0.102133864 | 2.29159447384 | 2.82895347577 | ns | YPD | KS.1 |
| FIM1_4177 | 3151.454551 | 3615.635692 | 6783.080192 | 0.179409805 | 0.163660835 | 0.246309702 | 0.31501048 | ns | YPD | KS.1 |
| FIM1_4178 | 2495.225595 | 3440.530481 | 6196.614027 | 0.433497688 | 0.150265926 | 0.002499898 | 0.005160577 | ns | YPD | KS.1 |
| FIM1_4180 | 23.4845806 | 19.52110017 | 24.9312206 | -0.210875976 | 0.304530745 | 0.42046835 | 0.498054772 | ns | YPD | KS.1 |

| | | | | | | | | | | |
|---|---|---|---|---|---|---|---|---|---|---|
| FIM1_4181 | 11656.59436 | 4029.764652 | 3689.600522 | -1.491177397 | 0.14933929 | 1.33505050734 | 2.51586222501 | Km<KS.1 | YPD | KS.1 |
| FIM1_4182 | 4886.569897 | 6968.822074 | 8869.187302 | 0.502476827 | 0.079957539 | 1.83163049557 | 1.01023965803 | ns | YPD | KS.1 |
| FIM1_4183 | 1192.953804 | 1446.524559 | 1556.276746 | 0.266484982 | 0.106023106 | 0.009794201 | 0.01795363 | ns | YPD | KS.1 |
| FIM1_4184 | 950.2716361 | 2455.781788 | 2060.641967 | 1.352347858 | 0.095652672 | 2.14711238126 | 1.66904439012 | Km>KS.1 | YPD | KS.1 |
| FIM1_4185 | 603.8564495 | 589.8805317 | 760.5632707 | -0.032097939 | 0.120786441 | 0.783543382 | 0.830379381 | ns | YPD | KS.1 |
| FIM1_4186 | 1237.462399 | 2144.65829 | 2235.521867 | 0.706731261 | 0.23393249 | 0.000918577 | 0.002046715 | ns | YPD | KS.1 |
| FIM1_4187 | 840.0335017 | 1109.06862 | 693.7005976 | 0.37798208 | 0.136371305 | 0.003908332 | 0.007765157 | ns | YPD | KS.1 |
| FIM1_4188 | 932.6884985 | 1523.812135 | 1297.856177 | 0.665562335 | 0.164439085 | 2.04901625306 | 5.95087907705 | ns | YPD | KS.1 |
| FIM1_4189 | 3809.7444 | 4114.31609 | 3247.958278 | 0.100022369 | 0.162932734 | 0.516775063 | 0.592522687 | ns | YPD | KS.1 |
| FIM1_4190 | 1098.143398 | 1428.478151 | 1947.08494 | 0.362359773 | 0.114877961 | 0.001156696 | 0.002533933 | ns | YPD | KS.1 |
| FIM1_4191 | 2007.030359 | 2406.102314 | 2019.30619 | 0.24418092 | 0.135618619 | 0.060515616 | 0.093095498 | ns | YPD | KS.1 |
| FIM1_4192 | 487.5095204 | 430.9491594 | 923.868796 | -0.149141487 | 0.222524065 | 0.462957971 | 0.540661011 | ns | YPD | KS.1 |
| FIM1_4193 | 134.0300821 | 286.2836586 | 265.2306088 | 1.006569225 | 0.224654381 | 1.40730205232 | 4.85192495517 | Km>KS.1 | YPD | KS.1 |
| FIM1_4194 | 868.8595104 | 1464.201805 | 1161.170013 | 0.701461524 | 0.179204208 | 3.31846743822 | 9.36967962835 | ns | YPD | KS.1 |
| FIM1_4195 | 2682.499208 | 2193.013982 | 2259.970138 | -0.27688794 | 0.11541285 | 0.013296962 | 0.023634293 | ns | YPD | KS.1 |
| FIM1_4196 | 10202.50016 | 5128.34601 | 3731.145569 | -0.933604622 | 0.185861084 | 1.11465243607 | 4.47209344311 | ns | YPD | KS.1 |
| FIM1_4197 | 665.98707 | 1573.561434 | 1354.845331 | 1.115318324 | 0.269173928 | 5.14642972700 | 1.62872060380 | Km>KS.1 | YPD | KS.1 |
| FIM1_4198 | 2604.630092 | 2725.842493 | 1718.026555 | 0.055448495 | 0.215312229 | 0.779160629 | 0.82686095 | ns | YPD | KS.1 |
| FIM1_4199 | 898.6557628 | 561.5155772 | 370.175666 | -0.613994486 | 0.205026993 | 0.0012255 | 0.002667043 | ns | YPD | KS.1 |
| FIM1_4201 | 725.3260734 | 584.7707433 | 544.8274497 | -0.304004168 | 0.073765884 | 2.94498733589 | 8.38175743482 | ns | YPD | KS.1 |
| FIM1_4203 | 268.0004143 | 256.114075 | 324.5406144 | -0.05713247 | 0.215377855 | 0.772690849 | 0.821923663 | ns | YPD | KS.1 |
| FIM1_4204 | 2370.199821 | 957.5732542 | 1148.355635 | -1.271956326 | 0.139784872 | 1.04595344192 | 1.44947587007 | Km<KS.1 | YPD | KS.1 |
| FIM1_4205 | 14653.9365 | 7700.087109 | 3643.20249 | -0.73731355 | 0.346766458 | 0.01141561 | 0.020629372 | ns | YPD | KS.1 |
| FIM1_4206 | 1366.041259 | 1791.012239 | 1485.64639 | 0.326931056 | 0.243953856 | 0.135826077 | 0.188858227 | ns | YPD | KS.1 |
| FIM1_4207 | 1624.951888 | 2379.043384 | 2188.568411 | 0.504147927 | 0.178986218 | 0.002713527 | 0.00555776 | ns | YPD | KS.1 |
| FIM1_4208 | 912.2570453 | 1800.820263 | 1001.676204 | 0.916070873 | 0.19550794 | 6.36128803572 | 2.32189346865 | ns | YPD | KS.1 |
| FIM1_4209 | 825.8746067 | 866.2341183 | 664.3409912 | 0.06303789 | 0.135409885 | 0.627943883 | 0.69575477 | ns | YPD | KS.1 |
| FIM1_4210 | 2309.751805 | 2291.015595 | 2048.586944 | -0.010524764 | 0.159346609 | 0.944537118 | 0.959973884 | ns | YPD | KS.1 |
| FIM1_4211 | 396.7088921 | 578.0905376 | 725.4670799 | 0.48123603 | 0.214103219 | 0.014362346 | 0.025346814 | ns | YPD | KS.1 |
| FIM1_4212 | 3882.434072 | 4254.302013 | 3234.851481 | 0.126952903 | 0.089697695 | 0.148211385 | 0.203632047 | ns | YPD | KS.1 |
| FIM1_4213 | 743.0639976 | 502.0944937 | 423.953216 | -0.530766263 | 0.156574281 | 0.00037092 | 0.000886324 | ns | YPD | KS.1 |
| FIM1_4214 | 1672.857617 | 1036.831141 | 1128.506859 | -0.665374441 | 0.124465897 | 3.51170614010 | 1.51524180806 | ns | YPD | KS.1 |
| FIM1_4215 | 11531.11255 | 8065.483804 | 5631.325258 | -0.427406103 | 0.264128557 | 0.067816057 | 0.102610975 | ns | YPD | KS.1 |
| FIM1_4216 | 1177.339759 | 977.8687848 | 743.7061317 | -0.23217286 | 0.205280989 | 0.220108271 | 0.286659332 | ns | YPD | KS.1 |
| FIM1_4217 | 1691.772021 | 1295.440132 | 889.1683139 | -0.336874392 | 0.208380411 | 0.078937567 | 0.117053471 | ns | YPD | KS.1 |
| FIM1_4218 | 64.17997741 | 44.66158425 | 40.80047592 | -0.461094765 | 0.213958727 | 0.018886329 | 0.032500688 | ns | YPD | KS.1 |
| FIM1_4219 | 567.4837004 | 435.2249496 | 472.1011329 | -0.350389865 | 0.168734756 | 0.027655824 | 0.045619271 | ns | YPD | KS.1 |
| FIM1_4220 | 638.3914171 | 680.6471463 | 752.1887105 | 0.08434802 | 0.157820324 | 0.573574275 | 0.646180258 | ns | YPD | KS.1 |
| FIM1_4221 | 74.99209809 | 63.68477918 | 215.5270907 | -0.203667756 | 0.202054397 | 0.275562021 | 0.34608905 | ns | YPD | KS.1 |
| FIM1_4222 | 714.8267573 | 1787.910355 | 1521.313095 | 1.185286945 | 0.28088746 | 3.18487045088 | 1.03695880190 | Km>KS.1 | YPD | KS.1 |
| FIM1_4223 | 1226.539464 | 2573.438234 | 2560.189644 | 1.02774751 | 0.153001295 | 3.34981721995 | 2.18132731273 | Km>KS.1 | YPD | KS.1 |
| FIM1_4224 | 1847.445 | 2532.770597 | 5080.741052 | 0.411727796 | 0.18395828 | 0.016688171 | 0.029043363 | ns | YPD | KS.1 |
| FIM1_4225 | 607.803868 | 1354.502627 | 581.9715583 | 0.913387681 | 0.381080128 | 0.003773377 | 0.007530105 | ns | YPD | KS.1 |
| FIM1_4226 | 99071.59371 | 46779.39684 | 38582.63717 | -0.991795982 | 0.231145301 | 3.46892184161 | 1.12429225811 | ns | YPD | KS.1 |
| FIM1_4227 | 6430.236481 | 9595.085898 | 2863.283207 | 0.418659377 | 0.357179335 | 0.154802095 | 0.211402806 | ns | YPD | KS.1 |

| | | | | | | | | | | |
|---|---|---|---|---|---|---|---|---|---|---|
| FIM1_4228 | 87.99677983 | 22.18643395 | 24.61517435 | -1.784909434 | 0.33726564 | 4.4097388036 | 2.1155690017 | Km<KS.1 | YPD | KS.1 |
| FIM1_4229 | 7930.805618 | 1301.770762 | 1314.53667 | -2.414750166 | 0.332892047 | 2.6292772312 | 2.2436797002 | Km<KS.1 | YPD | KS.1 |
| FIM1_4230 | 1862.707781 | 2350.514756 | 2387.443186 | 0.278915087 | 0.243975363 | 0.203341209 | 0.268619893 | ns | YPD | KS.1 |
| FIM1_4231 | 5067.058969 | 6163.508113 | 5772.270488 | 0.250870814 | 0.187211175 | 0.150679115 | 0.206509255 | ns | YPD | KS.1 |
| FIM1_4232 | 626.1579149 | 469.5032514 | 487.0612049 | -0.376136961 | 0.181704263 | 0.026989379 | 0.044668051 | ns | YPD | KS.1 |
| FIM1_4233 | 1895.245223 | 1185.63065 | 813.2059935 | -0.64562046 | 0.139809346 | 1.5944485570 | 5.4518086402 | ns | YPD | KS.1 |
| FIM1_4234 | 415.6204225 | 744.1080768 | 1164.257849 | 0.797321352 | 0.160351164 | 1.9350831799 | 7.5506186824 | ns | YPD | KS.1 |
| FIM1_4235 | 1052.89287 | 813.160279 | 998.3452731 | -0.35826327 | 0.109778917 | 0.000796368 | 0.001791918 | ns | YPD | KS.1 |
| FIM1_4236 | 3095.099472 | 2810.315654 | 2572.85615 | -0.131547495 | 0.113618273 | 0.232317229 | 0.300124179 | ns | YPD | KS.1 |
| FIM1_4237 | 5545.250627 | 7859.619562 | 7971.332114 | 0.488831091 | 0.099385539 | 5.0205734475 | 1.8474373447 | ns | YPD | KS.1 |
| FIM1_4238 | 768.9882494 | 2224.468813 | 2974.152138 | 1.50378993 | 0.123686447 | 3.7845802844 | 1.4595571251 | Km>KS.1 | YPD | KS.1 |
| FIM1_4239 | 1342.545777 | 2481.551029 | 2512.177134 | 0.85950801 | 0.126025024 | 2.2871300218 | 1.5191551213 | ns | YPD | KS.1 |
| FIM1_4240 | 1181.339866 | 616.2741427 | 1416.346129 | -0.820269136 | 0.269391651 | 0.000660102 | 0.001514764 | ns | YPD | KS.1 |
| FIM1_4241 | 2628.754009 | 3329.366246 | 2297.8411 | 0.282207873 | 0.24712476 | 0.202832249 | 0.268161158 | ns | YPD | KS.1 |
| FIM1_4242 | 720.1606061 | 397.8788081 | 256.0742388 | -0.784432487 | 0.209411204 | 5.4877471549 | 0.000150505 | ns | YPD | KS.1 |
| FIM1_4243 | 482.813925 | 464.3053296 | 683.6635556 | -0.051514018 | 0.114207962 | 0.641930878 | 0.707959681 | ns | YPD | KS.1 |
| FIM1_4244 | 893.9395257 | 945.3423956 | 691.9464541 | 0.069352181 | 0.196844261 | 0.704502052 | 0.764093679 | ns | YPD | KS.1 |
| FIM1_4245 | 184.6560739 | 64.40352693 | 45.78680657 | -1.397361567 | 0.262610061 | 8.9967282976 | 4.1674788902 | Km<KS.1 | YPD | KS.1 |
| FIM1_4246 | 777.1388847 | 773.7537077 | 598.2144239 | -0.005017985 | 0.113027016 | 0.963522425 | 0.97267456 | ns | YPD | KS.1 |
| FIM1_4247 | 746.9816797 | 677.1661745 | 496.5544882 | -0.122157052 | 0.199604364 | 0.50906677 | 0.585032844 | ns | YPD | KS.1 |
| FIM1_4248 | 967.4392783 | 774.2619645 | 1281.443992 | -0.312896293 | 0.09011834 | 0.000398743 | 0.000947348 | ns | YPD | KS.1 |
| FIM1_4249 | 7081.86646 | 7838.254481 | 5696.017768 | 0.136769871 | 0.127847266 | 0.266685409 | 0.336542656 | ns | YPD | KS.1 |
| FIM1_4250 | 1146.679969 | 1309.498898 | 1103.258929 | 0.188179072 | 0.063074496 | 0.002538526 | 0.005229468 | ns | YPD | KS.1 |
| FIM1_4251 | 1276.752015 | 2732.794499 | 3147.760199 | 1.083596123 | 0.088057115 | 1.3506425975 | 5.4189088086 | Km>KS.1 | YPD | KS.1 |
| FIM1_4252 | 983.6224609 | 1301.709522 | 1133.022365 | 0.38578594 | 0.118249123 | 0.00076558 | 0.001732042 | ns | YPD | KS.1 |
| FIM1_4253 | 1208.943148 | 5763.278382 | 2736.043885 | 2.172257817 | 0.208927857 | 3.3747638199 | 7.7370737346 | Km>KS.1 | YPD | KS.1 |
| FIM1_4254 | 1952.412403 | 1354.982987 | 1081.144263 | -0.503229762 | 0.127621104 | 4.5080188926 | 0.000125153 | ns | YPD | KS.1 |
| FIM1_4255 | 757.5948029 | 1812.888748 | 1779.909196 | 1.177446651 | 0.214440957 | 5.4183639371 | 2.5827462957 | Km>KS.1 | YPD | KS.1 |
| FIM1_4257 | 16281.59212 | 8227.490702 | 4226.826573 | -0.875283561 | 0.256988265 | 0.000165304 | 0.000419372 | ns | YPD | KS.1 |
| FIM1_4258 | 67.34295275 | 35.59924806 | 42.59403129 | -0.810725381 | 0.256814024 | 0.000461709 | 0.001084002 | ns | YPD | KS.1 |
| FIM1_4259 | 2034.513327 | 3626.589033 | 2675.666478 | 0.761752811 | 0.210972423 | 9.8154711551 | 0.00025674 | ns | YPD | KS.1 |
| FIM1_4260 | 743.2974594 | 1221.285816 | 767.4231294 | 0.628425287 | 0.240162687 | 0.003846392 | 0.007660449 | ns | YPD | KS.1 |
| FIM1_4261 | 3249.027772 | 937.8536374 | 920.3489585 | -1.704306598 | 0.218830367 | 3.2218354133 | 3.0299869907 | Km<KS.1 | YPD | KS.1 |
| FIM1_4262 | 2482.813053 | 3281.678845 | 2964.984714 | 0.368579361 | 0.168287972 | 0.020260952 | 0.034508125 | ns | YPD | KS.1 |
| FIM1_4263 | 672.3422334 | 607.0399842 | 569.9360588 | -0.139378299 | 0.111559994 | 0.197811345 | 0.262219942 | ns | YPD | KS.1 |
| FIM1_4265 | 313.2607796 | 664.3163225 | 891.4510669 | 1.053681467 | 0.132333491 | 2.8999290805 | 2.7428036455 | Km>KS.1 | YPD | KS.1 |
| FIM1_4266 | 9811.083795 | 12065.3101 | 9953.457295 | 0.25328665 | 0.224704757 | 0.215812367 | 0.282434192 | ns | YPD | KS.1 |
| FIM1_4267 | 1256.69948 | 1037.151271 | 1443.309574 | -0.256833898 | 0.144926461 | 0.06342898 | 0.096975776 | ns | YPD | KS.1 |
| FIM1_4268 | 3984.976512 | 4523.924261 | 3576.938436 | 0.159125396 | 0.197356242 | 0.384760554 | 0.461471494 | ns | YPD | KS.1 |
| FIM1_4269 | 403.5112446 | 416.7719099 | 394.092963 | 0.043623384 | 0.144645152 | 0.752522756 | 0.804599336 | ns | YPD | KS.1 |
| FIM1_4270 | 4850.082345 | 2951.694575 | 2359.182223 | -0.694136418 | 0.115938785 | 7.7525127244 | 4.0513393701 | ns | YPD | KS.1 |
| FIM1_4271 | 730.3232628 | 953.1918865 | 835.0517382 | 0.368783997 | 0.10918092 | 0.000520627 | 0.001216019 | ns | YPD | KS.1 |
| FIM1_4272 | 1355.43061 | 655.3711897 | 712.7728461 | -0.96451371 | 0.222274467 | 2.9484200374 | 9.6566094049 | ns | YPD | KS.1 |
| FIM1_4273 | 1415.756407 | 817.1133235 | 1940.035915 | -0.650789652 | 0.304899681 | 0.013549127 | 0.024048129 | ns | YPD | KS.1 |
| FIM1_4274 | 1997.529891 | 501.5843108 | 458.2280169 | -1.957626423 | 0.136761183 | 4.2373651257 | 3.5134819167 | Km<KS.1 | YPD | KS.1 |

| | | | | | | | | | | |
|---|---|---|---|---|---|---|---|---|---|---|
| FIM1_4275 | 176.003785 | 16.20859451 | 33.90945476 | -3.151537836 | 0.443077992 | 1.36263188635 | 1.20410188892 | Km<KS.1 | YPD | KS.1 |
| FIM1_4276 | 1746.744707 | 1873.270975 | 2355.293366 | 0.096101631 | 0.106362122 | 0.352955676 | 0.428273195 | ns | YPD | KS.1 |
| FIM1_4277 | 270.5187213 | 178.1360791 | 199.5390417 | -0.552826211 | 0.187693397 | 0.001636093 | 0.003496375 | ns | YPD | KS.1 |
| FIM1_4278 | 2390.150838 | 3210.414588 | 2990.652528 | 0.417126671 | 0.077429376 | 4.71922548724 | 1.99474484274 | ns | YPD | KS.1 |
| FIM1_4279 | 1279.77428 | 1375.263549 | 3477.358609 | 0.077265446 | 0.303809127 | 0.767649255 | 0.817784805 | ns | YPD | KS.1 |
| FIM1_4280 | 3687.683295 | 2241.988031 | 2995.245579 | -0.687942216 | 0.136177034 | 1.62875355278 | 6.41571569682 | ns | YPD | KS.1 |
| FIM1_4281 | 4776.177155 | 5691.95689 | 5184.495808 | 0.240064398 | 0.118051323 | 0.035543919 | 0.05713441 | ns | YPD | KS.1 |
| FIM1_4282 | 11872.3081 | 3354.360664 | 2506.198991 | -1.629837807 | 0.328337098 | 3.57660162488 | 1.53790778598 | Km<KS.1 | YPD | KS.1 |
| FIM1_4283 | 311.7977371 | 62.56709133 | 72.17510083 | -2.170479123 | 0.282029057 | 1.91716889920 | 1.84485788656 | Km<KS.1 | YPD | KS.1 |
| FIM1_4284 | 3254.22999 | 4037.127836 | 5208.636838 | 0.295056296 | 0.12079752 | 0.011526653 | 0.020807366 | ns | YPD | KS.1 |
| FIM1_4285 | 5145.645396 | 2342.760627 | 2583.026233 | -1.105597002 | 0.128934238 | 1.54688936804 | 1.76914358759 | Km<KS.1 | YPD | KS.1 |
| FIM1_4286 | 1806.750243 | 2709.635069 | 2558.176458 | 0.475489472 | 0.285855992 | 0.056608695 | 0.087598213 | ns | YPD | KS.1 |
| FIM1_4287 | 14.42328084 | 15.4997429 | 12.20518445 | 0.064784748 | 0.325274366 | 0.813804293 | 0.853970968 | ns | YPD | KS.1 |
| FIM1_4288 | 1103.683074 | 1694.910214 | 1283.172397 | 0.534550013 | 0.243307269 | 0.014748384 | 0.026000429 | ns | YPD | KS.1 |
| FIM1_4289 | 714.2402019 | 2302.415132 | 2144.079177 | 1.641621987 | 0.156427254 | 5.01089554180 | 1.13314569638 | Km>KS.1 | YPD | KS.1 |
| FIM1_4290 | 1450.94 | 1311.874555 | 1397.806259 | -0.135989071 | 0.124854984 | 0.258774823 | 0.328587225 | ns | YPD | KS.1 |
| FIM1_4291 | 613.4960005 | 692.7678748 | 822.1209317 | 0.170542508 | 0.084017944 | 0.038513139 | 0.061312918 | ns | YPD | KS.1 |
| FIM1_4292 | 280.6896082 | 617.8516219 | 926.2401936 | 1.11368688 | 0.123872967 | 3.77645311077 | 5.01009446028 | Km>KS.1 | YPD | KS.1 |
| FIM1_4293 | 681.3336735 | 565.2861891 | 651.6130055 | -0.25999537 | 0.094462405 | 0.004881834 | 0.009520628 | ns | YPD | KS.1 |
| FIM1_4294 | 970.679897 | 1103.343946 | 1014.23371 | 0.173364371 | 0.125787514 | 0.152703397 | 0.208995709 | ns | YPD | KS.1 |
| FIM1_4295 | 835.4142002 | 838.5219129 | 957.734905 | 0.005296422 | 0.083391689 | 0.948439687 | 0.963349825 | ns | YPD | KS.1 |
| FIM1_4296 | 1483.57519 | 918.076474 | 1234.676435 | -0.685226995 | 0.061001248 | 9.90988707240 | 2.65062839705 | ns | YPD | KS.1 |
| FIM1_4297 | 1500.128358 | 1335.275642 | 795.4135375 | -0.15044647 | 0.16976735 | 0.347203035 | 0.42233132 | ns | YPD | KS.1 |
| FIM1_4298 | 392.0646807 | 432.1868727 | 256.6677733 | 0.089622391 | 0.390782434 | 0.773535726 | 0.822646481 | ns | YPD | KS.1 |
| FIM1_4299 | 516.9430989 | 478.0284136 | 366.378426 | -0.10233893 | 0.149827589 | 0.473704763 | 0.55152848 | ns | YPD | KS.1 |
| FIM1_4300 | 340.4470015 | 284.4252782 | 223.9361699 | -0.242064475 | 0.145807616 | 0.081901342 | 0.121015497 | ns | YPD | KS.1 |
| FIM1_4303 | 114.5591467 | 42.72610322 | 82.48496482 | -1.321153502 | 0.23913706 | 3.37555339919 | 1.63997833603 | Km<KS.1 | YPD | KS.1 |
| FIM1_4304 | 51.80465592 | 65.67820019 | 30.91326256 | 0.238454935 | 0.352413302 | 0.412612628 | 0.490032901 | ns | YPD | KS.1 |
| FIM1_4307 | 776.7581087 | 337.538839 | 476.7071757 | -1.139456414 | 0.191398504 | 3.84234360551 | 2.07103569202 | Km<KS.1 | YPD | KS.1 |
| FIM1_4308 | 123.937361 | 102.3731972 | 116.7686975 | -0.239992854 | 0.21218732 | 0.218044213 | 0.284940888 | ns | YPD | KS.1 |
| FIM1_4314 | 298.7501334 | 418.5527785 | 646.7515623 | 0.427892684 | 0.212858111 | 0.028565648 | 0.046948826 | ns | YPD | KS.1 |
| FIM1_4780 | 36.36195436 | 73.70754726 | 82.75928701 | 0.805138761 | 0.36716341 | 0.008276182 | 0.015357705 | ns | YPD | KS.1 |
| FIM1_4781 | 390.2886433 | 68.98634146 | 87.57792087 | -2.424387476 | 0.201120426 | 1.03204455263 | 4.17432654419 | Km<KS.1 | YPD | KS.1 |
| FIM1_4782 | 700.46721 | 443.1751452 | 296.5047851 | -0.637962497 | 0.120004723 | 4.40087725041 | 1.86493733567 | ns | YPD | KS.1 |
| FIM1_4783 | 3656.233837 | 1385.92669 | 1729.24346 | -1.367370593 | 0.132842061 | 7.26440346444 | 1.51215093036 | Km<KS.1 | YPD | KS.1 |
| FIM1_4785 | 2857.330199 | 1251.897836 | 1037.162625 | -1.161109263 | 0.128234121 | 1.93135997830 | 2.63970216815 | Km<KS.1 | YPD | KS.1 |
| FIM1_4786 | 8848.683143 | 6254.952908 | 3276.771524 | -0.334152718 | 0.392803954 | 0.286038903 | 0.357638488 | ns | YPD | KS.1 |
| FIM1_4787 | 756.0845369 | 242.7261193 | 235.4642185 | -1.519544934 | 0.257502231 | 2.48653163750 | 1.35492824716 | Km<KS.1 | YPD | KS.1 |
| FIM1_4788 | 4330.189302 | 833.7668516 | 744.3069426 | -2.157105834 | 0.351267985 | 1.14438204219 | 7.10774114845 | Km<KS.1 | YPD | KS.1 |
| FIM1_4789 | 102.2443886 | 373.475493 | 264.372092 | 1.519478266 | 0.443069265 | 3.51757740270 | 9.89816039506 | Km>KS.1 | YPD | KS.1 |
| FIM1_4790 | 945.7671047 | 618.8736294 | 614.4716867 | -0.557888703 | 0.190231553 | 0.001686818 | 0.00359243 | ns | YPD | KS.1 |
| FIM1_4791 | 591.2843509 | 1592.292844 | 2073.260557 | 1.332798527 | 0.233136573 | 1.08912404651 | 5.62657542200 | Km>KS.1 | YPD | KS.1 |
| FIM1_4792 | 644.8950488 | 718.5679584 | 464.198878 | 0.124205407 | 0.260762522 | 0.592309344 | 0.663919748 | ns | YPD | KS.1 |
| FIM1_4793 | 3321.807969 | 3581.665613 | 2070.580399 | 0.098871095 | 0.152566227 | 0.496072968 | 0.573012077 | ns | YPD | KS.1 |
| FIM1_4794 | 5005.434969 | 5402.345302 | 2670.050722 | 0.092989385 | 0.216569709 | 0.639732047 | 0.705847624 | ns | YPD | KS.1 |

| | | | | | | | | | | |
|---|---|---|---|---|---|---|---|---|---|---|
| FIM1_4795 | 5621.563025 | 11655.50041 | 6331.766408 | 0.919929804 | 0.281406241 | 0.000245197 | 0.000606591 | ns | YPD | KS.1 |
| FIM1_4796 | 5918.72775 | 1989.458977 | 2521.152864 | -1.486972118 | 0.217902407 | 6.6529429595( | 4.6226803385: | Km<KS.1 | YPD | KS.1 |
| FIM1_4797 | 3326.659606 | 2550.06309 | 4622.760369 | -0.300824086 | 0.28808403 | 0.230135868 | 0.297537927 | ns | YPD | KS.1 |
| FIM1_4798 | 336.783179 | 463.5098596 | 907.3828427 | 0.398831078 | 0.226873035 | 0.053199151 | 0.08275978 | ns | YPD | KS.1 |
| FIM1_4799 | 700.0346511 | 870.0537378 | 915.778069 | 0.272613002 | 0.207864622 | 0.154249391 | 0.210821627 | ns | YPD | KS.1 |
| FIM1_4800 | 13.45119768 | 11.04515615 | 14.76985844 | -0.214064306 | 0.331539459 | 0.44250525 | 0.519580746 | ns | YPD | KS.1 |
| FIM1_4801 | 266.6042946 | 285.6744997 | 369.4995752 | 0.080039377 | 0.237496207 | 0.709506344 | 0.768268427 | ns | YPD | KS.1 |
| FIM1_4802 | 327.9408011 | 962.4068293 | 805.0057536 | 1.53225351 | 0.099519222 | 1.1604614080: | 1.3121126147 | Km>KS.1 | YPD | KS.1 |
| FIM1_4803 | 1951.405299 | 1960.694628 | 1781.19925 | 0.005735704 | 0.219356102 | 0.977230497 | 0.984049745 | ns | YPD | KS.1 |
| FIM1_4804 | 2107.366229 | 3659.507802 | 5526.854346 | 0.759011581 | 0.149118536 | 1.1373450785 | 4.5594615356! | ns | YPD | KS.1 |
| FIM1_4805 | 2625.04477 | 2558.755448 | 4310.284211 | -0.033491785 | 0.150614959 | 0.815627753 | 0.855523523 | ns | YPD | KS.1 |
| FIM1_4806 | 1945.097977 | 2276.34088 | 2008.96905 | 0.221792378 | 0.0716852 | 0.001698801 | 0.003616403 | ns | YPD | KS.1 |
| FIM1_4807 | 2159.697626 | 2818.863983 | 2654.061447 | 0.336351369 | 0.208058885 | 0.079002034 | 0.11711416 | ns | YPD | KS.1 |
| FIM1_4808 | 2462.770446 | 2988.655076 | 2096.452981 | 0.244250073 | 0.199172302 | 0.185272409 | 0.24784357 | ns | YPD | KS.1 |
| FIM1_4809 | 877.4735358 | 522.7958535 | 696.4791253 | -0.705803319 | 0.158782879 | 3.1655462690( | 1.0313420228: | ns | YPD | KS.1 |
| FIM1_4810 | 9902.726541 | 10859.45472 | 12749.87612 | 0.12776146 | 0.093073948 | 0.160277426 | 0.217922983 | ns | YPD | KS.1 |
| FIM1_4811 | 4053.022779 | 2543.79927 | 1936.960012 | -0.517334738 | 0.333210428 | 0.065177753 | 0.09928332 | ns | YPD | KS.1 |
| FIM1_4812 | 328.3797757 | 271.8844893 | 213.5856373 | -0.246517699 | 0.168917744 | 0.121475423 | 0.171638805 | ns | YPD | KS.1 |
| FIM1_4813 | 1613.012114 | 1059.981764 | 1787.613643 | -0.481160586 | 0.304340109 | 0.066164479 | 0.100448056 | ns | YPD | KS.1 |
| FIM1_4814 | 645.5751336 | 1168.031611 | 1173.57424 | 0.841512568 | 0.090477017 | 3.5605763394: | 5.0466858372( | ns | YPD | KS.1 |
| FIM1_4815 | 1066.674751 | 1013.809696 | 843.8898588 | -0.069231711 | 0.102377391 | 0.487776845 | 0.565135026 | ns | YPD | KS.1 |
| FIM1_4816 | 1392.261759 | 1082.311337 | 1091.397534 | -0.350267963 | 0.100043811 | 0.000339469 | 0.000817454 | ns | YPD | KS.1 |
| FIM1_4817 | 1447.215002 | 1118.632683 | 1649.15812 | -0.342637608 | 0.158292159 | 0.022494301 | 0.037807144 | ns | YPD | KS.1 |
| FIM1_4818 | 1173.253861 | 1095.738261 | 1041.875757 | -0.093909952 | 0.094036885 | 0.306982346 | 0.379909744 | ns | YPD | KS.1 |
| FIM1_4819 | 3073.151156 | 3908.092611 | 5748.894804 | 0.33539782 | 0.09746488 | 0.000433502 | 0.001021152 | ns | YPD | KS.1 |
| FIM1_4820 | 2284.8638 | 1733.675157 | 1825.771121 | -0.385006483 | 0.101597155 | 0.000104919 | 0.000273142 | ns | YPD | KS.1 |
| FIM1_4821 | 684.9736181 | 812.7059034 | 1098.858809 | 0.193874492 | 0.273171227 | 0.420342079 | 0.498023778 | ns | YPD | KS.1 |
| FIM1_4822 | 812.8035179 | 1415.64166 | 1621.314464 | 0.77425331 | 0.125575869 | 2.1285316090 | 1.1688128868( | ns | YPD | KS.1 |
| FIM1_4823 | 1234.091789 | 1183.249067 | 1166.873272 | -0.056885295 | 0.113720584 | 0.606129578 | 0.676271507 | ns | YPD | KS.1 |
| FIM1_4824 | 3425.826171 | 2133.009615 | 3418.530375 | -0.61727086 | 0.208436167 | 0.001356644 | 0.00293193 | ns | YPD | KS.1 |
| FIM1_4825 | 1468.410044 | 456.9438569 | 465.332088 | -1.587458382 | 0.230401869 | 3.3985419573( | 2.4153923196! | Km<KS.1 | YPD | KS.1 |
| FIM1_4826 | 946.9485849 | 1141.570198 | 1047.673213 | 0.258879365 | 0.104755971 | 0.011132042 | 0.020138876 | ns | YPD | KS.1 |
| FIM1_4827 | 820.6378469 | 1282.906952 | 1322.638848 | 0.626053618 | 0.109644354 | 4.8027776213! | 2.2993888164! | ns | YPD | KS.1 |
| FIM1_4828 | 1171.009279 | 852.553229 | 929.2585492 | -0.438615939 | 0.119894977 | 0.000160968 | 0.000408997 | ns | YPD | KS.1 |
| FIM1_4829 | 4004.156766 | 4824.492694 | 2408.544241 | 0.20678834 | 0.290973696 | 0.413348575 | 0.490789776 | ns | YPD | KS.1 |
| FIM1_4830 | 4060.895384 | 3929.48989 | 2396.706495 | -0.038084993 | 0.252088368 | 0.866053042 | 0.896880492 | ns | YPD | KS.1 |
| FIM1_4831 | 366.3973641 | 1262.862996 | 1966.919271 | 1.766364951 | 0.095460342 | 7.3364576353: | 3.0415730613: | Km>KS.1 | YPD | KS.1 |
| FIM1_4832 | 2045.685106 | 3263.398503 | 3576.343722 | 0.665644508 | 0.066959596 | 1.0291933817 | 1.7966404981: | ns | YPD | KS.1 |
| FIM1_4833 | 4904.320996 | 4656.398828 | 4511.192965 | -0.068438327 | 0.144907308 | 0.621329895 | 0.689981301 | ns | YPD | KS.1 |
| FIM1_4834 | 3274.352705 | 2634.27576 | 1340.639535 | -0.225395605 | 0.335751138 | 0.422936815 | 0.500425853 | ns | YPD | KS.1 |
| FIM1_4835 | 1776.345475 | 2273.733131 | 3751.275652 | 0.334642695 | 0.136099771 | 0.010431936 | 0.018975826 | ns | YPD | KS.1 |
| FIM1_4836 | 186.4899181 | 305.4638777 | 802.9952928 | 0.612941451 | 0.255433667 | 0.007351325 | 0.013769895 | ns | YPD | KS.1 |
| FIM1_4837 | 12.0607207 | 10.19806474 | 67.6932174 | -0.175054076 | 0.366000173 | 0.557715916 | 0.631029494 | ns | YPD | KS.1 |
| FIM1_4839 | 473.0625087 | 1031.733054 | 576.3763114 | 0.987575896 | 0.284382982 | 0.000100992 | 0.000263746 | ns | YPD | KS.1 |
| FIM1_4840 | 4050.919614 | 7170.503857 | 8620.223328 | 0.808406898 | 0.094450431 | 3.1317509068: | 3.4933768523: | ns | YPD | KS.1 |

| | | | | | | | | | | |
|---|---|---|---|---|---|---|---|---|---|---|
| FIM1_4841 | 845.9104933 | 278.6691775 | 248.0003937 | -1.564061493 | 0.14270439 | 3.83898669689 | 1.00520835879 | Km<KS.1 | YPD | KS.1 |
| FIM1_4842 | 75520.70811 | 41699.39184 | 58807.41069 | -0.832647576 | 0.119039979 | 7.05510209676 | 4.89527655947 | ns | YPD | KS.1 |
| FIM1_4844 | 7768.767821 | 583.7775632 | 1119.871416 | -3.413390381 | 0.481515999 | 1.35072554422 | 1.19570455204 | Km<KS.1 | YPD | KS.1 |
| FIM1_4845 | 1742.311186 | 804.7697023 | 600.696593 | -0.934165541 | 0.328459036 | 0.000977492 | 0.002171961 | ns | YPD | KS.1 |
| FIM1_4846 | 1344.66912 | 993.7722043 | 542.9534828 | -0.328232888 | 0.319606535 | 0.226389028 | 0.293456335 | ns | YPD | KS.1 |
| FIM1_4847 | 2648.283805 | 1773.057091 | 765.4735833 | -0.47336902 | 0.280931865 | 0.05434595 | 0.084411833 | ns | YPD | KS.1 |
| FIM1_4848 | 4737.785035 | 5711.655798 | 2313.99835 | 0.220143432 | 0.251246852 | 0.327371869 | 0.40125032 | ns | YPD | KS.1 |
| FIM1_4849 | 2442.116251 | 6004.73622 | 3562.945838 | 1.160828924 | 0.281583141 | 5.12442302076 | 1.62278832134 | Km>KS.1 | YPD | KS.1 |
| FIM1_4850 | 4294.493734 | 747.6450869 | 797.3107309 | -2.220314291 | 0.41649402 | 1.09981958096 | 5.67593611543 | Km<KS.1 | YPD | KS.1 |
| FIM1_4851 | 713.9216623 | 640.2884768 | 855.6509651 | -0.142580778 | 0.156227191 | 0.336561528 | 0.411500025 | ns | YPD | KS.1 |
| FIM1_4852 | 413.054123 | 290.8459678 | 468.2283045 | -0.466307154 | 0.171784833 | 0.003989889 | 0.007914553 | ns | YPD | KS.1 |
| FIM1_4853 | 1103.666351 | 791.3899107 | 464.3356436 | -0.427637874 | 0.200440888 | 0.021170409 | 0.035787559 | ns | YPD | KS.1 |
| FIM1_4854 | 5600.100191 | 4426.500842 | 3480.209368 | -0.257546256 | 0.304802096 | 0.325356882 | 0.399469519 | ns | YPD | KS.1 |
| FIM1_4855 | 1401.506791 | 1766.412793 | 915.6515042 | 0.23003246 | 0.360674166 | 0.436628998 | 0.513585314 | ns | YPD | KS.1 |
| FIM1_4856 | 165.9142976 | 108.1584856 | 90.94481753 | -0.532327686 | 0.24433253 | 0.01551904 | 0.027229385 | ns | YPD | KS.1 |
| FIM1_4857 | 607.9412719 | 68.1830039 | 97.34138546 | -2.911501011 | 0.398097541 | 4.50549890096 | 4.16633030339 | Km<KS.1 | YPD | KS.1 |
| FIM1_4858 | 1238.174875 | 624.2399809 | 1889.896656 | -0.830908856 | 0.31010047 | 0.0020426 | 0.004296802 | ns | YPD | KS.1 |
| FIM1_4859 | 2503.441798 | 2415.983231 | 1667.293549 | -0.049315068 | 0.082219234 | 0.541475546 | 0.6154537 | ns | YPD | KS.1 |
| FIM1_4860 | 326.4264578 | 857.6332616 | 551.1737234 | 1.266905311 | 0.267914736 | 2.52297218234 | 9.73761567662 | Km>KS.1 | YPD | KS.1 |
| FIM1_4861 | 522.8012065 | 817.0575401 | 825.2678814 | 0.625293603 | 0.106939532 | 2.12442937431 | 1.05690361372 | ns | YPD | KS.1 |
| FIM1_4862 | 580.8182794 | 1108.202719 | 1646.755377 | 0.886834849 | 0.164215168 | 1.60064979347 | 7.15474638142 | ns | YPD | KS.1 |
| FIM1_4863 | 2387.179467 | 2646.816194 | 5275.620652 | 0.117345568 | 0.268353178 | 0.621072117 | 0.689849025 | ns | YPD | KS.1 |
| FIM1_4864 | 45.46377785 | 37.33204208 | 64.56806349 | -0.234439061 | 0.251904876 | 0.297902674 | 0.370214275 | ns | YPD | KS.1 |
| FIM1_4865 | 1114.217773 | 990.1448385 | 766.4919362 | -0.144212781 | 0.217083371 | 0.468517796 | 0.546334373 | ns | YPD | KS.1 |
| FIM1_4866 | 1622.013271 | 2411.891672 | 1468.118296 | 0.489941599 | 0.24535638 | 0.026311326 | 0.043603844 | ns | YPD | KS.1 |
| FIM1_4867 | 1431.771351 | 1873.145159 | 1774.801931 | 0.345397965 | 0.193592331 | 0.054759967 | 0.08500182 | ns | YPD | KS.1 |
| FIM1_4868 | 836.1147747 | 1086.148888 | 1300.526989 | 0.355169286 | 0.134770732 | 0.006102044 | 0.011635748 | ns | YPD | KS.1 |
| FIM1_4869 | 377.9792447 | 1114.903574 | 2324.438527 | 1.508269783 | 0.168242547 | 2.26593324904 | 3.05501840487 | Km>KS.1 | YPD | KS.1 |
| FIM1_4870 | 1410.210209 | 608.2891444 | 695.124892 | -1.06786846 | 0.291479793 | 4.10393497899 | 0.000114767 | Km<KS.1 | YPD | KS.1 |
| FIM1_4871 | 1499.452392 | 641.6537943 | 519.1620767 | -1.167964742 | 0.179139048 | 9.77916402171 | 6.14284608687 | Km<KS.1 | YPD | KS.1 |
| FIM1_4872 | 3120.768625 | 598.5202416 | 580.6762812 | -2.302864808 | 0.208564582 | 2.07217693108 | 5.89090298979 | Km<KS.1 | YPD | KS.1 |
| FIM1_4873 | 1919.21178 | 896.5883323 | 1506.886808 | -1.037458052 | 0.187313294 | 5.41987363371 | 2.58274629576 | Km<KS.1 | YPD | KS.1 |
| FIM1_4874 | 1206.554293 | 2332.036722 | 2687.585499 | 0.86344753 | 0.229666222 | 4.36991633046 | 0.000121658 | ns | YPD | KS.1 |
| FIM1_4875 | 1606.125638 | 2465.077034 | 1454.363538 | 0.528108206 | 0.251847322 | 0.019189369 | 0.032919694 | ns | YPD | KS.1 |
| FIM1_4876 | 47.36614109 | 41.22844889 | 66.18190685 | -0.170852498 | 0.256410242 | 0.454981036 | 0.532470161 | ns | YPD | KS.1 |
| FIM1_4877 | 1903.279043 | 1935.631251 | 1003.914225 | 0.020614954 | 0.216344804 | 0.917322156 | 0.938063253 | ns | YPD | KS.1 |
| FIM1_4878 | 8377.596451 | 13071.58627 | 19245.88169 | 0.614240444 | 0.132524901 | 1.57452968532 | 5.39111162042 | ns | YPD | KS.1 |
| FIM1_4879 | 2439.388888 | 2437.895146 | 2685.028214 | -0.000619663 | 0.06836029 | 0.992673756 | 0.994302819 | ns | YPD | KS.1 |
| FIM1_4880 | 2189.581914 | 4871.067156 | 4901.205305 | 1.145487912 | 0.066581161 | 3.44337705462 | 1.22362863191 | Km>KS.1 | YPD | KS.1 |
| FIM1_4881 | 1792.466204 | 3057.716038 | 2085.045205 | 0.740701804 | 0.13403033 | 1.07520438697 | 4.93463268007 | ns | YPD | KS.1 |
| FIM1_4882 | 8485.306277 | 330.5047026 | 1008.366534 | -4.234148447 | 0.581846243 | 4.91131091718 | 4.51640883789 | Km<KS.1 | YPD | KS.1 |
| FIM1_4883 | 5976.196919 | 4651.061588 | 3633.9843 | -0.330491087 | 0.166921971 | 0.035917835 | 0.057647796 | ns | YPD | KS.1 |
| FIM1_4884 | 1915.496551 | 2526.891256 | 2324.54675 | 0.389721315 | 0.086572284 | 4.65734238325 | 1.48527425363 | ns | YPD | KS.1 |
| FIM1_4885 | 729.7083978 | 1254.687615 | 961.0478088 | 0.759384228 | 0.113647989 | 7.34372929469 | 4.65414690969 | ns | YPD | KS.1 |
| FIM1_4886 | 182.2934712 | 243.589807 | 262.1975986 | 0.382072587 | 0.165935441 | 0.014819062 | 0.026115775 | ns | YPD | KS.1 |

| | | | | | | | | | | |
|---|---|---|---|---|---|---|---|---|---|---|
| FIM1_4887 | 165.3074391 | 329.440095 | 366.9965181 | 0.971322818 | 0.125144461 | 1.6806934069 | 1.4694990684 | ns | YPD | KS.1 |
| FIM1_4888 | 626.9115548 | 1086.894538 | 846.6949265 | 0.731651686 | 0.195490846 | 6.2462324493 | 0.000169438 | ns | YPD | KS.1 |
| FIM1_4889 | 745.0529367 | 494.3314245 | 630.0680574 | -0.53394641 | 0.200455318 | 0.004074057 | 0.008071857 | ns | YPD | KS.1 |
| FIM1_4890 | 344.5112912 | 637.1248125 | 592.0101904 | 0.830780856 | 0.185950389 | 2.1636881511 | 7.2341052096 | ns | YPD | KS.1 |
| FIM1_4891 | 956.6375215 | 921.0330724 | 1119.446052 | -0.049770915 | 0.148514622 | 0.72553157 | 0.781619654 | ns | YPD | KS.1 |
| FIM1_4892 | 5240.836001 | 6857.910488 | 4920.135313 | 0.325793911 | 0.241081566 | 0.133344815 | 0.18576284 | ns | YPD | KS.1 |
| FIM1_4893 | 6903.656815 | 12209.8825 | 9611.487046 | 0.726799873 | 0.245446762 | 0.001069986 | 0.002358521 | ns | YPD | KS.1 |
| FIM1_4894 | 2106.162906 | 1452.247884 | 1464.534456 | -0.487905754 | 0.186683454 | 0.005169604 | 0.010007307 | ns | YPD | KS.1 |
| FIM1_4895 | 1485.199412 | 2329.37877 | 1695.610525 | 0.573725495 | 0.22627896 | 0.00545834 | 0.010496807 | ns | YPD | KS.1 |
| FIM1_4896 | 2681.264417 | 2723.297475 | 1266.705427 | 0.018950128 | 0.218216548 | 0.924536636 | 0.944276281 | ns | YPD | KS.1 |
| FIM1_4897 | 1825.556629 | 1615.684514 | 1069.82916 | -0.147558517 | 0.225835383 | 0.473313619 | 0.551202073 | ns | YPD | KS.1 |
| FIM1_4898 | 598.7271334 | 1298.544227 | 920.0857228 | 0.98943276 | 0.274288209 | 6.0461455974 | 0.00016491 | ns | YPD | KS.1 |
| FIM1_4899 | 4365.882113 | 3832.442473 | 7998.009529 | -0.175077801 | 0.135358821 | 0.177601389 | 0.238608402 | ns | YPD | KS.1 |
| FIM1_4900 | 2705.720324 | 3337.209681 | 4562.418037 | 0.278946407 | 0.153518652 | 0.055923314 | 0.086618458 | ns | YPD | KS.1 |
| FIM1_4901 | 535.267726 | 456.2452809 | 385.1794082 | -0.198196905 | 0.213056743 | 0.310976236 | 0.384088077 | ns | YPD | KS.1 |
| FIM1_4902 | 1449.166052 | 1205.840806 | 1002.088481 | -0.245291558 | 0.147242879 | 0.080626993 | 0.11923879 | ns | YPD | KS.1 |
| FIM1_4903 | 2263.104877 | 1578.485507 | 1339.826362 | -0.491444008 | 0.141301675 | 0.000289901 | 0.000707335 | ns | YPD | KS.1 |
| FIM1_4904 | 868.3955119 | 599.975169 | 1271.668189 | -0.478577595 | 0.200562813 | 0.009993532 | 0.018243938 | ns | YPD | KS.1 |
| FIM1_4905 | 572.0308863 | 842.5894093 | 445.754003 | 0.486678154 | 0.229716115 | 0.0197157 | 0.033683245 | ns | YPD | KS.1 |
| FIM1_4906 | 2419.260269 | 1501.937721 | 1602.179037 | -0.665325055 | 0.117234584 | 5.3863202256 | 2.5716836010 | ns | YPD | KS.1 |
| FIM1_4907 | 3078.171927 | 1390.991027 | 2107.583726 | -0.954719155 | 0.337743817 | 0.000984548 | 0.002185688 | ns | YPD | KS.1 |
| FIM1_4908 | 397.4086104 | 383.7430354 | 559.5296818 | -0.044011228 | 0.189004258 | 0.803030157 | 0.845518525 | ns | YPD | KS.1 |
| FIM1_4909 | 405.2721646 | 1961.322019 | 2808.153758 | 2.239880147 | 0.133228097 | 1.8595586894 | 5.1396135998 | Km>KS.1 | YPD | KS.1 |
| FIM1_4910 | 955.5302517 | 784.0249486 | 642.7546195 | -0.256603553 | 0.175366358 | 0.119128694 | 0.1688505 | ns | YPD | KS.1 |
| FIM1_4911 | 934.1074456 | 1338.56379 | 1207.163562 | 0.460420061 | 0.209741936 | 0.017087162 | 0.029641894 | ns | YPD | KS.1 |
| FIM1_4912 | 653.324077 | 947.8125607 | 1002.936123 | 0.486267934 | 0.191679072 | 0.006471775 | 0.012256217 | ns | YPD | KS.1 |
| FIM1_4913 | 1217.631888 | 790.9681578 | 1779.518337 | -0.537529507 | 0.243375381 | 0.01423206 | 0.025134717 | ns | YPD | KS.1 |
| FIM1_4914 | 3136.060704 | 2187.083019 | 2391.945927 | -0.494417583 | 0.13304782 | 0.000115441 | 0.000298193 | ns | YPD | KS.1 |
| FIM1_4915 | 3256.892193 | 3332.800863 | 2017.395644 | 0.026902778 | 0.246281317 | 0.903334706 | 0.926425513 | ns | YPD | KS.1 |
| FIM1_4916 | 1460.277791 | 357.3644516 | 1357.458541 | -1.744271822 | 0.400508389 | 4.9747442393 | 1.8358909399 | Km<KS.1 | YPD | KS.1 |
| FIM1_4917 | 3326.022112 | 1975.02912 | 1595.349315 | -0.725337341 | 0.126363826 | 3.2120524318 | 1.5666628282 | ns | YPD | KS.1 |
| FIM1_4918 | 135.3674847 | 164.873387 | 185.4433156 | 0.251001468 | 0.180231526 | 0.136900506 | 0.190139592 | ns | YPD | KS.1 |
| FIM1_4919 | 676.5366237 | 836.9226834 | 1474.08409 | 0.267205704 | 0.20576209 | 0.158901887 | 0.21640758 | ns | YPD | KS.1 |
| FIM1_4920 | 931.8133371 | 678.3013069 | 491.5382069 | -0.419418737 | 0.171795793 | 0.009562185 | 0.017554197 | ns | YPD | KS.1 |
| FIM1_4921 | 1905.734496 | 7106.150707 | 10694.03251 | 1.856126458 | 0.149784268 | 9.2704173913 | 3.9085022476 | Km>KS.1 | YPD | KS.1 |
| FIM1_4922 | 541.8665632 | 805.8163773 | 1043.283304 | 0.523371674 | 0.186193189 | 0.002659745 | 0.005452093 | ns | YPD | KS.1 |
| FIM1_4923 | 538.7446802 | 717.2552785 | 779.2656105 | 0.398652609 | 0.102488384 | 6.8241987654 | 0.000183219 | ns | YPD | KS.1 |
| FIM1_4924 | 445.4867829 | 818.2285895 | 2324.548194 | 0.75380317 | 0.277472857 | 0.002171126 | 0.004536477 | ns | YPD | KS.1 |
| FIM1_4925 | 854.5859613 | 224.8672532 | 244.9278878 | -1.664647426 | 0.382744357 | 6.2082089415 | 2.2676827814 | Km<KS.1 | YPD | KS.1 |
| FIM1_4926 | 1106.54276 | 1528.582188 | 1363.095632 | 0.455450059 | 0.085092788 | 5.3316254238 | 2.2383828256 | ns | YPD | KS.1 |
| FIM1_4927 | 967.8615392 | 548.1023613 | 1324.343166 | -0.674315414 | 0.307172934 | 0.011047475 | 0.020000433 | ns | YPD | KS.1 |
| FIM1_4928 | 805.9427296 | 367.5879234 | 1095.136634 | -0.951658246 | 0.328387359 | 0.000792755 | 0.001785403 | ns | YPD | KS.1 |
| FIM1_4929 | 237.3472844 | 576.4777722 | 701.8857728 | 1.257765562 | 0.119158292 | 5.6458119572 | 1.2709463568 | Km>KS.1 | YPD | KS.1 |
| FIM1_4931 | 3194.051745 | 4590.342234 | 3938.011143 | 0.480828235 | 0.17497296 | 0.003506935 | 0.007040759 | ns | YPD | KS.1 |
| FIM1_4932 | 9562.254347 | 8485.700641 | 4932.710896 | -0.1552712 | 0.165567566 | 0.32092294 | 0.394512386 | ns | YPD | KS.1 |

| | | | | | | | | | | |
|---|---|---|---|---|---|---|---|---|---|---|
| FIM1_4933 | 1076.496737 | 1590.944386 | 1783.902986 | 0.521422094 | 0.170138911 | 0.001177883 | 0.002574679 | ns | YPD | KS.1 |
| FIM1_4935 | 7859.174758 | 2590.558395 | 1921.499279 | -1.537604664 | 0.185988361 | 9.4003003577 | 1.0166291913 | Km<KS.1 | YPD | KS.1 |
| FIM1_4936 | 1959.716227 | 2106.437678 | 1229.813806 | 0.082016392 | 0.266150868 | 0.728073007 | 0.783337521 | ns | YPD | KS.1 |
| FIM1_4937 | 5542.868081 | 8987.81687 | 6854.844582 | 0.566037492 | 0.300908581 | 0.029567546 | 0.048387677 | ns | YPD | KS.1 |
| FIM1_4938 | 1419.692766 | 1063.384695 | 1253.715977 | -0.37998693 | 0.173818659 | 0.020101269 | 0.034271355 | ns | YPD | KS.1 |
| FIM1_4939 | 549.3818811 | 672.8133937 | 997.7523455 | 0.273741997 | 0.135009864 | 0.034637267 | 0.055930024 | ns | YPD | KS.1 |
| FIM1_4940 | 763.2414071 | 689.359412 | 1454.603785 | -0.120119434 | 0.240903176 | 0.580484995 | 0.653226159 | ns | YPD | KS.1 |
| FIM1_4941 | 4777.804103 | 6702.60003 | 8851.194038 | 0.470040986 | 0.113276498 | 1.9936563086 | 5.8070492597 | ns | YPD | KS.1 |
| FIM1_4942 | 1451.256609 | 1909.172722 | 1224.514551 | 0.293758547 | 0.32358418 | 0.283343476 | 0.354982068 | ns | YPD | KS.1 |
| FIM1_4943 | 3855.701177 | 2582.813491 | 2318.517883 | -0.541248124 | 0.157568005 | 0.000307611 | 0.000746884 | ns | YPD | KS.1 |
| FIM1_4944 | 3628.735293 | 2079.035524 | 1183.17116 | -0.662764435 | 0.302557663 | 0.011465289 | 0.020710058 | ns | YPD | KS.1 |
| FIM1_4945 | 5637.416846 | 8234.642774 | 7429.713428 | 0.497425512 | 0.18732806 | 0.004487674 | 0.008824577 | ns | YPD | KS.1 |
| FIM1_4946 | 2887.450954 | 4193.382846 | 3010.056622 | 0.500537923 | 0.163061631 | 0.001207532 | 0.002632546 | ns | YPD | KS.1 |
| FIM1_4947 | 1031.996878 | 1708.147132 | 1835.211868 | 0.699180637 | 0.130937715 | 3.3651114164 | 1.4570434549 | ns | YPD | KS.1 |
| FIM1_4948 | 717.5389726 | 553.7369714 | 502.0525132 | -0.350786851 | 0.138706669 | 0.008359033 | 0.015482572 | ns | YPD | KS.1 |
| FIM1_4949 | 1545.107563 | 1397.82762 | 1482.360849 | -0.141513353 | 0.065535021 | 0.028731253 | 0.047158688 | ns | YPD | KS.1 |
| FIM1_4950 | 1165.135294 | 2099.651714 | 3177.919928 | 0.834097108 | 0.095170438 | 4.8985875664 | 5.9008409547 | ns | YPD | KS.1 |
| FIM1_4951 | 2990.65122 | 3685.754893 | 2430.954616 | 0.27275446 | 0.171477924 | 0.090964661 | 0.132937568 | ns | YPD | KS.1 |
| FIM1_4952 | 2888.170463 | 3175.325415 | 6919.195298 | 0.108560041 | 0.262680059 | 0.641784454 | 0.707955135 | ns | YPD | KS.1 |
| FIM1_4953 | 1749.892304 | 987.4575526 | 2320.49064 | -0.662083093 | 0.325950013 | 0.017003683 | 0.029516163 | ns | YPD | KS.1 |
| FIM1_4954 | 107.9580207 | 93.8301921 | 227.721582 | -0.154788862 | 0.284828523 | 0.533760825 | 0.608976266 | ns | YPD | KS.1 |
| FIM1_4955 | 4532.809745 | 6974.526882 | 10080.4091 | 0.579011928 | 0.168072649 | 0.000273946 | 0.000672033 | ns | YPD | KS.1 |
| FIM1_4956 | 11408.2863 | 1636.381389 | 2186.763112 | -2.72634186 | 0.210437313 | 4.6565771224 | 2.5180946939 | Km<KS.1 | YPD | KS.1 |
| FIM1_4957 | 3291.716141 | 458.0423621 | 753.7054491 | -2.764720711 | 0.219396925 | 4.0381256300 | 1.9316995201 | Km<KS.1 | YPD | KS.1 |
| FIM1_4958 | 182.7148318 | 429.1298346 | 219.4088314 | 0.924829691 | 0.425956394 | 0.00657494 | 0.012437387 | ns | YPD | KS.1 |
| FIM1_4960 | 1127.817367 | 3598.999125 | 4081.875028 | 1.654433506 | 0.099723533 | 4.1701634073 | 9.4302558871 | Km>KS.1 | YPD | KS.1 |
| FIM1_4961 | 3029.088243 | 2755.942699 | 3685.306776 | -0.11697605 | 0.205883703 | 0.538324297 | 0.612711823 | ns | YPD | KS.1 |
| FIM1_4962 | 1405.045733 | 583.6279239 | 811.1420474 | -1.220777569 | 0.162015273 | 6.1766761797 | 5.0960139293 | Km<KS.1 | YPD | KS.1 |
| FIM1_4963 | 1280.297771 | 536.3019506 | 718.4865154 | -1.232779685 | 0.110236629 | 6.0699484667 | 1.6411953055 | Km<KS.1 | YPD | KS.1 |
| FIM1_4964 | 230.8570811 | 166.3822632 | 90.85514937 | -0.448575117 | 0.13378566 | 0.00049751 | 0.001164211 | ns | YPD | KS.1 |
| FIM1_4965 | 3604.736622 | 7059.94899 | 7135.825911 | 0.944280826 | 0.121534456 | 1.6608155513 | 1.4546755929 | ns | YPD | KS.1 |
| FIM1_4966 | 3512.559185 | 2545.227626 | 4089.50433 | -0.451770495 | 0.095921775 | 1.5358197884 | 5.2658190541 | ns | YPD | KS.1 |
| FIM1_4967 | 3263.998077 | 2132.228429 | 1829.677499 | -0.587852681 | 0.130040167 | 2.8853335640 | 9.4562150733 | ns | YPD | KS.1 |
| FIM1_4968 | 897.078933 | 1000.417621 | 1696.526117 | 0.118857652 | 0.296461766 | 0.643268039 | 0.708805868 | ns | YPD | KS.1 |
| FIM1_4969 | 11882.74095 | 9155.826505 | 13025.51777 | -0.369055602 | 0.07424056 | 4.7547092727 | 1.7587121659 | ns | YPD | KS.1 |
| FIM1_4970 | 8915.543541 | 13438.1878 | 13733.02455 | 0.576785178 | 0.098216277 | 2.0333719811 | 1.0146465001 | ns | YPD | KS.1 |
| FIM1_4971 | 6977.408194 | 8751.091121 | 5299.304992 | 0.29079904 | 0.188586258 | 0.097871439 | 0.14166727 | ns | YPD | KS.1 |
| FIM1_4972 | 2232.941837 | 3340.44384 | 1907.458323 | 0.493712557 | 0.252210195 | 0.028649074 | 0.04705485 | ns | YPD | KS.1 |
| FIM1_4973 | 1384.310668 | 1981.598617 | 1688.204938 | 0.462311997 | 0.201868382 | 0.013316836 | 0.023652716 | ns | YPD | KS.1 |
| FIM1_4974 | 501.5481989 | 434.4846352 | 495.5369414 | -0.195775105 | 0.115888686 | 0.080982499 | 0.119728954 | ns | YPD | KS.1 |
| FIM1_4976 | 4532.239333 | 9365.359679 | 7599.367736 | 1.02187213 | 0.11945421 | 2.1507680053 | 2.4373737645 | Km>KS.1 | YPD | KS.1 |
| FIM1_4977 | 911.4690369 | 695.8012436 | 1707.165664 | -0.274232526 | 0.353429014 | 0.34708941 | 0.422296359 | ns | YPD | KS.1 |
| FIM1_4978 | 549.3771569 | 558.8276574 | 947.8801932 | 0.022050682 | 0.138283697 | 0.868019204 | 0.89854256 | ns | YPD | KS.1 |
| FIM1_4979 | 14027.45778 | 14975.33126 | 16927.46391 | 0.090411898 | 0.09396824 | 0.325029526 | 0.399166105 | ns | YPD | KS.1 |
| FIM1_4980 | 1487.419729 | 593.8269582 | 556.3859116 | -1.268615103 | 0.177615179 | 1.0585026741 | 7.8131317564 | Km<KS.1 | YPD | KS.1 |

| | | | | | | | | | | |
|---|---|---|---|---|---|---|---|---|---|---|
| FIM1_4981 | 9443.898411 | 12741.15415 | 15859.94697 | 0.42551568 | 0.068152086 | 2.77236762675 | 1.50573460077 | ns | YPD | KS.1 |
| FIM1_4982 | 1282.382009 | 1051.446817 | 1010.171553 | -0.280611154 | 0.067484743 | 2.57977338988 | 7.40159897041 | ns | YPD | KS.1 |
| FIM1_4983 | 1310.40599 | 976.3836267 | 976.9536087 | -0.411734306 | 0.100700041 | 2.88641261851 | 8.22445748976 | ns | YPD | KS.1 |
| FIM1_4984 | 3683.667022 | 2792.165038 | 1549.734645 | -0.366042915 | 0.168195547 | 0.021070384 | 0.035642693 | ns | YPD | KS.1 |
| FIM1_4985 | 2694.755894 | 3648.557518 | 2672.857618 | 0.42547658 | 0.092199168 | 2.55804999703 | 8.44479013618 | ns | YPD | KS.1 |
| FIM1_4986 | 4631.238955 | 4969.809708 | 4455.986282 | 0.098039866 | 0.087578888 | 0.253293649 | 0.322450333 | ns | YPD | KS.1 |
| FIM1_4987 | 1905.022727 | 1895.048759 | 2820.403304 | -0.007044683 | 0.1777828 | 0.966369431 | 0.97499248 | ns | YPD | KS.1 |
| FIM1_4988 | 206.7323649 | 165.3220454 | 174.4108259 | -0.295610161 | 0.167823768 | 0.061787392 | 0.094874159 | ns | YPD | KS.1 |
| FIM1_4989 | 10103.24464 | 10166.99318 | 7408.666665 | 0.008155456 | 0.160535139 | 0.957300112 | 0.968592242 | ns | YPD | KS.1 |
| FIM1_4991 | 1713.67201 | 4469.842317 | 3383.67469 | 1.309534034 | 0.202980283 | 1.17631186329 | 7.26975344086 | Km>KS.1 | YPD | KS.1 |
| FIM1_4992 | 461.130819 | 839.7885698 | 1100.95135 | 0.802646122 | 0.191284228 | 7.92643145616 | 2.43720621102 | ns | YPD | KS.1 |
| FIM1_4993 | 3033.39707 | 3482.264782 | 2479.73763 | 0.178949543 | 0.170145578 | 0.264309444 | 0.334080153 | ns | YPD | KS.1 |
| FIM1_4995 | 3350.598269 | 1589.659441 | 1753.530347 | -1.001369096 | 0.208542493 | 3.00498807633 | 1.14910189698 | Km<KS.1 | YPD | KS.1 |
| FIM1_4996 | 669.9248783 | 715.5714305 | 533.7981596 | 0.08496172 | 0.163303053 | 0.582689622 | 0.655114321 | ns | YPD | KS.1 |
| FIM1_4997 | 1559.138545 | 1467.031776 | 1646.699176 | -0.081945305 | 0.122012904 | 0.487060922 | 0.56470009 | ns | YPD | KS.1 |
| FIM1_4998 | 3834.843731 | 3281.080655 | 2531.997194 | -0.203957105 | 0.164094676 | 0.188474319 | 0.251585655 | ns | YPD | KS.1 |
| FIM1_4999 | 2672.885061 | 2673.549011 | 3751.186881 | 0.000364746 | 0.14952682 | 0.997960462 | 0.998361814 | ns | YPD | KS.1 |
| FIM1_5000 | 1515.679763 | 1150.212885 | 621.3457797 | -0.33248808 | 0.245830371 | 0.131740998 | 0.184052644 | ns | YPD | KS.1 |
| FIM1_5001 | 4212.270865 | 1738.590223 | 1113.903104 | -1.15121231 | 0.269212528 | 2.66108308055 | 8.76747571242 | Km<KS.1 | YPD | KS.1 |
| FIM1_5002 | 2963.581065 | 3040.373044 | 1846.561683 | 0.032990259 | 0.165288795 | 0.832953266 | 0.871125184 | ns | YPD | KS.1 |
| FIM1_5003 | 911.8228134 | 1033.402034 | 1211.205547 | 0.166714231 | 0.140512318 | 0.215078308 | 0.281805263 | ns | YPD | KS.1 |
| FIM1_5004 | 3451.178562 | 2091.539441 | 3222.634587 | -0.62794541 | 0.249556769 | 0.005144956 | 0.009975119 | ns | YPD | KS.1 |
| FIM1_5005 | 10766.59791 | 13821.26665 | 20594.94691 | 0.344721148 | 0.114862703 | 0.001984737 | 0.004189253 | ns | YPD | KS.1 |
| FIM1_5006 | 228.0029812 | 182.4699618 | 269.8336058 | -0.290510073 | 0.169740491 | 0.069231983 | 0.104362163 | ns | YPD | KS.1 |
| FIM1_5007 | 4580.815544 | 4552.017408 | 2601.176588 | -0.006768737 | 0.300250692 | 0.979170149 | 0.985309767 | ns | YPD | KS.1 |
| FIM1_5008 | 3454.548358 | 4848.264221 | 4296.317491 | 0.464909336 | 0.130857665 | 0.000230189 | 0.000573744 | ns | YPD | KS.1 |
| FIM1_5009 | 302.865771 | 47.6867775 | 119.1624388 | -2.413490891 | 0.385243172 | 2.44654728344 | 1.61856020414 | Km<KS.1 | YPD | KS.1 |
| FIM1_5010 | 1549.834162 | 1212.697787 | 1071.331298 | -0.311721119 | 0.199309851 | 0.091028164 | 0.132960985 | ns | YPD | KS.1 |
| FIM1_5011 | 80.22689723 | 152.8889505 | 171.8248649 | 0.856319469 | 0.207665749 | 9.69861137986 | 2.94930266594 | ns | YPD | KS.1 |
| FIM1_5012 | 464.1987812 | 480.5887666 | 962.7448218 | 0.04275351 | 0.175056644 | 0.795250662 | 0.839281299 | ns | YPD | KS.1 |
| FIM1_5013 | 1183.520454 | 1291.412871 | 1207.073186 | 0.116840428 | 0.133625814 | 0.362779233 | 0.43785218 | ns | YPD | KS.1 |
| FIM1_5014 | 2506.72505 | 1656.733083 | 2375.999414 | -0.580564157 | 0.104980577 | 1.50777155380 | 6.77611877161 | ns | YPD | KS.1 |
| FIM1_5015 | 1301.731047 | 2761.81924 | 2910.350965 | 1.030408318 | 0.177001402 | 1.05136035373 | 5.44278643061 | Km>KS.1 | YPD | KS.1 |
| FIM1_5016 | 1499.522242 | 1412.978094 | 752.5604019 | -0.072239345 | 0.219407367 | 0.719210913 | 0.775986617 | ns | YPD | KS.1 |
| FIM1_5017 | 1012.82842 | 3487.570576 | 2331.695015 | 1.701689264 | 0.210734171 | 3.22411721232 | 3.34166315236 | Km>KS.1 | YPD | KS.1 |
| FIM1_5018 | 262.6473269 | 394.4081232 | 382.6164024 | 0.544225645 | 0.16678933 | 0.000569029 | 0.001319777 | ns | YPD | KS.1 |
| FIM1_5019 | 2084.038261 | 1930.117747 | 1121.73002 | -0.091368093 | 0.233020701 | 0.665645595 | 0.728782314 | ns | YPD | KS.1 |
| FIM1_5020 | 2811.322977 | 1792.248277 | 1100.992639 | -0.522017498 | 0.301760014 | 0.045002142 | 0.070805078 | ns | YPD | KS.1 |
| FIM1_5021 | 1881.897366 | 1016.829917 | 757.4105412 | -0.863881921 | 0.117029701 | 3.87497336427 | 2.97041486706 | ns | YPD | KS.1 |
| FIM1_5022 | 6053.01788 | 3646.572701 | 2655.25825 | -0.714852228 | 0.097802399 | 9.17577634408 | 6.82354070430 | ns | YPD | KS.1 |
| FIM1_5023 | 1584.834109 | 1660.770346 | 1964.965564 | 0.064397328 | 0.105316137 | 0.529934426 | 0.605656736 | ns | YPD | KS.1 |
| FIM1_5024 | 1263.95845 | 2630.391153 | 3413.721675 | 1.030397467 | 0.1236864 | 1.43851036856 | 1.54236833698 | Km>KS.1 | YPD | KS.1 |
| FIM1_5025 | 2404.451262 | 3625.377853 | 2924.38756 | 0.529447106 | 0.209746138 | 0.006166872 | 0.011732386 | ns | YPD | KS.1 |
| FIM1_5026 | 9546.093174 | 2496.739424 | 2627.883575 | -1.898208121 | 0.138522156 | 2.74265357107 | 1.81929353548 | Km<KS.1 | YPD | KS.1 |
| FIM1_5027 | 2986.98297 | 2229.246248 | 3523.227109 | -0.383864238 | 0.176821522 | 0.020758416 | 0.035173129 | ns | YPD | KS.1 |

| | | | | | | | | | | |
|---|---|---|---|---|---|---|---|---|---|---|
| FIM1_5028 | 5118.525935 | 4201.289538 | 3668.055305 | -0.274326357 | 0.101189834 | 0.005440849 | 0.010467218 | ns | YPD | KS.1 |
| FIM1_5029 | 2665.010242 | 2840.517172 | 2781.189831 | 0.089372891 | 0.082669077 | 0.27082837 | 0.341020284 | ns | YPD | KS.1 |
| FIM1_5030 | 1517.156859 | 2424.5285 | 3657.061808 | 0.666314696 | 0.076212698 | 8.5462153946 | 1.0027693770 | ns | YPD | KS.1 |
| FIM1_5031 | 3478.217409 | 3487.553976 | 5372.95306 | 0.003599195 | 0.126302932 | 0.976443423 | 0.983759828 | ns | YPD | KS.1 |
| FIM1_5032 | 1337.969832 | 1040.740723 | 2251.585274 | -0.285487165 | 0.283108023 | 0.24833387 | 0.317028741 | ns | YPD | KS.1 |
| FIM1_5033 | 1663.896904 | 1989.339526 | 1764.733732 | 0.252573255 | 0.071445677 | 0.000338593 | 0.000815739 | ns | YPD | KS.1 |
| FIM1_5034 | 6355.207741 | 3080.714933 | 4885.437389 | -1.014257029 | 0.131746657 | 2.5452086584 | 2.1756723498 | Km<KS.1 | YPD | KS.1 |
| FIM1_5035 | 1300.699517 | 1208.726592 | 1070.618224 | -0.090166754 | 0.208632703 | 0.639262795 | 0.705486336 | ns | YPD | KS.1 |
| FIM1_5036 | 1019.231699 | 882.6784843 | 1330.740806 | -0.187471098 | 0.167060465 | 0.234400687 | 0.302216453 | ns | YPD | KS.1 |
| FIM1_5037 | 1940.643547 | 1529.395397 | 1385.984308 | -0.337388321 | 0.074610914 | 4.5811853042 | 1.4628624446 | ns | YPD | KS.1 |
| FIM1_5038 | 2367.40933 | 3401.970565 | 3473.213098 | 0.505561088 | 0.108309084 | 1.6960070842 | 5.7673514998 | ns | YPD | KS.1 |
| FIM1_5039 | 1531.722729 | 1976.3156 | 1431.105986 | 0.347923782 | 0.12959579 | 0.005328982 | 0.010271865 | ns | YPD | KS.1 |
| FIM1_5040 | 7599.269848 | 10530.20573 | 15960.82951 | 0.445632875 | 0.135658824 | 0.000637944 | 0.001470018 | ns | YPD | KS.1 |
| FIM1_5041 | 564.1392318 | 575.9766365 | 428.8023825 | 0.026301665 | 0.151812845 | 0.855760337 | 0.888440667 | ns | YPD | KS.1 |
| FIM1_5042 | 15897.86236 | 22658.97951 | 18192.75871 | 0.481301331 | 0.145995511 | 0.000572378 | 0.001326924 | ns | YPD | KS.1 |
| FIM1_5043 | 862.7982571 | 853.9862292 | 914.906838 | -0.013244038 | 0.172678866 | 0.935144761 | 0.952935595 | ns | YPD | KS.1 |
| FIM1_5044 | 197.9752508 | 303.2352653 | 237.6434499 | 0.543877027 | 0.223554559 | 0.007739723 | 0.014415994 | ns | YPD | KS.1 |
| FIM1_5045 | 5046.483659 | 2833.895795 | 2635.336793 | -0.775682665 | 0.186221818 | 9.6289861015 | 2.9299208474 | ns | YPD | KS.1 |
| FIM1_5046 | 7968.716648 | 7557.740217 | 3462.930132 | -0.060785433 | 0.258709342 | 0.792078307 | 0.836643222 | ns | YPD | KS.1 |
| FIM1_5047 | 397.2309511 | 1573.980943 | 1684.120007 | 1.958027234 | 0.122510002 | 3.9197447952 | 6.5002434520 | Km>KS.1 | YPD | KS.1 |
| FIM1_5048 | 348.4711327 | 384.1864402 | 648.1284846 | 0.133135651 | 0.114806238 | 0.231322892 | 0.298917243 | ns | YPD | KS.1 |
| FIM1_5049 | 1865.947366 | 2371.031686 | 2315.334913 | 0.333199172 | 0.101310916 | 0.000755108 | 0.001711464 | ns | YPD | KS.1 |
| FIM1_5050 | 4247.053689 | 3422.367628 | 2158.68306 | -0.286136495 | 0.157459625 | 0.055381525 | 0.085888136 | ns | YPD | KS.1 |
| FIM1_5051 | 83.24462595 | 38.89464863 | 26.75475964 | -0.987862936 | 0.262602344 | 3.3098229797 | 9.3558916615 | ns | YPD | KS.1 |
| FIM1_5052 | 1012.476959 | 470.9225042 | 833.6461928 | -1.077514955 | 0.124402942 | 7.6398635102 | 9.0067111288 | Km<KS.1 | YPD | KS.1 |
| FIM1_5053 | 592.2893728 | 1058.244018 | 1307.8111 | 0.804022697 | 0.138848976 | 2.0022741599 | 1.0031534688 | ns | YPD | KS.1 |
| FIM1_5054 | 449.3998165 | 495.8417248 | 670.2714311 | 0.128752959 | 0.150929663 | 0.370453516 | 0.44646469 | ns | YPD | KS.1 |
| FIM1_5055 | 1219.15416 | 3040.934766 | 3470.755075 | 1.292208264 | 0.119092428 | 2.1904579397 | 5.4487641251 | Km>KS.1 | YPD | KS.1 |
| FIM1_5056 | 1606.948107 | 1615.651145 | 1420.427163 | 0.007111008 | 0.12430565 | 0.952774522 | 0.96666881 | ns | YPD | KS.1 |
| FIM1_5057 | 3327.052406 | 2445.157678 | 6219.004623 | -0.33627909 | 0.316728718 | 0.212132599 | 0.278385565 | ns | YPD | KS.1 |
| FIM1_5058 | 1473.621887 | 1107.293415 | 2622.191853 | -0.36304708 | 0.20488094 | 0.05471218 | 0.08495415 | ns | YPD | KS.1 |
| FIM1_5059 | 651.3788041 | 730.6353889 | 686.9090204 | 0.160387701 | 0.081874196 | 0.045984559 | 0.072213756 | ns | YPD | KS.1 |
| FIM1_5061 | 1786.996571 | 987.7089682 | 2153.880053 | -0.819862809 | 0.144328358 | 3.7108323618 | 1.7941099125 | ns | YPD | KS.1 |
| FIM1_5062 | 791.4882378 | 754.394728 | 633.3694877 | -0.066252921 | 0.11302773 | 0.545792282 | 0.619369663 | ns | YPD | KS.1 |
| FIM1_5063 | 923.0809141 | 1387.963996 | 1990.11932 | 0.558888169 | 0.139640799 | 3.1332111100 | 8.8717844463 | ns | YPD | KS.1 |
| FIM1_5064 | 408.5941534 | 659.8935234 | 645.2072038 | 0.618862224 | 0.218345231 | 0.002028688 | 0.004272957 | ns | YPD | KS.1 |
| FIM1_5065 | 6608.550481 | 4834.858014 | 3945.297633 | -0.403607909 | 0.194330092 | 0.025366247 | 0.042220501 | ns | YPD | KS.1 |
| FIM1_5066 | 761.3057944 | 543.5128846 | 475.1938222 | -0.470871459 | 0.101457672 | 2.0696411017 | 6.9336461153 | ns | YPD | KS.1 |
| FIM1_5067 | 1341.187234 | 3081.92229 | 3488.00408 | 1.175557191 | 0.117051132 | 1.3487340506 | 2.5320573215 | Km>KS.1 | YPD | KS.1 |
| FIM1_5068 | 456.5715261 | 750.5906169 | 1083.197022 | 0.662600148 | 0.183261653 | 0.000119849 | 0.000308616 | ns | YPD | KS.1 |
| FIM1_5069 | 6858.086463 | 1894.214232 | 1090.397409 | -1.630644201 | 0.3549252 | 2.1853336885 | 8.4673170564 | Km<KS.1 | YPD | KS.1 |
| FIM1_5070 | 103.7287608 | 47.19268909 | 66.93085334 | -1.072804476 | 0.197432844 | 9.1523317694 | 4.2277484264 | Km<KS.1 | YPD | KS.1 |
| FIM1_5071 | 3298.995003 | 4009.332406 | 7131.434308 | 0.267578719 | 0.11674343 | 0.017931675 | 0.030964971 | ns | YPD | KS.1 |
| FIM1_5072 | 76.314119 | 156.1610944 | 128.7444275 | 0.931044249 | 0.24132379 | 2.5336961357 | 7.2819978483 | ns | YPD | KS.1 |
| FIM1_5073 | 317.254182 | 471.2323327 | 611.9560529 | 0.523843098 | 0.176613014 | 0.001622647 | 0.003470623 | ns | YPD | KS.1 |

| | | | | | | | | | | |
|---|---|---|---|---|---|---|---|---|---|---|
| FIM1_5074 | 731.0584148 | 635.0490674 | 663.8664992 | -0.186045219 | 0.155742273 | 0.208349039 | 0.274143472 | ns | YPD | KS.1 |
| FIM1_5075 | 2917.469093 | 2833.643152 | 1277.11883 | -0.037860785 | 0.16104853 | 0.80423694 | 0.846251856 | ns | YPD | KS.1 |
| FIM1_5076 | 1511.944566 | 2294.859328 | 3651.790514 | 0.578948711 | 0.121512393 | 9.0127299767 | 3.2096157218 | ns | YPD | KS.1 |
| FIM1_5077 | 1917.409721 | 2865.657083 | 2209.852189 | 0.541892233 | 0.159814066 | 0.000361377 | 0.000865106 | ns | YPD | KS.1 |
| FIM1_5078 | 779.3146929 | 588.6733146 | 454.6684508 | -0.38197818 | 0.13639319 | 0.003552928 | 0.007121602 | ns | YPD | KS.1 |
| FIM1_5079 | 3958.958603 | 4839.533804 | 6011.899769 | 0.272910328 | 0.128692118 | 0.027624939 | 0.04558344 | ns | YPD | KS.1 |
| FIM1_5080 | 867.881653 | 1131.673003 | 1060.424088 | 0.37877572 | 0.061480993 | 5.0862833862 | 2.7057081101 | ns | YPD | KS.1 |
| FIM1_5081 | 1695.743931 | 1608.009468 | 1610.532415 | -0.072874213 | 0.110521282 | 0.497235263 | 0.574221317 | ns | YPD | KS.1 |
| FIM1_5082 | 278.4285485 | 394.6386598 | 557.9162857 | 0.474434096 | 0.137998399 | 0.00034753 | 0.000835247 | ns | YPD | KS.1 |
| FIM1_5083 | 3483.243899 | 5552.273234 | 4870.532015 | 0.592597231 | 0.231659644 | 0.004919366 | 0.009571312 | ns | YPD | KS.1 |
| FIM1_5084 | 975.6696637 | 401.8053976 | 802.1607 | -1.13179222 | 0.293033565 | 1.6225624338 | 4.7793061626 | Km<KS.1 | YPD | KS.1 |
| FIM1_5085 | 1720.997642 | 3154.389814 | 1763.991479 | 0.786385736 | 0.232154812 | 0.000215271 | 0.000538718 | ns | YPD | KS.1 |
| FIM1_5086 | 3090.312629 | 3374.173953 | 2732.781178 | 0.118370362 | 0.128572639 | 0.339076788 | 0.413964913 | ns | YPD | KS.1 |
| FIM1_5087 | 644.4621561 | 853.5739782 | 1469.97806 | 0.362959962 | 0.187250763 | 0.037679702 | 0.060159344 | ns | YPD | KS.1 |
| FIM1_5088 | 9097.225532 | 10684.23532 | 9428.848193 | 0.196909601 | 0.21959537 | 0.326724512 | 0.400753068 | ns | YPD | KS.1 |
| FIM1_5089 | 1132.31688 | 1678.389478 | 2468.934404 | 0.482043795 | 0.252071582 | 0.032518622 | 0.052697116 | ns | YPD | KS.1 |
| FIM1_5090 | 522.7952577 | 787.145796 | 387.9351323 | 0.478388206 | 0.288945863 | 0.057204861 | 0.088465708 | ns | YPD | KS.1 |
| FIM1_5091 | 275.5109867 | 393.8513332 | 492.4874872 | 0.429172293 | 0.256735389 | 0.060583228 | 0.093168952 | ns | YPD | KS.1 |
| FIM1_5093 | 1377.125512 | 1245.699971 | 1273.576786 | -0.138288784 | 0.096061578 | 0.140372023 | 0.193986337 | ns | YPD | KS.1 |
| FIM1_5094 | 56833.69205 | 61178.68436 | 89508.85285 | 0.101215491 | 0.103601491 | 0.3157208 | 0.388982412 | ns | YPD | KS.1 |
| FIM1_5095 | 229.8408188 | 393.1462586 | 503.4258247 | 0.737202256 | 0.143445299 | 9.1873188565 | 3.7403364411 | ns | YPD | KS.1 |
| FIM1_5096 | 788.6855543 | 951.2092624 | 1407.574219 | 0.243292643 | 0.174855976 | 0.138495642 | 0.192033394 | ns | YPD | KS.1 |
| FIM1_5097 | 7466.763015 | 10806.64309 | 9492.802646 | 0.524319192 | 0.076070234 | 2.8958174008 | 1.9006189405 | ns | YPD | KS.1 |
| FIM1_5098 | 348.8140136 | 464.7605475 | 297.09687 | 0.360873286 | 0.214743496 | 0.066701297 | 0.101139577 | ns | YPD | KS.1 |
| FIM1_5099 | 335.6572395 | 1056.606254 | 1756.661765 | 1.622871534 | 0.130899125 | 1.5139701500 | 6.3294130222 | Km>KS.1 | YPD | KS.1 |
| FIM1_5100 | 1092.835992 | 1227.418156 | 1058.239174 | 0.165015558 | 0.057696646 | 0.003859082 | 0.007679573 | ns | YPD | KS.1 |
| FIM1_5101 | 2624.211006 | 1245.455909 | 1189.433938 | -1.049036977 | 0.120913382 | 7.0871856908 | 8.3949401933 | Km<KS.1 | YPD | KS.1 |
| FIM1_5102 | 14325.67348 | 5299.847529 | 3568.104286 | -1.313248615 | 0.262249147 | 5.6477244943 | 2.3611285175 | Km<KS.1 | YPD | KS.1 |
| FIM1_5103 | 281.1486092 | 402.5135274 | 511.8376738 | 0.488193523 | 0.14164994 | 0.000327957 | 0.000793188 | ns | YPD | KS.1 |
| FIM1_5104 | 722.9794654 | 684.4674199 | 743.3096505 | -0.077289928 | 0.086260145 | 0.360865279 | 0.435859374 | ns | YPD | KS.1 |
| FIM1_5105 | 2009.146722 | 2164.95649 | 2556.219037 | 0.104500791 | 0.073670647 | 0.149741472 | 0.205450585 | ns | YPD | KS.1 |
| FIM1_5106 | 1160.337755 | 1552.889144 | 873.3045665 | 0.394396436 | 0.142659726 | 0.003889476 | 0.00773078 | ns | YPD | KS.1 |
| FIM1_5107 | 2615.644868 | 3173.019465 | 4318.641348 | 0.257996567 | 0.146214873 | 0.064402942 | 0.098163185 | ns | YPD | KS.1 |
| FIM1_5108 | 439.0635079 | 952.9103437 | 919.5715219 | 1.075613931 | 0.157259336 | 1.3111683304 | 8.9602506098 | Km>KS.1 | YPD | KS.1 |
| FIM1_5109 | 560.8604968 | 832.2511415 | 1055.894387 | 0.533723421 | 0.158603973 | 0.000403394 | 0.000957027 | ns | YPD | KS.1 |
| FIM1_5110 | 4672.047096 | 1926.591154 | 1771.777108 | -1.253485214 | 0.115498543 | 2.2920941302 | 5.6732180586 | Km<KS.1 | YPD | KS.1 |
| FIM1_5111 | 1385.353839 | 1862.259833 | 1833.436691 | 0.412094059 | 0.105422295 | 6.1616002072 | 0.000167546 | ns | YPD | KS.1 |
| FIM1_5112 | 1295.058093 | 874.7958115 | 912.3996043 | -0.543947467 | 0.118790123 | 2.4015807424 | 7.9599361718 | ns | YPD | KS.1 |
| FIM1_5113 | 311.9976098 | 146.7037519 | 185.8773579 | -1.022989815 | 0.194714105 | 2.7252827755 | 1.1908350619 | Km<KS.1 | YPD | KS.1 |
| FIM1_5114 | 619.2550592 | 285.9230797 | 473.4386546 | -1.031330898 | 0.21954914 | 4.7338678726 | 1.7523060019 | Km<KS.1 | YPD | KS.1 |
| FIM1_5115 | 770.3633683 | 1065.378393 | 1717.039464 | 0.445296654 | 0.126192235 | 0.000261506 | 0.000643737 | ns | YPD | KS.1 |
| FIM1_5116 | 661.0125117 | 1146.236346 | 1592.914278 | 0.766290173 | 0.125672062 | 3.3255131342 | 1.7963548146 | ns | YPD | KS.1 |
| FIM1_5117 | 635.6986438 | 640.5994269 | 553.3084487 | 0.00969703 | 0.229710412 | 0.962993678 | 0.972577301 | ns | YPD | KS.1 |
| FIM1_5118 | 1424.060252 | 1889.327773 | 2917.278492 | 0.382201068 | 0.143785402 | 0.005468444 | 0.010512176 | ns | YPD | KS.1 |
| FIM1_5119 | 1115.713789 | 1088.086916 | 1408.965236 | -0.034309771 | 0.1123614 | 0.753141974 | 0.80508838 | ns | YPD | KS.1 |

| | | | | | | | | | | |
|---|---|---|---|---|---|---|---|---|---|---|
| FIM1_5120 | 1095.705572 | 1897.626932 | 1218.891875 | 0.713738352 | 0.222361926 | 0.000476349 | 0.001116794 | ns | YPD | KS.1 |
| FIM1_5121 | 606.4086688 | 1113.291245 | 2267.146921 | 0.837709597 | 0.152062505 | 9.62173263343 | 4.42813319623 | ns | YPD | KS.1 |
| FIM1_5122 | 10240.08409 | 8653.661268 | 4430.879936 | -0.18271239 | 0.30288103 | 0.483455053 | 0.561566399 | ns | YPD | KS.1 |
| FIM1_5123 | 1747.136188 | 2808.249555 | 4067.119462 | 0.528404125 | 0.333017891 | 0.059601681 | 0.091858229 | ns | YPD | KS.1 |
| FIM1_5124 | 351.8673765 | 876.677179 | 969.6212758 | 1.296355974 | 0.106570502 | 5.24604151836 | 1.99229439342 | Km>KS.1 | YPD | KS.1 |
| FIM1_5125 | 1221.11897 | 1442.273648 | 2718.883313 | 0.190745968 | 0.268628502 | 0.421690904 | 0.499384015 | ns | YPD | KS.1 |
| FIM1_5126 | 1313.670999 | 1834.933081 | 1546.701462 | 0.474548938 | 0.069871139 | 6.51311187361 | 4.14357181217 | ns | YPD | KS.1 |
| FIM1_5127 | 6232.975589 | 5712.663732 | 4733.280554 | -0.115396898 | 0.146055744 | 0.408057788 | 0.48531855 | ns | YPD | KS.1 |
| FIM1_5128 | 3092.554732 | 3881.21377 | 3394.131703 | 0.317327652 | 0.095795407 | 0.000709397 | 0.001616539 | ns | YPD | KS.1 |
| FIM1_5129 | 2221.095066 | 1337.636181 | 1408.446859 | -0.701280636 | 0.136598662 | 1.02425949772 | 4.12606558798 | ns | YPD | KS.1 |
| FIM1_5130 | 2697.986384 | 2792.619976 | 2394.966011 | 0.044839158 | 0.157669881 | 0.764694333 | 0.815510034 | ns | YPD | KS.1 |
| FIM1_5131 | 2666.528637 | 3522.730262 | 2682.725242 | 0.276581706 | 0.366805702 | 0.35524636 | 0.430326428 | ns | YPD | KS.1 |
| FIM1_5132 | 368.9564921 | 274.4630155 | 193.6028101 | -0.31050037 | 0.33834248 | 0.27230881 | 0.342797654 | ns | YPD | KS.1 |
| FIM1_5133 | 2385.415786 | 2096.778029 | 1610.349592 | -0.171297577 | 0.145254387 | 0.216762811 | 0.283415239 | ns | YPD | KS.1 |
| FIM1_5134 | 174.0035879 | 311.8170666 | 596.8297995 | 0.789613964 | 0.181022802 | 3.86365343848 | 1.24573401532 | ns | YPD | KS.1 |
| FIM1_5135 | 548.0275319 | 1033.98718 | 1370.994416 | 0.872789594 | 0.159614522 | 1.12697812581 | 5.14849970243 | ns | YPD | KS.1 |
| FIM1_5136 | 1720.871479 | 2007.735593 | 2722.172896 | 0.206779099 | 0.136395091 | 0.11383176 | 0.162308223 | ns | YPD | KS.1 |
| FIM1_5137 | 657.8286387 | 938.0458381 | 955.2377765 | 0.467941914 | 0.177193733 | 0.004954803 | 0.009636492 | ns | YPD | KS.1 |
| FIM1_5138 | 1440.686255 | 2484.101541 | 2129.186013 | 0.674098481 | 0.268511708 | 0.00473436 | 0.009262069 | ns | YPD | KS.1 |
| FIM1_5139 | 1665.086097 | 4041.90966 | 3263.840028 | 1.142033088 | 0.281986777 | 7.27411594300 | 2.25615503843 | Km>KS.1 | YPD | KS.1 |
| FIM1_5140 | 1434.746673 | 2926.749924 | 3280.874518 | 0.887826658 | 0.291132018 | 0.000559035 | 0.001299018 | ns | YPD | KS.1 |
| FIM1_5141 | 6113.914788 | 4571.468525 | 2740.679032 | -0.388891826 | 0.156664063 | 0.008982036 | 0.016550233 | ns | YPD | KS.1 |
| FIM1_5142 | 1730.141822 | 2232.254356 | 1638.967762 | 0.315457379 | 0.222614845 | 0.120089844 | 0.170067456 | ns | YPD | KS.1 |
| FIM1_5143 | 2791.155537 | 2368.791753 | 2447.338046 | -0.229462009 | 0.085933164 | 0.006475463 | 0.012258535 | ns | YPD | KS.1 |
| FIM1_5144 | 1556.042693 | 1816.22247 | 1888.397872 | 0.211329245 | 0.117959575 | 0.063991722 | 0.097611467 | ns | YPD | KS.1 |
| FIM1_5145 | 1997.482622 | 1757.060526 | 2312.449822 | -0.172507596 | 0.131847893 | 0.173374863 | 0.233687332 | ns | YPD | KS.1 |
| FIM1_5146 | 825.9475845 | 375.3922338 | 627.1426176 | -1.035808067 | 0.244405934 | 4.00716492213 | 1.28700099984 | Km<KS.1 | YPD | KS.1 |
| FIM1_5147 | 13587.32828 | 18341.24065 | 29081.46093 | 0.413589901 | 0.120974959 | 0.000416494 | 0.000985287 | ns | YPD | KS.1 |
| FIM1_5148 | 236.6778963 | 243.4801217 | 298.8760442 | 0.033371848 | 0.238394866 | 0.876991695 | 0.90575746 | ns | YPD | KS.1 |
| FIM1_5149 | 5121.298901 | 2849.129837 | 4101.836441 | -0.632393803 | 0.374757861 | 0.038998977 | 0.062007002 | ns | YPD | KS.1 |
| FIM1_5150 | 8645.34198 | 11477.6996 | 9268.607682 | 0.357072511 | 0.212087474 | 0.066663749 | 0.10111346 | ns | YPD | KS.1 |
| FIM1_5151 | 171.4853324 | 232.728402 | 223.4158694 | 0.409050495 | 0.15227978 | 0.004811516 | 0.009405616 | ns | YPD | KS.1 |
| FIM1_5152 | 226.6980828 | 215.5995562 | 293.9752473 | -0.058019476 | 0.24422777 | 0.791948424 | 0.836643222 | ns | YPD | KS.1 |
| FIM1_5153 | 5593.845261 | 6669.372802 | 6107.404169 | 0.244753723 | 0.096718318 | 0.009568964 | 0.017560161 | ns | YPD | KS.1 |
| FIM1_5155 | 838.101241 | 870.8697788 | 1027.34733 | 0.051721519 | 0.132785351 | 0.685526333 | 0.747423517 | ns | YPD | KS.1 |
| FIM1_5156 | 452.5638832 | 831.8400406 | 907.6906643 | 0.789028447 | 0.23500112 | 0.00023852 | 0.000591543 | ns | YPD | KS.1 |
| FIM1_5157 | 1651.375116 | 2085.62624 | 2358.045634 | 0.330028215 | 0.071580721 | 3.03348317911 | 9.90911281424 | ns | YPD | KS.1 |
| FIM1_5158 | 2142.7607 | 1855.352424 | 1384.854333 | -0.179608829 | 0.203047884 | 0.338497016 | 0.413358531 | ns | YPD | KS.1 |
| FIM1_5159 | 913.1766797 | 871.1357765 | 963.2899021 | -0.064358722 | 0.123609184 | 0.58984459 | 0.661812547 | ns | YPD | KS.1 |
| FIM1_5160 | 4687.691082 | 8281.945093 | 11959.88436 | 0.72565977 | 0.245110008 | 0.001074717 | 0.002366851 | ns | YPD | KS.1 |
| FIM1_5161 | 1030.786344 | 3093.243277 | 1637.126416 | 1.441585738 | 0.284079567 | 3.06109638481 | 1.33003969558 | Km>KS.1 | YPD | KS.1 |
| FIM1_5162 | 1848.103776 | 2294.300722 | 2683.565014 | 0.300539032 | 0.102200639 | 0.002569152 | 0.005283807 | ns | YPD | KS.1 |
| FIM1_5163 | 4142.393033 | 3191.628264 | 3379.838754 | -0.341677159 | 0.173967835 | 0.036671609 | 0.058757248 | ns | YPD | KS.1 |
| FIM1_5164 | 1488.054055 | 257.6320266 | 390.2049425 | -2.501256892 | 0.1295248 | 1.56628121369 | 9.74031129767 | Km<KS.1 | YPD | KS.1 |
| FIM1_5165 | 2567.256736 | 1760.119728 | 1240.079075 | -0.517186346 | 0.136931797 | 8.65373703466 | 0.000228273 | ns | YPD | KS.1 |

| | | | | | | | | | | |
|---|---|---|---|---|---|---|---|---|---|---|
| FIM1_5166 | 2512.72118 | 3499.938395 | 2961.30963 | 0.419129353 | 0.215632264 | 0.033911552 | 0.05482937 | ns | YPD | KS.1 |
| FIM1_5167 | 301.1830712 | 724.0527333 | 637.241487 | 1.234716417 | 0.1277557 | 5.2455176021 | 8.6126897922 | Km>KS.1 | YPD | KS.1 |
| FIM1_5168 | 1582.314213 | 948.8764321 | 781.3317534 | -0.699871167 | 0.15207545 | 1.5238787029 | 5.2320887145 | ns | YPD | KS.1 |
| FIM1_5169 | 441.2314619 | 158.4762283 | 280.748438 | -1.400698685 | 0.20869126 | 1.7154957232 | 1.1580178050 | Km<KS.1 | YPD | KS.1 |
| FIM1_5170 | 3191.061202 | 2310.262636 | 2146.695084 | -0.45471696 | 0.087234779 | 1.1460366470 | 4.5906057318 | ns | YPD | KS.1 |
| FIM1_5171 | 2299.636156 | 1085.045639 | 1014.350572 | -1.034330003 | 0.168702382 | 1.5607103110 | 8.6948866710 | Km<KS.1 | YPD | KS.1 |
| FIM1_5172 | 3166.426503 | 5079.359498 | 7294.21892 | 0.657707264 | 0.121921716 | 2.7263540413 | 1.1908350619 | ns | YPD | KS.1 |
| FIM1_5173 | 2013.254109 | 1805.973724 | 1704.099513 | -0.145998686 | 0.133571617 | 0.255108529 | 0.32418006 | ns | YPD | KS.1 |
| FIM1_5174 | 5319.342667 | 4072.898055 | 4111.276212 | -0.375887165 | 0.084019134 | 5.4262848131 | 1.7118431798 | ns | YPD | KS.1 |
| FIM1_5175 | 1663.724553 | 2830.864386 | 3329.456953 | 0.731705232 | 0.146666853 | 2.0514485594 | 7.9858815207 | ns | YPD | KS.1 |
| FIM1_5176 | 601.5625798 | 217.826658 | 608.5618791 | -1.379194733 | 0.218263633 | 2.4384831585 | 1.4669230609 | Km<KS.1 | YPD | KS.1 |
| FIM1_5177 | 730.2525279 | 571.2270942 | 1527.121631 | -0.258753967 | 0.32909227 | 0.350625514 | 0.425661769 | ns | YPD | KS.1 |
| FIM1_5178 | 1318.569476 | 863.8583412 | 1130.801298 | -0.592775444 | 0.102757222 | 3.6561762796 | 1.7694043766 | ns | YPD | KS.1 |
| FIM1_5179 | 2686.759185 | 2303.622731 | 4531.64748 | -0.174787561 | 0.272143146 | 0.466286532 | 0.543909847 | ns | YPD | KS.1 |
| FIM1_5180 | 1181.550016 | 785.0552283 | 802.0512042 | -0.563119612 | 0.133852928 | 1.2798049419 | 3.8263398955 | ns | YPD | KS.1 |
| FIM1_5181 | 3248.58888 | 4389.771289 | 4942.464288 | 0.411478018 | 0.131672683 | 0.001180478 | 0.002578148 | ns | YPD | KS.1 |
| FIM1_5182 | 578.3082216 | 816.0161332 | 1281.7074 | 0.479715559 | 0.101348486 | 1.2958131710 | 4.4862007836 | ns | YPD | KS.1 |
| FIM1_5183 | 2691.657576 | 3787.12835 | 5023.530225 | 0.469013314 | 0.130114515 | 0.000187341 | 0.000472627 | ns | YPD | KS.1 |
| FIM1_5184 | 544.8942224 | 635.0082633 | 724.6511252 | 0.205289299 | 0.138810188 | 0.122506581 | 0.172997514 | ns | YPD | KS.1 |
| FIM1_5185 | 2066.170953 | 2989.188699 | 3451.371304 | 0.51518907 | 0.107678082 | 9.3536823172 | 3.3191561717 | ns | YPD | KS.1 |
| FIM1_5186 | 1541.345898 | 1613.252541 | 1683.684437 | 0.061857939 | 0.106847817 | 0.551811467 | 0.625202015 | ns | YPD | KS.1 |
| FIM1_5187 | 4178.434301 | 3290.051863 | 2096.002034 | -0.309430204 | 0.182912153 | 0.070344071 | 0.105806547 | ns | YPD | KS.1 |
| FIM1_5188 | 3555.597447 | 6295.547473 | 8105.092407 | 0.774096319 | 0.174514133 | 2.8430507324 | 9.3237820659 | ns | YPD | KS.1 |
| FIM1_5189 | 183.6082435 | 185.1161975 | 255.0617251 | 0.007030223 | 0.239037064 | 0.974047685 | 0.981743767 | ns | YPD | KS.1 |
| FIM1_5190 | 1101.107462 | 1045.600786 | 1079.103786 | -0.069915286 | 0.127915437 | 0.570690265 | 0.643368245 | ns | YPD | KS.1 |
| FIM1_5191 | 4339.041094 | 1322.703458 | 1573.276048 | -1.66032789 | 0.169311619 | 5.5611044001 | 1.0060543414 | Km<KS.1 | YPD | KS.1 |
| FIM1_5192 | 11872.9286 | 1792.994983 | 1933.053245 | -2.458330336 | 0.398776135 | 4.0595177748 | 2.6194683436 | Km<KS.1 | YPD | KS.1 |
| FIM1_5193 | 10824.51166 | 13223.16658 | 15741.76877 | 0.276730207 | 0.107881968 | 0.008343345 | 0.015470795 | ns | YPD | KS.1 |
| FIM1_5194 | 4171.397888 | 3058.772621 | 3900.873431 | -0.410482192 | 0.170351222 | 0.010597289 | 0.019234409 | ns | YPD | KS.1 |
| FIM1_5195 | 7187.917126 | 6163.909175 | 5536.513644 | -0.208685903 | 0.125654334 | 0.084846072 | 0.124773636 | ns | YPD | KS.1 |
| FIM1_5196 | 329.9007379 | 217.2023628 | 274.2174018 | -0.574418629 | 0.132959105 | 7.5170434301 | 2.3213712641 | ns | YPD | KS.1 |
| FIM1_5197 | 400.2668509 | 370.8235374 | 384.7279617 | -0.105184555 | 0.114139794 | 0.341802648 | 0.416883592 | ns | YPD | KS.1 |
| FIM1_5199 | 2676.063021 | 3215.131756 | 3164.931706 | 0.257121743 | 0.087642288 | 0.00277126 | 0.005666675 | ns | YPD | KS.1 |
| FIM1_5200 | 776.0664784 | 860.9967867 | 329.1609456 | 0.114262271 | 0.288317238 | 0.649204863 | 0.714398185 | ns | YPD | KS.1 |
| FIM1_5201 | 1424.504137 | 889.3019062 | 502.8709471 | -0.610008995 | 0.214325293 | 0.00199177 | 0.004198752 | ns | YPD | KS.1 |
| FIM1_5202 | 1095.548766 | 574.8735169 | 778.6488792 | -0.910530373 | 0.106813561 | 3.4403557513 | 3.8204843444 | ns | YPD | KS.1 |
| FIM1_5203 | 1324.292695 | 824.1360043 | 745.3949667 | -0.646719775 | 0.153819707 | 1.0784584613 | 3.2615992979 | ns | YPD | KS.1 |
| FIM1_5204 | 1259.525471 | 694.3992413 | 477.5816684 | -0.795913286 | 0.196098074 | 1.4636989578 | 4.3318871594 | ns | YPD | KS.1 |
| FIM1_5205 | 1241.758302 | 1637.677404 | 1564.827119 | 0.388486096 | 0.08873308 | 8.2842191167 | 2.5393709245 | ns | YPD | KS.1 |
| FIM1_5206 | 1466.897115 | 1067.44206 | 958.1753731 | -0.442751116 | 0.104503516 | 1.4286790218 | 4.2433899305 | ns | YPD | KS.1 |
| FIM1_5207 | 497.2825159 | 260.8573393 | 269.4435916 | -0.87957536 | 0.17472113 | 1.1771472122 | 4.7076425893 | ns | YPD | KS.1 |
| FIM1_5208 | 58.16171777 | 11.39301758 | 21.5559521 | -1.794657013 | 0.56518539 | 4.0755560740 | 0.000114038 | Km<KS.1 | YPD | KS.1 |
| FIM1_5209 | 71.98120359 | 47.03703431 | 37.8397702 | -0.462943889 | 0.342814407 | 0.105564717 | 0.151524659 | ns | YPD | KS.1 |
| FIM1_4316 | 5277.880815 | 2652.730751 | 4233.280858 | -0.920346463 | 0.206858362 | 1.9499063889 | 6.5545839762 | ns | YPD | KS.1 |
| FIM1_4318 | 1002.389709 | 307.5633106 | 297.7479064 | -1.656346188 | 0.159409127 | 1.4501395343 | 3.1781692437 | Km<KS.1 | YPD | KS.1 |

| | | | | | | | | | | |
|---|---|---|---|---|---|---|---|---|---|---|
| FIM1_4319 | 580.9416769 | 472.0873557 | 791.3821641 | -0.234821513 | 0.281827984 | 0.340799548 | 0.415864055 | ns | YPD | KS.1 |
| FIM1_4320 | 4872.187736 | 4730.028052 | 3853.218499 | -0.040140095 | 0.112141638 | 0.712392255 | 0.77063524 | ns | YPD | KS.1 |
| FIM1_4321 | 1325.294139 | 1209.262801 | 1574.930606 | -0.124678684 | 0.124371303 | 0.298790169 | 0.370971074 | ns | YPD | KS.1 |
| FIM1_4322 | 1972.616234 | 1115.866142 | 1448.110775 | -0.792547271 | 0.132154529 | 5.8640437866 | 3.1002781975 | ns | YPD | KS.1 |
| FIM1_4323 | 25517.03457 | 17016.74709 | 16724.97566 | -0.575876149 | 0.073139627 | 1.6085028645 | 1.4113406968 | ns | YPD | KS.1 |
| FIM1_4324 | 45.42733092 | 30.17729157 | 38.88673509 | -0.485573277 | 0.273435156 | 0.043783867 | 0.069019245 | ns | YPD | KS.1 |
| FIM1_4325 | 2063.892863 | 1894.622692 | 1983.336966 | -0.117357419 | 0.103537062 | 0.244343209 | 0.312817155 | ns | YPD | KS.1 |
| FIM1_4326 | 325.272492 | 1016.106285 | 1289.899005 | 1.588157838 | 0.169489359 | 4.4638842332 | 6.8122159694 | Km>KS.1 | YPD | KS.1 |
| FIM1_4327 | 2711.729266 | 2725.029905 | 2328.645034 | 0.006381076 | 0.13317688 | 0.96027539 | 0.970536119 | ns | YPD | KS.1 |
| FIM1_4328 | 783.4387342 | 507.4413428 | 872.2491588 | -0.569146244 | 0.196694047 | 0.001868068 | 0.003958108 | ns | YPD | KS.1 |
| FIM1_4329 | 1014.709133 | 544.4642289 | 704.8157214 | -0.85374568 | 0.161796724 | 3.3988851373 | 1.4703872659 | ns | YPD | KS.1 |
| FIM1_4330 | 1544.129957 | 3238.423915 | 2374.448184 | 1.033161333 | 0.14100506 | 4.2033061672 | 3.2072773285 | Km>KS.1 | YPD | KS.1 |
| FIM1_4331 | 9520.919493 | 4273.633014 | 5501.843841 | -1.082528164 | 0.205526242 | 2.2539454412 | 9.9321333660 | Km<KS.1 | YPD | KS.1 |
| FIM1_4332 | 2462.589087 | 1713.061203 | 1772.143234 | -0.514460868 | 0.076671412 | 1.0526119471 | 6.5705701848 | ns | YPD | KS.1 |
| FIM1_4333 | 1438.496743 | 2328.328813 | 2923.975554 | 0.680188682 | 0.092700062 | 8.0295328620 | 6.0161033115 | ns | YPD | KS.1 |
| FIM1_4334 | 1217.776791 | 677.3039864 | 898.2957246 | -0.799892409 | 0.167382477 | 5.1414636804 | 1.8905234153 | ns | YPD | KS.1 |
| FIM1_4335 | 2776.922183 | 2325.210977 | 2760.392648 | -0.251013404 | 0.069037728 | 0.000230546 | 0.000574345 | ns | YPD | KS.1 |
| FIM1_4336 | 852.9193803 | 769.9020146 | 963.1192203 | -0.134616214 | 0.150917737 | 0.348989897 | 0.424090068 | ns | YPD | KS.1 |
| FIM1_4337 | 1350.067095 | 1270.732387 | 1516.12613 | -0.082391071 | 0.114469204 | 0.457867379 | 0.53534435 | ns | YPD | KS.1 |
| FIM1_4338 | 2442.686811 | 4299.573956 | 4371.463693 | 0.807787018 | 0.06719026 | 7.2801724146 | 2.5506237861 | ns | YPD | KS.1 |
| FIM1_4339 | 156.5171199 | 316.2405669 | 495.8349819 | 0.962376573 | 0.171916946 | 4.4733985985 | 2.1440421991 | ns | YPD | KS.1 |
| FIM1_4340 | 2153.133565 | 4836.675453 | 10345.93131 | 1.14361958 | 0.114707294 | 2.9939551326 | 5.5166395499 | Km>KS.1 | YPD | KS.1 |
| FIM1_4341 | 266.9525645 | 236.7586018 | 351.1808717 | -0.142649831 | 0.239839989 | 0.509934204 | 0.585595583 | ns | YPD | KS.1 |
| FIM1_4342 | 604.8541086 | 370.83297 | 597.9498622 | -0.677784785 | 0.130753415 | 8.2702200794 | 3.3863658349 | ns | YPD | KS.1 |
| FIM1_4343 | 13175.72708 | 16099.99689 | 25916.70843 | 0.276722469 | 0.109849848 | 0.009518575 | 0.017480587 | ns | YPD | KS.1 |
| FIM1_4344 | 1744.878622 | 1027.043613 | 2485.044825 | -0.671446781 | 0.244775855 | 0.002401613 | 0.004967994 | ns | YPD | KS.1 |
| FIM1_4345 | 156.0590789 | 315.7360768 | 544.9502246 | 0.961609942 | 0.170032575 | 3.2092598268 | 1.5666628282 | ns | YPD | KS.1 |
| FIM1_4346 | 323.6253491 | 386.0212521 | 1382.390916 | 0.179592884 | 0.342187601 | 0.528766033 | 0.604479593 | ns | YPD | KS.1 |
| FIM1_4347 | 2775.463462 | 1935.914248 | 1730.147006 | -0.499672882 | 0.116975982 | 1.0952752409 | 3.3084361406 | ns | YPD | KS.1 |
| FIM1_4348 | 3434.393051 | 2770.572348 | 2477.390735 | -0.302497798 | 0.078679624 | 9.4486608295 | 0.000247797 | ns | YPD | KS.1 |
| FIM1_4349 | 3531.178465 | 248.9441963 | 484.6426657 | -3.671045279 | 0.335665237 | 1.0015989700 | 2.9140086995 | Km<KS.1 | YPD | KS.1 |
| FIM1_4350 | 2435.378156 | 2979.096908 | 2577.991275 | 0.285967123 | 0.067166786 | 1.6510965821 | 4.8547313808 | ns | YPD | KS.1 |
| FIM1_4351 | 1235.962092 | 4321.516087 | 4666.112367 | 1.774730902 | 0.12800262 | 4.0349770800 | 2.8273254892 | Km>KS.1 | YPD | KS.1 |
| FIM1_4352 | 687.9125427 | 674.962682 | 1263.868516 | -0.023603642 | 0.198240187 | 0.89796075 | 0.92224499 | ns | YPD | KS.1 |
| FIM1_4353 | 2275.382535 | 2100.349541 | 2362.725098 | -0.107192643 | 0.131532546 | 0.396839132 | 0.473674348 | ns | YPD | KS.1 |
| FIM1_4354 | 1086.841862 | 941.2134975 | 1627.87739 | -0.19028559 | 0.152996104 | 0.190855357 | 0.254437374 | ns | YPD | KS.1 |
| FIM1_4355 | 35.43526486 | 26.52588501 | 26.18476884 | -0.332774376 | 0.294676173 | 0.192097908 | 0.255804896 | ns | YPD | KS.1 |
| FIM1_4356 | 15927.56727 | 16419.95948 | 23485.72274 | 0.042743683 | 0.069110246 | 0.53081495 | 0.606523743 | ns | YPD | KS.1 |
| FIM1_4357 | 1140.173287 | 1267.487416 | 1923.814461 | 0.141691256 | 0.132259967 | 0.264995704 | 0.334777457 | ns | YPD | KS.1 |
| FIM1_4358 | 2504.202424 | 3043.258593 | 4355.850489 | 0.264059254 | 0.131666858 | 0.036953032 | 0.059151008 | ns | YPD | KS.1 |
| FIM1_4359 | 1502.531894 | 1682.537772 | 2335.321236 | 0.151824991 | 0.130521897 | 0.226500856 | 0.293524813 | ns | YPD | KS.1 |
| FIM1_4360 | 4588.104588 | 8192.236813 | 8671.074157 | 0.81998414 | 0.09688123 | 6.9581234295 | 7.5913736978 | ns | YPD | KS.1 |
| FIM1_4361 | 3276.331022 | 1920.745549 | 2926.548286 | -0.701207959 | 0.208662597 | 0.00028648 | 0.00070002 | ns | YPD | KS.1 |
| FIM1_4363 | 1245.670183 | 493.8808578 | 979.5832279 | -1.234214032 | 0.239360973 | 3.0358335249 | 1.3202160652 | Km<KS.1 | YPD | KS.1 |
| FIM1_4364 | 101.0049695 | 259.036793 | 393.1670421 | 1.281220317 | 0.202678783 | 2.9024804614 | 1.7355577278 | Km>KS.1 | YPD | KS.1 |

| | | | | | | | | | | |
|---|---|---|---|---|---|---|---|---|---|---|
| FIM1_4365 | 151.0599349 | 761.384021 | 1130.932375 | 2.29076736 | 0.146530989 | 3.4919691777 | 4.6952828808 | Km>KS.1 | YPD | KS.1 |
| FIM1_4366 | 9629.776902 | 8121.499174 | 9010.826394 | -0.238686228 | 0.085121511 | 0.00426573 | 0.00842812 | ns | YPD | KS.1 |
| FIM1_4367 | 1632.545748 | 1576.722998 | 1772.816264 | -0.046735295 | 0.134324303 | 0.717388559 | 0.774356278 | ns | YPD | KS.1 |
| FIM1_4368 | 1473.5825 | 1269.636551 | 1621.621862 | -0.202889073 | 0.122728183 | 0.086716917 | 0.127300915 | ns | YPD | KS.1 |
| FIM1_4369 | 1577.346318 | 1343.156121 | 3186.351165 | -0.184453969 | 0.265809925 | 0.433307551 | 0.510709563 | ns | YPD | KS.1 |
| FIM1_4370 | 793.9855037 | 460.0918051 | 886.3623511 | -0.728206482 | 0.190604568 | 4.5992237061 | 0.00012733 | ns | YPD | KS.1 |
| FIM1_4371 | 5928.764942 | 2932.604327 | 3441.943324 | -0.966632619 | 0.168540139 | 1.9886215554 | 9.9832414111 | ns | YPD | KS.1 |
| FIM1_4372 | 1430.886351 | 1783.952324 | 2499.143024 | 0.286365824 | 0.177594857 | 0.085502535 | 0.125701865 | ns | YPD | KS.1 |
| FIM1_4373 | 5635.92765 | 6053.368821 | 8223.638193 | 0.08798501 | 0.208660106 | 0.647437376 | 0.712926283 | ns | YPD | KS.1 |
| FIM1_4374 | 2122.644159 | 1124.058134 | 1187.237333 | -0.899973771 | 0.099778054 | 4.2981686139 | 5.6569811784 | ns | YPD | KS.1 |
| FIM1_4376 | 197.5911366 | 204.2353157 | 275.9932316 | 0.043502167 | 0.168493546 | 0.784601365 | 0.830508892 | ns | YPD | KS.1 |
| FIM1_4377 | 3349.631207 | 5254.427889 | 5376.730464 | 0.580412056 | 0.21579235 | 0.003427724 | 0.006895643 | ns | YPD | KS.1 |
| FIM1_4378 | 87.45859901 | 78.7341083 | 200.5203965 | -0.139927585 | 0.178238732 | 0.402823128 | 0.479666123 | ns | YPD | KS.1 |
| FIM1_4379 | 1625.62068 | 3258.289608 | 3467.010987 | 0.980473816 | 0.112884115 | 7.3817838568 | 8.7231293795 | ns | YPD | KS.1 |
| FIM1_4380 | 1644.228329 | 1827.07109 | 1529.100328 | 0.147135359 | 0.085841203 | 0.08035461 | 0.118906659 | ns | YPD | KS.1 |
| FIM1_4381 | 520.5694508 | 1333.571116 | 1501.693725 | 1.328403376 | 0.122425245 | 2.0418291732 | 5.1303536045 | Km>KS.1 | YPD | KS.1 |
| FIM1_4382 | 749.5881246 | 805.3614728 | 961.0533677 | 0.096593537 | 0.123945488 | 0.419491839 | 0.497371758 | ns | YPD | KS.1 |
| FIM1_4383 | 2542.558029 | 5345.413882 | 5562.631831 | 1.063032582 | 0.067649 | 1.9806281599 | 2.4634062738 | Km>KS.1 | YPD | KS.1 |
| FIM1_4384 | 4790.779879 | 1247.682236 | 1504.303601 | -1.839192548 | 0.234852256 | 1.6538745268 | 1.6007832239 | Km<KS.1 | YPD | KS.1 |
| FIM1_4385 | 1866.193445 | 2396.984449 | 1672.830195 | 0.347068398 | 0.108588121 | 0.001024515 | 0.002266693 | ns | YPD | KS.1 |
| FIM1_4386 | 6491.062759 | 11761.51 | 15426.11709 | 0.835057075 | 0.114071265 | 6.5042764260 | 4.9028447302 | ns | YPD | KS.1 |
| FIM1_4387 | 28.64660298 | 23.54253666 | 32.08724951 | -0.231766073 | 0.261645767 | 0.318441941 | 0.391946723 | ns | YPD | KS.1 |
| FIM1_4388 | 4279.421024 | 2168.601042 | 2828.08616 | -0.857616572 | 0.2732556 | 0.000443038 | 0.001042627 | ns | YPD | KS.1 |
| FIM1_4389 | 296.946734 | 358.6979465 | 261.3620366 | 0.243377035 | 0.175923222 | 0.140523976 | 0.194142399 | ns | YPD | KS.1 |
| FIM1_4390 | 167.396852 | 98.2931988 | 119.0014353 | -0.709043097 | 0.189392385 | 6.5365142523 | 0.000176831 | ns | YPD | KS.1 |
| FIM1_4391 | 6707.363018 | 6576.257912 | 11072.76012 | -0.024864804 | 0.186392288 | 0.886560495 | 0.913553948 | ns | YPD | KS.1 |
| FIM1_4392 | 4078.311412 | 2754.972922 | 2446.787469 | -0.513940366 | 0.191568891 | 0.004004582 | 0.007940533 | ns | YPD | KS.1 |
| FIM1_4393 | 93.04028384 | 146.1095331 | 102.1546705 | 0.56031307 | 0.247954388 | 0.011912095 | 0.021448669 | ns | YPD | KS.1 |
| FIM1_4394 | 400.1028743 | 623.2756086 | 574.9786187 | 0.61658157 | 0.125533407 | 3.9500793102 | 1.4820244772 | ns | YPD | KS.1 |
| FIM1_4396 | 1746.702563 | 2340.578985 | 3371.773329 | 0.392777158 | 0.152195624 | 0.006742715 | 0.012720898 | ns | YPD | KS.1 |
| FIM1_4397 | 1073.94973 | 1804.564535 | 1309.553351 | 0.725660688 | 0.114947008 | 9.2185437993 | 5.2354172833 | ns | YPD | KS.1 |
| FIM1_4398 | 504.4515229 | 497.9485336 | 199.3745941 | -0.01446492 | 0.291858868 | 0.954531352 | 0.967561833 | ns | YPD | KS.1 |
| FIM1_4399 | 6265.377361 | 13640.04029 | 14052.33915 | 1.042660551 | 0.215330007 | 2.2572775899 | 8.7324696810 | Km>KS.1 | YPD | KS.1 |
| FIM1_4400 | 802.1052707 | 1190.355799 | 1072.102269 | 0.554341133 | 0.098626343 | 9.4992709552 | 4.3798770159 | ns | YPD | KS.1 |
| FIM1_4401 | 848.7827356 | 1934.581364 | 2429.574297 | 1.161344564 | 0.123824855 | 9.3793393435 | 1.3846354075 | Km>KS.1 | YPD | KS.1 |
| FIM1_4402 | 133.0446889 | 107.4027176 | 172.6698895 | -0.279451253 | 0.171890538 | 0.083976044 | 0.123676975 | ns | YPD | KS.1 |
| FIM1_4403 | 12289.01304 | 9482.400286 | 5429.59483 | -0.296133589 | 0.281078202 | 0.228464249 | 0.29568409 | ns | YPD | KS.1 |
| FIM1_4404 | 394.5475256 | 396.7467428 | 460.0954742 | 0.007577981 | 0.154274242 | 0.958847975 | 0.969566804 | ns | YPD | KS.1 |
| FIM1_4405 | 243.5590578 | 536.7467432 | 609.2161831 | 1.085977752 | 0.171102137 | 3.5591703385 | 2.1129919372 | Km>KS.1 | YPD | KS.1 |
| FIM1_4406 | 1009.498243 | 1358.778142 | 1069.145318 | 0.408668411 | 0.121591505 | 0.00051851 | 0.001211643 | ns | YPD | KS.1 |
| FIM1_4407 | 673.2992512 | 65.66721001 | 122.619116 | -3.253261814 | 0.264344671 | 8.4461093190 | 3.6859117422 | Km<KS.1 | YPD | KS.1 |
| FIM1_4408 | 850.25371 | 588.0675974 | 448.4685894 | -0.426378752 | 0.289288479 | 0.090139694 | 0.1318568 | ns | YPD | KS.1 |
| FIM1_4409 | 1453.944555 | 1920.94006 | 6251.210809 | 0.316589037 | 0.286776821 | 0.204993509 | 0.270371874 | ns | YPD | KS.1 |
| FIM1_4410 | 825.5900199 | 865.0285676 | 823.5708034 | 0.062810555 | 0.129508638 | 0.614628803 | 0.683913732 | ns | YPD | KS.1 |
| FIM1_4412 | 648.8201691 | 335.5044035 | 240.9380461 | -0.803508242 | 0.302554266 | 0.002331632 | 0.004835293 | ns | YPD | KS.1 |

| | | | | | | | | | | |
|---|---|---|---|---|---|---|---|---|---|---|
| FIM1_4413 | 1210.58364 | 1264.682274 | 1434.163885 | 0.061434122 | 0.089788517 | 0.484847117 | 0.562920515 | ns | YPD | KS.1 |
| FIM1_4414 | 1456.01862 | 4012.281118 | 3221.545165 | 1.43211603 | 0.128156031 | 4.5781627681 | 1.2583624183 | Km>KS.1 | YPD | KS.1 |
| FIM1_4415 | 6548.051677 | 6381.171891 | 3335.323888 | -0.029234387 | 0.267024622 | 0.901652453 | 0.925463369 | ns | YPD | KS.1 |
| FIM1_4416 | 3060.425656 | 2599.885133 | 1896.95476 | -0.20516706 | 0.198428519 | 0.264435405 | 0.334154468 | ns | YPD | KS.1 |
| FIM1_4417 | 982.7602359 | 1567.677116 | 1298.851095 | 0.658732338 | 0.096022016 | 2.6778302942 | 1.7621965229 | ns | YPD | KS.1 |
| FIM1_4418 | 3402.457891 | 5124.731486 | 5851.324223 | 0.576444256 | 0.095885629 | 8.6928747385 | 4.5237501908 | ns | YPD | KS.1 |
| FIM1_4419 | 807.9813266 | 1014.29326 | 1039.942133 | 0.312612897 | 0.115091395 | 0.005097497 | 0.009894673 | ns | YPD | KS.1 |
| FIM1_4420 | 1266.836634 | 1281.041086 | 1050.982712 | 0.01462294 | 0.103667374 | 0.884914502 | 0.912613941 | ns | YPD | KS.1 |
| FIM1_4421 | 980.7403178 | 610.9833269 | 841.2736502 | -0.667179039 | 0.096859364 | 2.1591647805 | 1.4418583601 | ns | YPD | KS.1 |
| FIM1_4422 | 29.26781853 | 9.384052796 | 14.57156587 | -1.265066823 | 0.462050271 | 0.000616836 | 0.001424679 | Km<KS.1 | YPD | KS.1 |
| FIM1_4423 | 720.5967605 | 1433.955872 | 1754.733821 | 0.899508555 | 0.235211607 | 3.1129966945 | 8.8245917694 | ns | YPD | KS.1 |
| FIM1_4424 | 3109.891801 | 1060.174257 | 1053.413028 | -1.241465759 | 0.422432661 | 0.000355312 | 0.000852303 | Km<KS.1 | YPD | KS.1 |
| FIM1_4425 | 352.4570887 | 733.129641 | 643.1878699 | 1.045542961 | 0.07956206 | 3.2560797815 | 1.6873955117 | Km>KS.1 | YPD | KS.1 |
| FIM1_4426 | 347.8560336 | 745.0983312 | 728.6083259 | 1.066370243 | 0.131334762 | 7.8522786685 | 7.8601783452 | Km>KS.1 | YPD | KS.1 |
| FIM1_4427 | 4809.529459 | 5083.164423 | 5440.023767 | 0.075225089 | 0.113913471 | 0.495947421 | 0.573000098 | ns | YPD | KS.1 |
| FIM1_4428 | 5355.330784 | 9959.471872 | 6919.829657 | 0.801645198 | 0.239252152 | 0.000237852 | 0.000590346 | ns | YPD | KS.1 |
| FIM1_4429 | 1684.465243 | 2759.947738 | 2044.53562 | 0.667506416 | 0.167733365 | 2.7224692064 | 7.7840714379 | ns | YPD | KS.1 |
| FIM1_4430 | 234.1507047 | 227.0349501 | 343.7891576 | -0.038464568 | 0.16331464 | 0.803647614 | 0.845989607 | ns | YPD | KS.1 |
| FIM1_4431 | 5736.096694 | 3161.99774 | 2195.196975 | -0.842326483 | 0.098444081 | 2.9882273501 | 3.3407710263 | ns | YPD | KS.1 |
| FIM1_4432 | 4305.270207 | 5706.99759 | 5969.247125 | 0.395712225 | 0.08979109 | 7.1680058927 | 2.2260193081 | ns | YPD | KS.1 |
| FIM1_4433 | 346.7262695 | 454.500476 | 622.5379711 | 0.357366676 | 0.167885174 | 0.024068067 | 0.040261814 | ns | YPD | KS.1 |
| FIM1_4434 | 2840.716179 | 1952.756307 | 1987.486864 | -0.534815905 | 0.05798136 | 1.4558506409 | 2.0007892095 | ns | YPD | KS.1 |
| FIM1_4435 | 6419.559246 | 5326.18724 | 4237.227993 | -0.257758151 | 0.108027219 | 0.014103447 | 0.024942996 | ns | YPD | KS.1 |
| FIM1_4436 | 1862.911106 | 1873.772772 | 1632.471748 | 0.008193144 | 0.072223684 | 0.908399292 | 0.931235623 | ns | YPD | KS.1 |
| FIM1_4437 | 579.4995963 | 563.9992095 | 636.5513952 | -0.034401983 | 0.139849029 | 0.797514386 | 0.840956776 | ns | YPD | KS.1 |
| FIM1_4438 | 1671.147035 | 1918.254561 | 1274.270211 | 0.182096889 | 0.154751083 | 0.215487307 | 0.282180131 | ns | YPD | KS.1 |
| FIM1_4439 | 1043.75872 | 1187.921437 | 1607.302738 | 0.17641674 | 0.111244588 | 0.102101769 | 0.147191046 | ns | YPD | KS.1 |
| FIM1_4440 | 2064.448535 | 1872.322356 | 1509.242747 | -0.120120958 | 0.210997095 | 0.53618281 | 0.610693562 | ns | YPD | KS.1 |
| FIM1_4441 | 4821.719028 | 5126.551095 | 3396.226368 | 0.071224314 | 0.25084059 | 0.751403384 | 0.803575201 | ns | YPD | KS.1 |
| FIM1_4443 | 1630.078821 | 606.1340095 | 510.6500014 | -1.181321238 | 0.376862694 | 0.000216731 | 0.000541556 | Km<KS.1 | YPD | KS.1 |
| FIM1_4444 | 9969.686346 | 23733.74549 | 29229.62885 | 1.212205705 | 0.147785188 | 3.0269203601 | 3.1438264701 | Km>KS.1 | YPD | KS.1 |
| FIM1_4445 | 1235.481112 | 1370.542635 | 1353.071427 | 0.141821996 | 0.107064156 | 0.172933598 | 0.233155731 | ns | YPD | KS.1 |
| FIM1_4446 | 3483.875579 | 4891.79934 | 5081.720682 | 0.469035618 | 0.120810314 | 6.1997580702 | 0.000168453 | ns | YPD | KS.1 |
| FIM1_4447 | 2089.062632 | 1895.710544 | 1698.587544 | -0.135853455 | 0.075010324 | 0.065717098 | 0.099890792 | ns | YPD | KS.1 |
| FIM1_4448 | 447.731924 | 401.8093507 | 260.2087931 | -0.124184075 | 0.263338847 | 0.595305587 | 0.666887029 | ns | YPD | KS.1 |
| FIM1_4449 | 769.5670475 | 1172.368599 | 733.6709146 | 0.509475794 | 0.264825111 | 0.030046011 | 0.049089953 | ns | YPD | KS.1 |
| FIM1_4450 | 1704.198693 | 3195.160514 | 3259.486394 | 0.87266832 | 0.141186052 | 1.5683005695 | 8.7273997019 | ns | YPD | KS.1 |
| FIM1_4451 | 4069.456704 | 2014.776704 | 2267.005217 | -0.956463294 | 0.183573165 | 3.9423349659 | 1.6850474000 | ns | YPD | KS.1 |
| FIM1_4452 | 4881.024833 | 6465.465534 | 10157.37843 | 0.376122733 | 0.155346017 | 0.010866089 | 0.019693549 | ns | YPD | KS.1 |
| FIM1_4454 | 3738.536008 | 5851.29672 | 4077.397871 | 0.634321729 | 0.084556938 | 2.5784211501 | 2.0011927023 | ns | YPD | KS.1 |
| FIM1_4455 | 306.8911982 | 905.2711794 | 1386.220544 | 1.5025158 | 0.179405367 | 4.0719477007 | 4.4917826632 | Km>KS.1 | YPD | KS.1 |
| FIM1_4456 | 1956.393888 | 1571.43505 | 1521.687779 | -0.288083331 | 0.165664975 | 0.065506961 | 0.09975425 | ns | YPD | KS.1 |
| FIM1_4457 | 939.2618075 | 1572.487545 | 1899.474495 | 0.722962409 | 0.109079393 | 1.1500598549 | 7.1340994740 | ns | YPD | KS.1 |
| FIM1_4458 | 874.8240531 | 1247.573316 | 1523.401855 | 0.495111772 | 0.105545612 | 1.5469650833 | 5.3003796760 | ns | YPD | KS.1 |
| FIM1_4460 | 597.8771301 | 329.0729969 | 482.6566913 | -0.795397968 | 0.198767949 | 1.8695672763 | 5.4648044651 | ns | YPD | KS.1 |

| | | | | | | | | | | |
|---|---|---|---|---|---|---|---|---|---|---|
| FIM1_4461 | 265.7076936 | 101.6782596 | 235.8112276 | -1.304551917 | 0.20657108 | 2.88795720978 | 1.72895151849 | Km<KS.1 | YPD | KS.1 |
| FIM1_4462 | 881.6867826 | 1048.878501 | 1086.853569 | 0.239951697 | 0.106668208 | 0.020704111 | 0.035106664 | ns | YPD | KS.1 |
| FIM1_4463 | 4033.704269 | 2558.181079 | 1342.582568 | -0.629414926 | 0.131418084 | 7.11974646322 | 2.57792857747 | ns | YPD | KS.1 |
| FIM1_4464 | 1961.585428 | 1087.03986 | 1042.604224 | -0.815226136 | 0.146832061 | 7.88988677806 | 3.68219387625 | ns | YPD | KS.1 |
| FIM1_4465 | 975.6108283 | 955.0627145 | 1372.504425 | -0.02875754 | 0.125147352 | 0.811879724 | 0.85267081 | ns | YPD | KS.1 |
| FIM1_4466 | 729.0576866 | 1331.996848 | 1228.347793 | 0.848817258 | 0.106809432 | 4.86221340198 | 4.47953919905 | ns | YPD | KS.1 |
| FIM1_4467 | 1108.20763 | 2525.802013 | 2880.810584 | 1.176036951 | 0.078384094 | 9.31669820459 | 9.08834775840 | Km>KS.1 | YPD | KS.1 |
| FIM1_4468 | 408.5150065 | 230.2930276 | 290.415238 | -0.788390301 | 0.149748628 | 4.17187985418 | 1.77697793446 | ns | YPD | KS.1 |
| FIM1_4469 | 2312.512708 | 1960.765729 | 3092.130005 | -0.187495366 | 0.272922038 | 0.435491912 | 0.512559298 | ns | YPD | KS.1 |
| FIM1_4470 | 3366.298028 | 3625.348721 | 2972.149374 | 0.08896584 | 0.228752871 | 0.669105919 | 0.73128338 | ns | YPD | KS.1 |
| FIM1_4471 | 821.3447653 | 599.9194019 | 801.3701326 | -0.427345682 | 0.141178845 | 0.001594952 | 0.003418737 | ns | YPD | KS.1 |
| FIM1_4472 | 2394.925696 | 2838.467654 | 2324.054921 | 0.241046277 | 0.064301079 | 0.000149525 | 0.000380895 | ns | YPD | KS.1 |
| FIM1_4473 | 994.6255635 | 751.7794883 | 825.0001088 | -0.379091601 | 0.14145716 | 0.005151224 | 0.009979494 | ns | YPD | KS.1 |
| FIM1_4474 | 4678.93381 | 10984.08051 | 10933.34462 | 1.183838671 | 0.163238931 | 5.54782024705 | 4.19459053633 | Km>KS.1 | YPD | KS.1 |
| FIM1_4475 | 266.1439939 | 728.1022022 | 812.4556507 | 1.387921995 | 0.184096136 | 4.36364668195 | 3.65473775130 | Km>KS.1 | YPD | KS.1 |
| FIM1_4476 | 701.2326921 | 814.439007 | 1386.57011 | 0.191311598 | 0.184417978 | 0.266934134 | 0.336542656 | ns | YPD | KS.1 |
| FIM1_4477 | 1878.623943 | 2313.016526 | 2194.233526 | 0.289260633 | 0.096338411 | 0.002130881 | 0.004463635 | ns | YPD | KS.1 |
| FIM1_4478 | 2082.60693 | 1824.630736 | 1911.228193 | -0.177861788 | 0.1347962 | 0.169136742 | 0.228718481 | ns | YPD | KS.1 |
| FIM1_4479 | 1751.50127 | 2908.326922 | 2132.936426 | 0.653804444 | 0.223849806 | 0.001430512 | 0.003078201 | ns | YPD | KS.1 |
| FIM1_4480 | 435.9049367 | 445.0226788 | 436.876264 | 0.02666724 | 0.162899072 | 0.862837899 | 0.894481881 | ns | YPD | KS.1 |
| FIM1_4481 | 2032.590977 | 2451.322987 | 1967.364171 | 0.26387787 | 0.081662503 | 0.001011149 | 0.002239745 | ns | YPD | KS.1 |
| FIM1_4482 | 21595.80034 | 3611.628215 | 2864.161851 | -2.382323089 | 0.337027832 | 1.12333965107 | 8.98491119631 | Km<KS.1 | YPD | KS.1 |
| FIM1_4484 | 540.0119061 | 10.39248917 | 72.63694843 | -5.182418786 | 0.59501049 | 8.51261775213 | 1.18961441901 | Km<KS.1 | YPD | KS.1 |
| FIM1_4486 | 3829.845936 | 2430.816333 | 1769.97335 | -0.523584203 | 0.307264505 | 0.047378509 | 0.074238766 | ns | YPD | KS.1 |
| FIM1_4487 | 1981.656718 | 565.2260215 | 435.3429884 | -1.628083423 | 0.317530687 | 1.54265574259 | 6.92038982814 | Km<KS.1 | YPD | KS.1 |
| FIM1_4488 | 26.0163893 | 25.78983816 | 33.08812412 | -0.005056569 | 0.291721872 | 0.984092121 | 0.988218307 | ns | YPD | KS.1 |
| FIM1_4489 | 1058.646417 | 1372.187222 | 1176.216034 | 0.301444826 | 0.269820242 | 0.205341647 | 0.270687518 | ns | YPD | KS.1 |
| FIM1_4490 | 15256.70635 | 12703.36771 | 7550.038692 | -0.22128449 | 0.23219642 | 0.293090048 | 0.365078365 | ns | YPD | KS.1 |
| FIM1_4491 | 1275.160076 | 1475.583493 | 931.9821455 | 0.187536431 | 0.178717633 | 0.263048798 | 0.333039931 | ns | YPD | KS.1 |
| FIM1_4492 | 650.3699156 | 562.6334802 | 821.0718333 | -0.197115322 | 0.121247697 | 0.092296775 | 0.134616375 | ns | YPD | KS.1 |
| FIM1_4493 | 16928.48991 | 11610.7084 | 6962.732737 | -0.48311 | 0.210800341 | 0.012753914 | 0.022807593 | ns | YPD | KS.1 |
| FIM1_4494 | 44.204053 | 39.15246665 | 44.75258721 | -0.151750462 | 0.241118116 | 0.485203857 | 0.563203263 | ns | YPD | KS.1 |
| FIM1_4495 | 2961.153585 | 1535.283967 | 1200.271987 | -0.911877234 | 0.143899655 | 5.32307648228 | 3.10097254090 | ns | YPD | KS.1 |
| FIM1_4496 | 379.9682654 | 679.2631574 | 572.8180632 | 0.808074634 | 0.135142696 | 6.32531780898 | 3.33352289191 | ns | YPD | KS.1 |
| FIM1_4497 | 3825.232965 | 3316.059532 | 4143.687139 | -0.194973589 | 0.118090659 | 0.087791493 | 0.128704066 | ns | YPD | KS.1 |
| FIM1_4498 | 8775.202233 | 12371.76489 | 18755.82401 | 0.473354096 | 0.125555833 | 9.70168772988 | 0.000254031 | ns | YPD | KS.1 |
| FIM1_4499 | 3409.681303 | 2969.277565 | 3375.850452 | -0.187979465 | 0.122250769 | 0.11117403 | 0.158842848 | ns | YPD | KS.1 |
| FIM1_4500 | 3896.165341 | 3843.241422 | 2985.97264 | -0.019139308 | 0.080726736 | 0.809420252 | 0.850806202 | ns | YPD | KS.1 |
| FIM1_4501 | 240.1536783 | 839.8352832 | 1516.750278 | 1.739690525 | 0.189551919 | 1.92661407387 | 2.77823333841 | Km>KS.1 | YPD | KS.1 |
| FIM1_4502 | 122.8299504 | 180.1119564 | 211.4499236 | 0.481516059 | 0.225691984 | 0.019186281 | 0.032919694 | ns | YPD | KS.1 |
| FIM1_4503 | 581.5177333 | 682.1230352 | 692.7756756 | 0.219518533 | 0.103350222 | 0.029136424 | 0.047750151 | ns | YPD | KS.1 |
| FIM1_4504 | 1684.881534 | 1719.680126 | 1117.332412 | 0.026354559 | 0.17167727 | 0.870685578 | 0.900365984 | ns | YPD | KS.1 |
| FIM1_4505 | 2587.365515 | 1426.183177 | 2753.53033 | -0.682465003 | 0.337472171 | 0.01660825 | 0.028940821 | ns | YPD | KS.1 |
| FIM1_4506 | 495.0212248 | 849.8873586 | 1446.303108 | 0.726812817 | 0.180774479 | 2.00685209129 | 5.83864862817 | ns | YPD | KS.1 |
| FIM1_4507 | 2307.581229 | 2238.810938 | 2743.829087 | -0.04025721 | 0.140960755 | 0.765673542 | 0.816204386 | ns | YPD | KS.1 |

| | | | | | | | | | | |
|---|---|---|---|---|---|---|---|---|---|---|
| FIM1_4508 | 1207.839515 | 948.3877742 | 1156.549221 | -0.336884911 | 0.100975157 | 0.000634051 | 0.001463082 | ns | YPD | KS.1 |
| FIM1_4509 | 2452.8108 | 1821.556069 | 2958.750099 | -0.423866972 | 0.063759756 | 1.9318273018 | 1.1677813883 | ns | YPD | KS.1 |
| FIM1_4510 | 3246.069728 | 3762.435674 | 5040.076391 | 0.205790288 | 0.091224884 | 0.021127778 | 0.035727632 | ns | YPD | KS.1 |
| FIM1_4512 | 781.0353947 | 772.7340057 | 894.7769578 | -0.014325499 | 0.105432666 | 0.889035857 | 0.915882728 | ns | YPD | KS.1 |
| FIM1_4513 | 124.1947247 | 128.9361736 | 220.9323604 | 0.048988575 | 0.161961391 | 0.749655989 | 0.801906182 | ns | YPD | KS.1 |
| FIM1_4514 | 318.3737521 | 253.620801 | 316.521227 | -0.305291385 | 0.140403783 | 0.023146071 | 0.038863214 | ns | YPD | KS.1 |
| FIM1_4515 | 831.4984949 | 1301.70786 | 1229.686415 | 0.630268762 | 0.096777929 | 3.0830627680 | 1.8391171787 | ns | YPD | KS.1 |
| FIM1_4516 | 620.8400823 | 276.6884774 | 750.3798019 | -0.939320051 | 0.367670667 | 0.002291918 | 0.004762862 | ns | YPD | KS.1 |
| FIM1_4517 | 1319.888539 | 1836.092047 | 1121.894738 | 0.422147037 | 0.205758006 | 0.026125167 | 0.043338681 | ns | YPD | KS.1 |
| FIM1_4518 | 628.2176753 | 813.6823393 | 733.9843654 | 0.3454131 | 0.155887008 | 0.019667138 | 0.033628658 | ns | YPD | KS.1 |
| FIM1_4519 | 634.4177676 | 1072.459178 | 1030.65912 | 0.739320711 | 0.104899667 | 5.8876083548 | 4.1023601631 | ns | YPD | KS.1 |
| FIM1_4520 | 2391.813174 | 1815.576846 | 2052.447947 | -0.360137073 | 0.178558142 | 0.031462708 | 0.051085827 | ns | YPD | KS.1 |
| FIM1_4521 | 2364.739877 | 3230.115492 | 2375.04407 | 0.441918537 | 0.076150407 | 4.0953762822 | 1.9723617622 | ns | YPD | KS.1 |
| FIM1_4522 | 1684.502898 | 889.3330077 | 643.0435648 | -0.81456687 | 0.255787452 | 0.000416109 | 0.000984844 | ns | YPD | KS.1 |
| FIM1_4523 | 3621.296236 | 3374.848123 | 3284.485673 | -0.100023385 | 0.059513631 | 0.08935274 | 0.130744083 | ns | YPD | KS.1 |
| FIM1_4524 | 1693.115688 | 1111.962156 | 2028.069343 | -0.584185284 | 0.121505397 | 7.1764223240 | 2.5946730423 | ns | YPD | KS.1 |
| FIM1_4525 | 276.1410996 | 264.1027079 | 257.5943842 | -0.056197955 | 0.173008166 | 0.730260355 | 0.785183762 | ns | YPD | KS.1 |
| FIM1_4526 | 6142.953071 | 5576.857518 | 6663.423493 | -0.136691566 | 0.063449454 | 0.029213825 | 0.047856035 | ns | YPD | KS.1 |
| FIM1_4527 | 13422.89886 | 18259.65124 | 16071.36789 | 0.362679858 | 0.266688167 | 0.123985948 | 0.174739403 | ns | YPD | KS.1 |
| FIM1_4528 | 1135.574858 | 229.4187802 | 224.419495 | -2.119213668 | 0.323115495 | 8.5322314628 | 5.9037345657 | Km<KS.1 | YPD | KS.1 |
| FIM1_4529 | 1904.725328 | 689.2772126 | 548.8525816 | -1.302405828 | 0.30546557 | 2.0512225085 | 6.8765714150 | Km<KS.1 | YPD | KS.1 |
| FIM1_4530 | 2552.043915 | 2861.663388 | 2316.557567 | 0.158139835 | 0.103361724 | 0.11600026 | 0.164838987 | ns | YPD | KS.1 |
| FIM1_4531 | 1312.495596 | 1850.11259 | 3611.620623 | 0.480265131 | 0.098634751 | 6.5668723977 | 2.3934205259 | ns | YPD | KS.1 |
| FIM1_4532 | 128.3072815 | 1017.182754 | 1404.05236 | 2.943813454 | 0.158313468 | 4.1248685918 | 2.2801356938 | Km>KS.1 | YPD | KS.1 |
| FIM1_4533 | 4207.498365 | 5098.419478 | 8237.061385 | 0.221589942 | 0.266067293 | 0.34673785 | 0.422064461 | ns | YPD | KS.1 |
| FIM1_4534 | 531.3262882 | 475.3054854 | 795.2805864 | -0.1480745 | 0.142015734 | 0.275688624 | 0.34608905 | ns | YPD | KS.1 |
| FIM1_4535 | 848.8373724 | 1147.831197 | 1795.554662 | 0.398147448 | 0.172476731 | 0.014209845 | 0.025104395 | ns | YPD | KS.1 |
| FIM1_4536 | 1332.590419 | 2898.159307 | 2216.598367 | 1.017341912 | 0.24631193 | 6.6886543178 | 2.0849658666 | Km>KS.1 | YPD | KS.1 |
| FIM1_4537 | 614.0599789 | 824.5001048 | 926.008992 | 0.408411121 | 0.114918066 | 0.000253648 | 0.000625942 | ns | YPD | KS.1 |
| FIM1_4538 | 826.2614332 | 460.8292752 | 724.0558032 | -0.705196795 | 0.29587713 | 0.006293406 | 0.01194115 | ns | YPD | KS.1 |
| FIM1_4539 | 489.5135935 | 867.312759 | 1030.865874 | 0.807932459 | 0.101070304 | 3.6235357927 | 3.3821933524 | ns | YPD | KS.1 |
| FIM1_4540 | 245.3252081 | 285.1475754 | 575.5351737 | 0.198258706 | 0.16307723 | 0.198593544 | 0.263046561 | ns | YPD | KS.1 |
| FIM1_4542 | 2644.462227 | 4468.520844 | 4417.824229 | 0.716738673 | 0.156380072 | 1.6066342645 | 5.4821711015 | ns | YPD | KS.1 |
| FIM1_4543 | 2689.296638 | 1743.151706 | 3931.707626 | -0.477597787 | 0.332429391 | 0.087833259 | 0.128710004 | ns | YPD | KS.1 |
| FIM1_4544 | 1034.387699 | 929.0701795 | 678.0470273 | -0.139300349 | 0.168800493 | 0.381697028 | 0.45812852 | ns | YPD | KS.1 |
| FIM1_4545 | 2327.518843 | 1545.786447 | 2193.460746 | -0.510702209 | 0.238773042 | 0.01790013 | 0.030921231 | ns | YPD | KS.1 |
| FIM1_4546 | 4145.488956 | 2683.738001 | 5054.654355 | -0.517230079 | 0.280812735 | 0.035618213 | 0.057235339 | ns | YPD | KS.1 |
| FIM1_4547 | 51.48481531 | 36.80335438 | 93.14022584 | -0.390817034 | 0.285743228 | 0.116680586 | 0.165711081 | ns | YPD | KS.1 |
| FIM1_4548 | 945.1476445 | 1110.889362 | 1192.076338 | 0.217362472 | 0.138763975 | 0.1019044 | 0.146949099 | ns | YPD | KS.1 |
| FIM1_4549 | 2709.087251 | 2513.546432 | 2090.131992 | -0.097231328 | 0.164823792 | 0.532880647 | 0.608325199 | ns | YPD | KS.1 |
| FIM1_4550 | 69.51192438 | 62.77312468 | 58.11738184 | -0.114291005 | 0.272991397 | 0.634846996 | 0.701702689 | ns | YPD | KS.1 |
| FIM1_4551 | 184.5914508 | 118.1733042 | 155.7175167 | -0.602702267 | 0.171180751 | 0.000196122 | 0.00049328 | ns | YPD | KS.1 |
| FIM1_4552 | 681.4247515 | 1196.484238 | 991.3499352 | 0.771183205 | 0.156637155 | 2.6361310308 | 1.0150736747 | ns | YPD | KS.1 |
| FIM1_4553 | 778.3989486 | 522.3062739 | 556.4867473 | -0.526998497 | 0.18328749 | 0.00215782 | 0.004512465 | ns | YPD | KS.1 |
| FIM1_4554 | 2789.136597 | 2908.208852 | 2258.936361 | 0.054320271 | 0.160132265 | 0.720694861 | 0.777082127 | ns | YPD | KS.1 |

| | | | | | | | | | | |
|---|---|---|---|---|---|---|---|---|---|---|
| FIM1_4555 | 4412.135177 | 6235.084114 | 5241.801357 | 0.437874354 | 0.216852122 | 0.027469033 | 0.045341221 | ns | YPD | KS.1 |
| FIM1_4556 | 1731.872207 | 2659.786355 | 2797.311843 | 0.542939124 | 0.229354068 | 0.009276561 | 0.017055023 | ns | YPD | KS.1 |
| FIM1_4557 | 2799.420871 | 3743.210893 | 4748.96601 | 0.4062745 | 0.096954117 | 1.87084324001 | 5.46532302939 | ns | YPD | KS.1 |
| FIM1_4558 | 1719.802324 | 1826.498577 | 1668.102854 | 0.079076478 | 0.150654621 | 0.581883536 | 0.654503864 | ns | YPD | KS.1 |
| FIM1_4559 | 865.3876407 | 917.8377642 | 1020.512662 | 0.079496608 | 0.118360105 | 0.487754103 | 0.565135026 | ns | YPD | KS.1 |
| FIM1_4560 | 83.77953201 | 111.9512072 | 161.0635889 | 0.355591769 | 0.234250286 | 0.093307655 | 0.135885557 | ns | YPD | KS.1 |
| FIM1_4561 | 6269.872541 | 5953.064106 | 4062.928576 | -0.063468816 | 0.211903637 | 0.744846862 | 0.797592153 | ns | YPD | KS.1 |
| FIM1_4562 | 3241.919655 | 835.799857 | 665.2961242 | -1.680117776 | 0.392661039 | 8.16786731382 | 2.93121507837 | Km<KS.1 | YPD | KS.1 |
| FIM1_4563 | 971.6039916 | 1346.903287 | 2210.548623 | 0.406715376 | 0.228325004 | 0.049905144 | 0.077952305 | ns | YPD | KS.1 |
| FIM1_4564 | 187.6982571 | 144.3041037 | 338.2719058 | -0.334020134 | 0.208776373 | 0.082008238 | 0.121137465 | ns | YPD | KS.1 |
| FIM1_4565 | 6161.572546 | 6870.834795 | 4454.604135 | 0.127977354 | 0.245746282 | 0.562432523 | 0.635175773 | ns | YPD | KS.1 |
| FIM1_4566 | 2826.65112 | 14285.23596 | 10289.83615 | 2.16719899 | 0.307683773 | 2.57646353907 | 2.00119270231 | Km>KS.1 | YPD | KS.1 |
| FIM1_4567 | 2996.607535 | 5352.5299 | 6056.028043 | 0.813772257 | 0.115638775 | 5.41324227842 | 3.78242701336 | ns | YPD | KS.1 |
| FIM1_4568 | 4294.35245 | 5590.009976 | 5307.079298 | 0.375253052 | 0.060799525 | 4.76792035135 | 2.54784143372 | ns | YPD | KS.1 |
| FIM1_4569 | 1257.59137 | 1576.559209 | 2428.104162 | 0.302352303 | 0.147371158 | 0.031487425 | 0.051109279 | ns | YPD | KS.1 |
| FIM1_4570 | 2113.294845 | 1707.572591 | 1569.40606 | -0.294714848 | 0.107466976 | 0.004821848 | 0.009422111 | ns | YPD | KS.1 |
| FIM1_4571 | 785.0218492 | 927.0973696 | 1194.927305 | 0.225925934 | 0.123903581 | 0.058722155 | 0.09058689 | ns | YPD | KS.1 |
| FIM1_4572 | 5475.993598 | 5882.82442 | 4823.600807 | 0.081661444 | 0.264674426 | 0.728039326 | 0.783337521 | ns | YPD | KS.1 |
| FIM1_4573 | 3606.260906 | 3317.148197 | 1981.366376 | -0.107649418 | 0.17200355 | 0.50649034 | 0.582610275 | ns | YPD | KS.1 |
| FIM1_4574 | 903.9476727 | 645.7859004 | 667.0720463 | -0.472130671 | 0.095974868 | 5.16977601875 | 1.89952996257 | ns | YPD | KS.1 |
| FIM1_4575 | 5474.359171 | 5362.095085 | 8341.093234 | -0.028373768 | 0.105354861 | 0.782116991 | 0.829337621 | ns | YPD | KS.1 |
| FIM1_4576 | 559.5263245 | 1090.71997 | 1594.398959 | 0.931170575 | 0.136109409 | 1.70839873975 | 1.15479398509 | ns | YPD | KS.1 |
| FIM1_4577 | 203.7464774 | 562.5211174 | 482.5281874 | 1.381532294 | 0.21263177 | 7.55727726985 | 4.78339114726 | Km>KS.1 | YPD | KS.1 |
| FIM1_4578 | 430.0552948 | 348.6009477 | 446.1690164 | -0.280983943 | 0.144268988 | 0.041460045 | 0.065647271 | ns | YPD | KS.1 |
| FIM1_4579 | 271.334741 | 416.6118519 | 334.6155605 | 0.545119722 | 0.226122185 | 0.00820508 | 0.01523713 | ns | YPD | KS.1 |
| FIM1_4580 | 1230.135847 | 1638.566066 | 3068.994516 | 0.379854314 | 0.167015351 | 0.016015639 | 0.028006613 | ns | YPD | KS.1 |
| FIM1_4582 | 2528.738706 | 2611.775024 | 1719.339598 | 0.038906588 | 0.224143432 | 0.849242621 | 0.883333062 | ns | YPD | KS.1 |
| FIM1_4583 | 1879.253796 | 2121.741529 | 1529.542087 | 0.170477602 | 0.079423953 | 0.028927865 | 0.047450092 | ns | YPD | KS.1 |
| FIM1_4584 | 2805.129926 | 1925.847804 | 1894.691616 | -0.523176512 | 0.114312149 | 2.53725985862 | 8.38171832448 | ns | YPD | KS.1 |
| FIM1_4585 | 4006.449732 | 3079.362391 | 2464.633392 | -0.371519548 | 0.079000273 | 1.82556694446 | 6.15324901675 | ns | YPD | KS.1 |
| FIM1_4586 | 9100.296509 | 14500.17049 | 14634.50189 | 0.664274199 | 0.06845455 | 1.08756900182 | 1.72313878473 | ns | YPD | KS.1 |
| FIM1_4587 | 1746.817612 | 1280.276503 | 2336.525261 | -0.338812305 | 0.317891683 | 0.209958776 | 0.276042524 | ns | YPD | KS.1 |
| FIM1_4588 | 948.0190517 | 1926.57717 | 3198.471663 | 0.989791115 | 0.136572376 | 8.24433685896 | 6.15849487587 | ns | YPD | KS.1 |
| FIM1_4589 | 721.7248883 | 998.9781485 | 2506.601279 | 0.401863742 | 0.234253116 | 0.057946686 | 0.089501634 | ns | YPD | KS.1 |
| FIM1_4590 | 450.6579896 | 413.0392717 | 592.5620059 | -0.117133522 | 0.128145649 | 0.342645011 | 0.417501575 | ns | YPD | KS.1 |
| FIM1_4591 | 791.3599257 | 1740.7834 | 2635.107102 | 1.114009117 | 0.114496282 | 3.44948633037 | 5.75879009852 | Km>KS.1 | YPD | KS.1 |
| FIM1_4592 | 933.6783818 | 864.835295 | 1307.898843 | -0.10675393 | 0.087583922 | 0.213512594 | 0.279901227 | ns | YPD | KS.1 |
| FIM1_4593 | 713.2035543 | 615.5986113 | 726.3664995 | -0.194126398 | 0.150488188 | 0.175428453 | 0.235943918 | ns | YPD | KS.1 |
| FIM1_4594 | 836.1166616 | 1876.719243 | 742.0136083 | 1.068189791 | 0.238490889 | 1.24914359359 | 4.33669879842 | Km>KS.1 | YPD | KS.1 |
| FIM1_4595 | 2809.836929 | 4367.937452 | 4025.083491 | 0.610568994 | 0.128666816 | 9.24324423409 | 3.28700071941 | ns | YPD | KS.1 |
| FIM1_4596 | 2710.825811 | 1684.02396 | 3538.793639 | -0.60621254 | 0.231256262 | 0.003976644 | 0.007891426 | ns | YPD | KS.1 |
| FIM1_4597 | 449.0809384 | 2128.447759 | 1866.817906 | 2.220922973 | 0.106392482 | 8.43712764399 | 8.39494200577 | Km>KS.1 | YPD | KS.1 |
| FIM1_4598 | 507.5279734 | 412.5509123 | 478.6078264 | -0.27377437 | 0.16310227 | 0.075841627 | 0.113069252 | ns | YPD | KS.1 |
| FIM1_4599 | 270.666579 | 199.4839473 | 280.8019534 | -0.400160552 | 0.184737007 | 0.020452817 | 0.034763499 | ns | YPD | KS.1 |
| FIM1_4600 | 3861.80788 | 4383.341999 | 3711.087624 | 0.174604125 | 0.103954478 | 0.084345013 | 0.124183617 | ns | YPD | KS.1 |

| | | | | | | | | | | |
|---|---|---|---|---|---|---|---|---|---|---|
| FIM1_4601 | 674.9361181 | 811.8217789 | 781.6834496 | 0.25667235 | 0.092437826 | 0.004548345 | 0.008933288 | ns | YPD | KS.1 |
| FIM1_4602 | 117.7681075 | 252.6315609 | 450.4868022 | 0.970662484 | 0.277284084 | 9.44248411534 | 0.000247766 | ns | YPD | KS.1 |
| FIM1_4603 | 2798.816325 | 3835.039211 | 3864.028105 | 0.437434904 | 0.111526509 | 5.57740868595 | 0.000152795 | ns | YPD | KS.1 |
| FIM1_4604 | 225.8426094 | 142.006969 | 140.3676843 | -0.586728322 | 0.236966934 | 0.006286994 | 0.011933535 | ns | YPD | KS.1 |
| FIM1_4605 | 905.4489173 | 897.9709952 | 1105.088326 | -0.010708996 | 0.101808857 | 0.914111916 | 0.935549637 | ns | YPD | KS.1 |
| FIM1_4606 | 613.0814952 | 508.2057349 | 706.6727291 | -0.258872015 | 0.110170741 | 0.015539084 | 0.027239938 | ns | YPD | KS.1 |
| FIM1_4607 | 2895.677955 | 2300.633262 | 2344.433376 | -0.325389719 | 0.074232635 | 8.88113382825 | 2.71231680758 | ns | YPD | KS.1 |
| FIM1_4608 | 571.1028315 | 435.75257 | 390.933759 | -0.346772292 | 0.195625701 | 0.05603216 | 0.086760036 | ns | YPD | KS.1 |
| FIM1_4609 | 868.1791746 | 2624.767887 | 3251.156676 | 1.513187827 | 0.213785108 | 1.04434806642 | 7.72010643457 | Km>KS.1 | YPD | KS.1 |
| FIM1_4612 | 6618.653023 | 8370.602884 | 7107.770878 | 0.316630886 | 0.141731075 | 0.019564732 | 0.033482814 | ns | YPD | KS.1 |
| FIM1_4613 | 5289.750382 | 7750.539176 | 15088.26707 | 0.470566536 | 0.245222844 | 0.032728042 | 0.053019214 | ns | YPD | KS.1 |
| FIM1_4614 | 625.3907793 | 846.4245185 | 609.0939818 | 0.420790339 | 0.108209219 | 6.60025343486 | 0.000178226 | ns | YPD | KS.1 |
| FIM1_4615 | 1943.796795 | 1647.29838 | 1502.09062 | -0.230269398 | 0.093303675 | 0.011607376 | 0.020937889 | ns | YPD | KS.1 |
| FIM1_4616 | 762.580805 | 3169.023226 | 3201.066156 | 2.037382599 | 0.094022009 | 6.76231051885 | 8.41062370783 | Km>KS.1 | YPD | KS.1 |
| FIM1_4617 | 3100.308923 | 4847.478768 | 3570.518226 | 0.598962303 | 0.173368225 | 0.000253133 | 0.00062498 | ns | YPD | KS.1 |
| FIM1_4618 | 1609.511838 | 4153.862211 | 4057.374277 | 1.343576951 | 0.115172113 | 1.90502144530 | 6.19443247737 | Km>KS.1 | YPD | KS.1 |
| FIM1_4619 | 440.1077653 | 785.0613869 | 857.5893191 | 0.815922933 | 0.101598485 | 2.63453587496 | 2.51088428696 | ns | YPD | KS.1 |
| FIM1_4620 | 912.6818083 | 866.4350146 | 744.6249619 | -0.061861855 | 0.232629041 | 0.769582808 | 0.818967801 | ns | YPD | KS.1 |
| FIM1_4621 | 103.1040381 | 175.3656977 | 125.0342392 | 0.52531878 | 0.412144341 | 0.105821086 | 0.151848833 | ns | YPD | KS.1 |
| FIM1_4622 | 3324.23015 | 2124.702279 | 1327.016102 | -0.622018104 | 0.122003694 | 1.47797224617 | 5.84957193694 | ns | YPD | KS.1 |
| FIM1_4623 | 1346.014819 | 1113.882353 | 1412.356005 | -0.25909971 | 0.116715254 | 0.021858722 | 0.036838463 | ns | YPD | KS.1 |
| FIM1_4624 | 826.2689144 | 1326.791061 | 2292.836892 | 0.657505441 | 0.12474961 | 5.40540531148 | 2.26744447087 | ns | YPD | KS.1 |
| FIM1_4625 | 793.4931371 | 1305.331898 | 926.2415632 | 0.708130275 | 0.079537404 | 1.85309970421 | 2.31637463027 | ns | YPD | KS.1 |
| FIM1_4626 | 662.2151762 | 595.2010206 | 425.5798025 | -0.132732233 | 0.201524763 | 0.476739599 | 0.554672476 | ns | YPD | KS.1 |
| FIM1_4627 | 1473.367833 | 2913.717653 | 2357.628351 | 0.934799589 | 0.168463133 | 6.26422594560 | 2.96522588760 | ns | YPD | KS.1 |
| FIM1_4628 | 339.1779052 | 270.4007532 | 297.3288643 | -0.291389648 | 0.188261367 | 0.096681513 | 0.140067131 | ns | YPD | KS.1 |
| FIM1_4629 | 928.1750184 | 638.0060646 | 744.8255768 | -0.495848222 | 0.178234731 | 0.003070382 | 0.006234754 | ns | YPD | KS.1 |
| FIM1_4630 | 3101.7511 | 1481.661504 | 1317.615698 | -1.045852447 | 0.106211433 | 1.22270894097 | 2.11214478518 | Km<KS.1 | YPD | KS.1 |
| FIM1_4631 | 2079.479391 | 2055.131559 | 2080.043648 | -0.015700633 | 0.144199045 | 0.90940083 | 0.931878297 | ns | YPD | KS.1 |
| FIM1_4632 | 5597.122159 | 2930.703836 | 2347.234258 | -0.850292591 | 0.223931235 | 3.86671901253 | 0.000108377 | ns | YPD | KS.1 |
| FIM1_4633 | 180.2028118 | 56.56505267 | 76.16514234 | -1.52739985 | 0.282195963 | 4.13645769346 | 1.99022021518 | Km<KS.1 | YPD | KS.1 |
| FIM1_4634 | 6598.318466 | 3880.323942 | 2905.267577 | -0.61487829 | 0.317575489 | 0.023538614 | 0.03942916 | ns | YPD | KS.1 |
| FIM1_4635 | 1348.003637 | 1450.503745 | 1175.173052 | 0.0987262 | 0.125266965 | 0.413965847 | 0.491405414 | ns | YPD | KS.1 |
| FIM1_4636 | 453.8173833 | 489.1486865 | 580.978079 | 0.102953212 | 0.092241899 | 0.253842358 | 0.322983563 | ns | YPD | KS.1 |
| FIM1_4637 | 1193.199022 | 1018.729119 | 1091.158215 | -0.210325972 | 0.149159265 | 0.138813898 | 0.192367449 | ns | YPD | KS.1 |
| FIM1_4638 | 1111.577652 | 1291.788588 | 1228.748941 | 0.208038386 | 0.096842966 | 0.027760692 | 0.045746752 | ns | YPD | KS.1 |
| FIM1_4639 | 3388.289767 | 3782.279964 | 2308.383022 | 0.149555557 | 0.118744127 | 0.192765743 | 0.256351128 | ns | YPD | KS.1 |
| FIM1_4640 | 567.3814796 | 763.407982 | 889.2184359 | 0.393120721 | 0.167802196 | 0.013076729 | 0.023309469 | ns | YPD | KS.1 |
| FIM1_4641 | 5121.308059 | 4967.344445 | 6170.986705 | -0.041559703 | 0.1089048 | 0.694692556 | 0.755100604 | ns | YPD | KS.1 |
| FIM1_4642 | 733.2112868 | 3697.592567 | 3031.349814 | 2.288081998 | 0.155700973 | 5.75022311543 | 5.50141538448 | Km>KS.1 | YPD | KS.1 |
| FIM1_4643 | 445.9308094 | 330.1229586 | 499.9010469 | -0.404751012 | 0.150675378 | 0.004845698 | 0.009464996 | ns | YPD | KS.1 |
| FIM1_4644 | 718.2639174 | 782.0895428 | 877.6145571 | 0.116843773 | 0.116698346 | 0.30107905 | 0.373347027 | ns | YPD | KS.1 |
| FIM1_4645 | 681.9397727 | 984.6581552 | 1148.317039 | 0.501136408 | 0.143024707 | 0.000258183 | 0.000636188 | ns | YPD | KS.1 |
| FIM1_4646 | 491.9265246 | 1120.952313 | 1583.751112 | 1.16886057 | 0.107696698 | 2.65834835608 | 6.48298189779 | Km>KS.1 | YPD | KS.1 |
| FIM1_4647 | 261.7716214 | 1065.926911 | 1914.355097 | 1.988112917 | 0.138509795 | 2.19863232214 | 1.85393149197 | Km>KS.1 | YPD | KS.1 |

| | | | | | | | | | | |
|---|---|---|---|---|---|---|---|---|---|---|
| FIM1_4648 | 1645.32299 | 1593.121343 | 1698.51504 | -0.043775012 | 0.125065621 | 0.716940756 | 0.774208869 | ns | YPD | KS.1 |
| FIM1_4649 | 3074.5017 | 3194.934206 | 2547.851914 | 0.052749614 | 0.102424304 | 0.59726485 | 0.668479781 | ns | YPD | KS.1 |
| FIM1_4650 | 1242.531155 | 1266.630101 | 1549.062917 | 0.026425954 | 0.120822262 | 0.821130882 | 0.8609328 | ns | YPD | KS.1 |
| FIM1_4651 | 7456.072306 | 5063.133018 | 4372.04566 | -0.467267828 | 0.261699764 | 0.044377761 | 0.069933279 | ns | YPD | KS.1 |
| FIM1_4652 | 302.1969034 | 1533.203135 | 1412.47613 | 2.316472115 | 0.111490162 | 4.62957649859 | 3.83869051342 | Km>KS.1 | YPD | KS.1 |
| FIM1_4653 | 681.1862708 | 1422.607053 | 1046.675546 | 0.988824052 | 0.207033267 | 3.50214532791 | 1.32194028880 | ns | YPD | KS.1 |
| FIM1_4654 | 955.0384538 | 663.129769 | 1079.974788 | -0.483911361 | 0.175106502 | 0.003325033 | 0.006708046 | ns | YPD | KS.1 |
| FIM1_4655 | 4390.679411 | 2581.014648 | 1886.468117 | -0.67311405 | 0.245006126 | 0.002362246 | 0.004896739 | ns | YPD | KS.1 |
| FIM1_4656 | 3448.645418 | 3861.793402 | 2415.83052 | 0.129122074 | 0.266033513 | 0.583826827 | 0.655948163 | ns | YPD | KS.1 |
| FIM1_4657 | 933.6514949 | 1032.280556 | 980.8930235 | 0.136248853 | 0.125415211 | 0.259875134 | 0.329731903 | ns | YPD | KS.1 |
| FIM1_4658 | 3818.342253 | 15168.13912 | 20752.37853 | 1.929401174 | 0.17959538 | 1.71363444826 | 4.32757938076 | Km>KS.1 | YPD | KS.1 |
| FIM1_4660 | 872.1714854 | 250.4662881 | 183.5974283 | -1.706098181 | 0.22656125 | 2.38309917220 | 2.04412385891 | Km<KS.1 | YPD | KS.1 |
| FIM1_4661 | 356.4246405 | 303.7396353 | 320.9228849 | -0.193206488 | 0.229837333 | 0.354652578 | 0.429711781 | ns | YPD | KS.1 |
| FIM1_4662 | 1189.493767 | 1000.867524 | 3115.470066 | -0.191812753 | 0.288305735 | 0.444796495 | 0.521901548 | ns | YPD | KS.1 |
| FIM1_4664 | 45.22355892 | 25.45623725 | 39.5335887 | -0.722407975 | 0.259675744 | 0.001906493 | 0.004037804 | ns | YPD | KS.1 |
| FIM1_4665 | 2952.986751 | 3945.088903 | 3582.114891 | 0.340173267 | 0.266519088 | 0.148863371 | 0.204414924 | ns | YPD | KS.1 |
| FIM1_4666 | 607.6433222 | 1200.028364 | 786.747206 | 0.829833678 | 0.304682479 | 0.001793346 | 0.003807895 | ns | YPD | KS.1 |
| FIM1_4667 | 4472.967889 | 4098.709741 | 5171.262894 | -0.094036779 | 0.301984492 | 0.71791046 | 0.774751527 | ns | YPD | KS.1 |
| FIM1_4668 | 1147.249058 | 1169.873124 | 590.2764347 | 0.022894521 | 0.246952308 | 0.917866093 | 0.938426596 | ns | YPD | KS.1 |
| FIM1_4669 | 59.67629277 | 66.71340014 | 39.49713505 | 0.119098564 | 0.308306227 | 0.652340753 | 0.717055953 | ns | YPD | KS.1 |
| FIM1_4670 | 1984.015571 | 3585.465065 | 2421.800564 | 0.786718749 | 0.202175306 | 3.02446612547 | 8.59812512813 | ns | YPD | KS.1 |
| FIM1_4671 | 1629.716725 | 1680.954269 | 1252.57698 | 0.041709227 | 0.124022564 | 0.727708014 | 0.783337521 | ns | YPD | KS.1 |
| FIM1_4672 | 2152.28593 | 966.425274 | 1166.076171 | -1.131016725 | 0.115085911 | 1.27110704153 | 2.18815139502 | Km<KS.1 | YPD | KS.1 |
| FIM1_4673 | 639.6329403 | 552.9691655 | 398.5748486 | -0.197239802 | 0.129574142 | 0.113560529 | 0.161973519 | ns | YPD | KS.1 |
| FIM1_4674 | 600.7895671 | 246.1296406 | 279.8664622 | -1.26602302 | 0.107289448 | 4.50045529704 | 1.54412173122 | Km<KS.1 | YPD | KS.1 |
| FIM1_4675 | 6807.814966 | 4051.476931 | 4876.118353 | -0.724713151 | 0.120189276 | 5.56117315990 | 2.94641495958 | ns | YPD | KS.1 |
| FIM1_4676 | 1572.012245 | 2467.226085 | 2206.597425 | 0.641910845 | 0.07302581 | 5.93629433838 | 7.11640104420 | ns | YPD | KS.1 |
| FIM1_4678 | 1794.336829 | 2228.794261 | 2206.929083 | 0.233995217 | 0.311801931 | 0.379778643 | 0.456046041 | ns | YPD | KS.1 |
| FIM1_4679 | 2917.592871 | 1641.124631 | 1706.025796 | -0.721244067 | 0.262039327 | 0.002092961 | 0.004389747 | ns | YPD | KS.1 |
| FIM1_4680 | 1026.274252 | 766.5749126 | 650.5430871 | -0.406097937 | 0.106606762 | 9.35706381314 | 0.000245783 | ns | YPD | KS.1 |
| FIM1_4681 | 4171.636815 | 3115.803776 | 3003.550954 | -0.38624132 | 0.168236499 | 0.014982034 | 0.026356301 | ns | YPD | KS.1 |
| FIM1_4682 | 2797.137637 | 6056.646553 | 6334.836483 | 1.102219294 | 0.080644009 | 2.43196582986 | 1.45771445826 | Km>KS.1 | YPD | KS.1 |
| FIM1_4683 | 18.62532703 | 14.32551795 | 149.174951 | -0.271345008 | 0.331097345 | 0.329810502 | 0.40394073 | ns | YPD | KS.1 |
| FIM1_4684 | 1182.817 | 1410.050888 | 1653.177765 | 0.227760361 | 0.175582586 | 0.167105813 | 0.226402892 | ns | YPD | KS.1 |
| FIM1_4685 | 1941.471296 | 2429.392185 | 1895.683557 | 0.236536817 | 0.325737551 | 0.390014084 | 0.466872009 | ns | YPD | KS.1 |
| FIM1_4686 | 1458.656924 | 1056.00837 | 374.4480036 | -0.352473638 | 0.319704834 | 0.194065111 | 0.257803452 | ns | YPD | KS.1 |
| FIM1_4688 | 321.1228574 | 1036.376188 | 1030.77713 | 1.628867251 | 0.183497033 | 3.88547158594 | 5.14101620606 | Km>KS.1 | YPD | KS.1 |
| FIM1_4689 | 1455.958791 | 1222.796616 | 1260.925112 | -0.239198235 | 0.111757353 | 0.027376497 | 0.045233502 | ns | YPD | KS.1 |
| FIM1_4690 | 4568.095614 | 3288.314415 | 3273.689111 | -0.446575277 | 0.142624317 | 0.001086736 | 0.002388035 | ns | YPD | KS.1 |
| FIM1_4691 | 1600.471185 | 1500.472195 | 1392.037738 | -0.071359573 | 0.283797809 | 0.773724878 | 0.822671782 | ns | YPD | KS.1 |
| FIM1_4692 | 1282.963324 | 1066.885839 | 1443.260619 | -0.239809666 | 0.173915478 | 0.142295904 | 0.196372294 | ns | YPD | KS.1 |
| FIM1_4693 | 701.0265423 | 482.9883495 | 358.3793039 | -0.483527453 | 0.1971716 | 0.008255058 | 0.015324221 | ns | YPD | KS.1 |
| FIM1_4694 | 1569.346361 | 1255.008678 | 868.552117 | -0.258788656 | 0.26697686 | 0.272966916 | 0.343452303 | ns | YPD | KS.1 |
| FIM1_4695 | 581.1940977 | 1067.597223 | 611.148356 | 0.758565992 | 0.272884591 | 0.001771564 | 0.003766341 | ns | YPD | KS.1 |
| FIM1_4697 | 1362.83697 | 1353.088033 | 1508.15025 | -0.007027386 | 0.351078251 | 0.980687646 | 0.986039013 | ns | YPD | KS.1 |

| | | | | | | | | | | |
|---|---|---|---|---|---|---|---|---|---|---|
| FIM1_4698 | 1610.000496 | 1542.251922 | 1300.486597 | -0.056443552 | 0.149729207 | 0.692709276 | 0.753274021 | ns | YPD | KS.1 |
| FIM1_4699 | 1365.660906 | 1628.163126 | 876.9958142 | 0.201628506 | 0.268509868 | 0.395474126 | 0.472459474 | ns | YPD | KS.1 |
| FIM1_4700 | 10.31423118 | 30.27173604 | 31.51954424 | 1.338564548 | 0.339463717 | 7.46131022903 | 2.30559120431 | Km>KS.1 | YPD | KS.1 |
| FIM1_4701 | 74.5213052 | 328.1261993 | 949.344502 | 2.027758825 | 0.248508709 | 6.95221970141 | 7.59137369784 | Km>KS.1 | YPD | KS.1 |
| FIM1_4702 | 2988.487918 | 2781.138694 | 2349.804366 | -0.078613646 | 0.291408952 | 0.75632384 | 0.807795428 | ns | YPD | KS.1 |
| FIM1_4703 | 7958.525208 | 9785.280673 | 8353.968002 | 0.24210764 | 0.25699921 | 0.290242556 | 0.361984637 | ns | YPD | KS.1 |
| FIM1_4704 | 2174.365748 | 1443.928717 | 1487.561147 | -0.49498667 | 0.264216874 | 0.034659435 | 0.05594766 | ns | YPD | KS.1 |
| FIM1_4705 | 796.9264941 | 1351.426652 | 1309.422391 | 0.729609809 | 0.141141512 | 7.95436875358 | 3.26509773507 | ns | YPD | KS.1 |
| FIM1_4706 | 837.7376705 | 1221.946581 | 1323.038148 | 0.516984196 | 0.136909093 | 8.68688662155 | 0.000229026 | ns | YPD | KS.1 |
| FIM1_4707 | 994.2377275 | 613.6860215 | 651.8354446 | -0.621480079 | 0.221775224 | 0.002229325 | 0.004644426 | ns | YPD | KS.1 |
| FIM1_4708 | 1679.201638 | 1155.255783 | 1232.431846 | -0.419564179 | 0.310401978 | 0.114182651 | 0.16267431 | ns | YPD | KS.1 |
| FIM1_4709 | 1457.438774 | 3009.720654 | 1693.347632 | 0.932222877 | 0.260963708 | 7.83035202152 | 0.000207655 | ns | YPD | KS.1 |
| FIM1_4710 | 1235.703319 | 1371.908558 | 1338.096276 | 0.124714904 | 0.23557944 | 0.558767934 | 0.631788744 | ns | YPD | KS.1 |
| FIM1_4711 | 5829.98906 | 3204.404493 | 2530.178698 | -0.77796421 | 0.229425634 | 0.000216456 | 0.000541139 | ns | YPD | KS.1 |
| FIM1_4712 | 738.2891977 | 809.4481672 | 488.8438043 | 0.114100979 | 0.205433615 | 0.547607893 | 0.621005075 | ns | YPD | KS.1 |
| FIM1_4713 | 2540.299306 | 2654.056296 | 2362.926956 | 0.056367607 | 0.172092342 | 0.728228349 | 0.783337521 | ns | YPD | KS.1 |
| FIM1_4714 | 2909.113745 | 2197.164723 | 1656.996978 | -0.358482664 | 0.20028538 | 0.053003217 | 0.082480765 | ns | YPD | KS.1 |
| FIM1_4715 | 6747.313931 | 5947.084584 | 4836.093207 | -0.157285995 | 0.202652079 | 0.401267412 | 0.478042475 | ns | YPD | KS.1 |
| FIM1_4716 | 6869.652894 | 5614.167793 | 3401.173216 | -0.204302841 | 0.347307504 | 0.478232584 | 0.556279426 | ns | YPD | KS.1 |
| FIM1_4717 | 2094.622862 | 2104.355896 | 4812.049546 | 0.00528383 | 0.270995808 | 0.98239027 | 0.986983228 | ns | YPD | KS.1 |
| FIM1_4718 | 1755.996216 | 904.5736605 | 1948.392689 | -0.712475691 | 0.392787133 | 0.024923107 | 0.041510699 | ns | YPD | KS.1 |
| FIM1_4719 | 440.5001398 | 352.6054651 | 359.3621064 | -0.274173267 | 0.221687059 | 0.175238376 | 0.235752006 | ns | YPD | KS.1 |
| FIM1_4720 | 1992.082692 | 1869.934107 | 2687.385742 | -0.071630916 | 0.267995251 | 0.762677846 | 0.813708403 | ns | YPD | KS.1 |
| FIM1_4721 | 821.4722238 | 687.4355823 | 1004.264901 | -0.243244046 | 0.12662135 | 0.046170057 | 0.072459317 | ns | YPD | KS.1 |
| FIM1_4722 | 1821.056759 | 1209.218629 | 1807.401918 | -0.391763764 | 0.406123203 | 0.22133232 | 0.288102641 | ns | YPD | KS.1 |
| FIM1_4724 | 676.5238575 | 428.4791183 | 584.5339102 | -0.619578093 | 0.161786767 | 5.63116135188 | 0.000154183 | ns | YPD | KS.1 |
| FIM1_4725 | 211.6972207 | 191.5359952 | 211.8274937 | -0.049429306 | 0.636435162 | 0.894405988 | 0.919999071 | ns | YPD | KS.1 |
| FIM1_4726 | 51.66822034 | 42.06992932 | 61.86479955 | -0.122603739 | 0.57759539 | 0.740880486 | 0.794199616 | ns | YPD | KS.1 |
| FIM1_4727 | 511.3851998 | 240.311981 | 420.3880307 | -0.353725626 | 0.694920353 | 0.356081541 | 0.431181442 | ns | YPD | KS.1 |
| FIM1_4728 | 370.0142231 | 170.6439922 | 140.8377942 | -0.405974457 | 0.671045087 | 0.296061069 | 0.368318033 | ns | YPD | KS.1 |
| FIM1_4729 | 708.5558974 | 739.024224 | 766.2970696 | 0.019427528 | 0.656506487 | 0.958244529 | 0.969153595 | ns | YPD | KS.1 |
| FIM1_4730 | 134.0382383 | 216.1141118 | 481.3915826 | 0.571142614 | 0.281840512 | 0.020867371 | 0.035323297 | ns | YPD | KS.1 |
| FIM1_4731 | 1358.368886 | 973.3013409 | 1595.830275 | -0.3992086 | 0.258997649 | 0.083002542 | 0.122424442 | ns | YPD | KS.1 |
| FIM1_4732 | 4917.527907 | 3923.071909 | 3415.38404 | -0.264575804 | 0.259775941 | 0.251839586 | 0.321092245 | ns | YPD | KS.1 |
| FIM1_4733 | 3189.757039 | 1687.484808 | 1350.604209 | -0.857541589 | 0.19082574 | 1.80711312079 | 6.09931328084 | ns | YPD | KS.1 |
| FIM1_4734 | 1479.690958 | 585.1463556 | 494.3151773 | -1.252351658 | 0.220382237 | 1.55705297931 | 7.92059158700 | Km<KS.1 | YPD | KS.1 |
| FIM1_4736 | 530.6225542 | 390.8854531 | 252.8171641 | -0.370013956 | 0.246526857 | 0.094248743 | 0.136981448 | ns | YPD | KS.1 |
| FIM1_4737 | 1528.190163 | 976.7619326 | 807.1249009 | -0.571564527 | 0.224346101 | 0.005294434 | 0.010209228 | ns | YPD | KS.1 |
| FIM1_4738 | 195.5059591 | 256.2175092 | 122.0593403 | 0.311223149 | 0.273998397 | 0.19648107 | 0.260595394 | ns | YPD | KS.1 |
| FIM1_4739 | 566.4015078 | 675.712578 | 449.3723635 | 0.233235243 | 0.155000697 | 0.113069961 | 0.161320062 | ns | YPD | KS.1 |
| FIM1_4740 | 4003.942333 | 2939.335042 | 1494.907235 | -0.409639733 | 0.168495218 | 0.010002516 | 0.01824544 | ns | YPD | KS.1 |
| FIM1_4741 | 644.8712341 | 827.0962253 | 562.5953815 | 0.317854877 | 0.19389625 | 0.077498922 | 0.115091683 | ns | YPD | KS.1 |
| FIM1_4742 | 574.5836023 | 585.0041497 | 286.3233181 | 0.020695819 | 0.263662305 | 0.929579291 | 0.94845303 | ns | YPD | KS.1 |
| FIM1_4743 | 38.23175846 | 82.37433762 | 29.18473673 | 0.861875013 | 0.381337067 | 0.006107587 | 0.011640168 | ns | YPD | KS.1 |
| FIM1_4744 | 1585.465175 | 2601.542972 | 2880.341186 | 0.653623983 | 0.196743601 | 0.000365086 | 0.000873222 | ns | YPD | KS.1 |

| | | | | | | | | | | |
|---|---|---|---|---|---|---|---|---|---|---|
| FIM1_4745 | 485.0322835 | 795.0718572 | 978.5767618 | 0.681961983 | 0.134871333 | 1.6116361177( | 6.3533198776 | ns | YPD | KS.1 |
| FIM1_4746 | 2687.180233 | 2414.811118 | 2526.440432 | -0.128899125 | 0.227411743 | 0.533560416 | 0.608961475 | ns | YPD | KS.1 |
| FIM1_4747 | 629.3417779 | 567.7153296 | 669.4574419 | -0.140950591 | 0.118206444 | 0.217688167 | 0.28455035 | ns | YPD | KS.1 |
| FIM1_4748 | 2972.120388 | 3865.199985 | 4156.527977 | 0.366778893 | 0.097851441 | 0.000127333 | 0.000326704 | ns | YPD | KS.1 |
| FIM1_4749 | 683.1711935 | 438.7951368 | 434.7145254 | -0.617376327 | 0.116575712 | 5.1512787954! | 2.1644942573! | ns | YPD | KS.1 |
| FIM1_4750 | 3902.13221 | 5349.003954 | 4683.332238 | 0.44688918 | 0.075492837 | 2.0117214699( | 1.0058607349! | ns | YPD | KS.1 |
| FIM1_4751 | 248.0875252 | 189.6404043 | 204.7642249 | -0.345857172 | 0.194908882 | 0.055888513 | 0.086591514 | ns | YPD | KS.1 |
| FIM1_4753 | 1119.157491 | 1134.137741 | 2190.71695 | 0.015971081 | 0.232103761 | 0.939616211 | 0.956340149 | ns | YPD | KS.1 |
| FIM1_4754 | 1134.737155 | 1755.683112 | 1932.410155 | 0.606345074 | 0.120339104 | 2.0959556180! | 8.1463899999( | ns | YPD | KS.1 |
| FIM1_4755 | 379.5866866 | 407.9179865 | 420.6909983 | 0.091925166 | 0.157449513 | 0.538729536 | 0.612982309 | ns | YPD | KS.1 |
| FIM1_4756 | 1560.464699 | 1601.694881 | 1824.271366 | 0.034402872 | 0.151205615 | 0.811374643 | 0.852320281 | ns | YPD | KS.1 |
| FIM1_4757 | 1246.760121 | 776.4538593 | 1744.657445 | -0.577931481 | 0.267742664 | 0.01488493 | 0.026213284 | ns | YPD | KS.1 |
| FIM1_4758 | 1130.646157 | 1096.010775 | 689.9815069 | -0.042606681 | 0.11034145 | 0.691082192 | 0.752162307 | ns | YPD | KS.1 |
| FIM1_4759 | 7583.51891 | 6111.840614 | 6178.264648 | -0.283978198 | 0.164676774 | 0.067933802 | 0.102695432 | ns | YPD | KS.1 |
| FIM1_4760 | 10007.76856 | 5520.575125 | 4423.256503 | -0.717337793 | 0.299122223 | 0.005885839 | 0.011249346 | ns | YPD | KS.1 |
| FIM1_4761 | 658.7971293 | 345.2087652 | 374.2602876 | -0.880647538 | 0.173832839 | 9.9361433279! | 4.0188872403 | ns | YPD | KS.1 |
| FIM1_4762 | 148.6527193 | 166.1918507 | 127.7586392 | 0.146178782 | 0.168545168 | 0.357930523 | 0.432945382 | ns | YPD | KS.1 |
| FIM1_4763 | 180.1085989 | 214.7348375 | 172.8569932 | 0.213213066 | 0.227817164 | 0.303305564 | 0.375826944 | ns | YPD | KS.1 |
| FIM1_4764 | 295.8776158 | 302.2927572 | 328.2736609 | 0.023537358 | 0.249726706 | 0.916338408 | 0.937442644 | ns | YPD | KS.1 |
| FIM1_4765 | 621.4369563 | 237.9597223 | 314.2749395 | -1.268501667 | 0.25759188 | 9.4691870243 | 3.8456494241( | Km<KS.1 | YPD | KS.1 |
| FIM1_4766 | 296.0280836 | 140.1684741 | 195.8864339 | -1.006800418 | 0.206278006 | 1.9965058494( | 7.7841822892 | Km<KS.1 | YPD | KS.1 |
| FIM1_4767 | 1705.046828 | 610.85979 | 816.4770033 | -1.436385171 | 0.154616134 | 1.2995525642( | 1.9015511785! | Km<KS.1 | YPD | KS.1 |
| FIM1_4769 | 416.712122 | 90.54414922 | 118.1575199 | -1.989403076 | 0.344286556 | 1.6637977355 | 9.2381626498( | Km<KS.1 | YPD | KS.1 |
| FIM1_4770 | 643.865394 | 317.3025534 | 529.218885 | -0.757340849 | 0.404196936 | 0.019698975 | 0.033666232 | ns | YPD | KS.1 |
| FIM1_4771 | 473.0036961 | 862.7336384 | 746.2717156 | 0.795728115 | 0.206441161 | 3.4495749032( | 9.7123005906! | ns | YPD | KS.1 |
| FIM1_4772 | 278.2040493 | 79.12907903 | 75.1073442 | -1.730855258 | 0.210389809 | 8.5870487421 | 9.3073131791 | Km<KS.1 | YPD | KS.1 |
| FIM1_4773 | 2112.68373 | 974.3068269 | 2745.278261 | -1.064504508 | 0.173801579 | 1.5322152823 | 8.5457074323! | Km<KS.1 | YPD | KS.1 |
| FIM1_3 | 2711.794609 | 4134.440481 | 3037.529789 | 0.583648502 | 0.134220622 | 8.5379874008( | 2.7211074515! | ns | YPD | KS.3 |
| FIM1_5 | 570.5736754 | 763.0394869 | 633.8041188 | 0.348710475 | 0.282777829 | 0.174814156 | 0.223630863 | ns | YPD | KS.3 |
| FIM1_6 | 323.0090037 | 350.1681196 | 405.4975614 | 0.105288246 | 0.203853595 | 0.587150905 | 0.645113903 | ns | YPD | KS.3 |
| FIM1_7 | 74.65378534 | 64.94534338 | 60.8195808 | -0.176727713 | 0.226151944 | 0.406044607 | 0.469565765 | ns | YPD | KS.3 |
| FIM1_8 | 558.8011726 | 563.0297939 | 375.3172182 | 0.009792835 | 0.240091012 | 0.965152554 | 0.972384357 | ns | YPD | KS.3 |
| FIM1_9 | 3435.024734 | 3312.941117 | 2901.971283 | -0.040660072 | 0.324504991 | 0.887101756 | 0.906973127 | ns | YPD | KS.3 |
| FIM1_10 | 909.9154954 | 4944.338791 | 2943.370395 | 2.250406443 | 0.341560741 | 1.1710403072! | 1.8436473192 | Km>KS.3 | YPD | KS.3 |
| FIM1_11 | 404.8019085 | 350.1619611 | 340.6940978 | -0.194280099 | 0.170909759 | 0.238780039 | 0.293678787 | ns | YPD | KS.3 |
| FIM1_12 | 505.3741912 | 1523.123224 | 1729.337393 | 1.424293315 | 0.27570854 | 1.0688041371! | 6.2190650087( | Km>KS.3 | YPD | KS.3 |
| FIM1_13 | 28981.63037 | 21659.26469 | 35774.55391 | -0.292517926 | 0.394009037 | 0.371472708 | 0.434534851 | ns | YPD | KS.3 |
| FIM1_14 | 21.19068473 | 12.99318278 | 16.92066841 | -0.527699212 | 0.36750426 | 0.091633678 | 0.12398084 | ns | YPD | KS.3 |
| FIM1_15 | 12270.65671 | 7427.593607 | 14849.47076 | -0.434601093 | 0.47302364 | 0.229212391 | 0.283171504 | ns | YPD | KS.3 |
| FIM1_16 | 14815.25188 | 6312.094667 | 13864.7932 | -1.069757774 | 0.278065953 | 2.2009781008 | 6.4373110239 | Km<KS.3 | YPD | KS.3 |
| FIM1_17 | 18029.36399 | 11058.2523 | 21161.65642 | -0.453274942 | 0.442274077 | 0.195298855 | 0.246351878 | ns | YPD | KS.3 |
| FIM1_18 | 2981.084047 | 1292.096227 | 2232.947735 | -1.053408651 | 0.27162981 | 2.0529336398 | 6.0613322599! | Km<KS.3 | YPD | KS.3 |
| FIM1_19 | 1326.746002 | 1085.087411 | 965.1753006 | -0.283159276 | 0.093673866 | 0.002232715 | 0.004308672 | ns | YPD | KS.3 |
| FIM1_20 | 3813.024867 | 2515.191789 | 3333.389345 | -0.588119867 | 0.092304856 | 1.1548835125 | 9.9576178073 | ns | YPD | KS.3 |
| FIM1_21 | 944.0517827 | 403.9863633 | 585.7584931 | -1.182588417 | 0.135256603 | 2.8084249082( | 1.2818269649! | Km<KS.3 | YPD | KS.3 |

| | | | | | | | | | | |
|---|---|---|---|---|---|---|---|---|---|---|
| FIM1_22 | 1705.385523 | 1539.353995 | 1520.407378 | -0.145195133 | 0.072232532 | 0.042976617 | 0.063050208 | ns | YPD | KS.3 |
| FIM1_23 | 343.5796333 | 220.2223719 | 230.7427362 | -0.599987093 | 0.172120027 | 0.000295455 | 0.000685582 | ns | YPD | KS.3 |
| FIM1_24 | 1315.453015 | 670.9427168 | 1199.112786 | -0.836539612 | 0.272326978 | 0.00075843 | 0.001609036 | ns | YPD | KS.3 |
| FIM1_25 | 521.7253286 | 240.3485949 | 454.9892691 | -0.998041731 | 0.244580479 | 1.0567336638 | 3.3064465269 | ns | YPD | KS.3 |
| FIM1_26 | 375.4522088 | 178.6962462 | 284.9825584 | -0.97573899 | 0.218284057 | 1.9894089279 | 7.3096819915 | ns | YPD | KS.3 |
| FIM1_27 | 946.9028915 | 553.6774636 | 657.296323 | -0.731541015 | 0.161617801 | 2.8363108654 | 1.0071839083 | ns | YPD | KS.3 |
| FIM1_28 | 6722.884759 | 11598.73299 | 11506.37819 | 0.729251435 | 0.187650642 | 4.8073806755 | 0.000130479 | ns | YPD | KS.3 |
| FIM1_29 | 1731.605161 | 3778.58307 | 2774.21492 | 1.049165755 | 0.19177897 | 9.1662946303 | 5.4159519936 | Km>KS.3 | YPD | KS.3 |
| FIM1_30 | 2589.918052 | 1306.459053 | 2049.9388 | -0.954235735 | 0.1281634 | 2.6210892284 | 4.6405405379 | ns | YPD | KS.3 |
| FIM1_31 | 4429.81599 | 4982.144893 | 4258.824352 | 0.166168362 | 0.08875703 | 0.058584242 | 0.082964512 | ns | YPD | KS.3 |
| FIM1_32 | 2870.698448 | 2129.344564 | 2794.011073 | -0.413577842 | 0.131067237 | 0.001258694 | 0.002554877 | ns | YPD | KS.3 |
| FIM1_33 | 888.1424807 | 632.0316917 | 732.2157693 | -0.452698864 | 0.187176732 | 0.011544657 | 0.019286323 | ns | YPD | KS.3 |
| FIM1_34 | 2178.690158 | 1557.310613 | 1643.975651 | -0.477014166 | 0.076356277 | 3.0400894783 | 2.4083511392 | ns | YPD | KS.3 |
| FIM1_35 | 32.94541542 | 41.13337872 | 43.59138275 | 0.2666606 | 0.278961685 | 0.294244663 | 0.354618992 | ns | YPD | KS.3 |
| FIM1_36 | 6459.002237 | 3078.585804 | 6124.361802 | -0.750932036 | 0.418464626 | 0.027275886 | 0.042011621 | ns | YPD | KS.3 |
| FIM1_37 | 771.4804999 | 560.4785223 | 700.1965119 | -0.438246302 | 0.14677968 | 0.002154134 | 0.004171591 | ns | YPD | KS.3 |
| FIM1_38 | 1447.489309 | 552.2381282 | 1220.934355 | -1.315009662 | 0.182477126 | 4.1581368486 | 7.0603176867 | Km<KS.3 | YPD | KS.3 |
| FIM1_39 | 1456.074851 | 1202.720579 | 1622.369305 | -0.244196216 | 0.227573448 | 0.253466724 | 0.310055804 | ns | YPD | KS.3 |
| FIM1_40 | 1239.230681 | 719.2125459 | 1421.516646 | -0.7565097 | 0.127968896 | 1.5190118346 | 1.0554586421 | ns | YPD | KS.3 |
| FIM1_41 | 1277.726772 | 952.4846551 | 1004.945188 | -0.406080388 | 0.132996189 | 0.001791967 | 0.003529309 | ns | YPD | KS.3 |
| FIM1_42 | 1068.23933 | 650.8567564 | 1088.647994 | -0.67517539 | 0.161145747 | 1.4753156772 | 4.4781546639 | ns | YPD | KS.3 |
| FIM1_43 | 3004.757644 | 2339.777019 | 2347.041188 | -0.355509682 | 0.077218924 | 3.4739735017 | 1.2128433804 | ns | YPD | KS.3 |
| FIM1_44 | 250.1019274 | 117.9407358 | 223.7839843 | -0.995302971 | 0.210214118 | 5.2825716315 | 2.1846046439 | ns | YPD | KS.3 |
| FIM1_45 | 1052.140925 | 612.2322902 | 909.3249672 | -0.748703758 | 0.138375645 | 2.8666982631 | 1.5302386115 | ns | YPD | KS.3 |
| FIM1_46 | 4222.906642 | 4137.760307 | 3895.846385 | -0.027984453 | 0.136329803 | 0.833634469 | 0.861006491 | ns | YPD | KS.3 |
| FIM1_47 | 861.5000992 | 2309.26507 | 1453.094101 | 1.393966396 | 0.109663274 | 2.2146934974 | 6.8863125935 | Km>KS.3 | YPD | KS.3 |
| FIM1_48 | 2314.776774 | 1614.614981 | 2574.536415 | -0.512381243 | 0.074666441 | 4.6904380831 | 5.3891292063 | ns | YPD | KS.3 |
| FIM1_49 | 7108.364422 | 3476.932013 | 9518.429149 | -0.873794286 | 0.290025171 | 0.000830339 | 0.001750396 | ns | YPD | KS.3 |
| FIM1_50 | 4575.893552 | 1725.64094 | 4848.123799 | -1.290196222 | 0.229056674 | 1.4557422737 | 1.0143302257 | Km<KS.3 | YPD | KS.3 |
| FIM1_51 | 3892.605618 | 2076.197936 | 3374.321267 | -0.862565789 | 0.154391425 | 8.0930011570 | 4.8276595630 | ns | YPD | KS.3 |
| FIM1_52 | 3145.211667 | 2534.745798 | 3261.822567 | -0.297218088 | 0.137796429 | 0.027185146 | 0.041884826 | ns | YPD | KS.3 |
| FIM1_53 | 24783.91932 | 21185.86383 | 32534.50092 | -0.218822051 | 0.116482014 | 0.055995987 | 0.079662578 | ns | YPD | KS.3 |
| FIM1_54 | 1623.81942 | 5434.387673 | 3260.43686 | 1.624507378 | 0.233230694 | 5.0356653223 | 8.3508116596 | Km>KS.3 | YPD | KS.3 |
| FIM1_55 | 683.5114899 | 327.4623658 | 884.141548 | -0.85168 | 0.334512014 | 0.003547714 | 0.006588233 | ns | YPD | KS.3 |
| FIM1_56 | 36.99256598 | 27.04428853 | 110.1015889 | -0.353375154 | 0.320073958 | 0.212242095 | 0.264770417 | ns | YPD | KS.3 |
| FIM1_57 | 402.7935052 | 336.4064384 | 411.2940193 | -0.218781584 | 0.269974138 | 0.376740919 | 0.439766793 | ns | YPD | KS.3 |
| FIM1_58 | 3049.32834 | 2028.808259 | 3663.533176 | -0.540721571 | 0.192299061 | 0.003235137 | 0.006064359 | ns | YPD | KS.3 |
| FIM1_59 | 2635.806473 | 8563.48328 | 4886.402825 | 1.501872339 | 0.302699774 | 2.1723555091 | 1.1850294581 | Km>KS.3 | YPD | KS.3 |
| FIM1_60 | 2347.191755 | 1908.538004 | 2097.26308 | -0.288268469 | 0.119468957 | 0.01401155 | 0.023028563 | ns | YPD | KS.3 |
| FIM1_61 | 2232.438257 | 2073.944035 | 1514.364944 | -0.103689118 | 0.091876032 | 0.253928958 | 0.310498922 | ns | YPD | KS.3 |
| FIM1_62 | 1833.857671 | 743.5706172 | 1248.976291 | -1.24074271 | 0.165567596 | 6.6669738359 | 1.3161982076 | Km<KS.3 | YPD | KS.3 |
| FIM1_63 | 60.01519784 | 51.5932734 | 61.0388668 | -0.177525151 | 0.296353384 | 0.506109935 | 0.568374024 | ns | YPD | KS.3 |
| FIM1_64 | 1816.463524 | 1375.720175 | 2027.713438 | -0.392498769 | 0.09641374 | 3.7758548540 | 0.000104768 | ns | YPD | KS.3 |
| FIM1_65 | 249.2678714 | 337.6699113 | 253.6940693 | 0.422112675 | 0.122996412 | 0.000466791 | 0.001037197 | ns | YPD | KS.3 |
| FIM1_66 | 4354.084864 | 2934.197612 | 4634.515481 | -0.537847482 | 0.157837221 | 0.000436357 | 0.000975675 | ns | YPD | KS.3 |

| | | | | | | | | | | |
|---|---|---|---|---|---|---|---|---|---|---|
| FIM1_67 | 522.6805868 | 582.0396414 | 516.1440968 | 0.143682687 | 0.18197127 | 0.411098118 | 0.474968215 | ns | YPD | KS.3 |
| FIM1_68 | 10796.64229 | 10465.51643 | 15105.22591 | -0.0417827 | 0.173511899 | 0.802763721 | 0.835163009 | ns | YPD | KS.3 |
| FIM1_69 | 9956.158819 | 9133.372778 | 14277.90827 | -0.119236189 | 0.13177004 | 0.355163637 | 0.417716097 | ns | YPD | KS.3 |
| FIM1_70 | 1273.865833 | 791.5354286 | 1316.552668 | -0.634330484 | 0.189101445 | 0.000449564 | 0.001001156 | ns | YPD | KS.3 |
| FIM1_71 | 839.3958209 | 604.3997611 | 738.570201 | -0.449962463 | 0.148334488 | 0.001814933 | 0.003571714 | ns | YPD | KS.3 |
| FIM1_72 | 813.4493429 | 467.3562419 | 603.4281888 | -0.755790997 | 0.1606964 | 1.15015156624 | 4.44617037056 | ns | YPD | KS.3 |
| FIM1_73 | 10814.18629 | 11192.96914 | 13298.10691 | 0.04846043 | 0.09731995 | 0.614266942 | 0.670295437 | ns | YPD | KS.3 |
| FIM1_74 | 800.711874 | 680.3822034 | 775.8522912 | -0.229851555 | 0.093103836 | 0.012541176 | 0.020811324 | ns | YPD | KS.3 |
| FIM1_75 | 12279.06483 | 20240.17744 | 22825.45264 | 0.688399911 | 0.144104103 | 9.18313484146 | 3.63164513801 | ns | YPD | KS.3 |
| FIM1_76 | 272.5895237 | 580.358411 | 347.4029841 | 1.03473589 | 0.16307953 | 4.76381582473 | 4.45488415940 | Km>KS.3 | YPD | KS.3 |
| FIM1_77 | 829.2240989 | 1022.312154 | 859.8821874 | 0.294288446 | 0.105051279 | 0.004492442 | 0.008159876 | ns | YPD | KS.3 |
| FIM1_78 | 2857.136349 | 3728.374536 | 3222.916234 | 0.368763288 | 0.129333449 | 0.003586586 | 0.006648695 | ns | YPD | KS.3 |
| FIM1_79 | 2026.636044 | 6988.052883 | 3178.933275 | 1.673077052 | 0.229229169 | 3.40066926134 | 6.93374162917 | Km>KS.3 | YPD | KS.3 |
| FIM1_80 | 82.50116454 | 211.9830141 | 115.301577 | 1.27121797 | 0.199653904 | 1.70904583967 | 1.74885900857 | Km>KS.3 | YPD | KS.3 |
| FIM1_81 | 1347.271892 | 2309.586261 | 1455.405965 | 0.73575942 | 0.159193724 | 1.78313404834 | 6.63011352057 | ns | YPD | KS.3 |
| FIM1_82 | 2439.059877 | 1954.378665 | 2001.056325 | -0.312019297 | 0.097945101 | 0.001257726 | 0.002553953 | ns | YPD | KS.3 |
| FIM1_83 | 459.9262033 | 327.0989009 | 385.1962856 | -0.464465691 | 0.155072322 | 0.002019625 | 0.003932539 | ns | YPD | KS.3 |
| FIM1_84 | 904.2520351 | 905.1894011 | 813.4542519 | 0.001786695 | 0.145730859 | 0.989959147 | 0.99089732 | ns | YPD | KS.3 |
| FIM1_85 | 876.5805284 | 655.6841529 | 812.4313926 | -0.407712499 | 0.101767757 | 4.89141042961 | 0.00013247 | ns | YPD | KS.3 |
| FIM1_86 | 647.8586401 | 416.6530377 | 890.901293 | -0.616405215 | 0.11802967 | 1.04112436754 | 5.02394534837 | ns | YPD | KS.3 |
| FIM1_87 | 409.9825201 | 378.0873804 | 533.8379842 | -0.106532803 | 0.194847624 | 0.567067378 | 0.627203247 | ns | YPD | KS.3 |
| FIM1_88 | 751.8406388 | 868.393944 | 812.0837853 | 0.192998372 | 0.175797891 | 0.254288057 | 0.310831224 | ns | YPD | KS.3 |
| FIM1_89 | 667.2332567 | 1281.570918 | 613.585864 | 0.747356522 | 0.335680121 | 0.010764381 | 0.018128908 | ns | YPD | KS.3 |
| FIM1_90 | 3556.968411 | 3375.724576 | 2907.03043 | -0.074281333 | 0.078873131 | 0.342404273 | 0.404142647 | ns | YPD | KS.3 |
| FIM1_91 | 1534.562097 | 550.4821056 | 2110.39745 | -1.314236387 | 0.273480845 | 1.13422494637 | 5.40495125310 | Km<KS.3 | YPD | KS.3 |
| FIM1_92 | 375.6549846 | 329.8429015 | 320.0306234 | -0.17778025 | 0.141623901 | 0.198135721 | 0.249298233 | ns | YPD | KS.3 |
| FIM1_93 | 270.3674281 | 244.1331045 | 258.6469668 | -0.141962467 | 0.10793012 | 0.18192316 | 0.231550865 | ns | YPD | KS.3 |
| FIM1_94 | 498.7424481 | 372.8673838 | 357.0574842 | -0.398285755 | 0.146331413 | 0.005171879 | 0.009252104 | ns | YPD | KS.3 |
| FIM1_95 | 4965.09712 | 6631.442945 | 5406.626742 | 0.408417847 | 0.093929566 | 1.08837227127 | 3.39051474613 | ns | YPD | KS.3 |
| FIM1_96 | 399.9299698 | 916.5668963 | 692.0664714 | 1.050180133 | 0.265888863 | 1.54771407240 | 4.68077660195 | Km>KS.3 | YPD | KS.3 |
| FIM1_97 | 2750.065362 | 2432.687887 | 3895.884372 | -0.144426675 | 0.293120837 | 0.585312948 | 0.643378682 | ns | YPD | KS.3 |
| FIM1_98 | 243.2306447 | 121.21455 | 194.9117998 | -0.892791549 | 0.243852766 | 7.92134118886 | 0.000207196 | ns | YPD | KS.3 |
| FIM1_99 | 395.4228904 | 462.4020717 | 463.1981537 | 0.209039939 | 0.175323924 | 0.215758283 | 0.268617982 | ns | YPD | KS.3 |
| FIM1_100 | 2351.817545 | 4066.46233 | 3763.677311 | 0.75881649 | 0.135016516 | 8.51902473192 | 5.06357802166 | ns | YPD | KS.3 |
| FIM1_101 | 776.0143735 | 2099.312944 | 1500.413589 | 1.355738756 | 0.187405947 | 2.80452979958 | 4.89562657997 | Km>KS.3 | YPD | KS.3 |
| FIM1_102 | 322.4649545 | 203.8843009 | 221.6806448 | -0.618953267 | 0.171351086 | 0.000177161 | 0.000432472 | ns | YPD | KS.3 |
| FIM1_103 | 2732.244015 | 1469.509133 | 1767.344721 | -0.869406146 | 0.11537244 | 1.65766681438 | 3.05440459317 | ns | YPD | KS.3 |
| FIM1_104 | 877.87797 | 705.5163397 | 640.3642589 | -0.310681834 | 0.069079307 | 6.00286846242 | 2.00296918850 | ns | YPD | KS.3 |
| FIM1_105 | 3472.760365 | 2902.468069 | 3825.164874 | -0.238980058 | 0.182914127 | 0.173546771 | 0.222295362 | ns | YPD | KS.3 |
| FIM1_107 | 1180.787668 | 806.8111545 | 944.1502495 | -0.523681185 | 0.143679246 | 0.000182365 | 0.000443867 | ns | YPD | KS.3 |
| FIM1_108 | 2352.040456 | 1136.380394 | 2134.243348 | -0.994491526 | 0.164906724 | 3.96628272273 | 3.06878017815 | ns | YPD | KS.3 |
| FIM1_109 | 1063.093443 | 7670.872049 | 3009.743906 | 2.803156059 | 0.174789879 | 1.41946909357 | 2.35395291350 | Km>KS.3 | YPD | KS.3 |
| FIM1_110 | 748.5621509 | 1367.227637 | 762.545038 | 0.835416158 | 0.136698682 | 3.69453906788 | 2.87192685355 | ns | YPD | KS.3 |
| FIM1_111 | 6961.432054 | 5141.25739 | 5944.23296 | -0.426703357 | 0.099543637 | 1.40820962012 | 4.29016709133 | ns | YPD | KS.3 |
| FIM1_112 | 1203.478041 | 561.4800064 | 919.1620486 | -1.034713653 | 0.177626773 | 1.22286017474 | 8.58071843350 | Km<KS.3 | YPD | KS.3 |

| | | | | | | | | | | |
|---|---|---|---|---|---|---|---|---|---|---|
| FIM1_113 | 1625.147224 | 1887.810799 | 1147.542679 | 0.208213762 | 0.123991875 | 0.086900967 | 0.11825282 | ns | YPD | KS.3 |
| FIM1_114 | 1330.202728 | 841.0831378 | 985.3718464 | -0.636301371 | 0.129587045 | 5.1801479769 | 2.1476030154 | ns | YPD | KS.3 |
| FIM1_115 | 1349.046264 | 1759.383423 | 1055.82097 | 0.333190543 | 0.245293053 | 0.144209949 | 0.187186732 | ns | YPD | KS.3 |
| FIM1_116 | 409.8993133 | 411.4142746 | 432.800489 | 0.004716898 | 0.160217173 | 0.975765622 | 0.980693731 | ns | YPD | KS.3 |
| FIM1_117 | 11299.28234 | 8173.885227 | 11163.9285 | -0.427199141 | 0.196583284 | 0.022672766 | 0.035470096 | ns | YPD | KS.3 |
| FIM1_118 | 131.729086 | 154.7915164 | 186.7466403 | 0.208420762 | 0.219490275 | 0.314319043 | 0.375149209 | ns | YPD | KS.3 |
| FIM1_119 | 5823.155732 | 4691.96105 | 7368.886416 | -0.283979928 | 0.198345292 | 0.133132485 | 0.174344331 | ns | YPD | KS.3 |
| FIM1_120 | 1103.908842 | 945.9292213 | 1362.100667 | -0.216444621 | 0.111693354 | 0.04898296 | 0.070840182 | ns | YPD | KS.3 |
| FIM1_121 | 3950.606367 | 1914.838044 | 3572.133845 | -0.899111599 | 0.276604216 | 0.00034986 | 0.000798419 | ns | YPD | KS.3 |
| FIM1_122 | 4181.538398 | 2899.752655 | 3375.700208 | -0.516347945 | 0.096480284 | 6.0108159551 | 3.0297679206 | ns | YPD | KS.3 |
| FIM1_123 | 4771.752642 | 2256.05394 | 4792.686076 | -1.048824865 | 0.122385738 | 2.0829874054 | 7.6490925960 | Km<KS.3 | YPD | KS.3 |
| FIM1_124 | 726.9849447 | 1069.808266 | 724.0275505 | 0.538707889 | 0.119959572 | 4.7435996267 | 1.6097822744 | ns | YPD | KS.3 |
| FIM1_125 | 158.2659664 | 425.0830414 | 653.6319812 | 1.353744017 | 0.176075929 | 8.9278897264 | 1.9828683655 | Km>KS.3 | YPD | KS.3 |
| FIM1_126 | 1555.720376 | 1258.114467 | 1309.752804 | -0.299794001 | 0.092646447 | 0.001067182 | 0.00219935 | ns | YPD | KS.3 |
| FIM1_127 | 763.0762601 | 483.2104785 | 779.7362617 | -0.604810374 | 0.196631358 | 0.001244866 | 0.002530939 | ns | YPD | KS.3 |
| FIM1_128 | 11989.82468 | 11511.76441 | 15811.42649 | -0.056788102 | 0.11610171 | 0.6188637 | 0.674446202 | ns | YPD | KS.3 |
| FIM1_129 | 609.2973286 | 809.6710211 | 1036.64876 | 0.371882635 | 0.20438731 | 0.055534166 | 0.079073404 | ns | YPD | KS.3 |
| FIM1_130 | 7248.382196 | 7714.854907 | 9887.24238 | 0.086842217 | 0.119532301 | 0.459526268 | 0.523144893 | ns | YPD | KS.3 |
| FIM1_131 | 1709.169283 | 3409.52995 | 2248.066324 | 0.944055525 | 0.162657689 | 1.8233873241 | 1.2494975120 | ns | YPD | KS.3 |
| FIM1_132 | 14109.75302 | 14058.47575 | 21182.2828 | -0.005196617 | 0.089195737 | 0.953064723 | 0.963523064 | ns | YPD | KS.3 |
| FIM1_133 | 3622.876239 | 5414.202402 | 4961.324517 | 0.55121564 | 0.147407595 | 0.000120637 | 0.000305896 | ns | YPD | KS.3 |
| FIM1_134 | 3561.390904 | 4379.839363 | 4336.865324 | 0.288302661 | 0.118370987 | 0.013165248 | 0.021723751 | ns | YPD | KS.3 |
| FIM1_135 | 1587.922599 | 1320.464963 | 1364.517315 | -0.260845539 | 0.090960237 | 0.003748038 | 0.006918919 | ns | YPD | KS.3 |
| FIM1_136 | 63.55219979 | 330.4870471 | 244.4891628 | 2.222891621 | 0.305009656 | 7.9475515743 | 1.7972304128 | Km>KS.3 | YPD | KS.3 |
| FIM1_137 | 581.1536094 | 1368.526563 | 852.1868397 | 1.129635181 | 0.222458465 | 5.8987646913 | 2.9763036855 | Km>KS.3 | YPD | KS.3 |
| FIM1_138 | 7518.155101 | 6154.98843 | 11142.29983 | -0.271993012 | 0.157125242 | 0.074310057 | 0.102949745 | ns | YPD | KS.3 |
| FIM1_140 | 12041.21693 | 11064.6161 | 17517.12602 | -0.116649751 | 0.135442156 | 0.378282787 | 0.441256005 | ns | YPD | KS.3 |
| FIM1_141 | 770.668456 | 829.0550853 | 718.8621058 | 0.100735625 | 0.140304782 | 0.461940943 | 0.525533088 | ns | YPD | KS.3 |
| FIM1_142 | 10570.23018 | 9988.056968 | 12027.67333 | -0.080816079 | 0.065857741 | 0.217133167 | 0.270059377 | ns | YPD | KS.3 |
| FIM1_143 | 172.6243258 | 320.3346053 | 247.9532581 | 0.775249619 | 0.260263 | 0.00119798 | 0.002445609 | ns | YPD | KS.3 |
| FIM1_144 | 1250.469178 | 456.465688 | 911.1906525 | -1.157011677 | 0.372764858 | 0.000240094 | 0.000565294 | Km<KS.3 | YPD | KS.3 |
| FIM1_145 | 7695.902683 | 14942.18624 | 13754.70225 | 0.885877157 | 0.194825142 | 1.7847513781 | 6.6311241837 | ns | YPD | KS.3 |
| FIM1_146 | 1867.816528 | 2616.238942 | 2501.115526 | 0.476104914 | 0.090714526 | 1.1196324049 | 5.3507888709 | ns | YPD | KS.3 |
| FIM1_147 | 2258.00691 | 2055.110362 | 2010.46277 | -0.133427665 | 0.082698196 | 0.1035674 | 0.138805984 | ns | YPD | KS.3 |
| FIM1_148 | 952.2451533 | 899.6969456 | 787.1053614 | -0.079658242 | 0.085982385 | 0.34962365 | 0.411881994 | ns | YPD | KS.3 |
| FIM1_149 | 740.9169497 | 410.6816991 | 567.6685807 | -0.827793483 | 0.112904762 | 8.6010326418 | 1.3627432290 | ns | YPD | KS.3 |
| FIM1_150 | 1314.62813 | 966.3632629 | 1020.816971 | -0.435742373 | 0.086981028 | 4.1873754132 | 1.7669374623 | ns | YPD | KS.3 |
| FIM1_151 | 1311.812589 | 960.2194645 | 929.2554157 | -0.434063953 | 0.121784191 | 0.000280372 | 0.000653326 | ns | YPD | KS.3 |
| FIM1_153 | 806.8771745 | 242.6887445 | 573.2743016 | -1.652967758 | 0.191342082 | 6.7774109030 | 3.5492230781 | Km<KS.3 | YPD | KS.3 |
| FIM1_154 | 1102.202815 | 2986.613048 | 1999.653377 | 1.263145742 | 0.282645234 | 7.1269933100 | 2.8850115311 | Km>KS.3 | YPD | KS.3 |
| FIM1_155 | 1799.349692 | 4271.237087 | 3525.482726 | 1.148736325 | 0.213380634 | 1.0563196353 | 6.1536184844 | Km>KS.3 | YPD | KS.3 |
| FIM1_156 | 963.8674601 | 1407.025705 | 1443.315126 | 0.520839105 | 0.142329617 | 0.000173026 | 0.000423832 | ns | YPD | KS.3 |
| FIM1_157 | 1043.74788 | 3184.230465 | 2207.756938 | 1.370773619 | 0.331296143 | 1.9839854677 | 7.3005382413 | Km>KS.3 | YPD | KS.3 |
| FIM1_158 | 5736.596121 | 5896.697703 | 6922.7706 | 0.038463857 | 0.11352319 | 0.730584332 | 0.774154856 | ns | YPD | KS.3 |
| FIM1_159 | 5818.284004 | 6451.798472 | 6425.530611 | 0.14846957 | 0.032904889 | 6.2125615700 | 2.0604995873 | ns | YPD | KS.3 |

| | | | | | | | | | | |
|---|---|---|---|---|---|---|---|---|---|---|
| FIM1_160 | 5029.373837 | 5575.669114 | 5656.953157 | 0.146735432 | 0.073233996 | 0.043611441 | 0.063795037 | ns | YPD | KS.3 |
| FIM1_161 | 498.8993338 | 985.6707946 | 678.5940738 | 0.911852366 | 0.191486888 | 5.8789043089 | 2.4012765958 | ns | YPD | KS.3 |
| FIM1_162 | 1917.643653 | 1201.513074 | 1262.050445 | -0.637314088 | 0.159420105 | 3.6240516540 | 0.00010112 | ns | YPD | KS.3 |
| FIM1_163 | 559.3322249 | 313.7656568 | 516.5720716 | -0.74359754 | 0.232551115 | 0.000623798 | 0.001350476 | ns | YPD | KS.3 |
| FIM1_164 | 639.468745 | 1117.337087 | 893.1192894 | 0.790142029 | 0.092163501 | 4.1297444616 | 1.4367467620 | ns | YPD | KS.3 |
| FIM1_165 | 834.4444348 | 335.785821 | 587.4606674 | -1.266563228 | 0.144295921 | 1.4617537559 | 7.1296322900 | Km<KS.3 | YPD | KS.3 |
| FIM1_166 | 1269.029572 | 685.1402145 | 737.7645358 | -0.822959959 | 0.193110461 | 7.7817182632 | 2.5107001337 | ns | YPD | KS.3 |
| FIM1_167 | 656.3592079 | 467.1788089 | 515.6252124 | -0.45727348 | 0.173301474 | 0.006157382 | 0.010878187 | ns | YPD | KS.3 |
| FIM1_168 | 1890.756644 | 996.4003255 | 1298.756978 | -0.896147087 | 0.120120299 | 2.7461394098 | 4.8275772311 | ns | YPD | KS.3 |
| FIM1_169 | 1940.449257 | 2462.413638 | 2192.324479 | 0.33080997 | 0.126090365 | 0.007429403 | 0.012955233 | ns | YPD | KS.3 |
| FIM1_170 | 9675.924176 | 10035.32422 | 11208.52652 | 0.052213935 | 0.051960301 | 0.313188271 | 0.374096434 | ns | YPD | KS.3 |
| FIM1_171 | 2193.916178 | 1031.433784 | 1811.685558 | -1.046767927 | 0.141614664 | 2.9674707719 | 5.1619465351 | Km<KS.3 | YPD | KS.3 |
| FIM1_172 | 698.2716187 | 588.8644224 | 709.2605182 | -0.234289793 | 0.139215066 | 0.084700581 | 0.115606417 | ns | YPD | KS.3 |
| FIM1_173 | 911.2290143 | 596.9474906 | 894.9360852 | -0.59008395 | 0.121579963 | 7.4802032969 | 3.0132802755 | ns | YPD | KS.3 |
| FIM1_174 | 451.6764838 | 284.9532205 | 327.707724 | -0.497450119 | 0.361220142 | 0.107609563 | 0.143797361 | ns | YPD | KS.3 |
| FIM1_175 | 111.045513 | 70.87775178 | 88.16067366 | -0.554543881 | 0.267152262 | 0.023600439 | 0.036748728 | ns | YPD | KS.3 |
| FIM1_176 | 3018.908481 | 2044.077363 | 2120.157969 | -0.539125631 | 0.135112192 | 4.4039587016 | 0.000120317 | ns | YPD | KS.3 |
| FIM1_177 | 6934.926738 | 4471.753032 | 8182.819457 | -0.560840402 | 0.233480333 | 0.010220357 | 0.017312317 | ns | YPD | KS.3 |
| FIM1_178 | 1095.570823 | 693.5558728 | 898.2572507 | -0.647124277 | 0.085809931 | 2.5783714298 | 4.5976336428 | ns | YPD | KS.3 |
| FIM1_179 | 3531.66017 | 2395.718447 | 2568.32626 | -0.542511473 | 0.115313562 | 1.6901028789 | 6.3077733102 | ns | YPD | KS.3 |
| FIM1_180 | 1665.233961 | 1915.829243 | 1767.514357 | 0.196932837 | 0.101027387 | 0.048280916 | 0.069906158 | ns | YPD | KS.3 |
| FIM1_181 | 10885.45977 | 9127.008735 | 11963.46107 | -0.250790557 | 0.07255778 | 0.000500284 | 0.001102752 | ns | YPD | KS.3 |
| FIM1_182 | 7061.461845 | 5644.035703 | 5639.517076 | -0.283990859 | 0.235126678 | 0.19676015 | 0.247943704 | ns | YPD | KS.3 |
| FIM1_183 | 123.3984234 | 57.57469897 | 86.97005316 | -0.951246672 | 0.273464998 | 0.000133302 | 0.000334599 | ns | YPD | KS.3 |
| FIM1_184 | 3702.493665 | 3933.523593 | 4253.522987 | 0.083451138 | 0.135032105 | 0.527303192 | 0.589645624 | ns | YPD | KS.3 |
| FIM1_185 | 491.1331502 | 752.193175 | 1297.052696 | 0.56358352 | 0.197199569 | 0.002701317 | 0.005141183 | ns | YPD | KS.3 |
| FIM1_186 | 217.4680637 | 428.4114147 | 342.0256655 | 0.85861551 | 0.253984504 | 0.000247363 | 0.000580356 | ns | YPD | KS.3 |
| FIM1_187 | 4807.392442 | 5157.793995 | 7421.616443 | 0.085212966 | 0.27141538 | 0.731847227 | 0.775162861 | ns | YPD | KS.3 |
| FIM1_189 | 1246.17196 | 1235.122028 | 1349.127392 | -0.012498647 | 0.107959157 | 0.906474614 | 0.923742565 | ns | YPD | KS.3 |
| FIM1_190 | 202.6440757 | 108.3574684 | 166.9618692 | -0.831717161 | 0.197425682 | 9.4582864738 | 2.9781629878 | ns | YPD | KS.3 |
| FIM1_191 | 562.6177443 | 907.6995334 | 673.3985639 | 0.668086479 | 0.118150259 | 8.3873656755 | 4.9912851956 | ns | YPD | KS.3 |
| FIM1_192 | 1448.350753 | 6725.929863 | 3739.861374 | 1.967279686 | 0.37061852 | 5.0983684949 | 3.8372743210 | Km>KS.3 | YPD | KS.3 |
| FIM1_193 | 21.12783384 | 402.7995884 | 567.2888931 | 3.992912911 | 0.360895558 | 1.4819702521 | 2.3040006263 | Km>KS.3 | YPD | KS.3 |
| FIM1_194 | 1809.431386 | 3266.036909 | 2532.895346 | 0.779494963 | 0.205343941 | 6.2000594998 | 0.000165213 | ns | YPD | KS.3 |
| FIM1_195 | 1412.924644 | 721.5550002 | 1244.231713 | -0.915262703 | 0.168126181 | 1.5890690183 | 8.9734601204 | ns | YPD | KS.3 |
| FIM1_196 | 2395.447969 | 2336.447299 | 2924.485626 | -0.035014131 | 0.117377584 | 0.761512765 | 0.8009163 | ns | YPD | KS.3 |
| FIM1_197 | 2237.908344 | 2437.435346 | 2057.649556 | 0.110437771 | 0.214496822 | 0.586379528 | 0.644408693 | ns | YPD | KS.3 |
| FIM1_198 | 2824.616084 | 5380.113326 | 4654.542393 | 0.88789648 | 0.148014132 | 6.5188925452 | 4.8477564144 | ns | YPD | KS.3 |
| FIM1_199 | 616.2020598 | 1246.209152 | 1061.302746 | 0.953271165 | 0.177653239 | 2.2076413703 | 1.2029590161 | ns | YPD | KS.3 |
| FIM1_200 | 3677.117301 | 1855.767762 | 2473.384623 | -0.940370384 | 0.153165497 | 2.3519895788 | 1.9119523128 | ns | YPD | KS.3 |
| FIM1_201 | 2629.958229 | 1925.67517 | 2060.566516 | -0.435667822 | 0.114878086 | 0.000114466 | 0.000292334 | ns | YPD | KS.3 |
| FIM1_203 | 1872.192541 | 993.9797332 | 1224.968132 | -0.887287551 | 0.116273017 | 7.5784471322 | 1.4613478481 | ns | YPD | KS.3 |
| FIM1_204 | 182.8797336 | 148.0775875 | 149.289268 | -0.272961373 | 0.212636341 | 0.175157354 | 0.223954725 | ns | YPD | KS.3 |
| FIM1_205 | 832.7041878 | 1700.654105 | 1549.415675 | 0.883125511 | 0.279089332 | 0.000491175 | 0.001085561 | ns | YPD | KS.3 |
| FIM1_206 | 3028.166956 | 2949.20658 | 2888.196518 | -0.036926419 | 0.105342508 | 0.722223079 | 0.767201554 | ns | YPD | KS.3 |

| | | | | | | | | | | |
|---|---|---|---|---|---|---|---|---|---|---|
| FIM1_207 | 857.7070603 | 472.4258233 | 798.6830117 | -0.817802667 | 0.154148 | 4.3936526867 | 2.267471173 | ns | YPD | KS.3 |
| FIM1_208 | 3264.020913 | 2395.545282 | 3150.474754 | -0.429837356 | 0.125126941 | 0.000456932 | 0.001016199 | ns | YPD | KS.3 |
| FIM1_209 | 3980.0664 | 5922.792826 | 4312.461468 | 0.552601369 | 0.125939905 | 7.4909357128 | 2.427843985 | ns | YPD | KS.3 |
| FIM1_210 | 2701.321867 | 2332.706379 | 2926.221732 | -0.201461003 | 0.142621354 | 0.147439148 | 0.191067925 | ns | YPD | KS.3 |
| FIM1_211 | 2173.920531 | 1528.967784 | 1421.061327 | -0.490244997 | 0.120665261 | 3.4700764954 | 9.704120609 | ns | YPD | KS.3 |
| FIM1_212 | 1927.981957 | 1695.303485 | 1343.651911 | -0.17602015 | 0.145533896 | 0.214399386 | 0.267059826 | ns | YPD | KS.3 |
| FIM1_213 | 1989.472657 | 1193.698622 | 1407.992985 | -0.723487522 | 0.088337132 | 1.2427799059 | 3.271338641 | ns | YPD | KS.3 |
| FIM1_214 | 693.1308207 | 254.0867219 | 339.0121516 | -1.364947615 | 0.190485262 | 4.4457201355 | 7.472114079 | Km<KS.3 | YPD | KS.3 |
| FIM1_215 | 987.3761164 | 897.0879065 | 763.6384068 | -0.135812148 | 0.078691888 | 0.08188494 | 0.112132556 | ns | YPD | KS.3 |
| FIM1_216 | 3280.313725 | 5279.063854 | 3833.499387 | 0.635844468 | 0.185841752 | 0.000351574 | 0.000801229 | ns | YPD | KS.3 |
| FIM1_217 | 2261.7528 | 2244.690658 | 2626.542711 | -0.010579948 | 0.132226151 | 0.934831994 | 0.9468219 | ns | YPD | KS.3 |
| FIM1_218 | 883.6608223 | 2061.112334 | 1854.385565 | 1.061943438 | 0.277343471 | 2.4317448828 | 7.066548359 | Km>KS.3 | YPD | KS.3 |
| FIM1_219 | 1278.283364 | 1406.208435 | 1022.823757 | 0.134922008 | 0.090387884 | 0.131435332 | 0.17234865 | ns | YPD | KS.3 |
| FIM1_220 | 407.8160935 | 306.3119344 | 417.3276213 | -0.392051079 | 0.145675952 | 0.005708721 | 0.010143174 | ns | YPD | KS.3 |
| FIM1_221 | 1522.408704 | 815.3508122 | 1188.193571 | -0.859864157 | 0.148405067 | 2.4213284096 | 1.612598238 | ns | YPD | KS.3 |
| FIM1_222 | 1975.99612 | 2127.812973 | 1793.918807 | 0.10585579 | 0.065536695 | 0.104285304 | 0.139730511 | ns | YPD | KS.3 |
| FIM1_224 | 2241.529725 | 1236.656803 | 1990.069359 | -0.82493602 | 0.135391926 | 4.2522740001 | 3.274777577 | ns | YPD | KS.3 |
| FIM1_225 | 804.8966295 | 1357.779636 | 1133.049414 | 0.734775459 | 0.106932315 | 2.9757824796 | 3.550244085 | ns | YPD | KS.3 |
| FIM1_226 | 245.3696002 | 413.6915672 | 393.6260039 | 0.67398488 | 0.227835425 | 0.001602925 | 0.00318982 | ns | YPD | KS.3 |
| FIM1_227 | 503.7163801 | 3364.075943 | 1808.330015 | 2.695029429 | 0.168201436 | 1.2950218252 | 2.353952913 | Km>KS.3 | YPD | KS.3 |
| FIM1_228 | 60.68991196 | 57.22313552 | 100.5731306 | -0.079020752 | 0.140890456 | 0.565440709 | 0.625682279 | ns | YPD | KS.3 |
| FIM1_229 | 6276.89419 | 8019.635491 | 7860.361819 | 0.338750611 | 0.132621426 | 0.009012408 | 0.015407811 | ns | YPD | KS.3 |
| FIM1_230 | 2019.243259 | 1620.661268 | 1941.403259 | -0.309386122 | 0.09907102 | 0.001562912 | 0.003117677 | ns | YPD | KS.3 |
| FIM1_231 | 173.6959354 | 87.14600227 | 186.9610113 | -0.837876611 | 0.29245282 | 0.001459203 | 0.002926051 | ns | YPD | KS.3 |
| FIM1_233 | 4180.2798 | 5031.358985 | 4732.029427 | 0.252889483 | 0.151426576 | 0.085771585 | 0.116921183 | ns | YPD | KS.3 |
| FIM1_234 | 2120.794159 | 1931.363895 | 1866.763399 | -0.133460594 | 0.060805799 | 0.027403009 | 0.042168256 | ns | YPD | KS.3 |
| FIM1_235 | 1434.041518 | 782.6480396 | 816.0938578 | -0.840478786 | 0.133378881 | 1.0887103573 | 9.502340399 | ns | YPD | KS.3 |
| FIM1_236 | 1712.343526 | 3312.316013 | 2878.122841 | 0.916514917 | 0.134477811 | 2.8461083503 | 3.420142280 | ns | YPD | KS.3 |
| FIM1_237 | 393.9242974 | 527.0346874 | 509.869959 | 0.350125378 | 0.280697101 | 0.170492464 | 0.218946827 | ns | YPD | KS.3 |
| FIM1_238 | 476.9353876 | 439.0326101 | 439.7022536 | -0.114668257 | 0.136964875 | 0.391505593 | 0.454760758 | ns | YPD | KS.3 |
| FIM1_239 | 99.88566052 | 70.25345332 | 93.79866522 | -0.457608401 | 0.20710286 | 0.01985486 | 0.031548365 | ns | YPD | KS.3 |
| FIM1_240 | 699.8135087 | 859.986867 | 582.5653609 | 0.272906762 | 0.190620718 | 0.134535871 | 0.175904326 | ns | YPD | KS.3 |
| FIM1_241 | 4128.029345 | 3664.702981 | 3621.35305 | -0.169514916 | 0.066734067 | 0.010612608 | 0.017909675 | ns | YPD | KS.3 |
| FIM1_242 | 80.02552661 | 42.8293546 | 58.14148934 | -0.820895322 | 0.208281119 | 3.1064658535 | 8.796054423 | ns | YPD | KS.3 |
| FIM1_243 | 1278.403049 | 1532.99178 | 1207.709963 | 0.257891473 | 0.079714262 | 0.001105106 | 0.002270922 | ns | YPD | KS.3 |
| FIM1_245 | 3905.499767 | 2531.351929 | 4952.962543 | -0.551925227 | 0.237432549 | 0.012749236 | 0.021107304 | ns | YPD | KS.3 |
| FIM1_246 | 1858.668646 | 2771.677275 | 2323.651007 | 0.564528194 | 0.093477607 | 9.9350968077 | 7.071116826 | ns | YPD | KS.3 |
| FIM1_247 | 1455.27974 | 2720.437793 | 1343.159356 | 0.819625079 | 0.215725873 | 5.5598354466 | 0.000149031 | ns | YPD | KS.3 |
| FIM1_248 | 1005.923152 | 1006.789638 | 1115.187354 | 0.001480975 | 0.114171611 | 0.989479849 | 0.990874043 | ns | YPD | KS.3 |
| FIM1_249 | 811.8547952 | 380.2364369 | 995.5845502 | -0.880799706 | 0.334428501 | 0.002555111 | 0.004875979 | ns | YPD | KS.3 |
| FIM1_250 | 1278.740415 | 584.5973562 | 804.4931944 | -1.093994497 | 0.125904902 | 6.2974797648 | 2.655081511 | Km<KS.3 | YPD | KS.3 |
| FIM1_251 | 822.3944713 | 398.3019075 | 522.5479514 | -1.012936045 | 0.125360755 | 1.4648206117 | 3.775897691 | Km<KS.3 | YPD | KS.3 |
| FIM1_252 | 1997.330982 | 1999.713871 | 2061.09628 | 0.001347995 | 0.081416988 | 0.986676505 | 0.988663768 | ns | YPD | KS.3 |
| FIM1_253 | 4363.297501 | 4304.487521 | 4712.613198 | -0.019137922 | 0.101691898 | 0.848776952 | 0.873895971 | ns | YPD | KS.3 |
| FIM1_254 | 640.8730233 | 466.3968109 | 630.1023059 | -0.437019919 | 0.144817351 | 0.001942811 | 0.003797833 | ns | YPD | KS.3 |

| | | | | | | | | | | |
|---|---|---|---|---|---|---|---|---|---|---|
| FIM1_255 | 2869.706067 | 4037.21134 | 3916.679253 | 0.465624556 | 0.154699486 | 0.001921694 | 0.003760987 | ns | YPD | KS.3 |
| FIM1_256 | 1680.670516 | 2614.13579 | 2945.980669 | 0.593521175 | 0.177528255 | 0.000503321 | 0.001108956 | ns | YPD | KS.3 |
| FIM1_257 | 816.1612116 | 1001.633065 | 989.1991687 | 0.285744909 | 0.114076557 | 0.010875247 | 0.018303233 | ns | YPD | KS.3 |
| FIM1_258 | 1235.73111 | 1492.973125 | 1326.343152 | 0.264467873 | 0.11128521 | 0.015767475 | 0.025651795 | ns | YPD | KS.3 |
| FIM1_259 | 4300.547481 | 2303.730336 | 3182.099749 | -0.862193424 | 0.143619645 | 6.769602699 | 5.019191271 | ns | YPD | KS.3 |
| FIM1_260 | 716.4367011 | 477.4724951 | 531.1506979 | -0.553061238 | 0.155406194 | 0.000242603 | 0.000570932 | ns | YPD | KS.3 |
| FIM1_261 | 1287.743443 | 1672.923945 | 1417.32722 | 0.34064575 | 0.209386897 | 0.08617626 | 0.117362961 | ns | YPD | KS.3 |
| FIM1_262 | 1200.613251 | 677.8119183 | 1256.727153 | -0.756903547 | 0.201268958 | 7.517698549 | 0.00019726 | ns | YPD | KS.3 |
| FIM1_263 | 2890.477222 | 3718.377598 | 4356.538085 | 0.35170546 | 0.11563467 | 0.001974749 | 0.003849677 | ns | YPD | KS.3 |
| FIM1_264 | 798.5793777 | 1121.167717 | 998.4399574 | 0.477100503 | 0.100909914 | 1.653093960 | 6.197545181 | ns | YPD | KS.3 |
| FIM1_265 | 1889.730044 | 1885.957517 | 1462.899096 | -0.002952442 | 0.081054277 | 0.970695016 | 0.976979103 | ns | YPD | KS.3 |
| FIM1_266 | 842.6769589 | 1475.555528 | 972.3963985 | 0.774400311 | 0.139107959 | 1.119073800 | 6.503962799 | ns | YPD | KS.3 |
| FIM1_267 | 5244.113491 | 4575.271523 | 5139.402025 | -0.192762616 | 0.09182063 | 0.033814966 | 0.050900904 | ns | YPD | KS.3 |
| FIM1_268 | 3856.778075 | 5353.290899 | 5576.256847 | 0.450293429 | 0.144565313 | 0.001382067 | 0.002789365 | ns | YPD | KS.3 |
| FIM1_269 | 1313.234662 | 1228.726998 | 1362.757812 | -0.0928527 | 0.118786818 | 0.426256324 | 0.490204626 | ns | YPD | KS.3 |
| FIM1_270 | 152.158176 | 82.98523194 | 135.2654486 | -0.80863953 | 0.196611 | 1.547382451 | 4.680776601 | ns | YPD | KS.3 |
| FIM1_271 | 2773.126069 | 3540.609291 | 2806.474049 | 0.336762207 | 0.137396667 | 0.012076562 | 0.020087227 | ns | YPD | KS.3 |
| FIM1_272 | 533.1369735 | 772.9771612 | 690.4990834 | 0.511138723 | 0.140627912 | 0.000193149 | 0.000464885 | ns | YPD | KS.3 |
| FIM1_273 | 496.8905201 | 549.6631315 | 446.7911323 | 0.131604724 | 0.20380032 | 0.49722089 | 0.560161668 | ns | YPD | KS.3 |
| FIM1_274 | 1831.145517 | 997.6833822 | 1722.830084 | -0.791332725 | 0.220953688 | 0.000139438 | 0.000347895 | ns | YPD | KS.3 |
| FIM1_275 | 1585.148903 | 1074.990891 | 1602.25596 | -0.523958074 | 0.171375545 | 0.001512084 | 0.003022345 | ns | YPD | KS.3 |
| FIM1_276 | 837.9770101 | 1311.858175 | 906.7754852 | 0.621858328 | 0.129725207 | 9.565916988 | 3.771033044 | ns | YPD | KS.3 |
| FIM1_277 | 294.9724786 | 380.141939 | 311.8922662 | 0.356104083 | 0.106811528 | 0.000716033 | 0.00152683 | ns | YPD | KS.3 |
| FIM1_278 | 654.2113604 | 502.993005 | 637.3066224 | -0.359164662 | 0.15113757 | 0.014476499 | 0.023706577 | ns | YPD | KS.3 |
| FIM1_279 | 3143.590673 | 2500.994628 | 3531.57243 | -0.319545785 | 0.113708906 | 0.004276647 | 0.007794398 | ns | YPD | KS.3 |
| FIM1_280 | 30.41570845 | 17.79411007 | 26.86602388 | -0.601482438 | 0.337933707 | 0.041262193 | 0.060751528 | ns | YPD | KS.3 |
| FIM1_281 | 2752.93094 | 3846.308396 | 3055.708706 | 0.450884467 | 0.17072144 | 0.006165245 | 0.010888212 | ns | YPD | KS.3 |
| FIM1_282 | 146.0415509 | 643.2931873 | 392.1436837 | 1.866393989 | 0.381169984 | 6.508199304 | 3.924641399 | Km>KS.3 | YPD | KS.3 |
| FIM1_283 | 185.8477839 | 956.6347052 | 1789.244243 | 2.151574722 | 0.353708031 | 3.428117094 | 4.035029713 | Km>KS.3 | YPD | KS.3 |
| FIM1_284 | 801.5534429 | 881.7129912 | 1082.482432 | 0.130371296 | 0.146481441 | 0.360783532 | 0.423424881 | ns | YPD | KS.3 |
| FIM1_285 | 736.0936391 | 2117.409155 | 1445.590392 | 1.301832905 | 0.319375382 | 3.479210827 | 1.213047201 | Km>KS.3 | YPD | KS.3 |
| FIM1_286 | 2019.983306 | 4945.472245 | 3409.995974 | 1.218208649 | 0.181803588 | 2.288946806 | 2.796345770 | Km>KS.3 | YPD | KS.3 |
| FIM1_287 | 621.4366631 | 729.5544809 | 685.4018079 | 0.224654871 | 0.107750223 | 0.034345094 | 0.051590231 | ns | YPD | KS.3 |
| FIM1_288 | 119.5697989 | 103.7096613 | 132.9791323 | -0.192702611 | 0.179739689 | 0.264799346 | 0.322254586 | ns | YPD | KS.3 |
| FIM1_290 | 366.4679994 | 424.5192834 | 401.1793757 | 0.198689114 | 0.159262537 | 0.198077078 | 0.249287493 | ns | YPD | KS.3 |
| FIM1_291 | 1325.576642 | 1852.090593 | 1665.079634 | 0.454491683 | 0.160388016 | 0.003425891 | 0.006383448 | ns | YPD | KS.3 |
| FIM1_292 | 600.5593678 | 1531.860867 | 1255.979238 | 1.166842242 | 0.292780556 | 8.791799889 | 2.794837345 | Km>KS.3 | YPD | KS.3 |
| FIM1_293 | 2451.104516 | 4198.744448 | 3668.649036 | 0.747515366 | 0.131222254 | 5.593187083 | 3.414246103 | ns | YPD | KS.3 |
| FIM1_294 | 86.66060336 | 718.4367029 | 2209.438311 | 2.783703675 | 0.416517492 | 1.245144079 | 1.935809935 | Km>KS.3 | YPD | KS.3 |
| FIM1_295 | 21.81492811 | 42.61914981 | 54.48768277 | 0.77907444 | 0.323414256 | 0.00625094 | 0.01102781 | ns | YPD | KS.3 |
| FIM1_296 | 48.96570996 | 78.18672128 | 99.09863344 | 0.544768132 | 0.312282051 | 0.049854997 | 0.0719758 | ns | YPD | KS.3 |
| FIM1_297 | 21.1374218 | 45.49441513 | 31.93400973 | 0.887307548 | 0.334233732 | 0.002361818 | 0.004531448 | ns | YPD | KS.3 |
| FIM1_298 | 20.12228899 | 71.99454444 | 43.81990865 | 1.696743521 | 0.256198784 | 3.861451061 | 5.396269309 | Km>KS.3 | YPD | KS.3 |
| FIM1_299 | 323.0787971 | 779.8605833 | 625.4742574 | 1.122404243 | 0.264362449 | 3.392654587 | 1.187787232 | Km>KS.3 | YPD | KS.3 |
| FIM1_300 | 1350.896598 | 845.2799995 | 1750.706662 | -0.526548877 | 0.337046281 | 0.073525737 | 0.10200517 | ns | YPD | KS.3 |

| | | | | | | | | | | |
|---|---|---|---|---|---|---|---|---|---|---|
| FIM1_301 | 136.0686947 | 221.5919101 | 194.6034244 | 0.614092554 | 0.248511726 | 0.007687571 | 0.013349273 | ns | YPD | KS.3 |
| FIM1_302 | 393.9111398 | 727.9838759 | 737.3037931 | 0.835645601 | 0.165263978 | 1.5976702798 | 7.4353691697 | ns | YPD | KS.3 |
| FIM1_303 | 16446.54224 | 26552.72154 | 31660.2633 | 0.658818231 | 0.145902428 | 3.4497683463 | 1.2052791336 | ns | YPD | KS.3 |
| FIM1_304 | 127.3203026 | 1977.201623 | 807.8463057 | 3.864046569 | 0.216524854 | 3.4146753898 | 1.6988010064 | Km>KS.3 | YPD | KS.3 |
| FIM1_305 | 547.203784 | 1063.967734 | 843.7830533 | 0.775043114 | 0.32384319 | 0.006580501 | 0.011576376 | ns | YPD | KS.3 |
| FIM1_306 | 204.3787526 | 298.2550638 | 334.8470566 | 0.380672205 | 0.396423986 | 0.246695091 | 0.302218192 | ns | YPD | KS.3 |
| FIM1_307 | 1545.809653 | 898.6537694 | 1116.511687 | -0.757084943 | 0.12162067 | 2.1555890859 | 1.7667307582 | ns | YPD | KS.3 |
| FIM1_308 | 2591.794481 | 1680.509381 | 2450.767558 | -0.594200071 | 0.149184241 | 4.1575894807 | 0.00011415 | ns | YPD | KS.3 |
| FIM1_309 | 4623.334376 | 3916.551929 | 4752.360542 | -0.23513892 | 0.083643389 | 0.004554277 | 0.008257117 | ns | YPD | KS.3 |
| FIM1_310 | 3695.506843 | 3449.718469 | 4156.181608 | -0.093513903 | 0.157677007 | 0.540935957 | 0.602722595 | ns | YPD | KS.3 |
| FIM1_311 | 3784.020537 | 3081.680622 | 3860.09946 | -0.288359965 | 0.104161087 | 0.004996922 | 0.008977857 | ns | YPD | KS.3 |
| FIM1_312 | 3998.078759 | 3495.566578 | 3656.8811 | -0.177377966 | 0.19190169 | 0.333812026 | 0.395690929 | ns | YPD | KS.3 |
| FIM1_313 | 2545.506909 | 2725.043185 | 2693.26069 | 0.0907604 | 0.181253325 | 0.60231563 | 0.659300387 | ns | YPD | KS.3 |
| FIM1_314 | 2180.598021 | 2086.836153 | 2126.457402 | -0.057337655 | 0.206582636 | 0.77017151 | 0.807092422 | ns | YPD | KS.3 |
| FIM1_315 | 797.3977756 | 831.113858 | 659.2698316 | 0.058575821 | 0.086821762 | 0.495716211 | 0.558863091 | ns | YPD | KS.3 |
| FIM1_316 | 857.5232755 | 466.2663407 | 1101.901215 | -0.684201259 | 0.345850449 | 0.022399068 | 0.035097752 | ns | YPD | KS.3 |
| FIM1_317 | 145.8727657 | 60.21342172 | 101.9596112 | -1.122846452 | 0.268860466 | 4.5930017582 | 1.5640098389 | Km<KS.3 | YPD | KS.3 |
| FIM1_318 | 6878.609241 | 7714.668643 | 7843.167884 | 0.163922557 | 0.056691355 | 0.003683876 | 0.006808054 | ns | YPD | KS.3 |
| FIM1_319 | 266.7420757 | 311.5530188 | 301.5180789 | 0.211831557 | 0.149106303 | 0.144218441 | 0.187186732 | ns | YPD | KS.3 |
| FIM1_320 | 1476.12054 | 2050.468058 | 1810.685389 | 0.420459989 | 0.228971964 | 0.050337953 | 0.072483738 | ns | YPD | KS.3 |
| FIM1_321 | 95.97670466 | 38.12532614 | 107.0083954 | -1.057585624 | 0.363847512 | 0.000645068 | 0.001391076 | Km<KS.3 | YPD | KS.3 |
| FIM1_322 | 8970.227693 | 11782.9295 | 14723.54858 | 0.382960984 | 0.105002463 | 0.000215905 | 0.000513937 | ns | YPD | KS.3 |
| FIM1_323 | 550.3055256 | 535.5314942 | 571.029106 | -0.034710916 | 0.223026013 | 0.868806213 | 0.891014412 | ns | YPD | KS.3 |
| FIM1_324 | 2532.253507 | 3179.098381 | 5869.511829 | 0.289765626 | 0.230491215 | 0.180175528 | 0.229898244 | ns | YPD | KS.3 |
| FIM1_325 | 4.790638413 | 15.64920357 | 17.19711774 | 1.325544144 | 0.416475868 | 9.4258008445 | 0.000244364 | Km>KS.3 | YPD | KS.3 |
| FIM1_326 | 98.59837342 | 5271.617922 | 4974.569664 | 5.341133827 | 0.428010193 | 1.0219402038 | 3.9108865492 | Km>KS.3 | YPD | KS.3 |
| FIM1_327 | 1056.950918 | 1553.428477 | 1256.923338 | 0.539545808 | 0.111368501 | 8.4662332894 | 3.3668673553 | ns | YPD | KS.3 |
| FIM1_328 | 73.33950975 | 228.6456441 | 197.3835281 | 1.402548941 | 0.330940513 | 1.1098509104 | 4.3204290135 | Km>KS.3 | YPD | KS.3 |
| FIM1_329 | 566.9877637 | 1280.839033 | 674.5158635 | 1.047438844 | 0.248957368 | 5.2096602540 | 1.7571565941 | Km>KS.3 | YPD | KS.3 |
| FIM1_330 | 6210.90776 | 9076.359625 | 10680.29304 | 0.516208049 | 0.159084576 | 0.000806903 | 0.001704604 | ns | YPD | KS.3 |
| FIM1_331 | 366.854691 | 423.7255024 | 526.2211276 | 0.179412777 | 0.249861167 | 0.439168666 | 0.502961352 | ns | YPD | KS.3 |
| FIM1_332 | 667.82553 | 1320.845016 | 1159.272756 | 0.926434752 | 0.171456816 | 1.9340895454 | 1.0703109553 | ns | YPD | KS.3 |
| FIM1_333 | 638.6180434 | 1559.37928 | 1067.606057 | 1.232155059 | 0.156755982 | 3.9387998144 | 9.3758512329 | Km>KS.3 | YPD | KS.3 |
| FIM1_334 | 1929.762358 | 1813.893686 | 1901.909592 | -0.084557661 | 0.151534119 | 0.565993223 | 0.626154388 | ns | YPD | KS.3 |
| FIM1_335 | 906.9510419 | 621.1998216 | 1214.735243 | -0.444359144 | 0.302380362 | 0.101091576 | 0.135707121 | ns | YPD | KS.3 |
| FIM1_336 | 464.8083723 | 1060.512416 | 665.6980096 | 1.104588846 | 0.199953242 | 5.6133869473 | 3.4223774587 | Km>KS.3 | YPD | KS.3 |
| FIM1_337 | 1929.905164 | 3408.936383 | 3669.238901 | 0.785472712 | 0.142021053 | 1.3429331455 | 7.6794165508 | ns | YPD | KS.3 |
| FIM1_338 | 898.4683436 | 2235.344358 | 2051.336962 | 1.122858028 | 0.300748663 | 2.8610952488 | 8.1710383830 | Km>KS.3 | YPD | KS.3 |
| FIM1_339 | 3657.689006 | 3619.768365 | 3976.175577 | -0.014036818 | 0.163604356 | 0.929365518 | 0.942628635 | ns | YPD | KS.3 |
| FIM1_340 | 414.8114877 | 402.1846817 | 277.3983359 | -0.035326637 | 0.313484255 | 0.899241599 | 0.918064222 | ns | YPD | KS.3 |
| FIM1_341 | 4559.23042 | 4400.158339 | 4523.907596 | -0.049562403 | 0.117483165 | 0.667759925 | 0.719381902 | ns | YPD | KS.3 |
| FIM1_342 | 1150.372779 | 1015.968614 | 1112.983384 | -0.170512193 | 0.144498172 | 0.22593819 | 0.279751741 | ns | YPD | KS.3 |
| FIM1_343 | 1241.669189 | 3323.749721 | 2331.160854 | 1.283941243 | 0.248350601 | 1.9706861701 | 1.0857324137 | Km>KS.3 | YPD | KS.3 |
| FIM1_344 | 1402.315812 | 1434.742497 | 1776.822187 | 0.031812465 | 0.121493266 | 0.789687884 | 0.824836705 | ns | YPD | KS.3 |
| FIM1_345 | 3408.242901 | 4168.398533 | 3734.663347 | 0.257465437 | 0.226370607 | 0.226394719 | 0.280107866 | ns | YPD | KS.3 |

| | | | | | | | | | |
|---|---|---|---|---|---|---|---|---|---|
| FIM1_346 | 3844.227916 | 7254.895282 | 6352.248537 | 0.86871535 | 0.159567738 | 1.79362915589 | 1.00149327166 | ns | YPD | KS.3 |
| FIM1_347 | 1109.687606 | 1372.6274 | 1962.428707 | 0.296154466 | 0.118625287 | 0.011036273 | 0.01853036 | ns | YPD | KS.3 |
| FIM1_348 | 1446.156177 | 941.8284892 | 2060.195225 | -0.503380618 | 0.30459912 | 0.064708815 | 0.090888299 | ns | YPD | KS.3 |
| FIM1_349 | 2439.112717 | 3071.891352 | 2501.27156 | 0.321624654 | 0.117189573 | 0.005223492 | 0.009337719 | ns | YPD | KS.3 |
| FIM1_350 | 834.4927156 | 671.5719642 | 551.4005951 | -0.304259978 | 0.104146532 | 0.003051634 | 0.005748534 | ns | YPD | KS.3 |
| FIM1_351 | 1958.101982 | 2255.271181 | 1800.110609 | 0.201699215 | 0.068425129 | 0.003016078 | 0.005690586 | ns | YPD | KS.3 |
| FIM1_352 | 3032.631284 | 9572.926302 | 5972.361674 | 1.421334559 | 0.331026052 | 7.95812015987 | 3.18516876873 | Km>KS.3 | YPD | KS.3 |
| FIM1_354 | 296.8895912 | 810.7991623 | 493.2727055 | 1.235250697 | 0.313692138 | 8.18105309936 | 2.62078165932 | Km>KS.3 | YPD | KS.3 |
| FIM1_355 | 2402.057287 | 2169.416754 | 2709.148435 | -0.141884325 | 0.122476104 | 0.237831535 | 0.292584541 | ns | YPD | KS.3 |
| FIM1_356 | 1354.619505 | 2379.496765 | 2103.957352 | 0.728740146 | 0.226668313 | 0.000607221 | 0.001315734 | ns | YPD | KS.3 |
| FIM1_357 | 1863.677053 | 7107.99115 | 3991.281274 | 1.729055631 | 0.316125217 | 4.73714453189 | 3.60908025209 | Km>KS.3 | YPD | KS.3 |
| FIM1_358 | 2965.034282 | 3509.198448 | 6074.080717 | 0.199877771 | 0.288498408 | 0.444334258 | 0.507825163 | ns | YPD | KS.3 |
| FIM1_359 | 6345.755077 | 8600.225231 | 7789.665494 | 0.435788387 | 0.049642987 | 1.26351139769 | 4.79844977367 | ns | YPD | KS.3 |
| FIM1_360 | 1209.743926 | 1426.531545 | 1375.551515 | 0.231721852 | 0.095517331 | 0.014102742 | 0.023132588 | ns | YPD | KS.3 |
| FIM1_361 | 385.5158389 | 651.2820704 | 560.8953301 | 0.705426854 | 0.177539093 | 3.52018463700 | 9.83871829724 | ns | YPD | KS.3 |
| FIM1_362 | 2124.908627 | 3771.859156 | 2519.03007 | 0.765573339 | 0.191603138 | 2.81845789593 | 8.06777085390 | ns | YPD | KS.3 |
| FIM1_363 | 1061.76335 | 1212.891605 | 1212.245779 | 0.182039188 | 0.146720755 | 0.20260585 | 0.254023212 | ns | YPD | KS.3 |
| FIM1_364 | 2760.5364 | 2709.633455 | 2836.126551 | -0.026297171 | 0.065413036 | 0.685994728 | 0.736474702 | ns | YPD | KS.3 |
| FIM1_365 | 1597.939975 | 1151.950493 | 1520.484683 | -0.437921791 | 0.180169985 | 0.01139163 | 0.019065348 | ns | YPD | KS.3 |
| FIM1_366 | 2091.266654 | 2204.263561 | 1946.325552 | 0.071162573 | 0.161510708 | 0.649204871 | 0.70182404 | ns | YPD | KS.3 |
| FIM1_367 | 1108.844897 | 3497.157907 | 4689.156278 | 1.447740039 | 0.310480444 | 1.25335412554 | 5.92159237852 | Km>KS.3 | YPD | KS.3 |
| FIM1_368 | 215.4355326 | 541.6402373 | 489.1536025 | 1.122576005 | 0.312175334 | 4.85563124494 | 0.000131716 | Km>KS.3 | YPD | KS.3 |
| FIM1_369 | 13.35377939 | 29.21451799 | 35.23365655 | 0.8221426 | 0.397042119 | 0.012522187 | 0.020797131 | ns | YPD | KS.3 |
| FIM1_370 | 702.5808862 | 724.0524585 | 767.2498702 | 0.041328098 | 0.123276381 | 0.732605174 | 0.775296303 | ns | YPD | KS.3 |
| FIM1_371 | 26.21045839 | 203.5554891 | 98.26239274 | 2.817703759 | 0.29537164 | 1.54020891488 | 1.25615399205 | Km>KS.3 | YPD | KS.3 |
| FIM1_372 | 5724.523279 | 15924.52562 | 15260.98212 | 1.358744853 | 0.228795701 | 1.73129838476 | 1.45986601088 | Km>KS.3 | YPD | KS.3 |
| FIM1_373 | 591.9977584 | 2066.450788 | 1422.934025 | 1.717930319 | 0.199208981 | 5.48758346939 | 2.96747040872 | Km>KS.3 | YPD | KS.3 |
| FIM1_374 | 1006.171615 | 1574.49469 | 1849.472422 | 0.621193934 | 0.130428669 | 1.11698730257 | 4.34141549241 | ns | YPD | KS.3 |
| FIM1_375 | 540.8180382 | 2415.545298 | 1675.434603 | 2.051567728 | 0.242390795 | 6.74499240870 | 3.54922307815 | Km>KS.3 | YPD | KS.3 |
| FIM1_376 | 446.5461962 | 656.3232787 | 754.4049239 | 0.468999572 | 0.274359503 | 0.061245222 | 0.086487364 | ns | YPD | KS.3 |
| FIM1_377 | 432.0102275 | 605.2706712 | 809.5753069 | 0.442717411 | 0.201387192 | 0.020848703 | 0.0328443 | ns | YPD | KS.3 |
| FIM1_378 | 18397.50262 | 36888.20518 | 46593.75577 | 0.963666496 | 0.141126556 | 2.27696468943 | 2.79634577000 | ns | YPD | KS.3 |
| FIM1_379 | 266.0026071 | 1009.930324 | 566.4716419 | 1.654197958 | 0.364828233 | 9.17052280928 | 4.48166512536 | Km>KS.3 | YPD | KS.3 |
| FIM1_380 | 1369.714132 | 4571.087385 | 2248.476043 | 1.565672303 | 0.283426933 | 7.41740373790 | 5.45881414143 | Km>KS.3 | YPD | KS.3 |
| FIM1_381 | 1871.295516 | 3780.805812 | 2891.447937 | 0.864633662 | 0.283694165 | 0.000760669 | 0.00161241 | ns | YPD | KS.3 |
| FIM1_383 | 1254.120064 | 5882.485825 | 5227.277974 | 1.990472624 | 0.365414932 | 2.16913513070 | 1.77490909132 | Km>KS.3 | YPD | KS.3 |
| FIM1_384 | 35048.75476 | 38327.54768 | 47056.6348 | 0.115802911 | 0.212614032 | 0.565264931 | 0.625682279 | ns | YPD | KS.3 |
| FIM1_385 | 368.835069 | 1296.342784 | 715.9322575 | 1.737109554 | 0.187879163 | 1.71650968797 | 1.12363627600 | Km>KS.3 | YPD | KS.3 |
| FIM1_386 | 1890.997049 | 4555.373066 | 2795.587424 | 1.100859396 | 0.281881817 | 1.55874663750 | 4.70818155051 | Km>KS.3 | YPD | KS.3 |
| FIM1_387 | 1314.860453 | 725.697823 | 897.1630477 | -0.815442067 | 0.153092766 | 3.93381108818 | 2.04928902238 | ns | YPD | KS.3 |
| FIM1_388 | 464.7094162 | 904.0694341 | 624.7718793 | 0.918204284 | 0.145869719 | 9.30385145512 | 8.28026135764 | ns | YPD | KS.3 |
| FIM1_389 | 2406.953966 | 4013.631579 | 3232.834531 | 0.696815939 | 0.160690563 | 7.34355132092 | 2.38722065890 | ns | YPD | KS.3 |
| FIM1_390 | 387.4466745 | 337.9209936 | 386.9148257 | -0.177425619 | 0.211100417 | 0.375191918 | 0.438370078 | ns | YPD | KS.3 |
| FIM1_391 | 1151.351269 | 1496.483542 | 1297.739817 | 0.364346237 | 0.124710201 | 0.002883214 | 0.005464377 | ns | YPD | KS.3 |
| FIM1_392 | 4093.811862 | 2877.81742 | 4038.265816 | -0.487552548 | 0.133505618 | 0.000186723 | 0.000452043 | ns | YPD | KS.3 |

| | | | | | | | | | | |
|---|---|---|---|---|---|---|---|---|---|---|
| FIM1_393 | 1635.819039 | 2757.420728 | 1931.746616 | 0.739699215 | 0.088496 | 2.9181542882 | 8.961615792 | ns | YPD | KS.3 |
| FIM1_394 | 221.2245661 | 311.0247596 | 215.3585894 | 0.434872335 | 0.23147138 | 0.044946822 | 0.065574909 | ns | YPD | KS.3 |
| FIM1_395 | 1590.190894 | 1881.597774 | 1365.180795 | 0.230251851 | 0.147445828 | 0.108663058 | 0.144816157 | ns | YPD | KS.3 |
| FIM1_396 | 2777.825491 | 1925.973514 | 2343.481647 | -0.508236573 | 0.12889357 | 5.6140572088 | 0.000150404 | ns | YPD | KS.3 |
| FIM1_397 | 63.15580919 | 51.6575611 | 44.24618426 | -0.244628213 | 0.274348862 | 0.329372815 | 0.391174446 | ns | YPD | KS.3 |
| FIM1_398 | 3101.678057 | 2119.873361 | 2180.258315 | -0.528448628 | 0.127197487 | 2.2104247820 | 6.4611417688 | ns | YPD | KS.3 |
| FIM1_399 | 2735.519313 | 1548.607406 | 2774.821104 | -0.785313206 | 0.142615085 | 1.5413672014 | 8.7338289601 | ns | YPD | KS.3 |
| FIM1_401 | 2118.924137 | 1250.229733 | 1422.118783 | -0.732916784 | 0.130366071 | 8.9084447913 | 5.2698588391 | ns | YPD | KS.3 |
| FIM1_402 | 543.5235391 | 576.3867009 | 416.4323931 | 0.078054926 | 0.191014324 | 0.669353887 | 0.720786924 | ns | YPD | KS.3 |
| FIM1_403 | 779.6501875 | 1249.746177 | 967.7019184 | 0.637935474 | 0.170809539 | 0.000106082 | 0.000272461 | ns | YPD | KS.3 |
| FIM1_404 | 588.8053772 | 688.6345345 | 546.760275 | 0.201966624 | 0.218430625 | 0.327469015 | 0.3890992 | ns | YPD | KS.3 |
| FIM1_405 | 6416.913553 | 8536.484498 | 7626.635991 | 0.404080581 | 0.086228823 | 2.2175375770 | 8.0586190254 | ns | YPD | KS.3 |
| FIM1_406 | 7577.470635 | 10498.62089 | 9920.880166 | 0.462659365 | 0.081383606 | 9.7115590338 | 5.6975243152 | ns | YPD | KS.3 |
| FIM1_407 | 3896.899443 | 3839.502461 | 3459.852175 | -0.019823095 | 0.17859013 | 0.90815105 | 0.925071965 | ns | YPD | KS.3 |
| FIM1_408 | 1312.907228 | 1941.650538 | 1724.87704 | 0.555005607 | 0.081721211 | 7.2117629739 | 7.9730046212 | ns | YPD | KS.3 |
| FIM1_409 | 4160.602975 | 5433.726064 | 4668.864928 | 0.371489123 | 0.122116516 | 0.001931876 | 0.003777942 | ns | YPD | KS.3 |
| FIM1_410 | 162.5583287 | 175.0529496 | 198.6579872 | 0.097490455 | 0.199977294 | 0.608958477 | 0.665619816 | ns | YPD | KS.3 |
| FIM1_411 | 1041.031445 | 3435.535195 | 2144.197744 | 1.481671931 | 0.335276164 | 3.4135302103 | 1.4665209668 | Km>KS.3 | YPD | KS.3 |
| FIM1_412 | 644.4406224 | 855.9695325 | 728.237725 | 0.395521542 | 0.119056201 | 0.000716812 | 0.001527254 | ns | YPD | KS.3 |
| FIM1_413 | 4005.016807 | 4855.35031 | 4930.833769 | 0.271752164 | 0.092679027 | 0.003026761 | 0.005708163 | ns | YPD | KS.3 |
| FIM1_414 | 287.046743 | 818.2695689 | 811.4516496 | 1.327787445 | 0.288759177 | 2.9416456278 | 1.2814962345 | Km>KS.3 | YPD | KS.3 |
| FIM1_415 | 432.1979802 | 559.4598869 | 421.903523 | 0.349996505 | 0.161540629 | 0.025224348 | 0.039045156 | ns | YPD | KS.3 |
| FIM1_416 | 1043.155049 | 1490.420017 | 1370.58872 | 0.49791255 | 0.11803494 | 1.7437366332 | 5.2228114091 | ns | YPD | KS.3 |
| FIM1_417 | 4083.936862 | 5575.960397 | 6188.655595 | 0.426076156 | 0.149926236 | 0.003461383 | 0.006444753 | ns | YPD | KS.3 |
| FIM1_418 | 488.6138493 | 464.1823986 | 901.0270708 | -0.067638689 | 0.178465603 | 0.693638376 | 0.742917313 | ns | YPD | KS.3 |
| FIM1_419 | 749.7628439 | 1149.281182 | 844.6077708 | 0.587771419 | 0.142900256 | 2.4117296390 | 7.0124809785 | ns | YPD | KS.3 |
| FIM1_420 | 919.2081645 | 656.2663979 | 749.4564911 | -0.467057194 | 0.129489851 | 0.000228422 | 0.000540572 | ns | YPD | KS.3 |
| FIM1_421 | 1433.191509 | 1200.307083 | 1164.997138 | -0.248721059 | 0.102676917 | 0.014076767 | 0.023097598 | ns | YPD | KS.3 |
| FIM1_422 | 679.023368 | 817.4280055 | 623.0930477 | 0.257780397 | 0.121793395 | 0.031041956 | 0.04701179 | ns | YPD | KS.3 |
| FIM1_423 | 69.39527634 | 138.5757118 | 92.95811991 | 0.703368461 | 0.411991315 | 0.036811812 | 0.054864819 | ns | YPD | KS.3 |
| FIM1_425 | 3460.37764 | 1698.034242 | 3568.612381 | -0.871484329 | 0.288202191 | 0.000806498 | 0.001704473 | ns | YPD | KS.3 |
| FIM1_426 | 4392.342709 | 5099.531329 | 4407.258022 | 0.207796803 | 0.119987114 | 0.077816044 | 0.107239562 | ns | YPD | KS.3 |
| FIM1_427 | 395.8248471 | 263.5827905 | 348.6366388 | -0.557646047 | 0.146545902 | 9.1750039056 | 0.000238618 | ns | YPD | KS.3 |
| FIM1_428 | 329.1328057 | 355.5801363 | 345.022864 | 0.102104873 | 0.186389317 | 0.567783248 | 0.627715924 | ns | YPD | KS.3 |
| FIM1_429 | 1134.395563 | 1166.260226 | 1382.713337 | 0.038210097 | 0.111404169 | 0.727552182 | 0.771599255 | ns | YPD | KS.3 |
| FIM1_430 | 1114.224029 | 1087.300141 | 1029.537073 | -0.034084395 | 0.097075382 | 0.722326046 | 0.767201554 | ns | YPD | KS.3 |
| FIM1_431 | 741.6647344 | 763.9370075 | 613.3133998 | 0.039002456 | 0.196853493 | 0.835545881 | 0.862594056 | ns | YPD | KS.3 |
| FIM1_432 | 224.0041625 | 123.5098894 | 229.5730587 | -0.756638893 | 0.247027782 | 0.00095202 | 0.001976753 | ns | YPD | KS.3 |
| FIM1_433 | 2076.574139 | 1397.942759 | 1934.95982 | -0.549402517 | 0.12902165 | 1.3544662005 | 4.1416529487 | ns | YPD | KS.3 |
| FIM1_434 | 1167.388736 | 1215.243536 | 1025.508772 | 0.055315945 | 0.131021762 | 0.666264499 | 0.718081864 | ns | YPD | KS.3 |
| FIM1_435 | 1087.212536 | 805.1187244 | 1007.262228 | -0.422327992 | 0.103880248 | 3.7379648058 | 0.00010389 | ns | YPD | KS.3 |
| FIM1_436 | 487.9683076 | 247.598339 | 391.0584459 | -0.889926801 | 0.217139524 | 1.3400314729 | 4.1076134183 | ns | YPD | KS.3 |
| FIM1_437 | 566.158899 | 478.4770635 | 545.3349553 | -0.229336812 | 0.15598141 | 0.129757092 | 0.170417512 | ns | YPD | KS.3 |
| FIM1_438 | 14.05697531 | 51.8589638 | 26.92615975 | 1.481439853 | 0.439122123 | 2.5427300334 | 7.3546987886 | Km>KS.3 | YPD | KS.3 |
| FIM1_439 | 1252.732686 | 1473.412468 | 1895.423913 | 0.198657431 | 0.26374422 | 0.413169004 | 0.477139228 | ns | YPD | KS.3 |

| | | | | | | | | | | |
|---|---|---|---|---|---|---|---|---|---|---|
| FIM1_440 | 655.1905892 | 587.0914628 | 823.75812 | -0.15227723 | 0.121361041 | 0.201205265 | 0.252521743 | ns | YPD | KS.3 |
| FIM1_441 | 842.096798 | 1155.012958 | 966.3812468 | 0.443195839 | 0.1099031 | 4.1962759669 | 0.000115149 | ns | YPD | KS.3 |
| FIM1_442 | 881.6761398 | 1749.630755 | 1211.952098 | 0.9206752 | 0.187537297 | 2.739244474 | 1.2006820492 | ns | YPD | KS.3 |
| FIM1_443 | 115.814612 | 342.6881445 | 176.328668 | 1.345113826 | 0.317030793 | 1.4005148509 | 5.3309574471 | Km>KS.3 | YPD | KS.3 |
| FIM1_444 | 1223.809289 | 1069.733842 | 1310.064377 | -0.184528282 | 0.143653278 | 0.187578645 | 0.237819511 | ns | YPD | KS.3 |
| FIM1_445 | 1491.340783 | 2239.426914 | 1874.845332 | 0.564157874 | 0.128830082 | 7.657905032 | 2.4755086117 | ns | YPD | KS.3 |
| FIM1_446 | 475.2906524 | 177.7369418 | 323.4614719 | -1.346690535 | 0.176195061 | 1.3144396013 | 2.8308818255 | Km<KS.3 | YPD | KS.3 |
| FIM1_447 | 1011.430122 | 1331.487758 | 917.951041 | 0.390526737 | 0.079667613 | 7.6698936953 | 3.0773820689 | ns | YPD | KS.3 |
| FIM1_448 | 2907.344736 | 3321.664014 | 2575.742673 | 0.184317706 | 0.13039925 | 0.148794534 | 0.192673817 | ns | YPD | KS.3 |
| FIM1_449 | 3870.663752 | 5636.359182 | 4280.172542 | 0.495784429 | 0.197978877 | 0.008572432 | 0.014716304 | ns | YPD | KS.3 |
| FIM1_450 | 174.2481678 | 48.22547615 | 132.0429841 | -1.718098918 | 0.258869086 | 3.1624534183 | 4.4952016446 | Km<KS.3 | YPD | KS.3 |
| FIM1_451 | 1066.002466 | 2068.029029 | 2000.966981 | 0.851795575 | 0.23838152 | 0.000123746 | 0.000312664 | ns | YPD | KS.3 |
| FIM1_452 | 832.6176238 | 1290.437745 | 780.8864883 | 0.594221171 | 0.164720037 | 0.000188342 | 0.000455076 | ns | YPD | KS.3 |
| FIM1_453 | 1847.365069 | 1647.894051 | 2775.856282 | -0.154292921 | 0.166890163 | 0.338812938 | 0.400569003 | ns | YPD | KS.3 |
| FIM1_454 | 114.0504928 | 660.0282577 | 268.9446485 | 2.240500685 | 0.42423327 | 3.808981105 | 2.9562684865 | Km>KS.3 | YPD | KS.3 |
| FIM1_455 | 760.7451137 | 1197.221176 | 1245.188269 | 0.58278111 | 0.228184526 | 0.006465293 | 0.011385781 | ns | YPD | KS.3 |
| FIM1_456 | 647.2245487 | 1284.413455 | 1442.054858 | 0.893998719 | 0.223711532 | 2.0456660240 | 6.0470519725 | ns | YPD | KS.3 |
| FIM1_457 | 9975.980746 | 9799.66153 | 14320.3747 | -0.024581166 | 0.137207149 | 0.85450705 | 0.878886205 | ns | YPD | KS.3 |
| FIM1_458 | 13953.56768 | 13319.45607 | 20648.47836 | -0.064645844 | 0.122713851 | 0.591407876 | 0.649074384 | ns | YPD | KS.3 |
| FIM1_459 | 1298.616023 | 1236.3796 | 1542.097326 | -0.064781385 | 0.194957869 | 0.727944439 | 0.771850721 | ns | YPD | KS.3 |
| FIM1_460 | 498.3497418 | 264.1437347 | 413.5934671 | -0.858132917 | 0.176555869 | 4.1396752196 | 1.749777585 | ns | YPD | KS.3 |
| FIM1_461 | 646.5128312 | 1002.533257 | 842.2776394 | 0.607627677 | 0.132192809 | 2.571160088 | 9.2025334093 | ns | YPD | KS.3 |
| FIM1_462 | 617.8674311 | 687.1992629 | 709.7308589 | 0.150105612 | 0.077279466 | 0.050278848 | 0.072440564 | ns | YPD | KS.3 |
| FIM1_463 | 1622.692635 | 1011.792139 | 2168.265916 | -0.598905635 | 0.242714189 | 0.007989077 | 0.0138342 | ns | YPD | KS.3 |
| FIM1_464 | 4243.568671 | 7114.072049 | 5385.754652 | 0.702267383 | 0.164662763 | 1.0025204902 | 3.1467125797 | ns | YPD | KS.3 |
| FIM1_465 | 18.56014477 | 17.20838134 | 27.27999917 | -0.086546747 | 0.340322861 | 0.770428525 | 0.807092422 | ns | YPD | KS.3 |
| FIM1_466 | 5791.674175 | 7713.477005 | 7328.141002 | 0.399911404 | 0.116727742 | 0.000489313 | 0.001082406 | ns | YPD | KS.3 |
| FIM1_467 | 1809.85332 | 1581.3691 | 2370.570354 | -0.173546399 | 0.220600143 | 0.404266038 | 0.467944053 | ns | YPD | KS.3 |
| FIM1_468 | 46.68915495 | 54.80100785 | 101.8914025 | 0.202308899 | 0.21877342 | 0.327323358 | 0.389019041 | ns | YPD | KS.3 |
| FIM1_469 | 546.1549633 | 518.4983104 | 815.2713542 | -0.067110642 | 0.220587028 | 0.747086971 | 0.788450929 | ns | YPD | KS.3 |
| FIM1_470 | 565.7265003 | 2110.73632 | 1474.85134 | 1.784549654 | 0.236057035 | 2.693471657 | 6.6336740075 | Km>KS.3 | YPD | KS.3 |
| FIM1_471 | 555.0252836 | 1873.345164 | 924.8653515 | 1.516743536 | 0.334383087 | 1.680371468 | 7.776602840 | Km>KS.3 | YPD | KS.3 |
| FIM1_472 | 299.3149741 | 390.3574807 | 470.3324537 | 0.327026419 | 0.260133594 | 0.172760672 | 0.221460138 | ns | YPD | KS.3 |
| FIM1_473 | 1311.806141 | 2395.254308 | 1561.354367 | 0.798868734 | 0.199522151 | 2.5208678202 | 7.2957052970 | ns | YPD | KS.3 |
| FIM1_474 | 2057.530496 | 1120.448596 | 1664.142884 | -0.846783732 | 0.128290315 | 1.487372708 | 1.5383948494 | ns | YPD | KS.3 |
| FIM1_475 | 823.4011484 | 943.8108878 | 707.7649955 | 0.173295933 | 0.232139122 | 0.425876248 | 0.489994065 | ns | YPD | KS.3 |
| FIM1_476 | 8349.509347 | 6344.609919 | 9708.260431 | -0.371236463 | 0.165836887 | 0.020542669 | 0.032454678 | ns | YPD | KS.3 |
| FIM1_477 | 6444.162098 | 7137.082783 | 8762.515758 | 0.143303028 | 0.104732999 | 0.165303332 | 0.212557269 | ns | YPD | KS.3 |
| FIM1_478 | 2355.786845 | 3623.736782 | 3891.826898 | 0.512511711 | 0.294179585 | 0.053196586 | 0.076202999 | ns | YPD | KS.3 |
| FIM1_479 | 834.99492 | 2036.631911 | 1402.401075 | 1.161534613 | 0.240248651 | 1.825919250 | 8.3645932530 | Km>KS.3 | YPD | KS.3 |
| FIM1_480 | 1174.791901 | 718.2917962 | 1130.206243 | -0.685348322 | 0.124629367 | 1.983132868 | 1.091381197 | ns | YPD | KS.3 |
| FIM1_481 | 1640.769076 | 1915.546252 | 1343.131751 | 0.216418733 | 0.114517332 | 0.054675591 | 0.078029566 | ns | YPD | KS.3 |
| FIM1_482 | 1289.150664 | 1696.940982 | 1188.457089 | 0.375492872 | 0.150976584 | 0.010495125 | 0.017747534 | ns | YPD | KS.3 |
| FIM1_483 | 2523.351227 | 2122.085855 | 2054.337398 | -0.244952428 | 0.08898104 | 0.005414285 | 0.009642889 | ns | YPD | KS.3 |
| FIM1_485 | 53.33909256 | 121.4393688 | 116.1817359 | 1.096120582 | 0.202946553 | 1.1568250902 | 6.6920986325 | Km>KS.3 | YPD | KS.3 |

| | | | | | | | | | | |
|---|---|---|---|---|---|---|---|---|---|---|
| FIM1_486 | 1024.597533 | 1774.178468 | 1003.7575 | 0.740941482 | 0.175645579 | 1.13748475330 | 3.53245109094 | ns | YPD | KS.3 |
| FIM1_487 | 504.8721668 | 569.6346768 | 668.4113705 | 0.156539282 | 0.208638226 | 0.429152738 | 0.493079647 | ns | YPD | KS.3 |
| FIM1_488 | 3243.355767 | 4036.835686 | 5026.573853 | 0.290531093 | 0.187196091 | 0.105262625 | 0.140926146 | ns | YPD | KS.3 |
| FIM1_489 | 2154.469515 | 6505.977957 | 3569.253078 | 1.433291528 | 0.270212096 | 4.83490729263 | 2.99174922647 | Km>KS.3 | YPD | KS.3 |
| FIM1_490 | 1916.882714 | 2944.945915 | 1831.003238 | 0.597048276 | 0.126429825 | 1.42061317389 | 5.39919827358 | ns | YPD | KS.3 |
| FIM1_491 | 7265.452449 | 8992.756889 | 12395.74351 | 0.301031693 | 0.093208116 | 0.001089531 | 0.002241695 | ns | YPD | KS.3 |
| FIM1_492 | 1887.832343 | 2405.770328 | 2334.251937 | 0.342004433 | 0.093461764 | 0.00021456 | 0.000511158 | ns | YPD | KS.3 |
| FIM1_493 | 1351.106929 | 879.5618442 | 1321.065216 | -0.5931166 | 0.13777227 | 1.02326010604 | 3.20574246069 | ns | YPD | KS.3 |
| FIM1_494 | 2262.326439 | 1371.591933 | 1974.82989 | -0.700392795 | 0.116199893 | 8.38941838886 | 6.06647623322 | ns | YPD | KS.3 |
| FIM1_495 | 3806.687815 | 2846.315447 | 3267.866095 | -0.409863903 | 0.097988364 | 2.27867963552 | 6.65283520347 | ns | YPD | KS.3 |
| FIM1_496 | 5006.870495 | 5305.514006 | 5435.532296 | 0.08260373 | 0.063903558 | 0.193721334 | 0.244734291 | ns | YPD | KS.3 |
| FIM1_497 | 617.4998798 | 1423.854427 | 1598.935476 | 1.092959354 | 0.230390259 | 3.67109887554 | 1.56501430212 | Km>KS.3 | YPD | KS.3 |
| FIM1_498 | 471.816957 | 487.9050938 | 630.0746192 | 0.044532692 | 0.157290906 | 0.770423802 | 0.807092422 | ns | YPD | KS.3 |
| FIM1_499 | 2260.103465 | 2603.555047 | 3312.6156 | 0.197388063 | 0.114367131 | 0.07931508 | 0.109102112 | ns | YPD | KS.3 |
| FIM1_500 | 219.3247937 | 300.6012982 | 352.4480144 | 0.428328143 | 0.155766881 | 0.004585103 | 0.008300906 | ns | YPD | KS.3 |
| FIM1_501 | 423.3145996 | 570.2093059 | 642.575356 | 0.398195224 | 0.178835597 | 0.020536503 | 0.032454678 | ns | YPD | KS.3 |
| FIM1_502 | 2100.276407 | 2136.772478 | 2460.250386 | 0.023549564 | 0.137122016 | 0.860467204 | 0.883555075 | ns | YPD | KS.3 |
| FIM1_503 | 977.1597042 | 1597.356451 | 959.0798002 | 0.662618094 | 0.175060423 | 8.28878570521 | 0.000216579 | ns | YPD | KS.3 |
| FIM1_504 | 894.5214347 | 1239.889902 | 1291.733053 | 0.461012835 | 0.094716219 | 8.42031871991 | 3.35397002654 | ns | YPD | KS.3 |
| FIM1_505 | 667.3696483 | 803.1815069 | 1053.730224 | 0.24490191 | 0.189812888 | 0.177653927 | 0.226971825 | ns | YPD | KS.3 |
| FIM1_506 | 2333.322808 | 2042.847874 | 3207.087225 | -0.164163892 | 0.257090351 | 0.489644152 | 0.552786239 | ns | YPD | KS.3 |
| FIM1_507 | 1356.091095 | 1233.845834 | 1545.803788 | -0.130668524 | 0.135164737 | 0.32271219 | 0.384089269 | ns | YPD | KS.3 |
| FIM1_508 | 667.2638426 | 1384.296449 | 1038.605976 | 1.011335975 | 0.141298158 | 1.88519039672 | 2.79964842498 | Km>KS.3 | YPD | KS.3 |
| FIM1_509 | 672.688202 | 1567.363103 | 964.2370488 | 1.113054495 | 0.224504684 | 1.16778609238 | 5.55559605646 | Km>KS.3 | YPD | KS.3 |
| FIM1_510 | 1012.976731 | 794.3587464 | 854.0455006 | -0.335795579 | 0.13483428 | 0.010830739 | 0.018234492 | ns | YPD | KS.3 |
| FIM1_511 | 724.2621117 | 1279.290511 | 1599.389843 | 0.749323185 | 0.206819634 | 0.000130969 | 0.000329243 | ns | YPD | KS.3 |
| FIM1_512 | 191.5221552 | 200.5046651 | 324.3843601 | 0.059499389 | 0.209139759 | 0.7643064 | 0.802538646 | ns | YPD | KS.3 |
| FIM1_513 | 2732.602913 | 2426.556847 | 3189.871274 | -0.164885968 | 0.127051069 | 0.185426319 | 0.235270578 | ns | YPD | KS.3 |
| FIM1_514 | 1756.76587 | 2518.257084 | 1894.917338 | 0.484016675 | 0.174997374 | 0.004056785 | 0.007433704 | ns | YPD | KS.3 |
| FIM1_515 | 1496.736182 | 1733.020178 | 1777.12397 | 0.208054043 | 0.071964805 | 0.003604025 | 0.006677848 | ns | YPD | KS.3 |
| FIM1_516 | 1193.703188 | 1385.524621 | 1352.961096 | 0.208801009 | 0.107674198 | 0.049069619 | 0.070944886 | ns | YPD | KS.3 |
| FIM1_517 | 1062.364725 | 851.2094237 | 1368.583645 | -0.269454065 | 0.271191798 | 0.278025702 | 0.336294157 | ns | YPD | KS.3 |
| FIM1_518 | 16558.38902 | 21284.76236 | 24314.06549 | 0.356443096 | 0.079917712 | 6.85713104777 | 2.24731402916 | ns | YPD | KS.3 |
| FIM1_519 | 135.0034903 | 784.6238849 | 587.3415799 | 2.256249525 | 0.418145221 | 1.98193577507 | 1.64335508016 | Km>KS.3 | YPD | KS.3 |
| FIM1_522 | 2635.949671 | 1930.420779 | 2603.129516 | -0.390107544 | 0.248285141 | 0.090553237 | 0.122652424 | ns | YPD | KS.3 |
| FIM1_523 | 837.8728431 | 312.9783509 | 636.3143509 | -1.338775144 | 0.191179749 | 1.63524188615 | 2.45780313703 | Km<KS.3 | YPD | KS.3 |
| FIM1_524 | 2308.253237 | 2449.020384 | 1738.364328 | 0.08238565 | 0.119578673 | 0.483049346 | 0.546299272 | ns | YPD | KS.3 |
| FIM1_525 | 1466.748154 | 1629.417485 | 1775.306378 | 0.14973817 | 0.073212519 | 0.039427579 | 0.058413402 | ns | YPD | KS.3 |
| FIM1_526 | 302.6566908 | 158.8503919 | 136.6789371 | -0.74573536 | 0.327612504 | 0.009560292 | 0.016277362 | ns | YPD | KS.3 |
| FIM1_527 | 4179.384279 | 24421.6185 | 14481.84218 | 2.306845027 | 0.38698348 | 6.89062736097 | 7.66909868475 | Km>KS.3 | YPD | KS.3 |
| FIM1_528 | 1119.133368 | 6085.953545 | 3607.913838 | 2.242607076 | 0.349137489 | 3.55926571478 | 5.00207540425 | Km>KS.3 | YPD | KS.3 |
| FIM1_529 | 22.4501564 | 27.15763008 | 26.20148965 | 0.240850743 | 0.235418718 | 0.27421358 | 0.332329492 | ns | YPD | KS.3 |
| FIM1_530 | 3625.795681 | 2522.572471 | 3718.552015 | -0.49178139 | 0.164380111 | 0.001967251 | 0.003838068 | ns | YPD | KS.3 |
| FIM1_531 | 5402.428192 | 10247.51995 | 10265.85102 | 0.785156231 | 0.281445213 | 0.002104752 | 0.004087097 | ns | YPD | KS.3 |
| FIM1_532 | 1284.792924 | 3154.766366 | 4169.976391 | 0.99660026 | 0.381924517 | 0.001887876 | 0.003702083 | ns | YPD | KS.3 |

| | | | | | | | | | | |
|---|---|---|---|---|---|---|---|---|---|---|
| FIM1_533 | 884.8621104 | 1638.007497 | 1149.200053 | 0.866037903 | 0.10746731 | 2.6453983340 | 6.5476899064 | ns | YPD | KS.3 |
| FIM1_534 | 1177.283792 | 1217.108106 | 1090.027217 | 0.045903067 | 0.128696107 | 0.715786622 | 0.76237175 | ns | YPD | KS.3 |
| FIM1_535 | 763.8668784 | 2280.867974 | 1542.118228 | 1.297905353 | 0.359976971 | 2.3435627534 | 6.8262439685 | Km>KS.3 | YPD | KS.3 |
| FIM1_536 | 306.8754171 | 1279.668114 | 1058.344059 | 1.875771009 | 0.309631991 | 7.5147571887 | 8.2895603135 | Km>KS.3 | YPD | KS.3 |
| FIM1_537 | 645.3847335 | 2091.459743 | 995.1810894 | 1.491725164 | 0.307498597 | 4.0105050372 | 2.0870567531 | Km>KS.3 | YPD | KS.3 |
| FIM1_538 | 995.0302392 | 1076.119191 | 1018.384598 | 0.106273329 | 0.15529212 | 0.480885735 | 0.544457959 | ns | YPD | KS.3 |
| FIM1_539 | 670.2821106 | 453.210965 | 440.214433 | -0.544050789 | 0.123825745 | 7.3877856474 | 2.3959735069 | ns | YPD | KS.3 |
| FIM1_540 | 1733.435299 | 2969.704164 | 2288.724654 | 0.760074487 | 0.097263759 | 2.4333617024 | 5.0232259211 | ns | YPD | KS.3 |
| FIM1_541 | 724.6740198 | 466.8378831 | 552.7711887 | -0.625903864 | 0.071435631 | 1.1012823668 | 4.3140785631 | ns | YPD | KS.3 |
| FIM1_542 | 1184.039945 | 622.3218338 | 827.8097266 | -0.899056755 | 0.121864536 | 5.0982848415 | 8.4265671385 | ns | YPD | KS.3 |
| FIM1_543 | 346.8467129 | 879.648123 | 548.75533 | 1.234165386 | 0.221015437 | 2.4395181004 | 1.6203741721 | Km>KS.3 | YPD | KS.3 |
| FIM1_544 | 1643.538255 | 2354.934731 | 1915.488907 | 0.494196706 | 0.143958053 | 0.000423891 | 0.00095208 | ns | YPD | KS.3 |
| FIM1_545 | 1017.72119 | 1050.270542 | 1877.281489 | 0.042280566 | 0.171234967 | 0.798048545 | 0.831300568 | ns | YPD | KS.3 |
| FIM1_546 | 5197.016939 | 4121.29287 | 4994.01369 | -0.326284364 | 0.100053184 | 0.000954474 | 0.001981021 | ns | YPD | KS.3 |
| FIM1_547 | 1899.502586 | 2971.796379 | 2150.754405 | 0.605106571 | 0.169756022 | 0.000217781 | 0.000517906 | ns | YPD | KS.3 |
| FIM1_549 | 291.0892949 | 1679.540372 | 1034.980657 | 2.475590989 | 0.181471717 | 1.2100842124 | 7.5252111963 | Km>KS.3 | YPD | KS.3 |
| FIM1_550 | 8709.763955 | 9630.006046 | 11820.81753 | 0.139275358 | 0.125962849 | 0.259400148 | 0.316225371 | ns | YPD | KS.3 |
| FIM1_551 | 721.8231949 | 374.7406252 | 566.140344 | -0.886626075 | 0.17752506 | 1.9397219280 | 8.8128918648 | ns | YPD | KS.3 |
| FIM1_552 | 1901.603047 | 1061.722136 | 1160.952802 | -0.818423374 | 0.11072369 | 5.6174501771 | 9.2233711654 | ns | YPD | KS.3 |
| FIM1_553 | 2046.042045 | 1885.440282 | 1591.648014 | -0.115099967 | 0.093020813 | 0.210878668 | 0.263267597 | ns | YPD | KS.3 |
| FIM1_554 | 2337.862511 | 1534.249084 | 1887.284174 | -0.592873251 | 0.101587412 | 3.2758247544 | 2.1028681488 | ns | YPD | KS.3 |
| FIM1_555 | 935.4118508 | 413.8026653 | 575.5339994 | -1.130627024 | 0.144164532 | 6.7383977220 | 1.5520152160 | Km<KS.3 | YPD | KS.3 |
| FIM1_556 | 1375.798701 | 728.9525519 | 1202.338386 | -0.884793199 | 0.127920638 | 1.5212383720 | 1.9294216317 | ns | YPD | KS.3 |
| FIM1_557 | 808.9490945 | 938.0608079 | 994.4919767 | 0.208361334 | 0.101439709 | 0.03741231 | 0.05569307 | ns | YPD | KS.3 |
| FIM1_558 | 1419.085038 | 857.1748225 | 1298.567273 | -0.690607475 | 0.15368392 | 3.5958126420 | 1.2509907618 | ns | YPD | KS.3 |
| FIM1_559 | 1070.418355 | 668.5182727 | 786.9371466 | -0.658923525 | 0.115792934 | 6.9149257906 | 4.1447898564 | ns | YPD | KS.3 |
| FIM1_560 | 2476.095111 | 1483.677333 | 2200.518531 | -0.712550527 | 0.127685018 | 1.1800845606 | 6.8187232161 | ns | YPD | KS.3 |
| FIM1_561 | 1010.612601 | 1211.674294 | 945.6953352 | 0.249891856 | 0.139719165 | 0.066866055 | 0.093574859 | ns | YPD | KS.3 |
| FIM1_562 | 1349.030508 | 2162.769505 | 1953.332671 | 0.665739363 | 0.098655721 | 8.0483936634 | 8.7706650860 | ns | YPD | KS.3 |
| FIM1_563 | 248.1872247 | 132.9936166 | 220.2215981 | -0.793830618 | 0.246193735 | 0.000505554 | 0.001112891 | ns | YPD | KS.3 |
| FIM1_564 | 1744.49921 | 1049.221585 | 1556.202246 | -0.680423163 | 0.185459011 | 0.000126757 | 0.000319462 | ns | YPD | KS.3 |
| FIM1_565 | 1958.267549 | 1167.653235 | 1520.594745 | -0.685749802 | 0.196636428 | 0.000250299 | 0.000585994 | ns | YPD | KS.3 |
| FIM1_566 | 994.9217825 | 495.4362678 | 826.6143196 | -0.960943534 | 0.150324411 | 4.3719539402 | 4.1116201990 | ns | YPD | KS.3 |
| FIM1_567 | 2290.708418 | 1165.118793 | 1648.299936 | -0.948226771 | 0.116561734 | 1.1310267766 | 3.0051059325 | ns | YPD | KS.3 |
| FIM1_568 | 2431.714295 | 2789.139475 | 3256.95093 | 0.145777546 | 0.359276098 | 0.636116792 | 0.690224873 | ns | YPD | KS.3 |
| FIM1_569 | 2679.091165 | 1693.993532 | 2490.001199 | -0.550117756 | 0.289055312 | 0.03528717 | 0.052798096 | ns | YPD | KS.3 |
| FIM1_570 | 1051.032967 | 555.9736421 | 1243.278647 | -0.853028193 | 0.189189227 | 2.3263766666 | 8.4050282618 | ns | YPD | KS.3 |
| FIM1_571 | 1633.686885 | 2963.375808 | 2026.460833 | 0.700012973 | 0.312525534 | 0.01170775 | 0.01953519 | ns | YPD | KS.3 |
| FIM1_572 | 274.5882722 | 402.9712468 | 510.8466022 | 0.40395214 | 0.371889834 | 0.200234372 | 0.251366642 | ns | YPD | KS.3 |
| FIM1_573 | 3587.628472 | 4057.420743 | 4434.991654 | 0.169776263 | 0.135328539 | 0.199339637 | 0.250433003 | ns | YPD | KS.3 |
| FIM1_574 | 2633.794608 | 1820.936804 | 3334.420775 | -0.426221775 | 0.313600335 | 0.126355905 | 0.16638979 | ns | YPD | KS.3 |
| FIM1_575 | 2391.960586 | 1791.647022 | 2703.430966 | -0.403667348 | 0.115877337 | 0.000393686 | 0.000888248 | ns | YPD | KS.3 |
| FIM1_576 | 6311.239486 | 5052.20863 | 5823.24603 | -0.314216278 | 0.092460295 | 0.000589455 | 0.001280585 | ns | YPD | KS.3 |
| FIM1_577 | 1019.5233 | 483.0034956 | 905.1260683 | -1.03863132 | 0.135365493 | 3.5271713163 | 7.1623172647 | Km<KS.3 | YPD | KS.3 |
| FIM1_578 | 4619.121342 | 3853.593222 | 4422.84059 | -0.250956539 | 0.128477712 | 0.046129487 | 0.067083951 | ns | YPD | KS.3 |

| | | | | | | | | | | |
|---|---|---|---|---|---|---|---|---|---|---|
| FIM1_579 | 1282.68259 | 1223.968622 | 850.9124711 | -0.06475178 | 0.12378405 | 0.593897867 | 0.650945558 | ns | YPD | KS.3 |
| FIM1_580 | 338.2294325 | 232.3991229 | 260.9150765 | -0.523023855 | 0.125674015 | 2.1590341830 | 6.3444743416 | ns | YPD | KS.3 |
| FIM1_581 | 1314.077414 | 874.4993565 | 835.7897162 | -0.553901595 | 0.16051019 | 0.000363024 | 0.000824301 | ns | YPD | KS.3 |
| FIM1_582 | 2595.728511 | 1297.542894 | 3226.125734 | -0.808440636 | 0.325803329 | 0.004769543 | 0.008609751 | ns | YPD | KS.3 |
| FIM1_583 | 329.9952758 | 177.4926783 | 256.7458051 | -0.821034199 | 0.203146438 | 2.0381195726 | 6.0283263220 | ns | YPD | KS.3 |
| FIM1_584 | 1442.634913 | 764.8691452 | 916.6185616 | -0.887112249 | 0.121182352 | 8.0640917057 | 1.2941566527 | ns | YPD | KS.3 |
| FIM1_585 | 9816.850037 | 10106.0461 | 10000.39067 | 0.038438723 | 0.18808309 | 0.831099958 | 0.858895366 | ns | YPD | KS.3 |
| FIM1_586 | 1636.422272 | 1772.244212 | 2018.87692 | 0.104515146 | 0.200122688 | 0.583647403 | 0.642115398 | ns | YPD | KS.3 |
| FIM1_587 | 902.7264229 | 453.6108029 | 743.9692453 | -0.916357312 | 0.199137321 | 1.2673712813 | 4.8688587834 | ns | YPD | KS.3 |
| FIM1_588 | 877.6425977 | 390.698574 | 685.0246758 | -1.109410127 | 0.164302838 | 2.4224760145 | 2.9394678468 | Km<KS.3 | YPD | KS.3 |
| FIM1_589 | 1345.488237 | 1084.693258 | 965.9544655 | -0.302204533 | 0.104853568 | 0.003465046 | 0.006449161 | ns | YPD | KS.3 |
| FIM1_590 | 1224.101764 | 1063.353644 | 896.1242943 | -0.197651363 | 0.096135621 | 0.037479813 | 0.055760859 | ns | YPD | KS.3 |
| FIM1_591 | 1043.256289 | 691.4247278 | 799.1978164 | -0.571990732 | 0.122720038 | 1.9983832923 | 7.3372375493 | ns | YPD | KS.3 |
| FIM1_592 | 4926.209199 | 6500.02924 | 5793.999017 | 0.381697388 | 0.139516528 | 0.005049443 | 0.009055869 | ns | YPD | KS.3 |
| FIM1_593 | 1450.328577 | 1004.429783 | 1160.875021 | -0.498513374 | 0.163302248 | 0.001594274 | 0.003174073 | ns | YPD | KS.3 |
| FIM1_594 | 2844.905072 | 1412.106949 | 2091.221904 | -0.94427613 | 0.183848829 | 7.8782352764 | 3.8883155258 | ns | YPD | KS.3 |
| FIM1_595 | 2276.990439 | 1530.697557 | 2258.61927 | -0.54111444 | 0.157962135 | 0.000406488 | 0.000915058 | ns | YPD | KS.3 |
| FIM1_596 | 3191.988147 | 2672.844908 | 3810.382358 | -0.240718045 | 0.160718282 | 0.122066919 | 0.161339778 | ns | YPD | KS.3 |
| FIM1_597 | 1931.036213 | 1798.903003 | 1847.660826 | -0.098743383 | 0.11386159 | 0.378059612 | 0.441099102 | ns | YPD | KS.3 |
| FIM1_598 | 1121.664706 | 659.6650494 | 942.4573859 | -0.731318381 | 0.145341103 | 2.3019914524 | 1.0298927586 | ns | YPD | KS.3 |
| FIM1_599 | 802.391713 | 604.9789492 | 612.5166413 | -0.396217247 | 0.10552003 | 0.000139233 | 0.000347638 | ns | YPD | KS.3 |
| FIM1_600 | 2138.43192 | 1711.330427 | 1731.684023 | -0.297638414 | 0.179590859 | 0.084684805 | 0.115606417 | ns | YPD | KS.3 |
| FIM1_601 | 2186.103156 | 1386.563866 | 2786.680377 | -0.537188102 | 0.301967485 | 0.04712799 | 0.06837613 | ns | YPD | KS.3 |
| FIM1_602 | 2436.660351 | 2210.140202 | 2163.783996 | -0.138371195 | 0.081358383 | 0.086263257 | 0.117449289 | ns | YPD | KS.3 |
| FIM1_603 | 6469.579669 | 8640.828718 | 6877.477411 | 0.401165711 | 0.128801634 | 0.001467169 | 0.002939655 | ns | YPD | KS.3 |
| FIM1_604 | 2800.039926 | 1624.668989 | 2260.077787 | -0.758600751 | 0.125645023 | 6.9797283224 | 5.1519508018 | ns | YPD | KS.3 |
| FIM1_605 | 672.8785697 | 387.7841655 | 511.9090931 | -0.744899981 | 0.173470447 | 8.0452745577 | 2.5839406665 | ns | YPD | KS.3 |
| FIM1_606 | 1531.773759 | 1142.201429 | 1345.168045 | -0.40051667 | 0.152259459 | 0.00676742 | 0.011875808 | ns | YPD | KS.3 |
| FIM1_607 | 640.0192245 | 321.3565338 | 715.6332677 | -0.858877586 | 0.269995149 | 0.000496108 | 0.001095489 | ns | YPD | KS.3 |
| FIM1_608 | 602.474723 | 559.0351761 | 591.9919784 | -0.103559752 | 0.132405752 | 0.424087094 | 0.488161335 | ns | YPD | KS.3 |
| FIM1_609 | 5123.464479 | 11358.21959 | 7419.754934 | 1.046826964 | 0.221736859 | 4.7993006329 | 2.0064303066 | Km>KS.3 | YPD | KS.3 |
| FIM1_610 | 760.6062759 | 485.1908743 | 549.4681379 | -0.629776314 | 0.112946462 | 1.4179791074 | 8.0622240678 | ns | YPD | KS.3 |
| FIM1_611 | 180.7284987 | 372.90861 | 193.9310292 | 0.979625437 | 0.180480568 | 1.4477211148 | 8.2219321303 | ns | YPD | KS.3 |
| FIM1_612 | 2056.861793 | 2326.048614 | 1807.422097 | 0.175008998 | 0.077665775 | 0.023134691 | 0.0361026 | ns | YPD | KS.3 |
| FIM1_613 | 1412.101859 | 1553.443816 | 1297.621499 | 0.134755558 | 0.091794218 | 0.137811901 | 0.1794012 | ns | YPD | KS.3 |
| FIM1_615 | 1167.920961 | 1373.319724 | 988.5036954 | 0.225798454 | 0.120876788 | 0.057078902 | 0.081040964 | ns | YPD | KS.3 |
| FIM1_616 | 2968.345979 | 1819.454059 | 2442.610828 | -0.685461354 | 0.113607054 | 8.3170322928 | 6.0228872863 | ns | YPD | KS.3 |
| FIM1_617 | 487.9781065 | 213.2042887 | 457.5838486 | -0.976594084 | 0.329522076 | 0.000711837 | 0.001519258 | ns | YPD | KS.3 |
| FIM1_618 | 339.7889991 | 3105.056446 | 1419.095574 | 3.100559355 | 0.235419214 | 3.5601865923 | 1.6101752997 | Km>KS.3 | YPD | KS.3 |
| FIM1_619 | 7752.273731 | 12605.87122 | 13755.79097 | 0.661251629 | 0.162764849 | 2.6313591496 | 7.5758169962 | ns | YPD | KS.3 |
| FIM1_620 | 917.8067127 | 1003.379712 | 905.0081186 | 0.107682314 | 0.273495689 | 0.666964978 | 0.718681128 | ns | YPD | KS.3 |
| FIM1_621 | 234.4662377 | 659.1934452 | 628.3279031 | 1.311662512 | 0.28541543 | 3.1952395773 | 1.3834914619 | Km>KS.3 | YPD | KS.3 |
| FIM1_622 | 262.1520361 | 174.3306503 | 308.1685685 | -0.567383945 | 0.122261566 | 2.2202484841 | 8.0625811741 | ns | YPD | KS.3 |
| FIM1_623 | 3828.251033 | 2240.739487 | 3212.491914 | -0.750160384 | 0.114961447 | 3.0789581361 | 2.9917610795 | ns | YPD | KS.3 |
| FIM1_625 | 439.0672996 | 1817.134974 | 1665.134305 | 1.851795959 | 0.320007329 | 4.4237245016 | 4.1524583765 | Km>KS.3 | YPD | KS.3 |

| | | | | | | | | | | |
|---|---|---|---|---|---|---|---|---|---|---|
| FIM1_626 | 2307.58075 | 2240.286552 | 3460.275943 | -0.037847148 | 0.223206546 | 0.85717565 | 0.881084475 | ns | YPD | KS.3 |
| FIM1_627 | 975.6080192 | 1372.809286 | 1097.322672 | 0.472912659 | 0.13176602 | 0.000242765 | 0.000571042 | ns | YPD | KS.3 |
| FIM1_628 | 1712.398323 | 2888.22884 | 2869.052745 | 0.701712788 | 0.181712202 | 5.62899857956 | 0.000150642 | ns | YPD | KS.3 |
| FIM1_629 | 1291.280165 | 2755.323417 | 2629.388905 | 1.017463954 | 0.19252756 | 2.8452106384 | 1.5203998846 | Km>KS.3 | YPD | KS.3 |
| FIM1_630 | 707.1080134 | 889.7205497 | 786.9538102 | 0.31638355 | 0.136841929 | 0.017931221 | 0.028817994 | ns | YPD | KS.3 |
| FIM1_631 | 1256.300618 | 3436.424459 | 2014.126863 | 1.279074684 | 0.280432777 | 4.3286527200 | 1.8219160137 | Km>KS.3 | YPD | KS.3 |
| FIM1_632 | 314.9891753 | 430.5800197 | 338.091691 | 0.415045183 | 0.189638046 | 0.022150235 | 0.034740674 | ns | YPD | KS.3 |
| FIM1_633 | 5190.708071 | 4263.56279 | 5665.138552 | -0.276285267 | 0.104968155 | 0.007596172 | 0.013213621 | ns | YPD | KS.3 |
| FIM1_634 | 557.5440151 | 472.3863444 | 504.229274 | -0.232293443 | 0.106883085 | 0.027439046 | 0.042197605 | ns | YPD | KS.3 |
| FIM1_635 | 3310.581694 | 2752.416668 | 3749.836985 | -0.25882402 | 0.108442491 | 0.015400955 | 0.025088326 | ns | YPD | KS.3 |
| FIM1_636 | 490.5380984 | 682.6030582 | 505.1129773 | 0.467529702 | 0.089957375 | 1.4926409518 | 6.9726654794 | ns | YPD | KS.3 |
| FIM1_637 | 6597.978594 | 5518.0346 | 8546.879484 | -0.248498614 | 0.123173833 | 0.039744479 | 0.058812844 | ns | YPD | KS.3 |
| FIM1_638 | 1909.981343 | 1448.305831 | 1509.720033 | -0.388213664 | 0.106020462 | 0.00020289 | 0.000485278 | ns | YPD | KS.3 |
| FIM1_639 | 103.1836989 | 67.60167514 | 79.66084673 | -0.53617618 | 0.243490068 | 0.017918656 | 0.028812318 | ns | YPD | KS.3 |
| FIM1_640 | 1555.288798 | 1269.330705 | 1173.532448 | -0.288425889 | 0.078021762 | 0.000194116 | 0.000466986 | ns | YPD | KS.3 |
| FIM1_641 | 689.8673859 | 569.3843878 | 514.1290684 | -0.262528393 | 0.147298179 | 0.067105582 | 0.093857259 | ns | YPD | KS.3 |
| FIM1_642 | 3126.922734 | 3328.234952 | 3393.308776 | 0.080092854 | 0.221317053 | 0.701167336 | 0.74959406 | ns | YPD | KS.3 |
| FIM1_644 | 1867.458764 | 914.3304583 | 1767.328646 | -0.938881675 | 0.216764637 | 4.1882739351 | 1.4419835866 | ns | YPD | KS.3 |
| FIM1_645 | 258.3142357 | 2344.17339 | 1058.851261 | 3.107773501 | 0.211261964 | 1.6606936105 | 1.3769917853 | Km>KS.3 | YPD | KS.3 |
| FIM1_646 | 1918.456102 | 3736.928237 | 2889.920807 | 0.924493022 | 0.138241503 | 6.7270722754 | 7.5207156337 | ns | YPD | KS.3 |
| FIM1_647 | 1864.620528 | 1415.447047 | 1610.557098 | -0.364046486 | 0.194431747 | 0.049872043 | 0.071979522 | ns | YPD | KS.3 |
| FIM1_648 | 225.0724853 | 173.7376759 | 180.6262444 | -0.358366734 | 0.1230435 | 0.002985814 | 0.005641634 | ns | YPD | KS.3 |
| FIM1_650 | 566.7599569 | 787.0994267 | 655.9854887 | 0.455139995 | 0.129914008 | 0.000343693 | 0.000785066 | ns | YPD | KS.3 |
| FIM1_651 | 790.8865041 | 1671.575348 | 1317.559511 | 0.957367659 | 0.24896185 | 3.1782417381 | 8.9788487490 | ns | YPD | KS.3 |
| FIM1_652 | 2917.692267 | 2237.551839 | 2526.977574 | -0.373894203 | 0.09722757 | 9.8907451362 | 0.000256017 | ns | YPD | KS.3 |
| FIM1_653 | 3245.65248 | 1623.959655 | 3738.84694 | -0.829764349 | 0.304518996 | 0.002272444 | 0.004380688 | ns | YPD | KS.3 |
| FIM1_654 | 974.6648554 | 556.691956 | 618.8363942 | -0.783656308 | 0.117078309 | 9.1815851829 | 9.8657421781 | ns | YPD | KS.3 |
| FIM1_655 | 538.1057313 | 206.6802834 | 491.6011962 | -1.316572518 | 0.165003094 | 1.0454664767 | 2.8267368054 | Km<KS.3 | YPD | KS.3 |
| FIM1_656 | 2207.044014 | 1439.505998 | 1796.04403 | -0.594470404 | 0.125511964 | 1.3317242084 | 5.0964061055 | ns | YPD | KS.3 |
| FIM1_657 | 828.4474759 | 1001.033068 | 840.9049317 | 0.256173152 | 0.162189305 | 0.102790303 | 0.137875912 | ns | YPD | KS.3 |
| FIM1_658 | 596.6739446 | 879.6906496 | 678.8625973 | 0.522403215 | 0.17400924 | 0.001818097 | 0.003576526 | ns | YPD | KS.3 |
| FIM1_659 | 1792.601762 | 2054.350476 | 2457.652158 | 0.189983918 | 0.115643783 | 0.094763342 | 0.127971668 | ns | YPD | KS.3 |
| FIM1_660 | 759.6050402 | 784.3999922 | 935.7586285 | 0.045334358 | 0.093754598 | 0.624838651 | 0.680113584 | ns | YPD | KS.3 |
| FIM1_661 | 701.7699645 | 1152.322672 | 869.1390861 | 0.659729482 | 0.191808589 | 0.000314493 | 0.000724019 | ns | YPD | KS.3 |
| FIM1_662 | 9789.895332 | 8539.970432 | 13329.56743 | -0.183963667 | 0.169017328 | 0.25969963 | 0.316435381 | ns | YPD | KS.3 |
| FIM1_663 | 548.0726181 | 1411.288538 | 728.0410642 | 1.113240715 | 0.344591227 | 0.000185804 | 0.000450256 | Km>KS.3 | YPD | KS.3 |
| FIM1_664 | 2959.501921 | 2128.933683 | 3029.072366 | -0.453866162 | 0.139759268 | 0.000871163 | 0.001829479 | ns | YPD | KS.3 |
| FIM1_665 | 791.0351734 | 547.8535871 | 624.5284246 | -0.516230548 | 0.100379281 | 1.8695615894 | 8.5409264529 | ns | YPD | KS.3 |
| FIM1_666 | 285.9596945 | 306.0593382 | 221.6060011 | 0.085033644 | 0.245134872 | 0.709419141 | 0.757328314 | ns | YPD | KS.3 |
| FIM1_667 | 267.8983794 | 513.9088629 | 255.920656 | 0.871551066 | 0.192462021 | 2.0233432297 | 7.4234015989 | ns | YPD | KS.3 |
| FIM1_668 | 3648.431051 | 4330.984739 | 3917.240784 | 0.237156869 | 0.131001663 | 0.064364 | 0.090480616 | ns | YPD | KS.3 |
| FIM1_669 | 495.44601 | 2036.874055 | 1242.978932 | 1.897445286 | 0.270739861 | 1.0839246424 | 2.0426231424 | Km>KS.3 | YPD | KS.3 |
| FIM1_670 | 1903.347285 | 3519.788689 | 2971.716088 | 0.664482123 | 0.370429524 | 0.034597794 | 0.051860207 | ns | YPD | KS.3 |
| FIM1_671 | 675.070515 | 630.2024262 | 622.0532644 | -0.095318198 | 0.125059735 | 0.437043808 | 0.500989158 | ns | YPD | KS.3 |
| FIM1_672 | 837.6812035 | 978.3549659 | 662.1761727 | 0.21202788 | 0.150717941 | 0.147949064 | 0.191678801 | ns | YPD | KS.3 |

| | | | | | | | | | | |
|---|---|---|---|---|---|---|---|---|---|---|
| FIM1_673 | 174.7417262 | 433.6978126 | 349.8445058 | 1.118709482 | 0.301772734 | 3.2195070653 | 9.0851092741 | Km>KS.3 | YPD | KS.3 |
| FIM1_674 | 4699.836276 | 6986.790361 | 5215.570752 | 0.546998437 | 0.138973892 | 5.4603016341 | 0.000146679 | ns | YPD | KS.3 |
| FIM1_675 | 1029.436548 | 1685.956118 | 1473.001452 | 0.688809876 | 0.119488406 | 4.2199336030 | 2.6608580070 | ns | YPD | KS.3 |
| FIM1_676 | 1806.542866 | 2984.742775 | 3264.916752 | 0.640808956 | 0.237349383 | 0.003800547 | 0.007000268 | ns | YPD | KS.3 |
| FIM1_677 | 1434.675716 | 2208.55517 | 1919.82398 | 0.588147332 | 0.158112191 | 0.000122819 | 0.000310638 | ns | YPD | KS.3 |
| FIM1_678 | 1965.76185 | 4387.501771 | 2766.754028 | 1.104880867 | 0.157332766 | 3.6751710061 | 5.1504157058 | Km>KS.3 | YPD | KS.3 |
| FIM1_679 | 2760.334928 | 3525.929665 | 2876.776366 | 0.306301793 | 0.247205381 | 0.182405503 | 0.231910906 | ns | YPD | KS.3 |
| FIM1_680 | 3658.708392 | 4698.550563 | 5497.70073 | 0.307118733 | 0.263620224 | 0.205491569 | 0.257252278 | ns | YPD | KS.3 |
| FIM1_681 | 814.1481283 | 2765.355065 | 2011.579786 | 1.498251654 | 0.353866044 | 7.3575749666 | 2.9662832624 | Km>KS.3 | YPD | KS.3 |
| FIM1_682 | 656.4668448 | 2320.311088 | 1325.94449 | 1.610912706 | 0.316689204 | 6.7382002198 | 4.0486166780 | Km>KS.3 | YPD | KS.3 |
| FIM1_683 | 2254.090898 | 4414.514702 | 3487.012945 | 0.937652071 | 0.126800123 | 4.0310803012 | 6.8916235391 | ns | YPD | KS.3 |
| FIM1_684 | 1567.314941 | 2550.615572 | 1704.721099 | 0.686797628 | 0.099113309 | 2.1791923959 | 2.6901940867 | ns | YPD | KS.3 |
| FIM1_685 | 678.2756276 | 731.8623151 | 571.6491387 | 0.104470742 | 0.144115539 | 0.457049351 | 0.520801768 | ns | YPD | KS.3 |
| FIM1_686 | 3478.756245 | 3376.663178 | 3809.941275 | -0.042363383 | 0.071619287 | 0.551511634 | 0.612722282 | ns | YPD | KS.3 |
| FIM1_687 | 1351.989445 | 973.8592355 | 1055.214549 | -0.456955928 | 0.119735963 | 0.000101221 | 0.000261326 | ns | YPD | KS.3 |
| FIM1_688 | 1270.288031 | 859.6056367 | 1036.037415 | -0.548863805 | 0.102951227 | 6.4168173937 | 3.2148707486 | ns | YPD | KS.3 |
| FIM1_689 | 144.7592893 | 80.33626059 | 143.7030848 | -0.73821307 | 0.260384809 | 0.002052572 | 0.003990443 | ns | YPD | KS.3 |
| FIM1_690 | 2135.238196 | 1429.095224 | 1524.061999 | -0.562155307 | 0.11249684 | 3.7535036503 | 1.5960410820 | ns | YPD | KS.3 |
| FIM1_691 | 1875.766008 | 1249.614259 | 1611.399486 | -0.563230009 | 0.131074094 | 1.1136663189 | 3.4628062103 | ns | YPD | KS.3 |
| FIM1_692 | 1615.486379 | 1104.616955 | 1227.666889 | -0.524353362 | 0.138345292 | 0.000102571 | 0.000264399 | ns | YPD | KS.3 |
| FIM1_693 | 1977.694308 | 1374.417494 | 1255.218501 | -0.511979177 | 0.100143951 | 2.2100983245 | 9.9414458994 | ns | YPD | KS.3 |
| FIM1_694 | 936.4785955 | 590.0253628 | 543.4684132 | -0.642253762 | 0.125877626 | 1.8912879184 | 8.6164445004 | ns | YPD | KS.3 |
| FIM1_695 | 1157.247745 | 1170.515207 | 1000.654697 | 0.016069522 | 0.082554539 | 0.844316284 | 0.870564459 | ns | YPD | KS.3 |
| FIM1_696 | 2280.711242 | 1468.974212 | 1557.699312 | -0.615227818 | 0.115701609 | 6.2200371829 | 3.1225716432 | ns | YPD | KS.3 |
| FIM1_697 | 1113.023638 | 999.5629268 | 898.8205216 | -0.149726174 | 0.114344142 | 0.183129101 | 0.232711948 | ns | YPD | KS.3 |
| FIM1_698 | 811.1033101 | 797.2446253 | 812.8636411 | -0.024517668 | 0.138476742 | 0.856135629 | 0.880197304 | ns | YPD | KS.3 |
| FIM1_699 | 2362.37369 | 3816.865538 | 2938.740619 | 0.616629209 | 0.229399937 | 0.004123473 | 0.007542014 | ns | YPD | KS.3 |
| FIM1_700 | 7007.580684 | 3676.266259 | 6849.827476 | -0.91662813 | 0.083673943 | 1.8450496362 | 2.2947804850 | ns | YPD | KS.3 |
| FIM1_701 | 1757.069043 | 921.9876778 | 1751.349444 | -0.8440538 | 0.218080444 | 3.9253610266 | 0.000108432 | ns | YPD | KS.3 |
| FIM1_702 | 1538.755281 | 917.0463693 | 1186.573425 | -0.722636469 | 0.12080294 | 1.0620610070 | 7.4946858296 | ns | YPD | KS.3 |
| FIM1_703 | 3100.93563 | 2786.995868 | 2874.929683 | -0.145928458 | 0.149094307 | 0.314446674 | 0.375149209 | ns | YPD | KS.3 |
| FIM1_704 | 3834.866613 | 10192.44576 | 6127.943018 | 1.266977027 | 0.254994994 | 6.0815010496 | 3.0561078506 | Km>KS.3 | YPD | KS.3 |
| FIM1_705 | 1063.701388 | 1575.249306 | 1247.382072 | 0.482568876 | 0.266906095 | 0.048783487 | 0.070592743 | ns | YPD | KS.3 |
| FIM1_706 | 3171.836281 | 1802.744053 | 2196.0927 | -0.790547755 | 0.117812188 | 8.0566712448 | 8.7706650860 | ns | YPD | KS.3 |
| FIM1_707 | 1319.724125 | 1067.255055 | 994.1467093 | -0.300184248 | 0.089465472 | 0.000697627 | 0.00149213 | ns | YPD | KS.3 |
| FIM1_708 | 1099.055326 | 587.6301092 | 661.6962084 | -0.872448402 | 0.126643417 | 1.9093306257 | 2.3806816699 | ns | YPD | KS.3 |
| FIM1_709 | 1982.607049 | 1215.087688 | 1294.26645 | -0.687165327 | 0.1091594 | 1.5873735349 | 1.3476422075 | ns | YPD | KS.3 |
| FIM1_710 | 648.6916046 | 629.9078864 | 649.6720697 | -0.039811001 | 0.133093532 | 0.759788202 | 0.79965016 | ns | YPD | KS.3 |
| FIM1_711 | 497.0916879 | 387.8043734 | 444.1594199 | -0.333068389 | 0.17673615 | 0.050184487 | 0.072351665 | ns | YPD | KS.3 |
| FIM1_712 | 30.70321908 | 31.09884007 | 21.06410889 | 0.014414373 | 0.367723405 | 0.963269449 | 0.971273917 | ns | YPD | KS.3 |
| FIM1_713 | 1775.18098 | 865.7091334 | 1449.725859 | -0.870118908 | 0.297363727 | 0.001104645 | 0.002270912 | ns | YPD | KS.3 |
| FIM1_714 | 847.3347818 | 814.131209 | 955.3241447 | -0.054129465 | 0.155422997 | 0.719800428 | 0.765009 | ns | YPD | KS.3 |
| FIM1_715 | 1722.332119 | 1008.68623 | 1366.728225 | -0.688735283 | 0.230158393 | 0.00139246 | 0.002806925 | ns | YPD | KS.3 |
| FIM1_718 | 3594.346305 | 3526.389915 | 4056.178042 | -0.025618652 | 0.16914626 | 0.87538536 | 0.897206874 | ns | YPD | KS.3 |
| FIM1_719 | 7051.69541 | 4247.496299 | 7126.365091 | -0.699043252 | 0.142589644 | 4.7860631475 | 2.0025789873 | ns | YPD | KS.3 |

| | | | | | | | | | | |
|---|---|---|---|---|---|---|---|---|---|---|
| FIM1_720 | 693.0481928 | 650.7440082 | 588.1164666 | -0.086019757 | 0.138884135 | 0.525880023 | 0.588186401 | ns | YPD | KS.3 |
| FIM1_721 | 2444.579641 | 1787.103362 | 2180.455325 | -0.432953307 | 0.134576495 | 0.000994164 | 0.002059961 | ns | YPD | KS.3 |
| FIM1_722 | 5494.397352 | 4227.877245 | 4671.506458 | -0.370761502 | 0.088220002 | 2.1766208585 | 6.3810776494 | ns | YPD | KS.3 |
| FIM1_723 | 1158.468304 | 1136.311525 | 877.597854 | -0.026884888 | 0.122324673 | 0.82281557 | 0.852103968 | ns | YPD | KS.3 |
| FIM1_724 | 579.9992083 | 254.569499 | 388.5643602 | -1.099814931 | 0.204045439 | 1.2140469265 | 6.9825242306 | Km<KS.3 | YPD | KS.3 |
| FIM1_725 | 5658.431952 | 3201.214316 | 5829.750812 | -0.747240885 | 0.211677349 | 0.000187344 | 0.000453325 | ns | YPD | KS.3 |
| FIM1_726 | 1247.64598 | 817.3818561 | 898.0738828 | -0.588714973 | 0.12429793 | 1.3447675923 | 5.1384168753 | ns | YPD | KS.3 |
| FIM1_727 | 2227.382811 | 1155.955931 | 1229.378744 | -0.908466238 | 0.139331023 | 2.1753443540 | 2.1952004383 | ns | YPD | KS.3 |
| FIM1_728 | 2193.474553 | 2019.86649 | 2356.280893 | -0.110732458 | 0.172955857 | 0.506716069 | 0.568926302 | ns | YPD | KS.3 |
| FIM1_729 | 3607.215581 | 1974.080618 | 2429.537675 | -0.841751342 | 0.122943502 | 2.7752724581 | 3.3512088542 | ns | YPD | KS.3 |
| FIM1_730 | 2541.61214 | 1688.623192 | 1867.201485 | -0.576439689 | 0.098533193 | 3.0912575269 | 1.9946830346 | ns | YPD | KS.3 |
| FIM1_731 | 561.2839638 | 677.2480734 | 460.353051 | 0.264697208 | 0.099975628 | 0.007321304 | 0.012789146 | ns | YPD | KS.3 |
| FIM1_732 | 7737.190519 | 5536.505528 | 8995.403339 | -0.461315849 | 0.139154834 | 0.000679164 | 0.001457654 | ns | YPD | KS.3 |
| FIM1_733 | 2753.902048 | 1564.576957 | 2077.4532 | -0.723289267 | 0.238130361 | 0.001116897 | 0.002289478 | ns | YPD | KS.3 |
| FIM1_734 | 2066.664679 | 1856.048539 | 2175.091979 | -0.153609736 | 0.05945058 | 0.009429247 | 0.016081764 | ns | YPD | KS.3 |
| FIM1_735 | 697.6244138 | 871.8382932 | 672.6776497 | 0.31441224 | 0.094378318 | 0.000751195 | 0.001595729 | ns | YPD | KS.3 |
| FIM1_736 | 3823.19204 | 3027.483096 | 2714.657046 | -0.324852075 | 0.120300685 | 0.00595093 | 0.0105359 | ns | YPD | KS.3 |
| FIM1_737 | 2837.820458 | 2340.229432 | 2322.058371 | -0.272868255 | 0.085128487 | 0.00121275 | 0.002473887 | ns | YPD | KS.3 |
| FIM1_738 | 1750.458018 | 924.1816326 | 1834.221427 | -0.868183752 | 0.169494528 | 1.0415510813 | 5.0239453483 | ns | YPD | KS.3 |
| FIM1_739 | 604.3988062 | 312.781361 | 482.0774618 | -0.871523749 | 0.206059147 | 7.9423896820 | 2.5558466150 | ns | YPD | KS.3 |
| FIM1_740 | 1140.977711 | 676.7944626 | 827.4917073 | -0.723906975 | 0.133777515 | 3.0097516322 | 1.5946234686 | ns | YPD | KS.3 |
| FIM1_741 | 2242.216794 | 1347.900354 | 1896.080384 | -0.716591893 | 0.102865134 | 1.5806031797 | 1.9857325300 | ns | YPD | KS.3 |
| FIM1_742 | 2357.364035 | 2397.294305 | 1995.452123 | 0.023091403 | 0.125462865 | 0.851117817 | 0.875943554 | ns | YPD | KS.3 |
| FIM1_743 | 2481.203997 | 1601.849739 | 2245.619871 | -0.618057602 | 0.094093281 | 2.9776936415 | 2.8990265884 | ns | YPD | KS.3 |
| FIM1_744 | 640.5750091 | 383.3994683 | 472.0112613 | -0.69816653 | 0.164295895 | 1.0829371902 | 3.3756970686 | ns | YPD | KS.3 |
| FIM1_745 | 588.2921144 | 735.5158522 | 570.5548511 | 0.31461378 | 0.101625814 | 0.001704945 | 0.003371265 | ns | YPD | KS.3 |
| FIM1_746 | 1187.519497 | 1724.322989 | 1985.127197 | 0.50978909 | 0.152698317 | 0.000584328 | 0.001270555 | ns | YPD | KS.3 |
| FIM1_747 | 1424.853007 | 967.0979763 | 1178.845108 | -0.528776102 | 0.155055044 | 0.000438157 | 0.00097882 | ns | YPD | KS.3 |
| FIM1_748 | 1027.524497 | 990.1426358 | 794.7275218 | -0.046550899 | 0.240930956 | 0.83596789 | 0.862850674 | ns | YPD | KS.3 |
| FIM1_749 | 1354.093537 | 1056.660035 | 1109.475622 | -0.341270205 | 0.138676784 | 0.011698367 | 0.019529993 | ns | YPD | KS.3 |
| FIM1_750 | 1568.201093 | 1246.905772 | 1422.150315 | -0.321449404 | 0.106425302 | 0.002178758 | 0.004214356 | ns | YPD | KS.3 |
| FIM1_751 | 3857.106954 | 4049.084217 | 3565.154167 | 0.06904521 | 0.077350628 | 0.368330944 | 0.43116387 | ns | YPD | KS.3 |
| FIM1_752 | 4458.640618 | 5034.62963 | 5517.589437 | 0.171297581 | 0.095470811 | 0.069473002 | 0.096896043 | ns | YPD | KS.3 |
| FIM1_753 | 2132.005109 | 3599.144296 | 2091.90095 | 0.718637916 | 0.150600876 | 8.8731213405 | 3.5202375334 | ns | YPD | KS.3 |
| FIM1_754 | 1298.655553 | 3659.614983 | 2116.637094 | 1.41123107 | 0.191263988 | 7.3030999241 | 1.4137323783 | Km>KS.3 | YPD | KS.3 |
| FIM1_755 | 3053.333961 | 2338.18832 | 2519.886863 | -0.38074099 | 0.064762255 | 3.3663706627 | 2.1554303793 | ns | YPD | KS.3 |
| FIM1_756 | 985.0125737 | 496.7091204 | 822.4926349 | -0.959495913 | 0.11640738 | 4.4752528476 | 1.3252608879 | ns | YPD | KS.3 |
| FIM1_757 | 2560.144327 | 1699.83486 | 2005.582828 | -0.570720238 | 0.122286617 | 1.9436456309 | 7.1839799508 | ns | YPD | KS.3 |
| FIM1_758 | 510.2380208 | 448.8682845 | 378.9891539 | -0.179598841 | 0.106896128 | 0.088254612 | 0.11993081 | ns | YPD | KS.3 |
| FIM1_759 | 1082.959993 | 544.5051435 | 738.316749 | -0.967039157 | 0.110995378 | 7.6930794922 | 3.1371369241 | ns | YPD | KS.3 |
| FIM1_760 | 2538.745627 | 2987.418108 | 2381.124377 | 0.221359415 | 0.15545109 | 0.1423655 | 0.184974762 | ns | YPD | KS.3 |
| FIM1_761 | 2488.815926 | 2792.724321 | 2800.818831 | 0.154698844 | 0.171852298 | 0.350726672 | 0.412985844 | ns | YPD | KS.3 |
| FIM1_762 | 1116.133468 | 464.2605561 | 1122.594086 | -1.210662331 | 0.157478003 | 1.6842935702 | 3.5060085825 | Km<KS.3 | YPD | KS.3 |
| FIM1_763 | 757.5709905 | 763.4667976 | 775.0109219 | 0.010780419 | 0.122861423 | 0.928754432 | 0.942200917 | ns | YPD | KS.3 |
| FIM1_764 | 2293.746687 | 2205.101439 | 2454.664853 | -0.055677496 | 0.09937347 | 0.570467213 | 0.630263021 | ns | YPD | KS.3 |

| | | | | | | | | | | |
|---|---|---|---|---|---|---|---|---|---|---|
| FIM1_765 | 247.9801401 | 343.8878013 | 354.7709332 | 0.409815995 | 0.246614099 | 0.073699284 | 0.102217435 | ns | YPD | KS.3 |
| FIM1_767 | 534.2954681 | 1682.343081 | 1201.766306 | 1.435683632 | 0.317793793 | 2.65747811723 | 1.16896141761 | Km>KS.3 | YPD | KS.3 |
| FIM1_768 | 310.1522006 | 376.0937411 | 370.6882992 | 0.264556878 | 0.146803857 | 0.064167526 | 0.090257671 | ns | YPD | KS.3 |
| FIM1_769 | 3330.376385 | 2760.663782 | 2602.665713 | -0.267221841 | 0.069888618 | 0.000118854 | 0.000302299 | ns | YPD | KS.3 |
| FIM1_770 | 615.897999 | 2713.405737 | 2444.516323 | 2.0313905 | 0.241833697 | 1.21224233063 | 6.03090559492 | Km>KS.3 | YPD | KS.3 |
| FIM1_772 | 10.49512541 | 44.88320393 | 53.38291118 | 1.782059304 | 0.402775839 | 9.30755592192 | 4.53971477564 | Km>KS.3 | YPD | KS.3 |
| FIM1_773 | 2240.384582 | 1432.021372 | 2254.322242 | -0.589440148 | 0.202528126 | 0.002190851 | 0.00423281 | ns | YPD | KS.3 |
| FIM1_774 | 761.9163781 | 928.8811059 | 784.9324631 | 0.274793492 | 0.127223994 | 0.027497418 | 0.042274307 | ns | YPD | KS.3 |
| FIM1_776 | 1796.225155 | 2616.725958 | 2078.601108 | 0.519675434 | 0.136058619 | 9.17538947051 | 0.000238618 | ns | YPD | KS.3 |
| FIM1_777 | 2535.060133 | 2804.409265 | 2381.458233 | 0.141500792 | 0.107376463 | 0.181162885 | 0.230802908 | ns | YPD | KS.3 |
| FIM1_778 | 637.671842 | 899.3434543 | 665.8322517 | 0.470027998 | 0.151964913 | 0.001449584 | 0.002909108 | ns | YPD | KS.3 |
| FIM1_779 | 7685.341048 | 11760.82789 | 14674.89046 | 0.600122364 | 0.097351189 | 4.27824688547 | 3.28968752013 | ns | YPD | KS.3 |
| FIM1_780 | 119.9172476 | 559.8136579 | 453.4928227 | 2.097553718 | 0.264474649 | 5.56591454314 | 1.89632973250 | Km>KS.3 | YPD | KS.3 |
| FIM1_781 | 360.7266306 | 438.0751596 | 472.8245672 | 0.246360297 | 0.235393922 | 0.26333448 | 0.32070723 | ns | YPD | KS.3 |
| FIM1_782 | 1578.649776 | 3004.398038 | 1671.123472 | 0.82718053 | 0.237402768 | 0.00018386 | 0.000447068 | ns | YPD | KS.3 |
| FIM1_783 | 337.3352891 | 1129.824629 | 935.7181848 | 1.674232505 | 0.176895082 | 2.98525205246 | 2.21666103895 | Km>KS.3 | YPD | KS.3 |
| FIM1_784 | 271.6953212 | 702.7249425 | 488.4529459 | 1.258534452 | 0.224686084 | 1.99717011621 | 1.35182603104 | Km>KS.3 | YPD | KS.3 |
| FIM1_785 | 2887.279267 | 3874.266427 | 3305.208004 | 0.402648874 | 0.147719255 | 0.00508484 | 0.00911278 | ns | YPD | KS.3 |
| FIM1_786 | 2001.232429 | 2031.814571 | 2337.434031 | 0.02105432 | 0.114200103 | 0.851342292 | 0.875993362 | ns | YPD | KS.3 |
| FIM1_787 | 3567.07601 | 6118.064541 | 4819.528209 | 0.7265264 | 0.178290526 | 2.19067998813 | 6.41473392640 | ns | YPD | KS.3 |
| FIM1_788 | 198.7114772 | 220.7762667 | 200.3153645 | 0.129898069 | 0.256723471 | 0.584177972 | 0.642557023 | ns | YPD | KS.3 |
| FIM1_789 | 946.3349021 | 763.0486144 | 1008.009036 | -0.28289367 | 0.198804464 | 0.135443247 | 0.176811901 | ns | YPD | KS.3 |
| FIM1_790 | 250.1535956 | 602.8709685 | 368.2901065 | 1.164120074 | 0.220032278 | 1.66270989071 | 9.34687198454 | Km>KS.3 | YPD | KS.3 |
| FIM1_791 | 2217.616477 | 5856.602856 | 3356.780222 | 1.305892938 | 0.205859147 | 1.71194841643 | 1.74885900857 | Km>KS.3 | YPD | KS.3 |
| FIM1_792 | 1532.789213 | 1147.438558 | 1088.768169 | -0.4050344 | 0.11148899 | 0.000222716 | 0.000528884 | ns | YPD | KS.3 |
| FIM1_793 | 3866.696539 | 4373.230174 | 3888.038793 | 0.171321306 | 0.120385057 | 0.147269101 | 0.190897285 | ns | YPD | KS.3 |
| FIM1_794 | 785.0493658 | 1025.875553 | 1027.485245 | 0.376063076 | 0.102227278 | 0.000192288 | 0.000463261 | ns | YPD | KS.3 |
| FIM1_795 | 3414.652401 | 2025.766754 | 2824.487688 | -0.717941183 | 0.147757797 | 5.74571187036 | 2.35460597652 | ns | YPD | KS.3 |
| FIM1_796 | 1244.935543 | 1360.537492 | 1281.943094 | 0.125507014 | 0.093188781 | 0.173236009 | 0.221954454 | ns | YPD | KS.3 |
| FIM1_797 | 413.8526305 | 208.58357 | 336.0054251 | -0.907653273 | 0.206729513 | 3.49074324647 | 1.21613779070 | ns | YPD | KS.3 |
| FIM1_798 | 803.106942 | 543.8688433 | 648.7373743 | -0.523098796 | 0.178156923 | 0.002248833 | 0.004338094 | ns | YPD | KS.3 |
| FIM1_799 | 1622.613306 | 1842.537038 | 1734.643283 | 0.154174001 | 0.27143948 | 0.535253105 | 0.597059237 | ns | YPD | KS.3 |
| FIM1_800 | 1999.862377 | 1809.427258 | 2792.052274 | -0.136014139 | 0.156825515 | 0.371400578 | 0.434534851 | ns | YPD | KS.3 |
| FIM1_801 | 780.2382653 | 3832.363273 | 2056.706458 | 2.142752671 | 0.298259874 | 1.76840025707 | 3.66574636621 | Km>KS.3 | YPD | KS.3 |
| FIM1_802 | 663.7911683 | 1017.128044 | 852.4081707 | 0.567796849 | 0.189058871 | 0.001685573 | 0.003339596 | ns | YPD | KS.3 |
| FIM1_803 | 1395.800819 | 1326.590399 | 1145.42991 | -0.071670194 | 0.088026445 | 0.410881839 | 0.474938928 | ns | YPD | KS.3 |
| FIM1_804 | 3232.88113 | 5938.975093 | 5144.829841 | 0.803185066 | 0.205538361 | 3.72492686650 | 0.000103644 | ns | YPD | KS.3 |
| FIM1_805 | 1083.893924 | 1122.388797 | 1049.39095 | 0.048308193 | 0.133544321 | 0.711515854 | 0.758960414 | ns | YPD | KS.3 |
| FIM1_806 | 1410.527057 | 2436.050054 | 1397.046938 | 0.689004013 | 0.25126362 | 0.003035206 | 0.005721921 | ns | YPD | KS.3 |
| FIM1_807 | 236.5313079 | 203.0797604 | 604.1760257 | -0.204507997 | 0.170548637 | 0.214041021 | 0.26668021 | ns | YPD | KS.3 |
| FIM1_808 | 791.2577922 | 1302.748248 | 1277.977105 | 0.683261358 | 0.151709378 | 3.47943388193 | 1.21304720130 | ns | YPD | KS.3 |
| FIM1_809 | 11353.06118 | 9834.221527 | 20737.53121 | -0.199443027 | 0.124222703 | 0.101649551 | 0.136382556 | ns | YPD | KS.3 |
| FIM1_810 | 1982.351375 | 1493.235508 | 1437.83775 | -0.399097218 | 0.097305178 | 3.28709675813 | 9.25269574834 | ns | YPD | KS.3 |
| FIM1_811 | 2903.671183 | 2912.437654 | 2368.765038 | 0.004088001 | 0.171810775 | 0.980324305 | 0.983884087 | ns | YPD | KS.3 |
| FIM1_812 | 2266.344858 | 1014.968988 | 2025.255696 | -1.114973455 | 0.142106612 | 6.93742689812 | 1.59049303309 | Km<KS.3 | YPD | KS.3 |

| | | | | | | | | | | |
|---|---|---|---|---|---|---|---|---|---|---|
| FIM1_813 | 968.5493826 | 833.1840805 | 733.4134806 | -0.210545169 | 0.105615757 | 0.043173592 | 0.063247532 | ns | YPD | KS.3 |
| FIM1_814 | 56.56361444 | 60.70142868 | 59.6130047 | 0.086168148 | 0.276290032 | 0.732752405 | 0.775296303 | ns | YPD | KS.3 |
| FIM1_815 | 109.8010592 | 174.4707543 | 102.8411246 | 0.571832093 | 0.266066659 | 0.019164033 | 0.030577635 | ns | YPD | KS.3 |
| FIM1_816 | 365.3990369 | 311.8712775 | 391.056969 | -0.20345049 | 0.221317575 | 0.329560022 | 0.391303367 | ns | YPD | KS.3 |
| FIM1_817 | 1162.497311 | 1028.801533 | 1487.348653 | -0.163950478 | 0.172628635 | 0.324690289 | 0.386166433 | ns | YPD | KS.3 |
| FIM1_818 | 548.827047 | 82.87438035 | 327.2956797 | -2.667904556 | 0.197920686 | 2.2656698185 | 1.1271707347 | Km<KS.3 | YPD | KS.3 |
| FIM1_819 | 1212.32473 | 1336.877759 | 1264.18978 | 0.13517177 | 0.131119983 | 0.29225383 | 0.352390404 | ns | YPD | KS.3 |
| FIM1_820 | 4807.636157 | 6554.953797 | 6602.112733 | 0.424707872 | 0.148106106 | 0.003198652 | 0.006003313 | ns | YPD | KS.3 |
| FIM1_821 | 7397.30407 | 5768.29672 | 8074.419783 | -0.336357579 | 0.165120567 | 0.035122016 | 0.052582615 | ns | YPD | KS.3 |
| FIM1_822 | 501.4367032 | 785.3564833 | 963.4226905 | 0.603749212 | 0.175073818 | 0.000337098 | 0.000771063 | ns | YPD | KS.3 |
| FIM1_823 | 1180.145818 | 691.0244501 | 1053.613319 | -0.753317762 | 0.103218137 | 1.3084939172 | 2.0279617564 | ns | YPD | KS.3 |
| FIM1_824 | 4216.703694 | 2009.583519 | 4683.444448 | -0.859944458 | 0.333368683 | 0.003159199 | 0.00593767 | ns | YPD | KS.3 |
| FIM1_825 | 3595.937268 | 2111.729565 | 3341.939333 | -0.744678463 | 0.117303949 | 1.0003080460 | 8.8236392357 | ns | YPD | KS.3 |
| FIM1_826 | 116.0865363 | 77.76838125 | 94.20694577 | -0.470164013 | 0.300847326 | 0.081515182 | 0.111718466 | ns | YPD | KS.3 |
| FIM1_828 | 1322.317645 | 699.66782 | 1423.450385 | -0.878110729 | 0.145060545 | 4.7674624790 | 3.6266247451 | ns | YPD | KS.3 |
| FIM1_829 | 610.999597 | 223.382491 | 492.427216 | -1.397710062 | 0.153011747 | 3.0589634483 | 1.8788077969 | Km<KS.3 | YPD | KS.3 |
| FIM1_830 | 3177.105254 | 2515.999249 | 2874.1411 | -0.271349201 | 0.304235992 | 0.319359409 | 0.380554027 | ns | YPD | KS.3 |
| FIM1_831 | 3940.808316 | 2395.549874 | 4234.203468 | -0.68654778 | 0.142347023 | 7.3272634837 | 2.9577828360 | ns | YPD | KS.3 |
| FIM1_832 | 951.0927036 | 522.9122983 | 754.9707349 | -0.805929484 | 0.180028175 | 3.0346277889 | 1.0707285992 | ns | YPD | KS.3 |
| FIM1_833 | 3226.921431 | 2379.361761 | 3921.404092 | -0.390315572 | 0.227083485 | 0.067283731 | 0.094053544 | ns | YPD | KS.3 |
| FIM1_834 | 1393.545277 | 1545.711368 | 1221.750856 | 0.144777897 | 0.113294731 | 0.194036016 | 0.245007405 | ns | YPD | KS.3 |
| FIM1_835 | 1549.735192 | 1150.619179 | 953.2250199 | -0.41098323 | 0.135181799 | 0.001861039 | 0.003655219 | ns | YPD | KS.3 |
| FIM1_836 | 1798.943819 | 1339.215318 | 1089.792055 | -0.366899272 | 0.253727149 | 0.118175627 | 0.156362698 | ns | YPD | KS.3 |
| FIM1_840 | 403.8668129 | 450.7140428 | 485.8406089 | 0.149912926 | 0.15712552 | 0.325391719 | 0.386908174 | ns | YPD | KS.3 |
| FIM1_841 | 53.22113226 | 142.6329527 | 116.3551879 | 1.203681286 | 0.317893178 | 1.7089195553 | 5.1246984858 | Km>KS.3 | YPD | KS.3 |
| FIM1_842 | 161.1414275 | 264.4475286 | 255.2486176 | 0.633361642 | 0.23544717 | 0.003969877 | 0.007287873 | ns | YPD | KS.3 |
| FIM1_843 | 281.2631559 | 193.0691429 | 1017.35679 | -0.387760892 | 0.383711615 | 0.228385146 | 0.282359866 | ns | YPD | KS.3 |
| FIM1_844 | 62.89670755 | 126.3986806 | 112.3709184 | 0.919439255 | 0.211533481 | 4.1299218677 | 1.4238642614 | ns | YPD | KS.3 |
| FIM1_846 | 27.19047852 | 63.09457018 | 84.39783745 | 1.058769067 | 0.27212505 | 1.9150738702 | 5.7085035975 | Km>KS.3 | YPD | KS.3 |
| FIM1_847 | 9.187872849 | 20.40088299 | 15.97427327 | 0.986356559 | 0.281545318 | 0.000109168 | 0.000279954 | ns | YPD | KS.3 |
| FIM1_850 | 329.1378139 | 641.1723534 | 698.1875307 | 0.806783628 | 0.294487929 | 0.002312826 | 0.004451183 | ns | YPD | KS.3 |
| FIM1_851 | 297.6044434 | 358.178215 | 305.0748288 | 0.230543637 | 0.249679559 | 0.319823284 | 0.381015526 | ns | YPD | KS.3 |
| FIM1_852 | 55.10974449 | 37.96604856 | 63.46054071 | -0.487894865 | 0.201583873 | 0.010934154 | 0.018383715 | ns | YPD | KS.3 |
| FIM1_853 | 134.3760218 | 67.69094032 | 101.2122781 | -0.904480111 | 0.214487821 | 7.6879048964 | 2.4835926532 | ns | YPD | KS.3 |
| FIM1_854 | 2531.520655 | 943.6269962 | 1559.299267 | -1.264291833 | 0.269519529 | 2.4689287867 | 1.0996347998 | Km<KS.3 | YPD | KS.3 |
| FIM1_855 | 372.2967321 | 217.3061559 | 276.8693972 | -0.714470097 | 0.197004723 | 0.000139265 | 0.000347638 | ns | YPD | KS.3 |
| FIM1_856 | 345.154557 | 305.4317199 | 299.5086226 | -0.164850008 | 0.168794199 | 0.311916496 | 0.372742277 | ns | YPD | KS.3 |
| FIM1_857 | 198.0285661 | 202.2318897 | 194.7328761 | 0.030724072 | 0.134200407 | 0.814903947 | 0.845318419 | ns | YPD | KS.3 |
| FIM1_858 | 1496.852059 | 175.5648778 | 643.7882858 | -2.961013616 | 0.287668074 | 1.2650960553 | 1.3391176330 | Km<KS.3 | YPD | KS.3 |
| FIM1_859 | 353.2046027 | 338.7579533 | 460.6221006 | -0.059029401 | 0.097163623 | 0.53863613 | 0.600429027 | ns | YPD | KS.3 |
| FIM1_860 | 620.777353 | 458.0544038 | 530.8493055 | -0.4282686 | 0.096878011 | 7.6225069052 | 2.4661724048 | ns | YPD | KS.3 |
| FIM1_861 | 1148.838769 | 713.4690368 | 689.1671746 | -0.654186203 | 0.149158757 | 6.3607722026 | 2.1012511094 | ns | YPD | KS.3 |
| FIM1_862 | 197.5206309 | 166.2138263 | 158.4303416 | -0.230004893 | 0.183901517 | 0.192479786 | 0.243289363 | ns | YPD | KS.3 |
| FIM1_863 | 10.02408863 | 30.55986818 | 32.18866186 | 1.255057552 | 0.403509554 | 0.000159692 | 0.000394277 | Km>KS.3 | YPD | KS.3 |
| FIM1_864 | 284.9611597 | 703.0853926 | 574.4896196 | 1.143860421 | 0.272419261 | 3.8533599927 | 1.3368525776 | Km>KS.3 | YPD | KS.3 |

| | | | | | | | | | | |
|---|---|---|---|---|---|---|---|---|---|---|
| FIM1_865 | 916.1821843 | 708.9528812 | 947.753543 | -0.334403697 | 0.207642797 | 0.089670312 | 0.121622084 | ns | YPD | KS.3 |
| FIM1_866 | 613.0823119 | 594.4000097 | 514.4551748 | -0.041125022 | 0.174078818 | 0.806381858 | 0.837700928 | ns | YPD | KS.3 |
| FIM1_867 | 856.8851745 | 1701.292162 | 1299.862932 | 0.912107727 | 0.200721571 | 1.686021019[illegible] | 6.301994419[illegible] | ns | YPD | KS.3 |
| FIM1_868 | 1367.199395 | 998.2356928 | 901.8214115 | -0.427963531 | 0.157469418 | 0.005053973 | 0.009060726 | ns | YPD | KS.3 |
| FIM1_869 | 908.3274497 | 493.4099774 | 674.8613506 | -0.854796679 | 0.116181958 | 6.645844582[illegible] | 1.080492705[illegible] | ns | YPD | KS.3 |
| FIM1_870 | 1304.283239 | 906.0640356 | 1337.424702 | -0.47674342 | 0.20635323 | 0.014907581 | 0.024364395 | ns | YPD | KS.3 |
| FIM1_871 | 1263.293118 | 638.0534859 | 1085.450442 | -0.967851435 | 0.092585069 | 3.578047364[illegible] | 3.632813395[illegible] | ns | YPD | KS.3 |
| FIM1_872 | 3442.462623 | 2142.866545 | 2799.267011 | -0.667794047 | 0.102041545 | 3.201153977[illegible] | 3.104432950[illegible] | ns | YPD | KS.3 |
| FIM1_873 | 1086.116349 | 890.1929357 | 998.9041111 | -0.271936789 | 0.1470734 | 0.057515796 | 0.081568155 | ns | YPD | KS.3 |
| FIM1_874 | 2611.498335 | 8173.122793 | 4884.448618 | 1.53101711 | 0.227578146 | 4.361032462[illegible] | 6.043492061[illegible] | Km>KS.3 | YPD | KS.3 |
| FIM1_875 | 175.8052795 | 531.6407371 | 368.4667161 | 1.490221082 | 0.216259698 | 1.717676899[illegible] | 2.573928485[illegible] | Km>KS.3 | YPD | KS.3 |
| FIM1_876 | 1554.253333 | 2945.360384 | 2803.00508 | 0.798118673 | 0.265411191 | 0.001033597 | 0.002136329 | ns | YPD | KS.3 |
| FIM1_877 | 1493.876742 | 816.3062594 | 1254.645693 | -0.83459548 | 0.143268434 | 2.136897813[illegible] | 1.434691851[illegible] | ns | YPD | KS.3 |
| FIM1_878 | 3777.816171 | 3089.224025 | 3660.528863 | -0.281055638 | 0.114526359 | 0.012583473 | 0.020867593 | ns | YPD | KS.3 |
| FIM1_879 | 26.22476396 | 12.97120831 | 27.8276178 | -0.811242595 | 0.327567298 | 0.004794619 | 0.008645607 | ns | YPD | KS.3 |
| FIM1_880 | 1619.594215 | 1647.985045 | 1216.289069 | 0.024906291 | 0.101430205 | 0.803533873 | 0.835614761 | ns | YPD | KS.3 |
| FIM1_881 | 1861.759756 | 1503.877356 | 1551.362047 | -0.300981337 | 0.094648545 | 0.001294694 | 0.002621531 | ns | YPD | KS.3 |
| FIM1_882 | 715.9294071 | 482.6151194 | 510.0564056 | -0.545415241 | 0.132893601 | 2.682530451[illegible] | 7.696418106[illegible] | ns | YPD | KS.3 |
| FIM1_883 | 1810.335474 | 1879.324311 | 1562.433082 | 0.049355701 | 0.194231326 | 0.790293467 | 0.825228114 | ns | YPD | KS.3 |
| FIM1_884 | 607.6125774 | 360.6001647 | 330.5532116 | -0.660877323 | 0.245014506 | 0.003678376 | 0.006800416 | ns | YPD | KS.3 |
| FIM1_885 | 2168.471771 | 1782.880755 | 2050.872708 | -0.277605716 | 0.08283124 | 0.000720296 | 0.001533365 | ns | YPD | KS.3 |
| FIM1_886 | 3133.716369 | 5253.639298 | 4045.572158 | 0.670999512 | 0.220298608 | 0.001212826 | 0.002473887 | ns | YPD | KS.3 |
| FIM1_887 | 519.0679077 | 394.5195159 | 602.4879608 | -0.379804923 | 0.130120164 | 0.002861067 | 0.00542447 | ns | YPD | KS.3 |
| FIM1_888 | 1328.220713 | 1389.41957 | 1495.16734 | 0.062237535 | 0.137581767 | 0.643308156 | 0.697115677 | ns | YPD | KS.3 |
| FIM1_889 | 1135.083676 | 994.4368795 | 968.179931 | -0.186278213 | 0.097850217 | 0.053955827 | 0.077135127 | ns | YPD | KS.3 |
| FIM1_890 | 470.6929189 | 214.4262103 | 335.6733714 | -1.085949823 | 0.153151504 | 2.402687549[illegible] | 3.464745090[illegible] | Km<KS.3 | YPD | KS.3 |
| FIM1_891 | 4861.883827 | 4744.470283 | 4113.37712 | -0.033351755 | 0.149612262 | 0.818766868 | 0.848794576 | ns | YPD | KS.3 |
| FIM1_892 | 131.4492691 | 128.2124082 | 170.0596042 | -0.029314183 | 0.288087189 | 0.910600601 | 0.927187472 | ns | YPD | KS.3 |
| FIM1_893 | 20.23165744 | 47.37823328 | 28.89626081 | 1.012589182 | 0.325033792 | 0.000388625 | 0.000878423 | Km>KS.3 | YPD | KS.3 |
| FIM1_894 | 2974.5536 | 8435.155304 | 3726.986747 | 1.322852039 | 0.286879371 | 2.825431883[illegible] | 1.235195370[illegible] | Km>KS.3 | YPD | KS.3 |
| FIM1_895 | 626.2600447 | 983.5472564 | 707.5695422 | 0.627582301 | 0.125404688 | 3.239779850[illegible] | 1.397910212[illegible] | ns | YPD | KS.3 |
| FIM1_896 | 126.558497 | 132.9751846 | 140.4085589 | 0.067025822 | 0.151384306 | 0.648835915 | 0.701730147 | ns | YPD | KS.3 |
| FIM1_897 | 447.4674176 | 592.9227875 | 566.0714161 | 0.3641211 | 0.215655717 | 0.074002436 | 0.102580697 | ns | YPD | KS.3 |
| FIM1_898 | 4179.959656 | 6665.20276 | 5129.727209 | 0.62255057 | 0.187454314 | 0.000517668 | 0.001135536 | ns | YPD | KS.3 |
| FIM1_899 | 727.5054345 | 402.7483869 | 688.9489661 | -0.794164696 | 0.183260531 | 6.029535381[illegible] | 2.009024756[illegible] | ns | YPD | KS.3 |
| FIM1_900 | 2379.626827 | 2127.459216 | 1909.136672 | -0.149886296 | 0.17581233 | 0.376044517 | 0.43915997 | ns | YPD | KS.3 |
| FIM1_901 | 2239.136762 | 3949.804611 | 3456.279029 | 0.794120079 | 0.118207615 | 7.576098251[illegible] | 8.338736460[illegible] | ns | YPD | KS.3 |
| FIM1_902 | 680.7777257 | 502.7182781 | 486.7253884 | -0.424255739 | 0.115989748 | 0.000197873 | 0.000475105 | ns | YPD | KS.3 |
| FIM1_903 | 1370.10366 | 832.8169207 | 1651.287398 | -0.578859937 | 0.314850511 | 0.038340721 | 0.056938795 | ns | YPD | KS.3 |
| FIM1_904 | 1198.291848 | 807.3259349 | 1256.333312 | -0.475891989 | 0.28313308 | 0.064169122 | 0.090257671 | ns | YPD | KS.3 |
| FIM1_905 | 947.445608 | 1108.311813 | 879.7972504 | 0.216530256 | 0.134709644 | 0.100140037 | 0.134538667 | ns | YPD | KS.3 |
| FIM1_906 | 7580.23501 | 7237.935669 | 12090.97668 | -0.063551992 | 0.140013934 | 0.641919297 | 0.695913817 | ns | YPD | KS.3 |
| FIM1_907 | 10782.78615 | 12699.92252 | 16591.70845 | 0.227713004 | 0.120676801 | 0.054616786 | 0.077968008 | ns | YPD | KS.3 |
| FIM1_909 | 1205.511158 | 689.6047378 | 1077.98299 | -0.768686632 | 0.148263496 | 9.462542729[illegible] | 4.606276915[illegible] | ns | YPD | KS.3 |
| FIM1_910 | 247.9706666 | 550.999271 | 267.2848716 | 1.061827179 | 0.207983532 | 6.460711015[illegible] | 3.230355507[illegible] | Km>KS.3 | YPD | KS.3 |

| | | | | | | | | | |
|---|---|---|---|---|---|---|---|---|---|
| FIM1_912 | 70.69456787 | 108.0970424 | 111.3884135 | 0.537531728 | 0.244060294 | 0.017859155 | 0.028762733 | ns | YPD | KS.3 |
| FIM1_913 | 1643.556642 | 3060.55724 | 2001.170019 | 0.792655476 | 0.24442736 | 0.000475599 | 0.001055355 | ns | YPD | KS.3 |
| FIM1_914 | 478.0105651 | 647.9447586 | 592.0757581 | 0.42380808 | 0.119415189 | 0.000300682 | 0.000695764 | ns | YPD | KS.3 |
| FIM1_915 | 4930.498564 | 4815.065138 | 4187.559889 | -0.033182319 | 0.113104376 | 0.76560321 | 0.803560331 | ns | YPD | KS.3 |
| FIM1_916 | 979.7287721 | 1973.285955 | 1234.286868 | 0.960710592 | 0.157408563 | 2.78759742192 | 2.23322015685 | ns | YPD | KS.3 |
| FIM1_917 | 4155.341228 | 11065.20897 | 10820.5135 | 1.212884928 | 0.303988728 | 7.21281125009 | 2.35195414984 | Km>KS.3 | YPD | KS.3 |
| FIM1_918 | 75.16360862 | 325.2720823 | 235.7526735 | 1.985486557 | 0.259539514 | 6.46592631781 | 1.85942100758 | Km>KS.3 | YPD | KS.3 |
| FIM1_919 | 105.3217711 | 549.647938 | 300.1039835 | 2.238873461 | 0.294256044 | 6.91071287614 | 1.97590784820 | Km>KS.3 | YPD | KS.3 |
| FIM1_920 | 776.2563549 | 2455.106834 | 1881.743709 | 1.581397318 | 0.188235065 | 7.89900107353 | 3.19492116592 | Km>KS.3 | YPD | KS.3 |
| FIM1_921 | 1923.33438 | 1977.801112 | 2693.04147 | 0.038313512 | 0.142498279 | 0.782791714 | 0.818688965 | ns | YPD | KS.3 |
| FIM1_922 | 296.6661911 | 450.6736612 | 488.257024 | 0.561221591 | 0.178199092 | 0.001046453 | 0.002160196 | ns | YPD | KS.3 |
| FIM1_923 | 237.3004507 | 446.6180194 | 409.4636735 | 0.843077931 | 0.19380348 | 4.96905988640 | 1.68056240209 | ns | YPD | KS.3 |
| FIM1_924 | 637.547625 | 893.7355064 | 926.2492243 | 0.465895094 | 0.136055919 | 0.000454624 | 0.001011519 | ns | YPD | KS.3 |
| FIM1_925 | 203.8451114 | 252.5947142 | 227.6234873 | 0.284860212 | 0.184929565 | 0.108255155 | 0.144388579 | ns | YPD | KS.3 |
| FIM1_926 | 744.9748047 | 7071.783955 | 4723.754127 | 3.046025805 | 0.353853809 | 1.03675732867 | 5.23597221984 | Km>KS.3 | YPD | KS.3 |
| FIM1_927 | 5762.991664 | 5424.824215 | 6918.70648 | -0.085801185 | 0.083096532 | 0.297499174 | 0.35819419 | ns | YPD | KS.3 |
| FIM1_928 | 577.1620332 | 622.8359058 | 996.4788868 | 0.103069759 | 0.154890111 | 0.493153441 | 0.556335231 | ns | YPD | KS.3 |
| FIM1_929 | 783.8594065 | 2218.12446 | 1444.466614 | 1.373126649 | 0.238696132 | 4.96227948214 | 3.74050612480 | Km>KS.3 | YPD | KS.3 |
| FIM1_930 | 1178.710518 | 3124.705999 | 1751.919322 | 1.327589142 | 0.186355502 | 7.17780909174 | 1.15940260491 | Km>KS.3 | YPD | KS.3 |
| FIM1_931 | 1631.538592 | 1190.588016 | 1672.699267 | -0.438805991 | 0.12140722 | 0.000230046 | 0.000543696 | ns | YPD | KS.3 |
| FIM1_932 | 647.1806523 | 636.8274132 | 497.2585964 | -0.022404235 | 0.104825909 | 0.828427524 | 0.857380264 | ns | YPD | KS.3 |
| FIM1_933 | 2454.090301 | 2767.47892 | 2148.831949 | 0.17083648 | 0.078711477 | 0.0286667 | 0.043882103 | ns | YPD | KS.3 |
| FIM1_934 | 1273.501353 | 1210.862613 | 1323.273902 | -0.07033638 | 0.122129157 | 0.557374035 | 0.618268857 | ns | YPD | KS.3 |
| FIM1_935 | 997.7495532 | 2113.166175 | 1205.089316 | 0.860830051 | 0.342496557 | 0.00377463 | 0.006957682 | ns | YPD | KS.3 |
| FIM1_936 | 5014.717086 | 11655.14473 | 8369.136041 | 1.084610238 | 0.250910154 | 2.75004926932 | 9.78647719232 | Km>KS.3 | YPD | KS.3 |
| FIM1_937 | 2257.73883 | 1290.41521 | 2593.518284 | -0.783419495 | 0.115810161 | 5.62819068538 | 6.34926273464 | ns | YPD | KS.3 |
| FIM1_938 | 2078.304904 | 3905.71978 | 3427.445661 | 0.835872363 | 0.202713345 | 1.38359034741 | 4.22292145914 | ns | YPD | KS.3 |
| FIM1_939 | 2178.67275 | 1564.435415 | 1869.916936 | -0.470887569 | 0.077865401 | 1.07972365625 | 7.60853426329 | ns | YPD | KS.3 |
| FIM1_940 | 3722.586358 | 3392.647767 | 2972.43512 | -0.124399149 | 0.174461937 | 0.459340438 | 0.523053028 | ns | YPD | KS.3 |
| FIM1_941 | 552.9122312 | 521.0899965 | 506.1071318 | -0.07549457 | 0.227239617 | 0.72380486 | 0.768608149 | ns | YPD | KS.3 |
| FIM1_942 | 825.3274965 | 1598.851913 | 1787.929814 | 0.870381841 | 0.212321541 | 1.40559434992 | 4.28482346254 | ns | YPD | KS.3 |
| FIM1_943 | 6622.79566 | 5579.01326 | 8646.69721 | -0.235054439 | 0.145542542 | 0.097312499 | 0.131022918 | ns | YPD | KS.3 |
| FIM1_944 | 1015.721048 | 644.6816254 | 1283.016317 | -0.567032562 | 0.256924705 | 0.016764933 | 0.027106312 | ns | YPD | KS.3 |
| FIM1_945 | 2225.733598 | 2543.879344 | 2338.598098 | 0.187978055 | 0.101003039 | 0.059360342 | 0.083944771 | ns | YPD | KS.3 |
| FIM1_946 | 36.49749778 | 63.15846769 | 45.99085676 | 0.737722312 | 0.175835085 | 1.26692475973 | 3.89792868255 | ns | YPD | KS.3 |
| FIM1_947 | 7236.869897 | 5319.610655 | 5270.519313 | -0.426526686 | 0.129763051 | 0.000784051 | 0.001659852 | ns | YPD | KS.3 |
| FIM1_948 | 5202.560881 | 8319.42054 | 5590.866524 | 0.660749165 | 0.103198309 | 8.28931821080 | 7.45738844462 | ns | YPD | KS.3 |
| FIM1_949 | 992.0893663 | 3223.461806 | 1384.197803 | 1.565461763 | 0.247984767 | 5.91249478766 | 6.65490080738 | Km>KS.3 | YPD | KS.3 |
| FIM1_950 | 190.2686089 | 596.9506062 | 465.2174882 | 1.513130845 | 0.247857282 | 2.91754178614 | 2.85162483027 | Km>KS.3 | YPD | KS.3 |
| FIM1_951 | 10426.70278 | 20137.99036 | 20186.18416 | 0.908434789 | 0.1461553 | 1.58674062197 | 1.34764220754 | ns | YPD | KS.3 |
| FIM1_952 | 13.8082633 | 37.75023387 | 41.24879713 | 1.181934921 | 0.352380165 | 9.35507548236 | 0.000242858 | Km>KS.3 | YPD | KS.3 |
| FIM1_953 | 2101.756829 | 4970.552135 | 3895.405561 | 1.199035133 | 0.13765343 | 3.51611193827 | 1.52110059938 | Km>KS.3 | YPD | KS.3 |
| FIM1_954 | 1854.2672 | 1848.673165 | 1865.356409 | -0.003970864 | 0.221275637 | 0.984818954 | 0.987001268 | ns | YPD | KS.3 |
| FIM1_955 | 700.1868101 | 3154.749716 | 1946.284371 | 1.981653149 | 0.322338473 | 3.06370154250 | 3.64639118994 | Km>KS.3 | YPD | KS.3 |
| FIM1_957 | 11435.87696 | 11872.21532 | 11679.93381 | 0.04441879 | 0.287254096 | 0.864590329 | 0.887239457 | ns | YPD | KS.3 |

| | | | | | | | | | | |
|---|---|---|---|---|---|---|---|---|---|---|
| FIM1_958 | 3445.774016 | 3292.321982 | 4172.932193 | -0.060933283 | 0.175884432 | 0.719078017 | 0.764404516 | ns | YPD | KS.3 |
| FIM1_959 | 828.076605 | 958.9531241 | 1630.320037 | 0.185378832 | 0.236529585 | 0.402024062 | 0.465457228 | ns | YPD | KS.3 |
| FIM1_960 | 2481.212995 | 1897.685089 | 2383.03718 | -0.37293904 | 0.123315055 | 0.002046014 | 0.003979249 | ns | YPD | KS.3 |
| FIM1_961 | 5383.26324 | 4239.775969 | 4809.67305 | -0.337144753 | 0.092735059 | 0.000236376 | 0.000557596 | ns | YPD | KS.3 |
| FIM1_962 | 3669.305044 | 2205.31262 | 2784.4084 | -0.7182635 | 0.098091172 | 1.1773123158 | 1.8476746912 | ns | YPD | KS.3 |
| FIM1_963 | 859.7385666 | 909.4632279 | 705.9682752 | 0.076643188 | 0.154286179 | 0.609025407 | 0.665619816 | ns | YPD | KS.3 |
| FIM1_964 | 284.3292797 | 177.0044773 | 229.22586 | -0.65765966 | 0.133124201 | 4.274186713 | 1.8017508667 | ns | YPD | KS.3 |
| FIM1_965 | 1557.379058 | 1157.35601 | 1032.537501 | -0.419072999 | 0.090881904 | 3.1356185615 | 1.1040129047 | ns | YPD | KS.3 |
| FIM1_966 | 4038.720559 | 3094.687157 | 3950.198171 | -0.360470754 | 0.163437036 | 0.022612393 | 0.035398569 | ns | YPD | KS.3 |
| FIM1_967 | 1119.724011 | 891.7605454 | 1015.043269 | -0.317301571 | 0.116948124 | 0.005765013 | 0.01022858 | ns | YPD | KS.3 |
| FIM1_968 | 562.7697908 | 651.9151929 | 752.0076583 | 0.200374767 | 0.153617137 | 0.179315203 | 0.228859193 | ns | YPD | KS.3 |
| FIM1_969 | 417.7153743 | 525.0090973 | 440.0148115 | 0.310046559 | 0.163340693 | 0.049786685 | 0.071898043 | ns | YPD | KS.3 |
| FIM1_970 | 1310.046151 | 1051.899142 | 965.1903519 | -0.306619801 | 0.11092972 | 0.004986263 | 0.008961943 | ns | YPD | KS.3 |
| FIM1_971 | 198.2387275 | 139.3715225 | 131.9005992 | -0.470541261 | 0.179643839 | 0.006398209 | 0.011275626 | ns | YPD | KS.3 |
| FIM1_972 | 117.893671 | 105.5037826 | 157.1682748 | -0.148785743 | 0.201674333 | 0.438441274 | 0.50235959 | ns | YPD | KS.3 |
| FIM1_973 | 1269.502858 | 1880.452797 | 1188.232828 | 0.556500156 | 0.0874707 | 1.2912166882 | 1.1075522455 | ns | YPD | KS.3 |
| FIM1_974 | 957.6691406 | 1308.371521 | 1062.195313 | 0.421264471 | 0.168857067 | 0.009739048 | 0.016559044 | ns | YPD | KS.3 |
| FIM1_975 | 1129.998015 | 3152.727793 | 2546.823218 | 1.391980588 | 0.197029458 | 8.1221258442 | 1.2992789734 | Km>KS.3 | YPD | KS.3 |
| FIM1_976 | 30.67547045 | 171.0619351 | 101.3528942 | 2.298381342 | 0.3297243 | 8.2899653101 | 1.5862529776 | Km>KS.3 | YPD | KS.3 |
| FIM1_977 | 1453.639082 | 2891.251139 | 3415.972487 | 0.809017865 | 0.318367732 | 0.004049033 | 0.007423401 | ns | YPD | KS.3 |
| FIM1_978 | 1108.086575 | 2996.806321 | 4313.41991 | 1.081132603 | 0.409363506 | 0.001281045 | 0.002597484 | Km>KS.3 | YPD | KS.3 |
| FIM1_979 | 1077.218448 | 1341.073082 | 2201.409289 | 0.257378153 | 0.296392323 | 0.335002266 | 0.39681816 | ns | YPD | KS.3 |
| FIM1_980 | 1330.613928 | 3987.696016 | 2473.290594 | 1.449157669 | 0.245621061 | 1.4244558786 | 1.2176405491 | Km>KS.3 | YPD | KS.3 |
| FIM1_981 | 658.6962892 | 2200.991723 | 1418.350743 | 1.466082779 | 0.358873437 | 1.6297464853 | 6.1284873505 | Km>KS.3 | YPD | KS.3 |
| FIM1_982 | 1020.373302 | 1125.401252 | 1544.672182 | 0.130338531 | 0.183259402 | 0.458828425 | 0.522685169 | ns | YPD | KS.3 |
| FIM1_983 | 9161.796809 | 9654.306181 | 12436.09678 | 0.073434354 | 0.104928998 | 0.477909692 | 0.541594696 | ns | YPD | KS.3 |
| FIM1_984 | 20.25557608 | 50.37409379 | 91.15760918 | 0.796272783 | 0.509716447 | 0.034841332 | 0.052193805 | ns | YPD | KS.3 |
| FIM1_988 | 2224.049086 | 2385.531886 | 2634.034784 | 0.097035609 | 0.130417739 | 0.44729442 | 0.51085623 | ns | YPD | KS.3 |
| FIM1_989 | 90.03559774 | 125.7557573 | 71.54174204 | 0.439646106 | 0.206110837 | 0.024630135 | 0.038220499 | ns | YPD | KS.3 |
| FIM1_991 | 716.840555 | 1604.520262 | 1301.236003 | 1.049970986 | 0.232810677 | 1.3069343174 | 5.0086900956 | Km>KS.3 | YPD | KS.3 |
| FIM1_992 | 1345.974219 | 3207.502298 | 2183.789848 | 1.151942853 | 0.215890099 | 1.3589617423 | 7.7621523170 | Km>KS.3 | YPD | KS.3 |
| FIM1_993 | 1198.472844 | 829.0908377 | 1362.119294 | -0.463263654 | 0.24669686 | 0.04320814 | 0.063260888 | ns | YPD | KS.3 |
| FIM1_994 | 1715.633957 | 1559.801645 | 2129.332173 | -0.126257577 | 0.188475308 | 0.48436085 | 0.547533567 | ns | YPD | KS.3 |
| FIM1_995 | 1547.089139 | 1176.836212 | 2185.273744 | -0.338870438 | 0.256913296 | 0.153361351 | 0.198136533 | ns | YPD | KS.3 |
| FIM1_996 | 1524.652036 | 1221.088857 | 2898.353279 | -0.241431026 | 0.347823047 | 0.422800672 | 0.486793183 | ns | YPD | KS.3 |
| FIM1_997 | 176.3431367 | 229.1652431 | 234.4595379 | 0.329113405 | 0.246034813 | 0.150246047 | 0.194351036 | ns | YPD | KS.3 |
| FIM1_998 | 1024.872643 | 1399.099743 | 1120.239119 | 0.433303199 | 0.120996081 | 0.000263075 | 0.00061417 | ns | YPD | KS.3 |
| FIM1_999 | 1152.292762 | 838.5065848 | 1425.907567 | -0.439850982 | 0.133550125 | 0.000753694 | 0.00159967 | ns | YPD | KS.3 |
| FIM1_1000 | 518.7379753 | 2616.858682 | 961.7356256 | 2.129456448 | 0.346274703 | 2.293284571 | 2.7963457700 | Km>KS.3 | YPD | KS.3 |
| FIM1_1001 | 377.0971633 | 303.4379076 | 323.4595592 | -0.306105045 | 0.105039633 | 0.003118635 | 0.005868082 | ns | YPD | KS.3 |
| FIM1_1002 | 337.6079336 | 377.3161554 | 328.5117821 | 0.152936528 | 0.137481669 | 0.254786445 | 0.311287467 | ns | YPD | KS.3 |
| FIM1_1003 | 1802.021406 | 2315.946315 | 3583.500302 | 0.311789745 | 0.253179095 | 0.183544303 | 0.233060977 | ns | YPD | KS.3 |
| FIM1_1004 | 1373.314867 | 3331.969342 | 3041.230843 | 1.172233728 | 0.221246722 | 1.5465869095 | 8.7534355802 | Km>KS.3 | YPD | KS.3 |
| FIM1_1005 | 662.5293564 | 1520.351021 | 1178.634649 | 1.142970739 | 0.158595941 | 8.4147356065 | 1.3374859310 | Km>KS.3 | YPD | KS.3 |
| FIM1_1006 | 2779.236623 | 5078.773316 | 3500.720483 | 0.817578199 | 0.171014943 | 6.8151196722 | 2.7677730914 | ns | YPD | KS.3 |

| | | | | | | | | | | |
|---|---|---|---|---|---|---|---|---|---|---|
| FIM1_1007 | 734.707647 | 634.5610088 | 680.5163647 | -0.208452521 | 0.09248944 | 0.022679472 | 0.035470096 | ns | YPD | KS.3 |
| FIM1_1008 | 481.5805547 | 433.6454459 | 350.3893167 | -0.141689345 | 0.165247314 | 0.37533362 | 0.438432675 | ns | YPD | KS.3 |
| FIM1_1009 | 810.9565774 | 726.1198926 | 736.7478592 | -0.153101749 | 0.130547034 | 0.230942498 | 0.285025783 | ns | YPD | KS.3 |
| FIM1_1010 | 1696.639939 | 5897.020049 | 2615.082822 | 1.612074681 | 0.295754962 | 9.01869144075 | 6.48381357193 | Km>KS.3 | YPD | KS.3 |
| FIM1_1011 | 5785.090936 | 10431.27292 | 10635.097 | 0.769814397 | 0.217634166 | 0.00017333 | 0.000424214 | ns | YPD | KS.3 |
| FIM1_1012 | 950.6158713 | 3391.317456 | 2599.211892 | 1.532220084 | 0.380895281 | 1.59180317686 | 6.00851350902 | Km>KS.3 | YPD | KS.3 |
| FIM1_1013 | 2295.818317 | 2985.462321 | 2768.382107 | 0.337010661 | 0.224147084 | 0.110101799 | 0.146537306 | ns | YPD | KS.3 |
| FIM1_1014 | 2262.261764 | 2612.463114 | 2115.903474 | 0.15829843 | 0.339131997 | 0.592561051 | 0.650010924 | ns | YPD | KS.3 |
| FIM1_1015 | 1213.942392 | 1531.883422 | 2155.16098 | 0.280859499 | 0.276214154 | 0.265385978 | 0.322889518 | ns | YPD | KS.3 |
| FIM1_1016 | 123.8136411 | 50.02486185 | 196.0072282 | -1.153010554 | 0.266643272 | 2.13597348310 | 7.79066574667 | Km<KS.3 | YPD | KS.3 |
| FIM1_1017 | 4248.079061 | 2584.332823 | 4189.521793 | -0.68831373 | 0.135123761 | 1.81172864637 | 8.30723503754 | ns | YPD | KS.3 |
| FIM1_1018 | 1589.03908 | 926.366121 | 1304.181682 | -0.764026019 | 0.091708021 | 3.49360990527 | 1.05979934626 | ns | YPD | KS.3 |
| FIM1_1019 | 881.1207508 | 1203.561611 | 854.6122633 | 0.421930042 | 0.165317598 | 0.008272107 | 0.014259783 | ns | YPD | KS.3 |
| FIM1_1020 | 144.327879 | 209.7832121 | 134.9193748 | 0.436059511 | 0.306379567 | 0.111285435 | 0.147993863 | ns | YPD | KS.3 |
| FIM1_1021 | 376.3174671 | 1428.700347 | 651.045516 | 1.671663481 | 0.353209906 | 3.20351418389 | 1.69108840456 | Km>KS.3 | YPD | KS.3 |
| FIM1_1022 | 5334.743976 | 7345.069818 | 7715.193246 | 0.448273373 | 0.109038624 | 2.97668877745 | 8.47198321958 | ns | YPD | KS.3 |
| FIM1_1024 | 966.0885539 | 3542.417339 | 1785.986354 | 1.732188125 | 0.261949343 | 3.49525745291 | 4.94003006484 | Km>KS.3 | YPD | KS.3 |
| FIM1_1025 | 3388.321835 | 3528.462071 | 5296.423224 | 0.055125031 | 0.153047702 | 0.710880974 | 0.758445817 | ns | YPD | KS.3 |
| FIM1_1026 | 4444.419753 | 4674.859021 | 6893.761453 | 0.06867933 | 0.155464184 | 0.649080431 | 0.70182404 | ns | YPD | KS.3 |
| FIM1_1027 | 679.5358003 | 1088.81614 | 1025.936303 | 0.627245149 | 0.19096867 | 0.000584282 | 0.001270555 | ns | YPD | KS.3 |
| FIM1_1028 | 119.3586829 | 288.8496327 | 216.8554548 | 1.183095315 | 0.205197909 | 1.03490101933 | 7.31339853859 | Km>KS.3 | YPD | KS.3 |
| FIM1_1029 | 8965.500985 | 9988.486035 | 12682.87503 | 0.137062925 | 0.232582092 | 0.529553393 | 0.591629942 | ns | YPD | KS.3 |
| FIM1_1030 | 2151.008694 | 2548.795885 | 4336.793453 | 0.19019246 | 0.327572468 | 0.50954257 | 0.57158383 | ns | YPD | KS.3 |
| FIM1_1031 | 79.52156807 | 60.85962026 | 81.79274124 | -0.337142709 | 0.240259708 | 0.132507713 | 0.173663296 | ns | YPD | KS.3 |
| FIM1_1032 | 864.3706606 | 636.5253293 | 743.9069697 | -0.431111759 | 0.097747133 | 7.96444489152 | 2.56128722271 | ns | YPD | KS.3 |
| FIM1_1033 | 373.3817388 | 109.8824705 | 215.5946391 | -1.657873218 | 0.222938963 | 1.29674215828 | 2.80490966847 | Km<KS.3 | YPD | KS.3 |
| FIM1_1034 | 592.3569026 | 1023.764195 | 648.1549468 | 0.772616716 | 0.09566979 | 2.87080695784 | 7.00111010551 | ns | YPD | KS.3 |
| FIM1_1035 | 178.1954606 | 215.349436 | 164.1932233 | 0.255227561 | 0.160557727 | 0.100809047 | 0.135364374 | ns | YPD | KS.3 |
| FIM1_1036 | 849.6384719 | 350.8314588 | 581.8946502 | -1.23808051 | 0.127511573 | 2.67234921057 | 2.04537497270 | Km<KS.3 | YPD | KS.3 |
| FIM1_1037 | 1445.679416 | 1489.436661 | 1268.132034 | 0.042789342 | 0.066255303 | 0.515964835 | 0.578368883 | ns | YPD | KS.3 |
| FIM1_1038 | 3385.419262 | 3044.668109 | 3371.647619 | -0.150767851 | 0.078322247 | 0.052326929 | 0.075108619 | ns | YPD | KS.3 |
| FIM1_1039 | 3393.499149 | 3869.05344 | 3203.341312 | 0.18331387 | 0.113377468 | 0.100317866 | 0.134741194 | ns | YPD | KS.3 |
| FIM1_1040 | 1035.982346 | 2571.151445 | 1696.149232 | 1.2099777 | 0.214606132 | 1.96787843747 | 1.33381406355 | Km>KS.3 | YPD | KS.3 |
| FIM1_1041 | 960.4854757 | 2889.872056 | 2064.977982 | 1.392282653 | 0.299803579 | 1.77103901826 | 8.13565938674 | Km>KS.3 | YPD | KS.3 |
| FIM1_1042 | 49.4926891 | 88.83745533 | 77.10730827 | 0.785224769 | 0.185274512 | 9.44320433972 | 2.97529712413 | ns | YPD | KS.3 |
| FIM1_1043 | 145.34903 | 272.0861906 | 188.1524893 | 0.796504305 | 0.248118395 | 0.000527476 | 0.001155523 | ns | YPD | KS.3 |
| FIM1_1044 | 1544.315201 | 3369.705538 | 2014.807761 | 1.014510199 | 0.233682586 | 3.18721599547 | 1.11980223004 | Km>KS.3 | YPD | KS.3 |
| FIM1_1045 | 407.2469745 | 399.1480708 | 478.7153123 | -0.028251611 | 0.137716436 | 0.833661963 | 0.861006491 | ns | YPD | KS.3 |
| FIM1_1046 | 992.708542 | 1088.74338 | 813.5084914 | 0.128321722 | 0.124818264 | 0.294514454 | 0.354858176 | ns | YPD | KS.3 |
| FIM1_1047 | 1861.983306 | 1651.971452 | 1493.593789 | -0.170246514 | 0.069746081 | 0.014022794 | 0.023039433 | ns | YPD | KS.3 |
| FIM1_1048 | 2259.683961 | 1843.440578 | 1727.147317 | -0.285582233 | 0.106780273 | 0.006651276 | 0.011684356 | ns | YPD | KS.3 |
| FIM1_1049 | 2110.770455 | 1198.383798 | 2601.580577 | -0.658223989 | 0.318686078 | 0.019573233 | 0.031160587 | ns | YPD | KS.3 |
| FIM1_1050 | 54.24686384 | 90.25628926 | 107.7630703 | 0.618426004 | 0.280807385 | 0.015366512 | 0.025040419 | ns | YPD | KS.3 |
| FIM1_1051 | 527.4983795 | 494.7428289 | 591.553105 | -0.08644852 | 0.16641246 | 0.591048034 | 0.648822588 | ns | YPD | KS.3 |
| FIM1_1052 | 793.5597376 | 726.3790918 | 1052.997585 | -0.115177577 | 0.207044099 | 0.558064249 | 0.618896486 | ns | YPD | KS.3 |

| | | | | | | | | | | |
|---|---|---|---|---|---|---|---|---|---|---|
| FIM1_1053 | 91.90528059 | 102.5559335 | 89.13311434 | 0.14421426 | 0.188761067 | 0.425058329 | 0.489166132 | ns | YPD | KS.3 |
| FIM1_1054 | 2054.197459 | 2182.133317 | 2238.829363 | 0.084700377 | 0.097441234 | 0.378952134 | 0.441829592 | ns | YPD | KS.3 |
| FIM1_1055 | 280.3309626 | 142.3468066 | 204.8296976 | -0.921522651 | 0.170478024 | 1.9364295963 | 1.0704152491 | ns | YPD | KS.3 |
| FIM1_1056 | 785.3664063 | 395.8612665 | 564.8640782 | -0.943758541 | 0.150152637 | 9.2067623632 | 8.2209194867 | ns | YPD | KS.3 |
| FIM1_1057 | 1395.421791 | 2201.288262 | 1415.285574 | 0.641556727 | 0.102668231 | 2.3274121266 | 1.8950696121 | ns | YPD | KS.3 |
| FIM1_1058 | 2318.01423 | 1876.892885 | 2535.405671 | -0.296297637 | 0.106320617 | 0.004691007 | 0.00847721 | ns | YPD | KS.3 |
| FIM1_1060 | 4494.583317 | 3242.198586 | 4530.229662 | -0.428068758 | 0.204116604 | 0.02730612 | 0.042045171 | ns | YPD | KS.3 |
| FIM1_1061 | 785.3373962 | 732.803636 | 795.8899212 | -0.094891451 | 0.139392798 | 0.485617958 | 0.548705278 | ns | YPD | KS.3 |
| FIM1_1062 | 1125.940764 | 837.8946141 | 854.6353268 | -0.414977129 | 0.101360131 | 3.3341265955 | 9.3607673886 | ns | YPD | KS.3 |
| FIM1_1063 | 1042.870718 | 811.1304052 | 789.7390678 | -0.345676785 | 0.139767531 | 0.011254689 | 0.018852552 | ns | YPD | KS.3 |
| FIM1_1064 | 797.476152 | 1336.705794 | 1260.751839 | 0.705265855 | 0.158586796 | 4.3392914347 | 1.4877997855 | ns | YPD | KS.3 |
| FIM1_1065 | 1402.208762 | 1372.666742 | 1242.392912 | -0.02962269 | 0.097013323 | 0.757299849 | 0.797537415 | ns | YPD | KS.3 |
| FIM1_1066 | 631.3126488 | 349.696466 | 442.4206693 | -0.826499645 | 0.118301692 | 1.0737767702 | 1.4057998505 | ns | YPD | KS.3 |
| FIM1_1067 | 667.2449657 | 482.1486164 | 409.8270158 | -0.449172001 | 0.13214937 | 0.000509625 | 0.00112086 | ns | YPD | KS.3 |
| FIM1_1068 | 121.4577891 | 285.1337639 | 234.1648906 | 1.095571787 | 0.25284114 | 2.5266160965 | 9.0757509604 | Km>KS.3 | YPD | KS.3 |
| FIM1_1069 | 392.3834211 | 758.2749988 | 417.8969508 | 0.792598637 | 0.298640313 | 0.003079181 | 0.00579823 | ns | YPD | KS.3 |
| FIM1_1070 | 1044.548342 | 1589.352919 | 1189.605498 | 0.589832711 | 0.105480644 | 1.3847411242 | 7.8822506786 | ns | YPD | KS.3 |
| FIM1_1071 | 1708.13566 | 1630.986317 | 1522.70044 | -0.06601672 | 0.06551111 | 0.310844436 | 0.371564408 | ns | YPD | KS.3 |
| FIM1_1072 | 355.4114167 | 1094.441452 | 1457.758092 | 1.339675436 | 0.361836001 | 1.3554852243 | 4.1422229676 | Km>KS.3 | YPD | KS.3 |
| FIM1_1074 | 1356.875851 | 1023.20979 | 1221.662544 | -0.392751998 | 0.122834478 | 0.001115466 | 0.002287488 | ns | YPD | KS.3 |
| FIM1_1075 | 2197.513084 | 3158.365573 | 3065.067165 | 0.501748545 | 0.133157143 | 0.000115776 | 0.000295038 | ns | YPD | KS.3 |
| FIM1_1076 | 193.5342835 | 792.5810753 | 428.2304838 | 1.904777602 | 0.257228898 | 5.5017254774 | 1.2850274295 | Km>KS.3 | YPD | KS.3 |
| FIM1_1077 | 50.65546924 | 120.4085252 | 125.6029179 | 1.030088489 | 0.324085904 | 0.000297205 | 0.000688359 | Km>KS.3 | YPD | KS.3 |
| FIM1_1078 | 325.9228955 | 316.2533383 | 294.9642808 | -0.041434471 | 0.164112314 | 0.794166828 | 0.828471371 | ns | YPD | KS.3 |
| FIM1_1079 | 245.6193986 | 748.072623 | 387.3355343 | 1.515320691 | 0.201354025 | 1.3923156481 | 2.9350721819 | Km>KS.3 | YPD | KS.3 |
| FIM1_1080 | 885.089132 | 1250.686361 | 1084.702492 | 0.436798682 | 0.240982765 | 0.051802191 | 0.074448425 | ns | YPD | KS.3 |
| FIM1_1081 | 745.6292169 | 3275.778724 | 1316.689336 | 1.948531809 | 0.317132981 | 3.4307890826 | 4.0350297130 | Km>KS.3 | YPD | KS.3 |
| FIM1_1082 | 724.6837128 | 1699.983112 | 1101.869711 | 1.122439539 | 0.224297079 | 8.8898554613 | 4.3530542244 | Km>KS.3 | YPD | KS.3 |
| FIM1_1083 | 1514.202213 | 1934.195446 | 1554.803504 | 0.344072266 | 0.10214645 | 0.00063972 | 0.00138074 | ns | YPD | KS.3 |
| FIM1_1084 | 295.9408856 | 928.8065372 | 682.8854909 | 1.574764424 | 0.182291932 | 1.0419321603 | 5.2359722198 | Km>KS.3 | YPD | KS.3 |
| FIM1_1085 | 773.7602227 | 440.8535392 | 580.9697454 | -0.790720872 | 0.10673308 | 5.2789839052 | 8.6963393802 | ns | YPD | KS.3 |
| FIM1_1086 | 1010.730486 | 489.3865975 | 635.0051541 | -1.001169837 | 0.148171312 | 3.3464431018 | 3.9545259933 | Km<KS.3 | YPD | KS.3 |
| FIM1_1087 | 1029.935892 | 977.8072034 | 755.4464923 | -0.071689174 | 0.132563247 | 0.580462818 | 0.638894362 | ns | YPD | KS.3 |
| FIM1_1088 | 2763.95474 | 2263.206009 | 2300.985618 | -0.283157757 | 0.08343596 | 0.000615218 | 0.00133248 | ns | YPD | KS.3 |
| FIM1_1089 | 165.8459749 | 80.66082282 | 127.3354343 | -0.958027355 | 0.205312271 | 8.2505171871 | 3.2942474322 | ns | YPD | KS.3 |
| FIM1_1090 | 4228.357187 | 4528.270824 | 4998.7929 | 0.095613884 | 0.115653016 | 0.400497515 | 0.464121858 | ns | YPD | KS.3 |
| FIM1_1091 | 1615.930132 | 1467.500915 | 2368.850307 | -0.121999661 | 0.23489499 | 0.579051757 | 0.637482295 | ns | YPD | KS.3 |
| FIM1_1092 | 1094.119818 | 1120.436338 | 980.3221085 | 0.033662128 | 0.085959216 | 0.692584712 | 0.742137767 | ns | YPD | KS.3 |
| FIM1_1093 | 1046.510147 | 1005.095782 | 1159.130995 | -0.055264378 | 0.144779169 | 0.695284995 | 0.744520631 | ns | YPD | KS.3 |
| FIM1_1094 | 1751.357993 | 1799.66018 | 1676.028457 | 0.038315975 | 0.056863854 | 0.49856838 | 0.561298414 | ns | YPD | KS.3 |
| FIM1_1095 | 1418.143207 | 1663.736335 | 1197.186712 | 0.224569575 | 0.100598199 | 0.023730515 | 0.036916607 | ns | YPD | KS.3 |
| FIM1_1096 | 400.7997971 | 434.2669812 | 436.598392 | 0.107595455 | 0.168775428 | 0.509240953 | 0.571374321 | ns | YPD | KS.3 |
| FIM1_1097 | 6301.213817 | 11363.74178 | 8368.502534 | 0.793425875 | 0.181411512 | 5.0319007289 | 1.7006593835 | ns | YPD | KS.3 |
| FIM1_1098 | 3058.857686 | 2386.16359 | 3522.134248 | -0.333387465 | 0.174718427 | 0.047563997 | 0.068988596 | ns | YPD | KS.3 |
| FIM1_1099 | 538.2695343 | 1346.3165 | 822.1322244 | 1.266321936 | 0.156906252 | 6.2023900278 | 1.7940052551 | Km>KS.3 | YPD | KS.3 |

| | | | | | | | | | | |
|---|---|---|---|---|---|---|---|---|---|---|
| FIM1_1100 | 7007.612768 | 7126.88063 | 7850.830426 | 0.023811502 | 0.090156434 | 0.789561791 | 0.824836705 | ns | YPD | KS.3 |
| FIM1_1101 | 34.77126441 | 92.82807377 | 78.11312331 | 0.933506098 | 0.481970332 | 0.011393222 | 0.019065348 | ns | YPD | KS.3 |
| FIM1_1102 | 1118.536212 | 944.7951678 | 1220.438134 | -0.233806813 | 0.130841536 | 0.067902646 | 0.09489204 | ns | YPD | KS.3 |
| FIM1_1103 | 797.216765 | 704.4172775 | 973.7573959 | -0.156367333 | 0.23840405 | 0.482685348 | 0.546011734 | ns | YPD | KS.3 |
| FIM1_1104 | 794.5877218 | 1682.550782 | 1469.220352 | 1.013652074 | 0.182720879 | 6.63442128255 | 3.99591354487 | Km>KS.3 | YPD | KS.3 |
| FIM1_1105 | 3956.433785 | 3515.919674 | 4789.31649 | -0.168047977 | 0.072724359 | 0.019971606 | 0.03171361 | ns | YPD | KS.3 |
| FIM1_1106 | 6390.19833 | 7002.703749 | 6673.423361 | 0.124081296 | 0.159477138 | 0.422173369 | 0.486183452 | ns | YPD | KS.3 |
| FIM1_1107 | 711.0692986 | 780.7439686 | 984.9406452 | 0.126564336 | 0.15697808 | 0.405984809 | 0.469565765 | ns | YPD | KS.3 |
| FIM1_1108 | 1474.597148 | 992.444113 | 1788.027479 | -0.475930395 | 0.285360696 | 0.065852675 | 0.092286496 | ns | YPD | KS.3 |
| FIM1_1109 | 3368.877569 | 6625.078043 | 4698.985873 | 0.927731026 | 0.15675935 | 9.58897511628 | 6.84435454856 | ns | YPD | KS.3 |
| FIM1_1110 | 1653.412222 | 1022.482278 | 1348.385957 | -0.658539358 | 0.151822739 | 7.86599145935 | 2.53438451725 | ns | YPD | KS.3 |
| FIM1_1111 | 732.3702548 | 881.8983633 | 1127.910016 | 0.2590781 | 0.118005105 | 0.025448695 | 0.039367928 | ns | YPD | KS.3 |
| FIM1_1112 | 2885.2438 | 4507.781395 | 3978.287789 | 0.625320419 | 0.111847544 | 1.31138434206 | 7.50763763150 | ns | YPD | KS.3 |
| FIM1_1113 | 459.7755098 | 2560.667163 | 1419.349751 | 2.340944524 | 0.290473884 | 2.09100822726 | 7.64909259607 | Km>KS.3 | YPD | KS.3 |
| FIM1_1114 | 268.9777758 | 2080.040175 | 2368.019001 | 2.686988425 | 0.415340581 | 4.22810522411 | 5.87564901955 | Km>KS.3 | YPD | KS.3 |
| FIM1_1115 | 113.2444794 | 521.7735899 | 361.7384784 | 2.076968765 | 0.26541289 | 1.34080289377 | 4.33148986788 | Km>KS.3 | YPD | KS.3 |
| FIM1_1116 | 5065.193647 | 7290.842636 | 6624.83979 | 0.496974898 | 0.154785271 | 0.000934417 | 0.001945073 | ns | YPD | KS.3 |
| FIM1_1117 | 1452.769709 | 1830.825402 | 1887.62211 | 0.313894055 | 0.159771889 | 0.042586096 | 0.062534187 | ns | YPD | KS.3 |
| FIM1_1118 | 21.37675669 | 14.53058011 | 20.87135697 | -0.476861675 | 0.252670644 | 0.041440964 | 0.060960613 | ns | YPD | KS.3 |
| FIM1_1119 | 2356.209194 | 3651.171942 | 2937.395474 | 0.604577061 | 0.139038435 | 8.25032675814 | 2.63957399497 | ns | YPD | KS.3 |
| FIM1_1120 | 51.78620825 | 38.9355994 | 43.48615303 | -0.370362491 | 0.202265656 | 0.054243901 | 0.077502415 | ns | YPD | KS.3 |
| FIM1_1121 | 12450.02092 | 11939.38587 | 12996.19495 | -0.057310402 | 0.14616497 | 0.687289929 | 0.737706019 | ns | YPD | KS.3 |
| FIM1_1122 | 5087.926323 | 6550.717348 | 5340.92505 | 0.35899965 | 0.077749273 | 3.24467314435 | 1.13918481955 | ns | YPD | KS.3 |
| FIM1_1123 | 354.6057529 | 408.9848689 | 448.2279881 | 0.195463118 | 0.138857581 | 0.149366294 | 0.19336386 | ns | YPD | KS.3 |
| FIM1_1124 | 2669.603216 | 4900.703274 | 3235.534849 | 0.841804274 | 0.136994298 | 2.94770474350 | 2.33888853252 | ns | YPD | KS.3 |
| FIM1_1125 | 12170.32756 | 15545.08444 | 21740.66123 | 0.338211244 | 0.133323265 | 0.009478954 | 0.016155464 | ns | YPD | KS.3 |
| FIM1_1126 | 550.5573058 | 762.4454135 | 764.9335658 | 0.39254789 | 0.280410576 | 0.123985377 | 0.163658066 | ns | YPD | KS.3 |
| FIM1_1127 | 378.630273 | 525.390444 | 411.6442297 | 0.413153314 | 0.242319811 | 0.067172535 | 0.093924498 | ns | YPD | KS.3 |
| FIM1_1128 | 148.6246517 | 121.8638339 | 192.2800568 | -0.241344057 | 0.270658323 | 0.330505552 | 0.392238818 | ns | YPD | KS.3 |
| FIM1_1129 | 4009.418572 | 2958.090062 | 3127.035772 | -0.424871801 | 0.115056452 | 0.0001724 | 0.000422507 | ns | YPD | KS.3 |
| FIM1_1130 | 484.349561 | 453.1567594 | 445.2423528 | -0.090347086 | 0.157353788 | 0.553961597 | 0.614894901 | ns | YPD | KS.3 |
| FIM1_1131 | 1857.224017 | 1314.605564 | 1327.843283 | -0.484701614 | 0.108262678 | 5.46048170875 | 1.83553354736 | ns | YPD | KS.3 |
| FIM1_1132 | 461.1833896 | 1304.712159 | 990.3518956 | 1.39633023 | 0.214441053 | 3.72229736921 | 4.35727750867 | Km>KS.3 | YPD | KS.3 |
| FIM1_1133 | 206.6532348 | 1213.204699 | 653.9017944 | 2.441861576 | 0.264828326 | 1.03893291406 | 7.18737843369 | Km>KS.3 | YPD | KS.3 |
| FIM1_1134 | 87.2501349 | 417.2348294 | 307.6470545 | 2.11527172 | 0.283836698 | 2.39866186126 | 5.99665465315 | Km>KS.3 | YPD | KS.3 |
| FIM1_1135 | 13214.12324 | 11991.18845 | 14001.82884 | -0.122952849 | 0.234370814 | 0.575346307 | 0.634385611 | ns | YPD | KS.3 |
| FIM1_1136 | 734.3889777 | 976.9438545 | 1073.687292 | 0.393380749 | 0.137660935 | 0.003423476 | 0.006381338 | ns | YPD | KS.3 |
| FIM1_1137 | 2113.804661 | 4420.996614 | 3821.963831 | 0.928116267 | 0.265224819 | 0.00013219 | 0.000331977 | ns | YPD | KS.3 |
| FIM1_1138 | 735.9441092 | 452.9505055 | 592.9896694 | -0.675785426 | 0.122871224 | 2.01017737951 | 1.10504226111 | ns | YPD | KS.3 |
| FIM1_1139 | 8125.220897 | 8767.599594 | 9084.169046 | 0.10730385 | 0.093302587 | 0.244847377 | 0.300176367 | ns | YPD | KS.3 |
| FIM1_1140 | 3745.23848 | 1668.263694 | 3299.452156 | -0.894036064 | 0.373756603 | 0.004689254 | 0.008477121 | ns | YPD | KS.3 |
| FIM1_1141 | 3537.857921 | 1865.994982 | 3765.705268 | -0.755275616 | 0.312258285 | 0.006475714 | 0.011400099 | ns | YPD | KS.3 |
| FIM1_1142 | 690.1607891 | 1027.811456 | 869.6513995 | 0.559595187 | 0.102817635 | 3.39811079810 | 1.78517436331 | ns | YPD | KS.3 |
| FIM1_1143 | 1715.784805 | 1645.109099 | 1700.876473 | -0.059805345 | 0.09128618 | 0.507859909 | 0.570081915 | ns | YPD | KS.3 |
| FIM1_1144 | 1272.232032 | 712.5919412 | 867.3689057 | -0.753730692 | 0.221976143 | 0.000303306 | 0.000701509 | ns | YPD | KS.3 |

| | | | | | | | | | |
|---|---|---|---|---|---|---|---|---|---|
| FIM1_1145 | 7000.279847 | 8978.766635 | 9504.782481 | 0.351253092 | 0.093826665 | 0.000152539 | 0.00037944 | ns | YPD | KS.3 |
| FIM1_1146 | 769.4419271 | 573.4610468 | 629.5973332 | -0.40462201 | 0.13887293 | 0.002830501 | 0.005368564 | ns | YPD | KS.3 |
| FIM1_1147 | 3701.109482 | 3002.079957 | 3609.849336 | -0.294044541 | 0.104223268 | 0.00422398 | 0.00770884 | ns | YPD | KS.3 |
| FIM1_1148 | 574.2194618 | 522.4111578 | 672.4674812 | -0.129481515 | 0.15285718 | 0.383349512 | 0.446224572 | ns | YPD | KS.3 |
| FIM1_1149 | 383.1896868 | 225.9973659 | 286.7529395 | -0.717509506 | 0.166319783 | 7.7915425922 | 2.5121791572 | ns | YPD | KS.3 |
| FIM1_1150 | 1112.570866 | 657.1574469 | 929.2402497 | -0.719888542 | 0.156479492 | 2.0415365351 | 7.4846309966 | ns | YPD | KS.3 |
| FIM1_1151 | 5975.6959 | 5379.80644 | 6165.423123 | -0.14366545 | 0.147523318 | 0.317152064 | 0.378195475 | ns | YPD | KS.3 |
| FIM1_1152 | 389.6711075 | 1357.949767 | 803.2191854 | 1.671545758 | 0.246194041 | 1.3766428886 | 2.1269560158 | Km>KS.3 | YPD | KS.3 |
| FIM1_1153 | 86.12944278 | 104.558508 | 75.4474694 | 0.255342361 | 0.196441306 | 0.173141196 | 0.221890121 | ns | YPD | KS.3 |
| FIM1_1154 | 1603.151642 | 1143.955998 | 1354.118306 | -0.472602559 | 0.111888355 | 1.7604695004 | 5.2697567777 | ns | YPD | KS.3 |
| FIM1_1155 | 78.64483909 | 109.8564841 | 96.14274037 | 0.372088208 | 0.336388564 | 0.205585666 | 0.25730533 | ns | YPD | KS.3 |
| FIM1_1156 | 8861.675751 | 7445.386039 | 9766.387008 | -0.241842216 | 0.124517812 | 0.047649265 | 0.069071997 | ns | YPD | KS.3 |
| FIM1_1157 | 3918.005051 | 3622.432732 | 2947.350756 | -0.110433747 | 0.096127151 | 0.245044767 | 0.30034435 | ns | YPD | KS.3 |
| FIM1_1158 | 659.6295787 | 1537.93467 | 1034.973562 | 1.060475575 | 0.278095414 | 2.6017339956 | 7.5122615371 | Km>KS.3 | YPD | KS.3 |
| FIM1_1159 | 1942.659089 | 1833.611092 | 1818.015142 | -0.082106139 | 0.080471233 | 0.303515797 | 0.363765 | ns | YPD | KS.3 |
| FIM1_1160 | 11641.93894 | 9418.882019 | 13109.57996 | -0.300514261 | 0.082170433 | 0.000224416 | 0.00053216 | ns | YPD | KS.3 |
| FIM1_1161 | 85.52260436 | 107.61246 | 118.611153 | 0.259751635 | 0.325189639 | 0.365035693 | 0.427608329 | ns | YPD | KS.3 |
| FIM1_1162 | 6046.162408 | 9207.67677 | 6707.310668 | 0.494481197 | 0.302969593 | 0.068428796 | 0.095519994 | ns | YPD | KS.3 |
| FIM1_1163 | 1520.081566 | 1230.017641 | 1155.211628 | -0.298472076 | 0.09195682 | 0.001031943 | 0.002133798 | ns | YPD | KS.3 |
| FIM1_1164 | 39.87433863 | 30.59391991 | 38.21550992 | -0.325424678 | 0.265288176 | 0.18201907 | 0.231597155 | ns | YPD | KS.3 |
| FIM1_1165 | 1206.6744 | 758.3995947 | 916.3283181 | -0.629910057 | 0.165774099 | 8.3017848255 | 0.000216805 | ns | YPD | KS.3 |
| FIM1_1166 | 4667.73536 | 5019.702943 | 6055.591906 | 0.099462333 | 0.147829094 | 0.489467582 | 0.552786239 | ns | YPD | KS.3 |
| FIM1_1167 | 713.1333719 | 1315.760956 | 888.8186405 | 0.846596206 | 0.141705281 | 8.4106006032 | 6.0729663281 | ns | YPD | KS.3 |
| FIM1_1168 | 1598.397419 | 1834.032572 | 1564.529 | 0.192110931 | 0.114176654 | 0.087171161 | 0.118588057 | ns | YPD | KS.3 |
| FIM1_1169 | 1362.789244 | 576.3258722 | 944.2420511 | -1.184175463 | 0.159952356 | 1.6661405652 | 3.0586897830 | Km<KS.3 | YPD | KS.3 |
| FIM1_1170 | 1273.894533 | 946.6741192 | 1068.3569 | -0.408345554 | 0.141609753 | 0.003100647 | 0.005836443 | ns | YPD | KS.3 |
| FIM1_1171 | 22.1590683 | 60.04447654 | 54.07909024 | 1.331819699 | 0.216314424 | 5.0469935791 | 4.7020211716 | Km>KS.3 | YPD | KS.3 |
| FIM1_1172 | 2832.735807 | 2371.345571 | 2917.458834 | -0.252633187 | 0.075049597 | 0.000695524 | 0.001488912 | ns | YPD | KS.3 |
| FIM1_1173 | 851.0776579 | 281.675665 | 611.6997115 | -1.505569256 | 0.198270028 | 8.6772739964 | 1.9431024375 | Km<KS.3 | YPD | KS.3 |
| FIM1_1174 | 431.296393 | 721.9135957 | 552.607912 | 0.660259057 | 0.233886139 | 0.002520845 | 0.004814283 | ns | YPD | KS.3 |
| FIM1_1175 | 112.6870965 | 135.716609 | 154.079058 | 0.234518949 | 0.239022761 | 0.293401188 | 0.35368813 | ns | YPD | KS.3 |
| FIM1_1177 | 2690.97654 | 1623.249924 | 2264.533237 | -0.717069497 | 0.085989054 | 3.6088612967 | 1.0815713825 | ns | YPD | KS.3 |
| FIM1_1178 | 11828.0897 | 9503.614447 | 14048.49393 | -0.29925018 | 0.149040474 | 0.039006836 | 0.057841731 | ns | YPD | KS.3 |
| FIM1_1179 | 1407.126822 | 888.6889052 | 940.2419267 | -0.647488906 | 0.100027746 | 5.3421653251 | 4.9677144846 | ns | YPD | KS.3 |
| FIM1_1180 | 766.4608242 | 774.67973 | 1300.328035 | 0.015089866 | 0.147680322 | 0.916412191 | 0.932341646 | ns | YPD | KS.3 |
| FIM1_1181 | 1731.598831 | 2097.935806 | 2233.461848 | 0.255529876 | 0.18315722 | 0.146106417 | 0.189439516 | ns | YPD | KS.3 |
| FIM1_1182 | 61.81069173 | 139.8836894 | 106.0153218 | 1.094770223 | 0.195457522 | 3.7402640032 | 2.3785768916 | Km>KS.3 | YPD | KS.3 |
| FIM1_1183 | 625.2742709 | 700.5234003 | 1349.595068 | 0.152431317 | 0.174216015 | 0.363933561 | 0.426801816 | ns | YPD | KS.3 |
| FIM1_1184 | 701.3800414 | 448.4638348 | 592.4635588 | -0.610929225 | 0.155432025 | 5.0307521020 | 0.000135874 | ns | YPD | KS.3 |
| FIM1_1186 | 807.5042694 | 819.7537695 | 717.6328512 | 0.021472546 | 0.076932508 | 0.778506127 | 0.814696673 | ns | YPD | KS.3 |
| FIM1_1187 | 7187.192033 | 6424.696732 | 6179.320756 | -0.16026121 | 0.058576026 | 0.005985377 | 0.010593117 | ns | YPD | KS.3 |
| FIM1_1188 | 412.0570733 | 1021.493407 | 1370.245573 | 1.200295353 | 0.22350897 | 9.3409022167 | 5.5060412948 | Km>KS.3 | YPD | KS.3 |
| FIM1_1189 | 678.2192272 | 1679.572113 | 1684.525144 | 1.158788875 | 0.263668397 | 1.5167566680 | 5.7426669889 | Km>KS.3 | YPD | KS.3 |
| FIM1_1190 | 234.4815828 | 543.8240486 | 919.1149451 | 1.071780018 | 0.260784727 | 7.3285515959 | 2.3845352642 | Km>KS.3 | YPD | KS.3 |
| FIM1_1191 | 166.2250922 | 32.33762597 | 87.82668476 | -1.835337605 | 0.541463415 | 6.5834158252 | 2.1676038206 | Km<KS.3 | YPD | KS.3 |

| | | | | | | | | | | |
|---|---|---|---|---|---|---|---|---|---|---|
| FIM1_1192 | 1122.873488 | 704.4126549 | 1024.594112 | -0.625868449 | 0.180629462 | 0.000305059 | 0.000704908 | ns | YPD | KS.3 |
| FIM1_1193 | 4690.493818 | 4738.712618 | 4984.269302 | 0.014594689 | 0.065977883 | 0.823939839 | 0.853090676 | ns | YPD | KS.3 |
| FIM1_1194 | 3875.494561 | 2282.260658 | 3125.589537 | -0.741350347 | 0.11563411 | 6.6795817088 | 6.0862489014 | ns | YPD | KS.3 |
| FIM1_1195 | 347.3532805 | 178.8976419 | 301.4078431 | -0.906776125 | 0.162414892 | 7.3714416621 | 4.4131073729 | ns | YPD | KS.3 |
| FIM1_1196 | 11825.95505 | 11123.96933 | 16415.82363 | -0.084936639 | 0.124933616 | 0.488174283 | 0.551468451 | ns | YPD | KS.3 |
| FIM1_1197 | 4230.710678 | 3166.673232 | 4171.618958 | -0.403230845 | 0.121665673 | 0.000731007 | 0.0015555 | ns | YPD | KS.3 |
| FIM1_1198 | 928.9459838 | 1797.106754 | 1483.304579 | 0.877684954 | 0.199355439 | 3.5783644154 | 1.2457916701 | ns | YPD | KS.3 |
| FIM1_1199 | 1521.704077 | 1785.087391 | 1383.38834 | 0.224120516 | 0.105890998 | 0.031794108 | 0.048033917 | ns | YPD | KS.3 |
| FIM1_1200 | 2745.455697 | 6297.055383 | 4476.768721 | 1.095839956 | 0.219344549 | 1.0191926007 | 4.9323766426 | Km>KS.3 | YPD | KS.3 |
| FIM1_1201 | 120.5915878 | 189.881163 | 205.2568071 | 0.615236339 | 0.162912317 | 9.3579737461 | 0.000242858 | ns | YPD | KS.3 |
| FIM1_1202 | 166.4012511 | 373.2065154 | 329.0836406 | 1.051075811 | 0.233599567 | 1.3678550005 | 5.2146196380 | Km>KS.3 | YPD | KS.3 |
| FIM1_1203 | 896.7669153 | 997.0835861 | 1165.203984 | 0.128563534 | 0.271309855 | 0.605020293 | 0.661999825 | ns | YPD | KS.3 |
| FIM1_1204 | 2611.907749 | 5619.88014 | 4190.507741 | 1.049934663 | 0.162631759 | 2.1926549917 | 2.2037290068 | Km>KS.3 | YPD | KS.3 |
| FIM1_1205 | 622.4955847 | 847.5156794 | 847.5103943 | 0.42764329 | 0.128177442 | 0.000656514 | 0.001412086 | ns | YPD | KS.3 |
| FIM1_1206 | 843.6556529 | 760.959774 | 779.5023305 | -0.14358465 | 0.114268241 | 0.201463233 | 0.252781737 | ns | YPD | KS.3 |
| FIM1_1207 | 2116.634758 | 2392.39458 | 1871.703589 | 0.171427558 | 0.109212602 | 0.111034495 | 0.14769963 | ns | YPD | KS.3 |
| FIM1_1208 | 2782.41133 | 1532.599608 | 1630.311595 | -0.836472676 | 0.113493771 | 6.3162794530 | 1.0336674433 | ns | YPD | KS.3 |
| FIM1_1209 | 1213.396704 | 1044.870339 | 1031.406901 | -0.211349285 | 0.090833013 | 0.018686883 | 0.029893004 | ns | YPD | KS.3 |
| FIM1_1210 | 445.5020212 | 267.9653341 | 392.2183859 | -0.66854882 | 0.206168358 | 0.000625379 | 0.001352134 | ns | YPD | KS.3 |
| FIM1_1211 | 1367.176691 | 1354.579879 | 1425.858515 | -0.012441445 | 0.148313002 | 0.931320515 | 0.943877404 | ns | YPD | KS.3 |
| FIM1_1212 | 6919.387802 | 7656.428397 | 9933.849973 | 0.141698966 | 0.109039951 | 0.18710168 | 0.237275263 | ns | YPD | KS.3 |
| FIM1_1213 | 5641.86612 | 5972.353906 | 8591.439274 | 0.078370356 | 0.137043201 | 0.558340191 | 0.619064509 | ns | YPD | KS.3 |
| FIM1_1214 | 13671.78042 | 12635.3918 | 18090.46764 | -0.111127543 | 0.09568863 | 0.239989319 | 0.294874503 | ns | YPD | KS.3 |
| FIM1_1215 | 829.221339 | 1062.696412 | 930.5813569 | 0.350016434 | 0.093567151 | 0.000154453 | 0.000384009 | ns | YPD | KS.3 |
| FIM1_1216 | 894.6434099 | 1396.422909 | 932.4318863 | 0.607619704 | 0.156781983 | 6.3473564044 | 0.000168957 | ns | YPD | KS.3 |
| FIM1_1217 | 2683.238008 | 1901.317831 | 3883.21751 | -0.410708366 | 0.289545573 | 0.116807101 | 0.15467536 | ns | YPD | KS.3 |
| FIM1_1218 | 644.5156753 | 285.2306018 | 430.1686544 | -1.123536484 | 0.153759266 | 4.2997674982 | 7.2513028148 | Km<KS.3 | YPD | KS.3 |
| FIM1_1219 | 1483.226152 | 1404.16608 | 1348.642288 | -0.076878373 | 0.104646412 | 0.456328913 | 0.520219143 | ns | YPD | KS.3 |
| FIM1_1220 | 795.5200318 | 623.0038862 | 623.5037668 | -0.320690721 | 0.20068498 | 0.093291831 | 0.126018696 | ns | YPD | KS.3 |
| FIM1_1221 | 970.4840951 | 764.3920827 | 911.9478004 | -0.338002464 | 0.087222385 | 9.0754427602 | 0.000236265 | ns | YPD | KS.3 |
| FIM1_1222 | 6391.905284 | 6059.001404 | 5569.113365 | -0.076509384 | 0.056526938 | 0.174046248 | 0.222820403 | ns | YPD | KS.3 |
| FIM1_1223 | 459.1625833 | 593.9933444 | 358.2853896 | 0.350076716 | 0.15901603 | 0.023097754 | 0.036056268 | ns | YPD | KS.3 |
| FIM1_1224 | 384.9917967 | 218.0796891 | 283.1675351 | -0.783848344 | 0.140470368 | 1.0152381663 | 5.9281806072 | ns | YPD | KS.3 |
| FIM1_1225 | 2567.894029 | 3137.534234 | 2384.065711 | 0.273471644 | 0.151533998 | 0.06333818 | 0.089215018 | ns | YPD | KS.3 |
| FIM1_1227 | 2689.21384 | 7097.683232 | 4298.800342 | 1.307880614 | 0.202488236 | 7.9573403536 | 8.7006084086 | Km>KS.3 | YPD | KS.3 |
| FIM1_1228 | 1378.181652 | 1488.105213 | 1288.517643 | 0.104534718 | 0.153093795 | 0.482217901 | 0.545607018 | ns | YPD | KS.3 |
| FIM1_1230 | 2947.812307 | 1569.665654 | 2123.311934 | -0.864103156 | 0.155693045 | 9.9589194906 | 5.8288969959 | ns | YPD | KS.3 |
| FIM1_1231 | 3620.697446 | 3233.50911 | 3724.980557 | -0.152664614 | 0.165980928 | 0.341471211 | 0.403232679 | ns | YPD | KS.3 |
| FIM1_1232 | 3611.880341 | 1924.099179 | 3261.024632 | -0.863047567 | 0.156532112 | 1.2295593424 | 7.0635770537 | ns | YPD | KS.3 |
| FIM1_1233 | 1301.167479 | 657.4507141 | 1200.732231 | -0.900815998 | 0.210853088 | 6.0741119642 | 2.0199670469 | ns | YPD | KS.3 |
| FIM1_1234 | 8329.546694 | 7197.889622 | 8850.688557 | -0.205558223 | 0.099332738 | 0.036108734 | 0.053897676 | ns | YPD | KS.3 |
| FIM1_1235 | 1518.936028 | 1880.940377 | 2019.227918 | 0.296718787 | 0.126160986 | 0.016411281 | 0.026612165 | ns | YPD | KS.3 |
| FIM1_1236 | 286.7632124 | 325.2497613 | 389.0121877 | 0.169072364 | 0.170250784 | 0.30355459 | 0.363765 | ns | YPD | KS.3 |
| FIM1_1239 | 2585.47437 | 1971.499154 | 2726.193429 | -0.35822441 | 0.193563452 | 0.052679044 | 0.075548643 | ns | YPD | KS.3 |
| FIM1_1240 | 297.1364843 | 247.3971783 | 277.3851919 | -0.249386515 | 0.158977128 | 0.105588738 | 0.141286705 | ns | YPD | KS.3 |

| | | | | | | | | | | |
|---|---|---|---|---|---|---|---|---|---|---|
| FIM1_1242 | 913.3217639 | 775.9131653 | 774.0457167 | -0.224819745 | 0.135367434 | 0.089280248 | 0.121220411 | ns | YPD | KS.3 |
| FIM1_1243 | 3922.359701 | 3440.680707 | 3450.124753 | -0.186653964 | 0.065709139 | 0.004277127 | 0.007794398 | ns | YPD | KS.3 |
| FIM1_1244 | 1267.306823 | 656.7455025 | 884.1417876 | -0.921433311 | 0.115391017 | 4.15236810294 | 9.83715776768 | ns | YPD | KS.3 |
| FIM1_1245 | 1707.907016 | 1479.981128 | 1256.399244 | -0.197329744 | 0.134657183 | 0.133900594 | 0.175257947 | ns | YPD | KS.3 |
| FIM1_1246 | 645.6862649 | 558.3004965 | 571.5752836 | -0.204985759 | 0.098287946 | 0.034725434 | 0.052035854 | ns | YPD | KS.3 |
| FIM1_1247 | 670.566064 | 673.4948328 | 887.4453676 | 0.0061429 | 0.133749193 | 0.962547267 | 0.97113621 | ns | YPD | KS.3 |
| FIM1_1248 | 1134.080034 | 1145.711231 | 1548.077761 | 0.013649218 | 0.144359964 | 0.922721325 | 0.937227152 | ns | YPD | KS.3 |
| FIM1_1249 | 1602.221569 | 6120.392181 | 4249.76047 | 1.730934129 | 0.316537887 | 4.7310895445 | 3.6090802520 | Km>KS.3 | YPD | KS.3 |
| FIM1_1250 | 1542.058054 | 4480.674867 | 2503.527131 | 1.469483241 | 0.173869644 | 9.4036460057 | 3.7426511102 | Km>KS.3 | YPD | KS.3 |
| FIM1_1251 | 3026.540012 | 4745.145714 | 3902.077065 | 0.630264558 | 0.111755567 | 9.8000301317 | 5.7426560548 | ns | YPD | KS.3 |
| FIM1_1252 | 1611.47906 | 1355.068399 | 1817.874988 | -0.237093327 | 0.149497747 | 0.103011678 | 0.138098383 | ns | YPD | KS.3 |
| FIM1_1253 | 469.2038601 | 435.9617121 | 403.9005518 | -0.104007457 | 0.09527002 | 0.269398146 | 0.327131016 | ns | YPD | KS.3 |
| FIM1_1254 | 1633.626079 | 825.837172 | 1261.360215 | -0.913460136 | 0.192305364 | 6.2078346503 | 2.5294002772 | ns | YPD | KS.3 |
| FIM1_1255 | 2514.754493 | 1290.056774 | 2162.44669 | -0.807219977 | 0.29521787 | 0.002346601 | 0.004503988 | ns | YPD | KS.3 |
| FIM1_1256 | 2247.428694 | 1512.463762 | 1269.138431 | -0.552307588 | 0.119720728 | 2.5949854485 | 9.2744630792 | ns | YPD | KS.3 |
| FIM1_1257 | 1453.02184 | 2246.546368 | 1885.744787 | 0.602395388 | 0.136152904 | 5.8377498893 | 1.9544283781 | ns | YPD | KS.3 |
| FIM1_1258 | 1419.965528 | 698.5771339 | 1152.889444 | -0.990151821 | 0.126682838 | 1.3274244548 | 2.8465244236 | ns | YPD | KS.3 |
| FIM1_1259 | 387.1735796 | 337.8353178 | 347.9151025 | -0.179121691 | 0.195759366 | 0.337855248 | 0.399531699 | ns | YPD | KS.3 |
| FIM1_1261 | 2775.642293 | 1736.280064 | 2347.620967 | -0.628711385 | 0.182206806 | 0.00032007 | 0.000734478 | ns | YPD | KS.3 |
| FIM1_1262 | 1744.616743 | 1075.770197 | 2501.382779 | -0.635982056 | 0.204717495 | 0.00106084 | 0.002187186 | ns | YPD | KS.3 |
| FIM1_1263 | 970.6672561 | 1872.148613 | 2066.89919 | 0.894587522 | 0.16653699 | 2.5130603778 | 1.3537810067 | ns | YPD | KS.3 |
| FIM1_1264 | 2907.059599 | 2052.606074 | 2953.260636 | -0.461721637 | 0.190952315 | 0.011421135 | 0.019105631 | ns | YPD | KS.3 |
| FIM1_1265 | 3703.054513 | 6233.538021 | 5046.128415 | 0.704450046 | 0.171474727 | 1.9875437424 | 5.8997793072 | ns | YPD | KS.3 |
| FIM1_1266 | 849.8522406 | 563.6415863 | 805.668196 | -0.548901744 | 0.183978801 | 0.001854002 | 0.003642836 | ns | YPD | KS.3 |
| FIM1_1267 | 3151.231051 | 2265.453488 | 4172.094383 | -0.39330572 | 0.289386427 | 0.132991345 | 0.174205356 | ns | YPD | KS.3 |
| FIM1_1268 | 946.7760617 | 641.3556191 | 779.0367007 | -0.535998579 | 0.143401188 | 0.000124294 | 0.000313729 | ns | YPD | KS.3 |
| FIM1_1269 | 2959.479493 | 1907.793732 | 3666.06524 | -0.519296824 | 0.299369831 | 0.053363292 | 0.076419797 | ns | YPD | KS.3 |
| FIM1_1270 | 2265.250533 | 1650.587107 | 2826.111165 | -0.439598233 | 0.126965586 | 0.000408436 | 0.000919027 | ns | YPD | KS.3 |
| FIM1_1271 | 4550.247611 | 3886.390628 | 3309.465527 | -0.219533443 | 0.119488871 | 0.061395842 | 0.086650869 | ns | YPD | KS.3 |
| FIM1_1272 | 5472.302847 | 6677.542958 | 7662.356285 | 0.279772931 | 0.101899309 | 0.005394665 | 0.009612629 | ns | YPD | KS.3 |
| FIM1_1273 | 533.6810147 | 697.2375908 | 549.5565453 | 0.358455582 | 0.175581034 | 0.03394862 | 0.051056343 | ns | YPD | KS.3 |
| FIM1_1274 | 60.61819158 | 130.8949539 | 79.67623041 | 0.978243107 | 0.255518455 | 3.1951269953 | 9.0214283778 | ns | YPD | KS.3 |
| FIM1_1275 | 2715.273601 | 3038.496142 | 2704.232452 | 0.155548944 | 0.128807703 | 0.217629359 | 0.270541245 | ns | YPD | KS.3 |
| FIM1_1276 | 2754.924507 | 3043.997715 | 3207.325652 | 0.14332769 | 0.045123449 | 0.001446268 | 0.002904797 | ns | YPD | KS.3 |
| FIM1_1277 | 2373.349649 | 3634.235718 | 2437.463365 | 0.574832915 | 0.172258252 | 0.000531116 | 0.001162473 | ns | YPD | KS.3 |
| FIM1_1279 | 1286.355758 | 965.5138168 | 1120.380077 | -0.398783907 | 0.123214236 | 0.000967062 | 0.002005474 | ns | YPD | KS.3 |
| FIM1_1280 | 5725.834144 | 3239.839338 | 7070.403574 | -0.797900607 | 0.114920398 | 1.5624207654 | 1.9728536314 | ns | YPD | KS.3 |
| FIM1_1281 | 2323.196145 | 1793.825188 | 2186.258833 | -0.36216615 | 0.11094996 | 0.000912142 | 0.001906684 | ns | YPD | KS.3 |
| FIM1_1282 | 530.6293084 | 1100.088734 | 725.1315443 | 0.89501735 | 0.286835763 | 0.000550233 | 0.0012022 | ns | YPD | KS.3 |
| FIM1_1283 | 2635.883699 | 2651.79006 | 3946.865104 | 0.007324617 | 0.270831782 | 0.976461225 | 0.981194626 | ns | YPD | KS.3 |
| FIM1_1284 | 514.2759835 | 2134.965906 | 937.7489032 | 1.944387404 | 0.238478921 | 1.2131995913 | 4.6428215131 | Km>KS.3 | YPD | KS.3 |
| FIM1_1285 | 5276.788164 | 4900.266828 | 4577.267726 | -0.103943196 | 0.101991675 | 0.301749706 | 0.362347282 | ns | YPD | KS.3 |
| FIM1_1286 | 541.6346802 | 303.4331563 | 438.3162663 | -0.78604015 | 0.170449244 | 1.6768909780 | 6.2725809141 | ns | YPD | KS.3 |
| FIM1_1287 | 1780.499533 | 1646.936294 | 1377.703421 | -0.100083119 | 0.221292095 | 0.631524082 | 0.686289276 | ns | YPD | KS.3 |
| FIM1_1288 | 577.736409 | 793.2211736 | 608.8673932 | 0.437446335 | 0.135328136 | 0.000937248 | 0.00195015 | ns | YPD | KS.3 |

| | | | | | | | | | | |
|---|---|---|---|---|---|---|---|---|---|---|
| FIM1_1289 | 2580.437896 | 1681.904445 | 2209.892379 | -0.59942322 | 0.113692041 | 8.1750386627 | 4.0228305981 | ns | YPD | KS.3 |
| FIM1_1290 | 4373.840095 | 4700.945698 | 4253.796382 | 0.099616321 | 0.13276791 | 0.443140758 | 0.506942147 | ns | YPD | KS.3 |
| FIM1_1291 | 1464.434287 | 602.8824066 | 1057.93371 | -1.207573494 | 0.180853501 | 2.8074120870 | 3.3818099595 | Km<KS.3 | YPD | KS.3 |
| FIM1_1292 | 417.5112221 | 160.0333791 | 242.678051 | -1.312704823 | 0.175501762 | 5.4242332155 | 1.0751219221 | Km<KS.3 | YPD | KS.3 |
| FIM1_1293 | 1876.870449 | 1352.812503 | 2514.956573 | -0.421271123 | 0.223398686 | 0.045086235 | 0.065681997 | ns | YPD | KS.3 |
| FIM1_1294 | 1425.570686 | 671.2371137 | 1168.931978 | -0.919830958 | 0.293580305 | 0.000490535 | 0.001084626 | ns | YPD | KS.3 |
| FIM1_1295 | 1705.557 | 3929.596162 | 3331.000546 | 1.164547909 | 0.133047069 | 2.7659695442 | 1.2741387484 | Km>KS.3 | YPD | KS.3 |
| FIM1_1296 | 337.9947174 | 1753.079465 | 921.9107713 | 2.088937879 | 0.409312038 | 1.2164174297 | 8.5475659787 | Km>KS.3 | YPD | KS.3 |
| FIM1_1297 | 1798.463312 | 1314.506039 | 1960.959462 | -0.378305507 | 0.278798352 | 0.136395143 | 0.177914482 | ns | YPD | KS.3 |
| FIM1_1298 | 3237.194299 | 3110.58737 | 2757.110115 | -0.051439728 | 0.215888319 | 0.801077043 | 0.833931427 | ns | YPD | KS.3 |
| FIM1_1299 | 1437.966338 | 911.0939714 | 985.6162665 | -0.643728294 | 0.097524231 | 2.2914022025 | 2.2799451915 | ns | YPD | KS.3 |
| FIM1_1300 | 1810.34875 | 1334.122127 | 1455.409244 | -0.416722324 | 0.152686932 | 0.004948909 | 0.008904457 | ns | YPD | KS.3 |
| FIM1_1301 | 498.6003749 | 319.5256051 | 342.8589182 | -0.613348933 | 0.140312567 | 7.3156902259 | 2.3834681646 | ns | YPD | KS.3 |
| FIM1_1302 | 1045.016983 | 664.742357 | 736.6684656 | -0.632798959 | 0.112790379 | 1.1559654582 | 6.6920986325 | ns | YPD | KS.3 |
| FIM1_1303 | 1892.405078 | 1100.448583 | 2040.5456 | -0.71120551 | 0.210532467 | 0.000354512 | 0.000807553 | ns | YPD | KS.3 |
| FIM1_1304 | 922.9575948 | 2187.647447 | 1576.531809 | 1.084439356 | 0.276685531 | 1.5596147126 | 4.7081815505 | Km>KS.3 | YPD | KS.3 |
| FIM1_1305 | 4180.732116 | 5321.763948 | 4427.895953 | 0.328937042 | 0.154277138 | 0.028078796 | 0.043048385 | ns | YPD | KS.3 |
| FIM1_1306 | 191.5302975 | 85.49931091 | 117.1812301 | -1.071694714 | 0.210769831 | 6.9704433300 | 3.4747450467 | Km<KS.3 | YPD | KS.3 |
| FIM1_1307 | 1724.176465 | 1146.455965 | 1081.113627 | -0.566319953 | 0.128470833 | 6.6763526090 | 2.1952977019 | ns | YPD | KS.3 |
| FIM1_1308 | 1025.56452 | 916.4519792 | 883.5704345 | -0.156633509 | 0.117531699 | 0.175086722 | 0.223921965 | ns | YPD | KS.3 |
| FIM1_1309 | 986.4368145 | 1230.276865 | 1078.806146 | 0.299904992 | 0.159144317 | 0.051837054 | 0.074448425 | ns | YPD | KS.3 |
| FIM1_1310 | 2188.897866 | 2067.574006 | 2115.586315 | -0.080047089 | 0.10536679 | 0.441034458 | 0.504900495 | ns | YPD | KS.3 |
| FIM1_1311 | 781.0036291 | 2573.370462 | 2063.53469 | 1.57186646 | 0.261459997 | 3.8874090021 | 3.6978699399 | Km>KS.3 | YPD | KS.3 |
| FIM1_1312 | 1422.061658 | 1635.035268 | 1879.694692 | 0.178944993 | 0.222970779 | 0.394278725 | 0.457391304 | ns | YPD | KS.3 |
| FIM1_1313 | 23042.85952 | 23926.43424 | 32699.65459 | 0.051376736 | 0.149605153 | 0.724100649 | 0.768758158 | ns | YPD | KS.3 |
| FIM1_1314 | 1147.907888 | 886.6269292 | 895.8018927 | -0.336737745 | 0.208484104 | 0.088604989 | 0.120341201 | ns | YPD | KS.3 |
| FIM1_1315 | 3508.19784 | 2336.143938 | 3495.315727 | -0.547767185 | 0.173584991 | 0.001046881 | 0.002160196 | ns | YPD | KS.3 |
| FIM1_1316 | 1117.648627 | 769.4184352 | 918.0455375 | -0.489316992 | 0.205587175 | 0.01216381 | 0.020225586 | ns | YPD | KS.3 |
| FIM1_1317 | 488.1682046 | 544.4980068 | 631.2855543 | 0.144605076 | 0.182733415 | 0.409906231 | 0.473921333 | ns | YPD | KS.3 |
| FIM1_1318 | 1577.427876 | 940.6607147 | 1423.226802 | -0.715462497 | 0.136917259 | 8.4998439287 | 4.1744050883 | ns | YPD | KS.3 |
| FIM1_1319 | 2131.921854 | 6828.920029 | 3635.495891 | 1.555105609 | 0.237674274 | 1.3558872125 | 1.7296253545 | Km>KS.3 | YPD | KS.3 |
| FIM1_1320 | 10292.3668 | 14453.63088 | 13061.52587 | 0.445174926 | 0.203890371 | 0.02157397 | 0.03390713 | ns | YPD | KS.3 |
| FIM1_1321 | 967.1061533 | 762.3547574 | 788.6044429 | -0.331335764 | 0.11970522 | 0.004819315 | 0.008683843 | ns | YPD | KS.3 |
| FIM1_1322 | 7683.592779 | 7007.529018 | 7929.863146 | -0.118168953 | 0.221815409 | 0.572020138 | 0.631417836 | ns | YPD | KS.3 |
| FIM1_1323 | 674.1633095 | 794.3875707 | 817.2490221 | 0.220438216 | 0.174427209 | 0.189640427 | 0.240005374 | ns | YPD | KS.3 |
| FIM1_1324 | 1295.456056 | 508.5758566 | 1164.683031 | -1.185938658 | 0.274852378 | 1.9653214710 | 7.2533192273 | Km<KS.3 | YPD | KS.3 |
| FIM1_1325 | 1929.214777 | 2131.455358 | 1963.284815 | 0.135903181 | 0.153267769 | 0.361417881 | 0.423969336 | ns | YPD | KS.3 |
| FIM1_1326 | 1076.266358 | 1033.569224 | 1051.740856 | -0.056329258 | 0.099196021 | 0.565286262 | 0.625682279 | ns | YPD | KS.3 |
| FIM1_1327 | 2274.023496 | 2499.824871 | 2187.925136 | 0.132927803 | 0.10616495 | 0.204049776 | 0.255704695 | ns | YPD | KS.3 |
| FIM1_1328 | 1712.864094 | 2155.105142 | 1573.607562 | 0.327711159 | 0.070106217 | 2.5341031956 | 9.0960774880 | ns | YPD | KS.3 |
| FIM1_1329 | 773.6028497 | 2420.946964 | 1818.801545 | 1.516002255 | 0.242414177 | 1.1134972879 | 1.1836856854 | Km>KS.3 | YPD | KS.3 |
| FIM1_1331 | 233.3586625 | 235.6799649 | 153.9760707 | 0.012419078 | 0.274757878 | 0.960570894 | 0.969929003 | ns | YPD | KS.3 |
| FIM1_1332 | 575.3646452 | 478.1535624 | 565.8234931 | -0.25610257 | 0.129319841 | 0.043136008 | 0.063230714 | ns | YPD | KS.3 |
| FIM1_1333 | 6040.162673 | 4842.143422 | 7441.272709 | -0.284099685 | 0.222069545 | 0.174543757 | 0.223399844 | ns | YPD | KS.3 |
| FIM1_1334 | 2205.998631 | 2665.429511 | 3113.798748 | 0.24769547 | 0.202067597 | 0.197765408 | 0.248958225 | ns | YPD | KS.3 |

| | | | | | | | | | | |
|---|---|---|---|---|---|---|---|---|---|---|
| FIM1_1335 | 661.7281084 | 842.8971052 | 988.3723608 | 0.325226716 | 0.172254925 | 0.050188059 | 0.072351665 | ns | YPD | KS.3 |
| FIM1_1336 | 442.581293 | 2909.251341 | 1587.422816 | 2.582095792 | 0.294869036 | 9.9056716077 | 5.7303158428 | Km>KS.3 | YPD | KS.3 |
| FIM1_1337 | 2491.07232 | 2512.804267 | 3774.680074 | 0.010216086 | 0.291113704 | 0.969047073 | 0.975715278 | ns | YPD | KS.3 |
| FIM1_1338 | 1021.157096 | 858.1478999 | 857.7782986 | -0.228817027 | 0.195645188 | 0.220535744 | 0.27381216 | ns | YPD | KS.3 |
| FIM1_1339 | 4044.511596 | 6907.846454 | 5129.626516 | 0.716587005 | 0.185749924 | 5.5458000796 | 0.000148735 | ns | YPD | KS.3 |
| FIM1_1340 | 1928.56637 | 2726.351042 | 2228.020992 | 0.481968826 | 0.121345848 | 5.1609837824 | 0.000138939 | ns | YPD | KS.3 |
| FIM1_1341 | 882.5457501 | 664.164609 | 703.1858564 | -0.364814585 | 0.224759926 | 0.084421212 | 0.11528837 | ns | YPD | KS.3 |
| FIM1_1342 | 2569.518063 | 2575.766137 | 2388.158002 | 0.003219602 | 0.152061142 | 0.982622039 | 0.985349319 | ns | YPD | KS.3 |
| FIM1_1344 | 425.7185535 | 547.7786261 | 431.1494487 | 0.353420556 | 0.113824923 | 0.001594374 | 0.003174073 | ns | YPD | KS.3 |
| FIM1_1345 | 1249.504027 | 1479.883686 | 1529.858814 | 0.234715715 | 0.126153771 | 0.057681373 | 0.081779661 | ns | YPD | KS.3 |
| FIM1_1346 | 7525.736819 | 5590.814132 | 8008.702261 | -0.414457449 | 0.118825829 | 0.000382591 | 0.000865964 | ns | YPD | KS.3 |
| FIM1_1347 | 11488.99201 | 10100.32285 | 16702.93723 | -0.177513248 | 0.136879899 | 0.184378096 | 0.234000262 | ns | YPD | KS.3 |
| FIM1_1348 | 11061.5693 | 9616.969181 | 10458.08256 | -0.198163455 | 0.085027174 | 0.018644141 | 0.029843823 | ns | YPD | KS.3 |
| FIM1_1349 | 1839.77062 | 1072.167116 | 1253.590956 | -0.764350925 | 0.09108464 | 2.0880479019 | 6.6589989181 | ns | YPD | KS.3 |
| FIM1_1350 | 7247.880015 | 5760.463458 | 8388.563378 | -0.310437669 | 0.165525362 | 0.052380374 | 0.075150305 | ns | YPD | KS.3 |
| FIM1_1352 | 2227.192534 | 1460.389024 | 2375.419536 | -0.595587853 | 0.096711968 | 4.4814360872 | 3.4247533846 | ns | YPD | KS.3 |
| FIM1_1353 | 7447.249349 | 7334.052266 | 8281.703894 | -0.021542293 | 0.095765562 | 0.81996201 | 0.849856459 | ns | YPD | KS.3 |
| FIM1_1354 | 1432.121694 | 856.9420444 | 1056.633872 | -0.703239144 | 0.153418623 | 2.2881776627 | 8.2850683202 | ns | YPD | KS.3 |
| FIM1_1355 | 1796.873875 | 2137.621401 | 1650.334581 | 0.24832106 | 0.06242021 | 6.3585579661 | 0.000169165 | ns | YPD | KS.3 |
| FIM1_1356 | 1079.305354 | 651.4366472 | 718.3923749 | -0.709363105 | 0.107814975 | 2.3319269103 | 2.3156360038 | ns | YPD | KS.3 |
| FIM1_1357 | 1138.1371 | 992.6013116 | 1025.871029 | -0.186230046 | 0.15365657 | 0.212126348 | 0.264692395 | ns | YPD | KS.3 |
| FIM1_1358 | 1364.39352 | 1239.521947 | 1315.886861 | -0.133798131 | 0.122922194 | 0.267323243 | 0.32502911 | ns | YPD | KS.3 |
| FIM1_1359 | 367.8306686 | 686.9597701 | 495.7212686 | 0.863293184 | 0.141108836 | 3.3092002648 | 2.6090762785 | ns | YPD | KS.3 |
| FIM1_1360 | 3515.190385 | 5355.187198 | 2675.62504 | 0.537956923 | 0.23291357 | 0.013572061 | 0.022343151 | ns | YPD | KS.3 |
| FIM1_1361 | 338.3769228 | 181.7308694 | 279.5368154 | -0.848146793 | 0.163671889 | 7.9642993244 | 3.9268968423 | ns | YPD | KS.3 |
| FIM1_1362 | 468.6823264 | 320.3009724 | 484.6343487 | -0.519901225 | 0.152484157 | 0.000445133 | 0.000993066 | ns | YPD | KS.3 |
| FIM1_1363 | 4021.549977 | 1881.524005 | 3829.713585 | -0.936330998 | 0.285807713 | 0.000287278 | 0.000668167 | ns | YPD | KS.3 |
| FIM1_1364 | 429.281192 | 103.7442573 | 299.4874911 | -1.923631335 | 0.255424785 | 1.9563654235 | 4.9912399909 | Km<KS.3 | YPD | KS.3 |
| FIM1_1365 | 2522.443156 | 2922.653636 | 2515.753909 | 0.199833425 | 0.159500746 | 0.196133107 | 0.247278816 | ns | YPD | KS.3 |
| FIM1_1366 | 1774.412238 | 1050.396742 | 1390.123103 | -0.721819571 | 0.145655477 | 3.4815884882 | 1.4906112503 | ns | YPD | KS.3 |
| FIM1_1367 | 1250.712953 | 1253.485868 | 982.0969972 | 0.003075638 | 0.089609329 | 0.972336465 | 0.978433235 | ns | YPD | KS.3 |
| FIM1_1368 | 5470.21746 | 5875.837229 | 5774.293505 | 0.100926021 | 0.09465792 | 0.280808965 | 0.33949565 | ns | YPD | KS.3 |
| FIM1_1369 | 2336.392674 | 3108.424554 | 3750.884313 | 0.378293436 | 0.191023204 | 0.038352141 | 0.056938795 | ns | YPD | KS.3 |
| FIM1_1370 | 21.98806633 | 35.33106556 | 34.13613517 | 0.565294602 | 0.29451168 | 0.033077636 | 0.049851935 | ns | YPD | KS.3 |
| FIM1_1371 | 788.4718036 | 1313.561191 | 1244.944897 | 0.717728679 | 0.10498216 | 3.9311204457 | 4.5680883630 | ns | YPD | KS.3 |
| FIM1_1372 | 119.7498708 | 125.0838763 | 292.5230999 | 0.057139846 | 0.177286938 | 0.737758211 | 0.780231037 | ns | YPD | KS.3 |
| FIM1_1373 | 1417.024694 | 3060.782682 | 2413.523477 | 1.0834939 | 0.112631209 | 1.1652438319 | 7.8339027887 | Km>KS.3 | YPD | KS.3 |
| FIM1_1374 | 2727.646717 | 1681.672351 | 2208.060344 | -0.681112395 | 0.101745715 | 1.1313303421 | 1.1949826862 | ns | YPD | KS.3 |
| FIM1_1375 | 914.1432242 | 2595.274454 | 1570.463931 | 1.291238377 | 0.313255902 | 2.9912623151 | 1.0569268478 | Km>KS.3 | YPD | KS.3 |
| FIM1_1376 | 114.1564815 | 75.9635786 | 96.17648643 | -0.549586152 | 0.169557999 | 0.000776474 | 0.001644513 | ns | YPD | KS.3 |
| FIM1_1377 | 3848.686378 | 4955.407212 | 3826.541247 | 0.3544097 | 0.107370211 | 0.000807448 | 0.001705032 | ns | YPD | KS.3 |
| FIM1_1378 | 358.5957323 | 199.2265679 | 321.4348399 | -0.797132925 | 0.167778461 | 8.2819355746 | 3.3041402954 | ns | YPD | KS.3 |
| FIM1_1379 | 234.4312101 | 450.8992744 | 272.0880493 | 0.85762351 | 0.216370972 | 2.5807900887 | 7.4604478161 | ns | YPD | KS.3 |
| FIM1_1380 | 1393.893767 | 3376.037633 | 2315.659662 | 1.131730005 | 0.260251847 | 2.0695045647 | 7.5593136636 | Km>KS.3 | YPD | KS.3 |
| FIM1_1382 | 4389.331863 | 4493.554897 | 4035.366892 | 0.03366742 | 0.06480119 | 0.601362069 | 0.658401473 | ns | YPD | KS.3 |

| | | | | | | | | | | |
|---|---|---|---|---|---|---|---|---|---|---|
| FIM1_1383 | 2626.140354 | 2616.232453 | 2587.870545 | -0.005147598 | 0.119001416 | 0.964880898 | 0.972384357 | ns | YPD | KS.3 |
| FIM1_1384 | 1334.564225 | 846.9307618 | 1200.455473 | -0.621373862 | 0.155489656 | 3.75214953935 | 0.000104168 | ns | YPD | KS.3 |
| FIM1_1385 | 3627.815886 | 5447.905737 | 5001.549149 | 0.546366657 | 0.17681562 | 0.001309736 | 0.002649831 | ns | YPD | KS.3 |
| FIM1_1386 | 532.6587463 | 770.6605851 | 571.1931895 | 0.510899592 | 0.131366874 | 6.99368357168 | 0.000184581 | ns | YPD | KS.3 |
| FIM1_1387 | 3208.312511 | 2829.414219 | 2965.730192 | -0.180272641 | 0.044660209 | 5.17306884549 | 0.000139189 | ns | YPD | KS.3 |
| FIM1_1388 | 499.4490419 | 403.292316 | 383.3514424 | -0.300845941 | 0.097092339 | 0.001709309 | 0.00337721 | ns | YPD | KS.3 |
| FIM1_1389 | 731.1554404 | 371.088103 | 608.2360905 | -0.949687381 | 0.117115092 | 1.39538879363 | 3.63458599389 | ns | YPD | KS.3 |
| FIM1_1390 | 2054.201351 | 3748.910583 | 2724.645374 | 0.797580974 | 0.200501602 | 2.82007105827 | 8.06777085390 | ns | YPD | KS.3 |
| FIM1_1391 | 1217.344259 | 4825.296269 | 3696.141313 | 1.70580042 | 0.376059736 | 7.32257990356 | 3.64298350202 | Km>KS.3 | YPD | KS.3 |
| FIM1_1392 | 548.8064761 | 1079.94891 | 1101.82239 | 0.915124307 | 0.178242974 | 8.63602314749 | 4.23710208666 | ns | YPD | KS.3 |
| FIM1_1393 | 832.5303388 | 931.5701392 | 1062.710524 | 0.14775784 | 0.196217942 | 0.430189908 | 0.49415719 | ns | YPD | KS.3 |
| FIM1_1394 | 195.0944476 | 340.4695958 | 587.753791 | 0.696689157 | 0.258276195 | 0.003408886 | 0.006356524 | ns | YPD | KS.3 |
| FIM1_1395 | 681.0968765 | 1176.459625 | 753.8510585 | 0.687613134 | 0.252968324 | 0.003269759 | 0.006122338 | ns | YPD | KS.3 |
| FIM1_1396 | 1233.791222 | 1786.528104 | 1338.63058 | 0.4990665 | 0.171199769 | 0.00249271 | 0.004766039 | ns | YPD | KS.3 |
| FIM1_1397 | 3400.971844 | 3882.900024 | 4469.632152 | 0.181908393 | 0.142323934 | 0.18999176 | 0.240341865 | ns | YPD | KS.3 |
| FIM1_1399 | 1305.119899 | 2462.868637 | 2805.973654 | 0.835956482 | 0.210513818 | 2.64431904473 | 7.60253967288 | ns | YPD | KS.3 |
| FIM1_1400 | 1243.906388 | 3534.900547 | 1902.072314 | 1.386765818 | 0.231333322 | 1.07700597655 | 9.41670427652 | Km>KS.3 | YPD | KS.3 |
| FIM1_1401 | 2453.83142 | 5396.174526 | 3542.249151 | 0.98171966 | 0.278402943 | 0.000101763 | 0.000262589 | ns | YPD | KS.3 |
| FIM1_1402 | 1008.480147 | 1144.134824 | 872.8792913 | 0.17837288 | 0.093920472 | 0.054752535 | 0.07809457 | ns | YPD | KS.3 |
| FIM1_1403 | 627.8746162 | 717.3107775 | 747.890367 | 0.181677437 | 0.152670444 | 0.220705861 | 0.273955005 | ns | YPD | KS.3 |
| FIM1_1404 | 389.582225 | 332.3171588 | 894.6827909 | -0.184315656 | 0.305382323 | 0.50014822 | 0.562440641 | ns | YPD | KS.3 |
| FIM1_1405 | 840.4931698 | 643.1197755 | 849.0081097 | -0.348313953 | 0.210605189 | 0.080898081 | 0.110933835 | ns | YPD | KS.3 |
| FIM1_1406 | 3361.339355 | 6962.713459 | 6026.309982 | 0.939322431 | 0.238962213 | 2.36462230743 | 6.88355528349 | ns | YPD | KS.3 |
| FIM1_1407 | 1499.356748 | 1731.920645 | 1369.341169 | 0.201715334 | 0.109942534 | 0.062451376 | 0.08804069 | ns | YPD | KS.3 |
| FIM1_1408 | 5748.357817 | 6241.622622 | 7942.031075 | 0.114254559 | 0.125608314 | 0.353567334 | 0.41613378 | ns | YPD | KS.3 |
| FIM1_1409 | 641.2968042 | 1295.216413 | 2107.260258 | 0.81202415 | 0.333126143 | 0.00530581 | 0.00946447 | ns | YPD | KS.3 |
| FIM1_1410 | 2400.722898 | 3541.416619 | 3987.427979 | 0.530419599 | 0.155342384 | 0.000430295 | 0.000964287 | ns | YPD | KS.3 |
| FIM1_1411 | 506.6779698 | 1259.428123 | 1205.529616 | 1.113793561 | 0.307113571 | 4.46455325318 | 0.000121878 | Km>KS.3 | YPD | KS.3 |
| FIM1_1412 | 2765.884672 | 7384.404686 | 4924.602873 | 1.308916299 | 0.219495366 | 1.86619978259 | 1.56039393586 | Km>KS.3 | YPD | KS.3 |
| FIM1_1413 | 501.1821606 | 1204.342792 | 623.2557519 | 1.079338219 | 0.297673359 | 5.06269699066 | 0.000136588 | Km>KS.3 | YPD | KS.3 |
| FIM1_1414 | 148.3182472 | 410.1761602 | 305.0673911 | 1.383635778 | 0.190516866 | 1.98966821948 | 3.62586058313 | Km>KS.3 | YPD | KS.3 |
| FIM1_1415 | 1564.644625 | 1832.39616 | 1607.211688 | 0.223352875 | 0.086453292 | 0.009088331 | 0.015532273 | ns | YPD | KS.3 |
| FIM1_1416 | 904.1914705 | 862.5141319 | 1016.078387 | -0.065849752 | 0.102525829 | 0.515115666 | 0.577705238 | ns | YPD | KS.3 |
| FIM1_1418 | 1181.50354 | 939.1457553 | 1019.213384 | -0.324279623 | 0.091022602 | 0.000316559 | 0.000727429 | ns | YPD | KS.3 |
| FIM1_1420 | 1075.434297 | 1358.970752 | 1437.044234 | 0.326662966 | 0.114730684 | 0.00378578 | 0.006975651 | ns | YPD | KS.3 |
| FIM1_1421 | 339.4843505 | 756.1436511 | 553.3328384 | 1.033649643 | 0.243116574 | 4.48147184471 | 1.53232456545 | Km>KS.3 | YPD | KS.3 |
| FIM1_1422 | 268.9923633 | 492.9650528 | 486.6103039 | 0.805265966 | 0.195917025 | 1.57782263076 | 4.75737429580 | ns | YPD | KS.3 |
| FIM1_1423 | 944.4515871 | 1157.055526 | 793.3963286 | 0.28835276 | 0.081373279 | 0.000350825 | 0.000799888 | ns | YPD | KS.3 |
| FIM1_1424 | 1143.891334 | 1502.0617 | 1449.15332 | 0.386948921 | 0.079827527 | 1.01544831460 | 3.99356155347 | ns | YPD | KS.3 |
| FIM1_1425 | 907.8462874 | 1576.460968 | 1331.415899 | 0.752762514 | 0.160784539 | 1.28542546937 | 4.93440718373 | ns | YPD | KS.3 |
| FIM1_1426 | 1285.28451 | 6542.187659 | 3731.823932 | 2.082101113 | 0.393339311 | 4.29756059498 | 3.29943888272 | Km>KS.3 | YPD | KS.3 |
| FIM1_1427 | 33.31758018 | 76.97271293 | 38.6833852 | 1.046316297 | 0.278246359 | 3.36759164169 | 9.43874277039 | Km>KS.3 | YPD | KS.3 |
| FIM1_1428 | 2133.2399 | 2646.29214 | 2119.551811 | 0.293130227 | 0.156543982 | 0.05358022 | 0.076708372 | ns | YPD | KS.3 |
| FIM1_1429 | 2075.029603 | 3228.336794 | 2233.881656 | 0.598268469 | 0.168058113 | 0.000224316 | 0.00053216 | ns | YPD | KS.3 |
| FIM1_1430 | 765.8209183 | 1807.370667 | 1121.095545 | 1.168143899 | 0.17927493 | 9.71062210745 | 1.04117122811 | Km>KS.3 | YPD | KS.3 |

| | | | | | | | | | | |
|---|---|---|---|---|---|---|---|---|---|---|
| FIM1_1431 | 1251.40883 | 3041.925174 | 2212.056116 | 1.203023774 | 0.188266776 | 1.9493445543 | 1.9791814607 | Km>KS.3 | YPD | KS.3 |
| FIM1_1432 | 13358.35269 | 10338.75058 | 13282.27271 | -0.355712877 | 0.126173092 | 0.004017207 | 0.007372041 | ns | YPD | KS.3 |
| FIM1_1433 | 2039.923306 | 1715.748684 | 1941.906229 | -0.243423736 | 0.097108234 | 0.011170063 | 0.018723404 | ns | YPD | KS.3 |
| FIM1_1434 | 1038.743215 | 2956.436626 | 1918.794881 | 1.356750621 | 0.262007704 | 1.3668287281 | 7.7892015148 | Km>KS.3 | YPD | KS.3 |
| FIM1_1435 | 3070.869211 | 6249.95298 | 5628.031343 | 0.967164451 | 0.170460867 | 3.6789150646 | 2.3434830277 | ns | YPD | KS.3 |
| FIM1_1436 | 463.2298582 | 468.4672376 | 560.7691604 | 0.015579004 | 0.153124264 | 0.916609142 | 0.932351356 | ns | YPD | KS.3 |
| FIM1_1437 | 378.7951656 | 321.0676007 | 339.2170924 | -0.23396721 | 0.095685523 | 0.01335471 | 0.022014473 | ns | YPD | KS.3 |
| FIM1_1438 | 34061.49552 | 55519.2895 | 60172.18324 | 0.690935832 | 0.092727076 | 4.7173171511 | 7.8490477681 | ns | YPD | KS.3 |
| FIM1_1439 | 641.9688329 | 2112.539123 | 1220.359772 | 1.676662936 | 0.135986899 | 5.4068720254 | 1.4157467540 | Km>KS.3 | YPD | KS.3 |
| FIM1_1440 | 420.254696 | 311.664688 | 401.8067455 | -0.41672632 | 0.115857959 | 0.00025258 | 0.000591055 | ns | YPD | KS.3 |
| FIM1_1441 | 118.6551321 | 399.8183899 | 322.5394479 | 1.53706616 | 0.31722977 | 3.3408415450 | 1.7569436244 | Km>KS.3 | YPD | KS.3 |
| FIM1_1442 | 59.58715754 | 470.3928111 | 578.1003268 | 2.817358646 | 0.324765697 | 3.7685316175 | 2.1065668311 | Km>KS.3 | YPD | KS.3 |
| FIM1_1443 | 560.0260141 | 453.470997 | 494.6692785 | -0.288749272 | 0.149258658 | 0.046714691 | 0.067855646 | ns | YPD | KS.3 |
| FIM1_1444 | 8184.805017 | 6613.78194 | 12306.94573 | -0.292274651 | 0.145044748 | 0.038583209 | 0.057264756 | ns | YPD | KS.3 |
| FIM1_1445 | 1088.646587 | 591.5716432 | 1243.352178 | -0.745764419 | 0.282199445 | 0.003574011 | 0.006632116 | ns | YPD | KS.3 |
| FIM1_1446 | 875.3596966 | 563.9829512 | 640.2136342 | -0.600175012 | 0.156646674 | 7.6983528412 | 0.000201788 | ns | YPD | KS.3 |
| FIM1_1447 | 726.0077819 | 485.0223442 | 788.7167144 | -0.521208711 | 0.222032781 | 0.012633027 | 0.020928842 | ns | YPD | KS.3 |
| FIM1_1448 | 656.3695515 | 392.8845089 | 525.5591259 | -0.680334443 | 0.197419638 | 0.000294773 | 0.000684319 | ns | YPD | KS.3 |
| FIM1_1449 | 2195.073997 | 1547.458318 | 2542.263048 | -0.488153518 | 0.11679836 | 2.0981553930 | 6.1875062717 | ns | YPD | KS.3 |
| FIM1_1450 | 769.6678242 | 1389.782904 | 920.6746159 | 0.815702896 | 0.14306824 | 4.6611927550 | 2.8950604190 | ns | YPD | KS.3 |
| FIM1_1451 | 7615.602131 | 2947.160846 | 8214.938459 | -1.224890007 | 0.257411156 | 2.0838511334 | 9.4161302351 | Km<KS.3 | YPD | KS.3 |
| FIM1_1453 | 3712.607614 | 4354.90309 | 3848.212792 | 0.19543469 | 0.263821413 | 0.420906096 | 0.485060882 | ns | YPD | KS.3 |
| FIM1_1454 | 1292.466374 | 1230.329771 | 1125.145479 | -0.067528642 | 0.138455163 | 0.617482487 | 0.673235892 | ns | YPD | KS.3 |
| FIM1_1455 | 783.931865 | 583.6127408 | 1192.667858 | -0.387107202 | 0.203082795 | 0.044991693 | 0.065601898 | ns | YPD | KS.3 |
| FIM1_1456 | 642.5751825 | 532.5604867 | 560.0456439 | -0.255807749 | 0.155098445 | 0.089301067 | 0.121220411 | ns | YPD | KS.3 |
| FIM1_1457 | 2411.407281 | 3777.455625 | 2947.262511 | 0.613141237 | 0.155173939 | 4.5959530245 | 0.000125081 | ns | YPD | KS.3 |
| FIM1_1458 | 1400.935633 | 1184.951096 | 1399.980782 | -0.231673193 | 0.13179431 | 0.072423531 | 0.100588238 | ns | YPD | KS.3 |
| FIM1_1460 | 1305.473867 | 1037.595707 | 1103.395768 | -0.324977588 | 0.087481889 | 0.000175266 | 0.000428475 | ns | YPD | KS.3 |
| FIM1_1461 | 1210.979784 | 332.5079706 | 986.9451075 | -1.67970796 | 0.29870947 | 2.4199573243 | 1.9576077542 | Km<KS.3 | YPD | KS.3 |
| FIM1_1462 | 1913.194812 | 1209.203293 | 1477.699355 | -0.644712497 | 0.107249154 | 1.0303069920 | 7.2912905907 | ns | YPD | KS.3 |
| FIM1_1463 | 310.5816232 | 202.2934062 | 239.1789131 | -0.575147396 | 0.180560756 | 0.000904369 | 0.001892026 | ns | YPD | KS.3 |
| FIM1_1464 | 37.59404128 | 32.90270711 | 24.69833755 | -0.123465028 | 0.425394105 | 0.718136094 | 0.764254687 | ns | YPD | KS.3 |
| FIM1_1465 | 1455.431737 | 1489.299716 | 1340.776731 | 0.032443949 | 0.109742389 | 0.764057855 | 0.802538646 | ns | YPD | KS.3 |
| FIM1_1466 | 1499.182721 | 1060.253206 | 1195.574456 | -0.485070588 | 0.110868158 | 8.7498763337 | 2.7850694024 | ns | YPD | KS.3 |
| FIM1_1467 | 2694.923198 | 3338.780189 | 2349.248935 | 0.292507819 | 0.151419136 | 0.046858995 | 0.068045388 | ns | YPD | KS.3 |
| FIM1_1468 | 3710.757707 | 3212.167366 | 3250.144257 | -0.202856189 | 0.100582038 | 0.041044234 | 0.060502241 | ns | YPD | KS.3 |
| FIM1_1469 | 177.5342668 | 89.07415526 | 198.5580107 | -0.912895132 | 0.20332118 | 2.1715724360 | 7.9089113246 | ns | YPD | KS.3 |
| FIM1_1470 | 1687.283319 | 2466.067832 | 1860.483098 | 0.539394987 | 0.076940899 | 1.5787896060 | 1.9857325300 | ns | YPD | KS.3 |
| FIM1_1471 | 4183.010665 | 2221.174352 | 4424.68561 | -0.883306891 | 0.125370589 | 6.0986123742 | 8.3124922087 | ns | YPD | KS.3 |
| FIM1_1472 | 922.7119785 | 426.005138 | 1066.927055 | -1.062130623 | 0.159046845 | 4.7349173157 | 5.4276989967 | Km<KS.3 | YPD | KS.3 |
| FIM1_1473 | 1365.436859 | 1066.394367 | 1273.656244 | -0.342067854 | 0.130696554 | 0.007486724 | 0.013036909 | ns | YPD | KS.3 |
| FIM1_1474 | 3463.515885 | 2763.558682 | 2614.068586 | -0.307479572 | 0.154795195 | 0.040724287 | 0.060155383 | ns | YPD | KS.3 |
| FIM1_1475 | 1994.549319 | 1570.098147 | 1427.26863 | -0.336872059 | 0.096936491 | 0.00043504 | 0.000973168 | ns | YPD | KS.3 |
| FIM1_1476 | 159.5581151 | 391.3180762 | 1083.108515 | 1.156298252 | 0.253949807 | 7.3305451996 | 2.9577828360 | Km>KS.3 | YPD | KS.3 |
| FIM1_1477 | 922.7766191 | 2472.412327 | 1843.773248 | 1.264596158 | 0.267408409 | 2.0462888993 | 9.2548066130 | Km>KS.3 | YPD | KS.3 |

| | | | | | | | | | | |
|---|---|---|---|---|---|---|---|---|---|---|
| FIM1_1478 | 2272.273963 | 2425.28037 | 1710.692423 | 0.081651937 | 0.243430416 | 0.718869229 | 0.764345889 | ns | YPD | KS.3 |
| FIM1_1479 | 1371.667346 | 693.716909 | 809.3494458 | -0.96537234 | 0.091368336 | 1.09715672027 | 1.18659884421 | ns | YPD | KS.3 |
| FIM1_1480 | 1047.414199 | 480.2560847 | 905.3788425 | -1.019610896 | 0.227588781 | 1.65841318084 | 6.20813060549 | Km<KS.3 | YPD | KS.3 |
| FIM1_1481 | 716.6578267 | 255.383519 | 507.7294437 | -1.409496298 | 0.185698843 | 1.46108985786 | 3.06705571430 | Km<KS.3 | YPD | KS.3 |
| FIM1_1482 | 3812.372711 | 9005.39149 | 6288.323436 | 1.195160996 | 0.141527574 | 3.61236820368 | 1.27457672435 | Km>KS.3 | YPD | KS.3 |
| FIM1_1483 | 589.7420728 | 396.4160975 | 398.8084871 | -0.520132168 | 0.20773745 | 0.00826993 | 0.014259783 | ns | YPD | KS.3 |
| FIM1_1484 | 1074.804987 | 833.7293311 | 742.8195055 | -0.350693381 | 0.134119245 | 0.00747694 | 0.013024431 | ns | YPD | KS.3 |
| FIM1_1485 | 3651.250553 | 2736.351041 | 3134.913706 | -0.409496584 | 0.079840877 | 2.30663932304 | 1.03104498042 | ns | YPD | KS.3 |
| FIM1_1486 | 1863.824431 | 1108.06897 | 2021.714419 | -0.687376465 | 0.200446808 | 0.000309251 | 0.00071327 | ns | YPD | KS.3 |
| FIM1_1487 | 2145.45868 | 791.0203591 | 1944.335533 | -1.259666624 | 0.286894155 | 1.03714457990 | 4.07245010657 | Km<KS.3 | YPD | KS.3 |
| FIM1_1488 | 5731.68418 | 5608.627759 | 5118.618575 | -0.030906156 | 0.075397144 | 0.679635581 | 0.730435735 | ns | YPD | KS.3 |
| FIM1_1489 | 617.9153384 | 335.6791845 | 496.6185416 | -0.838999309 | 0.150753063 | 9.70875372639 | 5.69752431524 | ns | YPD | KS.3 |
| FIM1_1490 | 657.107514 | 479.622507 | 542.9241269 | -0.428110507 | 0.15776389 | 0.005116376 | 0.0091594 | ns | YPD | KS.3 |
| FIM1_1491 | 818.4814841 | 974.002563 | 1008.296104 | 0.246357944 | 0.084198512 | 0.003145916 | 0.00591494 | ns | YPD | KS.3 |
| FIM1_1492 | 1737.380228 | 1251.442699 | 1549.742596 | -0.456153664 | 0.123902379 | 0.000173352 | 0.000424214 | ns | YPD | KS.3 |
| FIM1_1493 | 733.9840169 | 662.3934597 | 639.7551531 | -0.139228999 | 0.155432783 | 0.356158835 | 0.418688612 | ns | YPD | KS.3 |
| FIM1_1494 | 180.1995645 | 159.4121936 | 226.628268 | -0.166535822 | 0.162694707 | 0.290360089 | 0.350351777 | ns | YPD | KS.3 |
| FIM1_1495 | 87.56981328 | 82.42058248 | 72.26210851 | -0.074384878 | 0.266139319 | 0.761098187 | 0.800690099 | ns | YPD | KS.3 |
| FIM1_1496 | 926.5162384 | 982.636974 | 888.9900235 | 0.082602916 | 0.109951048 | 0.445577969 | 0.50901272 | ns | YPD | KS.3 |
| FIM1_1497 | 594.7501042 | 720.3091092 | 541.8235652 | 0.268239902 | 0.112281749 | 0.015198083 | 0.024806582 | ns | YPD | KS.3 |
| FIM1_1498 | 873.1329857 | 455.0543389 | 589.1756182 | -0.896163846 | 0.150275573 | 7.92122347581 | 5.77831184635 | ns | YPD | KS.3 |
| FIM1_1499 | 993.8665733 | 2087.691165 | 1990.863723 | 0.988333085 | 0.202231868 | 2.52144029401 | 1.11901565233 | ns | YPD | KS.3 |
| FIM1_1500 | 1457.801166 | 1798.108378 | 1542.250736 | 0.289380713 | 0.13484864 | 0.028130238 | 0.043113966 | ns | YPD | KS.3 |
| FIM1_1501 | 2118.000339 | 1106.003316 | 1400.018085 | -0.900009188 | 0.139467319 | 3.47171994056 | 3.32790109909 | ns | YPD | KS.3 |
| FIM1_1503 | 2149.381458 | 2410.901237 | 2153.953742 | 0.161776897 | 0.092100659 | 0.075768597 | 0.104882796 | ns | YPD | KS.3 |
| FIM1_1504 | 1364.29965 | 1048.932459 | 912.8558115 | -0.372084184 | 0.085139538 | 1.02264464968 | 3.20574246069 | ns | YPD | KS.3 |
| FIM1_1505 | 732.9687531 | 638.796214 | 530.282422 | -0.189678301 | 0.128592827 | 0.132110035 | 0.173187727 | ns | YPD | KS.3 |
| FIM1_1506 | 238.1365588 | 148.0755924 | 147.5683477 | -0.601415288 | 0.245266189 | 0.008294789 | 0.014288981 | ns | YPD | KS.3 |
| FIM1_1507 | 4943.60918 | 3856.32395 | 4989.294237 | -0.342928798 | 0.134671448 | 0.009179588 | 0.015677464 | ns | YPD | KS.3 |
| FIM1_1508 | 1933.23547 | 1846.235557 | 1713.87884 | -0.06407706 | 0.118915976 | 0.583373013 | 0.641955483 | ns | YPD | KS.3 |
| FIM1_1509 | 2152.680766 | 1134.462683 | 1673.109911 | -0.887388967 | 0.139320992 | 6.22171534975 | 5.70037456077 | ns | YPD | KS.3 |
| FIM1_1510 | 2035.150804 | 842.9893131 | 1336.410245 | -1.237502008 | 0.121399758 | 2.04895077251 | 1.96029424870 | Km<KS.3 | YPD | KS.3 |
| FIM1_1511 | 344.1260243 | 182.9299123 | 223.070927 | -0.856455551 | 0.17291507 | 2.59408447542 | 1.14512602175 | ns | YPD | KS.3 |
| FIM1_1512 | 821.7082662 | 528.951126 | 639.4952849 | -0.6083093 | 0.138591875 | 6.79756357266 | 2.23073078984 | ns | YPD | KS.3 |
| FIM1_1513 | 4873.928068 | 3917.098439 | 4753.957716 | -0.309331944 | 0.086379579 | 0.000298912 | 0.000691991 | ns | YPD | KS.3 |
| FIM1_1514 | 7365.148697 | 9486.275861 | 11778.31794 | 0.351759073 | 0.124009159 | 0.003821269 | 0.007033228 | ns | YPD | KS.3 |
| FIM1_1515 | 255.5236328 | 350.690417 | 365.2917633 | 0.430496891 | 0.159439359 | 0.005312791 | 0.009473526 | ns | YPD | KS.3 |
| FIM1_1516 | 114.6788203 | 109.6042496 | 209.8896237 | -0.046595641 | 0.365603883 | 0.881204021 | 0.901869986 | ns | YPD | KS.3 |
| FIM1_1518 | 1744.012452 | 965.4838585 | 1559.237096 | -0.818378366 | 0.13942306 | 1.70165964090 | 1.17743486974 | ns | YPD | KS.3 |
| FIM1_1519 | 583.4106275 | 414.1546911 | 582.3236775 | -0.472441267 | 0.138395235 | 0.000468556 | 0.001040189 | ns | YPD | KS.3 |
| FIM1_1520 | 4358.013653 | 3983.556015 | 4137.183329 | -0.122528939 | 0.151347073 | 0.40501081 | 0.468588088 | ns | YPD | KS.3 |
| FIM1_1521 | 3200.165615 | 2406.607496 | 4002.233212 | -0.322644635 | 0.324086761 | 0.259242348 | 0.316110461 | ns | YPD | KS.3 |
| FIM1_1523 | 381.0208968 | 397.8919837 | 369.144513 | 0.059534392 | 0.127093935 | 0.632693717 | 0.687308713 | ns | YPD | KS.3 |
| FIM1_1524 | 1324.860862 | 740.363509 | 1103.512142 | -0.807508955 | 0.134703591 | 8.15085620390 | 5.91115300502 | ns | YPD | KS.3 |
| FIM1_1525 | 1881.134206 | 1654.310364 | 1670.902005 | -0.182408847 | 0.076797474 | 0.016683491 | 0.027009556 | ns | YPD | KS.3 |

| | | | | | | | | | | |
|---|---|---|---|---|---|---|---|---|---|---|
| FIM1_1526 | 3815.639983 | 3579.911074 | 7536.909485 | -0.088392409 | 0.124945505 | 0.470690879 | 0.534265828 | ns | YPD | KS.3 |
| FIM1_1527 | 585.0469026 | 506.4861095 | 531.9941973 | -0.202882606 | 0.101213976 | 0.04226443 | 0.062080171 | ns | YPD | KS.3 |
| FIM1_1528 | 326.0299222 | 614.9800307 | 668.6270175 | 0.738999215 | 0.321741433 | 0.00923574 | 0.015762541 | ns | YPD | KS.3 |
| FIM1_1529 | 1352.321594 | 1451.77252 | 1461.92079 | 0.098131799 | 0.133051844 | 0.450893416 | 0.514258309 | ns | YPD | KS.3 |
| FIM1_1530 | 2656.084058 | 3740.314422 | 2471.08405 | 0.469883564 | 0.145669605 | 0.000920715 | 0.001922988 | ns | YPD | KS.3 |
| FIM1_1531 | 566.829763 | 964.9428233 | 741.0917151 | 0.704654731 | 0.198571565 | 0.00019144 | 0.00046144 | ns | YPD | KS.3 |
| FIM1_1532 | 836.0515774 | 708.7488111 | 768.8674388 | -0.208486322 | 0.237356897 | 0.347386073 | 0.409439875 | ns | YPD | KS.3 |
| FIM1_1533 | 1102.188724 | 594.2313169 | 731.9239523 | -0.865769814 | 0.11671652 | 4.1060364932( | 6.9957299841! | ns | YPD | KS.3 |
| FIM1_1534 | 566.2722943 | 408.4843089 | 392.2872536 | -0.458007459 | 0.110112536 | 2.3829119065: | 6.9327407807( | ns | YPD | KS.3 |
| FIM1_1535 | 1013.879916 | 676.8728971 | 781.0826287 | -0.57429513 | 0.078508932 | 1.6187202492! | 2.4477608632! | ns | YPD | KS.3 |
| FIM1_1536 | 647.1935758 | 753.6977818 | 720.7602547 | 0.21151587 | 0.121793937 | 0.076831937 | 0.106089061 | ns | YPD | KS.3 |
| FIM1_1537 | 1837.053688 | 2110.167921 | 1969.124807 | 0.193622294 | 0.1114015 | 0.077465888 | 0.106786587 | ns | YPD | KS.3 |
| FIM1_1538 | 2017.311534 | 2690.593189 | 1902.472634 | 0.401046659 | 0.120667517 | 0.000708793 | 0.00151341 | ns | YPD | KS.3 |
| FIM1_1540 | 1268.415934 | 892.6778727 | 957.5712073 | -0.479598478 | 0.15309255 | 0.001251216 | 0.002541771 | ns | YPD | KS.3 |
| FIM1_1541 | 1487.424425 | 3597.753393 | 2516.562676 | 1.184760661 | 0.202028919 | 5.7007771655! | 4.2648671276: | Km>KS.3 | YPD | KS.3 |
| FIM1_1542 | 4113.559366 | 2184.86508 | 4261.464166 | -0.782379877 | 0.273875923 | 0.001728026 | 0.00341148 | ns | YPD | KS.3 |
| FIM1_1543 | 3471.369294 | 1610.169753 | 3473.59313 | -0.901662365 | 0.327176215 | 0.001689644 | 0.003344997 | ns | YPD | KS.3 |
| FIM1_1544 | 294.7239939 | 58.19420727 | 191.583174 | -2.279184307 | 0.193958409 | 1.8052605729. | 3.9048571088: | Km<KS.3 | YPD | KS.3 |
| FIM1_1545 | 2223.711687 | 918.1986947 | 1128.202135 | -1.231159936 | 0.140685035 | 2.1504123651: | 1.0092737279: | Km<KS.3 | YPD | KS.3 |
| FIM1_1546 | 455.2643469 | 171.7626837 | 298.4181561 | -1.352110346 | 0.153023449 | 5.8447543425: | 3.1266293391. | Km<KS.3 | YPD | KS.3 |
| FIM1_1547 | 1170.496873 | 643.8471452 | 968.3615341 | -0.82580236 | 0.141851058 | 2.2326692492. | 1.4969716327. | ns | YPD | KS.3 |
| FIM1_1548 | 1676.585246 | 917.4906204 | 1539.925275 | -0.799221498 | 0.201112271 | 2.8556488936: | 8.1601684352! | ns | YPD | KS.3 |
| FIM1_1549 | 1942.98755 | 1686.349073 | 1949.095125 | -0.195507437 | 0.133747997 | 0.134990135 | 0.176352272 | ns | YPD | KS.3 |
| FIM1_1550 | 753.1451813 | 927.4711505 | 1478.150675 | 0.276043813 | 0.188479441 | 0.126163971 | 0.166225042 | ns | YPD | KS.3 |
| FIM1_1551 | 10585.62864 | 9944.528745 | 21078.24352 | -0.078675409 | 0.238974734 | 0.724517337 | 0.768925588 | ns | YPD | KS.3 |
| FIM1_1552 | 555.8080619 | 216.3870526 | 398.9322626 | -1.261588353 | 0.211761623 | 2.3716625391! | 1.9247995322: | Km<KS.3 | YPD | KS.3 |
| FIM1_1553 | 3329.239393 | 1568.955451 | 3566.150704 | -0.880780643 | 0.32767773 | 0.00219011 | 0.00423281 | ns | YPD | KS.3 |
| FIM1_1554 | 1425.842827 | 975.9711036 | 1029.671585 | -0.521279546 | 0.143961905 | 0.0002007 | 0.000480503 | ns | YPD | KS.3 |
| FIM1_1556 | 666.5020272 | 267.2115936 | 412.6890585 | -1.232895706 | 0.19595881 | 3.2751323628: | 3.1601299955: | Km<KS.3 | YPD | KS.3 |
| FIM1_1557 | 547.4564152 | 183.499429 | 370.8202114 | -1.505242049 | 0.178607256 | 9.5393392730! | 3.7665248320: | Km<KS.3 | YPD | KS.3 |
| FIM1_1558 | 3021.195269 | 2734.625733 | 2132.708685 | -0.138839019 | 0.116318397 | 0.224713465 | 0.278443209 | ns | YPD | KS.3 |
| FIM1_1559 | 3470.954848 | 5793.022983 | 5562.228255 | 0.684220397 | 0.187594435 | 0.000136778 | 0.000341944 | ns | YPD | KS.3 |
| FIM1_1560 | 1590.668138 | 1325.039519 | 1399.809743 | -0.255735305 | 0.109638695 | 0.017859367 | 0.028762733 | ns | YPD | KS.3 |
| FIM1_1561 | 2506.115981 | 1942.094078 | 1903.390235 | -0.348110427 | 0.151545414 | 0.018085733 | 0.02903405 | ns | YPD | KS.3 |
| FIM1_1562 | 749.8351714 | 356.7386818 | 554.8368495 | -0.979894137 | 0.215046595 | 1.3077966701! | 5.0086900956( | ns | YPD | KS.3 |
| FIM1_1563 | 1148.013256 | 1433.727225 | 1104.718313 | 0.313109373 | 0.095729771 | 0.000933367 | 0.0019437 | ns | YPD | KS.3 |
| FIM1_1564 | 82.91948327 | 311.939237 | 328.3333144 | 1.699108555 | 0.323110799 | 1.7744444306! | 1.2210042935( | Km>KS.3 | YPD | KS.3 |
| FIM1_1566 | 2019.82036 | 3407.042507 | 2607.517748 | 0.699304157 | 0.186468765 | 8.8601356087: | 0.000231144 | ns | YPD | KS.3 |
| FIM1_1567 | 643.2583263 | 2254.748416 | 1116.362786 | 1.679027541 | 0.247668826 | 1.4247439229( | 2.1876854989( | Km>KS.3 | YPD | KS.3 |
| FIM1_1568 | 6906.860189 | 7477.416599 | 7098.304888 | 0.112391915 | 0.08585774 | 0.186313627 | 0.236336129 | ns | YPD | KS.3 |
| FIM1_1569 | 192.6355943 | 272.8936004 | 191.7269476 | 0.442311231 | 0.236002827 | 0.044926721 | 0.06556481 | ns | YPD | KS.3 |
| FIM1_1570 | 1135.809493 | 1266.059628 | 1038.993896 | 0.151648265 | 0.115091944 | 0.180313741 | 0.230015606 | ns | YPD | KS.3 |
| FIM1_1571 | 476.3899377 | 291.1758269 | 365.0476731 | -0.675029996 | 0.147839659 | 2.6333921030: | 9.3982250448! | ns | YPD | KS.3 |
| FIM1_1572 | 643.8901938 | 505.7716884 | 549.9838849 | -0.332538966 | 0.137029137 | 0.012964399 | 0.021427869 | ns | YPD | KS.3 |
| FIM1_1573 | 1403.722475 | 994.6152067 | 1173.957086 | -0.488993161 | 0.08271545 | 2.4253509275! | 1.6131177626( | ns | YPD | KS.3 |

| | | | | | | | | | | |
|---|---|---|---|---|---|---|---|---|---|---|
| FIM1_1574 | 3424.999599 | 3511.7532 | 3203.394133 | 0.03563853 | 0.070241646 | 0.609584612 | 0.666084657 | ns | YPD | KS.3 |
| FIM1_1575 | 987.2937419 | 1068.445777 | 851.4544938 | 0.110538999 | 0.109758664 | 0.306525557 | 0.36684259 | ns | YPD | KS.3 |
| FIM1_1576 | 23.23260162 | 34.06165671 | 38.68437699 | 0.482367052 | 0.2329278 | 0.026930092 | 0.041530442 | ns | YPD | KS.3 |
| FIM1_1577 | 372.3629628 | 549.3040594 | 674.6265987 | 0.53431519 | 0.144486781 | 0.000145793 | 0.000363023 | ns | YPD | KS.3 |
| FIM1_1578 | 7077.888588 | 8810.643279 | 9826.523273 | 0.309196511 | 0.09213376 | 0.000690728 | 0.001479919 | ns | YPD | KS.3 |
| FIM1_1579 | 1263.04017 | 958.6861295 | 1877.861067 | -0.331307648 | 0.281544809 | 0.195777024 | 0.246892444 | ns | YPD | KS.3 |
| FIM1_1580 | 2016.697867 | 1025.898584 | 1777.236402 | -0.898326969 | 0.201250357 | 2.5531570439 | 9.1485491929 | ns | YPD | KS.3 |
| FIM1_1581 | 5183.130833 | 3970.818872 | 4862.540858 | -0.377459259 | 0.084580511 | 6.6327941370 | 2.1824173830 | ns | YPD | KS.3 |
| FIM1_1582 | 93.83224021 | 72.20545629 | 111.7929124 | -0.316721306 | 0.271242444 | 0.202281827 | 0.25368089 | ns | YPD | KS.3 |
| FIM1_1583 | 674.6602518 | 1058.914718 | 887.9140318 | 0.628393815 | 0.12226649 | 1.5904248478 | 7.4115337106 | ns | YPD | KS.3 |
| FIM1_1584 | 4631.533732 | 6245.201282 | 5326.866002 | 0.413496519 | 0.132502011 | 0.001417096 | 0.002851963 | ns | YPD | KS.3 |
| FIM1_1585 | 542.5526644 | 720.9696507 | 919.5679531 | 0.391233099 | 0.140320344 | 0.004259288 | 0.007767579 | ns | YPD | KS.3 |
| FIM1_1586 | 691.2971837 | 1829.106973 | 2013.747848 | 1.269216629 | 0.246921707 | 2.4576224739 | 1.3275430844 | Km>KS.3 | YPD | KS.3 |
| FIM1_1587 | 5822.548151 | 7625.743181 | 7941.355191 | 0.377359429 | 0.112377432 | 0.000642597 | 0.00138635 | ns | YPD | KS.3 |
| FIM1_1588 | 4870.889228 | 6623.121634 | 6573.148533 | 0.410841172 | 0.180438135 | 0.017765709 | 0.028631164 | ns | YPD | KS.3 |
| FIM1_1589 | 194.2607811 | 118.0468307 | 174.21158 | -0.691552958 | 0.127473738 | 2.9736881571 | 1.5771960108 | ns | YPD | KS.3 |
| FIM1_1590 | 998.6085415 | 463.0373491 | 907.6081619 | -1.017949577 | 0.211229531 | 3.2397570288 | 1.3979102129 | Km<KS.3 | YPD | KS.3 |
| FIM1_1591 | 298.6317549 | 107.301179 | 193.798865 | -1.340583768 | 0.24783686 | 4.1584470109 | 2.6287514459 | Km<KS.3 | YPD | KS.3 |
| FIM1_1592 | 404.6287911 | 140.6844556 | 207.5354789 | -1.447425343 | 0.18215301 | 7.1827863594 | 2.0419635507 | Km<KS.3 | YPD | KS.3 |
| FIM1_1593 | 4637.019868 | 7387.03182 | 3711.678997 | 0.536181928 | 0.319934932 | 0.058128392 | 0.082389957 | ns | YPD | KS.3 |
| FIM1_1594 | 2104.686661 | 1914.379375 | 2701.158854 | -0.116637837 | 0.259197119 | 0.625938943 | 0.680963534 | ns | YPD | KS.3 |
| FIM1_1595 | 1499.60995 | 957.787747 | 1468.553781 | -0.609759145 | 0.162511003 | 0.000104138 | 0.000267884 | ns | YPD | KS.3 |
| FIM1_1596 | 1225.566908 | 2472.448252 | 1618.955995 | 0.924703627 | 0.213090039 | 4.2017796340 | 1.4439380160 | ns | YPD | KS.3 |
| FIM1_1597 | 990.2035849 | 932.6195337 | 1675.69388 | -0.07021575 | 0.297325697 | 0.793117382 | 0.827554396 | ns | YPD | KS.3 |
| FIM1_1598 | 2595.736999 | 1618.614292 | 3069.22103 | -0.548077579 | 0.314840679 | 0.049903508 | 0.072004047 | ns | YPD | KS.3 |
| FIM1_1599 | 2692.177431 | 2549.724106 | 2665.984911 | -0.076711116 | 0.088784512 | 0.382765725 | 0.445649306 | ns | YPD | KS.3 |
| FIM1_1601 | 14833.019 | 15833.46208 | 22763.71051 | 0.090046189 | 0.13422234 | 0.492697558 | 0.555947006 | ns | YPD | KS.3 |
| FIM1_1602 | 1764.179812 | 1325.613543 | 2256.884439 | -0.403509228 | 0.095516569 | 1.9085868659 | 5.6925777327 | ns | YPD | KS.3 |
| FIM1_1603 | 4870.846406 | 4018.539821 | 4657.555361 | -0.26913473 | 0.111730874 | 0.01439039 | 0.023573325 | ns | YPD | KS.3 |
| FIM1_1604 | 1399.056728 | 2769.218728 | 2197.326057 | 0.944239708 | 0.143401654 | 1.2803125645 | 1.3409589492 | ns | YPD | KS.3 |
| FIM1_1605 | 2010.604242 | 1468.385305 | 2906.898455 | -0.360398411 | 0.316466674 | 0.199400049 | 0.250445656 | ns | YPD | KS.3 |
| FIM1_1606 | 6376.566597 | 7546.060704 | 7623.839318 | 0.232211243 | 0.135701618 | 0.079952683 | 0.10981905 | ns | YPD | KS.3 |
| FIM1_1607 | 1647.157998 | 1329.165092 | 1263.574432 | -0.29939103 | 0.116314258 | 0.008827514 | 0.015107287 | ns | YPD | KS.3 |
| FIM1_1608 | 2504.061945 | 2516.448007 | 2705.623237 | 0.006676664 | 0.141747684 | 0.961490307 | 0.970266588 | ns | YPD | KS.3 |
| FIM1_1609 | 1030.632249 | 1126.364432 | 1307.238998 | 0.119423321 | 0.166810913 | 0.458912325 | 0.522685169 | ns | YPD | KS.3 |
| FIM1_1610 | 2405.461957 | 1669.320841 | 2102.081972 | -0.475025774 | 0.213556604 | 0.018654133 | 0.029850213 | ns | YPD | KS.3 |
| FIM1_1611 | 1905.749407 | 1655.45261 | 1829.231039 | -0.189846407 | 0.167488091 | 0.240769559 | 0.295760137 | ns | YPD | KS.3 |
| FIM1_1612 | 1115.648193 | 1068.241376 | 1563.798557 | -0.05633085 | 0.212967526 | 0.780024656 | 0.816114125 | ns | YPD | KS.3 |
| FIM1_1613 | 1119.073527 | 1749.029766 | 1328.538532 | 0.630004151 | 0.096837936 | 4.4378562117 | 4.1578784657 | ns | YPD | KS.3 |
| FIM1_1614 | 672.8409599 | 394.9026401 | 538.7302173 | -0.738097184 | 0.134301056 | 1.8155722945 | 1.0114750465 | ns | YPD | KS.3 |
| FIM1_1615 | 2505.293963 | 2948.694161 | 3138.809537 | 0.229151755 | 0.102055573 | 0.022876787 | 0.035767447 | ns | YPD | KS.3 |
| FIM1_1616 | 821.2904162 | 859.6609689 | 848.8679576 | 0.062950609 | 0.130733074 | 0.622827817 | 0.678320576 | ns | YPD | KS.3 |
| FIM1_1617 | 2939.521597 | 2741.61733 | 3551.417207 | -0.098187425 | 0.097116988 | 0.306185735 | 0.36652407 | ns | YPD | KS.3 |
| FIM1_1618 | 293.9770934 | 144.9190918 | 330.5806568 | -0.915934785 | 0.233503986 | 2.6155206158 | 7.5433130803 | ns | YPD | KS.3 |
| FIM1_1619 | 1178.035911 | 809.9624672 | 1644.712945 | -0.522923331 | 0.117528851 | 5.8929473400 | 1.9689330434 | ns | YPD | KS.3 |

| | | | | | | | | | | |
|---|---|---|---|---|---|---|---|---|---|---|
| FIM1_1621 | 3546.223163 | 3951.261507 | 3449.992768 | 0.152535989 | 0.095454802 | 0.105932494 | 0.141670472 | ns | YPD | KS.3 |
| FIM1_1622 | 627.2607391 | 690.4604855 | 889.5684964 | 0.131630118 | 0.150118692 | 0.367327018 | 0.430089883 | ns | YPD | KS.3 |
| FIM1_1623 | 282.1624846 | 398.1940643 | 369.2683476 | 0.465803468 | 0.165116935 | 0.003510222 | 0.00652348 | ns | YPD | KS.3 |
| FIM1_1624 | 741.8754024 | 270.4646039 | 674.7249446 | -1.393564903 | 0.164495386 | 1.170911653 | 4.551004276 | Km<KS.3 | YPD | KS.3 |
| FIM1_1625 | 1776.716074 | 1141.055319 | 1770.134835 | -0.609188476 | 0.144113427 | 1.411250268 | 4.291546507 | ns | YPD | KS.3 |
| FIM1_1627 | 1548.313874 | 1407.057345 | 2079.215773 | -0.129542423 | 0.162777273 | 0.411061011 | 0.474968215 | ns | YPD | KS.3 |
| FIM1_1628 | 274.2485095 | 363.8348729 | 650.5022275 | 0.297890213 | 0.36811461 | 0.341581423 | 0.403267105 | ns | YPD | KS.3 |
| FIM1_1629 | 480.6088591 | 305.1744473 | 370.7286386 | -0.629624757 | 0.132388508 | 1.138363233 | 4.414152055 | ns | YPD | KS.3 |
| FIM1_1630 | 2087.593538 | 3876.523237 | 2888.50703 | 0.86842503 | 0.113488926 | 6.758464263 | 1.318651259 | ns | YPD | KS.3 |
| FIM1_1631 | 894.0851211 | 723.2291901 | 737.8346959 | -0.28087133 | 0.188644615 | 0.119963825 | 0.158644346 | ns | YPD | KS.3 |
| FIM1_1632 | 795.7458236 | 2263.416985 | 1564.129185 | 1.281561307 | 0.322570073 | 5.847020463 | 1.956215655 | Km>KS.3 | YPD | KS.3 |
| FIM1_1633 | 1254.316058 | 234.1204127 | 1191.610824 | -2.213526615 | 0.354866626 | 1.213712937 | 1.560364177 | Km<KS.3 | YPD | KS.3 |
| FIM1_1634 | 2507.554017 | 2495.068736 | 2116.876201 | -0.00690665 | 0.144483881 | 0.960882467 | 0.970046728 | ns | YPD | KS.3 |
| FIM1_1635 | 5315.580598 | 6520.352976 | 6104.133034 | 0.290193849 | 0.077972277 | 0.000175608 | 0.000429102 | ns | YPD | KS.3 |
| FIM1_1636 | 526.7303942 | 731.2189392 | 829.9480188 | 0.452659774 | 0.136012671 | 0.000655697 | 0.001411537 | ns | YPD | KS.3 |
| FIM1_1637 | 149.722279 | 200.8677147 | 238.0021329 | 0.394094469 | 0.171817561 | 0.01735045 | 0.028016387 | ns | YPD | KS.3 |
| FIM1_1638 | 28.72089752 | 42.30168279 | 70.51401508 | 0.475850049 | 0.271478519 | 0.055396075 | 0.078921956 | ns | YPD | KS.3 |
| FIM1_1640 | 2880.924519 | 2497.143978 | 2570.685489 | -0.199602849 | 0.11438996 | 0.076059224 | 0.10522654 | ns | YPD | KS.3 |
| FIM1_1641 | 4564.957026 | 4169.049667 | 4617.052727 | -0.128411717 | 0.087390457 | 0.137823394 | 0.1794012 | ns | YPD | KS.3 |
| FIM1_1642 | 1718.050872 | 2399.889323 | 1800.68227 | 0.471595598 | 0.095370669 | 5.594326547 | 2.300146658 | ns | YPD | KS.3 |
| FIM1_1643 | 1730.045499 | 1696.874336 | 1577.718136 | -0.02669796 | 0.120207828 | 0.821088471 | 0.850846728 | ns | YPD | KS.3 |
| FIM1_1644 | 964.5837364 | 624.2344026 | 640.2083275 | -0.609727067 | 0.111184327 | 2.480372012 | 1.338378607 | ns | YPD | KS.3 |
| FIM1_1645 | 791.9112897 | 636.389102 | 638.4829664 | -0.302400846 | 0.131515087 | 0.018775765 | 0.029996606 | ns | YPD | KS.3 |
| FIM1_1646 | 2153.970018 | 4914.794031 | 3165.894573 | 1.129556963 | 0.166973738 | 2.065836497 | 2.562976702 | Km>KS.3 | YPD | KS.3 |
| FIM1_1647 | 85.48840014 | 69.63323903 | 78.19150405 | -0.258967491 | 0.237070336 | 0.242565096 | 0.297598361 | ns | YPD | KS.3 |
| FIM1_1648 | 3586.233822 | 1895.499364 | 3748.977745 | -0.867104494 | 0.168566396 | 9.302156057 | 4.539714775 | ns | YPD | KS.3 |
| FIM1_1649 | 2481.922364 | 1750.128889 | 2915.235222 | -0.464732327 | 0.187886272 | 0.0097645 | 0.01659098 | ns | YPD | KS.3 |
| FIM1_1650 | 1384.300522 | 1104.512891 | 1231.102463 | -0.282707519 | 0.245817601 | 0.216263896 | 0.269180106 | ns | YPD | KS.3 |
| FIM1_1651 | 650.3206718 | 376.0997088 | 475.7090115 | -0.740031884 | 0.173122716 | 8.873386848 | 2.817172914 | ns | YPD | KS.3 |
| FIM1_1652 | 3883.888775 | 2497.963421 | 2996.097767 | -0.61034763 | 0.136117519 | 4.363608492 | 1.495106904 | ns | YPD | KS.3 |
| FIM1_1653 | 786.2021563 | 787.9952417 | 589.1797007 | 0.003102017 | 0.13155304 | 0.980779647 | 0.984142547 | ns | YPD | KS.3 |
| FIM1_1654 | 537.9132682 | 648.0599735 | 472.4391557 | 0.242547033 | 0.207936349 | 0.219065387 | 0.272054494 | ns | YPD | KS.3 |
| FIM1_1655 | 1642.437892 | 1440.639315 | 1447.568048 | -0.183773502 | 0.107189589 | 0.08191782 | 0.112146713 | ns | YPD | KS.3 |
| FIM1_1656 | 1358.865025 | 2006.674759 | 1353.127567 | 0.541066795 | 0.127701164 | 1.508099739 | 4.572087876 | ns | YPD | KS.3 |
| FIM1_1657 | 17521.47698 | 25564.26808 | 30073.46668 | 0.531870365 | 0.100908401 | 9.168702624 | 4.481665125 | ns | YPD | KS.3 |
| FIM1_1658 | 1061.787288 | 705.0095419 | 1053.529141 | -0.564512537 | 0.139649427 | 3.394696013 | 9.504002626 | ns | YPD | KS.3 |
| FIM1_1659 | 2793.052391 | 4008.089814 | 2925.916133 | 0.507554487 | 0.104642311 | 8.618372190 | 3.421899573 | ns | YPD | KS.3 |
| FIM1_1660 | 3335.789316 | 1928.795025 | 2284.236219 | -0.771444161 | 0.103246075 | 3.407887307 | 5.866518808 | ns | YPD | KS.3 |
| FIM1_1661 | 2397.55068 | 1158.263526 | 1659.451143 | -1.008637584 | 0.140734595 | 1.773680379 | 2.641934098 | Km<KS.3 | YPD | KS.3 |
| FIM1_1664 | 1959.621977 | 1859.005852 | 2194.986819 | -0.075003657 | 0.080939767 | 0.350013932 | 0.412244156 | ns | YPD | KS.3 |
| FIM1_1665 | 455.5459832 | 346.976199 | 323.128888 | -0.355236619 | 0.206347328 | 0.069802337 | 0.097328091 | ns | YPD | KS.3 |
| FIM1_1666 | 1038.667603 | 830.4262484 | 919.0415446 | -0.317910215 | 0.077373827 | 3.450776933 | 9.655576627 | ns | YPD | KS.3 |
| FIM1_1667 | 1412.163728 | 609.114021 | 1963.193442 | -1.049521328 | 0.281239882 | 3.728126067 | 0.000103675 | Km<KS.3 | YPD | KS.3 |
| FIM1_1669 | 2327.0643 | 1956.689693 | 2080.923473 | -0.240981907 | 0.121287753 | 0.042964413 | 0.063050208 | ns | YPD | KS.3 |
| FIM1_1670 | 1826.552279 | 1065.677224 | 1932.306517 | -0.710763434 | 0.204258696 | 0.000244636 | 0.000574901 | ns | YPD | KS.3 |

| | | | | | | | | | | |
|---|---|---|---|---|---|---|---|---|---|---|
| FIM1_1671 | 5189.858194 | 2783.538236 | 5657.092212 | -0.761696642 | 0.28302231 | 0.002987095 | 0.005641913 | ns | YPD | KS.3 |
| FIM1_1672 | 4014.51578 | 4177.034778 | 4201.129198 | 0.05410867 | 0.152233768 | 0.714606166 | 0.761766805 | ns | YPD | KS.3 |
| FIM1_1673 | 4190.732414 | 4229.662872 | 3566.447103 | 0.013179726 | 0.084792195 | 0.875337062 | 0.897206874 | ns | YPD | KS.3 |
| FIM1_1674 | 1590.867075 | 1455.129964 | 1673.238799 | -0.123384849 | 0.128335149 | 0.326416635 | 0.388034112 | ns | YPD | KS.3 |
| FIM1_1675 | 13.19085123 | 11.29405136 | 18.27940201 | -0.171308109 | 0.332909077 | 0.557209546 | 0.618224239 | ns | YPD | KS.3 |
| FIM1_1676 | 678.7319406 | 558.6510084 | 574.8902043 | -0.271621549 | 0.115200271 | 0.016489665 | 0.026721852 | ns | YPD | KS.3 |
| FIM1_1677 | 670.66628 | 821.9229503 | 639.4036967 | 0.218765528 | 0.354014607 | 0.473162327 | 0.536826129 | ns | YPD | KS.3 |
| FIM1_1678 | 239.186439 | 246.3323357 | 212.1497853 | 0.037931657 | 0.228531553 | 0.859771741 | 0.883023206 | ns | YPD | KS.3 |
| FIM1_1679 | 878.3750217 | 833.0292717 | 646.7479553 | -0.067438644 | 0.225229259 | 0.750373379 | 0.791273727 | ns | YPD | KS.3 |
| FIM1_1680 | 370.3410652 | 403.4910245 | 512.5961361 | 0.115959331 | 0.164074917 | 0.465165474 | 0.52859713 | ns | YPD | KS.3 |
| FIM1_1681 | 91.62570571 | 42.33250084 | 59.03730574 | -0.90306349 | 0.328248845 | 0.001706123 | 0.003372254 | ns | YPD | KS.3 |
| FIM1_1682 | 647.238286 | 791.7438963 | 870.1066741 | 0.26816727 | 0.18247208 | 0.125860489 | 0.165869122 | ns | YPD | KS.3 |
| FIM1_1683 | 5427.903591 | 3700.175293 | 4383.462853 | -0.512422848 | 0.182069887 | 0.003353229 | 0.00626213 | ns | YPD | KS.3 |
| FIM1_1684 | 1727.516186 | 692.6406758 | 1764.440744 | -1.289289098 | 0.110319277 | 1.1106940934 | 2.0465567092 | Km<KS.3 | YPD | KS.3 |
| FIM1_1685 | 1865.527447 | 1305.090765 | 2269.185769 | -0.503660336 | 0.09685978 | 1.4018767948 | 6.5733619741 | ns | YPD | KS.3 |
| FIM1_1686 | 1216.741012 | 853.0955558 | 1051.0058 | -0.490672749 | 0.134325205 | 0.000185243 | 0.000449554 | ns | YPD | KS.3 |
| FIM1_1687 | 2230.537809 | 1606.893799 | 2583.761881 | -0.460102182 | 0.106724944 | 1.2103699083 | 3.7354778497 | ns | YPD | KS.3 |
| FIM1_1688 | 264.3965274 | 162.4758121 | 317.1652885 | -0.61512684 | 0.24531009 | 0.006937449 | 0.012152749 | ns | YPD | KS.3 |
| FIM1_1689 | 199.1327617 | 288.7384616 | 271.523357 | 0.491335101 | 0.194110643 | 0.007998072 | 0.013844958 | ns | YPD | KS.3 |
| FIM1_1690 | 4771.749924 | 4081.174266 | 3733.747297 | -0.221700581 | 0.081405991 | 0.00601302 | 0.010634474 | ns | YPD | KS.3 |
| FIM1_1691 | 1010.10583 | 4037.802793 | 2389.642711 | 1.592249064 | 0.450605778 | 8.9477540515 | 2.8389717096 | Km>KS.3 | YPD | KS.3 |
| FIM1_1692 | 979.3870821 | 1610.252141 | 1205.142974 | 0.665088794 | 0.185907726 | 0.000185703 | 0.000450229 | ns | YPD | KS.3 |
| FIM1_1693 | 286.7135686 | 584.4841552 | 768.7516542 | 0.866986805 | 0.293041382 | 0.001006157 | 0.002083944 | ns | YPD | KS.3 |
| FIM1_1694 | 29.19621981 | 64.63699426 | 53.63782133 | 0.984301713 | 0.282110136 | 0.000115821 | 0.000295038 | ns | YPD | KS.3 |
| FIM1_1695 | 50.54791771 | 299.5049742 | 186.2138148 | 2.325878669 | 0.387060265 | 5.1358205879 | 5.8468438044 | Km>KS.3 | YPD | KS.3 |
| FIM1_1697 | 154.6309862 | 188.0107905 | 205.3618799 | 0.24259011 | 0.253761407 | 0.301846332 | 0.362375845 | ns | YPD | KS.3 |
| FIM1_1698 | 942.4586192 | 1377.648445 | 2099.503148 | 0.472739409 | 0.255184412 | 0.045016618 | 0.065619008 | ns | YPD | KS.3 |
| FIM1_1701 | 767.3645979 | 400.6872957 | 548.8457104 | -0.858248537 | 0.207671546 | 1.2559275206 | 3.8688788949 | ns | YPD | KS.3 |
| FIM1_1703 | 131.4246129 | 125.3485578 | 84.05414166 | -0.056101597 | 0.272407106 | 0.822043224 | 0.851481374 | ns | YPD | KS.3 |
| FIM1_1705 | 37.58013656 | 20.79020019 | 22.09320857 | -0.709150119 | 0.297017015 | 0.007862098 | 0.0136333 | ns | YPD | KS.3 |
| FIM1_1707 | 405.7945665 | 495.1402842 | 374.05025 | 0.267817657 | 0.171779103 | 0.105972798 | 0.141686286 | ns | YPD | KS.3 |
| FIM1_1708 | 305.3717119 | 3252.120524 | 2951.506154 | 3.052228244 | 0.474315755 | 8.8491794495 | 1.1771301540 | Km>KS.3 | YPD | KS.3 |
| FIM1_1709 | 1009.278741 | 1140.08746 | 1248.502807 | 0.150250709 | 0.257512785 | 0.527761489 | 0.590025485 | ns | YPD | KS.3 |
| FIM1_1710 | 245.8345597 | 206.154865 | 440.0704167 | -0.20918739 | 0.287033537 | 0.421496924 | 0.485516832 | ns | YPD | KS.3 |
| FIM1_1711 | 782.9771321 | 1720.395631 | 1667.795688 | 0.889513307 | 0.356752255 | 0.003654815 | 0.006764399 | ns | YPD | KS.3 |
| FIM1_1712 | 1236.61496 | 1224.070625 | 2816.95815 | -0.011997108 | 0.296500424 | 0.964180569 | 0.971995609 | ns | YPD | KS.3 |
| FIM1_1713 | 61.42908305 | 57.05571338 | 91.60946069 | -0.095565069 | 0.219261965 | 0.644304919 | 0.697739872 | ns | YPD | KS.3 |
| FIM1_1714 | 221.308769 | 243.8436923 | 331.4930736 | 0.131621031 | 0.161962367 | 0.401378897 | 0.464926662 | ns | YPD | KS.3 |
| FIM1_1715 | 212.9808761 | 255.8395071 | 205.5618188 | 0.237967084 | 0.213818499 | 0.239582846 | 0.294593342 | ns | YPD | KS.3 |
| FIM1_1716 | 3703.50681 | 3640.648242 | 4759.1703 | -0.021258712 | 0.250004557 | 0.927016365 | 0.941013348 | ns | YPD | KS.3 |
| FIM1_1717 | 204.6584193 | 228.037934 | 180.8783533 | 0.149950899 | 0.125094799 | 0.221565034 | 0.274952867 | ns | YPD | KS.3 |
| FIM1_1718 | 232.1362913 | 160.4335291 | 178.6442181 | -0.511861234 | 0.127493112 | 4.1314607734 | 0.0001135 | ns | YPD | KS.3 |
| FIM1_1719 | 202.3852622 | 264.2333167 | 232.4189604 | 0.364932333 | 0.152972185 | 0.014035126 | 0.023052081 | ns | YPD | KS.3 |
| FIM1_1721 | 93.95964 | 26.07087615 | 37.01723315 | -1.549903126 | 0.380326633 | 1.1760633130 | 4.5391117007 | Km<KS.3 | YPD | KS.3 |
| FIM1_1724 | 8068.830976 | 12983.15563 | 10659.98046 | 0.607891224 | 0.234995342 | 0.005624787 | 0.010004761 | ns | YPD | KS.3 |

| | | | | | | | | | | | |
|---|---|---|---|---|---|---|---|---|---|---|---|
| FIM1_1725 | 16349.9828 | 12562.74025 | 18386.44368 | -0.341833117 | 0.213311826 | 0.090356575 | 0.122419379 | ns | | YPD | KS.3 |
| FIM1_1727 | 1363.724227 | 2718.937076 | 1983.948707 | 0.908498342 | 0.21336399 | 6.3388479567 | 2.0953999059 | ns | | YPD | KS.3 |
| FIM1_1728 | 4851.792413 | 6519.733609 | 6048.725339 | 0.407408899 | 0.137777287 | 0.002454027 | 0.004701111 | ns | | YPD | KS.3 |
| FIM1_1729 | 2784.882901 | 3550.945846 | 3532.289846 | 0.339139239 | 0.116081421 | 0.002955947 | 0.005589448 | ns | | YPD | KS.3 |
| FIM1_1730 | 4039.946533 | 5127.891883 | 4780.754088 | 0.339792435 | 0.069410899 | 8.3441043556 | 3.3262755744 | ns | | YPD | KS.3 |
| FIM1_1731 | 4505.970004 | 4321.811962 | 4615.337124 | -0.05940212 | 0.073726843 | 0.417107281 | 0.481240427 | ns | | YPD | KS.3 |
| FIM1_1732 | 668.8231277 | 1010.881276 | 1104.078117 | 0.564057961 | 0.153865311 | 0.000157858 | 0.000391109 | ns | | YPD | KS.3 |
| FIM1_1733 | 4813.450526 | 11364.20506 | 8127.148046 | 1.065987135 | 0.288606347 | 4.0901402786 | 0.000112609 | Km>KS.3 | | YPD | KS.3 |
| FIM1_1734 | 712.6198565 | 593.8582671 | 542.6370371 | -0.235261234 | 0.216330996 | 0.249897435 | 0.305990583 | ns | | YPD | KS.3 |
| FIM1_1735 | 2196.213502 | 1613.577916 | 2353.005583 | -0.418962211 | 0.159455433 | 0.006680127 | 0.011730897 | ns | | YPD | KS.3 |
| FIM1_1736 | 831.8796951 | 618.5689975 | 664.8273505 | -0.40622592 | 0.147056871 | 0.004531288 | 0.00822143 | ns | | YPD | KS.3 |
| FIM1_1737 | 443.7266482 | 630.916914 | 484.8333214 | 0.474532303 | 0.171340561 | 0.004064939 | 0.007445902 | ns | | YPD | KS.3 |
| FIM1_1738 | 1270.836181 | 1184.916996 | 1392.724953 | -0.095902013 | 0.142336232 | 0.489674162 | 0.552786239 | ns | | YPD | KS.3 |
| FIM1_1739 | 2808.269603 | 1935.017976 | 2986.10881 | -0.511480295 | 0.145925319 | 0.000316356 | 0.000727297 | ns | | YPD | KS.3 |
| FIM1_1740 | 866.5869286 | 876.2011216 | 944.6310137 | 0.015585487 | 0.113259966 | 0.888783971 | 0.908319691 | ns | | YPD | KS.3 |
| FIM1_1741 | 2334.844541 | 2999.077468 | 4603.886683 | 0.337374602 | 0.16931537 | 0.038980002 | 0.057819174 | ns | | YPD | KS.3 |
| FIM1_1742 | 886.6843187 | 1004.653308 | 1223.137733 | 0.169651266 | 0.154343603 | 0.257694833 | 0.314377586 | ns | | YPD | KS.3 |
| FIM1_1743 | 2351.355824 | 4116.138061 | 3528.491098 | 0.716647438 | 0.237247785 | 0.001196874 | 0.002444355 | ns | | YPD | KS.3 |
| FIM1_1744 | 1375.548482 | 4605.66367 | 2700.92026 | 1.555426039 | 0.295914833 | 3.5014119116 | 2.2361391861 | Km>KS.3 | | YPD | KS.3 |
| FIM1_1745 | 1247.569249 | 1480.350237 | 1396.955375 | 0.24099536 | 0.096202497 | 0.011240874 | 0.018835752 | ns | | YPD | KS.3 |
| FIM1_1746 | 602.751175 | 397.6036058 | 512.8983152 | -0.56103487 | 0.174460877 | 0.000833084 | 0.001754696 | ns | | YPD | KS.3 |
| FIM1_1747 | 2486.504747 | 4181.701847 | 3201.610679 | 0.703359802 | 0.170991475 | 1.9473841097 | 5.7909360106 | ns | | YPD | KS.3 |
| FIM1_1748 | 23.30135451 | 256.2501418 | 243.67275 | 3.333552009 | 0.266305054 | 1.5946653213 | 4.4074777632 | Km>KS.3 | | YPD | KS.3 |
| FIM1_1749 | 2788.229734 | 4819.364251 | 4469.165477 | 0.749977279 | 0.153295342 | 4.5342146151 | 1.9019998069 | ns | | YPD | KS.3 |
| FIM1_1750 | 2537.651205 | 4937.019857 | 4549.600814 | 0.835002332 | 0.262601581 | 0.000530987 | 0.001162473 | ns | | YPD | KS.3 |
| FIM1_1751 | 582.4503664 | 799.8779835 | 1262.995325 | 0.354521772 | 0.333127275 | 0.224503832 | 0.278252756 | ns | | YPD | KS.3 |
| FIM1_1752 | 473.6763778 | 1170.097607 | 1127.180991 | 1.114772908 | 0.299284386 | 3.0489694064 | 8.6480175581 | Km>KS.3 | | YPD | KS.3 |
| FIM1_1753 | 832.6076376 | 835.3731019 | 787.7175786 | 0.004935589 | 0.108103206 | 0.963043967 | 0.971243409 | ns | | YPD | KS.3 |
| FIM1_1754 | 202.3011227 | 940.4424089 | 560.3280302 | 2.094846532 | 0.260928439 | 2.5159064949 | 9.0700252263 | Km>KS.3 | | YPD | KS.3 |
| FIM1_1755 | 783.4753748 | 2538.370213 | 2207.912755 | 1.468872496 | 0.324448323 | 2.1810764762 | 9.8286734324 | Km>KS.3 | | YPD | KS.3 |
| FIM1_1756 | 554.0515239 | 2322.235008 | 1284.85607 | 1.957209574 | 0.240104283 | 1.1866382123 | 4.5763760515 | Km>KS.3 | | YPD | KS.3 |
| FIM1_1757 | 936.1237045 | 1443.947209 | 1650.215053 | 0.549190332 | 0.241413785 | 0.014569499 | 0.023843177 | ns | | YPD | KS.3 |
| FIM1_1758 | 214.2943985 | 194.6942824 | 332.026574 | -0.121983181 | 0.234559124 | 0.578629711 | 0.637158656 | ns | | YPD | KS.3 |
| FIM1_1759 | 100.7436023 | 565.7971153 | 363.4795938 | 2.364498892 | 0.278057887 | 5.2436215017 | 2.8667051616 | Km>KS.3 | | YPD | KS.3 |
| FIM1_1760 | 2268.169368 | 6578.802586 | 3933.361456 | 1.395328955 | 0.251686046 | 1.5085648903 | 1.0496657803 | Km>KS.3 | | YPD | KS.3 |
| FIM1_1761 | 1297.775355 | 4687.113185 | 2837.073276 | 1.676468754 | 0.290761265 | 1.0525415001 | 9.2515794407 | Km>KS.3 | | YPD | KS.3 |
| FIM1_1762 | 3089.801835 | 3052.129923 | 5193.355965 | -0.015029317 | 0.265723355 | 0.950954283 | 0.961975917 | ns | | YPD | KS.3 |
| FIM1_1763 | 78.27187436 | 53.76502269 | 106.1058246 | -0.45219845 | 0.286297134 | 0.081356847 | 0.111532189 | ns | | YPD | KS.3 |
| FIM1_1764 | 3442.308406 | 3269.32261 | 3825.694864 | -0.072638723 | 0.102116221 | 0.471028253 | 0.534526815 | ns | | YPD | KS.3 |
| FIM1_1765 | 1423.689578 | 2300.814953 | 2218.508472 | 0.67161581 | 0.116361489 | 4.1806830007 | 2.6394540518 | ns | | YPD | KS.3 |
| FIM1_1766 | 816.1802766 | 2686.507365 | 1836.890909 | 1.526380316 | 0.298623433 | 8.8051490328 | 5.2211700164 | Km>KS.3 | | YPD | KS.3 |
| FIM1_1767 | 1805.357223 | 1672.376967 | 1753.236508 | -0.10557967 | 0.132932886 | 0.416879621 | 0.481089334 | ns | | YPD | KS.3 |
| FIM1_1769 | 35.68084834 | 22.58498385 | 21.2872472 | -0.443584817 | 0.420703775 | 0.192715223 | 0.243525079 | ns | | YPD | KS.3 |
| FIM1_1770 | 1969.03013 | 2006.354951 | 2963.922448 | 0.026134189 | 0.104081217 | 0.799062925 | 0.832182971 | ns | | YPD | KS.3 |
| FIM1_1771 | 1126.716231 | 1216.403614 | 1673.299133 | 0.105298297 | 0.140954397 | 0.443914564 | 0.507578707 | ns | | YPD | KS.3 |

| | | | | | | | | | |
|---|---|---|---|---|---|---|---|---|---|
| FIM1_1772 | 2054.97987 | 991.1836915 | 1718.984209 | -0.998499966 | 0.161981575 | 1.6981920346 | 1.4343812177 | ns | YPD | KS.3 |
| FIM1_1774 | 1044.934282 | 655.7611122 | 925.4380031 | -0.647180335 | 0.127659847 | 2.2262367218 | 1.0004993397 | ns | YPD | KS.3 |
| FIM1_1775 | 6565.443712 | 6710.102414 | 6632.013889 | 0.029973038 | 0.140310915 | 0.826750774 | 0.85582295 | ns | YPD | KS.3 |
| FIM1_1776 | 2689.609237 | 3889.664825 | 2784.014127 | 0.501987947 | 0.158868353 | 0.001105841 | 0.002271495 | ns | YPD | KS.3 |
| FIM1_1777 | 3775.387865 | 752.4394542 | 3183.634497 | -2.274807021 | 0.176763888 | 1.7562325595 | 6.2408978455 | Km<KS.3 | YPD | KS.3 |
| FIM1_1778 | 2215.246931 | 4053.8783 | 2756.463574 | 0.828791622 | 0.154194619 | 2.9143472522 | 1.5490253824 | ns | YPD | KS.3 |
| FIM1_1779 | 6526.298898 | 5734.121704 | 5361.145866 | -0.163435118 | 0.236808127 | 0.46050125 | 0.524134916 | ns | YPD | KS.3 |
| FIM1_1780 | 1231.441377 | 813.8130279 | 982.3092643 | -0.576967959 | 0.122562657 | 1.5792116933 | 5.9655111423 | ns | YPD | KS.3 |
| FIM1_1781 | 845.5586019 | 719.122218 | 622.2993874 | -0.222198763 | 0.14159889 | 0.107662268 | 0.143829157 | ns | YPD | KS.3 |
| FIM1_1782 | 1262.081197 | 1226.088987 | 1110.647824 | -0.040746014 | 0.076751477 | 0.59267155 | 0.650010924 | ns | YPD | KS.3 |
| FIM1_1783 | 446.4093786 | 379.5070437 | 438.0035736 | -0.221025145 | 0.152332544 | 0.135407741 | 0.176811901 | ns | YPD | KS.3 |
| FIM1_1784 | 1829.659456 | 2269.320736 | 2015.541673 | 0.297793585 | 0.131734686 | 0.020864234 | 0.032858361 | ns | YPD | KS.3 |
| FIM1_1785 | 1476.10931 | 824.6042798 | 875.1746038 | -0.799871587 | 0.150095955 | 4.0215274155 | 2.0906059448 | ns | YPD | KS.3 |
| FIM1_1786 | 51.38625932 | 46.21951437 | 53.71768876 | -0.135734881 | 0.2119922 | 0.499164656 | 0.561730307 | ns | YPD | KS.3 |
| FIM1_1787 | 2158.35158 | 1212.130796 | 2637.023263 | -0.695864179 | 0.291565691 | 0.008126567 | 0.014038081 | ns | YPD | KS.3 |
| FIM1_1788 | 636.2515506 | 473.8146029 | 541.1282185 | -0.405139641 | 0.142334326 | 0.003498968 | 0.006504995 | ns | YPD | KS.3 |
| FIM1_1789 | 825.7586338 | 1979.009944 | 1310.35748 | 1.112916293 | 0.2641825 | 4.0604845448 | 1.4047921147 | Km>KS.3 | YPD | KS.3 |
| FIM1_1790 | 535.5924298 | 833.9300008 | 819.0189478 | 0.596695697 | 0.174218126 | 0.000372112 | 0.000844167 | ns | YPD | KS.3 |
| FIM1_1791 | 84.94868597 | 693.4742435 | 360.2382304 | 2.968109866 | 0.190396113 | 2.3750527637 | 2.9539718748 | Km>KS.3 | YPD | KS.3 |
| FIM1_1792 | 4310.080773 | 7448.915534 | 6424.741287 | 0.696579922 | 0.241880769 | 0.001959784 | 0.003826502 | ns | YPD | KS.3 |
| FIM1_1793 | 1869.826324 | 1697.354854 | 1819.898147 | -0.134471304 | 0.123620483 | 0.267533448 | 0.325154057 | ns | YPD | KS.3 |
| FIM1_1794 | 630.4780862 | 982.8000543 | 681.1152267 | 0.491924335 | 0.344193067 | 0.099688309 | 0.133967947 | ns | YPD | KS.3 |
| FIM1_1795 | 815.146058 | 1115.267025 | 820.1649257 | 0.448132383 | 0.058392614 | 1.2501664077 | 2.3470105202 | ns | YPD | KS.3 |
| FIM1_1796 | 74.43080719 | 52.74439317 | 82.68238282 | -0.441319381 | 0.230188932 | 0.040868617 | 0.06027909 | ns | YPD | KS.3 |
| FIM1_1797 | 1683.253843 | 1686.933782 | 1607.787515 | 0.002660177 | 0.233515695 | 0.990299794 | 0.99089732 | ns | YPD | KS.3 |
| FIM1_1798 | 799.4235054 | 3335.637511 | 1934.363292 | 1.883761765 | 0.303822606 | 2.9356694591 | 3.5108066248 | Km>KS.3 | YPD | KS.3 |
| FIM1_1799 | 1363.64334 | 1455.680175 | 1747.339532 | 0.090364838 | 0.121857729 | 0.449978437 | 0.513452879 | ns | YPD | KS.3 |
| FIM1_1800 | 472.1615354 | 941.5765772 | 846.4389477 | 0.895338323 | 0.230093983 | 3.1499167224 | 8.9038839170 | ns | YPD | KS.3 |
| FIM1_1801 | 926.8985056 | 699.5804225 | 2016.171127 | -0.328905928 | 0.302312269 | 0.225041566 | 0.278710926 | ns | YPD | KS.3 |
| FIM1_1802 | 752.9217172 | 5799.554942 | 3150.560927 | 2.815602284 | 0.289018003 | 2.2062971573 | 1.9956960650 | Km>KS.3 | YPD | KS.3 |
| FIM1_1803 | 824.8779529 | 4672.882288 | 4401.067168 | 2.212565252 | 0.420932213 | 4.4693086379 | 3.4207400729 | Km>KS.3 | YPD | KS.3 |
| FIM1_1804 | 293.9090374 | 934.7777886 | 737.2140736 | 1.51848365 | 0.261835284 | 1.8581268206 | 1.5562594162 | Km>KS.3 | YPD | KS.3 |
| FIM1_1805 | 70.20713266 | 39.86712325 | 87.50925761 | -0.676438833 | 0.298068155 | 0.011482528 | 0.01919791 | ns | YPD | KS.3 |
| FIM1_1806 | 765.8440403 | 1722.73539 | 1470.141121 | 1.00571951 | 0.284177232 | 8.9914382930 | 0.000234201 | Km>KS.3 | YPD | KS.3 |
| FIM1_1807 | 56.08063832 | 259.0537477 | 206.0439287 | 2.098937746 | 0.244770835 | 2.4131438144 | 1.3799299398 | Km>KS.3 | YPD | KS.3 |
| FIM1_1808 | 687.7541522 | 4693.936212 | 2362.410885 | 2.64624247 | 0.284234087 | 8.1724743470 | 5.8924724458 | Km>KS.3 | YPD | KS.3 |
| FIM1_1809 | 1181.573333 | 7051.350756 | 3503.93053 | 2.395582489 | 0.338835036 | 4.4575128964 | 8.9419865563 | Km>KS.3 | YPD | KS.3 |
| FIM1_1810 | 1115.755781 | 2095.921824 | 1536.574471 | 0.869633468 | 0.145595012 | 8.0062091763 | 5.8238549695 | ns | YPD | KS.3 |
| FIM1_1811 | 1643.70795 | 1364.734412 | 1497.327843 | -0.260390981 | 0.11133317 | 0.017500013 | 0.028221253 | ns | YPD | KS.3 |
| FIM1_1812 | 500.2203262 | 805.0272564 | 669.3733824 | 0.630155744 | 0.196279742 | 0.000752756 | 0.001598362 | ns | YPD | KS.3 |
| FIM1_1813 | 450.8594839 | 563.6855529 | 514.8656018 | 0.306204389 | 0.143623001 | 0.028695587 | 0.04391281 | ns | YPD | KS.3 |
| FIM1_1814 | 839.0677592 | 975.0853066 | 879.8759774 | 0.210311753 | 0.107231519 | 0.046570914 | 0.067666558 | ns | YPD | KS.3 |
| FIM1_1815 | 481.2499418 | 855.4132308 | 661.3282019 | 0.802812295 | 0.122510269 | 2.2745724628 | 2.2722887555 | ns | YPD | KS.3 |
| FIM1_1816 | 2362.301206 | 2743.509338 | 2257.114841 | 0.206892908 | 0.130366394 | 0.10500045 | 0.140650845 | ns | YPD | KS.3 |
| FIM1_1817 | 1700.027246 | 2317.236194 | 1715.826269 | 0.426746304 | 0.138643595 | 0.001610644 | 0.003201341 | ns | YPD | KS.3 |

| | | | | | | | | | |
|---|---|---|---|---|---|---|---|---|---|
| FIM1_1818 | 15007.26937 | 19281.94959 | 24018.39903 | 0.35564727 | 0.080853182 | 9.1245100007 | 2.8876868481 | ns | YPD | KS.3 |
| FIM1_1819 | 2126.234773 | 3331.780665 | 2817.807409 | 0.62598859 | 0.122157936 | 1.7308448943 | 7.9804942999 | ns | YPD | KS.3 |
| FIM1_1820 | 521.0419439 | 410.643781 | 505.4351108 | -0.319035443 | 0.178361082 | 0.062889568 | 0.088608214 | ns | YPD | KS.3 |
| FIM1_1821 | 1687.434361 | 1303.815824 | 1756.485633 | -0.356103926 | 0.136004455 | 0.007359677 | 0.012842649 | ns | YPD | KS.3 |
| FIM1_1822 | 502.3918781 | 561.9860292 | 470.8690079 | 0.155109023 | 0.127700801 | 0.215121663 | 0.267892434 | ns | YPD | KS.3 |
| FIM1_1823 | 236.2274571 | 157.0200615 | 249.6394333 | -0.460520961 | 0.333035899 | 0.114420316 | 0.151716704 | ns | YPD | KS.3 |
| FIM1_1824 | 618.5849802 | 1393.311303 | 1074.180864 | 1.098264097 | 0.184768614 | 4.8268149810 | 3.6494535761 | Km>KS.3 | YPD | KS.3 |
| FIM1_1825 | 4918.163798 | 2965.611746 | 4575.038093 | -0.686227556 | 0.167200769 | 2.0970294032 | 6.1875062717 | ns | YPD | KS.3 |
| FIM1_1826 | 5984.597198 | 9434.473663 | 6200.367689 | 0.588021669 | 0.223346255 | 0.005109065 | 0.009149603 | ns | YPD | KS.3 |
| FIM1_1827 | 4125.596056 | 4445.20732 | 5286.396663 | 0.1043658 | 0.10908622 | 0.331443773 | 0.393164704 | ns | YPD | KS.3 |
| FIM1_1828 | 3791.599011 | 5033.75597 | 3775.938771 | 0.39220653 | 0.131520277 | 0.002301259 | 0.004432351 | ns | YPD | KS.3 |
| FIM1_1829 | 181.8740802 | 267.659283 | 281.7080514 | 0.527908261 | 0.152982263 | 0.00037798 | 0.000856308 | ns | YPD | KS.3 |
| FIM1_1830 | 1867.743467 | 2009.551063 | 1821.326502 | 0.102347693 | 0.106062684 | 0.327673724 | 0.389249469 | ns | YPD | KS.3 |
| FIM1_1831 | 692.3945843 | 356.4340799 | 557.696713 | -0.920145688 | 0.137998423 | 7.7888059669 | 8.5539314978 | ns | YPD | KS.3 |
| FIM1_1832 | 588.024519 | 434.8990835 | 410.2668753 | -0.416375189 | 0.135197273 | 0.001621129 | 0.003219608 | ns | YPD | KS.3 |
| FIM1_1833 | 858.9372067 | 736.3294282 | 903.3919552 | -0.218947709 | 0.075484565 | 0.003473628 | 0.006460299 | ns | YPD | KS.3 |
| FIM1_1834 | 726.9661221 | 1387.767614 | 892.8793349 | 0.879289091 | 0.16900659 | 6.5850342378 | 3.2892113788 | ns | YPD | KS.3 |
| FIM1_1835 | 520.6027459 | 828.4709417 | 680.737761 | 0.650255708 | 0.113866905 | 6.2433616051 | 3.7786768838 | ns | YPD | KS.3 |
| FIM1_1836 | 1671.691114 | 2532.675947 | 2170.345231 | 0.555355395 | 0.183185748 | 0.001567434 | 0.003125444 | ns | YPD | KS.3 |
| FIM1_1837 | 3440.123905 | 4881.067068 | 5368.555356 | 0.453586783 | 0.215971659 | 0.026177658 | 0.040445295 | ns | YPD | KS.3 |
| FIM1_1838 | 107.1580142 | 155.0549751 | 170.4799956 | 0.473818917 | 0.223826859 | 0.024412326 | 0.037894328 | ns | YPD | KS.3 |
| FIM1_1839 | 2120.403307 | 2305.570801 | 3181.795781 | 0.115497792 | 0.132843479 | 0.374169191 | 0.437380574 | ns | YPD | KS.3 |
| FIM1_1840 | 4692.457436 | 7971.918927 | 7274.870358 | 0.711947694 | 0.181056591 | 4.1238500183 | 0.000113412 | ns | YPD | KS.3 |
| FIM1_1841 | 739.5941874 | 968.3307827 | 1025.017145 | 0.381029684 | 0.086970403 | 9.6404002033 | 3.0316682055 | ns | YPD | KS.3 |
| FIM1_1842 | 809.815007 | 1650.8862 | 1034.982679 | 0.900076909 | 0.25873388 | 0.000154532 | 0.000384013 | ns | YPD | KS.3 |
| FIM1_1843 | 7160.626902 | 9919.013227 | 8375.507969 | 0.457836139 | 0.104268052 | 8.4336081397 | 2.6912893197 | ns | YPD | KS.3 |
| FIM1_1844 | 76.73469086 | 58.53770431 | 62.51638684 | -0.342777845 | 0.238274818 | 0.123440166 | 0.162981642 | ns | YPD | KS.3 |
| FIM1_1845 | 560.5821294 | 515.5970646 | 543.7782126 | -0.116999804 | 0.115988053 | 0.304930615 | 0.365197354 | ns | YPD | KS.3 |
| FIM1_1846 | 3003.513269 | 3731.25973 | 3597.826933 | 0.300589751 | 0.127861754 | 0.016400687 | 0.026603658 | ns | YPD | KS.3 |
| FIM1_1847 | 3978.201818 | 5201.030763 | 4023.427754 | 0.359246179 | 0.176679437 | 0.034590649 | 0.051860207 | ns | YPD | KS.3 |
| FIM1_1848 | 4300.981533 | 7193.854954 | 7128.816569 | 0.684640579 | 0.192151038 | 0.000188812 | 0.000455769 | ns | YPD | KS.3 |
| FIM1_1849 | 1461.970683 | 1949.101291 | 3645.990782 | 0.375279214 | 0.207617559 | 0.056766795 | 0.080689944 | ns | YPD | KS.3 |
| FIM1_1850 | 422.266518 | 816.8549521 | 680.5337264 | 0.887256721 | 0.184742901 | 5.1302886589 | 2.1287060949 | ns | YPD | KS.3 |
| FIM1_1851 | 2883.016192 | 2566.209182 | 3843.198155 | -0.164391567 | 0.093082113 | 0.07412216 | 0.102718035 | ns | YPD | KS.3 |
| FIM1_1853 | 25378.78804 | 48156.95198 | 51825.76471 | 0.885213774 | 0.143167891 | 2.0743990741 | 1.7086316877 | ns | YPD | KS.3 |
| FIM1_1854 | 3237.452347 | 8070.888285 | 6421.667845 | 1.21499397 | 0.216156297 | 2.1314114947 | 1.4329421873 | Km>KS.3 | YPD | KS.3 |
| FIM1_1855 | 503.4072487 | 1369.864013 | 833.8984346 | 1.311072031 | 0.244587856 | 6.2196392558 | 3.7689044211 | Km>KS.3 | YPD | KS.3 |
| FIM1_1856 | 1099.707012 | 2419.392199 | 1627.702602 | 1.042229916 | 0.214606325 | 2.4861284898 | 1.1053162856 | Km>KS.3 | YPD | KS.3 |
| FIM1_1857 | 6777.278653 | 9056.700607 | 6646.68315 | 0.398393044 | 0.142880483 | 0.004223073 | 0.00770884 | ns | YPD | KS.3 |
| FIM1_1858 | 985.4500887 | 1049.57408 | 1277.778027 | 0.087479089 | 0.126684322 | 0.481139732 | 0.544510957 | ns | YPD | KS.3 |
| FIM1_1859 | 2160.172258 | 1196.612586 | 1973.780227 | -0.805868037 | 0.162180334 | 2.7024938008 | 1.1877126023 | ns | YPD | KS.3 |
| FIM1_1860 | 1194.112503 | 2519.084851 | 1463.585236 | 0.980283429 | 0.219870596 | 2.0710704761 | 7.5594832126 | ns | YPD | KS.3 |
| FIM1_1861 | 650.5045598 | 542.6587762 | 621.8867677 | -0.250087291 | 0.1342146 | 0.056677543 | 0.080586103 | ns | YPD | KS.3 |
| FIM1_1862 | 899.842326 | 1174.104699 | 889.5884481 | 0.36789408 | 0.131235026 | 0.004168135 | 0.007612509 | ns | YPD | KS.3 |
| FIM1_1863 | 965.822813 | 441.3627456 | 657.76852 | -1.113003953 | 0.085155471 | 7.4134075087 | 2.4587801570 | Km<KS.3 | YPD | KS.3 |

| | | | | | | | | | | |
|---|---|---|---|---|---|---|---|---|---|---|
| FIM1_1864 | 1602.605318 | 1190.832974 | 1432.97111 | -0.414326816 | 0.115295609 | 0.000256477 | 0.000599893 | ns | YPD | KS.3 |
| FIM1_1865 | 616.6510886 | 944.3595588 | 897.4371506 | 0.567310226 | 0.188519189 | 0.001654083 | 0.003281126 | ns | YPD | KS.3 |
| FIM1_1866 | 188.9604287 | 338.6901188 | 472.7962783 | 0.726464078 | 0.264928896 | 0.002794219 | 0.005301769 | ns | YPD | KS.3 |
| FIM1_1867 | 215.7241824 | 375.6443846 | 199.9969661 | 0.750108431 | 0.173940908 | 7.3242747769( | 2.38453526423 | ns | YPD | KS.3 |
| FIM1_1868 | 2265.731439 | 1761.061857 | 1780.688788 | -0.347995815 | 0.134341311 | 0.008048427 | 0.013922435 | ns | YPD | KS.3 |
| FIM1_1869 | 2654.871026 | 1497.064664 | 2045.223674 | -0.798987148 | 0.124604526 | 5.84169637198 | 5.38193323159 | ns | YPD | KS.3 |
| FIM1_1870 | 1722.081468 | 1171.190368 | 1245.736782 | -0.534033991 | 0.130907109 | 3.03510269469 | 8.61851364502 | ns | YPD | KS.3 |
| FIM1_1871 | 1809.783111 | 2509.757105 | 1710.552191 | 0.439801532 | 0.173508152 | 0.008500236 | 0.014607487 | ns | YPD | KS.3 |
| FIM1_1872 | 950.6856236 | 1219.399192 | 908.4532218 | 0.340007351 | 0.150598273 | 0.020205439 | 0.032013394 | ns | YPD | KS.3 |
| FIM1_1873 | 1605.101084 | 1032.814294 | 969.815206 | -0.615138147 | 0.119748071 | 1.65037582137 | 7.65916017849 | ns | YPD | KS.3 |
| FIM1_1874 | 3614.510706 | 2759.865991 | 1626.170728 | -0.365568443 | 0.162636956 | 0.020187052 | 0.031998682 | ns | YPD | KS.3 |
| FIM1_1875 | 111.5327061 | 85.77271418 | 58.7017236 | -0.341454209 | 0.212569052 | 0.08971147 | 0.121644744 | ns | YPD | KS.3 |
| FIM1_1876 | 304.5866157 | 816.3589675 | 644.4371434 | 1.147484204 | 0.358716447 | 0.000181952 | 0.000443078 | Km>KS.3 | YPD | KS.3 |
| FIM1_1877 | 822.9013609 | 620.7889568 | 1186.719207 | -0.316941788 | 0.328480423 | 0.272585845 | 0.330679 | ns | YPD | KS.3 |
| FIM1_1878 | 1465.39569 | 5336.723783 | 2956.918088 | 1.570434884 | 0.377062707 | 7.26282083053 | 2.93521800421 | Km>KS.3 | YPD | KS.3 |
| FIM1_1879 | 604.8265298 | 459.5680724 | 476.7881425 | -0.385815654 | 0.10516641 | 0.000197977 | 0.000475127 | ns | YPD | KS.3 |
| FIM1_1880 | 3820.820263 | 3072.632868 | 3492.685695 | -0.302023804 | 0.127948271 | 0.015962832 | 0.025952643 | ns | YPD | KS.3 |
| FIM1_1881 | 1044.997273 | 946.3490638 | 750.7455602 | -0.138332291 | 0.111366178 | 0.207044197 | 0.258935365 | ns | YPD | KS.3 |
| FIM1_1882 | 1801.587908 | 4152.761118 | 3247.16289 | 1.118185378 | 0.20098609 | 4.27597375949 | 2.69278094347 | Km>KS.3 | YPD | KS.3 |
| FIM1_1883 | 149.3271336 | 252.166824 | 133.49796 | 0.647246597 | 0.268054145 | 0.008385863 | 0.014425887 | ns | YPD | KS.3 |
| FIM1_1884 | 3966.268943 | 5030.201206 | 4422.520693 | 0.316951304 | 0.182099193 | 0.069869565 | 0.097394532 | ns | YPD | KS.3 |
| FIM1_1885 | 2056.547654 | 1206.195178 | 1568.879703 | -0.752467263 | 0.100117341 | 2.54054267158 | 4.54647474500( | ns | YPD | KS.3 |
| FIM1_1886 | 1089.226113 | 829.5793171 | 895.1978618 | -0.379224046 | 0.118196606 | 0.00108949 | 0.002241695 | ns | YPD | KS.3 |
| FIM1_1887 | 6969.165884 | 4579.717879 | 5765.02779 | -0.566111172 | 0.172754578 | 0.000667102 | 0.001433003 | ns | YPD | KS.3 |
| FIM1_1888 | 2336.523399 | 2037.99511 | 2288.547933 | -0.195705986 | 0.059508468 | 0.000951824 | 0.001976753 | ns | YPD | KS.3 |
| FIM1_1889 | 1630.243494 | 1606.43879 | 1442.223128 | -0.020629735 | 0.117020891 | 0.8576702 | 0.88141071 | ns | YPD | KS.3 |
| FIM1_1890 | 417.2624407 | 431.0212285 | 1155.395381 | 0.033720467 | 0.370472336 | 0.914632544 | 0.930911805 | ns | YPD | KS.3 |
| FIM1_1891 | 1263.493111 | 2625.681647 | 1454.40925 | 0.886053672 | 0.298624003 | 0.00092795 | 0.001935862 | ns | YPD | KS.3 |
| FIM1_1892 | 986.4820136 | 683.8802543 | 894.0127034 | -0.466463592 | 0.23459817 | 0.033493378 | 0.050447943 | ns | YPD | KS.3 |
| FIM1_1893 | 2037.628882 | 1437.03948 | 1753.076194 | -0.486579807 | 0.121589422 | 4.52106636902 | 0.000123313 | ns | YPD | KS.3 |
| FIM1_1894 | 1482.960196 | 1897.364958 | 1675.682039 | 0.336998322 | 0.149585781 | 0.020521212 | 0.032451695 | ns | YPD | KS.3 |
| FIM1_1895 | 1269.005425 | 1084.420754 | 1091.70638 | -0.219316885 | 0.11401968 | 0.050534352 | 0.072745486 | ns | YPD | KS.3 |
| FIM1_1896 | 1766.693198 | 934.9065197 | 1174.592518 | -0.89993595 | 0.094683296 | 6.22296009228 | 3.91888942520( | ns | YPD | KS.3 |
| FIM1_1897 | 2140.239816 | 1379.503471 | 1873.564261 | -0.619009564 | 0.100327172 | 4.00332593766 | 3.09263145029 | ns | YPD | KS.3 |
| FIM1_1898 | 1741.062479 | 1684.427793 | 1951.388759 | -0.046725298 | 0.093040015 | 0.611596237 | 0.667989304 | ns | YPD | KS.3 |
| FIM1_1899 | 6213.501834 | 7800.064091 | 8322.433107 | 0.31332917 | 0.138088165 | 0.020127259 | 0.031930202 | ns | YPD | KS.3 |
| FIM1_1900 | 578.6402513 | 369.5966741 | 403.9192299 | -0.625830927 | 0.117232285 | 5.43909212163 | 2.76965028711 | ns | YPD | KS.3 |
| FIM1_1901 | 3488.677084 | 5538.58459 | 4115.123878 | 0.626331925 | 0.167160322 | 0.000103252 | 0.000265879 | ns | YPD | KS.3 |
| FIM1_1902 | 951.2828573 | 678.6059509 | 880.7159253 | -0.461672686 | 0.152077248 | 0.00177374 | 0.003498952 | ns | YPD | KS.3 |
| FIM1_1903 | 1148.676719 | 963.7409899 | 936.9536444 | -0.247740915 | 0.089611962 | 0.005213364 | 0.009322964 | ns | YPD | KS.3 |
| FIM1_1904 | 4336.74985 | 4014.131295 | 5400.172419 | -0.106407216 | 0.136858934 | 0.42618326 | 0.490204626 | ns | YPD | KS.3 |
| FIM1_1905 | 1402.93265 | 898.8344421 | 1071.12881 | -0.618115111 | 0.1296006 | 1.08642904411 | 4.24586370341 | ns | YPD | KS.3 |
| FIM1_1907 | 1452.988366 | 815.2519857 | 1143.469546 | -0.779322365 | 0.178184183 | 5.19588365204 | 1.75369885813 | ns | YPD | KS.3 |
| FIM1_1908 | 4514.782161 | 2680.020201 | 4344.102142 | -0.715548411 | 0.150986168 | 1.04944011623 | 4.11423528625 | ns | YPD | KS.3 |
| FIM1_1909 | 4608.173488 | 5832.832482 | 4253.410281 | 0.331831379 | 0.098721702 | 0.000664054 | 0.001427688 | ns | YPD | KS.3 |

| | | | | | | | | | | |
|---|---|---|---|---|---|---|---|---|---|---|
| FIM1_1910 | 1693.109952 | 1328.416401 | 1422.179399 | -0.331556295 | 0.15009597 | 0.02309234 | 0.036056268 | ns | YPD | KS.3 |
| FIM1_1911 | 7377.493472 | 6544.661007 | 7619.23478 | -0.170121218 | 0.077567158 | 0.027068379 | 0.041717839 | ns | YPD | KS.3 |
| FIM1_1912 | 2397.373947 | 4125.83294 | 3167.10355 | 0.758759221 | 0.119274868 | 8.90848321907 | 7.98553225493 | ns | YPD | KS.3 |
| FIM1_1913 | 2541.426134 | 1129.015483 | 2600.344044 | -1.037846801 | 0.25421635 | 9.22816480485 | 2.91123144604 | Km<KS.3 | YPD | KS.3 |
| FIM1_1914 | 666.6967448 | 280.9622141 | 540.0841743 | -1.185565231 | 0.167662461 | 1.92349804989 | 2.83260058580 | Km<KS.3 | YPD | KS.3 |
| FIM1_1915 | 1660.177345 | 3302.620642 | 2001.753845 | 0.872868246 | 0.252980108 | 0.000184883 | 0.000449117 | ns | YPD | KS.3 |
| FIM1_1916 | 1041.597188 | 1191.377335 | 1040.49571 | 0.17550577 | 0.203213722 | 0.364004041 | 0.426801816 | ns | YPD | KS.3 |
| FIM1_1917 | 1141.460587 | 459.6713368 | 992.6185635 | -1.176152318 | 0.251067689 | 3.62284739384 | 1.54576893519 | Km<KS.3 | YPD | KS.3 |
| FIM1_1918 | 1095.18893 | 672.9644055 | 883.3691243 | -0.66389819 | 0.160384443 | 1.87971712584 | 5.61320091010 | ns | YPD | KS.3 |
| FIM1_1919 | 1270.508963 | 1655.092134 | 1482.241881 | 0.368999776 | 0.117625296 | 0.001407444 | 0.002833685 | ns | YPD | KS.3 |
| FIM1_1920 | 2326.715124 | 933.5068338 | 2593.445431 | -1.100018356 | 0.321337127 | 9.95201696539 | 0.000257335 | Km<KS.3 | YPD | KS.3 |
| FIM1_1921 | 5268.567023 | 1162.133752 | 2806.918951 | -2.131820747 | 0.165000986 | 7.20741909267 | 2.98807583217 | Km<KS.3 | YPD | KS.3 |
| FIM1_1922 | 2280.075378 | 2088.411738 | 2248.488215 | -0.125420575 | 0.060064348 | 0.035887999 | 0.053600359 | ns | YPD | KS.3 |
| FIM1_1923 | 2326.698466 | 2190.680732 | 2652.797944 | -0.085374822 | 0.089889548 | 0.337216407 | 0.398871047 | ns | YPD | KS.3 |
| FIM1_1924 | 896.44279 | 660.4551784 | 1067.849095 | -0.414365307 | 0.163593529 | 0.00882131 | 0.015101864 | ns | YPD | KS.3 |
| FIM1_1925 | 2836.364163 | 1932.178981 | 2161.135795 | -0.543111981 | 0.090450941 | 1.27247441824 | 8.91628201515 | ns | YPD | KS.3 |
| FIM1_1926 | 711.1246239 | 496.405714 | 614.4443236 | -0.480137221 | 0.1826397 | 0.00613914 | 0.010849812 | ns | YPD | KS.3 |
| FIM1_1927 | 3429.516231 | 2134.329405 | 3871.751739 | -0.569749131 | 0.289078455 | 0.029236905 | 0.044617669 | ns | YPD | KS.3 |
| FIM1_1928 | 2391.142645 | 1783.403619 | 1626.853045 | -0.409763215 | 0.114170091 | 0.000262353 | 0.00061306 | ns | YPD | KS.3 |
| FIM1_1929 | 1071.211605 | 795.9643204 | 577.4460461 | -0.371773452 | 0.248004844 | 0.106447634 | 0.142282907 | ns | YPD | KS.3 |
| FIM1_1930 | 988.9391084 | 619.6749994 | 719.1019424 | -0.642997311 | 0.14640794 | 6.31709797446 | 2.09011516699 | ns | YPD | KS.3 |
| FIM1_1931 | 9414.78394 | 6965.073773 | 12554.5319 | -0.428688626 | 0.075228844 | 9.35301647257 | 5.50665762734 | ns | YPD | KS.3 |
| FIM1_1932 | 587.656408 | 534.9642798 | 615.3969019 | -0.130398418 | 0.118886158 | 0.264227966 | 0.321637909 | ns | YPD | KS.3 |
| FIM1_1933 | 5029.645253 | 5926.330004 | 5025.61618 | 0.227853385 | 0.124150233 | 0.061316411 | 0.086563321 | ns | YPD | KS.3 |
| FIM1_1934 | 260.0404336 | 720.098898 | 592.1345006 | 1.393881516 | 0.180860265 | 6.35276906557 | 1.47000121400 | Km>KS.3 | YPD | KS.3 |
| FIM1_1935 | 1133.980554 | 2836.268204 | 2584.393175 | 1.262242802 | 0.163193848 | 9.40587820002 | 2.07054177190 | Km>KS.3 | YPD | KS.3 |
| FIM1_1936 | 4590.48227 | 6202.622277 | 5334.781665 | 0.430237539 | 0.0596224 | 4.12165824098 | 5.74376743667 | ns | YPD | KS.3 |
| FIM1_1937 | 2372.109774 | 1846.541234 | 2724.981043 | -0.351932903 | 0.103026438 | 0.000533846 | 0.001167934 | ns | YPD | KS.3 |
| FIM1_1938 | 1132.965117 | 768.8573708 | 1245.841524 | -0.524458877 | 0.168278968 | 0.001238987 | 0.002522078 | ns | YPD | KS.3 |
| FIM1_1939 | 219.0439276 | 207.2916126 | 307.9868761 | -0.069834346 | 0.225893096 | 0.742460812 | 0.784234084 | ns | YPD | KS.3 |
| FIM1_1940 | 2602.029771 | 3271.130658 | 3000.403003 | 0.294186766 | 0.221530887 | 0.158826375 | 0.204758025 | ns | YPD | KS.3 |
| FIM1_1941 | 994.112067 | 2474.946948 | 1548.488475 | 1.172638928 | 0.257508392 | 6.87425818992 | 2.78951341719 | Km>KS.3 | YPD | KS.3 |
| FIM1_1942 | 715.4342715 | 1173.911448 | 919.5417893 | 0.672164158 | 0.165980803 | 2.72386939843 | 7.80601973341 | ns | YPD | KS.3 |
| FIM1_1943 | 1304.28345 | 1034.213847 | 1103.740139 | -0.316181695 | 0.153911886 | 0.034372279 | 0.05161548 | ns | YPD | KS.3 |
| FIM1_1944 | 706.710664 | 616.7872795 | 568.8789386 | -0.19001575 | 0.111489747 | 0.083387768 | 0.114002238 | ns | YPD | KS.3 |
| FIM1_1945 | 510.726386 | 679.4837314 | 776.3111489 | 0.38894594 | 0.155096483 | 0.009756492 | 0.016583036 | ns | YPD | KS.3 |
| FIM1_1946 | 1746.408177 | 3150.521775 | 2199.170287 | 0.808488521 | 0.155068784 | 7.38884106035 | 3.67227615137 | ns | YPD | KS.3 |
| FIM1_1947 | 1110.281531 | 828.1368288 | 995.8216166 | -0.395469995 | 0.168955474 | 0.01530231 | 0.024952144 | ns | YPD | KS.3 |
| FIM1_1948 | 2142.590385 | 2353.997189 | 2939.991438 | 0.131684672 | 0.110501158 | 0.226195286 | 0.279930733 | ns | YPD | KS.3 |
| FIM1_1949 | 490.3748266 | 281.6964339 | 448.1257848 | -0.752622497 | 0.167991328 | 3.36959137255 | 1.18054345622 | ns | YPD | KS.3 |
| FIM1_1950 | 834.061609 | 1113.124814 | 1066.766058 | 0.40186623 | 0.119748284 | 0.000630356 | 0.001361711 | ns | YPD | KS.3 |
| FIM1_1951 | 1229.769401 | 1110.661946 | 1505.63966 | -0.13999331 | 0.140397586 | 0.306903896 | 0.367207043 | ns | YPD | KS.3 |
| FIM1_1952 | 1691.462892 | 2554.367027 | 2539.892516 | 0.580298909 | 0.100905458 | 5.55522237844 | 3.41200386824 | ns | YPD | KS.3 |
| FIM1_1953 | 1329.94258 | 4599.213718 | 3001.487266 | 1.675267354 | 0.231276992 | 5.07899408245 | 1.01071982240 | Km>KS.3 | YPD | KS.3 |
| FIM1_1954 | 786.7960887 | 1302.833978 | 2132.792289 | 0.517174605 | 0.392980171 | 0.113679056 | 0.150860663 | ns | YPD | KS.3 |

| | | | | | | | | | | |
|---|---|---|---|---|---|---|---|---|---|---|
| FIM1_1955 | 579.9166881 | 869.9790709 | 615.9159347 | 0.569582348 | 0.106930223 | 6.37352752525 | 3.19640115303 | ns | YPD | KS.3 |
| FIM1_1956 | 1132.44732 | 707.8495229 | 737.4129827 | -0.619235754 | 0.201824287 | 0.001244227 | 0.002530675 | ns | YPD | KS.3 |
| FIM1_1957 | 1779.659388 | 1203.758025 | 1281.058434 | -0.539357792 | 0.138541369 | 6.59687488978 | 0.000174851 | ns | YPD | KS.3 |
| FIM1_1958 | 1081.892311 | 1449.010903 | 1017.167306 | 0.397545774 | 0.157473541 | 0.009214545 | 0.015731765 | ns | YPD | KS.3 |
| FIM1_1959 | 792.3300996 | 916.4667474 | 825.9505333 | 0.202757402 | 0.117138477 | 0.078215553 | 0.107730447 | ns | YPD | KS.3 |
| FIM1_1960 | 2628.898498 | 2914.819552 | 2838.54034 | 0.146521615 | 0.075065728 | 0.049256294 | 0.071173413 | ns | YPD | KS.3 |
| FIM1_1961 | 2497.300511 | 6427.112087 | 3122.551629 | 1.277974333 | 0.195561191 | 5.47723171521 | 6.19300631436 | Km>KS.3 | YPD | KS.3 |
| FIM1_1962 | 19974.72145 | 30581.75794 | 36346.13897 | 0.593962611 | 0.12096514 | 5.57381772916 | 2.29550854326 | ns | YPD | KS.3 |
| FIM1_1963 | 441.5150612 | 601.4461422 | 534.5366959 | 0.419363956 | 0.160089759 | 0.006832794 | 0.011982076 | ns | YPD | KS.3 |
| FIM1_1964 | 12670.23562 | 11349.10109 | 12314.88059 | -0.140004927 | 0.230599816 | 0.517623994 | 0.579734212 | ns | YPD | KS.3 |
| FIM1_1965 | 922.895167 | 1879.484244 | 1120.310801 | 0.987303687 | 0.137361767 | 1.62676424422 | 2.45247033787 | ns | YPD | KS.3 |
| FIM1_1966 | 1579.376191 | 1559.79849 | 1545.494868 | -0.017151954 | 0.116519553 | 0.880979425 | 0.901825646 | ns | YPD | KS.3 |
| FIM1_1967 | 2303.627115 | 3256.944009 | 2172.736869 | 0.481683927 | 0.12407533 | 7.49267272271 | 0.000196707 | ns | YPD | KS.3 |
| FIM1_1968 | 1886.366113 | 2365.801213 | 2275.329364 | 0.291794554 | 0.219160026 | 0.158258482 | 0.204114117 | ns | YPD | KS.3 |
| FIM1_1970 | 530.4539193 | 357.7665304 | 570.7668803 | -0.533475717 | 0.168101812 | 0.001006784 | 0.002084373 | ns | YPD | KS.3 |
| FIM1_1971 | 486.8858277 | 373.4202946 | 411.3584587 | -0.361109102 | 0.159235573 | 0.019260641 | 0.030721926 | ns | YPD | KS.3 |
| FIM1_1972 | 886.3214346 | 3951.836387 | 3532.771367 | 1.954970801 | 0.330981107 | 1.51457773526 | 1.56328303588 | Km>KS.3 | YPD | KS.3 |
| FIM1_1973 | 3590.565178 | 8024.693175 | 7261.775774 | 1.028378876 | 0.253775643 | 1.08229763546 | 3.37569706860 | Km>KS.3 | YPD | KS.3 |
| FIM1_1975 | 1073.607656 | 1135.161968 | 1218.097813 | 0.078116117 | 0.103500358 | 0.444236422 | 0.507825163 | ns | YPD | KS.3 |
| FIM1_1976 | 2559.247287 | 4257.277149 | 2935.738703 | 0.701233038 | 0.143610249 | 5.26404850396 | 2.17875551640 | ns | YPD | KS.3 |
| FIM1_1977 | 753.7925236 | 1091.507188 | 1034.614862 | 0.523383361 | 0.090788788 | 5.58225014457 | 3.41424610340 | ns | YPD | KS.3 |
| FIM1_1978 | 1272.015251 | 3048.553565 | 2647.71356 | 1.132432281 | 0.245151654 | 5.84521877871 | 2.39144436053 | Km>KS.3 | YPD | KS.3 |
| FIM1_1979 | 885.4764057 | 1047.631883 | 814.5126033 | 0.238587159 | 0.078400627 | 0.002156155 | 0.00417388 | ns | YPD | KS.3 |
| FIM1_1980 | 10267.08658 | 7695.794701 | 8603.960782 | -0.384596683 | 0.182820845 | 0.028368723 | 0.043452709 | ns | YPD | KS.3 |
| FIM1_1981 | 8116.053737 | 5718.821346 | 10239.13225 | -0.460743761 | 0.200086015 | 0.015557098 | 0.0253261 | ns | YPD | KS.3 |
| FIM1_1982 | 15.59522173 | 129.1408558 | 64.93626001 | 2.735021816 | 0.450151302 | 5.22491541201 | 5.93469273396 | Km>KS.3 | YPD | KS.3 |
| FIM1_1983 | 4498.719262 | 4928.23449 | 4897.795103 | 0.127059146 | 0.117886212 | 0.272739165 | 0.330784336 | ns | YPD | KS.3 |
| FIM1_1984 | 695.9910333 | 462.3504503 | 569.3680288 | -0.572908521 | 0.108205655 | 7.55911386036 | 3.74567643977 | ns | YPD | KS.3 |
| FIM1_1985 | 3744.099602 | 3081.793071 | 2826.378344 | -0.271534009 | 0.117241151 | 0.018435856 | 0.029543139 | ns | YPD | KS.3 |
| FIM1_1986 | 2967.350172 | 2061.842963 | 2153.313448 | -0.516136444 | 0.083688881 | 4.78941030084 | 3.63775820560 | ns | YPD | KS.3 |
| FIM1_1987 | 989.8572695 | 833.4011343 | 767.2735327 | -0.226630254 | 0.195575351 | 0.224830135 | 0.278518407 | ns | YPD | KS.3 |
| FIM1_1988 | 3030.493308 | 2977.400178 | 2531.306992 | -0.023759775 | 0.17230664 | 0.886325058 | 0.906365295 | ns | YPD | KS.3 |
| FIM1_1989 | 788.160666 | 1574.338798 | 1456.817408 | 0.923450423 | 0.196668092 | 7.89432342067 | 3.16217866488 | ns | YPD | KS.3 |
| FIM1_1990 | 2075.38859 | 2184.90305 | 2173.972986 | 0.07191877 | 0.110358975 | 0.508126701 | 0.570252726 | ns | YPD | KS.3 |
| FIM1_1991 | 2727.780365 | 2785.828858 | 2246.902047 | 0.027876292 | 0.188980868 | 0.877604161 | 0.898925407 | ns | YPD | KS.3 |
| FIM1_1992 | 3200.620493 | 5693.173685 | 4555.279446 | 0.749809865 | 0.220230941 | 0.000296057 | 0.000686339 | ns | YPD | KS.3 |
| FIM1_1993 | 1184.661524 | 2038.470072 | 1575.670143 | 0.740036414 | 0.160948163 | 1.98119103566 | 7.30053824133 | ns | YPD | KS.3 |
| FIM1_1994 | 11413.39646 | 14808.59777 | 15695.74925 | 0.369776104 | 0.078984542 | 2.35104145282 | 8.48182104988 | ns | YPD | KS.3 |
| FIM1_1995 | 1868.406961 | 5070.106995 | 3370.377933 | 1.316179004 | 0.235848931 | 1.75483050301 | 1.21085738591 | Km>KS.3 | YPD | KS.3 |
| FIM1_1996 | 7778.128283 | 5235.928093 | 7937.596766 | -0.543023612 | 0.14757436 | 0.000154621 | 0.000384044 | ns | YPD | KS.3 |
| FIM1_1997 | 1897.218529 | 1275.255539 | 1520.024576 | -0.560456757 | 0.094436529 | 1.90019478916 | 1.29499576385 | ns | YPD | KS.3 |
| FIM1_1998 | 1368.529658 | 1533.930545 | 1350.747153 | 0.161597862 | 0.085075607 | 0.055193238 | 0.078678039 | ns | YPD | KS.3 |
| FIM1_1999 | 285.791857 | 849.4267994 | 411.8301405 | 1.472757705 | 0.209291404 | 6.62979729116 | 1.08049270573 | Km>KS.3 | YPD | KS.3 |
| FIM1_2000 | 5117.452264 | 4806.023105 | 5751.658379 | -0.08437485 | 0.171629444 | 0.61037322 | 0.666799905 | ns | YPD | KS.3 |
| FIM1_2001 | 2554.568108 | 2079.865286 | 3238.179242 | -0.250790402 | 0.267919659 | 0.307830171 | 0.368049772 | ns | YPD | KS.3 |

| | | | | | | | | | | |
|---|---|---|---|---|---|---|---|---|---|---|
| FIM1_2002 | 3573.666757 | 2782.682575 | 3293.617611 | -0.349699385 | 0.113791016 | 0.001780619 | 0.003509739 | ns | YPD | KS.3 |
| FIM1_2003 | 4527.687237 | 4652.727357 | 4467.052273 | 0.037130972 | 0.151063452 | 0.800442201 | 0.833444946 | ns | YPD | KS.3 |
| FIM1_2004 | 651.0573415 | 520.0034151 | 610.1034241 | -0.303566759 | 0.166022533 | 0.058529777 | 0.082911629 | ns | YPD | KS.3 |
| FIM1_2005 | 26625.20515 | 31570.60103 | 32739.38233 | 0.228708603 | 0.173176825 | 0.170685642 | 0.21911801 | ns | YPD | KS.3 |
| FIM1_2006 | 826.6697367 | 1533.28484 | 869.3747403 | 0.857325511 | 0.134807643 | 7.1603328680 | 6.4768465488 | ns | YPD | KS.3 |
| FIM1_2007 | 834.1428901 | 557.692284 | 634.0629093 | -0.559468698 | 0.124813839 | 4.7779881211 | 1.6181478569 | ns | YPD | KS.3 |
| FIM1_2008 | 811.950915 | 641.2980045 | 575.9856178 | -0.322086605 | 0.150458435 | 0.027670177 | 0.042500504 | ns | YPD | KS.3 |
| FIM1_2009 | 294.8402145 | 401.5404997 | 814.4961066 | 0.375528361 | 0.274151984 | 0.133890158 | 0.175257947 | ns | YPD | KS.3 |
| FIM1_2010 | 42.81954088 | 81.48547142 | 78.59222337 | 0.822924242 | 0.241610481 | 0.000247541 | 0.000580356 | ns | YPD | KS.3 |
| FIM1_2011 | 3369.354136 | 3312.016743 | 2751.075001 | -0.02410793 | 0.082335636 | 0.767705075 | 0.805087004 | ns | YPD | KS.3 |
| FIM1_2012 | 2809.704708 | 3419.358242 | 3168.947475 | 0.273827354 | 0.118116119 | 0.018292605 | 0.029329237 | ns | YPD | KS.3 |
| FIM1_2013 | 1222.578215 | 1406.970151 | 1113.463587 | 0.182616414 | 0.209300219 | 0.357686898 | 0.420286329 | ns | YPD | KS.3 |
| FIM1_2014 | 1504.654624 | 2030.799517 | 1707.956662 | 0.421494868 | 0.102915094 | 3.2881941875 | 9.2526957483 | ns | YPD | KS.3 |
| FIM1_2015 | 34.04717373 | 9.122806003 | 21.92099029 | -1.741137794 | 0.286000553 | 1.0692312325 | 1.1390632509 | Km<KS.3 | YPD | KS.3 |
| FIM1_2016 | 3498.722297 | 2818.813133 | 2361.274511 | -0.304513427 | 0.09640676 | 0.001389203 | 0.002801493 | ns | YPD | KS.3 |
| FIM1_2017 | 1060.645587 | 1454.577763 | 1226.622931 | 0.419108968 | 0.189638736 | 0.020876413 | 0.032867138 | ns | YPD | KS.3 |
| FIM1_2018 | 279.0261873 | 127.3277135 | 205.6604993 | -0.919372123 | 0.329813359 | 0.001458504 | 0.00292583 | ns | YPD | KS.3 |
| FIM1_2019 | 1486.39105 | 1374.664437 | 1473.133084 | -0.100094567 | 0.222482285 | 0.633101575 | 0.687402955 | ns | YPD | KS.3 |
| FIM1_2020 | 2477.493852 | 4248.759298 | 3206.016417 | 0.71427488 | 0.199261246 | 0.000163681 | 0.000402925 | ns | YPD | KS.3 |
| FIM1_2021 | 3910.341462 | 3834.654355 | 4196.697368 | -0.025922607 | 0.187280193 | 0.885179873 | 0.905508775 | ns | YPD | KS.3 |
| FIM1_2022 | 2048.872774 | 1130.994958 | 1584.664656 | -0.816724913 | 0.150716651 | 2.3478874757 | 1.2710272243 | ns | YPD | KS.3 |
| FIM1_2023 | 2872.083024 | 3521.800694 | 2191.806579 | 0.275845443 | 0.16377697 | 0.081705147 | 0.111917154 | ns | YPD | KS.3 |
| FIM1_2024 | 225.1186775 | 848.0968 | 535.9085137 | 1.255517448 | 0.555246898 | 0.001701622 | 0.003366032 | Km>KS.3 | YPD | KS.3 |
| FIM1_2025 | 305.1330829 | 171.6767325 | 215.5552265 | -0.765419421 | 0.194869716 | 3.7411368137 | 0.00010392 | ns | YPD | KS.3 |
| FIM1_2026 | 1796.785694 | 1190.389436 | 1235.45582 | -0.581378379 | 0.092455178 | 2.0036209033 | 1.6585713800 | ns | YPD | KS.3 |
| FIM1_2027 | 1606.566612 | 670.2337077 | 972.3894514 | -1.229369473 | 0.117768191 | 1.6431501690 | 1.7030566856 | Km<KS.3 | YPD | KS.3 |
| FIM1_2028 | 2133.493909 | 1634.079369 | 2123.234624 | -0.35717344 | 0.178140859 | 0.037073446 | 0.055238213 | ns | YPD | KS.3 |
| FIM1_2029 | 1977.594052 | 1772.928283 | 2108.789834 | -0.154068387 | 0.0934146 | 0.095341245 | 0.128682228 | ns | YPD | KS.3 |
| FIM1_2030 | 557.3356194 | 1415.877072 | 1023.682246 | 1.260440012 | 0.194526916 | 8.5400804656 | 9.2362826775 | Km>KS.3 | YPD | KS.3 |
| FIM1_2031 | 2408.16874 | 3545.869193 | 1632.790141 | 0.469042913 | 0.278120182 | 0.064158547 | 0.090257671 | ns | YPD | KS.3 |
| FIM1_2032 | 22.88226093 | 68.26911354 | 27.57214771 | 1.217967266 | 0.407838153 | 0.000275137 | 0.000641428 | Km>KS.3 | YPD | KS.3 |
| FIM1_2033 | 1869.003026 | 3019.317229 | 1731.269197 | 0.675984565 | 0.100270259 | 8.2538999628 | 8.9657537806 | ns | YPD | KS.3 |
| FIM1_2034 | 2204.044361 | 2220.415314 | 2139.809632 | 0.010586341 | 0.090429596 | 0.905829571 | 0.923463548 | ns | YPD | KS.3 |
| FIM1_2035 | 318.3706181 | 152.5748756 | 253.7108773 | -0.976244886 | 0.205085513 | 4.9351719628 | 2.0563216512 | ns | YPD | KS.3 |
| FIM1_2037 | 344.0939382 | 233.9907684 | 314.2585225 | -0.494557005 | 0.22685115 | 0.020247549 | 0.0320699 | ns | YPD | KS.3 |
| FIM1_2038 | 1555.870713 | 2930.164517 | 1938.2021 | 0.856119368 | 0.176154996 | 4.1685337678 | 1.7604800929 | ns | YPD | KS.3 |
| FIM1_2039 | 619.1528064 | 347.0063051 | 620.846494 | -0.778140981 | 0.183139268 | 9.1499811833 | 2.8920683854 | ns | YPD | KS.3 |
| FIM1_2040 | 498.9114076 | 277.112252 | 447.3371595 | -0.798938818 | 0.168829393 | 9.0520400714 | 3.5826491134 | ns | YPD | KS.3 |
| FIM1_2041 | 2223.127282 | 3659.794729 | 5070.218881 | 0.684055731 | 0.149881289 | 2.6123131475 | 9.3296898127 | ns | YPD | KS.3 |
| FIM1_2042 | 4092.920418 | 3092.341865 | 3704.298972 | -0.395402363 | 0.095118878 | 2.5935771101 | 7.4930581433 | ns | YPD | KS.3 |
| FIM1_2043 | 2329.407531 | 1552.227375 | 2435.732405 | -0.536241913 | 0.197687876 | 0.004406289 | 0.008012166 | ns | YPD | KS.3 |
| FIM1_2044 | 1266.071215 | 905.6583137 | 810.1224928 | -0.460640427 | 0.141972917 | 0.000872462 | 0.001831434 | ns | YPD | KS.3 |
| FIM1_2045 | 584.2575593 | 311.6018064 | 395.9114986 | -0.861016717 | 0.156159066 | 1.2347141378 | 7.0850090378 | ns | YPD | KS.3 |
| FIM1_2046 | 321.1632032 | 230.9793673 | 265.7848338 | -0.423563946 | 0.222940738 | 0.043546342 | 0.063718545 | ns | YPD | KS.3 |
| FIM1_2047 | 558.5752531 | 968.3478019 | 771.6282506 | 0.74441714 | 0.172215327 | 7.0820759551 | 2.3149361285 | ns | YPD | KS.3 |

| | | | | | | | | | |
|---|---|---|---|---|---|---|---|---|---|
| FIM1_2048 | 1470.132091 | 1512.689172 | 1837.727007 | 0.040135566 | 0.098290326 | 0.679323391 | 0.730257966 | ns | YPD | KS.3 |
| FIM1_2049 | 592.2806387 | 293.5053015 | 419.7850415 | -0.982228541 | 0.120419239 | 8.55402596721 | 2.39080220151 | ns | YPD | KS.3 |
| FIM1_2050 | 3781.129491 | 2664.963482 | 3690.857324 | -0.493184952 | 0.097635295 | 3.13031758121 | 1.35656184377 | ns | YPD | KS.3 |
| FIM1_2051 | 2051.923532 | 2038.082413 | 2042.719606 | -0.009387739 | 0.113038376 | 0.932744068 | 0.945092003 | ns | YPD | KS.3 |
| FIM1_2052 | 3323.76417 | 2173.925966 | 3036.137461 | -0.570237337 | 0.177902361 | 0.000850993 | 0.001790185 | ns | YPD | KS.3 |
| FIM1_2053 | 4822.515152 | 6276.762863 | 6991.329792 | 0.368698208 | 0.112046219 | 0.000825404 | 0.00174147 | ns | YPD | KS.3 |
| FIM1_2054 | 1122.050829 | 1075.623176 | 1495.778769 | -0.059033891 | 0.116320497 | 0.605719167 | 0.662442922 | ns | YPD | KS.3 |
| FIM1_2055 | 1116.890024 | 1308.914436 | 893.0585356 | 0.176502682 | 0.332119015 | 0.544612639 | 0.606547544 | ns | YPD | KS.3 |
| FIM1_2056 | 694.7369136 | 1236.829843 | 844.1085238 | 0.763480439 | 0.201006832 | 6.37337598429 | 0.000169386 | ns | YPD | KS.3 |
| FIM1_2057 | 469.312556 | 749.5467653 | 693.8576477 | 0.6538472 | 0.118059454 | 1.68398663212 | 9.44513359057 | ns | YPD | KS.3 |
| FIM1_2058 | 442.8909227 | 652.5446744 | 517.6529092 | 0.477144635 | 0.265378615 | 0.050274152 | 0.072440564 | ns | YPD | KS.3 |
| FIM1_2059 | 793.0021828 | 676.1460187 | 774.0261571 | -0.218906695 | 0.140110533 | 0.10938074 | 0.14569456 | ns | YPD | KS.3 |
| FIM1_2060 | 2892.266782 | 2875.027642 | 3723.71952 | -0.006886018 | 0.305177562 | 0.979896642 | 0.983851825 | ns | YPD | KS.3 |
| FIM1_2061 | 1847.681354 | 1671.509689 | 1823.673689 | -0.137105781 | 0.147367167 | 0.339271683 | 0.400920814 | ns | YPD | KS.3 |
| FIM1_2062 | 900.9945466 | 1859.083669 | 1165.197352 | 0.893247014 | 0.282644281 | 0.000483797 | 0.001071584 | ns | YPD | KS.3 |
| FIM1_2063 | 2346.70133 | 5335.446639 | 3667.742342 | 1.144196648 | 0.135949009 | 5.60322554127 | 1.89632973250 | Km>KS.3 | YPD | KS.3 |
| FIM1_2064 | 159.0978698 | 2369.872766 | 1152.213554 | 3.756270398 | 0.2715032 | 2.32073460178 | 1.64937923483 | Km>KS.3 | YPD | KS.3 |
| FIM1_2065 | 19.19775398 | 99.62845989 | 56.90405186 | 2.243370067 | 0.274969405 | 8.30567413838 | 3.33231684181 | Km>KS.3 | YPD | KS.3 |
| FIM1_2066 | 505.5549932 | 3421.722074 | 3256.199129 | 2.439597391 | 0.454177333 | 2.26123236869 | 1.84723005488 | Km>KS.3 | YPD | KS.3 |
| FIM1_2067 | 1526.355175 | 2693.88062 | 2073.776345 | 0.774369924 | 0.162001463 | 7.55821904607 | 3.03978494375 | ns | YPD | KS.3 |
| FIM1_2068 | 1298.442153 | 1693.866943 | 1436.187791 | 0.37220549 | 0.11053962 | 0.000624897 | 0.00135168 | ns | YPD | KS.3 |
| FIM1_2069 | 2291.697527 | 2074.158653 | 1555.198042 | -0.137949109 | 0.128620435 | 0.273572568 | 0.33171424 | ns | YPD | KS.3 |
| FIM1_2070 | 94.91199487 | 57.146728 | 80.28421507 | -0.644354804 | 0.246211763 | 0.004806006 | 0.008663001 | ns | YPD | KS.3 |
| FIM1_2071 | 2463.489033 | 2109.555745 | 2107.247229 | -0.22063726 | 0.068731007 | 0.001236383 | 0.002518839 | ns | YPD | KS.3 |
| FIM1_2072 | 1710.338752 | 1109.306659 | 1485.603666 | -0.597522496 | 0.13962637 | 1.13985883123 | 3.53761552425 | ns | YPD | KS.3 |
| FIM1_2073 | 3734.945171 | 2091.413116 | 3368.971596 | -0.761792131 | 0.210658792 | 0.000130748 | 0.000329187 | ns | YPD | KS.3 |
| FIM1_2074 | 1038.889178 | 734.729263 | 831.1518814 | -0.486563384 | 0.104160662 | 2.15483951679 | 7.85371911797 | ns | YPD | KS.3 |
| FIM1_2075 | 5246.308576 | 4687.64919 | 6104.843797 | -0.15469463 | 0.140931794 | 0.26059851 | 0.317452886 | ns | YPD | KS.3 |
| FIM1_2076 | 1469.169433 | 1350.655426 | 1345.166444 | -0.117367828 | 0.114643061 | 0.297954333 | 0.358464584 | ns | YPD | KS.3 |
| FIM1_2077 | 1689.080315 | 3052.423044 | 2543.597471 | 0.794882682 | 0.183488112 | 6.05950346691 | 2.01645683932 | ns | YPD | KS.3 |
| FIM1_2078 | 823.6674183 | 1866.942196 | 1355.849322 | 1.105339742 | 0.187105562 | 5.87815732540 | 4.39096586995 | Km>KS.3 | YPD | KS.3 |
| FIM1_2079 | 721.7004742 | 1022.348278 | 1965.955201 | 0.403588482 | 0.310308211 | 0.144523387 | 0.187533607 | ns | YPD | KS.3 |
| FIM1_2080 | 20.63659991 | 61.4525946 | 36.46740299 | 1.339056435 | 0.327959792 | 2.88118189612 | 1.02038042595 | Km>KS.3 | YPD | KS.3 |
| FIM1_2081 | 6578.49207 | 5759.993932 | 11195.6198 | -0.168686029 | 0.2323918 | 0.438728792 | 0.502573277 | ns | YPD | KS.3 |
| FIM1_2083 | 553.9906504 | 1028.864875 | 754.0544526 | 0.79806127 | 0.232806255 | 0.000242972 | 0.000571259 | ns | YPD | KS.3 |
| FIM1_2084 | 6185.487143 | 4975.454286 | 6222.755823 | -0.310286702 | 0.068237192 | 4.74756377863 | 1.61002929779 | ns | YPD | KS.3 |
| FIM1_2085 | 2970.091655 | 2801.941878 | 2721.631889 | -0.08263853 | 0.075564005 | 0.270547959 | 0.328286852 | ns | YPD | KS.3 |
| FIM1_2086 | 3245.797055 | 3236.793428 | 3151.420719 | -0.004076904 | 0.094299767 | 0.96512199 | 0.972384357 | ns | YPD | KS.3 |
| FIM1_2087 | 2056.073889 | 3782.4712 | 3960.918562 | 0.75154416 | 0.274928785 | 0.002702804 | 0.005142046 | ns | YPD | KS.3 |
| FIM1_2088 | 1198.394539 | 1610.23268 | 1507.92372 | 0.406135163 | 0.141925424 | 0.003331973 | 0.006229449 | ns | YPD | KS.3 |
| FIM1_2089 | 453.9399817 | 287.7296404 | 349.9156491 | -0.620541517 | 0.161098527 | 6.83491118838 | 0.000180678 | ns | YPD | KS.3 |
| FIM1_2090 | 1090.403319 | 881.3095788 | 845.9294823 | -0.299428265 | 0.097958934 | 0.001969316 | 0.003840591 | ns | YPD | KS.3 |
| FIM1_2091 | 1483.959409 | 662.7789672 | 1214.776412 | -1.121039605 | 0.138688611 | 9.95242468785 | 2.72051169352 | Km<KS.3 | YPD | KS.3 |
| FIM1_2092 | 817.9745475 | 295.7909632 | 677.1625973 | -1.285661412 | 0.288171519 | 6.71638512398 | 2.72990326730 | Km<KS.3 | YPD | KS.3 |
| FIM1_2093 | 2794.64633 | 1731.932358 | 3388.232517 | -0.566274248 | 0.300745024 | 0.035757957 | 0.053454278 | ns | YPD | KS.3 |

| | | | | | | | | | | |
|---|---|---|---|---|---|---|---|---|---|---|
| FIM1_2094 | 8509.505557 | 5270.819073 | 8107.293509 | -0.657589114 | 0.148834612 | 5.4439206623 | 1.8312038739 | ns | YPD | KS.3 |
| FIM1_2095 | 1153.798893 | 1522.39222 | 1015.029635 | 0.372719467 | 0.173423919 | 0.025706723 | 0.039742369 | ns | YPD | KS.3 |
| FIM1_2096 | 1006.851945 | 2603.837443 | 1200.383166 | 1.238787336 | 0.245087603 | 4.3800492106 | 2.2627980085 | Km>KS.3 | YPD | KS.3 |
| FIM1_2097 | 883.4513609 | 620.9209646 | 1071.318606 | -0.500081582 | 0.08515582 | 3.0288682926 | 1.9595084207 | ns | YPD | KS.3 |
| FIM1_2098 | 72.524036 | 607.2470381 | 1283.458349 | 2.75857948 | 0.447719727 | 3.2251444208 | 3.8202603556 | Km>KS.3 | YPD | KS.3 |
| FIM1_2099 | 9.508094395 | 8.56454576 | 15.19172552 | -0.110209287 | 0.38076752 | 0.730746833 | 0.774162158 | ns | YPD | KS.3 |
| FIM1_2100 | 1729.754674 | 1207.973061 | 1852.733756 | -0.48597141 | 0.166479626 | 0.002507443 | 0.004792366 | ns | YPD | KS.3 |
| FIM1_2101 | 896.6728881 | 686.2874063 | 761.0685894 | -0.368927556 | 0.137194274 | 0.005892379 | 0.010450832 | ns | YPD | KS.3 |
| FIM1_2102 | 1204.272324 | 1093.040698 | 1111.61358 | -0.134488561 | 0.123686042 | 0.267716433 | 0.325247681 | ns | YPD | KS.3 |
| FIM1_2103 | 3672.426568 | 8470.600758 | 5729.274012 | 1.076299285 | 0.248757212 | 2.7792011981 | 9.8760899720 | Km>KS.3 | YPD | KS.3 |
| FIM1_2104 | 1049.855073 | 424.8775449 | 766.9407679 | -1.201497462 | 0.217808425 | 4.0965674322 | 2.5929291317 | Km<KS.3 | YPD | KS.3 |
| FIM1_2105 | 672.9737618 | 321.2169779 | 459.8558034 | -1.012751817 | 0.162514051 | 1.0585744932 | 9.2881977138 | Km<KS.3 | YPD | KS.3 |
| FIM1_2106 | 1594.740457 | 1096.328107 | 901.2688859 | -0.502154062 | 0.178689951 | 0.003449451 | 0.006424941 | ns | YPD | KS.3 |
| FIM1_2107 | 908.5050853 | 963.796738 | 817.3913675 | 0.08196711 | 0.123386563 | 0.498370714 | 0.561202875 | ns | YPD | KS.3 |
| FIM1_2108 | 1065.232881 | 598.6009667 | 655.523916 | -0.793131453 | 0.147111894 | 2.8933669422 | 1.5411670811 | ns | YPD | KS.3 |
| FIM1_2109 | 298.3391943 | 155.9570422 | 233.2353216 | -0.872598918 | 0.186220302 | 9.4826909701 | 3.7441577441 | ns | YPD | KS.3 |
| FIM1_2110 | 704.0635224 | 405.841495 | 423.3050302 | -0.745371242 | 0.171995066 | 6.7213592161 | 2.2086368626 | ns | YPD | KS.3 |
| FIM1_2111 | 56.8969346 | 55.85733177 | 44.30100201 | -0.022123661 | 0.26656461 | 0.928045369 | 0.94167361 | ns | YPD | KS.3 |
| FIM1_2112 | 1356.971411 | 1027.123407 | 1126.871592 | -0.384473111 | 0.133818741 | 0.003292737 | 0.006163043 | ns | YPD | KS.3 |
| FIM1_2113 | 1372.964215 | 1677.591888 | 1409.47737 | 0.278438632 | 0.12239837 | 0.020431209 | 0.032329918 | ns | YPD | KS.3 |
| FIM1_2115 | 1517.250974 | 954.0497537 | 1230.235367 | -0.650513647 | 0.111152162 | 2.6849892463 | 1.7576080921 | ns | YPD | KS.3 |
| FIM1_2116 | 1680.246064 | 1301.339922 | 1207.977676 | -0.364114025 | 0.068674637 | 9.5109337618 | 4.6245579653 | ns | YPD | KS.3 |
| FIM1_2117 | 2839.64252 | 2570.369715 | 2470.402507 | -0.141833623 | 0.070192109 | 0.041970919 | 0.061685472 | ns | YPD | KS.3 |
| FIM1_2118 | 1662.394207 | 1740.897354 | 1670.173672 | 0.065854149 | 0.056269296 | 0.239866453 | 0.294796345 | ns | YPD | KS.3 |
| FIM1_2119 | 786.3722437 | 363.5399146 | 891.3692608 | -0.931981465 | 0.304519154 | 0.00060073 | 0.001302803 | ns | YPD | KS.3 |
| FIM1_2120 | 1601.268752 | 1105.549415 | 1712.28105 | -0.510935465 | 0.139021633 | 0.000165008 | 0.000405993 | ns | YPD | KS.3 |
| FIM1_2121 | 3573.683751 | 3889.629655 | 3712.965937 | 0.11928759 | 0.098251853 | 0.219019869 | 0.272054494 | ns | YPD | KS.3 |
| FIM1_2122 | 1799.737516 | 1106.232358 | 1269.252022 | -0.681752768 | 0.112789826 | 7.8326138241 | 5.7304784963 | ns | YPD | KS.3 |
| FIM1_2123 | 6345.066243 | 10275.02971 | 7668.136712 | 0.675682163 | 0.112191616 | 9.0745295410 | 6.5145432130 | ns | YPD | KS.3 |
| FIM1_2124 | 1833.15862 | 3426.273639 | 2574.161137 | 0.827275957 | 0.204653737 | 2.0009581503 | 5.9325189498 | ns | YPD | KS.3 |
| FIM1_2125 | 942.6949518 | 1572.232593 | 1307.919858 | 0.700560399 | 0.153255883 | 2.4432921996 | 8.7954983308 | ns | YPD | KS.3 |
| FIM1_2126 | 1172.658057 | 935.1696635 | 1140.724034 | -0.307524689 | 0.156765053 | 0.043136692 | 0.063230714 | ns | YPD | KS.3 |
| FIM1_2127 | 1158.862736 | 1806.343166 | 1172.311572 | 0.626564522 | 0.096550969 | 4.9755776469 | 4.6441836385 | ns | YPD | KS.3 |
| FIM1_2128 | 871.4775357 | 585.8950236 | 763.6469738 | -0.539431007 | 0.161438609 | 0.000553297 | 0.001208363 | ns | YPD | KS.3 |
| FIM1_2129 | 4041.947579 | 2081.566442 | 3207.282133 | -0.928894522 | 0.120093579 | 3.0124419695 | 6.1674480652 | ns | YPD | KS.3 |
| FIM1_2130 | 2362.165382 | 4160.797623 | 4591.296402 | 0.736708657 | 0.220477241 | 0.000382789 | 0.00086602 | ns | YPD | KS.3 |
| FIM1_2131 | 10159.23194 | 8165.558161 | 10641.61632 | -0.29958052 | 0.144997262 | 0.033921224 | 0.051030568 | ns | YPD | KS.3 |
| FIM1_2132 | 1717.829857 | 1001.85951 | 2400.954067 | -0.599470216 | 0.347577325 | 0.046271944 | 0.067271456 | ns | YPD | KS.3 |
| FIM1_2133 | 1499.075063 | 934.9046092 | 1195.454349 | -0.665458659 | 0.099451063 | 1.1892899238 | 1.2482526099 | ns | YPD | KS.3 |
| FIM1_2134 | 1309.582994 | 685.4286715 | 923.6418429 | -0.897471973 | 0.138320221 | 2.7688061286 | 2.7222945632 | ns | YPD | KS.3 |
| FIM1_2135 | 3035.983974 | 2034.108586 | 2432.676187 | -0.559124692 | 0.118281314 | 1.4758287604 | 5.5962256733 | ns | YPD | KS.3 |
| FIM1_2136 | 2823.070466 | 1581.495992 | 1894.50463 | -0.819065141 | 0.095283851 | 3.1645063169 | 1.1245299233 | ns | YPD | KS.3 |
| FIM1_2137 | 1086.668493 | 505.6713123 | 929.025873 | -1.061148034 | 0.142040867 | 1.5559702581 | 2.8864588464 | Km<KS.3 | YPD | KS.3 |
| FIM1_2138 | 1599.327673 | 1066.193075 | 1077.161676 | -0.53363555 | 0.201697169 | 0.005392814 | 0.009612629 | ns | YPD | KS.3 |
| FIM1_2139 | 2833.754671 | 1446.090128 | 2822.038722 | -0.930379383 | 0.143042932 | 2.2797368005 | 2.2728838842 | ns | YPD | KS.3 |

| | | | | | | | | | | |
|---|---|---|---|---|---|---|---|---|---|---|
| FIM1_2140 | 859.7685991 | 1394.787085 | 1049.093401 | 0.680409043 | 0.104736536 | 4.3030171165 | 4.0621461394 | ns | YPD | KS.3 |
| FIM1_2141 | 2040.375999 | 1016.346194 | 1365.113948 | -0.962029654 | 0.14733733 | 1.7615743783 | 1.7958673222 | ns | YPD | KS.3 |
| FIM1_2142 | 233.3566111 | 185.4069424 | 184.7391978 | -0.313001367 | 0.145272375 | 0.026959194 | 0.041562439 | ns | YPD | KS.3 |
| FIM1_2143 | 1116.66505 | 735.2323788 | 887.3095082 | -0.580131408 | 0.128913283 | 4.2537136063 | 1.4594638063 | ns | YPD | KS.3 |
| FIM1_2144 | 2148.10461 | 1381.978496 | 1324.008097 | -0.606899365 | 0.143622027 | 1.4261150612 | 4.3341004458 | ns | YPD | KS.3 |
| FIM1_2145 | 729.2277398 | 949.0784367 | 768.7166833 | 0.335029034 | 0.233327457 | 0.125188769 | 0.16507133 | ns | YPD | KS.3 |
| FIM1_2146 | 7882.759817 | 8826.024176 | 11469.12418 | 0.159196798 | 0.096621566 | 0.095427784 | 0.128694287 | ns | YPD | KS.3 |
| FIM1_2147 | 1051.486711 | 619.8347528 | 730.851883 | -0.716990934 | 0.167452014 | 9.0196829552 | 2.8563286252 | ns | YPD | KS.3 |
| FIM1_2148 | 936.192748 | 476.297486 | 554.9139861 | -0.904400263 | 0.191933946 | 7.6702588251 | 3.0773820689 | ns | YPD | KS.3 |
| FIM1_2149 | 1127.169808 | 919.2872294 | 947.8479098 | -0.286249057 | 0.103409022 | 0.005009772 | 0.008994447 | ns | YPD | KS.3 |
| FIM1_2150 | 1339.686764 | 1214.581825 | 1219.572483 | -0.130008982 | 0.18618454 | 0.466481437 | 0.529971489 | ns | YPD | KS.3 |
| FIM1_2151 | 778.1354562 | 806.3114691 | 1008.198779 | 0.049057682 | 0.142766856 | 0.724567468 | 0.768925588 | ns | YPD | KS.3 |
| FIM1_2152 | 255.0649987 | 293.3803532 | 256.6004841 | 0.195200721 | 0.122926519 | 0.105586628 | 0.141286705 | ns | YPD | KS.3 |
| FIM1_2153 | 487.8953968 | 445.8308125 | 445.3975149 | -0.124445303 | 0.121819149 | 0.298011964 | 0.358464584 | ns | YPD | KS.3 |
| FIM1_2154 | 1388.654638 | 464.6904448 | 1304.304847 | -1.26558098 | 0.381743458 | 7.6747351597 | 0.000201275 | Km<KS.3 | YPD | KS.3 |
| FIM1_2155 | 895.5831222 | 949.120791 | 789.1994816 | 0.081907659 | 0.098928389 | 0.401847716 | 0.465361356 | ns | YPD | KS.3 |
| FIM1_2156 | 1356.737383 | 723.4288026 | 1257.674473 | -0.859413907 | 0.161296522 | 3.5003140255 | 1.8369264005 | ns | YPD | KS.3 |
| FIM1_2157 | 3022.978639 | 3273.144941 | 3731.140676 | 0.112281708 | 0.090101588 | 0.207945063 | 0.259866035 | ns | YPD | KS.3 |
| FIM1_2158 | 2681.694922 | 2905.782947 | 2815.198628 | 0.111978127 | 0.114081899 | 0.318395308 | 0.379587025 | ns | YPD | KS.3 |
| FIM1_2159 | 133.1306595 | 99.73171128 | 86.34968685 | -0.341032301 | 0.294112283 | 0.198557405 | 0.2497025 | ns | YPD | KS.3 |
| FIM1_2160 | 1835.784919 | 1460.310014 | 1771.477552 | -0.314747546 | 0.140426353 | 0.021595077 | 0.033923431 | ns | YPD | KS.3 |
| FIM1_2161 | 1745.210783 | 1450.837704 | 1565.997904 | -0.264140107 | 0.055471427 | 1.7380195495 | 6.4720413613 | ns | YPD | KS.3 |
| FIM1_2162 | 840.2432959 | 501.1949478 | 767.1367039 | -0.72348325 | 0.113836955 | 9.9815073897 | 8.8202485371 | ns | YPD | KS.3 |
| FIM1_2163 | 1227.700069 | 693.4320239 | 761.4008034 | -0.76400691 | 0.188129822 | 2.1432163144 | 6.3054412564 | ns | YPD | KS.3 |
| FIM1_2164 | 810.7805861 | 732.0624401 | 769.3607597 | -0.144221288 | 0.1002645 | 0.145128744 | 0.188270014 | ns | YPD | KS.3 |
| FIM1_2165 | 2471.989975 | 2573.364036 | 2668.911091 | 0.055961954 | 0.113206703 | 0.615414263 | 0.671275149 | ns | YPD | KS.3 |
| FIM1_2166 | 741.3863195 | 340.7652447 | 398.1753113 | -1.011090897 | 0.232683002 | 3.1645965966 | 1.1126408528 | Km<KS.3 | YPD | KS.3 |
| FIM1_2167 | 4288.7884 | 5046.419192 | 4787.92655 | 0.226530982 | 0.119613047 | 0.053812533 | 0.076974512 | ns | YPD | KS.3 |
| FIM1_2168 | 4932.631973 | 3681.284095 | 4647.44812 | -0.416177765 | 0.075152606 | 2.4053407337 | 1.3007141467 | ns | YPD | KS.3 |
| FIM1_2169 | 2704.562441 | 2713.807807 | 2427.300803 | 0.004722125 | 0.103098345 | 0.962973234 | 0.971243409 | ns | YPD | KS.3 |
| FIM1_2170 | 2712.382616 | 3495.143847 | 3230.549092 | 0.349290802 | 0.138600925 | 0.009829717 | 0.016679005 | ns | YPD | KS.3 |
| FIM1_2171 | 643.2086962 | 1035.608383 | 1110.462969 | 0.639522237 | 0.179115314 | 0.00020022 | 0.000479814 | ns | YPD | KS.3 |
| FIM1_2172 | 1100.112596 | 2699.537456 | 2057.017079 | 1.264519148 | 0.114155111 | 1.3693034695 | 1.8411580434 | Km>KS.3 | YPD | KS.3 |
| FIM1_2173 | 825.6802708 | 585.3042579 | 774.6059075 | -0.458130775 | 0.186268548 | 0.010241686 | 0.01733664 | ns | YPD | KS.3 |
| FIM1_2174 | 1340.017113 | 1324.125225 | 1263.218158 | -0.015821471 | 0.170899411 | 0.923581979 | 0.937718438 | ns | YPD | KS.3 |
| FIM1_2175 | 2921.909322 | 4325.569016 | 3557.19596 | 0.545523181 | 0.125035834 | 8.4873802877 | 2.7067126238 | ns | YPD | KS.3 |
| FIM1_2176 | 1473.395067 | 739.1760371 | 925.9356295 | -0.942470194 | 0.163028924 | 2.0997585833 | 1.4135722533 | ns | YPD | KS.3 |
| FIM1_2177 | 98.11343363 | 71.94327174 | 79.54727981 | -0.402380999 | 0.206488636 | 0.0400767 | 0.059251584 | ns | YPD | KS.3 |
| FIM1_2178 | 4197.063443 | 3929.208992 | 4144.390667 | -0.085160909 | 0.215063169 | 0.675537692 | 0.726502382 | ns | YPD | KS.3 |
| FIM1_2179 | 8838.192457 | 9255.632956 | 9088.135023 | 0.06415597 | 0.122455339 | 0.593470713 | 0.650620714 | ns | YPD | KS.3 |
| FIM1_2180 | 2456.187447 | 1692.152274 | 2978.44097 | -0.443420343 | 0.292003423 | 0.092558785 | 0.12508365 | ns | YPD | KS.3 |
| FIM1_2181 | 2436.938476 | 2917.884436 | 2827.95635 | 0.229644167 | 0.228613342 | 0.284808984 | 0.343913761 | ns | YPD | KS.3 |
| FIM1_2182 | 606.3384384 | 972.2855701 | 1562.092515 | 0.535114057 | 0.331337463 | 0.06550339 | 0.091874644 | ns | YPD | KS.3 |
| FIM1_2183 | 1933.365299 | 3353.952393 | 2582.084018 | 0.773302834 | 0.110686444 | 1.2137908271 | 1.5603641770 | ns | YPD | KS.3 |
| FIM1_2184 | 1616.89895 | 3646.902923 | 2910.723947 | 1.130410152 | 0.139849629 | 9.6282889723 | 2.6611520909 | Km>KS.3 | YPD | KS.3 |

| | | | | | | | | | | |
|---|---|---|---|---|---|---|---|---|---|---|
| FIM1_2185 | 1582.989832 | 4100.641326 | 3145.59594 | 1.300162285 | 0.179574694 | 3.4653528639( | 5.9448725854{ | Km>KS.3 | YPD | KS.3 |
| FIM1_2186 | 1913.78396 | 1156.913757 | 1562.596842 | -0.703833556 | 0.11814847 | 1.2841122816: | 8.9851738411! | ns | YPD | KS.3 |
| FIM1_2187 | 1782.848143 | 1212.360696 | 1825.116442 | -0.491955485 | 0.233547301 | 0.02436003 | 0.037833662 | ns | YPD | KS.3 |
| FIM1_2188 | 5648.500369 | 4147.243775 | 4842.867628 | -0.435680808 | 0.095830762 | 4.1919921867{ | 1.4422656382( | ns | YPD | KS.3 |
| FIM1_2189 | 1297.644008 | 559.6851834 | 1001.661405 | -1.144230578 | 0.178526258 | 2.1498817468{ | 2.1739149778( | Km<KS.3 | YPD | KS.3 |
| FIM1_2190 | 1687.318345 | 2839.818599 | 2212.447532 | 0.7290121 | 0.114519226 | 9.2206493941! | 8.2209194867: | ns | YPD | KS.3 |
| FIM1_2191 | 7796.271595 | 6821.289327 | 7904.87394 | -0.189483052 | 0.081094262 | 0.018438481 | 0.029543139 | ns | YPD | KS.3 |
| FIM1_2192 | 1142.860549 | 588.6639424 | 907.4031976 | -0.900521869 | 0.171904831 | 5.1330809287: | 2.6191874482· | ns | YPD | KS.3 |
| FIM1_2193 | 3203.879241 | 2211.907242 | 4372.129069 | -0.440168722 | 0.293187908 | 0.096053161 | 0.129397367 | ns | YPD | KS.3 |
| FIM1_2194 | 603.6793941 | 420.5543684 | 674.7807856 | -0.458604599 | 0.238122164 | 0.039083674 | 0.057938402 | ns | YPD | KS.3 |
| FIM1_2195 | 683.4966343 | 675.5370561 | 827.4549321 | -0.016090872 | 0.133131573 | 0.901673415 | 0.920169279 | ns | YPD | KS.3 |
| FIM1_2196 | 470.5602731 | 404.6845713 | 346.3742841 | -0.202012513 | 0.17481889 | 0.230349003 | 0.284363843 | ns | YPD | KS.3 |
| FIM1_2197 | 1463.349858 | 1386.253763 | 1611.03317 | -0.075668813 | 0.113876977 | 0.499424609 | 0.561880921 | ns | YPD | KS.3 |
| FIM1_2198 | 2207.958582 | 2117.271641 | 2316.489743 | -0.053301894 | 0.230423574 | 0.805322787 | 0.836775452 | ns | YPD | KS.3 |
| FIM1_2199 | 82.65067219 | 122.7851426 | 102.5282884 | 0.49903149 | 0.244474309 | 0.028149322 | 0.043129928 | ns | YPD | KS.3 |
| FIM1_2200 | 954.2685986 | 1177.542048 | 1233.819232 | 0.286065467 | 0.15493537 | 0.057159715 | 0.081132549 | ns | YPD | KS.3 |
| FIM1_2201 | 1188.749804 | 1063.233779 | 1062.087388 | -0.157219524 | 0.08775755 | 0.070385186 | 0.097976022 | ns | YPD | KS.3 |
| FIM1_2202 | 3260.093699 | 4608.141235 | 4287.758579 | 0.477589871 | 0.137178691 | 0.000362195 | 0.000822796 | ns | YPD | KS.3 |
| FIM1_2203 | 951.3232579 | 977.5533131 | 970.0864016 | 0.038208764 | 0.123527789 | 0.752568347 | 0.793226171 | ns | YPD | KS.3 |
| FIM1_2204 | 1155.688301 | 1171.171399 | 982.1773121 | 0.018080517 | 0.156119733 | 0.905024139 | 0.922831542 | ns | YPD | KS.3 |
| FIM1_2205 | 1080.015025 | 801.6412094 | 972.1536766 | -0.409115097 | 0.143661714 | 0.003467923 | 0.006452101 | ns | YPD | KS.3 |
| FIM1_2206 | 1071.109506 | 557.6867163 | 1280.748421 | -0.774020483 | 0.309918216 | 0.005001048 | 0.008982026 | ns | YPD | KS.3 |
| FIM1_2207 | 343.3381642 | 1024.234086 | 462.4421567 | 1.311387547 | 0.349855534 | 1.2773466615: | 3.9251387530: | Km>KS.3 | YPD | KS.3 |
| FIM1_2209 | 1033.663841 | 1738.573773 | 1758.944291 | 0.666181834 | 0.23424162 | 0.002334438 | 0.004484104 | ns | YPD | KS.3 |
| FIM1_2210 | 4057.903983 | 2645.00623 | 4726.000485 | -0.539266192 | 0.246653609 | 0.018536708 | 0.02969096 | ns | YPD | KS.3 |
| FIM1_2211 | 5759.059278 | 3146.720704 | 4800.855103 | -0.853132869 | 0.100319804 | 6.4595597492: | 2.1713722805 | ns | YPD | KS.3 |
| FIM1_2212 | 1235.581646 | 850.2946387 | 1039.059064 | -0.518707635 | 0.127494597 | 3.2547036810: | 9.1688283199{ | ns | YPD | KS.3 |
| FIM1_2213 | 1215.884073 | 721.7301044 | 898.7242659 | -0.730900298 | 0.114032811 | 6.9004503558! | 6.2645511898! | ns | YPD | KS.3 |
| FIM1_2214 | 808.121349 | 566.1966284 | 624.8375672 | -0.499435623 | 0.107133332 | 2.2170949591· | 8.0586190254· | ns | YPD | KS.3 |
| FIM1_2215 | 6373.051708 | 3624.591017 | 6323.786572 | -0.750359267 | 0.195570397 | 5.6186353654: | 0.000150445 | ns | YPD | KS.3 |
| FIM1_2216 | 1302.234117 | 826.1942817 | 919.6030042 | -0.636297034 | 0.115856521 | 2.2606480393 | 1.2291501634{ | ns | YPD | KS.3 |
| FIM1_2217 | 758.7174035 | 1662.548465 | 851.7880038 | 1.011724248 | 0.243015157 | 7.0910860881( | 2.3163593754{ | Km>KS.3 | YPD | KS.3 |
| FIM1_2218 | 7010.089433 | 6839.465716 | 8006.289705 | -0.034754348 | 0.093853874 | 0.708037501 | 0.756060649 | ns | YPD | KS.3 |
| FIM1_2219 | 2729.217251 | 2135.712946 | 2623.452048 | -0.34634135 | 0.093433381 | 0.000177332 | 0.000432675 | ns | YPD | KS.3 |
| FIM1_2220 | 1743.235507 | 1681.398327 | 1521.415233 | -0.051123313 | 0.059547872 | 0.388360483 | 0.45152919 | ns | YPD | KS.3 |
| FIM1_2221 | 895.4552357 | 777.5611118 | 801.815568 | -0.199811924 | 0.087212402 | 0.020679652 | 0.032629645 | ns | YPD | KS.3 |
| FIM1_2222 | 696.6157115 | 612.8394274 | 622.0733073 | -0.172721116 | 0.164748012 | 0.278451871 | 0.336727773 | ns | YPD | KS.3 |
| FIM1_2223 | 1823.42618 | 1252.796147 | 1068.373156 | -0.519462323 | 0.132467729 | 6.0444195038( | 0.000161412 | ns | YPD | KS.3 |
| FIM1_2224 | 4878.175019 | 2395.685682 | 3951.563321 | -0.984807625 | 0.142368694 | 1.1463486389· | 1.4813206438: | ns | YPD | KS.3 |
| FIM1_2225 | 849.0735744 | 2473.754105 | 1245.852409 | 1.417362775 | 0.236664708 | 9.6418930993: | 8.5505201727· | Km>KS.3 | YPD | KS.3 |
| FIM1_2226 | 9462.206154 | 8461.806113 | 10280.44126 | -0.159630448 | 0.061234866 | 0.008794228 | 0.015065869 | ns | YPD | KS.3 |
| FIM1_2227 | 1392.049162 | 2512.974185 | 2047.478626 | 0.788701345 | 0.1912709 | 1.5491360871: | 4.6822308830( | ns | YPD | KS.3 |
| FIM1_2228 | 438.9501004 | 1433.300119 | 941.2334906 | 1.464112676 | 0.336178675 | 4.9657719543! | 2.0673402069: | Km>KS.3 | YPD | KS.3 |
| FIM1_2229 | 451.543911 | 1082.649273 | 888.5419249 | 1.164131068 | 0.211071139 | 4.7562443194( | 2.9467391642( | Km>KS.3 | YPD | KS.3 |
| FIM1_2230 | 347.7656686 | 644.5179089 | 429.8878394 | 0.779357574 | 0.253346531 | 0.000863455 | 0.001814057 | ns | YPD | KS.3 |

| | | | | | | | | | | |
|---|---|---|---|---|---|---|---|---|---|---|
| FIM1_2231 | 970.5219045 | 967.7516431 | 934.2259976 | -0.003494521 | 0.090004112 | 0.96870576 | 0.97556906 | ns | YPD | KS.3 |
| FIM1_2232 | 254.3967844 | 831.6928855 | 872.8070102 | 1.563152623 | 0.258603631 | 3.3115774473 | 3.186672688 | Km>KS.3 | YPD | KS.3 |
| FIM1_2233 | 2047.235992 | 1032.600481 | 1744.585935 | -0.947153773 | 0.14279001 | 9.141941309 | 9.844406496 | ns | YPD | KS.3 |
| FIM1_2234 | 583.6314062 | 803.6460135 | 525.7881748 | 0.449830583 | 0.100723933 | 6.016557494 | 2.006191255 | ns | YPD | KS.3 |
| FIM1_2235 | 700.9040292 | 399.3761833 | 542.9644863 | -0.749755128 | 0.193048828 | 4.645462087 | 0.00012636 | ns | YPD | KS.3 |
| FIM1_2236 | 2000.775477 | 776.7477127 | 1267.879751 | -1.323521261 | 0.133042004 | 1.693249503 | 1.559984496 | Km<KS.3 | YPD | KS.3 |
| FIM1_2237 | 40.74313171 | 36.52695401 | 40.526282 | -0.131348042 | 0.269717425 | 0.595431612 | 0.652482879 | ns | YPD | KS.3 |
| FIM1_2238 | 959.9962972 | 780.2462976 | 875.0395528 | -0.28470774 | 0.140250643 | 0.037480264 | 0.055760859 | ns | YPD | KS.3 |
| FIM1_2239 | 832.3804517 | 441.3631692 | 707.4400088 | -0.872079589 | 0.151565323 | 2.9808015547 | 1.930922882 | ns | YPD | KS.3 |
| FIM1_2240 | 862.2488814 | 577.3828753 | 670.9627868 | -0.547326496 | 0.155518682 | 0.000284147 | 0.000661811 | ns | YPD | KS.3 |
| FIM1_2241 | 3473.291678 | 2822.775087 | 2657.322158 | -0.274314034 | 0.191376011 | 0.133939497 | 0.175262756 | ns | YPD | KS.3 |
| FIM1_2243 | 1295.11708 | 978.6510456 | 1190.464514 | -0.387391312 | 0.134307302 | 0.003167237 | 0.00595053 | ns | YPD | KS.3 |
| FIM1_2244 | 5058.282711 | 5347.256781 | 5009.643501 | 0.077069686 | 0.124512482 | 0.528004836 | 0.59016492 | ns | YPD | KS.3 |
| FIM1_2245 | 1718.943911 | 1727.022919 | 1697.103408 | 0.006395041 | 0.173306965 | 0.969472045 | 0.975945654 | ns | YPD | KS.3 |
| FIM1_2246 | 9401.519715 | 9732.843799 | 11412.85161 | 0.048861839 | 0.091275049 | 0.588460103 | 0.64626689 | ns | YPD | KS.3 |
| FIM1_2247 | 1567.744024 | 1059.808686 | 1103.334595 | -0.54984552 | 0.105089542 | 1.100969183 | 5.276803165 | ns | YPD | KS.3 |
| FIM1_2248 | 2777.026122 | 965.9373466 | 2883.141521 | -1.299881342 | 0.320472546 | 3.819700373 | 1.326099746 | Km<KS.3 | YPD | KS.3 |
| FIM1_2249 | 3941.216844 | 2693.252774 | 4456.848732 | -0.465551551 | 0.270886859 | 0.060438534 | 0.085372433 | ns | YPD | KS.3 |
| FIM1_2250 | 200.5480351 | 81.90408684 | 393.1173166 | -1.106640241 | 0.296754146 | 3.086963060 | 8.745809353 | Km<KS.3 | YPD | KS.3 |
| FIM1_2251 | 491.403452 | 685.3052791 | 559.9824934 | 0.470281406 | 0.086969193 | 4.699570791 | 2.412834333 | ns | YPD | KS.3 |
| FIM1_2252 | 3053.778351 | 3999.490661 | 3579.467591 | 0.370691636 | 0.142716764 | 0.007704022 | 0.013368506 | ns | YPD | KS.3 |
| FIM1_2253 | 13490.65835 | 17300.53608 | 15903.47299 | 0.354006752 | 0.072757466 | 9.575303395 | 3.771744607 | ns | YPD | KS.3 |
| FIM1_2254 | 2887.458038 | 1793.013884 | 2607.848379 | -0.648831449 | 0.1606173 | 2.971085775 | 8.460876779 | ns | YPD | KS.3 |
| FIM1_2255 | 8925.101853 | 11290.98577 | 13851.80021 | 0.335448933 | 0.065608583 | 2.708679209 | 1.189380323 | ns | YPD | KS.3 |
| FIM1_2256 | 901.9728258 | 554.017992 | 1342.415325 | -0.647625317 | 0.192770387 | 0.000430506 | 0.000964325 | ns | YPD | KS.3 |
| FIM1_2257 | 567.8357748 | 1714.294711 | 1340.549922 | 1.269266494 | 0.388521089 | 8.932645089 | 0.000232792 | Km>KS.3 | YPD | KS.3 |
| FIM1_2258 | 2425.245528 | 1812.955485 | 2130.826195 | -0.394043219 | 0.16388016 | 0.012910966 | 0.021346645 | ns | YPD | KS.3 |
| FIM1_2259 | 615.7031632 | 313.9779932 | 421.2064794 | -0.933105405 | 0.139663858 | 6.864459583 | 7.657104580 | ns | YPD | KS.3 |
| FIM1_2260 | 2790.245282 | 1641.875379 | 1913.190993 | -0.745429145 | 0.106364887 | 1.103894253 | 1.430175498 | ns | YPD | KS.3 |
| FIM1_2261 | 4704.182101 | 2556.655084 | 3325.419316 | -0.855362598 | 0.113604751 | 1.803472779 | 3.298631279 | ns | YPD | KS.3 |
| FIM1_2262 | 2902.070657 | 1958.266953 | 2968.684737 | -0.517174125 | 0.202347612 | 0.007173216 | 0.012552497 | ns | YPD | KS.3 |
| FIM1_2263 | 3495.548174 | 2886.183122 | 4218.55243 | -0.261807076 | 0.149509547 | 0.071808505 | 0.099817634 | ns | YPD | KS.3 |
| FIM1_2264 | 1520.962297 | 1452.637875 | 1340.425027 | -0.064149148 | 0.116076012 | 0.574071214 | 0.633400818 | ns | YPD | KS.3 |
| FIM1_2265 | 946.4801444 | 1307.893709 | 992.9188256 | 0.455446353 | 0.099721357 | 3.705331666 | 1.288191826 | ns | YPD | KS.3 |
| FIM1_2266 | 934.1638217 | 1309.28599 | 1334.817771 | 0.46174965 | 0.149664373 | 0.001504301 | 0.003007998 | ns | YPD | KS.3 |
| FIM1_2267 | 1407.432295 | 1783.401147 | 1619.757706 | 0.326857384 | 0.135659244 | 0.013670115 | 0.022485 | ns | YPD | KS.3 |
| FIM1_2268 | 1888.520501 | 1073.72644 | 1336.216251 | -0.795792415 | 0.101036862 | 1.373039897 | 2.919176704 | ns | YPD | KS.3 |
| FIM1_2270 | 1092.584947 | 540.7232333 | 840.3921184 | -0.93767034 | 0.199264869 | 7.217999741 | 2.919474564 | ns | YPD | KS.3 |
| FIM1_2271 | 1972.533613 | 1363.680437 | 1494.176021 | -0.512429568 | 0.127464756 | 4.035503703 | 0.00011129 | ns | YPD | KS.3 |
| FIM1_2272 | 626.8970057 | 706.8994512 | 737.2469833 | 0.162103481 | 0.165807902 | 0.311979667 | 0.372742277 | ns | YPD | KS.3 |
| FIM1_2273 | 162.865961 | 140.0670524 | 169.1592428 | -0.191680863 | 0.229460571 | 0.373584931 | 0.436800242 | ns | YPD | KS.3 |
| FIM1_2274 | 1734.804967 | 2620.563266 | 2250.51838 | 0.523735495 | 0.238821992 | 0.018713985 | 0.029926735 | ns | YPD | KS.3 |
| FIM1_2275 | 2231.491532 | 1785.014876 | 2090.895444 | -0.298908145 | 0.176828177 | 0.07900287 | 0.108724558 | ns | YPD | KS.3 |
| FIM1_2276 | 697.8083809 | 990.4707255 | 871.5900199 | 0.481199893 | 0.143951925 | 0.000599043 | 0.001299711 | ns | YPD | KS.3 |
| FIM1_2278 | 1159.142899 | 2271.973648 | 1870.660216 | 0.888952934 | 0.208480428 | 6.507559574 | 2.144046946 | ns | YPD | KS.3 |

| | | | | | | | | | | |
|---|---|---|---|---|---|---|---|---|---|---|
| FIM1_2279 | 282.2504315 | 383.1977654 | 338.5058139 | 0.426632657 | 0.122040594 | 0.000366212 | 0.00083116 | ns | YPD | KS.3 |
| FIM1_2280 | 1969.855892 | 2148.8486 | 1950.211584 | 0.121901186 | 0.106551166 | 0.245794061 | 0.301188535 | ns | YPD | KS.3 |
| FIM1_2281 | 801.9518124 | 963.8042962 | 749.2942018 | 0.243483709 | 0.188673332 | 0.177782218 | 0.227077416 | ns | YPD | KS.3 |
| FIM1_2282 | 355.8596002 | 406.9323901 | 305.8777883 | 0.185961747 | 0.123999373 | 0.126293125 | 0.166351151 | ns | YPD | KS.3 |
| FIM1_2283 | 1151.978268 | 882.054195 | 1312.087633 | -0.370116311 | 0.130134041 | 0.003662621 | 0.006776203 | ns | YPD | KS.3 |
| FIM1_2284 | 1478.412745 | 3655.488054 | 2494.53559 | 1.194436651 | 0.225911214 | 1.50867405168 | 8.55832771624 | Km>KS.3 | YPD | KS.3 |
| FIM1_2285 | 4524.978208 | 4405.904222 | 4091.590869 | -0.035755637 | 0.173906352 | 0.83109937 | 0.858895366 | ns | YPD | KS.3 |
| FIM1_2286 | 1333.971705 | 795.2160923 | 1032.886163 | -0.718882122 | 0.129108383 | 1.25158818639 | 7.17356132176 | ns | YPD | KS.3 |
| FIM1_2287 | 1117.365654 | 1011.229386 | 927.4687669 | -0.137303262 | 0.136254649 | 0.302461565 | 0.362939287 | ns | YPD | KS.3 |
| FIM1_2288 | 1754.244158 | 1639.115046 | 1886.171791 | -0.094584142 | 0.119220482 | 0.419319253 | 0.483456149 | ns | YPD | KS.3 |
| FIM1_2289 | 3461.422342 | 1815.433306 | 3305.0036 | -0.888354272 | 0.149718159 | 9.70988226644 | 6.92072554091 | ns | YPD | KS.3 |
| FIM1_2290 | 2107.091965 | 3092.273072 | 2575.129183 | 0.513345505 | 0.181446647 | 0.003198951 | 0.006003313 | ns | YPD | KS.3 |
| FIM1_2291 | 2775.050048 | 2735.580385 | 3516.64279 | -0.019483671 | 0.155730578 | 0.897452821 | 0.916426064 | ns | YPD | KS.3 |
| FIM1_2292 | 1167.144254 | 621.9164592 | 872.7159229 | -0.853931807 | 0.172050089 | 2.47679789111 | 1.10215290772 | ns | YPD | KS.3 |
| FIM1_2293 | 824.9417244 | 396.1996571 | 613.4994378 | -0.975407437 | 0.204299353 | 4.61130512494 | 1.93270791883 | ns | YPD | KS.3 |
| FIM1_2294 | 9005.302121 | 8558.028981 | 9396.152219 | -0.072449204 | 0.076616304 | 0.340656056 | 0.402365593 | ns | YPD | KS.3 |
| FIM1_2295 | 1112.096745 | 611.2397091 | 746.0783644 | -0.817546804 | 0.159827651 | 1.22473987768 | 5.80293418236 | ns | YPD | KS.3 |
| FIM1_2297 | 1561.026435 | 1054.79095 | 1216.926502 | -0.552838185 | 0.097360274 | 8.89415727463 | 5.26767052872 | ns | YPD | KS.3 |
| FIM1_2298 | 1715.078265 | 3145.639927 | 1665.205528 | 0.82082724 | 0.174198454 | 9.49281584108 | 3.74518309353 | ns | YPD | KS.3 |
| FIM1_2299 | 1002.431501 | 1603.072231 | 1448.848662 | 0.64137832 | 0.155329846 | 2.04976641414 | 6.05557478861 | ns | YPD | KS.3 |
| FIM1_2300 | 812.547389 | 582.9627263 | 538.300696 | -0.453952763 | 0.150139148 | 0.001865502 | 0.003662465 | ns | YPD | KS.3 |
| FIM1_2301 | 3100.952248 | 2020.24528 | 2796.687825 | -0.566850912 | 0.196510055 | 0.002470922 | 0.004727582 | ns | YPD | KS.3 |
| FIM1_2302 | 3052.528956 | 1873.959265 | 3328.172383 | -0.684933454 | 0.108709525 | 1.53847836865 | 1.31060443219 | ns | YPD | KS.3 |
| FIM1_2303 | 166.4056259 | 501.0379407 | 302.1341551 | 1.465781111 | 0.235987521 | 1.88238558120 | 1.91510598496 | Km>KS.3 | YPD | KS.3 |
| FIM1_2304 | 173.4593467 | 381.0185339 | 326.5847066 | 1.073693874 | 0.170543359 | 5.76925644937 | 5.33495368692 | Km>KS.3 | YPD | KS.3 |
| FIM1_2305 | 823.3456886 | 885.4391037 | 762.5268359 | 0.103040695 | 0.086251494 | 0.227751338 | 0.281646261 | ns | YPD | KS.3 |
| FIM1_2306 | 388.2677258 | 404.237058 | 275.4655839 | 0.05565315 | 0.139972328 | 0.683762565 | 0.734236728 | ns | YPD | KS.3 |
| FIM1_2307 | 2716.80896 | 2178.09329 | 2506.191909 | -0.308921881 | 0.114072408 | 0.005897212 | 0.010455677 | ns | YPD | KS.3 |
| FIM1_2308 | 2239.500457 | 1158.54507 | 1907.014888 | -0.867666205 | 0.212112736 | 1.46992799371 | 4.46452488935 | ns | YPD | KS.3 |
| FIM1_2309 | 192.8514573 | 333.8406264 | 295.1554919 | 0.688618084 | 0.256340361 | 0.003584296 | 0.006648695 | ns | YPD | KS.3 |
| FIM1_2310 | 131.2691096 | 226.1537634 | 334.031678 | 0.708412155 | 0.215705552 | 0.000498544 | 0.001099404 | ns | YPD | KS.3 |
| FIM1_2311 | 450.3664168 | 2456.183863 | 1672.056037 | 2.298785905 | 0.301351972 | 6.16189995592 | 1.79271650764 | Km>KS.3 | YPD | KS.3 |
| FIM1_2312 | 5228.876136 | 4491.175741 | 4768.238598 | -0.213566847 | 0.102746691 | 0.035135601 | 0.052587129 | ns | YPD | KS.3 |
| FIM1_2313 | 849.9848891 | 1064.61535 | 1339.623606 | 0.305087785 | 0.162059476 | 0.051812263 | 0.074448425 | ns | YPD | KS.3 |
| FIM1_2314 | 1265.084433 | 932.0980735 | 1073.429431 | -0.431130292 | 0.095145592 | 4.52737161974 | 1.54695561869 | ns | YPD | KS.3 |
| FIM1_2315 | 931.8162863 | 1855.690033 | 1049.537091 | 0.907546137 | 0.212642166 | 6.07900606988 | 2.02024416818 | ns | YPD | KS.3 |
| FIM1_2316 | 657.6534694 | 1383.603195 | 886.4830386 | 0.96653113 | 0.231573212 | 7.78190875621 | 2.51070013373 | ns | YPD | KS.3 |
| FIM1_2317 | 1896.742705 | 3883.326995 | 2337.312787 | 0.998541511 | 0.130526217 | 4.77161587482 | 9.53365019167 | ns | YPD | KS.3 |
| FIM1_2318 | 11274.69844 | 5974.779484 | 13391.12303 | -0.847465951 | 0.194100613 | 4.57037811263 | 1.55843941812 | ns | YPD | KS.3 |
| FIM1_2319 | 159.679862 | 328.5863503 | 164.3750103 | 0.944102031 | 0.222486078 | 6.11714101251 | 2.03156051650 | ns | YPD | KS.3 |
| FIM1_2320 | 207.337889 | 128.7083598 | 214.5370624 | -0.660994557 | 0.132735064 | 3.46430066266 | 1.48576687903 | ns | YPD | KS.3 |
| FIM1_2321 | 1891.082802 | 1549.469557 | 1675.237695 | -0.284177321 | 0.063743495 | 7.37468717242 | 2.39328562836 | ns | YPD | KS.3 |
| FIM1_2322 | 911.6523006 | 623.9742978 | 811.9243081 | -0.528583603 | 0.120027286 | 7.21423524154 | 2.35195414984 | ns | YPD | KS.3 |
| FIM1_2323 | 3804.414316 | 2666.752752 | 3024.517997 | -0.495961469 | 0.118000768 | 1.87101984113 | 5.59058481061 | ns | YPD | KS.3 |
| FIM1_2324 | 3285.428084 | 5368.893634 | 3890.555805 | 0.665881251 | 0.167301507 | 3.69979671582 | 0.000103118 | ns | YPD | KS.3 |

| | | | | | | | | | | |
|---|---|---|---|---|---|---|---|---|---|---|
| FIM1_2325 | 39.96595767 | 41.49787689 | 40.96022622 | 0.050980243 | 0.175815541 | 0.763379626 | 0.802072574 | ns | YPD | KS.3 |
| FIM1_2326 | 635.2821207 | 416.3435132 | 442.7879753 | -0.589903538 | 0.119103315 | 4.5099517718 | 1.8934185709 | ns | YPD | KS.3 |
| FIM1_2327 | 3072.17231 | 3618.246521 | 3239.293127 | 0.230939586 | 0.093026163 | 0.012059131 | 0.020064943 | ns | YPD | KS.3 |
| FIM1_2328 | 6560.236715 | 8741.063122 | 7241.459513 | 0.406382698 | 0.086427265 | 2.0465913290 | 7.4976376009 | ns | YPD | KS.3 |
| FIM1_2329 | 520.3569112 | 424.8012942 | 371.0631883 | -0.26648687 | 0.199026891 | 0.160053221 | 0.206286211 | ns | YPD | KS.3 |
| FIM1_2330 | 422.1879048 | 338.6359558 | 471.7337333 | -0.288973958 | 0.200808664 | 0.130713889 | 0.171538275 | ns | YPD | KS.3 |
| FIM1_2331 | 2467.688752 | 2420.94891 | 2684.208701 | -0.025243599 | 0.191724938 | 0.890529644 | 0.909729976 | ns | YPD | KS.3 |
| FIM1_2332 | 7124.58443 | 6231.433883 | 8886.548113 | -0.179987771 | 0.171539216 | 0.276762829 | 0.334848024 | ns | YPD | KS.3 |
| FIM1_2333 | 590.5997994 | 391.0014776 | 387.8348301 | -0.558990795 | 0.164117145 | 0.000424567 | 0.000953168 | ns | YPD | KS.3 |
| FIM1_2334 | 619.9577925 | 780.4330161 | 624.1776974 | 0.313273654 | 0.158224117 | 0.041117986 | 0.06057506 | ns | YPD | KS.3 |
| FIM1_2335 | 3501.180195 | 6919.258288 | 6406.854433 | 0.924435273 | 0.173411511 | 2.9050311903 | 1.5457251520 | ns | YPD | KS.3 |
| FIM1_2336 | 636.5710998 | 3996.855288 | 1689.305228 | 2.558248762 | 0.243484555 | 4.5021585761 | 5.4629851016 | Km>KS.3 | YPD | KS.3 |
| FIM1_2337 | 154.7534679 | 301.4604749 | 216.0073424 | 0.862673391 | 0.231798435 | 6.7858219206 | 0.000179572 | ns | YPD | KS.3 |
| FIM1_2338 | 1618.131161 | 1703.951781 | 1538.303557 | 0.072825864 | 0.097383216 | 0.449098537 | 0.512681327 | ns | YPD | KS.3 |
| FIM1_2339 | 884.840033 | 1069.939271 | 640.8000692 | 0.263723682 | 0.127005466 | 0.034087103 | 0.051249121 | ns | YPD | KS.3 |
| FIM1_2340 | 1081.333489 | 840.5715929 | 1242.990799 | -0.329550236 | 0.204249511 | 0.08960651 | 0.12156869 | ns | YPD | KS.3 |
| FIM1_2341 | 389.8043938 | 688.7953218 | 716.9177179 | 0.762240731 | 0.186564988 | 1.9365617708 | 5.7690986886 | ns | YPD | KS.3 |
| FIM1_2342 | 1949.575184 | 1335.180858 | 1706.408292 | -0.520308188 | 0.144187937 | 0.000210834 | 0.000502586 | ns | YPD | KS.3 |
| FIM1_2343 | 60.69825299 | 52.33251942 | 62.07038615 | -0.174500147 | 0.282858306 | 0.497470236 | 0.560315695 | ns | YPD | KS.3 |
| FIM1_2344 | 531.8612509 | 1092.016615 | 771.224789 | 0.93397258 | 0.231576568 | 1.5663981075 | 4.7257917435 | ns | YPD | KS.3 |
| FIM1_2345 | 712.9486716 | 1193.547804 | 917.6130562 | 0.707339946 | 0.150529675 | 1.2997995108 | 4.9857382934 | ns | YPD | KS.3 |
| FIM1_2346 | 5716.921432 | 3651.735312 | 4009.822228 | -0.626308022 | 0.117626053 | 5.8642718576 | 2.9619038062 | ns | YPD | KS.3 |
| FIM1_2347 | 647.1175099 | 468.259116 | 620.2654514 | -0.435841926 | 0.172469979 | 0.008732019 | 0.014969605 | ns | YPD | KS.3 |
| FIM1_2348 | 3761.187599 | 2445.321111 | 2895.085466 | -0.60585891 | 0.102764425 | 2.2387002554 | 1.4989951239 | ns | YPD | KS.3 |
| FIM1_2349 | 3269.385086 | 2754.593198 | 2302.086355 | -0.242232506 | 0.085824892 | 0.00437742 | 0.007962583 | ns | YPD | KS.3 |
| FIM1_2350 | 2601.48999 | 2410.000121 | 2886.681087 | -0.106473351 | 0.11796055 | 0.358353901 | 0.42097064 | ns | YPD | KS.3 |
| FIM1_2351 | 1580.302592 | 1172.789725 | 1167.699938 | -0.393075495 | 0.196734253 | 0.036161097 | 0.053959645 | ns | YPD | KS.3 |
| FIM1_2352 | 937.7145823 | 1815.536286 | 1125.829973 | 0.860802371 | 0.223662849 | 4.1070717744 | 0.000113013 | ns | YPD | KS.3 |
| FIM1_2353 | 1129.694051 | 711.8570379 | 972.6931714 | -0.644180148 | 0.121020744 | 5.7328034252 | 2.9102752082 | ns | YPD | KS.3 |
| FIM1_2354 | 662.0029871 | 624.7732241 | 595.0128552 | -0.077252295 | 0.177421614 | 0.651005978 | 0.703465409 | ns | YPD | KS.3 |
| FIM1_2355 | 167.8328683 | 136.3877058 | 214.1355932 | -0.274472924 | 0.189838113 | 0.130858994 | 0.171683411 | ns | YPD | KS.3 |
| FIM1_2356 | 213.1033293 | 690.9203675 | 428.712158 | 1.523502458 | 0.282170263 | 1.8496120266 | 1.2657248738 | Km>KS.3 | YPD | KS.3 |
| FIM1_2357 | 2174.857203 | 1694.080906 | 2252.726222 | -0.327734326 | 0.201320696 | 0.087213225 | 0.118612847 | ns | YPD | KS.3 |
| FIM1_2358 | 1204.19219 | 564.9285889 | 1063.905746 | -1.064850433 | 0.111217026 | 1.9281767959 | 1.2458025402 | Km<KS.3 | YPD | KS.3 |
| FIM1_2359 | 3851.8141 | 3327.864729 | 4212.938516 | -0.2063127 | 0.095664314 | 0.029099015 | 0.0444345 | ns | YPD | KS.3 |
| FIM1_2360 | 2680.620413 | 2750.644253 | 2317.011115 | 0.034377335 | 0.183901052 | 0.845598025 | 0.871344278 | ns | YPD | KS.3 |
| FIM1_2361 | 3083.751886 | 2901.001442 | 3183.044409 | -0.086276262 | 0.090108434 | 0.333292225 | 0.395168927 | ns | YPD | KS.3 |
| FIM1_2362 | 2066.270871 | 1275.567328 | 1599.294081 | -0.68181136 | 0.093649004 | 1.7291885939 | 2.5833973738 | ns | YPD | KS.3 |
| FIM1_2363 | 2052.43135 | 2341.439361 | 2113.157998 | 0.179536572 | 0.152760254 | 0.226453205 | 0.280110565 | ns | YPD | KS.3 |
| FIM1_2364 | 2659.610552 | 3082.401174 | 2512.202957 | 0.209550489 | 0.07866842 | 0.007244563 | 0.012663985 | ns | YPD | KS.3 |
| FIM1_2365 | 14295.89831 | 12228.795 | 19621.02589 | -0.216077993 | 0.130669202 | 0.091160558 | 0.123374259 | ns | YPD | KS.3 |
| FIM1_2366 | 786.9000079 | 659.1478018 | 568.7596446 | -0.242320092 | 0.146580223 | 0.08954371 | 0.121516627 | ns | YPD | KS.3 |
| FIM1_2367 | 411.9281611 | 516.5916221 | 488.8553682 | 0.295936184 | 0.204202042 | 0.127390276 | 0.167441644 | ns | YPD | KS.3 |
| FIM1_2368 | 1443.519265 | 3314.814249 | 2404.573716 | 1.11670698 | 0.196299416 | 2.0806166152 | 1.4025836939 | Km>KS.3 | YPD | KS.3 |
| FIM1_2369 | 1326.801072 | 1062.501297 | 1138.461136 | -0.306895199 | 0.132554355 | 0.017933771 | 0.028817994 | ns | YPD | KS.3 |

| | | | | | | | | | | |
|---|---|---|---|---|---|---|---|---|---|---|
| FIM1_2370 | 618.9621862 | 229.6881528 | 394.0827039 | -1.227209328 | 0.306354719 | 6.3731991286 | 2.1039592345 | Km<KS.3 | YPD | KS.3 |
| FIM1_2371 | 1511.479323 | 8909.388075 | 4375.944249 | 2.393713172 | 0.323250858 | 3.7962886750 | 9.0800654608 | Km>KS.3 | YPD | KS.3 |
| FIM1_2372 | 3380.78386 | 5666.769087 | 4975.428215 | 0.71642829 | 0.132604655 | 3.2054198301 | 1.6910884045 | ns | YPD | KS.3 |
| FIM1_2373 | 322.8081515 | 475.6246671 | 648.7126715 | 0.45418677 | 0.304207163 | 0.095310979 | 0.128676287 | ns | YPD | KS.3 |
| FIM1_2374 | 1811.500815 | 1335.636088 | 1107.153194 | -0.426141419 | 0.113132017 | 0.000128342 | 0.000323291 | ns | YPD | KS.3 |
| FIM1_2375 | 1912.061424 | 2199.768971 | 1708.348738 | 0.194517484 | 0.125294112 | 0.113346578 | 0.150493522 | ns | YPD | KS.3 |
| FIM1_2376 | 1392.608248 | 1416.78261 | 1376.278378 | 0.023616574 | 0.135750988 | 0.8587379 | 0.882143517 | ns | YPD | KS.3 |
| FIM1_2377 | 7702.613652 | 9618.226492 | 9042.424857 | 0.30907132 | 0.1213531 | 0.009462098 | 0.01613226 | ns | YPD | KS.3 |
| FIM1_2378 | 703.1893724 | 813.1722172 | 689.8052412 | 0.197434319 | 0.15799077 | 0.19753751 | 0.248797243 | ns | YPD | KS.3 |
| FIM1_2379 | 2458.879178 | 2274.98578 | 1764.115152 | -0.107772919 | 0.125049733 | 0.379497885 | 0.442362226 | ns | YPD | KS.3 |
| FIM1_2380 | 997.8443043 | 521.2611709 | 815.8202954 | -0.880790188 | 0.173460237 | 1.2747718609 | 6.0170683189 | ns | YPD | KS.3 |
| FIM1_2381 | 1584.496822 | 1378.568852 | 1661.261604 | -0.19699042 | 0.084024525 | 0.017978873 | 0.028881141 | ns | YPD | KS.3 |
| FIM1_2382 | 813.5190194 | 734.4695349 | 593.4255858 | -0.140254706 | 0.140774861 | 0.30723861 | 0.367430789 | ns | YPD | KS.3 |
| FIM1_2383 | 750.4027534 | 1471.324785 | 893.42857 | 0.943430663 | 0.118902536 | 5.8914531172 | 1.3696251989 | ns | YPD | KS.3 |
| FIM1_2384 | 9083.130895 | 6183.158821 | 9984.682313 | -0.539300206 | 0.109682403 | 5.8714895922 | 2.4002186295 | ns | YPD | KS.3 |
| FIM1_2385 | 1095.662468 | 885.6472433 | 919.9138713 | -0.301630814 | 0.081188957 | 0.000178572 | 0.000435274 | ns | YPD | KS.3 |
| FIM1_2386 | 839.511127 | 758.4979647 | 686.0131183 | -0.128259994 | 0.234839132 | 0.559647616 | 0.620237667 | ns | YPD | KS.3 |
| FIM1_2387 | 637.7249244 | 465.5686306 | 480.6728324 | -0.442180093 | 0.097928327 | 4.8168909040 | 1.6302062753 | ns | YPD | KS.3 |
| FIM1_2388 | 5180.355555 | 3535.72872 | 6074.465123 | -0.539996149 | 0.091775859 | 2.6705295410 | 1.7504459112 | ns | YPD | KS.3 |
| FIM1_2389 | 1440.477504 | 2423.26542 | 1980.459844 | 0.708268136 | 0.162269482 | 6.3058751540 | 2.0894508362 | ns | YPD | KS.3 |
| FIM1_2390 | 1350.364879 | 1307.948193 | 1057.863545 | -0.044543816 | 0.085391548 | 0.598494195 | 0.655550115 | ns | YPD | KS.3 |
| FIM1_2391 | 1790.141525 | 811.133023 | 1136.747479 | -1.079678695 | 0.171679792 | 5.9080331384 | 5.4329879600 | Km<KS.3 | YPD | KS.3 |
| FIM1_2392 | 1324.001984 | 675.5481401 | 1043.287542 | -0.906792662 | 0.183552629 | 2.4292207097 | 1.0838899579 | ns | YPD | KS.3 |
| FIM1_2393 | 403.971632 | 146.2372063 | 366.9729904 | -1.354529954 | 0.224510973 | 9.8637934862 | 8.7317388957 | Km<KS.3 | YPD | KS.3 |
| FIM1_2394 | 1400.573125 | 1951.986097 | 1032.077971 | 0.444052992 | 0.180909941 | 0.010580529 | 0.017873728 | ns | YPD | KS.3 |
| FIM1_2395 | 13107.94562 | 13392.50718 | 17822.67553 | 0.03010715 | 0.103196064 | 0.767452626 | 0.804991949 | ns | YPD | KS.3 |
| FIM1_2396 | 2582.174444 | 2796.400448 | 2557.179755 | 0.112826329 | 0.090612515 | 0.208264506 | 0.260199879 | ns | YPD | KS.3 |
| FIM1_2397 | 4088.928718 | 4045.413605 | 3026.433657 | -0.01489918 | 0.107637596 | 0.888298192 | 0.908009761 | ns | YPD | KS.3 |
| FIM1_2398 | 1895.419345 | 859.7117958 | 2011.486547 | -1.093763762 | 0.147444544 | 2.0761768794 | 3.7697007209 | Km<KS.3 | YPD | KS.3 |
| FIM1_2399 | 666.1849526 | 348.5260201 | 828.3418785 | -0.891298012 | 0.150078849 | 9.3257259106 | 6.6756095547 | ns | YPD | KS.3 |
| FIM1_2400 | 913.529498 | 385.8088224 | 510.3101681 | -1.190650692 | 0.153006061 | 8.7097858003 | 1.9431024375 | Km<KS.3 | YPD | KS.3 |
| FIM1_2401 | 841.7297744 | 635.069909 | 714.9424682 | -0.388126696 | 0.138562444 | 0.004106636 | 0.007516745 | ns | YPD | KS.3 |
| FIM1_2402 | 2284.838378 | 1155.438707 | 1613.534404 | -0.950479961 | 0.12890494 | 4.5607085302 | 7.6139345429 | ns | YPD | KS.3 |
| FIM1_2403 | 3378.881751 | 2187.005066 | 2146.319139 | -0.609438816 | 0.111553779 | 2.7900503142 | 1.4925269154 | ns | YPD | KS.3 |
| FIM1_2404 | 4767.338439 | 5696.643875 | 5503.319899 | 0.246823195 | 0.127736124 | 0.048577069 | 0.070314495 | ns | YPD | KS.3 |
| FIM1_2405 | 2109.242562 | 1558.113226 | 1827.205237 | -0.424054437 | 0.111744835 | 0.00011494 | 0.000293395 | ns | YPD | KS.3 |
| FIM1_2406 | 7633.561656 | 11430.39609 | 8854.810501 | 0.490076907 | 0.277659957 | 0.052753716 | 0.07561214 | ns | YPD | KS.3 |
| FIM1_2407 | 1087.295078 | 864.7607744 | 1434.130017 | -0.304685152 | 0.185053764 | 0.0859966 | 0.117182439 | ns | YPD | KS.3 |
| FIM1_2408 | 696.5473374 | 324.8735696 | 317.70967 | -1.02124226 | 0.195984439 | 4.1977155784 | 2.1731149846 | Km<KS.3 | YPD | KS.3 |
| FIM1_2409 | 754.6577365 | 442.9728781 | 558.8150676 | -0.732929114 | 0.147638366 | 3.2532977401 | 1.4025265387 | ns | YPD | KS.3 |
| FIM1_2410 | 6795.30381 | 4250.320053 | 5763.511942 | -0.658618955 | 0.109309837 | 9.2170227494 | 6.6073037721 | ns | YPD | KS.3 |
| FIM1_2411 | 1037.514544 | 730.1008627 | 686.7980707 | -0.485774594 | 0.133486878 | 0.000196722 | 0.000472798 | ns | YPD | KS.3 |
| FIM1_2412 | 719.4677272 | 467.3388178 | 600.7828512 | -0.57721088 | 0.183308604 | 0.001021122 | 0.0021123 | ns | YPD | KS.3 |
| FIM1_2413 | 3043.243807 | 2543.17612 | 2181.1896 | -0.240931668 | 0.173387538 | 0.149405867 | 0.193364773 | ns | YPD | KS.3 |
| FIM1_2414 | 1776.742443 | 1573.105065 | 1461.678195 | -0.169021 | 0.124170473 | 0.165217753 | 0.212502151 | ns | YPD | KS.3 |

| | | | | | | | | | | |
|---|---|---|---|---|---|---|---|---|---|---|
| FIM1_2415 | 1162.467608 | 866.8578234 | 948.249549 | -0.407525653 | 0.124622673 | 0.000851013 | 0.001790185 | ns | YPD | KS.3 |
| FIM1_2416 | 5399.686774 | 3467.578805 | 4098.833739 | -0.627316456 | 0.087962742 | 5.7089134834 | 7.8241996088 | ns | YPD | KS.3 |
| FIM1_2417 | 1632.188841 | 1194.987279 | 1140.405553 | -0.43648285 | 0.111274962 | 6.7181348773 | 0.000177875 | ns | YPD | KS.3 |
| FIM1_2418 | 14641.9283 | 13623.52768 | 15167.9165 | -0.102502053 | 0.074579825 | 0.16622835 | 0.213691484 | ns | YPD | KS.3 |
| FIM1_2419 | 761.9972282 | 880.2350971 | 811.3745993 | 0.194163062 | 0.172642892 | 0.24348146 | 0.298582863 | ns | YPD | KS.3 |
| FIM1_2420 | 1959.582615 | 1460.648335 | 1741.522806 | -0.405996676 | 0.134127025 | 0.001955746 | 0.003820117 | ns | YPD | KS.3 |
| FIM1_2422 | 5544.329091 | 5950.610363 | 4611.739681 | 0.100700445 | 0.074225833 | 0.171785845 | 0.220323429 | ns | YPD | KS.3 |
| FIM1_2423 | 734.2893803 | 892.8858717 | 573.2406725 | 0.255373207 | 0.205326537 | 0.190584276 | 0.241015956 | ns | YPD | KS.3 |
| FIM1_2424 | 731.076974 | 1575.870564 | 1761.714842 | 1.045492124 | 0.173227115 | 3.2835987654 | 2.5930006123 | Km>KS.3 | YPD | KS.3 |
| FIM1_2425 | 515.5363367 | 398.5623686 | 549.0309316 | -0.338133805 | 0.200999551 | 0.077199309 | 0.106537188 | ns | YPD | KS.3 |
| FIM1_2426 | 2895.144488 | 3949.554506 | 3547.211572 | 0.431165255 | 0.126814177 | 0.000519157 | 0.001138301 | ns | YPD | KS.3 |
| FIM1_2427 | 408.7089094 | 207.1802868 | 261.1951881 | -0.927602323 | 0.16596383 | 6.7245008474 | 4.0452710660 | ns | YPD | KS.3 |
| FIM1_2428 | 318.3264697 | 512.5468744 | 668.4622509 | 0.594018734 | 0.258045495 | 0.012529294 | 0.020798544 | ns | YPD | KS.3 |
| FIM1_2429 | 1220.999494 | 1279.685206 | 1186.307702 | 0.066812642 | 0.077299703 | 0.383710397 | 0.44654017 | ns | YPD | KS.3 |
| FIM1_2430 | 2482.081357 | 1812.31136 | 1665.928015 | -0.440400504 | 0.110375606 | 5.0427926314 | 0.000136125 | ns | YPD | KS.3 |
| FIM1_2431 | 1696.289854 | 1736.466721 | 1976.13754 | 0.031827119 | 0.152188632 | 0.829648545 | 0.858108423 | ns | YPD | KS.3 |
| FIM1_2432 | 1490.723233 | 1789.071421 | 1147.157163 | 0.249999806 | 0.146499217 | 0.07967712 | 0.109501014 | ns | YPD | KS.3 |
| FIM1_2433 | 26473.38415 | 13550.12878 | 28152.16964 | -0.83232201 | 0.271765822 | 0.000790338 | 0.001671739 | ns | YPD | KS.3 |
| FIM1_2434 | 6795.928228 | 3800.433729 | 5279.855016 | -0.748257161 | 0.232438169 | 0.00057305 | 0.001249044 | ns | YPD | KS.3 |
| FIM1_2435 | 2057.695648 | 1923.89812 | 1608.339509 | -0.094091447 | 0.110385388 | 0.386717213 | 0.449828883 | ns | YPD | KS.3 |
| FIM1_2436 | 233.2691882 | 374.8631433 | 317.0982153 | 0.630874397 | 0.192576033 | 0.000598197 | 0.001298442 | ns | YPD | KS.3 |
| FIM1_2437 | 150.3135042 | 227.1933772 | 185.0976495 | 0.506364446 | 0.270341767 | 0.040751598 | 0.060169524 | ns | YPD | KS.3 |
| FIM1_2438 | 1306.463208 | 1973.632216 | 1531.349559 | 0.571267037 | 0.133399135 | 1.1744297914 | 3.6358358508 | ns | YPD | KS.3 |
| FIM1_2439 | 271.4694875 | 579.8500141 | 443.2228972 | 0.95260361 | 0.268826209 | 0.00010431 | 0.000268048 | ns | YPD | KS.3 |
| FIM1_2440 | 176.5134471 | 119.8740314 | 226.0552197 | -0.386559929 | 0.400591887 | 0.24249423 | 0.297584803 | ns | YPD | KS.3 |
| FIM1_2444 | 12850.80146 | 13142.5068 | 20677.52651 | 0.025540193 | 0.316350216 | 0.927565688 | 0.941378885 | ns | YPD | KS.3 |
| FIM1_2445 | 895.8949342 | 995.9653859 | 805.4567149 | 0.1273744 | 0.276931021 | 0.614381346 | 0.670295437 | ns | YPD | KS.3 |
| FIM1_2446 | 257.7589431 | 236.8382462 | 299.0757199 | -0.116518537 | 0.136283388 | 0.381639866 | 0.444650664 | ns | YPD | KS.3 |
| FIM1_2447 | 426.8596137 | 415.4832542 | 473.4177694 | -0.037072473 | 0.128832522 | 0.768946691 | 0.80604926 | ns | YPD | KS.3 |
| FIM1_2448 | 63.64567744 | 39.01325458 | 42.2117417 | -0.494331523 | 0.399255508 | 0.134350442 | 0.175708056 | ns | YPD | KS.3 |
| FIM1_2449 | 56.24333651 | 63.33022078 | 43.40792016 | 0.126757556 | 0.3603591 | 0.681400076 | 0.732015845 | ns | YPD | KS.3 |
| FIM1_2450 | 989.0245651 | 1570.359218 | 632.294383 | 0.604732137 | 0.210600168 | 0.002410835 | 0.004621929 | ns | YPD | KS.3 |
| FIM1_2451 | 2102.545953 | 2083.40517 | 931.4556968 | -0.007942679 | 0.45598578 | 0.9821012 | 0.985071264 | ns | YPD | KS.3 |
| FIM1_2452 | 71.6888623 | 78.83334775 | 52.95915738 | 0.115769799 | 0.263163022 | 0.632871577 | 0.687308713 | ns | YPD | KS.3 |
| FIM1_2453 | 317.7390595 | 292.2138066 | 209.9167577 | -0.112543242 | 0.164097156 | 0.478474149 | 0.541987452 | ns | YPD | KS.3 |
| FIM1_2454 | 692.6435827 | 615.1117398 | 378.7276646 | -0.152759341 | 0.217380205 | 0.457030928 | 0.520801768 | ns | YPD | KS.3 |
| FIM1_2455 | 1119.453396 | 3171.671293 | 1087.383669 | 1.1748013 | 0.391136922 | 0.000308902 | 0.000712795 | Km>KS.3 | YPD | KS.3 |
| FIM1_2456 | 419.2343276 | 268.9123343 | 256.7192958 | -0.569566588 | 0.229030749 | 0.007986936 | 0.0138342 | ns | YPD | KS.3 |
| FIM1_2457 | 336.4532742 | 743.2135515 | 402.8146455 | 1.077283873 | 0.176141774 | 1.7884200616 | 1.5054805087 | Km>KS.3 | YPD | KS.3 |
| FIM1_2458 | 2029.138541 | 2393.461461 | 2445.15704 | 0.219471529 | 0.185338824 | 0.216947206 | 0.269895561 | ns | YPD | KS.3 |
| FIM1_2459 | 1618.149033 | 1164.852363 | 2120.245717 | -0.41716159 | 0.236761817 | 0.059320209 | 0.083911868 | ns | YPD | KS.3 |
| FIM1_2460 | 2804.238047 | 1727.165403 | 4269.400428 | -0.584602837 | 0.286405783 | 0.024186943 | 0.03757965 | ns | YPD | KS.3 |
| FIM1_2461 | 3006.847904 | 3991.549882 | 2918.362486 | 0.400843234 | 0.088339479 | 4.5498925158 | 1.5525181938 | ns | YPD | KS.3 |
| FIM1_2462 | 5602.668673 | 5286.312222 | 5382.854023 | -0.082119128 | 0.089631894 | 0.354610612 | 0.41726296 | ns | YPD | KS.3 |
| FIM1_2463 | 947.6209771 | 1284.50557 | 944.635553 | 0.430555258 | 0.086780862 | 5.4069898006 | 2.2323464114 | ns | YPD | KS.3 |

| | | | | | | | | | | |
|---|---|---|---|---|---|---|---|---|---|---|
| FIM1_2464 | 2128.675221 | 2998.909477 | 1939.7883 | 0.464657773 | 0.163673047 | 0.003325312 | 0.006219334 | ns | YPD | KS.3 |
| FIM1_2465 | 650.8950172 | 395.2268051 | 596.242639 | -0.697212875 | 0.118479003 | 2.02245668895 | 1.36708179722 | ns | YPD | KS.3 |
| FIM1_2466 | 682.9917015 | 1614.572755 | 842.4749687 | 1.085855855 | 0.272035034 | 1.15127454086 | 3.57081723243 | Km>KS.3 | YPD | KS.3 |
| FIM1_2467 | 557.8419328 | 512.4577904 | 403.5880091 | -0.117533809 | 0.132997424 | 0.366337128 | 0.42903183 | ns | YPD | KS.3 |
| FIM1_2468 | 2920.094576 | 2194.575996 | 2249.658896 | -0.396322659 | 0.127015412 | 0.001447095 | 0.002905286 | ns | YPD | KS.3 |
| FIM1_2469 | 1695.487544 | 1907.351307 | 1894.960468 | 0.167592847 | 0.069343035 | 0.01500458 | 0.024506823 | ns | YPD | KS.3 |
| FIM1_2470 | 6350.858359 | 8206.543071 | 6846.182636 | 0.361120984 | 0.097845595 | 0.000186235 | 0.000451081 | ns | YPD | KS.3 |
| FIM1_2471 | 1465.126127 | 875.3948614 | 1109.614518 | -0.702056083 | 0.160576094 | 6.17476193085 | 2.04932892635 | ns | YPD | KS.3 |
| FIM1_2472 | 2460.484733 | 2450.692688 | 2006.773113 | -0.005324699 | 0.104530242 | 0.95880902 | 0.968543121 | ns | YPD | KS.3 |
| FIM1_2473 | 1411.805078 | 948.117284 | 1554.022244 | -0.525142889 | 0.198927773 | 0.005560087 | 0.009893217 | ns | YPD | KS.3 |
| FIM1_2474 | 3204.188849 | 3679.023118 | 3438.768156 | 0.195359596 | 0.090503747 | 0.029144976 | 0.044491026 | ns | YPD | KS.3 |
| FIM1_2475 | 1665.524084 | 1465.423113 | 1268.99965 | -0.179422066 | 0.10504181 | 0.083231466 | 0.113819831 | ns | YPD | KS.3 |
| FIM1_2476 | 6696.200508 | 6437.371946 | 9869.721191 | -0.055504928 | 0.101173726 | 0.578358407 | 0.637000902 | ns | YPD | KS.3 |
| FIM1_2477 | 1901.793855 | 2074.166437 | 2092.768011 | 0.120636159 | 0.119364409 | 0.303543072 | 0.363765 | ns | YPD | KS.3 |
| FIM1_2478 | 1476.339835 | 711.9159432 | 1118.601481 | -1.013812542 | 0.136265085 | 2.28686939784 | 4.10728348530 | Km<KS.3 | YPD | KS.3 |
| FIM1_2479 | 1131.071349 | 725.3629696 | 825.2483291 | -0.623666061 | 0.108407293 | 5.10096286024 | 3.14854717490 | ns | YPD | KS.3 |
| FIM1_2480 | 2606.554666 | 2310.593547 | 2053.053403 | -0.171452216 | 0.068152566 | 0.011366445 | 0.019033344 | ns | YPD | KS.3 |
| FIM1_2481 | 1904.879876 | 1448.663061 | 1282.433671 | -0.387070603 | 0.089075359 | 1.12785749655 | 3.50474143993 | ns | YPD | KS.3 |
| FIM1_2482 | 1533.641768 | 1433.281154 | 1389.641121 | -0.095618052 | 0.070914365 | 0.174687727 | 0.223526605 | ns | YPD | KS.3 |
| FIM1_2483 | 1912.770153 | 1536.485816 | 1291.08699 | -0.308859386 | 0.093466751 | 0.000831125 | 0.001751311 | ns | YPD | KS.3 |
| FIM1_2484 | 458.2880904 | 422.8008684 | 351.8027649 | -0.111050716 | 0.130571107 | 0.384999539 | 0.447935619 | ns | YPD | KS.3 |
| FIM1_2485 | 1928.383708 | 1912.02465 | 1569.017362 | -0.011640255 | 0.123310895 | 0.923358314 | 0.937682713 | ns | YPD | KS.3 |
| FIM1_2486 | 9365.650139 | 6599.899506 | 11891.10262 | -0.477985247 | 0.15341097 | 0.001328411 | 0.002686522 | ns | YPD | KS.3 |
| FIM1_2487 | 2185.980181 | 2725.062907 | 2468.994696 | 0.312465568 | 0.083513843 | 0.000159407 | 0.000393769 | ns | YPD | KS.3 |
| FIM1_2488 | 230.0738398 | 398.7325274 | 344.4512517 | 0.731947977 | 0.191649467 | 6.27882734167 | 0.000167223 | ns | YPD | KS.3 |
| FIM1_2489 | 310.5434815 | 460.1756527 | 585.8551589 | 0.528818516 | 0.172856686 | 0.001491405 | 0.002983409 | ns | YPD | KS.3 |
| FIM1_2490 | 692.3468469 | 1478.945807 | 1138.043493 | 1.044906844 | 0.153992395 | 2.39335101910 | 2.91122770661 | Km>KS.3 | YPD | KS.3 |
| FIM1_2491 | 204.2070573 | 541.6916768 | 811.3610947 | 1.114186838 | 0.372070091 | 0.000400079 | 0.000901854 | Km>KS.3 | YPD | KS.3 |
| FIM1_2492 | 1375.416743 | 2793.470032 | 2656.474913 | 0.927083288 | 0.221632392 | 8.38159941480 | 2.67640931249 | ns | YPD | KS.3 |
| FIM1_2493 | 1606.702186 | 1783.42271 | 1708.561039 | 0.148002279 | 0.068783909 | 0.030373554 | 0.046154072 | ns | YPD | KS.3 |
| FIM1_2494 | 764.1059703 | 1259.119111 | 863.5211927 | 0.693743236 | 0.128858818 | 3.72858107549 | 1.94645234528 | ns | YPD | KS.3 |
| FIM1_2495 | 5978.633995 | 3972.929857 | 5237.240303 | -0.549164037 | 0.176891404 | 0.001242013 | 0.002527205 | ns | YPD | KS.3 |
| FIM1_2496 | 1771.438868 | 1225.349579 | 1081.57116 | -0.508341749 | 0.138652362 | 0.000171495 | 0.00042084 | ns | YPD | KS.3 |
| FIM1_2497 | 970.8330584 | 512.7443283 | 585.9993221 | -0.864225305 | 0.174615608 | 2.59286606800 | 1.14512602175 | ns | YPD | KS.3 |
| FIM1_2498 | 153.7461747 | 88.11535909 | 134.5474941 | -0.708389063 | 0.241118687 | 0.001590788 | 0.003169471 | ns | YPD | KS.3 |
| FIM1_2499 | 1114.682173 | 1433.519789 | 1237.132266 | 0.353669748 | 0.102769333 | 0.00048528 | 0.001073962 | ns | YPD | KS.3 |
| FIM1_2500 | 1167.69747 | 2216.505025 | 1509.180097 | 0.840791602 | 0.214987847 | 3.35263327110 | 9.40741710306 | ns | YPD | KS.3 |
| FIM1_2502 | 1730.572759 | 2404.774136 | 3124.580927 | 0.447983007 | 0.15681436 | 0.003214464 | 0.006027876 | ns | YPD | KS.3 |
| FIM1_2503 | 277.4571207 | 456.6932976 | 574.3758891 | 0.647567614 | 0.217731447 | 0.001604938 | 0.00319255 | ns | YPD | KS.3 |
| FIM1_2504 | 820.7299833 | 917.9335211 | 853.0960671 | 0.139595635 | 0.24789988 | 0.544560483 | 0.606547544 | ns | YPD | KS.3 |
| FIM1_2505 | 447.7326399 | 398.4867203 | 596.8034414 | -0.155646888 | 0.177078677 | 0.361167614 | 0.423775679 | ns | YPD | KS.3 |
| FIM1_2506 | 731.1356677 | 907.9278073 | 537.1545035 | 0.287408946 | 0.188747936 | 0.111755624 | 0.148579431 | ns | YPD | KS.3 |
| FIM1_2507 | 1246.326639 | 774.8998331 | 1007.272581 | -0.660308062 | 0.127095961 | 1.11357603518 | 5.33208929262 | ns | YPD | KS.3 |
| FIM1_2508 | 25.91784144 | 23.28822557 | 19.23299098 | -0.114264366 | 0.363896241 | 0.713145157 | 0.760372301 | ns | YPD | KS.3 |
| FIM1_2510 | 359.7867808 | 173.2660648 | 247.8209449 | -0.974994334 | 0.199239827 | 2.53989892632 | 1.12419903545 | ns | YPD | KS.3 |

| | | | | | | | | | | |
|---|---|---|---|---|---|---|---|---|---|---|
| FIM1_2511 | 4052.102353 | 4175.765696 | 5768.840307 | 0.039623898 | 0.192104497 | 0.829260693 | 0.858064049 | ns | YPD | KS.3 |
| FIM1_2512 | 1619.732068 | 1149.364195 | 1212.02316 | -0.476471642 | 0.125694996 | 0.000109456 | 0.000280548 | ns | YPD | KS.3 |
| FIM1_2513 | 6308.01608 | 4406.729543 | 6384.272418 | -0.477082448 | 0.188131714 | 0.008056311 | 0.013926389 | ns | YPD | KS.3 |
| FIM1_2514 | 629.833885 | 622.5227449 | 481.8060497 | -0.01545563 | 0.139077025 | 0.9093841 | 0.926138361 | ns | YPD | KS.3 |
| FIM1_2515 | 98.30715342 | 108.9222316 | 94.90197683 | 0.136779306 | 0.182869133 | 0.436027142 | 0.499938934 | ns | YPD | KS.3 |
| FIM1_2516 | 5028.719443 | 4054.01997 | 4106.495846 | -0.301962754 | 0.108333647 | 0.00466131 | 0.008429669 | ns | YPD | KS.3 |
| FIM1_2517 | 2176.976375 | 1679.211879 | 2074.916218 | -0.350749335 | 0.166040148 | 0.028829151 | 0.044090079 | ns | YPD | KS.3 |
| FIM1_2518 | 2598.167362 | 1945.05972 | 2038.925988 | -0.41060639 | 0.081693575 | 3.95569816482 | 1.67771512105 | ns | YPD | KS.3 |
| FIM1_2519 | 1933.277167 | 1920.705933 | 2929.214204 | -0.007012473 | 0.355048226 | 0.981693646 | 0.984861039 | ns | YPD | KS.3 |
| FIM1_2520 | 717.6785008 | 574.8231546 | 689.008901 | -0.299560435 | 0.167825712 | 0.064605013 | 0.090768127 | ns | YPD | KS.3 |
| FIM1_2521 | 575.5325904 | 543.0336265 | 697.9728368 | -0.077865753 | 0.180674275 | 0.653893339 | 0.706125322 | ns | YPD | KS.3 |
| FIM1_2522 | 583.2870142 | 379.8582951 | 490.9830642 | -0.587253102 | 0.149853702 | 5.49725099975 | 0.000147513 | ns | YPD | KS.3 |
| FIM1_2523 | 4605.569602 | 5057.433326 | 4516.161231 | 0.133820238 | 0.05913389 | 0.022988614 | 0.035930994 | ns | YPD | KS.3 |
| FIM1_2524 | 546.4073554 | 3128.173135 | 1763.71944 | 2.385063581 | 0.287558397 | 3.21777834179 | 1.41667674782 | Km>KS.3 | YPD | KS.3 |
| FIM1_2525 | 532.8057845 | 494.1577453 | 706.1662017 | -0.103229739 | 0.150623872 | 0.480983463 | 0.544457959 | ns | YPD | KS.3 |
| FIM1_2526 | 3794.852718 | 2420.362649 | 5219.96843 | -0.619597059 | 0.142963555 | 8.58020370485 | 2.73281135926 | ns | YPD | KS.3 |
| FIM1_2527 | 91.8427516 | 137.0957021 | 199.4878147 | 0.4546224 | 0.325166449 | 0.112650121 | 0.149608743 | ns | YPD | KS.3 |
| FIM1_2528 | 14164.0983 | 18939.67743 | 23970.31229 | 0.408560404 | 0.102256053 | 5.11574025464 | 0.000137945 | ns | YPD | KS.3 |
| FIM1_2529 | 5509.985806 | 11355.49136 | 8482.783516 | 0.856547869 | 0.316136178 | 0.002207136 | 0.004262617 | ns | YPD | KS.3 |
| FIM1_2530 | 589.4823945 | 2522.253084 | 1352.405008 | 1.85905112 | 0.354268923 | 9.98883576569 | 7.08908101773 | Km>KS.3 | YPD | KS.3 |
| FIM1_2531 | 1485.539622 | 3824.185566 | 3138.28937 | 1.286696352 | 0.185087948 | 2.97354507748 | 4.26322385029 | Km>KS.3 | YPD | KS.3 |
| FIM1_2532 | 341.5048399 | 327.0192798 | 477.4910332 | -0.053517847 | 0.272391832 | 0.830103543 | 0.858400567 | ns | YPD | KS.3 |
| FIM1_2533 | 1315.760059 | 2638.045542 | 2574.912595 | 0.964934479 | 0.138057383 | 7.28706407713 | 9.82470021239 | ns | YPD | KS.3 |
| FIM1_2534 | 1329.286772 | 2292.920152 | 2106.468145 | 0.742459678 | 0.162680305 | 2.31924917083 | 8.38536673322 | ns | YPD | KS.3 |
| FIM1_2535 | 452.1048357 | 422.9675878 | 461.4695118 | -0.089743796 | 0.171675209 | 0.587934222 | 0.645831917 | ns | YPD | KS.3 |
| FIM1_2536 | 291.8372357 | 341.0857109 | 460.6005252 | 0.182327687 | 0.297374468 | 0.495843554 | 0.558863091 | ns | YPD | KS.3 |
| FIM1_2537 | 583.0875912 | 1098.812901 | 1077.293257 | 0.79141933 | 0.264753587 | 0.001113143 | 0.002283664 | ns | YPD | KS.3 |
| FIM1_2538 | 300.3233676 | 730.9231442 | 815.3711556 | 1.191248662 | 0.204193776 | 6.67373215332 | 4.95549514369 | Km>KS.3 | YPD | KS.3 |
| FIM1_2539 | 341.1393016 | 712.0932631 | 615.8128797 | 0.98429457 | 0.195329204 | 1.16807105026 | 5.55559605646 | ns | YPD | KS.3 |
| FIM1_2540 | 2323.278545 | 4688.071214 | 3341.158667 | 0.941117682 | 0.191444506 | 2.50174286368 | 1.11126524525 | ns | YPD | KS.3 |
| FIM1_2541 | 844.783553 | 1134.72884 | 1315.902532 | 0.404314472 | 0.14563801 | 0.004353785 | 0.007925386 | ns | YPD | KS.3 |
| FIM1_2542 | 384.4016241 | 263.5184139 | 401.8720866 | -0.507697289 | 0.179590948 | 0.003248276 | 0.006086694 | ns | YPD | KS.3 |
| FIM1_2543 | 806.6813025 | 1787.045306 | 1152.744984 | 1.091160149 | 0.161800888 | 2.73486302700 | 3.31044855458 | Km>KS.3 | YPD | KS.3 |
| FIM1_2544 | 599.1245376 | 983.2761565 | 859.9406074 | 0.69543159 | 0.1085431 | 7.54256487071 | 6.81021056838 | ns | YPD | KS.3 |
| FIM1_2546 | 831.0731075 | 1170.510522 | 1183.701345 | 0.473858011 | 0.130447075 | 0.000205034 | 0.000490171 | ns | YPD | KS.3 |
| FIM1_2547 | 607.716486 | 3524.934976 | 2021.126718 | 2.403054247 | 0.289219794 | 2.94862560817 | 1.32156868474 | Km>KS.3 | YPD | KS.3 |
| FIM1_2548 | 2453.774475 | 5952.165753 | 5162.908892 | 1.059434003 | 0.324424162 | 0.000199956 | 0.000479413 | Km>KS.3 | YPD | KS.3 |
| FIM1_2549 | 3488.130092 | 8824.9497 | 5865.03692 | 1.213719483 | 0.239349959 | 4.44694231835 | 2.28785295075 | Km>KS.3 | YPD | KS.3 |
| FIM1_2550 | 1833.923564 | 1348.389234 | 1720.231843 | -0.409873724 | 0.184674394 | 0.02062049 | 0.032567282 | ns | YPD | KS.3 |
| FIM1_2552 | 1114.123973 | 3154.429973 | 2587.608678 | 1.27781321 | 0.320610306 | 5.63641923216 | 1.89339538690 | Km>KS.3 | YPD | KS.3 |
| FIM1_2553 | 515.6817802 | 804.7585684 | 530.5381407 | 0.558669997 | 0.250633329 | 0.016103446 | 0.026155614 | ns | YPD | KS.3 |
| FIM1_2554 | 1274.398333 | 1898.701548 | 1249.548733 | 0.490404276 | 0.266452744 | 0.044920341 | 0.06556481 | ns | YPD | KS.3 |
| FIM1_2555 | 3660.182101 | 13413.33644 | 14232.97528 | 1.543898652 | 0.39952057 | 2.93960321000 | 1.04015120695 | Km>KS.3 | YPD | KS.3 |
| FIM1_2556 | 82.49983439 | 751.5237908 | 413.0204317 | 2.937591263 | 0.398301436 | 1.39034858606 | 2.93507218199 | Km>KS.3 | YPD | KS.3 |
| FIM1_2557 | 7474.692205 | 12407.8437 | 12207.00935 | 0.662795391 | 0.212326522 | 0.000957996 | 0.001987503 | ns | YPD | KS.3 |

| | | | | | | | | | | |
|---|---|---|---|---|---|---|---|---|---|---|
| FIM1_2558 | 1135.640834 | 572.1073135 | 1227.149074 | -0.874347357 | 0.248429663 | 0.000142641 | 0.000355708 | ns | YPD | KS.3 |
| FIM1_2559 | 384.6604325 | 891.2021394 | 458.2771054 | 1.018219049 | 0.308038494 | 0.000200348 | 0.000479891 | Km>KS.3 | YPD | KS.3 |
| FIM1_2560 | 4118.882984 | 3222.7356 | 3738.299824 | -0.337733091 | 0.139614796 | 0.013175948 | 0.021734197 | ns | YPD | KS.3 |
| FIM1_2561 | 414.5747064 | 483.224636 | 346.8324722 | 0.197833919 | 0.218253983 | 0.337122033 | 0.398854249 | ns | YPD | KS.3 |
| FIM1_2562 | 2510.174906 | 4195.820985 | 2830.694731 | 0.713855003 | 0.129532346 | 1.75136627847 | 9.80095302071 | ns | YPD | KS.3 |
| FIM1_2563 | 2296.250805 | 3463.387301 | 3001.103454 | 0.581232962 | 0.090435122 | 8.11509568282 | 7.31387699675 | ns | YPD | KS.3 |
| FIM1_2564 | 852.3512177 | 1778.133978 | 1739.693656 | 1.015209241 | 0.148351702 | 1.75823494814 | 2.19779368518 | Km>KS.3 | YPD | KS.3 |
| FIM1_2565 | 1059.021364 | 857.4023112 | 1138.975453 | -0.29095517 | 0.138752002 | 0.031708639 | 0.047948475 | ns | YPD | KS.3 |
| FIM1_2566 | 2315.463631 | 4485.024875 | 2318.819173 | 0.893103288 | 0.17903355 | 1.96597709472 | 8.91589429924 | ns | YPD | KS.3 |
| FIM1_2567 | 944.3286585 | 1153.997438 | 1056.417571 | 0.278336871 | 0.125816876 | 0.02401896 | 0.037341977 | ns | YPD | KS.3 |
| FIM1_2568 | 1079.206328 | 632.3998332 | 1217.730461 | -0.68257507 | 0.238660364 | 0.002147355 | 0.004161704 | ns | YPD | KS.3 |
| FIM1_2570 | 270.4450411 | 404.9700087 | 388.1636513 | 0.513154564 | 0.235500219 | 0.019715454 | 0.031366928 | ns | YPD | KS.3 |
| FIM1_2571 | 384.2737209 | 221.0216486 | 421.0415144 | -0.685676045 | 0.26663704 | 0.005020097 | 0.009006485 | ns | YPD | KS.3 |
| FIM1_2572 | 227.5669668 | 132.5385619 | 287.6389418 | -0.716012192 | 0.20019372 | 0.000168139 | 0.000413287 | ns | YPD | KS.3 |
| FIM1_2573 | 535.797629 | 453.7815019 | 484.8169296 | -0.225850785 | 0.155696399 | 0.135020262 | 0.176352272 | ns | YPD | KS.3 |
| FIM1_2574 | 1879.905137 | 1584.968729 | 1794.752805 | -0.237218159 | 0.124053768 | 0.051223954 | 0.073674233 | ns | YPD | KS.3 |
| FIM1_2575 | 978.9782741 | 884.8692886 | 933.2852499 | -0.140964269 | 0.120319008 | 0.232851676 | 0.287239545 | ns | YPD | KS.3 |
| FIM1_2576 | 11879.88181 | 14503.09516 | 14153.82679 | 0.285635121 | 0.053337499 | 7.60964724192 | 3.76696467946 | ns | YPD | KS.3 |
| FIM1_2577 | 3981.679317 | 5940.535255 | 5459.753975 | 0.543891024 | 0.161077389 | 0.00048399 | 0.001071584 | ns | YPD | KS.3 |
| FIM1_2578 | 26.56145073 | 77.47163407 | 73.6616089 | 1.43109009 | 0.215641424 | 1.35250225577 | 1.72962535450 | Km>KS.3 | YPD | KS.3 |
| FIM1_2579 | 112.6254153 | 335.7273819 | 269.9166729 | 1.509460524 | 0.167854202 | 6.24299659798 | 3.69748905654 | Km>KS.3 | YPD | KS.3 |
| FIM1_2580 | 469.3327705 | 2032.660642 | 1231.593787 | 2.004952386 | 0.24249798 | 3.94856315147 | 1.69345704125 | Km>KS.3 | YPD | KS.3 |
| FIM1_2581 | 12521.77438 | 16104.9351 | 20957.63236 | 0.34732882 | 0.13561133 | 0.008760554 | 0.01501335 | ns | YPD | KS.3 |
| FIM1_2582 | 653.7749833 | 523.0790068 | 655.2949581 | -0.299749507 | 0.172537114 | 0.071534918 | 0.099492932 | ns | YPD | KS.3 |
| FIM1_2583 | 676.1657361 | 176.7867537 | 481.033384 | -1.856985311 | 0.197013605 | 1.78458834143 | 1.43198822558 | Km<KS.3 | YPD | KS.3 |
| FIM1_2584 | 88.47789348 | 84.45564847 | 103.7048874 | -0.047076223 | 0.391736377 | 0.885192213 | 0.905508775 | ns | YPD | KS.3 |
| FIM1_2585 | 5830.289393 | 3661.138913 | 4837.266344 | -0.647073968 | 0.126805269 | 1.86800671024 | 8.54092645298 | ns | YPD | KS.3 |
| FIM1_2586 | 1357.906341 | 1104.097057 | 1096.603075 | -0.28958852 | 0.110269772 | 0.007646747 | 0.013287659 | ns | YPD | KS.3 |
| FIM1_2587 | 5257.042911 | 4590.370821 | 4513.054149 | -0.191235592 | 0.094624719 | 0.040916536 | 0.060331881 | ns | YPD | KS.3 |
| FIM1_2588 | 2214.036062 | 3492.848577 | 5707.663931 | 0.593265852 | 0.215635025 | 0.00356277 | 0.006613724 | ns | YPD | KS.3 |
| FIM1_2590 | 412.0548253 | 2402.272537 | 1955.598832 | 2.393503989 | 0.306998066 | 1.86758335432 | 5.99434012114 | Km>KS.3 | YPD | KS.3 |
| FIM1_2591 | 427.9592037 | 858.5974754 | 888.9165065 | 0.881076719 | 0.256108921 | 0.000187596 | 0.000453714 | ns | YPD | KS.3 |
| FIM1_2592 | 1945.020487 | 2011.450435 | 1822.145642 | 0.046702232 | 0.118946452 | 0.689418645 | 0.739352826 | ns | YPD | KS.3 |
| FIM1_2593 | 2029.923539 | 2121.053069 | 1588.488274 | 0.06226929 | 0.088079403 | 0.475193921 | 0.538885287 | ns | YPD | KS.3 |
| FIM1_2594 | 2160.177725 | 1979.033541 | 2182.106981 | -0.121908161 | 0.120377056 | 0.302415034 | 0.362939287 | ns | YPD | KS.3 |
| FIM1_2595 | 1609.042769 | 1209.723766 | 1316.272804 | -0.402371512 | 0.096540075 | 2.45281805566 | 7.11117122780 | ns | YPD | KS.3 |
| FIM1_2596 | 969.4414046 | 2627.062021 | 2024.028705 | 1.33649989 | 0.212656786 | 2.18326987891 | 2.19873839020 | Km>KS.3 | YPD | KS.3 |
| FIM1_2597 | 718.6398883 | 946.6037037 | 977.7203132 | 0.362875269 | 0.196760026 | 0.053152085 | 0.076161181 | ns | YPD | KS.3 |
| FIM1_2598 | 1337.355313 | 693.8025034 | 1013.907709 | -0.914986359 | 0.126558493 | 1.46700717463 | 2.23876094902 | ns | YPD | KS.3 |
| FIM1_2599 | 1158.126181 | 725.9863503 | 964.505576 | -0.604485804 | 0.222085207 | 0.003813631 | 0.007021766 | ns | YPD | KS.3 |
| FIM1_2600 | 1733.167378 | 953.589577 | 1325.314155 | -0.805432434 | 0.179415074 | 2.86520922156 | 1.01599542960 | ns | YPD | KS.3 |
| FIM1_2601 | 1955.655631 | 2080.661096 | 2074.916074 | 0.086142049 | 0.118949926 | 0.461041512 | 0.524629808 | ns | YPD | KS.3 |
| FIM1_2602 | 2054.499402 | 2571.545478 | 1905.859368 | 0.315260528 | 0.104762703 | 0.002272675 | 0.004380688 | ns | YPD | KS.3 |
| FIM1_2603 | 338.573342 | 460.1569583 | 286.3863057 | 0.360174523 | 0.299489545 | 0.180710296 | 0.230403312 | ns | YPD | KS.3 |
| FIM1_2604 | 357.3350989 | 508.1800675 | 543.7081307 | 0.460483813 | 0.206190965 | 0.01861934 | 0.029813717 | ns | YPD | KS.3 |

| | | | | | | | | | | |
|---|---|---|---|---|---|---|---|---|---|---|
| FIM1_2606 | 22.33986906 | 22.79164506 | 32.27504231 | 0.024286211 | 0.313955833 | 0.930713336 | 0.943610932 | ns | YPD | KS.3 |
| FIM1_2607 | 2823.745244 | 2523.229502 | 3002.133013 | -0.147324028 | 0.201771955 | 0.44315405 | 0.506942147 | ns | YPD | KS.3 |
| FIM1_2608 | 832.0055166 | 849.0282475 | 800.2735726 | 0.028569355 | 0.088425529 | 0.744159288 | 0.785694495 | ns | YPD | KS.3 |
| FIM1_2609 | 773.3614469 | 501.1806145 | 502.6738705 | -0.594209165 | 0.150621907 | 4.87334781622 | 0.000132125 | ns | YPD | KS.3 |
| FIM1_2610 | 1054.974047 | 1202.271733 | 1233.092503 | 0.181497338 | 0.121686503 | 0.128637279 | 0.168991408 | ns | YPD | KS.3 |
| FIM1_2611 | 2127.73674 | 1928.823911 | 1871.482649 | -0.127707007 | 0.208292721 | 0.518314391 | 0.580246196 | ns | YPD | KS.3 |
| FIM1_2612 | 1210.978566 | 5229.048096 | 3394.826906 | 1.801883033 | 0.402927476 | 6.89619287071 | 3.44117949165 | Km>KS.3 | YPD | KS.3 |
| FIM1_2613 | 1617.197017 | 5982.208864 | 3064.290298 | 1.699970594 | 0.301647649 | 2.05022436843 | 1.69432993902 | Km>KS.3 | YPD | KS.3 |
| FIM1_2614 | 923.5925047 | 660.7688404 | 709.2168211 | -0.462893101 | 0.136462206 | 0.000513463 | 0.001128303 | ns | YPD | KS.3 |
| FIM1_2615 | 53.33373372 | 49.95289793 | 64.47101098 | -0.081728713 | 0.26571249 | 0.737997635 | 0.780231037 | ns | YPD | KS.3 |
| FIM1_2616 | 893.3409155 | 802.2906971 | 737.3979886 | -0.151457806 | 0.088451644 | 0.083685897 | 0.114378389 | ns | YPD | KS.3 |
| FIM1_2617 | 1701.415985 | 932.0545903 | 1133.242136 | -0.854328016 | 0.083500325 | 4.98154776000 | 4.67607549170 | ns | YPD | KS.3 |
| FIM1_2618 | 415.2094523 | 362.2103269 | 394.8937208 | -0.186762129 | 0.151345403 | 0.204314329 | 0.255907298 | ns | YPD | KS.3 |
| FIM1_2619 | 418.5241831 | 234.7672492 | 276.3989874 | -0.78026979 | 0.177346582 | 4.60188533677 | 1.56596303354 | ns | YPD | KS.3 |
| FIM1_2620 | 823.6189593 | 335.1964376 | 670.0593794 | -1.247009973 | 0.147431548 | 2.59817078956 | 9.29920840150 | Km<KS.3 | YPD | KS.3 |
| FIM1_2621 | 6485.599961 | 10714.86335 | 8627.981714 | 0.678004566 | 0.173093871 | 4.70204502475 | 0.000127829 | ns | YPD | KS.3 |
| FIM1_2622 | 641.8675589 | 702.4698861 | 652.659282 | 0.124844457 | 0.132395502 | 0.335181226 | 0.396935635 | ns | YPD | KS.3 |
| FIM1_2623 | 2371.660575 | 1516.788771 | 1779.418525 | -0.625205554 | 0.114875471 | 3.05070764619 | 1.61460324891 | ns | YPD | KS.3 |
| FIM1_2624 | 5638.103067 | 7802.443531 | 7410.842857 | 0.451756321 | 0.124168479 | 0.000206379 | 0.000492676 | ns | YPD | KS.3 |
| FIM1_2625 | 468.1216326 | 628.3714453 | 576.4427825 | 0.398671032 | 0.163217709 | 0.011569478 | 0.0193213 | ns | YPD | KS.3 |
| FIM1_2626 | 1071.165654 | 1825.784896 | 1584.582197 | 0.742819024 | 0.124994268 | 1.29388730458 | 9.04085581502 | ns | YPD | KS.3 |
| FIM1_2627 | 822.9866544 | 434.3682024 | 1047.969986 | -0.86946757 | 0.167984915 | 7.79950769564 | 3.85586830512 | ns | YPD | KS.3 |
| FIM1_2628 | 999.3697203 | 680.5244928 | 833.5519857 | -0.53547338 | 0.122447206 | 8.22547726299 | 2.63331720613 | ns | YPD | KS.3 |
| FIM1_2629 | 6681.196769 | 10759.97948 | 7639.601927 | 0.592704602 | 0.260271598 | 0.01338489 | 0.022049612 | ns | YPD | KS.3 |
| FIM1_2630 | 626.5832432 | 293.5507671 | 544.7853279 | -1.027754392 | 0.178672612 | 1.93106692126 | 1.31243960837 | Km<KS.3 | YPD | KS.3 |
| FIM1_2631 | 242.8687178 | 288.71221 | 217.1279088 | 0.23774674 | 0.144862208 | 0.092084283 | 0.12452278 | ns | YPD | KS.3 |
| FIM1_2632 | 878.2062288 | 536.4749783 | 622.236406 | -0.678598938 | 0.1430787 | 1.10906132722 | 4.32042901350 | ns | YPD | KS.3 |
| FIM1_2633 | 907.0646955 | 1393.963095 | 808.6597244 | 0.585804118 | 0.158226688 | 0.000132109 | 0.00033194 | ns | YPD | KS.3 |
| FIM1_2634 | 2917.130983 | 4618.295178 | 5047.092429 | 0.566509956 | 0.267020561 | 0.020693286 | 0.032640805 | ns | YPD | KS.3 |
| FIM1_2635 | 17375.7195 | 19690.41363 | 31529.08503 | 0.174984887 | 0.110577932 | 0.108025223 | 0.144197876 | ns | YPD | KS.3 |
| FIM1_2636 | 3298.758877 | 4872.453765 | 3879.739344 | 0.55506074 | 0.074723213 | 7.13530853127 | 1.15629185482 | ns | YPD | KS.3 |
| FIM1_2637 | 4177.082326 | 3346.345359 | 4556.375614 | -0.297679948 | 0.173855163 | 0.075567572 | 0.10463364 | ns | YPD | KS.3 |
| FIM1_2638 | 1921.834891 | 1488.798507 | 1535.1718 | -0.355816838 | 0.118973616 | 0.002324845 | 0.004467402 | ns | YPD | KS.3 |
| FIM1_2639 | 2410.489115 | 3508.474355 | 3287.046383 | 0.496475611 | 0.19497733 | 0.007605115 | 0.013224553 | ns | YPD | KS.3 |
| FIM1_2640 | 2564.68978 | 3388.820626 | 3174.526742 | 0.38119381 | 0.149338212 | 0.008666799 | 0.014862917 | ns | YPD | KS.3 |
| FIM1_2641 | 1425.973463 | 774.6064705 | 974.5261912 | -0.859780566 | 0.105146951 | 1.01323177768 | 2.75455087103 | ns | YPD | KS.3 |
| FIM1_2642 | 380.9479338 | 519.647076 | 395.3580463 | 0.433636944 | 0.116943619 | 0.000160594 | 0.000396308 | ns | YPD | KS.3 |
| FIM1_2643 | 2293.193186 | 2858.272242 | 2442.68225 | 0.302299035 | 0.143803008 | 0.030993758 | 0.04695309 | ns | YPD | KS.3 |
| FIM1_2644 | 1458.887188 | 1511.133051 | 1769.190409 | 0.049091412 | 0.104191986 | 0.632876626 | 0.687308713 | ns | YPD | KS.3 |
| FIM1_2645 | 660.4515354 | 642.3659034 | 693.2951935 | -0.038391413 | 0.129655853 | 0.762386134 | 0.801536563 | ns | YPD | KS.3 |
| FIM1_2646 | 2399.244176 | 2601.676685 | 2678.289446 | 0.113955703 | 0.101200505 | 0.253953792 | 0.310498922 | ns | YPD | KS.3 |
| FIM1_2647 | 3550.891834 | 3827.780606 | 3353.209645 | 0.106131351 | 0.090776418 | 0.237370334 | 0.292161656 | ns | YPD | KS.3 |
| FIM1_2648 | 349.7226242 | 420.7258676 | 308.8158223 | 0.260355406 | 0.103075426 | 0.010452047 | 0.017680698 | ns | YPD | KS.3 |
| FIM1_2649 | 870.6796006 | 801.6437042 | 874.0597844 | -0.117263839 | 0.077121875 | 0.125438504 | 0.165356798 | ns | YPD | KS.3 |
| FIM1_2650 | 9938.420551 | 7332.141887 | 13759.61847 | -0.42053221 | 0.133592299 | 0.001281251 | 0.002597484 | ns | YPD | KS.3 |

| | | | | | | | | | | |
|---|---|---|---|---|---|---|---|---|---|---|
| FIM1_2651 | 224.6505874 | 442.5065039 | 381.7676173 | 0.894695199 | 0.209791984 | 6.3876864682 | 2.1073435132 | ns | YPD | KS.3 |
| FIM1_2652 | 6861.184147 | 7616.469913 | 7485.896065 | 0.147763747 | 0.08565988 | 0.08160414 | 0.111809583 | ns | YPD | KS.3 |
| FIM1_2653 | 2456.074509 | 1312.874557 | 1690.258184 | -0.878417738 | 0.114844436 | 6.7589160023 | 1.3186512592 | ns | YPD | KS.3 |
| FIM1_2654 | 1765.942922 | 2526.622772 | 2235.239421 | 0.498272271 | 0.123554545 | 3.9015009737 | 0.000107833 | ns | YPD | KS.3 |
| FIM1_2655 | 553.4901614 | 2021.384875 | 1444.349023 | 1.516676886 | 0.412103273 | 6.9176051575 | 2.2656409255 | Km>KS.3 | YPD | KS.3 |
| FIM1_2656 | 354.0444332 | 771.7113912 | 282.8426353 | 0.946832975 | 0.300106592 | 0.000420473 | 0.000944831 | ns | YPD | KS.3 |
| FIM1_2657 | 1111.078924 | 871.0615656 | 919.771808 | -0.336822014 | 0.130455972 | 0.008339785 | 0.01435655 | ns | YPD | KS.3 |
| FIM1_2659 | 2135.482088 | 1875.335947 | 1561.785365 | -0.179264538 | 0.132741056 | 0.1674472 | 0.215147164 | ns | YPD | KS.3 |
| FIM1_2660 | 1470.369289 | 732.7814153 | 1832.770411 | -0.867998252 | 0.271069651 | 0.000450547 | 0.001002896 | ns | YPD | KS.3 |
| FIM1_2661 | 912.4379138 | 2979.998306 | 1387.871653 | 1.456794717 | 0.341751095 | 7.7955252536 | 3.1251199143 | Km>KS.3 | YPD | KS.3 |
| FIM1_2662 | 4820.459561 | 7445.998347 | 5391.751476 | 0.60105441 | 0.13636265 | 6.3186597209 | 2.0901151669 | ns | YPD | KS.3 |
| FIM1_2663 | 207.444211 | 184.3471795 | 406.4431412 | -0.136999824 | 0.306261058 | 0.617071348 | 0.672935107 | ns | YPD | KS.3 |
| FIM1_2664 | 2809.414648 | 2151.08802 | 2489.737755 | -0.377201485 | 0.090751434 | 2.6500379592 | 7.6119739301 | ns | YPD | KS.3 |
| FIM1_2665 | 384.3689842 | 608.1418787 | 497.1695574 | 0.602109761 | 0.207294537 | 0.002182067 | 0.004219114 | ns | YPD | KS.3 |
| FIM1_2666 | 483.193902 | 2160.443263 | 1035.415032 | 2.089189597 | 0.198155836 | 1.2082275615 | 1.5412646458 | Km>KS.3 | YPD | KS.3 |
| FIM1_2667 | 3058.03228 | 3968.024559 | 3083.454991 | 0.370515488 | 0.074835598 | 6.0917876617 | 2.4841511161 | ns | YPD | KS.3 |
| FIM1_2668 | 1956.319421 | 4856.371168 | 3118.802518 | 1.148357007 | 0.27623945 | 4.5457928743 | 1.5521839086 | Km>KS.3 | YPD | KS.3 |
| FIM1_2669 | 20.37629302 | 59.49190756 | 47.98920621 | 1.351167555 | 0.297367252 | 3.4386257373 | 1.4760278725 | Km>KS.3 | YPD | KS.3 |
| FIM1_2671 | 38.46374282 | 17.86693252 | 54.40823865 | -0.933940041 | 0.29657138 | 0.000445913 | 0.00099436 | ns | YPD | KS.3 |
| FIM1_2672 | 1229.180857 | 961.1537726 | 872.2161792 | -0.345632193 | 0.102606759 | 0.000638553 | 0.001378821 | ns | YPD | KS.3 |
| FIM1_2674 | 4007.973131 | 4752.933119 | 5072.522202 | 0.23302491 | 0.148931867 | 0.107791069 | 0.143923931 | ns | YPD | KS.3 |
| FIM1_2675 | 837.2095422 | 20644.87172 | 10561.10044 | 3.744684433 | 0.676816317 | 1.1307807084 | 9.8202100828 | Km>KS.3 | YPD | KS.3 |
| FIM1_2676 | 9761.3561 | 30706.5406 | 23990.20896 | 1.439140487 | 0.314158998 | 1.9365359328 | 8.8124416950 | Km>KS.3 | YPD | KS.3 |
| FIM1_2677 | 3928.150026 | 3295.122148 | 3814.504978 | -0.2458327 | 0.112315892 | 0.02614097 | 0.040401157 | ns | YPD | KS.3 |
| FIM1_2678 | 603.3079123 | 462.5861645 | 579.2869035 | -0.364689925 | 0.14645911 | 0.010530075 | 0.017800586 | ns | YPD | KS.3 |
| FIM1_2679 | 1280.140391 | 2536.163738 | 1383.071188 | 0.936980933 | 0.158368708 | 9.4447115360 | 6.7510689499 | ns | YPD | KS.3 |
| FIM1_2680 | 4723.298025 | 3067.226968 | 4303.406281 | -0.594286625 | 0.143491987 | 2.1084690554 | 6.2142378856 | ns | YPD | KS.3 |
| FIM1_2681 | 2010.788382 | 4566.144465 | 4554.22403 | 1.111115753 | 0.183449949 | 2.3037494146 | 1.8788775963 | Km>KS.3 | YPD | KS.3 |
| FIM1_2682 | 275.7742016 | 1399.502688 | 945.3060971 | 2.170396639 | 0.318876725 | 2.6556999424 | 4.6851444019 | Km>KS.3 | YPD | KS.3 |
| FIM1_2683 | 1165.715223 | 1532.754539 | 2297.684861 | 0.367057701 | 0.175821962 | 0.030091996 | 0.045824205 | ns | YPD | KS.3 |
| FIM1_2684 | 4301.750732 | 2862.46827 | 5456.353642 | -0.517833167 | 0.237546161 | 0.019521245 | 0.031097724 | ns | YPD | KS.3 |
| FIM1_2685 | 1303.366855 | 777.399845 | 1047.779783 | -0.720319588 | 0.123373647 | 2.5499163638 | 1.6780203585 | ns | YPD | KS.3 |
| FIM1_2686 | 294.9319194 | 527.3641056 | 211.4828374 | 0.776430853 | 0.190473675 | 1.9523327365 | 5.8021836106 | ns | YPD | KS.3 |
| FIM1_2687 | 787.236702 | 1568.585451 | 777.3624947 | 0.841950263 | 0.288412593 | 0.001219644 | 0.002486775 | ns | YPD | KS.3 |
| FIM1_2688 | 1403.618066 | 2616.874999 | 1954.134537 | 0.866320232 | 0.130734415 | 1.1877141100 | 1.2482526099 | ns | YPD | KS.3 |
| FIM1_2689 | 1877.993928 | 1582.206453 | 1808.791337 | -0.240023131 | 0.108680284 | 0.02496201 | 0.03869566 | ns | YPD | KS.3 |
| FIM1_2690 | 6497.83934 | 5027.12779 | 6279.00711 | -0.363548072 | 0.085224585 | 1.6560443446 | 4.9811491018 | ns | YPD | KS.3 |
| FIM1_2691 | 1832.247972 | 1465.455358 | 1404.064091 | -0.317417593 | 0.074862834 | 1.9399645763 | 5.7757772396 | ns | YPD | KS.3 |
| FIM1_2692 | 1234.811935 | 647.572893 | 751.1524816 | -0.900159864 | 0.126218904 | 3.1289262632 | 4.4731057930 | ns | YPD | KS.3 |
| FIM1_2693 | 3038.129636 | 1976.514022 | 2325.402404 | -0.60822946 | 0.091023197 | 1.4032394723 | 1.4574355689 | ns | YPD | KS.3 |
| FIM1_2694 | 591.3457958 | 540.0110929 | 600.8518616 | -0.127181944 | 0.096716333 | 0.183252131 | 0.232749387 | ns | YPD | KS.3 |
| FIM1_2695 | 3269.644514 | 1846.603631 | 2827.478016 | -0.799428427 | 0.118488909 | 6.1367703823 | 6.8762235703 | ns | YPD | KS.3 |
| FIM1_2696 | 5271.572657 | 4213.161577 | 6745.90564 | -0.293095995 | 0.204303638 | 0.131260021 | 0.172164145 | ns | YPD | KS.3 |
| FIM1_2697 | 515.3614613 | 210.8437238 | 428.8738886 | -1.143476533 | 0.261572485 | 1.7860716394 | 6.6311241837 | Km<KS.3 | YPD | KS.3 |
| FIM1_2698 | 3717.571396 | 3199.597419 | 3296.22013 | -0.210874217 | 0.101585564 | 0.035431431 | 0.052982077 | ns | YPD | KS.3 |

| | | | | | | | | | |
|---|---|---|---|---|---|---|---|---|---|
| FIM1_2699 | 2330.861957 | 1086.718795 | 2342.557362 | -0.983623336 | 0.242221139 | 1.2020168304 | 3.7120010746 | ns | YPD | KS.3 |
| FIM1_2700 | 385.3483727 | 683.9826632 | 571.3223321 | 0.768718917 | 0.185842589 | 1.5330623318 | 4.6421090083 | ns | YPD | KS.3 |
| FIM1_2701 | 2057.080201 | 2486.125523 | 2051.847764 | 0.250288662 | 0.191990825 | 0.172761165 | 0.221460138 | ns | YPD | KS.3 |
| FIM1_2702 | 231.2358502 | 194.0049568 | 193.6273669 | -0.235575991 | 0.165133746 | 0.140170314 | 0.182360699 | ns | YPD | KS.3 |
| FIM1_2703 | 238.7265415 | 63.37269138 | 206.2115144 | -1.536635193 | 0.427317295 | 8.8314904619 | 2.8056618804 | Km<KS.3 | YPD | KS.3 |
| FIM1_2704 | 1497.225729 | 1671.532783 | 1576.395522 | 0.151938077 | 0.132726707 | 0.241958707 | 0.29700088 | ns | YPD | KS.3 |
| FIM1_2705 | 1928.64968 | 2005.427744 | 3068.511263 | 0.051245099 | 0.198875347 | 0.786977252 | 0.82235073 | ns | YPD | KS.3 |
| FIM1_2706 | 20.54054091 | 17.92416865 | 23.14872675 | -0.155126738 | 0.310809009 | 0.575646713 | 0.634576202 | ns | YPD | KS.3 |
| FIM1_2708 | 5166.142607 | 3252.455189 | 5778.449195 | -0.567670914 | 0.271510164 | 0.022230289 | 0.034844261 | ns | YPD | KS.3 |
| FIM1_2709 | 3350.933468 | 2461.612167 | 3182.780214 | -0.430616181 | 0.116762902 | 0.000174467 | 0.000426732 | ns | YPD | KS.3 |
| FIM1_2710 | 7999.156181 | 5764.260193 | 8876.301857 | -0.44067271 | 0.173671657 | 0.008427548 | 0.014492586 | ns | YPD | KS.3 |
| FIM1_2711 | 294.2878468 | 419.6450356 | 333.9744302 | 0.496247358 | 0.111369712 | 5.9360474594 | 1.9820024235 | ns | YPD | KS.3 |
| FIM1_2712 | 2772.489976 | 2535.741292 | 2749.901888 | -0.121663746 | 0.153270672 | 0.413921017 | 0.47778586 | ns | YPD | KS.3 |
| FIM1_2713 | 2815.815524 | 1874.028059 | 1923.715799 | -0.572653471 | 0.103960545 | 2.2957558343 | 1.2455163877 | ns | YPD | KS.3 |
| FIM1_2714 | 1496.763994 | 1541.073233 | 1521.290893 | 0.042085862 | 0.07145891 | 0.553243957 | 0.614372475 | ns | YPD | KS.3 |
| FIM1_2715 | 3003.893603 | 2575.228159 | 2877.232405 | -0.217439251 | 0.09283612 | 0.017867035 | 0.028762733 | ns | YPD | KS.3 |
| FIM1_2716 | 8882.569906 | 5293.860432 | 7226.098918 | -0.713512426 | 0.142997983 | 2.9715349312 | 1.2911254395 | ns | YPD | KS.3 |
| FIM1_2718 | 4476.241566 | 7320.551515 | 5181.679644 | 0.694272229 | 0.097318456 | 4.9491930036 | 6.8205637654 | ns | YPD | KS.3 |
| FIM1_2719 | 2303.661119 | 2008.493668 | 2146.318507 | -0.185807085 | 0.15952169 | 0.229464975 | 0.283413171 | ns | YPD | KS.3 |
| FIM1_2720 | 2538.198215 | 3663.446289 | 3393.652226 | 0.522230902 | 0.074303206 | 1.4250150145 | 1.8131584903 | ns | YPD | KS.3 |
| FIM1_2721 | 292.7405434 | 799.8382893 | 627.1013239 | 1.304464122 | 0.25630799 | 2.7635281883 | 1.4815250794 | Km>KS.3 | YPD | KS.3 |
| FIM1_2722 | 3375.49623 | 1490.333121 | 2572.884247 | -1.137760223 | 0.137869861 | 2.2982277060 | 7.2365081249 | Km<KS.3 | YPD | KS.3 |
| FIM1_2723 | 778.5668952 | 527.0096459 | 810.7871717 | -0.496775924 | 0.235814395 | 0.024166092 | 0.037558984 | ns | YPD | KS.3 |
| FIM1_2724 | 2514.483314 | 2143.839137 | 1911.980482 | -0.220714914 | 0.128974227 | 0.080580569 | 0.11057611 | ns | YPD | KS.3 |
| FIM1_2726 | 606.8482983 | 1077.527112 | 789.6774471 | 0.790350681 | 0.14765879 | 3.6110664315 | 1.8870856614 | ns | YPD | KS.3 |
| FIM1_2727 | 157.5767125 | 102.0090504 | 134.5850001 | -0.565377898 | 0.212699531 | 0.004935797 | 0.008884077 | ns | YPD | KS.3 |
| FIM1_2728 | 456.6996562 | 500.3948589 | 592.1077316 | 0.114600767 | 0.243854397 | 0.613914193 | 0.670079664 | ns | YPD | KS.3 |
| FIM1_2729 | 481.5484569 | 1179.984883 | 331.3030649 | 1.064375407 | 0.331109 | 0.00023384 | 0.000552137 | Km>KS.3 | YPD | KS.3 |
| FIM1_2730 | 62.60991537 | 146.0477801 | 150.8965523 | 1.070035988 | 0.269215648 | 1.3049510573 | 4.0050163543 | Km>KS.3 | YPD | KS.3 |
| FIM1_2731 | 3539.687648 | 5380.49334 | 4238.867516 | 0.589391896 | 0.101850166 | 4.4220496832 | 2.7637810520 | ns | YPD | KS.3 |
| FIM1_2732 | 8519.938858 | 9857.232712 | 8924.198354 | 0.205496569 | 0.095696416 | 0.029795667 | 0.045414657 | ns | YPD | KS.3 |
| FIM1_2733 | 432.8042789 | 324.0351655 | 314.6935133 | -0.394519455 | 0.153655346 | 0.008178795 | 0.014118497 | ns | YPD | KS.3 |
| FIM1_2734 | 282.5255931 | 210.5009785 | 239.54723 | -0.406303686 | 0.130282696 | 0.001438866 | 0.002891098 | ns | YPD | KS.3 |
| FIM1_2735 | 789.3182327 | 542.990162 | 651.349206 | -0.518707369 | 0.126117984 | 2.6878525746 | 7.7072429733 | ns | YPD | KS.3 |
| FIM1_2736 | 801.343261 | 312.7147327 | 595.3209154 | -1.307063061 | 0.148966208 | 1.2552881765 | 6.1832264142 | Km<KS.3 | YPD | KS.3 |
| FIM1_2737 | 2483.682226 | 1877.714391 | 2475.593407 | -0.390607957 | 0.116598135 | 0.000651955 | 0.001404102 | ns | YPD | KS.3 |
| FIM1_2738 | 607.5304371 | 550.0364457 | 482.4117522 | -0.125178259 | 0.239246663 | 0.575327993 | 0.634385611 | ns | YPD | KS.3 |
| FIM1_2739 | 2222.053071 | 2294.813826 | 2229.439946 | 0.038883799 | 0.274339992 | 0.87683559 | 0.898323118 | ns | YPD | KS.3 |
| FIM1_2740 | 5821.779288 | 4798.214126 | 5832.091995 | -0.274609874 | 0.078059215 | 0.000390636 | 0.000882168 | ns | YPD | KS.3 |
| FIM1_2741 | 1323.262046 | 673.3672114 | 950.3022383 | -0.9514792 | 0.10690029 | 1.5001993787 | 7.2461086495 | ns | YPD | KS.3 |
| FIM1_2742 | 7896.348286 | 7868.020522 | 6492.584788 | -0.004531806 | 0.237560882 | 0.983720925 | 0.986099456 | ns | YPD | KS.3 |
| FIM1_2743 | 500.3350429 | 692.1590163 | 767.070899 | 0.391660799 | 0.279054585 | 0.123349896 | 0.16290569 | ns | YPD | KS.3 |
| FIM1_2744 | 5064.262564 | 5230.309384 | 6478.222438 | 0.045616726 | 0.076313526 | 0.546970938 | 0.608356901 | ns | YPD | KS.3 |
| FIM1_2745 | 1628.734105 | 807.8911906 | 1426.642156 | -0.894734372 | 0.248571686 | 9.9818127742 | 0.000257972 | ns | YPD | KS.3 |
| FIM1_2746 | 1631.707607 | 1011.268492 | 1398.377701 | -0.664864761 | 0.128277163 | 1.1796291574 | 5.5998616970 | ns | YPD | KS.3 |

| | | | | | | | | | | |
|---|---|---|---|---|---|---|---|---|---|---|
| FIM1_2747 | 3458.58034 | 2282.291268 | 2989.429139 | -0.5671407 | 0.156513343 | 0.000185114 | 0.000449459 | ns | YPD | KS.3 |
| FIM1_2748 | 785.0451799 | 554.2540459 | 558.3447268 | -0.481852295 | 0.131703758 | 0.000183342 | 0.000446027 | ns | YPD | KS.3 |
| FIM1_2749 | 1184.833158 | 1360.944765 | 918.7957471 | 0.191792518 | 0.131721046 | 0.136774974 | 0.17836317 | ns | YPD | KS.3 |
| FIM1_2750 | 78.86393086 | 98.10158334 | 94.23558957 | 0.296955924 | 0.175501316 | 0.078876376 | 0.108580511 | ns | YPD | KS.3 |
| FIM1_2751 | 643.3822128 | 417.0678018 | 582.8865073 | -0.574908927 | 0.193354476 | 0.00183589 | 0.003608674 | ns | YPD | KS.3 |
| FIM1_2752 | 1872.250674 | 1472.651273 | 1326.891207 | -0.339960536 | 0.083872286 | 4.29405227963 | 0.000117692 | ns | YPD | KS.3 |
| FIM1_2753 | 1046.607399 | 1797.984473 | 1404.235879 | 0.739650685 | 0.157206805 | 1.18074812531 | 4.55013317075 | ns | YPD | KS.3 |
| FIM1_2754 | 1832.209033 | 4702.432363 | 3605.0053 | 1.245453239 | 0.227514922 | 4.35579789229 | 2.72923104712 | Km>KS.3 | YPD | KS.3 |
| FIM1_2755 | 1412.061225 | 950.8827383 | 1085.069687 | -0.551885237 | 0.118164195 | 1.96875054103 | 7.26058854088 | ns | YPD | KS.3 |
| FIM1_2756 | 1120.550089 | 734.8121962 | 909.1344341 | -0.514991461 | 0.2741131 | 0.039656332 | 0.058699867 | ns | YPD | KS.3 |
| FIM1_2757 | 4788.977329 | 3180.784229 | 5736.513145 | -0.487682782 | 0.292281116 | 0.064463798 | 0.090595309 | ns | YPD | KS.3 |
| FIM1_2758 | 387.819877 | 619.9025389 | 579.5831668 | 0.64190044 | 0.150891196 | 1.18142622072 | 3.65522104981 | ns | YPD | KS.3 |
| FIM1_2759 | 202.7472141 | 70.20159051 | 236.8115605 | -1.45111659 | 0.188242287 | 4.68897809462 | 1.10036160475 | Km<KS.3 | YPD | KS.3 |
| FIM1_2760 | 5750.669723 | 10559.85064 | 8012.268097 | 0.798901302 | 0.210995908 | 6.15912760615 | 0.000164376 | ns | YPD | KS.3 |
| FIM1_2761 | 1896.544131 | 1856.938881 | 1486.793316 | -0.029867633 | 0.061442098 | 0.62515629 | 0.680260836 | ns | YPD | KS.3 |
| FIM1_2762 | 2248.392297 | 2957.799143 | 2559.5786 | 0.361696359 | 0.195684166 | 0.052742107 | 0.07561214 | ns | YPD | KS.3 |
| FIM1_2763 | 68.66075006 | 1351.86538 | 643.8802253 | 3.519952833 | 0.65649899 | 3.40371462966 | 2.67511536849 | Km>KS.3 | YPD | KS.3 |
| FIM1_2764 | 664.2027746 | 1054.799786 | 1403.205605 | 0.623513563 | 0.174066072 | 0.000197233 | 0.000473797 | ns | YPD | KS.3 |
| FIM1_2765 | 1014.086319 | 625.4003634 | 1273.675358 | -0.673520507 | 0.124964307 | 3.75100814693 | 1.95610749800 | ns | YPD | KS.3 |
| FIM1_2766 | 2962.607886 | 3176.240986 | 3595.941157 | 0.097031946 | 0.118440096 | 0.404406071 | 0.467997256 | ns | YPD | KS.3 |
| FIM1_2767 | 1553.340514 | 2647.305843 | 2284.431879 | 0.703665078 | 0.202823266 | 0.000258014 | 0.000603206 | ns | YPD | KS.3 |
| FIM1_2768 | 1335.509445 | 1316.588219 | 1224.0717 | -0.018824556 | 0.18996915 | 0.917563543 | 0.933131362 | ns | YPD | KS.3 |
| FIM1_2769 | 2089.286519 | 1486.49305 | 2209.538832 | -0.480949548 | 0.09289359 | 1.63126811713 | 7.57755264496 | ns | YPD | KS.3 |
| FIM1_2770 | 1357.118087 | 1843.915533 | 1412.254494 | 0.415597284 | 0.162932058 | 0.008372067 | 0.014407137 | ns | YPD | KS.3 |
| FIM1_2771 | 440.4092681 | 415.7389682 | 445.9064314 | -0.075336673 | 0.200398554 | 0.693174315 | 0.74258015 | ns | YPD | KS.3 |
| FIM1_2772 | 2321.694672 | 2826.755913 | 1609.891903 | 0.278358611 | 0.09101273 | 0.001991186 | 0.00387868 | ns | YPD | KS.3 |
| FIM1_2773 | 412.7307965 | 381.4388939 | 405.4531913 | -0.106745883 | 0.164156381 | 0.501572406 | 0.563787329 | ns | YPD | KS.3 |
| FIM1_2774 | 640.5761676 | 1058.616863 | 928.7320496 | 0.631039056 | 0.2524392 | 0.006872509 | 0.012047475 | ns | YPD | KS.3 |
| FIM1_2775 | 4504.509232 | 3824.771424 | 4011.411568 | -0.231992074 | 0.081659508 | 0.004159098 | 0.007598793 | ns | YPD | KS.3 |
| FIM1_2776 | 1093.289361 | 883.3289303 | 718.1820608 | -0.296932943 | 0.119708383 | 0.011538068 | 0.01928179 | ns | YPD | KS.3 |
| FIM1_2777 | 1162.953196 | 850.9417793 | 817.4033035 | -0.42931982 | 0.14203533 | 0.001929928 | 0.003775616 | ns | YPD | KS.3 |
| FIM1_2778 | 807.0160966 | 586.94647 | 579.844977 | -0.446242187 | 0.108856805 | 3.14030771569 | 8.88682075403 | ns | YPD | KS.3 |
| FIM1_2779 | 1000.601166 | 745.7775267 | 941.1714081 | -0.404932193 | 0.139353187 | 0.00290094 | 0.005495879 | ns | YPD | KS.3 |
| FIM1_2780 | 115.8903983 | 262.5094509 | 312.6164471 | 1.041548686 | 0.259047792 | 1.18472738107 | 3.66315644551 | Km>KS.3 | YPD | KS.3 |
| FIM1_2781 | 1014.839543 | 3092.199585 | 1642.397071 | 1.430282323 | 0.283908991 | 2.04767483182 | 1.12317335042 | Km>KS.3 | YPD | KS.3 |
| FIM1_2782 | 1690.860212 | 844.1720184 | 1563.850147 | -0.95716268 | 0.150540934 | 5.52958100656 | 5.12284273886 | ns | YPD | KS.3 |
| FIM1_2783 | 5177.439367 | 3384.945068 | 6384.891685 | -0.508820629 | 0.289430482 | 0.051830054 | 0.074448425 | ns | YPD | KS.3 |
| FIM1_2784 | 1032.115295 | 960.4646065 | 907.651831 | -0.101551439 | 0.083983581 | 0.222366767 | 0.275741443 | ns | YPD | KS.3 |
| FIM1_2785 | 3204.719743 | 2032.066634 | 2810.77649 | -0.637299809 | 0.115383222 | 1.88988144905 | 1.04817839565 | ns | YPD | KS.3 |
| FIM1_2786 | 2236.473394 | 2919.847331 | 2411.717556 | 0.364383567 | 0.150388428 | 0.012685872 | 0.021009392 | ns | YPD | KS.3 |
| FIM1_2787 | 3179.390752 | 9227.872557 | 5756.818924 | 1.357047814 | 0.286163152 | 1.28517429775 | 6.05468004863 | Km>KS.3 | YPD | KS.3 |
| FIM1_2788 | 400.0304739 | 475.3884162 | 344.3151537 | 0.231196218 | 0.177482236 | 0.175825794 | 0.224751625 | ns | YPD | KS.3 |
| FIM1_2789 | 363.3393057 | 175.4929505 | 274.6503493 | -0.974054451 | 0.195970548 | 1.71971402723 | 7.94389720102 | ns | YPD | KS.3 |
| FIM1_2790 | 1448.433228 | 997.7937145 | 1088.731515 | -0.517459351 | 0.12574082 | 2.66472450759 | 7.64974288823 | ns | YPD | KS.3 |
| FIM1_2791 | 205.2971435 | 254.2014631 | 239.3295169 | 0.286489931 | 0.179199153 | 0.096343243 | 0.129753014 | ns | YPD | KS.3 |

| | | | | | | | | | | |
|---|---|---|---|---|---|---|---|---|---|---|
| FIM1_2792 | 727.6795435 | 469.287129 | 699.8149305 | -0.596531359 | 0.161215595 | 0.000130819 | 0.000329197 | ns | YPD | KS.3 |
| FIM1_2793 | 1803.364562 | 2071.872676 | 1821.019608 | 0.196632961 | 0.082950029 | 0.016765049 | 0.027106312 | ns | YPD | KS.3 |
| FIM1_2794 | 2859.747737 | 2033.902836 | 2224.607093 | -0.471912223 | 0.13170149 | 0.000248579 | 0.000582516 | ns | YPD | KS.3 |
| FIM1_2795 | 486.9285092 | 780.1097421 | 653.7490727 | 0.581328529 | 0.266810333 | 0.017509681 | 0.028227693 | ns | YPD | KS.3 |
| FIM1_2796 | 193.4637301 | 583.5456092 | 654.7569723 | 1.345324062 | 0.337641215 | 4.24887371602 | 1.45880929863 | Km>KS.3 | YPD | KS.3 |
| FIM1_2797 | 84.49919289 | 59.19037129 | 53.41088707 | -0.427462815 | 0.284823195 | 0.098082037 | 0.131951902 | ns | YPD | KS.3 |
| FIM1_2798 | 1651.64403 | 900.5567378 | 1657.686551 | -0.847494766 | 0.120531381 | 7.39759335346 | 9.91995335134 | ns | YPD | KS.3 |
| FIM1_2800 | 3787.105014 | 3162.268869 | 3329.2877 | -0.254047116 | 0.096809505 | 0.007905532 | 0.01370384 | ns | YPD | KS.3 |
| FIM1_2801 | 4900.612733 | 3497.92508 | 4352.614611 | -0.468791842 | 0.124651572 | 0.000124638 | 0.00031444 | ns | YPD | KS.3 |
| FIM1_2802 | 1046.9358 | 879.0176428 | 855.5880216 | -0.24161763 | 0.129545642 | 0.056810174 | 0.080721287 | ns | YPD | KS.3 |
| FIM1_2803 | 1752.258762 | 1545.928364 | 1290.444468 | -0.176169997 | 0.095599845 | 0.06225012 | 0.087781845 | ns | YPD | KS.3 |
| FIM1_2804 | 2339.740039 | 2192.545642 | 2299.026539 | -0.091893057 | 0.085230984 | 0.276453522 | 0.334636563 | ns | YPD | KS.3 |
| FIM1_2805 | 2015.694148 | 1227.561911 | 1868.510163 | -0.692113668 | 0.121174914 | 5.73251507698 | 3.49072980514 | ns | YPD | KS.3 |
| FIM1_2806 | 244.3164037 | 253.4416072 | 204.5735688 | 0.050850892 | 0.144918259 | 0.718785442 | 0.764345889 | ns | YPD | KS.3 |
| FIM1_2807 | 1283.867912 | 924.9949067 | 1116.620185 | -0.450647853 | 0.143195882 | 0.001238056 | 0.002521215 | ns | YPD | KS.3 |
| FIM1_2808 | 2144.413928 | 1563.486065 | 1536.198656 | -0.433727088 | 0.145312786 | 0.002173029 | 0.004204908 | ns | YPD | KS.3 |
| FIM1_2809 | 272.7778611 | 256.073446 | 172.0515214 | -0.07927909 | 0.23495868 | 0.718547544 | 0.764330561 | ns | YPD | KS.3 |
| FIM1_2810 | 340.5074745 | 67.03426035 | 237.534163 | -2.158484557 | 0.33127983 | 1.99242177887 | 2.91538187661 | Km<KS.3 | YPD | KS.3 |
| FIM1_2811 | 567.4671016 | 451.3256211 | 460.8452583 | -0.317954813 | 0.12760253 | 0.010973094 | 0.018436725 | ns | YPD | KS.3 |
| FIM1_2812 | 3048.107707 | 2407.650359 | 2959.470854 | -0.332535846 | 0.09487201 | 0.000390553 | 0.000882168 | ns | YPD | KS.3 |
| FIM1_2813 | 1029.537517 | 1095.392857 | 1060.003627 | 0.085341911 | 0.139133559 | 0.529851638 | 0.591830242 | ns | YPD | KS.3 |
| FIM1_2814 | 2249.220754 | 2303.887475 | 2193.081339 | 0.033105279 | 0.140661049 | 0.809423633 | 0.840509825 | ns | YPD | KS.3 |
| FIM1_2815 | 329.8642985 | 139.7657102 | 337.3905196 | -1.185227825 | 0.155911029 | 3.63543874988 | 7.35215763442 | Km<KS.3 | YPD | KS.3 |
| FIM1_2816 | 387.8514763 | 234.5042083 | 373.392304 | -0.690950191 | 0.151554245 | 2.63793754462 | 9.40769841182 | ns | YPD | KS.3 |
| FIM1_2817 | 484.2813851 | 324.5771366 | 534.8311899 | -0.548586051 | 0.151341055 | 0.000189722 | 0.000457522 | ns | YPD | KS.3 |
| FIM1_2818 | 643.0561794 | 526.6145987 | 428.9503715 | -0.279176024 | 0.111143602 | 0.010719186 | 0.018071145 | ns | YPD | KS.3 |
| FIM1_2819 | 2555.843706 | 1391.471205 | 2042.335681 | -0.864761478 | 0.079841669 | 8.26880089754 | 9.56681034076 | ns | YPD | KS.3 |
| FIM1_2820 | 1583.185759 | 9963.216539 | 2772.900424 | 2.54009445 | 0.270550333 | 2.85870830595 | 2.15485967001 | Km>KS.3 | YPD | KS.3 |
| FIM1_2821 | 1792.280384 | 2495.543577 | 1743.736036 | 0.460000763 | 0.125195718 | 0.000177507 | 0.000432892 | ns | YPD | KS.3 |
| FIM1_2822 | 663.4859047 | 1012.058664 | 639.5826493 | 0.580978294 | 0.143774282 | 3.32618717092 | 9.34374995786 | ns | YPD | KS.3 |
| FIM1_2823 | 3138.026173 | 2976.06935 | 3390.920902 | -0.075072726 | 0.08903579 | 0.39432187 | 0.457391304 | ns | YPD | KS.3 |
| FIM1_2824 | 1538.624927 | 1574.137944 | 1247.898865 | 0.032942708 | 0.082715398 | 0.687829102 | 0.73796631 | ns | YPD | KS.3 |
| FIM1_2825 | 1534.701478 | 1089.146091 | 1050.128606 | -0.47997184 | 0.11158397 | 1.23211879781 | 3.79788786811 | ns | YPD | KS.3 |
| FIM1_2826 | 821.1921969 | 1292.42783 | 888.972011 | 0.608979368 | 0.179169942 | 0.000400693 | 0.000902829 | ns | YPD | KS.3 |
| FIM1_2827 | 3571.847669 | 2046.329918 | 3463.262847 | -0.712831155 | 0.237451277 | 0.001282384 | 0.002598721 | ns | YPD | KS.3 |
| FIM1_2828 | 981.801201 | 982.9437768 | 768.7387174 | 0.002210793 | 0.103802016 | 0.982774537 | 0.985349319 | ns | YPD | KS.3 |
| FIM1_2829 | 2279.428711 | 2844.919644 | 2250.230989 | 0.306966747 | 0.128851853 | 0.014985454 | 0.024483623 | ns | YPD | KS.3 |
| FIM1_2830 | 1573.968022 | 2718.244342 | 2405.026122 | 0.752346525 | 0.14582933 | 1.12408242167 | 5.36690023784 | ns | YPD | KS.3 |
| FIM1_2831 | 76.28946515 | 309.6940551 | 339.8605135 | 1.633623544 | 0.441975091 | 4.02072630735 | 1.39200510640 | Km>KS.3 | YPD | KS.3 |
| FIM1_2832 | 1331.959825 | 2590.679347 | 1539.108163 | 0.933328171 | 0.11512335 | 1.48331918780 | 3.80387265943 | ns | YPD | KS.3 |
| FIM1_2833 | 1685.546013 | 1159.501855 | 1969.477907 | -0.442127483 | 0.297216572 | 0.098196223 | 0.132069806 | ns | YPD | KS.3 |
| FIM1_2834 | 726.6570754 | 4501.11582 | 1728.387067 | 2.420081587 | 0.366577426 | 1.18848558097 | 1.85934458029 | Km>KS.3 | YPD | KS.3 |
| FIM1_2835 | 493.2059422 | 1743.85764 | 1076.06063 | 1.615449017 | 0.313471237 | 4.61257854479 | 2.86844728254 | Km>KS.3 | YPD | KS.3 |
| FIM1_2836 | 722.731402 | 1185.100267 | 864.2023526 | 0.687620311 | 0.128140277 | 4.16065591457 | 2.15842160323 | ns | YPD | KS.3 |
| FIM1_2837 | 262.4459199 | 1107.688338 | 858.4962492 | 1.90745439 | 0.298342926 | 7.59717729150 | 1.01602035014 | Km>KS.3 | YPD | KS.3 |

| | | | | | | | | | | |
|---|---|---|---|---|---|---|---|---|---|---|
| FIM1_2838 | 2149.547872 | 3021.007675 | 2749.047553 | 0.47507795 | 0.116987182 | 3.57047136535 | 9.97366369603 | ns | YPD | KS.3 |
| FIM1_2839 | 1120.305091 | 940.7665874 | 734.8702793 | -0.245802818 | 0.093172679 | 0.007632606 | 0.01326772 | ns | YPD | KS.3 |
| FIM1_2840 | 143.7218292 | 124.818969 | 112.8839267 | -0.188523921 | 0.165883361 | 0.239826999 | 0.294796345 | ns | YPD | KS.3 |
| FIM1_2841 | 969.8798987 | 995.3422026 | 1116.009165 | 0.035833355 | 0.120287045 | 0.761635179 | 0.8009163 | ns | YPD | KS.3 |
| FIM1_2842 | 2934.408144 | 2131.422428 | 3122.825629 | -0.448908084 | 0.106677072 | 1.94577954494 | 5.78962514121 | ns | YPD | KS.3 |
| FIM1_2843 | 331.5995601 | 211.8583278 | 223.4923183 | -0.606187527 | 0.167160942 | 0.00017155 | 0.00042084 | ns | YPD | KS.3 |
| FIM1_2844 | 582.9977534 | 407.0823526 | 525.3170241 | -0.482078883 | 0.176794093 | 0.004579235 | 0.008293299 | ns | YPD | KS.3 |
| FIM1_2845 | 21687.91255 | 37649.92367 | 43296.34962 | 0.769373489 | 0.123593342 | 2.09748457637 | 1.72479103594 | ns | YPD | KS.3 |
| FIM1_2846 | 655.3763027 | 945.0436896 | 1216.290288 | 0.44422309 | 0.276179117 | 0.077900623 | 0.107326392 | ns | YPD | KS.3 |
| FIM1_2847 | 2780.437349 | 6113.877448 | 3194.810321 | 0.850985206 | 0.386474299 | 0.008516626 | 0.0146306 | ns | YPD | KS.3 |
| FIM1_2848 | 702.3866197 | 560.5771776 | 706.3170098 | -0.309479916 | 0.143182256 | 0.026586717 | 0.041039069 | ns | YPD | KS.3 |
| FIM1_2849 | 9221.604798 | 9846.525125 | 14759.95724 | 0.091150621 | 0.121469449 | 0.444646849 | 0.508065704 | ns | YPD | KS.3 |
| FIM1_2850 | 2210.641547 | 1398.207471 | 2126.038905 | -0.626879761 | 0.15336645 | 2.51940373073 | 7.29570529708 | ns | YPD | KS.3 |
| FIM1_2851 | 2433.353016 | 3544.770823 | 2482.05434 | 0.50437419 | 0.178711923 | 0.003312089 | 0.006196932 | ns | YPD | KS.3 |
| FIM1_2852 | 951.8570048 | 484.9322069 | 869.8070042 | -0.928593634 | 0.15015039 | 1.83051194547 | 1.53831029201 | ns | YPD | KS.3 |
| FIM1_2853 | 3976.812272 | 2108.373626 | 4109.292691 | -0.774855798 | 0.284879522 | 0.002658762 | 0.005069725 | ns | YPD | KS.3 |
| FIM1_2854 | 1896.05972 | 1464.440504 | 1743.33323 | -0.351200683 | 0.157742819 | 0.021652549 | 0.034002977 | ns | YPD | KS.3 |
| FIM1_2855 | 118.9646794 | 99.8084287 | 121.4704813 | -0.227295798 | 0.209434952 | 0.252505934 | 0.308956473 | ns | YPD | KS.3 |
| FIM1_2856 | 857.045581 | 877.328255 | 634.690912 | 0.031517093 | 0.175968528 | 0.852475091 | 0.876977581 | ns | YPD | KS.3 |
| FIM1_2857 | 705.8872707 | 478.8064052 | 751.6658429 | -0.526398292 | 0.164453155 | 0.0009261 | 0.001933422 | ns | YPD | KS.3 |
| FIM1_2858 | 825.0318921 | 1273.548286 | 1021.325247 | 0.604000756 | 0.124617235 | 7.55601496580 | 3.03978494375 | ns | YPD | KS.3 |
| FIM1_2859 | 4779.87096 | 2890.920803 | 3942.214549 | -0.682375091 | 0.166609169 | 2.19232818498 | 6.41578395311 | ns | YPD | KS.3 |
| FIM1_2860 | 7540.194675 | 7631.486614 | 11384.25993 | 0.014321113 | 0.284362003 | 0.955793946 | 0.965693518 | ns | YPD | KS.3 |
| FIM1_2862 | 250.162032 | 126.0408024 | 207.0002085 | -0.915616568 | 0.197695777 | 1.10051909049 | 4.29418233351 | ns | YPD | KS.3 |
| FIM1_2863 | 669.5511958 | 1667.075174 | 1263.715346 | 1.207189256 | 0.2226107 | 6.78231399596 | 4.07020652954 | Km>KS.3 | YPD | KS.3 |
| FIM1_2864 | 2530.647023 | 1775.647712 | 3071.00938 | -0.48881817 | 0.138445436 | 0.000296546 | 0.000687152 | ns | YPD | KS.3 |
| FIM1_2865 | 1313.112869 | 595.336561 | 1040.611865 | -1.078436931 | 0.172209582 | 7.03789624721 | 6.37769286519 | Km<KS.3 | YPD | KS.3 |
| FIM1_2866 | 876.3502024 | 771.7482988 | 682.8797352 | -0.173551926 | 0.145830918 | 0.221791979 | 0.275165859 | ns | YPD | KS.3 |
| FIM1_2867 | 1945.62016 | 1250.659792 | 1333.668172 | -0.608729616 | 0.1425586 | 1.16653213720 | 3.61588621969 | ns | YPD | KS.3 |
| FIM1_2868 | 50.45405136 | 73.60332361 | 53.04285322 | 0.471589136 | 0.254502058 | 0.045053496 | 0.065653527 | ns | YPD | KS.3 |
| FIM1_2869 | 209.4455219 | 257.3540757 | 263.2725796 | 0.275719155 | 0.182029302 | 0.114727058 | 0.152042385 | ns | YPD | KS.3 |
| FIM1_2870 | 5556.596617 | 4417.880089 | 4619.041052 | -0.31061627 | 0.162500562 | 0.048277202 | 0.069906158 | ns | YPD | KS.3 |
| FIM1_2871 | 1605.291437 | 1040.295939 | 1028.851863 | -0.608128213 | 0.11054636 | 2.25639739736 | 1.22818129670 | ns | YPD | KS.3 |
| FIM1_2872 | 1940.488485 | 5464.443596 | 3341.988258 | 1.442422523 | 0.148544557 | 9.89252472326 | 8.20255174970 | Km>KS.3 | YPD | KS.3 |
| FIM1_2873 | 151.2485131 | 565.9690608 | 379.4106986 | 1.82395013 | 0.194666066 | 3.50759792569 | 2.56622054122 | Km>KS.3 | YPD | KS.3 |
| FIM1_2874 | 2994.043107 | 5931.068415 | 5304.524092 | 0.932130136 | 0.166343555 | 6.10947479686 | 3.70666306273 | ns | YPD | KS.3 |
| FIM1_2875 | 1820.726337 | 2106.791339 | 2510.602835 | 0.206148742 | 0.089837593 | 0.020408873 | 0.032304849 | ns | YPD | KS.3 |
| FIM1_2876 | 2811.26757 | 2151.569375 | 2282.743622 | -0.375076117 | 0.108126069 | 0.000428904 | 0.000961982 | ns | YPD | KS.3 |
| FIM1_2877 | 2998.370506 | 1456.079363 | 3588.738224 | -0.896745268 | 0.276497878 | 0.000361214 | 0.00082094 | ns | YPD | KS.3 |
| FIM1_2878 | 2159.408328 | 1235.202201 | 1245.67938 | -0.77700879 | 0.127984599 | 5.44409814595 | 4.07897413796 | ns | YPD | KS.3 |
| FIM1_2879 | 1555.983539 | 2039.895092 | 1558.259213 | 0.368297778 | 0.158040301 | 0.016214209 | 0.026326923 | ns | YPD | KS.3 |
| FIM1_2880 | 1301.880422 | 1188.510293 | 1078.416442 | -0.124777625 | 0.144306676 | 0.374984868 | 0.438231083 | ns | YPD | KS.3 |
| FIM1_2881 | 1535.900045 | 768.0033755 | 1233.208042 | -0.933525438 | 0.184871966 | 1.28163976198 | 6.04375148425 | ns | YPD | KS.3 |
| FIM1_2882 | 2016.825918 | 1882.202807 | 1469.445907 | -0.095178962 | 0.132365752 | 0.462418746 | 0.525956393 | ns | YPD | KS.3 |
| FIM1_2883 | 3340.992187 | 2628.472657 | 2329.138013 | -0.338437275 | 0.093192274 | 0.00023975 | 0.000564751 | ns | YPD | KS.3 |

| | | | | | | | | | | |
|---|---|---|---|---|---|---|---|---|---|---|
| FIM1_2884 | 864.8045385 | 716.2517752 | 690.2018904 | -0.263977989 | 0.108014776 | 0.013116956 | 0.021658432 | ns | YPD | KS.3 |
| FIM1_2885 | 1331.714398 | 618.5816322 | 1656.05392 | -0.89711377 | 0.329560862 | 0.001890291 | 0.003705357 | ns | YPD | KS.3 |
| FIM1_2886 | 997.4096849 | 269.2787534 | 931.8734118 | -1.59732088 | 0.376928882 | 4.6691563465 | 1.9553074767 | Km<KS.3 | YPD | KS.3 |
| FIM1_2887 | 3940.92142 | 2006.114292 | 4640.995791 | -0.835124464 | 0.276467912 | 0.000899384 | 0.001882387 | ns | YPD | KS.3 |
| FIM1_2888 | 623.8484175 | 344.5212327 | 389.7551072 | -0.806029598 | 0.168141476 | 6.5646482964 | 2.6704108973 | ns | YPD | KS.3 |
| FIM1_2889 | 2540.470199 | 2049.133447 | 2031.263372 | -0.302910216 | 0.096813816 | 0.001540286 | 0.003074969 | ns | YPD | KS.3 |
| FIM1_2890 | 4442.170139 | 3503.954083 | 4347.024313 | -0.332186709 | 0.109952808 | 0.002152107 | 0.004169289 | ns | YPD | KS.3 |
| FIM1_2891 | 4181.475559 | 4081.946305 | 4319.216258 | -0.034012872 | 0.08849642 | 0.697847949 | 0.746782866 | ns | YPD | KS.3 |
| FIM1_2892 | 272.4993684 | 393.2054909 | 304.0822981 | 0.50347087 | 0.144966539 | 0.000360713 | 0.000820177 | ns | YPD | KS.3 |
| FIM1_2893 | 394.7129482 | 665.9637145 | 642.7355356 | 0.719202563 | 0.146055566 | 4.1137338610 | 1.7417724220 | ns | YPD | KS.3 |
| FIM1_2894 | 930.7088741 | 1431.360031 | 1040.38808 | 0.597389016 | 0.129805859 | 2.5461567302 | 9.1327539531 | ns | YPD | KS.3 |
| FIM1_2895 | 3957.309783 | 2578.047771 | 3125.276002 | -0.600448041 | 0.111525784 | 4.4146992484 | 2.2736158137 | ns | YPD | KS.3 |
| FIM1_2896 | 1465.286246 | 1637.468953 | 1441.378662 | 0.15763475 | 0.089528936 | 0.075240609 | 0.10420992 | ns | YPD | KS.3 |
| FIM1_2897 | 4075.97221 | 7787.172924 | 5776.34956 | 0.869382541 | 0.186414053 | 1.0648016908 | 4.1678901748 | ns | YPD | KS.3 |
| FIM1_2898 | 10818.5933 | 11210.70432 | 16191.39412 | 0.049569968 | 0.118497666 | 0.670299707 | 0.721634432 | ns | YPD | KS.3 |
| FIM1_2899 | 6132.983452 | 6868.957979 | 9340.909001 | 0.147818556 | 0.205137804 | 0.44842712 | 0.512032343 | ns | YPD | KS.3 |
| FIM1_2900 | 4504.670776 | 6225.260431 | 6646.927741 | 0.440050297 | 0.158336613 | 0.004130491 | 0.007552074 | ns | YPD | KS.3 |
| FIM1_2901 | 1550.447006 | 1424.006224 | 1677.89953 | -0.118714003 | 0.113585834 | 0.288117773 | 0.347740399 | ns | YPD | KS.3 |
| FIM1_2903 | 2009.960053 | 3439.615461 | 2443.122967 | 0.745375071 | 0.133005807 | 9.6243243284 | 5.6596942711 | ns | YPD | KS.3 |
| FIM1_2904 | 997.4334852 | 2220.616775 | 1819.270302 | 1.078070488 | 0.190265679 | 2.7152782177 | 1.7727702274 | Km>KS.3 | YPD | KS.3 |
| FIM1_2905 | 447.4296562 | 1481.884257 | 846.2092377 | 1.481242463 | 0.339183539 | 4.3505256321 | 1.8295743888 | Km>KS.3 | YPD | KS.3 |
| FIM1_2906 | 261.2889714 | 168.5893108 | 192.2999731 | -0.573271916 | 0.209758803 | 0.003898178 | 0.007164181 | ns | YPD | KS.3 |
| FIM1_2907 | 854.0850815 | 709.7667758 | 990.5456322 | -0.24375902 | 0.19689987 | 0.194428482 | 0.245378412 | ns | YPD | KS.3 |
| FIM1_2908 | 2872.642027 | 2632.52336 | 2481.130868 | -0.122283772 | 0.109609923 | 0.257347256 | 0.314030562 | ns | YPD | KS.3 |
| FIM1_2909 | 1392.001456 | 1196.690777 | 1242.605634 | -0.211428787 | 0.107134735 | 0.045229729 | 0.065871751 | ns | YPD | KS.3 |
| FIM1_2911 | 722.7084861 | 382.9387476 | 424.6130829 | -0.791752711 | 0.266406715 | 0.001175633 | 0.002404924 | ns | YPD | KS.3 |
| FIM1_2912 | 1651.341237 | 4348.36349 | 2982.181995 | 1.223240168 | 0.282420425 | 1.5755651842 | 5.9562589600 | Km>KS.3 | YPD | KS.3 |
| FIM1_2913 | 9.78854204 | 22.65472999 | 14.35777334 | 0.939840315 | 0.370429115 | 0.002774854 | 0.005269045 | ns | YPD | KS.3 |
| FIM1_2914 | 610.8751807 | 1907.572796 | 1880.422927 | 1.387733958 | 0.343333052 | 2.7736611820 | 9.8634484492 | Km>KS.3 | YPD | KS.3 |
| FIM1_2915 | 829.2476852 | 683.6704919 | 1489.707832 | -0.260381174 | 0.167791627 | 0.108180178 | 0.144365983 | ns | YPD | KS.3 |
| FIM1_2916 | 3197.346561 | 4144.333202 | 5132.835234 | 0.353061957 | 0.156173433 | 0.01978386 | 0.031445592 | ns | YPD | KS.3 |
| FIM1_2917 | 1680.748424 | 1705.205369 | 1446.996132 | 0.020765283 | 0.099027781 | 0.831856166 | 0.859498323 | ns | YPD | KS.3 |
| FIM1_2918 | 915.3526529 | 789.9485408 | 985.6773065 | -0.204835406 | 0.118834686 | 0.079321073 | 0.109102112 | ns | YPD | KS.3 |
| FIM1_2919 | 3605.762103 | 3619.205673 | 3139.017352 | 0.005252896 | 0.109622726 | 0.961199226 | 0.970169639 | ns | YPD | KS.3 |
| FIM1_2920 | 384.8103504 | 219.9556209 | 333.924461 | -0.71439889 | 0.24031322 | 0.001404496 | 0.00283004 | ns | YPD | KS.3 |
| FIM1_2921 | 2759.599468 | 1859.642615 | 2861.91268 | -0.480402101 | 0.275268423 | 0.055859529 | 0.079513922 | ns | YPD | KS.3 |
| FIM1_2922 | 1730.344191 | 1303.283675 | 940.8518398 | -0.382582786 | 0.167463942 | 0.01800583 | 0.028915108 | ns | YPD | KS.3 |
| FIM1_2923 | 4149.336625 | 3104.752331 | 3964.760454 | -0.403212955 | 0.123675749 | 0.000885355 | 0.001856149 | ns | YPD | KS.3 |
| FIM1_2924 | 6159.080902 | 4496.777926 | 6416.49765 | -0.436095304 | 0.129441866 | 0.000577517 | 0.001257946 | ns | YPD | KS.3 |
| FIM1_2925 | 53.38884484 | 65.6693922 | 55.21287527 | 0.23285162 | 0.328011831 | 0.419815593 | 0.48391626 | ns | YPD | KS.3 |
| FIM1_2926 | 2370.19039 | 2290.052219 | 2247.740019 | -0.045720633 | 0.184552834 | 0.796288215 | 0.829937816 | ns | YPD | KS.3 |
| FIM1_2927 | 4398.008709 | 4648.931658 | 5522.650867 | 0.078971703 | 0.069431189 | 0.252356649 | 0.308849774 | ns | YPD | KS.3 |
| FIM1_2928 | 414.1225506 | 2522.932654 | 1364.35551 | 2.458415056 | 0.307656483 | 4.4948409046 | 1.5529051041 | Km>KS.3 | YPD | KS.3 |
| FIM1_2929 | 1658.475204 | 3068.433559 | 1704.738101 | 0.803163463 | 0.219275263 | 9.9177778436 | 0.000256583 | ns | YPD | KS.3 |
| FIM1_2930 | 274.4034472 | 141.4817839 | 314.4429203 | -0.772310326 | 0.323422147 | 0.00672018 | 0.01179707 | ns | YPD | KS.3 |

| | | | | | | | | | |
|---|---|---|---|---|---|---|---|---|---|
| FIM1_2931 | 2994.8599 | 1984.48821 | 3084.728023 | -0.56430756 | 0.148921504 | 9.6815657779 | 0.000250863 | ns | YPD | KS.3 |
| FIM1_2932 | 1113.962992 | 811.4412432 | 921.3139034 | -0.436450201 | 0.139113889 | 0.001302718 | 0.002636705 | ns | YPD | KS.3 |
| FIM1_2933 | 2858.865552 | 3648.486265 | 2741.375958 | 0.344326605 | 0.092347632 | 0.000163015 | 0.000401485 | ns | YPD | KS.3 |
| FIM1_2934 | 2847.578215 | 2982.862846 | 3436.724809 | 0.0619896 | 0.178636474 | 0.718322596 | 0.764254687 | ns | YPD | KS.3 |
| FIM1_2935 | 11689.44432 | 12593.52936 | 14252.74012 | 0.100894432 | 0.160875521 | 0.517426437 | 0.579643442 | ns | YPD | KS.3 |
| FIM1_2936 | 4002.420863 | 5310.220742 | 4841.660457 | 0.37750778 | 0.181670804 | 0.030446224 | 0.046222143 | ns | YPD | KS.3 |
| FIM1_2937 | 598.8215361 | 898.1439469 | 929.7319652 | 0.537505361 | 0.192119638 | 0.003396178 | 0.00633758 | ns | YPD | KS.3 |
| FIM1_2939 | 740.0017944 | 739.5035953 | 579.7654092 | -0.000648477 | 0.168540672 | 0.996822134 | 0.997022541 | ns | YPD | KS.3 |
| FIM1_2941 | 1743.482419 | 1123.989429 | 1537.092147 | -0.608216415 | 0.133766282 | 3.2554622322 | 1.1421667563 | ns | YPD | KS.3 |
| FIM1_2942 | 5228.244636 | 5541.00267 | 7572.433439 | 0.080412192 | 0.129413936 | 0.525801418 | 0.588186401 | ns | YPD | KS.3 |
| FIM1_2943 | 1637.965555 | 1266.692486 | 1210.279347 | -0.361008951 | 0.104191656 | 0.000441662 | 0.000986206 | ns | YPD | KS.3 |
| FIM1_2944 | 1273.34299 | 691.1271863 | 950.1685742 | -0.842614351 | 0.145431774 | 2.5296682037 | 1.6691113148 | ns | YPD | KS.3 |
| FIM1_2945 | 929.8111391 | 489.5896888 | 787.5940644 | -0.85889651 | 0.190323603 | 2.2497383896 | 8.1577612888 | ns | YPD | KS.3 |
| FIM1_2946 | 2280.441349 | 2513.761094 | 1978.107633 | 0.137319325 | 0.096833351 | 0.151231371 | 0.195523927 | ns | YPD | KS.3 |
| FIM1_2947 | 4361.60316 | 2780.874493 | 3679.15989 | -0.626934328 | 0.123438963 | 2.1979492538 | 9.8957443783 | ns | YPD | KS.3 |
| FIM1_2948 | 1701.706701 | 1272.288508 | 1174.689626 | -0.40478606 | 0.121213557 | 0.000666728 | 0.001432818 | ns | YPD | KS.3 |
| FIM1_2949 | 3068.609273 | 3341.686985 | 3327.628688 | 0.117194017 | 0.140198839 | 0.391734582 | 0.454920528 | ns | YPD | KS.3 |
| FIM1_2950 | 65.7415653 | 168.5598633 | 91.11159101 | 1.261701431 | 0.207676169 | 1.1452649159 | 9.8918280500 | Km>KS.3 | YPD | KS.3 |
| FIM1_2951 | 3379.130573 | 2739.178796 | 2696.25655 | -0.290620967 | 0.130832927 | 0.023229001 | 0.036227046 | ns | YPD | KS.3 |
| FIM1_2952 | 1306.95453 | 953.8452654 | 991.1695776 | -0.438473727 | 0.120029475 | 0.000198083 | 0.000475152 | ns | YPD | KS.3 |
| FIM1_2953 | 1664.330284 | 2066.658268 | 1696.349832 | 0.289050422 | 0.18064118 | 0.095844024 | 0.129150601 | ns | YPD | KS.3 |
| FIM1_2954 | 840.1893484 | 713.5334915 | 669.8722657 | -0.231561176 | 0.077847152 | 0.002714696 | 0.005162696 | ns | YPD | KS.3 |
| FIM1_2955 | 1476.924513 | 687.5371798 | 1135.911101 | -1.048994604 | 0.161175837 | 1.5526464361 | 1.5992579750 | Km<KS.3 | YPD | KS.3 |
| FIM1_2956 | 495.6755998 | 301.3644956 | 356.0336572 | -0.67087917 | 0.175889359 | 7.2559104217 | 0.000191097 | ns | YPD | KS.3 |
| FIM1_2957 | 3645.092812 | 2391.523773 | 3906.709487 | -0.525156539 | 0.256384352 | 0.026515375 | 0.040941648 | ns | YPD | KS.3 |
| FIM1_2958 | 3125.006459 | 2267.640498 | 2248.188588 | -0.445770632 | 0.12371884 | 0.000238165 | 0.000561285 | ns | YPD | KS.3 |
| FIM1_2959 | 4029.997093 | 2917.55089 | 4410.205535 | -0.418135697 | 0.217113875 | 0.041366362 | 0.060882973 | ns | YPD | KS.3 |
| FIM1_2960 | 3136.225073 | 2107.977079 | 3342.8738 | -0.504204155 | 0.238812226 | 0.023615235 | 0.036760262 | ns | YPD | KS.3 |
| FIM1_2961 | 4381.433497 | 2814.081689 | 4279.333789 | -0.586549171 | 0.195318977 | 0.001634217 | 0.003243011 | ns | YPD | KS.3 |
| FIM1_2962 | 320.3916057 | 491.3121436 | 401.0509813 | 0.597226145 | 0.117981312 | 2.5271995937 | 1.1205720123 | ns | YPD | KS.3 |
| FIM1_2963 | 412.9317701 | 371.0533819 | 393.8680425 | -0.151374077 | 0.100405948 | 0.126731123 | 0.166759047 | ns | YPD | KS.3 |
| FIM1_2964 | 5770.080983 | 4957.952302 | 7234.558437 | -0.209477747 | 0.134034452 | 0.109982286 | 0.146417413 | ns | YPD | KS.3 |
| FIM1_2965 | 1581.7755 | 1152.543046 | 1402.398664 | -0.441221728 | 0.119002299 | 0.000159379 | 0.000393769 | ns | YPD | KS.3 |
| FIM1_2966 | 553.8237498 | 354.3630468 | 368.3976288 | -0.605853297 | 0.166039056 | 0.000157339 | 0.00039021 | ns | YPD | KS.3 |
| FIM1_2967 | 1519.136964 | 1617.004996 | 1546.006294 | 0.088306396 | 0.082881947 | 0.282392813 | 0.341244655 | ns | YPD | KS.3 |
| FIM1_2968 | 1908.600432 | 2170.180913 | 1890.686952 | 0.180899812 | 0.09544013 | 0.055148095 | 0.07863622 | ns | YPD | KS.3 |
| FIM1_2969 | 5377.490796 | 3806.138917 | 5685.816099 | -0.474141316 | 0.146622993 | 0.00089009 | 0.001864504 | ns | YPD | KS.3 |
| FIM1_2970 | 731.5040276 | 1025.689751 | 760.6778271 | 0.475182185 | 0.103611578 | 3.2991209048 | 1.1566685342 | ns | YPD | KS.3 |
| FIM1_2971 | 607.2105756 | 267.9397196 | 637.3109641 | -1.059005362 | 0.242387176 | 2.4322632429 | 8.7621358680 | Km<KS.3 | YPD | KS.3 |
| FIM1_2972 | 800.1988735 | 1235.24484 | 731.7172261 | 0.596993295 | 0.144359145 | 2.1556509878 | 6.3382763976 | ns | YPD | KS.3 |
| FIM1_2973 | 833.2938148 | 1363.824684 | 901.5278038 | 0.655218553 | 0.192309005 | 0.000357188 | 0.000812904 | ns | YPD | KS.3 |
| FIM1_2974 | 1844.652164 | 2679.604191 | 1991.706768 | 0.494402197 | 0.193544748 | 0.00745396 | 0.01298895 | ns | YPD | KS.3 |
| FIM1_2975 | 2490.364073 | 1834.594747 | 2788.527927 | -0.410894103 | 0.173707225 | 0.014061768 | 0.023088216 | ns | YPD | KS.3 |
| FIM1_2976 | 656.7977933 | 571.2737722 | 629.7327363 | -0.193836331 | 0.121843744 | 0.105059803 | 0.140692469 | ns | YPD | KS.3 |
| FIM1_2977 | 903.6630911 | 708.9235851 | 839.6745868 | -0.331442139 | 0.149390584 | 0.022532528 | 0.035284648 | ns | YPD | KS.3 |

| | | | | | | | | | | |
|---|---|---|---|---|---|---|---|---|---|---|
| FIM1_2978 | 2649.202323 | 2141.487127 | 2827.370299 | -0.291576764 | 0.14609682 | 0.040408387 | 0.05972422 | ns | YPD | KS.3 |
| FIM1_2979 | 375.5631018 | 239.4857312 | 219.4068092 | -0.610881012 | 0.163443714 | 0.00011012 | 0.000282105 | ns | YPD | KS.3 |
| FIM1_2980 | 1791.443437 | 1076.905858 | 926.1346837 | -0.699149863 | 0.148232892 | 1.2123808478 | 4.6684169644 | ns | YPD | KS.3 |
| FIM1_2981 | 4661.15717 | 2981.985018 | 5131.577162 | -0.566816369 | 0.240745729 | 0.011484016 | 0.01919791 | ns | YPD | KS.3 |
| FIM1_2982 | 2278.781493 | 4102.152072 | 3129.856232 | 0.825935881 | 0.109881386 | 2.1360479478 | 3.8510341968 | ns | YPD | KS.3 |
| FIM1_2983 | 5461.632206 | 3442.764285 | 6118.843466 | -0.620345778 | 0.17801046 | 0.000286106 | 0.000666063 | ns | YPD | KS.3 |
| FIM1_2984 | 546.0037609 | 171.0669161 | 607.7647513 | -1.484420245 | 0.296030858 | 1.7928676954 | 1.0014932716 | Km<KS.3 | YPD | KS.3 |
| FIM1_2985 | 12539.1614 | 13404.06524 | 15548.43375 | 0.077874734 | 0.298908518 | 0.772078735 | 0.808480679 | ns | YPD | KS.3 |
| FIM1_2986 | 1020.846799 | 1053.265752 | 930.6546089 | 0.044254946 | 0.088389954 | 0.613046912 | 0.669279874 | ns | YPD | KS.3 |
| FIM1_2987 | 1845.786236 | 2105.487875 | 1963.239033 | 0.186057321 | 0.087181802 | 0.031143981 | 0.047137604 | ns | YPD | KS.3 |
| FIM1_2988 | 832.8354677 | 1460.075028 | 931.4687696 | 0.77822929 | 0.135266484 | 3.7435692585 | 2.3785768916 | ns | YPD | KS.3 |
| FIM1_2989 | 873.7120105 | 545.6537928 | 992.0718388 | -0.630672301 | 0.183456768 | 0.000334348 | 0.000765476 | ns | YPD | KS.3 |
| FIM1_2990 | 6864.138581 | 12619.61083 | 12583.20332 | 0.866178864 | 0.079745736 | 5.8807288206 | 6.9658633053 | ns | YPD | KS.3 |
| FIM1_2991 | 231.1836693 | 150.9930551 | 416.6918034 | -0.556439549 | 0.213087287 | 0.005742719 | 0.01019266 | ns | YPD | KS.3 |
| FIM1_2992 | 73.75787445 | 66.09077297 | 85.80505748 | -0.140563919 | 0.231991771 | 0.518160008 | 0.580203925 | ns | YPD | KS.3 |
| FIM1_2993 | 1335.712264 | 1062.438674 | 1039.204649 | -0.32456052 | 0.079447106 | 3.7972289768 | 0.000105302 | ns | YPD | KS.3 |
| FIM1_2994 | 1163.721193 | 1021.38283 | 988.7851955 | -0.181156081 | 0.123457516 | 0.134741622 | 0.176080791 | ns | YPD | KS.3 |
| FIM1_2995 | 175.9743265 | 401.149368 | 250.7650778 | 1.052118815 | 0.256564599 | 8.1411936300 | 2.6096931900 | Km>KS.3 | YPD | KS.3 |
| FIM1_2996 | 358.9214281 | 890.8464776 | 702.7419252 | 1.063569189 | 0.344746537 | 0.000360112 | 0.000819184 | Km>KS.3 | YPD | KS.3 |
| FIM1_2997 | 1531.172965 | 2255.21245 | 1624.884602 | 0.542870868 | 0.110384894 | 5.8086368341 | 2.3784336008 | ns | YPD | KS.3 |
| FIM1_2998 | 2713.320245 | 3054.538093 | 3567.38224 | 0.16485214 | 0.12067857 | 0.164153195 | 0.211187522 | ns | YPD | KS.3 |
| FIM1_2999 | 647.2954179 | 981.6495522 | 807.5264055 | 0.570946695 | 0.148866517 | 7.9542949896 | 0.000207949 | ns | YPD | KS.3 |
| FIM1_3000 | 1313.771036 | 1561.155787 | 1759.153242 | 0.235978012 | 0.148713103 | 0.102919684 | 0.138012245 | ns | YPD | KS.3 |
| FIM1_3001 | 1284.951326 | 1018.636818 | 1124.216239 | -0.308295779 | 0.187791346 | 0.086528308 | 0.117777929 | ns | YPD | KS.3 |
| FIM1_3002 | 452.1683945 | 346.2079727 | 406.515444 | -0.36784726 | 0.131936958 | 0.00436947 | 0.007951029 | ns | YPD | KS.3 |
| FIM1_3003 | 1232.470032 | 602.7205202 | 974.3824504 | -0.906783567 | 0.256182165 | 0.000120988 | 0.00030663 | ns | YPD | KS.3 |
| FIM1_3004 | 7109.76622 | 5671.55839 | 5703.476767 | -0.319249347 | 0.091427832 | 0.000415431 | 0.000934344 | ns | YPD | KS.3 |
| FIM1_3005 | 2004.303074 | 695.1292618 | 2134.188148 | -1.316836636 | 0.31087349 | 1.6323848066 | 6.1337722154 | Km<KS.3 | YPD | KS.3 |
| FIM1_3006 | 4891.482946 | 4597.330498 | 4840.496114 | -0.085075982 | 0.144217006 | 0.545025445 | 0.606871439 | ns | YPD | KS.3 |
| FIM1_3007 | 4727.221909 | 2907.60024 | 3697.877283 | -0.678964193 | 0.118867868 | 5.8725743152 | 3.5716451367 | ns | YPD | KS.3 |
| FIM1_3008 | 3046.795535 | 2075.055468 | 2183.93518 | -0.544853272 | 0.082479535 | 2.6081668445 | 2.5694316933 | ns | YPD | KS.3 |
| FIM1_3009 | 1941.191872 | 1724.492229 | 1567.872644 | -0.167386607 | 0.088635453 | 0.056426067 | 0.080251481 | ns | YPD | KS.3 |
| FIM1_3010 | 927.5562163 | 504.6271864 | 653.7230729 | -0.853409788 | 0.115058904 | 4.2537532787 | 7.1981029121 | ns | YPD | KS.3 |
| FIM1_3011 | 422.6083121 | 777.6857583 | 622.8877379 | 0.832356705 | 0.162052239 | 1.0563498322 | 5.0797318475 | ns | YPD | KS.3 |
| FIM1_3012 | 2905.066702 | 1857.351866 | 2505.524487 | -0.630495159 | 0.099620651 | 1.4165440212 | 1.2129615328 | ns | YPD | KS.3 |
| FIM1_3013 | 2365.810201 | 2656.71592 | 2299.118373 | 0.163434775 | 0.094618147 | 0.080592759 | 0.11057611 | ns | YPD | KS.3 |
| FIM1_3014 | 4597.796735 | 3500.558737 | 6063.64154 | -0.330285417 | 0.275017714 | 0.188618764 | 0.238894692 | ns | YPD | KS.3 |
| FIM1_3015 | 4482.49692 | 2672.6441 | 4634.54022 | -0.722400303 | 0.119844113 | 8.0070689431 | 5.8238549695 | ns | YPD | KS.3 |
| FIM1_3016 | 1023.569693 | 1096.868545 | 759.6981876 | 0.094878355 | 0.144691988 | 0.501034035 | 0.563309452 | ns | YPD | KS.3 |
| FIM1_3017 | 12566.58778 | 16773.70496 | 21330.23418 | 0.402774434 | 0.118090097 | 0.000515888 | 0.001132131 | ns | YPD | KS.3 |
| FIM1_3018 | 654.8539247 | 272.7704215 | 535.4983651 | -1.193325425 | 0.178575102 | 2.8624931654 | 3.4315430116 | Km<KS.3 | YPD | KS.3 |
| FIM1_3019 | 3369.704616 | 4422.759083 | 4633.715476 | 0.379333647 | 0.11721677 | 0.000988903 | 0.002049914 | ns | YPD | KS.3 |
| FIM1_3020 | 90.61177386 | 163.1582375 | 96.90294568 | 0.779921374 | 0.200196573 | 4.1316421351 | 0.0001135 | ns | YPD | KS.3 |
| FIM1_3021 | 3021.129424 | 2585.566906 | 3834.209133 | -0.218149823 | 0.107746315 | 0.039899887 | 0.059007711 | ns | YPD | KS.3 |
| FIM1_3022 | 289.5466744 | 168.6261715 | 240.3611277 | -0.632144367 | 0.313084408 | 0.023065953 | 0.036040552 | ns | YPD | KS.3 |

| | | | | | | | | | | |
|---|---|---|---|---|---|---|---|---|---|---|
| FIM1_3023 | 632.4193669 | 1001.67759 | 752.7178282 | 0.589971232 | 0.230574128 | 0.00629764 | 0.011102324 | ns | YPD | KS.3 |
| FIM1_3024 | 135.8997976 | 49.61633659 | 74.4569741 | -1.044538678 | 0.440111066 | 0.002937766 | 0.005561194 | Km<KS.3 | YPD | KS.3 |
| FIM1_3025 | 1172.720015 | 1357.103319 | 931.7120277 | 0.189322473 | 0.21218913 | 0.346228991 | 0.40826955 | ns | YPD | KS.3 |
| FIM1_3026 | 3161.016134 | 2782.569034 | 2497.866158 | -0.170243604 | 0.179538947 | 0.324069098 | 0.385611998 | ns | YPD | KS.3 |
| FIM1_3028 | 3932.595324 | 5889.303416 | 4185.317805 | 0.560481031 | 0.128822044 | 8.765140150 | 2.788144005 | ns | YPD | KS.3 |
| FIM1_3029 | 1399.770835 | 1674.891364 | 1373.966389 | 0.24978345 | 0.121421831 | 0.036102206 | 0.053897676 | ns | YPD | KS.3 |
| FIM1_3030 | 3108.058661 | 2679.745078 | 3644.827954 | -0.210778781 | 0.072792164 | 0.003546244 | 0.006587963 | ns | YPD | KS.3 |
| FIM1_3031 | 1506.109913 | 1354.26194 | 1581.656361 | -0.147233104 | 0.126073632 | 0.23355521 | 0.287893253 | ns | YPD | KS.3 |
| FIM1_3032 | 1686.804262 | 2268.491513 | 2951.006234 | 0.400799316 | 0.164750099 | 0.011855953 | 0.019753305 | ns | YPD | KS.3 |
| FIM1_3033 | 961.1360248 | 1146.671623 | 1273.167036 | 0.244223366 | 0.126702006 | 0.049205234 | 0.07112029 | ns | YPD | KS.3 |
| FIM1_3034 | 1091.910098 | 4328.381408 | 2931.69041 | 1.780633422 | 0.322730687 | 2.891135058 | 2.305031570 | Km>KS.3 | YPD | KS.3 |
| FIM1_3035 | 7931.414536 | 8369.893539 | 11142.20316 | 0.072794648 | 0.162501742 | 0.643624824 | 0.697306946 | ns | YPD | KS.3 |
| FIM1_3036 | 2455.349281 | 4679.08057 | 3404.866561 | 0.891543613 | 0.142371323 | 1.232203609 | 1.060590476 | ns | YPD | KS.3 |
| FIM1_3037 | 1193.056897 | 684.0733955 | 1021.631874 | -0.641199345 | 0.324307989 | 0.024844289 | 0.038528784 | ns | YPD | KS.3 |
| FIM1_3038 | 12706.65734 | 10401.42504 | 17148.25544 | -0.275225749 | 0.140952851 | 0.045325821 | 0.06599238 | ns | YPD | KS.3 |
| FIM1_3039 | 3088.3543 | 1910.784122 | 2869.090948 | -0.666209038 | 0.131656515 | 2.257357170 | 1.011743416 | ns | YPD | KS.3 |
| FIM1_3040 | 2603.540055 | 979.6639662 | 2794.51789 | -1.178488678 | 0.328468173 | 4.055297717 | 0.000111773 | Km<KS.3 | YPD | KS.3 |
| FIM1_3041 | 2064.13562 | 1441.011893 | 1740.792045 | -0.505978616 | 0.099336542 | 2.462469369 | 1.097740601 | ns | YPD | KS.3 |
| FIM1_3043 | 213.1360245 | 224.1898038 | 207.6025605 | 0.066793472 | 0.199104291 | 0.724937565 | 0.769154273 | ns | YPD | KS.3 |
| FIM1_3044 | 795.207717 | 4993.571127 | 4677.982151 | 2.386679918 | 0.410302275 | 1.696160193 | 1.739875662 | Km>KS.3 | YPD | KS.3 |
| FIM1_3045 | 805.5426066 | 793.0541032 | 549.2223036 | -0.020271041 | 0.1944545 | 0.913119063 | 0.929561559 | ns | YPD | KS.3 |
| FIM1_3046 | 1582.734274 | 2706.05644 | 1642.342519 | 0.727411683 | 0.168301902 | 7.360443543 | 2.390222364 | ns | YPD | KS.3 |
| FIM1_3047 | 847.8974064 | 658.9816888 | 825.3510037 | -0.354597494 | 0.098398328 | 0.000262927 | 0.000614114 | ns | YPD | KS.3 |
| FIM1_3048 | 3524.416562 | 1526.221606 | 3099.753428 | -1.140459847 | 0.175874271 | 1.326938953 | 1.381071408 | Km<KS.3 | YPD | KS.3 |
| FIM1_3049 | 1051.764657 | 726.2930981 | 883.089918 | -0.514813922 | 0.123498006 | 2.120230461 | 6.241506830 | ns | YPD | KS.3 |
| FIM1_3050 | 738.5954564 | 589.0043313 | 792.229839 | -0.317180649 | 0.108290731 | 0.002945545 | 0.005571896 | ns | YPD | KS.3 |
| FIM1_3051 | 1277.502624 | 1039.023846 | 1014.141862 | -0.284629626 | 0.135964081 | 0.032156532 | 0.048522519 | ns | YPD | KS.3 |
| FIM1_3052 | 1048.140994 | 867.8103404 | 796.9924135 | -0.259789733 | 0.139220572 | 0.055914676 | 0.079569655 | ns | YPD | KS.3 |
| FIM1_3053 | 1981.044386 | 1289.683432 | 2143.005932 | -0.588920592 | 0.148188311 | 4.353462098 | 0.000119134 | ns | YPD | KS.3 |
| FIM1_3054 | 460.0157321 | 1685.205207 | 1456.614948 | 1.667889668 | 0.315054567 | 1.677437032 | 1.162290979 | Km>KS.3 | YPD | KS.3 |
| FIM1_3056 | 2462.125334 | 5109.273576 | 4038.32144 | 0.936395864 | 0.24500045 | 3.714977597 | 0.000103425 | ns | YPD | KS.3 |
| FIM1_3057 | 478.4796269 | 259.0504267 | 483.5889983 | -0.776580644 | 0.251485908 | 0.000836173 | 0.001760457 | ns | YPD | KS.3 |
| FIM1_3058 | 3803.567225 | 3420.563094 | 2914.185425 | -0.146353427 | 0.13503146 | 0.267574012 | 0.325154057 | ns | YPD | KS.3 |
| FIM1_3059 | 1175.358982 | 911.1744781 | 849.149499 | -0.356921954 | 0.104958464 | 0.000561943 | 0.001226699 | ns | YPD | KS.3 |
| FIM1_3060 | 865.5226995 | 613.6142867 | 717.4541639 | -0.478704037 | 0.123678664 | 7.890823825 | 0.000206615 | ns | YPD | KS.3 |
| FIM1_3061 | 3903.779273 | 3846.958313 | 3894.558609 | -0.020485633 | 0.105219012 | 0.843483839 | 0.869886421 | ns | YPD | KS.3 |
| FIM1_3062 | 1467.422433 | 1021.422021 | 1637.020043 | -0.482045077 | 0.188230651 | 0.00744995 | 0.012986511 | ns | YPD | KS.3 |
| FIM1_3063 | 1343.207717 | 1185.306625 | 1105.968508 | -0.176265766 | 0.086930729 | 0.04060122 | 0.059991408 | ns | YPD | KS.3 |
| FIM1_3064 | 4043.025465 | 5945.429525 | 6172.113327 | 0.5419772 | 0.104363213 | 1.374877962 | 6.465045238 | ns | YPD | KS.3 |
| FIM1_3065 | 2579.491725 | 2283.312715 | 2380.210882 | -0.171426629 | 0.103140808 | 0.092041249 | 0.124498426 | ns | YPD | KS.3 |
| FIM1_3066 | 4299.1155 | 3471.092703 | 4964.727518 | -0.300280685 | 0.105366175 | 0.003842768 | 0.007070182 | ns | YPD | KS.3 |
| FIM1_3067 | 717.5847012 | 565.037994 | 690.525503 | -0.330759329 | 0.131125319 | 0.009943179 | 0.016860026 | ns | YPD | KS.3 |
| FIM1_3068 | 3184.016224 | 6596.578975 | 5347.153281 | 0.968371744 | 0.203742401 | 5.236207587 | 2.169036865 | ns | YPD | KS.3 |
| FIM1_3069 | 1321.392764 | 712.842664 | 1141.648876 | -0.832409052 | 0.179732643 | 1.366232434 | 5.212428191 | ns | YPD | KS.3 |
| FIM1_3070 | 7796.447387 | 5794.133853 | 7452.029009 | -0.416285361 | 0.108198631 | 9.372482505 | 0.000243108 | ns | YPD | KS.3 |

| | | | | | | | | | | |
|---|---|---|---|---|---|---|---|---|---|---|
| FIM1_3071 | 4084.364426 | 2657.51774 | 3315.501244 | -0.595173958 | 0.133145995 | 4.7780084458 | 1.6181478569 | ns | YPD | KS.3 |
| FIM1_3072 | 1414.778833 | 1655.159903 | 1546.247319 | 0.209631959 | 0.180170922 | 0.226108124 | 0.279892489 | ns | YPD | KS.3 |
| FIM1_3074 | 1019.195353 | 833.9002187 | 871.4180666 | -0.280254109 | 0.111996342 | 0.011001073 | 0.018477495 | ns | YPD | KS.3 |
| FIM1_3075 | 1222.107473 | 863.6901188 | 1321.472421 | -0.470426023 | 0.16357808 | 0.00293877 | 0.005561194 | ns | YPD | KS.3 |
| FIM1_3076 | 1100.374445 | 1250.260488 | 1017.158479 | 0.180415822 | 0.093667304 | 0.05141792 | 0.073931835 | ns | YPD | KS.3 |
| FIM1_3077 | 3020.62296 | 2403.932409 | 2215.349619 | -0.316307166 | 0.128849122 | 0.012181209 | 0.02024775 | ns | YPD | KS.3 |
| FIM1_3078 | 962.203575 | 680.2487666 | 790.6237241 | -0.487421091 | 0.100153291 | 8.1572071237 | 3.2596068627 | ns | YPD | KS.3 |
| FIM1_3079 | 2496.198059 | 1167.087161 | 2045.88673 | -1.045878919 | 0.155919582 | 4.0748660901 | 4.7145253019 | Km<KS.3 | YPD | KS.3 |
| FIM1_3080 | 1640.455202 | 1473.272898 | 1517.297272 | -0.150472275 | 0.107863248 | 0.156852982 | 0.202476281 | ns | YPD | KS.3 |
| FIM1_3081 | 1902.661311 | 2403.841128 | 1776.031229 | 0.319308725 | 0.151560007 | 0.030164897 | 0.045907116 | ns | YPD | KS.3 |
| FIM1_3082 | 2131.689616 | 2194.492827 | 1874.539366 | 0.041246198 | 0.090692272 | 0.645794591 | 0.69920089 | ns | YPD | KS.3 |
| FIM1_3083 | 3680.548511 | 2181.300354 | 5074.020734 | -0.636754111 | 0.281022904 | 0.012624556 | 0.020928746 | ns | YPD | KS.3 |
| FIM1_3085 | 59.2990829 | 35.9622264 | 43.4414597 | -0.617834373 | 0.264868392 | 0.01105182 | 0.018550204 | ns | YPD | KS.3 |
| FIM1_3086 | 1586.407063 | 1418.893893 | 1403.542008 | -0.146932572 | 0.193645092 | 0.427224273 | 0.491124106 | ns | YPD | KS.3 |
| FIM1_3087 | 6733.789883 | 4146.937442 | 7192.444108 | -0.637137798 | 0.205965933 | 0.001106474 | 0.002271856 | ns | YPD | KS.3 |
| FIM1_3088 | 402.8832882 | 432.2878809 | 879.435673 | 0.088862872 | 0.240899935 | 0.692612191 | 0.742137767 | ns | YPD | KS.3 |
| FIM1_3089 | 456.1674969 | 577.9311752 | 526.3408947 | 0.313224816 | 0.191648553 | 0.087390868 | 0.118789499 | ns | YPD | KS.3 |
| FIM1_3090 | 380.1889413 | 363.7765568 | 396.6058563 | -0.060240987 | 0.132782033 | 0.642855667 | 0.696777112 | ns | YPD | KS.3 |
| FIM1_3092 | 377.7167976 | 499.2549956 | 500.4249974 | 0.374348632 | 0.176887751 | 0.027830722 | 0.04269437 | ns | YPD | KS.3 |
| FIM1_3098 | 1006.189412 | 1088.553291 | 1200.264754 | 0.103642885 | 0.194901307 | 0.577727474 | 0.636446897 | ns | YPD | KS.3 |
| FIM1_3099 | 2798.557991 | 2031.554101 | 2350.021231 | -0.381257592 | 0.289848306 | 0.145822916 | 0.189121222 | ns | YPD | KS.3 |
| FIM1_3100 | 1333.33732 | 771.3493943 | 982.5762384 | -0.751176206 | 0.150194644 | 2.5848196236 | 1.1430646780 | ns | YPD | KS.3 |
| FIM1_3101 | 1355.933799 | 874.5382025 | 1052.947167 | -0.602558759 | 0.145254454 | 2.0193833062 | 5.9764615994 | ns | YPD | KS.3 |
| FIM1_3102 | 11255.84999 | 8754.849601 | 16236.58602 | -0.345514736 | 0.141529382 | 0.012298893 | 0.020436538 | ns | YPD | KS.3 |
| FIM1_3103 | 200.1116851 | 127.3115004 | 162.3837864 | -0.52457041 | 0.314313176 | 0.060258836 | 0.085166963 | ns | YPD | KS.3 |
| FIM1_3104 | 2356.993576 | 2222.829016 | 1971.434572 | -0.081270914 | 0.12364067 | 0.502873576 | 0.565122214 | ns | YPD | KS.3 |
| FIM1_3105 | 20.94098704 | 8.561974445 | 14.7225111 | -0.994492995 | 0.377963408 | 0.001790865 | 0.003528536 | ns | YPD | KS.3 |
| FIM1_3106 | 1513.284746 | 1612.627881 | 1352.376741 | 0.088636637 | 0.114812066 | 0.432509259 | 0.496592099 | ns | YPD | KS.3 |
| FIM1_3107 | 1113.480999 | 458.1981111 | 1220.887682 | -0.962655636 | 0.395714835 | 0.003381948 | 0.006313393 | ns | YPD | KS.3 |
| FIM1_3108 | 2832.807719 | 2301.650955 | 2240.122344 | -0.28493206 | 0.142862441 | 0.040768696 | 0.060169524 | ns | YPD | KS.3 |
| FIM1_3109 | 3167.762897 | 1726.870687 | 3375.384685 | -0.802678977 | 0.203631363 | 3.2388312593 | 9.1344589090 | ns | YPD | KS.3 |
| FIM1_3110 | 1324.477989 | 5208.819603 | 4126.047925 | 1.491380261 | 0.487061491 | 7.1522732281 | 0.000188567 | Km>KS.3 | YPD | KS.3 |
| FIM1_3111 | 492.8237858 | 1989.620525 | 1294.786463 | 1.803046788 | 0.327738346 | 2.8992236171 | 2.3077819992 | Km>KS.3 | YPD | KS.3 |
| FIM1_3112 | 921.2121741 | 523.2923483 | 703.4952814 | -0.753457981 | 0.193109873 | 4.2802301236 | 0.000117388 | ns | YPD | KS.3 |
| FIM1_3113 | 742.5658194 | 3470.7287 | 1621.420472 | 2.161913363 | 0.18878893 | 4.9946305337 | 9.9393147621 | Km>KS.3 | YPD | KS.3 |
| FIM1_3114 | 1751.026299 | 1312.017004 | 1719.038682 | -0.389558423 | 0.168186524 | 0.016438693 | 0.02664793 | ns | YPD | KS.3 |
| FIM1_3115 | 399.5443271 | 277.2310175 | 260.7413305 | -0.496139929 | 0.161927318 | 0.001540884 | 0.003074969 | ns | YPD | KS.3 |
| FIM1_3116 | 1737.813686 | 2276.183516 | 1724.004441 | 0.35133253 | 0.209933348 | 0.077446892 | 0.106786587 | ns | YPD | KS.3 |
| FIM1_3117 | 18.40755857 | 10.84940698 | 19.37440892 | -0.658841216 | 0.280016788 | 0.009635168 | 0.016393625 | ns | YPD | KS.3 |
| FIM1_3118 | 3224.894968 | 5205.469536 | 4526.624002 | 0.640551708 | 0.184679421 | 0.000293095 | 0.000680741 | ns | YPD | KS.3 |
| FIM1_3119 | 1596.831396 | 961.7934929 | 946.5580616 | -0.676426373 | 0.18868901 | 0.000176616 | 0.000431353 | ns | YPD | KS.3 |
| FIM1_3120 | 3620.561408 | 2848.575912 | 2859.777166 | -0.337919452 | 0.096453958 | 0.000391055 | 0.000882713 | ns | YPD | KS.3 |
| FIM1_3121 | 1628.811509 | 1405.850056 | 1170.818666 | -0.206310863 | 0.104244318 | 0.044779791 | 0.065407945 | ns | YPD | KS.3 |
| FIM1_3122 | 2045.730373 | 1759.101051 | 1970.926694 | -0.205092626 | 0.158126269 | 0.18105237 | 0.230721194 | ns | YPD | KS.3 |
| FIM1_3123 | 2077.616331 | 2769.11973 | 2384.973826 | 0.402774212 | 0.108578902 | 0.000165514 | 0.000407036 | ns | YPD | KS.3 |

| | | | | | | | | | | |
|---|---|---|---|---|---|---|---|---|---|---|
| FIM1_3124 | 4916.479966 | 4861.669815 | 4368.791076 | -0.01612582 | 0.053994715 | 0.764307157 | 0.802538646 | ns | YPD | KS.3 |
| FIM1_3125 | 456.5764976 | 803.9106907 | 648.7975289 | 0.786315644 | 0.131163871 | 8.54705690597 | 6.16255189959 | ns | YPD | KS.3 |
| FIM1_3126 | 1536.271025 | 3339.633735 | 2225.153781 | 1.038634131 | 0.198579735 | 3.57565948547 | 1.87054741748 | Km>KS.3 | YPD | KS.3 |
| FIM1_3127 | 620.7633297 | 2012.494997 | 1641.556946 | 1.603314345 | 0.20522472 | 9.17007106331 | 2.54866500223 | Km>KS.3 | YPD | KS.3 |
| FIM1_3128 | 7714.708026 | 8138.579139 | 9997.73968 | 0.075586332 | 0.088574621 | 0.38868074 | 0.451795953 | ns | YPD | KS.3 |
| FIM1_3129 | 4652.281357 | 3380.125402 | 3927.025455 | -0.451834652 | 0.089489129 | 3.34284433318 | 1.43764562709 | ns | YPD | KS.3 |
| FIM1_3130 | 1655.223528 | 724.6499644 | 1143.616367 | -1.117943061 | 0.185409294 | 2.66170326465 | 2.13580221639 | Km<KS.3 | YPD | KS.3 |
| FIM1_3131 | 1522.672409 | 977.158752 | 1039.568712 | -0.610522938 | 0.143633557 | 1.26870251580 | 3.90098579490 | ns | YPD | KS.3 |
| FIM1_3132 | 4042.640044 | 2484.38832 | 3367.068562 | -0.689437542 | 0.089750866 | 8.02779539024 | 1.54201861260 | ns | YPD | KS.3 |
| FIM1_3133 | 546.4644315 | 729.3705869 | 594.2050343 | 0.389700925 | 0.169044731 | 0.016915308 | 0.02734037 | ns | YPD | KS.3 |
| FIM1_3134 | 9691.095984 | 12869.25698 | 11415.25866 | 0.373115929 | 0.198869678 | 0.048933275 | 0.070788905 | ns | YPD | KS.3 |
| FIM1_3135 | 1679.812894 | 899.9590429 | 1117.0395 | -0.876432263 | 0.111137189 | 1.04255530766 | 2.28489544301 | ns | YPD | KS.3 |
| FIM1_3136 | 4122.95348 | 2909.112954 | 4630.98272 | -0.465753082 | 0.182978529 | 0.007966885 | 0.013805383 | ns | YPD | KS.3 |
| FIM1_3137 | 40.65430486 | 26.21453289 | 24.44135902 | -0.464621719 | 0.373071227 | 0.141521426 | 0.184022242 | ns | YPD | KS.3 |
| FIM1_3138 | 5651.914736 | 3626.321176 | 6167.117837 | -0.5514902 | 0.259644727 | 0.021140543 | 0.033251408 | ns | YPD | KS.3 |
| FIM1_3139 | 465.3188263 | 430.677284 | 667.88201 | -0.102154609 | 0.188554938 | 0.571686489 | 0.631189588 | ns | YPD | KS.3 |
| FIM1_3140 | 686.7972791 | 447.68779 | 459.0135077 | -0.590904955 | 0.135634119 | 8.12736159338 | 2.60693900239 | ns | YPD | KS.3 |
| FIM1_3141 | 1067.558369 | 598.1919985 | 972.2596684 | -0.800775306 | 0.140367658 | 4.73277146238 | 2.93585262161 | ns | YPD | KS.3 |
| FIM1_3142 | 271.3551776 | 425.8556594 | 264.7518978 | 0.593618951 | 0.203223472 | 0.002102761 | 0.004084825 | ns | YPD | KS.3 |
| FIM1_3143 | 4576.014124 | 4086.218302 | 5562.129101 | -0.160346342 | 0.08603171 | 0.059871611 | 0.084643724 | ns | YPD | KS.3 |
| FIM1_3144 | 474.5212971 | 226.4231547 | 358.3171778 | -1.00156591 | 0.179843571 | 6.08670766441 | 3.69735905134 | Km<KS.3 | YPD | KS.3 |
| FIM1_3145 | 1822.664742 | 1257.161642 | 1300.969336 | -0.517704546 | 0.121252123 | 1.34907834377 | 4.12771510472 | ns | YPD | KS.3 |
| FIM1_3146 | 911.0441163 | 642.1225027 | 695.8146255 | -0.489059802 | 0.111497959 | 8.27205809706 | 2.64482577332 | ns | YPD | KS.3 |
| FIM1_3147 | 1407.737436 | 1592.324177 | 1134.359183 | 0.170181979 | 0.13549921 | 0.198808639 | 0.249892112 | ns | YPD | KS.3 |
| FIM1_3148 | 1196.607056 | 766.595052 | 864.5807955 | -0.623264219 | 0.112354341 | 1.68962015811 | 9.46605888135 | ns | YPD | KS.3 |
| FIM1_3149 | 1491.894232 | 950.2194395 | 1048.570728 | -0.623709685 | 0.135652018 | 2.48450094014 | 8.93091920319 | ns | YPD | KS.3 |
| FIM1_3150 | 4720.657247 | 2551.878347 | 5060.087817 | -0.765214758 | 0.267415464 | 0.001775317 | 0.003500674 | ns | YPD | KS.3 |
| FIM1_3151 | 965.1617824 | 639.9279673 | 787.651183 | -0.561362616 | 0.152981236 | 0.000156348 | 0.000387946 | ns | YPD | KS.3 |
| FIM1_3152 | 320.3757749 | 217.2856209 | 255.1933084 | -0.533385642 | 0.145278312 | 0.000162007 | 0.0003992 | ns | YPD | KS.3 |
| FIM1_3153 | 1230.250998 | 703.1697308 | 769.2644039 | -0.783367932 | 0.11461224 | 3.45534425689 | 4.05432492406 | ns | YPD | KS.3 |
| FIM1_3154 | 2223.062407 | 1587.059278 | 2146.446937 | -0.446099382 | 0.193319441 | 0.015671967 | 0.025504755 | ns | YPD | KS.3 |
| FIM1_3155 | 1910.580363 | 3390.755549 | 2866.78671 | 0.794358393 | 0.137394218 | 3.05151898746 | 1.97159830683 | ns | YPD | KS.3 |
| FIM1_3156 | 4585.047682 | 2850.350645 | 3973.487865 | -0.661151993 | 0.126881777 | 1.02324521269 | 4.94717680580 | ns | YPD | KS.3 |
| FIM1_3157 | 466.4639247 | 904.9543648 | 1292.04226 | 0.835086064 | 0.258339976 | 0.000443154 | 0.000989095 | ns | YPD | KS.3 |
| FIM1_3158 | 526.6947609 | 413.9380244 | 368.4361109 | -0.332062372 | 0.137926282 | 0.013671784 | 0.022485 | ns | YPD | KS.3 |
| FIM1_3159 | 471.6806312 | 280.4797405 | 404.0752316 | -0.717836118 | 0.141082653 | 1.76185302774 | 8.10843553471 | ns | YPD | KS.3 |
| FIM1_3160 | 363.5251929 | 504.984144 | 291.6823713 | 0.3892682 | 0.294029276 | 0.142066516 | 0.184682758 | ns | YPD | KS.3 |
| FIM1_3161 | 296.5950537 | 1830.251807 | 1401.267782 | 2.480088335 | 0.304991196 | 1.50444444383 | 5.67015993035 | Km>KS.3 | YPD | KS.3 |
| FIM1_3162 | 2484.327782 | 1973.300982 | 2181.720415 | -0.279292408 | 0.272987616 | 0.263441291 | 0.320758792 | ns | YPD | KS.3 |
| FIM1_3163 | 954.8767386 | 369.4597483 | 715.8975727 | -1.282561641 | 0.198011 | 7.88571901800 | 8.64128901202 | Km<KS.3 | YPD | KS.3 |
| FIM1_3164 | 3155.431955 | 3444.938496 | 2822.476065 | 0.125929189 | 0.05146976 | 0.014069985 | 0.023094086 | ns | YPD | KS.3 |
| FIM1_3165 | 869.5849052 | 561.1967905 | 793.2031306 | -0.600532521 | 0.150159852 | 3.84052285173 | 0.000106384 | ns | YPD | KS.3 |
| FIM1_3166 | 1430.849961 | 1126.691609 | 1168.677337 | -0.329033206 | 0.138224128 | 0.014759017 | 0.024129513 | ns | YPD | KS.3 |
| FIM1_3167 | 2426.593435 | 1924.50378 | 3498.114776 | -0.272855816 | 0.295780327 | 0.305934722 | 0.366311731 | ns | YPD | KS.3 |
| FIM1_3168 | 899.2734377 | 1115.072661 | 1110.963659 | 0.30064091 | 0.109004584 | 0.005105207 | 0.009145987 | ns | YPD | KS.3 |

| | | | | | | | | | | |
|---|---|---|---|---|---|---|---|---|---|---|
| FIM1_3169 | 1081.109112 | 905.7683889 | 1077.659817 | -0.240842161 | 0.156131898 | 0.111911468 | 0.148746875 | ns | YPD | KS.3 |
| FIM1_3170 | 66.12808158 | 82.96808913 | 121.4952025 | 0.288702091 | 0.234039575 | 0.187643281 | 0.237840846 | ns | YPD | KS.3 |
| FIM1_3171 | 1782.906389 | 1096.187187 | 1716.990805 | -0.669542926 | 0.145353761 | 2.19408721466 | 7.98506502775 | ns | YPD | KS.3 |
| FIM1_3172 | 1819.940477 | 1404.628083 | 1513.119302 | -0.339718099 | 0.201705822 | 0.076754236 | 0.106011194 | ns | YPD | KS.3 |
| FIM1_3173 | 1411.210846 | 2277.336449 | 1674.368048 | 0.659711838 | 0.142221471 | 1.91268934920 | 7.07481748126 | ns | YPD | KS.3 |
| FIM1_3174 | 265.5054355 | 660.1321639 | 492.5185698 | 1.20958354 | 0.21725871 | 2.96258792796 | 1.92561886619 | Km>KS.3 | YPD | KS.3 |
| FIM1_3175 | 980.472849 | 1402.893918 | 1395.309944 | 0.495884671 | 0.13133753 | 0.000113248 | 0.00028967 | ns | YPD | KS.3 |
| FIM1_3176 | 383.6501526 | 628.4445097 | 464.5748983 | 0.685944333 | 0.128016901 | 4.36142278886 | 2.25551750255 | ns | YPD | KS.3 |
| FIM1_3177 | 2496.871362 | 2406.00131 | 2248.049981 | -0.052615224 | 0.073352224 | 0.470073533 | 0.533686861 | ns | YPD | KS.3 |
| FIM1_3178 | 2207.664866 | 1575.61504 | 2050.784747 | -0.461196101 | 0.151151979 | 0.001686392 | 0.003339889 | ns | YPD | KS.3 |
| FIM1_3179 | 522.3136768 | 763.7765666 | 479.2921944 | 0.485688327 | 0.232113487 | 0.025441955 | 0.039367928 | ns | YPD | KS.3 |
| FIM1_3180 | 1542.532249 | 1934.800653 | 1588.096631 | 0.311984937 | 0.139251029 | 0.021701436 | 0.034055353 | ns | YPD | KS.3 |
| FIM1_3181 | 2299.379334 | 1676.170309 | 3024.698306 | -0.38549933 | 0.270961007 | 0.120241904 | 0.158969831 | ns | YPD | KS.3 |
| FIM1_3182 | 2442.610488 | 1183.245884 | 1881.758192 | -1.012226822 | 0.126638924 | 2.99125613003 | 7.25926792531 | Km<KS.3 | YPD | KS.3 |
| FIM1_3183 | 5191.801128 | 7234.746771 | 6908.476124 | 0.453769298 | 0.150839607 | 0.001961929 | 0.003829187 | ns | YPD | KS.3 |
| FIM1_3184 | 677.6487378 | 494.5617052 | 690.2581567 | -0.407970813 | 0.217005089 | 0.046475875 | 0.067548198 | ns | YPD | KS.3 |
| FIM1_3185 | 676.6869149 | 775.6363511 | 1068.012224 | 0.175912177 | 0.216014 | 0.38905502 | 0.452125373 | ns | YPD | KS.3 |
| FIM1_3186 | 2522.232811 | 4133.589959 | 3548.73503 | 0.687458345 | 0.12641916 | 2.78896695460 | 1.49252691541 | ns | YPD | KS.3 |
| FIM1_3187 | 988.3221967 | 402.7310889 | 663.5778738 | -1.242708683 | 0.151377261 | 2.17627848770 | 6.89616909320 | Km<KS.3 | YPD | KS.3 |
| FIM1_3188 | 1084.780639 | 625.1260745 | 690.8064976 | -0.766221049 | 0.128286429 | 1.02436437463 | 7.25956234162 | ns | YPD | KS.3 |
| FIM1_3189 | 542.6151118 | 362.0240401 | 411.7370225 | -0.543855973 | 0.177421189 | 0.001427984 | 0.002871552 | ns | YPD | KS.3 |
| FIM1_3190 | 1076.267739 | 584.9953154 | 680.2043265 | -0.843659672 | 0.139922365 | 6.03088099885 | 4.49829579749 | ns | YPD | KS.3 |
| FIM1_3191 | 1275.338076 | 649.1753886 | 883.2861558 | -0.910668887 | 0.182016124 | 1.73749544808 | 8.00374060575 | ns | YPD | KS.3 |
| FIM1_3192 | 798.5997536 | 932.9138253 | 729.3971384 | 0.21706882 | 0.112637571 | 0.050186636 | 0.072351665 | ns | YPD | KS.3 |
| FIM1_3193 | 46.97151758 | 46.14824947 | 43.75386283 | -0.026664666 | 0.246305104 | 0.907360658 | 0.924456128 | ns | YPD | KS.3 |
| FIM1_3194 | 1797.315701 | 2850.560208 | 2395.124519 | 0.607479293 | 0.202444556 | 0.001587008 | 0.003163207 | ns | YPD | KS.3 |
| FIM1_3195 | 3100.441653 | 1660.693211 | 3200.949999 | -0.800572431 | 0.238932632 | 0.000319329 | 0.000733115 | ns | YPD | KS.3 |
| FIM1_3196 | 779.4121372 | 397.3912684 | 477.2816158 | -0.921680326 | 0.161119508 | 3.18248267320 | 2.04823432072 | ns | YPD | KS.3 |
| FIM1_3197 | 291.1014225 | 1848.625611 | 1536.617768 | 2.505154569 | 0.32233094 | 2.81038395239 | 8.77368961879 | Km>KS.3 | YPD | KS.3 |
| FIM1_3198 | 288.472683 | 697.8069092 | 460.0031329 | 0.954336926 | 0.397220206 | 0.003754359 | 0.006928018 | ns | YPD | KS.3 |
| FIM1_3199 | 3315.567566 | 2343.607029 | 2359.023598 | -0.476598639 | 0.144146683 | 0.000686738 | 0.001472639 | ns | YPD | KS.3 |
| FIM1_3200 | 9030.939076 | 8008.855178 | 9302.489641 | -0.171904067 | 0.057032213 | 0.002466753 | 0.004721852 | ns | YPD | KS.3 |
| FIM1_3201 | 744.6562293 | 381.4859941 | 501.8279529 | -0.915636937 | 0.160536479 | 3.57268614786 | 2.27873251097 | ns | YPD | KS.3 |
| FIM1_3202 | 503.5678011 | 612.2220341 | 534.8867205 | 0.271489793 | 0.123203641 | 0.024690474 | 0.038302186 | ns | YPD | KS.3 |
| FIM1_3203 | 1370.193684 | 2645.095639 | 1449.933299 | 0.914135779 | 0.133497088 | 2.28735369310 | 2.79634577000 | ns | YPD | KS.3 |
| FIM1_3204 | 856.5081471 | 767.3213553 | 744.7715961 | -0.15635831 | 0.062408263 | 0.011786988 | 0.019651564 | ns | YPD | KS.3 |
| FIM1_3205 | 4323.462932 | 4367.550759 | 4777.253445 | 0.014217339 | 0.103837201 | 0.889607617 | 0.908974717 | ns | YPD | KS.3 |
| FIM1_3206 | 2304.169526 | 1489.304019 | 1815.968829 | -0.614144111 | 0.103544458 | 1.77784489268 | 1.22165446700 | ns | YPD | KS.3 |
| FIM1_3207 | 729.2361372 | 1441.430063 | 891.1447567 | 0.909124946 | 0.196442719 | 1.13931851058 | 4.41441556865 | ns | YPD | KS.3 |
| FIM1_3208 | 3419.116426 | 2355.260972 | 3247.202603 | -0.524823911 | 0.100131501 | 1.08737119532 | 5.22169082699 | ns | YPD | KS.3 |
| FIM1_3209 | 7309.637944 | 6921.32362 | 7110.124517 | -0.0756151 | 0.128230563 | 0.547263433 | 0.608410185 | ns | YPD | KS.3 |
| FIM1_3210 | 812.4710188 | 192.0120844 | 590.3318852 | -2.027583761 | 0.168661833 | 6.05885222428 | 1.50713940779 | Km<KS.3 | YPD | KS.3 |
| FIM1_3211 | 668.3641394 | 368.8894625 | 437.1034821 | -0.800344655 | 0.180471333 | 3.73721469865 | 1.29836893336 | ns | YPD | KS.3 |
| FIM1_3212 | 3768.403261 | 2577.230211 | 3159.226961 | -0.530893491 | 0.116326792 | 3.39688318519 | 1.18843135346 | ns | YPD | KS.3 |
| FIM1_3213 | 4254.988175 | 3951.450861 | 3801.11775 | -0.10402825 | 0.101256439 | 0.297945425 | 0.358464584 | ns | YPD | KS.3 |

| | | | | | | | | | | |
|---|---|---|---|---|---|---|---|---|---|---|
| FIM1_3214 | 3100.897047 | 1421.938475 | 3474.386174 | -0.86229017 | 0.370455408 | 0.006077532 | 0.010744749 | ns | YPD | KS.3 |
| FIM1_3215 | 2485.107321 | 3702.042375 | 2625.623474 | 0.564973761 | 0.085173689 | 2.1020225623( | 2.1298497449( | ns | YPD | KS.3 |
| FIM1_3216 | 63.37759952 | 67.12606017 | 80.9259995 | 0.075117159 | 0.234359143 | 0.732170085 | 0.775296303 | ns | YPD | KS.3 |
| FIM1_3217 | 1112.641709 | 2592.32793 | 2185.918002 | 1.083083422 | 0.255676032 | 4.0668917101 | 1.4060310116( | Km>KS.3 | YPD | KS.3 |
| FIM1_3218 | 143.4784578 | 150.8878672 | 239.287081 | 0.060544512 | 0.280532757 | 0.812696868 | 0.843556628 | ns | YPD | KS.3 |
| FIM1_3219 | 2860.731499 | 2711.569234 | 3158.916771 | -0.072043032 | 0.16905634 | 0.659044799 | 0.71107089 | ns | YPD | KS.3 |
| FIM1_3220 | 884.5996177 | 2902.151014 | 2141.848212 | 1.600643562 | 0.227445574 | 3.3888879577 | 5.8540686076 | Km>KS.3 | YPD | KS.3 |
| FIM1_3221 | 4948.357348 | 6452.761383 | 6394.062365 | 0.372496415 | 0.105867525 | 0.000357142 | 0.000812904 | ns | YPD | KS.3 |
| FIM1_3222 | 3045.972086 | 3115.77866 | 3571.471653 | 0.032344573 | 0.060777051 | 0.592783832 | 0.650010924 | ns | YPD | KS.3 |
| FIM1_3223 | 5032.580495 | 5126.299513 | 5186.784433 | 0.026062729 | 0.08505827 | 0.757107786 | 0.797503967 | ns | YPD | KS.3 |
| FIM1_3224 | 1200.34285 | 3340.225043 | 1877.403515 | 1.303154056 | 0.280701377 | 2.6472127269 | 1.1654763997( | Km>KS.3 | YPD | KS.3 |
| FIM1_3225 | 96.94402531 | 55.71425925 | 117.7906859 | -0.673827448 | 0.283981724 | 0.008853634 | 0.015146777 | ns | YPD | KS.3 |
| FIM1_3226 | 572.9867376 | 1033.086576 | 829.6867632 | 0.816562361 | 0.136260134 | 8.0800810947 | 5.8683800651 | ns | YPD | KS.3 |
| FIM1_3227 | 2964.87428 | 2196.903546 | 2360.123084 | -0.415428683 | 0.130113175 | 0.001104531 | 0.002270912 | ns | YPD | KS.3 |
| FIM1_3228 | 1910.095143 | 1321.85698 | 1687.076866 | -0.512241265 | 0.123797297 | 2.4352343677 | 7.0684311433 | ns | YPD | KS.3 |
| FIM1_3229 | 3118.99979 | 2537.401377 | 3769.560011 | -0.291445186 | 0.093799813 | 0.001673661 | 0.003318638 | ns | YPD | KS.3 |
| FIM1_3230 | 4606.269258 | 3538.026591 | 4397.562342 | -0.369777141 | 0.108972713 | 0.00056969 | 0.001242529 | ns | YPD | KS.3 |
| FIM1_3231 | 4562.231198 | 3967.013963 | 4952.197279 | -0.199074928 | 0.07253142 | 0.005713709 | 0.010148412 | ns | YPD | KS.3 |
| FIM1_3232 | 110.8886291 | 488.1606671 | 257.3779081 | 2.0270655 | 0.24483438 | 3.4241114636 | 1.4942942571 | Km>KS.3 | YPD | KS.3 |
| FIM1_3233 | 763.2064651 | 650.8129879 | 1744.519258 | -0.221172054 | 0.120463492 | 0.061500559 | 0.086774044 | ns | YPD | KS.3 |
| FIM1_3234 | 3124.867136 | 1817.089486 | 4010.201847 | -0.639691295 | 0.306494817 | 0.019393155 | 0.030913472 | ns | YPD | KS.3 |
| FIM1_3235 | 2208.800577 | 1303.7564 | 1430.647497 | -0.73987743 | 0.110479542 | 9.8621452288 | 1.0551434949 | ns | YPD | KS.3 |
| FIM1_3236 | 3739.074586 | 12666.72306 | 9277.944695 | 1.619677838 | 0.25560478 | 3.8446233035 | 4.4899063227 | Km>KS.3 | YPD | KS.3 |
| FIM1_3237 | 3995.045766 | 5453.036227 | 5182.316218 | 0.430433608 | 0.132324564 | 0.00088018 | 0.001846857 | ns | YPD | KS.3 |
| FIM1_3238 | 490.3808692 | 408.2298458 | 470.235462 | -0.256351093 | 0.119229712 | 0.028593009 | 0.04378277 | ns | YPD | KS.3 |
| FIM1_3239 | 2400.440743 | 2887.869656 | 3239.1185 | 0.258843882 | 0.109386234 | 0.01627486 | 0.02641678 | ns | YPD | KS.3 |
| FIM1_3240 | 2726.353695 | 2535.058851 | 2728.842769 | -0.100206129 | 0.137633881 | 0.456071152 | 0.52004446 | ns | YPD | KS.3 |
| FIM1_3241 | 1321.507572 | 1261.157579 | 1207.718789 | -0.06179397 | 0.188672081 | 0.73241948 | 0.775296303 | ns | YPD | KS.3 |
| FIM1_3242 | 886.2153294 | 995.4804481 | 739.1983989 | 0.161451073 | 0.129983002 | 0.204607296 | 0.25620974 | ns | YPD | KS.3 |
| FIM1_3243 | 2407.450899 | 2062.852971 | 2177.611347 | -0.214057971 | 0.128238591 | 0.088391353 | 0.120083829 | ns | YPD | KS.3 |
| FIM1_3244 | 581.1251475 | 504.7802725 | 1295.076085 | -0.181363052 | 0.218212454 | 0.378823182 | 0.441782778 | ns | YPD | KS.3 |
| FIM1_3245 | 1122.60925 | 1588.472496 | 1235.652559 | 0.477295221 | 0.142324303 | 0.000579259 | 0.001261187 | ns | YPD | KS.3 |
| FIM1_3246 | 1143.632964 | 1693.257015 | 1352.307553 | 0.528932488 | 0.172556913 | 0.001461803 | 0.002930084 | ns | YPD | KS.3 |
| FIM1_3247 | 4234.718741 | 9148.716118 | 5932.375847 | 0.962977438 | 0.273641884 | 0.000110725 | 0.000283508 | ns | YPD | KS.3 |
| FIM1_3248 | 267.2587337 | 300.0957179 | 270.241428 | 0.15932282 | 0.139465406 | 0.241899456 | 0.29700088 | ns | YPD | KS.3 |
| FIM1_3249 | 2661.365818 | 2742.692175 | 3260.562503 | 0.041463857 | 0.136279912 | 0.755553757 | 0.796204182 | ns | YPD | KS.3 |
| FIM1_3250 | 898.6639103 | 548.8591235 | 864.3547259 | -0.669949731 | 0.165113442 | 2.6452253615 | 7.6025396728 | ns | YPD | KS.3 |
| FIM1_3251 | 788.3450155 | 912.0560109 | 786.1267761 | 0.204803003 | 0.10813488 | 0.054567456 | 0.077919946 | ns | YPD | KS.3 |
| FIM1_3252 | 38.71422578 | 42.16223308 | 46.84830583 | 0.100139014 | 0.298943593 | 0.709562459 | 0.757328314 | ns | YPD | KS.3 |
| FIM1_3253 | 1480.248299 | 4129.633606 | 3578.117444 | 1.296436844 | 0.289469195 | 5.9220524204 | 2.4169163898( | Km>KS.3 | YPD | KS.3 |
| FIM1_3254 | 4470.716815 | 4836.870429 | 4440.083237 | 0.111081194 | 0.094074711 | 0.232389278 | 0.286740242 | ns | YPD | KS.3 |
| FIM1_3255 | 1071.030199 | 926.7846754 | 844.2191512 | -0.204626602 | 0.08195871 | 0.011780154 | 0.019646754 | ns | YPD | KS.3 |
| FIM1_3256 | 41.76902816 | 24.97383804 | 40.43753559 | -0.625551007 | 0.280825319 | 0.01422495 | 0.023317669 | ns | YPD | KS.3 |
| FIM1_3258 | 275.3164436 | 58.73618216 | 304.6439041 | -2.15438603 | 0.198564928 | 4.3498099609 | 6.1829441587 | Km<KS.3 | YPD | KS.3 |
| FIM1_3259 | 547.55544 | 134.1676432 | 420.9082459 | -1.910468951 | 0.245747094 | 3.0077558902 | 9.1801138367 | Km<KS.3 | YPD | KS.3 |

| | | | | | | | | | | | |
|---|---|---|---|---|---|---|---|---|---|---|---|
| FIM1_3260 | 1787.684962 | | 1958.740336 | 1532.399655 | 0.130080892 | 0.074767662 | 0.079674621 | 0.109501014 | ns | YPD | KS.3 |
| FIM1_3261 | 525.3504947 | | 607.0961479 | 583.1292843 | 0.196535818 | 0.151068889 | 0.180858141 | 0.230532731 | ns | YPD | KS.3 |
| FIM1_3262 | 22480.17316 | | 19807.54908 | 29873.94409 | -0.176442528 | 0.117510891 | 0.126555958 | 0.166609127 | ns | YPD | KS.3 |
| FIM1_3263 | 2779.749979 | | 2047.945252 | 1959.690367 | -0.429099297 | 0.104195344 | 2.9546463455 | 8.4237051970 | ns | YPD | KS.3 |
| FIM1_3264 | 2686.144827 | | 1257.755491 | 1677.565821 | -1.063118813 | 0.121124284 | 3.2087159959 | 1.4166767478 | Km<KS.3 | YPD | KS.3 |
| FIM1_3265 | 1545.319992 | | 1398.843927 | 1308.290788 | -0.140463107 | 0.093111465 | 0.127154527 | 0.16726435 | ns | YPD | KS.3 |
| FIM1_3266 | 650.7952861 | | 419.7084902 | 627.6223463 | -0.571393153 | 0.214366679 | 0.004780338 | 0.00862298 | ns | YPD | KS.3 |
| FIM1_3267 | 4092.416314 | | 2893.316894 | 3631.771014 | -0.485637304 | 0.111583411 | 9.7112948127 | 3.0520335876 | ns | YPD | KS.3 |
| FIM1_3268 | 1109.982101 | | 588.3674318 | 861.409376 | -0.861033486 | 0.172728378 | 2.1761336384 | 9.8152899829 | ns | YPD | KS.3 |
| FIM1_3269 | 762.2608386 | | 599.8694959 | 594.8101771 | -0.323436683 | 0.166258962 | 0.044118419 | 0.064460832 | ns | YPD | KS.3 |
| FIM1_3270 | 1236.462467 | | 6534.041553 | 3849.519547 | 2.161188924 | 0.378815645 | 3.5034915345 | 3.3518981508 | Km>KS.3 | YPD | KS.3 |
| FIM1_3271 | 3812.180087 | | 2604.711203 | 3839.313539 | -0.528708368 | 0.128007191 | 2.4559256701 | 7.1160339014 | ns | YPD | KS.3 |
| FIM1_3272 | 1631.172162 | | 1173.320322 | 1169.173293 | -0.454357636 | 0.13729014 | 0.000699255 | 0.00149497 | ns | YPD | KS.3 |
| FIM1_3273 | 1804.711348 | | 875.1800488 | 2005.529265 | -0.905339638 | 0.269687806 | 0.000236635 | 0.000557943 | ns | YPD | KS.3 |
| FIM1_3274 | 2766.351302 | | 2758.194582 | 2580.125895 | -0.004254598 | 0.048707199 | 0.930168668 | 0.943250943 | ns | YPD | KS.3 |
| FIM1_3275 | 2420.098657 | | 2698.153959 | 3593.207739 | 0.151271999 | 0.119722443 | 0.198284422 | 0.24942225 | ns | YPD | KS.3 |
| FIM1_3276 | 719.1691365 | | 1023.474609 | 476.724671 | 0.463909224 | 0.201324467 | 0.015425297 | 0.025119756 | ns | YPD | KS.3 |
| FIM1_3277 | 2667.811827 | | 2033.354187 | 2368.234626 | -0.381370739 | 0.105200184 | 0.000235502 | 0.000555798 | ns | YPD | KS.3 |
| FIM1_3278 | 5741.225956 | | 7366.075372 | 6815.733062 | 0.350938069 | 0.098654452 | 0.000315175 | 0.000725253 | ns | YPD | KS.3 |
| FIM1_3279 | 3826.868915 | | 3175.119537 | 3833.314864 | -0.258208675 | 0.132228825 | 0.045934808 | 0.066820372 | ns | YPD | KS.3 |
| FIM1_3280 | 1433.35233 | | 1969.286142 | 2105.880803 | 0.441501611 | 0.124778626 | 0.00030657 | 0.000708072 | ns | YPD | KS.3 |
| FIM1_3281 | 2849.332491 | | 2353.857325 | 4292.32629 | -0.26145848 | 0.147647299 | 0.0688487 | 0.096079182 | ns | YPD | KS.3 |
| FIM1_3282 | 2852.472894 | | 7395.82396 | 5569.056616 | 1.136972303 | 0.334027166 | 9.3141154063 | 0.000241972 | Km>KS.3 | YPD | KS.3 |
| FIM1_3283 | 1058.749006 | | 565.5866405 | 807.0213661 | -0.858319921 | 0.157662996 | 1.8431450069 | 1.0233980367 | ns | YPD | KS.3 |
| FIM1_3284 | 1627.146949 | | 1622.129686 | 1342.972478 | -0.004071409 | 0.15388431 | 0.978271417 | 0.982418308 | ns | YPD | KS.3 |
| FIM1_3286 | 2072.863256 | | 2242.191088 | 1644.190823 | 0.109774019 | 0.112480096 | 0.32138474 | 0.382784075 | ns | YPD | KS.3 |
| FIM1_3287 | 627.6892263 | | 409.4558032 | 460.2756596 | -0.574510876 | 0.176685877 | 0.000719327 | 0.001531958 | ns | YPD | KS.3 |
| FIM1_3288 | 14718.46548 | | 19619.91487 | 24758.36259 | 0.374651957 | 0.208930082 | 0.058600461 | 0.082964512 | ns | YPD | KS.3 |
| FIM1_3290 | 1155.787575 | | 1923.232424 | 1587.834695 | 0.707195918 | 0.129774267 | 2.5143590960 | 1.3537810067 | ns | YPD | KS.3 |
| FIM1_3291 | 13.30111772 | | 23.46581816 | 18.39346275 | 0.591399131 | 0.385371318 | 0.067015304 | 0.09375735 | ns | YPD | KS.3 |
| FIM1_3292 | 17005.95542 | | 20513.46849 | 28547.75295 | 0.260677775 | 0.122733349 | 0.030412104 | 0.046197251 | ns | YPD | KS.3 |
| FIM1_3293 | 445.0718628 | | 256.2320828 | 402.3163413 | -0.740447487 | 0.186295838 | 3.2538814373 | 9.1688283199 | ns | YPD | KS.3 |
| FIM1_3294 | 290.542128 | | 241.2777107 | 335.1507035 | -0.24685784 | 0.183276036 | 0.160549206 | 0.206871872 | ns | YPD | KS.3 |
| FIM1_3295 | 330.4368129 | | 3168.044988 | 1289.998448 | 3.185956389 | 0.211052169 | 5.4045657927 | 5.3775429637 | Km>KS.3 | YPD | KS.3 |
| FIM1_3297 | 1855.729042 | | 1631.323309 | 2133.083476 | -0.181829336 | 0.090020151 | 0.041240869 | 0.060738106 | ns | YPD | KS.3 |
| FIM1_3298 | 3430.942847 | | 2401.021361 | 3671.679717 | -0.448369968 | 0.246892511 | 0.050557775 | 0.072758152 | ns | YPD | KS.3 |
| FIM1_3299 | 4590.16717 | | 6849.597047 | 5009.030904 | 0.565707037 | 0.092738677 | 6.7890099902 | 5.0260899853 | ns | YPD | KS.3 |
| FIM1_3300 | 8421.521452 | | 6389.369519 | 12154.06664 | -0.377175838 | 0.151916052 | 0.010602633 | 0.017898913 | ns | YPD | KS.3 |
| FIM1_3301 | 2285.697758 | | 1505.493941 | 2184.731024 | -0.580855949 | 0.124863579 | 2.0581547617 | 7.5265017636 | ns | YPD | KS.3 |
| FIM1_3302 | 777.8902703 | | 173.7271428 | 695.5945618 | -1.86775234 | 0.39891664 | 1.9520962541 | 1.0766908733 | Km<KS.3 | YPD | KS.3 |
| FIM1_3303 | 780.3142322 | | 413.6920567 | 583.473781 | -0.863209642 | 0.167030365 | 8.2663703254 | 4.0637541866 | ns | YPD | KS.3 |
| FIM1_3304 | 6937.146295 | | 6659.733309 | 10610.29114 | -0.051489016 | 0.237707081 | 0.816765708 | 0.846896498 | ns | YPD | KS.3 |
| FIM1_3305 | 6644.040993 | | 11806.86105 | 11293.93362 | 0.800218849 | 0.128438989 | 1.8912716264 | 1.5760596887 | ns | YPD | KS.3 |
| FIM1_3306 | 274.1917048 | | 304.840488 | 385.9812727 | 0.137890453 | 0.199993581 | 0.469348682 | 0.53298555 | ns | YPD | KS.3 |
| FIM1_3307 | 916.9859644 | | 1312.180239 | 1497.973183 | 0.489789993 | 0.15205402 | 0.000910325 | 0.001903686 | ns | YPD | KS.3 |

| | | | | | | | | | | |
|---|---|---|---|---|---|---|---|---|---|---|
| FIM1_3308 | 254.0116295 | 798.570965 | 1459.473884 | 1.54088767 | 0.224155043 | 1.4863569616( | 2.2613534813: | Km>KS.3 | YPD | KS.3 |
| FIM1_3309 | 3696.585275 | 4559.915557 | 4207.45595 | 0.287555354 | 0.145902115 | 0.042988202 | 0.063050208 | ns | YPD | KS.3 |
| FIM1_3310 | 1758.884953 | 1843.945793 | 1586.122228 | 0.066187372 | 0.108111204 | 0.534393487 | 0.596234043 | ns | YPD | KS.3 |
| FIM1_3311 | 5587.701887 | 4842.798346 | 5799.004745 | -0.204436091 | 0.062054754 | 0.000928046 | 0.001935862 | ns | YPD | KS.3 |
| FIM1_3312 | 974.5727419 | 6357.04407 | 3030.972899 | 2.561605093 | 0.304943108 | 1.9870646950{ | 9.4149017695 | Km>KS.3 | YPD | KS.3 |
| FIM1_3313 | 202.6157241 | 1254.439054 | 449.383105 | 2.314608712 | 0.446269808 | 6.2011146746( | 4.6183451356( | Km>KS.3 | YPD | KS.3 |
| FIM1_3314 | 686.6309013 | 640.9944418 | 611.8314272 | -0.095237678 | 0.128692737 | 0.449980814 | 0.513452879 | ns | YPD | KS.3 |
| FIM1_3315 | 551.4342412 | 577.7632627 | 625.3048471 | 0.062490213 | 0.168519344 | 0.701081556 | 0.74959406 | ns | YPD | KS.3 |
| FIM1_3316 | 2185.771996 | 1650.534957 | 1921.22516 | -0.396949352 | 0.092054632 | 1.2981648234( | 3.9866481462 | ns | YPD | KS.3 |
| FIM1_3317 | 241.9857753 | 93.27567858 | 156.9880462 | -1.31094825 | 0.166367984 | 2.3901462534( | 5.9966546531! | Km<KS.3 | YPD | KS.3 |
| FIM1_3318 | 771.122087 | 330.9613435 | 902.019129 | -1.033779468 | 0.301554441 | 0.000123128 | 0.000311261 | Km<KS.3 | YPD | KS.3 |
| FIM1_3319 | 1337.517302 | 1225.904224 | 1424.631369 | -0.119523287 | 0.145378572 | 0.398749883 | 0.462311971 | ns | YPD | KS.3 |
| FIM1_3320 | 2431.993618 | 3329.138443 | 3754.133926 | 0.379081982 | 0.278846874 | 0.135646719 | 0.177031067 | ns | YPD | KS.3 |
| FIM1_3321 | 1009.01119 | 2067.201004 | 2821.384153 | 0.849162129 | 0.315885701 | 0.002396455 | 0.00459613 | ns | YPD | KS.3 |
| FIM1_3322 | 430.8584776 | 818.853801 | 576.8316103 | 0.80281447 | 0.264125351 | 0.000919473 | 0.0019212 | ns | YPD | KS.3 |
| FIM1_3323 | 1642.489889 | 1119.192527 | 1336.77278 | -0.530769251 | 0.132913769 | 4.4010694166 | 0.000120304 | ns | YPD | KS.3 |
| FIM1_3324 | 1435.148708 | 2393.024206 | 2165.389234 | 0.696211351 | 0.162210819 | 8.9850577944( | 2.8471759571 | ns | YPD | KS.3 |
| FIM1_3326 | 9784.985018 | 12308.12706 | 10761.74779 | 0.307673458 | 0.175223455 | 0.068243008 | 0.095287389 | ns | YPD | KS.3 |
| FIM1_3327 | 1355.230977 | 1543.937001 | 1547.231501 | 0.177650725 | 0.153301962 | 0.232931263 | 0.287266493 | ns | YPD | KS.3 |
| FIM1_3328 | 347.9970502 | 278.4922462 | 250.7530417 | -0.296403271 | 0.183597857 | 0.092496174 | 0.125045779 | ns | YPD | KS.3 |
| FIM1_3329 | 400.9306927 | 192.6317845 | 333.4106703 | -1.006295135 | 0.157296933 | 3.6978828471( | 3.5243232115( | Km<KS.3 | YPD | KS.3 |
| FIM1_3331 | 496.4111265 | 267.5864972 | 427.7347597 | -0.864710455 | 0.120852426 | 2.8938009438( | 4.1608843050: | ns | YPD | KS.3 |
| FIM1_3332 | 535.4490489 | 450.0653738 | 563.365356 | -0.243207007 | 0.109074306 | 0.023588372 | 0.036746131 | ns | YPD | KS.3 |
| FIM1_3333 | 3198.027499 | 1866.218872 | 3974.130456 | -0.714810076 | 0.196841774 | 0.000136651 | 0.0003418 | ns | YPD | KS.3 |
| FIM1_3334 | 16891.81764 | 30451.82234 | 33685.66399 | 0.821377282 | 0.12609322 | 2.8255259508: | 2.7671243317 | ns | YPD | KS.3 |
| FIM1_3335 | 1313.180478 | 1113.380906 | 1422.542747 | -0.217953256 | 0.193075993 | 0.237581922 | 0.292349756 | ns | YPD | KS.3 |
| FIM1_3336 | 1324.99519 | 700.3880214 | 1146.475696 | -0.855890418 | 0.186889007 | 1.6514759605{ | 6.1961484946 | ns | YPD | KS.3 |
| FIM1_3337 | 1980.976744 | 2337.360853 | 2308.848679 | 0.234607897 | 0.07984653 | 0.003047764 | 0.00574342 | ns | YPD | KS.3 |
| FIM1_3339 | 770.1805646 | 1041.20726 | 1165.07265 | 0.416383111 | 0.134399773 | 0.001524301 | 0.003044318 | ns | YPD | KS.3 |
| FIM1_3340 | 3316.404312 | 2840.429499 | 5574.560663 | -0.202297387 | 0.204642043 | 0.298431443 | 0.358882385 | ns | YPD | KS.3 |
| FIM1_3341 | 1304.792669 | 1332.80459 | 1121.771156 | 0.030083175 | 0.094093503 | 0.746429295 | 0.787923986 | ns | YPD | KS.3 |
| FIM1_3342 | 236.697477 | 173.0887913 | 158.9391856 | -0.40131529 | 0.224623627 | 0.057490621 | 0.081555699 | ns | YPD | KS.3 |
| FIM1_3343 | 3623.109064 | 10088.28767 | 8696.620167 | 1.307313923 | 0.277969039 | 1.9378515003{ | 8.8124416950( | Km>KS.3 | YPD | KS.3 |
| FIM1_3344 | 7106.50707 | 22120.68807 | 18575.06962 | 1.38724393 | 0.340547349 | 2.4299162822( | 8.7600242782: | Km>KS.3 | YPD | KS.3 |
| FIM1_3345 | 2725.314825 | 4605.67736 | 6481.108677 | 0.607914728 | 0.318715308 | 0.031142372 | 0.047137604 | ns | YPD | KS.3 |
| FIM1_3346 | 144.9901625 | 253.1786099 | 215.6425215 | 0.71213419 | 0.238156072 | 0.001334392 | 0.002697521 | ns | YPD | KS.3 |
| FIM1_3347 | 1231.692693 | 3106.18536 | 1658.200111 | 1.250350696 | 0.194025832 | 1.1253402902: | 1.1911846689( | Km>KS.3 | YPD | KS.3 |
| FIM1_3348 | 1259.598322 | 1635.5189 | 957.1650824 | 0.343185423 | 0.199975798 | 0.071557955 | 0.099497156 | ns | YPD | KS.3 |
| FIM1_3349 | 1622.404985 | 2549.287247 | 1789.716452 | 0.632620556 | 0.114510474 | 1.8923816103 | 1.0483962707 | ns | YPD | KS.3 |
| FIM1_3350 | 6074.375109 | 10145.1955 | 9211.614553 | 0.67895962 | 0.198802533 | 0.000331018 | 0.000758551 | ns | YPD | KS.3 |
| FIM1_3351 | 1593.754153 | 2054.831344 | 1694.361127 | 0.354905439 | 0.114405146 | 0.001606973 | 0.003195319 | ns | YPD | KS.3 |
| FIM1_3353 | 6195.542918 | 5080.21682 | 8404.574634 | -0.272114064 | 0.145145052 | 0.054283414 | 0.077536602 | ns | YPD | KS.3 |
| FIM1_3355 | 1761.381692 | 1316.079006 | 1440.681107 | -0.384352918 | 0.196049811 | 0.039874272 | 0.058987363 | ns | YPD | KS.3 |
| FIM1_3356 | 1610.422535 | 2157.276744 | 1282.228097 | 0.382070807 | 0.206369705 | 0.051154881 | 0.073596164 | ns | YPD | KS.3 |
| FIM1_3357 | 455.9593858 | 365.663932 | 631.5619316 | -0.299248457 | 0.157143529 | 0.049593596 | 0.071639994 | ns | YPD | KS.3 |

| | | | | | | | | | |
|---|---|---|---|---|---|---|---|---|---|
| FIM1_3358 | 19.55083047 | 4.315878622 | 16.18493289 | -1.598863663 | 0.546220137 | 7.3712368354 | 0.000193826 | Km<KS.3 | YPD | KS.3 |
| FIM1_3359 | 6367.955313 | 5172.332727 | 5452.627362 | -0.293350102 | 0.094572404 | 0.001700905 | 0.003365951 | ns | YPD | KS.3 |
| FIM1_3360 | 3591.359239 | 3079.476534 | 3710.643881 | -0.21456166 | 0.115255603 | 0.058339402 | 0.082665487 | ns | YPD | KS.3 |
| FIM1_3361 | 6839.116681 | 6193.547135 | 6897.756809 | -0.14021317 | 0.087524981 | 0.105686125 | 0.141378993 | ns | YPD | KS.3 |
| FIM1_3362 | 2888.330594 | 5243.689001 | 3212.704601 | 0.800716264 | 0.184392137 | 5.7019954117 | 1.9115516963 | ns | YPD | KS.3 |
| FIM1_3363 | 1273.195981 | 1339.642741 | 1497.899917 | 0.071163773 | 0.112918767 | 0.521908442 | 0.584138245 | ns | YPD | KS.3 |
| FIM1_3364 | 2213.760084 | 2508.153454 | 1841.479941 | 0.176746708 | 0.089319757 | 0.045577236 | 0.066339014 | ns | YPD | KS.3 |
| FIM1_3365 | 515.4834312 | 273.1121293 | 341.3008944 | -0.854542161 | 0.182824352 | 1.0513714058 | 4.1185612156 | ns | YPD | KS.3 |
| FIM1_3366 | 319.2388519 | 1086.070007 | 426.0664148 | 1.388633418 | 0.42207095 | 5.0296326431 | 0.000135874 | Km>KS.3 | YPD | KS.3 |
| FIM1_3367 | 617.1820435 | 679.2930117 | 713.2190129 | 0.129158482 | 0.170338181 | 0.432140768 | 0.496283546 | ns | YPD | KS.3 |
| FIM1_3368 | 652.342406 | 346.2689071 | 621.4381627 | -0.859287983 | 0.171417341 | 1.8911670009 | 8.6164445004 | ns | YPD | KS.3 |
| FIM1_3369 | 1102.022255 | 948.1638634 | 1156.707981 | -0.197176541 | 0.200249582 | 0.301347498 | 0.361951666 | ns | YPD | KS.3 |
| FIM1_3370 | 797.3310644 | 494.9472928 | 841.2917541 | -0.632827038 | 0.194617615 | 0.000649173 | 0.001398715 | ns | YPD | KS.3 |
| FIM1_3371 | 915.6811844 | 426.6253814 | 821.7171968 | -1.040056323 | 0.17272928 | 3.6457026415 | 2.8428480629 | Km<KS.3 | YPD | KS.3 |
| FIM1_3372 | 850.3731945 | 821.6743622 | 785.7021713 | -0.047973048 | 0.107915992 | 0.651902199 | 0.704280877 | ns | YPD | KS.3 |
| FIM1_3373 | 580.3547414 | 346.0853652 | 864.5272455 | -0.600257153 | 0.317040189 | 0.032630118 | 0.049222207 | ns | YPD | KS.3 |
| FIM1_3374 | 2614.216823 | 7308.220198 | 5453.42696 | 1.249033217 | 0.328587209 | 1.3434237469 | 4.1154760721 | Km>KS.3 | YPD | KS.3 |
| FIM1_3375 | 5343.144053 | 7538.107997 | 8288.966713 | 0.477311537 | 0.12905832 | 0.000157837 | 0.000391109 | ns | YPD | KS.3 |
| FIM1_3376 | 22.58421834 | 27.36280704 | 19.77240213 | 0.216436974 | 0.316442759 | 0.441064834 | 0.504900495 | ns | YPD | KS.3 |
| FIM1_3377 | 868.2185469 | 1247.800947 | 1614.212457 | 0.444796989 | 0.267517045 | 0.069920424 | 0.097438126 | ns | YPD | KS.3 |
| FIM1_3378 | 25.97323892 | 20.43283175 | 24.81555604 | -0.304001301 | 0.244646764 | 0.181800346 | 0.231496474 | ns | YPD | KS.3 |
| FIM1_3379 | 426.7791618 | 533.5606987 | 602.1434 | 0.30761228 | 0.136141493 | 0.02074028 | 0.032704562 | ns | YPD | KS.3 |
| FIM1_3380 | 251.5668083 | 402.278301 | 384.328297 | 0.619294774 | 0.200557101 | 0.001165135 | 0.002385411 | ns | YPD | KS.3 |
| FIM1_3381 | 12975.41409 | 10168.42414 | 18287.95914 | -0.336879379 | 0.133382243 | 0.009795625 | 0.016630137 | ns | YPD | KS.3 |
| FIM1_3382 | 401.1893442 | 426.9922601 | 540.4338346 | 0.084168816 | 0.154655037 | 0.575189735 | 0.634385611 | ns | YPD | KS.3 |
| FIM1_3383 | 2363.662782 | 2901.872513 | 2471.836453 | 0.28499007 | 0.123622328 | 0.018759881 | 0.029980857 | ns | YPD | KS.3 |
| FIM1_3384 | 9257.003787 | 4964.427175 | 10102.61494 | -0.761699302 | 0.28316447 | 0.002998801 | 0.005661873 | ns | YPD | KS.3 |
| FIM1_3385 | 93.47584307 | 80.35184585 | 83.71763438 | -0.188805589 | 0.251463048 | 0.418117863 | 0.482294545 | ns | YPD | KS.3 |
| FIM1_3386 | 812.142757 | 190.2531321 | 715.7074906 | -1.976783401 | 0.24944486 | 7.3174854488 | 2.4432543696 | Km<KS.3 | YPD | KS.3 |
| FIM1_3387 | 972.712105 | 2956.576377 | 1783.26767 | 1.460616223 | 0.25412851 | 3.3663609620 | 2.6499439535 | Km>KS.3 | YPD | KS.3 |
| FIM1_3388 | 1672.427888 | 3367.838819 | 3075.192619 | 0.885677694 | 0.256798645 | 0.000179464 | 0.000437234 | ns | YPD | KS.3 |
| FIM1_3389 | 1458.548975 | 985.0391982 | 1286.693949 | -0.467262125 | 0.292337248 | 0.076547296 | 0.105784111 | ns | YPD | KS.3 |
| FIM1_3390 | 3248.564921 | 4742.660828 | 3612.230617 | 0.524447614 | 0.130666908 | 4.0774100042 | 0.000112321 | ns | YPD | KS.3 |
| FIM1_3391 | 1696.687508 | 1844.853423 | 1527.696093 | 0.10943074 | 0.202331239 | 0.569918934 | 0.629797134 | ns | YPD | KS.3 |
| FIM1_3392 | 1197.79706 | 906.1499074 | 997.793671 | -0.384120436 | 0.139327396 | 0.004723192 | 0.008532273 | ns | YPD | KS.3 |
| FIM1_3393 | 117.7484714 | 91.50899234 | 168.986438 | -0.279297884 | 0.334857609 | 0.340535411 | 0.402318611 | ns | YPD | KS.3 |
| FIM1_3394 | 1735.733649 | 3246.982797 | 2718.688271 | 0.835955628 | 0.193295065 | 5.6754428027 | 1.9045830395 | ns | YPD | KS.3 |
| FIM1_3395 | 753.6208033 | 2965.18344 | 2191.632266 | 1.879873954 | 0.219642058 | 4.6748201606 | 2.5841366998 | Km>KS.3 | YPD | KS.3 |
| FIM1_3396 | 708.1995869 | 868.4111349 | 1060.624303 | 0.259012309 | 0.232104966 | 0.233826699 | 0.288156509 | ns | YPD | KS.3 |
| FIM1_3397 | 5280.871965 | 10464.96539 | 7705.802668 | 0.82581456 | 0.297641927 | 0.001982722 | 0.003863706 | ns | YPD | KS.3 |
| FIM1_3398 | 10.36907047 | 5.086057503 | 8.902902147 | -0.730314767 | 0.414794174 | 0.030878608 | 0.046792896 | ns | YPD | KS.3 |
| FIM1_3399 | 762.2244296 | 1297.751842 | 1046.559827 | 0.709451907 | 0.190589748 | 9.6888029674 | 0.00025092 | ns | YPD | KS.3 |
| FIM1_3400 | 1915.962546 | 1865.319513 | 1585.761323 | -0.037621939 | 0.105738002 | 0.718199046 | 0.764254687 | ns | YPD | KS.3 |
| FIM1_3401 | 1080.217261 | 2150.591639 | 1416.926159 | 0.949704511 | 0.148236521 | 4.1132873888 | 3.8978294779 | ns | YPD | KS.3 |
| FIM1_3402 | 3140.179545 | 2808.404678 | 3669.48146 | -0.158532601 | 0.080014566 | 0.045728862 | 0.066540242 | ns | YPD | KS.3 |

| | | | | | | | | | | |
|---|---|---|---|---|---|---|---|---|---|---|
| FIM1_3403 | 3438.942748 | 3774.675235 | 4961.124225 | 0.114864977 | 0.257100802 | 0.628853284 | 0.683834992 | ns | YPD | KS.3 |
| FIM1_3404 | 57.36978831 | 35.88215942 | 107.336423 | -0.50986343 | 0.355501599 | 0.095381751 | 0.128692193 | ns | YPD | KS.3 |
| FIM1_3405 | 7673.563054 | 8003.755084 | 10136.20419 | 0.057252423 | 0.1553685 | 0.704280939 | 0.752533864 | ns | YPD | KS.3 |
| FIM1_3407 | 1344.213656 | 1184.115536 | 1399.676544 | -0.172835249 | 0.15430854 | 0.248766092 | 0.304680282 | ns | YPD | KS.3 |
| FIM1_3408 | 857.602158 | 1549.293788 | 1106.331544 | 0.832519691 | 0.105800176 | 1.33615990252 | 2.85295944852 | ns | YPD | KS.3 |
| FIM1_3409 | 5838.54216 | 7437.833248 | 9335.100371 | 0.323259024 | 0.180709413 | 0.06260587 | 0.088233486 | ns | YPD | KS.3 |
| FIM1_3410 | 1342.781919 | 1436.146289 | 1501.152017 | 0.086134263 | 0.222001058 | 0.680656201 | 0.731374643 | ns | YPD | KS.3 |
| FIM1_3411 | 1589.249369 | 1189.287587 | 1892.610748 | -0.370537799 | 0.229075627 | 0.084773995 | 0.115674883 | ns | YPD | KS.3 |
| FIM1_3412 | 1283.827984 | 1281.59493 | 1428.368822 | -0.002712978 | 0.08950828 | 0.975570285 | 0.980693731 | ns | YPD | KS.3 |
| FIM1_3413 | 2034.199596 | 2443.06009 | 2766.208354 | 0.242673027 | 0.188498972 | 0.178863048 | 0.228340689 | ns | YPD | KS.3 |
| FIM1_3414 | 1161.315818 | 1380.957057 | 2553.054635 | 0.230535049 | 0.182679096 | 0.18873404 | 0.238979854 | ns | YPD | KS.3 |
| FIM1_3415 | 324.6330811 | 1932.842586 | 1188.797963 | 2.356444442 | 0.369684694 | 5.07794203321 | 6.97863429621 | Km>KS.3 | YPD | KS.3 |
| FIM1_3416 | 250.6332876 | 835.9216255 | 460.4770577 | 1.589659763 | 0.26110466 | 2.20534746426 | 2.21201686183 | Km>KS.3 | YPD | KS.3 |
| FIM1_3417 | 10522.35397 | 9274.663728 | 14646.90247 | -0.175691831 | 0.120412012 | 0.137304961 | 0.178913615 | ns | YPD | KS.3 |
| FIM1_3418 | 2965.389219 | 1442.003187 | 2580.809561 | -1.007562236 | 0.125653287 | 2.46465799730 | 6.13083676829 | Km<KS.3 | YPD | KS.3 |
| FIM1_3419 | 3182.00722 | 3925.433217 | 3343.998621 | 0.297407173 | 0.085523995 | 0.00044654 | 0.000995313 | ns | YPD | KS.3 |
| FIM1_3420 | 91.96456872 | 162.0455521 | 103.0118483 | 0.64388245 | 0.335187346 | 0.027924752 | 0.042825414 | ns | YPD | KS.3 |
| FIM1_3421 | 114.4827804 | 128.1945594 | 127.9908184 | 0.130792851 | 0.308660111 | 0.635167157 | 0.689344809 | ns | YPD | KS.3 |
| FIM1_3422 | 219.442294 | 189.627954 | 142.3187291 | -0.15937217 | 0.342220391 | 0.592477933 | 0.650010924 | ns | YPD | KS.3 |
| FIM1_3423 | 354.6028505 | 1110.905618 | 604.0889833 | 1.43584206 | 0.311950688 | 1.77050488375 | 8.13565938674 | Km>KS.3 | YPD | KS.3 |
| FIM1_3424 | 324.1558159 | 1317.822214 | 844.8276442 | 1.929313345 | 0.219004989 | 4.25428562946 | 2.58110622031 | Km>KS.3 | YPD | KS.3 |
| FIM1_3425 | 2314.144957 | 1074.239555 | 1240.169645 | -1.002075997 | 0.227960691 | 2.58641919972 | 9.25049282430 | Km<KS.3 | YPD | KS.3 |
| FIM1_3426 | 2244.963806 | 8884.936535 | 4545.849937 | 1.576400197 | 0.450295533 | 1.07960930975 | 3.36954599500 | Km>KS.3 | YPD | KS.3 |
| FIM1_3427 | 708.1803581 | 3149.910135 | 1391.919756 | 2.066433472 | 0.217755631 | 5.51570837789 | 4.73114641034 | Km>KS.3 | YPD | KS.3 |
| FIM1_3428 | 38.72308485 | 134.6849527 | 84.78845287 | 1.629787946 | 0.281445046 | 1.13105133215 | 9.82021008284 | Km>KS.3 | YPD | KS.3 |
| FIM1_3429 | 3228.202618 | 2453.051331 | 3127.203162 | -0.383711503 | 0.11522365 | 0.000705538 | 0.001507754 | ns | YPD | KS.3 |
| FIM1_3430 | 1592.251238 | 1347.054148 | 1770.705665 | -0.235279479 | 0.102493244 | 0.019985104 | 0.031724917 | ns | YPD | KS.3 |
| FIM1_3431 | 191.5553069 | 327.3204839 | 368.9373469 | 0.722491611 | 0.176278321 | 1.99598058727 | 5.92129005467 | ns | YPD | KS.3 |
| FIM1_3432 | 908.8780041 | 830.5050699 | 922.3058643 | -0.121629925 | 0.166819142 | 0.450674408 | 0.514126389 | ns | YPD | KS.3 |
| FIM1_3433 | 3598.435725 | 4998.451283 | 3764.933484 | 0.466725116 | 0.078902768 | 2.44605078434 | 1.62254702027 | ns | YPD | KS.3 |
| FIM1_3434 | 941.707965 | 878.6493623 | 717.5488816 | -0.097312788 | 0.088274292 | 0.265493968 | 0.322941929 | ns | YPD | KS.3 |
| FIM1_3435 | 3950.334365 | 4084.992185 | 3127.965965 | 0.045071551 | 0.16931045 | 0.782814755 | 0.818688965 | ns | YPD | KS.3 |
| FIM1_3437 | 13876.07369 | 16326.05492 | 16824.47536 | 0.216583072 | 0.182545114 | 0.216613025 | 0.269547224 | ns | YPD | KS.3 |
| FIM1_3438 | 1170.477141 | 2461.852939 | 2603.795388 | 0.989587268 | 0.202881714 | 2.63525909769 | 1.16124127644 | ns | YPD | KS.3 |
| FIM1_3439 | 1651.843588 | 3497.926985 | 1906.752553 | 0.949998051 | 0.259736263 | 6.82004749602 | 0.000180381 | ns | YPD | KS.3 |
| FIM1_3440 | 2763.940879 | 3657.383504 | 3000.803043 | 0.384730382 | 0.143524633 | 0.005941515 | 0.010529947 | ns | YPD | KS.3 |
| FIM1_3441 | 1432.091594 | 973.7157878 | 987.7667858 | -0.529085571 | 0.147284121 | 0.000221519 | 0.000526545 | ns | YPD | KS.3 |
| FIM1_3442 | 4315.027102 | 6347.895541 | 5052.080606 | 0.528022992 | 0.151711567 | 0.000338533 | 0.00077399 | ns | YPD | KS.3 |
| FIM1_3444 | 3046.167133 | 3173.32888 | 4131.371059 | 0.055959582 | 0.142682164 | 0.687564168 | 0.737841186 | ns | YPD | KS.3 |
| FIM1_3445 | 2807.477039 | 2533.456083 | 3482.366193 | -0.136452998 | 0.185662651 | 0.443502411 | 0.507224022 | ns | YPD | KS.3 |
| FIM1_3446 | 1435.466647 | 1528.802876 | 1375.250604 | 0.089492261 | 0.070117392 | 0.198933729 | 0.249986184 | ns | YPD | KS.3 |
| FIM1_3447 | 3083.662377 | 2092.691855 | 2612.233532 | -0.542061907 | 0.115269736 | 1.70883755114 | 6.37291365588 | ns | YPD | KS.3 |
| FIM1_3448 | 969.3628686 | 1080.433833 | 773.1970386 | 0.153670293 | 0.089820661 | 0.083857764 | 0.114581811 | ns | YPD | KS.3 |
| FIM1_3449 | 1160.514351 | 704.7511196 | 936.3167794 | -0.698651165 | 0.112336751 | 2.52357344354 | 2.03151745657 | ns | YPD | KS.3 |
| FIM1_3450 | 3105.073959 | 2530.137591 | 4650.317217 | -0.247955835 | 0.273773131 | 0.321994049 | 0.383417998 | ns | YPD | KS.3 |

| | | | | | | | | | | |
|---|---|---|---|---|---|---|---|---|---|---|
| FIM1_3452 | 1985.149057 | 1912.196888 | 1780.430393 | -0.052027566 | 0.121238238 | 0.662048204 | 0.714001694 | ns | YPD | KS.3 |
| FIM1_3454 | 9674.618771 | 21451.3118 | 12413.85407 | 1.035257159 | 0.235062667 | 2.2403759004( | 8.1297374941{ | Km>KS.3 | YPD | KS.3 |
| FIM1_3455 | 338.0753601 | 1238.167037 | 1212.410285 | 1.697021183 | 0.291420522 | 6.7587292994 | 6.1471075438 | Km>KS.3 | YPD | KS.3 |
| FIM1_3456 | 26.11081151 | 142.1955944 | 220.5758679 | 2.221882551 | 0.366896552 | 3.8657559639( | 4.5040130961 | Km>KS.3 | YPD | KS.3 |
| FIM1_3457 | 189.1876517 | 1784.799789 | 898.4169684 | 3.065629501 | 0.32631287 | 9.6965717706 | 6.8914920798 | Km>KS.3 | YPD | KS.3 |
| FIM1_3458 | 6996.623319 | 20775.73094 | 8765.855031 | 1.467691137 | 0.213414957 | 2.1235501114 | 3.0981412916 | Km>KS.3 | YPD | KS.3 |
| FIM1_3460 | 4604.919433 | 6948.146174 | 6233.295187 | 0.567916617 | 0.13789209 | 2.4333073716 | 7.0669609888 | ns | YPD | KS.3 |
| FIM1_3461 | 653.9228184 | 1596.774253 | 1201.563314 | 1.044708311 | 0.342387528 | 0.000429073 | 0.000961982 | Km>KS.3 | YPD | KS.3 |
| FIM1_3462 | 839.2173718 | 983.9921135 | 799.4370233 | 0.223375593 | 0.105457324 | 0.031681041 | 0.047921308 | ns | YPD | KS.3 |
| FIM1_3463 | 872.6210857 | 7253.78295 | 3452.321499 | 2.913989305 | 0.29997199 | 3.6331781335 | 3.1710633709 | Km>KS.3 | YPD | KS.3 |
| FIM1_3464 | 267.694297 | 269.0280004 | 332.051609 | 0.006065111 | 0.227080115 | 0.977338097 | 0.98187743 | ns | YPD | KS.3 |
| FIM1_3465 | 218.9587686 | 278.0373602 | 228.7605552 | 0.282695304 | 0.292050679 | 0.283936092 | 0.342942962 | ns | YPD | KS.3 |
| FIM1_3466 | 2246.162185 | 3390.964495 | 2895.335907 | 0.585599915 | 0.076084702 | 8.6120801807( | 1.6415746704( | ns | YPD | KS.3 |
| FIM1_3467 | 243.4170007 | 1723.87251 | 819.1413989 | 2.529297281 | 0.438019956 | 2.4209685863 | 2.3944967628 | Km>KS.3 | YPD | KS.3 |
| FIM1_3468 | 1543.739003 | 1785.166521 | 2493.683692 | 0.197301045 | 0.156939226 | 0.19502714 | 0.246071524 | ns | YPD | KS.3 |
| FIM1_3469 | 1031.595596 | 3878.809435 | 2023.438997 | 1.720464621 | 0.30534176 | 1.8816742289( | 1.5706928336 | Km>KS.3 | YPD | KS.3 |
| FIM1_3470 | 472.9611522 | 1179.131157 | 741.3694133 | 1.175724455 | 0.256133381 | 5.7131529725( | 2.3451267358 | Km>KS.3 | YPD | KS.3 |
| FIM1_3471 | 1999.457367 | 1560.568419 | 2447.582828 | -0.343904787 | 0.127625289 | 0.005944611 | 0.010529947 | ns | YPD | KS.3 |
| FIM1_3472 | 790.0634717 | 798.832884 | 647.6004953 | 0.015169462 | 0.131548641 | 0.906213589 | 0.923665767 | ns | YPD | KS.3 |
| FIM1_3473 | 1880.848901 | 2384.58539 | 3333.603423 | 0.271920669 | 0.314508526 | 0.330770368 | 0.392459476 | ns | YPD | KS.3 |
| FIM1_3474 | 251.7876096 | 1912.929182 | 705.3237644 | 2.736381803 | 0.350283847 | 3.5405120873( | 1.0675180384 | Km>KS.3 | YPD | KS.3 |
| FIM1_3475 | 3216.07658 | 3795.106832 | 3795.78794 | 0.232628512 | 0.101479342 | 0.020189721 | 0.031998682 | ns | YPD | KS.3 |
| FIM1_3476 | 1753.515831 | 1876.105724 | 1593.52545 | 0.095348943 | 0.09793656 | 0.324352938 | 0.385857453 | ns | YPD | KS.3 |
| FIM1_3477 | 1841.60607 | 1856.346203 | 1971.95525 | 0.010748538 | 0.149716423 | 0.941173503 | 0.952662905 | ns | YPD | KS.3 |
| FIM1_3478 | 139.3696842 | 197.9015474 | 204.791761 | 0.457931539 | 0.206668965 | 0.019536802 | 0.031112545 | ns | YPD | KS.3 |
| FIM1_3479 | 6021.356204 | 5793.73393 | 7511.9524 | -0.055144865 | 0.056895611 | 0.330334009 | 0.392128775 | ns | YPD | KS.3 |
| FIM1_3480 | 2783.571848 | 1177.5348 | 3057.765141 | -0.998430514 | 0.345234679 | 0.000827025 | 0.001744149 | ns | YPD | KS.3 |
| FIM1_3481 | 3255.186737 | 3956.412611 | 3386.072532 | 0.275874639 | 0.087903913 | 0.001523744 | 0.003044318 | ns | YPD | KS.3 |
| FIM1_3482 | 2441.954163 | 1676.56601 | 3183.553039 | -0.50181636 | 0.18445905 | 0.0045361 | 0.008227159 | ns | YPD | KS.3 |
| FIM1_3483 | 1429.019516 | 807.3864864 | 933.8487455 | -0.805707996 | 0.099348759 | 2.0122024692 | 5.1075037165 | ns | YPD | KS.3 |
| FIM1_3484 | 1773.472009 | 1602.15932 | 1506.444146 | -0.14358301 | 0.082884756 | 0.080494213 | 0.110532352 | ns | YPD | KS.3 |
| FIM1_3485 | 515.0029168 | 271.1450466 | 508.4485735 | -0.877267283 | 0.16316298 | 2.5585385131 | 1.3731099355 | ns | YPD | KS.3 |
| FIM1_3486 | 3566.005657 | 2774.832523 | 3483.811659 | -0.342699761 | 0.150855488 | 0.01943171 | 0.030965008 | ns | YPD | KS.3 |
| FIM1_3487 | 1039.30712 | 599.8552819 | 881.4953144 | -0.734124995 | 0.189851409 | 5.1503550854( | 0.000138728 | ns | YPD | KS.3 |
| FIM1_3488 | 468.8063104 | 538.9553553 | 422.2680436 | 0.195228406 | 0.107648989 | 0.065701444 | 0.092126462 | ns | YPD | KS.3 |
| FIM1_3489 | 938.3964749 | 661.4068922 | 767.3091327 | -0.49300203 | 0.10107658 | 7.6665179547 | 3.0773820689 | ns | YPD | KS.3 |
| FIM1_3490 | 1137.796337 | 970.0301308 | 1105.079606 | -0.224225693 | 0.107647581 | 0.034522938 | 0.051794818 | ns | YPD | KS.3 |
| FIM1_3491 | 2596.705033 | 2092.579839 | 2077.042796 | -0.29862025 | 0.130051822 | 0.01899715 | 0.030321085 | ns | YPD | KS.3 |
| FIM1_3492 | 886.4036748 | 859.3811856 | 782.9234932 | -0.041209152 | 0.185253986 | 0.816639287 | 0.846896498 | ns | YPD | KS.3 |
| FIM1_3493 | 35.54550221 | 199.2199309 | 111.3256679 | 2.333264707 | 0.304578084 | 4.9654221653 | 1.4617145131( | Km>KS.3 | YPD | KS.3 |
| FIM1_3494 | 7067.470109 | 9530.867488 | 9553.046138 | 0.419635359 | 0.106506036 | 6.3736897135 | 0.000169386 | ns | YPD | KS.3 |
| FIM1_3495 | 163.7880318 | 450.5319273 | 257.5342315 | 1.396451396 | 0.166451675 | 2.3353057315( | 8.4803985505 | Km>KS.3 | YPD | KS.3 |
| FIM1_3496 | 528.7801612 | 491.2103485 | 619.1826244 | -0.09919054 | 0.176423424 | 0.559297533 | 0.619987796 | ns | YPD | KS.3 |
| FIM1_3497 | 1433.466927 | 2880.875035 | 1782.456726 | 0.814125332 | 0.325819176 | 0.004482323 | 0.00814447 | ns | YPD | KS.3 |
| FIM1_3498 | 140.5275184 | 110.077283 | 150.7250461 | -0.311567869 | 0.231881449 | 0.151684294 | 0.196058552 | ns | YPD | KS.3 |

| | | | | | | | | | | |
|---|---|---|---|---|---|---|---|---|---|---|
| FIM1_3499 | 2324.720266 | 3159.807664 | 2687.064173 | 0.435014065 | 0.08427237 | 1.87734529502 | 8.56861728692 | ns | YPD | KS.3 |
| FIM1_3500 | 1481.059256 | 1716.445195 | 1777.103672 | 0.194694072 | 0.192628288 | 0.290422257 | 0.350351777 | ns | YPD | KS.3 |
| FIM1_3501 | 2022.149005 | 4069.199091 | 2906.741389 | 0.88895876 | 0.25207234 | 0.000133485 | 0.000334891 | ns | YPD | KS.3 |
| FIM1_3503 | 191.9587717 | 753.2036394 | 375.0021449 | 1.529928327 | 0.466511558 | 2.92059475945 | 8.33139846806 | Km>KS.3 | YPD | KS.3 |
| FIM1_3504 | 311.7291338 | 312.4294834 | 551.5915906 | 0.002456645 | 0.293645158 | 0.992606309 | 0.993005508 | ns | YPD | KS.3 |
| FIM1_3505 | 2406.874789 | 2780.775566 | 3680.734522 | 0.202661475 | 0.102375763 | 0.044829681 | 0.065461598 | ns | YPD | KS.3 |
| FIM1_3506 | 716.329148 | 731.1198102 | 702.3294603 | 0.027084663 | 0.181767404 | 0.876753452 | 0.898323118 | ns | YPD | KS.3 |
| FIM1_3507 | 1590.135926 | 1277.552016 | 1893.918144 | -0.30806439 | 0.100469131 | 0.00189338 | 0.00370995 | ns | YPD | KS.3 |
| FIM1_3508 | 1051.591776 | 769.071512 | 926.6437892 | -0.435149626 | 0.123245124 | 0.000317861 | 0.000730082 | ns | YPD | KS.3 |
| FIM1_3509 | 23.81652793 | 34.46312932 | 75.09163503 | 0.47223117 | 0.239629256 | 0.034585103 | 0.051860207 | ns | YPD | KS.3 |
| FIM1_3511 | 3813.106324 | 3904.213173 | 4938.838614 | 0.032555748 | 0.134784294 | 0.804883047 | 0.836493244 | ns | YPD | KS.3 |
| FIM1_3512 | 3219.121343 | 5928.165814 | 4645.559879 | 0.816085562 | 0.1909912 | 7.52275123460 | 2.43656818959 | ns | YPD | KS.3 |
| FIM1_3513 | 749.9331247 | 829.8550808 | 986.2109913 | 0.105327927 | 0.371125506 | 0.73804367 | 0.780231037 | ns | YPD | KS.3 |
| FIM1_3514 | 1400.901834 | 4142.69326 | 2916.016212 | 1.426227412 | 0.248944428 | 4.43579280618 | 3.40031883062 | Km>KS.3 | YPD | KS.3 |
| FIM1_3516 | 3060.484347 | 8409.863555 | 7410.048664 | 1.244886753 | 0.313433904 | 6.83720371822 | 2.24226028333 | Km>KS.3 | YPD | KS.3 |
| FIM1_3517 | 307.6308123 | 489.4105542 | 512.655349 | 0.648389865 | 0.115773694 | 1.19013060193 | 6.86083400302 | ns | YPD | KS.3 |
| FIM1_3518 | 958.34499 | 2651.625398 | 3047.35086 | 1.182408471 | 0.364990149 | 0.000138699 | 0.000346574 | Km>KS.3 | YPD | KS.3 |
| FIM1_3519 | 370.5070103 | 796.1890494 | 757.2062454 | 0.978247194 | 0.250197953 | 2.29542317822 | 6.69386301973 | ns | YPD | KS.3 |
| FIM1_3520 | 256.1391221 | 2148.594045 | 1141.632281 | 2.960632554 | 0.259720129 | 8.18661861839 | 1.31382024601 | Km>KS.3 | YPD | KS.3 |
| FIM1_3522 | 1804.750679 | 2367.385377 | 3616.222836 | 0.30240438 | 0.333729844 | 0.300870543 | 0.36155337 | ns | YPD | KS.3 |
| FIM1_3523 | 4637.487 | 9043.961976 | 7519.610076 | 0.898843711 | 0.184575223 | 3.55126925960 | 1.51652914734 | ns | YPD | KS.3 |
| FIM1_3524 | 1989.344696 | 1381.660243 | 1888.769054 | -0.492621854 | 0.16862353 | 0.002465614 | 0.004721489 | ns | YPD | KS.3 |
| FIM1_3525 | 557.1639112 | 567.1651989 | 478.5321531 | 0.022229296 | 0.239381473 | 0.920799978 | 0.935796952 | ns | YPD | KS.3 |
| FIM1_3526 | 2466.573813 | 2535.324368 | 3162.86158 | 0.038670908 | 0.093490877 | 0.675743012 | 0.72656613 | ns | YPD | KS.3 |
| FIM1_3527 | 830.2791071 | 1748.945663 | 1778.7921 | 0.963983188 | 0.236491656 | 1.19484994030 | 3.69216052983 | ns | YPD | KS.3 |
| FIM1_3528 | 1188.760268 | 4814.936384 | 3188.141061 | 1.760302953 | 0.362552552 | 1.18836240432 | 6.85858812239 | Km>KS.3 | YPD | KS.3 |
| FIM1_3529 | 866.1187194 | 500.7956876 | 616.9667557 | -0.726471947 | 0.19783069 | 0.00011382 | 0.000290983 | ns | YPD | KS.3 |
| FIM1_3530 | 606.476688 | 278.1519727 | 406.0561677 | -1.052549875 | 0.18638415 | 3.29986570961 | 2.11557112181 | Km<KS.3 | YPD | KS.3 |
| FIM1_3531 | 2245.452776 | 1408.263164 | 1385.990734 | -0.662086805 | 0.082841183 | 7.13042932976 | 1.61981214226 | ns | YPD | KS.3 |
| FIM1_3532 | 1407.070516 | 712.9991503 | 977.3404401 | -0.940324532 | 0.14287559 | 1.32232271198 | 1.37915209478 | ns | YPD | KS.3 |
| FIM1_3533 | 582.0488574 | 637.4728982 | 652.0218671 | 0.124801857 | 0.145936041 | 0.379981062 | 0.442821688 | ns | YPD | KS.3 |
| FIM1_3534 | 2663.072509 | 2722.910499 | 2703.360744 | 0.031726219 | 0.083439736 | 0.701228292 | 0.74959406 | ns | YPD | KS.3 |
| FIM1_3535 | 1725.560099 | 1305.03484 | 1527.632879 | -0.397696253 | 0.073944747 | 6.02863570196 | 3.03567435397 | ns | YPD | KS.3 |
| FIM1_3536 | 694.8309324 | 334.815763 | 460.5971985 | -1.017274834 | 0.131968052 | 2.85421539827 | 5.86765355637 | Km<KS.3 | YPD | KS.3 |
| FIM1_3537 | 7128.158642 | 4385.343327 | 8084.007475 | -0.598304444 | 0.269253611 | 0.015224736 | 0.024841935 | ns | YPD | KS.3 |
| FIM1_3538 | 1271.920234 | 925.3428985 | 903.5957502 | -0.443577969 | 0.117119968 | 0.000115695 | 0.00029502 | ns | YPD | KS.3 |
| FIM1_3539 | 614.1065683 | 408.310892 | 579.1258978 | -0.563714749 | 0.137453221 | 2.64137188417 | 7.60024587840 | ns | YPD | KS.3 |
| FIM1_3540 | 2543.009431 | 2849.307342 | 3361.474688 | 0.160411693 | 0.094440831 | 0.08578137 | 0.116921183 | ns | YPD | KS.3 |
| FIM1_3541 | 211.5264275 | 134.2867542 | 306.9328249 | -0.580981618 | 0.23149842 | 0.007346736 | 0.012829067 | ns | YPD | KS.3 |
| FIM1_3542 | 561.4195816 | 483.1469664 | 450.4154753 | -0.208705329 | 0.111376506 | 0.056966779 | 0.080904861 | ns | YPD | KS.3 |
| FIM1_3544 | 2687.324359 | 2491.479737 | 2705.31606 | -0.101826079 | 0.169208443 | 0.533196508 | 0.595165499 | ns | YPD | KS.3 |
| FIM1_3545 | 112.7216643 | 257.2890087 | 154.6222435 | 1.020677138 | 0.288647021 | 8.66034470011 | 0.00022605 | Km>KS.3 | YPD | KS.3 |
| FIM1_3546 | 2500.366225 | 1815.564973 | 1688.710571 | -0.435210839 | 0.15857884 | 0.004620567 | 0.008362066 | ns | YPD | KS.3 |
| FIM1_3547 | 969.8268458 | 422.7587211 | 926.7482602 | -1.16003542 | 0.130703047 | 9.41890778248 | 4.83083156885 | Km<KS.3 | YPD | KS.3 |
| FIM1_3548 | 454.5194161 | 731.6832195 | 437.2404573 | 0.564277622 | 0.299511364 | 0.035812893 | 0.053508036 | ns | YPD | KS.3 |

| | | | | | | | | | | |
|---|---|---|---|---|---|---|---|---|---|---|
| FIM1_3549 | 123.3392004 | 165.4598147 | 171.1134703 | 0.39439417 | 0.168338606 | 0.01523264 | 0.024846683 | ns | YPD | KS.3 |
| FIM1_3550 | 1192.012261 | 839.2635281 | 1459.170544 | -0.453810785 | 0.218706546 | 0.02781257 | 0.042679684 | ns | YPD | KS.3 |
| FIM1_3551 | 1166.31837 | 1999.101203 | 1689.147242 | 0.714871487 | 0.196837699 | 0.00013643 | 0.000341418 | ns | YPD | KS.3 |
| FIM1_3552 | 611.8619078 | 452.7679962 | 580.9714058 | -0.412833014 | 0.146968431 | 0.003895989 | 0.007162802 | ns | YPD | KS.3 |
| FIM1_3553 | 227.4081958 | 579.6629075 | 438.2540512 | 1.218768228 | 0.243964575 | 6.44680792492 | 3.22664682358 | Km>KS.3 | YPD | KS.3 |
| FIM1_3554 | 691.5304672 | 374.2415133 | 602.7025135 | -0.841608961 | 0.156675628 | 2.87715161309 | 1.53417248394 | ns | YPD | KS.3 |
| FIM1_3555 | 829.817082 | 560.8009773 | 575.0273797 | -0.536817889 | 0.149220766 | 0.000214635 | 0.000511158 | ns | YPD | KS.3 |
| FIM1_3556 | 1943.628766 | 2526.879979 | 2232.991527 | 0.36810439 | 0.106217064 | 0.000437316 | 0.000977379 | ns | YPD | KS.3 |
| FIM1_3557 | 3696.765353 | 2469.972793 | 4774.948837 | -0.523257643 | 0.217050554 | 0.010629089 | 0.017931406 | ns | YPD | KS.3 |
| FIM1_3558 | 1362.668222 | 1243.507576 | 1198.38517 | -0.125141133 | 0.145379184 | 0.376955132 | 0.439913624 | ns | YPD | KS.3 |
| FIM1_3559 | 199.5441346 | 228.1922651 | 243.8344568 | 0.180530818 | 0.157233134 | 0.236577615 | 0.291258014 | ns | YPD | KS.3 |
| FIM1_3560 | 1242.96914 | 1367.025413 | 1344.499968 | 0.133234998 | 0.110039502 | 0.218901927 | 0.271987284 | ns | YPD | KS.3 |
| FIM1_3561 | 1064.048462 | 1582.280256 | 872.0920139 | 0.51586558 | 0.215027005 | 0.011099322 | 0.018623652 | ns | YPD | KS.3 |
| FIM1_3562 | 2097.428861 | 2111.045596 | 1735.131374 | 0.009218665 | 0.108623963 | 0.931355613 | 0.943877404 | ns | YPD | KS.3 |
| FIM1_3563 | 2460.946798 | 1603.01559 | 1925.341098 | -0.599177344 | 0.115803825 | 1.39085723550 | 6.53400825931 | ns | YPD | KS.3 |
| FIM1_3565 | 110.1378796 | 278.7705799 | 110.8741976 | 1.112122315 | 0.32796335 | 0.000106882 | 0.000274232 | Km>KS.3 | YPD | KS.3 |
| FIM1_3566 | 379.8180711 | 239.0891592 | 242.8917698 | -0.603205801 | 0.215010679 | 0.002966939 | 0.0056081 | ns | YPD | KS.3 |
| FIM1_3567 | 6740.567535 | 6427.868636 | 7172.635201 | -0.064113046 | 0.1656159 | 0.688951675 | 0.739011337 | ns | YPD | KS.3 |
| FIM1_3568 | 3317.56863 | 6564.357065 | 4147.484486 | 0.892426558 | 0.22088082 | 1.70569747657 | 5.11812119780 | ns | YPD | KS.3 |
| FIM1_3569 | 691.570946 | 1074.664552 | 2003.273747 | 0.466827524 | 0.371327992 | 0.138399903 | 0.180104503 | ns | YPD | KS.3 |
| FIM1_3570 | 1049.037079 | 837.7619046 | 1301.032634 | -0.289761336 | 0.220203635 | 0.162949636 | 0.209747591 | ns | YPD | KS.3 |
| FIM1_3571 | 3536.180292 | 3636.215162 | 4121.434755 | 0.039010423 | 0.101486158 | 0.696945805 | 0.746138451 | ns | YPD | KS.3 |
| FIM1_3572 | 584.0150329 | 461.5158614 | 519.58149 | -0.327026067 | 0.126300531 | 0.008239768 | 0.014213886 | ns | YPD | KS.3 |
| FIM1_3573 | 4383.563333 | 3658.191195 | 4375.909811 | -0.254453183 | 0.101262689 | 0.010893899 | 0.018328424 | ns | YPD | KS.3 |
| FIM1_3574 | 1159.418259 | 7748.389748 | 3146.18648 | 2.480730431 | 0.40989341 | 4.35900750901 | 5.01992184198 | Km>KS.3 | YPD | KS.3 |
| FIM1_3575 | 1647.704188 | 2288.802736 | 1718.833637 | 0.419162618 | 0.231815715 | 0.053599599 | 0.076714041 | ns | YPD | KS.3 |
| FIM1_3576 | 3378.964544 | 3309.974141 | 3522.498511 | -0.028645406 | 0.124544581 | 0.814605107 | 0.845184652 | ns | YPD | KS.3 |
| FIM1_3577 | 558.5883387 | 403.9442473 | 487.2915457 | -0.446186772 | 0.139986482 | 0.001084354 | 0.002233101 | ns | YPD | KS.3 |
| FIM1_3579 | 1663.012626 | 3211.242204 | 2293.963276 | 0.881881334 | 0.189488822 | 1.08047995150 | 4.22593377258 | ns | YPD | KS.3 |
| FIM1_3580 | 4270.087169 | 2253.762247 | 5248.873732 | -0.768375775 | 0.297915044 | 0.00404967 | 0.007423401 | ns | YPD | KS.3 |
| FIM1_3581 | 6472.998032 | 2818.300267 | 4370.134313 | -1.087361698 | 0.230805918 | 4.39594146533 | 1.84711222888 | Km<KS.3 | YPD | KS.3 |
| FIM1_3582 | 2350.716979 | 2740.30653 | 2298.735857 | 0.216252876 | 0.093984075 | 0.019950332 | 0.031689944 | ns | YPD | KS.3 |
| FIM1_3583 | 1259.754972 | 3401.701884 | 1973.623488 | 1.285294271 | 0.258717365 | 5.64778289999 | 2.87004289350 | Km>KS.3 | YPD | KS.3 |
| FIM1_3584 | 494.7481017 | 468.0187992 | 589.4158153 | -0.075301385 | 0.167328055 | 0.641476698 | 0.695585565 | ns | YPD | KS.3 |
| FIM1_3585 | 1445.778041 | 1272.797181 | 1187.196258 | -0.168122126 | 0.193943873 | 0.364297575 | 0.426844191 | ns | YPD | KS.3 |
| FIM1_3586 | 1378.903845 | 4491.845319 | 2909.673499 | 1.614961937 | 0.200229413 | 1.10429288141 | 3.59075626474 | Km>KS.3 | YPD | KS.3 |
| FIM1_3587 | 51.33249919 | 178.6173008 | 85.38572007 | 1.57899464 | 0.322154939 | 2.06355183508 | 1.13063550435 | Km>KS.3 | YPD | KS.3 |
| FIM1_3588 | 606.0605759 | 1138.135003 | 1024.528335 | 0.77960274 | 0.27303025 | 0.001747033 | 0.003447636 | ns | YPD | KS.3 |
| FIM1_3589 | 1892.96571 | 6138.588115 | 3873.387259 | 1.597933494 | 0.212006245 | 8.20335037404 | 1.84668181497 | Km>KS.3 | YPD | KS.3 |
| FIM1_3591 | 665.5146327 | 683.470083 | 629.3839924 | 0.033243718 | 0.247966452 | 0.885305464 | 0.905508775 | ns | YPD | KS.3 |
| FIM1_3592 | 890.4884073 | 173.32326 | 1972.23227 | -2.140061632 | 0.36062748 | 8.84099809534 | 9.54099035235 | Km<KS.3 | YPD | KS.3 |
| FIM1_3594 | 1951.395811 | 3603.325246 | 3172.010947 | 0.858798984 | 0.116979191 | 7.41282988409 | 1.19348960107 | ns | YPD | KS.3 |
| FIM1_3595 | 3169.948825 | 5193.332312 | 5378.255139 | 0.672964817 | 0.159620671 | 1.31994972048 | 4.04855108472 | ns | YPD | KS.3 |
| FIM1_3596 | 22.81796884 | 55.84026335 | 37.01878554 | 1.012002265 | 0.366908728 | 0.001176824 | 0.002405382 | Km>KS.3 | YPD | KS.3 |
| FIM1_3597 | 6730.5912 | 6556.494562 | 10687.94615 | -0.036687012 | 0.112240141 | 0.739828955 | 0.78195221 | ns | YPD | KS.3 |

| | | | | | | | | | | |
|---|---|---|---|---|---|---|---|---|---|---|
| FIM1_3598 | 1592.652151 | 1836.220378 | 2566.044627 | 0.183076518 | 0.219159189 | 0.376170111 | 0.439203545 | ns | YPD | KS.3 |
| FIM1_3599 | 5225.097728 | 6398.743818 | 6212.493068 | 0.282838004 | 0.115623871 | 0.012842675 | 0.021240793 | ns | YPD | KS.3 |
| FIM1_3600 | 485.7725975 | 288.6025188 | 386.4645009 | -0.695979901 | 0.187380229 | 0.000102848 | 0.000264976 | ns | YPD | KS.3 |
| FIM1_3601 | 2726.65921 | 2995.091624 | 2242.782252 | 0.129311818 | 0.137389361 | 0.335307231 | 0.396990356 | ns | YPD | KS.3 |
| FIM1_3603 | 844.2763482 | 390.1774191 | 629.563039 | -1.078841872 | 0.124143451 | 6.56376070595 | 2.72122579267 | Km<KS.3 | YPD | KS.3 |
| FIM1_3604 | 3532.885517 | 2398.526135 | 3903.096472 | -0.5222019 | 0.171893176 | 0.001616856 | 0.003212404 | ns | YPD | KS.3 |
| FIM1_3605 | 1387.193133 | 1362.956002 | 1340.176429 | -0.025218187 | 0.066255702 | 0.701845544 | 0.750092713 | ns | YPD | KS.3 |
| FIM1_3606 | 885.6293535 | 564.2908842 | 573.6389977 | -0.59117819 | 0.206419379 | 0.002528515 | 0.004827076 | ns | YPD | KS.3 |
| FIM1_3607 | 2125.217907 | 1083.932824 | 1506.810348 | -0.936110518 | 0.133505253 | 6.73985669974 | 9.11162692424 | ns | YPD | KS.3 |
| FIM1_3608 | 542.1151455 | 635.3697989 | 542.6673427 | 0.218418322 | 0.139381285 | 0.108410712 | 0.144557302 | ns | YPD | KS.3 |
| FIM1_3609 | 889.5075465 | 1616.65375 | 1486.044669 | 0.752076063 | 0.255548594 | 0.001418715 | 0.002854066 | ns | YPD | KS.3 |
| FIM1_3610 | 2226.834469 | 1313.885468 | 1668.484706 | -0.738158337 | 0.117645216 | 1.63488102453 | 1.38325392807 | ns | YPD | KS.3 |
| FIM1_3611 | 1066.419543 | 961.8462321 | 868.238066 | -0.1450819 | 0.092301965 | 0.112035685 | 0.148872204 | ns | YPD | KS.3 |
| FIM1_3612 | 2030.40434 | 1749.651248 | 1620.530005 | -0.209272951 | 0.097847006 | 0.030372236 | 0.046154072 | ns | YPD | KS.3 |
| FIM1_3613 | 732.8762671 | 1046.015457 | 1052.78488 | 0.465621272 | 0.206113437 | 0.01730414 | 0.027950681 | ns | YPD | KS.3 |
| FIM1_3614 | 464.0676912 | 102.9001274 | 312.8522437 | -2.084960354 | 0.222820921 | 1.92742037808 | 1.52205021920 | Km<KS.3 | YPD | KS.3 |
| FIM1_3615 | 1752.435894 | 878.9390075 | 1455.642772 | -0.92847805 | 0.18619624 | 1.80359886412 | 8.27758703786 | ns | YPD | KS.3 |
| FIM1_3616 | 3905.082999 | 2381.052958 | 3554.016383 | -0.625288715 | 0.246578218 | 0.006285637 | 0.011085091 | ns | YPD | KS.3 |
| FIM1_3617 | 263.510227 | 129.1860778 | 224.0788788 | -0.934974118 | 0.221229235 | 6.77492622019 | 2.22476950135 | ns | YPD | KS.3 |
| FIM1_3618 | 3535.782452 | 2894.438325 | 2714.576116 | -0.281499942 | 0.09923809 | 0.00406796 | 0.007448694 | ns | YPD | KS.3 |
| FIM1_3619 | 369.7003461 | 242.1696037 | 359.8464444 | -0.563884333 | 0.189153494 | 0.001823608 | 0.003585949 | ns | YPD | KS.3 |
| FIM1_3620 | 1995.405395 | 1951.440971 | 1503.26998 | -0.030527058 | 0.138414301 | 0.821334146 | 0.850924069 | ns | YPD | KS.3 |
| FIM1_3621 | 1854.621794 | 1285.15434 | 1545.844025 | -0.505033772 | 0.141467587 | 0.000249852 | 0.000585222 | ns | YPD | KS.3 |
| FIM1_3622 | 2930.680205 | 2640.837343 | 2433.684599 | -0.147184284 | 0.08501299 | 0.080535921 | 0.110559108 | ns | YPD | KS.3 |
| FIM1_3623 | 336.470175 | 606.2648817 | 572.7992675 | 0.648743229 | 0.356841633 | 0.034254041 | 0.051484548 | ns | YPD | KS.3 |
| FIM1_3624 | 207.2198438 | 679.4778869 | 462.092434 | 1.445844728 | 0.35319702 | 1.72020992261 | 6.41052012359 | Km>KS.3 | YPD | KS.3 |
| FIM1_3625 | 11681.92688 | 18030.15591 | 18717.45703 | 0.619799851 | 0.064180538 | 2.64617548253 | 1.68778500328 | ns | YPD | KS.3 |
| FIM1_3626 | 223.8989338 | 181.9095927 | 201.9656237 | -0.287805807 | 0.135938479 | 0.03025234 | 0.04599798 | ns | YPD | KS.3 |
| FIM1_3627 | 381.5834971 | 1127.531166 | 1302.001326 | 1.461511963 | 0.212088719 | 1.98709664508 | 2.91538186761 | Km>KS.3 | YPD | KS.3 |
| FIM1_3628 | 4874.827569 | 12066.07359 | 11043.21588 | 1.079250559 | 0.330096643 | 0.000184003 | 0.000447198 | Km>KS.3 | YPD | KS.3 |
| FIM1_3629 | 2875.534656 | 6281.305749 | 4731.292686 | 1.037399125 | 0.20862899 | 1.39911993037 | 6.56662420150 | Km>KS.3 | YPD | KS.3 |
| FIM1_3630 | 157.5540823 | 361.141515 | 197.6624048 | 1.096225914 | 0.218802707 | 9.46007783797 | 4.60627691581 | Km>KS.3 | YPD | KS.3 |
| FIM1_3631 | 742.6949466 | 442.6026196 | 557.9884718 | -0.721510572 | 0.123786712 | 2.69870616595 | 1.76426585750 | ns | YPD | KS.3 |
| FIM1_3632 | 6749.009633 | 7895.143072 | 8567.738859 | 0.219951963 | 0.105746836 | 0.034872047 | 0.052224093 | ns | YPD | KS.3 |
| FIM1_3633 | 1791.581691 | 3155.339597 | 2506.69561 | 0.681110931 | 0.292621429 | 0.009797574 | 0.016630137 | ns | YPD | KS.3 |
| FIM1_3634 | 2893.160506 | 2737.414911 | 3704.855705 | -0.068432301 | 0.254833713 | 0.771734032 | 0.808289855 | ns | YPD | KS.3 |
| FIM1_3635 | 4253.711336 | 2742.093597 | 4861.765801 | -0.546489737 | 0.258070572 | 0.021662372 | 0.034007669 | ns | YPD | KS.3 |
| FIM1_3636 | 2535.975872 | 2414.088469 | 2761.731935 | -0.059316941 | 0.275884673 | 0.813946344 | 0.844677318 | ns | YPD | KS.3 |
| FIM1_3637 | 2745.21281 | 1764.212736 | 3180.97478 | -0.537491756 | 0.278097876 | 0.033824801 | 0.050900904 | ns | YPD | KS.3 |
| FIM1_3638 | 6505.822545 | 3960.77148 | 7351.407994 | -0.602202023 | 0.282105264 | 0.01872582 | 0.02993604 | ns | YPD | KS.3 |
| FIM1_3639 | 438.4538359 | 220.5507925 | 401.9788217 | -0.803138402 | 0.323080228 | 0.004780307 | 0.00862298 | ns | YPD | KS.3 |
| FIM1_3640 | 1177.726964 | 616.7373235 | 1134.694664 | -0.78480682 | 0.291078088 | 0.002780066 | 0.005276928 | ns | YPD | KS.3 |
| FIM1_3641 | 915.4679696 | 610.3795995 | 956.6364595 | -0.501302737 | 0.26208068 | 0.037648191 | 0.055993946 | ns | YPD | KS.3 |
| FIM1_3642 | 2858.013821 | 1820.306678 | 2394.135994 | -0.558496864 | 0.263125159 | 0.020950222 | 0.032972906 | ns | YPD | KS.3 |
| FIM1_3643 | 1296.108222 | 831.7128039 | 1267.188752 | -0.527312501 | 0.295745524 | 0.047600551 | 0.069021493 | ns | YPD | KS.3 |

| | | | | | | | | | | |
|---|---|---|---|---|---|---|---|---|---|---|
| FIM1_3644 | 2371.003377 | 1854.206401 | 2834.249379 | -0.305242196 | 0.253958954 | 0.19409419 | 0.245018675 | ns | YPD | KS.3 |
| FIM1_3645 | 3542.104182 | 2204.395796 | 3118.831835 | -0.585746959 | 0.266305645 | 0.01649689 | 0.026724854 | ns | YPD | KS.3 |
| FIM1_3646 | 991.9274569 | 771.7714732 | 1263.162388 | -0.305619079 | 0.270316638 | 0.217271046 | 0.270163323 | ns | YPD | KS.3 |
| FIM1_3647 | 14957.85759 | 12671.71661 | 25119.05881 | -0.213929587 | 0.217454082 | 0.297690535 | 0.358337869 | ns | YPD | KS.3 |
| FIM1_3648 | 1603.097902 | 882.9926365 | 2447.029702 | -0.61026647 | 0.400520265 | 0.06518602 | 0.091455287 | ns | YPD | KS.3 |
| FIM1_3649 | 1120.656129 | 772.4164332 | 1619.634937 | -0.450514368 | 0.278659143 | 0.075903251 | 0.105039965 | ns | YPD | KS.3 |
| FIM1_3650 | 1014.280895 | 1076.414484 | 1797.310892 | 0.071836831 | 0.27341638 | 0.774013807 | 0.810336424 | ns | YPD | KS.3 |
| FIM1_3651 | 1058.5833 | 812.7551437 | 1422.069128 | -0.31887959 | 0.277609888 | 0.207856635 | 0.259820794 | ns | YPD | KS.3 |
| FIM1_3652 | 267.3841058 | 199.7724442 | 400.6113962 | -0.310850832 | 0.360838603 | 0.314420523 | 0.375149209 | ns | YPD | KS.3 |
| FIM1_3653 | 3401.832261 | 3708.585151 | 5782.597739 | 0.104773504 | 0.270226776 | 0.672388061 | 0.723427899 | ns | YPD | KS.3 |
| FIM1_3654 | 6685.450661 | 4836.67161 | 7828.351318 | -0.412126431 | 0.233372867 | 0.059222877 | 0.083798013 | ns | YPD | KS.3 |
| FIM1_3655 | 4358.629523 | 2916.156935 | 4492.996544 | -0.503891764 | 0.250051173 | 0.029689009 | 0.045265957 | ns | YPD | KS.3 |
| FIM1_3656 | 4761.670331 | 3419.562094 | 5621.544174 | -0.410996678 | 0.256471753 | 0.082787261 | 0.113274649 | ns | YPD | KS.3 |
| FIM1_3657 | 732.2978351 | 681.7947354 | 773.3438998 | -0.084147106 | 0.292210918 | 0.74997814 | 0.791166507 | ns | YPD | KS.3 |
| FIM1_3658 | 1990.75379 | 753.55278 | 2222.721408 | -1.142100037 | 0.348968503 | 0.000144517 | 0.000360207 | Km<KS.3 | YPD | KS.3 |
| FIM1_3659 | 1105.029151 | 1618.887491 | 1990.138433 | 0.461297174 | 0.280643028 | 0.070754893 | 0.098435568 | ns | YPD | KS.3 |
| FIM1_3660 | 1316.088034 | 786.0866104 | 1551.995642 | -0.61935066 | 0.290782932 | 0.01829319 | 0.029329237 | ns | YPD | KS.3 |
| FIM1_3661 | 2049.687623 | 1287.363207 | 2798.114036 | -0.561830429 | 0.284178436 | 0.029273482 | 0.044659789 | ns | YPD | KS.3 |
| FIM1_3662 | 1769.359535 | 1052.221449 | 1358.714328 | -0.632753734 | 0.280479486 | 0.013051107 | 0.021556859 | ns | YPD | KS.3 |
| FIM1_3663 | 4993.866125 | 1537.309686 | 4056.103488 | -1.389513285 | 0.380755432 | 1.34445720446 | 4.11610744136 | Km<KS.3 | YPD | KS.3 |
| FIM1_3664 | 1541.324296 | 751.0678229 | 1737.776234 | -0.869896903 | 0.298504302 | 0.001147267 | 0.002349796 | ns | YPD | KS.3 |
| FIM1_3665 | 1525.145929 | 661.1791927 | 1234.791245 | -0.986556427 | 0.329394048 | 0.000624254 | 0.001350877 | ns | YPD | KS.3 |
| FIM1_3666 | 1732.149186 | 969.4796228 | 1461.323656 | -0.691544014 | 0.301195042 | 0.010373876 | 0.017554433 | ns | YPD | KS.3 |
| FIM1_3667 | 2143.314839 | 1266.782094 | 2133.783649 | -0.651187523 | 0.266490418 | 0.007694154 | 0.013356041 | ns | YPD | KS.3 |
| FIM1_3668 | 1513.142934 | 919.3288334 | 1477.657812 | -0.607240661 | 0.278722292 | 0.016658586 | 0.026978016 | ns | YPD | KS.3 |
| FIM1_3669 | 1239.781717 | 961.1653179 | 1245.2853 | -0.296813215 | 0.302936669 | 0.274410745 | 0.332487447 | ns | YPD | KS.3 |
| FIM1_3670 | 799.7329289 | 493.4855971 | 785.9254473 | -0.575837362 | 0.295150925 | 0.030214305 | 0.045954194 | ns | YPD | KS.3 |
| FIM1_3671 | 579.9017359 | 354.0360032 | 649.025862 | -0.559683814 | 0.331545308 | 0.054145196 | 0.07738361 | ns | YPD | KS.3 |
| FIM1_3672 | 1150.827671 | 1631.246173 | 2288.424868 | 0.424351951 | 0.27391897 | 0.090000589 | 0.121970288 | ns | YPD | KS.3 |
| FIM1_3673 | 2059.494197 | 1182.36581 | 2448.216564 | -0.635503241 | 0.328740492 | 0.027720333 | 0.042551267 | ns | YPD | KS.3 |
| FIM1_3674 | 46.82238565 | 43.74946248 | 71.71387351 | -0.076732507 | 0.317535111 | 0.7853497 | 0.820822428 | ns | YPD | KS.3 |
| FIM1_3675 | 3317.100487 | 2707.51418 | 4505.666315 | -0.24902905 | 0.263438233 | 0.304432538 | 0.364688629 | ns | YPD | KS.3 |
| FIM1_3676 | 104.1674909 | 119.0214875 | 187.0670644 | 0.158082398 | 0.290632986 | 0.547621338 | 0.608672063 | ns | YPD | KS.3 |
| FIM1_3677 | 1615.985746 | 1127.119734 | 1500.927836 | -0.439453068 | 0.272028936 | 0.077408302 | 0.106766372 | ns | YPD | KS.3 |
| FIM1_3678 | 2084.389473 | 1018.915051 | 2981.63063 | -0.7369345 | 0.407287893 | 0.027614299 | 0.042427775 | ns | YPD | KS.3 |
| FIM1_3679 | 27967.42267 | 42499.28293 | 61159.05026 | 0.542664056 | 0.218017097 | 0.008309063 | 0.014308614 | ns | YPD | KS.3 |
| FIM1_3680 | 218.9396508 | 114.2657536 | 178.89611 | -0.780750703 | 0.299869064 | 0.003663917 | 0.006776203 | ns | YPD | KS.3 |
| FIM1_3681 | 1099.731317 | 784.075002 | 1295.64304 | -0.418386644 | 0.259812258 | 0.080670443 | 0.110652179 | ns | YPD | KS.3 |
| FIM1_3682 | 1969.035742 | 1774.805013 | 3493.016974 | -0.1229018 | 0.289383656 | 0.639002812 | 0.693205188 | ns | YPD | KS.3 |
| FIM1_3683 | 1182.551793 | 612.7008429 | 1390.773785 | -0.827776199 | 0.259167146 | 0.0005154 | 0.001132059 | ns | YPD | KS.3 |
| FIM1_3684 | 630.4780625 | 493.1420353 | 675.516524 | -0.297827459 | 0.273173912 | 0.233293412 | 0.287641815 | ns | YPD | KS.3 |
| FIM1_3685 | 1030.467057 | 624.8836601 | 967.9350028 | -0.588272001 | 0.306425336 | 0.031579338 | 0.047781997 | ns | YPD | KS.3 |
| FIM1_3686 | 1145.679583 | 396.8795522 | 1007.071036 | -1.169760153 | 0.409930011 | 0.000498461 | 0.001099404 | Km<KS.3 | YPD | KS.3 |
| FIM1_3687 | 667.2185995 | 374.4225762 | 709.5175891 | -0.713105213 | 0.272165186 | 0.004114556 | 0.007528472 | ns | YPD | KS.3 |
| FIM1_3688 | 785.7059391 | 928.5162007 | 1342.19789 | 0.190097365 | 0.317724841 | 0.499976353 | 0.562374487 | ns | YPD | KS.3 |

| | | | | | | | | | | |
|---|---|---|---|---|---|---|---|---|---|---|
| FIM1_3689 | 1636.589248 | 1495.156946 | 1805.398193 | -0.111268407 | 0.258601001 | 0.641279546 | 0.69552338 | ns | YPD | KS.3 |
| FIM1_3690 | 1040.305876 | 752.6388762 | 1028.481125 | -0.394968844 | 0.270347478 | 0.110714767 | 0.147313711 | ns | YPD | KS.3 |
| FIM1_3691 | 3695.249238 | 2739.383235 | 3698.699931 | -0.37344993 | 0.251067611 | 0.108549912 | 0.144704129 | ns | YPD | KS.3 |
| FIM1_3692 | 2195.854963 | 1271.253092 | 2354.507026 | -0.67256087 | 0.273031824 | 0.006970933 | 0.012207108 | ns | YPD | KS.3 |
| FIM1_3693 | 2340.398485 | 1603.911111 | 2060.419245 | -0.459882935 | 0.274542398 | 0.066618042 | 0.093254012 | ns | YPD | KS.3 |
| FIM1_3694 | 1323.991991 | 762.2332406 | 1198.114767 | -0.682236467 | 0.269708634 | 0.005690069 | 0.010113645 | ns | YPD | KS.3 |
| FIM1_3695 | 3339.386398 | 4399.9337 | 5682.653673 | 0.332652456 | 0.278484932 | 0.190002926 | 0.240341865 | ns | YPD | KS.3 |
| FIM1_3696 | 3069.331439 | 2449.928957 | 4593.752869 | -0.27438582 | 0.269843723 | 0.267340526 | 0.32502911 | ns | YPD | KS.3 |
| FIM1_3697 | 7916.957879 | 7208.416722 | 11978.24542 | -0.12197377 | 0.207766493 | 0.536347266 | 0.598145628 | ns | YPD | KS.3 |
| FIM1_3698 | 2462.515196 | 2388.708128 | 3455.448409 | -0.03697542 | 0.268499949 | 0.880747724 | 0.901774012 | ns | YPD | KS.3 |
| FIM1_3699 | 225.5038922 | 467.0407607 | 1113.884538 | 0.77412313 | 0.390138009 | 0.017398529 | 0.028084907 | ns | YPD | KS.3 |
| FIM1_3700 | 743.7641483 | 898.5907215 | 866.2872135 | 0.217279788 | 0.312173162 | 0.434518509 | 0.498575123 | ns | YPD | KS.3 |
| FIM1_3701 | 1023.703534 | 424.4685114 | 989.2838633 | -1.063344372 | 0.315207757 | 0.000135357 | 0.000339075 | Km<KS.3 | YPD | KS.3 |
| FIM1_3702 | 701.9546428 | 478.4702279 | 650.5353628 | -0.457904978 | 0.289042886 | 0.079880157 | 0.109749732 | ns | YPD | KS.3 |
| FIM1_3703 | 3104.828935 | 2383.867579 | 2849.464825 | -0.324165438 | 0.264447818 | 0.182557177 | 0.232044445 | ns | YPD | KS.3 |
| FIM1_3704 | 2124.90245 | 601.0102044 | 1314.613492 | -1.601227197 | 0.324386308 | 1.5554981246 | 8.7938672389 | Km<KS.3 | YPD | KS.3 |
| FIM1_3705 | 3155.357558 | 1785.791301 | 2843.130502 | -0.680544134 | 0.297947448 | 0.010962607 | 0.018425327 | ns | YPD | KS.3 |
| FIM1_3706 | 4034.579239 | 3101.317516 | 5158.641043 | -0.32026982 | 0.270728674 | 0.196562643 | 0.247757575 | ns | YPD | KS.3 |
| FIM1_3707 | 714.773586 | 460.1058301 | 787.9552835 | -0.518857545 | 0.302458818 | 0.055521416 | 0.079073404 | ns | YPD | KS.3 |
| FIM1_3708 | 5675.538417 | 4420.060237 | 5903.164291 | -0.324240977 | 0.213322009 | 0.108232726 | 0.144388579 | ns | YPD | KS.3 |
| FIM1_3709 | 1762.022837 | 1071.939455 | 2293.754854 | -0.600804344 | 0.285257501 | 0.020104375 | 0.031904072 | ns | YPD | KS.3 |
| FIM1_3710 | 2558.928226 | 2449.143421 | 2787.408431 | -0.053157711 | 0.270637358 | 0.83033914 | 0.858465757 | ns | YPD | KS.3 |
| FIM1_3711 | 3207.506001 | 2842.820469 | 3620.269437 | -0.145288748 | 0.276809705 | 0.565349201 | 0.625682279 | ns | YPD | KS.3 |
| FIM1_3712 | 3570.468418 | 2286.177688 | 3898.949549 | -0.538544414 | 0.283282499 | 0.036213882 | 0.054022208 | ns | YPD | KS.3 |
| FIM1_3713 | 3519.153224 | 2634.44636 | 3243.832749 | -0.355796573 | 0.263774967 | 0.142566041 | 0.185186959 | ns | YPD | KS.3 |
| FIM1_3714 | 535.912052 | 565.3559854 | 1059.112359 | 0.058242358 | 0.344281587 | 0.845585833 | 0.871344278 | ns | YPD | KS.3 |
| FIM1_3715 | 330.9311724 | 301.2284024 | 396.9883538 | -0.104830949 | 0.330252701 | 0.717871274 | 0.764254687 | ns | YPD | KS.3 |
| FIM1_3716 | 3070.00447 | 3444.358806 | 5631.398114 | 0.143872244 | 0.245685599 | 0.529137482 | 0.591298062 | ns | YPD | KS.3 |
| FIM1_3717 | 8494.183552 | 5747.714533 | 8897.841964 | -0.499823009 | 0.23047763 | 0.020662306 | 0.032612618 | ns | YPD | KS.3 |
| FIM1_3718 | 2148.638507 | 797.4084115 | 1894.578112 | -1.204474826 | 0.323307055 | 2.1641680378 | 6.3520566302 | Km<KS.3 | YPD | KS.3 |
| FIM1_3719 | 1535.940867 | 963.506978 | 1434.633716 | -0.52539025 | 0.334636962 | 0.07264492 | 0.100867563 | ns | YPD | KS.3 |
| FIM1_3720 | 1438.368006 | 1171.820849 | 1754.689686 | -0.241844644 | 0.293114167 | 0.360765286 | 0.423424881 | ns | YPD | KS.3 |
| FIM1_3722 | 65.1327555 | 86.82257182 | 144.6627501 | 0.333788937 | 0.306488366 | 0.223174129 | 0.276673633 | ns | YPD | KS.3 |
| FIM1_3723 | 78.98255152 | 447.0287413 | 374.7185674 | 2.171494232 | 0.446722631 | 3.9233358012 | 2.4896167871 | Km>KS.3 | YPD | KS.3 |
| FIM1_3725 | 67.84837398 | 116.4095385 | 110.7036884 | 0.524523891 | 0.422601462 | 0.124683534 | 0.164492332 | ns | YPD | KS.3 |
| FIM1_3727 | 312.9398086 | 254.1102066 | 551.350287 | -0.223619392 | 0.355724662 | 0.464897209 | 0.52841298 | ns | YPD | KS.3 |
| FIM1_3731 | 67.25564055 | 10.04147328 | 113.1955211 | -2.45756852 | 0.42116407 | 1.5888388462 | 1.6331556322 | Km<KS.3 | YPD | KS.3 |
| FIM1_3732 | 217.3012128 | 151.865823 | 285.7986033 | -0.41994411 | 0.303579663 | 0.122461004 | 0.161774694 | ns | YPD | KS.3 |
| FIM1_3733 | 868.403712 | 982.2930022 | 1175.971791 | 0.168687008 | 0.148627434 | 0.243487573 | 0.298582863 | ns | YPD | KS.3 |
| FIM1_3734 | 778.7846957 | 533.6043884 | 827.4544815 | -0.521264514 | 0.138317205 | 0.000112344 | 0.000287506 | ns | YPD | KS.3 |
| FIM1_3735 | 1931.754993 | 1747.544814 | 1708.862647 | -0.142322031 | 0.071136301 | 0.044003085 | 0.064311206 | ns | YPD | KS.3 |
| FIM1_3736 | 1475.484607 | 1290.543403 | 1412.888503 | -0.181776126 | 0.157535015 | 0.234203269 | 0.288534371 | ns | YPD | KS.3 |
| FIM1_3737 | 582.8098213 | 1390.017254 | 793.9904804 | 1.11550365 | 0.255633184 | 2.0494292429 | 7.5025095537 | Km>KS.3 | YPD | KS.3 |
| FIM1_3738 | 429.6952426 | 648.988982 | 451.205716 | 0.5710552 | 0.135172823 | 1.5204503233 | 4.6067237263 | ns | YPD | KS.3 |
| FIM1_3739 | 1774.444639 | 1484.506045 | 1405.266701 | -0.230405251 | 0.215947941 | 0.25906074 | 0.315966458 | ns | YPD | KS.3 |

| | | | | | | | | | | |
|---|---|---|---|---|---|---|---|---|---|---|
| FIM1_3740 | 1958.444509 | 1629.430597 | 2403.374038 | -0.250791781 | 0.15274241 | 0.091027587 | 0.123227822 | ns | YPD | KS.3 |
| FIM1_3741 | 3652.562536 | 4672.998636 | 3060.206979 | 0.343513182 | 0.118661594 | 0.003205117 | 0.006012616 | ns | YPD | KS.3 |
| FIM1_3742 | 2909.620086 | 2659.794453 | 2198.695666 | -0.124904257 | 0.119084232 | 0.285660157 | 0.344857869 | ns | YPD | KS.3 |
| FIM1_3743 | 3750.273826 | 3809.705795 | 3417.466024 | 0.02225492 | 0.090327413 | 0.803383443 | 0.835614761 | ns | YPD | KS.3 |
| FIM1_3744 | 1522.368312 | 1839.692119 | 1593.700646 | 0.262904797 | 0.123862389 | 0.030463756 | 0.046234651 | ns | YPD | KS.3 |
| FIM1_3745 | 9346.478929 | 9519.782953 | 12179.15248 | 0.025816577 | 0.101292515 | 0.796244499 | 0.829937816 | ns | YPD | KS.3 |
| FIM1_3746 | 1892.092466 | 2048.548592 | 1963.194845 | 0.110465272 | 0.122589668 | 0.358521809 | 0.42106846 | ns | YPD | KS.3 |
| FIM1_3747 | 19203.02478 | 16530.91034 | 16972.88969 | -0.203610785 | 0.157153747 | 0.181684105 | 0.231407686 | ns | YPD | KS.3 |
| FIM1_3748 | 6845.679276 | 4306.945336 | 7906.555294 | -0.6377882 | 0.144392805 | 5.6773801963 | 1.9045830395 | ns | YPD | KS.3 |
| FIM1_3749 | 5763.012621 | 4002.067592 | 4416.571733 | -0.518150037 | 0.078496252 | 2.8058889843 | 2.7533131552 | ns | YPD | KS.3 |
| FIM1_3750 | 1526.925918 | 590.4111302 | 1375.601999 | -1.160811544 | 0.313231873 | 2.7654010124 | 7.9204778565 | Km<KS.3 | YPD | KS.3 |
| FIM1_3751 | 2709.953389 | 1876.744573 | 1850.963617 | -0.520684043 | 0.083853368 | 3.6419649085 | 2.8428480629 | ns | YPD | KS.3 |
| FIM1_3752 | 2685.761966 | 2816.580249 | 2341.916908 | 0.066830165 | 0.101593985 | 0.505098351 | 0.567494195 | ns | YPD | KS.3 |
| FIM1_3753 | 1616.337953 | 3320.995717 | 2476.245096 | 0.988171932 | 0.158166134 | 1.0304935706 | 9.0738150687 | ns | YPD | KS.3 |
| FIM1_3754 | 1536.00434 | 2923.03459 | 2359.349226 | 0.814949522 | 0.252114935 | 0.000466657 | 0.001037197 | ns | YPD | KS.3 |
| FIM1_3755 | 201.162511 | 131.6229343 | 213.6750868 | -0.55671332 | 0.206975269 | 0.004561621 | 0.008264407 | ns | YPD | KS.3 |
| FIM1_3756 | 15998.33417 | 14947.93472 | 22678.61819 | -0.094263294 | 0.124916077 | 0.44162983 | 0.505430965 | ns | YPD | KS.3 |
| FIM1_3757 | 818.8506587 | 645.499598 | 660.2333306 | -0.331731483 | 0.116469422 | 0.003756058 | 0.006928583 | ns | YPD | KS.3 |
| FIM1_3758 | 1341.831313 | 834.4470356 | 1542.398213 | -0.635962098 | 0.184240143 | 0.000314361 | 0.000724019 | ns | YPD | KS.3 |
| FIM1_3759 | 7063.041821 | 8570.395387 | 9864.711821 | 0.244442746 | 0.237279945 | 0.270393016 | 0.328178886 | ns | YPD | KS.3 |
| FIM1_3760 | 644.1311688 | 686.5638983 | 619.9495454 | 0.088849979 | 0.112224152 | 0.421183604 | 0.485268279 | ns | YPD | KS.3 |
| FIM1_3761 | 943.4006916 | 851.7284161 | 735.6103109 | -0.144574288 | 0.082938095 | 0.078614759 | 0.108250325 | ns | YPD | KS.3 |
| FIM1_3762 | 2048.866933 | 2387.774066 | 1837.459971 | 0.208791459 | 0.151785672 | 0.156997961 | 0.202610858 | ns | YPD | KS.3 |
| FIM1_3763 | 2065.819479 | 3225.343801 | 3123.577917 | 0.586500875 | 0.202455962 | 0.002298184 | 0.004428143 | ns | YPD | KS.3 |
| FIM1_3764 | 411.1477245 | 344.0089063 | 462.6643805 | -0.249187746 | 0.101234158 | 0.012628848 | 0.020928842 | ns | YPD | KS.3 |
| FIM1_3765 | 6461.077638 | 9899.227138 | 9241.215501 | 0.583993218 | 0.151776564 | 7.4026662351 | 0.00019455 | ns | YPD | KS.3 |
| FIM1_3766 | 107.8038013 | 114.495327 | 98.51668961 | 0.081355023 | 0.166036276 | 0.612345306 | 0.668660645 | ns | YPD | KS.3 |
| FIM1_3767 | 2938.800444 | 1961.563658 | 2270.164745 | -0.557745538 | 0.138598927 | 3.7070415307 | 0.000103262 | ns | YPD | KS.3 |
| FIM1_3768 | 24056.21807 | 11717.93451 | 26488.78546 | -0.793661397 | 0.366108111 | 0.010920777 | 0.018367432 | ns | YPD | KS.3 |
| FIM1_3769 | 1896.792743 | 2307.05451 | 2223.519758 | 0.277212194 | 0.085314777 | 0.001036812 | 0.002142084 | ns | YPD | KS.3 |
| FIM1_3770 | 5812.650652 | 6309.958164 | 6474.110678 | 0.117393889 | 0.05663926 | 0.03737806 | 0.055658739 | ns | YPD | KS.3 |
| FIM1_3771 | 1495.474731 | 1423.531125 | 1381.914983 | -0.067166065 | 0.154610116 | 0.65463533 | 0.70677317 | ns | YPD | KS.3 |
| FIM1_3772 | 2261.776012 | 2456.646687 | 2235.125208 | 0.111786234 | 0.163689451 | 0.480416127 | 0.544063335 | ns | YPD | KS.3 |
| FIM1_3773 | 8593.994579 | 9023.906613 | 9253.453081 | 0.065094387 | 0.180229431 | 0.707160863 | 0.755286666 | ns | YPD | KS.3 |
| FIM1_3774 | 5055.090089 | 3641.466025 | 4738.193851 | -0.44588867 | 0.159314293 | 0.003860153 | 0.007099541 | ns | YPD | KS.3 |
| FIM1_3775 | 323.5671372 | 239.0302173 | 237.9796987 | -0.420334169 | 0.123091304 | 0.000498094 | 0.001099387 | ns | YPD | KS.3 |
| FIM1_3776 | 430.3725502 | 374.89781 | 418.9431956 | -0.189794857 | 0.137732979 | 0.158285882 | 0.204114117 | ns | YPD | KS.3 |
| FIM1_3777 | 4444.072662 | 10685.40353 | 9059.563685 | 1.007554437 | 0.354954241 | 0.000942791 | 0.001960863 | Km>KS.3 | YPD | KS.3 |
| FIM1_3778 | 248.3034757 | 532.5812199 | 395.1968135 | 0.970254775 | 0.255998501 | 3.8176068419 | 0.000105808 | ns | YPD | KS.3 |
| FIM1_3779 | 1091.960736 | 959.5119986 | 1354.296373 | -0.178115434 | 0.14049843 | 0.193893868 | 0.244890072 | ns | YPD | KS.3 |
| FIM1_3780 | 9433.66111 | 6991.888398 | 10131.22759 | -0.42391587 | 0.088053153 | 1.1501952295 | 4.4461703705 | ns | YPD | KS.3 |
| FIM1_3782 | 3963.567125 | 3926.379089 | 3009.537331 | -0.012992945 | 0.12625502 | 0.916398091 | 0.932341646 | ns | YPD | KS.3 |
| FIM1_3783 | 1385.613445 | 2098.652696 | 1425.301828 | 0.583781307 | 0.104845723 | 1.6041798080 | 9.0485198924 | ns | YPD | KS.3 |
| FIM1_3784 | 3628.463398 | 2254.750979 | 3653.060382 | -0.66514121 | 0.117354021 | 7.8061267101 | 4.6621224949 | ns | YPD | KS.3 |
| FIM1_3785 | 520.9180398 | 296.1674048 | 374.948203 | -0.75218869 | 0.192896708 | 4.3324995023 | 0.000118625 | ns | YPD | KS.3 |

| | | | | | | | | | | |
|---|---|---|---|---|---|---|---|---|---|---|
| FIM1_3786 | 4983.474431 | 3985.514249 | 4454.690071 | -0.318306146 | 0.070458329 | 5.4224830001( | 1.8252268556( | ns | YPD | KS.3 |
| FIM1_3787 | 66.27368202 | 92.6028658 | 84.05890081 | 0.452382341 | 0.169436768 | 0.005651097 | 0.010047966 | ns | YPD | KS.3 |
| FIM1_3788 | 6724.729999 | 5279.560919 | 6370.812734 | -0.339231082 | 0.107534208 | 0.001365171 | 0.002756383 | ns | YPD | KS.3 |
| FIM1_3789 | 790.8499091 | 767.5700219 | 690.6040231 | -0.041058334 | 0.121396242 | 0.730470649 | 0.774154856 | ns | YPD | KS.3 |
| FIM1_3790 | 494.9363853 | 443.1001039 | 421.9346332 | -0.150048541 | 0.154926633 | 0.318526253 | 0.379652159 | ns | YPD | KS.3 |
| FIM1_3791 | 3786.210305 | 3683.910303 | 3342.107276 | -0.038749935 | 0.072608403 | 0.591006626 | 0.648822588 | ns | YPD | KS.3 |
| FIM1_3792 | 3597.382576 | 3838.76008 | 3126.315086 | 0.083294111 | 0.22208926 | 0.69073644 | 0.740446841 | ns | YPD | KS.3 |
| FIM1_3793 | 5441.0449 | 3854.887666 | 6434.595813 | -0.478281373 | 0.127968367 | 0.000135067 | 0.000338519 | ns | YPD | KS.3 |
| FIM1_3794 | 6191.396254 | 6052.720826 | 9539.717368 | -0.031526823 | 0.12325185 | 0.794336421 | 0.828474569 | ns | YPD | KS.3 |
| FIM1_3795 | 867.455556 | 1382.79074 | 881.351855 | 0.610769303 | 0.208978134 | 0.002028652 | 0.003947026 | ns | YPD | KS.3 |
| FIM1_3796 | 385.1268575 | 273.4735543 | 333.1994851 | -0.465153452 | 0.158620447 | 0.002471646 | 0.004727582 | ns | YPD | KS.3 |
| FIM1_3797 | 4339.607911 | 4417.952448 | 4135.845504 | 0.024688273 | 0.133018355 | 0.84953295 | 0.874493364 | ns | YPD | KS.3 |
| FIM1_3798 | 1859.752651 | 2336.816109 | 2050.789803 | 0.315982187 | 0.129610603 | 0.012768876 | 0.02112576 | ns | YPD | KS.3 |
| FIM1_3799 | 12273.03691 | 12358.64804 | 14279.63957 | 0.009355989 | 0.169710619 | 0.954470455 | 0.964552207 | ns | YPD | KS.3 |
| FIM1_3801 | 2087.242518 | 2932.877274 | 2398.246948 | 0.450755879 | 0.191930479 | 0.013977011 | 0.022979388 | ns | YPD | KS.3 |
| FIM1_3802 | 1991.559405 | 1782.7854 | 2210.137681 | -0.132418832 | 0.281414332 | 0.605182783 | 0.662001835 | ns | YPD | KS.3 |
| FIM1_3803 | 360.9507532 | 233.8957646 | 313.849329 | -0.580431552 | 0.184947732 | 0.001052476 | 0.002170842 | ns | YPD | KS.3 |
| FIM1_3804 | 388.610101 | 251.1482225 | 257.7622274 | -0.602479007 | 0.136899166 | 6.5061947600 | 2.1440469460 | ns | YPD | KS.3 |
| FIM1_3805 | 404.8676401 | 482.3543085 | 471.9418112 | 0.239308512 | 0.150321534 | 0.101603984 | 0.136358193 | ns | YPD | KS.3 |
| FIM1_3806 | 630.4737443 | 376.1324844 | 864.3943599 | -0.643638486 | 0.261028673 | 0.0073569 | 0.012842308 | ns | YPD | KS.3 |
| FIM1_3807 | 7870.891417 | 10799.12392 | 10336.56045 | 0.451412868 | 0.064614574 | 2.1197015970 | 2.6232625485 | ns | YPD | KS.3 |
| FIM1_3808 | 594.9798704 | 718.9886142 | 598.9609995 | 0.265220423 | 0.103440597 | 0.009338784 | 0.015932939 | ns | YPD | KS.3 |
| FIM1_3809 | 1894.171745 | 803.0291366 | 2060.132763 | -1.021537405 | 0.325046562 | 0.00034454 | 0.00078664 | Km<KS.3 | YPD | KS.3 |
| FIM1_3810 | 1996.788317 | 1290.268622 | 1956.994185 | -0.600723434 | 0.14478676 | 2.0193042482( | 5.9764615994 | ns | YPD | KS.3 |
| FIM1_3811 | 4192.558146 | 5744.705454 | 5184.452121 | 0.429278902 | 0.155661019 | 0.004469887 | 0.00812484 | ns | YPD | KS.3 |
| FIM1_3812 | 17.68598243 | 30.4474989 | 22.63589428 | 0.63762752 | 0.309831078 | 0.020847183 | 0.0328443 | ns | YPD | KS.3 |
| FIM1_3813 | 4853.128115 | 3357.497847 | 4199.466434 | -0.523303075 | 0.080522742 | 5.5201012363( | 5.1228427388( | ns | YPD | KS.3 |
| FIM1_3814 | 42.45181609 | 24.33145633 | 26.24877177 | -0.669071535 | 0.301664881 | 0.013271358 | 0.021884324 | ns | YPD | KS.3 |
| FIM1_3815 | 7987.888084 | 15281.58937 | 13481.12748 | 0.824999414 | 0.24857309 | 0.000336285 | 0.000769558 | ns | YPD | KS.3 |
| FIM1_3816 | 1830.885685 | 1299.938609 | 2349.079688 | -0.448083135 | 0.206479888 | 0.02221428 | 0.034830143 | ns | YPD | KS.3 |
| FIM1_3817 | 4184.398027 | 2439.54288 | 3144.887224 | -0.750840866 | 0.127594292 | 1.8119195696( | 1.2433517046 | ns | YPD | KS.3 |
| FIM1_3818 | 85.66559218 | 140.2857909 | 352.0551801 | 0.613973251 | 0.2610389 | 0.010584447 | 0.017874279 | ns | YPD | KS.3 |
| FIM1_3819 | 1737.317316 | 3413.562461 | 2295.003418 | 0.908772874 | 0.185162081 | 2.8487134652( | 1.2442800254( | ns | YPD | KS.3 |
| FIM1_3820 | 2107.296684 | 938.2888525 | 1699.424937 | -1.052794772 | 0.23505134 | 1.4966001323 | 5.6706669142 | Km<KS.3 | YPD | KS.3 |
| FIM1_3821 | 1047.375187 | 692.304093 | 934.0114174 | -0.557089787 | 0.175211177 | 0.00095031 | 0.001974851 | ns | YPD | KS.3 |
| FIM1_3822 | 3164.671496 | 6224.009231 | 5347.315536 | 0.944667672 | 0.12466683 | 9.8093280412 | 1.8626491223 | ns | YPD | KS.3 |
| FIM1_3823 | 74.0394267 | 171.0598332 | 134.5366798 | 1.150665578 | 0.160844184 | 1.2047725676 | 1.8789164652 | Km>KS.3 | YPD | KS.3 |
| FIM1_3824 | 9280.618514 | 31896.84338 | 20601.80896 | 1.662068301 | 0.235507828 | 2.1364531423 | 3.8510341968( | Km>KS.3 | YPD | KS.3 |
| FIM1_3825 | 330.2391365 | 127.6555875 | 253.0589104 | -1.151396208 | 0.320277283 | 4.3779015926 | 0.000119736 | Km<KS.3 | YPD | KS.3 |
| FIM1_3826 | 6431.176831 | 6027.360188 | 6236.632612 | -0.092676419 | 0.064874832 | 0.150852421 | 0.195084688 | ns | YPD | KS.3 |
| FIM1_3827 | 427.0008558 | 398.4352785 | 477.8659757 | -0.096204129 | 0.138975419 | 0.478344633 | 0.541964142 | ns | YPD | KS.3 |
| FIM1_3828 | 3222.658861 | 2096.930225 | 3076.068311 | -0.604848876 | 0.10295838 | 2.5449919772 | 1.6769980247 | ns | YPD | KS.3 |
| FIM1_3829 | 2312.080215 | 1313.700322 | 2219.783123 | -0.794406778 | 0.110245488 | 2.3655707355 | 3.4211379097 | ns | YPD | KS.3 |
| FIM1_3830 | 2531.025997 | 4285.74868 | 3249.216534 | 0.72639794 | 0.142602068 | 1.6786634122 | 7.7759315418( | ns | YPD | KS.3 |
| FIM1_3831 | 748.5657902 | 1032.171096 | 740.3250707 | 0.433485788 | 0.169539885 | 0.008047165 | 0.013922435 | ns | YPD | KS.3 |

| | | | | | | | | | | |
|---|---|---|---|---|---|---|---|---|---|---|
| FIM1_3832 | 630.08663 | 549.7376528 | 785.8791234 | -0.188923902 | 0.135541746 | 0.15384801 | 0.198648806 | ns | YPD | KS.3 |
| FIM1_3833 | 70.33121622 | 105.894542 | 255.9778003 | 0.479653803 | 0.306062576 | 0.079531958 | 0.109361937 | ns | YPD | KS.3 |
| FIM1_3834 | 2989.118331 | 3161.284367 | 3235.127828 | 0.079116786 | 0.095223584 | 0.400611729 | 0.464146099 | ns | YPD | KS.3 |
| FIM1_3835 | 421.8318373 | 563.4331189 | 381.0551814 | 0.404152999 | 0.116839553 | 0.000431075 | 0.000965166 | ns | YPD | KS.3 |
| FIM1_3836 | 4790.311139 | 3710.282185 | 4396.370425 | -0.349035826 | 0.151134765 | 0.017490301 | 0.028214736 | ns | YPD | KS.3 |
| FIM1_3837 | 2057.823123 | 1664.743015 | 2397.157586 | -0.296940834 | 0.109717694 | 0.005988944 | 0.01059566 | ns | YPD | KS.3 |
| FIM1_3838 | 1500.353075 | 2557.971104 | 1857.047638 | 0.74663079 | 0.116183487 | 5.979740776 | 5.488784199 | ns | YPD | KS.3 |
| FIM1_3839 | 3283.807011 | 2969.794151 | 2382.466326 | -0.127296628 | 0.234180602 | 0.561644142 | 0.622173148 | ns | YPD | KS.3 |
| FIM1_3840 | 903.9252479 | 632.5154996 | 898.5308475 | -0.474164358 | 0.18981229 | 0.008998519 | 0.015389354 | ns | YPD | KS.3 |
| FIM1_3841 | 1513.162014 | 3575.822865 | 2372.251608 | 1.148888763 | 0.205874256 | 3.469033560 | 2.218308735 | Km>KS.3 | YPD | KS.3 |
| FIM1_3842 | 184.244165 | 242.2388884 | 197.4601246 | 0.326453112 | 0.287121037 | 0.209623339 | 0.261766092 | ns | YPD | KS.3 |
| FIM1_3843 | 10179.18585 | 10361.36401 | 11982.34472 | 0.025156112 | 0.073238802 | 0.729419888 | 0.773250361 | ns | YPD | KS.3 |
| FIM1_3844 | 762.6511996 | 965.02132 | 1232.990209 | 0.293250234 | 0.250656299 | 0.207141631 | 0.258992113 | ns | YPD | KS.3 |
| FIM1_3845 | 1004.131981 | 357.2207269 | 1007.305287 | -1.345844496 | 0.25592765 | 9.303691840 | 5.490612919 | Km<KS.3 | YPD | KS.3 |
| FIM1_3846 | 792.7864491 | 770.5844492 | 975.6377112 | -0.039384727 | 0.118557312 | 0.735292103 | 0.777818033 | ns | YPD | KS.3 |
| FIM1_3847 | 22846.57857 | 23997.02213 | 31400.47508 | 0.069520542 | 0.086785298 | 0.418590106 | 0.482727348 | ns | YPD | KS.3 |
| FIM1_3848 | 1076.075255 | 712.7893201 | 750.7664885 | -0.52087051 | 0.242650415 | 0.021090017 | 0.033182428 | ns | YPD | KS.3 |
| FIM1_3849 | 21653.36305 | 25561.06261 | 30737.89021 | 0.227031473 | 0.147439379 | 0.113683743 | 0.150860663 | ns | YPD | KS.3 |
| FIM1_3850 | 1517.258171 | 3223.25939 | 2505.267773 | 1.010164053 | 0.194063822 | 4.476364470 | 2.300610871 | Km>KS.3 | YPD | KS.3 |
| FIM1_3851 | 1293.660289 | 1561.823328 | 1348.817359 | 0.259601914 | 0.136993509 | 0.052386182 | 0.075150305 | ns | YPD | KS.3 |
| FIM1_3852 | 1617.236314 | 719.5418763 | 1300.114399 | -1.113201111 | 0.160101853 | 5.859180006 | 8.008082563 | Km<KS.3 | YPD | KS.3 |
| FIM1_3853 | 2743.460285 | 1912.090711 | 1715.996936 | -0.503252915 | 0.119889382 | 1.896795476 | 5.660802336 | ns | YPD | KS.3 |
| FIM1_3854 | 6719.36434 | 4966.228222 | 5777.754768 | -0.413430358 | 0.150289423 | 0.004653017 | 0.008417731 | ns | YPD | KS.3 |
| FIM1_3855 | 4052.561939 | 3353.506506 | 3691.321226 | -0.263117238 | 0.122801824 | 0.02897893 | 0.044264715 | ns | YPD | KS.3 |
| FIM1_3856 | 958.4072816 | 820.061481 | 1475.589319 | -0.215754373 | 0.127844395 | 0.08497171 | 0.115912876 | ns | YPD | KS.3 |
| FIM1_3857 | 5889.250061 | 5498.170844 | 6700.010341 | -0.097454252 | 0.081899297 | 0.230017258 | 0.284024785 | ns | YPD | KS.3 |
| FIM1_3858 | 1126.444468 | 1028.589732 | 1106.416192 | -0.126496798 | 0.11285279 | 0.254689991 | 0.311246059 | ns | YPD | KS.3 |
| FIM1_3859 | 25.23576156 | 19.09640543 | 26.75156027 | -0.350699653 | 0.249919984 | 0.130347016 | 0.171147112 | ns | YPD | KS.3 |
| FIM1_3860 | 2824.382806 | 2880.629047 | 2591.085474 | 0.025283889 | 0.221789728 | 0.903763677 | 0.921735198 | ns | YPD | KS.3 |
| FIM1_3861 | 352.9836531 | 890.5125089 | 683.3851481 | 1.219979247 | 0.228322931 | 1.002911176 | 5.863082376 | Km>KS.3 | YPD | KS.3 |
| FIM1_3862 | 3705.172306 | 4510.944892 | 4239.368737 | 0.229244328 | 0.302304024 | 0.397905162 | 0.461440136 | ns | YPD | KS.3 |
| FIM1_3863 | 6121.955364 | 6621.678429 | 5834.400475 | 0.107111684 | 0.150347427 | 0.463864784 | 0.527359986 | ns | YPD | KS.3 |
| FIM1_3864 | 440.1962143 | 748.5822553 | 239.0951616 | 0.634504732 | 0.295932613 | 0.017130169 | 0.027678659 | ns | YPD | KS.3 |
| FIM1_3865 | 1117.129858 | 1194.605023 | 647.6237333 | 0.079194887 | 0.290210869 | 0.762984133 | 0.801826375 | ns | YPD | KS.3 |
| FIM1_3866 | 1764.056664 | 2317.403379 | 1486.262765 | 0.373972466 | 0.146237384 | 0.008617575 | 0.014783598 | ns | YPD | KS.3 |
| FIM1_3867 | 1731.742567 | 1427.295223 | 1536.73574 | -0.264645427 | 0.147781714 | 0.06576641 | 0.092191572 | ns | YPD | KS.3 |
| FIM1_3868 | 343.9335603 | 479.4262092 | 508.917387 | 0.44238471 | 0.186232376 | 0.013153269 | 0.021711186 | ns | YPD | KS.3 |
| FIM1_3869 | 13087.53375 | 14015.36714 | 15450.81622 | 0.089242703 | 0.206147503 | 0.648682796 | 0.701717092 | ns | YPD | KS.3 |
| FIM1_3870 | 1343.102282 | 1106.301391 | 1812.154242 | -0.265170434 | 0.147224224 | 0.064272628 | 0.090377706 | ns | YPD | KS.3 |
| FIM1_3871 | 2737.819676 | 1654.158436 | 2563.177529 | -0.651828051 | 0.22400686 | 0.001954736 | 0.003819642 | ns | YPD | KS.3 |
| FIM1_3872 | 5903.338236 | 5082.101657 | 5529.200178 | -0.21201562 | 0.085776988 | 0.012580116 | 0.020867593 | ns | YPD | KS.3 |
| FIM1_3873 | 1184.157368 | 1585.278219 | 1453.231919 | 0.39534184 | 0.162526929 | 0.011941587 | 0.019876011 | ns | YPD | KS.3 |
| FIM1_3874 | 670.0616725 | 539.6473476 | 510.1569601 | -0.300573292 | 0.120575313 | 0.011110453 | 0.018636043 | ns | YPD | KS.3 |
| FIM1_3875 | 436.4903087 | 331.63441 | 283.7631909 | -0.381573745 | 0.123447786 | 0.001623488 | 0.003223006 | ns | YPD | KS.3 |
| FIM1_3876 | 126.2567998 | 114.0598016 | 99.06440972 | -0.129883843 | 0.229269367 | 0.546322188 | 0.607771218 | ns | YPD | KS.3 |

| | | | | | | | | | | |
|---|---|---|---|---|---|---|---|---|---|---|
| FIM1_3877 | 155.5716228 | 364.0877995 | 174.706943 | 1.154657675 | 0.182247174 | 3.3450184435 | 3.2126383699 | Km>KS.3 | YPD | KS.3 |
| FIM1_3879 | 10153.79078 | 33159.89998 | 20269.80526 | 1.585525659 | 0.235987688 | 3.4946532305 | 4.9400300648 | Km>KS.3 | YPD | KS.3 |
| FIM1_3880 | 3619.629958 | 2485.491001 | 3181.816457 | -0.500908235 | 0.186248606 | 0.004983481 | 0.00896018 | ns | YPD | KS.3 |
| FIM1_3881 | 6156.607359 | 6152.57247 | 6875.92404 | -0.00079052 | 0.059505761 | 0.989350454 | 0.990874043 | ns | YPD | KS.3 |
| FIM1_3882 | 63.40610215 | 53.65078809 | 60.88332332 | -0.198282595 | 0.278492865 | 0.434908029 | 0.498770734 | ns | YPD | KS.3 |
| FIM1_3883 | 718.6958262 | 428.9844447 | 422.477108 | -0.710536905 | 0.144209538 | 4.1245278865 | 1.7448576731 | ns | YPD | KS.3 |
| FIM1_3884 | 461.971184 | 536.7896303 | 508.2338864 | 0.199187257 | 0.19014085 | 0.27372126 | 0.331813662 | ns | YPD | KS.3 |
| FIM1_3885 | 2568.609167 | 2771.917247 | 2432.3808 | 0.106600134 | 0.107344779 | 0.313628295 | 0.374442229 | ns | YPD | KS.3 |
| FIM1_3886 | 3562.964394 | 2972.311745 | 3862.400914 | -0.251557312 | 0.125416709 | 0.040770144 | 0.060169524 | ns | YPD | KS.3 |
| FIM1_3887 | 392.6034914 | 316.6394724 | 388.6433825 | -0.301257539 | 0.099358235 | 0.002133956 | 0.004137346 | ns | YPD | KS.3 |
| FIM1_3888 | 3452.694853 | 2056.096231 | 3161.601912 | -0.719618174 | 0.131761745 | 2.2906155163 | 1.2440843006 | ns | YPD | KS.3 |
| FIM1_3889 | 770.8009788 | 1019.494926 | 1106.781719 | 0.369954548 | 0.191396812 | 0.043203141 | 0.063260888 | ns | YPD | KS.3 |
| FIM1_3890 | 1698.836042 | 2282.214522 | 2590.63797 | 0.404881914 | 0.145396931 | 0.004238795 | 0.007733042 | ns | YPD | KS.3 |
| FIM1_3891 | 280.5574567 | 563.0426952 | 393.9541856 | 0.906648596 | 0.227146454 | 2.0156752540 | 5.9725934419 | ns | YPD | KS.3 |
| FIM1_3892 | 1405.129375 | 4234.111266 | 1681.82906 | 1.38599522 | 0.306479789 | 3.2572121693 | 1.4029983153 | Km>KS.3 | YPD | KS.3 |
| FIM1_3893 | 113.7432648 | 171.7222416 | 134.945167 | 0.561902051 | 0.15735147 | 0.000228389 | 0.000540572 | ns | YPD | KS.3 |
| FIM1_3894 | 830.222121 | 991.0279605 | 844.3097769 | 0.246706579 | 0.121611679 | 0.038749954 | 0.057495086 | ns | YPD | KS.3 |
| FIM1_3895 | 196.5835986 | 155.1676553 | 166.2505611 | -0.316217593 | 0.183148161 | 0.072017345 | 0.10007997 | ns | YPD | KS.3 |
| FIM1_3896 | 1703.023987 | 1574.858813 | 1224.149328 | -0.109022519 | 0.114497807 | 0.333041541 | 0.394965832 | ns | YPD | KS.3 |
| FIM1_3897 | 427.5310948 | 375.938269 | 509.3109975 | -0.176910273 | 0.135645583 | 0.182076714 | 0.231611264 | ns | YPD | KS.3 |
| FIM1_3898 | 3412.750681 | 2336.731104 | 4862.051882 | -0.429669186 | 0.326142976 | 0.134702267 | 0.176075612 | ns | YPD | KS.3 |
| FIM1_3899 | 1270.923087 | 1321.247739 | 974.2576244 | 0.055412867 | 0.092673673 | 0.545483638 | 0.606986445 | ns | YPD | KS.3 |
| FIM1_3900 | 577.6625218 | 298.5571499 | 381.9129481 | -0.913307562 | 0.141110326 | 2.9524242821 | 2.8800609418 | ns | YPD | KS.3 |
| FIM1_3901 | 1690.015549 | 1688.095803 | 1554.936371 | -0.001612083 | 0.133480655 | 0.990148965 | 0.99089732 | ns | YPD | KS.3 |
| FIM1_3902 | 613.4975887 | 1003.683672 | 1006.228109 | 0.654401712 | 0.192922277 | 0.0003779 | 0.000856308 | ns | YPD | KS.3 |
| FIM1_3903 | 1030.503114 | 435.3992745 | 596.3936974 | -1.190116219 | 0.15448943 | 1.6161055108 | 3.3782037464 | Km<KS.3 | YPD | KS.3 |
| FIM1_3904 | 6972.230814 | 5267.552358 | 7748.498376 | -0.391210442 | 0.117426761 | 0.000696386 | 0.001490116 | ns | YPD | KS.3 |
| FIM1_3905 | 5872.428673 | 11260.64327 | 6693.299249 | 0.878067355 | 0.180822646 | 4.0226449696 | 1.7046557686 | ns | YPD | KS.3 |
| FIM1_3906 | 1018.808059 | 595.3914986 | 747.9196432 | -0.734812298 | 0.156965114 | 1.3385861198 | 5.1187286289 | ns | YPD | KS.3 |
| FIM1_3907 | 1326.467916 | 1600.030725 | 1759.155748 | 0.262989928 | 0.107167676 | 0.012764562 | 0.021125647 | ns | YPD | KS.3 |
| FIM1_3908 | 1626.295394 | 1757.852973 | 1524.870191 | 0.110616605 | 0.074078446 | 0.132581719 | 0.173714525 | ns | YPD | KS.3 |
| FIM1_3909 | 1150.38246 | 487.6226762 | 1174.845268 | -1.088121321 | 0.267578307 | 8.3316937217 | 2.6621821622 | Km<KS.3 | YPD | KS.3 |
| FIM1_3910 | 5706.93021 | 5835.458202 | 6264.664992 | 0.031449857 | 0.08632708 | 0.713005917 | 0.760372301 | ns | YPD | KS.3 |
| FIM1_3911 | 302.6992926 | 176.4604776 | 182.5380654 | -0.708929262 | 0.209125779 | 0.000341285 | 0.000779923 | ns | YPD | KS.3 |
| FIM1_3912 | 3763.649683 | 10346.81547 | 4257.191824 | 1.343921323 | 0.226680406 | 1.9697098078 | 1.6359442894 | Km>KS.3 | YPD | KS.3 |
| FIM1_3913 | 288.8177664 | 1281.208663 | 592.3381723 | 2.086794449 | 0.184432593 | 2.3885421827 | 3.9609991197 | Km>KS.3 | YPD | KS.3 |
| FIM1_3914 | 512.1852652 | 627.7305144 | 465.4991353 | 0.277350725 | 0.154069858 | 0.063757036 | 0.08977958 | ns | YPD | KS.3 |
| FIM1_3915 | 2971.068909 | 3033.050498 | 2899.055429 | 0.028879525 | 0.102095215 | 0.774394847 | 0.810564773 | ns | YPD | KS.3 |
| FIM1_3916 | 1384.407282 | 1056.249039 | 852.7608378 | -0.368407362 | 0.155396062 | 0.014554533 | 0.023826523 | ns | YPD | KS.3 |
| FIM1_3917 | 1865.523693 | 2371.560637 | 1846.392987 | 0.328397246 | 0.148103473 | 0.022673276 | 0.035470096 | ns | YPD | KS.3 |
| FIM1_3918 | 1376.629959 | 530.0350155 | 1266.487463 | -1.259481429 | 0.230512688 | 4.3553830159 | 2.7292310471 | Km<KS.3 | YPD | KS.3 |
| FIM1_3919 | 357.5462514 | 290.9923543 | 331.3315058 | -0.278736117 | 0.161019567 | 0.073879039 | 0.102438188 | ns | YPD | KS.3 |
| FIM1_3920 | 266.1503445 | 205.6178812 | 231.2274604 | -0.351145944 | 0.149974863 | 0.01603584 | 0.026054312 | ns | YPD | KS.3 |
| FIM1_3921 | 3997.054182 | 3293.901989 | 2798.983204 | -0.26862999 | 0.124205651 | 0.027435253 | 0.042197605 | ns | YPD | KS.3 |
| FIM1_3922 | 807.983747 | 338.9057336 | 500.5854168 | -1.197686974 | 0.158140367 | 4.3102580800 | 8.6815926916 | Km<KS.3 | YPD | KS.3 |

| | | | | | | | | | | |
|---|---|---|---|---|---|---|---|---|---|---|
| FIM1_3923 | 3049.526949 | 1899.020174 | 3006.232563 | -0.660352828 | 0.123075201 | 4.3997796167 | 2.268280165 | ns | YPD | KS.3 |
| FIM1_3924 | 423.4464931 | 205.2521576 | 705.9763253 | -0.792487947 | 0.371928362 | 0.011939345 | 0.019876011 | ns | YPD | KS.3 |
| FIM1_3925 | 256.8595233 | 241.3260374 | 212.0267322 | -0.07940641 | 0.222702228 | 0.705162432 | 0.753313957 | ns | YPD | KS.3 |
| FIM1_3926 | 1591.940201 | 2601.879369 | 1880.232738 | 0.680031626 | 0.135826366 | 2.908028543 | 1.267961613 | ns | YPD | KS.3 |
| FIM1_3927 | 259.4854524 | 157.1293987 | 370.1716768 | -0.67101197 | 0.184883268 | 0.000150479 | 0.000374504 | ns | YPD | KS.3 |
| FIM1_3928 | 2162.49761 | 3207.272805 | 2189.577524 | 0.539667413 | 0.150557559 | 0.000224627 | 0.000532339 | ns | YPD | KS.3 |
| FIM1_3929 | 1541.067134 | 881.9248522 | 1378.074944 | -0.749887168 | 0.182367119 | 1.776839897 | 5.312366879 | ns | YPD | KS.3 |
| FIM1_3930 | 1309.170271 | 981.1035864 | 1070.974216 | -0.40691852 | 0.094096158 | 1.213870418 | 3.743958667 | ns | YPD | KS.3 |
| FIM1_3932 | 2378.048122 | 1388.82552 | 2228.120119 | -0.755606836 | 0.10784866 | 1.095930278 | 1.427291396 | ns | YPD | KS.3 |
| FIM1_3933 | 612.9727256 | 399.5288086 | 529.5501752 | -0.594518121 | 0.130284241 | 3.082003554 | 1.086673825 | ns | YPD | KS.3 |
| FIM1_3934 | 1343.437816 | 1258.814097 | 1094.394944 | -0.090706344 | 0.107999865 | 0.394025779 | 0.457367767 | ns | YPD | KS.3 |
| FIM1_3935 | 1853.616331 | 1451.770778 | 1625.570104 | -0.334610222 | 0.146475259 | 0.018950005 | 0.030257163 | ns | YPD | KS.3 |
| FIM1_3936 | 1366.611049 | 1065.605878 | 891.8872015 | -0.351646043 | 0.090048704 | 7.920893748 | 0.000207196 | ns | YPD | KS.3 |
| FIM1_3937 | 68.6537807 | 34.92549966 | 34.38314948 | -0.754786101 | 0.356371723 | 0.013648334 | 0.022461285 | ns | YPD | KS.3 |
| FIM1_3938 | 70.45928851 | 106.71288 | 80.08171728 | 0.513541136 | 0.261323954 | 0.032782549 | 0.049437157 | ns | YPD | KS.3 |
| FIM1_3939 | 301.6969999 | 897.0415487 | 588.5738225 | 1.422345719 | 0.260119841 | 2.044679992 | 1.380228353 | Km>KS.3 | YPD | KS.3 |
| FIM1_3940 | 1075.038653 | 4669.973277 | 2416.895088 | 1.917068616 | 0.328363733 | 2.586207621 | 2.552853753 | Km>KS.3 | YPD | KS.3 |
| FIM1_3942 | 719.6693769 | 680.9649738 | 686.7445039 | -0.077347465 | 0.110391861 | 0.47677128 | 0.540427687 | ns | YPD | KS.3 |
| FIM1_3943 | 4782.025607 | 5428.06726 | 4597.163711 | 0.179478067 | 0.084428075 | 0.031907761 | 0.048190987 | ns | YPD | KS.3 |
| FIM1_3944 | 1329.090276 | 675.9556709 | 950.884262 | -0.955633022 | 0.097215352 | 2.211349427 | 1.718978656 | ns | YPD | KS.3 |
| FIM1_3945 | 1162.691367 | 453.7329693 | 915.1070056 | -1.242800658 | 0.228418143 | 5.309753638 | 3.269309944 | Km<KS.3 | YPD | KS.3 |
| FIM1_3946 | 1691.538823 | 834.5127586 | 1684.428242 | -0.841226052 | 0.310765196 | 0.002320373 | 0.004462255 | ns | YPD | KS.3 |
| FIM1_3947 | 1649.322707 | 1562.487082 | 1402.901195 | -0.07484064 | 0.128768996 | 0.552974924 | 0.614210816 | ns | YPD | KS.3 |
| FIM1_3948 | 1960.387367 | 2159.206299 | 1624.501317 | 0.136936584 | 0.088490838 | 0.118014787 | 0.156191425 | ns | YPD | KS.3 |
| FIM1_3949 | 1994.94043 | 1895.955865 | 1524.536322 | -0.071403875 | 0.092089471 | 0.433133058 | 0.497193577 | ns | YPD | KS.3 |
| FIM1_3950 | 1602.713688 | 1434.768224 | 1387.801633 | -0.156785396 | 0.08943773 | 0.07653121 | 0.105784111 | ns | YPD | KS.3 |
| FIM1_3951 | 1919.858214 | 2344.804463 | 2008.560255 | 0.285025261 | 0.066803208 | 1.768960520 | 5.291989531 | ns | YPD | KS.3 |
| FIM1_3952 | 1000.873817 | 2955.774996 | 1720.543302 | 1.349723364 | 0.311826311 | 9.360747057 | 3.698944925 | Km>KS.3 | YPD | KS.3 |
| FIM1_3953 | 1523.26763 | 1820.059155 | 1806.123057 | 0.247942454 | 0.117493225 | 0.031757621 | 0.048007951 | ns | YPD | KS.3 |
| FIM1_3954 | 877.9331794 | 621.2133134 | 635.473512 | -0.47573635 | 0.140493079 | 0.000515799 | 0.001132131 | ns | YPD | KS.3 |
| FIM1_3955 | 1047.809309 | 1066.689069 | 717.4408983 | 0.023633958 | 0.184319701 | 0.893719412 | 0.912801083 | ns | YPD | KS.3 |
| FIM1_3956 | 3281.991838 | 1372.157644 | 3147.383048 | -1.136461781 | 0.238411514 | 2.809117697 | 1.229143407 | Km<KS.3 | YPD | KS.3 |
| FIM1_3957 | 2422.202027 | 3575.422935 | 2442.72559 | 0.556370403 | 0.06288155 | 5.741096873 | 2.441192901 | ns | YPD | KS.3 |
| FIM1_3958 | 4461.165582 | 2756.441739 | 3999.83523 | -0.580080938 | 0.287111723 | 0.025601453 | 0.039591926 | ns | YPD | KS.3 |
| FIM1_3959 | 1144.6207 | 950.1558987 | 783.6072442 | -0.260557586 | 0.109627795 | 0.015803106 | 0.025701358 | ns | YPD | KS.3 |
| FIM1_3960 | 272.647951 | 190.0586328 | 230.82045 | -0.455783189 | 0.243112119 | 0.043993674 | 0.064311206 | ns | YPD | KS.3 |
| FIM1_3961 | 1102.197899 | 1087.826993 | 941.5538678 | -0.015360124 | 0.296457942 | 0.954144331 | 0.964418538 | ns | YPD | KS.3 |
| FIM1_3962 | 1113.248239 | 920.8446269 | 1344.310423 | -0.263133345 | 0.127967593 | 0.035815429 | 0.053508036 | ns | YPD | KS.3 |
| FIM1_3963 | 625.2554457 | 1966.868911 | 1670.37721 | 1.496254009 | 0.2673587 | 6.861678142 | 5.072340083 | Km>KS.3 | YPD | KS.3 |
| FIM1_3964 | 1746.305203 | 3833.823906 | 2477.98014 | 0.952964509 | 0.302915476 | 0.00042523 | 0.000954226 | ns | YPD | KS.3 |
| FIM1_3965 | 543.2647802 | 490.7714369 | 578.3806888 | -0.141282696 | 0.132349567 | 0.275271061 | 0.333448631 | ns | YPD | KS.3 |
| FIM1_3966 | 442.716089 | 1356.522259 | 404.0608576 | 1.192635391 | 0.444110908 | 0.000740378 | 0.001573422 | Km>KS.3 | YPD | KS.3 |
| FIM1_3967 | 1254.573956 | 1082.465959 | 1412.758355 | -0.200107331 | 0.160743198 | 0.198786034 | 0.249892112 | ns | YPD | KS.3 |
| FIM1_3968 | 1450.83773 | 1504.271089 | 1315.968476 | 0.050328674 | 0.120573984 | 0.670791482 | 0.721866239 | ns | YPD | KS.3 |
| FIM1_3969 | 3074.573991 | 2919.943995 | 2221.040879 | -0.069595588 | 0.165740974 | 0.664141624 | 0.715948988 | ns | YPD | KS.3 |

| | | | | | | | | | | |
|---|---|---|---|---|---|---|---|---|---|---|
| FIM1_3970 | 928.6791005 | 1169.409646 | 906.8312738 | 0.299341582 | 0.210811512 | 0.134004222 | 0.175301343 | ns | YPD | KS.3 |
| FIM1_3971 | 305.1037605 | 532.7802049 | 324.094424 | 0.677995822 | 0.283187017 | 0.008292356 | 0.014288981 | ns | YPD | KS.3 |
| FIM1_3972 | 744.5475325 | 804.7642915 | 550.1518357 | 0.109776391 | 0.097862348 | 0.256154962 | 0.312805826 | ns | YPD | KS.3 |
| FIM1_3973 | 5471.798493 | 4141.534735 | 6210.686297 | -0.388392003 | 0.118958444 | 0.000885227 | 0.001856149 | ns | YPD | KS.3 |
| FIM1_3974 | 1369.215458 | 1067.355804 | 1268.56398 | -0.343222697 | 0.136818482 | 0.010216671 | 0.017311967 | ns | YPD | KS.3 |
| FIM1_3975 | 1026.347185 | 1373.056537 | 1007.307858 | 0.401313063 | 0.138216409 | 0.002934272 | 0.005556911 | ns | YPD | KS.3 |
| FIM1_3976 | 7951.481225 | 9736.523534 | 8974.615497 | 0.273968416 | 0.163891446 | 0.084007616 | 0.11475505 | ns | YPD | KS.3 |
| FIM1_3977 | 356.0235663 | 1870.322247 | 1321.053844 | 2.077452694 | 0.430694304 | 5.5149609174 | 3.3914623689 | Km>KS.3 | YPD | KS.3 |
| FIM1_3978 | 1009.578182 | 982.3603841 | 1124.950898 | -0.037344363 | 0.160336917 | 0.810067057 | 0.841002423 | ns | YPD | KS.3 |
| FIM1_3979 | 918.4182452 | 591.7233512 | 703.252064 | -0.618660042 | 0.101982495 | 7.6739290239 | 5.6392609887 | ns | YPD | KS.3 |
| FIM1_3980 | 11238.85687 | 9454.024748 | 15468.15665 | -0.24057456 | 0.121675739 | 0.04396559 | 0.064294182 | ns | YPD | KS.3 |
| FIM1_3981 | 346.2351635 | 710.1758445 | 506.7927928 | 0.863484185 | 0.304286674 | 0.00147804 | 0.002959055 | ns | YPD | KS.3 |
| FIM1_3982 | 4476.94675 | 3231.006793 | 3579.231815 | -0.455878221 | 0.114310327 | 4.9874741739 | 0.000134925 | ns | YPD | KS.3 |
| FIM1_3983 | 1941.967924 | 1587.2539 | 1834.252194 | -0.280329817 | 0.123993062 | 0.021157586 | 0.033267697 | ns | YPD | KS.3 |
| FIM1_3984 | 429.8000558 | 221.2744819 | 288.2195062 | -0.878342479 | 0.206682681 | 7.1318280317 | 2.3281394001 | ns | YPD | KS.3 |
| FIM1_3985 | 1012.986116 | 732.7709166 | 708.1430085 | -0.458590873 | 0.083789334 | 3.2988994506 | 1.7367221975 | ns | YPD | KS.3 |
| FIM1_3986 | 1369.450717 | 1827.683372 | 1607.990702 | 0.381278613 | 0.193932393 | 0.039509498 | 0.058517342 | ns | YPD | KS.3 |
| FIM1_3987 | 608.2519257 | 799.3259172 | 787.8348508 | 0.380705597 | 0.115012681 | 0.000760383 | 0.00161241 | ns | YPD | KS.3 |
| FIM1_3988 | 1466.97812 | 992.2181416 | 1013.687429 | -0.544583341 | 0.121912466 | 5.2546184291 | 1.7711196940 | ns | YPD | KS.3 |
| FIM1_3989 | 12511.01375 | 11449.98533 | 10098.21849 | -0.124554964 | 0.10140782 | 0.213341939 | 0.265875789 | ns | YPD | KS.3 |
| FIM1_3990 | 1686.936378 | 2491.784107 | 1567.976856 | 0.487947697 | 0.251680567 | 0.036295102 | 0.054127138 | ns | YPD | KS.3 |
| FIM1_3991 | 653.4559116 | 970.2636016 | 1565.42951 | 0.492228624 | 0.255553553 | 0.037087073 | 0.055241973 | ns | YPD | KS.3 |
| FIM1_3992 | 670.1620021 | 667.3353372 | 725.4346143 | -0.005729152 | 0.088316746 | 0.947757116 | 0.959117621 | ns | YPD | KS.3 |
| FIM1_3993 | 7370.297088 | 8044.104023 | 8151.152732 | 0.125280851 | 0.04752559 | 0.008187852 | 0.014129228 | ns | YPD | KS.3 |
| FIM1_3994 | 106.897072 | 72.65204414 | 83.69121574 | -0.506646265 | 0.201247523 | 0.008124965 | 0.014038081 | ns | YPD | KS.3 |
| FIM1_3995 | 161.8818749 | 221.350227 | 185.7070297 | 0.384080667 | 0.264720065 | 0.114488063 | 0.151766084 | ns | YPD | KS.3 |
| FIM1_3996 | 753.4909626 | 1253.573478 | 1615.367025 | 0.59627268 | 0.309807082 | 0.030735136 | 0.046603871 | ns | YPD | KS.3 |
| FIM1_3997 | 1728.462271 | 2271.804584 | 1901.564487 | 0.379264695 | 0.127114904 | 0.002322233 | 0.004464108 | ns | YPD | KS.3 |
| FIM1_3998 | 1726.532539 | 1166.166782 | 1173.428708 | -0.534223398 | 0.157999319 | 0.00048312 | 0.001070612 | ns | YPD | KS.3 |
| FIM1_3999 | 196.8971207 | 91.42665482 | 239.0508598 | -0.998255511 | 0.233153749 | 4.3899749151 | 1.5020718846 | ns | YPD | KS.3 |
| FIM1_4000 | 828.0249983 | 742.7708318 | 866.1570562 | -0.148072113 | 0.154052557 | 0.322360357 | 0.383762329 | ns | YPD | KS.3 |
| FIM1_4001 | 131.284787 | 100.0475792 | 118.8632034 | -0.352238757 | 0.217355132 | 0.086129127 | 0.117330889 | ns | YPD | KS.3 |
| FIM1_4002 | 4099.464355 | 4064.909119 | 4722.895656 | -0.012174902 | 0.080948731 | 0.879436116 | 0.900616442 | ns | YPD | KS.3 |
| FIM1_4003 | 3650.487294 | 2990.954976 | 4266.578467 | -0.279396663 | 0.107730545 | 0.008482132 | 0.014581412 | ns | YPD | KS.3 |
| FIM1_4004 | 1069.570897 | 867.8862871 | 1181.977227 | -0.294688007 | 0.097036971 | 0.002111618 | 0.00409723 | ns | YPD | KS.3 |
| FIM1_4005 | 1894.306811 | 3234.641548 | 2922.143101 | 0.729529421 | 0.160720461 | 2.6815254736 | 9.5562960110 | ns | YPD | KS.3 |
| FIM1_4006 | 3229.469566 | 2126.287126 | 2646.572787 | -0.59238634 | 0.085252974 | 2.2563379239 | 2.7785349435 | ns | YPD | KS.3 |
| FIM1_4007 | 1736.416964 | 1574.093806 | 1713.896372 | -0.1397077 | 0.07783455 | 0.070424184 | 0.098002885 | ns | YPD | KS.3 |
| FIM1_4008 | 628.1616122 | 331.819405 | 533.3020394 | -0.832639023 | 0.219972394 | 5.6977695239 | 0.0001524 | ns | YPD | KS.3 |
| FIM1_4009 | 2919.012005 | 2117.33358 | 3342.353643 | -0.438658769 | 0.152273127 | 0.003016297 | 0.005690586 | ns | YPD | KS.3 |
| FIM1_4010 | 1876.77404 | 2031.576119 | 1854.404496 | 0.112533733 | 0.08401816 | 0.176476857 | 0.22552591 | ns | YPD | KS.3 |
| FIM1_4011 | 2531.118946 | 1841.282987 | 1968.013867 | -0.454278577 | 0.061291539 | 9.3134755999 | 1.4709378130 | ns | YPD | KS.3 |
| FIM1_4012 | 2920.691807 | 2112.171385 | 2395.085632 | -0.454053776 | 0.110515382 | 2.9891436582 | 8.5025670095 | ns | YPD | KS.3 |
| FIM1_4013 | 1964.801081 | 1832.017624 | 1792.394935 | -0.100030069 | 0.07123567 | 0.157479218 | 0.20317923 | ns | YPD | KS.3 |
| FIM1_4014 | 456.4300233 | 203.2428182 | 366.1548634 | -1.092741165 | 0.186205661 | 7.7798758363 | 5.7002772143 | Km<KS.3 | YPD | KS.3 |

| | | | | | | | | | | |
|---|---|---|---|---|---|---|---|---|---|---|
| FIM1_4015 | 3999.823187 | 2082.952924 | 2846.782905 | -0.879460529 | 0.181592131 | 4.2771211530 | 1.8017508667 | ns | YPD | KS.3 |
| FIM1_4016 | 2618.035885 | 2950.281267 | 2731.732684 | 0.164403294 | 0.13989607 | 0.228577422 | 0.282527373 | ns | YPD | KS.3 |
| FIM1_4017 | 464.111461 | 267.333346 | 389.0880982 | -0.735585521 | 0.191536394 | 5.7165704298 | 0.000152821 | ns | YPD | KS.3 |
| FIM1_4018 | 3359.229885 | 2224.689898 | 4265.881815 | -0.471326703 | 0.322056342 | 0.097570128 | 0.131298725 | ns | YPD | KS.3 |
| FIM1_4019 | 1175.627255 | 781.9138016 | 802.5732301 | -0.570952339 | 0.110770923 | 1.6184419258 | 7.5249986738 | ns | YPD | KS.3 |
| FIM1_4020 | 4265.82637 | 2997.166295 | 3839.847189 | -0.50054181 | 0.083960696 | 1.7612471331 | 1.2136017295 | ns | YPD | KS.3 |
| FIM1_4021 | 1479.348773 | 852.4702141 | 1069.033636 | -0.764578342 | 0.134093964 | 5.2268779439 | 3.2222698601 | ns | YPD | KS.3 |
| FIM1_4022 | 39.94843995 | 17.30497026 | 31.43776702 | -0.984153628 | 0.333179304 | 0.000716306 | 0.00152683 | ns | YPD | KS.3 |
| FIM1_4023 | 7336.377776 | 6903.339784 | 7102.690264 | -0.087142847 | 0.056987176 | 0.1245779 | 0.164396566 | ns | YPD | KS.3 |
| FIM1_4024 | 2843.121627 | 2299.811931 | 2905.728058 | -0.241603868 | 0.317854025 | 0.391380096 | 0.454721153 | ns | YPD | KS.3 |
| FIM1_4025 | 402.6694858 | 171.6202757 | 257.0080608 | -1.153991526 | 0.187815894 | 1.1406847335 | 9.8694026947 | Km<KS.3 | YPD | KS.3 |
| FIM1_4026 | 318.4864951 | 172.8398535 | 370.4821554 | -0.695305325 | 0.337353758 | 0.018115219 | 0.029072004 | ns | YPD | KS.3 |
| FIM1_4027 | 404.4954251 | 256.1562082 | 621.9376762 | -0.502683807 | 0.34905545 | 0.095795689 | 0.129120443 | ns | YPD | KS.3 |
| FIM1_4028 | 1167.335016 | 997.5352089 | 1191.808455 | -0.222025659 | 0.088986008 | 0.011709334 | 0.01953519 | ns | YPD | KS.3 |
| FIM1_4029 | 91.67334701 | 191.9897176 | 104.9791132 | 0.905499999 | 0.289813164 | 0.000526621 | 0.001154159 | ns | YPD | KS.3 |
| FIM1_4030 | 5050.071394 | 3268.998802 | 5795.456378 | -0.529794065 | 0.276157626 | 0.035402995 | 0.052955472 | ns | YPD | KS.3 |
| FIM1_4031 | 3753.881003 | 2232.038759 | 3294.908715 | -0.723681745 | 0.126829489 | 5.5649401315 | 3.4137579722 | ns | YPD | KS.3 |
| FIM1_4033 | 1087.948033 | 596.2284313 | 824.2126454 | -0.835433442 | 0.131978399 | 9.1573111824 | 8.1938171102 | ns | YPD | KS.3 |
| FIM1_4034 | 655.8033104 | 336.0984165 | 554.5582895 | -0.915647327 | 0.159658161 | 2.9690417035 | 1.9258125782 | ns | YPD | KS.3 |
| FIM1_4035 | 470.7571842 | 217.1020202 | 436.1044789 | -1.047848446 | 0.183427974 | 2.2865867617 | 1.5269488778 | Km<KS.3 | YPD | KS.3 |
| FIM1_4036 | 2591.593415 | 1600.988442 | 1895.044124 | -0.664028754 | 0.142116929 | 1.6104063606 | 6.0695239728 | ns | YPD | KS.3 |
| FIM1_4037 | 2417.127083 | 1373.977545 | 1354.565124 | -0.793234656 | 0.109650536 | 1.9244602974 | 2.8326005858 | ns | YPD | KS.3 |
| FIM1_4038 | 1054.513035 | 713.2322834 | 775.7880711 | -0.54633511 | 0.114090574 | 1.1097973345 | 4.3204290135 | ns | YPD | KS.3 |
| FIM1_4039 | 864.206535 | 814.8430103 | 1193.073608 | -0.080460769 | 0.14592625 | 0.57137746 | 0.631061471 | ns | YPD | KS.3 |
| FIM1_4041 | 523.6074759 | 605.4144176 | 1151.064925 | 0.168844918 | 0.301610692 | 0.532841347 | 0.594902536 | ns | YPD | KS.3 |
| FIM1_4042 | 1937.043795 | 1625.682146 | 1666.45966 | -0.24519489 | 0.110047124 | 0.023654959 | 0.036810579 | ns | YPD | KS.3 |
| FIM1_4043 | 1229.523908 | 1736.913666 | 1573.467723 | 0.470335079 | 0.157205066 | 0.002025537 | 0.003942507 | ns | YPD | KS.3 |
| FIM1_4044 | 291.7278706 | 355.8779593 | 280.2021216 | 0.227852685 | 0.313011982 | 0.413447291 | 0.477349797 | ns | YPD | KS.3 |
| FIM1_4045 | 4325.749924 | 3482.957594 | 4361.899764 | -0.293131851 | 0.164332941 | 0.065154982 | 0.091437527 | ns | YPD | KS.3 |
| FIM1_4046 | 1403.480766 | 840.7868838 | 1270.611571 | -0.691406116 | 0.175015481 | 3.9835117008 | 0.000109917 | ns | YPD | KS.3 |
| FIM1_4047 | 8857.443179 | 9102.994122 | 7888.412633 | 0.037842435 | 0.129850543 | 0.766011199 | 0.803818966 | ns | YPD | KS.3 |
| FIM1_4048 | 376.9647996 | 333.9561147 | 405.4900098 | -0.159877551 | 0.196656628 | 0.394292361 | 0.457391304 | ns | YPD | KS.3 |
| FIM1_4049 | 1480.927356 | 510.9399597 | 1563.26928 | -1.254310498 | 0.360849495 | 4.5357507943 | 0.000123646 | Km<KS.3 | YPD | KS.3 |
| FIM1_4050 | 4267.360872 | 5160.043408 | 3856.464842 | 0.26968854 | 0.079188061 | 0.000595012 | 0.001292092 | ns | YPD | KS.3 |
| FIM1_4051 | 5510.2841 | 5301.8253 | 8292.48385 | -0.053963277 | 0.114308082 | 0.631302061 | 0.686203386 | ns | YPD | KS.3 |
| FIM1_4052 | 56.56934729 | 116.0727762 | 75.04869013 | 0.95493935 | 0.20022597 | 5.0220473019 | 2.0872752988 | ns | YPD | KS.3 |
| FIM1_4053 | 351.3796268 | 1890.879279 | 749.4921098 | 2.146856997 | 0.4098891 | 5.3196008386 | 3.9977362797 | Km>KS.3 | YPD | KS.3 |
| FIM1_4054 | 2108.458498 | 4047.19441 | 2907.120408 | 0.886543266 | 0.169353854 | 5.4321785177 | 2.7689639473 | ns | YPD | KS.3 |
| FIM1_4055 | 1692.018431 | 2204.459139 | 1475.51348 | 0.344040711 | 0.211079804 | 0.085326548 | 0.116365015 | ns | YPD | KS.3 |
| FIM1_4056 | 1238.883919 | 2514.996326 | 1623.584067 | 0.982942418 | 0.137484951 | 2.1783788773 | 3.1596020159 | ns | YPD | KS.3 |
| FIM1_4057 | 1364.244491 | 1201.692764 | 1189.612241 | -0.177877078 | 0.105497081 | 0.087246302 | 0.118625404 | ns | YPD | KS.3 |
| FIM1_4058 | 525.0599651 | 348.3913282 | 385.20337 | -0.560007919 | 0.15557807 | 0.000205408 | 0.000490663 | ns | YPD | KS.3 |
| FIM1_4059 | 1630.553462 | 6058.97723 | 4790.05668 | 1.70027904 | 0.307021745 | 3.6241024374 | 2.8348914507 | Km>KS.3 | YPD | KS.3 |
| FIM1_4060 | 1814.64342 | 1265.626602 | 2445.498945 | -0.449737797 | 0.252938039 | 0.054718904 | 0.078068984 | ns | YPD | KS.3 |
| FIM1_4061 | 4107.153915 | 2433.334649 | 5649.129174 | -0.643984357 | 0.272101359 | 0.009587033 | 0.016317306 | ns | YPD | KS.3 |

| | | | | | | | | | | |
|---|---|---|---|---|---|---|---|---|---|---|
| FIM1_4062 | 933.1691941 | 535.254817 | 1445.784866 | -0.651576799 | 0.31238851 | 0.018951019 | 0.030257163 | ns | YPD | KS.3 |
| FIM1_4063 | 2370.037323 | 2890.616044 | 2330.080871 | 0.27097623 | 0.151648021 | 0.065981591 | 0.092441119 | ns | YPD | KS.3 |
| FIM1_4064 | 4799.076602 | 2997.082094 | 5527.055317 | -0.628205956 | 0.187706717 | 0.000467418 | 0.001038127 | ns | YPD | KS.3 |
| FIM1_4065 | 2156.251181 | 2672.966079 | 4374.277099 | 0.254219562 | 0.29116808 | 0.334127512 | 0.395917581 | ns | YPD | KS.3 |
| FIM1_4066 | 1718.744923 | 4024.627488 | 3727.910755 | 1.196521157 | 0.116608364 | 1.2106069151( | 1.1809351769( | Km>KS.3 | YPD | KS.3 |
| FIM1_4067 | 716.5097156 | 1229.62728 | 1068.184694 | 0.760104006 | 0.104386086 | 1.4590513483( | 2.2334709101( | ns | YPD | KS.3 |
| FIM1_4068 | 286.7809739 | 293.3768372 | 406.8654862 | 0.030096129 | 0.174685445 | 0.858104825 | 0.881675239 | ns | YPD | KS.3 |
| FIM1_4069 | 2053.525824 | 4486.114749 | 2644.230048 | 0.991887999 | 0.259552311 | 3.1494508868< | 8.9038839170< | ns | YPD | KS.3 |
| FIM1_4070 | 2508.689494 | 3703.547932 | 2799.421974 | 0.483439526 | 0.258522215 | 0.04260255 | 0.06253989 | ns | YPD | KS.3 |
| FIM1_4071 | 67.29927998 | 127.6209435 | 102.4320302 | 0.810306806 | 0.25099741 | 0.00047886 | 0.0010617 | ns | YPD | KS.3 |
| FIM1_4072 | 2293.732916 | 1765.637168 | 1604.04108 | -0.36698733 | 0.107128495 | 0.00050743 | 0.001116526 | ns | YPD | KS.3 |
| FIM1_4073 | 1349.196669 | 779.9762181 | 1025.178863 | -0.765874986 | 0.119215939 | 5.8132249705( | 5.3656390034< | ns | YPD | KS.3 |
| FIM1_4074 | 1204.690982 | 1894.551666 | 1295.36021 | 0.535297072 | 0.300241481 | 0.046911631 | 0.068101945 | ns | YPD | KS.3 |
| FIM1_4075 | 689.5427909 | 1860.059437 | 728.1756904 | 1.276544694 | 0.265353958 | 1.2996378212! | 6.1170275880! | Km>KS.3 | YPD | KS.3 |
| FIM1_4076 | 7.413104574 | 21.9323961 | 18.61852671 | 1.240667371 | 0.387210439 | 0.000124106 | 0.000313415 | Km>KS.3 | YPD | KS.3 |
| FIM1_4077 | 13316.96082 | 39256.27338 | 36444.03178 | 1.361828243 | 0.300497665 | 3.4685939469( | 1.4863268635! | Km>KS.3 | YPD | KS.3 |
| FIM1_4078 | 112.4681455 | 591.1830993 | 253.8581896 | 1.995287956 | 0.480038675 | 1.7089035829! | 7.9012967707< | Km>KS.3 | YPD | KS.3 |
| FIM1_4080 | 3975.679216 | 9574.640049 | 3950.898547 | 1.076826318 | 0.302421598 | 6.5313186546( | 0.000173205 | Km>KS.3 | YPD | KS.3 |
| FIM1_4081 | 14.83676195 | 27.66445919 | 26.3944966 | 0.766606967 | 0.272547531 | 0.002054901 | 0.003993411 | ns | YPD | KS.3 |
| FIM1_4082 | 16.51807342 | 67.2344887 | 53.98004836 | 1.73112368 | 0.386710956 | 8.8090674455! | 4.3177448809( | Km>KS.3 | YPD | KS.3 |
| FIM1_4083 | 363.0049149 | 359.4183946 | 621.9285562 | -0.014218151 | 0.147112159 | 0.920937061 | 0.935796952 | ns | YPD | KS.3 |
| FIM1_4084 | 3133.320909 | 2643.568804 | 3451.86931 | -0.233297163 | 0.143611521 | 0.095585211 | 0.12887166 | ns | YPD | KS.3 |
| FIM1_4085 | 7865.763193 | 10488.75853 | 9779.713209 | 0.402289255 | 0.113763149 | 0.000323554 | 0.000742131 | ns | YPD | KS.3 |
| FIM1_4086 | 7222.547642 | 6848.24278 | 11459.64232 | -0.068415247 | 0.220042198 | 0.74179758 | 0.783699928 | ns | YPD | KS.3 |
| FIM1_4087 | 232.5289212 | 311.1483816 | 342.0848434 | 0.349864887 | 0.280636544 | 0.170735959 | 0.21911801 | ns | YPD | KS.3 |
| FIM1_4088 | 1494.144141 | 3918.137765 | 2166.353819 | 1.234660109 | 0.267226541 | 3.9235587018! | 1.6655038004! | Km>KS.3 | YPD | KS.3 |
| FIM1_4089 | 242.7787838 | 250.4051942 | 243.3551311 | 0.04285498 | 0.149450425 | 0.768205703 | 0.805442228 | ns | YPD | KS.3 |
| FIM1_4090 | 234.3941529 | 260.6770159 | 225.2102565 | 0.149741313 | 0.110837239 | 0.170022122 | 0.218399189 | ns | YPD | KS.3 |
| FIM1_4091 | 1975.64298 | 1130.043887 | 2847.784754 | -0.644637698 | 0.323711312 | 0.023881342 | 0.03713963 | ns | YPD | KS.3 |
| FIM1_4092 | 1467.651477 | 1094.556866 | 1310.075233 | -0.412666279 | 0.099876032 | 2.8392055200< | 8.1178433690( | ns | YPD | KS.3 |
| FIM1_4093 | 943.6673045 | 2870.326753 | 1490.982194 | 1.389912734 | 0.313904989 | 4.9741304492( | 2.0690885439! | Km>KS.3 | YPD | KS.3 |
| FIM1_4094 | 1457.421422 | 6189.265972 | 6504.902137 | 1.935592803 | 0.282119631 | 2.7655772020( | 4.8446290775! | Km>KS.3 | YPD | KS.3 |
| FIM1_4095 | 1174.12615 | 1165.34038 | 1717.111691 | -0.010281285 | 0.162609755 | 0.947935949 | 0.959117621 | ns | YPD | KS.3 |
| FIM1_4096 | 1223.197628 | 1041.960782 | 1803.497877 | -0.21764342 | 0.16026086 | 0.161027155 | 0.207380299 | ns | YPD | KS.3 |
| FIM1_4097 | 809.7266253 | 1204.460753 | 1921.574245 | 0.438450058 | 0.344163206 | 0.142300752 | 0.184938934 | ns | YPD | KS.3 |
| FIM1_4098 | 888.3195489 | 1252.393209 | 1806.243203 | 0.44235679 | 0.222123404 | 0.034427401 | 0.051682655 | ns | YPD | KS.3 |
| FIM1_4099 | 245.0494229 | 259.5703868 | 365.0170713 | 0.074667668 | 0.191092928 | 0.68301786 | 0.733595391 | ns | YPD | KS.3 |
| FIM1_4100 | 318.8810828 | 521.7710121 | 354.4049164 | 0.690468968 | 0.109726747 | 1.6021993218( | 1.3579116910! | ns | YPD | KS.3 |
| FIM1_4101 | 518.9617525 | 1042.843475 | 806.4622174 | 0.931923628 | 0.195955547 | 5.7333727004( | 2.3514863301! | ns | YPD | KS.3 |
| FIM1_4102 | 2086.740379 | 2679.089771 | 2062.510701 | 0.346027515 | 0.129773013 | 0.006453081 | 0.0113683 | ns | YPD | KS.3 |
| FIM1_4103 | 2373.653047 | 2484.655072 | 4905.46933 | 0.060020543 | 0.197775393 | 0.750397878 | 0.791273727 | ns | YPD | KS.3 |
| FIM1_4104 | 489.4652662 | 1118.996374 | 1512.017908 | 1.063001944 | 0.249616037 | 3.9442060207! | 1.3674163730! | Km>KS.3 | YPD | KS.3 |
| FIM1_4105 | 62.88150394 | 142.5103763 | 221.2919914 | 1.046179985 | 0.253356168 | 7.3464016658! | 2.3872206589( | Km>KS.3 | YPD | KS.3 |
| FIM1_4106 | 4510.919171 | 8173.866031 | 8180.424686 | 0.753808434 | 0.248227511 | 0.00104439 | 0.002156846 | ns | YPD | KS.3 |
| FIM1_4107 | 1290.210409 | 1037.312115 | 1193.939476 | -0.306690398 | 0.103831853 | 0.002744696 | 0.005217754 | ns | YPD | KS.3 |

| | | | | | | | | | | |
|---|---|---|---|---|---|---|---|---|---|---|
| FIM1_4108 | 3466.214333 | 2052.362389 | 2811.311319 | -0.744558548 | 0.081537281 | 3.05889770673 | 1.72932001033 | ns | YPD | KS.3 |
| FIM1_4109 | 386.8617425 | 257.3276437 | 326.6748471 | -0.536730615 | 0.202416927 | 0.005274482 | 0.0094221 | ns | YPD | KS.3 |
| FIM1_4110 | 682.7864752 | 858.7666248 | 929.8709923 | 0.315291698 | 0.139321386 | 0.020406751 | 0.032304849 | ns | YPD | KS.3 |
| FIM1_4111 | 2504.218086 | 2535.706391 | 2452.532085 | 0.017827247 | 0.05830837 | 0.758742548 | 0.798718615 | ns | YPD | KS.3 |
| FIM1_4112 | 1916.915614 | 3566.876589 | 2371.194331 | 0.804634615 | 0.227682955 | 0.000161539 | 0.000398244 | ns | YPD | KS.3 |
| FIM1_4113 | 1158.059002 | 1482.614249 | 1778.14431 | 0.294682358 | 0.286082792 | 0.256088374 | 0.312801291 | ns | YPD | KS.3 |
| FIM1_4114 | 599.1656539 | 1352.451956 | 2048.426331 | 0.890847723 | 0.380933814 | 0.005415524 | 0.009642889 | ns | YPD | KS.3 |
| FIM1_4115 | 833.0944396 | 1701.491941 | 1095.168122 | 0.95880327 | 0.189764262 | 1.17387120482 | 5.57785028079 | ns | YPD | KS.3 |
| FIM1_4116 | 112.0395159 | 292.3228448 | 254.7718731 | 1.136077282 | 0.339933705 | 0.000116266 | 0.000296019 | Km>KS.3 | YPD | KS.3 |
| FIM1_4117 | 56.2520811 | 163.6916362 | 180.9317383 | 1.266212998 | 0.354342477 | 3.01524417649 | 8.57190844459 | Km>KS.3 | YPD | KS.3 |
| FIM1_4118 | 1000.910511 | 2956.138272 | 2095.015767 | 1.400338158 | 0.270727755 | 1.15416006186 | 6.69209863250 | Km>KS.3 | YPD | KS.3 |
| FIM1_4119 | 1077.648126 | 1877.622829 | 1782.112594 | 0.766809042 | 0.140598046 | 2.16274751424 | 1.18108330223 | ns | YPD | KS.3 |
| FIM1_4120 | 1957.563845 | 1927.062015 | 2779.77814 | -0.017394015 | 0.335665102 | 0.952764253 | 0.963415073 | ns | YPD | KS.3 |
| FIM1_4121 | 838.3382967 | 578.5177201 | 988.6484663 | -0.454565012 | 0.268733575 | 0.064987467 | 0.091234919 | ns | YPD | KS.3 |
| FIM1_4122 | 739.6446164 | 834.9203901 | 644.6217503 | 0.166332054 | 0.143439562 | 0.234249312 | 0.288534371 | ns | YPD | KS.3 |
| FIM1_4123 | 684.615336 | 1407.662585 | 1041.902857 | 0.988826273 | 0.15797514 | 9.54196371730 | 8.47701240956 | ns | YPD | KS.3 |
| FIM1_4124 | 52.28013203 | 80.01080639 | 61.31069651 | 0.529207929 | 0.258520119 | 0.026409791 | 0.04079128 | ns | YPD | KS.3 |
| FIM1_4125 | 4687.097716 | 10249.22469 | 7126.221652 | 1.054571986 | 0.188497266 | 4.44385891349 | 2.77392698803 | Km>KS.3 | YPD | KS.3 |
| FIM1_4126 | 236.5645265 | 409.3486729 | 369.1405331 | 0.754485306 | 0.148148691 | 1.59105889506 | 7.41153371062 | ns | YPD | KS.3 |
| FIM1_4127 | 33.80436373 | 68.71418263 | 56.92117253 | 0.872108369 | 0.281435152 | 0.000630284 | 0.001361711 | ns | YPD | KS.3 |
| FIM1_4128 | 1166.170173 | 1078.007936 | 1315.209878 | -0.108202248 | 0.139704431 | 0.427530635 | 0.491329385 | ns | YPD | KS.3 |
| FIM1_4129 | 2414.920184 | 1284.262807 | 2444.613147 | -0.86516777 | 0.157463756 | 1.36342712379 | 7.77872699641 | ns | YPD | KS.3 |
| FIM1_4130 | 1474.788857 | 3182.171424 | 2108.407292 | 1.032712843 | 0.192770001 | 1.82125952204 | 1.01350851478 | Km>KS.3 | YPD | KS.3 |
| FIM1_4132 | 223.5357053 | 242.0459046 | 198.7253864 | 0.110137755 | 0.121321319 | 0.355126607 | 0.417716097 | ns | YPD | KS.3 |
| FIM1_4133 | 848.0535155 | 830.1654133 | 883.5089162 | -0.028930196 | 0.14443116 | 0.837170527 | 0.863912751 | ns | YPD | KS.3 |
| FIM1_4134 | 2010.136495 | 1340.939306 | 1743.048971 | -0.565924578 | 0.115051753 | 5.58023285047 | 2.29624966344 | ns | YPD | KS.3 |
| FIM1_4135 | 867.7421947 | 1697.605675 | 1187.428661 | 0.924771823 | 0.148725612 | 1.49452583593 | 1.27534580339 | ns | YPD | KS.3 |
| FIM1_4136 | 1860.340426 | 2159.488576 | 2361.138264 | 0.209259231 | 0.104374855 | 0.042060331 | 0.061798625 | ns | YPD | KS.3 |
| FIM1_4137 | 2561.064204 | 3926.802748 | 3197.217042 | 0.564870445 | 0.197450281 | 0.002671275 | 0.005089848 | ns | YPD | KS.3 |
| FIM1_4138 | 1402.363785 | 1462.547984 | 1404.198415 | 0.059184759 | 0.099134147 | 0.545494753 | 0.606986445 | ns | YPD | KS.3 |
| FIM1_4139 | 3727.477975 | 1941.343752 | 3528.830738 | -0.826416778 | 0.252662464 | 0.000396755 | 0.000894768 | ns | YPD | KS.3 |
| FIM1_4140 | 1380.567405 | 1158.077847 | 1085.210523 | -0.223976901 | 0.229545173 | 0.298595636 | 0.358993062 | ns | YPD | KS.3 |
| FIM1_4141 | 8104.903386 | 6268.034419 | 8667.610626 | -0.35336265 | 0.141684977 | 0.010537122 | 0.017806447 | ns | YPD | KS.3 |
| FIM1_4142 | 886.1157859 | 1107.423568 | 850.6829663 | 0.312463951 | 0.107534748 | 0.003186685 | 0.005984808 | ns | YPD | KS.3 |
| FIM1_4143 | 712.9562318 | 768.7100939 | 875.8348652 | 0.104596315 | 0.11110826 | 0.338994249 | 0.400688142 | ns | YPD | KS.3 |
| FIM1_4144 | 561.5913992 | 246.0611788 | 395.2838946 | -1.116270328 | 0.187426505 | 4.21677530147 | 3.25247397284 | Km<KS.3 | YPD | KS.3 |
| FIM1_4145 | 670.8040948 | 512.5559358 | 511.0689433 | -0.361950009 | 0.17297235 | 0.029905755 | 0.045559245 | ns | YPD | KS.3 |
| FIM1_4146 | 1210.138386 | 759.9051001 | 1621.817164 | -0.49061661 | 0.375405539 | 0.122114965 | 0.161360411 | ns | YPD | KS.3 |
| FIM1_4147 | 630.4970928 | 1018.094677 | 825.2580876 | 0.640186185 | 0.186191507 | 0.000327797 | 0.000751517 | ns | YPD | KS.3 |
| FIM1_4148 | 1626.184861 | 5779.240923 | 3486.418128 | 1.639822744 | 0.300662994 | 7.76045004498 | 5.69443052710 | Km>KS.3 | YPD | KS.3 |
| FIM1_4149 | 10465.57698 | 7688.257644 | 12080.15282 | -0.43218421 | 0.109564292 | 6.16203189896 | 0.000164376 | ns | YPD | KS.3 |
| FIM1_4150 | 1389.680341 | 2251.59496 | 2153.571342 | 0.661126874 | 0.151574099 | 7.01125048981 | 2.29480073597 | ns | YPD | KS.3 |
| FIM1_4151 | 2461.038083 | 4872.712501 | 3026.452796 | 0.948776491 | 0.135638278 | 7.33786098653 | 9.86644821838 | ns | YPD | KS.3 |
| FIM1_4152 | 1609.810996 | 676.9771188 | 1951.519265 | -1.013845372 | 0.339374848 | 0.000584242 | 0.001270555 | Km<KS.3 | YPD | KS.3 |
| FIM1_4153 | 1768.780862 | 2486.894129 | 1687.778482 | 0.462227006 | 0.162400651 | 0.003261176 | 0.006108567 | ns | YPD | KS.3 |

| | | | | | | | | | | |
|---|---|---|---|---|---|---|---|---|---|---|
| FIM1_4154 | 1186.129352 | 792.3653786 | 796.973279 | -0.529128264 | 0.205270666 | 0.006614434 | 0.011627848 | ns | YPD | KS.3 |
| FIM1_4155 | 3522.785162 | 2193.143148 | 3767.699817 | -0.617768242 | 0.214766185 | 0.002317451 | 0.004458359 | ns | YPD | KS.3 |
| FIM1_4156 | 2321.956501 | 2111.597573 | 3625.250613 | -0.114647855 | 0.27473381 | 0.648061825 | 0.70119782 | ns | YPD | KS.3 |
| FIM1_4157 | 1718.446648 | 2820.263003 | 3012.997044 | 0.657732487 | 0.194319344 | 0.000385492 | 0.000871738 | ns | YPD | KS.3 |
| FIM1_4158 | 2818.815606 | 5188.664035 | 4488.575145 | 0.81254593 | 0.195451474 | 1.2643930508 | 3.8925466755 | ns | YPD | KS.3 |
| FIM1_4159 | 910.4591725 | 1241.169636 | 1271.519633 | 0.433720966 | 0.110196618 | 6.3774623979 | 0.000169396 | ns | YPD | KS.3 |
| FIM1_4160 | 308.1553333 | 314.9498007 | 355.4111926 | 0.028582215 | 0.150505817 | 0.845189741 | 0.871284493 | ns | YPD | KS.3 |
| FIM1_4161 | 8923.481782 | 12341.78428 | 15530.23963 | 0.457144761 | 0.097344533 | 1.9830091000 | 7.3005382413 | ns | YPD | KS.3 |
| FIM1_4162 | 1231.373173 | 1161.312813 | 1415.571465 | -0.064032201 | 0.341881619 | 0.829619466 | 0.858108423 | ns | YPD | KS.3 |
| FIM1_4163 | 3852.636502 | 4687.908302 | 7457.914414 | 0.272158036 | 0.125990794 | 0.02754032 | 0.042327183 | ns | YPD | KS.3 |
| FIM1_4164 | 62.56432712 | 59.36251134 | 109.5779923 | -0.056806064 | 0.339428338 | 0.847822415 | 0.873093876 | ns | YPD | KS.3 |
| FIM1_4165 | 377.4848803 | 220.1237948 | 424.6697933 | -0.732067516 | 0.167565446 | 5.8915995608 | 1.9689330434 | ns | YPD | KS.3 |
| FIM1_4166 | 513.4448976 | 4057.511165 | 2248.026681 | 2.873363205 | 0.26365266 | 1.7844936152 | 2.3362778252 | Km>KS.3 | YPD | KS.3 |
| FIM1_4167 | 503.5191207 | 1724.239205 | 1085.647549 | 1.521425282 | 0.346429076 | 3.2368036347 | 1.3979102129 | Km>KS.3 | YPD | KS.3 |
| FIM1_4169 | 724.0289641 | 930.4484729 | 672.9160416 | 0.355074477 | 0.088326763 | 4.8776460672 | 0.000132169 | ns | YPD | KS.3 |
| FIM1_4170 | 1040.224364 | 559.3332399 | 738.4157999 | -0.848482395 | 0.159916167 | 4.0674810538 | 2.1122879168 | ns | YPD | KS.3 |
| FIM1_4171 | 62.29514032 | 24.02103982 | 117.2484734 | -1.167675557 | 0.316425462 | 2.8627919340 | 8.1711932711 | Km<KS.3 | YPD | KS.3 |
| FIM1_4172 | 482.1550311 | 1550.700812 | 1125.987572 | 1.549114496 | 0.248918988 | 1.1449942435 | 1.2068530426 | Km>KS.3 | YPD | KS.3 |
| FIM1_4173 | 764.0898659 | 867.2559454 | 1128.570396 | 0.17640604 | 0.11754369 | 0.126736876 | 0.166759047 | ns | YPD | KS.3 |
| FIM1_4174 | 4942.820139 | 4203.43632 | 5769.751469 | -0.200055326 | 0.257273309 | 0.400073763 | 0.463738809 | ns | YPD | KS.3 |
| FIM1_4175 | 2216.467609 | 1646.335083 | 1627.710598 | -0.413344925 | 0.124141612 | 0.000683938 | 0.001467266 | ns | YPD | KS.3 |
| FIM1_4176 | 1674.201085 | 1418.270151 | 1316.289622 | -0.233606085 | 0.098882114 | 0.016742642 | 0.027087689 | ns | YPD | KS.3 |
| FIM1_4177 | 5185.865889 | 3615.635692 | 6783.080192 | -0.488328968 | 0.16584931 | 0.002304393 | 0.004436669 | ns | YPD | KS.3 |
| FIM1_4178 | 5144.737197 | 3440.530481 | 6196.614027 | -0.55153293 | 0.149413292 | 0.000145779 | 0.000363023 | ns | YPD | KS.3 |
| FIM1_4180 | 34.32873026 | 19.52110017 | 24.9312206 | -0.674120488 | 0.303441201 | 0.013003755 | 0.021485779 | ns | YPD | KS.3 |
| FIM1_4181 | 2430.656237 | 4029.764652 | 3689.600522 | 0.695222341 | 0.14691886 | 1.1324688862 | 4.3981519977 | ns | YPD | KS.3 |
| FIM1_4182 | 9068.711838 | 6968.822074 | 8869.187302 | -0.374068065 | 0.079334054 | 1.9869537695 | 7.3060569130 | ns | YPD | KS.3 |
| FIM1_4183 | 1919.948759 | 1446.524559 | 1556.276746 | -0.397768702 | 0.105062377 | 0.000122706 | 0.00031051 | ns | YPD | KS.3 |
| FIM1_4184 | 2471.554419 | 2455.781788 | 2060.641967 | -0.009163042 | 0.092777677 | 0.920458385 | 0.93569278 | ns | YPD | KS.3 |
| FIM1_4185 | 1021.815388 | 589.8805317 | 760.5632707 | -0.766760407 | 0.121221579 | 1.1072225057 | 9.6469911841 | ns | YPD | KS.3 |
| FIM1_4186 | 1916.158792 | 2144.65829 | 2235.521867 | 0.144004932 | 0.224995276 | 0.496472888 | 0.559445666 | ns | YPD | KS.3 |
| FIM1_4187 | 464.5635294 | 1109.06862 | 693.7005976 | 1.212251127 | 0.137872433 | 1.6090718374 | 7.6972426838 | Km>KS.3 | YPD | KS.3 |
| FIM1_4188 | 987.4489884 | 1523.812135 | 1297.856177 | 0.589688251 | 0.161396341 | 0.000158753 | 0.000392543 | ns | YPD | KS.3 |
| FIM1_4189 | 3108.336037 | 4114.31609 | 3247.958278 | 0.379035399 | 0.165999344 | 0.018135378 | 0.029094971 | ns | YPD | KS.3 |
| FIM1_4190 | 2658.023786 | 1428.478151 | 1947.08494 | -0.871117402 | 0.113969841 | 7.1803102020 | 1.3953923146 | ns | YPD | KS.3 |
| FIM1_4191 | 2029.33288 | 2406.102314 | 2019.30619 | 0.234827955 | 0.135749925 | 0.076710979 | 0.105980872 | ns | YPD | KS.3 |
| FIM1_4192 | 942.5720123 | 430.9491594 | 923.868796 | -1.019060774 | 0.232296321 | 2.5542381565 | 9.1485491929 | Km<KS.3 | YPD | KS.3 |
| FIM1_4193 | 409.5013373 | 286.2836586 | 265.2306088 | -0.465964407 | 0.210355374 | 0.019347323 | 0.030850299 | ns | YPD | KS.3 |
| FIM1_4194 | 1151.883687 | 1464.201805 | 1161.170013 | 0.321972457 | 0.174407756 | 0.055301282 | 0.078809476 | ns | YPD | KS.3 |
| FIM1_4195 | 1766.2665 | 2193.013982 | 2259.970138 | 0.302046103 | 0.11539502 | 0.007761927 | 0.013464291 | ns | YPD | KS.3 |
| FIM1_4196 | 2530.008417 | 5128.34601 | 3731.145569 | 0.953326027 | 0.182899534 | 5.1085736943 | 2.6093587402 | ns | YPD | KS.3 |
| FIM1_4197 | 1416.918445 | 1573.561434 | 1354.845331 | 0.130243134 | 0.25161287 | 0.576712727 | 0.635469727 | ns | YPD | KS.3 |
| FIM1_4198 | 1334.396094 | 2725.842493 | 1718.026555 | 0.93148707 | 0.225958787 | 1.0772567353 | 3.3643140353 | ns | YPD | KS.3 |
| FIM1_4199 | 193.3396821 | 561.5155772 | 370.175666 | 1.437515584 | 0.210594923 | 3.5425754949 | 4.9927232541 | Km>KS.3 | YPD | KS.3 |
| FIM1_4201 | 573.5242772 | 584.7707433 | 544.8274497 | 0.027718532 | 0.072393204 | 0.699849658 | 0.748763881 | ns | YPD | KS.3 |

| | | | | | | | | | | |
|---|---|---|---|---|---|---|---|---|---|---|
| FIM1_4203 | 602.2034344 | 256.114075 | 324.5406144 | -1.124664483 | 0.225746021 | 9.8940373116 | 4.8022278659 | Km<KS.3 | YPD | KS.3 |
| FIM1_4204 | 1535.707387 | 957.5732542 | 1148.355635 | -0.652693823 | 0.137468633 | 1.1355591605 | 4.4067135909 | ns | YPD | KS.3 |
| FIM1_4205 | 861.140068 | 7700.087109 | 3643.20249 | 2.94991325 | 0.365767615 | 7.6676936742 | 2.5431184019 | Km>KS.3 | YPD | KS.3 |
| FIM1_4206 | 514.886764 | 1791.012239 | 1485.64639 | 1.648755928 | 0.264998499 | 7.0206492435 | 7.7963683006 | Km>KS.3 | YPD | KS.3 |
| FIM1_4207 | 1945.184531 | 2379.043384 | 2188.568411 | 0.269605206 | 0.176355233 | 0.112261115 | 0.149131922 | ns | YPD | KS.3 |
| FIM1_4208 | 1132.955994 | 1800.820263 | 1001.676204 | 0.617273395 | 0.189534464 | 0.000655975 | 0.001411537 | ns | YPD | KS.3 |
| FIM1_4209 | 771.0657795 | 866.2341183 | 664.3409912 | 0.16069267 | 0.136734535 | 0.229049732 | 0.283040839 | ns | YPD | KS.3 |
| FIM1_4210 | 1363.220907 | 2291.015595 | 2048.586944 | 0.705887807 | 0.163750644 | 8.0984095329 | 2.5993282210 | ns | YPD | KS.3 |
| FIM1_4211 | 520.703704 | 578.0905376 | 725.4670799 | 0.135342017 | 0.209468903 | 0.495796011 | 0.558863091 | ns | YPD | KS.3 |
| FIM1_4212 | 3069.61284 | 4254.302013 | 3234.851481 | 0.461386684 | 0.090084043 | 2.2517893848 | 1.0101579972 | ns | YPD | KS.3 |
| FIM1_4213 | 292.8655641 | 502.0944937 | 423.953216 | 0.736816145 | 0.156416574 | 1.1554371397 | 4.4629656602 | ns | YPD | KS.3 |
| FIM1_4214 | 775.0338717 | 1036.831141 | 1128.506859 | 0.404394855 | 0.123247711 | 0.00082124 | 0.00173342 | ns | YPD | KS.3 |
| FIM1_4215 | 3155.862517 | 8065.483804 | 5631.325258 | 1.191328522 | 0.273898177 | 1.6458977324 | 6.1798801650 | Km>KS.3 | YPD | KS.3 |
| FIM1_4216 | 633.783499 | 977.8687848 | 743.7061317 | 0.56761236 | 0.208980082 | 0.004140614 | 0.007567801 | ns | YPD | KS.3 |
| FIM1_4217 | 701.4253071 | 1295.440132 | 889.1683139 | 0.806349446 | 0.21150145 | 5.4473327674 | 0.00014641 | ns | YPD | KS.3 |
| FIM1_4218 | 27.65799424 | 44.66158425 | 40.80047592 | 0.620628864 | 0.216145822 | 0.002342516 | 0.004497883 | ns | YPD | KS.3 |
| FIM1_4219 | 319.6502334 | 435.2249496 | 472.1011329 | 0.415242941 | 0.168627185 | 0.010735463 | 0.018086329 | ns | YPD | KS.3 |
| FIM1_4220 | 489.1045366 | 680.6471463 | 752.1887105 | 0.448322255 | 0.160472944 | 0.003908328 | 0.007180181 | ns | YPD | KS.3 |
| FIM1_4221 | 253.9465243 | 63.68477918 | 215.5270907 | -1.910506241 | 0.207095739 | 9.8382720433 | 6.8937187909 | Km<KS.3 | YPD | KS.3 |
| FIM1_4222 | 881.3014863 | 1787.910355 | 1521.313095 | 0.884506918 | 0.268588598 | 0.000314318 | 0.000724019 | ns | YPD | KS.3 |
| FIM1_4223 | 1783.778629 | 2573.438234 | 2560.189644 | 0.501914742 | 0.14917734 | 0.000538097 | 0.001176717 | ns | YPD | KS.3 |
| FIM1_4224 | 3705.522258 | 2532.770597 | 5080.741052 | -0.508545664 | 0.18302318 | 0.003765741 | 0.00694387 | ns | YPD | KS.3 |
| FIM1_4225 | 238.9233974 | 1354.502627 | 581.9715583 | 2.216420551 | 0.41893897 | 3.6812981595 | 2.8661124168 | Km>KS.3 | YPD | KS.3 |
| FIM1_4226 | 15230.30954 | 46779.39684 | 38582.63717 | 1.498062919 | 0.233092848 | 3.9391153925 | 4.5680883630 | Km>KS.3 | YPD | KS.3 |
| FIM1_4227 | 888.3608481 | 9595.085898 | 2863.283207 | 3.170533932 | 0.39952899 | 2.8216891236 | 8.7736896187 | Km>KS.3 | YPD | KS.3 |
| FIM1_4228 | 9.280421193 | 22.18643395 | 24.61517435 | 1.004454896 | 0.342486679 | 0.000714996 | 0.001525345 | Km>KS.3 | YPD | KS.3 |
| FIM1_4229 | 255.1642664 | 1301.770762 | 1314.53667 | 2.149285665 | 0.344306522 | 1.2318160235 | 1.5794548240 | Km>KS.3 | YPD | KS.3 |
| FIM1_4230 | 888.1316731 | 2350.514756 | 2387.443186 | 1.25739778 | 0.258153053 | 1.0421530853 | 5.0239453483 | Km>KS.3 | YPD | KS.3 |
| FIM1_4231 | 3197.125771 | 6163.508113 | 5772.270488 | 0.878471079 | 0.191293802 | 1.4673624721 | 5.5683663608 | ns | YPD | KS.3 |
| FIM1_4232 | 348.5915225 | 469.5032514 | 487.0612049 | 0.397146545 | 0.181273525 | 0.022513602 | 0.035266111 | ns | YPD | KS.3 |
| FIM1_4233 | 671.0847438 | 1185.63065 | 813.2059935 | 0.786999021 | 0.139134415 | 6.4844467084 | 3.9150633949 | ns | YPD | KS.3 |
| FIM1_4234 | 1536.310441 | 744.1080768 | 1164.257849 | -0.996383128 | 0.156094627 | 4.1831051079 | 3.9564539756 | ns | YPD | KS.3 |
| FIM1_4235 | 1138.716277 | 813.160279 | 998.3452731 | -0.471660682 | 0.108799335 | 1.0693482190 | 3.3417131846 | ns | YPD | KS.3 |
| FIM1_4236 | 2495.969078 | 2810.315654 | 2572.85615 | 0.165641454 | 0.114356556 | 0.140873069 | 0.183227063 | ns | YPD | KS.3 |
| FIM1_4237 | 9970.872185 | 7859.619562 | 7971.332114 | -0.335033972 | 0.098710533 | 0.000587647 | 0.001277214 | ns | YPD | KS.3 |
| FIM1_4238 | 3644.852421 | 2224.468813 | 2974.152138 | -0.689626552 | 0.120168197 | 4.9087193204 | 3.0336495179 | ns | YPD | KS.3 |
| FIM1_4239 | 2446.275583 | 2481.551029 | 2512.177134 | 0.019818273 | 0.123112373 | 0.869692875 | 0.891739913 | ns | YPD | KS.3 |
| FIM1_4240 | 1067.883926 | 616.2741427 | 1416.346129 | -0.687516221 | 0.259180123 | 0.003964182 | 0.007280106 | ns | YPD | KS.3 |
| FIM1_4241 | 1216.43078 | 3329.366246 | 2297.8411 | 1.300157039 | 0.262626287 | 5.7923573375 | 2.9345191195 | Km>KS.3 | YPD | KS.3 |
| FIM1_4242 | 230.6191298 | 397.8788081 | 256.0742388 | 0.718413275 | 0.205364739 | 0.000224706 | 0.000532339 | ns | YPD | KS.3 |
| FIM1_4243 | 607.9082922 | 464.3053296 | 683.6635556 | -0.377175344 | 0.113619585 | 0.00073771 | 0.001568422 | ns | YPD | KS.3 |
| FIM1_4244 | 491.0957217 | 945.3423956 | 691.9464541 | 0.867022461 | 0.204767711 | 7.8706011641 | 2.5343845172 | ns | YPD | KS.3 |
| FIM1_4245 | 28.51904754 | 64.40352693 | 45.78680657 | 1.031720522 | 0.26417013 | 1.9730207004 | 5.8601659611 | Km>KS.3 | YPD | KS.3 |
| FIM1_4246 | 645.2465403 | 773.7537077 | 598.2144239 | 0.25383854 | 0.113761303 | 0.023311813 | 0.036344803 | ns | YPD | KS.3 |
| FIM1_4247 | 513.4255099 | 677.1661745 | 496.5544882 | 0.362340511 | 0.203987885 | 0.061655019 | 0.086967315 | ns | YPD | KS.3 |

| | | | | | | | | | | |
|---|---|---|---|---|---|---|---|---|---|---|
| FIM1_4248 | 1610.710613 | 774.2619645 | 1281.443992 | -1.039773774 | 0.088825098 | 2.3779415749 | 3.9609991197 | Km<KS.3 | YPD | KS.3 |
| FIM1_4249 | 4851.391658 | 7838.254481 | 5696.017768 | 0.666263733 | 0.129628775 | 1.4823964365 | 6.9313179245 | ns | YPD | KS.3 |
| FIM1_4250 | 938.3444692 | 1309.498898 | 1103.258929 | 0.476074926 | 0.062106591 | 1.2938133903 | 2.4198201566 | ns | YPD | KS.3 |
| FIM1_4251 | 3564.445552 | 2732.794499 | 3147.760199 | -0.376247746 | 0.085793146 | 9.4954961564 | 2.9879881959 | ns | YPD | KS.3 |
| FIM1_4252 | 1322.544132 | 1301.709522 | 1133.022365 | -0.022217065 | 0.116758434 | 0.846520583 | 0.872114288 | ns | YPD | KS.3 |
| FIM1_4253 | 3885.919212 | 5763.278382 | 2736.043885 | 0.519583059 | 0.199490684 | 0.006215082 | 0.01096844 | ns | YPD | KS.3 |
| FIM1_4254 | 770.75805 | 1354.982987 | 1081.144263 | 0.785391268 | 0.127666011 | 3.2187114051 | 2.5458011511 | ns | YPD | KS.3 |
| FIM1_4255 | 1940.765197 | 1812.888748 | 1779.909196 | -0.089063647 | 0.204119008 | 0.646412775 | 0.699717919 | ns | YPD | KS.3 |
| FIM1_4257 | 1643.948959 | 8227.490702 | 4226.826573 | 2.200369533 | 0.268927459 | 6.7160313002 | 2.7613434478 | Km>KS.3 | YPD | KS.3 |
| FIM1_4258 | 42.35834886 | 35.59924806 | 42.59403129 | -0.217570811 | 0.243056941 | 0.336696283 | 0.398445292 | ns | YPD | KS.3 |
| FIM1_4259 | 1760.427714 | 3626.589033 | 2675.666478 | 0.957451728 | 0.207777262 | 1.0938182386 | 4.2713859789 | ns | YPD | KS.3 |
| FIM1_4260 | 474.0350508 | 1221.285816 | 767.4231294 | 1.235785534 | 0.242400841 | 3.5264458461 | 1.8486899983 | Km>KS.3 | YPD | KS.3 |
| FIM1_4261 | 319.4855369 | 937.8536374 | 920.3489585 | 1.444492702 | 0.2200007 | 2.0406611159 | 2.5380722629 | Km>KS.3 | YPD | KS.3 |
| FIM1_4262 | 1894.192452 | 3281.678845 | 2964.984714 | 0.745208748 | 0.168961266 | 4.7309127835 | 1.6065727711 | ns | YPD | KS.3 |
| FIM1_4263 | 557.3037832 | 607.0399842 | 569.9360588 | 0.119561607 | 0.11139179 | 0.275594745 | 0.333759459 | ns | YPD | KS.3 |
| FIM1_4265 | 1473.294013 | 664.3163225 | 891.4510669 | -1.113832363 | 0.126782107 | 2.5000286821 | 1.1623965134 | Km<KS.3 | YPD | KS.3 |
| FIM1_4266 | 7028.844465 | 12065.3101 | 9953.457295 | 0.696020513 | 0.229839511 | 0.001220295 | 0.002487083 | ns | YPD | KS.3 |
| FIM1_4267 | 1461.383351 | 1037.151271 | 1443.309574 | -0.471019063 | 0.145163342 | 0.000860055 | 0.00180794 | ns | YPD | KS.3 |
| FIM1_4268 | 2439.957442 | 4523.924261 | 3576.938436 | 0.816947384 | 0.203622769 | 2.3325316050 | 6.7980929907 | ns | YPD | KS.3 |
| FIM1_4269 | 242.4509986 | 416.7719099 | 394.092963 | 0.743873645 | 0.147810139 | 2.2313835435 | 1.0019073221 | ns | YPD | KS.3 |
| FIM1_4270 | 1976.361717 | 2951.694575 | 2359.182223 | 0.561077559 | 0.115001918 | 6.8927999211 | 2.7947579142 | ns | YPD | KS.3 |
| FIM1_4271 | 803.9270301 | 953.1918865 | 835.0517382 | 0.238668252 | 0.107588165 | 0.024365639 | 0.037833662 | ns | YPD | KS.3 |
| FIM1_4272 | 725.2149956 | 655.3711897 | 712.7728461 | -0.130921048 | 0.2125403 | 0.515480558 | 0.577984173 | ns | YPD | KS.3 |
| FIM1_4273 | 1321.750556 | 817.1133235 | 1940.035915 | -0.57386058 | 0.294717524 | 0.030592308 | 0.046401443 | ns | YPD | KS.3 |
| FIM1_4274 | 436.1863937 | 501.5843108 | 458.2280169 | 0.193103848 | 0.134936509 | 0.143225916 | 0.185995544 | ns | YPD | KS.3 |
| FIM1_4275 | 24.35378223 | 16.20859451 | 33.90945476 | -0.424316891 | 0.376612395 | 0.18213274 | 0.231623308 | ns | YPD | KS.3 |
| FIM1_4276 | 2485.65173 | 1873.270975 | 2355.293366 | -0.396948407 | 0.106778727 | 0.000161415 | 0.000398137 | ns | YPD | KS.3 |
| FIM1_4277 | 291.0403181 | 178.1360791 | 199.5390417 | -0.656928088 | 0.183903968 | 0.00019255 | 0.000463667 | ns | YPD | KS.3 |
| FIM1_4278 | 2812.198507 | 3210.414588 | 2990.652528 | 0.188157468 | 0.076649171 | 0.013369981 | 0.022032346 | ns | YPD | KS.3 |
| FIM1_4279 | 2793.349484 | 1375.263549 | 3477.358609 | -0.829487982 | 0.324030101 | 0.003635969 | 0.006733717 | ns | YPD | KS.3 |
| FIM1_4280 | 1980.029573 | 2241.988031 | 2995.245579 | 0.171404609 | 0.134131089 | 0.191276312 | 0.241829645 | ns | YPD | KS.3 |
| FIM1_4281 | 3459.478406 | 5691.95689 | 5184.495808 | 0.695696781 | 0.118692131 | 2.3354708345 | 1.5575023326 | ns | YPD | KS.3 |
| FIM1_4282 | 761.3254662 | 3354.360664 | 2506.198991 | 1.920973634 | 0.342924937 | 1.0645251513 | 9.3239658946 | Km>KS.3 | YPD | KS.3 |
| FIM1_4283 | 32.20245603 | 62.56709133 | 72.17510083 | 0.822129896 | 0.274679515 | 0.001017239 | 0.002105143 | ns | YPD | KS.3 |
| FIM1_4284 | 3864.413632 | 4037.127836 | 5208.636838 | 0.060736728 | 0.120452772 | 0.607625626 | 0.664381866 | ns | YPD | KS.3 |
| FIM1_4285 | 1748.582173 | 2342.760627 | 2583.026233 | 0.405946653 | 0.127048692 | 0.001107523 | 0.002273072 | ns | YPD | KS.3 |
| FIM1_4286 | 1575.488066 | 2709.635069 | 2558.176458 | 0.657892971 | 0.284556627 | 0.010732668 | 0.018086329 | ns | YPD | KS.3 |
| FIM1_4287 | 12.25491135 | 15.4997429 | 12.20518445 | 0.263163343 | 0.326762668 | 0.360500037 | 0.423291878 | ns | YPD | KS.3 |
| FIM1_4288 | 652.7483188 | 1694.910214 | 1283.172397 | 1.240898907 | 0.248277212 | 5.8227395011 | 2.9469103782 | Km>KS.3 | YPD | KS.3 |
| FIM1_4289 | 2213.922468 | 2302.415132 | 2144.079177 | 0.053454507 | 0.150829041 | 0.715531445 | 0.762263156 | ns | YPD | KS.3 |
| FIM1_4290 | 2010.128538 | 1311.874555 | 1397.806259 | -0.593335932 | 0.125768727 | 1.4628025841 | 5.5552998900 | ns | YPD | KS.3 |
| FIM1_4291 | 880.4081031 | 692.7678748 | 822.1209317 | -0.340208078 | 0.081640789 | 2.6224569434 | 7.5589358596 | ns | YPD | KS.3 |
| FIM1_4292 | 1103.161315 | 617.8516219 | 926.2401936 | -0.811174425 | 0.117149826 | 1.7308114705 | 2.1689639964 | ns | YPD | KS.3 |
| FIM1_4293 | 729.437306 | 565.2861891 | 651.6130055 | -0.359879058 | 0.092879283 | 8.9044379762 | 0.000232178 | ns | YPD | KS.3 |
| FIM1_4294 | 1079.225816 | 1103.343946 | 1014.23371 | 0.030969232 | 0.125757999 | 0.801692709 | 0.834397746 | ns | YPD | KS.3 |

| | | | | | | | | | | |
|---|---|---|---|---|---|---|---|---|---|---|
| FIM1_4295 | 1262.46052 | 838.5219129 | 957.734905 | -0.58054732 | 0.082284194 | 1.0727290644 | 1.4057998505 | ns | YPD | KS.3 |
| FIM1_4296 | 1291.411042 | 918.076474 | 1234.676435 | -0.487462462 | 0.059872241 | 2.7829269846 | 6.8202274622 | ns | YPD | KS.3 |
| FIM1_4297 | 1307.684578 | 1335.275642 | 795.4135375 | 0.02827751 | 0.17185284 | 0.864572878 | 0.887239457 | ns | YPD | KS.3 |
| FIM1_4298 | 108.3218868 | 432.1868727 | 256.6677733 | 1.524032427 | 0.482488737 | 4.5471912054 | 0.00012389 | Km>KS.3 | YPD | KS.3 |
| FIM1_4299 | 393.2989752 | 478.0284136 | 366.378426 | 0.266642894 | 0.151474803 | 0.070127534 | 0.097672028 | ns | YPD | KS.3 |
| FIM1_4300 | 190.7064601 | 284.4252782 | 223.9361699 | 0.549214184 | 0.146778153 | 0.000119864 | 0.000304403 | ns | YPD | KS.3 |
| FIM1_4303 | 100.8688173 | 42.72610322 | 82.48496482 | -1.121076512 | 0.231217152 | 1.9760740003 | 8.9535229066 | Km<KS.3 | YPD | KS.3 |
| FIM1_4304 | 53.9067228 | 65.67820019 | 30.91326256 | 0.213365283 | 0.354937203 | 0.484948982 | 0.548073872 | ns | YPD | KS.3 |
| FIM1_4307 | 662.6559578 | 337.538839 | 476.7071757 | -0.906830756 | 0.186896611 | 3.8013522050 | 1.6150065943 | ns | YPD | KS.3 |
| FIM1_4308 | 144.0367275 | 102.3731972 | 116.7686975 | -0.443969271 | 0.211638976 | 0.026688875 | 0.04118398 | ns | YPD | KS.3 |
| FIM1_4314 | 595.9774072 | 418.5527785 | 646.7515623 | -0.46140049 | 0.209561373 | 0.020142975 | 0.031944948 | ns | YPD | KS.3 |
| FIM1_4780 | 57.7955232 | 73.70754726 | 82.75928701 | 0.270610145 | 0.332680477 | 0.353470563 | 0.416118327 | ns | YPD | KS.3 |
| FIM1_4781 | 61.89593669 | 68.98634146 | 87.57792087 | 0.146282623 | 0.197382342 | 0.437278229 | 0.501142407 | ns | YPD | KS.3 |
| FIM1_4782 | 260.1734656 | 443.1751452 | 296.5047851 | 0.744062787 | 0.120331127 | 2.8865553764 | 2.3050315570 | ns | YPD | KS.3 |
| FIM1_4783 | 1557.277188 | 1385.92669 | 1729.24346 | -0.16116374 | 0.130467397 | 0.207033485 | 0.258935365 | ns | YPD | KS.3 |
| FIM1_4785 | 578.6511599 | 1251.897836 | 1037.162625 | 1.077863691 | 0.128281252 | 7.9976863563 | 2.6349993127 | Km>KS.3 | YPD | KS.3 |
| FIM1_4786 | 817.2313131 | 6254.952908 | 3276.771524 | 2.573951754 | 0.487351403 | 3.9640919606 | 3.0687801781 | Km>KS.3 | YPD | KS.3 |
| FIM1_4787 | 126.0994396 | 242.7261193 | 235.4642185 | 0.832401825 | 0.249060316 | 0.000303889 | 0.000702533 | ns | YPD | KS.3 |
| FIM1_4788 | 212.6869394 | 833.7668516 | 744.3069426 | 1.714716692 | 0.358462943 | 2.0659241202 | 1.1306900438 | Km>KS.3 | YPD | KS.3 |
| FIM1_4789 | 270.6868402 | 373.475493 | 264.372092 | 0.332157857 | 0.380538604 | 0.299538816 | 0.360040012 | ns | YPD | KS.3 |
| FIM1_4790 | 493.250456 | 618.8736294 | 614.4716867 | 0.301390292 | 0.187012918 | 0.092574472 | 0.12508365 | ns | YPD | KS.3 |
| FIM1_4791 | 1571.123531 | 1592.292844 | 2073.260557 | 0.017027748 | 0.220070829 | 0.934654408 | 0.9468219 | ns | YPD | KS.3 |
| FIM1_4792 | 296.4590022 | 718.5679584 | 464.198878 | 1.110869131 | 0.280074633 | 1.1704837369 | 3.6258758352 | Km>KS.3 | YPD | KS.3 |
| FIM1_4793 | 2153.337909 | 3581.665613 | 2070.580399 | 0.695565428 | 0.155970854 | 4.1764436009 | 1.4389062960 | ns | YPD | KS.3 |
| FIM1_4794 | 1963.492864 | 5402.345302 | 2670.050722 | 1.339468765 | 0.232374996 | 5.4059227259 | 4.0564804768 | Km>KS.3 | YPD | KS.3 |
| FIM1_4795 | 4663.35924 | 11655.50041 | 6331.766408 | 1.156854988 | 0.277088359 | 4.0735640876 | 1.4073598150 | Km>KS.3 | YPD | KS.3 |
| FIM1_4796 | 1473.566743 | 1989.458977 | 2521.152864 | 0.391290247 | 0.208818381 | 0.048126233 | 0.069722775 | ns | YPD | KS.3 |
| FIM1_4797 | 2303.723793 | 2550.06309 | 4622.760369 | 0.120153407 | 0.289535014 | 0.646658565 | 0.699831708 | ns | YPD | KS.3 |
| FIM1_4798 | 493.6132324 | 463.5098596 | 907.3828427 | -0.081090903 | 0.223500542 | 0.700112947 | 0.748884522 | ns | YPD | KS.3 |
| FIM1_4799 | 609.5964202 | 870.0537378 | 915.778069 | 0.46368973 | 0.209703444 | 0.019605669 | 0.03120224 | ns | YPD | KS.3 |
| FIM1_4800 | 19.89124917 | 11.04515615 | 14.76985844 | -0.678551612 | 0.325667586 | 0.017907362 | 0.028803468 | ns | YPD | KS.3 |
| FIM1_4801 | 296.3490419 | 285.6744997 | 369.4995752 | -0.046392349 | 0.242520581 | 0.837431428 | 0.864002769 | ns | YPD | KS.3 |
| FIM1_4802 | 1259.937666 | 962.4068293 | 805.0057536 | -0.380200389 | 0.092699251 | 3.3552453484 | 9.4094394636 | ns | YPD | KS.3 |
| FIM1_4803 | 1244.602409 | 1960.694628 | 1781.19925 | 0.584533359 | 0.227510492 | 0.006171698 | 0.010895742 | ns | YPD | KS.3 |
| FIM1_4804 | 4631.041248 | 3659.507802 | 5526.854346 | -0.322773217 | 0.145894071 | 0.023086917 | 0.036056268 | ns | YPD | KS.3 |
| FIM1_4805 | 3144.428038 | 2558.755448 | 4310.284211 | -0.281048302 | 0.153284079 | 0.059066859 | 0.083601031 | ns | YPD | KS.3 |
| FIM1_4806 | 1941.054569 | 2276.34088 | 2008.96905 | 0.227113492 | 0.070996219 | 0.001280237 | 0.002597484 | ns | YPD | KS.3 |
| FIM1_4807 | 1457.712956 | 2818.863983 | 2654.061447 | 0.868350168 | 0.211419414 | 1.3659364518 | 4.1715984335 | ns | YPD | KS.3 |
| FIM1_4808 | 1136.488852 | 2988.655076 | 2096.452981 | 1.297537917 | 0.208258521 | 3.6857415392 | 3.5194940802 | Km>KS.3 | YPD | KS.3 |
| FIM1_4809 | 516.9516509 | 522.7958535 | 696.4791253 | 0.015208561 | 0.155411098 | 0.919712517 | 0.935125643 | ns | YPD | KS.3 |
| FIM1_4810 | 11277.2649 | 10859.45472 | 12749.87612 | -0.053291365 | 0.093475071 | 0.564266048 | 0.624938466 | ns | YPD | KS.3 |
| FIM1_4811 | 837.4227901 | 2543.79927 | 1936.960012 | 1.331896398 | 0.353914565 | 1.1040046310 | 3.4370607256 | Km>KS.3 | YPD | KS.3 |
| FIM1_4812 | 155.1599914 | 271.8844893 | 213.5856373 | 0.75840152 | 0.171933097 | 4.5865472929 | 1.5628816974 | ns | YPD | KS.3 |
| FIM1_4813 | 699.5691975 | 1059.981764 | 1787.613643 | 0.48995813 | 0.300370292 | 0.069102572 | 0.09640642 | ns | YPD | KS.3 |
| FIM1_4814 | 1493.19389 | 1168.031611 | 1173.57424 | -0.347524938 | 0.086869562 | 5.3356564664 | 0.000143486 | ns | YPD | KS.3 |

| | | | | | | | | | | |
|---|---|---|---|---|---|---|---|---|---|---|
| FIM1_4815 | 1278.077442 | 1013.809696 | 843.8898588 | -0.32514029 | 0.102427013 | 0.001292537 | 0.00261823 | ns | YPD | KS.3 |
| FIM1_4816 | 1393.466916 | 1082.311337 | 1091.397534 | -0.355155313 | 0.099235503 | 0.000288986 | 0.000671825 | ns | YPD | KS.3 |
| FIM1_4817 | 1656.606608 | 1118.632683 | 1649.15812 | -0.53502465 | 0.158138591 | 0.000478885 | 0.0010617 | ns | YPD | KS.3 |
| FIM1_4818 | 1531.070992 | 1095.738261 | 1041.875757 | -0.472049833 | 0.093754578 | 3.5070163981 | 1.5002069287 | ns | YPD | KS.3 |
| FIM1_4819 | 3373.016494 | 3908.092611 | 5748.894804 | 0.207404446 | 0.09698145 | 0.030420542 | 0.046197251 | ns | YPD | KS.3 |
| FIM1_4820 | 1853.78951 | 1733.675157 | 1825.771121 | -0.093770922 | 0.101022153 | 0.347044468 | 0.409134177 | ns | YPD | KS.3 |
| FIM1_4821 | 1575.095884 | 812.7059034 | 1098.858809 | -0.809329907 | 0.285194663 | 0.001710317 | 0.003377859 | ns | YPD | KS.3 |
| FIM1_4822 | 2358.20988 | 1415.64166 | 1621.314464 | -0.711751334 | 0.122598123 | 3.160781921 | 2.036902857 | ns | YPD | KS.3 |
| FIM1_4823 | 1655.79662 | 1183.249067 | 1166.873272 | -0.469390454 | 0.114456545 | 3.02738424129 | 8.6015057683 | ns | YPD | KS.3 |
| FIM1_4824 | 3655.465677 | 2133.009615 | 3418.530375 | -0.709756209 | 0.205318691 | 0.000266796 | 0.000622565 | ns | YPD | KS.3 |
| FIM1_4825 | 528.780424 | 456.9438569 | 465.332088 | -0.187662367 | 0.219040434 | 0.364095407 | 0.426808353 | ns | YPD | KS.3 |
| FIM1_4826 | 1388.217324 | 1141.570198 | 1047.673213 | -0.274247023 | 0.103507563 | 0.00722694 | 0.012642062 | ns | YPD | KS.3 |
| FIM1_4827 | 1632.306507 | 1282.906952 | 1322.638848 | -0.337769496 | 0.106987676 | 0.001355617 | 0.002738203 | ns | YPD | KS.3 |
| FIM1_4828 | 845.4465481 | 852.553229 | 929.2585492 | 0.011415406 | 0.118884945 | 0.922144274 | 0.936832298 | ns | YPD | KS.3 |
| FIM1_4829 | 1613.082984 | 4824.492694 | 2408.544241 | 1.35510676 | 0.321648014 | 1.5293214526 | 5.785835914 | Km>KS.3 | YPD | KS.3 |
| FIM1_4830 | 1645.306015 | 3929.48989 | 2396.706495 | 1.101108963 | 0.270916314 | 8.0332144716 | 2.581733979 | Km>KS.3 | YPD | KS.3 |
| FIM1_4831 | 2811.090317 | 1262.862996 | 1966.919271 | -1.136162894 | 0.089120589 | 4.4478758667 | 1.301657790 | Km<KS.3 | YPD | KS.3 |
| FIM1_4832 | 3889.510464 | 3263.398503 | 3576.343722 | -0.250477334 | 0.065536508 | 0.000121062 | 0.000306661 | ns | YPD | KS.3 |
| FIM1_4833 | 4057.277074 | 4656.398828 | 4511.192965 | 0.18845829 | 0.147231327 | 0.188589054 | 0.238894692 | ns | YPD | KS.3 |
| FIM1_4834 | 405.7833432 | 2634.27576 | 1340.639535 | 2.440727888 | 0.407111572 | 5.959520026 | 6.6926889683 | Km>KS.3 | YPD | KS.3 |
| FIM1_4835 | 2893.431059 | 2273.733131 | 3751.275652 | -0.332807116 | 0.135441026 | 0.011916704 | 0.019847875 | ns | YPD | KS.3 |
| FIM1_4836 | 906.5196225 | 305.4638777 | 802.9952928 | -1.42017945 | 0.25907612 | 1.9120394301 | 1.3012853850 | Km<KS.3 | YPD | KS.3 |
| FIM1_4837 | 54.81795877 | 10.19806474 | 67.6932174 | -2.125153035 | 0.424464314 | 1.9362780583 | 1.3141859946 | Km<KS.3 | YPD | KS.3 |
| FIM1_4839 | 378.7141238 | 1031.733054 | 576.3763114 | 1.27194299 | 0.281493919 | 5.4512183434 | 2.2487405687 | Km>KS.3 | YPD | KS.3 |
| FIM1_4840 | 6834.332841 | 7170.503857 | 8620.223328 | 0.067660142 | 0.09318265 | 0.462819725 | 0.526292144 | ns | YPD | KS.3 |
| FIM1_4841 | 143.9276767 | 278.6691775 | 248.0003937 | 0.911954041 | 0.14458009 | 8.7056892331 | 7.817834645 | ns | YPD | KS.3 |
| FIM1_4842 | 25182.64595 | 41699.39184 | 58807.41069 | 0.705115362 | 0.117861029 | 1.0960685387 | 7.7127878080 | ns | YPD | KS.3 |
| FIM1_4844 | 119.0436006 | 583.7775632 | 1119.871416 | 1.866220542 | 0.487223569 | 1.0209507314 | 4.0120299280 | Km>KS.3 | YPD | KS.3 |
| FIM1_4845 | 240.267753 | 804.7697023 | 600.696593 | 1.500785771 | 0.337282838 | 2.7161206484 | 1.1915961398 | Km>KS.3 | YPD | KS.3 |
| FIM1_4846 | 254.7372222 | 993.7722043 | 542.9534828 | 1.692753418 | 0.367742339 | 5.5725560816 | 2.8347102767 | Km>KS.3 | YPD | KS.3 |
| FIM1_4847 | 160.091989 | 1773.057091 | 765.4735833 | 3.316753958 | 0.298191352 | 2.3661430601 | 3.567139916 | Km>KS.3 | YPD | KS.3 |
| FIM1_4848 | 1203.014958 | 5711.655798 | 2313.99835 | 2.105059513 | 0.284303889 | 3.5049949905 | 8.4647330475 | Km>KS.3 | YPD | KS.3 |
| FIM1_4849 | 1411.759287 | 6004.73622 | 3562.945838 | 1.925922883 | 0.293154876 | 2.1774696497 | 3.1596020159 | Km>KS.3 | YPD | KS.3 |
| FIM1_4850 | 158.5098299 | 747.6450869 | 797.3107309 | 1.898591346 | 0.432051766 | 7.2826578487 | 3.6267490287 | Km>KS.3 | YPD | KS.3 |
| FIM1_4851 | 887.9855571 | 640.2884768 | 855.6509651 | -0.445272305 | 0.157743005 | 0.003586955 | 0.006648695 | ns | YPD | KS.3 |
| FIM1_4852 | 413.1436133 | 290.8459678 | 468.2283045 | -0.474118965 | 0.169298385 | 0.003698609 | 0.006830208 | ns | YPD | KS.3 |
| FIM1_4853 | 282.0615535 | 791.3899107 | 464.3356436 | 1.390500118 | 0.207651393 | 1.0957136566 | 1.4272913965 | Km>KS.3 | YPD | KS.3 |
| FIM1_4854 | 967.2907091 | 4426.500842 | 3480.209368 | 1.960338226 | 0.358474508 | 2.0729809253 | 1.7086316877 | Km>KS.3 | YPD | KS.3 |
| FIM1_4855 | 333.3086099 | 1766.412793 | 915.6515042 | 2.058989461 | 0.451322652 | 2.1521393015 | 1.176581651 | Km>KS.3 | YPD | KS.3 |
| FIM1_4856 | 49.49311949 | 108.1584856 | 90.94481753 | 0.999999742 | 0.249237837 | 1.403181476 | 4.2800906475 | ns | YPD | KS.3 |
| FIM1_4857 | 17.38342265 | 68.1830039 | 97.34138546 | 1.635754128 | 0.408617006 | 1.1152862968 | 4.338193375 | Km>KS.3 | YPD | KS.3 |
| FIM1_4858 | 1282.130771 | 624.2399809 | 1889.896656 | -0.870157212 | 0.299446392 | 0.001176422 | 0.002405382 | ns | YPD | KS.3 |
| FIM1_4859 | 1809.169188 | 2415.983231 | 1667.293549 | 0.410157336 | 0.082670097 | 5.536817713 | 2.2821597450 | ns | YPD | KS.3 |
| FIM1_4860 | 407.965407 | 857.6332616 | 551.1737234 | 0.942591315 | 0.256888776 | 6.6631848665 | 0.000176514 | ns | YPD | KS.3 |
| FIM1_4861 | 620.4388234 | 817.0575401 | 825.2678814 | 0.385693965 | 0.1039987 | 0.000169203 | 0.00041549 | ns | YPD | KS.3 |

| | | | | | | | | | | |
|---|---|---|---|---|---|---|---|---|---|---|
| FIM1_4862 | 1683.350731 | 1108.202719 | 1646.755377 | -0.569497307 | 0.159044987 | 0.000217393 | 0.000517231 | ns | YPD | KS.3 |
| FIM1_4863 | 3959.878914 | 2646.816194 | 5275.620652 | -0.488796885 | 0.278075504 | 0.053669672 | 0.07679224 | ns | YPD | KS.3 |
| FIM1_4864 | 90.29326689 | 37.33204208 | 64.56806349 | -1.138217607 | 0.254534271 | 1.1486311556 | 4.4461703705 | Km<KS.3 | YPD | KS.3 |
| FIM1_4865 | 538.8832883 | 990.1448385 | 766.4919362 | 0.789192655 | 0.225253113 | 0.000187846 | 0.000454099 | ns | YPD | KS.3 |
| FIM1_4866 | 945.9222462 | 2411.891672 | 1468.118296 | 1.212452326 | 0.251318821 | 1.5792534151 | 7.3703430958 | Km>KS.3 | YPD | KS.3 |
| FIM1_4867 | 1287.289574 | 1873.145159 | 1774.801931 | 0.496456712 | 0.193894513 | 0.007305654 | 0.01276629 | ns | YPD | KS.3 |
| FIM1_4868 | 1040.73749 | 1086.148888 | 1300.526989 | 0.058613999 | 0.133340125 | 0.653160072 | 0.705486617 | ns | YPD | KS.3 |
| FIM1_4869 | 2290.379315 | 1114.903574 | 2324.438527 | -0.985352633 | 0.162678402 | 3.4601957809 | 2.7152167208 | ns | YPD | KS.3 |
| FIM1_4870 | 254.4812572 | 608.2891444 | 695.124892 | 1.086954189 | 0.284375613 | 2.2912581917 | 6.6856360728 | Km>KS.3 | YPD | KS.3 |
| FIM1_4871 | 331.9689726 | 641.6537943 | 519.1620767 | 0.892168143 | 0.176815218 | 1.4632303174 | 6.8481381274 | ns | YPD | KS.3 |
| FIM1_4872 | 213.6556779 | 598.5202416 | 580.6762812 | 1.387831152 | 0.208698754 | 1.5241461332 | 1.9294216317 | Km>KS.3 | YPD | KS.3 |
| FIM1_4873 | 1300.554113 | 896.5883323 | 1506.886808 | -0.497618398 | 0.181699808 | 0.004320437 | 0.00787044 | ns | YPD | KS.3 |
| FIM1_4874 | 2520.691119 | 2332.036722 | 2687.585499 | -0.100128049 | 0.219201039 | 0.628488926 | 0.683588196 | ns | YPD | KS.3 |
| FIM1_4875 | 1691.176406 | 2465.077034 | 1454.363538 | 0.473493249 | 0.247236839 | 0.039152737 | 0.058023493 | ns | YPD | KS.3 |
| FIM1_4876 | 70.30566023 | 41.22844889 | 66.18190685 | -0.669452458 | 0.257243762 | 0.004761283 | 0.008597962 | ns | YPD | KS.3 |
| FIM1_4877 | 682.6670227 | 1935.631251 | 1003.914225 | 1.381184413 | 0.233673026 | 1.8486966286 | 1.5509722980 | Km>KS.3 | YPD | KS.3 |
| FIM1_4878 | 15000.19261 | 13071.58627 | 19245.88169 | -0.190365898 | 0.13078909 | 0.137034053 | 0.178654196 | ns | YPD | KS.3 |
| FIM1_4879 | 2930.344289 | 2437.895146 | 2685.028214 | -0.262296921 | 0.068116849 | 0.000106802 | 0.00027417 | ns | YPD | KS.3 |
| FIM1_4880 | 5219.839719 | 4871.067156 | 4901.205305 | -0.098685086 | 0.065069811 | 0.127226347 | 0.167314586 | ns | YPD | KS.3 |
| FIM1_4881 | 1863.927931 | 3057.716038 | 2085.045205 | 0.68647995 | 0.132121671 | 1.0567884346 | 5.0797318475 | ns | YPD | KS.3 |
| FIM1_4882 | 32.89925102 | 330.5047026 | 1008.366534 | 2.77516112 | 0.600926911 | 1.1544085928 | 6.6920986325 | Km>KS.3 | YPD | KS.3 |
| FIM1_4883 | 2924.956602 | 4651.061588 | 3633.9843 | 0.628212233 | 0.167785739 | 0.000104038 | 0.000267764 | ns | YPD | KS.3 |
| FIM1_4884 | 2838.855269 | 2526.891256 | 2324.54675 | -0.164695099 | 0.085582415 | 0.052047648 | 0.074729308 | ns | YPD | KS.3 |
| FIM1_4885 | 934.4727408 | 1254.687615 | 961.0478088 | 0.412334829 | 0.110903464 | 0.000158359 | 0.000391774 | ns | YPD | KS.3 |
| FIM1_4886 | 276.5339905 | 243.589807 | 262.1975986 | -0.172347593 | 0.160908575 | 0.268884987 | 0.326587601 | ns | YPD | KS.3 |
| FIM1_4887 | 380.0518703 | 329.440095 | 366.9965181 | -0.200039716 | 0.115051819 | 0.077060298 | 0.106374856 | ns | YPD | KS.3 |
| FIM1_4888 | 545.5309031 | 1086.894538 | 846.6949265 | 0.922552445 | 0.192349435 | 4.8154581255 | 2.0114948929 | ns | YPD | KS.3 |
| FIM1_4889 | 765.6067375 | 494.3314245 | 630.0680574 | -0.57808897 | 0.197574081 | 0.002126501 | 0.0041245 | ns | YPD | KS.3 |
| FIM1_4890 | 515.1748306 | 637.1248125 | 592.0101904 | 0.283976329 | 0.178588901 | 0.098238961 | 0.132091576 | ns | YPD | KS.3 |
| FIM1_4891 | 1161.706941 | 921.0330724 | 1119.446052 | -0.317291379 | 0.150568498 | 0.030187408 | 0.045927326 | ns | YPD | KS.3 |
| FIM1_4892 | 2356.216776 | 6857.910488 | 4920.135313 | 1.395823807 | 0.255870302 | 2.4913642817 | 1.6482097475 | Km>KS.3 | YPD | KS.3 |
| FIM1_4893 | 4671.786716 | 12209.8825 | 9611.487046 | 1.252353018 | 0.246311715 | 3.5402936435 | 1.8539958817 | Km>KS.3 | YPD | KS.3 |
| FIM1_4894 | 803.9983391 | 1452.247884 | 1464.534456 | 0.7921878 | 0.186553413 | 8.9570849324 | 2.8401209393 | ns | YPD | KS.3 |
| FIM1_4895 | 835.8667683 | 2329.37877 | 1695.610525 | 1.35872943 | 0.23111284 | 2.4876098657 | 2.0058118447 | Km>KS.3 | YPD | KS.3 |
| FIM1_4896 | 415.8451501 | 2723.297475 | 1266.705427 | 2.621548809 | 0.240254884 | 7.3590287628 | 1.0169768915 | Km>KS.3 | YPD | KS.3 |
| FIM1_4897 | 381.6372243 | 1615.684514 | 1069.82916 | 1.960465344 | 0.252677384 | 2.9157126600 | 8.9616157925 | Km>KS.3 | YPD | KS.3 |
| FIM1_4898 | 506.1863012 | 1298.544227 | 920.0857228 | 1.200670626 | 0.269911664 | 1.0096555592 | 3.9739212084 | Km>KS.3 | YPD | KS.3 |
| FIM1_4899 | 5914.219127 | 3832.442473 | 7998.009529 | -0.599740719 | 0.136781124 | 7.0426370378 | 2.3035581369 | ns | YPD | KS.3 |
| FIM1_4900 | 2401.819398 | 3337.209681 | 4562.418037 | 0.44864124 | 0.154057946 | 0.002695439 | 0.005133924 | ns | YPD | KS.3 |
| FIM1_4901 | 200.0326223 | 456.2452809 | 385.1794082 | 1.083786635 | 0.223197761 | 2.1725674913 | 9.8080973408 | Km>KS.3 | YPD | KS.3 |
| FIM1_4902 | 731.3108875 | 1205.840806 | 1002.088481 | 0.68628652 | 0.148778573 | 2.0590088241 | 7.5265017636 | ns | YPD | KS.3 |
| FIM1_4903 | 924.418375 | 1578.485507 | 1339.826362 | 0.738503396 | 0.141139129 | 7.8047424789 | 3.8558683051 | ns | YPD | KS.3 |
| FIM1_4904 | 560.1320178 | 599.975169 | 1271.668189 | 0.089582204 | 0.197488656 | 0.634478003 | 0.688747123 | ns | YPD | KS.3 |
| FIM1_4905 | 342.7143944 | 842.5894093 | 445.754003 | 1.178465585 | 0.233936768 | 6.0756029630 | 3.0561078506 | Km>KS.3 | YPD | KS.3 |
| FIM1_4906 | 1085.934678 | 1501.937721 | 1602.179037 | 0.452561932 | 0.116249972 | 7.4426203706 | 0.000195496 | ns | YPD | KS.3 |

| | | | | | | | | | | |
|---|---|---|---|---|---|---|---|---|---|---|
| FIM1_4907 | 963.3149227 | 1390.991027 | 2107.583726 | 0.424312002 | 0.313261867 | 0.127750137 | 0.167870294 | ns | YPD | KS.3 |
| FIM1_4908 | 549.6793292 | 383.7430354 | 559.5296818 | -0.476489627 | 0.192848453 | 0.009676513 | 0.016458343 | ns | YPD | KS.3 |
| FIM1_4909 | 3299.195993 | 1961.322019 | 2808.153758 | -0.723789322 | 0.127293603 | 6.2545218010 | 3.7808318298 | ns | YPD | KS.3 |
| FIM1_4910 | 846.0003986 | 784.0249486 | 642.7546195 | -0.10192711 | 0.175815727 | 0.547228311 | 0.608410185 | ns | YPD | KS.3 |
| FIM1_4911 | 944.9547085 | 1338.56379 | 1207.163562 | 0.454913581 | 0.207486523 | 0.020806008 | 0.03279781 | ns | YPD | KS.3 |
| FIM1_4912 | 1120.510092 | 947.8125607 | 1002.936123 | -0.221699411 | 0.188103044 | 0.218552148 | 0.271620518 | ns | YPD | KS.3 |
| FIM1_4913 | 1282.372502 | 790.9681578 | 1779.518337 | -0.614723889 | 0.239862876 | 0.005945452 | 0.010529947 | ns | YPD | KS.3 |
| FIM1_4914 | 1708.113328 | 2187.083019 | 2391.945927 | 0.341795731 | 0.132114121 | 0.008174711 | 0.014116345 | ns | YPD | KS.3 |
| FIM1_4915 | 1253.91506 | 3332.800863 | 2017.395644 | 1.253481975 | 0.267322517 | 2.6061007335 | 1.1494105628 | Km>KS.3 | YPD | KS.3 |
| FIM1_4916 | 506.3947446 | 357.3644516 | 1357.458541 | -0.378531903 | 0.352709051 | 0.213248165 | 0.265825513 | ns | YPD | KS.3 |
| FIM1_4917 | 1176.573201 | 1975.02912 | 1595.349315 | 0.72113624 | 0.125441864 | 4.3460574272 | 2.7292310471 | ns | YPD | KS.3 |
| FIM1_4918 | 235.6418769 | 164.873387 | 185.4433156 | -0.478854904 | 0.176635767 | 0.004823797 | 0.008688773 | ns | YPD | KS.3 |
| FIM1_4919 | 1685.349358 | 836.9226834 | 1474.08409 | -0.924901066 | 0.210105407 | 3.1563620993 | 1.1105305123 | ns | YPD | KS.3 |
| FIM1_4920 | 727.6251717 | 678.3013069 | 491.5382069 | -0.093849492 | 0.170069641 | 0.56759047 | 0.627642274 | ns | YPD | KS.3 |
| FIM1_4921 | 10993.40035 | 7106.150707 | 10694.03251 | -0.599624442 | 0.146082758 | 2.4524893286 | 7.1111712278 | ns | YPD | KS.3 |
| FIM1_4922 | 1072.981902 | 805.8163773 | 1043.283304 | -0.382282097 | 0.182333922 | 0.028940379 | 0.044219406 | ns | YPD | KS.3 |
| FIM1_4923 | 704.1828373 | 717.2552785 | 779.2656105 | 0.025943491 | 0.099901769 | 0.792548401 | 0.827303461 | ns | YPD | KS.3 |
| FIM1_4924 | 820.872152 | 818.2285895 | 2324.548194 | -0.003876496 | 0.262419098 | 0.987209087 | 0.98899823 | ns | YPD | KS.3 |
| FIM1_4925 | 94.89598344 | 224.8672532 | 244.9278878 | 0.98253017 | 0.358999983 | 0.001382867 | 0.002789848 | ns | YPD | KS.3 |
| FIM1_4926 | 1927.352825 | 1528.582188 | 1363.095632 | -0.328275765 | 0.083290261 | 6.9638068936 | 0.00018389 | ns | YPD | KS.3 |
| FIM1_4927 | 1174.409077 | 548.1023613 | 1324.343166 | -0.921747765 | 0.302525654 | 0.000647574 | 0.001395875 | ns | YPD | KS.3 |
| FIM1_4928 | 1004.579911 | 367.5879234 | 1095.136634 | -1.222994286 | 0.324259523 | 1.6778398398 | 5.0406118376 | Km<KS.3 | YPD | KS.3 |
| FIM1_4929 | 923.2845391 | 576.4777722 | 701.8857728 | -0.66017681 | 0.111093899 | 1.5281084959 | 1.0602984333 | ns | YPD | KS.3 |
| FIM1_4931 | 1972.494769 | 4590.342234 | 3938.011143 | 1.150428294 | 0.17658809 | 1.0460012636 | 1.1167073576 | Km>KS.3 | YPD | KS.3 |
| FIM1_4932 | 4208.261296 | 8485.700641 | 4932.710896 | 0.954490741 | 0.170201395 | 5.5929024759 | 3.4142461034 | ns | YPD | KS.3 |
| FIM1_4933 | 2661.05707 | 1590.944386 | 1783.902986 | -0.697255685 | 0.168278172 | 1.7304624020 | 5.1861749698 | ns | YPD | KS.3 |
| FIM1_4935 | 1299.061956 | 2590.558395 | 1921.499279 | 0.931040459 | 0.182576782 | 9.9316390703 | 4.8145144405 | ns | YPD | KS.3 |
| FIM1_4936 | 524.9094765 | 2106.437678 | 1229.813806 | 1.818671586 | 0.307436494 | 2.2727290142 | 2.2722887555 | Km>KS.3 | YPD | KS.3 |
| FIM1_4937 | 3533.122133 | 8987.81687 | 6854.844582 | 1.145885495 | 0.306948242 | 2.6313522024 | 7.5758169962 | Km>KS.3 | YPD | KS.3 |
| FIM1_4938 | 1456.499529 | 1063.384695 | 1253.715977 | -0.423212803 | 0.173103437 | 0.01116128 | 0.018714988 | ns | YPD | KS.3 |
| FIM1_4939 | 1465.889838 | 672.8133937 | 997.7523455 | -1.084244858 | 0.134555468 | 1.3921204432 | 3.6345859938 | Km<KS.3 | YPD | KS.3 |
| FIM1_4940 | 1007.976493 | 689.359412 | 1454.603785 | -0.477100896 | 0.248162255 | 0.038293865 | 0.056886228 | ns | YPD | KS.3 |
| FIM1_4941 | 8166.818401 | 6702.60003 | 8851.194038 | -0.276320019 | 0.11245626 | 0.012524262 | 0.020797131 | ns | YPD | KS.3 |
| FIM1_4942 | 644.261265 | 1909.172722 | 1224.514551 | 1.292912098 | 0.35595001 | 2.1619850927 | 6.3493954169 | Km>KS.3 | YPD | KS.3 |
| FIM1_4943 | 1390.670827 | 2582.813491 | 2318.517883 | 0.847706609 | 0.157218903 | 2.5330158678 | 1.3623517775 | ns | YPD | KS.3 |
| FIM1_4944 | 291.5917715 | 2079.035524 | 1183.17116 | 2.673670505 | 0.322326431 | 6.4047148640 | 2.6776013822 | Km>KS.3 | YPD | KS.3 |
| FIM1_4945 | 4633.011537 | 8234.642774 | 7429.713428 | 0.770350007 | 0.186691583 | 1.5956865291 | 4.8083225212 | ns | YPD | KS.3 |
| FIM1_4946 | 2419.333707 | 4193.382846 | 3010.056622 | 0.749285254 | 0.162358284 | 1.7897639349 | 6.6398773872 | ns | YPD | KS.3 |
| FIM1_4947 | 2650.648222 | 1708.147132 | 1835.211868 | -0.610204097 | 0.128307524 | 1.1781040184 | 4.5434631719 | ns | YPD | KS.3 |
| FIM1_4948 | 586.7292874 | 553.7369714 | 502.0525132 | -0.079650379 | 0.137611696 | 0.553501605 | 0.61452142 | ns | YPD | KS.3 |
| FIM1_4949 | 1310.32559 | 1397.82762 | 1482.360849 | 0.092117605 | 0.064881357 | 0.153371615 | 0.198136533 | ns | YPD | KS.3 |
| FIM1_4950 | 3066.070195 | 2099.651714 | 3177.919928 | -0.535099901 | 0.092866091 | 5.5784340120 | 3.4142461034 | ns | YPD | KS.3 |
| FIM1_4951 | 1773.138855 | 3685.754893 | 2430.954616 | 0.993618237 | 0.174821927 | 3.2121026101 | 2.0646266777 | ns | YPD | KS.3 |
| FIM1_4952 | 5982.743008 | 3175.325415 | 6919.195298 | -0.781457699 | 0.276009485 | 0.001872575 | 0.003673525 | ns | YPD | KS.3 |
| FIM1_4953 | 1663.322762 | 987.4575526 | 2320.49064 | -0.607626117 | 0.314482448 | 0.029524252 | 0.045028557 | ns | YPD | KS.3 |

| | | | | | | | | | | |
|---|---|---|---|---|---|---|---|---|---|---|
| FIM1_4954 | 228.0684299 | 93.8301921 | 227.721582 | -1.089599769 | 0.302992839 | 5.46608465443 | 0.000146755 | Km<KS.3 | YPD | KS.3 |
| FIM1_4955 | 11149.8856 | 6974.526882 | 10080.4091 | -0.636354603 | 0.16616991 | 7.27669587195 | 0.000191543 | ns | YPD | KS.3 |
| FIM1_4956 | 655.3233487 | 1636.381389 | 2186.763112 | 1.226239955 | 0.206224324 | 2.94425583845 | 2.33888853252 | Km>KS.3 | YPD | KS.3 |
| FIM1_4957 | 204.2965071 | 458.0423621 | 753.7054491 | 1.067831427 | 0.214225479 | 1.18910786062 | 5.63947722268 | Km>KS.3 | YPD | KS.3 |
| FIM1_4958 | 113.4439112 | 429.1298346 | 219.4088314 | 1.509125781 | 0.446697403 | 2.23033710529 | 6.51551796761 | Km>KS.3 | YPD | KS.3 |
| FIM1_4960 | 5202.716208 | 3598.999125 | 4081.875028 | -0.519868008 | 0.097062387 | 5.84417360257 | 2.95475240577 | ns | YPD | KS.3 |
| FIM1_4961 | 3766.452385 | 2755.942699 | 3685.306776 | -0.406673735 | 0.210978847 | 0.041828888 | 0.061494892 | ns | YPD | KS.3 |
| FIM1_4962 | 709.5321599 | 583.6279239 | 811.1420474 | -0.265302985 | 0.157868035 | 0.08308403 | 0.11364945 | ns | YPD | KS.3 |
| FIM1_4963 | 712.3941417 | 536.3019506 | 718.4865154 | -0.397687329 | 0.10884007 | 0.000207024 | 0.000493977 | ns | YPD | KS.3 |
| FIM1_4964 | 80.82177848 | 166.3822632 | 90.85514937 | 1.001917339 | 0.137771717 | 8.33776332117 | 1.32949911932 | Km>KS.3 | YPD | KS.3 |
| FIM1_4965 | 5594.801088 | 7059.94899 | 7135.825911 | 0.324016452 | 0.119443417 | 0.005737905 | 0.010187751 | ns | YPD | KS.3 |
| FIM1_4966 | 3536.566821 | 2545.227626 | 4089.50433 | -0.464011627 | 0.09540127 | 8.53872965661 | 3.39298562633 | ns | YPD | KS.3 |
| FIM1_4967 | 1814.789238 | 2132.228429 | 1829.677499 | 0.223452165 | 0.128624434 | 0.076125404 | 0.10528882 | ns | YPD | KS.3 |
| FIM1_4968 | 1019.360565 | 1000.417621 | 1696.526117 | -0.021822765 | 0.304448265 | 0.936231798 | 0.947853723 | ns | YPD | KS.3 |
| FIM1_4969 | 10601.62534 | 9155.826505 | 13025.51777 | -0.208577392 | 0.074061256 | 0.004558305 | 0.008261409 | ns | YPD | KS.3 |
| FIM1_4970 | 12525.75098 | 13438.1878 | 13733.02455 | 0.099001051 | 0.097403901 | 0.303588398 | 0.363765 | ns | YPD | KS.3 |
| FIM1_4971 | 5251.86905 | 8751.091121 | 5299.304992 | 0.680291145 | 0.190861108 | 0.000189525 | 0.000457268 | ns | YPD | KS.3 |
| FIM1_4972 | 1502.71047 | 3340.44384 | 1907.458323 | 1.019663148 | 0.255154307 | 1.40954777972 | 4.29154650743 | Km>KS.3 | YPD | KS.3 |
| FIM1_4973 | 1946.608631 | 1981.598617 | 1688.204938 | 0.023310312 | 0.198758692 | 0.902115468 | 0.920242865 | ns | YPD | KS.3 |
| FIM1_4974 | 619.293318 | 434.4846352 | 495.5369414 | -0.49495469 | 0.114588647 | 1.11324206186 | 3.46280621037 | ns | YPD | KS.3 |
| FIM1_4976 | 5998.535511 | 9365.359679 | 7599.367736 | 0.62239927 | 0.117801191 | 7.39664848368 | 3.67248764534 | ns | YPD | KS.3 |
| FIM1_4977 | 1413.350491 | 695.8012436 | 1707.165664 | -0.776178711 | 0.369977524 | 0.013474522 | 0.022189919 | ns | YPD | KS.3 |
| FIM1_4978 | 1202.129104 | 558.8276574 | 947.8801932 | -1.063791399 | 0.140365886 | 6.74330461992 | 1.31865125928 | Km<KS.3 | YPD | KS.3 |
| FIM1_4979 | 13809.7515 | 14975.33126 | 16927.46391 | 0.114253939 | 0.094634719 | 0.221999938 | 0.275355196 | ns | YPD | KS.3 |
| FIM1_4980 | 412.5733348 | 593.8269582 | 556.3859116 | 0.490109576 | 0.173410953 | 0.003339311 | 0.006240824 | ns | YPD | KS.3 |
| FIM1_4981 | 12229.84586 | 12741.15415 | 15859.94697 | 0.058281823 | 0.067839773 | 0.387404219 | 0.450522672 | ns | YPD | KS.3 |
| FIM1_4982 | 1158.232933 | 1051.446817 | 1010.171553 | -0.136907936 | 0.066428859 | 0.038167767 | 0.056715843 | ns | YPD | KS.3 |
| FIM1_4983 | 1246.118541 | 976.3836267 | 976.9536087 | -0.343248218 | 0.09976203 | 0.0004915 | 0.001085795 | ns | YPD | KS.3 |
| FIM1_4984 | 1639.572211 | 2792.165038 | 1549.734645 | 0.72171067 | 0.16901675 | 9.41337569540 | 2.96777845910 | ns | YPD | KS.3 |
| FIM1_4985 | 3167.728392 | 3648.557518 | 2672.857618 | 0.199647744 | 0.091481347 | 0.027379086 | 0.042144477 | ns | YPD | KS.3 |
| FIM1_4986 | 4647.896415 | 4969.809708 | 4455.986282 | 0.094764645 | 0.087910505 | 0.276267265 | 0.334492491 | ns | YPD | KS.3 |
| FIM1_4987 | 2168.356737 | 1895.048759 | 2820.403304 | -0.179561546 | 0.181582908 | 0.303336328 | 0.363765 | ns | YPD | KS.3 |
| FIM1_4988 | 255.391458 | 165.3220454 | 174.4108259 | -0.589623848 | 0.165780725 | 0.000230457 | 0.000544408 | ns | YPD | KS.3 |
| FIM1_4989 | 6613.591597 | 10166.99318 | 7408.666665 | 0.583393719 | 0.164657609 | 0.000245169 | 0.00057588 | ns | YPD | KS.3 |
| FIM1_4991 | 2885.648085 | 4469.842317 | 3383.67469 | 0.579288249 | 0.195863199 | 0.001917204 | 0.003753676 | ns | YPD | KS.3 |
| FIM1_4992 | 1182.699801 | 839.7885698 | 1100.95135 | -0.456603982 | 0.184388749 | 0.009796225 | 0.016630137 | ns | YPD | KS.3 |
| FIM1_4993 | 2128.644125 | 3482.264782 | 2479.73763 | 0.664330734 | 0.173377485 | 6.84597524868 | 0.000180875 | ns | YPD | KS.3 |
| FIM1_4995 | 1266.614694 | 1589.659441 | 1753.530347 | 0.298118922 | 0.200652783 | 0.118697697 | 0.15701171 | ns | YPD | KS.3 |
| FIM1_4996 | 726.4926383 | 715.5714305 | 533.7981596 | -0.020001063 | 0.165332231 | 0.900467216 | 0.919126877 | ns | YPD | KS.3 |
| FIM1_4997 | 2373.281016 | 1467.031776 | 1646.699176 | -0.670215949 | 0.123349916 | 2.95612476854 | 1.56955397262 | ns | YPD | KS.3 |
| FIM1_4998 | 2117.415919 | 3281.080655 | 2531.997194 | 0.593538197 | 0.166490791 | 0.000221795 | 0.000526948 | ns | YPD | KS.3 |
| FIM1_4999 | 3618.23685 | 2673.549011 | 3751.186881 | -0.413277509 | 0.152398983 | 0.005237503 | 0.009359402 | ns | YPD | KS.3 |
| FIM1_5000 | 398.2819002 | 1150.212885 | 621.3457797 | 1.378081694 | 0.261364934 | 7.44541554299 | 4.45203633730 | Km>KS.3 | YPD | KS.3 |
| FIM1_5001 | 729.5992332 | 1738.590223 | 1113.903104 | 1.106674716 | 0.262873518 | 4.20265778353 | 1.44393801609 | Km>KS.3 | YPD | KS.3 |
| FIM1_5002 | 1800.981807 | 3040.373044 | 1846.561683 | 0.709169322 | 0.170038824 | 1.50027782667 | 4.55114767544 | ns | YPD | KS.3 |

| | | | | | | | | | | |
|---|---|---|---|---|---|---|---|---|---|---|
| FIM1_5003 | 1384.773313 | 1033.402034 | 1211.205547 | -0.402923353 | 0.141064004 | 0.003400435 | 0.006343144 | ns | YPD | KS.3 |
| FIM1_5004 | 2830.908508 | 2091.539441 | 3222.634587 | -0.381596421 | 0.242085286 | 0.090672084 | 0.122779972 | ns | YPD | KS.3 |
| FIM1_5005 | 14502.55349 | 13821.26665 | 20594.94691 | -0.067177566 | 0.114517468 | 0.550938525 | 0.612222283 | ns | YPD | KS.3 |
| FIM1_5006 | 481.2858846 | 182.4699618 | 269.8336058 | -1.330445518 | 0.170787872 | 4.4707307826 | 1.0541178030 | Km<KS.3 | YPD | KS.3 |
| FIM1_5007 | 1068.294462 | 4552.017408 | 2601.176588 | 1.846147936 | 0.35889712 | 1.8523021108 | 1.2658245881 | Km>KS.3 | YPD | KS.3 |
| FIM1_5008 | 2765.657323 | 4848.264221 | 4296.317491 | 0.780054306 | 0.130616489 | 9.9730237917 | 7.0879704806 | ns | YPD | KS.3 |
| FIM1_5009 | 55.45174414 | 47.6867775 | 119.1624388 | -0.167772162 | 0.336648262 | 0.568545245 | 0.628418705 | ns | YPD | KS.3 |
| FIM1_5010 | 575.9746134 | 1212.697787 | 1071.331298 | 0.990059389 | 0.204043998 | 2.9922932777 | 1.2990103888 | ns | YPD | KS.3 |
| FIM1_5011 | 172.8778326 | 152.8889505 | 171.8248649 | -0.163523525 | 0.191225728 | 0.371412263 | 0.434534851 | ns | YPD | KS.3 |
| FIM1_5012 | 826.5326135 | 480.5887666 | 962.7448218 | -0.730301877 | 0.178925439 | 2.1122399187 | 6.2216658353 | ns | YPD | KS.3 |
| FIM1_5013 | 816.0227341 | 1291.412871 | 1207.073186 | 0.634717909 | 0.135538097 | 1.6143739440 | 6.0798715908 | ns | YPD | KS.3 |
| FIM1_5014 | 2436.002805 | 1656.733083 | 2375.999414 | -0.542359201 | 0.104038148 | 1.2344865713 | 5.8435496600 | ns | YPD | KS.3 |
| FIM1_5015 | 2890.773191 | 2761.81924 | 2910.350965 | -0.061395194 | 0.170874992 | 0.709681327 | 0.757328314 | ns | YPD | KS.3 |
| FIM1_5016 | 797.9350516 | 1412.978094 | 752.5604019 | 0.738170836 | 0.228446627 | 0.000562186 | 0.001226699 | ns | YPD | KS.3 |
| FIM1_5017 | 1761.006346 | 3487.570576 | 2331.695015 | 0.9060532 | 0.204382217 | 2.8816773838 | 1.0203804259 | ns | YPD | KS.3 |
| FIM1_5018 | 500.0093898 | 394.4081232 | 382.6164024 | -0.321405642 | 0.161332488 | 0.039642702 | 0.058697155 | ns | YPD | KS.3 |
| FIM1_5019 | 404.7695779 | 1930.117747 | 1121.73002 | 2.13130839 | 0.263822578 | 1.6113996605 | 6.0276039934 | Km>KS.3 | YPD | KS.3 |
| FIM1_5020 | 373.013309 | 1792.248277 | 1100.992639 | 2.063979509 | 0.337357633 | 3.0965057195 | 3.6766386527 | Km>KS.3 | YPD | KS.3 |
| FIM1_5021 | 776.5752981 | 1016.829917 | 757.4105412 | 0.376889691 | 0.115797206 | 0.000928942 | 0.001936918 | ns | YPD | KS.3 |
| FIM1_5022 | 2205.807215 | 3646.572701 | 2655.25825 | 0.709717562 | 0.097386499 | 1.5517734868 | 2.3536808223 | ns | YPD | KS.3 |
| FIM1_5023 | 1585.033573 | 1660.770346 | 1964.965564 | 0.065265143 | 0.105862591 | 0.531770215 | 0.593839914 | ns | YPD | KS.3 |
| FIM1_5024 | 4823.756797 | 2630.391153 | 3413.721675 | -0.847575849 | 0.12128077 | 1.0043657230 | 1.3253897803 | ns | YPD | KS.3 |
| FIM1_5025 | 3643.875494 | 3625.377853 | 2924.38756 | -0.006640054 | 0.205368354 | 0.972867317 | 0.978769444 | ns | YPD | KS.3 |
| FIM1_5026 | 1674.334044 | 2496.739424 | 2627.883575 | 0.552320409 | 0.136298719 | 3.3168037591 | 9.3226546338 | ns | YPD | KS.3 |
| FIM1_5027 | 2408.370227 | 2229.246248 | 3523.227109 | -0.103522202 | 0.175730658 | 0.540793301 | 0.602698628 | ns | YPD | KS.3 |
| FIM1_5028 | 4094.26614 | 4201.289538 | 3668.055305 | 0.036350781 | 0.101035005 | 0.7155034 | 0.762263156 | ns | YPD | KS.3 |
| FIM1_5029 | 3438.558378 | 2840.517172 | 2781.189831 | -0.270756579 | 0.082656531 | 0.000949314 | 0.001973606 | ns | YPD | KS.3 |
| FIM1_5030 | 3118.516553 | 2424.5285 | 3657.061808 | -0.35830903 | 0.074444785 | 1.2417387211 | 4.7740727492 | ns | YPD | KS.3 |
| FIM1_5031 | 4997.285495 | 3487.553976 | 5372.95306 | -0.499246466 | 0.128291959 | 7.0402900058 | 0.000185713 | ns | YPD | KS.3 |
| FIM1_5032 | 1633.217055 | 1040.740723 | 2251.585274 | -0.541037386 | 0.288468129 | 0.038117785 | 0.056658494 | ns | YPD | KS.3 |
| FIM1_5033 | 1792.748267 | 1989.339526 | 1764.733732 | 0.148145536 | 0.070516493 | 0.034457966 | 0.051712935 | ns | YPD | KS.3 |
| FIM1_5034 | 2727.250753 | 3080.714933 | 4885.437389 | 0.168708442 | 0.129431701 | 0.183193771 | 0.232734681 | ns | YPD | KS.3 |
| FIM1_5035 | 606.4416682 | 1208.726592 | 1070.618224 | 0.904255897 | 0.217947391 | 1.0359512585 | 3.2434597302 | ns | YPD | KS.3 |
| FIM1_5036 | 1143.275397 | 882.6784843 | 1330.740806 | -0.349197549 | 0.168589175 | 0.031920811 | 0.048196065 | ns | YPD | KS.3 |
| FIM1_5037 | 1431.378764 | 1529.395397 | 1385.984308 | 0.094365198 | 0.074127586 | 0.199775518 | 0.250853913 | ns | YPD | KS.3 |
| FIM1_5038 | 4635.877756 | 3401.970565 | 3473.213098 | -0.434151706 | 0.107212672 | 3.9486813210 | 0.000109016 | ns | YPD | KS.3 |
| FIM1_5039 | 1619.311036 | 1976.3156 | 1431.105986 | 0.275998047 | 0.128864842 | 0.02874518 | 0.043975176 | ns | YPD | KS.3 |
| FIM1_5040 | 11774.49293 | 10530.20573 | 15960.82951 | -0.154117342 | 0.134661159 | 0.241775007 | 0.296921911 | ns | YPD | KS.3 |
| FIM1_5041 | 323.4779495 | 575.9766365 | 428.8023825 | 0.789950736 | 0.15585405 | 1.6703403961 | 7.7445885098 | ns | YPD | KS.3 |
| FIM1_5042 | 13941.72911 | 22658.97951 | 18192.75871 | 0.668210045 | 0.145470948 | 2.3397791163 | 8.4473157502 | ns | YPD | KS.3 |
| FIM1_5043 | 630.3986928 | 853.9862292 | 914.906838 | 0.406706195 | 0.176641406 | 0.016703998 | 0.027033959 | ns | YPD | KS.3 |
| FIM1_5044 | 100.0806983 | 303.2352653 | 237.6434499 | 1.47794867 | 0.231025653 | 5.2990598681 | 6.0051988255 | Km>KS.3 | YPD | KS.3 |
| FIM1_5045 | 2013.348962 | 2833.895795 | 2635.336793 | 0.45681938 | 0.181674509 | 0.008801296 | 0.015072787 | ns | YPD | KS.3 |
| FIM1_5046 | 1779.479356 | 7557.740217 | 3462.930132 | 1.917458661 | 0.298654888 | 6.1474403974 | 8.3561519063 | Km>KS.3 | YPD | KS.3 |
| FIM1_5047 | 1531.742837 | 1573.980943 | 1684.120007 | 0.037780093 | 0.116553824 | 0.741620543 | 0.783679312 | ns | YPD | KS.3 |

| | | | | | | | | | | |
|---|---|---|---|---|---|---|---|---|---|---|
| FIM1_5048 | 737.7577472 | 384.1864402 | 648.1284846 | -0.916697832 | 0.112545488 | 1.13559781973 | 3.0051059325 | ns | YPD | KS.3 |
| FIM1_5049 | 2035.541118 | 2371.031686 | 2315.334913 | 0.214429331 | 0.100615565 | 0.030852908 | 0.046768195 | ns | YPD | KS.3 |
| FIM1_5050 | 1929.849281 | 3422.367628 | 2158.68306 | 0.782604936 | 0.159155298 | 3.71715798192 | 1.5819384910 | ns | YPD | KS.3 |
| FIM1_5051 | 23.16820103 | 38.89464863 | 26.75475964 | 0.639286088 | 0.258310821 | 0.007239809 | 0.012660122 | ns | YPD | KS.3 |
| FIM1_5052 | 722.2682897 | 470.9225042 | 833.6461928 | -0.596759193 | 0.122339657 | 6.53826260925 | 2.6618540491 | ns | YPD | KS.3 |
| FIM1_5053 | 1335.136235 | 1058.244018 | 1307.8111 | -0.32120086 | 0.13454467 | 0.014593872 | 0.023875211 | ns | YPD | KS.3 |
| FIM1_5054 | 979.4820085 | 495.8417248 | 670.2714311 | -0.936190023 | 0.152487782 | 2.38159045627 | 1.9297088794 | ns | YPD | KS.3 |
| FIM1_5055 | 4490.384164 | 3040.934766 | 3470.755075 | -0.544758585 | 0.11632205 | 1.87084770971 | 6.9251989254 | ns | YPD | KS.3 |
| FIM1_5056 | 2135.492384 | 1615.651145 | 1420.427163 | -0.387118669 | 0.125763262 | 0.00168474 | 0.003339276 | ns | YPD | KS.3 |
| FIM1_5057 | 5977.405871 | 2445.157678 | 6219.004623 | -1.056668523 | 0.334663264 | 0.00029096 | 0.000676098 | Km<KS.3 | YPD | KS.3 |
| FIM1_5058 | 2369.999072 | 1107.293415 | 2622.191853 | -1.00959702 | 0.208308909 | 2.89780474248 | 1.2646121573 | Km<KS.3 | YPD | KS.3 |
| FIM1_5059 | 716.9189911 | 730.6353889 | 686.9090204 | 0.027130086 | 0.08001689 | 0.732440999 | 0.775296303 | ns | YPD | KS.3 |
| FIM1_5061 | 2376.722629 | 987.7089682 | 2153.880053 | -1.220292482 | 0.143362223 | 1.82915554181 | 6.7910812093 | Km<KS.3 | YPD | KS.3 |
| FIM1_5062 | 849.4805704 | 754.394728 | 633.3694877 | -0.16559959 | 0.113223124 | 0.137160022 | 0.178771577 | ns | YPD | KS.3 |
| FIM1_5063 | 2015.680824 | 1387.963996 | 1990.11932 | -0.515210269 | 0.137265904 | 0.000120451 | 0.000305737 | ns | YPD | KS.3 |
| FIM1_5064 | 401.9548511 | 659.8935234 | 645.2072038 | 0.646729291 | 0.213584863 | 0.001349336 | 0.002726622 | ns | YPD | KS.3 |
| FIM1_5065 | 2728.915317 | 4834.858014 | 3945.297633 | 0.760785148 | 0.195230794 | 4.29604888651 | 0.000117692 | ns | YPD | KS.3 |
| FIM1_5066 | 347.6923941 | 543.5128846 | 475.1938222 | 0.62958723 | 0.101725458 | 3.48020344433 | 2.7266160843 | ns | YPD | KS.3 |
| FIM1_5067 | 4043.401501 | 3081.92229 | 3488.00408 | -0.3795871 | 0.11434661 | 0.000735876 | 0.001565192 | ns | YPD | KS.3 |
| FIM1_5068 | 1208.794552 | 750.5906169 | 1083.197022 | -0.640586902 | 0.178437116 | 0.000185479 | 0.000449907 | ns | YPD | KS.3 |
| FIM1_5069 | 194.4937369 | 1894.214232 | 1090.397409 | 3.079369545 | 0.355625899 | 7.02609871701 | 3.6411292830 | Km>KS.3 | YPD | KS.3 |
| FIM1_5070 | 37.77575051 | 47.19268909 | 66.93085334 | 0.292166229 | 0.194720274 | 0.116105043 | 0.153786631 | ns | YPD | KS.3 |
| FIM1_5071 | 3591.523957 | 4009.332406 | 7131.434308 | 0.153414698 | 0.116555419 | 0.180594909 | 0.230315219 | ns | YPD | KS.3 |
| FIM1_5072 | 95.20820842 | 156.1610944 | 128.7444275 | 0.637075033 | 0.226239852 | 0.002696758 | 0.00513447 | ns | YPD | KS.3 |
| FIM1_5073 | 686.293469 | 471.2323327 | 611.9560529 | -0.507362735 | 0.172142444 | 0.002221157 | 0.00428803 | ns | YPD | KS.3 |
| FIM1_5074 | 739.8388845 | 635.0490674 | 663.8664992 | -0.20754931 | 0.156481043 | 0.17164287 | 0.220196822 | ns | YPD | KS.3 |
| FIM1_5075 | 545.1434696 | 2833.643152 | 1277.11883 | 2.328889686 | 0.172181124 | 3.43289062053 | 1.8976256485 | Km>KS.3 | YPD | KS.3 |
| FIM1_5076 | 2511.893939 | 2294.859328 | 3651.790514 | -0.12605317 | 0.119566907 | 0.283131634 | 0.342054367 | ns | YPD | KS.3 |
| FIM1_5077 | 1334.677746 | 2865.657083 | 2209.852189 | 1.048529535 | 0.159911493 | 1.12406210958 | 1.1911846689 | Km>KS.3 | YPD | KS.3 |
| FIM1_5078 | 447.4959048 | 588.6733146 | 454.6684508 | 0.378810047 | 0.135932195 | 0.004336171 | 0.007896212 | ns | YPD | KS.3 |
| FIM1_5079 | 5552.540394 | 4839.533804 | 6011.899769 | -0.190413264 | 0.12859742 | 0.130631221 | 0.17147502 | ns | YPD | KS.3 |
| FIM1_5080 | 1169.797438 | 1131.673003 | 1060.424088 | -0.04655854 | 0.058917699 | 0.427253293 | 0.491124106 | ns | YPD | KS.3 |
| FIM1_5081 | 1469.691196 | 1608.009468 | 1610.532415 | 0.125920251 | 0.111333114 | 0.25062006 | 0.306799901 | ns | YPD | KS.3 |
| FIM1_5082 | 618.2429821 | 394.6386598 | 557.9162857 | -0.622247285 | 0.133387743 | 1.80160890738 | 6.6788407706 | ns | YPD | KS.3 |
| FIM1_5083 | 2877.220234 | 5552.273234 | 4870.532015 | 0.851845511 | 0.229589231 | 7.29792230957 | 0.000192 | ns | YPD | KS.3 |
| FIM1_5084 | 386.957811 | 401.8053976 | 802.1607 | 0.045341731 | 0.272094786 | 0.855621006 | 0.879850042 | ns | YPD | KS.3 |
| FIM1_5085 | 1653.510854 | 3154.389814 | 1763.991479 | 0.838704666 | 0.226673933 | 7.85689435820 | 0.000205835 | ns | YPD | KS.3 |
| FIM1_5086 | 2723.743704 | 3374.173953 | 2732.781178 | 0.296404691 | 0.129907925 | 0.019771911 | 0.031436644 | ns | YPD | KS.3 |
| FIM1_5087 | 1239.871514 | 853.5739782 | 1469.97806 | -0.497453862 | 0.186305268 | 0.005304871 | 0.00946447 | ns | YPD | KS.3 |
| FIM1_5088 | 5066.101728 | 10684.23532 | 9428.848193 | 0.97267335 | 0.228550731 | 5.32228977732 | 1.7927143968 | ns | YPD | KS.3 |
| FIM1_5089 | 1533.966008 | 1678.389478 | 2468.934404 | 0.112233171 | 0.246834328 | 0.62488426 | 0.680113584 | ns | YPD | KS.3 |
| FIM1_5090 | 177.4642471 | 787.145796 | 387.9351323 | 1.957045451 | 0.321927549 | 5.08107838333 | 5.7977901277 | Km>KS.3 | YPD | KS.3 |
| FIM1_5091 | 254.322428 | 393.8513332 | 492.4874872 | 0.545759805 | 0.254941177 | 0.020506488 | 0.032438721 | ns | YPD | KS.3 |
| FIM1_5093 | 912.7843413 | 1245.699971 | 1273.576786 | 0.438703034 | 0.096389502 | 4.07870782733 | 1.4081590174 | ns | YPD | KS.3 |
| FIM1_5094 | 47213.51525 | 61178.68436 | 89508.85285 | 0.363901793 | 0.10458076 | 0.000416821 | 0.000937046 | ns | YPD | KS.3 |

| | | | | | | | | | | |
|---|---|---|---|---|---|---|---|---|---|---|
| FIM1_5095 | 656.2130728 | 393.1462586 | 503.4258247 | -0.709667892 | 0.13650911 | 9.9387061919 | 4.8145144405 | ns | YPD | KS.3 |
| FIM1_5096 | 1920.506729 | 951.2092624 | 1407.574219 | -0.951539799 | 0.17758443 | 2.3148246585 | 1.2544937555 | ns | YPD | KS.3 |
| FIM1_5097 | 7694.218794 | 10806.64309 | 9492.802646 | 0.483072558 | 0.075688997 | 1.2584122446 | 1.0812782240 | ns | YPD | KS.3 |
| FIM1_5098 | 300.9506722 | 464.7605475 | 297.09687 | 0.565642524 | 0.214751154 | 0.005294723 | 0.009451469 | ns | YPD | KS.3 |
| FIM1_5099 | 1768.143088 | 1056.606254 | 1756.661765 | -0.717431413 | 0.12472077 | 4.2885366808 | 2.6972781273 | ns | YPD | KS.3 |
| FIM1_5100 | 1290.960346 | 1227.418156 | 1058.239174 | -0.071702939 | 0.055834979 | 0.197196191 | 0.248430248 | ns | YPD | KS.3 |
| FIM1_5101 | 802.1939663 | 1245.455909 | 1189.433938 | 0.613650521 | 0.119920702 | 1.8380475706 | 8.4124072347 | ns | YPD | KS.3 |
| FIM1_5102 | 1083.068141 | 5299.847529 | 3568.104286 | 2.163010611 | 0.272285271 | 4.7866677908 | 1.6423222247 | Km>KS.3 | YPD | KS.3 |
| FIM1_5103 | 380.7001972 | 402.5135274 | 511.8376738 | 0.075890207 | 0.137260723 | 0.571443604 | 0.631061471 | ns | YPD | KS.3 |
| FIM1_5104 | 768.7735133 | 684.4674199 | 743.3096505 | -0.165212867 | 0.085205087 | 0.050309582 | 0.072463859 | ns | YPD | KS.3 |
| FIM1_5105 | 2563.157388 | 2164.95649 | 2556.219037 | -0.240498881 | 0.073173555 | 0.000932843 | 0.001943423 | ns | YPD | KS.3 |
| FIM1_5106 | 1054.162915 | 1552.889144 | 873.3045665 | 0.533467316 | 0.141968958 | 0.000115278 | 0.000294107 | ns | YPD | KS.3 |
| FIM1_5107 | 3655.561812 | 3173.019465 | 4318.641348 | -0.193866344 | 0.14636564 | 0.173800097 | 0.222562544 | ns | YPD | KS.3 |
| FIM1_5108 | 910.5267746 | 952.9103437 | 919.5715219 | 0.061624039 | 0.151076333 | 0.674861113 | 0.725931683 | ns | YPD | KS.3 |
| FIM1_5109 | 1482.434598 | 832.2511415 | 1055.894387 | -0.790188117 | 0.156106487 | 1.7289523339 | 7.9791631367 | ns | YPD | KS.3 |
| FIM1_5110 | 2081.534455 | 1926.591154 | 1771.777108 | -0.107793798 | 0.1138982 | 0.336062022 | 0.397789331 | ns | YPD | KS.3 |
| FIM1_5111 | 2609.358323 | 1862.259833 | 1833.436691 | -0.47394051 | 0.104200966 | 3.9582937605 | 1.3713448091 | ns | YPD | KS.3 |
| FIM1_5112 | 901.840455 | 874.7958115 | 912.3996043 | -0.0421321 | 0.117474725 | 0.715185701 | 0.762221264 | ns | YPD | KS.3 |
| FIM1_5113 | 239.3968824 | 146.7037519 | 185.8773579 | -0.652973192 | 0.188296527 | 0.000287026 | 0.000667893 | ns | YPD | KS.3 |
| FIM1_5114 | 452.9688755 | 285.9230797 | 473.4386546 | -0.601085855 | 0.211340661 | 0.00264272 | 0.005041232 | ns | YPD | KS.3 |
| FIM1_5115 | 1781.47861 | 1065.378393 | 1717.039464 | -0.716832319 | 0.124430888 | 4.0808604076 | 2.5862777742 | ns | YPD | KS.3 |
| FIM1_5116 | 2217.657436 | 1146.236346 | 1592.914278 | -0.922569818 | 0.122598373 | 1.5607184516 | 2.8864588464 | ns | YPD | KS.3 |
| FIM1_5117 | 543.3963779 | 640.5994269 | 553.3084487 | 0.208130535 | 0.236118028 | 0.346017142 | 0.408116473 | ns | YPD | KS.3 |
| FIM1_5118 | 2388.037842 | 1889.327773 | 2917.278492 | -0.321879276 | 0.142674122 | 0.02065364 | 0.032609286 | ns | YPD | KS.3 |
| FIM1_5119 | 1343.190238 | 1088.086916 | 1408.965236 | -0.294530233 | 0.112972462 | 0.008052435 | 0.013924527 | ns | YPD | KS.3 |
| FIM1_5120 | 781.8337621 | 1897.626932 | 1218.891875 | 1.172241567 | 0.221745643 | 1.6502562097 | 9.2873581939 | Km>KS.3 | YPD | KS.3 |
| FIM1_5121 | 2019.724206 | 1113.291245 | 2267.146921 | -0.820199496 | 0.148127148 | 1.1941769225 | 6.8761923491 | ns | YPD | KS.3 |
| FIM1_5122 | 2228.467605 | 8653.661268 | 4430.879936 | 1.707217084 | 0.353096568 | 1.6391066604 | 9.2350573453 | Km>KS.3 | YPD | KS.3 |
| FIM1_5123 | 2480.969897 | 2808.249555 | 4067.119462 | 0.140017619 | 0.321613396 | 0.62255231 | 0.678168983 | ns | YPD | KS.3 |
| FIM1_5124 | 1240.632849 | 876.677179 | 969.6212758 | -0.488341767 | 0.100486653 | 8.4398016257 | 3.3590410470 | ns | YPD | KS.3 |
| FIM1_5125 | 2337.525005 | 1442.273648 | 2718.883313 | -0.588933831 | 0.277297463 | 0.019727848 | 0.031376612 | ns | YPD | KS.3 |
| FIM1_5126 | 1423.586112 | 1834.933081 | 1546.701462 | 0.361587645 | 0.068529892 | 1.0971405945 | 5.2635240675 | ns | YPD | KS.3 |
| FIM1_5127 | 3821.576886 | 5712.663732 | 4733.280554 | 0.551212313 | 0.148637569 | 0.000136361 | 0.000341417 | ns | YPD | KS.3 |
| FIM1_5128 | 3050.585761 | 3881.21377 | 3394.131703 | 0.339656815 | 0.095429033 | 0.000316153 | 0.000727167 | ns | YPD | KS.3 |
| FIM1_5129 | 1302.846175 | 1337.636181 | 1408.446859 | 0.03614143 | 0.134380549 | 0.783293685 | 0.818845573 | ns | YPD | KS.3 |
| FIM1_5130 | 1491.312827 | 2792.619976 | 2394.966011 | 0.856292298 | 0.162413467 | 4.7903116508 | 2.4543563813 | ns | YPD | KS.3 |
| FIM1_5131 | 3124.047507 | 3522.730262 | 2682.725242 | 0.125300221 | 0.369640371 | 0.689949393 | 0.73976255 | ns | YPD | KS.3 |
| FIM1_5132 | 67.24795117 | 274.4630155 | 193.6028101 | 1.713589881 | 0.399467042 | 2.2736283699 | 1.0181189145 | Km>KS.3 | YPD | KS.3 |
| FIM1_5133 | 1371.788189 | 2096.778029 | 1610.349592 | 0.582300543 | 0.147228189 | 4.7667763506 | 0.000129518 | ns | YPD | KS.3 |
| FIM1_5134 | 619.5607772 | 311.8170666 | 596.8297995 | -0.932550542 | 0.172979887 | 2.0354593470 | 1.1177053257 | ns | YPD | KS.3 |
| FIM1_5135 | 1748.918118 | 1033.98718 | 1370.994416 | -0.719475488 | 0.154922407 | 1.6563572349 | 6.2051033461 | ns | YPD | KS.3 |
| FIM1_5136 | 2576.412259 | 2007.735593 | 2722.172896 | -0.343894739 | 0.136795523 | 0.010057273 | 0.017047677 | ns | YPD | KS.3 |
| FIM1_5137 | 716.9056913 | 938.0458381 | 955.2377765 | 0.360721902 | 0.174737309 | 0.032060137 | 0.048391742 | ns | YPD | KS.3 |
| FIM1_5138 | 887.8215212 | 2484.101541 | 2129.186013 | 1.319843807 | 0.272934496 | 9.5186881537 | 4.6245579653 | Km>KS.3 | YPD | KS.3 |
| FIM1_5139 | 1792.943977 | 4041.90966 | 3263.840028 | 1.021043602 | 0.272551424 | 3.8704834521 | 0.000107035 | Km>KS.3 | YPD | KS.3 |

| | | | | | | | | | | |
|---|---|---|---|---|---|---|---|---|---|---|
| FIM1_5140 | 3231.245382 | 2926.749924 | 3280.874518 | -0.119770392 | 0.272628181 | 0.631307115 | 0.686203386 | ns | YPD | KS.3 |
| FIM1_5141 | 2489.80362 | 4571.468525 | 2740.679032 | 0.831701418 | 0.157442955 | 4.8128122350 | 2.4633478260 | ns | YPD | KS.3 |
| FIM1_5142 | 1393.691377 | 2232.254356 | 1638.967762 | 0.60781129 | 0.225074758 | 0.004039685 | 0.007410557 | ns | YPD | KS.3 |
| FIM1_5143 | 2526.924735 | 2368.791753 | 2447.338046 | -0.091340803 | 0.085618532 | 0.281493161 | 0.340240154 | ns | YPD | KS.3 |
| FIM1_5144 | 2268.101132 | 1816.22247 | 1888.397872 | -0.309899503 | 0.117759251 | 0.007396115 | 0.012901709 | ns | YPD | KS.3 |
| FIM1_5145 | 2037.573032 | 1757.060526 | 2312.449822 | -0.204463699 | 0.132532268 | 0.114810989 | 0.152113094 | ns | YPD | KS.3 |
| FIM1_5146 | 519.7342357 | 375.3922338 | 627.1426176 | -0.414854523 | 0.232581976 | 0.056821296 | 0.080721287 | ns | YPD | KS.3 |
| FIM1_5147 | 20194.94036 | 18341.24065 | 29081.46093 | -0.133992272 | 0.12037418 | 0.256991959 | 0.313673945 | ns | YPD | KS.3 |
| FIM1_5148 | 103.1584484 | 243.4801217 | 298.8760442 | 1.099288255 | 0.255460052 | 2.8524783288 | 1.0122025453 | Km>KS.3 | YPD | KS.3 |
| FIM1_5149 | 1594.693759 | 2849.129837 | 4101.836441 | 0.632820958 | 0.362471914 | 0.041099386 | 0.060565594 | ns | YPD | KS.3 |
| FIM1_5150 | 5231.546662 | 11477.6996 | 9268.607682 | 1.036680278 | 0.216807504 | 3.6802780904 | 1.5675842037 | Km>KS.3 | YPD | KS.3 |
| FIM1_5151 | 247.3543443 | 232.728402 | 223.4158694 | -0.083146884 | 0.146456716 | 0.559954158 | 0.620439184 | ns | YPD | KS.3 |
| FIM1_5152 | 500.9922047 | 215.5995562 | 293.9752473 | -1.075819793 | 0.258467896 | 5.7626962080 | 1.9306002447 | Km<KS.3 | YPD | KS.3 |
| FIM1_5153 | 6401.533953 | 6669.372802 | 6107.404169 | 0.057709077 | 0.096563645 | 0.545329927 | 0.606986445 | ns | YPD | KS.3 |
| FIM1_5155 | 1677.566448 | 870.8697788 | 1027.34733 | -0.910412429 | 0.134659093 | 4.2223352416 | 4.8738092406 | ns | YPD | KS.3 |
| FIM1_5156 | 916.4163059 | 831.8400406 | 907.6906643 | -0.123982663 | 0.223968267 | 0.556636827 | 0.61772657 | ns | YPD | KS.3 |
| FIM1_5157 | 2538.402528 | 2085.62624 | 2358.045634 | -0.279926893 | 0.070342841 | 6.1813553610 | 0.000164803 | ns | YPD | KS.3 |
| FIM1_5158 | 1233.607674 | 1855.352424 | 1384.854333 | 0.534220854 | 0.207574226 | 0.006637808 | 0.011664816 | ns | YPD | KS.3 |
| FIM1_5159 | 824.6052166 | 871.1357765 | 963.2899021 | 0.076081113 | 0.124529184 | 0.533363861 | 0.595218754 | ns | YPD | KS.3 |
| FIM1_5160 | 10286.78577 | 8281.945093 | 11959.88436 | -0.274621547 | 0.235551085 | 0.21267217 | 0.265240422 | ns | YPD | KS.3 |
| FIM1_5161 | 1364.726357 | 3093.243277 | 1637.126416 | 1.027361255 | 0.273445612 | 3.6328806857 | 0.000101309 | Km>KS.3 | YPD | KS.3 |
| FIM1_5162 | 3304.146848 | 2294.300722 | 2683.565014 | -0.513369807 | 0.101523734 | 2.9594352008 | 1.2869921437 | ns | YPD | KS.3 |
| FIM1_5163 | 2247.987772 | 3191.628264 | 3379.838754 | 0.471245981 | 0.174272522 | 0.00497674 | 0.008951294 | ns | YPD | KS.3 |
| FIM1_5164 | 312.1288606 | 257.6320266 | 390.2049425 | -0.266985725 | 0.128385134 | 0.033744828 | 0.050811295 | ns | YPD | KS.3 |
| FIM1_5165 | 1247.808905 | 1760.119728 | 1240.079075 | 0.474958736 | 0.136043291 | 0.000350527 | 0.000799575 | ns | YPD | KS.3 |
| FIM1_5166 | 2290.173744 | 3499.938395 | 2961.30963 | 0.551633943 | 0.215008082 | 0.006600204 | 0.011606934 | ns | YPD | KS.3 |
| FIM1_5167 | 632.2785882 | 724.0527333 | 637.241487 | 0.188484671 | 0.121450307 | 0.113885302 | 0.151087834 | ns | YPD | KS.3 |
| FIM1_5168 | 1020.140312 | 948.8764321 | 781.3317534 | -0.098802943 | 0.14909204 | 0.495834008 | 0.558863091 | ns | YPD | KS.3 |
| FIM1_5169 | 357.6494738 | 158.4762283 | 280.748438 | -1.087275303 | 0.204164651 | 1.8113638705 | 1.0102617999 | Km<KS.3 | YPD | KS.3 |
| FIM1_5170 | 2305.634714 | 2310.262636 | 2146.695084 | 0.002881915 | 0.086752597 | 0.973241489 | 0.978947919 | ns | YPD | KS.3 |
| FIM1_5171 | 971.7776555 | 1085.045639 | 1014.350572 | 0.149061621 | 0.164088826 | 0.347819604 | 0.409853749 | ns | YPD | KS.3 |
| FIM1_5172 | 6366.557272 | 5079.359498 | 7294.21892 | -0.31458473 | 0.12017488 | 0.007671815 | 0.013326564 | ns | YPD | KS.3 |
| FIM1_5173 | 1962.917157 | 1805.973724 | 1704.099513 | -0.114918215 | 0.134540411 | 0.382363507 | 0.445367579 | ns | YPD | KS.3 |
| FIM1_5174 | 2676.772734 | 4072.898055 | 4111.276212 | 0.59538813 | 0.084052567 | 8.5464089879 | 1.1399030754 | ns | YPD | KS.3 |
| FIM1_5175 | 4785.625752 | 2830.864386 | 3329.456953 | -0.723466276 | 0.14421041 | 2.5299731402 | 1.1208028827 | ns | YPD | KS.3 |
| FIM1_5176 | 592.827723 | 217.826658 | 608.5618791 | -1.339031353 | 0.216256266 | 3.9119991160 | 3.7141594661 | Km<KS.3 | YPD | KS.3 |
| FIM1_5177 | 1365.884895 | 571.2270942 | 1527.121631 | -1.004871295 | 0.351664422 | 0.000898815 | 0.001881989 | Km<KS.3 | YPD | KS.3 |
| FIM1_5178 | 1055.455377 | 863.8583412 | 1130.801298 | -0.281650502 | 0.101502024 | 0.004925953 | 0.008869569 | ns | YPD | KS.3 |
| FIM1_5179 | 4194.611448 | 2303.622731 | 4531.64748 | -0.730739945 | 0.283894598 | 0.004502736 | 0.008175588 | ns | YPD | KS.3 |
| FIM1_5180 | 959.5910667 | 785.0552283 | 802.0512042 | -0.277412177 | 0.132059437 | 0.031781535 | 0.048029507 | ns | YPD | KS.3 |
| FIM1_5181 | 5117.632561 | 4389.771289 | 4942.464288 | -0.212189721 | 0.130756895 | 0.097351363 | 0.131039781 | ns | YPD | KS.3 |
| FIM1_5182 | 1229.06996 | 816.0161332 | 1281.7074 | -0.578426101 | 0.098349625 | 2.5542250022 | 1.6786353218 | ns | YPD | KS.3 |
| FIM1_5183 | 5011.086514 | 3787.12835 | 5023.530225 | -0.388255584 | 0.128995226 | 0.002110676 | 0.004097001 | ns | YPD | KS.3 |
| FIM1_5184 | 1181.056992 | 635.0082633 | 724.6511252 | -0.858989935 | 0.138767924 | 2.1208827698 | 1.7411537590 | ns | YPD | KS.3 |
| FIM1_5185 | 4238.8327 | 2989.188699 | 3451.371304 | -0.490256669 | 0.106498977 | 2.9773588905 | 1.0527619389 | ns | YPD | KS.3 |

| | | | | | | | | | | |
|---|---|---|---|---|---|---|---|---|---|---|
| FIM1_5186 | 1910.799476 | 1613.252541 | 1683.684437 | -0.237155772 | 0.107358642 | 0.024981321 | 0.038705099 | ns | YPD | KS.3 |
| FIM1_5187 | 1940.978798 | 3290.051863 | 2096.002034 | 0.706856669 | 0.184848659 | 6.4918972976 | 0.000172344 | ns | YPD | KS.3 |
| FIM1_5188 | 10561.31485 | 6295.547473 | 8105.092407 | -0.700008169 | 0.171145601 | 2.1678191556 | 6.3590214027 | ns | YPD | KS.3 |
| FIM1_5189 | 355.6055681 | 185.1161975 | 255.0617251 | -0.830581631 | 0.24871341 | 0.000308209 | 0.000711526 | ns | YPD | KS.3 |
| FIM1_5190 | 1268.407572 | 1045.600786 | 1079.103786 | -0.267417346 | 0.129005732 | 0.03428381 | 0.051513729 | ns | YPD | KS.3 |
| FIM1_5191 | 770.232796 | 1322.703458 | 1573.276048 | 0.734593245 | 0.16618345 | 4.6243506601 | 1.5725320939 | ns | YPD | KS.3 |
| FIM1_5192 | 627.6653068 | 1792.994983 | 1933.053245 | 1.213706905 | 0.373847024 | 0.000119776 | 0.000304333 | Km>KS.3 | YPD | KS.3 |
| FIM1_5193 | 9445.601325 | 13223.16658 | 15741.76877 | 0.471877326 | 0.107996417 | 9.1447564953 | 2.8920683854 | ns | YPD | KS.3 |
| FIM1_5194 | 3960.327853 | 3058.772621 | 3900.873431 | -0.348094122 | 0.169367728 | 0.033217256 | 0.050047198 | ns | YPD | KS.3 |
| FIM1_5195 | 6259.370758 | 6163.909175 | 5536.513644 | -0.021359847 | 0.126102945 | 0.862832429 | 0.885800936 | ns | YPD | KS.3 |
| FIM1_5196 | 333.2618884 | 217.2023628 | 274.2174018 | -0.592516988 | 0.129726514 | 3.0311170904 | 1.0702489371 | ns | YPD | KS.3 |
| FIM1_5197 | 320.5417607 | 370.8235374 | 384.7279617 | 0.202760684 | 0.113786061 | 0.070085276 | 0.097640507 | ns | YPD | KS.3 |
| FIM1_5199 | 4385.062716 | 3215.131756 | 3164.931706 | -0.439248922 | 0.087119024 | 3.5233823464 | 1.5059129874 | ns | YPD | KS.3 |
| FIM1_5200 | 207.48912 | 860.9967867 | 329.1609456 | 1.830739305 | 0.339333435 | 4.8107458724 | 3.6483934017 | Km>KS.3 | YPD | KS.3 |
| FIM1_5201 | 327.0025444 | 889.3019062 | 502.8709471 | 1.336081405 | 0.218347557 | 6.2823644652 | 5.7453608850 | Km>KS.3 | YPD | KS.3 |
| FIM1_5202 | 983.6873465 | 574.8735169 | 778.6488792 | -0.755141739 | 0.105192009 | 3.1496407721 | 4.4898174331 | ns | YPD | KS.3 |
| FIM1_5203 | 628.1220544 | 824.1360043 | 745.3949667 | 0.37078846 | 0.151596959 | 0.011829064 | 0.019715106 | ns | YPD | KS.3 |
| FIM1_5204 | 244.1126812 | 694.3992413 | 477.5816684 | 1.417524691 | 0.198869041 | 4.5314613978 | 7.5905792774 | Km>KS.3 | YPD | KS.3 |
| FIM1_5205 | 1899.029139 | 1637.677404 | 1564.827119 | -0.209588166 | 0.087301451 | 0.015326189 | 0.024982893 | ns | YPD | KS.3 |
| FIM1_5206 | 826.3971761 | 1067.44206 | 958.1753731 | 0.360254684 | 0.10410097 | 0.000448988 | 0.00100032 | ns | YPD | KS.3 |
| FIM1_5207 | 340.3094859 | 260.8573393 | 269.4435916 | -0.357667004 | 0.169622811 | 0.028888211 | 0.044153257 | ns | YPD | KS.3 |
| FIM1_5208 | 19.73957143 | 11.39301758 | 21.5559521 | -0.490432451 | 0.462321483 | 0.170757894 | 0.21911801 | ns | YPD | KS.3 |
| FIM1_5209 | 50.91909849 | 47.03703431 | 37.8397702 | -0.086604262 | 0.33304427 | 0.766743174 | 0.804417396 | ns | YPD | KS.3 |
| FIM1_4316 | 2183.197584 | 2652.730751 | 4233.280858 | 0.255727456 | 0.199254995 | 0.17806811 | 0.227384201 | ns | YPD | KS.3 |
| FIM1_4318 | 377.4119193 | 307.5633106 | 297.7479064 | -0.277490604 | 0.155804303 | 0.066456383 | 0.093053899 | ns | YPD | KS.3 |
| FIM1_4319 | 1517.911107 | 472.0873557 | 791.3821641 | -1.474537525 | 0.31177235 | 7.9848211238 | 3.9331173357 | Km<KS.3 | YPD | KS.3 |
| FIM1_4320 | 4030.082571 | 4730.028052 | 3853.218499 | 0.223821992 | 0.113471042 | 0.044969755 | 0.065589133 | ns | YPD | KS.3 |
| FIM1_4321 | 1654.733375 | 1209.262801 | 1574.930606 | -0.43603451 | 0.125150801 | 0.000378351 | 0.000856758 | ns | YPD | KS.3 |
| FIM1_4322 | 1397.823881 | 1115.866142 | 1448.110775 | -0.311863198 | 0.129894076 | 0.014185776 | 0.02326112 | ns | YPD | KS.3 |
| FIM1_4323 | 14321.65651 | 17016.74709 | 16724.97566 | 0.245364532 | 0.072853508 | 0.000694736 | 0.001487865 | ns | YPD | KS.3 |
| FIM1_4324 | 63.06490861 | 30.17729157 | 38.88673509 | -0.918574877 | 0.266939179 | 0.000168272 | 0.000413409 | ns | YPD | KS.3 |
| FIM1_4325 | 2820.91707 | 1894.622692 | 1983.336966 | -0.559466751 | 0.103966525 | 4.7891962347 | 2.4543563813 | ns | YPD | KS.3 |
| FIM1_4326 | 1772.474553 | 1016.106285 | 1289.899005 | -0.758298294 | 0.162527033 | 1.3737650730 | 5.2331403052 | ns | YPD | KS.3 |
| FIM1_4327 | 2099.626911 | 2725.029905 | 2328.645034 | 0.359976811 | 0.135384101 | 0.006505503 | 0.011448489 | ns | YPD | KS.3 |
| FIM1_4328 | 886.7476422 | 507.4413428 | 872.2491588 | -0.743030561 | 0.194220211 | 5.9711095845 | 0.00015954 | ns | YPD | KS.3 |
| FIM1_4329 | 794.2032885 | 544.4642289 | 704.8157214 | -0.513887215 | 0.158061764 | 0.000791944 | 0.001674424 | ns | YPD | KS.3 |
| FIM1_4330 | 3142.19589 | 3238.423915 | 2374.448184 | 0.041594641 | 0.137377715 | 0.756620681 | 0.797159654 | ns | YPD | KS.3 |
| FIM1_4331 | 5633.387308 | 4273.633014 | 5501.843841 | -0.363639618 | 0.197846374 | 0.05384341 | 0.076996541 | ns | YPD | KS.3 |
| FIM1_4332 | 2002.585409 | 1713.061203 | 1772.143234 | -0.221331596 | 0.076026226 | 0.003352059 | 0.00626213 | ns | YPD | KS.3 |
| FIM1_4333 | 3908.331621 | 2328.328813 | 2923.975554 | -0.73308722 | 0.090946614 | 3.5438265288 | 8.5171676236 | ns | YPD | KS.3 |
| FIM1_4334 | 879.74213 | 677.3039864 | 898.2957246 | -0.354174182 | 0.163292805 | 0.024967451 | 0.03869566 | ns | YPD | KS.3 |
| FIM1_4335 | 3674.127606 | 2325.210977 | 2760.392648 | -0.652194242 | 0.068448851 | 8.7106235013 | 5.4169189899 | ns | YPD | KS.3 |
| FIM1_4336 | 1227.362361 | 769.9020146 | 963.1192203 | -0.638188157 | 0.152782435 | 1.6721481816 | 5.0265481592 | ns | YPD | KS.3 |
| FIM1_4337 | 2042.331088 | 1270.732387 | 1516.12613 | -0.664010572 | 0.115321498 | 4.6054351851 | 2.8675894926 | ns | YPD | KS.3 |
| FIM1_4338 | 5434.356212 | 4299.573956 | 4371.463693 | -0.334141403 | 0.065863841 | 3.3434291267 | 1.4376456270 | ns | YPD | KS.3 |

| | | | | | | | | | | |
|---|---|---|---|---|---|---|---|---|---|---|
| FIM1_4339 | 824.6385086 | 316.2405669 | 495.8349819 | -1.320494397 | 0.164146664 | 6.0183369372 | 1.7612486036 | Km<KS.3 | YPD | KS.3 |
| FIM1_4340 | 7021.025488 | 4836.675453 | 10345.93131 | -0.52177143 | 0.112608105 | 2.4638192922 | 8.8629797388 | ns | YPD | KS.3 |
| FIM1_4341 | 384.183077 | 236.7586018 | 351.1808717 | -0.611388372 | 0.246578387 | 0.007551 | 0.013144236 | ns | YPD | KS.3 |
| FIM1_4342 | 575.1216103 | 370.83297 | 597.9498622 | -0.608713536 | 0.128262422 | 1.2406718978 | 4.7736602410 | ns | YPD | KS.3 |
| FIM1_4343 | 17758.98256 | 16099.99689 | 25916.70843 | -0.137292699 | 0.109773238 | 0.204126134 | 0.255735965 | ns | YPD | KS.3 |
| FIM1_4344 | 2012.643758 | 1027.043613 | 2485.044825 | -0.863477691 | 0.240796308 | 0.000114378 | 0.000292261 | ns | YPD | KS.3 |
| FIM1_4345 | 747.5814331 | 315.7360768 | 544.9502246 | -1.185996662 | 0.161513864 | 2.6027405678 | 4.6245122589 | Km<KS.3 | YPD | KS.3 |
| FIM1_4346 | 1177.599152 | 386.0212521 | 1382.390916 | -1.288233573 | 0.386194667 | 6.5127748654 | 0.000172806 | Km<KS.3 | YPD | KS.3 |
| FIM1_4347 | 1601.727489 | 1935.914248 | 1730.147006 | 0.264600193 | 0.116206735 | 0.020541544 | 0.032454678 | ns | YPD | KS.3 |
| FIM1_4348 | 2847.585105 | 2770.572348 | 2477.390735 | -0.038785225 | 0.078337365 | 0.617731621 | 0.673359951 | ns | YPD | KS.3 |
| FIM1_4349 | 95.69336676 | 248.9441963 | 484.6426657 | 1.1604088 | 0.320650065 | 3.8654787981 | 0.000106974 | Km>KS.3 | YPD | KS.3 |
| FIM1_4350 | 3094.853524 | 2979.096908 | 2577.991275 | -0.054056739 | 0.066364338 | 0.412618563 | 0.476614198 | ns | YPD | KS.3 |
| FIM1_4351 | 4442.823994 | 4321.516087 | 4666.112367 | -0.038521652 | 0.124498484 | 0.752420954 | 0.793226171 | ns | YPD | KS.3 |
| FIM1_4352 | 1622.917905 | 674.962682 | 1263.868516 | -1.171332065 | 0.208277002 | 2.4833015861 | 1.6450633010 | Km<KS.3 | YPD | KS.3 |
| FIM1_4353 | 2942.918028 | 2100.349541 | 2362.725098 | -0.466643815 | 0.133067605 | 0.000334306 | 0.000765476 | ns | YPD | KS.3 |
| FIM1_4354 | 1768.192235 | 941.2134975 | 1627.87739 | -0.864642277 | 0.155107462 | 8.6457773560 | 5.1327854828 | ns | YPD | KS.3 |
| FIM1_4355 | 29.94805912 | 26.52588501 | 26.18476884 | -0.143275462 | 0.289526692 | 0.584613863 | 0.642752258 | ns | YPD | KS.3 |
| FIM1_4356 | 17644.09601 | 16419.95948 | 23485.72274 | -0.102425455 | 0.069442231 | 0.137711868 | 0.179349881 | ns | YPD | KS.3 |
| FIM1_4357 | 1854.372454 | 1267.487416 | 1923.814461 | -0.526657434 | 0.133083593 | 5.1502260240 | 0.000138728 | ns | YPD | KS.3 |
| FIM1_4358 | 4687.641427 | 3043.258593 | 4355.850489 | -0.598751351 | 0.131941279 | 3.4498843945 | 1.2052791336 | ns | YPD | KS.3 |
| FIM1_4359 | 2276.197503 | 1682.537772 | 2335.321236 | -0.418335769 | 0.131271762 | 0.001123956 | 0.002303 | ns | YPD | KS.3 |
| FIM1_4360 | 9396.077144 | 8192.236813 | 8671.074157 | -0.19325698 | 0.095590363 | 0.040797979 | 0.060192748 | ns | YPD | KS.3 |
| FIM1_4361 | 2423.561646 | 1920.745549 | 2926.548286 | -0.304561245 | 0.202591338 | 0.114058694 | 0.151277526 | ns | YPD | KS.3 |
| FIM1_4363 | 888.4190985 | 493.8808578 | 979.5832279 | -0.757911794 | 0.230163591 | 0.000433468 | 0.000970087 | ns | YPD | KS.3 |
| FIM1_4364 | 558.7590489 | 259.036793 | 393.1670421 | -1.034938584 | 0.189584737 | 1.0252252556 | 5.9794790702 | Km<KS.3 | YPD | KS.3 |
| FIM1_4365 | 1503.668674 | 761.384021 | 1130.932375 | -0.94585732 | 0.134780435 | 6.2830646727 | 8.5172334460 | ns | YPD | KS.3 |
| FIM1_4366 | 7417.445526 | 8121.499174 | 9010.826394 | 0.12848609 | 0.085073788 | 0.127386821 | 0.167441644 | ns | YPD | KS.3 |
| FIM1_4367 | 2111.840048 | 1576.722998 | 1772.816264 | -0.403239934 | 0.13615955 | 0.002430027 | 0.004656928 | ns | YPD | KS.3 |
| FIM1_4368 | 1752.700163 | 1269.636551 | 1621.621862 | -0.448846335 | 0.122893569 | 0.00019612 | 0.00047158 | ns | YPD | KS.3 |
| FIM1_4369 | 3587.423797 | 1343.156121 | 3186.351165 | -1.237957509 | 0.286925559 | 1.6005775604 | 6.0370533457 | Km<KS.3 | YPD | KS.3 |
| FIM1_4370 | 1066.587317 | 460.0918051 | 886.3623511 | -1.135680176 | 0.188917277 | 2.7923590427 | 2.2334383018 | Km<KS.3 | YPD | KS.3 |
| FIM1_4371 | 3214.182901 | 2932.604327 | 3441.943324 | -0.123901875 | 0.163862264 | 0.434596962 | 0.498575123 | ns | YPD | KS.3 |
| FIM1_4372 | 2075.522836 | 1783.952324 | 2499.143024 | -0.202488192 | 0.177691425 | 0.236310925 | 0.291001696 | ns | YPD | KS.3 |
| FIM1_4373 | 7587.205542 | 6053.368821 | 8223.638193 | -0.292613919 | 0.213948038 | 0.148299673 | 0.192083018 | ns | YPD | KS.3 |
| FIM1_4374 | 1348.369682 | 1124.058134 | 1187.237333 | -0.255975234 | 0.098581729 | 0.008557395 | 0.014695561 | ns | YPD | KS.3 |
| FIM1_4376 | 397.4996656 | 204.2353157 | 275.9932316 | -0.905361555 | 0.169931441 | 3.1119782611 | 1.6452807491 | ns | YPD | KS.3 |
| FIM1_4377 | 3956.323074 | 5254.427889 | 5376.730464 | 0.368996052 | 0.211113055 | 0.064992272 | 0.091234919 | ns | YPD | KS.3 |
| FIM1_4378 | 202.3684984 | 78.7341083 | 200.5203965 | -1.296155827 | 0.176294258 | 1.5326467160 | 2.8557743117 | Km<KS.3 | YPD | KS.3 |
| FIM1_4379 | 4010.490603 | 3258.289608 | 3467.010987 | -0.290603605 | 0.110504559 | 0.007559925 | 0.013155169 | ns | YPD | KS.3 |
| FIM1_4380 | 2037.774808 | 1827.07109 | 1529.100328 | -0.154224558 | 0.085314686 | 0.068027547 | 0.095019726 | ns | YPD | KS.3 |
| FIM1_4381 | 1967.723403 | 1333.571116 | 1501.693725 | -0.543235755 | 0.117901611 | 2.7051406166 | 9.6335537349 | ns | YPD | KS.3 |
| FIM1_4382 | 1379.377156 | 805.3614728 | 961.0533677 | -0.749737327 | 0.124510321 | 7.8631574262 | 5.7443771212 | ns | YPD | KS.3 |
| FIM1_4383 | 5662.761599 | 5345.413882 | 5562.631831 | -0.082321153 | 0.066321514 | 0.211856243 | 0.264421678 | ns | YPD | KS.3 |
| FIM1_4384 | 966.3679974 | 1247.682236 | 1504.303601 | 0.328378948 | 0.222918582 | 0.117739269 | 0.155868244 | ns | YPD | KS.3 |
| FIM1_4385 | 1670.118901 | 2396.984449 | 1672.830195 | 0.506867689 | 0.108171861 | 1.9526242656 | 7.2118082566 | ns | YPD | KS.3 |

| | | | | | | | | | | |
|---|---|---|---|---|---|---|---|---|---|---|
| FIM1_4386 | 10911.48143 | 11761.51 | 15426.11709 | 0.104830854 | 0.11236184 | 0.343166678 | 0.404890216 | ns | YPD | KS.3 |
| FIM1_4387 | 30.81420404 | 23.54253666 | 32.08724951 | -0.336697734 | 0.255317302 | 0.153841156 | 0.198648806 | ns | YPD | KS.3 |
| FIM1_4388 | 1109.726611 | 2168.601042 | 2828.08616 | 0.839052847 | 0.264564514 | 0.000540356 | 0.001181139 | ns | YPD | KS.3 |
| FIM1_4389 | 330.5349405 | 358.6979465 | 261.3620366 | 0.109485528 | 0.175157592 | 0.516459209 | 0.578690217 | ns | YPD | KS.3 |
| FIM1_4390 | 156.7304219 | 98.2931988 | 119.0014353 | -0.623858979 | 0.18300787 | 0.000376183 | 0.000853012 | ns | YPD | KS.3 |
| FIM1_4391 | 8493.206725 | 6576.257912 | 11072.76012 | -0.338686109 | 0.191130175 | 0.063880549 | 0.089928051 | ns | YPD | KS.3 |
| FIM1_4392 | 1308.969571 | 2754.972922 | 2446.787469 | 0.99863629 | 0.192329809 | 4.9736688803 | 2.5430629680 | ns | YPD | KS.3 |
| FIM1_4393 | 51.07126301 | 146.1095331 | 102.1546705 | 1.370093213 | 0.254909323 | 4.4013816519 | 2.7543237381 | Km>KS.3 | YPD | KS.3 |
| FIM1_4394 | 481.6994573 | 623.2756086 | 574.9786187 | 0.358254373 | 0.121918864 | 0.002747674 | 0.005221419 | ns | YPD | KS.3 |
| FIM1_4396 | 3969.714944 | 2340.578985 | 3371.773329 | -0.724438122 | 0.151907655 | 8.8931380260 | 3.5225606432 | ns | YPD | KS.3 |
| FIM1_4397 | 1841.600193 | 1804.564535 | 1309.553351 | -0.028533433 | 0.112363697 | 0.79640666 | 0.829937816 | ns | YPD | KS.3 |
| FIM1_4398 | 284.7832145 | 497.9485336 | 199.3745941 | 0.659104078 | 0.307693667 | 0.016315004 | 0.026473302 | ns | YPD | KS.3 |
| FIM1_4399 | 14158.06804 | 13640.04029 | 14052.33915 | -0.048571913 | 0.205926169 | 0.803982873 | 0.835760041 | ns | YPD | KS.3 |
| FIM1_4400 | 903.3475897 | 1190.355799 | 1072.102269 | 0.388273941 | 0.096677927 | 4.7925214303 | 0.000130146 | ns | YPD | KS.3 |
| FIM1_4401 | 2287.792979 | 1934.581364 | 2429.574297 | -0.233701305 | 0.12023433 | 0.047762552 | 0.069216049 | ns | YPD | KS.3 |
| FIM1_4402 | 134.9882086 | 107.4027176 | 172.6698895 | -0.309630498 | 0.168603848 | 0.057180262 | 0.081138563 | ns | YPD | KS.3 |
| FIM1_4403 | 2365.100897 | 9482.400286 | 5429.59483 | 1.803678631 | 0.31862565 | 1.1339636791 | 9.8283437345 | Km>KS.3 | YPD | KS.3 |
| FIM1_4404 | 772.1760408 | 396.7467428 | 460.0954742 | -0.913099535 | 0.156711749 | 1.7344575985 | 1.1984620212 | ns | YPD | KS.3 |
| FIM1_4405 | 939.5086305 | 536.7467432 | 609.2161831 | -0.762231462 | 0.163347297 | 1.3576672779 | 5.1837257928 | ns | YPD | KS.3 |
| FIM1_4406 | 1299.17005 | 1358.778142 | 1069.145318 | 0.06244809 | 0.120075761 | 0.596435119 | 0.653438607 | ns | YPD | KS.3 |
| FIM1_4407 | 55.51113034 | 65.66721001 | 122.619116 | 0.209983549 | 0.249293962 | 0.364277096 | 0.426844191 | ns | YPD | KS.3 |
| FIM1_4408 | 242.8717217 | 588.0675974 | 448.4685894 | 1.086673717 | 0.299986375 | 4.9999500703 | 0.000135189 | Km>KS.3 | YPD | KS.3 |
| FIM1_4409 | 7175.012108 | 1920.94006 | 6251.210809 | -1.689790319 | 0.321589142 | 1.8915446126 | 1.2908689229 | Km<KS.3 | YPD | KS.3 |
| FIM1_4410 | 661.1732147 | 865.0285676 | 823.5708034 | 0.371948643 | 0.130937954 | 0.003695462 | 0.00682693 | ns | YPD | KS.3 |
| FIM1_4412 | 213.1091011 | 335.5044035 | 240.9380461 | 0.54505284 | 0.28835753 | 0.036647536 | 0.054636348 | ns | YPD | KS.3 |
| FIM1_4413 | 1751.612646 | 1264.682274 | 1434.163885 | -0.460475715 | 0.089437488 | 1.9545075042 | 8.8719660892 | ns | YPD | KS.3 |
| FIM1_4414 | 3410.865032 | 4012.281118 | 3221.545165 | 0.225413241 | 0.125074402 | 0.066063668 | 0.092530054 | ns | YPD | KS.3 |
| FIM1_4415 | 2233.873094 | 6381.171891 | 3335.323888 | 1.323516692 | 0.294872247 | 5.0350215454 | 2.0909208838 | Km>KS.3 | YPD | KS.3 |
| FIM1_4416 | 1591.971848 | 2599.885133 | 1896.95476 | 0.646476769 | 0.202809738 | 0.000789956 | 0.001671642 | ns | YPD | KS.3 |
| FIM1_4417 | 1816.529757 | 1567.677116 | 1298.851095 | -0.207520744 | 0.093657354 | 0.02503177 | 0.038759121 | ns | YPD | KS.3 |
| FIM1_4418 | 5356.450937 | 5124.731486 | 5851.324223 | -0.062445323 | 0.094830331 | 0.505347055 | 0.567645428 | ns | YPD | KS.3 |
| FIM1_4419 | 887.3219044 | 1014.29326 | 1039.942133 | 0.186754558 | 0.113840006 | 0.095400363 | 0.128692193 | ns | YPD | KS.3 |
| FIM1_4420 | 1019.55468 | 1281.041086 | 1050.982712 | 0.320380567 | 0.104448633 | 0.001866201 | 0.003662465 | ns | YPD | KS.3 |
| FIM1_4421 | 869.323316 | 610.9833269 | 841.2736502 | -0.497817836 | 0.095310758 | 1.2469983683 | 5.8971643369 | ns | YPD | KS.3 |
| FIM1_4422 | 16.80668252 | 9.384052796 | 14.57156587 | -0.579443654 | 0.406597157 | 0.08277045 | 0.113274649 | ns | YPD | KS.3 |
| FIM1_4423 | 1785.639325 | 1433.955872 | 1754.733821 | -0.281383885 | 0.223941597 | 0.181936147 | 0.231550865 | ns | YPD | KS.3 |
| FIM1_4424 | 1211.740531 | 1060.174257 | 1053.413028 | -0.139332925 | 0.370004567 | 0.657541965 | 0.709603314 | ns | YPD | KS.3 |
| FIM1_4425 | 662.5607875 | 733.129641 | 643.1878699 | 0.144560031 | 0.073279632 | 0.046959144 | 0.068151034 | ns | YPD | KS.3 |
| FIM1_4426 | 1030.363002 | 745.0983312 | 728.6083259 | -0.450440449 | 0.125417055 | 0.000247518 | 0.000580356 | ns | YPD | KS.3 |
| FIM1_4427 | 7354.635361 | 5083.164423 | 5440.023767 | -0.51632981 | 0.115247612 | 5.1509653311 | 1.7397184333 | ns | YPD | KS.3 |
| FIM1_4428 | 8401.62823 | 9959.471872 | 6919.829657 | 0.216797885 | 0.22903026 | 0.313408296 | 0.374269388 | ns | YPD | KS.3 |
| FIM1_4429 | 1441.414531 | 2759.947738 | 2044.53562 | 0.884961794 | 0.165986979 | 3.2183690782 | 1.6961214157 | ns | YPD | KS.3 |
| FIM1_4430 | 375.3665898 | 227.0349501 | 343.7891576 | -0.683239408 | 0.164432065 | 1.6909040697 | 5.0767940535 | ns | YPD | KS.3 |
| FIM1_4431 | 1397.662748 | 3161.99774 | 2195.196975 | 1.155649918 | 0.098634059 | 1.3833266880 | 2.4578750974 | Km>KS.3 | YPD | KS.3 |
| FIM1_4432 | 5711.233223 | 5706.99759 | 5969.247125 | -0.001120471 | 0.089255929 | 0.989881151 | 0.99089732 | ns | YPD | KS.3 |

| | | | | | | | | | | |
|---|---|---|---|---|---|---|---|---|---|---|
| FIM1_4433 | 1213.061775 | 454.500476 | 622.5379711 | -1.3508604 | 0.168962331 | 7.81966679217 | 2.21038876653 | Km<KS.3 | YPD | KS.3 |
| FIM1_4434 | 2082.640098 | 1952.756307 | 1987.486864 | -0.092061941 | 0.057483743 | 0.107695802 | 0.143835333 | ns | YPD | KS.3 |
| FIM1_4435 | 4338.12794 | 5326.18724 | 4237.227993 | 0.287604923 | 0.108098678 | 0.006920476 | 0.012127288 | ns | YPD | KS.3 |
| FIM1_4436 | 1911.076617 | 1873.772772 | 1632.471748 | -0.027787446 | 0.07198019 | 0.697519832 | 0.746592333 | ns | YPD | KS.3 |
| FIM1_4437 | 838.6164224 | 563.9992095 | 636.5513952 | -0.546269724 | 0.141182091 | 7.19981454508 | 0.00018972 | ns | YPD | KS.3 |
| FIM1_4438 | 1344.160611 | 1918.254561 | 1274.270211 | 0.484675997 | 0.156533844 | 0.0014053 | 0.002830513 | ns | YPD | KS.3 |
| FIM1_4439 | 1756.76181 | 1187.921437 | 1607.302738 | -0.548548866 | 0.11083747 | 4.90952695376 | 2.04906850629 | ns | YPD | KS.3 |
| FIM1_4440 | 1182.855647 | 1872.322356 | 1509.242747 | 0.596590476 | 0.217539288 | 0.003636884 | 0.006733717 | ns | YPD | KS.3 |
| FIM1_4441 | 1773.505605 | 5126.551095 | 3396.226368 | 1.364247558 | 0.275108997 | 4.17917347979 | 2.16576958978 | Km>KS.3 | YPD | KS.3 |
| FIM1_4443 | 317.9457894 | 606.1340095 | 510.6500014 | 0.724869737 | 0.348007015 | 0.016001785 | 0.026007475 | ns | YPD | KS.3 |
| FIM1_4444 | 22868.20544 | 23733.74549 | 29229.62885 | 0.050913225 | 0.144298168 | 0.717354715 | 0.763878362 | ns | YPD | KS.3 |
| FIM1_4445 | 1532.958484 | 1370.542635 | 1353.071427 | -0.157047894 | 0.106883336 | 0.136019869 | 0.177471505 | ns | YPD | KS.3 |
| FIM1_4446 | 4715.126644 | 4891.79934 | 5081.720682 | 0.051156307 | 0.119761461 | 0.663648297 | 0.715572231 | ns | YPD | KS.3 |
| FIM1_4447 | 1947.905035 | 1895.710544 | 1698.587544 | -0.038332677 | 0.074663373 | 0.605047881 | 0.661999825 | ns | YPD | KS.3 |
| FIM1_4448 | 146.7007915 | 401.8093507 | 260.2087931 | 1.271221108 | 0.28800659 | 8.88987740426 | 3.52256064327 | Km>KS.3 | YPD | KS.3 |
| FIM1_4449 | 607.614057 | 1172.368599 | 733.6709146 | 0.822681594 | 0.264370351 | 0.000688766 | 0.001476351 | ns | YPD | KS.3 |
| FIM1_4450 | 2273.174084 | 3195.160514 | 3259.486394 | 0.469384363 | 0.138255584 | 0.000503666 | 0.001109224 | ns | YPD | KS.3 |
| FIM1_4451 | 1555.806834 | 2014.776704 | 2267.005217 | 0.346090678 | 0.178188018 | 0.043415477 | 0.063545748 | ns | YPD | KS.3 |
| FIM1_4452 | 7794.056743 | 6465.465534 | 10157.37843 | -0.254527386 | 0.154624964 | 0.089986447 | 0.121970288 | ns | YPD | KS.3 |
| FIM1_4454 | 3089.277953 | 5851.29672 | 4077.397871 | 0.907212764 | 0.084112704 | 1.21000707305 | 1.36813299736 | ns | YPD | KS.3 |
| FIM1_4455 | 1733.347536 | 905.2711794 | 1386.220544 | -0.881476859 | 0.172399232 | 1.05667909905 | 5.07973184755 | ns | YPD | KS.3 |
| FIM1_4456 | 1013.820133 | 1571.43505 | 1521.687779 | 0.593618323 | 0.166942778 | 0.000229625 | 0.000542958 | ns | YPD | KS.3 |
| FIM1_4457 | 1429.943157 | 1572.487545 | 1899.474495 | 0.132808485 | 0.106533694 | 0.205984097 | 0.257739155 | ns | YPD | KS.3 |
| FIM1_4458 | 2089.740933 | 1247.573316 | 1523.401855 | -0.726371686 | 0.103555705 | 1.10012500113 | 1.42901354586 | ns | YPD | KS.3 |
| FIM1_4460 | 414.1988497 | 329.0729969 | 482.6566913 | -0.30367167 | 0.192288482 | 0.09851387 | 0.132425427 | ns | YPD | KS.3 |
| FIM1_4461 | 184.706347 | 101.6782596 | 235.8112276 | -0.788520302 | 0.199849255 | 3.29686705299 | 9.27185618352 | ns | YPD | KS.3 |
| FIM1_4462 | 1346.093198 | 1048.878501 | 1086.853569 | -0.349591156 | 0.105444939 | 0.000770169 | 0.001631853 | ns | YPD | KS.3 |
| FIM1_4463 | 1361.772484 | 2558.181079 | 1342.582568 | 0.877187251 | 0.130978197 | 7.14036889703 | 7.91165596052 | ns | YPD | KS.3 |
| FIM1_4464 | 845.8533995 | 1087.03986 | 1042.604224 | 0.344288025 | 0.144241425 | 0.014288549 | 0.023414207 | ns | YPD | KS.3 |
| FIM1_4465 | 1507.119131 | 955.0627145 | 1372.504425 | -0.634661671 | 0.12643492 | 2.95605334016 | 1.28664613887 | ns | YPD | KS.3 |
| FIM1_4466 | 1171.07037 | 1331.996848 | 1228.347793 | 0.180778517 | 0.103725421 | 0.077245849 | 0.106571853 | ns | YPD | KS.3 |
| FIM1_4467 | 3254.818413 | 2525.802013 | 2880.810584 | -0.360764199 | 0.07581481 | 1.62574096117 | 6.11804938113 | ns | YPD | KS.3 |
| FIM1_4468 | 350.6737633 | 230.2930276 | 290.415238 | -0.576952036 | 0.146054707 | 4.90691508813 | 0.000132818 | ns | YPD | KS.3 |
| FIM1_4469 | 2242.159143 | 1960.765729 | 3092.130005 | -0.161181439 | 0.278514694 | 0.525673687 | 0.588186401 | ns | YPD | KS.3 |
| FIM1_4470 | 1569.00872 | 3625.348721 | 2972.149374 | 1.084460417 | 0.242726786 | 1.41653487709 | 5.38781423053 | Km>KS.3 | YPD | KS.3 |
| FIM1_4471 | 603.4782313 | 599.9194019 | 801.3701326 | -0.008165349 | 0.139799123 | 0.952288351 | 0.963129609 | ns | YPD | KS.3 |
| FIM1_4472 | 2657.870907 | 2838.467654 | 2324.054921 | 0.093913966 | 0.063559916 | 0.137409095 | 0.179002421 | ns | YPD | KS.3 |
| FIM1_4473 | 753.6669814 | 751.7794883 | 825.0001088 | -0.003849352 | 0.140429916 | 0.977593453 | 0.981935681 | ns | YPD | KS.3 |
| FIM1_4474 | 7104.08513 | 10984.08051 | 10933.34462 | 0.593436528 | 0.159482586 | 0.000121155 | 0.000306741 | ns | YPD | KS.3 |
| FIM1_4475 | 619.0909128 | 728.1022022 | 812.4556507 | 0.216965999 | 0.17483251 | 0.197678048 | 0.248911235 | ns | YPD | KS.3 |
| FIM1_4476 | 1184.929721 | 814.439007 | 1386.57011 | -0.499830675 | 0.186631267 | 0.005161555 | 0.009236955 | ns | YPD | KS.3 |
| FIM1_4477 | 1982.117067 | 2313.016526 | 2194.233526 | 0.217386495 | 0.09573135 | 0.021577884 | 0.03390713 | ns | YPD | KS.3 |
| FIM1_4478 | 1820.404638 | 1824.630736 | 1911.228193 | 0.003298077 | 0.1355146 | 0.980137419 | 0.983884087 | ns | YPD | KS.3 |
| FIM1_4479 | 1508.308664 | 2908.326922 | 2132.936426 | 0.857533526 | 0.220779171 | 3.58507627054 | 0.000100088 | ns | YPD | KS.3 |
| FIM1_4480 | 253.4985906 | 445.0226788 | 436.876264 | 0.763113491 | 0.167527076 | 2.31172289089 | 8.36423373251 | ns | YPD | KS.3 |

| | | | | | | | | | | |
|---|---|---|---|---|---|---|---|---|---|---|
| FIM1_4481 | 1663.177992 | 2451.322987 | 1967.364171 | 0.550660115 | 0.081309327 | 8.2838824073 | 8.9787178598 | ns | YPD | KS.3 |
| FIM1_4482 | 199.4084919 | 3611.628215 | 2864.161851 | 3.965844892 | 0.328813814 | 1.4669476459 | 3.4752688280 | Km>KS.3 | YPD | KS.3 |
| FIM1_4484 | 11.1319034 | 10.39248917 | 72.63694843 | -0.059986775 | 0.475011798 | 0.867775135 | 0.890140474 | ns | YPD | KS.3 |
| FIM1_4486 | 766.5550517 | 2430.816333 | 1769.97335 | 1.433782088 | 0.326812958 | 4.9211224760 | 2.0521864474 | Km>KS.3 | YPD | KS.3 |
| FIM1_4487 | 137.8101404 | 565.2260215 | 435.3429884 | 1.822653849 | 0.331153355 | 2.6496994313 | 2.1296049549 | Km>KS.3 | YPD | KS.3 |
| FIM1_4488 | 21.1568441 | 25.78983816 | 33.08812412 | 0.227472951 | 0.295349134 | 0.392892744 | 0.45615902 | ns | YPD | KS.3 |
| FIM1_4489 | 706.089237 | 1372.187222 | 1176.216034 | 0.81950463 | 0.277967483 | 0.001181626 | 0.002414205 | ns | YPD | KS.3 |
| FIM1_4490 | 4508.258577 | 12703.36771 | 7550.038692 | 1.357485781 | 0.248198969 | 2.7467796058 | 1.7909867023 | Km>KS.3 | YPD | KS.3 |
| FIM1_4491 | 676.7198652 | 1475.583493 | 931.9821455 | 1.05336539 | 0.1844623 | 2.2740432181 | 1.5206135766 | Km>KS.3 | YPD | KS.3 |
| FIM1_4492 | 822.9118891 | 562.6334802 | 821.0718333 | -0.529713436 | 0.120622415 | 7.6240666506 | 2.4661724048 | ns | YPD | KS.3 |
| FIM1_4493 | 4587.122451 | 11610.7084 | 6962.732737 | 1.237593342 | 0.214901645 | 8.6711745144 | 6.2429946757 | Km>KS.3 | YPD | KS.3 |
| FIM1_4494 | 41.27765019 | 39.15246665 | 44.75258721 | -0.067616493 | 0.239701875 | 0.762583479 | 0.801574648 | ns | YPD | KS.3 |
| FIM1_4495 | 1115.609635 | 1535.283967 | 1200.271987 | 0.439503866 | 0.141454056 | 0.001438228 | 0.002890984 | ns | YPD | KS.3 |
| FIM1_4496 | 451.6240885 | 679.2631574 | 572.8180632 | 0.56537363 | 0.130989363 | 1.0179694929 | 3.1931892984 | ns | YPD | KS.3 |
| FIM1_4497 | 3725.340213 | 3316.059532 | 4143.687139 | -0.162162593 | 0.118434056 | 0.163446255 | 0.210332416 | ns | YPD | KS.3 |
| FIM1_4498 | 14045.10559 | 12371.76489 | 18755.82401 | -0.176128431 | 0.124617954 | 0.149556148 | 0.19350893 | ns | YPD | KS.3 |
| FIM1_4499 | 2888.504649 | 2969.277565 | 3375.850452 | 0.038420931 | 0.122714184 | 0.749703299 | 0.791044308 | ns | YPD | KS.3 |
| FIM1_4500 | 2912.460012 | 3843.241422 | 2985.97264 | 0.393473029 | 0.081273358 | 1.0391037004 | 4.0769250078 | ns | YPD | KS.3 |
| FIM1_4501 | 1431.097889 | 839.8352832 | 1516.750278 | -0.716676018 | 0.180220175 | 3.3921785653 | 9.5023020059 | ns | YPD | KS.3 |
| FIM1_4502 | 252.7405204 | 180.1119564 | 211.4499236 | -0.439138238 | 0.218107412 | 0.032856477 | 0.049533628 | ns | YPD | KS.3 |
| FIM1_4503 | 885.5397157 | 682.1230352 | 692.7756756 | -0.366626109 | 0.10135965 | 0.000246732 | 0.000579278 | ns | YPD | KS.3 |
| FIM1_4504 | 1067.975636 | 1719.680126 | 1117.332412 | 0.641077727 | 0.176562471 | 0.000158363 | 0.000391774 | ns | YPD | KS.3 |
| FIM1_4505 | 1772.013712 | 1426.183177 | 2753.53033 | -0.246973625 | 0.319176931 | 0.382434231 | 0.445367579 | ns | YPD | KS.3 |
| FIM1_4506 | 1667.383333 | 849.8873586 | 1446.303108 | -0.912490433 | 0.176750376 | 7.4750528938 | 3.7077156676 | ns | YPD | KS.3 |
| FIM1_4507 | 3257.555279 | 2238.810938 | 2743.829087 | -0.515682091 | 0.14344983 | 0.000223715 | 0.000531004 | ns | YPD | KS.3 |
| FIM1_4508 | 1270.425131 | 948.3877742 | 1156.549221 | -0.411892013 | 0.100077198 | 3.0467325239 | 8.6466025708 | ns | YPD | KS.3 |
| FIM1_4509 | 2746.471124 | 1821.556069 | 2958.750099 | -0.58680878 | 0.063001636 | 7.5177750105 | 4.4001094914 | ns | YPD | KS.3 |
| FIM1_4510 | 4802.430266 | 3762.435674 | 5040.076391 | -0.34499181 | 0.090979258 | 0.000126454 | 0.000318859 | ns | YPD | KS.3 |
| FIM1_4512 | 977.7853304 | 772.7340057 | 894.7769578 | -0.330197156 | 0.105354385 | 0.001476429 | 0.002957019 | ns | YPD | KS.3 |
| FIM1_4513 | 273.7318489 | 128.9361736 | 220.9323604 | -1.03271522 | 0.160099718 | 2.4024568332 | 2.3809208656 | Km<KS.3 | YPD | KS.3 |
| FIM1_4514 | 452.5548188 | 253.620801 | 316.521227 | -0.801064313 | 0.138500233 | 2.9648724653 | 1.9256188661 | ns | YPD | KS.3 |
| FIM1_4515 | 1514.968869 | 1301.70786 | 1229.686415 | -0.213729014 | 0.094178621 | 0.021706437 | 0.034055353 | ns | YPD | KS.3 |
| FIM1_4516 | 472.5782862 | 276.6884774 | 750.3798019 | -0.600560456 | 0.341518001 | 0.043166685 | 0.063247532 | ns | YPD | KS.3 |
| FIM1_4517 | 1291.231067 | 1836.092047 | 1121.894738 | 0.461605686 | 0.204459316 | 0.017463803 | 0.028181129 | ns | YPD | KS.3 |
| FIM1_4518 | 776.2313661 | 813.6823393 | 733.9843654 | 0.063783297 | 0.154319541 | 0.670431024 | 0.721634432 | ns | YPD | KS.3 |
| FIM1_4519 | 858.5099881 | 1072.459178 | 1030.65912 | 0.31231252 | 0.101913057 | 0.001897544 | 0.003716647 | ns | YPD | KS.3 |
| FIM1_4520 | 1366.874998 | 1815.576846 | 2052.447947 | 0.379856218 | 0.178350749 | 0.026751647 | 0.041268045 | ns | YPD | KS.3 |
| FIM1_4521 | 2741.028694 | 3230.115492 | 2375.04407 | 0.233502522 | 0.075344942 | 0.001793982 | 0.00353188 | ns | YPD | KS.3 |
| FIM1_4522 | 841.7399821 | 889.3330077 | 643.0435648 | 0.068948247 | 0.243342043 | 0.761085124 | 0.800690099 | ns | YPD | KS.3 |
| FIM1_4523 | 2888.781868 | 3374.848123 | 3284.485673 | 0.221995234 | 0.059374039 | 0.000172154 | 0.000422112 | ns | YPD | KS.3 |
| FIM1_4524 | 1904.569009 | 1111.962156 | 2028.069343 | -0.751900944 | 0.120378918 | 1.9042145033 | 1.5841918318 | ns | YPD | KS.3 |
| FIM1_4525 | 198.1617404 | 264.1027079 | 257.5943842 | 0.384640462 | 0.176059431 | 0.023203467 | 0.036198573 | ns | YPD | KS.3 |
| FIM1_4526 | 5006.228142 | 5576.857518 | 6663.423493 | 0.153994911 | 0.063398344 | 0.014604572 | 0.02388486 | ns | YPD | KS.3 |
| FIM1_4527 | 6318.196667 | 18259.65124 | 16071.36789 | 1.353840201 | 0.283728041 | 1.1274996751 | 5.3780545388 | Km>KS.3 | YPD | KS.3 |
| FIM1_4528 | 43.82297032 | 229.4187802 | 224.419495 | 2.188616052 | 0.343173707 | 4.9138154388 | 6.7906199467 | Km>KS.3 | YPD | KS.3 |

| | | | | | | | | | | |
|---|---|---|---|---|---|---|---|---|---|---|
| FIM1_4529 | 312.5889818 | 689.2772126 | 548.8525816 | 0.97001897 | 0.292860161 | 0.000228507 | 0.000540572 | ns | YPD | KS.3 |
| FIM1_4530 | 2106.971908 | 2861.663388 | 2316.557567 | 0.430239029 | 0.103750284 | 2.6048302186 | 7.516838943 | ns | YPD | KS.3 |
| FIM1_4531 | 2499.327695 | 1850.11259 | 3611.620623 | -0.424072251 | 0.097117796 | 9.826601227 | 3.086322039 | ns | YPD | KS.3 |
| FIM1_4532 | 990.5791153 | 1017.182754 | 1404.05236 | 0.035892958 | 0.143307781 | 0.797273676 | 0.830667338 | ns | YPD | KS.3 |
| FIM1_4533 | 5228.215846 | 5098.419478 | 8237.061385 | -0.030534784 | 0.270031925 | 0.901864994 | 0.920176035 | ns | YPD | KS.3 |
| FIM1_4534 | 1269.800593 | 475.3054854 | 795.2805864 | -1.369147485 | 0.144496596 | 1.4194476388 | 9.415669337 | Km<KS.3 | YPD | KS.3 |
| FIM1_4535 | 1898.475466 | 1147.831197 | 1795.554662 | -0.680480191 | 0.171713232 | 3.866098143 | 0.000106974 | ns | YPD | KS.3 |
| FIM1_4536 | 2663.68444 | 2898.159307 | 2216.598367 | 0.107015742 | 0.232886679 | 0.623935095 | 0.679377785 | ns | YPD | KS.3 |
| FIM1_4537 | 1168.203149 | 824.5001048 | 926.008992 | -0.487506401 | 0.112556195 | 1.065881087 | 3.332971972 | ns | YPD | KS.3 |
| FIM1_4538 | 418.2972145 | 460.8292752 | 724.0558032 | 0.116066544 | 0.281011554 | 0.650088499 | 0.702626609 | ns | YPD | KS.3 |
| FIM1_4539 | 1405.600885 | 867.312759 | 1030.865874 | -0.681112037 | 0.096721241 | 9.875322801 | 1.306641780 | ns | YPD | KS.3 |
| FIM1_4540 | 840.0629315 | 285.1475754 | 575.5351737 | -1.49475125 | 0.165606071 | 4.989956731 | 2.990968041 | Km<KS.3 | YPD | KS.3 |
| FIM1_4542 | 5322.359495 | 4468.520844 | 4417.824229 | -0.23832108 | 0.152998499 | 0.108849502 | 0.145025783 | ns | YPD | KS.3 |
| FIM1_4543 | 2925.837715 | 1743.151706 | 3931.707626 | -0.590346032 | 0.329736183 | 0.041375946 | 0.060882973 | ns | YPD | KS.3 |
| FIM1_4544 | 970.1650346 | 929.0701795 | 678.0470273 | -0.057938995 | 0.170835442 | 0.725287034 | 0.76936098 | ns | YPD | KS.3 |
| FIM1_4545 | 1493.521065 | 1545.786447 | 2193.460746 | 0.043497161 | 0.233579599 | 0.842459469 | 0.869010131 | ns | YPD | KS.3 |
| FIM1_4546 | 4588.889273 | 2683.738001 | 5054.654355 | -0.655932731 | 0.277877537 | 0.009503119 | 0.016185559 | ns | YPD | KS.3 |
| FIM1_4547 | 75.33831296 | 36.80335438 | 93.14022584 | -0.883526238 | 0.284952123 | 0.000606818 | 0.001315434 | ns | YPD | KS.3 |
| FIM1_4548 | 1065.119802 | 1110.889362 | 1192.076338 | 0.057919351 | 0.138647422 | 0.668807596 | 0.720354577 | ns | YPD | KS.3 |
| FIM1_4549 | 1655.364181 | 2513.546432 | 2090.131992 | 0.564913838 | 0.168557761 | 0.000512579 | 0.001126857 | ns | YPD | KS.3 |
| FIM1_4550 | 39.50955877 | 62.77312468 | 58.11738184 | 0.558007739 | 0.283600129 | 0.030121678 | 0.04585537 | ns | YPD | KS.3 |
| FIM1_4551 | 175.0731019 | 118.1733042 | 155.7175167 | -0.534560766 | 0.166083779 | 0.00086018 | 0.00180794 | ns | YPD | KS.3 |
| FIM1_4552 | 805.612085 | 1196.484238 | 991.3499352 | 0.540457973 | 0.152844026 | 0.000269711 | 0.000629073 | ns | YPD | KS.3 |
| FIM1_4553 | 602.5535689 | 522.3062739 | 556.4867473 | -0.191051641 | 0.180060452 | 0.269705156 | 0.327423902 | ns | YPD | KS.3 |
| FIM1_4554 | 1842.288389 | 2908.208852 | 2258.936361 | 0.61984913 | 0.164106774 | 9.259045413 | 0.000240667 | ns | YPD | KS.3 |
| FIM1_4555 | 5712.409665 | 6235.084114 | 5241.801357 | 0.113200307 | 0.214166106 | 0.57652273 | 0.635401104 | ns | YPD | KS.3 |
| FIM1_4556 | 3270.989959 | 2659.786355 | 2797.311843 | -0.265144088 | 0.224108447 | 0.208795539 | 0.260797842 | ns | YPD | KS.3 |
| FIM1_4557 | 4665.756881 | 3743.210893 | 4748.96601 | -0.310632666 | 0.096210643 | 0.001084456 | 0.002233101 | ns | YPD | KS.3 |
| FIM1_4558 | 1622.381094 | 1826.498577 | 1668.102854 | 0.161611949 | 0.1528684 | 0.276581739 | 0.334710326 | ns | YPD | KS.3 |
| FIM1_4559 | 748.6824176 | 917.8377642 | 1020.512662 | 0.284160217 | 0.119091898 | 0.015126374 | 0.024697641 | ns | YPD | KS.3 |
| FIM1_4560 | 306.9342461 | 111.9512072 | 161.0635889 | -1.329879025 | 0.236706161 | 1.328368169 | 9.268768084 | Km<KS.3 | YPD | KS.3 |
| FIM1_4561 | 2803.031449 | 5953.064106 | 4062.928576 | 0.987190814 | 0.222657096 | 2.274825067 | 8.242720109 | ns | YPD | KS.3 |
| FIM1_4562 | 116.4779984 | 835.799857 | 665.2961242 | 2.585159056 | 0.410056222 | 1.029687584 | 1.355210511 | Km>KS.3 | YPD | KS.3 |
| FIM1_4563 | 2058.027481 | 1346.903287 | 2210.548623 | -0.544910274 | 0.227533177 | 0.010708221 | 0.01805878 | ns | YPD | KS.3 |
| FIM1_4564 | 324.7255225 | 144.3041037 | 338.2719058 | -1.078361439 | 0.210778061 | 5.774317898 | 2.928362033 | Km<KS.3 | YPD | KS.3 |
| FIM1_4565 | 2376.910302 | 6870.834795 | 4454.604135 | 1.372701323 | 0.267778056 | 1.677559509 | 9.419704923 | Km>KS.3 | YPD | KS.3 |
| FIM1_4566 | 14882.22723 | 14285.23596 | 10289.83615 | -0.049081051 | 0.279475136 | 0.847212981 | 0.872646911 | ns | YPD | KS.3 |
| FIM1_4567 | 5947.735357 | 5352.5299 | 6056.028043 | -0.147294452 | 0.113655105 | 0.187755012 | 0.237921851 | ns | YPD | KS.3 |
| FIM1_4568 | 5520.267219 | 5590.009976 | 5307.079298 | 0.017909804 | 0.060256294 | 0.765193689 | 0.80329998 | ns | YPD | KS.3 |
| FIM1_4569 | 2996.709475 | 1576.559209 | 2428.104162 | -0.885038137 | 0.147966055 | 7.308201190 | 5.386414951 | ns | YPD | KS.3 |
| FIM1_4570 | 2242.305792 | 1707.572591 | 1569.40606 | -0.381767597 | 0.107063888 | 0.000295696 | 0.000685823 | ns | YPD | KS.3 |
| FIM1_4571 | 1789.896978 | 927.0973696 | 1194.927305 | -0.919204381 | 0.123608188 | 3.095771491 | 5.366363474 | ns | YPD | KS.3 |
| FIM1_4572 | 2249.244194 | 5882.82442 | 4823.600807 | 1.207252832 | 0.287962612 | 3.109498965 | 1.095591880 | Km>KS.3 | YPD | KS.3 |
| FIM1_4573 | 1766.163274 | 3317.148197 | 1981.366376 | 0.851623602 | 0.17729349 | 5.599635494 | 2.300428289 | ns | YPD | KS.3 |
| FIM1_4574 | 657.3146985 | 645.7859004 | 667.0720463 | -0.024940359 | 0.094916694 | 0.790394364 | 0.825228114 | ns | YPD | KS.3 |

| | | | | | | | | | | |
|---|---|---|---|---|---|---|---|---|---|---|
| FIM1_4575 | 7323.673842 | 5362.095085 | 8341.093234 | -0.437580792 | 0.106555143 | 3.0756124382 | 8.7186164560 | ns | YPD | KS.3 |
| FIM1_4576 | 1671.472181 | 1090.71997 | 1594.398959 | -0.591653065 | 0.131780106 | 4.3843411618 | 1.5011766882 | ns | YPD | KS.3 |
| FIM1_4577 | 559.7804206 | 562.5211174 | 482.5281874 | 0.006317745 | 0.199686337 | 0.973520734 | 0.979030857 | ns | YPD | KS.3 |
| FIM1_4578 | 373.701571 | 348.6009477 | 446.1690164 | -0.095827024 | 0.143183417 | 0.492461387 | 0.555806579 | ns | YPD | KS.3 |
| FIM1_4579 | 289.2697504 | 416.6118519 | 334.6155605 | 0.470819115 | 0.220761329 | 0.023591385 | 0.036746131 | ns | YPD | KS.3 |
| FIM1_4580 | 2584.650611 | 1638.566066 | 3068.994516 | -0.618023106 | 0.166515499 | 0.000120548 | 0.000305827 | ns | YPD | KS.3 |
| FIM1_4582 | 1255.163501 | 2611.775024 | 1719.339598 | 0.947804681 | 0.236198553 | 1.6398714927 | 4.9354874025 | ns | YPD | KS.3 |
| FIM1_4583 | 1745.740935 | 2121.741529 | 1529.542087 | 0.277233198 | 0.079010321 | 0.000403506 | 0.000908755 | ns | YPD | KS.3 |
| FIM1_4584 | 1759.25814 | 1925.847804 | 1894.691616 | 0.126625042 | 0.113400979 | 0.25644062 | 0.313077812 | ns | YPD | KS.3 |
| FIM1_4585 | 2485.977266 | 3079.362391 | 2464.633392 | 0.304122306 | 0.078822966 | 0.000100211 | 0.000258851 | ns | YPD | KS.3 |
| FIM1_4586 | 12815.67794 | 14500.17049 | 14634.50189 | 0.175976837 | 0.067938363 | 0.009157278 | 0.015644732 | ns | YPD | KS.3 |
| FIM1_4587 | 1743.044079 | 1280.276503 | 2336.525261 | -0.352284924 | 0.319258934 | 0.212772412 | 0.265298935 | ns | YPD | KS.3 |
| FIM1_4588 | 3843.045233 | 1926.57717 | 3198.471663 | -0.960279365 | 0.133790171 | 1.8967058026 | 2.8083664786 | ns | YPD | KS.3 |
| FIM1_4589 | 2891.439316 | 998.9781485 | 2506.601279 | -1.400324207 | 0.244277665 | 4.9191549504 | 3.7136260816 | Km<KS.3 | YPD | KS.3 |
| FIM1_4590 | 864.4246374 | 413.0392717 | 592.5620059 | -1.029808045 | 0.128284199 | 2.1366899728 | 5.3959556420 | Km<KS.3 | YPD | KS.3 |
| FIM1_4591 | 3253.873974 | 1740.7834 | 2635.107102 | -0.878728704 | 0.111470705 | 1.0615465498 | 2.3163131953 | ns | YPD | KS.3 |
| FIM1_4592 | 1599.132273 | 864.835295 | 1307.898843 | -0.872050019 | 0.086640816 | 2.6190144945 | 2.3267137696 | ns | YPD | KS.3 |
| FIM1_4593 | 1087.883647 | 615.5986113 | 726.3664995 | -0.781245097 | 0.151745824 | 1.1166983164 | 5.3418981962 | ns | YPD | KS.3 |
| FIM1_4594 | 1180.501666 | 1876.719243 | 742.0136083 | 0.596356959 | 0.227995637 | 0.005288968 | 0.009444586 | ns | YPD | KS.3 |
| FIM1_4595 | 4093.432538 | 4367.937452 | 4025.083491 | 0.089926972 | 0.126792789 | 0.469318239 | 0.53298555 | ns | YPD | KS.3 |
| FIM1_4596 | 2915.441107 | 1684.02396 | 3538.793639 | -0.708883573 | 0.227537043 | 0.000889489 | 0.00186403 | ns | YPD | KS.3 |
| FIM1_4597 | 2350.635342 | 2128.447759 | 1866.817906 | -0.139509161 | 0.100776625 | 0.160807346 | 0.207150841 | ns | YPD | KS.3 |
| FIM1_4598 | 643.3394252 | 412.5509123 | 478.6078264 | -0.603115282 | 0.163127001 | 0.000130885 | 0.000329197 | ns | YPD | KS.3 |
| FIM1_4599 | 295.4488238 | 199.4839473 | 280.8019534 | -0.52588744 | 0.182468341 | 0.002659695 | 0.005069725 | ns | YPD | KS.3 |
| FIM1_4600 | 2914.563041 | 4383.341999 | 3711.087624 | 0.57362712 | 0.104491153 | 2.5480092249 | 1.3689358416 | ns | YPD | KS.3 |
| FIM1_4601 | 657.1239672 | 811.8217789 | 781.6834496 | 0.297894691 | 0.090947886 | 0.000929973 | 0.001938255 | ns | YPD | KS.3 |
| FIM1_4602 | 414.7811335 | 252.6315609 | 450.4868022 | -0.618928845 | 0.258980106 | 0.009484628 | 0.016159597 | ns | YPD | KS.3 |
| FIM1_4603 | 2640.219264 | 3835.039211 | 3864.028105 | 0.522817502 | 0.110987073 | 1.6900335338 | 6.3077733102 | ns | YPD | KS.3 |
| FIM1_4604 | 194.8684924 | 142.006969 | 140.3676843 | -0.405117626 | 0.230073233 | 0.060433405 | 0.085372433 | ns | YPD | KS.3 |
| FIM1_4605 | 1409.918809 | 897.9709952 | 1105.088326 | -0.63529917 | 0.101789618 | 2.4700634248 | 1.9948970030 | ns | YPD | KS.3 |
| FIM1_4606 | 708.2580427 | 508.2057349 | 706.6727291 | -0.465028281 | 0.108761091 | 1.4108314522 | 4.2915465074 | ns | YPD | KS.3 |
| FIM1_4607 | 1815.650925 | 2300.633262 | 2344.433376 | 0.336669442 | 0.074053272 | 4.6568496634 | 1.5825018494 | ns | YPD | KS.3 |
| FIM1_4608 | 433.3424886 | 435.75257 | 390.933759 | 0.007369346 | 0.194602886 | 0.968382922 | 0.975441392 | ns | YPD | KS.3 |
| FIM1_4609 | 2746.820858 | 2624.767887 | 3251.156676 | -0.059461993 | 0.203351655 | 0.758604699 | 0.798718615 | ns | YPD | KS.3 |
| FIM1_4612 | 6195.199476 | 8370.602884 | 7107.770878 | 0.413871573 | 0.141802871 | 0.002755587 | 0.005234457 | ns | YPD | KS.3 |
| FIM1_4613 | 9736.343472 | 7750.539176 | 15088.26707 | -0.287168479 | 0.2414309 | 0.2020668 | 0.253475122 | ns | YPD | KS.3 |
| FIM1_4614 | 632.8349847 | 846.4245185 | 609.0939818 | 0.40754615 | 0.106459887 | 0.000102376 | 0.000264034 | ns | YPD | KS.3 |
| FIM1_4615 | 1881.190196 | 1647.29838 | 1502.09062 | -0.187216264 | 0.092878674 | 0.04152924 | 0.061072411 | ns | YPD | KS.3 |
| FIM1_4616 | 2988.596779 | 3169.023226 | 3201.066156 | 0.082576238 | 0.090358876 | 0.355749266 | 0.418305979 | ns | YPD | KS.3 |
| FIM1_4617 | 3484.395356 | 4847.478768 | 3570.518226 | 0.445130957 | 0.170563274 | 0.006806078 | 0.011939435 | ns | YPD | KS.3 |
| FIM1_4618 | 4343.22286 | 4153.862211 | 4057.374277 | -0.062308417 | 0.112580546 | 0.573879184 | 0.633329401 | ns | YPD | KS.3 |
| FIM1_4619 | 856.2336326 | 785.0613869 | 857.5893191 | -0.122098434 | 0.097009674 | 0.20272131 | 0.254103935 | ns | YPD | KS.3 |
| FIM1_4620 | 443.1685192 | 866.4350146 | 744.6249619 | 0.856700129 | 0.244480083 | 0.000158284 | 0.000391774 | ns | YPD | KS.3 |
| FIM1_4621 | 86.27388666 | 175.3656977 | 125.0342392 | 0.72607397 | 0.409342687 | 0.030495989 | 0.046269455 | ns | YPD | KS.3 |
| FIM1_4622 | 1035.67174 | 2124.702279 | 1327.016102 | 1.005609505 | 0.122077085 | 4.0681319423 | 1.2119135576 | Km>KS.3 | YPD | KS.3 |

| | | | | | | | | | | |
|---|---|---|---|---|---|---|---|---|---|---|
| FIM1_4623 | 1671.422589 | 1113.882353 | 1412.356005 | -0.567256586 | 0.116404798 | 7.0267391613 | 2.8467448963 | ns | YPD | KS.3 |
| FIM1_4624 | 2576.984814 | 1326.791061 | 2292.836892 | -0.928095224 | 0.122441127 | 1.0087171285 | 1.9081246063 | ns | YPD | KS.3 |
| FIM1_4625 | 702.2709936 | 1305.331898 | 926.2415632 | 0.881999582 | 0.077910974 | 3.3259689980 | 4.8666752251 | ns | YPD | KS.3 |
| FIM1_4626 | 489.3336693 | 595.2010206 | 425.5798025 | 0.256063664 | 0.205389636 | 0.189509772 | 0.239901047 | ns | YPD | KS.3 |
| FIM1_4627 | 2625.817344 | 2913.717653 | 2357.628351 | 0.140705737 | 0.163295978 | 0.373347784 | 0.436625582 | ns | YPD | KS.3 |
| FIM1_4628 | 287.1516896 | 270.4007532 | 297.3288643 | -0.079865552 | 0.187664598 | 0.656983501 | 0.709154462 | ns | YPD | KS.3 |
| FIM1_4629 | 659.2192554 | 638.0060646 | 744.8255768 | -0.04388488 | 0.175548252 | 0.795240478 | 0.829243634 | ns | YPD | KS.3 |
| FIM1_4630 | 799.5797468 | 1481.661504 | 1317.615698 | 0.867998549 | 0.106132831 | 9.8020867819 | 2.6942199856 | ns | YPD | KS.3 |
| FIM1_4631 | 1281.300108 | 2055.131559 | 2080.043648 | 0.648910565 | 0.147497645 | 6.0331316291 | 2.0090247560 | ns | YPD | KS.3 |
| FIM1_4632 | 1150.875498 | 2930.703836 | 2347.234258 | 1.23844427 | 0.223080659 | 2.8885159518 | 1.8809380707 | Km>KS.3 | YPD | KS.3 |
| FIM1_4633 | 69.30865084 | 56.56505267 | 76.16514234 | -0.249821388 | 0.263859013 | 0.303541267 | 0.363765 | ns | YPD | KS.3 |
| FIM1_4634 | 1146.678188 | 3880.323942 | 2905.267577 | 1.516603575 | 0.337311009 | 2.0307523358 | 9.1928961519 | Km>KS.3 | YPD | KS.3 |
| FIM1_4635 | 1663.510894 | 1450.503745 | 1175.173052 | -0.189713374 | 0.126103935 | 0.124864279 | 0.164687112 | ns | YPD | KS.3 |
| FIM1_4636 | 746.7600497 | 489.1486865 | 580.978079 | -0.598758129 | 0.089604305 | 1.4248552524 | 1.4768031001 | ns | YPD | KS.3 |
| FIM1_4637 | 1470.318384 | 1018.729119 | 1091.158215 | -0.502202265 | 0.150048754 | 0.000573179 | 0.001249044 | ns | YPD | KS.3 |
| FIM1_4638 | 1798.77015 | 1291.788588 | 1228.748941 | -0.467057793 | 0.095845167 | 8.1162554165 | 3.2458497345 | ns | YPD | KS.3 |
| FIM1_4639 | 2136.60382 | 3782.279964 | 2308.383022 | 0.798240781 | 0.120234336 | 1.2866784514 | 1.3447952302 | ns | YPD | KS.3 |
| FIM1_4640 | 945.959396 | 763.407982 | 889.2184359 | -0.289877083 | 0.165640611 | 0.07026053 | 0.097829873 | ns | YPD | KS.3 |
| FIM1_4641 | 5162.031711 | 4967.344445 | 6170.986705 | -0.053820706 | 0.110045948 | 0.619475967 | 0.674965602 | ns | YPD | KS.3 |
| FIM1_4642 | 2166.904448 | 3697.592567 | 3031.349814 | 0.733384711 | 0.15071431 | 5.3669965393 | 2.2176750650 | ns | YPD | KS.3 |
| FIM1_4643 | 415.1012804 | 330.1229586 | 499.9010469 | -0.314145728 | 0.148724541 | 0.029908843 | 0.045559245 | ns | YPD | KS.3 |
| FIM1_4644 | 1301.71737 | 782.0895428 | 877.6145571 | -0.712534899 | 0.116569368 | 4.8197505717 | 3.6494535761 | ns | YPD | KS.3 |
| FIM1_4645 | 1530.435975 | 984.6581552 | 1148.317039 | -0.608160959 | 0.140695534 | 9.2171736001 | 2.9112314460 | ns | YPD | KS.3 |
| FIM1_4646 | 1912.122428 | 1120.952313 | 1583.751112 | -0.752245236 | 0.10320971 | 1.4105599990 | 2.1726117631 | ns | YPD | KS.3 |
| FIM1_4647 | 2432.058675 | 1065.926911 | 1914.355097 | -1.15188775 | 0.131598466 | 2.8985781325 | 1.3109478372 | Km<KS.3 | YPD | KS.3 |
| FIM1_4648 | 1802.743973 | 1593.121343 | 1698.51504 | -0.171355266 | 0.126397748 | 0.16665368 | 0.214182913 | ns | YPD | KS.3 |
| FIM1_4649 | 2690.74963 | 3194.934206 | 2547.851914 | 0.241355377 | 0.103312345 | 0.017870474 | 0.028762733 | ns | YPD | KS.3 |
| FIM1_4650 | 1477.780996 | 1266.630101 | 1549.062917 | -0.214549112 | 0.121848187 | 0.072816125 | 0.101077071 | ns | YPD | KS.3 |
| FIM1_4651 | 2309.757223 | 5063.133018 | 4372.04566 | 0.990426417 | 0.265683283 | 4.5872213979 | 0.000124912 | ns | YPD | KS.3 |
| FIM1_4652 | 1482.158489 | 1533.203135 | 1412.47613 | 0.047376521 | 0.103918653 | 0.64395527 | 0.697513056 | ns | YPD | KS.3 |
| FIM1_4653 | 983.2010219 | 1422.607053 | 1046.675546 | 0.486723196 | 0.199039781 | 0.010228028 | 0.017319414 | ns | YPD | KS.3 |
| FIM1_4654 | 1008.680956 | 663.129769 | 1079.974788 | -0.565272823 | 0.173374151 | 0.000708076 | 0.001512528 | ns | YPD | KS.3 |
| FIM1_4655 | 822.7427505 | 2581.014648 | 1886.468117 | 1.508904936 | 0.252303326 | 6.4703380997 | 5.9064095498 | Km>KS.3 | YPD | KS.3 |
| FIM1_4656 | 738.3183611 | 3861.793402 | 2415.83052 | 2.228358332 | 0.308275842 | 1.2383102616 | 2.6902155247 | Km>KS.3 | YPD | KS.3 |
| FIM1_4657 | 706.9063483 | 1032.280556 | 980.8930235 | 0.525775264 | 0.126431199 | 2.1803227605 | 6.3881659210 | ns | YPD | KS.3 |
| FIM1_4658 | 12168.16146 | 15168.13912 | 20752.37853 | 0.295945719 | 0.173339315 | 0.076473313 | 0.105740615 | ns | YPD | KS.3 |
| FIM1_4660 | 62.11682897 | 250.4662881 | 183.5974283 | 1.89703628 | 0.239685227 | 1.0133593434 | 3.3167517984 | Km>KS.3 | YPD | KS.3 |
| FIM1_4661 | 159.4264575 | 303.7396353 | 320.9228849 | 0.826078666 | 0.238734215 | 0.00020147 | 0.000482113 | ns | YPD | KS.3 |
| FIM1_4662 | 2770.845037 | 1000.867524 | 3115.470066 | -1.253733202 | 0.314814618 | 6.3082515698 | 2.0894508362 | Km<KS.3 | YPD | KS.3 |
| FIM1_4664 | 41.23816436 | 25.45623725 | 39.5335887 | -0.611832007 | 0.244891006 | 0.00716103 | 0.012535583 | ns | YPD | KS.3 |
| FIM1_4665 | 1873.129704 | 3945.088903 | 3582.114891 | 0.928064467 | 0.274722283 | 0.000205437 | 0.000490663 | ns | YPD | KS.3 |
| FIM1_4666 | 252.5805008 | 1200.028364 | 786.747206 | 2.059532484 | 0.32644446 | 9.0235174871 | 1.1971199866 | Km>KS.3 | YPD | KS.3 |
| FIM1_4667 | 2656.451801 | 4098.709741 | 5171.262894 | 0.501689741 | 0.315065599 | 0.07290091 | 0.101166535 | ns | YPD | KS.3 |
| FIM1_4668 | 117.425974 | 1169.873124 | 590.2764347 | 3.190390835 | 0.274193088 | 6.9147540177 | 1.3231115860 | Km>KS.3 | YPD | KS.3 |
| FIM1_4669 | 38.8888052 | 66.71340014 | 39.49713505 | 0.621295908 | 0.322030688 | 0.028847506 | 0.044104592 | ns | YPD | KS.3 |

| | | | | | | | | | |
|---|---|---|---|---|---|---|---|---|---|
| FIM1_4670 | 2010.825686 | 3585.465065 | 2421.800564 | 0.767863776 | 0.197723734 | 4.46600842543 | 0.000121878 | ns | YPD | KS.3 |
| FIM1_4671 | 1535.830624 | 1680.954269 | 1252.57698 | 0.125540043 | 0.125378233 | 0.307190935 | 0.367430789 | ns | YPD | KS.3 |
| FIM1_4672 | 1115.933911 | 966.425274 | 1166.076171 | -0.200588743 | 0.113485744 | 0.072407771 | 0.100588238 | ns | YPD | KS.3 |
| FIM1_4673 | 428.4952996 | 552.9691655 | 398.5748486 | 0.353136684 | 0.129965819 | 0.00550538 | 0.009799379 | ns | YPD | KS.3 |
| FIM1_4674 | 250.6880742 | 246.1296406 | 279.8664622 | -0.026129764 | 0.106821344 | 0.804009559 | 0.835760041 | ns | YPD | KS.3 |
| FIM1_4675 | 3837.010083 | 4051.476931 | 4876.118353 | 0.075676204 | 0.11859636 | 0.516056175 | 0.578368883 | ns | YPD | KS.3 |
| FIM1_4676 | 2382.61617 | 2467.226085 | 2206.597425 | 0.049645238 | 0.071443288 | 0.484205638 | 0.547482511 | ns | YPD | KS.3 |
| FIM1_4678 | 1105.688997 | 2228.794261 | 2206.929083 | 0.816062897 | 0.327364961 | 0.004529789 | 0.00822143 | ns | YPD | KS.3 |
| FIM1_4679 | 1723.670948 | 1641.124631 | 1706.025796 | -0.06094076 | 0.250370389 | 0.79312148 | 0.827554396 | ns | YPD | KS.3 |
| FIM1_4680 | 490.6639529 | 766.5749126 | 650.5430871 | 0.626777864 | 0.106784215 | 2.52735799648 | 1.66911131480 | ns | YPD | KS.3 |
| FIM1_4681 | 3179.469907 | 3115.803776 | 3003.550954 | -0.027227743 | 0.167214253 | 0.866214406 | 0.888722761 | ns | YPD | KS.3 |
| FIM1_4682 | 5853.119268 | 6056.646553 | 6334.836483 | 0.048405745 | 0.079327351 | 0.538418627 | 0.600321082 | ns | YPD | KS.3 |
| FIM1_4683 | 118.6493967 | 14.32551795 | 149.174951 | -2.861773561 | 0.351744597 | 3.66315270096 | 1.28339327878 | Km<KS.3 | YPD | KS.3 |
| FIM1_4684 | 1967.8573 | 1410.050888 | 1653.177765 | -0.44718442 | 0.176992521 | 0.008616327 | 0.014783598 | ns | YPD | KS.3 |
| FIM1_4685 | 1910.770614 | 2429.392185 | 1895.683557 | 0.268148317 | 0.331678222 | 0.356800165 | 0.419343449 | ns | YPD | KS.3 |
| FIM1_4686 | 84.24126095 | 1056.00837 | 374.4480036 | 3.427615611 | 0.35443516 | 7.96205677845 | 6.71376821573 | Km>KS.3 | YPD | KS.3 |
| FIM1_4688 | 909.0420317 | 1036.376188 | 1030.77713 | 0.175467069 | 0.174482667 | 0.29664476 | 0.357285646 | ns | YPD | KS.3 |
| FIM1_4689 | 1568.961723 | 1222.796616 | 1260.925112 | -0.348476809 | 0.11139834 | 0.001480758 | 0.002963303 | ns | YPD | KS.3 |
| FIM1_4690 | 3271.930286 | 3288.314415 | 3273.689111 | 0.006849919 | 0.141500767 | 0.960426197 | 0.969929003 | ns | YPD | KS.3 |
| FIM1_4691 | 337.4737729 | 1500.472195 | 1392.037738 | 1.94389887 | 0.335855807 | 3.24654989122 | 3.14233184996 | Km>KS.3 | YPD | KS.3 |
| FIM1_4692 | 1214.800034 | 1066.885839 | 1443.260619 | -0.174150439 | 0.174754815 | 0.300984553 | 0.361603031 | ns | YPD | KS.3 |
| FIM1_4693 | 229.1757441 | 482.9883495 | 358.3793039 | 0.99479188 | 0.199196383 | 1.43050964180 | 6.70130458379 | ns | YPD | KS.3 |
| FIM1_4694 | 490.2207568 | 1255.008678 | 868.552117 | 1.18126757 | 0.284521881 | 4.10636301879 | 1.41672371834 | Km>KS.3 | YPD | KS.3 |
| FIM1_4695 | 134.9338706 | 1067.597223 | 611.148356 | 2.859111967 | 0.281905265 | 4.72254843488 | 4.69893569270 | Km>KS.3 | YPD | KS.3 |
| FIM1_4697 | 379.8507451 | 1353.088033 | 1508.15025 | 1.467552296 | 0.417721356 | 1.61444809253 | 4.86191238521 | Km>KS.3 | YPD | KS.3 |
| FIM1_4698 | 815.1463877 | 1542.251922 | 1300.486597 | 0.874945469 | 0.15396842 | 4.49871550543 | 2.80465033076 | ns | YPD | KS.3 |
| FIM1_4699 | 211.9881741 | 1628.163126 | 876.9958142 | 2.800730095 | 0.300329408 | 1.09303980225 | 7.44914111807 | Km>KS.3 | YPD | KS.3 |
| FIM1_4700 | 31.33986345 | 30.27173604 | 31.51954424 | -0.051068288 | 0.265307647 | 0.834232026 | 0.861416423 | ns | YPD | KS.3 |
| FIM1_4701 | 872.4059318 | 328.1261993 | 949.344502 | -1.290218079 | 0.233143617 | 2.56726475096 | 1.68497917361 | Km<KS.3 | YPD | KS.3 |
| FIM1_4702 | 1265.25463 | 2781.138694 | 2349.804366 | 0.943093317 | 0.312999674 | 0.000676623 | 0.001452826 | ns | YPD | KS.3 |
| FIM1_4703 | 3604.978148 | 9785.280673 | 8353.968002 | 1.274184575 | 0.275335963 | 3.21478087679 | 1.39074216191 | Km>KS.3 | YPD | KS.3 |
| FIM1_4704 | 771.9854668 | 1443.928717 | 1487.561147 | 0.782134143 | 0.264115713 | 0.001245435 | 0.002531061 | ns | YPD | KS.3 |
| FIM1_4705 | 1225.395343 | 1351.426652 | 1309.422391 | 0.134720052 | 0.137521982 | 0.315930551 | 0.376829176 | ns | YPD | KS.3 |
| FIM1_4706 | 986.3820424 | 1221.946581 | 1323.038148 | 0.295183006 | 0.134759154 | 0.025030336 | 0.038759121 | ns | YPD | KS.3 |
| FIM1_4707 | 316.6459694 | 613.6860215 | 651.8354446 | 0.864209488 | 0.220123637 | 2.96608299380 | 8.45146786609 | ns | YPD | KS.3 |
| FIM1_4708 | 471.3786043 | 1155.255783 | 1232.431846 | 1.074366258 | 0.32332285 | 0.000155605 | 0.000386294 | Km>KS.3 | YPD | KS.3 |
| FIM1_4709 | 656.0019833 | 3009.720654 | 1693.347632 | 2.063753508 | 0.272978444 | 1.11496811966 | 2.98223997748 | Km>KS.3 | YPD | KS.3 |
| FIM1_4710 | 897.8650312 | 1371.908558 | 1338.096276 | 0.535804091 | 0.243233826 | 0.017894561 | 0.028792187 | ns | YPD | KS.3 |
| FIM1_4711 | 1093.92554 | 3204.404493 | 2530.178698 | 1.429528021 | 0.232361289 | 3.27764236722 | 3.16012995553 | Km>KS.3 | YPD | KS.3 |
| FIM1_4712 | 121.6380554 | 809.4481672 | 488.8438043 | 2.654382408 | 0.224944909 | 3.48211681142 | 7.21813797367 | Km>KS.3 | YPD | KS.3 |
| FIM1_4713 | 2349.456853 | 2654.056296 | 2362.926956 | 0.163271073 | 0.175520656 | 0.334162397 | 0.395917581 | ns | YPD | KS.3 |
| FIM1_4714 | 679.5467818 | 2197.164723 | 1656.996978 | 1.594343185 | 0.21092678 | 7.07862668608 | 1.61542053959 | Km>KS.3 | YPD | KS.3 |
| FIM1_4715 | 3391.91253 | 5947.084584 | 4836.093207 | 0.738084206 | 0.208875104 | 0.000188654 | 0.000455608 | ns | YPD | KS.3 |
| FIM1_4716 | 1875.932226 | 5614.167793 | 3401.173216 | 1.252805046 | 0.39081028 | 0.000117564 | 0.000299171 | Km>KS.3 | YPD | KS.3 |
| FIM1_4717 | 2632.123942 | 2104.355896 | 4812.049546 | -0.268881522 | 0.280331073 | 0.292167172 | 0.352371317 | ns | YPD | KS.3 |

| | | | | | | | | | | |
|---|---|---|---|---|---|---|---|---|---|---|
| FIM1_4718 | 610.5634919 | 904.5736605 | 1948.392689 | 0.41649978 | 0.368746926 | 0.184102306 | 0.233709868 | ns | YPD | KS.3 |
| FIM1_4719 | 187.0927053 | 352.6054651 | 359.3621064 | 0.82055804 | 0.227651422 | 0.000118986 | 0.00030248 | ns | YPD | KS.3 |
| FIM1_4720 | 1533.358356 | 1869.934107 | 2687.385742 | 0.239285143 | 0.276532963 | 0.34320041 | 0.404890216 | ns | YPD | KS.3 |
| FIM1_4721 | 778.456843 | 687.4355823 | 1004.264901 | -0.172730279 | 0.126089618 | 0.162289288 | 0.208951658 | ns | YPD | KS.3 |
| FIM1_4722 | 348.7806135 | 1209.218629 | 1807.401918 | 1.343442296 | 0.460957373 | 0.000207257 | 0.000494297 | Km>KS.3 | YPD | KS.3 |
| FIM1_4724 | 734.1226063 | 428.4791183 | 584.5339102 | -0.735392818 | 0.159296848 | 1.8305790408 | 6.7811844587 | ns | YPD | KS.3 |
| FIM1_4725 | 231.17188 | 191.5359952 | 211.8274937 | -0.092889038 | 0.636025956 | 0.802625698 | 0.835163009 | ns | YPD | KS.3 |
| FIM1_4726 | 84.29312864 | 42.06992932 | 61.86479955 | -0.433711992 | 0.598386749 | 0.259491758 | 0.316259553 | ns | YPD | KS.3 |
| FIM1_4727 | 417.7321127 | 240.311981 | 420.3880307 | -0.250166264 | 0.668066774 | 0.499177827 | 0.561730307 | ns | YPD | KS.3 |
| FIM1_4728 | 113.8058485 | 170.6439922 | 140.8377942 | 0.204612674 | 0.636428592 | 0.584524846 | 0.642752258 | ns | YPD | KS.3 |
| FIM1_4729 | 898.4734069 | 739.024224 | 766.2970696 | -0.089296563 | 0.654420637 | 0.808205546 | 0.839420166 | ns | YPD | KS.3 |
| FIM1_4730 | 609.0182615 | 216.1141118 | 481.3915826 | -1.31574347 | 0.28625317 | 3.1271681840 | 1.3563785279 | Km<KS.3 | YPD | KS.3 |
| FIM1_4731 | 1042.170504 | 973.3013409 | 1595.830275 | -0.084543555 | 0.256425118 | 0.721444055 | 0.766592092 | ns | YPD | KS.3 |
| FIM1_4732 | 1292.365287 | 3923.071909 | 3415.38404 | 1.427168236 | 0.281905463 | 1.8248048745 | 1.0143468436 | Km>KS.3 | YPD | KS.3 |
| FIM1_4733 | 318.7560146 | 1687.484808 | 1350.604209 | 2.340685258 | 0.196361882 | 2.8266475991 | 6.3920780936 | Km>KS.3 | YPD | KS.3 |
| FIM1_4734 | 200.605831 | 585.1463556 | 494.3151773 | 1.432658838 | 0.2224329 | 4.9846922284 | 5.7008836406 | Km>KS.3 | YPD | KS.3 |
| FIM1_4736 | 83.53530839 | 390.8854531 | 252.8171641 | 2.089857018 | 0.275220042 | 8.2889559203 | 2.3298054069 | Km>KS.3 | YPD | KS.3 |
| FIM1_4737 | 1029.590552 | 976.7619326 | 807.1249009 | -0.067725509 | 0.218595334 | 0.742860431 | 0.784489629 | ns | YPD | KS.3 |
| FIM1_4738 | 86.97591901 | 256.2175092 | 122.0593403 | 1.36535076 | 0.296080987 | 2.3390771112 | 1.0446058014 | Km>KS.3 | YPD | KS.3 |
| FIM1_4739 | 324.6112517 | 675.712578 | 449.3723635 | 1.005848295 | 0.157838641 | 4.3545055161 | 4.1029668452 | Km>KS.3 | YPD | KS.3 |
| FIM1_4740 | 804.2373897 | 2939.335042 | 1494.907235 | 1.804808005 | 0.175768075 | 4.8411304770 | 5.3521386940 | Km>KS.3 | YPD | KS.3 |
| FIM1_4741 | 817.8769043 | 827.0962253 | 562.5953815 | 0.014903523 | 0.193053525 | 0.935643708 | 0.947451139 | ns | YPD | KS.3 |
| FIM1_4742 | 105.362529 | 585.0041497 | 286.3233181 | 2.317754027 | 0.308266244 | 1.4607379538 | 3.7758976910 | Km>KS.3 | YPD | KS.3 |
| FIM1_4743 | 16.29228395 | 82.37433762 | 29.18473673 | 2.02333184 | 0.424677958 | 8.3616093393 | 4.9819169416 | Km>KS.3 | YPD | KS.3 |
| FIM1_4744 | 2613.122726 | 2601.542972 | 2880.341186 | -0.005907423 | 0.191236273 | 0.974245734 | 0.97956195 | ns | YPD | KS.3 |
| FIM1_4745 | 1235.654032 | 795.0718572 | 978.5767618 | -0.611359245 | 0.130882948 | 1.7823951003 | 6.6301135205 | ns | YPD | KS.3 |
| FIM1_4746 | 1834.091829 | 2414.811118 | 2526.440432 | 0.349739039 | 0.233336113 | 0.109438111 | 0.145731959 | ns | YPD | KS.3 |
| FIM1_4747 | 729.74677 | 567.7153296 | 669.4574419 | -0.349749035 | 0.117796075 | 0.002509579 | 0.004794607 | ns | YPD | KS.3 |
| FIM1_4748 | 4058.013966 | 3865.199985 | 4156.527977 | -0.068518164 | 0.097202569 | 0.475574455 | 0.539193919 | ns | YPD | KS.3 |
| FIM1_4749 | 457.9112229 | 438.7951368 | 434.7145254 | -0.059359017 | 0.114849734 | 0.599257232 | 0.65624141 | ns | YPD | KS.3 |
| FIM1_4750 | 5272.6631 | 5349.003954 | 4683.332238 | 0.020468043 | 0.074871578 | 0.783023162 | 0.818734811 | ns | YPD | KS.3 |
| FIM1_4751 | 178.2266218 | 189.6404043 | 204.7642249 | 0.081264624 | 0.193436228 | 0.660293655 | 0.712263862 | ns | YPD | KS.3 |
| FIM1_4753 | 2251.465688 | 1134.137741 | 2190.71695 | -0.878048881 | 0.243893923 | 0.000104228 | 0.000267976 | ns | YPD | KS.3 |
| FIM1_4754 | 2691.720414 | 1755.683112 | 1932.410155 | -0.596651267 | 0.11828274 | 2.7762637703 | 1.2158373466 | ns | YPD | KS.3 |
| FIM1_4755 | 588.3817386 | 407.9179865 | 420.6909983 | -0.498669516 | 0.158574543 | 0.001169765 | 0.002393904 | ns | YPD | KS.3 |
| FIM1_4756 | 2010.790968 | 1601.694881 | 1824.271366 | -0.310130102 | 0.153726757 | 0.037754491 | 0.056135264 | ns | YPD | KS.3 |
| FIM1_4757 | 1836.576648 | 776.4538593 | 1744.657445 | -1.089520266 | 0.269594637 | 9.2256121587 | 2.9112314460 | Km<KS.3 | YPD | KS.3 |
| FIM1_4758 | 826.5591145 | 1096.010775 | 689.9815069 | 0.395375048 | 0.111398246 | 0.000310368 | 0.000715515 | ns | YPD | KS.3 |
| FIM1_4759 | 4676.581142 | 6111.840614 | 6178.264648 | 0.361942197 | 0.165346901 | 0.02354433 | 0.036695816 | ns | YPD | KS.3 |
| FIM1_4760 | 918.2876968 | 5520.575125 | 4423.256503 | 2.426263267 | 0.320232336 | 1.0902274840 | 2.9318279637 | Km>KS.3 | YPD | KS.3 |
| FIM1_4761 | 216.6868184 | 345.2087652 | 374.2602876 | 0.6294573 | 0.171158926 | 0.000134906 | 0.000338285 | ns | YPD | KS.3 |
| FIM1_4762 | 83.43450223 | 166.1918507 | 127.7586392 | 0.933818733 | 0.1729945 | 1.9521108900 | 1.0766908733 | ns | YPD | KS.3 |
| FIM1_4763 | 147.843259 | 214.7348375 | 172.8569932 | 0.476663159 | 0.231090951 | 0.027702327 | 0.042536751 | ns | YPD | KS.3 |
| FIM1_4764 | 263.4668635 | 302.2927572 | 328.2736609 | 0.1699111 | 0.256629698 | 0.473894965 | 0.537534759 | ns | YPD | KS.3 |
| FIM1_4765 | 378.4387659 | 237.9597223 | 314.2749395 | -0.586054196 | 0.244635975 | 0.009941696 | 0.016860026 | ns | YPD | KS.3 |

| | | | | | | | | | | |
|---|---|---|---|---|---|---|---|---|---|---|
| FIM1_4766 | 209.6889983 | 140.1684741 | 195.8864339 | -0.530593683 | 0.198581958 | 0.005017137 | 0.009004422 | ns | YPD | KS.3 |
| FIM1_4767 | 839.6121942 | 610.85979 | 816.4770033 | -0.434145395 | 0.151220787 | 0.003128651 | 0.005884703 | ns | YPD | KS.3 |
| FIM1_4769 | 58.20552602 | 90.54414922 | 118.1575199 | 0.510776824 | 0.315339886 | 0.068032214 | 0.095019726 | ns | YPD | KS.3 |
| FIM1_4770 | 402.202463 | 317.3025534 | 529.218885 | -0.24686176 | 0.372963039 | 0.434637248 | 0.498575123 | ns | YPD | KS.3 |
| FIM1_4771 | 862.4244945 | 862.7336384 | 746.2717156 | 0.000407488 | 0.197878478 | 0.998277643 | 0.998277643 | ns | YPD | KS.3 |
| FIM1_4772 | 92.47484052 | 79.12907903 | 75.1073442 | -0.201696541 | 0.203147722 | 0.296672765 | 0.357285646 | ns | YPD | KS.3 |
| FIM1_4773 | 2570.248923 | 974.3068269 | 2745.278261 | -1.331311279 | 0.172950602 | 9.2299087954 | 2.0408353892 | Km<KS.3 | YPD | KS.3 |
| FIM1_3 | 2487.935177 | 4024.450814 | 3037.529789 | 0.635069549 | 0.138413238 | 5.6211645871 | 7.2448947723 | ns | NaCl | KS.1 |
| FIM1_5 | 364.5969707 | 476.8030815 | 633.8041188 | 0.235364411 | 0.255267286 | 0.219045596 | 0.343120856 | ns | NaCl | KS.1 |
| FIM1_6 | 315.5154852 | 394.5438021 | 405.4975614 | 0.234743556 | 0.197408258 | 0.150842722 | 0.256386247 | ns | NaCl | KS.1 |
| FIM1_7 | 35.87697542 | 74.93762594 | 60.8195808 | 0.774408007 | 0.298040317 | 0.000644968 | 0.002976545 | ns | NaCl | KS.1 |
| FIM1_8 | 164.5251811 | 328.3015477 | 375.3172182 | 0.776252339 | 0.26783785 | 0.000267382 | 0.001422702 | ns | NaCl | KS.1 |
| FIM1_9 | 1307.817626 | 5815.10095 | 2901.971283 | 1.803006307 | 0.400348137 | 5.3144628244 | 1.2711275265 | Km>KS.1 | NaCl | KS.1 |
| FIM1_10 | 573.4457868 | 2795.135043 | 2943.370395 | 2.032280794 | 0.357967121 | 5.8312820308 | 2.9303664751 | Km>KS.1 | NaCl | KS.1 |
| FIM1_11 | 199.7253284 | 349.6487519 | 340.6940978 | 0.693003255 | 0.194516475 | 3.7711084191 | 0.000271902 | ns | NaCl | KS.1 |
| FIM1_12 | 467.3904499 | 990.6146393 | 1729.337393 | 0.843712152 | 0.277561103 | 0.000129189 | 0.000774357 | ns | NaCl | KS.1 |
| FIM1_13 | 54237.11295 | 44428.48315 | 35774.55391 | -0.126128329 | 0.315522142 | 0.540451839 | 0.666914257 | ns | NaCl | KS.1 |
| FIM1_14 | 24.50401174 | 21.9617334 | 16.92066841 | -0.068401612 | 0.318350274 | 0.738740095 | 0.828688156 | ns | NaCl | KS.1 |
| FIM1_15 | 23753.75535 | 19494.81606 | 14849.47076 | -0.098417602 | 0.357117938 | 0.636049091 | 0.747025032 | ns | NaCl | KS.1 |
| FIM1_16 | 21074.14942 | 13427.54739 | 13864.7932 | -0.444277065 | 0.261046477 | 0.025052327 | 0.06009418 | ns | NaCl | KS.1 |
| FIM1_17 | 33482.14331 | 27568.25297 | 21161.65642 | -0.107103003 | 0.340068865 | 0.606654506 | 0.724291758 | ns | NaCl | KS.1 |
| FIM1_18 | 2398.097243 | 2951.512989 | 2232.947735 | 0.191575875 | 0.23243197 | 0.290456351 | 0.421534523 | ns | NaCl | KS.1 |
| FIM1_19 | 631.448533 | 571.9968294 | 965.1753006 | -0.126857512 | 0.104991408 | 0.19891021 | 0.318499612 | ns | NaCl | KS.1 |
| FIM1_20 | 3100.921672 | 3271.144042 | 3333.389345 | 0.070048012 | 0.091680173 | 0.423256459 | 0.559283104 | ns | NaCl | KS.1 |
| FIM1_21 | 397.4098377 | 535.2310194 | 585.7584931 | 0.372391762 | 0.144467686 | 0.003918749 | 0.013349278 | ns | NaCl | KS.1 |
| FIM1_22 | 1459.457925 | 1575.774396 | 1520.407378 | 0.104423274 | 0.077735004 | 0.164244693 | 0.274054561 | ns | NaCl | KS.1 |
| FIM1_23 | 196.7109415 | 165.945119 | 230.7427362 | -0.186878942 | 0.180269027 | 0.222650459 | 0.347128183 | ns | NaCl | KS.1 |
| FIM1_24 | 1681.435856 | 2092.246787 | 1199.112786 | 0.199237141 | 0.236235467 | 0.27629359 | 0.406049694 | ns | NaCl | KS.1 |
| FIM1_25 | 477.6366309 | 534.4969913 | 454.9892691 | 0.106326842 | 0.212244495 | 0.534433292 | 0.662218089 | ns | NaCl | KS.1 |
| FIM1_26 | 265.1625534 | 171.968155 | 284.9825584 | -0.473808168 | 0.228477941 | 0.009620778 | 0.027698709 | ns | NaCl | KS.1 |
| FIM1_27 | 670.9810982 | 556.1685608 | 657.296323 | -0.219447343 | 0.158831605 | 0.115184594 | 0.207174025 | ns | NaCl | KS.1 |
| FIM1_28 | 12180.75522 | 15262.90708 | 11506.37819 | 0.252190775 | 0.175523489 | 0.093223066 | 0.175410269 | ns | NaCl | KS.1 |
| FIM1_29 | 1936.230861 | 1547.363601 | 2774.21492 | -0.247162806 | 0.178957478 | 0.104841723 | 0.191834124 | ns | NaCl | KS.1 |
| FIM1_30 | 2005.34921 | 2095.350461 | 2049.9388 | 0.053740573 | 0.122587517 | 0.634757378 | 0.746199896 | ns | NaCl | KS.1 |
| FIM1_31 | 3513.568231 | 4830.093969 | 4258.824352 | 0.433253095 | 0.091408839 | 6.0859107352 | 7.7041745312 | ns | NaCl | KS.1 |
| FIM1_32 | 2865.77791 | 2475.969433 | 2794.011073 | -0.180459289 | 0.127050996 | 0.120866193 | 0.215368664 | ns | NaCl | KS.1 |
| FIM1_33 | 738.1585899 | 794.6297769 | 732.2157693 | 0.078447533 | 0.17311499 | 0.59836507 | 0.717663024 | ns | NaCl | KS.1 |
| FIM1_34 | 1479.142393 | 1890.084896 | 1643.975651 | 0.336139926 | 0.082222845 | 1.9843446492 | 0.000154493 | ns | NaCl | KS.1 |
| FIM1_35 | 21.27065811 | 30.09779015 | 43.59138275 | 0.252358568 | 0.305741267 | 0.225778576 | 0.350576908 | ns | NaCl | KS.1 |
| FIM1_36 | 6768.243779 | 8837.279451 | 6124.361802 | 0.16586945 | 0.325473672 | 0.427077213 | 0.563732856 | ns | NaCl | KS.1 |
| FIM1_37 | 883.9118786 | 1103.269273 | 700.1965119 | 0.265944689 | 0.145217945 | 0.040264197 | 0.088713188 | ns | NaCl | KS.1 |
| FIM1_38 | 1549.325408 | 2387.989086 | 1220.934355 | 0.524205342 | 0.18112381 | 0.000763423 | 0.003440245 | ns | NaCl | KS.1 |
| FIM1_39 | 2108.559913 | 2498.699382 | 1622.369305 | 0.168977156 | 0.205172372 | 0.313281697 | 0.446455584 | ns | NaCl | KS.1 |
| FIM1_40 | 817.4417338 | 608.800617 | 1421.516646 | -0.373566264 | 0.135304435 | 0.002255533 | 0.008437049 | ns | NaCl | KS.1 |
| FIM1_41 | 1049.155179 | 1152.470779 | 1004.945188 | 0.113652381 | 0.130172974 | 0.339168899 | 0.473048857 | ns | NaCl | KS.1 |

| | | | | | | | | | | |
|---|---|---|---|---|---|---|---|---|---|---|
| FIM1_42 | 1585.620445 | 1465.237835 | 1088.647994 | -0.09177244 | 0.15012821 | 0.492084936 | 0.62452106 | ns | NaCl | KS.1 |
| FIM1_43 | 2673.513434 | 2629.276338 | 2347.041188 | -0.022582237 | 0.078989365 | 0.767113007 | 0.849785618 | ns | NaCl | KS.1 |
| FIM1_44 | 231.3374412 | 182.4089105 | 223.7839843 | -0.246676616 | 0.203813544 | 0.139734639 | 0.240463448 | ns | NaCl | KS.1 |
| FIM1_45 | 663.0746616 | 544.6518981 | 909.3249672 | -0.240301177 | 0.141277741 | 0.058196468 | 0.120185732 | ns | NaCl | KS.1 |
| FIM1_46 | 3496.066844 | 4517.734938 | 3895.846385 | 0.320484115 | 0.135041643 | 0.008667466 | 0.025712965 | ns | NaCl | KS.1 |
| FIM1_47 | 1398.678715 | 1554.170423 | 1453.094101 | 0.135289134 | 0.108542751 | 0.183328501 | 0.298839874 | ns | NaCl | KS.1 |
| FIM1_48 | 2708.471931 | 2198.008846 | 2574.536415 | -0.284495313 | 0.077935474 | 0.000149264 | 0.000875696 | ns | NaCl | KS.1 |
| FIM1_49 | 9416.347424 | 22445.48988 | 9518.429149 | 0.964864452 | 0.304035176 | 4.9290413075 | 0.00034441 | ns | NaCl | KS.1 |
| FIM1_50 | 10257.96241 | 3270.612371 | 4848.123799 | -1.485288867 | 0.238839021 | 1.5246597501 | 9.1387738038 | Km<KS.1 | NaCl | KS.1 |
| FIM1_51 | 3613.443039 | 3498.277867 | 3374.321267 | -0.038241726 | 0.14282364 | 0.765906284 | 0.84901599 | ns | NaCl | KS.1 |
| FIM1_52 | 3229.555861 | 3905.934803 | 3261.822567 | 0.233966464 | 0.133750751 | 0.05373673 | 0.112057485 | ns | NaCl | KS.1 |
| FIM1_53 | 35778.7255 | 53127.16326 | 32534.50092 | 0.527124899 | 0.118052412 | 1.5033501450 | 1.6808365527 | ns | NaCl | KS.1 |
| FIM1_54 | 2188.874847 | 1899.32328 | 3260.43686 | -0.142960853 | 0.197676287 | 0.38176168 | 0.516947294 | ns | NaCl | KS.1 |
| FIM1_55 | 748.2435585 | 130.9360917 | 884.141548 | -2.216244212 | 0.408978735 | 2.4745190412 | 1.0258943525 | Km<KS.1 | NaCl | KS.1 |
| FIM1_56 | 26.80229349 | 18.01165942 | 110.1015889 | -0.276872775 | 0.335042265 | 0.198900864 | 0.318499612 | ns | NaCl | KS.1 |
| FIM1_57 | 450.8430951 | 419.3173799 | 411.2940193 | -0.062754099 | 0.233789358 | 0.729004464 | 0.820913809 | ns | NaCl | KS.1 |
| FIM1_58 | 4062.500624 | 6238.406292 | 3663.533176 | 0.505373669 | 0.193775791 | 0.00199798 | 0.007628511 | ns | NaCl | KS.1 |
| FIM1_59 | 3275.94494 | 4507.344638 | 4886.402825 | 0.290262527 | 0.254137257 | 0.130455507 | 0.228125183 | ns | NaCl | KS.1 |
| FIM1_60 | 2633.420583 | 2802.495963 | 2097.26308 | 0.077936397 | 0.11560572 | 0.468617838 | 0.602889513 | ns | NaCl | KS.1 |
| FIM1_61 | 1099.046906 | 971.3599079 | 1514.364944 | -0.165792498 | 0.098426183 | 0.0750668 | 0.147204309 | ns | NaCl | KS.1 |
| FIM1_62 | 1270.177817 | 1603.144493 | 1248.976291 | 0.272638915 | 0.157999062 | 0.049247551 | 0.104302498 | ns | NaCl | KS.1 |
| FIM1_63 | 43.91805327 | 46.09783121 | 61.0388668 | 0.042518272 | 0.274667519 | 0.828034417 | 0.894243309 | ns | NaCl | KS.1 |
| FIM1_64 | 2788.839837 | 1999.108888 | 2027.713438 | -0.450457529 | 0.099931551 | 1.7282880875 | 1.8938839726 | ns | NaCl | KS.1 |
| FIM1_65 | 206.626363 | 184.7163824 | 253.6940693 | -0.140027243 | 0.144990653 | 0.280778032 | 0.411086141 | ns | NaCl | KS.1 |
| FIM1_66 | 4167.283085 | 7383.285282 | 4634.515481 | 0.743724046 | 0.16203691 | 3.9740187702 | 5.3147159628 | ns | NaCl | KS.1 |
| FIM1_67 | 300.3340904 | 992.9267964 | 516.1440968 | 1.611613963 | 0.204161745 | 4.0968939919 | 7.0282922792 | Km>KS.1 | NaCl | KS.1 |
| FIM1_68 | 18044.0728 | 24016.67867 | 15105.22591 | 0.334183752 | 0.168812851 | 0.022042024 | 0.054099196 | ns | NaCl | KS.1 |
| FIM1_69 | 17404.96694 | 23182.6175 | 14277.90827 | 0.364215079 | 0.130671232 | 0.002167419 | 0.008157214 | ns | NaCl | KS.1 |
| FIM1_70 | 1477.344564 | 2317.06084 | 1316.552668 | 0.537498273 | 0.192378825 | 0.00099114 | 0.004258135 | ns | NaCl | KS.1 |
| FIM1_71 | 846.3894622 | 725.6850044 | 738.570201 | -0.182708524 | 0.145586872 | 0.160337226 | 0.269031264 | ns | NaCl | KS.1 |
| FIM1_72 | 740.0251226 | 681.8865974 | 603.4281888 | -0.092461901 | 0.152774268 | 0.494786113 | 0.627309101 | ns | NaCl | KS.1 |
| FIM1_73 | 16012.5016 | 18381.24466 | 13298.10691 | 0.181565017 | 0.095112725 | 0.044300325 | 0.095740277 | ns | NaCl | KS.1 |
| FIM1_74 | 931.0872121 | 993.715857 | 775.8522912 | 0.082009009 | 0.097316618 | 0.37441346 | 0.50941153 | ns | NaCl | KS.1 |
| FIM1_75 | 14605.87179 | 24531.78867 | 22825.45264 | 0.681683883 | 0.14680082 | 3.7110870240 | 5.0582624505 | ns | NaCl | KS.1 |
| FIM1_76 | 310.5544406 | 274.2395804 | 347.4029841 | -0.136856044 | 0.163340282 | 0.336905293 | 0.471036975 | ns | NaCl | KS.1 |
| FIM1_77 | 948.0569838 | 898.1796473 | 859.8821874 | -0.069311142 | 0.108069675 | 0.494239297 | 0.626775555 | ns | NaCl | KS.1 |
| FIM1_78 | 3168.847054 | 3047.690276 | 3222.916234 | -0.047841826 | 0.123652144 | 0.674648294 | 0.778922085 | ns | NaCl | KS.1 |
| FIM1_79 | 947.1561222 | 909.5718137 | 3178.933275 | -0.041185305 | 0.193382659 | 0.798144741 | 0.870903663 | ns | NaCl | KS.1 |
| FIM1_80 | 81.96324185 | 49.46063703 | 115.301577 | -0.536739294 | 0.254560285 | 0.006675654 | 0.020718266 | ns | NaCl | KS.1 |
| FIM1_81 | 1123.249504 | 1095.808383 | 1455.405965 | -0.029572705 | 0.149789534 | 0.824591729 | 0.892395878 | ns | NaCl | KS.1 |
| FIM1_82 | 1640.828999 | 2341.160039 | 2001.056325 | 0.479507946 | 0.103015179 | 7.4484677658 | 9.0823841017 | ns | NaCl | KS.1 |
| FIM1_83 | 404.7111912 | 318.4384564 | 385.1962856 | -0.286117999 | 0.161597201 | 0.042564818 | 0.092572596 | ns | NaCl | KS.1 |
| FIM1_84 | 696.7934355 | 357.0850063 | 813.4542519 | -0.884269879 | 0.162262709 | 3.0963410015 | 7.7799477184 | ns | NaCl | KS.1 |
| FIM1_85 | 547.5762637 | 562.4291907 | 812.4313926 | 0.034776169 | 0.110526443 | 0.736746063 | 0.827010755 | ns | NaCl | KS.1 |
| FIM1_86 | 646.1370909 | 351.5464372 | 890.901293 | -0.822700318 | 0.134603112 | 7.5529768776 | 2.8519866596 | ns | NaCl | KS.1 |

| | | | | | | | | | | |
|---|---|---|---|---|---|---|---|---|---|---|
| FIM1_87 | 459.0736985 | 733.7922917 | 533.8379842 | 0.552163089 | 0.204212923 | 0.001228663 | 0.005102337 | ns | NaCl | KS.1 |
| FIM1_88 | 705.4599681 | 388.108063 | 812.0837853 | -0.749459197 | 0.191371779 | 7.67673702618 | 6.80780155173 | ns | NaCl | KS.1 |
| FIM1_89 | 109.9736118 | 232.2440009 | 613.585864 | 0.675057906 | 0.353334871 | 0.005331657 | 0.017179399 | ns | NaCl | KS.1 |
| FIM1_90 | 2991.537001 | 3725.485333 | 2907.03043 | 0.298331063 | 0.081269378 | 0.000130288 | 0.000780003 | ns | NaCl | KS.1 |
| FIM1_91 | 1533.435533 | 7166.544965 | 2110.39745 | 2.047834826 | 0.295440127 | 1.60227310662 | 1.62698667912 | Km>KS.1 | NaCl | KS.1 |
| FIM1_92 | 249.6131517 | 220.3217237 | 320.0306234 | -0.14028914 | 0.153653649 | 0.302077544 | 0.434094679 | ns | NaCl | KS.1 |
| FIM1_93 | 237.0337384 | 165.6614996 | 258.6469668 | -0.455698636 | 0.143832122 | 0.000411268 | 0.002050158 | ns | NaCl | KS.1 |
| FIM1_94 | 251.3945887 | 128.6706213 | 357.0574842 | -0.873910698 | 0.185011796 | 1.38421629089 | 2.15202376475 | ns | NaCl | KS.1 |
| FIM1_95 | 6594.734389 | 6198.980706 | 5406.626742 | -0.080939342 | 0.092034104 | 0.356447712 | 0.491634979 | ns | NaCl | KS.1 |
| FIM1_96 | 289.2963629 | 142.7437537 | 692.0664714 | -0.765940913 | 0.287024228 | 0.000550631 | 0.002617606 | ns | NaCl | KS.1 |
| FIM1_97 | 6973.153083 | 1169.85175 | 3895.884372 | -2.373304187 | 0.343138649 | 2.60065506130 | 2.58765178599 | Km<KS.1 | NaCl | KS.1 |
| FIM1_98 | 224.4351675 | 140.3752101 | 194.9117998 | -0.490698942 | 0.252721416 | 0.012148638 | 0.033299986 | ns | NaCl | KS.1 |
| FIM1_99 | 596.2518358 | 365.6055214 | 463.1981537 | -0.597245359 | 0.189067273 | 0.000233823 | 0.001276917 | ns | NaCl | KS.1 |
| FIM1_100 | 5176.486415 | 4186.554212 | 3763.677311 | -0.263752622 | 0.13105886 | 0.026892256 | 0.063739387 | ns | NaCl | KS.1 |
| FIM1_101 | 852.5299027 | 769.5212603 | 1500.413589 | -0.111629627 | 0.170413221 | 0.448137669 | 0.583255256 | ns | NaCl | KS.1 |
| FIM1_102 | 168.0070586 | 162.5763321 | 221.6806448 | -0.037796003 | 0.176714964 | 0.802612866 | 0.873937443 | ns | NaCl | KS.1 |
| FIM1_103 | 1454.025089 | 1839.164561 | 1767.344721 | 0.302379757 | 0.116884587 | 0.005132162 | 0.016611911 | ns | NaCl | KS.1 |
| FIM1_104 | 525.1685237 | 491.9081625 | 640.3642589 | -0.092715888 | 0.088802468 | 0.275019649 | 0.404799631 | ns | NaCl | KS.1 |
| FIM1_105 | 4179.759143 | 6002.919284 | 3825.164874 | 0.424004926 | 0.182209028 | 0.006282769 | 0.019732814 | ns | NaCl | KS.1 |
| FIM1_107 | 937.9764424 | 959.4775604 | 944.1502495 | 0.02641907 | 0.138356946 | 0.832850444 | 0.897040692 | ns | NaCl | KS.1 |
| FIM1_108 | 3021.94185 | 3011.361766 | 2134.243348 | -0.004293216 | 0.150381385 | 0.974412342 | 0.988116877 | ns | NaCl | KS.1 |
| FIM1_109 | 1258.307191 | 7272.340601 | 3009.743906 | 2.476565161 | 0.179019155 | 2.96432299842 | 7.37375345857 | Km>KS.1 | NaCl | KS.1 |
| FIM1_110 | 430.8009047 | 1062.260051 | 762.545038 | 1.231815697 | 0.149620037 | 2.44658796854 | 4.68145197827 | Km>KS.1 | NaCl | KS.1 |
| FIM1_111 | 4017.944721 | 4360.148505 | 5944.23296 | 0.10637813 | 0.097969547 | 0.251943232 | 0.380630907 | ns | NaCl | KS.1 |
| FIM1_112 | 972.6644835 | 1336.255078 | 919.1620486 | 0.371316977 | 0.174185722 | 0.01307072 | 0.035264008 | ns | NaCl | KS.1 |
| FIM1_113 | 725.3985847 | 727.1094978 | 1147.542679 | 0.003396328 | 0.125916579 | 0.976593697 | 0.9889179 | ns | NaCl | KS.1 |
| FIM1_114 | 927.792876 | 943.919594 | 985.3718464 | 0.021142591 | 0.127282974 | 0.856371248 | 0.913083361 | ns | NaCl | KS.1 |
| FIM1_115 | 499.2925135 | 609.1156696 | 1055.82097 | 0.187808251 | 0.223214354 | 0.288271746 | 0.418892485 | ns | NaCl | KS.1 |
| FIM1_116 | 409.6167729 | 341.4041849 | 432.800489 | -0.214494281 | 0.162632598 | 0.130580545 | 0.228263602 | ns | NaCl | KS.1 |
| FIM1_117 | 11155.34199 | 16620.61041 | 11163.9285 | 0.460339881 | 0.196111716 | 0.005071771 | 0.016448539 | ns | NaCl | KS.1 |
| FIM1_118 | 99.43633649 | 93.68677399 | 186.7466403 | -0.062517407 | 0.2177965 | 0.719101846 | 0.813388223 | ns | NaCl | KS.1 |
| FIM1_119 | 8916.849505 | 13427.86581 | 7368.886416 | 0.471967318 | 0.198809916 | 0.004484413 | 0.014883226 | ns | NaCl | KS.1 |
| FIM1_120 | 1636.716133 | 1712.257687 | 1362.100667 | 0.057306008 | 0.109730842 | 0.57713613 | 0.699111821 | ns | NaCl | KS.1 |
| FIM1_121 | 2975.751741 | 6866.480874 | 3572.133845 | 0.946195532 | 0.288462937 | 3.78546524963 | 0.000272149 | ns | NaCl | KS.1 |
| FIM1_122 | 3229.852496 | 3959.331586 | 3375.700208 | 0.269471591 | 0.096962349 | 0.003287324 | 0.011492929 | ns | NaCl | KS.1 |
| FIM1_123 | 5617.002419 | 6321.947743 | 4792.686076 | 0.148931501 | 0.116734268 | 0.169105243 | 0.280578067 | ns | NaCl | KS.1 |
| FIM1_124 | 784.903111 | 854.0593712 | 724.0275505 | 0.107812915 | 0.121193361 | 0.335508681 | 0.469919957 | ns | NaCl | KS.1 |
| FIM1_125 | 772.0685914 | 267.8562824 | 653.6319812 | -1.411093434 | 0.189605279 | 4.50300909132 | 5.33392148318 | Km<KS.1 | NaCl | KS.1 |
| FIM1_126 | 1645.053205 | 1244.287821 | 1309.752804 | -0.375636452 | 0.098395548 | 5.14420785235 | 0.000357936 | ns | NaCl | KS.1 |
| FIM1_127 | 940.0227808 | 519.5424222 | 779.7362617 | -0.72014943 | 0.20781958 | 4.91371227891 | 0.000343822 | ns | NaCl | KS.1 |
| FIM1_128 | 18227.40603 | 20938.33604 | 15811.42649 | 0.176644244 | 0.112128385 | 0.091013012 | 0.17190565 | ns | NaCl | KS.1 |
| FIM1_129 | 1286.605321 | 1368.147084 | 1036.64876 | 0.063808318 | 0.183808634 | 0.68153182 | 0.784546705 | ns | NaCl | KS.1 |
| FIM1_130 | 8636.943024 | 8037.417925 | 9887.24238 | -0.090014214 | 0.114586344 | 0.398870039 | 0.535017105 | ns | NaCl | KS.1 |
| FIM1_131 | 2499.399455 | 2823.933414 | 2248.066324 | 0.141222442 | 0.151235791 | 0.292845224 | 0.424135368 | ns | NaCl | KS.1 |
| FIM1_132 | 23848.76544 | 34883.31944 | 21182.2828 | 0.522845206 | 0.090240246 | 1.26757107344 | 3.50342560578 | ns | NaCl | KS.1 |

| | | | | | | | | | | |
|---|---|---|---|---|---|---|---|---|---|---|
| FIM1_133 | 7214.268919 | 6823.118129 | 4961.324517 | -0.066266455 | 0.13754226 | 0.594658494 | 0.714595654 | ns | NaCl | KS.1 |
| FIM1_134 | 4053.726428 | 5148.667992 | 4336.865324 | 0.308337569 | 0.117636016 | 0.004535994 | 0.014994398 | ns | NaCl | KS.1 |
| FIM1_135 | 1477.992938 | 1582.533379 | 1364.517315 | 0.088514448 | 0.093318704 | 0.319296999 | 0.452564835 | ns | NaCl | KS.1 |
| FIM1_136 | 68.12362277 | 52.30016279 | 244.4891628 | -0.206789271 | 0.269190599 | 0.291932159 | 0.423429298 | ns | NaCl | KS.1 |
| FIM1_137 | 425.9560137 | 195.9135534 | 852.1868397 | -0.933998053 | 0.242129818 | 4.8995573290[illegible] | 4.6429138498[illegible] | ns | NaCl | KS.1 |
| FIM1_138 | 13707.71836 | 19140.92984 | 11142.29983 | 0.40959518 | 0.156347665 | 0.002939847 | 0.010537275 | ns | NaCl | KS.1 |
| FIM1_140 | 20506.30159 | 28387.47871 | 17517.12602 | 0.41440193 | 0.135465608 | 0.000719565 | 0.003275238 | ns | NaCl | KS.1 |
| FIM1_141 | 922.3941591 | 747.1665811 | 718.8621058 | -0.253792887 | 0.141309723 | 0.045423089 | 0.097604297 | ns | NaCl | KS.1 |
| FIM1_142 | 13715.27277 | 15637.05293 | 12027.67333 | 0.180685609 | 0.065702882 | 0.004750588 | 0.01561042 | ns | NaCl | KS.1 |
| FIM1_143 | 156.5224131 | 193.9680092 | 247.9532581 | 0.19816965 | 0.238792957 | 0.281821727 | 0.411781699 | ns | NaCl | KS.1 |
| FIM1_144 | 861.1057592 | 708.0704965 | 911.1906525 | -0.141713275 | 0.28725707 | 0.479605042 | 0.613692151 | ns | NaCl | KS.1 |
| FIM1_145 | 11768.31367 | 23255.77238 | 13754.70225 | 0.858470478 | 0.198983236 | 9.6914751994[illegible] | 1.1425376568[illegible] | ns | NaCl | KS.1 |
| FIM1_146 | 3031.331457 | 2913.338951 | 2501.115526 | -0.052071621 | 0.090246178 | 0.545961792 | 0.672316811 | ns | NaCl | KS.1 |
| FIM1_147 | 1838.192006 | 1890.544856 | 2010.46277 | 0.036323617 | 0.085455843 | 0.65789507 | 0.764726162 | ns | NaCl | KS.1 |
| FIM1_148 | 735.9797954 | 748.9611982 | 787.1053614 | 0.020726992 | 0.095075774 | 0.81874047 | 0.887886833 | ns | NaCl | KS.1 |
| FIM1_149 | 591.7980167 | 575.6570308 | 567.6685807 | -0.032285715 | 0.116958668 | 0.76639865 | 0.84918336 | ns | NaCl | KS.1 |
| FIM1_150 | 902.6773896 | 821.5074779 | 1020.816971 | -0.123954213 | 0.094845443 | 0.169136407 | 0.280578067 | ns | NaCl | KS.1 |
| FIM1_151 | 722.984064 | 722.0020004 | 929.2554157 | -0.003349113 | 0.123555831 | 0.976545928 | 0.9889179 | ns | NaCl | KS.1 |
| FIM1_153 | 513.2673686 | 634.1753681 | 573.2743016 | 0.233457677 | 0.176285579 | 0.121406241 | 0.215944243 | ns | NaCl | KS.1 |
| FIM1_154 | 1613.430506 | 1432.538509 | 1999.653377 | -0.105955905 | 0.231261305 | 0.556510595 | 0.680255579 | ns | NaCl | KS.1 |
| FIM1_155 | 3724.341449 | 3729.843508 | 3525.482726 | 0.001478408 | 0.184866673 | 0.992445035 | 0.996607017 | ns | NaCl | KS.1 |
| FIM1_156 | 1828.572016 | 980.5240517 | 1443.315126 | -0.82851363 | 0.149200619 | 2.0601609784[illegible] | 5.4517557807[illegible] | ns | NaCl | KS.1 |
| FIM1_157 | 1831.902879 | 734.9371896 | 2207.756938 | -0.968507012 | 0.334629146 | 0.000113807 | 0.000693861 | ns | NaCl | KS.1 |
| FIM1_158 | 7938.861561 | 7471.49171 | 6922.7706 | -0.076774402 | 0.109250809 | 0.453229778 | 0.588418096 | ns | NaCl | KS.1 |
| FIM1_159 | 6756.0706 | 7359.086709 | 6425.530611 | 0.122061223 | 0.036042102 | 0.000638496 | 0.002952155 | ns | NaCl | KS.1 |
| FIM1_160 | 6177.56556 | 6559.435276 | 5656.953157 | 0.08140572 | 0.073270114 | 0.251810728 | 0.380546285 | ns | NaCl | KS.1 |
| FIM1_161 | 472.5786702 | 506.0834038 | 678.5940738 | 0.0757477 | 0.177273579 | 0.617134523 | 0.732056331 | ns | NaCl | KS.1 |
| FIM1_162 | 1339.130722 | 1039.773066 | 1262.050445 | -0.301775883 | 0.1574341 | 0.029033731 | 0.067749912 | ns | NaCl | KS.1 |
| FIM1_163 | 220.7581442 | 219.4892289 | 516.5720716 | -0.003743721 | 0.210980159 | 0.982473909 | 0.991175946 | ns | NaCl | KS.1 |
| FIM1_164 | 641.2185362 | 825.9016882 | 893.1192894 | 0.332671491 | 0.102849536 | 0.000568774 | 0.002684679 | ns | NaCl | KS.1 |
| FIM1_165 | 667.9090626 | 737.2938999 | 587.4606674 | 0.116708571 | 0.138779558 | 0.351857855 | 0.486779562 | ns | NaCl | KS.1 |
| FIM1_166 | 571.2761282 | 659.1002739 | 737.7645358 | 0.150785266 | 0.179909236 | 0.324547008 | 0.457788875 | ns | NaCl | KS.1 |
| FIM1_167 | 194.8265959 | 332.4069929 | 515.6252124 | 0.652855779 | 0.197561984 | 0.000113031 | 0.000690824 | ns | NaCl | KS.1 |
| FIM1_168 | 1116.527146 | 1418.556483 | 1298.756978 | 0.30430421 | 0.12210482 | 0.00667369 | 0.020718266 | ns | NaCl | KS.1 |
| FIM1_169 | 1343.194461 | 1281.042499 | 2192.324479 | -0.058918734 | 0.123828478 | 0.605577246 | 0.723348571 | ns | NaCl | KS.1 |
| FIM1_170 | 10353.70329 | 14824.05569 | 11208.52652 | 0.508306638 | 0.053377969 | 3.0912211703[illegible] | 1.2815687768[illegible] | ns | NaCl | KS.1 |
| FIM1_171 | 2060.043806 | 2537.732624 | 1811.685558 | 0.255299247 | 0.136774448 | 0.038715323 | 0.086101356 | ns | NaCl | KS.1 |
| FIM1_172 | 696.2670529 | 668.1194718 | 709.2605182 | -0.050207553 | 0.136992498 | 0.685910953 | 0.787936089 | ns | NaCl | KS.1 |
| FIM1_173 | 852.2184821 | 1165.825888 | 894.9360852 | 0.404640697 | 0.126608769 | 0.000470737 | 0.00228703 | ns | NaCl | KS.1 |
| FIM1_174 | 337.6449833 | 283.5768762 | 327.707724 | -0.121219854 | 0.294838824 | 0.548282278 | 0.673840003 | ns | NaCl | KS.1 |
| FIM1_175 | 107.6057708 | 75.42042709 | 88.16067366 | -0.334789464 | 0.268141107 | 0.091768659 | 0.173197677 | ns | NaCl | KS.1 |
| FIM1_176 | 2234.49924 | 2416.863522 | 2120.157969 | 0.094899233 | 0.129217322 | 0.422106016 | 0.558336679 | ns | NaCl | KS.1 |
| FIM1_177 | 10294.05273 | 13070.70694 | 8182.819457 | 0.239767501 | 0.21252481 | 0.16261775 | 0.272306734 | ns | NaCl | KS.1 |
| FIM1_178 | 801.0874147 | 857.3026831 | 898.2572507 | 0.089828342 | 0.093299313 | 0.312066955 | 0.444851891 | ns | NaCl | KS.1 |
| FIM1_179 | 2118.340788 | 2708.115596 | 2568.32626 | 0.316883072 | 0.116234292 | 0.003202105 | 0.011266247 | ns | NaCl | KS.1 |

| | | | | | | | | | |
|---|---|---|---|---|---|---|---|---|---|
| FIM1_180 | 1629.835691 | 1677.37635 | 1767.514357 | 0.036712221 | 0.101554431 | 0.702067685 | 0.799981706 | ns | NaCl | KS.1 |
| FIM1_181 | 12985.38626 | 17256.15532 | 11963.46107 | 0.393305797 | 0.07322506 | 2.66855513131 | 5.2682784834 | ns | NaCl | KS.1 |
| FIM1_182 | 6715.088116 | 6931.275817 | 5639.517076 | 0.030407701 | 0.205053323 | 0.855842859 | 0.912715589 | ns | NaCl | KS.1 |
| FIM1_183 | 70.63363436 | 73.86630602 | 86.97005316 | 0.035885227 | 0.246994186 | 0.847362415 | 0.906976767 | ns | NaCl | KS.1 |
| FIM1_184 | 3639.755284 | 5231.492753 | 4253.522987 | 0.466313446 | 0.13733112 | 0.000173866 | 0.00099538 | ns | NaCl | KS.1 |
| FIM1_185 | 508.1615697 | 663.8046774 | 1297.052696 | 0.289960544 | 0.191601526 | 0.070236582 | 0.139324958 | ns | NaCl | KS.1 |
| FIM1_186 | 295.343141 | 167.2299603 | 342.0256655 | -0.593125916 | 0.267568991 | 0.003968196 | 0.013475071 | ns | NaCl | KS.1 |
| FIM1_187 | 8788.333058 | 9967.526424 | 7421.616443 | 0.111194529 | 0.233919909 | 0.539850286 | 0.666444387 | ns | NaCl | KS.1 |
| FIM1_189 | 1609.830949 | 1774.129656 | 1349.127392 | 0.124226752 | 0.107389694 | 0.217551505 | 0.341102659 | ns | NaCl | KS.1 |
| FIM1_190 | 178.8764124 | 178.9978306 | 166.9618692 | -0.001625512 | 0.188086281 | 0.991792322 | 0.996398788 | ns | NaCl | KS.1 |
| FIM1_191 | 811.4837633 | 342.306797 | 673.3985639 | -1.188193112 | 0.135626929 | 3.3435905349 | 9.7849193596 | Km<KS.1 | NaCl | KS.1 |
| FIM1_192 | 2617.419763 | 2492.670236 | 3739.861374 | -0.037768509 | 0.262126523 | 0.843926146 | 0.904271501 | ns | NaCl | KS.1 |
| FIM1_193 | 39.06402164 | 16.99305263 | 567.2888931 | -0.663391295 | 0.407975032 | 0.009195752 | 0.02678505 | ns | NaCl | KS.1 |
| FIM1_194 | 2967.85526 | 2590.911994 | 2532.895346 | -0.142935529 | 0.184840866 | 0.359676381 | 0.495401439 | ns | NaCl | KS.1 |
| FIM1_195 | 1589.68567 | 1386.171107 | 1244.231713 | -0.157339879 | 0.156833277 | 0.254527348 | 0.382734778 | ns | NaCl | KS.1 |
| FIM1_196 | 4350.907096 | 4108.977039 | 2924.485626 | -0.07244863 | 0.112957328 | 0.49185436 | 0.624387711 | ns | NaCl | KS.1 |
| FIM1_197 | 1634.591487 | 2056.263594 | 2057.649556 | 0.239626949 | 0.200252681 | 0.146492476 | 0.250017176 | ns | NaCl | KS.1 |
| FIM1_198 | 4213.415199 | 6255.925827 | 4654.542393 | 0.502431813 | 0.149333145 | 0.000166269 | 0.000958327 | ns | NaCl | KS.1 |
| FIM1_199 | 1112.626025 | 873.0573494 | 1061.302746 | -0.275625472 | 0.170780511 | 0.06111688 | 0.124982451 | ns | NaCl | KS.1 |
| FIM1_200 | 2094.182553 | 3291.194443 | 2473.384623 | 0.577132331 | 0.156488236 | 3.5644185157 | 0.000258122 | ns | NaCl | KS.1 |
| FIM1_201 | 2038.090699 | 2118.574511 | 2060.566516 | 0.04824942 | 0.112345963 | 0.64563791 | 0.754300314 | ns | NaCl | KS.1 |
| FIM1_203 | 901.2585407 | 855.9811148 | 1224.968132 | -0.065575693 | 0.117515849 | 0.547761878 | 0.673533203 | ns | NaCl | KS.1 |
| FIM1_204 | 155.2340052 | 80.04720941 | 149.289268 | -0.759261051 | 0.256310905 | 0.000235213 | 0.001278892 | ns | NaCl | KS.1 |
| FIM1_205 | 2155.424808 | 2219.845272 | 1549.415675 | 0.025791168 | 0.231040008 | 0.885983489 | 0.932071866 | ns | NaCl | KS.1 |
| FIM1_206 | 3277.98568 | 3004.455933 | 2888.196518 | -0.111585406 | 0.103623611 | 0.25306531 | 0.381284045 | ns | NaCl | KS.1 |
| FIM1_207 | 920.8471608 | 668.9619909 | 798.6830117 | -0.388626306 | 0.156978322 | 0.004882796 | 0.016002577 | ns | NaCl | KS.1 |
| FIM1_208 | 3138.093909 | 2924.629576 | 3150.474754 | -0.086724057 | 0.120309729 | 0.435806322 | 0.571765942 | ns | NaCl | KS.1 |
| FIM1_209 | 3664.418923 | 3557.564913 | 4312.461468 | -0.036746833 | 0.120420953 | 0.741576109 | 0.830184775 | ns | NaCl | KS.1 |
| FIM1_210 | 3521.878457 | 3740.497188 | 2926.221732 | 0.071477536 | 0.134596699 | 0.559029217 | 0.682663317 | ns | NaCl | KS.1 |
| FIM1_211 | 1010.516143 | 955.4843631 | 1421.061327 | -0.070714628 | 0.12098583 | 0.527218244 | 0.655891664 | ns | NaCl | KS.1 |
| FIM1_212 | 1226.075496 | 971.3418357 | 1343.651911 | -0.282692385 | 0.145969803 | 0.029885311 | 0.069476366 | ns | NaCl | KS.1 |
| FIM1_213 | 1386.311606 | 1587.301355 | 1407.992985 | 0.179367028 | 0.091752449 | 0.040099731 | 0.088426196 | ns | NaCl | KS.1 |
| FIM1_214 | 250.7864536 | 122.8750117 | 339.0121516 | -0.889209901 | 0.219797071 | 2.7369859158 | 2.8546131931 | ns | NaCl | KS.1 |
| FIM1_215 | 649.3302256 | 653.1230107 | 763.6384068 | 0.001883508 | 0.091456204 | 0.982788814 | 0.991175946 | ns | NaCl | KS.1 |
| FIM1_216 | 4946.097324 | 3933.01497 | 3833.499387 | -0.256984307 | 0.175195311 | 0.086725682 | 0.165691348 | ns | NaCl | KS.1 |
| FIM1_217 | 3385.412644 | 3061.770101 | 2626.542711 | -0.123461023 | 0.126761006 | 0.288278607 | 0.418892485 | ns | NaCl | KS.1 |
| FIM1_218 | 846.383076 | 359.6861221 | 1854.385565 | -0.967018926 | 0.292495383 | 3.160702420 | 0.000233648 | ns | NaCl | KS.1 |
| FIM1_219 | 639.7027761 | 629.6661225 | 1022.823757 | -0.025453343 | 0.101386824 | 0.790551886 | 0.865536011 | ns | NaCl | KS.1 |
| FIM1_220 | 352.3961504 | 319.0525704 | 417.3276213 | -0.117288651 | 0.149249602 | 0.377540773 | 0.51220762 | ns | NaCl | KS.1 |
| FIM1_221 | 1177.43701 | 1315.046275 | 1188.193571 | 0.130633686 | 0.14146774 | 0.304796953 | 0.437244764 | ns | NaCl | KS.1 |
| FIM1_222 | 1215.313231 | 1422.459844 | 1793.918807 | 0.216370136 | 0.074440201 | 0.002640397 | 0.009658807 | ns | NaCl | KS.1 |
| FIM1_224 | 2210.684209 | 2628.012634 | 1990.069359 | 0.212583618 | 0.130761133 | 0.07424993 | 0.145947611 | ns | NaCl | KS.1 |
| FIM1_225 | 1496.931305 | 682.8270003 | 1133.049414 | -1.093178686 | 0.116483981 | 1.8400632310 | 9.1543145746 | Km<KS.1 | NaCl | KS.1 |
| FIM1_226 | 269.624954 | 135.0540624 | 393.6260039 | -0.790478589 | 0.256133034 | 0.000139171 | 0.0008282 | ns | NaCl | KS.1 |
| FIM1_227 | 1425.149141 | 1579.258362 | 1808.330015 | 0.118239508 | 0.153282221 | 0.383691403 | 0.518995304 | ns | NaCl | KS.1 |

| | | | | | | | | | | |
|---|---|---|---|---|---|---|---|---|---|---|
| FIM1_228 | 111.422678 | 71.70348861 | 100.5731306 | -0.520229868 | 0.188728767 | 0.001198094 | 0.004992058 | ns | NaCl | KS.1 |
| FIM1_229 | 5819.818422 | 8384.203243 | 7860.361819 | 0.471774731 | 0.134413237 | 0.000110624 | 0.000681979 | ns | NaCl | KS.1 |
| FIM1_230 | 1895.608394 | 1653.490495 | 1941.403259 | -0.17686141 | 0.100490205 | 0.062273712 | 0.126712358 | ns | NaCl | KS.1 |
| FIM1_231 | 238.0426601 | 54.90711646 | 186.9610113 | -1.810819342 | 0.372071689 | 8.48497316299 | 2.45422915615 | Km<KS.1 | NaCl | KS.1 |
| FIM1_233 | 5269.97236 | 6206.70498 | 4732.029427 | 0.194656957 | 0.144109487 | 0.131434719 | 0.22943429 | ns | NaCl | KS.1 |
| FIM1_234 | 1737.19752 | 1761.686328 | 1866.763399 | 0.020407521 | 0.067180461 | 0.755213466 | 0.840535044 | ns | NaCl | KS.1 |
| FIM1_235 | 496.1127347 | 451.9703699 | 816.0938578 | -0.113785026 | 0.137212057 | 0.359615253 | 0.495401439 | ns | NaCl | KS.1 |
| FIM1_236 | 1815.86556 | 1900.119894 | 2878.122841 | 0.055043633 | 0.128385491 | 0.639845587 | 0.749480061 | ns | NaCl | KS.1 |
| FIM1_237 | 384.4055241 | 272.2036175 | 509.869959 | -0.308221571 | 0.263113311 | 0.115747524 | 0.207866226 | ns | NaCl | KS.1 |
| FIM1_238 | 457.2114765 | 258.9464303 | 439.7022536 | -0.74439226 | 0.158708445 | 2.44842474917 | 3.50026239285 | ns | NaCl | KS.1 |
| FIM1_239 | 128.1456584 | 49.68722804 | 93.79866522 | -1.132720104 | 0.268534402 | 4.57581430801 | 6.05094094062 | Km<KS.1 | NaCl | KS.1 |
| FIM1_240 | 469.07832 | 240.8406002 | 582.5653609 | -0.824792795 | 0.212583941 | 6.82948322693 | 6.08901058315 | ns | NaCl | KS.1 |
| FIM1_241 | 4177.696138 | 4309.372982 | 3621.35305 | 0.042540371 | 0.068160992 | 0.521483943 | 0.652118833 | ns | NaCl | KS.1 |
| FIM1_242 | 53.89579225 | 44.6082423 | 58.14148934 | -0.167893702 | 0.23240718 | 0.353736883 | 0.488708968 | ns | NaCl | KS.1 |
| FIM1_243 | 1394.031377 | 1296.132295 | 1207.709963 | -0.096121989 | 0.084803398 | 0.237699904 | 0.364312083 | ns | NaCl | KS.1 |
| FIM1_245 | 6009.773213 | 9252.529924 | 4952.962543 | 0.458834432 | 0.233537974 | 0.013302577 | 0.035792493 | ns | NaCl | KS.1 |
| FIM1_246 | 1897.019704 | 2038.502694 | 2323.651007 | 0.096452258 | 0.094561682 | 0.283556037 | 0.414056731 | ns | NaCl | KS.1 |
| FIM1_247 | 584.6016874 | 265.927436 | 1343.159356 | -0.956196949 | 0.234710963 | 1.86700080502 | 2.01482190997 | ns | NaCl | KS.1 |
| FIM1_248 | 1083.397461 | 1249.101125 | 1115.187354 | 0.181421069 | 0.115266333 | 0.090112851 | 0.170720271 | ns | NaCl | KS.1 |
| FIM1_249 | 1033.411392 | 2305.995531 | 995.5845502 | 0.781189313 | 0.344049599 | 0.001439742 | 0.005771729 | ns | NaCl | KS.1 |
| FIM1_250 | 796.2922655 | 698.8776928 | 804.4931944 | -0.161601311 | 0.126636751 | 0.163844962 | 0.273624936 | ns | NaCl | KS.1 |
| FIM1_251 | 509.1523061 | 413.5271931 | 522.5479514 | -0.261929308 | 0.133550047 | 0.030505156 | 0.070620359 | ns | NaCl | KS.1 |
| FIM1_252 | 2284.884887 | 2080.052294 | 2061.09628 | -0.12825116 | 0.083711635 | 0.110684061 | 0.200477721 | ns | NaCl | KS.1 |
| FIM1_253 | 4418.301952 | 5966.149488 | 4712.613198 | 0.400791171 | 0.103158881 | 3.46405654797 | 0.000251587 | ns | NaCl | KS.1 |
| FIM1_254 | 709.4377549 | 520.7140228 | 630.1023059 | -0.380131142 | 0.151236117 | 0.004554566 | 0.015025839 | ns | NaCl | KS.1 |
| FIM1_255 | 4498.907685 | 3412.420642 | 3916.679253 | -0.334467101 | 0.151999755 | 0.012859166 | 0.034712074 | ns | NaCl | KS.1 |
| FIM1_256 | 3692.40882 | 4764.532682 | 2945.980669 | 0.293560008 | 0.169890638 | 0.045249117 | 0.097367802 | ns | NaCl | KS.1 |
| FIM1_257 | 1167.084294 | 960.2158845 | 989.1991687 | -0.249677434 | 0.116942209 | 0.021031906 | 0.051875922 | ns | NaCl | KS.1 |
| FIM1_258 | 1071.205191 | 1009.807276 | 1326.343152 | -0.075828253 | 0.112891402 | 0.471633722 | 0.605516843 | ns | NaCl | KS.1 |
| FIM1_259 | 2927.291582 | 4209.814529 | 3182.099749 | 0.460648766 | 0.144722372 | 0.000382193 | 0.001928407 | ns | NaCl | KS.1 |
| FIM1_260 | 705.299048 | 500.5517801 | 531.1506979 | -0.417431456 | 0.162235203 | 0.003256057 | 0.011430866 | ns | NaCl | KS.1 |
| FIM1_261 | 1226.364112 | 1319.653572 | 1417.32722 | 0.074892922 | 0.1885425 | 0.636059907 | 0.747025032 | ns | NaCl | KS.1 |
| FIM1_262 | 1346.366752 | 2362.156289 | 1256.727153 | 0.67906234 | 0.207103847 | 0.000112084 | 0.000685879 | ns | NaCl | KS.1 |
| FIM1_263 | 3163.670665 | 4131.501639 | 4356.538085 | 0.347283234 | 0.11621014 | 0.001230085 | 0.00510398 | ns | NaCl | KS.1 |
| FIM1_264 | 878.1465102 | 1029.090707 | 998.4399574 | 0.205272912 | 0.105719112 | 0.038477368 | 0.085802289 | ns | NaCl | KS.1 |
| FIM1_265 | 1118.722137 | 1370.949176 | 1462.899096 | 0.272201791 | 0.088487037 | 0.00125348 | 0.00516658 | ns | NaCl | KS.1 |
| FIM1_266 | 986.6363665 | 868.2099622 | 972.3963985 | -0.154061312 | 0.136494711 | 0.212497489 | 0.335014135 | ns | NaCl | KS.1 |
| FIM1_267 | 6357.986845 | 7532.643398 | 5139.402025 | 0.225884832 | 0.091058472 | 0.009167964 | 0.026723864 | ns | NaCl | KS.1 |
| FIM1_268 | 4149.480988 | 5021.785997 | 5576.256847 | 0.231977403 | 0.139280764 | 0.064362198 | 0.129970864 | ns | NaCl | KS.1 |
| FIM1_269 | 1476.278013 | 1350.760651 | 1362.757812 | -0.112000561 | 0.117450411 | 0.304004955 | 0.436360257 | ns | NaCl | KS.1 |
| FIM1_270 | 157.6882818 | 108.4752797 | 135.2654486 | -0.412854435 | 0.214924577 | 0.018023845 | 0.045593295 | ns | NaCl | KS.1 |
| FIM1_271 | 2332.653473 | 2950.747793 | 2806.474049 | 0.290851788 | 0.135728746 | 0.017717225 | 0.044971018 | ns | NaCl | KS.1 |
| FIM1_272 | 689.2055509 | 704.9232594 | 690.4990834 | 0.025386564 | 0.137510021 | 0.838494957 | 0.901233357 | ns | NaCl | KS.1 |
| FIM1_273 | 248.014104 | 162.0053414 | 446.7911323 | -0.463020326 | 0.222926502 | 0.010013303 | 0.028597119 | ns | NaCl | KS.1 |
| FIM1_274 | 1873.108584 | 2852.397944 | 1722.830084 | 0.464443192 | 0.217869127 | 0.008701212 | 0.025785346 | ns | NaCl | KS.1 |

| | | | | | | | | | | |
|---|---|---|---|---|---|---|---|---|---|---|
| FIM1_275 | 1915.514604 | 1850.96218 | 1602.25596 | -0.039132106 | 0.157719384 | 0.778146348 | 0.85590937 | ns | NaCl | KS.1 |
| FIM1_276 | 1004.554471 | 820.324039 | 906.7754852 | -0.251294714 | 0.1315379 | 0.035589262 | 0.080297768 | ns | NaCl | KS.1 |
| FIM1_277 | 254.515883 | 232.839193 | 311.8922662 | -0.110468007 | 0.128531198 | 0.34778148 | 0.482760286 | ns | NaCl | KS.1 |
| FIM1_278 | 640.4261235 | 694.4677635 | 637.3066224 | 0.094578411 | 0.146578919 | 0.470534921 | 0.604574181 | ns | NaCl | KS.1 |
| FIM1_279 | 4347.423757 | 4767.431884 | 3531.57243 | 0.116925875 | 0.110006654 | 0.255911687 | 0.38425017 | ns | NaCl | KS.1 |
| FIM1_280 | 48.78495228 | 29.21883542 | 26.86602388 | -0.392571619 | 0.335445859 | 0.076725293 | 0.149755804 | ns | NaCl | KS.1 |
| FIM1_281 | 3776.466384 | 3204.349976 | 3055.708706 | -0.187309469 | 0.16029314 | 0.182059085 | 0.297584509 | ns | NaCl | KS.1 |
| FIM1_282 | 348.3170055 | 340.9856439 | 392.1436837 | -0.01626958 | 0.269715953 | 0.933226393 | 0.961658314 | ns | NaCl | KS.1 |
| FIM1_283 | 786.7095371 | 746.60866 | 1789.244243 | -0.041600034 | 0.254647468 | 0.826044245 | 0.893190637 | ns | NaCl | KS.1 |
| FIM1_284 | 1051.309291 | 1640.150284 | 1082.482432 | 0.571802726 | 0.152811972 | 2.9366843597( | 0.000217735 | ns | NaCl | KS.1 |
| FIM1_285 | 1530.455684 | 2259.911165 | 1445.590392 | 0.34416411 | 0.275671337 | 0.087950308 | 0.167365195 | ns | NaCl | KS.1 |
| FIM1_286 | 2929.243077 | 3142.818364 | 3409.995974 | 0.077930272 | 0.163113648 | 0.584376858 | 0.705649239 | ns | NaCl | KS.1 |
| FIM1_287 | 797.9487913 | 732.9746665 | 685.4018079 | -0.10962122 | 0.11166514 | 0.293201857 | 0.424528301 | ns | NaCl | KS.1 |
| FIM1_288 | 168.1614266 | 138.3168597 | 132.9791323 | -0.201865685 | 0.186050747 | 0.19778306 | 0.317000877 | ns | NaCl | KS.1 |
| FIM1_290 | 443.5442144 | 550.0647151 | 401.1793757 | 0.248838953 | 0.159717222 | 0.075188646 | 0.147362515 | ns | NaCl | KS.1 |
| FIM1_291 | 1837.940688 | 1396.083995 | 1665.079634 | -0.328175008 | 0.158625068 | 0.018283355 | 0.046149007 | ns | NaCl | KS.1 |
| FIM1_292 | 429.0088917 | 448.7216402 | 1255.979238 | 0.038996373 | 0.239191634 | 0.831564285 | 0.896043387 | ns | NaCl | KS.1 |
| FIM1_293 | 2627.844804 | 2693.848042 | 3668.649036 | 0.029966034 | 0.125010894 | 0.794499869 | 0.868711396 | ns | NaCl | KS.1 |
| FIM1_294 | 265.8183156 | 172.3624946 | 2209.438311 | -0.319903105 | 0.32160313 | 0.136679352 | 0.236844227 | ns | NaCl | KS.1 |
| FIM1_295 | 23.01235476 | 25.91180738 | 54.48768277 | 0.088988615 | 0.302504503 | 0.66099711 | 0.767256328 | ns | NaCl | KS.1 |
| FIM1_296 | 48.67289289 | 104.8115984 | 99.09863344 | 0.693574267 | 0.348250139 | 0.004120191 | 0.013896914 | ns | NaCl | KS.1 |
| FIM1_297 | 20.32424487 | 17.69312699 | 31.93400973 | -0.08794189 | 0.316997475 | 0.668385839 | 0.774207113 | ns | NaCl | KS.1 |
| FIM1_298 | 20.35569058 | 24.51290928 | 43.81990865 | 0.122611016 | 0.280203036 | 0.53612905 | 0.663658129 | ns | NaCl | KS.1 |
| FIM1_299 | 478.562073 | 579.0738004 | 625.4742574 | 0.17686937 | 0.228416941 | 0.323830608 | 0.457037524 | ns | NaCl | KS.1 |
| FIM1_300 | 2070.63462 | 3846.379916 | 1750.706662 | 0.538623458 | 0.330923705 | 0.018218481 | 0.0460086 | ns | NaCl | KS.1 |
| FIM1_301 | 172.7899041 | 134.8764327 | 194.6034244 | -0.228051635 | 0.236263863 | 0.213412693 | 0.33609628 | ns | NaCl | KS.1 |
| FIM1_302 | 779.680977 | 479.5075426 | 737.3037931 | -0.606509491 | 0.175291516 | 7.6482488910: | 0.000500658 | ns | NaCl | KS.1 |
| FIM1_303 | 24200.58633 | 43583.28177 | 31660.2633 | 0.779968572 | 0.14843767 | 1.2310458406( | 2.5732996037( | ns | NaCl | KS.1 |
| FIM1_304 | 408.8615114 | 348.940047 | 807.8463057 | -0.161487911 | 0.194815146 | 0.318389753 | 0.451536209 | ns | NaCl | KS.1 |
| FIM1_305 | 235.1723576 | 83.34116779 | 843.7830533 | -1.065279637 | 0.37382996 | 8.0546658766: | 0.000520362 | Km<KS.1 | NaCl | KS.1 |
| FIM1_306 | 121.72755 | 113.0272334 | 334.8470566 | -0.042312034 | 0.31672152 | 0.83608914 | 0.899555249 | ns | NaCl | KS.1 |
| FIM1_307 | 1065.570259 | 1261.045795 | 1116.511687 | 0.211790998 | 0.121831661 | 0.059042065 | 0.121528454 | ns | NaCl | KS.1 |
| FIM1_308 | 2733.845057 | 3858.827121 | 2450.767558 | 0.430044629 | 0.149813418 | 0.001243371 | 0.005137682 | ns | NaCl | KS.1 |
| FIM1_309 | 4321.582681 | 5578.761128 | 4752.360542 | 0.347168167 | 0.085130889 | 1.9712778121! | 0.000153958 | ns | NaCl | KS.1 |
| FIM1_310 | 3717.4146 | 7119.190198 | 4156.181608 | 0.854266691 | 0.162474124 | 9.7040844881 | 2.0899489319 | ns | NaCl | KS.1 |
| FIM1_311 | 3108.206132 | 3634.995989 | 3860.09946 | 0.203951663 | 0.103388102 | 0.035956432 | 0.081015964 | ns | NaCl | KS.1 |
| FIM1_312 | 3712.983253 | 2984.093668 | 3656.8811 | -0.23961575 | 0.180846944 | 0.118693405 | 0.212562884 | ns | NaCl | KS.1 |
| FIM1_313 | 3331.86861 | 3429.010247 | 2693.26069 | 0.031148914 | 0.165420253 | 0.828780596 | 0.894593939 | ns | NaCl | KS.1 |
| FIM1_314 | 2035.510336 | 1360.041611 | 2126.457402 | -0.453308939 | 0.208098031 | 0.0081328 | 0.024335617 | ns | NaCl | KS.1 |
| FIM1_315 | 585.9791749 | 399.9681661 | 659.2698316 | -0.513626082 | 0.10825838 | 4.1214868336 | 5.4971573719 | ns | NaCl | KS.1 |
| FIM1_316 | 1581.578351 | 2091.418083 | 1101.901215 | 0.211886129 | 0.288869509 | 0.295545095 | 0.42754779 | ns | NaCl | KS.1 |
| FIM1_317 | 67.84290279 | 22.16996088 | 101.9596112 | -1.177512891 | 0.375614406 | 1.9827350301 | 0.000154493 | Km<KS.1 | NaCl | KS.1 |
| FIM1_318 | 8685.83036 | 11622.68151 | 7843.167884 | 0.409245219 | 0.058076092 | 5.6076785046! | 3.7197600747 | ns | NaCl | KS.1 |
| FIM1_319 | 348.2616434 | 269.7855492 | 301.5180789 | -0.306148247 | 0.159616072 | 0.028458161 | 0.066656945 | ns | NaCl | KS.1 |
| FIM1_320 | 1439.902967 | 1364.106868 | 1810.685389 | -0.051933391 | 0.201813389 | 0.753919084 | 0.839505303 | ns | NaCl | KS.1 |

| | | | | | | | | | | |
|---|---|---|---|---|---|---|---|---|---|---|
| FIM1_321 | 136.119846 | 168.5275992 | 107.0083954 | 0.160587576 | 0.288308017 | 0.424655062 | 0.560982192 | ns | NaCl | KS.1 |
| FIM1_322 | 17267.4081 | 23126.02316 | 14723.54858 | 0.388309489 | 0.104969598 | 7.7968600901 | 0.000507714 | ns | NaCl | KS.1 |
| FIM1_323 | 901.6574386 | 251.7313247 | 571.029106 | -1.667565237 | 0.25686494 | 9.7254715581 | 8.064036833 | Km<KS.1 | NaCl | KS.1 |
| FIM1_324 | 13613.8611 | 5546.246708 | 5869.511829 | -1.107601776 | 0.244127894 | 1.2617667389 | 1.9739904170 | Km<KS.1 | NaCl | KS.1 |
| FIM1_325 | 26.46106418 | 14.28386115 | 17.19711774 | -0.418269618 | 0.374012428 | 0.069617515 | 0.138372807 | ns | NaCl | KS.1 |
| FIM1_326 | 206.639608 | 217.7980809 | 4974.569664 | 0.035237525 | 0.297611514 | 0.861036097 | 0.915702134 | ns | NaCl | KS.1 |
| FIM1_327 | 1365.103764 | 1134.477878 | 1256.923338 | -0.238772571 | 0.113662114 | 0.023751638 | 0.057500924 | ns | NaCl | KS.1 |
| FIM1_328 | 179.9528036 | 163.4961809 | 197.3835281 | -0.074806828 | 0.259596586 | 0.695599024 | 0.796273618 | ns | NaCl | KS.1 |
| FIM1_329 | 478.555123 | 331.2213248 | 674.5158635 | -0.36789886 | 0.238053476 | 0.047848572 | 0.101859926 | ns | NaCl | KS.1 |
| FIM1_330 | 13368.58986 | 12404.18374 | 10680.29304 | -0.086720119 | 0.147162451 | 0.509583245 | 0.640093083 | ns | NaCl | KS.1 |
| FIM1_331 | 501.6428595 | 277.2798299 | 526.2211276 | -0.632180344 | 0.265770849 | 0.002198022 | 0.008265425 | ns | NaCl | KS.1 |
| FIM1_332 | 1481.260139 | 839.4774272 | 1159.272756 | -0.719533224 | 0.178381936 | 5.2104225289 | 4.8817047234 | ns | NaCl | KS.1 |
| FIM1_333 | 968.8483518 | 722.2075784 | 1067.606057 | -0.354674959 | 0.156943496 | 0.010136299 | 0.02887271 | ns | NaCl | KS.1 |
| FIM1_334 | 1715.714618 | 2460.577654 | 1901.909592 | 0.448811856 | 0.15447155 | 0.001022265 | 0.004365468 | ns | NaCl | KS.1 |
| FIM1_335 | 1731.21869 | 2081.569035 | 1214.735243 | 0.152133274 | 0.256859496 | 0.425643486 | 0.561989475 | ns | NaCl | KS.1 |
| FIM1_336 | 416.1695274 | 228.3640249 | 665.6980096 | -0.718123528 | 0.217579013 | 8.955377572 | 0.000569004 | ns | NaCl | KS.1 |
| FIM1_337 | 4730.339095 | 3166.823183 | 3669.238901 | -0.51507266 | 0.144216443 | 7.234991814 | 0.000480291 | ns | NaCl | KS.1 |
| FIM1_338 | 2221.265485 | 3207.270746 | 2051.336962 | 0.332121114 | 0.265295398 | 0.092404623 | 0.174133712 | ns | NaCl | KS.1 |
| FIM1_339 | 2782.153488 | 5091.764613 | 3976.175577 | 0.783372798 | 0.169499761 | 3.0245027985 | 4.238563781 | ns | NaCl | KS.1 |
| FIM1_340 | 173.9404272 | 237.1486935 | 277.3983359 | 0.248237102 | 0.283667977 | 0.218998892 | 0.343120856 | ns | NaCl | KS.1 |
| FIM1_341 | 4852.513481 | 6470.106386 | 4523.907596 | 0.374251066 | 0.117950107 | 0.000585648 | 0.002756479 | ns | NaCl | KS.1 |
| FIM1_342 | 1200.250402 | 1173.183562 | 1112.983384 | -0.027125921 | 0.138374662 | 0.828464062 | 0.894446334 | ns | NaCl | KS.1 |
| FIM1_343 | 2156.219245 | 2164.46884 | 2331.160854 | 0.003719098 | 0.20701544 | 0.982379937 | 0.991175946 | ns | NaCl | KS.1 |
| FIM1_344 | 2231.818715 | 1639.036657 | 1776.822187 | -0.39915991 | 0.124348082 | 0.000459594 | 0.002246053 | ns | NaCl | KS.1 |
| FIM1_345 | 4626.605202 | 4694.93344 | 3734.663347 | 0.014533246 | 0.198779582 | 0.929397334 | 0.959682801 | ns | NaCl | KS.1 |
| FIM1_346 | 8305.683492 | 7086.536778 | 6352.248537 | -0.186109439 | 0.149256641 | 0.160913582 | 0.26963458 | ns | NaCl | KS.1 |
| FIM1_347 | 2339.891295 | 1965.961135 | 1962.428707 | -0.221044287 | 0.117057437 | 0.041387229 | 0.090544735 | ns | NaCl | KS.1 |
| FIM1_348 | 2612.570357 | 4798.003765 | 2060.195225 | 0.579667648 | 0.306158664 | 0.008902749 | 0.026171466 | ns | NaCl | KS.1 |
| FIM1_349 | 2157.324973 | 2334.873937 | 2501.27156 | 0.098469161 | 0.114624721 | 0.356144761 | 0.491626023 | ns | NaCl | KS.1 |
| FIM1_350 | 447.0846021 | 386.6074905 | 551.4005951 | -0.184452799 | 0.118693211 | 0.092915428 | 0.174897561 | ns | NaCl | KS.1 |
| FIM1_351 | 1601.663318 | 1641.449703 | 1800.110609 | 0.033145595 | 0.074285367 | 0.645191998 | 0.754300314 | ns | NaCl | KS.1 |
| FIM1_352 | 4306.268839 | 4806.00965 | 5972.361674 | 0.089135687 | 0.25396703 | 0.637723009 | 0.748471149 | ns | NaCl | KS.1 |
| FIM1_354 | 251.2869159 | 148.2622884 | 493.2727055 | -0.47731178 | 0.299500768 | 0.026344993 | 0.062711168 | ns | NaCl | KS.1 |
| FIM1_355 | 3174.81517 | 2741.946938 | 2709.148435 | -0.183941029 | 0.11951596 | 0.095789355 | 0.179020302 | ns | NaCl | KS.1 |
| FIM1_356 | 1235.746608 | 730.173099 | 2103.957352 | -0.593880579 | 0.233342343 | 0.001642036 | 0.006493744 | ns | NaCl | KS.1 |
| FIM1_357 | 2513.943238 | 3035.49763 | 3991.281274 | 0.163246354 | 0.245632492 | 0.381923184 | 0.517025263 | ns | NaCl | KS.1 |
| FIM1_358 | 7885.2717 | 13464.84102 | 6074.080717 | 0.514147981 | 0.287403311 | 0.015143827 | 0.039445308 | ns | NaCl | KS.1 |
| FIM1_359 | 7720.600429 | 9898.420436 | 7789.665494 | 0.351104392 | 0.051388837 | 3.443027775 | 1.9032292426 | ns | NaCl | KS.1 |
| FIM1_360 | 1557.179736 | 1518.397463 | 1375.551515 | -0.035454918 | 0.096866923 | 0.699895839 | 0.798938005 | ns | NaCl | KS.1 |
| FIM1_361 | 659.2702388 | 517.8966124 | 560.8953301 | -0.274164018 | 0.173950855 | 0.066186661 | 0.132613224 | ns | NaCl | KS.1 |
| FIM1_362 | 2180.3217 | 2091.499294 | 2519.03007 | -0.044650649 | 0.172502415 | 0.763769703 | 0.847402827 | ns | NaCl | KS.1 |
| FIM1_363 | 952.5314758 | 1223.523154 | 1212.245779 | 0.30421307 | 0.147287086 | 0.020313914 | 0.050304491 | ns | NaCl | KS.1 |
| FIM1_364 | 2698.640059 | 2639.444315 | 2836.126551 | -0.030483708 | 0.068674878 | 0.648325917 | 0.75625356 | ns | NaCl | KS.1 |
| FIM1_365 | 1663.238014 | 2055.363312 | 1520.484683 | 0.238399229 | 0.170988265 | 0.105653506 | 0.193173904 | ns | NaCl | KS.1 |
| FIM1_366 | 1277.299464 | 1484.909457 | 1946.325552 | 0.173142348 | 0.154675705 | 0.204859299 | 0.326031271 | ns | NaCl | KS.1 |

| | | | | | | | | | | |
|---|---|---|---|---|---|---|---|---|---|---|
| FIM1_367 | 6718.351103 | 4535.237337 | 4689.156278 | -0.359604211 | 0.267725275 | 0.070731821 | 0.139916823 | ns | NaCl | KS.1 |
| FIM1_368 | 370.0382833 | 511.6205547 | 489.1536025 | 0.277763924 | 0.269294173 | 0.160024953 | 0.268597889 | ns | NaCl | KS.1 |
| FIM1_369 | 35.32801507 | 35.42241762 | 35.23365655 | -0.003960922 | 0.315284152 | 0.984515867 | 0.992294659 | ns | NaCl | KS.1 |
| FIM1_370 | 782.1475417 | 653.7568871 | 767.2498702 | -0.224894315 | 0.127219895 | 0.053231164 | 0.111177599 | ns | NaCl | KS.1 |
| FIM1_371 | 92.77247414 | 79.31528669 | 98.26239274 | -0.134978437 | 0.24903413 | 0.472309632 | 0.60603485 | ns | NaCl | KS.1 |
| FIM1_372 | 16087.12031 | 19894.55145 | 15260.98212 | 0.218874753 | 0.200678901 | 0.185246337 | 0.301275098 | ns | NaCl | KS.1 |
| FIM1_373 | 1260.668454 | 1315.328229 | 1422.934025 | 0.045743946 | 0.172622402 | 0.758281889 | 0.843194545 | ns | NaCl | KS.1 |
| FIM1_374 | 2807.322947 | 1604.344204 | 1849.472422 | -0.75077328 | 0.134874701 | 2.36414761988 | 6.12503143894 | ns | NaCl | KS.1 |
| FIM1_375 | 1666.7317 | 1367.731737 | 1675.434603 | -0.200085539 | 0.204219154 | 0.231222313 | 0.357245655 | ns | NaCl | KS.1 |
| FIM1_376 | 657.0530642 | 278.7841543 | 754.4049239 | -0.952311697 | 0.300214445 | 5.24437305053 | 0.000361368 | ns | NaCl | KS.1 |
| FIM1_377 | 882.2365329 | 450.1464056 | 809.5753069 | -0.824963025 | 0.214798533 | 7.98214340728 | 7.04098642752 | ns | NaCl | KS.1 |
| FIM1_378 | 36880.17035 | 73465.2327 | 46593.75577 | 0.929191833 | 0.14232307 | 3.74354731352 | 2.02266172262 | ns | NaCl | KS.1 |
| FIM1_379 | 460.5409594 | 561.457316 | 566.4716419 | 0.154214735 | 0.272604431 | 0.432369874 | 0.568907729 | ns | NaCl | KS.1 |
| FIM1_380 | 1844.358069 | 1715.735529 | 2248.476043 | -0.064975158 | 0.225157777 | 0.714013634 | 0.809530043 | ns | NaCl | KS.1 |
| FIM1_381 | 2266.177784 | 4616.457694 | 2891.447937 | 0.759647396 | 0.291913754 | 0.000681656 | 0.003125567 | ns | NaCl | KS.1 |
| FIM1_383 | 1064.824365 | 3169.803234 | 5227.277974 | 1.200518542 | 0.355689125 | 8.11726475597 | 7.13487494009 | Km>KS.1 | NaCl | KS.1 |
| FIM1_384 | 56340.41158 | 78737.90822 | 47056.6348 | 0.363311014 | 0.205757324 | 0.031407011 | 0.072438517 | ns | NaCl | KS.1 |
| FIM1_385 | 579.9482256 | 674.8459189 | 715.9322575 | 0.165531897 | 0.170914254 | 0.261421992 | 0.389959587 | ns | NaCl | KS.1 |
| FIM1_386 | 2027.767182 | 1448.890874 | 2795.587424 | -0.313773336 | 0.25113944 | 0.100527855 | 0.185782347 | ns | NaCl | KS.1 |
| FIM1_387 | 798.8376464 | 809.4182826 | 897.1630477 | 0.014398486 | 0.145981567 | 0.91229963 | 0.948722964 | ns | NaCl | KS.1 |
| FIM1_388 | 535.3463738 | 455.1644679 | 624.7718793 | -0.191222738 | 0.147082046 | 0.144939899 | 0.247707317 | ns | NaCl | KS.1 |
| FIM1_389 | 3192.220472 | 3264.806827 | 3232.834531 | 0.026132027 | 0.148355194 | 0.843553865 | 0.904262115 | ns | NaCl | KS.1 |
| FIM1_390 | 397.1885795 | 290.3894268 | 386.9148257 | -0.332284313 | 0.21078393 | 0.052509471 | 0.110039856 | ns | NaCl | KS.1 |
| FIM1_391 | 1192.250947 | 1132.366839 | 1297.739817 | -0.063180672 | 0.122891174 | 0.577277644 | 0.699113019 | ns | NaCl | KS.1 |
| FIM1_392 | 4609.734124 | 4720.74753 | 4038.265816 | 0.028930293 | 0.12620641 | 0.8030112 | 0.874175212 | ns | NaCl | KS.1 |
| FIM1_393 | 1984.777307 | 2210.258583 | 1931.746616 | 0.143315439 | 0.090052733 | 0.095484562 | 0.178652011 | ns | NaCl | KS.1 |
| FIM1_394 | 204.5584849 | 246.8584855 | 215.3585894 | 0.180396676 | 0.217454868 | 0.299799111 | 0.432193734 | ns | NaCl | KS.1 |
| FIM1_395 | 1096.588247 | 1168.978677 | 1365.180795 | 0.075529653 | 0.141781095 | 0.554023886 | 0.678048914 | ns | NaCl | KS.1 |
| FIM1_396 | 2790.536935 | 3540.519518 | 2343.481647 | 0.299726891 | 0.127374299 | 0.009951166 | 0.028435986 | ns | NaCl | KS.1 |
| FIM1_397 | 52.92984378 | 40.18287092 | 44.24618426 | -0.227638507 | 0.278153323 | 0.25425277 | 0.382609659 | ns | NaCl | KS.1 |
| FIM1_398 | 1911.517473 | 1980.608304 | 2180.258315 | 0.042668617 | 0.122827005 | 0.706538649 | 0.802969348 | ns | NaCl | KS.1 |
| FIM1_399 | 2845.647582 | 2816.716227 | 2774.821104 | -0.012281776 | 0.133706534 | 0.919626349 | 0.954347327 | ns | NaCl | KS.1 |
| FIM1_401 | 1149.102528 | 1479.831618 | 1422.118783 | 0.317781646 | 0.131802913 | 0.007909077 | 0.023818196 | ns | NaCl | KS.1 |
| FIM1_402 | 222.2215937 | 152.4030617 | 416.4323931 | -0.416650665 | 0.211361913 | 0.015790422 | 0.04072439 | ns | NaCl | KS.1 |
| FIM1_403 | 644.139739 | 474.9115011 | 967.7019184 | -0.351378175 | 0.174208141 | 0.018789355 | 0.047115444 | ns | NaCl | KS.1 |
| FIM1_404 | 391.5598473 | 416.7531344 | 546.760275 | 0.06367468 | 0.199616692 | 0.698674456 | 0.797952697 | ns | NaCl | KS.1 |
| FIM1_405 | 8651.385935 | 10862.97729 | 7626.635991 | 0.307654881 | 0.086259168 | 0.000187579 | 0.001058055 | ns | NaCl | KS.1 |
| FIM1_406 | 10867.70623 | 13524.07394 | 9920.880166 | 0.297104694 | 0.081302516 | 0.000139714 | 0.000830439 | ns | NaCl | KS.1 |
| FIM1_407 | 3455.140217 | 2587.986165 | 3459.852175 | -0.332797977 | 0.174626745 | 0.026262571 | 0.062551756 | ns | NaCl | KS.1 |
| FIM1_408 | 1568.447881 | 1481.518302 | 1724.87704 | -0.07522245 | 0.085570075 | 0.359578171 | 0.495401439 | ns | NaCl | KS.1 |
| FIM1_409 | 4596.641173 | 4416.618707 | 4668.864928 | -0.049272382 | 0.117020341 | 0.650371571 | 0.758284173 | ns | NaCl | KS.1 |
| FIM1_410 | 172.6303036 | 83.64440136 | 198.6579872 | -0.847265747 | 0.243846975 | 2.93104817546 | 0.000217641 | ns | NaCl | KS.1 |
| FIM1_411 | 1220.329667 | 1132.143213 | 2144.197744 | -0.059331997 | 0.254477429 | 0.753951697 | 0.839505303 | ns | NaCl | KS.1 |
| FIM1_412 | 735.5242105 | 746.2788577 | 728.237725 | 0.013502297 | 0.120546541 | 0.903651997 | 0.942559044 | ns | NaCl | KS.1 |
| FIM1_413 | 7892.296145 | 5192.230419 | 4930.833769 | -0.576356555 | 0.094961386 | 1.92164229151 | 6.73251436639 | ns | NaCl | KS.1 |

| | | | | | | | | | |
|---|---|---|---|---|---|---|---|---|---|
| FIM1_414 | 447.8534909 | 263.624532 | 811.4516496 | -0.526762774 | 0.278851209 | 0.01156009 | 0.031986345 | ns | NaCl | KS.1 |
| FIM1_415 | 391.3511593 | 297.9224962 | 421.903523 | -0.314942699 | 0.169857828 | 0.031698928 | 0.072945429 | ns | NaCl | KS.1 |
| FIM1_416 | 1036.714008 | 1122.867614 | 1370.58872 | 0.097112996 | 0.118023942 | 0.374922144 | 0.509600707 | ns | NaCl | KS.1 |
| FIM1_417 | 5952.297117 | 4712.741488 | 6188.655595 | -0.282103123 | 0.145539196 | 0.029837416 | 0.069429909 | ns | NaCl | KS.1 |
| FIM1_418 | 569.5829296 | 613.5552273 | 901.0270708 | 0.083609661 | 0.168105083 | 0.566134501 | 0.688055153 | ns | NaCl | KS.1 |
| FIM1_419 | 640.6119217 | 609.697787 | 844.6077708 | -0.058575492 | 0.140945947 | 0.644779032 | 0.754061044 | ns | NaCl | KS.1 |
| FIM1_420 | 671.6678372 | 780.4971148 | 749.4564911 | 0.183330115 | 0.131005516 | 0.124553331 | 0.21981299 | ns | NaCl | KS.1 |
| FIM1_421 | 878.779116 | 1122.752573 | 1164.997138 | 0.318337607 | 0.109397612 | 0.001796457 | 0.006998727 | ns | NaCl | KS.1 |
| FIM1_422 | 547.972397 | 373.7692595 | 623.0930477 | -0.49647601 | 0.139025248 | 7.84932560488 | 0.000510463 | ns | NaCl | KS.1 |
| FIM1_423 | 23.18015928 | 48.42714655 | 92.95811991 | 0.474735503 | 0.407349647 | 0.046938467 | 0.100395044 | ns | NaCl | KS.1 |
| FIM1_425 | 4094.413447 | 7996.993365 | 3568.612381 | 0.696659075 | 0.29286539 | 0.001645814 | 0.006498354 | ns | NaCl | KS.1 |
| FIM1_426 | 4004.603157 | 5277.36944 | 4407.258022 | 0.356665537 | 0.120309165 | 0.00127588 | 0.005245872 | ns | NaCl | KS.1 |
| FIM1_427 | 379.4509563 | 356.5960759 | 348.6366388 | -0.07384444 | 0.14711025 | 0.574362602 | 0.696430403 | ns | NaCl | KS.1 |
| FIM1_428 | 405.8669468 | 457.5552507 | 345.022864 | 0.127428806 | 0.176987758 | 0.399615692 | 0.535584069 | ns | NaCl | KS.1 |
| FIM1_429 | 2042.090933 | 1697.528019 | 1382.713337 | -0.237320963 | 0.111450844 | 0.022247146 | 0.054521945 | ns | NaCl | KS.1 |
| FIM1_430 | 1133.983491 | 1104.503774 | 1029.537073 | -0.032869634 | 0.1001198 | 0.728682428 | 0.820736944 | ns | NaCl | KS.1 |
| FIM1_431 | 436.8919654 | 608.0162481 | 613.3133998 | 0.365619835 | 0.198606437 | 0.026440909 | 0.062879313 | ns | NaCl | KS.1 |
| FIM1_432 | 246.2844187 | 273.8992046 | 229.5730587 | 0.09742095 | 0.218186143 | 0.575725611 | 0.69791299 | ns | NaCl | KS.1 |
| FIM1_433 | 2469.417514 | 2578.878448 | 1934.95982 | 0.052957563 | 0.123233641 | 0.641196658 | 0.750753912 | ns | NaCl | KS.1 |
| FIM1_434 | 943.5028143 | 1034.997309 | 1025.508772 | 0.112146905 | 0.129555776 | 0.343701039 | 0.478431078 | ns | NaCl | KS.1 |
| FIM1_435 | 1080.213185 | 1080.010751 | 1007.262228 | -0.000250548 | 0.105467092 | 0.997986221 | 0.999191276 | ns | NaCl | KS.1 |
| FIM1_436 | 337.2145688 | 419.0240276 | 391.0584459 | 0.223298247 | 0.202893051 | 0.179763874 | 0.295059477 | ns | NaCl | KS.1 |
| FIM1_437 | 542.6500085 | 353.498798 | 545.3349553 | -0.535292113 | 0.170529086 | 0.000322329 | 0.001661747 | ns | NaCl | KS.1 |
| FIM1_438 | 19.76289557 | 10.70128086 | 26.92615975 | -0.359718706 | 0.399433169 | 0.113956055 | 0.205261179 | ns | NaCl | KS.1 |
| FIM1_439 | 2072.756254 | 3718.156768 | 1895.423913 | 0.613412766 | 0.271788001 | 0.003273694 | 0.011462115 | ns | NaCl | KS.1 |
| FIM1_440 | 991.2432345 | 555.0879231 | 823.75812 | -0.784788061 | 0.132328005 | 2.54381865508 | 8.58013464892 | ns | NaCl | KS.1 |
| FIM1_441 | 691.7532829 | 760.3862109 | 966.3812468 | 0.119908551 | 0.114764028 | 0.261488479 | 0.389959587 | ns | NaCl | KS.1 |
| FIM1_442 | 763.735074 | 685.7313689 | 1211.952098 | -0.115251268 | 0.173550792 | 0.439699427 | 0.575205009 | ns | NaCl | KS.1 |
| FIM1_443 | 93.71345325 | 94.16805248 | 176.328668 | 0.006621866 | 0.260702054 | 0.972386331 | 0.987204262 | ns | NaCl | KS.1 |
| FIM1_444 | 1275.133186 | 1344.712894 | 1310.064377 | 0.063767482 | 0.137558941 | 0.608674405 | 0.725132942 | ns | NaCl | KS.1 |
| FIM1_445 | 1582.352104 | 1460.756681 | 1874.845332 | -0.098174236 | 0.125365406 | 0.394240594 | 0.530237079 | ns | NaCl | KS.1 |
| FIM1_446 | 261.5354635 | 202.8430773 | 323.4614719 | -0.284603596 | 0.180484804 | 0.063513941 | 0.128604744 | ns | NaCl | KS.1 |
| FIM1_447 | 799.9099326 | 825.5547872 | 917.951041 | 0.043864265 | 0.089519333 | 0.608369796 | 0.724943649 | ns | NaCl | KS.1 |
| FIM1_448 | 1777.374027 | 2265.530381 | 2575.742673 | 0.304204824 | 0.130712924 | 0.010471912 | 0.029668428 | ns | NaCl | KS.1 |
| FIM1_449 | 3682.056124 | 3771.911806 | 4280.172542 | 0.025396074 | 0.177422684 | 0.866999868 | 0.92047041 | ns | NaCl | KS.1 |
| FIM1_450 | 143.5003778 | 117.4204651 | 132.0429841 | -0.19220613 | 0.22623571 | 0.281131322 | 0.411139031 | ns | NaCl | KS.1 |
| FIM1_451 | 2294.463183 | 2731.559019 | 2000.966981 | 0.17117823 | 0.20999825 | 0.314357285 | 0.447675071 | ns | NaCl | KS.1 |
| FIM1_452 | 677.0472093 | 600.3803968 | 780.8864883 | -0.133468887 | 0.158841382 | 0.338713548 | 0.472546243 | ns | NaCl | KS.1 |
| FIM1_453 | 2744.790009 | 3359.185704 | 2775.856282 | 0.233636726 | 0.15920011 | 0.094032318 | 0.176465779 | ns | NaCl | KS.1 |
| FIM1_454 | 200.8308685 | 187.6893071 | 268.9446485 | -0.045916928 | 0.288320922 | 0.817819774 | 0.887402992 | ns | NaCl | KS.1 |
| FIM1_455 | 1757.513613 | 1384.340054 | 1245.188269 | -0.242031908 | 0.209555584 | 0.154882677 | 0.261732784 | ns | NaCl | KS.1 |
| FIM1_456 | 1956.122314 | 1191.938276 | 1442.054858 | -0.559564897 | 0.225363198 | 0.002267594 | 0.008469428 | ns | NaCl | KS.1 |
| FIM1_457 | 16712.84778 | 22406.18646 | 14320.3747 | 0.369378904 | 0.13608062 | 0.002647887 | 0.009670084 | ns | NaCl | KS.1 |
| FIM1_458 | 23905.016 | 33602.33091 | 20648.47836 | 0.444790803 | 0.123455853 | 8.7497621982 | 0.000559512 | ns | NaCl | KS.1 |
| FIM1_459 | 2212.846503 | 1994.157845 | 1542.097326 | -0.111271812 | 0.177920088 | 0.463788986 | 0.598224061 | ns | NaCl | KS.1 |

| | | | | | | | | | | |
|---|---|---|---|---|---|---|---|---|---|---|
| FIM1_460 | 374.5330812 | 439.6966044 | 413.5934671 | 0.178065169 | 0.17028129 | 0.225669444 | 0.350516855 | ns | NaCl | KS.1 |
| FIM1_461 | 789.2922304 | 631.7772053 | 842.2776394 | -0.275408635 | 0.136032575 | 0.02503366 | 0.060078368 | ns | NaCl | KS.1 |
| FIM1_462 | 797.0663117 | 675.1964428 | 709.7308589 | -0.22155058 | 0.088595872 | 0.008786544 | 0.025896362 | ns | NaCl | KS.1 |
| FIM1_463 | 2314.379001 | 3619.769619 | 2168.265916 | 0.472450536 | 0.239320874 | 0.012247591 | 0.033460607 | ns | NaCl | KS.1 |
| FIM1_464 | 5945.33767 | 5175.671407 | 5385.754652 | -0.159694552 | 0.153386471 | 0.239293423 | 0.365965195 | ns | NaCl | KS.1 |
| FIM1_465 | 51.25103855 | 18.67557014 | 27.27999917 | -0.865211632 | 0.422825394 | 0.001499465 | 0.005982228 | ns | NaCl | KS.1 |
| FIM1_466 | 7831.223262 | 12145.41426 | 7328.141002 | 0.589120453 | 0.119039866 | 1.0909059209 | 1.7229387163 | ns | NaCl | KS.1 |
| FIM1_467 | 3397.441842 | 4091.227862 | 2370.570354 | 0.189014053 | 0.200729116 | 0.252527912 | 0.380935828 | ns | NaCl | KS.1 |
| FIM1_468 | 131.0746524 | 57.24574958 | 101.8914025 | -0.993615596 | 0.265811184 | 6.0185231997 | 5.4921500785 | ns | NaCl | KS.1 |
| FIM1_469 | 1076.26222 | 787.8236277 | 815.2713542 | -0.32783123 | 0.212542542 | 0.057081804 | 0.118079823 | ns | NaCl | KS.1 |
| FIM1_470 | 1357.947919 | 821.6136108 | 1474.85134 | -0.566707942 | 0.22695981 | 0.002114146 | 0.00797413 | ns | NaCl | KS.1 |
| FIM1_471 | 633.1263808 | 601.6177995 | 924.8653515 | -0.040474258 | 0.253810923 | 0.830411336 | 0.895382834 | ns | NaCl | KS.1 |
| FIM1_472 | 527.5344557 | 567.8251669 | 470.3324537 | 0.065923718 | 0.225702989 | 0.710419234 | 0.806005858 | ns | NaCl | KS.1 |
| FIM1_473 | 1346.677663 | 1224.850956 | 1561.354367 | -0.100344335 | 0.180307191 | 0.512899329 | 0.643711948 | ns | NaCl | KS.1 |
| FIM1_474 | 2041.822884 | 2119.531502 | 1664.142884 | 0.04569937 | 0.122584858 | 0.686256698 | 0.787936089 | ns | NaCl | KS.1 |
| FIM1_475 | 496.7254473 | 300.9909711 | 707.7649955 | -0.542411985 | 0.244511239 | 0.00495699 | 0.01620072 | ns | NaCl | KS.1 |
| FIM1_476 | 11467.21445 | 13550.42077 | 9708.260431 | 0.192828865 | 0.155730785 | 0.160019285 | 0.268597889 | ns | NaCl | KS.1 |
| FIM1_477 | 9728.652338 | 11654.44966 | 8762.515758 | 0.235917841 | 0.102830092 | 0.014696457 | 0.038633841 | ns | NaCl | KS.1 |
| FIM1_478 | 4313.691667 | 8217.67925 | 3891.826898 | 0.643279886 | 0.30133527 | 0.003842114 | 0.013128102 | ns | NaCl | KS.1 |
| FIM1_479 | 1273.999976 | 1403.113942 | 1402.401075 | 0.093730927 | 0.205519495 | 0.576125703 | 0.698057811 | ns | NaCl | KS.1 |
| FIM1_480 | 1292.469135 | 760.7791567 | 1130.206243 | -0.71072887 | 0.133036279 | 9.2335813665 | 2.0059854715 | ns | NaCl | KS.1 |
| FIM1_481 | 780.5215149 | 653.9441016 | 1343.131751 | -0.222843531 | 0.121363972 | 0.046207921 | 0.098960142 | ns | NaCl | KS.1 |
| FIM1_482 | 922.7518716 | 662.8914969 | 1188.457089 | -0.407297649 | 0.157305454 | 0.003248468 | 0.011413228 | ns | NaCl | KS.1 |
| FIM1_483 | 1951.633112 | 2336.195668 | 2054.337398 | 0.239980705 | 0.091537314 | 0.00585086 | 0.018675443 | ns | NaCl | KS.1 |
| FIM1_485 | 138.0434435 | 73.42717226 | 116.1817359 | -0.743709111 | 0.240738676 | 0.000166431 | 0.000958327 | ns | NaCl | KS.1 |
| FIM1_486 | 736.045542 | 454.8007261 | 1003.7575 | -0.588194114 | 0.187362729 | 0.000259319 | 0.001391708 | ns | NaCl | KS.1 |
| FIM1_487 | 821.2560685 | 762.3922633 | 668.4113705 | -0.076388466 | 0.189066297 | 0.629988785 | 0.742348225 | ns | NaCl | KS.1 |
| FIM1_488 | 5921.432844 | 4549.45162 | 5026.573853 | -0.296729055 | 0.17932277 | 0.051940602 | 0.109123519 | ns | NaCl | KS.1 |
| FIM1_489 | 3576.048636 | 3979.566962 | 3569.253078 | 0.098834745 | 0.220162542 | 0.572308497 | 0.69427817 | ns | NaCl | KS.1 |
| FIM1_490 | 1527.843408 | 1228.091929 | 1831.003238 | -0.273751982 | 0.127566105 | 0.018766306 | 0.047081377 | ns | NaCl | KS.1 |
| FIM1_491 | 15172.01548 | 20749.51608 | 12395.74351 | 0.424397981 | 0.093741687 | 1.7818649026 | 1.9483028331 | ns | NaCl | KS.1 |
| FIM1_492 | 3095.486876 | 1949.952048 | 2334.251937 | -0.637306626 | 0.098130657 | 1.0286132109 | 4.9205295428 | ns | NaCl | KS.1 |
| FIM1_493 | 1494.55726 | 1651.348529 | 1321.065216 | 0.119858008 | 0.13218155 | 0.319575478 | 0.452830533 | ns | NaCl | KS.1 |
| FIM1_494 | 2396.395229 | 2713.720674 | 1974.82989 | 0.15772561 | 0.113269932 | 0.134879439 | 0.234296511 | ns | NaCl | KS.1 |
| FIM1_495 | 2418.348994 | 3425.443854 | 3267.866095 | 0.47049557 | 0.101577449 | 8.6741861517 | 1.0348699305 | ns | NaCl | KS.1 |
| FIM1_496 | 6569.863691 | 6271.058131 | 5435.532296 | -0.064064984 | 0.064622067 | 0.309727234 | 0.442531013 | ns | NaCl | KS.1 |
| FIM1_497 | 2152.286088 | 1215.93318 | 1598.935476 | -0.656843967 | 0.233012577 | 0.000559647 | 0.002654187 | ns | NaCl | KS.1 |
| FIM1_498 | 720.6645348 | 694.6694515 | 630.0746192 | -0.044676945 | 0.150179928 | 0.738236292 | 0.828309777 | ns | NaCl | KS.1 |
| FIM1_499 | 3739.095257 | 4257.076576 | 3312.6156 | 0.165087119 | 0.111381047 | 0.112213713 | 0.202709958 | ns | NaCl | KS.1 |
| FIM1_500 | 498.9842816 | 402.2556088 | 352.4480144 | -0.256367457 | 0.156441726 | 0.062438258 | 0.126995231 | ns | NaCl | KS.1 |
| FIM1_501 | 798.2833319 | 820.4738443 | 642.575356 | 0.029476179 | 0.165418116 | 0.837843478 | 0.900858289 | ns | NaCl | KS.1 |
| FIM1_502 | 3296.178083 | 3153.546382 | 2460.250386 | -0.053319809 | 0.130042161 | 0.653855913 | 0.760920039 | ns | NaCl | KS.1 |
| FIM1_503 | 647.5192513 | 498.7275296 | 959.0798002 | -0.299002807 | 0.174947459 | 0.046065128 | 0.098696817 | ns | NaCl | KS.1 |
| FIM1_504 | 784.9471318 | 797.2627603 | 1291.733053 | 0.025300072 | 0.101344804 | 0.791708625 | 0.86642112 | ns | NaCl | KS.1 |
| FIM1_505 | 1461.484643 | 1486.114067 | 1053.730224 | 0.01817795 | 0.172350128 | 0.902565931 | 0.942343234 | ns | NaCl | KS.1 |

| | | | | | | | | | | |
|---|---|---|---|---|---|---|---|---|---|---|
| FIM1_506 | 4131.64499 | 6384.749485 | 3207.087225 | 0.437618587 | 0.251784411 | 0.023996607 | 0.05792674 | ns | NaCl | KS.1 |
| FIM1_507 | 1851.940859 | 1490.872964 | 1545.803788 | -0.269507907 | 0.134067829 | 0.02647055 | 0.062919725 | ns | NaCl | KS.1 |
| FIM1_508 | 916.6808293 | 676.4778043 | 1038.605976 | -0.376293092 | 0.145619076 | 0.00378119 | 0.012982348 | ns | NaCl | KS.1 |
| FIM1_509 | 567.9168104 | 394.5164122 | 964.2370488 | -0.384337525 | 0.218848403 | 0.029274908 | 0.068280668 | ns | NaCl | KS.1 |
| FIM1_510 | 1030.061985 | 907.3135892 | 854.0455006 | -0.152451487 | 0.132953482 | 0.207402952 | 0.328922438 | ns | NaCl | KS.1 |
| FIM1_511 | 1421.182228 | 1150.42999 | 1599.389843 | -0.222238702 | 0.191174972 | 0.164267477 | 0.274054561 | ns | NaCl | KS.1 |
| FIM1_512 | 394.4596972 | 169.2424141 | 324.3843601 | -1.052193789 | 0.234307218 | 2.02124948128 | 2.97506395543 | Km<KS.1 | NaCl | KS.1 |
| FIM1_513 | 4331.865699 | 4421.988482 | 3189.871274 | 0.025082948 | 0.120975813 | 0.822604395 | 0.890632614 | ns | NaCl | KS.1 |
| FIM1_514 | 1732.358249 | 1222.821958 | 1894.917338 | -0.412312845 | 0.176097394 | 0.006363905 | 0.019899704 | ns | NaCl | KS.1 |
| FIM1_515 | 2032.121205 | 1704.750046 | 1777.12397 | -0.24125148 | 0.076723426 | 0.001111691 | 0.004675116 | ns | NaCl | KS.1 |
| FIM1_516 | 1645.759787 | 1196.346807 | 1352.961096 | -0.422838738 | 0.113150546 | 5.70954497078 | 0.000389109 | ns | NaCl | KS.1 |
| FIM1_517 | 1852.724786 | 2534.246988 | 1368.583645 | 0.289694857 | 0.248682141 | 0.12650915 | 0.222632834 | ns | NaCl | KS.1 |
| FIM1_518 | 30194.12974 | 36628.45865 | 24314.06549 | 0.262593085 | 0.079149467 | 0.000562646 | 0.002663336 | ns | NaCl | KS.1 |
| FIM1_519 | 178.6688776 | 92.57622576 | 587.3415799 | -0.531241893 | 0.355125596 | 0.023414333 | 0.056739556 | ns | NaCl | KS.1 |
| FIM1_522 | 3106.464436 | 4369.690845 | 2603.129516 | 0.33953934 | 0.234321523 | 0.064523236 | 0.130118808 | ns | NaCl | KS.1 |
| FIM1_523 | 721.5536769 | 724.2027955 | 636.3143509 | 0.003139816 | 0.170659733 | 0.983007461 | 0.991175946 | ns | NaCl | KS.1 |
| FIM1_524 | 1259.285987 | 1326.207975 | 1738.364328 | 0.063552836 | 0.118768588 | 0.56376107 | 0.686419805 | ns | NaCl | KS.1 |
| FIM1_525 | 2021.442601 | 1452.317104 | 1775.306378 | -0.462035012 | 0.080007322 | 1.86073328473 | 5.00386383326 | ns | NaCl | KS.1 |
| FIM1_526 | 45.45544854 | 65.49211128 | 136.6789371 | 0.259477509 | 0.306755666 | 0.21408421 | 0.336833949 | ns | NaCl | KS.1 |
| FIM1_527 | 6065.973756 | 15045.68118 | 14481.84218 | 0.906395465 | 0.35894031 | 0.000419012 | 0.002082501 | ns | NaCl | KS.1 |
| FIM1_528 | 3097.313752 | 3140.604424 | 3607.913838 | 0.011313497 | 0.251708012 | 0.952052741 | 0.975182703 | ns | NaCl | KS.1 |
| FIM1_529 | 43.27999842 | 25.57922671 | 26.20148965 | -0.4772221 | 0.294558198 | 0.025115061 | 0.060215628 | ns | NaCl | KS.1 |
| FIM1_530 | 6730.92057 | 4360.198416 | 3718.552015 | -0.541461551 | 0.167357795 | 0.000224418 | 0.001228251 | ns | NaCl | KS.1 |
| FIM1_531 | 15986.23817 | 14116.85727 | 10265.85102 | -0.10882944 | 0.236073373 | 0.550525631 | 0.675698639 | ns | NaCl | KS.1 |
| FIM1_532 | 5618.862584 | 8940.004037 | 4169.976391 | 0.348500447 | 0.324355289 | 0.108186839 | 0.197009342 | ns | NaCl | KS.1 |
| FIM1_533 | 1163.641689 | 1009.14728 | 1149.200053 | -0.180611372 | 0.109946525 | 0.078718068 | 0.153156977 | ns | NaCl | KS.1 |
| FIM1_534 | 1225.524259 | 779.479778 | 1090.027217 | -0.591675453 | 0.138102978 | 2.69411679978 | 2.81580484851 | ns | NaCl | KS.1 |
| FIM1_535 | 1640.036436 | 1667.989272 | 1542.118228 | 0.012752122 | 0.269627097 | 0.947630239 | 0.972655341 | ns | NaCl | KS.1 |
| FIM1_536 | 619.5993056 | 1141.990809 | 1058.344059 | 0.626497506 | 0.286149088 | 0.003646367 | 0.012623993 | ns | NaCl | KS.1 |
| FIM1_537 | 777.3457093 | 631.9922494 | 995.1810894 | -0.179156443 | 0.248173613 | 0.340360785 | 0.474578168 | ns | NaCl | KS.1 |
| FIM1_538 | 995.5082731 | 989.2239912 | 1018.384598 | -0.007604116 | 0.147709113 | 0.954049057 | 0.976222554 | ns | NaCl | KS.1 |
| FIM1_539 | 366.7468024 | 409.6750416 | 440.214433 | 0.129451519 | 0.132315541 | 0.282690673 | 0.412914297 | ns | NaCl | KS.1 |
| FIM1_540 | 1759.502432 | 2094.854161 | 2288.724654 | 0.227994394 | 0.09907495 | 0.014807791 | 0.038773032 | ns | NaCl | KS.1 |
| FIM1_541 | 539.1976731 | 541.385349 | 552.7711887 | -0.000790764 | 0.087548341 | 0.992478594 | 0.996607017 | ns | NaCl | KS.1 |
| FIM1_542 | 862.9453987 | 861.5868798 | 827.8097266 | -0.000108293 | 0.120990779 | 0.999227815 | 0.999786056 | ns | NaCl | KS.1 |
| FIM1_543 | 457.0408968 | 536.2768679 | 548.75533 | 0.162225765 | 0.198205959 | 0.321854632 | 0.454893976 | ns | NaCl | KS.1 |
| FIM1_544 | 2524.097738 | 2517.809608 | 1915.488907 | -0.003737634 | 0.135567403 | 0.975780809 | 0.988899883 | ns | NaCl | KS.1 |
| FIM1_545 | 1593.886502 | 968.7237502 | 1877.281489 | -0.620284401 | 0.179438428 | 7.34353223267 | 0.00048518 | ns | NaCl | KS.1 |
| FIM1_546 | 4450.310806 | 6466.890212 | 4994.01369 | 0.506499607 | 0.102421791 | 1.53639735841 | 2.36643246381 | ns | NaCl | KS.1 |
| FIM1_547 | 1752.748039 | 1770.571697 | 2150.754405 | 0.011185681 | 0.156688427 | 0.93549562 | 0.962782522 | ns | NaCl | KS.1 |
| FIM1_549 | 1139.609559 | 1286.868033 | 1034.980657 | 0.137001457 | 0.162442876 | 0.334365372 | 0.468582458 | ns | NaCl | KS.1 |
| FIM1_550 | 11545.44063 | 20929.83107 | 11820.81753 | 0.80629475 | 0.128328618 | 2.68356844663 | 1.09432401819 | ns | NaCl | KS.1 |
| FIM1_551 | 656.205541 | 647.3221266 | 566.140344 | -0.015161261 | 0.16411446 | 0.915670842 | 0.951632011 | ns | NaCl | KS.1 |
| FIM1_552 | 889.1305823 | 1082.180384 | 1160.952802 | 0.252015037 | 0.114841171 | 0.017982435 | 0.045528049 | ns | NaCl | KS.1 |
| FIM1_553 | 948.8430157 | 1217.011044 | 1591.648014 | 0.329822213 | 0.100981983 | 0.000515191 | 0.002469244 | ns | NaCl | KS.1 |

| | | | | | | | | | |
|---|---|---|---|---|---|---|---|---|---|
| FIM1_554 | 1766.21366 | 1996.733132 | 1887.284174 | 0.158849823 | 0.102008848 | 0.098633827 | 0.182689237 | ns | NaCl | KS.1 |
| FIM1_555 | 497.8371827 | 488.3763575 | 575.5339994 | -0.023695565 | 0.141905041 | 0.852889357 | 0.910932708 | ns | NaCl | KS.1 |
| FIM1_556 | 1419.339345 | 1356.099762 | 1202.338386 | -0.055245874 | 0.123322622 | 0.627055365 | 0.739711913 | ns | NaCl | KS.1 |
| FIM1_557 | 1052.835159 | 1313.651984 | 994.4919767 | 0.290710131 | 0.105563204 | 0.003273619 | 0.011462115 | ns | NaCl | KS.1 |
| FIM1_558 | 1450.745353 | 1937.012138 | 1298.567273 | 0.351609417 | 0.152296987 | 0.009036096 | 0.026428323 | ns | NaCl | KS.1 |
| FIM1_559 | 662.432497 | 809.614691 | 786.9371466 | 0.254492996 | 0.121149031 | 0.022501192 | 0.054913959 | ns | NaCl | KS.1 |
| FIM1_560 | 1795.950774 | 1921.135056 | 2200.518531 | 0.082422231 | 0.123211772 | 0.468038152 | 0.602299484 | ns | NaCl | KS.1 |
| FIM1_561 | 585.6887249 | 804.415376 | 945.6953352 | 0.393183952 | 0.147436483 | 0.002749998 | 0.009957236 | ns | NaCl | KS.1 |
| FIM1_562 | 1596.73802 | 1607.869776 | 1953.332671 | 0.009971888 | 0.099398627 | 0.915651958 | 0.951632011 | ns | NaCl | KS.1 |
| FIM1_563 | 130.0376288 | 173.8789293 | 220.2215981 | 0.276816706 | 0.237862123 | 0.133627643 | 0.23236544 | ns | NaCl | KS.1 |
| FIM1_564 | 1688.209809 | 2154.149521 | 1556.202246 | 0.273857018 | 0.176330848 | 0.069276536 | 0.137860307 | ns | NaCl | KS.1 |
| FIM1_565 | 1404.321665 | 2704.637942 | 1520.594745 | 0.815965839 | 0.203905224 | 4.2897814100 | 4.2177198645 | ns | NaCl | KS.1 |
| FIM1_566 | 1009.510752 | 1185.22657 | 826.6143196 | 0.189898875 | 0.144056272 | 0.141052633 | 0.242312448 | ns | NaCl | KS.1 |
| FIM1_567 | 1709.98187 | 1835.479452 | 1648.299936 | 0.088643475 | 0.113728096 | 0.403027837 | 0.538179129 | ns | NaCl | KS.1 |
| FIM1_568 | 5087.514661 | 3422.531907 | 3256.95093 | -0.289767729 | 0.315644785 | 0.17210852 | 0.284465079 | ns | NaCl | KS.1 |
| FIM1_569 | 3570.101471 | 2625.881776 | 2490.001199 | -0.271793718 | 0.258389032 | 0.159826305 | 0.268536261 | ns | NaCl | KS.1 |
| FIM1_570 | 1050.087446 | 3185.278415 | 1243.278647 | 1.483848274 | 0.199760829 | 3.2380607214 | 4.0273380222 | Km>KS.1 | NaCl | KS.1 |
| FIM1_571 | 1341.709145 | 1249.629867 | 2026.460833 | -0.056553261 | 0.255772772 | 0.765667414 | 0.848940358 | ns | NaCl | KS.1 |
| FIM1_572 | 594.9763405 | 229.7659877 | 510.8466022 | -0.835789292 | 0.407747101 | 0.00174392 | 0.006842272 | ns | NaCl | KS.1 |
| FIM1_573 | 4636.006409 | 5882.892257 | 4434.991654 | 0.297176789 | 0.132939162 | 0.013685004 | 0.036562938 | ns | NaCl | KS.1 |
| FIM1_574 | 3714.038819 | 6742.843074 | 3334.420775 | 0.549221081 | 0.312468275 | 0.013593744 | 0.03641835 | ns | NaCl | KS.1 |
| FIM1_575 | 2996.52484 | 3477.594454 | 2703.430966 | 0.189487783 | 0.11327881 | 0.072261447 | 0.142602419 | ns | NaCl | KS.1 |
| FIM1_576 | 6433.533117 | 8036.705058 | 5823.24603 | 0.297346685 | 0.092421719 | 0.000701183 | 0.003206237 | ns | NaCl | KS.1 |
| FIM1_577 | 1331.545244 | 924.512769 | 905.1260683 | -0.469672939 | 0.139135634 | 0.000184278 | 0.001042984 | ns | NaCl | KS.1 |
| FIM1_578 | 4230.110688 | 3986.682502 | 4422.84059 | -0.072522128 | 0.122764433 | 0.521908426 | 0.652118833 | ns | NaCl | KS.1 |
| FIM1_579 | 630.7364267 | 703.6982392 | 850.9124711 | 0.132680795 | 0.127325652 | 0.255474395 | 0.383895345 | ns | NaCl | KS.1 |
| FIM1_580 | 244.92626 | 202.5662744 | 260.9150765 | -0.234746395 | 0.145539937 | 0.070885248 | 0.140164591 | ns | NaCl | KS.1 |
| FIM1_581 | 674.9744667 | 792.5187134 | 835.7897162 | 0.185461806 | 0.156254744 | 0.177782522 | 0.29248282 | ns | NaCl | KS.1 |
| FIM1_582 | 3638.459213 | 7308.890402 | 3226.125734 | 0.662096902 | 0.327878603 | 0.004549781 | 0.015025839 | ns | NaCl | KS.1 |
| FIM1_583 | 272.3200139 | 322.4552917 | 256.7458051 | 0.172186673 | 0.191574069 | 0.281832499 | 0.411781699 | ns | NaCl | KS.1 |
| FIM1_584 | 875.2595229 | 714.4743486 | 916.6185616 | -0.257574156 | 0.125195268 | 0.024712715 | 0.059422792 | ns | NaCl | KS.1 |
| FIM1_585 | 12685.92606 | 12667.22398 | 10000.39067 | -0.001660664 | 0.170000232 | 0.990986369 | 0.996192602 | ns | NaCl | KS.1 |
| FIM1_586 | 1559.928883 | 3249.645925 | 2018.87692 | 0.920091259 | 0.209822589 | 5.6449505842 | 7.2567517200 | ns | NaCl | KS.1 |
| FIM1_587 | 1008.74082 | 765.4747671 | 743.9692453 | -0.305696273 | 0.190114312 | 0.055028232 | 0.114498308 | ns | NaCl | KS.1 |
| FIM1_588 | 843.0909311 | 989.7051266 | 685.0246758 | 0.183893075 | 0.1550359 | 0.178828873 | 0.293912667 | ns | NaCl | KS.1 |
| FIM1_589 | 747.0138063 | 824.6948832 | 965.9544655 | 0.125107874 | 0.11034303 | 0.225304706 | 0.350059623 | ns | NaCl | KS.1 |
| FIM1_590 | 609.0306908 | 701.0764019 | 896.1242943 | 0.181689334 | 0.106193771 | 0.068205505 | 0.13594647 | ns | NaCl | KS.1 |
| FIM1_591 | 781.4959496 | 694.7410865 | 799.1978164 | -0.144652109 | 0.124747688 | 0.20706302 | 0.328488049 | ns | NaCl | KS.1 |
| FIM1_592 | 5140.142924 | 6118.992252 | 5793.999017 | 0.212879206 | 0.134225217 | 0.080273936 | 0.155636333 | ns | NaCl | KS.1 |
| FIM1_593 | 1076.063817 | 1261.75365 | 1160.875021 | 0.181764581 | 0.156069655 | 0.186207509 | 0.302541593 | ns | NaCl | KS.1 |
| FIM1_594 | 2367.604788 | 2946.798268 | 2091.221904 | 0.246216315 | 0.172046265 | 0.096220195 | 0.17942109 | ns | NaCl | KS.1 |
| FIM1_595 | 2346.010071 | 3420.162039 | 2258.61927 | 0.466852922 | 0.159698692 | 0.000890704 | 0.003897319 | ns | NaCl | KS.1 |
| FIM1_596 | 4843.910094 | 4258.587994 | 3810.382358 | -0.149503551 | 0.150769072 | 0.264219171 | 0.392502352 | ns | NaCl | KS.1 |
| FIM1_597 | 1636.392933 | 1743.526223 | 1847.660826 | 0.078819093 | 0.112543137 | 0.453095983 | 0.588397942 | ns | NaCl | KS.1 |
| FIM1_598 | 959.071676 | 1292.012515 | 942.4573859 | 0.367548222 | 0.146703629 | 0.004924113 | 0.016127361 | ns | NaCl | KS.1 |

| | | | | | | | | | | |
|---|---|---|---|---|---|---|---|---|---|---|
| FIM1_599 | 556.1364878 | 527.6930713 | 612.5166413 | -0.06729822 | 0.11370844 | 0.525570177 | 0.65433224 | ns | NaCl | KS.1 |
| FIM1_600 | 1348.184564 | 1179.937285 | 1731.684023 | -0.146059822 | 0.16832172 | 0.316319342 | 0.449194317 | ns | NaCl | KS.1 |
| FIM1_601 | 3202.612541 | 5603.205381 | 2786.680377 | 0.527162701 | 0.298128275 | 0.014833308 | 0.038819415 | ns | NaCl | KS.1 |
| FIM1_602 | 1993.409251 | 2076.563064 | 2163.783996 | 0.055026532 | 0.083909142 | 0.495031558 | 0.627375468 | ns | NaCl | KS.1 |
| FIM1_603 | 6909.465301 | 3704.914683 | 6877.477411 | -0.843923801 | 0.132181926 | 1.2601045877 | 5.8588974988 | ns | NaCl | KS.1 |
| FIM1_604 | 2398.309797 | 2654.342275 | 2260.077787 | 0.125236013 | 0.12103773 | 0.262580509 | 0.390885109 | ns | NaCl | KS.1 |
| FIM1_605 | 457.1331394 | 377.6264328 | 511.9090931 | -0.210780043 | 0.170547604 | 0.151849318 | 0.257657012 | ns | NaCl | KS.1 |
| FIM1_606 | 833.363891 | 472.7040184 | 1345.168045 | -0.735331343 | 0.165980929 | 8.6034121449 | 1.0288936399 | ns | NaCl | KS.1 |
| FIM1_607 | 690.885001 | 1469.212703 | 715.6332677 | 0.8418646 | 0.281038429 | 0.000149563 | 0.000876013 | ns | NaCl | KS.1 |
| FIM1_608 | 583.9833443 | 541.3948099 | 591.9919784 | -0.090969416 | 0.133836373 | 0.454878155 | 0.589942341 | ns | NaCl | KS.1 |
| FIM1_609 | 6660.879601 | 6568.870165 | 7419.754934 | -0.014049808 | 0.191217435 | 0.929945737 | 0.960049812 | ns | NaCl | KS.1 |
| FIM1_610 | 488.530002 | 419.6445429 | 549.4681379 | -0.18751312 | 0.123030896 | 0.097607699 | 0.181261032 | ns | NaCl | KS.1 |
| FIM1_611 | 154.6435016 | 101.2842518 | 193.9310292 | -0.46874791 | 0.2126977 | 0.007122218 | 0.021813375 | ns | NaCl | KS.1 |
| FIM1_612 | 1234.942915 | 1220.298606 | 1807.422097 | -0.017349089 | 0.084422563 | 0.830641055 | 0.895436456 | ns | NaCl | KS.1 |
| FIM1_613 | 1160.955488 | 1095.8351 | 1297.621499 | -0.075066461 | 0.096380866 | 0.412158797 | 0.547819935 | ns | NaCl | KS.1 |
| FIM1_615 | 789.6651017 | 514.0109417 | 988.5036954 | -0.559987166 | 0.134920674 | 5.5173942678 | 5.1306610248 | ns | NaCl | KS.1 |
| FIM1_616 | 2530.923003 | 2602.658212 | 2442.610828 | 0.03644085 | 0.110288034 | 0.724141841 | 0.816838783 | ns | NaCl | KS.1 |
| FIM1_617 | 635.8705239 | 597.3409828 | 457.5838486 | -0.048809601 | 0.261586327 | 0.799030922 | 0.871176602 | ns | NaCl | KS.1 |
| FIM1_618 | 1109.652989 | 1177.538836 | 1419.095574 | 0.058983594 | 0.198847801 | 0.719215034 | 0.813388223 | ns | NaCl | KS.1 |
| FIM1_619 | 12322.7829 | 12739.35544 | 13755.79097 | 0.038284458 | 0.149266123 | 0.773541953 | 0.853357329 | ns | NaCl | KS.1 |
| FIM1_620 | 1113.148485 | 674.4978911 | 905.0081186 | -0.491483662 | 0.27420404 | 0.01673327 | 0.042845093 | ns | NaCl | KS.1 |
| FIM1_621 | 535.3132111 | 359.0390328 | 628.3279031 | -0.380732711 | 0.260034883 | 0.052396801 | 0.109850014 | ns | NaCl | KS.1 |
| FIM1_622 | 352.6123623 | 227.1869874 | 308.1685685 | -0.578766118 | 0.144337838 | 9.4384309434 | 8.1239089868 | ns | NaCl | KS.1 |
| FIM1_623 | 3824.429334 | 3958.781702 | 3212.491914 | 0.043581348 | 0.110509967 | 0.673491614 | 0.778128374 | ns | NaCl | KS.1 |
| FIM1_625 | 232.10633 | 393.2360293 | 1665.134305 | 0.502772703 | 0.289118697 | 0.017681519 | 0.044903295 | ns | NaCl | KS.1 |
| FIM1_626 | 2246.370617 | 3460.738095 | 3460.275943 | 0.473134107 | 0.223981645 | 0.008783162 | 0.025896362 | ns | NaCl | KS.1 |
| FIM1_627 | 674.7021422 | 570.4931922 | 1097.322672 | -0.20502881 | 0.135689415 | 0.095068688 | 0.178141892 | ns | NaCl | KS.1 |
| FIM1_628 | 2603.559001 | 1527.348534 | 2869.052745 | -0.659835191 | 0.187884455 | 5.1547231063 | 0.000358167 | ns | NaCl | KS.1 |
| FIM1_629 | 2732.869786 | 3654.557531 | 2629.388905 | 0.328526902 | 0.182987862 | 0.034100373 | 0.07746546 | ns | NaCl | KS.1 |
| FIM1_630 | 603.904133 | 674.2533711 | 786.9538102 | 0.133841375 | 0.137248665 | 0.28114472 | 0.411139031 | ns | NaCl | KS.1 |
| FIM1_631 | 1477.956516 | 1814.113765 | 2014.126863 | 0.186459844 | 0.234976189 | 0.306491242 | 0.439295284 | ns | NaCl | KS.1 |
| FIM1_632 | 342.9465975 | 184.8266874 | 338.091691 | -0.757161542 | 0.214452133 | 3.3670153164 | 0.000244896 | ns | NaCl | KS.1 |
| FIM1_633 | 7490.695034 | 8070.378699 | 5665.138552 | 0.096305533 | 0.101656449 | 0.315766917 | 0.448840118 | ns | NaCl | KS.1 |
| FIM1_634 | 435.0674848 | 532.3479501 | 504.229274 | 0.257749617 | 0.118406692 | 0.018421846 | 0.046404396 | ns | NaCl | KS.1 |
| FIM1_635 | 4575.574556 | 4984.473975 | 3749.836985 | 0.109233642 | 0.105302878 | 0.270054341 | 0.399025942 | ns | NaCl | KS.1 |
| FIM1_636 | 474.1084509 | 501.6837565 | 505.1129773 | 0.072307424 | 0.103239349 | 0.457686417 | 0.592503233 | ns | NaCl | KS.1 |
| FIM1_637 | 10502.52097 | 13029.60627 | 8546.879484 | 0.273941243 | 0.120456554 | 0.01353799 | 0.036308085 | ns | NaCl | KS.1 |
| FIM1_638 | 1343.490526 | 1561.008717 | 1509.720033 | 0.193165532 | 0.107672053 | 0.055384008 | 0.115094169 | ns | NaCl | KS.1 |
| FIM1_639 | 67.88206392 | 62.89670461 | 79.66084673 | -0.06047966 | 0.237697013 | 0.740733947 | 0.829849233 | ns | NaCl | KS.1 |
| FIM1_640 | 1158.044765 | 1244.747523 | 1173.532448 | 0.095845256 | 0.084336672 | 0.236661847 | 0.36361726 | ns | NaCl | KS.1 |
| FIM1_641 | 456.4855508 | 348.7688186 | 514.1290684 | -0.323512402 | 0.15699667 | 0.019010528 | 0.04750245 | ns | NaCl | KS.1 |
| FIM1_642 | 2943.312306 | 3857.536923 | 3393.308776 | 0.281038631 | 0.208063772 | 0.097366028 | 0.180947324 | ns | NaCl | KS.1 |
| FIM1_644 | 2066.833069 | 2985.892196 | 1767.328646 | 0.403642406 | 0.208925278 | 0.018297738 | 0.046161889 | ns | NaCl | KS.1 |
| FIM1_645 | 899.5838191 | 950.2740839 | 1058.851261 | 0.057000476 | 0.18358241 | 0.713694273 | 0.809352406 | ns | NaCl | KS.1 |
| FIM1_646 | 3366.594421 | 3941.128836 | 2889.920807 | 0.192731659 | 0.132011778 | 0.108537114 | 0.197449218 | ns | NaCl | KS.1 |

| | | | | | | | | | | |
|---|---|---|---|---|---|---|---|---|---|---|
| FIM1_647 | 1612.093601 | 2598.333241 | 1610.557098 | 0.56655994 | 0.19989424 | 0.000773196 | 0.003474843 | ns | NaCl | KS.1 |
| FIM1_648 | 189.1809974 | 108.7373471 | 180.6262444 | -0.69943851 | 0.170268997 | 4.0585744585 | 4.0382815862 | ns | NaCl | KS.1 |
| FIM1_650 | 701.6905418 | 380.4370073 | 655.9854887 | -0.819272452 | 0.145514424 | 1.3623244313 | 3.7445105225 | ns | NaCl | KS.1 |
| FIM1_651 | 723.94276 | 667.041316 | 1317.559511 | -0.077548582 | 0.214400387 | 0.652431976 | 0.76014909 | ns | NaCl | KS.1 |
| FIM1_652 | 2417.593457 | 2910.835996 | 2526.977574 | 0.243244231 | 0.098126876 | 0.00870223 | 0.025785346 | ns | NaCl | KS.1 |
| FIM1_653 | 3867.213422 | 8066.823031 | 3738.84694 | 0.751735806 | 0.312962138 | 0.001203001 | 0.005005631 | ns | NaCl | KS.1 |
| FIM1_654 | 516.6763478 | 471.9768261 | 618.8363942 | -0.11259445 | 0.123919448 | 0.323646239 | 0.456964532 | ns | NaCl | KS.1 |
| FIM1_655 | 696.1861517 | 298.262872 | 491.6011962 | -1.11759982 | 0.176160528 | 5.5292120393 | 2.8069214179 | Km<KS.1 | NaCl | KS.1 |
| FIM1_656 | 1867.792974 | 2280.072014 | 1796.04403 | 0.250629865 | 0.124009368 | 0.027616038 | 0.065021196 | ns | NaCl | KS.1 |
| FIM1_657 | 701.4208553 | 670.1693399 | 840.9049317 | -0.052622468 | 0.155233323 | 0.701289885 | 0.799476897 | ns | NaCl | KS.1 |
| FIM1_658 | 559.079497 | 369.4969012 | 678.8625973 | -0.494001829 | 0.185570399 | 0.001787594 | 0.006970229 | ns | NaCl | KS.1 |
| FIM1_659 | 2560.280397 | 2518.800696 | 2457.652158 | -0.020841071 | 0.112257203 | 0.842476851 | 0.903496946 | ns | NaCl | KS.1 |
| FIM1_660 | 914.8117086 | 493.0349881 | 935.7586285 | -0.854669815 | 0.108553859 | 2.5916764523 | 3.3060488077 | ns | NaCl | KS.1 |
| FIM1_661 | 693.8094367 | 709.1319159 | 869.1390861 | 0.023954359 | 0.175784523 | 0.873650501 | 0.924571633 | ns | NaCl | KS.1 |
| FIM1_662 | 16369.51514 | 24777.34022 | 13329.56743 | 0.508979163 | 0.171320287 | 0.000634721 | 0.002948359 | ns | NaCl | KS.1 |
| FIM1_663 | 638.906213 | 501.9655813 | 728.0410642 | -0.186513996 | 0.278304405 | 0.3481735 | 0.482900241 | ns | NaCl | KS.1 |
| FIM1_664 | 4008.837769 | 3268.412774 | 3029.072366 | -0.250352142 | 0.135676964 | 0.041303851 | 0.090443071 | ns | NaCl | KS.1 |
| FIM1_665 | 552.3202404 | 577.7899377 | 624.5284246 | 0.05666305 | 0.108827852 | 0.578665521 | 0.700282405 | ns | NaCl | KS.1 |
| FIM1_666 | 138.0006612 | 140.4565831 | 221.6060011 | 0.01556598 | 0.227032174 | 0.930346709 | 0.960264497 | ns | NaCl | KS.1 |
| FIM1_667 | 224.0312281 | 167.0441202 | 255.920656 | -0.323799401 | 0.201755352 | 0.051449485 | 0.108137384 | ns | NaCl | KS.1 |
| FIM1_668 | 3644.487432 | 6144.48737 | 3917.240784 | 0.697583143 | 0.134783336 | 2.3650101388 | 4.7635325672 | ns | NaCl | KS.1 |
| FIM1_669 | 1136.848541 | 1339.392395 | 1242.978932 | 0.155099422 | 0.218960307 | 0.374455872 | 0.50941153 | ns | NaCl | KS.1 |
| FIM1_670 | 1416.83514 | 4023.338886 | 2971.716088 | 0.99202867 | 0.40851198 | 0.000344397 | 0.00176819 | ns | NaCl | KS.1 |
| FIM1_671 | 818.0366191 | 640.9984871 | 622.0532644 | -0.311292102 | 0.130661456 | 0.008778635 | 0.025896362 | ns | NaCl | KS.1 |
| FIM1_672 | 477.8080459 | 398.4992988 | 662.1761727 | -0.216705168 | 0.154915389 | 0.112706783 | 0.203496589 | ns | NaCl | KS.1 |
| FIM1_673 | 293.6608154 | 118.6350652 | 349.8445058 | -0.974133278 | 0.32766015 | 8.786363311 | 0.000560859 | ns | NaCl | KS.1 |
| FIM1_674 | 4225.366 | 3548.081138 | 5215.570752 | -0.213382219 | 0.134081777 | 0.079292423 | 0.153973746 | ns | NaCl | KS.1 |
| FIM1_675 | 1552.599766 | 1269.124136 | 1473.001452 | -0.255058677 | 0.120199213 | 0.02131444 | 0.052494723 | ns | NaCl | KS.1 |
| FIM1_676 | 2579.686599 | 2219.745388 | 3264.916752 | -0.146190495 | 0.209541827 | 0.389373235 | 0.524683598 | ns | NaCl | KS.1 |
| FIM1_677 | 783.5441916 | 1064.961467 | 1919.82398 | 0.371418882 | 0.160812971 | 0.008278985 | 0.024722659 | ns | NaCl | KS.1 |
| FIM1_678 | 2657.649484 | 2547.922458 | 2766.754028 | -0.048747841 | 0.14506951 | 0.707787842 | 0.803831036 | ns | NaCl | KS.1 |
| FIM1_679 | 2066.082072 | 3066.171571 | 2876.776366 | 0.401188227 | 0.239918089 | 0.032309894 | 0.074108679 | ns | NaCl | KS.1 |
| FIM1_680 | 5213.157201 | 3384.806417 | 5497.70073 | -0.426396987 | 0.256239352 | 0.029315001 | 0.068342141 | ns | NaCl | KS.1 |
| FIM1_681 | 2522.174163 | 1943.062163 | 2011.579786 | -0.208636089 | 0.274420268 | 0.292074451 | 0.423474094 | ns | NaCl | KS.1 |
| FIM1_682 | 1044.927769 | 587.9999158 | 1325.94449 | -0.562238568 | 0.29221141 | 0.009065434 | 0.026482965 | ns | NaCl | KS.1 |
| FIM1_683 | 3596.285526 | 4008.335976 | 3487.012945 | 0.134693666 | 0.12122234 | 0.228751483 | 0.354408521 | ns | NaCl | KS.1 |
| FIM1_684 | 1242.597886 | 1171.57879 | 1704.721099 | -0.075476677 | 0.102028054 | 0.433363553 | 0.569612596 | ns | NaCl | KS.1 |
| FIM1_685 | 455.8164332 | 323.5761363 | 571.6491387 | -0.425217518 | 0.159349116 | 0.002373181 | 0.008804305 | ns | NaCl | KS.1 |
| FIM1_686 | 3776.892283 | 4160.591748 | 3809.941275 | 0.132173266 | 0.072923496 | 0.0614818 | 0.125563199 | ns | NaCl | KS.1 |
| FIM1_687 | 961.0767427 | 936.6948844 | 1055.214549 | -0.032642927 | 0.119743228 | 0.76848896 | 0.851120342 | ns | NaCl | KS.1 |
| FIM1_688 | 793.2116584 | 836.5645955 | 1036.037415 | 0.069783185 | 0.107561617 | 0.489497333 | 0.622629484 | ns | NaCl | KS.1 |
| FIM1_689 | 198.4570247 | 192.2521014 | 143.7030848 | -0.030449273 | 0.225942636 | 0.863872735 | 0.918326252 | ns | NaCl | KS.1 |
| FIM1_690 | 1614.388618 | 1753.466125 | 1524.061999 | 0.10526484 | 0.111259603 | 0.311302124 | 0.44414341 | ns | NaCl | KS.1 |
| FIM1_691 | 1532.644561 | 2373.994969 | 1611.399486 | 0.573901415 | 0.135911113 | 3.7982172082 | 3.8235334126 | ns | NaCl | KS.1 |
| FIM1_692 | 1227.941062 | 1403.902374 | 1227.666889 | 0.16233483 | 0.13461582 | 0.183840581 | 0.299380325 | ns | NaCl | KS.1 |

| | | | | | | | | | | |
|---|---|---|---|---|---|---|---|---|---|---|
| FIM1_693 | 940.0100942 | 1176.116967 | 1255.218501 | 0.292912186 | 0.106412802 | 0.003258084 | 0.011430866 | ns | NaCl | KS.1 |
| FIM1_694 | 435.4049921 | 353.2119901 | 543.4684132 | -0.258136252 | 0.13764949 | 0.037586575 | 0.084193252 | ns | NaCl | KS.1 |
| FIM1_695 | 798.1875749 | 852.4683605 | 1000.654697 | 0.08510736 | 0.091765645 | 0.33091351 | 0.464399073 | ns | NaCl | KS.1 |
| FIM1_696 | 1387.485577 | 1710.483935 | 1557.699312 | 0.268184656 | 0.117002047 | 0.013197762 | 0.035548927 | ns | NaCl | KS.1 |
| FIM1_697 | 819.1217733 | 658.3801845 | 898.8205216 | -0.275897798 | 0.121407232 | 0.013498301 | 0.036240716 | ns | NaCl | KS.1 |
| FIM1_698 | 1073.646978 | 1194.078983 | 812.8636411 | 0.12667567 | 0.134479036 | 0.299658375 | 0.432116063 | ns | NaCl | KS.1 |
| FIM1_699 | 2761.170861 | 3051.909214 | 2938.740619 | 0.098588889 | 0.201609095 | 0.551563753 | 0.676035888 | ns | NaCl | KS.1 |
| FIM1_700 | 7499.311934 | 9830.462202 | 6849.827476 | 0.369194879 | 0.084032244 | 4.3492395659 | 4.2593730276 | ns | NaCl | KS.1 |
| FIM1_701 | 2044.231115 | 2963.663096 | 1751.349444 | 0.40602734 | 0.210965267 | 0.018415431 | 0.046404396 | ns | NaCl | KS.1 |
| FIM1_702 | 1138.118089 | 1436.480115 | 1186.573425 | 0.296853215 | 0.122476063 | 0.008311837 | 0.024790999 | ns | NaCl | KS.1 |
| FIM1_703 | 3106.845002 | 3905.532359 | 2874.929683 | 0.276535765 | 0.145301958 | 0.03300418 | 0.07549232 | ns | NaCl | KS.1 |
| FIM1_704 | 5546.175634 | 7619.405269 | 6127.943018 | 0.317308255 | 0.228579428 | 0.078858388 | 0.153370008 | ns | NaCl | KS.1 |
| FIM1_705 | 1233.700552 | 1477.958005 | 1247.382072 | 0.163534305 | 0.233480468 | 0.367572903 | 0.502797688 | ns | NaCl | KS.1 |
| FIM1_706 | 1786.922465 | 2462.815429 | 2196.0927 | 0.418128711 | 0.120608184 | 0.000166145 | 0.000958327 | ns | NaCl | KS.1 |
| FIM1_707 | 1003.900869 | 1201.956161 | 994.1467093 | 0.234012598 | 0.095529674 | 0.009734294 | 0.027944669 | ns | NaCl | KS.1 |
| FIM1_708 | 527.5925883 | 574.9473473 | 661.6962084 | 0.103466576 | 0.129592206 | 0.38252748 | 0.517702452 | ns | NaCl | KS.1 |
| FIM1_709 | 1086.154546 | 1198.618535 | 1294.26645 | 0.122779704 | 0.110661678 | 0.235258791 | 0.361908623 | ns | NaCl | KS.1 |
| FIM1_710 | 597.9316832 | 522.5233947 | 649.6720697 | -0.162459094 | 0.136018133 | 0.187262416 | 0.303774537 | ns | NaCl | KS.1 |
| FIM1_711 | 479.4205407 | 462.6092046 | 444.1594199 | -0.041536931 | 0.167447292 | 0.775041507 | 0.853721994 | ns | NaCl | KS.1 |
| FIM1_712 | 11.79661964 | 23.82688316 | 21.06410889 | 0.422832195 | 0.402248342 | 0.070458443 | 0.139542497 | ns | NaCl | KS.1 |
| FIM1_713 | 2039.693953 | 1149.259525 | 1449.725859 | -0.560299837 | 0.292796924 | 0.009358578 | 0.027132242 | ns | NaCl | KS.1 |
| FIM1_714 | 784.44048 | 959.2384439 | 955.3241447 | 0.235874847 | 0.153147075 | 0.081390673 | 0.157188896 | ns | NaCl | KS.1 |
| FIM1_715 | 590.1018413 | 735.9806901 | 1366.728225 | 0.220738347 | 0.211707378 | 0.197268083 | 0.316277381 | ns | NaCl | KS.1 |
| FIM1_718 | 3067.149864 | 3649.046017 | 4056.178042 | 0.199172968 | 0.160017292 | 0.155282559 | 0.262230391 | ns | NaCl | KS.1 |
| FIM1_719 | 7011.522531 | 10737.48725 | 7126.365091 | 0.55031325 | 0.144721355 | 2.5023903288 | 0.000187491 | ns | NaCl | KS.1 |
| FIM1_720 | 442.7653187 | 540.3062339 | 588.1164666 | 0.235950572 | 0.144218904 | 0.067410809 | 0.134524178 | ns | NaCl | KS.1 |
| FIM1_721 | 2333.542743 | 2632.527975 | 2180.455325 | 0.146118739 | 0.129401965 | 0.216538926 | 0.340158244 | ns | NaCl | KS.1 |
| FIM1_722 | 4288.386384 | 6341.349761 | 4671.506458 | 0.538035026 | 0.090769718 | 5.3182730409 | 1.6035399017 | ns | NaCl | KS.1 |
| FIM1_723 | 733.1210055 | 675.2899069 | 877.597854 | -0.104286652 | 0.125179321 | 0.364815674 | 0.5001262 | ns | NaCl | KS.1 |
| FIM1_724 | 370.723507 | 561.252277 | 388.5643602 | 0.468791797 | 0.206044241 | 0.005920183 | 0.018771772 | ns | NaCl | KS.1 |
| FIM1_725 | 6430.941149 | 11734.48515 | 5829.750812 | 0.721800928 | 0.217337572 | 8.2129991716 | 0.000528331 | ns | NaCl | KS.1 |
| FIM1_726 | 744.5785059 | 1006.469793 | 898.0738828 | 0.384814315 | 0.130560967 | 0.001189862 | 0.004966078 | ns | NaCl | KS.1 |
| FIM1_727 | 773.2301689 | 941.9309265 | 1229.378744 | 0.23988606 | 0.139508947 | 0.056071709 | 0.11628043 | ns | NaCl | KS.1 |
| FIM1_728 | 2887.107897 | 3641.889721 | 2356.280893 | 0.266921582 | 0.165939637 | 0.063807689 | 0.12909445 | ns | NaCl | KS.1 |
| FIM1_729 | 2593.6417 | 3340.137924 | 2429.537675 | 0.323319422 | 0.122275641 | 0.003984041 | 0.013483405 | ns | NaCl | KS.1 |
| FIM1_730 | 1746.852029 | 2004.152961 | 1867.201485 | 0.178546613 | 0.099705093 | 0.058033096 | 0.119898111 | ns | NaCl | KS.1 |
| FIM1_731 | 380.8069191 | 379.5551361 | 460.353051 | -0.008839654 | 0.115198542 | 0.934321169 | 0.962170941 | ns | NaCl | KS.1 |
| FIM1_732 | 10778.58895 | 13501.26388 | 8995.403339 | 0.27835339 | 0.135025164 | 0.022671039 | 0.055216151 | ns | NaCl | KS.1 |
| FIM1_733 | 1651.024854 | 2651.686818 | 2077.4532 | 0.512181937 | 0.237530716 | 0.006626249 | 0.020594638 | ns | NaCl | KS.1 |
| FIM1_734 | 2022.77294 | 1975.810016 | 2175.091979 | -0.033741351 | 0.064894038 | 0.594185249 | 0.714595654 | ns | NaCl | KS.1 |
| FIM1_735 | 691.0621622 | 452.906081 | 672.6776497 | -0.570867229 | 0.112031027 | 5.4091110630 | 9.2158655954 | ns | NaCl | KS.1 |
| FIM1_736 | 1967.458061 | 2260.199486 | 2714.657046 | 0.173205037 | 0.118593273 | 0.114458868 | 0.206090686 | ns | NaCl | KS.1 |
| FIM1_737 | 1983.194158 | 2384.894887 | 2322.058371 | 0.247389797 | 0.088159179 | 0.003273903 | 0.011462115 | ns | NaCl | KS.1 |
| FIM1_738 | 2039.767989 | 1881.591059 | 1834.221427 | -0.091387735 | 0.155860782 | 0.506476933 | 0.638227644 | ns | NaCl | KS.1 |
| FIM1_739 | 455.692267 | 432.7588703 | 482.0774618 | -0.054856613 | 0.186988338 | 0.727397348 | 0.819660658 | ns | NaCl | KS.1 |

| | | | | | | | | | |
|---|---|---|---|---|---|---|---|---|---|
| FIM1_740 | 724.9234848 | 851.4102585 | 827.4917073 | 0.193813649 | 0.133885429 | 0.11069594 | 0.200477721 | ns | NaCl | KS.1 |
| FIM1_741 | 1895.156213 | 2542.076751 | 1896.080384 | 0.38930645 | 0.105514896 | 8.06310846014 | 0.000520362 | ns | NaCl | KS.1 |
| FIM1_742 | 1776.495446 | 2159.397783 | 1995.452123 | 0.244179948 | 0.124713401 | 0.032727983 | 0.07492946 | ns | NaCl | KS.1 |
| FIM1_743 | 2435.032921 | 2457.776501 | 2245.619871 | 0.011183916 | 0.093604935 | 0.900196124 | 0.940658626 | ns | NaCl | KS.1 |
| FIM1_744 | 393.3752028 | 303.0341658 | 472.0112613 | -0.303972137 | 0.170471992 | 0.038652409 | 0.08605705 | ns | NaCl | KS.1 |
| FIM1_745 | 391.7941542 | 325.9544129 | 570.5548511 | -0.237556795 | 0.120693016 | 0.03268077 | 0.074855816 | ns | NaCl | KS.1 |
| FIM1_746 | 1077.818642 | 1297.919843 | 1985.127197 | 0.220560665 | 0.148793274 | 0.095554383 | 0.178715434 | ns | NaCl | KS.1 |
| FIM1_747 | 1052.528805 | 850.6338279 | 1178.845108 | -0.250474678 | 0.152803152 | 0.063704434 | 0.128937982 | ns | NaCl | KS.1 |
| FIM1_748 | 561.3012391 | 781.5479902 | 794.7275218 | 0.32995908 | 0.231680275 | 0.070227174 | 0.139324958 | ns | NaCl | KS.1 |
| FIM1_749 | 746.5570074 | 522.9476448 | 1109.475622 | -0.44517172 | 0.149218532 | 0.000806014 | 0.003594631 | ns | NaCl | KS.1 |
| FIM1_750 | 1435.914242 | 1269.576592 | 1422.150315 | -0.160284518 | 0.107874808 | 0.112771878 | 0.203496589 | ns | NaCl | KS.1 |
| FIM1_751 | 3060.780578 | 3405.324405 | 3565.154167 | 0.143182161 | 0.079039275 | 0.060267632 | 0.123641843 | ns | NaCl | KS.1 |
| FIM1_752 | 5660.283884 | 7376.540634 | 5517.589437 | 0.355389534 | 0.096143646 | 9.18210409762 | 0.000581923 | ns | NaCl | KS.1 |
| FIM1_753 | 1192.210819 | 3364.776494 | 2091.90095 | 1.420603443 | 0.158743092 | 1.46020212126 | 5.18893253806 | Km>KS.1 | NaCl | KS.1 |
| FIM1_754 | 1705.146124 | 1451.359446 | 2116.637094 | -0.177518676 | 0.17297806 | 0.232537604 | 0.358609602 | ns | NaCl | KS.1 |
| FIM1_755 | 1952.793355 | 2211.146273 | 2519.886863 | 0.16957262 | 0.069910823 | 0.012491996 | 0.033867944 | ns | NaCl | KS.1 |
| FIM1_756 | 715.6331548 | 680.6394424 | 822.4926349 | -0.063956313 | 0.118325725 | 0.560026486 | 0.683377917 | ns | NaCl | KS.1 |
| FIM1_757 | 1851.838968 | 2288.914527 | 2005.582828 | 0.267816183 | 0.121734111 | 0.016753096 | 0.042873792 | ns | NaCl | KS.1 |
| FIM1_758 | 359.3271047 | 244.9328275 | 378.9891539 | -0.508802096 | 0.134625919 | 3.26716312294 | 0.000239737 | ns | NaCl | KS.1 |
| FIM1_759 | 784.5427241 | 716.8065855 | 738.316749 | -0.11958213 | 0.114111576 | 0.260429541 | 0.389080171 | ns | NaCl | KS.1 |
| FIM1_760 | 1903.423111 | 2425.270964 | 2381.124377 | 0.289469256 | 0.152414899 | 0.031700104 | 0.072945429 | ns | NaCl | KS.1 |
| FIM1_761 | 2449.274485 | 2202.62382 | 2800.818831 | -0.119481546 | 0.160114686 | 0.394752418 | 0.530781967 | ns | NaCl | KS.1 |
| FIM1_762 | 1262.255301 | 1353.511825 | 1122.594086 | 0.079293456 | 0.145583226 | 0.543076483 | 0.669426537 | ns | NaCl | KS.1 |
| FIM1_763 | 791.9914949 | 641.2017121 | 775.0109219 | -0.266791294 | 0.128125585 | 0.022510494 | 0.054913959 | ns | NaCl | KS.1 |
| FIM1_764 | 2752.273035 | 2749.760223 | 2454.664853 | -0.003251104 | 0.098144505 | 0.972138079 | 0.987204262 | ns | NaCl | KS.1 |
| FIM1_765 | 330.6594215 | 419.7538693 | 354.7709332 | 0.227143041 | 0.227509132 | 0.20492458 | 0.326031271 | ns | NaCl | KS.1 |
| FIM1_767 | 600.1734544 | 578.0262617 | 1201.766306 | -0.030772111 | 0.246868586 | 0.86885592 | 0.921654201 | ns | NaCl | KS.1 |
| FIM1_768 | 329.1070792 | 243.2035779 | 370.6882992 | -0.364302624 | 0.162540665 | 0.010207249 | 0.029034342 | ns | NaCl | KS.1 |
| FIM1_769 | 2481.460597 | 2640.670175 | 2602.665713 | 0.082985203 | 0.072991291 | 0.241029731 | 0.367831355 | ns | NaCl | KS.1 |
| FIM1_770 | 2703.623479 | 3854.721828 | 2444.516323 | 0.378325065 | 0.215936491 | 0.03031163 | 0.070237707 | ns | NaCl | KS.1 |
| FIM1_772 | 81.53434324 | 51.89900671 | 53.38291118 | -0.346796841 | 0.315668268 | 0.106206571 | 0.194113773 | ns | NaCl | KS.1 |
| FIM1_773 | 2731.919289 | 3956.644905 | 2254.322242 | 0.417773202 | 0.199221892 | 0.011639041 | 0.032151154 | ns | NaCl | KS.1 |
| FIM1_774 | 701.5962798 | 816.688039 | 784.9324631 | 0.184078876 | 0.129186054 | 0.118668276 | 0.212562884 | ns | NaCl | KS.1 |
| FIM1_776 | 2140.169649 | 2481.020451 | 2078.601108 | 0.180370237 | 0.131500219 | 0.131992337 | 0.230084749 | ns | NaCl | KS.1 |
| FIM1_777 | 2351.223435 | 2193.704334 | 2381.458233 | -0.089589636 | 0.106191996 | 0.369397606 | 0.504073076 | ns | NaCl | KS.1 |
| FIM1_778 | 662.1322573 | 433.4334925 | 665.8322517 | -0.528249325 | 0.164494234 | 0.000258081 | 0.001386559 | ns | NaCl | KS.1 |
| FIM1_779 | 20160.51293 | 25595.80712 | 14674.89046 | 0.318408392 | 0.096365509 | 0.000472707 | 0.002292123 | ns | NaCl | KS.1 |
| FIM1_780 | 608.7400648 | 435.9614455 | 453.4928227 | -0.332889957 | 0.230037127 | 0.06660174 | 0.133391167 | ns | NaCl | KS.1 |
| FIM1_781 | 468.0698768 | 226.0746251 | 472.8245672 | -0.843610718 | 0.258766268 | 6.28685958915 | 0.000422093 | ns | NaCl | KS.1 |
| FIM1_782 | 1208.791054 | 480.3380801 | 1671.123472 | -1.131973378 | 0.253868497 | 1.59827158074 | 2.43989298426 | Km<KS.1 | NaCl | KS.1 |
| FIM1_783 | 1385.480882 | 631.3523919 | 935.7181848 | -1.032810393 | 0.179203088 | 2.94135464597 | 9.73976633508 | Km<KS.1 | NaCl | KS.1 |
| FIM1_784 | 418.3782362 | 173.180941 | 488.4529459 | -1.090487285 | 0.244562071 | 1.953263339e- | 2.88691424211 | Km<KS.1 | NaCl | KS.1 |
| FIM1_785 | 3059.71028 | 3466.061911 | 3305.208004 | 0.147990941 | 0.140230608 | 0.241484085 | 0.368296542 | ns | NaCl | KS.1 |
| FIM1_786 | 2940.952483 | 2405.114812 | 2337.434031 | -0.260020675 | 0.113772327 | 0.013835726 | 0.036845613 | ns | NaCl | KS.1 |
| FIM1_787 | 4576.860917 | 5522.695719 | 4819.528209 | 0.211741605 | 0.166360832 | 0.142432753 | 0.244261615 | ns | NaCl | KS.1 |

| | | | | | | | | | | | |
|---|---|---|---|---|---|---|---|---|---|---|---|
| FIM1_788 | 202.4934727 | 187.2569899 | 200.3153645 | -0.069878373 | 0.230410992 | 0.697347881 | 0.796991893 | ns | | NaCl | KS.1 |
| FIM1_789 | 900.5009885 | 917.7541964 | 1008.009036 | 0.020369976 | 0.180695586 | 0.89452407 | 0.936942838 | ns | | NaCl | KS.1 |
| FIM1_790 | 190.8864842 | 762.9924773 | 368.2901065 | 1.856539041 | 0.24770027 | 3.3987405158( | 5.1238588079; | Km>KS.1 | | NaCl | KS.1 |
| FIM1_791 | 3589.501259 | 4492.376985 | 3356.780222 | 0.242939504 | 0.186472269 | 0.121787594 | 0.216545133 | ns | | NaCl | KS.1 |
| FIM1_792 | 881.9574045 | 1144.609983 | 1088.768169 | 0.335852976 | 0.117352378 | 0.001942353 | 0.007467704 | ns | | NaCl | KS.1 |
| FIM1_793 | 3047.739229 | 3952.967535 | 3888.038793 | 0.335166232 | 0.120758013 | 0.002550304 | 0.009377502 | ns | | NaCl | KS.1 |
| FIM1_794 | 851.3163873 | 1030.833442 | 1027.485245 | 0.246065674 | 0.107530679 | 0.014423046 | 0.038126808 | ns | | NaCl | KS.1 |
| FIM1_795 | 2788.499066 | 3970.527324 | 2824.487688 | 0.443146372 | 0.148632635 | 0.00081837 | 0.003635943 | ns | | NaCl | KS.1 |
| FIM1_796 | 898.1471525 | 910.2885807 | 1281.943094 | 0.015453184 | 0.099189459 | 0.869382833 | 0.921820033 | ns | | NaCl | KS.1 |
| FIM1_797 | 347.3716452 | 389.2983184 | 336.0054251 | 0.115903858 | 0.189493946 | 0.465464106 | 0.599762737 | ns | | NaCl | KS.1 |
| FIM1_798 | 588.4642225 | 664.0048377 | 648.7373743 | 0.131430644 | 0.168581712 | 0.367864203 | 0.503057836 | ns | | NaCl | KS.1 |
| FIM1_799 | 1072.412709 | 1100.270944 | 1734.643283 | 0.023261621 | 0.232309528 | 0.897411996 | 0.938735214 | ns | | NaCl | KS.1 |
| FIM1_800 | 2812.139042 | 3686.94544 | 2792.052274 | 0.325706346 | 0.154011993 | 0.016514712 | 0.042329053 | ns | | NaCl | KS.1 |
| FIM1_801 | 2474.911646 | 3115.394204 | 2056.706458 | 0.210596154 | 0.237127295 | 0.251131568 | 0.379750623 | ns | | NaCl | KS.1 |
| FIM1_802 | 1023.42279 | 1148.911416 | 852.4081707 | 0.125045617 | 0.174117592 | 0.402904191 | 0.538179129 | ns | | NaCl | KS.1 |
| FIM1_803 | 1110.775874 | 980.116208 | 1145.42991 | -0.167150115 | 0.094517746 | 0.062670403 | 0.127363257 | ns | | NaCl | KS.1 |
| FIM1_804 | 5689.72665 | 9001.348496 | 5144.829841 | 0.532769506 | 0.20620392 | 0.001906924 | 0.007348664 | ns | | NaCl | KS.1 |
| FIM1_805 | 1015.580096 | 516.1658402 | 1049.39095 | -0.906063325 | 0.145832167 | 3.1202981358? | 1.2518938085‹ | ns | | NaCl | KS.1 |
| FIM1_806 | 502.5306516 | 316.316533 | 1397.046938 | -0.473530583 | 0.253603324 | 0.015498318 | 0.040137496 | ns | | NaCl | KS.1 |
| FIM1_807 | 242.3064703 | 235.3935574 | 604.1760257 | -0.037148023 | 0.169328524 | 0.799904095 | 0.87184705 | ns | | NaCl | KS.1 |
| FIM1_808 | 1272.430013 | 1135.381889 | 1277.977105 | -0.134185714 | 0.144758263 | 0.300765114 | 0.432636272 | ns | | NaCl | KS.1 |
| FIM1_809 | 19291.34501 | 14109.21342 | 20737.53121 | -0.405103725 | 0.12434831 | 0.000377959 | 0.00191092 | ns | | NaCl | KS.1 |
| FIM1_810 | 1334.238794 | 1471.827542 | 1437.83775 | 0.125903945 | 0.099726843 | 0.18191896 | 0.297517036 | ns | | NaCl | KS.1 |
| FIM1_811 | 2176.19938 | 2223.817314 | 2368.765038 | 0.024310572 | 0.158758838 | 0.861761095 | 0.916277291 | ns | | NaCl | KS.1 |
| FIM1_812 | 1956.838119 | 2167.214753 | 2025.255696 | 0.123267922 | 0.134093177 | 0.311730214 | 0.444553176 | ns | | NaCl | KS.1 |
| FIM1_813 | 501.1929802 | 722.1736439 | 733.4134806 | 0.484470036 | 0.119934663 | 1.2268652762( | 0.00010241 | ns | | NaCl | KS.1 |
| FIM1_814 | 58.94326234 | 43.93876567 | 59.6130047 | -0.237542949 | 0.277621514 | 0.23429809 | 0.360765397 | ns | | NaCl | KS.1 |
| FIM1_815 | 114.2781139 | 99.76656641 | 102.8411246 | -0.118405588 | 0.244122057 | 0.52375455 | 0.6532496 | ns | | NaCl | KS.1 |
| FIM1_816 | 420.4609712 | 413.7661615 | 391.056969 | -0.015472842 | 0.199707504 | 0.925075348 | 0.956808702 | ns | | NaCl | KS.1 |
| FIM1_817 | 1447.290726 | 2039.27258 | 1487.348653 | 0.407049757 | 0.173076484 | 0.006331869 | 0.01982445 | ns | | NaCl | KS.1 |
| FIM1_818 | 363.4349302 | 312.9412016 | 327.2956797 | -0.165997608 | 0.179024571 | 0.276358646 | 0.406049694 | ns | | NaCl | KS.1 |
| FIM1_819 | 1246.279578 | 1192.571006 | 1264.18978 | -0.054974238 | 0.127869732 | 0.639135898 | 0.749235812 | ns | | NaCl | KS.1 |
| FIM1_820 | 4543.042464 | 7347.015373 | 6602.112733 | 0.622830588 | 0.152047143 | 5.5358913624? | 5.1382573746( | ns | | NaCl | KS.1 |
| FIM1_821 | 8923.858134 | 11928.47326 | 8074.419783 | 0.345816975 | 0.161501511 | 0.01422754 | 0.037670044 | ns | | NaCl | KS.1 |
| FIM1_822 | 995.2769179 | 1015.868977 | 963.4226905 | 0.021384789 | 0.161440963 | 0.879797353 | 0.929031192 | ns | | NaCl | KS.1 |
| FIM1_823 | 1076.610484 | 831.1500934 | 1053.613319 | -0.342010544 | 0.109713197 | 0.000818544 | 0.003635943 | ns | | NaCl | KS.1 |
| FIM1_824 | 4148.510814 | 11332.33307 | 4683.444448 | 1.057066908 | 0.358596939 | 6.3338236800‹ | 0.000424102 | Km>KS.1 | | NaCl | KS.1 |
| FIM1_825 | 3238.900828 | 3831.3744 | 3341.939333 | 0.213843301 | 0.114617923 | 0.044524661 | 0.096099866 | ns | | NaCl | KS.1 |
| FIM1_826 | 64.46241811 | 87.30055955 | 94.20694577 | 0.242174217 | 0.282680184 | 0.22937378 | 0.354830396 | ns | | NaCl | KS.1 |
| FIM1_828 | 1343.028202 | 1236.781177 | 1423.450385 | -0.097730081 | 0.137830605 | 0.43319797 | 0.569545428 | ns | | NaCl | KS.1 |
| FIM1_829 | 474.4270881 | 444.0614675 | 492.427216 | -0.07855476 | 0.146947452 | 0.549823554 | 0.675233815 | ns | | NaCl | KS.1 |
| FIM1_830 | 2506.761298 | 2232.416176 | 2874.1411 | -0.093524193 | 0.255516439 | 0.622353694 | 0.737192769 | ns | | NaCl | KS.1 |
| FIM1_831 | 4536.010026 | 6264.083473 | 4234.203468 | 0.405928849 | 0.141812296 | 0.001421612 | 0.005708248 | ns | | NaCl | KS.1 |
| FIM1_832 | 551.1367455 | 1035.898937 | 754.9707349 | 0.79852548 | 0.189480094 | 1.8461690939( | 1.9993081763‹ | ns | | NaCl | KS.1 |
| FIM1_833 | 3969.299267 | 7351.993471 | 3921.404092 | 0.716784716 | 0.236168014 | 0.000221283 | 0.001213764 | ns | | NaCl | KS.1 |

| | | | | | | | | | | |
|---|---|---|---|---|---|---|---|---|---|---|
| FIM1_834 | 1141.891141 | 1222.847845 | 1221.750856 | 0.085373853 | 0.113825076 | 0.420965988 | 0.557144398 | ns | NaCl | KS.1 |
| FIM1_835 | 745.1697749 | 900.1111433 | 953.2250199 | 0.229303077 | 0.136747135 | 0.063453945 | 0.128535576 | ns | NaCl | KS.1 |
| FIM1_836 | 504.9857795 | 740.0221615 | 1089.792055 | 0.376650989 | 0.246276397 | 0.047437854 | 0.101158734 | ns | NaCl | KS.1 |
| FIM1_840 | 428.3049996 | 418.3858359 | 485.8406089 | -0.026869915 | 0.154156837 | 0.843909768 | 0.904271501 | ns | NaCl | KS.1 |
| FIM1_841 | 123.144405 | 57.16410452 | 116.3551879 | -0.720939981 | 0.339478219 | 0.00270785 | 0.009832141 | ns | NaCl | KS.1 |
| FIM1_842 | 469.6439551 | 173.498366 | 255.2486176 | -1.216701572 | 0.262033786 | 4.43111195646 | 7.86248065887 | Km<KS.1 | NaCl | KS.1 |
| FIM1_843 | 2597.531959 | 158.1320531 | 1017.35679 | -3.648682957 | 0.438367507 | 1.07501975398 | 2.32531446785 | Km<KS.1 | NaCl | KS.1 |
| FIM1_844 | 131.7315827 | 46.36865301 | 112.3709184 | -1.2755645 | 0.277173961 | 3.93250152956 | 7.19271879028 | Km<KS.1 | NaCl | KS.1 |
| FIM1_846 | 87.4329427 | 52.7131379 | 84.39783745 | -0.492166176 | 0.286255661 | 0.019338753 | 0.048129212 | ns | NaCl | KS.1 |
| FIM1_847 | 8.123504493 | 11.19312775 | 15.97427327 | 0.162921538 | 0.343510147 | 0.439088927 | 0.574632463 | ns | NaCl | KS.1 |
| FIM1_850 | 733.9591768 | 506.7254712 | 698.1875307 | -0.331026541 | 0.268138482 | 0.095326979 | 0.17855863 | ns | NaCl | KS.1 |
| FIM1_851 | 194.7847603 | 204.6222667 | 305.0748288 | 0.046021171 | 0.22550692 | 0.795338229 | 0.869436978 | ns | NaCl | KS.1 |
| FIM1_852 | 65.51374445 | 74.80952087 | 63.46054071 | 0.131600873 | 0.209549802 | 0.438360349 | 0.574359425 | ns | NaCl | KS.1 |
| FIM1_853 | 63.78824403 | 124.6738152 | 101.2122781 | 0.756712299 | 0.256963137 | 0.000251087 | 0.001354832 | ns | NaCl | KS.1 |
| FIM1_854 | 669.7412437 | 1335.992486 | 1559.299267 | 0.765007857 | 0.272322009 | 0.000367194 | 0.001865051 | ns | NaCl | KS.1 |
| FIM1_855 | 218.944783 | 277.268555 | 276.8693972 | 0.250943715 | 0.195310814 | 0.12191977 | 0.216702699 | ns | NaCl | KS.1 |
| FIM1_856 | 324.7235699 | 252.7603761 | 299.5086226 | -0.279034694 | 0.175405912 | 0.063143839 | 0.12816834 | ns | NaCl | KS.1 |
| FIM1_857 | 197.1419401 | 163.3191146 | 194.7328761 | -0.209321754 | 0.154912616 | 0.125540554 | 0.221084693 | ns | NaCl | KS.1 |
| FIM1_858 | 496.2916789 | 329.5167112 | 643.7882858 | -0.388122166 | 0.25947517 | 0.047902501 | 0.10191262 | ns | NaCl | KS.1 |
| FIM1_859 | 394.5494413 | 346.6413294 | 460.6221006 | -0.163138382 | 0.11268665 | 0.120277311 | 0.214626838 | ns | NaCl | KS.1 |
| FIM1_860 | 528.3040199 | 286.8829257 | 530.8493055 | -0.851813361 | 0.120930247 | 1.37771091812 | 1.05447874117 | ns | NaCl | KS.1 |
| FIM1_861 | 624.9840217 | 372.4176568 | 689.1671746 | -0.669182489 | 0.165004674 | 5.62086972783 | 5.19331254499 | ns | NaCl | KS.1 |
| FIM1_862 | 102.989959 | 120.6704647 | 158.4303416 | 0.156432301 | 0.197366681 | 0.338042014 | 0.471962694 | ns | NaCl | KS.1 |
| FIM1_863 | 12.69443404 | 15.39190813 | 32.18866186 | 0.101856922 | 0.343050725 | 0.624315094 | 0.738636764 | ns | NaCl | KS.1 |
| FIM1_864 | 585.8739748 | 394.9729115 | 574.4896196 | -0.3832434 | 0.253662242 | 0.047656185 | 0.101537269 | ns | NaCl | KS.1 |
| FIM1_865 | 694.4373242 | 792.8723213 | 947.753543 | 0.138326696 | 0.190834279 | 0.386004507 | 0.521132272 | ns | NaCl | KS.1 |
| FIM1_866 | 395.5532212 | 320.0718837 | 514.4551748 | -0.22852433 | 0.17538741 | 0.128042948 | 0.224616949 | ns | NaCl | KS.1 |
| FIM1_867 | 884.6905023 | 1020.723057 | 1299.862932 | 0.151503985 | 0.183213689 | 0.328548191 | 0.462145089 | ns | NaCl | KS.1 |
| FIM1_868 | 687.4503122 | 704.2653491 | 901.8214115 | 0.028934167 | 0.151335151 | 0.829681544 | 0.895199686 | ns | NaCl | KS.1 |
| FIM1_869 | 707.0675168 | 734.8445416 | 674.8613506 | 0.050697899 | 0.117736447 | 0.642708161 | 0.752346612 | ns | NaCl | KS.1 |
| FIM1_870 | 1054.950084 | 1990.808196 | 1337.424702 | 0.772588869 | 0.215903295 | 2.66197498359 | 0.000198551 | ns | NaCl | KS.1 |
| FIM1_871 | 1111.91973 | 1237.789487 | 1085.450442 | 0.139601956 | 0.095403034 | 0.123365663 | 0.218414297 | ns | NaCl | KS.1 |
| FIM1_872 | 2907.37881 | 3165.617738 | 2799.267011 | 0.109910624 | 0.100400626 | 0.246876399 | 0.374454295 | ns | NaCl | KS.1 |
| FIM1_873 | 655.226443 | 735.3530862 | 998.9041111 | 0.136851221 | 0.144982935 | 0.291841834 | 0.423421734 | ns | NaCl | KS.1 |
| FIM1_874 | 1983.209865 | 2592.711849 | 4884.448618 | 0.282225443 | 0.203470474 | 0.091014971 | 0.17190565 | ns | NaCl | KS.1 |
| FIM1_875 | 339.4170883 | 332.1461761 | 368.4667161 | -0.021841395 | 0.190370336 | 0.89096574 | 0.934546607 | ns | NaCl | KS.1 |
| FIM1_876 | 2609.755194 | 3272.510853 | 2803.00508 | 0.210290989 | 0.232656787 | 0.246543089 | 0.374062783 | ns | NaCl | KS.1 |
| FIM1_877 | 1299.715525 | 1461.926635 | 1254.645693 | 0.140357033 | 0.137133537 | 0.257993023 | 0.386598155 | ns | NaCl | KS.1 |
| FIM1_878 | 3582.500434 | 4234.821133 | 3660.528863 | 0.214088244 | 0.112509525 | 0.040935475 | 0.089873781 | ns | NaCl | KS.1 |
| FIM1_879 | 37.37997376 | 28.58983771 | 27.8276178 | -0.214909885 | 0.299699345 | 0.295793749 | 0.427658791 | ns | NaCl | KS.1 |
| FIM1_880 | 653.2226047 | 632.4425098 | 1216.289069 | -0.037156441 | 0.109850094 | 0.718009551 | 0.81294531 | ns | NaCl | KS.1 |
| FIM1_881 | 1335.516349 | 1473.969514 | 1551.362047 | 0.128872278 | 0.097495861 | 0.16316753 | 0.272676676 | ns | NaCl | KS.1 |
| FIM1_882 | 455.5579175 | 382.8241865 | 510.0564056 | -0.214609595 | 0.140331323 | 0.089118837 | 0.169310871 | ns | NaCl | KS.1 |
| FIM1_883 | 1335.756095 | 1264.042644 | 1562.433082 | -0.057832994 | 0.17739011 | 0.702858298 | 0.799981706 | ns | NaCl | KS.1 |
| FIM1_884 | 214.8721252 | 131.7299215 | 330.5532116 | -0.496817057 | 0.260196745 | 0.012699689 | 0.034300193 | ns | NaCl | KS.1 |

| | | | | | | | | | | |
|---|---|---|---|---|---|---|---|---|---|---|
| FIM1_885 | 1742.948213 | 2435.093428 | 2050.872708 | 0.458177793 | 0.087931488 | 4.6963717311 | 8.1980524079 | ns | NaCl | KS.1 |
| FIM1_886 | 3672.703101 | 3694.198275 | 4045.572158 | 0.005989668 | 0.192869398 | 0.970277285 | 0.986739472 | ns | NaCl | KS.1 |
| FIM1_887 | 567.6378932 | 499.5196885 | 602.4879608 | -0.154236459 | 0.132826922 | 0.20175775 | 0.322022716 | ns | NaCl | KS.1 |
| FIM1_888 | 1513.599758 | 1164.360229 | 1495.16734 | -0.325012464 | 0.139059536 | 0.009369214 | 0.027147258 | ns | NaCl | KS.1 |
| FIM1_889 | 982.1845573 | 940.8042246 | 968.179931 | -0.057341443 | 0.102070683 | 0.551930185 | 0.676085411 | ns | NaCl | KS.1 |
| FIM1_890 | 353.4884455 | 442.4615737 | 335.6733714 | 0.264456798 | 0.15492641 | 0.05275847 | 0.110329294 | ns | NaCl | KS.1 |
| FIM1_891 | 3457.226794 | 2152.88373 | 4113.37712 | -0.609512857 | 0.15532955 | 1.2083943651 | 0.000101208 | ns | NaCl | KS.1 |
| FIM1_892 | 167.8930955 | 57.15505877 | 170.0596042 | -1.186961495 | 0.354671362 | 9.5608679510 | 8.2009169062 | Km<KS.1 | NaCl | KS.1 |
| FIM1_893 | 23.71959125 | 25.79427072 | 28.89626081 | 0.049110339 | 0.298114009 | 0.807453895 | 0.877859075 | ns | NaCl | KS.1 |
| FIM1_894 | 1948.730175 | 1042.292389 | 3726.986747 | -0.65556274 | 0.281382697 | 0.002249032 | 0.008419061 | ns | NaCl | KS.1 |
| FIM1_895 | 761.9600435 | 592.4954046 | 707.5695422 | -0.316912093 | 0.131934657 | 0.0081347 | 0.024335617 | ns | NaCl | KS.1 |
| FIM1_896 | 150.8429869 | 126.2400467 | 140.4085589 | -0.196860561 | 0.170931955 | 0.181564066 | 0.297229756 | ns | NaCl | KS.1 |
| FIM1_897 | 619.4531571 | 559.0538593 | 566.0714161 | -0.103569706 | 0.195916026 | 0.523781438 | 0.6532496 | ns | NaCl | KS.1 |
| FIM1_898 | 4956.95311 | 4645.88268 | 5129.727209 | -0.070173128 | 0.169443822 | 0.632178056 | 0.744047634 | ns | NaCl | KS.1 |
| FIM1_899 | 583.4063524 | 897.6208908 | 688.9489661 | 0.514121949 | 0.188468593 | 0.00134279 | 0.00545265 | ns | NaCl | KS.1 |
| FIM1_900 | 1768.233785 | 1561.450056 | 1909.136672 | -0.138443668 | 0.164521864 | 0.333962027 | 0.468281027 | ns | NaCl | KS.1 |
| FIM1_901 | 3465.185847 | 3794.253097 | 3456.279029 | 0.114967839 | 0.113942607 | 0.278695838 | 0.408605338 | ns | NaCl | KS.1 |
| FIM1_902 | 345.2933773 | 463.48941 | 486.7253884 | 0.371053688 | 0.132166941 | 0.001975019 | 0.007558246 | ns | NaCl | KS.1 |
| FIM1_903 | 2236.691698 | 3316.023097 | 1651.287398 | 0.334918039 | 0.284881642 | 0.10195504 | 0.188070569 | ns | NaCl | KS.1 |
| FIM1_904 | 1214.828944 | 2039.030413 | 1256.333312 | 0.503597842 | 0.280292375 | 0.015668621 | 0.0404522 | ns | NaCl | KS.1 |
| FIM1_905 | 531.6984947 | 472.2246971 | 879.7972504 | -0.140110477 | 0.138806101 | 0.263565868 | 0.391882903 | ns | NaCl | KS.1 |
| FIM1_906 | 16535.83521 | 18744.91852 | 12090.97668 | 0.151983725 | 0.132750898 | 0.208229219 | 0.329812277 | ns | NaCl | KS.1 |
| FIM1_907 | 19609.42332 | 26658.8702 | 16591.70845 | 0.399571926 | 0.120610848 | 0.000318448 | 0.001648575 | ns | NaCl | KS.1 |
| FIM1_909 | 1499.739668 | 1326.73767 | 1077.98299 | -0.144929865 | 0.14108783 | 0.253737914 | 0.382066018 | ns | NaCl | KS.1 |
| FIM1_910 | 186.6596445 | 66.32001868 | 267.2848716 | -1.270406785 | 0.261825242 | 1.2121750239 | 2.5553265864 | Km<KS.1 | NaCl | KS.1 |
| FIM1_912 | 44.44843529 | 19.55702612 | 111.3884135 | -0.791338736 | 0.359889685 | 0.001611957 | 0.006390029 | ns | NaCl | KS.1 |
| FIM1_913 | 1704.625537 | 1805.875275 | 2001.170019 | 0.054083087 | 0.210141408 | 0.750407259 | 0.837493337 | ns | NaCl | KS.1 |
| FIM1_914 | 441.5475325 | 456.0990522 | 592.0757581 | 0.040750765 | 0.126082806 | 0.725070237 | 0.817408662 | ns | NaCl | KS.1 |
| FIM1_915 | 3502.4855 | 5061.406986 | 4187.559889 | 0.48928768 | 0.115992995 | 5.4857745221 | 5.1108105332 | ns | NaCl | KS.1 |
| FIM1_916 | 1347.611793 | 981.9573187 | 1234.286868 | -0.382209574 | 0.158095517 | 0.005902082 | 0.018762237 | ns | NaCl | KS.1 |
| FIM1_917 | 9805.731922 | 22660.26164 | 10820.5135 | 0.911055323 | 0.308411645 | 0.000124808 | 0.000751718 | ns | NaCl | KS.1 |
| FIM1_918 | 364.0067696 | 250.4453811 | 235.7526735 | -0.382830093 | 0.233648265 | 0.037468164 | 0.083965817 | ns | NaCl | KS.1 |
| FIM1_919 | 223.1983997 | 205.7635609 | 300.1039835 | -0.070433293 | 0.233406789 | 0.69716567 | 0.796991156 | ns | NaCl | KS.1 |
| FIM1_920 | 2547.704966 | 1812.779982 | 1881.743709 | -0.397443583 | 0.179407233 | 0.009455127 | 0.027300787 | ns | NaCl | KS.1 |
| FIM1_921 | 2987.501023 | 2239.421225 | 2693.04147 | -0.358061465 | 0.142474933 | 0.005019332 | 0.016342393 | ns | NaCl | KS.1 |
| FIM1_922 | 532.4605498 | 300.0481714 | 488.257024 | -0.714820594 | 0.194196358 | 2.2155471263 | 0.000170098 | ns | NaCl | KS.1 |
| FIM1_923 | 392.7508613 | 421.4184385 | 409.4636735 | 0.07464089 | 0.179243939 | 0.625046906 | 0.738774851 | ns | NaCl | KS.1 |
| FIM1_924 | 1354.784699 | 478.5593059 | 926.2492243 | -1.438935407 | 0.149384817 | 2.0654681929 | 1.2844630324 | Km<KS.1 | NaCl | KS.1 |
| FIM1_925 | 245.8916393 | 98.45320838 | 227.6234873 | -1.157577289 | 0.225653539 | 5.3816068369 | 1.2810284217 | Km<KS.1 | NaCl | KS.1 |
| FIM1_926 | 1754.197217 | 1387.153273 | 4723.754127 | -0.185915806 | 0.27286652 | 0.345461615 | 0.480434766 | ns | NaCl | KS.1 |
| FIM1_927 | 7971.025109 | 9964.857031 | 6918.70648 | 0.302559426 | 0.083250781 | 0.000147657 | 0.000867287 | ns | NaCl | KS.1 |
| FIM1_928 | 1697.638879 | 1014.095788 | 996.4788868 | -0.663376726 | 0.161574429 | 4.6203830532 | 4.4790016730 | ns | NaCl | KS.1 |
| FIM1_929 | 1222.956583 | 1314.421736 | 1444.466614 | 0.070994275 | 0.201644092 | 0.668106285 | 0.774063523 | ns | NaCl | KS.1 |
| FIM1_930 | 1542.481644 | 1470.934701 | 1751.919322 | -0.051388515 | 0.166526055 | 0.722553612 | 0.816421581 | ns | NaCl | KS.1 |
| FIM1_931 | 2277.856377 | 2047.518507 | 1672.699267 | -0.134156888 | 0.118018639 | 0.219957417 | 0.34411577 | ns | NaCl | KS.1 |

| | | | | | | | | | | |
|---|---|---|---|---|---|---|---|---|---|---|
| FIM1_932 | 413.8770574 | 388.1826195 | 497.2585964 | -0.075950335 | 0.118031285 | 0.487880365 | 0.621722545 | ns | NaCl | KS.1 |
| FIM1_933 | 1833.935171 | 1583.596799 | 2148.831949 | -0.195972783 | 0.083648495 | 0.014600022 | 0.038451619 | ns | NaCl | KS.1 |
| FIM1_934 | 1523.49147 | 1111.180696 | 1323.273902 | -0.407780184 | 0.126981762 | 0.000439815 | 0.00216427 | ns | NaCl | KS.1 |
| FIM1_935 | 1283.15222 | 974.7241411 | 1205.089316 | -0.211826292 | 0.28427116 | 0.292452616 | 0.423813506 | ns | NaCl | KS.1 |
| FIM1_936 | 7991.335837 | 7525.773537 | 8369.136041 | -0.056730354 | 0.21092163 | 0.7392143 | 0.828846325 | ns | NaCl | KS.1 |
| FIM1_937 | 3599.392639 | 2554.72192 | 2593.518284 | -0.452219815 | 0.117621862 | 3.2188739252 | 0.000236686 | ns | NaCl | KS.1 |
| FIM1_938 | 3779.429365 | 4167.234483 | 3427.445661 | 0.103064617 | 0.180807523 | 0.50237911 | 0.634671425 | ns | NaCl | KS.1 |
| FIM1_939 | 1974.972413 | 2251.787491 | 1869.916936 | 0.178265119 | 0.080764373 | 0.021786614 | 0.053604552 | ns | NaCl | KS.1 |
| FIM1_940 | 2702.836979 | 3335.200662 | 2972.43512 | 0.239355038 | 0.166303038 | 0.097114714 | 0.180682761 | ns | NaCl | KS.1 |
| FIM1_941 | 443.6775415 | 671.2724179 | 506.1071318 | 0.444127302 | 0.229866665 | 0.015304209 | 0.039758977 | ns | NaCl | KS.1 |
| FIM1_942 | 1060.784543 | 3294.222176 | 1787.929814 | 1.486501839 | 0.226744404 | 1.6545437964 | 1.2471750586 | Km>KS.1 | NaCl | KS.1 |
| FIM1_943 | 9796.842674 | 13321.68277 | 8646.69721 | 0.382472113 | 0.144504535 | 0.003060781 | 0.01086119 | ns | NaCl | KS.1 |
| FIM1_944 | 1525.707003 | 2100.912559 | 1283.016317 | 0.310082474 | 0.237627308 | 0.093473913 | 0.175749326 | ns | NaCl | KS.1 |
| FIM1_945 | 2784.576335 | 2079.221504 | 2338.598098 | -0.388802431 | 0.104194152 | 6.8428493887 | 0.000456225 | ns | NaCl | KS.1 |
| FIM1_946 | 37.33270945 | 38.05496652 | 45.99085676 | 0.019553584 | 0.22455412 | 0.912014608 | 0.948722964 | ns | NaCl | KS.1 |
| FIM1_947 | 4102.769185 | 6526.335186 | 5270.519313 | 0.614705285 | 0.133259467 | 5.3462339552 | 6.9084451759 | ns | NaCl | KS.1 |
| FIM1_948 | 4577.711336 | 6436.186603 | 5590.866524 | 0.457390996 | 0.105067874 | 3.4387175505 | 3.4913509824 | ns | NaCl | KS.1 |
| FIM1_949 | 1016.520405 | 511.8741204 | 1384.197803 | -0.799416109 | 0.248915835 | 8.8639348583 | 0.000563914 | ns | NaCl | KS.1 |
| FIM1_950 | 519.2149547 | 359.6121239 | 465.2174882 | -0.374775135 | 0.229565874 | 0.039269075 | 0.087007179 | ns | NaCl | KS.1 |
| FIM1_951 | 17318.99722 | 26712.89266 | 20186.18416 | 0.558451261 | 0.147575363 | 2.6101187792 | 0.000194975 | ns | NaCl | KS.1 |
| FIM1_952 | 32.1462115 | 27.14545819 | 41.24879713 | -0.116672376 | 0.30065964 | 0.565661386 | 0.687845561 | ns | NaCl | KS.1 |
| FIM1_953 | 4969.203147 | 3088.114363 | 3895.405561 | -0.625717382 | 0.139884681 | 1.0014534969 | 1.1695378280 | ns | NaCl | KS.1 |
| FIM1_954 | 2668.712073 | 1797.778317 | 1865.356409 | -0.428375309 | 0.219524376 | 0.015702393 | 0.040518365 | ns | NaCl | KS.1 |
| FIM1_955 | 1316.717039 | 1027.86108 | 1946.284371 | -0.216317424 | 0.251891488 | 0.254566709 | 0.382734778 | ns | NaCl | KS.1 |
| FIM1_957 | 13290.45429 | 8930.697584 | 11679.93381 | -0.36198026 | 0.270087432 | 0.070302185 | 0.139399509 | ns | NaCl | KS.1 |
| FIM1_958 | 3588.415188 | 4672.321834 | 4172.932193 | 0.304905706 | 0.170264901 | 0.037882858 | 0.08474575 | ns | NaCl | KS.1 |
| FIM1_959 | 1379.66252 | 3456.134975 | 1630.320037 | 1.125085278 | 0.25284887 | 1.7237823723 | 2.5952101534 | Km>KS.1 | NaCl | KS.1 |
| FIM1_960 | 3092.391866 | 3426.245598 | 2383.03718 | 0.128096087 | 0.118926402 | 0.244673683 | 0.372248188 | ns | NaCl | KS.1 |
| FIM1_961 | 4962.612484 | 4807.035035 | 4809.67305 | -0.042237407 | 0.091268182 | 0.627863048 | 0.740369439 | ns | NaCl | KS.1 |
| FIM1_962 | 2584.982182 | 3465.739531 | 2784.4084 | 0.392749504 | 0.100352511 | 3.1964152660 | 0.000235937 | ns | NaCl | KS.1 |
| FIM1_963 | 662.1967601 | 632.4146854 | 705.9682752 | -0.052765982 | 0.149715949 | 0.692335936 | 0.792720663 | ns | NaCl | KS.1 |
| FIM1_964 | 201.0408861 | 208.0486454 | 229.22586 | 0.047021471 | 0.146605034 | 0.719954612 | 0.81403959 | ns | NaCl | KS.1 |
| FIM1_965 | 741.3438178 | 741.8426459 | 1032.537501 | -0.001680449 | 0.099532757 | 0.985777311 | 0.993051943 | ns | NaCl | KS.1 |
| FIM1_966 | 3895.569715 | 3841.162223 | 3950.198171 | -0.015960022 | 0.150962335 | 0.905371173 | 0.943872872 | ns | NaCl | KS.1 |
| FIM1_967 | 842.4594379 | 1086.692403 | 1015.043269 | 0.325149985 | 0.12196426 | 0.003706763 | 0.012779728 | ns | NaCl | KS.1 |
| FIM1_968 | 619.3564277 | 648.7473071 | 752.0076583 | 0.053530986 | 0.148673075 | 0.686467963 | 0.787961288 | ns | NaCl | KS.1 |
| FIM1_969 | 300.8168352 | 320.026383 | 440.0148115 | 0.069294911 | 0.163017035 | 0.626563492 | 0.739711913 | ns | NaCl | KS.1 |
| FIM1_970 | 749.5573683 | 839.6522449 | 965.1903519 | 0.141549209 | 0.115282078 | 0.186519676 | 0.302850975 | ns | NaCl | KS.1 |
| FIM1_971 | 86.13052911 | 90.38156 | 131.9005992 | 0.04716882 | 0.19742165 | 0.772677751 | 0.852911429 | ns | NaCl | KS.1 |
| FIM1_972 | 199.6892091 | 201.366558 | 157.1682748 | -9.20E-05 | 0.189646362 | 0.999537872 | 0.999786056 | ns | NaCl | KS.1 |
| FIM1_973 | 795.5300445 | 927.4040486 | 1188.232828 | 0.201051998 | 0.096502525 | 0.027871273 | 0.065560086 | ns | NaCl | KS.1 |
| FIM1_974 | 1197.76611 | 1174.797342 | 1062.195313 | -0.022964948 | 0.157207118 | 0.868360462 | 0.921325079 | ns | NaCl | KS.1 |
| FIM1_975 | 2486.753251 | 2003.119342 | 2546.823218 | -0.238017933 | 0.179416065 | 0.119023499 | 0.212924095 | ns | NaCl | KS.1 |
| FIM1_976 | 51.76560197 | 38.93016938 | 101.3528942 | -0.204520844 | 0.284269526 | 0.308985594 | 0.441852064 | ns | NaCl | KS.1 |
| FIM1_977 | 4080.650414 | 5312.285294 | 3415.972487 | 0.216427344 | 0.269200032 | 0.27047619 | 0.399412005 | ns | NaCl | KS.1 |

| | | | | | | | | | | |
|---|---|---|---|---|---|---|---|---|---|---|
| FIM1_978 | 5619.33366 | 9158.667816 | 4313.41991 | 0.347772621 | 0.338141446 | 0.113828046 | 0.205179176 | ns | NaCl | KS.1 |
| FIM1_979 | 3280.803335 | 4851.387319 | 2201.409289 | 0.346795508 | 0.27494661 | 0.085231539 | 0.163590627 | ns | NaCl | KS.1 |
| FIM1_980 | 3373.508697 | 2276.395171 | 2473.290594 | -0.415770455 | 0.227691691 | 0.021969647 | 0.05394817 | ns | NaCl | KS.1 |
| FIM1_981 | 960.9432653 | 882.1739206 | 1418.350743 | -0.064506668 | 0.26696167 | 0.738924371 | 0.828708013 | ns | NaCl | KS.1 |
| FIM1_982 | 1907.714148 | 1222.644233 | 1544.672182 | -0.531603342 | 0.188714805 | 0.000945603 | 0.004097888 | ns | NaCl | KS.1 |
| FIM1_983 | 12940.71826 | 13734.9499 | 12436.09678 | 0.077163952 | 0.101311155 | 0.420217069 | 0.556301202 | ns | NaCl | KS.1 |
| FIM1_984 | 36.13225774 | 16.2163357 | 91.15760918 | -0.381804553 | 0.439979856 | 0.095416331 | 0.178652011 | ns | NaCl | KS.1 |
| FIM1_988 | 3276.351303 | 2387.879244 | 2634.034784 | -0.403083198 | 0.132154036 | 0.000780668 | 0.003498939 | ns | NaCl | KS.1 |
| FIM1_989 | 31.4847133 | 9.647761052 | 71.54174204 | -1.196589156 | 0.456978422 | 7.75267913928 | 0.000505499 | Km<KS.1 | NaCl | KS.1 |
| FIM1_991 | 1002.159464 | 1184.103925 | 1301.236003 | 0.166379691 | 0.205418503 | 0.321145394 | 0.454165739 | ns | NaCl | KS.1 |
| FIM1_992 | 2101.107001 | 1888.629489 | 2183.789848 | -0.109558285 | 0.189211513 | 0.489804129 | 0.622672274 | ns | NaCl | KS.1 |
| FIM1_993 | 1677.730709 | 1944.901717 | 1362.119294 | 0.140114182 | 0.21698623 | 0.419615776 | 0.555948998 | ns | NaCl | KS.1 |
| FIM1_994 | 2564.511422 | 3088.455847 | 2129.332173 | 0.204232551 | 0.17612876 | 0.175203931 | 0.288909366 | ns | NaCl | KS.1 |
| FIM1_995 | 2746.368856 | 4030.489693 | 2185.273744 | 0.378974841 | 0.245486716 | 0.045713969 | 0.098136935 | ns | NaCl | KS.1 |
| FIM1_996 | 3211.218668 | 6167.360924 | 2898.353279 | 0.548542381 | 0.345772238 | 0.01853446 | 0.046593704 | ns | NaCl | KS.1 |
| FIM1_997 | 104.2965677 | 101.292319 | 234.4595379 | -0.027655575 | 0.231738318 | 0.878010235 | 0.92819824 | ns | NaCl | KS.1 |
| FIM1_998 | 1296.435269 | 927.0518345 | 1120.239119 | -0.437784266 | 0.127484767 | 0.000171565 | 0.000983338 | ns | NaCl | KS.1 |
| FIM1_999 | 2178.881997 | 2179.196818 | 1425.907567 | -0.000355033 | 0.126683938 | 0.997565406 | 0.999172115 | ns | NaCl | KS.1 |
| FIM1_1000 | 386.8143344 | 277.5915265 | 961.7356256 | -0.280280884 | 0.2751074 | 0.160827052 | 0.269580385 | ns | NaCl | KS.1 |
| FIM1_1001 | 397.684172 | 338.0233069 | 323.4595592 | -0.204986089 | 0.119187957 | 0.062666087 | 0.127363257 | ns | NaCl | KS.1 |
| FIM1_1002 | 369.145903 | 332.6124277 | 328.5117821 | -0.125509944 | 0.143052852 | 0.328564257 | 0.462145089 | ns | NaCl | KS.1 |
| FIM1_1003 | 4089.561765 | 7200.255957 | 3583.500302 | 0.607314458 | 0.25981462 | 0.002734403 | 0.009907977 | ns | NaCl | KS.1 |
| FIM1_1004 | 2519.977699 | 2497.028173 | 3041.230843 | -0.00897883 | 0.190381357 | 0.955062263 | 0.976656682 | ns | NaCl | KS.1 |
| FIM1_1005 | 1157.402434 | 1210.328561 | 1178.634649 | 0.051431955 | 0.147465409 | 0.696286646 | 0.796514446 | ns | NaCl | KS.1 |
| FIM1_1006 | 3264.050267 | 3274.528977 | 3500.720483 | 0.0037858 | 0.155993838 | 0.978072332 | 0.989137805 | ns | NaCl | KS.1 |
| FIM1_1007 | 632.9348567 | 529.7159646 | 680.5163647 | -0.22818912 | 0.104601224 | 0.020131989 | 0.049878807 | ns | NaCl | KS.1 |
| FIM1_1008 | 272.7739774 | 196.6438104 | 350.3893167 | -0.37848544 | 0.183121034 | 0.014856732 | 0.038860273 | ns | NaCl | KS.1 |
| FIM1_1009 | 698.9495419 | 634.9228811 | 736.7478592 | -0.115826963 | 0.131640047 | 0.334508329 | 0.468650784 | ns | NaCl | KS.1 |
| FIM1_1010 | 1987.299304 | 1617.552139 | 2615.082822 | -0.184976768 | 0.238408868 | 0.314339972 | 0.447675071 | ns | NaCl | KS.1 |
| FIM1_1011 | 9615.05736 | 10368.71767 | 10635.097 | 0.076654527 | 0.190981939 | 0.631168072 | 0.743561724 | ns | NaCl | KS.1 |
| FIM1_1012 | 1582.137252 | 2016.586074 | 2599.211892 | 0.18340724 | 0.284622976 | 0.360654272 | 0.495924545 | ns | NaCl | KS.1 |
| FIM1_1013 | 2513.197162 | 2511.400734 | 2768.382107 | -0.000730508 | 0.197353952 | 0.996429583 | 0.998235437 | ns | NaCl | KS.1 |
| FIM1_1014 | 4486.351486 | 865.6599224 | 2115.903474 | -2.020146335 | 0.428537201 | 1.1546986873 | 3.2455513951 | Km<KS.1 | NaCl | KS.1 |
| FIM1_1015 | 2732.623584 | 3947.240447 | 2155.16098 | 0.342238297 | 0.258132866 | 0.078648968 | 0.1530824 | ns | NaCl | KS.1 |
| FIM1_1016 | 339.4233785 | 205.4173471 | 196.0072282 | -0.519163709 | 0.257991699 | 0.009079223 | 0.026492161 | ns | NaCl | KS.1 |
| FIM1_1017 | 5235.394536 | 7403.420479 | 4189.521793 | 0.444490732 | 0.135627453 | 0.000294632 | 0.001546198 | ns | NaCl | KS.1 |
| FIM1_1018 | 1262.429379 | 1596.323188 | 1304.181682 | 0.311886973 | 0.096030395 | 0.000593409 | 0.002783664 | ns | NaCl | KS.1 |
| FIM1_1019 | 544.1248846 | 775.448603 | 854.6122633 | 0.425124413 | 0.171841052 | 0.004177916 | 0.014072532 | ns | NaCl | KS.1 |
| FIM1_1020 | 119.8601611 | 50.9991872 | 134.9193748 | -0.813059506 | 0.358988191 | 0.001245455 | 0.005142026 | ns | NaCl | KS.1 |
| FIM1_1021 | 156.4948654 | 155.5917177 | 651.045516 | -0.004218627 | 0.266252744 | 0.982569837 | 0.991175946 | ns | NaCl | KS.1 |
| FIM1_1022 | 10079.11202 | 10002.91839 | 7715.193246 | -0.009750127 | 0.104686577 | 0.921242537 | 0.955028468 | ns | NaCl | KS.1 |
| FIM1_1024 | 1662.829692 | 1075.887343 | 1785.986354 | -0.455231933 | 0.240159084 | 0.015813614 | 0.040763074 | ns | NaCl | KS.1 |
| FIM1_1025 | 6835.031122 | 7161.575238 | 5296.423224 | 0.054633459 | 0.14250245 | 0.669976077 | 0.775687918 | ns | NaCl | KS.1 |
| FIM1_1026 | 7870.259909 | 8758.210519 | 6893.761453 | 0.125077534 | 0.145551708 | 0.336968964 | 0.471036975 | ns | NaCl | KS.1 |
| FIM1_1027 | 591.3221306 | 670.82589 | 1025.936303 | 0.136068626 | 0.178363639 | 0.371079201 | 0.505786035 | ns | NaCl | KS.1 |

| | | | | | | | | | | |
|---|---|---|---|---|---|---|---|---|---|---|
| FIM1_1028 | 139.5419646 | 103.4691846 | 216.8554548 | -0.308808513 | 0.216452002 | 0.076340133 | 0.149230713 | ns | NaCl | KS.1 |
| FIM1_1029 | 14075.87064 | 17477.01474 | 12682.87503 | 0.21542708 | 0.211673767 | 0.208161138 | 0.329809447 | ns | NaCl | KS.1 |
| FIM1_1030 | 5198.620832 | 10252.68204 | 4336.793453 | 0.616267102 | 0.335271667 | 0.008302512 | 0.024778042 | ns | NaCl | KS.1 |
| FIM1_1031 | 90.22548724 | 69.62954796 | 81.79274124 | -0.24199815 | 0.24306679 | 0.19437469 | 0.312242196 | ns | NaCl | KS.1 |
| FIM1_1032 | 716.560516 | 436.7651372 | 743.9069697 | -0.674322229 | 0.114943667 | 4.9709246884 | 1.5171994064 | ns | NaCl | KS.1 |
| FIM1_1033 | 244.5031731 | 207.4046209 | 215.5946391 | -0.171115334 | 0.200608263 | 0.299990133 | 0.432316514 | ns | NaCl | KS.1 |
| FIM1_1034 | 588.5921053 | 460.7955174 | 648.1549468 | -0.325520963 | 0.110858956 | 0.001600813 | 0.006361059 | ns | NaCl | KS.1 |
| FIM1_1035 | 117.9983019 | 130.0819018 | 164.1932233 | 0.093495683 | 0.178378385 | 0.538975363 | 0.665915914 | ns | NaCl | KS.1 |
| FIM1_1036 | 556.1602986 | 552.135448 | 581.8946502 | -0.015226618 | 0.127746052 | 0.896605741 | 0.938681305 | ns | NaCl | KS.1 |
| FIM1_1037 | 1094.432907 | 1263.152983 | 1268.132034 | 0.195242789 | 0.075513389 | 0.007451053 | 0.022644465 | ns | NaCl | KS.1 |
| FIM1_1038 | 3640.720333 | 4194.050015 | 3371.647619 | 0.191773263 | 0.079314562 | 0.012057283 | 0.033140875 | ns | NaCl | KS.1 |
| FIM1_1039 | 3113.560591 | 3539.548532 | 3203.341312 | 0.163782027 | 0.11117361 | 0.114499429 | 0.206090686 | ns | NaCl | KS.1 |
| FIM1_1040 | 1336.439303 | 1277.741743 | 1696.149232 | -0.045934813 | 0.187020034 | 0.770397148 | 0.851527619 | ns | NaCl | KS.1 |
| FIM1_1041 | 3069.895215 | 2976.50487 | 2064.977982 | -0.02643923 | 0.23560594 | 0.884390376 | 0.931380635 | ns | NaCl | KS.1 |
| FIM1_1042 | 99.11435545 | 108.0251623 | 77.10730827 | 0.074139259 | 0.188799453 | 0.639776419 | 0.749480061 | ns | NaCl | KS.1 |
| FIM1_1043 | 182.5838338 | 163.5943755 | 188.1524893 | -0.103226613 | 0.224359477 | 0.559832573 | 0.683308894 | ns | NaCl | KS.1 |
| FIM1_1044 | 1551.07984 | 1841.907687 | 2014.807761 | 0.170901706 | 0.205995898 | 0.309084832 | 0.441866965 | ns | NaCl | KS.1 |
| FIM1_1045 | 483.1574034 | 364.3234728 | 478.7153123 | -0.346499916 | 0.148009485 | 0.008473556 | 0.025182759 | ns | NaCl | KS.1 |
| FIM1_1046 | 638.6833739 | 520.4376351 | 813.5084914 | -0.252388193 | 0.132271102 | 0.035639358 | 0.080374347 | ns | NaCl | KS.1 |
| FIM1_1047 | 1196.161235 | 1257.704126 | 1493.593789 | 0.067179883 | 0.077703566 | 0.370863904 | 0.505631111 | ns | NaCl | KS.1 |
| FIM1_1048 | 1557.04946 | 1852.065515 | 1727.147317 | 0.223256806 | 0.108197187 | 0.027349808 | 0.064547103 | ns | NaCl | KS.1 |
| FIM1_1049 | 2705.153563 | 6030.022751 | 2601.580577 | 0.802160522 | 0.334586854 | 0.000958 | 0.004140789 | ns | NaCl | KS.1 |
| FIM1_1050 | 80.63043012 | 30.97674488 | 107.7630703 | -0.97988005 | 0.35993136 | 0.000174387 | 0.000997215 | ns | NaCl | KS.1 |
| FIM1_1051 | 597.9704721 | 439.9672973 | 591.553105 | -0.354808629 | 0.171170878 | 0.016201515 | 0.041655057 | ns | NaCl | KS.1 |
| FIM1_1052 | 621.8643442 | 353.3691702 | 1052.997585 | -0.661542646 | 0.223173696 | 0.000347054 | 0.001779996 | ns | NaCl | KS.1 |
| FIM1_1053 | 86.93905753 | 92.4250267 | 89.13311434 | 0.0507827 | 0.199938693 | 0.757769782 | 0.843001937 | ns | NaCl | KS.1 |
| FIM1_1054 | 2723.794716 | 3195.181952 | 2238.829363 | 0.209509799 | 0.097441353 | 0.023076955 | 0.056003831 | ns | NaCl | KS.1 |
| FIM1_1055 | 184.4556999 | 112.6422117 | 204.8296976 | -0.587333481 | 0.204291831 | 0.000623863 | 0.002906101 | ns | NaCl | KS.1 |
| FIM1_1056 | 546.5623911 | 641.0533006 | 564.8640782 | 0.189457648 | 0.147755239 | 0.150187246 | 0.255478488 | ns | NaCl | KS.1 |
| FIM1_1057 | 1216.436599 | 1197.246082 | 1415.285574 | -0.019166619 | 0.104609163 | 0.845795067 | 0.905688863 | ns | NaCl | KS.1 |
| FIM1_1058 | 2251.159072 | 2745.602423 | 2535.405671 | 0.259588505 | 0.106710016 | 0.00934072 | 0.027127894 | ns | NaCl | KS.1 |
| FIM1_1060 | 4234.283147 | 9150.694674 | 4530.229662 | 0.968988797 | 0.212294595 | 2.0607194241 | 3.0242121342 | ns | NaCl | KS.1 |
| FIM1_1061 | 602.3145028 | 934.610953 | 795.8899212 | 0.568452246 | 0.149604625 | 2.3558848900 | 0.000179213 | ns | NaCl | KS.1 |
| FIM1_1062 | 600.7093613 | 782.6177875 | 854.6353268 | 0.343490547 | 0.112136085 | 0.000976289 | 0.004208873 | ns | NaCl | KS.1 |
| FIM1_1063 | 468.6417012 | 436.0186028 | 789.7390678 | -0.087164529 | 0.142576792 | 0.496556672 | 0.628592734 | ns | NaCl | KS.1 |
| FIM1_1064 | 883.1833035 | 839.5807673 | 1260.751839 | -0.055979172 | 0.150435696 | 0.675795041 | 0.779703231 | ns | NaCl | KS.1 |
| FIM1_1065 | 1258.552532 | 1219.890906 | 1242.392912 | -0.042443056 | 0.099724332 | 0.653050673 | 0.760207849 | ns | NaCl | KS.1 |
| FIM1_1066 | 418.0962777 | 492.4811103 | 442.4206693 | 0.204585823 | 0.126165841 | 0.076729154 | 0.149755804 | ns | NaCl | KS.1 |
| FIM1_1067 | 294.1093297 | 329.7713793 | 409.8270158 | 0.138251895 | 0.142381078 | 0.279891921 | 0.410030126 | ns | NaCl | KS.1 |
| FIM1_1068 | 287.1828473 | 372.7777899 | 234.1648906 | 0.253138075 | 0.228140357 | 0.158935245 | 0.267309954 | ns | NaCl | KS.1 |
| FIM1_1069 | 168.1511274 | 236.6512807 | 417.8969508 | 0.293946624 | 0.273663289 | 0.140999664 | 0.242305122 | ns | NaCl | KS.1 |
| FIM1_1070 | 786.8414843 | 688.9282209 | 1189.605498 | -0.173125518 | 0.112221922 | 0.097911006 | 0.181756439 | ns | NaCl | KS.1 |
| FIM1_1071 | 1567.115714 | 1637.268179 | 1522.70044 | 0.061807398 | 0.071561496 | 0.373722123 | 0.50880979 | ns | NaCl | KS.1 |
| FIM1_1072 | 1288.933693 | 2557.88974 | 1457.758092 | 0.620834874 | 0.337239519 | 0.008067307 | 0.02421405 | ns | NaCl | KS.1 |
| FIM1_1074 | 1418.867206 | 1419.149894 | 1221.662544 | 0.001365757 | 0.11981511 | 0.990179896 | 0.995583061 | ns | NaCl | KS.1 |

| | | | | | | | | | | |
|---|---|---|---|---|---|---|---|---|---|---|
| FIM1_1075 | 3392.15828 | 3680.226956 | 3065.067165 | 0.099717817 | 0.126801011 | 0.391396049 | 0.526838566 | ns | NaCl | KS.1 |
| FIM1_1076 | 387.5069007 | 352.9125084 | 428.2304838 | -0.089561674 | 0.211910053 | 0.600350441 | 0.719177329 | ns | NaCl | KS.1 |
| FIM1_1077 | 176.2774978 | 179.4658963 | 125.6029179 | 0.01238867 | 0.259391567 | 0.948246405 | 0.973087018 | ns | NaCl | KS.1 |
| FIM1_1078 | 299.3723471 | 329.9975551 | 294.9642808 | 0.107941452 | 0.163331151 | 0.4489633 | 0.584098435 | ns | NaCl | KS.1 |
| FIM1_1079 | 554.9126888 | 265.3768755 | 387.3355343 | -0.930027037 | 0.21156435 | 5.1761336968 | 6.7235679220 | ns | NaCl | KS.1 |
| FIM1_1080 | 1477.932402 | 994.3842262 | 1084.702492 | -0.408667826 | 0.235404131 | 0.027441202 | 0.064732091 | ns | NaCl | KS.1 |
| FIM1_1081 | 1265.832748 | 1055.004906 | 1316.689336 | -0.157559438 | 0.244204822 | 0.397106839 | 0.533078153 | ns | NaCl | KS.1 |
| FIM1_1082 | 1368.714747 | 1365.053558 | 1101.869711 | -0.002757961 | 0.193529482 | 0.986343149 | 0.993331411 | ns | NaCl | KS.1 |
| FIM1_1083 | 1770.739158 | 1204.690528 | 1554.803504 | -0.520055477 | 0.109082515 | 3.5753662864 | 4.9001232163 | ns | NaCl | KS.1 |
| FIM1_1084 | 1007.459935 | 579.6407479 | 682.8854909 | -0.693658108 | 0.185558614 | 1.9067204346 | 0.00014962 | ns | NaCl | KS.1 |
| FIM1_1085 | 501.2835101 | 510.2788332 | 580.9697454 | 0.018015586 | 0.114471292 | 0.865888142 | 0.919776371 | ns | NaCl | KS.1 |
| FIM1_1086 | 585.2707948 | 344.2612387 | 635.0051541 | -0.68378397 | 0.163624721 | 3.1371349760 | 3.2246377078 | ns | NaCl | KS.1 |
| FIM1_1087 | 572.6441929 | 686.0859218 | 755.4464923 | 0.218920216 | 0.136556586 | 0.07613285 | 0.148942559 | ns | NaCl | KS.1 |
| FIM1_1088 | 2146.128777 | 2563.878254 | 2300.985618 | 0.23884789 | 0.086126297 | 0.003730401 | 0.012840939 | ns | NaCl | KS.1 |
| FIM1_1089 | 159.4299784 | 118.4833027 | 127.3354343 | -0.322696289 | 0.212320964 | 0.060839461 | 0.124558155 | ns | NaCl | KS.1 |
| FIM1_1090 | 4632.334627 | 6170.436344 | 4998.7929 | 0.374232646 | 0.116240874 | 0.000498182 | 0.002399281 | ns | NaCl | KS.1 |
| FIM1_1091 | 2812.439935 | 4735.700338 | 2368.850307 | 0.574230291 | 0.240579545 | 0.002772295 | 0.010023379 | ns | NaCl | KS.1 |
| FIM1_1092 | 793.445512 | 938.8716212 | 980.3221085 | 0.226144775 | 0.09496671 | 0.012034636 | 0.033096913 | ns | NaCl | KS.1 |
| FIM1_1093 | 1125.239155 | 722.7460335 | 1159.130995 | -0.565769414 | 0.154085784 | 3.9740563706 | 0.000284474 | ns | NaCl | KS.1 |
| FIM1_1094 | 1436.005336 | 1970.123189 | 1676.028457 | 0.441108605 | 0.065430253 | 4.1038452901 | 2.1491189809 | ns | NaCl | KS.1 |
| FIM1_1095 | 1003.554301 | 1053.433906 | 1197.186712 | 0.062096532 | 0.104335164 | 0.527549233 | 0.656139358 | ns | NaCl | KS.1 |
| FIM1_1096 | 368.3402351 | 463.6887349 | 436.598392 | 0.260965064 | 0.170130689 | 0.075385716 | 0.147655094 | ns | NaCl | KS.1 |
| FIM1_1097 | 5113.403757 | 11544.95255 | 8368.502534 | 1.063862252 | 0.186101333 | 3.3588395696 | 1.0921716901 | Km>KS.1 | NaCl | KS.1 |
| FIM1_1098 | 4656.288954 | 4402.963094 | 3522.134248 | -0.062518059 | 0.160049081 | 0.656217403 | 0.763132674 | ns | NaCl | KS.1 |
| FIM1_1099 | 682.6106734 | 432.2848235 | 822.1322244 | -0.576727792 | 0.1675004 | 9.1465854954 | 0.000580412 | ns | NaCl | KS.1 |
| FIM1_1100 | 8444.436844 | 11275.39589 | 7850.830426 | 0.391757964 | 0.090956699 | 5.7998602257 | 5.3218917233 | ns | NaCl | KS.1 |
| FIM1_1101 | 81.11182553 | 56.55722221 | 78.11312331 | -0.191305917 | 0.362806542 | 0.368915124 | 0.504044881 | ns | NaCl | KS.1 |
| FIM1_1102 | 1403.505416 | 1429.306601 | 1220.438134 | 0.021294312 | 0.126359756 | 0.854471448 | 0.911839437 | ns | NaCl | KS.1 |
| FIM1_1103 | 1409.739065 | 1183.992919 | 973.7573959 | -0.168975683 | 0.214172574 | 0.32669007 | 0.459899009 | ns | NaCl | KS.1 |
| FIM1_1104 | 1962.939009 | 1564.512841 | 1469.220352 | -0.256340696 | 0.171851679 | 0.08302482 | 0.160034281 | ns | NaCl | KS.1 |
| FIM1_1105 | 4592.810592 | 8456.208765 | 4789.31649 | 0.862866861 | 0.074593491 | 4.6198014887 | 4.5967024812 | ns | NaCl | KS.1 |
| FIM1_1106 | 6843.539216 | 7154.707025 | 6673.423361 | 0.051318064 | 0.147859852 | 0.697537275 | 0.797025251 | ns | NaCl | KS.1 |
| FIM1_1107 | 1260.13626 | 1109.850225 | 984.9406452 | -0.147976213 | 0.149795836 | 0.266692178 | 0.395122986 | ns | NaCl | KS.1 |
| FIM1_1108 | 2320.096462 | 4391.357721 | 1788.027479 | 0.649552865 | 0.293485618 | 0.003093742 | 0.010962511 | ns | NaCl | KS.1 |
| FIM1_1109 | 3290.177088 | 4364.276004 | 4698.985873 | 0.342660587 | 0.153110137 | 0.011287264 | 0.031423692 | ns | NaCl | KS.1 |
| FIM1_1110 | 1325.186518 | 1508.115844 | 1348.385957 | 0.151540629 | 0.144814951 | 0.242539791 | 0.369445505 | ns | NaCl | KS.1 |
| FIM1_1111 | 1571.255575 | 978.4964279 | 1127.910016 | -0.635833518 | 0.124917056 | 4.4720864222 | 7.8818660272 | ns | NaCl | KS.1 |
| FIM1_1112 | 3155.814262 | 3613.390054 | 3978.287789 | 0.173693829 | 0.109619715 | 0.090080365 | 0.170720271 | ns | NaCl | KS.1 |
| FIM1_1113 | 1629.292164 | 1669.505964 | 1419.349751 | 0.021905971 | 0.224316908 | 0.901421136 | 0.941543177 | ns | NaCl | KS.1 |
| FIM1_1114 | 551.4013845 | 480.8062838 | 2368.019001 | -0.095677437 | 0.289104257 | 0.632548895 | 0.744307179 | ns | NaCl | KS.1 |
| FIM1_1115 | 224.9180836 | 188.4313831 | 361.7384784 | -0.168886941 | 0.224551604 | 0.341090762 | 0.475462746 | ns | NaCl | KS.1 |
| FIM1_1116 | 7999.821287 | 6873.676447 | 6624.83979 | -0.179223488 | 0.146019376 | 0.16950902 | 0.280821636 | ns | NaCl | KS.1 |
| FIM1_1117 | 1357.393349 | 1008.561372 | 1887.62211 | -0.358050966 | 0.160949561 | 0.010944151 | 0.030726383 | ns | NaCl | KS.1 |
| FIM1_1118 | 17.69664429 | 21.01045139 | 20.87135697 | 0.150616473 | 0.28609187 | 0.452148868 | 0.587474698 | ns | NaCl | KS.1 |
| FIM1_1119 | 3373.221071 | 2923.246832 | 2937.395474 | -0.173381649 | 0.133206803 | 0.152072474 | 0.257947685 | ns | NaCl | KS.1 |

| | | | | | | | | | | |
|---|---|---|---|---|---|---|---|---|---|---|
| FIM1_1120 | 25.48082313 | 17.07567628 | 43.48615303 | -0.297985695 | 0.295948024 | 0.150436797 | 0.255783687 | ns | NaCl | KS.1 |
| FIM1_1121 | 9150.2916 | 13485.19329 | 12996.19495 | 0.492807779 | 0.148301215 | 0.000201611 | 0.001123199 | ns | NaCl | KS.1 |
| FIM1_1122 | 5049.588789 | 6197.285733 | 5340.92505 | 0.278708525 | 0.07879903 | 0.000236094 | 0.001282279 | ns | NaCl | KS.1 |
| FIM1_1123 | 495.5229917 | 372.2291866 | 448.2279881 | -0.34931458 | 0.148486315 | 0.008120042 | 0.024321017 | ns | NaCl | KS.1 |
| FIM1_1124 | 3490.984541 | 2460.443296 | 3235.534849 | -0.446450502 | 0.138668546 | 0.000358372 | 0.001826744 | ns | NaCl | KS.1 |
| FIM1_1125 | 27759.09062 | 35221.64283 | 21740.66123 | 0.298593689 | 0.130172222 | 0.011695681 | 0.032271778 | ns | NaCl | KS.1 |
| FIM1_1126 | 716.0053716 | 721.7115787 | 764.9335658 | 0.007641585 | 0.237675025 | 0.966633561 | 0.984442572 | ns | NaCl | KS.1 |
| FIM1_1127 | 298.2232486 | 183.85329 | 411.6442297 | -0.500423826 | 0.25467532 | 0.011025179 | 0.030866583 | ns | NaCl | KS.1 |
| FIM1_1128 | 221.1271026 | 99.89940194 | 192.2800568 | -0.851810861 | 0.306771985 | 0.000275383 | 0.001460587 | ns | NaCl | KS.1 |
| FIM1_1129 | 2867.364698 | 3152.254581 | 3127.035772 | 0.119972437 | 0.112177109 | 0.251793282 | 0.380546285 | ns | NaCl | KS.1 |
| FIM1_1130 | 461.4346903 | 427.0058206 | 445.2423528 | -0.088211393 | 0.154809491 | 0.519186787 | 0.650162398 | ns | NaCl | KS.1 |
| FIM1_1131 | 1241.982695 | 1322.95228 | 1327.843283 | 0.082115147 | 0.10896951 | 0.421342385 | 0.557494246 | ns | NaCl | KS.1 |
| FIM1_1132 | 249.0332241 | 370.3445659 | 990.3518956 | 0.440108825 | 0.216123182 | 0.012217814 | 0.033434338 | ns | NaCl | KS.1 |
| FIM1_1133 | 435.247099 | 729.7004654 | 653.9017944 | 0.552066315 | 0.250863466 | 0.004965488 | 0.01620072 | ns | NaCl | KS.1 |
| FIM1_1134 | 369.2614923 | 291.1539044 | 307.6470545 | -0.218746679 | 0.233144852 | 0.228816114 | 0.354408521 | ns | NaCl | KS.1 |
| FIM1_1135 | 12490.65027 | 21746.17882 | 14001.82884 | 0.619929669 | 0.241808831 | 0.001393504 | 0.005604434 | ns | NaCl | KS.1 |
| FIM1_1136 | 1387.181509 | 683.5384978 | 1073.687292 | -0.953266896 | 0.146356684 | 3.8002712722 | 2.0226617226 | ns | NaCl | KS.1 |
| FIM1_1137 | 5691.501495 | 2494.886973 | 3821.963831 | -0.951855145 | 0.275733814 | 2.0506657254 | 0.000158911 | ns | NaCl | KS.1 |
| FIM1_1138 | 576.7808053 | 316.4438027 | 592.9896694 | -0.805615008 | 0.141389143 | 9.5723943628 | 2.7369345951 | ns | NaCl | KS.1 |
| FIM1_1139 | 11382.50008 | 8763.455875 | 9084.169046 | -0.351760016 | 0.093664261 | 7.380013314 | 0.000486944 | ns | NaCl | KS.1 |
| FIM1_1140 | 4413.321423 | 2510.617746 | 3299.452156 | -0.443039754 | 0.337723195 | 0.048987966 | 0.103929694 | ns | NaCl | KS.1 |
| FIM1_1141 | 5989.793962 | 3096.349314 | 3765.705268 | -0.637102707 | 0.314407847 | 0.005062843 | 0.016441021 | ns | NaCl | KS.1 |
| FIM1_1142 | 982.552282 | 841.1820407 | 869.6513995 | -0.201405055 | 0.107455144 | 0.04533478 | 0.097509957 | ns | NaCl | KS.1 |
| FIM1_1143 | 1867.516439 | 1958.728774 | 1700.876473 | 0.061607692 | 0.092442394 | 0.484587412 | 0.618794244 | ns | NaCl | KS.1 |
| FIM1_1144 | 846.4405392 | 983.4742833 | 867.3689057 | 0.150312006 | 0.199439825 | 0.360646034 | 0.495924545 | ns | NaCl | KS.1 |
| FIM1_1145 | 9731.244047 | 15187.33537 | 9504.782481 | 0.613415965 | 0.095473096 | 1.743467711 | 7.6457018656 | ns | NaCl | KS.1 |
| FIM1_1146 | 634.9863471 | 487.6346967 | 629.5973332 | -0.318872964 | 0.145388284 | 0.013966495 | 0.037117154 | ns | NaCl | KS.1 |
| FIM1_1147 | 3309.20661 | 5536.237557 | 3609.849336 | 0.706455299 | 0.107237426 | 4.6351290788 | 2.4020590799 | ns | NaCl | KS.1 |
| FIM1_1148 | 668.4503075 | 715.2528745 | 672.4674812 | 0.078545503 | 0.147529111 | 0.551124613 | 0.675808574 | ns | NaCl | KS.1 |
| FIM1_1149 | 330.7205155 | 316.8339752 | 286.7529395 | -0.048671907 | 0.161959306 | 0.73129893 | 0.822602701 | ns | NaCl | KS.1 |
| FIM1_1150 | 1077.389769 | 1106.546682 | 929.2402497 | 0.029732059 | 0.146995733 | 0.821053773 | 0.889534521 | ns | NaCl | KS.1 |
| FIM1_1151 | 6586.997217 | 6993.541239 | 6165.423123 | 0.071127101 | 0.138255369 | 0.569527222 | 0.691578698 | ns | NaCl | KS.1 |
| FIM1_1152 | 693.067718 | 702.3125539 | 803.2191854 | 0.012731554 | 0.203318594 | 0.93903518 | 0.965227277 | ns | NaCl | KS.1 |
| FIM1_1153 | 38.93299615 | 36.81533464 | 75.4474694 | -0.077445268 | 0.240442099 | 0.673799691 | 0.778303567 | ns | NaCl | KS.1 |
| FIM1_1154 | 1283.726729 | 1137.253856 | 1354.118306 | -0.154672256 | 0.112986146 | 0.141752667 | 0.243256235 | ns | NaCl | KS.1 |
| FIM1_1155 | 106.3505207 | 94.04021944 | 96.14274037 | -0.091522517 | 0.285047634 | 0.645619165 | 0.754300314 | ns | NaCl | KS.1 |
| FIM1_1156 | 10144.67573 | 13455.23761 | 9766.387008 | 0.362971488 | 0.123845297 | 0.00138416 | 0.005571356 | ns | NaCl | KS.1 |
| FIM1_1157 | 2703.626843 | 2349.906043 | 2947.350756 | -0.184234539 | 0.097075569 | 0.045108735 | 0.097149765 | ns | NaCl | KS.1 |
| FIM1_1158 | 457.3585014 | 246.0954016 | 1034.973562 | -0.641294756 | 0.285405745 | 0.002957852 | 0.010571345 | ns | NaCl | KS.1 |
| FIM1_1159 | 1894.936773 | 2214.571835 | 1818.015142 | 0.210275244 | 0.083629876 | 0.00874924 | 0.025847666 | ns | NaCl | KS.1 |
| FIM1_1160 | 14450.20666 | 16298.18303 | 13109.57996 | 0.162148124 | 0.080919026 | 0.03731154 | 0.083652507 | ns | NaCl | KS.1 |
| FIM1_1161 | 86.53303315 | 75.05266136 | 118.611153 | -0.099165732 | 0.282906842 | 0.617496911 | 0.732311593 | ns | NaCl | KS.1 |
| FIM1_1162 | 2500.961804 | 4275.005434 | 6707.310668 | 0.497384423 | 0.296800748 | 0.02051693 | 0.050655944 | ns | NaCl | KS.1 |
| FIM1_1163 | 943.5105452 | 1151.823126 | 1155.211628 | 0.260746107 | 0.098771159 | 0.005158733 | 0.016676217 | ns | NaCl | KS.1 |
| FIM1_1164 | 34.88244179 | 41.22234249 | 38.21550992 | 0.119649084 | 0.265107364 | 0.536306712 | 0.663712909 | ns | NaCl | KS.1 |

| | | | | | | | | | | |
|---|---|---|---|---|---|---|---|---|---|---|
| FIM1_1165 | 948.4091822 | 1081.487127 | 916.3283181 | 0.149309965 | 0.157056025 | 0.280172859 | 0.410320864 | ns | NaCl | KS.1 |
| FIM1_1166 | 5885.436644 | 6154.512181 | 6055.591906 | 0.052887533 | 0.138417883 | 0.672723183 | 0.777782439 | ns | NaCl | KS.1 |
| FIM1_1167 | 811.4865385 | 781.7484368 | 888.8186405 | -0.04233282 | 0.137548761 | 0.733984917 | 0.824700809 | ns | NaCl | KS.1 |
| FIM1_1168 | 1428.206275 | 1726.371299 | 1564.529 | 0.242589594 | 0.115336236 | 0.023298541 | 0.056486472 | ns | NaCl | KS.1 |
| FIM1_1169 | 797.6881554 | 1105.841619 | 944.2420511 | 0.397769623 | 0.160330934 | 0.00462155 | 0.015216552 | ns | NaCl | KS.1 |
| FIM1_1170 | 1137.414373 | 1341.323218 | 1068.3569 | 0.199724686 | 0.138475903 | 0.109838963 | 0.199433883 | ns | NaCl | KS.1 |
| FIM1_1171 | 63.99730961 | 99.72608748 | 54.07909024 | 0.478731823 | 0.231018805 | 0.009446536 | 0.027291822 | ns | NaCl | KS.1 |
| FIM1_1172 | 3176.568928 | 3661.503852 | 2917.458834 | 0.194639652 | 0.076563259 | 0.008452435 | 0.025150037 | ns | NaCl | KS.1 |
| FIM1_1173 | 747.7828861 | 686.2960871 | 611.6997115 | -0.092239482 | 0.175365717 | 0.539468944 | 0.666300396 | ns | NaCl | KS.1 |
| FIM1_1174 | 482.5756419 | 355.3624251 | 552.607912 | -0.308055545 | 0.223894359 | 0.083687409 | 0.161124171 | ns | NaCl | KS.1 |
| FIM1_1175 | 160.2500296 | 136.3600405 | 154.079058 | -0.151420803 | 0.224776783 | 0.393406739 | 0.52940182 | ns | NaCl | KS.1 |
| FIM1_1177 | 1924.731335 | 2725.224539 | 2264.533237 | 0.475933268 | 0.090310798 | 3.13000380068 | 5.96619498405 | ns | NaCl | KS.1 |
| FIM1_1178 | 15236.66297 | 27243.04045 | 14048.49393 | 0.765844333 | 0.15254542 | 4.40455120380 | 7.85399363402 | ns | NaCl | KS.1 |
| FIM1_1179 | 689.5732553 | 888.5129593 | 940.2419267 | 0.329947577 | 0.109202957 | 0.001189731 | 0.004966078 | ns | NaCl | KS.1 |
| FIM1_1180 | 1628.601874 | 1414.773742 | 1300.328035 | -0.168271791 | 0.141671434 | 0.186381693 | 0.302725733 | ns | NaCl | KS.1 |
| FIM1_1181 | 2348.420759 | 2171.605264 | 2233.461848 | -0.084482333 | 0.167976921 | 0.561862982 | 0.685114788 | ns | NaCl | KS.1 |
| FIM1_1182 | 69.22735525 | 142.1132333 | 106.0153218 | 0.869327435 | 0.236771505 | 1.29940084435 | 0.000107668 | ns | NaCl | KS.1 |
| FIM1_1183 | 1260.846321 | 2475.319728 | 1349.595068 | 0.870899655 | 0.180802813 | 8.87614160108 | 1.43372741770 | ns | NaCl | KS.1 |
| FIM1_1184 | 615.1854238 | 636.5430763 | 592.4635588 | 0.038448807 | 0.14916816 | 0.772481396 | 0.852911429 | ns | NaCl | KS.1 |
| FIM1_1186 | 578.7471946 | 603.9080347 | 717.6328512 | 0.060300729 | 0.091390139 | 0.489384506 | 0.622629484 | ns | NaCl | KS.1 |
| FIM1_1187 | 6185.644886 | 7745.5566 | 6179.320756 | 0.313425501 | 0.060366497 | 1.04022831241 | 1.65869738918 | ns | NaCl | KS.1 |
| FIM1_1188 | 408.1438803 | 398.0203927 | 1370.245573 | -0.024599709 | 0.197247676 | 0.880177616 | 0.929031192 | ns | NaCl | KS.1 |
| FIM1_1189 | 710.9106116 | 460.6262462 | 1684.525144 | -0.435745482 | 0.25189653 | 0.024612634 | 0.059239408 | ns | NaCl | KS.1 |
| FIM1_1190 | 236.6143682 | 147.0342035 | 919.1149451 | -0.470533373 | 0.263626555 | 0.018734906 | 0.047026316 | ns | NaCl | KS.1 |
| FIM1_1191 | 38.03041755 | 36.74675923 | 87.82668476 | -0.025114145 | 0.343070873 | 0.903088556 | 0.942440089 | ns | NaCl | KS.1 |
| FIM1_1192 | 1368.10547 | 1258.842776 | 1024.594112 | -0.092426175 | 0.16599553 | 0.521957224 | 0.652118833 | ns | NaCl | KS.1 |
| FIM1_1193 | 5136.368105 | 6666.665735 | 4984.269302 | 0.361720799 | 0.067866914 | 3.91829691124 | 7.19271879028 | ns | NaCl | KS.1 |
| FIM1_1194 | 3742.349163 | 4844.497596 | 3125.589537 | 0.33492153 | 0.115100951 | 0.001674641 | 0.006596469 | ns | NaCl | KS.1 |
| FIM1_1195 | 367.7289573 | 200.1020374 | 301.4078431 | -0.774881662 | 0.182931068 | 1.81635769146 | 1.97300862773 | ns | NaCl | KS.1 |
| FIM1_1196 | 19193.07529 | 27196.74557 | 16415.82363 | 0.454450473 | 0.125879675 | 8.13611778839 | 0.000524316 | ns | NaCl | KS.1 |
| FIM1_1197 | 4495.115817 | 3704.516348 | 4171.618958 | -0.244838222 | 0.119400245 | 0.026265451 | 0.062551756 | ns | NaCl | KS.1 |
| FIM1_1198 | 1073.182438 | 1636.93059 | 1483.304579 | 0.489149042 | 0.200274966 | 0.003445701 | 0.011996054 | ns | NaCl | KS.1 |
| FIM1_1199 | 1369.070919 | 1418.91368 | 1383.38834 | 0.044053475 | 0.106539145 | 0.660021194 | 0.766481195 | ns | NaCl | KS.1 |
| FIM1_1200 | 4270.902424 | 5281.255836 | 4476.768721 | 0.221555406 | 0.19696323 | 0.17441012 | 0.287791159 | ns | NaCl | KS.1 |
| FIM1_1201 | 260.5785703 | 186.7149311 | 205.2568071 | -0.393904293 | 0.177212754 | 0.009419835 | 0.027252425 | ns | NaCl | KS.1 |
| FIM1_1202 | 341.1304765 | 295.4678491 | 329.0836406 | -0.141978971 | 0.208941019 | 0.402288 | 0.538105815 | ns | NaCl | KS.1 |
| FIM1_1203 | 943.0298299 | 645.7976389 | 1165.203984 | -0.353658599 | 0.258233422 | 0.0695814 | 0.1383563 | ns | NaCl | KS.1 |
| FIM1_1204 | 3933.249495 | 2813.24283 | 4190.507741 | -0.407904146 | 0.160628204 | 0.003742141 | 0.012866034 | ns | NaCl | KS.1 |
| FIM1_1205 | 1183.344473 | 613.9235615 | 847.5103943 | -0.887153152 | 0.137680638 | 7.54301288237 | 3.71549396928 | ns | NaCl | KS.1 |
| FIM1_1206 | 648.7480066 | 752.3912051 | 779.5023305 | 0.18594543 | 0.119326058 | 0.091766343 | 0.173197677 | ns | NaCl | KS.1 |
| FIM1_1207 | 1373.559034 | 1575.030096 | 1871.703589 | 0.174771935 | 0.110517337 | 0.090382242 | 0.171165456 | ns | NaCl | KS.1 |
| FIM1_1208 | 1421.921292 | 1429.246222 | 1630.311595 | 0.004170129 | 0.112319148 | 0.968298068 | 0.985532505 | ns | NaCl | KS.1 |
| FIM1_1209 | 1065.142806 | 1167.315353 | 1031.406901 | 0.122353979 | 0.095378664 | 0.176912364 | 0.291340288 | ns | NaCl | KS.1 |
| FIM1_1210 | 577.9545957 | 214.1262086 | 392.2183859 | -1.271290819 | 0.229040318 | 2.95619038512 | 9.73976633508 | Km<KS.1 | NaCl | KS.1 |
| FIM1_1211 | 1213.631494 | 825.4215314 | 1425.858515 | -0.481513648 | 0.155486703 | 0.000469016 | 0.002280896 | ns | NaCl | KS.1 |

| | | | | | | | | | | |
|---|---|---|---|---|---|---|---|---|---|---|
| FIM1_1212 | 9111.69804 | 16891.8509 | 9933.849973 | 0.851372064 | 0.110768933 | 1.142862580З | 1.236030725 | ns | NaCl | KS.1 |
| FIM1_1213 | 10442.99415 | 13059.73826 | 8591.439274 | 0.2771998 | 0.133391942 | 0.021876994 | 0.053800319 | ns | NaCl | KS.1 |
| FIM1_1214 | 19992.10535 | 29580.33655 | 18090.46764 | 0.535604025 | 0.096964921 | 5.849603585 | 1.366280649 | ns | NaCl | KS.1 |
| FIM1_1215 | 929.3865903 | 842.2777053 | 930.5813569 | -0.128480136 | 0.099595102 | 0.172610283 | 0.285010341 | ns | NaCl | KS.1 |
| FIM1_1216 | 748.6574486 | 916.0704334 | 932.4318863 | 0.237477042 | 0.154314441 | 0.081143935 | 0.156895094 | ns | NaCl | KS.1 |
| FIM1_1217 | 4293.351457 | 8525.508811 | 3883.21751 | 0.703198579 | 0.301667361 | 0.00180355 | 0.007020863 | ns | NaCl | KS.1 |
| FIM1_1218 | 287.1636998 | 198.286121 | 430.1686544 | -0.453547939 | 0.174293044 | 0.002548514 | 0.009377502 | ns | NaCl | KS.1 |
| FIM1_1219 | 1176.165075 | 1514.257143 | 1348.642288 | 0.332746505 | 0.10924393 | 0.001083722 | 0.004584625 | ns | NaCl | KS.1 |
| FIM1_1220 | 420.5153711 | 556.0241976 | 623.5037668 | 0.301927813 | 0.197530446 | 0.065114227 | 0.130833311 | ns | NaCl | KS.1 |
| FIM1_1221 | 893.7462181 | 799.0993791 | 911.9478004 | -0.150131047 | 0.094857602 | 0.095718794 | 0.178955656 | ns | NaCl | KS.1 |
| FIM1_1222 | 4840.514372 | 4962.162581 | 5569.113365 | 0.034372269 | 0.058781183 | 0.550706373 | 0.675698639 | ns | NaCl | KS.1 |
| FIM1_1223 | 226.6442539 | 194.4719472 | 358.2853896 | -0.17549597 | 0.170081625 | 0.232041049 | 0.357954796 | ns | NaCl | KS.1 |
| FIM1_1224 | 243.6879715 | 236.6699163 | 283.1675351 | -0.037146012 | 0.148904707 | 0.779690267 | 0.856660573 | ns | NaCl | KS.1 |
| FIM1_1225 | 1454.358291 | 2408.345102 | 2384.065711 | 0.650964166 | 0.158258728 | 4.663669999 | 4.496464776 | ns | NaCl | KS.1 |
| FIM1_1227 | 3550.73247 | 3628.779031 | 4298.800342 | 0.022953662 | 0.176397634 | 0.879189438 | 0.928852719 | ns | NaCl | KS.1 |
| FIM1_1228 | 1361.364886 | 1223.676291 | 1288.517643 | -0.124413945 | 0.146506604 | 0.342070778 | 0.476428365 | ns | NaCl | KS.1 |
| FIM1_1230 | 1819.080018 | 3249.066289 | 2123.311934 | 0.757234593 | 0.160064846 | 1.934711261 | 2.873190604 | ns | NaCl | KS.1 |
| FIM1_1231 | 3050.277568 | 5347.160044 | 3724.980557 | 0.718768619 | 0.17185523 | 2.756301564 | 2.868744829 | ns | NaCl | KS.1 |
| FIM1_1232 | 4525.959505 | 4660.498034 | 3261.024632 | 0.034580832 | 0.144110503 | 0.78924665 | 0.864684342 | ns | NaCl | KS.1 |
| FIM1_1233 | 1676.335436 | 1154.707303 | 1200.732231 | -0.414793404 | 0.205843165 | 0.0143643 | 0.037991702 | ns | NaCl | KS.1 |
| FIM1_1234 | 9473.475278 | 13058.3401 | 8850.688557 | 0.431911293 | 0.100303006 | 4.809008328 | 4.592095284 | ns | NaCl | KS.1 |
| FIM1_1235 | 1633.467842 | 1956.604674 | 2019.227918 | 0.227347831 | 0.124894269 | 0.047148396 | 0.100714155 | ns | NaCl | KS.1 |
| FIM1_1236 | 323.5975324 | 415.833849 | 389.0121877 | 0.288771473 | 0.173139379 | 0.052168543 | 0.109509916 | ns | NaCl | KS.1 |
| FIM1_1239 | 3958.85506 | 2570.406073 | 2726.193429 | -0.50649522 | 0.196296834 | 0.002156951 | 0.008129418 | ns | NaCl | KS.1 |
| FIM1_1240 | 281.2688224 | 223.5246999 | 277.3851919 | -0.267092471 | 0.168839292 | 0.067188232 | 0.134295482 | ns | NaCl | KS.1 |
| FIM1_1242 | 724.7574933 | 671.3601536 | 774.0457167 | -0.091266383 | 0.134778567 | 0.456020075 | 0.590961155 | ns | NaCl | KS.1 |
| FIM1_1243 | 3289.720732 | 3861.873312 | 3450.124753 | 0.218933469 | 0.06837088 | 0.000979454 | 0.004218862 | ns | NaCl | KS.1 |
| FIM1_1244 | 858.8144067 | 656.0675475 | 884.1417876 | -0.350909954 | 0.122758452 | 0.001838936 | 0.007130713 | ns | NaCl | KS.1 |
| FIM1_1245 | 993.2773445 | 1099.455806 | 1256.399244 | 0.123022661 | 0.132583858 | 0.308157189 | 0.44104776 | ns | NaCl | KS.1 |
| FIM1_1246 | 599.6799624 | 566.5587085 | 571.5752836 | -0.069074896 | 0.106783447 | 0.491143326 | 0.623941138 | ns | NaCl | KS.1 |
| FIM1_1247 | 862.3363115 | 401.1541063 | 887.4453676 | -1.033732213 | 0.148000638 | 1.083251686 | 8.692221194 | Km<KS.1 | NaCl | KS.1 |
| FIM1_1248 | 2083.038842 | 1852.491288 | 1548.077761 | -0.141005798 | 0.137946543 | 0.258097899 | 0.386598155 | ns | NaCl | KS.1 |
| FIM1_1249 | 5011.055585 | 6380.602345 | 4249.76047 | 0.211973761 | 0.249753338 | 0.261892256 | 0.390327733 | ns | NaCl | KS.1 |
| FIM1_1250 | 1339.665515 | 1352.400298 | 2503.527131 | 0.009316207 | 0.157077853 | 0.94636124 | 0.971553274 | ns | NaCl | KS.1 |
| FIM1_1251 | 3710.931295 | 3734.984405 | 3902.077065 | 0.008507585 | 0.108104958 | 0.933176414 | 0.961658314 | ns | NaCl | KS.1 |
| FIM1_1252 | 2697.261837 | 2420.849067 | 1817.874988 | -0.128035726 | 0.141441547 | 0.3144669 | 0.447675071 | ns | NaCl | KS.1 |
| FIM1_1253 | 468.2795014 | 305.7315382 | 403.9005518 | -0.572428631 | 0.119126029 | 2.407975303 | 3.479186395 | ns | NaCl | KS.1 |
| FIM1_1254 | 1468.321176 | 1577.572913 | 1261.360215 | 0.076622373 | 0.173079741 | 0.606855317 | 0.724353456 | ns | NaCl | KS.1 |
| FIM1_1255 | 2336.46399 | 3613.184476 | 2162.44669 | 0.402522708 | 0.274964211 | 0.047420855 | 0.101158734 | ns | NaCl | KS.1 |
| FIM1_1256 | 746.4748474 | 812.6756621 | 1269.138431 | 0.104512537 | 0.122348491 | 0.354439509 | 0.489543741 | ns | NaCl | KS.1 |
| FIM1_1257 | 1692.853872 | 1550.140771 | 1885.744787 | -0.107277459 | 0.131162571 | 0.370034266 | 0.504776658 | ns | NaCl | KS.1 |
| FIM1_1258 | 1173.5877 | 923.4520317 | 1152.889444 | -0.29993464 | 0.128372446 | 0.010395526 | 0.0294856 | ns | NaCl | KS.1 |
| FIM1_1259 | 289.3627794 | 188.859113 | 347.9151025 | -0.472211486 | 0.212883106 | 0.006761587 | 0.020893722 | ns | NaCl | KS.1 |
| FIM1_1261 | 2697.866895 | 3491.192829 | 2347.620967 | 0.293072992 | 0.17419934 | 0.049807767 | 0.105264927 | ns | NaCl | KS.1 |
| FIM1_1262 | 2641.441379 | 1663.558463 | 2501.382779 | -0.535830731 | 0.207608863 | 0.001897145 | 0.007322185 | ns | NaCl | KS.1 |

| | | | | | | | | | | |
|---|---|---|---|---|---|---|---|---|---|---|
| FIM1_1263 | 2810.930889 | 2191.963576 | 2066.89919 | -0.292623098 | 0.159856759 | 0.03647114 | 0.081879025 | ns | NaCl | KS.1 |
| FIM1_1264 | 4152.339704 | 3614.860973 | 2953.260636 | -0.150577895 | 0.17493695 | 0.315363808 | 0.448523426 | ns | NaCl | KS.1 |
| FIM1_1265 | 3895.212445 | 4940.181979 | 5046.128415 | 0.275828476 | 0.16372921 | 0.052959213 | 0.110702557 | ns | NaCl | KS.1 |
| FIM1_1266 | 845.8039042 | 1171.110204 | 805.668196 | 0.375279508 | 0.182482805 | 0.015443477 | 0.040058029 | ns | NaCl | KS.1 |
| FIM1_1267 | 4742.052033 | 8904.612589 | 4172.094383 | 0.631262253 | 0.297001614 | 0.004155049 | 0.014004994 | ns | NaCl | KS.1 |
| FIM1_1268 | 774.1284671 | 638.5921089 | 779.0367007 | -0.232928329 | 0.14400273 | 0.070671544 | 0.139906565 | ns | NaCl | KS.1 |
| FIM1_1269 | 3811.316088 | 8063.276733 | 3666.06524 | 0.769431056 | 0.314409826 | 0.000986606 | 0.004242321 | ns | NaCl | KS.1 |
| FIM1_1270 | 3206.804031 | 4483.320368 | 2826.111165 | 0.433851798 | 0.128217091 | 0.000210338 | 0.001166589 | ns | NaCl | KS.1 |
| FIM1_1271 | 2870.961949 | 3022.615851 | 3309.465527 | 0.064972406 | 0.115780813 | 0.546314912 | 0.672499733 | ns | NaCl | KS.1 |
| FIM1_1272 | 7379.0779 | 10768.90808 | 7662.356285 | 0.511940071 | 0.103679081 | 1.5601874832( | 2.3956582497( | ns | NaCl | KS.1 |
| FIM1_1273 | 418.301153 | 403.267399 | 549.5565453 | -0.041035165 | 0.168503254 | 0.778698296 | 0.856051837 | ns | NaCl | KS.1 |
| FIM1_1274 | 62.63381757 | 52.57348584 | 79.67623041 | -0.152313896 | 0.251487769 | 0.42012687 | 0.556301202 | ns | NaCl | KS.1 |
| FIM1_1275 | 3071.171699 | 2182.268382 | 2704.232452 | -0.440873429 | 0.13180252 | 0.000234701 | 0.001277946 | ns | NaCl | KS.1 |
| FIM1_1276 | 3149.323655 | 2805.749962 | 3207.325652 | -0.162611762 | 0.050592096 | 0.00109073 | 0.004602529 | ns | NaCl | KS.1 |
| FIM1_1277 | 1503.961989 | 1538.404373 | 2437.463365 | 0.025902106 | 0.15922721 | 0.85313877 | 0.910996874 | ns | NaCl | KS.1 |
| FIM1_1279 | 1059.497092 | 733.7848975 | 1120.380077 | -0.477682492 | 0.131992736 | 7.0812783635! | 0.000470981 | ns | NaCl | KS.1 |
| FIM1_1280 | 6121.614937 | 5055.378278 | 7070.403574 | -0.245619866 | 0.112419158 | 0.018837212 | 0.047211653 | ns | NaCl | KS.1 |
| FIM1_1281 | 2580.04918 | 3155.048353 | 2186.258833 | 0.260239377 | 0.11041432 | 0.011443718 | 0.031752647 | ns | NaCl | KS.1 |
| FIM1_1282 | 555.2943977 | 594.3737671 | 725.1315443 | 0.058472809 | 0.238803883 | 0.749642114 | 0.837108446 | ns | NaCl | KS.1 |
| FIM1_1283 | 2954.664908 | 2110.742201 | 3946.865104 | -0.31263393 | 0.25174216 | 0.102203452 | 0.188319323 | ns | NaCl | KS.1 |
| FIM1_1284 | 420.8472737 | 635.7092072 | 937.7489032 | 0.446013771 | 0.224470235 | 0.013374943 | 0.03594832 | ns | NaCl | KS.1 |
| FIM1_1285 | 3209.008222 | 5057.479594 | 4577.267726 | 0.621764186 | 0.10554234 | 4.9704482037! | 1.5171994064 | ns | NaCl | KS.1 |
| FIM1_1286 | 404.3128822 | 549.3237467 | 438.3162663 | 0.354613792 | 0.173044039 | 0.017162461 | 0.043786279 | ns | NaCl | KS.1 |
| FIM1_1287 | 1243.647591 | 954.8075259 | 1377.703421 | -0.271865589 | 0.209247762 | 0.110034701 | 0.199643559 | ns | NaCl | KS.1 |
| FIM1_1288 | 834.9662782 | 637.7012114 | 608.8673932 | -0.333867799 | 0.140106733 | 0.00798508 | 0.024003488 | ns | NaCl | KS.1 |
| FIM1_1289 | 2773.468338 | 3088.886474 | 2209.892379 | 0.137089185 | 0.110718491 | 0.185159945 | 0.301233069 | ns | NaCl | KS.1 |
| FIM1_1290 | 3374.625885 | 3428.088747 | 4253.796382 | 0.018801709 | 0.12650814 | 0.871475371 | 0.923251698 | ns | NaCl | KS.1 |
| FIM1_1291 | 1092.848715 | 1333.742152 | 1057.93371 | 0.224347009 | 0.168585583 | 0.123855586 | 0.218969986 | ns | NaCl | KS.1 |
| FIM1_1292 | 210.1868448 | 314.2413418 | 242.678051 | 0.470295486 | 0.187007493 | 0.003049939 | 0.010838177 | ns | NaCl | KS.1 |
| FIM1_1293 | 2327.297539 | 5224.90572 | 2514.956573 | 0.993302392 | 0.234447538 | 8.0421405460! | 9.7110799069! | ns | NaCl | KS.1 |
| FIM1_1294 | 1353.847396 | 1833.004503 | 1168.931978 | 0.269069987 | 0.256997587 | 0.162722755 | 0.272390884 | ns | NaCl | KS.1 |
| FIM1_1295 | 3610.372721 | 2721.812037 | 3331.000546 | -0.359335994 | 0.131643345 | 0.002642643 | 0.009659917 | ns | NaCl | KS.1 |
| FIM1_1296 | 635.8384984 | 319.4923913 | 921.9107713 | -0.592397515 | 0.349437838 | 0.012228472 | 0.03343686 | ns | NaCl | KS.1 |
| FIM1_1297 | 945.0215358 | 752.2352653 | 1960.959462 | -0.200229533 | 0.246723353 | 0.285683118 | 0.416795752 | ns | NaCl | KS.1 |
| FIM1_1298 | 1604.901335 | 1359.058099 | 2757.110115 | -0.169190118 | 0.197632747 | 0.300589251 | 0.432636272 | ns | NaCl | KS.1 |
| FIM1_1299 | 793.7728264 | 968.7621675 | 985.6162665 | 0.261351582 | 0.104698522 | 0.007779103 | 0.023483639 | ns | NaCl | KS.1 |
| FIM1_1300 | 1143.606715 | 1900.025737 | 1455.409244 | 0.654991409 | 0.1597235 | 4.8560133681: | 4.6260099507! | ns | NaCl | KS.1 |
| FIM1_1301 | 323.5660968 | 249.532764 | 342.8589182 | -0.316677911 | 0.155196188 | 0.020513062 | 0.050655944 | ns | NaCl | KS.1 |
| FIM1_1302 | 552.2381914 | 730.1017452 | 736.6684656 | 0.358177078 | 0.122646868 | 0.001462324 | 0.005857539 | ns | NaCl | KS.1 |
| FIM1_1303 | 2059.166766 | 4047.764389 | 2040.5456 | 0.827125431 | 0.217686727 | 9.3132536318! | 8.0300583741: | ns | NaCl | KS.1 |
| FIM1_1304 | 1293.195051 | 1640.988182 | 1576.531809 | 0.21813132 | 0.238292154 | 0.236081849 | 0.362950308 | ns | NaCl | KS.1 |
| FIM1_1305 | 3637.919186 | 4126.069637 | 4427.895953 | 0.148456587 | 0.145648871 | 0.254476832 | 0.382734778 | ns | NaCl | KS.1 |
| FIM1_1306 | 86.05677477 | 72.13274117 | 117.1812301 | -0.15525929 | 0.21851737 | 0.373367965 | 0.508626951 | ns | NaCl | KS.1 |
| FIM1_1307 | 864.7131875 | 1065.348121 | 1081.113627 | 0.258551435 | 0.130569412 | 0.029554597 | 0.068836198 | ns | NaCl | KS.1 |
| FIM1_1308 | 748.5960838 | 945.7414722 | 883.5704345 | 0.296047182 | 0.122966468 | 0.008716574 | 0.025797118 | ns | NaCl | KS.1 |

| | | | | | | | | | | |
|---|---|---|---|---|---|---|---|---|---|---|
| FIM1_1309 | 1054.465274 | 924.7413314 | 1078.806146 | -0.148879465 | 0.152804224 | 0.271279593 | 0.400276242 | ns | NaCl | KS.1 |
| FIM1_1310 | 2320.194424 | 2904.215864 | 2115.586315 | 0.294173305 | 0.106348825 | 0.003110683 | 0.010991229 | ns | NaCl | KS.1 |
| FIM1_1311 | 379.5102218 | 3617.860855 | 2063.53469 | 3.132954294 | 0.262538877 | 1.5860918452 | 2.6302689767 | Km>KS.1 | NaCl | KS.1 |
| FIM1_1312 | 2544.925289 | 1561.235318 | 1879.694692 | -0.545384971 | 0.2281093 | 0.003108855 | 0.010991229 | ns | NaCl | KS.1 |
| FIM1_1313 | 33623.55729 | 42552.966 | 32699.65459 | 0.285799265 | 0.144960126 | 0.027250913 | 0.064435976 | ns | NaCl | KS.1 |
| FIM1_1314 | 1400.3282 | 1152.028857 | 895.8018927 | -0.205129651 | 0.193224364 | 0.202525181 | 0.322956094 | ns | NaCl | KS.1 |
| FIM1_1315 | 3856.13154 | 4845.666558 | 3495.315727 | 0.262791711 | 0.165250535 | 0.067157251 | 0.134287509 | ns | NaCl | KS.1 |
| FIM1_1316 | 1405.072519 | 501.915078 | 918.0455375 | -1.332184045 | 0.223146992 | 1.760655712 | 7.6457018656 | Km<KS.1 | NaCl | KS.1 |
| FIM1_1317 | 1022.356124 | 685.7041079 | 631.2855543 | -0.470879044 | 0.187553166 | 0.003078889 | 0.010917658 | ns | NaCl | KS.1 |
| FIM1_1318 | 1678.842401 | 1365.300599 | 1423.226802 | -0.255367266 | 0.134764292 | 0.036330198 | 0.081710097 | ns | NaCl | KS.1 |
| FIM1_1319 | 3638.556515 | 2971.37335 | 3635.495891 | -0.205089009 | 0.204313387 | 0.219939074 | 0.34411577 | ns | NaCl | KS.1 |
| FIM1_1320 | 13944.309 | 15603.21349 | 13061.52587 | 0.11789057 | 0.183418629 | 0.44753808 | 0.582846118 | ns | NaCl | KS.1 |
| FIM1_1321 | 688.3178239 | 699.6862581 | 788.6044429 | 0.018097746 | 0.12197309 | 0.87238776 | 0.924021525 | ns | NaCl | KS.1 |
| FIM1_1322 | 8078.832429 | 8470.059273 | 7929.863146 | 0.047120195 | 0.195890246 | 0.771755786 | 0.852461154 | ns | NaCl | KS.1 |
| FIM1_1323 | 388.406911 | 259.1606388 | 817.2490221 | -0.472801151 | 0.190673785 | 0.003333869 | 0.011631134 | ns | NaCl | KS.1 |
| FIM1_1324 | 1424.349855 | 2113.715003 | 1164.683031 | 0.385120126 | 0.251310016 | 0.045422773 | 0.097604297 | ns | NaCl | KS.1 |
| FIM1_1325 | 1752.481995 | 1165.625041 | 1963.284815 | -0.510623815 | 0.159334762 | 0.000284656 | 0.001503358 | ns | NaCl | KS.1 |
| FIM1_1326 | 864.1501362 | 833.9181222 | 1051.740856 | -0.046094386 | 0.104333532 | 0.639147092 | 0.749235812 | ns | NaCl | KS.1 |
| FIM1_1327 | 1237.536391 | 1279.69354 | 2187.925136 | 0.041825122 | 0.107790922 | 0.679359351 | 0.782545212 | ns | NaCl | KS.1 |
| FIM1_1328 | 1173.533088 | 1106.922529 | 1573.607562 | -0.07537831 | 0.078974137 | 0.322649191 | 0.455887454 | ns | NaCl | KS.1 |
| FIM1_1329 | 682.5783566 | 547.3416186 | 1818.801545 | -0.219154552 | 0.212608165 | 0.201725645 | 0.322022716 | ns | NaCl | KS.1 |
| FIM1_1331 | 81.80582179 | 79.58652366 | 153.9760707 | -0.0186008 | 0.253682353 | 0.921550658 | 0.955142288 | ns | NaCl | KS.1 |
| FIM1_1332 | 484.3205923 | 422.0489588 | 565.8234931 | -0.169374407 | 0.135244104 | 0.167108397 | 0.27804234 | ns | NaCl | KS.1 |
| FIM1_1333 | 9601.705333 | 13743.98709 | 7441.272709 | 0.383882437 | 0.215445556 | 0.027782712 | 0.065382683 | ns | NaCl | KS.1 |
| FIM1_1334 | 4021.680053 | 2379.481784 | 3113.798748 | -0.623460784 | 0.209060632 | 0.000381293 | 0.001925819 | ns | NaCl | KS.1 |
| FIM1_1335 | 1222.182462 | 915.8934908 | 988.3723608 | -0.336209621 | 0.170662071 | 0.022350311 | 0.054693948 | ns | NaCl | KS.1 |
| FIM1_1336 | 694.0022959 | 2866.632782 | 1587.422816 | 1.825939001 | 0.311336866 | 2.8997715655 | 1.6393594930 | Km>KS.1 | NaCl | KS.1 |
| FIM1_1337 | 4136.004995 | 9114.998865 | 3774.680074 | 0.833970906 | 0.312201929 | 0.000402171 | 0.002014906 | ns | NaCl | KS.1 |
| FIM1_1338 | 578.828196 | 306.1254626 | 857.7782986 | -0.777573739 | 0.214527159 | 2.1957879117 | 0.000169103 | ns | NaCl | KS.1 |
| FIM1_1339 | 5609.125138 | 5562.232117 | 5129.626516 | -0.009093955 | 0.167072826 | 0.950031257 | 0.973834262 | ns | NaCl | KS.1 |
| FIM1_1340 | 2905.236442 | 2331.22768 | 2228.020992 | -0.280617106 | 0.120648625 | 0.011525403 | 0.031925878 | ns | NaCl | KS.1 |
| FIM1_1341 | 712.7839184 | 604.3891701 | 703.1858564 | -0.164026167 | 0.205298221 | 0.327839941 | 0.461387188 | ns | NaCl | KS.1 |
| FIM1_1342 | 2216.147195 | 2880.84034 | 2388.158002 | 0.317415912 | 0.150097661 | 0.017087533 | 0.04366229 | ns | NaCl | KS.1 |
| FIM1_1344 | 297.7676739 | 222.1732404 | 431.1494487 | -0.357846456 | 0.13993872 | 0.004430359 | 0.014753035 | ns | NaCl | KS.1 |
| FIM1_1345 | 1909.761367 | 1501.852752 | 1529.858814 | -0.303013794 | 0.126825443 | 0.00891145 | 0.026171466 | ns | NaCl | KS.1 |
| FIM1_1346 | 8907.718702 | 11918.77706 | 8008.702261 | 0.378673679 | 0.118572989 | 0.000534372 | 0.002551346 | ns | NaCl | KS.1 |
| FIM1_1347 | 19426.45148 | 26102.89447 | 16702.93723 | 0.372701504 | 0.135763969 | 0.002375893 | 0.008807799 | ns | NaCl | KS.1 |
| FIM1_1348 | 10626.36592 | 15035.02851 | 10458.08256 | 0.477086265 | 0.086291072 | 7.3072868093 | 1.6395928148 | ns | NaCl | KS.1 |
| FIM1_1349 | 1120.500612 | 905.7766906 | 1253.590956 | -0.286303846 | 0.098609936 | 0.002089187 | 0.007909975 | ns | NaCl | KS.1 |
| FIM1_1350 | 8385.564373 | 12113.71155 | 8388.563378 | 0.448136211 | 0.165987661 | 0.001950537 | 0.007487593 | ns | NaCl | KS.1 |
| FIM1_1352 | 2756.0812 | 2181.405408 | 2375.419536 | -0.311087879 | 0.0983817 | 0.000802185 | 0.003582468 | ns | NaCl | KS.1 |
| FIM1_1353 | 6714.354648 | 7696.740876 | 8281.703894 | 0.180156258 | 0.094390709 | 0.044496528 | 0.096080827 | ns | NaCl | KS.1 |
| FIM1_1354 | 1026.642713 | 1040.627671 | 1056.633872 | 0.014872942 | 0.145230923 | 0.909049325 | 0.946531862 | ns | NaCl | KS.1 |
| FIM1_1355 | 1388.981329 | 1567.811511 | 1650.334581 | 0.163777011 | 0.070358293 | 0.016491889 | 0.042314155 | ns | NaCl | KS.1 |
| FIM1_1356 | 663.2236862 | 604.5320106 | 718.3923749 | -0.119338098 | 0.114072129 | 0.261262402 | 0.389856163 | ns | NaCl | KS.1 |

| | | | | | | | | | | |
|---|---|---|---|---|---|---|---|---|---|---|
| FIM1_1357 | 1032.078198 | 938.088508 | 1025.871029 | -0.110738418 | 0.147582358 | 0.400610033 | 0.536483153 | ns | NaCl | KS.1 |
| FIM1_1358 | 1536.12059 | 1625.663598 | 1315.886861 | 0.070650829 | 0.119958959 | 0.524570641 | 0.653906024 | ns | NaCl | KS.1 |
| FIM1_1359 | 545.9718875 | 434.4516842 | 495.7212686 | -0.276636736 | 0.145920476 | 0.033544061 | 0.076436268 | ns | NaCl | KS.1 |
| FIM1_1360 | 1872.163534 | 1938.239254 | 2675.62504 | 0.033296531 | 0.203448467 | 0.841530144 | 0.903176394 | ns | NaCl | KS.1 |
| FIM1_1361 | 148.9948463 | 171.1285717 | 279.5368154 | 0.158817925 | 0.174889892 | 0.289477339 | 0.420236289 | ns | NaCl | KS.1 |
| FIM1_1362 | 383.3852664 | 375.2143456 | 484.6343487 | -0.02141662 | 0.151197908 | 0.873408966 | 0.924512682 | ns | NaCl | KS.1 |
| FIM1_1363 | 3799.095127 | 7877.034861 | 3829.713585 | 0.782966593 | 0.293085906 | 0.000506402 | 0.00242946 | ns | NaCl | KS.1 |
| FIM1_1364 | 304.9282819 | 356.5967631 | 299.4874911 | 0.149944501 | 0.212348483 | 0.381383122 | 0.516856724 | ns | NaCl | KS.1 |
| FIM1_1365 | 1984.57078 | 2756.115481 | 2515.753909 | 0.399183236 | 0.160234513 | 0.004461187 | 0.014834572 | ns | NaCl | KS.1 |
| FIM1_1366 | 1510.1663 | 1847.751571 | 1390.123103 | 0.243774925 | 0.14161988 | 0.055117633 | 0.114636382 | ns | NaCl | KS.1 |
| FIM1_1367 | 787.8206778 | 547.0091861 | 982.0969972 | -0.490175081 | 0.105737987 | 7.7888135757 | 9.4510603754 | ns | NaCl | KS.1 |
| FIM1_1368 | 4613.970067 | 6001.709553 | 5774.293505 | 0.353197334 | 0.095819879 | 9.6623371313 | 0.000606181 | ns | NaCl | KS.1 |
| FIM1_1369 | 1950.426048 | 6511.388263 | 3750.884313 | 1.623887484 | 0.205618465 | 3.7258094293 | 6.6199649682 | Km>KS.1 | NaCl | KS.1 |
| FIM1_1370 | 25.62194613 | 56.79911208 | 34.13613517 | 0.743791914 | 0.355302091 | 0.00257918 | 0.009476678 | ns | NaCl | KS.1 |
| FIM1_1371 | 1454.26344 | 1097.385469 | 1244.944897 | -0.371448055 | 0.109751268 | 0.000279497 | 0.00148083 | ns | NaCl | KS.1 |
| FIM1_1372 | 388.2968561 | 144.7576013 | 292.5230999 | -1.297157109 | 0.20273786 | 1.4392078871 | 8.8395793070 | Km<KS.1 | NaCl | KS.1 |
| FIM1_1373 | 2200.108407 | 2466.004404 | 2413.523477 | 0.146660779 | 0.110099027 | 0.154207473 | 0.260817937 | ns | NaCl | KS.1 |
| FIM1_1374 | 2647.338614 | 2141.772935 | 2208.060344 | -0.279090663 | 0.102811047 | 0.003857883 | 0.013163901 | ns | NaCl | KS.1 |
| FIM1_1375 | 814.3915093 | 1140.035661 | 1570.463931 | 0.294498149 | 0.267067195 | 0.135373565 | 0.234990748 | ns | NaCl | KS.1 |
| FIM1_1376 | 105.9658445 | 54.3607424 | 96.17648643 | -0.795252323 | 0.23397844 | 4.7111604880 | 0.000330579 | ns | NaCl | KS.1 |
| FIM1_1377 | 3402.763413 | 4145.603612 | 3826.541247 | 0.256509101 | 0.106915246 | 0.010348622 | 0.029402851 | ns | NaCl | KS.1 |
| FIM1_1378 | 352.9375767 | 332.8744758 | 321.4348399 | -0.074366607 | 0.162047838 | 0.599879114 | 0.718958947 | ns | NaCl | KS.1 |
| FIM1_1379 | 92.62506093 | 99.93056308 | 272.0880493 | 0.079296636 | 0.21647452 | 0.647086526 | 0.755235621 | ns | NaCl | KS.1 |
| FIM1_1380 | 1247.982576 | 1377.984437 | 2315.659662 | 0.092109064 | 0.218914608 | 0.597436147 | 0.716894557 | ns | NaCl | KS.1 |
| FIM1_1382 | 3970.041695 | 4402.243945 | 4035.366892 | 0.141202838 | 0.06679475 | 0.029929584 | 0.069546791 | ns | NaCl | KS.1 |
| FIM1_1383 | 2769.940284 | 2393.432807 | 2587.870545 | -0.184114072 | 0.116980369 | 0.089460191 | 0.16978992 | ns | NaCl | KS.1 |
| FIM1_1384 | 1301.08281 | 1499.185419 | 1200.455473 | 0.165292042 | 0.148053706 | 0.210315745 | 0.332693429 | ns | NaCl | KS.1 |
| FIM1_1385 | 3572.174388 | 4318.290983 | 5001.549149 | 0.215196829 | 0.166120266 | 0.135587502 | 0.235197986 | ns | NaCl | KS.1 |
| FIM1_1386 | 617.9758679 | 532.6770509 | 571.1931895 | -0.180447329 | 0.134494313 | 0.139131826 | 0.239508939 | ns | NaCl | KS.1 |
| FIM1_1387 | 2944.359422 | 3453.552122 | 2965.730192 | 0.223479969 | 0.049810855 | 5.1243950976 | 4.8192562591 | ns | NaCl | KS.1 |
| FIM1_1388 | 298.4555666 | 242.6406544 | 383.3514424 | -0.266117607 | 0.123804182 | 0.019134362 | 0.047716016 | ns | NaCl | KS.1 |
| FIM1_1389 | 660.2354219 | 396.0545278 | 608.2360905 | -0.690583512 | 0.131994891 | 1.7827693768 | 3.6649907643 | ns | NaCl | KS.1 |
| FIM1_1390 | 1930.766287 | 2292.522559 | 2724.645374 | 0.183498703 | 0.183376081 | 0.236884592 | 0.363847127 | ns | NaCl | KS.1 |
| FIM1_1391 | 3149.653391 | 2298.870676 | 3696.141313 | -0.248776491 | 0.285794041 | 0.219553066 | 0.343699341 | ns | NaCl | KS.1 |
| FIM1_1392 | 866.7558195 | 522.4893314 | 1101.82239 | -0.616745803 | 0.187450926 | 0.000136656 | 0.000815187 | ns | NaCl | KS.1 |
| FIM1_1393 | 1116.775399 | 871.0859573 | 1062.710524 | -0.268549882 | 0.188262051 | 0.089428079 | 0.16978992 | ns | NaCl | KS.1 |
| FIM1_1394 | 292.6292104 | 73.73263504 | 587.753791 | -1.721914986 | 0.322763344 | 9.4256978546 | 3.4735442094 | Km<KS.1 | NaCl | KS.1 |
| FIM1_1395 | 517.5878755 | 529.0317332 | 753.8510585 | 0.019235947 | 0.218957931 | 0.912142709 | 0.948722964 | ns | NaCl | KS.1 |
| FIM1_1396 | 1344.065307 | 1367.273003 | 1338.63058 | 0.01980748 | 0.158576518 | 0.887079182 | 0.932435862 | ns | NaCl | KS.1 |
| FIM1_1397 | 4736.793969 | 4231.273618 | 4469.632152 | -0.135343118 | 0.135192408 | 0.269812592 | 0.39878718 | ns | NaCl | KS.1 |
| FIM1_1399 | 2576.519871 | 2453.015795 | 2805.973654 | -0.050542258 | 0.185580942 | 0.74679834 | 0.835466998 | ns | NaCl | KS.1 |
| FIM1_1400 | 1872.184828 | 1764.670651 | 1902.072314 | -0.058787945 | 0.196039117 | 0.717558676 | 0.812808382 | ns | NaCl | KS.1 |
| FIM1_1401 | 1480.886497 | 4128.228044 | 3542.249151 | 1.21395095 | 0.295729703 | 4.8854001884 | 6.3792299048 | Km>KS.1 | NaCl | KS.1 |
| FIM1_1402 | 682.4826898 | 876.0249853 | 872.8792913 | 0.32942269 | 0.104377443 | 0.000757433 | 0.003419447 | ns | NaCl | KS.1 |
| FIM1_1403 | 613.8847224 | 637.7761831 | 747.890367 | 0.04606603 | 0.148181952 | 0.727648591 | 0.819758094 | ns | NaCl | KS.1 |

| | | | | | | | | | | |
|---|---|---|---|---|---|---|---|---|---|---|
| FIM1_1404 | 2476.290623 | 502.3927386 | 894.6827909 | -2.033897091 | 0.370293769 | 1.7122420457 | 7.6057180158 | Km<KS.1 | NaCl | KS.1 |
| FIM1_1405 | 785.7034306 | 974.7781242 | 849.0081097 | 0.227155601 | 0.197093882 | 0.163989348 | 0.273774164 | ns | NaCl | KS.1 |
| FIM1_1406 | 4347.333325 | 6042.828674 | 6026.309982 | 0.338669423 | 0.222240005 | 0.05648845 | 0.116998351 | ns | NaCl | KS.1 |
| FIM1_1407 | 978.8108258 | 1263.311394 | 1369.341169 | 0.329678368 | 0.115455315 | 0.002035462 | 0.007724199 | ns | NaCl | KS.1 |
| FIM1_1408 | 5974.622453 | 9165.628945 | 7942.031075 | 0.566667442 | 0.128499767 | 1.663696614 | 1.833908746 | ns | NaCl | KS.1 |
| FIM1_1409 | 2452.383878 | 3771.813378 | 2107.260258 | 0.358047105 | 0.297496063 | 0.087849872 | 0.167365195 | ns | NaCl | KS.1 |
| FIM1_1410 | 5067.596346 | 5080.975004 | 3987.427979 | 0.002997946 | 0.143802267 | 0.981482841 | 0.991175946 | ns | NaCl | KS.1 |
| FIM1_1411 | 1200.983428 | 932.1162279 | 1205.529616 | -0.216921342 | 0.257675481 | 0.258958854 | 0.387348256 | ns | NaCl | KS.1 |
| FIM1_1412 | 3883.290424 | 5090.173947 | 4924.602873 | 0.288494934 | 0.199606437 | 0.080074399 | 0.155309994 | ns | NaCl | KS.1 |
| FIM1_1413 | 428.1161231 | 428.1778557 | 623.2557519 | 0.001292219 | 0.242901196 | 0.994420621 | 0.997448369 | ns | NaCl | KS.1 |
| FIM1_1414 | 278.4988496 | 297.8728478 | 305.0673911 | 0.073728876 | 0.176794735 | 0.625877957 | 0.73939285 | ns | NaCl | KS.1 |
| FIM1_1415 | 1636.894629 | 1323.399047 | 1607.211688 | -0.286149762 | 0.091838682 | 0.001039449 | 0.004423661 | ns | NaCl | KS.1 |
| FIM1_1416 | 756.0581957 | 644.336256 | 1016.078387 | -0.207459359 | 0.11049287 | 0.044208164 | 0.095609381 | ns | NaCl | KS.1 |
| FIM1_1418 | 1249.947924 | 1123.074456 | 1019.213384 | -0.138464029 | 0.094986875 | 0.124964208 | 0.220303663 | ns | NaCl | KS.1 |
| FIM1_1420 | 1840.982789 | 1572.209635 | 1437.044234 | -0.199835913 | 0.114163754 | 0.059655444 | 0.122638774 | ns | NaCl | KS.1 |
| FIM1_1421 | 465.3719409 | 401.9380549 | 553.3328384 | -0.139628308 | 0.214264631 | 0.417567504 | 0.553555407 | ns | NaCl | KS.1 |
| FIM1_1422 | 568.5508718 | 361.0896075 | 486.6103039 | -0.530346955 | 0.204278531 | 0.00185778 | 0.007192572 | ns | NaCl | KS.1 |
| FIM1_1423 | 723.6337418 | 666.90502 | 793.3963286 | -0.108299523 | 0.093123133 | 0.222020513 | 0.346762078 | ns | NaCl | KS.1 |
| FIM1_1424 | 1268.070994 | 1105.552148 | 1449.15332 | -0.183330441 | 0.08645294 | 0.026715915 | 0.063442328 | ns | NaCl | KS.1 |
| FIM1_1425 | 1021.710736 | 789.004871 | 1331.415899 | -0.301390934 | 0.159565073 | 0.030973929 | 0.071605622 | ns | NaCl | KS.1 |
| FIM1_1426 | 4837.328299 | 4883.901569 | 3731.823932 | 0.007179044 | 0.270502733 | 0.970541938 | 0.986786069 | ns | NaCl | KS.1 |
| FIM1_1427 | 41.46302141 | 36.96778432 | 38.6833852 | -0.101534008 | 0.267465845 | 0.600804923 | 0.719548505 | ns | NaCl | KS.1 |
| FIM1_1428 | 1670.239761 | 1703.318197 | 2119.551811 | 0.02225075 | 0.147132723 | 0.865683339 | 0.919776371 | ns | NaCl | KS.1 |
| FIM1_1429 | 1746.076834 | 2131.760072 | 2233.881656 | 0.23067484 | 0.160445862 | 0.100329879 | 0.185485376 | ns | NaCl | KS.1 |
| FIM1_1430 | 1082.779621 | 681.4053189 | 1121.095545 | -0.56344445 | 0.184609144 | 0.000385579 | 0.001943523 | ns | NaCl | KS.1 |
| FIM1_1431 | 1389.952398 | 1152.603356 | 2212.056116 | -0.205257693 | 0.174004835 | 0.169215368 | 0.280615486 | ns | NaCl | KS.1 |
| FIM1_1432 | 15835.94138 | 17130.13039 | 13282.27271 | 0.097465822 | 0.119917643 | 0.379716587 | 0.514878719 | ns | NaCl | KS.1 |
| FIM1_1433 | 1676.833772 | 1976.889981 | 1941.906229 | 0.216974677 | 0.099111873 | 0.020456569 | 0.050582222 | ns | NaCl | KS.1 |
| FIM1_1434 | 1341.067397 | 1865.270295 | 1918.794881 | 0.326355265 | 0.234175357 | 0.075206719 | 0.147362515 | ns | NaCl | KS.1 |
| FIM1_1435 | 5540.42847 | 5338.259992 | 5628.031343 | -0.041814018 | 0.154715696 | 0.759937415 | 0.844776933 | ns | NaCl | KS.1 |
| FIM1_1436 | 521.3351328 | 342.4642492 | 560.7691604 | -0.52320792 | 0.167925234 | 0.000367387 | 0.001865051 | ns | NaCl | KS.1 |
| FIM1_1437 | 360.9022359 | 340.4305155 | 339.2170924 | -0.075044244 | 0.111776495 | 0.472402643 | 0.60603485 | ns | NaCl | KS.1 |
| FIM1_1438 | 62081.40425 | 97722.2473 | 60172.18324 | 0.627043417 | 0.093606129 | 2.6669315298 | 1.5250556736 | ns | NaCl | KS.1 |
| FIM1_1439 | 845.0906645 | 728.07637 | 1220.359772 | -0.179319074 | 0.133452612 | 0.139076237 | 0.239496116 | ns | NaCl | KS.1 |
| FIM1_1440 | 427.2652968 | 314.2448216 | 401.8067455 | -0.392686085 | 0.132722963 | 0.001106956 | 0.004659144 | ns | NaCl | KS.1 |
| FIM1_1441 | 168.4206069 | 181.2453577 | 322.5394479 | 0.057230774 | 0.250769711 | 0.760723051 | 0.845153456 | ns | NaCl | KS.1 |
| FIM1_1442 | 55.65311601 | 65.4827935 | 578.1003268 | 0.125642568 | 0.273274497 | 0.521885151 | 0.652118833 | ns | NaCl | KS.1 |
| FIM1_1443 | 557.9406982 | 589.2357958 | 494.6692785 | 0.061060199 | 0.145738002 | 0.639908072 | 0.749480061 | ns | NaCl | KS.1 |
| FIM1_1444 | 15690.17849 | 20478.60927 | 12306.94573 | 0.328566674 | 0.142079014 | 0.009856378 | 0.028213741 | ns | NaCl | KS.1 |
| FIM1_1445 | 1305.986563 | 2809.115213 | 1243.352178 | 0.832616384 | 0.295054588 | 0.000263322 | 0.001408629 | ns | NaCl | KS.1 |
| FIM1_1446 | 717.4534809 | 390.9197643 | 640.2136342 | -0.789052441 | 0.170812683 | 3.0039620703 | 4.2216698587 | ns | NaCl | KS.1 |
| FIM1_1447 | 865.0294683 | 1158.943082 | 788.7167144 | 0.304178057 | 0.210253854 | 0.075047194 | 0.147204309 | ns | NaCl | KS.1 |
| FIM1_1448 | 572.7944822 | 687.2006371 | 525.5591259 | 0.193659859 | 0.184736893 | 0.21431967 | 0.337097806 | ns | NaCl | KS.1 |
| FIM1_1449 | 3111.016394 | 3621.643265 | 2542.263048 | 0.19417628 | 0.113879789 | 0.066698938 | 0.133424694 | ns | NaCl | KS.1 |
| FIM1_1450 | 729.8607938 | 964.2095021 | 920.6746159 | 0.342014844 | 0.145950027 | 0.008583159 | 0.025478054 | ns | NaCl | KS.1 |

| | | | | | | | | | | |
|---|---|---|---|---|---|---|---|---|---|---|
| FIM1_1451 | 12051.04037 | 6237.158373 | 8214.938459 | -0.743927028 | 0.257898748 | 0.000319295 | 0.00165124 | ns | NaCl | KS.1 |
| FIM1_1453 | 2823.909996 | 3687.577751 | 3848.212792 | 0.248161075 | 0.239180179 | 0.179361106 | 0.294495545 | ns | NaCl | KS.1 |
| FIM1_1454 | 884.24302 | 773.9300822 | 1125.145479 | -0.158852804 | 0.13778453 | 0.202043942 | 0.322376077 | ns | NaCl | KS.1 |
| FIM1_1455 | 1255.15299 | 1689.70533 | 1192.667858 | 0.325223598 | 0.195405591 | 0.045437498 | 0.097604297 | ns | NaCl | KS.1 |
| FIM1_1456 | 476.3639408 | 730.593177 | 560.0456439 | 0.530968913 | 0.165831759 | 0.000263807 | 0.001409708 | ns | NaCl | KS.1 |
| FIM1_1457 | 2312.506561 | 2225.467529 | 2947.262511 | -0.045006111 | 0.144988494 | 0.729190201 | 0.820937147 | ns | NaCl | KS.1 |
| FIM1_1458 | 1572.950975 | 1484.373671 | 1399.980782 | -0.071490782 | 0.127427179 | 0.540750626 | 0.667055385 | ns | NaCl | KS.1 |
| FIM1_1460 | 1311.534037 | 1031.303173 | 1103.395768 | -0.323134462 | 0.094437215 | 0.000302459 | 0.001583929 | ns | NaCl | KS.1 |
| FIM1_1461 | 927.4749892 | 782.0947795 | 986.9451075 | -0.151797217 | 0.236945403 | 0.40668212 | 0.541698406 | ns | NaCl | KS.1 |
| FIM1_1462 | 1795.760095 | 1690.814504 | 1477.699355 | -0.077680563 | 0.106215251 | 0.436562993 | 0.572456745 | ns | NaCl | KS.1 |
| FIM1_1463 | 236.403399 | 285.2724634 | 239.1789131 | 0.200441996 | 0.179710452 | 0.189837722 | 0.307128936 | ns | NaCl | KS.1 |
| FIM1_1464 | 12.55674071 | 8.732642682 | 24.69833755 | -0.165955982 | 0.391666621 | 0.428078675 | 0.56475508 | ns | NaCl | KS.1 |
| FIM1_1465 | 1095.725797 | 978.7179971 | 1340.776731 | -0.146429064 | 0.1125409 | 0.162855876 | 0.272522026 | ns | NaCl | KS.1 |
| FIM1_1466 | 1033.716895 | 974.6139507 | 1195.574456 | -0.076360584 | 0.112530269 | 0.467280481 | 0.6017616 | ns | NaCl | KS.1 |
| FIM1_1467 | 1642.025734 | 1667.24689 | 2349.248935 | 0.018038272 | 0.143226185 | 0.888575714 | 0.933183795 | ns | NaCl | KS.1 |
| FIM1_1468 | 2975.607447 | 3329.858802 | 3250.144257 | 0.146618001 | 0.099773143 | 0.120074389 | 0.214341616 | ns | NaCl | KS.1 |
| FIM1_1469 | 125.9679168 | 59.36433619 | 198.5580107 | -0.872839276 | 0.25607504 | 3.34247918408 | 0.000243825 | ns | NaCl | KS.1 |
| FIM1_1470 | 1758.111513 | 1921.188056 | 1860.483098 | 0.120433135 | 0.080704339 | 0.12128334 | 0.215944243 | ns | NaCl | KS.1 |
| FIM1_1471 | 4507.334583 | 5432.751493 | 4424.68561 | 0.235238293 | 0.121408297 | 0.03534759 | 0.079861153 | ns | NaCl | KS.1 |
| FIM1_1472 | 898.8443875 | 890.8360216 | 1066.927055 | -0.01057769 | 0.148306677 | 0.936316804 | 0.963188249 | ns | NaCl | KS.1 |
| FIM1_1473 | 834.7585269 | 1633.302099 | 1273.656244 | 0.906559663 | 0.138233147 | 3.32556042968 | 1.85895091434 | ns | NaCl | KS.1 |
| FIM1_1474 | 2383.196404 | 3011.396006 | 2614.068586 | 0.279897526 | 0.150876392 | 0.036277512 | 0.081628503 | ns | NaCl | KS.1 |
| FIM1_1475 | 1182.835117 | 1343.26333 | 1427.26863 | 0.163326002 | 0.100584052 | 0.085486987 | 0.163891236 | ns | NaCl | KS.1 |
| FIM1_1476 | 71.3617406 | 82.08192175 | 1083.108515 | 0.121796404 | 0.241463117 | 0.50950713 | 0.640093083 | ns | NaCl | KS.1 |
| FIM1_1477 | 2005.32491 | 1517.869032 | 1843.773248 | -0.266265761 | 0.233541864 | 0.144373376 | 0.246823899 | ns | NaCl | KS.1 |
| FIM1_1478 | 2106.442755 | 924.7919106 | 1710.692423 | -0.975105089 | 0.261364667 | 6.70892494439 | 6.00737884057 | ns | NaCl | KS.1 |
| FIM1_1479 | 567.1729762 | 480.099398 | 809.3494458 | -0.216909455 | 0.105759605 | 0.028745968 | 0.067204506 | ns | NaCl | KS.1 |
| FIM1_1480 | 1066.481521 | 1360.5489 | 905.3788425 | 0.24954905 | 0.206679103 | 0.138892017 | 0.239344574 | ns | NaCl | KS.1 |
| FIM1_1481 | 619.4522909 | 419.9456711 | 507.7294437 | -0.459955628 | 0.186903782 | 0.003717606 | 0.012808232 | ns | NaCl | KS.1 |
| FIM1_1482 | 7774.476489 | 6462.686869 | 6288.323436 | -0.226267807 | 0.134405696 | 0.063223646 | 0.128225699 | ns | NaCl | KS.1 |
| FIM1_1483 | 424.0701575 | 394.6613388 | 398.8084871 | -0.073221156 | 0.190997375 | 0.646555649 | 0.755073792 | ns | NaCl | KS.1 |
| FIM1_1484 | 595.928701 | 457.3342256 | 742.8195055 | -0.32887763 | 0.143302892 | 0.010310347 | 0.029310845 | ns | NaCl | KS.1 |
| FIM1_1485 | 3063.526261 | 3118.039512 | 3134.913706 | 0.023519986 | 0.080804514 | 0.76266154 | 0.846739826 | ns | NaCl | KS.1 |
| FIM1_1486 | 1441.654299 | 1780.348403 | 2021.714419 | 0.227542689 | 0.186417286 | 0.147158333 | 0.250981388 | ns | NaCl | KS.1 |
| FIM1_1487 | 1755.931178 | 4107.147972 | 1944.335533 | 0.960976138 | 0.291257591 | 3.32176644741 | 0.000242669 | ns | NaCl | KS.1 |
| FIM1_1488 | 4376.387444 | 6004.451248 | 5118.618575 | 0.436808817 | 0.07787749 | 5.57670496920 | 1.30868430291 | ns | NaCl | KS.1 |
| FIM1_1489 | 597.3675748 | 551.1724901 | 496.6185416 | -0.094593227 | 0.145998798 | 0.46905819 | 0.603300024 | ns | NaCl | KS.1 |
| FIM1_1490 | 620.3600113 | 368.4945719 | 542.9241269 | -0.657812532 | 0.172762866 | 1.63894366166 | 0.000132151 | ns | NaCl | KS.1 |
| FIM1_1491 | 1103.437173 | 973.9812012 | 1008.296104 | -0.168364853 | 0.090438636 | 0.050954812 | 0.107324382 | ns | NaCl | KS.1 |
| FIM1_1492 | 1310.692451 | 1433.827537 | 1549.742596 | 0.111412205 | 0.121784577 | 0.321605186 | 0.454670588 | ns | NaCl | KS.1 |
| FIM1_1493 | 456.834415 | 288.0004336 | 639.7551531 | -0.573248411 | 0.174591522 | 0.000166189 | 0.000958327 | ns | NaCl | KS.1 |
| FIM1_1494 | 302.7001387 | 148.2391944 | 226.628268 | -0.923264399 | 0.189785359 | 5.82563378903 | 9.85800275525 | ns | NaCl | KS.1 |
| FIM1_1495 | 61.50349425 | 52.60839885 | 72.26210851 | -0.130845481 | 0.259827339 | 0.495090671 | 0.627375468 | ns | NaCl | KS.1 |
| FIM1_1496 | 828.4501107 | 963.8223942 | 888.9900235 | 0.191713371 | 0.113838921 | 0.070157513 | 0.139279181 | ns | NaCl | KS.1 |
| FIM1_1497 | 398.975536 | 532.9378677 | 541.8235652 | 0.374336358 | 0.127134655 | 0.001259267 | 0.005186138 | ns | NaCl | KS.1 |

| | | | | | | | | | | |
|---|---|---|---|---|---|---|---|---|---|---|
| FIM1_1498 | 631.670466 | 582.9159275 | 589.1756182 | -0.095799207 | 0.145853745 | 0.463046905 | 0.597421772 | ns | NaCl | KS.1 |
| FIM1_1499 | 2171.470376 | 1847.478935 | 1990.863723 | -0.172364859 | 0.182914285 | 0.265711936 | 0.394248995 | ns | NaCl | KS.1 |
| FIM1_1500 | 1231.78046 | 1245.215426 | 1542.250736 | 0.011062069 | 0.130763742 | 0.926237763 | 0.957612816 | ns | NaCl | KS.1 |
| FIM1_1501 | 1491.51114 | 1306.215003 | 1400.018085 | -0.161177556 | 0.134717119 | 0.187271659 | 0.303774537 | ns | NaCl | KS.1 |
| FIM1_1503 | 2113.350111 | 2413.433831 | 2153.953742 | 0.174945499 | 0.093423429 | 0.048912853 | 0.103814609 | ns | NaCl | KS.1 |
| FIM1_1504 | 741.0281317 | 766.651824 | 912.8558115 | 0.04317815 | 0.094744401 | 0.631967916 | 0.744047634 | ns | NaCl | KS.1 |
| FIM1_1505 | 381.2748881 | 246.8933985 | 530.282422 | -0.559979519 | 0.153700013 | 4.5479936282 | 0.000321396 | ns | NaCl | KS.1 |
| FIM1_1506 | 109.4123633 | 72.47964212 | 147.5683477 | -0.403023401 | 0.264540675 | 0.04257409 | 0.092572596 | ns | NaCl | KS.1 |
| FIM1_1507 | 4679.680233 | 4941.571178 | 4989.294237 | 0.066592295 | 0.127630159 | 0.569375912 | 0.69156376 | ns | NaCl | KS.1 |
| FIM1_1508 | 1755.003938 | 1588.477854 | 1713.87884 | -0.124660145 | 0.117356932 | 0.252156683 | 0.380833267 | ns | NaCl | KS.1 |
| FIM1_1509 | 1954.89764 | 2152.539727 | 1673.109911 | 0.115814617 | 0.132328397 | 0.336656172 | 0.47086434 | ns | NaCl | KS.1 |
| FIM1_1510 | 1366.81829 | 1537.921256 | 1336.410245 | 0.14812382 | 0.118501808 | 0.177016616 | 0.291415508 | ns | NaCl | KS.1 |
| FIM1_1511 | 208.7390225 | 183.30873 | 223.070927 | -0.141815757 | 0.176012904 | 0.346551225 | 0.481585583 | ns | NaCl | KS.1 |
| FIM1_1512 | 714.376782 | 707.3968335 | 639.4952849 | -0.011082249 | 0.135535762 | 0.928260441 | 0.958707846 | ns | NaCl | KS.1 |
| FIM1_1513 | 4726.94197 | 4242.069605 | 4753.957716 | -0.143764664 | 0.086186465 | 0.081668032 | 0.157663352 | ns | NaCl | KS.1 |
| FIM1_1514 | 12603.5751 | 10144.98843 | 11778.31794 | -0.275480692 | 0.12131468 | 0.013579486 | 0.036399753 | ns | NaCl | KS.1 |
| FIM1_1515 | 337.7189339 | 138.2729009 | 365.2917633 | -1.17240399 | 0.191875357 | 1.8288949679 | 7.8437521255 | Km<KS.1 | NaCl | KS.1 |
| FIM1_1516 | 213.1785537 | 136.417613 | 209.8896237 | -0.324874142 | 0.331050098 | 0.134860628 | 0.234296511 | ns | NaCl | KS.1 |
| FIM1_1518 | 1724.80868 | 2233.022461 | 1559.237096 | 0.320550465 | 0.137716361 | 0.009790434 | 0.028064898 | ns | NaCl | KS.1 |
| FIM1_1519 | 604.610035 | 431.7832436 | 582.3236775 | -0.425820402 | 0.148719216 | 0.001296851 | 0.005312038 | ns | NaCl | KS.1 |
| FIM1_1520 | 3601.527129 | 3350.10175 | 4137.183329 | -0.084646658 | 0.14229688 | 0.508417965 | 0.639487624 | ns | NaCl | KS.1 |
| FIM1_1521 | 4187.903192 | 8305.515787 | 4002.233212 | 0.632019837 | 0.332355511 | 0.006758315 | 0.020893722 | ns | NaCl | KS.1 |
| FIM1_1523 | 308.5482202 | 306.9340279 | 369.144513 | -0.008702435 | 0.136815556 | 0.944069872 | 0.969400953 | ns | NaCl | KS.1 |
| FIM1_1524 | 1147.772779 | 1283.131886 | 1103.512142 | 0.134437922 | 0.130561195 | 0.2592081 | 0.387604538 | ns | NaCl | KS.1 |
| FIM1_1525 | 1319.21434 | 1487.039066 | 1670.902005 | 0.162350037 | 0.082685824 | 0.040994039 | 0.089940451 | ns | NaCl | KS.1 |
| FIM1_1526 | 4784.631376 | 3278.443985 | 7536.909485 | -0.4943824 | 0.127947522 | 2.4604058181 | 0.000184902 | ns | NaCl | KS.1 |
| FIM1_1527 | 508.6909196 | 452.1492502 | 531.9941973 | -0.156044539 | 0.112644748 | 0.137201608 | 0.237418435 | ns | NaCl | KS.1 |
| FIM1_1528 | 754.9903808 | 946.4649126 | 668.6270175 | 0.180657892 | 0.269637035 | 0.356413489 | 0.491634979 | ns | NaCl | KS.1 |
| FIM1_1529 | 1078.809479 | 467.7143297 | 1461.92079 | -1.144769422 | 0.146991587 | 1.5066506920 | 2.0258343765 | Km<KS.1 | NaCl | KS.1 |
| FIM1_1530 | 1370.112897 | 1717.636842 | 2471.08405 | 0.274487989 | 0.144119291 | 0.033152717 | 0.075727626 | ns | NaCl | KS.1 |
| FIM1_1531 | 785.5243247 | 353.9607537 | 741.0917151 | -1.002595813 | 0.21359465 | 9.7584384660 | 1.5610363784 | Km<KS.1 | NaCl | KS.1 |
| FIM1_1532 | 853.2100848 | 468.1334484 | 768.8674388 | -0.668606503 | 0.251577876 | 0.000864502 | 0.003806103 | ns | NaCl | KS.1 |
| FIM1_1533 | 688.92472 | 798.2852479 | 731.9239523 | 0.183747929 | 0.119945892 | 0.097167135 | 0.180709814 | ns | NaCl | KS.1 |
| FIM1_1534 | 299.8329682 | 378.5724209 | 392.2872536 | 0.295136991 | 0.128201807 | 0.011600985 | 0.032063834 | ns | NaCl | KS.1 |
| FIM1_1535 | 846.4378648 | 756.7353272 | 781.0826287 | -0.153074278 | 0.088319267 | 0.069836298 | 0.13875223 | ns | NaCl | KS.1 |
| FIM1_1536 | 803.70521 | 801.7740727 | 720.7602547 | -0.003904228 | 0.121947493 | 0.972353116 | 0.987204262 | ns | NaCl | KS.1 |
| FIM1_1537 | 2451.343697 | 2414.871933 | 1969.124807 | -0.019257235 | 0.108797349 | 0.850300258 | 0.908752693 | ns | NaCl | KS.1 |
| FIM1_1538 | 1512.717838 | 1549.300066 | 1902.472634 | 0.028334159 | 0.118461656 | 0.796510026 | 0.870180595 | ns | NaCl | KS.1 |
| FIM1_1540 | 947.1858504 | 586.2719168 | 957.5712073 | -0.612395928 | 0.164000245 | 2.5952460906 | 0.000194156 | ns | NaCl | KS.1 |
| FIM1_1541 | 2265.013149 | 2144.726265 | 2516.562676 | -0.057527256 | 0.177832227 | 0.704859656 | 0.801892702 | ns | NaCl | KS.1 |
| FIM1_1542 | 3852.211329 | 9040.190108 | 4261.464166 | 0.97167114 | 0.28791374 | 2.4528574555 | 0.000184792 | ns | NaCl | KS.1 |
| FIM1_1543 | 3554.090056 | 7267.475179 | 3473.59313 | 0.686315485 | 0.328882465 | 0.0034852 | 0.012116612 | ns | NaCl | KS.1 |
| FIM1_1544 | 189.6020798 | 133.1110195 | 191.583174 | -0.382208669 | 0.197520604 | 0.019940962 | 0.049454778 | ns | NaCl | KS.1 |
| FIM1_1545 | 1169.964135 | 846.3985782 | 1128.202135 | -0.409985363 | 0.14406559 | 0.001471385 | 0.005889092 | ns | NaCl | KS.1 |
| FIM1_1546 | 345.1388231 | 226.6265651 | 298.4181561 | -0.527223221 | 0.169371477 | 0.000364812 | 0.001855766 | ns | NaCl | KS.1 |

| | | | | | | | | | | |
|---|---|---|---|---|---|---|---|---|---|---|
| FIM1_1547 | 1227.136622 | 825.8059755 | 968.3615341 | -0.507711234 | 0.147850598 | 0.000125768 | 0.000756587 | ns | NaCl | KS.1 |
| FIM1_1548 | 1692.293793 | 2342.264458 | 1539.925275 | 0.362414495 | 0.194078497 | 0.025283423 | 0.060473571 | ns | NaCl | KS.1 |
| FIM1_1549 | 1632.921047 | 1669.862207 | 1949.095125 | 0.026953149 | 0.128786851 | 0.819275017 | 0.887994163 | ns | NaCl | KS.1 |
| FIM1_1550 | 1450.205527 | 1217.982919 | 1478.150675 | -0.190445985 | 0.176395516 | 0.206724453 | 0.32826497 | ns | NaCl | KS.1 |
| FIM1_1551 | 15561.81324 | 11733.63978 | 21078.24352 | -0.281577181 | 0.222139022 | 0.111573661 | 0.201700205 | ns | NaCl | KS.1 |
| FIM1_1552 | 402.7529856 | 643.702399 | 398.9322626 | 0.53830285 | 0.213210776 | 0.002207446 | 0.008282085 | ns | NaCl | KS.1 |
| FIM1_1553 | 3838.923934 | 8183.878771 | 3566.150704 | 0.736973567 | 0.333824143 | 0.002069252 | 0.007840464 | ns | NaCl | KS.1 |
| FIM1_1554 | 900.3503216 | 1307.791279 | 1029.671585 | 0.471509019 | 0.149628284 | 0.000406671 | 0.0020296 | ns | NaCl | KS.1 |
| FIM1_1556 | 307.8331627 | 511.6049838 | 412.6890585 | 0.605796281 | 0.205069507 | 0.00044866 | 0.002199096 | ns | NaCl | KS.1 |
| FIM1_1557 | 374.1739285 | 415.1368199 | 370.8202114 | 0.111625776 | 0.165849974 | 0.438916221 | 0.574632463 | ns | NaCl | KS.1 |
| FIM1_1558 | 1661.49542 | 1779.404343 | 2132.708685 | 0.085764268 | 0.114918139 | 0.422701088 | 0.558846111 | ns | NaCl | KS.1 |
| FIM1_1559 | 6211.877023 | 7257.425116 | 5562.228255 | 0.171166771 | 0.172078902 | 0.24788537 | 0.37552671 | ns | NaCl | KS.1 |
| FIM1_1560 | 904.1982963 | 884.5385407 | 1399.809743 | -0.028662829 | 0.112600706 | 0.785168379 | 0.861384263 | ns | NaCl | KS.1 |
| FIM1_1561 | 1407.56169 | 1078.517954 | 1903.390235 | -0.3220491 | 0.152032849 | 0.01662171 | 0.042581363 | ns | NaCl | KS.1 |
| FIM1_1562 | 607.4040149 | 604.8113147 | 554.8368495 | -0.005137143 | 0.190296472 | 0.974272376 | 0.988116877 | ns | NaCl | KS.1 |
| FIM1_1563 | 1395.369411 | 1383.02072 | 1104.718313 | -0.012605185 | 0.097603787 | 0.891745285 | 0.93497003 | ns | NaCl | KS.1 |
| FIM1_1564 | 39.66062394 | 28.55242344 | 328.3333144 | -0.245047849 | 0.303560369 | 0.237687072 | 0.364312083 | ns | NaCl | KS.1 |
| FIM1_1566 | 1532.905704 | 2550.856446 | 2607.517748 | 0.622234459 | 0.192215344 | 0.000161386 | 0.000934689 | ns | NaCl | KS.1 |
| FIM1_1567 | 834.0837986 | 1002.043587 | 1116.362786 | 0.181079649 | 0.209721278 | 0.286850016 | 0.418007859 | ns | NaCl | KS.1 |
| FIM1_1568 | 6445.76276 | 7714.893338 | 7098.304888 | 0.242161476 | 0.08580582 | 0.003171194 | 0.011181213 | ns | NaCl | KS.1 |
| FIM1_1569 | 127.1192006 | 47.66531331 | 191.7269476 | -1.123445456 | 0.305373618 | 4.0828196035 | 4.0482841844 | Km<KS.1 | NaCl | KS.1 |
| FIM1_1570 | 942.1031326 | 740.0851516 | 1038.993896 | -0.30831792 | 0.121751663 | 0.005861138 | 0.018675443 | ns | NaCl | KS.1 |
| FIM1_1571 | 355.1296108 | 381.7128028 | 365.0476731 | 0.08191846 | 0.148376614 | 0.535998523 | 0.663658129 | ns | NaCl | KS.1 |
| FIM1_1572 | 439.81194 | 436.5619122 | 549.9838849 | -0.012214549 | 0.139406867 | 0.922749557 | 0.955595139 | ns | NaCl | KS.1 |
| FIM1_1573 | 1391.321546 | 1337.726012 | 1173.957086 | -0.054728026 | 0.086559461 | 0.509629889 | 0.640093083 | ns | NaCl | KS.1 |
| FIM1_1574 | 3268.765987 | 3455.020994 | 3203.394133 | 0.074693599 | 0.072165452 | 0.286160473 | 0.417238403 | ns | NaCl | KS.1 |
| FIM1_1575 | 793.7752085 | 514.0650537 | 851.4544938 | -0.577543247 | 0.124418448 | 5.2973879636 | 6.8631523748 | ns | NaCl | KS.1 |
| FIM1_1576 | 44.87685545 | 21.99664642 | 38.68437699 | -0.698198289 | 0.32345721 | 0.00279756 | 0.010092719 | ns | NaCl | KS.1 |
| FIM1_1577 | 793.8256209 | 381.1612831 | 674.6265987 | -0.985504052 | 0.157508265 | 1.7673481699 | 7.6457018656 | ns | NaCl | KS.1 |
| FIM1_1578 | 12367.58516 | 14655.39399 | 9826.523273 | 0.22581865 | 0.090757211 | 0.008979442 | 0.026309024 | ns | NaCl | KS.1 |
| FIM1_1579 | 2648.244361 | 3999.46549 | 1877.861067 | 0.384924461 | 0.266872684 | 0.053394596 | 0.111472142 | ns | NaCl | KS.1 |
| FIM1_1580 | 1540.160186 | 2726.517792 | 1777.236402 | 0.693345766 | 0.20643504 | 7.9958033609 | 0.00051931 | ns | NaCl | KS.1 |
| FIM1_1581 | 4522.00174 | 6432.772808 | 4862.540858 | 0.484996014 | 0.086816602 | 5.0592325706 | 1.2218292252 | ns | NaCl | KS.1 |
| FIM1_1582 | 111.5756208 | 36.82688163 | 111.7929124 | -1.235786501 | 0.353628246 | 4.5098194284 | 4.3820999329 | Km<KS.1 | NaCl | KS.1 |
| FIM1_1583 | 1035.606861 | 673.5825007 | 887.9140318 | -0.566707124 | 0.132006188 | 2.8438254102 | 2.9413786727 | ns | NaCl | KS.1 |
| FIM1_1584 | 6312.048923 | 6557.683366 | 5326.866002 | 0.046959389 | 0.125330521 | 0.68375256 | 0.786334506 | ns | NaCl | KS.1 |
| FIM1_1585 | 925.9619942 | 579.2455922 | 919.5679531 | -0.604845098 | 0.150697189 | 8.4236303027 | 7.3780916823 | ns | NaCl | KS.1 |
| FIM1_1586 | 966.0125418 | 810.568785 | 2013.747848 | -0.169149492 | 0.213502745 | 0.325233756 | 0.458237875 | ns | NaCl | KS.1 |
| FIM1_1587 | 8386.853901 | 9874.693303 | 7941.355191 | 0.20994662 | 0.10949953 | 0.040116166 | 0.088426196 | ns | NaCl | KS.1 |
| FIM1_1588 | 7133.410457 | 10178.65558 | 6573.148533 | 0.418741554 | 0.178981751 | 0.006217046 | 0.019600636 | ns | NaCl | KS.1 |
| FIM1_1589 | 193.0489763 | 208.6043373 | 174.21158 | 0.092244797 | 0.140991339 | 0.467752807 | 0.602088025 | ns | NaCl | KS.1 |
| FIM1_1590 | 1264.093434 | 1153.717771 | 907.6081619 | -0.094946785 | 0.186617501 | 0.545674408 | 0.672129284 | ns | NaCl | KS.1 |
| FIM1_1591 | 202.2559227 | 189.9699332 | 193.798865 | -0.062298556 | 0.215040242 | 0.717952137 | 0.81294531 | ns | NaCl | KS.1 |
| FIM1_1592 | 148.8381917 | 116.6296999 | 207.5354789 | -0.261573391 | 0.194768106 | 0.106274869 | 0.19416727 | ns | NaCl | KS.1 |
| FIM1_1593 | 2765.876901 | 1227.20938 | 3711.678997 | -0.805530274 | 0.339914889 | 0.001008785 | 0.004319023 | ns | NaCl | KS.1 |

| | | | | | | | | | | |
|---|---|---|---|---|---|---|---|---|---|---|
| FIM1_1594 | 2467.939433 | 5281.904492 | 2701.158854 | 0.859803042 | 0.27599643 | 9.4764728737 | 0.000596023 | ns | NaCl | KS.1 |
| FIM1_1595 | 1792.41355 | 1998.063579 | 1468.553781 | 0.124695703 | 0.151861991 | 0.354698602 | 0.489765624 | ns | NaCl | KS.1 |
| FIM1_1596 | 1404.039041 | 944.0639572 | 1618.955995 | -0.441461889 | 0.210508537 | 0.010486951 | 0.029677235 | ns | NaCl | KS.1 |
| FIM1_1597 | 2193.199108 | 3965.36424 | 1675.69388 | 0.567076154 | 0.301505359 | 0.009714105 | 0.027918932 | ns | NaCl | KS.1 |
| FIM1_1598 | 3512.352243 | 6330.421332 | 3069.22103 | 0.541805814 | 0.311101095 | 0.014548382 | 0.038364757 | ns | NaCl | KS.1 |
| FIM1_1599 | 2379.575625 | 2973.545282 | 2665.984911 | 0.300481861 | 0.09116924 | 0.000524818 | 0.002510547 | ns | NaCl | KS.1 |
| FIM1_1601 | 27027.42159 | 36992.05538 | 22763.71051 | 0.399714767 | 0.13388683 | 0.000982343 | 0.004227646 | ns | NaCl | KS.1 |
| FIM1_1602 | 3792.519655 | 2025.642882 | 2256.884439 | -0.874902898 | 0.098811872 | 6.0907934097 | 1.6834276229 | ns | NaCl | KS.1 |
| FIM1_1603 | 5475.543796 | 6263.300032 | 4657.555361 | 0.17230327 | 0.108788195 | 0.090524987 | 0.171370552 | ns | NaCl | KS.1 |
| FIM1_1604 | 1285.687574 | 1759.336033 | 2197.326057 | 0.393054829 | 0.144834209 | 0.002388355 | 0.008847405 | ns | NaCl | KS.1 |
| FIM1_1605 | 3253.785176 | 7289.870546 | 2906.898455 | 0.802997852 | 0.337477963 | 0.000997541 | 0.004278248 | ns | NaCl | KS.1 |
| FIM1_1606 | 8139.072499 | 8635.50065 | 7623.839318 | 0.072015254 | 0.12820271 | 0.539853443 | 0.666444387 | ns | NaCl | KS.1 |
| FIM1_1607 | 865.8266447 | 1039.128621 | 1263.574432 | 0.228094677 | 0.119954507 | 0.039280025 | 0.087007179 | ns | NaCl | KS.1 |
| FIM1_1608 | 3121.190528 | 3520.027922 | 2705.623237 | 0.144900751 | 0.135235496 | 0.237465123 | 0.364176629 | ns | NaCl | KS.1 |
| FIM1_1609 | 2014.922068 | 1613.391485 | 1307.238998 | -0.258502617 | 0.160975269 | 0.066152869 | 0.132598922 | ns | NaCl | KS.1 |
| FIM1_1610 | 2331.197876 | 2425.999559 | 2102.081972 | 0.040661309 | 0.189436638 | 0.797898919 | 0.870903663 | ns | NaCl | KS.1 |
| FIM1_1611 | 2103.183446 | 1747.353906 | 1829.231039 | -0.213471225 | 0.159877753 | 0.127382998 | 0.223853908 | ns | NaCl | KS.1 |
| FIM1_1612 | 1962.24646 | 2581.050855 | 1563.798557 | 0.290370275 | 0.201771793 | 0.080383973 | 0.155728297 | ns | NaCl | KS.1 |
| FIM1_1613 | 1357.030469 | 1185.804787 | 1328.538532 | -0.174542425 | 0.100177559 | 0.065046427 | 0.130821403 | ns | NaCl | KS.1 |
| FIM1_1614 | 492.9908713 | 329.3265365 | 538.7302173 | -0.520648586 | 0.150808932 | 0.000111118 | 0.000682481 | ns | NaCl | KS.1 |
| FIM1_1615 | 3378.167165 | 3629.193815 | 3138.809537 | 0.092473142 | 0.100183939 | 0.329120744 | 0.462666205 | ns | NaCl | KS.1 |
| FIM1_1616 | 809.2265516 | 869.7117854 | 848.8679576 | 0.086584556 | 0.12960589 | 0.465081194 | 0.599579927 | ns | NaCl | KS.1 |
| FIM1_1617 | 3251.81228 | 5968.57098 | 3551.417207 | 0.844676488 | 0.099664785 | 1.8339129336 | 3.6494867380 | ns | NaCl | KS.1 |
| FIM1_1618 | 341.1676386 | 668.4665159 | 330.5806568 | 0.786854409 | 0.242877102 | 8.4726730206 | 0.000543891 | ns | NaCl | KS.1 |
| FIM1_1619 | 2335.064686 | 2224.233224 | 1644.712945 | -0.060627191 | 0.113143384 | 0.565761615 | 0.687845561 | ns | NaCl | KS.1 |
| FIM1_1621 | 3278.104657 | 3372.250941 | 3449.992768 | 0.036720041 | 0.094552443 | 0.683232516 | 0.786096616 | ns | NaCl | KS.1 |
| FIM1_1622 | 786.1998989 | 914.9160695 | 889.5684964 | 0.18005265 | 0.146597299 | 0.168883107 | 0.280344831 | ns | NaCl | KS.1 |
| FIM1_1623 | 360.1721252 | 384.5630452 | 369.2683476 | 0.070841033 | 0.160867862 | 0.615331466 | 0.730440001 | ns | NaCl | KS.1 |
| FIM1_1624 | 1024.668804 | 709.8230928 | 674.7249446 | -0.449045998 | 0.163996947 | 0.001728519 | 0.006790996 | ns | NaCl | KS.1 |
| FIM1_1625 | 1677.9605 | 1459.872719 | 1770.134835 | -0.166669304 | 0.138706014 | 0.183033793 | 0.29865304 | ns | NaCl | KS.1 |
| FIM1_1627 | 2599.665996 | 2497.455592 | 2079.215773 | -0.045430945 | 0.151074084 | 0.735177642 | 0.825436418 | ns | NaCl | KS.1 |
| FIM1_1628 | 933.1734264 | 313.0621988 | 650.5022275 | -1.034896579 | 0.423282371 | 0.000266163 | 0.001419253 | Km<KS.1 | NaCl | KS.1 |
| FIM1_1629 | 387.8955858 | 339.00796 | 370.7286386 | -0.169187291 | 0.139393756 | 0.178233888 | 0.29312846 | ns | NaCl | KS.1 |
| FIM1_1630 | 2450.789287 | 2085.25717 | 2888.50703 | -0.20661263 | 0.112400575 | 0.048370559 | 0.102751295 | ns | NaCl | KS.1 |
| FIM1_1631 | 724.160221 | 590.8556094 | 737.8346959 | -0.217420589 | 0.180955098 | 0.157037976 | 0.264476618 | ns | NaCl | KS.1 |
| FIM1_1632 | 1566.293089 | 1269.347791 | 1564.129185 | -0.174185205 | 0.259173767 | 0.364544373 | 0.499892022 | ns | NaCl | KS.1 |
| FIM1_1633 | 1776.65558 | 1841.231442 | 1191.610824 | 0.028892951 | 0.253884631 | 0.878480142 | 0.92830049 | ns | NaCl | KS.1 |
| FIM1_1634 | 2148.166525 | 2227.275818 | 2116.876201 | 0.042517262 | 0.137015728 | 0.732051301 | 0.823198675 | ns | NaCl | KS.1 |
| FIM1_1635 | 6024.20897 | 6287.752447 | 6104.133034 | 0.058065515 | 0.077663108 | 0.439063619 | 0.574632463 | ns | NaCl | KS.1 |
| FIM1_1636 | 988.7810617 | 845.7847698 | 829.9480188 | -0.189534991 | 0.13432683 | 0.119837666 | 0.21407267 | ns | NaCl | KS.1 |
| FIM1_1637 | 303.7676613 | 218.6635508 | 238.0021329 | -0.376365233 | 0.181815174 | 0.014877762 | 0.038894833 | ns | NaCl | KS.1 |
| FIM1_1638 | 40.61466192 | 31.54091904 | 70.51401508 | -0.212252618 | 0.280122712 | 0.288438581 | 0.41897283 | ns | NaCl | KS.1 |
| FIM1_1640 | 2385.816657 | 2359.107809 | 2570.685489 | -0.014908167 | 0.111658407 | 0.886434041 | 0.9323487 | ns | NaCl | KS.1 |
| FIM1_1641 | 5329.391459 | 5392.623713 | 4617.052727 | 0.015298499 | 0.086258131 | 0.853315665 | 0.910996874 | ns | NaCl | KS.1 |
| FIM1_1642 | 1466.148006 | 1087.53909 | 1800.68227 | -0.397751482 | 0.102699422 | 3.6959870687 | 0.000266873 | ns | NaCl | KS.1 |

| | | | | | | | | | | |
|---|---|---|---|---|---|---|---|---|---|---|
| FIM1_1643 | 1380.967889 | 1363.654116 | 1577.718136 | -0.0152627 | 0.11848775 | 0.889552061 | 0.933524791 | ns | NaCl | KS.1 |
| FIM1_1644 | 539.1130559 | 505.1073427 | 640.2083275 | -0.085928932 | 0.118754519 | 0.434912534 | 0.570995045 | ns | NaCl | KS.1 |
| FIM1_1645 | 563.3831383 | 548.404313 | 638.4829664 | -0.035631677 | 0.13309953 | 0.768852363 | 0.851333297 | ns | NaCl | KS.1 |
| FIM1_1646 | 2286.061603 | 1361.933315 | 3165.894573 | -0.658433165 | 0.171604638 | 1.45204050814 | 0.000118619 | ns | NaCl | KS.1 |
| FIM1_1647 | 91.98728824 | 110.2113457 | 78.19150405 | 0.152005071 | 0.22802534 | 0.39562658 | 0.531526394 | ns | NaCl | KS.1 |
| FIM1_1648 | 4928.543279 | 4064.693223 | 3748.977745 | -0.223647324 | 0.158242126 | 0.107353029 | 0.195777609 | ns | NaCl | KS.1 |
| FIM1_1649 | 3280.959018 | 4902.845818 | 2915.235222 | 0.472603717 | 0.188605146 | 0.00309834 | 0.010970991 | ns | NaCl | KS.1 |
| FIM1_1650 | 863.6926626 | 1250.257045 | 1231.102463 | 0.372601382 | 0.237600179 | 0.044904387 | 0.096793469 | ns | NaCl | KS.1 |
| FIM1_1651 | 404.2596146 | 210.0549699 | 475.7090115 | -0.833080499 | 0.195341335 | 1.31698545558 | 1.49248351743 | ns | NaCl | KS.1 |
| FIM1_1652 | 3385.403677 | 3192.647112 | 2996.097767 | -0.070558226 | 0.129009637 | 0.550195303 | 0.675523601 | ns | NaCl | KS.1 |
| FIM1_1653 | 497.0693415 | 405.9067645 | 589.1797007 | -0.250653905 | 0.140441035 | 0.047053273 | 0.100554138 | ns | NaCl | KS.1 |
| FIM1_1654 | 485.1341718 | 178.5731653 | 472.4391557 | -1.283463136 | 0.238308881 | 6.93428239603 | 2.04130502486 | Km<KS.1 | NaCl | KS.1 |
| FIM1_1655 | 1588.113777 | 1334.850757 | 1447.568048 | -0.224023603 | 0.109196947 | 0.028072093 | 0.065938935 | ns | NaCl | KS.1 |
| FIM1_1656 | 610.5259684 | 307.54439 | 1353.127567 | -0.913776422 | 0.147966008 | 3.84860973172 | 1.50762467837 | ns | NaCl | KS.1 |
| FIM1_1657 | 37449.02319 | 49031.47375 | 30073.46668 | 0.359219283 | 0.100315685 | 0.00014152 | 0.000836202 | ns | NaCl | KS.1 |
| FIM1_1658 | 1272.631801 | 1214.731402 | 1053.529141 | -0.055957081 | 0.13364975 | 0.645506764 | 0.754300314 | ns | NaCl | KS.1 |
| FIM1_1659 | 2261.430085 | 2621.493307 | 2925.916133 | 0.191805466 | 0.104572558 | 0.050929469 | 0.107324382 | ns | NaCl | KS.1 |
| FIM1_1660 | 1629.067169 | 2256.47756 | 2284.236219 | 0.434484663 | 0.107540126 | 1.53762186678 | 0.000124587 | ns | NaCl | KS.1 |
| FIM1_1661 | 1641.844716 | 2118.891807 | 1659.451143 | 0.316118445 | 0.13875163 | 0.01135167 | 0.031567669 | ns | NaCl | KS.1 |
| FIM1_1664 | 2152.844966 | 2380.294931 | 2194.986819 | 0.133450194 | 0.083006955 | 0.094334615 | 0.176966331 | ns | NaCl | KS.1 |
| FIM1_1665 | 224.4281972 | 305.5923013 | 323.128888 | 0.329125555 | 0.209396208 | 0.053680218 | 0.112021429 | ns | NaCl | KS.1 |
| FIM1_1666 | 875.3958716 | 938.3856122 | 919.0415446 | 0.092593372 | 0.085842252 | 0.260736163 | 0.389304444 | ns | NaCl | KS.1 |
| FIM1_1667 | 1575.842676 | 5072.891336 | 1963.193442 | 1.427925254 | 0.30313165 | 1.02760481558 | 2.18475810150 | Km>KS.1 | NaCl | KS.1 |
| FIM1_1669 | 1783.375132 | 2502.695806 | 2080.923473 | 0.442362212 | 0.124861949 | 0.000111479 | 0.000683017 | ns | NaCl | KS.1 |
| FIM1_1670 | 2325.594763 | 3057.094467 | 1932.306517 | 0.297535374 | 0.192677111 | 0.064312302 | 0.129956824 | ns | NaCl | KS.1 |
| FIM1_1671 | 5380.301961 | 12046.72274 | 5657.092212 | 0.887542503 | 0.296661949 | 0.000123773 | 0.000746391 | ns | NaCl | KS.1 |
| FIM1_1672 | 3793.55124 | 4498.401166 | 4201.129198 | 0.202751862 | 0.145518493 | 0.118947653 | 0.212864955 | ns | NaCl | KS.1 |
| FIM1_1673 | 2885.098801 | 4043.506499 | 3566.447103 | 0.46243475 | 0.088297551 | 3.98401559409 | 7.21619628439 | ns | NaCl | KS.1 |
| FIM1_1674 | 735.7937753 | 782.3775828 | 1673.238799 | 0.076289612 | 0.129165998 | 0.51867402 | 0.649811949 | ns | NaCl | KS.1 |
| FIM1_1675 | 25.27070998 | 21.71052576 | 18.27940201 | -0.077762854 | 0.299933361 | 0.700589618 | 0.798938005 | ns | NaCl | KS.1 |
| FIM1_1676 | 469.0483854 | 542.586316 | 574.8902043 | 0.179838809 | 0.123256195 | 0.11274065 | 0.203496589 | ns | NaCl | KS.1 |
| FIM1_1677 | 301.5594256 | 303.1549103 | 639.4036967 | 0.004414197 | 0.288334325 | 0.982317212 | 0.991175946 | ns | NaCl | KS.1 |
| FIM1_1678 | 111.3378692 | 123.9772207 | 212.1497853 | 0.104937665 | 0.221896722 | 0.550745323 | 0.675698639 | ns | NaCl | KS.1 |
| FIM1_1679 | 411.2094396 | 550.7445024 | 646.7479553 | 0.300708774 | 0.218061139 | 0.085769403 | 0.16436933 | ns | NaCl | KS.1 |
| FIM1_1680 | 496.2469834 | 849.2217119 | 512.5961361 | 0.683320168 | 0.174666227 | 9.78357867504 | 8.35197056438 | ns | NaCl | KS.1 |
| FIM1_1681 | 35.32753662 | 22.80577512 | 59.03730574 | -0.302937975 | 0.329809401 | 0.160591304 | 0.269366736 | ns | NaCl | KS.1 |
| FIM1_1682 | 923.4676795 | 1055.846848 | 870.1066741 | 0.147594119 | 0.17072981 | 0.316377264 | 0.449194317 | ns | NaCl | KS.1 |
| FIM1_1683 | 4165.72285 | 5783.479648 | 4383.462853 | 0.381926351 | 0.178791721 | 0.012350725 | 0.033594783 | ns | NaCl | KS.1 |
| FIM1_1684 | 1089.726273 | 1039.953899 | 1764.440744 | -0.058557274 | 0.110179829 | 0.570228673 | 0.692261506 | ns | NaCl | KS.1 |
| FIM1_1685 | 2244.5699 | 1832.257711 | 2269.185769 | -0.26633514 | 0.098631086 | 0.004224478 | 0.014181363 | ns | NaCl | KS.1 |
| FIM1_1686 | 1043.771904 | 1029.777297 | 1051.0058 | -0.016791361 | 0.130530104 | 0.888075283 | 0.933183795 | ns | NaCl | KS.1 |
| FIM1_1687 | 1697.976834 | 1161.265199 | 2583.761881 | -0.50537347 | 0.113635066 | 1.79208954724 | 1.95498354703 | ns | NaCl | KS.1 |
| FIM1_1688 | 428.8426335 | 252.8420212 | 317.1652885 | -0.563390658 | 0.253939737 | 0.004512516 | 0.014946584 | ns | NaCl | KS.1 |
| FIM1_1689 | 305.9205241 | 361.1841739 | 271.523357 | 0.171175631 | 0.185286353 | 0.273422074 | 0.402925006 | ns | NaCl | KS.1 |
| FIM1_1690 | 3785.053412 | 2763.562152 | 3733.747297 | -0.432363737 | 0.08511838 | 1.07170511493 | 1.69800412318 | ns | NaCl | KS.1 |

| | | | | | | | | | | |
|---|---|---|---|---|---|---|---|---|---|---|
| FIM1_1691 | 286.1186338 | 1016.291323 | 2389.642711 | 1.231869423 | 0.463405527 | 5.89391066356 | 0.000400577 | Km>KS.1 | NaCl | KS.1 |
| FIM1_1692 | 471.0570786 | 632.334563 | 1205.142974 | 0.329731033 | 0.185734194 | 0.035422651 | 0.079994411 | ns | NaCl | KS.1 |
| FIM1_1693 | 678.6956415 | 349.6189845 | 768.7516542 | -0.669550256 | 0.300487966 | 0.002718503 | 0.009857546 | ns | NaCl | KS.1 |
| FIM1_1694 | 36.58792853 | 53.42600988 | 53.63782133 | 0.317747579 | 0.287389347 | 0.12137832 | 0.215944243 | ns | NaCl | KS.1 |
| FIM1_1695 | 167.7514941 | 97.96317553 | 186.2138148 | -0.439135438 | 0.323276626 | 0.046889771 | 0.100334026 | ns | NaCl | KS.1 |
| FIM1_1697 | 212.9668191 | 191.6804235 | 205.3618799 | -0.092362129 | 0.228150677 | 0.605337249 | 0.723235546 | ns | NaCl | KS.1 |
| FIM1_1698 | 2054.152155 | 3464.266607 | 2099.503148 | 0.54958461 | 0.257501997 | 0.005917664 | 0.018771772 | ns | NaCl | KS.1 |
| FIM1_1701 | 445.2274921 | 608.2126566 | 548.8457104 | 0.337866012 | 0.202289434 | 0.042617123 | 0.092625682 | ns | NaCl | KS.1 |
| FIM1_1703 | 55.07613961 | 52.93166174 | 84.05414166 | -0.0245129 | 0.260599226 | 0.898034552 | 0.938980214 | ns | NaCl | KS.1 |
| FIM1_1705 | 12.26839599 | 23.5071001 | 22.09320857 | 0.47878753 | 0.373016716 | 0.041399761 | 0.090544735 | ns | NaCl | KS.1 |
| FIM1_1707 | 304.7446486 | 413.0073929 | 374.05025 | 0.350088559 | 0.179175398 | 0.021894672 | 0.05381719 | ns | NaCl | KS.1 |
| FIM1_1708 | 582.8726159 | 5325.005068 | 2951.506154 | 2.790896155 | 0.499389025 | 2.59748268461 | 1.06797325255 | Km>KS.1 | NaCl | KS.1 |
| FIM1_1709 | 1004.726269 | 892.4694347 | 1248.502807 | -0.107940033 | 0.225521207 | 0.543316529 | 0.669556535 | ns | NaCl | KS.1 |
| FIM1_1710 | 328.3824832 | 278.5574201 | 440.0704167 | -0.139157112 | 0.250571831 | 0.460230162 | 0.594713002 | ns | NaCl | KS.1 |
| FIM1_1711 | 762.3508381 | 415.893937 | 1667.795688 | -0.502327803 | 0.336652358 | 0.027524656 | 0.06483891 | ns | NaCl | KS.1 |
| FIM1_1712 | 4854.683223 | 6435.751057 | 2816.95815 | 0.240770259 | 0.263011224 | 0.215993846 | 0.339409154 | ns | NaCl | KS.1 |
| FIM1_1713 | 72.3644681 | 82.81997379 | 91.60946069 | 0.128763191 | 0.220582345 | 0.46269307 | 0.597275044 | ns | NaCl | KS.1 |
| FIM1_1714 | 161.5219543 | 177.9626743 | 331.4930736 | 0.102229134 | 0.171814308 | 0.489947504 | 0.622672274 | ns | NaCl | KS.1 |
| FIM1_1715 | 132.7705201 | 96.52629975 | 205.5618188 | -0.319372285 | 0.229890766 | 0.078038907 | 0.152013925 | ns | NaCl | KS.1 |
| FIM1_1716 | 4023.600384 | 4272.948277 | 4759.1703 | 0.055899658 | 0.216829344 | 0.747093208 | 0.835608973 | ns | NaCl | KS.1 |
| FIM1_1717 | 165.9897838 | 166.9914071 | 180.8783533 | -3.76E-05 | 0.147697728 | 0.999772494 | 0.999786056 | ns | NaCl | KS.1 |
| FIM1_1718 | 126.0519602 | 174.9626317 | 178.6442181 | 0.385556453 | 0.163091957 | 0.006731074 | 0.020864233 | ns | NaCl | KS.1 |
| FIM1_1719 | 212.0261168 | 202.7187241 | 232.4189604 | -0.05259583 | 0.160381036 | 0.708499847 | 0.804378535 | ns | NaCl | KS.1 |
| FIM1_1721 | 33.62729251 | 33.41636517 | 37.01723315 | -0.002826794 | 0.299364376 | 0.988807857 | 0.995082984 | ns | NaCl | KS.1 |
| FIM1_1724 | 5279.153697 | 18313.71502 | 10659.98046 | 1.618633735 | 0.257574653 | 4.86908062766 | 3.27346974630 | Km>KS.1 | NaCl | KS.1 |
| FIM1_1725 | 19727.22999 | 28652.32998 | 18386.44368 | 0.410609695 | 0.20912292 | 0.016509029 | 0.042329053 | ns | NaCl | KS.1 |
| FIM1_1727 | 1709.017447 | 1085.782159 | 1983.948707 | -0.514538737 | 0.214424347 | 0.003471503 | 0.012077432 | ns | NaCl | KS.1 |
| FIM1_1728 | 5428.972471 | 6289.541813 | 6048.725339 | 0.180061524 | 0.131835073 | 0.133479776 | 0.23219245 | ns | NaCl | KS.1 |
| FIM1_1729 | 3057.027363 | 3358.000938 | 3532.289846 | 0.119615528 | 0.112924122 | 0.255960817 | 0.38425017 | ns | NaCl | KS.1 |
| FIM1_1730 | 5380.516678 | 5185.941636 | 4780.754088 | -0.049705556 | 0.070050554 | 0.46545884 | 0.599762737 | ns | NaCl | KS.1 |
| FIM1_1731 | 5254.711035 | 5220.021681 | 4615.337124 | -0.009211675 | 0.073968423 | 0.897772768 | 0.938915182 | ns | NaCl | KS.1 |
| FIM1_1732 | 1517.00886 | 1275.260102 | 1104.078117 | -0.205808156 | 0.147831741 | 0.118069751 | 0.211674599 | ns | NaCl | KS.1 |
| FIM1_1733 | 6971.883043 | 6520.556173 | 8127.148046 | -0.057932038 | 0.235014372 | 0.749779947 | 0.837108446 | ns | NaCl | KS.1 |
| FIM1_1734 | 466.130345 | 522.682933 | 542.6370371 | 0.113314527 | 0.198342387 | 0.489082389 | 0.622616398 | ns | NaCl | KS.1 |
| FIM1_1735 | 3292.723996 | 2424.024777 | 2353.005583 | -0.370507534 | 0.158107478 | 0.007596741 | 0.02303095 | ns | NaCl | KS.1 |
| FIM1_1736 | 559.1368541 | 593.1039527 | 664.8273505 | 0.06934769 | 0.144705822 | 0.593061923 | 0.713884119 | ns | NaCl | KS.1 |
| FIM1_1737 | 412.6833743 | 183.5257118 | 484.8333214 | -1.05474354 | 0.197254558 | 2.78973352869 | 7.08108382921 | Km<KS.1 | NaCl | KS.1 |
| FIM1_1738 | 1769.214147 | 1068.27723 | 1392.724953 | -0.658020604 | 0.149768316 | 1.30131836054 | 1.47809562642 | ns | NaCl | KS.1 |
| FIM1_1739 | 3180.462948 | 4726.490135 | 2986.10881 | 0.504421512 | 0.148353221 | 0.000144604 | 0.000851368 | ns | NaCl | KS.1 |
| FIM1_1740 | 984.7965409 | 761.9141926 | 944.6310137 | -0.328237218 | 0.119646311 | 0.00289814 | 0.010402776 | ns | NaCl | KS.1 |
| FIM1_1741 | 4898.402444 | 7700.753664 | 4603.886683 | 0.561358645 | 0.172930786 | 0.000199064 | 0.00111254 | ns | NaCl | KS.1 |
| FIM1_1742 | 1459.319492 | 1793.944148 | 1223.137733 | 0.244657197 | 0.149732293 | 0.065703138 | 0.131803675 | ns | NaCl | KS.1 |
| FIM1_1743 | 4031.388266 | 5908.448471 | 3528.491098 | 0.400114084 | 0.227835736 | 0.027326155 | 0.064521889 | ns | NaCl | KS.1 |
| FIM1_1744 | 2311.271063 | 2703.133222 | 2700.92026 | 0.13908224 | 0.235581136 | 0.445416469 | 0.580546747 | ns | NaCl | KS.1 |
| FIM1_1745 | 1726.337897 | 1509.759845 | 1396.955375 | -0.176321973 | 0.098152578 | 0.057657227 | 0.119171044 | ns | NaCl | KS.1 |

| | | | | | | | | | | |
|---|---|---|---|---|---|---|---|---|---|---|
| FIM1_1746 | 593.4443239 | 573.2719184 | 512.8983152 | -0.040597536 | 0.163776116 | 0.776404874 | 0.854749778 | ns | NaCl | KS.1 |
| FIM1_1747 | 2713.966979 | 2724.145572 | 3201.610679 | 0.004086665 | 0.156562556 | 0.976396708 | 0.9889179 | ns | NaCl | KS.1 |
| FIM1_1748 | 48.83726774 | 89.73517735 | 243.67275 | 0.628120796 | 0.291647304 | 0.003943991 | 0.013411726 | ns | NaCl | KS.1 |
| FIM1_1749 | 5055.585679 | 4375.741308 | 4469.165477 | -0.17064208 | 0.144281198 | 0.186697122 | 0.303040189 | ns | NaCl | KS.1 |
| FIM1_1750 | 4750.53606 | 5695.948081 | 4549.600814 | 0.168332914 | 0.226777597 | 0.345526752 | 0.480434766 | ns | NaCl | KS.1 |
| FIM1_1751 | 1632.130473 | 2699.51528 | 1262.995325 | 0.418099816 | 0.313530018 | 0.05417555 | 0.11287627 | ns | NaCl | KS.1 |
| FIM1_1752 | 1005.548515 | 1094.241206 | 1127.180991 | 0.07185029 | 0.242139453 | 0.697187037 | 0.796991156 | ns | NaCl | KS.1 |
| FIM1_1753 | 796.0014582 | 487.730662 | 787.7175786 | -0.660207779 | 0.12307105 | 9.41766325648 | 2.03708150874 | ns | NaCl | KS.1 |
| FIM1_1754 | 625.2328122 | 378.6319506 | 560.3280302 | -0.538599382 | 0.24600867 | 0.005424404 | 0.017455635 | ns | NaCl | KS.1 |
| FIM1_1755 | 2616.146884 | 2473.578507 | 2207.912755 | -0.046050604 | 0.247898755 | 0.805283744 | 0.875691066 | ns | NaCl | KS.1 |
| FIM1_1756 | 1744.214825 | 1204.40845 | 1284.85607 | -0.396823426 | 0.217593043 | 0.023997389 | 0.05792674 | ns | NaCl | KS.1 |
| FIM1_1757 | 1666.707236 | 1565.12264 | 1650.215053 | -0.059719212 | 0.209712555 | 0.725066895 | 0.817408662 | ns | NaCl | KS.1 |
| FIM1_1758 | 512.0929338 | 421.32233 | 332.026574 | -0.190492222 | 0.214281928 | 0.269225658 | 0.398220984 | ns | NaCl | KS.1 |
| FIM1_1759 | 366.344404 | 348.8876059 | 363.4795938 | -0.04468274 | 0.221141612 | 0.798914308 | 0.871176602 | ns | NaCl | KS.1 |
| FIM1_1760 | 3286.353209 | 3965.166008 | 3933.361456 | 0.18287068 | 0.213950795 | 0.288299069 | 0.418892485 | ns | NaCl | KS.1 |
| FIM1_1761 | 2319.312029 | 2189.940955 | 2837.073276 | -0.051175643 | 0.226955769 | 0.773839848 | 0.853357329 | ns | NaCl | KS.1 |
| FIM1_1762 | 6603.036396 | 11845.03178 | 5193.355965 | 0.610203279 | 0.273521407 | 0.003545845 | 0.012301659 | ns | NaCl | KS.1 |
| FIM1_1763 | 187.5742344 | 112.987862 | 106.1058246 | -0.483802631 | 0.288755393 | 0.021921545 | 0.053856636 | ns | NaCl | KS.1 |
| FIM1_1764 | 4637.526346 | 5579.562046 | 3825.694864 | 0.242344866 | 0.101143545 | 0.010963783 | 0.030729477 | ns | NaCl | KS.1 |
| FIM1_1765 | 1927.019423 | 1765.063058 | 2218.508472 | -0.110338251 | 0.114163333 | 0.299364172 | 0.431942215 | ns | NaCl | KS.1 |
| FIM1_1766 | 1466.852376 | 1353.982025 | 1836.890909 | -0.069276395 | 0.234444383 | 0.702593009 | 0.799981706 | ns | NaCl | KS.1 |
| FIM1_1767 | 1675.274193 | 1550.387134 | 1753.236508 | -0.093118875 | 0.128754176 | 0.429490075 | 0.566166699 | ns | NaCl | KS.1 |
| FIM1_1769 | 29.63867268 | 14.01985589 | 21.2872472 | -0.422074497 | 0.418932857 | 0.071689422 | 0.141585897 | ns | NaCl | KS.1 |
| FIM1_1770 | 3422.774375 | 4702.41391 | 2963.922448 | 0.423120876 | 0.105686469 | 1.89714960951 | 0.000149105 | ns | NaCl | KS.1 |
| FIM1_1771 | 1442.107249 | 2522.618674 | 1673.299133 | 0.741118416 | 0.14654502 | 3.90465626953 | 7.19271879028 | ns | NaCl | KS.1 |
| FIM1_1772 | 2229.009003 | 2305.505316 | 1718.984209 | 0.03852735 | 0.148687226 | 0.771461895 | 0.852325767 | ns | NaCl | KS.1 |
| FIM1_1774 | 924.5709646 | 791.9948699 | 925.4380031 | -0.190303127 | 0.128187927 | 0.104390043 | 0.191285621 | ns | NaCl | KS.1 |
| FIM1_1775 | 6374.978816 | 6848.768171 | 6632.013889 | 0.086637686 | 0.132515274 | 0.473033326 | 0.606667311 | ns | NaCl | KS.1 |
| FIM1_1776 | 2358.966333 | 3023.113145 | 2784.014127 | 0.29571268 | 0.15480013 | 0.030177822 | 0.069960236 | ns | NaCl | KS.1 |
| FIM1_1777 | 4088.623469 | 2691.752636 | 3183.634497 | -0.512431299 | 0.171750671 | 0.000599862 | 0.002807446 | ns | NaCl | KS.1 |
| FIM1_1778 | 2302.510832 | 2651.612802 | 2756.463574 | 0.16637771 | 0.145688674 | 0.20149871 | 0.321919101 | ns | NaCl | KS.1 |
| FIM1_1779 | 3161.590303 | 5499.175439 | 5361.145866 | 0.61460589 | 0.244622427 | 0.001647921 | 0.006501513 | ns | NaCl | KS.1 |
| FIM1_1780 | 916.1389054 | 1049.976118 | 982.3092643 | 0.169090224 | 0.122905638 | 0.135062944 | 0.234533384 | ns | NaCl | KS.1 |
| FIM1_1781 | 406.5059484 | 412.4669999 | 622.2993874 | 0.019831411 | 0.14431658 | 0.878320422 | 0.92830049 | ns | NaCl | KS.1 |
| FIM1_1782 | 878.083231 | 771.3150613 | 1110.647824 | -0.16842551 | 0.087936883 | 0.045141016 | 0.09717722 | ns | NaCl | KS.1 |
| FIM1_1783 | 267.462734 | 128.1853189 | 438.0035736 | -0.94386322 | 0.189341353 | 2.95440989030 | 5.69697255978 | ns | NaCl | KS.1 |
| FIM1_1784 | 1431.33606 | 1363.096195 | 2015.541673 | -0.058735371 | 0.128082131 | 0.616857743 | 0.731902522 | ns | NaCl | KS.1 |
| FIM1_1785 | 646.4829883 | 550.7606224 | 875.1746038 | -0.189626428 | 0.149392094 | 0.153415109 | 0.259870673 | ns | NaCl | KS.1 |
| FIM1_1786 | 60.65399173 | 48.54801991 | 53.71768876 | -0.219669468 | 0.235216475 | 0.229304755 | 0.354830396 | ns | NaCl | KS.1 |
| FIM1_1787 | 3270.49798 | 5435.291442 | 2637.023263 | 0.485731378 | 0.282451425 | 0.019924904 | 0.049439601 | ns | NaCl | KS.1 |
| FIM1_1788 | 530.9391497 | 695.0443201 | 541.1282185 | 0.328485888 | 0.147082065 | 0.01212026 | 0.033240516 | ns | NaCl | KS.1 |
| FIM1_1789 | 930.019402 | 864.9770755 | 1310.35748 | -0.066176197 | 0.221335142 | 0.70615371 | 0.802814147 | ns | NaCl | KS.1 |
| FIM1_1790 | 951.0776764 | 489.6221136 | 819.0189478 | -0.845775642 | 0.185678191 | 3.38788400285 | 4.65600080503 | ns | NaCl | KS.1 |
| FIM1_1791 | 303.1469457 | 340.1167135 | 360.2382304 | 0.125748059 | 0.175181864 | 0.402361614 | 0.538105815 | ns | NaCl | KS.1 |
| FIM1_1792 | 6594.857682 | 7455.855042 | 6424.741287 | 0.118035112 | 0.210375731 | 0.488033638 | 0.621758604 | ns | NaCl | KS.1 |

| | | | | | | | | | | |
|---|---|---|---|---|---|---|---|---|---|---|
| FIM1_1793 | 2133.927659 | 1597.334109 | 1819.898147 | -0.374636604 | 0.126230874 | 0.001165117 | 0.004875067 | ns | NaCl | KS.1 |
| FIM1_1794 | 410.0351803 | 308.2674152 | 681.1152267 | -0.211655378 | 0.294029827 | 0.299457048 | 0.431950947 | ns | NaCl | KS.1 |
| FIM1_1795 | 860.0809934 | 531.5386782 | 820.1649257 | -0.68639293 | 0.079679675 | 7.75284578345 | 1.75320035330 | ns | NaCl | KS.1 |
| FIM1_1796 | 129.6034875 | 69.51201129 | 82.68238282 | -0.69450926 | 0.266515779 | 0.000898109 | 0.003926269 | ns | NaCl | KS.1 |
| FIM1_1797 | 1956.496954 | 1353.363853 | 1607.787515 | -0.382665131 | 0.227545961 | 0.034390825 | 0.078035817 | ns | NaCl | KS.1 |
| FIM1_1798 | 1407.440289 | 1240.558079 | 1934.363292 | -0.111012407 | 0.234525798 | 0.541100672 | 0.667321726 | ns | NaCl | KS.1 |
| FIM1_1799 | 2074.132624 | 2111.465419 | 1747.339532 | 0.021817169 | 0.117840373 | 0.841909415 | 0.903176394 | ns | NaCl | KS.1 |
| FIM1_1800 | 701.1873191 | 317.7934726 | 846.4389477 | -0.954163031 | 0.24576831 | 4.08490183029 | 4.04828418440 | ns | NaCl | KS.1 |
| FIM1_1801 | 2036.501784 | 4627.663558 | 2016.171127 | 0.854022377 | 0.324205227 | 0.000405471 | 0.002027353 | ns | NaCl | KS.1 |
| FIM1_1802 | 2813.601548 | 2476.996665 | 3150.560927 | -0.113948268 | 0.23045842 | 0.526391535 | 0.655026985 | ns | NaCl | KS.1 |
| FIM1_1803 | 2083.423345 | 1787.844789 | 4401.067168 | -0.109373523 | 0.286148234 | 0.583673869 | 0.704971473 | ns | NaCl | KS.1 |
| FIM1_1804 | 543.6878077 | 482.1840502 | 737.2140736 | -0.111488367 | 0.217743164 | 0.521444995 | 0.652118833 | ns | NaCl | KS.1 |
| FIM1_1805 | 41.90040083 | 21.33273569 | 87.50925761 | -0.57888677 | 0.362138452 | 0.015431613 | 0.040048135 | ns | NaCl | KS.1 |
| FIM1_1806 | 1222.554588 | 1080.640727 | 1470.141121 | -0.107685229 | 0.236351514 | 0.554969675 | 0.678854977 | ns | NaCl | KS.1 |
| FIM1_1807 | 94.994141 | 122.0902276 | 206.0439287 | 0.23994049 | 0.227312519 | 0.180561638 | 0.295880814 | ns | NaCl | KS.1 |
| FIM1_1808 | 1953.518565 | 1528.218266 | 2362.410885 | -0.228541785 | 0.23514602 | 0.211168548 | 0.33361814 | ns | NaCl | KS.1 |
| FIM1_1809 | 2755.990854 | 2850.382564 | 3503.93053 | 0.027659583 | 0.248326844 | 0.882362058 | 0.930540801 | ns | NaCl | KS.1 |
| FIM1_1810 | 1374.063223 | 1299.23195 | 1536.574471 | -0.065231953 | 0.138337725 | 0.602143923 | 0.72064621 | ns | NaCl | KS.1 |
| FIM1_1811 | 1688.225536 | 1522.56564 | 1497.327843 | -0.132260469 | 0.110865203 | 0.201689193 | 0.322022716 | ns | NaCl | KS.1 |
| FIM1_1812 | 513.6684522 | 327.8874509 | 669.3733824 | -0.517093651 | 0.207328722 | 0.00263932 | 0.009658807 | ns | NaCl | KS.1 |
| FIM1_1813 | 565.0784321 | 459.7248847 | 514.8656018 | -0.250574201 | 0.147423932 | 0.05629983 | 0.116656249 | ns | NaCl | KS.1 |
| FIM1_1814 | 944.3138459 | 714.2577819 | 879.8759774 | -0.36353033 | 0.11566471 | 0.000681537 | 0.003125567 | ns | NaCl | KS.1 |
| FIM1_1815 | 740.7462687 | 420.2443173 | 661.3282019 | -0.76064589 | 0.136783064 | 2.37614284968 | 6.12503143894 | ns | NaCl | KS.1 |
| FIM1_1816 | 2123.357482 | 2261.092088 | 2257.114841 | 0.076809413 | 0.125609179 | 0.505894278 | 0.637968307 | ns | NaCl | KS.1 |
| FIM1_1817 | 1561.392353 | 1626.178267 | 1715.826269 | 0.049670886 | 0.132908474 | 0.681660937 | 0.784546705 | ns | NaCl | KS.1 |
| FIM1_1818 | 30033.16287 | 40411.32655 | 24018.39903 | 0.407728642 | 0.08112743 | 1.60370855447 | 2.43989298426 | ns | NaCl | KS.1 |
| FIM1_1819 | 2978.625534 | 2606.117657 | 2817.807409 | -0.16747028 | 0.118839625 | 0.127690833 | 0.224157337 | ns | NaCl | KS.1 |
| FIM1_1820 | 695.4021142 | 615.1345335 | 505.4351108 | -0.1358185 | 0.16844191 | 0.351764928 | 0.486779562 | ns | NaCl | KS.1 |
| FIM1_1821 | 2172.201646 | 2499.785461 | 1756.485633 | 0.171258079 | 0.13105888 | 0.151601054 | 0.25740662 | ns | NaCl | KS.1 |
| FIM1_1822 | 473.4944992 | 332.9907812 | 470.8690079 | -0.443151361 | 0.144361302 | 0.000609674 | 0.002845337 | ns | NaCl | KS.1 |
| FIM1_1823 | 281.3925053 | 298.3353282 | 249.6394333 | 0.041425624 | 0.273007398 | 0.831975681 | 0.896292554 | ns | NaCl | KS.1 |
| FIM1_1824 | 1063.360805 | 989.7412902 | 1074.180864 | -0.079019579 | 0.167876959 | 0.587279879 | 0.708638709 | ns | NaCl | KS.1 |
| FIM1_1825 | 5470.570907 | 5437.81022 | 4575.038093 | -0.007212642 | 0.152851772 | 0.957571546 | 0.978418246 | ns | NaCl | KS.1 |
| FIM1_1826 | 5090.803542 | 4865.04246 | 6200.367689 | -0.044918474 | 0.19570427 | 0.782009919 | 0.858845239 | ns | NaCl | KS.1 |
| FIM1_1827 | 6491.126623 | 5594.598909 | 5286.396663 | -0.192253266 | 0.106700751 | 0.054597408 | 0.113679606 | ns | NaCl | KS.1 |
| FIM1_1828 | 2811.760912 | 2982.833034 | 3775.938771 | 0.072585159 | 0.125910348 | 0.530427923 | 0.658567236 | ns | NaCl | KS.1 |
| FIM1_1829 | 251.4629143 | 225.8103478 | 281.7080514 | -0.117572681 | 0.158306802 | 0.398248225 | 0.534327108 | ns | NaCl | KS.1 |
| FIM1_1830 | 1862.934635 | 1829.388117 | 1821.326502 | -0.02448615 | 0.105358058 | 0.804985647 | 0.875558285 | ns | NaCl | KS.1 |
| FIM1_1831 | 631.771567 | 763.2945841 | 557.696713 | 0.22950429 | 0.137485892 | 0.064397624 | 0.129970864 | ns | NaCl | KS.1 |
| FIM1_1832 | 324.7987596 | 285.8587723 | 410.2668753 | -0.141825525 | 0.144858352 | 0.274277731 | 0.403946628 | ns | NaCl | KS.1 |
| FIM1_1833 | 973.3830661 | 700.1486097 | 903.3919552 | -0.453542777 | 0.088406568 | 7.39785488522 | 1.22273515129 | ns | NaCl | KS.1 |
| FIM1_1834 | 645.7087662 | 428.6602416 | 892.8793349 | -0.49422672 | 0.17863912 | 0.001298727 | 0.00531346 | ns | NaCl | KS.1 |
| FIM1_1835 | 582.1454063 | 439.1442493 | 680.737761 | -0.362817626 | 0.126691203 | 0.001716272 | 0.006755104 | ns | NaCl | KS.1 |
| FIM1_1836 | 2304.850637 | 2552.757583 | 2170.345231 | 0.112049482 | 0.167882312 | 0.441356901 | 0.576766636 | ns | NaCl | KS.1 |
| FIM1_1837 | 6317.615867 | 9447.45459 | 5368.555356 | 0.444952105 | 0.213507109 | 0.010662596 | 0.030088721 | ns | NaCl | KS.1 |

| | | | | | | | | | | |
|---|---|---|---|---|---|---|---|---|---|---|
| FIM1_1838 | 134.7166088 | 219.4405397 | 170.4799956 | 0.524470672 | 0.240528816 | 0.005905973 | 0.01876259 | ns | NaCl | KS.1 |
| FIM1_1839 | 4390.093945 | 3034.487467 | 3181.795781 | -0.477257852 | 0.135401094 | 0.000102015 | 0.000634405 | ns | NaCl | KS.1 |
| FIM1_1840 | 7354.011562 | 9606.346576 | 7274.870358 | 0.306490699 | 0.172964999 | 0.039189361 | 0.086904654 | ns | NaCl | KS.1 |
| FIM1_1841 | 1326.909358 | 741.9593471 | 1025.017145 | -0.807981353 | 0.097096622 | 7.4208225984 | 1.1909223363 | ns | NaCl | KS.1 |
| FIM1_1842 | 763.0724429 | 636.2914535 | 1034.982679 | -0.168998978 | 0.226244802 | 0.342952659 | 0.477522944 | ns | NaCl | KS.1 |
| FIM1_1843 | 7431.095243 | 11537.66401 | 8375.507969 | 0.599358079 | 0.10637985 | 2.4543179406 | 6.2939338943 | ns | NaCl | KS.1 |
| FIM1_1844 | 58.77984439 | 86.7818668 | 62.51638684 | 0.359315363 | 0.256696144 | 0.064485584 | 0.130095613 | ns | NaCl | KS.1 |
| FIM1_1845 | 670.9682308 | 607.5403924 | 543.7782126 | -0.129738732 | 0.119839835 | 0.241748872 | 0.368587386 | ns | NaCl | KS.1 |
| FIM1_1846 | 4913.156918 | 5239.069905 | 3597.826933 | 0.079451014 | 0.12184759 | 0.480092313 | 0.61415769 | ns | NaCl | KS.1 |
| FIM1_1847 | 3699.889044 | 3427.586872 | 4023.427754 | -0.084263993 | 0.162346042 | 0.552767184 | 0.676677348 | ns | NaCl | KS.1 |
| FIM1_1848 | 8314.267777 | 8428.257311 | 7128.816569 | 0.01466698 | 0.171795627 | 0.921131292 | 0.955028468 | ns | NaCl | KS.1 |
| FIM1_1849 | 2851.983662 | 2838.795011 | 3645.990782 | -0.00449625 | 0.184850748 | 0.977024687 | 0.988963812 | ns | NaCl | KS.1 |
| FIM1_1850 | 742.7012159 | 617.7708536 | 680.5337264 | -0.201318971 | 0.174506972 | 0.178484161 | 0.293443061 | ns | NaCl | KS.1 |
| FIM1_1851 | 5164.783075 | 4265.908302 | 3843.198155 | -0.254043844 | 0.092893624 | 0.003980437 | 0.013480376 | ns | NaCl | KS.1 |
| FIM1_1853 | 51235.22272 | 79520.17016 | 51825.76471 | 0.569955051 | 0.14460122 | 1.3031732015 | 0.000107696 | ns | NaCl | KS.1 |
| FIM1_1854 | 6076.563669 | 8345.846497 | 6421.667845 | 0.345187961 | 0.201565034 | 0.037902463 | 0.084748204 | ns | NaCl | KS.1 |
| FIM1_1855 | 718.617153 | 582.7136954 | 833.8984346 | -0.203388958 | 0.214749483 | 0.238933456 | 0.365639478 | ns | NaCl | KS.1 |
| FIM1_1856 | 1878.509112 | 1210.995952 | 1627.702602 | -0.496218499 | 0.213076749 | 0.00455437 | 0.015025839 | ns | NaCl | KS.1 |
| FIM1_1857 | 6658.618117 | 7288.801566 | 6646.68315 | 0.108444339 | 0.134707604 | 0.375428539 | 0.510037406 | ns | NaCl | KS.1 |
| FIM1_1858 | 1846.872792 | 1179.533756 | 1277.778027 | -0.591699193 | 0.132905946 | 1.2453795092 | 1.4210465729 | ns | NaCl | KS.1 |
| FIM1_1859 | 2578.916968 | 2706.530339 | 1973.780227 | 0.055224581 | 0.149391428 | 0.678256502 | 0.781636808 | ns | NaCl | KS.1 |
| FIM1_1860 | 912.0355512 | 916.366559 | 1463.585236 | 0.004628926 | 0.192981276 | 0.97703661 | 0.988963812 | ns | NaCl | KS.1 |
| FIM1_1861 | 727.1176994 | 733.7594312 | 621.8867677 | 0.005804427 | 0.132135479 | 0.961591037 | 0.981455879 | ns | NaCl | KS.1 |
| FIM1_1862 | 876.7959559 | 922.9073092 | 889.5884481 | 0.060475561 | 0.129421913 | 0.609585604 | 0.726044621 | ns | NaCl | KS.1 |
| FIM1_1863 | 606.6501685 | 684.6098177 | 657.76852 | 0.155576427 | 0.095251613 | 0.085410081 | 0.163806921 | ns | NaCl | KS.1 |
| FIM1_1864 | 1419.412243 | 1008.894802 | 1432.97111 | -0.447745182 | 0.121817111 | 6.4941783155 | 0.000434255 | ns | NaCl | KS.1 |
| FIM1_1865 | 754.7450045 | 1024.826831 | 897.4371506 | 0.34602474 | 0.185613586 | 0.027282237 | 0.064448778 | ns | NaCl | KS.1 |
| FIM1_1866 | 210.3772313 | 210.1637522 | 472.7962783 | -0.000242956 | 0.230879562 | 0.998921765 | 0.999786056 | ns | NaCl | KS.1 |
| FIM1_1867 | 75.0753514 | 151.2658108 | 199.9969661 | 0.874869179 | 0.225283907 | 5.5879744240 | 5.1769409236 | ns | NaCl | KS.1 |
| FIM1_1868 | 1355.627348 | 2027.015798 | 1780.688788 | 0.521516056 | 0.139820989 | 3.7784716966 | 0.000272039 | ns | NaCl | KS.1 |
| FIM1_1869 | 1967.622255 | 2873.503649 | 2045.223674 | 0.495986842 | 0.127714586 | 2.2526734366 | 0.000172416 | ns | NaCl | KS.1 |
| FIM1_1870 | 1013.464485 | 993.7976452 | 1245.736782 | -0.023977036 | 0.128413304 | 0.838554615 | 0.901233357 | ns | NaCl | KS.1 |
| FIM1_1871 | 1159.875077 | 1145.278228 | 1710.552191 | -0.013208653 | 0.161226865 | 0.925506551 | 0.957055725 | ns | NaCl | KS.1 |
| FIM1_1872 | 682.4550514 | 692.4351565 | 908.4532218 | 0.019354847 | 0.146247746 | 0.882470654 | 0.930540801 | ns | NaCl | KS.1 |
| FIM1_1873 | 688.6419102 | 869.328869 | 969.815206 | 0.294353004 | 0.12570199 | 0.010480306 | 0.029675312 | ns | NaCl | KS.1 |
| FIM1_1874 | 853.3873921 | 478.0703947 | 1626.170728 | -0.736589511 | 0.176862667 | 2.8086982815 | 2.9110987396 | ns | NaCl | KS.1 |
| FIM1_1875 | 27.5518386 | 10.57567704 | 58.7017236 | -0.820907229 | 0.406929279 | 0.002009186 | 0.007665413 | ns | NaCl | KS.1 |
| FIM1_1876 | 863.358675 | 742.2077622 | 644.4371434 | -0.112725039 | 0.277393575 | 0.567626391 | 0.689607153 | ns | NaCl | KS.1 |
| FIM1_1877 | 1393.405999 | 2626.458808 | 1186.719207 | 0.563908635 | 0.329392714 | 0.013763883 | 0.036696313 | ns | NaCl | KS.1 |
| FIM1_1878 | 1854.588049 | 2049.615179 | 2956.918088 | 0.074817823 | 0.272950397 | 0.701906856 | 0.799981706 | ns | NaCl | KS.1 |
| FIM1_1879 | 457.2013689 | 385.5630164 | 476.7881425 | -0.212931313 | 0.118704941 | 0.052275688 | 0.109688547 | ns | NaCl | KS.1 |
| FIM1_1880 | 3509.736317 | 3720.427928 | 3492.685695 | 0.072243344 | 0.122408595 | 0.522483371 | 0.65235834 | ns | NaCl | KS.1 |
| FIM1_1881 | 590.5102734 | 666.2171798 | 750.7455602 | 0.150091597 | 0.118082623 | 0.17001002 | 0.281370542 | ns | NaCl | KS.1 |
| FIM1_1882 | 3765.460613 | 2965.954281 | 3247.16289 | -0.261221301 | 0.185396291 | 0.094822957 | 0.177807728 | ns | NaCl | KS.1 |
| FIM1_1883 | 87.18070706 | 100.5402187 | 133.49796 | 0.116169371 | 0.247900715 | 0.53488895 | 0.662617661 | ns | NaCl | KS.1 |

| | | | | | | | | | | |
|---|---|---|---|---|---|---|---|---|---|---|
| FIM1_1884 | 4680.284645 | 3976.535682 | 4422.520693 | -0.180744812 | 0.169480046 | 0.217127023 | 0.340759286 | ns | NaCl | KS.1 |
| FIM1_1885 | 1339.93566 | 1416.182536 | 1568.879703 | 0.072701533 | 0.1014558 | 0.448197959 | 0.583255256 | ns | NaCl | KS.1 |
| FIM1_1886 | 787.3699626 | 587.4048718 | 895.1978618 | -0.377680026 | 0.127759126 | 0.001193948 | 0.004978952 | ns | NaCl | KS.1 |
| FIM1_1887 | 5604.371042 | 7568.784888 | 5765.02779 | 0.353716943 | 0.168561092 | 0.015292828 | 0.039754866 | ns | NaCl | KS.1 |
| FIM1_1888 | 2436.179732 | 2656.489665 | 2288.547933 | 0.118391291 | 0.063528964 | 0.056284165 | 0.116656249 | ns | NaCl | KS.1 |
| FIM1_1889 | 1316.839634 | 1806.200686 | 1442.223128 | 0.41304987 | 0.121501384 | 0.000218359 | 0.001202494 | ns | NaCl | KS.1 |
| FIM1_1890 | 1067.297276 | 2919.05115 | 1155.395381 | 0.909719036 | 0.414274818 | 0.000880374 | 0.003858908 | ns | NaCl | KS.1 |
| FIM1_1891 | 421.6389305 | 524.6202811 | 1454.40925 | 0.185699847 | 0.255497017 | 0.330603051 | 0.464094294 | ns | NaCl | KS.1 |
| FIM1_1892 | 1070.836893 | 835.5306654 | 894.0127034 | -0.246898622 | 0.216683079 | 0.155724127 | 0.262708556 | ns | NaCl | KS.1 |
| FIM1_1893 | 1561.276119 | 2272.588054 | 1753.076194 | 0.492531203 | 0.12595306 | 2.04493070132 | 0.000158713 | ns | NaCl | KS.1 |
| FIM1_1894 | 995.2037167 | 979.2896943 | 1675.682039 | -0.018977974 | 0.143589992 | 0.883048945 | 0.930953274 | ns | NaCl | KS.1 |
| FIM1_1895 | 1145.08553 | 1275.515081 | 1091.70638 | 0.136717497 | 0.114444482 | 0.199204153 | 0.318765089 | ns | NaCl | KS.1 |
| FIM1_1896 | 1034.075836 | 1150.703155 | 1174.592518 | 0.142628256 | 0.098726374 | 0.126904146 | 0.223248985 | ns | NaCl | KS.1 |
| FIM1_1897 | 1997.939222 | 2378.64791 | 1873.564261 | 0.228314291 | 0.100895977 | 0.016349529 | 0.042013897 | ns | NaCl | KS.1 |
| FIM1_1898 | 2523.706836 | 1862.555652 | 1951.388759 | -0.411074751 | 0.096961362 | 7.16843736802 | 6.37978102073 | ns | NaCl | KS.1 |
| FIM1_1899 | 7253.263057 | 9747.028858 | 8322.433107 | 0.371989421 | 0.137243728 | 0.002649311 | 0.009670084 | ns | NaCl | KS.1 |
| FIM1_1900 | 383.7205831 | 379.4402438 | 403.9192299 | -0.011380849 | 0.126021696 | 0.921736306 | 0.955142288 | ns | NaCl | KS.1 |
| FIM1_1901 | 3732.237712 | 4068.767914 | 4115.123878 | 0.09844856 | 0.154375949 | 0.470862034 | 0.604682143 | ns | NaCl | KS.1 |
| FIM1_1902 | 1023.642044 | 1099.579596 | 880.7159253 | 0.08335888 | 0.144689376 | 0.520519054 | 0.651467244 | ns | NaCl | KS.1 |
| FIM1_1903 | 951.5663941 | 946.2617019 | 936.9536444 | -0.001881027 | 0.095312143 | 0.983445155 | 0.991416342 | ns | NaCl | KS.1 |
| FIM1_1904 | 3381.31161 | 8198.118025 | 5400.172419 | 1.212819879 | 0.141375433 | 1.43311640568 | 3.75250216751 | Km>KS.1 | NaCl | KS.1 |
| FIM1_1905 | 1198.740821 | 1116.192214 | 1071.12881 | -0.088954281 | 0.126581673 | 0.443937734 | 0.579074522 | ns | NaCl | KS.1 |
| FIM1_1907 | 1228.989543 | 1530.796579 | 1143.469546 | 0.24867206 | 0.16936923 | 0.089130687 | 0.169310871 | ns | NaCl | KS.1 |
| FIM1_1908 | 4380.671082 | 6275.751621 | 4344.102142 | 0.44963504 | 0.151341373 | 0.00082289 | 0.003651989 | ns | NaCl | KS.1 |
| FIM1_1909 | 3206.782471 | 3153.131311 | 4253.410281 | -0.022267371 | 0.097633293 | 0.810052826 | 0.880240073 | ns | NaCl | KS.1 |
| FIM1_1910 | 900.3737542 | 779.8713249 | 1422.179399 | -0.167410638 | 0.147272843 | 0.202602205 | 0.322956094 | ns | NaCl | KS.1 |
| FIM1_1911 | 8039.299488 | 10351.7138 | 7619.23478 | 0.346385353 | 0.078321162 | 4.23029259695 | 4.16746646927 | ns | NaCl | KS.1 |
| FIM1_1912 | 2994.634782 | 3301.184828 | 3167.10355 | 0.122663531 | 0.115345577 | 0.252742336 | 0.381143717 | ns | NaCl | KS.1 |
| FIM1_1913 | 2936.534557 | 5326.810029 | 2600.344044 | 0.657126444 | 0.255428197 | 0.00115895 | 0.004853347 | ns | NaCl | KS.1 |
| FIM1_1914 | 589.7084182 | 624.7692249 | 540.0841743 | 0.064909909 | 0.156168343 | 0.637642525 | 0.748471149 | ns | NaCl | KS.1 |
| FIM1_1915 | 1574.823565 | 2309.230069 | 2001.753845 | 0.386275408 | 0.239248177 | 0.038678103 | 0.08605705 | ns | NaCl | KS.1 |
| FIM1_1916 | 999.8441148 | 971.8748488 | 1040.49571 | -0.02967455 | 0.184159881 | 0.848859186 | 0.907797604 | ns | NaCl | KS.1 |
| FIM1_1917 | 1077.97922 | 1575.72754 | 992.6185635 | 0.388834314 | 0.234087512 | 0.034892385 | 0.078904372 | ns | NaCl | KS.1 |
| FIM1_1918 | 941.9082695 | 913.8603205 | 883.3691243 | -0.034302186 | 0.150983866 | 0.798343373 | 0.870903663 | ns | NaCl | KS.1 |
| FIM1_1919 | 1565.238052 | 1559.287095 | 1482.241881 | -0.005227508 | 0.115387354 | 0.961186224 | 0.981305451 | ns | NaCl | KS.1 |
| FIM1_1920 | 2882.129725 | 6036.994968 | 2593.445431 | 0.739466846 | 0.321397257 | 0.001639453 | 0.006488685 | ns | NaCl | KS.1 |
| FIM1_1921 | 3120.585836 | 2641.105681 | 2806.918951 | -0.195646814 | 0.150402671 | 0.142863666 | 0.244831809 | ns | NaCl | KS.1 |
| FIM1_1922 | 2175.55914 | 2347.131731 | 2248.488215 | 0.104212907 | 0.064789496 | 0.099189675 | 0.183582081 | ns | NaCl | KS.1 |
| FIM1_1923 | 2567.212953 | 4328.393144 | 2652.797944 | 0.726326554 | 0.093166309 | 6.45064431363 | 7.46324545588 | ns | NaCl | KS.1 |
| FIM1_1924 | 1331.0409 | 1450.957182 | 1067.849095 | 0.098051975 | 0.152960794 | 0.469461472 | 0.603506673 | ns | NaCl | KS.1 |
| FIM1_1925 | 1928.030818 | 1814.961974 | 2161.135795 | -0.079792061 | 0.092168448 | 0.363949982 | 0.499352223 | ns | NaCl | KS.1 |
| FIM1_1926 | 773.2849726 | 547.9910659 | 614.4443236 | -0.397656361 | 0.185146334 | 0.011206757 | 0.031217031 | ns | NaCl | KS.1 |
| FIM1_1927 | 3678.80778 | 8766.474042 | 3871.751739 | 0.955078768 | 0.308909107 | 6.68048133738 | 0.000446113 | ns | NaCl | KS.1 |
| FIM1_1928 | 1477.538118 | 1421.035064 | 1626.853045 | -0.050387375 | 0.113344801 | 0.633703652 | 0.745151395 | ns | NaCl | KS.1 |
| FIM1_1929 | 398.1103909 | 441.0576731 | 577.4460461 | 0.094700441 | 0.220571547 | 0.588877713 | 0.709890983 | ns | NaCl | KS.1 |

| | | | | | | | | | | |
|---|---|---|---|---|---|---|---|---|---|---|
| FIM1_1930 | 803.7837136 | 730.4982998 | 719.1019424 | -0.117042438 | 0.14243179 | 0.360631309 | 0.495924545 | ns | NaCl | KS.1 |
| FIM1_1931 | 13923.24285 | 11963.89364 | 12554.5319 | -0.206630547 | 0.0747031 | 0.004217294 | 0.014166805 | ns | NaCl | KS.1 |
| FIM1_1932 | 724.2553188 | 416.215935 | 615.3969019 | -0.746770508 | 0.133854617 | 2.2244593400 | 5.8245711667 | ns | NaCl | KS.1 |
| FIM1_1933 | 3756.168176 | 6111.788464 | 5025.61618 | 0.652193243 | 0.127827162 | 3.9888522175 | 7.2161962843 | ns | NaCl | KS.1 |
| FIM1_1934 | 373.7062566 | 208.7527437 | 592.1345006 | -0.726054889 | 0.198837863 | 2.3856514375 | 0.0001812 | ns | NaCl | KS.1 |
| FIM1_1935 | 901.828733 | 765.1512515 | 2584.393175 | -0.190101724 | 0.156004714 | 0.166612831 | 0.277502121 | ns | NaCl | KS.1 |
| FIM1_1936 | 5628.719914 | 7503.268924 | 5334.781665 | 0.403322126 | 0.061676707 | 1.9658904581 | 8.3592350678 | ns | NaCl | KS.1 |
| FIM1_1937 | 2682.794437 | 3646.739929 | 2724.981043 | 0.409268452 | 0.105126483 | 3.2208331458 | 0.000236686 | ns | NaCl | KS.1 |
| FIM1_1938 | 1593.106852 | 1743.206316 | 1245.841524 | 0.101412629 | 0.156199237 | 0.461690522 | 0.596290329 | ns | NaCl | KS.1 |
| FIM1_1939 | 260.1017428 | 396.2375946 | 307.9868761 | 0.449214056 | 0.231210611 | 0.014598778 | 0.038451619 | ns | NaCl | KS.1 |
| FIM1_1940 | 3315.75637 | 2827.67969 | 3000.403003 | -0.160817713 | 0.199951223 | 0.328919321 | 0.462513742 | ns | NaCl | KS.1 |
| FIM1_1941 | 1803.880871 | 1183.140409 | 1548.488475 | -0.43138929 | 0.243377459 | 0.022975242 | 0.055801637 | ns | NaCl | KS.1 |
| FIM1_1942 | 824.9366164 | 621.4336801 | 919.5417893 | -0.330380708 | 0.166742346 | 0.022298467 | 0.054620815 | ns | NaCl | KS.1 |
| FIM1_1943 | 1151.719754 | 1222.790525 | 1103.740139 | 0.068971223 | 0.146137874 | 0.597966345 | 0.717357745 | ns | NaCl | KS.1 |
| FIM1_1944 | 674.5465352 | 440.4431388 | 568.8789386 | -0.563751392 | 0.127142485 | 1.5034618411 | 1.6808365527 | ns | NaCl | KS.1 |
| FIM1_1945 | 806.1135366 | 699.2147319 | 776.3111489 | -0.164446688 | 0.150588721 | 0.218852928 | 0.343035071 | ns | NaCl | KS.1 |
| FIM1_1946 | 2404.653928 | 1549.341089 | 2199.170287 | -0.55790763 | 0.159552073 | 8.0642997796 | 0.000520362 | ns | NaCl | KS.1 |
| FIM1_1947 | 1143.184339 | 1051.416454 | 995.8216166 | -0.094731645 | 0.158382823 | 0.496391258 | 0.628543271 | ns | NaCl | KS.1 |
| FIM1_1948 | 2479.132402 | 4939.880841 | 2939.991438 | 0.952857795 | 0.113633998 | 2.7851249108 | 5.1318505301 | ns | NaCl | KS.1 |
| FIM1_1949 | 479.7331742 | 422.4291263 | 448.1257848 | -0.1417327 | 0.162068547 | 0.317056469 | 0.449773577 | ns | NaCl | KS.1 |
| FIM1_1950 | 1129.696328 | 803.4150089 | 1066.766058 | -0.443407325 | 0.1272085 | 0.000137811 | 0.000821091 | ns | NaCl | KS.1 |
| FIM1_1951 | 1094.933379 | 382.6188948 | 1505.63966 | -1.433273763 | 0.157600521 | 3.5623168503 | 1.3632712561 | Km<KS.1 | NaCl | KS.1 |
| FIM1_1952 | 2917.067167 | 3410.639199 | 2539.892516 | 0.205246475 | 0.100083924 | 0.029789914 | 0.069351813 | ns | NaCl | KS.1 |
| FIM1_1953 | 2329.993495 | 2350.654782 | 3001.487266 | 0.008876943 | 0.192194711 | 0.955857273 | 0.977067995 | ns | NaCl | KS.1 |
| FIM1_1954 | 2514.486277 | 5094.517722 | 2132.792289 | 0.529831525 | 0.37793714 | 0.026791021 | 0.063590329 | ns | NaCl | KS.1 |
| FIM1_1955 | 503.5231861 | 502.0967317 | 615.9159347 | -0.006384219 | 0.115637093 | 0.952695539 | 0.975640244 | ns | NaCl | KS.1 |
| FIM1_1956 | 813.5838268 | 878.0237867 | 737.4129827 | 0.07844367 | 0.183080929 | 0.612898018 | 0.729108563 | ns | NaCl | KS.1 |
| FIM1_1957 | 1229.450154 | 1443.262469 | 1281.058434 | 0.19330823 | 0.135522388 | 0.115259306 | 0.207226626 | ns | NaCl | KS.1 |
| FIM1_1958 | 695.1728381 | 445.2752322 | 1017.167306 | -0.552340376 | 0.170665539 | 0.000213788 | 0.001183085 | ns | NaCl | KS.1 |
| FIM1_1959 | 946.8318604 | 961.249103 | 825.9505333 | 0.017868958 | 0.117252164 | 0.869682434 | 0.921941213 | ns | NaCl | KS.1 |
| FIM1_1960 | 3135.415423 | 3226.713701 | 2838.54034 | 0.03826648 | 0.076407017 | 0.604650027 | 0.722935324 | ns | NaCl | KS.1 |
| FIM1_1961 | 1259.028805 | 714.4619519 | 3122.551629 | -0.696802667 | 0.201558623 | 5.5185737550 | 0.000378688 | ns | NaCl | KS.1 |
| FIM1_1962 | 46456.19885 | 56692.7575 | 36346.13897 | 0.253820963 | 0.117390462 | 0.019377031 | 0.048200365 | ns | NaCl | KS.1 |
| FIM1_1963 | 624.2498221 | 520.5465462 | 534.5366959 | -0.213086651 | 0.158102278 | 0.124790385 | 0.22007521 | ns | NaCl | KS.1 |
| FIM1_1964 | 12056.60823 | 18535.79387 | 12314.88059 | 0.46263394 | 0.229356915 | 0.011575469 | 0.032011093 | ns | NaCl | KS.1 |
| FIM1_1965 | 1136.797932 | 1145.243587 | 1120.310801 | 0.009497778 | 0.131923073 | 0.937107969 | 0.963644512 | ns | NaCl | KS.1 |
| FIM1_1966 | 1716.228618 | 1330.037299 | 1545.494868 | -0.328516709 | 0.119460825 | 0.002837051 | 0.010215995 | ns | NaCl | KS.1 |
| FIM1_1967 | 1732.860706 | 1501.772753 | 2172.736869 | -0.177929258 | 0.122778327 | 0.115423524 | 0.207378849 | ns | NaCl | KS.1 |
| FIM1_1968 | 2514.556838 | 3909.609483 | 2275.329364 | 0.490145503 | 0.220462522 | 0.006167062 | 0.019467724 | ns | NaCl | KS.1 |
| FIM1_1970 | 851.3154304 | 557.3073537 | 570.7668803 | -0.521707513 | 0.175364507 | 0.000593169 | 0.002783664 | ns | NaCl | KS.1 |
| FIM1_1971 | 427.2037345 | 357.6181625 | 411.3584587 | -0.201410084 | 0.160510009 | 0.151650011 | 0.25740662 | ns | NaCl | KS.1 |
| FIM1_1972 | 684.1378805 | 867.1403417 | 3532.771367 | 0.20324251 | 0.256121368 | 0.288181707 | 0.418892485 | ns | NaCl | KS.1 |
| FIM1_1973 | 6025.465039 | 11616.21725 | 7261.775774 | 0.740771476 | 0.257734501 | 0.000334047 | 0.001718599 | ns | NaCl | KS.1 |
| FIM1_1975 | 1026.070123 | 1106.584383 | 1218.097813 | 0.097900876 | 0.106134904 | 0.326365517 | 0.459572162 | ns | NaCl | KS.1 |
| FIM1_1976 | 3114.348552 | 2960.956391 | 2935.738703 | -0.060393849 | 0.135103426 | 0.622654191 | 0.737197667 | ns | NaCl | KS.1 |

| | | | | | | | | | | |
|---|---|---|---|---|---|---|---|---|---|---|
| FIM1_1977 | 834.3614719 | 868.6381073 | 1034.614862 | 0.054954328 | 0.097117142 | 0.551077988 | 0.675808574 | ns | NaCl | KS.1 |
| FIM1_1978 | 3104.190691 | 3552.00355 | 2647.71356 | 0.130530336 | 0.209618663 | 0.44218516 | 0.577161997 | ns | NaCl | KS.1 |
| FIM1_1979 | 827.501693 | 548.2468608 | 814.5126033 | -0.572287138 | 0.095647129 | 3.32377167199 | 1.08787921501 | ns | NaCl | KS.1 |
| FIM1_1980 | 10921.44599 | 7942.021621 | 8603.960782 | -0.368914717 | 0.178864435 | 0.015641948 | 0.040406474 | ns | NaCl | KS.1 |
| FIM1_1981 | 12580.79596 | 14876.89162 | 10239.13225 | 0.179137385 | 0.182919154 | 0.24737933 | 0.374988473 | ns | NaCl | KS.1 |
| FIM1_1982 | 17.98394144 | 15.6559134 | 64.93626001 | -0.077955788 | 0.3425399 | 0.706774427 | 0.802969348 | ns | NaCl | KS.1 |
| FIM1_1983 | 5281.559894 | 6471.106095 | 4897.795103 | 0.25996558 | 0.115940561 | 0.015477975 | 0.04012659 | ns | NaCl | KS.1 |
| FIM1_1984 | 571.5027245 | 629.3498193 | 569.3680288 | 0.120473688 | 0.114038902 | 0.256620839 | 0.385008647 | ns | NaCl | KS.1 |
| FIM1_1985 | 1925.587293 | 3131.511743 | 2826.378344 | 0.655094464 | 0.122280151 | 9.94334629254 | 2.13224775023 | ns | NaCl | KS.1 |
| FIM1_1986 | 1821.341924 | 2237.63172 | 2153.313448 | 0.275646605 | 0.087412185 | 0.000952377 | 0.004120064 | ns | NaCl | KS.1 |
| FIM1_1987 | 834.9221566 | 979.0728556 | 767.2735327 | 0.17018936 | 0.182576434 | 0.271163963 | 0.400276242 | ns | NaCl | KS.1 |
| FIM1_1988 | 2456.119175 | 2542.251193 | 2531.306992 | 0.038236566 | 0.159012184 | 0.784419203 | 0.860905699 | ns | NaCl | KS.1 |
| FIM1_1989 | 965.9263883 | 1963.937998 | 1456.817408 | 0.894346144 | 0.203370616 | 5.87609696328 | 7.49482504281 | ns | NaCl | KS.1 |
| FIM1_1990 | 2006.827974 | 3193.191905 | 2173.972986 | 0.629227087 | 0.114879873 | 5.49153202961 | 1.29480435295 | ns | NaCl | KS.1 |
| FIM1_1991 | 2039.109858 | 1865.723276 | 2246.902047 | -0.095945277 | 0.173400268 | 0.51986099 | 0.650807354 | ns | NaCl | KS.1 |
| FIM1_1992 | 4227.162333 | 4619.910465 | 4555.279446 | 0.08964396 | 0.193659909 | 0.578113404 | 0.699784473 | ns | NaCl | KS.1 |
| FIM1_1993 | 1404.235536 | 835.1369086 | 1575.670143 | -0.659333529 | 0.169137834 | 1.12597004405 | 9.47834343349 | ns | NaCl | KS.1 |
| FIM1_1994 | 18989.74179 | 25671.4951 | 15695.74925 | 0.415267605 | 0.079478688 | 5.34914832656 | 9.14502162358 | ns | NaCl | KS.1 |
| FIM1_1995 | 2931.858894 | 3591.271412 | 3370.377933 | 0.204615356 | 0.205787619 | 0.223209942 | 0.347782481 | ns | NaCl | KS.1 |
| FIM1_1996 | 7197.363742 | 14052.7509 | 7937.596766 | 0.893214961 | 0.150518824 | 1.81837775756 | 6.41590733608 | ns | NaCl | KS.1 |
| FIM1_1997 | 1435.4939 | 1727.601613 | 1520.024576 | 0.24478079 | 0.097506778 | 0.00792404 | 0.023848818 | ns | NaCl | KS.1 |
| FIM1_1998 | 1327.46468 | 1554.088907 | 1350.747153 | 0.207949028 | 0.08963581 | 0.014992993 | 0.039131242 | ns | NaCl | KS.1 |
| FIM1_1999 | 381.5617975 | 375.8402786 | 411.8301405 | -0.013155312 | 0.186112839 | 0.933164842 | 0.961658314 | ns | NaCl | KS.1 |
| FIM1_2000 | 7108.097044 | 7654.984346 | 5751.658379 | 0.08325729 | 0.158007726 | 0.549304236 | 0.67492926 | ns | NaCl | KS.1 |
| FIM1_2001 | 3387.950312 | 6751.761414 | 3238.179242 | 0.746469816 | 0.281998281 | 0.000637532 | 0.00295044 | ns | NaCl | KS.1 |
| FIM1_2002 | 3928.218756 | 5126.211443 | 3293.617611 | 0.347176703 | 0.113940938 | 0.001013936 | 0.004333617 | ns | NaCl | KS.1 |
| FIM1_2003 | 3316.715747 | 5583.24666 | 4467.052273 | 0.676665855 | 0.156105924 | 1.60325495260 | 1.77643505327 | ns | NaCl | KS.1 |
| FIM1_2004 | 903.4273177 | 451.9524266 | 610.1034241 | -0.905086438 | 0.178356155 | 2.15140718115 | 4.36867376581 | ns | NaCl | KS.1 |
| FIM1_2005 | 39631.04788 | 42290.25548 | 32739.38233 | 0.072758169 | 0.158619313 | 0.60189349 | 0.72064621 | ns | NaCl | KS.1 |
| FIM1_2006 | 490.6740918 | 856.411079 | 869.3747403 | 0.736703662 | 0.147085793 | 5.08754923163 | 8.72777842323 | ns | NaCl | KS.1 |
| FIM1_2007 | 683.7230734 | 730.3978709 | 634.0629093 | 0.078116881 | 0.125399481 | 0.498065498 | 0.629861681 | ns | NaCl | KS.1 |
| FIM1_2008 | 461.2298043 | 437.3919057 | 575.9856178 | -0.06280869 | 0.149857312 | 0.637863435 | 0.748471149 | ns | NaCl | KS.1 |
| FIM1_2009 | 285.1391833 | 211.7246898 | 814.4961066 | -0.267859382 | 0.255390782 | 0.163142754 | 0.272676676 | ns | NaCl | KS.1 |
| FIM1_2010 | 26.32575295 | 37.77441177 | 78.59222337 | 0.306489881 | 0.285531502 | 0.13332173 | 0.232142575 | ns | NaCl | KS.1 |
| FIM1_2011 | 2315.729137 | 3124.165597 | 2751.075001 | 0.408206545 | 0.086356851 | 7.31739488196 | 8.96007986520 | ns | NaCl | KS.1 |
| FIM1_2012 | 2162.656657 | 3153.988322 | 3168.947475 | 0.499510857 | 0.122095604 | 9.18383326724 | 7.93221710148 | ns | NaCl | KS.1 |
| FIM1_2013 | 764.0576343 | 398.4371197 | 1113.463587 | -0.781785792 | 0.226337479 | 4.06786852836 | 0.000290771 | ns | NaCl | KS.1 |
| FIM1_2014 | 1629.75004 | 1140.510524 | 1707.956662 | -0.478375726 | 0.109935318 | 3.15064712350 | 3.23184936895 | ns | NaCl | KS.1 |
| FIM1_2015 | 18.09600083 | 12.21048392 | 21.92099029 | -0.26939074 | 0.321532838 | 0.205167427 | 0.326313283 | ns | NaCl | KS.1 |
| FIM1_2016 | 1201.390328 | 916.767572 | 2361.274511 | -0.361248647 | 0.105233044 | 0.000245044 | 0.001326542 | ns | NaCl | KS.1 |
| FIM1_2017 | 1303.361459 | 758.5433054 | 1226.622931 | -0.660409178 | 0.199580785 | 0.00010812 | 0.000668194 | ns | NaCl | KS.1 |
| FIM1_2018 | 205.3782741 | 50.73045148 | 205.6604993 | -1.611902412 | 0.418575448 | 1.95555798913 | 2.88691424211 | Km<KS.1 | NaCl | KS.1 |
| FIM1_2019 | 993.3039348 | 1282.545028 | 1473.133084 | 0.262300553 | 0.209244664 | 0.123006162 | 0.218010565 | ns | NaCl | KS.1 |
| FIM1_2020 | 3817.814808 | 2367.260946 | 3206.016417 | -0.565053995 | 0.202715521 | 0.000900864 | 0.003934853 | ns | NaCl | KS.1 |
| FIM1_2021 | 4182.041444 | 4937.41668 | 4196.697368 | 0.182315502 | 0.173995639 | 0.22213675 | 0.346762078 | ns | NaCl | KS.1 |

| | | | | | | | | | | |
|---|---|---|---|---|---|---|---|---|---|---|
| FIM1_2022 | 1664.943889 | 1745.371091 | 1584.664656 | 0.054941979 | 0.141399266 | 0.666257242 | 0.772640974 | ns | NaCl | KS.1 |
| FIM1_2023 | 1823.559766 | 1157.762708 | 2191.806579 | -0.567527907 | 0.170998692 | 0.000149671 | 0.000876013 | ns | NaCl | KS.1 |
| FIM1_2024 | 464.3235908 | 112.2311744 | 535.9085137 | -1.031788102 | 0.583311033 | 0.000938423 | 0.004072383 | Km<KS.1 | NaCl | KS.1 |
| FIM1_2025 | 157.8033878 | 117.9399933 | 215.5552265 | -0.315047389 | 0.209620121 | 0.064789906 | 0.130497887 | ns | NaCl | KS.1 |
| FIM1_2026 | 963.8544309 | 1051.120725 | 1235.45582 | 0.110967785 | 0.097885618 | 0.231665167 | 0.357596713 | ns | NaCl | KS.1 |
| FIM1_2027 | 855.5471298 | 980.287145 | 972.3894514 | 0.16847148 | 0.118491362 | 0.124467979 | 0.219740311 | ns | NaCl | KS.1 |
| FIM1_2028 | 2856.553997 | 3070.164978 | 2123.234624 | 0.079417434 | 0.163303003 | 0.577554915 | 0.699278584 | ns | NaCl | KS.1 |
| FIM1_2029 | 1806.576404 | 1335.0989 | 2108.789834 | -0.406365453 | 0.09940348 | 1.42901228415 | 0.00011693 | ns | NaCl | KS.1 |
| FIM1_2030 | 435.8794477 | 350.8830899 | 1023.682246 | -0.233810045 | 0.186367702 | 0.136270529 | 0.236218078 | ns | NaCl | KS.1 |
| FIM1_2031 | 733.4472211 | 465.0817808 | 1632.790141 | -0.434362065 | 0.272298372 | 0.032264605 | 0.074038936 | ns | NaCl | KS.1 |
| FIM1_2032 | 14.14883355 | 20.52110575 | 27.57214771 | 0.204507352 | 0.35569221 | 0.338243643 | 0.472023037 | ns | NaCl | KS.1 |
| FIM1_2033 | 1347.798457 | 1274.444868 | 1731.269197 | -0.072189888 | 0.102548325 | 0.455723036 | 0.590884051 | ns | NaCl | KS.1 |
| FIM1_2034 | 1678.245671 | 1967.952739 | 2139.809632 | 0.212268068 | 0.093408065 | 0.01676352 | 0.042878413 | ns | NaCl | KS.1 |
| FIM1_2035 | 290.9604979 | 195.988382 | 253.7108773 | -0.437399824 | 0.212684534 | 0.011780673 | 0.032488275 | ns | NaCl | KS.1 |
| FIM1_2037 | 408.1324585 | 205.080036 | 314.2585225 | -0.803394406 | 0.248350832 | 8.06242075826 | 0.000520362 | ns | NaCl | KS.1 |
| FIM1_2038 | 1461.127685 | 2027.768791 | 1938.2021 | 0.384543338 | 0.173908044 | 0.010106562 | 0.02883036 | ns | NaCl | KS.1 |
| FIM1_2039 | 508.7544607 | 1028.794013 | 620.846494 | 0.900064704 | 0.192084486 | 1.51389658102 | 2.33901723309 | ns | NaCl | KS.1 |
| FIM1_2040 | 441.2087814 | 838.6988768 | 447.3371595 | 0.826275144 | 0.178128228 | 2.43595768530 | 3.50026239285 | ns | NaCl | KS.1 |
| FIM1_2041 | 3185.880285 | 16967.89552 | 5070.218881 | 2.370093873 | 0.155333725 | 1.64547393186 | 8.18623281103 | Km>KS.1 | NaCl | KS.1 |
| FIM1_2042 | 4352.075416 | 4412.835336 | 3704.298972 | 0.017736259 | 0.093373759 | 0.841996405 | 0.903176394 | ns | NaCl | KS.1 |
| FIM1_2043 | 2506.566671 | 4830.968762 | 2435.732405 | 0.815731682 | 0.204799988 | 4.63655449574 | 4.47900167307 | ns | NaCl | KS.1 |
| FIM1_2044 | 583.7345378 | 600.4791354 | 810.1224928 | 0.03157195 | 0.140833624 | 0.80366934 | 0.874508961 | ns | NaCl | KS.1 |
| FIM1_2045 | 328.8781861 | 272.9597417 | 395.9114986 | -0.218696547 | 0.16160975 | 0.121378421 | 0.215944243 | ns | NaCl | KS.1 |
| FIM1_2046 | 254.4692886 | 218.5743826 | 265.7848338 | -0.150574565 | 0.209284217 | 0.374960642 | 0.509600707 | ns | NaCl | KS.1 |
| FIM1_2047 | 613.1111648 | 563.9133569 | 771.6282506 | -0.089612266 | 0.163114028 | 0.529299936 | 0.657659136 | ns | NaCl | KS.1 |
| FIM1_2048 | 2226.026721 | 2319.484589 | 1837.727007 | 0.053838525 | 0.097507317 | 0.56061613 | 0.683929683 | ns | NaCl | KS.1 |
| FIM1_2049 | 383.7775673 | 454.8986486 | 419.7850415 | 0.207766575 | 0.128425811 | 0.076567885 | 0.149617136 | ns | NaCl | KS.1 |
| FIM1_2050 | 4657.671254 | 4230.456646 | 3690.857324 | -0.125958731 | 0.09590118 | 0.166560874 | 0.277502121 | ns | NaCl | KS.1 |
| FIM1_2051 | 1852.979595 | 2272.023746 | 2042.719606 | 0.26326169 | 0.113688539 | 0.012632645 | 0.034174774 | ns | NaCl | KS.1 |
| FIM1_2052 | 1937.405391 | 2701.482399 | 3036.137461 | 0.390581296 | 0.176206071 | 0.009709254 | 0.027918932 | ns | NaCl | KS.1 |
| FIM1_2053 | 9217.320211 | 9421.108451 | 6991.329792 | 0.027913711 | 0.107417274 | 0.782023906 | 0.858845239 | ns | NaCl | KS.1 |
| FIM1_2054 | 1480.364314 | 1221.88207 | 1495.778769 | -0.243966152 | 0.117582913 | 0.024850368 | 0.05969608 | ns | NaCl | KS.1 |
| FIM1_2055 | 586.8823775 | 980.8623129 | 893.0585356 | 0.424174531 | 0.316796718 | 0.052106819 | 0.10942652 | ns | NaCl | KS.1 |
| FIM1_2056 | 696.0221347 | 845.6921076 | 844.1085238 | 0.204675038 | 0.187495265 | 0.194056088 | 0.312032657 | ns | NaCl | KS.1 |
| FIM1_2057 | 555.8275453 | 626.8787464 | 693.8576477 | 0.142897047 | 0.122495924 | 0.205596094 | 0.326786124 | ns | NaCl | KS.1 |
| FIM1_2058 | 275.5174243 | 360.5371041 | 517.6529092 | 0.244104923 | 0.244368668 | 0.191932188 | 0.309342198 | ns | NaCl | KS.1 |
| FIM1_2059 | 792.9155834 | 984.3981561 | 774.0261571 | 0.260591055 | 0.140549614 | 0.039053608 | 0.086659992 | ns | NaCl | KS.1 |
| FIM1_2060 | 3611.311555 | 4406.429821 | 3723.71952 | 0.162269908 | 0.261653224 | 0.400072684 | 0.536052142 | ns | NaCl | KS.1 |
| FIM1_2061 | 1910.616279 | 2317.664318 | 1823.673689 | 0.231721794 | 0.143094493 | 0.070698513 | 0.139906565 | ns | NaCl | KS.1 |
| FIM1_2062 | 623.1155645 | 675.3479139 | 1165.197352 | 0.069570915 | 0.236474639 | 0.702738341 | 0.799981706 | ns | NaCl | KS.1 |
| FIM1_2063 | 2576.39646 | 2047.310279 | 3667.742342 | -0.286337622 | 0.133280851 | 0.017785086 | 0.045120246 | ns | NaCl | KS.1 |
| FIM1_2064 | 481.1840539 | 232.6045349 | 1152.213554 | -0.806050393 | 0.279035741 | 0.000242387 | 0.001313591 | ns | NaCl | KS.1 |
| FIM1_2065 | 43.13839697 | 55.58757667 | 56.90405186 | 0.222283042 | 0.255855404 | 0.245823154 | 0.373405998 | ns | NaCl | KS.1 |
| FIM1_2066 | 912.6651252 | 753.1369182 | 3256.199129 | -0.128681234 | 0.302787735 | 0.5278059 | 0.656294515 | ns | NaCl | KS.1 |
| FIM1_2067 | 1409.605254 | 1582.509774 | 2073.776345 | 0.134608529 | 0.152445837 | 0.31912904 | 0.452455678 | ns | NaCl | KS.1 |

| | | | | | | | | | | |
|---|---|---|---|---|---|---|---|---|---|---|
| FIM1_2068 | 984.1337786 | 1280.88033 | 1436.187791 | 0.343465324 | 0.115990729 | 0.001367296 | 0.005523177 | ns | NaCl | KS.1 |
| FIM1_2069 | 864.3719528 | 754.4848837 | 1555.198042 | -0.166576203 | 0.130668776 | 0.16210718 | 0.271543171 | ns | NaCl | KS.1 |
| FIM1_2070 | 85.56050751 | 70.81048563 | 80.28421507 | -0.168286265 | 0.239223559 | 0.360549707 | 0.495924545 | ns | NaCl | KS.1 |
| FIM1_2071 | 1930.945721 | 2156.310816 | 2107.247229 | 0.150717823 | 0.073212906 | 0.033585504 | 0.076470427 | ns | NaCl | KS.1 |
| FIM1_2072 | 1587.943691 | 1824.194803 | 1485.603666 | 0.167249352 | 0.134622526 | 0.170888481 | 0.282636367 | ns | NaCl | KS.1 |
| FIM1_2073 | 3628.532516 | 5443.709376 | 3368.971596 | 0.456206717 | 0.207763494 | 0.007681979 | 0.023232732 | ns | NaCl | KS.1 |
| FIM1_2074 | 686.2539547 | 870.1983577 | 831.1518814 | 0.309674137 | 0.112097108 | 0.002953755 | 0.010571345 | ns | NaCl | KS.1 |
| FIM1_2075 | 6650.861857 | 5273.006277 | 6104.843797 | -0.285829744 | 0.137701703 | 0.021292735 | 0.05246724 | ns | NaCl | KS.1 |
| FIM1_2076 | 1085.343555 | 1193.983013 | 1345.166444 | 0.120360756 | 0.115463191 | 0.262201822 | 0.39055511 | ns | NaCl | KS.1 |
| FIM1_2077 | 2374.567894 | 3230.65241 | 2543.597471 | 0.355449559 | 0.178146694 | 0.019440454 | 0.048309819 | ns | NaCl | KS.1 |
| FIM1_2078 | 1667.0896 | 1269.526523 | 1355.849322 | -0.309237249 | 0.178316148 | 0.041981609 | 0.091564449 | ns | NaCl | KS.1 |
| FIM1_2079 | 1079.089966 | 216.2463179 | 1965.955201 | -2.035350628 | 0.383054608 | 4.72206874925 | 1.83533531465 | Km<KS.1 | NaCl | KS.1 |
| FIM1_2080 | 36.87522568 | 32.52211158 | 36.46740299 | -0.08466967 | 0.284614916 | 0.670211373 | 0.775779801 | ns | NaCl | KS.1 |
| FIM1_2081 | 12269.32757 | 26740.00997 | 11195.6198 | 0.935343338 | 0.244998499 | 5.68999371674 | 5.24217013718 | ns | NaCl | KS.1 |
| FIM1_2083 | 590.1903821 | 535.9710093 | 754.0544526 | -0.092952915 | 0.206081979 | 0.579962996 | 0.701511282 | ns | NaCl | KS.1 |
| FIM1_2084 | 6413.278764 | 8240.902755 | 6222.755823 | 0.347700501 | 0.069531527 | 2.44674956597 | 3.50026239285 | ns | NaCl | KS.1 |
| FIM1_2085 | 2329.325702 | 2439.391673 | 2721.631889 | 0.063248191 | 0.07827066 | 0.402732333 | 0.53816636 | ns | NaCl | KS.1 |
| FIM1_2086 | 3168.242259 | 2963.454893 | 3151.420719 | -0.087042269 | 0.093893714 | 0.330121139 | 0.463548593 | ns | NaCl | KS.1 |
| FIM1_2087 | 5898.765488 | 4521.973853 | 3960.918562 | -0.243377684 | 0.242709407 | 0.191509777 | 0.308936816 | ns | NaCl | KS.1 |
| FIM1_2088 | 1489.144316 | 2458.87472 | 1507.92372 | 0.65728457 | 0.147501057 | 9.76320829275 | 1.14827331575 | ns | NaCl | KS.1 |
| FIM1_2089 | 344.2197613 | 306.2429993 | 349.9156491 | -0.132590362 | 0.160944976 | 0.346783143 | 0.481643254 | ns | NaCl | KS.1 |
| FIM1_2090 | 625.7941567 | 833.110101 | 845.9294823 | 0.373816922 | 0.109167437 | 0.00023885 | 0.001295832 | ns | NaCl | KS.1 |
| FIM1_2091 | 1476.330239 | 1752.528184 | 1214.776412 | 0.20936109 | 0.133410414 | 0.083852901 | 0.161194408 | ns | NaCl | KS.1 |
| FIM1_2092 | 714.8354845 | 1071.732813 | 677.1625973 | 0.385539006 | 0.260108061 | 0.049655787 | 0.105032967 | ns | NaCl | KS.1 |
| FIM1_2093 | 3247.733966 | 7461.344153 | 3388.232517 | 0.878764245 | 0.320129486 | 0.000266107 | 0.001419253 | ns | NaCl | KS.1 |
| FIM1_2094 | 9352.547155 | 13386.52881 | 8107.293509 | 0.450529921 | 0.148972927 | 0.000687163 | 0.003147914 | ns | NaCl | KS.1 |
| FIM1_2095 | 700.8414062 | 358.2274029 | 1015.029635 | -0.858367114 | 0.189956662 | 3.84297786438 | 5.19533012915 | ns | NaCl | KS.1 |
| FIM1_2096 | 896.2598252 | 486.152177 | 1200.383166 | -0.691910212 | 0.250075635 | 0.000567484 | 0.002681131 | ns | NaCl | KS.1 |
| FIM1_2097 | 1051.785534 | 1014.625192 | 1071.318606 | -0.045671631 | 0.089750035 | 0.594543017 | 0.714595654 | ns | NaCl | KS.1 |
| FIM1_2098 | 93.53193541 | 8.856995882 | 1283.458349 | -2.830484564 | 0.62076898 | 3.44825136396 | 8.57752526787 | Km<KS.1 | NaCl | KS.1 |
| FIM1_2099 | 17.28013463 | 23.03736861 | 15.19172552 | 0.159992547 | 0.333598004 | 0.44524368 | 0.580546747 | ns | NaCl | KS.1 |
| FIM1_2100 | 2076.33341 | 2991.713796 | 1852.733756 | 0.443447843 | 0.167359779 | 0.002315447 | 0.008622267 | ns | NaCl | KS.1 |
| FIM1_2101 | 822.7149346 | 906.0241606 | 761.0685894 | 0.117140607 | 0.134650235 | 0.338105536 | 0.471962694 | ns | NaCl | KS.1 |
| FIM1_2102 | 860.155503 | 936.8043539 | 1111.61358 | 0.104453742 | 0.124101965 | 0.360529828 | 0.495924545 | ns | NaCl | KS.1 |
| FIM1_2103 | 4934.739454 | 5222.200094 | 5729.274012 | 0.053956644 | 0.209682162 | 0.750639759 | 0.837504553 | ns | NaCl | KS.1 |
| FIM1_2104 | 594.5941675 | 965.9611584 | 766.9407679 | 0.553597471 | 0.218726178 | 0.002030552 | 0.007711449 | ns | NaCl | KS.1 |
| FIM1_2105 | 377.6868639 | 516.4260742 | 459.8558034 | 0.368114161 | 0.167302492 | 0.011154883 | 0.03114228 | ns | NaCl | KS.1 |
| FIM1_2106 | 603.346591 | 439.0962666 | 901.2688859 | -0.363864664 | 0.182875744 | 0.019005898 | 0.04750245 | ns | NaCl | KS.1 |
| FIM1_2107 | 800.3972359 | 889.9392667 | 817.3913675 | 0.130977554 | 0.124406798 | 0.252321049 | 0.380833267 | ns | NaCl | KS.1 |
| FIM1_2108 | 521.5614421 | 486.7443239 | 655.523916 | -0.083468999 | 0.145789301 | 0.522495569 | 0.65235834 | ns | NaCl | KS.1 |
| FIM1_2109 | 275.4452163 | 309.9378223 | 233.2353216 | 0.122821309 | 0.177002852 | 0.416923656 | 0.553118717 | ns | NaCl | KS.1 |
| FIM1_2110 | 457.2374732 | 460.6178876 | 423.3050302 | 0.005633328 | 0.163381458 | 0.968510388 | 0.985546979 | ns | NaCl | KS.1 |
| FIM1_2111 | 36.97251404 | 43.13395226 | 44.30100201 | 0.117668579 | 0.268140005 | 0.545094792 | 0.671581622 | ns | NaCl | KS.1 |
| FIM1_2112 | 949.2907044 | 563.9075047 | 1126.871592 | -0.687580754 | 0.14599042 | 2.64173475542 | 3.76579667858 | ns | NaCl | KS.1 |
| FIM1_2113 | 1200.645256 | 1113.594034 | 1409.47737 | -0.093505235 | 0.121540429 | 0.404827125 | 0.539950388 | ns | NaCl | KS.1 |

| | | | | | | | | | | | |
|---|---|---|---|---|---|---|---|---|---|---|---|
| FIM1_2115 | 1170.638209 | 1374.418951 | 1230.235367 | 0.204257642 | 0.11236811 | 0.050941377 | 0.107324382 | ns | | NaCl | KS.1 |
| FIM1_2116 | 1050.397642 | 927.998114 | 1207.977676 | -0.174582898 | 0.079532147 | 0.022674693 | 0.055216151 | ns | | NaCl | KS.1 |
| FIM1_2117 | 2399.855551 | 2891.33712 | 2470.402507 | 0.253961329 | 0.073763213 | 0.00036807 | 0.001866616 | ns | | NaCl | KS.1 |
| FIM1_2118 | 1621.38257 | 1493.30856 | 1670.173672 | -0.117463867 | 0.06439462 | 0.061546013 | 0.125591228 | ns | | NaCl | KS.1 |
| FIM1_2119 | 948.3109268 | 1502.355038 | 891.3692608 | 0.421243605 | 0.282577534 | 0.041447029 | 0.090597086 | ns | | NaCl | KS.1 |
| FIM1_2120 | 1908.638065 | 1875.655382 | 1712.28105 | -0.021289566 | 0.131962654 | 0.859641742 | 0.91500164 | ns | | NaCl | KS.1 |
| FIM1_2121 | 3734.901906 | 3831.176301 | 3712.965937 | 0.03300345 | 0.096646839 | 0.719189185 | 0.813388223 | ns | | NaCl | KS.1 |
| FIM1_2122 | 1074.784831 | 1409.339536 | 1269.252022 | 0.352292269 | 0.117094886 | 0.001125911 | 0.0047304 | ns | | NaCl | KS.1 |
| FIM1_2123 | 5669.536453 | 7493.785561 | 7668.136712 | 0.365832714 | 0.112266981 | 0.000450912 | 0.002207844 | ns | | NaCl | KS.1 |
| FIM1_2124 | 2650.404898 | 997.2374894 | 2574.161137 | -1.265916413 | 0.216070278 | 5.0527714751 | 1.9486463634 | Km<KS.1 | | NaCl | KS.1 |
| FIM1_2125 | 1117.025785 | 1022.28441 | 1307.919858 | -0.103118839 | 0.146073662 | 0.430085597 | 0.566801548 | ns | | NaCl | KS.1 |
| FIM1_2126 | 1122.473851 | 737.5163188 | 1140.724034 | -0.521615365 | 0.164716207 | 0.000310169 | 0.001612423 | ns | | NaCl | KS.1 |
| FIM1_2127 | 891.2875549 | 955.1755637 | 1172.311572 | 0.087782954 | 0.101915716 | 0.361648862 | 0.496880168 | ns | | NaCl | KS.1 |
| FIM1_2128 | 943.3575016 | 707.5173961 | 763.6469738 | -0.346296389 | 0.162476103 | 0.014551617 | 0.038364757 | ns | | NaCl | KS.1 |
| FIM1_2129 | 4147.677761 | 4482.918964 | 3207.282133 | 0.097643809 | 0.114784018 | 0.360776233 | 0.49595517 | ns | | NaCl | KS.1 |
| FIM1_2130 | 2706.122357 | 2586.637435 | 4591.296402 | -0.045199112 | 0.19327282 | 0.778877595 | 0.856051837 | ns | | NaCl | KS.1 |
| FIM1_2131 | 11153.5268 | 14752.44962 | 10641.61632 | 0.346156322 | 0.142787724 | 0.006771729 | 0.020912073 | ns | | NaCl | KS.1 |
| FIM1_2132 | 2567.569209 | 5493.695651 | 2400.954067 | 0.684656052 | 0.356212801 | 0.004960286 | 0.01620072 | ns | | NaCl | KS.1 |
| FIM1_2133 | 1416.090731 | 1521.296122 | 1195.454349 | 0.091711684 | 0.100196298 | 0.333177581 | 0.467312789 | ns | | NaCl | KS.1 |
| FIM1_2134 | 822.7505753 | 888.8285753 | 923.6418429 | 0.091435682 | 0.13466064 | 0.454857878 | 0.589942341 | ns | | NaCl | KS.1 |
| FIM1_2135 | 2115.262484 | 3148.158329 | 2432.676187 | 0.527707093 | 0.122121176 | 2.9228946121 | 3.0106419659 | ns | | NaCl | KS.1 |
| FIM1_2136 | 1384.434582 | 1915.03217 | 1894.50463 | 0.43562614 | 0.100765158 | 4.3708688288 | 4.2721163897 | ns | | NaCl | KS.1 |
| FIM1_2137 | 1265.338179 | 1080.676728 | 929.025873 | -0.192746308 | 0.136897666 | 0.119379721 | 0.213351332 | ns | | NaCl | KS.1 |
| FIM1_2138 | 1155.41726 | 1151.010122 | 1077.161676 | -0.004212354 | 0.181574297 | 0.978202613 | 0.989137805 | ns | | NaCl | KS.1 |
| FIM1_2139 | 3027.392137 | 3143.651748 | 2822.038722 | 0.045372346 | 0.133679833 | 0.709227615 | 0.804837451 | ns | | NaCl | KS.1 |
| FIM1_2140 | 1067.927832 | 819.3022438 | 1049.093401 | -0.346173337 | 0.11187609 | 0.000869017 | 0.003819224 | ns | | NaCl | KS.1 |
| FIM1_2141 | 1553.679285 | 1720.274121 | 1365.113948 | 0.120867195 | 0.139365063 | 0.33658623 | 0.47086434 | ns | | NaCl | KS.1 |
| FIM1_2142 | 126.0255656 | 124.9842805 | 184.7391978 | -0.000688161 | 0.167755209 | 0.99622778 | 0.998234281 | ns | | NaCl | KS.1 |
| FIM1_2143 | 758.5582911 | 888.9467992 | 887.3095082 | 0.193495028 | 0.129829853 | 0.102308332 | 0.18837304 | ns | | NaCl | KS.1 |
| FIM1_2144 | 943.3765583 | 1213.659098 | 1324.008097 | 0.307829178 | 0.1445754 | 0.017140147 | 0.043751787 | ns | | NaCl | KS.1 |
| FIM1_2145 | 706.1213756 | 627.6019341 | 768.7166833 | -0.113470607 | 0.209037271 | 0.503262041 | 0.635141719 | ns | | NaCl | KS.1 |
| FIM1_2146 | 12235.16233 | 10636.7242 | 11469.12418 | -0.184629398 | 0.094723659 | 0.040080007 | 0.088424849 | ns | | NaCl | KS.1 |
| FIM1_2147 | 741.0021905 | 983.2340399 | 730.851883 | 0.329484678 | 0.166234545 | 0.022346338 | 0.054693948 | ns | | NaCl | KS.1 |
| FIM1_2148 | 462.1140969 | 515.5398878 | 554.9139861 | 0.114317494 | 0.178515793 | 0.452724548 | 0.588069093 | ns | | NaCl | KS.1 |
| FIM1_2149 | 892.9248121 | 948.946697 | 947.8479098 | 0.080496325 | 0.106937371 | 0.422887409 | 0.558943905 | ns | | NaCl | KS.1 |
| FIM1_2150 | 916.04065 | 983.0539619 | 1219.572483 | 0.07739262 | 0.172476248 | 0.602314137 | 0.720662056 | ns | | NaCl | KS.1 |
| FIM1_2151 | 760.9767253 | 1861.469186 | 1008.198779 | 1.217542664 | 0.151161821 | 1.1652742735 | 1.8116373470 | Km>KS.1 | | NaCl | KS.1 |
| FIM1_2152 | 184.8844977 | 312.7987763 | 256.6004841 | 0.686174405 | 0.152019707 | 6.7990242023 | 8.3933363292 | ns | | NaCl | KS.1 |
| FIM1_2153 | 300.6702835 | 287.4362594 | 445.3975149 | -0.052582671 | 0.13477105 | 0.667715965 | 0.773971791 | ns | | NaCl | KS.1 |
| FIM1_2154 | 1392.037738 | 2958.413006 | 1304.304847 | 0.662401695 | 0.360257839 | 0.006531997 | 0.020335848 | ns | | NaCl | KS.1 |
| FIM1_2155 | 715.1762401 | 738.9851718 | 789.1994816 | 0.04006234 | 0.105508828 | 0.686595414 | 0.787961288 | ns | | NaCl | KS.1 |
| FIM1_2156 | 1353.22996 | 1567.133824 | 1257.674473 | 0.169620817 | 0.151981044 | 0.207952579 | 0.329583969 | ns | | NaCl | KS.1 |
| FIM1_2157 | 3313.053345 | 3078.569236 | 3731.140676 | -0.096489325 | 0.090066654 | 0.262099521 | 0.390519652 | ns | | NaCl | KS.1 |
| FIM1_2158 | 2635.278104 | 4217.966443 | 2815.198628 | 0.635443614 | 0.11799098 | 9.0163425574 | 1.9673817641 | ns | | NaCl | KS.1 |
| FIM1_2159 | 63.66484328 | 65.95252499 | 86.34968685 | 0.028808355 | 0.266423847 | 0.881463399 | 0.930070077 | ns | | NaCl | KS.1 |

| | | | | | | | | | | |
|---|---|---|---|---|---|---|---|---|---|---|
| FIM1_2160 | 1332.473664 | 1616.245182 | 1771.477552 | 0.233799845 | 0.138209603 | 0.060641573 | 0.124306479 | ns | NaCl | KS.1 |
| FIM1_2161 | 1371.116874 | 1193.624402 | 1565.997904 | -0.195885321 | 0.06605866 | 0.002318926 | 0.008628763 | ns | NaCl | KS.1 |
| FIM1_2162 | 828.771513 | 906.6071186 | 767.1367039 | 0.112476439 | 0.11472434 | 0.292133285 | 0.423474094 | ns | NaCl | KS.1 |
| FIM1_2163 | 650.6290133 | 599.9192765 | 761.4008034 | -0.088362138 | 0.174410564 | 0.555091869 | 0.678854977 | ns | NaCl | KS.1 |
| FIM1_2164 | 878.523567 | 755.3036022 | 769.3607597 | -0.200662015 | 0.106373944 | 0.04422054 | 0.095609381 | ns | NaCl | KS.1 |
| FIM1_2165 | 3404.57547 | 3602.492389 | 2668.911091 | 0.071763553 | 0.109771611 | 0.485101433 | 0.619291668 | ns | NaCl | KS.1 |
| FIM1_2166 | 400.4403152 | 357.716085 | 398.1753113 | -0.10853217 | 0.206866404 | 0.519215117 | 0.650162398 | ns | NaCl | KS.1 |
| FIM1_2167 | 4916.376528 | 4342.101145 | 4787.92655 | -0.156049715 | 0.115983404 | 0.147309063 | 0.251152361 | ns | NaCl | KS.1 |
| FIM1_2168 | 4201.199784 | 6240.204933 | 4647.44812 | 0.551369771 | 0.077723738 | 2.0939619309 | 1.5548448666 | ns | NaCl | KS.1 |
| FIM1_2169 | 2358.844861 | 2884.303188 | 2427.300803 | 0.263776895 | 0.103994508 | 0.006882445 | 0.021227629 | ns | NaCl | KS.1 |
| FIM1_2170 | 3034.076178 | 2595.840392 | 3230.549092 | -0.189170069 | 0.133817903 | 0.119390976 | 0.213351332 | ns | NaCl | KS.1 |
| FIM1_2171 | 1264.237497 | 1171.994706 | 1110.462969 | -0.083249432 | 0.164818892 | 0.562052909 | 0.685178442 | ns | NaCl | KS.1 |
| FIM1_2172 | 2337.509696 | 2152.366021 | 2057.017079 | -0.105574404 | 0.110590265 | 0.307321866 | 0.440065552 | ns | NaCl | KS.1 |
| FIM1_2173 | 844.9415498 | 842.0660173 | 774.6059075 | -0.00321497 | 0.171029835 | 0.98262852 | 0.991175946 | ns | NaCl | KS.1 |
| FIM1_2174 | 1099.623504 | 948.9670177 | 1263.218158 | -0.163887694 | 0.162895729 | 0.248980795 | 0.377071371 | ns | NaCl | KS.1 |
| FIM1_2175 | 2963.559709 | 4253.149453 | 3557.19596 | 0.471275801 | 0.127498555 | 5.3755408446 | 0.000369383 | ns | NaCl | KS.1 |
| FIM1_2176 | 680.4097808 | 288.094189 | 925.9356295 | -1.136109225 | 0.180474478 | 6.8938266115 | 3.4296787392 | Km<KS.1 | NaCl | KS.1 |
| FIM1_2177 | 79.20509423 | 62.68040997 | 79.54727981 | -0.207375491 | 0.224247935 | 0.242608737 | 0.369445505 | ns | NaCl | KS.1 |
| FIM1_2178 | 3830.152709 | 3916.573336 | 4144.390667 | 0.022709474 | 0.190966925 | 0.886901158 | 0.932435862 | ns | NaCl | KS.1 |
| FIM1_2179 | 7867.415181 | 7898.6878 | 9088.135023 | 0.005113486 | 0.116801861 | 0.962430322 | 0.98169855 | ns | NaCl | KS.1 |
| FIM1_2180 | 3323.22119 | 6145.978916 | 2978.44097 | 0.608317109 | 0.297407942 | 0.005568076 | 0.017860204 | ns | NaCl | KS.1 |
| FIM1_2181 | 3581.189997 | 4221.746287 | 2827.95635 | 0.163412474 | 0.20542732 | 0.329849182 | 0.463297482 | ns | NaCl | KS.1 |
| FIM1_2182 | 762.2962312 | 1122.202521 | 1562.092515 | 0.311553633 | 0.295586452 | 0.133359493 | 0.232142575 | ns | NaCl | KS.1 |
| FIM1_2183 | 2981.427336 | 2334.779657 | 2582.084018 | -0.318472623 | 0.111412861 | 0.00211397 | 0.00797413 | ns | NaCl | KS.1 |
| FIM1_2184 | 2879.567917 | 2791.742093 | 2910.723947 | -0.037290378 | 0.130939835 | 0.755214401 | 0.840535044 | ns | NaCl | KS.1 |
| FIM1_2185 | 2012.144832 | 2404.348497 | 3145.59594 | 0.202244571 | 0.165272237 | 0.159180999 | 0.267542388 | ns | NaCl | KS.1 |
| FIM1_2186 | 1613.727705 | 1926.350229 | 1562.596842 | 0.225127437 | 0.117263934 | 0.038045787 | 0.085030454 | ns | NaCl | KS.1 |
| FIM1_2187 | 2059.173057 | 3150.06116 | 1825.116442 | 0.45402413 | 0.230652194 | 0.01347457 | 0.036196535 | ns | NaCl | KS.1 |
| FIM1_2188 | 6234.603038 | 4742.47336 | 4842.867628 | -0.367047516 | 0.096881435 | 6.0216702552 | 0.000407035 | ns | NaCl | KS.1 |
| FIM1_2189 | 1346.937539 | 1376.834518 | 1001.661405 | 0.024151627 | 0.161163643 | 0.864210105 | 0.91848863 | ns | NaCl | KS.1 |
| FIM1_2190 | 2041.378639 | 2018.963725 | 2212.447532 | -0.013846746 | 0.111777469 | 0.894568539 | 0.936942838 | ns | NaCl | KS.1 |
| FIM1_2191 | 6360.1419 | 9778.340141 | 7904.87394 | 0.599184811 | 0.083119793 | 7.7338451958 | 6.7501543595 | ns | NaCl | KS.1 |
| FIM1_2192 | 740.573972 | 1098.923623 | 907.4031976 | 0.477263514 | 0.175472631 | 0.001614388 | 0.006394572 | ns | NaCl | KS.1 |
| FIM1_2193 | 4390.076397 | 10119.57031 | 4372.129069 | 0.896565591 | 0.313844409 | 0.000177988 | 0.001014307 | ns | NaCl | KS.1 |
| FIM1_2194 | 688.8486796 | 1095.956847 | 674.7807856 | 0.496102861 | 0.239598301 | 0.008778333 | 0.025896362 | ns | NaCl | KS.1 |
| FIM1_2195 | 823.3322205 | 883.1094601 | 827.4549321 | 0.083270579 | 0.130950027 | 0.486125765 | 0.619963005 | ns | NaCl | KS.1 |
| FIM1_2196 | 169.1648187 | 297.1224133 | 346.3742841 | 0.692773006 | 0.202326601 | 6.3232055213 | 0.000423962 | ns | NaCl | KS.1 |
| FIM1_2197 | 1211.944092 | 1574.189534 | 1611.03317 | 0.338340272 | 0.117539945 | 0.001823273 | 0.007086549 | ns | NaCl | KS.1 |
| FIM1_2198 | 3186.179253 | 2254.157895 | 2316.489743 | -0.360137357 | 0.222247382 | 0.042842773 | 0.093027665 | ns | NaCl | KS.1 |
| FIM1_2199 | 68.73023191 | 184.6276488 | 102.5282884 | 1.179674425 | 0.290822802 | 7.1096664481 | 8.7550966781 | Km>KS.1 | NaCl | KS.1 |
| FIM1_2200 | 1170.916134 | 1595.941804 | 1233.819232 | 0.37774648 | 0.156074394 | 0.005973401 | 0.018916403 | ns | NaCl | KS.1 |
| FIM1_2201 | 1062.558461 | 1218.49786 | 1062.087388 | 0.18133557 | 0.093104926 | 0.040569142 | 0.089266467 | ns | NaCl | KS.1 |
| FIM1_2202 | 3223.821232 | 3097.087755 | 4287.758579 | -0.048450935 | 0.130091746 | 0.6837745 | 0.786334506 | ns | NaCl | KS.1 |
| FIM1_2203 | 826.6629681 | 500.0411545 | 970.0864016 | -0.670281853 | 0.137480409 | 1.2202548186 | 1.9150686822 | ns | NaCl | KS.1 |
| FIM1_2204 | 642.0584191 | 463.6401939 | 982.1773121 | -0.393600251 | 0.164760316 | 0.006073981 | 0.019186068 | ns | NaCl | KS.1 |

| | | | | | | | | | | | |
|---|---|---|---|---|---|---|---|---|---|---|---|
| FIM1_2205 | 805.307778 | 802.6668036 | 972.1536766 | -0.003335238 | 0.139455269 | 0.978882046 | 0.98962369 | ns | | NaCl | KS.1 |
| FIM1_2206 | 1700.591206 | 2678.781201 | 1280.748421 | 0.405474274 | 0.287227226 | 0.051063243 | 0.107507252 | ns | | NaCl | KS.1 |
| FIM1_2207 | 419.6733072 | 308.9067093 | 462.4421567 | -0.241917805 | 0.284306326 | 0.23103719 | 0.357070525 | ns | | NaCl | KS.1 |
| FIM1_2209 | 2335.046172 | 1728.916025 | 1758.944291 | -0.307571473 | 0.218749841 | 0.079574749 | 0.154461325 | ns | | NaCl | KS.1 |
| FIM1_2210 | 4487.512642 | 11297.5624 | 4726.000485 | 1.118114122 | 0.261767502 | 3.91224691925 | 5.24620712218 | Km>KS.1 | | NaCl | KS.1 |
| FIM1_2211 | 6015.492559 | 5877.821529 | 4800.855103 | -0.030553386 | 0.097175816 | 0.740485541 | 0.829849233 | ns | | NaCl | KS.1 |
| FIM1_2212 | 1154.444759 | 663.1349287 | 1039.059064 | -0.741745106 | 0.137487222 | 6.34833150485 | 1.45543544869 | ns | | NaCl | KS.1 |
| FIM1_2213 | 824.7012368 | 1017.772806 | 898.7242659 | 0.269910315 | 0.117963223 | 0.013253518 | 0.03567979 | ns | | NaCl | KS.1 |
| FIM1_2214 | 471.0223088 | 577.7334727 | 624.8375672 | 0.263096867 | 0.118152174 | 0.015922955 | 0.041002433 | ns | | NaCl | KS.1 |
| FIM1_2215 | 6902.704095 | 10983.74517 | 6323.786572 | 0.55146089 | 0.197409788 | 0.000935861 | 0.004066295 | ns | | NaCl | KS.1 |
| FIM1_2216 | 807.1732531 | 987.7406554 | 919.6030042 | 0.254792946 | 0.119762234 | 0.021046389 | 0.051885919 | ns | | NaCl | KS.1 |
| FIM1_2217 | 706.3928589 | 621.5954527 | 851.7880038 | -0.122735009 | 0.21206229 | 0.473139531 | 0.606667311 | ns | | NaCl | KS.1 |
| FIM1_2218 | 6876.466702 | 8109.943693 | 8006.289705 | 0.219024523 | 0.092942738 | 0.013153664 | 0.035449337 | ns | | NaCl | KS.1 |
| FIM1_2219 | 3296.578159 | 3781.299624 | 2623.452048 | 0.181046963 | 0.092991187 | 0.040667363 | 0.089443029 | ns | | NaCl | KS.1 |
| FIM1_2220 | 1177.450814 | 1444.073407 | 1521.415233 | 0.28137989 | 0.069627355 | 3.06494914972 | 0.000226907 | ns | | NaCl | KS.1 |
| FIM1_2221 | 804.4087299 | 767.6779539 | 801.815568 | -0.058436504 | 0.095087138 | 0.518169728 | 0.649343677 | ns | | NaCl | KS.1 |
| FIM1_2222 | 519.9522015 | 409.7337359 | 622.0733073 | -0.276081505 | 0.167480348 | 0.056947592 | 0.117851195 | ns | | NaCl | KS.1 |
| FIM1_2223 | 741.6060495 | 998.1236952 | 1068.373156 | 0.374051137 | 0.13837499 | 0.002683518 | 0.009759139 | ns | | NaCl | KS.1 |
| FIM1_2224 | 5133.42494 | 6021.29237 | 3951.563321 | 0.193718483 | 0.134866922 | 0.11303954 | 0.203831719 | ns | | NaCl | KS.1 |
| FIM1_2225 | 887.2427014 | 1175.190203 | 1245.852409 | 0.288771244 | 0.213168952 | 0.09381714 | 0.176194893 | ns | | NaCl | KS.1 |
| FIM1_2226 | 10817.18424 | 13495.44899 | 10280.44126 | 0.308424885 | 0.061958038 | 3.29088037688 | 4.56048185932 | ns | | NaCl | KS.1 |
| FIM1_2227 | 1682.731394 | 3605.972655 | 2047.478626 | 0.976260756 | 0.19735588 | 3.15832117269 | 5.99719382983 | ns | | NaCl | KS.1 |
| FIM1_2228 | 958.1338289 | 968.9653681 | 941.2334906 | 0.009042282 | 0.254239141 | 0.961854841 | 0.981455879 | ns | | NaCl | KS.1 |
| FIM1_2229 | 1136.119483 | 1154.391291 | 888.5419249 | 0.016166378 | 0.184356547 | 0.917373067 | 0.952803969 | ns | | NaCl | KS.1 |
| FIM1_2230 | 283.3866285 | 238.658999 | 429.8878394 | -0.155624255 | 0.227940699 | 0.384421078 | 0.519785037 | ns | | NaCl | KS.1 |
| FIM1_2231 | 848.4731435 | 818.9155318 | 934.2259976 | -0.049422884 | 0.097036377 | 0.59159939 | 0.712295974 | ns | | NaCl | KS.1 |
| FIM1_2232 | 365.9935353 | 413.5398425 | 872.8070102 | 0.114557854 | 0.216789539 | 0.508920736 | 0.639848537 | ns | | NaCl | KS.1 |
| FIM1_2233 | 2342.648009 | 2479.04376 | 1744.585935 | 0.068092327 | 0.133863664 | 0.576088555 | 0.698057811 | ns | | NaCl | KS.1 |
| FIM1_2234 | 444.0474466 | 337.2515925 | 525.7881748 | -0.355482064 | 0.12112453 | 0.00141245 | 0.005676042 | ns | | NaCl | KS.1 |
| FIM1_2235 | 515.1347527 | 631.1071653 | 542.9644863 | 0.2188672 | 0.183106857 | 0.157830047 | 0.265720637 | ns | | NaCl | KS.1 |
| FIM1_2236 | 1017.079192 | 1079.439345 | 1267.879751 | 0.072253424 | 0.128352945 | 0.53891184 | 0.665915914 | ns | | NaCl | KS.1 |
| FIM1_2237 | 31.24077235 | 18.11514785 | 40.526282 | -0.428636989 | 0.335968932 | 0.055496994 | 0.115280812 | ns | | NaCl | KS.1 |
| FIM1_2238 | 736.4274638 | 639.9252243 | 875.0395528 | -0.167299051 | 0.140259448 | 0.185360074 | 0.301361557 | ns | | NaCl | KS.1 |
| FIM1_2239 | 660.9884068 | 476.6695395 | 707.4400088 | -0.402101782 | 0.158067154 | 0.003790703 | 0.01299707 | ns | | NaCl | KS.1 |
| FIM1_2240 | 501.9905604 | 506.9549837 | 670.9627868 | 0.011137417 | 0.151563517 | 0.934091652 | 0.962133741 | ns | | NaCl | KS.1 |
| FIM1_2241 | 2613.26117 | 2434.849333 | 2657.322158 | -0.075702553 | 0.174100799 | 0.612712031 | 0.729070163 | ns | | NaCl | KS.1 |
| FIM1_2243 | 1429.573696 | 1396.821612 | 1190.464514 | -0.029328571 | 0.129224669 | 0.804203671 | 0.87489903 | ns | | NaCl | KS.1 |
| FIM1_2244 | 5000.024329 | 6175.557828 | 5009.643501 | 0.267255624 | 0.122302574 | 0.017427183 | 0.044325275 | ns | | NaCl | KS.1 |
| FIM1_2245 | 1120.456038 | 1834.20748 | 1697.103408 | 0.612191511 | 0.181191317 | 0.000101035 | 0.000629096 | ns | | NaCl | KS.1 |
| FIM1_2246 | 13404.71 | 16643.69206 | 11412.85161 | 0.290085547 | 0.090654648 | 0.000766948 | 0.003449879 | ns | | NaCl | KS.1 |
| FIM1_2247 | 869.3752208 | 1075.59011 | 1103.334595 | 0.277061206 | 0.110753092 | 0.00722944 | 0.022065316 | ns | | NaCl | KS.1 |
| FIM1_2248 | 2829.485713 | 6489.765045 | 2883.141521 | 0.875072411 | 0.320737825 | 0.000283699 | 0.001500369 | ns | | NaCl | KS.1 |
| FIM1_2249 | 4449.281444 | 9020.821151 | 4456.848732 | 0.767401748 | 0.28411241 | 0.000498712 | 0.00239951 | ns | | NaCl | KS.1 |
| FIM1_2250 | 365.7907653 | 833.2344684 | 393.1173166 | 0.900992244 | 0.302920064 | 0.000122864 | 0.000741808 | ns | | NaCl | KS.1 |
| FIM1_2251 | 484.4012059 | 502.4958017 | 559.9824934 | 0.048987434 | 0.100744393 | 0.607146523 | 0.724527213 | ns | | NaCl | KS.1 |

| | | | | | | | | | |
|---|---|---|---|---|---|---|---|---|---|
| FIM1_2252 | 3192.301387 | 3221.231359 | 3579.467591 | 0.011258478 | 0.134535638 | 0.926677685 | 0.957669606 | ns | NaCl | KS.1 |
| FIM1_2253 | 16088.10076 | 23395.47389 | 15903.47299 | 0.522835386 | 0.073774806 | 2.4600723169 | 1.7737477937 | ns | NaCl | KS.1 |
| FIM1_2254 | 2131.145974 | 2700.356538 | 2607.848379 | 0.28024228 | 0.155387278 | 0.040558213 | 0.089266467 | ns | NaCl | KS.1 |
| FIM1_2255 | 14562.79094 | 18643.55417 | 13851.80021 | 0.343437068 | 0.066036879 | 8.6529828516 | 1.4022341917 | ns | NaCl | KS.1 |
| FIM1_2256 | 605.3459318 | 747.7232515 | 1342.415325 | 0.230750795 | 0.183391054 | 0.136900733 | 0.23701727 | ns | NaCl | KS.1 |
| FIM1_2257 | 342.2457467 | 359.1314037 | 1340.549922 | 0.035027962 | 0.287361446 | 0.860301246 | 0.915507743 | ns | NaCl | KS.1 |
| FIM1_2258 | 3132.299277 | 2700.551634 | 2130.826195 | -0.172237599 | 0.154303523 | 0.206387314 | 0.327869075 | ns | NaCl | KS.1 |
| FIM1_2259 | 462.5613925 | 376.2707869 | 421.2064794 | -0.249984472 | 0.145080792 | 0.053742545 | 0.112057485 | ns | NaCl | KS.1 |
| FIM1_2260 | 1594.704614 | 1740.785222 | 1913.190993 | 0.112863135 | 0.106242348 | 0.258171733 | 0.386598155 | ns | NaCl | KS.1 |
| FIM1_2261 | 2515.054616 | 4120.223878 | 3325.419316 | 0.669421872 | 0.117035493 | 1.2062941252 | 3.3715243107 | ns | NaCl | KS.1 |
| FIM1_2262 | 3132.76995 | 2355.596507 | 2968.684737 | -0.3118136 | 0.193200664 | 0.05306672 | 0.110880693 | ns | NaCl | KS.1 |
| FIM1_2263 | 3857.71998 | 4377.306585 | 4218.55243 | 0.150183089 | 0.141550124 | 0.238127701 | 0.364743016 | ns | NaCl | KS.1 |
| FIM1_2264 | 1228.093325 | 1424.953848 | 1340.425027 | 0.187374881 | 0.116777421 | 0.083440507 | 0.160711003 | ns | NaCl | KS.1 |
| FIM1_2265 | 1064.263666 | 1060.134065 | 992.9188256 | -0.005216798 | 0.102608181 | 0.957052531 | 0.978088813 | ns | NaCl | KS.1 |
| FIM1_2266 | 1948.478948 | 1850.858749 | 1334.817771 | -0.060082692 | 0.140731688 | 0.635870067 | 0.747025032 | ns | NaCl | KS.1 |
| FIM1_2267 | 1499.839333 | 1827.346025 | 1619.757706 | 0.242103229 | 0.133795563 | 0.045951131 | 0.098494992 | ns | NaCl | KS.1 |
| FIM1_2268 | 1330.158037 | 1704.054014 | 1336.216251 | 0.325744547 | 0.104515215 | 0.000880187 | 0.003858908 | ns | NaCl | KS.1 |
| FIM1_2270 | 806.1812023 | 1126.1812 | 840.3921184 | 0.375257876 | 0.19424334 | 0.020693741 | 0.051067144 | ns | NaCl | KS.1 |
| FIM1_2271 | 1379.68775 | 1535.313438 | 1494.176021 | 0.129304218 | 0.12474941 | 0.259558676 | 0.388012143 | ns | NaCl | KS.1 |
| FIM1_2272 | 545.5482316 | 280.7646115 | 737.2469833 | -0.847937614 | 0.185107671 | 2.9818933593 | 4.2144657564 | ns | NaCl | KS.1 |
| FIM1_2273 | 141.8633019 | 78.91761045 | 169.1592428 | -0.613821583 | 0.266387358 | 0.002896313 | 0.010402776 | ns | NaCl | KS.1 |
| FIM1_2274 | 1837.65142 | 1163.276927 | 2250.51838 | -0.487220595 | 0.238940286 | 0.009846806 | 0.028202568 | ns | NaCl | KS.1 |
| FIM1_2275 | 2483.00594 | 1980.457004 | 2090.895444 | -0.256523251 | 0.169234042 | 0.07929261 | 0.153973746 | ns | NaCl | KS.1 |
| FIM1_2276 | 1253.206061 | 894.9685846 | 871.5900199 | -0.422452287 | 0.14816461 | 0.00136775 | 0.005523177 | ns | NaCl | KS.1 |
| FIM1_2278 | 1329.520249 | 1722.886512 | 1870.660216 | 0.278535203 | 0.195099761 | 0.085916612 | 0.16452469 | ns | NaCl | KS.1 |
| FIM1_2279 | 332.6959547 | 284.1320046 | 338.5058139 | -0.200098767 | 0.135406141 | 0.102736263 | 0.188881341 | ns | NaCl | KS.1 |
| FIM1_2280 | 1630.61857 | 1883.126984 | 1950.211584 | 0.185725941 | 0.10736285 | 0.064833493 | 0.130532833 | ns | NaCl | KS.1 |
| FIM1_2281 | 742.1714078 | 460.5142561 | 749.2942018 | -0.571374132 | 0.200491158 | 0.000721548 | 0.003281143 | ns | NaCl | KS.1 |
| FIM1_2282 | 279.3859019 | 113.0072041 | 305.8777883 | -1.206005236 | 0.169831254 | 1.9341064901 | 1.7494872342 | Km<KS.1 | NaCl | KS.1 |
| FIM1_2283 | 987.086815 | 707.221977 | 1312.087633 | -0.425245082 | 0.137672153 | 0.000619633 | 0.002889102 | ns | NaCl | KS.1 |
| FIM1_2284 | 2025.606171 | 1813.047591 | 2494.53559 | -0.111542026 | 0.195907091 | 0.49232332 | 0.624664248 | ns | NaCl | KS.1 |
| FIM1_2285 | 2847.495041 | 3706.259798 | 4091.590869 | 0.305480669 | 0.168946213 | 0.036405949 | 0.081769569 | ns | NaCl | KS.1 |
| FIM1_2286 | 955.8309751 | 1003.704824 | 1032.886163 | 0.05920908 | 0.1266195 | 0.610580223 | 0.726881218 | ns | NaCl | KS.1 |
| FIM1_2287 | 704.031505 | 664.5879388 | 927.4687669 | -0.070949505 | 0.135797721 | 0.564795671 | 0.687511247 | ns | NaCl | KS.1 |
| FIM1_2288 | 1712.382178 | 2322.034766 | 1886.171791 | 0.395223993 | 0.122044636 | 0.000423616 | 0.002101186 | ns | NaCl | KS.1 |
| FIM1_2289 | 3593.063764 | 5221.144918 | 3305.0036 | 0.471004643 | 0.150255702 | 0.000431476 | 0.002133789 | ns | NaCl | KS.1 |
| FIM1_2290 | 1396.883444 | 3028.37839 | 2575.129183 | 1.002916605 | 0.188935704 | 4.3134846261 | 1.0519404909 | Km>KS.1 | NaCl | KS.1 |
| FIM1_2291 | 3108.924641 | 5058.537539 | 3516.64279 | 0.623755983 | 0.160655638 | 1.3626264117 | 0.000112236 | ns | NaCl | KS.1 |
| FIM1_2292 | 1060.617711 | 1047.464443 | 872.7159229 | -0.014532411 | 0.158511141 | 0.916998428 | 0.952803969 | ns | NaCl | KS.1 |
| FIM1_2293 | 687.087357 | 635.207336 | 613.4994378 | -0.082456574 | 0.183774591 | 0.595823361 | 0.71564974 | ns | NaCl | KS.1 |
| FIM1_2294 | 10052.15009 | 13446.51787 | 9396.152219 | 0.40089538 | 0.077535357 | 7.7012094113 | 1.2620386998 | ns | NaCl | KS.1 |
| FIM1_2295 | 733.0272524 | 795.9695656 | 746.0783644 | 0.093095941 | 0.151956221 | 0.490002403 | 0.622672274 | ns | NaCl | KS.1 |
| FIM1_2297 | 1058.823018 | 1214.532764 | 1216.926502 | 0.176097076 | 0.101281157 | 0.065287597 | 0.131128703 | ns | NaCl | KS.1 |
| FIM1_2298 | 1125.006837 | 1182.135674 | 1665.205528 | 0.053515008 | 0.16105047 | 0.704538668 | 0.801710859 | ns | NaCl | KS.1 |
| FIM1_2299 | 1313.867374 | 1361.077059 | 1448.848662 | 0.040263952 | 0.146044047 | 0.758141653 | 0.843194545 | ns | NaCl | KS.1 |

| | | | | | | | | | | |
|---|---|---|---|---|---|---|---|---|---|---|
| FIM1_2300 | 393.1706998 | 375.2119732 | 538.300696 | -0.058350019 | 0.151184011 | 0.664122016 | 0.770524027 | ns | NaCl | KS.1 |
| FIM1_2301 | 2497.527711 | 3334.664608 | 2796.687825 | 0.321411967 | 0.188497149 | 0.042460762 | 0.092475236 | ns | NaCl | KS.1 |
| FIM1_2302 | 2631.814156 | 2679.248541 | 3328.172383 | 0.022143553 | 0.106018145 | 0.824279786 | 0.89225238 | ns | NaCl | KS.1 |
| FIM1_2303 | 227.5930391 | 247.9836454 | 302.1341551 | 0.081922461 | 0.206925041 | 0.626601324 | 0.739711913 | ns | NaCl | KS.1 |
| FIM1_2304 | 246.7052189 | 148.8264625 | 326.5847066 | -0.605326972 | 0.197037735 | 0.00030427 | 0.001590066 | ns | NaCl | KS.1 |
| FIM1_2305 | 674.4705801 | 561.5279775 | 762.5268359 | -0.246068224 | 0.099730623 | 0.008900316 | 0.026171466 | ns | NaCl | KS.1 |
| FIM1_2306 | 208.0427396 | 54.78276326 | 275.4655839 | -1.806235511 | 0.222111057 | 2.43469422145 | 5.93607347414 | Km<KS.1 | NaCl | KS.1 |
| FIM1_2307 | 3420.730812 | 3527.958544 | 2506.191909 | 0.039025146 | 0.110250662 | 0.705373923 | 0.802110918 | ns | NaCl | KS.1 |
| FIM1_2308 | 2251.214249 | 3083.915569 | 1907.014888 | 0.341867931 | 0.201358043 | 0.039617183 | 0.087597994 | ns | NaCl | KS.1 |
| FIM1_2309 | 255.1564106 | 131.5400575 | 295.1554919 | -0.707255633 | 0.280753837 | 0.001081306 | 0.004582195 | ns | NaCl | KS.1 |
| FIM1_2310 | 368.9951508 | 467.9128384 | 334.031678 | 0.249075894 | 0.201693798 | 0.133180325 | 0.231993039 | ns | NaCl | KS.1 |
| FIM1_2311 | 1147.762293 | 1408.788989 | 1672.056037 | 0.184729299 | 0.237988607 | 0.314497361 | 0.447675071 | ns | NaCl | KS.1 |
| FIM1_2312 | 5127.634647 | 5623.201908 | 4768.238598 | 0.119775873 | 0.100417788 | 0.206976345 | 0.328455284 | ns | NaCl | KS.1 |
| FIM1_2313 | 1262.328858 | 2278.892226 | 1339.623606 | 0.765427754 | 0.168581334 | 4.69536090534 | 6.16343548920 | ns | NaCl | KS.1 |
| FIM1_2314 | 1185.239848 | 738.9697298 | 1073.429431 | -0.653170463 | 0.106047432 | 8.64934106810 | 3.21122923983 | ns | NaCl | KS.1 |
| FIM1_2315 | 735.9890464 | 476.65299 | 1049.537091 | -0.487068302 | 0.216245522 | 0.005798419 | 0.018539288 | ns | NaCl | KS.1 |
| FIM1_2316 | 743.2119619 | 761.2907608 | 886.4830386 | 0.023771312 | 0.20136214 | 0.885736754 | 0.93200938 | ns | NaCl | KS.1 |
| FIM1_2317 | 1771.809329 | 2022.562712 | 2337.312787 | 0.162932244 | 0.126463717 | 0.159906458 | 0.268580226 | ns | NaCl | KS.1 |
| FIM1_2318 | 22776.83671 | 20110.38196 | 13391.12303 | -0.134551416 | 0.174732673 | 0.36929512 | 0.504073076 | ns | NaCl | KS.1 |
| FIM1_2319 | 114.0297864 | 129.9856451 | 164.3750103 | 0.120333785 | 0.215024052 | 0.485624848 | 0.619500146 | ns | NaCl | KS.1 |
| FIM1_2320 | 243.0767897 | 279.8528298 | 214.5370624 | 0.163666512 | 0.140404214 | 0.19549885 | 0.313946668 | ns | NaCl | KS.1 |
| FIM1_2321 | 1539.861658 | 1860.943643 | 1675.237695 | 0.259258962 | 0.070357908 | 0.000143593 | 0.000846417 | ns | NaCl | KS.1 |
| FIM1_2322 | 961.1977667 | 669.0685246 | 811.9243081 | -0.474504693 | 0.128889167 | 5.61913986175 | 0.000384 | ns | NaCl | KS.1 |
| FIM1_2323 | 3300.837759 | 4217.017104 | 3024.517997 | 0.315246403 | 0.117532538 | 0.003680387 | 0.012697591 | ns | NaCl | KS.1 |
| FIM1_2324 | 5003.836543 | 4571.616659 | 3890.555805 | -0.103148029 | 0.154243981 | 0.449618144 | 0.584610676 | ns | NaCl | KS.1 |
| FIM1_2325 | 38.13428307 | 25.10824459 | 40.96022622 | -0.397175525 | 0.268862382 | 0.047497019 | 0.101241504 | ns | NaCl | KS.1 |
| FIM1_2326 | 395.6012461 | 364.5828714 | 442.7879753 | -0.107411972 | 0.128698034 | 0.36182964 | 0.496985727 | ns | NaCl | KS.1 |
| FIM1_2327 | 2854.769471 | 2881.902919 | 3239.293127 | 0.012840641 | 0.092780631 | 0.88458783 | 0.931391419 | ns | NaCl | KS.1 |
| FIM1_2328 | 7003.730934 | 10238.96642 | 7241.459513 | 0.523428645 | 0.08822893 | 5.46497357656 | 1.63784599659 | ns | NaCl | KS.1 |
| FIM1_2329 | 299.7669341 | 347.9734661 | 371.0631883 | 0.157076546 | 0.190718757 | 0.324711951 | 0.457891711 | ns | NaCl | KS.1 |
| FIM1_2330 | 524.7556299 | 580.3392996 | 471.7337333 | 0.104616476 | 0.185022497 | 0.502963362 | 0.634925838 | ns | NaCl | KS.1 |
| FIM1_2331 | 2447.703665 | 3352.058187 | 2684.208701 | 0.355441485 | 0.187561209 | 0.02447651 | 0.058968832 | ns | NaCl | KS.1 |
| FIM1_2332 | 6758.598031 | 11707.58256 | 8886.548113 | 0.696419333 | 0.177042473 | 8.56002270972 | 7.47124789138 | ns | NaCl | KS.1 |
| FIM1_2333 | 306.9789007 | 382.9794288 | 387.8348301 | 0.254259726 | 0.168049002 | 0.080363514 | 0.155728297 | ns | NaCl | KS.1 |
| FIM1_2334 | 544.1772055 | 392.1506418 | 624.1776974 | -0.390759761 | 0.167383113 | 0.007123602 | 0.021813375 | ns | NaCl | KS.1 |
| FIM1_2335 | 7987.579849 | 6803.099802 | 6406.854433 | -0.182889488 | 0.160248394 | 0.192527341 | 0.309874966 | ns | NaCl | KS.1 |
| FIM1_2336 | 451.1676396 | 730.8214338 | 1689.305228 | 0.5263563 | 0.235645719 | 0.005118136 | 0.016577296 | ns | NaCl | KS.1 |
| FIM1_2337 | 138.3786695 | 187.2974688 | 216.0073424 | 0.298628874 | 0.229217509 | 0.098350086 | 0.182299432 | ns | NaCl | KS.1 |
| FIM1_2338 | 1978.571955 | 1753.32673 | 1538.303557 | -0.15940238 | 0.098540016 | 0.087370553 | 0.1667313 | ns | NaCl | KS.1 |
| FIM1_2339 | 413.0667208 | 350.646766 | 640.8000692 | -0.198700566 | 0.138169879 | 0.111033008 | 0.200888413 | ns | NaCl | KS.1 |
| FIM1_2340 | 1308.781438 | 901.7549766 | 1242.990799 | -0.41663665 | 0.204011668 | 0.013357606 | 0.035921129 | ns | NaCl | KS.1 |
| FIM1_2341 | 652.8123047 | 401.3557893 | 716.9177179 | -0.585563706 | 0.197313157 | 0.000465969 | 0.002270514 | ns | NaCl | KS.1 |
| FIM1_2342 | 2091.330987 | 1793.059246 | 1706.408292 | -0.184571249 | 0.138757323 | 0.140309989 | 0.241370053 | ns | NaCl | KS.1 |
| FIM1_2343 | 53.53264562 | 17.95588163 | 62.07038615 | -1.067415012 | 0.409467285 | 0.000152265 | 0.000887022 | Km<KS.1 | NaCl | KS.1 |
| FIM1_2344 | 213.8163216 | 214.8541074 | 771.224789 | 0.00995765 | 0.210233011 | 0.953303263 | 0.975932871 | ns | NaCl | KS.1 |

| | | | | | | | | | | |
|---|---|---|---|---|---|---|---|---|---|---|
| FIM1_2345 | 771.2249961 | 585.1827968 | 917.6130562 | -0.335425047 | 0.15421675 | 0.013657755 | 0.036550474 | ns | NaCl | KS.1 |
| FIM1_2346 | 3373.158427 | 4492.231852 | 4009.822228 | 0.372051621 | 0.118309077 | 0.000651893 | 0.003005717 | ns | NaCl | KS.1 |
| FIM1_2347 | 709.1532985 | 684.7006568 | 620.2654514 | -0.040651716 | 0.161642682 | 0.773939351 | 0.853357329 | ns | NaCl | KS.1 |
| FIM1_2348 | 3173.394311 | 3795.390848 | 2895.085466 | 0.233842892 | 0.102147976 | 0.014992108 | 0.039131242 | ns | NaCl | KS.1 |
| FIM1_2349 | 1607.155213 | 1588.986088 | 2302.086355 | -0.017164271 | 0.089397803 | 0.840908251 | 0.903028347 | ns | NaCl | KS.1 |
| FIM1_2350 | 2937.243334 | 2791.276543 | 2886.681087 | -0.063361526 | 0.114300074 | 0.55188879 | 0.676085411 | ns | NaCl | KS.1 |
| FIM1_2351 | 997.1656502 | 962.3962494 | 1167.699938 | -0.036679457 | 0.179159367 | 0.810174733 | 0.880240073 | ns | NaCl | KS.1 |
| FIM1_2352 | 740.7886627 | 894.5660297 | 1125.829973 | 0.19048693 | 0.202925465 | 0.252309746 | 0.380833267 | ns | NaCl | KS.1 |
| FIM1_2353 | 1193.484568 | 840.7220228 | 972.6931714 | -0.460563606 | 0.127740533 | 8.0605045067 | 0.000520362 | ns | NaCl | KS.1 |
| FIM1_2354 | 377.8917291 | 375.8430519 | 595.0128552 | -0.005346565 | 0.170811354 | 0.971087672 | 0.986958359 | ns | NaCl | KS.1 |
| FIM1_2355 | 286.6096883 | 168.504362 | 214.1355932 | -0.637814031 | 0.211400896 | 0.000321121 | 0.001657236 | ns | NaCl | KS.1 |
| FIM1_2356 | 323.674535 | 216.0842784 | 428.712158 | -0.385320101 | 0.258295155 | 0.048864684 | 0.103756638 | ns | NaCl | KS.1 |
| FIM1_2357 | 2159.345481 | 3964.754427 | 2252.726222 | 0.74076584 | 0.209639503 | 3.5381873403 | 0.000256596 | ns | NaCl | KS.1 |
| FIM1_2358 | 1304.794979 | 1032.71958 | 1063.905746 | -0.304861629 | 0.113577861 | 0.003818028 | 0.013081743 | ns | NaCl | KS.1 |
| FIM1_2359 | 5020.497315 | 6161.800469 | 4212.938516 | 0.272126609 | 0.095359627 | 0.002581888 | 0.009479625 | ns | NaCl | KS.1 |
| FIM1_2360 | 1927.413935 | 1894.61683 | 2317.011115 | -0.017652348 | 0.168340215 | 0.903719928 | 0.942559044 | ns | NaCl | KS.1 |
| FIM1_2361 | 3548.306545 | 3340.017757 | 3183.044409 | -0.079821988 | 0.089708393 | 0.351889643 | 0.486779562 | ns | NaCl | KS.1 |
| FIM1_2362 | 1485.011163 | 1788.511024 | 1599.294081 | 0.245540994 | 0.096579573 | 0.007218552 | 0.022045608 | ns | NaCl | KS.1 |
| FIM1_2363 | 2169.969931 | 2257.6323 | 2113.157998 | 0.045506279 | 0.143581915 | 0.724236356 | 0.816838783 | ns | NaCl | KS.1 |
| FIM1_2364 | 2494.515891 | 2646.422365 | 2512.202957 | 0.078007621 | 0.0806491 | 0.315510723 | 0.448604128 | ns | NaCl | KS.1 |
| FIM1_2365 | 22363.24397 | 32505.7884 | 19621.02589 | 0.486098296 | 0.132074628 | 5.3585878101 | 0.000368727 | ns | NaCl | KS.1 |
| FIM1_2366 | 452.4099417 | 397.4722046 | 568.7596446 | -0.15116698 | 0.149508908 | 0.255765197 | 0.384188361 | ns | NaCl | KS.1 |
| FIM1_2367 | 487.3111432 | 589.2760027 | 488.8553682 | 0.198244753 | 0.192417495 | 0.216741859 | 0.340262148 | ns | NaCl | KS.1 |
| FIM1_2368 | 1402.683717 | 1996.811708 | 2404.573716 | 0.403490166 | 0.191157838 | 0.01197678 | 0.032956018 | ns | NaCl | KS.1 |
| FIM1_2369 | 954.3926603 | 883.6525025 | 1138.461136 | -0.094898299 | 0.130933346 | 0.427217458 | 0.563768396 | ns | NaCl | KS.1 |
| FIM1_2370 | 325.0060122 | 385.0185608 | 394.0827039 | 0.14040598 | 0.250959488 | 0.456601428 | 0.591560444 | ns | NaCl | KS.1 |
| FIM1_2371 | 2621.887299 | 2259.003674 | 4375.944249 | -0.126808954 | 0.245261339 | 0.495932478 | 0.62812222 | ns | NaCl | KS.1 |
| FIM1_2372 | 6932.661062 | 6730.631554 | 4975.428215 | -0.036312524 | 0.124834557 | 0.751982489 | 0.838250702 | ns | NaCl | KS.1 |
| FIM1_2373 | 367.5478069 | 67.43488219 | 648.7126715 | -2.161983261 | 0.393637854 | 1.6114095092 | 7.2230838607 | Km<KS.1 | NaCl | KS.1 |
| FIM1_2374 | 1149.345533 | 887.7735329 | 1107.153194 | -0.332899897 | 0.119050046 | 0.002415147 | 0.00893335 | ns | NaCl | KS.1 |
| FIM1_2375 | 1700.119479 | 1805.655276 | 1708.348738 | 0.074112211 | 0.121990321 | 0.510538585 | 0.641072554 | ns | NaCl | KS.1 |
| FIM1_2376 | 1188.052209 | 1272.156862 | 1376.278378 | 0.082822252 | 0.13195836 | 0.491251746 | 0.623941138 | ns | NaCl | KS.1 |
| FIM1_2377 | 8048.429916 | 13523.56657 | 9042.424857 | 0.700951306 | 0.123947686 | 1.6168775336 | 4.4197613900 | ns | NaCl | KS.1 |
| FIM1_2378 | 732.7277626 | 577.4199234 | 689.8052412 | -0.282990084 | 0.159198412 | 0.042435507 | 0.092475236 | ns | NaCl | KS.1 |
| FIM1_2379 | 1536.078377 | 1845.519037 | 1764.115152 | 0.228331336 | 0.124575557 | 0.045724907 | 0.098136935 | ns | NaCl | KS.1 |
| FIM1_2380 | 882.5198068 | 884.814093 | 815.8202954 | 0.001921877 | 0.160002565 | 0.989082483 | 0.995082984 | ns | NaCl | KS.1 |
| FIM1_2381 | 1716.982389 | 1926.079264 | 1661.261604 | 0.153032969 | 0.086640994 | 0.065082004 | 0.130821403 | ns | NaCl | KS.1 |
| FIM1_2382 | 549.4696593 | 581.790515 | 593.4255858 | 0.069609626 | 0.140477434 | 0.582716912 | 0.704328629 | ns | NaCl | KS.1 |
| FIM1_2383 | 618.0857415 | 543.4725024 | 893.42857 | -0.157545366 | 0.124558125 | 0.168724733 | 0.280175416 | ns | NaCl | KS.1 |
| FIM1_2384 | 10861.39964 | 16202.92646 | 9984.682313 | 0.538489112 | 0.11124061 | 2.2943768033 | 3.3375802914 | ns | NaCl | KS.1 |
| FIM1_2385 | 933.2635284 | 829.6806769 | 919.9138713 | -0.155709089 | 0.089893911 | 0.069574291 | 0.1383563 | ns | NaCl | KS.1 |
| FIM1_2386 | 607.2502058 | 302.9693422 | 686.0131183 | -0.802941066 | 0.255802394 | 0.00011131 | 0.000682822 | ns | NaCl | KS.1 |
| FIM1_2387 | 390.5203258 | 412.0299522 | 480.6728324 | 0.0648726 | 0.112270045 | 0.536102532 | 0.663658129 | ns | NaCl | KS.1 |
| FIM1_2388 | 8006.074114 | 7175.980631 | 6074.465123 | -0.145212928 | 0.089964863 | 0.090876847 | 0.171775196 | ns | NaCl | KS.1 |
| FIM1_2389 | 1349.1927 | 2192.971791 | 1980.459844 | 0.615773951 | 0.167956924 | 3.3524113675 | 0.000244191 | ns | NaCl | KS.1 |

| | | | | | | | | | | |
|---|---|---|---|---|---|---|---|---|---|---|
| FIM1_2390 | 1011.328957 | 825.0710075 | 1057.863545 | -0.270782012 | 0.094883721 | 0.002593773 | 0.009509227 | ns | NaCl | KS.1 |
| FIM1_2391 | 1237.544202 | 1405.932914 | 1136.747479 | 0.145176848 | 0.159137186 | 0.298648055 | 0.431284201 | ns | NaCl | KS.1 |
| FIM1_2392 | 997.9356874 | 1107.826521 | 1043.287542 | 0.11419001 | 0.168536342 | 0.4340049 | 0.570154311 | ns | NaCl | KS.1 |
| FIM1_2393 | 341.8578227 | 336.2678987 | 366.9729904 | -0.017643643 | 0.195522385 | 0.913398113 | 0.949666794 | ns | NaCl | KS.1 |
| FIM1_2394 | 541.5826721 | 238.7862737 | 1032.077971 | -1.055818469 | 0.203229221 | 6.25729368058 | 1.44120537318 | Km<KS.1 | NaCl | KS.1 |
| FIM1_2395 | 16628.31226 | 31964.48794 | 17822.67553 | 0.907354269 | 0.104496074 | 2.50567925541 | 5.93607347414 | ns | NaCl | KS.1 |
| FIM1_2396 | 2418.858316 | 2570.88054 | 2557.179755 | 0.080744183 | 0.091220095 | 0.353710669 | 0.488708968 | ns | NaCl | KS.1 |
| FIM1_2397 | 2009.695305 | 2187.561829 | 3026.433657 | 0.106844374 | 0.107115637 | 0.287965745 | 0.418892485 | ns | NaCl | KS.1 |
| FIM1_2398 | 2007.656446 | 1359.786666 | 2011.486547 | -0.493425847 | 0.149899055 | 0.000225095 | 0.001230602 | ns | NaCl | KS.1 |
| FIM1_2399 | 724.2513152 | 470.0557748 | 828.3418785 | -0.544689461 | 0.159570124 | 0.0001162 | 0.000705856 | ns | NaCl | KS.1 |
| FIM1_2400 | 443.2303467 | 308.050142 | 510.3101681 | -0.448920965 | 0.165481682 | 0.00186856 | 0.007228681 | ns | NaCl | KS.1 |
| FIM1_2401 | 722.0020201 | 673.2461682 | 714.9424682 | -0.083673143 | 0.136764587 | 0.499583142 | 0.631620364 | ns | NaCl | KS.1 |
| FIM1_2402 | 1564.186827 | 1973.591712 | 1613.534404 | 0.292339845 | 0.127961567 | 0.0122804 | 0.033513433 | ns | NaCl | KS.1 |
| FIM1_2403 | 1442.321669 | 1913.020428 | 2146.319139 | 0.368009268 | 0.115264878 | 0.000562512 | 0.002663336 | ns | NaCl | KS.1 |
| FIM1_2404 | 6485.861476 | 6650.876938 | 5503.319899 | 0.030568871 | 0.121334717 | 0.785201534 | 0.861384263 | ns | NaCl | KS.1 |
| FIM1_2405 | 1976.164284 | 2174.071124 | 1827.205237 | 0.121172032 | 0.110032292 | 0.239108928 | 0.365795485 | ns | NaCl | KS.1 |
| FIM1_2406 | 5542.526352 | 8677.761891 | 8854.810501 | 0.431154089 | 0.267374926 | 0.031692898 | 0.072945429 | ns | NaCl | KS.1 |
| FIM1_2407 | 1320.883319 | 2514.491935 | 1434.130017 | 0.813349299 | 0.192826993 | 1.72790000447 | 1.89388397260 | ns | NaCl | KS.1 |
| FIM1_2408 | 291.8717488 | 188.6810681 | 317.70967 | -0.495665392 | 0.211397833 | 0.004375241 | 0.01459881 | ns | NaCl | KS.1 |
| FIM1_2409 | 727.8992796 | 481.2676619 | 558.8150676 | -0.52461327 | 0.15782103 | 0.000175251 | 0.001001002 | ns | NaCl | KS.1 |
| FIM1_2410 | 5285.198082 | 7161.125097 | 5763.511942 | 0.401708774 | 0.110020593 | 8.84883225525 | 0.000563674 | ns | NaCl | KS.1 |
| FIM1_2411 | 598.0202998 | 670.9677519 | 686.7980707 | 0.136799331 | 0.134665131 | 0.26313833 | 0.391598323 | ns | NaCl | KS.1 |
| FIM1_2412 | 668.9507651 | 711.3964128 | 600.7828512 | 0.065092537 | 0.169808304 | 0.657559488 | 0.764514712 | ns | NaCl | KS.1 |
| FIM1_2413 | 1483.258759 | 2207.215412 | 2181.1896 | 0.47945167 | 0.17716822 | 0.001667046 | 0.006571754 | ns | NaCl | KS.1 |
| FIM1_2414 | 1511.831668 | 1256.720723 | 1461.678195 | -0.231407968 | 0.12456878 | 0.042858172 | 0.093027665 | ns | NaCl | KS.1 |
| FIM1_2415 | 897.8654655 | 1097.81612 | 948.249549 | 0.251168216 | 0.126530084 | 0.029996615 | 0.06967001 | ns | NaCl | KS.1 |
| FIM1_2416 | 3905.912766 | 4452.83073 | 4098.833739 | 0.173843758 | 0.087848735 | 0.038460337 | 0.085802289 | ns | NaCl | KS.1 |
| FIM1_2417 | 886.8939529 | 843.0195092 | 1140.405553 | -0.063289555 | 0.114191298 | 0.552011848 | 0.676085411 | ns | NaCl | KS.1 |
| FIM1_2418 | 14227.14228 | 20843.49122 | 15167.9165 | 0.532594747 | 0.075743046 | 3.52637873940 | 2.40325126418 | ns | NaCl | KS.1 |
| FIM1_2419 | 773.9472159 | 860.0854525 | 811.3745993 | 0.117159048 | 0.16330185 | 0.411071311 | 0.546520516 | ns | NaCl | KS.1 |
| FIM1_2420 | 1327.119144 | 1908.839605 | 1741.522806 | 0.467268984 | 0.13861717 | 0.000189514 | 0.001067759 | ns | NaCl | KS.1 |
| FIM1_2422 | 3173.554078 | 4072.008405 | 4611.739681 | 0.340507486 | 0.077325458 | 4.74796190391 | 4.56005993667 | ns | NaCl | KS.1 |
| FIM1_2423 | 469.2532506 | 479.0686808 | 573.2406725 | 0.020267145 | 0.188814421 | 0.89821243 | 0.938980214 | ns | NaCl | KS.1 |
| FIM1_2424 | 1140.139644 | 870.3043474 | 1761.714842 | -0.313010266 | 0.168771859 | 0.031912374 | 0.073366018 | ns | NaCl | KS.1 |
| FIM1_2425 | 575.4179933 | 599.9539175 | 549.0309316 | 0.042769407 | 0.183608169 | 0.783117885 | 0.859667139 | ns | NaCl | KS.1 |
| FIM1_2426 | 2088.650571 | 1981.468331 | 3547.211572 | -0.065523036 | 0.122796646 | 0.562978293 | 0.685634518 | ns | NaCl | KS.1 |
| FIM1_2427 | 206.3097153 | 234.1707663 | 261.1951881 | 0.13732143 | 0.1691193 | 0.347896901 | 0.482785797 | ns | NaCl | KS.1 |
| FIM1_2428 | 342.3526744 | 165.3663332 | 668.4622509 | -0.800357116 | 0.283642938 | 0.000303713 | 0.001588822 | ns | NaCl | KS.1 |
| FIM1_2429 | 1286.60094 | 1060.362036 | 1186.307702 | -0.266644595 | 0.085156411 | 0.001062672 | 0.00450707 | ns | NaCl | KS.1 |
| FIM1_2430 | 1198.573968 | 1162.238434 | 1665.928015 | -0.040137253 | 0.111412635 | 0.700017937 | 0.798938005 | ns | NaCl | KS.1 |
| FIM1_2431 | 1513.843408 | 1555.54556 | 1976.13754 | 0.032318191 | 0.143992452 | 0.802617121 | 0.873937443 | ns | NaCl | KS.1 |
| FIM1_2432 | 592.116271 | 564.4206315 | 1147.157163 | -0.055840655 | 0.145329426 | 0.668085242 | 0.774063523 | ns | NaCl | KS.1 |
| FIM1_2433 | 26023.01683 | 41558.96125 | 28152.16964 | 0.467086365 | 0.261583316 | 0.019004328 | 0.04750245 | ns | NaCl | KS.1 |
| FIM1_2434 | 3844.445335 | 5809.064873 | 5279.855016 | 0.442867045 | 0.226749352 | 0.014668575 | 0.038591307 | ns | NaCl | KS.1 |
| FIM1_2435 | 1522.590287 | 1529.026999 | 1608.339509 | 0.004640446 | 0.109920953 | 0.96405201 | 0.98261806 | ns | NaCl | KS.1 |

| | | | | | | | | | | |
|---|---|---|---|---|---|---|---|---|---|---|
| FIM1_2436 | 240.233541 | 278.4860715 | 317.0982153 | 0.157652839 | 0.18682825 | 0.315974368 | 0.448962186 | ns | NaCl | KS.1 |
| FIM1_2437 | 89.47565996 | 50.02105933 | 185.0976495 | -0.534765881 | 0.307043945 | 0.015060143 | 0.039268453 | ns | NaCl | KS.1 |
| FIM1_2438 | 1290.515875 | 1279.238544 | 1531.349559 | -0.011240473 | 0.129109937 | 0.924252303 | 0.956237537 | ns | NaCl | KS.1 |
| FIM1_2439 | 278.9723381 | 298.0850992 | 443.2228972 | 0.057787637 | 0.230010042 | 0.747462491 | 0.835834096 | ns | NaCl | KS.1 |
| FIM1_2440 | 147.1757742 | 98.26751148 | 226.0552197 | -0.252844363 | 0.342132995 | 0.240120867 | 0.366892296 | ns | NaCl | KS.1 |
| FIM1_2444 | 17811.27738 | 33067.36073 | 20677.52651 | 0.565536067 | 0.319465522 | 0.012232178 | 0.03343686 | ns | NaCl | KS.1 |
| FIM1_2445 | 679.6450023 | 1016.740137 | 805.4567149 | 0.37491793 | 0.264941268 | 0.058559859 | 0.120735723 | ns | NaCl | KS.1 |
| FIM1_2446 | 243.8626354 | 278.4283558 | 299.0757199 | 0.163364577 | 0.146593199 | 0.212075034 | 0.334625213 | ns | NaCl | KS.1 |
| FIM1_2447 | 334.1098485 | 383.7098288 | 473.4177694 | 0.172934947 | 0.137753555 | 0.164668955 | 0.274540231 | ns | NaCl | KS.1 |
| FIM1_2448 | 28.08040765 | 18.4128155 | 42.2117417 | -0.23426471 | 0.362880685 | 0.277754105 | 0.407498282 | ns | NaCl | KS.1 |
| FIM1_2449 | 32.18908924 | 41.00479724 | 43.40792016 | 0.153641573 | 0.321810269 | 0.459847513 | 0.594527385 | ns | NaCl | KS.1 |
| FIM1_2450 | 214.9369101 | 309.8322479 | 632.294383 | 0.389412836 | 0.218101869 | 0.026912654 | 0.063757359 | ns | NaCl | KS.1 |
| FIM1_2451 | 52.39231546 | 743.8723843 | 931.4556968 | 3.188502512 | 0.607659005 | 2.55248226741 | 8.58013464892 | Km>KS.1 | NaCl | KS.1 |
| FIM1_2452 | 43.23035201 | 44.52686927 | 52.95915738 | 0.015617062 | 0.260595718 | 0.934926291 | 0.962594846 | ns | NaCl | KS.1 |
| FIM1_2453 | 119.9808928 | 136.4934395 | 209.9167577 | 0.140241205 | 0.182469625 | 0.364419855 | 0.499859052 | ns | NaCl | KS.1 |
| FIM1_2454 | 190.4389064 | 149.8169728 | 378.7276646 | -0.236241858 | 0.218444982 | 0.176440591 | 0.290755859 | ns | NaCl | KS.1 |
| FIM1_2455 | 331.2407797 | 446.3351482 | 1087.383669 | 0.210813392 | 0.306157614 | 0.30840966 | 0.441282157 | ns | NaCl | KS.1 |
| FIM1_2456 | 138.4334679 | 102.6294582 | 256.7192958 | -0.289462323 | 0.235741393 | 0.115110109 | 0.207114934 | ns | NaCl | KS.1 |
| FIM1_2457 | 291.5858816 | 245.8654739 | 402.8146455 | -0.185665456 | 0.17591525 | 0.217447527 | 0.341047115 | ns | NaCl | KS.1 |
| FIM1_2458 | 1307.532469 | 1493.229736 | 2445.15704 | 0.146218567 | 0.172694274 | 0.325012882 | 0.458122752 | ns | NaCl | KS.1 |
| FIM1_2459 | 2602.206204 | 3617.015342 | 2120.245717 | 0.336371207 | 0.224198612 | 0.059589451 | 0.122553749 | ns | NaCl | KS.1 |
| FIM1_2460 | 4724.662752 | 8120.194465 | 4269.400428 | 0.530459014 | 0.283638139 | 0.011830456 | 0.032571399 | ns | NaCl | KS.1 |
| FIM1_2461 | 2494.093187 | 3136.199889 | 2918.362486 | 0.308222778 | 0.09082349 | 0.000357173 | 0.00182437 | ns | NaCl | KS.1 |
| FIM1_2462 | 5318.515263 | 6777.198301 | 5382.854023 | 0.326509234 | 0.09040184 | 0.000145476 | 0.00085549 | ns | NaCl | KS.1 |
| FIM1_2463 | 744.7187776 | 708.9684827 | 944.635553 | -0.061251485 | 0.096448598 | 0.503716679 | 0.635554268 | ns | NaCl | KS.1 |
| FIM1_2464 | 1707.279941 | 1546.857683 | 1939.7883 | -0.114005335 | 0.153877584 | 0.402411872 | 0.538105815 | ns | NaCl | KS.1 |
| FIM1_2465 | 619.3258429 | 439.5687714 | 596.242639 | -0.446512951 | 0.130968379 | 0.000180678 | 0.001024244 | ns | NaCl | KS.1 |
| FIM1_2466 | 399.4332059 | 357.1564982 | 842.4749687 | -0.099132667 | 0.231129919 | 0.582024158 | 0.703662257 | ns | NaCl | KS.1 |
| FIM1_2467 | 294.1256268 | 333.5584352 | 403.5880091 | 0.153722558 | 0.143354122 | 0.231984636 | 0.357954796 | ns | NaCl | KS.1 |
| FIM1_2468 | 1899.530305 | 2336.340309 | 2249.658896 | 0.258672842 | 0.126005785 | 0.024883583 | 0.05974702 | ns | NaCl | KS.1 |
| FIM1_2469 | 2219.667021 | 1670.546271 | 1894.960468 | -0.393732734 | 0.075407855 | 6.06253043918 | 1.01895570726 | ns | NaCl | KS.1 |
| FIM1_2470 | 4953.160437 | 6671.203522 | 6846.182636 | 0.399603859 | 0.099310777 | 1.94545251569 | 0.00015218 | ns | NaCl | KS.1 |
| FIM1_2471 | 1012.856426 | 1053.193291 | 1109.614518 | 0.044345307 | 0.150916449 | 0.741073451 | 0.829852484 | ns | NaCl | KS.1 |
| FIM1_2472 | 1514.734181 | 1383.839414 | 2006.773113 | -0.117843822 | 0.106103963 | 0.237170317 | 0.363963368 | ns | NaCl | KS.1 |
| FIM1_2473 | 1551.590347 | 3128.143658 | 1554.022244 | 0.877130307 | 0.206729133 | 1.23952592457 | 1.41856803644 | ns | NaCl | KS.1 |
| FIM1_2474 | 3246.880798 | 4175.121424 | 3438.768156 | 0.338390891 | 0.092267282 | 0.000111111 | 0.000682481 | ns | NaCl | KS.1 |
| FIM1_2475 | 982.8724394 | 804.4838082 | 1268.99965 | -0.26261373 | 0.111903899 | 0.011658153 | 0.032186077 | ns | NaCl | KS.1 |
| FIM1_2476 | 13034.23435 | 14199.84321 | 9869.721191 | 0.11139841 | 0.098037965 | 0.23050857 | 0.356419896 | ns | NaCl | KS.1 |
| FIM1_2477 | 2122.653711 | 2208.776464 | 2092.768011 | 0.048531423 | 0.116071275 | 0.653011615 | 0.760207849 | ns | NaCl | KS.1 |
| FIM1_2478 | 1106.914085 | 1255.820884 | 1118.601481 | 0.152321364 | 0.131924333 | 0.204862175 | 0.326031271 | ns | NaCl | KS.1 |
| FIM1_2479 | 765.7986616 | 793.6481692 | 825.2483291 | 0.041817767 | 0.111854429 | 0.689123035 | 0.790133003 | ns | NaCl | KS.1 |
| FIM1_2480 | 1777.478276 | 1913.195551 | 2053.053403 | 0.100645884 | 0.073372536 | 0.157006527 | 0.264476618 | ns | NaCl | KS.1 |
| FIM1_2481 | 874.3455976 | 1067.010046 | 1282.433671 | 0.261343597 | 0.097386655 | 0.00452314 | 0.014971805 | ns | NaCl | KS.1 |
| FIM1_2482 | 1205.731705 | 1401.487766 | 1389.641121 | 0.20334952 | 0.078366707 | 0.007090123 | 0.021746831 | ns | NaCl | KS.1 |
| FIM1_2483 | 862.232431 | 957.9815753 | 1291.08699 | 0.134633063 | 0.100276312 | 0.155422589 | 0.2623778 | ns | NaCl | KS.1 |

| | | | | | | | | | | |
|---|---|---|---|---|---|---|---|---|---|---|
| FIM1_2484 | 322.9342314 | 284.490744 | 351.8027649 | -0.152112468 | 0.141807859 | 0.232821595 | 0.358825104 | ns | NaCl | KS.1 |
| FIM1_2485 | 1295.601107 | 817.8045877 | 1569.017362 | -0.608054798 | 0.133095691 | 6.7684439488 | 8.3763703097 | ns | NaCl | KS.1 |
| FIM1_2486 | 13643.82745 | 19374.23831 | 11891.10262 | 0.435618599 | 0.153263064 | 0.001326533 | 0.005413864 | ns | NaCl | KS.1 |
| FIM1_2487 | 2547.113199 | 2453.176293 | 2468.994696 | -0.050957678 | 0.084784703 | 0.531433741 | 0.65948687 | ns | NaCl | KS.1 |
| FIM1_2488 | 393.8376268 | 334.6374071 | 344.4512517 | -0.174204648 | 0.182673865 | 0.260190668 | 0.388840065 | ns | NaCl | KS.1 |
| FIM1_2489 | 732.2897389 | 406.7439896 | 585.8551589 | -0.744214462 | 0.185176219 | 5.1044715795 | 4.8096110053 | ns | NaCl | KS.1 |
| FIM1_2490 | 1285.099282 | 1107.535813 | 1138.043493 | -0.174832248 | 0.146696133 | 0.181861576 | 0.297517036 | ns | NaCl | KS.1 |
| FIM1_2491 | 270.2407187 | 101.1750543 | 811.3610947 | -0.934080224 | 0.398648163 | 0.00056376 | 0.00266607 | ns | NaCl | KS.1 |
| FIM1_2492 | 2524.236326 | 3272.341658 | 2656.474913 | 0.271296057 | 0.203666051 | 0.104368489 | 0.191285621 | ns | NaCl | KS.1 |
| FIM1_2493 | 1598.261495 | 1788.513402 | 1708.561039 | 0.152874997 | 0.074197633 | 0.033287908 | 0.075966671 | ns | NaCl | KS.1 |
| FIM1_2494 | 777.6776172 | 728.6167437 | 863.5211927 | -0.081430109 | 0.128713384 | 0.489593275 | 0.622629484 | ns | NaCl | KS.1 |
| FIM1_2495 | 5256.187416 | 4651.095833 | 5237.240303 | -0.136893731 | 0.16304481 | 0.336106985 | 0.470625457 | ns | NaCl | KS.1 |
| FIM1_2496 | 864.0036799 | 462.5274164 | 1081.57116 | -0.835482041 | 0.152405781 | 3.0109779623 | 7.6038656662 | ns | NaCl | KS.1 |
| FIM1_2497 | 545.591472 | 583.7391045 | 585.9993221 | 0.072525637 | 0.164257025 | 0.612604172 | 0.729070163 | ns | NaCl | KS.1 |
| FIM1_2498 | 179.1254099 | 126.0564359 | 134.5474941 | -0.35425101 | 0.240894127 | 0.058266833 | 0.12023123 | ns | NaCl | KS.1 |
| FIM1_2499 | 944.8034203 | 1105.593892 | 1237.132266 | 0.206966872 | 0.107264784 | 0.039368805 | 0.087126248 | ns | NaCl | KS.1 |
| FIM1_2500 | 1159.615141 | 353.5764191 | 1509.180097 | -1.557183707 | 0.238506276 | 1.3337346500 | 1.0367702943 | Km<KS.1 | NaCl | KS.1 |
| FIM1_2502 | 3030.036286 | 2169.35062 | 3124.580927 | -0.40857136 | 0.157308845 | 0.003155151 | 0.011132538 | ns | NaCl | KS.1 |
| FIM1_2503 | 666.9779943 | 589.0270242 | 574.3758891 | -0.125199375 | 0.196135898 | 0.441258319 | 0.576766636 | ns | NaCl | KS.1 |
| FIM1_2504 | 608.2629241 | 633.8206721 | 853.0960671 | 0.037868123 | 0.217637263 | 0.827440077 | 0.893922775 | ns | NaCl | KS.1 |
| FIM1_2505 | 579.9797518 | 568.6048002 | 596.8034414 | -0.022583114 | 0.166146044 | 0.875815907 | 0.926469091 | ns | NaCl | KS.1 |
| FIM1_2506 | 267.3745004 | 161.2660387 | 537.1545035 | -0.5834293 | 0.215997719 | 0.001086316 | 0.004587796 | ns | NaCl | KS.1 |
| FIM1_2507 | 932.9200029 | 1083.452843 | 1007.272581 | 0.185241136 | 0.126651562 | 0.110339132 | 0.200049993 | ns | NaCl | KS.1 |
| FIM1_2508 | 24.85658241 | 18.37915311 | 19.23299098 | -0.182758647 | 0.339663481 | 0.386915081 | 0.521795209 | ns | NaCl | KS.1 |
| FIM1_2510 | 173.6446546 | 306.3278521 | 247.8209449 | 0.674074817 | 0.218451786 | 0.000221664 | 0.001214514 | ns | NaCl | KS.1 |
| FIM1_2511 | 5984.116566 | 11860.32735 | 5768.840307 | 0.862029088 | 0.199545798 | 9.3331999955 | 1.1055397613 | ns | NaCl | KS.1 |
| FIM1_2512 | 1073.66613 | 869.1939281 | 1212.02316 | -0.265956449 | 0.128523478 | 0.023296567 | 0.056486472 | ns | NaCl | KS.1 |
| FIM1_2513 | 6242.874249 | 9649.454336 | 6384.272418 | 0.518420962 | 0.19033285 | 0.001335011 | 0.00543638 | ns | NaCl | KS.1 |
| FIM1_2514 | 301.0893667 | 302.4300761 | 481.8060497 | 0.004331458 | 0.146705296 | 0.973673284 | 0.987770104 | ns | NaCl | KS.1 |
| FIM1_2515 | 67.61668409 | 55.76520641 | 94.90197683 | -0.180571209 | 0.217085079 | 0.298818297 | 0.431404825 | ns | NaCl | KS.1 |
| FIM1_2516 | 3592.782344 | 4750.176412 | 4106.495846 | 0.367682407 | 0.109509435 | 0.000313637 | 0.001625802 | ns | NaCl | KS.1 |
| FIM1_2517 | 1917.131921 | 1452.903882 | 2074.916218 | -0.324887572 | 0.16370368 | 0.022636242 | 0.055176534 | ns | NaCl | KS.1 |
| FIM1_2518 | 1878.798867 | 2216.075601 | 2038.925988 | 0.221562575 | 0.08499086 | 0.006483742 | 0.020210912 | ns | NaCl | KS.1 |
| FIM1_2519 | 3149.314563 | 6291.301102 | 2929.214204 | 0.574552921 | 0.357428303 | 0.015578195 | 0.040323371 | ns | NaCl | KS.1 |
| FIM1_2520 | 680.6810481 | 939.8019146 | 689.008901 | 0.383047325 | 0.170131853 | 0.009169374 | 0.026723864 | ns | NaCl | KS.1 |
| FIM1_2521 | 636.6861026 | 961.9164935 | 697.9728368 | 0.490718454 | 0.186879591 | 0.002019887 | 0.007694441 | ns | NaCl | KS.1 |
| FIM1_2522 | 468.3244844 | 532.2853558 | 490.9830642 | 0.150472689 | 0.148535011 | 0.255492962 | 0.383895345 | ns | NaCl | KS.1 |
| FIM1_2523 | 3942.948473 | 5361.862713 | 4516.161231 | 0.431414604 | 0.062376075 | 1.2706318528 | 7.9017418350 | ns | NaCl | KS.1 |
| FIM1_2524 | 1268.458147 | 1686.022923 | 1763.71944 | 0.268627039 | 0.238418473 | 0.145856461 | 0.249102606 | ns | NaCl | KS.1 |
| FIM1_2525 | 978.5971177 | 658.0704064 | 706.1662017 | -0.497276267 | 0.157850941 | 0.000364444 | 0.001855766 | ns | NaCl | KS.1 |
| FIM1_2526 | 5385.216871 | 3693.902665 | 5219.96843 | -0.480481213 | 0.144685024 | 0.000214454 | 0.001185008 | ns | NaCl | KS.1 |
| FIM1_2527 | 204.79881 | 123.7147381 | 199.4878147 | -0.42139167 | 0.314113354 | 0.052654946 | 0.110298256 | ns | NaCl | KS.1 |
| FIM1_2528 | 32855.27498 | 34724.2605 | 23970.31229 | 0.071729552 | 0.098458689 | 0.442081275 | 0.577161997 | ns | NaCl | KS.1 |
| FIM1_2529 | 6440.528509 | 8307.719254 | 8482.783516 | 0.210277643 | 0.266296375 | 0.281516549 | 0.411561807 | ns | NaCl | KS.1 |
| FIM1_2530 | 1093.23941 | 619.6525335 | 1352.405008 | -0.518959205 | 0.307771835 | 0.01810017 | 0.045733035 | ns | NaCl | KS.1 |

| | | | | | | | | | | |
|---|---|---|---|---|---|---|---|---|---|---|
| FIM1_2531 | 3913.71158 | 3784.711306 | 3138.28937 | -0.036718175 | 0.16413053 | 0.797607378 | 0.870903663 | ns | NaCl | KS.1 |
| FIM1_2532 | 757.692315 | 415.9402202 | 477.4910332 | -0.618200402 | 0.284192334 | 0.00392026 | 0.013349278 | ns | NaCl | KS.1 |
| FIM1_2533 | 2907.928098 | 2248.032288 | 2574.912595 | -0.320968987 | 0.135873946 | 0.008909512 | 0.026171466 | ns | NaCl | KS.1 |
| FIM1_2534 | 2500.085621 | 1600.003562 | 2106.468145 | -0.558570346 | 0.166944469 | 0.000140749 | 0.000834593 | ns | NaCl | KS.1 |
| FIM1_2535 | 679.068116 | 476.6848383 | 461.4695118 | -0.420931809 | 0.177599688 | 0.005670242 | 0.018152801 | ns | NaCl | KS.1 |
| FIM1_2536 | 682.6763245 | 423.1604857 | 460.6005252 | -0.433118494 | 0.289158898 | 0.03867793 | 0.08605705 | ns | NaCl | KS.1 |
| FIM1_2537 | 1985.825174 | 1164.115097 | 1077.293257 | -0.556122402 | 0.264270385 | 0.006176617 | 0.019485523 | ns | NaCl | KS.1 |
| FIM1_2538 | 1200.380954 | 759.4465997 | 815.3711556 | -0.533927177 | 0.204152073 | 0.001729486 | 0.006790996 | ns | NaCl | KS.1 |
| FIM1_2539 | 712.2579283 | 716.809373 | 615.8128797 | 0.006692503 | 0.175885786 | 0.964578204 | 0.982953004 | ns | NaCl | KS.1 |
| FIM1_2540 | 2792.503025 | 3023.724447 | 3341.158667 | 0.085962613 | 0.171888433 | 0.561732907 | 0.685114788 | ns | NaCl | KS.1 |
| FIM1_2541 | 1957.62751 | 1646.575278 | 1315.902532 | -0.208980058 | 0.140554056 | 0.098305823 | 0.182285303 | ns | NaCl | KS.1 |
| FIM1_2542 | 483.428595 | 472.2579443 | 401.8720866 | -0.026482993 | 0.167998421 | 0.855775352 | 0.912715589 | ns | NaCl | KS.1 |
| FIM1_2543 | 1041.020524 | 983.1415738 | 1152.744984 | -0.065042488 | 0.15114558 | 0.62827174 | 0.740675807 | ns | NaCl | KS.1 |
| FIM1_2544 | 722.4435791 | 828.3810951 | 859.9406074 | 0.177345325 | 0.1130016 | 0.09191631 | 0.173410558 | ns | NaCl | KS.1 |
| FIM1_2546 | 1698.307523 | 1056.368289 | 1183.701345 | -0.627224973 | 0.137075125 | 6.12432066528 | 7.73312063700 | ns | NaCl | KS.1 |
| FIM1_2547 | 1433.158577 | 1190.118431 | 2021.126718 | -0.170759091 | 0.231291555 | 0.34428118 | 0.479104579 | ns | NaCl | KS.1 |
| FIM1_2548 | 6735.387529 | 7131.087422 | 5162.908892 | 0.045350135 | 0.255903156 | 0.811098691 | 0.880859199 | ns | NaCl | KS.1 |
| FIM1_2549 | 5400.878384 | 4570.043684 | 5865.03692 | -0.165284933 | 0.206672266 | 0.326260145 | 0.459553856 | ns | NaCl | KS.1 |
| FIM1_2550 | 2212.63211 | 1320.266264 | 1720.231843 | -0.635169257 | 0.191965299 | 0.000119116 | 0.000720924 | ns | NaCl | KS.1 |
| FIM1_2552 | 1402.99128 | 1642.01308 | 2587.608678 | 0.129790691 | 0.254471838 | 0.494152821 | 0.626775555 | ns | NaCl | KS.1 |
| FIM1_2553 | 307.4233256 | 337.4389552 | 530.5381407 | 0.085819077 | 0.222441376 | 0.625994509 | 0.73939285 | ns | NaCl | KS.1 |
| FIM1_2554 | 1277.601437 | 682.9110279 | 1249.548733 | -0.664604385 | 0.277865508 | 0.001838216 | 0.007130713 | ns | NaCl | KS.1 |
| FIM1_2555 | 5261.232792 | 20467.30714 | 14232.97528 | 1.506544488 | 0.425360547 | 1.02118636898 | 1.18978973904 | Km>KS.1 | NaCl | KS.1 |
| FIM1_2556 | 112.1855588 | 89.07153866 | 413.0204317 | -0.164731347 | 0.300426191 | 0.419931 | 0.55621851 | ns | NaCl | KS.1 |
| FIM1_2557 | 11412.90944 | 14455.11553 | 12207.00935 | 0.250117134 | 0.195924203 | 0.123967834 | 0.219012775 | ns | NaCl | KS.1 |
| FIM1_2558 | 1390.219139 | 2129.980324 | 1227.149074 | 0.443383971 | 0.239844732 | 0.018495232 | 0.046565677 | ns | NaCl | KS.1 |
| FIM1_2559 | 470.7083349 | 283.4967538 | 458.2771054 | -0.46478708 | 0.29334874 | 0.028476402 | 0.066668281 | ns | NaCl | KS.1 |
| FIM1_2560 | 2672.352299 | 3853.224249 | 3738.299824 | 0.467496374 | 0.142281828 | 0.000257823 | 0.001386559 | ns | NaCl | KS.1 |
| FIM1_2561 | 199.7099781 | 235.5500503 | 346.8324722 | 0.162686896 | 0.209351054 | 0.337943197 | 0.471962694 | ns | NaCl | KS.1 |
| FIM1_2562 | 2182.91167 | 1986.803981 | 2830.694731 | -0.115329565 | 0.125141549 | 0.316033281 | 0.448962186 | ns | NaCl | KS.1 |
| FIM1_2563 | 3148.799194 | 3520.42841 | 3001.103454 | 0.148417835 | 0.090280986 | 0.084946573 | 0.1631066 | ns | NaCl | KS.1 |
| FIM1_2564 | 1153.30947 | 874.2819595 | 1739.693656 | -0.338912953 | 0.149005632 | 0.010431458 | 0.029570657 | ns | NaCl | KS.1 |
| FIM1_2565 | 1046.327382 | 1017.708675 | 1138.975453 | -0.034604095 | 0.13428914 | 0.776942482 | 0.854963249 | ns | NaCl | KS.1 |
| FIM1_2566 | 1657.07154 | 1469.559837 | 2318.819173 | -0.133477947 | 0.165314317 | 0.35337184 | 0.488475939 | ns | NaCl | KS.1 |
| FIM1_2567 | 884.2664484 | 1079.149324 | 1056.417571 | 0.249294482 | 0.127854062 | 0.032758878 | 0.074965695 | ns | NaCl | KS.1 |
| FIM1_2568 | 1466.48972 | 1797.234478 | 1217.730461 | 0.199490093 | 0.213546375 | 0.246314973 | 0.373944763 | ns | NaCl | KS.1 |
| FIM1_2570 | 231.8764902 | 214.0099226 | 388.1636513 | -0.07601334 | 0.214997915 | 0.659416788 | 0.765958095 | ns | NaCl | KS.1 |
| FIM1_2571 | 335.2681777 | 637.4093959 | 421.0415144 | 0.687783631 | 0.27813646 | 0.001336421 | 0.00543638 | ns | NaCl | KS.1 |
| FIM1_2572 | 281.3532433 | 415.9470704 | 287.6389418 | 0.443756593 | 0.203648209 | 0.008466992 | 0.025178294 | ns | NaCl | KS.1 |
| FIM1_2573 | 465.1294358 | 303.9590171 | 484.8169296 | -0.528284814 | 0.17224889 | 0.000424443 | 0.002101421 | ns | NaCl | KS.1 |
| FIM1_2574 | 2286.972129 | 2303.858151 | 1794.752805 | 0.008853476 | 0.119513402 | 0.936269469 | 0.963188249 | ns | NaCl | KS.1 |
| FIM1_2575 | 1203.861653 | 1153.106593 | 933.2852499 | -0.053945589 | 0.118722342 | 0.624051985 | 0.7385011 | ns | NaCl | KS.1 |
| FIM1_2576 | 14658.93667 | 21231.73775 | 14153.82679 | 0.524513173 | 0.054366071 | 8.60882391424 | 4.75876655259 | ns | NaCl | KS.1 |
| FIM1_2577 | 3834.598424 | 6297.058935 | 5459.753975 | 0.63156684 | 0.165420499 | 1.72606499940 | 0.00013828 | ns | NaCl | KS.1 |
| FIM1_2578 | 95.73330165 | 78.62619591 | 73.6616089 | -0.206425237 | 0.209658784 | 0.224820174 | 0.349525114 | ns | NaCl | KS.1 |

| | | | | | | | | | | |
|---|---|---|---|---|---|---|---|---|---|---|
| FIM1_2579 | 270.1646682 | 220.4826557 | 269.9166729 | -0.225794052 | 0.167964896 | 0.120409716 | 0.214786065 | ns | NaCl | KS.1 |
| FIM1_2580 | 683.300193 | 505.8988144 | 1231.593787 | -0.309012948 | 0.215677643 | 0.075456595 | 0.147735757 | ns | NaCl | KS.1 |
| FIM1_2581 | 17197.36518 | 31405.36951 | 20957.63236 | 0.808822873 | 0.138171018 | 3.7863060169 | 1.2231735346 | ns | NaCl | KS.1 |
| FIM1_2582 | 947.9330444 | 366.5245399 | 655.2949581 | -1.26167657 | 0.188816399 | 2.6620242529 | 1.8919386655 | Km<KS.1 | NaCl | KS.1 |
| FIM1_2583 | 671.5316854 | 404.2819819 | 481.033384 | -0.612270625 | 0.196941203 | 0.0002614 | 0.001401365 | ns | NaCl | KS.1 |
| FIM1_2584 | 54.4661008 | 74.02084177 | 103.7048874 | 0.182893468 | 0.333945014 | 0.385240931 | 0.520629634 | ns | NaCl | KS.1 |
| FIM1_2585 | 5538.019325 | 5419.254917 | 4837.266344 | -0.026867932 | 0.120291345 | 0.809305072 | 0.879679426 | ns | NaCl | KS.1 |
| FIM1_2586 | 939.0663399 | 1222.108122 | 1096.603075 | 0.342709479 | 0.115820597 | 0.001381555 | 0.005565373 | ns | NaCl | KS.1 |
| FIM1_2587 | 5231.100672 | 5218.915835 | 4513.054149 | -0.00367338 | 0.092822376 | 0.966890981 | 0.984503199 | ns | NaCl | KS.1 |
| FIM1_2588 | 2769.604546 | 2833.588574 | 5707.663931 | 0.023332569 | 0.190179373 | 0.883493517 | 0.931027377 | ns | NaCl | KS.1 |
| FIM1_2590 | 1101.770298 | 357.3312064 | 1955.598832 | -1.341275089 | 0.311446133 | 1.0583924041 | 1.6822690769 | Km<KS.1 | NaCl | KS.1 |
| FIM1_2591 | 1002.117629 | 1043.256196 | 888.9165065 | 0.037271492 | 0.217133247 | 0.829881598 | 0.895199686 | ns | NaCl | KS.1 |
| FIM1_2592 | 1454.714337 | 1633.24868 | 1822.145642 | 0.144073231 | 0.118036696 | 0.187699327 | 0.304369019 | ns | NaCl | KS.1 |
| FIM1_2593 | 930.9230282 | 983.0837099 | 1588.488274 | 0.073790609 | 0.095319047 | 0.415488989 | 0.55150953 | ns | NaCl | KS.1 |
| FIM1_2594 | 2304.631092 | 2015.28991 | 2182.106981 | -0.167901591 | 0.118284575 | 0.125184124 | 0.220613183 | ns | NaCl | KS.1 |
| FIM1_2595 | 1243.788618 | 1533.817778 | 1316.272804 | 0.276615671 | 0.10061831 | 0.003507453 | 0.012176958 | ns | NaCl | KS.1 |
| FIM1_2596 | 2074.192503 | 1540.704994 | 2024.028705 | -0.32420647 | 0.197454574 | 0.04773188 | 0.101655009 | ns | NaCl | KS.1 |
| FIM1_2597 | 909.591186 | 1240.789253 | 977.7203132 | 0.345766889 | 0.192674425 | 0.031802274 | 0.0731467 | ns | NaCl | KS.1 |
| FIM1_2598 | 994.9131062 | 787.9691177 | 1013.907709 | -0.295949694 | 0.129519656 | 0.012095503 | 0.03320923 | ns | NaCl | KS.1 |
| FIM1_2599 | 1094.423091 | 1334.39326 | 964.505576 | 0.201721608 | 0.202453243 | 0.224748179 | 0.349522411 | ns | NaCl | KS.1 |
| FIM1_2600 | 1653.943877 | 1817.775885 | 1325.314155 | 0.104068092 | 0.164278599 | 0.467378255 | 0.6017616 | ns | NaCl | KS.1 |
| FIM1_2601 | 2577.198789 | 2951.895335 | 2074.916074 | 0.171021777 | 0.116277357 | 0.112916444 | 0.203683579 | ns | NaCl | KS.1 |
| FIM1_2602 | 1855.251164 | 1951.474192 | 1905.859368 | 0.065115018 | 0.104347565 | 0.507707789 | 0.639131136 | ns | NaCl | KS.1 |
| FIM1_2603 | 177.8442383 | 188.7122292 | 286.3863057 | 0.048589188 | 0.257348402 | 0.798430473 | 0.870903663 | ns | NaCl | KS.1 |
| FIM1_2604 | 552.9570663 | 419.0476656 | 543.7081307 | -0.297158981 | 0.200619329 | 0.072482082 | 0.14298111 | ns | NaCl | KS.1 |
| FIM1_2606 | 40.84297577 | 42.64335599 | 32.27504231 | 0.024451963 | 0.278356361 | 0.901060242 | 0.941363861 | ns | NaCl | KS.1 |
| FIM1_2607 | 3402.1912 | 4953.131031 | 3002.133013 | 0.423803668 | 0.200269197 | 0.010814401 | 0.030448016 | ns | NaCl | KS.1 |
| FIM1_2608 | 804.7956975 | 863.1983013 | 800.2735726 | 0.090145731 | 0.095534109 | 0.320750856 | 0.453978808 | ns | NaCl | KS.1 |
| FIM1_2609 | 408.1227338 | 292.890806 | 502.6738705 | -0.401778897 | 0.16472058 | 0.00510331 | 0.016540045 | ns | NaCl | KS.1 |
| FIM1_2610 | 1118.998959 | 788.146184 | 1233.092503 | -0.459190339 | 0.129888564 | 0.000106279 | 0.000657636 | ns | NaCl | KS.1 |
| FIM1_2611 | 1194.822425 | 1319.110263 | 1871.482649 | 0.101390219 | 0.189077262 | 0.522542309 | 0.65235834 | ns | NaCl | KS.1 |
| FIM1_2612 | 3054.048053 | 2886.929005 | 3394.826906 | -0.040754964 | 0.278005898 | 0.835827337 | 0.899468094 | ns | NaCl | KS.1 |
| FIM1_2613 | 2679.409715 | 2365.597117 | 3064.290298 | -0.109624585 | 0.234761691 | 0.546380688 | 0.672499733 | ns | NaCl | KS.1 |
| FIM1_2614 | 718.1952152 | 429.6442074 | 709.2168211 | -0.675291287 | 0.151008882 | 8.5621048716 | 1.0264210056 | ns | NaCl | KS.1 |
| FIM1_2615 | 51.57588013 | 71.69444286 | 64.47101098 | 0.275494355 | 0.26883193 | 0.162994417 | 0.272662147 | ns | NaCl | KS.1 |
| FIM1_2616 | 602.1884378 | 681.506435 | 737.3979886 | 0.156076499 | 0.099538071 | 0.097201902 | 0.180709814 | ns | NaCl | KS.1 |
| FIM1_2617 | 1072.662753 | 1032.413177 | 1133.242136 | -0.048103319 | 0.089437243 | 0.573837776 | 0.69596366 | ns | NaCl | KS.1 |
| FIM1_2618 | 475.5056753 | 321.2234549 | 394.8937208 | -0.488471705 | 0.165778761 | 0.000751267 | 0.003394692 | ns | NaCl | KS.1 |
| FIM1_2619 | 255.8546073 | 179.2184405 | 276.3989874 | -0.41129957 | 0.193562686 | 0.011178895 | 0.031174328 | ns | NaCl | KS.1 |
| FIM1_2620 | 1004.249739 | 631.926027 | 670.0593794 | -0.597742578 | 0.153460555 | 1.4234735837 | 0.000116669 | ns | NaCl | KS.1 |
| FIM1_2621 | 7233.83129 | 9894.570427 | 8627.981714 | 0.370373029 | 0.169169271 | 0.011358027 | 0.031567701 | ns | NaCl | KS.1 |
| FIM1_2622 | 600.1852742 | 456.887173 | 652.659282 | -0.341967077 | 0.141763384 | 0.007124972 | 0.021813375 | ns | NaCl | KS.1 |
| FIM1_2623 | 1638.137672 | 2005.01941 | 1779.418525 | 0.258321113 | 0.115301066 | 0.015642788 | 0.040406474 | ns | NaCl | KS.1 |
| FIM1_2624 | 6384.074483 | 7310.610396 | 7410.842857 | 0.170141103 | 0.119608336 | 0.123775628 | 0.218969986 | ns | NaCl | KS.1 |
| FIM1_2625 | 561.4720258 | 312.5435254 | 576.4427825 | -0.749544567 | 0.180558194 | 2.8540914069 | 2.9458723547 | ns | NaCl | KS.1 |

| | | | | | | | | | | |
|---|---|---|---|---|---|---|---|---|---|---|
| FIM1_2626 | 1320.532385 | 1386.436525 | 1584.582197 | 0.060939219 | 0.12191028 | 0.588325699 | 0.709556449 | ns | NaCl | KS.1 |
| FIM1_2627 | 811.9796582 | 652.5223815 | 1047.969986 | -0.255320437 | 0.163293182 | 0.072683827 | 0.143322251 | ns | NaCl | KS.1 |
| FIM1_2628 | 808.116242 | 922.2563393 | 833.5519857 | 0.163343048 | 0.123371191 | 0.150205945 | 0.255478488 | ns | NaCl | KS.1 |
| FIM1_2629 | 6878.733837 | 12794.12217 | 7639.601927 | 0.672085273 | 0.267860755 | 0.001297312 | 0.005312038 | ns | NaCl | KS.1 |
| FIM1_2630 | 491.0602086 | 735.9235328 | 544.7853279 | 0.485206374 | 0.182729537 | 0.001895171 | 0.007320245 | ns | NaCl | KS.1 |
| FIM1_2631 | 156.413305 | 188.1543082 | 217.1279088 | 0.210423719 | 0.16461642 | 0.141734201 | 0.243256235 | ns | NaCl | KS.1 |
| FIM1_2632 | 398.8531828 | 611.0499531 | 622.236406 | 0.543030426 | 0.156487772 | 9.47106748453 | 0.000596023 | ns | NaCl | KS.1 |
| FIM1_2633 | 702.5556559 | 636.4210812 | 808.6597244 | -0.115410746 | 0.153203372 | 0.395003271 | 0.530975756 | ns | NaCl | KS.1 |
| FIM1_2634 | 3429.55278 | 5564.785761 | 5047.092429 | 0.487103233 | 0.26277729 | 0.014999654 | 0.039131242 | ns | NaCl | KS.1 |
| FIM1_2635 | 35570.34045 | 50439.72112 | 31529.08503 | 0.465512712 | 0.111455816 | 7.34603661366 | 6.52616645588 | ns | NaCl | KS.1 |
| FIM1_2636 | 3827.603799 | 3960.847586 | 3879.739344 | 0.045634269 | 0.075624615 | 0.533008397 | 0.660946355 | ns | NaCl | KS.1 |
| FIM1_2637 | 4505.962055 | 3233.490355 | 4556.375614 | -0.39257855 | 0.172526963 | 0.008263673 | 0.024691757 | ns | NaCl | KS.1 |
| FIM1_2638 | 1124.547168 | 1555.009611 | 1535.1718 | 0.421841981 | 0.124318024 | 0.000214612 | 0.001185008 | ns | NaCl | KS.1 |
| FIM1_2639 | 5169.904495 | 2596.54124 | 3287.046383 | -0.865716544 | 0.202335347 | 1.10519036536 | 1.28166015564 | ns | NaCl | KS.1 |
| FIM1_2640 | 3938.481369 | 2256.881327 | 3174.526742 | -0.730066616 | 0.154597659 | 2.18634608293 | 3.19913875371 | ns | NaCl | KS.1 |
| FIM1_2641 | 996.9102718 | 807.7670551 | 974.5261912 | -0.278919391 | 0.110974023 | 0.006944525 | 0.021379339 | ns | NaCl | KS.1 |
| FIM1_2642 | 375.3739167 | 266.6701991 | 395.3580463 | -0.436251169 | 0.139136518 | 0.000502821 | 0.002416941 | ns | NaCl | KS.1 |
| FIM1_2643 | 2098.673695 | 1573.538096 | 2442.68225 | -0.358233785 | 0.144838889 | 0.005611328 | 0.017975761 | ns | NaCl | KS.1 |
| FIM1_2644 | 2261.013632 | 1416.549595 | 1769.190409 | -0.637827708 | 0.110036699 | 8.37263523979 | 2.43589826420 | ns | NaCl | KS.1 |
| FIM1_2645 | 833.3174076 | 590.6331712 | 693.2951935 | -0.438663827 | 0.138557965 | 0.000447658 | 0.002196348 | ns | NaCl | KS.1 |
| FIM1_2646 | 2575.395737 | 2532.974414 | 2678.289446 | -0.020689411 | 0.10009819 | 0.827140916 | 0.893922775 | ns | NaCl | KS.1 |
| FIM1_2647 | 2742.409163 | 2676.577905 | 3353.209645 | -0.031350079 | 0.091144418 | 0.718686971 | 0.813345696 | ns | NaCl | KS.1 |
| FIM1_2648 | 276.6240377 | 278.359489 | 308.8158223 | 0.003158215 | 0.122503307 | 0.977720848 | 0.989137805 | ns | NaCl | KS.1 |
| FIM1_2649 | 916.4783212 | 1079.743953 | 874.0597844 | 0.218868084 | 0.085060063 | 0.007210076 | 0.022033249 | ns | NaCl | KS.1 |
| FIM1_2650 | 15298.16596 | 22860.77063 | 13759.61847 | 0.523061473 | 0.135409601 | 2.18615940257 | 0.000168622 | ns | NaCl | KS.1 |
| FIM1_2651 | 364.3822801 | 358.0354529 | 381.7676173 | -0.018180911 | 0.190059566 | 0.909051102 | 0.946531862 | ns | NaCl | KS.1 |
| FIM1_2652 | 8969.451495 | 10717.7869 | 7485.896065 | 0.239232091 | 0.085117907 | 0.003320196 | 0.011599703 | ns | NaCl | KS.1 |
| FIM1_2653 | 1839.809673 | 1621.760529 | 1690.258184 | -0.160318485 | 0.113462564 | 0.129126098 | 0.226357412 | ns | NaCl | KS.1 |
| FIM1_2654 | 2009.893246 | 1950.975271 | 2235.239421 | -0.035358276 | 0.119650889 | 0.749685936 | 0.837108446 | ns | NaCl | KS.1 |
| FIM1_2655 | 1546.022888 | 1686.501933 | 1444.349023 | 0.060155914 | 0.289319614 | 0.76323653 | 0.847000164 | ns | NaCl | KS.1 |
| FIM1_2656 | 123.6062652 | 126.8448429 | 282.8426353 | 0.026945562 | 0.255598822 | 0.886972892 | 0.932435862 | ns | NaCl | KS.1 |
| FIM1_2657 | 836.6091707 | 930.5445543 | 919.771808 | 0.128071853 | 0.129677178 | 0.279777956 | 0.409983897 | ns | NaCl | KS.1 |
| FIM1_2659 | 1145.314423 | 1416.877321 | 1561.785365 | 0.263330898 | 0.133364756 | 0.029434581 | 0.068588777 | ns | NaCl | KS.1 |
| FIM1_2660 | 2132.319244 | 1396.485805 | 1832.770411 | -0.415431305 | 0.256533053 | 0.033620118 | 0.076514221 | ns | NaCl | KS.1 |
| FIM1_2661 | 945.8826613 | 901.5061464 | 1387.871653 | -0.038032999 | 0.257962363 | 0.841701694 | 0.903176394 | ns | NaCl | KS.1 |
| FIM1_2662 | 5467.962204 | 5764.937743 | 5391.751476 | 0.064044705 | 0.128616845 | 0.586675533 | 0.708088462 | ns | NaCl | KS.1 |
| FIM1_2663 | 618.5379826 | 588.5912272 | 406.4431412 | -0.039590736 | 0.255428848 | 0.834554863 | 0.898487436 | ns | NaCl | KS.1 |
| FIM1_2664 | 2336.190884 | 2135.26779 | 2489.737755 | -0.119900361 | 0.09188466 | 0.171058612 | 0.282823727 | ns | NaCl | KS.1 |
| FIM1_2665 | 224.5018662 | 107.3924127 | 497.1695574 | -0.885422251 | 0.244929467 | 1.50653149581 | 0.000122267 | ns | NaCl | KS.1 |
| FIM1_2666 | 448.8218215 | 499.4512562 | 1035.415032 | 0.116621029 | 0.177560252 | 0.441894512 | 0.577161997 | ns | NaCl | KS.1 |
| FIM1_2667 | 3143.198942 | 3185.919776 | 3083.454991 | 0.018393781 | 0.076403289 | 0.803474272 | 0.874487968 | ns | NaCl | KS.1 |
| FIM1_2668 | 2267.301604 | 3112.14462 | 3118.802518 | 0.298842035 | 0.244311618 | 0.111529937 | 0.201694452 | ns | NaCl | KS.1 |
| FIM1_2669 | 41.38717193 | 34.22367983 | 47.98920621 | -0.151109805 | 0.274096886 | 0.442667578 | 0.577569158 | ns | NaCl | KS.1 |
| FIM1_2671 | 48.84069741 | 35.88866927 | 54.40823865 | -0.288284798 | 0.287967054 | 0.158501527 | 0.266670644 | ns | NaCl | KS.1 |
| FIM1_2672 | 670.5774404 | 781.1145655 | 872.2161792 | 0.194752274 | 0.110126913 | 0.058256326 | 0.12023123 | ns | NaCl | KS.1 |

| | | | | | | | | | | |
|---|---|---|---|---|---|---|---|---|---|---|
| FIM1_2674 | 3201.788318 | 3648.532761 | 5072.522202 | 0.155537417 | 0.141625882 | 0.221955368 | 0.346762078 | ns | NaCl | KS.1 |
| FIM1_2675 | 397.9419516 | 1082.977979 | 10561.10044 | 0.482326073 | 0.480814113 | 0.044593139 | 0.096205927 | ns | NaCl | KS.1 |
| FIM1_2676 | 7867.242466 | 43907.17084 | 23990.20896 | 2.278705414 | 0.3367661 | 5.96939477999 | 5.60334698688 | Km>KS.1 | NaCl | KS.1 |
| FIM1_2677 | 4564.939931 | 5386.728367 | 3814.504978 | 0.212472303 | 0.110073918 | 0.038652892 | 0.08605705 | ns | NaCl | KS.1 |
| FIM1_2678 | 690.3816078 | 608.9249702 | 579.2869035 | -0.150122221 | 0.144159463 | 0.245266936 | 0.372808741 | ns | NaCl | KS.1 |
| FIM1_2679 | 910.3129113 | 809.1398139 | 1383.071188 | -0.136751811 | 0.151693368 | 0.30958327 | 0.44245239 | ns | NaCl | KS.1 |
| FIM1_2680 | 4208.569267 | 7637.051019 | 4303.406281 | 0.792122463 | 0.146747033 | 5.44311958472 | 1.28950094923 | ns | NaCl | KS.1 |
| FIM1_2681 | 2502.742749 | 4704.707897 | 4554.22403 | 0.802442025 | 0.18665418 | 1.25512986077 | 1.42889497879 | ns | NaCl | KS.1 |
| FIM1_2682 | 460.5680239 | 546.189489 | 945.3060971 | 0.147079554 | 0.245518773 | 0.430349776 | 0.566849387 | ns | NaCl | KS.1 |
| FIM1_2683 | 2949.656922 | 1407.494429 | 2297.684861 | -0.958821034 | 0.182908579 | 7.33646624815 | 1.63959281481 | ns | NaCl | KS.1 |
| FIM1_2684 | 4496.428 | 11689.93378 | 5456.353642 | 1.180523582 | 0.252255389 | 4.49939688794 | 7.88186602729 | Km>KS.1 | NaCl | KS.1 |
| FIM1_2685 | 1123.713872 | 854.3703662 | 1047.779783 | -0.354011171 | 0.127874859 | 0.002402401 | 0.008892818 | ns | NaCl | KS.1 |
| FIM1_2686 | 159.8978467 | 80.56798823 | 211.4828374 | -0.838402712 | 0.239441423 | 2.75419736127 | 0.000204815 | ns | NaCl | KS.1 |
| FIM1_2687 | 573.7582139 | 483.4987827 | 777.3624947 | -0.146763971 | 0.245735272 | 0.431549561 | 0.568279266 | ns | NaCl | KS.1 |
| FIM1_2688 | 1610.86206 | 2062.602614 | 1954.134537 | 0.310986798 | 0.130350212 | 0.008710264 | 0.025793787 | ns | NaCl | KS.1 |
| FIM1_2689 | 1592.152612 | 1456.13367 | 1808.791337 | -0.11396386 | 0.108871562 | 0.263820879 | 0.392144868 | ns | NaCl | KS.1 |
| FIM1_2690 | 5893.474052 | 8061.531989 | 6279.00711 | 0.428754137 | 0.086748741 | 2.23077074737 | 3.25457022528 | ns | NaCl | KS.1 |
| FIM1_2691 | 1265.7766 | 1452.900764 | 1404.064091 | 0.183487956 | 0.081314283 | 0.018941156 | 0.047448264 | ns | NaCl | KS.1 |
| FIM1_2692 | 708.4563558 | 665.9172829 | 751.1524816 | -0.079113314 | 0.126872369 | 0.496848974 | 0.628642839 | ns | NaCl | KS.1 |
| FIM1_2693 | 2217.017216 | 2603.883352 | 2325.402404 | 0.21251217 | 0.092404218 | 0.015596488 | 0.040349728 | ns | NaCl | KS.1 |
| FIM1_2694 | 590.7498331 | 602.6859116 | 600.8518616 | 0.021570204 | 0.104885939 | 0.827145248 | 0.893922775 | ns | NaCl | KS.1 |
| FIM1_2695 | 3549.636475 | 4178.140082 | 2827.478016 | 0.207090917 | 0.115187071 | 0.052744736 | 0.110329294 | ns | NaCl | KS.1 |
| FIM1_2696 | 7672.910193 | 13055.14327 | 6745.90564 | 0.630908845 | 0.210499384 | 0.000352366 | 0.001803241 | ns | NaCl | KS.1 |
| FIM1_2697 | 568.6564447 | 560.0069554 | 428.8738886 | -0.014428027 | 0.218416177 | 0.933945008 | 0.962133741 | ns | NaCl | KS.1 |
| FIM1_2698 | 2687.950692 | 2828.431972 | 3296.22013 | 0.066009894 | 0.100504162 | 0.487430595 | 0.621467763 | ns | NaCl | KS.1 |
| FIM1_2699 | 2187.814574 | 4822.116979 | 2342.557362 | 0.944777838 | 0.249466576 | 6.15804136758 | 5.57022832794 | ns | NaCl | KS.1 |
| FIM1_2700 | 440.4679939 | 389.097614 | 571.3223321 | -0.13089058 | 0.176445176 | 0.385861068 | 0.521080025 | ns | NaCl | KS.1 |
| FIM1_2701 | 2203.02348 | 1581.773035 | 2051.847764 | -0.37688671 | 0.189659234 | 0.01808242 | 0.045711402 | ns | NaCl | KS.1 |
| FIM1_2702 | 158.6697565 | 164.2723344 | 193.6273669 | 0.034977115 | 0.173254284 | 0.814467452 | 0.884324656 | ns | NaCl | KS.1 |
| FIM1_2703 | 179.521533 | 43.96088107 | 206.2115144 | -1.444170978 | 0.489573461 | 9.78733434982 | 8.35197056438 | Km<KS.1 | NaCl | KS.1 |
| FIM1_2704 | 1250.896704 | 1465.121 | 1576.395522 | 0.194319384 | 0.131057543 | 0.103545968 | 0.190018883 | ns | NaCl | KS.1 |
| FIM1_2705 | 2256.76211 | 6305.429292 | 3068.511263 | 1.345412324 | 0.21180104 | 1.46087110810 | 8.86321190585 | Km>KS.1 | NaCl | KS.1 |
| FIM1_2706 | 23.74627277 | 20.62334355 | 23.14872675 | -0.07904056 | 0.299914533 | 0.695936138 | 0.796476256 | ns | NaCl | KS.1 |
| FIM1_2708 | 5546.015871 | 13389.93926 | 5778.449195 | 1.010770193 | 0.289181081 | 1.35211400264 | 0.000111555 | Km>KS.1 | NaCl | KS.1 |
| FIM1_2709 | 3644.400043 | 4150.949267 | 3182.780214 | 0.164811766 | 0.113505587 | 0.118821614 | 0.212715916 | ns | NaCl | KS.1 |
| FIM1_2710 | 9604.944548 | 13700.48708 | 8876.301857 | 0.424394733 | 0.172593354 | 0.004371717 | 0.01459684 | ns | NaCl | KS.1 |
| FIM1_2711 | 430.2164044 | 348.3620966 | 333.9744302 | -0.270781356 | 0.124734059 | 0.017832374 | 0.045217156 | ns | NaCl | KS.1 |
| FIM1_2712 | 2846.313185 | 4211.675726 | 2749.901888 | 0.490658047 | 0.156225431 | 0.000387692 | 0.001952194 | ns | NaCl | KS.1 |
| FIM1_2713 | 1665.974027 | 1730.402726 | 1923.715799 | 0.047900991 | 0.103954605 | 0.624943864 | 0.738774851 | ns | NaCl | KS.1 |
| FIM1_2714 | 1204.354868 | 1170.874276 | 1521.290893 | -0.036566037 | 0.079149256 | 0.632178764 | 0.744047634 | ns | NaCl | KS.1 |
| FIM1_2715 | 3037.89952 | 4199.937652 | 2877.232405 | 0.438873507 | 0.095449313 | 1.18531436893 | 1.36187967331 | ns | NaCl | KS.1 |
| FIM1_2716 | 6055.507072 | 10508.30645 | 7226.098918 | 0.729422847 | 0.145941408 | 5.48680262352 | 9.31632868669 | ns | NaCl | KS.1 |
| FIM1_2718 | 3561.123421 | 4978.341829 | 5181.679644 | 0.452231425 | 0.099735631 | 1.51014043133 | 1.68451763360 | ns | NaCl | KS.1 |
| FIM1_2719 | 1987.303751 | 2985.854017 | 2146.318507 | 0.505811907 | 0.163422103 | 0.000424508 | 0.002101421 | ns | NaCl | KS.1 |
| FIM1_2720 | 2310.341793 | 2677.053479 | 3393.652226 | 0.199496252 | 0.077512741 | 0.007619181 | 0.02307086 | ns | NaCl | KS.1 |

| | | | | | | | | | | |
|---|---|---|---|---|---|---|---|---|---|---|
| FIM1_2721 | 510.7036009 | 1169.68793 | 627.1013239 | 0.982129613 | 0.261861354 | 6.0371654565 | 5.4921500785 | ns | NaCl | KS.1 |
| FIM1_2722 | 2947.721606 | 3738.68522 | 2572.884247 | 0.295742148 | 0.133914993 | 0.014745023 | 0.038671693 | ns | NaCl | KS.1 |
| FIM1_2723 | 753.7569627 | 510.1468281 | 810.7871717 | -0.405010221 | 0.2323927 | 0.027500555 | 0.06483891 | ns | NaCl | KS.1 |
| FIM1_2724 | 1036.588506 | 965.9006644 | 1911.980482 | -0.082233337 | 0.128025829 | 0.483350569 | 0.617531864 | ns | NaCl | KS.1 |
| FIM1_2726 | 591.4495355 | 327.3467859 | 789.6774471 | -0.761721018 | 0.164661468 | 3.1667848840 | 4.4130965820 | ns | NaCl | KS.1 |
| FIM1_2727 | 144.3631253 | 51.00975561 | 134.5850001 | -1.295211996 | 0.279448904 | 3.0305864444 | 5.7989106004 | Km<KS.1 | NaCl | KS.1 |
| FIM1_2728 | 300.947473 | 149.2568226 | 592.1077316 | -0.78888498 | 0.274157513 | 0.000269955 | 0.001434854 | ns | NaCl | KS.1 |
| FIM1_2729 | 24.73604221 | 51.4468033 | 331.3030649 | 0.609035989 | 0.375421941 | 0.012594165 | 0.03410777 | ns | NaCl | KS.1 |
| FIM1_2730 | 67.66194394 | 27.41071407 | 150.8965523 | -0.94069696 | 0.354089068 | 0.000251968 | 0.001356645 | ns | NaCl | KS.1 |
| FIM1_2731 | 3853.343787 | 4755.382267 | 4238.867516 | 0.277007394 | 0.101686729 | 0.003778465 | 0.012981951 | ns | NaCl | KS.1 |
| FIM1_2732 | 7726.213977 | 9756.674114 | 8924.198354 | 0.31143448 | 0.095636002 | 0.000576675 | 0.002716818 | ns | NaCl | KS.1 |
| FIM1_2733 | 223.0134426 | 361.2068721 | 314.6935133 | 0.601310323 | 0.175418097 | 8.793369092 | 0.000560859 | ns | NaCl | KS.1 |
| FIM1_2734 | 283.8154215 | 238.3112677 | 239.54723 | -0.206762245 | 0.143428948 | 0.107652071 | 0.196251028 | ns | NaCl | KS.1 |
| FIM1_2735 | 634.8250093 | 764.8442661 | 651.349206 | 0.229593662 | 0.129227398 | 0.051434379 | 0.108137384 | ns | NaCl | KS.1 |
| FIM1_2736 | 639.5094384 | 613.6989084 | 595.3209154 | -0.04990986 | 0.142043124 | 0.696289713 | 0.796514446 | ns | NaCl | KS.1 |
| FIM1_2737 | 3312.958741 | 3260.407285 | 2475.593407 | -0.020130102 | 0.11234183 | 0.847883066 | 0.907143711 | ns | NaCl | KS.1 |
| FIM1_2738 | 376.1211651 | 320.5296337 | 482.4117522 | -0.152871596 | 0.219221642 | 0.381683143 | 0.516947294 | ns | NaCl | KS.1 |
| FIM1_2739 | 2150.198216 | 982.093763 | 2229.439946 | -0.85709305 | 0.29534367 | 0.000186173 | 0.001052511 | ns | NaCl | KS.1 |
| FIM1_2740 | 5363.431321 | 7631.416093 | 5832.091995 | 0.488794458 | 0.080101067 | 2.2168254516 | 7.6906918004 | ns | NaCl | KS.1 |
| FIM1_2741 | 979.9096986 | 1033.855184 | 950.3022383 | 0.066169808 | 0.108136569 | 0.514258221 | 0.645091944 | ns | NaCl | KS.1 |
| FIM1_2742 | 4474.718713 | 6062.949599 | 6492.584788 | 0.306059337 | 0.223437109 | 0.085266267 | 0.163594169 | ns | NaCl | KS.1 |
| FIM1_2743 | 957.2373496 | 1321.380321 | 767.070899 | 0.292633838 | 0.255062445 | 0.128194939 | 0.224804309 | ns | NaCl | KS.1 |
| FIM1_2744 | 6794.480602 | 5365.781235 | 6478.222438 | -0.323209722 | 0.077596948 | 1.5028120890 | 0.000122165 | ns | NaCl | KS.1 |
| FIM1_2745 | 1752.567102 | 1942.731363 | 1426.642156 | 0.09723408 | 0.21308991 | 0.570873397 | 0.692875129 | ns | NaCl | KS.1 |
| FIM1_2746 | 1118.738253 | 1067.015755 | 1398.377701 | -0.058227582 | 0.125677801 | 0.614285861 | 0.729721146 | ns | NaCl | KS.1 |
| FIM1_2747 | 3438.758684 | 4236.061924 | 2989.429139 | 0.247324028 | 0.150012403 | 0.063195062 | 0.128219998 | ns | NaCl | KS.1 |
| FIM1_2748 | 457.6001414 | 379.0645398 | 558.3447268 | -0.23299997 | 0.140374852 | 0.064890329 | 0.130594412 | ns | NaCl | KS.1 |
| FIM1_2749 | 551.7349741 | 376.2704955 | 918.7957471 | -0.494345683 | 0.148701168 | 0.000199251 | 0.00111254 | ns | NaCl | KS.1 |
| FIM1_2750 | 92.58141777 | 91.14116402 | 94.23558957 | -0.027892961 | 0.190341606 | 0.861014899 | 0.915702134 | ns | NaCl | KS.1 |
| FIM1_2751 | 635.221037 | 672.5552827 | 582.8865073 | 0.06168466 | 0.177238621 | 0.683913449 | 0.786334506 | ns | NaCl | KS.1 |
| FIM1_2752 | 1253.886993 | 1165.17826 | 1326.891207 | -0.097745861 | 0.089444565 | 0.252915793 | 0.381174211 | ns | NaCl | KS.1 |
| FIM1_2753 | 1212.571209 | 1644.831741 | 1404.235879 | 0.370717911 | 0.157084094 | 0.00725849 | 0.022140396 | ns | NaCl | KS.1 |
| FIM1_2754 | 1415.648199 | 2225.079644 | 3605.0053 | 0.502997502 | 0.222415317 | 0.005279283 | 0.017032706 | ns | NaCl | KS.1 |
| FIM1_2755 | 1076.762556 | 972.9584222 | 1085.069687 | -0.126426729 | 0.118641745 | 0.249916354 | 0.378373055 | ns | NaCl | KS.1 |
| FIM1_2756 | 1001.624606 | 1021.849176 | 909.1344341 | 0.017127804 | 0.23212 | 0.924330917 | 0.956237537 | ns | NaCl | KS.1 |
| FIM1_2757 | 4932.622418 | 13438.38345 | 5736.513145 | 1.135238892 | 0.319347308 | 6.0386052119 | 5.4921500785 | Km>KS.1 | NaCl | KS.1 |
| FIM1_2758 | 779.2880067 | 685.2996151 | 579.5831668 | -0.150160591 | 0.14614866 | 0.250348715 | 0.378681927 | ns | NaCl | KS.1 |
| FIM1_2759 | 306.5861267 | 257.3381972 | 236.8115605 | -0.18667787 | 0.173601266 | 0.210486999 | 0.332752723 | ns | NaCl | KS.1 |
| FIM1_2760 | 6868.937472 | 9971.263505 | 8012.268097 | 0.414484588 | 0.205140541 | 0.014200898 | 0.037619526 | ns | NaCl | KS.1 |
| FIM1_2761 | 1389.503421 | 1411.692618 | 1486.793316 | 0.019362578 | 0.069693442 | 0.77512809 | 0.853721994 | ns | NaCl | KS.1 |
| FIM1_2762 | 2240.549702 | 2821.743937 | 2559.5786 | 0.252237913 | 0.184600898 | 0.10563205 | 0.193173904 | ns | NaCl | KS.1 |
| FIM1_2763 | 15.96581056 | 161.012473 | 643.8802253 | 2.157515404 | 0.867612295 | 5.2529441074 | 4.9122926569 | Km>KS.1 | NaCl | KS.1 |
| FIM1_2764 | 1393.591535 | 699.0002462 | 1403.205605 | -0.886123123 | 0.182932262 | 7.3706877627 | 1.2223057206 | ns | NaCl | KS.1 |
| FIM1_2765 | 1579.569678 | 1371.482643 | 1273.675358 | -0.174430979 | 0.122014051 | 0.120799077 | 0.215326195 | ns | NaCl | KS.1 |
| FIM1_2766 | 3982.863984 | 4506.48156 | 3595.941157 | 0.156244618 | 0.114932571 | 0.143630087 | 0.245637567 | ns | NaCl | KS.1 |

| | | | | | | | | | | |
|---|---|---|---|---|---|---|---|---|---|---|
| FIM1_2767 | 1913.323419 | 1467.649359 | 2284.431879 | -0.288450802 | 0.192123949 | 0.072251969 | 0.142602419 | ns | NaCl | KS.1 |
| FIM1_2768 | 1233.501107 | 855.5531271 | 1224.0717 | -0.417953795 | 0.192074217 | 0.009554914 | 0.027556925 | ns | NaCl | KS.1 |
| FIM1_2769 | 2107.130619 | 2054.631098 | 2209.538832 | -0.033445547 | 0.093246793 | 0.706628895 | 0.802969348 | ns | NaCl | KS.1 |
| FIM1_2770 | 943.6475782 | 588.153344 | 1412.254494 | -0.590608043 | 0.173994075 | 0.000103718 | 0.000644193 | ns | NaCl | KS.1 |
| FIM1_2771 | 495.3561339 | 447.5221959 | 445.9064314 | -0.105919244 | 0.186460528 | 0.499987536 | 0.631971034 | ns | NaCl | KS.1 |
| FIM1_2772 | 1110.923288 | 1255.67275 | 1609.891903 | 0.158954943 | 0.096532494 | 0.082473047 | 0.159093994 | ns | NaCl | KS.1 |
| FIM1_2773 | 385.9258676 | 386.1053472 | 405.4531913 | -0.002488749 | 0.160053782 | 0.985866039 | 0.993051943 | ns | NaCl | KS.1 |
| FIM1_2774 | 804.2294528 | 831.053994 | 928.7320496 | 0.031215908 | 0.217469617 | 0.857323683 | 0.913690395 | ns | NaCl | KS.1 |
| FIM1_2775 | 3389.446264 | 5023.092338 | 4011.411568 | 0.544773741 | 0.0848483 | 2.2632546011 | 9.461925748 | ns | NaCl | KS.1 |
| FIM1_2776 | 512.8588613 | 596.1085826 | 718.1820608 | 0.18528939 | 0.126708294 | 0.110380495 | 0.200052081 | ns | NaCl | KS.1 |
| FIM1_2777 | 717.9381998 | 394.0792213 | 817.4033035 | -0.792461564 | 0.157559221 | 3.887979080 | 7.190593280 | ns | NaCl | KS.1 |
| FIM1_2778 | 524.8337708 | 574.9384498 | 579.844977 | 0.110864331 | 0.116326005 | 0.304927558 | 0.437306025 | ns | NaCl | KS.1 |
| FIM1_2779 | 1030.179917 | 1231.099347 | 941.1714081 | 0.217459729 | 0.13717213 | 0.079228649 | 0.153969737 | ns | NaCl | KS.1 |
| FIM1_2780 | 557.1518965 | 82.12240068 | 312.6164471 | -2.617427399 | 0.299037322 | 3.892900789 | 1.291145428 | Km<KS.1 | NaCl | KS.1 |
| FIM1_2781 | 894.767518 | 672.819288 | 1642.397071 | -0.262976267 | 0.243158012 | 0.159144086 | 0.267542388 | ns | NaCl | KS.1 |
| FIM1_2782 | 2144.542135 | 2100.485664 | 1563.850147 | -0.024501718 | 0.139800727 | 0.846113513 | 0.905834888 | ns | NaCl | KS.1 |
| FIM1_2783 | 5603.344991 | 14550.53808 | 6384.891685 | 1.073813543 | 0.313485355 | 1.280940375 | 0.000106688 | Km>KS.1 | NaCl | KS.1 |
| FIM1_2784 | 856.9685677 | 707.5716706 | 907.651831 | -0.25571753 | 0.094903697 | 0.004467185 | 0.014835946 | ns | NaCl | KS.1 |
| FIM1_2785 | 2833.497555 | 2629.902128 | 2810.77649 | -0.094670966 | 0.112095224 | 0.365728889 | 0.501240007 | ns | NaCl | KS.1 |
| FIM1_2786 | 1945.548608 | 2330.511342 | 2411.717556 | 0.215483009 | 0.145189738 | 0.09679926 | 0.180163232 | ns | NaCl | KS.1 |
| FIM1_2787 | 5244.678497 | 6641.169744 | 5756.818924 | 0.215329251 | 0.238872605 | 0.242744097 | 0.36953852 | ns | NaCl | KS.1 |
| FIM1_2788 | 199.1934203 | 131.4842797 | 344.3151537 | -0.464768676 | 0.20610458 | 0.006352132 | 0.019875382 | ns | NaCl | KS.1 |
| FIM1_2789 | 274.5463543 | 232.0453632 | 274.6503493 | -0.178827443 | 0.188420189 | 0.258166181 | 0.386598155 | ns | NaCl | KS.1 |
| FIM1_2790 | 1006.430512 | 695.8861326 | 1088.731515 | -0.478011621 | 0.134838454 | 9.408970681 | 0.000593278 | ns | NaCl | KS.1 |
| FIM1_2791 | 192.3748022 | 131.6294977 | 239.3295169 | -0.423036506 | 0.203240089 | 0.011814292 | 0.032554957 | ns | NaCl | KS.1 |
| FIM1_2792 | 784.3515824 | 635.5305989 | 699.8149305 | -0.245476622 | 0.158360223 | 0.077248277 | 0.150650795 | ns | NaCl | KS.1 |
| FIM1_2793 | 1490.4092 | 1558.947011 | 1821.019608 | 0.059497371 | 0.086917779 | 0.475113652 | 0.608727895 | ns | NaCl | KS.1 |
| FIM1_2794 | 2292.518269 | 2397.397487 | 2224.607093 | 0.05525876 | 0.126057962 | 0.633335996 | 0.745057125 | ns | NaCl | KS.1 |
| FIM1_2795 | 550.7824621 | 648.5372656 | 653.7490727 | 0.146878706 | 0.2331454 | 0.417703513 | 0.553562861 | ns | NaCl | KS.1 |
| FIM1_2796 | 220.5606968 | 177.0802397 | 654.7569723 | -0.174956983 | 0.272438633 | 0.373809001 | 0.50880979 | ns | NaCl | KS.1 |
| FIM1_2797 | 34.8271751 | 28.6627284 | 53.41088707 | -0.156063957 | 0.290010367 | 0.438607881 | 0.57453244 | ns | NaCl | KS.1 |
| FIM1_2798 | 1397.755856 | 1797.326162 | 1657.686551 | 0.322689073 | 0.121475471 | 0.003852675 | 0.013155154 | ns | NaCl | KS.1 |
| FIM1_2800 | 2407.701287 | 5103.193765 | 3329.2877 | 1.051357247 | 0.100303806 | 3.773533887 | 2.681904441 | Km>KS.1 | NaCl | KS.1 |
| FIM1_2801 | 4840.352705 | 5244.585633 | 4352.614611 | 0.099661442 | 0.119243398 | 0.366740069 | 0.502214838 | ns | NaCl | KS.1 |
| FIM1_2802 | 624.8527946 | 774.5964941 | 855.5880216 | 0.266640009 | 0.134214289 | 0.028270135 | 0.066310194 | ns | NaCl | KS.1 |
| FIM1_2803 | 957.3383391 | 1263.939828 | 1290.444468 | 0.367475394 | 0.103348992 | 0.000150084 | 0.000877401 | ns | NaCl | KS.1 |
| FIM1_2804 | 2180.243348 | 2252.152197 | 2299.026539 | 0.042653691 | 0.086768796 | 0.608127254 | 0.724943649 | ns | NaCl | KS.1 |
| FIM1_2805 | 2494.673237 | 1990.721209 | 1868.510163 | -0.288455542 | 0.120493624 | 0.009318025 | 0.027077789 | ns | NaCl | KS.1 |
| FIM1_2806 | 180.6491771 | 165.1593563 | 204.5735688 | -0.109587103 | 0.161050617 | 0.437173358 | 0.573106048 | ns | NaCl | KS.1 |
| FIM1_2807 | 1369.010909 | 1047.114329 | 1116.620185 | -0.329564309 | 0.144146438 | 0.010517307 | 0.029712439 | ns | NaCl | KS.1 |
| FIM1_2808 | 1507.406696 | 1596.458668 | 1536.198656 | 0.067816628 | 0.13844736 | 0.588065704 | 0.709414859 | ns | NaCl | KS.1 |
| FIM1_2809 | 104.2813182 | 75.62115082 | 172.0515214 | -0.299033876 | 0.249018009 | 0.115297146 | 0.207226626 | ns | NaCl | KS.1 |
| FIM1_2810 | 189.6691615 | 256.9855679 | 237.534163 | 0.258073782 | 0.26377489 | 0.186167173 | 0.302541593 | ns | NaCl | KS.1 |
| FIM1_2811 | 353.3288348 | 473.1796072 | 460.8452583 | 0.367359784 | 0.140908042 | 0.003677157 | 0.012695249 | ns | NaCl | KS.1 |
| FIM1_2812 | 2897.256513 | 2673.452841 | 2959.470854 | -0.104396734 | 0.094599632 | 0.245885035 | 0.373405998 | ns | NaCl | KS.1 |

| | | | | | | | | | | | |
|---|---|---|---|---|---|---|---|---|---|---|---|
| FIM1_2813 | 802.5934752 | 671.5842434 | 1060.003627 | -0.213736452 | 0.140514187 | 0.090776084 | 0.17173184 | ns | | NaCl | KS.1 |
| FIM1_2814 | 2133.272633 | 1565.43059 | 2193.081339 | -0.387887739 | 0.142910731 | 0.002441128 | 0.009016043 | ns | | NaCl | KS.1 |
| FIM1_2815 | 491.1697953 | 354.5345177 | 337.3905196 | -0.401754944 | 0.159490934 | 0.004076907 | 0.013769595 | ns | | NaCl | KS.1 |
| FIM1_2816 | 403.0978355 | 404.3277354 | 373.392304 | 0.000651527 | 0.148438847 | 0.996076116 | 0.998234281 | ns | | NaCl | KS.1 |
| FIM1_2817 | 695.9656493 | 591.1666045 | 534.8311899 | -0.195959797 | 0.148294356 | 0.137773973 | 0.238077637 | ns | | NaCl | KS.1 |
| FIM1_2818 | 240.0077101 | 292.9967956 | 428.9503715 | 0.243267564 | 0.133924844 | 0.045082458 | 0.097135222 | ns | | NaCl | KS.1 |
| FIM1_2819 | 2343.871047 | 2009.055458 | 2042.335681 | -0.205774874 | 0.082406358 | 0.009307605 | 0.027063317 | ns | | NaCl | KS.1 |
| FIM1_2820 | 1523.141645 | 1417.433533 | 2772.900424 | -0.067081518 | 0.21687217 | 0.698830131 | 0.797952697 | ns | | NaCl | KS.1 |
| FIM1_2821 | 1175.558174 | 1432.859414 | 1743.736036 | 0.248003727 | 0.126138803 | 0.031693326 | 0.072945429 | ns | | NaCl | KS.1 |
| FIM1_2822 | 302.3836771 | 635.4563286 | 639.5826493 | 0.991785162 | 0.161084621 | 3.2336407638 | 1.2869890240 | ns | | NaCl | KS.1 |
| FIM1_2823 | 4333.953857 | 4337.258707 | 3390.920902 | 0.00054753 | 0.08796371 | 0.9948147 | 0.997622079 | ns | | NaCl | KS.1 |
| FIM1_2824 | 982.666995 | 1212.659399 | 1247.898865 | 0.280018651 | 0.090931429 | 0.001203364 | 0.005005631 | ns | | NaCl | KS.1 |
| FIM1_2825 | 954.8851352 | 899.3936646 | 1050.128606 | -0.078435309 | 0.113819538 | 0.459722364 | 0.594520083 | ns | | NaCl | KS.1 |
| FIM1_2826 | 691.4826706 | 472.5468576 | 888.972011 | -0.445101074 | 0.18597634 | 0.004785983 | 0.015705981 | ns | | NaCl | KS.1 |
| FIM1_2827 | 3416.979178 | 6889.823882 | 3463.262847 | 0.822568123 | 0.246396245 | 5.2345508512 | 0.000361191 | ns | | NaCl | KS.1 |
| FIM1_2828 | 485.2000097 | 426.564749 | 768.7387174 | -0.160822049 | 0.117099065 | 0.138532699 | 0.238973709 | ns | | NaCl | KS.1 |
| FIM1_2829 | 2028.814908 | 3111.605749 | 2250.230989 | 0.562437496 | 0.133430875 | 4.1029244323 | 4.0580614414 | ns | | NaCl | KS.1 |
| FIM1_2830 | 2393.239699 | 2194.016618 | 2405.026122 | -0.104302922 | 0.137731247 | 0.402578863 | 0.538105815 | ns | | NaCl | KS.1 |
| FIM1_2831 | 250.983518 | 152.9725104 | 339.8605135 | -0.356130947 | 0.340462634 | 0.106869014 | 0.19496639 | ns | | NaCl | KS.1 |
| FIM1_2832 | 951.8062354 | 1260.215677 | 1539.108163 | 0.363769246 | 0.120403185 | 0.001026118 | 0.004378161 | ns | | NaCl | KS.1 |
| FIM1_2833 | 2123.896447 | 3744.482924 | 1969.477907 | 0.541182488 | 0.296592966 | 0.012366357 | 0.033618123 | ns | | NaCl | KS.1 |
| FIM1_2834 | 937.0155338 | 850.9211991 | 1728.387067 | -0.07459287 | 0.263606787 | 0.698450886 | 0.797885914 | ns | | NaCl | KS.1 |
| FIM1_2835 | 1397.656684 | 1073.110703 | 1076.06063 | -0.232365246 | 0.25293356 | 0.222584865 | 0.347128183 | ns | | NaCl | KS.1 |
| FIM1_2836 | 735.1952843 | 719.5433293 | 864.2023526 | -0.02662955 | 0.127958964 | 0.820455283 | 0.889079728 | ns | | NaCl | KS.1 |
| FIM1_2837 | 577.7368525 | 582.935828 | 858.4962492 | 0.008558964 | 0.229753065 | 0.9619254 | 0.981455879 | ns | | NaCl | KS.1 |
| FIM1_2838 | 3696.659486 | 3008.119576 | 2749.047553 | -0.263710343 | 0.115726091 | 0.01389302 | 0.036941622 | ns | | NaCl | KS.1 |
| FIM1_2839 | 493.2451013 | 449.0295849 | 734.8702793 | -0.123082036 | 0.108273888 | 0.225177736 | 0.349971645 | ns | | NaCl | KS.1 |
| FIM1_2840 | 86.04762402 | 69.17136346 | 112.8839267 | -0.227065655 | 0.203092319 | 0.172841484 | 0.285297407 | ns | | NaCl | KS.1 |
| FIM1_2841 | 1184.470113 | 1219.777828 | 1116.009165 | 0.035842195 | 0.118607519 | 0.744526691 | 0.833300402 | ns | | NaCl | KS.1 |
| FIM1_2842 | 4126.341345 | 4617.806232 | 3122.825629 | 0.145318396 | 0.103985307 | 0.137597445 | 0.237893991 | ns | | NaCl | KS.1 |
| FIM1_2843 | 148.7210812 | 147.800624 | 223.4923183 | -0.008838116 | 0.176619768 | 0.953373618 | 0.975932871 | ns | | NaCl | KS.1 |
| FIM1_2844 | 535.1921018 | 364.3885683 | 525.3170241 | -0.453666461 | 0.185918515 | 0.00405024 | 0.013688819 | ns | | NaCl | KS.1 |
| FIM1_2845 | 36364.48506 | 72669.52633 | 43296.34962 | 0.948375986 | 0.124753104 | 1.6024592417 | 1.6269866791 | ns | | NaCl | KS.1 |
| FIM1_2846 | 1507.227001 | 2076.246365 | 1216.290288 | 0.294051125 | 0.252095118 | 0.123934521 | 0.219012775 | ns | | NaCl | KS.1 |
| FIM1_2847 | 1484.95371 | 580.0159888 | 3194.810321 | -0.830942888 | 0.404862749 | 0.00178052 | 0.006963906 | ns | | NaCl | KS.1 |
| FIM1_2848 | 836.6889283 | 575.2834078 | 706.3170098 | -0.470276727 | 0.151753421 | 0.000489229 | 0.002365319 | ns | | NaCl | KS.1 |
| FIM1_2849 | 18504.06835 | 23620.2018 | 14759.95724 | 0.313024073 | 0.119611948 | 0.004503744 | 0.014927467 | ns | | NaCl | KS.1 |
| FIM1_2850 | 2743.910868 | 2815.608339 | 2126.038905 | 0.029755713 | 0.142646315 | 0.816626331 | 0.886475234 | ns | | NaCl | KS.1 |
| FIM1_2851 | 2113.485235 | 2028.353482 | 2482.05434 | -0.044868781 | 0.16373998 | 0.75356118 | 0.839446232 | ns | | NaCl | KS.1 |
| FIM1_2852 | 928.6448615 | 669.2634104 | 869.8070042 | -0.402523062 | 0.153682152 | 0.003038727 | 0.01082024 | ns | | NaCl | KS.1 |
| FIM1_2853 | 4243.114288 | 9023.538659 | 4109.292691 | 0.813482394 | 0.296317984 | 0.000358195 | 0.001826744 | ns | | NaCl | KS.1 |
| FIM1_2854 | 2280.157696 | 1918.588625 | 1743.33323 | -0.202084151 | 0.150782217 | 0.130892905 | 0.22872926 | ns | | NaCl | KS.1 |
| FIM1_2855 | 156.7071846 | 67.94327857 | 121.4704813 | -1.00176953 | 0.258119987 | 3.3459862489 | 3.4111232763 | Km<KS.1 | | NaCl | KS.1 |
| FIM1_2856 | 415.0250976 | 322.1214604 | 634.690912 | -0.285996437 | 0.180228818 | 0.061960126 | 0.126332634 | ns | | NaCl | KS.1 |
| FIM1_2857 | 945.124757 | 1149.320243 | 751.6658429 | 0.227124197 | 0.158012523 | 0.1015948 | 0.187475567 | ns | | NaCl | KS.1 |

| | | | | | | | | | | |
|---|---|---|---|---|---|---|---|---|---|---|
| FIM1_2858 | 1098.506718 | 1211.660569 | 1021.325247 | 0.122662521 | 0.122874582 | 0.278708732 | 0.408605338 | ns | NaCl | KS.1 |
| FIM1_2859 | 3864.433177 | 4067.740071 | 3942.214549 | 0.058975511 | 0.153071681 | 0.663826803 | 0.770361172 | ns | NaCl | KS.1 |
| FIM1_2860 | 18778.73878 | 15626.46283 | 11384.25993 | -0.158256043 | 0.246299084 | 0.397210194 | 0.533078153 | ns | NaCl | KS.1 |
| FIM1_2862 | 227.906741 | 205.1835244 | 207.0002085 | -0.110904297 | 0.18768541 | 0.48202517 | 0.615996717 | ns | NaCl | KS.1 |
| FIM1_2863 | 784.2495042 | 1154.667836 | 1263.715346 | 0.423127527 | 0.215074062 | 0.015487501 | 0.040130373 | ns | NaCl | KS.1 |
| FIM1_2864 | 4099.655099 | 5326.156061 | 3071.00938 | 0.327019728 | 0.136138061 | 0.007799987 | 0.023518143 | ns | NaCl | KS.1 |
| FIM1_2865 | 1132.854668 | 1487.508655 | 1040.611865 | 0.318546983 | 0.166355659 | 0.027271854 | 0.064448778 | ns | NaCl | KS.1 |
| FIM1_2866 | 529.3136974 | 406.655528 | 682.8797352 | -0.315905444 | 0.154003492 | 0.020053351 | 0.04970873 | ns | NaCl | KS.1 |
| FIM1_2867 | 1266.74894 | 1209.485063 | 1333.668172 | -0.056414986 | 0.136680124 | 0.648902319 | 0.756748485 | ns | NaCl | KS.1 |
| FIM1_2868 | 51.41017075 | 16.8674488 | 53.04285322 | -1.145922332 | 0.395383076 | 4.6958284128 | 0.000329968 | Km<KS.1 | NaCl | KS.1 |
| FIM1_2869 | 65.55605308 | 19.73034058 | 263.2725796 | -1.405027437 | 0.341443764 | 1.7318789365 | 2.5952101534 | Km<KS.1 | NaCl | KS.1 |
| FIM1_2870 | 4680.095744 | 6085.428439 | 4619.041052 | 0.312034821 | 0.158157168 | 0.024503981 | 0.05900644 | ns | NaCl | KS.1 |
| FIM1_2871 | 925.2134913 | 1027.266036 | 1028.851863 | 0.131808124 | 0.112907986 | 0.210483048 | 0.332752723 | ns | NaCl | KS.1 |
| FIM1_2872 | 1278.393563 | 1470.542681 | 3341.988258 | 0.16553329 | 0.141019053 | 0.19200979 | 0.309342198 | ns | NaCl | KS.1 |
| FIM1_2873 | 453.5374952 | 477.0480167 | 379.4106986 | 0.052507073 | 0.171932275 | 0.723123588 | 0.816692332 | ns | NaCl | KS.1 |
| FIM1_2874 | 7326.803247 | 4269.918124 | 5304.524092 | -0.690551837 | 0.169591338 | 4.8786194859 | 4.6318954088 | ns | NaCl | KS.1 |
| FIM1_2875 | 2735.079915 | 3672.798511 | 2510.602835 | 0.400164672 | 0.092188371 | 4.7790758132 | 4.5722888789 | ns | NaCl | KS.1 |
| FIM1_2876 | 2401.406907 | 2252.543367 | 2282.743622 | -0.081127266 | 0.106504132 | 0.417586622 | 0.553555407 | ns | NaCl | KS.1 |
| FIM1_2877 | 4194.74392 | 8402.52423 | 3588.738224 | 0.751828675 | 0.283095811 | 0.000607613 | 0.002838378 | ns | NaCl | KS.1 |
| FIM1_2878 | 1197.601906 | 1019.752433 | 1245.67938 | -0.199613791 | 0.127676271 | 0.08735525 | 0.1667313 | ns | NaCl | KS.1 |
| FIM1_2879 | 1084.772939 | 1233.217232 | 1558.259213 | 0.147815916 | 0.151423881 | 0.271302811 | 0.400276242 | ns | NaCl | KS.1 |
| FIM1_2880 | 1058.94099 | 1369.062332 | 1078.416442 | 0.312881142 | 0.144987078 | 0.015636499 | 0.040406474 | ns | NaCl | KS.1 |
| FIM1_2881 | 1422.685901 | 1612.283022 | 1233.208042 | 0.137066102 | 0.169042246 | 0.348631691 | 0.483400966 | ns | NaCl | KS.1 |
| FIM1_2882 | 1048.286781 | 1260.367 | 1469.445907 | 0.226802917 | 0.132628237 | 0.059737612 | 0.122756968 | ns | NaCl | KS.1 |
| FIM1_2883 | 1849.546784 | 2046.675196 | 2329.138013 | 0.132934773 | 0.094932349 | 0.140585571 | 0.241676993 | ns | NaCl | KS.1 |
| FIM1_2884 | 532.058217 | 679.2223239 | 690.2018904 | 0.314933306 | 0.118539245 | 0.00397458 | 0.013478892 | ns | NaCl | KS.1 |
| FIM1_2885 | 1719.034418 | 3554.522471 | 1656.05392 | 0.69538356 | 0.332084124 | 0.003280704 | 0.011477848 | ns | NaCl | KS.1 |
| FIM1_2886 | 876.1571464 | 2086.807034 | 931.8734118 | 0.845160367 | 0.358357278 | 0.000853334 | 0.003770279 | ns | NaCl | KS.1 |
| FIM1_2887 | 5063.006369 | 9978.788837 | 4640.995791 | 0.728132076 | 0.283253355 | 0.000855963 | 0.00377854 | ns | NaCl | KS.1 |
| FIM1_2888 | 284.7282939 | 134.834288 | 389.7551072 | -0.955424857 | 0.200742883 | 8.6337090285 | 1.4022341917 | ns | NaCl | KS.1 |
| FIM1_2889 | 2046.294499 | 2061.973874 | 2031.263372 | 0.008359781 | 0.097047201 | 0.927710571 | 0.95833889 | ns | NaCl | KS.1 |
| FIM1_2890 | 4401.055318 | 4021.245604 | 4347.024313 | -0.114890589 | 0.107057546 | 0.252904637 | 0.381174211 | ns | NaCl | KS.1 |
| FIM1_2891 | 6646.150761 | 4461.771767 | 4319.216258 | -0.548335642 | 0.091016649 | 2.8111345582 | 9.3861707565 | ns | NaCl | KS.1 |
| FIM1_2892 | 304.7897325 | 181.1808256 | 304.0822981 | -0.659987725 | 0.173153964 | 1.6005538508 | 0.000129266 | ns | NaCl | KS.1 |
| FIM1_2893 | 859.8385832 | 472.1569571 | 642.7355356 | -0.789579598 | 0.157040388 | 3.9562952765 | 7.2097322347 | ns | NaCl | KS.1 |
| FIM1_2894 | 745.7211174 | 717.7632851 | 1040.38808 | -0.050195045 | 0.129772898 | 0.672402303 | 0.777592156 | ns | NaCl | KS.1 |
| FIM1_2895 | 2924.174881 | 3287.883119 | 3125.276002 | 0.149538298 | 0.109166959 | 0.143259937 | 0.245296269 | ns | NaCl | KS.1 |
| FIM1_2896 | 1369.237293 | 1658.52273 | 1441.378662 | 0.255426421 | 0.093802918 | 0.004104519 | 0.013853447 | ns | NaCl | KS.1 |
| FIM1_2897 | 4850.611439 | 6526.592575 | 5776.34956 | 0.339908742 | 0.178952398 | 0.025867245 | 0.061780865 | ns | NaCl | KS.1 |
| FIM1_2898 | 19874.75436 | 26003.6777 | 16191.39412 | 0.348169661 | 0.117519865 | 0.001332909 | 0.005435428 | ns | NaCl | KS.1 |
| FIM1_2899 | 9913.159121 | 20796.20102 | 9340.909001 | 0.924533255 | 0.213895315 | 7.3301557892 | 8.9600798652 | ns | NaCl | KS.1 |
| FIM1_2900 | 4925.065852 | 11457.17698 | 6646.927741 | 1.132069466 | 0.163025614 | 8.9174125254 | 7.5193436125 | Km>KS.1 | NaCl | KS.1 |
| FIM1_2901 | 1636.429994 | 2015.224666 | 1677.89953 | 0.268321106 | 0.114424089 | 0.011472586 | 0.031815003 | ns | NaCl | KS.1 |
| FIM1_2903 | 2212.609045 | 1792.133974 | 2443.122967 | -0.26060729 | 0.13116008 | 0.028878743 | 0.067483206 | ns | NaCl | KS.1 |
| FIM1_2904 | 1974.111207 | 1569.422803 | 1819.270302 | -0.25546981 | 0.177700843 | 0.091982657 | 0.173469946 | ns | NaCl | KS.1 |

| | | | | | | | | | | |
|---|---|---|---|---|---|---|---|---|---|---|
| FIM1_2905 | 830.8290002 | 797.6679403 | 846.2092377 | -0.032021134 | 0.255999113 | 0.865976386 | 0.919776371 | ns | NaCl | KS.1 |
| FIM1_2906 | 149.9439286 | 132.751865 | 192.2999731 | -0.127161108 | 0.206187146 | 0.449236676 | 0.584301298 | ns | NaCl | KS.1 |
| FIM1_2907 | 1453.123775 | 1243.319134 | 990.5456322 | -0.167101951 | 0.18193773 | 0.278755079 | 0.408605338 | ns | NaCl | KS.1 |
| FIM1_2908 | 2374.851943 | 3150.89324 | 2481.130868 | 0.371753905 | 0.11171092 | 0.000342822 | 0.001761919 | ns | NaCl | KS.1 |
| FIM1_2909 | 1190.32533 | 1420.259612 | 1242.605634 | 0.2287324 | 0.109600592 | 0.025387668 | 0.060693728 | ns | NaCl | KS.1 |
| FIM1_2911 | 360.22502 | 244.7860869 | 424.6130829 | -0.366945259 | 0.255776205 | 0.058703393 | 0.120981517 | ns | NaCl | KS.1 |
| FIM1_2912 | 1766.381256 | 2742.191175 | 2982.181995 | 0.429872684 | 0.261132988 | 0.029882016 | 0.069476366 | ns | NaCl | KS.1 |
| FIM1_2913 | 8.135414979 | 21.46863589 | 14.35777334 | 0.666665318 | 0.459083208 | 0.010956442 | 0.030729477 | ns | NaCl | KS.1 |
| FIM1_2914 | 2381.705488 | 3155.96318 | 1880.422927 | 0.229367353 | 0.273539869 | 0.247087904 | 0.374660873 | ns | NaCl | KS.1 |
| FIM1_2915 | 1608.709128 | 749.9475364 | 1489.707832 | -1.000471999 | 0.176620866 | 5.9900236454 | 1.7844531518 | Km<KS.1 | NaCl | KS.1 |
| FIM1_2916 | 4977.376719 | 5215.637882 | 5132.835234 | 0.054160646 | 0.145107978 | 0.677132565 | 0.780625275 | ns | NaCl | KS.1 |
| FIM1_2917 | 1565.388421 | 910.6090703 | 1446.996132 | -0.743382922 | 0.108103794 | 5.8756900157 | 3.8462576089 | ns | NaCl | KS.1 |
| FIM1_2918 | 1300.612468 | 1021.948492 | 985.6773065 | -0.307566109 | 0.12166524 | 0.005952731 | 0.018862954 | ns | NaCl | KS.1 |
| FIM1_2919 | 2368.299188 | 3001.956878 | 3139.017352 | 0.308281812 | 0.110877314 | 0.002812466 | 0.010139142 | ns | NaCl | KS.1 |
| FIM1_2920 | 366.3639344 | 425.0346812 | 333.924461 | 0.142081289 | 0.214556752 | 0.40985211 | 0.545045241 | ns | NaCl | KS.1 |
| FIM1_2921 | 3360.368256 | 5043.319366 | 2861.91268 | 0.386137827 | 0.26072876 | 0.049629503 | 0.105022023 | ns | NaCl | KS.1 |
| FIM1_2922 | 602.4800309 | 535.5417567 | 940.8518398 | -0.131771561 | 0.162486816 | 0.353280627 | 0.488475939 | ns | NaCl | KS.1 |
| FIM1_2923 | 3802.240852 | 5921.746952 | 3964.760454 | 0.589495567 | 0.126852243 | 4.9460698091 | 6.4415437959 | ns | NaCl | KS.1 |
| FIM1_2924 | 6877.517034 | 9572.184702 | 6416.49765 | 0.426013283 | 0.130000909 | 0.000320798 | 0.001657236 | ns | NaCl | KS.1 |
| FIM1_2925 | 58.71780904 | 40.22334985 | 55.21287527 | -0.283426329 | 0.315589952 | 0.181107613 | 0.296677765 | ns | NaCl | KS.1 |
| FIM1_2926 | 2667.32098 | 2309.848389 | 2247.740019 | -0.157521816 | 0.171385431 | 0.286238125 | 0.417238403 | ns | NaCl | KS.1 |
| FIM1_2927 | 5920.052167 | 6004.431409 | 5522.650867 | 0.019810779 | 0.069742877 | 0.770217561 | 0.851527619 | ns | NaCl | KS.1 |
| FIM1_2928 | 874.4878737 | 634.5380025 | 1364.35551 | -0.291242871 | 0.254466691 | 0.129471777 | 0.226817827 | ns | NaCl | KS.1 |
| FIM1_2929 | 1115.33374 | 898.3875834 | 1704.738101 | -0.222833774 | 0.201694491 | 0.178928543 | 0.29397936 | ns | NaCl | KS.1 |
| FIM1_2930 | 320.7642052 | 541.0906171 | 314.4429203 | 0.459299671 | 0.306674573 | 0.034032505 | 0.077346602 | ns | NaCl | KS.1 |
| FIM1_2931 | 3671.712981 | 5262.768383 | 3084.728023 | 0.451633181 | 0.149802058 | 0.000705475 | 0.003219944 | ns | NaCl | KS.1 |
| FIM1_2932 | 906.0275823 | 849.2175448 | 921.3139034 | -0.075235782 | 0.135752582 | 0.541388757 | 0.66751154 | ns | NaCl | KS.1 |
| FIM1_2933 | 2240.619681 | 2647.35516 | 2741.375958 | 0.219755769 | 0.094031526 | 0.01382544 | 0.036840687 | ns | NaCl | KS.1 |
| FIM1_2934 | 3395.404099 | 6035.050383 | 3436.724809 | 0.724043552 | 0.185224538 | 8.6246434192 | 7.5144660263 | ns | NaCl | KS.1 |
| FIM1_2935 | 14393.90869 | 20606.09906 | 14252.74012 | 0.440049038 | 0.161270022 | 0.001851877 | 0.007175303 | ns | NaCl | KS.1 |
| FIM1_2936 | 3649.656277 | 7794.054858 | 4841.660457 | 0.982096845 | 0.18797854 | 7.3493306070 | 1.6395928148 | ns | NaCl | KS.1 |
| FIM1_2937 | 1421.933107 | 1093.90917 | 929.7319652 | -0.290924096 | 0.18409704 | 0.061573053 | 0.125594891 | ns | NaCl | KS.1 |
| FIM1_2939 | 501.9332839 | 242.1023475 | 579.7654092 | -0.941357165 | 0.190310301 | 3.6232979541 | 6.8022291780 | ns | NaCl | KS.1 |
| FIM1_2941 | 1842.856724 | 1864.561013 | 1537.092147 | 0.013977779 | 0.127639608 | 0.904977896 | 0.943673241 | ns | NaCl | KS.1 |
| FIM1_2942 | 7334.163376 | 5900.617743 | 7572.433439 | -0.273131922 | 0.126754528 | 0.018400591 | 0.04639784 | ns | NaCl | KS.1 |
| FIM1_2943 | 1343.732095 | 1086.757914 | 1210.279347 | -0.277062726 | 0.108556914 | 0.006262365 | 0.019681154 | ns | NaCl | KS.1 |
| FIM1_2944 | 1012.347891 | 1255.181467 | 950.1685742 | 0.259677508 | 0.142906463 | 0.042490239 | 0.092475236 | ns | NaCl | KS.1 |
| FIM1_2945 | 891.8672964 | 998.3744877 | 787.5940644 | 0.121437756 | 0.174001289 | 0.416400749 | 0.552572346 | ns | NaCl | KS.1 |
| FIM1_2946 | 1399.147248 | 1352.392127 | 1978.107633 | -0.04577054 | 0.099506533 | 0.627155254 | 0.739711913 | ns | NaCl | KS.1 |
| FIM1_2947 | 3826.46 | 4625.029305 | 3679.15989 | 0.240061279 | 0.120532481 | 0.030688046 | 0.071010712 | ns | NaCl | KS.1 |
| FIM1_2948 | 860.400587 | 766.6323387 | 1174.689626 | -0.147319404 | 0.12375702 | 0.195753419 | 0.314254037 | ns | NaCl | KS.1 |
| FIM1_2949 | 3717.185713 | 2689.671263 | 3327.628688 | -0.407168183 | 0.141569896 | 0.001353618 | 0.005479454 | ns | NaCl | KS.1 |
| FIM1_2950 | 59.35396024 | 59.42693052 | 91.11159101 | -0.009122938 | 0.220434422 | 0.95843298 | 0.979097346 | ns | NaCl | KS.1 |
| FIM1_2951 | 2107.95151 | 2549.242366 | 2696.25655 | 0.23645734 | 0.128738427 | 0.044136968 | 0.095511707 | ns | NaCl | KS.1 |
| FIM1_2952 | 751.6093324 | 645.9346272 | 991.1695776 | -0.187676798 | 0.124579685 | 0.100903522 | 0.186407361 | ns | NaCl | KS.1 |

| | | | | | | | | | | | |
|---|---|---|---|---|---|---|---|---|---|---|---|
| FIM1_2953 | 1521.625954 | 1564.272636 | 1696.349832 | 0.030771902 | 0.166029178 | 0.831280157 | 0.895931279 | ns | | NaCl | KS.1 |
| FIM1_2954 | 589.1878413 | 602.029796 | 669.8722657 | 0.024625442 | 0.09168758 | 0.778396604 | 0.855995381 | ns | | NaCl | KS.1 |
| FIM1_2955 | 1188.564975 | 1516.028938 | 1135.911101 | 0.287994524 | 0.155892852 | 0.03578115 | 0.080657554 | ns | | NaCl | KS.1 |
| FIM1_2956 | 356.211043 | 398.9903103 | 356.0336572 | 0.12180094 | 0.169440911 | 0.405830225 | 0.540708455 | ns | | NaCl | KS.1 |
| FIM1_2957 | 3885.359537 | 8143.290462 | 3906.709487 | 0.841011252 | 0.269376777 | 0.000100416 | 0.000626025 | ns | | NaCl | KS.1 |
| FIM1_2958 | 1490.657236 | 2015.30215 | 2248.188588 | 0.387293344 | 0.12692345 | 0.000834948 | 0.0036989 | ns | | NaCl | KS.1 |
| FIM1_2959 | 5152.253551 | 8561.833101 | 4410.205535 | 0.58206469 | 0.221799757 | 0.001372991 | 0.005535359 | ns | | NaCl | KS.1 |
| FIM1_2960 | 3397.992952 | 6652.213728 | 3342.8738 | 0.776197914 | 0.24917992 | 0.000134237 | 0.000802679 | ns | | NaCl | KS.1 |
| FIM1_2961 | 3898.331916 | 8159.886029 | 4279.333789 | 0.936842882 | 0.201765591 | 1.616869063255 | 2.45241572855 | ns | | NaCl | KS.1 |
| FIM1_2962 | 320.2032437 | 286.304283 | 401.0509813 | -0.135836655 | 0.131869952 | 0.258224255 | 0.386598155 | ns | | NaCl | KS.1 |
| FIM1_2963 | 406.7929687 | 416.400233 | 393.8680425 | 0.027035346 | 0.112244171 | 0.796543202 | 0.870180595 | ns | | NaCl | KS.1 |
| FIM1_2964 | 8764.323457 | 10361.07874 | 7234.558437 | 0.20654746 | 0.128880206 | 0.07920826 | 0.153969737 | ns | | NaCl | KS.1 |
| FIM1_2965 | 1742.080823 | 1522.938867 | 1402.398664 | -0.169134902 | 0.117581163 | 0.120509574 | 0.214887144 | ns | | NaCl | KS.1 |
| FIM1_2966 | 447.2560277 | 382.8834441 | 368.3976288 | -0.177390328 | 0.163672681 | 0.213665834 | 0.336388457 | ns | | NaCl | KS.1 |
| FIM1_2967 | 1809.80735 | 1367.092022 | 1546.006294 | -0.381583454 | 0.089040492 | 6.71377414142 | 6.00737884057 | ns | | NaCl | KS.1 |
| FIM1_2968 | 1839.532372 | 1257.819018 | 1890.686952 | -0.517296082 | 0.102624595 | 8.92634169579 | 1.43716990086 | ns | | NaCl | KS.1 |
| FIM1_2969 | 6300.192935 | 8131.234995 | 5685.816099 | 0.312706898 | 0.143163202 | 0.01464949 | 0.038561488 | ns | | NaCl | KS.1 |
| FIM1_2970 | 867.3717703 | 670.4729886 | 760.6778271 | -0.337701822 | 0.112611168 | 0.001239482 | 0.00512587 | ns | | NaCl | KS.1 |
| FIM1_2971 | 917.1731741 | 1141.225773 | 637.3109641 | 0.215694834 | 0.214202929 | 0.211093743 | 0.333605899 | ns | | NaCl | KS.1 |
| FIM1_2972 | 847.8042222 | 556.4873846 | 731.7172261 | -0.53476242 | 0.154956425 | 0.000105261 | 0.000652149 | ns | | NaCl | KS.1 |
| FIM1_2973 | 196.0062834 | 155.7749133 | 901.5278038 | -0.238744338 | 0.199989882 | 0.147616291 | 0.251589944 | ns | | NaCl | KS.1 |
| FIM1_2974 | 1540.712671 | 1859.612361 | 1991.706768 | 0.204203562 | 0.18057651 | 0.183176333 | 0.298733842 | ns | | NaCl | KS.1 |
| FIM1_2975 | 3032.952578 | 5348.52602 | 2788.527927 | 0.719316687 | 0.17931265 | 5.72436424049 | 5.26408726367 | ns | | NaCl | KS.1 |
| FIM1_2976 | 670.9255344 | 695.9979795 | 629.7327363 | 0.044984569 | 0.123206941 | 0.692171535 | 0.792714868 | ns | | NaCl | KS.1 |
| FIM1_2977 | 813.4440437 | 501.4264196 | 839.6745868 | -0.617422653 | 0.161740665 | 1.82153490782 | 0.000144302 | ns | | NaCl | KS.1 |
| FIM1_2978 | 3250.798268 | 3895.777548 | 2827.370299 | 0.218418736 | 0.14049382 | 0.083853493 | 0.161194408 | ns | | NaCl | KS.1 |
| FIM1_2979 | 170.1400086 | 175.6657707 | 219.4068092 | 0.032638664 | 0.171222471 | 0.825187694 | 0.892652485 | ns | | NaCl | KS.1 |
| FIM1_2980 | 581.0048697 | 558.886526 | 926.1346837 | -0.044844779 | 0.145836358 | 0.731331004 | 0.822602701 | ns | | NaCl | KS.1 |
| FIM1_2981 | 4960.670041 | 10678.49219 | 5131.577162 | 0.906917657 | 0.252116444 | 1.47399395234 | 0.000120215 | ns | | NaCl | KS.1 |
| FIM1_2982 | 2879.321443 | 3197.774643 | 3129.856232 | 0.13503969 | 0.107414898 | 0.180115464 | 0.295246931 | ns | | NaCl | KS.1 |
| FIM1_2983 | 7667.957511 | 10767.55261 | 6118.843466 | 0.400507691 | 0.175079678 | 0.007732683 | 0.023357681 | ns | | NaCl | KS.1 |
| FIM1_2984 | 736.9038081 | 1036.504302 | 607.7647513 | 0.316929344 | 0.253094845 | 0.098737243 | 0.182812722 | ns | | NaCl | KS.1 |
| FIM1_2985 | 14231.92719 | 18572.90791 | 15548.43375 | 0.224609003 | 0.263069072 | 0.247768196 | 0.375463532 | ns | | NaCl | KS.1 |
| FIM1_2986 | 912.1345726 | 830.7817642 | 930.6546089 | -0.125634721 | 0.095830278 | 0.167372475 | 0.278301491 | ns | | NaCl | KS.1 |
| FIM1_2987 | 2404.390942 | 2134.859402 | 1963.239033 | -0.16019285 | 0.088787079 | 0.058975369 | 0.121441416 | ns | | NaCl | KS.1 |
| FIM1_2988 | 649.5198467 | 863.364384 | 931.4687696 | 0.353937017 | 0.140669579 | 0.005067224 | 0.016444513 | ns | | NaCl | KS.1 |
| FIM1_2989 | 1404.056587 | 1576.413261 | 992.0718388 | 0.126572787 | 0.168323811 | 0.385318254 | 0.520629634 | ns | | NaCl | KS.1 |
| FIM1_2990 | 14030.97017 | 16386.23909 | 12583.20332 | 0.210324901 | 0.078717286 | 0.00554119 | 0.017785432 | ns | | NaCl | KS.1 |
| FIM1_2991 | 162.5720416 | 72.36308404 | 416.6918034 | -0.940282422 | 0.260518629 | 1.22606295168 | 0.00010241 | ns | | NaCl | KS.1 |
| FIM1_2992 | 69.6818878 | 36.21068154 | 85.80505748 | -0.660414854 | 0.300016864 | 0.003027804 | 0.010798083 | ns | | NaCl | KS.1 |
| FIM1_2993 | 905.2262855 | 816.2956373 | 1039.204649 | -0.145913242 | 0.089218251 | 0.086902449 | 0.16596533 | ns | | NaCl | KS.1 |
| FIM1_2994 | 961.4817249 | 657.6895568 | 988.7851955 | -0.498388793 | 0.133741857 | 4.22819101304 | 0.000300075 | ns | | NaCl | KS.1 |
| FIM1_2995 | 224.304429 | 53.48985483 | 250.7650778 | -1.848519238 | 0.331517768 | 1.52593670589 | 6.90139556619 | Km<KS.1 | | NaCl | KS.1 |
| FIM1_2996 | 251.3585751 | 111.1723801 | 702.7419252 | -0.747644002 | 0.359076827 | 0.002590484 | 0.009504173 | ns | | NaCl | KS.1 |
| FIM1_2997 | 1368.942487 | 1381.210241 | 1624.884602 | 0.011417554 | 0.110222102 | 0.911913267 | 0.948722964 | ns | | NaCl | KS.1 |

| | | | | | | | | | | |
|---|---|---|---|---|---|---|---|---|---|---|
| FIM1_2998 | 3347.079555 | 2735.641256 | 3567.38224 | -0.255340996 | 0.119442524 | 0.020478979 | 0.05061248 | ns | NaCl | KS.1 |
| FIM1_2999 | 628.8148337 | 608.6887894 | 807.5264055 | -0.036887293 | 0.145150262 | 0.776808059 | 0.854963249 | ns | NaCl | KS.1 |
| FIM1_3000 | 1142.439256 | 2727.152895 | 1759.153242 | 1.177311891 | 0.155840669 | 7.7530829395 | 8.7662699146 | Km>KS.1 | NaCl | KS.1 |
| FIM1_3001 | 1005.083291 | 1300.224819 | 1124.216239 | 0.287427802 | 0.181630005 | 0.062145035 | 0.126554052 | ns | NaCl | KS.1 |
| FIM1_3002 | 386.2049607 | 278.9897374 | 406.515444 | -0.406145367 | 0.1492398 | 0.002201619 | 0.008266456 | ns | NaCl | KS.1 |
| FIM1_3003 | 886.4304665 | 1309.549317 | 974.3824504 | 0.391347242 | 0.242523274 | 0.037884331 | 0.08474575 | ns | NaCl | KS.1 |
| FIM1_3004 | 5063.421619 | 6338.370355 | 5703.476767 | 0.300841716 | 0.091952505 | 0.000570974 | 0.00269251 | ns | NaCl | KS.1 |
| FIM1_3005 | 2339.890521 | 4407.580836 | 2134.188148 | 0.632632385 | 0.298485736 | 0.004186556 | 0.014092094 | ns | NaCl | KS.1 |
| FIM1_3006 | 5146.263815 | 7971.640004 | 4840.496114 | 0.56430891 | 0.147713539 | 2.2017512730 | 0.0001693 | ns | NaCl | KS.1 |
| FIM1_3007 | 4424.403698 | 5338.390985 | 3697.877283 | 0.239608869 | 0.116136282 | 0.025940645 | 0.061926443 | ns | NaCl | KS.1 |
| FIM1_3008 | 1879.478178 | 2031.96233 | 2183.93518 | 0.102995683 | 0.085227719 | 0.207855462 | 0.32953503 | ns | NaCl | KS.1 |
| FIM1_3009 | 1249.682912 | 1296.105721 | 1567.872644 | 0.047158571 | 0.092925471 | 0.59443301 | 0.714595654 | ns | NaCl | KS.1 |
| FIM1_3010 | 606.961579 | 588.660366 | 653.7230729 | -0.038738883 | 0.118967697 | 0.725371022 | 0.81756249 | ns | NaCl | KS.1 |
| FIM1_3011 | 549.4726153 | 519.562017 | 622.8877379 | -0.064594654 | 0.155339591 | 0.637892999 | 0.748471149 | ns | NaCl | KS.1 |
| FIM1_3012 | 2613.006961 | 3605.522604 | 2505.524487 | 0.43308561 | 0.102073167 | 6.3604420718 | 5.7428673878 | ns | NaCl | KS.1 |
| FIM1_3013 | 1736.002477 | 2132.534125 | 2299.118373 | 0.271856428 | 0.09766691 | 0.003214704 | 0.011302581 | ns | NaCl | KS.1 |
| FIM1_3014 | 6135.953628 | 13781.15507 | 6063.64154 | 0.897601186 | 0.293657732 | 9.6782608983 | 0.000606415 | ns | NaCl | KS.1 |
| FIM1_3015 | 4937.023939 | 6370.582139 | 4634.54022 | 0.32851791 | 0.118557571 | 0.002660677 | 0.009690242 | ns | NaCl | KS.1 |
| FIM1_3016 | 541.2981299 | 430.3068675 | 759.6981876 | -0.275802767 | 0.151432652 | 0.039681646 | 0.087701549 | ns | NaCl | KS.1 |
| FIM1_3017 | 27607.10562 | 34237.55458 | 21330.23418 | 0.276364912 | 0.115434699 | 0.009754844 | 0.027987514 | ns | NaCl | KS.1 |
| FIM1_3018 | 696.8902407 | 738.239616 | 535.4983651 | 0.063472148 | 0.162790118 | 0.655501687 | 0.762656898 | ns | NaCl | KS.1 |
| FIM1_3019 | 4299.529382 | 4512.291165 | 4633.715476 | 0.060587655 | 0.112831389 | 0.565079163 | 0.687654109 | ns | NaCl | KS.1 |
| FIM1_3020 | 131.466772 | 102.7341972 | 96.90294568 | -0.256744188 | 0.20997958 | 0.131979211 | 0.230084749 | ns | NaCl | KS.1 |
| FIM1_3021 | 4131.942564 | 5366.44382 | 3834.209133 | 0.344346286 | 0.108008494 | 0.000635305 | 0.002948359 | ns | NaCl | KS.1 |
| FIM1_3022 | 196.0257077 | 315.3699815 | 240.3611277 | 0.413152453 | 0.298505859 | 0.051430601 | 0.108137384 | ns | NaCl | KS.1 |
| FIM1_3023 | 729.0363711 | 182.3275374 | 752.7178282 | -1.831330832 | 0.270630307 | 7.6990465811 | 7.0931031001 | Km<KS.1 | NaCl | KS.1 |
| FIM1_3024 | 35.37375336 | 35.52215417 | 74.4569741 | 0.006362067 | 0.334136872 | 0.975342644 | 0.988858703 | ns | NaCl | KS.1 |
| FIM1_3025 | 763.5016284 | 657.435991 | 931.7120277 | -0.152424927 | 0.195842374 | 0.348024219 | 0.4828278 | ns | NaCl | KS.1 |
| FIM1_3026 | 2522.912447 | 2608.731556 | 2497.866158 | 0.036686201 | 0.164504854 | 0.798119169 | 0.870903663 | ns | NaCl | KS.1 |
| FIM1_3028 | 3550.608119 | 4629.959912 | 4185.317805 | 0.336400524 | 0.12807664 | 0.003970745 | 0.013475071 | ns | NaCl | KS.1 |
| FIM1_3029 | 1224.640234 | 1123.421778 | 1373.966389 | -0.107382791 | 0.120834202 | 0.336217506 | 0.470647746 | ns | NaCl | KS.1 |
| FIM1_3030 | 3326.368208 | 3077.741171 | 3644.827954 | -0.106859266 | 0.074545325 | 0.138756844 | 0.23919449 | ns | NaCl | KS.1 |
| FIM1_3031 | 1753.786413 | 919.3478229 | 1581.656361 | -0.876114605 | 0.133720924 | 3.8217126015 | 2.0226617226 | ns | NaCl | KS.1 |
| FIM1_3032 | 4166.590727 | 3308.501584 | 2951.006234 | -0.270631205 | 0.158330278 | 0.051326648 | 0.108016106 | ns | NaCl | KS.1 |
| FIM1_3033 | 1471.394727 | 1091.099975 | 1273.167036 | -0.382058899 | 0.13050446 | 0.001284644 | 0.005273187 | ns | NaCl | KS.1 |
| FIM1_3034 | 1866.24844 | 1066.462984 | 2931.69041 | -0.543955277 | 0.289932018 | 0.010958662 | 0.030729477 | ns | NaCl | KS.1 |
| FIM1_3035 | 13864.27544 | 13813.99257 | 11142.20316 | -0.004082597 | 0.14975108 | 0.975586502 | 0.988899883 | ns | NaCl | KS.1 |
| FIM1_3036 | 3728.119518 | 3267.228356 | 3404.866561 | -0.159111747 | 0.135002458 | 0.193786702 | 0.311700241 | ns | NaCl | KS.1 |
| FIM1_3037 | 1028.642441 | 775.9684032 | 1021.631874 | -0.225360512 | 0.278021074 | 0.25880857 | 0.387348256 | ns | NaCl | KS.1 |
| FIM1_3038 | 18906.84106 | 28260.61478 | 17148.25544 | 0.517228907 | 0.142969214 | 5.9924644774 | 0.000405612 | ns | NaCl | KS.1 |
| FIM1_3039 | 3201.769323 | 4456.69725 | 2869.090948 | 0.4245764 | 0.13246016 | 0.000416024 | 0.00206972 | ns | NaCl | KS.1 |
| FIM1_3040 | 3063.773507 | 5723.816588 | 2794.51789 | 0.589117028 | 0.313199401 | 0.008723948 | 0.025802644 | ns | NaCl | KS.1 |
| FIM1_3041 | 1522.572484 | 1576.903852 | 1740.792045 | 0.04438831 | 0.100289801 | 0.63995805 | 0.749480061 | ns | NaCl | KS.1 |
| FIM1_3043 | 203.4189255 | 202.5011788 | 207.6025605 | -0.003800778 | 0.191092864 | 0.981017795 | 0.991175946 | ns | NaCl | KS.1 |
| FIM1_3044 | 408.1447418 | 439.4196726 | 4677.982151 | 0.053266069 | 0.282023607 | 0.787728923 | 0.863395328 | ns | NaCl | KS.1 |

| | | | | | | | | | | |
|---|---|---|---|---|---|---|---|---|---|---|
| FIM1_3045 | 321.9595098 | 286.4550669 | 549.2223036 | -0.121483597 | 0.187676265 | 0.441200453 | 0.576766636 | ns | NaCl | KS.1 |
| FIM1_3046 | 1125.866815 | 1095.566655 | 1642.342519 | -0.03014847 | 0.156799273 | 0.82740667 | 0.893922775 | ns | NaCl | KS.1 |
| FIM1_3047 | 899.045823 | 825.021898 | 825.3510037 | -0.110887239 | 0.102990414 | 0.253458922 | 0.381761471 | ns | NaCl | KS.1 |
| FIM1_3048 | 3403.258367 | 5420.720568 | 3099.753428 | 0.575893197 | 0.176789622 | 0.000180308 | 0.001024007 | ns | NaCl | KS.1 |
| FIM1_3049 | 707.8490666 | 748.3075839 | 883.089918 | 0.067177582 | 0.124579984 | 0.557996531 | 0.681569542 | ns | NaCl | KS.1 |
| FIM1_3050 | 1023.689883 | 909.2320155 | 792.229839 | -0.149713353 | 0.110171637 | 0.146018873 | 0.249294405 | ns | NaCl | KS.1 |
| FIM1_3051 | 852.9356294 | 425.5953755 | 1014.141862 | -0.932620203 | 0.150401324 | 3.0751140849 | 1.2437961441 | ns | NaCl | KS.1 |
| FIM1_3052 | 692.3022588 | 669.4136309 | 796.9924135 | -0.03996678 | 0.137623515 | 0.748446923 | 0.83655885 | ns | NaCl | KS.1 |
| FIM1_3053 | 3336.310614 | 1829.156104 | 2143.005932 | -0.794528995 | 0.152830486 | 1.5920607425 | 3.2865154331 | ns | NaCl | KS.1 |
| FIM1_3054 | 965.482915 | 1434.985686 | 1456.614948 | 0.364263567 | 0.267672034 | 0.067286448 | 0.134349815 | ns | NaCl | KS.1 |
| FIM1_3056 | 3376.862145 | 5445.57215 | 4038.32144 | 0.512670296 | 0.240911663 | 0.007089776 | 0.021746831 | ns | NaCl | KS.1 |
| FIM1_3057 | 559.6583627 | 401.1525643 | 483.5889983 | -0.329168505 | 0.236659328 | 0.074666277 | 0.146638982 | ns | NaCl | KS.1 |
| FIM1_3058 | 1927.087521 | 2195.510661 | 2914.185425 | 0.158860932 | 0.131133095 | 0.18381821 | 0.299380325 | ns | NaCl | KS.1 |
| FIM1_3059 | 527.7097895 | 526.2521716 | 849.149499 | -0.001415006 | 0.114310717 | 0.989403985 | 0.995205181 | ns | NaCl | KS.1 |
| FIM1_3060 | 634.7368664 | 623.9418809 | 717.4541639 | -0.021517359 | 0.125583339 | 0.85220556 | 0.910397823 | ns | NaCl | KS.1 |
| FIM1_3061 | 3169.475572 | 3138.560212 | 3894.558609 | -0.012005758 | 0.103202655 | 0.901870495 | 0.941814801 | ns | NaCl | KS.1 |
| FIM1_3062 | 1791.234819 | 2062.491621 | 1637.020043 | 0.153837247 | 0.173500193 | 0.302058716 | 0.434094679 | ns | NaCl | KS.1 |
| FIM1_3063 | 821.640734 | 604.0505839 | 1105.968508 | -0.418265334 | 0.101556585 | 1.1827335108 | 9.9225956432 | ns | NaCl | KS.1 |
| FIM1_3064 | 5951.590764 | 9490.643204 | 6172.113327 | 0.638123803 | 0.106484699 | 2.5483971755 | 8.5801346489 | ns | NaCl | KS.1 |
| FIM1_3065 | 2388.565946 | 2778.823196 | 2380.210882 | 0.197199328 | 0.103003558 | 0.041913448 | 0.0914865 | ns | NaCl | KS.1 |
| FIM1_3066 | 6345.831921 | 5385.73748 | 4964.727518 | -0.213419559 | 0.103524462 | 0.028327103 | 0.066412505 | ns | NaCl | KS.1 |
| FIM1_3067 | 835.0124043 | 834.9282466 | 690.525503 | -0.000710255 | 0.128705701 | 0.995193959 | 0.997801279 | ns | NaCl | KS.1 |
| FIM1_3068 | 5187.592868 | 5849.857954 | 5347.153281 | 0.127087519 | 0.18149686 | 0.409384022 | 0.544568319 | ns | NaCl | KS.1 |
| FIM1_3069 | 1510.405396 | 1574.335167 | 1141.648876 | 0.04553123 | 0.163403574 | 0.74966659 | 0.837108446 | ns | NaCl | KS.1 |
| FIM1_3070 | 7800.004222 | 10819.55207 | 7452.029009 | 0.435468352 | 0.109187769 | 1.9104066448 | 0.000149674 | ns | NaCl | KS.1 |
| FIM1_3071 | 3177.604839 | 2846.766469 | 3315.501244 | -0.13436402 | 0.127603072 | 0.250344941 | 0.378681927 | ns | NaCl | KS.1 |
| FIM1_3072 | 1012.584292 | 837.6446698 | 1546.247319 | -0.210589715 | 0.172712562 | 0.156096132 | 0.263246867 | ns | NaCl | KS.1 |
| FIM1_3074 | 681.0406846 | 858.1998531 | 871.4180666 | 0.296650108 | 0.11896977 | 0.006855258 | 0.021156892 | ns | NaCl | KS.1 |
| FIM1_3075 | 1212.598822 | 2058.047342 | 1321.472421 | 0.674580841 | 0.170050738 | 8.0273303457 | 7.0683130035 | ns | NaCl | KS.1 |
| FIM1_3076 | 829.1162723 | 979.9463733 | 1017.158479 | 0.217085469 | 0.100898918 | 0.022474523 | 0.054889912 | ns | NaCl | KS.1 |
| FIM1_3077 | 2014.226786 | 2569.689 | 2215.349619 | 0.306479405 | 0.128740351 | 0.0089957 | 0.026325651 | ns | NaCl | KS.1 |
| FIM1_3078 | 728.0472465 | 864.5031719 | 790.6237241 | 0.224620209 | 0.10672766 | 0.024642479 | 0.05928256 | ns | NaCl | KS.1 |
| FIM1_3079 | 1756.129296 | 3152.850084 | 2045.88673 | 0.765066682 | 0.159446966 | 1.3568054811 | 2.1160210874 | ns | NaCl | KS.1 |
| FIM1_3080 | 1451.891389 | 1757.183627 | 1517.297272 | 0.246406021 | 0.109592782 | 0.015979974 | 0.041127972 | ns | NaCl | KS.1 |
| FIM1_3081 | 1514.773696 | 1567.646793 | 1776.031229 | 0.037850728 | 0.14347536 | 0.769055837 | 0.851369112 | ns | NaCl | KS.1 |
| FIM1_3082 | 1718.503653 | 1595.872923 | 1874.539366 | -0.098215284 | 0.093275956 | 0.268876728 | 0.397995156 | ns | NaCl | KS.1 |
| FIM1_3083 | 5182.438563 | 11902.47036 | 5074.020734 | 0.923108147 | 0.297391743 | 7.4373652711 | 0.00048943 | ns | NaCl | KS.1 |
| FIM1_3085 | 24.1094194 | 41.09548804 | 43.4414597 | 0.447462407 | 0.316331138 | 0.041300551 | 0.090443071 | ns | NaCl | KS.1 |
| FIM1_3086 | 630.6772722 | 1310.991081 | 1403.542008 | 0.923151756 | 0.206144512 | 3.6650575652 | 5.0092476338 | ns | NaCl | KS.1 |
| FIM1_3087 | 9193.220638 | 13267.25395 | 7192.444108 | 0.410989215 | 0.201175934 | 0.013668143 | 0.036558609 | ns | NaCl | KS.1 |
| FIM1_3088 | 256.2800762 | 190.2043194 | 879.435673 | -0.28573383 | 0.235264796 | 0.119322588 | 0.213351332 | ns | NaCl | KS.1 |
| FIM1_3089 | 489.483073 | 454.3821851 | 526.3408947 | -0.078308657 | 0.179311984 | 0.608233969 | 0.724943649 | ns | NaCl | KS.1 |
| FIM1_3090 | 386.2869089 | 538.5562627 | 396.6058563 | 0.41937511 | 0.144181567 | 0.001154348 | 0.004838146 | ns | NaCl | KS.1 |
| FIM1_3092 | 345.9213298 | 494.0643258 | 500.4249974 | 0.420586083 | 0.184982503 | 0.007345738 | 0.022379085 | ns | NaCl | KS.1 |
| FIM1_3098 | 620.5535404 | 949.5685868 | 1200.264754 | 0.491699131 | 0.201573927 | 0.003436994 | 0.01197412 | ns | NaCl | KS.1 |

| | | | | | | | | | | |
|---|---|---|---|---|---|---|---|---|---|---|
| FIM1_3099 | 1614.550657 | 2425.961367 | 2350.021231 | 0.370474172 | 0.272415655 | 0.065493091 | 0.131463638 | ns | NaCl | KS.1 |
| FIM1_3100 | 786.157505 | 813.4946669 | 982.5762384 | 0.04063411 | 0.144145797 | 0.753500225 | 0.839446232 | ns | NaCl | KS.1 |
| FIM1_3101 | 950.9410103 | 1001.381218 | 1052.947167 | 0.061630622 | 0.139682732 | 0.625090153 | 0.738774851 | ns | NaCl | KS.1 |
| FIM1_3102 | 20700.82997 | 27125.55894 | 16236.58602 | 0.336149316 | 0.138933172 | 0.007158479 | 0.02190248 | ns | NaCl | KS.1 |
| FIM1_3103 | 127.3759033 | 196.0170419 | 162.3837864 | 0.359428552 | 0.296984842 | 0.086457562 | 0.165369615 | ns | NaCl | KS.1 |
| FIM1_3104 | 1639.152772 | 1618.564226 | 1971.434572 | -0.016123538 | 0.120745661 | 0.885227403 | 0.931670473 | ns | NaCl | KS.1 |
| FIM1_3105 | 12.07781807 | 19.66176517 | 14.7225111 | 0.285643179 | 0.358961773 | 0.192449339 | 0.309849663 | ns | NaCl | KS.1 |
| FIM1_3106 | 1306.901827 | 1184.536792 | 1352.376741 | -0.124219144 | 0.115266893 | 0.246481095 | 0.374062783 | ns | NaCl | KS.1 |
| FIM1_3107 | 4228.277472 | 381.9457707 | 1220.887682 | -3.141714854 | 0.435248243 | 3.1320459562 | 3.0552801240 | Km<KS.1 | NaCl | KS.1 |
| FIM1_3108 | 2909.693111 | 1400.142994 | 2240.122344 | -0.982461836 | 0.148887215 | 1.9239327773 | 1.1129727403 | ns | NaCl | KS.1 |
| FIM1_3109 | 4048.476477 | 6596.042126 | 3375.384685 | 0.574929441 | 0.205867845 | 0.000850448 | 0.00376087 | ns | NaCl | KS.1 |
| FIM1_3110 | 1517.931299 | 3286.322057 | 4126.047925 | 0.546962828 | 0.401631976 | 0.025281071 | 0.060473571 | ns | NaCl | KS.1 |
| FIM1_3111 | 362.1371902 | 773.615459 | 1294.786463 | 0.788784783 | 0.312981781 | 0.000745362 | 0.003374137 | ns | NaCl | KS.1 |
| FIM1_3112 | 533.5047197 | 633.0790113 | 703.4952814 | 0.184792228 | 0.181714053 | 0.230544568 | 0.356419896 | ns | NaCl | KS.1 |
| FIM1_3113 | 750.9318241 | 974.4029306 | 1621.420472 | 0.294246269 | 0.176125121 | 0.050780933 | 0.107094168 | ns | NaCl | KS.1 |
| FIM1_3114 | 1913.353706 | 1654.968857 | 1719.038682 | -0.162996833 | 0.158405703 | 0.241431817 | 0.368296542 | ns | NaCl | KS.1 |
| FIM1_3115 | 256.346105 | 129.4018568 | 260.7413305 | -0.86662272 | 0.196881573 | 6.3722160708 | 8.0257658107 | ns | NaCl | KS.1 |
| FIM1_3116 | 1070.177479 | 1273.036667 | 1724.004441 | 0.180640961 | 0.193561131 | 0.262283024 | 0.390559128 | ns | NaCl | KS.1 |
| FIM1_3117 | 23.69577028 | 15.03846269 | 19.37440892 | -0.335141595 | 0.321007171 | 0.119918134 | 0.214139525 | ns | NaCl | KS.1 |
| FIM1_3118 | 6849.215721 | 3076.291445 | 4526.624002 | -1.039089763 | 0.190669878 | 1.7058727899 | 4.6375503443 | Km<KS.1 | NaCl | KS.1 |
| FIM1_3119 | 892.9873259 | 807.53381 | 946.5580616 | -0.107944097 | 0.174302212 | 0.470678226 | 0.604602162 | ns | NaCl | KS.1 |
| FIM1_3120 | 2661.664801 | 3016.332695 | 2859.777166 | 0.163855983 | 0.096509919 | 0.073354424 | 0.144358488 | ns | NaCl | KS.1 |
| FIM1_3121 | 880.9944073 | 977.8807719 | 1170.818666 | 0.129100542 | 0.108661332 | 0.204678578 | 0.325952601 | ns | NaCl | KS.1 |
| FIM1_3122 | 2257.32981 | 1338.583577 | 1970.926694 | -0.669048268 | 0.165349136 | 5.8501908399 | 5.3501285714 | ns | NaCl | KS.1 |
| FIM1_3123 | 1677.108478 | 1568.972834 | 2384.973826 | -0.085135141 | 0.108474295 | 0.402563672 | 0.538105815 | ns | NaCl | KS.1 |
| FIM1_3124 | 4696.572048 | 4941.246884 | 4368.791076 | 0.07094582 | 0.056388599 | 0.199808034 | 0.319525865 | ns | NaCl | KS.1 |
| FIM1_3125 | 610.0455197 | 536.6354641 | 648.7975289 | -0.158260978 | 0.133348664 | 0.191643159 | 0.309051772 | ns | NaCl | KS.1 |
| FIM1_3126 | 1961.691651 | 1909.085925 | 2225.153781 | -0.028972969 | 0.176175452 | 0.847715957 | 0.907143711 | ns | NaCl | KS.1 |
| FIM1_3127 | 1321.745805 | 1374.665862 | 1641.556946 | 0.041730631 | 0.17716853 | 0.782944947 | 0.859666985 | ns | NaCl | KS.1 |
| FIM1_3128 | 10021.71288 | 12154.85836 | 9997.73968 | 0.259025918 | 0.087966301 | 0.002028411 | 0.0077092 | ns | NaCl | KS.1 |
| FIM1_3129 | 3815.798774 | 5201.204359 | 3927.025455 | 0.420456765 | 0.091571348 | 1.3352267326 | 1.5062931961 | ns | NaCl | KS.1 |
| FIM1_3130 | 1131.195207 | 1631.089361 | 1143.616367 | 0.429384764 | 0.182993737 | 0.005824501 | 0.018610722 | ns | NaCl | KS.1 |
| FIM1_3131 | 950.1518406 | 842.0380446 | 1039.568712 | -0.14308056 | 0.140422788 | 0.258035935 | 0.386598155 | ns | NaCl | KS.1 |
| FIM1_3132 | 4221.359962 | 4760.379874 | 3367.068562 | 0.160001779 | 0.08890083 | 0.059577539 | 0.122553749 | ns | NaCl | KS.1 |
| FIM1_3133 | 568.0017903 | 574.4383875 | 594.2050343 | 0.011802348 | 0.160671386 | 0.933243486 | 0.961658314 | ns | NaCl | KS.1 |
| FIM1_3134 | 7998.995604 | 15509.1451 | 11415.25866 | 0.822589015 | 0.206319795 | 4.4531209687 | 4.3354749157 | ns | NaCl | KS.1 |
| FIM1_3135 | 958.7248115 | 1201.405634 | 1117.0395 | 0.290749732 | 0.114912798 | 0.006326047 | 0.01982445 | ns | NaCl | KS.1 |
| FIM1_3136 | 4391.262052 | 9137.700045 | 4630.98272 | 0.943954682 | 0.188739778 | 2.7178599695 | 5.3340111765 | ns | NaCl | KS.1 |
| FIM1_3137 | 17.87292963 | 19.56857312 | 24.44135902 | 0.06834825 | 0.334265671 | 0.740775968 | 0.829849233 | ns | NaCl | KS.1 |
| FIM1_3138 | 5788.458025 | 13275.83562 | 6167.117837 | 0.961535912 | 0.274554266 | 1.6467394862 | 0.000132565 | ns | NaCl | KS.1 |
| FIM1_3139 | 814.7508391 | 1207.24758 | 667.88201 | 0.458747437 | 0.191148732 | 0.004426327 | 0.014749484 | ns | NaCl | KS.1 |
| FIM1_3140 | 364.1078605 | 394.156007 | 459.0135077 | 0.094305927 | 0.140631371 | 0.456918808 | 0.591817513 | ns | NaCl | KS.1 |
| FIM1_3141 | 805.3232035 | 1315.325579 | 972.2596684 | 0.641364647 | 0.147381436 | 1.6661844288 | 1.8339087463 | ns | NaCl | KS.1 |
| FIM1_3142 | 195.940053 | 141.9393555 | 264.7518978 | -0.336589037 | 0.21455143 | 0.052336338 | 0.109769511 | ns | NaCl | KS.1 |
| FIM1_3143 | 6148.493514 | 8858.993912 | 5562.129101 | 0.502805445 | 0.087737725 | 2.0091738612 | 5.3452620106 | ns | NaCl | KS.1 |

| | | | | | | | | | | |
|---|---|---|---|---|---|---|---|---|---|---|
| FIM1_3144 | 411.5871711 | 265.9372978 | 358.3171778 | -0.525853605 | 0.192015848 | 0.001235859 | 0.005118614 | ns | NaCl | KS.1 |
| FIM1_3145 | 1035.980659 | 1388.286357 | 1300.969336 | 0.375542516 | 0.125786742 | 0.001095117 | 0.004617125 | ns | NaCl | KS.1 |
| FIM1_3146 | 670.7335307 | 598.7443199 | 695.8146255 | -0.144279207 | 0.117287554 | 0.184652237 | 0.300505358 | ns | NaCl | KS.1 |
| FIM1_3147 | 600.9271039 | 533.5557336 | 1134.359183 | -0.139238633 | 0.138486541 | 0.265656608 | 0.394248995 | ns | NaCl | KS.1 |
| FIM1_3148 | 579.2265957 | 491.5167393 | 864.5807955 | -0.208602334 | 0.121694524 | 0.062730577 | 0.12740678 | ns | NaCl | KS.1 |
| FIM1_3149 | 812.4926855 | 921.9237586 | 1048.570728 | 0.152170581 | 0.134354708 | 0.212204749 | 0.334723724 | ns | NaCl | KS.1 |
| FIM1_3150 | 4252.467844 | 11642.69615 | 5060.087817 | 1.204317186 | 0.28553929 | 3.19508516003 | 4.44009739418 | Km>KS.1 | NaCl | KS.1 |
| FIM1_3151 | 1133.401266 | 1097.031737 | 787.651183 | -0.039242358 | 0.144473162 | 0.762105515 | 0.84631137 | ns | NaCl | KS.1 |
| FIM1_3152 | 265.2172462 | 248.1675282 | 255.1933084 | -0.083762099 | 0.151515067 | 0.533652677 | 0.66158038 | ns | NaCl | KS.1 |
| FIM1_3153 | 674.9802886 | 541.5902398 | 769.2644039 | -0.278243784 | 0.123149934 | 0.013842101 | 0.036845613 | ns | NaCl | KS.1 |
| FIM1_3154 | 2725.384427 | 3148.675895 | 2146.446937 | 0.15589067 | 0.17725865 | 0.303219887 | 0.435359001 | ns | NaCl | KS.1 |
| FIM1_3155 | 1827.110242 | 2544.851463 | 2866.78671 | 0.420743428 | 0.139177704 | 0.000788747 | 0.003528793 | ns | NaCl | KS.1 |
| FIM1_3156 | 3614.634126 | 4265.136568 | 3973.487865 | 0.206940887 | 0.122983183 | 0.06729643 | 0.134349815 | ns | NaCl | KS.1 |
| FIM1_3157 | 638.2132995 | 396.6241583 | 1292.04226 | -0.480669935 | 0.256273009 | 0.014728815 | 0.038667997 | ns | NaCl | KS.1 |
| FIM1_3158 | 366.1486741 | 261.8361679 | 368.4361109 | -0.410300169 | 0.156184105 | 0.002868553 | 0.010311455 | ns | NaCl | KS.1 |
| FIM1_3159 | 490.1568694 | 318.159276 | 404.0752316 | -0.561115154 | 0.156641259 | 5.74344498777 | 0.000390884 | ns | NaCl | KS.1 |
| FIM1_3160 | 148.5448913 | 165.6786125 | 291.6823713 | 0.089490433 | 0.25708933 | 0.638385394 | 0.748695742 | ns | NaCl | KS.1 |
| FIM1_3161 | 1922.85714 | 1155.744078 | 1401.267782 | -0.502320434 | 0.273344429 | 0.014465814 | 0.038219555 | ns | NaCl | KS.1 |
| FIM1_3162 | 2540.618541 | 1565.394933 | 2181.720415 | -0.474349901 | 0.269951607 | 0.01949039 | 0.048409731 | ns | NaCl | KS.1 |
| FIM1_3163 | 824.9965518 | 669.6196291 | 715.8975727 | -0.227461089 | 0.182772292 | 0.14161361 | 0.243192168 | ns | NaCl | KS.1 |
| FIM1_3164 | 2583.826698 | 2973.914803 | 2822.476065 | 0.195721408 | 0.05678789 | 0.000436321 | 0.002153526 | ns | NaCl | KS.1 |
| FIM1_3165 | 1001.33475 | 1039.968491 | 793.2031306 | 0.043420941 | 0.142425921 | 0.734737307 | 0.825128239 | ns | NaCl | KS.1 |
| FIM1_3166 | 1229.164649 | 1192.778566 | 1168.677337 | -0.035558302 | 0.133356998 | 0.769673602 | 0.851484583 | ns | NaCl | KS.1 |
| FIM1_3167 | 3631.838948 | 7617.489224 | 3498.114776 | 0.759886298 | 0.312929691 | 0.001082921 | 0.004584625 | ns | NaCl | KS.1 |
| FIM1_3168 | 1391.889946 | 1048.028884 | 1110.963659 | -0.371193897 | 0.114069848 | 0.000447153 | 0.002196039 | ns | NaCl | KS.1 |
| FIM1_3169 | 1265.955198 | 1193.53817 | 1077.659817 | -0.068482479 | 0.147655345 | 0.603555473 | 0.721973667 | ns | NaCl | KS.1 |
| FIM1_3170 | 64.3876162 | 48.26746516 | 121.4952025 | -0.259647499 | 0.254076672 | 0.174912799 | 0.288524925 | ns | NaCl | KS.1 |
| FIM1_3171 | 2291.442422 | 2415.202571 | 1716.990805 | 0.062540464 | 0.136516017 | 0.613381931 | 0.729117728 | ns | NaCl | KS.1 |
| FIM1_3172 | 1326.614236 | 1457.14631 | 1513.119302 | 0.098645241 | 0.18327882 | 0.524898471 | 0.65398695 | ns | NaCl | KS.1 |
| FIM1_3173 | 1090.936072 | 1201.197135 | 1674.368048 | 0.115993881 | 0.137634036 | 0.351557712 | 0.486779562 | ns | NaCl | KS.1 |
| FIM1_3174 | 601.3795264 | 640.7276847 | 492.5185698 | 0.064803731 | 0.190447045 | 0.6842764 | 0.78657003 | ns | NaCl | KS.1 |
| FIM1_3175 | 1987.329548 | 1353.230454 | 1395.309944 | -0.499130703 | 0.135898019 | 5.21706973923 | 0.000360601 | ns | NaCl | KS.1 |
| FIM1_3176 | 462.6149368 | 346.8043475 | 464.5748983 | -0.360010336 | 0.141254512 | 0.004497505 | 0.014916725 | ns | NaCl | KS.1 |
| FIM1_3177 | 2152.319524 | 2488.194075 | 2248.049981 | 0.195321488 | 0.076850475 | 0.008451794 | 0.025150037 | ns | NaCl | KS.1 |
| FIM1_3178 | 2002.929417 | 2353.474584 | 2050.784747 | 0.191861817 | 0.144688712 | 0.138361936 | 0.238844772 | ns | NaCl | KS.1 |
| FIM1_3179 | 308.4795223 | 208.8181306 | 479.2921944 | -0.399750189 | 0.236667742 | 0.031429472 | 0.072456729 | ns | NaCl | KS.1 |
| FIM1_3180 | 1027.29371 | 1011.575916 | 1588.096631 | -0.017064469 | 0.135353578 | 0.889616288 | 0.933524791 | ns | NaCl | KS.1 |
| FIM1_3181 | 3223.265882 | 6327.009244 | 3024.698306 | 0.722218789 | 0.28331204 | 0.00093203 | 0.004053191 | ns | NaCl | KS.1 |
| FIM1_3182 | 2169.718024 | 2487.994764 | 1881.758192 | 0.170414743 | 0.122164925 | 0.130035201 | 0.227550166 | ns | NaCl | KS.1 |
| FIM1_3183 | 9444.095716 | 8197.521672 | 6908.476124 | -0.168085009 | 0.142337466 | 0.188550466 | 0.30525173 | ns | NaCl | KS.1 |
| FIM1_3184 | 1001.490826 | 861.0268533 | 690.2581567 | -0.153787526 | 0.197098573 | 0.345824616 | 0.480714575 | ns | NaCl | KS.1 |
| FIM1_3185 | 1637.413408 | 1769.082934 | 1068.012224 | 0.078135589 | 0.192731275 | 0.626736327 | 0.739711913 | ns | NaCl | KS.1 |
| FIM1_3186 | 2889.331175 | 3089.97114 | 3548.73503 | 0.082931489 | 0.121190485 | 0.458974438 | 0.593861594 | ns | NaCl | KS.1 |
| FIM1_3187 | 583.2389309 | 713.8378472 | 663.5778738 | 0.238593147 | 0.14884534 | 0.071361783 | 0.141050803 | ns | NaCl | KS.1 |
| FIM1_3188 | 529.0364017 | 684.6630943 | 690.8064976 | 0.320182859 | 0.135294849 | 0.008835608 | 0.026025546 | ns | NaCl | KS.1 |

| | | | | | | | | | | |
|---|---|---|---|---|---|---|---|---|---|---|
| FIM1_3189 | 347.9201271 | 406.1844028 | 411.7370225 | 0.165835618 | 0.172868374 | 0.264567239 | 0.392784845 | ns | NaCl | KS.1 |
| FIM1_3190 | 640.064306 | 612.5056127 | 680.2043265 | -0.052237835 | 0.137795475 | 0.67539015 | 0.77941684 | ns | NaCl | KS.1 |
| FIM1_3191 | 1001.246975 | 1069.428381 | 883.2861558 | 0.072110516 | 0.166430582 | 0.618126047 | 0.732883003 | ns | NaCl | KS.1 |
| FIM1_3192 | 677.0338539 | 645.3024217 | 729.3971384 | -0.06388691 | 0.117326167 | 0.55757817 | 0.681393121 | ns | NaCl | KS.1 |
| FIM1_3193 | 43.2919089 | 74.27496584 | 43.75386283 | 0.504106073 | 0.284324483 | 0.016408751 | 0.042144315 | ns | NaCl | KS.1 |
| FIM1_3194 | 1756.914249 | 2416.839917 | 2395.124519 | 0.353147698 | 0.196001863 | 0.030357483 | 0.07031121 | ns | NaCl | KS.1 |
| FIM1_3195 | 3832.670171 | 6327.53578 | 3200.949999 | 0.54881527 | 0.238741178 | 0.003916086 | 0.013349278 | ns | NaCl | KS.1 |
| FIM1_3196 | 451.5625242 | 448.6168818 | 477.2816158 | -0.010774237 | 0.155364637 | 0.937454761 | 0.963801908 | ns | NaCl | KS.1 |
| FIM1_3197 | 459.2739098 | 773.0664405 | 1536.617768 | 0.496786216 | 0.287142661 | 0.018518208 | 0.046578947 | ns | NaCl | KS.1 |
| FIM1_3198 | 208.9180622 | 404.7688263 | 460.0031329 | 0.502301106 | 0.366206713 | 0.032604701 | 0.074715977 | ns | NaCl | KS.1 |
| FIM1_3199 | 2212.190223 | 2651.499155 | 2359.023598 | 0.219117856 | 0.139476227 | 0.081048239 | 0.156780129 | ns | NaCl | KS.1 |
| FIM1_3200 | 8218.582599 | 13448.94245 | 9302.489641 | 0.698972216 | 0.058586298 | 8.9247032644 | 1.1100099685 | ns | NaCl | KS.1 |
| FIM1_3201 | 459.6486806 | 565.1524111 | 501.8279529 | 0.236246604 | 0.158981636 | 0.090061908 | 0.170720271 | ns | NaCl | KS.1 |
| FIM1_3202 | 469.4872956 | 416.1267668 | 534.8867205 | -0.153165529 | 0.13076124 | 0.199032042 | 0.318592153 | ns | NaCl | KS.1 |
| FIM1_3203 | 1096.90271 | 1106.799283 | 1449.933299 | 0.01081952 | 0.129598346 | 0.927310547 | 0.958124605 | ns | NaCl | KS.1 |
| FIM1_3204 | 684.4511796 | 715.4408006 | 744.7715961 | 0.064527196 | 0.077766444 | 0.390420997 | 0.525953008 | ns | NaCl | KS.1 |
| FIM1_3205 | 4780.459776 | 5315.262916 | 4777.253445 | 0.137748108 | 0.101644476 | 0.150980677 | 0.256533084 | ns | NaCl | KS.1 |
| FIM1_3206 | 1553.398529 | 1879.646751 | 1815.968829 | 0.247017385 | 0.105330494 | 0.012372821 | 0.033618123 | ns | NaCl | KS.1 |
| FIM1_3207 | 647.1770705 | 643.1278046 | 891.1447567 | -0.005246969 | 0.178513208 | 0.972520219 | 0.987204262 | ns | NaCl | KS.1 |
| FIM1_3208 | 2595.360507 | 3696.372773 | 3247.202603 | 0.477149445 | 0.10326274 | 8.8832176073 | 1.0572729090 | ns | NaCl | KS.1 |
| FIM1_3209 | 6610.670279 | 8010.114509 | 7110.124517 | 0.240628353 | 0.12484235 | 0.035524666 | 0.08018839 | ns | NaCl | KS.1 |
| FIM1_3210 | 422.6841737 | 296.017703 | 590.3318852 | -0.426533448 | 0.172747792 | 0.00420879 | 0.014147789 | ns | NaCl | KS.1 |
| FIM1_3211 | 346.3012519 | 330.5327974 | 437.1034821 | -0.049733563 | 0.172435945 | 0.73773898 | 0.827938513 | ns | NaCl | KS.1 |
| FIM1_3212 | 3368.480385 | 3849.396207 | 3159.226961 | 0.169300337 | 0.113313154 | 0.108547459 | 0.197449218 | ns | NaCl | KS.1 |
| FIM1_3213 | 3273.999591 | 3743.872677 | 3801.11775 | 0.174723684 | 0.100486335 | 0.065507208 | 0.131463638 | ns | NaCl | KS.1 |
| FIM1_3214 | 3959.700932 | 7831.288518 | 3474.386174 | 0.567090426 | 0.354750359 | 0.016452929 | 0.042235976 | ns | NaCl | KS.1 |
| FIM1_3215 | 2449.290756 | 2655.537133 | 2625.623474 | 0.107107007 | 0.086459596 | 0.196143017 | 0.314777906 | ns | NaCl | KS.1 |
| FIM1_3216 | 81.24409487 | 36.35589951 | 80.9259995 | -0.878447732 | 0.309263066 | 0.000202822 | 0.001128681 | ns | NaCl | KS.1 |
| FIM1_3217 | 2529.17786 | 2637.908714 | 2185.918002 | 0.039475439 | 0.21402529 | 0.818426681 | 0.887848395 | ns | NaCl | KS.1 |
| FIM1_3218 | 168.6044263 | 131.7847207 | 239.287081 | -0.213958044 | 0.258753877 | 0.266438329 | 0.395006359 | ns | NaCl | KS.1 |
| FIM1_3219 | 4188.297232 | 1786.124953 | 3158.916771 | -1.129200765 | 0.176638534 | 3.8059944129 | 2.0226617226 | Km<KS.1 | NaCl | KS.1 |
| FIM1_3220 | 884.9799799 | 2132.242214 | 2141.848212 | 1.104965257 | 0.228151578 | 2.9842275669 | 5.7322517935 | Km>KS.1 | NaCl | KS.1 |
| FIM1_3221 | 6467.681676 | 8101.221391 | 6394.062365 | 0.295608146 | 0.105063561 | 0.002671501 | 0.009722544 | ns | NaCl | KS.1 |
| FIM1_3222 | 3303.868835 | 3298.812484 | 3571.471653 | -0.002210036 | 0.063605271 | 0.971625363 | 0.987114147 | ns | NaCl | KS.1 |
| FIM1_3223 | 5473.458367 | 7102.5425 | 5186.784433 | 0.353674602 | 0.086162946 | 1.7046265968 | 0.000136783 | ns | NaCl | KS.1 |
| FIM1_3224 | 1498.578427 | 1228.302496 | 1877.403515 | -0.181028597 | 0.234621497 | 0.320260033 | 0.453671317 | ns | NaCl | KS.1 |
| FIM1_3225 | 145.6117982 | 92.03291637 | 117.7906859 | -0.42215934 | 0.282489777 | 0.041001998 | 0.089940451 | ns | NaCl | KS.1 |
| FIM1_3226 | 861.5296921 | 671.343189 | 829.6867632 | -0.308467926 | 0.13937911 | 0.013850216 | 0.036847498 | ns | NaCl | KS.1 |
| FIM1_3227 | 2481.723404 | 3209.664837 | 2360.123084 | 0.323848423 | 0.129586836 | 0.00604113 | 0.019118717 | ns | NaCl | KS.1 |
| FIM1_3228 | 1684.653028 | 1624.480852 | 1687.076866 | -0.045651661 | 0.120283453 | 0.68172772 | 0.784546705 | ns | NaCl | KS.1 |
| FIM1_3229 | 4735.78151 | 5597.606183 | 3769.560011 | 0.221094897 | 0.092958504 | 0.012323098 | 0.033574707 | ns | NaCl | KS.1 |
| FIM1_3230 | 4495.42742 | 5589.565463 | 4397.562342 | 0.283366525 | 0.108168489 | 0.005030001 | 0.016366418 | ns | NaCl | KS.1 |
| FIM1_3231 | 6468.398392 | 7173.076072 | 4952.197279 | 0.14157254 | 0.072494236 | 0.043964636 | 0.095221621 | ns | NaCl | KS.1 |
| FIM1_3232 | 141.0590592 | 192.5821808 | 257.3779081 | 0.308927533 | 0.227707592 | 0.08616938 | 0.164945235 | ns | NaCl | KS.1 |
| FIM1_3233 | 1042.911543 | 378.450841 | 1744.519258 | -1.410685268 | 0.136933129 | 2.8468123008 | 2.3604818660 | Km<KS.1 | NaCl | KS.1 |

| | | | | | | | | | | |
|---|---|---|---|---|---|---|---|---|---|---|
| FIM1_3234 | 4134.175951 | 9026.309929 | 4010.201847 | 0.800446815 | 0.321585808 | 0.000765815 | 0.003447902 | ns | NaCl | KS.1 |
| FIM1_3235 | 1390.524359 | 1002.646555 | 1430.647497 | -0.433491779 | 0.117131307 | 6.2731120087 | 0.00042174 | ns | NaCl | KS.1 |
| FIM1_3236 | 7865.430798 | 14301.88906 | 9277.944695 | 0.674387627 | 0.246799738 | 0.000676564 | 0.003107947 | ns | NaCl | KS.1 |
| FIM1_3237 | 6204.255064 | 4282.242881 | 5182.316218 | -0.479321274 | 0.134447574 | 8.6616232376 | 0.000555304 | ns | NaCl | KS.1 |
| FIM1_3238 | 631.622064 | 352.78774 | 470.235462 | -0.792803415 | 0.135780147 | 4.3209612572 | 1.3510440821 | ns | NaCl | KS.1 |
| FIM1_3239 | 4614.869002 | 2927.095346 | 3239.1185 | -0.617355759 | 0.112892074 | 5.9859993647 | 1.3916049925 | ns | NaCl | KS.1 |
| FIM1_3240 | 3469.920973 | 3402.505353 | 2728.842769 | -0.02407198 | 0.130309903 | 0.839874534 | 0.902325017 | ns | NaCl | KS.1 |
| FIM1_3241 | 921.1575082 | 638.7113308 | 1207.718789 | -0.4175709 | 0.192392951 | 0.009707075 | 0.027918932 | ns | NaCl | KS.1 |
| FIM1_3242 | 572.2832471 | 589.0069949 | 739.1983989 | 0.035037214 | 0.132053996 | 0.771150586 | 0.852171072 | ns | NaCl | KS.1 |
| FIM1_3243 | 2131.818113 | 2982.977912 | 2177.611347 | 0.431958228 | 0.130843661 | 0.000286166 | 0.001506535 | ns | NaCl | KS.1 |
| FIM1_3244 | 1310.702018 | 866.9863358 | 1295.076085 | -0.452373896 | 0.218812972 | 0.010743121 | 0.030281601 | ns | NaCl | KS.1 |
| FIM1_3245 | 1163.250989 | 955.5304273 | 1235.652559 | -0.237230757 | 0.141290076 | 0.061516681 | 0.125582884 | ns | NaCl | KS.1 |
| FIM1_3246 | 957.2606814 | 699.4766369 | 1352.307553 | -0.368888417 | 0.174255498 | 0.013699509 | 0.036563872 | ns | NaCl | KS.1 |
| FIM1_3247 | 4741.1722 | 5648.574474 | 5932.375847 | 0.158704028 | 0.232239271 | 0.380342501 | 0.515586905 | ns | NaCl | KS.1 |
| FIM1_3248 | 261.0694034 | 87.66207216 | 270.241428 | -1.484886978 | 0.193859555 | 5.3983538001 | 7.8990618105 | Km<KS.1 | NaCl | KS.1 |
| FIM1_3249 | 3828.887762 | 3059.654168 | 3260.562503 | -0.277573132 | 0.133842095 | 0.022069115 | 0.054138977 | ns | NaCl | KS.1 |
| FIM1_3250 | 1086.594615 | 899.0514941 | 864.3547259 | -0.219932228 | 0.158591008 | 0.113955332 | 0.205261179 | ns | NaCl | KS.1 |
| FIM1_3251 | 569.5708377 | 744.11562 | 786.1267761 | 0.34465491 | 0.118495541 | 0.001611732 | 0.006390029 | ns | NaCl | KS.1 |
| FIM1_3252 | 26.89901272 | 26.1838798 | 46.84830583 | -0.035978121 | 0.293130217 | 0.857491549 | 0.913690395 | ns | NaCl | KS.1 |
| FIM1_3253 | 1548.822458 | 835.3872615 | 3578.117444 | -0.63872782 | 0.283656862 | 0.002957166 | 0.010571345 | ns | NaCl | KS.1 |
| FIM1_3254 | 4794.843177 | 5309.712686 | 4440.083237 | 0.134717565 | 0.092882833 | 0.12768345 | 0.224157337 | ns | NaCl | KS.1 |
| FIM1_3255 | 685.0286345 | 687.0665509 | 844.2191512 | 0.000925624 | 0.093225785 | 0.991687782 | 0.996398788 | ns | NaCl | KS.1 |
| FIM1_3256 | 59.32404055 | 46.97024144 | 40.43753559 | -0.212645775 | 0.267457467 | 0.277255152 | 0.407012879 | ns | NaCl | KS.1 |
| FIM1_3258 | 444.329028 | 335.5861879 | 304.6439041 | -0.319183835 | 0.180153123 | 0.037261505 | 0.083577993 | ns | NaCl | KS.1 |
| FIM1_3259 | 389.7218943 | 395.4259453 | 420.9082459 | 0.013917876 | 0.20219416 | 0.933127769 | 0.961658314 | ns | NaCl | KS.1 |
| FIM1_3260 | 1264.245017 | 1470.825341 | 1532.399655 | 0.199591376 | 0.08159073 | 0.010914452 | 0.030660303 | ns | NaCl | KS.1 |
| FIM1_3261 | 686.910981 | 545.3759019 | 583.1292843 | -0.27672812 | 0.153055344 | 0.040699658 | 0.089474502 | ns | NaCl | KS.1 |
| FIM1_3262 | 35298.10818 | 49371.80048 | 29873.94409 | 0.44150586 | 0.118120653 | 5.2187461709 | 0.000360601 | ns | NaCl | KS.1 |
| FIM1_3263 | 1771.978383 | 1941.883875 | 1959.690367 | 0.118304078 | 0.104284908 | 0.228181005 | 0.353755219 | ns | NaCl | KS.1 |
| FIM1_3264 | 1370.095936 | 1797.563574 | 1677.565821 | 0.348491809 | 0.12284111 | 0.001989284 | 0.007606984 | ns | NaCl | KS.1 |
| FIM1_3265 | 1396.421498 | 1444.879347 | 1308.290788 | 0.044649145 | 0.095535903 | 0.623030421 | 0.737467605 | ns | NaCl | KS.1 |
| FIM1_3266 | 888.237729 | 896.7930028 | 627.6223463 | 0.01012235 | 0.190461697 | 0.949360935 | 0.973834262 | ns | NaCl | KS.1 |
| FIM1_3267 | 4610.832416 | 4954.282859 | 3631.771014 | 0.091410149 | 0.107852107 | 0.366315334 | 0.501905476 | ns | NaCl | KS.1 |
| FIM1_3268 | 895.9598879 | 1099.800478 | 861.409376 | 0.233049941 | 0.164828862 | 0.103906901 | 0.190540667 | ns | NaCl | KS.1 |
| FIM1_3269 | 434.0389217 | 406.9640361 | 594.8101771 | -0.074336249 | 0.162203367 | 0.600293908 | 0.719177329 | ns | NaCl | KS.1 |
| FIM1_3270 | 6952.410238 | 3029.637463 | 3849.519547 | -0.818088339 | 0.346236979 | 0.000965103 | 0.004167872 | ns | NaCl | KS.1 |
| FIM1_3271 | 4281.624179 | 3746.966002 | 3839.313539 | -0.166233531 | 0.123259454 | 0.142786113 | 0.244783223 | ns | NaCl | KS.1 |
| FIM1_3272 | 876.1273162 | 713.208286 | 1169.173293 | -0.251838489 | 0.139551084 | 0.044893674 | 0.096793469 | ns | NaCl | KS.1 |
| FIM1_3273 | 2846.319001 | 3497.438122 | 2005.529265 | 0.189150979 | 0.232448434 | 0.296577051 | 0.428666713 | ns | NaCl | KS.1 |
| FIM1_3274 | 2163.280455 | 2057.744362 | 2580.125895 | -0.068747699 | 0.05609856 | 0.211827278 | 0.334340326 | ns | NaCl | KS.1 |
| FIM1_3275 | 3156.725201 | 5451.719466 | 3593.207739 | 0.741383147 | 0.123012682 | 1.5780510963 | 5.6480605785 | ns | NaCl | KS.1 |
| FIM1_3276 | 265.9663176 | 160.3399368 | 476.724671 | -0.570031493 | 0.226146383 | 0.001949285 | 0.007487593 | ns | NaCl | KS.1 |
| FIM1_3277 | 2071.599969 | 2862.802665 | 2368.234626 | 0.430261549 | 0.108515455 | 2.1601383501 | 0.000166874 | ns | NaCl | KS.1 |
| FIM1_3278 | 5555.91366 | 7832.883243 | 6815.733062 | 0.464044729 | 0.100490001 | 9.5674468733 | 1.1305949690 | ns | NaCl | KS.1 |
| FIM1_3279 | 5073.505784 | 5081.419303 | 3833.314864 | 0.001916155 | 0.125208479 | 0.986730124 | 0.99352001 | ns | NaCl | KS.1 |

| | | | | | | | | | | |
|---|---|---|---|---|---|---|---|---|---|---|
| FIM1_3280 | 2074.022054 | 1661.642123 | 2105.880803 | -0.279267275 | 0.124674039 | 0.014496236 | 0.038279605 | ns | NaCl | KS.1 |
| FIM1_3281 | 4317.664566 | 4742.027261 | 4292.32629 | 0.111210212 | 0.138973208 | 0.375598674 | 0.510129239 | ns | NaCl | KS.1 |
| FIM1_3282 | 4884.404861 | 6834.622085 | 5569.056616 | 0.277443926 | 0.28009065 | 0.168621782 | 0.280097951 | ns | NaCl | KS.1 |
| FIM1_3283 | 1028.664273 | 731.7020594 | 807.0213661 | -0.41742987 | 0.160767459 | 0.003040542 | 0.01082024 | ns | NaCl | KS.1 |
| FIM1_3284 | 1352.845736 | 682.6923558 | 1342.972478 | -0.907004421 | 0.164188667 | 1.89253592938 | 5.06202486487 | ns | NaCl | KS.1 |
| FIM1_3286 | 1636.103339 | 1740.956832 | 1644.190823 | 0.076994252 | 0.111512758 | 0.460034287 | 0.594614336 | ns | NaCl | KS.1 |
| FIM1_3287 | 491.0061757 | 603.1171071 | 460.2756596 | 0.226500121 | 0.17167508 | 0.125387835 | 0.220893937 | ns | NaCl | KS.1 |
| FIM1_3288 | 20563.56394 | 31939.78615 | 24758.36259 | 0.501868355 | 0.210146397 | 0.003785184 | 0.012987097 | ns | NaCl | KS.1 |
| FIM1_3290 | 1981.141913 | 1438.497428 | 1587.834695 | -0.410427044 | 0.132481859 | 0.000642439 | 0.002967628 | ns | NaCl | KS.1 |
| FIM1_3291 | 30.77747682 | 9.258151964 | 18.39346275 | -0.842701692 | 0.505552396 | 0.003031658 | 0.010804083 | ns | NaCl | KS.1 |
| FIM1_3292 | 35685.23253 | 44672.0797 | 28547.75295 | 0.286457762 | 0.120053969 | 0.009592939 | 0.027634552 | ns | NaCl | KS.1 |
| FIM1_3293 | 490.9175596 | 582.1924871 | 402.3163413 | 0.184677107 | 0.175662671 | 0.219423799 | 0.343605099 | ns | NaCl | KS.1 |
| FIM1_3294 | 438.6791337 | 461.6732215 | 335.1507035 | 0.053385268 | 0.171484286 | 0.718169776 | 0.81294531 | ns | NaCl | KS.1 |
| FIM1_3295 | 788.0938732 | 816.691662 | 1289.998448 | 0.037253745 | 0.18416562 | 0.810913885 | 0.880850781 | ns | NaCl | KS.1 |
| FIM1_3297 | 1914.708719 | 2446.269333 | 2133.083476 | 0.330522752 | 0.093090604 | 0.000180761 | 0.001024244 | ns | NaCl | KS.1 |
| FIM1_3298 | 3200.6316 | 7335.42221 | 3671.679717 | 0.982389989 | 0.261245034 | 5.80861749902 | 5.32189172332 | ns | NaCl | KS.1 |
| FIM1_3299 | 4497.365902 | 5944.379027 | 5009.030904 | 0.376028761 | 0.094095543 | 2.44508535874 | 0.000184792 | ns | NaCl | KS.1 |
| FIM1_3300 | 15150.1564 | 19214.99646 | 12154.06664 | 0.287239625 | 0.146874485 | 0.028114769 | 0.066008012 | ns | NaCl | KS.1 |
| FIM1_3301 | 2069.930275 | 3022.101616 | 2184.731024 | 0.496254176 | 0.128075519 | 2.33420073357 | 0.000178108 | ns | NaCl | KS.1 |
| FIM1_3302 | 706.7435313 | 1536.491364 | 695.5945618 | 0.712124947 | 0.35527644 | 0.003650703 | 0.012630213 | ns | NaCl | KS.1 |
| FIM1_3303 | 574.6734892 | 649.6430834 | 583.473781 | 0.137674655 | 0.159191599 | 0.32450119 | 0.457788875 | ns | NaCl | KS.1 |
| FIM1_3304 | 10659.90714 | 23670.93956 | 10610.29114 | 0.952033793 | 0.251627684 | 6.05838611508 | 5.50008593477 | ns | NaCl | KS.1 |
| FIM1_3305 | 8551.080438 | 7980.030399 | 11293.93362 | -0.085266259 | 0.121577264 | 0.44764925 | 0.582846118 | ns | NaCl | KS.1 |
| FIM1_3306 | 516.1419622 | 325.3518459 | 385.9812727 | -0.534074843 | 0.211146077 | 0.002218182 | 0.008316093 | ns | NaCl | KS.1 |
| FIM1_3307 | 1336.586646 | 908.4648884 | 1497.973183 | -0.481733469 | 0.157698621 | 0.000540289 | 0.002577122 | ns | NaCl | KS.1 |
| FIM1_3308 | 444.5823966 | 408.9470139 | 1459.473884 | -0.083658839 | 0.194770246 | 0.605151443 | 0.723187227 | ns | NaCl | KS.1 |
| FIM1_3309 | 4237.096333 | 5822.650985 | 4207.45595 | 0.396189543 | 0.145866405 | 0.002338978 | 0.008690377 | ns | NaCl | KS.1 |
| FIM1_3310 | 1350.498197 | 1442.714009 | 1586.122228 | 0.082629352 | 0.108711702 | 0.417466693 | 0.553555407 | ns | NaCl | KS.1 |
| FIM1_3311 | 6170.728605 | 8041.84239 | 5799.004745 | 0.369408084 | 0.063688532 | 2.52579918758 | 6.44402613242 | ns | NaCl | KS.1 |
| FIM1_3312 | 2229.855154 | 1867.297995 | 3030.972899 | -0.156584152 | 0.239749941 | 0.395249993 | 0.531163888 | ns | NaCl | KS.1 |
| FIM1_3313 | 208.0469293 | 205.8564809 | 449.383105 | -0.006011051 | 0.296018773 | 0.976124659 | 0.9889179 | ns | NaCl | KS.1 |
| FIM1_3314 | 683.3192449 | 488.2270962 | 611.8314272 | -0.432015393 | 0.139858385 | 0.000600688 | 0.002808667 | ns | NaCl | KS.1 |
| FIM1_3315 | 891.0120883 | 778.1888074 | 625.3048471 | -0.152869344 | 0.160534504 | 0.276943126 | 0.406788323 | ns | NaCl | KS.1 |
| FIM1_3316 | 2432.439712 | 2883.519038 | 1921.22516 | 0.224298776 | 0.092857941 | 0.011031314 | 0.030866583 | ns | NaCl | KS.1 |
| FIM1_3317 | 164.8461247 | 94.78800444 | 156.9880462 | -0.677744269 | 0.204480414 | 9.92981762329 | 0.000620614 | ns | NaCl | KS.1 |
| FIM1_3318 | 1025.703733 | 1935.224337 | 902.019129 | 0.635247064 | 0.29838898 | 0.004045018 | 0.013680466 | ns | NaCl | KS.1 |
| FIM1_3319 | 1658.62276 | 1970.11304 | 1424.631369 | 0.206703896 | 0.140793414 | 0.102517408 | 0.188618382 | ns | NaCl | KS.1 |
| FIM1_3320 | 1744.415868 | 4713.608644 | 3754.133926 | 1.150590888 | 0.304846372 | 2.49889258416 | 2.62832782372 | Km>KS.1 | NaCl | KS.1 |
| FIM1_3321 | 2917.223095 | 5620.270365 | 2821.384153 | 0.630252815 | 0.314990747 | 0.005534392 | 0.017775079 | ns | NaCl | KS.1 |
| FIM1_3322 | 582.5672391 | 448.1719152 | 576.8316103 | -0.24548322 | 0.238148108 | 0.182920953 | 0.298566844 | ns | NaCl | KS.1 |
| FIM1_3323 | 1148.9196 | 647.8363609 | 1336.77278 | -0.765001265 | 0.143361265 | 8.23803196248 | 1.82169525947 | ns | NaCl | KS.1 |
| FIM1_3324 | 1143.514966 | 1355.33209 | 2165.389234 | 0.197780279 | 0.155381257 | 0.148810745 | 0.253365317 | ns | NaCl | KS.1 |
| FIM1_3326 | 10358.67302 | 15299.58226 | 10761.74779 | 0.469509674 | 0.176212146 | 0.001974995 | 0.007558246 | ns | NaCl | KS.1 |
| FIM1_3327 | 1326.804592 | 1798.655363 | 1547.231501 | 0.370761632 | 0.154264877 | 0.006449899 | 0.020130644 | ns | NaCl | KS.1 |
| FIM1_3328 | 340.969733 | 202.0719263 | 250.7530417 | -0.628486377 | 0.205403037 | 0.000289238 | 0.001519493 | ns | NaCl | KS.1 |

| | | | | | | | | | | |
|---|---|---|---|---|---|---|---|---|---|---|
| FIM1_3329 | 371.5928317 | 445.6630272 | 333.4106703 | 0.211033857 | 0.15535633 | 0.123305436 | 0.218411246 | ns | NaCl | KS.1 |
| FIM1_3331 | 583.3468 | 431.6090937 | 427.7347597 | -0.38844957 | 0.13043324 | 0.001058734 | 0.004498036 | ns | NaCl | KS.1 |
| FIM1_3332 | 670.2419376 | 643.8410918 | 563.365356 | -0.04465473 | 0.112779788 | 0.6714706 | 0.776929074 | ns | NaCl | KS.1 |
| FIM1_3333 | 3723.976601 | 8119.273323 | 3974.130456 | 0.993928986 | 0.202698874 | 3.64777270318 | 6.82243202945 | ns | NaCl | KS.1 |
| FIM1_3334 | 40841.81154 | 52716.06119 | 33685.66399 | 0.32595311 | 0.123428017 | 0.003979405 | 0.013480376 | ns | NaCl | KS.1 |
| FIM1_3335 | 1593.632811 | 2193.736656 | 1422.542747 | 0.360442978 | 0.189255704 | 0.023458256 | 0.056818316 | ns | NaCl | KS.1 |
| FIM1_3336 | 1449.121528 | 1602.823875 | 1146.475696 | 0.109505359 | 0.169933883 | 0.455892253 | 0.590949442 | ns | NaCl | KS.1 |
| FIM1_3337 | 1438.926987 | 1142.748752 | 2308.848679 | -0.309423544 | 0.087792062 | 0.000218503 | 0.001202494 | ns | NaCl | KS.1 |
| FIM1_3339 | 1179.460262 | 933.5083095 | 1165.07265 | -0.289940317 | 0.135655383 | 0.018026535 | 0.045593295 | ns | NaCl | KS.1 |
| FIM1_3340 | 4151.774383 | 3050.872311 | 5574.560663 | -0.336830992 | 0.197539304 | 0.039852001 | 0.087960828 | ns | NaCl | KS.1 |
| FIM1_3341 | 999.0097716 | 1052.661069 | 1121.771156 | 0.062870073 | 0.098857731 | 0.502072157 | 0.63444475 | ns | NaCl | KS.1 |
| FIM1_3342 | 131.6940382 | 95.94751392 | 158.9391856 | -0.314522552 | 0.236055967 | 0.087688799 | 0.167210339 | ns | NaCl | KS.1 |
| FIM1_3343 | 7889.050577 | 9640.513907 | 8696.620167 | 0.184199612 | 0.231826746 | 0.308545279 | 0.441349271 | ns | NaCl | KS.1 |
| FIM1_3344 | 11835.16057 | 13392.83865 | 18575.06962 | 0.098293379 | 0.260045854 | 0.607818027 | 0.724943649 | ns | NaCl | KS.1 |
| FIM1_3345 | 6244.648819 | 7033.460914 | 6481.108677 | 0.093816296 | 0.261629353 | 0.625174296 | 0.738774851 | ns | NaCl | KS.1 |
| FIM1_3346 | 182.6584442 | 199.1137563 | 215.6425215 | 0.08525326 | 0.216406934 | 0.622530132 | 0.737197667 | ns | NaCl | KS.1 |
| FIM1_3347 | 1288.158031 | 915.8406488 | 1658.200111 | -0.38782246 | 0.188880439 | 0.014745769 | 0.038671693 | ns | NaCl | KS.1 |
| FIM1_3348 | 763.1215154 | 300.5969036 | 957.1650824 | -1.201127269 | 0.221203383 | 8.54312729641 | 2.45676637570 | Km<KS.1 | NaCl | KS.1 |
| FIM1_3349 | 1296.782114 | 1306.022237 | 1789.716452 | 0.010650977 | 0.11392499 | 0.92013791 | 0.954679062 | ns | NaCl | KS.1 |
| FIM1_3350 | 7168.469498 | 12361.077 | 9211.614553 | 0.658928656 | 0.203682646 | 0.000141121 | 0.000835807 | ns | NaCl | KS.1 |
| FIM1_3351 | 1241.905411 | 1257.068603 | 1694.361127 | 0.01560176 | 0.114198911 | 0.883480946 | 0.931027377 | ns | NaCl | KS.1 |
| FIM1_3353 | 10148.51985 | 13879.92894 | 8404.574634 | 0.390681465 | 0.144291737 | 0.002459803 | 0.009071549 | ns | NaCl | KS.1 |
| FIM1_3355 | 1707.814519 | 1453.334079 | 1440.681107 | -0.172702629 | 0.181354221 | 0.261844239 | 0.390327733 | ns | NaCl | KS.1 |
| FIM1_3356 | 854.4518632 | 587.075771 | 1282.228097 | -0.4148556 | 0.208230594 | 0.015153113 | 0.039448842 | ns | NaCl | KS.1 |
| FIM1_3357 | 534.5419247 | 447.9851108 | 631.5619316 | -0.204794072 | 0.156395437 | 0.136869417 | 0.23701727 | ns | NaCl | KS.1 |
| FIM1_3358 | 16.17172863 | 9.155914163 | 16.18493289 | -0.30225058 | 0.406806059 | 0.17242406 | 0.284797376 | ns | NaCl | KS.1 |
| FIM1_3359 | 5799.658778 | 7375.468541 | 5452.627362 | 0.321044709 | 0.094886736 | 0.000351101 | 0.001798893 | ns | NaCl | KS.1 |
| FIM1_3360 | 3012.428363 | 2881.744321 | 3710.643881 | -0.056177886 | 0.112016626 | 0.591403229 | 0.712295974 | ns | NaCl | KS.1 |
| FIM1_3361 | 7190.657434 | 10242.93834 | 6897.756809 | 0.485505282 | 0.089156189 | 1.12878922768 | 2.38967081179 | ns | NaCl | KS.1 |
| FIM1_3362 | 2022.224598 | 2543.851368 | 3212.704601 | 0.258155153 | 0.17432775 | 0.084149337 | 0.161638205 | ns | NaCl | KS.1 |
| FIM1_3363 | 1569.97032 | 1569.193582 | 1497.899917 | 0.000288095 | 0.111406874 | 0.997793613 | 0.999191276 | ns | NaCl | KS.1 |
| FIM1_3364 | 1444.277583 | 1373.893881 | 1841.479941 | -0.065410437 | 0.093116039 | 0.46098743 | 0.595536865 | ns | NaCl | KS.1 |
| FIM1_3365 | 296.4265776 | 299.3755063 | 341.3008944 | 0.010592868 | 0.174376468 | 0.94361136 | 0.969330274 | ns | NaCl | KS.1 |
| FIM1_3366 | 126.5299307 | 107.9588161 | 426.0664148 | -0.100910977 | 0.307928156 | 0.621106862 | 0.735891079 | ns | NaCl | KS.1 |
| FIM1_3367 | 948.0007555 | 529.2396692 | 713.2190129 | -0.743956729 | 0.182345705 | 3.94809804576 | 3.94413409189 | ns | NaCl | KS.1 |
| FIM1_3368 | 571.3113866 | 1119.902054 | 621.4381627 | 0.870398711 | 0.178852168 | 6.95296174805 | 1.16077129854 | ns | NaCl | KS.1 |
| FIM1_3369 | 1591.252646 | 1320.360769 | 1156.707981 | -0.198214412 | 0.18627765 | 0.206409235 | 0.327869075 | ns | NaCl | KS.1 |
| FIM1_3370 | 723.8390913 | 901.5911617 | 841.2917541 | 0.239866254 | 0.184246556 | 0.123320038 | 0.218411246 | ns | NaCl | KS.1 |
| FIM1_3371 | 960.7067528 | 1044.308446 | 821.7171968 | 0.092554905 | 0.15893345 | 0.507484163 | 0.639011317 | ns | NaCl | KS.1 |
| FIM1_3372 | 856.2103503 | 663.1361794 | 785.7021713 | -0.332703678 | 0.116433405 | 0.001996704 | 0.007628511 | ns | NaCl | KS.1 |
| FIM1_3373 | 1064.402891 | 1776.06826 | 864.5272455 | 0.453644036 | 0.301910728 | 0.034713283 | 0.078606547 | ns | NaCl | KS.1 |
| FIM1_3374 | 3652.008261 | 5941.221551 | 5453.42696 | 0.435049323 | 0.294206832 | 0.039593517 | 0.087584593 | ns | NaCl | KS.1 |
| FIM1_3375 | 9331.582281 | 12334.11497 | 8288.966713 | 0.355786618 | 0.127802234 | 0.002272856 | 0.008482714 | ns | NaCl | KS.1 |
| FIM1_3376 | 31.63193503 | 15.86163962 | 19.77240213 | -0.475748786 | 0.367455193 | 0.041632304 | 0.090962104 | ns | NaCl | KS.1 |
| FIM1_3377 | 1834.357306 | 2727.775878 | 1614.212457 | 0.383498802 | 0.254496979 | 0.04794438 | 0.101933029 | ns | NaCl | KS.1 |

| | | | | | | | | | | |
|---|---|---|---|---|---|---|---|---|---|---|
| FIM1_3378 | 26.82421081 | 21.01420326 | 24.81555604 | -0.207120122 | 0.283949281 | 0.302798442 | 0.435004981 | ns | NaCl | KS.1 |
| FIM1_3379 | 488.8698968 | 423.7537349 | 602.1434 | -0.168371378 | 0.140127069 | 0.182211443 | 0.297604048 | ns | NaCl | KS.1 |
| FIM1_3380 | 271.3078434 | 305.9919348 | 384.328297 | 0.130120962 | 0.190051311 | 0.413487988 | 0.549293122 | ns | NaCl | KS.1 |
| FIM1_3381 | 22412.95882 | 30949.36426 | 18287.95914 | 0.412605133 | 0.133209094 | 0.000637328 | 0.00295044 | ns | NaCl | KS.1 |
| FIM1_3382 | 527.5261664 | 452.7352871 | 540.4338346 | -0.171958481 | 0.153556987 | 0.205401381 | 0.326580974 | ns | NaCl | KS.1 |
| FIM1_3383 | 2282.703589 | 2396.575002 | 2471.836453 | 0.060810926 | 0.119613238 | 0.583012662 | 0.704514936 | ns | NaCl | KS.1 |
| FIM1_3384 | 8151.670047 | 24434.72406 | 10102.61494 | 1.305154833 | 0.307989795 | 1.7303233656 | 2.5952101534 | Km>KS.1 | NaCl | KS.1 |
| FIM1_3385 | 115.2117606 | 90.37376487 | 83.71763438 | -0.231216632 | 0.244932265 | 0.216669222 | 0.340255486 | ns | NaCl | KS.1 |
| FIM1_3386 | 756.6505074 | 1255.433252 | 715.7074906 | 0.563149289 | 0.234073611 | 0.002763072 | 0.009997298 | ns | NaCl | KS.1 |
| FIM1_3387 | 1584.25522 | 1556.473728 | 1783.26767 | -0.016839853 | 0.20883755 | 0.92078366 | 0.954950742 | ns | NaCl | KS.1 |
| FIM1_3388 | 2449.812666 | 3261.241064 | 3075.192619 | 0.277050673 | 0.231453518 | 0.126958431 | 0.223265533 | ns | NaCl | KS.1 |
| FIM1_3389 | 967.5676694 | 1084.870573 | 1286.693949 | 0.096608843 | 0.247131948 | 0.604956787 | 0.723128308 | ns | NaCl | KS.1 |
| FIM1_3390 | 3470.065026 | 4046.827533 | 3612.230617 | 0.191425776 | 0.126295675 | 0.098096166 | 0.181964365 | ns | NaCl | KS.1 |
| FIM1_3391 | 1661.307521 | 2079.450954 | 1527.696093 | 0.239983577 | 0.190225396 | 0.131807687 | 0.229923999 | ns | NaCl | KS.1 |
| FIM1_3392 | 1019.783502 | 641.7045426 | 997.793671 | -0.601783048 | 0.149795781 | 8.3851066936 | 7.3573026103 | ns | NaCl | KS.1 |
| FIM1_3393 | 263.3456772 | 136.973448 | 168.986438 | -0.568472972 | 0.342271546 | 0.014722823 | 0.038667997 | ns | NaCl | KS.1 |
| FIM1_3394 | 2305.666484 | 1580.833639 | 2718.688271 | -0.435176171 | 0.191728043 | 0.006960865 | 0.021403154 | ns | NaCl | KS.1 |
| FIM1_3395 | 2363.429103 | 3130.802332 | 2191.632266 | 0.304653519 | 0.195682122 | 0.061024757 | 0.124886123 | ns | NaCl | KS.1 |
| FIM1_3396 | 1415.888279 | 991.1503415 | 1060.624303 | -0.37081202 | 0.22515151 | 0.038825959 | 0.086308823 | ns | NaCl | KS.1 |
| FIM1_3397 | 7811.043214 | 5962.556472 | 7705.802668 | -0.234433867 | 0.256938385 | 0.222436362 | 0.347120735 | ns | NaCl | KS.1 |
| FIM1_3398 | 11.72896402 | 11.13734996 | 8.902902147 | -0.019590108 | 0.348077444 | 0.92432625 | 0.956237537 | ns | NaCl | KS.1 |
| FIM1_3399 | 986.822472 | 780.3904186 | 1046.559827 | -0.259669079 | 0.182168402 | 0.09260047 | 0.174370681 | ns | NaCl | KS.1 |
| FIM1_3400 | 1292.833527 | 1317.947273 | 1585.761323 | 0.022997327 | 0.106945612 | 0.818997523 | 0.887886833 | ns | NaCl | KS.1 |
| FIM1_3401 | 1574.975667 | 1420.097158 | 1416.926159 | -0.121759119 | 0.140617329 | 0.33654361 | 0.47086434 | ns | NaCl | KS.1 |
| FIM1_3402 | 4228.714683 | 5215.288561 | 3669.48146 | 0.28586481 | 0.080958491 | 0.000234782 | 0.001277946 | ns | NaCl | KS.1 |
| FIM1_3403 | 5087.630638 | 10925.82775 | 4961.124225 | 0.869046572 | 0.273376572 | 7.4062720801 | 0.000488029 | ns | NaCl | KS.1 |
| FIM1_3404 | 112.4403728 | 18.64315838 | 107.336423 | -2.110258166 | 0.535174798 | 3.6956602935 | 6.8861085993 | Km<KS.1 | NaCl | KS.1 |
| FIM1_3405 | 11081.92038 | 19123.87664 | 10136.20419 | 0.708382494 | 0.159670084 | 9.0902497376 | 1.0793315619 | ns | NaCl | KS.1 |
| FIM1_3407 | 1600.802027 | 1680.852377 | 1399.676544 | 0.056283148 | 0.145324761 | 0.665604809 | 0.772064333 | ns | NaCl | KS.1 |
| FIM1_3408 | 1141.845333 | 915.550762 | 1106.331544 | -0.288197092 | 0.110833004 | 0.005223743 | 0.016875405 | ns | NaCl | KS.1 |
| FIM1_3409 | 10336.42168 | 12582.06534 | 9335.100371 | 0.221026165 | 0.169486269 | 0.131070645 | 0.22895943 | ns | NaCl | KS.1 |
| FIM1_3410 | 1669.10706 | 2135.637967 | 1501.152017 | 0.253328215 | 0.207026816 | 0.133481489 | 0.23219245 | ns | NaCl | KS.1 |
| FIM1_3411 | 2706.581709 | 2898.23398 | 1892.610748 | 0.066950771 | 0.20128641 | 0.685622518 | 0.787935326 | ns | NaCl | KS.1 |
| FIM1_3412 | 1669.247338 | 1466.485944 | 1428.368822 | -0.171900727 | 0.092380589 | 0.050608376 | 0.106820818 | ns | NaCl | KS.1 |
| FIM1_3413 | 3977.664389 | 3573.716142 | 2766.208354 | -0.116046382 | 0.172330245 | 0.434165882 | 0.570215223 | ns | NaCl | KS.1 |
| FIM1_3414 | 2993.172188 | 2661.400902 | 2553.054635 | -0.129359282 | 0.168191505 | 0.374646906 | 0.509532082 | ns | NaCl | KS.1 |
| FIM1_3415 | 1052.092174 | 1215.375985 | 1188.797963 | 0.112938957 | 0.265932796 | 0.559801818 | 0.683308894 | ns | NaCl | KS.1 |
| FIM1_3416 | 518.5747039 | 462.4646932 | 460.4770577 | -0.106519183 | 0.216218861 | 0.538399337 | 0.665640333 | ns | NaCl | KS.1 |
| FIM1_3417 | 16806.48035 | 23206.61588 | 14646.90247 | 0.421349626 | 0.120800433 | 0.000151195 | 0.000882857 | ns | NaCl | KS.1 |
| FIM1_3418 | 3381.234069 | 3911.532601 | 2580.809561 | 0.18227 | 0.120610774 | 0.101511403 | 0.18739118 | ns | NaCl | KS.1 |
| FIM1_3419 | 2793.696507 | 2989.459304 | 3343.998621 | 0.090499322 | 0.086391669 | 0.274504961 | 0.404161639 | ns | NaCl | KS.1 |
| FIM1_3420 | 52.87352952 | 27.93622336 | 103.0118483 | -0.485818291 | 0.361423804 | 0.037061729 | 0.083167389 | ns | NaCl | KS.1 |
| FIM1_3421 | 142.1816671 | 109.0085841 | 127.9908184 | -0.205779772 | 0.278037892 | 0.301444572 | 0.433435476 | ns | NaCl | KS.1 |
| FIM1_3422 | 44.13102696 | 71.5399071 | 142.3187291 | 0.350850038 | 0.335119009 | 0.109934729 | 0.199534943 | ns | NaCl | KS.1 |
| FIM1_3423 | 322.7007703 | 178.2079009 | 604.0889833 | -0.573627385 | 0.301505227 | 0.008990529 | 0.026325651 | ns | NaCl | KS.1 |

| | | | | | | | | | | |
|---|---|---|---|---|---|---|---|---|---|---|
| FIM1_3424 | 685.8216264 | 758.6189836 | 844.8276442 | 0.104582926 | 0.187075536 | 0.506391618 | 0.638227644 | ns | NaCl | KS.1 |
| FIM1_3425 | 850.8584192 | 494.9569136 | 1240.169645 | -0.608513945 | 0.234517253 | 0.001338016 | 0.005438421 | ns | NaCl | KS.1 |
| FIM1_3426 | 1890.383091 | 3867.897191 | 4545.849937 | 0.549799642 | 0.376792202 | 0.022202104 | 0.054438376 | ns | NaCl | KS.1 |
| FIM1_3427 | 884.8758268 | 860.0487254 | 1391.919756 | -0.029179598 | 0.184542089 | 0.851563859 | 0.909907688 | ns | NaCl | KS.1 |
| FIM1_3428 | 97.9961389 | 46.0513712 | 84.78845287 | -0.783013088 | 0.313140653 | 0.000806353 | 0.003594631 | ns | NaCl | KS.1 |
| FIM1_3429 | 3554.863111 | 4285.265296 | 3127.203162 | 0.239601553 | 0.113388129 | 0.022978705 | 0.055801637 | ns | NaCl | KS.1 |
| FIM1_3430 | 2361.990199 | 2044.494277 | 1770.705665 | -0.186708183 | 0.102324922 | 0.052741153 | 0.110329294 | ns | NaCl | KS.1 |
| FIM1_3431 | 182.4313694 | 567.4320922 | 368.9373469 | 1.525626741 | 0.200572412 | 6.44328200022 | 9.15866512889 | Km>KS.1 | NaCl | KS.1 |
| FIM1_3432 | 1413.753304 | 940.6336783 | 922.3058643 | -0.49846806 | 0.172395044 | 0.000860018 | 0.003789718 | ns | NaCl | KS.1 |
| FIM1_3433 | 3826.826923 | 4164.21462 | 3764.933484 | 0.113635739 | 0.079541552 | 0.138418316 | 0.238859217 | ns | NaCl | KS.1 |
| FIM1_3434 | 585.6378448 | 465.37015 | 717.5488816 | -0.300745706 | 0.105138624 | 0.00226004 | 0.008447557 | ns | NaCl | KS.1 |
| FIM1_3435 | 2893.036895 | 3966.663478 | 3127.965965 | 0.374478487 | 0.167919468 | 0.010056007 | 0.028702601 | ns | NaCl | KS.1 |
| FIM1_3437 | 17895.0138 | 16001.57772 | 16824.47536 | -0.123242299 | 0.167381105 | 0.395981436 | 0.531859515 | ns | NaCl | KS.1 |
| FIM1_3438 | 4570.27935 | 2696.750454 | 2603.795388 | -0.631670093 | 0.205585438 | 0.00027367 | 0.001453051 | ns | NaCl | KS.1 |
| FIM1_3439 | 1028.853703 | 892.8437734 | 1906.752553 | -0.129589984 | 0.223174422 | 0.462989861 | 0.597421772 | ns | NaCl | KS.1 |
| FIM1_3440 | 2806.131408 | 3456.066891 | 3000.803043 | 0.254292582 | 0.139639114 | 0.042945157 | 0.093147792 | ns | NaCl | KS.1 |
| FIM1_3441 | 604.26461 | 452.183204 | 987.7667858 | -0.348430427 | 0.155158596 | 0.010800443 | 0.030425937 | ns | NaCl | KS.1 |
| FIM1_3442 | 5126.567003 | 5378.386917 | 5052.080606 | 0.055857738 | 0.141302681 | 0.660860443 | 0.767256328 | ns | NaCl | KS.1 |
| FIM1_3444 | 5836.142669 | 5913.270716 | 4131.371059 | 0.015744989 | 0.133840246 | 0.897158574 | 0.938735214 | ns | NaCl | KS.1 |
| FIM1_3445 | 4384.640334 | 4822.195459 | 3482.366193 | 0.103494036 | 0.169742145 | 0.480659176 | 0.614609043 | ns | NaCl | KS.1 |
| FIM1_3446 | 1601.619574 | 1342.563399 | 1375.250604 | -0.242288462 | 0.07691005 | 0.001085858 | 0.004587796 | ns | NaCl | KS.1 |
| FIM1_3447 | 2348.128824 | 3085.776952 | 2612.233532 | 0.354233887 | 0.116446147 | 0.000999158 | 0.00428149 | ns | NaCl | KS.1 |
| FIM1_3448 | 642.9517568 | 553.9334161 | 773.1970386 | -0.198959247 | 0.102792787 | 0.039747008 | 0.087807001 | ns | NaCl | KS.1 |
| FIM1_3449 | 1136.560947 | 811.2805159 | 936.3167794 | -0.445154751 | 0.119698085 | 5.53159699513 | 0.000379059 | ns | NaCl | KS.1 |
| FIM1_3450 | 5276.867844 | 10338.76567 | 4650.317217 | 0.71297772 | 0.28678562 | 0.001152687 | 0.004835258 | ns | NaCl | KS.1 |
| FIM1_3452 | 1529.315041 | 1450.086146 | 1780.430393 | -0.066385672 | 0.119280565 | 0.547980536 | 0.673635573 | ns | NaCl | KS.1 |
| FIM1_3454 | 8281.527148 | 2616.754844 | 12413.85407 | -1.485222605 | 0.250414927 | 9.23134485917 | 4.50254320337 | Km<KS.1 | NaCl | KS.1 |
| FIM1_3455 | 147.9144612 | 69.49796305 | 1212.410285 | -0.793810682 | 0.313363865 | 0.000703647 | 0.003214547 | ns | NaCl | KS.1 |
| FIM1_3456 | 41.76642412 | 36.89545704 | 220.5758679 | -0.082852813 | 0.28890221 | 0.678480606 | 0.781713991 | ns | NaCl | KS.1 |
| FIM1_3457 | 479.7490888 | 567.2888029 | 898.4169684 | 0.136546473 | 0.255832365 | 0.473290303 | 0.606704266 | ns | NaCl | KS.1 |
| FIM1_3458 | 3853.043229 | 5809.579872 | 8765.855031 | 0.46691723 | 0.204578703 | 0.005858308 | 0.018675443 | ns | NaCl | KS.1 |
| FIM1_3460 | 5685.19461 | 8178.673985 | 6233.295187 | 0.466066428 | 0.139168871 | 0.000206483 | 0.00114649 | ns | NaCl | KS.1 |
| FIM1_3461 | 1105.476647 | 1574.768735 | 1201.563314 | 0.284573222 | 0.289162479 | 0.164487303 | 0.27432931 | ns | NaCl | KS.1 |
| FIM1_3462 | 694.7912945 | 642.8373493 | 799.4370233 | -0.099465192 | 0.111970849 | 0.341451225 | 0.47583189 | ns | NaCl | KS.1 |
| FIM1_3463 | 2087.843869 | 2839.760169 | 3452.321499 | 0.280246633 | 0.251747501 | 0.141797604 | 0.243256235 | ns | NaCl | KS.1 |
| FIM1_3464 | 290.7785923 | 284.8283271 | 332.051609 | -0.021395846 | 0.20642388 | 0.898714205 | 0.939307388 | ns | NaCl | KS.1 |
| FIM1_3465 | 168.8247377 | 69.60618194 | 228.7605552 | -0.904694241 | 0.342694425 | 0.000313723 | 0.001625802 | ns | NaCl | KS.1 |
| FIM1_3466 | 2810.49718 | 2944.68065 | 2895.335907 | 0.062223039 | 0.077683776 | 0.40710663 | 0.541973637 | ns | NaCl | KS.1 |
| FIM1_3467 | 536.4324848 | 552.5524806 | 819.1413989 | 0.020448058 | 0.292170618 | 0.918593802 | 0.953673658 | ns | NaCl | KS.1 |
| FIM1_3468 | 3185.533899 | 3586.772558 | 2493.683692 | 0.137879384 | 0.147452007 | 0.29474111 | 0.426632826 | ns | NaCl | KS.1 |
| FIM1_3469 | 1860.837326 | 1538.45636 | 2023.438997 | -0.168109456 | 0.240709728 | 0.362708891 | 0.497924044 | ns | NaCl | KS.1 |
| FIM1_3470 | 748.3925574 | 749.2865472 | 741.3694133 | 0.000935071 | 0.214790533 | 0.99567033 | 0.998077754 | ns | NaCl | KS.1 |
| FIM1_3471 | 2502.061982 | 4062.451582 | 2447.582828 | 0.645048799 | 0.131418051 | 1.11580618706 | 1.75668853817 | ns | NaCl | KS.1 |
| FIM1_3472 | 461.5436923 | 474.7390221 | 647.6004953 | 0.036763801 | 0.135446348 | 0.765040216 | 0.848552421 | ns | NaCl | KS.1 |
| FIM1_3473 | 3316.067577 | 7639.194823 | 3333.603423 | 0.84217898 | 0.338645624 | 0.000631339 | 0.002938179 | ns | NaCl | KS.1 |

| | | | | | | | | | | |
|---|---|---|---|---|---|---|---|---|---|---|
| FIM1_3474 | 145.464672 | 117.5797313 | 705.3237644 | -0.164500076 | 0.276145205 | 0.405393492 | 0.540416029 | ns | NaCl | KS.1 |
| FIM1_3475 | 3707.197043 | 3677.506891 | 3795.78794 | -0.010802962 | 0.099441972 | 0.908687366 | 0.946531862 | ns | NaCl | KS.1 |
| FIM1_3476 | 1133.94231 | 1188.968812 | 1593.52545 | 0.058646972 | 0.101393434 | 0.540502168 | 0.666914257 | ns | NaCl | KS.1 |
| FIM1_3477 | 2287.301675 | 1906.288849 | 1971.95525 | -0.217120611 | 0.14479874 | 0.093564206 | 0.175786225 | ns | NaCl | KS.1 |
| FIM1_3478 | 193.3009196 | 175.1091003 | 204.791761 | -0.102186813 | 0.197921971 | 0.532145664 | 0.660041056 | ns | NaCl | KS.1 |
| FIM1_3479 | 6478.472548 | 7770.046913 | 7511.9524 | 0.253560299 | 0.058402462 | 9.0489860341 | 7.8429800557 | ns | NaCl | KS.1 |
| FIM1_3480 | 3425.011307 | 6959.850728 | 3057.765141 | 0.651502638 | 0.339088998 | 0.005886505 | 0.018736636 | ns | NaCl | KS.1 |
| FIM1_3481 | 3461.679047 | 3866.136115 | 3386.072532 | 0.147050034 | 0.088021535 | 0.080667138 | 0.156169423 | ns | NaCl | KS.1 |
| FIM1_3482 | 3096.778261 | 2237.759711 | 3183.553039 | -0.374635861 | 0.181206517 | 0.015082962 | 0.039307352 | ns | NaCl | KS.1 |
| FIM1_3483 | 862.1313354 | 719.9996998 | 933.8487455 | -0.23558754 | 0.106842657 | 0.018519228 | 0.046578947 | ns | NaCl | KS.1 |
| FIM1_3484 | 1235.492364 | 959.9452303 | 1506.444146 | -0.336123261 | 0.091990143 | 0.000118374 | 0.00071731 | ns | NaCl | KS.1 |
| FIM1_3485 | 470.9448379 | 465.1268613 | 508.4485735 | -0.013437753 | 0.155515644 | 0.92209543 | 0.955315444 | ns | NaCl | KS.1 |
| FIM1_3486 | 3544.590727 | 3258.892678 | 3483.811659 | -0.098749165 | 0.141842758 | 0.439146256 | 0.574632463 | ns | NaCl | KS.1 |
| FIM1_3487 | 994.0784707 | 928.6550844 | 881.4953144 | -0.073885162 | 0.173110341 | 0.61981584 | 0.734711414 | ns | NaCl | KS.1 |
| FIM1_3488 | 415.4211306 | 342.1700614 | 422.2680436 | -0.244149598 | 0.123379413 | 0.031182493 | 0.071987427 | ns | NaCl | KS.1 |
| FIM1_3489 | 704.9331261 | 804.1910335 | 767.3091327 | 0.173837896 | 0.107274019 | 0.083811214 | 0.161194408 | ns | NaCl | KS.1 |
| FIM1_3490 | 1034.816433 | 1250.052956 | 1105.079606 | 0.241438771 | 0.110844315 | 0.01943143 | 0.048309819 | ns | NaCl | KS.1 |
| FIM1_3491 | 1839.7398 | 1679.033525 | 2077.042796 | -0.113099759 | 0.126408847 | 0.329620083 | 0.463227461 | ns | NaCl | KS.1 |
| FIM1_3492 | 843.5153519 | 448.2678999 | 782.9234932 | -0.790757451 | 0.199822149 | 5.6265235412 | 5.1933125449 | ns | NaCl | KS.1 |
| FIM1_3493 | 129.1312327 | 115.2086864 | 111.3256679 | -0.099225542 | 0.242464549 | 0.591550251 | 0.712295974 | ns | NaCl | KS.1 |
| FIM1_3494 | 7547.755845 | 15151.35571 | 9553.046138 | 0.966688561 | 0.108194767 | 2.1473958649 | 6.6770590175 | ns | NaCl | KS.1 |
| FIM1_3495 | 143.1890682 | 135.0459953 | 257.5342315 | -0.0533661 | 0.175950405 | 0.723285711 | 0.816692332 | ns | NaCl | KS.1 |
| FIM1_3496 | 642.0495612 | 877.7890048 | 619.1826244 | 0.360869106 | 0.176982174 | 0.017128736 | 0.043745102 | ns | NaCl | KS.1 |
| FIM1_3497 | 1422.876337 | 499.4254985 | 1782.456726 | -1.126670549 | 0.357895137 | 2.4522423547 | 0.000184792 | Km<KS.1 | NaCl | KS.1 |
| FIM1_3498 | 223.5980486 | 181.4401004 | 150.7250461 | -0.205286671 | 0.218706656 | 0.240027968 | 0.366863023 | ns | NaCl | KS.1 |
| FIM1_3499 | 2451.119558 | 2406.758264 | 2687.064173 | -0.025095445 | 0.085608063 | 0.760044533 | 0.844776933 | ns | NaCl | KS.1 |
| FIM1_3500 | 1958.972604 | 1844.238138 | 1777.103672 | -0.06367585 | 0.175174 | 0.671672753 | 0.776929074 | ns | NaCl | KS.1 |
| FIM1_3501 | 2772.068845 | 1739.188068 | 2906.741389 | -0.488028707 | 0.246741802 | 0.011320457 | 0.031498476 | ns | NaCl | KS.1 |
| FIM1_3503 | 216.4817434 | 257.4775772 | 375.0021449 | 0.108729236 | 0.317662504 | 0.597433236 | 0.716894557 | ns | NaCl | KS.1 |
| FIM1_3504 | 505.5853527 | 931.0246351 | 551.5915906 | 0.594494587 | 0.302088077 | 0.00706555 | 0.021698217 | ns | NaCl | KS.1 |
| FIM1_3505 | 5628.151807 | 5246.157312 | 3680.734522 | -0.09138617 | 0.099492611 | 0.331841081 | 0.465569481 | ns | NaCl | KS.1 |
| FIM1_3506 | 789.8726418 | 664.1314201 | 702.3294603 | -0.191469285 | 0.173774833 | 0.199324425 | 0.318854988 | ns | NaCl | KS.1 |
| FIM1_3507 | 2636.78906 | 1866.79685 | 1893.918144 | -0.466145287 | 0.104207535 | 1.8904413239 | 2.0313057423 | ns | NaCl | KS.1 |
| FIM1_3508 | 1014.854941 | 1215.271226 | 926.6437892 | 0.226020861 | 0.123739497 | 0.046708249 | 0.099988615 | ns | NaCl | KS.1 |
| FIM1_3509 | 23.86204878 | 10.9640355 | 75.09163503 | -0.63774144 | 0.388167107 | 0.010391776 | 0.0294856 | ns | NaCl | KS.1 |
| FIM1_3511 | 6990.770803 | 5044.716335 | 4938.838614 | -0.41591688 | 0.135556699 | 0.000693037 | 0.003171904 | ns | NaCl | KS.1 |
| FIM1_3512 | 4305.139797 | 3695.190055 | 4645.559879 | -0.16646249 | 0.174434277 | 0.26593679 | 0.394464976 | ns | NaCl | KS.1 |
| FIM1_3513 | 730.5431319 | 575.7440936 | 986.2109913 | -0.160489043 | 0.307213991 | 0.43498919 | 0.570995045 | ns | NaCl | KS.1 |
| FIM1_3514 | 4203.032997 | 2553.339008 | 2916.016212 | -0.543886188 | 0.239284536 | 0.004280677 | 0.014340987 | ns | NaCl | KS.1 |
| FIM1_3516 | 8697.073981 | 12229.03416 | 7410.048664 | 0.299166285 | 0.267058336 | 0.129479925 | 0.226817827 | ns | NaCl | KS.1 |
| FIM1_3517 | 598.7724233 | 357.938509 | 512.655349 | -0.68518034 | 0.131953974 | 2.2369431587 | 4.5238992742 | ns | NaCl | KS.1 |
| FIM1_3518 | 3828.588127 | 6177.517306 | 3047.35086 | 0.386480767 | 0.313653651 | 0.073337038 | 0.144358488 | ns | NaCl | KS.1 |
| FIM1_3519 | 998.8789526 | 858.7018533 | 757.2062454 | -0.143788555 | 0.216567444 | 0.406995708 | 0.541970998 | ns | NaCl | KS.1 |
| FIM1_3520 | 1356.201292 | 1029.18988 | 1141.632281 | -0.270438126 | 0.225906459 | 0.130311833 | 0.227954068 | ns | NaCl | KS.1 |
| FIM1_3522 | 4352.220397 | 7865.180724 | 3616.222836 | 0.508470389 | 0.327055172 | 0.024101616 | 0.058121929 | ns | NaCl | KS.1 |

| | | | | | | | | | | |
|---|---|---|---|---|---|---|---|---|---|---|
| FIM1_3523 | 7500.96929 | 10479.56603 | 7519.610076 | 0.389464851 | 0.179471958 | 0.010981834 | 0.030762738 | ns | NaCl | KS.1 |
| FIM1_3524 | 2020.678774 | 2710.558163 | 1888.769054 | 0.346883298 | 0.165611949 | 0.015890764 | 0.040940729 | ns | NaCl | KS.1 |
| FIM1_3525 | 549.8543504 | 480.4994273 | 478.5321531 | -0.128398414 | 0.215529639 | 0.457576489 | 0.592503233 | ns | NaCl | KS.1 |
| FIM1_3526 | 4458.729562 | 3330.125338 | 3162.86158 | -0.393406783 | 0.095397597 | 1.30067762043 | 0.000107668 | ns | NaCl | KS.1 |
| FIM1_3527 | 2477.094881 | 1522.065187 | 1778.7921 | -0.534411408 | 0.234867988 | 0.004435291 | 0.014759579 | ns | NaCl | KS.1 |
| FIM1_3528 | 1347.593289 | 2082.97199 | 3188.141061 | 0.369805791 | 0.29431053 | 0.077013713 | 0.150252244 | ns | NaCl | KS.1 |
| FIM1_3529 | 615.422027 | 675.6265308 | 616.9667557 | 0.097814032 | 0.181187198 | 0.525031845 | 0.653989342 | ns | NaCl | KS.1 |
| FIM1_3530 | 425.9037084 | 390.9182416 | 406.0561677 | -0.092058612 | 0.173891737 | 0.537767698 | 0.665024682 | ns | NaCl | KS.1 |
| FIM1_3531 | 1086.461479 | 1168.128071 | 1385.990734 | 0.091893574 | 0.089138773 | 0.280971002 | 0.411126687 | ns | NaCl | KS.1 |
| FIM1_3532 | 999.6096121 | 1096.929104 | 977.3404401 | 0.109645818 | 0.137219335 | 0.377414544 | 0.51220762 | ns | NaCl | KS.1 |
| FIM1_3533 | 352.8023609 | 309.4095398 | 652.0218671 | -0.149050404 | 0.152213291 | 0.269269425 | 0.398220984 | ns | NaCl | KS.1 |
| FIM1_3534 | 1993.890364 | 2359.401764 | 2703.360744 | 0.226307156 | 0.086554511 | 0.006245587 | 0.019665694 | ns | NaCl | KS.1 |
| FIM1_3535 | 1560.422225 | 1797.796877 | 1527.632879 | 0.192869758 | 0.078603758 | 0.01089256 | 0.030616094 | ns | NaCl | KS.1 |
| FIM1_3536 | 466.7910879 | 481.7102755 | 460.5971985 | 0.036219313 | 0.13309699 | 0.765146967 | 0.848552421 | ns | NaCl | KS.1 |
| FIM1_3537 | 7683.255322 | 17642.8986 | 8084.007475 | 0.945592366 | 0.284807946 | 3.31955247145 | 0.000242669 | ns | NaCl | KS.1 |
| FIM1_3538 | 836.9493423 | 767.0182153 | 903.5957502 | -0.109893493 | 0.119582126 | 0.32070099 | 0.453978808 | ns | NaCl | KS.1 |
| FIM1_3539 | 557.671884 | 611.2306475 | 579.1258978 | 0.110144137 | 0.136701146 | 0.37376316 | 0.50880979 | ns | NaCl | KS.1 |
| FIM1_3540 | 4109.161672 | 3485.683531 | 3361.474688 | -0.217151183 | 0.094290415 | 0.015249729 | 0.039679605 | ns | NaCl | KS.1 |
| FIM1_3541 | 215.0548869 | 554.456878 | 306.9328249 | 1.174138267 | 0.251649281 | 4.97075801243 | 8.55692772037 | Km>KS.1 | NaCl | KS.1 |
| FIM1_3542 | 370.0153184 | 363.553281 | 450.4154753 | -0.029472444 | 0.123445881 | 0.795686218 | 0.869626304 | ns | NaCl | KS.1 |
| FIM1_3544 | 3415.218711 | 3427.205892 | 2705.31606 | 0.003777318 | 0.15583907 | 0.978104628 | 0.989137805 | ns | NaCl | KS.1 |
| FIM1_3545 | 123.5385189 | 42.6383535 | 154.6222435 | -1.139811654 | 0.35360948 | 1.81034719678 | 0.000143644 | Km<KS.1 | NaCl | KS.1 |
| FIM1_3546 | 1586.457688 | 2456.022067 | 1688.710571 | 0.548949532 | 0.163616467 | 0.000141524 | 0.000836202 | ns | NaCl | KS.1 |
| FIM1_3547 | 1021.535478 | 1259.757765 | 926.7482602 | 0.261709919 | 0.128648457 | 0.025727727 | 0.061477157 | ns | NaCl | KS.1 |
| FIM1_3548 | 160.1210187 | 579.4490892 | 437.2404573 | 1.526044258 | 0.355561025 | 4.37879844633 | 7.83615909011 | Km>KS.1 | NaCl | KS.1 |
| FIM1_3549 | 196.0215228 | 148.3494993 | 171.1134703 | -0.318881023 | 0.183856407 | 0.040367026 | 0.088900378 | ns | NaCl | KS.1 |
| FIM1_3550 | 2083.362223 | 823.9238418 | 1459.170544 | -1.170331334 | 0.233112715 | 8.86231755702 | 1.94229206370 | Km<KS.1 | NaCl | KS.1 |
| FIM1_3551 | 2351.594536 | 1965.08787 | 1689.147242 | -0.194076098 | 0.181154684 | 0.206931247 | 0.328455284 | ns | NaCl | KS.1 |
| FIM1_3552 | 635.8393753 | 546.8472755 | 580.9714058 | -0.180554178 | 0.146173344 | 0.166709901 | 0.277558423 | ns | NaCl | KS.1 |
| FIM1_3553 | 520.6973649 | 480.4266848 | 438.2540512 | -0.077525287 | 0.208835351 | 0.647149638 | 0.755235621 | ns | NaCl | KS.1 |
| FIM1_3554 | 652.8800462 | 578.9054884 | 602.7025135 | -0.14046146 | 0.151210807 | 0.295397588 | 0.427458696 | ns | NaCl | KS.1 |
| FIM1_3555 | 530.9111337 | 405.268449 | 575.0273797 | -0.326363654 | 0.15641718 | 0.017650318 | 0.044869868 | ns | NaCl | KS.1 |
| FIM1_3556 | 1944.117737 | 2401.444505 | 2232.991527 | 0.276578218 | 0.107503926 | 0.005915554 | 0.018771772 | ns | NaCl | KS.1 |
| FIM1_3557 | 5585.232553 | 9219.718452 | 4774.948837 | 0.574208834 | 0.220772059 | 0.001519895 | 0.006058878 | ns | NaCl | KS.1 |
| FIM1_3558 | 1076.270218 | 1119.239833 | 1198.38517 | 0.047071859 | 0.139804283 | 0.709220459 | 0.804837451 | ns | NaCl | KS.1 |
| FIM1_3559 | 243.1858979 | 173.5843263 | 243.8344568 | -0.391884164 | 0.177239091 | 0.009793098 | 0.028064898 | ns | NaCl | KS.1 |
| FIM1_3560 | 1504.391786 | 1215.191376 | 1344.499968 | -0.276764678 | 0.112922754 | 0.00833678 | 0.024850498 | ns | NaCl | KS.1 |
| FIM1_3561 | 392.1019334 | 260.3852439 | 872.0920139 | -0.44348516 | 0.223730041 | 0.013675528 | 0.036558707 | ns | NaCl | KS.1 |
| FIM1_3562 | 1209.213195 | 1649.069167 | 1735.131374 | 0.409979421 | 0.114354546 | 0.000110055 | 0.000679308 | ns | NaCl | KS.1 |
| FIM1_3563 | 1731.721237 | 2155.842588 | 1925.341098 | 0.280408117 | 0.116373064 | 0.009211408 | 0.026814953 | ns | NaCl | KS.1 |
| FIM1_3565 | 101.3344963 | 100.3784512 | 110.8741976 | -0.009100992 | 0.268731312 | 0.962557895 | 0.98169855 | ns | NaCl | KS.1 |
| FIM1_3566 | 242.3416278 | 192.6698069 | 242.8917698 | -0.233783617 | 0.209912469 | 0.169826408 | 0.281253788 | ns | NaCl | KS.1 |
| FIM1_3567 | 7713.045939 | 10675.20988 | 7172.635201 | 0.390622336 | 0.164226573 | 0.006329839 | 0.01982445 | ns | NaCl | KS.1 |
| FIM1_3568 | 3688.047147 | 3099.375522 | 4147.484486 | -0.177838094 | 0.197369663 | 0.276147305 | 0.406049694 | ns | NaCl | KS.1 |
| FIM1_3569 | 2844.912838 | 4374.458987 | 2003.273747 | 0.311187956 | 0.324066061 | 0.147983298 | 0.252129077 | ns | NaCl | KS.1 |

| | | | | | | | | | | |
|---|---|---|---|---|---|---|---|---|---|---|
| FIM1_3570 | 1847.296568 | 1650.624124 | 1301.032634 | -0.112907508 | 0.196938324 | 0.48857837 | 0.622293238 | ns | NaCl | KS.1 |
| FIM1_3571 | 5394.090761 | 5062.136841 | 4121.434755 | -0.082743111 | 0.098983186 | 0.377455105 | 0.51220762 | ns | NaCl | KS.1 |
| FIM1_3572 | 519.2320022 | 471.5089087 | 519.58149 | -0.119405596 | 0.130608295 | 0.316582893 | 0.449358029 | ns | NaCl | KS.1 |
| FIM1_3573 | 5321.640189 | 6000.532014 | 4375.909811 | 0.156430433 | 0.099211846 | 0.095464618 | 0.178652011 | ns | NaCl | KS.1 |
| FIM1_3574 | 1801.787705 | 1673.786132 | 3146.18648 | -0.052761819 | 0.280675487 | 0.789317103 | 0.864684342 | ns | NaCl | KS.1 |
| FIM1_3575 | 1909.173204 | 1525.167863 | 1718.833637 | -0.224286528 | 0.212133909 | 0.190780894 | 0.307960723 | ns | NaCl | KS.1 |
| FIM1_3576 | 4609.11162 | 3371.150446 | 3522.498511 | -0.403404663 | 0.125907086 | 0.000460375 | 0.002247658 | ns | NaCl | KS.1 |
| FIM1_3577 | 481.4772631 | 333.5527262 | 487.2915457 | -0.460132996 | 0.154727081 | 0.000778049 | 0.003493498 | ns | NaCl | KS.1 |
| FIM1_3579 | 2197.084123 | 2247.825311 | 2293.963276 | 0.024682154 | 0.17011214 | 0.866715651 | 0.920365073 | ns | NaCl | KS.1 |
| FIM1_3580 | 5426.122504 | 11517.16552 | 5248.873732 | 0.785384817 | 0.309071087 | 0.000716316 | 0.003263434 | ns | NaCl | KS.1 |
| FIM1_3581 | 4358.868834 | 2539.713401 | 4370.134313 | -0.614318707 | 0.231530007 | 0.001100479 | 0.004635805 | ns | NaCl | KS.1 |
| FIM1_3582 | 2302.017024 | 2238.491891 | 2298.735857 | -0.036947843 | 0.094356088 | 0.680805981 | 0.784030035 | ns | NaCl | KS.1 |
| FIM1_3583 | 2684.25454 | 2693.634081 | 1973.623488 | 0.003181385 | 0.213199163 | 0.985202492 | 0.992785578 | ns | NaCl | KS.1 |
| FIM1_3584 | 976.9138299 | 719.8552979 | 589.4158153 | -0.364561325 | 0.168273082 | 0.012336652 | 0.033593236 | ns | NaCl | KS.1 |
| FIM1_3585 | 1327.468205 | 785.7317244 | 1187.196258 | -0.632790134 | 0.204077494 | 0.000247811 | 0.001340063 | ns | NaCl | KS.1 |
| FIM1_3586 | 2789.875243 | 3863.026836 | 2909.673499 | 0.370080295 | 0.187003206 | 0.018976456 | 0.04750245 | ns | NaCl | KS.1 |
| FIM1_3587 | 65.87298297 | 95.66904524 | 85.38572007 | 0.299997774 | 0.284028269 | 0.140392812 | 0.241429049 | ns | NaCl | KS.1 |
| FIM1_3588 | 1075.509033 | 1032.615008 | 1024.528335 | -0.035804629 | 0.229273176 | 0.841551336 | 0.903176394 | ns | NaCl | KS.1 |
| FIM1_3589 | 2654.170779 | 3045.61666 | 3873.387259 | 0.145462928 | 0.183680029 | 0.34908804 | 0.483764066 | ns | NaCl | KS.1 |
| FIM1_3591 | 672.2808171 | 238.0058049 | 629.3839924 | -1.263145575 | 0.280135408 | 6.46770904724 | 1.08339570740 | Km<KS.1 | NaCl | KS.1 |
| FIM1_3592 | 2841.297869 | 2549.916773 | 1972.23227 | -0.086054986 | 0.257662603 | 0.651630903 | 0.759396519 | ns | NaCl | KS.1 |
| FIM1_3594 | 3467.376044 | 4260.571129 | 3172.010947 | 0.264469789 | 0.11495529 | 0.013084012 | 0.035280736 | ns | NaCl | KS.1 |
| FIM1_3595 | 4587.49623 | 4825.949854 | 5378.255139 | 0.058675107 | 0.147425794 | 0.656006884 | 0.763066225 | ns | NaCl | KS.1 |
| FIM1_3596 | 36.09099668 | 36.39081253 | 37.01878554 | -0.001873807 | 0.30223068 | 0.992600557 | 0.996607017 | ns | NaCl | KS.1 |
| FIM1_3597 | 12317.54692 | 14736.1516 | 10687.94615 | 0.231057953 | 0.109531548 | 0.023848578 | 0.057623445 | ns | NaCl | KS.1 |
| FIM1_3598 | 3060.058713 | 4911.287203 | 2566.044627 | 0.531691361 | 0.222662016 | 0.003326729 | 0.011614371 | ns | NaCl | KS.1 |
| FIM1_3599 | 5572.800075 | 7671.449722 | 6212.493068 | 0.420013028 | 0.116756978 | 0.000100218 | 0.000625575 | ns | NaCl | KS.1 |
| FIM1_3600 | 484.5008764 | 410.0272365 | 386.4645009 | -0.183129504 | 0.178502526 | 0.228753683 | 0.354408521 | ns | NaCl | KS.1 |
| FIM1_3601 | 1666.817707 | 2027.876647 | 2242.782252 | 0.240471461 | 0.135436334 | 0.04971528 | 0.105114117 | ns | NaCl | KS.1 |
| FIM1_3603 | 541.2017883 | 545.9996057 | 629.563039 | 0.006684019 | 0.125978144 | 0.953973921 | 0.976222554 | ns | NaCl | KS.1 |
| FIM1_3604 | 4336.003112 | 7353.169415 | 3903.096472 | 0.666199511 | 0.176575678 | 1.82958435080 | 0.000144479 | ns | NaCl | KS.1 |
| FIM1_3605 | 1105.79816 | 961.1890099 | 1340.176429 | -0.191136757 | 0.077035608 | 0.010138819 | 0.02887271 | ns | NaCl | KS.1 |
| FIM1_3606 | 432.0771254 | 336.1400657 | 573.6389977 | -0.266090278 | 0.200830786 | 0.10774755 | 0.196281237 | ns | NaCl | KS.1 |
| FIM1_3607 | 1348.263838 | 2095.992276 | 1506.810348 | 0.57857655 | 0.13814467 | 4.34926934284 | 4.25937302769 | ns | NaCl | KS.1 |
| FIM1_3608 | 329.7211115 | 525.5789431 | 542.6673427 | 0.601455831 | 0.156880738 | 1.80672681376 | 0.000143586 | ns | NaCl | KS.1 |
| FIM1_3609 | 1617.606058 | 632.4081266 | 1486.044669 | -1.12601391 | 0.273402198 | 7.19044080040 | 8.83270197086 | Km<KS.1 | NaCl | KS.1 |
| FIM1_3610 | 1685.795624 | 1508.367739 | 1668.484706 | -0.140646059 | 0.116063605 | 0.191949562 | 0.309342198 | ns | NaCl | KS.1 |
| FIM1_3611 | 680.1264723 | 863.577614 | 868.238066 | 0.313066631 | 0.102707982 | 0.001169637 | 0.004889868 | ns | NaCl | KS.1 |
| FIM1_3612 | 1483.98948 | 1402.875829 | 1620.530005 | -0.073382384 | 0.099872354 | 0.437514157 | 0.573401721 | ns | NaCl | KS.1 |
| FIM1_3613 | 507.9392692 | 388.1862475 | 1052.78488 | -0.283036079 | 0.201219895 | 0.087662485 | 0.167210339 | ns | NaCl | KS.1 |
| FIM1_3614 | 284.0484995 | 288.5443064 | 312.8522437 | 0.015546283 | 0.190709876 | 0.922366285 | 0.955397099 | ns | NaCl | KS.1 |
| FIM1_3615 | 1602.179695 | 2497.692761 | 1455.642772 | 0.533239143 | 0.187912427 | 0.000879692 | 0.003858908 | ns | NaCl | KS.1 |
| FIM1_3616 | 3948.831441 | 6157.086876 | 3554.016383 | 0.464419628 | 0.241877072 | 0.014363028 | 0.037991702 | ns | NaCl | KS.1 |
| FIM1_3617 | 270.4217533 | 355.8388299 | 224.0788788 | 0.280290714 | 0.209338446 | 0.099633538 | 0.184266488 | ns | NaCl | KS.1 |
| FIM1_3618 | 1885.442041 | 1961.89657 | 2714.576116 | 0.050072984 | 0.099796119 | 0.596094022 | 0.715802018 | ns | NaCl | KS.1 |

| | | | | | | | | | | |
|---|---|---|---|---|---|---|---|---|---|---|
| FIM1_3619 | 422.597648 | 322.2807266 | 359.8464444 | -0.3029604 | 0.188340762 | 0.055563127 | 0.115370016 | ns | NaCl | KS.1 |
| FIM1_3620 | 1048.355206 | 922.466529 | 1503.26998 | -0.153606643 | 0.136889874 | 0.214943608 | 0.337972329 | ns | NaCl | KS.1 |
| FIM1_3621 | 1630.436031 | 1364.947425 | 1545.844025 | -0.216425312 | 0.138266345 | 0.082664758 | 0.159402004 | ns | NaCl | KS.1 |
| FIM1_3622 | 2258.917717 | 2379.723397 | 2433.684599 | 0.06944233 | 0.086632582 | 0.403066419 | 0.538179129 | ns | NaCl | KS.1 |
| FIM1_3623 | 106.7106254 | 67.97944221 | 572.7992675 | -0.327294327 | 0.328609823 | 0.131276211 | 0.22923803 | ns | NaCl | KS.1 |
| FIM1_3624 | 201.8021604 | 52.00499638 | 462.092434 | -1.519382986 | 0.417579851 | 6.71685038732 | 8.35408266923 | Km<KS.1 | NaCl | KS.1 |
| FIM1_3625 | 22095.23541 | 30723.67499 | 18717.45703 | 0.462064456 | 0.06474628 | 2.24782976548 | 1.64455192401 | ns | NaCl | KS.1 |
| FIM1_3626 | 203.8829864 | 105.3318331 | 201.9656237 | -0.852681114 | 0.180597208 | 1.50203671318 | 2.32792294333 | ns | NaCl | KS.1 |
| FIM1_3627 | 709.2264885 | 705.8340819 | 1302.001326 | -0.004065568 | 0.184292034 | 0.979180869 | 0.98972467 | ns | NaCl | KS.1 |
| FIM1_3628 | 10763.36143 | 15318.23373 | 11043.21588 | 0.294499419 | 0.280990846 | 0.145449012 | 0.248492045 | ns | NaCl | KS.1 |
| FIM1_3629 | 4137.544127 | 4394.292931 | 4731.292686 | 0.062638075 | 0.18294662 | 0.686078957 | 0.787936089 | ns | NaCl | KS.1 |
| FIM1_3630 | 171.5287569 | 107.7390416 | 197.6624048 | -0.502105475 | 0.239781871 | 0.008075779 | 0.024217602 | ns | NaCl | KS.1 |
| FIM1_3631 | 524.8101413 | 514.7776343 | 557.9884718 | -0.026273644 | 0.126941381 | 0.821640692 | 0.889782857 | ns | NaCl | KS.1 |
| FIM1_3632 | 8240.251877 | 11609.07328 | 8567.738859 | 0.459069961 | 0.107102263 | 4.63194040246 | 4.47900167307 | ns | NaCl | KS.1 |
| FIM1_3633 | 1875.030352 | 415.1754848 | 2506.69561 | -1.925116769 | 0.345110025 | 1.24002355718 | 5.84965915046 | Km<KS.1 | NaCl | KS.1 |
| FIM1_3634 | 5552.533378 | 3055.003991 | 3704.855705 | -0.644666951 | 0.264644124 | 0.001785124 | 0.006970229 | ns | NaCl | KS.1 |
| FIM1_3635 | 8775.417408 | 5044.033656 | 4861.765801 | -0.586939236 | 0.261738803 | 0.00382426 | 0.013094076 | ns | NaCl | KS.1 |
| FIM1_3636 | 3515.435607 | 1727.436346 | 2761.731935 | -0.756770885 | 0.2926491 | 0.000722374 | 0.003281143 | ns | NaCl | KS.1 |
| FIM1_3637 | 5241.383354 | 2265.723744 | 3180.97478 | -0.935419075 | 0.296355661 | 5.95446115729 | 0.00040414 | ns | NaCl | KS.1 |
| FIM1_3638 | 11925.12993 | 6123.383951 | 7351.407994 | -0.695930778 | 0.291188219 | 0.001604142 | 0.006369199 | ns | NaCl | KS.1 |
| FIM1_3639 | 590.6660016 | 249.7023265 | 401.9788217 | -0.868096983 | 0.34488138 | 0.000515807 | 0.002469817 | ns | NaCl | KS.1 |
| FIM1_3640 | 1667.930692 | 776.104012 | 1134.694664 | -0.813233787 | 0.304589649 | 0.000443801 | 0.002181731 | ns | NaCl | KS.1 |
| FIM1_3641 | 1244.736879 | 813.2498606 | 956.6364595 | -0.420499556 | 0.254427005 | 0.030791308 | 0.07121653 | ns | NaCl | KS.1 |
| FIM1_3642 | 1918.671492 | 1370.879661 | 2394.135994 | -0.321141657 | 0.244421163 | 0.087904133 | 0.167365195 | ns | NaCl | KS.1 |
| FIM1_3643 | 1772.442817 | 788.8203911 | 1267.188752 | -0.85514291 | 0.315996799 | 0.000331034 | 0.001704862 | ns | NaCl | KS.1 |
| FIM1_3644 | 4807.293889 | 1861.512696 | 2834.249379 | -1.136290224 | 0.273804127 | 5.99493133979 | 7.60836311619 | Km<KS.1 | NaCl | KS.1 |
| FIM1_3645 | 4229.795049 | 1963.223141 | 3118.831835 | -0.860751561 | 0.280878037 | 0.000110997 | 0.000682481 | ns | NaCl | KS.1 |
| FIM1_3646 | 2119.809021 | 1176.190693 | 1263.162388 | -0.608572378 | 0.278111153 | 0.003969853 | 0.013475071 | ns | NaCl | KS.1 |
| FIM1_3647 | 38470.36758 | 36901.33875 | 25119.05881 | -0.041935461 | 0.19218487 | 0.793702356 | 0.868221025 | ns | NaCl | KS.1 |
| FIM1_3648 | 3255.980168 | 4221.164044 | 2447.029702 | 0.166843635 | 0.318733083 | 0.422203535 | 0.558336679 | ns | NaCl | KS.1 |
| FIM1_3649 | 2459.886641 | 1543.395994 | 1619.634937 | -0.448627532 | 0.270787788 | 0.026842006 | 0.063680964 | ns | NaCl | KS.1 |
| FIM1_3650 | 2976.056756 | 1307.537135 | 1797.310892 | -0.915173515 | 0.294589582 | 7.64417768008 | 0.000500658 | ns | NaCl | KS.1 |
| FIM1_3651 | 1893.236594 | 1213.284082 | 1422.069128 | -0.424390091 | 0.268750798 | 0.034825935 | 0.078789916 | ns | NaCl | KS.1 |
| FIM1_3652 | 415.1529106 | 186.1730155 | 400.6113962 | -0.675546463 | 0.379389292 | 0.006753607 | 0.020893722 | ns | NaCl | KS.1 |
| FIM1_3653 | 8850.639971 | 6948.595317 | 5782.597739 | -0.219519097 | 0.241172959 | 0.236375505 | 0.363289508 | ns | NaCl | KS.1 |
| FIM1_3654 | 11998.52272 | 7677.287299 | 7828.351318 | -0.481271247 | 0.232278932 | 0.009347363 | 0.027131349 | ns | NaCl | KS.1 |
| FIM1_3655 | 6079.527047 | 3355.805676 | 4492.996544 | -0.651542387 | 0.257710686 | 0.00134519 | 0.005454215 | ns | NaCl | KS.1 |
| FIM1_3656 | 8652.072613 | 5695.074401 | 5621.544174 | -0.419901067 | 0.24842378 | 0.028654481 | 0.067022118 | ns | NaCl | KS.1 |
| FIM1_3657 | 1037.724893 | 462.4396614 | 773.3438998 | -0.848459182 | 0.317581791 | 0.000375739 | 0.00190163 | ns | NaCl | KS.1 |
| FIM1_3658 | 3610.310354 | 3082.871505 | 2222.721408 | -0.120792215 | 0.271873151 | 0.537037753 | 0.664287126 | ns | NaCl | KS.1 |
| FIM1_3659 | 2750.369107 | 1311.898805 | 1990.138433 | -0.791662217 | 0.296719726 | 0.000491783 | 0.002375362 | ns | NaCl | KS.1 |
| FIM1_3660 | 2705.4772 | 1405.420317 | 1551.995642 | -0.664590143 | 0.298515482 | 0.002794856 | 0.010090282 | ns | NaCl | KS.1 |
| FIM1_3661 | 4245.814404 | 1616.655241 | 2798.114036 | -1.101985796 | 0.307664113 | 6.45688372648 | 5.81938343102 | Km<KS.1 | NaCl | KS.1 |
| FIM1_3662 | 1450.099158 | 608.6574851 | 1358.714328 | -0.969788391 | 0.301338509 | 4.17035966940 | 0.000296817 | ns | NaCl | KS.1 |
| FIM1_3663 | 6090.781475 | 5212.446774 | 4056.103488 | -0.11400293 | 0.281385291 | 0.56550889 | 0.687845561 | ns | NaCl | KS.1 |

| | | | | | | | | | | |
|---|---|---|---|---|---|---|---|---|---|---|
| FIM1_3664 | 2683.530827 | 1806.168131 | 1737.776234 | -0.357898212 | 0.2713006 | 0.074070633 | 0.145652727 | ns | NaCl | KS.1 |
| FIM1_3665 | 1618.204669 | 610.9986195 | 1234.791245 | -1.028075806 | 0.350440119 | 7.6089372717 | 0.000499399 | Km<KS.1 | NaCl | KS.1 |
| FIM1_3666 | 2360.304065 | 1268.378561 | 1461.323656 | -0.605172003 | 0.303231856 | 0.006316268 | 0.01982445 | ns | NaCl | KS.1 |
| FIM1_3667 | 3106.615239 | 1573.636744 | 2133.783649 | -0.741847299 | 0.277185042 | 0.000598482 | 0.002803624 | ns | NaCl | KS.1 |
| FIM1_3668 | 2647.138451 | 1187.160562 | 1477.657812 | -0.883618828 | 0.295512727 | 0.0001266 | 0.000759901 | ns | NaCl | KS.1 |
| FIM1_3669 | 1770.287183 | 595.336725 | 1245.2853 | -1.226639379 | 0.342691116 | 3.4965067531 | 3.5427945207 | Km<KS.1 | NaCl | KS.1 |
| FIM1_3670 | 1202.610668 | 409.9475292 | 785.9254473 | -1.23192983 | 0.329242981 | 1.8742321601 | 2.0182478348 | Km<KS.1 | NaCl | KS.1 |
| FIM1_3671 | 984.9906792 | 317.6813536 | 649.025862 | -1.216455302 | 0.378799029 | 1.2827459228 | 0.000106688 | Km<KS.1 | NaCl | KS.1 |
| FIM1_3672 | 3231.26447 | 1399.559063 | 2288.424868 | -0.937862655 | 0.293406742 | 5.1748523994 | 0.000358564 | ns | NaCl | KS.1 |
| FIM1_3673 | 2719.495763 | 2874.675594 | 2448.216564 | 0.042494686 | 0.265536804 | 0.825703909 | 0.893016728 | ns | NaCl | KS.1 |
| FIM1_3674 | 111.853868 | 43.57002136 | 71.71387351 | -0.909987962 | 0.376044105 | 0.000533053 | 0.002547493 | ns | NaCl | KS.1 |
| FIM1_3675 | 7555.117192 | 3999.997802 | 4505.666315 | -0.682519303 | 0.274601864 | 0.001321829 | 0.005399095 | ns | NaCl | KS.1 |
| FIM1_3676 | 315.6172711 | 104.1564614 | 187.0670644 | -1.263315268 | 0.341198203 | 1.8486065550 | 1.9993081763 | Km<KS.1 | NaCl | KS.1 |
| FIM1_3677 | 2016.614954 | 1160.940667 | 1500.927836 | -0.56034709 | 0.275617863 | 0.00716601 | 0.021912046 | ns | NaCl | KS.1 |
| FIM1_3678 | 6351.819009 | 4315.37407 | 2981.63063 | -0.25443785 | 0.33234383 | 0.234254893 | 0.360765397 | ns | NaCl | KS.1 |
| FIM1_3679 | 99602.09887 | 81232.77276 | 61159.05026 | -0.210745095 | 0.198377283 | 0.198536729 | 0.318106353 | ns | NaCl | KS.1 |
| FIM1_3680 | 223.9179398 | 214.7674405 | 178.89611 | -0.03360096 | 0.250160618 | 0.857867789 | 0.913895557 | ns | NaCl | KS.1 |
| FIM1_3681 | 1714.811476 | 802.8746159 | 1295.64304 | -0.855360455 | 0.276671826 | 0.000104211 | 0.000646449 | ns | NaCl | KS.1 |
| FIM1_3682 | 3746.877947 | 2906.557853 | 3493.016974 | -0.219072101 | 0.25564194 | 0.252383373 | 0.380833267 | ns | NaCl | KS.1 |
| FIM1_3683 | 2075.984733 | 1237.91259 | 1390.773785 | -0.541755009 | 0.257573895 | 0.006608746 | 0.020561922 | ns | NaCl | KS.1 |
| FIM1_3684 | 843.3937839 | 432.9157784 | 675.516524 | -0.698740991 | 0.289212025 | 0.001479911 | 0.005913702 | ns | NaCl | KS.1 |
| FIM1_3685 | 1380.510475 | 455.6672984 | 967.9350028 | -1.252126374 | 0.344322553 | 2.4982027368 | 2.6283278237 | Km<KS.1 | NaCl | KS.1 |
| FIM1_3686 | 1366.096922 | 1209.544206 | 1007.071036 | -0.080542951 | 0.300115883 | 0.690545301 | 0.791184588 | ns | NaCl | KS.1 |
| FIM1_3687 | 1144.561116 | 372.32488 | 709.5175891 | -1.356640743 | 0.301892152 | 4.2002553137 | 7.5437798505 | Km<KS.1 | NaCl | KS.1 |
| FIM1_3688 | 1883.263452 | 740.6395928 | 1342.19789 | -0.962961856 | 0.351230357 | 0.000179763 | 0.001022082 | ns | NaCl | KS.1 |
| FIM1_3689 | 2119.78015 | 1061.492217 | 1805.398193 | -0.762901766 | 0.274485432 | 0.000404836 | 0.002026218 | ns | NaCl | KS.1 |
| FIM1_3690 | 647.9667635 | 255.9259381 | 1028.481125 | -1.059712077 | 0.300065414 | 9.4619963075 | 8.1301263609 | Km<KS.1 | NaCl | KS.1 |
| FIM1_3691 | 4925.884614 | 2947.736495 | 3698.699931 | -0.542639373 | 0.253881735 | 0.0060597 | 0.019165293 | ns | NaCl | KS.1 |
| FIM1_3692 | 4453.240465 | 2270.822072 | 2354.507026 | -0.722239949 | 0.282457641 | 0.000912199 | 0.003980868 | ns | NaCl | KS.1 |
| FIM1_3693 | 2833.827458 | 1211.259245 | 2060.419245 | -0.955160758 | 0.294559803 | 4.1157598372 | 0.000293351 | ns | NaCl | KS.1 |
| FIM1_3694 | 1809.863749 | 925.587445 | 1198.114767 | -0.722407347 | 0.280213572 | 0.000859905 | 0.003789718 | ns | NaCl | KS.1 |
| FIM1_3695 | 4275.389478 | 2826.174015 | 5682.653673 | -0.389538522 | 0.265170517 | 0.049910234 | 0.105405281 | ns | NaCl | KS.1 |
| FIM1_3696 | 6793.97829 | 2990.273985 | 4593.752869 | -0.922968393 | 0.288919232 | 5.5652087593 | 0.000380838 | ns | NaCl | KS.1 |
| FIM1_3697 | 13774.68363 | 10788.84195 | 11978.24542 | -0.260403933 | 0.19466771 | 0.107714093 | 0.196281237 | ns | NaCl | KS.1 |
| FIM1_3698 | 4475.971983 | 3091.181874 | 3455.448409 | -0.351124746 | 0.253572987 | 0.068602432 | 0.136682858 | ns | NaCl | KS.1 |
| FIM1_3699 | 288.4980477 | 99.05536023 | 1113.884538 | -0.948648488 | 0.435988345 | 0.000747239 | 0.00337956 | ns | NaCl | KS.1 |
| FIM1_3700 | 634.1255229 | 177.2404845 | 866.2872135 | -1.463635603 | 0.376823571 | 3.3418812842 | 4.6182942747 | Km<KS.1 | NaCl | KS.1 |
| FIM1_3701 | 1875.480137 | 850.0650328 | 989.2838633 | -0.815920344 | 0.321989626 | 0.000633349 | 0.002944777 | ns | NaCl | KS.1 |
| FIM1_3702 | 645.8159856 | 187.1226317 | 650.5353628 | -1.472033657 | 0.33971359 | 4.7567156404 | 8.2743567522 | Km<KS.1 | NaCl | KS.1 |
| FIM1_3703 | 3146.787659 | 1696.085855 | 2849.464825 | -0.65640961 | 0.274783834 | 0.001921757 | 0.007399955 | ns | NaCl | KS.1 |
| FIM1_3704 | 1672.267974 | 728.3513446 | 1314.613492 | -0.886284319 | 0.316471118 | 0.00021956 | 0.001206971 | ns | NaCl | KS.1 |
| FIM1_3705 | 3834.06069 | 2081.956395 | 2843.130502 | -0.598059465 | 0.299399901 | 0.006508602 | 0.020275702 | ns | NaCl | KS.1 |
| FIM1_3706 | 8288.443658 | 4626.306842 | 5158.641043 | -0.601874911 | 0.277088722 | 0.004257184 | 0.014271893 | ns | NaCl | KS.1 |
| FIM1_3707 | 945.5645585 | 447.7386336 | 787.9552835 | -0.753662235 | 0.320114532 | 0.001343708 | 0.00545265 | ns | NaCl | KS.1 |
| FIM1_3708 | 8508.945705 | 4255.546394 | 5903.164291 | -0.844388096 | 0.222823929 | 9.0878243140 | 7.8629436456 | ns | NaCl | KS.1 |

| | | | | | | | | | | |
|---|---|---|---|---|---|---|---|---|---|---|
| FIM1_3709 | 4333.953845 | 2218.40686 | 2293.754854 | -0.693761847 | 0.294579975 | 0.001772202 | 0.006942288 | ns | NaCl | KS.1 |
| FIM1_3710 | 3488.958342 | 1502.555995 | 2787.408431 | -0.948548542 | 0.292218794 | 4.19385246983 | 0.000298063 | ns | NaCl | KS.1 |
| FIM1_3711 | 4432.416143 | 1915.724053 | 3620.269437 | -0.931397198 | 0.298700156 | 6.85025999310 | 0.000456225 | ns | NaCl | KS.1 |
| FIM1_3712 | 4547.168774 | 2549.912391 | 3898.949549 | -0.5784288 | 0.285868626 | 0.006736318 | 0.020867484 | ns | NaCl | KS.1 |
| FIM1_3713 | 3395.447235 | 2064.425327 | 3243.832749 | -0.504479096 | 0.26348759 | 0.012119823 | 0.033240516 | ns | NaCl | KS.1 |
| FIM1_3714 | 1504.054141 | 319.6605744 | 1059.112359 | -1.824473849 | 0.439561992 | 2.55825820150 | 5.07065121613 | Km<KS.1 | NaCl | KS.1 |
| FIM1_3715 | 458.3648543 | 236.0524656 | 396.9883538 | -0.582127674 | 0.33940013 | 0.012458419 | 0.033813766 | ns | NaCl | KS.1 |
| FIM1_3716 | 7236.78844 | 3313.110546 | 5631.398114 | -0.913896357 | 0.260927329 | 2.01937155873 | 0.000156975 | ns | NaCl | KS.1 |
| FIM1_3717 | 11650.02613 | 6572.059311 | 8897.841964 | -0.65212056 | 0.237377921 | 0.000710315 | 0.003239064 | ns | NaCl | KS.1 |
| FIM1_3718 | 2954.376568 | 1079.480783 | 1894.578112 | -1.107117281 | 0.336959493 | 1.73986988123 | 0.000139162 | Km<KS.1 | NaCl | KS.1 |
| FIM1_3719 | 1421.153339 | 502.2258153 | 1434.633716 | -1.074994714 | 0.374805285 | 7.28527007613 | 0.000481971 | Km<KS.1 | NaCl | KS.1 |
| FIM1_3720 | 2444.886539 | 980.4318382 | 1754.689686 | -0.998405235 | 0.320803363 | 5.11856345914 | 0.000356651 | ns | NaCl | KS.1 |
| FIM1_3722 | 216.5431243 | 85.32719091 | 144.6627501 | -0.97278267 | 0.349812787 | 0.000153838 | 0.00089459 | ns | NaCl | KS.1 |
| FIM1_3723 | 401.3568285 | 171.7008131 | 374.7185674 | -0.722859166 | 0.389484091 | 0.004581051 | 0.015093197 | ns | NaCl | KS.1 |
| FIM1_3725 | 69.94364168 | 101.6346326 | 110.7036884 | 0.21866605 | 0.348889 | 0.306813543 | 0.439590707 | ns | NaCl | KS.1 |
| FIM1_3727 | 1031.702824 | 526.1461626 | 551.350287 | -0.554280992 | 0.354921295 | 0.018691218 | 0.046940338 | ns | NaCl | KS.1 |
| FIM1_3731 | 150.0284504 | 213.224303 | 113.1955211 | 0.276329849 | 0.294313258 | 0.179853505 | 0.29510923 | ns | NaCl | KS.1 |
| FIM1_3732 | 419.2123138 | 330.9553623 | 285.7986033 | -0.196895045 | 0.264102277 | 0.310742659 | 0.443472383 | ns | NaCl | KS.1 |
| FIM1_3733 | 1015.396377 | 1020.761302 | 1175.971791 | 0.007178063 | 0.142172308 | 0.955273959 | 0.976672409 | ns | NaCl | KS.1 |
| FIM1_3734 | 640.0036004 | 664.9413843 | 827.4544815 | 0.043796432 | 0.136286042 | 0.723149979 | 0.816692332 | ns | NaCl | KS.1 |
| FIM1_3735 | 1198.449313 | 1463.450865 | 1708.862647 | 0.27235741 | 0.079262158 | 0.000352674 | 0.001803241 | ns | NaCl | KS.1 |
| FIM1_3736 | 1248.262813 | 1788.304196 | 1412.888503 | 0.441441055 | 0.160826891 | 0.001751487 | 0.006866547 | ns | NaCl | KS.1 |
| FIM1_3737 | 235.089931 | 88.02149869 | 793.9904804 | -1.149329395 | 0.295352187 | 1.60196678389 | 1.77643505327 | Km<KS.1 | NaCl | KS.1 |
| FIM1_3738 | 418.664481 | 238.0663132 | 451.205716 | -0.736017345 | 0.159011389 | 3.37937892558 | 4.65600080503 | ns | NaCl | KS.1 |
| FIM1_3739 | 1455.639876 | 677.8299897 | 1405.266701 | -0.933252135 | 0.23037291 | 2.25298768155 | 2.39683371226 | ns | NaCl | KS.1 |
| FIM1_3740 | 2259.888169 | 2894.569615 | 2403.374038 | 0.298745636 | 0.149592937 | 0.024420104 | 0.058861443 | ns | NaCl | KS.1 |
| FIM1_3741 | 1973.884681 | 1579.108671 | 3060.206979 | -0.285661167 | 0.120296542 | 0.009924241 | 0.028375343 | ns | NaCl | KS.1 |
| FIM1_3742 | 1576.541637 | 1623.49353 | 2198.695666 | 0.036214126 | 0.117231288 | 0.739454372 | 0.828928684 | ns | NaCl | KS.1 |
| FIM1_3743 | 2998.763083 | 3875.620418 | 3417.466024 | 0.345607219 | 0.092518028 | 8.21966145729 | 0.000528331 | ns | NaCl | KS.1 |
| FIM1_3744 | 1341.875468 | 1177.709214 | 1593.700646 | -0.162466563 | 0.123365148 | 0.152397552 | 0.258322938 | ns | NaCl | KS.1 |
| FIM1_3745 | 12657.20019 | 18304.4461 | 12179.15248 | 0.499405311 | 0.102691647 | 2.41270212337 | 3.47918639530 | ns | NaCl | KS.1 |
| FIM1_3746 | 1933.071344 | 1769.117723 | 1963.194845 | -0.110074393 | 0.119889758 | 0.32097013 | 0.454159954 | ns | NaCl | KS.1 |
| FIM1_3747 | 17830.4231 | 21600.55098 | 16972.88969 | 0.226859016 | 0.149657768 | 0.087877571 | 0.167365195 | ns | NaCl | KS.1 |
| FIM1_3748 | 7226.763529 | 12862.37976 | 7906.555294 | 0.764133627 | 0.147251432 | 1.82522176410 | 3.73682233597 | ns | NaCl | KS.1 |
| FIM1_3749 | 3218.052621 | 4798.968527 | 4416.571733 | 0.555512819 | 0.081921384 | 1.89834762094 | 1.11091695578 | ns | NaCl | KS.1 |
| FIM1_3750 | 1489.691872 | 2660.15349 | 1375.601999 | 0.555414557 | 0.298504423 | 0.010718262 | 0.030228659 | ns | NaCl | KS.1 |
| FIM1_3751 | 1391.36317 | 1766.950304 | 1850.963617 | 0.320730032 | 0.089581755 | 0.000168371 | 0.00096726 | ns | NaCl | KS.1 |
| FIM1_3752 | 1618.371618 | 1782.666143 | 2341.916908 | 0.122957563 | 0.102814127 | 0.204619562 | 0.325952601 | ns | NaCl | KS.1 |
| FIM1_3753 | 1399.478274 | 3400.671273 | 2476.245096 | 1.195331795 | 0.163517689 | 4.44229021435 | 4.25007573392 | Km>KS.1 | NaCl | KS.1 |
| FIM1_3754 | 2508.786504 | 3637.187604 | 2359.349226 | 0.373959489 | 0.237424867 | 0.044057448 | 0.095381116 | ns | NaCl | KS.1 |
| FIM1_3755 | 276.0170407 | 314.2470508 | 213.6750868 | 0.134455317 | 0.192138625 | 0.401650746 | 0.537732095 | ns | NaCl | KS.1 |
| FIM1_3756 | 27282.95428 | 36464.43918 | 22678.61819 | 0.373416045 | 0.124206329 | 0.001032205 | 0.004396591 | ns | NaCl | KS.1 |
| FIM1_3757 | 644.4353207 | 596.4841434 | 660.2333306 | -0.097733808 | 0.120850533 | 0.381568476 | 0.516947294 | ns | NaCl | KS.1 |
| FIM1_3758 | 1746.829654 | 2177.592204 | 1542.398213 | 0.246440652 | 0.17377995 | 0.098421082 | 0.182363084 | ns | NaCl | KS.1 |
| FIM1_3759 | 11874.18776 | 12849.51872 | 9864.711821 | 0.075828537 | 0.207550721 | 0.653091125 | 0.760207849 | ns | NaCl | KS.1 |

| | | | | | | | | | | |
|---|---|---|---|---|---|---|---|---|---|---|
| FIM1_3760 | 916.3794167 | 588.9247864 | 619.9495454 | -0.590284306 | 0.123782283 | 2.70920571604 | 3.85094241066 | ns | NaCl | KS.1 |
| FIM1_3761 | 677.0918904 | 719.4462372 | 735.6103109 | 0.08127033 | 0.093613206 | 0.361925485 | 0.496985727 | ns | NaCl | KS.1 |
| FIM1_3762 | 1908.161771 | 2091.108596 | 1837.459971 | 0.107705868 | 0.143858046 | 0.404071036 | 0.539231064 | ns | NaCl | KS.1 |
| FIM1_3763 | 2865.275144 | 3585.58468 | 3123.577917 | 0.241213833 | 0.188388943 | 0.12721593 | 0.223639312 | ns | NaCl | KS.1 |
| FIM1_3764 | 395.1624415 | 378.0272975 | 462.6643805 | -0.057150339 | 0.113936363 | 0.590530312 | 0.711525382 | ns | NaCl | KS.1 |
| FIM1_3765 | 8197.427341 | 9064.597425 | 9241.215501 | 0.118763502 | 0.141830181 | 0.351948961 | 0.486779562 | ns | NaCl | KS.1 |
| FIM1_3766 | 103.2783037 | 65.93166022 | 98.51668961 | -0.48438375 | 0.217448895 | 0.006255834 | 0.019681154 | ns | NaCl | KS.1 |
| FIM1_3767 | 2266.954142 | 1929.973574 | 2270.164745 | -0.194887888 | 0.134301459 | 0.109634366 | 0.199207806 | ns | NaCl | KS.1 |
| FIM1_3768 | 42439.18365 | 38069.94359 | 26488.78546 | -0.076568688 | 0.286396214 | 0.700656586 | 0.798938005 | ns | NaCl | KS.1 |
| FIM1_3769 | 1571.263391 | 2638.277301 | 2223.519758 | 0.721430799 | 0.090918686 | 2.15225634619 | 2.81775666376 | ns | NaCl | KS.1 |
| FIM1_3770 | 6301.797778 | 8007.698881 | 6474.110678 | 0.336229651 | 0.05845977 | 3.96417987998 | 9.71516990291 | ns | NaCl | KS.1 |
| FIM1_3771 | 1553.502123 | 1546.332306 | 1381.914983 | -0.005575171 | 0.145660398 | 0.965924205 | 0.983921564 | ns | NaCl | KS.1 |
| FIM1_3772 | 1875.492021 | 2145.140667 | 2235.125208 | 0.15521736 | 0.154795448 | 0.256160182 | 0.384433456 | ns | NaCl | KS.1 |
| FIM1_3773 | 7835.292237 | 11088.77732 | 9253.453081 | 0.407653274 | 0.178863771 | 0.007663094 | 0.023189717 | ns | NaCl | KS.1 |
| FIM1_3774 | 5282.740216 | 2839.901196 | 4738.193851 | -0.811038761 | 0.164380677 | 5.99698927218 | 1.01135666539 | ns | NaCl | KS.1 |
| FIM1_3775 | 195.5825266 | 181.6100781 | 237.9796987 | -0.077469686 | 0.144296375 | 0.549565965 | 0.675084118 | ns | NaCl | KS.1 |
| FIM1_3776 | 258.9241552 | 188.7184823 | 418.9431956 | -0.386614664 | 0.16262244 | 0.006462383 | 0.020156962 | ns | NaCl | KS.1 |
| FIM1_3777 | 7086.092257 | 9076.032007 | 9059.563685 | 0.187208662 | 0.284788459 | 0.351111489 | 0.486432654 | ns | NaCl | KS.1 |
| FIM1_3778 | 272.5740471 | 301.7643513 | 395.1968135 | 0.092553844 | 0.223130961 | 0.599860299 | 0.718958947 | ns | NaCl | KS.1 |
| FIM1_3779 | 1799.186404 | 1389.534619 | 1354.296373 | -0.319559973 | 0.140257653 | 0.011190836 | 0.031190145 | ns | NaCl | KS.1 |
| FIM1_3780 | 9257.064289 | 13812.69932 | 10131.22759 | 0.552079218 | 0.089574159 | 1.16071367853 | 4.18445692082 | ns | NaCl | KS.1 |
| FIM1_3782 | 2567.678971 | 2305.839854 | 3009.537331 | -0.132848918 | 0.12281497 | 0.240432403 | 0.367255513 | ns | NaCl | KS.1 |
| FIM1_3783 | 980.9963884 | 909.8963414 | 1425.301828 | -0.09185067 | 0.108655954 | 0.367144399 | 0.502487864 | ns | NaCl | KS.1 |
| FIM1_3784 | 2855.100096 | 3026.859549 | 3653.060382 | 0.073765441 | 0.113441521 | 0.485638305 | 0.619500146 | ns | NaCl | KS.1 |
| FIM1_3785 | 345.0473465 | 236.1102006 | 374.948203 | -0.418561097 | 0.202078103 | 0.012349761 | 0.033594783 | ns | NaCl | KS.1 |
| FIM1_3786 | 4636.697919 | 3971.338548 | 4454.690071 | -0.212043001 | 0.07217259 | 0.002419027 | 0.008941054 | ns | NaCl | KS.1 |
| FIM1_3787 | 90.69859812 | 76.84923571 | 84.05890081 | -0.171427316 | 0.193895166 | 0.287963826 | 0.418892485 | ns | NaCl | KS.1 |
| FIM1_3788 | 6113.053924 | 6860.129363 | 6370.812734 | 0.149193981 | 0.104700972 | 0.129706019 | 0.227133912 | ns | NaCl | KS.1 |
| FIM1_3789 | 676.6282274 | 626.1952409 | 690.6040231 | -0.095676919 | 0.124448735 | 0.403479057 | 0.53858554 | ns | NaCl | KS.1 |
| FIM1_3790 | 505.9109501 | 332.6597232 | 421.9346332 | -0.520965049 | 0.169914243 | 0.000439177 | 0.002163274 | ns | NaCl | KS.1 |
| FIM1_3791 | 3006.19499 | 3308.978379 | 3342.107276 | 0.129616511 | 0.074775312 | 0.073286448 | 0.144338906 | ns | NaCl | KS.1 |
| FIM1_3792 | 2692.69408 | 2732.580782 | 3126.315086 | 0.015008035 | 0.196403823 | 0.926515083 | 0.957669606 | ns | NaCl | KS.1 |
| FIM1_3793 | 6332.15425 | 6917.746769 | 6434.595813 | 0.109646584 | 0.121889999 | 0.329706219 | 0.463227461 | ns | NaCl | KS.1 |
| FIM1_3794 | 11923.07663 | 16053.0366 | 9539.717368 | 0.384342018 | 0.122974682 | 0.00065888 | 0.00303512 | ns | NaCl | KS.1 |
| FIM1_3795 | 420.3378568 | 811.8622324 | 881.351855 | 0.798198037 | 0.22323263 | 2.44579589025 | 0.000184792 | ns | NaCl | KS.1 |
| FIM1_3796 | 264.4209098 | 342.5669022 | 333.1994851 | 0.302988382 | 0.167087739 | 0.036355417 | 0.081729869 | ns | NaCl | KS.1 |
| FIM1_3797 | 4579.472312 | 4249.353987 | 4135.845504 | -0.091066563 | 0.12678344 | 0.433795768 | 0.570030097 | ns | NaCl | KS.1 |
| FIM1_3798 | 1984.244547 | 1700.310249 | 2050.789803 | -0.190095601 | 0.127169907 | 0.102453103 | 0.188569806 | ns | NaCl | KS.1 |
| FIM1_3799 | 13018.70685 | 11643.73397 | 14279.63957 | -0.126479267 | 0.157319255 | 0.361098842 | 0.49626153 | ns | NaCl | KS.1 |
| FIM1_3801 | 2331.426175 | 2424.612662 | 2398.246948 | 0.042399779 | 0.173581663 | 0.776346907 | 0.854749778 | ns | NaCl | KS.1 |
| FIM1_3802 | 1738.018357 | 2092.92297 | 2210.137681 | 0.161417021 | 0.245016886 | 0.38659356 | 0.521682192 | ns | NaCl | KS.1 |
| FIM1_3803 | 381.4821461 | 297.8285976 | 313.849329 | -0.279570755 | 0.184507265 | 0.07285302 | 0.143598959 | ns | NaCl | KS.1 |
| FIM1_3804 | 181.5839669 | 84.12595187 | 257.7622274 | -0.997492984 | 0.192595354 | 8.76744625276 | 1.93000199590 | ns | NaCl | KS.1 |
| FIM1_3805 | 374.6097921 | 311.2827567 | 471.9418112 | -0.215648307 | 0.156424666 | 0.117243035 | 0.210344067 | ns | NaCl | KS.1 |
| FIM1_3806 | 1081.033613 | 1221.183957 | 864.3943599 | 0.111370353 | 0.224454919 | 0.529456607 | 0.657689543 | ns | NaCl | KS.1 |

| | | | | | | | | | | |
|---|---|---|---|---|---|---|---|---|---|---|
| FIM1_3807 | 10946.60761 | 14564.16292 | 10336.56045 | 0.398162078 | 0.065549494 | 4.0988649810 | 1.2988441580 | ns | NaCl | KS.1 |
| FIM1_3808 | 551.3965099 | 363.6758395 | 598.9609995 | -0.546756488 | 0.123415644 | 1.6354742981 | 1.8081076963 | ns | NaCl | KS.1 |
| FIM1_3809 | 2340.829209 | 4614.226439 | 2060.132763 | 0.651069303 | 0.320635067 | 0.004687084 | 0.015422119 | ns | NaCl | KS.1 |
| FIM1_3810 | 2523.122709 | 2717.364164 | 1956.994185 | 0.088004828 | 0.136360478 | 0.476545759 | 0.610091393 | ns | NaCl | KS.1 |
| FIM1_3811 | 4757.640037 | 6030.470262 | 5184.452121 | 0.284364941 | 0.150716968 | 0.033245354 | 0.075904376 | ns | NaCl | KS.1 |
| FIM1_3812 | 17.18579749 | 13.61744918 | 22.63589428 | -0.145644937 | 0.32520698 | 0.483905993 | 0.618082751 | ns | NaCl | KS.1 |
| FIM1_3813 | 4845.632995 | 5961.129082 | 4199.466434 | 0.281486731 | 0.081219084 | 0.000305094 | 0.001592699 | ns | NaCl | KS.1 |
| FIM1_3814 | 15.1290746 | 19.80016661 | 26.24877177 | 0.170424927 | 0.319671157 | 0.412945065 | 0.548718403 | ns | NaCl | KS.1 |
| FIM1_3815 | 12730.63889 | 12829.1705 | 13481.12748 | 0.00728866 | 0.211030185 | 0.965891165 | 0.983921564 | ns | NaCl | KS.1 |
| FIM1_3816 | 2296.995843 | 3898.168608 | 2349.079688 | 0.624528544 | 0.212614757 | 0.000439117 | 0.002163274 | ns | NaCl | KS.1 |
| FIM1_3817 | 1654.121395 | 1365.683787 | 3144.887224 | -0.2401174 | 0.127185391 | 0.038912838 | 0.086463318 | ns | NaCl | KS.1 |
| FIM1_3818 | 79.36127522 | 72.53291866 | 352.0551801 | -0.07992586 | 0.245260744 | 0.667290243 | 0.773658578 | ns | NaCl | KS.1 |
| FIM1_3819 | 1827.278898 | 1388.945843 | 2295.003418 | -0.310444366 | 0.177995644 | 0.040920146 | 0.089873781 | ns | NaCl | KS.1 |
| FIM1_3820 | 1651.512167 | 3501.637084 | 1699.424937 | 0.902040972 | 0.240929653 | 8.6629908498 | 7.5346817269 | ns | NaCl | KS.1 |
| FIM1_3821 | 821.222693 | 1182.722832 | 934.0114174 | 0.433161679 | 0.177699202 | 0.00446379 | 0.014834572 | ns | NaCl | KS.1 |
| FIM1_3822 | 5160.896145 | 5099.928637 | 5347.315536 | -0.014608708 | 0.118130064 | 0.893982348 | 0.93672329 | ns | NaCl | KS.1 |
| FIM1_3823 | 80.87961372 | 207.8160733 | 134.5366798 | 1.230275071 | 0.201433307 | 1.3523928587 | 6.2297726594 | Km>KS.1 | NaCl | KS.1 |
| FIM1_3824 | 9734.253902 | 21591.74273 | 20601.80896 | 0.980490615 | 0.23128381 | 8.3971513613 | 1.0115212596 | ns | NaCl | KS.1 |
| FIM1_3825 | 222.6585853 | 145.4230346 | 253.0589104 | -0.369588481 | 0.291026978 | 0.075600746 | 0.14795976 | ns | NaCl | KS.1 |
| FIM1_3826 | 6845.579272 | 8398.673935 | 6236.632612 | 0.283139388 | 0.065985888 | 1.0174535134 | 8.6232218555 | ns | NaCl | KS.1 |
| FIM1_3827 | 705.6245595 | 488.9139578 | 477.8659757 | -0.469487779 | 0.148494003 | 0.000395733 | 0.001988657 | ns | NaCl | KS.1 |
| FIM1_3828 | 4220.192201 | 4269.966413 | 3076.068311 | 0.014969475 | 0.099951972 | 0.874297026 | 0.925059061 | ns | NaCl | KS.1 |
| FIM1_3829 | 3110.962694 | 2737.372978 | 2219.783123 | -0.164932379 | 0.107663672 | 0.102083275 | 0.188237322 | ns | NaCl | KS.1 |
| FIM1_3830 | 3544.019134 | 2584.758257 | 3249.216534 | -0.394763678 | 0.142764883 | 0.002026189 | 0.007706642 | ns | NaCl | KS.1 |
| FIM1_3831 | 545.5910186 | 216.5499666 | 740.3250707 | -1.216369811 | 0.193919349 | 5.1611194648 | 2.6470690038 | Km<KS.1 | NaCl | KS.1 |
| FIM1_3832 | 945.911182 | 879.9345804 | 785.8791234 | -0.087676231 | 0.132090574 | 0.466549845 | 0.60100608 | ns | NaCl | KS.1 |
| FIM1_3833 | 67.59714886 | 37.07683865 | 255.9778003 | -0.511838715 | 0.337445328 | 0.025181546 | 0.060316894 | ns | NaCl | KS.1 |
| FIM1_3834 | 3867.018141 | 3158.580716 | 3235.127828 | -0.267798099 | 0.095972133 | 0.003195384 | 0.011250555 | ns | NaCl | KS.1 |
| FIM1_3835 | 348.7263641 | 332.8794977 | 381.0551814 | -0.060024016 | 0.128574305 | 0.610297709 | 0.726718789 | ns | NaCl | KS.1 |
| FIM1_3836 | 5628.596244 | 6331.239644 | 4396.370425 | 0.138865575 | 0.142230585 | 0.277345916 | 0.407019449 | ns | NaCl | KS.1 |
| FIM1_3837 | 2766.521555 | 2471.064034 | 2397.157586 | -0.144646159 | 0.107826274 | 0.15237128 | 0.258322938 | ns | NaCl | KS.1 |
| FIM1_3838 | 1602.1513 | 2100.74923 | 1857.047638 | 0.351070988 | 0.118295068 | 0.001294285 | 0.005308382 | ns | NaCl | KS.1 |
| FIM1_3839 | 2292.250026 | 1412.136305 | 2382.466326 | -0.526418047 | 0.238616919 | 0.005488899 | 0.017640358 | ns | NaCl | KS.1 |
| FIM1_3840 | 705.937737 | 1534.643723 | 898.5308475 | 0.993934835 | 0.199578723 | 2.4829890172 | 4.9809961133 | ns | NaCl | KS.1 |
| FIM1_3841 | 1955.72331 | 2081.143119 | 2372.251608 | 0.065205355 | 0.181055885 | 0.67166869 | 0.776929074 | ns | NaCl | KS.1 |
| FIM1_3842 | 163.7751576 | 156.3727499 | 197.4601246 | -0.038926426 | 0.250163993 | 0.835654242 | 0.899468094 | ns | NaCl | KS.1 |
| FIM1_3843 | 13113.12437 | 18597.6836 | 11982.34472 | 0.48635726 | 0.074288142 | 1.2463595375 | 5.8496591504 | ns | NaCl | KS.1 |
| FIM1_3844 | 869.6803207 | 1631.433064 | 1232.990209 | 0.693346032 | 0.262992092 | 0.000827436 | 0.003668889 | ns | NaCl | KS.1 |
| FIM1_3845 | 1005.033554 | 1969.531782 | 1007.305287 | 0.768273983 | 0.255425524 | 0.000196184 | 0.001100353 | ns | NaCl | KS.1 |
| FIM1_3846 | 1083.942226 | 1293.370662 | 975.6377112 | 0.222933954 | 0.11915078 | 0.042764499 | 0.092905408 | ns | NaCl | KS.1 |
| FIM1_3847 | 33611.82392 | 52931.01964 | 31400.47508 | 0.630825544 | 0.087839727 | 8.6325266224 | 7.4046241287 | ns | NaCl | KS.1 |
| FIM1_3848 | 951.4071509 | 780.2205839 | 750.7664885 | -0.191616531 | 0.218601602 | 0.272456118 | 0.401739534 | ns | NaCl | KS.1 |
| FIM1_3849 | 32798.42887 | 46431.90577 | 30737.89021 | 0.436195549 | 0.147714453 | 0.000927064 | 0.004035123 | ns | NaCl | KS.1 |
| FIM1_3850 | 2933.093331 | 2303.7544 | 2505.267773 | -0.267147093 | 0.181204631 | 0.082403039 | 0.159020606 | ns | NaCl | KS.1 |
| FIM1_3851 | 897.9174886 | 886.6238852 | 1348.817359 | -0.017038083 | 0.134277217 | 0.889062294 | 0.933337184 | ns | NaCl | KS.1 |

| | | | | | | | | | | |
|---|---|---|---|---|---|---|---|---|---|---|
| FIM1_3852 | 1703.605296 | 1643.367986 | 1300.114399 | -0.042148666 | 0.147352871 | 0.74892655 | 0.836906915 | ns | NaCl | KS.1 |
| FIM1_3853 | 1295.818829 | 1640.828753 | 1715.996936 | 0.299747729 | 0.122059323 | 0.007523096 | 0.02283551 | ns | NaCl | KS.1 |
| FIM1_3854 | 5266.698909 | 5218.013491 | 5777.754768 | -0.010687556 | 0.140100501 | 0.932654181 | 0.961658314 | ns | NaCl | KS.1 |
| FIM1_3855 | 3805.46791 | 3252.612038 | 3691.321226 | -0.197024218 | 0.119868812 | 0.075048891 | 0.147204309 | ns | NaCl | KS.1 |
| FIM1_3856 | 1446.219779 | 665.922992 | 1475.589319 | -1.057875695 | 0.137045583 | 4.04890868749 | 4.91300505373 | Km<KS.1 | NaCl | KS.1 |
| FIM1_3857 | 6645.616472 | 9559.365298 | 6700.010341 | 0.501953943 | 0.083636409 | 3.91507919496 | 1.25661412870 | ns | NaCl | KS.1 |
| FIM1_3858 | 925.4857251 | 775.1265907 | 1106.416192 | -0.225451527 | 0.117671767 | 0.03834443 | 0.085659425 | ns | NaCl | KS.1 |
| FIM1_3859 | 33.50284419 | 19.78486775 | 26.75156027 | -0.447510364 | 0.314565392 | 0.040775251 | 0.089601093 | ns | NaCl | KS.1 |
| FIM1_3860 | 2187.491819 | 1974.196618 | 2591.085474 | -0.102366835 | 0.198376006 | 0.532050803 | 0.660041056 | ns | NaCl | KS.1 |
| FIM1_3861 | 603.341167 | 501.2966487 | 683.3851481 | -0.186514438 | 0.204119438 | 0.264177266 | 0.392502352 | ns | NaCl | KS.1 |
| FIM1_3862 | 2774.093847 | 3684.158951 | 4239.368737 | 0.238490198 | 0.26698464 | 0.223767827 | 0.348542561 | ns | NaCl | KS.1 |
| FIM1_3863 | 5096.266628 | 7107.78839 | 5834.400475 | 0.412812871 | 0.150711878 | 0.002024981 | 0.007706642 | ns | NaCl | KS.1 |
| FIM1_3864 | 24.34383196 | 16.65115417 | 239.0951616 | -0.216935297 | 0.323676878 | 0.303198273 | 0.435359001 | ns | NaCl | KS.1 |
| FIM1_3865 | 279.6140242 | 89.10492902 | 647.6237333 | -1.283787203 | 0.3520323 | 2.04756028537 | 2.19066933757 | Km<KS.1 | NaCl | KS.1 |
| FIM1_3866 | 1553.144985 | 1003.248019 | 1486.262765 | -0.560042322 | 0.153754189 | 4.56247459064 | 0.000321962 | ns | NaCl | KS.1 |
| FIM1_3867 | 1886.192393 | 1841.327631 | 1536.73574 | -0.029045537 | 0.139613835 | 0.817837732 | 0.887402992 | ns | NaCl | KS.1 |
| FIM1_3868 | 356.1535052 | 201.5152558 | 508.917387 | -0.693537604 | 0.20934719 | 9.40896939565 | 0.000593278 | ns | NaCl | KS.1 |
| FIM1_3869 | 17762.49846 | 21350.08723 | 15450.81622 | 0.194039339 | 0.189330571 | 0.221413726 | 0.346067636 | ns | NaCl | KS.1 |
| FIM1_3870 | 1873.136603 | 1041.445411 | 1812.154242 | -0.774837037 | 0.15454081 | 4.44091872390 | 7.86248065887 | ns | NaCl | KS.1 |
| FIM1_3871 | 2692.951019 | 4542.026813 | 2563.177529 | 0.594760936 | 0.228102267 | 0.001366973 | 0.005523177 | ns | NaCl | KS.1 |
| FIM1_3872 | 5547.165334 | 6342.32986 | 5529.200178 | 0.179502117 | 0.085385964 | 0.028225253 | 0.066236148 | ns | NaCl | KS.1 |
| FIM1_3873 | 1089.201234 | 853.3058578 | 1453.231919 | -0.286901425 | 0.160990989 | 0.041404879 | 0.090544735 | ns | NaCl | KS.1 |
| FIM1_3874 | 393.2734271 | 230.6150319 | 510.1569601 | -0.698993258 | 0.147858787 | 2.34037668647 | 3.39456968372 | ns | NaCl | KS.1 |
| FIM1_3875 | 254.0122833 | 138.1497983 | 283.7631909 | -0.801096566 | 0.163823878 | 7.71175406561 | 1.26203869988 | ns | NaCl | KS.1 |
| FIM1_3876 | 111.0370449 | 27.20498785 | 99.06440972 | -1.828435518 | 0.357818147 | 2.22604948595 | 7.69069180042 | Km<KS.1 | NaCl | KS.1 |
| FIM1_3877 | 64.31805694 | 21.8373802 | 174.706943 | -1.239494919 | 0.322809419 | 1.24035597156 | 1.41856803644 | Km<KS.1 | NaCl | KS.1 |
| FIM1_3879 | 12021.08755 | 10745.92491 | 20269.80526 | -0.112373675 | 0.197450341 | 0.49139093 | 0.623958621 | ns | NaCl | KS.1 |
| FIM1_3880 | 3244.988561 | 2821.841835 | 3181.816457 | -0.152971279 | 0.171571773 | 0.300801782 | 0.432636272 | ns | NaCl | KS.1 |
| FIM1_3881 | 5605.110368 | 6368.59489 | 6875.92404 | 0.177345275 | 0.061107272 | 0.002981318 | 0.010647565 | ns | NaCl | KS.1 |
| FIM1_3882 | 31.00531221 | 27.12334279 | 60.88332332 | -0.079026959 | 0.283479353 | 0.690478777 | 0.791184588 | ns | NaCl | KS.1 |
| FIM1_3883 | 293.7746776 | 175.3961996 | 422.477108 | -0.648743841 | 0.174645028 | 2.45523297308 | 0.000184792 | ns | NaCl | KS.1 |
| FIM1_3884 | 428.018654 | 281.648717 | 508.2338864 | -0.480836341 | 0.203016802 | 0.00437855 | 0.014600059 | ns | NaCl | KS.1 |
| FIM1_3885 | 2428.909652 | 2134.515003 | 2432.3808 | -0.166781887 | 0.106933182 | 0.096272223 | 0.179450847 | ns | NaCl | KS.1 |
| FIM1_3886 | 3726.81352 | 3559.774337 | 3862.400914 | -0.056238612 | 0.120149809 | 0.613037411 | 0.729108563 | ns | NaCl | KS.1 |
| FIM1_3887 | 388.3828986 | 260.2124928 | 388.6433825 | -0.530774294 | 0.125273671 | 4.22569713926 | 4.16746646927 | ns | NaCl | KS.1 |
| FIM1_3888 | 3942.656453 | 3438.497071 | 3161.601912 | -0.169439438 | 0.126328188 | 0.143450113 | 0.245414138 | ns | NaCl | KS.1 |
| FIM1_3889 | 1123.034853 | 721.8820863 | 1106.781719 | -0.519969356 | 0.19794952 | 0.001775165 | 0.006948425 | ns | NaCl | KS.1 |
| FIM1_3890 | 2864.895365 | 2522.600163 | 2590.63797 | -0.151405145 | 0.138467718 | 0.225961499 | 0.350642064 | ns | NaCl | KS.1 |
| FIM1_3891 | 326.6875873 | 142.0717758 | 393.9541856 | -1.001487503 | 0.254348129 | 2.67977988605 | 2.80671682802 | Km<KS.1 | NaCl | KS.1 |
| FIM1_3892 | 1213.896002 | 679.4258067 | 1681.82906 | -0.572975706 | 0.291031772 | 0.007812952 | 0.023542967 | ns | NaCl | KS.1 |
| FIM1_3893 | 90.20862107 | 146.8156796 | 134.945167 | 0.581643427 | 0.200714748 | 0.000593619 | 0.002783664 | ns | NaCl | KS.1 |
| FIM1_3894 | 645.0966328 | 511.9034869 | 844.3097769 | -0.289453152 | 0.130427466 | 0.014700274 | 0.038633841 | ns | NaCl | KS.1 |
| FIM1_3895 | 162.2439618 | 154.9351869 | 166.2505611 | -0.050109067 | 0.184342573 | 0.747756462 | 0.835974921 | ns | NaCl | KS.1 |
| FIM1_3896 | 945.2087896 | 859.4074935 | 1224.149328 | -0.119518338 | 0.11731847 | 0.272163012 | 0.401426322 | ns | NaCl | KS.1 |
| FIM1_3897 | 514.647067 | 631.5346232 | 509.3109975 | 0.252553054 | 0.138736779 | 0.043275882 | 0.093811553 | ns | NaCl | KS.1 |

| | | | | | | | | | | |
|---|---|---|---|---|---|---|---|---|---|---|
| FIM1_3898 | 5565.680308 | 11481.97772 | 4862.051882 | 0.678797885 | 0.337374698 | 0.004253734 | 0.014269944 | ns | NaCl | KS.1 |
| FIM1_3899 | 671.821923 | 751.0750544 | 974.2576244 | 0.143194887 | 0.102221571 | 0.137412343 | 0.237700419 | ns | NaCl | KS.1 |
| FIM1_3900 | 369.4223372 | 449.4269891 | 381.9129481 | 0.233131552 | 0.14562265 | 0.072987215 | 0.143806493 | ns | NaCl | KS.1 |
| FIM1_3901 | 1172.637549 | 1657.687882 | 1554.936371 | 0.442442891 | 0.138415473 | 0.000395663 | 0.001988657 | ns | NaCl | KS.1 |
| FIM1_3902 | 490.8403756 | 571.3685382 | 1006.228109 | 0.163462399 | 0.181859334 | 0.289220032 | 0.419985307 | ns | NaCl | KS.1 |
| FIM1_3903 | 620.5449588 | 502.0559807 | 596.3936974 | -0.249187329 | 0.153502182 | 0.066056853 | 0.132459833 | ns | NaCl | KS.1 |
| FIM1_3904 | 8150.582092 | 11633.16939 | 7748.498376 | 0.469745763 | 0.118751119 | 1.8718915159 | 0.000147352 | ns | NaCl | KS.1 |
| FIM1_3905 | 5005.528654 | 6662.865562 | 6693.299249 | 0.330557042 | 0.173607369 | 0.026609219 | 0.063219132 | ns | NaCl | KS.1 |
| FIM1_3906 | 584.1831482 | 432.7520537 | 747.9196432 | -0.360054156 | 0.163179193 | 0.011365648 | 0.031571245 | ns | NaCl | KS.1 |
| FIM1_3907 | 1587.472101 | 1061.969744 | 1759.155748 | -0.539016585 | 0.11472191 | 4.6591324803 | 6.1320592829 | ns | NaCl | KS.1 |
| FIM1_3908 | 1388.174315 | 1525.688661 | 1524.870191 | 0.127308374 | 0.079838083 | 0.098014526 | 0.181880742 | ns | NaCl | KS.1 |
| FIM1_3909 | 1260.556274 | 2079.942004 | 1174.845268 | 0.516485553 | 0.259042726 | 0.00959222 | 0.027634552 | ns | NaCl | KS.1 |
| FIM1_3910 | 6210.681402 | 8366.183626 | 6264.664992 | 0.406336252 | 0.087663941 | 1.1579055622 | 1.3334676324 | ns | NaCl | KS.1 |
| FIM1_3911 | 133.4311622 | 74.92955879 | 182.5380654 | -0.632075562 | 0.252143162 | 0.001552918 | 0.006180616 | ns | NaCl | KS.1 |
| FIM1_3912 | 720.2941304 | 1721.390956 | 4257.191824 | 1.086977585 | 0.232583935 | 7.3675102453 | 1.2223057206 | Km>KS.1 | NaCl | KS.1 |
| FIM1_3913 | 260.5043322 | 506.6356158 | 592.3381723 | 0.845346659 | 0.194179841 | 8.5247549199 | 1.0244119740 | ns | NaCl | KS.1 |
| FIM1_3914 | 432.5771898 | 454.3394769 | 465.4991353 | 0.054185753 | 0.152167845 | 0.688290488 | 0.789542351 | ns | NaCl | KS.1 |
| FIM1_3915 | 3168.261044 | 3454.570047 | 2899.055429 | 0.111798333 | 0.100613685 | 0.239730965 | 0.366521681 | ns | NaCl | KS.1 |
| FIM1_3916 | 628.2551817 | 715.3748553 | 852.7608378 | 0.147843989 | 0.152180647 | 0.273093974 | 0.402560747 | ns | NaCl | KS.1 |
| FIM1_3917 | 1568.711428 | 1915.595805 | 1846.392987 | 0.239551373 | 0.144569828 | 0.063836218 | 0.129099668 | ns | NaCl | KS.1 |
| FIM1_3918 | 1345.199543 | 988.7155948 | 1266.487463 | -0.322366775 | 0.21234871 | 0.061122071 | 0.124982451 | ns | NaCl | KS.1 |
| FIM1_3919 | 190.3113208 | 115.5309707 | 331.3315058 | -0.615981512 | 0.201027189 | 0.000299522 | 0.001570204 | ns | NaCl | KS.1 |
| FIM1_3920 | 182.9279237 | 170.2529537 | 231.2274604 | -0.081700789 | 0.162493084 | 0.565189478 | 0.687654109 | ns | NaCl | KS.1 |
| FIM1_3921 | 2073.179879 | 2630.221095 | 2798.983204 | 0.302184885 | 0.124519738 | 0.008069728 | 0.02421405 | ns | NaCl | KS.1 |
| FIM1_3922 | 571.3967794 | 392.3169966 | 500.5854168 | -0.464380179 | 0.166364525 | 0.001370259 | 0.005528825 | ns | NaCl | KS.1 |
| FIM1_3923 | 4359.020846 | 3932.085976 | 3006.232563 | -0.129266829 | 0.118021842 | 0.237291698 | 0.364022879 | ns | NaCl | KS.1 |
| FIM1_3924 | 781.3907292 | 1831.220736 | 705.9763253 | 0.754076914 | 0.381751497 | 0.003108948 | 0.010991229 | ns | NaCl | KS.1 |
| FIM1_3925 | 102.0043846 | 43.3967069 | 212.0267322 | -0.954493328 | 0.297684503 | 4.6479052700 | 0.000327063 | ns | NaCl | KS.1 |
| FIM1_3926 | 1617.812565 | 1410.207353 | 1880.232738 | -0.165307241 | 0.132200425 | 0.169466344 | 0.280821636 | ns | NaCl | KS.1 |
| FIM1_3927 | 331.9829065 | 128.6507351 | 370.1716768 | -1.211846555 | 0.214898505 | 2.4776922395 | 8.5010475113 | Km<KS.1 | NaCl | KS.1 |
| FIM1_3928 | 2139.734555 | 2690.789138 | 2189.577524 | 0.275861921 | 0.146798926 | 0.03491806 | 0.078926555 | ns | NaCl | KS.1 |
| FIM1_3929 | 1431.341771 | 1978.846079 | 1378.074944 | 0.37532952 | 0.178965533 | 0.013987573 | 0.037153325 | ns | NaCl | KS.1 |
| FIM1_3930 | 975.193068 | 1103.312545 | 1070.974216 | 0.159763044 | 0.099003959 | 0.087989329 | 0.167365195 | ns | NaCl | KS.1 |
| FIM1_3932 | 2587.711538 | 1743.306888 | 2228.120119 | -0.532109635 | 0.112165055 | 3.8250649906 | 5.1852039042 | ns | NaCl | KS.1 |
| FIM1_3933 | 531.7506294 | 485.7382426 | 529.5501752 | -0.111447953 | 0.132717925 | 0.356403532 | 0.491634979 | ns | NaCl | KS.1 |
| FIM1_3934 | 865.5098108 | 994.70178 | 1094.394944 | 0.175903818 | 0.112154817 | 0.092449145 | 0.174151646 | ns | NaCl | KS.1 |
| FIM1_3935 | 1370.600519 | 1527.693478 | 1625.570104 | 0.129466672 | 0.140650631 | 0.306874075 | 0.439590707 | ns | NaCl | KS.1 |
| FIM1_3936 | 840.2433007 | 600.2634042 | 891.8872015 | -0.455122232 | 0.104382707 | 3.3676003642 | 3.4261373849 | ns | NaCl | KS.1 |
| FIM1_3937 | 18.71785945 | 11.05472631 | 34.38314948 | -0.335664042 | 0.384200711 | 0.135439338 | 0.235022917 | ns | NaCl | KS.1 |
| FIM1_3938 | 104.8299009 | 108.7014555 | 80.08171728 | 0.023580884 | 0.236017199 | 0.896910325 | 0.938735214 | ns | NaCl | KS.1 |
| FIM1_3939 | 421.4865777 | 537.3426606 | 588.5738225 | 0.235299404 | 0.226806927 | 0.18844533 | 0.305180832 | ns | NaCl | KS.1 |
| FIM1_3940 | 1337.03313 | 3029.234365 | 2416.895088 | 0.876939201 | 0.31112472 | 0.000217598 | 0.001200164 | ns | NaCl | KS.1 |
| FIM1_3942 | 656.3208434 | 466.2186308 | 686.7445039 | -0.448598559 | 0.124651728 | 8.7335865322 | 0.000559197 | ns | NaCl | KS.1 |
| FIM1_3943 | 3659.816724 | 5745.222883 | 4597.163711 | 0.626418677 | 0.087505554 | 1.0329349164 | 8.4243462448 | ns | NaCl | KS.1 |
| FIM1_3944 | 878.9422619 | 923.1731285 | 950.884262 | 0.060761749 | 0.10132443 | 0.525739485 | 0.654379268 | ns | NaCl | KS.1 |

| | | | | | | | | | | |
|---|---|---|---|---|---|---|---|---|---|---|
| FIM1_3945 | 1043.566751 | 1273.622235 | 915.1070056 | 0.20244672 | 0.202077168 | 0.222513464 | 0.347128183 | ns | NaCl | KS.1 |
| FIM1_3946 | 1592.654378 | 2280.740221 | 1684.428242 | 0.309435574 | 0.274950018 | 0.122816079 | 0.217828874 | ns | NaCl | KS.1 |
| FIM1_3947 | 1158.084026 | 1118.45366 | 1402.901195 | -0.04218189 | 0.126517118 | 0.716611981 | 0.812105833 | ns | NaCl | KS.1 |
| FIM1_3948 | 1087.597518 | 1158.9328 | 1624.501317 | 0.083967309 | 0.094160569 | 0.348752147 | 0.483433249 | ns | NaCl | KS.1 |
| FIM1_3949 | 1030.980362 | 1035.353547 | 1524.536322 | 0.005761849 | 0.097566255 | 0.950379088 | 0.973869405 | ns | NaCl | KS.1 |
| FIM1_3950 | 1370.894101 | 1171.612309 | 1387.801633 | -0.21020676 | 0.094563262 | 0.019166142 | 0.047771321 | ns | NaCl | KS.1 |
| FIM1_3951 | 2116.023279 | 1867.556468 | 2008.560255 | -0.169641536 | 0.071769484 | 0.014776352 | 0.038731481 | ns | NaCl | KS.1 |
| FIM1_3952 | 1289.745571 | 1575.001431 | 1720.543302 | 0.170134892 | 0.251290473 | 0.368139297 | 0.503295686 | ns | NaCl | KS.1 |
| FIM1_3953 | 1646.017023 | 1457.300516 | 1806.123057 | -0.154279455 | 0.116741273 | 0.154284339 | 0.260817937 | ns | NaCl | KS.1 |
| FIM1_3954 | 658.1723181 | 593.3454467 | 635.473512 | -0.118595435 | 0.139676672 | 0.346644819 | 0.481585583 | ns | NaCl | KS.1 |
| FIM1_3955 | 585.2355317 | 466.7933734 | 717.4408983 | -0.25043916 | 0.181502444 | 0.103855541 | 0.19051671 | ns | NaCl | KS.1 |
| FIM1_3956 | 3369.864681 | 6602.688154 | 3147.383048 | 0.788970259 | 0.241564319 | 7.67787249375 | 0.000501937 | ns | NaCl | KS.1 |
| FIM1_3957 | 2271.862761 | 2404.598649 | 2442.72559 | 0.078064251 | 0.06735465 | 0.234043932 | 0.360597263 | ns | NaCl | KS.1 |
| FIM1_3958 | 2295.175602 | 8683.244277 | 3999.83523 | 1.64993184 | 0.329392923 | 7.49445249542 | 2.19322948028 | Km>KS.1 | NaCl | KS.1 |
| FIM1_3959 | 765.2800033 | 564.3433017 | 783.6072442 | -0.393199019 | 0.12128062 | 0.000422011 | 0.002095316 | ns | NaCl | KS.1 |
| FIM1_3960 | 274.3696759 | 363.7185283 | 230.82045 | 0.273302799 | 0.22924535 | 0.129826026 | 0.227264067 | ns | NaCl | KS.1 |
| FIM1_3961 | 796.6451882 | 852.5819828 | 941.5538678 | 0.054865587 | 0.250452944 | 0.770175566 | 0.851527619 | ns | NaCl | KS.1 |
| FIM1_3962 | 2071.269291 | 1236.239113 | 1344.310423 | -0.688182284 | 0.133827732 | 2.90044983142 | 5.61468401219 | ns | NaCl | KS.1 |
| FIM1_3963 | 1782.006237 | 1471.846488 | 1670.37721 | -0.180663157 | 0.22281012 | 0.30636826 | 0.43924556 | ns | NaCl | KS.1 |
| FIM1_3964 | 1961.483707 | 2888.025849 | 2477.98014 | 0.346279608 | 0.271963577 | 0.083897212 | 0.161216156 | ns | NaCl | KS.1 |
| FIM1_3965 | 725.4396649 | 534.5490172 | 578.3806888 | -0.385709509 | 0.140182699 | 0.002200047 | 0.008266456 | ns | NaCl | KS.1 |
| FIM1_3966 | 150.7143435 | 226.8418954 | 404.0608576 | 0.261260702 | 0.342418866 | 0.225941403 | 0.350642064 | ns | NaCl | KS.1 |
| FIM1_3967 | 1703.924442 | 1736.291676 | 1412.758355 | 0.020813704 | 0.149949392 | 0.87614429 | 0.926619439 | ns | NaCl | KS.1 |
| FIM1_3968 | 1124.068623 | 1303.282429 | 1315.968476 | 0.185620234 | 0.121114658 | 0.096538807 | 0.179880362 | ns | NaCl | KS.1 |
| FIM1_3969 | 1961.581188 | 2263.783499 | 2221.040879 | 0.164054224 | 0.156751921 | 0.234571804 | 0.361075101 | ns | NaCl | KS.1 |
| FIM1_3970 | 920.8766719 | 1054.975526 | 906.8312738 | 0.138964242 | 0.192475775 | 0.386621556 | 0.521682192 | ns | NaCl | KS.1 |
| FIM1_3971 | 160.2963418 | 225.6488524 | 324.094424 | 0.303054566 | 0.265826273 | 0.123819753 | 0.218969986 | ns | NaCl | KS.1 |
| FIM1_3972 | 422.2129664 | 290.1607497 | 550.1518357 | -0.492058303 | 0.124352552 | 1.68566061337 | 0.000135479 | ns | NaCl | KS.1 |
| FIM1_3973 | 7831.402373 | 10108.19788 | 6210.686297 | 0.329221313 | 0.117732566 | 0.002452963 | 0.009053034 | ns | NaCl | KS.1 |
| FIM1_3974 | 1135.8926 | 1200.957054 | 1268.56398 | 0.068427472 | 0.132564881 | 0.571108831 | 0.692991813 | ns | NaCl | KS.1 |
| FIM1_3975 | 867.5757746 | 447.8467094 | 1007.307858 | -0.880944138 | 0.152041913 | 4.36260277588 | 1.35104408210 | ns | NaCl | KS.1 |
| FIM1_3976 | 5972.421508 | 7582.297202 | 8974.615497 | 0.281345409 | 0.158068483 | 0.042492139 | 0.092475236 | ns | NaCl | KS.1 |
| FIM1_3977 | 452.3859293 | 111.7916255 | 1321.053844 | -1.520499916 | 0.454175291 | 1.92660481194 | 2.06570235763 | Km<KS.1 | NaCl | KS.1 |
| FIM1_3978 | 1606.15174 | 1457.826049 | 1124.950898 | -0.111949596 | 0.15109694 | 0.404254501 | 0.539331226 | ns | NaCl | KS.1 |
| FIM1_3979 | 793.8354261 | 743.6554977 | 703.252064 | -0.084458282 | 0.10664765 | 0.399267046 | 0.53540527 | ns | NaCl | KS.1 |
| FIM1_3980 | 17680.92604 | 26179.91081 | 15468.15665 | 0.519152922 | 0.123377517 | 5.05107930644 | 4.76833388037 | ns | NaCl | KS.1 |
| FIM1_3981 | 196.2402175 | 117.383466 | 506.7927928 | -0.452717732 | 0.300415868 | 0.034612637 | 0.078423106 | ns | NaCl | KS.1 |
| FIM1_3982 | 3840.022439 | 2920.296008 | 3579.231815 | -0.356418233 | 0.115359615 | 0.000843549 | 0.003733679 | ns | NaCl | KS.1 |
| FIM1_3983 | 2269.889023 | 1798.887921 | 1834.252194 | -0.296111009 | 0.124195987 | 0.009284201 | 0.027011052 | ns | NaCl | KS.1 |
| FIM1_3984 | 342.9971053 | 252.3080296 | 288.2195062 | -0.330951256 | 0.205328384 | 0.049282324 | 0.104331728 | ns | NaCl | KS.1 |
| FIM1_3985 | 624.4986126 | 553.2389076 | 708.1430085 | -0.166369717 | 0.097457292 | 0.07150663 | 0.14128097 | ns | NaCl | KS.1 |
| FIM1_3986 | 2200.392954 | 1748.903316 | 1607.990702 | -0.251837833 | 0.183325403 | 0.104441211 | 0.191308919 | ns | NaCl | KS.1 |
| FIM1_3987 | 904.8402646 | 731.2256545 | 787.8348508 | -0.271073722 | 0.11983588 | 0.014121727 | 0.037429725 | ns | NaCl | KS.1 |
| FIM1_3988 | 984.2614854 | 672.3039126 | 1013.687429 | -0.50028966 | 0.131896529 | 3.20636423408 | 0.000236321 | ns | NaCl | KS.1 |
| FIM1_3989 | 8891.406638 | 12050.34154 | 10098.21849 | 0.406755556 | 0.102283846 | 2.29654952999 | 0.000175504 | ns | NaCl | KS.1 |

| | | | | | | | | | | |
|---|---|---|---|---|---|---|---|---|---|---|
| FIM1_3990 | 1453.866082 | 821.3067542 | 1567.976856 | -0.614648725 | 0.260152479 | 0.002477683 | 0.009130721 | ns | NaCl | KS.1 |
| FIM1_3991 | 2381.694877 | 2725.32039 | 1565.42951 | 0.125003324 | 0.221804842 | 0.477302738 | 0.610903298 | ns | NaCl | KS.1 |
| FIM1_3992 | 768.7200511 | 702.5373309 | 725.4346143 | -0.119245623 | 0.096551371 | 0.193090623 | 0.310681064 | ns | NaCl | KS.1 |
| FIM1_3993 | 8732.436784 | 11722.04039 | 8151.152732 | 0.416842452 | 0.049174863 | 6.6740919728 | 1.1067869188 | ns | NaCl | KS.1 |
| FIM1_3994 | 45.92951631 | 47.05161447 | 83.69121574 | 0.011297914 | 0.229248824 | 0.949693211 | 0.973834262 | ns | NaCl | KS.1 |
| FIM1_3995 | 277.7968509 | 159.036732 | 185.7070297 | -0.567450513 | 0.279323955 | 0.006959094 | 0.021403154 | ns | NaCl | KS.1 |
| FIM1_3996 | 1844.420015 | 2606.160255 | 1615.367025 | 0.293618375 | 0.275739446 | 0.142927356 | 0.24485661 | ns | NaCl | KS.1 |
| FIM1_3997 | 1776.201218 | 2058.412316 | 1901.564487 | 0.183687612 | 0.124612172 | 0.108470393 | 0.197449218 | ns | NaCl | KS.1 |
| FIM1_3998 | 853.7959847 | 920.1024245 | 1173.428708 | 0.085448754 | 0.150854873 | 0.524012043 | 0.653373412 | ns | NaCl | KS.1 |
| FIM1_3999 | 364.3108327 | 253.7966399 | 239.0508598 | -0.382366368 | 0.224777226 | 0.033103595 | 0.075650154 | ns | NaCl | KS.1 |
| FIM1_4000 | 1043.44611 | 1141.976752 | 866.1570562 | 0.103501786 | 0.146813957 | 0.430235405 | 0.566848819 | ns | NaCl | KS.1 |
| FIM1_4001 | 153.9160653 | 118.2909375 | 118.8632034 | -0.269960802 | 0.220083144 | 0.124682353 | 0.219962661 | ns | NaCl | KS.1 |
| FIM1_4002 | 5140.002014 | 6752.303391 | 4722.895656 | 0.372791125 | 0.082310009 | 2.2654248151 | 2.3979762670 | ns | NaCl | KS.1 |
| FIM1_4003 | 4805.628147 | 4599.908882 | 4266.578467 | -0.056433332 | 0.104430022 | 0.566210612 | 0.688055153 | ns | NaCl | KS.1 |
| FIM1_4004 | 1299.68073 | 1215.217807 | 1181.977227 | -0.089186934 | 0.09905948 | 0.341731273 | 0.476088794 | ns | NaCl | KS.1 |
| FIM1_4005 | 2427.230469 | 2207.684748 | 2922.143101 | -0.108852589 | 0.149885443 | 0.414440744 | 0.550411826 | ns | NaCl | KS.1 |
| FIM1_4006 | 2656.101417 | 2839.635534 | 2646.572787 | 0.090162331 | 0.08593444 | 0.273950464 | 0.403584115 | ns | NaCl | KS.1 |
| FIM1_4007 | 1831.539022 | 1597.724855 | 1713.896372 | -0.184634541 | 0.0821228 | 0.019291884 | 0.048036599 | ns | NaCl | KS.1 |
| FIM1_4008 | 382.7157503 | 843.1839404 | 533.3020394 | 0.971086915 | 0.232641425 | 1.1534663704 | 1.3314373997 | ns | NaCl | KS.1 |
| FIM1_4009 | 2924.441157 | 5230.034312 | 3342.353643 | 0.762761096 | 0.156705987 | 9.6783717064 | 1.5532225561 | ns | NaCl | KS.1 |
| FIM1_4010 | 1420.659465 | 1560.832338 | 1854.404496 | 0.124954369 | 0.08829076 | 0.13897097 | 0.239397707 | ns | NaCl | KS.1 |
| FIM1_4011 | 1523.827905 | 1625.342422 | 1968.013867 | 0.087905251 | 0.068653328 | 0.188157778 | 0.304913663 | ns | NaCl | KS.1 |
| FIM1_4012 | 2642.601603 | 2824.422473 | 2395.085632 | 0.084326875 | 0.107979203 | 0.405157772 | 0.54024656 | ns | NaCl | KS.1 |
| FIM1_4013 | 1599.339312 | 2224.803365 | 1792.394935 | 0.456374983 | 0.077592959 | 1.0059662794 | 2.8598184230 | ns | NaCl | KS.1 |
| FIM1_4014 | 313.0349202 | 300.9212882 | 366.1548634 | -0.04005498 | 0.174790751 | 0.789456078 | 0.864684342 | ns | NaCl | KS.1 |
| FIM1_4015 | 2143.112103 | 2244.502402 | 2846.782905 | 0.050735911 | 0.164607371 | 0.723638321 | 0.816751773 | ns | NaCl | KS.1 |
| FIM1_4016 | 2137.063806 | 2552.193748 | 2731.732684 | 0.217084584 | 0.136185677 | 0.077971467 | 0.151942049 | ns | NaCl | KS.1 |
| FIM1_4017 | 398.9533269 | 573.2577412 | 389.0880982 | 0.413343736 | 0.193840598 | 0.010875302 | 0.030584865 | ns | NaCl | KS.1 |
| FIM1_4018 | 4563.240129 | 9591.468331 | 4265.881815 | 0.711869888 | 0.335406463 | 0.002839944 | 0.010215995 | ns | NaCl | KS.1 |
| FIM1_4019 | 552.2123761 | 672.5913032 | 802.5732301 | 0.250985989 | 0.119437855 | 0.022739855 | 0.055320674 | ns | NaCl | KS.1 |
| FIM1_4020 | 5025.708317 | 5166.492746 | 3839.847189 | 0.036723395 | 0.08304927 | 0.645741716 | 0.754300314 | ns | NaCl | KS.1 |
| FIM1_4021 | 1074.519068 | 986.3257712 | 1069.033636 | -0.105072466 | 0.130854708 | 0.37903675 | 0.51409701 | ns | NaCl | KS.1 |
| FIM1_4022 | 34.75665894 | 37.91531446 | 31.43776702 | 0.059080464 | 0.285456721 | 0.766268689 | 0.84918336 | ns | NaCl | KS.1 |
| FIM1_4023 | 7299.505548 | 8127.24568 | 7102.690264 | 0.149301378 | 0.058115932 | 0.008728814 | 0.025802644 | ns | NaCl | KS.1 |
| FIM1_4024 | 2314.508224 | 2894.329247 | 2905.728058 | 0.177192368 | 0.271435161 | 0.367030413 | 0.50247009 | ns | NaCl | KS.1 |
| FIM1_4025 | 176.3564948 | 185.7128738 | 257.0080608 | 0.0503272 | 0.183218333 | 0.745642177 | 0.834361186 | ns | NaCl | KS.1 |
| FIM1_4026 | 118.8590503 | 163.9116573 | 370.4821554 | 0.242041284 | 0.295557336 | 0.238434938 | 0.365101205 | ns | NaCl | KS.1 |
| FIM1_4027 | 295.0544085 | 209.8194763 | 621.9376762 | -0.252255925 | 0.303557951 | 0.224729169 | 0.349522411 | ns | NaCl | KS.1 |
| FIM1_4028 | 1058.910411 | 1637.572792 | 1191.808455 | 0.598647899 | 0.096471861 | 7.5670801823 | 2.8519866596 | ns | NaCl | KS.1 |
| FIM1_4029 | 77.4685394 | 116.9560275 | 104.9791132 | 0.363439207 | 0.283241493 | 0.076327162 | 0.149230713 | ns | NaCl | KS.1 |
| FIM1_4030 | 5285.850253 | 13283.34294 | 5795.456378 | 1.056906408 | 0.296317371 | 8.4580689428 | 7.3952360264 | Km>KS.1 | NaCl | KS.1 |
| FIM1_4031 | 3890.916013 | 4363.492155 | 3294.908715 | 0.142399743 | 0.121465167 | 0.20391514 | 0.32494485 | ns | NaCl | KS.1 |
| FIM1_4033 | 965.7422062 | 742.5117023 | 824.2126454 | -0.327916815 | 0.135260475 | 0.007312415 | 0.022291217 | ns | NaCl | KS.1 |
| FIM1_4034 | 492.658964 | 1002.647281 | 554.5582895 | 0.936769624 | 0.167776819 | 1.2192556358 | 3.3887132896 | ns | NaCl | KS.1 |
| FIM1_4035 | 506.8703167 | 767.7430494 | 436.1044789 | 0.494591273 | 0.185853331 | 0.001787741 | 0.006970229 | ns | NaCl | KS.1 |

| | | | | | | | | | | |
|---|---|---|---|---|---|---|---|---|---|---|
| FIM1_4036 | 1715.791743 | 2176.476514 | 1895.044124 | 0.291898934 | 0.140078171 | 0.020395062 | 0.050480315 | ns | NaCl | KS.1 |
| FIM1_4037 | 1035.44447 | 1127.079063 | 1354.565124 | 0.1078371 | 0.11138923 | 0.300097665 | 0.432316514 | ns | NaCl | KS.1 |
| FIM1_4038 | 627.9730466 | 785.8294136 | 775.7880711 | 0.286488233 | 0.121079743 | 0.01012232 | 0.028858763 | ns | NaCl | KS.1 |
| FIM1_4039 | 1081.711579 | 1688.425926 | 1193.073608 | 0.571646045 | 0.152101174 | 2.750484205 | 0.000204815 | ns | NaCl | KS.1 |
| FIM1_4041 | 1073.56303 | 2165.204138 | 1151.064925 | 0.694731152 | 0.31640866 | 0.002609243 | 0.009558898 | ns | NaCl | KS.1 |
| FIM1_4042 | 1315.735414 | 1766.308924 | 1666.45966 | 0.384944817 | 0.114524593 | 0.000285952 | 0.001506535 | ns | NaCl | KS.1 |
| FIM1_4043 | 1269.81753 | 1536.620285 | 1573.467723 | 0.224123604 | 0.152069181 | 0.095955009 | 0.179128018 | ns | NaCl | KS.1 |
| FIM1_4044 | 200.4454167 | 347.6770491 | 280.2021216 | 0.485699765 | 0.312666793 | 0.027128239 | 0.06417641 | ns | NaCl | KS.1 |
| FIM1_4045 | 4160.722996 | 7055.569638 | 4361.899764 | 0.673737349 | 0.169435394 | 7.738619398 | 6.850468239 | ns | NaCl | KS.1 |
| FIM1_4046 | 1331.970442 | 1577.73691 | 1270.611571 | 0.190635617 | 0.164388718 | 0.182785758 | 0.298444091 | ns | NaCl | KS.1 |
| FIM1_4047 | 6864.449302 | 10586.43556 | 7888.412633 | 0.570652815 | 0.132868933 | 2.778218856 | 2.885519584 | ns | NaCl | KS.1 |
| FIM1_4048 | 572.954279 | 640.8364425 | 405.4900098 | 0.118565541 | 0.181894447 | 0.442239504 | 0.577161997 | ns | NaCl | KS.1 |
| FIM1_4049 | 1799.264636 | 3066.488652 | 1563.26928 | 0.448638991 | 0.317785511 | 0.041228136 | 0.09039664 | ns | NaCl | KS.1 |
| FIM1_4050 | 3319.668049 | 4056.953817 | 3856.464842 | 0.272558935 | 0.081215508 | 0.000472318 | 0.002292123 | ns | NaCl | KS.1 |
| FIM1_4051 | 8544.398526 | 5890.932448 | 8292.48385 | -0.494138943 | 0.116419251 | 4.774426109 | 4.572288789 | ns | NaCl | KS.1 |
| FIM1_4052 | 97.34340369 | 77.39811101 | 75.04869013 | -0.236352988 | 0.213717548 | 0.170197394 | 0.281586975 | ns | NaCl | KS.1 |
| FIM1_4053 | 773.8728269 | 334.7478552 | 749.4921098 | -0.769166232 | 0.368757173 | 0.00230202 | 0.008578689 | ns | NaCl | KS.1 |
| FIM1_4054 | 2279.305502 | 2335.818694 | 2907.120408 | 0.027514346 | 0.155086058 | 0.84095082 | 0.903028347 | ns | NaCl | KS.1 |
| FIM1_4055 | 1408.730314 | 1085.215646 | 1475.51348 | -0.274386903 | 0.200249103 | 0.096603429 | 0.179917915 | ns | NaCl | KS.1 |
| FIM1_4056 | 1195.017959 | 1062.557408 | 1623.584067 | -0.142591182 | 0.133831501 | 0.240984977 | 0.367831355 | ns | NaCl | KS.1 |
| FIM1_4057 | 1170.164051 | 1028.340836 | 1189.612241 | -0.165610061 | 0.108549191 | 0.103148297 | 0.189428858 | ns | NaCl | KS.1 |
| FIM1_4058 | 441.0626072 | 370.5488983 | 385.20337 | -0.199293894 | 0.156792881 | 0.148569984 | 0.253041996 | ns | NaCl | KS.1 |
| FIM1_4059 | 3326.110425 | 5662.494473 | 4790.05668 | 0.528150429 | 0.277728886 | 0.011174851 | 0.031174328 | ns | NaCl | KS.1 |
| FIM1_4060 | 3202.910461 | 4877.710861 | 2445.498945 | 0.426878707 | 0.245697804 | 0.025224031 | 0.060389584 | ns | NaCl | KS.1 |
| FIM1_4061 | 4329.166199 | 13983.66999 | 5649.129174 | 1.437974967 | 0.300486644 | 6.747368520 | 1.525825381 | Km>KS.1 | NaCl | KS.1 |
| FIM1_4062 | 1342.026715 | 3593.068216 | 1445.784866 | 1.067837963 | 0.339559144 | 3.310633001 | 0.000242568 | Km>KS.1 | NaCl | KS.1 |
| FIM1_4063 | 1853.101791 | 1908.737372 | 2330.080871 | 0.034849889 | 0.143057246 | 0.786419036 | 0.862339587 | ns | NaCl | KS.1 |
| FIM1_4064 | 5599.47719 | 8477.506635 | 5527.055317 | 0.491400421 | 0.188292779 | 0.00211193 | 0.00797413 | ns | NaCl | KS.1 |
| FIM1_4065 | 5553.244789 | 9821.570761 | 4374.277099 | 0.552335229 | 0.293288315 | 0.010372456 | 0.02945375 | ns | NaCl | KS.1 |
| FIM1_4066 | 2883.721289 | 3260.179644 | 3727.910755 | 0.156656553 | 0.112779802 | 0.136070448 | 0.23595346 | ns | NaCl | KS.1 |
| FIM1_4067 | 744.0319223 | 906.8157419 | 1068.184694 | 0.25407724 | 0.11065259 | 0.013741031 | 0.03665503 | ns | NaCl | KS.1 |
| FIM1_4068 | 409.1818764 | 419.1465667 | 406.8654862 | 0.025958863 | 0.166168236 | 0.857451254 | 0.913690395 | ns | NaCl | KS.1 |
| FIM1_4069 | 2052.958659 | 2432.365315 | 2644.230048 | 0.158608106 | 0.222961842 | 0.369092563 | 0.504044881 | ns | NaCl | KS.1 |
| FIM1_4070 | 2234.499833 | 2033.597042 | 2799.421974 | -0.085609533 | 0.222789342 | 0.627156753 | 0.739711913 | ns | NaCl | KS.1 |
| FIM1_4071 | 73.66821621 | 94.9784318 | 102.4320302 | 0.228420061 | 0.245819269 | 0.223131448 | 0.347769096 | ns | NaCl | KS.1 |
| FIM1_4072 | 1166.740184 | 1309.832769 | 1604.04108 | 0.145874813 | 0.109241375 | 0.153600045 | 0.259918443 | ns | NaCl | KS.1 |
| FIM1_4073 | 710.9392021 | 863.6007274 | 1025.178863 | 0.247553619 | 0.123413713 | 0.028930338 | 0.067572033 | ns | NaCl | KS.1 |
| FIM1_4074 | 820.8802604 | 1569.549186 | 1295.36021 | 0.635234062 | 0.308100449 | 0.004720776 | 0.015522711 | ns | NaCl | KS.1 |
| FIM1_4075 | 564.6295347 | 218.5588117 | 728.1756904 | -1.119055034 | 0.28277736 | 1.433137769 | 1.613090588 | Km<KS.1 | NaCl | KS.1 |
| FIM1_4076 | 24.22519541 | 12.66866842 | 18.61852671 | -0.438786802 | 0.384859783 | 0.060242666 | 0.123641609 | ns | NaCl | KS.1 |
| FIM1_4077 | 26863.8146 | 61582.52176 | 36444.03178 | 0.914926816 | 0.300286036 | 9.231422912 | 0.000583725 | ns | NaCl | KS.1 |
| FIM1_4078 | 79.49667713 | 182.361615 | 253.8581896 | 0.634363761 | 0.405245294 | 0.01181794 | 0.032554957 | ns | NaCl | KS.1 |
| FIM1_4080 | 1085.265427 | 1899.438374 | 3950.898547 | 0.541770801 | 0.291588322 | 0.011506055 | 0.031890042 | ns | NaCl | KS.1 |
| FIM1_4081 | 25.66768442 | 26.05577473 | 26.3944966 | 0.016527944 | 0.271661722 | 0.932370851 | 0.961658314 | ns | NaCl | KS.1 |
| FIM1_4082 | 45.67270798 | 73.88758597 | 53.98004836 | 0.377410663 | 0.322565749 | 0.08321919 | 0.160346813 | ns | NaCl | KS.1 |

| | | | | | | | | | | |
|---|---|---|---|---|---|---|---|---|---|---|
| FIM1_4083 | 709.9124675 | 750.0222411 | 621.9285562 | 0.065538447 | 0.1412898 | 0.606658041 | 0.724291758 | ns | NaCl | KS.1 |
| FIM1_4084 | 3911.07005 | 5107.318002 | 3451.86931 | 0.329656748 | 0.141615431 | 0.009422666 | 0.027252425 | ns | NaCl | KS.1 |
| FIM1_4085 | 9480.065652 | 14439.81465 | 9779.713209 | 0.56549027 | 0.115831028 | 1.67541387173 | 2.53349058111 | ns | NaCl | KS.1 |
| FIM1_4086 | 11533.38979 | 25447.57935 | 11459.64232 | 0.973033929 | 0.231162939 | 9.85349199367 | 1.15343818043 | ns | NaCl | KS.1 |
| FIM1_4087 | 439.8927248 | 330.0003284 | 342.0848434 | -0.256044887 | 0.255408154 | 0.182079848 | 0.297584509 | ns | NaCl | KS.1 |
| FIM1_4088 | 2009.089849 | 2463.165962 | 2166.353819 | 0.190849981 | 0.227223587 | 0.28578219 | 0.416818057 | ns | NaCl | KS.1 |
| FIM1_4089 | 249.8174584 | 137.2798893 | 243.3551311 | -0.752503851 | 0.183903543 | 3.64768427311 | 3.68846123144 | ns | NaCl | KS.1 |
| FIM1_4090 | 205.7948697 | 179.1564096 | 225.2102565 | -0.162687459 | 0.137426832 | 0.190383561 | 0.307718719 | ns | NaCl | KS.1 |
| FIM1_4091 | 3074.130432 | 6953.989002 | 2847.784754 | 0.809902364 | 0.340706864 | 0.00097011 | 0.004185859 | ns | NaCl | KS.1 |
| FIM1_4092 | 1284.057962 | 1064.021006 | 1310.075233 | -0.245119071 | 0.104265049 | 0.012263883 | 0.033486727 | ns | NaCl | KS.1 |
| FIM1_4093 | 1021.559873 | 1099.411303 | 1490.982194 | 0.061411873 | 0.245356869 | 0.74111248 | 0.829852484 | ns | NaCl | KS.1 |
| FIM1_4094 | 5419.677342 | 10879.06843 | 6504.902137 | 0.781410828 | 0.268024346 | 0.000248122 | 0.001340289 | ns | NaCl | KS.1 |
| FIM1_4095 | 1971.908034 | 2306.581366 | 1717.111691 | 0.181184569 | 0.154076296 | 0.183203408 | 0.298733842 | ns | NaCl | KS.1 |
| FIM1_4096 | 2197.858876 | 2168.272851 | 1803.497877 | -0.016182804 | 0.148889132 | 0.903008845 | 0.942440089 | ns | NaCl | KS.1 |
| FIM1_4097 | 3143.87562 | 3931.299249 | 1921.574245 | 0.165679256 | 0.285972288 | 0.408899319 | 0.544214583 | ns | NaCl | KS.1 |
| FIM1_4098 | 2868.114349 | 1582.188149 | 1806.243203 | -0.693955182 | 0.230912596 | 0.000267019 | 0.001422292 | ns | NaCl | KS.1 |
| FIM1_4099 | 555.2644034 | 284.6512608 | 365.0170713 | -0.831505467 | 0.207556608 | 3.98512775104 | 3.97314840910 | ns | NaCl | KS.1 |
| FIM1_4100 | 375.5158958 | 260.0489113 | 354.4049164 | -0.475635258 | 0.133879058 | 9.23399874118 | 0.000583725 | ns | NaCl | KS.1 |
| FIM1_4101 | 683.1458094 | 524.2917244 | 806.4622174 | -0.290303368 | 0.189070571 | 0.06727918 | 0.134349815 | ns | NaCl | KS.1 |
| FIM1_4102 | 1469.789745 | 1216.013866 | 2062.510701 | -0.23482949 | 0.129858795 | 0.047213904 | 0.100810803 | ns | NaCl | KS.1 |
| FIM1_4103 | 3272.049831 | 2293.074242 | 4905.46933 | -0.401244091 | 0.195862223 | 0.014068688 | 0.03732892 | ns | NaCl | KS.1 |
| FIM1_4104 | 1367.49685 | 1937.279953 | 1512.017908 | 0.352941696 | 0.230950827 | 0.052728487 | 0.110329294 | ns | NaCl | KS.1 |
| FIM1_4105 | 495.5491089 | 279.8908075 | 221.2919914 | -0.625792394 | 0.253733694 | 0.001784652 | 0.006970229 | ns | NaCl | KS.1 |
| FIM1_4106 | 11977.55784 | 17249.03431 | 8180.424686 | 0.369766714 | 0.233880477 | 0.044365481 | 0.095839456 | ns | NaCl | KS.1 |
| FIM1_4107 | 1187.720423 | 1408.110181 | 1193.939476 | 0.220683108 | 0.10639257 | 0.026863168 | 0.063700791 | ns | NaCl | KS.1 |
| FIM1_4108 | 3216.543848 | 3623.068126 | 2811.311319 | 0.159418674 | 0.082076675 | 0.043344077 | 0.09391846 | ns | NaCl | KS.1 |
| FIM1_4109 | 359.3982908 | 311.8702969 | 326.6748471 | -0.148678152 | 0.190825356 | 0.351440434 | 0.486752828 | ns | NaCl | KS.1 |
| FIM1_4110 | 1267.443807 | 1182.357004 | 929.8709923 | -0.082759439 | 0.133912909 | 0.496846344 | 0.628642839 | ns | NaCl | KS.1 |
| FIM1_4111 | 2249.758599 | 2813.682986 | 2452.532085 | 0.312635879 | 0.063509766 | 4.29488127719 | 5.70155963588 | ns | NaCl | KS.1 |
| FIM1_4112 | 2109.844206 | 2663.443071 | 2371.194331 | 0.238139457 | 0.207224351 | 0.158491379 | 0.266670644 | ns | NaCl | KS.1 |
| FIM1_4113 | 1967.631913 | 982.4026333 | 1778.14431 | -0.717596141 | 0.301496775 | 0.001490218 | 0.005950106 | ns | NaCl | KS.1 |
| FIM1_4114 | 2741.940197 | 4887.522968 | 2048.426331 | 0.447681793 | 0.343434457 | 0.048282068 | 0.102607129 | ns | NaCl | KS.1 |
| FIM1_4115 | 1150.682156 | 1122.639347 | 1095.168122 | -0.026215642 | 0.171274688 | 0.859219874 | 0.914748315 | ns | NaCl | KS.1 |
| FIM1_4116 | 314.4492924 | 381.7975125 | 254.7718731 | 0.150761623 | 0.271604246 | 0.441872343 | 0.577161997 | ns | NaCl | KS.1 |
| FIM1_4117 | 217.5568961 | 215.2231238 | 180.9317383 | -0.007949674 | 0.269665965 | 0.967339305 | 0.984758142 | ns | NaCl | KS.1 |
| FIM1_4118 | 1914.882169 | 1987.064249 | 2095.015767 | 0.034041343 | 0.219379524 | 0.845374956 | 0.905628855 | ns | NaCl | KS.1 |
| FIM1_4119 | 2087.44669 | 2617.211157 | 1782.112594 | 0.277850442 | 0.137214134 | 0.024783883 | 0.059565128 | ns | NaCl | KS.1 |
| FIM1_4120 | 3556.268498 | 5584.438948 | 2779.77814 | 0.362001232 | 0.309207982 | 0.089988093 | 0.170678902 | ns | NaCl | KS.1 |
| FIM1_4121 | 897.1626961 | 1452.814551 | 988.6484663 | 0.479508579 | 0.265701029 | 0.017242654 | 0.0439458 | ns | NaCl | KS.1 |
| FIM1_4122 | 585.8538705 | 429.9722008 | 644.6217503 | -0.378126469 | 0.15336856 | 0.00526078 | 0.016984024 | ns | NaCl | KS.1 |
| FIM1_4123 | 1014.774055 | 799.684146 | 1041.902857 | -0.282483331 | 0.15542555 | 0.039014835 | 0.086612586 | ns | NaCl | KS.1 |
| FIM1_4124 | 61.67568533 | 48.43521371 | 61.31069651 | -0.197824548 | 0.259757627 | 0.304512212 | 0.436962289 | ns | NaCl | KS.1 |
| FIM1_4125 | 7419.045741 | 6735.311011 | 7126.221652 | -0.105794213 | 0.168957871 | 0.469422069 | 0.603506673 | ns | NaCl | KS.1 |
| FIM1_4126 | 387.1213583 | 370.4192513 | 369.1405331 | -0.050305765 | 0.147273409 | 0.7023254 | 0.799981706 | ns | NaCl | KS.1 |
| FIM1_4127 | 44.75240713 | 29.01185857 | 56.92117253 | -0.372352499 | 0.307865282 | 0.08119971 | 0.156941941 | ns | NaCl | KS.1 |

| | | | | | | | | | | |
|---|---|---|---|---|---|---|---|---|---|---|
| FIM1_4128 | 1783.875641 | 1853.281633 | 1315.209878 | 0.045392221 | 0.132956695 | 0.707856034 | 0.803831036 | ns | NaCl | KS.1 |
| FIM1_4129 | 3168.202108 | 3452.215092 | 2444.613147 | 0.099609989 | 0.145806917 | 0.445302451 | 0.580546747 | ns | NaCl | KS.1 |
| FIM1_4130 | 1603.184336 | 1611.319215 | 2108.407292 | 0.004950653 | 0.172326728 | 0.973400846 | 0.987695127 | ns | NaCl | KS.1 |
| FIM1_4132 | 188.2206692 | 133.4167543 | 198.7253864 | -0.4074333 | 0.159465538 | 0.003585604 | 0.012430927 | ns | NaCl | KS.1 |
| FIM1_4133 | 799.374132 | 414.9240051 | 883.5089162 | -0.866297074 | 0.158832819 | 3.19933117318 | 7.99832793296 | ns | NaCl | KS.1 |
| FIM1_4134 | 2176.073729 | 1465.3117 | 1743.048971 | -0.526067824 | 0.120145777 | 2.26083262622 | 2.39797626709 | ns | NaCl | KS.1 |
| FIM1_4135 | 1030.321538 | 996.5443748 | 1187.428661 | -0.03969721 | 0.141517228 | 0.755518018 | 0.840684889 | ns | NaCl | KS.1 |
| FIM1_4136 | 2717.565792 | 2368.5766 | 2361.138264 | -0.177373644 | 0.103756064 | 0.069189706 | 0.137750403 | ns | NaCl | KS.1 |
| FIM1_4137 | 2434.117256 | 3510.481153 | 3197.217042 | 0.416751245 | 0.195090824 | 0.010613688 | 0.029967707 | ns | NaCl | KS.1 |
| FIM1_4138 | 1534.214086 | 1634.283435 | 1404.198415 | 0.082348148 | 0.100088229 | 0.384484208 | 0.519785037 | ns | NaCl | KS.1 |
| FIM1_4139 | 3069.001557 | 7341.275555 | 3528.830738 | 1.039440407 | 0.264214213 | 2.29118645213 | 2.42009609328 | Km>KS.1 | NaCl | KS.1 |
| FIM1_4140 | 1073.725879 | 1228.54482 | 1085.210523 | 0.131944239 | 0.205952006 | 0.43204255 | 0.568777901 | ns | NaCl | KS.1 |
| FIM1_4141 | 9214.932582 | 11994.09498 | 8667.610626 | 0.326995338 | 0.139081075 | 0.008953748 | 0.026280174 | ns | NaCl | KS.1 |
| FIM1_4142 | 829.6993527 | 760.3134684 | 850.6829663 | -0.111046837 | 0.11197821 | 0.288136981 | 0.418892485 | ns | NaCl | KS.1 |
| FIM1_4143 | 1007.887313 | 1007.647375 | 875.8348652 | -0.001573435 | 0.111607251 | 0.987969143 | 0.994566266 | ns | NaCl | KS.1 |
| FIM1_4144 | 430.2112626 | 299.2046931 | 395.2838946 | -0.418787899 | 0.192116044 | 0.009427422 | 0.027252425 | ns | NaCl | KS.1 |
| FIM1_4145 | 473.7555026 | 262.5719907 | 511.0689433 | -0.740033247 | 0.19303604 | 1.10871829638 | 9.34893817716 | ns | NaCl | KS.1 |
| FIM1_4146 | 2016.760947 | 3379.217913 | 1621.817164 | 0.379965748 | 0.338381136 | 0.086644256 | 0.165599376 | ns | NaCl | KS.1 |
| FIM1_4147 | 913.7507733 | 718.0115906 | 825.2580876 | -0.267735473 | 0.179494362 | 0.079624125 | 0.154496889 | ns | NaCl | KS.1 |
| FIM1_4148 | 2534.606092 | 2763.39942 | 3486.418128 | 0.075661131 | 0.233473353 | 0.676000459 | 0.779747207 | ns | NaCl | KS.1 |
| FIM1_4149 | 11900.507 | 13758.03392 | 12080.15282 | 0.187099771 | 0.1063727 | 0.06073716 | 0.124399906 | ns | NaCl | KS.1 |
| FIM1_4150 | 2267.716268 | 2253.508807 | 2153.571342 | -0.007779197 | 0.141571223 | 0.95136662 | 0.974680587 | ns | NaCl | KS.1 |
| FIM1_4151 | 1792.908664 | 2068.770868 | 3026.452796 | 0.174788778 | 0.131106545 | 0.143429304 | 0.245414138 | ns | NaCl | KS.1 |
| FIM1_4152 | 2305.156544 | 4370.382628 | 1951.519265 | 0.580456749 | 0.326125782 | 0.011140902 | 0.031120711 | ns | NaCl | KS.1 |
| FIM1_4153 | 1359.335025 | 1335.860481 | 1687.778482 | -0.018459714 | 0.152116972 | 0.891284704 | 0.934684106 | ns | NaCl | KS.1 |
| FIM1_4154 | 879.5139151 | 454.2080352 | 796.973279 | -0.802072438 | 0.219355283 | 1.77741006070 | 0.000141482 | ns | NaCl | KS.1 |
| FIM1_4155 | 4662.213726 | 5414.222654 | 3767.699817 | 0.15388437 | 0.192700992 | 0.3380917 | 0.471962694 | ns | NaCl | KS.1 |
| FIM1_4156 | 4568.84837 | 7678.887227 | 3625.250613 | 0.514843989 | 0.275033105 | 0.012681856 | 0.034270631 | ns | NaCl | KS.1 |
| FIM1_4157 | 2997.21047 | 3836.495015 | 3012.997044 | 0.273099527 | 0.183127633 | 0.078082689 | 0.152039679 | ns | NaCl | KS.1 |
| FIM1_4158 | 6148.380012 | 5356.097291 | 4488.575145 | -0.148764675 | 0.176841761 | 0.325059962 | 0.458122752 | ns | NaCl | KS.1 |
| FIM1_4159 | 1711.048355 | 1121.512397 | 1271.519633 | -0.567337749 | 0.116687939 | 1.84226469318 | 2.74409187083 | ns | NaCl | KS.1 |
| FIM1_4160 | 417.6386126 | 324.1431031 | 355.4111926 | -0.300367211 | 0.157839559 | 0.030124369 | 0.069868875 | ns | NaCl | KS.1 |
| FIM1_4161 | 17691.37002 | 20231.96679 | 15530.23963 | 0.176654226 | 0.094895537 | 0.049916551 | 0.105405281 | ns | NaCl | KS.1 |
| FIM1_4162 | 1023.856801 | 1256.145165 | 1415.571465 | 0.149983105 | 0.286299642 | 0.45414992 | 0.589305126 | ns | NaCl | KS.1 |
| FIM1_4163 | 9646.089546 | 9597.493327 | 7457.914414 | -0.006279974 | 0.119370306 | 0.95472419 | 0.976656682 | ns | NaCl | KS.1 |
| FIM1_4164 | 51.7441631 | 39.81412661 | 109.5779923 | -0.179217169 | 0.309137917 | 0.385579963 | 0.52084179 | ns | NaCl | KS.1 |
| FIM1_4165 | 512.238821 | 555.5338683 | 424.6697933 | 0.092449405 | 0.157986418 | 0.506014661 | 0.637968307 | ns | NaCl | KS.1 |
| FIM1_4166 | 742.5420831 | 1088.102591 | 2248.026681 | 0.385723193 | 0.239986544 | 0.039323446 | 0.087064594 | ns | NaCl | KS.1 |
| FIM1_4167 | 1272.950095 | 802.0791203 | 1085.647549 | -0.400864692 | 0.295194459 | 0.056889934 | 0.117780866 | ns | NaCl | KS.1 |
| FIM1_4169 | 582.6413765 | 560.6441634 | 672.9160416 | -0.044626332 | 0.100362614 | 0.638383261 | 0.748695742 | ns | NaCl | KS.1 |
| FIM1_4170 | 836.2636401 | 336.1774799 | 738.4157999 | -1.221539753 | 0.175524348 | 4.88399633795 | 4.33890746094 | Km<KS.1 | NaCl | KS.1 |
| FIM1_4171 | 66.30131245 | 20.2118911 | 117.2484734 | -1.308025641 | 0.426561668 | 1.28454086150 | 0.000106688 | Km<KS.1 | NaCl | KS.1 |
| FIM1_4172 | 999.6954834 | 747.1222071 | 1125.987572 | -0.293495311 | 0.220585628 | 0.095905511 | 0.179128018 | ns | NaCl | KS.1 |
| FIM1_4173 | 1376.249169 | 844.0876406 | 1128.570396 | -0.656081057 | 0.125675882 | 2.09557287735 | 4.27273568231 | ns | NaCl | KS.1 |
| FIM1_4174 | 5110.33259 | 11977.59677 | 5769.751469 | 0.99176378 | 0.275205755 | 9.99271197866 | 8.48357373615 | ns | NaCl | KS.1 |

| | | | | | | | | | | |
|---|---|---|---|---|---|---|---|---|---|---|
| FIM1_4175 | 1390.798412 | 1425.117158 | 1627.710598 | 0.029965163 | 0.121512278 | 0.789600194 | 0.864684342 | ns | NaCl | KS.1 |
| FIM1_4176 | 1271.91353 | 1028.658662 | 1316.289622 | -0.279958512 | 0.104560884 | 0.004297333 | 0.014377426 | ns | NaCl | KS.1 |
| FIM1_4177 | 7582.887068 | 9943.380694 | 6783.080192 | 0.320980361 | 0.160768062 | 0.022366754 | 0.054707278 | ns | NaCl | KS.1 |
| FIM1_4178 | 6918.439253 | 9062.711561 | 6196.614027 | 0.330644488 | 0.146079905 | 0.011113561 | 0.03106178 | ns | NaCl | KS.1 |
| FIM1_4180 | 35.40281698 | 23.61183915 | 24.9312206 | -0.301913703 | 0.312443388 | 0.154288882 | 0.260817937 | ns | NaCl | KS.1 |
| FIM1_4181 | 1774.48361 | 2809.156073 | 3689.600522 | 0.592802844 | 0.151778053 | 1.3873919992 | 0.000113899 | ns | NaCl | KS.1 |
| FIM1_4182 | 8915.709364 | 12137.39246 | 8869.187302 | 0.424404265 | 0.080408164 | 3.8263203539 | 7.1029640899 | ns | NaCl | KS.1 |
| FIM1_4183 | 1507.005557 | 1650.452208 | 1556.276746 | 0.117020789 | 0.105474631 | 0.237993075 | 0.364649075 | ns | NaCl | KS.1 |
| FIM1_4184 | 1952.353795 | 2516.401153 | 2060.641967 | 0.3407145 | 0.096142957 | 0.000177258 | 0.001011307 | ns | NaCl | KS.1 |
| FIM1_4185 | 792.1815896 | 760.65424 | 760.5632707 | -0.051425331 | 0.121547066 | 0.647035908 | 0.755235621 | ns | NaCl | KS.1 |
| FIM1_4186 | 1859.29708 | 1090.713248 | 2235.521867 | -0.603300345 | 0.233819913 | 0.001426632 | 0.005723787 | ns | NaCl | KS.1 |
| FIM1_4187 | 753.8140472 | 607.4410904 | 693.7005976 | -0.262115979 | 0.13918132 | 0.036399131 | 0.081769569 | ns | NaCl | KS.1 |
| FIM1_4188 | 1613.415246 | 1710.817747 | 1297.856177 | 0.067287289 | 0.150409037 | 0.61509568 | 0.730440001 | ns | NaCl | KS.1 |
| FIM1_4189 | 2594.189863 | 2485.789224 | 3247.958278 | -0.048766973 | 0.153843602 | 0.720371517 | 0.814325902 | ns | NaCl | KS.1 |
| FIM1_4190 | 1627.806969 | 1778.176674 | 1947.08494 | 0.111409267 | 0.112269885 | 0.28765842 | 0.418892485 | ns | NaCl | KS.1 |
| FIM1_4191 | 1786.727137 | 1860.329835 | 2019.30619 | 0.049586116 | 0.130319282 | 0.677221847 | 0.780625275 | ns | NaCl | KS.1 |
| FIM1_4192 | 1053.045076 | 1682.236826 | 923.868796 | 0.516066051 | 0.22944062 | 0.005145579 | 0.016644511 | ns | NaCl | KS.1 |
| FIM1_4193 | 251.8306288 | 216.1101457 | 265.2306088 | -0.155438884 | 0.201406866 | 0.34767192 | 0.482742897 | ns | NaCl | KS.1 |
| FIM1_4194 | 1366.542722 | 815.670015 | 1161.170013 | -0.640483637 | 0.183742158 | 6.1084483255 | 0.000411782 | ns | NaCl | KS.1 |
| FIM1_4195 | 2401.512022 | 2370.77571 | 2259.970138 | -0.016281644 | 0.112208419 | 0.876578968 | 0.926882119 | ns | NaCl | KS.1 |
| FIM1_4196 | 2118.850305 | 2098.654988 | 3731.145569 | -0.01036439 | 0.165142877 | 0.942565006 | 0.968655423 | ns | NaCl | KS.1 |
| FIM1_4197 | 1357.262807 | 562.8941775 | 1354.845331 | -1.038869451 | 0.273600473 | 3.9201094976 | 3.9240532698 | Km<KS.1 | NaCl | KS.1 |
| FIM1_4198 | 1593.517532 | 949.3473992 | 1718.026555 | -0.586634912 | 0.229547157 | 0.001643418 | 0.006494048 | ns | NaCl | KS.1 |
| FIM1_4199 | 296.0119716 | 280.0635585 | 370.175666 | -0.057544692 | 0.189640739 | 0.717252486 | 0.812646576 | ns | NaCl | KS.1 |
| FIM1_4201 | 508.0459953 | 391.0263174 | 544.8274497 | -0.359907643 | 0.095986332 | 7.2815478951 | 0.000481971 | ns | NaCl | KS.1 |
| FIM1_4203 | 275.2570151 | 274.7572946 | 324.5406144 | -0.002283584 | 0.201551236 | 0.988992545 | 0.995082984 | ns | NaCl | KS.1 |
| FIM1_4204 | 909.2747495 | 600.3008185 | 1148.355635 | -0.534108921 | 0.147825694 | 5.6858952886 | 0.000388029 | ns | NaCl | KS.1 |
| FIM1_4205 | 1103.221423 | 2897.738844 | 3643.20249 | 0.990259444 | 0.361167707 | 0.000157215 | 0.000911588 | ns | NaCl | KS.1 |
| FIM1_4206 | 1696.926571 | 2069.719806 | 1485.64639 | 0.190814858 | 0.219651254 | 0.275891786 | 0.40584318 | ns | NaCl | KS.1 |
| FIM1_4207 | 2269.711025 | 1789.808811 | 2188.568411 | -0.271113856 | 0.169865039 | 0.064387604 | 0.129970864 | ns | NaCl | KS.1 |
| FIM1_4208 | 796.9336642 | 651.1187672 | 1001.676204 | -0.219121214 | 0.180796538 | 0.153555122 | 0.259918443 | ns | NaCl | KS.1 |
| FIM1_4209 | 512.8033125 | 385.5845684 | 664.3409912 | -0.349775275 | 0.148326921 | 0.007982348 | 0.024003488 | ns | NaCl | KS.1 |
| FIM1_4210 | 2022.040719 | 2393.974717 | 2048.586944 | 0.196053913 | 0.154729527 | 0.151055432 | 0.256572473 | ns | NaCl | KS.1 |
| FIM1_4211 | 1148.81288 | 1046.706089 | 725.4670799 | -0.095581526 | 0.189128221 | 0.546723954 | 0.672755793 | ns | NaCl | KS.1 |
| FIM1_4212 | 2751.366946 | 3396.523789 | 3234.851481 | 0.28176881 | 0.091803189 | 0.001236698 | 0.005118614 | ns | NaCl | KS.1 |
| FIM1_4213 | 367.3456007 | 347.5773125 | 423.953216 | -0.063323139 | 0.154873004 | 0.64375386 | 0.753349903 | ns | NaCl | KS.1 |
| FIM1_4214 | 1137.931525 | 1148.758298 | 1128.506859 | 0.01327316 | 0.120950292 | 0.905548788 | 0.943872872 | ns | NaCl | KS.1 |
| FIM1_4215 | 3499.06883 | 5665.283744 | 5631.325258 | 0.488848282 | 0.25997523 | 0.014041462 | 0.037276559 | ns | NaCl | KS.1 |
| FIM1_4216 | 632.6433746 | 467.1546333 | 743.7061317 | -0.322230953 | 0.204255131 | 0.054611911 | 0.113679606 | ns | NaCl | KS.1 |
| FIM1_4217 | 668.7933546 | 352.7551994 | 889.1683139 | -0.761348787 | 0.226057606 | 5.9878840763 | 0.000405612 | ns | NaCl | KS.1 |
| FIM1_4218 | 44.93021848 | 24.29786527 | 40.80047592 | -0.599623874 | 0.3031595 | 0.006745474 | 0.020882844 | ns | NaCl | KS.1 |
| FIM1_4219 | 536.752079 | 479.5798505 | 472.1011329 | -0.130506077 | 0.162123619 | 0.357111908 | 0.492414563 | ns | NaCl | KS.1 |
| FIM1_4220 | 981.9735881 | 848.6244245 | 752.1887105 | -0.169683261 | 0.153704931 | 0.211825959 | 0.334340326 | ns | NaCl | KS.1 |
| FIM1_4221 | 263.4711731 | 173.4961368 | 215.5270907 | -0.47444782 | 0.205735681 | 0.005322895 | 0.017162283 | ns | NaCl | KS.1 |
| FIM1_4222 | 2199.168149 | 1506.451562 | 1521.313095 | -0.366750766 | 0.24953654 | 0.055236707 | 0.114836028 | ns | NaCl | KS.1 |

| | | | | | | | | | | |
|---|---|---|---|---|---|---|---|---|---|---|
| FIM1_4223 | 3098.760832 | 2807.954147 | 2560.189644 | -0.11606828 | 0.140600601 | 0.359617442 | 0.495401439 | ns | NaCl | KS.1 |
| FIM1_4224 | 6085.027223 | 7141.007047 | 5080.741052 | 0.177639388 | 0.169156758 | 0.224485579 | 0.349441725 | ns | NaCl | KS.1 |
| FIM1_4225 | 409.0388336 | 574.3034659 | 581.9715583 | 0.252129176 | 0.302946063 | 0.224601085 | 0.349512168 | ns | NaCl | KS.1 |
| FIM1_4226 | 18811.56231 | 69046.88101 | 38582.63717 | 1.727433612 | 0.242677918 | 9.1795216952 | 1.0148471207 | Km>KS.1 | NaCl | KS.1 |
| FIM1_4227 | 1945.287282 | 1442.252554 | 2863.283207 | -0.218499461 | 0.300081858 | 0.28829184 | 0.418892485 | ns | NaCl | KS.1 |
| FIM1_4228 | 16.22747376 | 15.28716908 | 24.61517435 | -0.037221602 | 0.316622309 | 0.855531435 | 0.912715589 | ns | NaCl | KS.1 |
| FIM1_4229 | 461.0958479 | 311.5520365 | 1314.53667 | -0.341467236 | 0.280613198 | 0.093284457 | 0.175459422 | ns | NaCl | KS.1 |
| FIM1_4230 | 3818.44245 | 3638.815442 | 2387.443186 | -0.045142692 | 0.21326529 | 0.792475766 | 0.867069923 | ns | NaCl | KS.1 |
| FIM1_4231 | 7043.966077 | 9286.986882 | 5772.270488 | 0.311505943 | 0.181222847 | 0.042950962 | 0.093147792 | ns | NaCl | KS.1 |
| FIM1_4232 | 522.7089138 | 409.270964 | 487.0612049 | -0.272121629 | 0.179315892 | 0.07461028 | 0.146598003 | ns | NaCl | KS.1 |
| FIM1_4233 | 422.4205967 | 600.5836024 | 813.2059935 | 0.440246755 | 0.150712817 | 0.001009794 | 0.004319626 | ns | NaCl | KS.1 |
| FIM1_4234 | 1321.448512 | 1573.18552 | 1164.257849 | 0.204904382 | 0.148582432 | 0.121199927 | 0.215886013 | ns | NaCl | KS.1 |
| FIM1_4235 | 874.5367398 | 966.8330246 | 998.3452731 | 0.128477225 | 0.111527275 | 0.217317008 | 0.340949895 | ns | NaCl | KS.1 |
| FIM1_4236 | 2372.049855 | 2203.276016 | 2572.85615 | -0.094067106 | 0.112172986 | 0.369089213 | 0.504044881 | ns | NaCl | KS.1 |
| FIM1_4237 | 6782.777009 | 9775.334027 | 7971.332114 | 0.495573172 | 0.100574394 | 1.7661948174 | 2.6386844495 | ns | NaCl | KS.1 |
| FIM1_4238 | 3379.238722 | 4338.472397 | 2974.152138 | 0.321042575 | 0.119302853 | 0.003499582 | 0.012158116 | ns | NaCl | KS.1 |
| FIM1_4239 | 3333.750795 | 3411.413859 | 2512.177134 | 0.029064491 | 0.118135207 | 0.790898992 | 0.86572552 | ns | NaCl | KS.1 |
| FIM1_4240 | 1581.041201 | 2744.885216 | 1416.346129 | 0.584400958 | 0.261478017 | 0.0039417 | 0.013411726 | ns | NaCl | KS.1 |
| FIM1_4241 | 2604.798938 | 2573.874893 | 2297.8411 | -0.010927882 | 0.215445384 | 0.949528377 | 0.973834262 | ns | NaCl | KS.1 |
| FIM1_4242 | 179.6502671 | 123.7752464 | 256.0742388 | -0.394551316 | 0.222453901 | 0.026986987 | 0.063903027 | ns | NaCl | KS.1 |
| FIM1_4243 | 875.6890962 | 810.4332806 | 683.6635556 | -0.095056259 | 0.114428882 | 0.37237672 | 0.507415553 | ns | NaCl | KS.1 |
| FIM1_4244 | 549.7662176 | 306.6024258 | 691.9464541 | -0.692534468 | 0.219432394 | 0.00016278 | 0.000941661 | ns | NaCl | KS.1 |
| FIM1_4245 | 30.19182811 | 8.925571291 | 45.78680657 | -1.083810419 | 0.470297111 | 0.000283788 | 0.001500369 | Km<KS.1 | NaCl | KS.1 |
| FIM1_4246 | 584.951861 | 407.4215334 | 598.2144239 | -0.475503482 | 0.130147928 | 6.2456482392 | 0.000420461 | ns | NaCl | KS.1 |
| FIM1_4247 | 421.2172743 | 504.5061694 | 496.5544882 | 0.188544547 | 0.192991656 | 0.241031199 | 0.367831355 | ns | NaCl | KS.1 |
| FIM1_4248 | 1644.081903 | 1245.794938 | 1281.443992 | -0.37444736 | 0.093756652 | 2.4887476548 | 0.00018675 | ns | NaCl | KS.1 |
| FIM1_4249 | 5379.251485 | 6445.620094 | 5696.017768 | 0.226066592 | 0.125420034 | 0.04921724 | 0.104282695 | ns | NaCl | KS.1 |
| FIM1_4250 | 1227.995568 | 1380.309294 | 1103.258929 | 0.159219928 | 0.07000665 | 0.01917921 | 0.047779955 | ns | NaCl | KS.1 |
| FIM1_4251 | 3230.214631 | 3356.665297 | 3147.760199 | 0.051961972 | 0.085882542 | 0.528288158 | 0.656565971 | ns | NaCl | KS.1 |
| FIM1_4252 | 1036.866291 | 831.7931393 | 1133.022365 | -0.282751457 | 0.121777064 | 0.011554404 | 0.031986345 | ns | NaCl | KS.1 |
| FIM1_4253 | 1470.62475 | 2190.197322 | 2736.043885 | 0.455245317 | 0.200345415 | 0.006331139 | 0.01982445 | ns | NaCl | KS.1 |
| FIM1_4254 | 861.6544071 | 1090.842657 | 1081.144263 | 0.29716668 | 0.130130428 | 0.012080862 | 0.03318735 | ns | NaCl | KS.1 |
| FIM1_4255 | 2060.540351 | 2035.165564 | 1779.909196 | -0.013445841 | 0.183304488 | 0.930964032 | 0.960702356 | ns | NaCl | KS.1 |
| FIM1_4257 | 1758.814166 | 1773.358791 | 4226.826573 | 0.007959055 | 0.212476923 | 0.962913807 | 0.981860256 | ns | NaCl | KS.1 |
| FIM1_4258 | 36.69998792 | 50.53752287 | 42.59403129 | 0.255207341 | 0.263048534 | 0.190349089 | 0.307718719 | ns | NaCl | KS.1 |
| FIM1_4259 | 2194.553428 | 2383.743044 | 2675.666478 | 0.086281557 | 0.183955543 | 0.579136825 | 0.700682321 | ns | NaCl | KS.1 |
| FIM1_4260 | 774.2856098 | 591.182882 | 767.4231294 | -0.270407745 | 0.219113822 | 0.123003681 | 0.218010565 | ns | NaCl | KS.1 |
| FIM1_4261 | 477.589548 | 977.9253277 | 920.3489585 | 0.87814169 | 0.22389963 | 4.7276048004 | 4.5492908863 | ns | NaCl | KS.1 |
| FIM1_4262 | 3183.16448 | 4701.420098 | 2964.984714 | 0.476398096 | 0.16966929 | 0.001234595 | 0.005118427 | ns | NaCl | KS.1 |
| FIM1_4263 | 664.4210053 | 323.6625118 | 569.9360588 | -0.990520256 | 0.130458819 | 1.4605784470 | 1.5460378242 | ns | NaCl | KS.1 |
| FIM1_4265 | 862.5295435 | 1116.96115 | 891.4510669 | 0.32775612 | 0.129222679 | 0.005347303 | 0.017218661 | ns | NaCl | KS.1 |
| FIM1_4266 | 6824.763779 | 15670.37298 | 9953.457295 | 1.018205487 | 0.239462149 | 6.7611351416 | 8.3763703097 | Km>KS.1 | NaCl | KS.1 |
| FIM1_4267 | 1457.762229 | 1929.348877 | 1443.309574 | 0.345531973 | 0.144956244 | 0.007585637 | 0.023011307 | ns | NaCl | KS.1 |
| FIM1_4268 | 3496.201736 | 4402.638738 | 3576.938436 | 0.248865025 | 0.188510634 | 0.115778354 | 0.207866226 | ns | NaCl | KS.1 |
| FIM1_4269 | 477.029538 | 539.7178583 | 394.092963 | 0.145292456 | 0.14518196 | 0.263540984 | 0.391882903 | ns | NaCl | KS.1 |

| | | | | | | | | | | |
|---|---|---|---|---|---|---|---|---|---|---|
| FIM1_4270 | 2014.12695 | 1962.877777 | 2359.182223 | -0.032049126 | 0.112548023 | 0.760423729 | 0.845009617 | ns | NaCl | KS.1 |
| FIM1_4271 | 870.6884877 | 902.8386983 | 835.0517382 | 0.043514865 | 0.110247853 | 0.673320402 | 0.778111266 | ns | NaCl | KS.1 |
| FIM1_4272 | 458.6476754 | 594.5608771 | 712.7728461 | 0.273561355 | 0.20489353 | 0.102926739 | 0.189091775 | ns | NaCl | KS.1 |
| FIM1_4273 | 2103.307583 | 4359.789288 | 1940.035915 | 0.753347822 | 0.307804383 | 0.001062369 | 0.00450707 | ns | NaCl | KS.1 |
| FIM1_4274 | 188.5238757 | 210.0933628 | 458.2280169 | 0.121985625 | 0.153199973 | 0.368574835 | 0.503752693 | ns | NaCl | KS.1 |
| FIM1_4275 | 12.26601389 | 15.39315876 | 33.90945476 | 0.108640542 | 0.349649996 | 0.602156039 | 0.72064621 | ns | NaCl | KS.1 |
| FIM1_4276 | 2455.615988 | 2633.547426 | 2355.293366 | 0.089142324 | 0.104935315 | 0.366743017 | 0.502214838 | ns | NaCl | KS.1 |
| FIM1_4277 | 162.8076932 | 91.11348271 | 199.5390417 | -0.683384763 | 0.226098983 | 0.000263283 | 0.001408629 | ns | NaCl | KS.1 |
| FIM1_4278 | 2822.262345 | 3055.949864 | 2990.652528 | 0.108041201 | 0.07840105 | 0.153359635 | 0.259865185 | ns | NaCl | KS.1 |
| FIM1_4279 | 4311.729943 | 7519.933062 | 3477.358609 | 0.499964713 | 0.308864257 | 0.022517483 | 0.054913959 | ns | NaCl | KS.1 |
| FIM1_4280 | 2949.463535 | 2722.294376 | 2995.245579 | -0.096922308 | 0.128202019 | 0.409169462 | 0.544428477 | ns | NaCl | KS.1 |
| FIM1_4281 | 5445.780484 | 6227.207134 | 5184.495808 | 0.169750942 | 0.114583988 | 0.111043846 | 0.200888413 | ns | NaCl | KS.1 |
| FIM1_4282 | 972.9387463 | 1805.541034 | 2506.198991 | 0.59575673 | 0.306270454 | 0.007377506 | 0.022462112 | ns | NaCl | KS.1 |
| FIM1_4283 | 47.79699581 | 27.91535858 | 72.17510083 | -0.451322456 | 0.314417745 | 0.039198803 | 0.086904654 | ns | NaCl | KS.1 |
| FIM1_4284 | 5010.049017 | 4651.518666 | 5208.636838 | -0.092358925 | 0.115859699 | 0.391229752 | 0.526757244 | ns | NaCl | KS.1 |
| FIM1_4285 | 2003.806567 | 2754.084402 | 2583.026233 | 0.409482865 | 0.129213692 | 0.000505666 | 0.002428271 | ns | NaCl | KS.1 |
| FIM1_4286 | 2089.620385 | 3509.245899 | 2558.176458 | 0.506253625 | 0.279339639 | 0.014979919 | 0.039131242 | ns | NaCl | KS.1 |
| FIM1_4287 | 11.49693355 | 10.20818335 | 12.20518445 | -0.070815008 | 0.339603 | 0.732191786 | 0.823198675 | ns | NaCl | KS.1 |
| FIM1_4288 | 1469.754417 | 1652.537567 | 1283.172397 | 0.112933236 | 0.210240938 | 0.506837228 | 0.638519931 | ns | NaCl | KS.1 |
| FIM1_4289 | 2395.917083 | 2234.74646 | 2144.079177 | -0.081733252 | 0.142388678 | 0.523353635 | 0.653091685 | ns | NaCl | KS.1 |
| FIM1_4290 | 1162.049862 | 1374.678096 | 1397.806259 | 0.210241256 | 0.125451941 | 0.067561228 | 0.134770292 | ns | NaCl | KS.1 |
| FIM1_4291 | 808.9586889 | 971.0183008 | 822.1209317 | 0.241302325 | 0.090146655 | 0.004951057 | 0.01620072 | ns | NaCl | KS.1 |
| FIM1_4292 | 1120.985927 | 1220.483048 | 926.2401936 | 0.106653286 | 0.11562176 | 0.321156798 | 0.454165739 | ns | NaCl | KS.1 |
| FIM1_4293 | 648.4438486 | 727.0975356 | 651.6130055 | 0.145255446 | 0.100968042 | 0.127598766 | 0.224153906 | ns | NaCl | KS.1 |
| FIM1_4294 | 1030.350411 | 665.0950846 | 1014.23371 | -0.576141443 | 0.136514717 | 3.80431967694 | 3.82353341268 | ns | NaCl | KS.1 |
| FIM1_4295 | 874.7509675 | 805.3110284 | 957.734905 | -0.11367803 | 0.091050117 | 0.19070684 | 0.3079411 | ns | NaCl | KS.1 |
| FIM1_4296 | 1384.55123 | 1160.467598 | 1234.676435 | -0.242024999 | 0.068768546 | 0.000288391 | 0.001516644 | ns | NaCl | KS.1 |
| FIM1_4297 | 258.9247189 | 326.7919295 | 795.4135375 | 0.263616005 | 0.179020985 | 0.083758565 | 0.161194408 | ns | NaCl | KS.1 |
| FIM1_4298 | 218.3984816 | 259.6751645 | 256.6677733 | 0.104338482 | 0.323132012 | 0.613341808 | 0.729117728 | ns | NaCl | KS.1 |
| FIM1_4299 | 300.8100618 | 435.5816983 | 366.378426 | 0.450707424 | 0.165884678 | 0.001830416 | 0.007108759 | ns | NaCl | KS.1 |
| FIM1_4300 | 247.9889485 | 216.060621 | 223.9361699 | -0.164340102 | 0.156272733 | 0.232642588 | 0.358660328 | ns | NaCl | KS.1 |
| FIM1_4303 | 70.53215093 | 143.8266206 | 82.48496482 | 0.809845298 | 0.260194322 | 0.000117989 | 0.000715846 | ns | NaCl | KS.1 |
| FIM1_4304 | 13.31867476 | 10.5395134 | 30.91326256 | -0.094950382 | 0.352973611 | 0.647871868 | 0.75590116 | ns | NaCl | KS.1 |
| FIM1_4307 | 399.9080247 | 480.9097774 | 476.7071757 | 0.199529995 | 0.178617543 | 0.189895198 | 0.307128936 | ns | NaCl | KS.1 |
| FIM1_4308 | 73.95179674 | 181.2295148 | 116.7686975 | 1.096843836 | 0.254827316 | 3.75986703808 | 5.11074822799 | Km>KS.1 | NaCl | KS.1 |
| FIM1_4314 | 508.6378232 | 1390.722847 | 646.7515623 | 1.29581778 | 0.225555497 | 8.36819567585 | 3.13020853288 | Km>KS.1 | NaCl | KS.1 |
| FIM1_4780 | 48.73102012 | 250.2768164 | 82.75928701 | 1.988314269 | 0.442263099 | 3.54347685308 | 8.77054594233 | Km>KS.1 | NaCl | KS.1 |
| FIM1_4781 | 51.54596044 | 63.27824654 | 87.57792087 | 0.206820554 | 0.225788596 | 0.245838232 | 0.373405998 | ns | NaCl | KS.1 |
| FIM1_4782 | 202.0555241 | 169.193868 | 296.5047851 | -0.209844547 | 0.147193732 | 0.109768669 | 0.199379018 | ns | NaCl | KS.1 |
| FIM1_4783 | 630.3972019 | 506.8233937 | 1729.24346 | -0.262228272 | 0.137800841 | 0.034811141 | 0.078789916 | ns | NaCl | KS.1 |
| FIM1_4785 | 678.7573498 | 1040.522116 | 1037.162625 | 0.56070343 | 0.136495663 | 6.63165862471 | 5.95700684055 | ns | NaCl | KS.1 |
| FIM1_4786 | 2065.054596 | 6318.212105 | 3276.771524 | 0.987179169 | 0.450881719 | 0.000593662 | 0.002783664 | ns | NaCl | KS.1 |
| FIM1_4787 | 189.4009158 | 257.5964934 | 235.4642185 | 0.296742109 | 0.236030272 | 0.10668366 | 0.194699637 | ns | NaCl | KS.1 |
| FIM1_4788 | 189.5799662 | 155.3541431 | 744.3069426 | -0.154475419 | 0.274251885 | 0.432820819 | 0.569350495 | ns | NaCl | KS.1 |
| FIM1_4789 | 436.6636719 | 228.7842175 | 264.372092 | -0.484311317 | 0.366753419 | 0.038458779 | 0.085802289 | ns | NaCl | KS.1 |

| | | | | | | | | | | |
|---|---|---|---|---|---|---|---|---|---|---|
| FIM1_4790 | 472.654812 | 715.383085 | 614.4716867 | 0.485352623 | 0.194560789 | 0.003025039 | 0.010795961 | ns | NaCl | KS.1 |
| FIM1_4791 | 3526.029427 | 4294.139338 | 2073.260557 | 0.201078753 | 0.201142607 | 0.224202084 | 0.349109662 | ns | NaCl | KS.1 |
| FIM1_4792 | 454.4545525 | 402.1294609 | 464.198878 | -0.106499402 | 0.234727451 | 0.55781204 | 0.681511517 | ns | NaCl | KS.1 |
| FIM1_4793 | 1257.121469 | 1496.836212 | 2070.580399 | 0.205594136 | 0.150403568 | 0.123560387 | 0.218681225 | ns | NaCl | KS.1 |
| FIM1_4794 | 2065.407374 | 1187.499 | 2670.050722 | -0.633169307 | 0.231666155 | 0.000801743 | 0.003582468 | ns | NaCl | KS.1 |
| FIM1_4795 | 8627.388078 | 4851.341675 | 6331.766408 | -0.601648827 | 0.271558307 | 0.00382968 | 0.013103617 | ns | NaCl | KS.1 |
| FIM1_4796 | 3402.34135 | 1337.977387 | 2521.152864 | -1.192231208 | 0.221305589 | 1.13350478938 | 3.20408314043 | Km<KS.1 | NaCl | KS.1 |
| FIM1_4797 | 7112.47374 | 9558.386828 | 4622.760369 | 0.258291857 | 0.25962878 | 0.182176315 | 0.297604048 | ns | NaCl | KS.1 |
| FIM1_4798 | 1419.554716 | 1050.858825 | 907.3828427 | -0.31306472 | 0.212734561 | 0.069193619 | 0.137750403 | ns | NaCl | KS.1 |
| FIM1_4799 | 988.0112101 | 848.122133 | 915.778069 | -0.158268775 | 0.192001759 | 0.323297283 | 0.456673476 | ns | NaCl | KS.1 |
| FIM1_4800 | 10.63294696 | 18.50475693 | 14.76985844 | 0.329968859 | 0.36101264 | 0.138000876 | 0.238386931 | ns | NaCl | KS.1 |
| FIM1_4801 | 479.2055674 | 474.5933696 | 369.4995752 | -0.010736089 | 0.213585447 | 0.950149046 | 0.973834262 | ns | NaCl | KS.1 |
| FIM1_4802 | 683.6746607 | 814.072059 | 805.0057536 | 0.228200177 | 0.102195109 | 0.0176648 | 0.044883748 | ns | NaCl | KS.1 |
| FIM1_4803 | 1750.750501 | 1686.844271 | 1781.19925 | -0.036378997 | 0.199330595 | 0.824795648 | 0.892422434 | ns | NaCl | KS.1 |
| FIM1_4804 | 6220.51956 | 8068.230119 | 5526.854346 | 0.319583376 | 0.142924213 | 0.01248668 | 0.033867944 | ns | NaCl | KS.1 |
| FIM1_4805 | 3641.549953 | 9581.479631 | 4310.284211 | 1.315312516 | 0.159202561 | 1.15232666730 | 2.38867715409 | Km>KS.1 | NaCl | KS.1 |
| FIM1_4806 | 2064.592244 | 2256.073897 | 2008.96905 | 0.121563986 | 0.074617226 | 0.092387784 | 0.174133712 | ns | NaCl | KS.1 |
| FIM1_4807 | 3389.748278 | 3285.598163 | 2654.061447 | -0.032211886 | 0.185519346 | 0.836940053 | 0.900081444 | ns | NaCl | KS.1 |
| FIM1_4808 | 2258.88247 | 1719.729141 | 2096.452981 | -0.296859819 | 0.192490455 | 0.064731655 | 0.130433368 | ns | NaCl | KS.1 |
| FIM1_4809 | 900.7529366 | 760.1822987 | 696.4791253 | -0.198613418 | 0.151271725 | 0.138675594 | 0.239137289 | ns | NaCl | KS.1 |
| FIM1_4810 | 14683.86902 | 13902.7077 | 12749.87612 | -0.072214601 | 0.090980293 | 0.405766518 | 0.540708455 | ns | NaCl | KS.1 |
| FIM1_4811 | 1599.421174 | 1499.322035 | 1936.960012 | -0.048980285 | 0.26656629 | 0.799996338 | 0.87184705 | ns | NaCl | KS.1 |
| FIM1_4812 | 139.0664716 | 143.1215438 | 213.5856373 | 0.031583352 | 0.179672501 | 0.836475888 | 0.899776766 | ns | NaCl | KS.1 |
| FIM1_4813 | 2554.530731 | 3992.681886 | 1787.613643 | 0.402749552 | 0.283045447 | 0.050780177 | 0.107094168 | ns | NaCl | KS.1 |
| FIM1_4814 | 1051.232374 | 1219.889942 | 1173.57424 | 0.199130555 | 0.092835287 | 0.024089331 | 0.058120475 | ns | NaCl | KS.1 |
| FIM1_4815 | 607.2102889 | 609.5129256 | 843.8898588 | 0.003541897 | 0.11066531 | 0.972720249 | 0.987205884 | ns | NaCl | KS.1 |
| FIM1_4816 | 674.6963203 | 768.5002811 | 1091.397534 | 0.16229957 | 0.10726319 | 0.106540575 | 0.194581263 | ns | NaCl | KS.1 |
| FIM1_4817 | 1943.032045 | 1979.924502 | 1649.15812 | 0.021522262 | 0.147161646 | 0.870061585 | 0.922146652 | ns | NaCl | KS.1 |
| FIM1_4818 | 791.9567961 | 708.5045891 | 1041.875757 | -0.14554378 | 0.102347152 | 0.131481909 | 0.229436163 | ns | NaCl | KS.1 |
| FIM1_4819 | 4604.650856 | 5173.848091 | 5748.894804 | 0.153389526 | 0.095559165 | 0.090850109 | 0.171775196 | ns | NaCl | KS.1 |
| FIM1_4820 | 1926.832854 | 1553.161134 | 1825.771121 | -0.282688471 | 0.103891586 | 0.003732261 | 0.012840939 | ns | NaCl | KS.1 |
| FIM1_4821 | 1003.937739 | 944.3111409 | 1098.858809 | -0.05252153 | 0.237404876 | 0.773674416 | 0.853357329 | ns | NaCl | KS.1 |
| FIM1_4822 | 1524.002589 | 1126.573855 | 1621.314464 | -0.389313651 | 0.126849651 | 0.000779625 | 0.003497415 | ns | NaCl | KS.1 |
| FIM1_4823 | 701.0316971 | 960.4651543 | 1166.873272 | 0.408821553 | 0.12342981 | 0.000305992 | 0.001595714 | ns | NaCl | KS.1 |
| FIM1_4824 | 3218.34631 | 5876.386257 | 3418.530375 | 0.730638094 | 0.211456235 | 4.90363689484 | 0.0003436 | ns | NaCl | KS.1 |
| FIM1_4825 | 301.0661047 | 175.9576004 | 465.332088 | -0.588013101 | 0.240778748 | 0.002246461 | 0.00841577 | ns | NaCl | KS.1 |
| FIM1_4826 | 989.4272043 | 1131.216623 | 1047.673213 | 0.17130662 | 0.107107542 | 0.087999277 | 0.167365195 | ns | NaCl | KS.1 |
| FIM1_4827 | 1351.700743 | 1607.933458 | 1322.638848 | 0.222725428 | 0.108696977 | 0.028357692 | 0.066452905 | ns | NaCl | KS.1 |
| FIM1_4828 | 1023.358645 | 767.4708532 | 929.2585492 | -0.367409054 | 0.125307425 | 0.001351795 | 0.005476532 | ns | NaCl | KS.1 |
| FIM1_4829 | 1729.682014 | 835.9333635 | 2408.544241 | -0.734390516 | 0.315231666 | 0.001564459 | 0.006221571 | ns | NaCl | KS.1 |
| FIM1_4830 | 1623.942978 | 1047.962394 | 2396.706495 | -0.435357984 | 0.255872995 | 0.026119613 | 0.062323777 | ns | NaCl | KS.1 |
| FIM1_4831 | 2105.677433 | 2451.084137 | 1966.919271 | 0.201329536 | 0.090076154 | 0.019065224 | 0.047591315 | ns | NaCl | KS.1 |
| FIM1_4832 | 3785.424954 | 3105.485192 | 3576.343722 | -0.273249789 | 0.068669022 | 4.11462373919 | 0.000293351 | ns | NaCl | KS.1 |
| FIM1_4833 | 4904.965049 | 4946.751853 | 4511.192965 | 0.010152372 | 0.137844185 | 0.935168457 | 0.962644956 | ns | NaCl | KS.1 |
| FIM1_4834 | 801.1024667 | 1086.835154 | 1340.639535 | 0.22905514 | 0.295340424 | 0.263504747 | 0.391882903 | ns | NaCl | KS.1 |

| | | | | | | | | | | |
|---|---|---|---|---|---|---|---|---|---|---|
| FIM1_4835 | 4421.342614 | 5975.362521 | 3751.275652 | 0.380886908 | 0.135043213 | 0.001815202 | 0.007060696 | ns | NaCl | KS.1 |
| FIM1_4836 | 559.7817832 | 433.8879058 | 802.9952928 | -0.247375407 | 0.226957277 | 0.167160731 | 0.27804234 | ns | NaCl | KS.1 |
| FIM1_4837 | 72.54561845 | 25.24789666 | 67.6932174 | -0.984275805 | 0.423014589 | 0.000451332 | 0.002207844 | ns | NaCl | KS.1 |
| FIM1_4839 | 515.7054436 | 307.9011722 | 576.3763114 | -0.515079916 | 0.272591526 | 0.0121936 | 0.033386439 | ns | NaCl | KS.1 |
| FIM1_4840 | 9797.700988 | 12951.43417 | 8620.223328 | 0.376109932 | 0.093608252 | 2.2291489563 | 0.000170879 | ns | NaCl | KS.1 |
| FIM1_4841 | 222.425039 | 104.0155587 | 248.0003937 | -0.991924847 | 0.185911632 | 3.8962797625 | 9.5960355537 | ns | NaCl | KS.1 |
| FIM1_4842 | 54213.9611 | 90342.75792 | 58807.41069 | 0.692451978 | 0.11932166 | 6.9231566339 | 2.0413050248 | ns | NaCl | KS.1 |
| FIM1_4844 | 236.6696349 | 200.4923286 | 1119.871416 | -0.101792029 | 0.316215076 | 0.620372479 | 0.735196065 | ns | NaCl | KS.1 |
| FIM1_4845 | 406.6529941 | 426.8980362 | 600.696593 | 0.039660713 | 0.256241721 | 0.834513721 | 0.898487436 | ns | NaCl | KS.1 |
| FIM1_4846 | 207.4665233 | 199.7566592 | 542.9534828 | -0.028160918 | 0.270095389 | 0.884777454 | 0.931393955 | ns | NaCl | KS.1 |
| FIM1_4847 | 343.0618897 | 353.1690099 | 765.4735833 | 0.025111506 | 0.238694818 | 0.89092375 | 0.934546607 | ns | NaCl | KS.1 |
| FIM1_4848 | 766.0237116 | 2543.666941 | 2313.99835 | 1.503443043 | 0.287989172 | 5.0061268702 | 1.5186269011 | Km>KS.1 | NaCl | KS.1 |
| FIM1_4849 | 4037.451957 | 5020.789139 | 3562.945838 | 0.200745242 | 0.233777437 | 0.269792964 | 0.39878718 | ns | NaCl | KS.1 |
| FIM1_4850 | 418.7506392 | 116.4626386 | 797.3107309 | -1.321850816 | 0.443156588 | 1.5482578505 | 0.000125245 | Km<KS.1 | NaCl | KS.1 |
| FIM1_4851 | 981.2351028 | 757.8609023 | 855.6509651 | -0.307473521 | 0.157512609 | 0.026189978 | 0.062461718 | ns | NaCl | KS.1 |
| FIM1_4852 | 579.1228358 | 310.4108905 | 468.2283045 | -0.800376494 | 0.184670902 | 1.0813234840 | 1.2569122273 | ns | NaCl | KS.1 |
| FIM1_4853 | 323.581054 | 318.2371886 | 464.3356436 | -0.016004819 | 0.187260795 | 0.919078354 | 0.953977636 | ns | NaCl | KS.1 |
| FIM1_4854 | 4848.899948 | 4467.828748 | 3480.209368 | -0.065053316 | 0.2572836 | 0.732500195 | 0.823359348 | ns | NaCl | KS.1 |
| FIM1_4855 | 688.651135 | 1044.822156 | 915.6515042 | 0.298827676 | 0.323287835 | 0.163285783 | 0.272782663 | ns | NaCl | KS.1 |
| FIM1_4856 | 75.44078945 | 39.70813693 | 90.94481753 | -0.631802414 | 0.300481158 | 0.004370719 | 0.01459684 | ns | NaCl | KS.1 |
| FIM1_4857 | 37.29230447 | 45.20482824 | 97.34138546 | 0.128085403 | 0.30690799 | 0.531321605 | 0.65948687 | ns | NaCl | KS.1 |
| FIM1_4858 | 1826.016623 | 4171.92828 | 1889.896656 | 0.886127888 | 0.312537517 | 0.000198778 | 0.001112398 | ns | NaCl | KS.1 |
| FIM1_4859 | 1351.486535 | 1408.025328 | 1667.293549 | 0.053755939 | 0.087487619 | 0.521282852 | 0.652118833 | ns | NaCl | KS.1 |
| FIM1_4860 | 522.1199309 | 453.9991011 | 551.1737234 | -0.129877429 | 0.22310894 | 0.461922212 | 0.596434727 | ns | NaCl | KS.1 |
| FIM1_4861 | 1110.417689 | 805.8857904 | 825.2678814 | -0.424800525 | 0.111584703 | 4.2650407023 | 0.000302259 | ns | NaCl | KS.1 |
| FIM1_4862 | 2290.245042 | 2311.857305 | 1646.755377 | 0.010500346 | 0.147391801 | 0.936470666 | 0.963188249 | ns | NaCl | KS.1 |
| FIM1_4863 | 6395.636074 | 11982.61232 | 5275.620652 | 0.650047709 | 0.285807864 | 0.002653733 | 0.00967912 | ns | NaCl | KS.1 |
| FIM1_4864 | 85.77452877 | 62.24809275 | 64.56806349 | -0.295144673 | 0.251633234 | 0.122191786 | 0.217031108 | ns | NaCl | KS.1 |
| FIM1_4865 | 858.162351 | 616.3788959 | 766.4919362 | -0.346572075 | 0.216242636 | 0.046996006 | 0.100474916 | ns | NaCl | KS.1 |
| FIM1_4866 | 980.7930487 | 538.9966471 | 1468.118296 | -0.662000934 | 0.254540931 | 0.001047742 | 0.004455144 | ns | NaCl | KS.1 |
| FIM1_4867 | 2300.064575 | 2776.585349 | 1774.801931 | 0.204411547 | 0.179895341 | 0.181532383 | 0.297229756 | ns | NaCl | KS.1 |
| FIM1_4868 | 1605.334364 | 1610.22661 | 1300.526989 | 0.002762707 | 0.127984652 | 0.981218999 | 0.991175946 | ns | NaCl | KS.1 |
| FIM1_4869 | 3498.654212 | 2472.996346 | 2324.438527 | -0.42412464 | 0.161158562 | 0.002660627 | 0.009690242 | ns | NaCl | KS.1 |
| FIM1_4870 | 933.5922781 | 858.5983842 | 695.124892 | -0.072931637 | 0.233406476 | 0.686995966 | 0.788239144 | ns | NaCl | KS.1 |
| FIM1_4871 | 516.0885944 | 389.6197673 | 519.1620767 | -0.322108195 | 0.177476696 | 0.033555137 | 0.076436268 | ns | NaCl | KS.1 |
| FIM1_4872 | 315.495487 | 185.6009031 | 580.6762812 | -0.619808896 | 0.223094196 | 0.000738589 | 0.003349572 | ns | NaCl | KS.1 |
| FIM1_4873 | 1539.095162 | 2351.490212 | 1506.886808 | 0.508288396 | 0.184727911 | 0.001280429 | 0.005260228 | ns | NaCl | KS.1 |
| FIM1_4874 | 2915.126657 | 4367.967433 | 2687.585499 | 0.442264495 | 0.217877124 | 0.012290947 | 0.033523828 | ns | NaCl | KS.1 |
| FIM1_4875 | 1053.674533 | 443.0068402 | 1454.363538 | -1.023660455 | 0.267730883 | 3.7959180764 | 3.8235334126 | Km<KS.1 | NaCl | KS.1 |
| FIM1_4876 | 74.2681575 | 34.29657053 | 66.18190685 | -0.786382311 | 0.321814336 | 0.000919727 | 0.004010205 | ns | NaCl | KS.1 |
| FIM1_4877 | 754.6658256 | 791.6098481 | 1003.914225 | 0.04746379 | 0.199246048 | 0.772646297 | 0.852911429 | ns | NaCl | KS.1 |
| FIM1_4878 | 21095.6462 | 31776.44246 | 19245.88169 | 0.536547551 | 0.132899115 | 9.9143537800 | 8.4314376164 | ns | NaCl | KS.1 |
| FIM1_4879 | 2540.547666 | 3115.042405 | 2685.028214 | 0.280975094 | 0.07164618 | 5.0784526079 | 0.000354352 | ns | NaCl | KS.1 |
| FIM1_4880 | 4985.945637 | 6500.484721 | 4901.205305 | 0.368762945 | 0.0671491 | 1.5298449872 | 3.1712411715 | ns | NaCl | KS.1 |
| FIM1_4881 | 1870.595977 | 2373.905237 | 2085.045205 | 0.298091 | 0.131197317 | 0.012403866 | 0.033684079 | ns | NaCl | KS.1 |

| | | | | | | | | | | |
|---|---|---|---|---|---|---|---|---|---|---|
| FIM1_4882 | 65.39958993 | 38.71664803 | 1008.366534 | -0.276030879 | 0.390294796 | 0.20854806 | 0.330212157 | ns | NaCl | KS.1 |
| FIM1_4883 | 2991.536809 | 3116.629062 | 3633.9843 | 0.046414095 | 0.154372362 | 0.734025163 | 0.824700809 | ns | NaCl | KS.1 |
| FIM1_4884 | 1885.564207 | 2176.298636 | 2324.54675 | 0.19254859 | 0.088508404 | 0.022735163 | 0.055320674 | ns | NaCl | KS.1 |
| FIM1_4885 | 1141.256536 | 643.5966813 | 961.0478088 | -0.778851656 | 0.121887647 | 1.4440291775 | 6.5908671179 | ns | NaCl | KS.1 |
| FIM1_4886 | 322.1784062 | 189.6149651 | 262.1975986 | -0.652076812 | 0.185303852 | 5.1655583271 | 0.000358419 | ns | NaCl | KS.1 |
| FIM1_4887 | 409.8136458 | 340.0715041 | 366.9965181 | -0.230328064 | 0.12736311 | 0.047914295 | 0.10191262 | ns | NaCl | KS.1 |
| FIM1_4888 | 1000.571869 | 580.7920664 | 846.6949265 | -0.661528387 | 0.200857067 | 0.000113742 | 0.000693861 | ns | NaCl | KS.1 |
| FIM1_4889 | 535.0476446 | 235.3508686 | 630.0680574 | -1.028938735 | 0.218249221 | 7.8970154616 | 1.2881197351 | Km<KS.1 | NaCl | KS.1 |
| FIM1_4890 | 814.7780247 | 510.968191 | 592.0101904 | -0.565083428 | 0.188778617 | 0.000465719 | 0.002270514 | ns | NaCl | KS.1 |
| FIM1_4891 | 1498.227434 | 1253.838346 | 1119.446052 | -0.212664538 | 0.146039532 | 0.102872637 | 0.189062198 | ns | NaCl | KS.1 |
| FIM1_4892 | 5629.166805 | 6144.95609 | 4920.135313 | 0.083323472 | 0.211138301 | 0.625162466 | 0.738774851 | ns | NaCl | KS.1 |
| FIM1_4893 | 11462.50911 | 13674.19532 | 9611.487046 | 0.17266741 | 0.21107396 | 0.311768046 | 0.444553176 | ns | NaCl | KS.1 |
| FIM1_4894 | 1569.737735 | 1133.471258 | 1464.534456 | -0.373969458 | 0.183228434 | 0.016120031 | 0.041466989 | ns | NaCl | KS.1 |
| FIM1_4895 | 2241.910864 | 1789.211371 | 1695.610525 | -0.231365793 | 0.204126369 | 0.16632606 | 0.277210101 | ns | NaCl | KS.1 |
| FIM1_4896 | 862.2198757 | 809.0176899 | 1266.705427 | -0.062464478 | 0.200851333 | 0.705269685 | 0.802110918 | ns | NaCl | KS.1 |
| FIM1_4897 | 1094.060735 | 891.9792536 | 1069.82916 | -0.202200396 | 0.211574649 | 0.237179746 | 0.363963368 | ns | NaCl | KS.1 |
| FIM1_4898 | 1257.307802 | 817.6013627 | 920.0857228 | -0.429446304 | 0.252935412 | 0.027027478 | 0.06396846 | ns | NaCl | KS.1 |
| FIM1_4899 | 10061.26647 | 11041.27856 | 7998.009529 | 0.113169208 | 0.128914846 | 0.337304353 | 0.471373358 | ns | NaCl | KS.1 |
| FIM1_4900 | 5434.072677 | 6550.167367 | 4562.418037 | 0.222215993 | 0.146741706 | 0.08948526 | 0.16978992 | ns | NaCl | KS.1 |
| FIM1_4901 | 348.6706291 | 321.2565733 | 385.1794082 | -0.078280527 | 0.199232066 | 0.633715689 | 0.745151395 | ns | NaCl | KS.1 |
| FIM1_4902 | 1190.388247 | 734.9427412 | 1002.088481 | -0.621529104 | 0.157302093 | 1.0324530333 | 8.7354657161 | ns | NaCl | KS.1 |
| FIM1_4903 | 1794.889919 | 845.485417 | 1339.826362 | -1.016850578 | 0.148243933 | 2.8142010110 | 1.9719225394 | Km<KS.1 | NaCl | KS.1 |
| FIM1_4904 | 1373.325216 | 1268.361868 | 1271.668189 | -0.084416917 | 0.179330782 | 0.580566186 | 0.702070193 | ns | NaCl | KS.1 |
| FIM1_4905 | 337.2498266 | 277.906307 | 445.754003 | -0.190771007 | 0.212837332 | 0.266460571 | 0.395006359 | ns | NaCl | KS.1 |
| FIM1_4906 | 1866.996551 | 1501.135812 | 1602.179037 | -0.27924316 | 0.117124906 | 0.00991338 | 0.028360588 | ns | NaCl | KS.1 |
| FIM1_4907 | 2254.608813 | 4151.851616 | 2107.583726 | 0.566203286 | 0.314082987 | 0.011441436 | 0.031752647 | ns | NaCl | KS.1 |
| FIM1_4908 | 630.4436203 | 604.5596919 | 559.5296818 | -0.045668177 | 0.177076013 | 0.762960955 | 0.846883256 | ns | NaCl | KS.1 |
| FIM1_4909 | 3565.217912 | 4120.213581 | 2808.153758 | 0.180211766 | 0.122551925 | 0.110250145 | 0.199961529 | ns | NaCl | KS.1 |
| FIM1_4910 | 465.0275694 | 679.3241465 | 642.7546195 | 0.446633235 | 0.18372593 | 0.004285127 | 0.014346235 | ns | NaCl | KS.1 |
| FIM1_4911 | 1248.651131 | 1324.430451 | 1207.163562 | 0.060438623 | 0.186282563 | 0.700179819 | 0.798938005 | ns | NaCl | KS.1 |
| FIM1_4912 | 860.8699162 | 1115.568272 | 1002.936123 | 0.290348306 | 0.182756832 | 0.060704626 | 0.124384479 | ns | NaCl | KS.1 |
| FIM1_4913 | 1954.581445 | 4231.069395 | 1779.518337 | 0.916803331 | 0.250910509 | 1.1468774599 | 9.6380327081 | ns | NaCl | KS.1 |
| FIM1_4914 | 2206.773976 | 3739.201879 | 2391.945927 | 0.703956287 | 0.136461147 | 2.5423184140 | 5.0706512161 | ns | NaCl | KS.1 |
| FIM1_4915 | 1554.718977 | 1596.687216 | 2017.395644 | 0.024478245 | 0.219620899 | 0.888541877 | 0.933183795 | ns | NaCl | KS.1 |
| FIM1_4916 | 1386.314285 | 3038.815255 | 1357.458541 | 0.692747383 | 0.366578456 | 0.005059672 | 0.016441021 | ns | NaCl | KS.1 |
| FIM1_4917 | 1287.182575 | 1381.150128 | 1595.349315 | 0.086206225 | 0.122691461 | 0.446196884 | 0.581411603 | ns | NaCl | KS.1 |
| FIM1_4918 | 208.0879995 | 221.9858777 | 185.4433156 | 0.071091603 | 0.174654654 | 0.635314562 | 0.746678466 | ns | NaCl | KS.1 |
| FIM1_4919 | 1693.790596 | 2415.338462 | 1474.08409 | 0.392648104 | 0.202863025 | 0.019039271 | 0.047550389 | ns | NaCl | KS.1 |
| FIM1_4920 | 296.2952651 | 129.2800048 | 491.5382069 | -1.06827167 | 0.20644743 | 6.5892436038 | 1.5030550796 | Km<KS.1 | NaCl | KS.1 |
| FIM1_4921 | 12690.03112 | 14602.30751 | 10694.03251 | 0.168576108 | 0.137875936 | 0.175929127 | 0.290009082 | ns | NaCl | KS.1 |
| FIM1_4922 | 1384.251292 | 1417.604657 | 1043.283304 | 0.025853585 | 0.166822771 | 0.858435648 | 0.914122436 | ns | NaCl | KS.1 |
| FIM1_4923 | 1014.204115 | 800.7108199 | 779.2656105 | -0.312433236 | 0.106600599 | 0.001721501 | 0.006770332 | ns | NaCl | KS.1 |
| FIM1_4924 | 4950.028317 | 1361.891395 | 2324.548194 | -1.634851168 | 0.299265553 | 6.7047215312 | 2.5658453552 | Km<KS.1 | NaCl | KS.1 |
| FIM1_4925 | 210.2114211 | 242.4090609 | 244.9278878 | 0.10301524 | 0.283321888 | 0.604212713 | 0.722586117 | ns | NaCl | KS.1 |
| FIM1_4926 | 1133.090128 | 1275.579341 | 1363.095632 | 0.158471958 | 0.089387652 | 0.063377319 | 0.128432653 | ns | NaCl | KS.1 |

| | | | | | | | | | | |
|---|---|---|---|---|---|---|---|---|---|---|
| FIM1_4927 | 1398.278593 | 2512.375979 | 1324.343166 | 0.568822652 | 0.296363081 | 0.008875916 | 0.026128805 | ns | NaCl | KS.1 |
| FIM1_4928 | 1147.481478 | 2114.954817 | 1095.136634 | 0.581120847 | 0.308019694 | 0.008965293 | 0.026298545 | ns | NaCl | KS.1 |
| FIM1_4929 | 656.6921887 | 519.3688164 | 701.8857728 | -0.297521258 | 0.120756437 | 0.00742212 | 0.022570323 | ns | NaCl | KS.1 |
| FIM1_4931 | 5497.427353 | 5060.88192 | 3938.011143 | -0.092791342 | 0.159242952 | 0.507034104 | 0.638606245 | ns | NaCl | KS.1 |
| FIM1_4932 | 3792.73662 | 4369.835256 | 4932.710896 | 0.161963095 | 0.157426394 | 0.242175461 | 0.369011613 | ns | NaCl | KS.1 |
| FIM1_4933 | 2002.878802 | 1594.401549 | 1783.902986 | -0.265168114 | 0.161915705 | 0.060601186 | 0.124274897 | ns | NaCl | KS.1 |
| FIM1_4935 | 974.8149738 | 787.23498 | 1921.499279 | -0.237642698 | 0.174150901 | 0.111614407 | 0.201700571 | ns | NaCl | KS.1 |
| FIM1_4936 | 977.9945814 | 838.3994145 | 1229.813806 | -0.134292802 | 0.238962337 | 0.464722427 | 0.599272699 | ns | NaCl | KS.1 |
| FIM1_4937 | 5877.870933 | 6147.793692 | 6854.844582 | 0.037431423 | 0.2441298 | 0.839933096 | 0.902325017 | ns | NaCl | KS.1 |
| FIM1_4938 | 1283.310975 | 1593.658519 | 1253.715977 | 0.247282889 | 0.16619193 | 0.086355669 | 0.165238251 | ns | NaCl | KS.1 |
| FIM1_4939 | 787.1202099 | 972.3529388 | 997.7523455 | 0.259830016 | 0.134896647 | 0.033316428 | 0.075996896 | ns | NaCl | KS.1 |
| FIM1_4940 | 1881.324133 | 2895.498812 | 1454.603785 | 0.445268753 | 0.242968548 | 0.018988549 | 0.04750245 | ns | NaCl | KS.1 |
| FIM1_4941 | 8965.469904 | 13602.0614 | 8851.194038 | 0.560620131 | 0.114543173 | 1.60205572676 | 2.43989298426 | ns | NaCl | KS.1 |
| FIM1_4942 | 1211.338243 | 1241.885738 | 1224.514551 | 0.018709265 | 0.268021905 | 0.923039863 | 0.955696841 | ns | NaCl | KS.1 |
| FIM1_4943 | 1906.540251 | 1470.422133 | 2318.517883 | -0.310215194 | 0.154076646 | 0.022453924 | 0.054866539 | ns | NaCl | KS.1 |
| FIM1_4944 | 702.2297715 | 891.4626612 | 1183.17116 | 0.204795912 | 0.255265759 | 0.283730479 | 0.414189887 | ns | NaCl | KS.1 |
| FIM1_4945 | 8295.62073 | 8474.421506 | 7429.713428 | 0.023158226 | 0.16742174 | 0.873399131 | 0.924512682 | ns | NaCl | KS.1 |
| FIM1_4946 | 2726.334632 | 3777.715799 | 3010.056622 | 0.395056061 | 0.160938863 | 0.005033831 | 0.016368177 | ns | NaCl | KS.1 |
| FIM1_4947 | 1601.220253 | 1458.994896 | 1835.211868 | -0.114239669 | 0.12514264 | 0.320641757 | 0.453978808 | ns | NaCl | KS.1 |
| FIM1_4948 | 402.22281 | 347.4903736 | 502.0525132 | -0.174810232 | 0.144757346 | 0.177271661 | 0.291738841 | ns | NaCl | KS.1 |
| FIM1_4949 | 1176.542902 | 1220.48109 | 1482.360849 | 0.047739647 | 0.073495746 | 0.50289729 | 0.634925838 | ns | NaCl | KS.1 |
| FIM1_4950 | 3355.057955 | 2899.89853 | 3177.919928 | -0.193562842 | 0.09297149 | 0.02856495 | 0.066844133 | ns | NaCl | KS.1 |
| FIM1_4951 | 2255.325588 | 2144.270562 | 2430.954616 | -0.056507896 | 0.159141171 | 0.6861501 | 0.787936089 | ns | NaCl | KS.1 |
| FIM1_4952 | 6597.187225 | 15275.06849 | 6919.195298 | 0.948861842 | 0.289740386 | 3.80662010136 | 0.000273275 | ns | NaCl | KS.1 |
| FIM1_4953 | 2422.805202 | 5549.297944 | 2320.49064 | 0.845472287 | 0.333422472 | 0.000548165 | 0.002609684 | ns | NaCl | KS.1 |
| FIM1_4954 | 269.9519814 | 286.3499126 | 227.721582 | 0.04780016 | 0.24590638 | 0.797235964 | 0.870746195 | ns | NaCl | KS.1 |
| FIM1_4955 | 8682.011594 | 15784.4184 | 10080.4091 | 0.772447324 | 0.170112075 | 4.58533614998 | 6.05094094062 | ns | NaCl | KS.1 |
| FIM1_4956 | 833.7116826 | 1774.914073 | 2186.763112 | 0.950517043 | 0.211024774 | 2.92582555989 | 4.14700346452 | ns | NaCl | KS.1 |
| FIM1_4957 | 425.2395358 | 900.2608339 | 753.7054491 | 0.92905599 | 0.222198012 | 1.32738497227 | 1.50085005387 | ns | NaCl | KS.1 |
| FIM1_4958 | 239.2958102 | 38.23634814 | 219.4088314 | -2.134524596 | 0.551737146 | 4.93070192894 | 8.51744517240 | Km<KS.1 | NaCl | KS.1 |
| FIM1_4960 | 3988.924649 | 4704.998034 | 4081.875028 | 0.217687511 | 0.096497256 | 0.017214061 | 0.043895414 | ns | NaCl | KS.1 |
| FIM1_4961 | 3184.783882 | 4625.987461 | 3685.306776 | 0.412927585 | 0.207539524 | 0.015376933 | 0.039927058 | ns | NaCl | KS.1 |
| FIM1_4962 | 684.0261945 | 1218.580503 | 811.1420474 | 0.747512773 | 0.166840337 | 6.56806032706 | 8.21007540882 | ns | NaCl | KS.1 |
| FIM1_4963 | 722.4976067 | 622.5586969 | 718.4865154 | -0.187801763 | 0.114646624 | 0.07791561 | 0.151892695 | ns | NaCl | KS.1 |
| FIM1_4964 | 65.61999207 | 66.61893696 | 90.85514937 | 0.025785104 | 0.185573413 | 0.869170818 | 0.92179169 | ns | NaCl | KS.1 |
| FIM1_4965 | 8861.924552 | 10881.86337 | 7135.825911 | 0.262201778 | 0.116859878 | 0.015294636 | 0.039754866 | ns | NaCl | KS.1 |
| FIM1_4966 | 5017.396709 | 3087.307316 | 4089.50433 | -0.669525819 | 0.09848815 | 1.20454226110 | 7.58556677085 | ns | NaCl | KS.1 |
| FIM1_4967 | 1516.549441 | 1339.985807 | 1829.677499 | -0.150683046 | 0.126649888 | 0.194372473 | 0.312242196 | ns | NaCl | KS.1 |
| FIM1_4968 | 2380.630401 | 2802.299946 | 1696.526117 | 0.132321688 | 0.258644559 | 0.489317914 | 0.622629484 | ns | NaCl | KS.1 |
| FIM1_4969 | 14181.75021 | 14753.93034 | 13025.51777 | 0.053848158 | 0.073042946 | 0.44719588 | 0.58256075 | ns | NaCl | KS.1 |
| FIM1_4970 | 12442.03625 | 20378.92051 | 13733.02455 | 0.68121946 | 0.099064581 | 6.77558978614 | 4.37773495923 | ns | NaCl | KS.1 |
| FIM1_4971 | 4092.402891 | 4104.745718 | 5299.304992 | 0.003256788 | 0.171331105 | 0.982425362 | 0.991175946 | ns | NaCl | KS.1 |
| FIM1_4972 | 1876.432134 | 1396.455512 | 1907.458323 | -0.288767125 | 0.230542141 | 0.111040215 | 0.200888413 | ns | NaCl | KS.1 |
| FIM1_4973 | 1380.373628 | 1242.84356 | 1688.204938 | -0.110220393 | 0.182130015 | 0.475450479 | 0.608845852 | ns | NaCl | KS.1 |
| FIM1_4974 | 425.6136461 | 446.8513448 | 495.5369414 | 0.061532242 | 0.122668947 | 0.586681536 | 0.708088462 | ns | NaCl | KS.1 |

| | | | | | | | | | | |
|---|---|---|---|---|---|---|---|---|---|---|
| FIM1_4976 | 7650.960401 | 10093.01704 | 7599.367736 | 0.359862897 | 0.117199309 | 0.000885908 | 0.003879748 | ns | NaCl | KS.1 |
| FIM1_4977 | 2425.947683 | 3801.37078 | 1707.165664 | 0.337468785 | 0.321008305 | 0.117550822 | 0.210820238 | ns | NaCl | KS.1 |
| FIM1_4978 | 729.5392966 | 563.201029 | 947.8801932 | -0.317556575 | 0.144133737 | 0.01369181 | 0.036562938 | ns | NaCl | KS.1 |
| FIM1_4979 | 19249.50852 | 20087.67168 | 16927.46391 | 0.056113596 | 0.09182136 | 0.521843056 | 0.652118833 | ns | NaCl | KS.1 |
| FIM1_4980 | 270.2045846 | 263.653607 | 556.3859116 | -0.02821014 | 0.171431227 | 0.848733629 | 0.907797604 | ns | NaCl | KS.1 |
| FIM1_4981 | 19397.95133 | 22447.52089 | 15859.94697 | 0.201023741 | 0.067419468 | 0.002167605 | 0.008157214 | ns | NaCl | KS.1 |
| FIM1_4982 | 883.928845 | 937.9804129 | 1010.171553 | 0.077898248 | 0.077680065 | 0.299262709 | 0.431921084 | ns | NaCl | KS.1 |
| FIM1_4983 | 901.3655436 | 792.3320379 | 976.9536087 | -0.166622471 | 0.105827354 | 0.093553041 | 0.175786225 | ns | NaCl | KS.1 |
| FIM1_4984 | 733.6246202 | 560.0125213 | 1549.734645 | -0.313404407 | 0.16993605 | 0.032599886 | 0.074715977 | ns | NaCl | KS.1 |
| FIM1_4985 | 2321.287096 | 2092.322614 | 2672.857618 | -0.136217369 | 0.093054092 | 0.124116097 | 0.219196871 | ns | NaCl | KS.1 |
| FIM1_4986 | 3677.846999 | 4032.032396 | 4455.986282 | 0.122045817 | 0.087935958 | 0.146925874 | 0.250670859 | ns | NaCl | KS.1 |
| FIM1_4987 | 3693.137542 | 1876.770513 | 2820.403304 | -0.864061451 | 0.189142006 | 2.99264479525 | 4.21767927942 | ns | NaCl | KS.1 |
| FIM1_4988 | 159.1998516 | 121.9297108 | 174.4108259 | -0.293509141 | 0.187504 | 0.062743037 | 0.12740678 | ns | NaCl | KS.1 |
| FIM1_4989 | 5777.215818 | 9724.403763 | 7408.666665 | 0.663648619 | 0.168892391 | 9.74698969346 | 8.34617447934 | ns | NaCl | KS.1 |
| FIM1_4991 | 3399.345863 | 4031.384391 | 3383.67469 | 0.184141582 | 0.18003161 | 0.229063892 | 0.354580092 | ns | NaCl | KS.1 |
| FIM1_4992 | 1096.066457 | 955.1415055 | 1100.95135 | -0.1503741 | 0.172159515 | 0.310321756 | 0.443125929 | ns | NaCl | KS.1 |
| FIM1_4993 | 2152.334662 | 2707.320669 | 2479.73763 | 0.263754042 | 0.16567076 | 0.066688468 | 0.133424694 | ns | NaCl | KS.1 |
| FIM1_4995 | 1358.080043 | 2233.087738 | 1753.530347 | 0.586572206 | 0.207317615 | 0.000722841 | 0.003281143 | ns | NaCl | KS.1 |
| FIM1_4996 | 472.8905489 | 332.9369412 | 533.7981596 | -0.414571067 | 0.176872161 | 0.006258767 | 0.019681154 | ns | NaCl | KS.1 |
| FIM1_4997 | 1638.470148 | 1297.682793 | 1646.699176 | -0.296924677 | 0.12475173 | 0.009357513 | 0.027132242 | ns | NaCl | KS.1 |
| FIM1_4998 | 1998.523345 | 2111.258347 | 2531.997194 | 0.063110159 | 0.154272042 | 0.643868098 | 0.753349903 | ns | NaCl | KS.1 |
| FIM1_4999 | 4213.867014 | 5690.952955 | 3751.186881 | 0.368329178 | 0.150807814 | 0.005854965 | 0.018675443 | ns | NaCl | KS.1 |
| FIM1_5000 | 455.5226543 | 439.8103892 | 621.3457797 | -0.032059622 | 0.217720237 | 0.853611733 | 0.911117436 | ns | NaCl | KS.1 |
| FIM1_5001 | 507.4925475 | 704.1072047 | 1113.903104 | 0.317738636 | 0.240342551 | 0.088005497 | 0.167365195 | ns | NaCl | KS.1 |
| FIM1_5002 | 1338.751102 | 1270.120608 | 1846.561683 | -0.059396042 | 0.15790802 | 0.669164363 | 0.77492847 | ns | NaCl | KS.1 |
| FIM1_5003 | 1645.315719 | 900.4615047 | 1211.205547 | -0.803697661 | 0.14875109 | 5.12202579838 | 1.23101827763 | ns | NaCl | KS.1 |
| FIM1_5004 | 3968.097682 | 4557.535384 | 3222.634587 | 0.132570963 | 0.213167985 | 0.439972526 | 0.575410967 | ns | NaCl | KS.1 |
| FIM1_5005 | 24037.20441 | 33410.83841 | 20594.94691 | 0.43457561 | 0.115110107 | 4.63343209707 | 0.000326506 | ns | NaCl | KS.1 |
| FIM1_5006 | 209.5930023 | 167.1003132 | 269.8336058 | -0.254194952 | 0.179583737 | 0.096175383 | 0.179404774 | ns | NaCl | KS.1 |
| FIM1_5007 | 2107.345359 | 3049.802332 | 2601.176588 | 0.310888411 | 0.282588061 | 0.126204358 | 0.222175047 | ns | NaCl | KS.1 |
| FIM1_5008 | 6402.542154 | 4751.124644 | 4296.317491 | -0.380939761 | 0.129930412 | 0.001273204 | 0.005239199 | ns | NaCl | KS.1 |
| FIM1_5009 | 276.2620394 | 21.94893578 | 119.1624388 | -3.34788814 | 0.431561524 | 8.74816883167 | 1.20894833159 | Km<KS.1 | NaCl | KS.1 |
| FIM1_5010 | 1348.286345 | 983.4236512 | 1071.331298 | -0.349015878 | 0.195798664 | 0.032192794 | 0.073908236 | ns | NaCl | KS.1 |
| FIM1_5011 | 185.624892 | 196.9311816 | 171.8248649 | 0.05787283 | 0.186064401 | 0.712106414 | 0.807735844 | ns | NaCl | KS.1 |
| FIM1_5012 | 960.0648704 | 908.9781635 | 962.7448218 | -0.060288026 | 0.164562912 | 0.674276011 | 0.77867297 | ns | NaCl | KS.1 |
| FIM1_5013 | 1352.470695 | 1070.501248 | 1207.073186 | -0.290460816 | 0.135747361 | 0.017884245 | 0.045302503 | ns | NaCl | KS.1 |
| FIM1_5014 | 1985.237077 | 1633.220454 | 2375.999414 | -0.253675578 | 0.105817695 | 0.010506909 | 0.029699928 | ns | NaCl | KS.1 |
| FIM1_5015 | 3819.194836 | 3133.634606 | 2910.350965 | -0.227166127 | 0.162475681 | 0.10902592 | 0.198247059 | ns | NaCl | KS.1 |
| FIM1_5016 | 464.5800722 | 342.8428696 | 752.5604019 | -0.307238938 | 0.220038483 | 0.081052963 | 0.156780129 | ns | NaCl | KS.1 |
| FIM1_5017 | 2542.701939 | 2598.393492 | 2331.695015 | 0.022709495 | 0.180576421 | 0.88243832 | 0.930540801 | ns | NaCl | KS.1 |
| FIM1_5018 | 377.9371959 | 299.2524038 | 382.6164024 | -0.270325914 | 0.167414854 | 0.062243967 | 0.126703656 | ns | NaCl | KS.1 |
| FIM1_5019 | 907.6510194 | 1155.21211 | 1121.73002 | 0.236254941 | 0.220649942 | 0.17917579 | 0.294288398 | ns | NaCl | KS.1 |
| FIM1_5020 | 712.3974238 | 1027.120689 | 1100.992639 | 0.322108357 | 0.271713802 | 0.106641203 | 0.194693573 | ns | NaCl | KS.1 |
| FIM1_5021 | 527.0346687 | 324.7502574 | 757.4105412 | -0.645602769 | 0.137272408 | 3.11122319425 | 4.34784702005 | ns | NaCl | KS.1 |
| FIM1_5022 | 1768.483441 | 2202.101494 | 2655.25825 | 0.289949156 | 0.100133001 | 0.002111004 | 0.00797413 | ns | NaCl | KS.1 |

| | | | | | | | | | | |
|---|---|---|---|---|---|---|---|---|---|---|
| FIM1_5023 | 2359.695714 | 2691.700232 | 1964.965564 | 0.169097214 | 0.104746925 | 0.085888439 | 0.16452469 | ns | NaCl | KS.1 |
| FIM1_5024 | 3590.003534 | 4947.230711 | 3413.721675 | 0.417910021 | 0.122062647 | 0.000195278 | 0.00109651 | ns | NaCl | KS.1 |
| FIM1_5025 | 2748.54085 | 2636.643941 | 2924.38756 | -0.043735745 | 0.184362506 | 0.778964154 | 0.856051837 | ns | NaCl | KS.1 |
| FIM1_5026 | 1446.675626 | 2999.010318 | 2627.883575 | 0.988403653 | 0.141284786 | 1.21514394476 | 9.59577956384 | ns | NaCl | KS.1 |
| FIM1_5027 | 4693.449445 | 6549.837411 | 3523.227109 | 0.393168077 | 0.173901042 | 0.008562782 | 0.025432739 | ns | NaCl | KS.1 |
| FIM1_5028 | 2755.21195 | 3985.197096 | 3668.055305 | 0.498811026 | 0.104613549 | 3.90438210142 | 5.24620712218 | ns | NaCl | KS.1 |
| FIM1_5029 | 2074.917406 | 2595.228889 | 2781.189831 | 0.303200064 | 0.086321348 | 0.000234693 | 0.001277946 | ns | NaCl | KS.1 |
| FIM1_5030 | 3883.170127 | 3676.131807 | 3657.061808 | -0.073811011 | 0.075197551 | 0.310622235 | 0.443427725 | ns | NaCl | KS.1 |
| FIM1_5031 | 5846.542694 | 10325.12803 | 5372.95306 | 0.766886211 | 0.130813074 | 4.01664926102 | 1.28095064574 | ns | NaCl | KS.1 |
| FIM1_5032 | 2476.466581 | 5110.751422 | 2251.585274 | 0.759042733 | 0.301892929 | 0.000868177 | 0.003818906 | ns | NaCl | KS.1 |
| FIM1_5033 | 1496.082839 | 1490.551114 | 1764.733732 | -0.004661783 | 0.076555241 | 0.949804967 | 0.973834262 | ns | NaCl | KS.1 |
| FIM1_5034 | 3253.765202 | 3272.112361 | 4885.437389 | 0.008017526 | 0.123564204 | 0.943893901 | 0.969400953 | ns | NaCl | KS.1 |
| FIM1_5035 | 1302.240776 | 1044.414164 | 1070.618224 | -0.229442604 | 0.199383157 | 0.163115523 | 0.272676676 | ns | NaCl | KS.1 |
| FIM1_5036 | 1620.678417 | 1582.868342 | 1330.740806 | -0.027306056 | 0.156003618 | 0.842850232 | 0.903702566 | ns | NaCl | KS.1 |
| FIM1_5037 | 1208.748666 | 1264.01997 | 1385.984308 | 0.061606412 | 0.080896977 | 0.429340964 | 0.566166699 | ns | NaCl | KS.1 |
| FIM1_5038 | 2960.257249 | 4095.528317 | 3473.213098 | 0.431415161 | 0.109656828 | 2.44502543205 | 0.000184792 | ns | NaCl | KS.1 |
| FIM1_5039 | 1177.18981 | 1107.845547 | 1431.105986 | -0.074291117 | 0.126843 | 0.523392271 | 0.653091685 | ns | NaCl | KS.1 |
| FIM1_5040 | 18345.13319 | 25623.51719 | 15960.82951 | 0.427269883 | 0.135010549 | 0.000473357 | 0.002293041 | ns | NaCl | KS.1 |
| FIM1_5041 | 439.7402754 | 414.7641949 | 428.8023825 | -0.060885392 | 0.152438312 | 0.652582264 | 0.76014909 | ns | NaCl | KS.1 |
| FIM1_5042 | 17182.05417 | 22817.9607 | 18192.75871 | 0.35177433 | 0.142695774 | 0.005902091 | 0.018762237 | ns | NaCl | KS.1 |
| FIM1_5043 | 1099.005288 | 919.4596699 | 914.906838 | -0.200633775 | 0.167923654 | 0.167626245 | 0.278630327 | ns | NaCl | KS.1 |
| FIM1_5044 | 316.8041855 | 244.9695546 | 237.6434499 | -0.264127435 | 0.211124887 | 0.122649342 | 0.217610726 | ns | NaCl | KS.1 |
| FIM1_5045 | 2057.143271 | 1673.251238 | 2635.336793 | -0.232533933 | 0.172302438 | 0.116611596 | 0.209286684 | ns | NaCl | KS.1 |
| FIM1_5046 | 2462.723801 | 2624.792831 | 3462.930132 | 0.056564492 | 0.228759897 | 0.751884167 | 0.838250702 | ns | NaCl | KS.1 |
| FIM1_5047 | 2252.832644 | 2727.233202 | 1684.120007 | 0.243409862 | 0.115482161 | 0.022982423 | 0.055801637 | ns | NaCl | KS.1 |
| FIM1_5048 | 577.970681 | 556.3243665 | 648.1284846 | -0.047911881 | 0.116864576 | 0.659043933 | 0.765882637 | ns | NaCl | KS.1 |
| FIM1_5049 | 2296.055271 | 2584.352628 | 2315.334913 | 0.153789695 | 0.100386243 | 0.104843615 | 0.191834124 | ns | NaCl | KS.1 |
| FIM1_5050 | 1536.583891 | 1435.989356 | 2158.68306 | -0.078512359 | 0.149227675 | 0.554899002 | 0.678854977 | ns | NaCl | KS.1 |
| FIM1_5051 | 17.1786512 | 13.09068928 | 26.75475964 | -0.183867798 | 0.317477069 | 0.37737232 | 0.51220762 | ns | NaCl | KS.1 |
| FIM1_5052 | 941.2740065 | 997.720181 | 833.6461928 | 0.073563563 | 0.120256576 | 0.508505134 | 0.639487624 | ns | NaCl | KS.1 |
| FIM1_5053 | 1489.067646 | 1673.790433 | 1307.8111 | 0.142301463 | 0.13034845 | 0.231662998 | 0.357596713 | ns | NaCl | KS.1 |
| FIM1_5054 | 667.2725385 | 644.9971265 | 670.2714311 | -0.03980937 | 0.146242657 | 0.761040681 | 0.845317569 | ns | NaCl | KS.1 |
| FIM1_5055 | 3936.470456 | 4475.098213 | 3470.755075 | 0.162540148 | 0.112981336 | 0.122498158 | 0.217420028 | ns | NaCl | KS.1 |
| FIM1_5056 | 1037.122223 | 980.882791 | 1420.427163 | -0.06662188 | 0.124960248 | 0.562270838 | 0.685276193 | ns | NaCl | KS.1 |
| FIM1_5057 | 5709.734109 | 14742.80817 | 6219.004623 | 0.990533671 | 0.349430515 | 0.000121372 | 0.000733688 | ns | NaCl | KS.1 |
| FIM1_5058 | 2945.092497 | 2127.287486 | 2622.191853 | -0.359409563 | 0.198436167 | 0.028966183 | 0.067624007 | ns | NaCl | KS.1 |
| FIM1_5059 | 755.0042104 | 662.1970078 | 686.9090204 | -0.173649561 | 0.091251571 | 0.045827433 | 0.09827219 | ns | NaCl | KS.1 |
| FIM1_5061 | 2564.909317 | 2821.486724 | 2153.880053 | 0.115053932 | 0.133954298 | 0.344733663 | 0.479600104 | ns | NaCl | KS.1 |
| FIM1_5062 | 466.3456101 | 362.1274081 | 633.3694877 | -0.313928085 | 0.129121478 | 0.007601509 | 0.023031368 | ns | NaCl | KS.1 |
| FIM1_5063 | 2197.290748 | 2451.158966 | 1990.11932 | 0.132001563 | 0.131290964 | 0.270190622 | 0.399108772 | ns | NaCl | KS.1 |
| FIM1_5064 | 657.4512472 | 905.2455059 | 645.2072038 | 0.344425444 | 0.206499136 | 0.041656593 | 0.09097522 | ns | NaCl | KS.1 |
| FIM1_5065 | 2830.474821 | 4342.150053 | 3945.297633 | 0.501484929 | 0.195969806 | 0.002344787 | 0.00870546 | ns | NaCl | KS.1 |
| FIM1_5066 | 487.8863168 | 410.3701135 | 475.1938222 | -0.22514039 | 0.114370117 | 0.034019686 | 0.077346602 | ns | NaCl | KS.1 |
| FIM1_5067 | 4338.342589 | 4651.386165 | 3488.00408 | 0.088094899 | 0.110437779 | 0.393860616 | 0.529869271 | ns | NaCl | KS.1 |
| FIM1_5068 | 1590.225006 | 1064.76627 | 1083.197022 | -0.480537439 | 0.181299749 | 0.001962161 | 0.007526407 | ns | NaCl | KS.1 |

| | | | | | | | | | | |
|---|---|---|---|---|---|---|---|---|---|---|
| FIM1_5069 | 123.3826249 | 131.0099609 | 1090.397409 | 0.049305263 | 0.273745943 | 0.800895756 | 0.87263609 | ns | NaCl | KS.1 |
| FIM1_5070 | 55.35019175 | 39.59783198 | 66.93085334 | -0.308837776 | 0.239115266 | 0.096018008 | 0.179178391 | ns | NaCl | KS.1 |
| FIM1_5071 | 10152.56107 | 7458.488841 | 7131.434308 | -0.403851422 | 0.116949121 | 0.000186748 | 0.001054567 | ns | NaCl | KS.1 |
| FIM1_5072 | 145.3462269 | 65.63955847 | 128.7444275 | -0.902795304 | 0.273725953 | 4.30990070756 | 0.000305004 | ns | NaCl | KS.1 |
| FIM1_5073 | 760.8506859 | 505.169937 | 611.9560529 | -0.496261458 | 0.18015705 | 0.001336033 | 0.00543638 | ns | NaCl | KS.1 |
| FIM1_5074 | 831.9769664 | 534.2984968 | 663.8664992 | -0.555719845 | 0.167165923 | 0.000153931 | 0.00089459 | ns | NaCl | KS.1 |
| FIM1_5075 | 883.1143939 | 606.3308142 | 1277.11883 | -0.454172931 | 0.172290054 | 0.002299408 | 0.008575379 | ns | NaCl | KS.1 |
| FIM1_5076 | 4050.022998 | 4495.618893 | 3651.790514 | 0.131260418 | 0.115393708 | 0.221088785 | 0.345668355 | ns | NaCl | KS.1 |
| FIM1_5077 | 2853.458148 | 1853.345478 | 2209.852189 | -0.542354652 | 0.162422398 | 0.000154103 | 0.00089459 | ns | NaCl | KS.1 |
| FIM1_5078 | 375.5308482 | 297.4293987 | 454.6684508 | -0.279005953 | 0.148408671 | 0.034445814 | 0.078107531 | ns | NaCl | KS.1 |
| FIM1_5079 | 8353.812508 | 7455.560426 | 6011.899769 | -0.141186438 | 0.122993799 | 0.212670074 | 0.335138935 | ns | NaCl | KS.1 |
| FIM1_5080 | 1250.78466 | 939.1600999 | 1060.424088 | -0.400399747 | 0.071366194 | 6.61646356663 | 1.50305507963 | ns | NaCl | KS.1 |
| FIM1_5081 | 1758.080934 | 1586.842899 | 1610.532415 | -0.129714596 | 0.110733302 | 0.210052917 | 0.332383352 | ns | NaCl | KS.1 |
| FIM1_5082 | 522.885013 | 743.2647618 | 557.9162857 | 0.445457413 | 0.1416819 | 0.000468219 | 0.002279247 | ns | NaCl | KS.1 |
| FIM1_5083 | 4530.981239 | 6238.763523 | 4870.532015 | 0.334479155 | 0.215056 | 0.05418061 | 0.11287627 | ns | NaCl | KS.1 |
| FIM1_5084 | 809.4841051 | 1170.241126 | 802.1607 | 0.345831465 | 0.256602878 | 0.074690087 | 0.146638982 | ns | NaCl | KS.1 |
| FIM1_5085 | 1255.39541 | 1430.131449 | 1763.991479 | 0.129664856 | 0.200690612 | 0.432239064 | 0.56888607 | ns | NaCl | KS.1 |
| FIM1_5086 | 2116.014883 | 2765.977422 | 2732.781178 | 0.338596276 | 0.130477936 | 0.004313351 | 0.014421319 | ns | NaCl | KS.1 |
| FIM1_5087 | 1145.612004 | 1285.107212 | 1469.97806 | 0.124866528 | 0.17201683 | 0.399378266 | 0.535410098 | ns | NaCl | KS.1 |
| FIM1_5088 | 8342.366252 | 11718.2715 | 9428.848193 | 0.36000827 | 0.214927152 | 0.038441092 | 0.085802289 | ns | NaCl | KS.1 |
| FIM1_5089 | 3637.951417 | 2407.173468 | 2468.934404 | -0.422528171 | 0.241709738 | 0.025159465 | 0.060293034 | ns | NaCl | KS.1 |
| FIM1_5090 | 319.9312861 | 232.4156495 | 387.9351323 | -0.277344964 | 0.263084873 | 0.155626577 | 0.262633047 | ns | NaCl | KS.1 |
| FIM1_5091 | 971.4919625 | 235.9848688 | 492.4874872 | -1.839053237 | 0.296537642 | 3.34982998348 | 2.31463946775 | Km<KS.1 | NaCl | KS.1 |
| FIM1_5093 | 1426.277748 | 1009.613675 | 1273.576786 | -0.460708108 | 0.103738841 | 2.25470990419 | 2.39683371226 | ns | NaCl | KS.1 |
| FIM1_5094 | 101066.831 | 141388.2164 | 89508.85285 | 0.450268813 | 0.10509134 | 4.86312201858 | 4.62600995075 | ns | NaCl | KS.1 |
| FIM1_5095 | 520.9079469 | 577.7023116 | 503.4258247 | 0.123903945 | 0.136637616 | 0.316731971 | 0.449441402 | ns | NaCl | KS.1 |
| FIM1_5096 | 1489.277845 | 2181.282487 | 1407.574219 | 0.45689814 | 0.177350607 | 0.002709521 | 0.009832141 | ns | NaCl | KS.1 |
| FIM1_5097 | 9558.070656 | 13343.86594 | 9492.802646 | 0.46242471 | 0.076905524 | 4.37222305968 | 1.35104408210 | ns | NaCl | KS.1 |
| FIM1_5098 | 257.7941135 | 272.8223188 | 297.09687 | 0.056552323 | 0.198863821 | 0.730341144 | 0.822046876 | ns | NaCl | KS.1 |
| FIM1_5099 | 2257.525838 | 2553.18031 | 1756.661765 | 0.153194368 | 0.120387241 | 0.168333821 | 0.279713012 | ns | NaCl | KS.1 |
| FIM1_5100 | 862.7865434 | 947.288653 | 1058.239174 | 0.127366243 | 0.07028324 | 0.062083514 | 0.126480541 | ns | NaCl | KS.1 |
| FIM1_5101 | 1038.170422 | 768.3675887 | 1189.433938 | -0.388502893 | 0.126641132 | 0.000786051 | 0.003519897 | ns | NaCl | KS.1 |
| FIM1_5102 | 2022.212154 | 2131.404812 | 3568.104286 | 0.049209754 | 0.214379368 | 0.774974417 | 0.853721994 | ns | NaCl | KS.1 |
| FIM1_5103 | 684.0140018 | 438.109093 | 511.8376738 | -0.577491903 | 0.149539013 | 1.76384987265 | 0.000140627 | ns | NaCl | KS.1 |
| FIM1_5104 | 798.3695814 | 736.2606815 | 743.3096505 | -0.107587853 | 0.093673106 | 0.22756299 | 0.352907069 | ns | NaCl | KS.1 |
| FIM1_5105 | 2804.956204 | 2998.547212 | 2556.219037 | 0.089944842 | 0.074986342 | 0.215348533 | 0.338502039 | ns | NaCl | KS.1 |
| FIM1_5106 | 594.4471469 | 397.2134932 | 873.3045665 | -0.503412385 | 0.156699942 | 0.000285598 | 0.001506535 | ns | NaCl | KS.1 |
| FIM1_5107 | 4533.112663 | 6398.557261 | 4318.641348 | 0.432358983 | 0.14726902 | 0.000997255 | 0.004278248 | ns | NaCl | KS.1 |
| FIM1_5108 | 1131.950015 | 1147.130852 | 919.5715219 | 0.015688181 | 0.143616142 | 0.903227004 | 0.942440089 | ns | NaCl | KS.1 |
| FIM1_5109 | 1038.229481 | 1452.926923 | 1055.894387 | 0.410998282 | 0.157736363 | 0.003045953 | 0.010831749 | ns | NaCl | KS.1 |
| FIM1_5110 | 1022.330394 | 1331.385985 | 1771.777108 | 0.340499993 | 0.119129873 | 0.001926114 | 0.007410996 | ns | NaCl | KS.1 |
| FIM1_5111 | 1542.950837 | 2184.425342 | 1833.436691 | 0.463725339 | 0.1091585 | 5.38373401896 | 5.02515511150 | ns | NaCl | KS.1 |
| FIM1_5112 | 871.873715 | 884.5709525 | 912.3996043 | 0.016527134 | 0.118284297 | 0.880292068 | 0.929031192 | ns | NaCl | KS.1 |
| FIM1_5113 | 179.7185825 | 196.84272 | 185.8773579 | 0.095804009 | 0.184635176 | 0.539023796 | 0.665915914 | ns | NaCl | KS.1 |
| FIM1_5114 | 479.5460259 | 725.1905274 | 473.4386546 | 0.462445091 | 0.213368519 | 0.008031547 | 0.024128591 | ns | NaCl | KS.1 |

| | | | | | | | | | | |
|---|---|---|---|---|---|---|---|---|---|---|
| FIM1_5115 | 1949.88504 | 2700.836003 | 1717.039464 | 0.42230483 | 0.126106746 | 0.000251953 | 0.001356645 | ns | NaCl | KS.1 |
| FIM1_5116 | 1820.744193 | 1739.464069 | 1592.914278 | -0.056431081 | 0.118484256 | 0.607534504 | 0.724816345 | ns | NaCl | KS.1 |
| FIM1_5117 | 545.7697917 | 566.9025448 | 553.3084487 | 0.036337884 | 0.209456239 | 0.830404332 | 0.895382834 | ns | NaCl | KS.1 |
| FIM1_5118 | 3893.91929 | 3848.821465 | 2917.278492 | -0.013928332 | 0.133965077 | 0.909039714 | 0.946531862 | ns | NaCl | KS.1 |
| FIM1_5119 | 1760.758474 | 1203.616434 | 1408.965236 | -0.505545182 | 0.11888454 | 4.3840592347 | 4.2766068025 | ns | NaCl | KS.1 |
| FIM1_5120 | 1183.502441 | 1182.664152 | 1218.891875 | 0.000143554 | 0.191824509 | 0.999284842 | 0.999786056 | ns | NaCl | KS.1 |
| FIM1_5121 | 2576.464993 | 2614.020712 | 2267.146921 | 0.016430842 | 0.137977804 | 0.89534294 | 0.937556541 | ns | NaCl | KS.1 |
| FIM1_5122 | 2629.98491 | 3758.404486 | 4430.879936 | 0.299866056 | 0.280417501 | 0.13814835 | 0.238558849 | ns | NaCl | KS.1 |
| FIM1_5123 | 5138.349804 | 6023.391997 | 4067.119462 | 0.122960843 | 0.2693398 | 0.528125987 | 0.656528432 | ns | NaCl | KS.1 |
| FIM1_5124 | 904.2223141 | 1045.83491 | 969.6212758 | 0.185586833 | 0.105005827 | 0.059891901 | 0.122972433 | ns | NaCl | KS.1 |
| FIM1_5125 | 3399.921516 | 4025.551467 | 2718.883313 | 0.149374345 | 0.237987128 | 0.415316092 | 0.551427157 | ns | NaCl | KS.1 |
| FIM1_5126 | 1777.931871 | 1775.004295 | 1546.701462 | -0.001965481 | 0.073159821 | 0.977903198 | 0.989137805 | ns | NaCl | KS.1 |
| FIM1_5127 | 4054.227076 | 5062.147896 | 4733.280554 | 0.269505321 | 0.143908983 | 0.036270184 | 0.081628503 | ns | NaCl | KS.1 |
| FIM1_5128 | 2987.200544 | 3398.823667 | 3394.131703 | 0.171233864 | 0.095343627 | 0.058462817 | 0.120585619 | ns | NaCl | KS.1 |
| FIM1_5129 | 1383.424649 | 1551.230746 | 1408.446859 | 0.138750918 | 0.130765736 | 0.244810808 | 0.372326961 | ns | NaCl | KS.1 |
| FIM1_5130 | 2750.591319 | 1958.573876 | 2394.966011 | -0.411686804 | 0.161777234 | 0.00362395 | 0.012555118 | ns | NaCl | KS.1 |
| FIM1_5131 | 1761.181386 | 752.2531042 | 2682.725242 | -0.716277297 | 0.392215814 | 0.005000367 | 0.016291306 | ns | NaCl | KS.1 |
| FIM1_5132 | 173.977977 | 206.6693615 | 193.6028101 | 0.124307418 | 0.286744567 | 0.534226649 | 0.66212695 | ns | NaCl | KS.1 |
| FIM1_5133 | 1344.472213 | 1042.795095 | 1610.349592 | -0.308798413 | 0.147365121 | 0.018546284 | 0.046599881 | ns | NaCl | KS.1 |
| FIM1_5134 | 785.4044347 | 796.8015163 | 596.8297995 | 0.015859341 | 0.159034071 | 0.909678274 | 0.946986695 | ns | NaCl | KS.1 |
| FIM1_5135 | 1423.104995 | 1652.224419 | 1370.994416 | 0.174615781 | 0.147543912 | 0.184423099 | 0.300230666 | ns | NaCl | KS.1 |
| FIM1_5136 | 3091.893907 | 2618.274468 | 2722.172896 | -0.203544082 | 0.132387432 | 0.090784872 | 0.17173184 | ns | NaCl | KS.1 |
| FIM1_5137 | 1257.861607 | 929.0807089 | 955.2377765 | -0.353372009 | 0.173689972 | 0.017873195 | 0.045297578 | ns | NaCl | KS.1 |
| FIM1_5138 | 2938.90764 | 2619.045204 | 2129.186013 | -0.105333305 | 0.223767277 | 0.551242451 | 0.675808574 | ns | NaCl | KS.1 |
| FIM1_5139 | 3120.302881 | 3004.878627 | 3263.840028 | -0.033717892 | 0.225349887 | 0.849202715 | 0.907969806 | ns | NaCl | KS.1 |
| FIM1_5140 | 3756.54874 | 3353.076951 | 3280.874518 | -0.099528102 | 0.234415675 | 0.583529635 | 0.704968416 | ns | NaCl | KS.1 |
| FIM1_5141 | 1922.795801 | 2005.344725 | 2740.679032 | 0.047628214 | 0.146668969 | 0.716589134 | 0.812105833 | ns | NaCl | KS.1 |
| FIM1_5142 | 1518.190318 | 1245.462461 | 1638.967762 | -0.200277479 | 0.204764192 | 0.231602718 | 0.357596713 | ns | NaCl | KS.1 |
| FIM1_5143 | 2519.238231 | 2684.475122 | 2447.338046 | 0.083443557 | 0.08656855 | 0.31462071 | 0.447722549 | ns | NaCl | KS.1 |
| FIM1_5144 | 2164.469974 | 1328.933281 | 1888.397872 | -0.658810648 | 0.124192916 | 1.313966712 | 2.735139914 | ns | NaCl | KS.1 |
| FIM1_5145 | 1904.134846 | 2063.184206 | 2312.449822 | 0.09912253 | 0.127749369 | 0.397198281 | 0.533078153 | ns | NaCl | KS.1 |
| FIM1_5146 | 528.3407034 | 1157.007992 | 627.1426176 | 0.938840956 | 0.247436595 | 6.142508985 | 5.566299126 | ns | NaCl | KS.1 |
| FIM1_5147 | 33415.37487 | 47877.84654 | 29081.46093 | 0.473482073 | 0.121475353 | 2.346333783 | 0.00017876 | ns | NaCl | KS.1 |
| FIM1_5148 | 349.1083862 | 206.2885067 | 298.8760442 | -0.555924157 | 0.25813142 | 0.005485063 | 0.017639423 | ns | NaCl | KS.1 |
| FIM1_5149 | 6051.919136 | 917.7863128 | 4101.836441 | -2.392886606 | 0.444896866 | 3.675072912 | 1.451070455 | Km<KS.1 | NaCl | KS.1 |
| FIM1_5150 | 9056.591461 | 13529.72371 | 9268.607682 | 0.447461616 | 0.210336403 | 0.009480547 | 0.027358308 | ns | NaCl | KS.1 |
| FIM1_5151 | 262.8036669 | 233.0315582 | 223.4158694 | -0.143654317 | 0.154526439 | 0.292737689 | 0.424103088 | ns | NaCl | KS.1 |
| FIM1_5152 | 220.0971143 | 110.9064176 | 293.9752473 | -0.744950505 | 0.283051301 | 0.000669957 | 0.003082155 | ns | NaCl | KS.1 |
| FIM1_5153 | 5020.963298 | 7594.508216 | 6107.404169 | 0.566000074 | 0.099136846 | 1.773281909 | 4.794607335 | ns | NaCl | KS.1 |
| FIM1_5155 | 810.9069783 | 955.5397258 | 1027.34733 | 0.198379967 | 0.133986956 | 0.102720335 | 0.188881341 | ns | NaCl | KS.1 |
| FIM1_5156 | 1042.572903 | 1255.047108 | 907.6906643 | 0.185892837 | 0.204719396 | 0.266698088 | 0.395122986 | ns | NaCl | KS.1 |
| FIM1_5157 | 2335.49568 | 2841.355996 | 2358.045634 | 0.267457794 | 0.073855508 | 0.000178414 | 0.001015569 | ns | NaCl | KS.1 |
| FIM1_5158 | 1150.467269 | 944.1216728 | 1384.854333 | -0.207686418 | 0.192890575 | 0.196423943 | 0.315025505 | ns | NaCl | KS.1 |
| FIM1_5159 | 1438.198039 | 808.4772969 | 963.2899021 | -0.775723501 | 0.132637496 | 4.268893235 | 1.344161003 | ns | NaCl | KS.1 |
| FIM1_5160 | 12433.01608 | 26089.11488 | 11959.88436 | 0.877053325 | 0.247836463 | 2.061750592 | 0.000159521 | ns | NaCl | KS.1 |

| | | | | | | | | | | |
|---|---|---|---|---|---|---|---|---|---|---|
| FIM1_5161 | 1471.537168 | 1655.088457 | 1637.126416 | 0.105632779 | 0.228889029 | 0.555376211 | 0.679035795 | ns | NaCl | KS.1 |
| FIM1_5162 | 2696.56145 | 3235.760289 | 2683.565014 | 0.238380651 | 0.101561316 | 0.01267349 | 0.034266638 | ns | NaCl | KS.1 |
| FIM1_5163 | 3081.385516 | 2946.127006 | 3379.838754 | -0.049873016 | 0.159760164 | 0.722167953 | 0.816171188 | ns | NaCl | KS.1 |
| FIM1_5164 | 319.0370881 | 221.008437 | 390.2049425 | -0.465121959 | 0.151498888 | 0.000551965 | 0.002620255 | ns | NaCl | KS.1 |
| FIM1_5165 | 866.7383687 | 858.8064492 | 1240.079075 | -0.012638014 | 0.13358631 | 0.917241892 | 0.952803969 | ns | NaCl | KS.1 |
| FIM1_5166 | 2435.986181 | 3007.056576 | 2961.30963 | 0.219077912 | 0.197450965 | 0.180005482 | 0.295182273 | ns | NaCl | KS.1 |
| FIM1_5167 | 754.2973516 | 555.5822712 | 637.241487 | -0.394004567 | 0.131028776 | 0.000938899 | 0.004072383 | ns | NaCl | KS.1 |
| FIM1_5168 | 387.6926136 | 450.7121024 | 781.3317534 | 0.174733857 | 0.152741961 | 0.196284665 | 0.314903647 | ns | NaCl | KS.1 |
| FIM1_5169 | 244.9248246 | 241.3967072 | 280.748438 | -0.017641397 | 0.188426686 | 0.911216353 | 0.948389405 | ns | NaCl | KS.1 |
| FIM1_5170 | 1791.124839 | 1639.728229 | 2146.695084 | -0.117972724 | 0.08987408 | 0.1693654 | 0.280770698 | ns | NaCl | KS.1 |
| FIM1_5171 | 747.7900126 | 897.7108944 | 1014.350572 | 0.209902297 | 0.159913994 | 0.133958825 | 0.232859942 | ns | NaCl | KS.1 |
| FIM1_5172 | 9475.771486 | 11447.56788 | 7294.21892 | 0.240413699 | 0.117010157 | 0.026405594 | 0.062825361 | ns | NaCl | KS.1 |
| FIM1_5173 | 1722.853979 | 1392.805756 | 1704.099513 | -0.263479423 | 0.134041114 | 0.030028695 | 0.069711974 | ns | NaCl | KS.1 |
| FIM1_5174 | 4186.162009 | 4034.664811 | 4111.276212 | -0.048403624 | 0.083638386 | 0.547189964 | 0.673162728 | ns | NaCl | KS.1 |
| FIM1_5175 | 2616.659982 | 3233.786157 | 3329.456953 | 0.258774605 | 0.139978665 | 0.039782387 | 0.08784615 | ns | NaCl | KS.1 |
| FIM1_5176 | 889.1373515 | 1094.22353 | 608.5618791 | 0.219239786 | 0.192620327 | 0.172281767 | 0.284656856 | ns | NaCl | KS.1 |
| FIM1_5177 | 1526.004669 | 3275.793699 | 1527.121631 | 0.706430631 | 0.350154993 | 0.003658408 | 0.012639293 | ns | NaCl | KS.1 |
| FIM1_5178 | 1036.201273 | 1431.008189 | 1130.801298 | 0.430642984 | 0.107642846 | 1.85375541813 | 0.000146156 | ns | NaCl | KS.1 |
| FIM1_5179 | 4294.687397 | 9904.776697 | 4531.64748 | 0.925679285 | 0.299239542 | 7.6010269998 | 0.000499399 | ns | NaCl | KS.1 |
| FIM1_5180 | 680.519433 | 717.7288066 | 802.0512042 | 0.064059399 | 0.131842286 | 0.594276551 | 0.714595654 | ns | NaCl | KS.1 |
| FIM1_5181 | 4840.687799 | 4786.556259 | 4942.464288 | -0.013356913 | 0.124205939 | 0.90707285 | 0.945263391 | ns | NaCl | KS.1 |
| FIM1_5182 | 1492.720465 | 1231.370718 | 1281.7074 | -0.255100372 | 0.101327502 | 0.00749078 | 0.0227513 | ns | NaCl | KS.1 |
| FIM1_5183 | 4904.077033 | 7451.367923 | 5023.530225 | 0.549835068 | 0.131752932 | 5.1948398126 | 4.8762883146 | ns | NaCl | KS.1 |
| FIM1_5184 | 607.2425912 | 930.1918205 | 724.6511252 | 0.546256798 | 0.147715935 | 3.8734093539 | 0.000277669 | ns | NaCl | KS.1 |
| FIM1_5185 | 3317.257885 | 4301.637853 | 3451.371304 | 0.342281301 | 0.107290027 | 0.000636459 | 0.00295044 | ns | NaCl | KS.1 |
| FIM1_5186 | 1487.474409 | 1485.235408 | 1683.684437 | -0.001286062 | 0.107357924 | 0.989824915 | 0.99542732 | ns | NaCl | KS.1 |
| FIM1_5187 | 1268.853783 | 1012.539161 | 2096.002034 | -0.25069264 | 0.176558965 | 0.09663129 | 0.179917915 | ns | NaCl | KS.1 |
| FIM1_5188 | 8681.141863 | 9408.364967 | 8105.092407 | 0.090617087 | 0.156583199 | 0.511587282 | 0.642227285 | ns | NaCl | KS.1 |
| FIM1_5189 | 261.2950381 | 232.0612255 | 255.0617251 | -0.110260795 | 0.221168102 | 0.529943574 | 0.658130125 | ns | NaCl | KS.1 |
| FIM1_5190 | 962.0697562 | 977.7103932 | 1079.103786 | 0.018978707 | 0.127114044 | 0.870792839 | 0.922725106 | ns | NaCl | KS.1 |
| FIM1_5191 | 1510.611842 | 1441.351184 | 1573.276048 | -0.052003208 | 0.153756791 | 0.702539503 | 0.799981706 | ns | NaCl | KS.1 |
| FIM1_5192 | 775.9294287 | 535.2792548 | 1933.053245 | -0.282432822 | 0.304344881 | 0.176729652 | 0.291135767 | ns | NaCl | KS.1 |
| FIM1_5193 | 18207.12769 | 23582.97705 | 15741.76877 | 0.340306276 | 0.107089468 | 0.00067033 | 0.003082155 | ns | NaCl | KS.1 |
| FIM1_5194 | 4433.447764 | 4757.536468 | 3900.873431 | 0.079753126 | 0.156127707 | 0.562651511 | 0.685404326 | ns | NaCl | KS.1 |
| FIM1_5195 | 5309.851476 | 5481.351157 | 5536.513644 | 0.039374089 | 0.120336498 | 0.723666696 | 0.816751773 | ns | NaCl | KS.1 |
| FIM1_5196 | 250.4429328 | 173.1183467 | 274.2174018 | -0.462950666 | 0.158672573 | 0.000923046 | 0.004021152 | ns | NaCl | KS.1 |
| FIM1_5197 | 411.0927019 | 489.0805989 | 384.7279617 | 0.215647734 | 0.122542767 | 0.055757708 | 0.11572574 | ns | NaCl | KS.1 |
| FIM1_5199 | 2842.33101 | 3889.776303 | 3164.931706 | 0.42689999 | 0.090204176 | 6.5005679860 | 8.1556656036 | ns | NaCl | KS.1 |
| FIM1_5200 | 236.9221372 | 194.3531599 | 329.1609456 | -0.162436318 | 0.262704518 | 0.400508122 | 0.536483153 | ns | NaCl | KS.1 |
| FIM1_5201 | 251.2631802 | 377.082832 | 502.8709471 | 0.444335787 | 0.21934506 | 0.012299442 | 0.033528616 | ns | NaCl | KS.1 |
| FIM1_5202 | 914.0081454 | 620.1265803 | 778.6488792 | -0.519798235 | 0.116156733 | 1.4801037205 | 1.6621932302 | ns | NaCl | KS.1 |
| FIM1_5203 | 603.6480746 | 693.4427604 | 745.3949667 | 0.162105789 | 0.149214132 | 0.222136833 | 0.346762078 | ns | NaCl | KS.1 |
| FIM1_5204 | 377.3878525 | 377.3020431 | 477.5816684 | -4.10E-05 | 0.179651647 | 0.999786056 | 0.999786056 | ns | NaCl | KS.1 |
| FIM1_5205 | 1329.449939 | 1849.4982 | 1564.827119 | 0.449636772 | 0.093794954 | 4.2976580305 | 5.7015596538 | ns | NaCl | KS.1 |
| FIM1_5206 | 853.9461833 | 1036.569813 | 958.1753731 | 0.251978623 | 0.109303383 | 0.013514581 | 0.036264854 | ns | NaCl | KS.1 |

| | | | | | | | | | | |
|---|---|---|---|---|---|---|---|---|---|---|
| FIM1_5207 | 225.0703618 | 174.6521664 | 269.4435916 | -0.289283197 | 0.183090992 | 0.062000402 | 0.126362965 | ns | NaCl | KS.1 |
| FIM1_5208 | 17.12290607 | 20.8745512 | 21.5559521 | 0.087360371 | 0.367743624 | 0.673041407 | 0.777969563 | ns | NaCl | KS.1 |
| FIM1_5209 | 24.17945711 | 34.80246566 | 37.8397702 | 0.247312639 | 0.32709087 | 0.244875139 | 0.372326961 | ns | NaCl | KS.1 |
| FIM1_4316 | 4422.265608 | 6602.240727 | 4233.280858 | 0.459433235 | 0.199327045 | 0.005703428 | 0.0182473 | ns | NaCl | KS.1 |
| FIM1_4318 | 144.3208167 | 101.8140764 | 297.7479064 | -0.385186819 | 0.193761897 | 0.017416638 | 0.044321113 | ns | NaCl | KS.1 |
| FIM1_4319 | 595.3597979 | 624.2876744 | 791.3821641 | 0.039753234 | 0.242939104 | 0.829730665 | 0.895199686 | ns | NaCl | KS.1 |
| FIM1_4320 | 2569.382822 | 4043.447819 | 3853.218499 | 0.611380376 | 0.117690956 | 2.76582408336 | 5.37499016200 | ns | NaCl | KS.1 |
| FIM1_4321 | 1231.774634 | 1304.788987 | 1574.930606 | 0.069924676 | 0.122774168 | 0.536965502 | 0.664287126 | ns | NaCl | KS.1 |
| FIM1_4322 | 1182.896865 | 2160.562739 | 1448.110775 | 0.812320483 | 0.135694654 | 1.68497585437 | 5.98768205392 | ns | NaCl | KS.1 |
| FIM1_4323 | 12949.26406 | 21328.71356 | 16724.97566 | 0.702689631 | 0.074009996 | 2.37399798894 | 1.07369450881 | ns | NaCl | KS.1 |
| FIM1_4324 | 51.65697225 | 18.71173378 | 38.88673509 | -1.029019534 | 0.379480117 | 0.000140702 | 0.000834593 | Km<KS.1 | NaCl | KS.1 |
| FIM1_4325 | 1762.343027 | 1700.064564 | 1983.336966 | -0.046883291 | 0.103885433 | 0.632116957 | 0.744047634 | ns | NaCl | KS.1 |
| FIM1_4326 | 1205.714548 | 1531.851622 | 1289.899005 | 0.281369246 | 0.158084705 | 0.042491189 | 0.092475236 | ns | NaCl | KS.1 |
| FIM1_4327 | 2618.532633 | 2841.472034 | 2328.645034 | 0.099151734 | 0.129306065 | 0.401850314 | 0.537854536 | ns | NaCl | KS.1 |
| FIM1_4328 | 1012.623933 | 1219.244518 | 872.2491588 | 0.201468197 | 0.18015733 | 0.188373429 | 0.305163728 | ns | NaCl | KS.1 |
| FIM1_4329 | 549.827951 | 345.0648015 | 704.8157214 | -0.581394438 | 0.173562849 | 0.000126625 | 0.000759901 | ns | NaCl | KS.1 |
| FIM1_4330 | 2390.041341 | 2379.312889 | 2374.448184 | -0.00613435 | 0.13097048 | 0.959109662 | 0.97958747 | ns | NaCl | KS.1 |
| FIM1_4331 | 4760.486906 | 8237.888969 | 5501.843841 | 0.663039383 | 0.204414801 | 0.000134839 | 0.00080531 | ns | NaCl | KS.1 |
| FIM1_4332 | 1368.157009 | 1800.18028 | 1772.143234 | 0.37564039 | 0.08286268 | 2.19080859977 | 2.33889973902 | ns | NaCl | KS.1 |
| FIM1_4333 | 2678.318212 | 3257.065654 | 2923.975554 | 0.261328287 | 0.092142274 | 0.002838813 | 0.010215995 | ns | NaCl | KS.1 |
| FIM1_4334 | 837.9414037 | 786.5734274 | 898.2957246 | -0.073041676 | 0.154886573 | 0.593723798 | 0.714507957 | ns | NaCl | KS.1 |
| FIM1_4335 | 1791.978058 | 1858.225277 | 2760.392648 | 0.049838697 | 0.073680649 | 0.485334085 | 0.619429727 | ns | NaCl | KS.1 |
| FIM1_4336 | 1064.610334 | 759.7011491 | 963.1192203 | -0.417559966 | 0.157145752 | 0.002539836 | 0.009352839 | ns | NaCl | KS.1 |
| FIM1_4337 | 1348.796905 | 1552.443532 | 1516.12613 | 0.177157165 | 0.115114154 | 0.09758148 | 0.181261032 | ns | NaCl | KS.1 |
| FIM1_4338 | 4377.872836 | 4956.134892 | 4371.463693 | 0.171623461 | 0.067516115 | 0.008973516 | 0.026307155 | ns | NaCl | KS.1 |
| FIM1_4339 | 495.5149893 | 248.1687789 | 495.8349819 | -0.902232923 | 0.180454949 | 3.18990647825 | 6.03413868034 | ns | NaCl | KS.1 |
| FIM1_4340 | 9104.79391 | 11194.616 | 10345.93131 | 0.267250958 | 0.110409881 | 0.009390981 | 0.027194489 | ns | NaCl | KS.1 |
| FIM1_4341 | 243.293369 | 362.3259027 | 351.1808717 | 0.401327097 | 0.244209395 | 0.034191858 | 0.077637833 | ns | NaCl | KS.1 |
| FIM1_4342 | 488.6308107 | 541.782066 | 597.9498622 | 0.124426001 | 0.13089037 | 0.297487171 | 0.429824108 | ns | NaCl | KS.1 |
| FIM1_4343 | 32563.88191 | 40725.17372 | 25916.70843 | 0.291626982 | 0.108012662 | 0.003840142 | 0.013128102 | ns | NaCl | KS.1 |
| FIM1_4344 | 2744.061042 | 5150.64506 | 2485.044825 | 0.720334531 | 0.246463912 | 0.000312538 | 0.001623045 | ns | NaCl | KS.1 |
| FIM1_4345 | 535.0090377 | 401.0301683 | 544.9502246 | -0.340161625 | 0.164207088 | 0.017284818 | 0.044030705 | ns | NaCl | KS.1 |
| FIM1_4346 | 1963.711989 | 3004.054595 | 1382.390916 | 0.324356783 | 0.313378 | 0.128005053 | 0.224616949 | ns | NaCl | KS.1 |
| FIM1_4347 | 1175.754075 | 1115.589661 | 1730.147006 | -0.064271359 | 0.116445933 | 0.55262889 | 0.676674558 | ns | NaCl | KS.1 |
| FIM1_4348 | 1935.472332 | 2233.654528 | 2477.390735 | 0.19209721 | 0.082030113 | 0.014796456 | 0.038763754 | ns | NaCl | KS.1 |
| FIM1_4349 | 74.50066642 | 45.67706098 | 484.6426657 | -0.391492613 | 0.314768399 | 0.070383218 | 0.139449028 | ns | NaCl | KS.1 |
| FIM1_4350 | 2898.558463 | 2695.828256 | 2577.991275 | -0.096045366 | 0.069496027 | 0.155174493 | 0.262136877 | ns | NaCl | KS.1 |
| FIM1_4351 | 5355.996827 | 6467.838014 | 4666.112367 | 0.237875267 | 0.121494045 | 0.033437096 | 0.076237192 | ns | NaCl | KS.1 |
| FIM1_4352 | 1289.702704 | 1890.196537 | 1263.868516 | 0.430421709 | 0.202064838 | 0.010148569 | 0.028883942 | ns | NaCl | KS.1 |
| FIM1_4353 | 1917.713541 | 3106.078856 | 2362.725098 | 0.637132807 | 0.137960464 | 4.83228151927 | 6.32647383115 | ns | NaCl | KS.1 |
| FIM1_4354 | 1565.465435 | 1174.248357 | 1627.87739 | -0.347795271 | 0.154063533 | 0.010502594 | 0.029699928 | ns | NaCl | KS.1 |
| FIM1_4355 | 33.5800282 | 13.51396076 | 26.18476884 | -0.793229359 | 0.41311788 | 0.002787326 | 0.010070403 | ns | NaCl | KS.1 |
| FIM1_4356 | 23170.11993 | 32270.66227 | 23485.72274 | 0.462277785 | 0.07011163 | 1.01893602579 | 4.92052954287 | ns | NaCl | KS.1 |
| FIM1_4357 | 2190.798114 | 1292.074274 | 1923.814461 | -0.702575245 | 0.139088354 | 4.49476451402 | 7.88186602729 | ns | NaCl | KS.1 |
| FIM1_4358 | 4602.218368 | 4622.019056 | 4355.850489 | 0.005250054 | 0.124763833 | 0.963543231 | 0.982300732 | ns | NaCl | KS.1 |

| | | | | | | | | | | |
|---|---|---|---|---|---|---|---|---|---|---|
| FIM1_4359 | 3259.667656 | 2035.428061 | 2335.321236 | -0.623173974 | 0.135864287 | 5.89040520952 | 7.49482504281 | ns | NaCl | KS.1 |
| FIM1_4360 | 10267.54589 | 10468.67422 | 8671.074157 | 0.025353219 | 0.09295517 | 0.774783859 | 0.853721994 | ns | NaCl | KS.1 |
| FIM1_4361 | 2414.195874 | 5734.640313 | 2926.548286 | 1.104932973 | 0.212349761 | 4.75910349815 | 1.15495316601 | Km>KS.1 | NaCl | KS.1 |
| FIM1_4363 | 866.4350673 | 1963.991423 | 979.5832279 | 0.998428762 | 0.24063627 | 1.14174672121 | 1.32097440419 | ns | NaCl | KS.1 |
| FIM1_4364 | 346.2899207 | 381.2192901 | 393.1670421 | 0.100189055 | 0.177310336 | 0.508462352 | 0.639487624 | ns | NaCl | KS.1 |
| FIM1_4365 | 1283.356139 | 1302.19927 | 1130.932375 | 0.016573369 | 0.128974704 | 0.888413015 | 0.933183795 | ns | NaCl | KS.1 |
| FIM1_4366 | 7690.685992 | 10748.37622 | 9010.826394 | 0.459559443 | 0.086559548 | 2.72329414840 | 5.33401117650 | ns | NaCl | KS.1 |
| FIM1_4367 | 1808.378499 | 1927.3502 | 1772.816264 | 0.077413924 | 0.13049607 | 0.516128357 | 0.646948494 | ns | NaCl | KS.1 |
| FIM1_4368 | 1612.581564 | 1749.108603 | 1621.621862 | 0.099559351 | 0.119878983 | 0.369417173 | 0.504073076 | ns | NaCl | KS.1 |
| FIM1_4369 | 3120.159483 | 6939.598722 | 3186.351165 | 0.890025324 | 0.290079116 | 9.65151143486 | 0.000606181 | ns | NaCl | KS.1 |
| FIM1_4370 | 720.6732877 | 1583.652877 | 886.3623511 | 1.01617447 | 0.194352852 | 6.24794329465 | 1.44120537318 | Km>KS.1 | NaCl | KS.1 |
| FIM1_4371 | 2650.151035 | 3410.338343 | 3441.943324 | 0.297777832 | 0.159617137 | 0.033089601 | 0.075650154 | ns | NaCl | KS.1 |
| FIM1_4372 | 2724.148633 | 3578.366649 | 2499.143024 | 0.313491623 | 0.172291467 | 0.034398502 | 0.078035817 | ns | NaCl | KS.1 |
| FIM1_4373 | 8694.21474 | 15991.54975 | 8223.638193 | 0.726406539 | 0.222407884 | 9.78731888990 | 0.000612477 | ns | NaCl | KS.1 |
| FIM1_4374 | 848.488106 | 963.8933227 | 1187.237333 | 0.163857376 | 0.10430548 | 0.094854615 | 0.177807728 | ns | NaCl | KS.1 |
| FIM1_4376 | 301.7800192 | 298.2478452 | 275.9932316 | -0.014831184 | 0.164485006 | 0.917639755 | 0.952882025 | ns | NaCl | KS.1 |
| FIM1_4377 | 6048.5547 | 9189.593171 | 5376.730464 | 0.470096253 | 0.210845044 | 0.006627541 | 0.020594638 | ns | NaCl | KS.1 |
| FIM1_4378 | 254.2784237 | 214.4451562 | 200.5203965 | -0.18706492 | 0.170050892 | 0.202575498 | 0.322956094 | ns | NaCl | KS.1 |
| FIM1_4379 | 2362.424687 | 3117.505845 | 3467.010987 | 0.363997525 | 0.11257034 | 0.000496704 | 0.002396488 | ns | NaCl | KS.1 |
| FIM1_4380 | 1162.742897 | 1523.125523 | 1529.100328 | 0.362772244 | 0.092591467 | 3.61376946127 | 0.000261315 | ns | NaCl | KS.1 |
| FIM1_4381 | 1458.785645 | 1929.060112 | 1501.693725 | 0.361049814 | 0.120486568 | 0.001126739 | 0.0047304 | ns | NaCl | KS.1 |
| FIM1_4382 | 892.3028527 | 1005.391514 | 961.0533677 | 0.146355745 | 0.124217971 | 0.200121766 | 0.319924738 | ns | NaCl | KS.1 |
| FIM1_4383 | 5848.029722 | 7442.047447 | 5562.631831 | 0.334016909 | 0.067861179 | 3.91180801218 | 5.24620712218 | ns | NaCl | KS.1 |
| FIM1_4384 | 835.7124838 | 1426.146381 | 1504.303601 | 0.608319056 | 0.231715243 | 0.001225354 | 0.005092846 | ns | NaCl | KS.1 |
| FIM1_4385 | 1341.733192 | 1427.346737 | 1672.830195 | 0.079330026 | 0.108716216 | 0.436341344 | 0.572316948 | ns | NaCl | KS.1 |
| FIM1_4386 | 17653.11002 | 20945.26305 | 15426.11709 | 0.220254662 | 0.109395031 | 0.031107814 | 0.071848364 | ns | NaCl | KS.1 |
| FIM1_4387 | 46.30933762 | 38.49785215 | 32.08724951 | -0.164858825 | 0.255929021 | 0.387488696 | 0.522427171 | ns | NaCl | KS.1 |
| FIM1_4388 | 2997.699193 | 7178.744536 | 2828.08616 | 1.019775076 | 0.277059162 | 6.63353123551 | 5.95700684055 | Km>KS.1 | NaCl | KS.1 |
| FIM1_4389 | 196.3586578 | 162.3557222 | 261.3620366 | -0.204030958 | 0.185075939 | 0.191352286 | 0.308782881 | ns | NaCl | KS.1 |
| FIM1_4390 | 97.10127571 | 77.46418518 | 119.0014353 | -0.224973814 | 0.206617652 | 0.18181975 | 0.297517036 | ns | NaCl | KS.1 |
| FIM1_4391 | 12919.32788 | 21004.00097 | 11072.76012 | 0.58442745 | 0.19551785 | 0.000436332 | 0.002153526 | ns | NaCl | KS.1 |
| FIM1_4392 | 2854.313843 | 2745.434692 | 2446.787469 | -0.042613911 | 0.171347173 | 0.773162279 | 0.853256951 | ns | NaCl | KS.1 |
| FIM1_4393 | 148.2899013 | 66.98267878 | 102.1546705 | -0.867567786 | 0.284991524 | 0.000114887 | 0.000699588 | ns | NaCl | KS.1 |
| FIM1_4394 | 867.1181148 | 518.2905368 | 574.9786187 | -0.690690797 | 0.133734191 | 2.55666435891 | 5.07065121613 | ns | NaCl | KS.1 |
| FIM1_4396 | 3260.226558 | 3164.747467 | 3371.773329 | -0.034468563 | 0.1413624 | 0.7867058 | 0.862463939 | ns | NaCl | KS.1 |
| FIM1_4397 | 1261.883184 | 712.458415 | 1309.553351 | -0.779255401 | 0.123169717 | 2.17257118481 | 9.15978105463 | ns | NaCl | KS.1 |
| FIM1_4398 | 86.39829104 | 73.48920317 | 199.3745941 | -0.134250839 | 0.275393844 | 0.495480282 | 0.627709295 | ns | NaCl | KS.1 |
| FIM1_4399 | 13307.95264 | 20339.47679 | 14052.33915 | 0.483034688 | 0.207089376 | 0.00477802 | 0.015690197 | ns | NaCl | KS.1 |
| FIM1_4400 | 1229.612448 | 991.5590207 | 1072.102269 | -0.284009968 | 0.102254183 | 0.003126386 | 0.011038871 | ns | NaCl | KS.1 |
| FIM1_4401 | 3345.615619 | 3546.137655 | 2429.574297 | 0.072930868 | 0.115666348 | 0.497878128 | 0.629784817 | ns | NaCl | KS.1 |
| FIM1_4402 | 298.6700663 | 263.7352521 | 172.6698895 | -0.145714638 | 0.163732043 | 0.307382471 | 0.440065552 | ns | NaCl | KS.1 |
| FIM1_4403 | 4009.892539 | 4543.201378 | 5429.59483 | 0.106974002 | 0.24185832 | 0.562506665 | 0.685395704 | ns | NaCl | KS.1 |
| FIM1_4404 | 417.8487156 | 370.2530254 | 460.0954742 | -0.143230375 | 0.155921309 | 0.297550598 | 0.429824108 | ns | NaCl | KS.1 |
| FIM1_4405 | 668.0066427 | 511.875099 | 609.2161831 | -0.314533877 | 0.164681949 | 0.028027411 | 0.065865078 | ns | NaCl | KS.1 |
| FIM1_4406 | 916.1289833 | 731.0129686 | 1069.145318 | -0.287280806 | 0.12590483 | 0.012618078 | 0.034153938 | ns | NaCl | KS.1 |

| | | | | | | | | | | |
|---|---|---|---|---|---|---|---|---|---|---|
| FIM1_4407 | 64.92943027 | 29.63835503 | 122.619116 | -0.767194286 | 0.32778224 | 0.001303411 | 0.005328241 | ns | NaCl | KS.1 |
| FIM1_4408 | 410.7856924 | 131.0107964 | 448.4685894 | -1.316376422 | 0.337255474 | 6.5081391852 | 8.1556656036 | Km<KS.1 | NaCl | KS.1 |
| FIM1_4409 | 8799.94486 | 14116.02779 | 6251.210809 | 0.443682664 | 0.279139107 | 0.031212413 | 0.072023076 | ns | NaCl | KS.1 |
| FIM1_4410 | 921.8519829 | 650.2765395 | 823.5708034 | -0.443186919 | 0.139033923 | 0.000406736 | 0.0020296 | ns | NaCl | KS.1 |
| FIM1_4412 | 167.7981791 | 167.3867252 | 240.9380461 | 0.001303636 | 0.248904321 | 0.994440987 | 0.997448369 | ns | NaCl | KS.1 |
| FIM1_4413 | 1426.762089 | 1416.416492 | 1434.163885 | -0.011355128 | 0.092289664 | 0.897369717 | 0.938735214 | ns | NaCl | KS.1 |
| FIM1_4414 | 2979.421638 | 4543.890497 | 3221.545165 | 0.557461435 | 0.128805402 | 2.5145134112 | 2.6391781056 | ns | NaCl | KS.1 |
| FIM1_4415 | 1937.409732 | 2032.792739 | 3335.323888 | 0.041555154 | 0.234815505 | 0.818925406 | 0.887886833 | ns | NaCl | KS.1 |
| FIM1_4416 | 1322.718678 | 1368.077802 | 1896.95476 | 0.03487981 | 0.181959591 | 0.821271835 | 0.889577048 | ns | NaCl | KS.1 |
| FIM1_4417 | 1012.475184 | 1322.069411 | 1298.851095 | 0.353872779 | 0.100985942 | 0.000194131 | 0.001091301 | ns | NaCl | KS.1 |
| FIM1_4418 | 6073.690564 | 8827.335151 | 5851.324223 | 0.509724829 | 0.096776913 | 2.7377893332 | 5.3413733069 | ns | NaCl | KS.1 |
| FIM1_4419 | 1145.763718 | 960.7096886 | 1039.942133 | -0.223082919 | 0.116406381 | 0.038594217 | 0.086024297 | ns | NaCl | KS.1 |
| FIM1_4420 | 875.5079507 | 734.3015042 | 1050.982712 | -0.223489921 | 0.111223455 | 0.031084203 | 0.071827176 | ns | NaCl | KS.1 |
| FIM1_4421 | 1051.798306 | 703.3190503 | 841.2736502 | -0.549125124 | 0.105730785 | 3.4750866383 | 6.5486954643 | ns | NaCl | KS.1 |
| FIM1_4422 | 8.624344458 | 11.75355005 | 14.57156587 | 0.150059963 | 0.373131879 | 0.474813514 | 0.60850006 | ns | NaCl | KS.1 |
| FIM1_4423 | 2188.141541 | 2042.951507 | 1754.733821 | -0.068408262 | 0.198288418 | 0.676146624 | 0.779747207 | ns | NaCl | KS.1 |
| FIM1_4424 | 668.9728883 | 340.9843597 | 1053.413028 | -0.526029166 | 0.366210887 | 0.026203298 | 0.062463539 | ns | NaCl | KS.1 |
| FIM1_4425 | 750.0943329 | 759.7598627 | 643.1878699 | 0.015310503 | 0.084111077 | 0.849784323 | 0.908396435 | ns | NaCl | KS.1 |
| FIM1_4426 | 787.4897327 | 786.2089984 | 728.6083259 | -0.004096338 | 0.125215599 | 0.971637785 | 0.987114147 | ns | NaCl | KS.1 |
| FIM1_4427 | 4945.543828 | 5416.222516 | 5440.023767 | 0.115433666 | 0.111350778 | 0.267126138 | 0.395639338 | ns | NaCl | KS.1 |
| FIM1_4428 | 6938.470804 | 5293.283859 | 6919.829657 | -0.276211157 | 0.213567913 | 0.109264087 | 0.198607538 | ns | NaCl | KS.1 |
| FIM1_4429 | 2355.571415 | 2393.813351 | 2044.53562 | 0.018297439 | 0.152237233 | 0.89230091 | 0.935355463 | ns | NaCl | KS.1 |
| FIM1_4430 | 366.1366681 | 465.3851575 | 343.7891576 | 0.273912588 | 0.164811307 | 0.055864629 | 0.115899304 | ns | NaCl | KS.1 |
| FIM1_4431 | 1735.842222 | 1710.473223 | 2195.196975 | -0.018232774 | 0.098793814 | 0.845611208 | 0.905686924 | ns | NaCl | KS.1 |
| FIM1_4432 | 6578.755497 | 7888.352107 | 5969.247125 | 0.242628083 | 0.088868677 | 0.004199071 | 0.014124663 | ns | NaCl | KS.1 |
| FIM1_4433 | 539.4755379 | 662.9534233 | 622.5379711 | 0.236349761 | 0.162814417 | 0.095934088 | 0.179128018 | ns | NaCl | KS.1 |
| FIM1_4434 | 1406.507585 | 1739.073481 | 1987.486864 | 0.294345875 | 0.066304156 | 4.9256279908 | 4.6587451054 | ns | NaCl | KS.1 |
| FIM1_4435 | 3007.808505 | 3741.517934 | 4237.227993 | 0.284365098 | 0.108423194 | 0.00496605 | 0.01620072 | ns | NaCl | KS.1 |
| FIM1_4436 | 1356.047176 | 1325.005337 | 1632.471748 | -0.030259433 | 0.07875721 | 0.690676315 | 0.791184588 | ns | NaCl | KS.1 |
| FIM1_4437 | 531.3405054 | 454.955949 | 636.5513952 | -0.188999596 | 0.144203708 | 0.143282605 | 0.245296269 | ns | NaCl | KS.1 |
| FIM1_4438 | 1045.547896 | 774.0014115 | 1274.270211 | -0.364662627 | 0.159517091 | 0.009070748 | 0.026482965 | ns | NaCl | KS.1 |
| FIM1_4439 | 1968.099859 | 1536.727921 | 1607.302738 | -0.323219919 | 0.112943774 | 0.002050798 | 0.007776464 | ns | NaCl | KS.1 |
| FIM1_4440 | 1122.770404 | 506.9058742 | 1509.242747 | -0.976375164 | 0.232091348 | 9.8188946767 | 1.1520990805 | ns | NaCl | KS.1 |
| FIM1_4441 | 2841.392241 | 2896.87041 | 3396.226368 | 0.017492065 | 0.222046262 | 0.920740098 | 0.954950742 | ns | NaCl | KS.1 |
| FIM1_4443 | 493.1786841 | 151.8139795 | 510.6500014 | -1.238164114 | 0.404246648 | 1.8285169386 | 0.000144479 | Km<KS.1 | NaCl | KS.1 |
| FIM1_4444 | 31203.91068 | 44672.64526 | 29229.62885 | 0.454105372 | 0.145178057 | 0.000476123 | 0.002304195 | ns | NaCl | KS.1 |
| FIM1_4445 | 1448.511282 | 1316.748121 | 1353.071427 | -0.123849696 | 0.10783189 | 0.22062604 | 0.3450533 | ns | NaCl | KS.1 |
| FIM1_4446 | 4459.91746 | 4767.755907 | 5081.720682 | 0.084119709 | 0.115339099 | 0.433121428 | 0.569545428 | ns | NaCl | KS.1 |
| FIM1_4447 | 1456.051525 | 1518.518207 | 1698.587544 | 0.05446541 | 0.080118921 | 0.480815758 | 0.614609043 | ns | NaCl | KS.1 |
| FIM1_4448 | 205.2971671 | 261.8273942 | 260.2087931 | 0.212569284 | 0.248134669 | 0.258887942 | 0.387348256 | ns | NaCl | KS.1 |
| FIM1_4449 | 762.9452301 | 482.9573016 | 733.6709146 | -0.453468255 | 0.258262897 | 0.021521031 | 0.052977304 | ns | NaCl | KS.1 |
| FIM1_4450 | 3426.163531 | 3107.697515 | 3259.486394 | -0.118383323 | 0.131561598 | 0.323687037 | 0.456964532 | ns | NaCl | KS.1 |
| FIM1_4451 | 1590.407107 | 1246.93815 | 2267.005217 | -0.277382273 | 0.172523744 | 0.061452682 | 0.125555275 | ns | NaCl | KS.1 |
| FIM1_4452 | 12193.60582 | 15653.09685 | 10157.37843 | 0.300855506 | 0.150072869 | 0.023785729 | 0.057543505 | ns | NaCl | KS.1 |
| FIM1_4454 | 4472.055986 | 4843.577624 | 4077.397871 | 0.10680883 | 0.083566859 | 0.183467392 | 0.298968318 | ns | NaCl | KS.1 |

| | | | | | | | | | | |
|---|---|---|---|---|---|---|---|---|---|---|
| FIM1_4455 | 1928.971451 | 1906.863281 | 1386.220544 | -0.013304671 | 0.157183202 | 0.923496306 | 0.955970479 | ns | NaCl | KS.1 |
| FIM1_4456 | 1880.59719 | 854.9897935 | 1521.687779 | -1.0379231 | 0.175046795 | 1.0730932353 | 3.925469739 | Km<KS.1 | NaCl | KS.1 |
| FIM1_4457 | 2509.867388 | 2046.64066 | 1899.474495 | -0.266961719 | 0.107114626 | 0.007706045 | 0.023291357 | ns | NaCl | KS.1 |
| FIM1_4458 | 1346.643468 | 1258.099158 | 1523.401855 | -0.089714116 | 0.10488439 | 0.363466585 | 0.498826555 | ns | NaCl | KS.1 |
| FIM1_4460 | 364.0649774 | 396.3515032 | 482.6566913 | 0.09255858 | 0.181238504 | 0.547628279 | 0.673533203 | ns | NaCl | KS.1 |
| FIM1_4461 | 228.8645863 | 226.455895 | 235.8112276 | -0.007775762 | 0.185502584 | 0.960375043 | 0.980678539 | ns | NaCl | KS.1 |
| FIM1_4462 | 1122.682917 | 1152.287855 | 1086.853569 | 0.031953351 | 0.106920217 | 0.750461366 | 0.837493337 | ns | NaCl | KS.1 |
| FIM1_4463 | 601.6882624 | 640.5327938 | 1342.582568 | 0.076766376 | 0.132455098 | 0.524809662 | 0.65398695 | ns | NaCl | KS.1 |
| FIM1_4464 | 1175.6412 | 714.6907864 | 1042.604224 | -0.645609289 | 0.153810342 | 3.2858963126 | 3.363649003 | ns | NaCl | KS.1 |
| FIM1_4465 | 1425.695423 | 1252.187405 | 1372.504425 | -0.161760118 | 0.124342319 | 0.156965427 | 0.264476618 | ns | NaCl | KS.1 |
| FIM1_4466 | 1308.324509 | 1105.948588 | 1228.347793 | -0.212787425 | 0.106989969 | 0.033723602 | 0.076714641 | ns | NaCl | KS.1 |
| FIM1_4467 | 3581.856255 | 3711.973077 | 2880.810584 | 0.048963351 | 0.076543336 | 0.50837011 | 0.639487624 | ns | NaCl | KS.1 |
| FIM1_4468 | 312.3294013 | 202.4359401 | 290.415238 | -0.551097237 | 0.169627261 | 0.000205589 | 0.0011428 | ns | NaCl | KS.1 |
| FIM1_4469 | 3599.151134 | 5835.096685 | 3092.130005 | 0.466931487 | 0.273936982 | 0.022436076 | 0.054849866 | ns | NaCl | KS.1 |
| FIM1_4470 | 2928.346543 | 4431.250181 | 2972.149374 | 0.437805787 | 0.231849534 | 0.017372223 | 0.044230711 | ns | NaCl | KS.1 |
| FIM1_4471 | 689.8505659 | 883.8439034 | 801.3701326 | 0.304444803 | 0.142307675 | 0.016942917 | 0.043315011 | ns | NaCl | KS.1 |
| FIM1_4472 | 2066.457154 | 1908.640805 | 2324.054921 | -0.109447868 | 0.069140096 | 0.10353183 | 0.190018883 | ns | NaCl | KS.1 |
| FIM1_4473 | 1012.015036 | 714.4750358 | 825.0001088 | -0.439684084 | 0.14717034 | 0.000814622 | 0.003628239 | ns | NaCl | KS.1 |
| FIM1_4474 | 11774.99709 | 14533.075 | 10933.34462 | 0.24909911 | 0.151896747 | 0.06392101 | 0.129218621 | ns | NaCl | KS.1 |
| FIM1_4475 | 783.3464728 | 482.8944016 | 812.4556507 | -0.593600239 | 0.186644458 | 0.000220248 | 0.001209418 | ns | NaCl | KS.1 |
| FIM1_4476 | 1782.621624 | 1635.855925 | 1386.57011 | -0.093511371 | 0.170388855 | 0.525192192 | 0.654025321 | ns | NaCl | KS.1 |
| FIM1_4477 | 2113.552252 | 2043.712543 | 2194.233526 | -0.044000685 | 0.096110562 | 0.629950703 | 0.742348225 | ns | NaCl | KS.1 |
| FIM1_4478 | 1985.840273 | 1705.404386 | 1911.228193 | -0.185710466 | 0.13214643 | 0.122402483 | 0.21732775 | ns | NaCl | KS.1 |
| FIM1_4479 | 1674.380195 | 2608.422957 | 2132.936426 | 0.494532495 | 0.219448004 | 0.005594533 | 0.017933505 | ns | NaCl | KS.1 |
| FIM1_4480 | 543.2063067 | 383.6420889 | 436.876264 | -0.411632827 | 0.173478376 | 0.005863526 | 0.018675443 | ns | NaCl | KS.1 |
| FIM1_4481 | 1669.2944 | 1761.595766 | 1967.364171 | 0.070408448 | 0.08470932 | 0.386741861 | 0.521703025 | ns | NaCl | KS.1 |
| FIM1_4482 | 72.02266967 | 30.29572125 | 2864.161851 | -0.822788827 | 0.383128257 | 0.001533417 | 0.006107887 | ns | NaCl | KS.1 |
| FIM1_4484 | 14.23602439 | 23.80851963 | 72.63694843 | 0.227578167 | 0.399770207 | 0.287975483 | 0.418892485 | ns | NaCl | KS.1 |
| FIM1_4486 | 1560.72666 | 2005.076467 | 1769.97335 | 0.210679203 | 0.261565565 | 0.276344926 | 0.406049694 | ns | NaCl | KS.1 |
| FIM1_4487 | 255.2165471 | 134.1052817 | 435.3429884 | -0.616638396 | 0.311982794 | 0.006233195 | 0.019639103 | ns | NaCl | KS.1 |
| FIM1_4488 | 32.95340539 | 31.42936346 | 33.08812412 | -0.031458119 | 0.277827382 | 0.872850593 | 0.924314963 | ns | NaCl | KS.1 |
| FIM1_4489 | 1436.433313 | 1294.46216 | 1176.216034 | -0.091463341 | 0.23349372 | 0.613542909 | 0.729117728 | ns | NaCl | KS.1 |
| FIM1_4490 | 3907.455531 | 6263.738078 | 7550.038692 | 0.50954755 | 0.237548822 | 0.006894218 | 0.021250765 | ns | NaCl | KS.1 |
| FIM1_4491 | 822.0698187 | 873.3792379 | 931.9821455 | 0.067355905 | 0.16855206 | 0.644626196 | 0.754059564 | ns | NaCl | KS.1 |
| FIM1_4492 | 1136.770974 | 1091.738285 | 821.0718333 | -0.049578618 | 0.118275691 | 0.651350973 | 0.759248147 | ns | NaCl | KS.1 |
| FIM1_4493 | 4570.164194 | 7132.758237 | 6962.732737 | 0.508625212 | 0.210321428 | 0.003383661 | 0.011796577 | ns | NaCl | KS.1 |
| FIM1_4494 | 44.6051854 | 91.97213608 | 44.75258721 | 0.794139721 | 0.289823652 | 0.000396263 | 0.001989314 | ns | NaCl | KS.1 |
| FIM1_4495 | 669.1610638 | 990.5708827 | 1200.271987 | 0.499058748 | 0.150334745 | 0.000198623 | 0.001112398 | ns | NaCl | KS.1 |
| FIM1_4496 | 738.5097354 | 688.2814424 | 572.8180632 | -0.084746678 | 0.129863861 | 0.475355643 | 0.608845852 | ns | NaCl | KS.1 |
| FIM1_4497 | 3477.439948 | 5660.50721 | 4143.687139 | 0.657019145 | 0.121891402 | 8.2388227815 | 1.821695259 | ns | NaCl | KS.1 |
| FIM1_4498 | 19957.38937 | 31010.76879 | 18755.82401 | 0.585941395 | 0.126915613 | 5.8043218516 | 7.442397219 | ns | NaCl | KS.1 |
| FIM1_4499 | 2600.782459 | 2355.135807 | 3375.850452 | -0.123349655 | 0.119461689 | 0.264483071 | 0.392777098 | ns | NaCl | KS.1 |
| FIM1_4500 | 2846.162889 | 2548.10647 | 2985.97264 | -0.148369555 | 0.08317351 | 0.063310075 | 0.128348665 | ns | NaCl | KS.1 |
| FIM1_4501 | 1775.799274 | 1817.328283 | 1516.750278 | 0.025494172 | 0.163760925 | 0.858448245 | 0.914122436 | ns | NaCl | KS.1 |
| FIM1_4502 | 219.5098543 | 208.4662078 | 211.4499236 | -0.056303209 | 0.202705545 | 0.734704019 | 0.825128239 | ns | NaCl | KS.1 |

| | | | | | | | | | | |
|---|---|---|---|---|---|---|---|---|---|---|
| FIM1_4503 | 591.4788861 | 526.6925366 | 692.7756756 | -0.15459146 | 0.111349526 | 0.136922137 | 0.23701727 | ns | NaCl | KS.1 |
| FIM1_4504 | 768.2048023 | 824.0455842 | 1117.332412 | 0.077468553 | 0.165099532 | 0.590015881 | 0.711077764 | ns | NaCl | KS.1 |
| FIM1_4505 | 3217.623925 | 5763.597492 | 2753.53033 | 0.521508161 | 0.316198162 | 0.019129961 | 0.047716016 | ns | NaCl | KS.1 |
| FIM1_4506 | 1800.454925 | 2045.405338 | 1446.303108 | 0.142646963 | 0.162567005 | 0.315071368 | 0.448235646 | ns | NaCl | KS.1 |
| FIM1_4507 | 3015.507604 | 2919.120914 | 2743.829087 | -0.038825048 | 0.135055013 | 0.751711689 | 0.838250702 | ns | NaCl | KS.1 |
| FIM1_4508 | 1155.626809 | 1062.113835 | 1156.549221 | -0.109206392 | 0.102943328 | 0.260544663 | 0.389135304 | ns | NaCl | KS.1 |
| FIM1_4509 | 3413.808609 | 2789.30747 | 2958.750099 | -0.278416808 | 0.066137206 | 1.49037537809 | 0.000121352 | ns | NaCl | KS.1 |
| FIM1_4510 | 5027.539633 | 7933.645125 | 5040.076391 | 0.630675437 | 0.093166008 | 1.62385987895 | 9.61750344977 | ns | NaCl | KS.1 |
| FIM1_4512 | 832.4549268 | 602.3654413 | 894.7769578 | -0.426874498 | 0.11672531 | 7.69037637124 | 0.000502095 | ns | NaCl | KS.1 |
| FIM1_4513 | 228.1134245 | 259.0848317 | 220.9323604 | 0.142983715 | 0.160812577 | 0.309916142 | 0.442673788 | ns | NaCl | KS.1 |
| FIM1_4514 | 298.9166763 | 213.9174177 | 316.521227 | -0.414097589 | 0.159258739 | 0.003052759 | 0.010840455 | ns | NaCl | KS.1 |
| FIM1_4515 | 1256.699174 | 876.495132 | 1229.686415 | -0.48513161 | 0.103828384 | 6.65961608922 | 8.30365665260 | ns | NaCl | KS.1 |
| FIM1_4516 | 819.2118044 | 1514.930787 | 750.3798019 | 0.528661562 | 0.332310303 | 0.020429146 | 0.050539532 | ns | NaCl | KS.1 |
| FIM1_4517 | 903.8526756 | 456.6276261 | 1121.894738 | -0.83425829 | 0.219373426 | 8.95387375478 | 7.77408759686 | ns | NaCl | KS.1 |
| FIM1_4518 | 555.7038624 | 634.3485394 | 733.9843654 | 0.149292395 | 0.152125733 | 0.268273962 | 0.39722112 | ns | NaCl | KS.1 |
| FIM1_4519 | 1095.21552 | 1191.386045 | 1030.65912 | 0.10643662 | 0.104010556 | 0.277259628 | 0.407012879 | ns | NaCl | KS.1 |
| FIM1_4520 | 2190.566712 | 2675.414071 | 2052.447947 | 0.225400891 | 0.168558884 | 0.12205173 | 0.216859771 | ns | NaCl | KS.1 |
| FIM1_4521 | 1876.158771 | 2057.906692 | 2375.04407 | 0.12517446 | 0.079435279 | 0.102163168 | 0.188314843 | ns | NaCl | KS.1 |
| FIM1_4522 | 506.3109563 | 176.3331613 | 643.0435648 | -1.281677632 | 0.279520417 | 4.11583936023 | 7.41894957144 | Km<KS.1 | NaCl | KS.1 |
| FIM1_4523 | 3547.958828 | 4010.562776 | 3284.485673 | 0.169988771 | 0.061990506 | 0.004994428 | 0.016282621 | ns | NaCl | KS.1 |
| FIM1_4524 | 2773.680007 | 2108.483043 | 2028.069343 | -0.353017582 | 0.120722566 | 0.001477742 | 0.005909779 | ns | NaCl | KS.1 |
| FIM1_4525 | 305.4670541 | 231.4535077 | 257.5943842 | -0.308825946 | 0.182425039 | 0.045811506 | 0.09827219 | ns | NaCl | KS.1 |
| FIM1_4526 | 7089.082326 | 7743.092049 | 6663.423493 | 0.122741094 | 0.063934473 | 0.04917572 | 0.104239118 | ns | NaCl | KS.1 |
| FIM1_4527 | 15058.72958 | 19938.07694 | 16071.36789 | 0.260820645 | 0.241446385 | 0.16081025 | 0.269580385 | ns | NaCl | KS.1 |
| FIM1_4528 | 130.2571695 | 59.5909272 | 224.419495 | -0.748852695 | 0.339812164 | 0.001970045 | 0.007550827 | ns | NaCl | KS.1 |
| FIM1_4529 | 329.0139758 | 178.23141 | 548.8525816 | -0.59930905 | 0.299243921 | 0.006395472 | 0.019985851 | ns | NaCl | KS.1 |
| FIM1_4530 | 2123.469839 | 2149.760165 | 2316.557567 | 0.016141207 | 0.102838946 | 0.867925507 | 0.921060025 | ns | NaCl | KS.1 |
| FIM1_4531 | 3847.219972 | 2842.952845 | 3611.620623 | -0.406487607 | 0.099228151 | 1.37677682959 | 0.000113214 | ns | NaCl | KS.1 |
| FIM1_4532 | 1617.838003 | 1676.088879 | 1404.05236 | 0.042516333 | 0.136105143 | 0.730643135 | 0.822200768 | ns | NaCl | KS.1 |
| FIM1_4533 | 8520.366724 | 18465.50719 | 8237.061385 | 0.85606821 | 0.288184489 | 0.000151882 | 0.000885828 | ns | NaCl | KS.1 |
| FIM1_4534 | 820.7419776 | 512.5825533 | 795.2805864 | -0.61003152 | 0.153581292 | 9.85941708119 | 8.39907533886 | ns | NaCl | KS.1 |
| FIM1_4535 | 2348.243216 | 2508.575405 | 1795.554662 | 0.073923295 | 0.157537198 | 0.594143556 | 0.714595654 | ns | NaCl | KS.1 |
| FIM1_4536 | 1735.747613 | 1581.019928 | 2216.598367 | -0.089353789 | 0.206356764 | 0.595027685 | 0.714866635 | ns | NaCl | KS.1 |
| FIM1_4537 | 874.4149606 | 1095.477344 | 926.008992 | 0.287567608 | 0.116883569 | 0.007793806 | 0.023513757 | ns | NaCl | KS.1 |
| FIM1_4538 | 1232.54959 | 302.7640312 | 724.0558032 | -1.766316362 | 0.334882688 | 1.11365544481 | 4.04411375034 | Km<KS.1 | NaCl | KS.1 |
| FIM1_4539 | 1134.811252 | 1084.561978 | 1030.865874 | -0.062241288 | 0.099676692 | 0.509506227 | 0.640093083 | ns | NaCl | KS.1 |
| FIM1_4540 | 557.563163 | 424.5941536 | 575.5351737 | -0.323366483 | 0.164491224 | 0.02379236 | 0.057543505 | ns | NaCl | KS.1 |
| FIM1_4542 | 3972.295211 | 4067.05633 | 4417.824229 | 0.027894592 | 0.142839192 | 0.828096267 | 0.894243309 | ns | NaCl | KS.1 |
| FIM1_4543 | 3813.251177 | 9177.493397 | 3931.707626 | 0.875159471 | 0.352752308 | 0.000545969 | 0.002601719 | ns | NaCl | KS.1 |
| FIM1_4544 | 468.1731529 | 263.0710693 | 678.0470273 | -0.718488473 | 0.192222503 | 1.75047198135 | 0.000139785 | ns | NaCl | KS.1 |
| FIM1_4545 | 2169.762769 | 4001.42642 | 2193.460746 | 0.698765286 | 0.244258157 | 0.000414107 | 0.002062245 | ns | NaCl | KS.1 |
| FIM1_4546 | 5202.48094 | 10460.41927 | 5054.654355 | 0.747409676 | 0.288382416 | 0.000741234 | 0.003358504 | ns | NaCl | KS.1 |
| FIM1_4547 | 147.7497039 | 149.7938789 | 93.14022584 | 0.006919615 | 0.237435381 | 0.969767929 | 0.986423113 | ns | NaCl | KS.1 |
| FIM1_4548 | 1410.159945 | 1170.18963 | 1192.076338 | -0.228128602 | 0.136903202 | 0.065079815 | 0.130821403 | ns | NaCl | KS.1 |
| FIM1_4549 | 1897.668289 | 785.5628826 | 2090.131992 | -1.169523985 | 0.178494879 | 1.07600899017 | 6.86300605912 | Km<KS.1 | NaCl | KS.1 |

| | | | | | | | | | | |
|---|---|---|---|---|---|---|---|---|---|---|
| FIM1_4550 | 94.6481475 | 52.74429905 | 58.11738184 | -0.541031767 | 0.30742616 | 0.014093973 | 0.037376074 | ns | NaCl | KS.1 |
| FIM1_4551 | 162.9091767 | 97.07392443 | 155.7175167 | -0.627261638 | 0.206258128 | 0.000309251 | 0.001610607 | ns | NaCl | KS.1 |
| FIM1_4552 | 1094.598616 | 1006.796536 | 991.3499352 | -0.096539109 | 0.145795755 | 0.459451625 | 0.594324451 | ns | NaCl | KS.1 |
| FIM1_4553 | 576.4077526 | 602.795238 | 556.4867473 | 0.046064788 | 0.168912472 | 0.752837706 | 0.839016037 | ns | NaCl | KS.1 |
| FIM1_4554 | 1828.043984 | 1587.950802 | 2258.936361 | -0.161745371 | 0.154978938 | 0.237080486 | 0.363963368 | ns | NaCl | KS.1 |
| FIM1_4555 | 4271.624807 | 6817.743411 | 5241.801357 | 0.530334105 | 0.217892726 | 0.002949299 | 0.010563545 | ns | NaCl | KS.1 |
| FIM1_4556 | 2436.472182 | 3022.817691 | 2797.311843 | 0.219111174 | 0.20572471 | 0.192140923 | 0.309453251 | ns | NaCl | KS.1 |
| FIM1_4557 | 5057.89348 | 5445.929284 | 4748.96601 | 0.096806073 | 0.094360504 | 0.280861745 | 0.411087726 | ns | NaCl | KS.1 |
| FIM1_4558 | 1115.008324 | 725.4560553 | 1668.102854 | -0.538267021 | 0.162143208 | 0.000168189 | 0.00096726 | ns | NaCl | KS.1 |
| FIM1_4559 | 700.1136931 | 448.5591662 | 1020.512662 | -0.588555569 | 0.134506391 | 1.79583413265 | 1.95498354703 | ns | NaCl | KS.1 |
| FIM1_4560 | 125.0063549 | 71.51431187 | 161.0635889 | -0.59797581 | 0.266854095 | 0.003654672 | 0.012635158 | ns | NaCl | KS.1 |
| FIM1_4561 | 3549.854738 | 3633.417226 | 4062.928576 | 0.023464443 | 0.192664622 | 0.883863546 | 0.931220064 | ns | NaCl | KS.1 |
| FIM1_4562 | 180.9788784 | 420.7513342 | 665.2961242 | 0.761804716 | 0.375993509 | 0.002701836 | 0.009818577 | ns | NaCl | KS.1 |
| FIM1_4563 | 2390.267975 | 3971.923707 | 2210.548623 | 0.56866864 | 0.231267837 | 0.002331599 | 0.008669434 | ns | NaCl | KS.1 |
| FIM1_4564 | 382.253417 | 236.5604468 | 338.2719058 | -0.552700166 | 0.219267948 | 0.002099398 | 0.007942589 | ns | NaCl | KS.1 |
| FIM1_4565 | 3559.838396 | 3024.247598 | 4454.604135 | -0.152248206 | 0.222587354 | 0.388040708 | 0.523029673 | ns | NaCl | KS.1 |
| FIM1_4566 | 4743.154117 | 5610.617406 | 10289.83615 | 0.146281094 | 0.242174005 | 0.429430289 | 0.566166699 | ns | NaCl | KS.1 |
| FIM1_4567 | 7262.721768 | 7698.449054 | 6056.028043 | 0.073951717 | 0.109343186 | 0.470353842 | 0.60449764 | ns | NaCl | KS.1 |
| FIM1_4568 | 5177.098018 | 5765.734666 | 5307.079298 | 0.149226908 | 0.061920153 | 0.013642301 | 0.036528766 | ns | NaCl | KS.1 |
| FIM1_4569 | 2772.446181 | 2629.210679 | 2428.104162 | -0.062835607 | 0.138208717 | 0.615288594 | 0.730440001 | ns | NaCl | KS.1 |
| FIM1_4570 | 1074.799596 | 1212.813787 | 1569.40606 | 0.152422261 | 0.10978416 | 0.13761988 | 0.237893991 | ns | NaCl | KS.1 |
| FIM1_4571 | 1179.974353 | 1238.488583 | 1194.927305 | 0.058719425 | 0.120925069 | 0.599468903 | 0.71881364 | ns | NaCl | KS.1 |
| FIM1_4572 | 5490.618595 | 7944.279856 | 4823.600807 | 0.346978704 | 0.256100292 | 0.073444202 | 0.144478016 | ns | NaCl | KS.1 |
| FIM1_4573 | 1145.61427 | 1103.696588 | 1981.366376 | -0.041629588 | 0.163225843 | 0.770309282 | 0.851527619 | ns | NaCl | KS.1 |
| FIM1_4574 | 630.3287958 | 518.4783147 | 667.0720463 | -0.254493329 | 0.107093305 | 0.011086566 | 0.031003746 | ns | NaCl | KS.1 |
| FIM1_4575 | 9131.68829 | 10589.66798 | 8341.093234 | 0.192048498 | 0.103808803 | 0.049146144 | 0.104220829 | ns | NaCl | KS.1 |
| FIM1_4576 | 1848.119656 | 2119.763076 | 1594.398959 | 0.168014099 | 0.127471983 | 0.149945077 | 0.255209291 | ns | NaCl | KS.1 |
| FIM1_4577 | 514.0871531 | 297.6582189 | 482.5281874 | -0.639284603 | 0.216891603 | 0.000400527 | 0.00200869 | ns | NaCl | KS.1 |
| FIM1_4578 | 596.9357297 | 437.0323503 | 446.1690164 | -0.379335827 | 0.150717188 | 0.004526796 | 0.014973941 | ns | NaCl | KS.1 |
| FIM1_4579 | 328.9117216 | 404.2028381 | 334.6155605 | 0.209846942 | 0.207371848 | 0.213957575 | 0.336741201 | ns | NaCl | KS.1 |
| FIM1_4580 | 3827.792388 | 5947.269051 | 3068.994516 | 0.548517492 | 0.169161602 | 0.000212478 | 0.001177149 | ns | NaCl | KS.1 |
| FIM1_4582 | 1476.004812 | 1654.864075 | 1719.339598 | 0.111616709 | 0.205322728 | 0.50536338 | 0.637470288 | ns | NaCl | KS.1 |
| FIM1_4583 | 1159.355659 | 1401.475112 | 1529.542087 | 0.256291179 | 0.086306944 | 0.001889561 | 0.007304248 | ns | NaCl | KS.1 |
| FIM1_4584 | 1859.457246 | 1920.607327 | 1894.691616 | 0.041714667 | 0.111445321 | 0.688905983 | 0.79006622 | ns | NaCl | KS.1 |
| FIM1_4585 | 1926.292561 | 2131.236372 | 2464.633392 | 0.138124223 | 0.082232916 | 0.080674456 | 0.156169423 | ns | NaCl | KS.1 |
| FIM1_4586 | 14883.77291 | 18565.15551 | 14634.50189 | 0.305887283 | 0.068235115 | 3.82826399446 | 3.83984140573 | ns | NaCl | KS.1 |
| FIM1_4587 | 2923.953123 | 4849.531411 | 2336.525261 | 0.439433942 | 0.304976962 | 0.041253152 | 0.090411643 | ns | NaCl | KS.1 |
| FIM1_4588 | 2754.693192 | 3415.440705 | 3198.471663 | 0.268076788 | 0.130320087 | 0.023807543 | 0.057552249 | ns | NaCl | KS.1 |
| FIM1_4589 | 2489.833392 | 5351.654583 | 2506.601279 | 0.916843894 | 0.244189039 | 7.83274962924 | 6.92147946811 | ns | NaCl | KS.1 |
| FIM1_4590 | 514.5612006 | 629.6206604 | 592.5620059 | 0.252304278 | 0.132600686 | 0.036085216 | 0.08126933 | ns | NaCl | KS.1 |
| FIM1_4591 | 3057.508447 | 3366.605008 | 2635.107102 | 0.12263553 | 0.108194894 | 0.22656995 | 0.351476614 | ns | NaCl | KS.1 |
| FIM1_4592 | 1711.262432 | 1542.111411 | 1307.898843 | -0.138550591 | 0.088713885 | 0.102294404 | 0.18837304 | ns | NaCl | KS.1 |
| FIM1_4593 | 618.3018632 | 458.0064949 | 726.3664995 | -0.362336212 | 0.158168805 | 0.009047584 | 0.026446374 | ns | NaCl | KS.1 |
| FIM1_4594 | 356.2815693 | 392.9605866 | 742.0136083 | 0.090976908 | 0.206813649 | 0.58888846 | 0.709890983 | ns | NaCl | KS.1 |
| FIM1_4595 | 3658.674318 | 5723.169451 | 4025.083491 | 0.593074386 | 0.130452406 | 7.93732530525 | 9.60783294248 | ns | NaCl | KS.1 |

| | | | | | | | | | | |
|---|---|---|---|---|---|---|---|---|---|---|
| FIM1_4596 | 3108.935176 | 7671.921483 | 3538.793639 | 1.124834749 | 0.238053982 | 4.80066682709 | 8.32171340237 | Km>KS.1 | NaCl | KS.1 |
| FIM1_4597 | 1642.294075 | 2135.061974 | 1866.817906 | 0.348442523 | 0.104431661 | 0.000369065 | 0.001869752 | ns | NaCl | KS.1 |
| FIM1_4598 | 434.0002444 | 401.1952724 | 478.6078264 | -0.091094768 | 0.15925046 | 0.51485576 | 0.645678701 | ns | NaCl | KS.1 |
| FIM1_4599 | 425.6596666 | 293.4921224 | 280.8019534 | -0.42856293 | 0.189899481 | 0.007412842 | 0.022555897 | ns | NaCl | KS.1 |
| FIM1_4600 | 3713.828685 | 4813.158528 | 3711.087624 | 0.342625018 | 0.105087354 | 0.000497121 | 0.002396488 | ns | NaCl | KS.1 |
| FIM1_4601 | 968.2560353 | 688.8917852 | 781.6834496 | -0.460863409 | 0.10197779 | 1.55824672335 | 1.73429025698 | ns | NaCl | KS.1 |
| FIM1_4602 | 561.5152865 | 569.6602823 | 450.4868022 | 0.012475665 | 0.22186332 | 0.94339638 | 0.96930958 | ns | NaCl | KS.1 |
| FIM1_4603 | 4709.72236 | 4605.105801 | 3864.028105 | -0.028815032 | 0.107025765 | 0.774463938 | 0.853721994 | ns | NaCl | KS.1 |
| FIM1_4604 | 84.6288653 | 96.35479933 | 140.3676843 | 0.11967239 | 0.227727563 | 0.502944572 | 0.634925838 | ns | NaCl | KS.1 |
| FIM1_4605 | 1147.82309 | 1247.629069 | 1105.088326 | 0.107884081 | 0.103588622 | 0.26903484 | 0.398110747 | ns | NaCl | KS.1 |
| FIM1_4606 | 753.3334935 | 874.8160943 | 706.6727291 | 0.191116465 | 0.112256237 | 0.067643236 | 0.134879798 | ns | NaCl | KS.1 |
| FIM1_4607 | 2409.760045 | 2587.261021 | 2344.433376 | 0.09681146 | 0.076333391 | 0.189661197 | 0.306950051 | ns | NaCl | KS.1 |
| FIM1_4608 | 372.6343474 | 146.3428835 | 390.933759 | -1.194221758 | 0.226769622 | 2.16210625525 | 5.69125852904 | Km<KS.1 | NaCl | KS.1 |
| FIM1_4609 | 4100.914745 | 4605.33736 | 3251.156676 | 0.1213802 | 0.184219588 | 0.435616499 | 0.571667655 | ns | NaCl | KS.1 |
| FIM1_4612 | 6161.775277 | 7700.180563 | 7107.770878 | 0.273778319 | 0.137798419 | 0.027525584 | 0.06483891 | ns | NaCl | KS.1 |
| FIM1_4613 | 27866.73035 | 19838.55773 | 15088.26707 | -0.344060896 | 0.22899831 | 0.057450564 | 0.118793249 | ns | NaCl | KS.1 |
| FIM1_4614 | 619.7646475 | 560.0189175 | 609.0939818 | -0.12509986 | 0.114031534 | 0.238737971 | 0.365452741 | ns | NaCl | KS.1 |
| FIM1_4615 | 1092.766677 | 1394.148871 | 1502.09062 | 0.323422503 | 0.099361613 | 0.000550881 | 0.002617606 | ns | NaCl | KS.1 |
| FIM1_4616 | 4285.385957 | 4620.967182 | 3201.066156 | 0.099916583 | 0.08933507 | 0.241988202 | 0.368839247 | ns | NaCl | KS.1 |
| FIM1_4617 | 3289.997673 | 2842.397993 | 3570.518226 | -0.166189145 | 0.15969587 | 0.235239498 | 0.361908623 | ns | NaCl | KS.1 |
| FIM1_4618 | 4017.074097 | 4023.05857 | 4057.374277 | 0.002188269 | 0.109019576 | 0.982920574 | 0.991175946 | ns | NaCl | KS.1 |
| FIM1_4619 | 902.189609 | 893.031671 | 857.5893191 | -0.016011037 | 0.101346121 | 0.867273496 | 0.920564464 | ns | NaCl | KS.1 |
| FIM1_4620 | 817.4071358 | 819.3222782 | 744.6249619 | 0.001866489 | 0.210008843 | 0.991236321 | 0.996242565 | ns | NaCl | KS.1 |
| FIM1_4621 | 116.8220488 | 168.5018608 | 125.0342392 | 0.232457839 | 0.335273878 | 0.275632967 | 0.405582375 | ns | NaCl | KS.1 |
| FIM1_4622 | 976.5723029 | 644.9861185 | 1327.016102 | -0.545528898 | 0.132558217 | 6.81489000649 | 6.08690803991 | ns | NaCl | KS.1 |
| FIM1_4623 | 1104.920812 | 1422.172597 | 1412.356005 | 0.325237048 | 0.119750951 | 0.003189894 | 0.011239179 | ns | NaCl | KS.1 |
| FIM1_4624 | 2348.346325 | 1989.141087 | 2292.836892 | -0.208204267 | 0.119811408 | 0.059770342 | 0.122773515 | ns | NaCl | KS.1 |
| FIM1_4625 | 1114.144162 | 974.836933 | 926.2415632 | -0.177052196 | 0.085158349 | 0.030050967 | 0.069731139 | ns | NaCl | KS.1 |
| FIM1_4626 | 292.2279353 | 272.9459848 | 425.5798025 | -0.067618662 | 0.193655788 | 0.674851262 | 0.778975645 | ns | NaCl | KS.1 |
| FIM1_4627 | 2214.783059 | 1777.179094 | 2357.628351 | -0.256502512 | 0.158269403 | 0.064702862 | 0.130428175 | ns | NaCl | KS.1 |
| FIM1_4628 | 313.9515092 | 147.9636421 | 297.3288643 | -0.944824498 | 0.216923212 | 5.84737343291 | 7.47832463463 | ns | NaCl | KS.1 |
| FIM1_4629 | 973.0286073 | 580.2042348 | 744.8255768 | -0.641452349 | 0.186517632 | 7.24056381680 | 0.000480291 | ns | NaCl | KS.1 |
| FIM1_4630 | 1176.728917 | 793.5126842 | 1317.615698 | -0.5302727 | 0.115256162 | 7.77313667586 | 9.45106037546 | ns | NaCl | KS.1 |
| FIM1_4631 | 2493.43451 | 1678.900303 | 2080.043648 | -0.50034198 | 0.150964806 | 0.000201385 | 0.001123196 | ns | NaCl | KS.1 |
| FIM1_4632 | 1814.369526 | 2045.039964 | 2347.234258 | 0.121935541 | 0.193881102 | 0.44970956 | 0.584610676 | ns | NaCl | KS.1 |
| FIM1_4633 | 78.49728862 | 35.11543218 | 76.16514234 | -0.79059408 | 0.330558202 | 0.001027299 | 0.004379445 | ns | NaCl | KS.1 |
| FIM1_4634 | 1917.997434 | 1465.256494 | 2905.267577 | -0.223346119 | 0.267094213 | 0.253964107 | 0.382290902 | ns | NaCl | KS.1 |
| FIM1_4635 | 774.171531 | 785.4595425 | 1175.173052 | 0.017585176 | 0.126838744 | 0.879988728 | 0.929031192 | ns | NaCl | KS.1 |
| FIM1_4636 | 572.4732511 | 538.0287963 | 580.978079 | -0.085934658 | 0.100828835 | 0.367266889 | 0.502517264 | ns | NaCl | KS.1 |
| FIM1_4637 | 952.3504417 | 923.9750398 | 1091.158215 | -0.033681531 | 0.143744731 | 0.794193589 | 0.868567401 | ns | NaCl | KS.1 |
| FIM1_4638 | 1113.890123 | 1014.338929 | 1228.748941 | -0.121483707 | 0.100427635 | 0.200600415 | 0.320586915 | ns | NaCl | KS.1 |
| FIM1_4639 | 1818.898499 | 1862.363282 | 2308.383022 | 0.029504394 | 0.117025716 | 0.786116405 | 0.862197777 | ns | NaCl | KS.1 |
| FIM1_4640 | 1110.547385 | 1020.860208 | 889.2184359 | -0.095530045 | 0.155960519 | 0.487595737 | 0.621519035 | ns | NaCl | KS.1 |
| FIM1_4641 | 6555.75861 | 8485.663886 | 6170.986705 | 0.338007653 | 0.109776744 | 0.00094806 | 0.004104958 | ns | NaCl | KS.1 |
| FIM1_4642 | 4253.458345 | 3719.706875 | 3031.349814 | -0.159082456 | 0.141993205 | 0.21252354 | 0.335014135 | ns | NaCl | KS.1 |

| | | | | | | | | | | |
|---|---|---|---|---|---|---|---|---|---|---|
| FIM1_4643 | 651.4422307 | 592.191899 | 499.9010469 | -0.111179399 | 0.144446478 | 0.390611852 | 0.526067667 | ns | NaCl | KS.1 |
| FIM1_4644 | 714.7732624 | 650.6054972 | 877.6145571 | -0.11923953 | 0.12043566 | 0.28412143 | 0.414638931 | ns | NaCl | KS.1 |
| FIM1_4645 | 1127.56859 | 1381.580235 | 1148.317039 | 0.248119944 | 0.138875889 | 0.047282785 | 0.100914568 | ns | NaCl | KS.1 |
| FIM1_4646 | 2063.505664 | 1909.217109 | 1583.751112 | -0.099758522 | 0.102144508 | 0.300771701 | 0.432636272 | ns | NaCl | KS.1 |
| FIM1_4647 | 2449.254482 | 2523.278365 | 1914.355097 | 0.036197301 | 0.124139981 | 0.751609876 | 0.838250702 | ns | NaCl | KS.1 |
| FIM1_4648 | 1752.678325 | 1774.429701 | 1698.51504 | 0.015146739 | 0.122408257 | 0.893382214 | 0.936291661 | ns | NaCl | KS.1 |
| FIM1_4649 | 1853.058657 | 2341.526277 | 2547.851914 | 0.307473767 | 0.105829778 | 0.00190696 | 0.007348664 | ns | NaCl | KS.1 |
| FIM1_4650 | 1907.866401 | 1821.345539 | 1549.062917 | -0.058031212 | 0.118321849 | 0.596949996 | 0.716656909 | ns | NaCl | KS.1 |
| FIM1_4651 | 4573.413325 | 5434.938307 | 4372.04566 | 0.159586175 | 0.226590296 | 0.370805463 | 0.505631111 | ns | NaCl | KS.1 |
| FIM1_4652 | 1618.855864 | 1802.389018 | 1412.47613 | 0.138907426 | 0.104290097 | 0.156945422 | 0.264476618 | ns | NaCl | KS.1 |
| FIM1_4653 | 868.2313602 | 1132.592304 | 1046.675546 | 0.289728126 | 0.191287574 | 0.070132948 | 0.139279181 | ns | NaCl | KS.1 |
| FIM1_4654 | 1537.149008 | 942.7774837 | 1079.974788 | -0.60350457 | 0.180059971 | 0.000115292 | 0.000701196 | ns | NaCl | KS.1 |
| FIM1_4655 | 1984.218838 | 1328.799462 | 1886.468117 | -0.418998223 | 0.232792691 | 0.022913064 | 0.055714807 | ns | NaCl | KS.1 |
| FIM1_4656 | 2424.422604 | 2501.145561 | 2415.83052 | 0.027254076 | 0.232526088 | 0.879993065 | 0.929031192 | ns | NaCl | KS.1 |
| FIM1_4657 | 1275.224642 | 1156.610321 | 980.8930235 | -0.11920506 | 0.123830825 | 0.295696984 | 0.427643168 | ns | NaCl | KS.1 |
| FIM1_4658 | 28016.06133 | 34923.32705 | 20752.37853 | 0.253375518 | 0.164542655 | 0.076607007 | 0.149634809 | ns | NaCl | KS.1 |
| FIM1_4660 | 100.3070917 | 80.36073935 | 183.5974283 | -0.205984708 | 0.22765241 | 0.250097475 | 0.378532077 | ns | NaCl | KS.1 |
| FIM1_4661 | 476.3490689 | 600.1839884 | 320.9228849 | 0.228418166 | 0.216003034 | 0.187914893 | 0.304619287 | ns | NaCl | KS.1 |
| FIM1_4662 | 3706.145372 | 6852.29231 | 3115.470066 | 0.60070779 | 0.300875611 | 0.00644116 | 0.020115988 | ns | NaCl | KS.1 |
| FIM1_4664 | 43.1098118 | 21.01170202 | 39.5335887 | -0.661023731 | 0.331861554 | 0.004850252 | 0.015906397 | ns | NaCl | KS.1 |
| FIM1_4665 | 3500.190863 | 7448.094189 | 3582.114891 | 0.838264756 | 0.283474141 | 0.000171095 | 0.000981774 | ns | NaCl | KS.1 |
| FIM1_4666 | 479.7017285 | 453.8594491 | 786.747206 | -0.046690447 | 0.244100692 | 0.801101511 | 0.872669152 | ns | NaCl | KS.1 |
| FIM1_4667 | 5772.434505 | 11390.19326 | 5171.262894 | 0.648321167 | 0.322469412 | 0.004962547 | 0.01620072 | ns | NaCl | KS.1 |
| FIM1_4668 | 546.6962723 | 767.4875123 | 590.2764347 | 0.32641668 | 0.242210283 | 0.081288695 | 0.157052916 | ns | NaCl | KS.1 |
| FIM1_4669 | 29.82391728 | 23.15240403 | 39.49713505 | -0.147854522 | 0.309363855 | 0.472096846 | 0.605955059 | ns | NaCl | KS.1 |
| FIM1_4670 | 2316.261714 | 2363.345279 | 2421.800564 | 0.021146083 | 0.176596074 | 0.888728607 | 0.933183795 | ns | NaCl | KS.1 |
| FIM1_4671 | 1096.445523 | 797.8800679 | 1252.57698 | -0.405240345 | 0.132692869 | 0.000763101 | 0.003440245 | ns | NaCl | KS.1 |
| FIM1_4672 | 1230.7787 | 996.1158288 | 1166.076171 | -0.270875339 | 0.116802035 | 0.012172649 | 0.033347428 | ns | NaCl | KS.1 |
| FIM1_4673 | 288.8010525 | 286.3385088 | 398.5748486 | -0.004655277 | 0.140585726 | 0.970719804 | 0.986786069 | ns | NaCl | KS.1 |
| FIM1_4674 | 274.8559464 | 214.2328855 | 279.8664622 | -0.323572739 | 0.132335011 | 0.007031587 | 0.021607254 | ns | NaCl | KS.1 |
| FIM1_4675 | 4733.845509 | 6281.884151 | 4876.118353 | 0.366741332 | 0.118853839 | 0.000816393 | 0.003632876 | ns | NaCl | KS.1 |
| FIM1_4676 | 2445.174092 | 2609.712218 | 2206.597425 | 0.089798565 | 0.074209464 | 0.211613315 | 0.33421468 | ns | NaCl | KS.1 |
| FIM1_4678 | 3196.532826 | 3515.230079 | 2206.929083 | 0.073958804 | 0.263103074 | 0.700636214 | 0.798938005 | ns | NaCl | KS.1 |
| FIM1_4679 | 1139.185551 | 1911.058057 | 1706.025796 | 0.54525067 | 0.255710799 | 0.006063744 | 0.019165901 | ns | NaCl | KS.1 |
| FIM1_4680 | 568.3913518 | 678.8131007 | 650.5430871 | 0.227851779 | 0.114420122 | 0.031976919 | 0.07348045 | ns | NaCl | KS.1 |
| FIM1_4681 | 2672.802299 | 3678.042214 | 3003.550954 | 0.381609149 | 0.166318828 | 0.008245797 | 0.024653149 | ns | NaCl | KS.1 |
| FIM1_4682 | 7131.164704 | 7925.148707 | 6334.836483 | 0.142318692 | 0.078848693 | 0.061250522 | 0.125193652 | ns | NaCl | KS.1 |
| FIM1_4683 | 261.692219 | 185.3674955 | 149.174951 | -0.299452625 | 0.272312693 | 0.132992921 | 0.231747735 | ns | NaCl | KS.1 |
| FIM1_4684 | 1865.112866 | 2206.568743 | 1653.177765 | 0.188771156 | 0.166089876 | 0.190569377 | 0.307819042 | ns | NaCl | KS.1 |
| FIM1_4685 | 2122.831423 | 1091.018243 | 1895.683557 | -0.593012802 | 0.33713922 | 0.010870887 | 0.030584865 | ns | NaCl | KS.1 |
| FIM1_4686 | 178.6607745 | 62.86777267 | 374.4480036 | -1.036455445 | 0.384715917 | 0.000142037 | 0.000838237 | Km<KS.1 | NaCl | KS.1 |
| FIM1_4688 | 1613.954699 | 1456.683266 | 1030.77713 | -0.113072418 | 0.162334508 | 0.425518401 | 0.561973466 | ns | NaCl | KS.1 |
| FIM1_4689 | 1097.134735 | 1230.52459 | 1260.925112 | 0.144258666 | 0.112779357 | 0.169986995 | 0.281370542 | ns | NaCl | KS.1 |
| FIM1_4690 | 2680.967295 | 3190.280601 | 3273.689111 | 0.211018073 | 0.136994781 | 0.088241416 | 0.16774973 | ns | NaCl | KS.1 |
| FIM1_4691 | 1736.3608 | 3150.147288 | 1392.037738 | 0.578752324 | 0.298275211 | 0.008089864 | 0.024245226 | ns | NaCl | KS.1 |

| | | | | | | | | | | |
|---|---|---|---|---|---|---|---|---|---|---|
| FIM1_4692 | 969.6876464 | 1931.824919 | 1443.260619 | 0.887244401 | 0.183261519 | 7.43648354069 | 1.2250498548( | ns | NaCl | KS.1 |
| FIM1_4693 | 286.0744514 | 307.8408071 | 358.3793039 | 0.078303545 | 0.185490096 | 0.616780882 | 0.731902522 | ns | NaCl | KS.1 |
| FIM1_4694 | 773.402561 | 602.9008318 | 868.552117 | -0.224539329 | 0.24403097 | 0.22906943 | 0.354580092 | ns | NaCl | KS.1 |
| FIM1_4695 | 759.9625807 | 733.9401398 | 611.148356 | -0.030994868 | 0.224443283 | 0.860927386 | 0.915702134 | ns | NaCl | KS.1 |
| FIM1_4697 | 2112.170378 | 3459.106285 | 1508.15025 | 0.367797299 | 0.331469129 | 0.093926972 | 0.176334598 | ns | NaCl | KS.1 |
| FIM1_4698 | 1525.370175 | 1112.126412 | 1300.486597 | -0.386726885 | 0.154586237 | 0.004575291 | 0.015084211 | ns | NaCl | KS.1 |
| FIM1_4699 | 1000.760065 | 873.1512531 | 876.9958142 | -0.119025168 | 0.237402616 | 0.51526003 | 0.646022845 | ns | NaCl | KS.1 |
| FIM1_4700 | 34.88958808 | 34.45583674 | 31.51954424 | -0.010849912 | 0.263510157 | 0.95498853 | 0.976656682 | ns | NaCl | KS.1 |
| FIM1_4701 | 1379.191428 | 919.7369073 | 949.344502 | -0.437129818 | 0.222058905 | 0.014520462 | 0.038323235 | ns | NaCl | KS.1 |
| FIM1_4702 | 2433.256087 | 2273.141764 | 2349.804366 | -0.055222236 | 0.251893838 | 0.769362919 | 0.851376188 | ns | NaCl | KS.1 |
| FIM1_4703 | 8689.151781 | 7871.799611 | 8353.968002 | -0.089496557 | 0.224931096 | 0.613631342 | 0.729117728 | ns | NaCl | KS.1 |
| FIM1_4704 | 1588.437016 | 1543.541756 | 1487.561147 | -0.025659678 | 0.222740353 | 0.884166465 | 0.931341978 | ns | NaCl | KS.1 |
| FIM1_4705 | 1461.629413 | 1560.791859 | 1309.422391 | 0.07873536 | 0.132309246 | 0.513837084 | 0.644726228 | ns | NaCl | KS.1 |
| FIM1_4706 | 1769.842588 | 1497.878314 | 1323.038148 | -0.205448206 | 0.131758167 | 0.086536399 | 0.165456797 | ns | NaCl | KS.1 |
| FIM1_4707 | 821.64504 | 536.6535556 | 651.8354446 | -0.469940101 | 0.219930263 | 0.008387442 | 0.024986542 | ns | NaCl | KS.1 |
| FIM1_4708 | 1895.05686 | 1481.690366 | 1232.431846 | -0.202054465 | 0.266249539 | 0.300358289 | 0.43249855 | ns | NaCl | KS.1 |
| FIM1_4709 | 2053.215115 | 2256.841697 | 1693.347632 | 0.088553185 | 0.215586542 | 0.608302321 | 0.724943649 | ns | NaCl | KS.1 |
| FIM1_4710 | 1571.874759 | 1234.904174 | 1338.096276 | -0.236258731 | 0.221333752 | 0.180016686 | 0.295182273 | ns | NaCl | KS.1 |
| FIM1_4711 | 2129.897869 | 1834.031464 | 2530.178698 | -0.150525968 | 0.19944954 | 0.359978971 | 0.49568098 | ns | NaCl | KS.1 |
| FIM1_4712 | 486.0559182 | 486.51969 | 488.8438043 | 0.001199751 | 0.191551808 | 0.994017362 | 0.99742565 | ns | NaCl | KS.1 |
| FIM1_4713 | 1943.944488 | 2177.269785 | 2362.926956 | 0.12650652 | 0.16343957 | 0.375004661 | 0.509600707 | ns | NaCl | KS.1 |
| FIM1_4714 | 1758.8619 | 1465.310039 | 1656.996978 | -0.195026235 | 0.185832435 | 0.21302591 | 0.335593383 | ns | NaCl | KS.1 |
| FIM1_4715 | 4075.705385 | 4874.583152 | 4836.093207 | 0.188308238 | 0.189559011 | 0.235754658 | 0.362559327 | ns | NaCl | KS.1 |
| FIM1_4716 | 1870.522675 | 3341.396931 | 3401.173216 | 0.460703935 | 0.339617858 | 0.041927482 | 0.0914865 | ns | NaCl | KS.1 |
| FIM1_4717 | 5470.650688 | 9711.478604 | 4812.049546 | 0.574513028 | 0.284676199 | 0.00694359 | 0.021379339 | ns | NaCl | KS.1 |
| FIM1_4718 | 2604.072737 | 4208.645038 | 1948.392689 | 0.354777645 | 0.330229758 | 0.104641943 | 0.19160606 | ns | NaCl | KS.1 |
| FIM1_4719 | 407.3175444 | 457.9672666 | 359.3621064 | 0.115089414 | 0.203027286 | 0.489041556 | 0.622616398 | ns | NaCl | KS.1 |
| FIM1_4720 | 2781.487402 | 6079.822642 | 2687.385742 | 0.85538882 | 0.29525103 | 0.000190345 | 0.001071229 | ns | NaCl | KS.1 |
| FIM1_4721 | 870.2858173 | 679.6827376 | 1004.264901 | -0.308002896 | 0.131098532 | 0.009722985 | 0.02792832 | ns | NaCl | KS.1 |
| FIM1_4722 | 2213.664597 | 4419.458524 | 1807.401918 | 0.489557852 | 0.382733711 | 0.038994077 | 0.086605148 | ns | NaCl | KS.1 |
| FIM1_4724 | 732.2271142 | 574.8530335 | 584.5339102 | -0.289119882 | 0.15882918 | 0.037801704 | 0.084637028 | ns | NaCl | KS.1 |
| FIM1_4725 | 226.9948404 | 225.5951607 | 211.8274937 | -0.001538707 | 0.446431209 | 0.993291306 | 0.996978696 | ns | NaCl | KS.1 |
| FIM1_4726 | 45.53435485 | 72.86299809 | 61.86479955 | 0.148790827 | 0.433410338 | 0.453472337 | 0.58857941 | ns | NaCl | KS.1 |
| FIM1_4727 | 542.8398664 | 406.4541171 | 420.3880307 | -0.062904678 | 0.460791366 | 0.723903093 | 0.81683327 | ns | NaCl | KS.1 |
| FIM1_4728 | 63.61263343 | 65.36442138 | 140.8377942 | 0.007045417 | 0.445719692 | 0.969350805 | 0.986200462 | ns | NaCl | KS.1 |
| FIM1_4729 | 803.2415972 | 634.2810858 | 766.2970696 | -0.052483253 | 0.457573831 | 0.769404491 | 0.851376188 | ns | NaCl | KS.1 |
| FIM1_4730 | 743.4055972 | 702.7325982 | 481.3915826 | -0.05077094 | 0.23056748 | 0.777538699 | 0.855430126 | ns | NaCl | KS.1 |
| FIM1_4731 | 2266.235575 | 2813.91845 | 1595.830275 | 0.202273376 | 0.229495985 | 0.260973218 | 0.389541482 | ns | NaCl | KS.1 |
| FIM1_4732 | 3950.48432 | 2718.338716 | 3415.38404 | -0.361097713 | 0.249458045 | 0.058907103 | 0.121351072 | ns | NaCl | KS.1 |
| FIM1_4733 | 1566.626916 | 1009.477755 | 1350.604209 | -0.523178644 | 0.191019317 | 0.001248967 | 0.00515225 | ns | NaCl | KS.1 |
| FIM1_4734 | 497.4975749 | 388.557622 | 494.3151773 | -0.260449804 | 0.203480389 | 0.118644463 | 0.212562884 | ns | NaCl | KS.1 |
| FIM1_4736 | 334.7574418 | 293.5531747 | 252.8171641 | -0.124011792 | 0.22180403 | 0.480794159 | 0.614609043 | ns | NaCl | KS.1 |
| FIM1_4737 | 575.6288409 | 508.2286931 | 807.1249009 | -0.122182655 | 0.200486196 | 0.458953633 | 0.593861594 | ns | NaCl | KS.1 |
| FIM1_4738 | 74.07367147 | 91.03392373 | 122.0593403 | 0.167861633 | 0.260478374 | 0.383180111 | 0.518444671 | ns | NaCl | KS.1 |
| FIM1_4739 | 451.8886096 | 525.5266451 | 449.3723635 | 0.177297996 | 0.154739445 | 0.194308194 | 0.312242196 | ns | NaCl | KS.1 |

| | | | | | | | | | | |
|---|---|---|---|---|---|---|---|---|---|---|
| FIM1_4740 | 653.5946098 | 566.6418762 | 1494.907235 | -0.161577935 | 0.163438518 | 0.256935251 | 0.385364146 | ns | NaCl | KS.1 |
| FIM1_4741 | 392.8641597 | 282.1103813 | 562.5953815 | -0.363865962 | 0.200005787 | 0.027929618 | 0.06566628 | ns | NaCl | KS.1 |
| FIM1_4742 | 227.470127 | 206.397289 | 286.3233181 | -0.081207972 | 0.240850844 | 0.659250158 | 0.765943375 | ns | NaCl | KS.1 |
| FIM1_4743 | 19.2282278 | 16.93852546 | 29.18473673 | -0.082579538 | 0.330555131 | 0.689460309 | 0.790337566 | ns | NaCl | KS.1 |
| FIM1_4744 | 3314.15999 | 3498.430527 | 2880.341186 | 0.058109636 | 0.173469932 | 0.696866545 | 0.796991049 | ns | NaCl | KS.1 |
| FIM1_4745 | 1086.694562 | 622.9756959 | 978.5767618 | -0.744600632 | 0.141203564 | 1.2274405465 | 2.5732996037 | ns | NaCl | KS.1 |
| FIM1_4746 | 2748.472445 | 4227.475072 | 2526.440432 | 0.460211478 | 0.231526055 | 0.012534597 | 0.033964934 | ns | NaCl | KS.1 |
| FIM1_4747 | 666.1744114 | 749.4725355 | 669.4574419 | 0.145938012 | 0.120724183 | 0.190542558 | 0.307819042 | ns | NaCl | KS.1 |
| FIM1_4748 | 4406.64509 | 3645.897177 | 4156.527977 | -0.249655708 | 0.097363316 | 0.006686562 | 0.020739179 | ns | NaCl | KS.1 |
| FIM1_4749 | 344.9207983 | 301.8035796 | 434.7145254 | -0.169032455 | 0.129721509 | 0.153472685 | 0.259879716 | ns | NaCl | KS.1 |
| FIM1_4750 | 4145.645844 | 5646.943781 | 4683.332238 | 0.426798674 | 0.077440019 | 1.0266455038 | 2.1847581015 | ns | NaCl | KS.1 |
| FIM1_4751 | 207.3130167 | 255.8526322 | 204.7642249 | 0.221620227 | 0.192024508 | 0.166758216 | 0.277558423 | ns | NaCl | KS.1 |
| FIM1_4753 | 2155.805305 | 4913.855372 | 2190.71695 | 0.987127383 | 0.25317416 | 3.3235483125 | 3.3952059250 | ns | NaCl | KS.1 |
| FIM1_4754 | 1686.733955 | 2377.683973 | 1932.410155 | 0.450237073 | 0.121973704 | 6.0335923552 | 0.000407288 | ns | NaCl | KS.1 |
| FIM1_4755 | 400.4030679 | 320.5795735 | 420.6909983 | -0.257894786 | 0.163691127 | 0.070364545 | 0.139449028 | ns | NaCl | KS.1 |
| FIM1_4756 | 1796.471416 | 2928.219107 | 1824.271366 | 0.626974993 | 0.15928833 | 1.0784309138 | 9.1089877695 | ns | NaCl | KS.1 |
| FIM1_4757 | 1650.60634 | 3011.786484 | 1744.657445 | 0.643191946 | 0.268806438 | 0.00201678 | 0.007688492 | ns | NaCl | KS.1 |
| FIM1_4758 | 611.712702 | 354.4772172 | 689.9815069 | -0.731640489 | 0.131219999 | 2.3588264550 | 6.1250314389 | ns | NaCl | KS.1 |
| FIM1_4759 | 5522.653363 | 7860.3764 | 6178.264648 | 0.428471366 | 0.165284054 | 0.002930765 | 0.010512296 | ns | NaCl | KS.1 |
| FIM1_4760 | 6562.135598 | 5001.820666 | 4423.256503 | -0.239321409 | 0.253261996 | 0.209609616 | 0.331787413 | ns | NaCl | KS.1 |
| FIM1_4761 | 418.7993239 | 365.2505533 | 374.2602876 | -0.149244229 | 0.165693702 | 0.300144973 | 0.432316514 | ns | NaCl | KS.1 |
| FIM1_4762 | 200.9142472 | 121.3577415 | 127.7586392 | -0.613596748 | 0.200703957 | 0.000309496 | 0.001610607 | ns | NaCl | KS.1 |
| FIM1_4763 | 234.2584323 | 189.6996749 | 172.8569932 | -0.203196884 | 0.218439642 | 0.244456868 | 0.372032094 | ns | NaCl | KS.1 |
| FIM1_4764 | 466.0511517 | 338.1112051 | 328.2736609 | -0.307127025 | 0.24317804 | 0.101236251 | 0.186952616 | ns | NaCl | KS.1 |
| FIM1_4765 | 184.5405993 | 152.3813615 | 314.2749395 | -0.173363512 | 0.229132188 | 0.334351562 | 0.468582458 | ns | NaCl | KS.1 |
| FIM1_4766 | 172.6671733 | 172.299213 | 195.8864339 | 0.001328726 | 0.191437263 | 0.993371537 | 0.996978696 | ns | NaCl | KS.1 |
| FIM1_4767 | 571.4123862 | 742.316979 | 816.4770033 | 0.31683994 | 0.154210814 | 0.019820019 | 0.04920389 | ns | NaCl | KS.1 |
| FIM1_4769 | 69.25403677 | 120.326065 | 118.1575199 | 0.474634916 | 0.319501593 | 0.032129712 | 0.073797469 | ns | NaCl | KS.1 |
| FIM1_4770 | 481.5896601 | 842.7317227 | 529.218885 | 0.413538318 | 0.346966826 | 0.066684668 | 0.133424694 | ns | NaCl | KS.1 |
| FIM1_4771 | 679.7521758 | 928.906292 | 746.2717156 | 0.343726806 | 0.195431008 | 0.034616511 | 0.078423106 | ns | NaCl | KS.1 |
| FIM1_4772 | 26.58160439 | 21.21992949 | 75.1073442 | -0.147812805 | 0.272165851 | 0.451158436 | 0.586340966 | ns | NaCl | KS.1 |
| FIM1_4773 | 2019.731112 | 1649.215917 | 2745.278261 | -0.232118834 | 0.161293113 | 0.09958609 | 0.184247229 | ns | NaCl | KS.1 |
| FIM1_3 | 3972.683628 | 4024.450814 | 3037.529789 | 0.006689878 | 0.082023731 | 0.890535362 | 0.977268396 | ns | NaCl | KS.3 |
| FIM1_5 | 541.1120197 | 476.8030815 | 633.8041188 | -0.023054474 | 0.114826297 | 0.564437785 | 0.867532568 | ns | NaCl | KS.3 |
| FIM1_6 | 493.6296728 | 394.5438021 | 405.4975614 | -0.070178608 | 0.116922825 | 0.147509851 | 0.562037426 | ns | NaCl | KS.3 |
| FIM1_7 | 52.21799034 | 74.93762594 | 60.8195808 | 0.087205906 | 0.15694481 | 0.08533778 | 0.453584891 | ns | NaCl | KS.3 |
| FIM1_8 | 352.2877542 | 328.3015477 | 375.3172182 | -0.014992925 | 0.106250909 | 0.717264988 | 0.93000992 | ns | NaCl | KS.3 |
| FIM1_9 | 5072.061404 | 5815.10095 | 2901.971283 | 0.020499888 | 0.121255112 | 0.591787368 | 0.878786986 | ns | NaCl | KS.3 |
| FIM1_10 | 961.0099126 | 2795.135043 | 2943.370395 | 1.254704556 | 0.366860209 | 3.2802727034 | 0.00050998 | Km>KS.3 | NaCl | KS.3 |
| FIM1_11 | 341.199521 | 349.6487519 | 340.6940978 | 0.008872893 | 0.093610338 | 0.845459613 | 0.967147129 | ns | NaCl | KS.3 |
| FIM1_12 | 510.9969533 | 990.6146393 | 1729.337393 | 0.516178288 | 0.38345423 | 0.000743487 | 0.030071932 | ns | NaCl | KS.3 |
| FIM1_13 | 42702.44797 | 44428.48315 | 35774.55391 | 0.004260356 | 0.128509536 | 0.903214742 | 0.979395371 | ns | NaCl | KS.3 |
| FIM1_14 | 15.29048541 | 21.9617334 | 16.92066841 | 0.037525866 | 0.154753113 | 0.317087925 | 0.73382286 | ns | NaCl | KS.3 |
| FIM1_15 | 17895.52351 | 19494.81606 | 14849.47076 | 0.006755358 | 0.141449426 | 0.837566751 | 0.962829707 | ns | NaCl | KS.3 |
| FIM1_16 | 14224.48292 | 13427.54739 | 13864.7932 | -0.011552574 | 0.108728645 | 0.774593942 | 0.950214521 | ns | NaCl | KS.3 |

| | | | | | | | | | | |
|---|---|---|---|---|---|---|---|---|---|---|
| FIM1_17 | 26452.11556 | 27568.25297 | 21161.65642 | 0.003692618 | 0.135672778 | 0.912812118 | 0.98221021 | ns | NaCl | KS.3 |
| FIM1_18 | 3005.91678 | 2951.512989 | 2232.947735 | -0.003636813 | 0.107227585 | 0.928336465 | 0.984238275 | ns | NaCl | KS.3 |
| FIM1_19 | 586.5392585 | 571.9968294 | 965.1753006 | -0.01658363 | 0.074582148 | 0.736269887 | 0.938734671 | ns | NaCl | KS.3 |
| FIM1_20 | 3467.268442 | 3271.144042 | 3333.389345 | -0.04299969 | 0.069235624 | 0.380714657 | 0.778493059 | ns | NaCl | KS.3 |
| FIM1_21 | 520.684461 | 535.2310194 | 585.7584931 | 0.013291856 | 0.08482541 | 0.782575683 | 0.951646805 | ns | NaCl | KS.3 |
| FIM1_22 | 1555.536925 | 1575.774396 | 1520.407378 | 0.010968824 | 0.061610578 | 0.815626073 | 0.958816322 | ns | NaCl | KS.3 |
| FIM1_23 | 188.0059098 | 165.945119 | 230.7427362 | -0.040223055 | 0.100678091 | 0.380412325 | 0.778493059 | ns | NaCl | KS.3 |
| FIM1_24 | 1894.43911 | 2092.246787 | 1199.112786 | 0.020168469 | 0.110233885 | 0.620781928 | 0.892479944 | ns | NaCl | KS.3 |
| FIM1_25 | 504.9700485 | 534.4969913 | 454.9892691 | 0.013095001 | 0.104675128 | 0.753906053 | 0.943805388 | ns | NaCl | KS.3 |
| FIM1_26 | 220.1461398 | 171.968155 | 284.9825584 | -0.070681444 | 0.125086913 | 0.141067975 | 0.552608798 | ns | NaCl | KS.3 |
| FIM1_27 | 588.6296305 | 556.1685608 | 657.296323 | -0.02259844 | 0.091266445 | 0.629366312 | 0.896016329 | ns | NaCl | KS.3 |
| FIM1_28 | 15267.50163 | 15262.90708 | 11506.37819 | -0.000109479 | 0.093755122 | 0.998073985 | 0.999279146 | ns | NaCl | KS.3 |
| FIM1_29 | 1597.340142 | 1547.363601 | 2774.21492 | -0.01013662 | 0.094950896 | 0.822113446 | 0.960240659 | ns | NaCl | KS.3 |
| FIM1_30 | 2007.844809 | 2095.350461 | 2049.9388 | 0.022675596 | 0.081024035 | 0.643521339 | 0.898798052 | ns | NaCl | KS.3 |
| FIM1_31 | 4133.70466 | 4830.093969 | 4258.824352 | 0.127370885 | 0.075554715 | 0.014645668 | 0.175571561 | ns | NaCl | KS.3 |
| FIM1_32 | 2775.195563 | 2475.969433 | 2794.011073 | -0.060552968 | 0.085461877 | 0.227716898 | 0.651504817 | ns | NaCl | KS.3 |
| FIM1_33 | 724.7950449 | 794.6297769 | 732.2157693 | 0.030084174 | 0.097212328 | 0.509719569 | 0.84584885 | ns | NaCl | KS.3 |
| FIM1_34 | 1964.818807 | 1890.084896 | 1643.975651 | -0.030749892 | 0.062971588 | 0.517506673 | 0.850188733 | ns | NaCl | KS.3 |
| FIM1_35 | 25.38475763 | 30.09779015 | 43.59138275 | 0.022669609 | 0.127446861 | 0.545095708 | 0.857313452 | ns | NaCl | KS.3 |
| FIM1_36 | 7821.741848 | 8837.279451 | 6124.361802 | 0.012766499 | 0.132007103 | 0.716311005 | 0.929477573 | ns | NaCl | KS.3 |
| FIM1_37 | 838.8148372 | 1103.269273 | 700.1965119 | 0.157342496 | 0.133291218 | 0.011524973 | 0.151684497 | ns | NaCl | KS.3 |
| FIM1_38 | 1952.177696 | 2387.989086 | 1220.934355 | 0.079283598 | 0.107869547 | 0.116716904 | 0.519941265 | ns | NaCl | KS.3 |
| FIM1_39 | 2095.194457 | 2498.699382 | 1622.369305 | 0.046948247 | 0.110839047 | 0.295638744 | 0.714058464 | ns | NaCl | KS.3 |
| FIM1_40 | 671.9905791 | 608.800617 | 1421.516646 | -0.050671188 | 0.085795849 | 0.30663847 | 0.726095377 | ns | NaCl | KS.3 |
| FIM1_41 | 1078.626621 | 1152.470779 | 1004.945188 | 0.033093292 | 0.084068153 | 0.498620047 | 0.837486406 | ns | NaCl | KS.3 |
| FIM1_42 | 1537.546239 | 1465.237835 | 1088.647994 | -0.019782266 | 0.089445919 | 0.675184971 | 0.912806589 | ns | NaCl | KS.3 |
| FIM1_43 | 2997.042352 | 2629.276338 | 2347.041188 | -0.114672931 | 0.067495656 | 0.021111577 | 0.221497506 | ns | NaCl | KS.3 |
| FIM1_44 | 217.2130415 | 182.4089105 | 223.7839843 | -0.048638923 | 0.109785814 | 0.283544925 | 0.704471025 | ns | NaCl | KS.3 |
| FIM1_45 | 628.0855824 | 544.6518981 | 909.3249672 | -0.068508721 | 0.09241888 | 0.173172608 | 0.600790605 | ns | NaCl | KS.3 |
| FIM1_46 | 4086.407441 | 4517.734938 | 3895.846385 | 0.051090217 | 0.0857034 | 0.302995585 | 0.72182705 | ns | NaCl | KS.3 |
| FIM1_47 | 642.1716736 | 1554.170423 | 1453.094101 | 1.238441811 | 0.120907657 | 1.6595724743 | 4.1281865299 | Km>KS.3 | NaCl | KS.3 |
| FIM1_48 | 2010.91032 | 2198.008846 | 2574.536415 | 0.076831179 | 0.063772564 | 0.110256754 | 0.505654054 | ns | NaCl | KS.3 |
| FIM1_49 | 17216.2429 | 22445.48988 | 9518.429149 | 0.05464544 | 0.130997701 | 0.2150845 | 0.64036229 | ns | NaCl | KS.3 |
| FIM1_50 | 3712.304279 | 3270.612371 | 4848.123799 | -0.034455042 | 0.104569261 | 0.435367651 | 0.808096352 | ns | NaCl | KS.3 |
| FIM1_51 | 3850.223374 | 3498.277867 | 3374.321267 | -0.042415338 | 0.089531665 | 0.381630426 | 0.778609403 | ns | NaCl | KS.3 |
| FIM1_52 | 3269.796325 | 3905.934803 | 3261.822567 | 0.095041267 | 0.095778591 | 0.071532549 | 0.418675804 | ns | NaCl | KS.3 |
| FIM1_53 | 46294.02904 | 53127.16326 | 32534.50092 | 0.086653917 | 0.082860835 | 0.093876093 | 0.470201108 | ns | NaCl | KS.3 |
| FIM1_54 | 1676.485081 | 1899.32328 | 3260.43686 | 0.034052054 | 0.104518714 | 0.440311051 | 0.808917498 | ns | NaCl | KS.3 |
| FIM1_55 | 362.9196306 | 130.9360917 | 884.141548 | -0.956457713 | 0.511377393 | 0.000105754 | 0.007852613 | ns | NaCl | KS.3 |
| FIM1_56 | 22.80613455 | 18.01165942 | 110.1015889 | -0.027965786 | 0.140186557 | 0.446058173 | 0.812914509 | ns | NaCl | KS.3 |
| FIM1_57 | 410.2485078 | 419.3173799 | 411.2940193 | 0.004260521 | 0.10943688 | 0.914597622 | 0.982343702 | ns | NaCl | KS.3 |
| FIM1_58 | 4976.887642 | 6238.406292 | 3663.533176 | 0.082581307 | 0.115421173 | 0.103470326 | 0.491042668 | ns | NaCl | KS.3 |
| FIM1_59 | 3697.665199 | 4507.344638 | 4886.402825 | 0.03938447 | 0.120132227 | 0.348044059 | 0.756654396 | ns | NaCl | KS.3 |
| FIM1_60 | 2495.985765 | 2802.495963 | 2097.26308 | 0.068383464 | 0.08232091 | 0.177749991 | 0.606104322 | ns | NaCl | KS.3 |
| FIM1_61 | 981.0850293 | 971.3599079 | 1514.364944 | -0.009086351 | 0.071504189 | 0.852974783 | 0.969345976 | ns | NaCl | KS.3 |

| | | | | | | | | | | |
|---|---|---|---|---|---|---|---|---|---|---|
| FIM1_62 | 1549.251386 | 1603.144493 | 1248.976291 | 0.013384342 | 0.089577743 | 0.775590645 | 0.950385088 | ns | NaCl | KS.3 |
| FIM1_63 | 44.40935838 | 46.09783121 | 61.0388668 | 0.00640789 | 0.120070496 | 0.862188145 | 0.969856868 | ns | NaCl | KS.3 |
| FIM1_64 | 1877.312243 | 1999.108888 | 2027.713438 | 0.043477972 | 0.071713512 | 0.378962775 | 0.778493059 | ns | NaCl | KS.3 |
| FIM1_65 | 179.3756087 | 184.7163824 | 253.6940693 | 0.008842518 | 0.08826441 | 0.851479387 | 0.968696536 | ns | NaCl | KS.3 |
| FIM1_66 | 6447.102203 | 7383.285282 | 4634.515481 | 0.059341633 | 0.093593629 | 0.228667677 | 0.652306017 | ns | NaCl | KS.3 |
| FIM1_67 | 664.3731964 | 992.9267964 | 516.1440968 | 0.265392667 | 0.224670674 | 0.002782807 | 0.065304085 | ns | NaCl | KS.3 |
| FIM1_68 | 20577.48682 | 24016.67867 | 15105.22591 | 0.060243853 | 0.099109075 | 0.21590273 | 0.640629323 | ns | NaCl | KS.3 |
| FIM1_69 | 19879.62591 | 23182.6175 | 14277.90827 | 0.085212186 | 0.089614255 | 0.100184542 | 0.483636126 | ns | NaCl | KS.3 |
| FIM1_70 | 1852.197206 | 2317.06084 | 1316.552668 | 0.083649213 | 0.114801219 | 0.10042009 | 0.483636126 | ns | NaCl | KS.3 |
| FIM1_71 | 674.9072817 | 725.6850044 | 738.570201 | 0.03101611 | 0.088680554 | 0.517623675 | 0.850188733 | ns | NaCl | KS.3 |
| FIM1_72 | 663.4710775 | 681.8865974 | 603.4281888 | 0.011200249 | 0.089917713 | 0.810823447 | 0.958816322 | ns | NaCl | KS.3 |
| FIM1_73 | 18657.99189 | 18381.24466 | 13298.10691 | -0.010677724 | 0.069529202 | 0.826861062 | 0.960240659 | ns | NaCl | KS.3 |
| FIM1_74 | 937.3053238 | 993.715857 | 775.8522912 | 0.039257103 | 0.071946715 | 0.426565925 | 0.802854185 | ns | NaCl | KS.3 |
| FIM1_75 | 19085.21291 | 24531.78867 | 22825.45264 | 0.147594693 | 0.121290078 | 0.013816039 | 0.170372387 | ns | NaCl | KS.3 |
| FIM1_76 | 278.8985135 | 274.2395804 | 347.4029841 | -0.005565379 | 0.092517543 | 0.903352372 | 0.979395371 | ns | NaCl | KS.3 |
| FIM1_77 | 869.9485338 | 898.1796473 | 859.8821874 | 0.019387155 | 0.075835532 | 0.693995987 | 0.920525145 | ns | NaCl | KS.3 |
| FIM1_78 | 3281.796135 | 3047.690276 | 3222.916234 | -0.039382966 | 0.08241098 | 0.425012625 | 0.802854185 | ns | NaCl | KS.3 |
| FIM1_79 | 835.7512763 | 909.5718137 | 3178.933275 | 0.022667162 | 0.101826365 | 0.602568205 | 0.881535538 | ns | NaCl | KS.3 |
| FIM1_80 | 60.28533029 | 49.46063703 | 115.301577 | -0.044471495 | 0.118868924 | 0.303229629 | 0.72182705 | ns | NaCl | KS.3 |
| FIM1_81 | 1190.800721 | 1095.808383 | 1455.405965 | -0.034438113 | 0.090879003 | 0.469781042 | 0.821848579 | ns | NaCl | KS.3 |
| FIM1_82 | 2324.094829 | 2341.160039 | 2001.056325 | 0.004812893 | 0.071024537 | 0.921712713 | 0.983890455 | ns | NaCl | KS.3 |
| FIM1_83 | 372.4983448 | 318.4384564 | 385.1962856 | -0.065356403 | 0.099350433 | 0.18487629 | 0.615853002 | ns | NaCl | KS.3 |
| FIM1_84 | 505.7972787 | 357.0850063 | 813.4542519 | -0.248290948 | 0.189000556 | 0.002270079 | 0.058108556 | ns | NaCl | KS.3 |
| FIM1_85 | 612.3902798 | 562.4291907 | 812.4313926 | -0.051170117 | 0.078292363 | 0.306106605 | 0.725181123 | ns | NaCl | KS.3 |
| FIM1_86 | 391.6403385 | 351.5464372 | 890.901293 | -0.055643791 | 0.086161175 | 0.26419202 | 0.688347233 | ns | NaCl | KS.3 |
| FIM1_87 | 583.1132782 | 733.7922917 | 533.8379842 | 0.080274 | 0.11782252 | 0.109964877 | 0.505147982 | ns | NaCl | KS.3 |
| FIM1_88 | 498.2873108 | 388.108063 | 812.0837853 | -0.098519174 | 0.122607095 | 0.064310662 | 0.398841896 | ns | NaCl | KS.3 |
| FIM1_89 | 190.8860004 | 232.2440009 | 613.585864 | 0.028761152 | 0.127418409 | 0.456795337 | 0.817337042 | ns | NaCl | KS.3 |
| FIM1_90 | 3555.657132 | 3725.485333 | 2907.03043 | 0.038678457 | 0.063096705 | 0.416386943 | 0.800387073 | ns | NaCl | KS.3 |
| FIM1_91 | 4529.580097 | 7166.544965 | 2110.39745 | 0.166804984 | 0.223034917 | 0.017403674 | 0.197678715 | ns | NaCl | KS.3 |
| FIM1_92 | 264.9113828 | 220.3217237 | 320.0306234 | -0.07843433 | 0.102405984 | 0.122015341 | 0.528318345 | ns | NaCl | KS.3 |
| FIM1_93 | 197.3287503 | 165.6614996 | 258.6469668 | -0.086530144 | 0.097436787 | 0.095299348 | 0.471286535 | ns | NaCl | KS.3 |
| FIM1_94 | 169.5348731 | 128.6706213 | 357.0574842 | -0.122331749 | 0.137815225 | 0.033537726 | 0.292771172 | ns | NaCl | KS.3 |
| FIM1_95 | 6523.89677 | 6198.980706 | 5406.626742 | -0.037214495 | 0.069176744 | 0.447317539 | 0.812914509 | ns | NaCl | KS.3 |
| FIM1_96 | 180.8348596 | 142.7437537 | 692.0664714 | -0.050047634 | 0.126006224 | 0.250037612 | 0.673319638 | ns | NaCl | KS.3 |
| FIM1_97 | 2669.490579 | 1169.85175 | 3895.884372 | -0.729445279 | 0.447857189 | 0.000262146 | 0.01526215 | ns | NaCl | KS.3 |
| FIM1_98 | 198.2217573 | 140.3752101 | 194.9117998 | -0.097294199 | 0.15784243 | 0.066522228 | 0.404582009 | ns | NaCl | KS.3 |
| FIM1_99 | 457.8984697 | 365.6055214 | 463.1981537 | -0.085941264 | 0.114847382 | 0.093823335 | 0.470201108 | ns | NaCl | KS.3 |
| FIM1_100 | 4345.108458 | 4186.554212 | 3763.677311 | -0.018796955 | 0.082421528 | 0.699641633 | 0.922863518 | ns | NaCl | KS.3 |
| FIM1_101 | 817.7826821 | 769.5212603 | 1500.413589 | -0.020336258 | 0.095010615 | 0.6552628 | 0.901110473 | ns | NaCl | KS.3 |
| FIM1_102 | 157.7291768 | 162.5763321 | 221.6806448 | 0.008517395 | 0.096648921 | 0.847941139 | 0.967208916 | ns | NaCl | KS.3 |
| FIM1_103 | 1886.106968 | 1839.164561 | 1767.344721 | -0.014706797 | 0.077202405 | 0.764939656 | 0.947367385 | ns | NaCl | KS.3 |
| FIM1_104 | 503.0191616 | 491.9081625 | 640.3642589 | -0.019500051 | 0.068012546 | 0.688405482 | 0.918490387 | ns | NaCl | KS.3 |
| FIM1_105 | 5049.910861 | 6002.919284 | 3825.164874 | 0.063373337 | 0.103405144 | 0.19200607 | 0.618952432 | ns | NaCl | KS.3 |
| FIM1_107 | 913.5070582 | 959.4775604 | 944.1502495 | 0.022162575 | 0.086338098 | 0.64477034 | 0.898899754 | ns | NaCl | KS.3 |

| | | | | | | | | | |
|---|---|---|---|---|---|---|---|---|---|
| FIM1_108 | 2819.203008 | 3011.361766 | 2134.243348 | 0.026550866 | 0.090255144 | 0.574663422 | 0.872163065 | ns | NaCl | KS.3 |
| FIM1_109 | 1246.281018 | 7272.340601 | 3009.743906 | 2.453722302 | 0.174491778 | 1.50748568039 | 7.49974125996 | Km>KS.3 | NaCl | KS.3 |
| FIM1_110 | 437.8864582 | 1062.260051 | 762.545038 | 1.22218938 | 0.152602412 | 1.72026437936 | 2.85277176244 | Km>KS.3 | NaCl | KS.3 |
| FIM1_111 | 5067.228651 | 4360.148505 | 5944.23296 | -0.110770006 | 0.079213384 | 0.034528399 | 0.294141756 | ns | NaCl | KS.3 |
| FIM1_112 | 1216.542786 | 1336.255078 | 919.1620486 | 0.034285389 | 0.094458889 | 0.463352426 | 0.819880744 | ns | NaCl | KS.3 |
| FIM1_113 | 762.840905 | 727.1094978 | 1147.542679 | -0.024674339 | 0.082389197 | 0.613631398 | 0.889162818 | ns | NaCl | KS.3 |
| FIM1_114 | 992.3926136 | 943.919594 | 985.3718464 | -0.025007139 | 0.082811227 | 0.60847278 | 0.88569286 | ns | NaCl | KS.3 |
| FIM1_115 | 506.7828316 | 609.1156696 | 1055.82097 | 0.042474087 | 0.11478213 | 0.328040151 | 0.740319311 | ns | NaCl | KS.3 |
| FIM1_116 | 384.7608924 | 341.4041849 | 432.800489 | -0.047273338 | 0.096040124 | 0.321255906 | 0.737180727 | ns | NaCl | KS.3 |
| FIM1_117 | 16547.97373 | 16620.61041 | 11163.9285 | 0.00136594 | 0.095712662 | 0.975589492 | 0.996164232 | ns | NaCl | KS.3 |
| FIM1_118 | 91.71049558 | 93.68677399 | 186.7466403 | 0.002470309 | 0.106166056 | 0.951693567 | 0.987831315 | ns | NaCl | KS.3 |
| FIM1_119 | 11446.45208 | 13427.86581 | 7368.886416 | 0.051850012 | 0.104113205 | 0.267826386 | 0.689648447 | ns | NaCl | KS.3 |
| FIM1_120 | 1527.561897 | 1712.257687 | 1362.100667 | 0.071272811 | 0.080161797 | 0.160893858 | 0.582143233 | ns | NaCl | KS.3 |
| FIM1_121 | 5086.046854 | 6866.480874 | 3572.133845 | 0.069683772 | 0.138789292 | 0.138846617 | 0.549532156 | ns | NaCl | KS.3 |
| FIM1_122 | 3800.642515 | 3959.331586 | 3375.700208 | 0.02893248 | 0.070264548 | 0.555222188 | 0.863466829 | ns | NaCl | KS.3 |
| FIM1_123 | 6029.248077 | 6321.947743 | 4792.686076 | 0.027268823 | 0.078706599 | 0.58024258 | 0.8749481 | ns | NaCl | KS.3 |
| FIM1_124 | 757.8973748 | 854.0593712 | 724.0275505 | 0.068074151 | 0.084894108 | 0.179286321 | 0.608009168 | ns | NaCl | KS.3 |
| FIM1_125 | 202.8348518 | 267.8562824 | 653.6319812 | 0.115791675 | 0.136080725 | 0.040041588 | 0.315700318 | ns | NaCl | KS.3 |
| FIM1_126 | 1388.042518 | 1244.287821 | 1309.752804 | -0.079977698 | 0.07410966 | 0.113767539 | 0.513140081 | ns | NaCl | KS.3 |
| FIM1_127 | 622.2348232 | 519.5424222 | 779.7362617 | -0.057320185 | 0.107328999 | 0.224260179 | 0.648059715 | ns | NaCl | KS.3 |
| FIM1_128 | 21630.07461 | 20938.33604 | 15811.42649 | -0.019510961 | 0.076552849 | 0.692082994 | 0.91988055 | ns | NaCl | KS.3 |
| FIM1_129 | 1158.325738 | 1368.147084 | 1036.64876 | 0.051404934 | 0.106276326 | 0.267659154 | 0.689593109 | ns | NaCl | KS.3 |
| FIM1_130 | 8230.156457 | 8037.417925 | 9887.24238 | -0.013544653 | 0.077723126 | 0.782862214 | 0.951646805 | ns | NaCl | KS.3 |
| FIM1_131 | 2659.78588 | 2823.933414 | 2248.066324 | 0.024413404 | 0.089729725 | 0.606048001 | 0.883689315 | ns | NaCl | KS.3 |
| FIM1_132 | 31535.30278 | 34883.31944 | 21182.2828 | 0.079836323 | 0.068943623 | 0.107317969 | 0.498909937 | ns | NaCl | KS.3 |
| FIM1_133 | 5087.86089 | 6823.118129 | 4961.324517 | 0.188707141 | 0.146714302 | 0.005284205 | 0.095944957 | ns | NaCl | KS.3 |
| FIM1_134 | 4767.813837 | 5148.667992 | 4336.865324 | 0.045652474 | 0.079053362 | 0.359355274 | 0.763021562 | ns | NaCl | KS.3 |
| FIM1_135 | 1344.875142 | 1582.533379 | 1364.517315 | 0.12524719 | 0.079429679 | 0.018045993 | 0.202175238 | ns | NaCl | KS.3 |
| FIM1_136 | 42.87963849 | 52.30016279 | 244.4891628 | 0.031680911 | 0.126322912 | 0.421630707 | 0.801533346 | ns | NaCl | KS.3 |
| FIM1_137 | 240.7356656 | 195.9135534 | 852.1868397 | -0.053719276 | 0.115753788 | 0.237196878 | 0.659616806 | ns | NaCl | KS.3 |
| FIM1_138 | 15423.66859 | 19140.92984 | 11142.29983 | 0.103643685 | 0.108654536 | 0.054455399 | 0.363645114 | ns | NaCl | KS.3 |
| FIM1_140 | 24938.87485 | 28387.47871 | 17517.12602 | 0.068127804 | 0.087552949 | 0.178071497 | 0.606369404 | ns | NaCl | KS.3 |
| FIM1_141 | 808.4992444 | 747.1665811 | 718.8621058 | -0.035735027 | 0.08702482 | 0.461534365 | 0.819880744 | ns | NaCl | KS.3 |
| FIM1_142 | 15275.32662 | 15637.05293 | 12027.67333 | 0.022408407 | 0.054606927 | 0.613445201 | 0.889162818 | ns | NaCl | KS.3 |
| FIM1_143 | 174.9189202 | 193.9680092 | 247.9532581 | 0.021497361 | 0.111285324 | 0.596966047 | 0.879972174 | ns | NaCl | KS.3 |
| FIM1_144 | 963.1564523 | 708.0704965 | 911.1906525 | -0.045359704 | 0.142227701 | 0.266305966 | 0.689593109 | ns | NaCl | KS.3 |
| FIM1_145 | 17583.45689 | 23255.77238 | 13754.70225 | 0.111984619 | 0.134210133 | 0.044339617 | 0.331713673 | ns | NaCl | KS.3 |
| FIM1_146 | 2752.871726 | 2913.338951 | 2501.115526 | 0.042568229 | 0.068744731 | 0.384628979 | 0.778665419 | ns | NaCl | KS.3 |
| FIM1_147 | 1920.087196 | 1890.544856 | 2010.46277 | -0.012834139 | 0.065611575 | 0.789692353 | 0.953804189 | ns | NaCl | KS.3 |
| FIM1_148 | 719.9495017 | 748.9611982 | 787.1053614 | 0.025972708 | 0.070723037 | 0.596588775 | 0.879797482 | ns | NaCl | KS.3 |
| FIM1_149 | 614.9759475 | 575.6570308 | 567.6685807 | -0.036539366 | 0.079927079 | 0.460178149 | 0.819880744 | ns | NaCl | KS.3 |
| FIM1_150 | 790.5174019 | 821.5074779 | 1020.816971 | 0.026110772 | 0.070655312 | 0.594588963 | 0.87962202 | ns | NaCl | KS.3 |
| FIM1_151 | 760.3355978 | 722.0020004 | 929.2554157 | -0.027613839 | 0.081733891 | 0.573367503 | 0.871789624 | ns | NaCl | KS.3 |
| FIM1_153 | 622.4080427 | 634.1753681 | 573.2743016 | 0.005936261 | 0.094197432 | 0.895623494 | 0.977776362 | ns | NaCl | KS.3 |
| FIM1_154 | 1376.955301 | 1432.538509 | 1999.653377 | 0.007803325 | 0.108594504 | 0.845876187 | 0.967147129 | ns | NaCl | KS.3 |

| | | | | | | | | | | |
|---|---|---|---|---|---|---|---|---|---|---|
| FIM1_155 | 3448.578322 | 3729.843508 | 3525.482726 | 0.022844388 | 0.099551096 | 0.605817791 | 0.883689315 | ns | NaCl | KS.3 |
| FIM1_156 | 1027.663578 | 980.5240517 | 1443.315126 | -0.021255673 | 0.08590998 | 0.658785463 | 0.903627703 | ns | NaCl | KS.3 |
| FIM1_157 | 1164.455039 | 734.9371896 | 2207.756938 | -0.107703517 | 0.194096072 | 0.052461566 | 0.358511389 | ns | NaCl | KS.3 |
| FIM1_158 | 7700.099517 | 7471.49171 | 6922.7706 | -0.018261062 | 0.075870061 | 0.710869305 | 0.925563973 | ns | NaCl | KS.3 |
| FIM1_159 | 6891.463361 | 7359.086709 | 6425.530611 | 0.081844298 | 0.034259304 | 0.009464609 | 0.13642083 | ns | NaCl | KS.3 |
| FIM1_160 | 6149.05538 | 6559.435276 | 5656.953157 | 0.057610475 | 0.0601553 | 0.217631737 | 0.641504077 | ns | NaCl | KS.3 |
| FIM1_161 | 436.5281765 | 506.0834038 | 678.5940738 | 0.048732689 | 0.103034922 | 0.294878318 | 0.713530949 | ns | NaCl | KS.3 |
| FIM1_162 | 1419.729207 | 1039.773066 | 1262.050445 | -0.183079472 | 0.158328399 | 0.00734091 | 0.117365229 | ns | NaCl | KS.3 |
| FIM1_163 | 477.6175905 | 219.4892289 | 516.5720716 | -0.912447613 | 0.297950557 | 1.31840334626 | 0.001490695 | ns | NaCl | KS.3 |
| FIM1_164 | 660.0510788 | 825.9016882 | 893.1192894 | 0.179937147 | 0.100039144 | 0.002497085 | 0.061197035 | ns | NaCl | KS.3 |
| FIM1_165 | 739.4192782 | 737.2938999 | 587.4606674 | -0.001289294 | 0.085417432 | 0.978527559 | 0.996164232 | ns | NaCl | KS.3 |
| FIM1_166 | 641.7645627 | 659.1002739 | 737.7645358 | 0.007678854 | 0.096096961 | 0.86329765 | 0.969856868 | ns | NaCl | KS.3 |
| FIM1_167 | 310.0781144 | 332.4069929 | 515.6252124 | 0.024018375 | 0.095569004 | 0.598541634 | 0.880167289 | ns | NaCl | KS.3 |
| FIM1_168 | 1389.469512 | 1418.556483 | 1298.756978 | 0.01121437 | 0.078885967 | 0.819119802 | 0.959981393 | ns | NaCl | KS.3 |
| FIM1_169 | 1211.096782 | 1281.042499 | 2192.324479 | 0.029061207 | 0.08191815 | 0.553601958 | 0.862564904 | ns | NaCl | KS.3 |
| FIM1_170 | 14456.90777 | 14824.05569 | 11208.52652 | 0.02708548 | 0.046112703 | 0.496543398 | 0.836358994 | ns | NaCl | KS.3 |
| FIM1_171 | 2310.652956 | 2537.732624 | 1811.685558 | 0.045773894 | 0.08632325 | 0.352289013 | 0.756912594 | ns | NaCl | KS.3 |
| FIM1_172 | 668.5330509 | 668.1194718 | 709.2605182 | -0.00027458 | 0.085322056 | 0.995428521 | 0.99924489 | ns | NaCl | KS.3 |
| FIM1_173 | 1120.397227 | 1165.825888 | 894.9360852 | 0.022161638 | 0.079825665 | 0.651831608 | 0.900060184 | ns | NaCl | KS.3 |
| FIM1_174 | 398.9207506 | 283.5768762 | 327.707724 | -0.048049137 | 0.14874082 | 0.240758007 | 0.665058903 | ns | NaCl | KS.3 |
| FIM1_175 | 112.9460853 | 75.42042709 | 88.16067366 | -0.102562706 | 0.180648841 | 0.058708482 | 0.378457602 | ns | NaCl | KS.3 |
| FIM1_176 | 2400.097431 | 2416.863522 | 2120.157969 | 0.003337774 | 0.08255817 | 0.945131721 | 0.98719931 | ns | NaCl | KS.3 |
| FIM1_177 | 12676.66569 | 13070.70694 | 8182.819457 | 0.007564232 | 0.102300937 | 0.858077045 | 0.969856868 | ns | NaCl | KS.3 |
| FIM1_178 | 922.3087821 | 857.3026831 | 898.2572507 | -0.053027999 | 0.07046791 | 0.283134441 | 0.704025235 | ns | NaCl | KS.3 |
| FIM1_179 | 2759.597707 | 2708.115596 | 2568.32626 | -0.011426403 | 0.076837724 | 0.816242192 | 0.958816322 | ns | NaCl | KS.3 |
| FIM1_180 | 1566.385797 | 1677.37635 | 1767.514357 | 0.045309833 | 0.074136005 | 0.361770809 | 0.763449629 | ns | NaCl | KS.3 |
| FIM1_181 | 16383.07251 | 17256.15532 | 11963.46107 | 0.046791054 | 0.05873941 | 0.310615839 | 0.730644821 | ns | NaCl | KS.3 |
| FIM1_182 | 6103.968873 | 6931.275817 | 5639.517076 | 0.031501439 | 0.107000559 | 0.465417198 | 0.820834625 | ns | NaCl | KS.3 |
| FIM1_183 | 88.2933132 | 73.86630602 | 86.97005316 | -0.03225422 | 0.120084181 | 0.426377682 | 0.802854185 | ns | NaCl | KS.3 |
| FIM1_184 | 4633.22974 | 5231.492753 | 4253.522987 | 0.063249996 | 0.086944896 | 0.208400178 | 0.633002483 | ns | NaCl | KS.3 |
| FIM1_185 | 571.7206719 | 663.8046774 | 1297.052696 | 0.046420635 | 0.10350343 | 0.314009037 | 0.732050122 | ns | NaCl | KS.3 |
| FIM1_186 | 209.3555816 | 167.2299603 | 342.0256655 | -0.048586313 | 0.121768736 | 0.265912158 | 0.689593109 | ns | NaCl | KS.3 |
| FIM1_187 | 7745.752581 | 9967.526424 | 7421.616443 | 0.054390502 | 0.127818911 | 0.219575962 | 0.642906874 | ns | NaCl | KS.3 |
| FIM1_189 | 1544.677289 | 1774.129656 | 1349.127392 | 0.090623623 | 0.081569589 | 0.080392803 | 0.438546266 | ns | NaCl | KS.3 |
| FIM1_190 | 192.6523055 | 178.9978306 | 166.9618692 | -0.021237035 | 0.100320656 | 0.62839918 | 0.896016329 | ns | NaCl | KS.3 |
| FIM1_191 | 411.8913056 | 342.306797 | 673.3985639 | -0.102715933 | 0.097812853 | 0.054990037 | 0.364767243 | ns | NaCl | KS.3 |
| FIM1_192 | 2014.128156 | 2492.670236 | 3739.861374 | 0.034358108 | 0.126679136 | 0.389676767 | 0.779482559 | ns | NaCl | KS.3 |
| FIM1_193 | 17.4465321 | 16.99305263 | 567.2888931 | -0.004965636 | 0.133126795 | 0.884644024 | 0.975081831 | ns | NaCl | KS.3 |
| FIM1_194 | 2362.140612 | 2590.911994 | 2532.895346 | 0.027760311 | 0.099503519 | 0.53497972 | 0.855153333 | ns | NaCl | KS.3 |
| FIM1_195 | 1282.394226 | 1386.171107 | 1244.231713 | 0.029941505 | 0.091908505 | 0.52482937 | 0.854940852 | ns | NaCl | KS.3 |
| FIM1_196 | 3889.382413 | 4108.977039 | 2924.485626 | 0.032296085 | 0.077912782 | 0.513763518 | 0.848315135 | ns | NaCl | KS.3 |
| FIM1_197 | 2005.088071 | 2056.263594 | 2057.649556 | 0.007085903 | 0.099599039 | 0.869952233 | 0.971243188 | ns | NaCl | KS.3 |
| FIM1_198 | 5189.903378 | 6255.925827 | 4654.542393 | 0.093728271 | 0.099432664 | 0.074934459 | 0.426055924 | ns | NaCl | KS.3 |
| FIM1_199 | 765.5036198 | 873.0573494 | 1061.302746 | 0.048153735 | 0.098000172 | 0.309519045 | 0.729444457 | ns | NaCl | KS.3 |
| FIM1_200 | 3137.308929 | 3291.194443 | 2473.384623 | 0.02091766 | 0.087172252 | 0.661907153 | 0.904388188 | ns | NaCl | KS.3 |

| | | | | | | | | | | |
|---|---|---|---|---|---|---|---|---|---|---|
| FIM1_201 | 1925.835039 | 2118.574511 | 2060.566516 | 0.057056426 | 0.079659953 | 0.255901492 | 0.678991959 | ns | NaCl | KS.3 |
| FIM1_203 | 952.3953135 | 855.9811148 | 1224.968132 | -0.061810246 | 0.082363687 | 0.22008444 | 0.642906874 | ns | NaCl | KS.3 |
| FIM1_204 | 108.9903557 | 80.04720941 | 149.289268 | -0.084082481 | 0.143688633 | 0.093958987 | 0.470201108 | ns | NaCl | KS.3 |
| FIM1_205 | 1919.089517 | 2219.845272 | 1549.415675 | 0.02913204 | 0.114084508 | 0.478930177 | 0.826171162 | ns | NaCl | KS.3 |
| FIM1_206 | 2904.360887 | 3004.455933 | 2888.196518 | 0.022323652 | 0.073764707 | 0.650512879 | 0.899642849 | ns | NaCl | KS.3 |
| FIM1_207 | 873.2412786 | 668.9619909 | 798.6830117 | -0.138744664 | 0.128907302 | 0.019716528 | 0.21299357 | ns | NaCl | KS.3 |
| FIM1_208 | 3415.754491 | 2924.629576 | 3150.474754 | -0.089896442 | 0.088411841 | 0.084834967 | 0.452362228 | ns | NaCl | KS.3 |
| FIM1_209 | 3775.837058 | 3557.564913 | 4312.461468 | -0.032642182 | 0.08075907 | 0.507820223 | 0.844423736 | ns | NaCl | KS.3 |
| FIM1_210 | 3804.179111 | 3740.497188 | 2926.221732 | -0.008101755 | 0.084383016 | 0.866462666 | 0.970371341 | ns | NaCl | KS.3 |
| FIM1_211 | 1044.495533 | 955.4843631 | 1421.061327 | -0.049453119 | 0.082370658 | 0.32073736 | 0.737029267 | ns | NaCl | KS.3 |
| FIM1_212 | 1139.200576 | 971.3418357 | 1343.651911 | -0.075372351 | 0.095410919 | 0.137665831 | 0.546598172 | ns | NaCl | KS.3 |
| FIM1_213 | 1686.060138 | 1587.301355 | 1407.992985 | -0.044664939 | 0.068903753 | 0.362189467 | 0.763449629 | ns | NaCl | KS.3 |
| FIM1_214 | 176.3692379 | 122.8750117 | 339.0121516 | -0.152438349 | 0.181296699 | 0.01923873 | 0.209721297 | ns | NaCl | KS.3 |
| FIM1_215 | 684.2681742 | 653.1230107 | 763.6384068 | -0.037147908 | 0.069269465 | 0.448263255 | 0.812914509 | ns | NaCl | KS.3 |
| FIM1_216 | 3926.256268 | 3933.01497 | 3833.499387 | 0.000560455 | 0.093669861 | 0.990147128 | 0.999134858 | ns | NaCl | KS.3 |
| FIM1_217 | 3114.552568 | 3061.770101 | 2626.542711 | -0.008805788 | 0.081748687 | 0.856425793 | 0.969856868 | ns | NaCl | KS.3 |
| FIM1_218 | 843.7275286 | 359.6861221 | 1854.385565 | -0.938642804 | 0.367573639 | 3.42999638945 | 0.003281583 | ns | NaCl | KS.3 |
| FIM1_219 | 777.8937703 | 629.6661225 | 1022.823757 | -0.171356833 | 0.097807662 | 0.003451029 | 0.075302063 | ns | NaCl | KS.3 |
| FIM1_220 | 282.506272 | 319.0525704 | 417.3276213 | 0.049020168 | 0.093464987 | 0.310268159 | 0.7303513 | ns | NaCl | KS.3 |
| FIM1_221 | 1387.622426 | 1315.046275 | 1188.193571 | -0.023908269 | 0.086825797 | 0.618611181 | 0.892055254 | ns | NaCl | KS.3 |
| FIM1_222 | 1755.922821 | 1422.459844 | 1793.918807 | -0.219861557 | 0.077839115 | 5.67192668816 | 0.004702973 | ns | NaCl | KS.3 |
| FIM1_224 | 2605.455918 | 2628.012634 | 1990.069359 | 0.004248731 | 0.082308875 | 0.930258636 | 0.984325045 | ns | NaCl | KS.3 |
| FIM1_225 | 838.0059678 | 682.8270003 | 1133.049414 | -0.142790028 | 0.098670259 | 0.012177696 | 0.157397666 | ns | NaCl | KS.3 |
| FIM1_226 | 181.396442 | 135.0540624 | 393.6260039 | -0.077875967 | 0.137105531 | 0.112043115 | 0.509054336 | ns | NaCl | KS.3 |
| FIM1_227 | 1700.619718 | 1579.258362 | 1808.330015 | -0.029330979 | 0.090911184 | 0.535212821 | 0.855153333 | ns | NaCl | KS.3 |
| FIM1_228 | 83.23609063 | 71.70348861 | 100.5731306 | -0.046405074 | 0.102192211 | 0.316783256 | 0.73382286 | ns | NaCl | KS.3 |
| FIM1_229 | 7599.179585 | 8384.203243 | 7860.361819 | 0.051913543 | 0.084303069 | 0.297020178 | 0.715581302 | ns | NaCl | KS.3 |
| FIM1_230 | 1829.579221 | 1653.490495 | 1941.403259 | -0.068970243 | 0.07498026 | 0.170906821 | 0.597342667 | ns | NaCl | KS.3 |
| FIM1_231 | 68.10662671 | 54.90711646 | 186.9610113 | -0.034482481 | 0.127811394 | 0.386376196 | 0.779278134 | ns | NaCl | KS.3 |
| FIM1_233 | 6183.942898 | 6206.70498 | 4732.029427 | 0.001630142 | 0.08636629 | 0.97272077 | 0.995313014 | ns | NaCl | KS.3 |
| FIM1_234 | 1811.06775 | 1761.686328 | 1866.763399 | -0.025508524 | 0.055993313 | 0.570609763 | 0.870665206 | ns | NaCl | KS.3 |
| FIM1_235 | 470.5831299 | 451.9703699 | 816.0938578 | -0.019548119 | 0.085752838 | 0.684542653 | 0.91584821 | ns | NaCl | KS.3 |
| FIM1_236 | 2034.380505 | 1900.119894 | 2878.122841 | -0.03451837 | 0.083681302 | 0.48132437 | 0.828577419 | ns | NaCl | KS.3 |
| FIM1_237 | 280.3549608 | 272.2036175 | 509.869959 | -0.005183215 | 0.111669841 | 0.894461467 | 0.977268396 | ns | NaCl | KS.3 |
| FIM1_238 | 332.7015864 | 258.9464303 | 439.7022536 | -0.128771626 | 0.122698661 | 0.025777076 | 0.252442825 | ns | NaCl | KS.3 |
| FIM1_239 | 82.03316458 | 49.68722804 | 93.79866522 | -0.200666539 | 0.252000852 | 0.010967048 | 0.149455888 | ns | NaCl | KS.3 |
| FIM1_240 | 378.5420505 | 240.8406002 | 582.5653609 | -0.299883803 | 0.260048939 | 0.002423061 | 0.060255275 | ns | NaCl | KS.3 |
| FIM1_241 | 4287.510035 | 4309.372982 | 3621.35305 | 0.004703827 | 0.05625747 | 0.916878976 | 0.982343702 | ns | NaCl | KS.3 |
| FIM1_242 | 60.61588602 | 44.6082423 | 58.14148934 | -0.070213186 | 0.137945626 | 0.137228517 | 0.545297021 | ns | NaCl | KS.3 |
| FIM1_243 | 1356.477131 | 1296.132295 | 1207.709963 | -0.034829418 | 0.065864156 | 0.470796245 | 0.822127061 | ns | NaCl | KS.3 |
| FIM1_245 | 7800.714822 | 9252.529924 | 4952.962543 | 0.04327026 | 0.11119322 | 0.327397929 | 0.740319311 | ns | NaCl | KS.3 |
| FIM1_246 | 1868.060399 | 2038.502694 | 2323.651007 | 0.064220289 | 0.072006813 | 0.197756001 | 0.620924909 | ns | NaCl | KS.3 |
| FIM1_247 | 456.3340418 | 265.927436 | 1343.159356 | -0.398781435 | 0.312831286 | 0.00120479 | 0.041316566 | ns | NaCl | KS.3 |
| FIM1_248 | 1248.576144 | 1249.101125 | 1115.187354 | 0.001120432 | 0.077383456 | 0.981793603 | 0.996363429 | ns | NaCl | KS.3 |
| FIM1_249 | 1654.013065 | 2305.995531 | 995.5845502 | 0.057727384 | 0.146528374 | 0.186428506 | 0.618044911 | ns | NaCl | KS.3 |

| | | | | | | | | | | |
|---|---|---|---|---|---|---|---|---|---|---|
| FIM1_250 | 769.7813924 | 698.8776928 | 804.4931944 | -0.051209688 | 0.084425733 | 0.30314259 | 0.72182705 | ns | NaCl | KS.3 |
| FIM1_251 | 496.702646 | 413.5271931 | 522.5479514 | -0.101249702 | 0.09784997 | 0.05788239 | 0.377713236 | ns | NaCl | KS.3 |
| FIM1_252 | 2199.940657 | 2080.052294 | 2061.09628 | -0.046246502 | 0.065511564 | 0.338381957 | 0.747703619 | ns | NaCl | KS.3 |
| FIM1_253 | 5943.360144 | 5966.149488 | 4712.613198 | 0.002482361 | 0.071537762 | 0.959609296 | 0.991702586 | ns | NaCl | KS.3 |
| FIM1_254 | 576.290543 | 520.7140228 | 630.1023059 | -0.044810046 | 0.090019318 | 0.356190774 | 0.759883833 | ns | NaCl | KS.3 |
| FIM1_255 | 3714.789754 | 3412.420642 | 3916.679253 | -0.036619896 | 0.089233624 | 0.446926055 | 0.812914509 | ns | NaCl | KS.3 |
| FIM1_256 | 3690.399569 | 4764.532682 | 2945.980669 | 0.110755839 | 0.124350464 | 0.044966338 | 0.335229179 | ns | NaCl | KS.3 |
| FIM1_257 | 871.7491749 | 960.2158845 | 989.1991687 | 0.055973931 | 0.080893596 | 0.264478596 | 0.688347233 | ns | NaCl | KS.3 |
| FIM1_258 | 1076.120671 | 1009.807276 | 1326.343152 | -0.038519246 | 0.078342829 | 0.437447439 | 0.808096352 | ns | NaCl | KS.3 |
| FIM1_259 | 4192.212532 | 4209.814529 | 3182.099749 | 0.001955629 | 0.084229697 | 0.967616167 | 0.99408528 | ns | NaCl | KS.3 |
| FIM1_260 | 490.4471896 | 500.5517801 | 531.1506979 | 0.007841479 | 0.089591169 | 0.866995097 | 0.970371341 | ns | NaCl | KS.3 |
| FIM1_261 | 1457.876618 | 1319.653572 | 1417.32722 | -0.028633454 | 0.101249314 | 0.518316066 | 0.850188733 | ns | NaCl | KS.3 |
| FIM1_262 | 1853.230506 | 2362.156289 | 1256.727153 | 0.085185886 | 0.120761496 | 0.094773118 | 0.470201108 | ns | NaCl | KS.3 |
| FIM1_263 | 3428.24099 | 4131.501639 | 4356.538085 | 0.123920012 | 0.09217895 | 0.023843388 | 0.239908881 | ns | NaCl | KS.3 |
| FIM1_264 | 1025.190644 | 1029.090707 | 998.4399574 | 0.002221519 | 0.073688113 | 0.963943118 | 0.993691273 | ns | NaCl | KS.3 |
| FIM1_265 | 1369.565051 | 1370.949176 | 1462.899096 | -7.22E-05 | 0.065982913 | 0.998803726 | 0.999406383 | ns | NaCl | KS.3 |
| FIM1_266 | 829.510401 | 868.2099622 | 972.3963985 | 0.021068965 | 0.085323255 | 0.662373483 | 0.904388188 | ns | NaCl | KS.3 |
| FIM1_267 | 6995.280488 | 7532.643398 | 5139.402025 | 0.056041734 | 0.069043275 | 0.254718048 | 0.678747879 | ns | NaCl | KS.3 |
| FIM1_268 | 5092.445104 | 5021.785997 | 5576.256847 | -0.006295683 | 0.08471471 | 0.895844771 | 0.977803364 | ns | NaCl | KS.3 |
| FIM1_269 | 1405.792551 | 1350.760651 | 1362.757812 | -0.022643749 | 0.079107243 | 0.645250271 | 0.898899754 | ns | NaCl | KS.3 |
| FIM1_270 | 102.65133 | 108.4752797 | 135.2654486 | 0.012978772 | 0.102161653 | 0.76077285 | 0.945911573 | ns | NaCl | KS.3 |
| FIM1_271 | 2774.521247 | 2950.747793 | 2806.474049 | 0.030236021 | 0.084159906 | 0.535370125 | 0.855153333 | ns | NaCl | KS.3 |
| FIM1_272 | 725.4068493 | 704.9232594 | 690.4990834 | -0.013388079 | 0.085626477 | 0.780262766 | 0.951646805 | ns | NaCl | KS.3 |
| FIM1_273 | 194.9435656 | 162.0053414 | 446.7911323 | -0.049498674 | 0.111677909 | 0.273616843 | 0.694794376 | ns | NaCl | KS.3 |
| FIM1_274 | 2597.066084 | 2852.397944 | 1722.830084 | 0.025630425 | 0.102096605 | 0.558008728 | 0.864019117 | ns | NaCl | KS.3 |
| FIM1_275 | 1827.031634 | 1850.96218 | 1602.25596 | 0.004715952 | 0.09110294 | 0.918866308 | 0.982665495 | ns | NaCl | KS.3 |
| FIM1_276 | 858.1489824 | 820.324039 | 906.7754852 | -0.02263039 | 0.083121747 | 0.642175731 | 0.898798052 | ns | NaCl | KS.3 |
| FIM1_277 | 257.3948034 | 232.839193 | 311.8922662 | -0.051699169 | 0.085884044 | 0.29744033 | 0.715635559 | ns | NaCl | KS.3 |
| FIM1_278 | 649.2225273 | 694.4677635 | 637.3066224 | 0.028710231 | 0.089147589 | 0.547284825 | 0.859115631 | ns | NaCl | KS.3 |
| FIM1_279 | 4397.857056 | 4767.431884 | 3531.57243 | 0.049534034 | 0.077665454 | 0.321256584 | 0.737180727 | ns | NaCl | KS.3 |
| FIM1_280 | 27.93285243 | 29.21883542 | 26.86602388 | 0.004065786 | 0.126384594 | 0.908668657 | 0.980825899 | ns | NaCl | KS.3 |
| FIM1_281 | 3988.235207 | 3204.349976 | 3055.708706 | -0.093333414 | 0.111362683 | 0.075429656 | 0.427519783 | ns | NaCl | KS.3 |
| FIM1_282 | 233.5660836 | 340.9856439 | 392.1436837 | 0.065026037 | 0.157393237 | 0.148615521 | 0.563109076 | ns | NaCl | KS.3 |
| FIM1_283 | 643.2352039 | 746.60866 | 1789.244243 | 0.024717126 | 0.119393934 | 0.529335537 | 0.855153333 | ns | NaCl | KS.3 |
| FIM1_284 | 1138.598436 | 1640.150284 | 1082.482432 | 0.318399324 | 0.198502707 | 0.000606601 | 0.025793499 | ns | NaCl | KS.3 |
| FIM1_285 | 1569.779165 | 2259.911165 | 1445.590392 | 0.076723625 | 0.156677612 | 0.110777159 | 0.506448597 | ns | NaCl | KS.3 |
| FIM1_286 | 2918.300986 | 3142.818364 | 3409.995974 | 0.026635437 | 0.093782943 | 0.565444312 | 0.868073805 | ns | NaCl | KS.3 |
| FIM1_287 | 668.9326228 | 732.9746665 | 685.4018079 | 0.054031589 | 0.079461674 | 0.280841979 | 0.701610837 | ns | NaCl | KS.3 |
| FIM1_288 | 152.529217 | 138.3168597 | 132.9791323 | -0.027191054 | 0.099657825 | 0.542462637 | 0.857313452 | ns | NaCl | KS.3 |
| FIM1_290 | 471.9691236 | 550.0647151 | 401.1793757 | 0.061530498 | 0.097707857 | 0.209372544 | 0.633579568 | ns | NaCl | KS.3 |
| FIM1_291 | 1511.712605 | 1396.083995 | 1665.079634 | -0.032342329 | 0.090774754 | 0.496191151 | 0.836358994 | ns | NaCl | KS.3 |
| FIM1_292 | 687.2384149 | 448.7216402 | 1255.979238 | -0.112002244 | 0.186413157 | 0.047795394 | 0.343442 | ns | NaCl | KS.3 |
| FIM1_293 | 2327.866876 | 2693.848042 | 3668.649036 | 0.0792655 | 0.088840625 | 0.122877576 | 0.529277872 | ns | NaCl | KS.3 |
| FIM1_294 | 137.0152441 | 172.3624946 | 2209.438311 | 0.029247946 | 0.133860978 | 0.439339834 | 0.808917498 | ns | NaCl | KS.3 |
| FIM1_295 | 20.44725242 | 25.91180738 | 54.48768277 | 0.028839274 | 0.138052218 | 0.437881557 | 0.808096352 | ns | NaCl | KS.3 |

| | | | | | | | | | | |
|---|---|---|---|---|---|---|---|---|---|---|
| FIM1_296 | 46.30669709 | 104.8115984 | 99.09863344 | 0.388483575 | 0.436821669 | 0.002618861 | 0.063246767 | ns | NaCl | KS.3 |
| FIM1_297 | 17.51977398 | 17.69312699 | 31.93400973 | 0.000167038 | 0.131280842 | 0.996128443 | 0.99924489 | ns | NaCl | KS.3 |
| FIM1_298 | 24.64150173 | 24.51290928 | 43.81990865 | -0.003874312 | 0.119214802 | 0.916603931 | 0.982343702 | ns | NaCl | KS.3 |
| FIM1_299 | 537.9713504 | 579.0738004 | 625.4742574 | 0.015528879 | 0.107842298 | 0.704549734 | 0.924593755 | ns | NaCl | KS.3 |
| FIM1_300 | 2842.64518 | 3846.379916 | 1750.706662 | 0.048003985 | 0.140744007 | 0.248714238 | 0.67096552 | ns | NaCl | KS.3 |
| FIM1_301 | 133.7046355 | 134.8764327 | 194.6034244 | 0.001804416 | 0.108009723 | 0.96414639 | 0.993691273 | ns | NaCl | KS.3 |
| FIM1_302 | 510.5451815 | 479.5075426 | 737.3037931 | -0.023094639 | 0.092242671 | 0.619941212 | 0.892479944 | ns | NaCl | KS.3 |
| FIM1_303 | 32511.22932 | 43583.28177 | 31660.2633 | 0.193761414 | 0.146782342 | 0.004539747 | 0.086695706 | ns | NaCl | KS.3 |
| FIM1_304 | 414.3756449 | 348.940047 | 807.8463057 | -0.049297845 | 0.108359293 | 0.280679344 | 0.701610837 | ns | NaCl | KS.3 |
| FIM1_305 | 197.7668253 | 83.34116779 | 843.7830533 | -0.531254588 | 0.488184799 | 0.001094632 | 0.039178381 | ns | NaCl | KS.3 |
| FIM1_306 | 133.9530468 | 113.0272334 | 334.8470566 | -0.017357518 | 0.134499543 | 0.623818013 | 0.893754273 | ns | NaCl | KS.3 |
| FIM1_307 | 1312.067698 | 1261.045795 | 1116.511687 | -0.02181867 | 0.079799567 | 0.656843549 | 0.901977674 | ns | NaCl | KS.3 |
| FIM1_308 | 3252.477913 | 3858.827121 | 2450.767558 | 0.082439006 | 0.096727113 | 0.109135022 | 0.503194378 | ns | NaCl | KS.3 |
| FIM1_309 | 4945.614926 | 5578.761128 | 4752.360542 | 0.100331041 | 0.068670695 | 0.044182695 | 0.331037512 | ns | NaCl | KS.3 |
| FIM1_310 | 5597.389606 | 7119.190198 | 4156.181608 | 0.11962887 | 0.11745302 | 0.033480701 | 0.292771172 | ns | NaCl | KS.3 |
| FIM1_311 | 3239.020777 | 3634.995989 | 3860.09946 | 0.078453257 | 0.077088449 | 0.123248667 | 0.529611582 | ns | NaCl | KS.3 |
| FIM1_312 | 3053.605365 | 2984.093668 | 3656.8811 | -0.007282174 | 0.09531584 | 0.871083366 | 0.971243188 | ns | NaCl | KS.3 |
| FIM1_313 | 3216.202949 | 3429.010247 | 2693.26069 | 0.022282363 | 0.094145302 | 0.627685583 | 0.896016329 | ns | NaCl | KS.3 |
| FIM1_314 | 1400.834618 | 1360.041611 | 2126.457402 | -0.008619104 | 0.098457031 | 0.843843653 | 0.966641072 | ns | NaCl | KS.3 |
| FIM1_315 | 366.3288227 | 399.9681661 | 659.2698316 | 0.055719922 | 0.077671678 | 0.26645417 | 0.689593109 | ns | NaCl | KS.3 |
| FIM1_316 | 1601.597665 | 2091.418083 | 1101.901215 | 0.040112356 | 0.135458435 | 0.317934655 | 0.733979075 | ns | NaCl | KS.3 |
| FIM1_317 | 26.60646247 | 22.16996088 | 101.9596112 | -0.021563506 | 0.127459985 | 0.562794201 | 0.867434473 | ns | NaCl | KS.3 |
| FIM1_318 | 10769.60232 | 11622.68151 | 7843.167884 | 0.079877585 | 0.050406038 | 0.058967541 | 0.379022628 | ns | NaCl | KS.3 |
| FIM1_319 | 273.3646999 | 269.7855492 | 301.5180789 | -0.006161727 | 0.090256101 | 0.894780999 | 0.977285504 | ns | NaCl | KS.3 |
| FIM1_320 | 1324.602353 | 1364.106868 | 1810.685389 | 0.007544371 | 0.102110861 | 0.858672212 | 0.969856868 | ns | NaCl | KS.3 |
| FIM1_321 | 113.5419425 | 168.5275992 | 107.0083954 | 0.064481159 | 0.162277863 | 0.14847628 | 0.563010285 | ns | NaCl | KS.3 |
| FIM1_322 | 20648.67799 | 23126.02316 | 14723.54858 | 0.077810232 | 0.076369682 | 0.125609458 | 0.534982546 | ns | NaCl | KS.3 |
| FIM1_323 | 349.1133326 | 251.7313247 | 571.029106 | -0.099957468 | 0.149873311 | 0.062194238 | 0.390677191 | ns | NaCl | KS.3 |
| FIM1_324 | 4824.977409 | 5546.246708 | 5869.511829 | 0.03591292 | 0.107288608 | 0.411662127 | 0.798759393 | ns | NaCl | KS.3 |
| FIM1_325 | 16.54974783 | 14.28386115 | 17.19711774 | -0.014087008 | 0.137002295 | 0.682954017 | 0.915673519 | ns | NaCl | KS.3 |
| FIM1_326 | 98.27303941 | 217.7980809 | 4974.569664 | 0.22183203 | 0.355880998 | 0.010241619 | 0.142722848 | ns | NaCl | KS.3 |
| FIM1_327 | 1019.305187 | 1134.477878 | 1256.923338 | 0.064137304 | 0.080473328 | 0.204282851 | 0.627737607 | ns | NaCl | KS.3 |
| FIM1_328 | 127.6468431 | 163.4961809 | 197.3835281 | 0.041317772 | 0.13080294 | 0.313303829 | 0.731871738 | ns | NaCl | KS.3 |
| FIM1_329 | 381.4390441 | 331.2213248 | 674.5158635 | -0.032020344 | 0.110511516 | 0.450204318 | 0.813595672 | ns | NaCl | KS.3 |
| FIM1_330 | 15705.79854 | 12404.18374 | 10680.29304 | -0.1157975 | 0.115660151 | 0.037547928 | 0.305729858 | ns | NaCl | KS.3 |
| FIM1_331 | 460.211822 | 277.2798299 | 526.2211276 | -0.214762497 | 0.261154678 | 0.009163113 | 0.133293825 | ns | NaCl | KS.3 |
| FIM1_332 | 1155.666905 | 839.4774272 | 1159.272756 | -0.169082433 | 0.159760286 | 0.010881679 | 0.148997467 | ns | NaCl | KS.3 |
| FIM1_333 | 904.4702019 | 722.2075784 | 1067.606057 | -0.106448855 | 0.112013801 | 0.050053528 | 0.349544574 | ns | NaCl | KS.3 |
| FIM1_334 | 2535.931798 | 2460.577654 | 1901.909592 | -0.013170039 | 0.086948303 | 0.78232234 | 0.951646805 | ns | NaCl | KS.3 |
| FIM1_335 | 1797.227656 | 2081.569035 | 1214.735243 | 0.025130284 | 0.118311213 | 0.525509372 | 0.854940852 | ns | NaCl | KS.3 |
| FIM1_336 | 292.917987 | 228.3640249 | 665.6980096 | -0.079809788 | 0.123149994 | 0.109856362 | 0.505115899 | ns | NaCl | KS.3 |
| FIM1_337 | 3554.051462 | 3166.823183 | 3669.238901 | -0.056484589 | 0.088062144 | 0.255789784 | 0.678991959 | ns | NaCl | KS.3 |
| FIM1_338 | 2700.070284 | 3207.270746 | 2051.336962 | 0.032225387 | 0.118400481 | 0.430125872 | 0.805047964 | ns | NaCl | KS.3 |
| FIM1_339 | 5094.341362 | 5091.764613 | 3976.175577 | 2.09674361929113 | 0.089252355 | 0.999642978 | 0.999642978 | ns | NaCl | KS.3 |
| FIM1_340 | 259.1057484 | 237.1486935 | 277.3983359 | -0.013222818 | 0.11846499 | 0.726891996 | 0.934304934 | ns | NaCl | KS.3 |

| | | | | | | | | | | |
|---|---|---|---|---|---|---|---|---|---|---|
| FIM1_341 | 5905.171949 | 6470.106386 | 4523.907596 | 0.054731377 | 0.079447744 | 0.27491632 | 0.695399585 | ns | NaCl | KS.3 |
| FIM1_342 | 1223.381453 | 1173.183562 | 1112.983384 | -0.018895541 | 0.086093567 | 0.693961398 | 0.920525145 | ns | NaCl | KS.3 |
| FIM1_343 | 2061.839088 | 2164.46884 | 2331.160854 | 0.011563195 | 0.103780387 | 0.782946832 | 0.951646805 | ns | NaCl | KS.3 |
| FIM1_344 | 1673.128924 | 1639.036657 | 1776.822187 | -0.011167327 | 0.079289978 | 0.819730005 | 0.960016822 | ns | NaCl | KS.3 |
| FIM1_345 | 5865.894583 | 4694.93344 | 3734.663347 | -0.062542123 | 0.118082337 | 0.182473987 | 0.611396712 | ns | NaCl | KS.3 |
| FIM1_346 | 7276.386399 | 7086.536778 | 6352.248537 | -0.011030992 | 0.08806574 | 0.815690563 | 0.958816322 | ns | NaCl | KS.3 |
| FIM1_347 | 1729.510679 | 1965.961135 | 1962.428707 | 0.076097105 | 0.08349344 | 0.136969407 | 0.545297021 | ns | NaCl | KS.3 |
| FIM1_348 | 3292.421586 | 4798.003765 | 2060.195225 | 0.078573236 | 0.160239693 | 0.105052262 | 0.495197752 | ns | NaCl | KS.3 |
| FIM1_349 | 2315.798366 | 2334.873937 | 2501.27156 | 0.004317715 | 0.077592907 | 0.92991579 | 0.984325045 | ns | NaCl | KS.3 |
| FIM1_350 | 413.9168438 | 386.6074905 | 551.4005951 | -0.038879799 | 0.080592939 | 0.432238078 | 0.805772482 | ns | NaCl | KS.3 |
| FIM1_351 | 1845.135954 | 1641.449703 | 1800.110609 | -0.106274447 | 0.063509393 | 0.027782026 | 0.260899256 | ns | NaCl | KS.3 |
| FIM1_352 | 4655.591313 | 4806.00965 | 5972.361674 | 0.005240685 | 0.113877421 | 0.891544532 | 0.977268396 | ns | NaCl | KS.3 |
| FIM1_354 | 140.5131922 | 148.2622884 | 493.2727055 | 0.008767768 | 0.115657569 | 0.818241734 | 0.959981393 | ns | NaCl | KS.3 |
| FIM1_355 | 3201.956024 | 2741.946938 | 2709.148435 | -0.092041192 | 0.087784115 | 0.07840326 | 0.433395797 | ns | NaCl | KS.3 |
| FIM1_356 | 702.0982264 | 730.173099 | 2103.957352 | 0.009498469 | 0.101968398 | 0.823167818 | 0.960240659 | ns | NaCl | KS.3 |
| FIM1_357 | 2900.429707 | 3035.49763 | 3991.281274 | 0.008326265 | 0.111050231 | 0.832683949 | 0.961831996 | ns | NaCl | KS.3 |
| FIM1_358 | 9825.549636 | 13464.84102 | 6074.080717 | 0.065712725 | 0.142596168 | 0.152597905 | 0.568642049 | ns | NaCl | KS.3 |
| FIM1_359 | 9208.783788 | 9898.420436 | 7789.665494 | 0.080574126 | 0.045840271 | 0.042066288 | 0.323956287 | ns | NaCl | KS.3 |
| FIM1_360 | 1502.805818 | 1518.397463 | 1375.551515 | 0.006018926 | 0.07089696 | 0.902164085 | 0.979395371 | ns | NaCl | KS.3 |
| FIM1_361 | 581.4088527 | 517.8966124 | 560.8953301 | -0.041001995 | 0.097298599 | 0.380147915 | 0.778493059 | ns | NaCl | KS.3 |
| FIM1_362 | 2379.58298 | 2091.499294 | 2519.03007 | -0.043289994 | 0.099729152 | 0.351342891 | 0.756912594 | ns | NaCl | KS.3 |
| FIM1_363 | 1195.358858 | 1223.523154 | 1212.245779 | 0.010298317 | 0.086306681 | 0.829316294 | 0.961288108 | ns | NaCl | KS.3 |
| FIM1_364 | 2609.457226 | 2639.444315 | 2836.126551 | 0.010184293 | 0.05675488 | 0.822044252 | 0.960240659 | ns | NaCl | KS.3 |
| FIM1_365 | 1844.526425 | 2055.363312 | 1520.484683 | 0.038679858 | 0.096218228 | 0.407795583 | 0.79494601 | ns | NaCl | KS.3 |
| FIM1_366 | 1674.083559 | 1484.909457 | 1946.325552 | -0.049252067 | 0.093757691 | 0.307690678 | 0.727796074 | ns | NaCl | KS.3 |
| FIM1_367 | 2406.803698 | 4535.237337 | 4689.156278 | 0.321842812 | 0.351047842 | 0.003628551 | 0.076817191 | ns | NaCl | KS.3 |
| FIM1_368 | 461.4730164 | 511.6205547 | 489.1536025 | 0.017514621 | 0.115526767 | 0.653913334 | 0.900747416 | ns | NaCl | KS.3 |
| FIM1_369 | 36.20115453 | 35.42241762 | 35.23365655 | -0.003163424 | 0.130479866 | 0.927202715 | 0.984176127 | ns | NaCl | KS.3 |
| FIM1_370 | 639.0770134 | 653.7568871 | 767.2498702 | 0.010755262 | 0.081762411 | 0.825174846 | 0.960240659 | ns | NaCl | KS.3 |
| FIM1_371 | 72.7328648 | 79.31528669 | 98.26239274 | 0.01420682 | 0.114380442 | 0.715433146 | 0.929315901 | ns | NaCl | KS.3 |
| FIM1_372 | 15905.32297 | 19894.55145 | 15260.98212 | 0.066622933 | 0.117554207 | 0.162860386 | 0.586304673 | ns | NaCl | KS.3 |
| FIM1_373 | 1262.550985 | 1315.328229 | 1422.934025 | 0.013315832 | 0.095318568 | 0.767687102 | 0.948794469 | ns | NaCl | KS.3 |
| FIM1_374 | 1562.063303 | 1604.344204 | 1849.472422 | 0.013675937 | 0.08175285 | 0.778982399 | 0.951027591 | ns | NaCl | KS.3 |
| FIM1_375 | 1283.866874 | 1367.731737 | 1675.434603 | 0.016377548 | 0.101980169 | 0.70283166 | 0.923421278 | ns | NaCl | KS.3 |
| FIM1_376 | 503.8671793 | 278.7841543 | 754.4049239 | -0.271861873 | 0.317201234 | 0.005473952 | 0.09831376 | ns | NaCl | KS.3 |
| FIM1_377 | 722.044176 | 450.1464056 | 809.5753069 | -0.328474691 | 0.271673967 | 0.001828322 | 0.051389285 | ns | NaCl | KS.3 |
| FIM1_378 | 49556.78388 | 73465.2327 | 46593.75577 | 0.430440644 | 0.193074491 | 7.22006529660 | 0.005612473 | ns | NaCl | KS.3 |
| FIM1_379 | 531.7638656 | 561.457316 | 566.4716419 | 0.008086552 | 0.117541627 | 0.829832215 | 0.961438115 | ns | NaCl | KS.3 |
| FIM1_380 | 1550.887351 | 1715.735529 | 2248.476043 | 0.021007661 | 0.109708154 | 0.60850215 | 0.88569286 | ns | NaCl | KS.3 |
| FIM1_381 | 3450.01187 | 4616.457694 | 2891.447937 | 0.063732166 | 0.136512192 | 0.164135187 | 0.589056105 | ns | NaCl | KS.3 |
| FIM1_383 | 1516.323268 | 3169.803234 | 5227.277974 | 0.384014976 | 0.410450069 | 0.002537289 | 0.06187751 | ns | NaCl | KS.3 |
| FIM1_384 | 62885.66661 | 78737.90822 | 47056.6348 | 0.069934944 | 0.117180869 | 0.148388988 | 0.563010285 | ns | NaCl | KS.3 |
| FIM1_385 | 449.8597695 | 674.8459189 | 715.9322575 | 0.265353498 | 0.225185351 | 0.002804183 | 0.065496772 | ns | NaCl | KS.3 |
| FIM1_386 | 1735.150949 | 1448.890874 | 2795.587424 | -0.036956455 | 0.117273405 | 0.378340059 | 0.778493059 | ns | NaCl | KS.3 |
| FIM1_387 | 799.2429757 | 809.4182826 | 897.1630477 | 0.004991365 | 0.087961144 | 0.915891879 | 0.982343702 | ns | NaCl | KS.3 |

| | | | | | | | | | | |
|---|---|---|---|---|---|---|---|---|---|---|
| FIM1_388 | 550.5644691 | 455.1644679 | 624.7718793 | -0.088822847 | 0.101964635 | 0.087876495 | 0.459228534 | ns | NaCl | KS.3 |
| FIM1_389 | 3433.49162 | 3264.806827 | 3232.834531 | -0.020604491 | 0.089131974 | 0.663295612 | 0.904576664 | ns | NaCl | KS.3 |
| FIM1_390 | 300.861817 | 290.3894268 | 386.9148257 | -0.009758179 | 0.101027229 | 0.819887385 | 0.960016822 | ns | NaCl | KS.3 |
| FIM1_391 | 1322.168521 | 1132.366839 | 1297.739817 | -0.087412652 | 0.08936556 | 0.092767663 | 0.470201108 | ns | NaCl | KS.3 |
| FIM1_392 | 4584.746301 | 4720.74753 | 4038.265816 | 0.014967622 | 0.081967338 | 0.758670517 | 0.945250644 | ns | NaCl | KS.3 |
| FIM1_393 | 2034.773886 | 2210.258583 | 1931.746616 | 0.063069242 | 0.069471645 | 0.201709403 | 0.624846998 | ns | NaCl | KS.3 |
| FIM1_394 | 248.1729322 | 246.8584855 | 215.3585894 | -0.001613576 | 0.10401751 | 0.96900682 | 0.99411987 | ns | NaCl | KS.3 |
| FIM1_395 | 1193.118735 | 1168.978677 | 1365.180795 | -0.00898644 | 0.086584953 | 0.850505676 | 0.967955712 | ns | NaCl | KS.3 |
| FIM1_396 | 3187.573647 | 3540.519518 | 2343.481647 | 0.056901186 | 0.083896694 | 0.255893079 | 0.678991959 | ns | NaCl | KS.3 |
| FIM1_397 | 33.2023075 | 40.18287092 | 44.24618426 | 0.026128093 | 0.125801562 | 0.496085573 | 0.836358994 | ns | NaCl | KS.3 |
| FIM1_398 | 1981.581508 | 1980.608304 | 2180.258315 | -0.000645066 | 0.080619912 | 0.989451665 | 0.999134858 | ns | NaCl | KS.3 |
| FIM1_399 | 2608.083341 | 2816.716227 | 2774.821104 | 0.036810488 | 0.085864116 | 0.450570755 | 0.813595672 | ns | NaCl | KS.3 |
| FIM1_401 | 1493.184286 | 1479.831618 | 1422.118783 | -0.004715722 | 0.08168602 | 0.922780475 | 0.983890455 | ns | NaCl | KS.3 |
| FIM1_402 | 179.5009612 | 152.4030617 | 416.4323931 | -0.047121541 | 0.10822055 | 0.298866902 | 0.717010926 | ns | NaCl | KS.3 |
| FIM1_403 | 674.6101192 | 474.9115011 | 967.7019184 | -0.196703628 | 0.181384225 | 0.006850208 | 0.111371848 | ns | NaCl | KS.3 |
| FIM1_404 | 517.0499668 | 416.7531344 | 546.760275 | -0.059721531 | 0.116362042 | 0.199346402 | 0.622566448 | ns | NaCl | KS.3 |
| FIM1_405 | 10199.31735 | 10862.97729 | 7626.635991 | 0.05006979 | 0.065975996 | 0.301483386 | 0.720057535 | ns | NaCl | KS.3 |
| FIM1_406 | 13376.81168 | 13524.07394 | 9920.880166 | 0.008913721 | 0.062597228 | 0.850633643 | 0.967955712 | ns | NaCl | KS.3 |
| FIM1_407 | 4398.611125 | 2587.986165 | 3459.852175 | -0.611195289 | 0.240881838 | 4.42810641761 | 0.003933898 | ns | NaCl | KS.3 |
| FIM1_408 | 1767.510635 | 1481.518302 | 1724.87704 | -0.152723728 | 0.078080871 | 0.004275509 | 0.08508263 | ns | NaCl | KS.3 |
| FIM1_409 | 4587.133054 | 4416.618707 | 4668.864928 | -0.021103615 | 0.078941725 | 0.667782407 | 0.907957769 | ns | NaCl | KS.3 |
| FIM1_410 | 109.4425174 | 83.64440136 | 198.6579872 | -0.069897353 | 0.127885806 | 0.142803384 | 0.555038853 | ns | NaCl | KS.3 |
| FIM1_411 | 1189.320103 | 1132.143213 | 2144.197744 | -0.007844359 | 0.114616103 | 0.838000529 | 0.962829707 | ns | NaCl | KS.3 |
| FIM1_412 | 673.1381424 | 746.2788577 | 728.237725 | 0.055000658 | 0.083232631 | 0.271525671 | 0.693144962 | ns | NaCl | KS.3 |
| FIM1_413 | 5032.366492 | 5192.230419 | 4930.833769 | 0.022621681 | 0.068380108 | 0.642338231 | 0.898798052 | ns | NaCl | KS.3 |
| FIM1_414 | 333.3509177 | 263.624532 | 811.4516496 | -0.048243243 | 0.125903361 | 0.263204003 | 0.688222424 | ns | NaCl | KS.3 |
| FIM1_415 | 349.7702183 | 297.9224962 | 421.903523 | -0.060472269 | 0.100543391 | 0.212686906 | 0.636270209 | ns | NaCl | KS.3 |
| FIM1_416 | 1283.771453 | 1122.867614 | 1370.58872 | -0.079828727 | 0.084962538 | 0.120505047 | 0.526812484 | ns | NaCl | KS.3 |
| FIM1_417 | 4027.40185 | 4712.741488 | 6188.655595 | 0.074358728 | 0.09485005 | 0.142476635 | 0.555038853 | ns | NaCl | KS.3 |
| FIM1_418 | 557.9543172 | 613.5552273 | 901.0270708 | 0.033470369 | 0.096301918 | 0.468768047 | 0.821764392 | ns | NaCl | KS.3 |
| FIM1_419 | 647.6660103 | 609.697787 | 844.6077708 | -0.027027816 | 0.087422591 | 0.573555935 | 0.871789624 | ns | NaCl | KS.3 |
| FIM1_420 | 722.0107242 | 780.4971148 | 749.4564911 | 0.038758865 | 0.084371651 | 0.429960424 | 0.805047964 | ns | NaCl | KS.3 |
| FIM1_421 | 1075.406175 | 1122.752573 | 1164.997138 | 0.026693196 | 0.074773771 | 0.588705205 | 0.878638624 | ns | NaCl | KS.3 |
| FIM1_422 | 380.7145363 | 373.7692595 | 623.0930477 | -0.010170692 | 0.083826895 | 0.833288728 | 0.962305344 | ns | NaCl | KS.3 |
| FIM1_423 | 66.48491682 | 48.42714655 | 92.95811991 | -0.031730704 | 0.147675401 | 0.387395342 | 0.779278134 | ns | NaCl | KS.3 |
| FIM1_425 | 5829.833009 | 7996.993365 | 3568.612381 | 0.069704457 | 0.143040183 | 0.137107742 | 0.545297021 | ns | NaCl | KS.3 |
| FIM1_426 | 4994.288239 | 5277.36944 | 4407.258022 | 0.03185059 | 0.078705466 | 0.519214918 | 0.850629872 | ns | NaCl | KS.3 |
| FIM1_427 | 379.3294422 | 356.5960759 | 348.6366388 | -0.026241344 | 0.089329242 | 0.580895191 | 0.8749481 | ns | NaCl | KS.3 |
| FIM1_428 | 479.3181161 | 457.5552507 | 345.022864 | -0.015211893 | 0.095994371 | 0.734940877 | 0.938375543 | ns | NaCl | KS.3 |
| FIM1_429 | 1851.249094 | 1697.528019 | 1382.713337 | -0.053121796 | 0.078024383 | 0.288632223 | 0.708650347 | ns | NaCl | KS.3 |
| FIM1_430 | 978.7130188 | 1104.503774 | 1029.537073 | 0.083925606 | 0.077378377 | 0.100717357 | 0.484124495 | ns | NaCl | KS.3 |
| FIM1_431 | 601.1644961 | 608.0162481 | 613.3133998 | 0.003422444 | 0.097193687 | 0.938146689 | 0.986790704 | ns | NaCl | KS.3 |
| FIM1_432 | 291.1441402 | 273.8992046 | 229.5730587 | -0.013647408 | 0.10618811 | 0.741138966 | 0.941221011 | ns | NaCl | KS.3 |
| FIM1_433 | 2553.500966 | 2578.878448 | 1934.95982 | 0.00514303 | 0.080748664 | 0.916045657 | 0.982343702 | ns | NaCl | KS.3 |
| FIM1_434 | 1100.444386 | 1034.997309 | 1025.508772 | -0.03102788 | 0.083585008 | 0.525736893 | 0.855027345 | ns | NaCl | KS.3 |

| | | | | | | | | | | |
|---|---|---|---|---|---|---|---|---|---|---|
| FIM1_435 | 1165.294404 | 1080.010751 | 1007.262228 | -0.048508298 | 0.075959991 | 0.330635352 | 0.742254041 | ns | NaCl | KS.3 |
| FIM1_436 | 399.9354377 | 419.0240276 | 391.0584459 | 0.012644543 | 0.100796676 | 0.769184145 | 0.948835638 | ns | NaCl | KS.3 |
| FIM1_437 | 436.5112403 | 353.498798 | 545.3349553 | -0.092253159 | 0.110025354 | 0.078082009 | 0.432946334 | ns | NaCl | KS.3 |
| FIM1_438 | 10.5802296 | 10.70128086 | 26.92615975 | -0.00123447 | 0.143643007 | 0.96954702 | 0.99411987 | ns | NaCl | KS.3 |
| FIM1_439 | 2814.886059 | 3718.156768 | 1895.423913 | 0.064873454 | 0.133071388 | 0.161019964 | 0.582176107 | ns | NaCl | KS.3 |
| FIM1_440 | 653.782043 | 555.0879231 | 823.75812 | -0.09239379 | 0.091835039 | 0.078082936 | 0.432946334 | ns | NaCl | KS.3 |
| FIM1_441 | 863.3212021 | 760.3862109 | 966.3812468 | -0.07685811 | 0.082670904 | 0.133258716 | 0.540311419 | ns | NaCl | KS.3 |
| FIM1_442 | 762.5893003 | 685.7313689 | 1211.952098 | -0.034966781 | 0.097887309 | 0.446491248 | 0.812914509 | ns | NaCl | KS.3 |
| FIM1_443 | 89.15301906 | 94.16805248 | 176.328668 | 0.008609936 | 0.116906091 | 0.819921905 | 0.960016822 | ns | NaCl | KS.3 |
| FIM1_444 | 1292.825319 | 1344.712894 | 1310.064377 | 0.01844421 | 0.085737049 | 0.701308762 | 0.923421278 | ns | NaCl | KS.3 |
| FIM1_445 | 1580.100493 | 1460.756681 | 1874.845332 | -0.04135014 | 0.083118337 | 0.402335925 | 0.789854857 | ns | NaCl | KS.3 |
| FIM1_446 | 230.7582918 | 202.8430773 | 323.4614719 | -0.043631623 | 0.100095799 | 0.347187211 | 0.756654396 | ns | NaCl | KS.3 |
| FIM1_447 | 829.0441527 | 825.5547872 | 917.951041 | -0.002765465 | 0.067656331 | 0.954538611 | 0.989477692 | ns | NaCl | KS.3 |
| FIM1_448 | 2395.992343 | 2265.530381 | 2575.742673 | -0.029206151 | 0.082328978 | 0.551241744 | 0.860945181 | ns | NaCl | KS.3 |
| FIM1_449 | 4548.614596 | 3771.911806 | 4280.172542 | -0.062426534 | 0.108053371 | 0.192515871 | 0.618952432 | ns | NaCl | KS.3 |
| FIM1_450 | 263.9793934 | 117.4204651 | 132.0429841 | -0.950777268 | 0.311631488 | 1.2753177692[illegible] | 0.001490695 | ns | NaCl | KS.3 |
| FIM1_451 | 2284.821254 | 2731.559019 | 2000.966981 | 0.045823323 | 0.112018862 | 0.303111737 | 0.72182705 | ns | NaCl | KS.3 |
| FIM1_452 | 603.1040232 | 600.3803968 | 780.8864883 | -0.001227525 | 0.09108354 | 0.978838784 | 0.996164232 | ns | NaCl | KS.3 |
| FIM1_453 | 3438.330822 | 3359.185704 | 2775.856282 | -0.009092197 | 0.090143723 | 0.845550723 | 0.967147129 | ns | NaCl | KS.3 |
| FIM1_454 | 158.9964575 | 187.6893071 | 268.9446485 | 0.021365425 | 0.128143537 | 0.564811656 | 0.867532568 | ns | NaCl | KS.3 |
| FIM1_455 | 1452.197865 | 1384.340054 | 1245.188269 | -0.012099682 | 0.102041883 | 0.77647028 | 0.950410881 | ns | NaCl | KS.3 |
| FIM1_456 | 1164.998138 | 1191.938276 | 1442.054858 | 0.00600252 | 0.100531907 | 0.888725846 | 0.977268396 | ns | NaCl | KS.3 |
| FIM1_457 | 19365.55806 | 22406.18646 | 14320.3747 | 0.076445425 | 0.089954573 | 0.134870644 | 0.542426399 | ns | NaCl | KS.3 |
| FIM1_458 | 29312.88633 | 33602.33091 | 20648.47836 | 0.080984185 | 0.084712279 | 0.115689359 | 0.518050912 | ns | NaCl | KS.3 |
| FIM1_459 | 1849.868179 | 1994.157845 | 1542.097326 | 0.023520068 | 0.097328166 | 0.601450863 | 0.880762097 | ns | NaCl | KS.3 |
| FIM1_460 | 407.2073957 | 439.6966044 | 413.5934671 | 0.026556086 | 0.095001235 | 0.56354818 | 0.867434473 | ns | NaCl | KS.3 |
| FIM1_461 | 620.1407021 | 631.7772053 | 842.2776394 | 0.008516993 | 0.083983231 | 0.859939423 | 0.969856868 | ns | NaCl | KS.3 |
| FIM1_462 | 741.9495092 | 675.1964428 | 709.7308589 | -0.073048986 | 0.069740168 | 0.141066519 | 0.552608798 | ns | NaCl | KS.3 |
| FIM1_463 | 2843.503147 | 3619.769619 | 2168.265916 | 0.062956427 | 0.123267623 | 0.176276522 | 0.605047427 | ns | NaCl | KS.3 |
| FIM1_464 | 5601.664943 | 5175.671407 | 5385.754652 | -0.031812614 | 0.090924273 | 0.502635175 | 0.842535794 | ns | NaCl | KS.3 |
| FIM1_465 | 36.03448255 | 18.67557014 | 27.27999917 | -0.109525487 | 0.236543764 | 0.048355769 | 0.344656093 | ns | NaCl | KS.3 |
| FIM1_466 | 10843.03385 | 12145.41426 | 7328.141002 | 0.069756201 | 0.080554732 | 0.169528959 | 0.594663296 | ns | NaCl | KS.3 |
| FIM1_467 | 3311.379452 | 4091.227862 | 2370.570354 | 0.061044834 | 0.115360953 | 0.192919272 | 0.618952432 | ns | NaCl | KS.3 |
| FIM1_468 | 88.63701255 | 57.24574958 | 101.8914025 | -0.15726562 | 0.215084086 | 0.020221736 | 0.215832354 | ns | NaCl | KS.3 |
| FIM1_469 | 724.905829 | 787.8236277 | 815.2713542 | 0.021646172 | 0.102645754 | 0.616108167 | 0.890538796 | ns | NaCl | KS.3 |
| FIM1_470 | 886.1475683 | 821.6136108 | 1474.85134 | -0.019781027 | 0.102095489 | 0.646834128 | 0.898973628 | ns | NaCl | KS.3 |
| FIM1_471 | 603.8459897 | 601.6177995 | 924.8653515 | -0.000538301 | 0.114133775 | 0.988772962 | 0.999134858 | ns | NaCl | KS.3 |
| FIM1_472 | 560.0699414 | 567.8251669 | 470.3324537 | 0.002734175 | 0.107372086 | 0.946001414 | 0.987278589 | ns | NaCl | KS.3 |
| FIM1_473 | 1252.918556 | 1224.850956 | 1561.354367 | -0.007037185 | 0.096652062 | 0.874001808 | 0.971243188 | ns | NaCl | KS.3 |
| FIM1_474 | 2121.43982 | 2119.531502 | 1664.142884 | -0.000481803 | 0.080527044 | 0.99212325 | 0.999134858 | ns | NaCl | KS.3 |
| FIM1_475 | 364.3051579 | 300.9909711 | 707.7649955 | -0.047534072 | 0.115168138 | 0.283629239 | 0.704471025 | ns | NaCl | KS.3 |
| FIM1_476 | 14830.60801 | 13550.42077 | 9708.260431 | -0.036171042 | 0.091808569 | 0.446853533 | 0.812914509 | ns | NaCl | KS.3 |
| FIM1_477 | 10590.34102 | 11654.44966 | 8762.515758 | 0.065039402 | 0.075152813 | 0.195754607 | 0.618952432 | ns | NaCl | KS.3 |
| FIM1_478 | 5754.110201 | 8217.67925 | 3891.826898 | 0.077220678 | 0.154684419 | 0.109852613 | 0.505115899 | ns | NaCl | KS.3 |
| FIM1_479 | 1221.2668 | 1403.113942 | 1402.401075 | 0.034922159 | 0.107744065 | 0.421972573 | 0.801548921 | ns | NaCl | KS.3 |

| | | | | | | | | | | |
|---|---|---|---|---|---|---|---|---|---|---|
| FIM1_480 | 824.8737922 | 760.7791567 | 1130.206243 | -0.04316178 | 0.083234979 | 0.382697007 | 0.778665419 | ns | NaCl | KS.3 |
| FIM1_481 | 757.6202625 | 653.9441016 | 1343.131751 | -0.085938769 | 0.0873666 | 0.097490945 | 0.476631548 | ns | NaCl | KS.3 |
| FIM1_482 | 767.1450388 | 662.8914969 | 1188.457089 | -0.064546455 | 0.095439951 | 0.193311137 | 0.618952432 | ns | NaCl | KS.3 |
| FIM1_483 | 2152.797738 | 2336.195668 | 2054.337398 | 0.061772539 | 0.069594529 | 0.211152439 | 0.634491393 | ns | NaCl | KS.3 |
| FIM1_485 | 59.90594664 | 73.42717226 | 116.1817359 | 0.044735266 | 0.115833048 | 0.305915836 | 0.725074457 | ns | NaCl | KS.3 |
| FIM1_486 | 653.2758693 | 454.8007261 | 1003.7575 | -0.205972533 | 0.191363436 | 0.006194418 | 0.106434152 | ns | NaCl | KS.3 |
| FIM1_487 | 800.8004663 | 762.3922633 | 668.4113705 | -0.013781896 | 0.099388741 | 0.752243643 | 0.943523399 | ns | NaCl | KS.3 |
| FIM1_488 | 5061.912268 | 4549.45162 | 5026.573853 | -0.036332707 | 0.097312526 | 0.431822366 | 0.805772482 | ns | NaCl | KS.3 |
| FIM1_489 | 3679.5222 | 3979.566962 | 3569.253078 | 0.017088451 | 0.10723039 | 0.678983158 | 0.913941356 | ns | NaCl | KS.3 |
| FIM1_490 | 1318.285195 | 1228.091929 | 1831.003238 | -0.037392669 | 0.082470744 | 0.448009463 | 0.812914509 | ns | NaCl | KS.3 |
| FIM1_491 | 17038.71566 | 20749.51608 | 12395.74351 | 0.167959375 | 0.085958267 | 0.002650853 | 0.063710103 | ns | NaCl | KS.3 |
| FIM1_492 | 2198.017154 | 1949.952048 | 2334.251937 | -0.088996022 | 0.074210669 | 0.080040736 | 0.438066737 | ns | NaCl | KS.3 |
| FIM1_493 | 1709.06533 | 1651.348529 | 1321.065216 | -0.016923554 | 0.083664177 | 0.726973749 | 0.934304934 | ns | NaCl | KS.3 |
| FIM1_494 | 2665.097162 | 2713.720674 | 1974.82989 | 0.010733814 | 0.076897534 | 0.827171769 | 0.960240659 | ns | NaCl | KS.3 |
| FIM1_495 | 4045.047505 | 3425.443854 | 3267.866095 | -0.126076819 | 0.081391112 | 0.018168437 | 0.202663614 | ns | NaCl | KS.3 |
| FIM1_496 | 6137.842678 | 6271.058131 | 5435.532296 | 0.020593051 | 0.054260296 | 0.641269595 | 0.898798052 | ns | NaCl | KS.3 |
| FIM1_497 | 1085.350607 | 1215.93318 | 1598.935476 | 0.029665859 | 0.104565888 | 0.495560024 | 0.836358994 | ns | NaCl | KS.3 |
| FIM1_498 | 726.3942601 | 694.6694515 | 630.0746192 | -0.018596297 | 0.089621566 | 0.693050872 | 0.920513182 | ns | NaCl | KS.3 |
| FIM1_499 | 4147.019198 | 4257.076576 | 3312.6156 | 0.015787035 | 0.076256529 | 0.748449194 | 0.943274417 | ns | NaCl | KS.3 |
| FIM1_500 | 450.7222636 | 402.2556088 | 352.4480144 | -0.047426125 | 0.093508596 | 0.324676981 | 0.739131368 | ns | NaCl | KS.3 |
| FIM1_501 | 932.5274612 | 820.4738443 | 642.575356 | -0.046165203 | 0.098001007 | 0.327541326 | 0.740319311 | ns | NaCl | KS.3 |
| FIM1_502 | 2982.008729 | 3153.546382 | 2460.250386 | 0.02775343 | 0.083843122 | 0.569096636 | 0.869550295 | ns | NaCl | KS.3 |
| FIM1_503 | 471.008915 | 498.7275296 | 959.0798002 | 0.01872063 | 0.094288322 | 0.682006939 | 0.915044369 | ns | NaCl | KS.3 |
| FIM1_504 | 805.7469201 | 797.2627603 | 1291.733053 | -0.004280373 | 0.072824566 | 0.930542364 | 0.984325045 | ns | NaCl | KS.3 |
| FIM1_505 | 1218.417734 | 1486.114067 | 1053.730224 | 0.070145704 | 0.109322332 | 0.152514429 | 0.568642049 | ns | NaCl | KS.3 |
| FIM1_506 | 5319.062576 | 6384.749485 | 3207.087225 | 0.04067642 | 0.115606083 | 0.343585425 | 0.752349247 | ns | NaCl | KS.3 |
| FIM1_507 | 1392.237784 | 1490.872964 | 1545.803788 | 0.033289908 | 0.084363322 | 0.495747542 | 0.836358994 | ns | NaCl | KS.3 |
| FIM1_508 | 637.4205775 | 676.4778043 | 1038.605976 | 0.026936171 | 0.086669316 | 0.576125887 | 0.873849478 | ns | NaCl | KS.3 |
| FIM1_509 | 533.2034255 | 394.5164122 | 964.2370488 | -0.093992957 | 0.140067503 | 0.072675105 | 0.421203992 | ns | NaCl | KS.3 |
| FIM1_510 | 839.2318397 | 907.3135892 | 854.0455006 | 0.038365587 | 0.085515598 | 0.432925114 | 0.806063788 | ns | NaCl | KS.3 |
| FIM1_511 | 1131.450875 | 1150.42999 | 1599.389843 | 0.005345743 | 0.097895999 | 0.903021204 | 0.979395371 | ns | NaCl | KS.3 |
| FIM1_512 | 211.3585638 | 169.2424141 | 324.3843601 | -0.062962951 | 0.118782077 | 0.179731862 | 0.608447876 | ns | NaCl | KS.3 |
| FIM1_513 | 4325.151567 | 4421.988482 | 3189.871274 | 0.011835069 | 0.080112609 | 0.80884773 | 0.95855585 | ns | NaCl | KS.3 |
| FIM1_514 | 1347.226948 | 1222.821958 | 1894.917338 | -0.035542505 | 0.094874896 | 0.447165993 | 0.812914509 | ns | NaCl | KS.3 |
| FIM1_515 | 2004.952582 | 1704.750046 | 1777.12397 | -0.153107089 | 0.070236463 | 0.002741686 | 0.064671708 | ns | NaCl | KS.3 |
| FIM1_516 | 1262.55692 | 1196.346807 | 1352.961096 | -0.033773273 | 0.076386845 | 0.495193294 | 0.836358994 | ns | NaCl | KS.3 |
| FIM1_517 | 2059.940875 | 2534.246988 | 1368.583645 | 0.043164177 | 0.120489728 | 0.312057956 | 0.731871738 | ns | NaCl | KS.3 |
| FIM1_518 | 31752.30867 | 36628.45865 | 24314.06549 | 0.127867813 | 0.06825551 | 0.010774916 | 0.148080678 | ns | NaCl | KS.3 |
| FIM1_519 | 77.09942126 | 92.57622576 | 587.3415799 | 0.022321737 | 0.130230486 | 0.54569581 | 0.857742326 | ns | NaCl | KS.3 |
| FIM1_522 | 3644.274826 | 4369.690845 | 2603.129516 | 0.04287622 | 0.114190675 | 0.325404897 | 0.739131368 | ns | NaCl | KS.3 |
| FIM1_523 | 788.4214535 | 724.2027955 | 636.3143509 | -0.028502544 | 0.096422197 | 0.532992629 | 0.855153333 | ns | NaCl | KS.3 |
| FIM1_524 | 1731.540027 | 1326.207975 | 1738.364328 | -0.200626136 | 0.128693223 | 0.002469491 | 0.060820392 | ns | NaCl | KS.3 |
| FIM1_525 | 1366.922027 | 1452.317104 | 1775.306378 | 0.047284907 | 0.063310905 | 0.321662445 | 0.737180727 | ns | NaCl | KS.3 |
| FIM1_526 | 80.89736608 | 65.49211128 | 136.6789371 | -0.03083503 | 0.131002066 | 0.423955136 | 0.802854185 | ns | NaCl | KS.3 |
| FIM1_527 | 10576.74377 | 15045.68118 | 14481.84218 | 0.060391852 | 0.151122883 | 0.170978082 | 0.597342667 | ns | NaCl | KS.3 |

| | | | | | | | | | | |
|---|---|---|---|---|---|---|---|---|---|---|
| FIM1_528 | 2614.508258 | 3140.604424 | 3607.913838 | 0.031539394 | 0.121691731 | 0.431964967 | 0.805772482 | ns | NaCl | KS.3 |
| FIM1_529 | 21.77037498 | 25.57922671 | 26.20148965 | 0.025633127 | 0.125003284 | 0.504727651 | 0.844040357 | ns | NaCl | KS.3 |
| FIM1_530 | 5347.94075 | 4360.198416 | 3718.552015 | -0.090475014 | 0.106600337 | 0.082859063 | 0.446129696 | ns | NaCl | KS.3 |
| FIM1_531 | 14168.20889 | 14116.85727 | 10265.85102 | -0.000687559 | 0.10925918 | 0.986190321 | 0.999134858 | ns | NaCl | KS.3 |
| FIM1_532 | 5960.582185 | 8940.004037 | 4169.976391 | 0.061311848 | 0.162422006 | 0.160867892 | 0.582143233 | ns | NaCl | KS.3 |
| FIM1_533 | 1002.392977 | 1009.14728 | 1149.200053 | 0.005133879 | 0.075891676 | 0.916831914 | 0.982343702 | ns | NaCl | KS.3 |
| FIM1_534 | 997.8794291 | 779.479778 | 1090.027217 | -0.152032099 | 0.11787478 | 0.011382932 | 0.151642712 | ns | NaCl | KS.3 |
| FIM1_535 | 1489.855676 | 1667.989272 | 1542.118228 | 0.016454348 | 0.120674531 | 0.662995023 | 0.904422872 | ns | NaCl | KS.3 |
| FIM1_536 | 661.1531557 | 1141.990809 | 1058.344059 | 0.218460383 | 0.279295842 | 0.009411351 | 0.136108932 | ns | NaCl | KS.3 |
| FIM1_537 | 563.4307747 | 631.9922494 | 995.1810894 | 0.020934386 | 0.114064911 | 0.599620777 | 0.880219278 | ns | NaCl | KS.3 |
| FIM1_538 | 1211.160225 | 989.2239912 | 1018.384598 | -0.093029578 | 0.105487229 | 0.076494434 | 0.429525743 | ns | NaCl | KS.3 |
| FIM1_539 | 388.7447825 | 409.6750416 | 440.214433 | 0.023651107 | 0.084194032 | 0.62615065 | 0.89540925 | ns | NaCl | KS.3 |
| FIM1_540 | 2253.408587 | 2094.854161 | 2288.724654 | -0.051694209 | 0.072368149 | 0.297761931 | 0.715635559 | ns | NaCl | KS.3 |
| FIM1_541 | 607.4063144 | 541.385349 | 552.7711887 | -0.094550501 | 0.070988931 | 0.060311416 | 0.383943936 | ns | NaCl | KS.3 |
| FIM1_542 | 890.5746715 | 861.5868798 | 827.8097266 | -0.016804567 | 0.080378916 | 0.731427039 | 0.936960396 | ns | NaCl | KS.3 |
| FIM1_543 | 465.6376371 | 536.2768679 | 548.75533 | 0.038708689 | 0.105747611 | 0.384482394 | 0.778665419 | ns | NaCl | KS.3 |
| FIM1_544 | 2341.784465 | 2517.809608 | 1915.488907 | 0.033738536 | 0.086222523 | 0.48734897 | 0.831183108 | ns | NaCl | KS.3 |
| FIM1_545 | 980.2572765 | 968.7237502 | 1877.281489 | -0.004441859 | 0.09181041 | 0.923172026 | 0.983924964 | ns | NaCl | KS.3 |
| FIM1_546 | 5360.379941 | 6466.890212 | 4994.01369 | 0.145509826 | 0.086765656 | 0.008377253 | 0.12706352 | ns | NaCl | KS.3 |
| FIM1_547 | 1689.799799 | 1770.571697 | 2150.754405 | 0.01748704 | 0.091389895 | 0.707164913 | 0.925190614 | ns | NaCl | KS.3 |
| FIM1_549 | 979.1094127 | 1286.868033 | 1034.980657 | 0.124158538 | 0.132597983 | 0.031177455 | 0.277971041 | ns | NaCl | KS.3 |
| FIM1_550 | 16290.4992 | 20929.83107 | 11820.81753 | 0.175539899 | 0.119276403 | 0.00498441 | 0.092527753 | ns | NaCl | KS.3 |
| FIM1_551 | 560.4897169 | 647.3221266 | 566.140344 | 0.052193089 | 0.099478263 | 0.272994437 | 0.694349346 | ns | NaCl | KS.3 |
| FIM1_552 | 1090.630343 | 1082.180384 | 1160.952802 | -0.004670861 | 0.076708502 | 0.924269867 | 0.984002266 | ns | NaCl | KS.3 |
| FIM1_553 | 1228.279527 | 1217.011044 | 1591.648014 | -0.006592186 | 0.070937046 | 0.892911509 | 0.977268396 | ns | NaCl | KS.3 |
| FIM1_554 | 2027.993833 | 1996.733132 | 1887.284174 | -0.010375377 | 0.072531504 | 0.832685509 | 0.961831996 | ns | NaCl | KS.3 |
| FIM1_555 | 487.9981881 | 488.3763575 | 575.5339994 | 0.000143489 | 0.086865692 | 0.997591694 | 0.99924489 | ns | NaCl | KS.3 |
| FIM1_556 | 1271.414112 | 1356.099762 | 1202.338386 | 0.034460621 | 0.08209288 | 0.483806404 | 0.829184191 | ns | NaCl | KS.3 |
| FIM1_557 | 1112.088671 | 1313.651984 | 994.4919767 | 0.118777841 | 0.084627203 | 0.026642676 | 0.254519471 | ns | NaCl | KS.3 |
| FIM1_558 | 2000.509077 | 1937.012138 | 1298.567273 | -0.013849409 | 0.08726556 | 0.771094638 | 0.949083579 | ns | NaCl | KS.3 |
| FIM1_559 | 847.2249641 | 809.614691 | 786.9371466 | -0.025647221 | 0.07930411 | 0.602404349 | 0.881535538 | ns | NaCl | KS.3 |
| FIM1_560 | 1898.410758 | 1921.135056 | 2200.518531 | 0.006127092 | 0.080677748 | 0.900090097 | 0.978786499 | ns | NaCl | KS.3 |
| FIM1_561 | 705.592953 | 804.415376 | 945.6953352 | 0.061326034 | 0.090826591 | 0.21812437 | 0.64211168 | ns | NaCl | KS.3 |
| FIM1_562 | 1547.64294 | 1607.869776 | 1953.332671 | 0.02647437 | 0.072366003 | 0.590782791 | 0.878786986 | ns | NaCl | KS.3 |
| FIM1_563 | 168.8573506 | 173.8789293 | 220.2215981 | 0.006467148 | 0.106711625 | 0.873757244 | 0.971243188 | ns | NaCl | KS.3 |
| FIM1_564 | 2079.055506 | 2154.149521 | 1556.202246 | 0.012056379 | 0.093987475 | 0.790981901 | 0.953972111 | ns | NaCl | KS.3 |
| FIM1_565 | 2117.478993 | 2704.637942 | 1520.594745 | 0.088841038 | 0.12117801 | 0.085240143 | 0.45357575 | ns | NaCl | KS.3 |
| FIM1_566 | 972.2514723 | 1185.22657 | 826.6143196 | 0.095985685 | 0.103016983 | 0.06955995 | 0.413948267 | ns | NaCl | KS.3 |
| FIM1_567 | 1704.989751 | 1835.479452 | 1648.299936 | 0.043169439 | 0.078884623 | 0.385349503 | 0.778665419 | ns | NaCl | KS.3 |
| FIM1_568 | 2277.339835 | 3422.531907 | 3256.95093 | 0.061358069 | 0.162873614 | 0.160438141 | 0.58211082 | ns | NaCl | KS.3 |
| FIM1_569 | 2143.174568 | 2625.881776 | 2490.001199 | 0.038702878 | 0.121818972 | 0.35190249 | 0.756912594 | ns | NaCl | KS.3 |
| FIM1_570 | 1843.644686 | 3185.278415 | 1243.278647 | 0.622597703 | 0.252449934 | 5.2957754109[illegible] | 0.004542497 | ns | NaCl | KS.3 |
| FIM1_571 | 696.5344557 | 1249.629867 | 2026.460833 | 0.188760915 | 0.277092838 | 0.014301258 | 0.173179792 | ns | NaCl | KS.3 |
| FIM1_572 | 104.6895327 | 229.7659877 | 510.8466022 | 0.210793244 | 0.34617259 | 0.011485816 | 0.151642712 | ns | NaCl | KS.3 |
| FIM1_573 | 5316.916315 | 5882.892257 | 4434.991654 | 0.052118601 | 0.085338844 | 0.294315566 | 0.712862678 | ns | NaCl | KS.3 |

| | | | | | | | | | | |
|---|---|---|---|---|---|---|---|---|---|---|
| FIM1_574 | 5534.232729 | 6742.843074 | 3334.420775 | 0.032673617 | 0.124203985 | 0.413507546 | 0.8001568 | ns | NaCl | KS.3 |
| FIM1_575 | 3342.553917 | 3477.594454 | 2703.430966 | 0.023712747 | 0.077064199 | 0.630627402 | 0.897160803 | ns | NaCl | KS.3 |
| FIM1_576 | 7312.765656 | 8036.705058 | 5823.24603 | 0.071338514 | 0.070333371 | 0.151331598 | 0.566863767 | ns | NaCl | KS.3 |
| FIM1_577 | 1035.489579 | 924.512769 | 905.1260683 | -0.057968443 | 0.087180493 | 0.245326916 | 0.668767895 | ns | NaCl | KS.3 |
| FIM1_578 | 4238.85303 | 3986.682502 | 4422.84059 | -0.032596299 | 0.081527489 | 0.507755957 | 0.844423736 | ns | NaCl | KS.3 |
| FIM1_579 | 713.7429165 | 703.6982392 | 850.9124711 | -0.007808081 | 0.081827693 | 0.872489129 | 0.971243188 | ns | NaCl | KS.3 |
| FIM1_580 | 210.0086748 | 202.5662744 | 260.9150765 | -0.01786442 | 0.087917528 | 0.707124809 | 0.925190614 | ns | NaCl | KS.3 |
| FIM1_581 | 793.1613099 | 792.5187134 | 835.7897162 | -0.000240724 | 0.089714532 | 0.995888623 | 0.99924489 | ns | NaCl | KS.3 |
| FIM1_582 | 5705.578983 | 7308.890402 | 3226.125734 | 0.041287647 | 0.130781733 | 0.313602859 | 0.731871738 | ns | NaCl | KS.3 |
| FIM1_583 | 239.5674935 | 322.4552917 | 256.7458051 | 0.102191505 | 0.139352952 | 0.058392194 | 0.377936577 | ns | NaCl | KS.3 |
| FIM1_584 | 728.4849632 | 714.4743486 | 916.6185616 | -0.011347196 | 0.080814522 | 0.816245652 | 0.958816322 | ns | NaCl | KS.3 |
| FIM1_585 | 13656.15984 | 12667.22398 | 10000.39067 | -0.025203904 | 0.095754232 | 0.580871549 | 0.8749481 | ns | NaCl | KS.3 |
| FIM1_586 | 2582.210618 | 3249.645925 | 2018.87692 | 0.078456195 | 0.117376129 | 0.116051805 | 0.519206593 | ns | NaCl | KS.3 |
| FIM1_587 | 668.0394651 | 765.4747671 | 743.9692453 | 0.042069382 | 0.101889166 | 0.358623451 | 0.762783954 | ns | NaCl | KS.3 |
| FIM1_588 | 870.0569194 | 989.7051266 | 685.0246758 | 0.052230767 | 0.094565235 | 0.280926389 | 0.701610837 | ns | NaCl | KS.3 |
| FIM1_589 | 945.5262227 | 824.6948832 | 965.9544655 | -0.087602346 | 0.082131108 | 0.090343947 | 0.463573758 | ns | NaCl | KS.3 |
| FIM1_590 | 670.35478 | 701.0764019 | 896.1242943 | 0.028831282 | 0.074702049 | 0.559450233 | 0.865174047 | ns | NaCl | KS.3 |
| FIM1_591 | 624.1706472 | 694.7410865 | 799.1978164 | 0.0571022 | 0.084971279 | 0.253651601 | 0.676969706 | ns | NaCl | KS.3 |
| FIM1_592 | 5770.34144 | 6118.992252 | 5793.999017 | 0.028627654 | 0.084288275 | 0.556646545 | 0.863699344 | ns | NaCl | KS.3 |
| FIM1_593 | 1237.815049 | 1261.75365 | 1160.875021 | 0.007164834 | 0.089802853 | 0.878182707 | 0.972834689 | ns | NaCl | KS.3 |
| FIM1_594 | 2545.653652 | 2946.798268 | 2091.221904 | 0.052924354 | 0.099728479 | 0.266844912 | 0.689593109 | ns | NaCl | KS.3 |
| FIM1_595 | 3078.210652 | 3420.162039 | 2258.61927 | 0.044962951 | 0.091136904 | 0.352668019 | 0.756912594 | ns | NaCl | KS.3 |
| FIM1_596 | 5473.619814 | 4258.587994 | 3810.382358 | -0.124130549 | 0.122051562 | 0.029701094 | 0.271163489 | ns | NaCl | KS.3 |
| FIM1_597 | 1464.325933 | 1743.526223 | 1847.660826 | 0.11107426 | 0.089389262 | 0.038585053 | 0.311585109 | ns | NaCl | KS.3 |
| FIM1_598 | 1185.046267 | 1292.012515 | 942.4573859 | 0.039614073 | 0.087537612 | 0.415566723 | 0.800387073 | ns | NaCl | KS.3 |
| FIM1_599 | 551.2668847 | 527.6930713 | 612.5166413 | -0.025759986 | 0.078217115 | 0.601343423 | 0.880762097 | ns | NaCl | KS.3 |
| FIM1_600 | 1733.444518 | 1179.937285 | 1731.684023 | -0.241330922 | 0.209230761 | 0.003614868 | 0.076817191 | ns | NaCl | KS.3 |
| FIM1_601 | 4647.529185 | 5603.205381 | 2786.680377 | 0.032966545 | 0.121491069 | 0.415099711 | 0.800387073 | ns | NaCl | KS.3 |
| FIM1_602 | 2035.792245 | 2076.563064 | 2163.783996 | 0.015932679 | 0.064896715 | 0.739748778 | 0.940888181 | ns | NaCl | KS.3 |
| FIM1_603 | 4590.016194 | 3704.914683 | 6877.477411 | -0.131534977 | 0.103785889 | 0.020207194 | 0.215832354 | ns | NaCl | KS.3 |
| FIM1_604 | 3046.558869 | 2654.342275 | 2260.077787 | -0.079111039 | 0.086194078 | 0.123636738 | 0.530709897 | ns | NaCl | KS.3 |
| FIM1_605 | 475.9901534 | 377.6264328 | 511.9090931 | -0.089959932 | 0.115425506 | 0.083169237 | 0.447068602 | ns | NaCl | KS.3 |
| FIM1_606 | 533.8175056 | 472.7040184 | 1345.168045 | -0.050887258 | 0.093875502 | 0.293289519 | 0.71211096 | ns | NaCl | KS.3 |
| FIM1_607 | 1261.618504 | 1469.212703 | 715.6332677 | 0.032180846 | 0.113461648 | 0.441332578 | 0.809513863 | ns | NaCl | KS.3 |
| FIM1_608 | 523.5297644 | 541.3948099 | 591.9919784 | 0.015929877 | 0.084715964 | 0.741247019 | 0.941221011 | ns | NaCl | KS.3 |
| FIM1_609 | 6892.276571 | 6568.870165 | 7419.754934 | -0.013165291 | 0.100026587 | 0.761597847 | 0.946054754 | ns | NaCl | KS.3 |
| FIM1_610 | 442.7727677 | 419.6445429 | 549.4681379 | -0.027766784 | 0.081366308 | 0.571620861 | 0.870809186 | ns | NaCl | KS.3 |
| FIM1_611 | 112.0769977 | 101.2842518 | 193.9310292 | -0.026691888 | 0.102719013 | 0.541207214 | 0.857165292 | ns | NaCl | KS.3 |
| FIM1_612 | 1282.637079 | 1220.298606 | 1807.422097 | -0.040408591 | 0.06583493 | 0.403130371 | 0.789854857 | ns | NaCl | KS.3 |
| FIM1_613 | 1045.506399 | 1095.8351 | 1297.621499 | 0.032576974 | 0.071438152 | 0.508094985 | 0.844423736 | ns | NaCl | KS.3 |
| FIM1_615 | 613.0969237 | 514.0109417 | 988.5036954 | -0.096533432 | 0.094035041 | 0.067733942 | 0.407961695 | ns | NaCl | KS.3 |
| FIM1_616 | 2870.685661 | 2602.658212 | 2442.610828 | -0.059896219 | 0.07890965 | 0.233951319 | 0.656462386 | ns | NaCl | KS.3 |
| FIM1_617 | 545.587487 | 597.3409828 | 457.5838486 | 0.01396221 | 0.11769964 | 0.714162244 | 0.928149729 | ns | NaCl | KS.3 |
| FIM1_618 | 886.7927479 | 1177.538836 | 1419.095574 | 0.086643273 | 0.134455164 | 0.088946769 | 0.461910414 | ns | NaCl | KS.3 |
| FIM1_619 | 11169.05053 | 12739.35544 | 13755.79097 | 0.055416711 | 0.09413803 | 0.256417898 | 0.679999489 | ns | NaCl | KS.3 |

| | | | | | | | | | | |
|---|---|---|---|---|---|---|---|---|---|---|
| FIM1_620 | 857.7573701 | 674.4978911 | 905.0081186 | -0.049771254 | 0.126194945 | 0.251770166 | 0.675897894 | ns | NaCl | KS.3 |
| FIM1_621 | 420.0309418 | 359.0390328 | 628.3279031 | -0.03141831 | 0.115487367 | 0.446273571 | 0.812914509 | ns | NaCl | KS.3 |
| FIM1_622 | 266.7646138 | 227.1869874 | 308.1685685 | -0.081846968 | 0.09517472 | 0.111634459 | 0.50812574 | ns | NaCl | KS.3 |
| FIM1_623 | 3910.506214 | 3958.781702 | 3212.491914 | 0.00736792 | 0.076228583 | 0.880873438 | 0.973854523 | ns | NaCl | KS.3 |
| FIM1_625 | 335.6934655 | 393.2360293 | 1665.134305 | 0.02910523 | 0.117914287 | 0.470807161 | 0.822127061 | ns | NaCl | KS.3 |
| FIM1_626 | 2452.859202 | 3460.738095 | 3460.275943 | 0.12168177 | 0.161982305 | 0.036806146 | 0.303163208 | ns | NaCl | KS.3 |
| FIM1_627 | 737.5979543 | 570.4931922 | 1097.322672 | -0.15585737 | 0.124664187 | 0.010901523 | 0.148997467 | ns | NaCl | KS.3 |
| FIM1_628 | 1684.624855 | 1527.348534 | 2869.052745 | -0.034062482 | 0.095796634 | 0.462758208 | 0.819880744 | ns | NaCl | KS.3 |
| FIM1_629 | 3154.87786 | 3654.557531 | 2629.388905 | 0.050548552 | 0.101215561 | 0.283166733 | 0.704025235 | ns | NaCl | KS.3 |
| FIM1_630 | 684.1048003 | 674.2533711 | 786.9538102 | -0.005830539 | 0.084878888 | 0.903420086 | 0.979395371 | ns | NaCl | KS.3 |
| FIM1_631 | 1739.345887 | 1814.113765 | 2014.126863 | 0.008457947 | 0.108216647 | 0.83376936 | 0.962365393 | ns | NaCl | KS.3 |
| FIM1_632 | 226.4339698 | 184.8266874 | 338.091691 | -0.064981551 | 0.113773337 | 0.173656801 | 0.601631326 | ns | NaCl | KS.3 |
| FIM1_633 | 7969.941627 | 8070.378699 | 5665.138552 | 0.008291056 | 0.072598957 | 0.865925714 | 0.970184124 | ns | NaCl | KS.3 |
| FIM1_634 | 458.8891633 | 532.3479501 | 504.229274 | 0.089961293 | 0.086324059 | 0.084197382 | 0.450895561 | ns | NaCl | KS.3 |
| FIM1_635 | 5251.06322 | 4984.473975 | 3749.836985 | -0.033623259 | 0.074724338 | 0.4967995 | 0.836358994 | ns | NaCl | KS.3 |
| FIM1_636 | 445.8308287 | 501.6837565 | 505.1129773 | 0.077942375 | 0.07851005 | 0.126548024 | 0.534982546 | ns | NaCl | KS.3 |
| FIM1_637 | 12015.18312 | 13029.60627 | 8546.879484 | 0.046154848 | 0.08042141 | 0.353808551 | 0.757528931 | ns | NaCl | KS.3 |
| FIM1_638 | 1361.452194 | 1561.008717 | 1509.720033 | 0.090203917 | 0.081157196 | 0.081472319 | 0.440570421 | ns | NaCl | KS.3 |
| FIM1_639 | 46.22782005 | 62.89670461 | 79.66084673 | 0.06304846 | 0.142010406 | 0.164144049 | 0.589056105 | ns | NaCl | KS.3 |
| FIM1_640 | 1260.875622 | 1244.747523 | 1173.532448 | -0.010820399 | 0.064951498 | 0.821509601 | 0.960240659 | ns | NaCl | KS.3 |
| FIM1_641 | 400.9575591 | 348.7688186 | 514.1290684 | -0.05920305 | 0.095410684 | 0.22740989 | 0.651504817 | ns | NaCl | KS.3 |
| FIM1_642 | 3855.999164 | 3857.536923 | 3393.308776 | 0.000190564 | 0.10046663 | 0.996450846 | 0.99924489 | ns | NaCl | KS.3 |
| FIM1_644 | 2322.193827 | 2985.892196 | 1767.328646 | 0.080742229 | 0.124023358 | 0.106770687 | 0.498296595 | ns | NaCl | KS.3 |
| FIM1_645 | 632.56721 | 950.2740839 | 1058.851261 | 0.212155168 | 0.216396127 | 0.007022241 | 0.11342743 | ns | NaCl | KS.3 |
| FIM1_646 | 3035.214999 | 3941.128836 | 2889.920807 | 0.166000593 | 0.12642949 | 0.007938973 | 0.123041715 | ns | NaCl | KS.3 |
| FIM1_647 | 2128.592943 | 2598.333241 | 1610.557098 | 0.068472159 | 0.109836917 | 0.15983858 | 0.58211082 | ns | NaCl | KS.3 |
| FIM1_648 | 135.3845857 | 108.7373471 | 180.6262444 | -0.087012484 | 0.10973783 | 0.091786592 | 0.467388224 | ns | NaCl | KS.3 |
| FIM1_650 | 493.0497474 | 380.4370073 | 655.9854887 | -0.154604292 | 0.126738672 | 0.011657347 | 0.152920569 | ns | NaCl | KS.3 |
| FIM1_651 | 756.3957695 | 667.041316 | 1317.559511 | -0.02919537 | 0.108808943 | 0.490812363 | 0.835079661 | ns | NaCl | KS.3 |
| FIM1_652 | 2909.141802 | 2910.835996 | 2526.977574 | -0.000274228 | 0.070421143 | 0.995526051 | 0.99924489 | ns | NaCl | KS.3 |
| FIM1_653 | 6043.773296 | 8066.823031 | 3738.84694 | 0.055886126 | 0.136509511 | 0.203297703 | 0.626494461 | ns | NaCl | KS.3 |
| FIM1_654 | 464.8253497 | 471.9768261 | 618.8363942 | 0.007364011 | 0.081344739 | 0.879854651 | 0.973483611 | ns | NaCl | KS.3 |
| FIM1_655 | 348.5911811 | 298.262872 | 491.6011962 | -0.058399821 | 0.099192166 | 0.227966015 | 0.651799382 | ns | NaCl | KS.3 |
| FIM1_656 | 2206.42434 | 2280.072014 | 1796.04403 | 0.017839108 | 0.080154872 | 0.715823694 | 0.929477573 | ns | NaCl | KS.3 |
| FIM1_657 | 735.966832 | 670.1693399 | 840.9049317 | -0.037031417 | 0.092936129 | 0.434224956 | 0.807589234 | ns | NaCl | KS.3 |
| FIM1_658 | 446.8674573 | 369.4969012 | 678.8625973 | -0.070148086 | 0.106927182 | 0.154171805 | 0.571538546 | ns | NaCl | KS.3 |
| FIM1_659 | 2698.440142 | 2518.800696 | 2457.652158 | -0.041124834 | 0.078213396 | 0.407814381 | 0.79494601 | ns | NaCl | KS.3 |
| FIM1_660 | 555.3615462 | 493.0349881 | 935.7586285 | -0.077837165 | 0.07939243 | 0.127532236 | 0.535324115 | ns | NaCl | KS.3 |
| FIM1_661 | 701.6776739 | 709.1319159 | 869.1390861 | 0.003543811 | 0.095795906 | 0.936708305 | 0.986375069 | ns | NaCl | KS.3 |
| FIM1_662 | 21558.78296 | 24777.34022 | 13329.56743 | 0.055638937 | 0.096382376 | 0.251491857 | 0.675578827 | ns | NaCl | KS.3 |
| FIM1_663 | 472.7743817 | 501.9655813 | 728.0410642 | 0.008839781 | 0.118397724 | 0.813505481 | 0.958816322 | ns | NaCl | KS.3 |
| FIM1_664 | 3071.786653 | 3268.412774 | 3029.072366 | 0.030267662 | 0.084650983 | 0.534276272 | 0.855153333 | ns | NaCl | KS.3 |
| FIM1_665 | 666.6028008 | 577.7899377 | 624.5284246 | -0.092858275 | 0.082596386 | 0.074313523 | 0.423493444 | ns | NaCl | KS.3 |
| FIM1_666 | 150.6281164 | 140.4565831 | 221.6060011 | -0.014569756 | 0.109167957 | 0.718780821 | 0.930263939 | ns | NaCl | KS.3 |
| FIM1_667 | 203.3935565 | 167.0441202 | 255.920656 | -0.061406367 | 0.113017174 | 0.193194692 | 0.618952432 | ns | NaCl | KS.3 |

| | | | | | | | | | | |
|---|---|---|---|---|---|---|---|---|---|---|
| FIM1_668 | 4905.392728 | 6144.48737 | 3917.240784 | 0.139386319 | 0.108263059 | 0.015892965 | 0.185169786 | ns | NaCl | KS.3 |
| FIM1_669 | 760.8093112 | 1339.392395 | 1242.978932 | 0.351474419 | 0.326108597 | 0.002281281 | 0.058108556 | ns | NaCl | KS.3 |
| FIM1_670 | 2624.190336 | 4023.338886 | 2971.716088 | 0.065463878 | 0.167966377 | 0.14233687 | 0.554957625 | ns | NaCl | KS.3 |
| FIM1_671 | 667.4309588 | 640.9984871 | 622.0532644 | -0.022730377 | 0.082521578 | 0.641362363 | 0.898798052 | ns | NaCl | KS.3 |
| FIM1_672 | 505.6019518 | 398.4992988 | 662.1761727 | -0.115422609 | 0.119040034 | 0.038455334 | 0.311585109 | ns | NaCl | KS.3 |
| FIM1_673 | 133.6487629 | 118.6350652 | 349.8445058 | -0.019632441 | 0.117422142 | 0.613870105 | 0.889162818 | ns | NaCl | KS.3 |
| FIM1_674 | 4001.51577 | 3548.081138 | 5215.570752 | -0.060334181 | 0.087890957 | 0.227553866 | 0.651504817 | ns | NaCl | KS.3 |
| FIM1_675 | 1257.379591 | 1269.124136 | 1473.001452 | 0.005422477 | 0.078952123 | 0.911870984 | 0.982011252 | ns | NaCl | KS.3 |
| FIM1_676 | 2107.824009 | 2219.745388 | 3264.916752 | 0.012484797 | 0.103487366 | 0.766995413 | 0.948794469 | ns | NaCl | KS.3 |
| FIM1_677 | 1026.394095 | 1064.961467 | 1919.82398 | 0.015625515 | 0.089274186 | 0.740111162 | 0.940888181 | ns | NaCl | KS.3 |
| FIM1_678 | 2696.507002 | 2547.922458 | 2766.754028 | -0.024011611 | 0.0883596 | 0.614460657 | 0.889162818 | ns | NaCl | KS.3 |
| FIM1_679 | 4720.118982 | 3066.171571 | 2876.776366 | -0.158639215 | 0.208943294 | 0.019229154 | 0.209721297 | ns | NaCl | KS.3 |
| FIM1_680 | 4506.02728 | 3384.806417 | 5497.70073 | -0.067499844 | 0.134980791 | 0.148969865 | 0.563164954 | ns | NaCl | KS.3 |
| FIM1_681 | 1656.305243 | 1943.062163 | 2011.579786 | 0.02507745 | 0.121727978 | 0.519232229 | 0.850629872 | ns | NaCl | KS.3 |
| FIM1_682 | 665.9337451 | 587.9999158 | 1325.94449 | -0.022514719 | 0.115260876 | 0.571845952 | 0.870809186 | ns | NaCl | KS.3 |
| FIM1_683 | 3548.560254 | 4008.335976 | 3487.012945 | 0.068482991 | 0.084513772 | 0.176979802 | 0.605137125 | ns | NaCl | KS.3 |
| FIM1_684 | 1144.424529 | 1171.57879 | 1704.721099 | 0.015111783 | 0.073234671 | 0.758638912 | 0.945250644 | ns | NaCl | KS.3 |
| FIM1_685 | 411.7194929 | 323.5761363 | 571.6491387 | -0.119246182 | 0.119303883 | 0.034162134 | 0.292771172 | ns | NaCl | KS.3 |
| FIM1_686 | 4217.786044 | 4160.591748 | 3809.941275 | -0.011731165 | 0.058806404 | 0.799065341 | 0.95648403 | ns | NaCl | KS.3 |
| FIM1_687 | 1051.103517 | 936.6948844 | 1055.214549 | -0.065664264 | 0.083864292 | 0.194162565 | 0.618952432 | ns | NaCl | KS.3 |
| FIM1_688 | 801.5540342 | 836.5645955 | 1036.037415 | 0.027273276 | 0.075762612 | 0.580815598 | 0.8749481 | ns | NaCl | KS.3 |
| FIM1_689 | 180.0676536 | 192.2521014 | 143.7030848 | 0.012850583 | 0.108823353 | 0.750508958 | 0.943274417 | ns | NaCl | KS.3 |
| FIM1_690 | 1496.032451 | 1753.466125 | 1524.061999 | 0.101960067 | 0.086152714 | 0.053634827 | 0.362470256 | ns | NaCl | KS.3 |
| FIM1_691 | 2167.599119 | 2373.994969 | 1611.399486 | 0.047844578 | 0.08378574 | 0.334878319 | 0.745307602 | ns | NaCl | KS.3 |
| FIM1_692 | 1277.039016 | 1403.902374 | 1227.666889 | 0.046350237 | 0.086568703 | 0.34627116 | 0.755569745 | ns | NaCl | KS.3 |
| FIM1_693 | 1085.301748 | 1176.116967 | 1255.218501 | 0.05260015 | 0.075299065 | 0.292184854 | 0.71186075 | ns | NaCl | KS.3 |
| FIM1_694 | 419.3891496 | 353.2119901 | 543.4684132 | -0.088356605 | 0.09607105 | 0.089808044 | 0.463545584 | ns | NaCl | KS.3 |
| FIM1_695 | 927.5970007 | 852.4683605 | 1000.654697 | -0.064179879 | 0.07040575 | 0.195643918 | 0.618952432 | ns | NaCl | KS.3 |
| FIM1_696 | 1795.426625 | 1710.483935 | 1557.699312 | -0.027909786 | 0.077936996 | 0.571787723 | 0.870809186 | ns | NaCl | KS.3 |
| FIM1_697 | 655.8455745 | 658.3801845 | 898.8205216 | 0.002673257 | 0.079486894 | 0.956429837 | 0.989856135 | ns | NaCl | KS.3 |
| FIM1_698 | 1256.47293 | 1194.078983 | 812.8636411 | -0.024753771 | 0.084722509 | 0.609695119 | 0.887169704 | ns | NaCl | KS.3 |
| FIM1_699 | 3097.959204 | 3051.909214 | 2938.740619 | -0.003804194 | 0.101531768 | 0.928635471 | 0.984238275 | ns | NaCl | KS.3 |
| FIM1_700 | 9608.511963 | 9830.462202 | 6849.827476 | 0.018484241 | 0.063705386 | 0.698181945 | 0.922074642 | ns | NaCl | KS.3 |
| FIM1_701 | 2422.589102 | 2963.663096 | 1751.349444 | 0.060153414 | 0.112871737 | 0.200427637 | 0.624375388 | ns | NaCl | KS.3 |
| FIM1_702 | 1383.61742 | 1436.480115 | 1186.573425 | 0.021303739 | 0.07937441 | 0.664622482 | 0.905641427 | ns | NaCl | KS.3 |
| FIM1_703 | 3214.584868 | 3905.532359 | 2874.929683 | 0.095789919 | 0.101779741 | 0.069992593 | 0.415528819 | ns | NaCl | KS.3 |
| FIM1_704 | 5505.044542 | 7619.405269 | 6127.943018 | 0.094133649 | 0.148806744 | 0.072157987 | 0.420358296 | ns | NaCl | KS.3 |
| FIM1_705 | 1591.368198 | 1477.958005 | 1247.382072 | -0.014900235 | 0.109136494 | 0.712956577 | 0.927067164 | ns | NaCl | KS.3 |
| FIM1_706 | 2680.699246 | 2462.815429 | 2196.0927 | -0.050403271 | 0.079551421 | 0.313076684 | 0.731871738 | ns | NaCl | KS.3 |
| FIM1_707 | 1125.186781 | 1201.956161 | 994.1467093 | 0.0453554 | 0.070573759 | 0.357556585 | 0.761817563 | ns | NaCl | KS.3 |
| FIM1_708 | 533.9588487 | 574.9473473 | 661.6962084 | 0.03665845 | 0.084363649 | 0.454509791 | 0.816788914 | ns | NaCl | KS.3 |
| FIM1_709 | 1171.085934 | 1198.618535 | 1294.26645 | 0.012806292 | 0.076196572 | 0.794648395 | 0.955299664 | ns | NaCl | KS.3 |
| FIM1_710 | 606.3250096 | 522.5233947 | 649.6720697 | -0.073894958 | 0.092206104 | 0.145848345 | 0.560211748 | ns | NaCl | KS.3 |
| FIM1_711 | 402.6168726 | 462.6092046 | 444.1594199 | 0.047934933 | 0.099657595 | 0.308224569 | 0.728391584 | ns | NaCl | KS.3 |
| FIM1_712 | 20.54303497 | 23.82688316 | 21.06410889 | 0.012891846 | 0.136587068 | 0.707759939 | 0.925190614 | ns | NaCl | KS.3 |

| | | | | | | | | | | |
|---|---|---|---|---|---|---|---|---|---|---|
| FIM1_713 | 1560.21825 | 1149.259525 | 1449.725859 | -0.062860797 | 0.140554057 | 0.165767229 | 0.589908414 | ns | NaCl | KS.3 |
| FIM1_714 | 814.1100744 | 959.2384439 | 955.3241447 | 0.071016935 | 0.097951844 | 0.15641544 | 0.57556717 | ns | NaCl | KS.3 |
| FIM1_715 | 754.7923285 | 735.9806901 | 1366.728225 | -0.006258637 | 0.10230795 | 0.882241183 | 0.974110648 | ns | NaCl | KS.3 |
| FIM1_718 | 3426.203628 | 3649.046017 | 4056.178042 | 0.024135244 | 0.09164007 | 0.606059888 | 0.883689315 | ns | NaCl | KS.3 |
| FIM1_719 | 10060.77088 | 10737.48725 | 7126.365091 | 0.031311826 | 0.085004983 | 0.520114066 | 0.851453596 | ns | NaCl | KS.3 |
| FIM1_720 | 501.1805271 | 540.3062339 | 588.1164666 | 0.03267628 | 0.087538272 | 0.49844484 | 0.837486406 | ns | NaCl | KS.3 |
| FIM1_721 | 2250.110164 | 2632.527975 | 2180.455325 | 0.083175855 | 0.091260639 | 0.107387883 | 0.498909937 | ns | NaCl | KS.3 |
| FIM1_722 | 5125.835405 | 6341.349761 | 4671.506458 | 0.194721442 | 0.089260239 | 0.000726505 | 0.029870765 | ns | NaCl | KS.3 |
| FIM1_723 | 624.640614 | 675.2899069 | 877.597854 | 0.039205724 | 0.083275179 | 0.426148388 | 0.802854185 | ns | NaCl | KS.3 |
| FIM1_724 | 470.8756335 | 561.252277 | 388.5643602 | 0.054582624 | 0.107266429 | 0.242817021 | 0.666657537 | ns | NaCl | KS.3 |
| FIM1_725 | 9195.659835 | 11734.48515 | 5829.750812 | 0.079815764 | 0.12161985 | 0.110278321 | 0.505654054 | ns | NaCl | KS.3 |
| FIM1_726 | 874.1103355 | 1006.469793 | 898.0738828 | 0.077830249 | 0.087690994 | 0.129122843 | 0.536663444 | ns | NaCl | KS.3 |
| FIM1_727 | 1007.111177 | 941.9309265 | 1229.378744 | -0.031489229 | 0.085779452 | 0.516636596 | 0.849956041 | ns | NaCl | KS.3 |
| FIM1_728 | 2757.029044 | 3641.889721 | 2356.280893 | 0.130129447 | 0.134679043 | 0.026480068 | 0.254519471 | ns | NaCl | KS.3 |
| FIM1_729 | 2954.380519 | 3340.137924 | 2429.537675 | 0.071110002 | 0.083644591 | 0.162296431 | 0.585090395 | ns | NaCl | KS.3 |
| FIM1_730 | 1857.271635 | 2004.152961 | 1867.201485 | 0.051969394 | 0.073146965 | 0.296101027 | 0.714058464 | ns | NaCl | KS.3 |
| FIM1_731 | 350.0922613 | 379.5551361 | 460.353051 | 0.043123614 | 0.0803982 | 0.38524953 | 0.778665419 | ns | NaCl | KS.3 |
| FIM1_732 | 12908.97528 | 13501.26388 | 8995.403339 | 0.02202136 | 0.083576234 | 0.650539994 | 0.899642849 | ns | NaCl | KS.3 |
| FIM1_733 | 2606.077696 | 2651.686818 | 2077.4532 | 0.004242506 | 0.102838603 | 0.919543504 | 0.983178365 | ns | NaCl | KS.3 |
| FIM1_734 | 1737.719364 | 1975.810016 | 2175.091979 | 0.126215236 | 0.059230738 | 0.006886785 | 0.111601803 | ns | NaCl | KS.3 |
| FIM1_735 | 349.5658994 | 452.906081 | 672.6776497 | 0.198950767 | 0.121159936 | 0.00214582 | 0.057394915 | ns | NaCl | KS.3 |
| FIM1_736 | 2438.072822 | 2260.199486 | 2714.657046 | -0.04370347 | 0.080224574 | 0.379179198 | 0.778493059 | ns | NaCl | KS.3 |
| FIM1_737 | 2076.380246 | 2384.894887 | 2322.058371 | 0.112745111 | 0.072687989 | 0.027460577 | 0.259726943 | ns | NaCl | KS.3 |
| FIM1_738 | 2203.505234 | 1881.591059 | 1834.221427 | -0.063792015 | 0.098698164 | 0.194513572 | 0.618952432 | ns | NaCl | KS.3 |
| FIM1_739 | 432.4583341 | 432.7588703 | 482.0774618 | -0.000123748 | 0.098491616 | 0.997734864 | 0.99924489 | ns | NaCl | KS.3 |
| FIM1_740 | 853.8683016 | 851.4102585 | 827.4917073 | -0.00207862 | 0.083376291 | 0.965704163 | 0.993691273 | ns | NaCl | KS.3 |
| FIM1_741 | 2471.7208 | 2542.076751 | 1896.080384 | 0.018182691 | 0.072636598 | 0.711544781 | 0.926199708 | ns | NaCl | KS.3 |
| FIM1_742 | 2481.733363 | 2159.397783 | 1995.452123 | -0.078906245 | 0.086740915 | 0.124516233 | 0.532646824 | ns | NaCl | KS.3 |
| FIM1_743 | 2298.666817 | 2457.776501 | 2245.619871 | 0.047793433 | 0.070624986 | 0.33275332 | 0.743353286 | ns | NaCl | KS.3 |
| FIM1_744 | 309.7231294 | 303.0341658 | 472.0112613 | -0.008723642 | 0.092754217 | 0.848983704 | 0.96736917 | ns | NaCl | KS.3 |
| FIM1_745 | 290.4482747 | 325.9544129 | 570.5548511 | 0.060409052 | 0.084297261 | 0.229363401 | 0.652793432 | ns | NaCl | KS.3 |
| FIM1_746 | 1195.959917 | 1297.919843 | 1985.127197 | 0.035429254 | 0.089356886 | 0.460896749 | 0.819880744 | ns | NaCl | KS.3 |
| FIM1_747 | 918.0370499 | 850.6338279 | 1178.845108 | -0.031161708 | 0.089965822 | 0.51311139 | 0.847552971 | ns | NaCl | KS.3 |
| FIM1_748 | 794.9703745 | 781.5479902 | 794.7275218 | -0.004015658 | 0.10443015 | 0.922765251 | 0.983890455 | ns | NaCl | KS.3 |
| FIM1_749 | 622.8198152 | 522.9476448 | 1109.475622 | -0.083714324 | 0.097410956 | 0.104535111 | 0.493886208 | ns | NaCl | KS.3 |
| FIM1_750 | 1237.871732 | 1269.576592 | 1422.150315 | 0.014254107 | 0.075264576 | 0.77214385 | 0.94938462 | ns | NaCl | KS.3 |
| FIM1_751 | 3369.164565 | 3405.324405 | 3565.154167 | 0.0082931 | 0.062056729 | 0.860454063 | 0.969856868 | ns | NaCl | KS.3 |
| FIM1_752 | 6166.73394 | 7376.540634 | 5517.589437 | 0.143539672 | 0.08271202 | 0.008194217 | 0.125927317 | ns | NaCl | KS.3 |
| FIM1_753 | 2936.825737 | 3364.776494 | 2091.90095 | 0.063222597 | 0.092271125 | 0.204413539 | 0.627751455 | ns | NaCl | KS.3 |
| FIM1_754 | 1494.689635 | 1451.359446 | 2116.637094 | -0.009957167 | 0.094309415 | 0.826051568 | 0.960240659 | ns | NaCl | KS.3 |
| FIM1_755 | 2608.713503 | 2211.146273 | 2519.886863 | -0.167337367 | 0.065715642 | 0.000737113 | 0.030058491 | ns | NaCl | KS.3 |
| FIM1_756 | 710.3050974 | 680.6394424 | 822.4926349 | -0.024319625 | 0.07975019 | 0.62085645 | 0.892479944 | ns | NaCl | KS.3 |
| FIM1_757 | 2355.955699 | 2288.914527 | 2005.582828 | -0.016258286 | 0.079177178 | 0.740417534 | 0.940888181 | ns | NaCl | KS.3 |
| FIM1_758 | 246.0389317 | 244.9328275 | 378.9891539 | -0.00681779 | 0.083250531 | 0.887965505 | 0.976703159 | ns | NaCl | KS.3 |
| FIM1_759 | 672.741838 | 716.8065855 | 738.316749 | 0.034190026 | 0.078812148 | 0.48948241 | 0.833678532 | ns | NaCl | KS.3 |

| | | | | | | | | | | |
|---|---|---|---|---|---|---|---|---|---|---|
| FIM1_760 | 2600.512285 | 2425.270964 | 2381.124377 | -0.0297301 | 0.088996052 | 0.533921559 | 0.855153333 | ns | NaCl | KS.3 |
| FIM1_761 | 2732.159506 | 2202.62382 | 2800.818831 | -0.08969862 | 0.110740706 | 0.084396827 | 0.451260889 | ns | NaCl | KS.3 |
| FIM1_762 | 1447.583256 | 1353.511825 | 1122.594086 | -0.029256905 | 0.088802845 | 0.540619374 | 0.857165292 | ns | NaCl | KS.3 |
| FIM1_763 | 578.6512984 | 641.2017121 | 775.0109219 | 0.052007833 | 0.084778235 | 0.295790644 | 0.714058464 | ns | NaCl | KS.3 |
| FIM1_764 | 2910.483196 | 2749.760223 | 2454.664853 | -0.040267332 | 0.072206812 | 0.415173735 | 0.800387073 | ns | NaCl | KS.3 |
| FIM1_765 | 433.5682505 | 419.7538693 | 354.7709332 | -0.007251116 | 0.105638381 | 0.85996627 | 0.969856868 | ns | NaCl | KS.3 |
| FIM1_767 | 641.9952021 | 578.0262617 | 1201.766306 | -0.018188084 | 0.114963684 | 0.64342153 | 0.898798052 | ns | NaCl | KS.3 |
| FIM1_768 | 280.4709316 | 243.2035779 | 370.6882992 | -0.058572213 | 0.096932072 | 0.229784671 | 0.653618489 | ns | NaCl | KS.3 |
| FIM1_769 | 3096.969011 | 2640.670175 | 2602.665713 | -0.155056082 | 0.067364279 | 0.001979494 | 0.054109796 | ns | NaCl | KS.3 |
| FIM1_770 | 1538.584547 | 3854.721828 | 2444.516323 | 1.180651803 | 0.243345632 | 2.3042922829 | 6.743443592 | Km>KS.3 | NaCl | KS.3 |
| FIM1_772 | 8.780546721 | 51.89900671 | 53.38291118 | 1.859834791 | 0.569585944 | 1.603189173 | 0.000266045 | Km>KS.3 | NaCl | KS.3 |
| FIM1_773 | 3016.843563 | 3956.644905 | 2254.322242 | 0.098320475 | 0.130352974 | 0.064715306 | 0.399942992 | ns | NaCl | KS.3 |
| FIM1_774 | 824.7522476 | 816.688039 | 784.9324631 | -0.006134975 | 0.082028524 | 0.899554893 | 0.978786499 | ns | NaCl | KS.3 |
| FIM1_776 | 2347.532563 | 2481.020451 | 2078.601108 | 0.02749454 | 0.083612613 | 0.572927679 | 0.871656025 | ns | NaCl | KS.3 |
| FIM1_777 | 2232.491281 | 2193.704334 | 2381.458233 | -0.011623994 | 0.074612418 | 0.813230282 | 0.958816322 | ns | NaCl | KS.3 |
| FIM1_778 | 518.5138919 | 433.4334925 | 665.8322517 | -0.077029078 | 0.100955077 | 0.12812922 | 0.536116795 | ns | NaCl | KS.3 |
| FIM1_779 | 17617.06438 | 25595.80712 | 14674.89046 | 0.502052234 | 0.110790519 | 4.600628687 | 1.204638301 | ns | NaCl | KS.3 |
| FIM1_780 | 303.2104331 | 435.9614455 | 453.4928227 | 0.112114923 | 0.166964074 | 0.046783737 | 0.341725286 | ns | NaCl | KS.3 |
| FIM1_781 | 216.4622772 | 226.0746251 | 472.8245672 | 0.00913392 | 0.105625073 | 0.824543369 | 0.960240659 | ns | NaCl | KS.3 |
| FIM1_782 | 716.2618185 | 480.3380801 | 1671.123472 | -0.145162613 | 0.191917647 | 0.023719949 | 0.239851113 | ns | NaCl | KS.3 |
| FIM1_783 | 680.7922877 | 631.3523919 | 935.7181848 | -0.028217342 | 0.093411174 | 0.544467354 | 0.857313452 | ns | NaCl | KS.3 |
| FIM1_784 | 238.8495819 | 173.180941 | 488.4529459 | -0.10114994 | 0.150617601 | 0.060350585 | 0.383943936 | ns | NaCl | KS.3 |
| FIM1_785 | 3608.158362 | 3466.061911 | 3305.208004 | -0.018369939 | 0.086031309 | 0.702054277 | 0.923421278 | ns | NaCl | KS.3 |
| FIM1_786 | 2400.036823 | 2405.114812 | 2337.434031 | 0.000605823 | 0.076546174 | 0.990162775 | 0.999134858 | ns | NaCl | KS.3 |
| FIM1_787 | 4222.216408 | 5522.695719 | 4819.528209 | 0.118762762 | 0.12962413 | 0.036061917 | 0.301915458 | ns | NaCl | KS.3 |
| FIM1_788 | 158.3146786 | 187.2569899 | 200.3153645 | 0.03368371 | 0.116080756 | 0.417511502 | 0.800387073 | ns | NaCl | KS.3 |
| FIM1_789 | 894.5083115 | 917.7541964 | 1008.009036 | 0.007919124 | 0.09713929 | 0.857875413 | 0.969856868 | ns | NaCl | KS.3 |
| FIM1_790 | 540.1401816 | 762.9924773 | 368.2901065 | 0.130018948 | 0.164518701 | 0.03026512 | 0.272769874 | ns | NaCl | KS.3 |
| FIM1_791 | 2716.408138 | 4492.376985 | 3356.780222 | 0.443775588 | 0.286312276 | 0.000533611 | 0.02370281 | ns | NaCl | KS.3 |
| FIM1_792 | 1108.752104 | 1144.609983 | 1088.768169 | 0.018422589 | 0.077198352 | 0.708251231 | 0.925190614 | ns | NaCl | KS.3 |
| FIM1_793 | 4316.654212 | 3952.967535 | 3888.038793 | -0.050679896 | 0.080347587 | 0.310344251 | 0.7303513 | ns | NaCl | KS.3 |
| FIM1_794 | 923.3000288 | 1030.833442 | 1027.485245 | 0.071634287 | 0.078049012 | 0.157956879 | 0.57787696 | ns | NaCl | KS.3 |
| FIM1_795 | 3650.501778 | 3970.527324 | 2824.487688 | 0.038671483 | 0.08734638 | 0.426547478 | 0.802854185 | ns | NaCl | KS.3 |
| FIM1_796 | 921.4847299 | 910.2885807 | 1281.943094 | -0.009071081 | 0.071945109 | 0.853325453 | 0.969345976 | ns | NaCl | KS.3 |
| FIM1_797 | 349.8289071 | 389.2983184 | 336.0054251 | 0.030538707 | 0.10174313 | 0.49163299 | 0.835238267 | ns | NaCl | KS.3 |
| FIM1_798 | 644.5521815 | 664.0048377 | 648.7373743 | 0.009930218 | 0.093594833 | 0.827447579 | 0.960240659 | ns | NaCl | KS.3 |
| FIM1_799 | 1159.16781 | 1100.270944 | 1734.643283 | -0.009835125 | 0.10966411 | 0.805438852 | 0.957859861 | ns | NaCl | KS.3 |
| FIM1_800 | 3146.053624 | 3686.94544 | 2792.052274 | 0.070454282 | 0.096522578 | 0.160168182 | 0.58211082 | ns | NaCl | KS.3 |
| FIM1_801 | 2076.515119 | 3115.394204 | 2056.706458 | 0.116371051 | 0.182034507 | 0.043267633 | 0.328487717 | ns | NaCl | KS.3 |
| FIM1_802 | 931.1607561 | 1148.911416 | 852.4081707 | 0.075293182 | 0.111629398 | 0.129781107 | 0.537155579 | ns | NaCl | KS.3 |
| FIM1_803 | 1056.55054 | 980.116208 | 1145.42991 | -0.055654198 | 0.071371898 | 0.261757754 | 0.686421476 | ns | NaCl | KS.3 |
| FIM1_804 | 6874.138606 | 9001.348496 | 5144.829841 | 0.096535243 | 0.12976649 | 0.068029708 | 0.40875338 | ns | NaCl | KS.3 |
| FIM1_805 | 622.6419036 | 516.1658402 | 1049.39095 | -0.094459475 | 0.098927241 | 0.073113943 | 0.421631393 | ns | NaCl | KS.3 |
| FIM1_806 | 492.8375916 | 316.316533 | 1397.046938 | -0.155902762 | 0.212815671 | 0.020592954 | 0.217054975 | ns | NaCl | KS.3 |
| FIM1_807 | 213.9742806 | 235.3935574 | 604.1760257 | 0.029677138 | 0.096793513 | 0.516216678 | 0.849827258 | ns | NaCl | KS.3 |

| | | | | | | | | | | |
|---|---|---|---|---|---|---|---|---|---|---|
| FIM1_808 | 988.94201 | 1135.381889 | 1277.977105 | 0.061220345 | 0.093410107 | 0.21619375 | 0.640629323 | ns | NaCl | KS.3 |
| FIM1_809 | 14407.24677 | 14109.21342 | 20737.53121 | -0.011604299 | 0.079003823 | 0.812920406 | 0.958816322 | ns | NaCl | KS.3 |
| FIM1_810 | 1644.140575 | 1471.827542 | 1437.83775 | -0.078312944 | 0.075474649 | 0.122518481 | 0.528318345 | ns | NaCl | KS.3 |
| FIM1_811 | 2096.341022 | 2223.817314 | 2368.765038 | 0.021921323 | 0.092277574 | 0.637285609 | 0.898332475 | ns | NaCl | KS.3 |
| FIM1_812 | 2406.070291 | 2167.214753 | 2025.255696 | -0.050885336 | 0.08711999 | 0.303239906 | 0.72182705 | ns | NaCl | KS.3 |
| FIM1_813 | 606.1231478 | 722.1736439 | 733.4134806 | 0.114225985 | 0.090020679 | 0.03428642 | 0.292771172 | ns | NaCl | KS.3 |
| FIM1_814 | 35.97438429 | 43.93876567 | 59.6130047 | 0.030301138 | 0.127562628 | 0.43657089 | 0.808096352 | ns | NaCl | KS.3 |
| FIM1_815 | 74.46492564 | 99.76656641 | 102.8411246 | 0.056692594 | 0.136706881 | 0.198695971 | 0.620924909 | ns | NaCl | KS.3 |
| FIM1_816 | 363.4920015 | 413.7661615 | 391.056969 | 0.033665686 | 0.106350166 | 0.440311273 | 0.808917498 | ns | NaCl | KS.3 |
| FIM1_817 | 1683.122918 | 2039.27258 | 1487.348653 | 0.077040876 | 0.105383262 | 0.12604038 | 0.534982546 | ns | NaCl | KS.3 |
| FIM1_818 | 338.0935904 | 312.9412016 | 327.2956797 | -0.025850106 | 0.097628593 | 0.566784303 | 0.868965626 | ns | NaCl | KS.3 |
| FIM1_819 | 1019.995254 | 1192.571006 | 1264.18978 | 0.082051039 | 0.09117677 | 0.111568847 | 0.50812574 | ns | NaCl | KS.3 |
| FIM1_820 | 5501.659173 | 7347.015373 | 6602.112733 | 0.1819523 | 0.14406842 | 0.00621945 | 0.106434152 | ns | NaCl | KS.3 |
| FIM1_821 | 11571.54704 | 11928.47326 | 8074.419783 | 0.012009662 | 0.0896303 | 0.797890538 | 0.955924408 | ns | NaCl | KS.3 |
| FIM1_822 | 1009.807193 | 1015.868977 | 963.4226905 | 0.001812255 | 0.092100572 | 0.96853201 | 0.99408528 | ns | NaCl | KS.3 |
| FIM1_823 | 867.3877005 | 831.1500934 | 1053.613319 | -0.028393118 | 0.075541965 | 0.565515818 | 0.868073805 | ns | NaCl | KS.3 |
| FIM1_824 | 7680.834238 | 11332.33307 | 4683.444448 | 0.072579817 | 0.161664389 | 0.121439301 | 0.527999953 | ns | NaCl | KS.3 |
| FIM1_825 | 4243.831278 | 3831.3744 | 3341.939333 | -0.061757641 | 0.080081017 | 0.220591363 | 0.642906874 | ns | NaCl | KS.3 |
| FIM1_826 | 100.3769601 | 87.30055955 | 94.20694577 | -0.02142752 | 0.120490461 | 0.578118692 | 0.874933697 | ns | NaCl | KS.3 |
| FIM1_828 | 1080.586367 | 1236.781177 | 1423.450385 | 0.063835842 | 0.091368511 | 0.201361915 | 0.624846998 | ns | NaCl | KS.3 |
| FIM1_829 | 394.9899142 | 444.0614675 | 492.427216 | 0.048845852 | 0.09251182 | 0.313493994 | 0.731871738 | ns | NaCl | KS.3 |
| FIM1_830 | 2886.912681 | 2232.416176 | 2874.1411 | -0.045928852 | 0.131192727 | 0.274344377 | 0.694940567 | ns | NaCl | KS.3 |
| FIM1_831 | 6032.163668 | 6264.083473 | 4234.203468 | 0.017969663 | 0.08423231 | 0.710358244 | 0.925563973 | ns | NaCl | KS.3 |
| FIM1_832 | 923.3282047 | 1035.898937 | 754.9707349 | 0.041306698 | 0.096829511 | 0.378030822 | 0.778493059 | ns | NaCl | KS.3 |
| FIM1_833 | 5785.240079 | 7351.993471 | 3921.404092 | 0.068741453 | 0.121783887 | 0.150749626 | 0.565595316 | ns | NaCl | KS.3 |
| FIM1_834 | 1266.166224 | 1222.847845 | 1221.750856 | -0.020630778 | 0.077667992 | 0.675239685 | 0.912806589 | ns | NaCl | KS.3 |
| FIM1_835 | 787.9064359 | 900.1111433 | 953.2250199 | 0.065446338 | 0.089796999 | 0.192650089 | 0.618952432 | ns | NaCl | KS.3 |
| FIM1_836 | 1796.220572 | 740.0221615 | 1089.792055 | -1.066414562 | 0.302732323 | 3.59407055553 | 0.000525897 | Km<KS.3 | NaCl | KS.3 |
| FIM1_840 | 362.9651451 | 418.3858359 | 485.8406089 | 0.056134668 | 0.096954965 | 0.247005024 | 0.669893726 | ns | NaCl | KS.3 |
| FIM1_841 | 62.20709828 | 57.16410452 | 116.3551879 | -0.010365103 | 0.11953933 | 0.781261186 | 0.951646805 | ns | NaCl | KS.3 |
| FIM1_842 | 205.799424 | 173.498366 | 255.2486176 | -0.038340463 | 0.112547876 | 0.373250113 | 0.775331654 | ns | NaCl | KS.3 |
| FIM1_843 | 182.8777978 | 158.1320531 | 1017.35679 | -0.015736723 | 0.130919851 | 0.659563883 | 0.903691294 | ns | NaCl | KS.3 |
| FIM1_844 | 77.97977919 | 46.36865301 | 112.3709184 | -0.218537335 | 0.269490253 | 0.009009322 | 0.132216457 | ns | NaCl | KS.3 |
| FIM1_846 | 42.34158579 | 52.7131379 | 84.39783745 | 0.035452959 | 0.125168041 | 0.38013666 | 0.778493059 | ns | NaCl | KS.3 |
| FIM1_847 | 9.403606118 | 11.19312775 | 15.97427327 | 0.01304041 | 0.137332512 | 0.703736619 | 0.924258099 | ns | NaCl | KS.3 |
| FIM1_850 | 463.1820141 | 506.7254712 | 698.1875307 | 0.015975799 | 0.113720236 | 0.684631054 | 0.91584821 | ns | NaCl | KS.3 |
| FIM1_851 | 190.6671718 | 204.6222667 | 305.0748288 | 0.014662997 | 0.108818963 | 0.717816538 | 0.93000992 | ns | NaCl | KS.3 |
| FIM1_852 | 80.65370566 | 74.80952087 | 63.46054071 | -0.017250031 | 0.103948093 | 0.683739774 | 0.91584821 | ns | NaCl | KS.3 |
| FIM1_853 | 124.4254669 | 124.6738152 | 101.2122781 | 0.000640951 | 0.102510733 | 0.987846028 | 0.999134858 | ns | NaCl | KS.3 |
| FIM1_854 | 1977.210369 | 1335.992486 | 1559.299267 | -0.117469466 | 0.177811565 | 0.041976428 | 0.323956287 | ns | NaCl | KS.3 |
| FIM1_855 | 363.8769616 | 277.268555 | 276.8693972 | -0.095228815 | 0.130772471 | 0.07053292 | 0.417739615 | ns | NaCl | KS.3 |
| FIM1_856 | 294.4528475 | 252.7603761 | 299.5086226 | -0.052780104 | 0.100861372 | 0.2660808 | 0.689593109 | ns | NaCl | KS.3 |
| FIM1_857 | 181.9880428 | 163.3191146 | 194.7328761 | -0.039363386 | 0.092780887 | 0.408098917 | 0.79494601 | ns | NaCl | KS.3 |
| FIM1_858 | 838.4393941 | 329.5167112 | 643.7882858 | -1.099823921 | 0.330154263 | 5.34462943908 | 0.000718636 | Km<KS.3 | NaCl | KS.3 |
| FIM1_859 | 469.9985584 | 346.6413294 | 460.6221006 | -0.289153884 | 0.147487459 | 0.000232537 | 0.014282363 | ns | NaCl | KS.3 |

| | | | | | | | | | | |
|---|---|---|---|---|---|---|---|---|---|---|
| FIM1_860 | 316.5368574 | 286.8829257 | 530.8493055 | -0.062600059 | 0.083220195 | 0.214419008 | 0.639529117 | ns | NaCl | KS.3 |
| FIM1_861 | 480.2162792 | 372.4176568 | 689.1671746 | -0.12824033 | 0.125534729 | 0.026646949 | 0.254519471 | ns | NaCl | KS.3 |
| FIM1_862 | 163.6193541 | 120.6704647 | 158.4303416 | -0.109367828 | 0.1454711 | 0.048579362 | 0.345754403 | ns | NaCl | KS.3 |
| FIM1_863 | 14.60732776 | 15.39190813 | 32.18866186 | 0.003934574 | 0.136710904 | 0.906667357 | 0.979989779 | ns | NaCl | KS.3 |
| FIM1_864 | 405.8884709 | 394.9729115 | 574.4896196 | -0.005744728 | 0.108159848 | 0.886343417 | 0.97621397 | ns | NaCl | KS.3 |
| FIM1_865 | 877.6553888 | 792.8723213 | 947.753543 | -0.0285682 | 0.101224557 | 0.519268929 | 0.850629872 | ns | NaCl | KS.3 |
| FIM1_866 | 419.6868246 | 320.0718837 | 514.4551748 | -0.107315346 | 0.128521757 | 0.050095534 | 0.349544574 | ns | NaCl | KS.3 |
| FIM1_867 | 975.6222252 | 1020.723057 | 1299.862932 | 0.013734969 | 0.097240699 | 0.757158633 | 0.945250644 | ns | NaCl | KS.3 |
| FIM1_868 | 841.9846892 | 704.2653491 | 901.8214115 | -0.075963764 | 0.100775618 | 0.132548988 | 0.539053419 | ns | NaCl | KS.3 |
| FIM1_869 | 698.7736358 | 734.8445416 | 674.8613506 | 0.029339335 | 0.079821276 | 0.551593389 | 0.860945181 | ns | NaCl | KS.3 |
| FIM1_870 | 1765.989909 | 1990.808196 | 1337.424702 | 0.035979639 | 0.101789732 | 0.42456423 | 0.802854185 | ns | NaCl | KS.3 |
| FIM1_871 | 1330.9327 | 1237.789487 | 1085.450442 | -0.052427394 | 0.071201096 | 0.28960387 | 0.709394019 | ns | NaCl | KS.3 |
| FIM1_872 | 3218.44952 | 3165.617738 | 2799.267011 | -0.011207514 | 0.072081543 | 0.81939236 | 0.960016822 | ns | NaCl | KS.3 |
| FIM1_873 | 857.7339145 | 735.3530862 | 998.9041111 | -0.069625782 | 0.094929607 | 0.165493347 | 0.589355333 | ns | NaCl | KS.3 |
| FIM1_874 | 2041.432755 | 2592.711849 | 4884.448618 | 0.073539633 | 0.120987004 | 0.132028977 | 0.539053419 | ns | NaCl | KS.3 |
| FIM1_875 | 297.1620225 | 332.1461761 | 368.4667161 | 0.031034342 | 0.102946056 | 0.481987379 | 0.828664253 | ns | NaCl | KS.3 |
| FIM1_876 | 2531.640474 | 3272.510853 | 2803.00508 | 0.059754951 | 0.127962877 | 0.189006768 | 0.618952432 | ns | NaCl | KS.3 |
| FIM1_877 | 1583.74864 | 1461.926635 | 1254.645693 | -0.037837463 | 0.086516458 | 0.437509574 | 0.808096352 | ns | NaCl | KS.3 |
| FIM1_878 | 4333.79476 | 4234.821133 | 3660.528863 | -0.013752714 | 0.07631623 | 0.779874198 | 0.951646805 | ns | NaCl | KS.3 |
| FIM1_879 | 18.26679177 | 28.58983771 | 27.8276178 | 0.057507365 | 0.165321259 | 0.175603263 | 0.605047427 | ns | NaCl | KS.3 |
| FIM1_880 | 715.0655599 | 632.4425098 | 1216.289069 | -0.075225404 | 0.0806782 | 0.140491985 | 0.551655583 | ns | NaCl | KS.3 |
| FIM1_881 | 1580.027776 | 1473.969514 | 1551.362047 | -0.048479039 | 0.072009272 | 0.327798054 | 0.740319311 | ns | NaCl | KS.3 |
| FIM1_882 | 414.6300506 | 382.8241865 | 510.0564056 | -0.038402316 | 0.087593005 | 0.429206339 | 0.804596805 | ns | NaCl | KS.3 |
| FIM1_883 | 1265.761047 | 1264.042644 | 1562.433082 | -0.000254944 | 0.096041954 | 0.995430361 | 0.99924489 | ns | NaCl | KS.3 |
| FIM1_884 | 213.9869451 | 131.7299215 | 330.5532116 | -0.18285975 | 0.239627771 | 0.013978795 | 0.171291879 | ns | NaCl | KS.3 |
| FIM1_885 | 2279.241913 | 2435.093428 | 2050.872708 | 0.052944979 | 0.066120868 | 0.275224479 | 0.695399585 | ns | NaCl | KS.3 |
| FIM1_886 | 4325.53139 | 3694.198275 | 4045.572158 | -0.044315893 | 0.107183817 | 0.325662581 | 0.739131368 | ns | NaCl | KS.3 |
| FIM1_887 | 558.7796869 | 499.5196885 | 602.4879608 | -0.055557785 | 0.087471737 | 0.263641436 | 0.688243886 | ns | NaCl | KS.3 |
| FIM1_888 | 1162.792754 | 1164.360229 | 1495.16734 | 0.000675867 | 0.0840882 | 0.988811785 | 0.999134858 | ns | NaCl | KS.3 |
| FIM1_889 | 1064.041265 | 940.8042246 | 968.179931 | -0.085124059 | 0.078007687 | 0.096699964 | 0.474908511 | ns | NaCl | KS.3 |
| FIM1_890 | 443.364863 | 442.4615737 | 335.6733714 | -0.001202042 | 0.088472218 | 0.979636289 | 0.996363429 | ns | NaCl | KS.3 |
| FIM1_891 | 2852.078486 | 2152.88373 | 4113.37712 | -0.165504274 | 0.138456142 | 0.009515182 | 0.13642083 | ns | NaCl | KS.3 |
| FIM1_892 | 135.1723532 | 57.15505877 | 170.0596042 | -0.670026951 | 0.4870903 | 0.000460723 | 0.022039378 | ns | NaCl | KS.3 |
| FIM1_893 | 13.47202375 | 25.79427072 | 28.89626081 | 0.099593973 | 0.228852434 | 0.058418739 | 0.377936577 | ns | NaCl | KS.3 |
| FIM1_894 | 2077.59285 | 1042.292389 | 3726.986747 | -0.526870946 | 0.3988294 | 0.000759069 | 0.030210955 | ns | NaCl | KS.3 |
| FIM1_895 | 737.424253 | 592.4954046 | 707.5695422 | -0.129638431 | 0.106636944 | 0.022259018 | 0.229180264 | ns | NaCl | KS.3 |
| FIM1_896 | 136.7692006 | 126.2400467 | 140.4085589 | -0.027934195 | 0.096020828 | 0.541801916 | 0.857187397 | ns | NaCl | KS.3 |
| FIM1_897 | 555.6644052 | 559.0538593 | 566.0714161 | 0.001485463 | 0.100276166 | 0.972389198 | 0.995313014 | ns | NaCl | KS.3 |
| FIM1_898 | 5577.345031 | 4645.88268 | 5129.727209 | -0.065848167 | 0.10557797 | 0.176357465 | 0.605047427 | ns | NaCl | KS.3 |
| FIM1_899 | 962.9056426 | 897.6208908 | 688.9489661 | -0.024097971 | 0.094820978 | 0.599218577 | 0.880219278 | ns | NaCl | KS.3 |
| FIM1_900 | 1910.376115 | 1561.450056 | 1909.136672 | -0.079149036 | 0.10798105 | 0.117150938 | 0.52010139 | ns | NaCl | KS.3 |
| FIM1_901 | 3290.65902 | 3794.253097 | 3456.279029 | 0.087837422 | 0.084490086 | 0.090442228 | 0.463573758 | ns | NaCl | KS.3 |
| FIM1_902 | 418.2674684 | 463.48941 | 486.7253884 | 0.052686045 | 0.084261173 | 0.290330666 | 0.709776444 | ns | NaCl | KS.3 |
| FIM1_903 | 2726.147397 | 3316.023097 | 1651.287398 | 0.032524515 | 0.123904994 | 0.415830638 | 0.800387073 | ns | NaCl | KS.3 |
| FIM1_904 | 1754.39932 | 2039.030413 | 1256.333312 | 0.028586414 | 0.115859152 | 0.482217421 | 0.828681061 | ns | NaCl | KS.3 |

| | | | | | | | | | | |
|---|---|---|---|---|---|---|---|---|---|---|
| FIM1_905 | 432.0902514 | 472.2246971 | 879.7972504 | 0.040772163 | 0.08816116 | 0.401685759 | 0.789189152 | ns | NaCl | KS.3 |
| FIM1_906 | 16902.57675 | 18744.91852 | 12090.97668 | 0.051420574 | 0.086533185 | 0.299192595 | 0.717010926 | ns | NaCl | KS.3 |
| FIM1_907 | 22783.70047 | 26658.8702 | 16591.70845 | 0.096778786 | 0.086960239 | 0.065630835 | 0.402110102 | ns | NaCl | KS.3 |
| FIM1_909 | 1308.845932 | 1326.73767 | 1077.98299 | 0.006342264 | 0.086091869 | 0.894377921 | 0.977268396 | ns | NaCl | KS.3 |
| FIM1_910 | 111.2473409 | 66.32001868 | 267.2848716 | -0.24295786 | 0.273794281 | 0.006413919 | 0.107078006 | ns | NaCl | KS.3 |
| FIM1_912 | 38.67835989 | 19.55702612 | 111.3884135 | -0.218240342 | 0.326983307 | 0.010570008 | 0.145864059 | ns | NaCl | KS.3 |
| FIM1_913 | 1798.256281 | 1805.875275 | 2001.170019 | 0.000815984 | 0.103779493 | 0.984354332 | 0.998198696 | ns | NaCl | KS.3 |
| FIM1_914 | 417.6678646 | 456.0990522 | 592.0757581 | 0.045109115 | 0.084381389 | 0.361313101 | 0.763371684 | ns | NaCl | KS.3 |
| FIM1_915 | 4957.35153 | 5061.406986 | 4187.559889 | 0.012485412 | 0.075805134 | 0.799700447 | 0.956602482 | ns | NaCl | KS.3 |
| FIM1_916 | 1003.916134 | 981.9573187 | 1234.286868 | -0.008389017 | 0.0886075 | 0.858701939 | 0.969856868 | ns | NaCl | KS.3 |
| FIM1_917 | 14035.35246 | 22660.26164 | 10820.5135 | 0.139987248 | 0.216862848 | 0.028215569 | 0.262869768 | ns | NaCl | KS.3 |
| FIM1_918 | 213.9006029 | 250.4453811 | 235.7526735 | 0.036943049 | 0.111458648 | 0.390556044 | 0.779482559 | ns | NaCl | KS.3 |
| FIM1_919 | 196.5432599 | 205.7635609 | 300.1039835 | 0.008723203 | 0.109828792 | 0.826544922 | 0.960240659 | ns | NaCl | KS.3 |
| FIM1_920 | 1678.000801 | 1812.779982 | 1881.743709 | 0.027094206 | 0.094681235 | 0.55682356 | 0.863699344 | ns | NaCl | KS.3 |
| FIM1_921 | 2310.392273 | 2239.421225 | 2693.04147 | -0.014522765 | 0.084842882 | 0.76311687 | 0.946287744 | ns | NaCl | KS.3 |
| FIM1_922 | 330.2159576 | 300.0481714 | 488.257024 | -0.032421155 | 0.097534781 | 0.478695075 | 0.826171162 | ns | NaCl | KS.3 |
| FIM1_923 | 409.3071214 | 421.4184385 | 409.4636735 | 0.009085492 | 0.096684887 | 0.837953396 | 0.962829707 | ns | NaCl | KS.3 |
| FIM1_924 | 591.1880219 | 478.5593059 | 926.2492243 | -0.112454681 | 0.107373799 | 0.040605252 | 0.317627561 | ns | NaCl | KS.3 |
| FIM1_925 | 129.7080656 | 98.45320838 | 227.6234873 | -0.083973347 | 0.130933937 | 0.096192919 | 0.473352892 | ns | NaCl | KS.3 |
| FIM1_926 | 1319.874159 | 1387.153273 | 4723.754127 | 0.007546239 | 0.116838052 | 0.841629851 | 0.965884316 | ns | NaCl | KS.3 |
| FIM1_927 | 10285.89718 | 9964.857031 | 6918.70648 | -0.025918297 | 0.063736451 | 0.586921492 | 0.878311243 | ns | NaCl | KS.3 |
| FIM1_928 | 945.602686 | 1014.095788 | 996.4788868 | 0.029172535 | 0.089408046 | 0.540533893 | 0.857165292 | ns | NaCl | KS.3 |
| FIM1_929 | 1201.908317 | 1314.421736 | 1444.466614 | 0.022758788 | 0.103800648 | 0.596021272 | 0.87962202 | ns | NaCl | KS.3 |
| FIM1_930 | 1441.582894 | 1470.934701 | 1751.919322 | 0.007052034 | 0.093453513 | 0.876921338 | 0.972783118 | ns | NaCl | KS.3 |
| FIM1_931 | 2027.615377 | 2047.518507 | 1672.699267 | 0.005189933 | 0.078777904 | 0.915661143 | 0.982343702 | ns | NaCl | KS.3 |
| FIM1_932 | 456.5556688 | 388.1826195 | 497.2585964 | -0.094709771 | 0.088505549 | 0.071298718 | 0.418675804 | ns | NaCl | KS.3 |
| FIM1_933 | 1680.535693 | 1583.596799 | 2148.831949 | -0.047177524 | 0.065453222 | 0.328685695 | 0.741064881 | ns | NaCl | KS.3 |
| FIM1_934 | 1335.891726 | 1111.180696 | 1323.273902 | -0.110167483 | 0.094428751 | 0.041426767 | 0.322888102 | ns | NaCl | KS.3 |
| FIM1_935 | 1065.172497 | 974.7241411 | 1205.089316 | -0.01267798 | 0.11983606 | 0.735215104 | 0.938375543 | ns | NaCl | KS.3 |
| FIM1_936 | 7514.861723 | 7525.773537 | 8369.136041 | 0.000344213 | 0.103935803 | 0.993390453 | 0.999215023 | ns | NaCl | KS.3 |
| FIM1_937 | 2954.322702 | 2554.72192 | 2593.518284 | -0.09133297 | 0.084136869 | 0.079378651 | 0.436363304 | ns | NaCl | KS.3 |
| FIM1_938 | 4228.043254 | 4167.234483 | 3427.445661 | -0.004374998 | 0.096600614 | 0.921401289 | 0.983890455 | ns | NaCl | KS.3 |
| FIM1_939 | 2310.177224 | 2251.787491 | 1869.916936 | -0.020479726 | 0.062904133 | 0.666035276 | 0.906573323 | ns | NaCl | KS.3 |
| FIM1_940 | 3734.951048 | 3335.200662 | 2972.43512 | -0.042404097 | 0.095432587 | 0.36978343 | 0.770382145 | ns | NaCl | KS.3 |
| FIM1_941 | 604.1254544 | 671.2724179 | 506.1071318 | 0.027220099 | 0.105257448 | 0.527002769 | 0.855133325 | ns | NaCl | KS.3 |
| FIM1_942 | 1728.943329 | 3294.222176 | 1787.929814 | 0.754844308 | 0.268533816 | 2.56807348163 | 0.002555233 | ns | NaCl | KS.3 |
| FIM1_943 | 12383.22689 | 13321.68277 | 8646.69721 | 0.034166666 | 0.086252494 | 0.481979275 | 0.828664253 | ns | NaCl | KS.3 |
| FIM1_944 | 1919.498577 | 2100.912559 | 1283.016317 | 0.019597919 | 0.107990907 | 0.635375554 | 0.898009483 | ns | NaCl | KS.3 |
| FIM1_945 | 1886.033186 | 2079.221504 | 2338.598098 | 0.066290968 | 0.075365691 | 0.18779566 | 0.618637711 | ns | NaCl | KS.3 |
| FIM1_946 | 46.95229717 | 38.05496652 | 45.99085676 | -0.048089889 | 0.12000731 | 0.272081164 | 0.693444564 | ns | NaCl | KS.3 |
| FIM1_947 | 6163.60151 | 6526.335186 | 5270.519313 | 0.030325214 | 0.081540161 | 0.537170302 | 0.855981197 | ns | NaCl | KS.3 |
| FIM1_948 | 5481.299258 | 6436.186603 | 5590.866524 | 0.116081937 | 0.081976674 | 0.028695654 | 0.265110548 | ns | NaCl | KS.3 |
| FIM1_949 | 901.1562545 | 511.8741204 | 1384.197803 | -0.412016264 | 0.331664177 | 0.001251524 | 0.041599773 | ns | NaCl | KS.3 |
| FIM1_950 | 407.8952762 | 359.6121239 | 465.2174882 | -0.029564408 | 0.107516365 | 0.4891953 | 0.833474869 | ns | NaCl | KS.3 |
| FIM1_951 | 19868.92224 | 26712.89266 | 20186.18416 | 0.198558727 | 0.148859626 | 0.004073917 | 0.082277904 | ns | NaCl | KS.3 |

| | | | | | | | | | | |
|---|---|---|---|---|---|---|---|---|---|---|
| FIM1_952 | 36.48238786 | 27.14545819 | 41.24879713 | -0.038999934 | 0.14284314 | 0.318593611 | 0.73515919 | ns | NaCl | KS.3 |
| FIM1_953 | 3436.708458 | 3088.114363 | 3895.405561 | -0.054442715 | 0.086058836 | 0.274003168 | 0.694804019 | ns | NaCl | KS.3 |
| FIM1_954 | 2021.611059 | 1797.778317 | 1865.356409 | -0.031520569 | 0.104398131 | 0.471917303 | 0.822127061 | ns | NaCl | KS.3 |
| FIM1_955 | 809.8458507 | 1027.86108 | 1946.284371 | 0.046562704 | 0.126610675 | 0.275089309 | 0.695399585 | ns | NaCl | KS.3 |
| FIM1_957 | 14048.17244 | 8930.697584 | 11679.93381 | -0.121967049 | 0.19952874 | 0.039185275 | 0.313586284 | ns | NaCl | KS.3 |
| FIM1_958 | 4118.109838 | 4672.321834 | 4172.932193 | 0.047263664 | 0.096735641 | 0.319980789 | 0.736735876 | ns | NaCl | KS.3 |
| FIM1_959 | 2333.647661 | 3456.134975 | 1630.320037 | 0.141912076 | 0.18743564 | 0.025021912 | 0.247483127 | ns | NaCl | KS.3 |
| FIM1_960 | 3137.222661 | 3426.245598 | 2383.03718 | 0.049914604 | 0.081273177 | 0.317050933 | 0.73382286 | ns | NaCl | KS.3 |
| FIM1_961 | 5085.690513 | 4807.035035 | 4809.67305 | -0.041837883 | 0.069062458 | 0.393289205 | 0.781740625 | ns | NaCl | KS.3 |
| FIM1_962 | 3288.651106 | 3465.739531 | 2784.4084 | 0.036682128 | 0.071226485 | 0.456434321 | 0.817337042 | ns | NaCl | KS.3 |
| FIM1_963 | 708.1420061 | 632.4146854 | 705.9682752 | -0.046801016 | 0.092649541 | 0.33210668 | 0.742667065 | ns | NaCl | KS.3 |
| FIM1_964 | 252.3483874 | 208.0486454 | 229.22586 | -0.086533016 | 0.101605607 | 0.094652213 | 0.470201108 | ns | NaCl | KS.3 |
| FIM1_965 | 848.7663875 | 741.8426459 | 1032.537501 | -0.097019946 | 0.078236081 | 0.060722032 | 0.385321569 | ns | NaCl | KS.3 |
| FIM1_966 | 3719.5768 | 3841.162223 | 3950.198171 | 0.01286737 | 0.089497801 | 0.784080453 | 0.951646805 | ns | NaCl | KS.3 |
| FIM1_967 | 1024.624385 | 1086.692403 | 1015.043269 | 0.032954818 | 0.079518649 | 0.504676347 | 0.844040357 | ns | NaCl | KS.3 |
| FIM1_968 | 565.353501 | 648.7473071 | 752.0076583 | 0.058158408 | 0.094848434 | 0.235373666 | 0.659616806 | ns | NaCl | KS.3 |
| FIM1_969 | 285.8549327 | 320.026383 | 440.0148115 | 0.040547678 | 0.09632241 | 0.387213303 | 0.779278134 | ns | NaCl | KS.3 |
| FIM1_970 | 884.2731447 | 839.6522449 | 965.1903519 | -0.03127672 | 0.078349691 | 0.526796712 | 0.855077861 | ns | NaCl | KS.3 |
| FIM1_971 | 89.43425238 | 90.38156 | 131.9005992 | 0.002203125 | 0.101261196 | 0.958709031 | 0.991553507 | ns | NaCl | KS.3 |
| FIM1_972 | 189.2726653 | 201.366558 | 157.1682748 | 0.014896044 | 0.100209403 | 0.73164084 | 0.936960396 | ns | NaCl | KS.3 |
| FIM1_973 | 832.8637539 | 927.4040486 | 1188.232828 | 0.077156869 | 0.073757038 | 0.126101153 | 0.534982546 | ns | NaCl | KS.3 |
| FIM1_974 | 755.1266002 | 1174.797342 | 1062.195313 | 0.405247701 | 0.245243682 | 0.000421675 | 0.021012238 | ns | NaCl | KS.3 |
| FIM1_975 | 1587.39228 | 2003.119342 | 2546.823218 | 0.085496423 | 0.117321958 | 0.094578307 | 0.470201108 | ns | NaCl | KS.3 |
| FIM1_976 | 51.91373718 | 38.93016938 | 101.3528942 | -0.041497106 | 0.135900452 | 0.30499141 | 0.724521545 | ns | NaCl | KS.3 |
| FIM1_977 | 4653.613164 | 5312.285294 | 3415.972487 | 0.021541472 | 0.118585379 | 0.580146019 | 0.8749481 | ns | NaCl | KS.3 |
| FIM1_978 | 6297.968865 | 9158.667816 | 4313.41991 | 0.049560024 | 0.154993261 | 0.225444514 | 0.648597616 | ns | NaCl | KS.3 |
| FIM1_979 | 3375.081034 | 4851.387319 | 2201.409289 | 0.076879068 | 0.156285594 | 0.110436503 | 0.505913076 | ns | NaCl | KS.3 |
| FIM1_980 | 2565.285148 | 2276.395171 | 2473.290594 | -0.030115227 | 0.106156382 | 0.485456274 | 0.829376704 | ns | NaCl | KS.3 |
| FIM1_981 | 819.5115088 | 882.1739206 | 1418.350743 | 0.011101205 | 0.11825716 | 0.768377494 | 0.948835638 | ns | NaCl | KS.3 |
| FIM1_982 | 1243.37151 | 1222.644233 | 1544.672182 | -0.005107136 | 0.093950253 | 0.910281957 | 0.981715313 | ns | NaCl | KS.3 |
| FIM1_983 | 15775.56817 | 13734.9499 | 12436.09678 | -0.096788799 | 0.079071423 | 0.061862616 | 0.390501742 | ns | NaCl | KS.3 |
| FIM1_984 | 22.19247221 | 16.2163357 | 91.15760918 | -0.020806009 | 0.158038673 | 0.524615919 | 0.854940852 | ns | NaCl | KS.3 |
| FIM1_988 | 2491.673376 | 2387.879244 | 2634.034784 | -0.021478132 | 0.081834465 | 0.660284995 | 0.903691294 | ns | NaCl | KS.3 |
| FIM1_989 | 24.65277206 | 9.647761052 | 71.54174204 | -0.4228964 | 0.515013417 | 0.002244197 | 0.058108556 | ns | NaCl | KS.3 |
| FIM1_991 | 1162.769839 | 1184.103925 | 1301.236003 | 0.004693306 | 0.101686021 | 0.911938087 | 0.982011252 | ns | NaCl | KS.3 |
| FIM1_992 | 1834.328008 | 1888.629489 | 2183.789848 | 0.008248137 | 0.098763776 | 0.850054928 | 0.967955712 | ns | NaCl | KS.3 |
| FIM1_993 | 1922.331606 | 1944.901717 | 1362.119294 | 0.002646989 | 0.104564608 | 0.9489526 | 0.987736646 | ns | NaCl | KS.3 |
| FIM1_994 | 2831.649974 | 3088.455847 | 2129.332173 | 0.028968626 | 0.096513817 | 0.526441409 | 0.855027345 | ns | NaCl | KS.3 |
| FIM1_995 | 3412.002489 | 4030.489693 | 2185.273744 | 0.036971387 | 0.113728283 | 0.385366396 | 0.778665419 | ns | NaCl | KS.3 |
| FIM1_996 | 4617.746097 | 6167.360924 | 2898.353279 | 0.041990576 | 0.139482654 | 0.29631415 | 0.714226208 | ns | NaCl | KS.3 |
| FIM1_997 | 132.1077727 | 101.292319 | 234.4595379 | -0.058933649 | 0.131123547 | 0.190906765 | 0.618952432 | ns | NaCl | KS.3 |
| FIM1_998 | 1022.707294 | 927.0518345 | 1120.239119 | -0.055595185 | 0.083083803 | 0.266750697 | 0.689593109 | ns | NaCl | KS.3 |
| FIM1_999 | 1921.512363 | 2179.196818 | 1425.907567 | 0.066128207 | 0.087124794 | 0.190099484 | 0.618952432 | ns | NaCl | KS.3 |
| FIM1_1000 | 264.3695529 | 277.5915265 | 961.7356256 | 0.007574202 | 0.1151509 | 0.842858587 | 0.966485679 | ns | NaCl | KS.3 |
| FIM1_1001 | 393.9954894 | 338.0233069 | 323.4595592 | -0.089611641 | 0.087569188 | 0.085551171 | 0.454233803 | ns | NaCl | KS.3 |

| | | | | | | | | | | |
|---|---|---|---|---|---|---|---|---|---|---|
| FIM1_1002 | 352.9807727 | 332.6124277 | 328.5117821 | -0.026412203 | 0.087980233 | 0.581097087 | 0.874951829 | ns | NaCl | KS.3 |
| FIM1_1003 | 5546.201922 | 7200.255957 | 3583.500302 | 0.064229672 | 0.128457125 | 0.166658169 | 0.590711667 | ns | NaCl | KS.3 |
| FIM1_1004 | 2340.370554 | 2497.028173 | 3041.230843 | 0.017984185 | 0.100359486 | 0.679999216 | 0.913941356 | ns | NaCl | KS.3 |
| FIM1_1005 | 1305.61035 | 1210.328561 | 1178.634649 | -0.031778759 | 0.089750327 | 0.505588475 | 0.84439816 | ns | NaCl | KS.3 |
| FIM1_1006 | 3771.812132 | 3274.528977 | 3500.720483 | -0.055899919 | 0.096919201 | 0.248776461 | 0.67096552 | ns | NaCl | KS.3 |
| FIM1_1007 | 668.2771618 | 529.7159646 | 680.5163647 | -0.183855136 | 0.10463301 | 0.002434434 | 0.060255275 | ns | NaCl | KS.3 |
| FIM1_1008 | 261.9430641 | 196.6438104 | 350.3893167 | -0.121682247 | 0.138852877 | 0.034279621 | 0.292771172 | ns | NaCl | KS.3 |
| FIM1_1009 | 713.7959575 | 634.9228811 | 736.7478592 | -0.058739886 | 0.087642353 | 0.239209287 | 0.662163805 | ns | NaCl | KS.3 |
| FIM1_1010 | 1754.234045 | 1617.552139 | 2615.082822 | -0.015978455 | 0.110173383 | 0.691783654 | 0.91988055 | ns | NaCl | KS.3 |
| FIM1_1011 | 11409.52874 | 10368.71767 | 10635.097 | -0.026786318 | 0.101574285 | 0.542963819 | 0.857313452 | ns | NaCl | KS.3 |
| FIM1_1012 | 2078.807536 | 2016.586074 | 2599.211892 | -0.004183464 | 0.119233063 | 0.909992927 | 0.981715313 | ns | NaCl | KS.3 |
| FIM1_1013 | 2989.664717 | 2511.400734 | 2768.382107 | -0.047587242 | 0.110087063 | 0.291625645 | 0.711543689 | ns | NaCl | KS.3 |
| FIM1_1014 | 3612.916485 | 865.6599224 | 2115.903474 | -1.70844552 | 0.396597667 | 1.2682929322[illegible] | 3.0046463513[illegible] | Km<KS.3 | NaCl | KS.3 |
| FIM1_1015 | 3082.327563 | 3947.240447 | 2155.16098 | 0.051771529 | 0.127245386 | 0.236873677 | 0.659616806 | ns | NaCl | KS.3 |
| FIM1_1016 | 229.0412785 | 205.4173471 | 196.0072282 | -0.022848247 | 0.109670914 | 0.579743617 | 0.8749481 | ns | NaCl | KS.3 |
| FIM1_1017 | 6689.896759 | 7403.420479 | 4189.521793 | 0.052803437 | 0.084998746 | 0.288710065 | 0.708650347 | ns | NaCl | KS.3 |
| FIM1_1018 | 1738.464868 | 1596.323188 | 1304.181682 | -0.063519296 | 0.071096279 | 0.201253303 | 0.624846998 | ns | NaCl | KS.3 |
| FIM1_1019 | 730.7809895 | 775.448603 | 854.6122633 | 0.022515636 | 0.09179919 | 0.629462728 | 0.896016329 | ns | NaCl | KS.3 |
| FIM1_1020 | 82.41206907 | 50.9991872 | 134.9193748 | -0.089981613 | 0.188594303 | 0.076829539 | 0.430435761 | ns | NaCl | KS.3 |
| FIM1_1021 | 114.5673467 | 155.5917177 | 651.045516 | 0.049836947 | 0.141195186 | 0.235907883 | 0.659616806 | ns | NaCl | KS.3 |
| FIM1_1022 | 8764.333945 | 10002.91839 | 7715.193246 | 0.088398475 | 0.079866585 | 0.086521833 | 0.45668467 | ns | NaCl | KS.3 |
| FIM1_1024 | 1183.873977 | 1075.887343 | 1785.986354 | -0.02231993 | 0.106427147 | 0.595991609 | 0.87962202 | ns | NaCl | KS.3 |
| FIM1_1025 | 4804.947992 | 7161.575238 | 5296.423224 | 0.385491696 | 0.214211364 | 0.000275762 | 0.015769121 | ns | NaCl | KS.3 |
| FIM1_1026 | 7314.195215 | 8758.210519 | 6893.761453 | 0.083201997 | 0.099739591 | 0.105749843 | 0.497264148 | ns | NaCl | KS.3 |
| FIM1_1027 | 755.9685775 | 670.82589 | 1025.936303 | -0.038404577 | 0.099642577 | 0.40252935 | 0.789854857 | ns | NaCl | KS.3 |
| FIM1_1028 | 110.1411711 | 103.4691846 | 216.8554548 | -0.01536386 | 0.103768054 | 0.716108917 | 0.929477573 | ns | NaCl | KS.3 |
| FIM1_1029 | 20258.86205 | 17477.01474 | 12682.87503 | -0.037599036 | 0.108359969 | 0.390377799 | 0.779482559 | ns | NaCl | KS.3 |
| FIM1_1030 | 7588.539927 | 10252.68204 | 4336.793453 | 0.049038552 | 0.140230877 | 0.242126411 | 0.666322592 | ns | NaCl | KS.3 |
| FIM1_1031 | 83.78401955 | 69.62954796 | 81.79274124 | -0.038351612 | 0.118601645 | 0.361199004 | 0.763371684 | ns | NaCl | KS.3 |
| FIM1_1032 | 438.0733031 | 436.7651372 | 743.9069697 | -0.003332191 | 0.076966172 | 0.945917972 | 0.987278589 | ns | NaCl | KS.3 |
| FIM1_1033 | 274.2239266 | 207.4046209 | 215.5946391 | -0.089108281 | 0.134320064 | 0.083213171 | 0.447068602 | ns | NaCl | KS.3 |
| FIM1_1034 | 415.0934826 | 460.7955174 | 648.1549468 | 0.061073962 | 0.079474363 | 0.225409585 | 0.648597616 | ns | NaCl | KS.3 |
| FIM1_1035 | 129.9052659 | 130.0819018 | 164.1932233 | -0.002596646 | 0.096220242 | 0.953426501 | 0.988804845 | ns | NaCl | KS.3 |
| FIM1_1036 | 608.4796162 | 552.135448 | 581.8946502 | -0.052708746 | 0.085478208 | 0.289089996 | 0.708833284 | ns | NaCl | KS.3 |
| FIM1_1037 | 1215.855869 | 1263.152983 | 1268.132034 | 0.032520572 | 0.060455671 | 0.486392113 | 0.830120331 | ns | NaCl | KS.3 |
| FIM1_1038 | 4010.673131 | 4194.050015 | 3371.647619 | 0.037755712 | 0.062544132 | 0.425933287 | 0.802854185 | ns | NaCl | KS.3 |
| FIM1_1039 | 3066.443999 | 3539.548532 | 3203.341312 | 0.09167116 | 0.083398229 | 0.078090485 | 0.432946334 | ns | NaCl | KS.3 |
| FIM1_1040 | 1324.308172 | 1277.741743 | 1696.149232 | -0.010169806 | 0.098724151 | 0.816053503 | 0.958816322 | ns | NaCl | KS.3 |
| FIM1_1041 | 1746.640926 | 2976.50487 | 2064.977982 | 0.199514656 | 0.265517629 | 0.011812652 | 0.15408178 | ns | NaCl | KS.3 |
| FIM1_1042 | 65.98815293 | 108.0251623 | 77.10730827 | 0.27298116 | 0.268636584 | 0.003957087 | 0.081686757 | ns | NaCl | KS.3 |
| FIM1_1043 | 132.7251969 | 163.5943755 | 188.1524893 | 0.044727818 | 0.119357368 | 0.300268755 | 0.717845774 | ns | NaCl | KS.3 |
| FIM1_1044 | 2040.464928 | 1841.907687 | 2014.807761 | -0.026155793 | 0.104387339 | 0.544343201 | 0.857313452 | ns | NaCl | KS.3 |
| FIM1_1045 | 399.3447058 | 364.3234728 | 478.7153123 | -0.040656722 | 0.089053419 | 0.401369351 | 0.789189152 | ns | NaCl | KS.3 |
| FIM1_1046 | 618.9556118 | 520.4376351 | 813.5084914 | -0.093019799 | 0.094517999 | 0.076617985 | 0.429734472 | ns | NaCl | KS.3 |
| FIM1_1047 | 1258.891431 | 1257.704126 | 1493.593789 | -0.001478505 | 0.061685239 | 0.974934598 | 0.995954749 | ns | NaCl | KS.3 |

| | | | | | | | | | | |
|---|---|---|---|---|---|---|---|---|---|---|
| FIM1_1048 | 1808.659576 | 1852.065515 | 1727.147317 | 0.014483256 | 0.074684169 | 0.768573224 | 0.948835638 | ns | NaCl | KS.3 |
| FIM1_1049 | 4265.503995 | 6030.022751 | 2601.580577 | 0.06460895 | 0.150121307 | 0.153543537 | 0.570975106 | ns | NaCl | KS.3 |
| FIM1_1050 | 43.91400394 | 30.97674488 | 107.7630703 | -0.055774924 | 0.150039922 | 0.194050726 | 0.618952432 | ns | NaCl | KS.3 |
| FIM1_1051 | 509.6624864 | 439.9672973 | 591.553105 | -0.053940942 | 0.09831659 | 0.261316616 | 0.686178949 | ns | NaCl | KS.3 |
| FIM1_1052 | 440.6871466 | 353.3691702 | 1052.997585 | -0.065952321 | 0.116761683 | 0.166584533 | 0.590711667 | ns | NaCl | KS.3 |
| FIM1_1053 | 72.7647872 | 92.4250267 | 89.13311434 | 0.060127965 | 0.119699055 | 0.193922421 | 0.618952432 | ns | NaCl | KS.3 |
| FIM1_1054 | 3003.293116 | 3195.181952 | 2238.829363 | 0.04349041 | 0.07143173 | 0.378498768 | 0.778493059 | ns | NaCl | KS.3 |
| FIM1_1055 | 125.8776732 | 112.6422117 | 204.8296976 | -0.033512348 | 0.101344558 | 0.455099873 | 0.816788914 | ns | NaCl | KS.3 |
| FIM1_1056 | 619.5970295 | 641.0533006 | 564.8640782 | 0.015042071 | 0.087774648 | 0.751490354 | 0.943392508 | ns | NaCl | KS.3 |
| FIM1_1057 | 1280.115287 | 1197.246082 | 1415.285574 | -0.04276973 | 0.075236663 | 0.389376463 | 0.779482559 | ns | NaCl | KS.3 |
| FIM1_1058 | 2939.308906 | 2745.602423 | 2535.405671 | -0.043656015 | 0.074952753 | 0.379690176 | 0.778493059 | ns | NaCl | KS.3 |
| FIM1_1060 | 7178.672964 | 9150.694674 | 4530.229662 | 0.082774583 | 0.120993811 | 0.10151018 | 0.485353528 | ns | NaCl | KS.3 |
| FIM1_1061 | 943.3697889 | 934.610953 | 795.8899212 | -0.003189681 | 0.08468772 | 0.947101434 | 0.987736646 | ns | NaCl | KS.3 |
| FIM1_1062 | 763.9973403 | 782.6177875 | 854.6353268 | 0.013815165 | 0.075116015 | 0.778957538 | 0.951027591 | ns | NaCl | KS.3 |
| FIM1_1063 | 475.0130673 | 436.0186028 | 789.7390678 | -0.038564946 | 0.08902005 | 0.424775502 | 0.802854185 | ns | NaCl | KS.3 |
| FIM1_1064 | 759.8703385 | 839.5807673 | 1260.751839 | 0.041251919 | 0.09217971 | 0.388825356 | 0.779482559 | ns | NaCl | KS.3 |
| FIM1_1065 | 1308.904099 | 1219.890906 | 1242.392912 | -0.048905737 | 0.073471475 | 0.325157798 | 0.739131368 | ns | NaCl | KS.3 |
| FIM1_1066 | 451.9064108 | 492.4811103 | 442.4206693 | 0.046567546 | 0.083325391 | 0.347743909 | 0.756654396 | ns | NaCl | KS.3 |
| FIM1_1067 | 339.4665145 | 329.7713793 | 409.8270158 | -0.0113324 | 0.086468252 | 0.812399051 | 0.958816322 | ns | NaCl | KS.3 |
| FIM1_1068 | 197.7870109 | 372.7777899 | 234.1648906 | 0.494030466 | 0.368713278 | 0.000798441 | 0.031277519 | ns | NaCl | KS.3 |
| FIM1_1069 | 249.1974061 | 236.6512807 | 417.8969508 | -0.008459984 | 0.114007401 | 0.826398074 | 0.960240659 | ns | NaCl | KS.3 |
| FIM1_1070 | 704.9482526 | 688.9282209 | 1189.605498 | -0.015656416 | 0.077021248 | 0.750340236 | 0.943274417 | ns | NaCl | KS.3 |
| FIM1_1071 | 1625.647044 | 1637.268179 | 1522.70044 | 0.007576266 | 0.058361595 | 0.868936822 | 0.971243188 | ns | NaCl | KS.3 |
| FIM1_1072 | 2082.096486 | 2557.88974 | 1457.758092 | 0.031219367 | 0.127235641 | 0.425670866 | 0.802854185 | ns | NaCl | KS.3 |
| FIM1_1074 | 1341.406343 | 1419.149894 | 1221.662544 | 0.031404865 | 0.08064007 | 0.523820652 | 0.854940852 | ns | NaCl | KS.3 |
| FIM1_1075 | 3486.70859 | 3680.226956 | 3065.067165 | 0.027747996 | 0.082542352 | 0.570711417 | 0.870665206 | ns | NaCl | KS.3 |
| FIM1_1076 | 440.2246729 | 352.9125084 | 428.2304838 | -0.056080184 | 0.119659094 | 0.217456032 | 0.641504077 | ns | NaCl | KS.3 |
| FIM1_1077 | 159.8096138 | 179.4658963 | 125.6029179 | 0.017786346 | 0.118568637 | 0.643193315 | 0.898798052 | ns | NaCl | KS.3 |
| FIM1_1078 | 328.1275177 | 329.9975551 | 294.9642808 | 0.002165595 | 0.092278894 | 0.962352232 | 0.993505365 | ns | NaCl | KS.3 |
| FIM1_1079 | 311.6275127 | 265.3768755 | 387.3355343 | -0.050659154 | 0.106327282 | 0.273356572 | 0.694726351 | ns | NaCl | KS.3 |
| FIM1_1080 | 1263.91429 | 994.3842262 | 1084.702492 | -0.061727934 | 0.123065866 | 0.1826066 | 0.611396712 | ns | NaCl | KS.3 |
| FIM1_1081 | 1002.60566 | 1055.004906 | 1316.689336 | 0.009464636 | 0.111039112 | 0.810576802 | 0.958816322 | ns | NaCl | KS.3 |
| FIM1_1082 | 1139.606321 | 1365.053558 | 1101.869711 | 0.05103783 | 0.110220575 | 0.264121666 | 0.688347233 | ns | NaCl | KS.3 |
| FIM1_1083 | 1121.146959 | 1204.690528 | 1554.803504 | 0.045355697 | 0.075490331 | 0.362053519 | 0.763449629 | ns | NaCl | KS.3 |
| FIM1_1084 | 205.4674435 | 579.6407479 | 682.8854909 | 1.398968803 | 0.203347094 | 7.3596107457[illegible] | 6.1023439100[illegible] | Km>KS.3 | NaCl | KS.3 |
| FIM1_1085 | 515.6131646 | 510.2788332 | 580.9697454 | -0.007717052 | 0.078008661 | 0.875063856 | 0.971808705 | ns | NaCl | KS.3 |
| FIM1_1086 | 380.6727377 | 344.2612387 | 635.0051541 | -0.041981374 | 0.091983823 | 0.3814915 | 0.778609403 | ns | NaCl | KS.3 |
| FIM1_1087 | 664.8355502 | 686.0859218 | 755.4464923 | 0.014562905 | 0.084235744 | 0.763081302 | 0.946287744 | ns | NaCl | KS.3 |
| FIM1_1088 | 2395.990016 | 2563.878254 | 2300.985618 | 0.053619403 | 0.066449117 | 0.270016109 | 0.6917251 | ns | NaCl | KS.3 |
| FIM1_1089 | 143.5115447 | 118.4833027 | 127.3354343 | -0.056006311 | 0.11462508 | 0.223593468 | 0.648059715 | ns | NaCl | KS.3 |
| FIM1_1090 | 5973.881096 | 6170.436344 | 4998.7929 | 0.019522985 | 0.076683226 | 0.691884056 | 0.91988055 | ns | NaCl | KS.3 |
| FIM1_1091 | 3664.649252 | 4735.700338 | 2368.850307 | 0.070598844 | 0.126451716 | 0.140735262 | 0.552175021 | ns | NaCl | KS.3 |
| FIM1_1092 | 907.0905649 | 938.8716212 | 980.3221085 | 0.026023038 | 0.069584434 | 0.594770132 | 0.87962202 | ns | NaCl | KS.3 |
| FIM1_1093 | 785.0963071 | 722.7460335 | 1159.130995 | -0.036690539 | 0.088487022 | 0.447519571 | 0.812914509 | ns | NaCl | KS.3 |
| FIM1_1094 | 2036.258567 | 1970.123189 | 1676.028457 | -0.032550378 | 0.053312607 | 0.456990867 | 0.817337042 | ns | NaCl | KS.3 |

| | | | | | | | | | | |
|---|---|---|---|---|---|---|---|---|---|---|
| FIM1_1095 | 1027.873458 | 1053.433906 | 1197.186712 | 0.015215888 | 0.074063028 | 0.757188948 | 0.945250644 | ns | NaCl | KS.3 |
| FIM1_1096 | 449.9227606 | 463.6887349 | 436.598392 | 0.011010316 | 0.092563298 | 0.81058756 | 0.958816322 | ns | NaCl | KS.3 |
| FIM1_1097 | 8924.094642 | 11544.95255 | 8368.502534 | 0.109496336 | 0.125142039 | 0.046747556 | 0.341725286 | ns | NaCl | KS.3 |
| FIM1_1098 | 4776.587275 | 4402.963094 | 3522.134248 | -0.030219551 | 0.093375402 | 0.517682247 | 0.850188733 | ns | NaCl | KS.3 |
| FIM1_1099 | 511.9443188 | 432.2848235 | 822.1322244 | -0.072279793 | 0.100198419 | 0.148954393 | 0.563164954 | ns | NaCl | KS.3 |
| FIM1_1100 | 10570.71022 | 11275.39589 | 7850.830426 | 0.049419762 | 0.067771721 | 0.31178316 | 0.731871738 | ns | NaCl | KS.3 |
| FIM1_1101 | 32.68442771 | 56.55722221 | 78.11312331 | 0.051731689 | 0.182658 | 0.189314314 | 0.618952432 | ns | NaCl | KS.3 |
| FIM1_1102 | 1496.635233 | 1429.306601 | 1220.438134 | -0.023793986 | 0.082406271 | 0.626156539 | 0.89540925 | ns | NaCl | KS.3 |
| FIM1_1103 | 1107.323323 | 1183.992919 | 973.7573959 | 0.015701973 | 0.104708592 | 0.708269546 | 0.925190614 | ns | NaCl | KS.3 |
| FIM1_1104 | 1545.962075 | 1564.512841 | 1469.220352 | 0.004176492 | 0.093008627 | 0.927093552 | 0.984176127 | ns | NaCl | KS.3 |
| FIM1_1105 | 7065.388935 | 8456.208765 | 4789.31649 | 0.17966114 | 0.070404741 | 0.000478749 | 0.022683569 | ns | NaCl | KS.3 |
| FIM1_1106 | 7977.594516 | 7154.707025 | 6673.423361 | -0.045975705 | 0.091757401 | 0.341623768 | 0.750617902 | ns | NaCl | KS.3 |
| FIM1_1107 | 1189.92844 | 1109.850225 | 984.9406452 | -0.028832271 | 0.089745726 | 0.544413804 | 0.857313452 | ns | NaCl | KS.3 |
| FIM1_1108 | 2797.039995 | 4391.357721 | 1788.027479 | 0.125561594 | 0.20063351 | 0.036470751 | 0.302547392 | ns | NaCl | KS.3 |
| FIM1_1109 | 4471.014071 | 4364.276004 | 4698.985873 | -0.010145985 | 0.087482552 | 0.830739395 | 0.961620266 | ns | NaCl | KS.3 |
| FIM1_1110 | 1540.437176 | 1508.115844 | 1348.385957 | -0.009212406 | 0.087009392 | 0.846457155 | 0.967147129 | ns | NaCl | KS.3 |
| FIM1_1111 | 1033.171074 | 978.4964279 | 1127.910016 | -0.030844815 | 0.079726387 | 0.531750638 | 0.855153333 | ns | NaCl | KS.3 |
| FIM1_1112 | 3196.601588 | 3613.390054 | 3978.287789 | 0.078210323 | 0.080322483 | 0.126289515 | 0.534982546 | ns | NaCl | KS.3 |
| FIM1_1113 | 1319.725259 | 1669.505964 | 1419.349751 | 0.052812814 | 0.123913806 | 0.233600249 | 0.656217525 | ns | NaCl | KS.3 |
| FIM1_1114 | 458.1515885 | 480.8062838 | 2368.019001 | 0.006196941 | 0.122649922 | 0.864533856 | 0.970128052 | ns | NaCl | KS.3 |
| FIM1_1115 | 188.4735675 | 188.4313831 | 361.7384784 | -0.0008876 | 0.106240737 | 0.982615769 | 0.996842057 | ns | NaCl | KS.3 |
| FIM1_1116 | 8171.058622 | 6873.676447 | 6624.83979 | -0.080019038 | 0.098190521 | 0.117596661 | 0.520518737 | ns | NaCl | KS.3 |
| FIM1_1117 | 1173.353664 | 1008.561372 | 1887.62211 | -0.064089503 | 0.097167051 | 0.194323832 | 0.618952432 | ns | NaCl | KS.3 |
| FIM1_1118 | 10.5445453 | 21.01045139 | 20.87135697 | 0.153870414 | 0.289232965 | 0.023090268 | 0.23491633 | ns | NaCl | KS.3 |
| FIM1_1119 | 2655.976263 | 2923.246832 | 2937.395474 | 0.047596856 | 0.08618329 | 0.334681711 | 0.745307602 | ns | NaCl | KS.3 |
| FIM1_1120 | 25.16501668 | 17.07567628 | 43.48615303 | -0.06060255 | 0.154634097 | 0.168086483 | 0.592944764 | ns | NaCl | KS.3 |
| FIM1_1121 | 13732.39852 | 13485.19329 | 12996.19495 | -0.008299395 | 0.085076933 | 0.862816426 | 0.969856868 | ns | NaCl | KS.3 |
| FIM1_1122 | 6093.444287 | 6197.285733 | 5340.92505 | 0.014228132 | 0.0614487 | 0.762079479 | 0.946287744 | ns | NaCl | KS.3 |
| FIM1_1123 | 411.5227872 | 372.2291866 | 448.2279881 | -0.044048776 | 0.089585138 | 0.364624929 | 0.765404651 | ns | NaCl | KS.3 |
| FIM1_1124 | 2686.491393 | 2460.443296 | 3235.534849 | -0.044033642 | 0.085119662 | 0.371481215 | 0.773001053 | ns | NaCl | KS.3 |
| FIM1_1125 | 30887.00897 | 35221.64283 | 21740.66123 | 0.070610218 | 0.087118829 | 0.164408727 | 0.589224032 | ns | NaCl | KS.3 |
| FIM1_1126 | 643.2925752 | 721.7115787 | 764.9335658 | 0.021695123 | 0.113609522 | 0.588461037 | 0.878629549 | ns | NaCl | KS.3 |
| FIM1_1127 | 222.8069227 | 183.85329 | 411.6442297 | -0.043506897 | 0.117055263 | 0.314647927 | 0.732852732 | ns | NaCl | KS.3 |
| FIM1_1128 | 164.4132959 | 99.89940194 | 192.2800568 | -0.149051738 | 0.227844915 | 0.024390025 | 0.242680752 | ns | NaCl | KS.3 |
| FIM1_1129 | 2985.456291 | 3152.254581 | 3127.035772 | 0.032497058 | 0.077421975 | 0.511303729 | 0.846104012 | ns | NaCl | KS.3 |
| FIM1_1130 | 378.233569 | 427.0058206 | 445.2423528 | 0.047449882 | 0.094977225 | 0.321692333 | 0.737180727 | ns | NaCl | KS.3 |
| FIM1_1131 | 1444.133209 | 1322.95228 | 1327.843283 | -0.054144753 | 0.077670808 | 0.279704763 | 0.700652348 | ns | NaCl | KS.3 |
| FIM1_1132 | 366.9725802 | 370.3445659 | 990.3518956 | 0.003518147 | 0.09951853 | 0.935133964 | 0.985446192 | ns | NaCl | KS.3 |
| FIM1_1133 | 496.81538 | 729.7004654 | 653.9017944 | 0.123488624 | 0.1774893 | 0.036613105 | 0.302575075 | ns | NaCl | KS.3 |
| FIM1_1134 | 310.6307942 | 291.1539044 | 307.6470545 | -0.012852381 | 0.107985869 | 0.752018521 | 0.943523399 | ns | NaCl | KS.3 |
| FIM1_1135 | 23078.72259 | 21746.17882 | 14001.82884 | -0.014497816 | 0.103452789 | 0.731717091 | 0.936960396 | ns | NaCl | KS.3 |
| FIM1_1136 | 725.7810183 | 683.5384978 | 1073.687292 | -0.028390851 | 0.085960237 | 0.557375784 | 0.863845647 | ns | NaCl | KS.3 |
| FIM1_1137 | 2727.611508 | 2494.886973 | 3821.963831 | -0.018894756 | 0.10853218 | 0.645667125 | 0.898899754 | ns | NaCl | KS.3 |
| FIM1_1138 | 324.5760215 | 316.4438027 | 592.9896694 | -0.012636239 | 0.084738577 | 0.79307674 | 0.955175866 | ns | NaCl | KS.3 |
| FIM1_1139 | 10570.83638 | 8763.455875 | 9084.169046 | -0.156004013 | 0.083701425 | 0.004520508 | 0.086695706 | ns | NaCl | KS.3 |

| | | | | | | | | | | |
|---|---|---|---|---|---|---|---|---|---|---|
| FIM1_1140 | 3097.35416 | 2510.617746 | 3299.452156 | -0.027748034 | 0.131021479 | 0.464077221 | 0.819880744 | ns | NaCl | KS.3 |
| FIM1_1141 | 3771.523508 | 3096.349314 | 3765.705268 | -0.033726467 | 0.123330113 | 0.402775695 | 0.789854857 | ns | NaCl | KS.3 |
| FIM1_1142 | 782.1650974 | 841.1820407 | 869.6513995 | 0.045189877 | 0.076574283 | 0.364120718 | 0.764991795 | ns | NaCl | KS.3 |
| FIM1_1143 | 1857.187617 | 1958.728774 | 1700.876473 | 0.038312584 | 0.069583755 | 0.434760644 | 0.807880183 | ns | NaCl | KS.3 |
| FIM1_1144 | 1019.86575 | 983.4742833 | 867.3689057 | -0.009723562 | 0.100775289 | 0.820885965 | 0.960240659 | ns | NaCl | KS.3 |
| FIM1_1145 | 12960.64135 | 15187.33537 | 9504.782481 | 0.126024466 | 0.077480762 | 0.016545754 | 0.190104212 | ns | NaCl | KS.3 |
| FIM1_1146 | 528.0964294 | 487.6346967 | 629.5973332 | -0.034420056 | 0.087693038 | 0.476371523 | 0.825190921 | ns | NaCl | KS.3 |
| FIM1_1147 | 5092.628029 | 5536.237557 | 3609.849336 | 0.056413049 | 0.074490733 | 0.258963498 | 0.682746901 | ns | NaCl | KS.3 |
| FIM1_1148 | 648.0637523 | 715.2528745 | 672.4674812 | 0.041842682 | 0.091127621 | 0.384666782 | 0.778665419 | ns | NaCl | KS.3 |
| FIM1_1149 | 273.1764022 | 316.8339752 | 286.7529395 | 0.054194901 | 0.099631397 | 0.257304583 | 0.680871076 | ns | NaCl | KS.3 |
| FIM1_1150 | 1090.762784 | 1106.546682 | 929.2402497 | 0.00570495 | 0.088200071 | 0.903799828 | 0.979395371 | ns | NaCl | KS.3 |
| FIM1_1151 | 7023.882343 | 6993.541239 | 6165.423123 | -0.001909348 | 0.085417672 | 0.96820646 | 0.99408528 | ns | NaCl | KS.3 |
| FIM1_1152 | 641.2316966 | 702.3125539 | 803.2191854 | 0.022469775 | 0.104519651 | 0.598573777 | 0.880167289 | ns | NaCl | KS.3 |
| FIM1_1153 | 63.69500422 | 36.81533464 | 75.4474694 | -0.283902178 | 0.318801361 | 0.004771142 | 0.089910726 | ns | NaCl | KS.3 |
| FIM1_1154 | 1173.061191 | 1137.253856 | 1354.118306 | -0.018868977 | 0.077254802 | 0.701538727 | 0.923421278 | ns | NaCl | KS.3 |
| FIM1_1155 | 80.86781144 | 94.04021944 | 96.14274037 | 0.019155474 | 0.125947034 | 0.605844417 | 0.883689315 | ns | NaCl | KS.3 |
| FIM1_1156 | 14363.302 | 13455.23761 | 9766.387008 | -0.036657635 | 0.080148241 | 0.458627826 | 0.818684405 | ns | NaCl | KS.3 |
| FIM1_1157 | 2557.656462 | 2349.906043 | 2947.350756 | -0.060284288 | 0.072558754 | 0.226570479 | 0.650801464 | ns | NaCl | KS.3 |
| FIM1_1158 | 326.2052705 | 246.0954016 | 1034.973562 | -0.060977675 | 0.134765297 | 0.178002079 | 0.606369404 | ns | NaCl | KS.3 |
| FIM1_1159 | 1977.612798 | 2214.571835 | 1818.015142 | 0.094429721 | 0.068131209 | 0.05700555 | 0.374640173 | ns | NaCl | KS.3 |
| FIM1_1160 | 19923.444 | 16298.18303 | 13109.57996 | -0.19190159 | 0.081173508 | 0.00051059 | 0.023520242 | ns | NaCl | KS.3 |
| FIM1_1161 | 94.85348926 | 75.05266136 | 118.611153 | -0.032740101 | 0.131941178 | 0.399489723 | 0.787737366 | ns | NaCl | KS.3 |
| FIM1_1162 | 6421.142303 | 4275.005434 | 6707.310668 | -0.089923007 | 0.171774448 | 0.078675748 | 0.434419361 | ns | NaCl | KS.3 |
| FIM1_1163 | 1088.088873 | 1151.823126 | 1155.211628 | 0.03871924 | 0.071476621 | 0.43236443 | 0.805772482 | ns | NaCl | KS.3 |
| FIM1_1164 | 43.49943049 | 41.22234249 | 38.21550992 | -0.010340938 | 0.116417335 | 0.785958508 | 0.95289761 | ns | NaCl | KS.3 |
| FIM1_1165 | 1034.917252 | 1081.487127 | 916.3283181 | 0.016896332 | 0.090784029 | 0.717550702 | 0.93000992 | ns | NaCl | KS.3 |
| FIM1_1166 | 5513.431834 | 6154.512181 | 6055.591906 | 0.051045557 | 0.089177369 | 0.299179345 | 0.717010926 | ns | NaCl | KS.3 |
| FIM1_1167 | 678.4836583 | 781.7484368 | 888.8186405 | 0.067695306 | 0.092526331 | 0.177599857 | 0.606007743 | ns | NaCl | KS.3 |
| FIM1_1168 | 1586.569591 | 1726.371299 | 1564.529 | 0.050359532 | 0.079126523 | 0.313540474 | 0.731871738 | ns | NaCl | KS.3 |
| FIM1_1169 | 1210.493014 | 1105.841619 | 944.2420511 | -0.037170193 | 0.090668439 | 0.437567821 | 0.808096352 | ns | NaCl | KS.3 |
| FIM1_1170 | 1274.329346 | 1341.323218 | 1068.3569 | 0.024255471 | 0.085460675 | 0.615748132 | 0.890507836 | ns | NaCl | KS.3 |
| FIM1_1171 | 45.55217085 | 99.72608748 | 54.07909024 | 0.891190824 | 0.320702578 | 2.40957618864 | 0.002497425 | ns | NaCl | KS.3 |
| FIM1_1172 | 3102.358288 | 3661.503852 | 2917.458834 | 0.15675104 | 0.070746167 | 0.002259917 | 0.058108556 | ns | NaCl | KS.3 |
| FIM1_1173 | 641.320809 | 686.2960871 | 611.6997115 | 0.021313438 | 0.096665778 | 0.636341887 | 0.898332475 | ns | NaCl | KS.3 |
| FIM1_1174 | 367.4815268 | 355.3624251 | 552.607912 | -0.007865619 | 0.103858666 | 0.850571985 | 0.967955712 | ns | NaCl | KS.3 |
| FIM1_1175 | 99.94359255 | 136.3600405 | 154.079058 | 0.075513783 | 0.141802096 | 0.11801825 | 0.520642976 | ns | NaCl | KS.3 |
| FIM1_1177 | 2403.885981 | 2725.224539 | 2264.533237 | 0.100873032 | 0.070985548 | 0.045617725 | 0.338223816 | ns | NaCl | KS.3 |
| FIM1_1178 | 20724.64333 | 27243.04045 | 14048.49393 | 0.162084555 | 0.133704477 | 0.009976319 | 0.140601095 | ns | NaCl | KS.3 |
| FIM1_1179 | 857.5892733 | 888.5129593 | 940.2419267 | 0.022133189 | 0.074412443 | 0.653436669 | 0.900747416 | ns | NaCl | KS.3 |
| FIM1_1180 | 1253.820555 | 1414.773742 | 1300.328035 | 0.054956901 | 0.090569586 | 0.264835787 | 0.68837933 | ns | NaCl | KS.3 |
| FIM1_1181 | 2536.739626 | 2171.605264 | 2233.461848 | -0.055321164 | 0.10115641 | 0.246659249 | 0.669893726 | ns | NaCl | KS.3 |
| FIM1_1182 | 106.6789227 | 142.1132333 | 106.0153218 | 0.097554579 | 0.138978633 | 0.066110574 | 0.403405574 | ns | NaCl | KS.3 |
| FIM1_1183 | 1798.943951 | 2475.319728 | 1349.595068 | 0.168602582 | 0.160405678 | 0.011090948 | 0.149938773 | ns | NaCl | KS.3 |
| FIM1_1184 | 671.5356598 | 636.5430763 | 592.4635588 | -0.021840275 | 0.089484707 | 0.644156232 | 0.898899754 | ns | NaCl | KS.3 |
| FIM1_1186 | 636.7966032 | 603.9080347 | 717.6328512 | -0.036027812 | 0.069014344 | 0.461568683 | 0.819880744 | ns | NaCl | KS.3 |

| | | | | | | | | | |
|---|---|---|---|---|---|---|---|---|---|
| FIM1_1187 | 7373.766487 | 7745.5566 | 6179.320756 | 0.049608377 | 0.051371582 | 0.246439087 | 0.669893726 | ns | NaCl | KS.3 |
| FIM1_1188 | 388.435404 | 398.0203927 | 1370.245573 | 0.006338388 | 0.101172462 | 0.881919726 | 0.974110648 | ns | NaCl | KS.3 |
| FIM1_1189 | 647.9210178 | 460.6262462 | 1684.525144 | -0.091051483 | 0.153341348 | 0.077810482 | 0.432946334 | ns | NaCl | KS.3 |
| FIM1_1190 | 190.0229965 | 147.0342035 | 919.1149451 | -0.055648844 | 0.12804851 | 0.211791057 | 0.635060483 | ns | NaCl | KS.3 |
| FIM1_1191 | 69.00389014 | 36.74675923 | 87.82668476 | -0.075525483 | 0.210451505 | 0.097529832 | 0.476631548 | ns | NaCl | KS.3 |
| FIM1_1192 | 1306.833971 | 1258.842776 | 1024.594112 | -0.013195827 | 0.093395074 | 0.772934721 | 0.94938462 | ns | NaCl | KS.3 |
| FIM1_1193 | 6007.535544 | 6666.665735 | 4984.269302 | 0.100971852 | 0.057954732 | 0.028341516 | 0.263328375 | ns | NaCl | KS.3 |
| FIM1_1194 | 4197.471181 | 4844.497596 | 3125.589537 | 0.090812972 | 0.083594938 | 0.080725104 | 0.438915185 | ns | NaCl | KS.3 |
| FIM1_1195 | 232.8824161 | 200.1020374 | 301.4078431 | -0.055344417 | 0.100617725 | 0.247304716 | 0.669893726 | ns | NaCl | KS.3 |
| FIM1_1196 | 22688.19547 | 27196.74557 | 16415.82363 | 0.110532444 | 0.092905653 | 0.040469107 | 0.317627561 | ns | NaCl | KS.3 |
| FIM1_1197 | 4011.651267 | 3704.516348 | 4171.618958 | -0.045299515 | 0.080279706 | 0.362553926 | 0.763634962 | ns | NaCl | KS.3 |
| FIM1_1198 | 1522.359677 | 1636.93059 | 1483.304579 | 0.022733868 | 0.097587803 | 0.612518746 | 0.889162818 | ns | NaCl | KS.3 |
| FIM1_1199 | 1206.515641 | 1418.91368 | 1383.38834 | 0.107791455 | 0.084960973 | 0.042181272 | 0.323956287 | ns | NaCl | KS.3 |
| FIM1_1200 | 5273.214934 | 5281.255836 | 4476.768721 | 0.000464032 | 0.098977046 | 0.991470324 | 0.999134858 | ns | NaCl | KS.3 |
| FIM1_1201 | 196.574224 | 186.7149311 | 205.2568071 | -0.018501784 | 0.094272382 | 0.685480971 | 0.916492296 | ns | NaCl | KS.3 |
| FIM1_1202 | 295.3135117 | 295.4678491 | 329.0836406 | -0.000361813 | 0.102988746 | 0.993109924 | 0.999134858 | ns | NaCl | KS.3 |
| FIM1_1203 | 880.5185613 | 645.7976389 | 1165.203984 | -0.069606224 | 0.1410556 | 0.138232637 | 0.547974001 | ns | NaCl | KS.3 |
| FIM1_1204 | 3122.40856 | 2813.24283 | 4190.507741 | -0.043357296 | 0.091849907 | 0.367481099 | 0.768177319 | ns | NaCl | KS.3 |
| FIM1_1205 | 639.2841052 | 613.9235615 | 847.5103943 | -0.020170774 | 0.083377286 | 0.678095415 | 0.913295325 | ns | NaCl | KS.3 |
| FIM1_1206 | 778.8515958 | 752.3912051 | 779.5023305 | -0.019759451 | 0.079029703 | 0.687609375 | 0.918490387 | ns | NaCl | KS.3 |
| FIM1_1207 | 1595.474996 | 1575.030096 | 1871.703589 | -0.008155602 | 0.075729952 | 0.868286612 | 0.971161397 | ns | NaCl | KS.3 |
| FIM1_1208 | 1596.12566 | 1429.246222 | 1630.311595 | -0.068392335 | 0.080850602 | 0.177551995 | 0.606007743 | ns | NaCl | KS.3 |
| FIM1_1209 | 1101.535598 | 1167.315353 | 1031.406901 | 0.042312129 | 0.071012392 | 0.390859539 | 0.779482559 | ns | NaCl | KS.3 |
| FIM1_1210 | 295.0965283 | 214.1262086 | 392.2183859 | -0.111787521 | 0.150808273 | 0.046018808 | 0.34018361 | ns | NaCl | KS.3 |
| FIM1_1211 | 847.832107 | 825.4215314 | 1425.858515 | -0.011054015 | 0.087399232 | 0.816002006 | 0.958816322 | ns | NaCl | KS.3 |
| FIM1_1212 | 14761.97413 | 16891.8509 | 9933.849973 | 0.090592632 | 0.08009066 | 0.07967823 | 0.437357005 | ns | NaCl | KS.3 |
| FIM1_1213 | 12084.37625 | 13059.73826 | 8591.439274 | 0.039061011 | 0.084264195 | 0.426640358 | 0.802854185 | ns | NaCl | KS.3 |
| FIM1_1214 | 25759.95738 | 29580.33655 | 18090.46764 | 0.105850569 | 0.07526256 | 0.039930045 | 0.315700318 | ns | NaCl | KS.3 |
| FIM1_1215 | 898.610089 | 842.2777053 | 930.5813569 | -0.043670757 | 0.073130072 | 0.378301181 | 0.778493059 | ns | NaCl | KS.3 |
| FIM1_1216 | 861.4699031 | 916.0704334 | 932.4318863 | 0.025153497 | 0.089751699 | 0.595412115 | 0.87962202 | ns | NaCl | KS.3 |
| FIM1_1217 | 6382.532815 | 8525.508811 | 3883.21751 | 0.058821704 | 0.136397791 | 0.187665793 | 0.618637711 | ns | NaCl | KS.3 |
| FIM1_1218 | 213.6047579 | 198.286121 | 430.1686544 | -0.029438741 | 0.094151455 | 0.526149279 | 0.855027345 | ns | NaCl | KS.3 |
| FIM1_1219 | 1410.579737 | 1514.257143 | 1348.642288 | 0.046290818 | 0.07567137 | 0.352542844 | 0.756912594 | ns | NaCl | KS.3 |
| FIM1_1220 | 593.9121705 | 556.0241976 | 623.5037668 | -0.019281459 | 0.098917553 | 0.662564061 | 0.904388188 | ns | NaCl | KS.3 |
| FIM1_1221 | 870.9072589 | 799.0993791 | 911.9478004 | -0.063419475 | 0.072176769 | 0.20347467 | 0.626494461 | ns | NaCl | KS.3 |
| FIM1_1222 | 5139.326929 | 4962.162581 | 5569.113365 | -0.036115187 | 0.050686324 | 0.394721446 | 0.782366212 | ns | NaCl | KS.3 |
| FIM1_1223 | 227.2885503 | 194.4719472 | 358.2853896 | -0.057934678 | 0.102193994 | 0.227049145 | 0.651424162 | ns | NaCl | KS.3 |
| FIM1_1224 | 243.419453 | 236.6699163 | 283.1675351 | -0.013079659 | 0.089280147 | 0.780923862 | 0.951646805 | ns | NaCl | KS.3 |
| FIM1_1225 | 2249.985385 | 2408.345102 | 2384.065711 | 0.029828913 | 0.088060869 | 0.534426235 | 0.855153333 | ns | NaCl | KS.3 |
| FIM1_1227 | 3298.625857 | 3628.779031 | 4298.800342 | 0.030246843 | 0.098330184 | 0.504613985 | 0.844040357 | ns | NaCl | KS.3 |
| FIM1_1228 | 1083.774952 | 1223.676291 | 1288.517643 | 0.052504669 | 0.092302706 | 0.282227091 | 0.703799386 | ns | NaCl | KS.3 |
| FIM1_1230 | 2807.24428 | 3249.066289 | 2123.311934 | 0.065576608 | 0.094354117 | 0.188245363 | 0.618637711 | ns | NaCl | KS.3 |
| FIM1_1231 | 4654.457312 | 5347.160044 | 3724.980557 | 0.056498831 | 0.095883233 | 0.245880347 | 0.669176546 | ns | NaCl | KS.3 |
| FIM1_1232 | 4563.215429 | 4660.498034 | 3261.024632 | 0.009139955 | 0.087366663 | 0.84734762 | 0.967147129 | ns | NaCl | KS.3 |
| FIM1_1233 | 1608.07144 | 1154.707303 | 1200.732231 | -0.12963506 | 0.158658099 | 0.029901068 | 0.271952126 | ns | NaCl | KS.3 |

| | | | | | | | | | | |
|---|---|---|---|---|---|---|---|---|---|---|
| FIM1_1234 | 11732.71094 | 13058.3401 | 8850.688557 | 0.07696581 | 0.073848505 | 0.127105517 | 0.534982546 | ns | NaCl | KS.3 |
| FIM1_1235 | 1791.705686 | 1956.604674 | 2019.227918 | 0.048504568 | 0.08270627 | 0.329449837 | 0.741299385 | ns | NaCl | KS.3 |
| FIM1_1236 | 493.0671474 | 415.833849 | 389.0121877 | -0.063331596 | 0.101933112 | 0.193798452 | 0.618952432 | ns | NaCl | KS.3 |
| FIM1_1239 | 2497.227179 | 2570.406073 | 2726.193429 | 0.009213897 | 0.09565472 | 0.837065918 | 0.962829707 | ns | NaCl | KS.3 |
| FIM1_1240 | 191.0052206 | 223.5246999 | 277.3851919 | 0.055977093 | 0.101033582 | 0.242153014 | 0.666322592 | ns | NaCl | KS.3 |
| FIM1_1242 | 650.2427602 | 671.3601536 | 774.0457167 | 0.015076339 | 0.084886192 | 0.754385343 | 0.944041568 | ns | NaCl | KS.3 |
| FIM1_1243 | 3597.674667 | 3861.873312 | 3450.124753 | 0.065619651 | 0.057185977 | 0.150020466 | 0.564839452 | ns | NaCl | KS.3 |
| FIM1_1244 | 677.4933239 | 656.0675475 | 884.1417876 | -0.020187381 | 0.079725604 | 0.680883548 | 0.914523664 | ns | NaCl | KS.3 |
| FIM1_1245 | 1170.443122 | 1099.455806 | 1256.399244 | -0.030621541 | 0.08450544 | 0.52982989 | 0.855153333 | ns | NaCl | KS.3 |
| FIM1_1246 | 549.8901987 | 566.5587085 | 571.5752836 | 0.019707452 | 0.075592044 | 0.689232407 | 0.918545734 | ns | NaCl | KS.3 |
| FIM1_1247 | 487.076452 | 401.1541063 | 887.4453676 | -0.097963853 | 0.101730947 | 0.065114634 | 0.400922405 | ns | NaCl | KS.3 |
| FIM1_1248 | 1789.276722 | 1852.491288 | 1548.077761 | 0.015765767 | 0.085419095 | 0.743027652 | 0.941460666 | ns | NaCl | KS.3 |
| FIM1_1249 | 4650.694736 | 6380.602345 | 4249.76047 | 0.06834901 | 0.143021201 | 0.142114096 | 0.554849885 | ns | NaCl | KS.3 |
| FIM1_1250 | 978.3509459 | 1352.400298 | 2503.527131 | 0.175134615 | 0.162780377 | 0.009566395 | 0.136760965 | ns | NaCl | KS.3 |
| FIM1_1251 | 3591.742156 | 3734.984405 | 3902.077065 | 0.024354014 | 0.075748811 | 0.621623149 | 0.892479944 | ns | NaCl | KS.3 |
| FIM1_1252 | 2286.247532 | 2420.849067 | 1817.874988 | 0.0254816 | 0.087063171 | 0.595784208 | 0.87962202 | ns | NaCl | KS.3 |
| FIM1_1253 | 347.4417752 | 305.7315382 | 403.9005518 | -0.077355224 | 0.083799892 | 0.131175564 | 0.538132312 | ns | NaCl | KS.3 |
| FIM1_1254 | 1584.659616 | 1577.572913 | 1261.360215 | -0.001527058 | 0.094845772 | 0.972905967 | 0.995313014 | ns | NaCl | KS.3 |
| FIM1_1255 | 3200.690908 | 3613.184476 | 2162.44669 | 0.021988898 | 0.114748201 | 0.581249406 | 0.874951829 | ns | NaCl | KS.3 |
| FIM1_1256 | 912.2271131 | 812.6756621 | 1269.138431 | -0.06446119 | 0.084397729 | 0.201690254 | 0.624846998 | ns | NaCl | KS.3 |
| FIM1_1257 | 1622.007783 | 1550.140771 | 1885.744787 | -0.02253384 | 0.08375742 | 0.64285041 | 0.898798052 | ns | NaCl | KS.3 |
| FIM1_1258 | 1145.671547 | 923.4520317 | 1152.889444 | -0.129616633 | 0.104040213 | 0.021746075 | 0.225389007 | ns | NaCl | KS.3 |
| FIM1_1259 | 188.8068962 | 188.859113 | 347.9151025 | 0.001059596 | 0.099743521 | 0.980393962 | 0.996363429 | ns | NaCl | KS.3 |
| FIM1_1261 | 3364.053712 | 3491.192829 | 2347.620967 | 0.012859781 | 0.093297174 | 0.778661621 | 0.951027591 | ns | NaCl | KS.3 |
| FIM1_1262 | 1703.852125 | 1663.558463 | 2501.382779 | -0.007018493 | 0.097513125 | 0.873428483 | 0.971243188 | ns | NaCl | KS.3 |
| FIM1_1263 | 2076.325753 | 2191.963576 | 2066.89919 | 0.021391468 | 0.090464433 | 0.648966691 | 0.899642849 | ns | NaCl | KS.3 |
| FIM1_1264 | 3475.950884 | 3614.860973 | 2953.260636 | 0.012735148 | 0.095162449 | 0.777689972 | 0.950849745 | ns | NaCl | KS.3 |
| FIM1_1265 | 5091.209159 | 4940.181979 | 5046.128415 | -0.011368265 | 0.090905088 | 0.80682018 | 0.957859861 | ns | NaCl | KS.3 |
| FIM1_1266 | 1124.661887 | 1171.110204 | 805.668196 | 0.013944622 | 0.094186841 | 0.7592146 | 0.945454978 | ns | NaCl | KS.3 |
| FIM1_1267 | 7005.236438 | 8904.612589 | 4172.094383 | 0.046505692 | 0.127032938 | 0.274995103 | 0.695399585 | ns | NaCl | KS.3 |
| FIM1_1268 | 666.7441286 | 638.5921089 | 779.0367007 | -0.019528672 | 0.086861492 | 0.683263446 | 0.915742361 | ns | NaCl | KS.3 |
| FIM1_1269 | 6066.664745 | 8063.276733 | 3666.06524 | 0.054690165 | 0.135657676 | 0.21080279 | 0.634379635 | ns | NaCl | KS.3 |
| FIM1_1270 | 3855.243844 | 4483.320368 | 2826.111165 | 0.086722136 | 0.08806894 | 0.094916231 | 0.470201108 | ns | NaCl | KS.3 |
| FIM1_1271 | 3778.346627 | 3022.615851 | 3309.465527 | -0.151706638 | 0.105155001 | 0.009508471 | 0.13642083 | ns | NaCl | KS.3 |
| FIM1_1272 | 9316.175553 | 10768.90808 | 7662.356285 | 0.104927423 | 0.078858129 | 0.044106339 | 0.330963856 | ns | NaCl | KS.3 |
| FIM1_1273 | 438.4461519 | 403.267399 | 549.5565453 | -0.028949531 | 0.095888356 | 0.528321716 | 0.855153333 | ns | NaCl | KS.3 |
| FIM1_1274 | 49.87037855 | 52.57348584 | 79.67623041 | 0.007120644 | 0.113878911 | 0.853367994 | 0.969345976 | ns | NaCl | KS.3 |
| FIM1_1275 | 2592.52947 | 2182.268382 | 2704.232452 | -0.097940704 | 0.092833429 | 0.064257931 | 0.398841896 | ns | NaCl | KS.3 |
| FIM1_1276 | 2654.489613 | 2805.749962 | 3207.325652 | 0.060303628 | 0.046019862 | 0.129379293 | 0.536832346 | ns | NaCl | KS.3 |
| FIM1_1277 | 1573.781732 | 1538.404373 | 2437.463365 | -0.00820526 | 0.091593266 | 0.859011163 | 0.969856868 | ns | NaCl | KS.3 |
| FIM1_1279 | 920.6254905 | 733.7848975 | 1120.380077 | -0.141800724 | 0.10928301 | 0.014726259 | 0.176113313 | ns | NaCl | KS.3 |
| FIM1_1280 | 5723.118242 | 5055.378278 | 7070.403574 | -0.077762456 | 0.081031125 | 0.128611346 | 0.536136415 | ns | NaCl | KS.3 |
| FIM1_1281 | 2757.631185 | 3155.048353 | 2186.258833 | 0.087479551 | 0.08141403 | 0.090425491 | 0.463573758 | ns | NaCl | KS.3 |
| FIM1_1282 | 596.3744483 | 594.3737671 | 725.1315443 | -0.000594388 | 0.110424851 | 0.98794922 | 0.999134858 | ns | NaCl | KS.3 |
| FIM1_1283 | 2870.630464 | 2110.742201 | 3946.865104 | -0.070432956 | 0.140715183 | 0.135360961 | 0.543081274 | ns | NaCl | KS.3 |

| | | | | | | | | | | |
|---|---|---|---|---|---|---|---|---|---|---|
| FIM1_1284 | 602.3472339 | 635.7092072 | 937.7489032 | 0.014006387 | 0.101851855 | 0.743495452 | 0.941460666 | ns | NaCl | KS.3 |
| FIM1_1285 | 4866.792995 | 5057.479594 | 4577.267726 | 0.025974036 | 0.072214358 | 0.59768333 | 0.880167289 | ns | NaCl | KS.3 |
| FIM1_1286 | 455.6597535 | 549.3237467 | 438.3162663 | 0.07195891 | 0.104445699 | 0.147654858 | 0.562037426 | ns | NaCl | KS.3 |
| FIM1_1287 | 1176.390207 | 954.8075259 | 1377.703421 | -0.058477406 | 0.115259359 | 0.207692331 | 0.633002483 | ns | NaCl | KS.3 |
| FIM1_1288 | 682.8056126 | 637.7012114 | 608.8673932 | -0.032015013 | 0.085696497 | 0.509994167 | 0.845956462 | ns | NaCl | KS.3 |
| FIM1_1289 | 2915.758511 | 3088.886474 | 2209.892379 | 0.035201215 | 0.076919961 | 0.477409799 | 0.825839273 | ns | NaCl | KS.3 |
| FIM1_1290 | 3879.885695 | 3428.088747 | 4253.796382 | -0.065596005 | 0.08675944 | 0.193542924 | 0.618952432 | ns | NaCl | KS.3 |
| FIM1_1291 | 1245.239487 | 1333.742152 | 1057.93371 | 0.024372558 | 0.093679927 | 0.597965389 | 0.880167289 | ns | NaCl | KS.3 |
| FIM1_1292 | 289.1012327 | 314.2413418 | 242.678051 | 0.027918483 | 0.09520726 | 0.544100916 | 0.857313452 | ns | NaCl | KS.3 |
| FIM1_1293 | 3820.337502 | 5224.90572 | 2514.956573 | 0.10415687 | 0.146358884 | 0.055719291 | 0.369084797 | ns | NaCl | KS.3 |
| FIM1_1294 | 1672.987714 | 1833.004503 | 1168.931978 | 0.01670007 | 0.112723923 | 0.673846058 | 0.91246166 | ns | NaCl | KS.3 |
| FIM1_1295 | 2404.812259 | 2721.812037 | 3331.000546 | 0.065286254 | 0.086407772 | 0.19563735 | 0.618952432 | ns | NaCl | KS.3 |
| FIM1_1296 | 286.5071569 | 319.4923913 | 921.9107713 | 0.014531097 | 0.12312828 | 0.694510177 | 0.920646984 | ns | NaCl | KS.3 |
| FIM1_1297 | 1347.355013 | 752.2352653 | 1960.959462 | -0.258356348 | 0.309118048 | 0.00624087 | 0.106434152 | ns | NaCl | KS.3 |
| FIM1_1298 | 1734.293297 | 1359.058099 | 2757.110115 | -0.074245916 | 0.122159041 | 0.128913705 | 0.536563223 | ns | NaCl | KS.3 |
| FIM1_1299 | 896.0319298 | 968.7621675 | 985.6162665 | 0.052167404 | 0.074814466 | 0.29574258 | 0.714058464 | ns | NaCl | KS.3 |
| FIM1_1300 | 1676.172972 | 1900.025737 | 1455.409244 | 0.055884848 | 0.092065586 | 0.255695869 | 0.678991959 | ns | NaCl | KS.3 |
| FIM1_1301 | 259.8565127 | 249.532764 | 342.8589182 | -0.018146677 | 0.089578578 | 0.700077772 | 0.922863518 | ns | NaCl | KS.3 |
| FIM1_1302 | 626.9046872 | 730.1017452 | 736.6684656 | 0.090701439 | 0.087138971 | 0.082158602 | 0.443317834 | ns | NaCl | KS.3 |
| FIM1_1303 | 3039.001352 | 4047.764389 | 2040.5456 | 0.101274391 | 0.136244334 | 0.059753809 | 0.382593564 | ns | NaCl | KS.3 |
| FIM1_1304 | 1342.059989 | 1640.988182 | 1576.531809 | 0.042801192 | 0.119176201 | 0.317424481 | 0.73382286 | ns | NaCl | KS.3 |
| FIM1_1305 | 4315.58837 | 4126.069637 | 4427.895953 | -0.019091408 | 0.087666484 | 0.688681601 | 0.918490387 | ns | NaCl | KS.3 |
| FIM1_1306 | 79.22777126 | 72.13274117 | 117.1812301 | -0.017174976 | 0.106954954 | 0.678133252 | 0.913295325 | ns | NaCl | KS.3 |
| FIM1_1307 | 1084.633159 | 1065.348121 | 1081.113627 | -0.009494506 | 0.081878192 | 0.845291961 | 0.967147129 | ns | NaCl | KS.3 |
| FIM1_1308 | 889.918716 | 945.7414722 | 883.5704345 | 0.033682317 | 0.080165763 | 0.495097579 | 0.836358994 | ns | NaCl | KS.3 |
| FIM1_1309 | 1176.468199 | 924.7413314 | 1078.806146 | -0.112195163 | 0.117054924 | 0.04226083 | 0.323956287 | ns | NaCl | KS.3 |
| FIM1_1310 | 2616.6166 | 2904.215864 | 2115.586315 | 0.069614242 | 0.076796191 | 0.16859452 | 0.592944764 | ns | NaCl | KS.3 |
| FIM1_1311 | 1599.428476 | 3617.860855 | 2063.53469 | 0.976384654 | 0.293261387 | 6.69106329044 | 0.000876001 | ns | NaCl | KS.3 |
| FIM1_1312 | 1475.093281 | 1561.235318 | 1879.694692 | 0.01485174 | 0.101788876 | 0.729092372 | 0.935336397 | ns | NaCl | KS.3 |
| FIM1_1313 | 42503.43833 | 42552.966 | 32699.65459 | 0.000514369 | 0.085776876 | 0.991416965 | 0.999134858 | ns | NaCl | KS.3 |
| FIM1_1314 | 1035.033628 | 1152.028857 | 895.8018927 | 0.03062153 | 0.101585662 | 0.490976484 | 0.835079661 | ns | NaCl | KS.3 |
| FIM1_1315 | 4917.893932 | 4845.666558 | 3495.315727 | -0.005517033 | 0.091246011 | 0.905069388 | 0.979880338 | ns | NaCl | KS.3 |
| FIM1_1316 | 1029.905475 | 501.915078 | 918.0455375 | -0.891732456 | 0.239711992 | 2.93299319708 | 0.000470698 | ns | NaCl | KS.3 |
| FIM1_1317 | 788.2247783 | 685.7041079 | 631.2855543 | -0.048157603 | 0.100044759 | 0.305519877 | 0.724521545 | ns | NaCl | KS.3 |
| FIM1_1318 | 1505.149625 | 1365.300599 | 1423.226802 | -0.048640065 | 0.086151139 | 0.324846674 | 0.739131368 | ns | NaCl | KS.3 |
| FIM1_1319 | 3153.37205 | 2971.37335 | 3635.495891 | -0.015427833 | 0.101740593 | 0.719382063 | 0.930557921 | ns | NaCl | KS.3 |
| FIM1_1320 | 18674.55546 | 15603.21349 | 13061.52587 | -0.057249867 | 0.107639528 | 0.224313333 | 0.648059715 | ns | NaCl | KS.3 |
| FIM1_1321 | 809.840636 | 699.6862581 | 788.6044429 | -0.083552319 | 0.088020512 | 0.106070603 | 0.49734665 | ns | NaCl | KS.3 |
| FIM1_1322 | 10168.32225 | 8470.059273 | 7929.863146 | -0.051028455 | 0.110826275 | 0.263253723 | 0.688222424 | ns | NaCl | KS.3 |
| FIM1_1323 | 336.138957 | 259.1606388 | 817.2490221 | -0.099767432 | 0.125930067 | 0.062067797 | 0.390501742 | ns | NaCl | KS.3 |
| FIM1_1324 | 1608.434818 | 2113.715003 | 1164.683031 | 0.06390232 | 0.131749494 | 0.166131712 | 0.590360903 | ns | NaCl | KS.3 |
| FIM1_1325 | 1181.903755 | 1165.625041 | 1963.284815 | -0.005745121 | 0.087907163 | 0.903295339 | 0.979395371 | ns | NaCl | KS.3 |
| FIM1_1326 | 754.0509766 | 833.9181222 | 1051.740856 | 0.064847823 | 0.077369968 | 0.198577861 | 0.620924909 | ns | NaCl | KS.3 |
| FIM1_1327 | 1610.88998 | 1279.69354 | 2187.925136 | -0.176119485 | 0.106262565 | 0.003577733 | 0.076720786 | ns | NaCl | KS.3 |
| FIM1_1328 | 1116.217893 | 1106.922529 | 1573.607562 | -0.005760404 | 0.062646073 | 0.903163049 | 0.979395371 | ns | NaCl | KS.3 |

| | | | | | | | | | | |
|---|---|---|---|---|---|---|---|---|---|---|
| FIM1_1329 | 588.4826982 | 547.3416186 | 1818.801545 | -0.017623938 | 0.103867454 | 0.677693162 | 0.913287355 | ns | NaCl | KS.3 |
| FIM1_1331 | 85.2381298 | 79.58652366 | 153.9760707 | -0.010910593 | 0.115344125 | 0.776324497 | 0.950410881 | ns | NaCl | KS.3 |
| FIM1_1332 | 441.8661226 | 422.0489588 | 565.8234931 | -0.023079861 | 0.08509842 | 0.633192297 | 0.898009483 | ns | NaCl | KS.3 |
| FIM1_1333 | 11550.24013 | 13743.98709 | 7441.272709 | 0.048370704 | 0.109598586 | 0.286038254 | 0.705523209 | ns | NaCl | KS.3 |
| FIM1_1334 | 3046.370311 | 2379.481784 | 3113.798748 | -0.085062497 | 0.122265385 | 0.09482793 | 0.470201108 | ns | NaCl | KS.3 |
| FIM1_1335 | 934.4416774 | 915.8934908 | 988.3723608 | -0.007324909 | 0.091981244 | 0.873605907 | 0.971243188 | ns | NaCl | KS.3 |
| FIM1_1336 | 1789.441012 | 2866.632782 | 1587.422816 | 0.160440211 | 0.224707002 | 0.01968522 | 0.21299357 | ns | NaCl | KS.3 |
| FIM1_1337 | 5935.550745 | 9114.998865 | 3774.680074 | 0.105801061 | 0.184620522 | 0.054658329 | 0.36402301 | ns | NaCl | KS.3 |
| FIM1_1338 | 464.2154212 | 306.1254626 | 857.7782986 | -0.226089467 | 0.225461382 | 0.005849707 | 0.102113302 | ns | NaCl | KS.3 |
| FIM1_1339 | 5879.522346 | 5562.232117 | 5129.626516 | -0.019011525 | 0.094305291 | 0.677411759 | 0.913287355 | ns | NaCl | KS.3 |
| FIM1_1340 | 2269.476486 | 2331.22768 | 2228.020992 | 0.0147988 | 0.078942624 | 0.762991513 | 0.946287744 | ns | NaCl | KS.3 |
| FIM1_1341 | 634.657388 | 604.3891701 | 703.1858564 | -0.012326914 | 0.102331867 | 0.771925915 | 0.94938462 | ns | NaCl | KS.3 |
| FIM1_1342 | 2784.540883 | 2880.84034 | 2388.158002 | 0.014666511 | 0.087112653 | 0.758290946 | 0.945250644 | ns | NaCl | KS.3 |
| FIM1_1344 | 248.4015604 | 222.1732404 | 431.1494487 | -0.050802258 | 0.088309143 | 0.302483624 | 0.72182705 | ns | NaCl | KS.3 |
| FIM1_1345 | 1125.758647 | 1501.852752 | 1529.858814 | 0.2186791 | 0.143438759 | 0.001935064 | 0.05348302 | ns | NaCl | KS.3 |
| FIM1_1346 | 11317.44868 | 11918.77706 | 8008.702261 | 0.030416197 | 0.077895346 | 0.538220651 | 0.857036524 | ns | NaCl | KS.3 |
| FIM1_1347 | 23766.44359 | 26102.89447 | 16702.93723 | 0.047707168 | 0.085033487 | 0.334954925 | 0.745307602 | ns | NaCl | KS.3 |
| FIM1_1348 | 12971.12589 | 15035.02851 | 10458.08256 | 0.125956019 | 0.071772853 | 0.013835265 | 0.170372387 | ns | NaCl | KS.3 |
| FIM1_1349 | 1093.579837 | 905.7766906 | 1253.590956 | -0.151310423 | 0.0879505 | 0.006574291 | 0.108271704 | ns | NaCl | KS.3 |
| FIM1_1350 | 11261.10175 | 12113.71155 | 8388.563378 | 0.02903704 | 0.090936186 | 0.539101939 | 0.857036524 | ns | NaCl | KS.3 |
| FIM1_1352 | 2224.807925 | 2181.405408 | 2375.419536 | -0.013958563 | 0.070601413 | 0.775580346 | 0.950385088 | ns | NaCl | KS.3 |
| FIM1_1353 | 7463.356039 | 7696.740876 | 8281.703894 | 0.022173933 | 0.06942921 | 0.650006914 | 0.899642849 | ns | NaCl | KS.3 |
| FIM1_1354 | 1075.391533 | 1040.627671 | 1056.633872 | -0.014104321 | 0.087936562 | 0.766192839 | 0.948447219 | ns | NaCl | KS.3 |
| FIM1_1355 | 1409.643506 | 1567.811511 | 1650.334581 | 0.098528933 | 0.06056406 | 0.036567808 | 0.302575075 | ns | NaCl | KS.3 |
| FIM1_1356 | 596.740673 | 604.5320106 | 718.3923749 | 0.006564058 | 0.077849614 | 0.893618724 | 0.977268396 | ns | NaCl | KS.3 |
| FIM1_1357 | 1004.254603 | 938.088508 | 1025.871029 | -0.028459375 | 0.089310133 | 0.550351622 | 0.860945181 | ns | NaCl | KS.3 |
| FIM1_1358 | 1915.125336 | 1625.663598 | 1315.886861 | -0.096285806 | 0.089707846 | 0.067583542 | 0.407549238 | ns | NaCl | KS.3 |
| FIM1_1359 | 435.5431028 | 434.4516842 | 495.7212686 | -0.001323571 | 0.086748178 | 0.977803294 | 0.996164232 | ns | NaCl | KS.3 |
| FIM1_1360 | 2991.218081 | 1938.239254 | 2675.62504 | -0.188711579 | 0.22192341 | 0.011491317 | 0.151642712 | ns | NaCl | KS.3 |
| FIM1_1361 | 252.2624964 | 171.1285717 | 279.5368154 | -0.224438256 | 0.203235093 | 0.004759446 | 0.089910726 | ns | NaCl | KS.3 |
| FIM1_1362 | 419.8319142 | 375.2143456 | 484.6343487 | -0.044382438 | 0.092846689 | 0.355176381 | 0.75934787 | ns | NaCl | KS.3 |
| FIM1_1363 | 6089.919341 | 7877.034861 | 3829.713585 | 0.054190986 | 0.129139985 | 0.219541417 | 0.642906874 | ns | NaCl | KS.3 |
| FIM1_1364 | 291.5789172 | 356.5967631 | 299.4874911 | 0.050304903 | 0.115788107 | 0.261296268 | 0.686178949 | ns | NaCl | KS.3 |
| FIM1_1365 | 2357.863082 | 2756.115481 | 2515.753909 | 0.067408014 | 0.096950992 | 0.175719918 | 0.605047427 | ns | NaCl | KS.3 |
| FIM1_1366 | 1758.406663 | 1847.751571 | 1390.123103 | 0.022809443 | 0.085898301 | 0.635971854 | 0.898332475 | ns | NaCl | KS.3 |
| FIM1_1367 | 650.5606081 | 547.0091861 | 982.0969972 | -0.124245678 | 0.085542963 | 0.021372044 | 0.222905493 | ns | NaCl | KS.3 |
| FIM1_1368 | 5746.498845 | 6001.709553 | 5774.293505 | 0.031898978 | 0.069393756 | 0.514612751 | 0.849077129 | ns | NaCl | KS.3 |
| FIM1_1369 | 4116.250911 | 6511.388263 | 3750.884313 | 0.371467629 | 0.26368264 | 0.000911427 | 0.033587767 | ns | NaCl | KS.3 |
| FIM1_1370 | 41.14054836 | 56.79911208 | 34.13613517 | 0.056051833 | 0.145703063 | 0.195610615 | 0.618952432 | ns | NaCl | KS.3 |
| FIM1_1371 | 993.8876228 | 1097.385469 | 1244.944897 | 0.063216694 | 0.077906024 | 0.209841775 | 0.633800632 | ns | NaCl | KS.3 |
| FIM1_1372 | 116.3519973 | 144.7576013 | 292.5230999 | 0.068817575 | 0.115014438 | 0.154615604 | 0.572502577 | ns | NaCl | KS.3 |
| FIM1_1373 | 1375.370821 | 2466.004404 | 2413.523477 | 0.804811485 | 0.117504058 | 9.35125302282 | 5.16916486539 | ns | NaCl | KS.3 |
| FIM1_1374 | 2183.498429 | 2141.772935 | 2208.060344 | -0.013248434 | 0.072516867 | 0.787409462 | 0.95289761 | ns | NaCl | KS.3 |
| FIM1_1375 | 1012.712656 | 1140.035661 | 1570.463931 | 0.020797479 | 0.115510197 | 0.598670593 | 0.880167289 | ns | NaCl | KS.3 |
| FIM1_1376 | 72.10040834 | 54.3607424 | 96.17648643 | -0.081288806 | 0.134442948 | 0.102802095 | 0.48941667 | ns | NaCl | KS.3 |

| | | | | | | | | | | |
|---|---|---|---|---|---|---|---|---|---|---|
| FIM1_1377 | 3788.397739 | 4145.603612 | 3826.541247 | 0.058578866 | 0.076260059 | 0.242944747 | 0.666657537 | ns | NaCl | KS.3 |
| FIM1_1378 | 373.0918189 | 332.8744758 | 321.4348399 | -0.044932309 | 0.09656133 | 0.342191236 | 0.750617902 | ns | NaCl | KS.3 |
| FIM1_1379 | 109.4701987 | 99.93056308 | 272.0880493 | -0.019625787 | 0.10687154 | 0.637604779 | 0.898332475 | ns | NaCl | KS.3 |
| FIM1_1380 | 1287.697643 | 1377.984437 | 2315.659662 | 0.014877549 | 0.106646437 | 0.718462426 | 0.930194994 | ns | NaCl | KS.3 |
| FIM1_1382 | 4038.454912 | 4402.243945 | 4035.366892 | 0.08224121 | 0.057135076 | 0.071449096 | 0.418675804 | ns | NaCl | KS.3 |
| FIM1_1383 | 2445.79002 | 2393.432807 | 2587.870545 | -0.012136352 | 0.078227466 | 0.804775988 | 0.957859861 | ns | NaCl | KS.3 |
| FIM1_1384 | 1397.260864 | 1499.185419 | 1200.455473 | 0.029790191 | 0.089087261 | 0.532940206 | 0.855153333 | ns | NaCl | KS.3 |
| FIM1_1385 | 3972.327688 | 4318.290983 | 5001.549149 | 0.030703838 | 0.093865699 | 0.5101245 | 0.845956462 | ns | NaCl | KS.3 |
| FIM1_1386 | 607.5551812 | 532.6770509 | 571.1931895 | -0.06543071 | 0.089588116 | 0.192889224 | 0.618952432 | ns | NaCl | KS.3 |
| FIM1_1387 | 3300.447571 | 3453.552122 | 2965.730192 | 0.049635552 | 0.044448701 | 0.200716769 | 0.624846998 | ns | NaCl | KS.3 |
| FIM1_1388 | 268.2098215 | 242.6406544 | 383.3514424 | -0.05494206 | 0.083893143 | 0.271607115 | 0.693144962 | ns | NaCl | KS.3 |
| FIM1_1389 | 472.0155838 | 396.0545278 | 608.2360905 | -0.102498178 | 0.09484282 | 0.054914873 | 0.364754995 | ns | NaCl | KS.3 |
| FIM1_1390 | 2306.758092 | 2292.522559 | 2724.645374 | -0.00195753 | 0.096394369 | 0.964825416 | 0.993691273 | ns | NaCl | KS.3 |
| FIM1_1391 | 2152.690461 | 2298.870676 | 3696.141313 | 0.009622204 | 0.118442039 | 0.797612077 | 0.955924408 | ns | NaCl | KS.3 |
| FIM1_1392 | 844.212673 | 522.4893314 | 1101.82239 | -0.454554591 | 0.261841908 | 0.000317943 | 0.017192375 | ns | NaCl | KS.3 |
| FIM1_1393 | 1028.7455 | 871.0859573 | 1062.710524 | -0.052765407 | 0.105050171 | 0.259343935 | 0.68302598 | ns | NaCl | KS.3 |
| FIM1_1394 | 149.4773164 | 73.73263504 | 587.753791 | -0.464833326 | 0.40370556 | 0.001270598 | 0.041599773 | ns | NaCl | KS.3 |
| FIM1_1395 | 512.7824423 | 529.0317332 | 753.8510585 | 0.006504642 | 0.106243176 | 0.873526735 | 0.971243188 | ns | NaCl | KS.3 |
| FIM1_1396 | 1482.757646 | 1367.273003 | 1338.63058 | -0.030163916 | 0.09305359 | 0.519192693 | 0.850629872 | ns | NaCl | KS.3 |
| FIM1_1397 | 4947.378079 | 4231.273618 | 4469.632152 | -0.078254061 | 0.092887277 | 0.126268894 | 0.534982546 | ns | NaCl | KS.3 |
| FIM1_1399 | 2665.687101 | 2453.015795 | 2805.973654 | -0.024196736 | 0.099824985 | 0.58495495 | 0.877826895 | ns | NaCl | KS.3 |
| FIM1_1400 | 1697.568192 | 1764.670651 | 1902.072314 | 0.010224143 | 0.100867784 | 0.811788888 | 0.958816322 | ns | NaCl | KS.3 |
| FIM1_1401 | 2613.725905 | 4128.228044 | 3542.249151 | 0.146495808 | 0.212891078 | 0.024705868 | 0.244844015 | ns | NaCl | KS.3 |
| FIM1_1402 | 794.261019 | 876.0249853 | 872.8792913 | 0.066503343 | 0.075312912 | 0.186395572 | 0.618044911 | ns | NaCl | KS.3 |
| FIM1_1403 | 623.7584135 | 637.7761831 | 747.890367 | 0.009817257 | 0.088663009 | 0.835000738 | 0.962365393 | ns | NaCl | KS.3 |
| FIM1_1404 | 769.5282736 | 502.3927386 | 894.6827909 | -0.092996704 | 0.178591379 | 0.072811142 | 0.421203992 | ns | NaCl | KS.3 |
| FIM1_1405 | 715.3848913 | 974.7781242 | 849.0081097 | 0.110642717 | 0.146687828 | 0.047074449 | 0.34239091 | ns | NaCl | KS.3 |
| FIM1_1406 | 5218.0025 | 6042.828674 | 6026.309982 | 0.03714759 | 0.10842154 | 0.395171629 | 0.782558381 | ns | NaCl | KS.3 |
| FIM1_1407 | 1387.215287 | 1263.311394 | 1369.341169 | -0.058130098 | 0.078867358 | 0.247390172 | 0.669893726 | ns | NaCl | KS.3 |
| FIM1_1408 | 7782.335591 | 9165.628945 | 7942.031075 | 0.096640765 | 0.089719414 | 0.066716519 | 0.404774002 | ns | NaCl | KS.3 |
| FIM1_1409 | 2544.908457 | 3771.813378 | 2107.260258 | 0.073344787 | 0.162942769 | 0.118794894 | 0.522550484 | ns | NaCl | KS.3 |
| FIM1_1410 | 4660.423142 | 5080.975004 | 3987.427979 | 0.037479854 | 0.089304447 | 0.436714462 | 0.808096352 | ns | NaCl | KS.3 |
| FIM1_1411 | 894.8721965 | 932.1162279 | 1205.529616 | 0.007024934 | 0.112610487 | 0.856617538 | 0.969856868 | ns | NaCl | KS.3 |
| FIM1_1412 | 4584.644965 | 5090.173947 | 4924.602873 | 0.030048293 | 0.101550513 | 0.498581005 | 0.837486406 | ns | NaCl | KS.3 |
| FIM1_1413 | 415.8579004 | 428.1778557 | 623.2557519 | 0.005295937 | 0.111871682 | 0.892027409 | 0.977268396 | ns | NaCl | KS.3 |
| FIM1_1414 | 251.7338101 | 297.8728478 | 305.0673911 | 0.055830276 | 0.105605314 | 0.236552188 | 0.659616806 | ns | NaCl | KS.3 |
| FIM1_1415 | 1147.307374 | 1323.399047 | 1607.211688 | 0.10970246 | 0.075392563 | 0.033707532 | 0.292771172 | ns | NaCl | KS.3 |
| FIM1_1416 | 687.5641865 | 644.336256 | 1016.078387 | -0.040805853 | 0.077272582 | 0.411468536 | 0.798695266 | ns | NaCl | KS.3 |
| FIM1_1418 | 1000.945926 | 1123.074456 | 1019.213384 | 0.084802819 | 0.074635382 | 0.09498535 | 0.470201108 | ns | NaCl | KS.3 |
| FIM1_1420 | 1396.108117 | 1572.209635 | 1437.044234 | 0.072431077 | 0.081873135 | 0.15507018 | 0.572735074 | ns | NaCl | KS.3 |
| FIM1_1421 | 402.8112259 | 401.9380549 | 553.3328384 | -0.00051713 | 0.104126804 | 0.990053785 | 0.999134858 | ns | NaCl | KS.3 |
| FIM1_1422 | 369.7911018 | 361.0896075 | 486.6103039 | -0.007441652 | 0.097383817 | 0.866047778 | 0.970184124 | ns | NaCl | KS.3 |
| FIM1_1423 | 821.4007986 | 666.90502 | 793.3963286 | -0.179550718 | 0.091704633 | 0.00185213 | 0.051765993 | ns | NaCl | KS.3 |
| FIM1_1424 | 1223.824187 | 1105.552148 | 1449.15332 | -0.08042282 | 0.068957312 | 0.104855161 | 0.494928296 | ns | NaCl | KS.3 |
| FIM1_1425 | 885.537204 | 789.004871 | 1331.415899 | -0.045260692 | 0.093315993 | 0.345607323 | 0.754462295 | ns | NaCl | KS.3 |

| | | | | | | | | | | |
|---|---|---|---|---|---|---|---|---|---|---|
| FIM1_1426 | 4240.40722 | 4883.901569 | 3731.823932 | 0.02059233 | 0.122401638 | 0.588053555 | 0.878311243 | ns | NaCl | KS.3 |
| FIM1_1427 | 27.9741719 | 36.96778432 | 38.6833852 | 0.039542271 | 0.135237841 | 0.32348907 | 0.739131368 | ns | NaCl | KS.3 |
| FIM1_1428 | 1892.327084 | 1703.318197 | 2119.551811 | -0.044779175 | 0.091391856 | 0.354021058 | 0.757528931 | ns | NaCl | KS.3 |
| FIM1_1429 | 2045.139935 | 2131.760072 | 2233.881656 | 0.015989403 | 0.090811463 | 0.731877418 | 0.936960396 | ns | NaCl | KS.3 |
| FIM1_1430 | 773.8676458 | 681.4053189 | 1121.095545 | -0.045495073 | 0.097796328 | 0.334239498 | 0.745307602 | ns | NaCl | KS.3 |
| FIM1_1431 | 1123.55052 | 1152.603356 | 2212.056116 | 0.008967614 | 0.094275677 | 0.843043836 | 0.966485679 | ns | NaCl | KS.3 |
| FIM1_1432 | 19981.77185 | 17130.13039 | 13282.27271 | -0.090001136 | 0.088117266 | 0.084465715 | 0.451260889 | ns | NaCl | KS.3 |
| FIM1_1433 | 1844.238076 | 1976.889981 | 1941.906229 | 0.048569271 | 0.072433102 | 0.327397027 | 0.740319311 | ns | NaCl | KS.3 |
| FIM1_1434 | 1672.145426 | 1865.270295 | 1918.794881 | 0.024762922 | 0.108031258 | 0.555024653 | 0.863429533 | ns | NaCl | KS.3 |
| FIM1_1435 | 5531.096538 | 5338.259992 | 5628.031343 | -0.01360368 | 0.090522605 | 0.770698282 | 0.949065335 | ns | NaCl | KS.3 |
| FIM1_1436 | 408.6055748 | 342.4642492 | 560.7691604 | -0.075017712 | 0.101844116 | 0.135941329 | 0.544093414 | ns | NaCl | KS.3 |
| FIM1_1437 | 372.5608108 | 340.4305155 | 339.2170924 | -0.054204163 | 0.079426501 | 0.279373349 | 0.700652348 | ns | NaCl | KS.3 |
| FIM1_1438 | 82193.80094 | 97722.2473 | 60172.18324 | 0.142819355 | 0.079445474 | 0.00763143 | 0.119762487 | ns | NaCl | KS.3 |
| FIM1_1439 | 722.840906 | 728.07637 | 1220.359772 | 0.004195145 | 0.083762085 | 0.930751904 | 0.984325045 | ns | NaCl | KS.3 |
| FIM1_1440 | 293.374359 | 314.2448216 | 401.8067455 | 0.034069702 | 0.08432359 | 0.486050697 | 0.83010718 | ns | NaCl | KS.3 |
| FIM1_1441 | 175.1118641 | 181.2453577 | 322.5394479 | 0.005166437 | 0.113558938 | 0.893310287 | 0.977268396 | ns | NaCl | KS.3 |
| FIM1_1442 | 42.17254604 | 65.4827935 | 578.1003268 | 0.079838054 | 0.175881829 | 0.098777468 | 0.480369407 | ns | NaCl | KS.3 |
| FIM1_1443 | 467.2048992 | 589.2357958 | 494.6692785 | 0.111048772 | 0.113635621 | 0.043396468 | 0.328611002 | ns | NaCl | KS.3 |
| FIM1_1444 | 17282.77519 | 20478.60927 | 12306.94573 | 0.085093282 | 0.095466419 | 0.100215832 | 0.483636126 | ns | NaCl | KS.3 |
| FIM1_1445 | 2326.942844 | 2809.115213 | 1243.352178 | 0.037627393 | 0.11882521 | 0.368277438 | 0.768249278 | ns | NaCl | KS.3 |
| FIM1_1446 | 401.5796449 | 390.9197643 | 640.2136342 | -0.011446404 | 0.090733057 | 0.805749197 | 0.957859861 | ns | NaCl | KS.3 |
| FIM1_1447 | 1040.576025 | 1158.943082 | 788.7167144 | 0.02857958 | 0.103613319 | 0.512673233 | 0.847552971 | ns | NaCl | KS.3 |
| FIM1_1448 | 570.674631 | 687.2006371 | 525.5591259 | 0.06021127 | 0.108256664 | 0.205125213 | 0.628759992 | ns | NaCl | KS.3 |
| FIM1_1449 | 3561.153873 | 3621.643265 | 2542.263048 | 0.010234488 | 0.076892033 | 0.835081688 | 0.962365393 | ns | NaCl | KS.3 |
| FIM1_1450 | 734.4273686 | 964.2095021 | 920.6746159 | 0.160330821 | 0.133035558 | 0.010458513 | 0.14533828 | ns | NaCl | KS.3 |
| FIM1_1451 | 9817.258892 | 6237.158373 | 8214.938459 | -0.183131142 | 0.22805173 | 0.013187674 | 0.165678479 | ns | NaCl | KS.3 |
| FIM1_1453 | 3918.703666 | 3687.577751 | 3848.212792 | -0.012325676 | 0.108386868 | 0.76091285 | 0.945911573 | ns | NaCl | KS.3 |
| FIM1_1454 | 726.2800733 | 773.9300822 | 1125.145479 | 0.0295024 | 0.086304602 | 0.541879074 | 0.857187397 | ns | NaCl | KS.3 |
| FIM1_1455 | 1217.918529 | 1689.70533 | 1192.667858 | 0.132257243 | 0.157284182 | 0.027829382 | 0.260899256 | ns | NaCl | KS.3 |
| FIM1_1456 | 500.4908193 | 730.593177 | 560.0456439 | 0.29315711 | 0.208985224 | 0.001265621 | 0.041599773 | ns | NaCl | KS.3 |
| FIM1_1457 | 2327.485558 | 2225.467529 | 2947.262511 | -0.01922571 | 0.088043154 | 0.686039329 | 0.916746082 | ns | NaCl | KS.3 |
| FIM1_1458 | 1548.521842 | 1484.373671 | 1399.980782 | -0.02163631 | 0.082614644 | 0.657219913 | 0.901977674 | ns | NaCl | KS.3 |
| FIM1_1460 | 1089.08096 | 1031.303173 | 1103.395768 | -0.040070958 | 0.069959891 | 0.414735921 | 0.800387073 | ns | NaCl | KS.3 |
| FIM1_1461 | 924.8610133 | 782.0947795 | 986.9451075 | -0.033885059 | 0.116146247 | 0.415012709 | 0.800387073 | ns | NaCl | KS.3 |
| FIM1_1462 | 1649.609598 | 1690.814504 | 1477.699355 | 0.015178804 | 0.074797637 | 0.757834633 | 0.945250644 | ns | NaCl | KS.3 |
| FIM1_1463 | 265.9922825 | 285.2724634 | 239.1789131 | 0.02179394 | 0.096613893 | 0.629143385 | 0.896016329 | ns | NaCl | KS.3 |
| FIM1_1464 | 14.51905366 | 8.732642682 | 24.69833755 | -0.039000088 | 0.177076613 | 0.274108327 | 0.694804019 | ns | NaCl | KS.3 |
| FIM1_1465 | 974.1881985 | 978.7179971 | 1340.776731 | 0.001309735 | 0.076951553 | 0.978727818 | 0.996164232 | ns | NaCl | KS.3 |
| FIM1_1466 | 955.7171618 | 974.6139507 | 1195.574456 | 0.010513916 | 0.077242752 | 0.830599903 | 0.961620266 | ns | NaCl | KS.3 |
| FIM1_1467 | 1731.055974 | 1667.24689 | 2349.248935 | -0.016156707 | 0.087401654 | 0.734436511 | 0.938375543 | ns | NaCl | KS.3 |
| FIM1_1468 | 3395.278553 | 3329.858802 | 3250.144257 | -0.012983488 | 0.071697616 | 0.791301622 | 0.954126411 | ns | NaCl | KS.3 |
| FIM1_1469 | 74.2264059 | 59.36433619 | 198.5580107 | -0.051318818 | 0.120577144 | 0.247625038 | 0.669893726 | ns | NaCl | KS.3 |
| FIM1_1470 | 1725.810392 | 1921.188056 | 1860.483098 | 0.09094491 | 0.066576201 | 0.064130535 | 0.398841896 | ns | NaCl | KS.3 |
| FIM1_1471 | 5846.4652 | 5432.751493 | 4424.68561 | -0.041108766 | 0.080634742 | 0.407026016 | 0.79441131 | ns | NaCl | KS.3 |
| FIM1_1472 | 1047.48744 | 890.8360216 | 1066.927055 | -0.070242159 | 0.097570508 | 0.160497619 | 0.58211082 | ns | NaCl | KS.3 |

| | | | | | | | | | | |
|---|---|---|---|---|---|---|---|---|---|---|
| FIM1_1473 | 1254.506155 | 1633.302099 | 1273.656244 | 0.176098317 | 0.127867797 | 0.0057587 | 0.100878634 | ns | NaCl | KS.3 |
| FIM1_1474 | 2875.863648 | 3011.396006 | 2614.068586 | 0.019657256 | 0.087961709 | 0.67960902 | 0.913941356 | ns | NaCl | KS.3 |
| FIM1_1475 | 1440.281249 | 1343.26333 | 1427.26863 | -0.048885162 | 0.073231986 | 0.325136878 | 0.739131368 | ns | NaCl | KS.3 |
| FIM1_1476 | 116.1365127 | 82.08192175 | 1083.108515 | -0.081536351 | 0.152567998 | 0.098918575 | 0.480585849 | ns | NaCl | KS.3 |
| FIM1_1477 | 1257.262241 | 1517.869032 | 1843.773248 | 0.04250945 | 0.116097389 | 0.325346764 | 0.739131368 | ns | NaCl | KS.3 |
| FIM1_1478 | 961.5362991 | 924.7919106 | 1710.692423 | -0.009082309 | 0.105096994 | 0.826258713 | 0.960240659 | ns | NaCl | KS.3 |
| FIM1_1479 | 547.2721447 | 480.099398 | 809.3494458 | -0.088153999 | 0.080182742 | 0.087514865 | 0.45830153 | ns | NaCl | KS.3 |
| FIM1_1480 | 1174.806424 | 1360.5489 | 905.3788425 | 0.039605591 | 0.106814267 | 0.372569326 | 0.77466206 | ns | NaCl | KS.3 |
| FIM1_1481 | 452.1561856 | 419.9456711 | 507.7294437 | -0.025424241 | 0.095867852 | 0.577421054 | 0.874645237 | ns | NaCl | KS.3 |
| FIM1_1482 | 4710.940456 | 6462.686869 | 6288.323436 | 0.243105066 | 0.163572187 | 0.00152546 | 0.046559274 | ns | NaCl | KS.3 |
| FIM1_1483 | 375.6258705 | 394.6613388 | 398.8084871 | 0.013639987 | 0.100052434 | 0.753377971 | 0.943523399 | ns | NaCl | KS.3 |
| FIM1_1484 | 557.8612462 | 457.3342256 | 742.8195055 | -0.103551177 | 0.103273245 | 0.054119628 | 0.363353775 | ns | NaCl | KS.3 |
| FIM1_1485 | 3209.141548 | 3118.039512 | 3134.913706 | -0.023926798 | 0.063393863 | 0.615217568 | 0.889999244 | ns | NaCl | KS.3 |
| FIM1_1486 | 1561.975207 | 1780.348403 | 2021.714419 | 0.041238618 | 0.101587123 | 0.367732427 | 0.768177319 | ns | NaCl | KS.3 |
| FIM1_1487 | 3223.291556 | 4107.147972 | 1944.335533 | 0.052143651 | 0.125926479 | 0.235820932 | 0.659616806 | ns | NaCl | KS.3 |
| FIM1_1488 | 6039.275726 | 6004.451248 | 5118.618575 | -0.005198931 | 0.060280749 | 0.91117274 | 0.982011252 | ns | NaCl | KS.3 |
| FIM1_1489 | 661.6953447 | 551.1724901 | 496.6185416 | -0.083736627 | 0.100539306 | 0.103759119 | 0.491152821 | ns | NaCl | KS.3 |
| FIM1_1490 | 583.8231589 | 368.4945719 | 542.9241269 | -0.476118842 | 0.23944963 | 0.00015706 | 0.011005282 | ns | NaCl | KS.3 |
| FIM1_1491 | 959.3090191 | 973.9812012 | 1008.296104 | 0.009415211 | 0.068166895 | 0.846416369 | 0.967147129 | ns | NaCl | KS.3 |
| FIM1_1492 | 1336.262253 | 1433.827537 | 1549.742596 | 0.038530124 | 0.081523136 | 0.435589191 | 0.808096352 | ns | NaCl | KS.3 |
| FIM1_1493 | 450.0644502 | 288.0004336 | 639.7551531 | -0.433895952 | 0.244112977 | 0.000285721 | 0.016152987 | ns | NaCl | KS.3 |
| FIM1_1494 | 185.3728083 | 148.2391944 | 226.628268 | -0.084408171 | 0.116021067 | 0.097920012 | 0.478068751 | ns | NaCl | KS.3 |
| FIM1_1495 | 54.98582609 | 52.60839885 | 72.26210851 | -0.008547202 | 0.115869244 | 0.822414253 | 0.960240659 | ns | NaCl | KS.3 |
| FIM1_1496 | 890.8310424 | 963.8223942 | 888.9900235 | 0.046581161 | 0.078675669 | 0.350029543 | 0.756912594 | ns | NaCl | KS.3 |
| FIM1_1497 | 467.353692 | 532.9378677 | 541.8235652 | 0.074204466 | 0.085629645 | 0.146161779 | 0.560211748 | ns | NaCl | KS.3 |
| FIM1_1498 | 588.1346812 | 582.9159275 | 589.1756182 | -0.004839666 | 0.087822538 | 0.91850343 | 0.982488618 | ns | NaCl | KS.3 |
| FIM1_1499 | 1540.317897 | 1847.478935 | 1990.863723 | 0.059232653 | 0.107534986 | 0.211899578 | 0.635060483 | ns | NaCl | KS.3 |
| FIM1_1500 | 1295.22729 | 1245.215426 | 1542.250736 | -0.020205105 | 0.083729285 | 0.677196932 | 0.913287355 | ns | NaCl | KS.3 |
| FIM1_1501 | 1347.20243 | 1306.215003 | 1400.018085 | -0.015094869 | 0.084304787 | 0.754684782 | 0.944067586 | ns | NaCl | KS.3 |
| FIM1_1503 | 2582.464304 | 2413.433831 | 2153.953742 | -0.04988901 | 0.069991986 | 0.311349597 | 0.731871738 | ns | NaCl | KS.3 |
| FIM1_1504 | 777.8519997 | 766.651824 | 912.8558115 | -0.011350654 | 0.070040826 | 0.816391144 | 0.958816322 | ns | NaCl | KS.3 |
| FIM1_1505 | 287.0104249 | 246.8933985 | 530.282422 | -0.070511382 | 0.095454604 | 0.160601486 | 0.58211082 | ns | NaCl | KS.3 |
| FIM1_1506 | 90.84752232 | 72.47964212 | 147.5683477 | -0.047451083 | 0.125635468 | 0.269494053 | 0.691198798 | ns | NaCl | KS.3 |
| FIM1_1507 | 4839.234342 | 4941.571178 | 4989.294237 | 0.010727264 | 0.082240164 | 0.825355024 | 0.960240659 | ns | NaCl | KS.3 |
| FIM1_1508 | 1609.502113 | 1588.477854 | 1713.87884 | -0.00714872 | 0.078632029 | 0.884077671 | 0.975013614 | ns | NaCl | KS.3 |
| FIM1_1509 | 2237.339236 | 2152.539727 | 1673.109911 | -0.018888791 | 0.083809113 | 0.69686897 | 0.921808861 | ns | NaCl | KS.3 |
| FIM1_1510 | 1612.85257 | 1537.921256 | 1336.410245 | -0.026415751 | 0.079276017 | 0.591716975 | 0.878786986 | ns | NaCl | KS.3 |
| FIM1_1511 | 199.0368916 | 183.30873 | 223.070927 | -0.026464959 | 0.097350025 | 0.558792742 | 0.864443132 | ns | NaCl | KS.3 |
| FIM1_1512 | 736.0771956 | 707.3968335 | 639.4952849 | -0.017988092 | 0.085234359 | 0.708909874 | 0.925190614 | ns | NaCl | KS.3 |
| FIM1_1513 | 4836.071596 | 4242.069605 | 4753.957716 | -0.106631685 | 0.071128837 | 0.03515108 | 0.295899532 | ns | NaCl | KS.3 |
| FIM1_1514 | 9429.987239 | 10144.98843 | 11778.31794 | 0.041236828 | 0.080377189 | 0.405760756 | 0.792875005 | ns | NaCl | KS.3 |
| FIM1_1515 | 178.3393393 | 138.2729009 | 365.2917633 | -0.097749907 | 0.12506333 | 0.065762283 | 0.402419872 | ns | NaCl | KS.3 |
| FIM1_1516 | 76.85574551 | 136.417613 | 209.8896237 | 0.091928806 | 0.208921175 | 0.070888347 | 0.418675804 | ns | NaCl | KS.3 |
| FIM1_1518 | 2149.605573 | 2233.022461 | 1559.237096 | 0.018503515 | 0.083801318 | 0.702708796 | 0.923421278 | ns | NaCl | KS.3 |
| FIM1_1519 | 491.2796779 | 431.7832436 | 582.3236775 | -0.061271505 | 0.091753318 | 0.217580191 | 0.641504077 | ns | NaCl | KS.3 |

| | | | | | | | | | | |
|---|---|---|---|---|---|---|---|---|---|---|
| FIM1_1520 | 3818.072521 | 3350.10175 | 4137.183329 | -0.059198734 | 0.091986698 | 0.231522852 | 0.654993872 | ns | NaCl | KS.3 |
| FIM1_1521 | 6692.838605 | 8305.515787 | 4002.233212 | 0.033934382 | 0.127568822 | 0.392953774 | 0.781740625 | ns | NaCl | KS.3 |
| FIM1_1523 | 423.6265917 | 306.9340279 | 369.144513 | -0.246601409 | 0.170540997 | 0.001631826 | 0.04775491 | ns | NaCl | KS.3 |
| FIM1_1524 | 1296.033985 | 1283.131886 | 1103.512142 | -0.005253315 | 0.082802747 | 0.91368399 | 0.982311132 | ns | NaCl | KS.3 |
| FIM1_1525 | 1628.608828 | 1487.039066 | 1670.902005 | -0.074090945 | 0.06589059 | 0.128252839 | 0.536136415 | ns | NaCl | KS.3 |
| FIM1_1526 | 3483.460195 | 3278.443985 | 7536.909485 | -0.03305813 | 0.080621615 | 0.50264246 | 0.842535794 | ns | NaCl | KS.3 |
| FIM1_1527 | 509.8727997 | 452.1492502 | 531.9941973 | -0.076069716 | 0.082047518 | 0.13682201 | 0.545297021 | ns | NaCl | KS.3 |
| FIM1_1528 | 877.6590894 | 946.4649126 | 668.6270175 | 0.011729973 | 0.116853076 | 0.758144613 | 0.945250644 | ns | NaCl | KS.3 |
| FIM1_1529 | 571.0661539 | 467.7143297 | 1461.92079 | -0.106743266 | 0.104319691 | 0.04872438 | 0.34611662 | ns | NaCl | KS.3 |
| FIM1_1530 | 1621.826963 | 1717.636842 | 2471.08405 | 0.025970976 | 0.086525728 | 0.58980237 | 0.87877872 | ns | NaCl | KS.3 |
| FIM1_1531 | 476.1342541 | 353.9607537 | 741.0917151 | -0.107390787 | 0.140026996 | 0.050807108 | 0.352025696 | ns | NaCl | KS.3 |
| FIM1_1532 | 750.7141267 | 468.1334484 | 768.8674388 | -0.210828109 | 0.246051286 | 0.008979966 | 0.132175538 | ns | NaCl | KS.3 |
| FIM1_1533 | 886.3184506 | 798.2852479 | 731.9239523 | -0.060327843 | 0.082208073 | 0.230695396 | 0.654338423 | ns | NaCl | KS.3 |
| FIM1_1534 | 362.9787832 | 378.5724209 | 392.2872536 | 0.02243411 | 0.081174961 | 0.646899616 | 0.898973628 | ns | NaCl | KS.3 |
| FIM1_1535 | 820.7655232 | 756.7353272 | 781.0826287 | -0.063782453 | 0.069036455 | 0.195974481 | 0.618952432 | ns | NaCl | KS.3 |
| FIM1_1536 | 763.3776529 | 801.7740727 | 720.7602547 | 0.025848054 | 0.081225947 | 0.598224092 | 0.880167289 | ns | NaCl | KS.3 |
| FIM1_1537 | 2364.20185 | 2414.871933 | 1969.124807 | 0.012762266 | 0.075747276 | 0.795368664 | 0.955325714 | ns | NaCl | KS.3 |
| FIM1_1538 | 1698.91318 | 1549.300066 | 1902.472634 | -0.052730832 | 0.081664939 | 0.291529933 | 0.711543689 | ns | NaCl | KS.3 |
| FIM1_1540 | 741.1047742 | 586.2719168 | 957.5712073 | -0.113480202 | 0.116242875 | 0.040491614 | 0.317627561 | ns | NaCl | KS.3 |
| FIM1_1541 | 2089.264882 | 2144.726265 | 2516.562676 | 0.008127617 | 0.096303168 | 0.855191796 | 0.969815177 | ns | NaCl | KS.3 |
| FIM1_1542 | 7053.639611 | 9040.190108 | 4261.464166 | 0.054591033 | 0.12675644 | 0.219372566 | 0.642906874 | ns | NaCl | KS.3 |
| FIM1_1543 | 5761.586549 | 7267.475179 | 3473.59313 | 0.038520859 | 0.128595008 | 0.342768056 | 0.750889951 | ns | NaCl | KS.3 |
| FIM1_1544 | 201.592974 | 133.1110195 | 191.583174 | -0.21732718 | 0.216062221 | 0.006324773 | 0.106776808 | ns | NaCl | KS.3 |
| FIM1_1545 | 1106.746619 | 846.3985782 | 1128.202135 | -0.166694661 | 0.132686314 | 0.00849661 | 0.127329254 | ns | NaCl | KS.3 |
| FIM1_1546 | 244.8721537 | 226.6265651 | 298.4181561 | -0.031462146 | 0.092688686 | 0.503111477 | 0.842754074 | ns | NaCl | KS.3 |
| FIM1_1547 | 750.814043 | 825.8059755 | 968.3615341 | 0.042993468 | 0.087912927 | 0.378327465 | 0.778493059 | ns | NaCl | KS.3 |
| FIM1_1548 | 2072.282213 | 2342.264458 | 1539.925275 | 0.03849213 | 0.100678703 | 0.399024313 | 0.787737366 | ns | NaCl | KS.3 |
| FIM1_1549 | 1520.562149 | 1669.862207 | 1949.095125 | 0.047201281 | 0.085238731 | 0.339605859 | 0.748909198 | ns | NaCl | KS.3 |
| FIM1_1550 | 1199.49671 | 1217.982919 | 1478.150675 | 0.004962613 | 0.094859673 | 0.912172631 | 0.982051252 | ns | NaCl | KS.3 |
| FIM1_1551 | 13898.21136 | 11733.63978 | 21078.24352 | -0.041836908 | 0.111579928 | 0.340142864 | 0.749761075 | ns | NaCl | KS.3 |
| FIM1_1552 | 627.2099294 | 643.702399 | 398.9322626 | 0.007767168 | 0.098199467 | 0.859321901 | 0.969856868 | ns | NaCl | KS.3 |
| FIM1_1553 | 5988.231057 | 8183.878771 | 3566.150704 | 0.055072698 | 0.142080967 | 0.20355335 | 0.626494461 | ns | NaCl | KS.3 |
| FIM1_1554 | 1236.179685 | 1307.791279 | 1029.671585 | 0.026138615 | 0.086200952 | 0.587983767 | 0.878311243 | ns | NaCl | KS.3 |
| FIM1_1556 | 533.4324868 | 511.6049838 | 412.6890585 | -0.012781166 | 0.096351549 | 0.774878455 | 0.950214521 | ns | NaCl | KS.3 |
| FIM1_1557 | 484.0786093 | 415.1368199 | 370.8202114 | -0.057743798 | 0.10055357 | 0.230568447 | 0.654338423 | ns | NaCl | KS.3 |
| FIM1_1558 | 1896.933662 | 1779.404343 | 2132.708685 | -0.03739888 | 0.078841661 | 0.450447978 | 0.813595672 | ns | NaCl | KS.3 |
| FIM1_1559 | 6484.814143 | 7257.425116 | 5562.228255 | 0.038829695 | 0.09747439 | 0.403122586 | 0.789854857 | ns | NaCl | KS.3 |
| FIM1_1560 | 954.8105166 | 884.5385407 | 1399.809743 | -0.046367642 | 0.078826079 | 0.352152269 | 0.756912594 | ns | NaCl | KS.3 |
| FIM1_1561 | 1417.141694 | 1078.517954 | 1903.390235 | -0.150729638 | 0.133540749 | 0.014189115 | 0.172593755 | ns | NaCl | KS.3 |
| FIM1_1562 | 540.3866276 | 604.8113147 | 554.8368495 | 0.031307885 | 0.102814609 | 0.478822456 | 0.826171162 | ns | NaCl | KS.3 |
| FIM1_1563 | 1394.608836 | 1383.02072 | 1104.718313 | -0.006347889 | 0.071199214 | 0.896909053 | 0.977971161 | ns | NaCl | KS.3 |
| FIM1_1564 | 25.76366212 | 28.55242344 | 328.3333144 | 0.011636067 | 0.125945992 | 0.747204001 | 0.943274417 | ns | NaCl | KS.3 |
| FIM1_1566 | 2551.284362 | 2550.856446 | 2607.517748 | 3.36079993269654 | 0.093850672 | 0.9994083 | 0.999642978 | ns | NaCl | KS.3 |
| FIM1_1567 | 1032.219439 | 1002.043587 | 1116.362786 | -0.007251964 | 0.102548356 | 0.863551991 | 0.969856868 | ns | NaCl | KS.3 |
| FIM1_1568 | 8070.938793 | 7714.893338 | 7098.304888 | -0.03533692 | 0.065488309 | 0.463567607 | 0.819880744 | ns | NaCl | KS.3 |

| | | | | | | | | | | |
|---|---|---|---|---|---|---|---|---|---|---|
| FIM1_1569 | 66.74080589 | 47.66531331 | 191.7269476 | -0.069969272 | 0.147688714 | 0.13444791 | 0.54183277 | ns | NaCl | KS.3 |
| FIM1_1570 | 726.4195431 | 740.0851516 | 1038.993896 | 0.010018288 | 0.079429558 | 0.837926529 | 0.962829707 | ns | NaCl | KS.3 |
| FIM1_1571 | 397.2009184 | 381.7128028 | 365.0476731 | -0.016410133 | 0.088779843 | 0.728472905 | 0.935336397 | ns | NaCl | KS.3 |
| FIM1_1572 | 421.956487 | 436.5619122 | 549.9838849 | 0.014035062 | 0.086491683 | 0.769046913 | 0.948835638 | ns | NaCl | KS.3 |
| FIM1_1573 | 1272.065212 | 1337.726012 | 1173.957086 | 0.036764272 | 0.066990808 | 0.448997555 | 0.81346061 | ns | NaCl | KS.3 |
| FIM1_1574 | 3132.164709 | 3455.020994 | 3203.394133 | 0.089061371 | 0.061156143 | 0.059701064 | 0.382593564 | ns | NaCl | KS.3 |
| FIM1_1575 | 618.8456967 | 514.0650537 | 851.4544938 | -0.114589943 | 0.093964282 | 0.034970223 | 0.295879009 | ns | NaCl | KS.3 |
| FIM1_1576 | 32.8797547 | 21.99664642 | 38.68437699 | -0.075144737 | 0.16501233 | 0.113116439 | 0.511380136 | ns | NaCl | KS.3 |
| FIM1_1577 | 436.9774453 | 381.1612831 | 674.6265987 | -0.062028363 | 0.094355626 | 0.209933244 | 0.633800632 | ns | NaCl | KS.3 |
| FIM1_1578 | 13868.18607 | 14655.39399 | 9826.523273 | 0.041453828 | 0.068178901 | 0.396061132 | 0.783460887 | ns | NaCl | KS.3 |
| FIM1_1579 | 3024.518855 | 3999.46549 | 1877.861067 | 0.058633445 | 0.133909063 | 0.190414652 | 0.618952432 | ns | NaCl | KS.3 |
| FIM1_1580 | 2437.124257 | 2726.517792 | 1777.236402 | 0.035450548 | 0.099820943 | 0.435780859 | 0.808096352 | ns | NaCl | KS.3 |
| FIM1_1581 | 5771.693294 | 6432.772808 | 4862.540858 | 0.089257555 | 0.067972697 | 0.071301364 | 0.418675804 | ns | NaCl | KS.3 |
| FIM1_1582 | 54.30643033 | 36.82688163 | 111.7929124 | -0.071700971 | 0.162067489 | 0.123991095 | 0.531066893 | ns | NaCl | KS.3 |
| FIM1_1583 | 661.2525954 | 673.5825007 | 887.9140318 | 0.00931492 | 0.081349923 | 0.848421733 | 0.967357303 | ns | NaCl | KS.3 |
| FIM1_1584 | 6382.631944 | 6557.683366 | 5326.866002 | 0.014092135 | 0.081606772 | 0.772564968 | 0.94938462 | ns | NaCl | KS.3 |
| FIM1_1585 | 559.5549112 | 579.2455922 | 919.5679531 | 0.016223508 | 0.086347034 | 0.734776681 | 0.938375543 | ns | NaCl | KS.3 |
| FIM1_1586 | 1010.308231 | 810.568785 | 2013.747848 | -0.056370824 | 0.118795091 | 0.216596926 | 0.640647865 | ns | NaCl | KS.3 |
| FIM1_1587 | 9999.636909 | 9874.693303 | 7941.355191 | -0.007838795 | 0.075149907 | 0.873366137 | 0.971243188 | ns | NaCl | KS.3 |
| FIM1_1588 | 8617.011141 | 10178.65558 | 6573.148533 | 0.062299329 | 0.101961994 | 0.199823502 | 0.623535394 | ns | NaCl | KS.3 |
| FIM1_1589 | 225.2489382 | 208.6043373 | 174.21158 | -0.03402783 | 0.087694047 | 0.481215175 | 0.828577419 | ns | NaCl | KS.3 |
| FIM1_1590 | 1115.986489 | 1153.717771 | 907.6081619 | 0.009454531 | 0.098357108 | 0.829191351 | 0.961288108 | ns | NaCl | KS.3 |
| FIM1_1591 | 250.2594376 | 189.9699332 | 193.798865 | -0.072699944 | 0.132862749 | 0.130218183 | 0.537622789 | ns | NaCl | KS.3 |
| FIM1_1592 | 132.7308321 | 116.6296999 | 207.5354789 | -0.038710037 | 0.103753946 | 0.389191186 | 0.779482559 | ns | NaCl | KS.3 |
| FIM1_1593 | 2124.750248 | 1227.20938 | 3711.678997 | -0.14534776 | 0.240558354 | 0.026479808 | 0.254519471 | ns | NaCl | KS.3 |
| FIM1_1594 | 3650.789473 | 5281.904492 | 2701.158854 | 0.103056859 | 0.16542177 | 0.057928683 | 0.377713236 | ns | NaCl | KS.3 |
| FIM1_1595 | 1860.675342 | 1998.063579 | 1468.553781 | 0.028818298 | 0.090450134 | 0.54310976 | 0.857313452 | ns | NaCl | KS.3 |
| FIM1_1596 | 1046.567503 | 944.0639572 | 1618.955995 | -0.029396503 | 0.101718196 | 0.506776858 | 0.844423736 | ns | NaCl | KS.3 |
| FIM1_1597 | 2701.37464 | 3965.36424 | 1675.69388 | 0.083050321 | 0.163416633 | 0.093647669 | 0.470201108 | ns | NaCl | KS.3 |
| FIM1_1598 | 4945.251952 | 6330.421332 | 3069.22103 | 0.042286555 | 0.130223225 | 0.305536925 | 0.724521545 | ns | NaCl | KS.3 |
| FIM1_1599 | 2760.931214 | 2973.545282 | 2665.984911 | 0.057650201 | 0.068777042 | 0.241069289 | 0.665549231 | ns | NaCl | KS.3 |
| FIM1_1601 | 32137.38829 | 36992.05538 | 22763.71051 | 0.075471136 | 0.088488747 | 0.139660421 | 0.550332947 | ns | NaCl | KS.3 |
| FIM1_1602 | 2150.184002 | 2025.642882 | 2256.884439 | -0.042934417 | 0.071130598 | 0.384170894 | 0.778665419 | ns | NaCl | KS.3 |
| FIM1_1603 | 5670.932243 | 6263.300032 | 4657.555361 | 0.063072096 | 0.078071756 | 0.21090732 | 0.634379635 | ns | NaCl | KS.3 |
| FIM1_1604 | 1410.153939 | 1759.336033 | 2197.326057 | 0.119786487 | 0.10933598 | 0.031997729 | 0.283253921 | ns | NaCl | KS.3 |
| FIM1_1605 | 4790.245354 | 7289.870546 | 2906.898455 | 0.085708872 | 0.173567213 | 0.086563546 | 0.45668467 | ns | NaCl | KS.3 |
| FIM1_1606 | 9734.261488 | 8635.50065 | 7623.839318 | -0.062227745 | 0.086787249 | 0.215305085 | 0.640629323 | ns | NaCl | KS.3 |
| FIM1_1607 | 1184.421755 | 1039.128621 | 1263.574432 | -0.078398073 | 0.084447037 | 0.126594887 | 0.534982546 | ns | NaCl | KS.3 |
| FIM1_1608 | 3190.509144 | 3520.027922 | 2705.623237 | 0.047461431 | 0.087008934 | 0.334919647 | 0.745307602 | ns | NaCl | KS.3 |
| FIM1_1609 | 1589.903189 | 1613.391485 | 1307.238998 | 0.005508074 | 0.090392393 | 0.905783264 | 0.979880338 | ns | NaCl | KS.3 |
| FIM1_1610 | 2648.030674 | 2425.999559 | 2102.081972 | -0.02456811 | 0.100957769 | 0.576491044 | 0.873870489 | ns | NaCl | KS.3 |
| FIM1_1611 | 1589.711504 | 1747.353906 | 1829.231039 | 0.036341316 | 0.093027484 | 0.442087889 | 0.809513863 | ns | NaCl | KS.3 |
| FIM1_1612 | 2156.787771 | 2581.050855 | 1563.798557 | 0.052885217 | 0.10921741 | 0.25201821 | 0.675897894 | ns | NaCl | KS.3 |
| FIM1_1613 | 1307.891344 | 1185.804787 | 1328.538532 | -0.066970013 | 0.074747947 | 0.182931356 | 0.611615252 | ns | NaCl | KS.3 |
| FIM1_1614 | 350.5706796 | 329.3265365 | 538.7302173 | -0.030546553 | 0.087787129 | 0.525467399 | 0.854940852 | ns | NaCl | KS.3 |

| | | | | | | | | | | |
|---|---|---|---|---|---|---|---|---|---|---|
| FIM1_1615 | 3455.049266 | 3629.193815 | 3138.809537 | 0.033053629 | 0.072665551 | 0.502977681 | 0.842754074 | ns | NaCl | KS.3 |
| FIM1_1616 | 765.3909318 | 869.7117854 | 848.8679576 | 0.065168678 | 0.088174314 | 0.195413295 | 0.618952432 | ns | NaCl | KS.3 |
| FIM1_1617 | 4684.959329 | 5968.57098 | 3551.417207 | 0.222109349 | 0.106361237 | 0.000443367 | 0.021482506 | ns | NaCl | KS.3 |
| FIM1_1618 | 505.8433771 | 668.4665159 | 330.5806568 | 0.082227194 | 0.132897269 | 0.100523449 | 0.483659728 | ns | NaCl | KS.3 |
| FIM1_1619 | 2009.356291 | 2224.233224 | 1644.712945 | 0.061053753 | 0.080462894 | 0.225637134 | 0.648597616 | ns | NaCl | KS.3 |
| FIM1_1621 | 3242.852084 | 3372.250941 | 3449.992768 | 0.0275069 | 0.070124665 | 0.574621309 | 0.872163065 | ns | NaCl | KS.3 |
| FIM1_1622 | 709.590991 | 914.9160695 | 889.5684964 | 0.13345188 | 0.123849528 | 0.022410687 | 0.229732317 | ns | NaCl | KS.3 |
| FIM1_1623 | 423.2453347 | 384.5630452 | 369.2683476 | -0.03625507 | 0.094391767 | 0.43989932 | 0.808917498 | ns | NaCl | KS.3 |
| FIM1_1624 | 686.2847511 | 709.8230928 | 674.7249446 | 0.013388138 | 0.089788968 | 0.77523846 | 0.950385088 | ns | NaCl | KS.3 |
| FIM1_1625 | 1460.956755 | 1459.872719 | 1770.134835 | -0.000169184 | 0.085187353 | 0.997185094 | 0.99924489 | ns | NaCl | KS.3 |
| FIM1_1627 | 2540.500813 | 2497.455592 | 2079.215773 | -0.006658063 | 0.089290895 | 0.887094799 | 0.976547586 | ns | NaCl | KS.3 |
| FIM1_1628 | 259.9387465 | 313.0621988 | 650.5022275 | 0.022397705 | 0.131012757 | 0.543130942 | 0.857313452 | ns | NaCl | KS.3 |
| FIM1_1629 | 364.0032571 | 339.00796 | 370.7286386 | -0.035400708 | 0.087375016 | 0.464965169 | 0.820575281 | ns | NaCl | KS.3 |
| FIM1_1630 | 2064.814795 | 2085.25717 | 2888.50703 | 0.005667586 | 0.076436683 | 0.908199661 | 0.980825899 | ns | NaCl | KS.3 |
| FIM1_1631 | 681.0646951 | 590.8556094 | 737.8346959 | -0.045275125 | 0.101281365 | 0.328991167 | 0.741064881 | ns | NaCl | KS.3 |
| FIM1_1632 | 1067.07767 | 1269.347791 | 1564.129185 | 0.029855378 | 0.120595895 | 0.455420232 | 0.816788914 | ns | NaCl | KS.3 |
| FIM1_1633 | 1648.17193 | 1841.231442 | 1191.610824 | 0.018269318 | 0.116883933 | 0.638116188 | 0.898332475 | ns | NaCl | KS.3 |
| FIM1_1634 | 2659.013803 | 2227.275818 | 2116.876201 | -0.088366223 | 0.097353077 | 0.089684352 | 0.463545584 | ns | NaCl | KS.3 |
| FIM1_1635 | 6075.882077 | 6287.752447 | 6104.133034 | 0.029208493 | 0.06179279 | 0.535590307 | 0.855153333 | ns | NaCl | KS.3 |
| FIM1_1636 | 848.204446 | 845.7847698 | 829.9480188 | -0.000955855 | 0.083874065 | 0.98419206 | 0.998198696 | ns | NaCl | KS.3 |
| FIM1_1637 | 238.0791238 | 218.6635508 | 238.0021329 | -0.027149508 | 0.096190789 | 0.552216296 | 0.860945181 | ns | NaCl | KS.3 |
| FIM1_1638 | 24.03534784 | 31.54091904 | 70.51401508 | 0.0358916 | 0.134594709 | 0.36050147 | 0.763371684 | ns | NaCl | KS.3 |
| FIM1_1640 | 2654.425197 | 2359.107809 | 2570.685489 | -0.072948236 | 0.081134078 | 0.152184847 | 0.568361587 | ns | NaCl | KS.3 |
| FIM1_1641 | 5252.798968 | 5392.623713 | 4617.052727 | 0.020207336 | 0.066052973 | 0.675333577 | 0.912806589 | ns | NaCl | KS.3 |
| FIM1_1642 | 1255.64978 | 1087.53909 | 1800.68227 | -0.102470942 | 0.079184492 | 0.049102265 | 0.34611662 | ns | NaCl | KS.3 |
| FIM1_1643 | 1356.991205 | 1363.654116 | 1577.718136 | 0.002909809 | 0.079242852 | 0.952603618 | 0.988569671 | ns | NaCl | KS.3 |
| FIM1_1644 | 506.6969099 | 505.1073427 | 640.2083275 | -0.003627876 | 0.079557419 | 0.940889958 | 0.986790704 | ns | NaCl | KS.3 |
| FIM1_1645 | 582.6370924 | 548.404313 | 638.4829664 | -0.030376805 | 0.085199363 | 0.532040873 | 0.855153333 | ns | NaCl | KS.3 |
| FIM1_1646 | 1496.819754 | 1361.933315 | 3165.894573 | -0.037698047 | 0.092589315 | 0.42730917 | 0.803728968 | ns | NaCl | KS.3 |
| FIM1_1647 | 105.5013102 | 110.2113457 | 78.19150405 | 0.006098447 | 0.107061423 | 0.880512903 | 0.973672304 | ns | NaCl | KS.3 |
| FIM1_1648 | 4696.593102 | 4064.693223 | 3748.977745 | -0.058994733 | 0.096715528 | 0.227180747 | 0.65142606 | ns | NaCl | KS.3 |
| FIM1_1649 | 3742.491118 | 4902.845818 | 2915.235222 | 0.10985338 | 0.130210632 | 0.046717771 | 0.341725286 | ns | NaCl | KS.3 |
| FIM1_1650 | 1230.955987 | 1250.257045 | 1231.102463 | 0.003669849 | 0.104843667 | 0.929133324 | 0.984238275 | ns | NaCl | KS.3 |
| FIM1_1651 | 235.5117404 | 210.0549699 | 475.7090115 | -0.039188339 | 0.099522836 | 0.39424467 | 0.781740625 | ns | NaCl | KS.3 |
| FIM1_1652 | 3643.789361 | 3192.647112 | 2996.097767 | -0.068446764 | 0.088265017 | 0.175926838 | 0.605047427 | ns | NaCl | KS.3 |
| FIM1_1653 | 370.6733177 | 405.9067645 | 589.1797007 | 0.040297632 | 0.087965951 | 0.407235793 | 0.794509048 | ns | NaCl | KS.3 |
| FIM1_1654 | 318.7374756 | 178.5731653 | 472.4391557 | -0.540353395 | 0.330307941 | 0.000375613 | 0.019264677 | ns | NaCl | KS.3 |
| FIM1_1655 | 1386.639677 | 1334.850757 | 1447.568048 | -0.023428731 | 0.075612184 | 0.634835162 | 0.898009483 | ns | NaCl | KS.3 |
| FIM1_1656 | 397.7286311 | 307.54439 | 1353.127567 | -0.143017741 | 0.123427383 | 0.016431605 | 0.189229716 | ns | NaCl | KS.3 |
| FIM1_1657 | 40793.29251 | 49031.47375 | 30073.46668 | 0.142400946 | 0.08557106 | 0.009396423 | 0.136108932 | ns | NaCl | KS.3 |
| FIM1_1658 | 1163.535964 | 1214.731402 | 1053.529141 | 0.020232347 | 0.084706016 | 0.675632543 | 0.912892966 | ns | NaCl | KS.3 |
| FIM1_1659 | 2635.706486 | 2621.493307 | 2925.916133 | -0.003338013 | 0.073327672 | 0.945831748 | 0.987278589 | ns | NaCl | KS.3 |
| FIM1_1660 | 2097.730308 | 2256.47756 | 2284.236219 | 0.048356894 | 0.074470109 | 0.331289722 | 0.742417282 | ns | NaCl | KS.3 |
| FIM1_1661 | 1979.500679 | 2118.891807 | 1659.451143 | 0.033131265 | 0.085057855 | 0.496765199 | 0.836358994 | ns | NaCl | KS.3 |
| FIM1_1664 | 2385.474697 | 2380.294931 | 2194.986819 | -0.002113842 | 0.064076091 | 0.964677885 | 0.993691273 | ns | NaCl | KS.3 |

| | | | | | | | | | | |
|---|---|---|---|---|---|---|---|---|---|---|
| FIM1_1665 | 312.2505874 | 305.5923013 | 323.128888 | -0.005517944 | 0.099856113 | 0.898223365 | 0.977971161 | ns | NaCl | KS.3 |
| FIM1_1666 | 978.3378916 | 938.3856122 | 919.0415446 | -0.032162386 | 0.066054258 | 0.505645677 | 0.84439816 | ns | NaCl | KS.3 |
| FIM1_1667 | 3462.748187 | 5072.891336 | 1963.193442 | 0.100986632 | 0.168704939 | 0.060873503 | 0.385790671 | ns | NaCl | KS.3 |
| FIM1_1669 | 2155.22294 | 2502.695806 | 2080.923473 | 0.089016621 | 0.086912243 | 0.087267693 | 0.457488698 | ns | NaCl | KS.3 |
| FIM1_1670 | 2908.96543 | 3057.094467 | 1932.306517 | 0.014871317 | 0.09760149 | 0.737395081 | 0.939206485 | ns | NaCl | KS.3 |
| FIM1_1671 | 9688.06055 | 12046.72274 | 5657.092212 | 0.044273249 | 0.12260784 | 0.299198931 | 0.717010926 | ns | NaCl | KS.3 |
| FIM1_1672 | 4133.261182 | 4498.401166 | 4201.129198 | 0.037487989 | 0.088735038 | 0.437707691 | 0.808096352 | ns | NaCl | KS.3 |
| FIM1_1673 | 4052.761782 | 4043.506499 | 3566.447103 | -0.001920074 | 0.065089972 | 0.968080346 | 0.99408528 | ns | NaCl | KS.3 |
| FIM1_1674 | 1074.053888 | 782.3775828 | 1673.238799 | -0.25279442 | 0.162247424 | 0.001126138 | 0.039503502 | ns | NaCl | KS.3 |
| FIM1_1675 | 15.78395124 | 21.71052576 | 18.27940201 | 0.041305607 | 0.151787473 | 0.287655849 | 0.707758579 | ns | NaCl | KS.3 |
| FIM1_1676 | 609.6126725 | 542.586316 | 574.8902043 | -0.064995805 | 0.084135987 | 0.198331641 | 0.620924909 | ns | NaCl | KS.3 |
| FIM1_1677 | 356.2580522 | 303.1549103 | 639.4036967 | -0.020564709 | 0.127623508 | 0.578964573 | 0.8749481 | ns | NaCl | KS.3 |
| FIM1_1678 | 156.2502041 | 123.9772207 | 212.1497853 | -0.053237727 | 0.121204535 | 0.233841632 | 0.656462386 | ns | NaCl | KS.3 |
| FIM1_1679 | 598.9535685 | 550.7445024 | 646.7479553 | -0.020305689 | 0.103746278 | 0.634360936 | 0.898009483 | ns | NaCl | KS.3 |
| FIM1_1680 | 745.4489407 | 849.2217119 | 512.5961361 | 0.052801854 | 0.095535172 | 0.27471931 | 0.695399585 | ns | NaCl | KS.3 |
| FIM1_1681 | 37.61255119 | 22.80577512 | 59.03730574 | -0.07343754 | 0.18390997 | 0.112147558 | 0.509063959 | ns | NaCl | KS.3 |
| FIM1_1682 | 1020.757919 | 1055.846848 | 870.1066741 | 0.011601558 | 0.094055698 | 0.798522383 | 0.956343008 | ns | NaCl | KS.3 |
| FIM1_1683 | 5220.280622 | 5783.479648 | 4383.462853 | 0.036464366 | 0.095918225 | 0.433747586 | 0.80729302 | ns | NaCl | KS.3 |
| FIM1_1684 | 1040.685344 | 1039.953899 | 1764.440744 | 3.06036882443008 | 0.076264276 | 0.999503143 | 0.999642978 | ns | NaCl | KS.3 |
| FIM1_1685 | 1657.619809 | 1832.257711 | 2269.185769 | 0.071481227 | 0.07412899 | 0.155599259 | 0.573412083 | ns | NaCl | KS.3 |
| FIM1_1686 | 1023.028483 | 1029.777297 | 1051.0058 | 0.003158971 | 0.083291318 | 0.947921614 | 0.987736646 | ns | NaCl | KS.3 |
| FIM1_1687 | 1240.686599 | 1161.265199 | 2583.761881 | -0.039956377 | 0.076608905 | 0.421042513 | 0.80133378 | ns | NaCl | KS.3 |
| FIM1_1688 | 213.8066503 | 252.8420212 | 317.1652885 | 0.037610909 | 0.113488688 | 0.3789958 | 0.778493059 | ns | NaCl | KS.3 |
| FIM1_1689 | 293.3043539 | 361.1841739 | 271.523357 | 0.06587777 | 0.111851625 | 0.170775053 | 0.597342667 | ns | NaCl | KS.3 |
| FIM1_1690 | 2792.335686 | 2763.562152 | 3733.747297 | -0.009188565 | 0.064341138 | 0.847590248 | 0.967147129 | ns | NaCl | KS.3 |
| FIM1_1691 | 664.3120225 | 1016.291323 | 2389.642711 | 0.057504387 | 0.165395896 | 0.175569263 | 0.605047427 | ns | NaCl | KS.3 |
| FIM1_1692 | 659.8937886 | 632.334563 | 1205.142974 | -0.013993098 | 0.095384366 | 0.75627385 | 0.944867505 | ns | NaCl | KS.3 |
| FIM1_1693 | 343.8260137 | 349.6189845 | 768.7516542 | 0.003200479 | 0.111909903 | 0.934490272 | 0.985446192 | ns | NaCl | KS.3 |
| FIM1_1694 | 38.08911151 | 53.42600988 | 53.63782133 | 0.060306519 | 0.148373701 | 0.172929062 | 0.600364328 | ns | NaCl | KS.3 |
| FIM1_1695 | 83.15205758 | 97.96317553 | 186.2138148 | 0.022800899 | 0.126373557 | 0.544979837 | 0.857313452 | ns | NaCl | KS.3 |
| FIM1_1697 | 201.3929268 | 191.6804235 | 205.3618799 | -0.009555296 | 0.108394647 | 0.812615874 | 0.958816322 | ns | NaCl | KS.3 |
| FIM1_1698 | 3118.027236 | 3464.266607 | 2099.503148 | 0.023068764 | 0.108550316 | 0.579136444 | 0.8749481 | ns | NaCl | KS.3 |
| FIM1_1701 | 640.6701603 | 608.2126566 | 548.8457104 | -0.014682582 | 0.098940167 | 0.737858026 | 0.939315169 | ns | NaCl | KS.3 |
| FIM1_1703 | 47.37063251 | 52.93166174 | 84.05414166 | 0.018034388 | 0.119825911 | 0.6364645 | 0.898332475 | ns | NaCl | KS.3 |
| FIM1_1705 | 17.98882585 | 23.5071001 | 22.09320857 | 0.032585197 | 0.139194572 | 0.390043473 | 0.779482559 | ns | NaCl | KS.3 |
| FIM1_1707 | 419.9403459 | 413.0073929 | 374.05025 | -0.005689695 | 0.093207127 | 0.900692635 | 0.979013733 | ns | NaCl | KS.3 |
| FIM1_1708 | 1761.64137 | 5325.005068 | 2951.506154 | 0.613899807 | 0.603390284 | 0.000830444 | 0.031433557 | ns | NaCl | KS.3 |
| FIM1_1709 | 979.0646778 | 892.4694347 | 1248.502807 | -0.01950091 | 0.108995562 | 0.634588364 | 0.898009483 | ns | NaCl | KS.3 |
| FIM1_1710 | 289.0481484 | 278.5574201 | 440.0704167 | -0.006590916 | 0.112633925 | 0.865288285 | 0.970128052 | ns | NaCl | KS.3 |
| FIM1_1711 | 448.6184195 | 415.893937 | 1667.795688 | -0.009932174 | 0.121642137 | 0.787339576 | 0.95289761 | ns | NaCl | KS.3 |
| FIM1_1712 | 3119.244818 | 6435.751057 | 2816.95815 | 0.451480669 | 0.416547638 | 0.001501647 | 0.046346385 | ns | NaCl | KS.3 |
| FIM1_1713 | 60.33792009 | 82.81997379 | 91.60946069 | 0.082237526 | 0.146208294 | 0.098179279 | 0.47886462 | ns | NaCl | KS.3 |
| FIM1_1714 | 193.5026143 | 177.9626743 | 331.4930736 | -0.02852656 | 0.096207751 | 0.533223374 | 0.855153333 | ns | NaCl | KS.3 |
| FIM1_1715 | 109.3918009 | 96.52629975 | 205.5618188 | -0.028940755 | 0.109319153 | 0.492936761 | 0.835912774 | ns | NaCl | KS.3 |
| FIM1_1716 | 4464.777333 | 4272.948277 | 4759.1703 | -0.009602139 | 0.105759398 | 0.815620348 | 0.958816322 | ns | NaCl | KS.3 |

| | | | | | | | | | | |
|---|---|---|---|---|---|---|---|---|---|---|
| FIM1_1717 | 152.8198474 | 166.9914071 | 180.8783533 | 0.032634497 | 0.091095896 | 0.491776105 | 0.835238267 | ns | NaCl | KS.3 |
| FIM1_1718 | 163.5860223 | 174.9626317 | 178.6442181 | 0.025426126 | 0.089898763 | 0.591238059 | 0.878786986 | ns | NaCl | KS.3 |
| FIM1_1719 | 182.0415322 | 202.7187241 | 232.4189604 | 0.03746477 | 0.095588377 | 0.423001382 | 0.80229961 | ns | NaCl | KS.3 |
| FIM1_1721 | 54.74167118 | 33.41636517 | 37.01723315 | -0.076826155 | 0.185167971 | 0.103479843 | 0.491042668 | ns | NaCl | KS.3 |
| FIM1_1724 | 14535.79284 | 18313.71502 | 10659.98046 | 0.062876304 | 0.120414722 | 0.178860422 | 0.607807787 | ns | NaCl | KS.3 |
| FIM1_1725 | 30147.71567 | 28652.32998 | 18386.44368 | -0.01427151 | 0.099457872 | 0.743769343 | 0.941460666 | ns | NaCl | KS.3 |
| FIM1_1727 | 1286.240701 | 1085.782159 | 1983.948707 | -0.049912055 | 0.107626221 | 0.277038608 | 0.698564154 | ns | NaCl | KS.3 |
| FIM1_1728 | 6426.970867 | 6289.541813 | 6048.725339 | -0.010540991 | 0.082981267 | 0.82788918 | 0.960529074 | ns | NaCl | KS.3 |
| FIM1_1729 | 3724.371871 | 3358.000938 | 3532.289846 | -0.062394658 | 0.080042389 | 0.216110675 | 0.640629323 | ns | NaCl | KS.3 |
| FIM1_1730 | 5353.931283 | 5185.941636 | 4780.754088 | -0.028712213 | 0.057574106 | 0.529264064 | 0.855153333 | ns | NaCl | KS.3 |
| FIM1_1731 | 5627.510644 | 5220.021681 | 4615.337124 | -0.066997768 | 0.060965964 | 0.154707154 | 0.572502577 | ns | NaCl | KS.3 |
| FIM1_1732 | 1062.383405 | 1275.260102 | 1104.078117 | 0.083522426 | 0.100429686 | 0.104499852 | 0.493886208 | ns | NaCl | KS.3 |
| FIM1_1733 | 6865.19095 | 6520.556173 | 8127.148046 | -0.009761861 | 0.110121274 | 0.806170377 | 0.957859861 | ns | NaCl | KS.3 |
| FIM1_1734 | 522.040375 | 522.682933 | 542.6370371 | -6.63E-05 | 0.100610776 | 0.998763342 | 0.999406383 | ns | NaCl | KS.3 |
| FIM1_1735 | 2649.162863 | 2424.024777 | 2353.005583 | -0.036953907 | 0.09064557 | 0.440140945 | 0.808917498 | ns | NaCl | KS.3 |
| FIM1_1736 | 547.6707863 | 593.1039527 | 664.8273505 | 0.034370733 | 0.089325102 | 0.473856552 | 0.82307921 | ns | NaCl | KS.3 |
| FIM1_1737 | 224.2277164 | 183.5257118 | 484.8333214 | -0.071542971 | 0.111648886 | 0.144902331 | 0.558828756 | ns | NaCl | KS.3 |
| FIM1_1738 | 1068.982968 | 1068.27723 | 1392.724953 | -0.000174844 | 0.085355451 | 0.997088533 | 0.99924489 | ns | NaCl | KS.3 |
| FIM1_1739 | 4090.701598 | 4726.490135 | 2986.10881 | 0.07000405 | 0.0920163 | 0.165388798 | 0.589355333 | ns | NaCl | KS.3 |
| FIM1_1740 | 748.0260496 | 761.9141926 | 944.6310137 | 0.011147738 | 0.078642727 | 0.820247637 | 0.960172234 | ns | NaCl | KS.3 |
| FIM1_1741 | 5488.109293 | 7700.753664 | 4603.886683 | 0.204464121 | 0.176621042 | 0.0053459 | 0.096361786 | ns | NaCl | KS.3 |
| FIM1_1742 | 1672.380556 | 1793.944148 | 1223.137733 | 0.029851206 | 0.088788293 | 0.532725954 | 0.855153333 | ns | NaCl | KS.3 |
| FIM1_1743 | 3926.714659 | 5908.448471 | 3528.491098 | 0.1588749 | 0.199967825 | 0.018389509 | 0.20421386 | ns | NaCl | KS.3 |
| FIM1_1744 | 2372.671833 | 2703.133222 | 2700.92026 | 0.025976914 | 0.112921846 | 0.525309521 | 0.854940852 | ns | NaCl | KS.3 |
| FIM1_1745 | 1460.997593 | 1509.759845 | 1396.955375 | 0.02226389 | 0.071478778 | 0.650324608 | 0.899642849 | ns | NaCl | KS.3 |
| FIM1_1746 | 575.8163504 | 573.2719184 | 512.8983152 | -0.002226191 | 0.092768671 | 0.961156305 | 0.992888832 | ns | NaCl | KS.3 |
| FIM1_1747 | 3151.217606 | 2724.145572 | 3201.610679 | -0.057544439 | 0.097569981 | 0.23605263 | 0.659616806 | ns | NaCl | KS.3 |
| FIM1_1748 | 28.12817942 | 89.73517735 | 243.67275 | 1.399945513 | 0.367113034 | 9.01913434494 | 0.000160251 | Km>KS.3 | NaCl | KS.3 |
| FIM1_1749 | 4691.629217 | 4375.741308 | 4469.165477 | -0.030548646 | 0.087959179 | 0.525119366 | 0.854940852 | ns | NaCl | KS.3 |
| FIM1_1750 | 5643.439265 | 5695.948081 | 4549.600814 | 0.001962219 | 0.106326602 | 0.961561489 | 0.993101185 | ns | NaCl | KS.3 |
| FIM1_1751 | 2035.388154 | 2699.51528 | 1262.995325 | 0.04432259 | 0.137305877 | 0.279762513 | 0.700652348 | ns | NaCl | KS.3 |
| FIM1_1752 | 929.7498259 | 1094.241206 | 1127.180991 | 0.030335967 | 0.117637945 | 0.455340505 | 0.816788914 | ns | NaCl | KS.3 |
| FIM1_1753 | 521.461528 | 487.730662 | 787.7175786 | -0.038352106 | 0.080246672 | 0.438566516 | 0.808710033 | ns | NaCl | KS.3 |
| FIM1_1754 | 332.1916942 | 378.6319506 | 560.3280302 | 0.030261661 | 0.108968779 | 0.476360947 | 0.825190921 | ns | NaCl | KS.3 |
| FIM1_1755 | 2151.028375 | 2473.578507 | 2207.912755 | 0.024393559 | 0.117175836 | 0.538885564 | 0.857036524 | ns | NaCl | KS.3 |
| FIM1_1756 | 1140.966086 | 1204.40845 | 1284.85607 | 0.014103906 | 0.101503996 | 0.742514623 | 0.941460666 | ns | NaCl | KS.3 |
| FIM1_1757 | 1315.093837 | 1565.12264 | 1650.215053 | 0.042656316 | 0.112414894 | 0.330762453 | 0.742254041 | ns | NaCl | KS.3 |
| FIM1_1758 | 414.3201622 | 421.32233 | 332.026574 | 0.00393967 | 0.103531649 | 0.924805572 | 0.984176127 | ns | NaCl | KS.3 |
| FIM1_1759 | 343.1550694 | 348.8876059 | 363.4795938 | 0.003391173 | 0.106582806 | 0.933515892 | 0.985410898 | ns | NaCl | KS.3 |
| FIM1_1760 | 3633.293811 | 3965.166008 | 3933.361456 | 0.020905554 | 0.105193841 | 0.621216863 | 0.892479944 | ns | NaCl | KS.3 |
| FIM1_1761 | 2067.052806 | 2189.940955 | 2837.073276 | 0.011677023 | 0.108489654 | 0.772692615 | 0.94938462 | ns | NaCl | KS.3 |
| FIM1_1762 | 8280.718732 | 11845.03178 | 5193.355965 | 0.092918647 | 0.158889527 | 0.074029312 | 0.422357599 | ns | NaCl | KS.3 |
| FIM1_1763 | 116.012539 | 112.987862 | 106.1058246 | -0.005730726 | 0.113341259 | 0.881999588 | 0.974110648 | ns | NaCl | KS.3 |
| FIM1_1764 | 5348.51603 | 5579.562046 | 3825.694864 | 0.028635149 | 0.072174939 | 0.560925068 | 0.865840156 | ns | NaCl | KS.3 |
| FIM1_1765 | 1517.342603 | 1765.063058 | 2218.508472 | 0.092919573 | 0.086007898 | 0.075449722 | 0.427519783 | ns | NaCl | KS.3 |

| | | | | | | | | | | |
|---|---|---|---|---|---|---|---|---|---|---|
| FIM1_1766 | 1232.451312 | 1353.982025 | 1836.890909 | 0.018183669 | 0.111444755 | 0.650997841 | 0.899642849 | ns | NaCl | KS.3 |
| FIM1_1767 | 1578.458671 | 1550.387134 | 1753.236508 | -0.008432178 | 0.082577891 | 0.862064274 | 0.969856868 | ns | NaCl | KS.3 |
| FIM1_1769 | 17.20285635 | 14.01985589 | 21.2872472 | -0.014980678 | 0.147027592 | 0.650937608 | 0.899642849 | ns | NaCl | KS.3 |
| FIM1_1770 | 4373.147703 | 4702.41391 | 2963.922448 | 0.048343629 | 0.073852852 | 0.330955941 | 0.742254041 | ns | NaCl | KS.3 |
| FIM1_1771 | 2167.453568 | 2522.618674 | 1673.299133 | 0.076741297 | 0.092226318 | 0.132910742 | 0.539340082 | ns | NaCl | KS.3 |
| FIM1_1772 | 2298.452349 | 2305.505316 | 1718.984209 | 0.001242219 | 0.088561987 | 0.978944106 | 0.996164232 | ns | NaCl | KS.3 |
| FIM1_1774 | 897.9992303 | 791.9948699 | 925.4380031 | -0.065889752 | 0.086871001 | 0.19167958 | 0.618952432 | ns | NaCl | KS.3 |
| FIM1_1775 | 7165.277685 | 6848.768171 | 6632.013889 | -0.021740138 | 0.084132353 | 0.654002558 | 0.900747416 | ns | NaCl | KS.3 |
| FIM1_1776 | 3039.147322 | 3023.113145 | 2784.014127 | -0.002112571 | 0.088241012 | 0.964272275 | 0.993691273 | ns | NaCl | KS.3 |
| FIM1_1777 | 4719.294448 | 2691.752636 | 3183.634497 | -0.708538983 | 0.1969211 | 3.98616718746 | 0.000566605 | ns | NaCl | KS.3 |
| FIM1_1778 | 2487.023007 | 2651.612802 | 2756.463574 | 0.027689943 | 0.088160893 | 0.563089606 | 0.867434473 | ns | NaCl | KS.3 |
| FIM1_1779 | 6559.150888 | 5499.175439 | 5361.145866 | -0.044275037 | 0.112082503 | 0.316577894 | 0.73382286 | ns | NaCl | KS.3 |
| FIM1_1780 | 1021.768696 | 1049.976118 | 982.3092643 | 0.014664934 | 0.080319642 | 0.764419069 | 0.947366775 | ns | NaCl | KS.3 |
| FIM1_1781 | 487.7155479 | 412.4669999 | 622.2993874 | -0.075513597 | 0.097135042 | 0.136239325 | 0.544735099 | ns | NaCl | KS.3 |
| FIM1_1782 | 840.593045 | 771.3150613 | 1110.647824 | -0.063777883 | 0.068809286 | 0.195579213 | 0.618952432 | ns | NaCl | KS.3 |
| FIM1_1783 | 232.1772123 | 128.1853189 | 438.0035736 | -0.720448075 | 0.227699385 | 1.16768066772 | 0.001416881 | ns | NaCl | KS.3 |
| FIM1_1784 | 1450.886897 | 1363.096195 | 2015.541673 | -0.031393473 | 0.083420125 | 0.52118976 | 0.852373129 | ns | NaCl | KS.3 |
| FIM1_1785 | 641.1715677 | 550.7606224 | 875.1746038 | -0.067459214 | 0.096120043 | 0.176129285 | 0.605047427 | ns | NaCl | KS.3 |
| FIM1_1786 | 52.06398282 | 48.54801991 | 53.71768876 | -0.016681538 | 0.110278389 | 0.67929469 | 0.913941356 | ns | NaCl | KS.3 |
| FIM1_1787 | 4308.37537 | 5435.291442 | 2637.023263 | 0.045239994 | 0.125697283 | 0.286806306 | 0.706716876 | ns | NaCl | KS.3 |
| FIM1_1788 | 590.5696827 | 695.0443201 | 541.1282185 | 0.077452892 | 0.095623135 | 0.12867274 | 0.536136415 | ns | NaCl | KS.3 |
| FIM1_1789 | 912.4201769 | 864.9770755 | 1310.35748 | -0.011347893 | 0.107004535 | 0.781303268 | 0.951646805 | ns | NaCl | KS.3 |
| FIM1_1790 | 497.3722269 | 489.6221136 | 819.0189478 | -0.004439939 | 0.093221677 | 0.922387521 | 0.983890455 | ns | NaCl | KS.3 |
| FIM1_1791 | 223.0107624 | 340.1167135 | 360.2382304 | 0.272553818 | 0.23833614 | 0.002917249 | 0.067450821 | ns | NaCl | KS.3 |
| FIM1_1792 | 8109.58377 | 7455.855042 | 6424.741287 | -0.020064241 | 0.105067419 | 0.634931421 | 0.898009483 | ns | NaCl | KS.3 |
| FIM1_1793 | 1928.609246 | 1597.334109 | 1819.898147 | -0.115495514 | 0.095965473 | 0.034308662 | 0.292771172 | ns | NaCl | KS.3 |
| FIM1_1794 | 386.8070546 | 308.2674152 | 681.1152267 | -0.032022277 | 0.131416229 | 0.408817676 | 0.795392815 | ns | NaCl | KS.3 |
| FIM1_1795 | 622.1136658 | 531.5386782 | 820.1649257 | -0.146295086 | 0.073204036 | 0.004864364 | 0.090978232 | ns | NaCl | KS.3 |
| FIM1_1796 | 94.37926605 | 69.51201129 | 82.68238282 | -0.078867782 | 0.144080144 | 0.107561211 | 0.498909937 | ns | NaCl | KS.3 |
| FIM1_1797 | 1848.724411 | 1353.363853 | 1607.787515 | -0.093425677 | 0.144358608 | 0.073625724 | 0.421631393 | ns | NaCl | KS.3 |
| FIM1_1798 | 1226.462969 | 1240.558079 | 1934.363292 | 0.002232847 | 0.108952961 | 0.955303428 | 0.989719816 | ns | NaCl | KS.3 |
| FIM1_1799 | 2070.718264 | 2111.465419 | 1747.339532 | 0.010816283 | 0.079042628 | 0.825373285 | 0.960240659 | ns | NaCl | KS.3 |
| FIM1_1800 | 365.4104128 | 317.7934726 | 846.4389477 | -0.035207771 | 0.108325489 | 0.417259096 | 0.800387073 | ns | NaCl | KS.3 |
| FIM1_1801 | 4099.68818 | 4627.663558 | 2016.171127 | 0.020491172 | 0.116723365 | 0.601085003 | 0.880762097 | ns | NaCl | KS.3 |
| FIM1_1802 | 2228.356575 | 2476.996665 | 3150.560927 | 0.021493784 | 0.110663472 | 0.598491989 | 0.880167289 | ns | NaCl | KS.3 |
| FIM1_1803 | 2338.038752 | 1787.844789 | 4401.067168 | -0.038003373 | 0.136606679 | 0.33628155 | 0.746209059 | ns | NaCl | KS.3 |
| FIM1_1804 | 543.8128019 | 482.1840502 | 737.2140736 | -0.026913761 | 0.108756731 | 0.522206813 | 0.853429187 | ns | NaCl | KS.3 |
| FIM1_1805 | 58.73356421 | 21.33273569 | 87.50925761 | -0.808324581 | 0.569650515 | 0.000291494 | 0.016232695 | ns | NaCl | KS.3 |
| FIM1_1806 | 1248.316478 | 1080.640727 | 1470.141121 | -0.028211687 | 0.114482574 | 0.490398295 | 0.834666958 | ns | NaCl | KS.3 |
| FIM1_1807 | 139.8976412 | 122.0902276 | 206.0439287 | -0.031731467 | 0.109552369 | 0.456136892 | 0.817337042 | ns | NaCl | KS.3 |
| FIM1_1808 | 1835.145454 | 1528.218266 | 2362.410885 | -0.039747449 | 0.116272204 | 0.351463659 | 0.756912594 | ns | NaCl | KS.3 |
| FIM1_1809 | 2475.388178 | 2850.382564 | 3503.93053 | 0.024496723 | 0.117442732 | 0.536764442 | 0.855981197 | ns | NaCl | KS.3 |
| FIM1_1810 | 1430.333596 | 1299.23195 | 1536.574471 | -0.043982501 | 0.088145429 | 0.367709454 | 0.768177319 | ns | NaCl | KS.3 |
| FIM1_1811 | 1587.906293 | 1522.56564 | 1497.327843 | -0.025696199 | 0.076704043 | 0.602633641 | 0.881535538 | ns | NaCl | KS.3 |
| FIM1_1812 | 446.9135276 | 327.8874509 | 669.3733824 | -0.117665577 | 0.147249752 | 0.039206165 | 0.313586284 | ns | NaCl | KS.3 |

| | | | | | | | | | | |
|---|---|---|---|---|---|---|---|---|---|---|
| FIM1_1813 | 477.6799645 | 459.7248847 | 514.8656018 | -0.01757744 | 0.087696402 | 0.711893563 | 0.926245097 | ns | NaCl | KS.3 |
| FIM1_1814 | 792.1710649 | 714.2577819 | 879.8759774 | -0.062755744 | 0.080136282 | 0.213615383 | 0.637897078 | ns | NaCl | KS.3 |
| FIM1_1815 | 494.6292689 | 420.2443173 | 661.3282019 | -0.087000106 | 0.092671617 | 0.094194845 | 0.470201108 | ns | NaCl | KS.3 |
| FIM1_1816 | 2756.685037 | 2261.092088 | 2257.114841 | -0.11544671 | 0.099270708 | 0.035148532 | 0.295899532 | ns | NaCl | KS.3 |
| FIM1_1817 | 1913.986346 | 1626.178267 | 1715.826269 | -0.083021159 | 0.093488929 | 0.107682481 | 0.498909937 | ns | NaCl | KS.3 |
| FIM1_1818 | 32926.00525 | 40411.32655 | 24018.39903 | 0.200502349 | 0.081330633 | 0.000299346 | 0.016365368 | ns | NaCl | KS.3 |
| FIM1_1819 | 2702.244337 | 2606.117657 | 2817.807409 | -0.020266591 | 0.079171408 | 0.680012466 | 0.913941356 | ns | NaCl | KS.3 |
| FIM1_1820 | 531.1614736 | 615.1345335 | 505.4351108 | 0.051954155 | 0.100335438 | 0.273421949 | 0.694726351 | ns | NaCl | KS.3 |
| FIM1_1821 | 2275.332564 | 2499.785461 | 1756.485633 | 0.047496221 | 0.085317294 | 0.336668717 | 0.746466327 | ns | NaCl | KS.3 |
| FIM1_1822 | 346.3106969 | 332.9907812 | 470.8690079 | -0.017134378 | 0.085692393 | 0.721462885 | 0.931688886 | ns | NaCl | KS.3 |
| FIM1_1823 | 379.8084136 | 298.3353282 | 249.6394333 | -0.03673334 | 0.131961789 | 0.355671125 | 0.759483959 | ns | NaCl | KS.3 |
| FIM1_1824 | 872.0157965 | 989.7412902 | 1074.180864 | 0.043970154 | 0.098366779 | 0.347667192 | 0.756654396 | ns | NaCl | KS.3 |
| FIM1_1825 | 6292.729908 | 5437.81022 | 4575.038093 | -0.059991565 | 0.096704478 | 0.22050802 | 0.642906874 | ns | NaCl | KS.3 |
| FIM1_1826 | 7229.51589 | 4865.04246 | 6200.367689 | -0.167384022 | 0.19757661 | 0.015190563 | 0.180365277 | ns | NaCl | KS.3 |
| FIM1_1827 | 6368.149056 | 5594.598909 | 5286.396663 | -0.08613334 | 0.079849296 | 0.094252651 | 0.470201108 | ns | NaCl | KS.3 |
| FIM1_1828 | 3304.314114 | 2982.833034 | 3775.938771 | -0.053737223 | 0.084632018 | 0.281085142 | 0.701655084 | ns | NaCl | KS.3 |
| FIM1_1829 | 190.0403505 | 225.8103478 | 281.7080514 | 0.068547687 | 0.103059248 | 0.164745855 | 0.589224032 | ns | NaCl | KS.3 |
| FIM1_1830 | 1982.896161 | 1829.388117 | 1821.326502 | -0.052579704 | 0.076167695 | 0.292841044 | 0.711969309 | ns | NaCl | KS.3 |
| FIM1_1831 | 684.3714652 | 763.2945841 | 557.696713 | 0.052749727 | 0.087865141 | 0.286213697 | 0.705606117 | ns | NaCl | KS.3 |
| FIM1_1832 | 300.6055075 | 285.8587723 | 410.2668753 | -0.018131615 | 0.088005511 | 0.702867581 | 0.923421278 | ns | NaCl | KS.3 |
| FIM1_1833 | 734.3836073 | 700.1486097 | 903.3919552 | -0.038198696 | 0.067357868 | 0.432347381 | 0.805772482 | ns | NaCl | KS.3 |
| FIM1_1834 | 500.9147464 | 428.6602416 | 892.8793349 | -0.058529226 | 0.100261051 | 0.225672859 | 0.648597616 | ns | NaCl | KS.3 |
| FIM1_1835 | 441.6755311 | 439.1442493 | 680.737761 | -0.004403235 | 0.080870721 | 0.928056831 | 0.984238275 | ns | NaCl | KS.3 |
| FIM1_1836 | 2666.754513 | 2552.757583 | 2170.345231 | -0.015111824 | 0.093797849 | 0.740757167 | 0.941079393 | ns | NaCl | KS.3 |
| FIM1_1837 | 7804.495388 | 9447.45459 | 5368.555356 | 0.056401988 | 0.111089382 | 0.22538944 | 0.648597616 | ns | NaCl | KS.3 |
| FIM1_1838 | 189.8562939 | 219.4405397 | 170.4799956 | 0.035286534 | 0.108326717 | 0.416347532 | 0.800387073 | ns | NaCl | KS.3 |
| FIM1_1839 | 3251.096541 | 3034.487467 | 3181.795781 | -0.035541296 | 0.083145001 | 0.469438071 | 0.821764392 | ns | NaCl | KS.3 |
| FIM1_1840 | 9069.445392 | 9606.346576 | 7274.870358 | 0.020452957 | 0.093382697 | 0.657109241 | 0.901977674 | ns | NaCl | KS.3 |
| FIM1_1841 | 803.075272 | 741.9593471 | 1025.017145 | -0.056336122 | 0.072067087 | 0.257021092 | 0.680871076 | ns | NaCl | KS.3 |
| FIM1_1842 | 710.8774403 | 636.2914535 | 1034.982679 | -0.023991061 | 0.109340994 | 0.563200907 | 0.867434473 | ns | NaCl | KS.3 |
| FIM1_1843 | 10364.25658 | 11537.66401 | 8375.507969 | 0.073644441 | 0.075764092 | 0.145536617 | 0.559788274 | ns | NaCl | KS.3 |
| FIM1_1844 | 72.06659732 | 86.7818668 | 62.51638684 | 0.036813667 | 0.117043103 | 0.380324185 | 0.778493059 | ns | NaCl | KS.3 |
| FIM1_1845 | 576.7469304 | 607.5403924 | 543.7782126 | 0.026051388 | 0.080465389 | 0.596002502 | 0.87962202 | ns | NaCl | KS.3 |
| FIM1_1846 | 4550.809951 | 5239.069905 | 3597.826933 | 0.07943363 | 0.087050999 | 0.122279833 | 0.528318345 | ns | NaCl | KS.3 |
| FIM1_1847 | 4826.034943 | 3427.586872 | 4023.427754 | -0.190515768 | 0.176182732 | 0.00749829 | 0.11809776 | ns | NaCl | KS.3 |
| FIM1_1848 | 9675.413437 | 8428.257311 | 7128.816569 | -0.046690732 | 0.100421889 | 0.317792405 | 0.733979075 | ns | NaCl | KS.3 |
| FIM1_1849 | 2928.464861 | 2838.795011 | 3645.990782 | -0.008892169 | 0.098170706 | 0.839381438 | 0.963748593 | ns | NaCl | KS.3 |
| FIM1_1850 | 556.78051 | 617.7708536 | 680.5337264 | 0.035493361 | 0.097576788 | 0.441013664 | 0.809513863 | ns | NaCl | KS.3 |
| FIM1_1851 | 3978.878701 | 4265.908302 | 3843.198155 | 0.051817471 | 0.069749 | 0.292910902 | 0.711969309 | ns | NaCl | KS.3 |
| FIM1_1853 | 69757.51944 | 79520.17016 | 51825.76471 | 0.065286155 | 0.089360654 | 0.193926406 | 0.618952432 | ns | NaCl | KS.3 |
| FIM1_1854 | 7832.566203 | 8345.846497 | 6421.667845 | 0.018291003 | 0.099212902 | 0.677716199 | 0.913287355 | ns | NaCl | KS.3 |
| FIM1_1855 | 671.1211324 | 582.7136954 | 833.8984346 | -0.034069368 | 0.108489006 | 0.430195238 | 0.805047964 | ns | NaCl | KS.3 |
| FIM1_1856 | 1423.441906 | 1210.995952 | 1627.702602 | -0.047598822 | 0.106625046 | 0.297757652 | 0.715635559 | ns | NaCl | KS.3 |
| FIM1_1857 | 7027.857296 | 7288.801566 | 6646.68315 | 0.01721193 | 0.084534954 | 0.721625529 | 0.931688886 | ns | NaCl | KS.3 |
| FIM1_1858 | 1210.497299 | 1179.533756 | 1277.778027 | -0.013084774 | 0.081246656 | 0.788576305 | 0.953379471 | ns | NaCl | KS.3 |

| | | | | | | | | | | |
|---|---|---|---|---|---|---|---|---|---|---|
| FIM1_1859 | 2563.619173 | 2706.530339 | 1973.780227 | 0.022080197 | 0.08949893 | 0.640569261 | 0.898798052 | ns | NaCl | KS.3 |
| FIM1_1860 | 1084.377329 | 916.366559 | 1463.585236 | -0.047698232 | 0.108471424 | 0.293558296 | 0.712220852 | ns | NaCl | KS.3 |
| FIM1_1861 | 711.4154317 | 733.7594312 | 621.8867677 | 0.012690753 | 0.084043899 | 0.792809297 | 0.955175866 | ns | NaCl | KS.3 |
| FIM1_1862 | 967.4426023 | 922.9073092 | 889.5884481 | -0.024180694 | 0.083380367 | 0.61956027 | 0.892479944 | ns | NaCl | KS.3 |
| FIM1_1863 | 769.4766754 | 684.6098177 | 657.76852 | -0.089507472 | 0.073957794 | 0.078147901 | 0.432946334 | ns | NaCl | KS.3 |
| FIM1_1864 | 898.2602907 | 1008.894802 | 1432.97111 | 0.067332288 | 0.082803097 | 0.18402234 | 0.614437006 | ns | NaCl | KS.3 |
| FIM1_1865 | 966.8233355 | 1024.826831 | 897.4371506 | 0.019069752 | 0.09566957 | 0.673487471 | 0.91246166 | ns | NaCl | KS.3 |
| FIM1_1866 | 228.932337 | 210.1637522 | 472.7962783 | -0.016632819 | 0.11051629 | 0.67963427 | 0.913941356 | ns | NaCl | KS.3 |
| FIM1_1867 | 126.1946062 | 151.2658108 | 199.9969661 | 0.061343481 | 0.110671661 | 0.195919629 | 0.618952432 | ns | NaCl | KS.3 |
| FIM1_1868 | 2426.030192 | 2027.015798 | 1780.688788 | -0.097542492 | 0.095383388 | 0.065541207 | 0.402056108 | ns | NaCl | KS.3 |
| FIM1_1869 | 2510.603973 | 2873.503649 | 2045.223674 | 0.077607389 | 0.085508659 | 0.130182268 | 0.537622789 | ns | NaCl | KS.3 |
| FIM1_1870 | 986.0295638 | 993.7976452 | 1245.736782 | 0.003665592 | 0.082626278 | 0.939739486 | 0.986790704 | ns | NaCl | KS.3 |
| FIM1_1871 | 1268.159376 | 1145.278228 | 1710.552191 | -0.037308803 | 0.094912911 | 0.426407085 | 0.802854185 | ns | NaCl | KS.3 |
| FIM1_1872 | 774.9443044 | 692.4351565 | 908.4532218 | -0.047343017 | 0.091511131 | 0.329047703 | 0.741064881 | ns | NaCl | KS.3 |
| FIM1_1873 | 829.8244564 | 869.328869 | 969.815206 | 0.025167129 | 0.080602285 | 0.608279145 | 0.88569286 | ns | NaCl | KS.3 |
| FIM1_1874 | 1273.331863 | 478.0703947 | 1626.170728 | -1.334749476 | 0.180189582 | 1.60828573365 | 2.00030538123 | Km<KS.3 | NaCl | KS.3 |
| FIM1_1875 | 71.78488484 | 10.57567704 | 58.7017236 | -2.309469795 | 0.387351845 | 3.23479302298 | 1.34109127411 | Km<KS.3 | NaCl | KS.3 |
| FIM1_1876 | 533.6010908 | 742.2077622 | 644.4371434 | 0.05257747 | 0.146048382 | 0.214718533 | 0.63965551 | ns | NaCl | KS.3 |
| FIM1_1877 | 1913.09912 | 2626.458808 | 1186.719207 | 0.052442854 | 0.143311276 | 0.217741096 | 0.641504077 | ns | NaCl | KS.3 |
| FIM1_1878 | 2091.424452 | 2049.615179 | 2956.918088 | -0.002840829 | 0.11848815 | 0.939037754 | 0.986790704 | ns | NaCl | KS.3 |
| FIM1_1879 | 472.4626048 | 385.5630164 | 476.7881425 | -0.128756335 | 0.098191491 | 0.02113444 | 0.221497506 | ns | NaCl | KS.3 |
| FIM1_1880 | 4055.358899 | 3720.427928 | 3492.685695 | -0.04648417 | 0.082471784 | 0.349274659 | 0.756912594 | ns | NaCl | KS.3 |
| FIM1_1881 | 701.8428905 | 666.2171798 | 750.7455602 | -0.030095433 | 0.079255628 | 0.541880274 | 0.857187397 | ns | NaCl | KS.3 |
| FIM1_1882 | 3071.691272 | 2965.954281 | 3247.16289 | -0.010982814 | 0.096249638 | 0.805706157 | 0.957859861 | ns | NaCl | KS.3 |
| FIM1_1883 | 96.74754524 | 100.5402187 | 133.49796 | 0.00571613 | 0.11220481 | 0.88327962 | 0.97456556 | ns | NaCl | KS.3 |
| FIM1_1884 | 4244.55272 | 3976.535682 | 4422.520693 | -0.022620644 | 0.094256528 | 0.622371785 | 0.893077483 | ns | NaCl | KS.3 |
| FIM1_1885 | 1498.549524 | 1416.182536 | 1568.879703 | -0.036935136 | 0.073405166 | 0.455263229 | 0.816788914 | ns | NaCl | KS.3 |
| FIM1_1886 | 627.4948199 | 587.4048718 | 895.1978618 | -0.036170597 | 0.081996561 | 0.463027143 | 0.819880744 | ns | NaCl | KS.3 |
| FIM1_1887 | 7938.752609 | 7568.784888 | 5765.02779 | -0.017942803 | 0.091552684 | 0.699711913 | 0.922863518 | ns | NaCl | KS.3 |
| FIM1_1888 | 2720.489964 | 2656.489665 | 2288.547933 | -0.023871299 | 0.053465028 | 0.585982536 | 0.877826895 | ns | NaCl | KS.3 |
| FIM1_1889 | 1618.46862 | 1806.200686 | 1442.223128 | 0.065364619 | 0.08166766 | 0.196323003 | 0.618952432 | ns | NaCl | KS.3 |
| FIM1_1890 | 2106.453831 | 2919.05115 | 1155.395381 | 0.04296468 | 0.146566955 | 0.280120105 | 0.700652348 | ns | NaCl | KS.3 |
| FIM1_1891 | 536.6825334 | 524.6202811 | 1454.40925 | -0.003850833 | 0.112421112 | 0.920947626 | 0.983890455 | ns | NaCl | KS.3 |
| FIM1_1892 | 813.0057039 | 835.5306654 | 894.0127034 | 0.006582607 | 0.103165701 | 0.875311519 | 0.971808705 | ns | NaCl | KS.3 |
| FIM1_1893 | 2452.130496 | 2272.588054 | 1753.076194 | -0.043103537 | 0.08033207 | 0.385498076 | 0.778665419 | ns | NaCl | KS.3 |
| FIM1_1894 | 1013.758681 | 979.2896943 | 1675.682039 | -0.015025831 | 0.087514807 | 0.752089825 | 0.943523399 | ns | NaCl | KS.3 |
| FIM1_1895 | 1252.242459 | 1275.515081 | 1091.70638 | 0.010812535 | 0.077471454 | 0.825831706 | 0.960240659 | ns | NaCl | KS.3 |
| FIM1_1896 | 1275.571267 | 1150.703155 | 1174.592518 | -0.071119175 | 0.074134912 | 0.157628088 | 0.57787696 | ns | NaCl | KS.3 |
| FIM1_1897 | 2272.627557 | 2378.64791 | 1873.564261 | 0.030990066 | 0.072250164 | 0.529470134 | 0.855153333 | ns | NaCl | KS.3 |
| FIM1_1898 | 1837.848377 | 1862.555652 | 1951.388759 | 0.008748914 | 0.069630352 | 0.857786112 | 0.969856868 | ns | NaCl | KS.3 |
| FIM1_1899 | 9193.222201 | 9747.028858 | 8322.433107 | 0.029087648 | 0.083816263 | 0.551085968 | 0.860945181 | ns | NaCl | KS.3 |
| FIM1_1900 | 421.2611007 | 379.4402438 | 403.9192299 | -0.05406957 | 0.085021294 | 0.27799133 | 0.700256641 | ns | NaCl | KS.3 |
| FIM1_1901 | 4436.026363 | 4068.767914 | 4115.123878 | -0.034079322 | 0.091931491 | 0.47185423 | 0.822127061 | ns | NaCl | KS.3 |
| FIM1_1902 | 1057.110218 | 1099.579596 | 880.7159253 | 0.016757254 | 0.087733466 | 0.724584252 | 0.933222742 | ns | NaCl | KS.3 |
| FIM1_1903 | 1077.103423 | 946.2617019 | 936.9536444 | -0.09210614 | 0.075145221 | 0.071418536 | 0.418675804 | ns | NaCl | KS.3 |

| | | | | | | | | | | |
|---|---|---|---|---|---|---|---|---|---|---|
| FIM1_1904 | 6267.538218 | 8198.118025 | 5400.172419 | 0.176953163 | 0.130861931 | 0.00589461 | 0.102180094 | ns | NaCl | KS.3 |
| FIM1_1905 | 1206.144755 | 1116.192214 | 1071.12881 | -0.040724047 | 0.083568994 | 0.40875972 | 0.795392815 | ns | NaCl | KS.3 |
| FIM1_1907 | 1458.573472 | 1530.796579 | 1143.469546 | 0.017251425 | 0.092947493 | 0.707944261 | 0.925190614 | ns | NaCl | KS.3 |
| FIM1_1908 | 6571.291991 | 6275.751621 | 4344.102142 | -0.020629851 | 0.086554171 | 0.667156358 | 0.907354532 | ns | NaCl | KS.3 |
| FIM1_1909 | 3882.826904 | 3153.131311 | 4253.410281 | -0.168299547 | 0.093262911 | 0.003438345 | 0.075302063 | ns | NaCl | KS.3 |
| FIM1_1910 | 1104.429671 | 779.8713249 | 1422.179399 | -0.25385666 | 0.186377354 | 0.001898435 | 0.052763765 | ns | NaCl | KS.3 |
| FIM1_1911 | 10099.98315 | 10351.7138 | 7619.23478 | 0.020968882 | 0.061021902 | 0.654627903 | 0.900905621 | ns | NaCl | KS.3 |
| FIM1_1912 | 2943.74872 | 3301.184828 | 3167.10355 | 0.068063081 | 0.081923706 | 0.179635823 | 0.608447876 | ns | NaCl | KS.3 |
| FIM1_1913 | 3847.199203 | 5326.810029 | 2600.344044 | 0.092247652 | 0.148552724 | 0.075728608 | 0.427638847 | ns | NaCl | KS.3 |
| FIM1_1914 | 606.5123924 | 624.7692249 | 540.0841743 | 0.011441241 | 0.090863051 | 0.805667275 | 0.957859861 | ns | NaCl | KS.3 |
| FIM1_1915 | 2109.908841 | 2309.230069 | 2001.753845 | 0.020365672 | 0.106918173 | 0.625538856 | 0.895040497 | ns | NaCl | KS.3 |
| FIM1_1916 | 887.2222489 | 971.8748488 | 1040.49571 | 0.026619532 | 0.100013325 | 0.549522294 | 0.860945181 | ns | NaCl | KS.3 |
| FIM1_1917 | 1432.197616 | 1575.72754 | 992.6185635 | 0.0224595 | 0.106177944 | 0.594473282 | 0.87962202 | ns | NaCl | KS.3 |
| FIM1_1918 | 1012.961936 | 913.8603205 | 883.3691243 | -0.041660191 | 0.092253448 | 0.384331247 | 0.778665419 | ns | NaCl | KS.3 |
| FIM1_1919 | 1368.00101 | 1559.287095 | 1482.241881 | 0.077294268 | 0.083922246 | 0.131469849 | 0.538323043 | ns | NaCl | KS.3 |
| FIM1_1920 | 4454.170968 | 6036.994968 | 2593.445431 | 0.056902865 | 0.140051408 | 0.195101028 | 0.618952432 | ns | NaCl | KS.3 |
| FIM1_1921 | 3067.233712 | 2641.105681 | 2806.918951 | -0.065271282 | 0.095664431 | 0.188844484 | 0.618907317 | ns | NaCl | KS.3 |
| FIM1_1922 | 2136.206253 | 2347.131731 | 2248.488215 | 0.091741618 | 0.056548005 | 0.043272057 | 0.328487717 | ns | NaCl | KS.3 |
| FIM1_1923 | 3413.744846 | 4328.393144 | 2652.797944 | 0.228584948 | 0.100381885 | 0.000226293 | 0.014250729 | ns | NaCl | KS.3 |
| FIM1_1924 | 1442.918304 | 1450.957182 | 1067.849095 | 0.002195524 | 0.089525458 | 0.962566818 | 0.99352073 | ns | NaCl | KS.3 |
| FIM1_1925 | 2255.223231 | 1814.961974 | 2161.135795 | -0.191141885 | 0.093930432 | 0.00112558 | 0.039503502 | ns | NaCl | KS.3 |
| FIM1_1926 | 561.1479814 | 547.9910659 | 614.4443236 | -0.007805778 | 0.094638917 | 0.862683467 | 0.969856868 | ns | NaCl | KS.3 |
| FIM1_1927 | 6597.713214 | 8766.474042 | 3871.751739 | 0.058231641 | 0.135143827 | 0.191621065 | 0.618952432 | ns | NaCl | KS.3 |
| FIM1_1928 | 1113.269296 | 1421.035064 | 1626.853045 | 0.176019982 | 0.1141311 | 0.004375868 | 0.085706644 | ns | NaCl | KS.3 |
| FIM1_1929 | 412.2110702 | 441.0576731 | 577.4460461 | 0.014529979 | 0.107025846 | 0.723804398 | 0.93308324 | ns | NaCl | KS.3 |
| FIM1_1930 | 777.3430072 | 730.4982998 | 719.1019424 | -0.028934089 | 0.087788985 | 0.546889905 | 0.858831212 | ns | NaCl | KS.3 |
| FIM1_1931 | 12590.90912 | 11963.89364 | 12554.5319 | -0.044941698 | 0.060234172 | 0.335821829 | 0.745521464 | ns | NaCl | KS.3 |
| FIM1_1932 | 488.0197704 | 416.215935 | 615.3969019 | -0.087663513 | 0.091595908 | 0.092074386 | 0.467895885 | ns | NaCl | KS.3 |
| FIM1_1933 | 5355.369584 | 6111.788464 | 5025.61618 | 0.076487897 | 0.084959585 | 0.135261895 | 0.543081274 | ns | NaCl | KS.3 |
| FIM1_1934 | 235.6502672 | 208.7527437 | 592.1345006 | -0.040754392 | 0.100588244 | 0.375058715 | 0.778113889 | ns | NaCl | KS.3 |
| FIM1_1935 | 688.550639 | 765.1512515 | 2584.393175 | 0.041500747 | 0.093135581 | 0.384173563 | 0.778665419 | ns | NaCl | KS.3 |
| FIM1_1936 | 6269.994376 | 7503.268924 | 5334.781665 | 0.199988749 | 0.061278387 | 2.7651066017[illegible] | 0.002697334 | ns | NaCl | KS.3 |
| FIM1_1937 | 3405.46766 | 3646.739929 | 2724.981043 | 0.046178973 | 0.073556502 | 0.352374209 | 0.756912594 | ns | NaCl | KS.3 |
| FIM1_1938 | 1649.049543 | 1743.206316 | 1245.841524 | 0.021231995 | 0.091190576 | 0.649909855 | 0.899642849 | ns | NaCl | KS.3 |
| FIM1_1939 | 278.8754783 | 396.2375946 | 307.9868761 | 0.116743124 | 0.16379113 | 0.041668626 | 0.323298895 | ns | NaCl | KS.3 |
| FIM1_1940 | 2602.702528 | 2827.67969 | 3000.403003 | 0.021931299 | 0.102254094 | 0.612712894 | 0.889162818 | ns | NaCl | KS.3 |
| FIM1_1941 | 1624.348129 | 1183.140409 | 1548.488475 | -0.086539394 | 0.144860375 | 0.08800697 | 0.45942778 | ns | NaCl | KS.3 |
| FIM1_1942 | 664.3796139 | 621.4336801 | 919.5417893 | -0.024631362 | 0.092256092 | 0.597519028 | 0.880167289 | ns | NaCl | KS.3 |
| FIM1_1943 | 1215.618277 | 1222.790525 | 1103.740139 | 0.00217343 | 0.087779642 | 0.963345701 | 0.993691273 | ns | NaCl | KS.3 |
| FIM1_1944 | 565.1957623 | 440.4431388 | 568.8789386 | -0.171728978 | 0.118615196 | 0.005665242 | 0.099945317 | ns | NaCl | KS.3 |
| FIM1_1945 | 680.0781073 | 699.2147319 | 776.3111489 | 0.011567093 | 0.088910904 | 0.805996097 | 0.957859861 | ns | NaCl | KS.3 |
| FIM1_1946 | 1421.558055 | 1549.341089 | 2199.170287 | 0.03617064 | 0.089713047 | 0.451298984 | 0.814367952 | ns | NaCl | KS.3 |
| FIM1_1947 | 991.0586105 | 1051.416454 | 995.8216166 | 0.0219715 | 0.092034162 | 0.637059042 | 0.898332475 | ns | NaCl | KS.3 |
| FIM1_1948 | 3596.899864 | 4939.880841 | 2939.991438 | 0.3391689 | 0.147792332 | 5.5491139951[illegible] | 0.004679126 | ns | NaCl | KS.3 |
| FIM1_1949 | 670.560419 | 422.4291263 | 448.1257848 | -0.459409031 | 0.248562344 | 0.000231916 | 0.014282363 | ns | NaCl | KS.3 |

| | | | | | | | | | | |
|---|---|---|---|---|---|---|---|---|---|---|
| FIM1_1950 | 1055.949542 | 803.4150089 | 1066.766058 | -0.204978954 | 0.133322073 | 0.002378822 | 0.059470559 | ns | NaCl | KS.3 |
| FIM1_1951 | 735.1306124 | 382.6188948 | 1505.63966 | -0.865755576 | 0.15796238 | 3.3037826346[illegible] | 1.0272699129[illegible] | ns | NaCl | KS.3 |
| FIM1_1952 | 2979.929357 | 3410.639199 | 2539.892516 | 0.096968506 | 0.077853178 | 0.060576662 | 0.384890029 | ns | NaCl | KS.3 |
| FIM1_1953 | 1958.860933 | 2350.654782 | 3001.487266 | 0.052391462 | 0.110195317 | 0.254139239 | 0.677568443 | ns | NaCl | KS.3 |
| FIM1_1954 | 3416.945986 | 5094.517722 | 2132.792289 | 0.051477308 | 0.159671731 | 0.210211648 | 0.634204334 | ns | NaCl | KS.3 |
| FIM1_1955 | 475.6304762 | 502.0967317 | 615.9159347 | 0.029315725 | 0.079360481 | 0.552186612 | 0.860945181 | ns | NaCl | KS.3 |
| FIM1_1956 | 778.9952913 | 878.0237867 | 737.4129827 | 0.035827142 | 0.101282458 | 0.427626369 | 0.803933114 | ns | NaCl | KS.3 |
| FIM1_1957 | 1386.431655 | 1443.262469 | 1281.058434 | 0.018880516 | 0.084321335 | 0.696420605 | 0.921625594 | ns | NaCl | KS.3 |
| FIM1_1958 | 643.4173593 | 445.2752322 | 1017.167306 | -0.255556558 | 0.200436555 | 0.002353853 | 0.059143519 | ns | NaCl | KS.3 |
| FIM1_1959 | 821.5702715 | 961.249103 | 825.9505333 | 0.092817103 | 0.088232951 | 0.076324385 | 0.429055159 | ns | NaCl | KS.3 |
| FIM1_1960 | 3514.122097 | 3226.713701 | 2838.54034 | -0.074792378 | 0.062766322 | 0.117355543 | 0.520359917 | ns | NaCl | KS.3 |
| FIM1_1961 | 664.3218068 | 714.4619519 | 3122.551629 | 0.021811293 | 0.097139186 | 0.627585607 | 0.896016329 | ns | NaCl | KS.3 |
| FIM1_1962 | 47479.42916 | 56692.7575 | 36346.13897 | 0.112229399 | 0.090778355 | 0.037323053 | 0.304670827 | ns | NaCl | KS.3 |
| FIM1_1963 | 569.0523653 | 520.5465462 | 534.5366959 | -0.03575714 | 0.092550141 | 0.450104899 | 0.813595672 | ns | NaCl | KS.3 |
| FIM1_1964 | 18630.16167 | 18535.79387 | 12314.88059 | -0.001262071 | 0.101941936 | 0.976192147 | 0.996164232 | ns | NaCl | KS.3 |
| FIM1_1965 | 938.8998494 | 1145.243587 | 1120.310801 | 0.10626993 | 0.101339918 | 0.049097622 | 0.34611662 | ns | NaCl | KS.3 |
| FIM1_1966 | 1384.816075 | 1330.037299 | 1545.494868 | -0.023244778 | 0.078579226 | 0.636840094 | 0.898332475 | ns | NaCl | KS.3 |
| FIM1_1967 | 1594.719152 | 1501.772753 | 2172.736869 | -0.03237845 | 0.08113323 | 0.510882452 | 0.846104012 | ns | NaCl | KS.3 |
| FIM1_1968 | 3237.853652 | 3909.609483 | 2275.329364 | 0.053914358 | 0.111319091 | 0.241730125 | 0.666264473 | ns | NaCl | KS.3 |
| FIM1_1970 | 678.1666053 | 557.3073537 | 570.7668803 | -0.080782615 | 0.10676243 | 0.111897563 | 0.508857748 | ns | NaCl | KS.3 |
| FIM1_1971 | 414.2131093 | 357.6181625 | 411.3584587 | -0.057215925 | 0.097780615 | 0.238070001 | 0.660936525 | ns | NaCl | KS.3 |
| FIM1_1972 | 627.0991632 | 867.1403417 | 3532.771367 | 0.066578447 | 0.145187134 | 0.147976341 | 0.562830502 | ns | NaCl | KS.3 |
| FIM1_1973 | 7729.961695 | 11616.21725 | 7261.775774 | 0.140965441 | 0.192557591 | 0.026002644 | 0.253653239 | ns | NaCl | KS.3 |
| FIM1_1975 | 1022.801698 | 1106.584383 | 1218.097813 | 0.050940912 | 0.076416794 | 0.307795566 | 0.727796074 | ns | NaCl | KS.3 |
| FIM1_1976 | 2682.145878 | 2960.956391 | 2935.738703 | 0.047060244 | 0.087431702 | 0.338193547 | 0.747703619 | ns | NaCl | KS.3 |
| FIM1_1977 | 799.2745236 | 868.6381073 | 1034.614862 | 0.059792063 | 0.073333077 | 0.231184787 | 0.654979679 | ns | NaCl | KS.3 |
| FIM1_1978 | 2918.406882 | 3552.00355 | 2647.71356 | 0.050286278 | 0.114821118 | 0.262825909 | 0.688188894 | ns | NaCl | KS.3 |
| FIM1_1979 | 622.862019 | 548.2468608 | 814.5126033 | -0.100416675 | 0.075633042 | 0.050875673 | 0.352025696 | ns | NaCl | KS.3 |
| FIM1_1980 | 10292.05982 | 7942.021621 | 8603.960782 | -0.107761711 | 0.125862662 | 0.049248232 | 0.34611662 | ns | NaCl | KS.3 |
| FIM1_1981 | 15465.14614 | 14876.89162 | 10239.13225 | -0.011941787 | 0.096714629 | 0.788533261 | 0.953379471 | ns | NaCl | KS.3 |
| FIM1_1982 | 10.47881188 | 15.6559134 | 64.93626001 | 0.032417102 | 0.159510811 | 0.362004511 | 0.763449629 | ns | NaCl | KS.3 |
| FIM1_1983 | 5787.470224 | 6471.106095 | 4897.795103 | 0.067465694 | 0.081032022 | 0.183152141 | 0.611942177 | ns | NaCl | KS.3 |
| FIM1_1984 | 670.5735473 | 629.3498193 | 569.3680288 | -0.038354421 | 0.078384044 | 0.439357612 | 0.808917498 | ns | NaCl | KS.3 |
| FIM1_1985 | 3028.894721 | 3131.511743 | 2826.378344 | 0.019359957 | 0.077625338 | 0.694101783 | 0.920525145 | ns | NaCl | KS.3 |
| FIM1_1986 | 2338.561234 | 2237.63172 | 2153.313448 | -0.035249331 | 0.065994334 | 0.46577209 | 0.820834625 | ns | NaCl | KS.3 |
| FIM1_1987 | 905.6412489 | 979.0728556 | 767.2735327 | 0.024309432 | 0.098016314 | 0.58804532 | 0.878311243 | ns | NaCl | KS.3 |
| FIM1_1988 | 3015.036631 | 2542.251193 | 2531.306992 | -0.067295774 | 0.101493009 | 0.172428302 | 0.599043855 | ns | NaCl | KS.3 |
| FIM1_1989 | 1495.470123 | 1963.937998 | 1456.817408 | 0.104884918 | 0.131313164 | 0.053867037 | 0.362636686 | ns | NaCl | KS.3 |
| FIM1_1990 | 2761.073334 | 3193.191905 | 2173.972986 | 0.09568422 | 0.082646544 | 0.066693836 | 0.404774002 | ns | NaCl | KS.3 |
| FIM1_1991 | 1776.165184 | 1865.723276 | 2246.902047 | 0.015860333 | 0.095469906 | 0.725346796 | 0.933661141 | ns | NaCl | KS.3 |
| FIM1_1992 | 4757.086511 | 4619.910465 | 4555.279446 | -0.007926987 | 0.099909167 | 0.854388362 | 0.969345976 | ns | NaCl | KS.3 |
| FIM1_1993 | 1125.63385 | 835.1369086 | 1575.670143 | -0.159870669 | 0.14675564 | 0.012325965 | 0.158864447 | ns | NaCl | KS.3 |
| FIM1_1994 | 21983.65777 | 25671.4951 | 15695.74925 | 0.141857083 | 0.069436192 | 0.005152105 | 0.094234281 | ns | NaCl | KS.3 |
| FIM1_1995 | 3314.175199 | 3591.271412 | 3370.377933 | 0.020748366 | 0.102807252 | 0.629718599 | 0.89612415 | ns | NaCl | KS.3 |
| FIM1_1996 | 11580.3382 | 14052.7509 | 7937.596766 | 0.098034637 | 0.100940256 | 0.064917404 | 0.400203329 | ns | NaCl | KS.3 |

| | | | | | | | | | | |
|---|---|---|---|---|---|---|---|---|---|---|
| FIM1_1997 | 1817.576715 | 1727.601613 | 1520.024576 | -0.035651452 | 0.070759996 | 0.4685416 | 0.821764392 | ns | NaCl | KS.3 |
| FIM1_1998 | 1428.522566 | 1554.088907 | 1350.747153 | 0.063891172 | 0.069030326 | 0.195218005 | 0.618952432 | ns | NaCl | KS.3 |
| FIM1_1999 | 354.5508538 | 375.8402786 | 411.8301405 | 0.017173386 | 0.099381559 | 0.695496299 | 0.921340753 | ns | NaCl | KS.3 |
| FIM1_2000 | 8031.13709 | 7654.984346 | 5751.658379 | -0.018065929 | 0.09152178 | 0.697843624 | 0.922074642 | ns | NaCl | KS.3 |
| FIM1_2001 | 4792.255235 | 6751.761414 | 3238.179242 | 0.085543818 | 0.152648058 | 0.089407924 | 0.463337937 | ns | NaCl | KS.3 |
| FIM1_2002 | 4382.819096 | 5126.211443 | 3293.617611 | 0.102120231 | 0.085135656 | 0.05290131 | 0.360032854 | ns | NaCl | KS.3 |
| FIM1_2003 | 5531.066152 | 5583.24666 | 4467.052273 | 0.004233207 | 0.086366316 | 0.929258364 | 0.984238275 | ns | NaCl | KS.3 |
| FIM1_2004 | 495.1475385 | 451.9524266 | 610.1034241 | -0.035564257 | 0.09443098 | 0.447982969 | 0.812914509 | ns | NaCl | KS.3 |
| FIM1_2005 | 49427.12301 | 42290.25548 | 32739.38233 | -0.06116372 | 0.099191568 | 0.209950441 | 0.633800632 | ns | NaCl | KS.3 |
| FIM1_2006 | 656.7770975 | 856.411079 | 869.3747403 | 0.163020975 | 0.128794506 | 0.009075809 | 0.132411004 | ns | NaCl | KS.3 |
| FIM1_2007 | 635.2771646 | 730.3978709 | 634.0629093 | 0.073688716 | 0.088078368 | 0.14828547 | 0.563010285 | ns | NaCl | KS.3 |
| FIM1_2008 | 446.1167082 | 437.3919057 | 575.9856178 | -0.008949422 | 0.089185694 | 0.848867507 | 0.96736917 | ns | NaCl | KS.3 |
| FIM1_2009 | 232.8030599 | 211.7246898 | 814.4961066 | -0.017905175 | 0.1130203 | 0.652345283 | 0.90050438 | ns | NaCl | KS.3 |
| FIM1_2010 | 37.38528646 | 37.77441177 | 78.59222337 | 0.002743026 | 0.114656381 | 0.942664283 | 0.986790704 | ns | NaCl | KS.3 |
| FIM1_2011 | 2787.574036 | 3124.165597 | 2751.075001 | 0.093458378 | 0.06836533 | 0.05985066 | 0.382721124 | ns | NaCl | KS.3 |
| FIM1_2012 | 2373.042271 | 3153.988322 | 3168.947475 | 0.240596121 | 0.139221609 | 0.000824503 | 0.031433557 | ns | NaCl | KS.3 |
| FIM1_2013 | 616.7000207 | 398.4371197 | 1113.463587 | -0.229316457 | 0.236693874 | 0.006103084 | 0.105426535 | ns | NaCl | KS.3 |
| FIM1_2014 | 1205.391564 | 1140.510524 | 1707.956662 | -0.036072334 | 0.075184164 | 0.466548163 | 0.821323144 | ns | NaCl | KS.3 |
| FIM1_2015 | 14.51905366 | 12.21048392 | 21.92099029 | -0.018468835 | 0.13247596 | 0.606920931 | 0.884426372 | ns | NaCl | KS.3 |
| FIM1_2016 | 921.3903108 | 916.767572 | 2361.274511 | -0.005347533 | 0.073324697 | 0.913333909 | 0.982238694 | ns | NaCl | KS.3 |
| FIM1_2017 | 844.9604244 | 758.5433054 | 1226.622931 | -0.035433804 | 0.098809948 | 0.438572798 | 0.808710033 | ns | NaCl | KS.3 |
| FIM1_2018 | 108.4367916 | 50.73045148 | 205.6604993 | -0.294557972 | 0.38984864 | 0.005261975 | 0.095891295 | ns | NaCl | KS.3 |
| FIM1_2019 | 1288.494299 | 1282.545028 | 1473.133084 | -0.001198101 | 0.100991796 | 0.977587019 | 0.996164232 | ns | NaCl | KS.3 |
| FIM1_2020 | 2734.029446 | 2367.260946 | 3206.016417 | -0.046356855 | 0.102524078 | 0.316546177 | 0.73382286 | ns | NaCl | KS.3 |
| FIM1_2021 | 4356.395031 | 4937.41668 | 4196.697368 | 0.042849178 | 0.098797583 | 0.35779182 | 0.761992425 | ns | NaCl | KS.3 |
| FIM1_2022 | 1801.542553 | 1745.371091 | 1584.664656 | -0.013977937 | 0.086682625 | 0.769735157 | 0.94905388 | ns | NaCl | KS.3 |
| FIM1_2023 | 1606.28237 | 1157.762708 | 2191.806579 | -0.193811172 | 0.169161549 | 0.006341568 | 0.106776808 | ns | NaCl | KS.3 |
| FIM1_2024 | 108.6152131 | 112.2311744 | 535.9085137 | 0.002446715 | 0.142772642 | 0.939996727 | 0.986790704 | ns | NaCl | KS.3 |
| FIM1_2025 | 167.1172715 | 117.9399933 | 215.5552265 | -0.134074245 | 0.170962591 | 0.028112764 | 0.262403381 | ns | NaCl | KS.3 |
| FIM1_2026 | 985.02198 | 1051.120725 | 1235.45582 | 0.043580543 | 0.072286587 | 0.378477187 | 0.778493059 | ns | NaCl | KS.3 |
| FIM1_2027 | 1102.83648 | 980.287145 | 972.3894514 | -0.070063592 | 0.083044724 | 0.168085267 | 0.592944764 | ns | NaCl | KS.3 |
| FIM1_2028 | 3156.743798 | 3070.164978 | 2123.234624 | -0.010010394 | 0.092534875 | 0.827435813 | 0.960240659 | ns | NaCl | KS.3 |
| FIM1_2029 | 1546.323041 | 1335.0989 | 2108.789834 | -0.108946719 | 0.078246745 | 0.036776044 | 0.303163208 | ns | NaCl | KS.3 |
| FIM1_2030 | 406.2941724 | 350.8830899 | 1023.682246 | -0.046339738 | 0.103188535 | 0.315361867 | 0.73382286 | ns | NaCl | KS.3 |
| FIM1_2031 | 686.7612808 | 465.0817808 | 1632.790141 | -0.096864507 | 0.170129788 | 0.067016284 | 0.405603424 | ns | NaCl | KS.3 |
| FIM1_2032 | 27.0999283 | 20.52110575 | 27.57214771 | -0.027660567 | 0.145407137 | 0.441929873 | 0.809513863 | ns | NaCl | KS.3 |
| FIM1_2033 | 1268.198297 | 1274.444868 | 1731.269197 | 0.002777293 | 0.073251951 | 0.95491651 | 0.989525023 | ns | NaCl | KS.3 |
| FIM1_2034 | 1893.591159 | 1967.952739 | 2139.809632 | 0.028772494 | 0.06904176 | 0.555980413 | 0.863699344 | ns | NaCl | KS.3 |
| FIM1_2035 | 188.4880899 | 195.988382 | 253.7108773 | 0.010127656 | 0.100449907 | 0.814158934 | 0.958816322 | ns | NaCl | KS.3 |
| FIM1_2037 | 208.1901978 | 205.080036 | 314.2585225 | -0.003991594 | 0.1038548 | 0.923606008 | 0.983924964 | ns | NaCl | KS.3 |
| FIM1_2038 | 1499.345247 | 2027.768791 | 1938.2021 | 0.146478534 | 0.147378211 | 0.018051136 | 0.202175238 | ns | NaCl | KS.3 |
| FIM1_2039 | 833.151596 | 1028.794013 | 620.846494 | 0.080516906 | 0.111111055 | 0.111323751 | 0.507640384 | ns | NaCl | KS.3 |
| FIM1_2040 | 627.6948978 | 838.6988768 | 447.3371595 | 0.143361674 | 0.141430027 | 0.018871333 | 0.207709917 | ns | NaCl | KS.3 |
| FIM1_2041 | 9420.722877 | 16967.89552 | 5070.218881 | 0.783493892 | 0.154250457 | 2.9325024250[illegible] | 8.1051108691[illegible] | ns | NaCl | KS.3 |
| FIM1_2042 | 3970.573424 | 4412.835336 | 3704.298972 | 0.077654295 | 0.072598774 | 0.122396389 | 0.528318345 | ns | NaCl | KS.3 |

| | | | | | | | | | | |
|---|---|---|---|---|---|---|---|---|---|---|
| FIM1_2043 | 3613.113848 | 4830.968762 | 2435.732405 | 0.114032517 | 0.138844053 | 0.042330415 | 0.323990486 | ns | NaCl | KS.3 |
| FIM1_2044 | 734.6807572 | 600.4791354 | 810.1224928 | -0.101402978 | 0.104417479 | 0.058249181 | 0.37782226 | ns | NaCl | KS.3 |
| FIM1_2045 | 336.9648212 | 272.9597417 | 395.9114986 | -0.091239892 | 0.110871026 | 0.080486074 | 0.438574171 | ns | NaCl | KS.3 |
| FIM1_2046 | 247.3362462 | 218.5743826 | 265.7848338 | -0.030917149 | 0.10709822 | 0.472621922 | 0.822130792 | ns | NaCl | KS.3 |
| FIM1_2047 | 670.7346301 | 563.9133569 | 771.6282506 | -0.064966908 | 0.102302744 | 0.184180442 | 0.614552448 | ns | NaCl | KS.3 |
| FIM1_2048 | 2170.92148 | 2319.484589 | 1837.727007 | 0.046321662 | 0.072291582 | 0.349710702 | 0.756912594 | ns | NaCl | KS.3 |
| FIM1_2049 | 424.0388127 | 454.8986486 | 419.7850415 | 0.035574943 | 0.083053103 | 0.469133683 | 0.821764392 | ns | NaCl | KS.3 |
| FIM1_2050 | 4174.397352 | 4230.456646 | 3690.857324 | 0.009369884 | 0.070193028 | 0.848033225 | 0.967208916 | ns | NaCl | KS.3 |
| FIM1_2051 | 2036.194305 | 2272.023746 | 2042.719606 | 0.06816665 | 0.080150403 | 0.178747399 | 0.607807787 | ns | NaCl | KS.3 |
| FIM1_2052 | 2258.397263 | 2701.482399 | 3036.137461 | 0.068704912 | 0.103703432 | 0.163454458 | 0.587986932 | ns | NaCl | KS.3 |
| FIM1_2053 | 7824.42825 | 9421.108451 | 6991.329792 | 0.128488682 | 0.090335365 | 0.019264851 | 0.209721297 | ns | NaCl | KS.3 |
| FIM1_2054 | 1251.285143 | 1221.88207 | 1495.778769 | -0.013656354 | 0.078255029 | 0.780974353 | 0.951646805 | ns | NaCl | KS.3 |
| FIM1_2055 | 747.2311189 | 980.8623129 | 893.0585356 | 0.041627208 | 0.135836414 | 0.303939205 | 0.723145646 | ns | NaCl | KS.3 |
| FIM1_2056 | 662.4334013 | 845.6921076 | 844.1085238 | 0.082101077 | 0.120778147 | 0.103538444 | 0.491042668 | ns | NaCl | KS.3 |
| FIM1_2057 | 641.936406 | 626.8787464 | 693.8576477 | -0.015425946 | 0.080308637 | 0.752643677 | 0.943523399 | ns | NaCl | KS.3 |
| FIM1_2058 | 372.1870624 | 360.5371041 | 517.6529092 | -0.006405166 | 0.108842507 | 0.872771521 | 0.971243188 | ns | NaCl | KS.3 |
| FIM1_2059 | 1122.950435 | 984.3981561 | 774.0261571 | -0.065116325 | 0.090455096 | 0.194183477 | 0.618952432 | ns | NaCl | KS.3 |
| FIM1_2060 | 2886.380029 | 4406.429821 | 3723.71952 | 0.092436406 | 0.17716079 | 0.073844706 | 0.421788075 | ns | NaCl | KS.3 |
| FIM1_2061 | 2224.433471 | 2317.664318 | 1823.673689 | 0.018361906 | 0.086199188 | 0.701951436 | 0.923421278 | ns | NaCl | KS.3 |
| FIM1_2062 | 690.0292092 | 675.3479139 | 1165.197352 | -0.004095076 | 0.109883739 | 0.917599341 | 0.982366413 | ns | NaCl | KS.3 |
| FIM1_2063 | 2116.929092 | 2047.310279 | 3667.742342 | -0.016865063 | 0.082917581 | 0.728582102 | 0.935336397 | ns | NaCl | KS.3 |
| FIM1_2064 | 211.109704 | 232.6045349 | 1152.213554 | 0.019308651 | 0.110406322 | 0.634484646 | 0.898009483 | ns | NaCl | KS.3 |
| FIM1_2065 | 35.57245997 | 55.58757667 | 56.90405186 | 0.10612091 | 0.189693435 | 0.054274329 | 0.363411554 | ns | NaCl | KS.3 |
| FIM1_2066 | 1017.237616 | 753.1369182 | 3256.199129 | -0.038577905 | 0.142769441 | 0.322609951 | 0.738943142 | ns | NaCl | KS.3 |
| FIM1_2067 | 1355.491759 | 1582.509774 | 2073.776345 | 0.06574901 | 0.097496317 | 0.184416156 | 0.614926525 | ns | NaCl | KS.3 |
| FIM1_2068 | 1222.584575 | 1280.88033 | 1436.187791 | 0.028446509 | 0.077062159 | 0.564724969 | 0.867532568 | ns | NaCl | KS.3 |
| FIM1_2069 | 914.7057739 | 754.4848837 | 1555.198042 | -0.106765042 | 0.098992412 | 0.047882513 | 0.343442 | ns | NaCl | KS.3 |
| FIM1_2070 | 103.7307768 | 70.81048563 | 80.28421507 | -0.098606024 | 0.16791987 | 0.064375888 | 0.398841896 | ns | NaCl | KS.3 |
| FIM1_2071 | 2070.621013 | 2156.310816 | 2107.247229 | 0.036459541 | 0.059412249 | 0.43164739 | 0.805772482 | ns | NaCl | KS.3 |
| FIM1_2072 | 1678.865647 | 1824.194803 | 1485.603666 | 0.040094759 | 0.085798441 | 0.412912677 | 0.799937916 | ns | NaCl | KS.3 |
| FIM1_2073 | 4776.419983 | 5443.709376 | 3368.971596 | 0.038815043 | 0.102869304 | 0.390183855 | 0.779482559 | ns | NaCl | KS.3 |
| FIM1_2074 | 913.3342872 | 870.1983577 | 831.1518814 | -0.029383447 | 0.075746587 | 0.552190095 | 0.860945181 | ns | NaCl | KS.3 |
| FIM1_2075 | 5793.559291 | 5273.006277 | 6104.843797 | -0.045840282 | 0.086367523 | 0.351570969 | 0.756912594 | ns | NaCl | KS.3 |
| FIM1_2076 | 1350.462936 | 1193.983013 | 1345.166444 | -0.073204121 | 0.082516737 | 0.151133152 | 0.566606957 | ns | NaCl | KS.3 |
| FIM1_2077 | 2878.97292 | 3230.65241 | 2543.597471 | 0.040983806 | 0.096957984 | 0.381125257 | 0.778609403 | ns | NaCl | KS.3 |
| FIM1_2078 | 1455.319082 | 1269.526523 | 1355.849322 | -0.047959113 | 0.099489433 | 0.308340236 | 0.728391584 | ns | NaCl | KS.3 |
| FIM1_2079 | 230.5512153 | 216.2463179 | 1965.955201 | -0.010077727 | 0.117181129 | 0.790133278 | 0.953873589 | ns | NaCl | KS.3 |
| FIM1_2080 | 38.77790434 | 32.52211158 | 36.46740299 | -0.022985165 | 0.127350717 | 0.540310519 | 0.857165292 | ns | NaCl | KS.3 |
| FIM1_2081 | 19349.15563 | 26740.00997 | 11195.6198 | 0.101706918 | 0.149613969 | 0.059473996 | 0.381784685 | ns | NaCl | KS.3 |
| FIM1_2083 | 520.0881834 | 535.9710093 | 754.0544526 | 0.007305191 | 0.102931681 | 0.862117519 | 0.969856868 | ns | NaCl | KS.3 |
| FIM1_2084 | 7192.394911 | 8240.902755 | 6222.755823 | 0.133756335 | 0.061521366 | 0.005060215 | 0.093585763 | ns | NaCl | KS.3 |
| FIM1_2085 | 2649.982238 | 2439.391673 | 2721.631889 | -0.070109345 | 0.063564459 | 0.143945744 | 0.558168415 | ns | NaCl | KS.3 |
| FIM1_2086 | 3034.521486 | 2963.454893 | 3151.420719 | -0.016631093 | 0.069525091 | 0.733488248 | 0.938375543 | ns | NaCl | KS.3 |
| FIM1_2087 | 3646.917133 | 4521.973853 | 3960.918562 | 0.045458579 | 0.121418545 | 0.291031659 | 0.711059783 | ns | NaCl | KS.3 |
| FIM1_2088 | 1906.25193 | 2458.87472 | 1507.92372 | 0.150944644 | 0.123435218 | 0.012655133 | 0.161434064 | ns | NaCl | KS.3 |

| | | | | | | | | | | |
|---|---|---|---|---|---|---|---|---|---|---|
| FIM1_2089 | 319.3765562 | 306.2429993 | 349.9156491 | -0.015430397 | 0.092242528 | 0.738379278 | 0.939738272 | ns | NaCl | KS.3 |
| FIM1_2090 | 741.4170763 | 833.110101 | 845.9294823 | 0.075149337 | 0.077997321 | 0.13954798 | 0.550332947 | ns | NaCl | KS.3 |
| FIM1_2091 | 1689.069251 | 1752.528184 | 1214.776412 | 0.018056251 | 0.083499246 | 0.709817994 | 0.925563973 | ns | NaCl | KS.3 |
| FIM1_2092 | 885.1877928 | 1071.732813 | 677.1625973 | 0.039093646 | 0.118617121 | 0.353647576 | 0.757528931 | ns | NaCl | KS.3 |
| FIM1_2093 | 5715.933575 | 7461.344153 | 3388.232517 | 0.050162231 | 0.13224627 | 0.242439296 | 0.666657537 | ns | NaCl | KS.3 |
| FIM1_2094 | 11835.99103 | 13386.52881 | 8107.293509 | 0.05777299 | 0.090168096 | 0.243822077 | 0.667638752 | ns | NaCl | KS.3 |
| FIM1_2095 | 442.1543852 | 358.2274029 | 1015.029635 | -0.079949411 | 0.111848971 | 0.112971117 | 0.511380136 | ns | NaCl | KS.3 |
| FIM1_2096 | 599.0393982 | 486.152177 | 1200.383166 | -0.053090411 | 0.117592564 | 0.239138606 | 0.662163805 | ns | NaCl | KS.3 |
| FIM1_2097 | 994.8730863 | 1014.625192 | 1071.318606 | 0.015682704 | 0.068030096 | 0.746978036 | 0.943274417 | ns | NaCl | KS.3 |
| FIM1_2098 | 26.15571023 | 8.856995882 | 1283.458349 | -0.225215594 | 0.423226028 | 0.00860854 | 0.12856213 | ns | NaCl | KS.3 |
| FIM1_2099 | 19.34199753 | 23.03736861 | 15.19172552 | 0.015247794 | 0.135737952 | 0.661836338 | 0.904388188 | ns | NaCl | KS.3 |
| FIM1_2100 | 2498.412254 | 2991.713796 | 1852.733756 | 0.075816711 | 0.101838419 | 0.132623394 | 0.539053419 | ns | NaCl | KS.3 |
| FIM1_2101 | 843.2030807 | 906.0241606 | 761.0685894 | 0.034870253 | 0.085734151 | 0.474136307 | 0.82307921 | ns | NaCl | KS.3 |
| FIM1_2102 | 882.6543404 | 936.8043539 | 1111.61358 | 0.03126223 | 0.081926058 | 0.524544244 | 0.854940852 | ns | NaCl | KS.3 |
| FIM1_2103 | 5711.522555 | 5222.200094 | 5729.274012 | -0.021084018 | 0.10566426 | 0.617217022 | 0.891518014 | ns | NaCl | KS.3 |
| FIM1_2104 | 1450.093463 | 965.9611584 | 766.9407679 | -0.197250028 | 0.211820446 | 0.009071152 | 0.132411004 | ns | NaCl | KS.3 |
| FIM1_2105 | 471.119294 | 516.4260742 | 459.8558034 | 0.034813876 | 0.092673929 | 0.461168302 | 0.819880744 | ns | NaCl | KS.3 |
| FIM1_2106 | 575.7843693 | 439.0962666 | 901.2688859 | -0.110189794 | 0.13071595 | 0.046316141 | 0.340862131 | ns | NaCl | KS.3 |
| FIM1_2107 | 854.0429612 | 889.9392667 | 817.3913675 | 0.021927639 | 0.081362951 | 0.654116179 | 0.900747416 | ns | NaCl | KS.3 |
| FIM1_2108 | 590.3048451 | 486.7443239 | 655.523916 | -0.091568133 | 0.103457903 | 0.080295132 | 0.438494274 | ns | NaCl | KS.3 |
| FIM1_2109 | 273.578821 | 309.9378223 | 233.2353216 | 0.039665825 | 0.10007924 | 0.387758564 | 0.779433881 | ns | NaCl | KS.3 |
| FIM1_2110 | 375.2845235 | 460.6178876 | 423.3050302 | 0.07787143 | 0.108904758 | 0.121375612 | 0.527999953 | ns | NaCl | KS.3 |
| FIM1_2111 | 32.15527758 | 43.13395226 | 44.30100201 | 0.046257872 | 0.138465848 | 0.263612959 | 0.688243886 | ns | NaCl | KS.3 |
| FIM1_2112 | 578.8414413 | 563.9075047 | 1126.871592 | -0.012993697 | 0.085018928 | 0.787130056 | 0.95289761 | ns | NaCl | KS.3 |
| FIM1_2113 | 1216.89918 | 1113.594034 | 1409.47737 | -0.048665252 | 0.082424146 | 0.328121423 | 0.740319311 | ns | NaCl | KS.3 |
| FIM1_2115 | 1328.142219 | 1374.418951 | 1230.235367 | 0.020279319 | 0.076531461 | 0.680679857 | 0.914496972 | ns | NaCl | KS.3 |
| FIM1_2116 | 1093.702447 | 927.998114 | 1207.977676 | -0.155152885 | 0.073360389 | 0.002928518 | 0.067450821 | ns | NaCl | KS.3 |
| FIM1_2117 | 2537.287697 | 2891.33712 | 2470.402507 | 0.120201756 | 0.063898715 | 0.01331363 | 0.166477316 | ns | NaCl | KS.3 |
| FIM1_2118 | 1420.992321 | 1493.30856 | 1670.173672 | 0.044680279 | 0.055182103 | 0.316997638 | 0.73382286 | ns | NaCl | KS.3 |
| FIM1_2119 | 1428.881823 | 1502.355038 | 891.3692608 | 0.008693216 | 0.112864782 | 0.823196392 | 0.960240659 | ns | NaCl | KS.3 |
| FIM1_2120 | 1994.640083 | 1875.655382 | 1712.28105 | -0.029837713 | 0.084655259 | 0.539953395 | 0.857165292 | ns | NaCl | KS.3 |
| FIM1_2121 | 3509.629365 | 3831.176301 | 3712.965937 | 0.061739109 | 0.072884743 | 0.216258466 | 0.640629323 | ns | NaCl | KS.3 |
| FIM1_2122 | 1414.833664 | 1409.339536 | 1269.252022 | -0.001615115 | 0.076708281 | 0.973775302 | 0.995588189 | ns | NaCl | KS.3 |
| FIM1_2123 | 6869.36868 | 7493.785561 | 7668.136712 | 0.054836159 | 0.077320721 | 0.273728036 | 0.694794376 | ns | NaCl | KS.3 |
| FIM1_2124 | 1124.778747 | 997.2374894 | 2574.161137 | -0.036542007 | 0.101682407 | 0.418367262 | 0.800387073 | ns | NaCl | KS.3 |
| FIM1_2125 | 1083.357455 | 1022.28441 | 1307.919858 | -0.024627343 | 0.088574282 | 0.605217509 | 0.883689315 | ns | NaCl | KS.3 |
| FIM1_2126 | 925.4091082 | 737.5163188 | 1140.724034 | -0.104264369 | 0.112444357 | 0.053676559 | 0.362470256 | ns | NaCl | KS.3 |
| FIM1_2127 | 944.5374178 | 955.1755637 | 1172.311572 | 0.006535169 | 0.072869218 | 0.894149957 | 0.977268396 | ns | NaCl | KS.3 |
| FIM1_2128 | 847.0779833 | 707.5173961 | 763.6469738 | -0.077452815 | 0.102399403 | 0.125799972 | 0.534982546 | ns | NaCl | KS.3 |
| FIM1_2129 | 4204.50925 | 4482.918964 | 3207.282133 | 0.037378768 | 0.078784142 | 0.450707472 | 0.813595672 | ns | NaCl | KS.3 |
| FIM1_2130 | 2400.348228 | 2586.637435 | 4591.296402 | 0.020267216 | 0.101327537 | 0.641029858 | 0.898798052 | ns | NaCl | KS.3 |
| FIM1_2131 | 14758.60353 | 14752.44962 | 10641.61632 | -0.000195874 | 0.084626626 | 0.996749617 | 0.99924489 | ns | NaCl | KS.3 |
| FIM1_2132 | 3940.663529 | 5493.695651 | 2400.954067 | 0.051691369 | 0.146614521 | 0.219525078 | 0.642906874 | ns | NaCl | KS.3 |
| FIM1_2133 | 1537.335864 | 1521.296122 | 1195.454349 | -0.00753579 | 0.072062494 | 0.877943423 | 0.972834689 | ns | NaCl | KS.3 |
| FIM1_2134 | 967.9534375 | 888.8285753 | 923.6418429 | -0.041242817 | 0.086326333 | 0.399485962 | 0.787737366 | ns | NaCl | KS.3 |

| | | | | | | | | | | |
|---|---|---|---|---|---|---|---|---|---|---|
| FIM1_2135 | 2891.34144 | 3148.158329 | 2432.676187 | 0.050243867 | 0.079630095 | 0.314537715 | 0.732852732 | ns | NaCl | KS.3 |
| FIM1_2136 | 1950.307562 | 1915.03217 | 1894.50463 | -0.013464305 | 0.070360609 | 0.783195969 | 0.951646805 | ns | NaCl | KS.3 |
| FIM1_2137 | 1029.425267 | 1080.676728 | 929.025873 | 0.022196461 | 0.085185767 | 0.645943039 | 0.898899754 | ns | NaCl | KS.3 |
| FIM1_2138 | 1148.670484 | 1151.010122 | 1077.161676 | 0.000566206 | 0.097177409 | 0.989753385 | 0.999134858 | ns | NaCl | KS.3 |
| FIM1_2139 | 3102.731208 | 3143.651748 | 2822.038722 | 0.00640539 | 0.084158578 | 0.894302477 | 0.977268396 | ns | NaCl | KS.3 |
| FIM1_2140 | 835.6789648 | 819.3022438 | 1049.093401 | -0.011991577 | 0.075864662 | 0.807445867 | 0.957859861 | ns | NaCl | KS.3 |
| FIM1_2141 | 1703.856004 | 1720.274121 | 1365.113948 | 0.004419402 | 0.0855642 | 0.926460664 | 0.984176127 | ns | NaCl | KS.3 |
| FIM1_2142 | 144.8759867 | 124.9842805 | 184.7391978 | -0.050493234 | 0.100013537 | 0.285779159 | 0.705523209 | ns | NaCl | KS.3 |
| FIM1_2143 | 870.2926816 | 888.9467992 | 887.3095082 | 0.010278377 | 0.082269521 | 0.832517724 | 0.961831996 | ns | NaCl | KS.3 |
| FIM1_2144 | 1201.984332 | 1213.659098 | 1324.008097 | 0.004488588 | 0.08541513 | 0.925368241 | 0.984176127 | ns | NaCl | KS.3 |
| FIM1_2145 | 506.1898827 | 627.6019341 | 768.7166833 | 0.055339566 | 0.118011519 | 0.223892051 | 0.648059715 | ns | NaCl | KS.3 |
| FIM1_2146 | 12735.53114 | 10636.7242 | 11469.12418 | -0.14354041 | 0.083188744 | 0.008313853 | 0.126487513 | ns | NaCl | KS.3 |
| FIM1_2147 | 919.3502936 | 983.2340399 | 730.851883 | 0.024892045 | 0.091781655 | 0.594833951 | 0.87962202 | ns | NaCl | KS.3 |
| FIM1_2148 | 503.275981 | 515.5398878 | 554.9139861 | 0.006845139 | 0.096086178 | 0.877969339 | 0.972834689 | ns | NaCl | KS.3 |
| FIM1_2149 | 892.8175395 | 948.946697 | 947.8479098 | 0.039628938 | 0.076076866 | 0.424709966 | 0.802854185 | ns | NaCl | KS.3 |
| FIM1_2150 | 1081.877644 | 983.0539619 | 1219.572483 | -0.031268844 | 0.097084347 | 0.494583115 | 0.836358994 | ns | NaCl | KS.3 |
| FIM1_2151 | 1451.041501 | 1861.469186 | 1008.198779 | 0.144118478 | 0.120975061 | 0.0154687 | 0.182493537 | ns | NaCl | KS.3 |
| FIM1_2152 | 291.5300893 | 312.7987763 | 256.6004841 | 0.033258173 | 0.085614679 | 0.494329345 | 0.836358994 | ns | NaCl | KS.3 |
| FIM1_2153 | 291.4943897 | 287.4362594 | 445.3975149 | -0.006038632 | 0.084980907 | 0.899947292 | 0.978786499 | ns | NaCl | KS.3 |
| FIM1_2154 | 2020.381826 | 2958.413006 | 1304.304847 | 0.060151892 | 0.157306448 | 0.168583612 | 0.592944764 | ns | NaCl | KS.3 |
| FIM1_2155 | 705.5835713 | 738.9851718 | 789.1994816 | 0.028308099 | 0.075133991 | 0.566619252 | 0.868965626 | ns | NaCl | KS.3 |
| FIM1_2156 | 1525.084271 | 1567.133824 | 1257.674473 | 0.010896336 | 0.088931909 | 0.816959249 | 0.959030737 | ns | NaCl | KS.3 |
| FIM1_2157 | 2829.326868 | 3078.569236 | 3731.140676 | 0.063960289 | 0.069785194 | 0.19609975 | 0.618952432 | ns | NaCl | KS.3 |
| FIM1_2158 | 3947.759583 | 4217.966443 | 2815.198628 | 0.040339804 | 0.077280638 | 0.416714767 | 0.800387073 | ns | NaCl | KS.3 |
| FIM1_2159 | 83.41214435 | 65.95252499 | 86.34968685 | -0.036634083 | 0.12926271 | 0.36083327 | 0.763371684 | ns | NaCl | KS.3 |
| FIM1_2160 | 1688.847639 | 1616.245182 | 1771.477552 | -0.021036469 | 0.084825547 | 0.663514081 | 0.904585408 | ns | NaCl | KS.3 |
| FIM1_2161 | 1164.99574 | 1193.624402 | 1565.997904 | 0.018935722 | 0.055872008 | 0.67333178 | 0.91246166 | ns | NaCl | KS.3 |
| FIM1_2162 | 979.5585768 | 906.6071186 | 767.1367039 | -0.045429666 | 0.079192504 | 0.361599834 | 0.763449629 | ns | NaCl | KS.3 |
| FIM1_2163 | 813.7958404 | 599.9192765 | 761.4008034 | -0.130280471 | 0.147640814 | 0.028092315 | 0.262403381 | ns | NaCl | KS.3 |
| FIM1_2164 | 762.4549559 | 755.3036022 | 769.3607597 | -0.008351845 | 0.074625136 | 0.865153366 | 0.970128052 | ns | NaCl | KS.3 |
| FIM1_2165 | 3572.980726 | 3602.492389 | 2668.911091 | 0.005056813 | 0.075867664 | 0.918076284 | 0.982488618 | ns | NaCl | KS.3 |
| FIM1_2166 | 382.4728997 | 357.716085 | 398.1753113 | -0.015996184 | 0.103827679 | 0.705168795 | 0.924674422 | ns | NaCl | KS.3 |
| FIM1_2167 | 4797.35433 | 4342.101145 | 4787.92655 | -0.058357164 | 0.080788203 | 0.245673687 | 0.668980073 | ns | NaCl | KS.3 |
| FIM1_2168 | 5934.025582 | 6240.204933 | 4647.44812 | 0.043896386 | 0.060635283 | 0.348585948 | 0.756654396 | ns | NaCl | KS.3 |
| FIM1_2169 | 2957.477399 | 2884.303188 | 2427.300803 | -0.016335934 | 0.072802419 | 0.739696035 | 0.940888181 | ns | NaCl | KS.3 |
| FIM1_2170 | 2617.238284 | 2595.840392 | 3230.549092 | -0.00366128 | 0.083523794 | 0.93960187 | 0.986790704 | ns | NaCl | KS.3 |
| FIM1_2171 | 862.1477847 | 1171.994706 | 1110.462969 | 0.144490186 | 0.150076586 | 0.019421927 | 0.210969626 | ns | NaCl | KS.3 |
| FIM1_2172 | 1908.802749 | 2152.366021 | 2057.017079 | 0.074426231 | 0.080884995 | 0.144533085 | 0.558679104 | ns | NaCl | KS.3 |
| FIM1_2173 | 628.3226187 | 842.0660173 | 774.6059075 | 0.122166313 | 0.140987556 | 0.034081565 | 0.292771172 | ns | NaCl | KS.3 |
| FIM1_2174 | 1129.906077 | 948.9670177 | 1263.218158 | -0.067267026 | 0.102104998 | 0.172048254 | 0.5990025 | ns | NaCl | KS.3 |
| FIM1_2175 | 4028.431634 | 4253.149453 | 3557.19596 | 0.02989603 | 0.080191113 | 0.543938294 | 0.857313452 | ns | NaCl | KS.3 |
| FIM1_2176 | 407.501714 | 288.094189 | 925.9356295 | -0.198487986 | 0.181370273 | 0.006563462 | 0.108271704 | ns | NaCl | KS.3 |
| FIM1_2177 | 84.66953348 | 62.68040997 | 79.54727981 | -0.07267089 | 0.134362338 | 0.129766905 | 0.537155579 | ns | NaCl | KS.3 |
| FIM1_2178 | 4072.147919 | 3916.573336 | 4144.390667 | -0.010633777 | 0.099750214 | 0.806281403 | 0.957859861 | ns | NaCl | KS.3 |
| FIM1_2179 | 8581.747563 | 7898.6878 | 9088.135023 | -0.047362718 | 0.080477805 | 0.341777043 | 0.750617902 | ns | NaCl | KS.3 |

| | | | | | | | | | | |
|---|---|---|---|---|---|---|---|---|---|---|
| FIM1_2180 | 4716.34215 | 6145.978916 | 2978.44097 | 0.051797019 | 0.13147209 | 0.23237471 | 0.654993872 | ns | NaCl | KS.3 |
| FIM1_2181 | 3675.474332 | 4221.746287 | 2827.95635 | 0.035948667 | 0.106972233 | 0.412017787 | 0.799137814 | ns | NaCl | KS.3 |
| FIM1_2182 | 487.8424706 | 1122.202521 | 1562.092515 | 0.500010142 | 0.472996427 | 0.001287712 | 0.041599773 | ns | NaCl | KS.3 |
| FIM1_2183 | 2360.486906 | 2334.779657 | 2582.084018 | -0.006839982 | 0.075379994 | 0.889378438 | 0.977268396 | ns | NaCl | KS.3 |
| FIM1_2184 | 2752.721468 | 2791.742093 | 2910.723947 | 0.006898929 | 0.083346429 | 0.886600443 | 0.976280921 | ns | NaCl | KS.3 |
| FIM1_2185 | 1520.360441 | 2404.348497 | 3145.59594 | 0.438418379 | 0.252592428 | 0.000321385 | 0.017192375 | ns | NaCl | KS.3 |
| FIM1_2186 | 1905.507682 | 1926.350229 | 1562.596842 | 0.006573536 | 0.077823683 | 0.893469808 | 0.977268396 | ns | NaCl | KS.3 |
| FIM1_2187 | 2409.317939 | 3150.06116 | 1825.116442 | 0.076323764 | 0.129503496 | 0.119049625 | 0.522746589 | ns | NaCl | KS.3 |
| FIM1_2188 | 5283.899295 | 4742.47336 | 4842.867628 | -0.079655336 | 0.072986686 | 0.113883872 | 0.513199513 | ns | NaCl | KS.3 |
| FIM1_2189 | 1316.728734 | 1376.834518 | 1001.661405 | 0.016179334 | 0.092535179 | 0.725820257 | 0.933787374 | ns | NaCl | KS.3 |
| FIM1_2190 | 2136.384149 | 2018.963725 | 2212.447532 | -0.033766953 | 0.077601462 | 0.495157738 | 0.836358994 | ns | NaCl | KS.3 |
| FIM1_2191 | 9230.695672 | 9778.340141 | 7904.87394 | 0.048222156 | 0.063515918 | 0.312846142 | 0.731871738 | ns | NaCl | KS.3 |
| FIM1_2192 | 939.626753 | 1098.923623 | 907.4031976 | 0.061424992 | 0.09953764 | 0.207910869 | 0.633002483 | ns | NaCl | KS.3 |
| FIM1_2193 | 7649.492532 | 10119.57031 | 4372.129069 | 0.055190351 | 0.134464016 | 0.20889629 | 0.633579568 | ns | NaCl | KS.3 |
| FIM1_2194 | 952.5452126 | 1095.956847 | 674.7807856 | 0.03439939 | 0.108689633 | 0.425809378 | 0.802854185 | ns | NaCl | KS.3 |
| FIM1_2195 | 781.037781 | 883.1094601 | 827.4549321 | 0.061373272 | 0.088071508 | 0.220204324 | 0.642906874 | ns | NaCl | KS.3 |
| FIM1_2196 | 257.3802963 | 297.1224133 | 346.3742841 | 0.049344962 | 0.101602882 | 0.292405628 | 0.711969309 | ns | NaCl | KS.3 |
| FIM1_2197 | 1410.921112 | 1574.189534 | 1611.03317 | 0.066091899 | 0.080826127 | 0.191644587 | 0.618952432 | ns | NaCl | KS.3 |
| FIM1_2198 | 1858.464667 | 2254.157895 | 2316.489743 | 0.050761718 | 0.113620486 | 0.261041138 | 0.686178949 | ns | NaCl | KS.3 |
| FIM1_2199 | 112.2277378 | 184.6276488 | 102.5282884 | 0.210281567 | 0.259913604 | 0.009768259 | 0.138844204 | ns | NaCl | KS.3 |
| FIM1_2200 | 1504.305566 | 1595.941804 | 1233.819232 | 0.025352221 | 0.088680666 | 0.594692482 | 0.87962202 | ns | NaCl | KS.3 |
| FIM1_2201 | 1152.296616 | 1218.49786 | 1062.087388 | 0.041470494 | 0.069611769 | 0.398287328 | 0.786613521 | ns | NaCl | KS.3 |
| FIM1_2202 | 3352.736602 | 3097.087755 | 4287.758579 | -0.039554294 | 0.084749287 | 0.420421729 | 0.80133378 | ns | NaCl | KS.3 |
| FIM1_2203 | 717.1140746 | 500.0411545 | 970.0864016 | -0.364583238 | 0.18616822 | 0.000169232 | 0.011693431 | ns | NaCl | KS.3 |
| FIM1_2204 | 515.4403782 | 463.6401939 | 982.1773121 | -0.042941253 | 0.093552422 | 0.368296387 | 0.768249278 | ns | NaCl | KS.3 |
| FIM1_2205 | 881.4555814 | 802.6668036 | 972.1536766 | -0.042292362 | 0.088432827 | 0.384852829 | 0.778665419 | ns | NaCl | KS.3 |
| FIM1_2206 | 2055.115579 | 2678.781201 | 1280.748421 | 0.048348288 | 0.132362347 | 0.254963968 | 0.678991959 | ns | NaCl | KS.3 |
| FIM1_2207 | 316.527491 | 308.9067093 | 462.4421567 | -0.003502018 | 0.117720162 | 0.925306212 | 0.984176127 | ns | NaCl | KS.3 |
| FIM1_2209 | 1532.294932 | 1728.916025 | 1758.944291 | 0.030044995 | 0.106171626 | 0.48633728 | 0.830120331 | ns | NaCl | KS.3 |
| FIM1_2210 | 7614.406488 | 11297.5624 | 4726.000485 | 0.133134609 | 0.184733549 | 0.030064547 | 0.27272008 | ns | NaCl | KS.3 |
| FIM1_2211 | 6634.434226 | 5877.821529 | 4800.855103 | -0.087302737 | 0.075541226 | 0.086954269 | 0.456808328 | ns | NaCl | KS.3 |
| FIM1_2212 | 739.7774413 | 663.1349287 | 1039.059064 | -0.056697769 | 0.08629451 | 0.255780235 | 0.678991959 | ns | NaCl | KS.3 |
| FIM1_2213 | 1018.371281 | 1017.772806 | 898.7242659 | 0.00021877 | 0.077634596 | 0.996443712 | 0.99924489 | ns | NaCl | KS.3 |
| FIM1_2214 | 558.8961368 | 577.7334727 | 624.8375672 | 0.019906457 | 0.078096915 | 0.685816413 | 0.916694426 | ns | NaCl | KS.3 |
| FIM1_2215 | 9560.638676 | 10983.74517 | 6323.786572 | 0.046003295 | 0.101013142 | 0.322832564 | 0.739010075 | ns | NaCl | KS.3 |
| FIM1_2216 | 932.3908537 | 987.7406554 | 919.6030042 | 0.032161481 | 0.079334393 | 0.514907678 | 0.849077129 | ns | NaCl | KS.3 |
| FIM1_2217 | 586.3162502 | 621.5954527 | 851.7880038 | 0.013504949 | 0.104605563 | 0.746774118 | 0.943274417 | ns | NaCl | KS.3 |
| FIM1_2218 | 8576.569619 | 8109.943693 | 8006.289705 | -0.041116931 | 0.069181708 | 0.401601674 | 0.789189152 | ns | NaCl | KS.3 |
| FIM1_2219 | 3393.929558 | 3781.299624 | 2623.452048 | 0.080933351 | 0.072183603 | 0.107432253 | 0.498909937 | ns | NaCl | KS.3 |
| FIM1_2220 | 1342.524216 | 1444.073407 | 1521.415233 | 0.0677245 | 0.057809448 | 0.139712666 | 0.550332947 | ns | NaCl | KS.3 |
| FIM1_2221 | 772.817843 | 767.6779539 | 801.815568 | -0.002997866 | 0.070338608 | 0.951113535 | 0.987736646 | ns | NaCl | KS.3 |
| FIM1_2222 | 365.4897843 | 409.7337359 | 622.0733073 | 0.041133174 | 0.09591151 | 0.381944395 | 0.778609403 | ns | NaCl | KS.3 |
| FIM1_2223 | 845.1666626 | 998.1236952 | 1068.373156 | 0.087184853 | 0.09366012 | 0.093568875 | 0.470201108 | ns | NaCl | KS.3 |
| FIM1_2224 | 5752.335558 | 6021.29237 | 3951.563321 | 0.021973045 | 0.084250186 | 0.650440317 | 0.899642849 | ns | NaCl | KS.3 |
| FIM1_2225 | 1063.201376 | 1175.190203 | 1245.852409 | 0.025796236 | 0.104023181 | 0.550463783 | 0.860945181 | ns | NaCl | KS.3 |

| | | | | | | | | | | |
|---|---|---|---|---|---|---|---|---|---|---|
| FIM1_2226 | 12864.58605 | 13495.44899 | 10280.44126 | 0.048049605 | 0.052376804 | 0.267301028 | 0.689593109 | ns | NaCl | KS.3 |
| FIM1_2227 | 3386.187499 | 3605.972655 | 2047.478626 | 0.02091001 | 0.095568091 | 0.645146768 | 0.898899754 | ns | NaCl | KS.3 |
| FIM1_2228 | 887.3560835 | 968.9653681 | 941.2334906 | 0.014315559 | 0.116007902 | 0.710505575 | 0.925563973 | ns | NaCl | KS.3 |
| FIM1_2229 | 1049.0415 | 1154.391291 | 888.5419249 | 0.02809719 | 0.100319944 | 0.528143633 | 0.855153333 | ns | NaCl | KS.3 |
| FIM1_2230 | 311.1758762 | 238.658999 | 429.8878394 | -0.061955957 | 0.129381939 | 0.176831247 | 0.605137125 | ns | NaCl | KS.3 |
| FIM1_2231 | 913.5089161 | 818.9155318 | 934.2259976 | -0.079999759 | 0.074791706 | 0.11438399 | 0.514859894 | ns | NaCl | KS.3 |
| FIM1_2232 | 348.8348722 | 413.5398425 | 872.8070102 | 0.039471782 | 0.112924408 | 0.360792518 | 0.763371684 | ns | NaCl | KS.3 |
| FIM1_2233 | 2220.465528 | 2479.04376 | 1744.585935 | 0.053753634 | 0.087867252 | 0.27782165 | 0.700183744 | ns | NaCl | KS.3 |
| FIM1_2234 | 347.4558771 | 337.2515925 | 525.7881748 | -0.017001603 | 0.079716999 | 0.728795412 | 0.935336397 | ns | NaCl | KS.3 |
| FIM1_2235 | 562.5884969 | 631.1071653 | 542.9644863 | 0.036459518 | 0.099649183 | 0.424458869 | 0.802854185 | ns | NaCl | KS.3 |
| FIM1_2236 | 1096.689109 | 1079.439345 | 1267.879751 | -0.007851009 | 0.082448895 | 0.871528894 | 0.971243188 | ns | NaCl | KS.3 |
| FIM1_2237 | 23.68461919 | 18.11514785 | 40.526282 | -0.033466919 | 0.138202856 | 0.381482208 | 0.778609403 | ns | NaCl | KS.3 |
| FIM1_2238 | 582.7168997 | 639.9252243 | 875.0395528 | 0.042602477 | 0.088507244 | 0.381450617 | 0.778609403 | ns | NaCl | KS.3 |
| FIM1_2239 | 514.3651725 | 476.6695395 | 707.4400088 | -0.03240104 | 0.090229565 | 0.496620818 | 0.836358994 | ns | NaCl | KS.3 |
| FIM1_2240 | 515.3483576 | 506.9549837 | 670.9627868 | -0.006255165 | 0.08958735 | 0.893669776 | 0.977268396 | ns | NaCl | KS.3 |
| FIM1_2241 | 3177.582411 | 2434.849333 | 2657.322158 | -0.103082937 | 0.128554514 | 0.056537919 | 0.372058397 | ns | NaCl | KS.3 |
| FIM1_2243 | 1318.666839 | 1396.821612 | 1190.464514 | 0.028144055 | 0.083730671 | 0.563922934 | 0.867434473 | ns | NaCl | KS.3 |
| FIM1_2244 | 5923.35314 | 6175.557828 | 5009.643501 | 0.02302487 | 0.07965868 | 0.639438651 | 0.898798052 | ns | NaCl | KS.3 |
| FIM1_2245 | 1820.173895 | 1834.20748 | 1697.103408 | 0.003223344 | 0.091724548 | 0.944227392 | 0.987155352 | ns | NaCl | KS.3 |
| FIM1_2246 | 15084.73279 | 16643.69206 | 11412.85161 | 0.07596117 | 0.069842727 | 0.126468887 | 0.534982546 | ns | NaCl | KS.3 |
| FIM1_2247 | 1034.526134 | 1075.59011 | 1103.334595 | 0.024448845 | 0.075523652 | 0.620271208 | 0.892479944 | ns | NaCl | KS.3 |
| FIM1_2248 | 5108.003205 | 6489.765045 | 2883.141521 | 0.043918884 | 0.127863801 | 0.294780603 | 0.713530949 | ns | NaCl | KS.3 |
| FIM1_2249 | 7220.435339 | 9020.821151 | 4456.848732 | 0.047589799 | 0.122467703 | 0.272564687 | 0.693610905 | ns | NaCl | KS.3 |
| FIM1_2250 | 631.030025 | 833.2344684 | 393.1173166 | 0.058454649 | 0.133749132 | 0.191469775 | 0.618952432 | ns | NaCl | KS.3 |
| FIM1_2251 | 534.5631446 | 502.4958017 | 559.9824934 | -0.040317944 | 0.073445194 | 0.415642402 | 0.800387073 | ns | NaCl | KS.3 |
| FIM1_2252 | 3243.791361 | 3221.231359 | 3579.467591 | -0.003101495 | 0.084432487 | 0.94862093 | 0.987736646 | ns | NaCl | KS.3 |
| FIM1_2253 | 20946.19586 | 23395.47389 | 15903.47299 | 0.102760056 | 0.061449177 | 0.030526872 | 0.274523423 | ns | NaCl | KS.3 |
| FIM1_2254 | 2672.829662 | 2700.356538 | 2607.848379 | 0.004213966 | 0.088574733 | 0.928678249 | 0.984238275 | ns | NaCl | KS.3 |
| FIM1_2255 | 16914.64434 | 18643.55417 | 13851.80021 | 0.095743886 | 0.056539084 | 0.034955289 | 0.295879009 | ns | NaCl | KS.3 |
| FIM1_2256 | 1024.621673 | 747.7232515 | 1342.415325 | -0.132691146 | 0.151499462 | 0.026820503 | 0.255616863 | ns | NaCl | KS.3 |
| FIM1_2257 | 293.3931992 | 359.1314037 | 1340.549922 | 0.026779267 | 0.130695461 | 0.478041946 | 0.826171162 | ns | NaCl | KS.3 |
| FIM1_2258 | 2661.877478 | 2700.551634 | 2130.826195 | 0.005395934 | 0.089450444 | 0.908266488 | 0.980825899 | ns | NaCl | KS.3 |
| FIM1_2259 | 360.2029292 | 376.2707869 | 421.2064794 | 0.017896616 | 0.087380153 | 0.707428999 | 0.925190614 | ns | NaCl | KS.3 |
| FIM1_2260 | 1808.714085 | 1740.785222 | 1913.190993 | -0.023926575 | 0.074729819 | 0.627628568 | 0.896016329 | ns | NaCl | KS.3 |
| FIM1_2261 | 3867.860044 | 4120.223878 | 3325.419316 | 0.038811043 | 0.076846424 | 0.434231397 | 0.807589234 | ns | NaCl | KS.3 |
| FIM1_2262 | 2996.423712 | 2355.596507 | 2968.684737 | -0.082324115 | 0.120236954 | 0.103012125 | 0.489947725 | ns | NaCl | KS.3 |
| FIM1_2263 | 4004.493629 | 4377.306585 | 4218.55243 | 0.040403813 | 0.088263396 | 0.405551297 | 0.792777094 | ns | NaCl | KS.3 |
| FIM1_2264 | 1594.270706 | 1424.953848 | 1340.425027 | -0.066698653 | 0.081725546 | 0.18791784 | 0.618637711 | ns | NaCl | KS.3 |
| FIM1_2265 | 846.0423157 | 1060.134065 | 992.9188256 | 0.17562001 | 0.102791813 | 0.00331245 | 0.073568922 | ns | NaCl | KS.3 |
| FIM1_2266 | 1688.335002 | 1850.858749 | 1334.817771 | 0.041467115 | 0.088757281 | 0.393107864 | 0.781740625 | ns | NaCl | KS.3 |
| FIM1_2267 | 1571.204411 | 1827.346025 | 1619.757706 | 0.077858858 | 0.090987543 | 0.128437316 | 0.536136415 | ns | NaCl | KS.3 |
| FIM1_2268 | 1530.187806 | 1704.054014 | 1336.216251 | 0.072800329 | 0.076053314 | 0.15023779 | 0.564839452 | ns | NaCl | KS.3 |
| FIM1_2270 | 1015.67907 | 1126.1812 | 840.3921184 | 0.032539563 | 0.099209117 | 0.472799337 | 0.822151941 | ns | NaCl | KS.3 |
| FIM1_2271 | 1578.455643 | 1535.313438 | 1494.176021 | -0.015649181 | 0.081150328 | 0.748682419 | 0.943274417 | ns | NaCl | KS.3 |
| FIM1_2272 | 426.5643668 | 280.7646115 | 737.2469833 | -0.311812386 | 0.236304014 | 0.001456156 | 0.045562107 | ns | NaCl | KS.3 |

| | | | | | | | | | |
|---|---|---|---|---|---|---|---|---|---|
| FIM1_2273 | 117.9939261 | 78.91761045 | 169.1592428 | -0.113616998 | 0.177367327 | 0.045762435 | 0.338791839 | ns | NaCl | KS.3 |
| FIM1_2274 | 1597.839378 | 1163.276927 | 2250.51838 | -0.093516893 | 0.146298883 | 0.073386356 | 0.421631393 | ns | NaCl | KS.3 |
| FIM1_2275 | 2133.436175 | 1980.457004 | 2090.895444 | -0.026532934 | 0.093762345 | 0.566931712 | 0.868965626 | ns | NaCl | KS.3 |
| FIM1_2276 | 1131.003701 | 894.9685846 | 871.5900199 | -0.126154496 | 0.114516911 | 0.0266542 | 0.254519471 | ns | NaCl | KS.3 |
| FIM1_2278 | 1429.300699 | 1722.886512 | 1870.660216 | 0.058257674 | 0.109373884 | 0.215725271 | 0.640629323 | ns | NaCl | KS.3 |
| FIM1_2279 | 230.8240406 | 284.1320046 | 338.5058139 | 0.101736827 | 0.103062885 | 0.057493442 | 0.375860543 | ns | NaCl | KS.3 |
| FIM1_2280 | 2272.699991 | 1883.126984 | 1950.211584 | -0.132859025 | 0.090075631 | 0.015935964 | 0.185236963 | ns | NaCl | KS.3 |
| FIM1_2281 | 570.5759555 | 460.5142561 | 749.2942018 | -0.075467826 | 0.114158892 | 0.127930523 | 0.536116795 | ns | NaCl | KS.3 |
| FIM1_2282 | 145.5816697 | 113.0072041 | 305.8777883 | -0.107685475 | 0.121778449 | 0.049024283 | 0.34611662 | ns | NaCl | KS.3 |
| FIM1_2283 | 718.9067128 | 707.221977 | 1312.087633 | -0.008510121 | 0.08325971 | 0.860454899 | 0.969856868 | ns | NaCl | KS.3 |
| FIM1_2284 | 1828.66312 | 1813.047591 | 2494.53559 | -0.002279367 | 0.100105265 | 0.95771451 | 0.990773484 | ns | NaCl | KS.3 |
| FIM1_2285 | 4433.140299 | 3706.259798 | 4091.590869 | -0.070364822 | 0.102970836 | 0.155999432 | 0.57446127 | ns | NaCl | KS.3 |
| FIM1_2286 | 1149.592176 | 1003.704824 | 1032.886163 | -0.07234812 | 0.087885941 | 0.155055866 | 0.572735074 | ns | NaCl | KS.3 |
| FIM1_2287 | 705.8352193 | 664.5879388 | 927.4687669 | -0.028631903 | 0.085867994 | 0.554263247 | 0.862821173 | ns | NaCl | KS.3 |
| FIM1_2288 | 2048.274518 | 2322.034766 | 1886.171791 | 0.074073597 | 0.083307307 | 0.146802259 | 0.562037426 | ns | NaCl | KS.3 |
| FIM1_2289 | 4665.449194 | 5221.144918 | 3305.0036 | 0.052218092 | 0.089440154 | 0.288730503 | 0.708650347 | ns | NaCl | KS.3 |
| FIM1_2290 | 2690.686471 | 3028.37839 | 2575.129183 | 0.042444036 | 0.097098182 | 0.365684274 | 0.766012322 | ns | NaCl | KS.3 |
| FIM1_2291 | 4658.588996 | 5058.537539 | 3516.64279 | 0.035627127 | 0.089362893 | 0.458501298 | 0.818684405 | ns | NaCl | KS.3 |
| FIM1_2292 | 1092.233784 | 1047.464443 | 872.7159229 | -0.015578407 | 0.091774033 | 0.736811528 | 0.938943994 | ns | NaCl | KS.3 |
| FIM1_2293 | 665.6764619 | 635.207336 | 613.4994378 | -0.013905129 | 0.098073445 | 0.752700982 | 0.943523399 | ns | NaCl | KS.3 |
| FIM1_2294 | 12075.6911 | 13446.51787 | 9396.152219 | 0.095738478 | 0.063538543 | 0.047060578 | 0.34239091 | ns | NaCl | KS.3 |
| FIM1_2295 | 740.4836188 | 795.9695656 | 746.0783644 | 0.028442248 | 0.090731615 | 0.547589078 | 0.859115631 | ns | NaCl | KS.3 |
| FIM1_2297 | 1268.101059 | 1214.532764 | 1216.926502 | -0.030318194 | 0.072580907 | 0.538589176 | 0.857036524 | ns | NaCl | KS.3 |
| FIM1_2298 | 1234.634334 | 1182.135674 | 1665.205528 | -0.016356091 | 0.092394233 | 0.723250007 | 0.932893126 | ns | NaCl | KS.3 |
| FIM1_2299 | 1437.75661 | 1361.077059 | 1448.848662 | -0.023394418 | 0.088623433 | 0.622914143 | 0.893340404 | ns | NaCl | KS.3 |
| FIM1_2300 | 471.5225971 | 375.2119732 | 538.300696 | -0.108534863 | 0.115013767 | 0.047147943 | 0.342424841 | ns | NaCl | KS.3 |
| FIM1_2301 | 3428.439177 | 3334.664608 | 2796.687825 | -0.008651235 | 0.09593896 | 0.846481346 | 0.967147129 | ns | NaCl | KS.3 |
| FIM1_2302 | 2757.76613 | 2679.248541 | 3328.172383 | -0.018538347 | 0.074719564 | 0.706673636 | 0.925190614 | ns | NaCl | KS.3 |
| FIM1_2303 | 217.3501435 | 247.9836454 | 302.1341551 | 0.032014296 | 0.107695122 | 0.457202839 | 0.817337042 | ns | NaCl | KS.3 |
| FIM1_2304 | 149.9825317 | 148.8264625 | 326.5847066 | -0.001739943 | 0.09648306 | 0.968708631 | 0.99408528 | ns | NaCl | KS.3 |
| FIM1_2305 | 709.7031692 | 561.5279775 | 762.5268359 | -0.204855277 | 0.105902217 | 0.000981334 | 0.035377818 | ns | NaCl | KS.3 |
| FIM1_2306 | 99.35573829 | 54.78276326 | 275.4655839 | -0.602475852 | 0.31354457 | 0.000183548 | 0.01250889 | ns | NaCl | KS.3 |
| FIM1_2307 | 3207.69433 | 3527.958544 | 2506.191909 | 0.058588992 | 0.078857242 | 0.243839118 | 0.667638752 | ns | NaCl | KS.3 |
| FIM1_2308 | 2792.42656 | 3083.915569 | 1907.014888 | 0.02888926 | 0.100826818 | 0.515996239 | 0.849801744 | ns | NaCl | KS.3 |
| FIM1_2309 | 150.6558131 | 131.5400575 | 295.1554919 | -0.026586004 | 0.113444868 | 0.515383804 | 0.849359886 | ns | NaCl | KS.3 |
| FIM1_2310 | 358.4090015 | 467.9128384 | 334.031678 | 0.085263881 | 0.129534944 | 0.093061212 | 0.470201108 | ns | NaCl | KS.3 |
| FIM1_2311 | 950.1332616 | 1408.788989 | 1672.056037 | 0.106256695 | 0.174397219 | 0.053963345 | 0.362794107 | ns | NaCl | KS.3 |
| FIM1_2312 | 5182.340927 | 5623.201908 | 4768.238598 | 0.055661625 | 0.073931945 | 0.264685077 | 0.688347233 | ns | NaCl | KS.3 |
| FIM1_2313 | 1707.54474 | 2278.892226 | 1339.623606 | 0.155502329 | 0.142506287 | 0.013414161 | 0.166838629 | ns | NaCl | KS.3 |
| FIM1_2314 | 843.3284178 | 738.9697298 | 1073.429431 | -0.094084343 | 0.079506949 | 0.069203031 | 0.413804182 | ns | NaCl | KS.3 |
| FIM1_2315 | 603.4595487 | 476.65299 | 1049.537091 | -0.07281126 | 0.120407829 | 0.135054251 | 0.54272609 | ns | NaCl | KS.3 |
| FIM1_2316 | 953.3048362 | 761.2907608 | 886.4830386 | -0.061877693 | 0.118827149 | 0.185306053 | 0.616653921 | ns | NaCl | KS.3 |
| FIM1_2317 | 1778.826943 | 2022.562712 | 2337.312787 | 0.068767805 | 0.086706493 | 0.174746358 | 0.604143941 | ns | NaCl | KS.3 |
| FIM1_2318 | 18716.28643 | 20110.38196 | 13391.12303 | 0.023539743 | 0.096198257 | 0.60401495 | 0.883036843 | ns | NaCl | KS.3 |
| FIM1_2319 | 127.7726748 | 129.9856451 | 164.3750103 | 0.002680286 | 0.104434294 | 0.948370253 | 0.987736646 | ns | NaCl | KS.3 |

| | | | | | | | | | | |
|---|---|---|---|---|---|---|---|---|---|---|
| FIM1_2320 | 265.0701106 | 279.8528298 | 214.5370624 | 0.023294614 | 0.086458864 | 0.628065222 | 0.896016329 | ns | NaCl | KS.3 |
| FIM1_2321 | 1787.057221 | 1860.943643 | 1675.237695 | 0.036063946 | 0.057316568 | 0.428516233 | 0.804596805 | ns | NaCl | KS.3 |
| FIM1_2322 | 708.014691 | 669.0685246 | 811.9243081 | -0.031425086 | 0.081450144 | 0.522880269 | 0.854014884 | ns | NaCl | KS.3 |
| FIM1_2323 | 3696.796812 | 4217.017104 | 3024.517997 | 0.080242433 | 0.083098755 | 0.118412536 | 0.52179129 | ns | NaCl | KS.3 |
| FIM1_2324 | 4349.796889 | 4571.616659 | 3890.555805 | 0.019367178 | 0.090490091 | 0.679622484 | 0.913941356 | ns | NaCl | KS.3 |
| FIM1_2325 | 39.81366391 | 25.10824459 | 40.96022622 | -0.135981856 | 0.212204578 | 0.030204777 | 0.27272008 | ns | NaCl | KS.3 |
| FIM1_2326 | 397.6375534 | 364.5828714 | 442.7879753 | -0.046958471 | 0.085176562 | 0.342035158 | 0.750617902 | ns | NaCl | KS.3 |
| FIM1_2327 | 2781.389079 | 2881.902919 | 3239.293127 | 0.025878803 | 0.069322226 | 0.596517685 | 0.879797482 | ns | NaCl | KS.3 |
| FIM1_2328 | 9155.52335 | 10238.96642 | 7241.459513 | 0.091009593 | 0.06881688 | 0.067287554 | 0.406565836 | ns | NaCl | KS.3 |
| FIM1_2329 | 318.7816574 | 347.9734661 | 371.0631883 | 0.025493439 | 0.100580716 | 0.564050356 | 0.867434473 | ns | NaCl | KS.3 |
| FIM1_2330 | 550.4593048 | 580.3392996 | 471.7337333 | 0.015439356 | 0.098323417 | 0.726448809 | 0.934112904 | ns | NaCl | KS.3 |
| FIM1_2331 | 2804.711776 | 3352.058187 | 2684.208701 | 0.06124473 | 0.105910054 | 0.201644101 | 0.624846998 | ns | NaCl | KS.3 |
| FIM1_2332 | 12086.19766 | 11707.58256 | 8886.548113 | -0.011910542 | 0.091097912 | 0.79760478 | 0.955924408 | ns | NaCl | KS.3 |
| FIM1_2333 | 334.0772835 | 382.9794288 | 387.8348301 | 0.051144166 | 0.097878638 | 0.284226521 | 0.704898775 | ns | NaCl | KS.3 |
| FIM1_2334 | 402.2867373 | 392.1506418 | 624.1776974 | -0.009698344 | 0.09109882 | 0.834387721 | 0.962365393 | ns | NaCl | KS.3 |
| FIM1_2335 | 6794.235267 | 6803.099802 | 6406.854433 | 0.000499084 | 0.090809876 | 0.991411889 | 0.999134858 | ns | NaCl | KS.3 |
| FIM1_2336 | 748.6792655 | 730.8214338 | 1689.305228 | -0.005989641 | 0.102271284 | 0.887281022 | 0.976547586 | ns | NaCl | KS.3 |
| FIM1_2337 | 189.1712629 | 187.2974688 | 216.0073424 | -0.002698282 | 0.104247394 | 0.948108541 | 0.987736646 | ns | NaCl | KS.3 |
| FIM1_2338 | 1596.790579 | 1753.32673 | 1538.303557 | 0.063774084 | 0.073925589 | 0.203101621 | 0.626494461 | ns | NaCl | KS.3 |
| FIM1_2339 | 359.0718816 | 350.646766 | 640.8000692 | -0.011694836 | 0.085460446 | 0.807490303 | 0.957859861 | ns | NaCl | KS.3 |
| FIM1_2340 | 1059.335684 | 901.7549766 | 1242.990799 | -0.049313092 | 0.105944094 | 0.284762734 | 0.70510214 | ns | NaCl | KS.3 |
| FIM1_2341 | 508.3003228 | 401.3557893 | 716.9177179 | -0.086923245 | 0.119141385 | 0.090406799 | 0.463573758 | ns | NaCl | KS.3 |
| FIM1_2342 | 1992.751557 | 1793.059246 | 1706.408292 | -0.04952515 | 0.088281548 | 0.313954269 | 0.732050122 | ns | NaCl | KS.3 |
| FIM1_2343 | 15.92105326 | 17.95588163 | 62.07038615 | 0.013597643 | 0.131752606 | 0.699943603 | 0.922863518 | ns | NaCl | KS.3 |
| FIM1_2344 | 215.5166803 | 214.8541074 | 771.224789 | 0.000479909 | 0.10411485 | 0.990770577 | 0.999134858 | ns | NaCl | KS.3 |
| FIM1_2345 | 635.3706101 | 585.1827968 | 917.6130562 | -0.035449971 | 0.090061094 | 0.459216559 | 0.819441312 | ns | NaCl | KS.3 |
| FIM1_2346 | 4215.624568 | 4492.231852 | 4009.822228 | 0.037405645 | 0.078218688 | 0.45056435 | 0.813595672 | ns | NaCl | KS.3 |
| FIM1_2347 | 672.3221085 | 684.7006568 | 620.2654514 | 0.006154088 | 0.092275256 | 0.893400561 | 0.977268396 | ns | NaCl | KS.3 |
| FIM1_2348 | 3600.945964 | 3795.390848 | 2895.085466 | 0.035345561 | 0.072910618 | 0.474325245 | 0.82307921 | ns | NaCl | KS.3 |
| FIM1_2349 | 1844.622195 | 1588.986088 | 2302.086355 | -0.121131067 | 0.07547625 | 0.019828026 | 0.213516076 | ns | NaCl | KS.3 |
| FIM1_2350 | 2858.400662 | 2791.276543 | 2886.681087 | -0.013408755 | 0.077739209 | 0.78497708 | 0.952502676 | ns | NaCl | KS.3 |
| FIM1_2351 | 1181.752103 | 962.3962494 | 1167.699938 | -0.068331815 | 0.111513248 | 0.159260061 | 0.581305064 | ns | NaCl | KS.3 |
| FIM1_2352 | 904.6763048 | 894.5660297 | 1125.829973 | -0.003030589 | 0.100735924 | 0.943498749 | 0.986944129 | ns | NaCl | KS.3 |
| FIM1_2353 | 857.0591189 | 840.7220228 | 972.6931714 | -0.012037169 | 0.080283204 | 0.805579949 | 0.957859861 | ns | NaCl | KS.3 |
| FIM1_2354 | 367.5355107 | 375.8430519 | 595.0128552 | 0.007267564 | 0.094802601 | 0.871864776 | 0.971243188 | ns | NaCl | KS.3 |
| FIM1_2355 | 171.8975018 | 168.504362 | 214.1355932 | -0.007162491 | 0.098749779 | 0.869512774 | 0.971243188 | ns | NaCl | KS.3 |
| FIM1_2356 | 283.8618421 | 216.0842784 | 428.712158 | -0.059679693 | 0.131791404 | 0.186531656 | 0.618044911 | ns | NaCl | KS.3 |
| FIM1_2357 | 3157.937669 | 3964.754427 | 2252.726222 | 0.077092137 | 0.116792155 | 0.120942893 | 0.527799029 | ns | NaCl | KS.3 |
| FIM1_2358 | 997.9371416 | 1032.71958 | 1063.905746 | 0.019438973 | 0.076811204 | 0.693114046 | 0.920513182 | ns | NaCl | KS.3 |
| FIM1_2359 | 5665.405879 | 6161.800469 | 4212.938516 | 0.061413983 | 0.071178124 | 0.216333118 | 0.640629323 | ns | NaCl | KS.3 |
| FIM1_2360 | 1715.44117 | 1894.61683 | 2317.011115 | 0.034332478 | 0.096684327 | 0.457215877 | 0.817337042 | ns | NaCl | KS.3 |
| FIM1_2361 | 3548.051087 | 3340.017757 | 3183.044409 | -0.045263583 | 0.06844823 | 0.355043254 | 0.75934787 | ns | NaCl | KS.3 |
| FIM1_2362 | 1739.386592 | 1788.511024 | 1599.294081 | 0.019533814 | 0.069978733 | 0.689691832 | 0.918911854 | ns | NaCl | KS.3 |
| FIM1_2363 | 2393.152221 | 2257.6323 | 2113.157998 | -0.025421519 | 0.087994523 | 0.594989889 | 0.87962202 | ns | NaCl | KS.3 |
| FIM1_2364 | 2435.045843 | 2646.422365 | 2512.202957 | 0.068250289 | 0.065004694 | 0.15844522 | 0.578755484 | ns | NaCl | KS.3 |

| | | | | | | | | | | |
|---|---|---|---|---|---|---|---|---|---|---|
| FIM1_2365 | 27924.68958 | 32505.7884 | 19621.02589 | 0.085006568 | 0.089022628 | 0.100881905 | 0.484447374 | ns | NaCl | KS.3 |
| FIM1_2366 | 469.2294944 | 397.4722046 | 568.7596446 | -0.072834242 | 0.098236916 | 0.147618724 | 0.562037426 | ns | NaCl | KS.3 |
| FIM1_2367 | 417.0696983 | 589.2760027 | 488.8553682 | 0.137446603 | 0.166301927 | 0.025438418 | 0.250606195 | ns | NaCl | KS.3 |
| FIM1_2368 | 1823.666488 | 1996.811708 | 2404.573716 | 0.029399604 | 0.097521943 | 0.518002536 | 0.850188733 | ns | NaCl | KS.3 |
| FIM1_2369 | 999.6527175 | 883.6525025 | 1138.461136 | -0.063417002 | 0.088169889 | 0.206519003 | 0.631463595 | ns | NaCl | KS.3 |
| FIM1_2370 | 346.1590264 | 385.0185608 | 394.0827039 | 0.018218335 | 0.114701029 | 0.643447759 | 0.898798052 | ns | NaCl | KS.3 |
| FIM1_2371 | 1685.819459 | 2259.003674 | 4375.944249 | 0.060002488 | 0.13709823 | 0.18125297 | 0.611396712 | ns | NaCl | KS.3 |
| FIM1_2372 | 6457.356609 | 6730.631554 | 4975.428215 | 0.021501183 | 0.081733827 | 0.660036531 | 0.903691294 | ns | NaCl | KS.3 |
| FIM1_2373 | 151.9169642 | 67.43488219 | 648.7126715 | -0.467417047 | 0.458638792 | 0.001547134 | 0.04693289 | ns | NaCl | KS.3 |
| FIM1_2374 | 1135.177224 | 887.7735329 | 1107.153194 | -0.179185863 | 0.115978151 | 0.004037205 | 0.082277904 | ns | NaCl | KS.3 |
| FIM1_2375 | 1908.794313 | 1805.655276 | 1708.348738 | -0.029899737 | 0.081059253 | 0.543216803 | 0.857313452 | ns | NaCl | KS.3 |
| FIM1_2376 | 1182.081667 | 1272.156862 | 1376.278378 | 0.035803378 | 0.085052615 | 0.463808891 | 0.819880744 | ns | NaCl | KS.3 |
| FIM1_2377 | 12911.54038 | 13523.56657 | 9042.424857 | 0.026561142 | 0.078567835 | 0.59005718 | 0.87877872 | ns | NaCl | KS.3 |
| FIM1_2378 | 584.102473 | 577.4199234 | 689.8052412 | -0.0055806 | 0.089930752 | 0.904846504 | 0.979880338 | ns | NaCl | KS.3 |
| FIM1_2379 | 1549.863685 | 1845.519037 | 1764.115152 | 0.100647066 | 0.092412144 | 0.058208088 | 0.37782226 | ns | NaCl | KS.3 |
| FIM1_2380 | 949.6106002 | 884.814093 | 815.8202954 | -0.026115743 | 0.093052947 | 0.574502147 | 0.872163065 | ns | NaCl | KS.3 |
| FIM1_2381 | 1755.2255 | 1926.079264 | 1661.261604 | 0.073407013 | 0.068410786 | 0.136845883 | 0.545297021 | ns | NaCl | KS.3 |
| FIM1_2382 | 530.3243085 | 581.790515 | 593.4255858 | 0.042765029 | 0.088971831 | 0.37894745 | 0.778493059 | ns | NaCl | KS.3 |
| FIM1_2383 | 388.7630821 | 543.4725024 | 893.42857 | 0.298627431 | 0.174820693 | 0.000486511 | 0.022833891 | ns | NaCl | KS.3 |
| FIM1_2384 | 13637.46599 | 16202.92646 | 9984.682313 | 0.120167685 | 0.08624114 | 0.025725458 | 0.252434228 | ns | NaCl | KS.3 |
| FIM1_2385 | 794.1087647 | 829.6806769 | 919.9138713 | 0.032343797 | 0.06863374 | 0.507740838 | 0.844423736 | ns | NaCl | KS.3 |
| FIM1_2386 | 344.6739547 | 302.9693422 | 686.0131183 | -0.030576863 | 0.109029577 | 0.472124023 | 0.822127061 | ns | NaCl | KS.3 |
| FIM1_2387 | 414.6258095 | 412.0299522 | 480.6728324 | -0.005184529 | 0.07709077 | 0.915945523 | 0.982343702 | ns | NaCl | KS.3 |
| FIM1_2388 | 7140.733545 | 7175.980631 | 6074.465123 | 0.003564924 | 0.067399963 | 0.941362853 | 0.986790704 | ns | NaCl | KS.3 |
| FIM1_2389 | 1924.88868 | 2192.971791 | 1980.459844 | 0.054653577 | 0.094408702 | 0.261874967 | 0.686421476 | ns | NaCl | KS.3 |
| FIM1_2390 | 958.7911853 | 825.0710075 | 1057.863545 | -0.114953326 | 0.077626197 | 0.027846735 | 0.260899256 | ns | NaCl | KS.3 |
| FIM1_2391 | 1531.706913 | 1405.932914 | 1136.747479 | -0.032374985 | 0.092772169 | 0.491225872 | 0.83512061 | ns | NaCl | KS.3 |
| FIM1_2392 | 1244.02547 | 1107.826521 | 1043.287542 | -0.040687674 | 0.097300143 | 0.383459996 | 0.778665419 | ns | NaCl | KS.3 |
| FIM1_2393 | 350.8885862 | 336.2678987 | 366.9729904 | -0.011217998 | 0.101036747 | 0.793834482 | 0.955299664 | ns | NaCl | KS.3 |
| FIM1_2394 | 337.4301419 | 238.7862737 | 1032.077971 | -0.161181188 | 0.175025851 | 0.0150916 | 0.180049669 | ns | NaCl | KS.3 |
| FIM1_2395 | 26546.50301 | 31964.48794 | 17822.67553 | 0.140644706 | 0.087032788 | 0.010584307 | 0.145864059 | ns | NaCl | KS.3 |
| FIM1_2396 | 2113.359862 | 2570.88054 | 2557.179755 | 0.164970047 | 0.08652535 | 0.003172443 | 0.071740481 | ns | NaCl | KS.3 |
| FIM1_2397 | 1953.326069 | 2187.561829 | 3026.433657 | 0.072153126 | 0.078780741 | 0.155523031 | 0.573412083 | ns | NaCl | KS.3 |
| FIM1_2398 | 1321.882685 | 1359.786666 | 2011.486547 | 0.012939685 | 0.085923534 | 0.787104764 | 0.95289761 | ns | NaCl | KS.3 |
| FIM1_2399 | 450.6852005 | 470.0557748 | 828.3418785 | 0.017961192 | 0.088784165 | 0.70427033 | 0.924470947 | ns | NaCl | KS.3 |
| FIM1_2400 | 336.1023271 | 308.050142 | 510.3101681 | -0.035849572 | 0.092449848 | 0.449232182 | 0.813589409 | ns | NaCl | KS.3 |
| FIM1_2401 | 706.9916699 | 673.2461682 | 714.9424682 | -0.022419369 | 0.08574756 | 0.641886185 | 0.898798052 | ns | NaCl | KS.3 |
| FIM1_2402 | 1801.152826 | 1973.591712 | 1613.534404 | 0.048977945 | 0.08325149 | 0.324556402 | 0.739131368 | ns | NaCl | KS.3 |
| FIM1_2403 | 1845.71384 | 1913.020428 | 2146.319139 | 0.021436891 | 0.076381706 | 0.663665676 | 0.904585408 | ns | NaCl | KS.3 |
| FIM1_2404 | 6739.830885 | 6650.876938 | 5503.319899 | -0.007296663 | 0.080129695 | 0.881333586 | 0.974110648 | ns | NaCl | KS.3 |
| FIM1_2405 | 2073.546063 | 2174.071124 | 1827.205237 | 0.028799617 | 0.076433694 | 0.560032114 | 0.865535809 | ns | NaCl | KS.3 |
| FIM1_2406 | 10038.83601 | 8677.761891 | 8854.810501 | -0.028564629 | 0.114575744 | 0.485404017 | 0.829376704 | ns | NaCl | KS.3 |
| FIM1_2407 | 2071.625496 | 2514.491935 | 1434.130017 | 0.071401693 | 0.107519477 | 0.148121617 | 0.562952669 | ns | NaCl | KS.3 |
| FIM1_2408 | 226.414276 | 188.6810681 | 317.70967 | -0.054127906 | 0.109532905 | 0.242744555 | 0.666657537 | ns | NaCl | KS.3 |
| FIM1_2409 | 525.9083989 | 481.2676619 | 558.8150676 | -0.03879423 | 0.090084163 | 0.42011573 | 0.80133378 | ns | NaCl | KS.3 |

| | | | | | | | | | | |
|---|---|---|---|---|---|---|---|---|---|---|
| FIM1_2410 | 7059.107514 | 7161.125097 | 5763.511942 | 0.009056265 | 0.074129263 | 0.853886537 | 0.969345976 | ns | NaCl | KS.3 |
| FIM1_2411 | 631.4073567 | 670.9677519 | 686.7980707 | 0.028518592 | 0.085123927 | 0.55693417 | 0.863699344 | ns | NaCl | KS.3 |
| FIM1_2412 | 666.6807476 | 711.3964128 | 600.7828512 | 0.021521055 | 0.095250099 | 0.63657311 | 0.898332475 | ns | NaCl | KS.3 |
| FIM1_2413 | 1915.741712 | 2207.215412 | 2181.1896 | 0.053905896 | 0.097988251 | 0.262107934 | 0.686670339 | ns | NaCl | KS.3 |
| FIM1_2414 | 1415.261847 | 1256.720723 | 1461.678195 | -0.065278844 | 0.084826497 | 0.196322073 | 0.618952432 | ns | NaCl | KS.3 |
| FIM1_2415 | 1082.2531 | 1097.81612 | 948.249549 | 0.007364565 | 0.08068329 | 0.880070155 | 0.973483611 | ns | NaCl | KS.3 |
| FIM1_2416 | 4657.80993 | 4452.83073 | 4098.833739 | -0.035112981 | 0.066762095 | 0.469053465 | 0.821764392 | ns | NaCl | KS.3 |
| FIM1_2417 | 875.4718999 | 843.0195092 | 1140.405553 | -0.022047976 | 0.078115963 | 0.654331892 | 0.900747416 | ns | NaCl | KS.3 |
| FIM1_2418 | 20114.60372 | 20843.49122 | 15167.9165 | 0.031395399 | 0.059423677 | 0.498197203 | 0.837486406 | ns | NaCl | KS.3 |
| FIM1_2419 | 749.4463795 | 860.0854525 | 811.3745993 | 0.051053658 | 0.098075858 | 0.284624005 | 0.70510214 | ns | NaCl | KS.3 |
| FIM1_2420 | 1819.665784 | 1908.839605 | 1741.522806 | 0.0243808 | 0.08314983 | 0.616974421 | 0.891503846 | ns | NaCl | KS.3 |
| FIM1_2422 | 4143.527173 | 4072.008405 | 4611.739681 | -0.015775997 | 0.060381375 | 0.735225988 | 0.938375543 | ns | NaCl | KS.3 |
| FIM1_2423 | 433.7936419 | 479.0686808 | 573.2406725 | 0.027622918 | 0.101613554 | 0.531180656 | 0.855153333 | ns | NaCl | KS.3 |
| FIM1_2424 | 958.2371268 | 870.3043474 | 1761.714842 | -0.035614969 | 0.094231917 | 0.447857552 | 0.812914509 | ns | NaCl | KS.3 |
| FIM1_2425 | 579.2734581 | 599.9539175 | 549.0309316 | 0.010244006 | 0.097954473 | 0.81591384 | 0.958816322 | ns | NaCl | KS.3 |
| FIM1_2426 | 2235.790171 | 1981.468331 | 3547.211572 | -0.066737005 | 0.085262678 | 0.1871565 | 0.618483916 | ns | NaCl | KS.3 |
| FIM1_2427 | 350.0340363 | 234.1707663 | 261.1951881 | -0.285920434 | 0.226061642 | 0.001947969 | 0.053542241 | ns | NaCl | KS.3 |
| FIM1_2428 | 196.0018811 | 165.3663332 | 668.4622509 | -0.034877585 | 0.116428271 | 0.402980923 | 0.789854857 | ns | NaCl | KS.3 |
| FIM1_2429 | 1212.358385 | 1060.362036 | 1186.307702 | -0.114950897 | 0.07138213 | 0.023786233 | 0.239908881 | ns | NaCl | KS.3 |
| FIM1_2430 | 1194.219239 | 1162.238434 | 1665.928015 | -0.017104327 | 0.076891748 | 0.728242135 | 0.935336397 | ns | NaCl | KS.3 |
| FIM1_2431 | 1725.887368 | 1555.54556 | 1976.13754 | -0.045187246 | 0.090333082 | 0.35188968 | 0.756912594 | ns | NaCl | KS.3 |
| FIM1_2432 | 693.4445029 | 564.4206315 | 1147.157163 | -0.09798036 | 0.105982536 | 0.06523454 | 0.401164198 | ns | NaCl | KS.3 |
| FIM1_2433 | 45630.70205 | 41558.96125 | 28152.16964 | -0.019087309 | 0.109755704 | 0.639643759 | 0.898798052 | ns | NaCl | KS.3 |
| FIM1_2434 | 7121.865158 | 5809.064873 | 5279.855016 | -0.055108841 | 0.114924399 | 0.229017671 | 0.652325274 | ns | NaCl | KS.3 |
| FIM1_2435 | 1823.146676 | 1529.026999 | 1608.339509 | -0.11710843 | 0.088891458 | 0.030201744 | 0.27272008 | ns | NaCl | KS.3 |
| FIM1_2436 | 251.5281124 | 278.4860715 | 317.0982153 | 0.030730682 | 0.100548125 | 0.492342923 | 0.83568954 | ns | NaCl | KS.3 |
| FIM1_2437 | 71.39471001 | 50.02105933 | 185.0976495 | -0.066723553 | 0.152394727 | 0.14436052 | 0.558679104 | ns | NaCl | KS.3 |
| FIM1_2438 | 1269.163182 | 1279.238544 | 1531.349559 | 0.003564372 | 0.082811367 | 0.941359918 | 0.986790704 | ns | NaCl | KS.3 |
| FIM1_2439 | 258.5484377 | 298.0850992 | 443.2228972 | 0.028311229 | 0.113491269 | 0.491336287 | 0.83512061 | ns | NaCl | KS.3 |
| FIM1_2440 | 103.1490369 | 98.26751148 | 226.0552197 | -0.004574838 | 0.131361972 | 0.8945689 | 0.977268396 | ns | NaCl | KS.3 |
| FIM1_2444 | 34628.65862 | 33067.36073 | 20677.52651 | -0.007071655 | 0.116409123 | 0.851782764 | 0.96882013 | ns | NaCl | KS.3 |
| FIM1_2445 | 1303.794136 | 1016.740137 | 805.4567149 | -0.0512848 | 0.127814126 | 0.239516852 | 0.662365947 | ns | NaCl | KS.3 |
| FIM1_2446 | 305.3917934 | 278.4283558 | 299.0757199 | -0.037217452 | 0.089139391 | 0.440090933 | 0.808917498 | ns | NaCl | KS.3 |
| FIM1_2447 | 363.7215753 | 383.7098288 | 473.4177694 | 0.027148329 | 0.085777538 | 0.57464097 | 0.872163065 | ns | NaCl | KS.3 |
| FIM1_2448 | 39.62773398 | 18.4128155 | 42.2117417 | -0.108484 | 0.256988152 | 0.046278677 | 0.340862131 | ns | NaCl | KS.3 |
| FIM1_2449 | 34.43528267 | 41.00479724 | 43.40792016 | 0.019089803 | 0.134517192 | 0.592669472 | 0.87962202 | ns | NaCl | KS.3 |
| FIM1_2450 | 390.5848725 | 309.8322479 | 632.294383 | -0.068400147 | 0.119534706 | 0.153560742 | 0.570975106 | ns | NaCl | KS.3 |
| FIM1_2451 | 1702.006276 | 743.8723843 | 931.4556968 | -0.1107296 | 0.268679223 | 0.042789055 | 0.325996249 | ns | NaCl | KS.3 |
| FIM1_2452 | 45.64044495 | 44.52686927 | 52.95915738 | -0.005551489 | 0.116458227 | 0.883100368 | 0.97456556 | ns | NaCl | KS.3 |
| FIM1_2453 | 158.6503904 | 136.4934395 | 209.9167577 | -0.046667486 | 0.102588527 | 0.313638301 | 0.731871738 | ns | NaCl | KS.3 |
| FIM1_2454 | 207.3385103 | 149.8169728 | 378.7276646 | -0.096837406 | 0.149398262 | 0.067338743 | 0.406565836 | ns | NaCl | KS.3 |
| FIM1_2455 | 690.4611853 | 446.3351482 | 1087.383669 | -0.067131439 | 0.170167962 | 0.135682829 | 0.543933984 | ns | NaCl | KS.3 |
| FIM1_2456 | 162.0145038 | 102.6294582 | 256.7192958 | -0.173491567 | 0.226846871 | 0.015632991 | 0.182568377 | ns | NaCl | KS.3 |
| FIM1_2457 | 293.6818796 | 245.8654739 | 402.8146455 | -0.06184153 | 0.105616123 | 0.198465979 | 0.620924909 | ns | NaCl | KS.3 |
| FIM1_2458 | 1629.842405 | 1493.229736 | 2445.15704 | -0.029047985 | 0.096216298 | 0.526146063 | 0.855027345 | ns | NaCl | KS.3 |

| | | | | | | | | | | |
|---|---|---|---|---|---|---|---|---|---|---|
| FIM1_2459 | 3007.753742 | 3617.015342 | 2120.245717 | 0.047041967 | 0.113010419 | 0.291141463 | 0.711059783 | ns | NaCl | KS.3 |
| FIM1_2460 | 7120.540633 | 8120.194465 | 4269.400428 | 0.024619316 | 0.11460657 | 0.54124004 | 0.857165292 | ns | NaCl | KS.3 |
| FIM1_2461 | 2882.415266 | 3136.199889 | 2918.362486 | 0.064853114 | 0.069015519 | 0.188691213 | 0.618812647 | ns | NaCl | KS.3 |
| FIM1_2462 | 6643.45656 | 6777.198301 | 5382.854023 | 0.015057565 | 0.066779151 | 0.755598233 | 0.944735162 | ns | NaCl | KS.3 |
| FIM1_2463 | 709.3632471 | 708.9684827 | 944.635553 | 0.000847699 | 0.070971833 | 0.986186941 | 0.999134858 | ns | NaCl | KS.3 |
| FIM1_2464 | 1436.626898 | 1546.857683 | 1939.7883 | 0.02865184 | 0.091209491 | 0.543692698 | 0.857313452 | ns | NaCl | KS.3 |
| FIM1_2465 | 427.6451924 | 439.5687714 | 596.242639 | 0.011999492 | 0.081874787 | 0.805329003 | 0.957859861 | ns | NaCl | KS.3 |
| FIM1_2466 | 419.2863531 | 357.1564982 | 842.4749687 | -0.032924934 | 0.114935499 | 0.428996676 | 0.804596805 | ns | NaCl | KS.3 |
| FIM1_2467 | 672.9639773 | 333.5584352 | 403.5880091 | -0.942354421 | 0.151278766 | 2.80618373058 | 1.26916036906 | ns | NaCl | KS.3 |
| FIM1_2468 | 2182.810159 | 2336.340309 | 2249.658896 | 0.036141763 | 0.081723594 | 0.463625864 | 0.819880744 | ns | NaCl | KS.3 |
| FIM1_2469 | 1611.686827 | 1670.546271 | 1894.960468 | 0.030921254 | 0.060491954 | 0.508143033 | 0.844423736 | ns | NaCl | KS.3 |
| FIM1_2470 | 6274.004792 | 6671.203522 | 6846.182636 | 0.04341451 | 0.071102423 | 0.378914068 | 0.778493059 | ns | NaCl | KS.3 |
| FIM1_2471 | 1248.745049 | 1053.193291 | 1109.614518 | -0.072995935 | 0.0995269 | 0.14609836 | 0.560211748 | ns | NaCl | KS.3 |
| FIM1_2472 | 1403.833541 | 1383.839414 | 2006.773113 | -0.010182482 | 0.074578968 | 0.836008568 | 0.962829707 | ns | NaCl | KS.3 |
| FIM1_2473 | 2181.114542 | 3128.143658 | 1554.022244 | 0.171702574 | 0.182384434 | 0.012447264 | 0.160013282 | ns | NaCl | KS.3 |
| FIM1_2474 | 3643.656655 | 4175.121424 | 3438.768156 | 0.107369748 | 0.07375385 | 0.036169168 | 0.301915458 | ns | NaCl | KS.3 |
| FIM1_2475 | 849.6181058 | 804.4838082 | 1268.99965 | -0.035649355 | 0.077114269 | 0.471900382 | 0.822127061 | ns | NaCl | KS.3 |
| FIM1_2476 | 13036.81373 | 14199.84321 | 9869.721191 | 0.059670714 | 0.073117674 | 0.231865768 | 0.654993872 | ns | NaCl | KS.3 |
| FIM1_2477 | 2294.736296 | 2208.776464 | 2092.768011 | -0.022377095 | 0.078641343 | 0.649337238 | 0.899642849 | ns | NaCl | KS.3 |
| FIM1_2478 | 1288.773282 | 1255.820884 | 1118.601481 | -0.012986366 | 0.083254104 | 0.788759854 | 0.953379471 | ns | NaCl | KS.3 |
| FIM1_2479 | 805.7853466 | 793.6481692 | 825.2483291 | -0.010503931 | 0.07690989 | 0.830808967 | 0.961620266 | ns | NaCl | KS.3 |
| FIM1_2480 | 2069.702109 | 1913.195551 | 2053.053403 | -0.070268626 | 0.060600237 | 0.134614131 | 0.54183277 | ns | NaCl | KS.3 |
| FIM1_2481 | 1151.443414 | 1067.010046 | 1282.433671 | -0.056114452 | 0.071503315 | 0.258101846 | 0.681920702 | ns | NaCl | KS.3 |
| FIM1_2482 | 1314.81632 | 1401.487766 | 1389.641121 | 0.053893177 | 0.062663183 | 0.257161721 | 0.680871076 | ns | NaCl | KS.3 |
| FIM1_2483 | 916.5763895 | 957.9815753 | 1291.08699 | 0.028813731 | 0.072528477 | 0.558737105 | 0.864443132 | ns | NaCl | KS.3 |
| FIM1_2484 | 240.8915311 | 284.490744 | 351.8027649 | 0.074859913 | 0.097027551 | 0.139147537 | 0.550000428 | ns | NaCl | KS.3 |
| FIM1_2485 | 929.0834845 | 817.8045877 | 1569.017362 | -0.068895224 | 0.086464034 | 0.174105137 | 0.602764827 | ns | NaCl | KS.3 |
| FIM1_2486 | 16954.97358 | 19374.23831 | 11891.10262 | 0.060508778 | 0.092304822 | 0.222135836 | 0.645894671 | ns | NaCl | KS.3 |
| FIM1_2487 | 2505.527928 | 2453.176293 | 2468.994696 | -0.016909615 | 0.065351166 | 0.725033375 | 0.933499234 | ns | NaCl | KS.3 |
| FIM1_2488 | 328.2209478 | 334.6374071 | 344.4512517 | 0.005674746 | 0.096821938 | 0.898029908 | 0.977971161 | ns | NaCl | KS.3 |
| FIM1_2489 | 468.1481436 | 406.7439896 | 585.8551589 | -0.051376788 | 0.099249778 | 0.27992314 | 0.700652348 | ns | NaCl | KS.3 |
| FIM1_2490 | 1160.847066 | 1107.535813 | 1138.043493 | -0.019859358 | 0.087976469 | 0.676522928 | 0.913287355 | ns | NaCl | KS.3 |
| FIM1_2491 | 132.0204876 | 101.1750543 | 811.3610947 | -0.036783969 | 0.138722656 | 0.345433485 | 0.754462295 | ns | NaCl | KS.3 |
| FIM1_2492 | 2897.662107 | 3272.341658 | 2656.474913 | 0.033668676 | 0.103767925 | 0.446869536 | 0.812914509 | ns | NaCl | KS.3 |
| FIM1_2493 | 1917.918454 | 1788.513402 | 1708.561039 | -0.062089888 | 0.060548632 | 0.18552422 | 0.616967242 | ns | NaCl | KS.3 |
| FIM1_2494 | 685.6104807 | 728.6167437 | 863.5211927 | 0.029240371 | 0.083724786 | 0.549158724 | 0.86076391 | ns | NaCl | KS.3 |
| FIM1_2495 | 7138.532266 | 4651.095833 | 5237.240303 | -0.360453228 | 0.242609928 | 0.000752441 | 0.030188656 | ns | NaCl | KS.3 |
| FIM1_2496 | 589.5521795 | 462.5274164 | 1081.57116 | -0.133788803 | 0.120362447 | 0.021612286 | 0.224469984 | ns | NaCl | KS.3 |
| FIM1_2497 | 603.7847978 | 583.7391045 | 585.9993221 | -0.012588354 | 0.092973502 | 0.783672532 | 0.951646805 | ns | NaCl | KS.3 |
| FIM1_2498 | 157.5920748 | 126.0564359 | 134.5474941 | -0.053358989 | 0.122391804 | 0.231706664 | 0.654993872 | ns | NaCl | KS.3 |
| FIM1_2499 | 1137.808993 | 1105.593892 | 1237.132266 | -0.016184091 | 0.074395667 | 0.742357348 | 0.941460666 | ns | NaCl | KS.3 |
| FIM1_2500 | 794.9932923 | 353.5764191 | 1509.180097 | -1.022730416 | 0.24210145 | 5.01699363849 | 9.59982436596 | Km<KS.3 | NaCl | KS.3 |
| FIM1_2502 | 3034.860096 | 2169.35062 | 3124.580927 | -0.227062119 | 0.176715595 | 0.003046809 | 0.06921403 | ns | NaCl | KS.3 |
| FIM1_2503 | 645.7757443 | 589.0270242 | 574.3758891 | -0.024715548 | 0.102252385 | 0.57087707 | 0.870665206 | ns | NaCl | KS.3 |
| FIM1_2504 | 742.8955852 | 633.8206721 | 853.0960671 | -0.036060448 | 0.112406546 | 0.398218075 | 0.786613521 | ns | NaCl | KS.3 |

| | | | | | | | | | | |
|---|---|---|---|---|---|---|---|---|---|---|
| FIM1_2505 | 637.2477759 | 568.6048002 | 596.8034414 | -0.040409652 | 0.097079071 | 0.386928806 | 0.779278134 | ns | NaCl | KS.3 |
| FIM1_2506 | 198.4710836 | 161.2660387 | 537.1545035 | -0.062164565 | 0.113822736 | 0.188264823 | 0.618637711 | ns | NaCl | KS.3 |
| FIM1_2507 | 937.536381 | 1083.452843 | 1007.272581 | 0.079346294 | 0.088768991 | 0.122548617 | 0.528318345 | ns | NaCl | KS.3 |
| FIM1_2508 | 15.91917998 | 18.37915311 | 19.23299098 | 0.012371596 | 0.138378529 | 0.716245309 | 0.929477573 | ns | NaCl | KS.3 |
| FIM1_2510 | 265.2335152 | 306.3278521 | 247.8209449 | 0.044202683 | 0.104340743 | 0.33257032 | 0.74327823 | ns | NaCl | KS.3 |
| FIM1_2511 | 8930.41192 | 11860.32735 | 5768.840307 | 0.114264027 | 0.136095049 | 0.041785164 | 0.323298895 | ns | NaCl | KS.3 |
| FIM1_2512 | 1070.893466 | 869.1939281 | 1212.02316 | -0.12498156 | 0.102914021 | 0.025509023 | 0.250805116 | ns | NaCl | KS.3 |
| FIM1_2513 | 9079.156756 | 9649.454336 | 6384.272418 | 0.020442574 | 0.095056372 | 0.653519416 | 0.900747416 | ns | NaCl | KS.3 |
| FIM1_2514 | 304.9133598 | 302.4300761 | 481.8060497 | -0.003528276 | 0.088340458 | 0.940339245 | 0.986790704 | ns | NaCl | KS.3 |
| FIM1_2515 | 57.18179812 | 55.76520641 | 94.90197683 | -0.003168907 | 0.105525356 | 0.938412351 | 0.986790704 | ns | NaCl | KS.3 |
| FIM1_2516 | 4903.807198 | 4750.176412 | 4106.495846 | -0.020317208 | 0.074330299 | 0.680082593 | 0.913941356 | ns | NaCl | KS.3 |
| FIM1_2517 | 1659.001188 | 1452.903882 | 2074.916218 | -0.052183346 | 0.095567084 | 0.27967022 | 0.700652348 | ns | NaCl | KS.3 |
| FIM1_2518 | 2294.15565 | 2216.075601 | 2038.925988 | -0.027702723 | 0.064931593 | 0.564042031 | 0.867434473 | ns | NaCl | KS.3 |
| FIM1_2519 | 4957.985647 | 6291.301102 | 2929.214204 | 0.032026978 | 0.133754875 | 0.404981598 | 0.792597737 | ns | NaCl | KS.3 |
| FIM1_2520 | 889.0539588 | 939.8019146 | 689.008901 | 0.021372482 | 0.091782171 | 0.64657611 | 0.898973628 | ns | NaCl | KS.3 |
| FIM1_2521 | 975.5348507 | 961.9164935 | 697.9728368 | -0.004718312 | 0.093479415 | 0.917380671 | 0.982343702 | ns | NaCl | KS.3 |
| FIM1_2522 | 519.9952477 | 532.2853558 | 490.9830642 | 0.010334941 | 0.088162541 | 0.826983205 | 0.960240659 | ns | NaCl | KS.3 |
| FIM1_2523 | 4769.4635 | 5361.862713 | 4516.161231 | 0.121104651 | 0.055085473 | 0.006816991 | 0.111371848 | ns | NaCl | KS.3 |
| FIM1_2524 | 1313.534695 | 1686.022923 | 1763.71944 | 0.057536036 | 0.126600824 | 0.202194905 | 0.625219184 | ns | NaCl | KS.3 |
| FIM1_2525 | 663.2583965 | 658.0704064 | 706.1662017 | -0.003304043 | 0.08782457 | 0.944295944 | 0.987155352 | ns | NaCl | KS.3 |
| FIM1_2526 | 5694.301792 | 3693.902665 | 5219.96843 | -0.515906054 | 0.189780675 | 2.16533474278 | 0.00229203 | ns | NaCl | KS.3 |
| FIM1_2527 | 211.3379117 | 123.7147381 | 199.4878147 | -0.121370097 | 0.222601718 | 0.03998736 | 0.315700318 | ns | NaCl | KS.3 |
| FIM1_2528 | 32589.01621 | 34724.2605 | 23970.31229 | 0.043649441 | 0.072386229 | 0.377836972 | 0.778493059 | ns | NaCl | KS.3 |
| FIM1_2529 | 7685.464161 | 8307.719254 | 8482.783516 | 0.012698434 | 0.115518547 | 0.741673593 | 0.941460666 | ns | NaCl | KS.3 |
| FIM1_2530 | 632.4609955 | 619.6525335 | 1352.405008 | -0.003261743 | 0.115109897 | 0.931647569 | 0.984483147 | ns | NaCl | KS.3 |
| FIM1_2531 | 3334.364704 | 3784.711306 | 3138.28937 | 0.045913943 | 0.097543044 | 0.3308246 | 0.742254041 | ns | NaCl | KS.3 |
| FIM1_2532 | 456.879585 | 415.9402202 | 477.4910332 | -0.018630163 | 0.111944533 | 0.642353852 | 0.898798052 | ns | NaCl | KS.3 |
| FIM1_2533 | 2149.709705 | 2248.032288 | 2574.912595 | 0.021927512 | 0.083637796 | 0.651842581 | 0.900060184 | ns | NaCl | KS.3 |
| FIM1_2534 | 1943.644156 | 1600.003562 | 2106.468145 | -0.085146787 | 0.104169801 | 0.098402074 | 0.479012053 | ns | NaCl | KS.3 |
| FIM1_2535 | 512.5884941 | 476.6848383 | 461.4695118 | -0.026353121 | 0.094184446 | 0.568430042 | 0.869550295 | ns | NaCl | KS.3 |
| FIM1_2536 | 673.1310978 | 423.1604857 | 460.6005252 | -0.115153081 | 0.199256925 | 0.045011677 | 0.335229179 | ns | NaCl | KS.3 |
| FIM1_2537 | 1854.152367 | 1164.115097 | 1077.293257 | -0.168637574 | 0.226706301 | 0.017069073 | 0.194321832 | ns | NaCl | KS.3 |
| FIM1_2538 | 727.3844592 | 759.4465997 | 815.3711556 | 0.012995468 | 0.097408816 | 0.76936743 | 0.948835638 | ns | NaCl | KS.3 |
| FIM1_2539 | 775.8369063 | 716.809373 | 615.8128797 | -0.024942663 | 0.09740137 | 0.580419923 | 0.8749481 | ns | NaCl | KS.3 |
| FIM1_2540 | 3131.195986 | 3023.724447 | 3341.158667 | -0.011492973 | 0.094786179 | 0.79924936 | 0.95648403 | ns | NaCl | KS.3 |
| FIM1_2541 | 1670.006582 | 1646.575278 | 1315.902532 | -0.00664524 | 0.085440346 | 0.889730166 | 0.977268396 | ns | NaCl | KS.3 |
| FIM1_2542 | 529.7533291 | 472.2579443 | 401.8720866 | -0.039894044 | 0.097585528 | 0.391261755 | 0.779858666 | ns | NaCl | KS.3 |
| FIM1_2543 | 1062.24446 | 983.1415738 | 1152.744984 | -0.031169732 | 0.090856021 | 0.511087805 | 0.846104012 | ns | NaCl | KS.3 |
| FIM1_2544 | 782.6594935 | 828.3810951 | 859.9406074 | 0.034963351 | 0.077663553 | 0.480245766 | 0.827579731 | ns | NaCl | KS.3 |
| FIM1_2546 | 1063.053016 | 1056.368289 | 1183.701345 | -0.003188473 | 0.082243962 | 0.947640089 | 0.987736646 | ns | NaCl | KS.3 |
| FIM1_2547 | 1054.59268 | 1190.118431 | 2021.126718 | 0.024980615 | 0.110935377 | 0.544571817 | 0.857313452 | ns | NaCl | KS.3 |
| FIM1_2548 | 5881.846965 | 7131.087422 | 5162.908892 | 0.032309114 | 0.123270674 | 0.419586222 | 0.80133378 | ns | NaCl | KS.3 |
| FIM1_2549 | 4813.475866 | 4570.043684 | 5865.03692 | -0.012951103 | 0.10257934 | 0.760462129 | 0.945911573 | ns | NaCl | KS.3 |
| FIM1_2550 | 1547.720542 | 1320.266264 | 1720.231843 | -0.056950263 | 0.102445516 | 0.233348568 | 0.655880863 | ns | NaCl | KS.3 |
| FIM1_2552 | 1632.23784 | 1642.01308 | 2587.608678 | 0.001035206 | 0.113189791 | 0.978563234 | 0.996164232 | ns | NaCl | KS.3 |

| | | | | | | | | | | |
|---|---|---|---|---|---|---|---|---|---|---|
| FIM1_2553 | 380.3891519 | 337.4389552 | 530.5381407 | -0.025947925 | 0.109951679 | 0.532935 | 0.855153333 | ns | NaCl | KS.3 |
| FIM1_2554 | 870.6240476 | 682.9110279 | 1249.548733 | -0.053506432 | 0.126041505 | 0.226859501 | 0.651255637 | ns | NaCl | KS.3 |
| FIM1_2555 | 11347.15426 | 20467.30714 | 14232.97528 | 0.130738936 | 0.241583126 | 0.03376373 | 0.292771172 | ns | NaCl | KS.3 |
| FIM1_2556 | 100.6882579 | 89.07153866 | 413.0204317 | -0.015901129 | 0.12671439 | 0.663001048 | 0.904422872 | ns | NaCl | KS.3 |
| FIM1_2557 | 13854.71618 | 14455.11553 | 12207.00935 | 0.012099539 | 0.098806861 | 0.78224327 | 0.951646805 | ns | NaCl | KS.3 |
| FIM1_2558 | 1744.351911 | 2129.980324 | 1227.149074 | 0.049114875 | 0.116101812 | 0.269806195 | 0.691543442 | ns | NaCl | KS.3 |
| FIM1_2559 | 298.1578414 | 283.4967538 | 458.2771054 | -0.008456076 | 0.114293253 | 0.826129477 | 0.960240659 | ns | NaCl | KS.3 |
| FIM1_2560 | 5086.993917 | 3853.224249 | 3738.299824 | -0.181479589 | 0.136169948 | 0.005559534 | 0.098781009 | ns | NaCl | KS.3 |
| FIM1_2561 | 230.5160408 | 235.5500503 | 346.8324722 | 0.005367171 | 0.102649272 | 0.898554006 | 0.977971161 | ns | NaCl | KS.3 |
| FIM1_2562 | 2241.743834 | 1986.803981 | 2830.694731 | -0.064917844 | 0.085788908 | 0.198236554 | 0.620924909 | ns | NaCl | KS.3 |
| FIM1_2563 | 3365.016039 | 3520.42841 | 3001.103454 | 0.034102971 | 0.068059883 | 0.484176899 | 0.829184191 | ns | NaCl | KS.3 |
| FIM1_2564 | 793.3777375 | 874.2819595 | 1739.693656 | 0.042572935 | 0.089471607 | 0.380081216 | 0.778493059 | ns | NaCl | KS.3 |
| FIM1_2565 | 1128.287976 | 1017.708675 | 1138.975453 | -0.050156295 | 0.087576018 | 0.309184357 | 0.729346692 | ns | NaCl | KS.3 |
| FIM1_2566 | 1429.047972 | 1469.559837 | 2318.819173 | 0.009756164 | 0.09279734 | 0.831405971 | 0.961620266 | ns | NaCl | KS.3 |
| FIM1_2567 | 990.7671418 | 1079.149324 | 1056.417571 | 0.045600725 | 0.08329625 | 0.357426461 | 0.761817563 | ns | NaCl | KS.3 |
| FIM1_2568 | 1762.529421 | 1797.234478 | 1217.730461 | 0.004643315 | 0.103009796 | 0.911860965 | 0.982011252 | ns | NaCl | KS.3 |
| FIM1_2570 | 220.8048988 | 214.0099226 | 388.1636513 | -0.00731477 | 0.105247767 | 0.859217363 | 0.969856868 | ns | NaCl | KS.3 |
| FIM1_2571 | 537.166243 | 637.4093959 | 421.0415144 | 0.036419386 | 0.115584881 | 0.387524443 | 0.779278134 | ns | NaCl | KS.3 |
| FIM1_2572 | 398.4113836 | 415.9470704 | 287.6389418 | 0.013724162 | 0.097637082 | 0.756593654 | 0.945029734 | ns | NaCl | KS.3 |
| FIM1_2573 | 358.5427468 | 303.9590171 | 484.8169296 | -0.066855585 | 0.100726709 | 0.175418397 | 0.605047427 | ns | NaCl | KS.3 |
| FIM1_2574 | 2103.043914 | 2303.858151 | 1794.752805 | 0.050732056 | 0.082022025 | 0.309156886 | 0.729346692 | ns | NaCl | KS.3 |
| FIM1_2575 | 1016.914278 | 1153.106593 | 933.2852499 | 0.071413743 | 0.084615458 | 0.160599628 | 0.58211082 | ns | NaCl | KS.3 |
| FIM1_2576 | 18683.09208 | 21231.73775 | 14153.82679 | 0.14277109 | 0.0500537 | 0.000687318 | 0.028734494 | ns | NaCl | KS.3 |
| FIM1_2577 | 4852.451409 | 6297.058935 | 5459.753975 | 0.131914299 | 0.1263942 | 0.023918554 | 0.239908881 | ns | NaCl | KS.3 |
| FIM1_2578 | 63.77764315 | 78.62619591 | 73.6616089 | 0.049113807 | 0.116293867 | 0.269532798 | 0.691198798 | ns | NaCl | KS.3 |
| FIM1_2579 | 161.1858207 | 220.4826557 | 269.9166729 | 0.147239592 | 0.155935016 | 0.018766179 | 0.207470533 | ns | NaCl | KS.3 |
| FIM1_2580 | 585.7242595 | 505.8988144 | 1231.593787 | -0.03780817 | 0.107716643 | 0.389590665 | 0.779482559 | ns | NaCl | KS.3 |
| FIM1_2581 | 23999.66605 | 31405.36951 | 20957.63236 | 0.18101826 | 0.130925546 | 0.005145657 | 0.094234281 | ns | NaCl | KS.3 |
| FIM1_2582 | 428.777229 | 366.5245399 | 655.2949581 | -0.057255146 | 0.101798152 | 0.232173237 | 0.654993872 | ns | NaCl | KS.3 |
| FIM1_2583 | 444.3432973 | 404.2819819 | 481.033384 | -0.030230797 | 0.09773992 | 0.506392838 | 0.844423736 | ns | NaCl | KS.3 |
| FIM1_2584 | 91.23582387 | 74.02084177 | 103.7048874 | -0.022489281 | 0.138027994 | 0.529955672 | 0.855153333 | ns | NaCl | KS.3 |
| FIM1_2585 | 6057.397341 | 5419.254917 | 4837.266344 | -0.062347171 | 0.083574025 | 0.216046124 | 0.640629323 | ns | NaCl | KS.3 |
| FIM1_2586 | 1200.321751 | 1222.108122 | 1096.603075 | 0.010980266 | 0.076441313 | 0.82333172 | 0.960240659 | ns | NaCl | KS.3 |
| FIM1_2587 | 4880.772359 | 5218.915835 | 4513.054149 | 0.048419032 | 0.070183623 | 0.325899829 | 0.739131368 | ns | NaCl | KS.3 |
| FIM1_2588 | 3359.624864 | 2833.588574 | 5707.663931 | -0.049525517 | 0.108091905 | 0.279316972 | 0.700652348 | ns | NaCl | KS.3 |
| FIM1_2590 | 556.5950618 | 357.3312064 | 1955.598832 | -0.123330307 | 0.196682929 | 0.037981749 | 0.308756863 | ns | NaCl | KS.3 |
| FIM1_2591 | 985.7662096 | 1043.256196 | 888.9165065 | 0.012413956 | 0.106281956 | 0.763110057 | 0.946287744 | ns | NaCl | KS.3 |
| FIM1_2592 | 1767.444781 | 1633.24868 | 1822.145642 | -0.045495807 | 0.080333536 | 0.360524121 | 0.763371684 | ns | NaCl | KS.3 |
| FIM1_2593 | 1204.167117 | 983.0837099 | 1588.488274 | -0.165973767 | 0.089664412 | 0.003396788 | 0.074774419 | ns | NaCl | KS.3 |
| FIM1_2594 | 2121.786774 | 2015.28991 | 2182.106981 | -0.028822027 | 0.079362396 | 0.558803842 | 0.864443132 | ns | NaCl | KS.3 |
| FIM1_2595 | 1527.340163 | 1533.817778 | 1316.272804 | 0.003068831 | 0.071203495 | 0.950048639 | 0.987736646 | ns | NaCl | KS.3 |
| FIM1_2596 | 1525.696763 | 1540.704994 | 2024.028705 | 0.002699712 | 0.097756463 | 0.950944645 | 0.987736646 | ns | NaCl | KS.3 |
| FIM1_2597 | 1071.265956 | 1240.789253 | 977.7203132 | 0.047151098 | 0.102916082 | 0.308711771 | 0.728923143 | ns | NaCl | KS.3 |
| FIM1_2598 | 867.3393057 | 787.9691177 | 1013.907709 | -0.051813517 | 0.084488879 | 0.297743652 | 0.715635559 | ns | NaCl | KS.3 |
| FIM1_2599 | 1311.258304 | 1334.39326 | 964.505576 | 0.004600449 | 0.100531449 | 0.91453376 | 0.982343702 | ns | NaCl | KS.3 |

| | | | | | | | | | | |
|---|---|---|---|---|---|---|---|---|---|---|
| FIM1_2600 | 1624.495415 | 1817.775885 | 1325.314155 | 0.040678312 | 0.096127025 | 0.386271353 | 0.779278134 | ns | NaCl | KS.3 |
| FIM1_2601 | 2503.45882 | 2951.895335 | 2074.916074 | 0.101872198 | 0.088461618 | 0.054614642 | 0.36402301 | ns | NaCl | KS.3 |
| FIM1_2602 | 1734.708916 | 1951.474192 | 1905.859368 | 0.078219522 | 0.078383854 | 0.125207511 | 0.534684433 | ns | NaCl | KS.3 |
| FIM1_2603 | 197.8992198 | 188.7122292 | 286.3863057 | -0.007412464 | 0.115364368 | 0.845906826 | 0.967147129 | ns | NaCl | KS.3 |
| FIM1_2604 | 515.1135844 | 419.0476656 | 543.7081307 | -0.063286597 | 0.113705082 | 0.182398148 | 0.611396712 | ns | NaCl | KS.3 |
| FIM1_2606 | 22.79486421 | 42.64335599 | 32.27504231 | 0.124889775 | 0.243659528 | 0.037093694 | 0.304670827 | ns | NaCl | KS.3 |
| FIM1_2607 | 4206.502088 | 4953.131031 | 3002.133013 | 0.05162342 | 0.105268932 | 0.267646441 | 0.689593109 | ns | NaCl | KS.3 |
| FIM1_2608 | 878.5365697 | 863.1983013 | 800.2735726 | -0.012518886 | 0.070195993 | 0.797980717 | 0.955924408 | ns | NaCl | KS.3 |
| FIM1_2609 | 320.5639402 | 292.890806 | 502.6738705 | -0.035768431 | 0.092673753 | 0.449687498 | 0.813595672 | ns | NaCl | KS.3 |
| FIM1_2610 | 1005.142003 | 788.146184 | 1233.092503 | -0.16368212 | 0.117494909 | 0.007459833 | 0.11809776 | ns | NaCl | KS.3 |
| FIM1_2611 | 1262.179469 | 1319.110263 | 1871.482649 | 0.012412108 | 0.099041951 | 0.776422029 | 0.950410881 | ns | NaCl | KS.3 |
| FIM1_2612 | 2981.537832 | 2886.929005 | 3394.826906 | -0.004367771 | 0.120206502 | 0.905478729 | 0.979880338 | ns | NaCl | KS.3 |
| FIM1_2613 | 2748.353938 | 2365.597117 | 3064.290298 | -0.029890131 | 0.114600338 | 0.467771231 | 0.821764392 | ns | NaCl | KS.3 |
| FIM1_2614 | 527.074667 | 429.6442074 | 709.2168211 | -0.104658165 | 0.105111043 | 0.052347326 | 0.358438293 | ns | NaCl | KS.3 |
| FIM1_2615 | 76.9984827 | 71.69444286 | 64.47101098 | -0.014158783 | 0.113782883 | 0.71740515 | 0.93000992 | ns | NaCl | KS.3 |
| FIM1_2616 | 808.4433566 | 681.506435 | 737.3979886 | -0.132441336 | 0.083779164 | 0.014306913 | 0.173179792 | ns | NaCl | KS.3 |
| FIM1_2617 | 1213.606917 | 1032.413177 | 1133.242136 | -0.130056374 | 0.077019111 | 0.013409141 | 0.166838629 | ns | NaCl | KS.3 |
| FIM1_2618 | 362.2115772 | 321.2234549 | 394.8937208 | -0.049450177 | 0.094436133 | 0.304741989 | 0.724521545 | ns | NaCl | KS.3 |
| FIM1_2619 | 189.1726417 | 179.2184405 | 276.3989874 | -0.018220408 | 0.097560648 | 0.682675271 | 0.915673519 | ns | NaCl | KS.3 |
| FIM1_2620 | 745.0004209 | 631.926027 | 670.0593794 | -0.078017392 | 0.096317053 | 0.126092822 | 0.534982546 | ns | NaCl | KS.3 |
| FIM1_2621 | 9827.146194 | 9894.570427 | 8627.981714 | 0.002570026 | 0.090939616 | 0.955764248 | 0.989784998 | ns | NaCl | KS.3 |
| FIM1_2622 | 511.1616167 | 456.887173 | 652.659282 | -0.05521991 | 0.089035859 | 0.264625665 | 0.688347233 | ns | NaCl | KS.3 |
| FIM1_2623 | 1954.53234 | 2005.01941 | 1779.418525 | 0.014933333 | 0.07704372 | 0.761471088 | 0.946054754 | ns | NaCl | KS.3 |
| FIM1_2624 | 7445.526322 | 7310.610396 | 7410.842857 | -0.010023262 | 0.079041101 | 0.837969118 | 0.962829707 | ns | NaCl | KS.3 |
| FIM1_2625 | 420.3291113 | 312.5435254 | 576.4427825 | -0.144373858 | 0.146582352 | 0.019025693 | 0.208486389 | ns | NaCl | KS.3 |
| FIM1_2626 | 1468.952742 | 1386.436525 | 1584.582197 | -0.030827459 | 0.081135754 | 0.530939052 | 0.855153333 | ns | NaCl | KS.3 |
| FIM1_2627 | 730.9922917 | 652.5223815 | 1047.969986 | -0.044379967 | 0.094947823 | 0.350940223 | 0.756912594 | ns | NaCl | KS.3 |
| FIM1_2628 | 818.6752406 | 922.2563393 | 833.5519857 | 0.065478538 | 0.084965219 | 0.195017929 | 0.618952432 | ns | NaCl | KS.3 |
| FIM1_2629 | 10400.87086 | 12794.12217 | 7639.601927 | 0.046905146 | 0.118829184 | 0.283085165 | 0.704025235 | ns | NaCl | KS.3 |
| FIM1_2630 | 660.0324047 | 735.9235328 | 544.7853279 | 0.039934416 | 0.09621132 | 0.394057102 | 0.781740625 | ns | NaCl | KS.3 |
| FIM1_2631 | 203.2513292 | 188.1543082 | 217.1279088 | -0.029189333 | 0.092868528 | 0.532612749 | 0.855153333 | ns | NaCl | KS.3 |
| FIM1_2632 | 544.8879337 | 611.0499531 | 622.236406 | 0.052083459 | 0.090535336 | 0.288353407 | 0.708650347 | ns | NaCl | KS.3 |
| FIM1_2633 | 642.5510266 | 636.4210812 | 808.6597244 | -0.004630996 | 0.089721361 | 0.921071094 | 0.983890455 | ns | NaCl | KS.3 |
| FIM1_2634 | 3337.462793 | 5564.785761 | 5047.092429 | 0.207797324 | 0.261168004 | 0.010215213 | 0.142722848 | ns | NaCl | KS.3 |
| FIM1_2635 | 44130.57236 | 50439.72112 | 31529.08503 | 0.08854887 | 0.080388238 | 0.086329445 | 0.45668467 | ns | NaCl | KS.3 |
| FIM1_2636 | 3840.561628 | 3960.847586 | 3879.739344 | 0.026057649 | 0.060695103 | 0.577515061 | 0.874645237 | ns | NaCl | KS.3 |
| FIM1_2637 | 3990.033014 | 3233.490355 | 4556.375614 | -0.086040601 | 0.109508446 | 0.094599225 | 0.470201108 | ns | NaCl | KS.3 |
| FIM1_2638 | 1588.446757 | 1555.009611 | 1535.1718 | -0.011832889 | 0.078664988 | 0.809424398 | 0.958816322 | ns | NaCl | KS.3 |
| FIM1_2639 | 2958.697947 | 2596.54124 | 3287.046383 | -0.042669361 | 0.100637951 | 0.355456208 | 0.759483959 | ns | NaCl | KS.3 |
| FIM1_2640 | 2349.354549 | 2256.881327 | 3174.526742 | -0.018045847 | 0.086687379 | 0.706122643 | 0.925190614 | ns | NaCl | KS.3 |
| FIM1_2641 | 896.77096 | 807.7670551 | 974.5261912 | -0.068494366 | 0.079341553 | 0.176540781 | 0.605047427 | ns | NaCl | KS.3 |
| FIM1_2642 | 265.2682105 | 266.6701991 | 395.3580463 | 0.001321938 | 0.084460728 | 0.978083198 | 0.996164232 | ns | NaCl | KS.3 |
| FIM1_2643 | 1803.035016 | 1573.538096 | 2442.68225 | -0.06600687 | 0.091521499 | 0.188040305 | 0.618637711 | ns | NaCl | KS.3 |
| FIM1_2644 | 1670.399592 | 1416.549595 | 1769.190409 | -0.114522304 | 0.084778129 | 0.031947776 | 0.283253921 | ns | NaCl | KS.3 |
| FIM1_2645 | 568.1560321 | 590.6331712 | 693.2951935 | 0.018480479 | 0.083934865 | 0.702913845 | 0.923421278 | ns | NaCl | KS.3 |

| | | | | | | | | | | |
|---|---|---|---|---|---|---|---|---|---|---|
| FIM1_2646 | 2571.098182 | 2532.974414 | 2678.289446 | -0.009513712 | 0.07218163 | 0.846315218 | 0.967147129 | ns | NaCl | KS.3 |
| FIM1_2647 | 3149.128251 | 2676.577905 | 3353.209645 | -0.129679689 | 0.078182142 | 0.014134951 | 0.172593755 | ns | NaCl | KS.3 |
| FIM1_2648 | 266.4810281 | 278.359489 | 308.8158223 | 0.021391048 | 0.081709713 | 0.661674216 | 0.904388188 | ns | NaCl | KS.3 |
| FIM1_2649 | 989.7533977 | 1079.743953 | 874.0597844 | 0.069086917 | 0.067019924 | 0.158088551 | 0.57787696 | ns | NaCl | KS.3 |
| FIM1_2650 | 19874.8605 | 22860.77063 | 13759.61847 | 0.07558701 | 0.08816498 | 0.139186038 | 0.550000428 | ns | NaCl | KS.3 |
| FIM1_2651 | 322.1744258 | 358.0354529 | 381.7676173 | 0.029221434 | 0.102422304 | 0.507220189 | 0.844423736 | ns | NaCl | KS.3 |
| FIM1_2652 | 9790.882565 | 10717.7869 | 7485.896065 | 0.07296036 | 0.066997782 | 0.136430137 | 0.544735099 | ns | NaCl | KS.3 |
| FIM1_2653 | 1662.606904 | 1621.760529 | 1690.258184 | -0.014732135 | 0.077216922 | 0.764544874 | 0.947366775 | ns | NaCl | KS.3 |
| FIM1_2654 | 2009.855468 | 1950.975271 | 2235.239421 | -0.015553044 | 0.079778011 | 0.750966296 | 0.943274417 | ns | NaCl | KS.3 |
| FIM1_2655 | 1440.019888 | 1686.501933 | 1444.349023 | 0.020265337 | 0.127411189 | 0.584355874 | 0.877414869 | ns | NaCl | KS.3 |
| FIM1_2656 | 222.3581204 | 126.8448429 | 282.8426353 | -0.168089434 | 0.255736391 | 0.018697305 | 0.207169468 | ns | NaCl | KS.3 |
| FIM1_2657 | 911.0270646 | 930.5445543 | 919.771808 | 0.010027122 | 0.082685623 | 0.836309886 | 0.962829707 | ns | NaCl | KS.3 |
| FIM1_2659 | 1381.188459 | 1416.877321 | 1561.785365 | 0.012478774 | 0.082812701 | 0.797120794 | 0.955924408 | ns | NaCl | KS.3 |
| FIM1_2660 | 1842.658822 | 1396.485805 | 1832.770411 | -0.064205429 | 0.132971627 | 0.164040819 | 0.589056105 | ns | NaCl | KS.3 |
| FIM1_2661 | 1050.304029 | 901.5061464 | 1387.871653 | -0.024772909 | 0.120318697 | 0.526593726 | 0.855027345 | ns | NaCl | KS.3 |
| FIM1_2662 | 5611.889425 | 5764.937743 | 5391.751476 | 0.013370276 | 0.082634283 | 0.783170273 | 0.951646805 | ns | NaCl | KS.3 |
| FIM1_2663 | 563.8491688 | 588.5912272 | 406.4431412 | 0.006857184 | 0.1149284 | 0.857632971 | 0.969856868 | ns | NaCl | KS.3 |
| FIM1_2664 | 2239.928412 | 2135.26779 | 2489.737755 | -0.035999199 | 0.069106834 | 0.462061005 | 0.819880744 | ns | NaCl | KS.3 |
| FIM1_2665 | 149.0608543 | 107.3924127 | 497.1695574 | -0.10266543 | 0.153718477 | 0.058170824 | 0.37782226 | ns | NaCl | KS.3 |
| FIM1_2666 | 427.2442877 | 499.4512562 | 1035.415032 | 0.051829433 | 0.103693065 | 0.268689314 | 0.690146084 | ns | NaCl | KS.3 |
| FIM1_2667 | 3112.893313 | 3185.919776 | 3083.454991 | 0.019761452 | 0.061077097 | 0.673426859 | 0.91246166 | ns | NaCl | KS.3 |
| FIM1_2668 | 2235.813731 | 3112.14462 | 3118.802518 | 0.082619523 | 0.148571665 | 0.096853365 | 0.475192793 | ns | NaCl | KS.3 |
| FIM1_2669 | 12.34986335 | 34.22367983 | 47.98920621 | 0.5164107 | 0.551047113 | 0.001332486 | 0.041956436 | ns | NaCl | KS.3 |
| FIM1_2671 | 29.31607324 | 35.88866927 | 54.40823865 | 0.02191585 | 0.124836547 | 0.561882109 | 0.866779377 | ns | NaCl | KS.3 |
| FIM1_2672 | 810.8092987 | 781.1145655 | 872.2161792 | -0.023677706 | 0.075735603 | 0.631267933 | 0.897558722 | ns | NaCl | KS.3 |
| FIM1_2674 | 3433.022089 | 3648.532761 | 5072.522202 | 0.027455754 | 0.086986897 | 0.568422759 | 0.869550295 | ns | NaCl | KS.3 |
| FIM1_2675 | 660.3390669 | 1082.977979 | 10561.10044 | 0.034312588 | 0.173324852 | 0.320612027 | 0.737029267 | ns | NaCl | KS.3 |
| FIM1_2676 | 26931.75611 | 43907.17084 | 23990.20896 | 0.143991617 | 0.222070455 | 0.026440504 | 0.254519471 | ns | NaCl | KS.3 |
| FIM1_2677 | 5261.570281 | 5386.728367 | 3814.504978 | 0.014439424 | 0.075459431 | 0.769270976 | 0.948835638 | ns | NaCl | KS.3 |
| FIM1_2678 | 616.2037513 | 608.9249702 | 579.2869035 | -0.005609074 | 0.087076574 | 0.906020011 | 0.979880338 | ns | NaCl | KS.3 |
| FIM1_2679 | 839.2177811 | 809.1398139 | 1383.071188 | -0.014985117 | 0.089445924 | 0.750044453 | 0.943274417 | ns | NaCl | KS.3 |
| FIM1_2680 | 6430.133908 | 7637.051019 | 4303.406281 | 0.087730383 | 0.095543347 | 0.091751825 | 0.467388224 | ns | NaCl | KS.3 |
| FIM1_2681 | 3047.240516 | 4704.707897 | 4554.22403 | 0.358073449 | 0.247347115 | 0.000846653 | 0.031433557 | ns | NaCl | KS.3 |
| FIM1_2682 | 481.7746694 | 546.189489 | 945.3060971 | 0.023205792 | 0.114625998 | 0.562560078 | 0.867434473 | ns | NaCl | KS.3 |
| FIM1_2683 | 1639.841791 | 1407.494429 | 2297.684861 | -0.056774895 | 0.099597799 | 0.238604847 | 0.661799251 | ns | NaCl | KS.3 |
| FIM1_2684 | 8280.387264 | 11689.93378 | 5456.353642 | 0.110469856 | 0.159154582 | 0.048175095 | 0.344355024 | ns | NaCl | KS.3 |
| FIM1_2685 | 912.595257 | 854.3703662 | 1047.779783 | -0.036995772 | 0.082049441 | 0.453112701 | 0.816716052 | ns | NaCl | KS.3 |
| FIM1_2686 | 155.9078682 | 80.56798823 | 211.4828374 | -0.751444692 | 0.313371392 | 5.99485331936 | 0.004889245 | ns | NaCl | KS.3 |
| FIM1_2687 | 605.9015336 | 483.4987827 | 777.3624947 | -0.043874991 | 0.124647178 | 0.299617206 | 0.717667597 | ns | NaCl | KS.3 |
| FIM1_2688 | 1925.097584 | 2062.602614 | 1954.134537 | 0.036154846 | 0.082695967 | 0.462516017 | 0.819880744 | ns | NaCl | KS.3 |
| FIM1_2689 | 1487.421859 | 1456.13367 | 1808.791337 | -0.012880937 | 0.07567583 | 0.793515951 | 0.955175866 | ns | NaCl | KS.3 |
| FIM1_2690 | 7812.544171 | 8061.531989 | 6279.00711 | 0.025079579 | 0.064852987 | 0.601259157 | 0.880762097 | ns | NaCl | KS.3 |
| FIM1_2691 | 1486.335368 | 1452.900764 | 1404.064091 | -0.019900129 | 0.063140861 | 0.675359513 | 0.912806589 | ns | NaCl | KS.3 |
| FIM1_2692 | 728.4704408 | 665.9172829 | 751.1524816 | -0.047703651 | 0.084478291 | 0.335569237 | 0.745307602 | ns | NaCl | KS.3 |
| FIM1_2693 | 2626.306998 | 2603.883352 | 2325.402404 | -0.006611058 | 0.068145446 | 0.891794526 | 0.977268396 | ns | NaCl | KS.3 |

| | | | | | | | | | | |
|---|---|---|---|---|---|---|---|---|---|---|
| FIM1_2694 | 606.1470673 | 602.6859116 | 600.8518616 | -0.005483074 | 0.074339509 | 0.911201914 | 0.982011252 | ns | NaCl | KS.3 |
| FIM1_2695 | 3938.671052 | 4178.140082 | 2827.478016 | 0.034749942 | 0.078175631 | 0.482755831 | 0.829184191 | ns | NaCl | KS.3 |
| FIM1_2696 | 10746.39502 | 13055.14327 | 6745.90564 | 0.062377206 | 0.110169468 | 0.190641249 | 0.618952432 | ns | NaCl | KS.3 |
| FIM1_2697 | 542.0707449 | 560.0069554 | 428.8738886 | 0.006933236 | 0.10616743 | 0.865412722 | 0.970128052 | ns | NaCl | KS.3 |
| FIM1_2698 | 3293.413595 | 2828.431972 | 3296.22013 | -0.108907475 | 0.080649434 | 0.038237711 | 0.310330524 | ns | NaCl | KS.3 |
| FIM1_2699 | 3756.118771 | 4822.116979 | 2342.557362 | 0.067964047 | 0.12487144 | 0.152286632 | 0.568361587 | ns | NaCl | KS.3 |
| FIM1_2700 | 408.0431388 | 389.097614 | 571.3223321 | -0.014359865 | 0.096041956 | 0.748955109 | 0.943274417 | ns | NaCl | KS.3 |
| FIM1_2701 | 1607.913941 | 1581.773035 | 2051.847764 | -0.00560533 | 0.095515516 | 0.900349745 | 0.978854891 | ns | NaCl | KS.3 |
| FIM1_2702 | 179.2929698 | 164.2723344 | 193.6273669 | -0.029302236 | 0.097276687 | 0.51995431 | 0.851453596 | ns | NaCl | KS.3 |
| FIM1_2703 | 106.5469614 | 43.96088107 | 206.2115144 | -0.282181603 | 0.4181953 | 0.005891517 | 0.102180094 | ns | NaCl | KS.3 |
| FIM1_2704 | 1387.424867 | 1465.121 | 1576.395522 | 0.027516809 | 0.083375978 | 0.572913631 | 0.871656025 | ns | NaCl | KS.3 |
| FIM1_2705 | 3934.256321 | 6305.429292 | 3068.511263 | 0.365307522 | 0.272906793 | 0.001127537 | 0.039503502 | ns | NaCl | KS.3 |
| FIM1_2706 | 17.69632218 | 20.62334355 | 23.14872675 | 0.020028832 | 0.133384208 | 0.578247637 | 0.874933697 | ns | NaCl | KS.3 |
| FIM1_2708 | 9544.642909 | 13389.93926 | 5778.449195 | 0.083428126 | 0.150975134 | 0.09453233 | 0.470201108 | ns | NaCl | KS.3 |
| FIM1_2709 | 4079.41758 | 4150.949267 | 3182.780214 | 0.010263678 | 0.076929041 | 0.834613734 | 0.962365393 | ns | NaCl | KS.3 |
| FIM1_2710 | 11823.33329 | 13700.48708 | 8876.301857 | 0.057298498 | 0.098163746 | 0.23695454 | 0.659616806 | ns | NaCl | KS.3 |
| FIM1_2711 | 356.483367 | 348.3620966 | 333.9744302 | -0.014091498 | 0.08115171 | 0.772864293 | 0.94938462 | ns | NaCl | KS.3 |
| FIM1_2712 | 3374.104325 | 4211.675726 | 2749.901888 | 0.109854754 | 0.110079459 | 0.044649747 | 0.333532266 | ns | NaCl | KS.3 |
| FIM1_2713 | 1732.56011 | 1730.402726 | 1923.715799 | -0.001350558 | 0.073685518 | 0.978074332 | 0.996164232 | ns | NaCl | KS.3 |
| FIM1_2714 | 1150.19923 | 1170.874276 | 1521.290893 | 0.015110249 | 0.06282323 | 0.749955894 | 0.943274417 | ns | NaCl | KS.3 |
| FIM1_2715 | 3660.001077 | 4199.937652 | 2877.232405 | 0.106120152 | 0.074762707 | 0.03907388 | 0.313536379 | ns | NaCl | KS.3 |
| FIM1_2716 | 10396.8434 | 10508.30645 | 7226.098918 | 0.005157015 | 0.083990706 | 0.914858176 | 0.982343702 | ns | NaCl | KS.3 |
| FIM1_2718 | 4841.068712 | 4978.341829 | 5181.679644 | 0.01958935 | 0.070173501 | 0.689006256 | 0.918490387 | ns | NaCl | KS.3 |
| FIM1_2719 | 2068.630573 | 2985.854017 | 2146.318507 | 0.27672855 | 0.200772483 | 0.001509169 | 0.046346385 | ns | NaCl | KS.3 |
| FIM1_2720 | 2440.171169 | 2677.053479 | 3393.652226 | 0.080115372 | 0.063476518 | 0.095407003 | 0.471350386 | ns | NaCl | KS.3 |
| FIM1_2721 | 630.0833805 | 1169.68793 | 627.1013239 | 0.496347909 | 0.358531948 | 0.000721027 | 0.029870765 | ns | NaCl | KS.3 |
| FIM1_2722 | 3312.472511 | 3738.68522 | 2572.884247 | 0.06236687 | 0.087222208 | 0.214086406 | 0.638920137 | ns | NaCl | KS.3 |
| FIM1_2723 | 582.8628737 | 510.1468281 | 810.7871717 | -0.03219541 | 0.108267161 | 0.453514128 | 0.816716052 | ns | NaCl | KS.3 |
| FIM1_2724 | 1188.174216 | 965.9006644 | 1911.980482 | -0.116957804 | 0.101709136 | 0.033790973 | 0.292771172 | ns | NaCl | KS.3 |
| FIM1_2726 | 426.9869739 | 327.3467859 | 789.6774471 | -0.129382193 | 0.126380365 | 0.025854397 | 0.2527026 | ns | NaCl | KS.3 |
| FIM1_2727 | 82.0463082 | 51.00975561 | 134.5850001 | -0.186401139 | 0.245635706 | 0.013493042 | 0.167275542 | ns | NaCl | KS.3 |
| FIM1_2728 | 212.1778498 | 149.2568226 | 592.1077316 | -0.096437399 | 0.160642122 | 0.067895939 | 0.408442922 | ns | NaCl | KS.3 |
| FIM1_2729 | 141.4926307 | 51.4468033 | 331.3030649 | -0.808481422 | 0.540933343 | 0.000263828 | 0.01526215 | ns | NaCl | KS.3 |
| FIM1_2730 | 37.92383365 | 27.41071407 | 150.8965523 | -0.055351213 | 0.149276014 | 0.196810109 | 0.619310748 | ns | NaCl | KS.3 |
| FIM1_2731 | 4468.477873 | 4755.382267 | 4238.867516 | 0.04249161 | 0.072790606 | 0.390802567 | 0.779482559 | ns | NaCl | KS.3 |
| FIM1_2732 | 9658.249488 | 9756.674114 | 8924.198354 | 0.007349263 | 0.069052299 | 0.880146589 | 0.973483611 | ns | NaCl | KS.3 |
| FIM1_2733 | 279.3074518 | 361.2068721 | 314.6935133 | 0.120249876 | 0.125730926 | 0.034009755 | 0.292771172 | ns | NaCl | KS.3 |
| FIM1_2734 | 284.7022593 | 238.3112677 | 239.54723 | -0.083337936 | 0.098279413 | 0.10563386 | 0.497188698 | ns | NaCl | KS.3 |
| FIM1_2735 | 825.1433347 | 764.8442661 | 651.349206 | -0.03967373 | 0.083155575 | 0.420964117 | 0.80133378 | ns | NaCl | KS.3 |
| FIM1_2736 | 582.7671651 | 613.6989084 | 595.3209154 | 0.021923594 | 0.087579551 | 0.646308468 | 0.898905404 | ns | NaCl | KS.3 |
| FIM1_2737 | 3297.885356 | 3260.407285 | 2475.593407 | -0.006732144 | 0.076954424 | 0.891022042 | 0.977268396 | ns | NaCl | KS.3 |
| FIM1_2738 | 375.2290868 | 320.5296337 | 482.4117522 | -0.037066824 | 0.111977884 | 0.388071526 | 0.779482559 | ns | NaCl | KS.3 |
| FIM1_2739 | 1273.568273 | 982.093763 | 2229.439946 | -0.054613134 | 0.129638177 | 0.216513199 | 0.640647865 | ns | NaCl | KS.3 |
| FIM1_2740 | 7127.429256 | 7631.416093 | 5832.091995 | 0.058700471 | 0.062535257 | 0.21707511 | 0.641504077 | ns | NaCl | KS.3 |
| FIM1_2741 | 1028.602551 | 1033.855184 | 950.3022383 | 0.002016568 | 0.075349297 | 0.967281773 | 0.99408528 | ns | NaCl | KS.3 |

| | | | | | | | | | | |
|---|---|---|---|---|---|---|---|---|---|---|
| FIM1_2742 | 8555.694371 | 6062.949599 | 6492.584788 | -0.107983688 | 0.15827848 | 0.051158371 | 0.352999854 | ns | NaCl | KS.3 |
| FIM1_2743 | 996.0300007 | 1321.380321 | 767.070899 | 0.060107447 | 0.134537283 | 0.182440811 | 0.611396712 | ns | NaCl | KS.3 |
| FIM1_2744 | 5708.736257 | 5365.781235 | 6478.222438 | -0.053412297 | 0.061789822 | 0.258558893 | 0.682403445 | ns | NaCl | KS.3 |
| FIM1_2745 | 1789.080914 | 1942.731363 | 1426.642156 | 0.018997978 | 0.105822088 | 0.650378494 | 0.899642849 | ns | NaCl | KS.3 |
| FIM1_2746 | 1140.127063 | 1067.015755 | 1398.377701 | -0.034313578 | 0.082816923 | 0.484871059 | 0.829376704 | ns | NaCl | KS.3 |
| FIM1_2747 | 4075.308617 | 4236.061924 | 2989.429139 | 0.016303721 | 0.087885773 | 0.731458152 | 0.936960396 | ns | NaCl | KS.3 |
| FIM1_2748 | 399.6231662 | 379.0645398 | 558.3447268 | -0.02554969 | 0.086408905 | 0.595912211 | 0.87962202 | ns | NaCl | KS.3 |
| FIM1_2749 | 451.7049695 | 376.2704955 | 918.7957471 | -0.094180848 | 0.100586762 | 0.073815235 | 0.421788075 | ns | NaCl | KS.3 |
| FIM1_2750 | 76.88203275 | 91.14116402 | 94.23558957 | 0.043527995 | 0.10812333 | 0.331067682 | 0.742254041 | ns | NaCl | KS.3 |
| FIM1_2751 | 648.8220711 | 672.5552827 | 582.8865073 | 0.011703684 | 0.096366272 | 0.793174164 | 0.955175866 | ns | NaCl | KS.3 |
| FIM1_2752 | 1192.654419 | 1165.17826 | 1326.891207 | -0.018089303 | 0.067752098 | 0.709581988 | 0.925563973 | ns | NaCl | KS.3 |
| FIM1_2753 | 1766.128974 | 1644.831741 | 1404.235879 | -0.029437431 | 0.089307942 | 0.537192463 | 0.855981197 | ns | NaCl | KS.3 |
| FIM1_2754 | 1774.095704 | 2225.079644 | 3605.0053 | 0.06668093 | 0.118373682 | 0.16202105 | 0.584521192 | ns | NaCl | KS.3 |
| FIM1_2755 | 1022.78064 | 972.9584222 | 1085.069687 | -0.02758075 | 0.079758851 | 0.575393429 | 0.87300467 | ns | NaCl | KS.3 |
| FIM1_2756 | 1022.380064 | 1021.849176 | 909.1344341 | -0.000130408 | 0.109051751 | 0.997384931 | 0.99924489 | ns | NaCl | KS.3 |
| FIM1_2757 | 9771.301824 | 13438.38345 | 5736.513145 | 0.066246381 | 0.14352413 | 0.150025785 | 0.564839452 | ns | NaCl | KS.3 |
| FIM1_2758 | 740.0587639 | 685.2996151 | 579.5831668 | -0.032456266 | 0.08906179 | 0.498331569 | 0.837486406 | ns | NaCl | KS.3 |
| FIM1_2759 | 342.0230481 | 257.3381972 | 236.8115605 | -0.115796358 | 0.13420939 | 0.039832353 | 0.315550884 | ns | NaCl | KS.3 |
| FIM1_2760 | 8965.209739 | 9971.263505 | 8012.268097 | 0.031273367 | 0.10106223 | 0.483946853 | 0.829184191 | ns | NaCl | KS.3 |
| FIM1_2761 | 1374.634428 | 1411.692618 | 1486.793316 | 0.022037297 | 0.05764187 | 0.629295089 | 0.896016329 | ns | NaCl | KS.3 |
| FIM1_2762 | 2690.184956 | 2821.743937 | 2559.5786 | 0.015025368 | 0.09640715 | 0.737209001 | 0.939206485 | ns | NaCl | KS.3 |
| FIM1_2763 | 88.91968599 | 161.012473 | 643.8802253 | 0.045550451 | 0.186185253 | 0.217789022 | 0.641504077 | ns | NaCl | KS.3 |
| FIM1_2764 | 712.5992658 | 699.0002462 | 1403.205605 | -0.005849453 | 0.09257288 | 0.898413088 | 0.977971161 | ns | NaCl | KS.3 |
| FIM1_2765 | 1401.179908 | 1371.482643 | 1273.675358 | -0.010794083 | 0.080069708 | 0.825354857 | 0.960240659 | ns | NaCl | KS.3 |
| FIM1_2766 | 4055.98939 | 4506.48156 | 3595.941157 | 0.062953765 | 0.080862101 | 0.212300395 | 0.635496068 | ns | NaCl | KS.3 |
| FIM1_2767 | 1634.322859 | 1467.649359 | 2284.431879 | -0.032883155 | 0.100281761 | 0.465667926 | 0.820834625 | ns | NaCl | KS.3 |
| FIM1_2768 | 1032.051404 | 855.5531271 | 1224.0717 | -0.063458271 | 0.107412792 | 0.187429289 | 0.618483916 | ns | NaCl | KS.3 |
| FIM1_2769 | 2155.973735 | 2054.631098 | 2209.538832 | -0.034728789 | 0.069796301 | 0.478845017 | 0.826171162 | ns | NaCl | KS.3 |
| FIM1_2770 | 760.1707251 | 588.153344 | 1412.254494 | -0.118326704 | 0.125178082 | 0.035958257 | 0.3016734 | ns | NaCl | KS.3 |
| FIM1_2771 | 462.4640998 | 447.5221959 | 445.9064314 | -0.009698452 | 0.098340947 | 0.824915648 | 0.960240659 | ns | NaCl | KS.3 |
| FIM1_2772 | 1246.996472 | 1255.67275 | 1609.891903 | 0.003644527 | 0.070236541 | 0.940572971 | 0.986790704 | ns | NaCl | KS.3 |
| FIM1_2773 | 353.9019556 | 386.1053472 | 405.4531913 | 0.031103501 | 0.09402462 | 0.504525675 | 0.844040357 | ns | NaCl | KS.3 |
| FIM1_2774 | 840.0333666 | 831.053994 | 928.7320496 | -0.002091055 | 0.105620155 | 0.959287449 | 0.991590145 | ns | NaCl | KS.3 |
| FIM1_2775 | 4560.96113 | 5023.092338 | 4011.411568 | 0.080347493 | 0.066070979 | 0.099948971 | 0.483636126 | ns | NaCl | KS.3 |
| FIM1_2776 | 603.2190944 | 596.1085826 | 718.1820608 | -0.00672478 | 0.081226542 | 0.890242329 | 0.977268396 | ns | NaCl | KS.3 |
| FIM1_2777 | 529.3152566 | 394.0792213 | 817.4033035 | -0.178900228 | 0.149609123 | 0.007369996 | 0.117365229 | ns | NaCl | KS.3 |
| FIM1_2778 | 611.4357803 | 574.9384498 | 579.844977 | -0.036966142 | 0.079232433 | 0.455431648 | 0.816788914 | ns | NaCl | KS.3 |
| FIM1_2779 | 1139.129235 | 1231.099347 | 941.1714081 | 0.037523963 | 0.085955615 | 0.442037652 | 0.809513863 | ns | NaCl | KS.3 |
| FIM1_2780 | 144.1064129 | 82.12240068 | 312.6164471 | -0.240821585 | 0.295402442 | 0.007383983 | 0.117365229 | ns | NaCl | KS.3 |
| FIM1_2781 | 594.6888782 | 672.819288 | 1642.397071 | 0.025077086 | 0.112054867 | 0.540447425 | 0.857165292 | ns | NaCl | KS.3 |
| FIM1_2782 | 2063.513357 | 2100.485664 | 1563.850147 | 0.007953594 | 0.086129042 | 0.867791343 | 0.97082571 | ns | NaCl | KS.3 |
| FIM1_2783 | 10900.06037 | 14550.53808 | 6384.891685 | 0.059101199 | 0.136212635 | 0.186366284 | 0.618044911 | ns | NaCl | KS.3 |
| FIM1_2784 | 740.0469685 | 707.5716706 | 907.651831 | -0.032605199 | 0.070421128 | 0.506679354 | 0.844423736 | ns | NaCl | KS.3 |
| FIM1_2785 | 2791.090658 | 2629.902128 | 2810.77649 | -0.035651897 | 0.077688068 | 0.471789672 | 0.822127061 | ns | NaCl | KS.3 |
| FIM1_2786 | 2385.09909 | 2330.511342 | 2411.717556 | -0.010079602 | 0.086604448 | 0.832618438 | 0.961831996 | ns | NaCl | KS.3 |

| | | | | | | | | | | |
|---|---|---|---|---|---|---|---|---|---|---|
| FIM1_2787 | 5754.942296 | 6641.169744 | 5756.818924 | 0.029299608 | 0.113298649 | 0.478524048 | 0.826171162 | ns | NaCl | KS.3 |
| FIM1_2788 | 196.8122646 | 131.4842797 | 344.3151537 | -0.19936295 | 0.209037452 | 0.008497148 | 0.127329254 | ns | NaCl | KS.3 |
| FIM1_2789 | 249.3969894 | 232.0453632 | 274.6503493 | -0.02122784 | 0.099373979 | 0.630937762 | 0.897345731 | ns | NaCl | KS.3 |
| FIM1_2790 | 751.9169607 | 695.8861326 | 1088.731515 | -0.040336659 | 0.083831076 | 0.412771076 | 0.799937916 | ns | NaCl | KS.3 |
| FIM1_2791 | 190.469477 | 131.6294977 | 239.3295169 | -0.163368364 | 0.183425188 | 0.015186558 | 0.180365277 | ns | NaCl | KS.3 |
| FIM1_2792 | 637.2228674 | 635.5305989 | 699.8149305 | -0.00111077 | 0.090015618 | 0.980992692 | 0.996363429 | ns | NaCl | KS.3 |
| FIM1_2793 | 1434.660166 | 1558.947011 | 1821.019608 | 0.064433766 | 0.068348464 | 0.190226234 | 0.618952432 | ns | NaCl | KS.3 |
| FIM1_2794 | 3117.781334 | 2397.397487 | 2224.607093 | -0.175968836 | 0.126651418 | 0.005659472 | 0.099945317 | ns | NaCl | KS.3 |
| FIM1_2795 | 734.7080641 | 648.5372656 | 653.7490727 | -0.025431077 | 0.111847195 | 0.535771245 | 0.855153333 | ns | NaCl | KS.3 |
| FIM1_2796 | 215.0208161 | 177.0802397 | 654.7569723 | -0.031034097 | 0.125345537 | 0.431354747 | 0.805772482 | ns | NaCl | KS.3 |
| FIM1_2797 | 36.90121769 | 28.6627284 | 53.41088707 | -0.038329564 | 0.137322243 | 0.332147582 | 0.742667065 | ns | NaCl | KS.3 |
| FIM1_2798 | 1606.720565 | 1797.326162 | 1657.686551 | 0.065333053 | 0.082487192 | 0.196485501 | 0.619072431 | ns | NaCl | KS.3 |
| FIM1_2800 | 4371.024018 | 5103.193765 | 3329.2877 | 0.118535565 | 0.07884431 | 0.024277086 | 0.242041091 | ns | NaCl | KS.3 |
| FIM1_2801 | 4751.754152 | 5244.585633 | 4352.614611 | 0.05564159 | 0.082126126 | 0.26681064 | 0.689593109 | ns | NaCl | KS.3 |
| FIM1_2802 | 804.6411554 | 774.5964941 | 855.5880216 | -0.018589595 | 0.083124685 | 0.702103979 | 0.923421278 | ns | NaCl | KS.3 |
| FIM1_2803 | 1290.055147 | 1263.939828 | 1290.444468 | -0.014689749 | 0.071693213 | 0.764683094 | 0.947366775 | ns | NaCl | KS.3 |
| FIM1_2804 | 2168.492472 | 2252.152197 | 2299.026539 | 0.028812056 | 0.066577209 | 0.551724621 | 0.860945181 | ns | NaCl | KS.3 |
| FIM1_2805 | 1896.465417 | 1990.721209 | 1868.510163 | 0.026961198 | 0.07932575 | 0.584152609 | 0.877414869 | ns | NaCl | KS.3 |
| FIM1_2806 | 138.5768711 | 165.1593563 | 204.5735688 | 0.062536199 | 0.102779571 | 0.197520484 | 0.620924909 | ns | NaCl | KS.3 |
| FIM1_2807 | 1069.691588 | 1047.114329 | 1116.620185 | -0.009488623 | 0.08554938 | 0.843100161 | 0.966485679 | ns | NaCl | KS.3 |
| FIM1_2808 | 1995.037121 | 1596.458668 | 1536.198656 | -0.117513185 | 0.109772927 | 0.034623368 | 0.294446588 | ns | NaCl | KS.3 |
| FIM1_2809 | 104.7773276 | 75.62115082 | 172.0515214 | -0.076793425 | 0.146549188 | 0.112942149 | 0.511380136 | ns | NaCl | KS.3 |
| FIM1_2810 | 287.5912346 | 256.9855679 | 237.534163 | -0.020318436 | 0.115330621 | 0.606883094 | 0.884426372 | ns | NaCl | KS.3 |
| FIM1_2811 | 496.9074007 | 473.1796072 | 460.8452583 | -0.023086055 | 0.084101064 | 0.634418095 | 0.898009483 | ns | NaCl | KS.3 |
| FIM1_2812 | 3103.352697 | 2673.452841 | 2959.470854 | -0.11252867 | 0.077481552 | 0.030967288 | 0.276592922 | ns | NaCl | KS.3 |
| FIM1_2813 | 728.6385529 | 671.5842434 | 1060.003627 | -0.037143868 | 0.087297369 | 0.444314024 | 0.812278382 | ns | NaCl | KS.3 |
| FIM1_2814 | 1646.52669 | 1565.43059 | 2193.081339 | -0.024095485 | 0.085223239 | 0.618381468 | 0.891982547 | ns | NaCl | KS.3 |
| FIM1_2815 | 315.3846326 | 354.5345177 | 337.3905196 | 0.045984885 | 0.09323865 | 0.338761442 | 0.748041799 | ns | NaCl | KS.3 |
| FIM1_2816 | 410.4817822 | 404.3277354 | 373.392304 | -0.007535815 | 0.08880288 | 0.872755051 | 0.971243188 | ns | NaCl | KS.3 |
| FIM1_2817 | 626.2055237 | 591.1666045 | 534.8311899 | -0.025416695 | 0.088774855 | 0.59359902 | 0.87962202 | ns | NaCl | KS.3 |
| FIM1_2818 | 284.2191489 | 292.9967956 | 428.9503715 | 0.014141586 | 0.083143231 | 0.770655536 | 0.949065335 | ns | NaCl | KS.3 |
| FIM1_2819 | 2160.264259 | 2009.055458 | 2042.335681 | -0.057957093 | 0.065159732 | 0.230433588 | 0.654338423 | ns | NaCl | KS.3 |
| FIM1_2820 | 1439.3295 | 1417.433533 | 2772.900424 | -0.00347153 | 0.105337771 | 0.932662253 | 0.985136881 | ns | NaCl | KS.3 |
| FIM1_2821 | 1478.211228 | 1432.859414 | 1743.736036 | -0.016571586 | 0.080765222 | 0.734750021 | 0.938375543 | ns | NaCl | KS.3 |
| FIM1_2822 | 463.2566934 | 635.4563286 | 639.5826493 | 0.21091751 | 0.165242141 | 0.003811922 | 0.079348592 | ns | NaCl | KS.3 |
| FIM1_2823 | 4563.894369 | 4337.258707 | 3390.920902 | -0.039054599 | 0.06736928 | 0.42220735 | 0.801548921 | ns | NaCl | KS.3 |
| FIM1_2824 | 1193.700967 | 1212.659399 | 1247.898865 | 0.010752043 | 0.067110643 | 0.824286936 | 0.960240659 | ns | NaCl | KS.3 |
| FIM1_2825 | 925.3484235 | 899.3936646 | 1050.128606 | -0.018248345 | 0.07783014 | 0.71070344 | 0.925563973 | ns | NaCl | KS.3 |
| FIM1_2826 | 580.4214292 | 472.5468576 | 888.972011 | -0.0753133 | 0.109981589 | 0.130509914 | 0.5379344 | ns | NaCl | KS.3 |
| FIM1_2827 | 5130.56352 | 6889.823882 | 3463.262847 | 0.08652988 | 0.138010959 | 0.088791928 | 0.46158813 | ns | NaCl | KS.3 |
| FIM1_2828 | 450.529335 | 426.564749 | 768.7387174 | -0.029856252 | 0.079536045 | 0.544889136 | 0.857313452 | ns | NaCl | KS.3 |
| FIM1_2829 | 2608.532176 | 3111.605749 | 2250.230989 | 0.101286384 | 0.093429674 | 0.057082619 | 0.374651756 | ns | NaCl | KS.3 |
| FIM1_2830 | 2450.710871 | 2194.016618 | 2405.026122 | -0.052405099 | 0.088910551 | 0.287841095 | 0.707864285 | ns | NaCl | KS.3 |
| FIM1_2831 | 78.83008798 | 152.9725104 | 339.8605135 | 0.121296868 | 0.249469688 | 0.038942311 | 0.312985452 | ns | NaCl | KS.3 |
| FIM1_2832 | 1164.778892 | 1260.215677 | 1539.108163 | 0.046188436 | 0.079602061 | 0.353776241 | 0.757528931 | ns | NaCl | KS.3 |

| | | | | | | | | | | |
|---|---|---|---|---|---|---|---|---|---|---|
| FIM1_2833 | 2952.260845 | 3744.482924 | 1969.477907 | 0.04408344 | 0.127407453 | 0.294020669 | 0.71284251 | ns | NaCl | KS.3 |
| FIM1_2834 | 814.5735218 | 850.9211991 | 1728.387067 | 0.006679075 | 0.116589124 | 0.859671041 | 0.969856868 | ns | NaCl | KS.3 |
| FIM1_2835 | 975.868238 | 1073.110703 | 1076.06063 | 0.01732166 | 0.112951741 | 0.662545616 | 0.904388188 | ns | NaCl | KS.3 |
| FIM1_2836 | 643.696732 | 719.5433293 | 864.2023526 | 0.056676158 | 0.086491522 | 0.255780451 | 0.678991959 | ns | NaCl | KS.3 |
| FIM1_2837 | 553.0755008 | 582.935828 | 858.4962492 | 0.010459181 | 0.109220672 | 0.79427038 | 0.955299664 | ns | NaCl | KS.3 |
| FIM1_2838 | 3065.726926 | 3008.119576 | 2749.047553 | -0.010997836 | 0.077245205 | 0.822942554 | 0.960240659 | ns | NaCl | KS.3 |
| FIM1_2839 | 462.3157427 | 449.0295849 | 734.8702793 | -0.019764972 | 0.076055229 | 0.688357627 | 0.918490387 | ns | NaCl | KS.3 |
| FIM1_2840 | 119.552301 | 69.17136346 | 112.8839267 | -0.470189661 | 0.313955325 | 0.000584528 | 0.025069192 | ns | NaCl | KS.3 |
| FIM1_2841 | 1240.766057 | 1219.777828 | 1116.009165 | -0.00944413 | 0.079288251 | 0.847124921 | 0.967147129 | ns | NaCl | KS.3 |
| FIM1_2842 | 4493.772422 | 4617.806232 | 3122.825629 | 0.017709864 | 0.073534584 | 0.718970457 | 0.930267366 | ns | NaCl | KS.3 |
| FIM1_2843 | 161.0905173 | 147.800624 | 223.4923183 | -0.027558623 | 0.098165885 | 0.541349417 | 0.857165292 | ns | NaCl | KS.3 |
| FIM1_2844 | 400.257496 | 364.3885683 | 525.3170241 | -0.032063461 | 0.096703106 | 0.485392146 | 0.829376704 | ns | NaCl | KS.3 |
| FIM1_2845 | 49319.27839 | 72669.52633 | 43296.34962 | 0.474663411 | 0.15969866 | 8.11553045678 | 0.001009369 | ns | NaCl | KS.3 |
| FIM1_2846 | 1550.021559 | 2076.246365 | 1216.290288 | 0.064537776 | 0.136784522 | 0.160447299 | 0.58211082 | ns | NaCl | KS.3 |
| FIM1_2847 | 1038.370235 | 580.0159888 | 3194.810321 | -0.106033072 | 0.222828067 | 0.052799816 | 0.360032854 | ns | NaCl | KS.3 |
| FIM1_2848 | 520.3314233 | 575.2834078 | 706.3170098 | 0.04490204 | 0.08977372 | 0.355698015 | 0.759483959 | ns | NaCl | KS.3 |
| FIM1_2849 | 20708.78927 | 23620.2018 | 14759.95724 | 0.078523566 | 0.083768715 | 0.125942977 | 0.534982546 | ns | NaCl | KS.3 |
| FIM1_2850 | 2653.154422 | 2815.608339 | 2126.038905 | 0.025789165 | 0.087802966 | 0.590150694 | 0.87877872 | ns | NaCl | KS.3 |
| FIM1_2851 | 2193.112456 | 2028.353482 | 2482.05434 | -0.027768558 | 0.094236794 | 0.548595801 | 0.860423742 | ns | NaCl | KS.3 |
| FIM1_2852 | 867.35966 | 669.2634104 | 869.8070042 | -0.137560764 | 0.125277206 | 0.019891902 | 0.213741282 | ns | NaCl | KS.3 |
| FIM1_2853 | 7071.77716 | 9023.538659 | 4109.292691 | 0.050102687 | 0.126817185 | 0.248720989 | 0.67096552 | ns | NaCl | KS.3 |
| FIM1_2854 | 1942.066057 | 1918.588625 | 1743.33323 | -0.004840851 | 0.088273265 | 0.918263818 | 0.982488618 | ns | NaCl | KS.3 |
| FIM1_2855 | 121.4693181 | 67.94327857 | 121.4704813 | -0.394930515 | 0.340213909 | 0.001596829 | 0.047109988 | ns | NaCl | KS.3 |
| FIM1_2856 | 444.1541306 | 322.1214604 | 634.690912 | -0.149595509 | 0.159572397 | 0.018084016 | 0.202175238 | ns | NaCl | KS.3 |
| FIM1_2857 | 905.430905 | 1149.320243 | 751.6658429 | 0.109210453 | 0.11756433 | 0.046423813 | 0.341149881 | ns | NaCl | KS.3 |
| FIM1_2858 | 1159.901949 | 1211.660569 | 1021.325247 | 0.024015543 | 0.08095354 | 0.624320686 | 0.894068915 | ns | NaCl | KS.3 |
| FIM1_2859 | 5911.223804 | 4067.740071 | 3942.214549 | -0.273876094 | 0.205332099 | 0.001743399 | 0.050135331 | ns | NaCl | KS.3 |
| FIM1_2860 | 19707.80884 | 15626.46283 | 11384.25993 | -0.045491571 | 0.125512926 | 0.285017167 | 0.70510214 | ns | NaCl | KS.3 |
| FIM1_2862 | 237.3804394 | 205.1835244 | 207.0002085 | -0.043901373 | 0.104784651 | 0.334469097 | 0.745307602 | ns | NaCl | KS.3 |
| FIM1_2863 | 951.6751514 | 1154.667836 | 1263.715346 | 0.056307983 | 0.112031443 | 0.22481693 | 0.648385059 | ns | NaCl | KS.3 |
| FIM1_2864 | 4813.830706 | 5326.156061 | 3071.00938 | 0.051097066 | 0.086015481 | 0.302608819 | 0.72182705 | ns | NaCl | KS.3 |
| FIM1_2865 | 1404.724823 | 1487.508655 | 1040.611865 | 0.021724961 | 0.091681989 | 0.641494333 | 0.898798052 | ns | NaCl | KS.3 |
| FIM1_2866 | 479.2016968 | 406.655528 | 682.8797352 | -0.071584292 | 0.097585313 | 0.153914614 | 0.571010593 | ns | NaCl | KS.3 |
| FIM1_2867 | 1250.064814 | 1209.485063 | 1333.668172 | -0.015705723 | 0.085389532 | 0.743998161 | 0.941460666 | ns | NaCl | KS.3 |
| FIM1_2868 | 40.90577518 | 16.8674488 | 53.04285322 | -0.464864653 | 0.488605541 | 0.001685066 | 0.049024579 | ns | NaCl | KS.3 |
| FIM1_2869 | 36.1185156 | 19.73034058 | 263.2725796 | -0.192100381 | 0.28596381 | 0.013877464 | 0.17047008 | ns | NaCl | KS.3 |
| FIM1_2870 | 5743.028008 | 6085.428439 | 4619.041052 | 0.023465112 | 0.089822556 | 0.619591441 | 0.892479944 | ns | NaCl | KS.3 |
| FIM1_2871 | 1024.92113 | 1027.266036 | 1028.851863 | 0.001272322 | 0.076843137 | 0.979337342 | 0.996360588 | ns | NaCl | KS.3 |
| FIM1_2872 | 1278.456812 | 1470.542681 | 3341.988258 | 0.064879291 | 0.092122974 | 0.194301372 | 0.618952432 | ns | NaCl | KS.3 |
| FIM1_2873 | 302.727264 | 477.0480167 | 379.4106986 | 0.343473478 | 0.260807537 | 0.001287311 | 0.041599773 | ns | NaCl | KS.3 |
| FIM1_2874 | 4434.554945 | 4269.918124 | 5304.524092 | -0.015043458 | 0.089804923 | 0.74857431 | 0.943274417 | ns | NaCl | KS.3 |
| FIM1_2875 | 3138.855736 | 3672.798511 | 2510.602835 | 0.128102863 | 0.076421707 | 0.014503961 | 0.174292767 | ns | NaCl | KS.3 |
| FIM1_2876 | 2734.439419 | 2252.543367 | 2282.743622 | -0.137484724 | 0.091956173 | 0.013516536 | 0.167275542 | ns | NaCl | KS.3 |
| FIM1_2877 | 6143.308845 | 8402.52423 | 3588.738224 | 0.074156286 | 0.142571585 | 0.122086591 | 0.528318345 | ns | NaCl | KS.3 |
| FIM1_2878 | 1099.87762 | 1019.752433 | 1245.67938 | -0.039611527 | 0.083293296 | 0.421519703 | 0.801533346 | ns | NaCl | KS.3 |

| | | | | | | | | | | |
|---|---|---|---|---|---|---|---|---|---|---|
| FIM1_2879 | 1360.771781 | 1233.217232 | 1558.259213 | -0.040926587 | 0.091514859 | 0.393682188 | 0.781740625 | ns | NaCl | KS.3 |
| FIM1_2880 | 1103.873244 | 1369.062332 | 1078.416442 | 0.110495758 | 0.107078698 | 0.043314159 | 0.328487717 | ns | NaCl | KS.3 |
| FIM1_2881 | 1537.688497 | 1612.283022 | 1233.208042 | 0.016140374 | 0.093999112 | 0.723691901 | 0.93308324 | ns | NaCl | KS.3 |
| FIM1_2882 | 1386.06185 | 1260.367 | 1469.445907 | -0.047975368 | 0.085274462 | 0.332127101 | 0.742667065 | ns | NaCl | KS.3 |
| FIM1_2883 | 2125.238726 | 2046.675196 | 2329.138013 | -0.026790883 | 0.069963584 | 0.584402663 | 0.877414869 | ns | NaCl | KS.3 |
| FIM1_2884 | 621.2967195 | 679.2223239 | 690.2018904 | 0.053021242 | 0.079963729 | 0.289503475 | 0.709394019 | ns | NaCl | KS.3 |
| FIM1_2885 | 2748.335277 | 3554.522471 | 1656.05392 | 0.042917113 | 0.132298193 | 0.297385316 | 0.715635559 | ns | NaCl | KS.3 |
| FIM1_2886 | 1559.093753 | 2086.807034 | 931.8734118 | 0.04648333 | 0.138743721 | 0.261649048 | 0.686421476 | ns | NaCl | KS.3 |
| FIM1_2887 | 7688.348622 | 9978.788837 | 4640.995791 | 0.05752621 | 0.129362816 | 0.199907327 | 0.623535394 | ns | NaCl | KS.3 |
| FIM1_2888 | 157.1939123 | 134.834288 | 389.7551072 | -0.049821034 | 0.104541516 | 0.283131051 | 0.704025235 | ns | NaCl | KS.3 |
| FIM1_2889 | 2100.084799 | 2061.973874 | 2031.263372 | -0.013563904 | 0.070946612 | 0.781912658 | 0.951646805 | ns | NaCl | KS.3 |
| FIM1_2890 | 4790.206662 | 4021.245604 | 4347.024313 | -0.119921406 | 0.087319822 | 0.026388749 | 0.254519471 | ns | NaCl | KS.3 |
| FIM1_2891 | 5325.617401 | 4461.771767 | 4319.216258 | -0.149548106 | 0.079230696 | 0.005328686 | 0.096361786 | ns | NaCl | KS.3 |
| FIM1_2892 | 187.4279164 | 181.1808256 | 304.0822981 | -0.012469761 | 0.092065556 | 0.787004341 | 0.95289761 | ns | NaCl | KS.3 |
| FIM1_2893 | 404.7216929 | 472.1569571 | 642.7355356 | 0.067356763 | 0.095814755 | 0.176939803 | 0.605137125 | ns | NaCl | KS.3 |
| FIM1_2894 | 744.0717295 | 717.7632851 | 1040.38808 | -0.019447642 | 0.08344507 | 0.688842673 | 0.918490387 | ns | NaCl | KS.3 |
| FIM1_2895 | 3216.09677 | 3287.883119 | 3125.276002 | 0.013510321 | 0.075458804 | 0.783700996 | 0.951646805 | ns | NaCl | KS.3 |
| FIM1_2896 | 1384.912861 | 1658.52273 | 1441.378662 | 0.145971463 | 0.082762757 | 0.007275859 | 0.117086974 | ns | NaCl | KS.3 |
| FIM1_2897 | 5705.229094 | 6526.592575 | 5776.34956 | 0.047631897 | 0.098951339 | 0.312292851 | 0.731871738 | ns | NaCl | KS.3 |
| FIM1_2898 | 21769.72128 | 26003.6777 | 16191.39412 | 0.114578414 | 0.090273039 | 0.033894325 | 0.292771172 | ns | NaCl | KS.3 |
| FIM1_2899 | 15972.03156 | 20796.20102 | 9340.909001 | 0.09186945 | 0.127709999 | 0.077613535 | 0.432393435 | ns | NaCl | KS.3 |
| FIM1_2900 | 8147.36813 | 11457.17698 | 6646.927741 | 0.237303828 | 0.181383255 | 0.002568446 | 0.062331789 | ns | NaCl | KS.3 |
| FIM1_2901 | 1817.899867 | 2015.224666 | 1677.89953 | 0.06335077 | 0.079848121 | 0.209495154 | 0.633579568 | ns | NaCl | KS.3 |
| FIM1_2903 | 1850.981037 | 1792.133974 | 2443.122967 | -0.015844987 | 0.082471726 | 0.744680331 | 0.94197423 | ns | NaCl | KS.3 |
| FIM1_2904 | 1443.233172 | 1569.422803 | 1819.270302 | 0.028004549 | 0.09624873 | 0.540246358 | 0.857165292 | ns | NaCl | KS.3 |
| FIM1_2905 | 729.4058023 | 797.6679403 | 846.2092377 | 0.014410419 | 0.116435597 | 0.70799104 | 0.925190614 | ns | NaCl | KS.3 |
| FIM1_2906 | 130.2855491 | 132.751865 | 192.2999731 | 0.002300919 | 0.102922966 | 0.956244465 | 0.989856135 | ns | NaCl | KS.3 |
| FIM1_2907 | 1030.00272 | 1243.319134 | 990.5456322 | 0.06174943 | 0.108392167 | 0.195998774 | 0.618952432 | ns | NaCl | KS.3 |
| FIM1_2908 | 2629.624188 | 3150.89324 | 2481.130868 | 0.124726739 | 0.089058919 | 0.02212698 | 0.228385325 | ns | NaCl | KS.3 |
| FIM1_2909 | 1416.791057 | 1420.259612 | 1242.605634 | 0.001983039 | 0.075005228 | 0.967826111 | 0.99408528 | ns | NaCl | KS.3 |
| FIM1_2911 | 250.7219853 | 244.7860869 | 424.6130829 | -0.004355116 | 0.108960615 | 0.913047996 | 0.98221021 | ns | NaCl | KS.3 |
| FIM1_2912 | 2397.574291 | 2742.191175 | 2982.181995 | 0.027311535 | 0.112699471 | 0.506950007 | 0.844423736 | ns | NaCl | KS.3 |
| FIM1_2913 | 14.031223 | 21.46863589 | 14.35777334 | 0.048066326 | 0.165538834 | 0.225135274 | 0.648597616 | ns | NaCl | KS.3 |
| FIM1_2914 | 2390.381939 | 3155.96318 | 1880.422927 | 0.048134542 | 0.135332484 | 0.253307513 | 0.676801759 | ns | NaCl | KS.3 |
| FIM1_2915 | 799.3722101 | 749.9475364 | 1489.707832 | -0.023579905 | 0.092325732 | 0.612632775 | 0.889162818 | ns | NaCl | KS.3 |
| FIM1_2916 | 4274.015748 | 5215.637882 | 5132.835234 | 0.093291029 | 0.104008606 | 0.075923259 | 0.427766945 | ns | NaCl | KS.3 |
| FIM1_2917 | 881.1874177 | 910.6090703 | 1446.996132 | 0.021348288 | 0.074337349 | 0.664890024 | 0.905757905 | ns | NaCl | KS.3 |
| FIM1_2918 | 889.5393324 | 1021.948492 | 985.6773065 | 0.08031294 | 0.085971249 | 0.118588709 | 0.522105158 | ns | NaCl | KS.3 |
| FIM1_2919 | 3009.606695 | 3001.956878 | 3139.017352 | -0.001869708 | 0.074936507 | 0.969663777 | 0.99411987 | ns | NaCl | KS.3 |
| FIM1_2920 | 362.8181516 | 425.0346812 | 333.924461 | 0.037822989 | 0.111008178 | 0.382027247 | 0.778609403 | ns | NaCl | KS.3 |
| FIM1_2921 | 4206.918815 | 5043.319366 | 2861.91268 | 0.036690153 | 0.117598638 | 0.380565405 | 0.778493059 | ns | NaCl | KS.3 |
| FIM1_2922 | 571.3145219 | 535.5417567 | 940.8518398 | -0.023641896 | 0.093107936 | 0.609952779 | 0.88728511 | ns | NaCl | KS.3 |
| FIM1_2923 | 5389.056128 | 5921.746952 | 3964.760454 | 0.053617684 | 0.081487727 | 0.283944299 | 0.704550068 | ns | NaCl | KS.3 |
| FIM1_2924 | 9414.339564 | 9572.184702 | 6416.49765 | 0.00879904 | 0.080479599 | 0.857050901 | 0.969856868 | ns | NaCl | KS.3 |
| FIM1_2925 | 52.93495901 | 40.22334985 | 55.21287527 | -0.037625592 | 0.14022183 | 0.335053168 | 0.745307602 | ns | NaCl | KS.3 |

| | | | | | | | | | | |
|---|---|---|---|---|---|---|---|---|---|---|
| FIM1_2926 | 2581.970018 | 2309.848389 | 2247.740019 | -0.038230909 | 0.097341344 | 0.41009224 | 0.7969566 | ns | NaCl | KS.3 |
| FIM1_2927 | 5913.799136 | 6004.431409 | 5522.650867 | 0.014435743 | 0.057222108 | 0.750903711 | 0.943274417 | ns | NaCl | KS.3 |
| FIM1_2928 | 605.8616543 | 634.5380025 | 1364.35551 | 0.008691937 | 0.110451566 | 0.826330092 | 0.960240659 | ns | NaCl | KS.3 |
| FIM1_2929 | 1072.284269 | 898.3875834 | 1704.738101 | -0.050121997 | 0.10977967 | 0.271824286 | 0.693144962 | ns | NaCl | KS.3 |
| FIM1_2930 | 432.0653583 | 541.0906171 | 314.4429203 | 0.036923583 | 0.127957713 | 0.359886719 | 0.763371684 | ns | NaCl | KS.3 |
| FIM1_2931 | 4281.108115 | 5262.768383 | 3084.728023 | 0.104824786 | 0.104653846 | 0.052007203 | 0.356877014 | ns | NaCl | KS.3 |
| FIM1_2932 | 860.9651823 | 849.2175448 | 921.3139034 | -0.005518764 | 0.084868623 | 0.90856063 | 0.980825899 | ns | NaCl | KS.3 |
| FIM1_2933 | 2494.263432 | 2647.35516 | 2741.375958 | 0.042755176 | 0.069811846 | 0.38430908 | 0.778665419 | ns | NaCl | KS.3 |
| FIM1_2934 | 5026.754932 | 6035.050383 | 3436.724809 | 0.070001637 | 0.104409798 | 0.156667879 | 0.57585691 | ns | NaCl | KS.3 |
| FIM1_2935 | 22209.28864 | 20606.09906 | 14252.74012 | -0.030976968 | 0.090010453 | 0.515420602 | 0.849359886 | ns | NaCl | KS.3 |
| FIM1_2936 | 6141.349645 | 7794.054858 | 4841.660457 | 0.096222007 | 0.11827144 | 0.068847006 | 0.412170702 | ns | NaCl | KS.3 |
| FIM1_2937 | 958.0197689 | 1093.90917 | 929.7319652 | 0.043574582 | 0.100639834 | 0.346547292 | 0.755840763 | ns | NaCl | KS.3 |
| FIM1_2939 | 287.0874287 | 242.1023475 | 579.7654092 | -0.061282311 | 0.104571231 | 0.202938853 | 0.626494461 | ns | NaCl | KS.3 |
| FIM1_2941 | 1797.401372 | 1864.561013 | 1537.092147 | 0.018429497 | 0.082644368 | 0.704987228 | 0.924674422 | ns | NaCl | KS.3 |
| FIM1_2942 | 5962.613495 | 5900.617743 | 7572.433439 | -0.005506467 | 0.080670032 | 0.910160056 | 0.981715313 | ns | NaCl | KS.3 |
| FIM1_2943 | 1096.843552 | 1086.757914 | 1210.279347 | -0.006322378 | 0.074854931 | 0.897697791 | 0.977971161 | ns | NaCl | KS.3 |
| FIM1_2944 | 1054.128231 | 1255.181467 | 950.1685742 | 0.084956414 | 0.097102104 | 0.10042133 | 0.483636126 | ns | NaCl | KS.3 |
| FIM1_2945 | 969.787859 | 998.3744877 | 787.5940644 | 0.009378593 | 0.094996109 | 0.83507322 | 0.962365393 | ns | NaCl | KS.3 |
| FIM1_2946 | 1494.613021 | 1352.392127 | 1978.107633 | -0.070051867 | 0.074970667 | 0.164551878 | 0.589224032 | ns | NaCl | KS.3 |
| FIM1_2947 | 4656.528597 | 4625.029305 | 3679.15989 | -0.003530194 | 0.078807572 | 0.9425648 | 0.986790704 | ns | NaCl | KS.3 |
| FIM1_2948 | 916.7439305 | 766.6323387 | 1174.689626 | -0.106556278 | 0.094673529 | 0.047382924 | 0.342900262 | ns | NaCl | KS.3 |
| FIM1_2949 | 2887.868839 | 2689.671263 | 3327.628688 | -0.034108031 | 0.085315814 | 0.48416672 | 0.829184191 | ns | NaCl | KS.3 |
| FIM1_2950 | 57.53816194 | 59.42693052 | 91.11159101 | 0.004369925 | 0.107211648 | 0.913993614 | 0.982311132 | ns | NaCl | KS.3 |
| FIM1_2951 | 3945.305395 | 2549.242366 | 2696.25655 | -0.555576789 | 0.162128837 | 3.47145946103 | 0.000523349 | ns | NaCl | KS.3 |
| FIM1_2952 | 683.7038063 | 645.9346272 | 991.1695776 | -0.029812976 | 0.081771397 | 0.543722002 | 0.857313452 | ns | NaCl | KS.3 |
| FIM1_2953 | 1712.831491 | 1564.272636 | 1696.349832 | -0.031587228 | 0.095443878 | 0.494718361 | 0.836358994 | ns | NaCl | KS.3 |
| FIM1_2954 | 658.3814534 | 602.029796 | 669.8722657 | -0.068519646 | 0.070938158 | 0.168646601 | 0.592944764 | ns | NaCl | KS.3 |
| FIM1_2955 | 1503.455703 | 1516.028938 | 1135.911101 | 0.003375077 | 0.088590048 | 0.94283603 | 0.986790704 | ns | NaCl | KS.3 |
| FIM1_2956 | 406.1086756 | 398.9903103 | 356.0336572 | -0.006765053 | 0.093743108 | 0.881614149 | 0.974110648 | ns | NaCl | KS.3 |
| FIM1_2957 | 5957.767877 | 8143.290462 | 3906.709487 | 0.081637873 | 0.143075321 | 0.100246455 | 0.483636126 | ns | NaCl | KS.3 |
| FIM1_2958 | 2031.790025 | 2015.30215 | 2248.188588 | -0.0044516 | 0.079665196 | 0.927489625 | 0.984238275 | ns | NaCl | KS.3 |
| FIM1_2959 | 7097.254763 | 8561.833101 | 4410.205535 | 0.054703124 | 0.110727177 | 0.237131366 | 0.659616806 | ns | NaCl | KS.3 |
| FIM1_2960 | 4922.295121 | 6652.213728 | 3342.8738 | 0.087331732 | 0.140111498 | 0.08669297 | 0.456808328 | ns | NaCl | KS.3 |
| FIM1_2961 | 6341.904231 | 8159.886029 | 4279.333789 | 0.093884248 | 0.123431117 | 0.073534417 | 0.421631393 | ns | NaCl | KS.3 |
| FIM1_2962 | 238.4655367 | 286.304283 | 401.0509813 | 0.09342853 | 0.098286551 | 0.075683702 | 0.427638847 | ns | NaCl | KS.3 |
| FIM1_2963 | 339.4030902 | 416.400233 | 393.8680425 | 0.131277681 | 0.097620555 | 0.019023208 | 0.208486389 | ns | NaCl | KS.3 |
| FIM1_2964 | 10107.07084 | 10361.07874 | 7234.558437 | 0.012658228 | 0.081919831 | 0.794883197 | 0.955299664 | ns | NaCl | KS.3 |
| FIM1_2965 | 1455.924352 | 1522.938867 | 1402.398664 | 0.025492953 | 0.079086042 | 0.60468319 | 0.883494529 | ns | NaCl | KS.3 |
| FIM1_2966 | 397.3042248 | 382.8834441 | 368.3976288 | -0.013683267 | 0.092439511 | 0.766452497 | 0.94853263 | ns | NaCl | KS.3 |
| FIM1_2967 | 1502.173319 | 1367.092022 | 1546.006294 | -0.074252244 | 0.069012084 | 0.133568538 | 0.541126611 | ns | NaCl | KS.3 |
| FIM1_2968 | 1457.871403 | 1257.819018 | 1890.686952 | -0.108241442 | 0.07954727 | 0.038705447 | 0.311585109 | ns | NaCl | KS.3 |
| FIM1_2969 | 8029.165833 | 8131.234995 | 5685.816099 | 0.005744267 | 0.08506749 | 0.904757039 | 0.979880338 | ns | NaCl | KS.3 |
| FIM1_2970 | 693.7228867 | 670.4729886 | 760.6778271 | -0.021959413 | 0.076513838 | 0.656022061 | 0.90157728 | ns | NaCl | KS.3 |
| FIM1_2971 | 932.3204281 | 1141.225773 | 637.3109641 | 0.052265755 | 0.115569919 | 0.247440987 | 0.669893726 | ns | NaCl | KS.3 |
| FIM1_2972 | 533.7311201 | 556.4873846 | 731.7172261 | 0.017651947 | 0.087608775 | 0.71087034 | 0.925563973 | ns | NaCl | KS.3 |

| | | | | | | | | | | |
|---|---|---|---|---|---|---|---|---|---|---|
| FIM1_2973 | 149.4627786 | 155.7749133 | 901.5278038 | 0.010932919 | 0.100723842 | 0.799409165 | 0.95648403 | ns | NaCl | KS.3 |
| FIM1_2974 | 1808.33281 | 1859.612361 | 1991.706768 | 0.008861539 | 0.095696105 | 0.843125196 | 0.966485679 | ns | NaCl | KS.3 |
| FIM1_2975 | 4111.224313 | 5348.52602 | 2788.527927 | 0.119393253 | 0.127572901 | 0.03513901 | 0.295899532 | ns | NaCl | KS.3 |
| FIM1_2976 | 755.1083619 | 695.9979795 | 629.7327363 | -0.043282805 | 0.082596248 | 0.382010787 | 0.778609403 | ns | NaCl | KS.3 |
| FIM1_2977 | 626.3656022 | 501.4264196 | 839.6745868 | -0.105927114 | 0.111243017 | 0.05082036 | 0.352025696 | ns | NaCl | KS.3 |
| FIM1_2978 | 3513.945393 | 3895.777548 | 2827.370299 | 0.048453858 | 0.088327937 | 0.32373683 | 0.739131368 | ns | NaCl | KS.3 |
| FIM1_2979 | 227.1669443 | 175.6657707 | 219.4068092 | -0.102438107 | 0.125466821 | 0.05745807 | 0.375860543 | ns | NaCl | KS.3 |
| FIM1_2980 | 660.2576608 | 558.886526 | 926.1346837 | -0.074909693 | 0.097768056 | 0.138552216 | 0.548803562 | ns | NaCl | KS.3 |
| FIM1_2981 | 8185.782234 | 10678.49219 | 5131.577162 | 0.0723927 | 0.129038799 | 0.132816529 | 0.539340082 | ns | NaCl | KS.3 |
| FIM1_2982 | 2929.611451 | 3197.774643 | 3129.856232 | 0.056111481 | 0.077054363 | 0.263020941 | 0.688222424 | ns | NaCl | KS.3 |
| FIM1_2983 | 8638.710983 | 10767.55261 | 6118.843466 | 0.089796558 | 0.112387849 | 0.083942299 | 0.450499393 | ns | NaCl | KS.3 |
| FIM1_2984 | 962.1825136 | 1036.504302 | 607.7647513 | 0.01449097 | 0.110213394 | 0.718126618 | 0.930143173 | ns | NaCl | KS.3 |
| FIM1_2985 | 18404.18707 | 18572.90791 | 15548.43375 | 0.001531227 | 0.113176681 | 0.968303535 | 0.99408528 | ns | NaCl | KS.3 |
| FIM1_2986 | 768.9442579 | 830.7817642 | 930.6546089 | 0.052537745 | 0.072392808 | 0.290142376 | 0.709776444 | ns | NaCl | KS.3 |
| FIM1_2987 | 2005.19155 | 2134.859402 | 1963.239033 | 0.04654892 | 0.068124115 | 0.341084988 | 0.750290367 | ns | NaCl | KS.3 |
| FIM1_2988 | 621.2065261 | 863.364384 | 931.4687696 | 0.257063921 | 0.172677311 | 0.001305503 | 0.041708911 | ns | NaCl | KS.3 |
| FIM1_2989 | 1316.387344 | 1576.413261 | 992.0718388 | 0.066039963 | 0.104655295 | 0.176189524 | 0.605047427 | ns | NaCl | KS.3 |
| FIM1_2990 | 11052.62225 | 16386.23909 | 12583.20332 | 0.550412404 | 0.082189135 | 2.2320052447 | 1.1104226092 | ns | NaCl | KS.3 |
| FIM1_2991 | 129.4789276 | 72.36308404 | 416.6918034 | -0.359594911 | 0.330769048 | 0.002157992 | 0.057411816 | ns | NaCl | KS.3 |
| FIM1_2992 | 92.2227402 | 36.21068154 | 85.80505748 | -1.053173534 | 0.377075389 | 1.7914391646 | 0.00193748 | Km<KS.3 | NaCl | KS.3 |
| FIM1_2993 | 946.2717571 | 816.2956373 | 1039.204649 | -0.124978647 | 0.076010727 | 0.016704296 | 0.191483583 | ns | NaCl | KS.3 |
| FIM1_2994 | 828.3215335 | 657.6895568 | 988.7851955 | -0.1459449 | 0.11237301 | 0.01316385 | 0.165678479 | ns | NaCl | KS.3 |
| FIM1_2995 | 135.6132292 | 53.48985483 | 250.7650778 | -1.068822598 | 0.376945608 | 1.6051311531 | 0.001774562 | Km<KS.3 | NaCl | KS.3 |
| FIM1_2996 | 199.0171699 | 111.1723801 | 702.7419252 | -0.131736513 | 0.238273668 | 0.033244763 | 0.292213245 | ns | NaCl | KS.3 |
| FIM1_2997 | 1378.710369 | 1381.210241 | 1624.884602 | 0.001161484 | 0.076215396 | 0.981145547 | 0.996363429 | ns | NaCl | KS.3 |
| FIM1_2998 | 2777.915066 | 2735.641256 | 3567.38224 | -0.008320846 | 0.078540668 | 0.865280749 | 0.970128052 | ns | NaCl | KS.3 |
| FIM1_2999 | 665.8605492 | 608.6887894 | 807.5264055 | -0.038112006 | 0.089865059 | 0.428317962 | 0.804596805 | ns | NaCl | KS.3 |
| FIM1_3000 | 2077.901742 | 2727.152895 | 1759.153242 | 0.15873221 | 0.133230756 | 0.011025188 | 0.149455888 | ns | NaCl | KS.3 |
| FIM1_3001 | 1251.279579 | 1300.224819 | 1124.216239 | 0.012578293 | 0.095056792 | 0.78050263 | 0.951646805 | ns | NaCl | KS.3 |
| FIM1_3002 | 356.5120373 | 278.9897374 | 406.515444 | -0.131798489 | 0.119553054 | 0.022918042 | 0.233641926 | ns | NaCl | KS.3 |
| FIM1_3003 | 1132.316991 | 1309.549317 | 974.3824504 | 0.032854544 | 0.111016859 | 0.438743919 | 0.808725824 | ns | NaCl | KS.3 |
| FIM1_3004 | 5650.358893 | 6338.370355 | 5703.476767 | 0.088644225 | 0.071620897 | 0.0782668 | 0.433122723 | ns | NaCl | KS.3 |
| FIM1_3005 | 3546.144394 | 4407.580836 | 2134.188148 | 0.041131796 | 0.123900921 | 0.325082993 | 0.739131368 | ns | NaCl | KS.3 |
| FIM1_3006 | 6517.860421 | 7971.640004 | 4840.496114 | 0.105369258 | 0.102490206 | 0.050799999 | 0.352025696 | ns | NaCl | KS.3 |
| FIM1_3007 | 4702.301688 | 5338.390985 | 3697.877283 | 0.077016416 | 0.082669581 | 0.132524853 | 0.539053419 | ns | NaCl | KS.3 |
| FIM1_3008 | 2203.411329 | 2031.96233 | 2183.93518 | -0.06525107 | 0.06703051 | 0.182040299 | 0.611396712 | ns | NaCl | KS.3 |
| FIM1_3009 | 1545.549749 | 1296.105721 | 1567.872644 | -0.141523082 | 0.081740123 | 0.008784298 | 0.130065127 | ns | NaCl | KS.3 |
| FIM1_3010 | 636.4411543 | 588.660366 | 653.7230729 | -0.043519069 | 0.081178651 | 0.380592848 | 0.778493059 | ns | NaCl | KS.3 |
| FIM1_3011 | 502.1538206 | 519.562017 | 622.8877379 | 0.01244534 | 0.090842846 | 0.789152329 | 0.953383399 | ns | NaCl | KS.3 |
| FIM1_3012 | 3473.402563 | 3605.522604 | 2505.524487 | 0.026133268 | 0.071323825 | 0.594830588 | 0.87962202 | ns | NaCl | KS.3 |
| FIM1_3013 | 2103.268273 | 2132.534125 | 2299.118373 | 0.009412719 | 0.070102556 | 0.847314237 | 0.967147129 | ns | NaCl | KS.3 |
| FIM1_3014 | 10251.78551 | 13781.15507 | 6063.64154 | 0.065980214 | 0.137640367 | 0.153889124 | 0.571010593 | ns | NaCl | KS.3 |
| FIM1_3015 | 5876.35513 | 6370.582139 | 4634.54022 | 0.047514529 | 0.07938453 | 0.340640109 | 0.750043208 | ns | NaCl | KS.3 |
| FIM1_3016 | 457.4829226 | 430.3068675 | 759.6981876 | -0.026507509 | 0.089150166 | 0.577529569 | 0.874645237 | ns | NaCl | KS.3 |
| FIM1_3017 | 27839.89731 | 34237.55458 | 21330.23418 | 0.140964724 | 0.098203565 | 0.013017457 | 0.165208794 | ns | NaCl | KS.3 |

| | | | | | | | | | | |
|---|---|---|---|---|---|---|---|---|---|---|
| FIM1_3018 | 666.6338083 | 738.239616 | 535.4983651 | 0.036927777 | 0.095357719 | 0.429731839 | 0.805047964 | ns | NaCl | KS.3 |
| FIM1_3019 | 4426.83572 | 4512.291165 | 4633.715476 | 0.011132445 | 0.077112801 | 0.820838266 | 0.960240659 | ns | NaCl | KS.3 |
| FIM1_3020 | 53.38147021 | 102.7341972 | 96.90294568 | 0.551521874 | 0.370462793 | 0.000508636 | 0.023520242 | ns | NaCl | KS.3 |
| FIM1_3021 | 4465.435653 | 5366.44382 | 3834.209133 | 0.131849326 | 0.088744637 | 0.016255399 | 0.18807119 | ns | NaCl | KS.3 |
| FIM1_3022 | 291.9043323 | 315.3699815 | 240.3611277 | 0.012226189 | 0.11664435 | 0.748884516 | 0.943274417 | ns | NaCl | KS.3 |
| FIM1_3023 | 327.9678469 | 182.3275374 | 752.7178282 | -0.430729866 | 0.344564929 | 0.001160875 | 0.040387089 | ns | NaCl | KS.3 |
| FIM1_3024 | 56.84935378 | 35.52215417 | 74.4569741 | -0.049595512 | 0.169886103 | 0.212137063 | 0.635389456 | ns | NaCl | KS.3 |
| FIM1_3025 | 728.847472 | 657.435991 | 931.7120277 | -0.028401333 | 0.102551532 | 0.517954166 | 0.850188733 | ns | NaCl | KS.3 |
| FIM1_3026 | 2714.462913 | 2608.731556 | 2497.866158 | -0.014021542 | 0.093217715 | 0.759616574 | 0.945718832 | ns | NaCl | KS.3 |
| FIM1_3028 | 4484.931234 | 4629.959912 | 4185.317805 | 0.016749419 | 0.080726194 | 0.732053278 | 0.936960396 | ns | NaCl | KS.3 |
| FIM1_3029 | 1373.762088 | 1123.421778 | 1373.966389 | -0.124332681 | 0.098889581 | 0.025212849 | 0.248876828 | ns | NaCl | KS.3 |
| FIM1_3030 | 2931.476578 | 3077.741171 | 3644.827954 | 0.041392925 | 0.060556302 | 0.3763641 | 0.778493059 | ns | NaCl | KS.3 |
| FIM1_3031 | 1164.521504 | 919.3478229 | 1581.656361 | -0.149767753 | 0.113780282 | 0.011680365 | 0.152920569 | ns | NaCl | KS.3 |
| FIM1_3032 | 3539.500947 | 3308.501584 | 2951.006234 | -0.026886302 | 0.090696634 | 0.569046637 | 0.869550295 | ns | NaCl | KS.3 |
| FIM1_3033 | 1189.246707 | 1091.099975 | 1273.167036 | -0.045589509 | 0.083332846 | 0.357505992 | 0.761817563 | ns | NaCl | KS.3 |
| FIM1_3034 | 1152.447807 | 1066.462984 | 2931.69041 | -0.013863261 | 0.112803239 | 0.724631046 | 0.933222742 | ns | NaCl | KS.3 |
| FIM1_3035 | 13859.71356 | 13813.99257 | 11142.20316 | -0.001302029 | 0.088872202 | 0.977887519 | 0.996164232 | ns | NaCl | KS.3 |
| FIM1_3036 | 3092.60066 | 3267.228356 | 3404.866561 | 0.026304744 | 0.084886942 | 0.58765478 | 0.878311243 | ns | NaCl | KS.3 |
| FIM1_3037 | 869.1958195 | 775.9684032 | 1021.631874 | -0.017350303 | 0.119224742 | 0.649615622 | 0.899642849 | ns | NaCl | KS.3 |
| FIM1_3038 | 25018.53975 | 28260.61478 | 17148.25544 | 0.060833069 | 0.088216064 | 0.223807838 | 0.648059715 | ns | NaCl | KS.3 |
| FIM1_3039 | 4021.465036 | 4456.69725 | 2869.090948 | 0.054898879 | 0.084323395 | 0.271670728 | 0.693144962 | ns | NaCl | KS.3 |
| FIM1_3040 | 4726.138274 | 5723.816588 | 2794.51789 | 0.032188175 | 0.123142772 | 0.421282646 | 0.801484192 | ns | NaCl | KS.3 |
| FIM1_3041 | 1525.135288 | 1576.903852 | 1740.792045 | 0.021705774 | 0.072540706 | 0.65906997 | 0.903691294 | ns | NaCl | KS.3 |
| FIM1_3043 | 158.8706105 | 202.5011788 | 207.6025605 | 0.073537901 | 0.122629374 | 0.131308848 | 0.538132312 | ns | NaCl | KS.3 |
| FIM1_3044 | 743.7675863 | 439.4196726 | 4677.982151 | -0.102784816 | 0.207693901 | 0.057416109 | 0.375860543 | ns | NaCl | KS.3 |
| FIM1_3045 | 386.8390077 | 286.4550669 | 549.2223036 | -0.110117225 | 0.14274291 | 0.047420177 | 0.342900262 | ns | NaCl | KS.3 |
| FIM1_3046 | 1199.755989 | 1095.566655 | 1642.342519 | -0.03461948 | 0.093100965 | 0.46253522 | 0.819880744 | ns | NaCl | KS.3 |
| FIM1_3047 | 799.8017698 | 825.021898 | 825.3510037 | 0.020189683 | 0.073857733 | 0.681874864 | 0.915044369 | ns | NaCl | KS.3 |
| FIM1_3048 | 4815.521462 | 5420.720568 | 3099.753428 | 0.04537814 | 0.095193208 | 0.340723146 | 0.750043208 | ns | NaCl | KS.3 |
| FIM1_3049 | 760.6013478 | 748.3075839 | 883.089918 | -0.008517472 | 0.081274802 | 0.861269811 | 0.969856868 | ns | NaCl | KS.3 |
| FIM1_3050 | 955.5651169 | 909.2320155 | 792.229839 | -0.028995901 | 0.076642332 | 0.557373687 | 0.863845647 | ns | NaCl | KS.3 |
| FIM1_3051 | 631.5665356 | 425.5953755 | 1014.141862 | -0.398001774 | 0.208457047 | 0.000193787 | 0.012853871 | ns | NaCl | KS.3 |
| FIM1_3052 | 753.1044341 | 669.4136309 | 796.9924135 | -0.055134375 | 0.089630376 | 0.264600729 | 0.688347233 | ns | NaCl | KS.3 |
| FIM1_3053 | 2120.136333 | 1829.156104 | 2143.005932 | -0.069613624 | 0.09334474 | 0.1666234 | 0.590711667 | ns | NaCl | KS.3 |
| FIM1_3054 | 1221.846728 | 1434.985686 | 1456.614948 | 0.030302619 | 0.117160428 | 0.456788201 | 0.817337042 | ns | NaCl | KS.3 |
| FIM1_3056 | 4743.929662 | 5445.57215 | 4038.32144 | 0.033516338 | 0.108520955 | 0.43672805 | 0.808096352 | ns | NaCl | KS.3 |
| FIM1_3057 | 439.3602902 | 401.1525643 | 483.5889983 | -0.020285768 | 0.107771844 | 0.624744945 | 0.894161709 | ns | NaCl | KS.3 |
| FIM1_3058 | 2215.53862 | 2195.510661 | 2914.185425 | -0.004664522 | 0.082814209 | 0.923317717 | 0.983924964 | ns | NaCl | KS.3 |
| FIM1_3059 | 530.3721627 | 526.2521716 | 849.149499 | -0.003545824 | 0.077965241 | 0.942389671 | 0.986790704 | ns | NaCl | KS.3 |
| FIM1_3060 | 614.3924083 | 623.9418809 | 717.4541639 | 0.00792605 | 0.081882766 | 0.870559348 | 0.971243188 | ns | NaCl | KS.3 |
| FIM1_3061 | 3209.302376 | 3138.560212 | 3894.558609 | -0.014163598 | 0.073504894 | 0.77337894 | 0.949545959 | ns | NaCl | KS.3 |
| FIM1_3062 | 1963.997902 | 2062.491621 | 1637.020043 | 0.01626403 | 0.094987632 | 0.719746647 | 0.930667987 | ns | NaCl | KS.3 |
| FIM1_3063 | 635.3307587 | 604.0505839 | 1105.968508 | -0.03611814 | 0.072804744 | 0.464753168 | 0.820492197 | ns | NaCl | KS.3 |
| FIM1_3064 | 7758.296231 | 9490.643204 | 6172.113327 | 0.154994262 | 0.092279518 | 0.006246989 | 0.106434152 | ns | NaCl | KS.3 |
| FIM1_3065 | 2805.206108 | 2778.823196 | 2380.210882 | -0.005971549 | 0.072688619 | 0.903208842 | 0.979395371 | ns | NaCl | KS.3 |

| | | | | | | | | | | |
|---|---|---|---|---|---|---|---|---|---|---|
| FIM1_3066 | 5141.415096 | 5385.73748 | 4964.727518 | 0.030598524 | 0.073503493 | 0.535407598 | 0.855153333 | ns | NaCl | KS.3 |
| FIM1_3067 | 846.5268176 | 834.9282466 | 690.525503 | -0.006932101 | 0.08275331 | 0.88631468 | 0.97621397 | ns | NaCl | KS.3 |
| FIM1_3068 | 5452.159343 | 5849.857954 | 5347.153281 | 0.021608546 | 0.097820031 | 0.628985537 | 0.896016329 | ns | NaCl | KS.3 |
| FIM1_3069 | 1481.947693 | 1574.335167 | 1141.648876 | 0.021522279 | 0.093508437 | 0.640571369 | 0.898798052 | ns | NaCl | KS.3 |
| FIM1_3070 | 10333.88061 | 10819.55207 | 7452.029009 | 0.029611652 | 0.074197628 | 0.548817868 | 0.860500754 | ns | NaCl | KS.3 |
| FIM1_3071 | 2956.973726 | 2846.766469 | 3315.501244 | -0.019182368 | 0.082264588 | 0.693984455 | 0.920525145 | ns | NaCl | KS.3 |
| FIM1_3072 | 1037.620361 | 837.6446698 | 1546.247319 | -0.080948437 | 0.111771377 | 0.109683736 | 0.505115899 | ns | NaCl | KS.3 |
| FIM1_3074 | 835.663512 | 858.1998531 | 871.4180666 | 0.015842073 | 0.077975018 | 0.747252599 | 0.943274417 | ns | NaCl | KS.3 |
| FIM1_3075 | 1523.794001 | 2058.047342 | 1321.472421 | 0.164833652 | 0.149553762 | 0.011015709 | 0.149455888 | ns | NaCl | KS.3 |
| FIM1_3076 | 998.1869317 | 979.9463733 | 1017.158479 | -0.013320223 | 0.071745407 | 0.786044091 | 0.95289761 | ns | NaCl | KS.3 |
| FIM1_3077 | 2238.239618 | 2569.689 | 2215.349619 | 0.075473766 | 0.087365124 | 0.139852007 | 0.550445992 | ns | NaCl | KS.3 |
| FIM1_3078 | 858.6081274 | 864.5031719 | 790.6237241 | 0.005261971 | 0.073956805 | 0.914752731 | 0.982343702 | ns | NaCl | KS.3 |
| FIM1_3079 | 2636.909917 | 3152.850084 | 2045.88673 | 0.083149197 | 0.099320485 | 0.106032895 | 0.49734665 | ns | NaCl | KS.3 |
| FIM1_3080 | 1463.877159 | 1757.183627 | 1517.297272 | 0.125818234 | 0.089680388 | 0.021332578 | 0.222905493 | ns | NaCl | KS.3 |
| FIM1_3081 | 1316.576541 | 1567.646793 | 1776.031229 | 0.079540069 | 0.098313281 | 0.119373045 | 0.523704495 | ns | NaCl | KS.3 |
| FIM1_3082 | 1620.939066 | 1595.872923 | 1874.539366 | -0.012031744 | 0.06929574 | 0.805217265 | 0.957859861 | ns | NaCl | KS.3 |
| FIM1_3083 | 8744.57135 | 11902.47036 | 5074.020734 | 0.068700947 | 0.140950863 | 0.141640357 | 0.553543421 | ns | NaCl | KS.3 |
| FIM1_3085 | 40.35408437 | 41.09548804 | 43.4414597 | 0.002816786 | 0.116059364 | 0.94053964 | 0.986790704 | ns | NaCl | KS.3 |
| FIM1_3086 | 1206.513531 | 1310.991081 | 1403.542008 | 0.026597447 | 0.097790869 | 0.55576505 | 0.863699344 | ns | NaCl | KS.3 |
| FIM1_3087 | 11182.99476 | 13267.25395 | 7192.444108 | 0.053653489 | 0.106553573 | 0.250478473 | 0.673948297 | ns | NaCl | KS.3 |
| FIM1_3088 | 242.8227115 | 190.2043194 | 879.435673 | -0.057367129 | 0.124911492 | 0.204655957 | 0.627757602 | ns | NaCl | KS.3 |
| FIM1_3089 | 467.2208157 | 454.3821851 | 526.3408947 | -0.008477849 | 0.096715424 | 0.848552215 | 0.967357303 | ns | NaCl | KS.3 |
| FIM1_3090 | 440.772212 | 538.5562627 | 396.6058563 | 0.106993864 | 0.102879359 | 0.048115169 | 0.344355024 | ns | NaCl | KS.3 |
| FIM1_3092 | 443.876254 | 494.0643258 | 500.4249974 | 0.039003068 | 0.09735502 | 0.401501581 | 0.789189152 | ns | NaCl | KS.3 |
| FIM1_3098 | 1462.84329 | 949.5685868 | 1200.264754 | -0.28012718 | 0.243388157 | 0.002728026 | 0.064671708 | ns | NaCl | KS.3 |
| FIM1_3099 | 2401.172941 | 2425.961367 | 2350.021231 | 0.001864704 | 0.111561511 | 0.961880681 | 0.993224655 | ns | NaCl | KS.3 |
| FIM1_3100 | 880.5264109 | 813.4946669 | 982.5762384 | -0.033785189 | 0.088949409 | 0.481845179 | 0.828664253 | ns | NaCl | KS.3 |
| FIM1_3101 | 1028.949186 | 1001.381218 | 1052.947167 | -0.011670562 | 0.086094714 | 0.807295449 | 0.957859861 | ns | NaCl | KS.3 |
| FIM1_3102 | 23132.58099 | 27125.55894 | 16236.58602 | 0.081559639 | 0.092884215 | 0.113171765 | 0.511380136 | ns | NaCl | KS.3 |
| FIM1_3103 | 220.0245645 | 196.0170419 | 162.3837864 | -0.017587707 | 0.119037639 | 0.645782069 | 0.898899754 | ns | NaCl | KS.3 |
| FIM1_3104 | 1687.523497 | 1618.564226 | 1971.434572 | -0.022972548 | 0.080441277 | 0.63967425 | 0.898798052 | ns | NaCl | KS.3 |
| FIM1_3105 | 20.63130907 | 19.66176517 | 14.7225111 | -0.005843156 | 0.131152406 | 0.865999427 | 0.970184124 | ns | NaCl | KS.3 |
| FIM1_3106 | 1307.182752 | 1184.536792 | 1352.376741 | -0.058203309 | 0.080771236 | 0.246859091 | 0.669893726 | ns | NaCl | KS.3 |
| FIM1_3107 | 586.9227939 | 381.9457707 | 1220.887682 | -0.060843761 | 0.167145605 | 0.160224382 | 0.58211082 | ns | NaCl | KS.3 |
| FIM1_3108 | 2122.819782 | 1400.142994 | 2240.122344 | -0.460001448 | 0.202636033 | 6.76448148232 | 0.005427951 | ns | NaCl | KS.3 |
| FIM1_3109 | 4910.225394 | 6596.042126 | 3375.384685 | 0.113080862 | 0.14065966 | 0.043618494 | 0.329289841 | ns | NaCl | KS.3 |
| FIM1_3110 | 1429.853673 | 3286.322057 | 4126.047925 | 0.179855113 | 0.332772747 | 0.015986229 | 0.185388092 | ns | NaCl | KS.3 |
| FIM1_3111 | 646.4735623 | 773.615459 | 1294.786463 | 0.032755291 | 0.120146325 | 0.420223335 | 0.80133378 | ns | NaCl | KS.3 |
| FIM1_3112 | 539.4311699 | 633.0790113 | 703.4952814 | 0.052362675 | 0.104248889 | 0.263676954 | 0.688243886 | ns | NaCl | KS.3 |
| FIM1_3113 | 923.5158752 | 974.4029306 | 1621.420472 | 0.018063883 | 0.094063049 | 0.692828516 | 0.920513182 | ns | NaCl | KS.3 |
| FIM1_3114 | 1728.031885 | 1654.968857 | 1719.038682 | -0.01567841 | 0.091011274 | 0.736503765 | 0.93879227 | ns | NaCl | KS.3 |
| FIM1_3115 | 203.0742712 | 129.4018568 | 260.7413305 | -0.346357695 | 0.261641671 | 0.00125022 | 0.041599773 | ns | NaCl | KS.3 |
| FIM1_3116 | 1423.381897 | 1273.036667 | 1724.004441 | -0.031780544 | 0.102072384 | 0.474769755 | 0.823563295 | ns | NaCl | KS.3 |
| FIM1_3117 | 16.555383 | 15.03846269 | 19.37440892 | -0.0090126 | 0.127171181 | 0.800173788 | 0.956938604 | ns | NaCl | KS.3 |
| FIM1_3118 | 2452.862884 | 3076.291445 | 4526.624002 | 0.086977884 | 0.114738713 | 0.090997703 | 0.464798331 | ns | NaCl | KS.3 |

| | | | | | | | | | | |
|---|---|---|---|---|---|---|---|---|---|---|
| FIM1_3119 | 927.7533326 | 807.53381 | 946.5580616 | -0.046370947 | 0.100751813 | 0.320017332 | 0.736735876 | ns | NaCl | KS.3 |
| FIM1_3120 | 2770.539169 | 3016.332695 | 2859.777166 | 0.06054969 | 0.072318255 | 0.224284286 | 0.648059715 | ns | NaCl | KS.3 |
| FIM1_3121 | 1016.026906 | 977.8807719 | 1170.818666 | -0.025470061 | 0.07566356 | 0.605859739 | 0.883689315 | ns | NaCl | KS.3 |
| FIM1_3122 | 1622.743607 | 1338.583577 | 1970.926694 | -0.085560925 | 0.103234951 | 0.097320799 | 0.476631548 | ns | NaCl | KS.3 |
| FIM1_3123 | 1827.294932 | 1568.972834 | 2384.973826 | -0.100418619 | 0.08399154 | 0.05603604 | 0.370224834 | ns | NaCl | KS.3 |
| FIM1_3124 | 5506.448017 | 4941.246884 | 4368.791076 | -0.116178583 | 0.051190973 | 0.006545304 | 0.108271704 | ns | NaCl | KS.3 |
| FIM1_3125 | 529.5636219 | 536.6354641 | 648.7975289 | 0.004813691 | 0.083988061 | 0.920504041 | 0.983890455 | ns | NaCl | KS.3 |
| FIM1_3126 | 1867.64395 | 1909.085925 | 2225.153781 | 0.006801113 | 0.095928088 | 0.878901972 | 0.973188807 | ns | NaCl | KS.3 |
| FIM1_3127 | 1240.27451 | 1374.665862 | 1641.556946 | 0.032605626 | 0.098983076 | 0.472567686 | 0.822130792 | ns | NaCl | KS.3 |
| FIM1_3128 | 12125.36695 | 12154.85836 | 9997.73968 | 0.001898406 | 0.065918288 | 0.968562425 | 0.99408528 | ns | NaCl | KS.3 |
| FIM1_3129 | 4994.232593 | 5201.204359 | 3927.025455 | 0.030600805 | 0.067169831 | 0.528456839 | 0.855153333 | ns | NaCl | KS.3 |
| FIM1_3130 | 1469.59272 | 1631.089361 | 1143.616367 | 0.036621111 | 0.096390748 | 0.430761343 | 0.805047964 | ns | NaCl | KS.3 |
| FIM1_3131 | 1087.360621 | 842.0380446 | 1039.568712 | -0.14312921 | 0.123689321 | 0.016412756 | 0.189229716 | ns | NaCl | KS.3 |
| FIM1_3132 | 4559.728382 | 4760.379874 | 3367.068562 | 0.032985195 | 0.067309809 | 0.497254669 | 0.836605336 | ns | NaCl | KS.3 |
| FIM1_3133 | 555.2286979 | 574.4383875 | 594.2050343 | 0.012144279 | 0.092290717 | 0.792063876 | 0.954759779 | ns | NaCl | KS.3 |
| FIM1_3134 | 13297.91428 | 15509.1451 | 11415.25866 | 0.049697025 | 0.103493348 | 0.286021808 | 0.705523209 | ns | NaCl | KS.3 |
| FIM1_3135 | 1150.255536 | 1201.405634 | 1117.0395 | 0.026303316 | 0.077011673 | 0.594120378 | 0.87962202 | ns | NaCl | KS.3 |
| FIM1_3136 | 7266.45126 | 9137.700045 | 4630.98272 | 0.09050324 | 0.115465727 | 0.081827251 | 0.442009309 | ns | NaCl | KS.3 |
| FIM1_3137 | 22.211251 | 19.56857312 | 24.44135902 | -0.009143013 | 0.135630417 | 0.788682129 | 0.953379471 | ns | NaCl | KS.3 |
| FIM1_3138 | 9766.529958 | 13275.83562 | 6167.117837 | 0.077917561 | 0.14109278 | 0.110951567 | 0.506448597 | ns | NaCl | KS.3 |
| FIM1_3139 | 597.1454009 | 1207.24758 | 667.88201 | 0.902037236 | 0.207868694 | 4.50155533255 | 8.95809511179 | ns | NaCl | KS.3 |
| FIM1_3140 | 463.0622507 | 394.156007 | 459.0135077 | -0.076612032 | 0.095086379 | 0.132424159 | 0.539053419 | ns | NaCl | KS.3 |
| FIM1_3141 | 1177.520772 | 1315.325579 | 972.2596684 | 0.053905653 | 0.087874432 | 0.276560357 | 0.697711854 | ns | NaCl | KS.3 |
| FIM1_3142 | 171.0528435 | 141.9393555 | 264.7518978 | -0.050054071 | 0.112075468 | 0.268706124 | 0.690146084 | ns | NaCl | KS.3 |
| FIM1_3143 | 7286.477488 | 8858.993912 | 5562.129101 | 0.177255761 | 0.082244214 | 0.001320402 | 0.04184076 | ns | NaCl | KS.3 |
| FIM1_3144 | 290.8281222 | 265.9372978 | 358.3171778 | -0.030743019 | 0.097502052 | 0.50030922 | 0.839756535 | ns | NaCl | KS.3 |
| FIM1_3145 | 1445.777054 | 1388.286357 | 1300.969336 | -0.022617368 | 0.079756382 | 0.645263459 | 0.898899754 | ns | NaCl | KS.3 |
| FIM1_3146 | 618.6072356 | 598.7443199 | 695.8146255 | -0.019530662 | 0.079079391 | 0.690992201 | 0.919659229 | ns | NaCl | KS.3 |
| FIM1_3147 | 576.4210668 | 533.5557336 | 1134.359183 | -0.034915319 | 0.087026853 | 0.471474482 | 0.822127061 | ns | NaCl | KS.3 |
| FIM1_3148 | 487.9432612 | 491.5167393 | 864.5807955 | 0.002968165 | 0.080160668 | 0.951553976 | 0.987831315 | ns | NaCl | KS.3 |
| FIM1_3149 | 930.0080981 | 921.9237586 | 1048.570728 | -0.004173692 | 0.083854673 | 0.931078311 | 0.984325045 | ns | NaCl | KS.3 |
| FIM1_3150 | 8650.144856 | 11642.69615 | 5060.087817 | 0.071854695 | 0.137991087 | 0.131315101 | 0.538132312 | ns | NaCl | KS.3 |
| FIM1_3151 | 937.3315677 | 1097.031737 | 787.651183 | 0.069499236 | 0.095932663 | 0.16544084 | 0.589355333 | ns | NaCl | KS.3 |
| FIM1_3152 | 240.5145152 | 248.1675282 | 255.1933084 | 0.009933399 | 0.089975904 | 0.831713732 | 0.961620266 | ns | NaCl | KS.3 |
| FIM1_3153 | 588.2887347 | 541.5902398 | 769.2644039 | -0.045324411 | 0.081912177 | 0.361354837 | 0.763371684 | ns | NaCl | KS.3 |
| FIM1_3154 | 2636.493307 | 3148.675895 | 2146.446937 | 0.060532489 | 0.105855602 | 0.205960002 | 0.630415219 | ns | NaCl | KS.3 |
| FIM1_3155 | 2463.613642 | 2544.851463 | 2866.78671 | 0.016078633 | 0.083275272 | 0.740371604 | 0.940888181 | ns | NaCl | KS.3 |
| FIM1_3156 | 4873.694156 | 4265.136568 | 3973.487865 | -0.07544373 | 0.085763554 | 0.140156501 | 0.550917434 | ns | NaCl | KS.3 |
| FIM1_3157 | 551.2359385 | 396.6241583 | 1292.04226 | -0.084024393 | 0.147250968 | 0.093612053 | 0.470201108 | ns | NaCl | KS.3 |
| FIM1_3158 | 270.6940562 | 261.8361679 | 368.4361109 | -0.013247976 | 0.088837716 | 0.778780286 | 0.951027591 | ns | NaCl | KS.3 |
| FIM1_3159 | 424.0669732 | 318.159276 | 404.0752316 | -0.176626257 | 0.147898855 | 0.007697139 | 0.120041586 | ns | NaCl | KS.3 |
| FIM1_3160 | 176.3805082 | 165.6786125 | 291.6823713 | -0.010012502 | 0.115177199 | 0.794143853 | 0.955299664 | ns | NaCl | KS.3 |
| FIM1_3161 | 553.8652015 | 1155.744078 | 1401.267782 | 0.615474896 | 0.415201629 | 0.000444763 | 0.021482506 | ns | NaCl | KS.3 |
| FIM1_3162 | 2832.172865 | 1565.394933 | 2181.720415 | -0.297344317 | 0.326546336 | 0.004232694 | 0.084568893 | ns | NaCl | KS.3 |
| FIM1_3163 | 773.0210963 | 669.6196291 | 715.8975727 | -0.046785642 | 0.102103802 | 0.313555295 | 0.731871738 | ns | NaCl | KS.3 |

| | | | | | | | | | | |
|---|---|---|---|---|---|---|---|---|---|---|
| FIM1_3164 | 2918.587345 | 2973.914803 | 2822.476065 | 0.01938979 | 0.04896986 | 0.640324371 | 0.898798052 | ns | NaCl | KS.3 |
| FIM1_3165 | 987.2973083 | 1039.968491 | 793.2031306 | 0.0224194 | 0.087552349 | 0.639052255 | 0.898798052 | ns | NaCl | KS.3 |
| FIM1_3166 | 1236.38483 | 1192.778566 | 1168.677337 | -0.017066996 | 0.084438694 | 0.723960061 | 0.93308324 | ns | NaCl | KS.3 |
| FIM1_3167 | 5671.791001 | 7617.489224 | 3498.114776 | 0.057612172 | 0.137829211 | 0.192940937 | 0.618952432 | ns | NaCl | KS.3 |
| FIM1_3168 | 968.4511597 | 1048.028884 | 1110.963659 | 0.048078573 | 0.078192417 | 0.335264029 | 0.745307602 | ns | NaCl | KS.3 |
| FIM1_3169 | 1117.363147 | 1193.53817 | 1077.659817 | 0.027312458 | 0.08948423 | 0.565687493 | 0.868073805 | ns | NaCl | KS.3 |
| FIM1_3170 | 63.26163665 | 48.26746516 | 121.4952025 | -0.055495533 | 0.133064194 | 0.208304384 | 0.633002483 | ns | NaCl | KS.3 |
| FIM1_3171 | 2186.047164 | 2415.202571 | 1716.990805 | 0.046999371 | 0.087912031 | 0.338096335 | 0.747703619 | ns | NaCl | KS.3 |
| FIM1_3172 | 1604.79003 | 1457.14631 | 1513.119302 | -0.028977285 | 0.099638332 | 0.51801661 | 0.850188733 | ns | NaCl | KS.3 |
| FIM1_3173 | 1266.342367 | 1201.197135 | 1674.368048 | -0.024064047 | 0.08574598 | 0.618066784 | 0.891787196 | ns | NaCl | KS.3 |
| FIM1_3174 | 517.7678325 | 640.7276847 | 492.5185698 | 0.064221443 | 0.115102835 | 0.176377947 | 0.605047427 | ns | NaCl | KS.3 |
| FIM1_3175 | 1371.851603 | 1353.230454 | 1395.309944 | -0.007196258 | 0.08211964 | 0.882279916 | 0.974110648 | ns | NaCl | KS.3 |
| FIM1_3176 | 302.7836003 | 346.8043475 | 464.5748983 | 0.06369568 | 0.091408263 | 0.202206566 | 0.625219184 | ns | NaCl | KS.3 |
| FIM1_3177 | 2303.995007 | 2488.194075 | 2248.049981 | 0.065869241 | 0.062330208 | 0.165911633 | 0.590000266 | ns | NaCl | KS.3 |
| FIM1_3178 | 2331.256602 | 2353.474584 | 2050.784747 | 0.004393551 | 0.086574138 | 0.926506496 | 0.984176127 | ns | NaCl | KS.3 |
| FIM1_3179 | 255.7885896 | 208.8181306 | 479.2921944 | -0.049121617 | 0.116888926 | 0.268608294 | 0.690146084 | ns | NaCl | KS.3 |
| FIM1_3180 | 1198.149283 | 1011.575916 | 1588.096631 | -0.084407303 | 0.095304582 | 0.10255071 | 0.488687531 | ns | NaCl | KS.3 |
| FIM1_3181 | 4949.233566 | 6327.009244 | 3024.698306 | 0.053306681 | 0.126430922 | 0.227732036 | 0.651504817 | ns | NaCl | KS.3 |
| FIM1_3182 | 2379.07872 | 2487.994764 | 1881.758192 | 0.024324039 | 0.080390502 | 0.620360524 | 0.892479944 | ns | NaCl | KS.3 |
| FIM1_3183 | 7813.636905 | 8197.521672 | 6908.476124 | 0.021422164 | 0.086716925 | 0.655114268 | 0.901110473 | ns | NaCl | KS.3 |
| FIM1_3184 | 815.4698116 | 861.0268533 | 690.2581567 | 0.014567004 | 0.10076554 | 0.736082487 | 0.938734671 | ns | NaCl | KS.3 |
| FIM1_3185 | 1290.267407 | 1769.082934 | 1068.012224 | 0.110176701 | 0.14830456 | 0.047772603 | 0.343442 | ns | NaCl | KS.3 |
| FIM1_3186 | 2940.606564 | 3089.97114 | 3548.73503 | 0.026766799 | 0.080611363 | 0.585971478 | 0.877826895 | ns | NaCl | KS.3 |
| FIM1_3187 | 698.7754477 | 713.8378472 | 663.5778738 | 0.008735308 | 0.087341974 | 0.854028553 | 0.969345976 | ns | NaCl | KS.3 |
| FIM1_3188 | 642.9576278 | 684.6630943 | 690.8064976 | 0.030636097 | 0.08370565 | 0.530704769 | 0.855153333 | ns | NaCl | KS.3 |
| FIM1_3189 | 408.5351951 | 406.1844028 | 411.7370225 | -0.002853931 | 0.094084024 | 0.949707785 | 0.987736646 | ns | NaCl | KS.3 |
| FIM1_3190 | 590.8603285 | 612.5056127 | 680.2043265 | 0.016063831 | 0.085967942 | 0.737734641 | 0.939315169 | ns | NaCl | KS.3 |
| FIM1_3191 | 1014.798682 | 1069.428381 | 883.2861558 | 0.018137106 | 0.093948792 | 0.691909535 | 0.91988055 | ns | NaCl | KS.3 |
| FIM1_3192 | 738.5816339 | 645.3024217 | 729.3971384 | -0.08111531 | 0.085250646 | 0.115239242 | 0.51696594 | ns | NaCl | KS.3 |
| FIM1_3193 | 60.11817914 | 74.27496584 | 43.75386283 | 0.038346665 | 0.120320528 | 0.358126151 | 0.76205201 | ns | NaCl | KS.3 |
| FIM1_3194 | 2523.631423 | 2416.839917 | 2395.124519 | -0.012934326 | 0.097358131 | 0.77049831 | 0.949065335 | ns | NaCl | KS.3 |
| FIM1_3195 | 5162.873263 | 6327.53578 | 3200.949999 | 0.053099626 | 0.1155931 | 0.241618024 | 0.666264473 | ns | NaCl | KS.3 |
| FIM1_3196 | 501.142475 | 448.6168818 | 477.2816158 | -0.044230167 | 0.094223142 | 0.353916611 | 0.757528931 | ns | NaCl | KS.3 |
| FIM1_3197 | 525.5422509 | 773.0664405 | 1536.617768 | 0.08893802 | 0.166107803 | 0.081013956 | 0.439771952 | ns | NaCl | KS.3 |
| FIM1_3198 | 265.9585021 | 404.7688263 | 460.0031329 | 0.057880451 | 0.164591054 | 0.174379594 | 0.603295189 | ns | NaCl | KS.3 |
| FIM1_3199 | 2572.961738 | 2651.499155 | 2359.023598 | 0.01400874 | 0.085081746 | 0.770977804 | 0.949083579 | ns | NaCl | KS.3 |
| FIM1_3200 | 12978.99906 | 13448.94245 | 9302.489641 | 0.036544106 | 0.049548928 | 0.382206677 | 0.778656108 | ns | NaCl | KS.3 |
| FIM1_3201 | 486.5267238 | 565.1524111 | 501.8279529 | 0.059802517 | 0.097162097 | 0.22119633 | 0.643915588 | ns | NaCl | KS.3 |
| FIM1_3202 | 385.8802212 | 416.1267668 | 534.8867205 | 0.033811281 | 0.084836101 | 0.488558046 | 0.832674298 | ns | NaCl | KS.3 |
| FIM1_3203 | 1116.772477 | 1106.799283 | 1449.933299 | -0.004565306 | 0.082964925 | 0.924900582 | 0.984176127 | ns | NaCl | KS.3 |
| FIM1_3204 | 813.0245286 | 715.4408006 | 744.7715961 | -0.109024676 | 0.065881851 | 0.026512514 | 0.254519471 | ns | NaCl | KS.3 |
| FIM1_3205 | 5643.404111 | 5315.262916 | 4777.253445 | -0.039854122 | 0.073371431 | 0.420893727 | 0.80133378 | ns | NaCl | KS.3 |
| FIM1_3206 | 2065.221903 | 1879.646751 | 1815.968829 | -0.06332233 | 0.075875093 | 0.207938133 | 0.633002483 | ns | NaCl | KS.3 |
| FIM1_3207 | 704.9792907 | 643.1278046 | 891.1447567 | -0.028146559 | 0.098622456 | 0.53203452 | 0.855153333 | ns | NaCl | KS.3 |
| FIM1_3208 | 3852.871238 | 3696.372773 | 3247.202603 | -0.028324752 | 0.071623758 | 0.564744632 | 0.867532568 | ns | NaCl | KS.3 |

| | | | | | | | | | | |
|---|---|---|---|---|---|---|---|---|---|---|
| FIM1_3209 | 7946.793231 | 8010.114509 | 7110.124517 | 0.004215149 | 0.080231941 | 0.931237714 | 0.984325045 | ns | NaCl | KS.3 |
| FIM1_3210 | 304.4273717 | 296.017703 | 590.3318852 | -0.010642988 | 0.09219747 | 0.817228147 | 0.959120082 | ns | NaCl | KS.3 |
| FIM1_3211 | 371.5264454 | 330.5327974 | 437.1034821 | -0.038679387 | 0.098747224 | 0.401690773 | 0.789189152 | ns | NaCl | KS.3 |
| FIM1_3212 | 3711.528583 | 3849.396207 | 3159.226961 | 0.021660275 | 0.077122307 | 0.66027582 | 0.903691294 | ns | NaCl | KS.3 |
| FIM1_3213 | 3591.306747 | 3743.872677 | 3801.11775 | 0.028040264 | 0.072263469 | 0.569094612 | 0.869550295 | ns | NaCl | KS.3 |
| FIM1_3214 | 6007.873453 | 7831.288518 | 3474.386174 | 0.036509247 | 0.136699707 | 0.351058818 | 0.756912594 | ns | NaCl | KS.3 |
| FIM1_3215 | 2566.602045 | 2655.537133 | 2625.623474 | 0.026163587 | 0.06617451 | 0.588071307 | 0.878311243 | ns | NaCl | KS.3 |
| FIM1_3216 | 37.69282236 | 36.35589951 | 80.9259995 | -0.008765542 | 0.11541505 | 0.818582871 | 0.959981393 | ns | NaCl | KS.3 |
| FIM1_3217 | 2280.504502 | 2637.908714 | 2185.918002 | 0.033756279 | 0.110451432 | 0.429226115 | 0.804596805 | ns | NaCl | KS.3 |
| FIM1_3218 | 163.8076365 | 131.7847207 | 239.287081 | -0.040670454 | 0.125443329 | 0.326985741 | 0.740319311 | ns | NaCl | KS.3 |
| FIM1_3219 | 2888.739124 | 1786.124953 | 3158.916771 | -0.525264279 | 0.23913418 | 9.32070051793 | 0.007025831 | ns | NaCl | KS.3 |
| FIM1_3220 | 1257.047374 | 2132.242214 | 2141.848212 | 0.426450058 | 0.307057648 | 0.000840099 | 0.031433557 | ns | NaCl | KS.3 |
| FIM1_3221 | 7371.877185 | 8101.221391 | 6394.062365 | 0.063287883 | 0.075579624 | 0.207953168 | 0.633002483 | ns | NaCl | KS.3 |
| FIM1_3222 | 3627.308532 | 3298.812484 | 3571.471653 | -0.094852501 | 0.055628894 | 0.034944936 | 0.295879009 | ns | NaCl | KS.3 |
| FIM1_3223 | 6171.064285 | 7102.5425 | 5186.784433 | 0.118065637 | 0.071267852 | 0.020317975 | 0.215832354 | ns | NaCl | KS.3 |
| FIM1_3224 | 1245.497326 | 1228.302496 | 1877.403515 | -0.002873064 | 0.107934594 | 0.943001078 | 0.986790704 | ns | NaCl | KS.3 |
| FIM1_3225 | 107.3141367 | 92.03291637 | 117.7906859 | -0.027097475 | 0.118783218 | 0.496296029 | 0.836358994 | ns | NaCl | KS.3 |
| FIM1_3226 | 639.5662535 | 671.343189 | 829.6867632 | 0.022469405 | 0.085200303 | 0.641931842 | 0.898798052 | ns | NaCl | KS.3 |
| FIM1_3227 | 2898.951284 | 3209.664837 | 2360.123084 | 0.054090208 | 0.084116331 | 0.278511754 | 0.700652348 | ns | NaCl | KS.3 |
| FIM1_3228 | 1521.235925 | 1624.480852 | 1687.076866 | 0.035663387 | 0.081052468 | 0.46998175 | 0.821848579 | ns | NaCl | KS.3 |
| FIM1_3229 | 5074.429893 | 5597.606183 | 3769.560011 | 0.073161017 | 0.071215172 | 0.1427845 | 0.555038853 | ns | NaCl | KS.3 |
| FIM1_3230 | 5801.973024 | 5589.565463 | 4397.562342 | -0.023824103 | 0.074503929 | 0.629053037 | 0.896016329 | ns | NaCl | KS.3 |
| FIM1_3231 | 6943.576307 | 7173.076072 | 4952.197279 | 0.029496716 | 0.058780583 | 0.522349652 | 0.853429187 | ns | NaCl | KS.3 |
| FIM1_3232 | 132.2923238 | 192.5821808 | 257.3779081 | 0.119455409 | 0.173108866 | 0.039794652 | 0.315550884 | ns | NaCl | KS.3 |
| FIM1_3233 | 498.1750865 | 378.450841 | 1744.519258 | -0.191653733 | 0.138186726 | 0.004144494 | 0.083140565 | ns | NaCl | KS.3 |
| FIM1_3234 | 6808.331888 | 9026.309929 | 4010.201847 | 0.052517534 | 0.135397775 | 0.224094177 | 0.648059715 | ns | NaCl | KS.3 |
| FIM1_3235 | 1042.63803 | 1002.646555 | 1430.647497 | -0.024641226 | 0.077390487 | 0.617342625 | 0.891518014 | ns | NaCl | KS.3 |
| FIM1_3236 | 10825.9208 | 14301.88906 | 9277.944695 | 0.077824764 | 0.132596704 | 0.113373379 | 0.511826282 | ns | NaCl | KS.3 |
| FIM1_3237 | 5055.291878 | 4282.242881 | 5182.316218 | -0.091790035 | 0.091974433 | 0.079771962 | 0.437357005 | ns | NaCl | KS.3 |
| FIM1_3238 | 359.1977133 | 352.78774 | 470.235462 | -0.01202928 | 0.083202547 | 0.803976419 | 0.957859861 | ns | NaCl | KS.3 |
| FIM1_3239 | 3193.065649 | 2927.095346 | 3239.1185 | -0.05537069 | 0.07695254 | 0.269110095 | 0.690826998 | ns | NaCl | KS.3 |
| FIM1_3240 | 3250.489112 | 3402.505353 | 2728.842769 | 0.022306151 | 0.083658258 | 0.646284603 | 0.898905404 | ns | NaCl | KS.3 |
| FIM1_3241 | 868.677104 | 638.7113308 | 1207.718789 | -0.127023089 | 0.14702241 | 0.030570046 | 0.274523423 | ns | NaCl | KS.3 |
| FIM1_3242 | 559.4980497 | 589.0069949 | 739.1983989 | 0.024460286 | 0.084565305 | 0.614082867 | 0.889162818 | ns | NaCl | KS.3 |
| FIM1_3243 | 2498.173583 | 2982.977912 | 2177.611347 | 0.101910875 | 0.093337524 | 0.0557893 | 0.369084797 | ns | NaCl | KS.3 |
| FIM1_3244 | 696.0359078 | 866.9863358 | 1295.076085 | 0.063743939 | 0.117146541 | 0.177106373 | 0.605153989 | ns | NaCl | KS.3 |
| FIM1_3245 | 1093.801513 | 955.5304273 | 1235.652559 | -0.063902398 | 0.091287488 | 0.201014516 | 0.624846998 | ns | NaCl | KS.3 |
| FIM1_3246 | 838.4134406 | 699.4766369 | 1352.307553 | -0.070607634 | 0.104393737 | 0.153843914 | 0.571010593 | ns | NaCl | KS.3 |
| FIM1_3247 | 5560.234279 | 5648.574474 | 5932.375847 | 0.003181759 | 0.107727406 | 0.937006749 | 0.986375069 | ns | NaCl | KS.3 |
| FIM1_3248 | 142.3335271 | 87.66207216 | 270.241428 | -0.465836751 | 0.277828931 | 0.000370428 | 0.019196669 | ns | NaCl | KS.3 |
| FIM1_3249 | 3217.151069 | 3059.654168 | 3260.562503 | -0.024696778 | 0.083461317 | 0.612121185 | 0.889139531 | ns | NaCl | KS.3 |
| FIM1_3250 | 978.9275144 | 899.0514941 | 864.3547259 | -0.03359558 | 0.092209814 | 0.477203926 | 0.825770272 | ns | NaCl | KS.3 |
| FIM1_3251 | 644.4202508 | 744.11562 | 786.1267761 | 0.087405148 | 0.084587517 | 0.091909305 | 0.467534553 | ns | NaCl | KS.3 |
| FIM1_3252 | 31.44394408 | 26.1838798 | 46.84830583 | -0.027026368 | 0.132467574 | 0.471542325 | 0.822127061 | ns | NaCl | KS.3 |
| FIM1_3253 | 960.0898506 | 835.3872615 | 3578.117444 | -0.02726123 | 0.114020437 | 0.50455312 | 0.844040357 | ns | NaCl | KS.3 |

| | | | | | | | | | | |
|---|---|---|---|---|---|---|---|---|---|---|
| FIM1_3254 | 4903.691786 | 5309.712686 | 4440.083237 | 0.058771819 | 0.070534236 | 0.235277666 | 0.659616806 | ns | NaCl | KS.3 |
| FIM1_3255 | 673.016042 | 687.0665509 | 844.2191512 | 0.013427562 | 0.069678679 | 0.783407504 | 0.951646805 | ns | NaCl | KS.3 |
| FIM1_3256 | 44.67416536 | 46.97024144 | 40.43753559 | 0.004461314 | 0.116191819 | 0.905994935 | 0.979880338 | ns | NaCl | KS.3 |
| FIM1_3258 | 319.3182853 | 335.5861879 | 304.6439041 | 0.016112218 | 0.095244526 | 0.721695594 | 0.931688886 | ns | NaCl | KS.3 |
| FIM1_3259 | 350.4040076 | 395.4259453 | 420.9082459 | 0.030132026 | 0.106085982 | 0.485419628 | 0.829376704 | ns | NaCl | KS.3 |
| FIM1_3260 | 1414.904059 | 1470.825341 | 1532.399655 | 0.029296467 | 0.063516374 | 0.538760294 | 0.857036524 | ns | NaCl | KS.3 |
| FIM1_3261 | 545.1410065 | 545.3759019 | 583.1292843 | -0.000421505 | 0.088458433 | 0.992859059 | 0.999134858 | ns | NaCl | KS.3 |
| FIM1_3262 | 41957.77753 | 49371.80048 | 29873.94409 | 0.104018791 | 0.086928796 | 0.049804086 | 0.349460758 | ns | NaCl | KS.3 |
| FIM1_3263 | 1794.391255 | 1941.883875 | 1959.690367 | 0.051736885 | 0.07547276 | 0.30005911 | 0.717845774 | ns | NaCl | KS.3 |
| FIM1_3264 | 1702.863379 | 1797.563574 | 1677.565821 | 0.030269531 | 0.079494456 | 0.539447004 | 0.857153895 | ns | NaCl | KS.3 |
| FIM1_3265 | 1539.708356 | 1444.879347 | 1308.290788 | -0.044886893 | 0.071225557 | 0.36334586 | 0.764655523 | ns | NaCl | KS.3 |
| FIM1_3266 | 836.5053668 | 896.7930028 | 627.6223463 | 0.019382385 | 0.100618578 | 0.656833868 | 0.901977674 | ns | NaCl | KS.3 |
| FIM1_3267 | 4412.697246 | 4954.282859 | 3631.771014 | 0.074094559 | 0.079340611 | 0.145601211 | 0.559788274 | ns | NaCl | KS.3 |
| FIM1_3268 | 1022.985708 | 1099.800478 | 861.409376 | 0.026672223 | 0.092821609 | 0.567228996 | 0.868965626 | ns | NaCl | KS.3 |
| FIM1_3269 | 521.1816888 | 406.9640361 | 594.8101771 | -0.107128907 | 0.122250583 | 0.049884633 | 0.349460758 | ns | NaCl | KS.3 |
| FIM1_3270 | 1768.508639 | 3029.637463 | 3849.519547 | 0.132603312 | 0.229825653 | 0.032729079 | 0.28818968 | ns | NaCl | KS.3 |
| FIM1_3271 | 3176.695217 | 3746.966002 | 3839.313539 | 0.094408194 | 0.090746488 | 0.072542988 | 0.421203992 | ns | NaCl | KS.3 |
| FIM1_3272 | 869.2183908 | 713.208286 | 1169.173293 | -0.103430335 | 0.102105946 | 0.054209937 | 0.363411554 | ns | NaCl | KS.3 |
| FIM1_3273 | 2980.825384 | 3497.438122 | 2005.529265 | 0.034030777 | 0.113912542 | 0.418108325 | 0.800387073 | ns | NaCl | KS.3 |
| FIM1_3274 | 2103.624204 | 2057.744362 | 2580.125895 | -0.022972833 | 0.049144258 | 0.580746745 | 0.8749481 | ns | NaCl | KS.3 |
| FIM1_3275 | 4696.282445 | 5451.719466 | 3593.207739 | 0.091366571 | 0.085756291 | 0.079823148 | 0.437357005 | ns | NaCl | KS.3 |
| FIM1_3276 | 191.1714288 | 160.3399368 | 476.724671 | -0.047427343 | 0.110253989 | 0.292661264 | 0.711969309 | ns | NaCl | KS.3 |
| FIM1_3277 | 2850.815889 | 2862.802665 | 2368.234626 | 0.002533811 | 0.07331175 | 0.95886712 | 0.991553507 | ns | NaCl | KS.3 |
| FIM1_3278 | 7838.223904 | 7832.883243 | 6815.733062 | -0.000534594 | 0.070241389 | 0.991274757 | 0.999134858 | ns | NaCl | KS.3 |
| FIM1_3279 | 4873.282291 | 5081.419303 | 3833.314864 | 0.02169357 | 0.081903617 | 0.657064406 | 0.901977674 | ns | NaCl | KS.3 |
| FIM1_3280 | 1843.706125 | 1661.642123 | 2105.880803 | -0.057272894 | 0.083302885 | 0.253301908 | 0.676801759 | ns | NaCl | KS.3 |
| FIM1_3281 | 4480.008478 | 4742.027261 | 4292.32629 | 0.025918679 | 0.086337999 | 0.590841478 | 0.878786986 | ns | NaCl | KS.3 |
| FIM1_3282 | 5678.731387 | 6834.622085 | 5569.056616 | 0.03003569 | 0.123331992 | 0.447732712 | 0.812914509 | ns | NaCl | KS.3 |
| FIM1_3283 | 772.588705 | 731.7020594 | 807.0213661 | -0.02264085 | 0.089722238 | 0.631860372 | 0.89774822 | ns | NaCl | KS.3 |
| FIM1_3284 | 916.2879329 | 682.6923558 | 1342.972478 | -0.16880492 | 0.148816913 | 0.009760625 | 0.138844204 | ns | NaCl | KS.3 |
| FIM1_3286 | 1280.991806 | 1740.956832 | 1644.190823 | 0.290221504 | 0.149400372 | 0.00024432 | 0.014626776 | ns | NaCl | KS.3 |
| FIM1_3287 | 465.1023419 | 603.1171071 | 460.2756596 | 0.104897021 | 0.124783956 | 0.053462474 | 0.362364862 | ns | NaCl | KS.3 |
| FIM1_3288 | 30593.60671 | 31939.78615 | 24758.36259 | 0.012438308 | 0.098465898 | 0.776999653 | 0.950472898 | ns | NaCl | KS.3 |
| FIM1_3290 | 1439.884017 | 1438.497428 | 1587.834695 | -0.00082376 | 0.081562957 | 0.986492513 | 0.999134858 | ns | NaCl | KS.3 |
| FIM1_3291 | 14.20400931 | 9.258151964 | 18.39346275 | -0.033456954 | 0.161417021 | 0.347956783 | 0.756654396 | ns | NaCl | KS.3 |
| FIM1_3292 | 39683.26631 | 44672.0797 | 28547.75295 | 0.069348533 | 0.082788809 | 0.172142138 | 0.5990025 | ns | NaCl | KS.3 |
| FIM1_3293 | 535.4505012 | 582.1924871 | 402.3163413 | 0.027262929 | 0.096443275 | 0.549984364 | 0.860945181 | ns | NaCl | KS.3 |
| FIM1_3294 | 434.7063861 | 461.6732215 | 335.1507035 | 0.019389473 | 0.095605782 | 0.668616483 | 0.908843443 | ns | NaCl | KS.3 |
| FIM1_3295 | 778.6080554 | 816.691662 | 1289.998448 | 0.013960443 | 0.098362138 | 0.751206682 | 0.943274417 | ns | NaCl | KS.3 |
| FIM1_3297 | 2237.162779 | 2446.269333 | 2133.083476 | 0.068661295 | 0.070250696 | 0.166705363 | 0.590711667 | ns | NaCl | KS.3 |
| FIM1_3298 | 5566.210783 | 7335.42221 | 3671.679717 | 0.072561165 | 0.13200061 | 0.131037364 | 0.538132312 | ns | NaCl | KS.3 |
| FIM1_3299 | 5080.891695 | 5944.379027 | 5009.030904 | 0.124634424 | 0.077267571 | 0.017587287 | 0.198856255 | ns | NaCl | KS.3 |
| FIM1_3300 | 17405.65473 | 19214.99646 | 12154.06664 | 0.044474089 | 0.089123268 | 0.361102757 | 0.763371684 | ns | NaCl | KS.3 |
| FIM1_3301 | 2641.843319 | 3022.101616 | 2184.731024 | 0.07744577 | 0.085630974 | 0.130909367 | 0.538132312 | ns | NaCl | KS.3 |
| FIM1_3302 | 1134.487664 | 1536.491364 | 695.5945618 | 0.047003842 | 0.141143266 | 0.255372952 | 0.678991959 | ns | NaCl | KS.3 |

| | | | | | | | | | | |
|---|---|---|---|---|---|---|---|---|---|---|
| FIM1_3303 | 603.1618431 | 649.6430834 | 583.473781 | 0.02780188 | 0.092457421 | 0.552426553 | 0.861003165 | ns | NaCl | KS.3 |
| FIM1_3304 | 17604.79159 | 23670.93956 | 10610.29114 | 0.084762524 | 0.13827039 | 0.093038746 | 0.470201108 | ns | NaCl | KS.3 |
| FIM1_3305 | 8709.518642 | 7980.030399 | 11293.93362 | -0.047782931 | 0.082252608 | 0.336697575 | 0.746466327 | ns | NaCl | KS.3 |
| FIM1_3306 | 329.0552507 | 325.3518459 | 385.9812727 | -0.00398522 | 0.098550989 | 0.927172147 | 0.984176127 | ns | NaCl | KS.3 |
| FIM1_3307 | 1250.201934 | 908.4648884 | 1497.973183 | -0.205869305 | 0.164709724 | 0.004329353 | 0.085470362 | ns | NaCl | KS.3 |
| FIM1_3308 | 379.9412312 | 408.9470139 | 1459.473884 | 0.019696234 | 0.10165722 | 0.649271784 | 0.899642849 | ns | NaCl | KS.3 |
| FIM1_3309 | 5128.861691 | 5822.650985 | 4207.45595 | 0.060471423 | 0.090135168 | 0.224601021 | 0.648385059 | ns | NaCl | KS.3 |
| FIM1_3310 | 1529.325386 | 1442.714009 | 1586.122228 | -0.036437391 | 0.076359244 | 0.462331299 | 0.819880744 | ns | NaCl | KS.3 |
| FIM1_3311 | 7185.948023 | 8041.84239 | 5799.004745 | 0.114238526 | 0.055841391 | 0.011342006 | 0.151642712 | ns | NaCl | KS.3 |
| FIM1_3312 | 1657.596351 | 1867.297995 | 3030.972899 | 0.022995028 | 0.112830735 | 0.56992483 | 0.870548366 | ns | NaCl | KS.3 |
| FIM1_3313 | 200.3938252 | 205.8564809 | 449.383105 | 0.003277676 | 0.125029768 | 0.926822654 | 0.984176127 | ns | NaCl | KS.3 |
| FIM1_3314 | 565.298049 | 488.2270962 | 611.8314272 | -0.076396447 | 0.091645055 | 0.134611299 | 0.54183277 | ns | NaCl | KS.3 |
| FIM1_3315 | 772.5521057 | 778.1888074 | 625.3048471 | 0.002904388 | 0.091345414 | 0.949873807 | 0.987736646 | ns | NaCl | KS.3 |
| FIM1_3316 | 2606.64831 | 2883.519038 | 1921.22516 | 0.074803214 | 0.071324645 | 0.134461394 | 0.54183277 | ns | NaCl | KS.3 |
| FIM1_3317 | 108.5588614 | 94.78800444 | 156.9880462 | -0.042312241 | 0.103668828 | 0.352215397 | 0.756912594 | ns | NaCl | KS.3 |
| FIM1_3318 | 1479.56546 | 1935.224337 | 902.019129 | 0.053114328 | 0.132106131 | 0.223402608 | 0.648059715 | ns | NaCl | KS.3 |
| FIM1_3319 | 1873.187053 | 1970.11304 | 1424.631369 | 0.022971877 | 0.086011331 | 0.633439261 | 0.898009483 | ns | NaCl | KS.3 |
| FIM1_3320 | 5412.167476 | 4713.608644 | 3754.133926 | -0.026598508 | 0.114338066 | 0.513131568 | 0.847552971 | ns | NaCl | KS.3 |
| FIM1_3321 | 4696.697213 | 5620.270365 | 2821.384153 | 0.030316186 | 0.121720177 | 0.447260156 | 0.812914509 | ns | NaCl | KS.3 |
| FIM1_3322 | 615.5084083 | 448.1719152 | 576.8316103 | -0.078740088 | 0.14460541 | 0.107807404 | 0.498909937 | ns | NaCl | KS.3 |
| FIM1_3323 | 711.5985607 | 647.8363609 | 1336.77278 | -0.046504366 | 0.086756361 | 0.344511313 | 0.754044779 | ns | NaCl | KS.3 |
| FIM1_3324 | 991.6968374 | 1355.33209 | 2165.389234 | 0.177671618 | 0.15790083 | 0.008465379 | 0.127329254 | ns | NaCl | KS.3 |
| FIM1_3326 | 14923.90565 | 15299.58226 | 10761.74779 | 0.009113167 | 0.09172568 | 0.843522716 | 0.966543148 | ns | NaCl | KS.3 |
| FIM1_3327 | 1858.676978 | 1798.655363 | 1547.231501 | -0.013993681 | 0.087579108 | 0.768409746 | 0.948835638 | ns | NaCl | KS.3 |
| FIM1_3328 | 192.0498828 | 202.0719263 | 250.7530417 | 0.014421427 | 0.098187335 | 0.743767414 | 0.941460666 | ns | NaCl | KS.3 |
| FIM1_3329 | 454.9851392 | 445.6630272 | 333.4106703 | -0.008056487 | 0.089269748 | 0.863673677 | 0.969856868 | ns | NaCl | KS.3 |
| FIM1_3331 | 533.617038 | 431.6090937 | 427.7347597 | -0.1288344 | 0.104340322 | 0.022442192 | 0.229732317 | ns | NaCl | KS.3 |
| FIM1_3332 | 722.1685242 | 643.8410918 | 563.365356 | -0.066196252 | 0.08088852 | 0.190993657 | 0.618952432 | ns | NaCl | KS.3 |
| FIM1_3333 | 7071.91032 | 8119.273323 | 3974.130456 | 0.045397927 | 0.101215257 | 0.327989209 | 0.740319311 | ns | NaCl | KS.3 |
| FIM1_3334 | 43337.54731 | 52716.06119 | 33685.66399 | 0.121917498 | 0.096726893 | 0.027080574 | 0.256889329 | ns | NaCl | KS.3 |
| FIM1_3335 | 1839.652375 | 2193.736656 | 1422.542747 | 0.059631479 | 0.105795618 | 0.211533601 | 0.635060483 | ns | NaCl | KS.3 |
| FIM1_3336 | 1487.297524 | 1602.823875 | 1146.475696 | 0.02533657 | 0.095360185 | 0.57995809 | 0.8749481 | ns | NaCl | KS.3 |
| FIM1_3337 | 1176.497244 | 1142.748752 | 2308.848679 | -0.0216619 | 0.066543965 | 0.654189932 | 0.900747416 | ns | NaCl | KS.3 |
| FIM1_3339 | 871.3246513 | 933.5083095 | 1165.07265 | 0.033321922 | 0.084858899 | 0.494653144 | 0.836358994 | ns | NaCl | KS.3 |
| FIM1_3340 | 3015.937435 | 3050.872311 | 5574.560663 | 0.003460056 | 0.097562008 | 0.93727215 | 0.986445726 | ns | NaCl | KS.3 |
| FIM1_3341 | 1099.377558 | 1052.661069 | 1121.771156 | -0.032108191 | 0.072145454 | 0.514770479 | 0.849077129 | ns | NaCl | KS.3 |
| FIM1_3342 | 99.05948806 | 95.94751392 | 158.9391856 | -0.007845059 | 0.106960486 | 0.847081855 | 0.967147129 | ns | NaCl | KS.3 |
| FIM1_3343 | 8200.360069 | 9640.513907 | 8696.620167 | 0.034590822 | 0.114064657 | 0.411265969 | 0.798613659 | ns | NaCl | KS.3 |
| FIM1_3344 | 11713.33639 | 13392.83865 | 18575.06962 | 0.021619844 | 0.118932705 | 0.57817221 | 0.874933697 | ns | NaCl | KS.3 |
| FIM1_3345 | 6774.176751 | 7033.460914 | 6481.108677 | 0.005864175 | 0.115712908 | 0.877243789 | 0.972783118 | ns | NaCl | KS.3 |
| FIM1_3346 | 207.3277344 | 199.1137563 | 215.6425215 | -0.007931782 | 0.105324056 | 0.847486755 | 0.967147129 | ns | NaCl | KS.3 |
| FIM1_3347 | 941.9573879 | 915.8406488 | 1658.200111 | -0.008693367 | 0.095416929 | 0.846390202 | 0.967147129 | ns | NaCl | KS.3 |
| FIM1_3348 | 438.8883174 | 300.5969036 | 957.1650824 | -0.175006061 | 0.195806821 | 0.012764985 | 0.16241893 | ns | NaCl | KS.3 |
| FIM1_3349 | 1251.331226 | 1306.022237 | 1789.716452 | 0.025468255 | 0.078167396 | 0.605445528 | 0.883689315 | ns | NaCl | KS.3 |
| FIM1_3350 | 8887.340341 | 12361.077 | 9211.614553 | 0.143808051 | 0.160990457 | 0.021148003 | 0.221497506 | ns | NaCl | KS.3 |

| | | | | | | | | | |
|---|---|---|---|---|---|---|---|---|---|
| FIM1_3351 | 1521.555736 | 1257.068603 | 1694.361127 | -0.123632922 | 0.093843504 | 0.024594168 | 0.244223524 | ns | NaCl | KS.3 |
| FIM1_3353 | 11803.87925 | 13879.92894 | 8404.574634 | 0.08043049 | 0.094227127 | 0.117192695 | 0.52010139 | ns | NaCl | KS.3 |
| FIM1_3355 | 1355.953183 | 1453.334079 | 1440.681107 | 0.021695816 | 0.097420277 | 0.628647516 | 0.896016329 | ns | NaCl | KS.3 |
| FIM1_3356 | 883.3238039 | 587.075771 | 1282.228097 | -0.203032075 | 0.214819063 | 0.008286929 | 0.126464636 | ns | NaCl | KS.3 |
| FIM1_3357 | 435.4379232 | 447.9851108 | 631.5619316 | 0.010955381 | 0.090272532 | 0.814437245 | 0.958816322 | ns | NaCl | KS.3 |
| FIM1_3358 | 13.48516736 | 9.155914163 | 16.18493289 | -0.032412772 | 0.165977227 | 0.351936339 | 0.756912594 | ns | NaCl | KS.3 |
| FIM1_3359 | 6716.004555 | 7375.468541 | 5452.627362 | 0.069335221 | 0.071237723 | 0.16422529 | 0.589056105 | ns | NaCl | KS.3 |
| FIM1_3360 | 3120.339408 | 2881.744321 | 3710.643881 | -0.0478419 | 0.078562408 | 0.337558288 | 0.747375382 | ns | NaCl | KS.3 |
| FIM1_3361 | 8696.191726 | 10242.93834 | 6897.756809 | 0.138706663 | 0.075676537 | 0.008226408 | 0.125927317 | ns | NaCl | KS.3 |
| FIM1_3362 | 2534.935485 | 2543.851368 | 3212.704601 | 0.001234373 | 0.093393664 | 0.978349523 | 0.996164232 | ns | NaCl | KS.3 |
| FIM1_3363 | 1520.88208 | 1569.193582 | 1497.899917 | 0.019207099 | 0.076961995 | 0.696545173 | 0.921625594 | ns | NaCl | KS.3 |
| FIM1_3364 | 1341.421839 | 1373.893881 | 1841.479941 | 0.017136985 | 0.06944901 | 0.725662855 | 0.933787374 | ns | NaCl | KS.3 |
| FIM1_3365 | 319.7586821 | 299.3755063 | 341.3008944 | -0.021398644 | 0.096619795 | 0.635150965 | 0.898009483 | ns | NaCl | KS.3 |
| FIM1_3366 | 102.6851411 | 107.9588161 | 426.0664148 | 0.005679287 | 0.127678346 | 0.872061705 | 0.971243188 | ns | NaCl | KS.3 |
| FIM1_3367 | 585.7771778 | 529.2396692 | 713.2190129 | -0.038369638 | 0.095554002 | 0.412839216 | 0.799937916 | ns | NaCl | KS.3 |
| FIM1_3368 | 821.7341273 | 1119.902054 | 621.4381627 | 0.161935834 | 0.154178403 | 0.012496466 | 0.160231753 | ns | NaCl | KS.3 |
| FIM1_3369 | 1138.149455 | 1320.360769 | 1156.707981 | 0.046443708 | 0.103741153 | 0.313329715 | 0.731871738 | ns | NaCl | KS.3 |
| FIM1_3370 | 914.0739446 | 901.5911617 | 841.2917541 | -0.003863826 | 0.095895653 | 0.930955873 | 0.984325045 | ns | NaCl | KS.3 |
| FIM1_3371 | 886.0455361 | 1044.308446 | 821.7171968 | 0.064112853 | 0.100321569 | 0.190973708 | 0.618952432 | ns | NaCl | KS.3 |
| FIM1_3372 | 697.6480727 | 663.1361794 | 785.7021713 | -0.030777286 | 0.078183461 | 0.533380986 | 0.855153333 | ns | NaCl | KS.3 |
| FIM1_3373 | 1445.363615 | 1776.06826 | 864.5272455 | 0.034122105 | 0.125201018 | 0.394897191 | 0.782402837 | ns | NaCl | KS.3 |
| FIM1_3374 | 6335.305283 | 5941.221551 | 5453.42696 | -0.010417435 | 0.115079606 | 0.786333395 | 0.95289761 | ns | NaCl | KS.3 |
| FIM1_3375 | 9501.104249 | 12334.11497 | 8288.966713 | 0.183065499 | 0.125606498 | 0.004319314 | 0.085470362 | ns | NaCl | KS.3 |
| FIM1_3376 | 18.73208175 | 15.86163962 | 19.77240213 | -0.013023326 | 0.132867705 | 0.709965828 | 0.925563973 | ns | NaCl | KS.3 |
| FIM1_3377 | 1998.804234 | 2727.775878 | 1614.212457 | 0.074483489 | 0.141987321 | 0.12121108 | 0.527999953 | ns | NaCl | KS.3 |
| FIM1_3378 | 23.9001191 | 21.01420326 | 24.81555604 | -0.021801967 | 0.124899051 | 0.563597681 | 0.867434473 | ns | NaCl | KS.3 |
| FIM1_3379 | 389.4636551 | 423.7537349 | 602.1434 | 0.039245532 | 0.088314498 | 0.418360818 | 0.800387073 | ns | NaCl | KS.3 |
| FIM1_3380 | 245.7055724 | 305.9919348 | 384.328297 | 0.069604256 | 0.117206188 | 0.149784961 | 0.564839452 | ns | NaCl | KS.3 |
| FIM1_3381 | 26753.87351 | 30949.36426 | 18287.95914 | 0.079182837 | 0.088854668 | 0.123220838 | 0.529611582 | ns | NaCl | KS.3 |
| FIM1_3382 | 489.0490565 | 452.7352871 | 540.4338346 | -0.029040837 | 0.090868146 | 0.539200868 | 0.857036524 | ns | NaCl | KS.3 |
| FIM1_3383 | 2226.060837 | 2396.575002 | 2471.836453 | 0.041000999 | 0.081059203 | 0.407944 | 0.79494601 | ns | NaCl | KS.3 |
| FIM1_3384 | 18466.96389 | 24434.72406 | 10102.61494 | 0.060216141 | 0.133888952 | 0.182339549 | 0.611396712 | ns | NaCl | KS.3 |
| FIM1_3385 | 80.73118858 | 90.37376487 | 83.71763438 | 0.019139877 | 0.112789518 | 0.631942566 | 0.89774822 | ns | NaCl | KS.3 |
| FIM1_3386 | 1128.112011 | 1255.433252 | 715.7074906 | 0.027501967 | 0.104659607 | 0.524666877 | 0.854940852 | ns | NaCl | KS.3 |
| FIM1_3387 | 1491.618986 | 1556.473728 | 1783.26767 | 0.009948201 | 0.104067079 | 0.811799579 | 0.958816322 | ns | NaCl | KS.3 |
| FIM1_3388 | 2973.047643 | 3261.241064 | 3075.192619 | 0.020544339 | 0.10749729 | 0.621258379 | 0.892479944 | ns | NaCl | KS.3 |
| FIM1_3389 | 1401.367984 | 1084.870573 | 1286.693949 | -0.048910905 | 0.129795946 | 0.253778593 | 0.676969706 | ns | NaCl | KS.3 |
| FIM1_3390 | 4531.138067 | 4046.827533 | 3612.230617 | -0.060406811 | 0.084912874 | 0.22906779 | 0.652325274 | ns | NaCl | KS.3 |
| FIM1_3391 | 1792.73383 | 2079.450954 | 1527.696093 | 0.045683823 | 0.10382304 | 0.319986035 | 0.736735876 | ns | NaCl | KS.3 |
| FIM1_3392 | 756.2257561 | 641.7045426 | 997.793671 | -0.081024744 | 0.095726453 | 0.114549864 | 0.514859894 | ns | NaCl | KS.3 |
| FIM1_3393 | 110.8632804 | 136.973448 | 168.986438 | 0.028490164 | 0.129450185 | 0.456795903 | 0.817337042 | ns | NaCl | KS.3 |
| FIM1_3394 | 1744.109183 | 1580.833639 | 2718.688271 | -0.031877088 | 0.097827201 | 0.484828654 | 0.829376704 | ns | NaCl | KS.3 |
| FIM1_3395 | 2578.725795 | 3130.802332 | 2191.632266 | 0.061371693 | 0.110297792 | 0.19614791 | 0.618952432 | ns | NaCl | KS.3 |
| FIM1_3396 | 979.5759461 | 991.1503415 | 1060.624303 | 0.002692461 | 0.10264643 | 0.948938431 | 0.987736646 | ns | NaCl | KS.3 |
| FIM1_3397 | 9422.720428 | 5962.556472 | 7705.802668 | -0.122669746 | 0.201265248 | 0.03869712 | 0.311585109 | ns | NaCl | KS.3 |

| | | | | | | | | | | |
|---|---|---|---|---|---|---|---|---|---|---|
| FIM1_3398 | 8.733592085 | 11.13734996 | 8.902902147 | 0.019135985 | 0.149584816 | 0.568014286 | 0.869550295 | ns | NaCl | KS.3 |
| FIM1_3399 | 870.4001243 | 780.3904186 | 1046.559827 | -0.035968377 | 0.098749269 | 0.432444729 | 0.805772482 | ns | NaCl | KS.3 |
| FIM1_3400 | 1399.584424 | 1317.947273 | 1585.761323 | -0.038534023 | 0.075918915 | 0.437355629 | 0.808096352 | ns | NaCl | KS.3 |
| FIM1_3401 | 1120.472404 | 1420.097158 | 1416.926159 | 0.125379635 | 0.11575149 | 0.027556545 | 0.260140056 | ns | NaCl | KS.3 |
| FIM1_3402 | 4415.791891 | 5215.288561 | 3669.48146 | 0.151630746 | 0.072901042 | 0.003515001 | 0.076045768 | ns | NaCl | KS.3 |
| FIM1_3403 | 7965.094415 | 10925.82775 | 4961.124225 | 0.081235045 | 0.144073241 | 0.101129321 | 0.484750818 | ns | NaCl | KS.3 |
| FIM1_3404 | 56.73291917 | 18.64315838 | 107.336423 | -0.614248456 | 0.612140444 | 0.000832777 | 0.031433557 | ns | NaCl | KS.3 |
| FIM1_3405 | 15667.43634 | 19123.87664 | 10136.20419 | 0.094624304 | 0.103875065 | 0.072718569 | 0.421203992 | ns | NaCl | KS.3 |
| FIM1_3407 | 1520.540103 | 1680.852377 | 1399.676544 | 0.043046687 | 0.090523309 | 0.373216609 | 0.775331654 | ns | NaCl | KS.3 |
| FIM1_3408 | 875.1672723 | 915.550762 | 1106.331544 | 0.026901053 | 0.076293723 | 0.585934189 | 0.877826895 | ns | NaCl | KS.3 |
| FIM1_3409 | 13453.66699 | 12582.06534 | 9335.100371 | -0.023710835 | 0.093873237 | 0.607158828 | 0.884513959 | ns | NaCl | KS.3 |
| FIM1_3410 | 1893.636673 | 2135.637967 | 1501.152017 | 0.032111399 | 0.104631804 | 0.463840405 | 0.819880744 | ns | NaCl | KS.3 |
| FIM1_3411 | 2670.511645 | 2898.23398 | 1892.610748 | 0.020761297 | 0.103352617 | 0.628135804 | 0.896016329 | ns | NaCl | KS.3 |
| FIM1_3412 | 1434.106092 | 1466.485944 | 1428.368822 | 0.015996478 | 0.068889326 | 0.742808815 | 0.941460666 | ns | NaCl | KS.3 |
| FIM1_3413 | 3401.72782 | 3573.716142 | 2766.208354 | 0.016330435 | 0.095052466 | 0.718540575 | 0.930194994 | ns | NaCl | KS.3 |
| FIM1_3414 | 2173.018011 | 2661.400902 | 2553.054635 | 0.076347192 | 0.108934866 | 0.127041417 | 0.534982546 | ns | NaCl | KS.3 |
| FIM1_3415 | 946.1850711 | 1215.375985 | 1188.797963 | 0.041273637 | 0.131448288 | 0.312816839 | 0.731871738 | ns | NaCl | KS.3 |
| FIM1_3416 | 421.6795254 | 462.4646932 | 460.4770577 | 0.021035987 | 0.107140251 | 0.614299315 | 0.889162818 | ns | NaCl | KS.3 |
| FIM1_3417 | 21480.54476 | 23206.61588 | 14646.90247 | 0.045155647 | 0.07933518 | 0.364375185 | 0.765203269 | ns | NaCl | KS.3 |
| FIM1_3418 | 3416.255442 | 3911.532601 | 2580.809561 | 0.07807034 | 0.085489528 | 0.128124877 | 0.536116795 | ns | NaCl | KS.3 |
| FIM1_3419 | 2952.605327 | 2989.459304 | 3343.998621 | 0.009828185 | 0.065896924 | 0.838346879 | 0.963005246 | ns | NaCl | KS.3 |
| FIM1_3420 | 37.99896413 | 27.93622336 | 103.0118483 | -0.038082432 | 0.145352134 | 0.323944874 | 0.739131368 | ns | NaCl | KS.3 |
| FIM1_3421 | 114.6499856 | 109.0085841 | 127.9908184 | -0.006774187 | 0.117895604 | 0.856479089 | 0.969856868 | ns | NaCl | KS.3 |
| FIM1_3422 | 153.4049008 | 71.5399071 | 142.3187291 | -0.236508518 | 0.356256697 | 0.008860377 | 0.130802304 | ns | NaCl | KS.3 |
| FIM1_3423 | 190.0154421 | 178.2079009 | 604.0889833 | -0.011305541 | 0.114392121 | 0.770108052 | 0.949065335 | ns | NaCl | KS.3 |
| FIM1_3424 | 800.4934556 | 758.6189836 | 844.8276442 | -0.015403559 | 0.098747918 | 0.72615903 | 0.93398169 | ns | NaCl | KS.3 |
| FIM1_3425 | 1688.527936 | 494.9569136 | 1240.169645 | -1.635341227 | 0.242833289 | 2.4526375428 | 1.7431245393 | Km<KS.3 | NaCl | KS.3 |
| FIM1_3426 | 6397.706919 | 3867.897191 | 4545.849937 | -0.074790133 | 0.186564586 | 0.107904943 | 0.498909937 | ns | NaCl | KS.3 |
| FIM1_3427 | 808.1457276 | 860.0487254 | 1391.919756 | 0.018075025 | 0.098916247 | 0.681893516 | 0.915044369 | ns | NaCl | KS.3 |
| FIM1_3428 | 34.2817543 | 46.0513712 | 84.78845287 | 0.049831962 | 0.139075684 | 0.237923498 | 0.660898607 | ns | NaCl | KS.3 |
| FIM1_3429 | 4243.153276 | 4285.265296 | 3127.203162 | 0.005986195 | 0.076362409 | 0.903070888 | 0.979395371 | ns | NaCl | KS.3 |
| FIM1_3430 | 1939.549869 | 2044.494277 | 1770.705665 | 0.035387271 | 0.073544392 | 0.474223982 | 0.82307921 | ns | NaCl | KS.3 |
| FIM1_3431 | 531.2817442 | 567.4320922 | 368.9373469 | 0.024943371 | 0.093907733 | 0.589173558 | 0.878638624 | ns | NaCl | KS.3 |
| FIM1_3432 | 1003.746569 | 940.6336783 | 922.3058643 | -0.024738198 | 0.091949029 | 0.596670694 | 0.879797482 | ns | NaCl | KS.3 |
| FIM1_3433 | 3696.091927 | 4164.21462 | 3764.933484 | 0.102747789 | 0.06670841 | 0.037241228 | 0.304670827 | ns | NaCl | KS.3 |
| FIM1_3434 | 572.7677018 | 465.37015 | 717.5488816 | -0.156450944 | 0.095447664 | 0.006352952 | 0.106776808 | ns | NaCl | KS.3 |
| FIM1_3435 | 3655.41508 | 3966.663478 | 3127.965965 | 0.031304386 | 0.092417202 | 0.505790255 | 0.84439816 | ns | NaCl | KS.3 |
| FIM1_3437 | 19797.98052 | 16001.57772 | 16824.47536 | -0.08209796 | 0.111136012 | 0.106167265 | 0.49734665 | ns | NaCl | KS.3 |
| FIM1_3438 | 3221.633397 | 2696.750454 | 2603.795388 | -0.057991023 | 0.106848113 | 0.220550991 | 0.642906874 | ns | NaCl | KS.3 |
| FIM1_3439 | 1529.039017 | 892.8437734 | 1906.752553 | -0.270744939 | 0.294557835 | 0.004932785 | 0.091912385 | ns | NaCl | KS.3 |
| FIM1_3440 | 3320.748073 | 3456.066891 | 3000.803043 | 0.018888728 | 0.08501162 | 0.695472373 | 0.921340753 | ns | NaCl | KS.3 |
| FIM1_3441 | 609.8347225 | 452.183204 | 987.7667858 | -0.172853275 | 0.148717725 | 0.008678601 | 0.128883701 | ns | NaCl | KS.3 |
| FIM1_3442 | 5750.607519 | 5378.386917 | 5052.080606 | -0.029868306 | 0.087596648 | 0.534765381 | 0.855153333 | ns | NaCl | KS.3 |
| FIM1_3444 | 5722.930462 | 5913.270716 | 4131.371059 | 0.015518651 | 0.084474965 | 0.747900055 | 0.943274417 | ns | NaCl | KS.3 |
| FIM1_3445 | 5173.461983 | 4822.195459 | 3482.366193 | -0.023931514 | 0.095139083 | 0.600875274 | 0.880762097 | ns | NaCl | KS.3 |

| | | | | | | | | | | |
|---|---|---|---|---|---|---|---|---|---|---|
| FIM1_3446 | 1504.789978 | 1342.563399 | 1375.250604 | -0.102051678 | 0.064893051 | 0.036261969 | 0.302183074 | ns | NaCl | KS.3 |
| FIM1_3447 | 2911.095569 | 3085.776952 | 2612.233532 | 0.034550639 | 0.077508677 | 0.485384378 | 0.829376704 | ns | NaCl | KS.3 |
| FIM1_3448 | 618.1645017 | 553.9334161 | 773.1970386 | -0.076938428 | 0.077241712 | 0.13036117 | 0.537766849 | ns | NaCl | KS.3 |
| FIM1_3449 | 879.2290658 | 811.2805159 | 936.3167794 | -0.048025697 | 0.079680312 | 0.335575684 | 0.745307602 | ns | NaCl | KS.3 |
| FIM1_3450 | 7757.411373 | 10338.76567 | 4650.317217 | 0.063683219 | 0.135483382 | 0.164935294 | 0.589355333 | ns | NaCl | KS.3 |
| FIM1_3452 | 1527.855211 | 1450.086146 | 1780.430393 | -0.029120741 | 0.080164299 | 0.554286727 | 0.862821173 | ns | NaCl | KS.3 |
| FIM1_3454 | 4211.967398 | 2616.754844 | 12413.85407 | -0.256441991 | 0.263001769 | 0.004852532 | 0.090978232 | ns | NaCl | KS.3 |
| FIM1_3455 | 62.57473243 | 69.49796305 | 1212.410285 | 0.015062427 | 0.116406685 | 0.696033799 | 0.921625594 | ns | NaCl | KS.3 |
| FIM1_3456 | 29.89781317 | 36.89545704 | 220.5758679 | 0.026828464 | 0.132864398 | 0.473631259 | 0.82307921 | ns | NaCl | KS.3 |
| FIM1_3457 | 412.9651759 | 567.2888029 | 898.4169684 | 0.060891386 | 0.142829243 | 0.173439308 | 0.601296555 | ns | NaCl | KS.3 |
| FIM1_3458 | 4966.633368 | 5809.579872 | 8765.855031 | 0.048574275 | 0.104789522 | 0.29294496 | 0.711969309 | ns | NaCl | KS.3 |
| FIM1_3460 | 7431.256834 | 8178.673985 | 6233.295187 | 0.048414923 | 0.085462916 | 0.327751057 | 0.740319311 | ns | NaCl | KS.3 |
| FIM1_3461 | 1291.704133 | 1574.768735 | 1201.563314 | 0.03055485 | 0.125960077 | 0.436237546 | 0.808096352 | ns | NaCl | KS.3 |
| FIM1_3462 | 680.7237074 | 642.8373493 | 799.4370233 | -0.034814538 | 0.077860068 | 0.482041075 | 0.828664253 | ns | NaCl | KS.3 |
| FIM1_3463 | 2221.111998 | 2839.760169 | 3452.321499 | 0.051247703 | 0.127032814 | 0.240611239 | 0.665022731 | ns | NaCl | KS.3 |
| FIM1_3464 | 272.8989308 | 284.8283271 | 332.051609 | 0.009817334 | 0.103663575 | 0.814814831 | 0.958816322 | ns | NaCl | KS.3 |
| FIM1_3465 | 111.2440735 | 69.60618194 | 228.7605552 | -0.099758061 | 0.19147278 | 0.062087815 | 0.390501742 | ns | NaCl | KS.3 |
| FIM1_3466 | 3009.475751 | 2944.68065 | 2895.335907 | -0.01877932 | 0.061635085 | 0.689887555 | 0.91892653 | ns | NaCl | KS.3 |
| FIM1_3467 | 488.7420304 | 552.5524806 | 819.1413989 | 0.015181756 | 0.126541119 | 0.676799967 | 0.913287355 | ns | NaCl | KS.3 |
| FIM1_3468 | 3266.501421 | 3586.772558 | 2493.683692 | 0.039754354 | 0.090229424 | 0.409077037 | 0.795392815 | ns | NaCl | KS.3 |
| FIM1_3469 | 1656.053039 | 1538.45636 | 2023.438997 | -0.014082908 | 0.110690962 | 0.724570469 | 0.933222742 | ns | NaCl | KS.3 |
| FIM1_3470 | 841.0959512 | 749.2865472 | 741.3694133 | -0.026230514 | 0.108427555 | 0.532725098 | 0.855153333 | ns | NaCl | KS.3 |
| FIM1_3471 | 3064.306695 | 4062.451582 | 2447.582828 | 0.213882621 | 0.138566792 | 0.002003334 | 0.054462224 | ns | NaCl | KS.3 |
| FIM1_3472 | 530.9750646 | 474.7390221 | 647.6004953 | -0.052742342 | 0.08832606 | 0.285723264 | 0.705523209 | ns | NaCl | KS.3 |
| FIM1_3473 | 5376.959106 | 7639.194823 | 3333.603423 | 0.065312545 | 0.151365567 | 0.150190437 | 0.564839452 | ns | NaCl | KS.3 |
| FIM1_3474 | 99.70931385 | 117.5797313 | 705.3237644 | 0.024117599 | 0.12396634 | 0.529186878 | 0.855153333 | ns | NaCl | KS.3 |
| FIM1_3475 | 4200.286741 | 3677.506891 | 3795.78794 | -0.094500093 | 0.077784311 | 0.066894644 | 0.405360357 | ns | NaCl | KS.3 |
| FIM1_3476 | 1162.844908 | 1188.968812 | 1593.52545 | 0.013277111 | 0.072816958 | 0.787039181 | 0.95289761 | ns | NaCl | KS.3 |
| FIM1_3477 | 2034.841462 | 1906.288849 | 1971.95525 | -0.028636091 | 0.08760588 | 0.551242986 | 0.860945181 | ns | NaCl | KS.3 |
| FIM1_3478 | 198.6274435 | 175.1091003 | 204.791761 | -0.034820092 | 0.105498102 | 0.428706513 | 0.804596805 | ns | NaCl | KS.3 |
| FIM1_3479 | 7669.072121 | 7770.046913 | 7511.9524 | 0.013408457 | 0.04980118 | 0.749276258 | 0.943274417 | ns | NaCl | KS.3 |
| FIM1_3480 | 5188.184387 | 6959.850728 | 3057.765141 | 0.047549485 | 0.138963517 | 0.253702805 | 0.676969706 | ns | NaCl | KS.3 |
| FIM1_3481 | 3846.562817 | 3866.136115 | 3386.072532 | 0.003952182 | 0.066434941 | 0.934756861 | 0.985446192 | ns | NaCl | KS.3 |
| FIM1_3482 | 2070.485905 | 2237.759711 | 3183.553039 | 0.026725689 | 0.095305723 | 0.560400937 | 0.865836851 | ns | NaCl | KS.3 |
| FIM1_3483 | 825.6334591 | 719.9996998 | 933.8487455 | -0.092325086 | 0.081033246 | 0.075052577 | 0.426240375 | ns | NaCl | KS.3 |
| FIM1_3484 | 1189.436774 | 959.9452303 | 1506.444146 | -0.187028543 | 0.090989172 | 0.001212506 | 0.041316566 | ns | NaCl | KS.3 |
| FIM1_3485 | 602.3900062 | 465.1268613 | 508.4485735 | -0.120048413 | 0.125027554 | 0.034121097 | 0.292771172 | ns | NaCl | KS.3 |
| FIM1_3486 | 3215.2574 | 3258.892678 | 3483.811659 | 0.006003529 | 0.086459387 | 0.899789481 | 0.978786499 | ns | NaCl | KS.3 |
| FIM1_3487 | 950.6618712 | 928.6550844 | 881.4953144 | -0.00771102 | 0.09507837 | 0.863854437 | 0.969856868 | ns | NaCl | KS.3 |
| FIM1_3488 | 372.8857314 | 342.1700614 | 422.2680436 | -0.046389831 | 0.082634693 | 0.350086032 | 0.756912594 | ns | NaCl | KS.3 |
| FIM1_3489 | 942.8330383 | 804.1910335 | 767.3091327 | -0.106388157 | 0.083382417 | 0.043916091 | 0.33003407 | ns | NaCl | KS.3 |
| FIM1_3490 | 1218.650993 | 1250.052956 | 1105.079606 | 0.014967722 | 0.075551235 | 0.761102317 | 0.945911573 | ns | NaCl | KS.3 |
| FIM1_3491 | 1671.102876 | 1679.033525 | 2077.042796 | 0.001845041 | 0.081682029 | 0.969741453 | 0.99411987 | ns | NaCl | KS.3 |
| FIM1_3492 | 597.1119337 | 448.2678999 | 782.9234932 | -0.115723886 | 0.13843941 | 0.040368689 | 0.317627561 | ns | NaCl | KS.3 |
| FIM1_3493 | 97.33492034 | 115.2086864 | 111.3256679 | 0.030356941 | 0.118412508 | 0.453438856 | 0.816716052 | ns | NaCl | KS.3 |

| | | | | | | | | | | |
|---|---|---|---|---|---|---|---|---|---|---|
| FIM1_3494 | 13368.74421 | 15151.35571 | 9553.046138 | 0.085311919 | 0.078088181 | 0.096070238 | 0.473217264 | ns | NaCl | KS.3 |
| FIM1_3495 | 120.130279 | 135.0459953 | 257.5342315 | 0.039795555 | 0.101451952 | 0.383092378 | 0.778665419 | ns | NaCl | KS.3 |
| FIM1_3496 | 768.6959639 | 877.7890048 | 619.1826244 | 0.047847847 | 0.09803161 | 0.312173927 | 0.731871738 | ns | NaCl | KS.3 |
| FIM1_3497 | 1020.301325 | 499.4254985 | 1782.456726 | -0.322479745 | 0.38315815 | 0.004027609 | 0.082277904 | ns | NaCl | KS.3 |
| FIM1_3498 | 151.6211624 | 181.4401004 | 150.7250461 | 0.041515773 | 0.113616609 | 0.339255451 | 0.748468235 | ns | NaCl | KS.3 |
| FIM1_3499 | 2425.775128 | 2406.758264 | 2687.064173 | -0.006710217 | 0.065667725 | 0.889094272 | 0.977268396 | ns | NaCl | KS.3 |
| FIM1_3500 | 1664.202554 | 1844.238138 | 1777.103672 | 0.03338447 | 0.098440163 | 0.464299938 | 0.819983028 | ns | NaCl | KS.3 |
| FIM1_3501 | 1893.500381 | 1739.188068 | 2906.741389 | -0.019309374 | 0.106596065 | 0.643425509 | 0.898798052 | ns | NaCl | KS.3 |
| FIM1_3503 | 293.2485734 | 257.4775772 | 375.0021449 | -0.013620597 | 0.131783548 | 0.699443645 | 0.922863518 | ns | NaCl | KS.3 |
| FIM1_3504 | 705.8114351 | 931.0246351 | 551.5915906 | 0.053340866 | 0.134371238 | 0.219945785 | 0.642906874 | ns | NaCl | KS.3 |
| FIM1_3505 | 4232.301096 | 5246.157312 | 3680.734522 | 0.171658303 | 0.096432879 | 0.003256683 | 0.072981982 | ns | NaCl | KS.3 |
| FIM1_3506 | 644.231428 | 664.1314201 | 702.3294603 | 0.009934837 | 0.094517567 | 0.826155622 | 0.960240659 | ns | NaCl | KS.3 |
| FIM1_3507 | 1826.57363 | 1866.79685 | 1893.918144 | 0.013694517 | 0.072283086 | 0.780374843 | 0.951646805 | ns | NaCl | KS.3 |
| FIM1_3508 | 1205.396374 | 1215.271226 | 926.6437892 | 0.004565702 | 0.079962392 | 0.925586241 | 0.984176127 | ns | NaCl | KS.3 |
| FIM1_3509 | 19.95942176 | 10.9640355 | 75.09163503 | -0.102056036 | 0.226149839 | 0.056324004 | 0.371633849 | ns | NaCl | KS.3 |
| FIM1_3511 | 5142.316991 | 5044.716335 | 4938.838614 | -0.009632639 | 0.082271223 | 0.842873536 | 0.966485679 | ns | NaCl | KS.3 |
| FIM1_3512 | 4146.57578 | 3695.190055 | 4645.559879 | -0.038775808 | 0.098261035 | 0.401812286 | 0.789189152 | ns | NaCl | KS.3 |
| FIM1_3513 | 770.7613407 | 575.7440936 | 986.2109913 | -0.036986948 | 0.141809565 | 0.339192635 | 0.748468235 | ns | NaCl | KS.3 |
| FIM1_3514 | 2218.196809 | 2553.339008 | 2916.016212 | 0.034898818 | 0.108278624 | 0.420953195 | 0.80133378 | ns | NaCl | KS.3 |
| FIM1_3516 | 9346.383495 | 12229.03416 | 7410.048664 | 0.051744884 | 0.132438924 | 0.231765323 | 0.654993872 | ns | NaCl | KS.3 |
| FIM1_3517 | 367.5012205 | 357.938509 | 512.655349 | -0.012248417 | 0.082032228 | 0.80128997 | 0.957352931 | ns | NaCl | KS.3 |
| FIM1_3518 | 4224.064475 | 6177.517306 | 3047.35086 | 0.063419092 | 0.157595577 | 0.154777078 | 0.572502577 | ns | NaCl | KS.3 |
| FIM1_3519 | 862.5155669 | 858.7018533 | 757.2062454 | -0.000999558 | 0.104486221 | 0.980717119 | 0.996363429 | ns | NaCl | KS.3 |
| FIM1_3520 | 875.8265845 | 1029.18988 | 1141.632281 | 0.038064013 | 0.111716196 | 0.377928782 | 0.778493059 | ns | NaCl | KS.3 |
| FIM1_3522 | 6050.463239 | 7865.180724 | 3616.222836 | 0.040338754 | 0.134392044 | 0.317287963 | 0.73382286 | ns | NaCl | KS.3 |
| FIM1_3523 | 9783.130408 | 10479.56603 | 7519.610076 | 0.024083609 | 0.09427049 | 0.600760904 | 0.880762097 | ns | NaCl | KS.3 |
| FIM1_3524 | 2441.245365 | 2710.558163 | 1888.769054 | 0.040832837 | 0.09349268 | 0.390522124 | 0.779482559 | ns | NaCl | KS.3 |
| FIM1_3525 | 482.5376037 | 480.4994273 | 478.5321531 | -0.001113872 | 0.104510714 | 0.978508146 | 0.996164232 | ns | NaCl | KS.3 |
| FIM1_3526 | 3292.490595 | 3330.125338 | 3162.86158 | 0.008363491 | 0.068848949 | 0.863682855 | 0.969856868 | ns | NaCl | KS.3 |
| FIM1_3527 | 1506.060106 | 1522.065187 | 1778.7921 | 0.002586886 | 0.102252349 | 0.95110572 | 0.987736646 | ns | NaCl | KS.3 |
| FIM1_3528 | 1570.991565 | 2082.97199 | 3188.141061 | 0.048183853 | 0.136304882 | 0.251943053 | 0.675897894 | ns | NaCl | KS.3 |
| FIM1_3529 | 707.1423206 | 675.6265308 | 616.9667557 | -0.013959589 | 0.097216465 | 0.753422581 | 0.943523399 | ns | NaCl | KS.3 |
| FIM1_3530 | 471.256396 | 390.9182416 | 406.0561677 | -0.064828144 | 0.107064492 | 0.180363896 | 0.60960532 | ns | NaCl | KS.3 |
| FIM1_3531 | 1395.528161 | 1168.128071 | 1385.990734 | -0.153230266 | 0.080760583 | 0.004661112 | 0.088507754 | ns | NaCl | KS.3 |
| FIM1_3532 | 1078.155517 | 1096.929104 | 977.3404401 | 0.007411026 | 0.085057514 | 0.877362382 | 0.972783118 | ns | NaCl | KS.3 |
| FIM1_3533 | 320.5155915 | 309.4095398 | 652.0218671 | -0.013173672 | 0.089733028 | 0.7787926 | 0.951027591 | ns | NaCl | KS.3 |
| FIM1_3534 | 2355.701713 | 2359.401764 | 2703.360744 | 0.001512912 | 0.065387349 | 0.974882406 | 0.995954749 | ns | NaCl | KS.3 |
| FIM1_3535 | 1905.956564 | 1797.796877 | 1527.632879 | -0.048660614 | 0.062353935 | 0.304897049 | 0.724521545 | ns | NaCl | KS.3 |
| FIM1_3536 | 472.7376931 | 481.7102755 | 460.5971985 | 0.008356525 | 0.084257062 | 0.862387695 | 0.969856868 | ns | NaCl | KS.3 |
| FIM1_3537 | 12722.93304 | 17642.8986 | 8084.007475 | 0.080489887 | 0.147138824 | 0.102542525 | 0.488687531 | ns | NaCl | KS.3 |
| FIM1_3538 | 788.3219244 | 767.0182153 | 903.5957502 | -0.015541151 | 0.079777642 | 0.751149779 | 0.943274417 | ns | NaCl | KS.3 |
| FIM1_3539 | 578.0451751 | 611.2306475 | 579.1258978 | 0.02643334 | 0.085821668 | 0.58447294 | 0.877414869 | ns | NaCl | KS.3 |
| FIM1_3540 | 3589.083007 | 3485.683531 | 3361.474688 | -0.020672298 | 0.069341072 | 0.672129512 | 0.912198685 | ns | NaCl | KS.3 |
| FIM1_3541 | 503.7512979 | 554.456878 | 306.9328249 | 0.024977977 | 0.104613019 | 0.560757504 | 0.865840156 | ns | NaCl | KS.3 |
| FIM1_3542 | 472.7382029 | 363.553281 | 450.4154753 | -0.193390979 | 0.131052252 | 0.003374307 | 0.074609683 | ns | NaCl | KS.3 |

| | | | | | | | | | | |
|---|---|---|---|---|---|---|---|---|---|---|
| FIM1_3544 | 3030.993863 | 3427.205892 | 2705.31606 | 0.047992506 | 0.095121164 | 0.316476243 | 0.73382286 | ns | NaCl | KS.3 |
| FIM1_3545 | 89.54508249 | 42.6383535 | 154.6222435 | -0.340780754 | 0.394629028 | 0.003515684 | 0.076045768 | ns | NaCl | KS.3 |
| FIM1_3546 | 2145.972832 | 2456.022067 | 1688.710571 | 0.057740295 | 0.094101666 | 0.239310658 | 0.662163805 | ns | NaCl | KS.3 |
| FIM1_3547 | 1288.712661 | 1259.757765 | 926.7482602 | -0.011894656 | 0.08131419 | 0.807312096 | 0.957859861 | ns | NaCl | KS.3 |
| FIM1_3548 | 700.6611443 | 579.4490892 | 437.2404573 | -0.033422812 | 0.121808876 | 0.409127681 | 0.795392815 | ns | NaCl | KS.3 |
| FIM1_3549 | 155.4919006 | 148.3494993 | 171.1134703 | -0.016967001 | 0.096574248 | 0.705061098 | 0.924674422 | ns | NaCl | KS.3 |
| FIM1_3550 | 909.5432517 | 823.9238418 | 1459.170544 | -0.027013329 | 0.103055396 | 0.535781495 | 0.855153333 | ns | NaCl | KS.3 |
| FIM1_3551 | 2266.150619 | 1965.08787 | 1689.147242 | -0.046429021 | 0.101916544 | 0.31712537 | 0.73382286 | ns | NaCl | KS.3 |
| FIM1_3552 | 552.9322384 | 546.8472755 | 580.9714058 | -0.005565882 | 0.087472331 | 0.906530955 | 0.979989779 | ns | NaCl | KS.3 |
| FIM1_3553 | 403.1816917 | 480.4266848 | 438.2540512 | 0.042963427 | 0.112348848 | 0.328030344 | 0.740319311 | ns | NaCl | KS.3 |
| FIM1_3554 | 605.7855475 | 578.9054884 | 602.7025135 | -0.018531574 | 0.089546115 | 0.694180296 | 0.920525145 | ns | NaCl | KS.3 |
| FIM1_3555 | 412.3382806 | 405.268449 | 575.0273797 | -0.007747244 | 0.08905694 | 0.869019373 | 0.971243188 | ns | NaCl | KS.3 |
| FIM1_3556 | 2294.900333 | 2401.444505 | 2232.991527 | 0.029537866 | 0.074519555 | 0.549907048 | 0.860945181 | ns | NaCl | KS.3 |
| FIM1_3557 | 8167.150629 | 9219.718452 | 4774.948837 | 0.034082024 | 0.103381023 | 0.442875551 | 0.810336839 | ns | NaCl | KS.3 |
| FIM1_3558 | 958.995894 | 1119.239833 | 1198.38517 | 0.073020579 | 0.094626073 | 0.148815874 | 0.563164954 | ns | NaCl | KS.3 |
| FIM1_3559 | 229.6802545 | 173.5843263 | 243.8344568 | -0.121549247 | 0.134429695 | 0.033857311 | 0.292771172 | ns | NaCl | KS.3 |
| FIM1_3560 | 1118.671253 | 1215.191376 | 1344.499968 | 0.049758757 | 0.078405066 | 0.319183821 | 0.736179653 | ns | NaCl | KS.3 |
| FIM1_3561 | 351.4632535 | 260.3852439 | 872.0920139 | -0.094299625 | 0.140930938 | 0.072061976 | 0.420358296 | ns | NaCl | KS.3 |
| FIM1_3562 | 2075.512092 | 1649.069167 | 1735.131374 | -0.172686409 | 0.10559206 | 0.004065567 | 0.082277904 | ns | NaCl | KS.3 |
| FIM1_3563 | 2183.54204 | 2155.842588 | 1925.341098 | -0.007846662 | 0.077098898 | 0.873120627 | 0.971243188 | ns | NaCl | KS.3 |
| FIM1_3565 | 63.82507694 | 100.3784512 | 110.8741976 | 0.084084814 | 0.18077506 | 0.088747798 | 0.46158813 | ns | NaCl | KS.3 |
| FIM1_3566 | 252.6741922 | 192.6698069 | 242.8917698 | -0.080179173 | 0.130979252 | 0.106719761 | 0.498296595 | ns | NaCl | KS.3 |
| FIM1_3567 | 10406.91144 | 10675.20988 | 7172.635201 | 0.00999193 | 0.089754859 | 0.830981653 | 0.961620266 | ns | NaCl | KS.3 |
| FIM1_3568 | 3948.137842 | 3099.375522 | 4147.484486 | -0.074141598 | 0.121686318 | 0.129494899 | 0.53686427 | ns | NaCl | KS.3 |
| FIM1_3569 | 3368.224881 | 4374.458987 | 2003.273747 | 0.03375376 | 0.137465625 | 0.379368813 | 0.778493059 | ns | NaCl | KS.3 |
| FIM1_3570 | 1540.263404 | 1650.624124 | 1301.032634 | 0.018426174 | 0.101558958 | 0.669966801 | 0.90993307 | ns | NaCl | KS.3 |
| FIM1_3571 | 5201.081534 | 5062.136841 | 4121.434755 | -0.018553037 | 0.071726413 | 0.705560104 | 0.924943746 | ns | NaCl | KS.3 |
| FIM1_3572 | 471.6479649 | 471.5089087 | 519.58149 | -0.00087837 | 0.083335967 | 0.985505461 | 0.998958775 | ns | NaCl | KS.3 |
| FIM1_3573 | 5693.229553 | 6000.532014 | 4375.909811 | 0.035903173 | 0.072098678 | 0.466806017 | 0.821323144 | ns | NaCl | KS.3 |
| FIM1_3574 | 1658.619543 | 1673.786132 | 3146.18648 | 0.001228269 | 0.12060678 | 0.973235853 | 0.9954458 | ns | NaCl | KS.3 |
| FIM1_3575 | 1714.669934 | 1525.167863 | 1718.833637 | -0.029443963 | 0.105935274 | 0.494894708 | 0.836358994 | ns | NaCl | KS.3 |
| FIM1_3576 | 3660.720001 | 3371.150446 | 3522.498511 | -0.045540679 | 0.081391013 | 0.359502019 | 0.763021562 | ns | NaCl | KS.3 |
| FIM1_3577 | 361.805891 | 333.5527262 | 487.2915457 | -0.035956148 | 0.089567178 | 0.454149578 | 0.816788914 | ns | NaCl | KS.3 |
| FIM1_3579 | 2145.67784 | 2247.825311 | 2293.963276 | 0.015426614 | 0.094845658 | 0.733652578 | 0.938375543 | ns | NaCl | KS.3 |
| FIM1_3580 | 9000.498468 | 11517.16552 | 5248.873732 | 0.047265463 | 0.128355412 | 0.267534567 | 0.689593109 | ns | NaCl | KS.3 |
| FIM1_3581 | 4793.142796 | 2539.713401 | 4370.134313 | -0.662672427 | 0.318982394 | 0.000124581 | 0.009114558 | ns | NaCl | KS.3 |
| FIM1_3582 | 2291.688219 | 2238.491891 | 2298.735857 | -0.017022964 | 0.069826108 | 0.727683667 | 0.934975786 | ns | NaCl | KS.3 |
| FIM1_3583 | 1422.220588 | 2693.634081 | 1973.623488 | 0.539717496 | 0.364385773 | 0.000530079 | 0.02370281 | ns | NaCl | KS.3 |
| FIM1_3584 | 540.5090286 | 719.8552979 | 589.4158153 | 0.133760829 | 0.137237938 | 0.024182987 | 0.242041091 | ns | NaCl | KS.3 |
| FIM1_3585 | 1079.198726 | 785.7317244 | 1187.196258 | -0.133529002 | 0.154418029 | 0.026597044 | 0.254519471 | ns | NaCl | KS.3 |
| FIM1_3586 | 3810.71964 | 3863.026836 | 2909.673499 | 0.004397342 | 0.094867327 | 0.922123134 | 0.983890455 | ns | NaCl | KS.3 |
| FIM1_3587 | 99.89525912 | 95.66904524 | 85.38572007 | -0.008378801 | 0.115670286 | 0.826037291 | 0.960240659 | ns | NaCl | KS.3 |
| FIM1_3588 | 1017.805532 | 1032.615008 | 1024.528335 | 0.002854552 | 0.108391834 | 0.943153728 | 0.986790704 | ns | NaCl | KS.3 |
| FIM1_3589 | 3006.636702 | 3045.61666 | 3873.387259 | 0.003927201 | 0.096911822 | 0.929227531 | 0.984238275 | ns | NaCl | KS.3 |
| FIM1_3591 | 353.4469777 | 238.0058049 | 629.3839924 | -0.118169843 | 0.17984115 | 0.041447872 | 0.322888102 | ns | NaCl | KS.3 |

| | | | | | | | | | | |
|---|---|---|---|---|---|---|---|---|---|---|
| FIM1_3592 | 2556.538214 | 2549.916773 | 1972.23227 | -0.000364917 | 0.114591856 | 0.992363426 | 0.999134858 | ns | NaCl | KS.3 |
| FIM1_3594 | 3782.215239 | 4260.571129 | 3172.010947 | 0.073102453 | 0.081327093 | 0.151429537 | 0.566863767 | ns | NaCl | KS.3 |
| FIM1_3595 | 4681.790474 | 4825.949854 | 5378.255139 | 0.012588537 | 0.08837456 | 0.790018576 | 0.953873589 | ns | NaCl | KS.3 |
| FIM1_3596 | 34.70385154 | 36.39081253 | 37.01878554 | 0.004501761 | 0.127371583 | 0.898423504 | 0.977971161 | ns | NaCl | KS.3 |
| FIM1_3597 | 13135.91987 | 14736.1516 | 10687.94615 | 0.073710532 | 0.079194345 | 0.147502096 | 0.562037426 | ns | NaCl | KS.3 |
| FIM1_3598 | 4018.529054 | 4911.287203 | 2566.044627 | 0.057984041 | 0.113238398 | 0.212879594 | 0.636463931 | ns | NaCl | KS.3 |
| FIM1_3599 | 7848.737116 | 7671.449722 | 6212.493068 | -0.01363321 | 0.076424705 | 0.78172139 | 0.951646805 | ns | NaCl | KS.3 |
| FIM1_3600 | 409.7394846 | 410.0272365 | 386.4645009 | -0.000417958 | 0.095585547 | 0.99253709 | 0.999134858 | ns | NaCl | KS.3 |
| FIM1_3601 | 2064.876797 | 2027.876647 | 2242.782252 | -0.008832874 | 0.08353809 | 0.855073745 | 0.969815177 | ns | NaCl | KS.3 |
| FIM1_3603 | 611.556382 | 545.9996057 | 629.563039 | -0.062019922 | 0.08601545 | 0.217183037 | 0.641504077 | ns | NaCl | KS.3 |
| FIM1_3604 | 5780.279028 | 7353.169415 | 3903.096472 | 0.106387664 | 0.11837794 | 0.050699299 | 0.352025696 | ns | NaCl | KS.3 |
| FIM1_3605 | 1083.302003 | 961.1890099 | 1340.176429 | -0.1062947 | 0.065611222 | 0.030129859 | 0.27272008 | ns | NaCl | KS.3 |
| FIM1_3606 | 395.3143249 | 336.1400657 | 573.6389977 | -0.047145689 | 0.107539685 | 0.299914151 | 0.717845774 | ns | NaCl | KS.3 |
| FIM1_3607 | 2015.03371 | 2095.992276 | 1506.810348 | 0.020272789 | 0.082345055 | 0.677617403 | 0.913287355 | ns | NaCl | KS.3 |
| FIM1_3608 | 537.3102977 | 525.5789431 | 542.6673427 | -0.009064562 | 0.086119872 | 0.849585762 | 0.967648619 | ns | NaCl | KS.3 |
| FIM1_3609 | 548.0449399 | 632.4081266 | 1486.044669 | 0.030838348 | 0.111362975 | 0.46306633 | 0.819880744 | ns | NaCl | KS.3 |
| FIM1_3610 | 1598.830176 | 1508.367739 | 1668.484706 | -0.033934807 | 0.079003108 | 0.492586896 | 0.835818489 | ns | NaCl | KS.3 |
| FIM1_3611 | 850.9374818 | 863.577614 | 868.238066 | 0.008629202 | 0.071742144 | 0.860345233 | 0.969856868 | ns | NaCl | KS.3 |
| FIM1_3612 | 1480.214631 | 1402.875829 | 1620.530005 | -0.036539665 | 0.072854468 | 0.459657897 | 0.819640873 | ns | NaCl | KS.3 |
| FIM1_3613 | 528.5217633 | 388.1862475 | 1052.78488 | -0.107983831 | 0.14469628 | 0.050323592 | 0.350644077 | ns | NaCl | KS.3 |
| FIM1_3614 | 304.1334893 | 288.5443064 | 312.8522437 | -0.014273698 | 0.100099609 | 0.742441898 | 0.941460666 | ns | NaCl | KS.3 |
| FIM1_3615 | 2361.607277 | 2497.692761 | 1455.642772 | 0.019246449 | 0.094283811 | 0.673788746 | 0.91246166 | ns | NaCl | KS.3 |
| FIM1_3616 | 5295.987037 | 6157.086876 | 3554.016383 | 0.035757009 | 0.110353952 | 0.406228635 | 0.793477604 | ns | NaCl | KS.3 |
| FIM1_3617 | 325.2915807 | 355.8388299 | 224.0788788 | 0.022753902 | 0.102641434 | 0.599120512 | 0.880219278 | ns | NaCl | KS.3 |
| FIM1_3618 | 2070.825574 | 1961.89657 | 2714.576116 | -0.037354022 | 0.072751582 | 0.44975786 | 0.813595672 | ns | NaCl | KS.3 |
| FIM1_3619 | 407.4998994 | 322.2807266 | 359.8464444 | -0.083487571 | 0.11909979 | 0.099882252 | 0.483636126 | ns | NaCl | KS.3 |
| FIM1_3620 | 1131.597162 | 922.466529 | 1503.26998 | -0.107485187 | 0.103496858 | 0.047403702 | 0.342900262 | ns | NaCl | KS.3 |
| FIM1_3621 | 1504.618989 | 1364.947425 | 1545.844025 | -0.046645075 | 0.087468021 | 0.342158127 | 0.750617902 | ns | NaCl | KS.3 |
| FIM1_3622 | 2378.51017 | 2379.723397 | 2433.684599 | 0.00049536 | 0.0660311 | 0.991798907 | 0.999134858 | ns | NaCl | KS.3 |
| FIM1_3623 | 99.55622129 | 67.97944221 | 572.7992675 | -0.053203311 | 0.157115044 | 0.202644198 | 0.626183159 | ns | NaCl | KS.3 |
| FIM1_3624 | 123.7737525 | 52.00499638 | 462.092434 | -0.459859399 | 0.480730072 | 0.00171684 | 0.049658604 | ns | NaCl | KS.3 |
| FIM1_3625 | 24591.96826 | 30723.67499 | 18717.45703 | 0.26205599 | 0.072025767 | 7.41166956725 | 0.000136567 | ns | NaCl | KS.3 |
| FIM1_3626 | 127.6562249 | 105.3318331 | 201.9656237 | -0.072875848 | 0.108060424 | 0.141420371 | 0.553543421 | ns | NaCl | KS.3 |
| FIM1_3627 | 514.0059924 | 705.8340819 | 1302.001326 | 0.120074932 | 0.150875249 | 0.037182517 | 0.304670827 | ns | NaCl | KS.3 |
| FIM1_3628 | 12562.4242 | 15318.23373 | 11043.21588 | 0.032667778 | 0.124393231 | 0.413232224 | 0.8001568 | ns | NaCl | KS.3 |
| FIM1_3629 | 4679.525723 | 4394.292931 | 4731.292686 | -0.018697476 | 0.098325548 | 0.673271842 | 0.91246166 | ns | NaCl | KS.3 |
| FIM1_3630 | 120.2011531 | 107.7390416 | 197.6624048 | -0.02605029 | 0.107635138 | 0.537332518 | 0.855981197 | ns | NaCl | KS.3 |
| FIM1_3631 | 501.2584151 | 514.7776343 | 557.9884718 | 0.012153002 | 0.082500041 | 0.802499163 | 0.957859861 | ns | NaCl | KS.3 |
| FIM1_3632 | 10759.98147 | 11609.07328 | 8567.738859 | 0.050596425 | 0.074424992 | 0.309867721 | 0.729920413 | ns | NaCl | KS.3 |
| FIM1_3633 | 1061.657508 | 415.1754848 | 2506.69561 | -1.033658065 | 0.394353703 | 2.55952108000 | 0.002555233 | Km<KS.3 | NaCl | KS.3 |
| FIM1_3634 | 2527.861667 | 3055.003991 | 3704.855705 | 0.042723231 | 0.116267104 | 0.323084807 | 0.739010075 | ns | NaCl | KS.3 |
| FIM1_3635 | 5275.594281 | 5044.033656 | 4861.765801 | -0.009582195 | 0.106628296 | 0.814701935 | 0.958816322 | ns | NaCl | KS.3 |
| FIM1_3636 | 1667.357042 | 1727.436346 | 2761.731935 | 0.006642141 | 0.109965847 | 0.866972118 | 0.970371341 | ns | NaCl | KS.3 |
| FIM1_3637 | 2249.083261 | 2265.723744 | 3180.97478 | 0.001402205 | 0.109448462 | 0.971801609 | 0.995313014 | ns | NaCl | KS.3 |
| FIM1_3638 | 8623.471121 | 6123.383951 | 7351.407994 | -0.079224006 | 0.151329655 | 0.105111322 | 0.495197752 | ns | NaCl | KS.3 |

| | | | | | | | | | | |
|---|---|---|---|---|---|---|---|---|---|---|
| FIM1_3639 | 275.8108825 | 249.7023265 | 401.9788217 | -0.015165722 | 0.118658549 | 0.6901098 | 0.918976514 | ns | NaCl | KS.3 |
| FIM1_3640 | 796.8249843 | 776.104012 | 1134.694664 | -0.004800284 | 0.1113199 | 0.902443335 | 0.979395371 | ns | NaCl | KS.3 |
| FIM1_3641 | 829.1605873 | 813.2498606 | 956.6364595 | -0.003925616 | 0.107380972 | 0.92257773 | 0.983890455 | ns | NaCl | KS.3 |
| FIM1_3642 | 1418.971997 | 1370.879661 | 2394.135994 | -0.007033129 | 0.107481064 | 0.862031648 | 0.969856868 | ns | NaCl | KS.3 |
| FIM1_3643 | 744.0116159 | 788.8203911 | 1267.188752 | 0.009954334 | 0.113270905 | 0.797940266 | 0.955924408 | ns | NaCl | KS.3 |
| FIM1_3644 | 1870.098958 | 1861.512696 | 2834.249379 | -0.00100122 | 0.105970076 | 0.980436411 | 0.996363429 | ns | NaCl | KS.3 |
| FIM1_3645 | 1999.547342 | 1963.223141 | 3118.831835 | -0.003737822 | 0.107630697 | 0.926109332 | 0.984176127 | ns | NaCl | KS.3 |
| FIM1_3646 | 919.870977 | 1176.190693 | 1263.162388 | 0.052651877 | 0.126605055 | 0.231767023 | 0.654993872 | ns | NaCl | KS.3 |
| FIM1_3647 | 31789.32647 | 36901.33875 | 25119.05881 | 0.042065941 | 0.106076051 | 0.349354785 | 0.756912594 | ns | NaCl | KS.3 |
| FIM1_3648 | 3408.848378 | 4221.164044 | 2447.029702 | 0.024095257 | 0.135693921 | 0.508765948 | 0.844548078 | ns | NaCl | KS.3 |
| FIM1_3649 | 1191.21726 | 1543.395994 | 1619.634937 | 0.054180811 | 0.129633235 | 0.219140448 | 0.642906874 | ns | NaCl | KS.3 |
| FIM1_3650 | 1202.394664 | 1307.537135 | 1797.310892 | 0.016096327 | 0.110966639 | 0.688065935 | 0.918490387 | ns | NaCl | KS.3 |
| FIM1_3651 | 1188.852172 | 1213.284082 | 1422.069128 | 0.003778764 | 0.109910192 | 0.923911025 | 0.983924964 | ns | NaCl | KS.3 |
| FIM1_3652 | 215.0138328 | 186.1730155 | 400.6113962 | -0.017330774 | 0.127642153 | 0.635213136 | 0.898009483 | ns | NaCl | KS.3 |
| FIM1_3653 | 5358.143959 | 6948.595317 | 5782.597739 | 0.0568778 | 0.129257768 | 0.203626441 | 0.626494461 | ns | NaCl | KS.3 |
| FIM1_3654 | 7465.766578 | 7677.287299 | 7828.351318 | 0.00686147 | 0.102451252 | 0.8708968 | 0.971243188 | ns | NaCl | KS.3 |
| FIM1_3655 | 3176.57459 | 3355.805676 | 4492.996544 | 0.012232115 | 0.105721247 | 0.767421343 | 0.948794469 | ns | NaCl | KS.3 |
| FIM1_3656 | 5222.831165 | 5695.074401 | 5621.544174 | 0.018672416 | 0.107911868 | 0.650800036 | 0.899642849 | ns | NaCl | KS.3 |
| FIM1_3657 | 489.0193819 | 462.4396614 | 773.3438998 | -0.009400728 | 0.113532331 | 0.808407067 | 0.958451332 | ns | NaCl | KS.3 |
| FIM1_3658 | 2473.499976 | 3082.871505 | 2222.721408 | 0.03410937 | 0.12843755 | 0.389519085 | 0.779482559 | ns | NaCl | KS.3 |
| FIM1_3659 | 1292.100842 | 1311.898805 | 1990.138433 | 0.002851781 | 0.110153048 | 0.942396149 | 0.986790704 | ns | NaCl | KS.3 |
| FIM1_3660 | 1373.354511 | 1405.420317 | 1551.995642 | 0.004071844 | 0.11141406 | 0.917078731 | 0.982343702 | ns | NaCl | KS.3 |
| FIM1_3661 | 1819.175794 | 1616.655241 | 2798.114036 | -0.022113599 | 0.113684316 | 0.581678679 | 0.875038346 | ns | NaCl | KS.3 |
| FIM1_3662 | 629.4512986 | 608.6574851 | 1358.714328 | -0.006341034 | 0.110426916 | 0.872450936 | 0.971243188 | ns | NaCl | KS.3 |
| FIM1_3663 | 4654.597982 | 5212.446774 | 4056.103488 | 0.015590098 | 0.122396912 | 0.675844994 | 0.912932078 | ns | NaCl | KS.3 |
| FIM1_3664 | 1685.866131 | 1806.168131 | 1737.776234 | 0.012181036 | 0.112762471 | 0.756034799 | 0.944867505 | ns | NaCl | KS.3 |
| FIM1_3665 | 645.8813137 | 610.9986195 | 1234.791245 | -0.008582854 | 0.116715411 | 0.820693868 | 0.960240659 | ns | NaCl | KS.3 |
| FIM1_3666 | 1186.693816 | 1268.378561 | 1461.323656 | 0.011332558 | 0.113727934 | 0.77061572 | 0.949065335 | ns | NaCl | KS.3 |
| FIM1_3667 | 1521.838271 | 1573.636744 | 2133.783649 | 0.006744217 | 0.107761719 | 0.867302713 | 0.9704973 | ns | NaCl | KS.3 |
| FIM1_3668 | 1124.345374 | 1187.160562 | 1477.657812 | 0.01029386 | 0.110237839 | 0.795842845 | 0.955664531 | ns | NaCl | KS.3 |
| FIM1_3669 | 637.9172954 | 595.336725 | 1245.2853 | -0.011368374 | 0.115240679 | 0.767597854 | 0.948794469 | ns | NaCl | KS.3 |
| FIM1_3670 | 468.4030744 | 409.9475292 | 785.9254473 | -0.023297705 | 0.116837964 | 0.556245604 | 0.863699344 | ns | NaCl | KS.3 |
| FIM1_3671 | 355.9171384 | 317.6813536 | 649.025862 | -0.016494789 | 0.121061896 | 0.661575417 | 0.904388188 | ns | NaCl | KS.3 |
| FIM1_3672 | 1392.771731 | 1399.559063 | 2288.424868 | 0.000954919 | 0.109085118 | 0.980849099 | 0.996363429 | ns | NaCl | KS.3 |
| FIM1_3673 | 2317.609544 | 2874.675594 | 2448.216564 | 0.03388027 | 0.127511744 | 0.393665272 | 0.781740625 | ns | NaCl | KS.3 |
| FIM1_3674 | 47.40444353 | 43.57002136 | 71.71387351 | -0.010370714 | 0.123326648 | 0.776301753 | 0.950410881 | ns | NaCl | KS.3 |
| FIM1_3675 | 3783.169471 | 3999.997802 | 4505.666315 | 0.011378385 | 0.108137205 | 0.778825283 | 0.951027591 | ns | NaCl | KS.3 |
| FIM1_3676 | 73.96018947 | 104.1564614 | 187.0670644 | 0.061555645 | 0.148406411 | 0.167279234 | 0.591901982 | ns | NaCl | KS.3 |
| FIM1_3677 | 1076.074706 | 1160.940667 | 1500.927836 | 0.014801949 | 0.110193012 | 0.712590215 | 0.926833025 | ns | NaCl | KS.3 |
| FIM1_3678 | 3360.779476 | 4315.37407 | 2981.63063 | 0.028425054 | 0.138964001 | 0.441879749 | 0.809513863 | ns | NaCl | KS.3 |
| FIM1_3679 | 71906.29906 | 81232.77276 | 61159.05026 | 0.034130069 | 0.103549399 | 0.441863509 | 0.809513863 | ns | NaCl | KS.3 |
| FIM1_3680 | 148.723254 | 214.7674405 | 178.89611 | 0.07456973 | 0.156971462 | 0.116843082 | 0.519941265 | ns | NaCl | KS.3 |
| FIM1_3681 | 900.3379088 | 802.8746159 | 1295.64304 | -0.024056706 | 0.110316588 | 0.559805702 | 0.865454744 | ns | NaCl | KS.3 |
| FIM1_3682 | 2215.649196 | 2906.557853 | 3493.016974 | 0.053322152 | 0.132760981 | 0.221513104 | 0.64446064 | ns | NaCl | KS.3 |
| FIM1_3683 | 1547.667351 | 1237.91259 | 1390.773785 | -0.052093505 | 0.121122437 | 0.241611038 | 0.666264473 | ns | NaCl | KS.3 |

| | | | | | | | | | | |
|---|---|---|---|---|---|---|---|---|---|---|
| FIM1_3684 | 431.7605647 | 432.9157784 | 675.516524 | 0.000594362 | 0.109804519 | 0.988009246 | 0.999134858 | ns | NaCl | KS.3 |
| FIM1_3685 | 491.5623053 | 455.6672984 | 967.9350028 | -0.012313942 | 0.115561539 | 0.748924197 | 0.943274417 | ns | NaCl | KS.3 |
| FIM1_3686 | 938.1397467 | 1209.544206 | 1007.071036 | 0.031526665 | 0.137460663 | 0.405143261 | 0.79260233 | ns | NaCl | KS.3 |
| FIM1_3687 | 432.3658342 | 372.32488 | 709.5175891 | -0.029757915 | 0.114668659 | 0.469343075 | 0.821764392 | ns | NaCl | KS.3 |
| FIM1_3688 | 790.6465163 | 740.6395928 | 1342.19789 | -0.009900596 | 0.117378546 | 0.793400159 | 0.955175866 | ns | NaCl | KS.3 |
| FIM1_3689 | 1015.926898 | 1061.492217 | 1805.398193 | 0.009035342 | 0.107512228 | 0.823728907 | 0.960240659 | ns | NaCl | KS.3 |
| FIM1_3690 | 319.9639313 | 255.9259381 | 1028.481125 | -0.044775015 | 0.123544654 | 0.293621099 | 0.712220852 | ns | NaCl | KS.3 |
| FIM1_3691 | 2815.13251 | 2947.736495 | 3698.699931 | 0.010104086 | 0.105843488 | 0.806173132 | 0.957859861 | ns | NaCl | KS.3 |
| FIM1_3692 | 1959.331333 | 2270.822072 | 2354.507026 | 0.029974212 | 0.113887064 | 0.468301406 | 0.821764392 | ns | NaCl | KS.3 |
| FIM1_3693 | 1334.700104 | 1211.259245 | 2060.419245 | -0.018805854 | 0.11140553 | 0.640596773 | 0.898798052 | ns | NaCl | KS.3 |
| FIM1_3694 | 998.4591578 | 925.587445 | 1198.114767 | -0.015211397 | 0.109464274 | 0.706770513 | 0.925190614 | ns | NaCl | KS.3 |
| FIM1_3695 | 2347.817441 | 2826.174015 | 5682.653673 | 0.036529544 | 0.118737725 | 0.380097743 | 0.778493059 | ns | NaCl | KS.3 |
| FIM1_3696 | 3016.325927 | 2990.273985 | 4593.752869 | -0.001711553 | 0.108472793 | 0.965859357 | 0.993691273 | ns | NaCl | KS.3 |
| FIM1_3697 | 10565.79362 | 10788.84195 | 11978.24542 | 0.00607139 | 0.098060978 | 0.889822981 | 0.977268396 | ns | NaCl | KS.3 |
| FIM1_3698 | 2805.024404 | 3091.181874 | 3455.448409 | 0.0193858 | 0.110628897 | 0.63269954 | 0.897798121 | ns | NaCl | KS.3 |
| FIM1_3699 | 81.79464485 | 99.05536023 | 1113.884538 | 0.021160751 | 0.134276321 | 0.557734408 | 0.863863225 | ns | NaCl | KS.3 |
| FIM1_3700 | 295.6393472 | 177.2404845 | 866.2872135 | -0.118296806 | 0.213934287 | 0.042398469 | 0.32401288 | ns | NaCl | KS.3 |
| FIM1_3701 | 858.0140975 | 850.0650328 | 989.2838633 | -0.001641235 | 0.113576192 | 0.965927838 | 0.993691273 | ns | NaCl | KS.3 |
| FIM1_3702 | 228.9337158 | 187.1226317 | 650.5353628 | -0.035269634 | 0.122921382 | 0.386094335 | 0.779278134 | ns | NaCl | KS.3 |
| FIM1_3703 | 1780.940335 | 1696.085855 | 2849.464825 | -0.009891454 | 0.108227606 | 0.806503863 | 0.957859861 | ns | NaCl | KS.3 |
| FIM1_3704 | 910.5960341 | 728.3513446 | 1314.613492 | -0.041300771 | 0.125361686 | 0.321285031 | 0.737180727 | ns | NaCl | KS.3 |
| FIM1_3705 | 2120.792258 | 2081.956395 | 2843.130502 | -0.003202551 | 0.112259415 | 0.934273434 | 0.985446192 | ns | NaCl | KS.3 |
| FIM1_3706 | 4220.575517 | 4626.306842 | 5158.641043 | 0.018185172 | 0.110503551 | 0.653052079 | 0.900747416 | ns | NaCl | KS.3 |
| FIM1_3707 | 454.6630069 | 447.7386336 | 787.9552835 | -0.002442512 | 0.113935271 | 0.949195112 | 0.987736646 | ns | NaCl | KS.3 |
| FIM1_3708 | 4728.493472 | 4255.546394 | 5903.164291 | -0.029823743 | 0.101939233 | 0.500500037 | 0.839793485 | ns | NaCl | KS.3 |
| FIM1_3709 | 2042.851848 | 2218.40686 | 2293.754854 | 0.015241941 | 0.111957038 | 0.701220519 | 0.923421278 | ns | NaCl | KS.3 |
| FIM1_3710 | 1454.12143 | 1502.555995 | 2787.408431 | 0.006326464 | 0.109132603 | 0.874021257 | 0.971243188 | ns | NaCl | KS.3 |
| FIM1_3711 | 1818.262209 | 1915.724053 | 3620.269437 | 0.009804838 | 0.110398681 | 0.80494446 | 0.957859861 | ns | NaCl | KS.3 |
| FIM1_3712 | 3055.465651 | 2549.912391 | 3898.949549 | -0.034973623 | 0.118544701 | 0.397629464 | 0.78625063 | ns | NaCl | KS.3 |
| FIM1_3713 | 2050.051571 | 2064.425327 | 3243.832749 | 0.001410794 | 0.107494031 | 0.972081023 | 0.995313014 | ns | NaCl | KS.3 |
| FIM1_3714 | 396.0247614 | 319.6605744 | 1059.112359 | -0.029018474 | 0.130429992 | 0.448104423 | 0.812914509 | ns | NaCl | KS.3 |
| FIM1_3715 | 269.4005036 | 236.0524656 | 396.9883538 | -0.018688893 | 0.122887246 | 0.619393018 | 0.892479944 | ns | NaCl | KS.3 |
| FIM1_3716 | 3311.028237 | 3313.110546 | 5631.398114 | 0.000130637 | 0.104600693 | 0.997476905 | 0.99924489 | ns | NaCl | KS.3 |
| FIM1_3717 | 7356.65212 | 6572.059311 | 8897.841964 | -0.028828241 | 0.105058708 | 0.505453397 | 0.84439816 | ns | NaCl | KS.3 |
| FIM1_3718 | 1226.595155 | 1079.480783 | 1894.578112 | -0.021288328 | 0.117619222 | 0.586232034 | 0.877936294 | ns | NaCl | KS.3 |
| FIM1_3719 | 767.0149967 | 502.2258153 | 1434.633716 | -0.076771798 | 0.170986071 | 0.10730354 | 0.498909937 | ns | NaCl | KS.3 |
| FIM1_3720 | 1091.816935 | 980.4318382 | 1754.689686 | -0.01865678 | 0.115054027 | 0.635231613 | 0.898009483 | ns | NaCl | KS.3 |
| FIM1_3722 | 90.16249139 | 85.32719091 | 144.6627501 | -0.009033437 | 0.118232599 | 0.80976913 | 0.958816322 | ns | NaCl | KS.3 |
| FIM1_3723 | 159.5133789 | 171.7008131 | 374.7185674 | 0.008767923 | 0.125837221 | 0.8067609 | 0.957859861 | ns | NaCl | KS.3 |
| FIM1_3725 | 80.00810559 | 101.6346326 | 110.7036884 | 0.023151125 | 0.142121883 | 0.512844938 | 0.847552971 | ns | NaCl | KS.3 |
| FIM1_3727 | 437.3812122 | 526.1461626 | 551.350287 | 0.023944058 | 0.129105641 | 0.522350125 | 0.853429187 | ns | NaCl | KS.3 |
| FIM1_3731 | 221.7687953 | 213.224303 | 113.1955211 | -0.005391763 | 0.118658452 | 0.884746355 | 0.975081831 | ns | NaCl | KS.3 |
| FIM1_3732 | 350.2171627 | 330.9553623 | 285.7986033 | -0.009455243 | 0.114926377 | 0.805479948 | 0.957859861 | ns | NaCl | KS.3 |
| FIM1_3733 | 921.6819455 | 1020.761302 | 1175.971791 | 0.045310187 | 0.089970171 | 0.351304723 | 0.756912594 | ns | NaCl | KS.3 |
| FIM1_3734 | 744.8182375 | 664.9413843 | 827.4544815 | -0.054747676 | 0.088929333 | 0.268511079 | 0.690146084 | ns | NaCl | KS.3 |

| | | | | | | | | | | |
|---|---|---|---|---|---|---|---|---|---|---|
| FIM1_3735 | 1502.000112 | 1463.450865 | 1708.862647 | -0.021669082 | 0.061627669 | 0.645264047 | 0.898899754 | ns | NaCl | KS.3 |
| FIM1_3736 | 1646.721145 | 1788.304196 | 1412.888503 | 0.034349445 | 0.090190747 | 0.472338946 | 0.822130792 | ns | NaCl | KS.3 |
| FIM1_3737 | 148.6702003 | 88.02149869 | 793.9904804 | -0.197434563 | 0.261563756 | 0.012020019 | 0.156134712 | ns | NaCl | KS.3 |
| FIM1_3738 | 245.9281169 | 238.0663132 | 451.205716 | -0.013593979 | 0.088947298 | 0.773056703 | 0.94938462 | ns | NaCl | KS.3 |
| FIM1_3739 | 1142.006601 | 677.8299897 | 1405.266701 | -0.382584302 | 0.30317374 | 0.001307857 | 0.041708911 | ns | NaCl | KS.3 |
| FIM1_3740 | 2608.476185 | 2894.569615 | 2403.374038 | 0.046181486 | 0.090104035 | 0.342545322 | 0.75073259 | ns | NaCl | KS.3 |
| FIM1_3741 | 2075.980407 | 1579.108671 | 3060.206979 | -0.216137479 | 0.133155072 | 0.001600319 | 0.047109988 | ns | NaCl | KS.3 |
| FIM1_3742 | 1754.348904 | 1623.49353 | 2198.695666 | -0.044321238 | 0.080385427 | 0.372616343 | 0.77466206 | ns | NaCl | KS.3 |
| FIM1_3743 | 3599.709156 | 3875.620418 | 3417.466024 | 0.055738071 | 0.069037457 | 0.257245814 | 0.680871076 | ns | NaCl | KS.3 |
| FIM1_3744 | 1224.899257 | 1177.709214 | 1593.700646 | -0.0213716 | 0.080917555 | 0.662612049 | 0.904388188 | ns | NaCl | KS.3 |
| FIM1_3745 | 14720.88093 | 18304.4461 | 12179.15248 | 0.179443387 | 0.096922459 | 0.002289302 | 0.058108556 | ns | NaCl | KS.3 |
| FIM1_3746 | 1776.246597 | 1769.117723 | 1963.194845 | -0.00205434 | 0.07951043 | 0.966507836 | 0.994082382 | ns | NaCl | KS.3 |
| FIM1_3747 | 24870.23662 | 21600.55098 | 16972.88969 | -0.062143502 | 0.094094478 | 0.209471523 | 0.633579568 | ns | NaCl | KS.3 |
| FIM1_3748 | 10039.38934 | 12862.37976 | 7906.555294 | 0.14367612 | 0.11967989 | 0.015492245 | 0.182493537 | ns | NaCl | KS.3 |
| FIM1_3749 | 4686.325693 | 4798.968527 | 4416.571733 | 0.019812801 | 0.062126255 | 0.674784897 | 0.912806589 | ns | NaCl | KS.3 |
| FIM1_3750 | 1996.243997 | 2660.15349 | 1375.601999 | 0.055444573 | 0.136183516 | 0.206041235 | 0.630415219 | ns | NaCl | KS.3 |
| FIM1_3751 | 1582.407249 | 1766.950304 | 1850.963617 | 0.086250489 | 0.070013346 | 0.08416222 | 0.450895561 | ns | NaCl | KS.3 |
| FIM1_3752 | 2005.789191 | 1782.666143 | 2341.916908 | -0.081358562 | 0.077443612 | 0.110960597 | 0.506448597 | ns | NaCl | KS.3 |
| FIM1_3753 | 2031.310406 | 3400.671273 | 2476.245096 | 0.651524005 | 0.188166153 | 4.93291042062 | 0.000681701 | ns | NaCl | KS.3 |
| FIM1_3754 | 2942.114898 | 3637.187604 | 2359.349226 | 0.051341014 | 0.118495189 | 0.250109486 | 0.673319638 | ns | NaCl | KS.3 |
| FIM1_3755 | 281.0714503 | 314.2470508 | 213.6750868 | 0.032083233 | 0.102652223 | 0.469401257 | 0.821764392 | ns | NaCl | KS.3 |
| FIM1_3756 | 32078.56295 | 36464.43918 | 22678.61819 | 0.074008026 | 0.084430407 | 0.147183855 | 0.562037426 | ns | NaCl | KS.3 |
| FIM1_3757 | 595.001073 | 596.4841434 | 660.2333306 | 0.00074385 | 0.080172608 | 0.987850625 | 0.999134858 | ns | NaCl | KS.3 |
| FIM1_3758 | 2272.375559 | 2177.592204 | 1542.398213 | -0.014759753 | 0.093865257 | 0.746384296 | 0.943274417 | ns | NaCl | KS.3 |
| FIM1_3759 | 14718.08382 | 12849.51872 | 9864.711821 | -0.03338507 | 0.108004596 | 0.439577968 | 0.808917498 | ns | NaCl | KS.3 |
| FIM1_3760 | 649.9207018 | 588.9247864 | 619.9495454 | -0.055623286 | 0.081696537 | 0.267115489 | 0.689593109 | ns | NaCl | KS.3 |
| FIM1_3761 | 729.463023 | 719.4462372 | 735.6103109 | -0.009006904 | 0.069402071 | 0.853550948 | 0.969345976 | ns | NaCl | KS.3 |
| FIM1_3762 | 2239.652096 | 2091.108596 | 1837.459971 | -0.029749877 | 0.088155222 | 0.535297283 | 0.855153333 | ns | NaCl | KS.3 |
| FIM1_3763 | 3202.012414 | 3585.58468 | 3123.577917 | 0.034796342 | 0.100378186 | 0.442100335 | 0.809513863 | ns | NaCl | KS.3 |
| FIM1_3764 | 391.6750645 | 378.0272975 | 462.6643805 | -0.020823452 | 0.07829392 | 0.672185001 | 0.912198685 | ns | NaCl | KS.3 |
| FIM1_3765 | 9218.118004 | 9064.597425 | 9241.215501 | -0.007395687 | 0.086320487 | 0.876849306 | 0.972783118 | ns | NaCl | KS.3 |
| FIM1_3766 | 77.53840868 | 65.93166022 | 98.51668961 | -0.039252578 | 0.10748537 | 0.374734334 | 0.777765254 | ns | NaCl | KS.3 |
| FIM1_3767 | 2322.21684 | 1929.973574 | 2270.164745 | -0.097402098 | 0.097470491 | 0.066076124 | 0.403405574 | ns | NaCl | KS.3 |
| FIM1_3768 | 34662.18935 | 38069.94359 | 26488.78546 | 0.012167368 | 0.123376382 | 0.740040993 | 0.940888181 | ns | NaCl | KS.3 |
| FIM1_3769 | 2478.756637 | 2638.277301 | 2223.519758 | 0.048691686 | 0.066894179 | 0.316992574 | 0.73382286 | ns | NaCl | KS.3 |
| FIM1_3770 | 7697.03108 | 8007.698881 | 6474.110678 | 0.041467811 | 0.049921888 | 0.323771915 | 0.739131368 | ns | NaCl | KS.3 |
| FIM1_3771 | 1510.16836 | 1546.332306 | 1381.914983 | 0.00987475 | 0.087930614 | 0.834764168 | 0.962365393 | ns | NaCl | KS.3 |
| FIM1_3772 | 2177.482635 | 2145.140667 | 2235.125208 | -0.005565115 | 0.08964993 | 0.905284615 | 0.979880338 | ns | NaCl | KS.3 |
| FIM1_3773 | 11030.26362 | 11088.77732 | 9253.453081 | 0.001863119 | 0.092695756 | 0.967504114 | 0.99408528 | ns | NaCl | KS.3 |
| FIM1_3774 | 3999.724806 | 2839.901196 | 4738.193851 | -0.233735052 | 0.182160674 | 0.002839438 | 0.066010301 | ns | NaCl | KS.3 |
| FIM1_3775 | 222.1406859 | 181.6100781 | 237.9796987 | -0.093397008 | 0.103402758 | 0.07568617 | 0.427638847 | ns | NaCl | KS.3 |
| FIM1_3776 | 207.7291489 | 188.7184823 | 418.9431956 | -0.038444477 | 0.093065573 | 0.417748793 | 0.800387073 | ns | NaCl | KS.3 |
| FIM1_3777 | 8160.83999 | 9076.032007 | 9059.563685 | 0.014995377 | 0.121258155 | 0.688817627 | 0.918490387 | ns | NaCl | KS.3 |
| FIM1_3778 | 259.0700794 | 301.7643513 | 395.1968135 | 0.032809411 | 0.11267709 | 0.435438114 | 0.808096352 | ns | NaCl | KS.3 |
| FIM1_3779 | 1525.808075 | 1389.534619 | 1354.296373 | -0.044980041 | 0.086909308 | 0.359435627 | 0.763021562 | ns | NaCl | KS.3 |

| | | | | | | | | | | |
|---|---|---|---|---|---|---|---|---|---|---|
| FIM1_3780 | 13605.28075 | 13812.69932 | 10131.22759 | 0.011733481 | 0.065651776 | 0.807380303 | 0.957859861 | ns | NaCl | KS.3 |
| FIM1_3782 | 2288.716199 | 2305.839854 | 3009.537331 | 0.003991029 | 0.080386699 | 0.934858636 | 0.985446192 | ns | NaCl | KS.3 |
| FIM1_3783 | 904.3363673 | 909.8963414 | 1425.301828 | 0.005213916 | 0.075726201 | 0.915545569 | 0.982343702 | ns | NaCl | KS.3 |
| FIM1_3784 | 3893.110429 | 3026.859549 | 3653.060382 | -0.190626628 | 0.118327688 | 0.002742744 | 0.064671708 | ns | NaCl | KS.3 |
| FIM1_3785 | 260.3875515 | 236.1102006 | 374.948203 | -0.026893663 | 0.100173774 | 0.545234123 | 0.857313452 | ns | NaCl | KS.3 |
| FIM1_3786 | 4384.352646 | 3971.338548 | 4454.690071 | -0.091404816 | 0.060909625 | 0.052900602 | 0.360032854 | ns | NaCl | KS.3 |
| FIM1_3787 | 56.46295618 | 76.84923571 | 84.05890081 | 0.095479086 | 0.143970072 | 0.069798458 | 0.414871362 | ns | NaCl | KS.3 |
| FIM1_3788 | 7590.177865 | 6860.129363 | 6370.812734 | -0.066530952 | 0.076631735 | 0.187232615 | 0.618483916 | ns | NaCl | KS.3 |
| FIM1_3789 | 576.1276635 | 626.1952409 | 690.6040231 | 0.043428129 | 0.083483324 | 0.379637391 | 0.778493059 | ns | NaCl | KS.3 |
| FIM1_3790 | 341.5741231 | 332.6597232 | 421.9346332 | -0.011196836 | 0.090945138 | 0.809607617 | 0.958816322 | ns | NaCl | KS.3 |
| FIM1_3791 | 3338.15066 | 3308.978379 | 3342.107276 | -0.008092556 | 0.059819982 | 0.86167984 | 0.969856868 | ns | NaCl | KS.3 |
| FIM1_3792 | 3687.191695 | 2732.580782 | 3126.315086 | -0.096276646 | 0.140306304 | 0.068395272 | 0.410454137 | ns | NaCl | KS.3 |
| FIM1_3793 | 7476.229161 | 6917.746769 | 6434.595813 | -0.042205886 | 0.081736352 | 0.394248689 | 0.781740625 | ns | NaCl | KS.3 |
| FIM1_3794 | 13621.04879 | 16053.0366 | 9539.717368 | 0.099445442 | 0.08899445 | 0.059986946 | 0.383100198 | ns | NaCl | KS.3 |
| FIM1_3795 | 696.157919 | 811.8622324 | 881.351855 | 0.045466248 | 0.105949232 | 0.317672548 | 0.733979075 | ns | NaCl | KS.3 |
| FIM1_3796 | 315.6395786 | 342.5669022 | 333.1994851 | 0.030793774 | 0.092950159 | 0.511168599 | 0.846104012 | ns | NaCl | KS.3 |
| FIM1_3797 | 3975.542182 | 4249.353987 | 4135.845504 | 0.034403253 | 0.083052118 | 0.483496358 | 0.829184191 | ns | NaCl | KS.3 |
| FIM1_3798 | 1787.780482 | 1700.310249 | 2050.789803 | -0.025699358 | 0.082145625 | 0.599441219 | 0.880219278 | ns | NaCl | KS.3 |
| FIM1_3799 | 15188.06192 | 11643.73397 | 14279.63957 | -0.125336176 | 0.12873348 | 0.029589347 | 0.271099454 | ns | NaCl | KS.3 |
| FIM1_3801 | 2706.806269 | 2424.612662 | 2398.246948 | -0.036011938 | 0.098457915 | 0.43267582 | 0.805901236 | ns | NaCl | KS.3 |
| FIM1_3802 | 1711.996334 | 2092.92297 | 2210.137681 | 0.039067875 | 0.121103384 | 0.349492435 | 0.756912594 | ns | NaCl | KS.3 |
| FIM1_3803 | 348.093984 | 297.8285976 | 313.849329 | -0.053108494 | 0.103953895 | 0.258535105 | 0.682403445 | ns | NaCl | KS.3 |
| FIM1_3804 | 122.4454739 | 84.12595187 | 257.7622274 | -0.202672026 | 0.199927456 | 0.007295872 | 0.117086974 | ns | NaCl | KS.3 |
| FIM1_3805 | 330.36433 | 311.2827567 | 471.9418112 | -0.023178065 | 0.090940353 | 0.621512561 | 0.892479944 | ns | NaCl | KS.3 |
| FIM1_3806 | 1155.058307 | 1221.183957 | 864.3943599 | 0.011714982 | 0.107323758 | 0.774001817 | 0.95007625 | ns | NaCl | KS.3 |
| FIM1_3807 | 12747.02912 | 14564.16292 | 10336.56045 | 0.135354349 | 0.058492601 | 0.003534546 | 0.0761228 | ns | NaCl | KS.3 |
| FIM1_3808 | 472.9622156 | 363.6758395 | 598.9609995 | -0.188956762 | 0.123023763 | 0.003290693 | 0.073413451 | ns | NaCl | KS.3 |
| FIM1_3809 | 3256.006869 | 4614.226439 | 2060.132763 | 0.066420616 | 0.150909688 | 0.146122239 | 0.560211748 | ns | NaCl | KS.3 |
| FIM1_3810 | 2567.306333 | 2717.364164 | 1956.994185 | 0.026381989 | 0.085678496 | 0.585408888 | 0.877826895 | ns | NaCl | KS.3 |
| FIM1_3811 | 4723.707575 | 6030.470262 | 5184.452121 | 0.12501595 | 0.118933625 | 0.02840528 | 0.263328375 | ns | NaCl | KS.3 |
| FIM1_3812 | 18.85416687 | 13.61744918 | 22.63589428 | -0.035916381 | 0.148670759 | 0.34066143 | 0.750043208 | ns | NaCl | KS.3 |
| FIM1_3813 | 5634.077174 | 5961.129082 | 4199.466434 | 0.046980394 | 0.063437613 | 0.325145289 | 0.739131368 | ns | NaCl | KS.3 |
| FIM1_3814 | 22.35962336 | 19.80016661 | 26.24877177 | -0.013750802 | 0.128842695 | 0.700991196 | 0.923421278 | ns | NaCl | KS.3 |
| FIM1_3815 | 11322.30206 | 12829.1705 | 13481.12748 | 0.029356239 | 0.108206308 | 0.490195671 | 0.834607619 | ns | NaCl | KS.3 |
| FIM1_3816 | 3171.442589 | 3898.168608 | 2349.079688 | 0.065964514 | 0.112618657 | 0.169724142 | 0.594663296 | ns | NaCl | KS.3 |
| FIM1_3817 | 2829.320541 | 1365.683787 | 3144.887224 | -1.002253825 | 0.134332091 | 3.5529077641[illegible] | 3.5351432253[illegible] | Km<KS.3 | NaCl | KS.3 |
| FIM1_3818 | 89.37415413 | 72.53291866 | 352.0551801 | -0.040534661 | 0.123938861 | 0.330599877 | 0.742254041 | ns | NaCl | KS.3 |
| FIM1_3819 | 1499.504176 | 1388.945843 | 2295.003418 | -0.025926166 | 0.095259513 | 0.571797303 | 0.870809186 | ns | NaCl | KS.3 |
| FIM1_3820 | 2791.724639 | 3501.637084 | 1699.424937 | 0.063012375 | 0.11910144 | 0.179221336 | 0.608009168 | ns | NaCl | KS.3 |
| FIM1_3821 | 1160.705738 | 1182.722832 | 934.0114174 | 0.006764401 | 0.092177427 | 0.883012302 | 0.97456556 | ns | NaCl | KS.3 |
| FIM1_3822 | 5074.237587 | 5099.928637 | 5347.315536 | 0.002881768 | 0.07900585 | 0.953081809 | 0.988653461 | ns | NaCl | KS.3 |
| FIM1_3823 | 131.8007466 | 207.8160733 | 134.5366798 | 0.399159114 | 0.258497761 | 0.000582717 | 0.025069192 | ns | NaCl | KS.3 |
| FIM1_3824 | 15578.45104 | 21591.74273 | 20601.80896 | 0.114180476 | 0.153161554 | 0.043481842 | 0.328757089 | ns | NaCl | KS.3 |
| FIM1_3825 | 204.1687502 | 145.4230346 | 253.0589104 | -0.063637554 | 0.149579654 | 0.157761589 | 0.57787696 | ns | NaCl | KS.3 |
| FIM1_3826 | 8874.144333 | 8398.673935 | 6236.632612 | -0.0531614 | 0.05500296 | 0.233135695 | 0.655652958 | ns | NaCl | KS.3 |

| | | | | | | | | | | |
|---|---|---|---|---|---|---|---|---|---|---|
| FIM1_3827 | 453.632342 | 488.9139578 | 477.8659757 | 0.031286357 | 0.087422966 | 0.516481301 | 0.849956041 | ns | NaCl | KS.3 |
| FIM1_3828 | 3800.175831 | 4269.966413 | 3076.068311 | 0.080846776 | 0.076403716 | 0.112266456 | 0.509139122 | ns | NaCl | KS.3 |
| FIM1_3829 | 2576.71637 | 2737.372978 | 2219.783123 | 0.037597725 | 0.075849972 | 0.448369329 | 0.812914509 | ns | NaCl | KS.3 |
| FIM1_3830 | 2811.260654 | 2584.758257 | 3249.216534 | -0.039839389 | 0.086311668 | 0.415005757 | 0.800387073 | ns | NaCl | KS.3 |
| FIM1_3831 | 339.6913682 | 216.5499666 | 740.3250707 | -0.364322673 | 0.260067568 | 0.000950843 | 0.034528782 | ns | NaCl | KS.3 |
| FIM1_3832 | 802.7902768 | 879.9345804 | 785.8791234 | 0.044623577 | 0.086221799 | 0.364001865 | 0.764991795 | ns | NaCl | KS.3 |
| FIM1_3833 | 51.5517382 | 37.07683865 | 255.9778003 | -0.05058335 | 0.148375663 | 0.224802604 | 0.648385059 | ns | NaCl | KS.3 |
| FIM1_3834 | 2803.019416 | 3158.580716 | 3235.127828 | 0.088324451 | 0.07433102 | 0.082352694 | 0.443883698 | ns | NaCl | KS.3 |
| FIM1_3835 | 353.5634276 | 332.8794977 | 381.0551814 | -0.031414622 | 0.083898381 | 0.520312314 | 0.851497948 | ns | NaCl | KS.3 |
| FIM1_3836 | 5828.319815 | 6331.239644 | 4396.370425 | 0.03696071 | 0.088171116 | 0.44491923 | 0.812581927 | ns | NaCl | KS.3 |
| FIM1_3837 | 2734.538783 | 2471.064034 | 2397.157586 | -0.064630919 | 0.078027589 | 0.200337167 | 0.624375388 | ns | NaCl | KS.3 |
| FIM1_3838 | 1849.281734 | 2100.74923 | 1857.047638 | 0.077409934 | 0.082633361 | 0.130707486 | 0.538035296 | ns | NaCl | KS.3 |
| FIM1_3839 | 1490.106992 | 1412.136305 | 2382.466326 | -0.013133817 | 0.103680067 | 0.755132571 | 0.944390281 | ns | NaCl | KS.3 |
| FIM1_3840 | 1483.566709 | 1534.643723 | 898.5308475 | 0.011124545 | 0.095021257 | 0.805147313 | 0.957859861 | ns | NaCl | KS.3 |
| FIM1_3841 | 1794.08561 | 2081.143119 | 2372.251608 | 0.04648438 | 0.103528565 | 0.313391378 | 0.731871738 | ns | NaCl | KS.3 |
| FIM1_3842 | 157.3878452 | 156.3727499 | 197.4601246 | -0.001122891 | 0.113452129 | 0.976702913 | 0.996164232 | ns | NaCl | KS.3 |
| FIM1_3843 | 16503.51628 | 18597.6836 | 11982.34472 | 0.111045173 | 0.062436246 | 0.020543131 | 0.216989551 | ns | NaCl | KS.3 |
| FIM1_3844 | 1413.080069 | 1631.433064 | 1232.990209 | 0.03251145 | 0.110905932 | 0.443197455 | 0.810627697 | ns | NaCl | KS.3 |
| FIM1_3845 | 1588.95458 | 1969.531782 | 1007.305287 | 0.053862684 | 0.118445017 | 0.232939495 | 0.655471713 | ns | NaCl | KS.3 |
| FIM1_3846 | 1007.890639 | 1293.370662 | 975.6377112 | 0.178498016 | 0.118545616 | 0.004393004 | 0.085706644 | ns | NaCl | KS.3 |
| FIM1_3847 | 45010.43521 | 52931.01964 | 31400.47508 | 0.139055623 | 0.074619316 | 0.007741403 | 0.120354622 | ns | NaCl | KS.3 |
| FIM1_3848 | 785.7611651 | 780.2205839 | 750.7664885 | -0.001729419 | 0.104278886 | 0.96670821 | 0.994082957 | ns | NaCl | KS.3 |
| FIM1_3849 | 47786.1023 | 46431.90577 | 30737.89021 | -0.013161533 | 0.085421812 | 0.784071695 | 0.951646805 | ns | NaCl | KS.3 |
| FIM1_3850 | 2745.36641 | 2303.7544 | 2505.267773 | -0.060333272 | 0.105152393 | 0.207993111 | 0.633002483 | ns | NaCl | KS.3 |
| FIM1_3851 | 848.7837722 | 886.6238852 | 1348.817359 | 0.019616742 | 0.084976805 | 0.684522855 | 0.91584821 | ns | NaCl | KS.3 |
| FIM1_3852 | 1497.218515 | 1643.367986 | 1300.114399 | 0.038955475 | 0.090717812 | 0.417022345 | 0.800387073 | ns | NaCl | KS.3 |
| FIM1_3853 | 1533.142035 | 1640.828753 | 1715.996936 | 0.037629326 | 0.080102717 | 0.447115222 | 0.812914509 | ns | NaCl | KS.3 |
| FIM1_3854 | 5858.979615 | 5218.013491 | 5777.754768 | -0.052852973 | 0.090156728 | 0.28242833 | 0.703948368 | ns | NaCl | KS.3 |
| FIM1_3855 | 3495.761679 | 3252.612038 | 3691.321226 | -0.040347515 | 0.080416266 | 0.415648663 | 0.800387073 | ns | NaCl | KS.3 |
| FIM1_3856 | 783.3877325 | 665.922992 | 1475.589319 | -0.086973466 | 0.092487898 | 0.094284993 | 0.470201108 | ns | NaCl | KS.3 |
| FIM1_3857 | 8917.988967 | 9559.365298 | 6700.010341 | 0.057414657 | 0.064263854 | 0.232348386 | 0.654993872 | ns | NaCl | KS.3 |
| FIM1_3858 | 761.9474035 | 775.1265907 | 1106.416192 | 0.009099692 | 0.078558433 | 0.85282107 | 0.969345976 | ns | NaCl | KS.3 |
| FIM1_3859 | 24.46871541 | 19.78486775 | 26.75156027 | -0.030758853 | 0.13032684 | 0.426038305 | 0.802854185 | ns | NaCl | KS.3 |
| FIM1_3860 | 1946.288663 | 1974.196618 | 2591.085474 | 0.003614158 | 0.100785003 | 0.932628284 | 0.985136881 | ns | NaCl | KS.3 |
| FIM1_3861 | 500.6396121 | 501.2966487 | 683.3851481 | 0.000287603 | 0.10125348 | 0.994606244 | 0.99924489 | ns | NaCl | KS.3 |
| FIM1_3862 | 3497.138471 | 3684.158951 | 4239.368737 | 0.00863711 | 0.114261027 | 0.822563276 | 0.960240659 | ns | NaCl | KS.3 |
| FIM1_3863 | 6879.41002 | 7107.78839 | 5834.400475 | 0.014449612 | 0.086399165 | 0.762581977 | 0.946287744 | ns | NaCl | KS.3 |
| FIM1_3864 | 82.27309378 | 16.65115417 | 239.0951616 | -1.803125264 | 0.434581357 | 2.0223546337( | 4.3744410011( | Km<KS.3 | NaCl | KS.3 |
| FIM1_3865 | 132.3364316 | 89.10492902 | 647.6237333 | -0.076128567 | 0.16302367 | 0.110919232 | 0.506448597 | ns | NaCl | KS.3 |
| FIM1_3866 | 1073.486581 | 1003.248019 | 1486.262765 | -0.030656802 | 0.08769744 | 0.524187495 | 0.854940852 | ns | NaCl | KS.3 |
| FIM1_3867 | 1777.084598 | 1841.327631 | 1536.73574 | 0.01562961 | 0.086327257 | 0.744085093 | 0.941460666 | ns | NaCl | KS.3 |
| FIM1_3868 | 199.2021542 | 201.5152558 | 508.917387 | 0.002233916 | 0.098027124 | 0.959301179 | 0.991590145 | ns | NaCl | KS.3 |
| FIM1_3869 | 21582.7421 | 21350.08723 | 15450.81622 | -0.003161853 | 0.097662175 | 0.942614235 | 0.986790704 | ns | NaCl | KS.3 |
| FIM1_3870 | 1214.97545 | 1041.445411 | 1812.154242 | -0.072647091 | 0.094918665 | 0.150434628 | 0.564839452 | ns | NaCl | KS.3 |
| FIM1_3871 | 3663.190738 | 4542.026813 | 2563.177529 | 0.061912036 | 0.116156809 | 0.187472109 | 0.618483916 | ns | NaCl | KS.3 |

| | | | | | | | | | | |
|---|---|---|---|---|---|---|---|---|---|---|
| FIM1_3872 | 5910.209488 | 6342.32986 | 5529.200178 | 0.056120848 | 0.066439467 | 0.248557001 | 0.67096552 | ns | NaCl | KS.3 |
| FIM1_3873 | 968.4360381 | 853.3058578 | 1453.231919 | -0.05101195 | 0.095057092 | 0.290232099 | 0.709776444 | ns | NaCl | KS.3 |
| FIM1_3874 | 323.8623202 | 230.6150319 | 510.1569601 | -0.266161296 | 0.180796233 | 0.001257648 | 0.041599773 | ns | NaCl | KS.3 |
| FIM1_3875 | 228.541683 | 138.1497983 | 283.7631909 | -0.623226859 | 0.204283318 | 1.2964421381 | 0.001490695 | ns | NaCl | KS.3 |
| FIM1_3876 | 47.89790936 | 27.20498785 | 99.06440972 | -0.210473189 | 0.301881169 | 0.011203661 | 0.150643816 | ns | NaCl | KS.3 |
| FIM1_3877 | 60.21021513 | 21.8373802 | 174.706943 | -1.162565096 | 0.372998478 | 7.6788562361 | 0.000979546 | Km<KS.3 | NaCl | KS.3 |
| FIM1_3879 | 11424.02088 | 10745.92491 | 20269.80526 | -0.016196421 | 0.101340382 | 0.707293882 | 0.925190614 | ns | NaCl | KS.3 |
| FIM1_3880 | 2942.356716 | 2821.841835 | 3181.816457 | -0.013990477 | 0.094373933 | 0.758128329 | 0.945250644 | ns | NaCl | KS.3 |
| FIM1_3881 | 6691.974099 | 6368.59489 | 6875.92404 | -0.049436687 | 0.052150727 | 0.252494909 | 0.676266095 | ns | NaCl | KS.3 |
| FIM1_3882 | 45.31976543 | 27.12334279 | 60.88332332 | -0.09325089 | 0.19395523 | 0.071040889 | 0.418675804 | ns | NaCl | KS.3 |
| FIM1_3883 | 228.1087946 | 175.3961996 | 422.477108 | -0.118063894 | 0.127087185 | 0.036488128 | 0.302547392 | ns | NaCl | KS.3 |
| FIM1_3884 | 283.9820233 | 281.648717 | 508.2338864 | -0.002719873 | 0.097562645 | 0.950661224 | 0.987736646 | ns | NaCl | KS.3 |
| FIM1_3885 | 2228.852423 | 2134.515003 | 2432.3808 | -0.0277863 | 0.075064533 | 0.573716428 | 0.871789624 | ns | NaCl | KS.3 |
| FIM1_3886 | 4236.043469 | 3559.774337 | 3862.400914 | -0.102899567 | 0.091753218 | 0.053455922 | 0.362364862 | ns | NaCl | KS.3 |
| FIM1_3887 | 270.4386309 | 260.2124928 | 388.6433825 | -0.021534114 | 0.080958643 | 0.660197636 | 0.903691294 | ns | NaCl | KS.3 |
| FIM1_3888 | 4209.732879 | 3438.497071 | 3161.601912 | -0.119329365 | 0.100423513 | 0.030750866 | 0.275153886 | ns | NaCl | KS.3 |
| FIM1_3889 | 971.6124223 | 721.8820863 | 1106.781719 | -0.120238796 | 0.142640462 | 0.036120519 | 0.301915458 | ns | NaCl | KS.3 |
| FIM1_3890 | 3236.439564 | 2522.600163 | 2590.63797 | -0.140180635 | 0.120148207 | 0.01749122 | 0.198220547 | ns | NaCl | KS.3 |
| FIM1_3891 | 273.4121057 | 142.0717758 | 393.9541856 | -0.62755951 | 0.354753619 | 0.000246965 | 0.014626776 | ns | NaCl | KS.3 |
| FIM1_3892 | 1095.066713 | 679.4258067 | 1681.82906 | -0.138983328 | 0.216227309 | 0.02872253 | 0.265110548 | ns | NaCl | KS.3 |
| FIM1_3893 | 143.031717 | 146.8156796 | 134.945167 | 0.008002649 | 0.094496922 | 0.859427669 | 0.969856868 | ns | NaCl | KS.3 |
| FIM1_3894 | 553.6013528 | 511.9034869 | 844.3097769 | -0.039659908 | 0.083820483 | 0.42037884 | 0.80133378 | ns | NaCl | KS.3 |
| FIM1_3895 | 181.3574596 | 154.9351869 | 166.2505611 | -0.048080518 | 0.105390812 | 0.29597694 | 0.714058464 | ns | NaCl | KS.3 |
| FIM1_3896 | 968.9713485 | 859.4074935 | 1224.149328 | -0.070281279 | 0.083266268 | 0.166865949 | 0.590859855 | ns | NaCl | KS.3 |
| FIM1_3897 | 613.0255985 | 631.5346232 | 509.3109975 | 0.015377212 | 0.084576995 | 0.750001485 | 0.943274417 | ns | NaCl | KS.3 |
| FIM1_3898 | 8354.322235 | 11481.97772 | 4862.051882 | 0.053879397 | 0.143414569 | 0.209256456 | 0.633579568 | ns | NaCl | KS.3 |
| FIM1_3899 | 703.5584969 | 751.0750544 | 974.2576244 | 0.041906395 | 0.073993762 | 0.398181908 | 0.786613521 | ns | NaCl | KS.3 |
| FIM1_3900 | 404.3850382 | 449.4269891 | 381.9129481 | 0.046764752 | 0.089794638 | 0.337437786 | 0.747375382 | ns | NaCl | KS.3 |
| FIM1_3901 | 1596.055462 | 1657.687882 | 1554.936371 | 0.019030687 | 0.082980416 | 0.695589119 | 0.921340753 | ns | NaCl | KS.3 |
| FIM1_3902 | 565.510822 | 571.3685382 | 1006.228109 | 0.00357891 | 0.096292497 | 0.9358183 | 0.985958501 | ns | NaCl | KS.3 |
| FIM1_3903 | 745.7043072 | 502.0559807 | 596.3936974 | -0.345658852 | 0.218260519 | 0.0005844 | 0.025069192 | ns | NaCl | KS.3 |
| FIM1_3904 | 10826.31897 | 11633.16939 | 7748.498376 | 0.042995457 | 0.07818013 | 0.38735257 | 0.779278134 | ns | NaCl | KS.3 |
| FIM1_3905 | 5033.718419 | 6662.865562 | 6693.299249 | 0.126167714 | 0.135580922 | 0.029813878 | 0.271655755 | ns | NaCl | KS.3 |
| FIM1_3906 | 374.8351654 | 432.7520537 | 747.9196432 | 0.056474655 | 0.096918154 | 0.244586131 | 0.66821307 | ns | NaCl | KS.3 |
| FIM1_3907 | 1069.13185 | 1061.969744 | 1759.155748 | -0.004019618 | 0.075883091 | 0.934832846 | 0.985446192 | ns | NaCl | KS.3 |
| FIM1_3908 | 1423.600572 | 1525.688661 | 1524.870191 | 0.057794017 | 0.063986461 | 0.228529241 | 0.652285125 | ns | NaCl | KS.3 |
| FIM1_3909 | 1869.132363 | 2079.942004 | 1174.845268 | 0.022899856 | 0.109274617 | 0.579925338 | 0.8749481 | ns | NaCl | KS.3 |
| FIM1_3910 | 8144.115926 | 8366.183626 | 6264.664992 | 0.021273311 | 0.065284062 | 0.658136049 | 0.902985891 | ns | NaCl | KS.3 |
| FIM1_3911 | 141.4700747 | 74.92955879 | 182.5380654 | -0.535227476 | 0.360725296 | 0.000532185 | 0.02370281 | ns | NaCl | KS.3 |
| FIM1_3912 | 1479.410653 | 1721.390956 | 4257.191824 | 0.042341906 | 0.106715817 | 0.34528701 | 0.754462295 | ns | NaCl | KS.3 |
| FIM1_3913 | 359.9833083 | 506.6356158 | 592.3381723 | 0.176093416 | 0.174798816 | 0.010515572 | 0.145724155 | ns | NaCl | KS.3 |
| FIM1_3914 | 437.6409398 | 454.3394769 | 465.4991353 | 0.014317905 | 0.090003398 | 0.759914999 | 0.945853671 | ns | NaCl | KS.3 |
| FIM1_3915 | 3766.432491 | 3454.570047 | 2899.055429 | -0.059194266 | 0.074244609 | 0.236670358 | 0.659616806 | ns | NaCl | KS.3 |
| FIM1_3916 | 732.0716365 | 715.3748553 | 852.7608378 | -0.01010527 | 0.089110884 | 0.829790741 | 0.961438115 | ns | NaCl | KS.3 |
| FIM1_3917 | 2099.750616 | 1915.595805 | 1846.392987 | -0.041724837 | 0.088473683 | 0.390832656 | 0.779482559 | ns | NaCl | KS.3 |

| | | | | | | | | | | |
|---|---|---|---|---|---|---|---|---|---|---|
| FIM1_3918 | 885.8464625 | 988.7155948 | 1266.487463 | 0.02886817 | 0.104089514 | 0.507493761 | 0.844423736 | ns | NaCl | KS.3 |
| FIM1_3919 | 111.5065128 | 115.5309707 | 331.3315058 | 0.005965432 | 0.097777796 | 0.89198733 | 0.977268396 | ns | NaCl | KS.3 |
| FIM1_3920 | 163.5634816 | 170.2529537 | 231.2274604 | 0.013697723 | 0.093332667 | 0.764748738 | 0.947366775 | ns | NaCl | KS.3 |
| FIM1_3921 | 2640.853767 | 2630.221095 | 2798.983204 | -0.002123709 | 0.079599402 | 0.965370935 | 0.993691273 | ns | NaCl | KS.3 |
| FIM1_3922 | 497.1214605 | 392.3169966 | 500.5854168 | -0.110119021 | 0.11780191 | 0.045163625 | 0.335858052 | ns | NaCl | KS.3 |
| FIM1_3923 | 4598.65863 | 3932.085976 | 3006.232563 | -0.093913793 | 0.087809922 | 0.073232152 | 0.421631393 | ns | NaCl | KS.3 |
| FIM1_3924 | 1342.265297 | 1831.220736 | 705.9763253 | 0.043412674 | 0.143416854 | 0.28006489 | 0.700652348 | ns | NaCl | KS.3 |
| FIM1_3925 | 73.70289095 | 43.3967069 | 212.0267322 | -0.184691172 | 0.256971707 | 0.014441523 | 0.1741395 | ns | NaCl | KS.3 |
| FIM1_3926 | 1329.276297 | 1410.207353 | 1880.232738 | 0.029628753 | 0.084307973 | 0.543210736 | 0.857313452 | ns | NaCl | KS.3 |
| FIM1_3927 | 198.7016157 | 128.6507351 | 370.1716768 | -0.238069115 | 0.234367074 | 0.005116685 | 0.094234281 | ns | NaCl | KS.3 |
| FIM1_3928 | 2241.513087 | 2690.789138 | 2189.577524 | 0.087099691 | 0.099416053 | 0.093269905 | 0.470201108 | ns | NaCl | KS.3 |
| FIM1_3929 | 1680.088731 | 1978.846079 | 1378.074944 | 0.059912152 | 0.101904852 | 0.214558508 | 0.639561759 | ns | NaCl | KS.3 |
| FIM1_3930 | 1129.458485 | 1103.312545 | 1070.974216 | -0.017002415 | 0.071380166 | 0.728928206 | 0.935336397 | ns | NaCl | KS.3 |
| FIM1_3932 | 1714.375447 | 1743.306888 | 2228.120119 | 0.009707141 | 0.07487494 | 0.843563889 | 0.966543148 | ns | NaCl | KS.3 |
| FIM1_3933 | 462.6148552 | 485.7382426 | 529.5501752 | 0.022775997 | 0.08470846 | 0.638132054 | 0.898332475 | ns | NaCl | KS.3 |
| FIM1_3934 | 952.3004739 | 994.70178 | 1094.394944 | 0.025770395 | 0.076869167 | 0.601570853 | 0.880762097 | ns | NaCl | KS.3 |
| FIM1_3935 | 1568.424961 | 1527.693478 | 1625.570104 | -0.011376 | 0.086014301 | 0.812120518 | 0.958816322 | ns | NaCl | KS.3 |
| FIM1_3936 | 590.2757416 | 600.2634042 | 891.8872015 | 0.008625238 | 0.073206142 | 0.860670922 | 0.969856868 | ns | NaCl | KS.3 |
| FIM1_3937 | 9.998489668 | 11.05472631 | 34.38314948 | 0.003785127 | 0.14212109 | 0.90773503 | 0.980825899 | ns | NaCl | KS.3 |
| FIM1_3938 | 63.69875078 | 108.7014555 | 80.08171728 | 0.169876753 | 0.247937827 | 0.017865363 | 0.201086384 | ns | NaCl | KS.3 |
| FIM1_3939 | 344.1519232 | 537.3426606 | 588.5738225 | 0.167151165 | 0.21915855 | 0.017020035 | 0.194207964 | ns | NaCl | KS.3 |
| FIM1_3940 | 2087.114588 | 3029.234365 | 2416.895088 | 0.084048117 | 0.160691677 | 0.091779635 | 0.467388224 | ns | NaCl | KS.3 |
| FIM1_3942 | 363.7577081 | 466.2186308 | 686.7445039 | 0.162965742 | 0.117479175 | 0.00765517 | 0.119762487 | ns | NaCl | KS.3 |
| FIM1_3943 | 4937.056593 | 5745.222883 | 4597.163711 | 0.129059878 | 0.072687459 | 0.012180523 | 0.157397666 | ns | NaCl | KS.3 |
| FIM1_3944 | 927.0602527 | 923.1731285 | 950.884262 | -0.004701857 | 0.072683768 | 0.923712144 | 0.983924964 | ns | NaCl | KS.3 |
| FIM1_3945 | 1279.937852 | 1273.622235 | 915.1070056 | -0.001396342 | 0.100362278 | 0.974025057 | 0.995638927 | ns | NaCl | KS.3 |
| FIM1_3946 | 2025.721426 | 2280.740221 | 1684.428242 | 0.01993407 | 0.116691072 | 0.610348889 | 0.887342408 | ns | NaCl | KS.3 |
| FIM1_3947 | 1229.314236 | 1118.45366 | 1402.901195 | -0.049189439 | 0.084503883 | 0.321481446 | 0.737180727 | ns | NaCl | KS.3 |
| FIM1_3948 | 1057.189052 | 1158.9328 | 1624.501317 | 0.066194381 | 0.072182498 | 0.184941585 | 0.615853002 | ns | NaCl | KS.3 |
| FIM1_3949 | 1002.413075 | 1035.353547 | 1524.536322 | 0.021913803 | 0.071571013 | 0.655501267 | 0.901110473 | ns | NaCl | KS.3 |
| FIM1_3950 | 1235.254667 | 1171.612309 | 1387.801633 | -0.039213216 | 0.070356684 | 0.425231021 | 0.802854185 | ns | NaCl | KS.3 |
| FIM1_3951 | 2053.553432 | 1867.556468 | 2008.560255 | -0.086537025 | 0.06068083 | 0.066166623 | 0.403405574 | ns | NaCl | KS.3 |
| FIM1_3952 | 1447.654251 | 1575.001431 | 1720.543302 | 0.014928635 | 0.113330411 | 0.704199879 | 0.924470947 | ns | NaCl | KS.3 |
| FIM1_3953 | 1374.898814 | 1457.300516 | 1806.123057 | 0.032654615 | 0.079244793 | 0.508643438 | 0.844548078 | ns | NaCl | KS.3 |
| FIM1_3954 | 735.3921827 | 593.3454467 | 635.473512 | -0.108061354 | 0.106106367 | 0.046845557 | 0.341725286 | ns | NaCl | KS.3 |
| FIM1_3955 | 470.8567501 | 466.7933734 | 717.4408983 | -0.003542804 | 0.095509401 | 0.936875915 | 0.986375069 | ns | NaCl | KS.3 |
| FIM1_3956 | 5532.646099 | 6602.688154 | 3147.383048 | 0.046579217 | 0.111202687 | 0.298108804 | 0.716123274 | ns | NaCl | KS.3 |
| FIM1_3957 | 2150.623948 | 2404.598649 | 2442.72559 | 0.108282278 | 0.05932719 | 0.020346809 | 0.215832354 | ns | NaCl | KS.3 |
| FIM1_3958 | 6960.67793 | 8683.244277 | 3999.83523 | 0.043330865 | 0.123792979 | 0.305534931 | 0.724521545 | ns | NaCl | KS.3 |
| FIM1_3959 | 504.9343055 | 564.3433017 | 783.6072442 | 0.063520476 | 0.083045412 | 0.208243864 | 0.633002483 | ns | NaCl | KS.3 |
| FIM1_3960 | 298.8912364 | 363.7185283 | 230.82045 | 0.046043935 | 0.116139186 | 0.294286358 | 0.712862678 | ns | NaCl | KS.3 |
| FIM1_3961 | 857.8784049 | 852.5819828 | 941.5538678 | -0.001161511 | 0.113197278 | 0.975947832 | 0.996164232 | ns | NaCl | KS.3 |
| FIM1_3962 | 1170.683186 | 1236.239113 | 1344.310423 | 0.028741314 | 0.082211755 | 0.55760138 | 0.863863225 | ns | NaCl | KS.3 |
| FIM1_3963 | 1202.879722 | 1471.846488 | 1670.37721 | 0.047400987 | 0.117333154 | 0.281358318 | 0.701984771 | ns | NaCl | KS.3 |
| FIM1_3964 | 2490.961082 | 2888.025849 | 2477.98014 | 0.026472414 | 0.117211605 | 0.508479166 | 0.844548078 | ns | NaCl | KS.3 |

| | | | | | | | | | | |
|---|---|---|---|---|---|---|---|---|---|---|
| FIM1_3965 | 633.9240083 | 534.5490172 | 578.3806888 | -0.088316584 | 0.095363901 | 0.089970333 | 0.463573758 | ns | NaCl | KS.3 |
| FIM1_3966 | 312.1483034 | 226.8418954 | 404.0608576 | -0.035340754 | 0.147232589 | 0.348465768 | 0.756654396 | ns | NaCl | KS.3 |
| FIM1_3967 | 1553.297832 | 1736.291676 | 1412.758355 | 0.045711622 | 0.092650203 | 0.342513809 | 0.75073259 | ns | NaCl | KS.3 |
| FIM1_3968 | 1268.068731 | 1303.282429 | 1315.968476 | 0.015075547 | 0.079601679 | 0.758423442 | 0.945250644 | ns | NaCl | KS.3 |
| FIM1_3969 | 2459.331926 | 2263.783499 | 2221.040879 | -0.032479014 | 0.091911307 | 0.49190917 | 0.835238267 | ns | NaCl | KS.3 |
| FIM1_3970 | 993.3424361 | 1054.975526 | 906.8312738 | 0.016660529 | 0.099917936 | 0.702733091 | 0.923421278 | ns | NaCl | KS.3 |
| FIM1_3971 | 174.0235147 | 225.6488524 | 324.094424 | 0.049967442 | 0.130589345 | 0.245507135 | 0.66889266 | ns | NaCl | KS.3 |
| FIM1_3972 | 338.3616801 | 290.1607497 | 550.1518357 | -0.089388841 | 0.088374713 | 0.086381563 | 0.45668467 | ns | NaCl | KS.3 |
| FIM1_3973 | 8710.354958 | 10108.19788 | 6210.686297 | 0.092136468 | 0.085232791 | 0.077420442 | 0.432155137 | ns | NaCl | KS.3 |
| FIM1_3974 | 1159.53475 | 1200.957054 | 1268.56398 | 0.0175758 | 0.084126028 | 0.716491757 | 0.929477573 | ns | NaCl | KS.3 |
| FIM1_3975 | 492.0046061 | 447.8467094 | 1007.307858 | -0.042396004 | 0.089082816 | 0.382633797 | 0.778665419 | ns | NaCl | KS.3 |
| FIM1_3976 | 8384.438698 | 7582.297202 | 8974.615497 | -0.040786446 | 0.092109482 | 0.393968533 | 0.781740625 | ns | NaCl | KS.3 |
| FIM1_3977 | 170.4123841 | 111.7916255 | 1321.053844 | -0.062763432 | 0.167054145 | 0.152704679 | 0.568642049 | ns | NaCl | KS.3 |
| FIM1_3978 | 1366.752582 | 1457.826049 | 1124.950898 | 0.025909961 | 0.090148025 | 0.583881004 | 0.877414869 | ns | NaCl | KS.3 |
| FIM1_3979 | 710.0168348 | 743.6554977 | 703.252064 | 0.028733441 | 0.075717589 | 0.560925191 | 0.865840156 | ns | NaCl | KS.3 |
| FIM1_3980 | 20130.52646 | 26179.91081 | 15468.15665 | 0.200787083 | 0.125482507 | 0.002256607 | 0.058108556 | ns | NaCl | KS.3 |
| FIM1_3981 | 149.7994487 | 117.383466 | 506.7927928 | -0.039972271 | 0.129298099 | 0.327769306 | 0.740319311 | ns | NaCl | KS.3 |
| FIM1_3982 | 3534.326693 | 2920.296008 | 3579.231815 | -0.128091233 | 0.092866561 | 0.020313749 | 0.215832354 | ns | NaCl | KS.3 |
| FIM1_3983 | 1914.652043 | 1798.887921 | 1834.252194 | -0.034554885 | 0.08088256 | 0.48370764 | 0.829184191 | ns | NaCl | KS.3 |
| FIM1_3984 | 312.9116836 | 252.3080296 | 288.2195062 | -0.065056065 | 0.115833793 | 0.171626464 | 0.5990025 | ns | NaCl | KS.3 |
| FIM1_3985 | 615.0749974 | 553.2389076 | 708.1430085 | -0.079537464 | 0.075010717 | 0.116611616 | 0.519941265 | ns | NaCl | KS.3 |
| FIM1_3986 | 1814.226706 | 1748.903316 | 1607.990702 | -0.011657238 | 0.095971205 | 0.794603674 | 0.955299664 | ns | NaCl | KS.3 |
| FIM1_3987 | 772.386318 | 731.2256545 | 787.8348508 | -0.030750934 | 0.07965774 | 0.533021735 | 0.855153333 | ns | NaCl | KS.3 |
| FIM1_3988 | 720.8246883 | 672.3039126 | 1013.687429 | -0.037940296 | 0.082689903 | 0.441365283 | 0.809513863 | ns | NaCl | KS.3 |
| FIM1_3989 | 11220.75451 | 12050.34154 | 10098.21849 | 0.049290416 | 0.072643027 | 0.320631288 | 0.737029267 | ns | NaCl | KS.3 |
| FIM1_3990 | 967.7039939 | 821.3067542 | 1567.976856 | -0.037102255 | 0.113092919 | 0.385291826 | 0.778665419 | ns | NaCl | KS.3 |
| FIM1_3991 | 1936.671054 | 2725.32039 | 1565.42951 | 0.094085967 | 0.154147296 | 0.072092447 | 0.420358296 | ns | NaCl | KS.3 |
| FIM1_3992 | 614.5707557 | 702.5373309 | 725.4346143 | 0.094911191 | 0.07740787 | 0.065515514 | 0.402056108 | ns | NaCl | KS.3 |
| FIM1_3993 | 10620.25719 | 11722.04039 | 8151.152732 | 0.112758914 | 0.044911514 | 0.003843815 | 0.079679077 | ns | NaCl | KS.3 |
| FIM1_3994 | 54.02330839 | 47.05161447 | 83.69121574 | -0.02982832 | 0.113715567 | 0.470598929 | 0.822127061 | ns | NaCl | KS.3 |
| FIM1_3995 | 155.1388042 | 159.036732 | 185.7070297 | 0.003766508 | 0.110171296 | 0.923999484 | 0.983924964 | ns | NaCl | KS.3 |
| FIM1_3996 | 2089.495083 | 2606.160255 | 1615.367025 | 0.038268781 | 0.126183519 | 0.349024571 | 0.756912594 | ns | NaCl | KS.3 |
| FIM1_3997 | 1612.205592 | 2058.412316 | 1901.564487 | 0.16073468 | 0.116826909 | 0.00821351 | 0.125927317 | ns | NaCl | KS.3 |
| FIM1_3998 | 848.5626678 | 920.1024245 | 1173.428708 | 0.032510131 | 0.090869372 | 0.493851574 | 0.836358994 | ns | NaCl | KS.3 |
| FIM1_3999 | 252.8727713 | 253.7966399 | 239.0508598 | -0.000197257 | 0.102767629 | 0.996250851 | 0.99924489 | ns | NaCl | KS.3 |
| FIM1_4000 | 712.7066471 | 1141.976752 | 866.1570562 | 0.546184464 | 0.215005079 | 3.89754509616 | 0.003658545 | ns | NaCl | KS.3 |
| FIM1_4001 | 145.7267747 | 118.2909375 | 118.8632034 | -0.054272114 | 0.117947447 | 0.230841054 | 0.654378488 | ns | NaCl | KS.3 |
| FIM1_4002 | 5984.041968 | 6752.303391 | 4722.895656 | 0.10351248 | 0.067158846 | 0.036405602 | 0.302547392 | ns | NaCl | KS.3 |
| FIM1_4003 | 5137.102524 | 4599.908882 | 4266.578467 | -0.073076542 | 0.07759828 | 0.149912992 | 0.564839452 | ns | NaCl | KS.3 |
| FIM1_4004 | 1225.68253 | 1215.217807 | 1181.977227 | -0.006581325 | 0.071820833 | 0.893257476 | 0.977268396 | ns | NaCl | KS.3 |
| FIM1_4005 | 2067.93955 | 2207.684748 | 2922.143101 | 0.026907464 | 0.089845087 | 0.570553327 | 0.870665206 | ns | NaCl | KS.3 |
| FIM1_4006 | 2894.620055 | 2839.635534 | 2646.572787 | -0.01426195 | 0.065696565 | 0.767048669 | 0.948794469 | ns | NaCl | KS.3 |
| FIM1_4007 | 1696.046788 | 1597.724855 | 1713.896372 | -0.049163145 | 0.064780761 | 0.307157269 | 0.726977836 | ns | NaCl | KS.3 |
| FIM1_4008 | 803.4161985 | 843.1839404 | 533.3020394 | 0.013296235 | 0.100564563 | 0.758307896 | 0.945250644 | ns | NaCl | KS.3 |
| FIM1_4009 | 4615.002291 | 5230.034312 | 3342.353643 | 0.056900545 | 0.091431539 | 0.248830429 | 0.67096552 | ns | NaCl | KS.3 |

| | | | | | | | | | | |
|---|---|---|---|---|---|---|---|---|---|---|
| FIM1_4010 | 1476.551494 | 1560.832338 | 1854.404496 | 0.042315017 | 0.067614529 | 0.385334971 | 0.778665419 | ns | NaCl | KS.3 |
| FIM1_4011 | 1690.946852 | 1625.342422 | 1968.013867 | -0.03685022 | 0.056860308 | 0.416609061 | 0.800387073 | ns | NaCl | KS.3 |
| FIM1_4012 | 2678.141797 | 2824.422473 | 2395.085632 | 0.032886725 | 0.075936908 | 0.50640318 | 0.844423736 | ns | NaCl | KS.3 |
| FIM1_4013 | 2153.381922 | 2224.803365 | 1792.394935 | 0.028031815 | 0.060166007 | 0.547529481 | 0.859115631 | ns | NaCl | KS.3 |
| FIM1_4014 | 310.3260005 | 300.9212882 | 366.1548634 | -0.00915122 | 0.09583553 | 0.837918332 | 0.962829707 | ns | NaCl | KS.3 |
| FIM1_4015 | 2198.295505 | 2244.502402 | 2846.782905 | 0.00721107 | 0.092916639 | 0.874687601 | 0.971766596 | ns | NaCl | KS.3 |
| FIM1_4016 | 2517.493418 | 2552.193748 | 2731.732684 | 0.006984583 | 0.083949207 | 0.884923511 | 0.975081831 | ns | NaCl | KS.3 |
| FIM1_4017 | 529.406393 | 573.2577412 | 389.0880982 | 0.02560492 | 0.097336872 | 0.57105137 | 0.870665206 | ns | NaCl | KS.3 |
| FIM1_4018 | 7127.970725 | 9591.468331 | 4265.881815 | 0.05060276 | 0.139088867 | 0.232867073 | 0.655471713 | ns | NaCl | KS.3 |
| FIM1_4019 | 739.230486 | 672.5913032 | 802.5732301 | -0.054617162 | 0.08067313 | 0.27572207 | 0.695949924 | ns | NaCl | KS.3 |
| FIM1_4020 | 4752.983365 | 5166.492746 | 3839.847189 | 0.06763384 | 0.066250798 | 0.165144639 | 0.589355333 | ns | NaCl | KS.3 |
| FIM1_4021 | 1025.603719 | 986.3257712 | 1069.033636 | -0.019675734 | 0.083603394 | 0.685250666 | 0.916430662 | ns | NaCl | KS.3 |
| FIM1_4022 | 42.01290334 | 37.91531446 | 31.43776702 | -0.013756064 | 0.123625832 | 0.708569671 | 0.925190614 | ns | NaCl | KS.3 |
| FIM1_4023 | 8579.760118 | 8127.24568 | 7102.690264 | -0.055879552 | 0.050376299 | 0.186238862 | 0.618044911 | ns | NaCl | KS.3 |
| FIM1_4024 | 2482.64831 | 2894.329247 | 2905.728058 | 0.024031132 | 0.121312318 | 0.535705008 | 0.855153333 | ns | NaCl | KS.3 |
| FIM1_4025 | 206.6478442 | 185.7128738 | 257.0080608 | -0.033019692 | 0.100713563 | 0.462831311 | 0.819880744 | ns | NaCl | KS.3 |
| FIM1_4026 | 171.6613345 | 163.9116573 | 370.4821554 | -0.006494272 | 0.120110423 | 0.860333308 | 0.969856868 | ns | NaCl | KS.3 |
| FIM1_4027 | 255.2232913 | 209.8194763 | 621.9376762 | -0.026322445 | 0.129212207 | 0.487104638 | 0.831051295 | ns | NaCl | KS.3 |
| FIM1_4028 | 1412.579872 | 1637.572792 | 1191.808455 | 0.115117632 | 0.07590943 | 0.026619959 | 0.254519471 | ns | NaCl | KS.3 |
| FIM1_4029 | 90.47614161 | 116.9560275 | 104.9791132 | 0.047003843 | 0.131764245 | 0.265480937 | 0.689593109 | ns | NaCl | KS.3 |
| FIM1_4030 | 9775.635028 | 13283.34294 | 5795.456378 | 0.069668475 | 0.140527568 | 0.138211436 | 0.547974001 | ns | NaCl | KS.3 |
| FIM1_4031 | 4533.507311 | 4363.492155 | 3294.908715 | -0.020782437 | 0.080129626 | 0.671836228 | 0.912198685 | ns | NaCl | KS.3 |
| FIM1_4033 | 749.3078834 | 742.5117023 | 824.2126454 | -0.004171564 | 0.083183918 | 0.931300097 | 0.984325045 | ns | NaCl | KS.3 |
| FIM1_4034 | 878.9841006 | 1002.647281 | 554.5582895 | 0.056168368 | 0.094116854 | 0.250787509 | 0.674415058 | ns | NaCl | KS.3 |
| FIM1_4035 | 715.2684847 | 767.7430494 | 436.1044789 | 0.024837023 | 0.094565648 | 0.589112065 | 0.878638624 | ns | NaCl | KS.3 |
| FIM1_4036 | 2114.695198 | 2176.476514 | 1895.044124 | 0.013673534 | 0.084565281 | 0.776775818 | 0.950432783 | ns | NaCl | KS.3 |
| FIM1_4037 | 1211.477081 | 1127.079063 | 1354.565124 | -0.04390836 | 0.077779217 | 0.377646618 | 0.778493059 | ns | NaCl | KS.3 |
| FIM1_4038 | 885.514602 | 785.8294136 | 775.7880711 | -0.06942304 | 0.082782148 | 0.171715339 | 0.5990025 | ns | NaCl | KS.3 |
| FIM1_4039 | 1585.535269 | 1688.425926 | 1193.073608 | 0.02873598 | 0.086552042 | 0.551741326 | 0.860945181 | ns | NaCl | KS.3 |
| FIM1_4041 | 1899.425176 | 2165.204138 | 1151.064925 | 0.022129574 | 0.117448211 | 0.573151159 | 0.871729446 | ns | NaCl | KS.3 |
| FIM1_4042 | 1763.667068 | 1766.308924 | 1666.45966 | 0.000597846 | 0.075630199 | 0.990296833 | 0.999134858 | ns | NaCl | KS.3 |
| FIM1_4043 | 1592.141036 | 1536.620285 | 1573.467723 | -0.014682941 | 0.088559107 | 0.75613307 | 0.944867505 | ns | NaCl | KS.3 |
| FIM1_4044 | 285.8324073 | 347.6770491 | 280.2021216 | 0.031166232 | 0.12501847 | 0.430311682 | 0.805047964 | ns | NaCl | KS.3 |
| FIM1_4045 | 5882.709201 | 7055.569638 | 4361.899764 | 0.078181369 | 0.101706246 | 0.123274414 | 0.529611582 | ns | NaCl | KS.3 |
| FIM1_4046 | 1646.707581 | 1577.73691 | 1270.611571 | -0.015681522 | 0.092208395 | 0.734397059 | 0.938375543 | ns | NaCl | KS.3 |
| FIM1_4047 | 10520.69695 | 10586.43556 | 7888.412633 | 0.00330489 | 0.080636939 | 0.946000411 | 0.987278589 | ns | NaCl | KS.3 |
| FIM1_4048 | 491.1984637 | 640.8364425 | 405.4900098 | 0.095266423 | 0.128650356 | 0.070513673 | 0.417739615 | ns | NaCl | KS.3 |
| FIM1_4049 | 2733.82785 | 3066.488652 | 1563.26928 | 0.016590264 | 0.120971454 | 0.660009166 | 0.903691294 | ns | NaCl | KS.3 |
| FIM1_4050 | 3569.36489 | 4056.953817 | 3856.464842 | 0.110682582 | 0.067781068 | 0.026128629 | 0.254383425 | ns | NaCl | KS.3 |
| FIM1_4051 | 4497.852681 | 5890.932448 | 8292.48385 | 0.226565596 | 0.128410358 | 0.000940092 | 0.034389391 | ns | NaCl | KS.3 |
| FIM1_4052 | 73.47375294 | 77.39811101 | 75.04869013 | 0.009697564 | 0.104379255 | 0.815925632 | 0.958816322 | ns | NaCl | KS.3 |
| FIM1_4053 | 422.3772515 | 334.7478552 | 749.4921098 | -0.032576415 | 0.131950913 | 0.401388618 | 0.789189152 | ns | NaCl | KS.3 |
| FIM1_4054 | 2328.748687 | 2335.818694 | 2907.120408 | 0.001065201 | 0.090348374 | 0.981730799 | 0.996363429 | ns | NaCl | KS.3 |
| FIM1_4055 | 1243.382215 | 1085.215646 | 1475.51348 | -0.038630952 | 0.104190956 | 0.388998381 | 0.779482559 | ns | NaCl | KS.3 |
| FIM1_4056 | 1018.722461 | 1062.557408 | 1623.584067 | 0.019760091 | 0.084440829 | 0.683028026 | 0.915673519 | ns | NaCl | KS.3 |

| | | | | | | | | | | |
|---|---|---|---|---|---|---|---|---|---|---|
| FIM1_4057 | 1061.118464 | 1028.340836 | 1189.612241 | -0.019371703 | 0.075611913 | 0.694233235 | 0.920525145 | ns | NaCl | KS.3 |
| FIM1_4058 | 459.7869365 | 370.5488983 | 385.20337 | -0.093199094 | 0.109250708 | 0.075872494 | 0.427766945 | ns | NaCl | KS.3 |
| FIM1_4059 | 3918.417164 | 5662.494473 | 4790.05668 | 0.090117311 | 0.161168447 | 0.079128217 | 0.435950031 | ns | NaCl | KS.3 |
| FIM1_4060 | 4164.117241 | 4877.710861 | 2445.498945 | 0.035950815 | 0.112186283 | 0.399927231 | 0.787975436 | ns | NaCl | KS.3 |
| FIM1_4061 | 9785.269093 | 13983.66999 | 5649.129174 | 0.091174368 | 0.158161495 | 0.077300683 | 0.43210213 | ns | NaCl | KS.3 |
| FIM1_4062 | 2637.164977 | 3593.068216 | 1445.784866 | 0.057421944 | 0.141278382 | 0.191506553 | 0.618952432 | ns | NaCl | KS.3 |
| FIM1_4063 | 2047.037769 | 1908.737372 | 2330.080871 | -0.030346782 | 0.088264272 | 0.527194923 | 0.855153333 | ns | NaCl | KS.3 |
| FIM1_4064 | 7883.203176 | 8477.506635 | 5527.055317 | 0.02464453 | 0.095283731 | 0.590126416 | 0.87877872 | ns | NaCl | KS.3 |
| FIM1_4065 | 7490.386621 | 9821.570761 | 4374.277099 | 0.053072767 | 0.132685593 | 0.223123291 | 0.648059715 | ns | NaCl | KS.3 |
| FIM1_4066 | 2618.319573 | 3260.179644 | 3727.910755 | 0.154662055 | 0.103019222 | 0.008117316 | 0.125415053 | ns | NaCl | KS.3 |
| FIM1_4067 | 877.8323115 | 906.8157419 | 1068.184694 | 0.019114983 | 0.075483439 | 0.698067986 | 0.922074642 | ns | NaCl | KS.3 |
| FIM1_4068 | 441.534216 | 419.1465667 | 406.8654862 | -0.017545328 | 0.094004096 | 0.701170518 | 0.923421278 | ns | NaCl | KS.3 |
| FIM1_4069 | 2314.281894 | 2432.365315 | 2644.230048 | 0.010828446 | 0.106204437 | 0.792210727 | 0.954759779 | ns | NaCl | KS.3 |
| FIM1_4070 | 3283.948123 | 2033.597042 | 2799.421974 | -0.186533311 | 0.239424395 | 0.013111181 | 0.165678479 | ns | NaCl | KS.3 |
| FIM1_4071 | 116.5229103 | 94.9784318 | 102.4320302 | -0.042398408 | 0.119874235 | 0.319917025 | 0.736735876 | ns | NaCl | KS.3 |
| FIM1_4072 | 1480.698248 | 1309.832769 | 1604.04108 | -0.079664625 | 0.080284382 | 0.119852627 | 0.524421123 | ns | NaCl | KS.3 |
| FIM1_4073 | 938.4415887 | 863.6007274 | 1025.178863 | -0.04496844 | 0.081392929 | 0.365347044 | 0.765950925 | ns | NaCl | KS.3 |
| FIM1_4074 | 1779.485718 | 1569.549186 | 1295.36021 | -0.021480606 | 0.116468572 | 0.585578295 | 0.877826895 | ns | NaCl | KS.3 |
| FIM1_4075 | 335.1275961 | 218.5588117 | 728.1756904 | -0.131510411 | 0.195328753 | 0.031975714 | 0.283253921 | ns | NaCl | KS.3 |
| FIM1_4076 | 11.20892417 | 12.66866842 | 18.61852671 | 0.009682292 | 0.139367166 | 0.772946188 | 0.94938462 | ns | NaCl | KS.3 |
| FIM1_4077 | 43327.03326 | 61582.52176 | 36444.03178 | 0.082003744 | 0.154400956 | 0.097471557 | 0.476631548 | ns | NaCl | KS.3 |
| FIM1_4078 | 99.12055996 | 182.361615 | 253.8581896 | 0.096317777 | 0.2209219 | 0.063289218 | 0.396553977 | ns | NaCl | KS.3 |
| FIM1_4080 | 1794.82835 | 1899.438374 | 3950.898547 | 0.010095071 | 0.112333402 | 0.79655013 | 0.955821731 | ns | NaCl | KS.3 |
| FIM1_4081 | 25.21573321 | 26.05577473 | 26.3944966 | 0.006055388 | 0.119998796 | 0.869689017 | 0.971243188 | ns | NaCl | KS.3 |
| FIM1_4082 | 48.65619749 | 73.88758597 | 53.98004836 | 0.068482896 | 0.168418886 | 0.131946623 | 0.539053419 | ns | NaCl | KS.3 |
| FIM1_4083 | 769.2865631 | 750.0222411 | 621.9285562 | -0.010714349 | 0.086549952 | 0.82233926 | 0.960240659 | ns | NaCl | KS.3 |
| FIM1_4084 | 4570.250101 | 5107.318002 | 3451.86931 | 0.053476969 | 0.088206946 | 0.279720488 | 0.700652348 | ns | NaCl | KS.3 |
| FIM1_4085 | 13756.9449 | 14439.81465 | 9779.713209 | 0.0298097 | 0.076137659 | 0.546514944 | 0.858513371 | ns | NaCl | KS.3 |
| FIM1_4086 | 19361.7058 | 25447.57935 | 11459.64232 | 0.086057377 | 0.130953259 | 0.090841809 | 0.464798331 | ns | NaCl | KS.3 |
| FIM1_4087 | 379.4848132 | 330.0003284 | 342.0848434 | -0.02617228 | 0.115732928 | 0.516030366 | 0.849801744 | ns | NaCl | KS.3 |
| FIM1_4088 | 2363.011902 | 2463.165962 | 2166.353819 | 0.008919453 | 0.106448836 | 0.827401171 | 0.960240659 | ns | NaCl | KS.3 |
| FIM1_4089 | 166.2280264 | 137.2798893 | 243.3551311 | -0.06808325 | 0.105989894 | 0.164682234 | 0.589224032 | ns | NaCl | KS.3 |
| FIM1_4090 | 173.9995799 | 179.1564096 | 225.2102565 | 0.015029035 | 0.086421188 | 0.753407667 | 0.943523399 | ns | NaCl | KS.3 |
| FIM1_4091 | 4848.044956 | 6953.989002 | 2847.784754 | 0.066484785 | 0.153773035 | 0.144678569 | 0.558679104 | ns | NaCl | KS.3 |
| FIM1_4092 | 1227.789765 | 1064.021006 | 1310.075233 | -0.098420703 | 0.080412111 | 0.058744056 | 0.378457602 | ns | NaCl | KS.3 |
| FIM1_4093 | 899.1810531 | 1099.411303 | 1490.982194 | 0.03707233 | 0.122207929 | 0.367953076 | 0.768177319 | ns | NaCl | KS.3 |
| FIM1_4094 | 5407.598359 | 10879.06843 | 6504.902137 | 0.660814776 | 0.37528266 | 0.000237578 | 0.014414029 | ns | NaCl | KS.3 |
| FIM1_4095 | 1734.267963 | 2306.581366 | 1717.111691 | 0.149404093 | 0.13960966 | 0.015614314 | 0.182568377 | ns | NaCl | KS.3 |
| FIM1_4096 | 2124.180528 | 2168.272851 | 1803.497877 | 0.008212649 | 0.088777729 | 0.8614901 | 0.969856868 | ns | NaCl | KS.3 |
| FIM1_4097 | 3089.761563 | 3931.299249 | 1921.574245 | 0.034976909 | 0.132439637 | 0.373515906 | 0.775559947 | ns | NaCl | KS.3 |
| FIM1_4098 | 1561.280091 | 1582.188149 | 1806.243203 | 0.003422939 | 0.100619392 | 0.936272765 | 0.986228458 | ns | NaCl | KS.3 |
| FIM1_4099 | 392.5897126 | 284.6512608 | 365.0170713 | -0.131047743 | 0.154967816 | 0.028423585 | 0.263328375 | ns | NaCl | KS.3 |
| FIM1_4100 | 288.0584592 | 260.0489113 | 354.4049164 | -0.052197863 | 0.085767066 | 0.2932153 | 0.71211096 | ns | NaCl | KS.3 |
| FIM1_4101 | 521.2803029 | 524.2917244 | 806.4622174 | 0.001604393 | 0.0966274 | 0.971109457 | 0.995112162 | ns | NaCl | KS.3 |
| FIM1_4102 | 1446.722697 | 1216.013866 | 2062.510701 | -0.095789696 | 0.093958937 | 0.069519478 | 0.413948267 | ns | NaCl | KS.3 |

| | | | | | | | | | | |
|---|---|---|---|---|---|---|---|---|---|---|
| FIM1_4103 | 2623.471151 | 2293.074242 | 4905.46933 | -0.042626967 | 0.101661519 | 0.353581405 | 0.757528931 | ns | NaCl | KS.3 |
| FIM1_4104 | 1603.322863 | 1937.279953 | 1512.017908 | 0.046627482 | 0.114519429 | 0.292070231 | 0.71186075 | ns | NaCl | KS.3 |
| FIM1_4105 | 197.5296971 | 279.8908075 | 221.2919914 | 0.098453987 | 0.157110867 | 0.064652128 | 0.399942992 | ns | NaCl | KS.3 |
| FIM1_4106 | 12131.50063 | 17249.03431 | 8180.424686 | 0.107121405 | 0.160768247 | 0.052378822 | 0.358438293 | ns | NaCl | KS.3 |
| FIM1_4107 | 1431.75142 | 1408.110181 | 1193.939476 | -0.010668373 | 0.073894442 | 0.828259078 | 0.960734183 | ns | NaCl | KS.3 |
| FIM1_4108 | 3485.878933 | 3623.068126 | 2811.311319 | 0.031144513 | 0.063921358 | 0.514524134 | 0.849077129 | ns | NaCl | KS.3 |
| FIM1_4109 | 316.4302683 | 311.8702969 | 326.6748471 | -0.004521501 | 0.098882166 | 0.917186578 | 0.982343702 | ns | NaCl | KS.3 |
| FIM1_4110 | 1133.048528 | 1182.357004 | 929.8709923 | 0.020381925 | 0.084732302 | 0.673366306 | 0.91246166 | ns | NaCl | KS.3 |
| FIM1_4111 | 2697.449087 | 2813.682986 | 2452.532085 | 0.042176168 | 0.053150296 | 0.334400215 | 0.745307602 | ns | NaCl | KS.3 |
| FIM1_4112 | 3024.373119 | 2663.443071 | 2371.194331 | -0.03351508 | 0.105314864 | 0.444756346 | 0.812581927 | ns | NaCl | KS.3 |
| FIM1_4113 | 2098.183201 | 982.4026333 | 1778.14431 | -0.607327155 | 0.431907997 | 0.000521216 | 0.02370281 | ns | NaCl | KS.3 |
| FIM1_4114 | 3132.383164 | 4887.522968 | 2048.426331 | 0.069063874 | 0.172684096 | 0.128482547 | 0.536136415 | ns | NaCl | KS.3 |
| FIM1_4115 | 1025.543815 | 1122.639347 | 1095.168122 | 0.030035961 | 0.096896473 | 0.51118644 | 0.846104012 | ns | NaCl | KS.3 |
| FIM1_4116 | 260.3922717 | 381.7975125 | 254.7718731 | 0.069845297 | 0.159460245 | 0.130750688 | 0.538035296 | ns | NaCl | KS.3 |
| FIM1_4117 | 163.096303 | 215.2231238 | 180.9317383 | 0.043284062 | 0.136653471 | 0.288919214 | 0.708763851 | ns | NaCl | KS.3 |
| FIM1_4118 | 1954.318882 | 1987.064249 | 2095.015767 | 0.003605254 | 0.106082656 | 0.929635308 | 0.984238275 | ns | NaCl | KS.3 |
| FIM1_4119 | 2297.061444 | 2617.211157 | 1782.112594 | 0.06486589 | 0.08917407 | 0.19664195 | 0.619173229 | ns | NaCl | KS.3 |
| FIM1_4120 | 4228.088073 | 5584.438948 | 2779.77814 | 0.042401273 | 0.137020776 | 0.295830263 | 0.714058464 | ns | NaCl | KS.3 |
| FIM1_4121 | 1315.088786 | 1452.814551 | 988.6484663 | 0.020086787 | 0.110528739 | 0.621424475 | 0.892479944 | ns | NaCl | KS.3 |
| FIM1_4122 | 493.5348052 | 429.9722008 | 644.6217503 | -0.060603054 | 0.093677618 | 0.219997935 | 0.642906874 | ns | NaCl | KS.3 |
| FIM1_4123 | 890.5346415 | 799.684146 | 1041.902857 | -0.044827255 | 0.092194925 | 0.35202991 | 0.756912594 | ns | NaCl | KS.3 |
| FIM1_4124 | 55.8023392 | 48.43521371 | 61.31069651 | -0.022432572 | 0.117947487 | 0.567316654 | 0.868965626 | ns | NaCl | KS.3 |
| FIM1_4125 | 9103.744791 | 6735.311011 | 7126.221652 | -0.135440392 | 0.146163649 | 0.024256969 | 0.242041091 | ns | NaCl | KS.3 |
| FIM1_4126 | 207.7812442 | 370.4192513 | 369.1405331 | 0.741334081 | 0.187491147 | 1.60429385868 | 0.000266045 | ns | NaCl | KS.3 |
| FIM1_4127 | 36.03774995 | 29.01185857 | 56.92117253 | -0.035751603 | 0.133214458 | 0.364031663 | 0.764991795 | ns | NaCl | KS.3 |
| FIM1_4128 | 1682.817645 | 1853.281633 | 1315.209878 | 0.046815716 | 0.086676648 | 0.341583978 | 0.750617902 | ns | NaCl | KS.3 |
| FIM1_4129 | 3330.810804 | 3452.215092 | 2444.613147 | 0.015076749 | 0.087842452 | 0.750850601 | 0.943274417 | ns | NaCl | KS.3 |
| FIM1_4130 | 1437.633159 | 1611.319215 | 2108.407292 | 0.037655806 | 0.098577909 | 0.413508167 | 0.8001568 | ns | NaCl | KS.3 |
| FIM1_4132 | 130.6086118 | 133.4167543 | 198.7253864 | 0.011479316 | 0.090768932 | 0.805159325 | 0.957859861 | ns | NaCl | KS.3 |
| FIM1_4133 | 544.7743026 | 414.9240051 | 883.5089162 | -0.147631791 | 0.132937464 | 0.015516536 | 0.182493537 | ns | NaCl | KS.3 |
| FIM1_4134 | 1645.636935 | 1465.3117 | 1743.048971 | -0.069948078 | 0.081604848 | 0.168637531 | 0.592944764 | ns | NaCl | KS.3 |
| FIM1_4135 | 929.7962707 | 996.5443748 | 1187.428661 | 0.030071013 | 0.087940565 | 0.53144542 | 0.855153333 | ns | NaCl | KS.3 |
| FIM1_4136 | 2228.503765 | 2368.5766 | 2361.138264 | 0.04031737 | 0.074387216 | 0.416239435 | 0.800387073 | ns | NaCl | KS.3 |
| FIM1_4137 | 3153.089238 | 3510.481153 | 3197.217042 | 0.034100915 | 0.099151323 | 0.45366526 | 0.816716052 | ns | NaCl | KS.3 |
| FIM1_4138 | 1443.601344 | 1634.283435 | 1404.198415 | 0.086905359 | 0.077306909 | 0.089820692 | 0.463545584 | ns | NaCl | KS.3 |
| FIM1_4139 | 5581.693516 | 7341.275555 | 3528.830738 | 0.070929446 | 0.131494239 | 0.137208797 | 0.545297021 | ns | NaCl | KS.3 |
| FIM1_4140 | 1229.65281 | 1228.54482 | 1085.210523 | -0.000254201 | 0.102142256 | 0.995195116 | 0.99924489 | ns | NaCl | KS.3 |
| FIM1_4141 | 12755.54002 | 11994.09498 | 8667.610626 | -0.029730982 | 0.084801484 | 0.541161192 | 0.857165292 | ns | NaCl | KS.3 |
| FIM1_4142 | 783.3698814 | 760.3134684 | 850.6829663 | -0.017787744 | 0.077131042 | 0.717836803 | 0.93000992 | ns | NaCl | KS.3 |
| FIM1_4143 | 859.1264864 | 1007.647375 | 875.8348652 | 0.099689876 | 0.08658619 | 0.05868209 | 0.378457602 | ns | NaCl | KS.3 |
| FIM1_4144 | 400.3697049 | 299.2046931 | 395.2838946 | -0.117493183 | 0.140295018 | 0.038650384 | 0.311585109 | ns | NaCl | KS.3 |
| FIM1_4145 | 314.1310206 | 262.5719907 | 511.0689433 | -0.063861563 | 0.10602373 | 0.186593659 | 0.618044911 | ns | NaCl | KS.3 |
| FIM1_4146 | 2570.620459 | 3379.217913 | 1621.817164 | 0.034942026 | 0.139226874 | 0.363821253 | 0.764991795 | ns | NaCl | KS.3 |
| FIM1_4147 | 815.0242154 | 718.0115906 | 825.2580876 | -0.042517303 | 0.099286327 | 0.360020843 | 0.763371684 | ns | NaCl | KS.3 |
| FIM1_4148 | 2498.892727 | 2763.39942 | 3486.418128 | 0.019626012 | 0.111429898 | 0.626908526 | 0.895712211 | ns | NaCl | KS.3 |

| | | | | | | | | | | |
|---|---|---|---|---|---|---|---|---|---|---|
| FIM1_4149 | 16057.33994 | 13758.03392 | 12080.15282 | -0.105382893 | 0.082971755 | 0.045544743 | 0.338186712 | ns | NaCl | KS.3 |
| FIM1_4150 | 2565.677941 | 2253.508807 | 2153.571342 | -0.059053863 | 0.091891784 | 0.232649707 | 0.655397674 | ns | NaCl | KS.3 |
| FIM1_4151 | 2507.952982 | 2068.770868 | 3026.452796 | -0.106455309 | 0.098647126 | 0.048347286 | 0.344656093 | ns | NaCl | KS.3 |
| FIM1_4152 | 3565.340351 | 4370.382628 | 1951.519265 | 0.031929036 | 0.126163891 | 0.418901605 | 0.800936005 | ns | NaCl | KS.3 |
| FIM1_4153 | 1364.881097 | 1335.860481 | 1687.778482 | -0.008075423 | 0.089688772 | 0.862991113 | 0.969856868 | ns | NaCl | KS.3 |
| FIM1_4154 | 647.6199427 | 454.2080352 | 796.973279 | -0.144424175 | 0.172083425 | 0.022102549 | 0.228385325 | ns | NaCl | KS.3 |
| FIM1_4155 | 5738.509745 | 5414.222654 | 3767.699817 | -0.016321134 | 0.099732156 | 0.708768244 | 0.925190614 | ns | NaCl | KS.3 |
| FIM1_4156 | 5589.921776 | 7678.887227 | 3625.250613 | 0.072438374 | 0.143619495 | 0.127390862 | 0.535278326 | ns | NaCl | KS.3 |
| FIM1_4157 | 3471.957506 | 3836.495015 | 3012.997044 | 0.032592156 | 0.097991266 | 0.475346981 | 0.82427718 | ns | NaCl | KS.3 |
| FIM1_4158 | 5070.827354 | 5356.097291 | 4488.575145 | 0.017572047 | 0.096009671 | 0.696522708 | 0.921625594 | ns | NaCl | KS.3 |
| FIM1_4159 | 1119.359057 | 1121.512397 | 1271.519633 | 0.001112149 | 0.076440997 | 0.981943697 | 0.996363429 | ns | NaCl | KS.3 |
| FIM1_4160 | 324.650237 | 324.1431031 | 355.4111926 | -0.000324302 | 0.089714155 | 0.994461201 | 0.99924489 | ns | NaCl | KS.3 |
| FIM1_4161 | 19462.11305 | 20231.96679 | 15530.23963 | 0.027773937 | 0.069807531 | 0.570594954 | 0.870665206 | ns | NaCl | KS.3 |
| FIM1_4162 | 1521.309303 | 1256.145165 | 1415.571465 | -0.026719871 | 0.127762152 | 0.484076397 | 0.829184191 | ns | NaCl | KS.3 |
| FIM1_4163 | 7298.727248 | 9597.493327 | 7457.914414 | 0.207065206 | 0.133162901 | 0.002203011 | 0.057845441 | ns | NaCl | KS.3 |
| FIM1_4164 | 69.59878902 | 39.81412661 | 109.5779923 | -0.097442094 | 0.211985536 | 0.063256149 | 0.396553977 | ns | NaCl | KS.3 |
| FIM1_4165 | 378.2640233 | 555.5338683 | 424.6697933 | 0.268476793 | 0.212922962 | 0.002209173 | 0.057845441 | ns | NaCl | KS.3 |
| FIM1_4166 | 926.6694029 | 1088.102591 | 2248.026681 | 0.037340028 | 0.112180036 | 0.384675103 | 0.778665419 | ns | NaCl | KS.3 |
| FIM1_4167 | 832.239285 | 802.0791203 | 1085.647549 | -0.005771879 | 0.115629295 | 0.879216726 | 0.973320697 | ns | NaCl | KS.3 |
| FIM1_4169 | 648.4844121 | 560.6441634 | 672.9160416 | -0.100350789 | 0.07934714 | 0.053696599 | 0.362470256 | ns | NaCl | KS.3 |
| FIM1_4170 | 636.9082896 | 336.1774799 | 738.4157999 | -0.839909579 | 0.180419056 | 1.58761032582 | 3.59016425954 | ns | NaCl | KS.3 |
| FIM1_4171 | 40.23012597 | 20.2118911 | 117.2484734 | -0.21218843 | 0.339526424 | 0.011348605 | 0.151642712 | ns | NaCl | KS.3 |
| FIM1_4172 | 755.1675478 | 747.1222071 | 1125.987572 | -0.002563761 | 0.103041822 | 0.951205281 | 0.987736646 | ns | NaCl | KS.3 |
| FIM1_4173 | 928.1641288 | 844.0876406 | 1128.570396 | -0.053543563 | 0.081827757 | 0.284442571 | 0.705083104 | ns | NaCl | KS.3 |
| FIM1_4174 | 9608.660814 | 11977.59677 | 5769.751469 | 0.050529916 | 0.120869272 | 0.252835163 | 0.676266095 | ns | NaCl | KS.3 |
| FIM1_4175 | 1520.904006 | 1425.117158 | 1627.710598 | -0.035058437 | 0.081335787 | 0.477135369 | 0.825770272 | ns | NaCl | KS.3 |
| FIM1_4176 | 1181.784922 | 1028.658662 | 1316.289622 | -0.096971378 | 0.08010485 | 0.062061276 | 0.390501742 | ns | NaCl | KS.3 |
| FIM1_4177 | 7788.674633 | 9943.380694 | 6783.080192 | 0.114515199 | 0.119334263 | 0.039591418 | 0.315147688 | ns | NaCl | KS.3 |
| FIM1_4178 | 6925.143148 | 9062.711561 | 6196.614027 | 0.15674996 | 0.131047815 | 0.011443136 | 0.151642712 | ns | NaCl | KS.3 |
| FIM1_4180 | 23.57193114 | 23.61183915 | 24.9312206 | 0.000897172 | 0.124583216 | 0.979968397 | 0.996363429 | ns | NaCl | KS.3 |
| FIM1_4181 | 2631.93868 | 2809.156073 | 3689.600522 | 0.030104073 | 0.086580738 | 0.533432681 | 0.855153333 | ns | NaCl | KS.3 |
| FIM1_4182 | 13073.90598 | 12137.39246 | 8869.187302 | -0.063658396 | 0.063080927 | 0.182658165 | 0.611396712 | ns | NaCl | KS.3 |
| FIM1_4183 | 1546.676556 | 1650.452208 | 1556.276746 | 0.041825226 | 0.075335676 | 0.399767586 | 0.787972955 | ns | NaCl | KS.3 |
| FIM1_4184 | 2202.935414 | 2516.401153 | 2060.641967 | 0.101106258 | 0.075145456 | 0.049002588 | 0.34611662 | ns | NaCl | KS.3 |
| FIM1_4185 | 692.187547 | 760.65424 | 760.5632707 | 0.049952358 | 0.083060209 | 0.315630844 | 0.73382286 | ns | NaCl | KS.3 |
| FIM1_4186 | 1827.039246 | 1090.713248 | 2235.521867 | -0.341733558 | 0.299321066 | 0.002050464 | 0.055440543 | ns | NaCl | KS.3 |
| FIM1_4187 | 559.0839834 | 607.4410904 | 693.7005976 | 0.039431629 | 0.087030426 | 0.418453624 | 0.800387073 | ns | NaCl | KS.3 |
| FIM1_4188 | 1634.742199 | 1710.817747 | 1297.856177 | 0.018321256 | 0.089520781 | 0.697473473 | 0.922074642 | ns | NaCl | KS.3 |
| FIM1_4189 | 3102.081998 | 2485.789224 | 3247.958278 | -0.098237835 | 0.111767295 | 0.064794784 | 0.399942992 | ns | NaCl | KS.3 |
| FIM1_4190 | 1772.363538 | 1778.176674 | 1947.08494 | 0.00179598 | 0.076678158 | 0.970840672 | 0.995041685 | ns | NaCl | KS.3 |
| FIM1_4191 | 1857.399042 | 1860.329835 | 2019.30619 | 0.001129427 | 0.083065901 | 0.981383179 | 0.996363429 | ns | NaCl | KS.3 |
| FIM1_4192 | 1307.938805 | 1682.236826 | 923.868796 | 0.073037404 | 0.124875304 | 0.132188841 | 0.539053419 | ns | NaCl | KS.3 |
| FIM1_4193 | 250.0533654 | 216.1101457 | 265.2306088 | -0.039067718 | 0.10714278 | 0.377451425 | 0.778493059 | ns | NaCl | KS.3 |
| FIM1_4194 | 946.1567319 | 815.670015 | 1161.170013 | -0.05485447 | 0.099519336 | 0.252564887 | 0.676266095 | ns | NaCl | KS.3 |
| FIM1_4195 | 2301.352585 | 2370.77571 | 2259.970138 | 0.01758852 | 0.077161377 | 0.720843765 | 0.931688886 | ns | NaCl | KS.3 |

| | | | | | | | | | | |
|---|---|---|---|---|---|---|---|---|---|---|
| FIM1_4196 | 2329.746996 | 2098.654988 | 3731.145569 | -0.037036964 | 0.096093997 | 0.426682706 | 0.802854185 | ns | NaCl | KS.3 |
| FIM1_4197 | 853.2634679 | 562.8941775 | 1354.845331 | -0.13553331 | 0.194364226 | 0.029324711 | 0.269170553 | ns | NaCl | KS.3 |
| FIM1_4198 | 1083.308102 | 949.3473992 | 1718.026555 | -0.035329184 | 0.105783667 | 0.422048386 | 0.801548921 | ns | NaCl | KS.3 |
| FIM1_4199 | 276.4400716 | 280.0635585 | 370.175666 | 0.003057822 | 0.099251905 | 0.943728382 | 0.986976813 | ns | NaCl | KS.3 |
| FIM1_4201 | 363.2323199 | 391.0263174 | 544.8274497 | 0.046508449 | 0.07264302 | 0.348173474 | 0.756654396 | ns | NaCl | KS.3 |
| FIM1_4203 | 327.8147824 | 274.7572946 | 324.5406144 | -0.046873063 | 0.111325819 | 0.295419656 | 0.714058464 | ns | NaCl | KS.3 |
| FIM1_4204 | 678.2314391 | 600.3008185 | 1148.355635 | -0.058736634 | 0.090057181 | 0.236912263 | 0.659616806 | ns | NaCl | KS.3 |
| FIM1_4205 | 2033.941877 | 2897.738844 | 3643.20249 | 0.061602759 | 0.151605723 | 0.165354715 | 0.589355333 | ns | NaCl | KS.3 |
| FIM1_4206 | 2079.988755 | 2069.719806 | 1485.64639 | -0.0010435 | 0.104386606 | 0.979887141 | 0.996363429 | ns | NaCl | KS.3 |
| FIM1_4207 | 2004.238414 | 1789.808811 | 2188.568411 | -0.041353427 | 0.095990116 | 0.379449791 | 0.778493059 | ns | NaCl | KS.3 |
| FIM1_4208 | 668.6194084 | 651.1187672 | 1001.676204 | -0.008210572 | 0.095752933 | 0.854388001 | 0.969345976 | ns | NaCl | KS.3 |
| FIM1_4209 | 517.4677463 | 385.5845684 | 664.3409912 | -0.183534323 | 0.147985256 | 0.006289515 | 0.106776808 | ns | NaCl | KS.3 |
| FIM1_4210 | 2152.574865 | 2393.974717 | 2048.586944 | 0.043370997 | 0.092570641 | 0.365913095 | 0.766169043 | ns | NaCl | KS.3 |
| FIM1_4211 | 834.5920222 | 1046.706089 | 725.4670799 | 0.07060378 | 0.117547404 | 0.145363062 | 0.559737797 | ns | NaCl | KS.3 |
| FIM1_4212 | 3349.810234 | 3396.523789 | 3234.851481 | 0.010306855 | 0.067557396 | 0.831729046 | 0.961620266 | ns | NaCl | KS.3 |
| FIM1_4213 | 333.3753623 | 347.5773125 | 423.953216 | 0.015724674 | 0.091046048 | 0.735706333 | 0.938575431 | ns | NaCl | KS.3 |
| FIM1_4214 | 1137.097703 | 1148.758298 | 1128.506859 | 0.006257066 | 0.08013529 | 0.898125445 | 0.977971161 | ns | NaCl | KS.3 |
| FIM1_4215 | 5766.12325 | 5665.283744 | 5631.325258 | -0.003598902 | 0.107085381 | 0.929164011 | 0.984238275 | ns | NaCl | KS.3 |
| FIM1_4216 | 638.6737869 | 467.1546333 | 743.7061317 | -0.109969182 | 0.146453281 | 0.047892371 | 0.343442 | ns | NaCl | KS.3 |
| FIM1_4217 | 556.1095403 | 352.7551994 | 889.1683139 | -0.253975946 | 0.250605043 | 0.004538721 | 0.086695706 | ns | NaCl | KS.3 |
| FIM1_4218 | 33.35631502 | 24.29786527 | 40.80047592 | -0.058045606 | 0.143320094 | 0.186930297 | 0.618483916 | ns | NaCl | KS.3 |
| FIM1_4219 | 516.993118 | 479.5798505 | 472.1011329 | -0.028771315 | 0.09358433 | 0.536519769 | 0.855981197 | ns | NaCl | KS.3 |
| FIM1_4220 | 846.8343535 | 848.6244245 | 752.1887105 | 0.000498156 | 0.089412053 | 0.991509039 | 0.999134858 | ns | NaCl | KS.3 |
| FIM1_4221 | 192.2597629 | 173.4961368 | 215.5270907 | -0.029550904 | 0.101076612 | 0.506449027 | 0.844423736 | ns | NaCl | KS.3 |
| FIM1_4222 | 1829.427133 | 1506.451562 | 1521.313095 | -0.042155116 | 0.117743913 | 0.325731621 | 0.739131368 | ns | NaCl | KS.3 |
| FIM1_4223 | 3240.663651 | 2807.954147 | 2560.189644 | -0.066799061 | 0.092735154 | 0.182514861 | 0.611396712 | ns | NaCl | KS.3 |
| FIM1_4224 | 6445.420606 | 7141.007047 | 5080.741052 | 0.036192534 | 0.096022616 | 0.436668374 | 0.808096352 | ns | NaCl | KS.3 |
| FIM1_4225 | 470.4596813 | 574.3034659 | 581.9715583 | 0.027117126 | 0.129298272 | 0.47577058 | 0.824724263 | ns | NaCl | KS.3 |
| FIM1_4226 | 39053.18915 | 69046.88101 | 38582.63717 | 0.513455416 | 0.321175421 | 0.000426309 | 0.021012238 | ns | NaCl | KS.3 |
| FIM1_4227 | 1166.816864 | 1442.252554 | 2863.283207 | 0.028448791 | 0.130709302 | 0.45516302 | 0.816788914 | ns | NaCl | KS.3 |
| FIM1_4228 | 24.42176078 | 15.28716908 | 24.61517435 | -0.062959039 | 0.176064033 | 0.147346877 | 0.562037426 | ns | NaCl | KS.3 |
| FIM1_4229 | 407.0454006 | 311.5520365 | 1314.53667 | -0.048528415 | 0.133256942 | 0.252712622 | 0.676266095 | ns | NaCl | KS.3 |
| FIM1_4230 | 2750.774366 | 3638.815442 | 2387.443186 | 0.07354016 | 0.133051587 | 0.127212935 | 0.534982546 | ns | NaCl | KS.3 |
| FIM1_4231 | 7121.874497 | 9286.986882 | 5772.270488 | 0.106002711 | 0.128292924 | 0.052000764 | 0.356877014 | ns | NaCl | KS.3 |
| FIM1_4232 | 398.8465938 | 409.270964 | 487.0612049 | 0.008655409 | 0.095014515 | 0.847535636 | 0.967147129 | ns | NaCl | KS.3 |
| FIM1_4233 | 537.5587089 | 600.5836024 | 813.2059935 | 0.05089078 | 0.089442979 | 0.300162134 | 0.717845774 | ns | NaCl | KS.3 |
| FIM1_4234 | 1472.135133 | 1573.18552 | 1164.257849 | 0.028114462 | 0.088713376 | 0.556228528 | 0.863699344 | ns | NaCl | KS.3 |
| FIM1_4235 | 1112.068304 | 966.8330246 | 998.3452731 | -0.088009215 | 0.082835473 | 0.089285033 | 0.463183565 | ns | NaCl | KS.3 |
| FIM1_4236 | 2256.137474 | 2203.276016 | 2572.85615 | -0.014456069 | 0.076930305 | 0.768871551 | 0.948835638 | ns | NaCl | KS.3 |
| FIM1_4237 | 10339.49506 | 9775.334027 | 7971.332114 | -0.039684134 | 0.071043204 | 0.420621399 | 0.80133378 | ns | NaCl | KS.3 |
| FIM1_4238 | 3985.172161 | 4338.472397 | 2974.152138 | 0.049613309 | 0.079902276 | 0.320359196 | 0.737029267 | ns | NaCl | KS.3 |
| FIM1_4239 | 3164.483177 | 3411.413859 | 2512.177134 | 0.042448174 | 0.080648103 | 0.392365362 | 0.781432215 | ns | NaCl | KS.3 |
| FIM1_4240 | 2160.933404 | 2744.885216 | 1416.346129 | 0.056448937 | 0.124050772 | 0.210798676 | 0.634379635 | ns | NaCl | KS.3 |
| FIM1_4241 | 2040.857124 | 2573.874893 | 2297.8411 | 0.056311455 | 0.122211811 | 0.213416471 | 0.637685853 | ns | NaCl | KS.3 |
| FIM1_4242 | 98.27489736 | 123.7752464 | 256.0742388 | 0.05958145 | 0.120797735 | 0.195934647 | 0.618952432 | ns | NaCl | KS.3 |

| | | | | | | | | | | |
|---|---|---|---|---|---|---|---|---|---|---|
| FIM1_4243 | 697.3782143 | 810.4332806 | 683.6635556 | 0.092262737 | 0.086633973 | 0.077483896 | 0.432155137 | ns | NaCl | KS.3 |
| FIM1_4244 | 353.3136375 | 306.6024258 | 691.9464541 | -0.040544184 | 0.104851992 | 0.367339004 | 0.768177319 | ns | NaCl | KS.3 |
| FIM1_4245 | 19.83170147 | 8.925571291 | 45.78680657 | -0.150234056 | 0.299848901 | 0.023680553 | 0.239851113 | ns | NaCl | KS.3 |
| FIM1_4246 | 459.8897331 | 407.4215334 | 598.2144239 | -0.067427451 | 0.086085151 | 0.182743098 | 0.611396712 | ns | NaCl | KS.3 |
| FIM1_4247 | 526.6371478 | 504.5061694 | 496.5544882 | -0.012119191 | 0.098925113 | 0.781695723 | 0.951646805 | ns | NaCl | KS.3 |
| FIM1_4248 | 1225.500811 | 1245.794938 | 1281.443992 | 0.011676591 | 0.068745157 | 0.810543836 | 0.958816322 | ns | NaCl | KS.3 |
| FIM1_4249 | 6080.433504 | 6445.620094 | 5696.017768 | 0.031255962 | 0.081459955 | 0.525066684 | 0.854940852 | ns | NaCl | KS.3 |
| FIM1_4250 | 1301.093563 | 1380.309294 | 1103.258929 | 0.054107226 | 0.058171401 | 0.239009343 | 0.662163805 | ns | NaCl | KS.3 |
| FIM1_4251 | 3602.078559 | 3356.665297 | 3147.760199 | -0.054743686 | 0.066941937 | 0.261368665 | 0.686178949 | ns | NaCl | KS.3 |
| FIM1_4252 | 1139.14193 | 831.7931393 | 1133.022365 | -0.291853509 | 0.15889775 | 0.000339291 | 0.017957182 | ns | NaCl | KS.3 |
| FIM1_4253 | 2454.6983 | 2190.197322 | 2736.043885 | -0.035482062 | 0.100132317 | 0.434609893 | 0.807880183 | ns | NaCl | KS.3 |
| FIM1_4254 | 1030.199997 | 1090.842657 | 1081.144263 | 0.030424987 | 0.082325493 | 0.535080644 | 0.855153333 | ns | NaCl | KS.3 |
| FIM1_4255 | 1928.656932 | 2035.165564 | 1779.909196 | 0.015647917 | 0.098302455 | 0.723024387 | 0.932843964 | ns | NaCl | KS.3 |
| FIM1_4257 | 1583.402321 | 1773.358791 | 4226.826573 | 0.026173602 | 0.107795699 | 0.535154906 | 0.855153333 | ns | NaCl | KS.3 |
| FIM1_4258 | 38.65581922 | 50.53752287 | 42.59403129 | 0.045549787 | 0.129191019 | 0.279772219 | 0.700652348 | ns | NaCl | KS.3 |
| FIM1_4259 | 2406.769367 | 2383.743044 | 2675.666478 | -0.002771297 | 0.097448646 | 0.949779352 | 0.987736646 | ns | NaCl | KS.3 |
| FIM1_4260 | 612.8960202 | 591.182882 | 767.4231294 | -0.008696499 | 0.103448374 | 0.835714437 | 0.962829707 | ns | NaCl | KS.3 |
| FIM1_4261 | 787.8947937 | 977.9253277 | 920.3489585 | 0.066917809 | 0.115616182 | 0.162868956 | 0.586304673 | ns | NaCl | KS.3 |
| FIM1_4262 | 3681.741179 | 4701.420098 | 2964.984714 | 0.112103867 | 0.11973532 | 0.042694067 | 0.325771447 | ns | NaCl | KS.3 |
| FIM1_4263 | 369.8812492 | 323.6625118 | 569.9360588 | -0.074668689 | 0.088353446 | 0.143473524 | 0.557205918 | ns | NaCl | KS.3 |
| FIM1_4265 | 971.4246344 | 1116.96115 | 891.4510669 | 0.07710939 | 0.087672267 | 0.132314824 | 0.539053419 | ns | NaCl | KS.3 |
| FIM1_4266 | 11152.54823 | 15670.37298 | 9953.457295 | 0.116007025 | 0.158772452 | 0.042004159 | 0.323956287 | ns | NaCl | KS.3 |
| FIM1_4267 | 1560.621783 | 1929.348877 | 1443.309574 | 0.110133342 | 0.106090851 | 0.043699636 | 0.32940256 | ns | NaCl | KS.3 |
| FIM1_4268 | 3852.212942 | 4402.638738 | 3576.938436 | 0.041689881 | 0.101947156 | 0.362312477 | 0.763449629 | ns | NaCl | KS.3 |
| FIM1_4269 | 461.1668006 | 539.7178583 | 394.092963 | 0.070616089 | 0.095868926 | 0.159799176 | 0.58211082 | ns | NaCl | KS.3 |
| FIM1_4270 | 1845.155227 | 1962.877777 | 2359.182223 | 0.036750987 | 0.078169986 | 0.45838317 | 0.818684405 | ns | NaCl | KS.3 |
| FIM1_4271 | 958.8517474 | 902.8386983 | 835.0517382 | -0.037293584 | 0.077173969 | 0.452065707 | 0.815160163 | ns | NaCl | KS.3 |
| FIM1_4272 | 645.7039578 | 594.5608771 | 712.7728461 | -0.02172636 | 0.101445692 | 0.61795659 | 0.891787196 | ns | NaCl | KS.3 |
| FIM1_4273 | 3236.589461 | 4359.789288 | 1940.035915 | 0.059969272 | 0.138394998 | 0.180572747 | 0.609877403 | ns | NaCl | KS.3 |
| FIM1_4274 | 201.6793748 | 210.0933628 | 458.2280169 | 0.016162028 | 0.090127002 | 0.730266791 | 0.936118857 | ns | NaCl | KS.3 |
| FIM1_4275 | 8.816231018 | 15.39315876 | 33.90945476 | 0.047950058 | 0.179719518 | 0.211746985 | 0.635060483 | ns | NaCl | KS.3 |
| FIM1_4276 | 2639.334731 | 2633.547426 | 2355.293366 | -0.001680674 | 0.073932946 | 0.972721213 | 0.995313014 | ns | NaCl | KS.3 |
| FIM1_4277 | 144.2416723 | 91.11348271 | 199.5390417 | -0.256472905 | 0.255349365 | 0.004548257 | 0.086695706 | ns | NaCl | KS.3 |
| FIM1_4278 | 3151.892473 | 3055.949864 | 2990.652528 | -0.025318114 | 0.061997088 | 0.591664366 | 0.878786986 | ns | NaCl | KS.3 |
| FIM1_4279 | 5927.093413 | 7519.933062 | 3477.358609 | 0.039869226 | 0.129243823 | 0.328817421 | 0.741064881 | ns | NaCl | KS.3 |
| FIM1_4280 | 2988.116774 | 2722.294376 | 2995.245579 | -0.047378595 | 0.084717219 | 0.338458 | 0.747703619 | ns | NaCl | KS.3 |
| FIM1_4281 | 6244.116035 | 6227.207134 | 5184.495808 | -0.001559678 | 0.077202554 | 0.974663423 | 0.995954749 | ns | NaCl | KS.3 |
| FIM1_4282 | 1122.909914 | 1805.541034 | 2506.198991 | 0.121753364 | 0.206033569 | 0.039565634 | 0.315147688 | ns | NaCl | KS.3 |
| FIM1_4283 | 46.79452775 | 27.91535858 | 72.17510083 | -0.109875487 | 0.208583597 | 0.049943035 | 0.349460758 | ns | NaCl | KS.3 |
| FIM1_4284 | 4411.160168 | 4651.518666 | 5208.636838 | 0.030777471 | 0.078865738 | 0.533099601 | 0.855153333 | ns | NaCl | KS.3 |
| FIM1_4285 | 2227.230382 | 2754.084402 | 2583.026233 | 0.13137375 | 0.103005527 | 0.020163624 | 0.215832354 | ns | NaCl | KS.3 |
| FIM1_4286 | 3642.675012 | 3509.245899 | 2558.176458 | -0.006895369 | 0.110415589 | 0.861512086 | 0.969856868 | ns | NaCl | KS.3 |
| FIM1_4287 | 13.48516736 | 10.20818335 | 12.20518445 | -0.026041003 | 0.147114735 | 0.461630544 | 0.819880744 | ns | NaCl | KS.3 |
| FIM1_4288 | 1446.533515 | 1652.537567 | 1283.172397 | 0.032381346 | 0.108127534 | 0.451554257 | 0.81453315 | ns | NaCl | KS.3 |
| FIM1_4289 | 2460.238617 | 2234.74646 | 2144.079177 | -0.042657236 | 0.089285977 | 0.379527071 | 0.778493059 | ns | NaCl | KS.3 |

| | | | | | | | | | | |
|---|---|---|---|---|---|---|---|---|---|---|
| FIM1_4290 | 1253.082965 | 1374.678096 | 1397.806259 | 0.050017428 | 0.083250165 | 0.314890991 | 0.733075657 | ns | NaCl | KS.3 |
| FIM1_4291 | 929.7802495 | 971.0183008 | 822.1209317 | 0.03140153 | 0.067484549 | 0.518315098 | 0.850188733 | ns | NaCl | KS.3 |
| FIM1_4292 | 1226.546972 | 1220.483048 | 926.2401936 | -0.00260357 | 0.077932999 | 0.957681813 | 0.990773484 | ns | NaCl | KS.3 |
| FIM1_4293 | 561.9118894 | 727.0975356 | 651.6130055 | 0.224041389 | 0.117003551 | 0.000669754 | 0.028237509 | ns | NaCl | KS.3 |
| FIM1_4294 | 872.7252441 | 665.0950846 | 1014.23371 | -0.187577004 | 0.133802925 | 0.00435623 | 0.085661036 | ns | NaCl | KS.3 |
| FIM1_4295 | 834.9887625 | 805.3110284 | 957.734905 | -0.029108237 | 0.068859832 | 0.551168151 | 0.860945181 | ns | NaCl | KS.3 |
| FIM1_4296 | 1157.63131 | 1160.467598 | 1234.676435 | 0.003324954 | 0.05722178 | 0.94170365 | 0.986790704 | ns | NaCl | KS.3 |
| FIM1_4297 | 308.1263279 | 326.7919295 | 795.4135375 | 0.020703184 | 0.095387635 | 0.648743809 | 0.899642849 | ns | NaCl | KS.3 |
| FIM1_4298 | 205.1322415 | 259.6751645 | 256.6677733 | 0.024699135 | 0.139773582 | 0.492978508 | 0.835912774 | ns | NaCl | KS.3 |
| FIM1_4299 | 398.8376172 | 435.5816983 | 366.378426 | 0.035202503 | 0.091306971 | 0.459545928 | 0.819640873 | ns | NaCl | KS.3 |
| FIM1_4300 | 203.9039279 | 216.060621 | 223.9361699 | 0.019645135 | 0.091591847 | 0.673231785 | 0.91246166 | ns | NaCl | KS.3 |
| FIM1_4303 | 105.5562678 | 143.8266206 | 82.48496482 | 0.087005859 | 0.141820913 | 0.087261877 | 0.457488698 | ns | NaCl | KS.3 |
| FIM1_4304 | 17.52917103 | 10.5395134 | 30.91326256 | -0.044335993 | 0.169966569 | 0.244526334 | 0.66821307 | ns | NaCl | KS.3 |
| FIM1_4307 | 515.740421 | 480.9097774 | 476.7071757 | -0.022922482 | 0.096293208 | 0.612943972 | 0.889162818 | ns | NaCl | KS.3 |
| FIM1_4308 | 117.3600909 | 181.2295148 | 116.7686975 | 0.197686367 | 0.22879425 | 0.010155263 | 0.142316716 | ns | NaCl | KS.3 |
| FIM1_4314 | 1142.912928 | 1390.722847 | 646.7515623 | 0.06075368 | 0.111459537 | 0.198458966 | 0.620924909 | ns | NaCl | KS.3 |
| FIM1_4780 | 146.4254744 | 250.2768164 | 82.75928701 | 0.118369219 | 0.221239199 | 0.042256495 | 0.323956287 | ns | NaCl | KS.3 |
| FIM1_4781 | 44.64787812 | 63.27824654 | 87.57792087 | 0.099030448 | 0.162640536 | 0.063755112 | 0.398469449 | ns | NaCl | KS.3 |
| FIM1_4782 | 181.8203972 | 169.193868 | 296.5047851 | -0.030449794 | 0.089519531 | 0.523344768 | 0.854493017 | ns | NaCl | KS.3 |
| FIM1_4783 | 633.0084912 | 506.8233937 | 1729.24346 | -0.118921005 | 0.107359065 | 0.032598407 | 0.287548008 | ns | NaCl | KS.3 |
| FIM1_4785 | 785.559334 | 1040.522116 | 1037.162625 | 0.202803342 | 0.13961015 | 0.002965852 | 0.06799591 | ns | NaCl | KS.3 |
| FIM1_4786 | 2599.754279 | 6318.212105 | 3276.771524 | 0.272475071 | 0.411497552 | 0.006379472 | 0.106861534 | ns | NaCl | KS.3 |
| FIM1_4787 | 189.0510511 | 257.5964934 | 235.4642185 | 0.07900589 | 0.141513197 | 0.107723208 | 0.498909937 | ns | NaCl | KS.3 |
| FIM1_4788 | 240.5797848 | 155.3541431 | 744.3069426 | -0.086060004 | 0.178193861 | 0.08524489 | 0.45357575 | ns | NaCl | KS.3 |
| FIM1_4789 | 241.1891448 | 228.7842175 | 264.372092 | -0.00650647 | 0.127488508 | 0.85398096 | 0.969345976 | ns | NaCl | KS.3 |
| FIM1_4790 | 595.5874745 | 715.383085 | 614.4716867 | 0.063642211 | 0.106809672 | 0.187018067 | 0.618483916 | ns | NaCl | KS.3 |
| FIM1_4791 | 3236.204136 | 4294.139338 | 2073.260557 | 0.090039068 | 0.134186666 | 0.08114787 | 0.439771952 | ns | NaCl | KS.3 |
| FIM1_4792 | 368.9131423 | 402.1294609 | 464.198878 | 0.016884802 | 0.110969194 | 0.674298217 | 0.912806589 | ns | NaCl | KS.3 |
| FIM1_4793 | 1628.041134 | 1496.836212 | 2070.580399 | -0.035398929 | 0.089900815 | 0.460162133 | 0.819880744 | ns | NaCl | KS.3 |
| FIM1_4794 | 1874.253456 | 1187.499 | 2670.050722 | -0.238211055 | 0.248005295 | 0.005757804 | 0.100878634 | ns | NaCl | KS.3 |
| FIM1_4795 | 4677.603885 | 4851.341675 | 6331.766408 | 0.007320888 | 0.107911152 | 0.855995833 | 0.969856868 | ns | NaCl | KS.3 |
| FIM1_4796 | 1689.963682 | 1337.977387 | 2521.152864 | -0.074586492 | 0.119079928 | 0.128990877 | 0.536563223 | ns | NaCl | KS.3 |
| FIM1_4797 | 7147.189767 | 9558.386828 | 4622.760369 | 0.0584193 | 0.136719818 | 0.189513579 | 0.618952432 | ns | NaCl | KS.3 |
| FIM1_4798 | 1063.042179 | 1050.858825 | 907.3828427 | -0.002947707 | 0.101092785 | 0.944867426 | 0.987187877 | ns | NaCl | KS.3 |
| FIM1_4799 | 860.6627257 | 848.122133 | 915.778069 | -0.00426203 | 0.098843286 | 0.921940802 | 0.983890455 | ns | NaCl | KS.3 |
| FIM1_4800 | 21.86052236 | 18.50475693 | 14.76985844 | -0.022466738 | 0.133918827 | 0.537098113 | 0.855981197 | ns | NaCl | KS.3 |
| FIM1_4801 | 462.1161874 | 474.5933696 | 369.4995752 | 0.005590732 | 0.104965113 | 0.892251631 | 0.977268396 | ns | NaCl | KS.3 |
| FIM1_4802 | 873.1158954 | 814.072059 | 805.0057536 | -0.047601392 | 0.073328815 | 0.337831792 | 0.747648205 | ns | NaCl | KS.3 |
| FIM1_4803 | 1818.551102 | 1686.844271 | 1781.19925 | -0.019249211 | 0.102762111 | 0.653723891 | 0.900747416 | ns | NaCl | KS.3 |
| FIM1_4804 | 8031.969353 | 8068.230119 | 5526.854346 | 0.002056085 | 0.084902249 | 0.965849621 | 0.993691273 | ns | NaCl | KS.3 |
| FIM1_4805 | 6516.090908 | 9581.479631 | 4310.284211 | 0.352033928 | 0.209621239 | 0.00042658 | 0.021012238 | ns | NaCl | KS.3 |
| FIM1_4806 | 2168.8438 | 2256.073897 | 2008.96905 | 0.035265521 | 0.060256389 | 0.449791219 | 0.813595672 | ns | NaCl | KS.3 |
| FIM1_4807 | 3213.518725 | 3285.598163 | 2654.061447 | 0.006352707 | 0.098184494 | 0.884650421 | 0.975081831 | ns | NaCl | KS.3 |
| FIM1_4808 | 1902.063226 | 1719.729141 | 2096.452981 | -0.030586723 | 0.099775942 | 0.496291953 | 0.836358994 | ns | NaCl | KS.3 |
| FIM1_4809 | 809.3260083 | 760.1822987 | 696.4791253 | -0.02579714 | 0.089562158 | 0.586672861 | 0.878311243 | ns | NaCl | KS.3 |

| | | | | | | | | | | |
|---|---|---|---|---|---|---|---|---|---|---|
| FIM1_4810 | 16935.2698 | 13902.7077 | 12749.87612 | -0.167646552 | 0.086376328 | 0.00274286 | 0.064671708 | ns | NaCl | KS.3 |
| FIM1_4811 | 1566.899648 | 1499.322035 | 1936.960012 | -0.006467784 | 0.117506558 | 0.863191389 | 0.969856868 | ns | NaCl | KS.3 |
| FIM1_4812 | 130.2034047 | 143.1215438 | 213.5856373 | 0.028729069 | 0.099754431 | 0.521057942 | 0.852373129 | ns | NaCl | KS.3 |
| FIM1_4813 | 2917.078601 | 3992.681886 | 1787.613643 | 0.061770406 | 0.142240101 | 0.169733041 | 0.594663296 | ns | NaCl | KS.3 |
| FIM1_4814 | 1421.99851 | 1219.889942 | 1173.57424 | -0.120000607 | 0.076494968 | 0.021490316 | 0.223670134 | ns | NaCl | KS.3 |
| FIM1_4815 | 572.2254234 | 609.5129256 | 843.8898588 | 0.036287368 | 0.077774144 | 0.4640542 | 0.819880744 | ns | NaCl | KS.3 |
| FIM1_4816 | 896.5802792 | 768.5002811 | 1091.397534 | -0.107727084 | 0.083745743 | 0.041752864 | 0.323298895 | ns | NaCl | KS.3 |
| FIM1_4817 | 2005.265645 | 1979.924502 | 1649.15812 | -0.005291598 | 0.088203763 | 0.910729457 | 0.981985056 | ns | NaCl | KS.3 |
| FIM1_4818 | 820.2719834 | 708.5045891 | 1041.875757 | -0.103576178 | 0.080899192 | 0.047909296 | 0.343442 | ns | NaCl | KS.3 |
| FIM1_4819 | 4541.238628 | 5173.848091 | 5748.894804 | 0.096957181 | 0.075559669 | 0.058803563 | 0.378457602 | ns | NaCl | KS.3 |
| FIM1_4820 | 1643.665949 | 1553.161134 | 1825.771121 | -0.037492442 | 0.073750168 | 0.448796595 | 0.813392736 | ns | NaCl | KS.3 |
| FIM1_4821 | 1120.119759 | 944.3111409 | 1098.858809 | -0.032981335 | 0.117467449 | 0.422934377 | 0.80229961 | ns | NaCl | KS.3 |
| FIM1_4822 | 1251.723559 | 1126.573855 | 1621.314464 | -0.058643053 | 0.083467998 | 0.24273118 | 0.666657537 | ns | NaCl | KS.3 |
| FIM1_4823 | 946.0896784 | 960.4651543 | 1166.873272 | 0.008353402 | 0.078351226 | 0.864800104 | 0.970128052 | ns | NaCl | KS.3 |
| FIM1_4824 | 4659.093543 | 5876.386257 | 3418.530375 | 0.078051085 | 0.118121506 | 0.117151176 | 0.52010139 | ns | NaCl | KS.3 |
| FIM1_4825 | 189.6572356 | 175.9576004 | 465.332088 | -0.016214971 | 0.104618471 | 0.699693365 | 0.922863518 | ns | NaCl | KS.3 |
| FIM1_4826 | 1032.884264 | 1131.216623 | 1047.673213 | 0.058087434 | 0.077019661 | 0.247166183 | 0.669893726 | ns | NaCl | KS.3 |
| FIM1_4827 | 1372.13705 | 1607.933458 | 1322.638848 | 0.10534468 | 0.084329579 | 0.04624526 | 0.340862131 | ns | NaCl | KS.3 |
| FIM1_4828 | 826.8392194 | 767.4708532 | 929.2585492 | -0.03983048 | 0.081383064 | 0.420818577 | 0.80133378 | ns | NaCl | KS.3 |
| FIM1_4829 | 1512.717587 | 835.9333635 | 2408.544241 | -0.21833324 | 0.295386533 | 0.009957813 | 0.140601095 | ns | NaCl | KS.3 |
| FIM1_4830 | 1401.60199 | 1047.962394 | 2396.706495 | -0.06960232 | 0.136254262 | 0.140193261 | 0.550917434 | ns | NaCl | KS.3 |
| FIM1_4831 | 2360.138978 | 2451.084137 | 1966.919271 | 0.027801726 | 0.067547717 | 0.567262 | 0.868965626 | ns | NaCl | KS.3 |
| FIM1_4832 | 3451.773701 | 3105.485192 | 3576.343722 | -0.101635542 | 0.059171794 | 0.029108411 | 0.268174713 | ns | NaCl | KS.3 |
| FIM1_4833 | 5645.826221 | 4946.751853 | 4511.192965 | -0.062421174 | 0.091027463 | 0.210696411 | 0.634379635 | ns | NaCl | KS.3 |
| FIM1_4834 | 858.3731151 | 1086.835154 | 1340.639535 | 0.033778645 | 0.13222318 | 0.387121744 | 0.779278134 | ns | NaCl | KS.3 |
| FIM1_4835 | 5252.044903 | 5975.362521 | 3751.275652 | 0.067495815 | 0.087599238 | 0.181714849 | 0.611396712 | ns | NaCl | KS.3 |
| FIM1_4836 | 1632.511427 | 433.8879058 | 802.9952928 | -1.755121422 | 0.26461466 | 5.7095159460[illegible] | 3.5506052289[illegible] | Km<KS.3 | NaCl | KS.3 |
| FIM1_4837 | 118.1253776 | 25.24789666 | 67.6932174 | -1.797894697 | 0.438993856 | 2.4141190117[illegible] | 5.0042675347[illegible] | Km<KS.3 | NaCl | KS.3 |
| FIM1_4839 | 355.7931953 | 307.9011722 | 576.3763114 | -0.028715977 | 0.114380399 | 0.483813248 | 0.829184191 | ns | NaCl | KS.3 |
| FIM1_4840 | 12239.24553 | 12951.43417 | 8620.223328 | 0.041997404 | 0.068714656 | 0.390916379 | 0.779482559 | ns | NaCl | KS.3 |
| FIM1_4841 | 127.9454611 | 104.0155587 | 248.0003937 | -0.076766108 | 0.112117049 | 0.124040147 | 0.531066893 | ns | NaCl | KS.3 |
| FIM1_4842 | 70053.87948 | 90342.75792 | 58807.41069 | 0.198098946 | 0.119667541 | 0.002125004 | 0.057145368 | ns | NaCl | KS.3 |
| FIM1_4844 | 129.6334296 | 200.4923286 | 1119.871416 | 0.054157232 | 0.166936426 | 0.190389724 | 0.618952432 | ns | NaCl | KS.3 |
| FIM1_4845 | 558.1712852 | 426.8980362 | 600.696593 | -0.046999864 | 0.133081364 | 0.264010854 | 0.688347233 | ns | NaCl | KS.3 |
| FIM1_4846 | 252.0389629 | 199.7566592 | 542.9534828 | -0.035704754 | 0.130355898 | 0.368917049 | 0.769221425 | ns | NaCl | KS.3 |
| FIM1_4847 | 349.8673797 | 353.1690099 | 765.4735833 | 0.001691744 | 0.110647279 | 0.965658086 | 0.993691273 | ns | NaCl | KS.3 |
| FIM1_4848 | 1772.576775 | 2543.666941 | 2313.99835 | 0.101492301 | 0.162309429 | 0.060081548 | 0.383212437 | ns | NaCl | KS.3 |
| FIM1_4849 | 4516.44967 | 5020.789139 | 3562.945838 | 0.022014285 | 0.11004333 | 0.591777908 | 0.878786986 | ns | NaCl | KS.3 |
| FIM1_4850 | 191.6029078 | 116.4626386 | 797.3107309 | -0.080524274 | 0.18948936 | 0.094235404 | 0.470201108 | ns | NaCl | KS.3 |
| FIM1_4851 | 945.6201473 | 757.8609023 | 855.6509651 | -0.101730405 | 0.111134415 | 0.058056447 | 0.37782226 | ns | NaCl | KS.3 |
| FIM1_4852 | 443.4948771 | 310.4108905 | 468.2283045 | -0.211283545 | 0.191778487 | 0.005521463 | 0.098456203 | ns | NaCl | KS.3 |
| FIM1_4853 | 365.0607037 | 318.2371886 | 464.3356436 | -0.039868411 | 0.103867731 | 0.376614764 | 0.778493059 | ns | NaCl | KS.3 |
| FIM1_4854 | 3446.331002 | 4467.828748 | 3480.209368 | 0.045266935 | 0.132005598 | 0.27854255 | 0.700652348 | ns | NaCl | KS.3 |
| FIM1_4855 | 842.6702426 | 1044.822156 | 915.6515042 | 0.026923271 | 0.132992204 | 0.472089642 | 0.822127061 | ns | NaCl | KS.3 |
| FIM1_4856 | 53.99889443 | 39.70813693 | 90.94481753 | -0.057449517 | 0.139599356 | 0.192536415 | 0.618952432 | ns | NaCl | KS.3 |

| | | | | | | | | | | |
|---|---|---|---|---|---|---|---|---|---|---|
| FIM1_4857 | 20.88625516 | 45.20482824 | 97.34138546 | 0.161658659 | 0.305328215 | 0.020483364 | 0.216818586 | ns | NaCl | KS.3 |
| FIM1_4858 | 3234.780704 | 4171.92828 | 1889.896656 | 0.049272971 | 0.12962736 | 0.251396238 | 0.675578827 | ns | NaCl | KS.3 |
| FIM1_4859 | 1404.915776 | 1408.025328 | 1667.293549 | 0.000727426 | 0.066554457 | 0.987984156 | 0.999134858 | ns | NaCl | KS.3 |
| FIM1_4860 | 431.5413182 | 453.9991011 | 551.1737234 | 0.010559416 | 0.106815611 | 0.796194185 | 0.955756014 | ns | NaCl | KS.3 |
| FIM1_4861 | 902.3681392 | 805.8857904 | 825.2678814 | -0.072234147 | 0.079161294 | 0.155267952 | 0.573040105 | ns | NaCl | KS.3 |
| FIM1_4862 | 2218.564412 | 2311.857305 | 1646.755377 | 0.016969414 | 0.088693073 | 0.71984129 | 0.930667987 | ns | NaCl | KS.3 |
| FIM1_4863 | 8643.115268 | 11982.61232 | 5275.620652 | 0.075179199 | 0.146383472 | 0.117809869 | 0.520518737 | ns | NaCl | KS.3 |
| FIM1_4864 | 61.93476753 | 62.24809275 | 64.56806349 | 0.000537973 | 0.110639208 | 0.989074329 | 0.999134858 | ns | NaCl | KS.3 |
| FIM1_4865 | 621.1699422 | 616.3788959 | 766.4919362 | -0.00187615 | 0.10134864 | 0.964803048 | 0.993691273 | ns | NaCl | KS.3 |
| FIM1_4866 | 704.7013707 | 538.9966471 | 1468.118296 | -0.069251326 | 0.129727565 | 0.144475985 | 0.558679104 | ns | NaCl | KS.3 |
| FIM1_4867 | 2194.919626 | 2776.585349 | 1774.801931 | 0.08510589 | 0.118090219 | 0.095510763 | 0.471394887 | ns | NaCl | KS.3 |
| FIM1_4868 | 1585.272845 | 1610.22661 | 1300.526989 | 0.007626926 | 0.08247079 | 0.875150438 | 0.971808705 | ns | NaCl | KS.3 |
| FIM1_4869 | 2826.713519 | 2472.996346 | 2324.438527 | -0.056530791 | 0.094336159 | 0.247796943 | 0.669994451 | ns | NaCl | KS.3 |
| FIM1_4870 | 640.4687623 | 858.5983842 | 695.124892 | 0.06507975 | 0.137046984 | 0.157976729 | 0.57787696 | ns | NaCl | KS.3 |
| FIM1_4871 | 262.1759793 | 389.6197673 | 519.1620767 | 0.240894098 | 0.216295986 | 0.003989961 | 0.082025025 | ns | NaCl | KS.3 |
| FIM1_4872 | 139.9802648 | 185.6009031 | 580.6762812 | 0.084016072 | 0.132273126 | 0.095854063 | 0.472620379 | ns | NaCl | KS.3 |
| FIM1_4873 | 1826.678104 | 2351.490212 | 1506.886808 | 0.104035258 | 0.123305109 | 0.054741351 | 0.36408853 | ns | NaCl | KS.3 |
| FIM1_4874 | 3723.564235 | 4367.967433 | 2687.585499 | 0.044518537 | 0.107601306 | 0.322970088 | 0.739010075 | ns | NaCl | KS.3 |
| FIM1_4875 | 771.2013198 | 443.0068402 | 1454.363538 | -0.313544258 | 0.31336401 | 0.003236617 | 0.072860489 | ns | NaCl | KS.3 |
| FIM1_4876 | 50.13659498 | 34.29657053 | 66.18190685 | -0.073990895 | 0.159948022 | 0.117739687 | 0.520518737 | ns | NaCl | KS.3 |
| FIM1_4877 | 714.6214012 | 791.6098481 | 1003.914225 | 0.026434048 | 0.104086605 | 0.541185682 | 0.857165292 | ns | NaCl | KS.3 |
| FIM1_4878 | 27454.67906 | 31776.44246 | 19245.88169 | 0.081262164 | 0.088259914 | 0.114817129 | 0.515537198 | ns | NaCl | KS.3 |
| FIM1_4879 | 3058.033362 | 3115.042405 | 2685.028214 | 0.017051478 | 0.057632193 | 0.708703039 | 0.925190614 | ns | NaCl | KS.3 |
| FIM1_4880 | 7086.710415 | 6500.484721 | 4901.205305 | -0.083852255 | 0.056442994 | 0.06428516 | 0.398841896 | ns | NaCl | KS.3 |
| FIM1_4881 | 2269.178439 | 2373.905237 | 2085.045205 | 0.022983788 | 0.082255202 | 0.637962536 | 0.898332475 | ns | NaCl | KS.3 |
| FIM1_4882 | 23.35219018 | 38.71664803 | 1008.366534 | 0.040028016 | 0.172566668 | 0.272465428 | 0.693610905 | ns | NaCl | KS.3 |
| FIM1_4883 | 3269.909867 | 3116.629062 | 3633.9843 | -0.018611467 | 0.090693521 | 0.69089744 | 0.919659229 | ns | NaCl | KS.3 |
| FIM1_4884 | 2302.884537 | 2176.298636 | 2324.54675 | -0.04288372 | 0.067234706 | 0.378270028 | 0.778493059 | ns | NaCl | KS.3 |
| FIM1_4885 | 795.8700364 | 643.5966813 | 961.0478088 | -0.139999475 | 0.101152712 | 0.014155803 | 0.172593755 | ns | NaCl | KS.3 |
| FIM1_4886 | 243.3504675 | 189.6149651 | 262.1975986 | -0.097406615 | 0.120203491 | 0.066445853 | 0.404582009 | ns | NaCl | KS.3 |
| FIM1_4887 | 344.4025823 | 340.0715041 | 366.9965181 | -0.005974665 | 0.082073895 | 0.902149157 | 0.979395371 | ns | NaCl | KS.3 |
| FIM1_4888 | 722.0032157 | 580.7920664 | 846.6949265 | -0.076368559 | 0.113937651 | 0.124703648 | 0.532990249 | ns | NaCl | KS.3 |
| FIM1_4889 | 594.1550232 | 235.3508686 | 630.0680574 | -1.208410382 | 0.224909545 | 1.4060406226[illegible] | 4.9964657841[illegible] | Km<KS.3 | NaCl | KS.3 |
| FIM1_4890 | 679.4733475 | 510.968191 | 592.0101904 | -0.122140767 | 0.13739859 | 0.03366561 | 0.292771172 | ns | NaCl | KS.3 |
| FIM1_4891 | 1364.218156 | 1253.838346 | 1119.446052 | -0.03723948 | 0.088910122 | 0.440273241 | 0.808917498 | ns | NaCl | KS.3 |
| FIM1_4892 | 5574.247585 | 6144.95609 | 4920.135313 | 0.022933566 | 0.106214521 | 0.587113972 | 0.878311243 | ns | NaCl | KS.3 |
| FIM1_4893 | 12929.24974 | 13674.19532 | 9611.487046 | 0.013549217 | 0.103549007 | 0.748073753 | 0.943274417 | ns | NaCl | KS.3 |
| FIM1_4894 | 1189.561227 | 1133.471258 | 1464.534456 | -0.016146987 | 0.094760389 | 0.722096015 | 0.931887853 | ns | NaCl | KS.3 |
| FIM1_4895 | 1647.222599 | 1789.211371 | 1695.610525 | 0.021904664 | 0.102246517 | 0.613142808 | 0.889162818 | ns | NaCl | KS.3 |
| FIM1_4896 | 725.2312901 | 809.0176899 | 1266.705427 | 0.027948167 | 0.104839441 | 0.518023352 | 0.850188733 | ns | NaCl | KS.3 |
| FIM1_4897 | 870.8064557 | 891.9792536 | 1069.82916 | 0.005814143 | 0.102719085 | 0.890125135 | 0.977268396 | ns | NaCl | KS.3 |
| FIM1_4898 | 822.281577 | 817.6013627 | 920.0857228 | -0.001203216 | 0.106881005 | 0.976308772 | 0.996164232 | ns | NaCl | KS.3 |
| FIM1_4899 | 9227.907326 | 11041.27856 | 7998.009529 | 0.098174347 | 0.095421447 | 0.064111703 | 0.398841896 | ns | NaCl | KS.3 |
| FIM1_4900 | 5135.762057 | 6550.167367 | 4562.418037 | 0.126354631 | 0.118293976 | 0.027108924 | 0.256889329 | ns | NaCl | KS.3 |
| FIM1_4901 | 315.9711233 | 321.2565733 | 385.1794082 | 0.005080699 | 0.101427577 | 0.904937686 | 0.979880338 | ns | NaCl | KS.3 |

| | | | | | | | | | | |
|---|---|---|---|---|---|---|---|---|---|---|
| FIM1_4902 | 759.2016419 | 734.9427412 | 1002.088481 | -0.014338525 | 0.087529628 | 0.762976594 | 0.946287744 | ns | NaCl | KS.3 |
| FIM1_4903 | 867.4308626 | 845.485417 | 1339.826362 | -0.012475494 | 0.085475584 | 0.794962937 | 0.955299664 | ns | NaCl | KS.3 |
| FIM1_4904 | 954.7959176 | 1268.361868 | 1271.668189 | 0.106631251 | 0.135411428 | 0.051557999 | 0.354773227 | ns | NaCl | KS.3 |
| FIM1_4905 | 248.0790076 | 277.906307 | 445.754003 | 0.026823593 | 0.106632821 | 0.528945379 | 0.855153333 | ns | NaCl | KS.3 |
| FIM1_4906 | 1432.303172 | 1501.135812 | 1602.179037 | 0.027123894 | 0.078332911 | 0.582394284 | 0.875615462 | ns | NaCl | KS.3 |
| FIM1_4907 | 3058.263321 | 4151.851616 | 2107.583726 | 0.054973986 | 0.140581567 | 0.205247483 | 0.628759992 | ns | NaCl | KS.3 |
| FIM1_4908 | 603.9567585 | 604.5596919 | 559.5296818 | 7.7950892551940 | 0.096079171 | 0.998602354 | 0.999406383 | ns | NaCl | KS.3 |
| FIM1_4909 | 3838.89789 | 4120.213581 | 2808.153758 | 0.038606221 | 0.081316278 | 0.434875001 | 0.807880183 | ns | NaCl | KS.3 |
| FIM1_4910 | 434.0331353 | 679.3241465 | 642.7546195 | 0.372585482 | 0.256142643 | 0.000797804 | 0.031277519 | ns | NaCl | KS.3 |
| FIM1_4911 | 1414.069728 | 1324.430451 | 1207.163562 | -0.018951535 | 0.099234367 | 0.667061656 | 0.907354532 | ns | NaCl | KS.3 |
| FIM1_4912 | 1176.776245 | 1115.568272 | 1002.936123 | -0.016887942 | 0.095538179 | 0.708531604 | 0.925190614 | ns | NaCl | KS.3 |
| FIM1_4913 | 3055.020709 | 4231.069395 | 1779.518337 | 0.098893073 | 0.149881488 | 0.063903607 | 0.398841896 | ns | NaCl | KS.3 |
| FIM1_4914 | 2918.200677 | 3739.201879 | 2391.945927 | 0.160372647 | 0.119127898 | 0.008631106 | 0.12856213 | ns | NaCl | KS.3 |
| FIM1_4915 | 1605.631854 | 1596.687216 | 2017.395644 | -0.001159275 | 0.106111299 | 0.977322101 | 0.996164232 | ns | NaCl | KS.3 |
| FIM1_4916 | 2246.270793 | 3038.815255 | 1357.458541 | 0.04367537 | 0.141645541 | 0.27996238 | 0.700652348 | ns | NaCl | KS.3 |
| FIM1_4917 | 1328.459751 | 1381.150128 | 1595.349315 | 0.020414977 | 0.080893742 | 0.67673359 | 0.913287355 | ns | NaCl | KS.3 |
| FIM1_4918 | 206.7938334 | 221.9858777 | 185.4433156 | 0.022910268 | 0.096942014 | 0.6115055 | 0.888504633 | ns | NaCl | KS.3 |
| FIM1_4919 | 2131.461643 | 2415.338462 | 1474.08409 | 0.037259427 | 0.102193156 | 0.408961345 | 0.795392815 | ns | NaCl | KS.3 |
| FIM1_4920 | 205.0110714 | 129.2800048 | 491.5382069 | -0.337826054 | 0.269810503 | 0.001569613 | 0.046934852 | ns | NaCl | KS.3 |
| FIM1_4921 | 17841.16701 | 14602.30751 | 10694.03251 | -0.104046471 | 0.10238228 | 0.053121309 | 0.361036217 | ns | NaCl | KS.3 |
| FIM1_4922 | 1240.685712 | 1417.604657 | 1043.283304 | 0.0470943 | 0.098879673 | 0.317255186 | 0.73382286 | ns | NaCl | KS.3 |
| FIM1_4923 | 1089.766022 | 800.7108199 | 779.2656105 | -0.330823516 | 0.141728507 | 4.9433412332 | 0.004314583 | ns | NaCl | KS.3 |
| FIM1_4924 | 1269.514552 | 1361.891395 | 2324.548194 | 0.014008463 | 0.108632509 | 0.729891175 | 0.935878504 | ns | NaCl | KS.3 |
| FIM1_4925 | 167.6042025 | 242.4090609 | 244.9278878 | 0.057518112 | 0.154477715 | 0.182383395 | 0.611396712 | ns | NaCl | KS.3 |
| FIM1_4926 | 1236.254757 | 1275.579341 | 1363.095632 | 0.024017824 | 0.067391875 | 0.620743359 | 0.892479944 | ns | NaCl | KS.3 |
| FIM1_4927 | 2074.690988 | 2512.375979 | 1324.343166 | 0.035024626 | 0.121156418 | 0.392048398 | 0.781113649 | ns | NaCl | KS.3 |
| FIM1_4928 | 1847.101017 | 2114.954817 | 1095.136634 | 0.022788216 | 0.117898348 | 0.561855316 | 0.866779377 | ns | NaCl | KS.3 |
| FIM1_4929 | 531.5675343 | 519.3688164 | 701.8857728 | -0.012601195 | 0.07938694 | 0.797083622 | 0.955924408 | ns | NaCl | KS.3 |
| FIM1_4931 | 4365.577816 | 5060.88192 | 3938.011143 | 0.057361901 | 0.098283957 | 0.23634285 | 0.659616806 | ns | NaCl | KS.3 |
| FIM1_4932 | 3955.323558 | 4369.835256 | 4932.710896 | 0.039046961 | 0.093091081 | 0.410933353 | 0.798279355 | ns | NaCl | KS.3 |
| FIM1_4933 | 1559.155069 | 1594.401549 | 1783.902986 | 0.008386556 | 0.090629862 | 0.856897573 | 0.969856868 | ns | NaCl | KS.3 |
| FIM1_4935 | 847.6123814 | 787.23498 | 1921.499279 | -0.025004213 | 0.095207498 | 0.585117271 | 0.877826895 | ns | NaCl | KS.3 |
| FIM1_4936 | 693.6122072 | 838.3994145 | 1229.813806 | 0.037667183 | 0.11914773 | 0.367261876 | 0.768177319 | ns | NaCl | KS.3 |
| FIM1_4937 | 6033.935148 | 6147.793692 | 6854.844582 | 0.003277643 | 0.111860459 | 0.93294311 | 0.985224363 | ns | NaCl | KS.3 |
| FIM1_4938 | 1458.02839 | 1593.658519 | 1253.715977 | 0.032994831 | 0.093798347 | 0.480943233 | 0.828494663 | ns | NaCl | KS.3 |
| FIM1_4939 | 862.3577235 | 972.3529388 | 997.7523455 | 0.060261788 | 0.087721754 | 0.228200194 | 0.652094179 | ns | NaCl | KS.3 |
| FIM1_4940 | 2170.394741 | 2895.498812 | 1454.603785 | 0.076476422 | 0.135564192 | 0.116745504 | 0.519941265 | ns | NaCl | KS.3 |
| FIM1_4941 | 13346.66119 | 13602.0614 | 8851.194038 | 0.011712746 | 0.07516899 | 0.811849113 | 0.958816322 | ns | NaCl | KS.3 |
| FIM1_4942 | 1275.308408 | 1241.885738 | 1224.514551 | -0.003874856 | 0.117782206 | 0.917373266 | 0.982343702 | ns | NaCl | KS.3 |
| FIM1_4943 | 1580.323455 | 1470.422133 | 2318.517883 | -0.02985036 | 0.089459729 | 0.531384582 | 0.855153333 | ns | NaCl | KS.3 |
| FIM1_4944 | 655.6273604 | 891.4626612 | 1183.17116 | 0.06224331 | 0.140729696 | 0.168442405 | 0.592944764 | ns | NaCl | KS.3 |
| FIM1_4945 | 9042.87889 | 8474.421506 | 7429.713428 | -0.022253164 | 0.094684223 | 0.626860084 | 0.895712211 | ns | NaCl | KS.3 |
| FIM1_4946 | 3187.25595 | 3777.715799 | 3010.056622 | 0.073525128 | 0.099161383 | 0.143925994 | 0.558168415 | ns | NaCl | KS.3 |
| FIM1_4947 | 2114.227935 | 1458.994896 | 1835.211868 | -0.402107749 | 0.181001847 | 7.0738003401 | 0.005586057 | ns | NaCl | KS.3 |
| FIM1_4948 | 365.7264307 | 347.4903736 | 502.0525132 | -0.022249507 | 0.087914216 | 0.64093476 | 0.898798052 | ns | NaCl | KS.3 |

| | | | | | | | | | | |
|---|---|---|---|---|---|---|---|---|---|---|
| FIM1_4949 | 1221.889726 | 1220.48109 | 1482.360849 | -0.002891561 | 0.059507377 | 0.950237451 | 0.987736646 | ns | NaCl | KS.3 |
| FIM1_4950 | 3425.243567 | 2899.89853 | 3177.919928 | -0.132946247 | 0.079599847 | 0.012553673 | 0.160551468 | ns | NaCl | KS.3 |
| FIM1_4951 | 2141.613335 | 2144.270562 | 2430.954616 | 0.000409425 | 0.091382191 | 0.992925795 | 0.999134858 | ns | NaCl | KS.3 |
| FIM1_4952 | 12671.3961 | 15275.06849 | 6919.195298 | 0.038713699 | 0.117774246 | 0.35904026 | 0.763018066 | ns | NaCl | KS.3 |
| FIM1_4953 | 4135.222761 | 5549.297944 | 2320.49064 | 0.052849087 | 0.138158582 | 0.219625644 | 0.642906874 | ns | NaCl | KS.3 |
| FIM1_4954 | 275.3793882 | 286.3499126 | 227.721582 | 0.006514966 | 0.112549014 | 0.866896418 | 0.970371341 | ns | NaCl | KS.3 |
| FIM1_4955 | 15691.65531 | 15784.4184 | 10080.4091 | 0.002319194 | 0.089440751 | 0.960482442 | 0.992398785 | ns | NaCl | KS.3 |
| FIM1_4956 | 1327.98616 | 1774.914073 | 2186.763112 | 0.109715421 | 0.138666732 | 0.047609839 | 0.343442 | ns | NaCl | KS.3 |
| FIM1_4957 | 627.2719009 | 900.2608339 | 753.7054491 | 0.152571368 | 0.178315859 | 0.01886811 | 0.207709917 | ns | NaCl | KS.3 |
| FIM1_4958 | 86.17812215 | 38.23634814 | 219.4088314 | -0.175262238 | 0.327297049 | 0.016937561 | 0.193711183 | ns | NaCl | KS.3 |
| FIM1_4960 | 6183.194116 | 4704.998034 | 4081.875028 | -0.279210097 | 0.120189805 | 7.69299277966 | 0.005888098 | ns | NaCl | KS.3 |
| FIM1_4961 | 6563.345858 | 4625.987461 | 3685.306776 | -0.141918907 | 0.169351334 | 0.023154135 | 0.235085349 | ns | NaCl | KS.3 |
| FIM1_4962 | 877.6379862 | 1218.580503 | 811.1420474 | 0.208995901 | 0.171575348 | 0.004436511 | 0.086217362 | ns | NaCl | KS.3 |
| FIM1_4963 | 668.416647 | 622.5586969 | 718.4865154 | -0.041323494 | 0.078984005 | 0.405417616 | 0.792777094 | ns | NaCl | KS.3 |
| FIM1_4964 | 53.07345515 | 66.61893696 | 90.85514937 | 0.069734796 | 0.120541959 | 0.147280574 | 0.562037426 | ns | NaCl | KS.3 |
| FIM1_4965 | 9499.710004 | 10881.86337 | 7135.825911 | 0.08267113 | 0.083714445 | 0.108762222 | 0.502406736 | ns | NaCl | KS.3 |
| FIM1_4966 | 3672.142162 | 3087.307316 | 4089.50433 | -0.135445 | 0.081814374 | 0.011831003 | 0.15408178 | ns | NaCl | KS.3 |
| FIM1_4967 | 1398.543662 | 1339.985807 | 1829.677499 | -0.021311116 | 0.082042914 | 0.662548315 | 0.904388188 | ns | NaCl | KS.3 |
| FIM1_4968 | 2437.365006 | 2802.299946 | 1696.526117 | 0.023200688 | 0.118498158 | 0.554191875 | 0.862821173 | ns | NaCl | KS.3 |
| FIM1_4969 | 14342.84981 | 14753.93034 | 13025.51777 | 0.02507188 | 0.059142043 | 0.587604723 | 0.878311243 | ns | NaCl | KS.3 |
| FIM1_4970 | 18311.09464 | 20378.92051 | 13733.02455 | 0.078525096 | 0.073006492 | 0.118918729 | 0.522633106 | ns | NaCl | KS.3 |
| FIM1_4971 | 3943.677071 | 4104.745718 | 5299.304992 | 0.013117637 | 0.095011531 | 0.771533958 | 0.94938462 | ns | NaCl | KS.3 |
| FIM1_4972 | 1378.385449 | 1396.455512 | 1907.458323 | 0.002849821 | 0.105091897 | 0.944800932 | 0.987187877 | ns | NaCl | KS.3 |
| FIM1_4973 | 1173.85798 | 1242.84356 | 1688.204938 | 0.017247239 | 0.097753316 | 0.697941931 | 0.922074642 | ns | NaCl | KS.3 |
| FIM1_4974 | 440.0617781 | 446.8513448 | 495.5369414 | 0.008473533 | 0.080830172 | 0.862150191 | 0.969856868 | ns | NaCl | KS.3 |
| FIM1_4976 | 9336.241503 | 10093.01704 | 7599.367736 | 0.046622579 | 0.078571725 | 0.349626985 | 0.756912594 | ns | NaCl | KS.3 |
| FIM1_4977 | 2797.847023 | 3801.37078 | 1707.165664 | 0.042846664 | 0.142711203 | 0.285416996 | 0.705523209 | ns | NaCl | KS.3 |
| FIM1_4978 | 1054.819317 | 563.201029 | 947.8801932 | -0.842297655 | 0.150309022 | 1.86444312121 | 6.18373635203 | ns | NaCl | KS.3 |
| FIM1_4979 | 20099.756 | 20087.67168 | 16927.46391 | -0.000420384 | 0.068379444 | 0.993102721 | 0.999134858 | ns | NaCl | KS.3 |
| FIM1_4980 | 233.346312 | 263.653607 | 556.3859116 | 0.03993203 | 0.099370929 | 0.386607826 | 0.779278134 | ns | NaCl | KS.3 |
| FIM1_4981 | 21682.17148 | 22447.52089 | 15859.94697 | 0.03268976 | 0.055723483 | 0.466285778 | 0.821323144 | ns | NaCl | KS.3 |
| FIM1_4982 | 875.7071983 | 937.9804129 | 1010.171553 | 0.056246839 | 0.063154979 | 0.238646805 | 0.661799251 | ns | NaCl | KS.3 |
| FIM1_4983 | 739.1746288 | 792.3320379 | 976.9536087 | 0.043816021 | 0.076010089 | 0.378400046 | 0.778493059 | ns | NaCl | KS.3 |
| FIM1_4984 | 682.7858908 | 560.0125213 | 1549.734645 | -0.079058242 | 0.10719872 | 0.117778204 | 0.520518737 | ns | NaCl | KS.3 |
| FIM1_4985 | 2347.722663 | 2092.322614 | 2672.857618 | -0.086092711 | 0.073000344 | 0.088546479 | 0.46158813 | ns | NaCl | KS.3 |
| FIM1_4986 | 4494.421802 | 4032.032396 | 4455.986282 | -0.085598842 | 0.069947543 | 0.086358027 | 0.45668467 | ns | NaCl | KS.3 |
| FIM1_4987 | 1956.670538 | 1876.770513 | 2820.403304 | -0.014334942 | 0.093784801 | 0.753491149 | 0.943523399 | ns | NaCl | KS.3 |
| FIM1_4988 | 121.8167794 | 121.9297108 | 174.4108259 | 0.00083587 | 0.097243116 | 0.984865514 | 0.998513538 | ns | NaCl | KS.3 |
| FIM1_4989 | 8244.969799 | 9724.403763 | 7408.666665 | 0.070037833 | 0.098769897 | 0.160650886 | 0.58211082 | ns | NaCl | KS.3 |
| FIM1_4991 | 4181.93611 | 4031.384391 | 3383.67469 | -0.011560869 | 0.095935845 | 0.796303252 | 0.955756014 | ns | NaCl | KS.3 |
| FIM1_4992 | 1059.776099 | 955.1415055 | 1100.95135 | -0.035294123 | 0.097163969 | 0.444426483 | 0.812278382 | ns | NaCl | KS.3 |
| FIM1_4993 | 2428.027001 | 2707.320669 | 2479.73763 | 0.04110134 | 0.094709813 | 0.384981279 | 0.778665419 | ns | NaCl | KS.3 |
| FIM1_4995 | 1842.58894 | 2233.087738 | 1753.530347 | 0.062582283 | 0.109527971 | 0.19014793 | 0.618952432 | ns | NaCl | KS.3 |
| FIM1_4996 | 442.4704032 | 332.9369412 | 533.7981596 | -0.12904781 | 0.137716916 | 0.027741663 | 0.260899256 | ns | NaCl | KS.3 |
| FIM1_4997 | 1246.961761 | 1297.682793 | 1646.699176 | 0.020907756 | 0.080522112 | 0.669723521 | 0.909851042 | ns | NaCl | KS.3 |

| | | | | | | | | | | |
|---|---|---|---|---|---|---|---|---|---|---|
| FIM1_4998 | 2312.178038 | 2111.258347 | 2531.997194 | -0.035456843 | 0.092268407 | 0.454338126 | 0.816788914 | ns | NaCl | KS.3 |
| FIM1_4999 | 5081.356411 | 5690.952955 | 3751.186881 | 0.050906941 | 0.090412038 | 0.298620522 | 0.717006321 | ns | NaCl | KS.3 |
| FIM1_5000 | 373.9928596 | 439.8103892 | 621.3457797 | 0.036453273 | 0.112981031 | 0.392616229 | 0.781618943 | ns | NaCl | KS.3 |
| FIM1_5001 | 625.4642637 | 704.1072047 | 1113.903104 | 0.025875593 | 0.109982487 | 0.533899012 | 0.855153333 | ns | NaCl | KS.3 |
| FIM1_5002 | 1256.396752 | 1270.120608 | 1846.561683 | 0.003862711 | 0.091116005 | 0.933488726 | 0.985410898 | ns | NaCl | KS.3 |
| FIM1_5003 | 1066.919811 | 900.4615047 | 1211.205547 | -0.085250548 | 0.096450511 | 0.099554451 | 0.482907447 | ns | NaCl | KS.3 |
| FIM1_5004 | 4350.400849 | 4557.535384 | 3222.634587 | 0.010833884 | 0.104294078 | 0.795265705 | 0.955325714 | ns | NaCl | KS.3 |
| FIM1_5005 | 28313.64629 | 33410.83841 | 20594.94691 | 0.109123465 | 0.086512543 | 0.040603518 | 0.317627561 | ns | NaCl | KS.3 |
| FIM1_5006 | 196.154 | 167.1003132 | 269.8336058 | -0.055007761 | 0.103676098 | 0.245112534 | 0.668549812 | ns | NaCl | KS.3 |
| FIM1_5007 | 2233.594526 | 3049.802332 | 2601.176588 | 0.056190688 | 0.141812176 | 0.197640556 | 0.620924909 | ns | NaCl | KS.3 |
| FIM1_5008 | 4380.130683 | 4751.124644 | 4296.317491 | 0.04339452 | 0.082722084 | 0.380718314 | 0.778493059 | ns | NaCl | KS.3 |
| FIM1_5009 | 69.88376891 | 21.94893578 | 119.1624388 | -0.925656151 | 0.631014834 | 0.000196361 | 0.012853871 | ns | NaCl | KS.3 |
| FIM1_5010 | 918.8309763 | 983.4236512 | 1071.331298 | 0.020227624 | 0.098334375 | 0.649066084 | 0.899642849 | ns | NaCl | KS.3 |
| FIM1_5011 | 202.5385862 | 196.9311816 | 171.8248649 | -0.009016496 | 0.098447827 | 0.836808144 | 0.962829707 | ns | NaCl | KS.3 |
| FIM1_5012 | 925.4137671 | 908.9781635 | 962.7448218 | -0.006185036 | 0.09291911 | 0.892353102 | 0.977268396 | ns | NaCl | KS.3 |
| FIM1_5013 | 1175.340312 | 1070.501248 | 1207.073186 | -0.046498841 | 0.085952068 | 0.345606543 | 0.754462295 | ns | NaCl | KS.3 |
| FIM1_5014 | 1588.998343 | 1633.220454 | 2375.999414 | 0.017580247 | 0.074028333 | 0.721037603 | 0.931688886 | ns | NaCl | KS.3 |
| FIM1_5015 | 3299.562911 | 3133.634606 | 2910.350965 | -0.019495895 | 0.091663133 | 0.675385137 | 0.912806589 | ns | NaCl | KS.3 |
| FIM1_5016 | 376.6940342 | 342.8428696 | 752.5604019 | -0.022617294 | 0.104977631 | 0.595130256 | 0.87962202 | ns | NaCl | KS.3 |
| FIM1_5017 | 2834.033436 | 2598.393492 | 2331.695015 | -0.026407483 | 0.098866816 | 0.555589451 | 0.863699344 | ns | NaCl | KS.3 |
| FIM1_5018 | 320.6841521 | 299.2524038 | 382.6164024 | -0.025525821 | 0.093488728 | 0.581834542 | 0.875038346 | ns | NaCl | KS.3 |
| FIM1_5019 | 1050.406961 | 1155.21211 | 1121.73002 | 0.022373827 | 0.10615542 | 0.595850798 | 0.87962202 | ns | NaCl | KS.3 |
| FIM1_5020 | 800.2841006 | 1027.120689 | 1100.992639 | 0.046651662 | 0.129340112 | 0.271008568 | 0.693144962 | ns | NaCl | KS.3 |
| FIM1_5021 | 418.4787426 | 324.7502574 | 757.4105412 | -0.163025696 | 0.124016353 | 0.00847351 | 0.127329254 | ns | NaCl | KS.3 |
| FIM1_5022 | 1792.413166 | 2202.101494 | 2655.25825 | 0.16422053 | 0.092557705 | 0.004084953 | 0.082277904 | ns | NaCl | KS.3 |
| FIM1_5023 | 2508.561274 | 2691.700232 | 1964.965564 | 0.045432578 | 0.074940176 | 0.360992438 | 0.763371684 | ns | NaCl | KS.3 |
| FIM1_5024 | 4480.971009 | 4947.230711 | 3413.721675 | 0.057883052 | 0.080992158 | 0.249305082 | 0.671881247 | ns | NaCl | KS.3 |
| FIM1_5025 | 2367.705246 | 2636.643941 | 2924.38756 | 0.03163957 | 0.100886137 | 0.479728212 | 0.827260954 | ns | NaCl | KS.3 |
| FIM1_5026 | 2117.25569 | 2999.010318 | 2627.883575 | 0.321881448 | 0.18328534 | 0.000369222 | 0.019196669 | ns | NaCl | KS.3 |
| FIM1_5027 | 4752.585845 | 6549.837411 | 3523.227109 | 0.167581346 | 0.160763841 | 0.011432918 | 0.151642712 | ns | NaCl | KS.3 |
| FIM1_5028 | 4001.488315 | 3985.197096 | 3668.055305 | -0.002724901 | 0.0716331 | 0.955674231 | 0.989784998 | ns | NaCl | KS.3 |
| FIM1_5029 | 2726.938321 | 2595.228889 | 2781.189831 | -0.039217219 | 0.0654562 | 0.416160677 | 0.800387073 | ns | NaCl | KS.3 |
| FIM1_5030 | 3586.376602 | 3676.131807 | 3657.061808 | 0.021525081 | 0.060446842 | 0.644741654 | 0.898899754 | ns | NaCl | KS.3 |
| FIM1_5031 | 7817.601413 | 10325.12803 | 5372.95306 | 0.209275502 | 0.136215043 | 0.002205806 | 0.057845441 | ns | NaCl | KS.3 |
| FIM1_5032 | 3938.245884 | 5110.751422 | 2251.585274 | 0.052240913 | 0.130387063 | 0.230578143 | 0.654338423 | ns | NaCl | KS.3 |
| FIM1_5033 | 1596.769958 | 1490.551114 | 1764.733732 | -0.059625625 | 0.062285204 | 0.209201761 | 0.633579568 | ns | NaCl | KS.3 |
| FIM1_5034 | 3717.540504 | 3272.112361 | 4885.437389 | -0.069257219 | 0.086082608 | 0.172175593 | 0.5990025 | ns | NaCl | KS.3 |
| FIM1_5035 | 1019.309385 | 1044.414164 | 1070.618224 | 0.006543342 | 0.099672656 | 0.879710128 | 0.973483611 | ns | NaCl | KS.3 |
| FIM1_5036 | 1567.621948 | 1582.868342 | 1330.740806 | 0.003488588 | 0.090653057 | 0.940108782 | 0.986790704 | ns | NaCl | KS.3 |
| FIM1_5037 | 1197.170816 | 1264.01997 | 1385.984308 | 0.045298932 | 0.064300615 | 0.345105402 | 0.754462295 | ns | NaCl | KS.3 |
| FIM1_5038 | 3949.778497 | 4095.528317 | 3473.213098 | 0.023203314 | 0.074116553 | 0.637877996 | 0.898332475 | ns | NaCl | KS.3 |
| FIM1_5039 | 1142.148706 | 1107.845547 | 1431.105986 | -0.015914358 | 0.082239562 | 0.743795497 | 0.941460666 | ns | NaCl | KS.3 |
| FIM1_5040 | 23785.32734 | 25623.51719 | 15960.82951 | 0.038301113 | 0.08341336 | 0.436388823 | 0.808096352 | ns | NaCl | KS.3 |
| FIM1_5041 | 393.0160815 | 414.7641949 | 428.8023825 | 0.02317292 | 0.090850085 | 0.621775977 | 0.892479944 | ns | NaCl | KS.3 |
| FIM1_5042 | 22642.59036 | 22817.9607 | 18192.75871 | 0.003614578 | 0.084571717 | 0.940101571 | 0.986790704 | ns | NaCl | KS.3 |

| | | | | | | | | | | | |
|---|---|---|---|---|---|---|---|---|---|---|---|
| FIM1_5043 | 1138.527832 | 919.4596699 | 914.906838 | -0.084231709 | 0.111043698 | 0.099590561 | 0.482907447 | ns | | NaCl | KS.3 |
| FIM1_5044 | 190.3540007 | 244.9695546 | 237.6434499 | 0.069691834 | 0.124850524 | 0.1451346 | 0.559290966 | ns | | NaCl | KS.3 |
| FIM1_5045 | 1424.458746 | 1673.251238 | 2635.336793 | 0.057763583 | 0.101742379 | 0.228808004 | 0.652325274 | ns | | NaCl | KS.3 |
| FIM1_5046 | 2066.193584 | 2624.792831 | 3462.930132 | 0.052239899 | 0.125126215 | 0.236052626 | 0.659616806 | ns | | NaCl | KS.3 |
| FIM1_5047 | 2391.641356 | 2727.233202 | 1684.120007 | 0.079913858 | 0.082826545 | 0.119726215 | 0.524421123 | ns | | NaCl | KS.3 |
| FIM1_5048 | 533.4348546 | 556.3243665 | 648.1284846 | 0.023464101 | 0.07948003 | 0.633230947 | 0.898009483 | ns | | NaCl | KS.3 |
| FIM1_5049 | 2492.795773 | 2584.352628 | 2315.334913 | 0.024322794 | 0.072218767 | 0.621048101 | 0.892479944 | ns | | NaCl | KS.3 |
| FIM1_5050 | 1357.780442 | 1435.989356 | 2158.68306 | 0.022615047 | 0.089497164 | 0.63267409 | 0.897798121 | ns | | NaCl | KS.3 |
| FIM1_5051 | 14.01995266 | 13.09068928 | 26.75475964 | -0.008076428 | 0.130699341 | 0.81683697 | 0.959030737 | ns | | NaCl | KS.3 |
| FIM1_5052 | 859.2288344 | 997.720181 | 833.6461928 | 0.086247424 | 0.088090611 | 0.096521994 | 0.474502885 | ns | | NaCl | KS.3 |
| FIM1_5053 | 1637.023657 | 1673.790433 | 1307.8111 | 0.011108432 | 0.082806898 | 0.81892187 | 0.959981393 | ns | | NaCl | KS.3 |
| FIM1_5054 | 730.965436 | 644.9971265 | 670.2714311 | -0.054235883 | 0.092778586 | 0.267521827 | 0.689593109 | ns | | NaCl | KS.3 |
| FIM1_5055 | 4213.129122 | 4475.098213 | 3470.755075 | 0.035947094 | 0.077689507 | 0.468193931 | 0.821764392 | ns | | NaCl | KS.3 |
| FIM1_5056 | 1088.228719 | 980.882791 | 1420.427163 | -0.054580904 | 0.084510564 | 0.274150812 | 0.694804019 | ns | | NaCl | KS.3 |
| FIM1_5057 | 11057.61872 | 14742.80817 | 6219.004623 | 0.049230876 | 0.137279119 | 0.243696709 | 0.667638752 | ns | | NaCl | KS.3 |
| FIM1_5058 | 2180.991065 | 2127.287486 | 2622.191853 | -0.007543863 | 0.097589935 | 0.864001134 | 0.969856868 | ns | | NaCl | KS.3 |
| FIM1_5059 | 596.3157135 | 662.1970078 | 686.9090204 | 0.077803752 | 0.072501289 | 0.121582404 | 0.527999953 | ns | | NaCl | KS.3 |
| FIM1_5061 | 2516.538648 | 2821.486724 | 2153.880053 | 0.056241933 | 0.088022541 | 0.2577107 | 0.681249061 | ns | | NaCl | KS.3 |
| FIM1_5062 | 391.2248067 | 362.1274081 | 633.3694877 | -0.038270625 | 0.083365064 | 0.436796637 | 0.808096352 | ns | | NaCl | KS.3 |
| FIM1_5063 | 2654.687277 | 2451.158966 | 1990.11932 | -0.039903545 | 0.084792725 | 0.41643398 | 0.800387073 | ns | | NaCl | KS.3 |
| FIM1_5064 | 734.0352618 | 905.2455059 | 645.2072038 | 0.063327297 | 0.114476621 | 0.181501605 | 0.611396712 | ns | | NaCl | KS.3 |
| FIM1_5065 | 3930.265966 | 4342.150053 | 3945.297633 | 0.032346575 | 0.098033894 | 0.478328343 | 0.826171162 | ns | | NaCl | KS.3 |
| FIM1_5066 | 372.4147909 | 410.3701135 | 475.1938222 | 0.055120136 | 0.080857196 | 0.271488771 | 0.693144962 | ns | | NaCl | KS.3 |
| FIM1_5067 | 4499.344374 | 4651.386165 | 3488.00408 | 0.02002904 | 0.07634838 | 0.684409804 | 0.91584821 | ns | | NaCl | KS.3 |
| FIM1_5068 | 974.3830157 | 1064.76627 | 1083.197022 | 0.031255078 | 0.094976854 | 0.50018527 | 0.839756535 | ns | | NaCl | KS.3 |
| FIM1_5069 | 149.5393186 | 131.0099609 | 1090.397409 | -0.018082242 | 0.122057715 | 0.631484996 | 0.897610816 | ns | | NaCl | KS.3 |
| FIM1_5070 | 45.12256516 | 39.59783198 | 66.93085334 | -0.022677956 | 0.111229137 | 0.578599364 | 0.8749481 | ns | | NaCl | KS.3 |
| FIM1_5071 | 7759.92297 | 7458.488841 | 7131.434308 | -0.023588905 | 0.076978042 | 0.632416749 | 0.897798121 | ns | | NaCl | KS.3 |
| FIM1_5072 | 79.16203784 | 65.63955847 | 128.7444275 | -0.03766 | 0.116949988 | 0.371506637 | 0.773001053 | ns | | NaCl | KS.3 |
| FIM1_5073 | 554.4431642 | 505.169937 | 611.9560529 | -0.034126929 | 0.095129986 | 0.463636599 | 0.819880744 | ns | | NaCl | KS.3 |
| FIM1_5074 | 623.4429486 | 534.2984968 | 663.8664992 | -0.064499582 | 0.097648161 | 0.191435316 | 0.618952432 | ns | | NaCl | KS.3 |
| FIM1_5075 | 916.0810504 | 606.3308142 | 1277.11883 | -0.341352512 | 0.232692226 | 0.000842742 | 0.031433557 | ns | | NaCl | KS.3 |
| FIM1_5076 | 4870.183992 | 4495.618893 | 3651.790514 | -0.046791481 | 0.079504651 | 0.347761487 | 0.756654396 | ns | | NaCl | KS.3 |
| FIM1_5077 | 2034.496878 | 1853.345478 | 2209.852189 | -0.039161462 | 0.090759454 | 0.414624851 | 0.800387073 | ns | | NaCl | KS.3 |
| FIM1_5078 | 296.2060549 | 297.4293987 | 454.6684508 | 0.002151539 | 0.087666618 | 0.963738725 | 0.993691273 | ns | | NaCl | KS.3 |
| FIM1_5079 | 8071.122907 | 7455.560426 | 6011.899769 | -0.043033631 | 0.082088559 | 0.385083798 | 0.778665419 | ns | | NaCl | KS.3 |
| FIM1_5080 | 882.369735 | 939.1600999 | 1060.424088 | 0.054269598 | 0.059931021 | 0.244490426 | 0.66821307 | ns | | NaCl | KS.3 |
| FIM1_5081 | 1592.154167 | 1586.842899 | 1610.532415 | -0.001695218 | 0.076205528 | 0.972484742 | 0.995313014 | ns | | NaCl | KS.3 |
| FIM1_5082 | 720.6829401 | 743.2647618 | 557.9162857 | 0.014734594 | 0.083540732 | 0.76103076 | 0.945911573 | ns | | NaCl | KS.3 |
| FIM1_5083 | 5690.524796 | 6238.763523 | 4870.532015 | 0.023949192 | 0.103257022 | 0.579523242 | 0.8749481 | ns | | NaCl | KS.3 |
| FIM1_5084 | 1157.150195 | 1170.241126 | 802.1607 | 0.002263457 | 0.109007093 | 0.954671944 | 0.989477692 | ns | | NaCl | KS.3 |
| FIM1_5085 | 1683.487606 | 1430.131449 | 1763.991479 | -0.044223825 | 0.108797458 | 0.323295896 | 0.739131368 | ns | | NaCl | KS.3 |
| FIM1_5086 | 2682.940824 | 2765.977422 | 2732.781178 | 0.015769424 | 0.081396763 | 0.746659505 | 0.943274417 | ns | | NaCl | KS.3 |
| FIM1_5087 | 1224.382023 | 1285.107212 | 1469.97806 | 0.016153406 | 0.094880059 | 0.721754566 | 0.931688886 | ns | | NaCl | KS.3 |
| FIM1_5088 | 10779.76312 | 11718.2715 | 9428.848193 | 0.022018493 | 0.102439215 | 0.610891943 | 0.887872455 | ns | | NaCl | KS.3 |

| | | | | | | | | | | |
|---|---|---|---|---|---|---|---|---|---|---|
| FIM1_5089 | 2209.584685 | 2407.173468 | 2468.934404 | 0.019308502 | 0.10665322 | 0.643302529 | 0.898798052 | ns | NaCl | KS.3 |
| FIM1_5090 | 304.6147265 | 232.4156495 | 387.9351323 | -0.052007123 | 0.132334555 | 0.230177784 | 0.654338423 | ns | NaCl | KS.3 |
| FIM1_5091 | 407.6806453 | 235.9848688 | 492.4874872 | -0.26305278 | 0.296054027 | 0.005511276 | 0.098456203 | ns | NaCl | KS.3 |
| FIM1_5093 | 915.0687618 | 1009.613675 | 1273.576786 | 0.068837199 | 0.075690005 | 0.172329583 | 0.599043855 | ns | NaCl | KS.3 |
| FIM1_5094 | 122508.3526 | 141388.2164 | 89508.85285 | 0.101596869 | 0.079373697 | 0.051032097 | 0.352617612 | ns | NaCl | KS.3 |
| FIM1_5095 | 531.9051347 | 577.7023116 | 503.4258247 | 0.038865864 | 0.086838615 | 0.425169002 | 0.802854185 | ns | NaCl | KS.3 |
| FIM1_5096 | 1804.059383 | 2181.282487 | 1407.574219 | 0.074882991 | 0.105397459 | 0.134591753 | 0.54183277 | ns | NaCl | KS.3 |
| FIM1_5097 | 12109.73132 | 13343.86594 | 9492.802646 | 0.086618437 | 0.0623582 | 0.069498329 | 0.413948267 | ns | NaCl | KS.3 |
| FIM1_5098 | 272.2270435 | 272.8223188 | 297.09687 | 0.00040111 | 0.10123563 | 0.992478804 | 0.999134858 | ns | NaCl | KS.3 |
| FIM1_5099 | 2374.863008 | 2553.18031 | 1756.661765 | 0.040155751 | 0.080843169 | 0.417535771 | 0.800387073 | ns | NaCl | KS.3 |
| FIM1_5100 | 910.0814524 | 947.288653 | 1058.239174 | 0.036511011 | 0.058077406 | 0.425916333 | 0.802854185 | ns | NaCl | KS.3 |
| FIM1_5101 | 894.7951953 | 768.3675887 | 1189.433938 | -0.087937626 | 0.088650584 | 0.090993455 | 0.464798331 | ns | NaCl | KS.3 |
| FIM1_5102 | 1618.400624 | 2131.404812 | 3568.104286 | 0.07122915 | 0.131888369 | 0.135927178 | 0.544093414 | ns | NaCl | KS.3 |
| FIM1_5103 | 520.5399092 | 438.109093 | 511.8376738 | -0.084613326 | 0.09711118 | 0.101558397 | 0.485353528 | ns | NaCl | KS.3 |
| FIM1_5104 | 779.6944855 | 736.2606815 | 743.3096505 | -0.041808845 | 0.070514819 | 0.39579893 | 0.783253649 | ns | NaCl | KS.3 |
| FIM1_5105 | 3302.69937 | 2998.547212 | 2556.219037 | -0.086168411 | 0.062444439 | 0.071118178 | 0.418675804 | ns | NaCl | KS.3 |
| FIM1_5106 | 553.9600564 | 397.2134932 | 873.3045665 | -0.2255849 | 0.173519484 | 0.002990994 | 0.068257782 | ns | NaCl | KS.3 |
| FIM1_5107 | 6307.810218 | 6398.557261 | 4318.641348 | 0.006518413 | 0.085145639 | 0.891971312 | 0.977268396 | ns | NaCl | KS.3 |
| FIM1_5108 | 970.4949541 | 1147.130852 | 919.5715219 | 0.076060361 | 0.09744733 | 0.13374807 | 0.541152807 | ns | NaCl | KS.3 |
| FIM1_5109 | 1361.818765 | 1452.926923 | 1055.894387 | 0.027187088 | 0.088927297 | 0.568533246 | 0.869550295 | ns | NaCl | KS.3 |
| FIM1_5110 | 1206.531678 | 1331.385985 | 1771.777108 | 0.058131606 | 0.080498017 | 0.247439478 | 0.669893726 | ns | NaCl | KS.3 |
| FIM1_5111 | 1902.339706 | 2184.425342 | 1833.436691 | 0.09353856 | 0.079986758 | 0.071006155 | 0.418675804 | ns | NaCl | KS.3 |
| FIM1_5112 | 798.981031 | 884.5709525 | 912.3996043 | 0.056558131 | 0.082415422 | 0.25932016 | 0.68302598 | ns | NaCl | KS.3 |
| FIM1_5113 | 182.0439 | 196.84272 | 185.8773579 | 0.023588232 | 0.099519993 | 0.594764434 | 0.87962202 | ns | NaCl | KS.3 |
| FIM1_5114 | 568.0283858 | 725.1905274 | 473.4386546 | 0.077751037 | 0.12237031 | 0.116685714 | 0.519941265 | ns | NaCl | KS.3 |
| FIM1_5115 | 2675.457217 | 2700.836003 | 1717.039464 | 0.005343103 | 0.079220208 | 0.913109995 | 0.98221021 | ns | NaCl | KS.3 |
| FIM1_5116 | 2011.491175 | 1739.464069 | 1592.914278 | -0.084837432 | 0.086478642 | 0.101237407 | 0.484750818 | ns | NaCl | KS.3 |
| FIM1_5117 | 397.5713995 | 566.9025448 | 553.3084487 | 0.109905818 | 0.162639838 | 0.04905689 | 0.34611662 | ns | NaCl | KS.3 |
| FIM1_5118 | 3395.368392 | 3848.821465 | 2917.278492 | 0.061316006 | 0.089286903 | 0.219596035 | 0.642906874 | ns | NaCl | KS.3 |
| FIM1_5119 | 1309.520985 | 1203.616434 | 1408.965236 | -0.050487277 | 0.079166459 | 0.312354732 | 0.731871738 | ns | NaCl | KS.3 |
| FIM1_5120 | 1138.828053 | 1182.664152 | 1218.891875 | 0.010504163 | 0.1000255 | 0.808135054 | 0.958396161 | ns | NaCl | KS.3 |
| FIM1_5121 | 2577.68791 | 2614.020712 | 2267.146921 | 0.006135184 | 0.085529991 | 0.898090765 | 0.977971161 | ns | NaCl | KS.3 |
| FIM1_5122 | 2798.756462 | 3758.404486 | 4430.879936 | 0.052619466 | 0.138351068 | 0.220856766 | 0.64330352 | ns | NaCl | KS.3 |
| FIM1_5123 | 4394.75645 | 6023.391997 | 4067.119462 | 0.053684908 | 0.142813098 | 0.21084614 | 0.634379635 | ns | NaCl | KS.3 |
| FIM1_5124 | 1196.237779 | 1045.83491 | 969.6212758 | -0.093366416 | 0.079735397 | 0.071331872 | 0.418675804 | ns | NaCl | KS.3 |
| FIM1_5125 | 3681.9535 | 4025.551467 | 2718.883313 | 0.017298025 | 0.110979608 | 0.66713916 | 0.907354532 | ns | NaCl | KS.3 |
| FIM1_5126 | 1502.593976 | 1775.004295 | 1546.701462 | 0.159272102 | 0.069486817 | 0.001763039 | 0.050408731 | ns | NaCl | KS.3 |
| FIM1_5127 | 4687.350845 | 5062.147896 | 4733.280554 | 0.035252885 | 0.087308222 | 0.466874744 | 0.821323144 | ns | NaCl | KS.3 |
| FIM1_5128 | 4017.425133 | 3398.823667 | 3394.131703 | -0.129106495 | 0.080431014 | 0.015426547 | 0.182493537 | ns | NaCl | KS.3 |
| FIM1_5129 | 1419.004458 | 1551.230746 | 1408.446859 | 0.044643895 | 0.085168532 | 0.365145538 | 0.765851202 | ns | NaCl | KS.3 |
| FIM1_5130 | 1827.870229 | 1958.573876 | 2394.966011 | 0.02831627 | 0.090334712 | 0.550158903 | 0.860945181 | ns | NaCl | KS.3 |
| FIM1_5131 | 2325.166018 | 752.2531042 | 2682.725242 | -0.910362993 | 0.601500782 | 0.000194666 | 0.012853871 | ns | NaCl | KS.3 |
| FIM1_5132 | 127.7417107 | 206.6693615 | 193.6028101 | 0.084850047 | 0.187600391 | 0.086081688 | 0.456563326 | ns | NaCl | KS.3 |
| FIM1_5133 | 1153.521619 | 1042.795095 | 1610.349592 | -0.045286686 | 0.089381321 | 0.352535762 | 0.756912594 | ns | NaCl | KS.3 |
| FIM1_5134 | 688.2764783 | 796.8015163 | 596.8297995 | 0.056111601 | 0.098498345 | 0.244920608 | 0.668549812 | ns | NaCl | KS.3 |

| | | | | | | | | | | |
|---|---|---|---|---|---|---|---|---|---|---|
| FIM1_5135 | 1701.442628 | 1652.224419 | 1370.994416 | -0.012589398 | 0.08769677 | 0.790802357 | 0.953972111 | ns | NaCl | KS.3 |
| FIM1_5136 | 2871.959986 | 2618.274468 | 2722.172896 | -0.046293965 | 0.085421724 | 0.348267209 | 0.756654396 | ns | NaCl | KS.3 |
| FIM1_5137 | 855.3735336 | 929.0807089 | 955.2377765 | 0.029396125 | 0.094258972 | 0.526452392 | 0.855027345 | ns | NaCl | KS.3 |
| FIM1_5138 | 2396.209907 | 2619.045204 | 2129.186013 | 0.018905748 | 0.108441916 | 0.645695307 | 0.898899754 | ns | NaCl | KS.3 |
| FIM1_5139 | 3622.397551 | 3004.878627 | 3263.840028 | -0.0402141 | 0.116942666 | 0.345611974 | 0.754462295 | ns | NaCl | KS.3 |
| FIM1_5140 | 4461.215371 | 3353.076951 | 3280.874518 | -0.063131603 | 0.134995358 | 0.167712565 | 0.592592338 | ns | NaCl | KS.3 |
| FIM1_5141 | 1826.472646 | 2005.344725 | 2740.679032 | 0.03907815 | 0.090435761 | 0.416214004 | 0.800387073 | ns | NaCl | KS.3 |
| FIM1_5142 | 1264.215304 | 1245.462461 | 1638.967762 | -0.003960581 | 0.101135716 | 0.925972278 | 0.984176127 | ns | NaCl | KS.3 |
| FIM1_5143 | 2582.907489 | 2684.475122 | 2447.338046 | 0.029213796 | 0.066373129 | 0.5458517 | 0.857742326 | ns | NaCl | KS.3 |
| FIM1_5144 | 1283.87323 | 1328.933281 | 1888.397872 | 0.018235366 | 0.078913249 | 0.710488254 | 0.925563973 | ns | NaCl | KS.3 |
| FIM1_5145 | 2002.509384 | 2063.184206 | 2312.449822 | 0.015701554 | 0.082330032 | 0.747012792 | 0.943274417 | ns | NaCl | KS.3 |
| FIM1_5146 | 856.8156678 | 1157.007992 | 627.1426176 | 0.08985032 | 0.140309583 | 0.081090846 | 0.439771952 | ns | NaCl | KS.3 |
| FIM1_5147 | 41775.23242 | 47877.84654 | 29081.46093 | 0.082656651 | 0.083950581 | 0.108879491 | 0.502481882 | ns | NaCl | KS.3 |
| FIM1_5148 | 187.3570423 | 206.2885067 | 298.8760442 | 0.020130972 | 0.108892063 | 0.624556566 | 0.894149328 | ns | NaCl | KS.3 |
| FIM1_5149 | 1985.446473 | 917.7863128 | 4101.836441 | -0.264417472 | 0.37537767 | 0.006835483 | 0.111371848 | ns | NaCl | KS.3 |
| FIM1_5150 | 11676.95482 | 13529.72371 | 9268.607682 | 0.043160161 | 0.104964148 | 0.341137047 | 0.750290367 | ns | NaCl | KS.3 |
| FIM1_5151 | 236.8949765 | 233.0315582 | 223.4158694 | -0.007506636 | 0.090371428 | 0.871953519 | 0.971243188 | ns | NaCl | KS.3 |
| FIM1_5152 | 163.928748 | 110.9064176 | 293.9752473 | -0.105947787 | 0.174883025 | 0.054356505 | 0.363472596 | ns | NaCl | KS.3 |
| FIM1_5153 | 7415.21316 | 7594.508216 | 6107.404169 | 0.016898194 | 0.069629543 | 0.72945853 | 0.935564884 | ns | NaCl | KS.3 |
| FIM1_5155 | 938.9176673 | 955.5397258 | 1027.34733 | 0.007905845 | 0.083458672 | 0.870155188 | 0.971243188 | ns | NaCl | KS.3 |
| FIM1_5156 | 1152.587126 | 1255.047108 | 907.6906643 | 0.02195907 | 0.103055384 | 0.610214826 | 0.887342408 | ns | NaCl | KS.3 |
| FIM1_5157 | 2681.318543 | 2841.355996 | 2358.045634 | 0.051191323 | 0.059688196 | 0.271207464 | 0.693144962 | ns | NaCl | KS.3 |
| FIM1_5158 | 1084.95709 | 944.1216728 | 1384.854333 | -0.041130266 | 0.103983689 | 0.363338059 | 0.764655523 | ns | NaCl | KS.3 |
| FIM1_5159 | 780.9809655 | 808.4772969 | 963.2899021 | 0.017861208 | 0.08172467 | 0.714393024 | 0.92820718 | ns | NaCl | KS.3 |
| FIM1_5160 | 19083.66988 | 26089.11488 | 11959.88436 | 0.094073791 | 0.144647279 | 0.072385773 | 0.421192069 | ns | NaCl | KS.3 |
| FIM1_5161 | 1764.922716 | 1655.088457 | 1637.126416 | -0.013022089 | 0.108609867 | 0.747774041 | 0.943274417 | ns | NaCl | KS.3 |
| FIM1_5162 | 3021.385113 | 3235.760289 | 2683.565014 | 0.046521396 | 0.07321993 | 0.348593342 | 0.756654396 | ns | NaCl | KS.3 |
| FIM1_5163 | 2783.252441 | 2946.127006 | 3379.838754 | 0.020960349 | 0.092416414 | 0.651462295 | 0.90003469 | ns | NaCl | KS.3 |
| FIM1_5164 | 268.4436211 | 221.008437 | 390.2049425 | -0.094990163 | 0.103282033 | 0.071864388 | 0.42012377 | ns | NaCl | KS.3 |
| FIM1_5165 | 875.8160658 | 858.8064492 | 1240.079075 | -0.01027317 | 0.084356134 | 0.831269319 | 0.961620266 | ns | NaCl | KS.3 |
| FIM1_5166 | 3382.25969 | 3007.056576 | 2961.30963 | -0.033289396 | 0.102901722 | 0.453749381 | 0.816716052 | ns | NaCl | KS.3 |
| FIM1_5167 | 638.2825618 | 555.5822712 | 637.241487 | -0.076759163 | 0.088345214 | 0.133792553 | 0.541152807 | ns | NaCl | KS.3 |
| FIM1_5168 | 460.7935466 | 450.7121024 | 781.3317534 | -0.00943069 | 0.088988294 | 0.841070817 | 0.965465463 | ns | NaCl | KS.3 |
| FIM1_5169 | 262.9980816 | 241.3967072 | 280.748438 | -0.024865396 | 0.100922024 | 0.572239958 | 0.871142531 | ns | NaCl | KS.3 |
| FIM1_5170 | 1792.371862 | 1639.728229 | 2146.695084 | -0.0681208 | 0.06986799 | 0.169343029 | 0.594552976 | ns | NaCl | KS.3 |
| FIM1_5171 | 851.6150349 | 897.7108944 | 1014.350572 | 0.020224807 | 0.091185705 | 0.665169722 | 0.905890875 | ns | NaCl | KS.3 |
| FIM1_5172 | 10321.38002 | 11447.56788 | 7294.21892 | 0.061272575 | 0.080884721 | 0.224063807 | 0.648059715 | ns | NaCl | KS.3 |
| FIM1_5173 | 1424.010818 | 1392.805756 | 1704.099513 | -0.011283712 | 0.083236268 | 0.815837076 | 0.958816322 | ns | NaCl | KS.3 |
| FIM1_5174 | 3642.293564 | 4034.664811 | 4111.276212 | 0.083733823 | 0.067782358 | 0.089769065 | 0.463545584 | ns | NaCl | KS.3 |
| FIM1_5175 | 4938.285293 | 3233.786157 | 3329.456953 | -0.485821416 | 0.196787716 | 4.00919756105 | 0.003693659 | ns | NaCl | KS.3 |
| FIM1_5176 | 771.8942923 | 1094.22353 | 608.5618791 | 0.14356623 | 0.169733468 | 0.02229613 | 0.229180264 | ns | NaCl | KS.3 |
| FIM1_5177 | 2573.373747 | 3275.793699 | 1527.121631 | 0.036547666 | 0.131673377 | 0.358022732 | 0.76205201 | ns | NaCl | KS.3 |
| FIM1_5178 | 1252.654902 | 1431.008189 | 1130.801298 | 0.09203779 | 0.07918655 | 0.074784707 | 0.425690982 | ns | NaCl | KS.3 |
| FIM1_5179 | 7535.625516 | 9904.776697 | 4531.64748 | 0.057914381 | 0.132488621 | 0.195294531 | 0.618952432 | ns | NaCl | KS.3 |
| FIM1_5180 | 711.7864532 | 717.7288066 | 802.0512042 | 0.004269297 | 0.083614574 | 0.929574507 | 0.984238275 | ns | NaCl | KS.3 |

| | | | | | | | | | | |
|---|---|---|---|---|---|---|---|---|---|---|
| FIM1_5181 | 5349.438373 | 4786.556259 | 4942.464288 | -0.059481876 | 0.084850201 | 0.235795323 | 0.659616806 | ns | NaCl | KS.3 |
| FIM1_5182 | 1555.034121 | 1231.370718 | 1281.7074 | -0.198596607 | 0.105190668 | 0.0012736 | 0.041599773 | ns | NaCl | KS.3 |
| FIM1_5183 | 7279.92615 | 7451.367923 | 5023.530225 | 0.012348365 | 0.080490977 | 0.800572997 | 0.957185932 | ns | NaCl | KS.3 |
| FIM1_5184 | 728.4347412 | 930.1918205 | 724.6511252 | 0.138292854 | 0.117726041 | 0.018217042 | 0.202751192 | ns | NaCl | KS.3 |
| FIM1_5185 | 3953.199462 | 4301.637853 | 3451.371304 | 0.055693896 | 0.075540492 | 0.26568331 | 0.689593109 | ns | NaCl | KS.3 |
| FIM1_5186 | 1619.29438 | 1485.235408 | 1683.684437 | -0.054255282 | 0.077181322 | 0.278618387 | 0.700652348 | ns | NaCl | KS.3 |
| FIM1_5187 | 1322.800605 | 1012.539161 | 2096.002034 | -0.107944181 | 0.128764282 | 0.049218138 | 0.34611662 | ns | NaCl | KS.3 |
| FIM1_5188 | 10331.29578 | 9408.364967 | 8105.092407 | -0.036333444 | 0.092975834 | 0.442299444 | 0.809580476 | ns | NaCl | KS.3 |
| FIM1_5189 | 231.2963598 | 232.0612255 | 255.0617251 | 0.00065543 | 0.106135259 | 0.987173695 | 0.999134858 | ns | NaCl | KS.3 |
| FIM1_5190 | 909.8161356 | 977.7103932 | 1079.103786 | 0.036563541 | 0.083686342 | 0.456495362 | 0.817337042 | ns | NaCl | KS.3 |
| FIM1_5191 | 1309.113632 | 1441.351184 | 1573.276048 | 0.038461302 | 0.092730828 | 0.418304465 | 0.800387073 | ns | NaCl | KS.3 |
| FIM1_5192 | 633.4759983 | 535.2792548 | 1933.053245 | -0.023167564 | 0.126428505 | 0.539146016 | 0.857036524 | ns | NaCl | KS.3 |
| FIM1_5193 | 21400.75797 | 23582.97705 | 15741.76877 | 0.064226763 | 0.076225043 | 0.202018261 | 0.625219184 | ns | NaCl | KS.3 |
| FIM1_5194 | 3822.874169 | 4757.536468 | 3900.873431 | 0.094382874 | 0.111190897 | 0.073035071 | 0.421631393 | ns | NaCl | KS.3 |
| FIM1_5195 | 5857.656255 | 5481.351157 | 5536.513644 | -0.036385412 | 0.080940966 | 0.461344755 | 0.819880744 | ns | NaCl | KS.3 |
| FIM1_5196 | 203.3954452 | 173.1183467 | 274.2174018 | -0.071842181 | 0.098716284 | 0.15195111 | 0.567961513 | ns | NaCl | KS.3 |
| FIM1_5197 | 391.8083893 | 489.0805989 | 384.7279617 | 0.139139566 | 0.105954789 | 0.015599815 | 0.182568377 | ns | NaCl | KS.3 |
| FIM1_5199 | 3580.478975 | 3889.776303 | 3164.931706 | 0.064601425 | 0.068049336 | 0.188501411 | 0.618812647 | ns | NaCl | KS.3 |
| FIM1_5200 | 165.2875549 | 194.3531599 | 329.1609456 | 0.025980236 | 0.120688918 | 0.508181842 | 0.844423736 | ns | NaCl | KS.3 |
| FIM1_5201 | 294.3692783 | 377.082832 | 502.8709471 | 0.074461275 | 0.123416954 | 0.127616965 | 0.535324115 | ns | NaCl | KS.3 |
| FIM1_5202 | 657.351329 | 620.1265803 | 778.6488792 | -0.035896742 | 0.077665661 | 0.468810467 | 0.821764392 | ns | NaCl | KS.3 |
| FIM1_5203 | 648.2125298 | 693.4427604 | 745.3949667 | 0.027986793 | 0.089343847 | 0.556714926 | 0.863699344 | ns | NaCl | KS.3 |
| FIM1_5204 | 215.7096089 | 377.3020431 | 477.5816684 | 0.558226232 | 0.297978634 | 0.000211613 | 0.013672404 | ns | NaCl | KS.3 |
| FIM1_5205 | 1831.452582 | 1849.4982 | 1564.827119 | 0.007518121 | 0.06740369 | 0.876739521 | 0.972783118 | ns | NaCl | KS.3 |
| FIM1_5206 | 813.8963738 | 1036.569813 | 958.1753731 | 0.19045135 | 0.112185735 | 0.002320102 | 0.05859141 | ns | NaCl | KS.3 |
| FIM1_5207 | 141.6217144 | 174.6521664 | 269.4435916 | 0.066695122 | 0.110969263 | 0.167469546 | 0.592154224 | ns | NaCl | KS.3 |
| FIM1_5208 | 18.53488148 | 20.8745512 | 21.5559521 | 0.007735262 | 0.145546407 | 0.811212177 | 0.958816322 | ns | NaCl | KS.3 |
| FIM1_5209 | 41.18937627 | 34.80246566 | 37.8397702 | -0.01883486 | 0.130013545 | 0.604481022 | 0.883458603 | ns | NaCl | KS.3 |
| FIM1_4316 | 5980.971318 | 6602.240727 | 4233.280858 | 0.03081452 | 0.099000986 | 0.495398178 | 0.836358994 | ns | NaCl | KS.3 |
| FIM1_4318 | 138.3435074 | 101.8140764 | 297.7479064 | -0.117038681 | 0.143545936 | 0.039478073 | 0.315147688 | ns | NaCl | KS.3 |
| FIM1_4319 | 1240.561205 | 624.2876744 | 791.3821641 | -0.417731685 | 0.394481079 | 0.00179354 | 0.050964538 | ns | NaCl | KS.3 |
| FIM1_4320 | 4000.130207 | 4043.447819 | 3853.218499 | 0.006606745 | 0.075974489 | 0.893107626 | 0.977268396 | ns | NaCl | KS.3 |
| FIM1_4321 | 1295.299168 | 1304.788987 | 1574.930606 | 0.003566758 | 0.080575593 | 0.941739712 | 0.986790704 | ns | NaCl | KS.3 |
| FIM1_4322 | 1578.344212 | 2160.562739 | 1448.110775 | 0.263593335 | 0.16157752 | 0.000816728 | 0.031433557 | ns | NaCl | KS.3 |
| FIM1_4323 | 19888.59725 | 21328.71356 | 16724.97566 | 0.063505564 | 0.059446796 | 0.171975409 | 0.5990025 | ns | NaCl | KS.3 |
| FIM1_4324 | 35.08462931 | 18.71173378 | 38.88673509 | -0.152599196 | 0.267438616 | 0.023877787 | 0.239908881 | ns | NaCl | KS.3 |
| FIM1_4325 | 1662.824262 | 1700.064564 | 1983.336966 | 0.013892611 | 0.073882747 | 0.777653463 | 0.950849745 | ns | NaCl | KS.3 |
| FIM1_4326 | 1574.303807 | 1531.851622 | 1289.899005 | -0.011069885 | 0.089378454 | 0.813603961 | 0.958816322 | ns | NaCl | KS.3 |
| FIM1_4327 | 2718.075898 | 2841.472034 | 2328.645034 | 0.022230127 | 0.083096184 | 0.648028934 | 0.899642849 | ns | NaCl | KS.3 |
| FIM1_4328 | 1137.822613 | 1219.244518 | 872.2491588 | 0.022145944 | 0.096708969 | 0.623575745 | 0.893754273 | ns | NaCl | KS.3 |
| FIM1_4329 | 330.5079822 | 345.0648015 | 704.8157214 | 0.015639224 | 0.091813124 | 0.735767674 | 0.938575431 | ns | NaCl | KS.3 |
| FIM1_4330 | 2354.730606 | 2379.312889 | 2374.448184 | 0.004706792 | 0.083357866 | 0.922463676 | 0.983890455 | ns | NaCl | KS.3 |
| FIM1_4331 | 7161.950673 | 8237.888969 | 5501.843841 | 0.045102128 | 0.101887339 | 0.329328616 | 0.741299385 | ns | NaCl | KS.3 |
| FIM1_4332 | 1690.071724 | 1800.18028 | 1772.143234 | 0.052845235 | 0.063968872 | 0.270398946 | 0.692349335 | ns | NaCl | KS.3 |
| FIM1_4333 | 3375.281599 | 3257.065654 | 2923.975554 | -0.026093805 | 0.068058346 | 0.591922651 | 0.878786986 | ns | NaCl | KS.3 |

| | | | | | | | | | | |
|---|---|---|---|---|---|---|---|---|---|---|
| FIM1_4334 | 895.3591735 | 786.5734274 | 898.2957246 | -0.051816806 | 0.09539526 | 0.282958964 | 0.704025235 | ns | NaCl | KS.3 |
| FIM1_4335 | 2469.434158 | 1858.225277 | 2760.392648 | -0.363665601 | 0.091049756 | 5.33636385693 | 1.32742050941 | ns | NaCl | KS.3 |
| FIM1_4336 | 733.5445229 | 759.7011491 | 963.1192203 | 0.013420383 | 0.088397543 | 0.776566881 | 0.950410881 | ns | NaCl | KS.3 |
| FIM1_4337 | 1553.140452 | 1552.443532 | 1516.12613 | -0.00045644 | 0.077352215 | 0.992582768 | 0.999134858 | ns | NaCl | KS.3 |
| FIM1_4338 | 5259.866009 | 4956.134892 | 4371.463693 | -0.055406151 | 0.056335301 | 0.220150083 | 0.642906874 | ns | NaCl | KS.3 |
| FIM1_4339 | 339.6585001 | 248.1687789 | 495.8349819 | -0.161572082 | 0.158960116 | 0.013174616 | 0.165678479 | ns | NaCl | KS.3 |
| FIM1_4340 | 10449.56264 | 11194.616 | 10345.93131 | 0.043042586 | 0.076426697 | 0.386789214 | 0.779278134 | ns | NaCl | KS.3 |
| FIM1_4341 | 366.7345243 | 362.3259027 | 351.1808717 | -0.002417976 | 0.105370384 | 0.953035013 | 0.988653461 | ns | NaCl | KS.3 |
| FIM1_4342 | 572.2475436 | 541.782066 | 597.9498622 | -0.026865604 | 0.083684001 | 0.581487232 | 0.875038346 | ns | NaCl | KS.3 |
| FIM1_4343 | 36891.80374 | 40725.17372 | 25916.70843 | 0.064389416 | 0.076997517 | 0.201418991 | 0.624846998 | ns | NaCl | KS.3 |
| FIM1_4344 | 3797.097608 | 5150.64506 | 2485.044825 | 0.089998666 | 0.14159275 | 0.080681884 | 0.438915185 | ns | NaCl | KS.3 |
| FIM1_4345 | 453.5389732 | 401.0301683 | 544.9502246 | -0.047927092 | 0.094998234 | 0.317300453 | 0.73382286 | ns | NaCl | KS.3 |
| FIM1_4346 | 2228.431445 | 3004.054595 | 1382.390916 | 0.043144657 | 0.141042109 | 0.284910696 | 0.70510214 | ns | NaCl | KS.3 |
| FIM1_4347 | 1135.328099 | 1115.589661 | 1730.147006 | -0.009329843 | 0.078567021 | 0.849145963 | 0.96736917 | ns | NaCl | KS.3 |
| FIM1_4348 | 2274.992464 | 2233.654528 | 2477.390735 | -0.015575411 | 0.063422881 | 0.743411429 | 0.941460666 | ns | NaCl | KS.3 |
| FIM1_4349 | 69.86361133 | 45.67706098 | 484.6426657 | -0.072061583 | 0.168455414 | 0.120904647 | 0.527799029 | ns | NaCl | KS.3 |
| FIM1_4350 | 2638.439955 | 2695.828256 | 2577.991275 | 0.021447567 | 0.057349663 | 0.637629786 | 0.898332475 | ns | NaCl | KS.3 |
| FIM1_4351 | 6856.13279 | 6467.838014 | 4666.112367 | -0.032460258 | 0.080062488 | 0.510643299 | 0.846104012 | ns | NaCl | KS.3 |
| FIM1_4352 | 1723.929521 | 1890.196537 | 1263.868516 | 0.02772428 | 0.099417794 | 0.535707729 | 0.855153333 | ns | NaCl | KS.3 |
| FIM1_4353 | 3085.848312 | 3106.078856 | 2362.725098 | 0.003358944 | 0.08183175 | 0.944922345 | 0.987187877 | ns | NaCl | KS.3 |
| FIM1_4354 | 1144.344288 | 1174.248357 | 1627.87739 | 0.01101527 | 0.087996163 | 0.816020377 | 0.958816322 | ns | NaCl | KS.3 |
| FIM1_4355 | 15.84781139 | 13.51396076 | 26.18476884 | -0.01910813 | 0.136598381 | 0.589029651 | 0.878638624 | ns | NaCl | KS.3 |
| FIM1_4356 | 30415.15672 | 32270.66227 | 23485.72274 | 0.055393528 | 0.057088571 | 0.223548057 | 0.648059715 | ns | NaCl | KS.3 |
| FIM1_4357 | 1348.354833 | 1292.074274 | 1923.814461 | -0.021403168 | 0.083139677 | 0.660096924 | 0.903691294 | ns | NaCl | KS.3 |
| FIM1_4358 | 4672.717079 | 4622.019056 | 4355.850489 | -0.005593942 | 0.081304632 | 0.908575019 | 0.980825899 | ns | NaCl | KS.3 |
| FIM1_4359 | 2141.011333 | 2035.428061 | 2335.321236 | -0.0262642 | 0.082357795 | 0.591343785 | 0.878786986 | ns | NaCl | KS.3 |
| FIM1_4360 | 11297.70111 | 10468.67422 | 8671.074157 | -0.055862256 | 0.070567979 | 0.258822387 | 0.682736678 | ns | NaCl | KS.3 |
| FIM1_4361 | 4461.865074 | 5734.640313 | 2926.548286 | 0.087356227 | 0.123455596 | 0.088638315 | 0.46158813 | ns | NaCl | KS.3 |
| FIM1_4363 | 1377.382359 | 1963.991423 | 979.5832279 | 0.125348419 | 0.167159197 | 0.034163352 | 0.292771172 | ns | NaCl | KS.3 |
| FIM1_4364 | 393.0155717 | 381.2192901 | 393.1670421 | -0.010010062 | 0.096000165 | 0.822826786 | 0.960240659 | ns | NaCl | KS.3 |
| FIM1_4365 | 1406.267717 | 1302.19927 | 1130.932375 | -0.038855254 | 0.084346309 | 0.428888272 | 0.804596805 | ns | NaCl | KS.3 |
| FIM1_4366 | 10132.20124 | 10748.37622 | 9010.826394 | 0.047424433 | 0.065365685 | 0.325960648 | 0.739131368 | ns | NaCl | KS.3 |
| FIM1_4367 | 2018.107688 | 1927.3502 | 1772.816264 | -0.022445251 | 0.083559267 | 0.644372762 | 0.898899754 | ns | NaCl | KS.3 |
| FIM1_4368 | 1833.907664 | 1749.108603 | 1621.621862 | -0.02651299 | 0.08002893 | 0.589903929 | 0.87877872 | ns | NaCl | KS.3 |
| FIM1_4369 | 5316.490068 | 6939.598722 | 3186.351165 | 0.058533264 | 0.130649258 | 0.193394541 | 0.618952432 | ns | NaCl | KS.3 |
| FIM1_4370 | 1197.708101 | 1583.652877 | 886.3623511 | 0.118163896 | 0.135106067 | 0.037356624 | 0.304670827 | ns | NaCl | KS.3 |
| FIM1_4371 | 3009.194625 | 3410.338343 | 3441.943324 | 0.051218495 | 0.094268447 | 0.289827323 | 0.709591995 | ns | NaCl | KS.3 |
| FIM1_4372 | 3435.063776 | 3578.366649 | 2499.143024 | 0.014492071 | 0.092690609 | 0.752814866 | 0.943523399 | ns | NaCl | KS.3 |
| FIM1_4373 | 12780.80926 | 15991.54975 | 8223.638193 | 0.069143932 | 0.117131952 | 0.151817117 | 0.567887336 | ns | NaCl | KS.3 |
| FIM1_4374 | 847.5842182 | 963.8933227 | 1187.237333 | 0.086179291 | 0.0791469 | 0.093663515 | 0.470201108 | ns | NaCl | KS.3 |
| FIM1_4376 | 303.6846103 | 298.2478452 | 275.9932316 | -0.0064584 | 0.093161992 | 0.887431484 | 0.976547586 | ns | NaCl | KS.3 |
| FIM1_4377 | 7558.620345 | 9189.593171 | 5376.730464 | 0.059732001 | 0.111207067 | 0.204667906 | 0.627757602 | ns | NaCl | KS.3 |
| FIM1_4378 | 213.4521751 | 214.4451562 | 200.5203965 | 0.002498945 | 0.093925314 | 0.956009233 | 0.989832661 | ns | NaCl | KS.3 |
| FIM1_4379 | 3219.511704 | 3117.505845 | 3467.010987 | -0.01955858 | 0.075405376 | 0.691447795 | 0.91988055 | ns | NaCl | KS.3 |
| FIM1_4380 | 1552.12157 | 1523.125523 | 1529.100328 | -0.014787383 | 0.067330414 | 0.760352953 | 0.945911573 | ns | NaCl | KS.3 |

| | | | | | | | | | | |
|---|---|---|---|---|---|---|---|---|---|---|
| FIM1_4381 | 1998.607531 | 1929.060112 | 1501.693725 | -0.020449821 | 0.078121385 | 0.677760184 | 0.913287355 | ns | NaCl | KS.3 |
| FIM1_4382 | 1050.995101 | 1005.391514 | 961.0533677 | -0.024030882 | 0.081172327 | 0.623921324 | 0.893754273 | ns | NaCl | KS.3 |
| FIM1_4383 | 7066.221257 | 7442.047447 | 5562.631831 | 0.049037921 | 0.055911063 | 0.275680024 | 0.695949924 | ns | NaCl | KS.3 |
| FIM1_4384 | 1104.81772 | 1426.146381 | 1504.303601 | 0.07606205 | 0.125996202 | 0.121084042 | 0.527951891 | ns | NaCl | KS.3 |
| FIM1_4385 | 1442.562182 | 1427.346737 | 1672.830195 | -0.006372943 | 0.075496043 | 0.896882052 | 0.977971161 | ns | NaCl | KS.3 |
| FIM1_4386 | 19668.28569 | 20945.26305 | 15426.11709 | 0.03936592 | 0.076119819 | 0.427740895 | 0.803933114 | ns | NaCl | KS.3 |
| FIM1_4387 | 34.49725421 | 38.49785215 | 32.08724951 | 0.015661318 | 0.11715103 | 0.683750861 | 0.91584821 | ns | NaCl | KS.3 |
| FIM1_4388 | 4234.376474 | 7178.744536 | 2828.08616 | 0.246280356 | 0.282829343 | 0.006443448 | 0.107211218 | ns | NaCl | KS.3 |
| FIM1_4389 | 166.4815783 | 162.3557222 | 261.3620366 | -0.007217526 | 0.097435574 | 0.869971074 | 0.971243188 | ns | NaCl | KS.3 |
| FIM1_4390 | 93.06555588 | 77.46418518 | 119.0014353 | -0.048213453 | 0.111655336 | 0.283838191 | 0.704550068 | ns | NaCl | KS.3 |
| FIM1_4391 | 17640.98843 | 21004.00097 | 11072.76012 | 0.060602291 | 0.104913487 | 0.206637735 | 0.631463595 | ns | NaCl | KS.3 |
| FIM1_4392 | 2195.911341 | 2745.434692 | 2446.787469 | 0.081877933 | 0.114433092 | 0.105908531 | 0.49734665 | ns | NaCl | KS.3 |
| FIM1_4393 | 94.00601206 | 66.98267878 | 102.1546705 | -0.073369646 | 0.144537639 | 0.12403482 | 0.531066893 | ns | NaCl | KS.3 |
| FIM1_4394 | 424.819147 | 518.2905368 | 574.9786187 | 0.109314986 | 0.099326434 | 0.043815131 | 0.329773488 | ns | NaCl | KS.3 |
| FIM1_4396 | 3403.660591 | 3164.747467 | 3371.773329 | -0.03206472 | 0.087887045 | 0.505598084 | 0.84439816 | ns | NaCl | KS.3 |
| FIM1_4397 | 1013.266102 | 712.458415 | 1309.553351 | -0.392888535 | 0.166394599 | 4.35738087956 | 0.003933898 | ns | NaCl | KS.3 |
| FIM1_4398 | 59.94587198 | 73.48920317 | 199.3745941 | 0.027229775 | 0.126714131 | 0.478858355 | 0.826171162 | ns | NaCl | KS.3 |
| FIM1_4399 | 20011.64537 | 20339.47679 | 14052.33915 | 0.004751046 | 0.097666916 | 0.913909918 | 0.982311132 | ns | NaCl | KS.3 |
| FIM1_4400 | 1098.245521 | 991.5590207 | 1072.102269 | -0.070683287 | 0.075585951 | 0.161495462 | 0.58304784 | ns | NaCl | KS.3 |
| FIM1_4401 | 3003.809181 | 3546.137655 | 2429.574297 | 0.102318303 | 0.088922325 | 0.053843036 | 0.362636686 | ns | NaCl | KS.3 |
| FIM1_4402 | 156.7103227 | 263.7352521 | 172.6698895 | 0.54389092 | 0.265178013 | 0.000137463 | 0.009911306 | ns | NaCl | KS.3 |
| FIM1_4403 | 4079.821816 | 4543.201378 | 5429.59483 | 0.019919713 | 0.113315495 | 0.617839465 | 0.891787196 | ns | NaCl | KS.3 |
| FIM1_4404 | 402.7360955 | 370.2530254 | 460.0954742 | -0.034490995 | 0.0924611 | 0.465591815 | 0.820834625 | ns | NaCl | KS.3 |
| FIM1_4405 | 581.3599355 | 511.875099 | 609.2161831 | -0.050628318 | 0.095941803 | 0.291932671 | 0.71186075 | ns | NaCl | KS.3 |
| FIM1_4406 | 825.6780767 | 731.0129686 | 1069.145318 | -0.068509262 | 0.0855522 | 0.176588716 | 0.605047427 | ns | NaCl | KS.3 |
| FIM1_4407 | 43.81071294 | 29.63835503 | 122.619116 | -0.069486272 | 0.157962035 | 0.132465019 | 0.539053419 | ns | NaCl | KS.3 |
| FIM1_4408 | 117.509842 | 131.0107964 | 448.4685894 | 0.018198389 | 0.11701059 | 0.639085733 | 0.898798052 | ns | NaCl | KS.3 |
| FIM1_4409 | 10702.1684 | 14116.02779 | 6251.210809 | 0.055664421 | 0.133692224 | 0.206813413 | 0.631612481 | ns | NaCl | KS.3 |
| FIM1_4410 | 645.749429 | 650.2765395 | 823.5708034 | 0.003951998 | 0.08354651 | 0.934812269 | 0.985446192 | ns | NaCl | KS.3 |
| FIM1_4412 | 139.4375072 | 167.3867252 | 240.9380461 | 0.032203545 | 0.121699075 | 0.423836538 | 0.802854185 | ns | NaCl | KS.3 |
| FIM1_4413 | 1428.956339 | 1416.416492 | 1434.163885 | -0.007466275 | 0.068879288 | 0.878191073 | 0.972834689 | ns | NaCl | KS.3 |
| FIM1_4414 | 3420.97916 | 4543.890497 | 3221.545165 | 0.222002581 | 0.139470246 | 0.00156183 | 0.046934852 | ns | NaCl | KS.3 |
| FIM1_4415 | 1933.163981 | 2032.792739 | 3335.323888 | 0.009500964 | 0.110157852 | 0.811118512 | 0.958816322 | ns | NaCl | KS.3 |
| FIM1_4416 | 1397.807312 | 1368.077802 | 1896.95476 | -0.006459031 | 0.097315404 | 0.883597601 | 0.974700236 | ns | NaCl | KS.3 |
| FIM1_4417 | 1429.199056 | 1322.069411 | 1298.851095 | -0.055936195 | 0.072459453 | 0.260822731 | 0.686178949 | ns | NaCl | KS.3 |
| FIM1_4418 | 7764.672903 | 8827.335151 | 5851.324223 | 0.097364895 | 0.074061775 | 0.056478481 | 0.372058397 | ns | NaCl | KS.3 |
| FIM1_4419 | 888.3678649 | 960.7096886 | 1039.942133 | 0.045677816 | 0.079859393 | 0.3588618 | 0.762964725 | ns | NaCl | KS.3 |
| FIM1_4420 | 750.3248795 | 734.3015042 | 1050.982712 | -0.012393564 | 0.076351426 | 0.801096637 | 0.957352931 | ns | NaCl | KS.3 |
| FIM1_4421 | 665.2670291 | 703.3190503 | 841.2736502 | 0.034952846 | 0.074220032 | 0.47995078 | 0.827357981 | ns | NaCl | KS.3 |
| FIM1_4422 | 12.92033294 | 11.75355005 | 14.57156587 | -0.006830268 | 0.142522541 | 0.83497067 | 0.962365393 | ns | NaCl | KS.3 |
| FIM1_4423 | 2031.087217 | 2042.951507 | 1754.733821 | 0.001415071 | 0.100962418 | 0.973537231 | 0.995549378 | ns | NaCl | KS.3 |
| FIM1_4424 | 530.1710626 | 340.9843597 | 1053.413028 | -0.064370498 | 0.17115433 | 0.144751229 | 0.558679104 | ns | NaCl | KS.3 |
| FIM1_4425 | 692.8486584 | 759.7598627 | 643.1878699 | 0.071791758 | 0.067955009 | 0.144613866 | 0.558679104 | ns | NaCl | KS.3 |
| FIM1_4426 | 803.7857212 | 786.2089984 | 728.6083259 | -0.01187563 | 0.081708925 | 0.807390213 | 0.957859861 | ns | NaCl | KS.3 |
| FIM1_4427 | 5526.774117 | 5416.222516 | 5440.023767 | -0.012028513 | 0.076383749 | 0.806822762 | 0.957859861 | ns | NaCl | KS.3 |

| | | | | | | | | | | |
|---|---|---|---|---|---|---|---|---|---|---|
| FIM1_4428 | 7905.133497 | 5293.283859 | 6919.829657 | -0.160194101 | 0.197467682 | 0.017669793 | 0.199336097 | ns | NaCl | KS.3 |
| FIM1_4429 | 2175.627448 | 2393.813351 | 2044.53562 | 0.038213501 | 0.092198972 | 0.422283361 | 0.801548921 | ns | NaCl | KS.3 |
| FIM1_4430 | 388.3658933 | 465.3851575 | 343.7891576 | 0.071621964 | 0.102643592 | 0.150371635 | 0.564839452 | ns | NaCl | KS.3 |
| FIM1_4431 | 1448.872673 | 1710.473223 | 2195.196975 | 0.121954909 | 0.082918246 | 0.02255805 | 0.230444142 | ns | NaCl | KS.3 |
| FIM1_4432 | 7623.305658 | 7888.352107 | 5969.247125 | 0.026044356 | 0.066722692 | 0.590664335 | 0.878786986 | ns | NaCl | KS.3 |
| FIM1_4433 | 609.3839396 | 662.9534233 | 622.5379711 | 0.032119523 | 0.092889262 | 0.494199927 | 0.836358994 | ns | NaCl | KS.3 |
| FIM1_4434 | 1846.806912 | 1739.073481 | 1987.486864 | -0.057886093 | 0.054975012 | 0.194157579 | 0.618952432 | ns | NaCl | KS.3 |
| FIM1_4435 | 3589.194255 | 3741.517934 | 4237.227993 | 0.026363058 | 0.074701255 | 0.593272757 | 0.87962202 | ns | NaCl | KS.3 |
| FIM1_4436 | 1359.237369 | 1325.005337 | 1632.471748 | -0.021450559 | 0.06252525 | 0.650492047 | 0.899642849 | ns | NaCl | KS.3 |
| FIM1_4437 | 555.5747063 | 454.955949 | 636.5513952 | -0.099522039 | 0.104574232 | 0.061984967 | 0.390501742 | ns | NaCl | KS.3 |
| FIM1_4438 | 913.6548901 | 774.0014115 | 1274.270211 | -0.072303948 | 0.099026801 | 0.149587951 | 0.564839452 | ns | NaCl | KS.3 |
| FIM1_4439 | 1706.226967 | 1536.727921 | 1607.302738 | -0.06587172 | 0.079314188 | 0.192730001 | 0.618952432 | ns | NaCl | KS.3 |
| FIM1_4440 | 814.8128825 | 506.9058742 | 1509.242747 | -0.277101174 | 0.26644465 | 0.003649089 | 0.076924644 | ns | NaCl | KS.3 |
| FIM1_4441 | 2735.216652 | 2896.87041 | 3396.226368 | 0.012039034 | 0.10746509 | 0.767808348 | 0.948794469 | ns | NaCl | KS.3 |
| FIM1_4443 | 216.9951739 | 151.8139795 | 510.6500014 | -0.055117372 | 0.152447406 | 0.195815345 | 0.618952432 | ns | NaCl | KS.3 |
| FIM1_4444 | 44322.4811 | 44672.64526 | 29229.62885 | 0.003687341 | 0.084451045 | 0.938931803 | 0.986790704 | ns | NaCl | KS.3 |
| FIM1_4445 | 1318.534893 | 1316.748121 | 1353.071427 | -0.001766363 | 0.075185873 | 0.971339727 | 0.995143151 | ns | NaCl | KS.3 |
| FIM1_4446 | 5597.00037 | 4767.755907 | 5081.720682 | -0.098724617 | 0.087578268 | 0.0612007 | 0.387370843 | ns | NaCl | KS.3 |
| FIM1_4447 | 1541.473267 | 1518.518207 | 1698.587544 | -0.013185129 | 0.062954461 | 0.781088627 | 0.951646805 | ns | NaCl | KS.3 |
| FIM1_4448 | 201.6544664 | 261.8273942 | 260.2087931 | 0.052682834 | 0.129684403 | 0.228440179 | 0.652285125 | ns | NaCl | KS.3 |
| FIM1_4449 | 662.2701627 | 482.9573016 | 733.6709146 | -0.076572959 | 0.142627553 | 0.114562794 | 0.514859894 | ns | NaCl | KS.3 |
| FIM1_4450 | 2939.295104 | 3107.697515 | 3259.486394 | 0.027196464 | 0.08407799 | 0.576422227 | 0.873870489 | ns | NaCl | KS.3 |
| FIM1_4451 | 1195.659739 | 1246.93815 | 2267.005217 | 0.014319571 | 0.093366151 | 0.754474226 | 0.944041568 | ns | NaCl | KS.3 |
| FIM1_4452 | 14263.46621 | 15653.09685 | 10157.37843 | 0.040685858 | 0.08946073 | 0.400301779 | 0.788401167 | ns | NaCl | KS.3 |
| FIM1_4454 | 4903.868795 | 4843.577624 | 4077.397871 | -0.009780577 | 0.064430498 | 0.837981407 | 0.962829707 | ns | NaCl | KS.3 |
| FIM1_4455 | 1662.088199 | 1906.863281 | 1386.220544 | 0.053377823 | 0.096891947 | 0.267957879 | 0.689648447 | ns | NaCl | KS.3 |
| FIM1_4456 | 931.6639808 | 854.9897935 | 1521.687779 | -0.032630038 | 0.092896375 | 0.487700735 | 0.831497997 | ns | NaCl | KS.3 |
| FIM1_4457 | 2110.536439 | 2046.64066 | 1899.474495 | -0.020217045 | 0.074382656 | 0.681575027 | 0.915044369 | ns | NaCl | KS.3 |
| FIM1_4458 | 1569.152119 | 1258.099158 | 1523.401855 | -0.171918723 | 0.101709399 | 0.003779988 | 0.07901446 | ns | NaCl | KS.3 |
| FIM1_4460 | 394.6138439 | 396.3515032 | 482.6566913 | 0.001950598 | 0.096806241 | 0.96482745 | 0.993691273 | ns | NaCl | KS.3 |
| FIM1_4461 | 218.1027965 | 226.455895 | 235.8112276 | 0.011705574 | 0.098848298 | 0.789002224 | 0.953383399 | ns | NaCl | KS.3 |
| FIM1_4462 | 1071.809972 | 1152.287855 | 1086.853569 | 0.044824806 | 0.076506823 | 0.367899877 | 0.768177319 | ns | NaCl | KS.3 |
| FIM1_4463 | 534.0330055 | 640.5327938 | 1342.582568 | 0.094583319 | 0.097467503 | 0.072762405 | 0.421203992 | ns | NaCl | KS.3 |
| FIM1_4464 | 730.7682357 | 714.6907864 | 1042.604224 | -0.009852252 | 0.086591939 | 0.836337596 | 0.962829707 | ns | NaCl | KS.3 |
| FIM1_4465 | 1510.884834 | 1252.187405 | 1372.504425 | -0.110353361 | 0.096047856 | 0.041530913 | 0.322888102 | ns | NaCl | KS.3 |
| FIM1_4466 | 1249.646586 | 1105.948588 | 1228.347793 | -0.07786669 | 0.078964482 | 0.12716281 | 0.534982546 | ns | NaCl | KS.3 |
| FIM1_4467 | 3686.308173 | 3711.973077 | 2880.810584 | 0.006598076 | 0.060953895 | 0.887946373 | 0.976703159 | ns | NaCl | KS.3 |
| FIM1_4468 | 197.4658369 | 202.4359401 | 290.415238 | 0.006031263 | 0.091300826 | 0.896245156 | 0.977971161 | ns | NaCl | KS.3 |
| FIM1_4469 | 4390.77058 | 5835.096685 | 3092.130005 | 0.060744022 | 0.134946452 | 0.179022732 | 0.607944091 | ns | NaCl | KS.3 |
| FIM1_4470 | 3254.066985 | 4431.250181 | 2972.149374 | 0.092133617 | 0.143136699 | 0.076200509 | 0.428843365 | ns | NaCl | KS.3 |
| FIM1_4471 | 835.459777 | 883.8439034 | 801.3701326 | 0.026753989 | 0.085718389 | 0.580181271 | 0.8749481 | ns | NaCl | KS.3 |
| FIM1_4472 | 1975.999361 | 1908.640805 | 2324.054921 | -0.032920999 | 0.057334447 | 0.469842191 | 0.821848579 | ns | NaCl | KS.3 |
| FIM1_4473 | 786.053159 | 714.4750358 | 825.0001088 | -0.045115509 | 0.088207131 | 0.35613845 | 0.759883833 | ns | NaCl | KS.3 |
| FIM1_4474 | 15324.8032 | 14533.075 | 10933.34462 | -0.022175748 | 0.088724181 | 0.640591833 | 0.898798052 | ns | NaCl | KS.3 |
| FIM1_4475 | 531.6858882 | 482.8944016 | 812.4556507 | -0.033925519 | 0.096067037 | 0.463754623 | 0.819880744 | ns | NaCl | KS.3 |

| | | | | | | | | | | |
|---|---|---|---|---|---|---|---|---|---|---|
| FIM1_4476 | 1539.072197 | 1635.855925 | 1386.57011 | 0.020396784 | 0.095110332 | 0.654101047 | 0.900747416 | ns | NaCl | KS.3 |
| FIM1_4477 | 2186.361268 | 2043.712543 | 2194.233526 | -0.047661266 | 0.07168295 | 0.335485548 | 0.745307602 | ns | NaCl | KS.3 |
| FIM1_4478 | 1596.315069 | 1705.404386 | 1911.228193 | 0.032820644 | 0.084346723 | 0.501696402 | 0.84151706 | ns | NaCl | KS.3 |
| FIM1_4479 | 2699.602975 | 2608.422957 | 2132.936426 | -0.009224351 | 0.100009367 | 0.830941027 | 0.961620266 | ns | NaCl | KS.3 |
| FIM1_4480 | 404.891737 | 383.6420889 | 436.876264 | -0.018759004 | 0.092745495 | 0.684627052 | 0.91584821 | ns | NaCl | KS.3 |
| FIM1_4481 | 1808.103613 | 1761.595766 | 1967.364171 | -0.02145135 | 0.065288149 | 0.655470987 | 0.901110473 | ns | NaCl | KS.3 |
| FIM1_4482 | 78.19577358 | 30.29572125 | 2864.161851 | -0.671165542 | 0.549761996 | 0.000565863 | 0.024912979 | ns | NaCl | KS.3 |
| FIM1_4484 | 10.62718423 | 23.80851963 | 72.63694843 | 0.067545473 | 0.222528754 | 0.10642089 | 0.497597675 | ns | NaCl | KS.3 |
| FIM1_4486 | 1796.45692 | 2005.076467 | 1769.97335 | 0.018708568 | 0.115864845 | 0.632668056 | 0.897798121 | ns | NaCl | KS.3 |
| FIM1_4487 | 159.8195054 | 134.1052817 | 435.3429884 | -0.029077903 | 0.12154754 | 0.463690833 | 0.819880744 | ns | NaCl | KS.3 |
| FIM1_4488 | 45.90526725 | 31.42936346 | 33.08812412 | -0.06199794 | 0.15598543 | 0.161380405 | 0.58304784 | ns | NaCl | KS.3 |
| FIM1_4489 | 1212.423264 | 1294.46216 | 1176.216034 | 0.012712771 | 0.109911911 | 0.751062822 | 0.943274417 | ns | NaCl | KS.3 |
| FIM1_4490 | 6223.095591 | 6263.738078 | 7550.038692 | 0.001612248 | 0.102834087 | 0.969351419 | 0.99411987 | ns | NaCl | KS.3 |
| FIM1_4491 | 893.5435614 | 873.3792379 | 931.9821455 | -0.007225762 | 0.093879001 | 0.873511499 | 0.971243188 | ns | NaCl | KS.3 |
| FIM1_4492 | 960.1541872 | 1091.738285 | 821.0718333 | 0.074047515 | 0.084885982 | 0.146980381 | 0.562037426 | ns | NaCl | KS.3 |
| FIM1_4493 | 5516.041708 | 7132.758237 | 6962.732737 | 0.086510429 | 0.125553877 | 0.090478416 | 0.463573758 | ns | NaCl | KS.3 |
| FIM1_4494 | 66.76898174 | 91.97213608 | 44.75258721 | 0.0809561 | 0.147680587 | 0.101223339 | 0.484750818 | ns | NaCl | KS.3 |
| FIM1_4495 | 894.9928901 | 990.5708827 | 1200.271987 | 0.047691234 | 0.088419733 | 0.330755902 | 0.742254041 | ns | NaCl | KS.3 |
| FIM1_4496 | 554.6690655 | 688.2814424 | 572.8180632 | 0.121470009 | 0.10657299 | 0.029705347 | 0.271163489 | ns | NaCl | KS.3 |
| FIM1_4497 | 7037.762742 | 5660.50721 | 4143.687139 | -0.149748648 | 0.102407224 | 0.009795842 | 0.138844204 | ns | NaCl | KS.3 |
| FIM1_4498 | 27003.5331 | 31010.76879 | 18755.82401 | 0.080771011 | 0.085494312 | 0.11666691 | 0.519941265 | ns | NaCl | KS.3 |
| FIM1_4499 | 2407.801335 | 2355.135807 | 3375.850452 | -0.012128701 | 0.079387955 | 0.80451438 | 0.957859861 | ns | NaCl | KS.3 |
| FIM1_4500 | 2743.338215 | 2548.10647 | 2985.97264 | -0.06001583 | 0.065749373 | 0.215991214 | 0.640629323 | ns | NaCl | KS.3 |
| FIM1_4501 | 1787.474761 | 1817.328283 | 1516.750278 | 0.005868086 | 0.092733842 | 0.897968983 | 0.977971161 | ns | NaCl | KS.3 |
| FIM1_4502 | 221.0852172 | 208.4662078 | 211.4499236 | -0.015675233 | 0.103227646 | 0.71195201 | 0.926245097 | ns | NaCl | KS.3 |
| FIM1_4503 | 561.8775839 | 526.6925366 | 692.7756756 | -0.042372692 | 0.078015437 | 0.394110781 | 0.781740625 | ns | NaCl | KS.3 |
| FIM1_4504 | 926.5250623 | 824.0455842 | 1117.332412 | -0.042164759 | 0.096772842 | 0.369267738 | 0.769630079 | ns | NaCl | KS.3 |
| FIM1_4505 | 4316.416681 | 5763.597492 | 2753.53033 | 0.048978115 | 0.137671109 | 0.245034267 | 0.668549812 | ns | NaCl | KS.3 |
| FIM1_4506 | 1837.875681 | 2045.405338 | 1446.303108 | 0.039701502 | 0.095036785 | 0.39930868 | 0.787737366 | ns | NaCl | KS.3 |
| FIM1_4507 | 2774.527046 | 2919.120914 | 2743.829087 | 0.02378518 | 0.085274384 | 0.622761731 | 0.893340404 | ns | NaCl | KS.3 |
| FIM1_4508 | 1148.282182 | 1062.113835 | 1156.549221 | -0.051597831 | 0.07502202 | 0.301047795 | 0.719362527 | ns | NaCl | KS.3 |
| FIM1_4509 | 3092.008034 | 2789.30747 | 2958.750099 | -0.099881734 | 0.05743552 | 0.029263052 | 0.269101083 | ns | NaCl | KS.3 |
| FIM1_4510 | 6901.415954 | 7933.645125 | 5040.076391 | 0.110935798 | 0.073664278 | 0.030645541 | 0.274705523 | ns | NaCl | KS.3 |
| FIM1_4512 | 688.79812 | 602.3654413 | 894.7769578 | -0.083274926 | 0.083149381 | 0.106268951 | 0.497354685 | ns | NaCl | KS.3 |
| FIM1_4513 | 274.4286353 | 259.0848317 | 220.9323604 | -0.021699718 | 0.092068047 | 0.641055658 | 0.898798052 | ns | NaCl | KS.3 |
| FIM1_4514 | 212.9385057 | 213.9174177 | 316.521227 | 0.000266391 | 0.089569618 | 0.995454619 | 0.99924489 | ns | NaCl | KS.3 |
| FIM1_4515 | 1084.456626 | 876.495132 | 1229.686415 | -0.168139795 | 0.095753994 | 0.003756959 | 0.07886443 | ns | NaCl | KS.3 |
| FIM1_4516 | 1412.125898 | 1514.930787 | 750.3798019 | 0.00988959 | 0.119881998 | 0.790397653 | 0.953961262 | ns | NaCl | KS.3 |
| FIM1_4517 | 509.1689619 | 456.6276261 | 1121.894738 | -0.031100796 | 0.102007383 | 0.48363775 | 0.829184191 | ns | NaCl | KS.3 |
| FIM1_4518 | 754.9942643 | 634.3485394 | 733.9843654 | -0.077349255 | 0.100398078 | 0.127100163 | 0.534982546 | ns | NaCl | KS.3 |
| FIM1_4519 | 1160.589737 | 1191.386045 | 1030.65912 | 0.016105148 | 0.073766978 | 0.743471086 | 0.941460666 | ns | NaCl | KS.3 |
| FIM1_4520 | 2633.060118 | 2675.414071 | 2052.447947 | 0.005584137 | 0.092471792 | 0.90306192 | 0.979395371 | ns | NaCl | KS.3 |
| FIM1_4521 | 1886.865631 | 2057.906692 | 2375.04407 | 0.073272913 | 0.064534381 | 0.129327805 | 0.536832346 | ns | NaCl | KS.3 |
| FIM1_4522 | 326.2357861 | 176.3331613 | 643.0435648 | -0.407982031 | 0.356228989 | 0.001575502 | 0.046934852 | ns | NaCl | KS.3 |
| FIM1_4523 | 3640.221622 | 4010.562776 | 3284.485673 | 0.098400975 | 0.05459425 | 0.026966594 | 0.256517796 | ns | NaCl | KS.3 |

| | | | | | | | | | | | |
|---|---|---|---|---|---|---|---|---|---|---|---|
| FIM1_4524 | 2232.930014 | 2108.483043 | 2028.069343 | -0.032316671 | 0.079130141 | 0.513022378 | 0.847552971 | ns | | NaCl | KS.3 |
| FIM1_4525 | 206.8957456 | 231.4535077 | 257.5943842 | 0.036833132 | 0.099328538 | 0.420976478 | 0.80133378 | ns | | NaCl | KS.3 |
| FIM1_4526 | 7809.038208 | 7743.092049 | 6663.423493 | -0.007513714 | 0.053552119 | 0.863979982 | 0.969856868 | ns | | NaCl | KS.3 |
| FIM1_4527 | 15438.37905 | 19938.07694 | 16071.36789 | 0.057470295 | 0.12809117 | 0.201283344 | 0.624846998 | ns | | NaCl | KS.3 |
| FIM1_4528 | 45.7362275 | 59.5909272 | 224.419495 | 0.04094273 | 0.136181258 | 0.309512094 | 0.729444457 | ns | | NaCl | KS.3 |
| FIM1_4529 | 205.4430448 | 178.23141 | 548.8525816 | -0.024620632 | 0.117186708 | 0.535471882 | 0.855153333 | ns | | NaCl | KS.3 |
| FIM1_4530 | 2112.217964 | 2149.760165 | 2316.557567 | 0.011456472 | 0.073364171 | 0.815736105 | 0.958816322 | ns | | NaCl | KS.3 |
| FIM1_4531 | 2826.913043 | 2842.952845 | 3611.620623 | 0.003985997 | 0.070341126 | 0.935033338 | 0.985446192 | ns | | NaCl | KS.3 |
| FIM1_4532 | 1888.052176 | 1676.088879 | 1404.05236 | -0.056756759 | 0.089200893 | 0.252503366 | 0.676266095 | ns | | NaCl | KS.3 |
| FIM1_4533 | 13068.64336 | 18465.50719 | 8237.061385 | 0.08507418 | 0.153378677 | 0.090383765 | 0.463573758 | ns | | NaCl | KS.3 |
| FIM1_4534 | 631.2861559 | 512.5825533 | 795.2805864 | -0.105262475 | 0.106541597 | 0.051460806 | 0.35459489 | ns | | NaCl | KS.3 |
| FIM1_4535 | 2379.356235 | 2508.575405 | 1795.554662 | 0.019906074 | 0.091600648 | 0.669212065 | 0.90940454 | ns | | NaCl | KS.3 |
| FIM1_4536 | 2071.019436 | 1581.019928 | 2216.598367 | -0.075516431 | 0.129786812 | 0.121625718 | 0.527999953 | ns | | NaCl | KS.3 |
| FIM1_4537 | 1094.379055 | 1095.477344 | 926.008992 | -0.000133019 | 0.077124749 | 0.997838917 | 0.99924489 | ns | | NaCl | KS.3 |
| FIM1_4538 | 484.0476758 | 302.7640312 | 724.0558032 | -0.132436698 | 0.210153335 | 0.032242064 | 0.284909887 | ns | | NaCl | KS.3 |
| FIM1_4539 | 1082.2 | 1084.561978 | 1030.865874 | -0.00067944 | 0.072129238 | 0.988950439 | 0.999134858 | ns | | NaCl | KS.3 |
| FIM1_4540 | 503.4473732 | 424.5941536 | 575.5351737 | -0.070353987 | 0.101349933 | 0.157225885 | 0.57726847 | ns | | NaCl | KS.3 |
| FIM1_4542 | 4528.867652 | 4067.05633 | 4417.824229 | -0.047487698 | 0.090275641 | 0.329781557 | 0.741710329 | ns | | NaCl | KS.3 |
| FIM1_4543 | 7173.419029 | 9177.493397 | 3931.707626 | 0.039211618 | 0.131418941 | 0.33190783 | 0.742667065 | ns | | NaCl | KS.3 |
| FIM1_4544 | 315.5926547 | 263.0710693 | 678.0470273 | -0.06471823 | 0.10609996 | 0.181862182 | 0.611396712 | ns | | NaCl | KS.3 |
| FIM1_4545 | 3189.492553 | 4001.42642 | 2193.460746 | 0.060834733 | 0.119735259 | 0.190056555 | 0.618952432 | ns | | NaCl | KS.3 |
| FIM1_4546 | 8672.591849 | 10460.41927 | 5054.654355 | 0.038032724 | 0.118309236 | 0.365027143 | 0.765851202 | ns | | NaCl | KS.3 |
| FIM1_4547 | 115.6317612 | 149.7938789 | 93.14022584 | 0.05065965 | 0.129339406 | 0.242107936 | 0.666322592 | ns | | NaCl | KS.3 |
| FIM1_4548 | 1107.904479 | 1170.18963 | 1192.076338 | 0.02529213 | 0.085079703 | 0.60157379 | 0.880762097 | ns | | NaCl | KS.3 |
| FIM1_4549 | 875.2253773 | 785.5628826 | 2090.131992 | -0.040651543 | 0.094761436 | 0.389660148 | 0.779482559 | ns | | NaCl | KS.3 |
| FIM1_4550 | 49.15717177 | 52.74429905 | 58.11738184 | 0.010838112 | 0.11769417 | 0.774363377 | 0.950214521 | ns | | NaCl | KS.3 |
| FIM1_4551 | 162.069446 | 97.07392443 | 155.7175167 | -0.473691479 | 0.292152879 | 0.000424897 | 0.021012238 | ns | | NaCl | KS.3 |
| FIM1_4552 | 1033.721926 | 1006.796536 | 991.3499352 | -0.010847678 | 0.087817211 | 0.818941432 | 0.959981393 | ns | | NaCl | KS.3 |
| FIM1_4553 | 554.1811888 | 602.795238 | 556.4867473 | 0.027791801 | 0.095885636 | 0.544108739 | 0.857313452 | ns | | NaCl | KS.3 |
| FIM1_4554 | 1780.31062 | 1587.950802 | 2258.936361 | -0.046085865 | 0.093541731 | 0.337213756 | 0.7472777 | ns | | NaCl | KS.3 |
| FIM1_4555 | 6298.434383 | 6817.743411 | 5241.801357 | 0.022081278 | 0.100772795 | 0.614239265 | 0.889162818 | ns | | NaCl | KS.3 |
| FIM1_4556 | 3418.809328 | 3022.817691 | 2797.311843 | -0.032529867 | 0.105010598 | 0.457632044 | 0.817787148 | ns | | NaCl | KS.3 |
| FIM1_4557 | 5718.081923 | 5445.929284 | 4748.96601 | -0.034921438 | 0.070082442 | 0.476800138 | 0.825645905 | ns | | NaCl | KS.3 |
| FIM1_4558 | 865.1584119 | 725.4560553 | 1668.102854 | -0.077412935 | 0.099735713 | 0.127136718 | 0.534982546 | ns | | NaCl | KS.3 |
| FIM1_4559 | 604.9586943 | 448.5591662 | 1020.512662 | -0.233787433 | 0.152209597 | 0.001506749 | 0.046346385 | ns | | NaCl | KS.3 |
| FIM1_4560 | 89.85446101 | 71.51431187 | 161.0635889 | -0.052245291 | 0.124959861 | 0.236198996 | 0.659616806 | ns | | NaCl | KS.3 |
| FIM1_4561 | 3425.809781 | 3633.417226 | 4062.928576 | 0.015935959 | 0.100664293 | 0.713147699 | 0.927073374 | ns | | NaCl | KS.3 |
| FIM1_4562 | 331.104859 | 420.7513342 | 665.2961242 | 0.033399634 | 0.133436262 | 0.389511054 | 0.779482559 | ns | | NaCl | KS.3 |
| FIM1_4563 | 3293.035656 | 3971.923707 | 2210.548623 | 0.051083951 | 0.112053296 | 0.260973379 | 0.686178949 | ns | | NaCl | KS.3 |
| FIM1_4564 | 231.1559751 | 236.5604468 | 338.2719058 | 0.00487554 | 0.100133822 | 0.909769941 | 0.981715313 | ns | | NaCl | KS.3 |
| FIM1_4565 | 2933.582977 | 3024.247598 | 4454.604135 | 0.006610792 | 0.105867923 | 0.871893765 | 0.971243188 | ns | | NaCl | KS.3 |
| FIM1_4566 | 10005.87264 | 5610.617406 | 10289.83615 | -0.252768233 | 0.305456822 | 0.006594236 | 0.108271704 | ns | | NaCl | KS.3 |
| FIM1_4567 | 7744.501091 | 7698.449054 | 6056.028043 | -0.003671533 | 0.075664004 | 0.940468564 | 0.986790704 | ns | | NaCl | KS.3 |
| FIM1_4568 | 6228.714892 | 5765.734666 | 5307.079298 | -0.077682528 | 0.053515202 | 0.076939037 | 0.430564353 | ns | | NaCl | KS.3 |
| FIM1_4569 | 2505.722295 | 2629.210679 | 2428.104162 | 0.021909104 | 0.086114386 | 0.648833939 | 0.899642849 | ns | | NaCl | KS.3 |

| | | | | | | | | | | |
|---|---|---|---|---|---|---|---|---|---|---|
| FIM1_4570 | 1400.707484 | 1212.813787 | 1569.40606 | -0.094673394 | 0.082932249 | 0.069479445 | 0.413948267 | ns | NaCl | KS.3 |
| FIM1_4571 | 1293.72203 | 1238.488583 | 1194.927305 | -0.02412122 | 0.080458648 | 0.623196234 | 0.893487396 | ns | NaCl | KS.3 |
| FIM1_4572 | 5615.932787 | 7944.279856 | 4823.600807 | 0.083993785 | 0.15348372 | 0.092854963 | 0.470201108 | ns | NaCl | KS.3 |
| FIM1_4573 | 1390.620284 | 1103.696588 | 1981.366376 | -0.094200001 | 0.115877378 | 0.073222656 | 0.421631393 | ns | NaCl | KS.3 |
| FIM1_4574 | 520.927701 | 518.4783147 | 667.0720463 | -0.003315764 | 0.074938796 | 0.946231612 | 0.987311717 | ns | NaCl | KS.3 |
| FIM1_4575 | 11474.49484 | 10589.66798 | 8341.093234 | -0.05326437 | 0.074863858 | 0.286009007 | 0.705523209 | ns | NaCl | KS.3 |
| FIM1_4576 | 1897.283531 | 2119.763076 | 1594.398959 | 0.058275736 | 0.085402078 | 0.244398079 | 0.66821307 | ns | NaCl | KS.3 |
| FIM1_4577 | 526.8629597 | 297.6582189 | 482.5281874 | -0.561824119 | 0.304242728 | 0.000224833 | 0.014250729 | ns | NaCl | KS.3 |
| FIM1_4578 | 537.7892717 | 437.0323503 | 446.1690164 | -0.098849651 | 0.104715975 | 0.063373282 | 0.396581229 | ns | NaCl | KS.3 |
| FIM1_4579 | 343.851386 | 404.2028381 | 334.6155605 | 0.043206449 | 0.109314357 | 0.331677827 | 0.742667065 | ns | NaCl | KS.3 |
| FIM1_4580 | 4851.832624 | 5947.269051 | 3068.994516 | 0.088984462 | 0.106604492 | 0.086829932 | 0.456808328 | ns | NaCl | KS.3 |
| FIM1_4582 | 1637.074855 | 1654.864075 | 1719.339598 | 0.002640529 | 0.102245145 | 0.950097149 | 0.987736646 | ns | NaCl | KS.3 |
| FIM1_4583 | 1410.54822 | 1401.475112 | 1529.542087 | -0.004765254 | 0.065065456 | 0.920884401 | 0.983890455 | ns | NaCl | KS.3 |
| FIM1_4584 | 1715.864103 | 1920.607327 | 1894.691616 | 0.069591501 | 0.080759547 | 0.170522765 | 0.597009679 | ns | NaCl | KS.3 |
| FIM1_4585 | 2312.282121 | 2131.236372 | 2464.633392 | -0.065696595 | 0.065308646 | 0.175120765 | 0.60501792 | ns | NaCl | KS.3 |
| FIM1_4586 | 19829.85608 | 18565.15551 | 14634.50189 | -0.062354125 | 0.056606547 | 0.168921326 | 0.59349124 | ns | NaCl | KS.3 |
| FIM1_4587 | 3682.124798 | 4849.531411 | 2336.525261 | 0.046344427 | 0.135231609 | 0.266502203 | 0.689593109 | ns | NaCl | KS.3 |
| FIM1_4588 | 3582.960946 | 3415.440705 | 3198.471663 | -0.024505197 | 0.082295036 | 0.616127545 | 0.890538796 | ns | NaCl | KS.3 |
| FIM1_4589 | 4373.532987 | 5351.654583 | 2506.601279 | 0.053802491 | 0.114956215 | 0.237652051 | 0.660513382 | ns | NaCl | KS.3 |
| FIM1_4590 | 712.8452326 | 629.6206604 | 592.5620059 | -0.063244306 | 0.086916776 | 0.208454458 | 0.633002483 | ns | NaCl | KS.3 |
| FIM1_4591 | 3689.655724 | 3366.605008 | 2635.107102 | -0.058064005 | 0.07751885 | 0.247552808 | 0.669893726 | ns | NaCl | KS.3 |
| FIM1_4592 | 1585.605126 | 1542.111411 | 1307.898843 | -0.021288944 | 0.067304945 | 0.66074909 | 0.904077757 | ns | NaCl | KS.3 |
| FIM1_4593 | 512.4852031 | 458.0064949 | 726.3664995 | -0.046465828 | 0.092583502 | 0.335404406 | 0.745307602 | ns | NaCl | KS.3 |
| FIM1_4594 | 523.8959151 | 392.9605866 | 742.0136083 | -0.085174676 | 0.13697081 | 0.092199804 | 0.468055125 | ns | NaCl | KS.3 |
| FIM1_4595 | 4547.734357 | 5723.169451 | 4025.083491 | 0.149285152 | 0.109412979 | 0.011176169 | 0.150643816 | ns | NaCl | KS.3 |
| FIM1_4596 | 6074.106827 | 7671.921483 | 3538.793639 | 0.067398733 | 0.120254036 | 0.157519761 | 0.57787696 | ns | NaCl | KS.3 |
| FIM1_4597 | 2191.177152 | 2135.061974 | 1866.817906 | -0.016647734 | 0.072381176 | 0.734782271 | 0.938375543 | ns | NaCl | KS.3 |
| FIM1_4598 | 396.5674883 | 401.1952724 | 478.6078264 | 0.003432894 | 0.091568963 | 0.940677644 | 0.986790704 | ns | NaCl | KS.3 |
| FIM1_4599 | 294.237317 | 293.4921224 | 280.8019534 | -0.00051156 | 0.095492598 | 0.990872901 | 0.999134858 | ns | NaCl | KS.3 |
| FIM1_4600 | 4315.13397 | 4813.158528 | 3711.087624 | 0.07438835 | 0.076354728 | 0.142197709 | 0.554849885 | ns | NaCl | KS.3 |
| FIM1_4601 | 654.3600057 | 688.8917852 | 781.6834496 | 0.033541526 | 0.07292729 | 0.496940138 | 0.836358994 | ns | NaCl | KS.3 |
| FIM1_4602 | 598.1379804 | 569.6602823 | 450.4868022 | -0.010254322 | 0.1072354 | 0.801210084 | 0.957352931 | ns | NaCl | KS.3 |
| FIM1_4603 | 4460.290155 | 4605.105801 | 3864.028105 | 0.019786303 | 0.075158813 | 0.688051223 | 0.918490387 | ns | NaCl | KS.3 |
| FIM1_4604 | 95.02626024 | 96.35479933 | 140.3676843 | 0.002555136 | 0.107325196 | 0.949548328 | 0.987736646 | ns | NaCl | KS.3 |
| FIM1_4605 | 1299.184039 | 1247.629069 | 1105.088326 | -0.025877101 | 0.07374871 | 0.599787609 | 0.880219278 | ns | NaCl | KS.3 |
| FIM1_4606 | 691.9772924 | 874.8160943 | 706.6727291 | 0.173013569 | 0.110094262 | 0.004487823 | 0.086695706 | ns | NaCl | KS.3 |
| FIM1_4607 | 2119.980062 | 2587.261021 | 2344.433376 | 0.197272099 | 0.078530831 | 0.000293657 | 0.016232695 | ns | NaCl | KS.3 |
| FIM1_4608 | 193.1884849 | 146.3428835 | 390.933759 | -0.090432293 | 0.134558545 | 0.080257674 | 0.438494274 | ns | NaCl | KS.3 |
| FIM1_4609 | 4710.412875 | 4605.33736 | 3251.156676 | -0.006738275 | 0.097345271 | 0.878592314 | 0.973062502 | ns | NaCl | KS.3 |
| FIM1_4612 | 7316.864513 | 7700.180563 | 7107.770878 | 0.024477235 | 0.084584171 | 0.613816028 | 0.889162818 | ns | NaCl | KS.3 |
| FIM1_4613 | 16301.1384 | 19838.55773 | 15088.26707 | 0.048941248 | 0.115296551 | 0.272337821 | 0.693610905 | ns | NaCl | KS.3 |
| FIM1_4614 | 571.9108275 | 560.0189175 | 609.0939818 | -0.011233174 | 0.077848796 | 0.819104022 | 0.959981393 | ns | NaCl | KS.3 |
| FIM1_4615 | 1441.649558 | 1394.148871 | 1502.09062 | -0.023130918 | 0.070586429 | 0.637065535 | 0.898332475 | ns | NaCl | KS.3 |
| FIM1_4616 | 4182.255362 | 4620.967182 | 3201.066156 | 0.07692283 | 0.070197948 | 0.122434702 | 0.528318345 | ns | NaCl | KS.3 |
| FIM1_4617 | 3442.786978 | 2842.397993 | 3570.518226 | -0.078766923 | 0.104834709 | 0.119776845 | 0.524421123 | ns | NaCl | KS.3 |

| | | | | | | | | | | |
|---|---|---|---|---|---|---|---|---|---|---|
| FIM1_4618 | 4420.614134 | 4023.05857 | 4057.374277 | -0.058668244 | 0.078260306 | 0.243098734 | 0.66671235 | ns | NaCl | KS.3 |
| FIM1_4619 | 940.9483487 | 893.031671 | 857.5893191 | -0.035607983 | 0.073546009 | 0.471506235 | 0.822127061 | ns | NaCl | KS.3 |
| FIM1_4620 | 750.1896201 | 819.3222782 | 744.6249619 | 0.020469997 | 0.10589405 | 0.626391145 | 0.895487341 | ns | NaCl | KS.3 |
| FIM1_4621 | 153.7434134 | 168.5018608 | 125.0342392 | 0.009480879 | 0.130894146 | 0.78624598 | 0.95289761 | ns | NaCl | KS.3 |
| FIM1_4622 | 784.151146 | 644.9861185 | 1327.016102 | -0.115178843 | 0.098662175 | 0.035357096 | 0.297131001 | ns | NaCl | KS.3 |
| FIM1_4623 | 1552.927366 | 1422.172597 | 1412.356005 | -0.051248417 | 0.079961525 | 0.305262175 | 0.724521545 | ns | NaCl | KS.3 |
| FIM1_4624 | 2495.540156 | 1989.141087 | 2292.836892 | -0.150895053 | 0.107089222 | 0.01013917 | 0.142316716 | ns | NaCl | KS.3 |
| FIM1_4625 | 982.9087363 | 974.836933 | 926.2415632 | -0.005333483 | 0.06567305 | 0.911742396 | 0.982011252 | ns | NaCl | KS.3 |
| FIM1_4626 | 317.2369206 | 272.9459848 | 425.5798025 | -0.041794034 | 0.106423702 | 0.351274135 | 0.756912594 | ns | NaCl | KS.3 |
| FIM1_4627 | 2034.590793 | 1777.179094 | 2357.628351 | -0.055053529 | 0.095339322 | 0.257430853 | 0.680871076 | ns | NaCl | KS.3 |
| FIM1_4628 | 189.5388817 | 147.9636421 | 297.3288643 | -0.081720481 | 0.124982227 | 0.103639339 | 0.491053061 | ns | NaCl | KS.3 |
| FIM1_4629 | 629.6869994 | 580.2042348 | 744.8255768 | -0.029043287 | 0.095091852 | 0.529096326 | 0.855153333 | ns | NaCl | KS.3 |
| FIM1_4630 | 808.8269813 | 793.5126842 | 1317.615698 | -0.012522316 | 0.076358366 | 0.799077913 | 0.95648403 | ns | NaCl | KS.3 |
| FIM1_4631 | 1969.285764 | 1678.900303 | 2080.043648 | -0.07571943 | 0.094766671 | 0.136423796 | 0.544735099 | ns | NaCl | KS.3 |
| FIM1_4632 | 1764.873455 | 2045.039964 | 2347.234258 | 0.041762313 | 0.105838365 | 0.352863547 | 0.757005668 | ns | NaCl | KS.3 |
| FIM1_4633 | 64.87351631 | 35.11543218 | 76.16514234 | -0.188595416 | 0.281563604 | 0.014456204 | 0.1741395 | ns | NaCl | KS.3 |
| FIM1_4634 | 1403.134123 | 1465.256494 | 2905.267577 | 0.00709878 | 0.114535095 | 0.853129107 | 0.969345976 | ns | NaCl | KS.3 |
| FIM1_4635 | 883.3251954 | 785.4595425 | 1175.173052 | -0.061599035 | 0.086274198 | 0.219912209 | 0.642906874 | ns | NaCl | KS.3 |
| FIM1_4636 | 608.2847404 | 538.0287963 | 580.978079 | -0.088612459 | 0.077934869 | 0.084537718 | 0.451260889 | ns | NaCl | KS.3 |
| FIM1_4637 | 1074.13422 | 923.9750398 | 1091.158215 | -0.067290479 | 0.094676062 | 0.178226765 | 0.606483006 | ns | NaCl | KS.3 |
| FIM1_4638 | 1037.140426 | 1014.338929 | 1228.748941 | -0.015080334 | 0.072477808 | 0.758915284 | 0.945318863 | ns | NaCl | KS.3 |
| FIM1_4639 | 1721.782379 | 1862.363282 | 2308.383022 | 0.044495076 | 0.080458415 | 0.370769926 | 0.772114015 | ns | NaCl | KS.3 |
| FIM1_4640 | 988.1227086 | 1020.860208 | 889.2184359 | 0.012477179 | 0.090744506 | 0.788765809 | 0.953379471 | ns | NaCl | KS.3 |
| FIM1_4641 | 7072.771883 | 8485.663886 | 6170.986705 | 0.127358183 | 0.088829123 | 0.019736691 | 0.21299357 | ns | NaCl | KS.3 |
| FIM1_4642 | 3774.996847 | 3719.706875 | 3031.349814 | -0.006464447 | 0.086146104 | 0.892328197 | 0.977268396 | ns | NaCl | KS.3 |
| FIM1_4643 | 584.0428692 | 592.191899 | 499.9010469 | 0.006256536 | 0.087417262 | 0.895045414 | 0.977359731 | ns | NaCl | KS.3 |
| FIM1_4644 | 785.6883871 | 650.6054972 | 877.6145571 | -0.115897562 | 0.095678451 | 0.033722595 | 0.292771172 | ns | NaCl | KS.3 |
| FIM1_4645 | 1274.4557 | 1381.580235 | 1148.317039 | 0.038474389 | 0.086288178 | 0.430515468 | 0.805047964 | ns | NaCl | KS.3 |
| FIM1_4646 | 1869.592423 | 1909.217109 | 1583.751112 | 0.014070639 | 0.073080066 | 0.774733702 | 0.950214521 | ns | NaCl | KS.3 |
| FIM1_4647 | 2432.872427 | 2523.278365 | 1914.355097 | 0.019014442 | 0.081430969 | 0.697211028 | 0.922016178 | ns | NaCl | KS.3 |
| FIM1_4648 | 1731.834358 | 1774.429701 | 1698.51504 | 0.012957377 | 0.080707275 | 0.790879353 | 0.953972111 | ns | NaCl | KS.3 |
| FIM1_4649 | 2436.424509 | 2341.526277 | 2547.851914 | -0.025899722 | 0.073446766 | 0.599355838 | 0.880219278 | ns | NaCl | KS.3 |
| FIM1_4650 | 1718.21543 | 1821.345539 | 1549.062917 | 0.032400045 | 0.08007643 | 0.511403973 | 0.846104012 | ns | NaCl | KS.3 |
| FIM1_4651 | 4723.250051 | 5434.938307 | 4372.04566 | 0.030524573 | 0.111509239 | 0.466743486 | 0.821323144 | ns | NaCl | KS.3 |
| FIM1_4652 | 1695.268432 | 1802.389018 | 1412.47613 | 0.040142377 | 0.074626127 | 0.418352995 | 0.800387073 | ns | NaCl | KS.3 |
| FIM1_4653 | 1279.888575 | 1132.592304 | 1046.675546 | -0.038005414 | 0.100947531 | 0.403730002 | 0.790459174 | ns | NaCl | KS.3 |
| FIM1_4654 | 1107.506615 | 942.7774837 | 1079.974788 | -0.060767487 | 0.100233731 | 0.211199547 | 0.634491393 | ns | NaCl | KS.3 |
| FIM1_4655 | 1199.131459 | 1328.799462 | 1886.468117 | 0.024549144 | 0.10599387 | 0.563431473 | 0.867434473 | ns | NaCl | KS.3 |
| FIM1_4656 | 1894.12849 | 2501.145561 | 2415.83052 | 0.060685125 | 0.13335807 | 0.180369655 | 0.60960532 | ns | NaCl | KS.3 |
| FIM1_4657 | 1021.522247 | 1156.610321 | 980.8930235 | 0.067930421 | 0.086095138 | 0.179782588 | 0.608447876 | ns | NaCl | KS.3 |
| FIM1_4658 | 31709.01032 | 34923.32705 | 20752.37853 | 0.036551845 | 0.093759383 | 0.437915128 | 0.808096352 | ns | NaCl | KS.3 |
| FIM1_4660 | 92.77679877 | 80.36073935 | 183.5974283 | -0.029970767 | 0.111451896 | 0.474064294 | 0.82307921 | ns | NaCl | KS.3 |
| FIM1_4661 | 481.1755448 | 600.1839884 | 320.9228849 | 0.057680913 | 0.118965958 | 0.208540918 | 0.633002483 | ns | NaCl | KS.3 |
| FIM1_4662 | 5818.209848 | 6852.29231 | 3115.470066 | 0.029179383 | 0.118716032 | 0.46813001 | 0.821764392 | ns | NaCl | KS.3 |
| FIM1_4664 | 31.55099697 | 21.01170202 | 39.5335887 | -0.068520634 | 0.161665001 | 0.134363657 | 0.54183277 | ns | NaCl | KS.3 |

| | | | | | | | | | | |
|---|---|---|---|---|---|---|---|---|---|---|
| FIM1_4665 | 5281.233871 | 7448.094189 | 3582.114891 | 0.086640913 | 0.153161399 | 0.086922356 | 0.456808328 | ns | NaCl | KS.3 |
| FIM1_4666 | 634.4142067 | 453.8594491 | 786.747206 | -0.071151392 | 0.148271559 | 0.130183052 | 0.537622789 | ns | NaCl | KS.3 |
| FIM1_4667 | 8805.215511 | 11390.19326 | 5171.262894 | 0.044208899 | 0.131862788 | 0.287202091 | 0.706991787 | ns | NaCl | KS.3 |
| FIM1_4668 | 523.208695 | 767.4875123 | 590.2764347 | 0.110154374 | 0.171977023 | 0.049256395 | 0.34611662 | ns | NaCl | KS.3 |
| FIM1_4669 | 44.27036776 | 23.15240403 | 39.49713505 | -0.12062838 | 0.24242617 | 0.039820414 | 0.315550884 | ns | NaCl | KS.3 |
| FIM1_4670 | 2337.81285 | 2363.345279 | 2421.800564 | 0.003322799 | 0.095935986 | 0.940575332 | 0.986790704 | ns | NaCl | KS.3 |
| FIM1_4671 | 902.1817301 | 797.8800679 | 1252.57698 | -0.06485121 | 0.086518781 | 0.198317835 | 0.620924909 | ns | NaCl | KS.3 |
| FIM1_4672 | 936.0568679 | 996.1158288 | 1166.076171 | 0.035931968 | 0.078951793 | 0.467995066 | 0.821764392 | ns | NaCl | KS.3 |
| FIM1_4673 | 323.3937629 | 286.3385088 | 398.5748486 | -0.054078983 | 0.090657473 | 0.271718159 | 0.693144962 | ns | NaCl | KS.3 |
| FIM1_4674 | 228.1439998 | 214.2328855 | 279.8664622 | -0.035474243 | 0.084682386 | 0.468311707 | 0.821764392 | ns | NaCl | KS.3 |
| FIM1_4675 | 6343.074222 | 6281.884151 | 4876.118353 | -0.005755343 | 0.077334823 | 0.906712151 | 0.979989779 | ns | NaCl | KS.3 |
| FIM1_4676 | 2650.307128 | 2609.712218 | 2206.597425 | -0.012530733 | 0.059659167 | 0.78708953 | 0.95289761 | ns | NaCl | KS.3 |
| FIM1_4678 | 2858.837378 | 3515.230079 | 2206.929083 | 0.033287219 | 0.125899868 | 0.40326258 | 0.789854857 | ns | NaCl | KS.3 |
| FIM1_4679 | 2392.680473 | 1911.058057 | 1706.025796 | -0.053525294 | 0.121016576 | 0.232166294 | 0.654993872 | ns | NaCl | KS.3 |
| FIM1_4680 | 691.383803 | 678.8131007 | 650.5430871 | -0.010008024 | 0.076751571 | 0.83869283 | 0.963180246 | ns | NaCl | KS.3 |
| FIM1_4681 | 3876.688214 | 3678.042214 | 3003.550954 | -0.020216778 | 0.09084329 | 0.665964245 | 0.906573323 | ns | NaCl | KS.3 |
| FIM1_4682 | 7852.30069 | 7925.148707 | 6334.836483 | 0.007610211 | 0.061926965 | 0.871732876 | 0.971243188 | ns | NaCl | KS.3 |
| FIM1_4683 | 165.6340858 | 185.3674955 | 149.174951 | 0.018483665 | 0.116601379 | 0.635003235 | 0.898009483 | ns | NaCl | KS.3 |
| FIM1_4684 | 2000.757341 | 2206.568743 | 1653.177765 | 0.035625513 | 0.094904775 | 0.446105257 | 0.812914509 | ns | NaCl | KS.3 |
| FIM1_4685 | 1671.795122 | 1091.018243 | 1895.683557 | -0.078603038 | 0.172716411 | 0.102360292 | 0.488687531 | ns | NaCl | KS.3 |
| FIM1_4686 | 110.1238018 | 62.86777267 | 374.4480036 | -0.112885951 | 0.219470186 | 0.046789045 | 0.341725286 | ns | NaCl | KS.3 |
| FIM1_4688 | 1160.033041 | 1456.683266 | 1030.77713 | 0.094137385 | 0.114981863 | 0.073405073 | 0.421631393 | ns | NaCl | KS.3 |
| FIM1_4689 | 1199.20139 | 1230.52459 | 1260.925112 | 0.014660842 | 0.076909381 | 0.765706108 | 0.948080609 | ns | NaCl | KS.3 |
| FIM1_4690 | 3389.441847 | 3190.280601 | 3273.689111 | -0.028953645 | 0.085164937 | 0.551011833 | 0.860945181 | ns | NaCl | KS.3 |
| FIM1_4691 | 1943.491926 | 3150.147288 | 1392.037738 | 0.133563498 | 0.214668483 | 0.031750701 | 0.28257556 | ns | NaCl | KS.3 |
| FIM1_4692 | 2432.679506 | 1931.824919 | 1443.260619 | -0.096270185 | 0.115411368 | 0.068807389 | 0.412170702 | ns | NaCl | KS.3 |
| FIM1_4693 | 326.9292838 | 307.8408071 | 358.3793039 | -0.016987709 | 0.09871989 | 0.700023945 | 0.922863518 | ns | NaCl | KS.3 |
| FIM1_4694 | 577.0770376 | 602.9008318 | 868.552117 | 0.00835016 | 0.109636321 | 0.834006624 | 0.962365393 | ns | NaCl | KS.3 |
| FIM1_4695 | 538.6376919 | 733.9401398 | 611.148356 | 0.07551401 | 0.141693862 | 0.118047292 | 0.520642976 | ns | NaCl | KS.3 |
| FIM1_4697 | 2194.239329 | 3459.106285 | 1508.15025 | 0.070810371 | 0.175432183 | 0.122231561 | 0.528318345 | ns | NaCl | KS.3 |
| FIM1_4698 | 1192.336976 | 1112.126412 | 1300.486597 | -0.029845654 | 0.088885919 | 0.532606258 | 0.855153333 | ns | NaCl | KS.3 |
| FIM1_4699 | 633.8142871 | 873.1512531 | 876.9958142 | 0.072471189 | 0.144500824 | 0.126996983 | 0.534982546 | ns | NaCl | KS.3 |
| FIM1_4700 | 52.797857 | 34.45583674 | 31.51954424 | -0.089346213 | 0.176902881 | 0.07924527 | 0.436111968 | ns | NaCl | KS.3 |
| FIM1_4701 | 930.3075851 | 919.7369073 | 949.344502 | -0.002717655 | 0.101119496 | 0.949148419 | 0.987736646 | ns | NaCl | KS.3 |
| FIM1_4702 | 2271.407203 | 2273.141764 | 2349.804366 | 0.000113221 | 0.113528147 | 0.997649002 | 0.99924489 | ns | NaCl | KS.3 |
| FIM1_4703 | 7666.211439 | 7871.799611 | 8353.968002 | 0.0054869 | 0.107082969 | 0.892339118 | 0.977268396 | ns | NaCl | KS.3 |
| FIM1_4704 | 1506.012808 | 1543.541756 | 1487.561147 | 0.005175384 | 0.106987284 | 0.898471259 | 0.977971161 | ns | NaCl | KS.3 |
| FIM1_4705 | 1644.330761 | 1560.791859 | 1309.422391 | -0.025278186 | 0.084303192 | 0.602840421 | 0.881578805 | ns | NaCl | KS.3 |
| FIM1_4706 | 1636.220708 | 1497.878314 | 1323.038148 | -0.044609604 | 0.085065131 | 0.365620951 | 0.766012322 | ns | NaCl | KS.3 |
| FIM1_4707 | 527.3808242 | 536.6535556 | 651.8354446 | 0.004477612 | 0.10052853 | 0.916804447 | 0.982343702 | ns | NaCl | KS.3 |
| FIM1_4708 | 1307.308194 | 1481.690366 | 1232.431846 | 0.020484958 | 0.117923081 | 0.598690674 | 0.880167289 | ns | NaCl | KS.3 |
| FIM1_4709 | 1621.61768 | 2256.841697 | 1693.347632 | 0.093303747 | 0.150389652 | 0.073647775 | 0.421631393 | ns | NaCl | KS.3 |
| FIM1_4710 | 1593.638609 | 1234.904174 | 1338.096276 | -0.066686515 | 0.126705809 | 0.15672568 | 0.57585691 | ns | NaCl | KS.3 |
| FIM1_4711 | 1909.429619 | 1834.031464 | 2530.178698 | -0.010559516 | 0.100898221 | 0.805759781 | 0.957859861 | ns | NaCl | KS.3 |
| FIM1_4712 | 366.8979289 | 486.51969 | 488.8438043 | 0.091115198 | 0.134126386 | 0.078812648 | 0.434692818 | ns | NaCl | KS.3 |

| | | | | | | | | | | |
|---|---|---|---|---|---|---|---|---|---|---|
| FIM1_4713 | 2420.449127 | 2177.269785 | 2362.926956 | -0.038975688 | 0.095262856 | 0.406768933 | 0.794221131 | ns | NaCl | KS.3 |
| FIM1_4714 | 1491.208098 | 1465.310039 | 1656.996978 | -0.005346651 | 0.096986121 | 0.903748872 | 0.979395371 | ns | NaCl | KS.3 |
| FIM1_4715 | 5129.898089 | 4874.583152 | 4836.093207 | -0.014902744 | 0.098410852 | 0.735233742 | 0.938375543 | ns | NaCl | KS.3 |
| FIM1_4716 | 2840.458997 | 3341.396931 | 3401.173216 | 0.021226016 | 0.127186668 | 0.568793981 | 0.869550295 | ns | NaCl | KS.3 |
| FIM1_4717 | 8045.760948 | 9711.478604 | 4812.049546 | 0.036831581 | 0.119201078 | 0.375978388 | 0.778493059 | ns | NaCl | KS.3 |
| FIM1_4718 | 3333.558753 | 4208.645038 | 1948.392689 | 0.029873686 | 0.134286395 | 0.430604929 | 0.805047964 | ns | NaCl | KS.3 |
| FIM1_4719 | 457.6670533 | 457.9672666 | 359.3621064 | 0.000240899 | 0.101671895 | 0.995465397 | 0.99924489 | ns | NaCl | KS.3 |
| FIM1_4720 | 4468.907963 | 6079.822642 | 2687.385742 | 0.06873685 | 0.140837367 | 0.141551925 | 0.553543421 | ns | NaCl | KS.3 |
| FIM1_4721 | 676.6323746 | 679.6827376 | 1004.264901 | 0.003254767 | 0.082200875 | 0.946561294 | 0.987448613 | ns | NaCl | KS.3 |
| FIM1_4722 | 2525.174802 | 4419.458524 | 1807.401918 | 0.078994641 | 0.199351822 | 0.094672708 | 0.470201108 | ns | NaCl | KS.3 |
| FIM1_4724 | 631.2115199 | 574.8530335 | 584.5339102 | -0.038865119 | 0.092320682 | 0.414658971 | 0.800387073 | ns | NaCl | KS.3 |
| FIM1_4725 | 207.680321 | 225.5951607 | 211.8274937 | 0.002987468 | 0.172742689 | 0.912963407 | 0.98221021 | ns | NaCl | KS.3 |
| FIM1_4726 | 73.36530592 | 72.86299809 | 61.86479955 | -0.000321167 | 0.16235604 | 0.991229759 | 0.999134858 | ns | NaCl | KS.3 |
| FIM1_4727 | 416.3696352 | 406.4541171 | 420.3880307 | -0.000777357 | 0.176746966 | 0.9765308 | 0.996164232 | ns | NaCl | KS.3 |
| FIM1_4728 | 83.37271349 | 65.36442138 | 140.8377942 | -0.008911998 | 0.174353927 | 0.746870125 | 0.943274417 | ns | NaCl | KS.3 |
| FIM1_4729 | 688.8342247 | 634.2810858 | 766.2970696 | -0.002727331 | 0.176081003 | 0.918423464 | 0.982488618 | ns | NaCl | KS.3 |
| FIM1_4730 | 737.2054704 | 702.7325982 | 481.3915826 | -0.009632108 | 0.109130121 | 0.81008716 | 0.958816322 | ns | NaCl | KS.3 |
| FIM1_4731 | 2431.133484 | 2813.91845 | 1595.830275 | 0.031960254 | 0.111935621 | 0.447575321 | 0.812914509 | ns | NaCl | KS.3 |
| FIM1_4732 | 2768.747821 | 2718.338716 | 3415.38404 | -0.00371576 | 0.107444387 | 0.926656527 | 0.984176127 | ns | NaCl | KS.3 |
| FIM1_4733 | 898.3715846 | 1009.477755 | 1350.604209 | 0.039338709 | 0.098413241 | 0.395290294 | 0.782558381 | ns | NaCl | KS.3 |
| FIM1_4734 | 359.689785 | 388.557622 | 494.3151773 | 0.019796284 | 0.101642165 | 0.647714375 | 0.899642849 | ns | NaCl | KS.3 |
| FIM1_4736 | 159.0274062 | 293.5531747 | 252.8171641 | 0.392420805 | 0.353877407 | 0.001802967 | 0.050964538 | ns | NaCl | KS.3 |
| FIM1_4737 | 823.2431797 | 508.2286931 | 807.1249009 | -0.280535254 | 0.270716221 | 0.003603285 | 0.076817191 | ns | NaCl | KS.3 |
| FIM1_4738 | 60.19705618 | 91.03392373 | 122.0593403 | 0.085296103 | 0.171053 | 0.087734679 | 0.458969535 | ns | NaCl | KS.3 |
| FIM1_4739 | 441.85211 | 525.5266451 | 449.3723635 | 0.074319982 | 0.100967607 | 0.139393804 | 0.550332947 | ns | NaCl | KS.3 |
| FIM1_4740 | 479.6463194 | 566.6418762 | 1494.907235 | 0.062667377 | 0.101494381 | 0.198157585 | 0.620924909 | ns | NaCl | KS.3 |
| FIM1_4741 | 367.6646404 | 282.1103813 | 562.5953815 | -0.090210291 | 0.127715892 | 0.081308518 | 0.440163087 | ns | NaCl | KS.3 |
| FIM1_4742 | 196.2760698 | 206.397289 | 286.3233181 | 0.009408994 | 0.111590634 | 0.810887577 | 0.958816322 | ns | NaCl | KS.3 |
| FIM1_4743 | 29.77385477 | 16.93852546 | 29.18473673 | -0.076790471 | 0.201851908 | 0.098310756 | 0.479012053 | ns | NaCl | KS.3 |
| FIM1_4744 | 3521.567066 | 3498.430527 | 2880.341186 | -0.002061819 | 0.095022638 | 0.963373389 | 0.993691273 | ns | NaCl | KS.3 |
| FIM1_4745 | 904.0278731 | 622.9756959 | 978.5767618 | -0.383207842 | 0.192635366 | 0.0001506 | 0.010703359 | ns | NaCl | KS.3 |
| FIM1_4746 | 2416.493622 | 4227.475072 | 2526.440432 | 0.412321171 | 0.326736009 | 0.001193523 | 0.04123457 | ns | NaCl | KS.3 |
| FIM1_4747 | 670.859037 | 749.4725355 | 669.4574419 | 0.061895494 | 0.083527049 | 0.219179372 | 0.642906874 | ns | NaCl | KS.3 |
| FIM1_4748 | 3873.314341 | 3645.897177 | 4156.527977 | -0.042337057 | 0.071237487 | 0.390862696 | 0.779482559 | ns | NaCl | KS.3 |
| FIM1_4749 | 314.6583128 | 301.8035796 | 434.7145254 | -0.022400235 | 0.083576419 | 0.645008524 | 0.898899754 | ns | NaCl | KS.3 |
| FIM1_4750 | 5259.234169 | 5646.943781 | 4683.332238 | 0.062069591 | 0.061425235 | 0.188579265 | 0.618812647 | ns | NaCl | KS.3 |
| FIM1_4751 | 232.2053728 | 255.8526322 | 204.7642249 | 0.028028855 | 0.100394315 | 0.528880406 | 0.855153333 | ns | NaCl | KS.3 |
| FIM1_4753 | 3425.996437 | 4913.855372 | 2190.71695 | 0.117112627 | 0.166766669 | 0.041537364 | 0.322888102 | ns | NaCl | KS.3 |
| FIM1_4754 | 2133.348239 | 2377.683973 | 1932.410155 | 0.064260649 | 0.081453538 | 0.203510632 | 0.626494461 | ns | NaCl | KS.3 |
| FIM1_4755 | 317.7322597 | 320.5795735 | 420.6909983 | 0.002674096 | 0.091530655 | 0.953780286 | 0.988965594 | ns | NaCl | KS.3 |
| FIM1_4756 | 2176.776959 | 2928.219107 | 1824.271366 | 0.177640792 | 0.148190565 | 0.007501285 | 0.11809776 | ns | NaCl | KS.3 |
| FIM1_4757 | 2264.589937 | 3011.786484 | 1744.657445 | 0.069091599 | 0.134722364 | 0.142803966 | 0.555038853 | ns | NaCl | KS.3 |
| FIM1_4758 | 411.4161547 | 354.4772172 | 689.9815069 | -0.081095375 | 0.089212929 | 0.115432833 | 0.517367878 | ns | NaCl | KS.3 |
| FIM1_4759 | 7195.14916 | 7860.3764 | 6178.264648 | 0.035488296 | 0.091685782 | 0.455268454 | 0.816788914 | ns | NaCl | KS.3 |
| FIM1_4760 | 1280.097946 | 5001.820666 | 4423.256503 | 1.743471126 | 0.313142066 | 3.5088953277 | 1.3428272504 | Km>KS.3 | NaCl | KS.3 |

| | | | | | | | | | | |
|---|---|---|---|---|---|---|---|---|---|---|
| FIM1_4761 | 321.7861396 | 365.2505533 | 374.2602876 | 0.046387 | 0.098110018 | 0.325265731 | 0.739131368 | ns | NaCl | KS.3 |
| FIM1_4762 | 124.3366983 | 121.3577415 | 127.7586392 | -0.009525746 | 0.097583401 | 0.829027696 | 0.961288108 | ns | NaCl | KS.3 |
| FIM1_4763 | 170.0973398 | 189.6996749 | 172.8569932 | 0.024851556 | 0.107617492 | 0.554777481 | 0.863314973 | ns | NaCl | KS.3 |
| FIM1_4764 | 331.5662517 | 338.1112051 | 328.2736609 | 0.003602801 | 0.107564894 | 0.928804285 | 0.984238275 | ns | NaCl | KS.3 |
| FIM1_4765 | 229.3187653 | 152.3813615 | 314.2749395 | -0.123471161 | 0.184623692 | 0.037158874 | 0.304670827 | ns | NaCl | KS.3 |
| FIM1_4766 | 211.5590621 | 172.299213 | 195.8864339 | -0.059529005 | 0.11288217 | 0.204041993 | 0.62738499 | ns | NaCl | KS.3 |
| FIM1_4767 | 731.1669386 | 742.316979 | 816.4770033 | 0.007305244 | 0.087862832 | 0.877301737 | 0.972783118 | ns | NaCl | KS.3 |
| FIM1_4769 | 90.62027292 | 120.326065 | 118.1575199 | 0.046001034 | 0.137639527 | 0.266426255 | 0.689593109 | ns | NaCl | KS.3 |
| FIM1_4770 | 782.7482315 | 842.7317227 | 529.218885 | 0.008672842 | 0.126037181 | 0.808566883 | 0.958451332 | ns | NaCl | KS.3 |
| FIM1_4771 | 1175.427508 | 928.906292 | 746.2717156 | -0.081013761 | 0.118995596 | 0.107301242 | 0.498909937 | ns | NaCl | KS.3 |
| FIM1_4772 | 28.51459236 | 21.21992949 | 75.1073442 | -0.043706048 | 0.135243123 | 0.28715817 | 0.706991787 | ns | NaCl | KS.3 |
| FIM1_4773 | 2235.929414 | 1649.215917 | 2745.278261 | -0.158190205 | 0.149916964 | 0.013318185 | 0.166477316 | ns | NaCl | KS.3 |
| FIM1_3 | 1894.308797 | 2973.550777 | 3037.529789 | 0.558115161 | 0.164424797 | 2.96646310611 | 0.000162178 | ns | TM | KS.1 |
| FIM1_5 | 579.2685586 | 625.0760985 | 633.8041188 | 0.022344546 | 0.142502775 | 0.725361785 | 0.907961357 | ns | TM | KS.1 |
| FIM1_6 | 390.106426 | 405.6170494 | 405.4975614 | 0.01574171 | 0.118465312 | 0.801829259 | 0.936908579 | ns | TM | KS.1 |
| FIM1_7 | 18.54391998 | 39.42728799 | 60.8195808 | 0.571788952 | 0.439130778 | 0.00222932 | 0.02814941 | ns | TM | KS.1 |
| FIM1_8 | 244.3924337 | 337.7417933 | 375.3172182 | 0.142817942 | 0.18543166 | 0.081861634 | 0.304774234 | ns | TM | KS.1 |
| FIM1_9 | 1058.008074 | 1078.663684 | 2901.971283 | 0.004721123 | 0.151996433 | 0.93969698 | 0.983546982 | ns | TM | KS.1 |
| FIM1_10 | 982.053837 | 953.0646011 | 2943.370395 | -0.008261641 | 0.144598021 | 0.895458433 | 0.967972723 | ns | TM | KS.1 |
| FIM1_11 | 149.9026969 | 212.377455 | 340.6940978 | 0.234100752 | 0.186849691 | 0.014874749 | 0.110285958 | ns | TM | KS.1 |
| FIM1_12 | 1030.547373 | 571.9435466 | 1729.337393 | -0.464927549 | 0.340590138 | 0.002540772 | 0.030888701 | ns | TM | KS.1 |
| FIM1_13 | 45346.45834 | 34960.31201 | 35774.55391 | -0.052354248 | 0.18752612 | 0.424753687 | 0.740157476 | ns | TM | KS.1 |
| FIM1_14 | 13.35328137 | 14.37730315 | 16.92066841 | 0.013841973 | 0.180649017 | 0.820866686 | 0.943851992 | ns | TM | KS.1 |
| FIM1_15 | 18757.59105 | 15788.12038 | 14849.47076 | -0.025325294 | 0.195914533 | 0.679789495 | 0.887492141 | ns | TM | KS.1 |
| FIM1_16 | 21175.36724 | 14366.7213 | 13864.7932 | -0.171877426 | 0.21393634 | 0.053742518 | 0.24174415 | ns | TM | KS.1 |
| FIM1_17 | 27050.44122 | 21703.81208 | 21161.65642 | -0.036740366 | 0.192151727 | 0.559633417 | 0.826391733 | ns | TM | KS.1 |
| FIM1_18 | 2329.561898 | 1928.59211 | 2232.947735 | -0.065386372 | 0.149869112 | 0.337546411 | 0.665879925 | ns | TM | KS.1 |
| FIM1_19 | 1146.851215 | 1062.324111 | 965.1753006 | -0.058280647 | 0.080075878 | 0.272668607 | 0.599172403 | ns | TM | KS.1 |
| FIM1_20 | 3412.488334 | 3740.908132 | 3333.389345 | 0.075595029 | 0.081204729 | 0.163559464 | 0.45366779 | ns | TM | KS.1 |
| FIM1_21 | 611.2869441 | 612.0671481 | 585.7584931 | 0.000851605 | 0.089926348 | 0.987923151 | 0.996167262 | ns | TM | KS.1 |
| FIM1_22 | 1373.575174 | 1416.33382 | 1520.407378 | 0.024167497 | 0.059869033 | 0.57401347 | 0.833542619 | ns | TM | KS.1 |
| FIM1_23 | 208.4000849 | 210.5399599 | 230.7427362 | 0.005818344 | 0.109674815 | 0.924442686 | 0.976662213 | ns | TM | KS.1 |
| FIM1_24 | 703.2911926 | 981.6143765 | 1199.112786 | 0.133400604 | 0.19015679 | 0.098325033 | 0.338336381 | ns | TM | KS.1 |
| FIM1_25 | 437.5087892 | 497.8944151 | 454.9892691 | 0.047510503 | 0.135931674 | 0.470236712 | 0.771448038 | ns | TM | KS.1 |
| FIM1_26 | 395.7902032 | 284.9720257 | 284.9825584 | -0.179464106 | 0.18578909 | 0.041589099 | 0.208364318 | ns | TM | KS.1 |
| FIM1_27 | 741.8621899 | 702.1323353 | 657.296323 | -0.028127346 | 0.103617262 | 0.642513362 | 0.872815354 | ns | TM | KS.1 |
| FIM1_28 | 10849.16284 | 12848.41434 | 11506.37819 | 0.087704304 | 0.129829881 | 0.208064178 | 0.515497652 | ns | TM | KS.1 |
| FIM1_29 | 3418.749935 | 3252.353971 | 2774.21492 | -0.022705701 | 0.111759698 | 0.714469686 | 0.905084971 | ns | TM | KS.1 |
| FIM1_30 | 2132.108501 | 2109.040388 | 2049.9388 | -0.006361212 | 0.084799516 | 0.906957035 | 0.97188159 | ns | TM | KS.1 |
| FIM1_31 | 3845.805266 | 4136.74939 | 4258.824352 | 0.058447678 | 0.074860265 | 0.249286762 | 0.574965989 | ns | TM | KS.1 |
| FIM1_32 | 3382.182571 | 3151.366704 | 2794.011073 | -0.043643266 | 0.09266476 | 0.450318223 | 0.754323807 | ns | TM | KS.1 |
| FIM1_33 | 795.179714 | 803.145219 | 732.2157693 | 0.004498339 | 0.110504115 | 0.941670826 | 0.98379092 | ns | TM | KS.1 |
| FIM1_34 | 1493.954708 | 1566.308475 | 1643.975651 | 0.037556708 | 0.064473442 | 0.407621165 | 0.728957685 | ns | TM | KS.1 |
| FIM1_35 | 28.48249875 | 36.01776224 | 43.59138275 | 0.058894216 | 0.169325711 | 0.37983458 | 0.703660035 | ns | TM | KS.1 |
| FIM1_36 | 6118.472479 | 8764.592362 | 6124.361802 | 0.07435008 | 0.206830648 | 0.284419032 | 0.611419177 | ns | TM | KS.1 |

| | | | | | | | | | | |
|---|---|---|---|---|---|---|---|---|---|---|
| FIM1_37 | 631.130347 | 598.713297 | 700.1965119 | -0.028234682 | 0.098529309 | 0.633745084 | 0.869547581 | ns | TM | KS.1 |
| FIM1_38 | 706.8561185 | 1005.565519 | 1220.934355 | 0.274468458 | 0.1876175 | 0.006830744 | 0.065101441 | ns | TM | KS.1 |
| FIM1_39 | 1299.778154 | 1504.707474 | 1622.369305 | 0.058156563 | 0.135341872 | 0.384655235 | 0.708500478 | ns | TM | KS.1 |
| FIM1_40 | 2500.609603 | 2376.341907 | 1421.516646 | -0.031280364 | 0.087672675 | 0.574571066 | 0.833865535 | ns | TM | KS.1 |
| FIM1_41 | 861.3886694 | 895.0522114 | 1004.945188 | 0.022164367 | 0.090877058 | 0.696058987 | 0.894950919 | ns | TM | KS.1 |
| FIM1_42 | 996.9752528 | 1059.286967 | 1088.647994 | 0.031371405 | 0.103442848 | 0.605032613 | 0.851547247 | ns | TM | KS.1 |
| FIM1_43 | 2081.207104 | 2115.22232 | 2347.041188 | 0.012320748 | 0.059816235 | 0.7744949 | 0.926451582 | ns | TM | KS.1 |
| FIM1_44 | 309.938807 | 281.623735 | 223.7839843 | -0.039895739 | 0.123491675 | 0.535802902 | 0.812908026 | ns | TM | KS.1 |
| FIM1_45 | 1354.656295 | 1259.362577 | 909.3249672 | -0.043598208 | 0.096270232 | 0.460171851 | 0.763054265 | ns | TM | KS.1 |
| FIM1_46 | 2902.496939 | 3130.415645 | 3895.846385 | 0.045608419 | 0.09595969 | 0.439491975 | 0.747832613 | ns | TM | KS.1 |
| FIM1_47 | 1156.269003 | 1232.982256 | 1453.094101 | 0.043741314 | 0.083396353 | 0.419998944 | 0.737778527 | ns | TM | KS.1 |
| FIM1_48 | 3846.030739 | 3367.076708 | 2574.536415 | -0.145248114 | 0.078278698 | 0.012309283 | 0.097517084 | ns | TM | KS.1 |
| FIM1_49 | 5223.928178 | 8055.8092 | 9518.429149 | 0.190410885 | 0.236190162 | 0.043088996 | 0.212459177 | ns | TM | KS.1 |
| FIM1_50 | 9116.169427 | 4484.855769 | 4848.123799 | -0.887937757 | 0.266693895 | 1.0978004838[illegible] | 0.000444029 | ns | TM | KS.1 |
| FIM1_51 | 3261.928173 | 3065.936114 | 3374.321267 | -0.033756386 | 0.100094067 | 0.57280089 | 0.832094673 | ns | TM | KS.1 |
| FIM1_52 | 3648.004161 | 3524.569887 | 3261.822567 | -0.019771881 | 0.090707751 | 0.727067631 | 0.907961357 | ns | TM | KS.1 |
| FIM1_53 | 28977.55307 | 34161.16121 | 32534.50092 | 0.139732686 | 0.108677019 | 0.045176423 | 0.219529062 | ns | TM | KS.1 |
| FIM1_54 | 2677.280524 | 3606.686675 | 3260.43686 | 0.153438253 | 0.17354849 | 0.064294842 | 0.266080163 | ns | TM | KS.1 |
| FIM1_55 | 2409.083491 | 1168.64038 | 884.141548 | -0.481091885 | 0.422840747 | 0.00411923 | 0.044550371 | ns | TM | KS.1 |
| FIM1_56 | 41.99975282 | 15.8974698 | 110.1015889 | -0.751505817 | 0.538480404 | 0.001203254 | 0.017351278 | ns | TM | KS.1 |
| FIM1_57 | 498.9368985 | 361.9690452 | 411.2940193 | -0.121681094 | 0.18573847 | 0.120583775 | 0.381550647 | ns | TM | KS.1 |
| FIM1_58 | 2903.020542 | 3687.691403 | 3663.533176 | 0.134022331 | 0.149925684 | 0.085130773 | 0.31075147 | ns | TM | KS.1 |
| FIM1_59 | 3760.003249 | 4572.198914 | 4886.402825 | 0.062906017 | 0.154260677 | 0.353836067 | 0.679784721 | ns | TM | KS.1 |
| FIM1_60 | 1904.929985 | 2011.939452 | 2097.26308 | 0.035231304 | 0.085769963 | 0.522233692 | 0.807870838 | ns | TM | KS.1 |
| FIM1_61 | 1363.229337 | 1329.293114 | 1514.364944 | -0.017099108 | 0.069567793 | 0.720900818 | 0.906676397 | ns | TM | KS.1 |
| FIM1_62 | 1022.812144 | 1236.183497 | 1248.976291 | 0.116812696 | 0.130378458 | 0.10938191 | 0.35800987 | ns | TM | KS.1 |
| FIM1_63 | 67.70959211 | 84.13967784 | 61.0388668 | 0.060015005 | 0.16499668 | 0.37260383 | 0.695052139 | ns | TM | KS.1 |
| FIM1_64 | 2696.502343 | 2177.572136 | 2027.713438 | -0.232055384 | 0.099057715 | 0.002005096 | 0.02591001 | ns | TM | KS.1 |
| FIM1_65 | 252.852792 | 198.9679854 | 253.6940693 | -0.188728897 | 0.136166796 | 0.021625094 | 0.143551826 | ns | TM | KS.1 |
| FIM1_66 | 4250.045667 | 4537.989004 | 4634.515481 | 0.035114195 | 0.102075064 | 0.56126419 | 0.826391733 | ns | TM | KS.1 |
| FIM1_67 | 253.9480482 | 294.702463 | 516.1440968 | 0.070735314 | 0.127496141 | 0.295242384 | 0.622913851 | ns | TM | KS.1 |
| FIM1_68 | 13126.77094 | 16931.07465 | 15105.22591 | 0.167935039 | 0.150966819 | 0.041553911 | 0.208364318 | ns | TM | KS.1 |
| FIM1_69 | 13133.19171 | 15149.17464 | 14277.90827 | 0.101720731 | 0.109343602 | 0.126426094 | 0.392125822 | ns | TM | KS.1 |
| FIM1_70 | 801.2069018 | 1035.713776 | 1316.552668 | 0.150066726 | 0.155470657 | 0.062798129 | 0.262637093 | ns | TM | KS.1 |
| FIM1_71 | 793.2859953 | 726.2915504 | 738.570201 | -0.049477338 | 0.103673286 | 0.420867429 | 0.737778527 | ns | TM | KS.1 |
| FIM1_72 | 536.8267643 | 572.8221206 | 603.4281888 | 0.032699292 | 0.104891358 | 0.592256877 | 0.846446108 | ns | TM | KS.1 |
| FIM1_73 | 12096.08054 | 12936.36281 | 13298.10691 | 0.050482417 | 0.076899392 | 0.327058811 | 0.654510064 | ns | TM | KS.1 |
| FIM1_74 | 855.2844183 | 785.7038402 | 775.8522912 | -0.064405472 | 0.082267943 | 0.235514388 | 0.554774659 | ns | TM | KS.1 |
| FIM1_75 | 22922.46598 | 21876.02587 | 22825.45264 | -0.026481896 | 0.093958061 | 0.647018607 | 0.873595603 | ns | TM | KS.1 |
| FIM1_76 | 279.7224354 | 273.9535694 | 347.4029841 | -0.010299059 | 0.104967349 | 0.864812559 | 0.959975149 | ns | TM | KS.1 |
| FIM1_77 | 665.070475 | 624.17644 | 859.8821874 | -0.042337838 | 0.083933148 | 0.436731558 | 0.747832613 | ns | TM | KS.1 |
| FIM1_78 | 3218.401445 | 3097.778633 | 3222.916234 | -0.022868631 | 0.087413809 | 0.680380909 | 0.887492141 | ns | TM | KS.1 |
| FIM1_79 | 1108.275062 | 1046.791296 | 3178.933275 | -0.022756249 | 0.121992699 | 0.719212196 | 0.905913973 | ns | TM | KS.1 |
| FIM1_80 | 69.36401548 | 74.04577491 | 115.301577 | 0.020955054 | 0.127211051 | 0.741503504 | 0.914738314 | ns | TM | KS.1 |
| FIM1_81 | 1233.963202 | 1105.738059 | 1455.405965 | -0.060328173 | 0.110515441 | 0.344335632 | 0.670129201 | ns | TM | KS.1 |

| | | | | | | | | | | | |
|---|---|---|---|---|---|---|---|---|---|---|---|
| FIM1_82 | 1914.786933 | 1957.057294 | 2001.056325 | 0.015388703 | 0.071360163 | 0.752070838 | 0.918662314 | ns | | TM | KS.1 |
| FIM1_83 | 383.5886774 | 402.9913881 | 385.1962856 | 0.023793472 | 0.102303326 | 0.692419351 | 0.893131011 | ns | | TM | KS.1 |
| FIM1_84 | 1116.128489 | 894.5121432 | 813.4542519 | -0.164101083 | 0.134535245 | 0.037965952 | 0.196574108 | ns | | TM | KS.1 |
| FIM1_85 | 1056.190007 | 1173.645841 | 812.4313926 | 0.082803143 | 0.090220653 | 0.156866889 | 0.442160212 | ns | | TM | KS.1 |
| FIM1_86 | 1783.575537 | 1438.877322 | 890.901293 | -0.202759955 | 0.116608271 | 0.009848366 | 0.083624294 | ns | | TM | KS.1 |
| FIM1_87 | 453.7486595 | 549.810017 | 533.8379842 | 0.094589161 | 0.138353624 | 0.185638128 | 0.48817438 | ns | | TM | KS.1 |
| FIM1_88 | 907.2407245 | 780.8128172 | 812.0837853 | -0.077967869 | 0.123714959 | 0.249966243 | 0.575730011 | ns | | TM | KS.1 |
| FIM1_89 | 541.4546175 | 443.6438489 | 613.585864 | -0.050523573 | 0.165561176 | 0.44301717 | 0.748899226 | ns | | TM | KS.1 |
| FIM1_90 | 2650.668949 | 2638.766652 | 2907.03043 | -0.003216621 | 0.059728303 | 0.940340403 | 0.983546982 | ns | | TM | KS.1 |
| FIM1_91 | 525.9872436 | 1070.871715 | 2110.39745 | 0.74998271 | 0.363216647 | 0.000276533 | 0.005661525 | ns | | TM | KS.1 |
| FIM1_92 | 402.9154715 | 385.1960404 | 320.0306234 | -0.025451 | 0.097331207 | 0.665377425 | 0.880679852 | ns | | TM | KS.1 |
| FIM1_93 | 348.8631408 | 319.6621741 | 258.6469668 | -0.059930294 | 0.092390132 | 0.303679309 | 0.63187142 | ns | | TM | KS.1 |
| FIM1_94 | 492.2939508 | 406.8557693 | 357.0574842 | -0.127333337 | 0.128195992 | 0.084033442 | 0.308763938 | ns | | TM | KS.1 |
| FIM1_95 | 4733.170747 | 4727.480924 | 5406.626742 | -0.000902824 | 0.067130457 | 0.984577661 | 0.996034691 | ns | | TM | KS.1 |
| FIM1_96 | 375.0023261 | 313.983054 | 692.0664714 | -0.060987266 | 0.148149414 | 0.366727214 | 0.690301888 | ns | | TM | KS.1 |
| FIM1_97 | 7658.082636 | 2035.626225 | 3895.884372 | -1.629726736 | 0.302556369 | 3.9333910584 | 6.988793041 | Km<KS.1 | | TM | KS.1 |
| FIM1_98 | 215.8502007 | 195.1583209 | 194.9117998 | -0.035702331 | 0.134304925 | 0.58092046 | 0.838424819 | ns | | TM | KS.1 |
| FIM1_99 | 548.4064823 | 515.7010415 | 463.1981537 | -0.02972377 | 0.110069011 | 0.631810197 | 0.86950366 | ns | | TM | KS.1 |
| FIM1_100 | 3972.694555 | 3425.045914 | 3763.677311 | -0.104356682 | 0.111759459 | 0.121993907 | 0.384126384 | ns | | TM | KS.1 |
| FIM1_101 | 931.5571767 | 804.7445107 | 1500.413589 | -0.074021151 | 0.124731795 | 0.273126172 | 0.599198132 | ns | | TM | KS.1 |
| FIM1_102 | 232.9424685 | 205.4882559 | 221.6806448 | -0.062361694 | 0.12084627 | 0.344057722 | 0.670129201 | ns | | TM | KS.1 |
| FIM1_103 | 1649.556951 | 1878.887383 | 1767.344721 | 0.100578763 | 0.101365867 | 0.115366095 | 0.370766358 | ns | | TM | KS.1 |
| FIM1_104 | 686.4867393 | 675.9740681 | 640.3642589 | -0.011588527 | 0.06021844 | 0.788575629 | 0.933067989 | ns | | TM | KS.1 |
| FIM1_105 | 2517.383686 | 3581.472871 | 3825.164874 | 0.268452883 | 0.188870634 | 0.00781311 | 0.070772899 | ns | | TM | KS.1 |
| FIM1_107 | 1022.439037 | 1020.278166 | 944.1502495 | -0.001117688 | 0.093021613 | 0.984430473 | 0.996034691 | ns | | TM | KS.1 |
| FIM1_108 | 2128.716847 | 2166.353715 | 2134.243348 | 0.00896763 | 0.10052821 | 0.88008287 | 0.965258439 | ns | | TM | KS.1 |
| FIM1_109 | 1217.157895 | 4978.030273 | 3009.743906 | 1.928584094 | 0.173700539 | 2.0803021572 | 5.174751616 | Km>KS.1 | | TM | KS.1 |
| FIM1_110 | 441.6932437 | 866.7297866 | 762.545038 | 0.933113007 | 0.148766832 | 5.5792251366 | 2.775664505 | ns | | TM | KS.1 |
| FIM1_111 | 7427.087438 | 7857.43515 | 5944.23296 | 0.040669571 | 0.075665031 | 0.423372582 | 0.739045122 | ns | | TM | KS.1 |
| FIM1_112 | 707.3856145 | 941.9513615 | 919.1620486 | 0.195742137 | 0.162634248 | 0.025547432 | 0.156140634 | ns | | TM | KS.1 |
| FIM1_113 | 1099.677632 | 953.7478862 | 1147.542679 | -0.104025255 | 0.108617352 | 0.117902023 | 0.376242827 | ns | | TM | KS.1 |
| FIM1_114 | 985.0578984 | 1013.181312 | 985.3718464 | 0.016486379 | 0.087934361 | 0.766997789 | 0.924883104 | ns | | TM | KS.1 |
| FIM1_115 | 832.4431934 | 873.4886779 | 1055.82097 | 0.016522994 | 0.130584884 | 0.794478186 | 0.934404013 | ns | | TM | KS.1 |
| FIM1_116 | 491.3101829 | 392.4098367 | 432.800489 | -0.146679614 | 0.141212777 | 0.060926771 | 0.26041916 | ns | | TM | KS.1 |
| FIM1_117 | 9604.452268 | 11218.60407 | 11163.9285 | 0.074404784 | 0.129093986 | 0.27489909 | 0.600258288 | ns | | TM | KS.1 |
| FIM1_118 | 307.7998137 | 185.5820915 | 186.7466403 | -0.416441983 | 0.283918541 | 0.002377698 | 0.02935247 | ns | | TM | KS.1 |
| FIM1_119 | 5806.475657 | 7062.973654 | 7368.886416 | 0.097797498 | 0.13892432 | 0.173902182 | 0.469686946 | ns | | TM | KS.1 |
| FIM1_120 | 1511.079088 | 1351.89305 | 1362.100667 | -0.083017697 | 0.095183368 | 0.169627999 | 0.463676862 | ns | | TM | KS.1 |
| FIM1_121 | 1983.733356 | 3133.694408 | 3572.133845 | 0.232249126 | 0.253017096 | 0.024429064 | 0.153646767 | ns | | TM | KS.1 |
| FIM1_122 | 3121.470749 | 3198.817483 | 3375.700208 | 0.01708269 | 0.07000888 | 0.722327665 | 0.907011644 | ns | | TM | KS.1 |
| FIM1_123 | 5330.614162 | 5022.621797 | 4792.686076 | -0.038263697 | 0.086407303 | 0.489447801 | 0.788598425 | ns | | TM | KS.1 |
| FIM1_124 | 671.4423737 | 771.5669356 | 724.0275505 | 0.100538508 | 0.10651041 | 0.125318182 | 0.389417837 | ns | | TM | KS.1 |
| FIM1_125 | 801.5794448 | 342.9734285 | 653.6319812 | -1.155728343 | 0.177166145 | 1.5878162078 | 8.777095149 | Km<KS.1 | | TM | KS.1 |
| FIM1_126 | 1431.31382 | 1346.988423 | 1309.752804 | -0.044520185 | 0.075120853 | 0.378882421 | 0.703083325 | ns | | TM | KS.1 |
| FIM1_127 | 1118.770367 | 729.969962 | 779.7362617 | -0.352832744 | 0.228663865 | 0.002894066 | 0.03410525 | ns | | TM | KS.1 |

| | | | | | | | | | | |
|---|---|---|---|---|---|---|---|---|---|---|
| FIM1_128 | 16503.30276 | 17340.85182 | 15811.42649 | 0.032329245 | 0.082239905 | 0.546340668 | 0.818934867 | ns | TM | KS.1 |
| FIM1_129 | 799.7640533 | 968.1790301 | 1036.64876 | 0.09038109 | 0.139411442 | 0.203256141 | 0.51148169 | ns | TM | KS.1 |
| FIM1_130 | 13306.50694 | 11452.71749 | 9887.24238 | -0.119354288 | 0.106850824 | 0.076064207 | 0.294489829 | ns | TM | KS.1 |
| FIM1_131 | 1924.056734 | 2019.308942 | 2248.066324 | 0.025034355 | 0.101959278 | 0.676991026 | 0.887002016 | ns | TM | KS.1 |
| FIM1_132 | 19177.20273 | 21994.36638 | 21182.2828 | 0.135401255 | 0.088837862 | 0.028643288 | 0.168042873 | ns | TM | KS.1 |
| FIM1_133 | 4879.284668 | 4211.798991 | 4961.324517 | -0.094390989 | 0.115055129 | 0.162008524 | 0.451789467 | ns | TM | KS.1 |
| FIM1_134 | 4310.348879 | 4720.07677 | 4336.865324 | 0.062676064 | 0.091821008 | 0.281189989 | 0.606295641 | ns | TM | KS.1 |
| FIM1_135 | 1462.862562 | 1470.877417 | 1364.517315 | 0.003358458 | 0.067283169 | 0.942765197 | 0.984319543 | ns | TM | KS.1 |
| FIM1_136 | 95.77065664 | 84.25389719 | 244.4891628 | -0.036464834 | 0.148093613 | 0.572551861 | 0.832094673 | ns | TM | KS.1 |
| FIM1_137 | 691.8825218 | 452.2108762 | 852.1868397 | -0.291722813 | 0.235923833 | 0.008604878 | 0.07658187 | ns | TM | KS.1 |
| FIM1_138 | 9729.735748 | 12445.94793 | 11142.29983 | 0.181645683 | 0.143565171 | 0.028329645 | 0.166668193 | ns | TM | KS.1 |
| FIM1_140 | 16361.35096 | 18865.49261 | 17517.12602 | 0.098685002 | 0.110369337 | 0.138441618 | 0.412176572 | ns | TM | KS.1 |
| FIM1_141 | 806.7513835 | 780.1169253 | 718.8621058 | -0.019330828 | 0.093792108 | 0.737419112 | 0.913453034 | ns | TM | KS.1 |
| FIM1_142 | 11505.89077 | 12763.33607 | 12027.67333 | 0.111760749 | 0.067880589 | 0.025466375 | 0.156140634 | ns | TM | KS.1 |
| FIM1_143 | 165.6576437 | 133.6053606 | 247.9532581 | -0.072830611 | 0.155962655 | 0.293423673 | 0.620392168 | ns | TM | KS.1 |
| FIM1_144 | 967.0699129 | 822.2272707 | 911.1906525 | -0.037657702 | 0.164934903 | 0.557119097 | 0.825232602 | ns | TM | KS.1 |
| FIM1_145 | 9800.964757 | 12623.94068 | 13754.70225 | 0.145327722 | 0.154269193 | 0.068825794 | 0.276450087 | ns | TM | KS.1 |
| FIM1_146 | 2280.114872 | 2168.436172 | 2501.115526 | -0.036953076 | 0.071183926 | 0.447642455 | 0.752446492 | ns | TM | KS.1 |
| FIM1_147 | 2000.266375 | 1976.851442 | 2010.46277 | -0.008287844 | 0.062510497 | 0.851918958 | 0.95392681 | ns | TM | KS.1 |
| FIM1_148 | 737.02257 | 808.6424116 | 787.1053614 | 0.074256199 | 0.082008505 | 0.173522146 | 0.469348363 | ns | TM | KS.1 |
| FIM1_149 | 659.9305367 | 640.2786626 | 567.6685807 | -0.018554413 | 0.081367496 | 0.726881367 | 0.907961357 | ns | TM | KS.1 |
| FIM1_150 | 1115.844924 | 1203.162866 | 1020.816971 | 0.060923824 | 0.076868599 | 0.238992837 | 0.559787835 | ns | TM | KS.1 |
| FIM1_151 | 877.572727 | 839.9831491 | 929.2554157 | -0.026743605 | 0.086757217 | 0.628759377 | 0.867709349 | ns | TM | KS.1 |
| FIM1_153 | 660.9097259 | 803.2441415 | 573.2743016 | 0.106678084 | 0.136321692 | 0.142319038 | 0.418709174 | ns | TM | KS.1 |
| FIM1_154 | 1881.618143 | 1908.786819 | 1999.653377 | 0.004415002 | 0.135605928 | 0.944215171 | 0.984794649 | ns | TM | KS.1 |
| FIM1_155 | 3665.017536 | 3612.844804 | 3525.482726 | -0.005963858 | 0.116862092 | 0.923804319 | 0.976662213 | ns | TM | KS.1 |
| FIM1_156 | 1617.162534 | 1064.471081 | 1443.315126 | -0.475003493 | 0.173936379 | 4.8254859636 | 0.001356316 | ns | TM | KS.1 |
| FIM1_157 | 1970.860399 | 1349.395521 | 2207.756938 | -0.130288998 | 0.207939693 | 0.107425257 | 0.355110069 | ns | TM | KS.1 |
| FIM1_158 | 8759.406689 | 7605.860048 | 6922.7706 | -0.114866812 | 0.102457135 | 0.079035794 | 0.299697464 | ns | TM | KS.1 |
| FIM1_159 | 6520.865513 | 6735.447938 | 6425.530611 | 0.035903438 | 0.032438066 | 0.189877718 | 0.49354318 | ns | TM | KS.1 |
| FIM1_160 | 5702.225333 | 5852.079874 | 5656.953157 | 0.020853629 | 0.056927773 | 0.614975831 | 0.858612877 | ns | TM | KS.1 |
| FIM1_161 | 753.5892634 | 612.253163 | 678.5940738 | -0.109864415 | 0.141112933 | 0.135944575 | 0.407922054 | ns | TM | KS.1 |
| FIM1_162 | 1260.643712 | 862.3874805 | 1262.050445 | -0.360279185 | 0.183971214 | 0.001078979 | 0.016290797 | ns | TM | KS.1 |
| FIM1_163 | 264.1673969 | 297.5781382 | 516.5720716 | 0.043859988 | 0.133660853 | 0.502270965 | 0.796049077 | ns | TM | KS.1 |
| FIM1_164 | 941.5709596 | 825.6056512 | 893.1192894 | -0.117400731 | 0.093482069 | 0.059216257 | 0.255563352 | ns | TM | KS.1 |
| FIM1_165 | 626.2879576 | 603.099 | 587.4606674 | -0.020754784 | 0.094842503 | 0.720496074 | 0.906676397 | ns | TM | KS.1 |
| FIM1_166 | 673.8632592 | 875.0449489 | 737.7645358 | 0.149899184 | 0.157476558 | 0.063763713 | 0.265060417 | ns | TM | KS.1 |
| FIM1_167 | 348.2368913 | 363.9628548 | 515.6252124 | 0.02138245 | 0.109479196 | 0.728605086 | 0.908701505 | ns | TM | KS.1 |
| FIM1_168 | 1286.625914 | 1377.967691 | 1298.756978 | 0.044368744 | 0.088547151 | 0.430806555 | 0.743536592 | ns | TM | KS.1 |
| FIM1_169 | 1767.730793 | 1632.013647 | 2192.324479 | -0.051022669 | 0.093256641 | 0.381121667 | 0.70494176 | ns | TM | KS.1 |
| FIM1_170 | 8925.36727 | 11290.70089 | 11208.52652 | 0.315817516 | 0.0528659 | 1.9836084831 | 5.8049718843 | ns | TM | KS.1 |
| FIM1_171 | 1844.154954 | 1972.062597 | 1811.685558 | 0.039291068 | 0.096051819 | 0.504164243 | 0.796765282 | ns | TM | KS.1 |
| FIM1_172 | 837.3478265 | 808.8152968 | 709.2605182 | -0.01907296 | 0.092880546 | 0.739509246 | 0.914051801 | ns | TM | KS.1 |
| FIM1_173 | 799.2539006 | 867.3609117 | 894.9360852 | 0.053430331 | 0.092849391 | 0.358538234 | 0.683292083 | ns | TM | KS.1 |
| FIM1_174 | 340.7343211 | 391.7010551 | 327.707724 | 0.030440974 | 0.166172171 | 0.630335143 | 0.868799252 | ns | TM | KS.1 |

| | | | | | | | | | | |
|---|---|---|---|---|---|---|---|---|---|---|
| FIM1_175 | 96.80462958 | 92.77669077 | 88.16067366 | -0.012146744 | 0.140197879 | 0.847508118 | 0.952348261 | ns | TM | KS.1 |
| FIM1_176 | 1625.920131 | 2010.843329 | 2120.157969 | 0.170725109 | 0.127737097 | 0.029202529 | 0.16972335 | ns | TM | KS.1 |
| FIM1_177 | 7357.374939 | 8916.825206 | 8182.819457 | 0.078482523 | 0.144380818 | 0.260971813 | 0.588210523 | ns | TM | KS.1 |
| FIM1_178 | 1023.293547 | 1008.453085 | 898.2572507 | -0.009867061 | 0.065713315 | 0.830157828 | 0.946661434 | ns | TM | KS.1 |
| FIM1_179 | 2183.452324 | 2189.205192 | 2568.32626 | 0.0017293 | 0.07899822 | 0.973522611 | 0.993872484 | ns | TM | KS.1 |
| FIM1_180 | 1654.790075 | 1565.233204 | 1767.514357 | -0.0389014 | 0.077853247 | 0.452420899 | 0.756061126 | ns | TM | KS.1 |
| FIM1_181 | 9864.894885 | 11389.90328 | 11963.46107 | 0.165402005 | 0.075924153 | 0.004902178 | 0.051235998 | ns | TM | KS.1 |
| FIM1_182 | 6352.085672 | 4834.003445 | 5639.517076 | -0.121533284 | 0.166344233 | 0.116335267 | 0.372917495 | ns | TM | KS.1 |
| FIM1_183 | 86.2082825 | 95.46490628 | 86.97005316 | 0.029843459 | 0.143660806 | 0.641660479 | 0.872440799 | ns | TM | KS.1 |
| FIM1_184 | 4324.115422 | 4383.969867 | 4253.522987 | 0.007933884 | 0.088186005 | 0.886681975 | 0.967028314 | ns | TM | KS.1 |
| FIM1_185 | 473.8484017 | 424.1783717 | 1297.052696 | -0.049107917 | 0.123095439 | 0.450623187 | 0.754323807 | ns | TM | KS.1 |
| FIM1_186 | 350.6953162 | 265.5884053 | 342.0256655 | -0.110563997 | 0.169884886 | 0.144578545 | 0.422359521 | ns | TM | KS.1 |
| FIM1_187 | 9228.407248 | 8242.911449 | 7421.616443 | -0.035532737 | 0.141909599 | 0.582660551 | 0.839483417 | ns | TM | KS.1 |
| FIM1_189 | 1295.36441 | 1217.801291 | 1349.127392 | -0.041743495 | 0.082514222 | 0.438113567 | 0.747832613 | ns | TM | KS.1 |
| FIM1_190 | 208.2628471 | 198.7729324 | 166.9618692 | -0.019122841 | 0.117326467 | 0.760399062 | 0.923255837 | ns | TM | KS.1 |
| FIM1_191 | 692.120008 | 519.5514212 | 673.3985639 | -0.290127248 | 0.130322142 | 0.00132148 | 0.018660515 | ns | TM | KS.1 |
| FIM1_192 | 3522.647546 | 3612.944116 | 3739.861374 | 0.006393198 | 0.149565785 | 0.9186246 | 0.975695428 | ns | TM | KS.1 |
| FIM1_193 | 27.7684911 | 16.84274193 | 567.2888931 | -0.131379802 | 0.252005952 | 0.11166926 | 0.363820936 | ns | TM | KS.1 |
| FIM1_194 | 2424.302656 | 2437.283669 | 2532.895346 | 0.002181998 | 0.115341191 | 0.971994315 | 0.99362472 | ns | TM | KS.1 |
| FIM1_195 | 1488.052008 | 1387.100409 | 1244.231713 | -0.035837898 | 0.10633625 | 0.560161453 | 0.826391733 | ns | TM | KS.1 |
| FIM1_196 | 2798.036418 | 2848.682913 | 2924.485626 | 0.011335733 | 0.080228147 | 0.829455637 | 0.946484629 | ns | TM | KS.1 |
| FIM1_197 | 2107.745524 | 1606.872402 | 2057.649556 | -0.136412634 | 0.163711606 | 0.086226546 | 0.312966141 | ns | TM | KS.1 |
| FIM1_198 | 3144.629759 | 3139.809456 | 4654.542393 | -0.000801089 | 0.093397126 | 0.988863202 | 0.996167262 | ns | TM | KS.1 |
| FIM1_199 | 1143.648651 | 1137.781231 | 1061.302746 | -0.002411873 | 0.105683044 | 0.968241398 | 0.993374633 | ns | TM | KS.1 |
| FIM1_200 | 1991.039461 | 2300.951095 | 2473.384623 | 0.089017996 | 0.116036329 | 0.185766406 | 0.48821335 | ns | TM | KS.1 |
| FIM1_201 | 2202.889572 | 2174.606386 | 2060.566516 | -0.008320148 | 0.078994009 | 0.87313545 | 0.962759265 | ns | TM | KS.1 |
| FIM1_203 | 1354.395971 | 1473.023084 | 1224.968132 | 0.057524068 | 0.090220874 | 0.315860945 | 0.64270274 | ns | TM | KS.1 |
| FIM1_204 | 197.7007199 | 145.240084 | 149.289268 | -0.152698242 | 0.177383288 | 0.066245892 | 0.270807979 | ns | TM | KS.1 |
| FIM1_205 | 1235.486485 | 1370.523793 | 1549.415675 | 0.032462139 | 0.14127806 | 0.614207427 | 0.858462917 | ns | TM | KS.1 |
| FIM1_206 | 2952.513293 | 3019.147645 | 2888.196518 | 0.014932835 | 0.074528345 | 0.765966496 | 0.924600573 | ns | TM | KS.1 |
| FIM1_207 | 971.4435624 | 946.2695922 | 798.6830117 | -0.013779827 | 0.097524967 | 0.814387214 | 0.940696014 | ns | TM | KS.1 |
| FIM1_208 | 3514.757166 | 3442.387064 | 3150.474754 | -0.01257562 | 0.083837474 | 0.816117058 | 0.941382417 | ns | TM | KS.1 |
| FIM1_209 | 3616.807802 | 3500.349752 | 4312.461468 | -0.019959973 | 0.08521736 | 0.714980227 | 0.905084971 | ns | TM | KS.1 |
| FIM1_210 | 2926.766478 | 3084.44636 | 2926.221732 | 0.03000127 | 0.094855644 | 0.606177424 | 0.851596861 | ns | TM | KS.1 |
| FIM1_211 | 1457.46279 | 1453.736149 | 1421.061327 | -0.001554539 | 0.082221284 | 0.976783781 | 0.993872484 | ns | TM | KS.1 |
| FIM1_212 | 1312.073837 | 1320.182369 | 1343.651911 | 0.003137807 | 0.093865148 | 0.95651071 | 0.989353426 | ns | TM | KS.1 |
| FIM1_213 | 1400.164518 | 1361.484535 | 1407.992985 | -0.019403151 | 0.06750407 | 0.678832097 | 0.887492141 | ns | TM | KS.1 |
| FIM1_214 | 461.1953487 | 359.0967915 | 339.0121516 | -0.14355741 | 0.15306337 | 0.070923292 | 0.282027284 | ns | TM | KS.1 |
| FIM1_215 | 686.8697036 | 720.5216289 | 763.6384068 | 0.034988307 | 0.06896356 | 0.462119985 | 0.764495222 | ns | TM | KS.1 |
| FIM1_216 | 3588.500099 | 3503.005224 | 3833.499387 | -0.011049434 | 0.109114641 | 0.856994936 | 0.955916258 | ns | TM | KS.1 |
| FIM1_217 | 2752.563337 | 2636.927716 | 2626.542711 | -0.025526576 | 0.089373805 | 0.650028673 | 0.874068749 | ns | TM | KS.1 |
| FIM1_218 | 1076.774796 | 860.0241218 | 1854.385565 | -0.077999694 | 0.15816471 | 0.266113171 | 0.592885367 | ns | TM | KS.1 |
| FIM1_219 | 1097.85156 | 971.1090283 | 1022.823757 | -0.109239009 | 0.090905366 | 0.07111807 | 0.282372224 | ns | TM | KS.1 |
| FIM1_220 | 421.0029822 | 437.2276632 | 417.3276213 | 0.02050937 | 0.097554416 | 0.72728161 | 0.907961357 | ns | TM | KS.1 |
| FIM1_221 | 1196.270128 | 1279.480893 | 1188.193571 | 0.037273293 | 0.099220492 | 0.532576251 | 0.809787573 | ns | TM | KS.1 |

| | | | | | | | | | | |
|---|---|---|---|---|---|---|---|---|---|---|
| FIM1_222 | 1362.754625 | 1596.178937 | 1793.918807 | 0.187233209 | 0.07467547 | 0.001862146 | 0.024508402 | ns | TM | KS.1 |
| FIM1_224 | 2202.014355 | 2174.501847 | 1990.069359 | -0.006992282 | 0.088179819 | 0.900044711 | 0.968452273 | ns | TM | KS.1 |
| FIM1_225 | 1584.506114 | 1367.478589 | 1133.049414 | -0.12599022 | 0.101933766 | 0.056943152 | 0.251815274 | ns | TM | KS.1 |
| FIM1_226 | 720.1850192 | 444.6020756 | 393.6260039 | -0.366457234 | 0.271016397 | 0.004022191 | 0.043978899 | ns | TM | KS.1 |
| FIM1_227 | 1675.26739 | 2121.166621 | 1808.330015 | 0.160709827 | 0.142840841 | 0.045168769 | 0.219529062 | ns | TM | KS.1 |
| FIM1_228 | 130.8253592 | 159.2512226 | 100.5731306 | 0.13170331 | 0.128606686 | 0.076127473 | 0.294505582 | ns | TM | KS.1 |
| FIM1_229 | 9915.638497 | 10839.21336 | 7860.361819 | 0.056436445 | 0.096589227 | 0.343843776 | 0.670129201 | ns | TM | KS.1 |
| FIM1_230 | 2103.345753 | 2143.30711 | 1941.403259 | 0.012807105 | 0.071307039 | 0.792521 | 0.933303839 | ns | TM | KS.1 |
| FIM1_231 | 459.0287034 | 252.1306551 | 186.9610113 | -0.392305219 | 0.348303963 | 0.005863943 | 0.058229772 | ns | TM | KS.1 |
| FIM1_233 | 4562.355729 | 4969.296608 | 4732.029427 | 0.048200585 | 0.102954707 | 0.431034028 | 0.743536592 | ns | TM | KS.1 |
| FIM1_234 | 1862.314253 | 1812.547766 | 1866.763399 | -0.022407172 | 0.052185286 | 0.564730541 | 0.828150948 | ns | TM | KS.1 |
| FIM1_235 | 937.4447838 | 909.6309757 | 816.0938578 | -0.017581168 | 0.0897934 | 0.754865901 | 0.920022902 | ns | TM | KS.1 |
| FIM1_236 | 2302.55188 | 2389.351015 | 2878.122841 | 0.021574488 | 0.089370717 | 0.701092215 | 0.897235589 | ns | TM | KS.1 |
| FIM1_237 | 738.5917448 | 730.2500658 | 509.869959 | -0.003368128 | 0.139441723 | 0.957356901 | 0.989353426 | ns | TM | KS.1 |
| FIM1_238 | 612.020927 | 530.1737527 | 439.7022536 | -0.09408 | 0.113070264 | 0.160064671 | 0.447371762 | ns | TM | KS.1 |
| FIM1_239 | 127.92654 | 97.0166118 | 93.79866522 | -0.131073106 | 0.165981236 | 0.096531696 | 0.336392346 | ns | TM | KS.1 |
| FIM1_240 | 710.4786226 | 543.718753 | 582.5653609 | -0.15497772 | 0.159901623 | 0.058253527 | 0.253775214 | ns | TM | KS.1 |
| FIM1_241 | 2832.480254 | 2214.943592 | 3621.35305 | -0.313683513 | 0.070398178 | 6.7316856741 | 5.4016348755 | ns | TM | KS.1 |
| FIM1_242 | 69.45602368 | 66.17263008 | 58.14148934 | -0.018234393 | 0.127497708 | 0.773708433 | 0.925733395 | ns | TM | KS.1 |
| FIM1_243 | 1169.029554 | 1137.039336 | 1207.709963 | -0.021143028 | 0.064133151 | 0.639829217 | 0.870968656 | ns | TM | KS.1 |
| FIM1_245 | 3707.045167 | 4618.770728 | 4952.962543 | 0.09054041 | 0.151433454 | 0.207545158 | 0.514724407 | ns | TM | KS.1 |
| FIM1_246 | 2252.238182 | 2092.303652 | 2323.651007 | -0.056499189 | 0.077728409 | 0.277432833 | 0.602049362 | ns | TM | KS.1 |
| FIM1_247 | 1570.764154 | 1243.074093 | 1343.159356 | -0.111916574 | 0.151838504 | 0.135438029 | 0.407659726 | ns | TM | KS.1 |
| FIM1_248 | 910.067869 | 848.9781407 | 1115.187354 | -0.044902611 | 0.087424927 | 0.42179273 | 0.738117813 | ns | TM | KS.1 |
| FIM1_249 | 594.786416 | 884.5898796 | 995.5845502 | 0.126150024 | 0.214541815 | 0.116444724 | 0.37302801 | ns | TM | KS.1 |
| FIM1_250 | 817.5612264 | 738.4672766 | 804.4931944 | -0.066496737 | 0.098559459 | 0.273707197 | 0.599198132 | ns | TM | KS.1 |
| FIM1_251 | 589.6243214 | 649.0428452 | 522.5479514 | 0.062304987 | 0.097668372 | 0.301054831 | 0.630048398 | ns | TM | KS.1 |
| FIM1_252 | 2235.382645 | 2048.964485 | 2061.09628 | -0.075184332 | 0.075465736 | 0.144973379 | 0.423016165 | ns | TM | KS.1 |
| FIM1_253 | 3807.739909 | 4133.595082 | 4712.613198 | 0.061628881 | 0.082991782 | 0.258280234 | 0.587268814 | ns | TM | KS.1 |
| FIM1_254 | 664.8152195 | 667.5303304 | 630.1023059 | 0.002679874 | 0.094317034 | 0.962940891 | 0.992465493 | ns | TM | KS.1 |
| FIM1_255 | 4644.646343 | 4338.535522 | 3916.679253 | -0.037212576 | 0.101313639 | 0.537236898 | 0.81337601 | ns | TM | KS.1 |
| FIM1_256 | 2838.740783 | 2907.333028 | 2945.980669 | 0.01125882 | 0.106096622 | 0.852953781 | 0.954011929 | ns | TM | KS.1 |
| FIM1_257 | 957.0782293 | 950.623006 | 989.1991687 | -0.003909645 | 0.079972311 | 0.94064654 | 0.983546982 | ns | TM | KS.1 |
| FIM1_258 | 1303.189181 | 1305.273476 | 1326.343152 | 0.000977445 | 0.077899487 | 0.984897775 | 0.996110272 | ns | TM | KS.1 |
| FIM1_259 | 2769.300448 | 3208.557055 | 3182.099749 | 0.097306573 | 0.114037406 | 0.149388731 | 0.430313147 | ns | TM | KS.1 |
| FIM1_260 | 632.0125091 | 371.0793312 | 531.1506979 | -0.6699069 | 0.20109914 | 4.9228740267 | 0.000235493 | ns | TM | KS.1 |
| FIM1_261 | 1525.806215 | 1609.417823 | 1417.32722 | 0.021827306 | 0.119414254 | 0.729111162 | 0.908876981 | ns | TM | KS.1 |
| FIM1_262 | 846.7231546 | 1093.035728 | 1256.727153 | 0.137921977 | 0.156611559 | 0.081185274 | 0.304368302 | ns | TM | KS.1 |
| FIM1_263 | 2692.750724 | 2562.493577 | 4356.538085 | -0.031981921 | 0.082846492 | 0.552483228 | 0.822689033 | ns | TM | KS.1 |
| FIM1_264 | 921.034083 | 922.0570034 | 998.4399574 | 0.000144023 | 0.073670473 | 0.997691538 | 0.998293524 | ns | TM | KS.1 |
| FIM1_265 | 1387.226853 | 1520.328785 | 1462.899096 | 0.078347263 | 0.077462863 | 0.136853202 | 0.409362716 | ns | TM | KS.1 |
| FIM1_266 | 821.6118695 | 857.9054438 | 972.3963985 | 0.024485705 | 0.093786176 | 0.671513154 | 0.883336315 | ns | TM | KS.1 |
| FIM1_267 | 4498.020314 | 4806.871193 | 5139.402025 | 0.051393043 | 0.074577145 | 0.308310537 | 0.636779799 | ns | TM | KS.1 |
| FIM1_268 | 5963.980164 | 6054.155049 | 5576.256847 | 0.008302112 | 0.092240325 | 0.884297092 | 0.966744674 | ns | TM | KS.1 |
| FIM1_269 | 1343.093033 | 1387.733751 | 1362.757812 | 0.020458116 | 0.083036626 | 0.70379687 | 0.898252804 | ns | TM | KS.1 |

| | | | | | | | | | | |
|---|---|---|---|---|---|---|---|---|---|---|
| FIM1_270 | 224.7629484 | 185.3172846 | 135.2654486 | -0.093517205 | 0.138887716 | 0.190165924 | 0.494034189 | ns | TM | KS.1 |
| FIM1_271 | 2392.15515 | 2743.168176 | 2806.474049 | 0.091178169 | 0.109939373 | 0.166353034 | 0.456988595 | ns | TM | KS.1 |
| FIM1_272 | 732.5708649 | 664.3920651 | 690.4990834 | -0.057667575 | 0.103014978 | 0.349924158 | 0.675804614 | ns | TM | KS.1 |
| FIM1_273 | 386.8397108 | 539.9554574 | 446.7911323 | 0.198968596 | 0.187775947 | 0.029212649 | 0.16972335 | ns | TM | KS.1 |
| FIM1_274 | 1180.898296 | 1665.319891 | 1722.830084 | 0.194991455 | 0.192861116 | 0.032471743 | 0.180382188 | ns | TM | KS.1 |
| FIM1_275 | 1642.653841 | 1799.619941 | 1602.25596 | 0.046701626 | 0.11088368 | 0.458753849 | 0.762486824 | ns | TM | KS.1 |
| FIM1_276 | 955.8263485 | 978.7887596 | 906.7754852 | 0.013870675 | 0.087839276 | 0.802943653 | 0.936908579 | ns | TM | KS.1 |
| FIM1_277 | 331.214407 | 315.948904 | 311.8922662 | -0.031713906 | 0.085745797 | 0.564158285 | 0.827931406 | ns | TM | KS.1 |
| FIM1_278 | 768.4998862 | 756.3340178 | 637.3066224 | -0.008505322 | 0.096858721 | 0.884309844 | 0.966744674 | ns | TM | KS.1 |
| FIM1_279 | 3365.308481 | 3447.992157 | 3531.57243 | 0.015674939 | 0.078639354 | 0.762952564 | 0.924230468 | ns | TM | KS.1 |
| FIM1_280 | 32.62937521 | 16.80196182 | 26.86602388 | -0.209264461 | 0.317879978 | 0.042058608 | 0.209871189 | ns | TM | KS.1 |
| FIM1_281 | 3395.603745 | 2289.612652 | 3055.708706 | -0.362846727 | 0.196610906 | 0.001366452 | 0.019149579 | ns | TM | KS.1 |
| FIM1_282 | 326.3658215 | 301.9677255 | 392.1436837 | -0.018986991 | 0.154524887 | 0.762580197 | 0.924230468 | ns | TM | KS.1 |
| FIM1_283 | 469.7166184 | 367.2576377 | 1789.244243 | -0.071664432 | 0.169529795 | 0.300120238 | 0.628445786 | ns | TM | KS.1 |
| FIM1_284 | 810.1258754 | 929.8788768 | 1082.482432 | 0.085216321 | 0.113836752 | 0.199557469 | 0.50730629 | ns | TM | KS.1 |
| FIM1_285 | 1361.666507 | 1528.15169 | 1445.590392 | 0.032349867 | 0.148697703 | 0.614313471 | 0.858462917 | ns | TM | KS.1 |
| FIM1_286 | 3136.986629 | 3080.723374 | 3409.995974 | -0.008669834 | 0.106273702 | 0.886507235 | 0.967028314 | ns | TM | KS.1 |
| FIM1_287 | 675.949798 | 645.6871601 | 685.4018079 | -0.02990329 | 0.081673894 | 0.574996216 | 0.833922256 | ns | TM | KS.1 |
| FIM1_288 | 147.5238197 | 144.5182461 | 132.9791323 | -0.007277264 | 0.113365333 | 0.906467979 | 0.971703985 | ns | TM | KS.1 |
| FIM1_290 | 359.5359376 | 373.5515522 | 401.1793757 | 0.019105587 | 0.103786984 | 0.751720541 | 0.918662314 | ns | TM | KS.1 |
| FIM1_291 | 1707.750861 | 1288.789305 | 1665.079634 | -0.215580537 | 0.155195956 | 0.015318535 | 0.112569734 | ns | TM | KS.1 |
| FIM1_292 | 994.5436475 | 306.8901533 | 1255.979238 | -1.463229075 | 0.291081374 | 4.03591168704 | 4.89723430318 | Km<KS.1 | TM | KS.1 |
| FIM1_293 | 2518.339812 | 2217.918888 | 3668.649036 | -0.086998657 | 0.105529775 | 0.175794152 | 0.473511588 | ns | TM | KS.1 |
| FIM1_294 | 130.5110013 | 158.6628089 | 2209.438311 | 0.043707041 | 0.173072943 | 0.49899521 | 0.793892283 | ns | TM | KS.1 |
| FIM1_295 | 26.30787336 | 26.10651341 | 54.48768277 | -0.00186158 | 0.161426808 | 0.975972168 | 0.993872484 | ns | TM | KS.1 |
| FIM1_296 | 56.01195463 | 64.28412741 | 99.09863344 | 0.032591426 | 0.158639071 | 0.609584151 | 0.854035807 | ns | TM | KS.1 |
| FIM1_297 | 20.95758059 | 18.66882436 | 31.93400973 | -0.024126384 | 0.171007232 | 0.698997537 | 0.896699645 | ns | TM | KS.1 |
| FIM1_298 | 37.51758279 | 33.34960804 | 43.81990865 | -0.035103772 | 0.146657656 | 0.586409998 | 0.841613393 | ns | TM | KS.1 |
| FIM1_299 | 596.8910093 | 644.6249458 | 625.4742574 | 0.025335474 | 0.134799959 | 0.69183327 | 0.893131011 | ns | TM | KS.1 |
| FIM1_300 | 865.228041 | 1569.200965 | 1750.706662 | 0.2576456 | 0.322292263 | 0.023643169 | 0.150569038 | ns | TM | KS.1 |
| FIM1_301 | 220.2843716 | 238.9704967 | 194.6034244 | 0.027431105 | 0.13427607 | 0.668467638 | 0.882180579 | ns | TM | KS.1 |
| FIM1_302 | 882.0525244 | 789.5507942 | 737.3037931 | -0.059333297 | 0.112391233 | 0.354978261 | 0.679784721 | ns | TM | KS.1 |
| FIM1_303 | 27988.51131 | 26346.44688 | 31660.2633 | -0.034405432 | 0.096419379 | 0.558440486 | 0.82614557 | ns | TM | KS.1 |
| FIM1_304 | 528.7086257 | 516.1451871 | 807.8463057 | -0.009997166 | 0.118149686 | 0.873018173 | 0.962759265 | ns | TM | KS.1 |
| FIM1_305 | 430.9068758 | 200.6799825 | 843.7830533 | -0.575782303 | 0.445405589 | 0.002254131 | 0.028318942 | ns | TM | KS.1 |
| FIM1_306 | 216.6647168 | 103.3387086 | 334.8470566 | -0.258992801 | 0.373405437 | 0.026642022 | 0.160541482 | ns | TM | KS.1 |
| FIM1_307 | 1243.511165 | 1284.814122 | 1116.511687 | 0.020504994 | 0.084179098 | 0.705506946 | 0.899257553 | ns | TM | KS.1 |
| FIM1_308 | 1746.468068 | 2264.504289 | 2450.767558 | 0.208783062 | 0.144130901 | 0.015250951 | 0.112405156 | ns | TM | KS.1 |
| FIM1_309 | 4985.44045 | 5732.402846 | 4752.360542 | 0.14440597 | 0.085888047 | 0.018735266 | 0.130718431 | ns | TM | KS.1 |
| FIM1_310 | 2821.458893 | 3648.471355 | 4156.181608 | 0.191656124 | 0.147081707 | 0.023574091 | 0.150553405 | ns | TM | KS.1 |
| FIM1_311 | 3040.442312 | 2651.270222 | 3860.09946 | -0.117375938 | 0.098135301 | 0.066817835 | 0.272267737 | ns | TM | KS.1 |
| FIM1_312 | 4518.926485 | 4052.403223 | 3656.8811 | -0.050643444 | 0.11978607 | 0.434559963 | 0.746009599 | ns | TM | KS.1 |
| FIM1_313 | 2520.419413 | 2750.071928 | 2693.26069 | 0.041952551 | 0.113586788 | 0.507272944 | 0.799198286 | ns | TM | KS.1 |
| FIM1_314 | 3428.711352 | 2993.177275 | 2126.457402 | -0.059775589 | 0.128175793 | 0.36914996 | 0.69186398 | ns | TM | KS.1 |
| FIM1_315 | 691.0723242 | 635.8770052 | 659.2698316 | -0.064635359 | 0.080800959 | 0.228287876 | 0.545762703 | ns | TM | KS.1 |

| | | | | | | | | | | |
|---|---|---|---|---|---|---|---|---|---|---|
| FIM1_316 | 717.0485656 | 1028.114984 | 1101.901215 | 0.100629979 | 0.201895035 | 0.178724482 | 0.477526476 | ns | TM | KS.1 |
| FIM1_317 | 217.2659173 | 143.5367356 | 101.9596112 | -0.200229175 | 0.229002499 | 0.035755436 | 0.188635517 | ns | TM | KS.1 |
| FIM1_318 | 5623.114065 | 7227.978082 | 7843.167884 | 0.333227809 | 0.058529297 | 7.43806111133 | 1.84058836967 | ns | TM | KS.1 |
| FIM1_319 | 301.7222976 | 260.4189306 | 301.5180789 | -0.086306463 | 0.119406746 | 0.202848667 | 0.510973224 | ns | TM | KS.1 |
| FIM1_320 | 1700.283829 | 1390.14957 | 1810.685389 | -0.084204276 | 0.146020335 | 0.233493437 | 0.551843159 | ns | TM | KS.1 |
| FIM1_321 | 154.2319723 | 106.0907285 | 107.0083954 | -0.103177178 | 0.206928218 | 0.171439823 | 0.465692069 | ns | TM | KS.1 |
| FIM1_322 | 12890.66217 | 15261.79794 | 14723.54858 | 0.159379829 | 0.103195807 | 0.022403976 | 0.146622513 | ns | TM | KS.1 |
| FIM1_323 | 958.5291377 | 656.666813 | 571.029106 | -0.222016336 | 0.210519329 | 0.022428087 | 0.146622513 | ns | TM | KS.1 |
| FIM1_324 | 9380.253221 | 5331.450822 | 5869.511829 | -0.528705836 | 0.30971092 | 0.000878457 | 0.013786502 | ns | TM | KS.1 |
| FIM1_325 | 18.07171709 | 17.69504118 | 17.19711774 | -0.003769225 | 0.170607876 | 0.950965015 | 0.987490021 | ns | TM | KS.1 |
| FIM1_326 | 137.7329249 | 139.3628141 | 4974.569664 | 0.002286506 | 0.165092011 | 0.970386897 | 0.99362472 | ns | TM | KS.1 |
| FIM1_327 | 1177.010362 | 1236.688326 | 1256.923338 | 0.032507172 | 0.082011251 | 0.543434985 | 0.816794276 | ns | TM | KS.1 |
| FIM1_328 | 154.281192 | 136.188598 | 197.3835281 | -0.033258927 | 0.153152784 | 0.603968986 | 0.851547247 | ns | TM | KS.1 |
| FIM1_329 | 663.7337676 | 685.3760932 | 674.5158635 | 0.011022604 | 0.129042845 | 0.861579845 | 0.958274028 | ns | TM | KS.1 |
| FIM1_330 | 8645.762686 | 9357.130501 | 10680.29304 | 0.042708394 | 0.104320827 | 0.486166926 | 0.786100234 | ns | TM | KS.1 |
| FIM1_331 | 372.932336 | 319.5214853 | 526.2211276 | -0.053907422 | 0.142965487 | 0.418471879 | 0.736954903 | ns | TM | KS.1 |
| FIM1_332 | 1190.391902 | 1058.651884 | 1159.272756 | -0.061278938 | 0.115084498 | 0.344560904 | 0.670129201 | ns | TM | KS.1 |
| FIM1_333 | 896.3750005 | 913.0951985 | 1067.606057 | 0.009563987 | 0.098107591 | 0.870813657 | 0.96208467 | ns | TM | KS.1 |
| FIM1_334 | 1799.39114 | 1862.635994 | 1901.909592 | 0.018524943 | 0.097205425 | 0.752211283 | 0.918662314 | ns | TM | KS.1 |
| FIM1_335 | 934.8674241 | 1185.10772 | 1214.735243 | 0.070583634 | 0.166427261 | 0.306280388 | 0.634850461 | ns | TM | KS.1 |
| FIM1_336 | 691.8240765 | 528.7872742 | 665.6980096 | -0.150201987 | 0.160673592 | 0.064494558 | 0.266495371 | ns | TM | KS.1 |
| FIM1_337 | 4408.170907 | 3938.279261 | 3669.238901 | -0.070486554 | 0.104950257 | 0.263105919 | 0.589544539 | ns | TM | KS.1 |
| FIM1_338 | 947.539467 | 1333.315425 | 2051.336962 | 0.123081647 | 0.193635914 | 0.119092672 | 0.378827394 | ns | TM | KS.1 |
| FIM1_339 | 3009.428085 | 3275.224274 | 3976.175577 | 0.044606145 | 0.107320334 | 0.472892417 | 0.773385856 | ns | TM | KS.1 |
| FIM1_340 | 223.391501 | 316.5980941 | 277.3983359 | 0.105314052 | 0.197020584 | 0.163990722 | 0.45366779 | ns | TM | KS.1 |
| FIM1_341 | 3375.128195 | 4029.140281 | 4523.907596 | 0.151914927 | 0.112225373 | 0.03507658 | 0.187206368 | ns | TM | KS.1 |
| FIM1_342 | 1067.510151 | 1112.767067 | 1112.983384 | 0.022947712 | 0.095489315 | 0.693588629 | 0.893671231 | ns | TM | KS.1 |
| FIM1_343 | 2419.552863 | 2330.789759 | 2331.160854 | -0.013486639 | 0.12720099 | 0.831111567 | 0.946802573 | ns | TM | KS.1 |
| FIM1_344 | 1952.161808 | 1823.788434 | 1776.822187 | -0.043701617 | 0.088733001 | 0.438214965 | 0.747832613 | ns | TM | KS.1 |
| FIM1_345 | 1840.553426 | 993.2536828 | 3734.663347 | -0.659085206 | 0.312025227 | 0.000246249 | 0.005227779 | ns | TM | KS.1 |
| FIM1_346 | 5406.750335 | 4636.305498 | 6352.248537 | -0.092802041 | 0.119651437 | 0.174983577 | 0.471583585 | ns | TM | KS.1 |
| FIM1_347 | 2327.022602 | 1949.29265 | 1962.428707 | -0.150798057 | 0.112882355 | 0.036677937 | 0.191874592 | ns | TM | KS.1 |
| FIM1_348 | 1119.338296 | 1756.292777 | 2060.195225 | 0.178254895 | 0.242054405 | 0.053175613 | 0.240936862 | ns | TM | KS.1 |
| FIM1_349 | 2337.133419 | 2564.178605 | 2501.27156 | 0.063895118 | 0.092154009 | 0.273392535 | 0.599198132 | ns | TM | KS.1 |
| FIM1_350 | 551.5173417 | 580.6811948 | 551.4005951 | 0.033864701 | 0.082001679 | 0.52682137 | 0.808531286 | ns | TM | KS.1 |
| FIM1_351 | 1617.267448 | 1651.366538 | 1800.110609 | 0.016864176 | 0.056275539 | 0.681787454 | 0.888627871 | ns | TM | KS.1 |
| FIM1_352 | 4949.659293 | 5923.381633 | 5972.361674 | 0.050913452 | 0.1569997 | 0.441337143 | 0.747837973 | ns | TM | KS.1 |
| FIM1_354 | 438.4316957 | 624.7641271 | 493.2727055 | 0.122750028 | 0.198221825 | 0.120639029 | 0.381550647 | ns | TM | KS.1 |
| FIM1_355 | 3182.092608 | 3008.625763 | 2709.148435 | -0.035516471 | 0.086698737 | 0.521643921 | 0.807711954 | ns | TM | KS.1 |
| FIM1_356 | 935.1063774 | 685.024702 | 2103.957352 | -0.156349881 | 0.179576104 | 0.062362984 | 0.261819279 | ns | TM | KS.1 |
| FIM1_357 | 3949.799253 | 4235.981531 | 3991.281274 | 0.020631011 | 0.141708381 | 0.745347486 | 0.915807296 | ns | TM | KS.1 |
| FIM1_358 | 3443.255939 | 5044.330479 | 6074.080717 | 0.145319943 | 0.209781705 | 0.083341977 | 0.306449621 | ns | TM | KS.1 |
| FIM1_359 | 6220.558784 | 7178.769763 | 7789.665494 | 0.188194267 | 0.052278986 | 6.38495918343 | 0.001671851 | ns | TM | KS.1 |
| FIM1_360 | 1332.947458 | 1279.887749 | 1375.551515 | -0.028800659 | 0.073137195 | 0.561098562 | 0.826391733 | ns | TM | KS.1 |
| FIM1_361 | 503.0600697 | 547.2599944 | 560.8953301 | 0.039770168 | 0.112861923 | 0.5279809 | 0.808531286 | ns | TM | KS.1 |

| | | | | | | | | | | |
|---|---|---|---|---|---|---|---|---|---|---|
| FIM1_362 | 2537.392032 | 2551.463513 | 2519.03007 | 0.002443651 | 0.110650826 | 0.968300268 | 0.993374633 | ns | TM | KS.1 |
| FIM1_363 | 1176.615662 | 1322.767778 | 1212.245779 | 0.070280167 | 0.108563544 | 0.271969943 | 0.598783418 | ns | TM | KS.1 |
| FIM1_364 | 3288.24759 | 2935.254093 | 2836.126551 | -0.12373024 | 0.070352188 | 0.01783335 | 0.125845271 | ns | TM | KS.1 |
| FIM1_365 | 1272.052903 | 1524.964043 | 1520.484683 | 0.098504378 | 0.13195478 | 0.166292905 | 0.456988595 | ns | TM | KS.1 |
| FIM1_366 | 2062.899641 | 1975.585194 | 1946.325552 | -0.022238653 | 0.101825625 | 0.710862803 | 0.901949106 | ns | TM | KS.1 |
| FIM1_367 | 3764.488834 | 2094.849405 | 4689.156278 | -0.368216641 | 0.338759538 | 0.007135697 | 0.066581689 | ns | TM | KS.1 |
| FIM1_368 | 309.1807219 | 415.9913199 | 489.1536025 | 0.093884118 | 0.179997019 | 0.199381268 | 0.50711749 | ns | TM | KS.1 |
| FIM1_369 | 43.53975163 | 42.29881508 | 35.23365655 | -0.00607498 | 0.169280029 | 0.921203564 | 0.975881018 | ns | TM | KS.1 |
| FIM1_370 | 1022.760018 | 973.5985144 | 767.2498702 | -0.030714403 | 0.087371417 | 0.58068299 | 0.838424819 | ns | TM | KS.1 |
| FIM1_371 | 70.45429615 | 93.3641182 | 98.26239274 | 0.09306017 | 0.17436798 | 0.201812954 | 0.509840799 | ns | TM | KS.1 |
| FIM1_372 | 13359.12177 | 15703.04195 | 15260.98212 | 0.068617049 | 0.135516825 | 0.313553466 | 0.640101968 | ns | TM | KS.1 |
| FIM1_373 | 1312.965459 | 1463.735129 | 1422.934025 | 0.050789243 | 0.119125445 | 0.432660109 | 0.74505982 | ns | TM | KS.1 |
| FIM1_374 | 2093.374597 | 1457.79129 | 1849.472422 | -0.397232262 | 0.147867414 | 0.000109852 | 0.00259011 | ns | TM | KS.1 |
| FIM1_375 | 1489.079771 | 1575.095467 | 1675.434603 | 0.021474273 | 0.123917981 | 0.734740938 | 0.91155681 | ns | TM | KS.1 |
| FIM1_376 | 1250.499344 | 906.9672096 | 754.4049239 | -0.122040271 | 0.185771934 | 0.119803653 | 0.379844802 | ns | TM | KS.1 |
| FIM1_377 | 1384.187958 | 1088.921106 | 809.5753069 | -0.126131384 | 0.151732211 | 0.101214708 | 0.343715475 | ns | TM | KS.1 |
| FIM1_378 | 41170.52437 | 42262.87679 | 46593.75577 | 0.014929265 | 0.090530663 | 0.791725707 | 0.933303839 | ns | TM | KS.1 |
| FIM1_379 | 398.1327229 | 498.3150779 | 566.4716419 | 0.059693501 | 0.168134852 | 0.374454792 | 0.69745885 | ns | TM | KS.1 |
| FIM1_380 | 2036.549098 | 2168.053067 | 2248.476043 | 0.020205559 | 0.135423267 | 0.750786574 | 0.918418758 | ns | TM | KS.1 |
| FIM1_381 | 2439.71729 | 2844.294576 | 2891.447937 | 0.048279303 | 0.147580072 | 0.464138929 | 0.765867719 | ns | TM | KS.1 |
| FIM1_383 | 2053.815399 | 1773.676978 | 5227.277974 | -0.038781995 | 0.155768065 | 0.548421814 | 0.820817847 | ns | TM | KS.1 |
| FIM1_384 | 41564.57141 | 45479.24745 | 47056.6348 | 0.037165119 | 0.123314456 | 0.562825075 | 0.827194903 | ns | TM | KS.1 |
| FIM1_385 | 604.5634198 | 755.3687493 | 715.9322575 | 0.127717769 | 0.143210582 | 0.093758481 | 0.330902114 | ns | TM | KS.1 |
| FIM1_386 | 3393.713943 | 3071.260406 | 2795.587424 | -0.03149651 | 0.140329712 | 0.624421564 | 0.864597072 | ns | TM | KS.1 |
| FIM1_387 | 975.5358468 | 1059.028041 | 897.1630477 | 0.045463648 | 0.10361348 | 0.458042916 | 0.761873457 | ns | TM | KS.1 |
| FIM1_388 | 622.4281155 | 640.6473965 | 624.7718793 | 0.01544544 | 0.095608619 | 0.790576644 | 0.933303839 | ns | TM | KS.1 |
| FIM1_389 | 3709.888426 | 3445.979569 | 3232.834531 | -0.039389604 | 0.104172377 | 0.51944754 | 0.805989904 | ns | TM | KS.1 |
| FIM1_390 | 497.8818964 | 467.0824308 | 386.9148257 | -0.025487117 | 0.121764233 | 0.687713105 | 0.891826565 | ns | TM | KS.1 |
| FIM1_391 | 1358.861524 | 1486.33631 | 1297.739817 | 0.058603394 | 0.094879219 | 0.321542244 | 0.649745193 | ns | TM | KS.1 |
| FIM1_392 | 4460.872441 | 4368.546749 | 4038.265816 | -0.012172101 | 0.087592074 | 0.82635584 | 0.94595497 | ns | TM | KS.1 |
| FIM1_393 | 1506.662285 | 1574.666902 | 1931.746616 | 0.032083427 | 0.069880732 | 0.504019768 | 0.796765282 | ns | TM | KS.1 |
| FIM1_394 | 147.4037877 | 173.2783155 | 215.3585894 | 0.058599274 | 0.141507534 | 0.382964732 | 0.707482192 | ns | TM | KS.1 |
| FIM1_395 | 1319.853942 | 1242.84309 | 1365.180795 | -0.033563986 | 0.098697845 | 0.572403251 | 0.832094673 | ns | TM | KS.1 |
| FIM1_396 | 1612.033761 | 2197.036376 | 2343.481647 | 0.314440082 | 0.137804195 | 0.0008606 | 0.013548997 | ns | TM | KS.1 |
| FIM1_397 | 30.55100817 | 37.33731997 | 44.24618426 | 0.055048178 | 0.166859762 | 0.407850397 | 0.728957685 | ns | TM | KS.1 |
| FIM1_398 | 2128.82289 | 2254.61457 | 2180.258315 | 0.035497039 | 0.089258141 | 0.52902863 | 0.808531286 | ns | TM | KS.1 |
| FIM1_399 | 3895.217488 | 3589.608717 | 2774.821104 | -0.04844542 | 0.09867468 | 0.419196564 | 0.737776855 | ns | TM | KS.1 |
| FIM1_401 | 1301.064462 | 1504.728716 | 1422.118783 | 0.102858711 | 0.110241156 | 0.124260342 | 0.387735747 | ns | TM | KS.1 |
| FIM1_402 | 317.3282964 | 221.0010137 | 416.4323931 | -0.243052383 | 0.200226491 | 0.014243532 | 0.106880197 | ns | TM | KS.1 |
| FIM1_403 | 638.5859584 | 635.2177215 | 967.7019184 | -0.003440329 | 0.105015679 | 0.95461021 | 0.988678089 | ns | TM | KS.1 |
| FIM1_404 | 422.6456021 | 453.1448006 | 546.760275 | 0.026520875 | 0.124954519 | 0.677128474 | 0.887002016 | ns | TM | KS.1 |
| FIM1_405 | 5523.152826 | 6831.822405 | 7626.635991 | 0.247381628 | 0.087446183 | 0.000510302 | 0.009198371 | ns | TM | KS.1 |
| FIM1_406 | 7410.108573 | 9143.275254 | 9920.880166 | 0.250774755 | 0.081926673 | 0.000257983 | 0.005370152 | ns | TM | KS.1 |
| FIM1_407 | 3096.059096 | 3008.424395 | 3459.852175 | -0.01371491 | 0.107253962 | 0.822154464 | 0.944618544 | ns | TM | KS.1 |
| FIM1_408 | 1554.801567 | 1459.752903 | 1724.87704 | -0.050402278 | 0.070478607 | 0.298333511 | 0.627042339 | ns | TM | KS.1 |

| | | | | | | | | | | |
|---|---|---|---|---|---|---|---|---|---|---|
| FIM1_409 | 4796.964181 | 4708.35798 | 4668.864928 | -0.011547351 | 0.082199228 | 0.828889203 | 0.94647133 | ns | TM | KS.1 |
| FIM1_410 | 240.3612023 | 185.0743817 | 198.6579872 | -0.134564174 | 0.160409987 | 0.088332418 | 0.318214179 | ns | TM | KS.1 |
| FIM1_411 | 1735.338023 | 1277.679061 | 2144.197744 | -0.095890307 | 0.183348941 | 0.192537097 | 0.496564053 | ns | TM | KS.1 |
| FIM1_412 | 643.50237 | 516.6622031 | 728.237725 | -0.189231946 | 0.123797035 | 0.016935314 | 0.121227606 | ns | TM | KS.1 |
| FIM1_413 | 5545.311202 | 3109.635095 | 4930.833769 | -0.819004179 | 0.095834427 | 2.24536612207 | 2.79267411432 | ns | TM | KS.1 |
| FIM1_414 | 423.9905747 | 442.7160081 | 811.4516496 | 0.012955302 | 0.138235186 | 0.837706448 | 0.949337033 | ns | TM | KS.1 |
| FIM1_415 | 477.0006923 | 375.6496412 | 421.903523 | -0.158550729 | 0.145868918 | 0.048768989 | 0.230174256 | ns | TM | KS.1 |
| FIM1_416 | 1157.752601 | 1198.287083 | 1370.58872 | 0.021508133 | 0.083124518 | 0.689600348 | 0.892497849 | ns | TM | KS.1 |
| FIM1_417 | 7746.811272 | 5576.180398 | 6188.655595 | -0.304853323 | 0.159094324 | 0.002105412 | 0.026995939 | ns | TM | KS.1 |
| FIM1_418 | 1151.821356 | 1211.323972 | 901.0270708 | 0.02420689 | 0.108616943 | 0.69411563 | 0.893729024 | ns | TM | KS.1 |
| FIM1_419 | 927.9843439 | 900.6056066 | 844.6077708 | -0.016457376 | 0.093953912 | 0.775354626 | 0.926727011 | ns | TM | KS.1 |
| FIM1_420 | 717.6936383 | 766.4134803 | 749.4564911 | 0.039802518 | 0.093310572 | 0.492040069 | 0.790867874 | ns | TM | KS.1 |
| FIM1_421 | 957.7625706 | 918.6580381 | 1164.997138 | -0.027584245 | 0.077800022 | 0.593353526 | 0.846854399 | ns | TM | KS.1 |
| FIM1_422 | 716.0601073 | 588.3949045 | 623.0930477 | -0.162967011 | 0.121131803 | 0.031529055 | 0.177624525 | ns | TM | KS.1 |
| FIM1_423 | 42.2796991 | 41.82198661 | 92.95811991 | -0.001094446 | 0.177492296 | 0.985659139 | 0.996167262 | ns | TM | KS.1 |
| FIM1_425 | 2049.982536 | 3073.185883 | 3568.612381 | 0.169518729 | 0.221338235 | 0.05731254 | 0.25255083 | ns | TM | KS.1 |
| FIM1_426 | 3569.593655 | 4005.652095 | 4407.258022 | 0.083012826 | 0.098680708 | 0.178657426 | 0.477526476 | ns | TM | KS.1 |
| FIM1_427 | 381.6608073 | 370.9317065 | 348.6366388 | -0.01444882 | 0.097523254 | 0.80557918 | 0.937486882 | ns | TM | KS.1 |
| FIM1_428 | 306.7796884 | 309.2417015 | 345.022864 | 0.00324543 | 0.112784953 | 0.958131619 | 0.989353426 | ns | TM | KS.1 |
| FIM1_429 | 1452.9729 | 1501.248519 | 1382.713337 | 0.020602916 | 0.07917814 | 0.693172229 | 0.893633543 | ns | TM | KS.1 |
| FIM1_430 | 1159.452347 | 950.3024508 | 1029.537073 | -0.204572872 | 0.10373341 | 0.005988123 | 0.059108955 | ns | TM | KS.1 |
| FIM1_431 | 495.8762876 | 570.558517 | 613.3133998 | 0.063768266 | 0.127907561 | 0.340687301 | 0.668341572 | ns | TM | KS.1 |
| FIM1_432 | 247.8579049 | 291.9956352 | 229.5730587 | 0.059982584 | 0.142758446 | 0.373233532 | 0.695965825 | ns | TM | KS.1 |
| FIM1_433 | 1862.017071 | 1942.11079 | 1934.95982 | 0.025462812 | 0.088009008 | 0.647953322 | 0.873595603 | ns | TM | KS.1 |
| FIM1_434 | 816.7525094 | 1010.389414 | 1025.508772 | 0.172862288 | 0.127580467 | 0.027618959 | 0.163966971 | ns | TM | KS.1 |
| FIM1_435 | 890.2326764 | 870.1599251 | 1007.262228 | -0.014162527 | 0.076115516 | 0.780721039 | 0.929207457 | ns | TM | KS.1 |
| FIM1_436 | 358.2658608 | 447.9321476 | 391.0584459 | 0.102264166 | 0.149598248 | 0.163668992 | 0.45366779 | ns | TM | KS.1 |
| FIM1_437 | 766.0559957 | 600.6774341 | 545.3349553 | -0.173490673 | 0.143960938 | 0.034261666 | 0.185199792 | ns | TM | KS.1 |
| FIM1_438 | 20.28603301 | 26.48231293 | 26.92615975 | 0.047533427 | 0.19211065 | 0.462231483 | 0.764495222 | ns | TM | KS.1 |
| FIM1_439 | 1123.297643 | 1744.05252 | 1895.423913 | 0.221384107 | 0.242977871 | 0.027370277 | 0.163074407 | ns | TM | KS.1 |
| FIM1_440 | 1053.171087 | 720.7893945 | 823.75812 | -0.439930688 | 0.144149827 | 2.09518643403 | 0.000742966 | ns | TM | KS.1 |
| FIM1_441 | 696.9088592 | 713.4408632 | 966.3812468 | 0.013997978 | 0.080162799 | 0.790173174 | 0.933303839 | ns | TM | KS.1 |
| FIM1_442 | 1393.895947 | 1256.257032 | 1211.952098 | -0.049644074 | 0.117372532 | 0.440675756 | 0.747832613 | ns | TM | KS.1 |
| FIM1_443 | 157.9459396 | 180.956801 | 176.328668 | 0.037487264 | 0.151882455 | 0.561780269 | 0.826391733 | ns | TM | KS.1 |
| FIM1_444 | 1399.186946 | 1440.295673 | 1310.064377 | 0.016196539 | 0.093638407 | 0.778438856 | 0.928304554 | ns | TM | KS.1 |
| FIM1_445 | 1793.94079 | 1969.206533 | 1874.845332 | 0.060098792 | 0.096760853 | 0.315196367 | 0.641836948 | ns | TM | KS.1 |
| FIM1_446 | 436.3843571 | 452.831681 | 323.4614719 | 0.018268734 | 0.10643411 | 0.764374889 | 0.924230468 | ns | TM | KS.1 |
| FIM1_447 | 728.0127403 | 802.5185945 | 917.951041 | 0.083807312 | 0.081328786 | 0.125608072 | 0.389831665 | ns | TM | KS.1 |
| FIM1_448 | 2605.267641 | 2548.004254 | 2575.742673 | -0.013158731 | 0.086866869 | 0.811674553 | 0.939971031 | ns | TM | KS.1 |
| FIM1_449 | 4294.80537 | 4526.755583 | 4280.172542 | 0.022925816 | 0.115072065 | 0.714039557 | 0.905084971 | ns | TM | KS.1 |
| FIM1_450 | 90.09274694 | 123.6826991 | 132.0429841 | 0.129958197 | 0.181021733 | 0.102618127 | 0.346588717 | ns | TM | KS.1 |
| FIM1_451 | 1325.164514 | 1773.188389 | 2000.966981 | 0.133483701 | 0.172449853 | 0.09395012 | 0.331213943 | ns | TM | KS.1 |
| FIM1_452 | 871.4584816 | 815.5587139 | 780.8864883 | -0.033806087 | 0.10605447 | 0.581711124 | 0.838601229 | ns | TM | KS.1 |
| FIM1_453 | 4158.058482 | 3692.735283 | 2775.856282 | -0.0641921 | 0.114119639 | 0.322275878 | 0.650699064 | ns | TM | KS.1 |
| FIM1_454 | 260.4786155 | 268.3445969 | 268.9446485 | 0.006364396 | 0.159583184 | 0.918224731 | 0.975634767 | ns | TM | KS.1 |

| | | | | | | | | | | |
|---|---|---|---|---|---|---|---|---|---|---|
| FIM1_455 | 1464.888283 | 1282.960806 | 1245.188269 | -0.052360927 | 0.132827703 | 0.428700436 | 0.742835117 | ns | TM | KS.1 |
| FIM1_456 | 1955.598169 | 1125.545171 | 1442.054858 | -0.537506225 | 0.29575813 | 0.000637459 | 0.010714057 | ns | TM | KS.1 |
| FIM1_457 | 12338.50355 | 15577.51038 | 14320.3747 | 0.195826447 | 0.131712352 | 0.016835986 | 0.120864403 | ns | TM | KS.1 |
| FIM1_458 | 18905.38471 | 21676.01053 | 20648.47836 | 0.102746953 | 0.104730494 | 0.114999212 | 0.37039344 | ns | TM | KS.1 |
| FIM1_459 | 1999.815765 | 1292.6779 | 1542.097326 | -0.371518014 | 0.23223399 | 0.002235761 | 0.028159262 | ns | TM | KS.1 |
| FIM1_460 | 344.1631998 | 439.1724655 | 413.5934671 | 0.147563399 | 0.149605522 | 0.063758264 | 0.265060417 | ns | TM | KS.1 |
| FIM1_461 | 978.6431678 | 845.8877861 | 842.2776394 | -0.101407755 | 0.111524279 | 0.131074021 | 0.400376242 | ns | TM | KS.1 |
| FIM1_462 | 739.871184 | 697.6475151 | 709.7308589 | -0.046104237 | 0.069925458 | 0.338143951 | 0.666507986 | ns | TM | KS.1 |
| FIM1_463 | 2235.130109 | 2747.767241 | 2168.265916 | 0.081512589 | 0.149182843 | 0.247192687 | 0.57146079 | ns | TM | KS.1 |
| FIM1_464 | 5442.880378 | 5525.073754 | 5385.754652 | 0.007559554 | 0.100501139 | 0.898768444 | 0.968452273 | ns | TM | KS.1 |
| FIM1_465 | 31.34871543 | 28.3368415 | 27.27999917 | -0.021191389 | 0.165660112 | 0.734211122 | 0.911352378 | ns | TM | KS.1 |
| FIM1_466 | 5238.745871 | 6759.306594 | 7328.141002 | 0.259553414 | 0.118762804 | 0.002101812 | 0.026995939 | ns | TM | KS.1 |
| FIM1_467 | 1778.102431 | 1949.177773 | 2370.570354 | 0.036396463 | 0.126216486 | 0.571947967 | 0.832094673 | ns | TM | KS.1 |
| FIM1_468 | 100.5937277 | 66.60986361 | 101.8914025 | -0.220282287 | 0.221733744 | 0.024789186 | 0.154932414 | ns | TM | KS.1 |
| FIM1_469 | 931.588602 | 902.2424927 | 815.2713542 | -0.012233397 | 0.122254483 | 0.84590329 | 0.952348261 | ns | TM | KS.1 |
| FIM1_470 | 1501.182226 | 1299.476297 | 1474.85134 | -0.059768221 | 0.132968239 | 0.371901202 | 0.694536971 | ns | TM | KS.1 |
| FIM1_471 | 860.827091 | 902.6871248 | 924.8653515 | 0.012735129 | 0.14631825 | 0.839659505 | 0.949982862 | ns | TM | KS.1 |
| FIM1_472 | 348.1072367 | 347.1484941 | 470.3324537 | -0.000764319 | 0.134642878 | 0.990333116 | 0.996341203 | ns | TM | KS.1 |
| FIM1_473 | 1480.664191 | 1420.085148 | 1561.354367 | -0.018048904 | 0.114836289 | 0.77227282 | 0.925733395 | ns | TM | KS.1 |
| FIM1_474 | 1600.302325 | 1727.655591 | 1664.142884 | 0.048266117 | 0.092983256 | 0.405858296 | 0.728407296 | ns | TM | KS.1 |
| FIM1_475 | 783.7883608 | 708.3496292 | 707.7649955 | -0.037684989 | 0.131004912 | 0.560522431 | 0.826391733 | ns | TM | KS.1 |
| FIM1_476 | 8849.323416 | 9494.535986 | 9708.260431 | 0.036380144 | 0.105633202 | 0.553341615 | 0.822981923 | ns | TM | KS.1 |
| FIM1_477 | 7329.985595 | 8634.796969 | 8762.515758 | 0.152435815 | 0.102644441 | 0.026978013 | 0.161316847 | ns | TM | KS.1 |
| FIM1_478 | 1915.500322 | 3119.32629 | 3891.826898 | 0.225023288 | 0.266852931 | 0.029019561 | 0.169342179 | ns | TM | KS.1 |
| FIM1_479 | 1328.313242 | 1435.868861 | 1402.401075 | 0.028842052 | 0.128174967 | 0.652223179 | 0.87555594 | ns | TM | KS.1 |
| FIM1_480 | 2026.22852 | 1296.04072 | 1130.206243 | -0.570140697 | 0.149908939 | 5.20111886528 | 4.70464842814 | ns | TM | KS.1 |
| FIM1_481 | 1207.098231 | 1000.443988 | 1343.131751 | -0.163715548 | 0.114545522 | 0.026907035 | 0.161086041 | ns | TM | KS.1 |
| FIM1_482 | 1027.111803 | 1080.900795 | 1188.457089 | 0.026956436 | 0.099071361 | 0.64984971 | 0.874068749 | ns | TM | KS.1 |
| FIM1_483 | 1906.957732 | 2036.327893 | 2054.337398 | 0.0502497 | 0.073952027 | 0.316162667 | 0.642990763 | ns | TM | KS.1 |
| FIM1_485 | 138.5594796 | 136.2300356 | 116.1817359 | -0.006205246 | 0.118183896 | 0.920907127 | 0.975881018 | ns | TM | KS.1 |
| FIM1_486 | 1080.127038 | 1103.839191 | 1003.7575 | 0.010288654 | 0.105970866 | 0.865411396 | 0.959975149 | ns | TM | KS.1 |
| FIM1_487 | 835.5491456 | 781.8582619 | 668.4113705 | -0.027076338 | 0.120567056 | 0.669221588 | 0.882180579 | ns | TM | KS.1 |
| FIM1_488 | 7033.714206 | 6039.867952 | 5026.573853 | -0.076531967 | 0.126343129 | 0.260521058 | 0.588210523 | ns | TM | KS.1 |
| FIM1_489 | 2510.831399 | 3087.087761 | 3569.253078 | 0.075971913 | 0.151538268 | 0.275813478 | 0.600775854 | ns | TM | KS.1 |
| FIM1_490 | 1654.350102 | 1663.556488 | 1831.003238 | 0.003104977 | 0.084755498 | 0.954486922 | 0.988678089 | ns | TM | KS.1 |
| FIM1_491 | 10745.5445 | 12797.02562 | 12395.74351 | 0.183814886 | 0.094827645 | 0.007738768 | 0.070256154 | ns | TM | KS.1 |
| FIM1_492 | 3003.689084 | 1959.143974 | 2334.251937 | -0.57645974 | 0.108134954 | 2.25901813838 | 6.24367513249 | ns | TM | KS.1 |
| FIM1_493 | 1325.710671 | 1356.792518 | 1321.065216 | 0.01349646 | 0.090551196 | 0.811298276 | 0.939964817 | ns | TM | KS.1 |
| FIM1_494 | 1832.595421 | 2134.970474 | 1974.82989 | 0.125112091 | 0.107087999 | 0.065282952 | 0.268193795 | ns | TM | KS.1 |
| FIM1_495 | 3791.735579 | 4096.634279 | 3267.866095 | 0.058890837 | 0.080034974 | 0.267666273 | 0.594482013 | ns | TM | KS.1 |
| FIM1_496 | 5613.461393 | 5211.297983 | 5435.532296 | -0.071925768 | 0.061387944 | 0.105291188 | 0.351323716 | ns | TM | KS.1 |
| FIM1_497 | 2551.626435 | 1918.384754 | 1598.935476 | -0.139309654 | 0.169314222 | 0.083139364 | 0.306383953 | ns | TM | KS.1 |
| FIM1_498 | 702.2038731 | 701.3171762 | 630.0746192 | -0.000530313 | 0.099292251 | 0.992838351 | 0.997449676 | ns | TM | KS.1 |
| FIM1_499 | 3517.008195 | 3787.946388 | 3312.6156 | 0.051063799 | 0.086571303 | 0.359359776 | 0.683664672 | ns | TM | KS.1 |
| FIM1_500 | 312.3170956 | 310.5117006 | 352.4480144 | -0.003003635 | 0.101211072 | 0.959787904 | 0.990303759 | ns | TM | KS.1 |

| | | | | | | | | | | |
|---|---|---|---|---|---|---|---|---|---|---|
| FIM1_501 | 498.3246883 | 592.5344583 | 642.575356 | 0.090500256 | 0.130039103 | 0.196070314 | 0.50256523 | ns | TM | KS.1 |
| FIM1_502 | 2486.038185 | 2090.829427 | 2460.250386 | -0.125439422 | 0.119217808 | 0.079148723 | 0.299835289 | ns | TM | KS.1 |
| FIM1_503 | 1014.057104 | 830.8989843 | 959.0798002 | -0.114734629 | 0.135806384 | 0.119017455 | 0.378827394 | ns | TM | KS.1 |
| FIM1_504 | 1208.717679 | 1113.614825 | 1291.733053 | -0.062727958 | 0.081431515 | 0.244222902 | 0.567522712 | ns | TM | KS.1 |
| FIM1_505 | 779.3728898 | 987.6703925 | 1053.730224 | 0.131620373 | 0.149487547 | 0.089332792 | 0.320425841 | ns | TM | KS.1 |
| FIM1_506 | 2042.881598 | 2703.063269 | 3207.087225 | 0.109671936 | 0.170904699 | 0.147272339 | 0.426043493 | ns | TM | KS.1 |
| FIM1_507 | 2004.831446 | 1786.272539 | 1545.803788 | -0.074779106 | 0.104063821 | 0.235159702 | 0.554201572 | ns | TM | KS.1 |
| FIM1_508 | 936.7460746 | 881.6238657 | 1038.605976 | -0.034650522 | 0.096136995 | 0.555094904 | 0.824357358 | ns | TM | KS.1 |
| FIM1_509 | 791.5938138 | 562.5432993 | 964.2370488 | -0.189251489 | 0.192117051 | 0.035919257 | 0.189299049 | ns | TM | KS.1 |
| FIM1_510 | 836.7678902 | 888.7957616 | 854.0455006 | 0.035412703 | 0.094258046 | 0.542411355 | 0.815959873 | ns | TM | KS.1 |
| FIM1_511 | 2077.782724 | 1513.934016 | 1599.389843 | -0.187489888 | 0.179252395 | 0.034362123 | 0.185199792 | ns | TM | KS.1 |
| FIM1_512 | 302.1108008 | 201.3568922 | 324.3843601 | -0.268828488 | 0.22445938 | 0.011269547 | 0.091959747 | ns | TM | KS.1 |
| FIM1_513 | 3007.772045 | 3071.957775 | 3189.871274 | 0.01272909 | 0.084995976 | 0.815409832 | 0.940791141 | ns | TM | KS.1 |
| FIM1_514 | 2162.162367 | 1816.267643 | 1894.917338 | -0.097242636 | 0.128928317 | 0.168233657 | 0.461390543 | ns | TM | KS.1 |
| FIM1_515 | 1780.620837 | 1746.32347 | 1777.12397 | -0.015520349 | 0.057434024 | 0.710065886 | 0.901619224 | ns | TM | KS.1 |
| FIM1_516 | 1367.93198 | 1319.87994 | 1352.961096 | -0.023387308 | 0.078140464 | 0.651540118 | 0.875111255 | ns | TM | KS.1 |
| FIM1_517 | 835.2369004 | 1199.78699 | 1368.583645 | 0.147790757 | 0.201943235 | 0.078335177 | 0.298177127 | ns | TM | KS.1 |
| FIM1_518 | 20323.59518 | 24363.33976 | 24314.06549 | 0.21219366 | 0.081236952 | 0.001215165 | 0.01742204 | ns | TM | KS.1 |
| FIM1_519 | 305.0046695 | 206.4540479 | 587.3415799 | -0.105749212 | 0.211895932 | 0.164485943 | 0.454116296 | ns | TM | KS.1 |
| FIM1_522 | 2015.290927 | 2358.049191 | 2603.129516 | 0.056914698 | 0.141509865 | 0.395368217 | 0.718896697 | ns | TM | KS.1 |
| FIM1_523 | 810.104116 | 724.60455 | 636.3143509 | -0.053201506 | 0.118776881 | 0.412070228 | 0.732007978 | ns | TM | KS.1 |
| FIM1_524 | 1671.64967 | 1646.198045 | 1738.364328 | -0.009262247 | 0.082026814 | 0.862176701 | 0.958723533 | ns | TM | KS.1 |
| FIM1_525 | 2412.667071 | 1932.373413 | 1775.306378 | -0.274217489 | 0.07552988 | 3.26845639943 | 0.00102915 | ns | TM | KS.1 |
| FIM1_526 | 56.90443962 | 130.9837371 | 136.6789371 | 0.626579148 | 0.479977313 | 0.001939802 | 0.02513155 | ns | TM | KS.1 |
| FIM1_527 | 6242.314435 | 6557.514136 | 14481.84218 | 0.011961401 | 0.152968621 | 0.84837387 | 0.952348261 | ns | TM | KS.1 |
| FIM1_528 | 2597.645981 | 2956.613382 | 3607.913838 | 0.03572383 | 0.151254864 | 0.579346116 | 0.837375633 | ns | TM | KS.1 |
| FIM1_529 | 23.69486806 | 15.96015389 | 26.20148965 | -0.116848951 | 0.20921653 | 0.135587847 | 0.407805228 | ns | TM | KS.1 |
| FIM1_530 | 3624.983548 | 3438.108145 | 3718.552015 | -0.02722609 | 0.103221863 | 0.652543041 | 0.875599148 | ns | TM | KS.1 |
| FIM1_531 | 9198.655232 | 8950.412551 | 10265.85102 | -0.008299977 | 0.137932851 | 0.89537811 | 0.967972723 | ns | TM | KS.1 |
| FIM1_532 | 1853.71897 | 2940.089478 | 4169.976391 | 0.130750296 | 0.23737533 | 0.111150469 | 0.362605627 | ns | TM | KS.1 |
| FIM1_533 | 1032.295657 | 949.9188591 | 1149.200053 | -0.059436603 | 0.087627622 | 0.291884242 | 0.618715 | ns | TM | KS.1 |
| FIM1_534 | 1291.946801 | 985.6778051 | 1090.027217 | -0.254033346 | 0.134197156 | 0.004048216 | 0.044069751 | ns | TM | KS.1 |
| FIM1_535 | 1411.167404 | 1427.762318 | 1542.118228 | 0.002764613 | 0.152775724 | 0.964625062 | 0.993216478 | ns | TM | KS.1 |
| FIM1_536 | 710.4339267 | 862.0463126 | 1058.344059 | 0.062347755 | 0.15372 | 0.357563491 | 0.682799636 | ns | TM | KS.1 |
| FIM1_537 | 1004.625822 | 860.4925655 | 995.1810894 | -0.047351182 | 0.149486554 | 0.471843233 | 0.772376515 | ns | TM | KS.1 |
| FIM1_538 | 1102.852476 | 1162.362184 | 1018.384598 | 0.027496589 | 0.100735348 | 0.645909696 | 0.873206722 | ns | TM | KS.1 |
| FIM1_539 | 406.7599854 | 457.7991299 | 440.214433 | 0.077049618 | 0.103973207 | 0.222100927 | 0.536805479 | ns | TM | KS.1 |
| FIM1_540 | 2191.89728 | 2332.64501 | 2288.724654 | 0.045474992 | 0.076697356 | 0.37568897 | 0.699234053 | ns | TM | KS.1 |
| FIM1_541 | 561.2731245 | 568.251458 | 552.7711887 | 0.008559795 | 0.062028341 | 0.846277853 | 0.952348261 | ns | TM | KS.1 |
| FIM1_542 | 895.5235634 | 895.699332 | 827.8097266 | 0.000814308 | 0.08328603 | 0.987932076 | 0.996167262 | ns | TM | KS.1 |
| FIM1_543 | 412.093004 | 466.5903508 | 548.75533 | 0.050752487 | 0.12948293 | 0.44054958 | 0.747832613 | ns | TM | KS.1 |
| FIM1_544 | 1453.880193 | 1714.968816 | 1915.488907 | 0.111070993 | 0.119330306 | 0.11295377 | 0.366160178 | ns | TM | KS.1 |
| FIM1_545 | 3719.742338 | 3535.640454 | 1877.281489 | -0.0251646 | 0.105511705 | 0.679650694 | 0.887492141 | ns | TM | KS.1 |
| FIM1_546 | 3534.901353 | 3128.122656 | 4994.01369 | -0.103589433 | 0.092695441 | 0.08860736 | 0.318837658 | ns | TM | KS.1 |
| FIM1_547 | 1993.858856 | 2001.988909 | 2150.754405 | 0.001946074 | 0.102946525 | 0.974117547 | 0.993872484 | ns | TM | KS.1 |

| | | | | | | | | | | |
|---|---|---|---|---|---|---|---|---|---|---|
| FIM1_549 | 755.0746403 | 869.0121365 | 1034.980657 | 0.073590186 | 0.121272997 | 0.271801356 | 0.598783418 | ns | TM | KS.1 |
| FIM1_550 | 8258.025172 | 11322.84086 | 11820.81753 | 0.333060455 | 0.133585064 | 0.000411109 | 0.007660186 | ns | TM | KS.1 |
| FIM1_551 | 643.8834818 | 694.9960008 | 566.140344 | 0.036963786 | 0.111109682 | 0.554574445 | 0.824127468 | ns | TM | KS.1 |
| FIM1_552 | 1110.417952 | 1213.996562 | 1160.952802 | 0.063458946 | 0.089910708 | 0.269344645 | 0.596877332 | ns | TM | KS.1 |
| FIM1_553 | 1659.234722 | 1727.439357 | 1591.648014 | 0.028357196 | 0.071253576 | 0.560100843 | 0.826391733 | ns | TM | KS.1 |
| FIM1_554 | 1939.618699 | 2071.435895 | 1887.284174 | 0.047558017 | 0.079707178 | 0.367038934 | 0.690366236 | ns | TM | KS.1 |
| FIM1_555 | 670.8209378 | 685.0866595 | 575.5339994 | 0.011632923 | 0.093791989 | 0.839854443 | 0.949982862 | ns | TM | KS.1 |
| FIM1_556 | 1412.720706 | 1066.545026 | 1202.338386 | -0.274034094 | 0.133143119 | 0.002293458 | 0.028740431 | ns | TM | KS.1 |
| FIM1_557 | 854.1753145 | 839.830654 | 994.4919767 | -0.010568156 | 0.07475907 | 0.833488369 | 0.948012034 | ns | TM | KS.1 |
| FIM1_558 | 1098.588821 | 1281.545289 | 1298.567273 | 0.095503148 | 0.119216749 | 0.163723507 | 0.45366779 | ns | TM | KS.1 |
| FIM1_559 | 755.2020205 | 792.3225424 | 786.9371466 | 0.03036587 | 0.084776792 | 0.578077679 | 0.836271141 | ns | TM | KS.1 |
| FIM1_560 | 2838.708045 | 3166.543076 | 2200.518531 | 0.074328657 | 0.099648511 | 0.227136235 | 0.54408821 | ns | TM | KS.1 |
| FIM1_561 | 648.4368909 | 767.48188 | 945.6953352 | 0.114183421 | 0.120319069 | 0.105865763 | 0.352295767 | ns | TM | KS.1 |
| FIM1_562 | 1479.498656 | 1703.547522 | 1953.332671 | 0.126488015 | 0.097196665 | 0.049747322 | 0.232359781 | ns | TM | KS.1 |
| FIM1_563 | 178.5079288 | 160.1569647 | 220.2215981 | -0.037695406 | 0.136867105 | 0.561218234 | 0.826391733 | ns | TM | KS.1 |
| FIM1_564 | 1461.6402 | 1690.997486 | 1556.202246 | 0.073470671 | 0.124632663 | 0.276228349 | 0.601096377 | ns | TM | KS.1 |
| FIM1_565 | 1080.6692 | 1394.350156 | 1520.594745 | 0.141597389 | 0.155709017 | 0.074947389 | 0.291527179 | ns | TM | KS.1 |
| FIM1_566 | 858.6830673 | 866.8383028 | 826.6143196 | 0.005369961 | 0.095528779 | 0.926294517 | 0.976735299 | ns | TM | KS.1 |
| FIM1_567 | 1762.333926 | 1740.389336 | 1648.299936 | -0.007747423 | 0.07966833 | 0.882431296 | 0.966130215 | ns | TM | KS.1 |
| FIM1_568 | 6277.54715 | 2751.667434 | 3256.95093 | -0.501547886 | 0.472399524 | 0.00445922 | 0.047708855 | ns | TM | KS.1 |
| FIM1_569 | 4080.378132 | 2310.629326 | 2490.001199 | -0.339314135 | 0.326215426 | 0.009129328 | 0.079541868 | ns | TM | KS.1 |
| FIM1_570 | 767.2454845 | 1070.166026 | 1243.278647 | 0.232243966 | 0.183337301 | 0.014899505 | 0.110305115 | ns | TM | KS.1 |
| FIM1_571 | 2090.248856 | 2705.281135 | 2026.460833 | 0.076240241 | 0.171593239 | 0.275675538 | 0.600738415 | ns | TM | KS.1 |
| FIM1_572 | 950.5383862 | 998.9973296 | 510.8466022 | 0.01033575 | 0.163984892 | 0.867296444 | 0.960282219 | ns | TM | KS.1 |
| FIM1_573 | 3491.144854 | 3596.544202 | 4434.991654 | 0.017353673 | 0.089383271 | 0.757331433 | 0.920753636 | ns | TM | KS.1 |
| FIM1_574 | 2185.790905 | 3027.964001 | 3334.420775 | 0.100476669 | 0.189708013 | 0.177860076 | 0.476606421 | ns | TM | KS.1 |
| FIM1_575 | 2388.697976 | 2505.752149 | 2703.430966 | 0.031161746 | 0.082672061 | 0.562130608 | 0.826662659 | ns | TM | KS.1 |
| FIM1_576 | 4925.18088 | 5180.728694 | 5823.24603 | 0.037514645 | 0.071372731 | 0.441651945 | 0.747895515 | ns | TM | KS.1 |
| FIM1_577 | 1191.561123 | 839.7674422 | 905.1260683 | -0.366445854 | 0.151161582 | 0.000324308 | 0.006402512 | ns | TM | KS.1 |
| FIM1_578 | 5172.910518 | 5189.849545 | 4422.84059 | 0.00197918 | 0.084718798 | 0.970971604 | 0.99362472 | ns | TM | KS.1 |
| FIM1_579 | 753.5508618 | 871.3923904 | 850.9124711 | 0.105107902 | 0.109563844 | 0.116178016 | 0.372653532 | ns | TM | KS.1 |
| FIM1_580 | 288.7793723 | 279.3798878 | 260.9150765 | -0.022001467 | 0.092123824 | 0.700282485 | 0.897138176 | ns | TM | KS.1 |
| FIM1_581 | 688.314993 | 734.2146886 | 835.7897162 | 0.033245594 | 0.104807922 | 0.586090938 | 0.841613393 | ns | TM | KS.1 |
| FIM1_582 | 1892.362249 | 2902.153078 | 3226.125734 | 0.148445135 | 0.228147961 | 0.082366752 | 0.304891809 | ns | TM | KS.1 |
| FIM1_583 | 288.1509167 | 294.9654273 | 256.7458051 | 0.009944032 | 0.1173094 | 0.873510943 | 0.962759265 | ns | TM | KS.1 |
| FIM1_584 | 991.4333354 | 1054.709124 | 916.6185616 | 0.039814119 | 0.088414126 | 0.478392833 | 0.779543161 | ns | TM | KS.1 |
| FIM1_585 | 8725.039307 | 9955.921277 | 10000.39067 | 0.064219103 | 0.122588437 | 0.332916427 | 0.659601443 | ns | TM | KS.1 |
| FIM1_586 | 1130.732052 | 1568.633428 | 2018.87692 | 0.203466581 | 0.183675147 | 0.026066558 | 0.158641625 | ns | TM | KS.1 |
| FIM1_587 | 1050.052688 | 735.7842192 | 743.9692453 | -0.243175059 | 0.194952093 | 0.013534455 | 0.103590633 | ns | TM | KS.1 |
| FIM1_588 | 687.6825474 | 701.5148222 | 685.0246758 | 0.010239213 | 0.101450625 | 0.863787391 | 0.959694581 | ns | TM | KS.1 |
| FIM1_589 | 731.9405568 | 878.723235 | 965.9544655 | 0.167355429 | 0.109878231 | 0.021662414 | 0.143551826 | ns | TM | KS.1 |
| FIM1_590 | 687.8055406 | 735.9854856 | 896.1242943 | 0.048606305 | 0.081063274 | 0.362151533 | 0.685319085 | ns | TM | KS.1 |
| FIM1_591 | 837.6401947 | 611.4874892 | 799.1978164 | -0.325799018 | 0.136659754 | 0.000588368 | 0.010163652 | ns | TM | KS.1 |
| FIM1_592 | 6204.829643 | 6053.004126 | 5793.999017 | -0.014135962 | 0.090544896 | 0.802549423 | 0.936908579 | ns | TM | KS.1 |
| FIM1_593 | 1273.196325 | 1241.56885 | 1160.875021 | -0.01247094 | 0.101433593 | 0.83455507 | 0.948141464 | ns | TM | KS.1 |

| | | | | | | | | | | |
|---|---|---|---|---|---|---|---|---|---|---|
| FIM1_594 | 2307.95522 | 1781.422503 | 2091.221904 | -0.161552093 | 0.154382186 | 0.048979568 | 0.230707449 | ns | TM | KS.1 |
| FIM1_595 | 1968.254984 | 2097.26828 | 2258.61927 | 0.033793984 | 0.102243519 | 0.576063915 | 0.834571338 | ns | TM | KS.1 |
| FIM1_596 | 4151.632259 | 4211.036424 | 3810.382358 | 0.007351127 | 0.099351021 | 0.901052084 | 0.96882086 | ns | TM | KS.1 |
| FIM1_597 | 1651.037829 | 1338.100979 | 1847.660826 | -0.197291073 | 0.116088857 | 0.011272696 | 0.091959747 | ns | TM | KS.1 |
| FIM1_598 | 868.249785 | 997.09616 | 942.4573859 | 0.086854998 | 0.113166428 | 0.190829255 | 0.495240242 | ns | TM | KS.1 |
| FIM1_599 | 608.7379916 | 620.7332647 | 612.5166413 | 0.013012503 | 0.078332938 | 0.801783875 | 0.936908579 | ns | TM | KS.1 |
| FIM1_600 | 1946.476344 | 1753.310232 | 1731.684023 | -0.051431346 | 0.115681232 | 0.42331351 | 0.739045122 | ns | TM | KS.1 |
| FIM1_601 | 1777.927615 | 2434.626181 | 2786.680377 | 0.102293691 | 0.185215398 | 0.171533385 | 0.465692069 | ns | TM | KS.1 |
| FIM1_602 | 1991.883824 | 1882.289158 | 2163.783996 | -0.04358072 | 0.068063897 | 0.355750381 | 0.680714672 | ns | TM | KS.1 |
| FIM1_603 | 8665.64981 | 5519.133923 | 6877.477411 | -0.572017914 | 0.154552944 | 8.16908440475 | 6.11475692855 | ns | TM | KS.1 |
| FIM1_604 | 2171.748745 | 2142.859995 | 2260.077787 | -0.008091125 | 0.083958915 | 0.881132842 | 0.9656535 | ns | TM | KS.1 |
| FIM1_605 | 807.4362836 | 613.1049995 | 511.9090931 | -0.187680713 | 0.158164036 | 0.029034951 | 0.169342179 | ns | TM | KS.1 |
| FIM1_606 | 2372.08501 | 1747.203444 | 1345.168045 | -0.264186244 | 0.156300466 | 0.005184432 | 0.053400721 | ns | TM | KS.1 |
| FIM1_607 | 392.2057972 | 572.8269261 | 715.6332677 | 0.165652888 | 0.210030091 | 0.05893905 | 0.254891426 | ns | TM | KS.1 |
| FIM1_608 | 692.2942352 | 636.8570018 | 591.9919784 | -0.050910541 | 0.097779459 | 0.394469936 | 0.718333796 | ns | TM | KS.1 |
| FIM1_609 | 5416.705376 | 6039.913856 | 7419.754934 | 0.044674021 | 0.126606385 | 0.492313708 | 0.790867874 | ns | TM | KS.1 |
| FIM1_610 | 633.2828162 | 591.7950592 | 549.4681379 | -0.044291134 | 0.087531051 | 0.42836703 | 0.742665244 | ns | TM | KS.1 |
| FIM1_611 | 200.7631149 | 150.4649371 | 193.9310292 | -0.169185764 | 0.166423584 | 0.045414262 | 0.220040594 | ns | TM | KS.1 |
| FIM1_612 | 1651.623168 | 1491.225575 | 1807.422097 | -0.095902798 | 0.078735811 | 0.076994645 | 0.294813117 | ns | TM | KS.1 |
| FIM1_613 | 1331.181144 | 1343.526307 | 1297.621499 | 0.005968867 | 0.068215835 | 0.899427792 | 0.968452273 | ns | TM | KS.1 |
| FIM1_615 | 1034.83482 | 802.3354128 | 988.5036954 | -0.242665858 | 0.127335368 | 0.004530307 | 0.048319252 | ns | TM | KS.1 |
| FIM1_616 | 2382.678303 | 2406.033932 | 2442.610828 | 0.006198144 | 0.077962911 | 0.904512296 | 0.971072221 | ns | TM | KS.1 |
| FIM1_617 | 459.9351045 | 435.4591305 | 457.5838486 | -0.013717547 | 0.150200361 | 0.827064684 | 0.946035952 | ns | TM | KS.1 |
| FIM1_618 | 1057.019616 | 1070.764362 | 1419.095574 | 0.004744552 | 0.122744322 | 0.939819546 | 0.983546982 | ns | TM | KS.1 |
| FIM1_619 | 14230.15271 | 10998.2416 | 13755.79097 | -0.187665315 | 0.148311046 | 0.026178672 | 0.159021846 | ns | TM | KS.1 |
| FIM1_620 | 1084.781484 | 1029.241286 | 905.0081186 | -0.016004948 | 0.138778046 | 0.800578461 | 0.936908579 | ns | TM | KS.1 |
| FIM1_621 | 977.8037645 | 1155.893513 | 628.3279031 | 0.055238711 | 0.147732849 | 0.408379153 | 0.729248487 | ns | TM | KS.1 |
| FIM1_622 | 415.6701303 | 438.0049444 | 308.1685685 | 0.032850896 | 0.088342809 | 0.557470479 | 0.825421319 | ns | TM | KS.1 |
| FIM1_623 | 3440.528413 | 2961.161892 | 3212.491914 | -0.122994589 | 0.105286456 | 0.066729065 | 0.272267737 | ns | TM | KS.1 |
| FIM1_625 | 266.4795886 | 218.4612143 | 1665.134305 | -0.060167962 | 0.157703866 | 0.372221273 | 0.694598963 | ns | TM | KS.1 |
| FIM1_626 | 3939.369028 | 3348.244661 | 3460.275943 | -0.067148561 | 0.13686049 | 0.323404462 | 0.650941236 | ns | TM | KS.1 |
| FIM1_627 | 857.5952527 | 872.0727021 | 1097.322672 | 0.009436822 | 0.08887973 | 0.866003712 | 0.959975149 | ns | TM | KS.1 |
| FIM1_628 | 4005.363733 | 3307.52408 | 2869.052745 | -0.10634689 | 0.13426912 | 0.141818278 | 0.4174828 | ns | TM | KS.1 |
| FIM1_629 | 2553.687083 | 2155.556788 | 2629.388905 | -0.085930496 | 0.130813321 | 0.21696701 | 0.528087512 | ns | TM | KS.1 |
| FIM1_630 | 738.9146164 | 869.6461316 | 786.9538102 | 0.113052264 | 0.117731621 | 0.105652953 | 0.351822919 | ns | TM | KS.1 |
| FIM1_631 | 1541.317995 | 1627.773471 | 2014.126863 | 0.017251627 | 0.136213002 | 0.785754303 | 0.931631949 | ns | TM | KS.1 |
| FIM1_632 | 382.1114175 | 264.1335806 | 338.091691 | -0.258329774 | 0.200643663 | 0.010811351 | 0.089198126 | ns | TM | KS.1 |
| FIM1_633 | 5232.01496 | 5546.892572 | 5665.138552 | 0.040991731 | 0.07889002 | 0.432808619 | 0.74505982 | ns | TM | KS.1 |
| FIM1_634 | 428.3069629 | 468.5489361 | 504.229274 | 0.061257923 | 0.091654109 | 0.291221948 | 0.618364998 | ns | TM | KS.1 |
| FIM1_635 | 3423.03522 | 3801.728471 | 3749.836985 | 0.080168328 | 0.091669899 | 0.17368248 | 0.469348363 | ns | TM | KS.1 |
| FIM1_636 | 384.1738243 | 497.0751686 | 505.1129773 | 0.280559235 | 0.107535597 | 0.000617717 | 0.010560622 | ns | TM | KS.1 |
| FIM1_637 | 7501.080411 | 8586.201668 | 8546.879484 | 0.10059151 | 0.104580622 | 0.121564977 | 0.38326094 | ns | TM | KS.1 |
| FIM1_638 | 1386.706435 | 1556.363691 | 1509.720033 | 0.090797135 | 0.094544604 | 0.134589865 | 0.406322117 | ns | TM | KS.1 |
| FIM1_639 | 69.94362325 | 74.13874779 | 79.66084673 | 0.020550207 | 0.137861866 | 0.746643298 | 0.916442434 | ns | TM | KS.1 |
| FIM1_640 | 1127.843704 | 1104.477496 | 1173.532448 | -0.014287463 | 0.062467645 | 0.74739487 | 0.916738037 | ns | TM | KS.1 |

| | | | | | | | | | | |
|---|---|---|---|---|---|---|---|---|---|---|
| FIM1_641 | 603.5332452 | 550.5897984 | 514.1290684 | -0.050777663 | 0.10459316 | 0.411118301 | 0.731735607 | ns | TM | KS.1 |
| FIM1_642 | 2891.92299 | 3214.909867 | 3393.308776 | 0.042089438 | 0.127874215 | 0.516905176 | 0.804981347 | ns | TM | KS.1 |
| FIM1_644 | 1353.03914 | 1784.178779 | 1767.328646 | 0.142667014 | 0.165310647 | 0.076665169 | 0.294813117 | ns | TM | KS.1 |
| FIM1_645 | 1009.322542 | 970.1028909 | 1058.851261 | -0.016803254 | 0.116624372 | 0.78821477 | 0.933067989 | ns | TM | KS.1 |
| FIM1_646 | 3015.67836 | 2508.004839 | 2889.920807 | -0.137612277 | 0.121825727 | 0.06059674 | 0.259663032 | ns | TM | KS.1 |
| FIM1_647 | 1047.107555 | 1446.006924 | 1610.557098 | 0.207437731 | 0.180499705 | 0.023559743 | 0.150553405 | ns | TM | KS.1 |
| FIM1_648 | 200.8726573 | 158.3587358 | 180.6262444 | -0.191731877 | 0.139447291 | 0.021183491 | 0.141841004 | ns | TM | KS.1 |
| FIM1_650 | 638.0591086 | 553.06652 | 655.9854887 | -0.097171673 | 0.110973445 | 0.144878647 | 0.422987833 | ns | TM | KS.1 |
| FIM1_651 | 858.4230529 | 871.1214392 | 1317.559511 | 0.004955183 | 0.128815757 | 0.937401789 | 0.98242551 | ns | TM | KS.1 |
| FIM1_652 | 2227.575741 | 2311.96738 | 2526.977574 | 0.026291515 | 0.072430381 | 0.593206002 | 0.846854399 | ns | TM | KS.1 |
| FIM1_653 | 1925.717764 | 3173.826408 | 3738.84694 | 0.229489023 | 0.273313182 | 0.02801399 | 0.165718907 | ns | TM | KS.1 |
| FIM1_654 | 690.3773314 | 635.85543 | 618.8363942 | -0.05561249 | 0.092271772 | 0.338135254 | 0.666507986 | ns | TM | KS.1 |
| FIM1_655 | 880.0830429 | 457.8357619 | 491.6011962 | -0.886526861 | 0.176636937 | 2.5780021612[illegible] | 3.2063901880[illegible] | ns | TM | KS.1 |
| FIM1_656 | 1592.510088 | 1779.777228 | 1796.04403 | 0.075726688 | 0.099950788 | 0.219882607 | 0.533357373 | ns | TM | KS.1 |
| FIM1_657 | 1039.357901 | 859.2209875 | 840.9049317 | -0.117310848 | 0.130882123 | 0.108574919 | 0.357485256 | ns | TM | KS.1 |
| FIM1_658 | 757.7019884 | 675.6721015 | 678.8625973 | -0.057877407 | 0.116374948 | 0.371906913 | 0.694536971 | ns | TM | KS.1 |
| FIM1_659 | 2788.321037 | 2801.0304 | 2457.652158 | 0.002831039 | 0.07876387 | 0.956585611 | 0.989353426 | ns | TM | KS.1 |
| FIM1_660 | 1434.781826 | 866.3344081 | 935.7586285 | -0.703801421 | 0.105142722 | 3.5821568987[illegible] | 2.9702050951[illegible] | ns | TM | KS.1 |
| FIM1_661 | 756.3518052 | 711.8837965 | 869.1390861 | -0.026788152 | 0.11459815 | 0.668974068 | 0.882180579 | ns | TM | KS.1 |
| FIM1_662 | 9156.47671 | 12475.58426 | 13329.56743 | 0.238303658 | 0.166521006 | 0.010856726 | 0.08927638 | ns | TM | KS.1 |
| FIM1_663 | 583.2449378 | 542.7692442 | 728.0410642 | -0.017631891 | 0.153300812 | 0.779001506 | 0.928304554 | ns | TM | KS.1 |
| FIM1_664 | 3842.224125 | 2937.285529 | 3029.072366 | -0.236976607 | 0.139670849 | 0.007131969 | 0.066581689 | ns | TM | KS.1 |
| FIM1_665 | 694.6234708 | 659.9767545 | 624.5284246 | -0.034718159 | 0.078914178 | 0.505979776 | 0.798222804 | ns | TM | KS.1 |
| FIM1_666 | 233.3762014 | 228.2770962 | 221.6060011 | -0.007227912 | 0.1319363 | 0.908897747 | 0.972251892 | ns | TM | KS.1 |
| FIM1_667 | 237.9177087 | 235.1009121 | 255.920656 | -0.005361309 | 0.115320416 | 0.931280021 | 0.978943629 | ns | TM | KS.1 |
| FIM1_668 | 2873.232223 | 3502.834465 | 3917.240784 | 0.160187858 | 0.122609412 | 0.034783195 | 0.186153968 | ns | TM | KS.1 |
| FIM1_669 | 1692.768249 | 1815.206045 | 1242.978932 | 0.024270524 | 0.131507148 | 0.703738767 | 0.898252804 | ns | TM | KS.1 |
| FIM1_670 | 2132.482947 | 2299.356337 | 2971.716088 | 0.016080095 | 0.162790995 | 0.796044892 | 0.935498955 | ns | TM | KS.1 |
| FIM1_671 | 633.6894857 | 606.1276331 | 622.0532644 | -0.026143642 | 0.089064916 | 0.641550483 | 0.872440799 | ns | TM | KS.1 |
| FIM1_672 | 673.7582269 | 625.4730384 | 662.1761727 | -0.040467052 | 0.103106086 | 0.506496492 | 0.798422068 | ns | TM | KS.1 |
| FIM1_673 | 262.4625333 | 162.9054939 | 349.8445058 | -0.210414548 | 0.259554145 | 0.034798631 | 0.186153968 | ns | TM | KS.1 |
| FIM1_674 | 5894.743938 | 4832.931834 | 5215.570752 | -0.152212914 | 0.125449711 | 0.044508833 | 0.217302694 | ns | TM | KS.1 |
| FIM1_675 | 1674.825622 | 1554.691751 | 1473.001452 | -0.049409437 | 0.089313429 | 0.383908222 | 0.708354209 | ns | TM | KS.1 |
| FIM1_676 | 4875.146892 | 4455.136606 | 3264.916752 | -0.03340127 | 0.129776929 | 0.603831614 | 0.851547247 | ns | TM | KS.1 |
| FIM1_677 | 1502.350754 | 1268.052611 | 1919.82398 | -0.103978745 | 0.123973549 | 0.139656991 | 0.413618606 | ns | TM | KS.1 |
| FIM1_678 | 2772.014402 | 2226.041191 | 2766.754028 | -0.153375064 | 0.13618536 | 0.049788045 | 0.232359781 | ns | TM | KS.1 |
| FIM1_679 | 2270.710916 | 2414.680932 | 2876.776366 | 0.021103689 | 0.131297155 | 0.740213731 | 0.914311536 | ns | TM | KS.1 |
| FIM1_680 | 6265.404198 | 6386.762088 | 5497.70073 | 0.006120148 | 0.134334176 | 0.922778446 | 0.976369825 | ns | TM | KS.1 |
| FIM1_681 | 1595.000223 | 1418.387295 | 2011.579786 | -0.030237094 | 0.154038346 | 0.635466044 | 0.869849417 | ns | TM | KS.1 |
| FIM1_682 | 1198.427103 | 1144.558756 | 1325.94449 | -0.01335304 | 0.141791041 | 0.832556536 | 0.947385354 | ns | TM | KS.1 |
| FIM1_683 | 2735.128893 | 2966.576986 | 3487.012945 | 0.052498861 | 0.092945016 | 0.366995328 | 0.690366236 | ns | TM | KS.1 |
| FIM1_684 | 1447.881828 | 1513.976858 | 1704.721099 | 0.031250001 | 0.075294391 | 0.536498369 | 0.813136987 | ns | TM | KS.1 |
| FIM1_685 | 616.0010613 | 492.9389107 | 571.6491387 | -0.162428519 | 0.135735232 | 0.040103158 | 0.204001238 | ns | TM | KS.1 |
| FIM1_686 | 3613.469797 | 3637.952261 | 3809.941275 | 0.005639153 | 0.05504249 | 0.889283457 | 0.967028314 | ns | TM | KS.1 |
| FIM1_687 | 1024.994024 | 928.0518081 | 1055.214549 | -0.067328674 | 0.0959684 | 0.260966743 | 0.588210523 | ns | TM | KS.1 |

| | | | | | | | | | | |
|---|---|---|---|---|---|---|---|---|---|---|
| FIM1_688 | 1266.845814 | 1295.162169 | 1036.037415 | 0.014999128 | 0.074320942 | 0.764532003 | 0.924230468 | ns | TM | KS.1 |
| FIM1_689 | 133.1670553 | 121.4994295 | 143.7030848 | -0.029548438 | 0.139420902 | 0.645270888 | 0.873039206 | ns | TM | KS.1 |
| FIM1_690 | 1451.16565 | 1264.625816 | 1524.061999 | -0.109078161 | 0.102640775 | 0.093265665 | 0.330011865 | ns | TM | KS.1 |
| FIM1_691 | 1209.568032 | 1626.807675 | 1611.399486 | 0.288622384 | 0.138621254 | 0.001824789 | 0.024090145 | ns | TM | KS.1 |
| FIM1_692 | 1148.439585 | 1327.973954 | 1227.666889 | 0.09750695 | 0.112990867 | 0.146995709 | 0.425671509 | ns | TM | KS.1 |
| FIM1_693 | 1071.07587 | 1155.657745 | 1255.218501 | 0.055224355 | 0.082809055 | 0.308691059 | 0.636779799 | ns | TM | KS.1 |
| FIM1_694 | 580.999265 | 560.2094658 | 543.4684132 | -0.021974998 | 0.088989138 | 0.695144997 | 0.894091614 | ns | TM | KS.1 |
| FIM1_695 | 935.6548085 | 918.4673475 | 1000.654697 | -0.012780448 | 0.065476694 | 0.78056678 | 0.929207457 | ns | TM | KS.1 |
| FIM1_696 | 1367.14014 | 1411.308021 | 1557.699312 | 0.019956214 | 0.081472376 | 0.7074314 | 0.900120515 | ns | TM | KS.1 |
| FIM1_697 | 849.3947969 | 824.2466283 | 898.8205216 | -0.019205018 | 0.082206036 | 0.719450474 | 0.905913973 | ns | TM | KS.1 |
| FIM1_698 | 803.9401558 | 716.5867672 | 812.8636411 | -0.07156184 | 0.106435626 | 0.259615462 | 0.588210523 | ns | TM | KS.1 |
| FIM1_699 | 2780.411729 | 2788.879166 | 2938.740619 | 0.001116545 | 0.123415312 | 0.985828758 | 0.996167262 | ns | TM | KS.1 |
| FIM1_700 | 6778.734965 | 6380.797816 | 6849.827476 | -0.048560777 | 0.068526808 | 0.306387127 | 0.634850461 | ns | TM | KS.1 |
| FIM1_701 | 1252.419663 | 1812.924802 | 1751.349444 | 0.228476208 | 0.205079987 | 0.019269372 | 0.132643057 | ns | TM | KS.1 |
| FIM1_702 | 1049.554971 | 1124.236235 | 1186.573425 | 0.044287721 | 0.089452705 | 0.434435267 | 0.746009599 | ns | TM | KS.1 |
| FIM1_703 | 2822.405683 | 2581.06457 | 2874.929683 | -0.051266613 | 0.103154934 | 0.403838558 | 0.725739808 | ns | TM | KS.1 |
| FIM1_704 | 5110.293214 | 5235.165198 | 6127.943018 | 0.008432851 | 0.128535283 | 0.893759505 | 0.967972723 | ns | TM | KS.1 |
| FIM1_705 | 1076.171619 | 1079.598932 | 1247.382072 | 0.000878368 | 0.135188653 | 0.988889475 | 0.996167262 | ns | TM | KS.1 |
| FIM1_706 | 2142.268303 | 2084.393139 | 2196.0927 | -0.017377194 | 0.081222925 | 0.74327666 | 0.91508234 | ns | TM | KS.1 |
| FIM1_707 | 930.8631877 | 881.4309841 | 994.1467093 | -0.039619756 | 0.074015669 | 0.428359405 | 0.742665244 | ns | TM | KS.1 |
| FIM1_708 | 596.8991262 | 666.4133583 | 661.6962084 | 0.071869362 | 0.101628905 | 0.246777062 | 0.571296362 | ns | TM | KS.1 |
| FIM1_709 | 1259.548524 | 1275.720041 | 1294.26645 | 0.008237362 | 0.077110011 | 0.872436429 | 0.962759265 | ns | TM | KS.1 |
| FIM1_710 | 660.7752826 | 696.6961224 | 649.6720697 | 0.030178416 | 0.092963466 | 0.600211355 | 0.850484618 | ns | TM | KS.1 |
| FIM1_711 | 530.0389021 | 550.1664407 | 444.1594199 | 0.017495781 | 0.108447683 | 0.775423744 | 0.926727011 | ns | TM | KS.1 |
| FIM1_712 | 17.19650628 | 33.39130948 | 21.06410889 | 0.180473271 | 0.31330459 | 0.059588161 | 0.256462934 | ns | TM | KS.1 |
| FIM1_713 | 1704.783901 | 1434.175069 | 1449.725859 | -0.051954631 | 0.153145097 | 0.433529642 | 0.745784913 | ns | TM | KS.1 |
| FIM1_714 | 742.4506264 | 650.2822959 | 955.3241447 | -0.076741316 | 0.115584083 | 0.245943863 | 0.569897865 | ns | TM | KS.1 |
| FIM1_715 | 1967.254816 | 2289.242145 | 1366.728225 | 0.060577951 | 0.135883988 | 0.367287571 | 0.690572814 | ns | TM | KS.1 |
| FIM1_718 | 4668.907318 | 5051.657643 | 4056.178042 | 0.040157064 | 0.108162273 | 0.517950748 | 0.804981347 | ns | TM | KS.1 |
| FIM1_719 | 6237.884295 | 7672.373175 | 7126.365091 | 0.157959477 | 0.12850859 | 0.0404612 | 0.204775657 | ns | TM | KS.1 |
| FIM1_720 | 466.8452346 | 511.912137 | 588.1164666 | 0.053437984 | 0.102878582 | 0.38444173 | 0.708394027 | ns | TM | KS.1 |
| FIM1_721 | 1968.12509 | 2133.985484 | 2180.455325 | 0.049547789 | 0.096349763 | 0.40315344 | 0.72512233 | ns | TM | KS.1 |
| FIM1_722 | 4219.740727 | 4636.412325 | 4671.506458 | 0.080965201 | 0.079967209 | 0.133431877 | 0.405017444 | ns | TM | KS.1 |
| FIM1_723 | 673.9249442 | 732.0988026 | 877.597854 | 0.054150638 | 0.094816917 | 0.358065994 | 0.683043834 | ns | TM | KS.1 |
| FIM1_724 | 310.7979642 | 375.3980484 | 388.5643602 | 0.088498267 | 0.13885197 | 0.210987195 | 0.51871941 | ns | TM | KS.1 |
| FIM1_725 | 3439.873745 | 4847.582603 | 5829.750812 | 0.208865352 | 0.19145073 | 0.024907544 | 0.154938568 | ns | TM | KS.1 |
| FIM1_726 | 737.5676978 | 856.4708341 | 898.0738828 | 0.109717159 | 0.110644016 | 0.104689648 | 0.350588067 | ns | TM | KS.1 |
| FIM1_727 | 1282.958824 | 1357.631586 | 1229.378744 | 0.032452 | 0.09432182 | 0.576334673 | 0.834720524 | ns | TM | KS.1 |
| FIM1_728 | 2169.643098 | 2005.200118 | 2356.280893 | -0.03912053 | 0.109794345 | 0.53073611 | 0.808531286 | ns | TM | KS.1 |
| FIM1_729 | 2174.67622 | 2425.658567 | 2429.537675 | 0.076053908 | 0.098246374 | 0.213662265 | 0.521776083 | ns | TM | KS.1 |
| FIM1_730 | 1631.196087 | 1925.265174 | 1867.201485 | 0.159951714 | 0.100832628 | 0.020427763 | 0.138457929 | ns | TM | KS.1 |
| FIM1_731 | 413.0530665 | 447.3731663 | 460.353051 | 0.0541048 | 0.087295853 | 0.334728469 | 0.661873662 | ns | TM | KS.1 |
| FIM1_732 | 8017.771849 | 9089.056098 | 8995.403339 | 0.081776724 | 0.10720733 | 0.204104094 | 0.512578428 | ns | TM | KS.1 |
| FIM1_733 | 1580.054682 | 1890.220298 | 2077.4532 | 0.070634207 | 0.142670939 | 0.303949348 | 0.632168898 | ns | TM | KS.1 |
| FIM1_734 | 2865.118006 | 2833.295279 | 2175.091979 | -0.009145198 | 0.04829791 | 0.803881471 | 0.936908579 | ns | TM | KS.1 |

| | | | | | | | | | | |
|---|---|---|---|---|---|---|---|---|---|---|
| FIM1_735 | 812.1202394 | 721.167852 | 672.6776497 | -0.099439001 | 0.092264337 | 0.099493487 | 0.340998887 | ns | TM | KS.1 |
| FIM1_736 | 2248.173951 | 2588.82802 | 2714.657046 | 0.10715167 | 0.105632972 | 0.103646527 | 0.348223325 | ns | TM | KS.1 |
| FIM1_737 | 2284.423775 | 2357.977536 | 2322.058371 | 0.02373115 | 0.065435918 | 0.604600104 | 0.851547247 | ns | TM | KS.1 |
| FIM1_738 | 2354.842389 | 2126.791554 | 1834.221427 | -0.053333266 | 0.111256727 | 0.401050386 | 0.723750973 | ns | TM | KS.1 |
| FIM1_739 | 506.6576116 | 542.5369861 | 482.0774618 | 0.029437921 | 0.119644906 | 0.642456414 | 0.872815354 | ns | TM | KS.1 |
| FIM1_740 | 743.2480736 | 873.7828761 | 827.4917073 | 0.113964325 | 0.116152208 | 0.101286094 | 0.343723273 | ns | TM | KS.1 |
| FIM1_741 | 1550.830702 | 1740.069661 | 1896.080384 | 0.092809051 | 0.092810321 | 0.122745298 | 0.38575986 | ns | TM | KS.1 |
| FIM1_742 | 1949.957869 | 2053.821688 | 1995.452123 | 0.031947177 | 0.087945891 | 0.56723003 | 0.829990999 | ns | TM | KS.1 |
| FIM1_743 | 2621.262293 | 2108.434437 | 2245.619871 | -0.240776465 | 0.096938085 | 0.001315135 | 0.018640447 | ns | TM | KS.1 |
| FIM1_744 | 595.7858656 | 655.1586062 | 472.0112613 | 0.049126192 | 0.109857538 | 0.435209958 | 0.746610187 | ns | TM | KS.1 |
| FIM1_745 | 537.143935 | 466.3834224 | 570.5548511 | -0.113193849 | 0.102242135 | 0.082485024 | 0.305102598 | ns | TM | KS.1 |
| FIM1_746 | 1088.290406 | 1031.706948 | 1985.127197 | -0.028759992 | 0.099979599 | 0.629863402 | 0.868506215 | ns | TM | KS.1 |
| FIM1_747 | 1471.711659 | 1514.321341 | 1178.845108 | 0.015066146 | 0.098042842 | 0.797967208 | 0.936276987 | ns | TM | KS.1 |
| FIM1_748 | 742.3820634 | 826.9285006 | 794.7275218 | 0.038590444 | 0.134104219 | 0.552179164 | 0.822689033 | ns | TM | KS.1 |
| FIM1_749 | 1377.969723 | 1170.002121 | 1109.475622 | -0.113121654 | 0.11750797 | 0.10518947 | 0.351219876 | ns | TM | KS.1 |
| FIM1_750 | 1421.555199 | 1345.989922 | 1422.150315 | -0.036734692 | 0.080256486 | 0.486623453 | 0.786534009 | ns | TM | KS.1 |
| FIM1_751 | 3766.363842 | 3880.982442 | 3565.154167 | 0.023309932 | 0.060204761 | 0.589209909 | 0.844030895 | ns | TM | KS.1 |
| FIM1_752 | 4093.966029 | 3918.551841 | 5517.589437 | -0.031283197 | 0.071860197 | 0.522860703 | 0.807872226 | ns | TM | KS.1 |
| FIM1_753 | 1285.745493 | 1493.554668 | 2091.90095 | 0.09305563 | 0.117165913 | 0.170480552 | 0.464989445 | ns | TM | KS.1 |
| FIM1_754 | 1995.88094 | 2013.828284 | 2116.637094 | 0.003961395 | 0.108975757 | 0.948402368 | 0.986388143 | ns | TM | KS.1 |
| FIM1_755 | 2839.575443 | 2804.095578 | 2519.886863 | -0.009425407 | 0.051716966 | 0.807603364 | 0.937868053 | ns | TM | KS.1 |
| FIM1_756 | 1068.894583 | 880.0004575 | 822.4926349 | -0.173810373 | 0.115602733 | 0.02103733 | 0.141433402 | ns | TM | KS.1 |
| FIM1_757 | 2098.131149 | 2370.458678 | 2005.582828 | 0.087564552 | 0.101333409 | 0.16414111 | 0.45366779 | ns | TM | KS.1 |
| FIM1_758 | 460.7302339 | 363.3405968 | 378.9891539 | -0.22708392 | 0.12071972 | 0.005699502 | 0.057052359 | ns | TM | KS.1 |
| FIM1_759 | 866.8254313 | 724.4634913 | 738.316749 | -0.157781829 | 0.111470664 | 0.029384417 | 0.169985436 | ns | TM | KS.1 |
| FIM1_760 | 2130.690398 | 2417.785423 | 2381.124377 | 0.073382684 | 0.112769466 | 0.260688621 | 0.588210523 | ns | TM | KS.1 |
| FIM1_761 | 3114.725813 | 3406.430353 | 2800.818831 | 0.045452766 | 0.110737309 | 0.470143166 | 0.771448038 | ns | TM | KS.1 |
| FIM1_762 | 1328.730718 | 1330.024022 | 1122.594086 | 0.001052389 | 0.096927087 | 0.985632291 | 0.996167262 | ns | TM | KS.1 |
| FIM1_763 | 953.2058833 | 709.9218588 | 775.0109219 | -0.299324036 | 0.133281613 | 0.001127863 | 0.016554684 | ns | TM | KS.1 |
| FIM1_764 | 2494.940535 | 2159.202778 | 2454.664853 | -0.130459439 | 0.097455679 | 0.044472432 | 0.217302694 | ns | TM | KS.1 |
| FIM1_765 | 341.690697 | 371.9951709 | 354.7709332 | 0.029325244 | 0.133861829 | 0.647727994 | 0.873595603 | ns | TM | KS.1 |
| FIM1_767 | 852.9169592 | 468.5648494 | 1201.766306 | -0.364060496 | 0.346027416 | 0.007838365 | 0.070772899 | ns | TM | KS.1 |
| FIM1_768 | 492.0154851 | 385.5271755 | 370.6882992 | -0.1842234 | 0.142425079 | 0.026317539 | 0.159323047 | ns | TM | KS.1 |
| FIM1_769 | 2350.591683 | 2343.602579 | 2602.665713 | -0.002175402 | 0.054955402 | 0.957127287 | 0.989353426 | ns | TM | KS.1 |
| FIM1_770 | 3166.020999 | 3023.787599 | 2444.516323 | -0.017740779 | 0.122910201 | 0.778801198 | 0.928304554 | ns | TM | KS.1 |
| FIM1_772 | 80.56083768 | 122.108497 | 53.38291118 | 0.12920637 | 0.222528513 | 0.111854183 | 0.363946737 | ns | TM | KS.1 |
| FIM1_773 | 1729.139957 | 2249.681387 | 2254.322242 | 0.143340894 | 0.159223242 | 0.07360365 | 0.288556468 | ns | TM | KS.1 |
| FIM1_774 | 752.6538916 | 732.8729869 | 784.9324631 | -0.015823721 | 0.087875054 | 0.776002454 | 0.926727011 | ns | TM | KS.1 |
| FIM1_776 | 2256.773283 | 2272.76805 | 2078.601108 | 0.003869494 | 0.088726291 | 0.944765611 | 0.984955766 | ns | TM | KS.1 |
| FIM1_777 | 2644.428024 | 2597.734263 | 2381.458233 | -0.011549085 | 0.075356712 | 0.819234794 | 0.94310261 | ns | TM | KS.1 |
| FIM1_778 | 763.1732303 | 651.6828171 | 665.8322517 | -0.097096652 | 0.120331042 | 0.159172588 | 0.44660975 | ns | TM | KS.1 |
| FIM1_779 | 16774.20811 | 18282.99214 | 14674.89046 | 0.068169547 | 0.081367093 | 0.207040001 | 0.513839538 | ns | TM | KS.1 |
| FIM1_780 | 431.6998378 | 333.1220586 | 453.4928227 | -0.107551718 | 0.163396935 | 0.151569577 | 0.433367037 | ns | TM | KS.1 |
| FIM1_781 | 441.6198534 | 274.5021243 | 472.8245672 | -0.317402685 | 0.269083223 | 0.007881809 | 0.070983926 | ns | TM | KS.1 |
| FIM1_782 | 1961.23277 | 1279.538289 | 1671.123472 | -0.263561307 | 0.237917351 | 0.013708898 | 0.104063892 | ns | TM | KS.1 |

| | | | | | | | | | | |
|---|---|---|---|---|---|---|---|---|---|---|
| FIM1_783 | 1085.956783 | 918.3876341 | 935.7181848 | -0.095297528 | 0.12641596 | 0.173121556 | 0.468851247 | ns | TM | KS.1 |
| FIM1_784 | 549.8613009 | 316.5373176 | 488.4529459 | -0.529859595 | 0.297564265 | 0.000716961 | 0.011733155 | ns | TM | KS.1 |
| FIM1_785 | 3592.82388 | 4201.351547 | 3305.208004 | 0.101786451 | 0.11761611 | 0.138950068 | 0.412702442 | ns | TM | KS.1 |
| FIM1_786 | 2688.959587 | 2370.151216 | 2337.434031 | -0.096836783 | 0.099483478 | 0.123886737 | 0.387735747 | ns | TM | KS.1 |
| FIM1_787 | 3656.087338 | 3467.702687 | 4819.528209 | -0.025368235 | 0.108044821 | 0.679873615 | 0.887492141 | ns | TM | KS.1 |
| FIM1_788 | 154.024059 | 153.3453118 | 200.3153645 | -0.001375781 | 0.13625228 | 0.982593858 | 0.995804531 | ns | TM | KS.1 |
| FIM1_789 | 1446.840315 | 1473.028774 | 1008.009036 | 0.007784608 | 0.114195294 | 0.900171376 | 0.968452273 | ns | TM | KS.1 |
| FIM1_790 | 182.8366407 | 181.5026568 | 368.2901065 | -0.004111671 | 0.123856264 | 0.94788325 | 0.986388143 | ns | TM | KS.1 |
| FIM1_791 | 2960.890609 | 2095.533534 | 3356.780222 | -0.227835019 | 0.190840388 | 0.017383669 | 0.12351946 | ns | TM | KS.1 |
| FIM1_792 | 1031.454009 | 1110.078088 | 1088.768169 | 0.049449033 | 0.086805352 | 0.375199883 | 0.698585112 | ns | TM | KS.1 |
| FIM1_793 | 2788.850712 | 3173.513623 | 3888.038793 | 0.095063777 | 0.10241363 | 0.136407321 | 0.408811098 | ns | TM | KS.1 |
| FIM1_794 | 782.4032709 | 865.7541154 | 1027.485245 | 0.077023429 | 0.090742326 | 0.187146753 | 0.489513719 | ns | TM | KS.1 |
| FIM1_795 | 2452.889655 | 2815.84286 | 2824.487688 | 0.086903866 | 0.113048708 | 0.190407699 | 0.494404125 | ns | TM | KS.1 |
| FIM1_796 | 1346.307332 | 1574.842899 | 1281.943094 | 0.152731355 | 0.096760235 | 0.022109354 | 0.145110867 | ns | TM | KS.1 |
| FIM1_797 | 381.3968463 | 450.6503469 | 336.0054251 | 0.075197998 | 0.134073459 | 0.273935141 | 0.599308411 | ns | TM | KS.1 |
| FIM1_798 | 716.4286353 | 767.9894838 | 648.7373743 | 0.032933495 | 0.110975903 | 0.597277045 | 0.849348181 | ns | TM | KS.1 |
| FIM1_799 | 1675.613477 | 1139.581854 | 1734.643283 | -0.163425361 | 0.212370853 | 0.061681592 | 0.260865744 | ns | TM | KS.1 |
| FIM1_800 | 3199.57872 | 3171.880224 | 2792.052274 | -0.00439503 | 0.097683774 | 0.940274247 | 0.983546982 | ns | TM | KS.1 |
| FIM1_801 | 898.4977302 | 740.4917274 | 2056.706458 | -0.065276358 | 0.151966358 | 0.338367908 | 0.666685284 | ns | TM | KS.1 |
| FIM1_802 | 643.9228438 | 785.2213455 | 852.4081707 | 0.10341095 | 0.138440095 | 0.154152964 | 0.437933732 | ns | TM | KS.1 |
| FIM1_803 | 1162.709576 | 1116.973695 | 1145.42991 | -0.02877564 | 0.069872022 | 0.548891178 | 0.82102634 | ns | TM | KS.1 |
| FIM1_804 | 3189.420264 | 4602.963636 | 5144.829841 | 0.247668579 | 0.201406606 | 0.013235938 | 0.101775569 | ns | TM | KS.1 |
| FIM1_805 | 1176.533703 | 784.4758259 | 1049.39095 | -0.46335064 | 0.163617721 | 3.69248031588 | 0.001112321 | ns | TM | KS.1 |
| FIM1_806 | 1190.426437 | 779.2363097 | 1397.046938 | -0.229589193 | 0.235147116 | 0.023061917 | 0.148299176 | ns | TM | KS.1 |
| FIM1_807 | 259.7579167 | 244.6117346 | 604.1760257 | -0.029196917 | 0.110660075 | 0.638334068 | 0.870535085 | ns | TM | KS.1 |
| FIM1_808 | 1572.4656 | 1467.764849 | 1277.977105 | -0.037901684 | 0.100566134 | 0.528484089 | 0.808531286 | ns | TM | KS.1 |
| FIM1_809 | 35646.54667 | 33559.46611 | 20737.53121 | -0.038357711 | 0.087545348 | 0.491842108 | 0.790857947 | ns | TM | KS.1 |
| FIM1_810 | 1240.721065 | 1248.273405 | 1437.83775 | 0.004176238 | 0.071290078 | 0.931642328 | 0.979070676 | ns | TM | KS.1 |
| FIM1_811 | 2664.132268 | 2101.880716 | 2368.765038 | -0.151253528 | 0.145627247 | 0.057141338 | 0.252243261 | ns | TM | KS.1 |
| FIM1_812 | 2383.396437 | 2455.086699 | 2025.255696 | 0.01688371 | 0.091519942 | 0.76662752 | 0.924883104 | ns | TM | KS.1 |
| FIM1_813 | 558.1875878 | 610.1835808 | 733.4134806 | 0.062882621 | 0.090501594 | 0.275459773 | 0.600738415 | ns | TM | KS.1 |
| FIM1_814 | 39.99895776 | 27.95239331 | 59.6130047 | -0.09437382 | 0.192714704 | 0.198823255 | 0.506475009 | ns | TM | KS.1 |
| FIM1_815 | 92.96816426 | 66.57713958 | 102.8411246 | -0.115764161 | 0.189461839 | 0.134983055 | 0.406994362 | ns | TM | KS.1 |
| FIM1_816 | 417.7365271 | 329.2153793 | 391.056969 | -0.105686611 | 0.154898517 | 0.154686347 | 0.438498334 | ns | TM | KS.1 |
| FIM1_817 | 990.3089716 | 1207.583534 | 1487.348653 | 0.115941551 | 0.135272069 | 0.11550098 | 0.370960217 | ns | TM | KS.1 |
| FIM1_818 | 408.9032288 | 409.4956113 | 327.2956797 | 0.000770195 | 0.11059506 | 0.990004294 | 0.996341203 | ns | TM | KS.1 |
| FIM1_819 | 1251.313657 | 1343.751737 | 1264.18978 | 0.043865915 | 0.093962699 | 0.451579718 | 0.755416643 | ns | TM | KS.1 |
| FIM1_820 | 4862.163689 | 5089.963251 | 6602.112733 | 0.025340361 | 0.096060058 | 0.664706224 | 0.880679852 | ns | TM | KS.1 |
| FIM1_821 | 7249.008415 | 8179.501741 | 8074.419783 | 0.066311626 | 0.113948593 | 0.307460939 | 0.636056189 | ns | TM | KS.1 |
| FIM1_822 | 1314.736037 | 1100.591016 | 963.4226905 | -0.099865114 | 0.12971444 | 0.159454614 | 0.44660975 | ns | TM | KS.1 |
| FIM1_823 | 1494.840997 | 1354.981951 | 1053.613319 | -0.075237827 | 0.088132859 | 0.188718891 | 0.491815863 | ns | TM | KS.1 |
| FIM1_824 | 2613.194937 | 3890.42595 | 4683.444448 | 0.127705531 | 0.21493588 | 0.113483142 | 0.367564213 | ns | TM | KS.1 |
| FIM1_825 | 3058.353588 | 3316.205227 | 3341.939333 | 0.055601683 | 0.089232282 | 0.328503827 | 0.655821244 | ns | TM | KS.1 |
| FIM1_826 | 72.78425715 | 92.82710579 | 94.20694577 | 0.068108849 | 0.170705178 | 0.320489955 | 0.648936722 | ns | TM | KS.1 |
| FIM1_828 | 2073.709813 | 1942.594265 | 1423.450385 | -0.037398181 | 0.096906375 | 0.526409688 | 0.808531286 | ns | TM | KS.1 |

| | | | | | | | | | | | |
|---|---|---|---|---|---|---|---|---|---|---|---|
| FIM1_829 | 602.0407802 | 701.8299792 | 492.427216 | 0.0961588 | 0.118050624 | 0.159619065 | 0.44660975 | ns | | TM | KS.1 |
| FIM1_830 | 3953.031347 | 2822.606051 | 2874.1411 | -0.109633485 | 0.192587716 | 0.151244064 | 0.433183201 | ns | | TM | KS.1 |
| FIM1_831 | 4244.433024 | 4660.043152 | 4234.203468 | 0.056493184 | 0.100986362 | 0.354657692 | 0.679784721 | ns | | TM | KS.1 |
| FIM1_832 | 651.8557204 | 784.6304966 | 754.9707349 | 0.101698845 | 0.133348969 | 0.156207072 | 0.441088748 | ns | | TM | KS.1 |
| FIM1_833 | 2592.614987 | 3677.387832 | 3921.404092 | 0.189658234 | 0.196011304 | 0.03642536 | 0.190954861 | ns | | TM | KS.1 |
| FIM1_834 | 1364.950775 | 1363.951816 | 1221.750856 | -0.000723166 | 0.078833593 | 0.988911152 | 0.996167262 | ns | | TM | KS.1 |
| FIM1_835 | 875.2345258 | 842.8261198 | 953.2250199 | -0.021245812 | 0.091865133 | 0.709631531 | 0.901619224 | ns | | TM | KS.1 |
| FIM1_836 | 476.4217415 | 316.1250396 | 1089.792055 | -0.201946369 | 0.226897372 | 0.034438457 | 0.185223053 | ns | | TM | KS.1 |
| FIM1_840 | 558.0435904 | 565.7904549 | 485.8406089 | 0.006583428 | 0.099971766 | 0.911571994 | 0.973608989 | ns | | TM | KS.1 |
| FIM1_841 | 124.107609 | 44.47093591 | 116.3551879 | -1.15521506 | 0.400699212 | 3.88591638139 | 0.001157631 | Km<KS.1 | | TM | KS.1 |
| FIM1_842 | 433.5141754 | 134.4360197 | 255.2486176 | -1.519421084 | 0.247126292 | 7.52584774065 | 2.88008403921 | Km<KS.1 | | TM | KS.1 |
| FIM1_843 | 4371.690431 | 195.975701 | 1017.35679 | -4.38004797 | 0.458303636 | 4.65366242524 | 7.71732352186 | Km<KS.1 | | TM | KS.1 |
| FIM1_844 | 123.9562634 | 81.33109866 | 112.3709184 | -0.269014111 | 0.236521195 | 0.012435588 | 0.098138201 | ns | | TM | KS.1 |
| FIM1_846 | 90.279136 | 47.60364952 | 84.39783745 | -0.498163155 | 0.378156467 | 0.00258962 | 0.031270285 | ns | | TM | KS.1 |
| FIM1_847 | 11.05005527 | 10.09508809 | 15.97427327 | -0.027186503 | 0.174387244 | 0.663744414 | 0.880619203 | ns | | TM | KS.1 |
| FIM1_850 | 643.0180352 | 605.1253389 | 698.1875307 | -0.017325064 | 0.142898903 | 0.784315129 | 0.930812921 | ns | | TM | KS.1 |
| FIM1_851 | 285.9531496 | 319.0074971 | 305.0748288 | 0.037265975 | 0.137612052 | 0.56548663 | 0.828581174 | ns | | TM | KS.1 |
| FIM1_852 | 65.77677148 | 67.70447322 | 63.46054071 | 0.012191817 | 0.124787399 | 0.846780999 | 0.952348261 | ns | | TM | KS.1 |
| FIM1_853 | 56.66662492 | 57.72123416 | 101.2122781 | 0.009105942 | 0.132147989 | 0.885445183 | 0.967028314 | ns | | TM | KS.1 |
| FIM1_854 | 747.9364597 | 970.5857297 | 1559.299267 | 0.099680577 | 0.165507562 | 0.176840631 | 0.474852995 | ns | | TM | KS.1 |
| FIM1_855 | 233.1202204 | 254.7286618 | 276.8693972 | 0.036927629 | 0.121185007 | 0.563988266 | 0.827926121 | ns | | TM | KS.1 |
| FIM1_856 | 286.8136207 | 283.5729493 | 299.5086226 | -0.00610463 | 0.10713268 | 0.920139215 | 0.975881018 | ns | | TM | KS.1 |
| FIM1_857 | 170.9209038 | 124.1496288 | 194.7328761 | -0.266128274 | 0.168226828 | 0.006116534 | 0.060019243 | ns | | TM | KS.1 |
| FIM1_858 | 809.5275629 | 403.5727166 | 643.7882858 | -0.678965751 | 0.378910224 | 0.000572923 | 0.010001027 | ns | | TM | KS.1 |
| FIM1_859 | 471.7904502 | 452.595917 | 460.6221006 | -0.024120182 | 0.076635035 | 0.637096764 | 0.870169358 | ns | | TM | KS.1 |
| FIM1_860 | 781.6774847 | 507.7808382 | 530.8493055 | -0.576438357 | 0.121366497 | 5.44094731505 | 9.02290429746 | ns | | TM | KS.1 |
| FIM1_861 | 776.5477759 | 757.4479579 | 689.1671746 | -0.012930158 | 0.096550504 | 0.824796398 | 0.94482203 | ns | | TM | KS.1 |
| FIM1_862 | 114.0417184 | 129.8593664 | 158.4303416 | 0.058123213 | 0.128729767 | 0.381735732 | 0.705735886 | ns | | TM | KS.1 |
| FIM1_863 | 41.84009866 | 56.93010508 | 32.18866186 | 0.070611746 | 0.191627137 | 0.30500066 | 0.633687915 | ns | | TM | KS.1 |
| FIM1_864 | 692.6574842 | 722.5471762 | 574.4896196 | 0.013520126 | 0.134725405 | 0.830940691 | 0.946802573 | ns | | TM | KS.1 |
| FIM1_865 | 1178.843386 | 990.1465823 | 947.753543 | -0.07937946 | 0.136087027 | 0.252683571 | 0.579042268 | ns | | TM | KS.1 |
| FIM1_866 | 575.5943801 | 473.7854656 | 514.4551748 | -0.109309927 | 0.135301564 | 0.133674231 | 0.405258563 | ns | | TM | KS.1 |
| FIM1_867 | 1066.006044 | 1079.492436 | 1299.862932 | 0.005221959 | 0.114119347 | 0.932896268 | 0.979974437 | ns | | TM | KS.1 |
| FIM1_868 | 807.6623632 | 832.6668791 | 901.8214115 | 0.015365066 | 0.100358778 | 0.796161322 | 0.935498955 | ns | | TM | KS.1 |
| FIM1_869 | 707.6073072 | 775.4575028 | 674.8613506 | 0.062926487 | 0.093255953 | 0.283796859 | 0.610678794 | ns | | TM | KS.1 |
| FIM1_870 | 936.848504 | 1297.847634 | 1337.424702 | 0.193835604 | 0.183759236 | 0.031340644 | 0.176980365 | ns | | TM | KS.1 |
| FIM1_871 | 1160.111311 | 1323.563672 | 1085.450442 | 0.12200765 | 0.092366061 | 0.049683058 | 0.232305651 | ns | | TM | KS.1 |
| FIM1_872 | 2382.88398 | 2686.148478 | 2799.267011 | 0.099416354 | 0.093124574 | 0.101497686 | 0.343972064 | ns | | TM | KS.1 |
| FIM1_873 | 750.6022595 | 786.72051 | 998.9041111 | 0.02500622 | 0.097753842 | 0.671470967 | 0.883336315 | ns | | TM | KS.1 |
| FIM1_874 | 3212.626397 | 2803.267862 | 4884.448618 | -0.057312864 | 0.130484697 | 0.388844652 | 0.713088169 | ns | | TM | KS.1 |
| FIM1_875 | 296.5171543 | 325.2564185 | 368.4667161 | 0.037942066 | 0.123949776 | 0.555357671 | 0.824501467 | ns | | TM | KS.1 |
| FIM1_876 | 2479.445664 | 2611.842735 | 2803.00508 | 0.016877235 | 0.134530127 | 0.790287161 | 0.933303839 | ns | | TM | KS.1 |
| FIM1_877 | 1221.668609 | 1371.714075 | 1254.645693 | 0.071301653 | 0.106710229 | 0.261804212 | 0.588461463 | ns | | TM | KS.1 |
| FIM1_878 | 3501.479903 | 3668.062996 | 3660.528863 | 0.03036354 | 0.081594408 | 0.568918742 | 0.831239571 | ns | | TM | KS.1 |
| FIM1_879 | 30.19200932 | 11.63715871 | 27.8276178 | -0.648044807 | 0.535669798 | 0.00215836 | 0.02746251 | ns | | TM | KS.1 |

| | | | | | | | | | | |
|---|---|---|---|---|---|---|---|---|---|---|
| FIM1_880 | 1165.463041 | 1240.28202 | 1216.289069 | 0.042876194 | 0.079959946 | 0.416379519 | 0.735551151 | ns | TM | KS.1 |
| FIM1_881 | 1326.145488 | 1389.362938 | 1551.362047 | 0.033570608 | 0.073786628 | 0.500886767 | 0.795351314 | ns | TM | KS.1 |
| FIM1_882 | 451.6778757 | 428.2495103 | 510.0564056 | -0.030977024 | 0.094847895 | 0.594643002 | 0.847499664 | ns | TM | KS.1 |
| FIM1_883 | 1582.502648 | 1613.033749 | 1562.433082 | 0.008345685 | 0.112926926 | 0.892730762 | 0.967822084 | ns | TM | KS.1 |
| FIM1_884 | 379.6694045 | 321.4163312 | 330.5532116 | -0.060676668 | 0.142999208 | 0.368377377 | 0.691630801 | ns | TM | KS.1 |
| FIM1_885 | 1907.325249 | 1877.925715 | 2050.872708 | -0.010704166 | 0.062821068 | 0.810119167 | 0.939473859 | ns | TM | KS.1 |
| FIM1_886 | 4322.62463 | 4191.53619 | 4045.572158 | -0.012093762 | 0.120852752 | 0.84738275 | 0.952348261 | ns | TM | KS.1 |
| FIM1_887 | 831.3015305 | 846.1430468 | 602.4879608 | 0.011262975 | 0.087252665 | 0.838773439 | 0.949982862 | ns | TM | KS.1 |
| FIM1_888 | 2280.081859 | 1665.479979 | 1495.16734 | -0.304875242 | 0.146514488 | 0.001500142 | 0.020447135 | ns | TM | KS.1 |
| FIM1_889 | 834.4336107 | 964.6300614 | 968.179931 | 0.129213925 | 0.099511148 | 0.048810822 | 0.230174256 | ns | TM | KS.1 |
| FIM1_890 | 263.2725392 | 329.8592774 | 335.6733714 | 0.150331349 | 0.139747839 | 0.055379243 | 0.2466533 | ns | TM | KS.1 |
| FIM1_891 | 4246.502624 | 3225.672429 | 4113.37712 | -0.227021386 | 0.147546629 | 0.010526592 | 0.088016463 | ns | TM | KS.1 |
| FIM1_892 | 219.2411296 | 172.0540598 | 170.0596042 | -0.07582704 | 0.167032738 | 0.277742835 | 0.602049362 | ns | TM | KS.1 |
| FIM1_893 | 14.81666865 | 13.16992318 | 28.89626081 | -0.020033276 | 0.175048657 | 0.745987815 | 0.91614154 | ns | TM | KS.1 |
| FIM1_894 | 1388.820885 | 1700.612766 | 3726.986747 | 0.067932196 | 0.15400595 | 0.321873726 | 0.650150949 | ns | TM | KS.1 |
| FIM1_895 | 570.8275185 | 676.4678652 | 707.5695422 | 0.126536312 | 0.116327124 | 0.073866696 | 0.288677781 | ns | TM | KS.1 |
| FIM1_896 | 141.3647415 | 158.4925537 | 140.4085589 | 0.06056817 | 0.115839977 | 0.350860166 | 0.676561755 | ns | TM | KS.1 |
| FIM1_897 | 644.8551426 | 643.6089984 | 566.0714161 | -0.001074842 | 0.120355461 | 0.98631247 | 0.996167262 | ns | TM | KS.1 |
| FIM1_898 | 5172.054178 | 5396.514697 | 5129.727209 | 0.019403956 | 0.110523105 | 0.753344283 | 0.919275891 | ns | TM | KS.1 |
| FIM1_899 | 526.3553997 | 694.4139952 | 688.9489661 | 0.175654517 | 0.161525946 | 0.038462281 | 0.197675463 | ns | TM | KS.1 |
| FIM1_900 | 1916.285953 | 1822.27149 | 1909.136672 | -0.024123718 | 0.107838981 | 0.694381859 | 0.893803299 | ns | TM | KS.1 |
| FIM1_901 | 4103.196876 | 3159.423007 | 3456.279029 | -0.266517971 | 0.120475059 | 0.001825525 | 0.024090145 | ns | TM | KS.1 |
| FIM1_902 | 495.9175272 | 547.6247058 | 486.7253884 | 0.067655861 | 0.096636448 | 0.260662119 | 0.588210523 | ns | TM | KS.1 |
| FIM1_903 | 1057.60933 | 1420.546351 | 1651.287398 | 0.088215478 | 0.181091613 | 0.221357137 | 0.535714496 | ns | TM | KS.1 |
| FIM1_904 | 956.4435777 | 1168.070513 | 1256.333312 | 0.063280836 | 0.155936588 | 0.351330079 | 0.677205403 | ns | TM | KS.1 |
| FIM1_905 | 735.3684751 | 627.7884131 | 879.7972504 | -0.108334568 | 0.116305341 | 0.116743056 | 0.373743053 | ns | TM | KS.1 |
| FIM1_906 | 11307.41895 | 13251.15864 | 12090.97668 | 0.110160643 | 0.115929769 | 0.111071853 | 0.362586921 | ns | TM | KS.1 |
| FIM1_907 | 13997.38485 | 16970.57943 | 16591.70845 | 0.169170998 | 0.115853129 | 0.023965829 | 0.151692113 | ns | TM | KS.1 |
| FIM1_909 | 1197.332659 | 1196.082299 | 1077.98299 | -0.000437036 | 0.094286043 | 0.993952964 | 0.99786059 | ns | TM | KS.1 |
| FIM1_910 | 247.3511537 | 189.5797022 | 267.2848716 | -0.131184417 | 0.161748613 | 0.094971278 | 0.333005827 | ns | TM | KS.1 |
| FIM1_912 | 54.24465571 | 53.59230573 | 111.3884135 | -0.002236324 | 0.139592475 | 0.971672469 | 0.99362472 | ns | TM | KS.1 |
| FIM1_913 | 1094.70194 | 1857.875914 | 2001.170019 | 0.409177675 | 0.301552535 | 0.003199596 | 0.036847194 | ns | TM | KS.1 |
| FIM1_914 | 510.5733211 | 590.1005542 | 592.0757581 | 0.107146898 | 0.108922626 | 0.109149957 | 0.35800987 | ns | TM | KS.1 |
| FIM1_915 | 3557.341125 | 3697.242498 | 4187.559889 | 0.025016946 | 0.079948159 | 0.634149927 | 0.869547581 | ns | TM | KS.1 |
| FIM1_916 | 1108.471612 | 1020.59668 | 1234.286868 | -0.04465122 | 0.105036517 | 0.4685066 | 0.771134367 | ns | TM | KS.1 |
| FIM1_917 | 6304.410065 | 5408.883683 | 10820.5135 | -0.046030087 | 0.149938971 | 0.483179102 | 0.784023495 | ns | TM | KS.1 |
| FIM1_918 | 256.5886804 | 295.971277 | 235.7526735 | 0.053033263 | 0.137715666 | 0.424659124 | 0.740157476 | ns | TM | KS.1 |
| FIM1_919 | 231.8958684 | 271.3974632 | 300.1039835 | 0.050871424 | 0.146414594 | 0.442722263 | 0.748839657 | ns | TM | KS.1 |
| FIM1_920 | 1816.5665 | 1798.997505 | 1881.743709 | -0.004658532 | 0.107374559 | 0.939050091 | 0.983546982 | ns | TM | KS.1 |
| FIM1_921 | 3384.561478 | 2917.020571 | 2693.04147 | -0.098599517 | 0.1142908 | 0.145112145 | 0.423172872 | ns | TM | KS.1 |
| FIM1_922 | 659.4907012 | 590.9291068 | 488.257024 | -0.054506479 | 0.116534482 | 0.398794829 | 0.722244002 | ns | TM | KS.1 |
| FIM1_923 | 331.0218219 | 379.0186507 | 409.4636735 | 0.061847887 | 0.125798653 | 0.352448008 | 0.678105331 | ns | TM | KS.1 |
| FIM1_924 | 1248.147872 | 695.616282 | 926.2492243 | -0.800114993 | 0.152114221 | 4.20358932332 | 7.2113299598 | ns | TM | KS.1 |
| FIM1_925 | 325.1863706 | 199.788283 | 227.6234873 | -0.474145446 | 0.250787451 | 0.000622328 | 0.010603017 | ns | TM | KS.1 |
| FIM1_926 | 2196.629669 | 1627.57108 | 4723.754127 | -0.08685157 | 0.183147773 | 0.227086775 | 0.54408821 | ns | TM | KS.1 |

| | | | | | | | | | | |
|---|---|---|---|---|---|---|---|---|---|---|
| FIM1_927 | 5582.281453 | 6618.878663 | 6918.70648 | 0.190842654 | 0.086108514 | 0.003823268 | 0.042174626 | ns | TM | KS.1 |
| FIM1_928 | 1538.081651 | 1125.594179 | 996.4788868 | -0.266951559 | 0.159929437 | 0.005206944 | 0.053411437 | ns | TM | KS.1 |
| FIM1_929 | 1327.127595 | 1370.033487 | 1444.466614 | 0.011839911 | 0.124189246 | 0.851056383 | 0.953390116 | ns | TM | KS.1 |
| FIM1_930 | 1501.288454 | 1629.620309 | 1751.919322 | 0.039132825 | 0.112995837 | 0.534539794 | 0.811762966 | ns | TM | KS.1 |
| FIM1_931 | 1739.1318 | 1683.981117 | 1672.699267 | -0.019761468 | 0.083602119 | 0.714543895 | 0.905084971 | ns | TM | KS.1 |
| FIM1_932 | 487.5520448 | 492.2285504 | 497.2585964 | 0.006298345 | 0.079041681 | 0.903821736 | 0.970540284 | ns | TM | KS.1 |
| FIM1_933 | 2122.87898 | 1942.430837 | 2148.831949 | -0.078480796 | 0.07493647 | 0.127365883 | 0.393568488 | ns | TM | KS.1 |
| FIM1_934 | 1454.21952 | 1467.366964 | 1323.273902 | 0.005931494 | 0.08294687 | 0.91206365 | 0.973674801 | ns | TM | KS.1 |
| FIM1_935 | 1259.064146 | 777.4232878 | 1205.089316 | -0.167861427 | 0.252499037 | 0.064130578 | 0.265874687 | ns | TM | KS.1 |
| FIM1_936 | 8513.123897 | 8966.507202 | 8369.136041 | 0.018288242 | 0.129278507 | 0.773256069 | 0.925733395 | ns | TM | KS.1 |
| FIM1_937 | 3373.998626 | 2631.066924 | 2593.518284 | -0.252690251 | 0.117358227 | 0.002441123 | 0.02998663 | ns | TM | KS.1 |
| FIM1_938 | 2559.094493 | 2918.885655 | 3427.445661 | 0.059660534 | 0.125703222 | 0.368185916 | 0.691630801 | ns | TM | KS.1 |
| FIM1_939 | 1701.204685 | 1983.917651 | 1869.916936 | 0.169853997 | 0.084313812 | 0.007171644 | 0.066581689 | ns | TM | KS.1 |
| FIM1_940 | 2534.719924 | 2407.583641 | 2972.43512 | -0.024940213 | 0.107320565 | 0.68420316 | 0.890505759 | ns | TM | KS.1 |
| FIM1_941 | 508.6210454 | 390.6375258 | 506.1071318 | -0.117982823 | 0.163432014 | 0.123776861 | 0.387735747 | ns | TM | KS.1 |
| FIM1_942 | 495.9098505 | 794.4379936 | 1787.929814 | 0.388886706 | 0.259455704 | 0.002532881 | 0.030888701 | ns | TM | KS.1 |
| FIM1_943 | 8143.512661 | 8937.827657 | 8646.69721 | 0.055137433 | 0.102187069 | 0.368605967 | 0.691744506 | ns | TM | KS.1 |
| FIM1_944 | 1118.337638 | 1430.196348 | 1283.016317 | 0.093575602 | 0.161035775 | 0.197852106 | 0.505976981 | ns | TM | KS.1 |
| FIM1_945 | 2257.325047 | 1785.685005 | 2338.598098 | -0.25251362 | 0.104864261 | 0.001389982 | 0.019268155 | ns | TM | KS.1 |
| FIM1_946 | 51.97525258 | 47.10820873 | 45.99085676 | -0.042410834 | 0.129930118 | 0.514653636 | 0.804256034 | ns | TM | KS.1 |
| FIM1_947 | 4462.978298 | 4823.708472 | 5270.519313 | 0.049005498 | 0.093443622 | 0.400220812 | 0.723750973 | ns | TM | KS.1 |
| FIM1_948 | 4368.301933 | 4769.789856 | 5590.866524 | 0.066563151 | 0.084941711 | 0.23063983 | 0.548995638 | ns | TM | KS.1 |
| FIM1_949 | 1431.392811 | 1211.098565 | 1384.197803 | -0.065977412 | 0.139971084 | 0.332019506 | 0.65912278 | ns | TM | KS.1 |
| FIM1_950 | 607.5356156 | 467.393684 | 465.2174882 | -0.115265315 | 0.163373942 | 0.130492586 | 0.39950807 | ns | TM | KS.1 |
| FIM1_951 | 15120.80026 | 14368.93313 | 20186.18416 | -0.028802384 | 0.095111388 | 0.620967149 | 0.862695216 | ns | TM | KS.1 |
| FIM1_952 | 21.43687156 | 19.16663548 | 41.24879713 | -0.021870373 | 0.16980989 | 0.72536549 | 0.907961357 | ns | TM | KS.1 |
| FIM1_953 | 3599.841967 | 3447.977096 | 3895.405561 | -0.025177478 | 0.090708076 | 0.657145431 | 0.877664032 | ns | TM | KS.1 |
| FIM1_954 | 1997.536721 | 1909.408335 | 1865.356409 | -0.017470781 | 0.122853375 | 0.782036084 | 0.929216508 | ns | TM | KS.1 |
| FIM1_955 | 1406.494928 | 1076.375149 | 1946.284371 | -0.088959245 | 0.170651738 | 0.217439274 | 0.528978185 | ns | TM | KS.1 |
| FIM1_957 | 10826.78452 | 12824.12218 | 11679.93381 | 0.051048786 | 0.152334286 | 0.440857686 | 0.747832613 | ns | TM | KS.1 |
| FIM1_958 | 4370.53984 | 3687.169095 | 4172.932193 | -0.093792225 | 0.128030607 | 0.180664003 | 0.480643537 | ns | TM | KS.1 |
| FIM1_959 | 646.9625623 | 978.8175799 | 1630.320037 | 0.23775286 | 0.230213785 | 0.01962428 | 0.134663165 | ns | TM | KS.1 |
| FIM1_960 | 2144.825979 | 2262.435494 | 2383.03718 | 0.033817956 | 0.087059314 | 0.542551811 | 0.815959873 | ns | TM | KS.1 |
| FIM1_961 | 5210.619132 | 5432.482615 | 4809.67305 | 0.030439034 | 0.069830697 | 0.525884683 | 0.808531286 | ns | TM | KS.1 |
| FIM1_962 | 2481.120948 | 2743.209678 | 2784.4084 | 0.081329304 | 0.086475791 | 0.152547223 | 0.434912569 | ns | TM | KS.1 |
| FIM1_963 | 681.7276526 | 693.8586343 | 705.9682752 | 0.008626979 | 0.099012096 | 0.883837003 | 0.966744674 | ns | TM | KS.1 |
| FIM1_964 | 297.311175 | 287.086069 | 229.22586 | -0.015579836 | 0.093116412 | 0.786040156 | 0.931748814 | ns | TM | KS.1 |
| FIM1_965 | 860.7823267 | 899.5683282 | 1032.537501 | 0.031086522 | 0.07305164 | 0.530169783 | 0.808531286 | ns | TM | KS.1 |
| FIM1_966 | 5104.43633 | 4761.675234 | 3950.198171 | -0.036221631 | 0.104736049 | 0.55365116 | 0.823196211 | ns | TM | KS.1 |
| FIM1_967 | 879.5978843 | 1117.471617 | 1015.043269 | 0.228854027 | 0.122009954 | 0.005644185 | 0.056841744 | ns | TM | KS.1 |
| FIM1_968 | 673.888185 | 634.4372785 | 752.0076583 | -0.031183879 | 0.101750082 | 0.604595277 | 0.851547247 | ns | TM | KS.1 |
| FIM1_969 | 236.4980021 | 312.7012486 | 440.0148115 | 0.187141793 | 0.159851181 | 0.029831471 | 0.171016808 | ns | TM | KS.1 |
| FIM1_970 | 933.7145029 | 913.9131258 | 965.1903519 | -0.013474222 | 0.079510187 | 0.796816637 | 0.935826905 | ns | TM | KS.1 |
| FIM1_971 | 109.866558 | 124.0965796 | 131.9005992 | 0.052022157 | 0.125453148 | 0.427659085 | 0.742555695 | ns | TM | KS.1 |
| FIM1_972 | 172.1228395 | 168.6166805 | 157.1682748 | -0.008713307 | 0.119066996 | 0.889327104 | 0.967028314 | ns | TM | KS.1 |

| | | | | | | | | | | |
|---|---|---|---|---|---|---|---|---|---|---|
| FIM1_973 | 987.8890716 | 1022.80622 | 1188.232828 | 0.024940762 | 0.069545502 | 0.602177768 | 0.851547247 | ns | TM | KS.1 |
| FIM1_974 | 1172.557064 | 1032.778222 | 1062.195313 | -0.067821584 | 0.117019972 | 0.302014227 | 0.630781183 | ns | TM | KS.1 |
| FIM1_975 | 3689.503179 | 3472.714923 | 2546.823218 | -0.027485455 | 0.113228543 | 0.659995137 | 0.878874681 | ns | TM | KS.1 |
| FIM1_976 | 33.79188046 | 37.67293196 | 101.3528942 | 0.025864947 | 0.157504491 | 0.682461605 | 0.889273568 | ns | TM | KS.1 |
| FIM1_977 | 2518.771879 | 3067.107874 | 3415.972487 | 0.05470215 | 0.161006678 | 0.411296426 | 0.731735607 | ns | TM | KS.1 |
| FIM1_978 | 2361.770128 | 3661.484263 | 4313.41991 | 0.109339071 | 0.227347357 | 0.156220779 | 0.441088748 | ns | TM | KS.1 |
| FIM1_979 | 1217.687038 | 1835.282709 | 2201.409289 | 0.155900936 | 0.222204808 | 0.071920692 | 0.284298304 | ns | TM | KS.1 |
| FIM1_980 | 2110.598401 | 1965.917509 | 2473.290594 | -0.026355489 | 0.127613566 | 0.679651703 | 0.887492141 | ns | TM | KS.1 |
| FIM1_981 | 1212.56692 | 573.8107379 | 1418.350743 | -0.525526903 | 0.437434811 | 0.003138939 | 0.036399552 | ns | TM | KS.1 |
| FIM1_982 | 1894.599159 | 1205.20209 | 1544.672182 | -0.436710327 | 0.229075685 | 0.000706593 | 0.011628895 | ns | TM | KS.1 |
| FIM1_983 | 13071.20842 | 12281.83772 | 12436.09678 | -0.044200958 | 0.079377815 | 0.400017112 | 0.723667321 | ns | TM | KS.1 |
| FIM1_984 | 53.94776375 | 14.58001856 | 91.15760918 | -0.504730401 | 0.607659187 | 0.004636382 | 0.04900527 | ns | TM | KS.1 |
| FIM1_988 | 3101.051822 | 2266.786324 | 2634.034784 | -0.318378639 | 0.139092886 | 0.000805917 | 0.012933671 | ns | TM | KS.1 |
| FIM1_989 | 54.08500155 | 25.52714206 | 71.54174204 | -0.760134703 | 0.393606509 | 0.00038313 | 0.007359345 | ns | TM | KS.1 |
| FIM1_991 | 1467.113826 | 1315.991155 | 1301.236003 | -0.042335181 | 0.12974006 | 0.515298299 | 0.804775792 | ns | TM | KS.1 |
| FIM1_992 | 2213.808027 | 2141.104458 | 2183.789848 | -0.013648526 | 0.118382541 | 0.827556576 | 0.946035952 | ns | TM | KS.1 |
| FIM1_993 | 1277.032722 | 1508.545061 | 1362.119294 | 0.061343339 | 0.142619724 | 0.363667436 | 0.687524631 | ns | TM | KS.1 |
| FIM1_994 | 1802.600012 | 2052.848015 | 2129.332173 | 0.062681888 | 0.122641185 | 0.343675194 | 0.670129201 | ns | TM | KS.1 |
| FIM1_995 | 1280.201976 | 1897.319081 | 2185.273744 | 0.188758678 | 0.217607338 | 0.041052983 | 0.206928663 | ns | TM | KS.1 |
| FIM1_996 | 1883.653443 | 2846.806104 | 2898.353279 | 0.117080817 | 0.219626023 | 0.136490573 | 0.40881433 | ns | TM | KS.1 |
| FIM1_997 | 90.33422321 | 117.9294823 | 234.4595379 | 0.096130655 | 0.167907549 | 0.189638273 | 0.493436406 | ns | TM | KS.1 |
| FIM1_998 | 1008.503118 | 1047.087443 | 1120.239119 | 0.023309037 | 0.08523683 | 0.669996289 | 0.882274097 | ns | TM | KS.1 |
| FIM1_999 | 1409.694687 | 1389.805768 | 1425.907567 | -0.007766903 | 0.088123116 | 0.889005755 | 0.967028314 | ns | TM | KS.1 |
| FIM1_1000 | 591.7358111 | 552.4861624 | 961.7356256 | -0.01869584 | 0.147015409 | 0.767392231 | 0.924883104 | ns | TM | KS.1 |
| FIM1_1001 | 337.2953246 | 339.7412927 | 323.4595592 | 0.003901504 | 0.080827121 | 0.941161939 | 0.983672404 | ns | TM | KS.1 |
| FIM1_1002 | 353.2890715 | 334.8308701 | 328.5117821 | -0.029687175 | 0.09746153 | 0.614760541 | 0.858612877 | ns | TM | KS.1 |
| FIM1_1003 | 2080.337941 | 3166.890016 | 3583.500302 | 0.220372688 | 0.23277045 | 0.026356355 | 0.159323047 | ns | TM | KS.1 |
| FIM1_1004 | 3893.444687 | 3632.334097 | 3041.230843 | -0.028124873 | 0.121913035 | 0.658106284 | 0.878592731 | ns | TM | KS.1 |
| FIM1_1005 | 1137.919384 | 1134.505791 | 1178.634649 | -0.001557977 | 0.098161653 | 0.978854916 | 0.994649348 | ns | TM | KS.1 |
| FIM1_1006 | 3555.54364 | 3427.144243 | 3500.720483 | -0.018168333 | 0.103998821 | 0.763633227 | 0.924230468 | ns | TM | KS.1 |
| FIM1_1007 | 676.9267378 | 856.0793852 | 680.5163647 | 0.255932525 | 0.101377365 | 0.001015492 | 0.015592815 | ns | TM | KS.1 |
| FIM1_1008 | 305.9536288 | 290.1293003 | 350.3893167 | -0.026376529 | 0.108431097 | 0.668460079 | 0.882180579 | ns | TM | KS.1 |
| FIM1_1009 | 720.9104769 | 685.7242381 | 736.7478592 | -0.02940531 | 0.091720099 | 0.606857644 | 0.851659878 | ns | TM | KS.1 |
| FIM1_1010 | 2117.31752 | 1797.103994 | 2615.082822 | -0.053494239 | 0.147685538 | 0.421799409 | 0.738117813 | ns | TM | KS.1 |
| FIM1_1011 | 12182.53646 | 13244.13004 | 10635.097 | 0.034041793 | 0.123219224 | 0.594868106 | 0.847499664 | ns | TM | KS.1 |
| FIM1_1012 | 2383.227605 | 1844.524243 | 2599.211892 | -0.065758819 | 0.176217168 | 0.334262864 | 0.661742041 | ns | TM | KS.1 |
| FIM1_1013 | 3298.180567 | 3558.618417 | 2768.382107 | 0.029424016 | 0.125394336 | 0.645108778 | 0.873039206 | ns | TM | KS.1 |
| FIM1_1014 | 2009.63594 | 755.6273036 | 2115.903474 | -0.950451171 | 0.484130169 | 0.000294013 | 0.005945992 | ns | TM | KS.1 |
| FIM1_1015 | 1423.27968 | 2039.893442 | 2155.16098 | 0.142354305 | 0.200750929 | 0.085894579 | 0.312600975 | ns | TM | KS.1 |
| FIM1_1016 | 274.9030943 | 200.2337067 | 196.0072282 | -0.132858758 | 0.182979432 | 0.097701914 | 0.3377271 | ns | TM | KS.1 |
| FIM1_1017 | 3174.114848 | 3726.957617 | 4189.521793 | 0.115854028 | 0.115002147 | 0.095115835 | 0.333005827 | ns | TM | KS.1 |
| FIM1_1018 | 1332.412397 | 1372.858641 | 1304.181682 | 0.021687266 | 0.069407371 | 0.649908133 | 0.874068749 | ns | TM | KS.1 |
| FIM1_1019 | 730.9708283 | 789.7121607 | 854.6122633 | 0.038903269 | 0.107929259 | 0.530373579 | 0.808531286 | ns | TM | KS.1 |
| FIM1_1020 | 219.0285692 | 144.1835555 | 134.9193748 | -0.147324785 | 0.224834064 | 0.083308643 | 0.306449621 | ns | TM | KS.1 |
| FIM1_1021 | 356.5542811 | 261.261396 | 651.045516 | -0.091470871 | 0.185499345 | 0.208905908 | 0.515893552 | ns | TM | KS.1 |

| | | | | | | | | | | |
|---|---|---|---|---|---|---|---|---|---|---|
| FIM1_1022 | 6211.31265 | 6770.390994 | 7715.193246 | 0.062745587 | 0.086804691 | 0.263970422 | 0.590340989 | ns | TM | KS.1 |
| FIM1_1024 | 1548.222777 | 1232.945559 | 1785.986354 | -0.091375538 | 0.154504667 | 0.204994464 | 0.512865935 | ns | TM | KS.1 |
| FIM1_1025 | 6852.523702 | 5232.142144 | 5296.423224 | -0.215314941 | 0.147765364 | 0.01386311 | 0.104657014 | ns | TM | KS.1 |
| FIM1_1026 | 8032.300057 | 7148.932591 | 6893.761453 | -0.067219456 | 0.110237231 | 0.295031391 | 0.622741988 | ns | TM | KS.1 |
| FIM1_1027 | 1031.554518 | 1104.168399 | 1025.936303 | 0.030556109 | 0.114455253 | 0.626585344 | 0.866840608 | ns | TM | KS.1 |
| FIM1_1028 | 172.1369472 | 125.5091103 | 216.8554548 | -0.171938208 | 0.181442419 | 0.046841569 | 0.225156332 | ns | TM | KS.1 |
| FIM1_1029 | 11198.49808 | 12492.49297 | 12682.87503 | 0.041356417 | 0.131198292 | 0.524970498 | 0.808531286 | ns | TM | KS.1 |
| FIM1_1030 | 2147.502542 | 3548.00551 | 4336.793453 | 0.184890713 | 0.264246484 | 0.051331171 | 0.235847526 | ns | TM | KS.1 |
| FIM1_1031 | 85.90437086 | 66.24396283 | 81.79274124 | -0.095350219 | 0.165860965 | 0.192120244 | 0.496260756 | ns | TM | KS.1 |
| FIM1_1032 | 1052.210132 | 680.5177336 | 743.9069697 | -0.582848359 | 0.119414144 | 2.8171944450[illegible] | 5.3905932169[illegible] | ns | TM | KS.1 |
| FIM1_1033 | 231.6030349 | 209.1581117 | 215.5946391 | -0.040741437 | 0.126503351 | 0.529151864 | 0.808531286 | ns | TM | KS.1 |
| FIM1_1034 | 690.0438219 | 666.7477851 | 648.1549468 | -0.02318425 | 0.074576553 | 0.644204521 | 0.872978256 | ns | TM | KS.1 |
| FIM1_1035 | 169.7245071 | 192.8764576 | 164.1932233 | 0.063252017 | 0.119018083 | 0.335637156 | 0.663406774 | ns | TM | KS.1 |
| FIM1_1036 | 683.5977094 | 712.9414084 | 581.8946502 | 0.024543304 | 0.088032422 | 0.659831991 | 0.878874681 | ns | TM | KS.1 |
| FIM1_1037 | 1196.183604 | 1337.914501 | 1268.132034 | 0.119383492 | 0.074927393 | 0.027438705 | 0.163286551 | ns | TM | KS.1 |
| FIM1_1038 | 3428.336553 | 3669.128474 | 3371.647619 | 0.057714685 | 0.068432345 | 0.225259063 | 0.541907079 | ns | TM | KS.1 |
| FIM1_1039 | 2772.89885 | 2203.339397 | 3203.341312 | -0.227521072 | 0.11596812 | 0.004827525 | 0.050668642 | ns | TM | KS.1 |
| FIM1_1040 | 1983.691987 | 1514.498554 | 1696.149232 | -0.14186256 | 0.162345233 | 0.076929508 | 0.294813117 | ns | TM | KS.1 |
| FIM1_1041 | 2044.863168 | 1565.229013 | 2064.977982 | -0.091626122 | 0.169777661 | 0.206732644 | 0.513839538 | ns | TM | KS.1 |
| FIM1_1042 | 74.21814 | 63.24078923 | 77.10730827 | -0.067065368 | 0.136153664 | 0.323635307 | 0.650941236 | ns | TM | KS.1 |
| FIM1_1043 | 254.7503091 | 202.7504169 | 188.1524893 | -0.086289015 | 0.155834852 | 0.226636305 | 0.54408821 | ns | TM | KS.1 |
| FIM1_1044 | 1593.096882 | 1767.129191 | 2014.807761 | 0.039845712 | 0.129478426 | 0.538933812 | 0.814399145 | ns | TM | KS.1 |
| FIM1_1045 | 571.3824356 | 534.1998096 | 478.7153123 | -0.03854096 | 0.09701535 | 0.514341937 | 0.804256034 | ns | TM | KS.1 |
| FIM1_1046 | 914.182825 | 897.2765208 | 813.5084914 | -0.01105707 | 0.085778161 | 0.840130129 | 0.949982862 | ns | TM | KS.1 |
| FIM1_1047 | 1485.762561 | 1466.34716 | 1493.593789 | -0.010190295 | 0.056567459 | 0.805147776 | 0.937204068 | ns | TM | KS.1 |
| FIM1_1048 | 1588.968172 | 1773.723513 | 1727.147317 | 0.084585873 | 0.093159683 | 0.15696335 | 0.442181577 | ns | TM | KS.1 |
| FIM1_1049 | 1424.078608 | 2307.856707 | 2601.580577 | 0.189408213 | 0.25757851 | 0.047108615 | 0.226003239 | ns | TM | KS.1 |
| FIM1_1050 | 206.5309202 | 151.2358101 | 107.7630703 | -0.111967972 | 0.182942613 | 0.143421204 | 0.420432957 | ns | TM | KS.1 |
| FIM1_1051 | 841.7318173 | 638.3946367 | 591.553105 | -0.196761033 | 0.15679959 | 0.023571154 | 0.150553405 | ns | TM | KS.1 |
| FIM1_1052 | 2355.841157 | 1894.825637 | 1052.997585 | -0.105892055 | 0.1461317 | 0.150531843 | 0.431640299 | ns | TM | KS.1 |
| FIM1_1053 | 105.5228463 | 104.4125994 | 89.13311434 | -0.003301022 | 0.118956191 | 0.95790662 | 0.989353426 | ns | TM | KS.1 |
| FIM1_1054 | 1897.524658 | 2039.134695 | 2238.829363 | 0.053701203 | 0.079447746 | 0.308512419 | 0.636779799 | ns | TM | KS.1 |
| FIM1_1055 | 285.4034431 | 252.1016813 | 204.8296976 | -0.06244456 | 0.118006669 | 0.340099525 | 0.668193429 | ns | TM | KS.1 |
| FIM1_1056 | 451.1038754 | 549.5133425 | 564.8640782 | 0.132193622 | 0.130610446 | 0.076833234 | 0.294813117 | ns | TM | KS.1 |
| FIM1_1057 | 1271.788476 | 1284.954421 | 1415.285574 | 0.006432482 | 0.073932912 | 0.897426027 | 0.967972723 | ns | TM | KS.1 |
| FIM1_1058 | 2957.57714 | 3215.964315 | 2535.405671 | 0.061380767 | 0.085372594 | 0.268968795 | 0.596620808 | ns | TM | KS.1 |
| FIM1_1060 | 2437.761467 | 3765.23557 | 4530.229662 | 0.347934894 | 0.235860968 | 0.003469638 | 0.039377778 | ns | TM | KS.1 |
| FIM1_1061 | 585.7925031 | 683.2209821 | 795.8899212 | 0.101281117 | 0.116616738 | 0.139251677 | 0.412895859 | ns | TM | KS.1 |
| FIM1_1062 | 760.710982 | 745.775908 | 854.6353268 | -0.012388215 | 0.075565087 | 0.806702009 | 0.937659684 | ns | TM | KS.1 |
| FIM1_1063 | 816.3566146 | 714.2705968 | 789.7390678 | -0.084896759 | 0.111095178 | 0.196327516 | 0.502950252 | ns | TM | KS.1 |
| FIM1_1064 | 1266.814952 | 1154.405872 | 1260.751839 | -0.050618111 | 0.107108335 | 0.41732367 | 0.735551151 | ns | TM | KS.1 |
| FIM1_1065 | 1292.713083 | 1183.572386 | 1242.392912 | -0.069355996 | 0.084564352 | 0.211229422 | 0.51871941 | ns | TM | KS.1 |
| FIM1_1066 | 406.6612176 | 496.6631307 | 442.4206693 | 0.165472008 | 0.120997876 | 0.029485841 | 0.170326428 | ns | TM | KS.1 |
| FIM1_1067 | 414.4067447 | 438.1885198 | 409.8270158 | 0.031806729 | 0.094969106 | 0.585187094 | 0.841613393 | ns | TM | KS.1 |
| FIM1_1068 | 246.155414 | 153.7640628 | 234.1648906 | -0.279408071 | 0.264566244 | 0.013103625 | 0.101458781 | ns | TM | KS.1 |

| | | | | | | | | | | |
|---|---|---|---|---|---|---|---|---|---|---|
| FIM1_1069 | 374.5427503 | 417.1080973 | 417.8969508 | 0.030304968 | 0.14750195 | 0.636106431 | 0.869849417 | ns | TM | KS.1 |
| FIM1_1070 | 1392.05441 | 1274.863371 | 1189.605498 | -0.064770331 | 0.087270747 | 0.251114963 | 0.576776057 | ns | TM | KS.1 |
| FIM1_1071 | 1543.336602 | 1439.629782 | 1522.70044 | -0.063535894 | 0.064225434 | 0.163634261 | 0.45366779 | ns | TM | KS.1 |
| FIM1_1072 | 609.1932488 | 965.5635963 | 1457.758092 | 0.154853987 | 0.241610418 | 0.076469436 | 0.294813117 | ns | TM | KS.1 |
| FIM1_1074 | 1307.328474 | 1213.800693 | 1221.662544 | -0.047528064 | 0.091027588 | 0.407144339 | 0.728957685 | ns | TM | KS.1 |
| FIM1_1075 | 2689.229167 | 2489.160606 | 3065.067165 | -0.047710986 | 0.094940926 | 0.416571144 | 0.735551151 | ns | TM | KS.1 |
| FIM1_1076 | 441.2153649 | 493.0969618 | 428.2304838 | 0.04036076 | 0.133020897 | 0.534876473 | 0.812026382 | ns | TM | KS.1 |
| FIM1_1077 | 87.01152882 | 104.1885077 | 125.6029179 | 0.046428349 | 0.161165619 | 0.477818171 | 0.77913648 | ns | TM | KS.1 |
| FIM1_1078 | 240.7854744 | 291.3888731 | 294.9642808 | 0.10712271 | 0.133378049 | 0.138729215 | 0.412292618 | ns | TM | KS.1 |
| FIM1_1079 | 349.686034 | 365.3808115 | 387.3355343 | 0.01822288 | 0.115519939 | 0.770476559 | 0.925705893 | ns | TM | KS.1 |
| FIM1_1080 | 1028.623288 | 1105.618292 | 1084.702492 | 0.025508651 | 0.130250013 | 0.689598018 | 0.892497849 | ns | TM | KS.1 |
| FIM1_1081 | 1010.759126 | 1462.323519 | 1316.689336 | 0.143721151 | 0.204488617 | 0.084615933 | 0.310216851 | ns | TM | KS.1 |
| FIM1_1082 | 988.8768295 | 760.6991772 | 1101.869711 | -0.1259835 | 0.161213614 | 0.105111665 | 0.351195791 | ns | TM | KS.1 |
| FIM1_1083 | 1779.889858 | 1408.564756 | 1554.803504 | -0.249210899 | 0.106686371 | 0.001701536 | 0.022634069 | ns | TM | KS.1 |
| FIM1_1084 | 1005.48967 | 697.1605241 | 682.8854909 | -0.301565505 | 0.19100274 | 0.004195512 | 0.045081369 | ns | TM | KS.1 |
| FIM1_1085 | 582.8338814 | 618.412334 | 580.9697454 | 0.038913946 | 0.083282385 | 0.47205002 | 0.772376515 | ns | TM | KS.1 |
| FIM1_1086 | 940.789988 | 832.9009993 | 635.0051541 | -0.073874204 | 0.109845595 | 0.252501961 | 0.579042268 | ns | TM | KS.1 |
| FIM1_1087 | 614.7917774 | 691.4407537 | 755.4464923 | 0.076048347 | 0.106067034 | 0.232308417 | 0.549826059 | ns | TM | KS.1 |
| FIM1_1088 | 2052.059489 | 2178.681121 | 2300.985618 | 0.047491461 | 0.069986332 | 0.324282306 | 0.651040335 | ns | TM | KS.1 |
| FIM1_1089 | 168.0331628 | 143.6173745 | 127.3354343 | -0.067305922 | 0.133605931 | 0.320863389 | 0.649347037 | ns | TM | KS.1 |
| FIM1_1090 | 4121.288203 | 4358.876269 | 4998.7929 | 0.036988378 | 0.08361273 | 0.495101614 | 0.792222775 | ns | TM | KS.1 |
| FIM1_1091 | 1345.270701 | 2058.33429 | 2368.850307 | 0.257429538 | 0.236893032 | 0.014993091 | 0.11083303 | ns | TM | KS.1 |
| FIM1_1092 | 863.4924824 | 1036.330039 | 980.3221085 | 0.194144953 | 0.095301322 | 0.005685903 | 0.057030983 | ns | TM | KS.1 |
| FIM1_1093 | 1540.571389 | 1006.806735 | 1159.130995 | -0.481624365 | 0.178763624 | 5.25895144406 | 0.001437543 | ns | TM | KS.1 |
| FIM1_1094 | 1302.820957 | 1459.180667 | 1676.028457 | 0.128453745 | 0.067924221 | 0.012420822 | 0.098138201 | ns | TM | KS.1 |
| FIM1_1095 | 1036.881874 | 1072.11651 | 1197.186712 | 0.022833785 | 0.075569537 | 0.65211409 | 0.87555594 | ns | TM | KS.1 |
| FIM1_1096 | 375.7423134 | 417.2129003 | 436.598392 | 0.052508138 | 0.114638279 | 0.412867067 | 0.732913544 | ns | TM | KS.1 |
| FIM1_1097 | 4728.554882 | 5526.157987 | 8368.502534 | 0.082242276 | 0.125441782 | 0.229644609 | 0.547690282 | ns | TM | KS.1 |
| FIM1_1098 | 3441.896826 | 3395.988068 | 3522.134248 | -0.006397248 | 0.104944814 | 0.915715184 | 0.975605516 | ns | TM | KS.1 |
| FIM1_1099 | 806.8191663 | 769.8373124 | 822.1322244 | -0.024598733 | 0.100211836 | 0.679963863 | 0.887492141 | ns | TM | KS.1 |
| FIM1_1100 | 5874.946275 | 7295.592749 | 7850.830426 | 0.247559817 | 0.091446927 | 0.000702807 | 0.01161617 | ns | TM | KS.1 |
| FIM1_1101 | 73.98161423 | 72.80718482 | 78.11312331 | -0.001506156 | 0.186186758 | 0.980115924 | 0.995320825 | ns | TM | KS.1 |
| FIM1_1102 | 1222.704106 | 1434.222076 | 1220.438134 | 0.116763178 | 0.114262767 | 0.091986024 | 0.327580865 | ns | TM | KS.1 |
| FIM1_1103 | 1062.556341 | 1066.378395 | 973.7573959 | 0.001287434 | 0.127257009 | 0.983703198 | 0.996034691 | ns | TM | KS.1 |
| FIM1_1104 | 1615.001074 | 1701.876415 | 1469.220352 | 0.024742573 | 0.10925168 | 0.688364625 | 0.891826565 | ns | TM | KS.1 |
| FIM1_1105 | 3409.841525 | 4379.741756 | 4789.31649 | 0.31564956 | 0.074496079 | 1.69571571272 | 0.000109382 | ns | TM | KS.1 |
| FIM1_1106 | 6529.131681 | 6877.60752 | 6673.423361 | 0.02724125 | 0.101303696 | 0.649752363 | 0.874068749 | ns | TM | KS.1 |
| FIM1_1107 | 982.5526949 | 1132.358009 | 984.9406452 | 0.083401382 | 0.117235482 | 0.21360895 | 0.521776083 | ns | TM | KS.1 |
| FIM1_1108 | 927.9197845 | 1547.902266 | 1788.027479 | 0.265914867 | 0.28588422 | 0.01804024 | 0.127124924 | ns | TM | KS.1 |
| FIM1_1109 | 3895.741334 | 4127.585698 | 4698.985873 | 0.031062039 | 0.100349534 | 0.603736781 | 0.851547247 | ns | TM | KS.1 |
| FIM1_1110 | 1197.307776 | 1444.675098 | 1348.385957 | 0.125940468 | 0.127400607 | 0.086137283 | 0.312966141 | ns | TM | KS.1 |
| FIM1_1111 | 1706.951648 | 1242.166046 | 1127.910016 | -0.347487547 | 0.127298636 | 0.000182949 | 0.004027314 | ns | TM | KS.1 |
| FIM1_1112 | 3020.917839 | 2936.108633 | 3978.287789 | -0.018358522 | 0.078190604 | 0.722966582 | 0.907127048 | ns | TM | KS.1 |
| FIM1_1113 | 1204.787446 | 1341.695622 | 1419.349751 | 0.035206053 | 0.138961343 | 0.586115402 | 0.841613393 | ns | TM | KS.1 |
| FIM1_1114 | 531.2987494 | 252.3323152 | 2368.019001 | -0.385329285 | 0.41841359 | 0.009150377 | 0.079585886 | ns | TM | KS.1 |

| | | | | | | | | | | |
|---|---|---|---|---|---|---|---|---|---|---|
| FIM1_1115 | 208.6458113 | 204.658717 | 361.7384784 | -0.006273769 | 0.130708308 | 0.920851561 | 0.975881018 | ns | TM | KS.1 |
| FIM1_1116 | 5800.50297 | 7688.946828 | 6624.83979 | 0.228812868 | 0.151404072 | 0.01075834 | 0.088908211 | ns | TM | KS.1 |
| FIM1_1117 | 2522.556138 | 2513.576797 | 1887.62211 | -0.001870985 | 0.098902088 | 0.974693752 | 0.993872484 | ns | TM | KS.1 |
| FIM1_1118 | 9.546977461 | 16.77167302 | 20.87135697 | 0.160998624 | 0.273559126 | 0.073821032 | 0.288677781 | ns | TM | KS.1 |
| FIM1_1119 | 2790.600627 | 2501.273603 | 2937.395474 | -0.068788411 | 0.103693451 | 0.271151292 | 0.598713828 | ns | TM | KS.1 |
| FIM1_1120 | 35.85916937 | 26.04475066 | 43.48615303 | -0.118748933 | 0.186433763 | 0.127336216 | 0.393568488 | ns | TM | KS.1 |
| FIM1_1121 | 12496.2386 | 12652.31126 | 12996.19495 | 0.006806799 | 0.092919113 | 0.905371581 | 0.971575413 | ns | TM | KS.1 |
| FIM1_1122 | 4725.476552 | 5195.334511 | 5340.92505 | 0.088363305 | 0.074589356 | 0.088121685 | 0.317685061 | ns | TM | KS.1 |
| FIM1_1123 | 426.0730002 | 446.6773804 | 448.2279881 | 0.026984678 | 0.096105559 | 0.644683621 | 0.873039206 | ns | TM | KS.1 |
| FIM1_1124 | 2932.13931 | 2902.18902 | 3235.534849 | -0.005759917 | 0.088798246 | 0.917905259 | 0.975634767 | ns | TM | KS.1 |
| FIM1_1125 | 18682.52866 | 23133.37669 | 21740.66123 | 0.177884021 | 0.125990091 | 0.023661756 | 0.150569038 | ns | TM | KS.1 |
| FIM1_1126 | 939.9045227 | 544.5726311 | 764.9335658 | -0.3185138 | 0.311175063 | 0.010637059 | 0.088198944 | ns | TM | KS.1 |
| FIM1_1127 | 502.1208061 | 426.1860529 | 411.6442297 | -0.060704025 | 0.142106807 | 0.368038884 | 0.691630801 | ns | TM | KS.1 |
| FIM1_1128 | 252.2680421 | 169.3888685 | 192.2800568 | -0.16900349 | 0.218656528 | 0.057305736 | 0.25255083 | ns | TM | KS.1 |
| FIM1_1129 | 3229.464976 | 3079.230566 | 3127.035772 | -0.030996048 | 0.082080179 | 0.562422375 | 0.826847315 | ns | TM | KS.1 |
| FIM1_1130 | 429.6269848 | 490.354752 | 445.2423528 | 0.074586732 | 0.116746105 | 0.259944461 | 0.588210523 | ns | TM | KS.1 |
| FIM1_1131 | 1260.608575 | 1359.329988 | 1327.843283 | 0.052964757 | 0.085625 | 0.338840571 | 0.667250632 | ns | TM | KS.1 |
| FIM1_1132 | 575.4302408 | 440.9737222 | 990.3518956 | -0.137154096 | 0.161450673 | 0.084215918 | 0.309206046 | ns | TM | KS.1 |
| FIM1_1133 | 423.173289 | 555.3175669 | 653.9017944 | 0.112289427 | 0.167180469 | 0.139264171 | 0.412895859 | ns | TM | KS.1 |
| FIM1_1134 | 306.9765236 | 336.8305643 | 307.6470545 | 0.030211938 | 0.136473831 | 0.638239687 | 0.870535085 | ns | TM | KS.1 |
| FIM1_1135 | 10151.67109 | 12985.56765 | 14001.82884 | 0.10560068 | 0.158146951 | 0.155973527 | 0.441000098 | ns | TM | KS.1 |
| FIM1_1136 | 1595.218366 | 1165.852876 | 1073.687292 | -0.303141514 | 0.146778903 | 0.001582582 | 0.021279312 | ns | TM | KS.1 |
| FIM1_1137 | 4954.805697 | 4425.379962 | 3821.963831 | -0.037937757 | 0.138319119 | 0.558882957 | 0.82614557 | ns | TM | KS.1 |
| FIM1_1138 | 882.8159434 | 607.6655556 | 592.9896694 | -0.426578338 | 0.145593022 | 3.6714641528[illegible] | 0.001112321 | ns | TM | KS.1 |
| FIM1_1139 | 10132.65936 | 8606.233105 | 9084.169046 | -0.16666048 | 0.094658456 | 0.013230095 | 0.101775569 | ns | TM | KS.1 |
| FIM1_1140 | 5310.18481 | 3397.012278 | 3299.452156 | -0.126394289 | 0.231872545 | 0.11849592 | 0.377411781 | ns | TM | KS.1 |
| FIM1_1141 | 6056.717014 | 3802.289531 | 3765.705268 | -0.1869435 | 0.249844462 | 0.04769465 | 0.22749845 | ns | TM | KS.1 |
| FIM1_1142 | 904.8814073 | 817.6858416 | 869.6513995 | -0.077323525 | 0.090671545 | 0.185336679 | 0.488144311 | ns | TM | KS.1 |
| FIM1_1143 | 1543.676376 | 1669.242414 | 1700.876473 | 0.060606936 | 0.078367073 | 0.247623187 | 0.572190133 | ns | TM | KS.1 |
| FIM1_1144 | 857.7703719 | 976.0083628 | 867.3689057 | 0.052534232 | 0.130793078 | 0.426470165 | 0.741071977 | ns | TM | KS.1 |
| FIM1_1145 | 6312.903991 | 8048.359456 | 9504.782481 | 0.279878113 | 0.094705063 | 0.000251972 | 0.005293345 | ns | TM | KS.1 |
| FIM1_1146 | 792.4583184 | 665.7261562 | 629.5973332 | -0.121582251 | 0.121053851 | 0.089135333 | 0.320179266 | ns | TM | KS.1 |
| FIM1_1147 | 2475.018477 | 3042.691445 | 3609.849336 | 0.210099493 | 0.106880851 | 0.005753338 | 0.057381231 | ns | TM | KS.1 |
| FIM1_1148 | 575.6112276 | 680.61942 | 672.4674812 | 0.103647858 | 0.123146053 | 0.139790264 | 0.413618606 | ns | TM | KS.1 |
| FIM1_1149 | 298.864061 | 278.9470661 | 286.7529395 | -0.033458845 | 0.109245386 | 0.589613496 | 0.844122919 | ns | TM | KS.1 |
| FIM1_1150 | 902.5111672 | 976.257743 | 929.2402497 | 0.042385 | 0.104362973 | 0.48944879 | 0.788598425 | ns | TM | KS.1 |
| FIM1_1151 | 5313.528104 | 4972.809732 | 6165.423123 | -0.037286028 | 0.09846019 | 0.530902231 | 0.808531286 | ns | TM | KS.1 |
| FIM1_1152 | 697.4243458 | 725.982648 | 803.2191854 | 0.014635427 | 0.12551493 | 0.816878698 | 0.942042541 | ns | TM | KS.1 |
| FIM1_1153 | 59.47310736 | 76.9682447 | 75.4474694 | 0.116594497 | 0.16543902 | 0.127741986 | 0.393996515 | ns | TM | KS.1 |
| FIM1_1154 | 1630.067596 | 1523.791494 | 1354.118306 | -0.045501076 | 0.084735527 | 0.406624501 | 0.728526463 | ns | TM | KS.1 |
| FIM1_1155 | 89.95524906 | 77.69008261 | 96.14274037 | -0.032944733 | 0.164050453 | 0.604584714 | 0.851547247 | ns | TM | KS.1 |
| FIM1_1156 | 8136.662783 | 9044.600881 | 9766.387008 | 0.072790982 | 0.097591462 | 0.230824274 | 0.548995638 | ns | TM | KS.1 |
| FIM1_1157 | 3014.211589 | 2689.694355 | 2947.350756 | -0.096644905 | 0.089060487 | 0.100584007 | 0.342978366 | ns | TM | KS.1 |
| FIM1_1158 | 1260.859332 | 1270.412082 | 1034.973562 | 0.002298338 | 0.135710828 | 0.97093271 | 0.99362472 | ns | TM | KS.1 |
| FIM1_1159 | 1750.87978 | 1941.4519 | 1818.015142 | 0.096072863 | 0.07978701 | 0.079192677 | 0.299835289 | ns | TM | KS.1 |

| | | | | | | | | | | |
|---|---|---|---|---|---|---|---|---|---|---|
| FIM1_1160 | 10865.13048 | 8741.625416 | 13109.57996 | -0.260176661 | 0.082353574 | 0.000175696 | 0.003884834 | ns | TM | KS.1 |
| FIM1_1161 | 67.70405304 | 49.91915823 | 118.611153 | -0.077356943 | 0.188634954 | 0.270049505 | 0.597663125 | ns | TM | KS.1 |
| FIM1_1162 | 6575.633478 | 7173.980331 | 6707.310668 | 0.023958913 | 0.147326761 | 0.706127974 | 0.899352156 | ns | TM | KS.1 |
| FIM1_1163 | 964.7447038 | 907.8688316 | 1155.211628 | -0.044011899 | 0.076358939 | 0.389620484 | 0.713295089 | ns | TM | KS.1 |
| FIM1_1164 | 36.93649441 | 41.85135407 | 38.21550992 | 0.037057668 | 0.152218991 | 0.565923791 | 0.828754274 | ns | TM | KS.1 |
| FIM1_1165 | 871.1632754 | 975.7102392 | 916.3283181 | 0.060408453 | 0.113454468 | 0.348557562 | 0.674195903 | ns | TM | KS.1 |
| FIM1_1166 | 3960.055872 | 3212.519982 | 6055.591906 | -0.151589004 | 0.131319903 | 0.049016638 | 0.230707449 | ns | TM | KS.1 |
| FIM1_1167 | 840.4500786 | 828.2955104 | 888.8186405 | -0.007805344 | 0.092699712 | 0.891448877 | 0.967215407 | ns | TM | KS.1 |
| FIM1_1168 | 1435.626572 | 1553.588294 | 1564.529 | 0.053747567 | 0.088563613 | 0.342161304 | 0.669124405 | ns | TM | KS.1 |
| FIM1_1169 | 1000.414416 | 963.8963522 | 944.2420511 | -0.019340483 | 0.100130108 | 0.745114478 | 0.915747166 | ns | TM | KS.1 |
| FIM1_1170 | 1091.494097 | 1098.299384 | 1068.3569 | 0.003412104 | 0.092180785 | 0.952274522 | 0.987816044 | ns | TM | KS.1 |
| FIM1_1171 | 44.88993486 | 38.12844905 | 54.07909024 | -0.056867846 | 0.144497651 | 0.396039846 | 0.719349482 | ns | TM | KS.1 |
| FIM1_1172 | 3136.677637 | 2701.491526 | 2917.458834 | -0.167394933 | 0.080141682 | 0.00607514 | 0.059730875 | ns | TM | KS.1 |
| FIM1_1173 | 860.5164882 | 702.8618161 | 611.6997115 | -0.106424097 | 0.139504652 | 0.145237416 | 0.423239382 | ns | TM | KS.1 |
| FIM1_1174 | 464.4780925 | 311.2403445 | 552.607912 | -0.224871792 | 0.222128018 | 0.023024079 | 0.148299176 | ns | TM | KS.1 |
| FIM1_1175 | 194.7933321 | 156.574091 | 154.079058 | -0.081054157 | 0.152138334 | 0.250080915 | 0.575730011 | ns | TM | KS.1 |
| FIM1_1177 | 2504.536125 | 2292.539688 | 2264.533237 | -0.075016269 | 0.077754385 | 0.154240921 | 0.437933732 | ns | TM | KS.1 |
| FIM1_1178 | 9545.309194 | 12736.0968 | 14048.49393 | 0.248092337 | 0.149529971 | 0.006645186 | 0.064069383 | ns | TM | KS.1 |
| FIM1_1179 | 818.6122134 | 910.404284 | 940.2419267 | 0.084562653 | 0.091464139 | 0.15241405 | 0.434782052 | ns | TM | KS.1 |
| FIM1_1180 | 1775.004445 | 954.5925272 | 1300.328035 | -0.84796254 | 0.161126086 | 6.40678734606 | 1.02818603376 | ns | TM | KS.1 |
| FIM1_1181 | 2502.564121 | 2175.592812 | 2233.461848 | -0.070647004 | 0.123076238 | 0.29155078 | 0.618644347 | ns | TM | KS.1 |
| FIM1_1182 | 67.48473305 | 55.42703682 | 106.0153218 | -0.074514803 | 0.144841775 | 0.282306276 | 0.608260599 | ns | TM | KS.1 |
| FIM1_1183 | 812.1486351 | 550.9966133 | 1349.595068 | -0.331016856 | 0.199730932 | 0.002717449 | 0.032342849 | ns | TM | KS.1 |
| FIM1_1184 | 633.7923369 | 654.9967769 | 592.4635588 | 0.017379113 | 0.09972976 | 0.769660018 | 0.925705893 | ns | TM | KS.1 |
| FIM1_1186 | 622.8132607 | 671.5014717 | 717.6328512 | 0.06478282 | 0.076068027 | 0.208258183 | 0.515579507 | ns | TM | KS.1 |
| FIM1_1187 | 5185.524024 | 5472.425358 | 6179.320756 | 0.050191893 | 0.053459815 | 0.205815089 | 0.512865935 | ns | TM | KS.1 |
| FIM1_1188 | 321.8012408 | 366.8636844 | 1370.245573 | 0.052605038 | 0.131811421 | 0.426330243 | 0.741071977 | ns | TM | KS.1 |
| FIM1_1189 | 882.7236067 | 789.7637371 | 1684.525144 | -0.037899403 | 0.137370726 | 0.559233978 | 0.826310971 | ns | TM | KS.1 |
| FIM1_1190 | 372.7643733 | 236.9946831 | 919.1149451 | -0.248262159 | 0.252529174 | 0.019161803 | 0.132384765 | ns | TM | KS.1 |
| FIM1_1191 | 22.5877101 | 32.89257704 | 87.82668476 | 0.063977863 | 0.217847511 | 0.340222958 | 0.668193429 | ns | TM | KS.1 |
| FIM1_1192 | 1073.39212 | 1036.074278 | 1024.594112 | -0.01643085 | 0.108415751 | 0.788653446 | 0.933067989 | ns | TM | KS.1 |
| FIM1_1193 | 4071.149935 | 4796.553199 | 4984.269302 | 0.202103355 | 0.069280039 | 0.000554427 | 0.009746554 | ns | TM | KS.1 |
| FIM1_1194 | 2861.175438 | 2840.089708 | 3125.589537 | -0.004383162 | 0.078700645 | 0.932796769 | 0.979974437 | ns | TM | KS.1 |
| FIM1_1195 | 499.9341855 | 415.8351353 | 301.4078431 | -0.112333008 | 0.129001669 | 0.119870621 | 0.379844802 | ns | TM | KS.1 |
| FIM1_1196 | 14129.53432 | 17318.69138 | 16415.82363 | 0.176996572 | 0.119953654 | 0.021388674 | 0.14253455 | ns | TM | KS.1 |
| FIM1_1197 | 5406.721451 | 4846.04499 | 4171.618958 | -0.077158716 | 0.097467277 | 0.205506083 | 0.512865935 | ns | TM | KS.1 |
| FIM1_1198 | 1051.297177 | 1264.372621 | 1483.304579 | 0.090250315 | 0.136128445 | 0.201810216 | 0.509840799 | ns | TM | KS.1 |
| FIM1_1199 | 1208.394616 | 1341.160253 | 1383.38834 | 0.078866463 | 0.091896975 | 0.180921167 | 0.480985595 | ns | TM | KS.1 |
| FIM1_1200 | 3952.49942 | 3551.825158 | 4476.768721 | -0.044403826 | 0.125659054 | 0.494287375 | 0.792222775 | ns | TM | KS.1 |
| FIM1_1201 | 233.0811869 | 225.7603211 | 205.2568071 | -0.017488651 | 0.105419622 | 0.773319391 | 0.925733395 | ns | TM | KS.1 |
| FIM1_1202 | 369.2125013 | 351.2363785 | 329.0836406 | -0.01867502 | 0.126036967 | 0.768244553 | 0.925179691 | ns | TM | KS.1 |
| FIM1_1203 | 2206.789203 | 1120.833686 | 1165.203984 | -0.61297536 | 0.38218864 | 0.00099683 | 0.015497584 | ns | TM | KS.1 |
| FIM1_1204 | 3906.26703 | 3330.371614 | 4190.507741 | -0.095687943 | 0.121731368 | 0.166272353 | 0.456988595 | ns | TM | KS.1 |
| FIM1_1205 | 1161.384442 | 825.2221893 | 847.5103943 | -0.363735371 | 0.1439644 | 0.000255891 | 0.005348977 | ns | TM | KS.1 |
| FIM1_1206 | 811.3872169 | 780.7989086 | 779.5023305 | -0.024012775 | 0.082868195 | 0.655164772 | 0.87699044 | ns | TM | KS.1 |

| | | | | | | | | | | |
|---|---|---|---|---|---|---|---|---|---|---|
| FIM1_1207 | 1629.962658 | 1808.574587 | 1871.703589 | 0.077005902 | 0.092449629 | 0.192321461 | 0.496423026 | ns | TM | KS.1 |
| FIM1_1208 | 1740.692363 | 1578.027726 | 1630.311595 | -0.07057881 | 0.092505564 | 0.229759269 | 0.547701181 | ns | TM | KS.1 |
| FIM1_1209 | 1017.453542 | 1075.386532 | 1031.406901 | 0.040450529 | 0.074145914 | 0.419377064 | 0.737776855 | ns | TM | KS.1 |
| FIM1_1210 | 605.8301385 | 468.6612026 | 392.2183859 | -0.132950097 | 0.157806812 | 0.090303071 | 0.323439726 | ns | TM | KS.1 |
| FIM1_1211 | 1939.475055 | 1883.372975 | 1425.858515 | -0.015989465 | 0.095312461 | 0.783084199 | 0.930239707 | ns | TM | KS.1 |
| FIM1_1212 | 7521.046002 | 8939.514156 | 9933.849973 | 0.158899785 | 0.106484772 | 0.025020881 | 0.155210576 | ns | TM | KS.1 |
| FIM1_1213 | 7946.9316 | 9034.632634 | 8591.439274 | 0.085117203 | 0.107362109 | 0.188174236 | 0.491426155 | ns | TM | KS.1 |
| FIM1_1214 | 15548.30828 | 18675.26191 | 18090.46764 | 0.192658336 | 0.097074149 | 0.00648235 | 0.062742592 | ns | TM | KS.1 |
| FIM1_1215 | 951.1876224 | 892.8846972 | 930.5813569 | -0.046741537 | 0.077507961 | 0.366158018 | 0.690275157 | ns | TM | KS.1 |
| FIM1_1216 | 748.4054528 | 811.6559467 | 932.4318863 | 0.043315071 | 0.105545225 | 0.482371965 | 0.783685372 | ns | TM | KS.1 |
| FIM1_1217 | 2027.531226 | 3431.068663 | 3883.21751 | 0.274322915 | 0.294317411 | 0.016886555 | 0.121052755 | ns | TM | KS.1 |
| FIM1_1218 | 559.0175691 | 583.3605627 | 430.1686544 | 0.022745293 | 0.09913646 | 0.701290252 | 0.897235589 | ns | TM | KS.1 |
| FIM1_1219 | 1145.65423 | 1129.009215 | 1348.642288 | -0.009267539 | 0.075504394 | 0.854679933 | 0.955083708 | ns | TM | KS.1 |
| FIM1_1220 | 508.3586767 | 475.9737635 | 623.5037668 | -0.027550416 | 0.119014994 | 0.663112887 | 0.880619203 | ns | TM | KS.1 |
| FIM1_1221 | 803.8542502 | 806.0217095 | 911.9478004 | 0.001725732 | 0.067503042 | 0.970638981 | 0.99362472 | ns | TM | KS.1 |
| FIM1_1222 | 5591.363265 | 5506.943646 | 5569.113365 | -0.013567475 | 0.046230946 | 0.703611714 | 0.898252804 | ns | TM | KS.1 |
| FIM1_1223 | 338.4597073 | 304.3300288 | 358.2853896 | -0.056753801 | 0.113513228 | 0.376687893 | 0.700569072 | ns | TM | KS.1 |
| FIM1_1224 | 339.7120832 | 292.4685795 | 283.1675351 | -0.094431796 | 0.116940755 | 0.164682149 | 0.454405819 | ns | TM | KS.1 |
| FIM1_1225 | 1747.905334 | 1840.364327 | 2384.065711 | 0.027950662 | 0.098854965 | 0.637645147 | 0.870169358 | ns | TM | KS.1 |
| FIM1_1227 | 3690.930297 | 3422.133213 | 4298.800342 | -0.03359374 | 0.11672844 | 0.595504648 | 0.848163648 | ns | TM | KS.1 |
| FIM1_1228 | 1350.452497 | 1275.909209 | 1288.517643 | -0.030336435 | 0.100280978 | 0.611978282 | 0.855989876 | ns | TM | KS.1 |
| FIM1_1230 | 1544.106521 | 1880.001863 | 2123.311934 | 0.131124383 | 0.130275509 | 0.078512548 | 0.298395664 | ns | TM | KS.1 |
| FIM1_1231 | 2938.978461 | 3330.487241 | 3724.980557 | 0.068263162 | 0.115356222 | 0.296716211 | 0.624434497 | ns | TM | KS.1 |
| FIM1_1232 | 2747.324891 | 2566.217152 | 3261.024632 | -0.037037131 | 0.101847423 | 0.540082596 | 0.814399145 | ns | TM | KS.1 |
| FIM1_1233 | 1161.095736 | 1428.386179 | 1200.732231 | 0.09867395 | 0.143661086 | 0.173363455 | 0.469250918 | ns | TM | KS.1 |
| FIM1_1234 | 8156.863959 | 9032.325348 | 8850.688557 | 0.082919261 | 0.086710074 | 0.145989041 | 0.424167573 | ns | TM | KS.1 |
| FIM1_1235 | 1811.485422 | 1816.232812 | 2019.227918 | 0.001644333 | 0.084440564 | 0.975834001 | 0.993872484 | ns | TM | KS.1 |
| FIM1_1236 | 330.0792686 | 355.9339232 | 389.0121877 | 0.037993667 | 0.111456845 | 0.544346895 | 0.817435092 | ns | TM | KS.1 |
| FIM1_1239 | 3550.922628 | 2890.48451 | 2726.193429 | -0.10732893 | 0.140587416 | 0.143133702 | 0.419864485 | ns | TM | KS.1 |
| FIM1_1240 | 295.5174012 | 309.1406725 | 277.3851919 | 0.023475428 | 0.104773312 | 0.699065073 | 0.896699645 | ns | TM | KS.1 |
| FIM1_1242 | 719.6612424 | 803.469812 | 774.0457167 | 0.070054714 | 0.104647643 | 0.26516604 | 0.591039896 | ns | TM | KS.1 |
| FIM1_1243 | 3194.308682 | 3328.681257 | 3450.124753 | 0.035620277 | 0.055903052 | 0.383695844 | 0.7083068 | ns | TM | KS.1 |
| FIM1_1244 | 1131.062023 | 939.4366886 | 884.1417876 | -0.162290277 | 0.113816025 | 0.027488233 | 0.163385854 | ns | TM | KS.1 |
| FIM1_1245 | 987.9843445 | 1103.824885 | 1256.399244 | 0.070183446 | 0.103821094 | 0.262478071 | 0.589006948 | ns | TM | KS.1 |
| FIM1_1246 | 652.3131152 | 593.2537269 | 571.5752836 | -0.071141624 | 0.088143835 | 0.212464424 | 0.520438458 | ns | TM | KS.1 |
| FIM1_1247 | 1230.900132 | 894.4679416 | 887.4453676 | -0.317936553 | 0.14516624 | 0.001011719 | 0.015592815 | ns | TM | KS.1 |
| FIM1_1248 | 1780.269131 | 1745.418807 | 1548.077761 | -0.010862132 | 0.093075073 | 0.849738252 | 0.95298643 | ns | TM | KS.1 |
| FIM1_1249 | 3707.735022 | 3140.359959 | 4249.76047 | -0.051069752 | 0.150872062 | 0.440836211 | 0.747832613 | ns | TM | KS.1 |
| FIM1_1250 | 1606.671523 | 1499.233171 | 2503.527131 | -0.034863281 | 0.107261365 | 0.571906712 | 0.832094673 | ns | TM | KS.1 |
| FIM1_1251 | 3715.346583 | 3412.41412 | 3902.077065 | -0.060269824 | 0.087834038 | 0.286129737 | 0.612585758 | ns | TM | KS.1 |
| FIM1_1252 | 1825.276235 | 1699.570366 | 1817.874988 | -0.039542111 | 0.100383927 | 0.510848023 | 0.802123385 | ns | TM | KS.1 |
| FIM1_1253 | 522.8345128 | 418.3146858 | 403.9005518 | -0.226018474 | 0.108793648 | 0.00385615 | 0.04244324 | ns | TM | KS.1 |
| FIM1_1254 | 1192.161475 | 1249.544894 | 1261.360215 | 0.021208229 | 0.112599088 | 0.732898128 | 0.911086503 | ns | TM | KS.1 |
| FIM1_1255 | 1384.893317 | 1979.624311 | 2162.44669 | 0.13089382 | 0.199598132 | 0.104766768 | 0.350588067 | ns | TM | KS.1 |
| FIM1_1256 | 1170.003303 | 1195.937726 | 1269.138431 | 0.013387391 | 0.083008157 | 0.803335138 | 0.936908579 | ns | TM | KS.1 |

| | | | | | | | | | | |
|---|---|---|---|---|---|---|---|---|---|---|
| FIM1_1257 | 1946.999384 | 1663.042788 | 1885.744787 | -0.11083496 | 0.114950349 | 0.10789054 | 0.35617481 | ns | TM | KS.1 |
| FIM1_1258 | 1567.900485 | 1424.414445 | 1152.889444 | -0.063452937 | 0.096296706 | 0.288877998 | 0.616017163 | ns | TM | KS.1 |
| FIM1_1259 | 504.0831996 | 360.821979 | 347.9151025 | -0.21216995 | 0.186466383 | 0.022553386 | 0.146759546 | ns | TM | KS.1 |
| FIM1_1261 | 1748.991507 | 2184.057909 | 2347.620967 | 0.129058002 | 0.143241693 | 0.091100665 | 0.32535952 | ns | TM | KS.1 |
| FIM1_1262 | 5300.27626 | 3682.671461 | 2501.382779 | -0.245208746 | 0.199990766 | 0.013664071 | 0.104063892 | ns | TM | KS.1 |
| FIM1_1263 | 2336.891893 | 2073.760751 | 2066.89919 | -0.065288184 | 0.113809987 | 0.314202199 | 0.640900344 | ns | TM | KS.1 |
| FIM1_1264 | 3803.530141 | 2853.510632 | 2953.260636 | -0.17800923 | 0.16558591 | 0.037768892 | 0.196103802 | ns | TM | KS.1 |
| FIM1_1265 | 4473.383071 | 4924.421375 | 5046.128415 | 0.049555694 | 0.111097912 | 0.433308451 | 0.745662242 | ns | TM | KS.1 |
| FIM1_1266 | 642.1764123 | 839.4411775 | 805.668196 | 0.164476825 | 0.15830006 | 0.047382118 | 0.226713798 | ns | TM | KS.1 |
| FIM1_1267 | 2228.68305 | 3371.772297 | 4172.094383 | 0.167391669 | 0.225036262 | 0.060034419 | 0.257475201 | ns | TM | KS.1 |
| FIM1_1268 | 876.1432311 | 839.8602811 | 779.0367007 | -0.023142198 | 0.095169851 | 0.690649534 | 0.892989583 | ns | TM | KS.1 |
| FIM1_1269 | 1867.561772 | 2820.462045 | 3666.06524 | 0.157330714 | 0.223165859 | 0.070421152 | 0.280725344 | ns | TM | KS.1 |
| FIM1_1270 | 2662.177731 | 2730.870127 | 2826.111165 | 0.015424089 | 0.085147171 | 0.777559641 | 0.928020669 | ns | TM | KS.1 |
| FIM1_1271 | 2921.741129 | 3186.656567 | 3309.465527 | 0.05845768 | 0.091586842 | 0.31273369 | 0.639671112 | ns | TM | KS.1 |
| FIM1_1272 | 5572.055288 | 6932.762856 | 7662.356285 | 0.232514846 | 0.103517131 | 0.002490019 | 0.030511938 | ns | TM | KS.1 |
| FIM1_1273 | 653.5818048 | 617.5555991 | 549.5565453 | -0.026907872 | 0.109330309 | 0.663179902 | 0.880619203 | ns | TM | KS.1 |
| FIM1_1274 | 79.84096248 | 84.32470227 | 79.67623041 | 0.019009262 | 0.1382278 | 0.764736838 | 0.924230468 | ns | TM | KS.1 |
| FIM1_1275 | 3523.563378 | 2979.988869 | 2704.232452 | -0.128435605 | 0.114897038 | 0.068788582 | 0.276450087 | ns | TM | KS.1 |
| FIM1_1276 | 3608.758872 | 3300.124601 | 3207.325652 | -0.105290881 | 0.051040555 | 0.009071661 | 0.079456889 | ns | TM | KS.1 |
| FIM1_1277 | 2190.736187 | 2269.995788 | 2437.463365 | 0.017315868 | 0.104994366 | 0.775153145 | 0.926727011 | ns | TM | KS.1 |
| FIM1_1279 | 1524.459266 | 1179.424869 | 1120.380077 | -0.246322854 | 0.127035916 | 0.004066159 | 0.044168425 | ns | TM | KS.1 |
| FIM1_1280 | 12024.02609 | 10384.19372 | 7070.403574 | -0.120661596 | 0.103867126 | 0.069148807 | 0.27676212 | ns | TM | KS.1 |
| FIM1_1281 | 2197.023508 | 2257.898241 | 2186.258833 | 0.017835773 | 0.077984318 | 0.730090578 | 0.909157553 | ns | TM | KS.1 |
| FIM1_1282 | 607.7011123 | 616.4258302 | 725.1315443 | 0.004139812 | 0.139400665 | 0.947607695 | 0.986388143 | ns | TM | KS.1 |
| FIM1_1283 | 4858.813384 | 3332.141434 | 3946.865104 | -0.158670536 | 0.208403062 | 0.06607939 | 0.270807979 | ns | TM | KS.1 |
| FIM1_1284 | 556.5783257 | 472.5861418 | 937.7489032 | -0.067118328 | 0.137452317 | 0.323835102 | 0.650941236 | ns | TM | KS.1 |
| FIM1_1285 | 3320.601387 | 3833.479912 | 4577.267726 | 0.127755528 | 0.098349579 | 0.049417129 | 0.231715567 | ns | TM | KS.1 |
| FIM1_1286 | 440.676148 | 601.5977095 | 438.3162663 | 0.230760158 | 0.169477541 | 0.013211596 | 0.101775569 | ns | TM | KS.1 |
| FIM1_1287 | 1663.660461 | 1557.910984 | 1377.703421 | -0.025476841 | 0.124201092 | 0.688681152 | 0.892004357 | ns | TM | KS.1 |
| FIM1_1288 | 564.75587 | 547.2253586 | 608.8673932 | -0.017708144 | 0.092584944 | 0.757075495 | 0.920753636 | ns | TM | KS.1 |
| FIM1_1289 | 2087.027202 | 2145.945127 | 2209.892379 | 0.018049021 | 0.079365807 | 0.729952677 | 0.909157553 | ns | TM | KS.1 |
| FIM1_1290 | 4688.566001 | 4400.506953 | 4253.796382 | -0.038339553 | 0.092130432 | 0.504802712 | 0.797520957 | ns | TM | KS.1 |
| FIM1_1291 | 1341.460791 | 1134.354546 | 1057.93371 | -0.091037592 | 0.127882675 | 0.191797675 | 0.496260756 | ns | TM | KS.1 |
| FIM1_1292 | 161.6735878 | 239.1002356 | 242.678051 | 0.316698706 | 0.207869733 | 0.004042092 | 0.044069751 | ns | TM | KS.1 |
| FIM1_1293 | 1539.704006 | 2315.896073 | 2514.956573 | 0.261115445 | 0.226320506 | 0.013014396 | 0.101166593 | ns | TM | KS.1 |
| FIM1_1294 | 739.7467102 | 1089.341597 | 1168.931978 | 0.151490074 | 0.212296647 | 0.075496744 | 0.292976834 | ns | TM | KS.1 |
| FIM1_1295 | 3331.97751 | 3072.410593 | 3331.000546 | -0.050798465 | 0.094864437 | 0.387728484 | 0.712272102 | ns | TM | KS.1 |
| FIM1_1296 | 872.3475651 | 790.9812796 | 921.9107713 | -0.022676222 | 0.158999757 | 0.718382812 | 0.905913973 | ns | TM | KS.1 |
| FIM1_1297 | 1543.679714 | 603.6479486 | 1960.959462 | -1.137238015 | 0.325172777 | 9.7947693788[illegible] | 0.000414015 | Km<KS.1 | TM | KS.1 |
| FIM1_1298 | 3497.464055 | 3385.457219 | 2757.110115 | -0.012887476 | 0.120377036 | 0.837452937 | 0.949337033 | ns | TM | KS.1 |
| FIM1_1299 | 995.8984913 | 1028.472414 | 985.6162665 | 0.022285544 | 0.073662391 | 0.654395299 | 0.87699044 | ns | TM | KS.1 |
| FIM1_1300 | 1021.64255 | 1207.336329 | 1455.409244 | 0.10523858 | 0.1224589 | 0.13389879 | 0.40527648 | ns | TM | KS.1 |
| FIM1_1301 | 391.7688294 | 336.0100678 | 342.8589182 | -0.097148926 | 0.11759286 | 0.15526839 | 0.439647263 | ns | TM | KS.1 |
| FIM1_1302 | 540.1910017 | 656.8969192 | 736.6684656 | 0.171853252 | 0.117299042 | 0.023072073 | 0.148299176 | ns | TM | KS.1 |
| FIM1_1303 | 1224.975524 | 1825.286011 | 2040.5456 | 0.277908762 | 0.219104095 | 0.009193618 | 0.07968336 | ns | TM | KS.1 |

| | | | | | | | | | | |
|---|---|---|---|---|---|---|---|---|---|---|
| FIM1_1304 | 1248.728618 | 1409.555577 | 1576.531809 | 0.039071003 | 0.141411699 | 0.547811899 | 0.820398615 | ns | TM | KS.1 |
| FIM1_1305 | 3628.452166 | 3602.381585 | 4427.895953 | -0.003824883 | 0.09661573 | 0.947742035 | 0.986388143 | ns | TM | KS.1 |
| FIM1_1306 | 111.9735375 | 110.5217533 | 117.1812301 | -0.006097229 | 0.123562202 | 0.922788628 | 0.976369825 | ns | TM | KS.1 |
| FIM1_1307 | 1049.73061 | 1164.613425 | 1081.113627 | 0.06767812 | 0.099836938 | 0.269085967 | 0.596620808 | ns | TM | KS.1 |
| FIM1_1308 | 705.420123 | 789.4272038 | 883.5704345 | 0.079510276 | 0.099579742 | 0.198413411 | 0.505976981 | ns | TM | KS.1 |
| FIM1_1309 | 1200.682113 | 1094.082487 | 1078.806146 | -0.050365444 | 0.107786512 | 0.420759389 | 0.737778527 | ns | TM | KS.1 |
| FIM1_1310 | 1733.566766 | 2079.566489 | 2115.586315 | 0.172370586 | 0.106937252 | 0.017276245 | 0.122960396 | ns | TM | KS.1 |
| FIM1_1311 | 493.9991428 | 471.8352651 | 2063.53469 | -0.015776765 | 0.130738361 | 0.803483749 | 0.936908579 | ns | TM | KS.1 |
| FIM1_1312 | 2582.635267 | 1589.612529 | 1879.694692 | -0.38196064 | 0.27072304 | 0.003226838 | 0.037075097 | ns | TM | KS.1 |
| FIM1_1313 | 38052.09271 | 40081.23572 | 32699.65459 | 0.028689299 | 0.097172011 | 0.626053458 | 0.866615457 | ns | TM | KS.1 |
| FIM1_1314 | 800.4551793 | 751.906836 | 895.8018927 | -0.025496878 | 0.120420014 | 0.687040391 | 0.891826565 | ns | TM | KS.1 |
| FIM1_1315 | 3100.585902 | 3456.452244 | 3495.315727 | 0.056004642 | 0.114138456 | 0.383542279 | 0.708286132 | ns | TM | KS.1 |
| FIM1_1316 | 1500.666027 | 917.2807881 | 918.0455375 | -0.451713713 | 0.263063183 | 0.00108387 | 0.016290797 | ns | TM | KS.1 |
| FIM1_1317 | 722.3794716 | 635.6234162 | 631.2855543 | -0.06252744 | 0.121339371 | 0.34341676 | 0.670129201 | ns | TM | KS.1 |
| FIM1_1318 | 1760.343509 | 1585.346362 | 1423.226802 | -0.065994487 | 0.101984392 | 0.285821366 | 0.612477571 | ns | TM | KS.1 |
| FIM1_1319 | 3084.926495 | 3317.284585 | 3635.495891 | 0.028071646 | 0.12511179 | 0.659889113 | 0.878874681 | ns | TM | KS.1 |
| FIM1_1320 | 11160.94453 | 12960.62696 | 13061.52587 | 0.068265087 | 0.12961926 | 0.312269795 | 0.63931779 | ns | TM | KS.1 |
| FIM1_1321 | 729.3071793 | 729.9815736 | 788.6044429 | 0.000754583 | 0.083561454 | 0.988839311 | 0.996167262 | ns | TM | KS.1 |
| FIM1_1322 | 8088.352903 | 8417.265241 | 7929.863146 | 0.015396371 | 0.122441419 | 0.807078 | 0.937695714 | ns | TM | KS.1 |
| FIM1_1323 | 493.3729915 | 352.8211368 | 817.2490221 | -0.244645607 | 0.181406611 | 0.011430183 | 0.092454675 | ns | TM | KS.1 |
| FIM1_1324 | 892.2606717 | 1076.820795 | 1164.683031 | 0.064741033 | 0.150001822 | 0.341711632 | 0.668903606 | ns | TM | KS.1 |
| FIM1_1325 | 2329.98346 | 1352.128619 | 1963.284815 | -0.707636759 | 0.185554825 | 9.53700909671 | 6.77808860802 | ns | TM | KS.1 |
| FIM1_1326 | 1033.745265 | 972.6061427 | 1051.740856 | -0.043408384 | 0.079253893 | 0.407923255 | 0.728957685 | ns | TM | KS.1 |
| FIM1_1327 | 1683.182374 | 1824.272527 | 2187.925136 | 0.057511918 | 0.085413418 | 0.299380259 | 0.628303452 | ns | TM | KS.1 |
| FIM1_1328 | 1589.646833 | 1543.644215 | 1573.607562 | -0.024310293 | 0.058359782 | 0.564588138 | 0.828150948 | ns | TM | KS.1 |
| FIM1_1329 | 786.255653 | 588.8555148 | 1818.801545 | -0.135035217 | 0.171593679 | 0.090967998 | 0.325119102 | ns | TM | KS.1 |
| FIM1_1331 | 159.0762393 | 177.0506493 | 153.9760707 | 0.031201179 | 0.145095383 | 0.626990956 | 0.866840608 | ns | TM | KS.1 |
| FIM1_1332 | 784.9980416 | 684.3550806 | 565.8234931 | -0.095785743 | 0.108978296 | 0.146492489 | 0.424708703 | ns | TM | KS.1 |
| FIM1_1333 | 4816.749008 | 6572.617591 | 7441.272709 | 0.161104295 | 0.178830448 | 0.056812128 | 0.251459372 | ns | TM | KS.1 |
| FIM1_1334 | 3693.115545 | 3462.077242 | 3113.798748 | -0.02769828 | 0.117654205 | 0.660675032 | 0.879309333 | ns | TM | KS.1 |
| FIM1_1335 | 1120.051987 | 1047.196856 | 988.3723608 | -0.033215115 | 0.108388244 | 0.591229004 | 0.845462574 | ns | TM | KS.1 |
| FIM1_1336 | 619.4225485 | 913.0924108 | 1587.422816 | 0.167938329 | 0.213575917 | 0.057369175 | 0.252576678 | ns | TM | KS.1 |
| FIM1_1337 | 1774.364102 | 3032.495151 | 3774.680074 | 0.277894509 | 0.299954953 | 0.016616341 | 0.120154503 | ns | TM | KS.1 |
| FIM1_1338 | 776.1708025 | 575.7720223 | 857.7782986 | -0.177651015 | 0.171760553 | 0.039679305 | 0.202466195 | ns | TM | KS.1 |
| FIM1_1339 | 4278.927277 | 4346.720621 | 5129.626516 | 0.007212972 | 0.108628456 | 0.9061476 | 0.971578849 | ns | TM | KS.1 |
| FIM1_1340 | 2254.956975 | 2218.701365 | 2228.020992 | -0.00987972 | 0.082233837 | 0.853319468 | 0.954152748 | ns | TM | KS.1 |
| FIM1_1341 | 921.1110231 | 859.6536102 | 703.1858564 | -0.026486578 | 0.12556289 | 0.677687369 | 0.887002016 | ns | TM | KS.1 |
| FIM1_1342 | 2378.24446 | 2380.712866 | 2388.158002 | 0.000526216 | 0.09594587 | 0.992780839 | 0.997449676 | ns | TM | KS.1 |
| FIM1_1344 | 343.6272726 | 303.0833177 | 431.1494487 | -0.087866643 | 0.105164857 | 0.171179361 | 0.465692069 | ns | TM | KS.1 |
| FIM1_1345 | 1563.097006 | 1075.901571 | 1529.858814 | -0.420656618 | 0.147344596 | 5.01444597455 | 0.001401509 | ns | TM | KS.1 |
| FIM1_1346 | 6802.703422 | 7580.333901 | 8008.702261 | 0.077822661 | 0.096122273 | 0.198309956 | 0.505976981 | ns | TM | KS.1 |
| FIM1_1347 | 16349.07222 | 17768.75977 | 16702.93723 | 0.051069221 | 0.097022298 | 0.391098077 | 0.71457503 | ns | TM | KS.1 |
| FIM1_1348 | 8611.047125 | 10074.08091 | 10458.08256 | 0.168763963 | 0.087630582 | 0.008921978 | 0.078450448 | ns | TM | KS.1 |
| FIM1_1349 | 1436.668457 | 1356.42001 | 1253.590956 | -0.042925568 | 0.07368723 | 0.389695689 | 0.713295089 | ns | TM | KS.1 |
| FIM1_1350 | 7348.189588 | 9025.011838 | 8388.563378 | 0.130424895 | 0.134925364 | 0.083263107 | 0.306449621 | ns | TM | KS.1 |

| | | | | | | | | | | |
|---|---|---|---|---|---|---|---|---|---|---|
| FIM1_1352 | 3428.820373 | 2638.805797 | 2375.419536 | -0.300936211 | 0.098480147 | 0.000143368 | 0.003292826 | ns | TM | KS.1 |
| FIM1_1353 | 10451.2463 | 11003.33574 | 8281.703894 | 0.037728033 | 0.072803362 | 0.445398261 | 0.751385662 | ns | TM | KS.1 |
| FIM1_1354 | 1054.408135 | 1106.151282 | 1056.633872 | 0.025483814 | 0.099256716 | 0.667876792 | 0.882180579 | ns | TM | KS.1 |
| FIM1_1355 | 1440.141413 | 1419.197162 | 1650.334581 | -0.01133474 | 0.053040266 | 0.773683761 | 0.925733395 | ns | TM | KS.1 |
| FIM1_1356 | 717.1251087 | 734.2407645 | 718.3923749 | 0.015431238 | 0.07881304 | 0.766848756 | 0.924883104 | ns | TM | KS.1 |
| FIM1_1357 | 1180.919262 | 1035.854659 | 1025.871029 | -0.076430074 | 0.113660148 | 0.244588556 | 0.567815242 | ns | TM | KS.1 |
| FIM1_1358 | 1244.786943 | 1329.526581 | 1315.886861 | 0.041892231 | 0.08971223 | 0.459996757 | 0.763054265 | ns | TM | KS.1 |
| FIM1_1359 | 547.2364921 | 490.3108792 | 495.7212686 | -0.066770748 | 0.106587769 | 0.290881859 | 0.618193599 | ns | TM | KS.1 |
| FIM1_1360 | 1628.486383 | 1620.18187 | 2675.62504 | -0.001870991 | 0.124981516 | 0.9762867 | 0.993872484 | ns | TM | KS.1 |
| FIM1_1361 | 216.5150571 | 227.596631 | 279.5368154 | 0.024753408 | 0.107954489 | 0.687031614 | 0.891826565 | ns | TM | KS.1 |
| FIM1_1362 | 493.7708572 | 456.7026776 | 484.6343487 | -0.042076331 | 0.104262556 | 0.492341195 | 0.790867874 | ns | TM | KS.1 |
| FIM1_1363 | 2180.266317 | 3141.458927 | 3829.713585 | 0.145311155 | 0.202986354 | 0.082013706 | 0.304774234 | ns | TM | KS.1 |
| FIM1_1364 | 316.450688 | 380.8358206 | 299.4874911 | 0.073562854 | 0.14430077 | 0.287468745 | 0.614064838 | ns | TM | KS.1 |
| FIM1_1365 | 2360.450218 | 2519.821624 | 2515.753909 | 0.034466663 | 0.103222533 | 0.570243414 | 0.832094673 | ns | TM | KS.1 |
| FIM1_1366 | 1288.867077 | 1307.438948 | 1390.123103 | 0.008150587 | 0.093611642 | 0.887245595 | 0.967028314 | ns | TM | KS.1 |
| FIM1_1367 | 1078.075964 | 904.1802294 | 982.0969972 | -0.182486245 | 0.097858232 | 0.009222527 | 0.079794909 | ns | TM | KS.1 |
| FIM1_1368 | 5273.223637 | 5987.557917 | 5774.293505 | 0.115309281 | 0.090835967 | 0.058629581 | 0.254317312 | ns | TM | KS.1 |
| FIM1_1369 | 1320.066299 | 1737.688568 | 3750.884313 | 0.164294506 | 0.161849741 | 0.048712876 | 0.230174256 | ns | TM | KS.1 |
| FIM1_1370 | 12.56576022 | 14.37638182 | 34.13613517 | 0.026951137 | 0.175629541 | 0.666072494 | 0.880910621 | ns | TM | KS.1 |
| FIM1_1371 | 1563.18575 | 1607.150886 | 1244.944897 | 0.019053935 | 0.075182127 | 0.705762207 | 0.899257553 | ns | TM | KS.1 |
| FIM1_1372 | 866.2097261 | 422.4007276 | 292.5230999 | -0.973845347 | 0.182825203 | 1.01449331034 | 1.44202977685 | ns | TM | KS.1 |
| FIM1_1373 | 2824.36009 | 2860.380094 | 2413.523477 | 0.008405609 | 0.077032062 | 0.869769486 | 0.96136485 | ns | TM | KS.1 |
| FIM1_1374 | 2522.02448 | 2224.169916 | 2208.060344 | -0.106209464 | 0.094356822 | 0.085136534 | 0.31075147 | ns | TM | KS.1 |
| FIM1_1375 | 1102.024966 | 1393.903852 | 1570.463931 | 0.072454815 | 0.16450122 | 0.295789141 | 0.623274449 | ns | TM | KS.1 |
| FIM1_1376 | 92.0267881 | 59.3433511 | 96.17648643 | -0.355463372 | 0.247051192 | 0.003561061 | 0.0399916 | ns | TM | KS.1 |
| FIM1_1377 | 2862.169702 | 3087.735096 | 3826.541247 | 0.054086406 | 0.084255413 | 0.32396593 | 0.650941236 | ns | TM | KS.1 |
| FIM1_1378 | 306.4078023 | 301.4451815 | 321.4348399 | -0.007994646 | 0.105451577 | 0.894989671 | 0.967972723 | ns | TM | KS.1 |
| FIM1_1379 | 210.4642074 | 237.7663435 | 272.0880493 | 0.048649388 | 0.130792261 | 0.458944087 | 0.762486824 | ns | TM | KS.1 |
| FIM1_1380 | 1517.927111 | 1533.322767 | 2315.659662 | 0.00338157 | 0.130725233 | 0.957265445 | 0.989353426 | ns | TM | KS.1 |
| FIM1_1382 | 3539.317898 | 3813.951868 | 4035.366892 | 0.071028064 | 0.062669742 | 0.11482494 | 0.37039344 | ns | TM | KS.1 |
| FIM1_1383 | 2530.909135 | 2634.768851 | 2587.870545 | 0.025308035 | 0.083157225 | 0.638658473 | 0.870738806 | ns | TM | KS.1 |
| FIM1_1384 | 1076.557248 | 1091.300179 | 1200.455473 | 0.007645381 | 0.097997423 | 0.896484719 | 0.967972723 | ns | TM | KS.1 |
| FIM1_1385 | 4935.604659 | 4082.89287 | 5001.549149 | -0.108102698 | 0.132986729 | 0.135449553 | 0.407659726 | ns | TM | KS.1 |
| FIM1_1386 | 610.4535614 | 631.4022373 | 571.1931895 | 0.01843661 | 0.090385568 | 0.744285069 | 0.91508234 | ns | TM | KS.1 |
| FIM1_1387 | 2811.605196 | 3004.752452 | 2965.730192 | 0.073215992 | 0.048592538 | 0.050585441 | 0.23366998 | ns | TM | KS.1 |
| FIM1_1388 | 387.667251 | 373.424735 | 383.3514424 | -0.024250709 | 0.079214626 | 0.642566809 | 0.872815354 | ns | TM | KS.1 |
| FIM1_1389 | 887.790992 | 704.137034 | 608.2360905 | -0.219473555 | 0.120403607 | 0.006958622 | 0.066067072 | ns | TM | KS.1 |
| FIM1_1390 | 2562.21394 | 2599.14076 | 2724.645374 | 0.006119574 | 0.113776843 | 0.921349375 | 0.975881018 | ns | TM | KS.1 |
| FIM1_1391 | 3910.092057 | 5000.060479 | 3696.141313 | 0.065380468 | 0.172541464 | 0.336888829 | 0.665088066 | ns | TM | KS.1 |
| FIM1_1392 | 1390.757504 | 1528.933578 | 1101.82239 | 0.047153158 | 0.112934818 | 0.457472556 | 0.761179252 | ns | TM | KS.1 |
| FIM1_1393 | 1385.406251 | 1322.140626 | 1062.710524 | -0.020382475 | 0.11477431 | 0.744067624 | 0.91508234 | ns | TM | KS.1 |
| FIM1_1394 | 782.2913422 | 405.3251762 | 587.753791 | -0.640263517 | 0.358238571 | 0.00060071 | 0.01030528 | ns | TM | KS.1 |
| FIM1_1395 | 770.963355 | 905.347024 | 753.8510585 | 0.057235218 | 0.142844685 | 0.393158435 | 0.716470042 | ns | TM | KS.1 |
| FIM1_1396 | 1177.326175 | 1310.908963 | 1338.63058 | 0.055381001 | 0.113863639 | 0.388150967 | 0.712302124 | ns | TM | KS.1 |
| FIM1_1397 | 5615.742055 | 5007.190515 | 4469.632152 | -0.071383383 | 0.105768886 | 0.259292854 | 0.588210523 | ns | TM | KS.1 |

| | | | | | | | | | |
|---|---|---|---|---|---|---|---|---|---|
| FIM1_1399 | 2501.628954 | 2509.683821 | 2805.973654 | 0.001293471 | 0.116831572 | 0.983441104 | 0.996034691 | ns | TM | KS.1 |
| FIM1_1400 | 1533.829149 | 1704.413933 | 1902.072314 | 0.041963205 | 0.1277533 | 0.518046851 | 0.804981347 | ns | TM | KS.1 |
| FIM1_1401 | 1887.262045 | 3138.185931 | 3542.249151 | 0.294319879 | 0.287548021 | 0.012447277 | 0.098138201 | ns | TM | KS.1 |
| FIM1_1402 | 703.9024722 | 773.679664 | 872.8792913 | 0.073774845 | 0.086572111 | 0.191806827 | 0.496260756 | ns | TM | KS.1 |
| FIM1_1403 | 743.5988323 | 608.5101924 | 747.890367 | -0.132587168 | 0.131637511 | 0.076909263 | 0.294813117 | ns | TM | KS.1 |
| FIM1_1404 | 1777.538816 | 520.5324969 | 894.6827909 | -1.484198442 | 0.320640817 | 2.08688815622 | 2.16297262024 | Km<KS.1 | TM | KS.1 |
| FIM1_1405 | 1338.657493 | 1182.977296 | 849.0081097 | -0.052378775 | 0.127228515 | 0.42586912 | 0.741071977 | ns | TM | KS.1 |
| FIM1_1406 | 5590.82797 | 6307.328519 | 6026.309982 | 0.045779688 | 0.132732215 | 0.484712703 | 0.785178463 | ns | TM | KS.1 |
| FIM1_1407 | 1164.183081 | 1361.919106 | 1369.341169 | 0.132481755 | 0.106315943 | 0.052395066 | 0.239582218 | ns | TM | KS.1 |
| FIM1_1408 | 4971.244745 | 4809.395168 | 7942.031075 | -0.020091022 | 0.085045368 | 0.712865209 | 0.904028655 | ns | TM | KS.1 |
| FIM1_1409 | 1470.278403 | 2155.996874 | 2107.260258 | 0.119471799 | 0.209085811 | 0.129588009 | 0.39771767 | ns | TM | KS.1 |
| FIM1_1410 | 4651.759426 | 4325.137147 | 3987.427979 | -0.039729296 | 0.102105311 | 0.512231177 | 0.802566276 | ns | TM | KS.1 |
| FIM1_1411 | 1189.866309 | 1357.678402 | 1205.529616 | 0.038115345 | 0.148955518 | 0.556202673 | 0.825176412 | ns | TM | KS.1 |
| FIM1_1412 | 3562.857775 | 3454.658378 | 4924.602873 | -0.012535367 | 0.118659708 | 0.841425576 | 0.95052049 | ns | TM | KS.1 |
| FIM1_1413 | 618.9456301 | 576.5225053 | 623.2557519 | -0.020718603 | 0.142903129 | 0.744176484 | 0.91508234 | ns | TM | KS.1 |
| FIM1_1414 | 297.6849385 | 326.7744137 | 305.0673911 | 0.042986731 | 0.117600462 | 0.501552991 | 0.795417956 | ns | TM | KS.1 |
| FIM1_1415 | 2004.464702 | 1852.281456 | 1607.211688 | -0.064336431 | 0.075984422 | 0.210935804 | 0.51871941 | ns | TM | KS.1 |
| FIM1_1416 | 1483.314322 | 1457.708804 | 1016.078387 | -0.011218866 | 0.073189602 | 0.820914085 | 0.943851992 | ns | TM | KS.1 |
| FIM1_1418 | 1110.100329 | 1116.917002 | 1019.213384 | 0.005339004 | 0.068098987 | 0.909886889 | 0.972251892 | ns | TM | KS.1 |
| FIM1_1420 | 1374.146307 | 1490.230744 | 1437.044234 | 0.055310561 | 0.089090437 | 0.330490571 | 0.657690769 | ns | TM | KS.1 |
| FIM1_1421 | 615.3569056 | 593.8257235 | 553.3328384 | -0.012765136 | 0.127577478 | 0.839997842 | 0.949982862 | ns | TM | KS.1 |
| FIM1_1422 | 621.0406828 | 619.5016516 | 486.6103039 | -0.000970002 | 0.11283275 | 0.987480819 | 0.996167262 | ns | TM | KS.1 |
| FIM1_1423 | 770.0866636 | 786.2568265 | 793.3963286 | 0.014267917 | 0.066027689 | 0.757178188 | 0.920753636 | ns | TM | KS.1 |
| FIM1_1424 | 1279.267576 | 1317.580085 | 1449.15332 | 0.022068996 | 0.06367685 | 0.623454472 | 0.863738792 | ns | TM | KS.1 |
| FIM1_1425 | 1240.034052 | 1097.553025 | 1331.415899 | -0.068181831 | 0.113433799 | 0.294226483 | 0.621825298 | ns | TM | KS.1 |
| FIM1_1426 | 2960.807119 | 2930.988883 | 3731.823932 | -0.002442982 | 0.153202124 | 0.968724196 | 0.993374633 | ns | TM | KS.1 |
| FIM1_1427 | 26.23281084 | 38.94174594 | 38.6833852 | 0.132130077 | 0.220207809 | 0.106189253 | 0.353136053 | ns | TM | KS.1 |
| FIM1_1428 | 2298.613877 | 2338.028457 | 2119.551811 | 0.008769718 | 0.098049613 | 0.881413576 | 0.9656535 | ns | TM | KS.1 |
| FIM1_1429 | 2031.92591 | 2039.599953 | 2233.881656 | 0.001751496 | 0.102290166 | 0.976645041 | 0.993872484 | ns | TM | KS.1 |
| FIM1_1430 | 1250.647468 | 1187.869699 | 1121.095545 | -0.024887029 | 0.108054061 | 0.685549516 | 0.890505759 | ns | TM | KS.1 |
| FIM1_1431 | 1901.631076 | 2083.783996 | 2212.056116 | 0.0435299 | 0.114942361 | 0.49364602 | 0.791966768 | ns | TM | KS.1 |
| FIM1_1432 | 11884.21594 | 12732.15517 | 13282.27271 | 0.043845495 | 0.090018716 | 0.440654267 | 0.747832613 | ns | TM | KS.1 |
| FIM1_1433 | 1801.562172 | 1850.834335 | 1941.906229 | 0.018690925 | 0.071429009 | 0.70138017 | 0.897235589 | ns | TM | KS.1 |
| FIM1_1434 | 1523.02086 | 1785.891776 | 1918.794881 | 0.057578108 | 0.141921159 | 0.390504749 | 0.713987918 | ns | TM | KS.1 |
| FIM1_1435 | 5091.531411 | 4902.349331 | 5628.031343 | -0.01889699 | 0.103475128 | 0.754040285 | 0.919674042 | ns | TM | KS.1 |
| FIM1_1436 | 639.4799674 | 494.5953305 | 560.7691604 | -0.190650449 | 0.147145236 | 0.024135568 | 0.152378744 | ns | TM | KS.1 |
| FIM1_1437 | 360.6749428 | 362.9967989 | 339.2170924 | 0.007097057 | 0.076358453 | 0.889271432 | 0.967028314 | ns | TM | KS.1 |
| FIM1_1438 | 47668.12714 | 59973.43748 | 60172.18324 | 0.263970241 | 0.092474984 | 0.000400496 | 0.007633977 | ns | TM | KS.1 |
| FIM1_1439 | 945.2590107 | 906.6128751 | 1220.359772 | -0.024301905 | 0.090855216 | 0.668548694 | 0.882180579 | ns | TM | KS.1 |
| FIM1_1440 | 416.7594505 | 356.3825693 | 401.8067455 | -0.120466133 | 0.112009955 | 0.080781257 | 0.303082015 | ns | TM | KS.1 |
| FIM1_1441 | 137.6787297 | 134.8519418 | 322.5394479 | -0.005933764 | 0.144980383 | 0.924726692 | 0.976735299 | ns | TM | KS.1 |
| FIM1_1442 | 51.98234066 | 48.274216 | 578.1003268 | -0.018508594 | 0.153941793 | 0.768390495 | 0.925179691 | ns | TM | KS.1 |
| FIM1_1443 | 399.9440993 | 474.0875621 | 494.6692785 | 0.106975071 | 0.124148546 | 0.130445821 | 0.39950807 | ns | TM | KS.1 |
| FIM1_1444 | 11031.73867 | 13398.22951 | 12306.94573 | 0.140821033 | 0.126428201 | 0.059643559 | 0.256462934 | ns | TM | KS.1 |
| FIM1_1445 | 637.0190765 | 912.5741447 | 1243.352178 | 0.140557637 | 0.20071522 | 0.088633665 | 0.318837658 | ns | TM | KS.1 |

| | | | | | | | | | | |
|---|---|---|---|---|---|---|---|---|---|---|
| FIM1_1446 | 990.9121729 | 675.3665145 | 640.2136342 | -0.371640266 | 0.182677605 | 0.000812631 | 0.01299948 | ns | TM | KS.1 |
| FIM1_1447 | 784.193358 | 945.6162777 | 788.7167144 | 0.080180469 | 0.141683061 | 0.251333939 | 0.577012619 | ns | TM | KS.1 |
| FIM1_1448 | 569.9839931 | 581.2771369 | 525.5591259 | 0.008607594 | 0.114304145 | 0.889727554 | 0.967028314 | ns | TM | KS.1 |
| FIM1_1449 | 2570.758958 | 2407.019697 | 2542.263048 | -0.043568056 | 0.085970964 | 0.430709494 | 0.743536592 | ns | TM | KS.1 |
| FIM1_1450 | 713.5912661 | 791.2716389 | 920.6746159 | 0.060770246 | 0.104885499 | 0.330053873 | 0.657690769 | ns | TM | KS.1 |
| FIM1_1451 | 11975.32322 | 10806.83993 | 8214.938459 | -0.036331469 | 0.134038347 | 0.574551169 | 0.833865535 | ns | TM | KS.1 |
| FIM1_1453 | 3660.450361 | 3763.594417 | 3848.212792 | 0.008824211 | 0.134813139 | 0.888933511 | 0.967028314 | ns | TM | KS.1 |
| FIM1_1454 | 926.8054815 | 869.6586234 | 1125.145479 | -0.036940753 | 0.096381322 | 0.530244019 | 0.808531286 | ns | TM | KS.1 |
| FIM1_1455 | 1441.457488 | 1571.333385 | 1192.667858 | 0.037390945 | 0.12008893 | 0.558561088 | 0.82614557 | ns | TM | KS.1 |
| FIM1_1456 | 471.8916841 | 533.6486205 | 560.0456439 | 0.06947953 | 0.113977737 | 0.286735995 | 0.612762705 | ns | TM | KS.1 |
| FIM1_1457 | 2933.941212 | 3204.968403 | 2947.262511 | 0.04908328 | 0.104688252 | 0.426459778 | 0.741071977 | ns | TM | KS.1 |
| FIM1_1458 | 1482.14621 | 1434.102718 | 1399.980782 | -0.019166775 | 0.088741618 | 0.731945903 | 0.910813123 | ns | TM | KS.1 |
| FIM1_1460 | 1224.336373 | 1267.914187 | 1103.395768 | 0.02546217 | 0.068319459 | 0.589983038 | 0.844166125 | ns | TM | KS.1 |
| FIM1_1461 | 1512.500921 | 1558.62835 | 986.9451075 | 0.00935418 | 0.136689253 | 0.882281136 | 0.966130215 | ns | TM | KS.1 |
| FIM1_1462 | 1464.82648 | 1496.405744 | 1477.699355 | 0.014181094 | 0.076256717 | 0.780702183 | 0.929207457 | ns | TM | KS.1 |
| FIM1_1463 | 201.4507161 | 242.7512153 | 239.1789131 | 0.094701407 | 0.135805115 | 0.183533529 | 0.48542228 | ns | TM | KS.1 |
| FIM1_1464 | 19.89565223 | 15.30007873 | 24.69833755 | -0.038276782 | 0.203676975 | 0.541802652 | 0.815819671 | ns | TM | KS.1 |
| FIM1_1465 | 1072.696377 | 1014.100578 | 1340.776731 | -0.036824827 | 0.082950125 | 0.494817595 | 0.792222775 | ns | TM | KS.1 |
| FIM1_1466 | 1455.351423 | 1314.542405 | 1195.574456 | -0.074712495 | 0.092675088 | 0.205668858 | 0.512865935 | ns | TM | KS.1 |
| FIM1_1467 | 2430.678985 | 2347.226497 | 2349.248935 | -0.018852116 | 0.09695624 | 0.74770661 | 0.916894352 | ns | TM | KS.1 |
| FIM1_1468 | 3099.375311 | 3182.294788 | 3250.144257 | 0.017982265 | 0.072402312 | 0.714728291 | 0.905084971 | ns | TM | KS.1 |
| FIM1_1469 | 326.0142392 | 323.053745 | 198.5580107 | -0.004683587 | 0.114797096 | 0.939880847 | 0.983546982 | ns | TM | KS.1 |
| FIM1_1470 | 1723.920948 | 2000.393526 | 1860.483098 | 0.161225726 | 0.083561495 | 0.009308747 | 0.079846584 | ns | TM | KS.1 |
| FIM1_1471 | 4491.694263 | 5032.888146 | 4424.68561 | 0.079550844 | 0.099654238 | 0.19838816 | 0.505976981 | ns | TM | KS.1 |
| FIM1_1472 | 1499.416866 | 1666.938517 | 1066.927055 | 0.059935519 | 0.108597831 | 0.343830445 | 0.670129201 | ns | TM | KS.1 |
| FIM1_1473 | 657.0453052 | 933.9471216 | 1273.656244 | 0.371432878 | 0.151995614 | 0.000291065 | 0.00591353 | ns | TM | KS.1 |
| FIM1_1474 | 2455.917954 | 2581.393924 | 2614.068586 | 0.026607215 | 0.099587039 | 0.654815397 | 0.87699044 | ns | TM | KS.1 |
| FIM1_1475 | 1062.891635 | 1125.089261 | 1427.26863 | 0.040129106 | 0.077392335 | 0.436570374 | 0.747832613 | ns | TM | KS.1 |
| FIM1_1476 | 92.05082187 | 58.95890292 | 1083.108515 | -0.206917211 | 0.243354148 | 0.034287109 | 0.185199792 | ns | TM | KS.1 |
| FIM1_1477 | 2067.138713 | 1990.652333 | 1843.773248 | -0.012565045 | 0.132279984 | 0.842656883 | 0.951443988 | ns | TM | KS.1 |
| FIM1_1478 | 2585.436384 | 1439.39144 | 1710.692423 | -0.518224152 | 0.329965703 | 0.001294895 | 0.018406006 | ns | TM | KS.1 |
| FIM1_1479 | 1021.120731 | 863.0066415 | 809.3494458 | -0.168448402 | 0.09802779 | 0.014307821 | 0.107039717 | ns | TM | KS.1 |
| FIM1_1480 | 824.4445667 | 972.1835074 | 905.3788425 | 0.068469446 | 0.137394999 | 0.315306432 | 0.641836948 | ns | TM | KS.1 |
| FIM1_1481 | 649.7622168 | 505.5947397 | 507.7294437 | -0.152850815 | 0.152106569 | 0.057952188 | 0.253530772 | ns | TM | KS.1 |
| FIM1_1482 | 6285.739935 | 5099.32711 | 6288.323436 | -0.16308628 | 0.128114814 | 0.035478074 | 0.188204876 | ns | TM | KS.1 |
| FIM1_1483 | 417.0277301 | 419.5881302 | 398.8084871 | 0.002661048 | 0.118520951 | 0.966037794 | 0.993374633 | ns | TM | KS.1 |
| FIM1_1484 | 947.2028411 | 856.7801518 | 742.8195055 | -0.062826226 | 0.101336878 | 0.306675304 | 0.635183029 | ns | TM | KS.1 |
| FIM1_1485 | 3533.301763 | 3505.658573 | 3134.913706 | -0.005948553 | 0.059668378 | 0.88983015 | 0.967028314 | ns | TM | KS.1 |
| FIM1_1486 | 2946.890804 | 3230.937307 | 2021.714419 | 0.04081137 | 0.119357993 | 0.524031793 | 0.808390131 | ns | TM | KS.1 |
| FIM1_1487 | 1050.290708 | 1749.355512 | 1944.335533 | 0.291489656 | 0.288191964 | 0.012980315 | 0.101166593 | ns | TM | KS.1 |
| FIM1_1488 | 4674.37803 | 5130.658749 | 5118.618575 | 0.087909053 | 0.072904602 | 0.084589862 | 0.310216851 | ns | TM | KS.1 |
| FIM1_1489 | 540.1180767 | 516.2123717 | 496.6185416 | -0.024393589 | 0.099107456 | 0.680945588 | 0.887949093 | ns | TM | KS.1 |
| FIM1_1490 | 691.2900952 | 629.6448535 | 542.9241269 | -0.050406113 | 0.107944355 | 0.420677718 | 0.737778527 | ns | TM | KS.1 |
| FIM1_1491 | 1184.76284 | 1082.230738 | 1008.296104 | -0.07692333 | 0.079110577 | 0.149463541 | 0.430313147 | ns | TM | KS.1 |
| FIM1_1492 | 1874.029272 | 1897.719721 | 1549.742596 | 0.007906136 | 0.083358644 | 0.883330929 | 0.966744674 | ns | TM | KS.1 |

| | | | | | | | | | | |
|---|---|---|---|---|---|---|---|---|---|---|
| FIM1_1493 | 905.1704858 | 725.2436425 | 639.7551531 | -0.152158351 | 0.137853171 | 0.052128595 | 0.238705802 | ns | TM | KS.1 |
| FIM1_1494 | 330.3130248 | 254.5856675 | 226.628268 | -0.17744564 | 0.152067492 | 0.034144123 | 0.184838966 | ns | TM | KS.1 |
| FIM1_1495 | 69.21853747 | 72.08804598 | 72.26210851 | 0.010538779 | 0.142880614 | 0.867178148 | 0.960282219 | ns | TM | KS.1 |
| FIM1_1496 | 736.9473158 | 781.7969619 | 888.9900235 | 0.03985383 | 0.084783895 | 0.466589417 | 0.768382108 | ns | TM | KS.1 |
| FIM1_1497 | 639.8322751 | 581.248362 | 541.8235652 | -0.066034692 | 0.093724404 | 0.263191643 | 0.589544539 | ns | TM | KS.1 |
| FIM1_1498 | 675.5558539 | 525.3506452 | 589.1756182 | -0.191810476 | 0.144394576 | 0.02268574 | 0.147024228 | ns | TM | KS.1 |
| FIM1_1499 | 3065.315851 | 3017.918625 | 1990.863723 | -0.006691486 | 0.113702487 | 0.914020904 | 0.974699214 | ns | TM | KS.1 |
| FIM1_1500 | 1698.190668 | 1631.036419 | 1542.250736 | -0.023644551 | 0.090710029 | 0.676678763 | 0.887002016 | ns | TM | KS.1 |
| FIM1_1501 | 1536.011524 | 1585.209622 | 1400.018085 | 0.017914024 | 0.09143495 | 0.752768073 | 0.918798126 | ns | TM | KS.1 |
| FIM1_1503 | 2223.176144 | 2516.415784 | 2153.953742 | 0.112228833 | 0.08970965 | 0.062564128 | 0.262221175 | ns | TM | KS.1 |
| FIM1_1504 | 837.7027669 | 867.3264099 | 912.8558115 | 0.025114858 | 0.06906599 | 0.59787283 | 0.849348181 | ns | TM | KS.1 |
| FIM1_1505 | 641.5881674 | 514.6748017 | 530.282422 | -0.181126627 | 0.129168153 | 0.023132662 | 0.148496766 | ns | TM | KS.1 |
| FIM1_1506 | 182.1974355 | 134.7282174 | 147.5683477 | -0.123303774 | 0.177819856 | 0.115350022 | 0.370766358 | ns | TM | KS.1 |
| FIM1_1507 | 6212.930264 | 6112.255272 | 4989.294237 | -0.009426687 | 0.087854162 | 0.865280685 | 0.959975149 | ns | TM | KS.1 |
| FIM1_1508 | 1598.962068 | 1842.734492 | 1713.87884 | 0.108550082 | 0.105699272 | 0.099919044 | 0.341335565 | ns | TM | KS.1 |
| FIM1_1509 | 1469.683702 | 1625.716644 | 1673.109911 | 0.062345411 | 0.102078833 | 0.311980514 | 0.639251671 | ns | TM | KS.1 |
| FIM1_1510 | 1278.174075 | 1337.432806 | 1336.410245 | 0.028872207 | 0.08509818 | 0.597643213 | 0.849348181 | ns | TM | KS.1 |
| FIM1_1511 | 237.1710986 | 223.4048886 | 223.070927 | -0.028344433 | 0.111381623 | 0.648658994 | 0.874068749 | ns | TM | KS.1 |
| FIM1_1512 | 592.8383904 | 615.1757626 | 639.4952849 | 0.020007885 | 0.093690151 | 0.728478465 | 0.908701505 | ns | TM | KS.1 |
| FIM1_1513 | 5573.788782 | 5373.060334 | 4753.957716 | -0.027273759 | 0.065578176 | 0.552354371 | 0.822689033 | ns | TM | KS.1 |
| FIM1_1514 | 16603.63545 | 13651.04367 | 11778.31794 | -0.168662377 | 0.118063567 | 0.025535171 | 0.156140634 | ns | TM | KS.1 |
| FIM1_1515 | 409.4541558 | 275.3342973 | 365.2917633 | -0.370772241 | 0.195109863 | 0.001125569 | 0.016554684 | ns | TM | KS.1 |
| FIM1_1516 | 242.8671312 | 49.34097205 | 209.8896237 | -1.74751113 | 0.409615141 | 2.75419371795 | 2.63502187438 | Km<KS.1 | TM | KS.1 |
| FIM1_1518 | 1404.719966 | 1505.897053 | 1559.237096 | 0.041030685 | 0.096187776 | 0.486197069 | 0.786100234 | ns | TM | KS.1 |
| FIM1_1519 | 729.6001496 | 683.5164066 | 582.3236775 | -0.037031355 | 0.096019158 | 0.528454227 | 0.808531286 | ns | TM | KS.1 |
| FIM1_1520 | 4998.146457 | 4613.476859 | 4137.183329 | -0.044871358 | 0.102035316 | 0.460593358 | 0.763054265 | ns | TM | KS.1 |
| FIM1_1521 | 2060.374476 | 3310.53979 | 4002.233212 | 0.170774003 | 0.250095715 | 0.061004595 | 0.26041916 | ns | TM | KS.1 |
| FIM1_1523 | 345.3180284 | 367.2373774 | 369.144513 | 0.037195013 | 0.094957944 | 0.524256196 | 0.808485609 | ns | TM | KS.1 |
| FIM1_1524 | 1129.137251 | 1247.736438 | 1103.512142 | 0.063033974 | 0.100576058 | 0.303286004 | 0.631432423 | ns | TM | KS.1 |
| FIM1_1525 | 1684.81518 | 1853.304627 | 1670.902005 | 0.088112036 | 0.075920855 | 0.092896691 | 0.32940915 | ns | TM | KS.1 |
| FIM1_1526 | 14774.0388 | 11450.31636 | 7536.909485 | -0.24633915 | 0.125632256 | 0.003895538 | 0.042782125 | ns | TM | KS.1 |
| FIM1_1527 | 573.1824054 | 518.7733527 | 531.9941973 | -0.072954136 | 0.090847533 | 0.210284251 | 0.518673352 | ns | TM | KS.1 |
| FIM1_1528 | 508.3673138 | 729.2409245 | 668.6270175 | 0.113838086 | 0.201209071 | 0.141643294 | 0.417214559 | ns | TM | KS.1 |
| FIM1_1529 | 1374.941411 | 1193.962765 | 1461.92079 | -0.096496695 | 0.110295083 | 0.14618601 | 0.424167573 | ns | TM | KS.1 |
| FIM1_1530 | 1783.751274 | 1916.873009 | 2471.08405 | 0.040868112 | 0.099147653 | 0.494547934 | 0.792222775 | ns | TM | KS.1 |
| FIM1_1531 | 1314.623074 | 869.9647101 | 741.0917151 | -0.326052752 | 0.222510692 | 0.004187919 | 0.045081369 | ns | TM | KS.1 |
| FIM1_1532 | 1238.89141 | 880.9732307 | 768.8674388 | -0.167023781 | 0.192614754 | 0.053972187 | 0.242339016 | ns | TM | KS.1 |
| FIM1_1533 | 737.3977892 | 719.3655853 | 731.9239523 | -0.015495291 | 0.082721413 | 0.772732628 | 0.925733395 | ns | TM | KS.1 |
| FIM1_1534 | 362.6627822 | 426.024974 | 392.2872536 | 0.128568318 | 0.111609217 | 0.064800032 | 0.267310992 | ns | TM | KS.1 |
| FIM1_1535 | 737.8593108 | 810.6634821 | 781.0826287 | 0.080995792 | 0.07915217 | 0.130603822 | 0.399602716 | ns | TM | KS.1 |
| FIM1_1536 | 600.8552951 | 612.9652969 | 720.7602547 | 0.012212626 | 0.085822654 | 0.823755267 | 0.94482203 | ns | TM | KS.1 |
| FIM1_1537 | 1852.613818 | 1815.926544 | 1969.124807 | -0.012849963 | 0.078024294 | 0.803718632 | 0.936908579 | ns | TM | KS.1 |
| FIM1_1538 | 1660.269395 | 1819.974593 | 1902.472634 | 0.061899323 | 0.093785633 | 0.292977263 | 0.620392168 | ns | TM | KS.1 |
| FIM1_1540 | 1320.579331 | 939.1884524 | 957.5712073 | -0.310583237 | 0.167049468 | 0.002214022 | 0.028027381 | ns | TM | KS.1 |
| FIM1_1541 | 2244.513616 | 1745.230926 | 2516.562676 | -0.138300745 | 0.154719485 | 0.079820503 | 0.300838638 | ns | TM | KS.1 |

| | | | | | | | | | | |
|---|---|---|---|---|---|---|---|---|---|---|
| FIM1_1542 | 2209.1902 | 3400.561762 | 4261.464166 | 0.207292884 | 0.237303125 | 0.033177775 | 0.181488253 | ns | TM | KS.1 |
| FIM1_1543 | 1923.224188 | 2977.811047 | 3473.59313 | 0.155245843 | 0.232816985 | 0.074565861 | 0.290497381 | ns | TM | KS.1 |
| FIM1_1544 | 289.2270212 | 257.5767019 | 191.583174 | -0.054863816 | 0.118515841 | 0.3982497 | 0.721782243 | ns | TM | KS.1 |
| FIM1_1545 | 1164.485964 | 1104.164649 | 1128.202135 | -0.0310398 | 0.094395111 | 0.592982515 | 0.846854399 | ns | TM | KS.1 |
| FIM1_1546 | 362.0509005 | 358.6138837 | 298.4181561 | -0.00515 | 0.097896115 | 0.930112586 | 0.97843743 | ns | TM | KS.1 |
| FIM1_1547 | 1325.854488 | 983.6370907 | 968.3615341 | -0.273223644 | 0.147565346 | 0.003466595 | 0.039377778 | ns | TM | KS.1 |
| FIM1_1548 | 1288.629331 | 1594.224528 | 1539.925275 | 0.107994508 | 0.143739461 | 0.142921156 | 0.419584366 | ns | TM | KS.1 |
| FIM1_1549 | 2561.187917 | 2833.089383 | 1949.095125 | 0.064447445 | 0.100012725 | 0.29172656 | 0.618644347 | ns | TM | KS.1 |
| FIM1_1550 | 2776.442746 | 2157.213973 | 1478.150675 | -0.148493146 | 0.153431841 | 0.064093092 | 0.265874687 | ns | TM | KS.1 |
| FIM1_1551 | 42087.52681 | 39016.55492 | 21078.24352 | -0.027338811 | 0.129797949 | 0.669083265 | 0.882180579 | ns | TM | KS.1 |
| FIM1_1552 | 258.1407178 | 265.1158826 | 398.9322626 | 0.012026729 | 0.119726747 | 0.84798835 | 0.952348261 | ns | TM | KS.1 |
| FIM1_1553 | 1813.949183 | 3063.996098 | 3566.150704 | 0.219851166 | 0.282138788 | 0.03317291 | 0.181488253 | ns | TM | KS.1 |
| FIM1_1554 | 749.799385 | 907.0363947 | 1029.671585 | 0.133335915 | 0.126826854 | 0.071788972 | 0.284128986 | ns | TM | KS.1 |
| FIM1_1556 | 342.5052815 | 471.1224046 | 412.6890585 | 0.201280638 | 0.178971593 | 0.026244186 | 0.159225397 | ns | TM | KS.1 |
| FIM1_1557 | 356.9335153 | 391.4613378 | 370.8202114 | 0.045706413 | 0.11347321 | 0.471462909 | 0.772376515 | ns | TM | KS.1 |
| FIM1_1558 | 2041.46373 | 2062.932737 | 2132.708685 | 0.006483993 | 0.079915549 | 0.90168493 | 0.968873116 | ns | TM | KS.1 |
| FIM1_1559 | 5508.899223 | 6120.959459 | 5562.228255 | 0.050417017 | 0.117415939 | 0.434034099 | 0.745878978 | ns | TM | KS.1 |
| FIM1_1560 | 1916.990623 | 1990.832353 | 1399.809743 | 0.024728661 | 0.078499732 | 0.634048081 | 0.869547581 | ns | TM | KS.1 |
| FIM1_1561 | 1981.523724 | 1654.756952 | 1903.390235 | -0.118874873 | 0.125223343 | 0.099592238 | 0.340998887 | ns | TM | KS.1 |
| FIM1_1562 | 591.0004159 | 582.3992713 | 554.8368495 | -0.00583233 | 0.119049586 | 0.925738264 | 0.976735299 | ns | TM | KS.1 |
| FIM1_1563 | 937.766695 | 845.5779861 | 1104.718313 | -0.082488071 | 0.08876831 | 0.154083759 | 0.437933732 | ns | TM | KS.1 |
| FIM1_1564 | 56.57339619 | 40.35065615 | 328.3333144 | -0.099713539 | 0.19561335 | 0.180985936 | 0.480985595 | ns | TM | KS.1 |
| FIM1_1566 | 1685.931495 | 1974.171982 | 2607.517748 | 0.080157952 | 0.127395722 | 0.242154779 | 0.564800763 | ns | TM | KS.1 |
| FIM1_1567 | 748.0741743 | 684.2614223 | 1116.362786 | -0.033298545 | 0.129319963 | 0.604799771 | 0.851547247 | ns | TM | KS.1 |
| FIM1_1568 | 6859.326398 | 7345.035613 | 7098.304888 | 0.055284923 | 0.071791091 | 0.261059062 | 0.588210523 | ns | TM | KS.1 |
| FIM1_1569 | 242.1048643 | 187.3401329 | 191.7269476 | -0.105537203 | 0.162791251 | 0.157435195 | 0.443009104 | ns | TM | KS.1 |
| FIM1_1570 | 1032.636626 | 1080.380552 | 1038.993896 | 0.028866086 | 0.083521284 | 0.593483595 | 0.846854399 | ns | TM | KS.1 |
| FIM1_1571 | 381.5864018 | 371.2946444 | 365.0476731 | -0.014094997 | 0.098134494 | 0.810771958 | 0.939792751 | ns | TM | KS.1 |
| FIM1_1572 | 448.4102685 | 502.2321146 | 549.9838849 | 0.069498508 | 0.106838603 | 0.273450159 | 0.599198132 | ns | TM | KS.1 |
| FIM1_1573 | 1143.882872 | 1013.40512 | 1173.957086 | -0.114109322 | 0.086306605 | 0.052936671 | 0.240511361 | ns | TM | KS.1 |
| FIM1_1574 | 3015.686848 | 2931.592996 | 3203.394133 | -0.0224808 | 0.056427052 | 0.58520372 | 0.841613393 | ns | TM | KS.1 |
| FIM1_1575 | 1046.625878 | 899.4536461 | 851.4544938 | -0.126617929 | 0.105737102 | 0.060909292 | 0.26041916 | ns | TM | KS.1 |
| FIM1_1576 | 60.22835454 | 43.67921425 | 38.68437699 | -0.129625935 | 0.182812288 | 0.103662059 | 0.348223325 | ns | TM | KS.1 |
| FIM1_1577 | 1159.373666 | 897.0342146 | 674.6265987 | -0.211150691 | 0.140809202 | 0.013656342 | 0.104063892 | ns | TM | KS.1 |
| FIM1_1578 | 7613.580971 | 9143.04587 | 9826.523273 | 0.196905401 | 0.094192286 | 0.004919987 | 0.051314325 | ns | TM | KS.1 |
| FIM1_1579 | 1133.206501 | 1600.854956 | 1877.861067 | 0.129468917 | 0.195024083 | 0.106536511 | 0.353581815 | ns | TM | KS.1 |
| FIM1_1580 | 1110.73275 | 1383.241718 | 1777.236402 | 0.112478005 | 0.145564035 | 0.131054424 | 0.400376242 | ns | TM | KS.1 |
| FIM1_1581 | 3693.034172 | 4052.449384 | 4862.540858 | 0.081475753 | 0.07798112 | 0.124643296 | 0.388047807 | ns | TM | KS.1 |
| FIM1_1582 | 149.6959356 | 74.16870784 | 111.7929124 | -0.617114303 | 0.401952528 | 0.001175158 | 0.017044925 | ns | TM | KS.1 |
| FIM1_1583 | 830.521054 | 756.9559708 | 887.9140318 | -0.061404157 | 0.095828885 | 0.302655784 | 0.631432423 | ns | TM | KS.1 |
| FIM1_1584 | 5072.026815 | 4720.524411 | 5326.866002 | -0.044191206 | 0.093365225 | 0.446700612 | 0.752446492 | ns | TM | KS.1 |
| FIM1_1585 | 1049.671013 | 950.3745102 | 919.5679531 | -0.059938063 | 0.102477594 | 0.330975105 | 0.658113969 | ns | TM | KS.1 |
| FIM1_1586 | 1232.999695 | 1075.794597 | 2013.747848 | -0.051346533 | 0.135944436 | 0.437904167 | 0.747832613 | ns | TM | KS.1 |
| FIM1_1587 | 7209.054493 | 7361.253434 | 7941.355191 | 0.013520518 | 0.077312888 | 0.792504491 | 0.933303839 | ns | TM | KS.1 |
| FIM1_1588 | 5078.917401 | 5854.990648 | 6573.148533 | 0.07339611 | 0.122534272 | 0.274411258 | 0.599699421 | ns | TM | KS.1 |

| | | | | | | | | | | |
|---|---|---|---|---|---|---|---|---|---|---|
| FIM1_1589 | 205.398508 | 180.6106328 | 174.21158 | -0.083229949 | 0.11111473 | 0.204403763 | 0.512813274 | ns | TM | KS.1 |
| FIM1_1590 | 935.2310016 | 1004.021005 | 907.6081619 | 0.029735751 | 0.120158215 | 0.639459475 | 0.870968656 | ns | TM | KS.1 |
| FIM1_1591 | 228.6977847 | 173.9116811 | 193.798865 | -0.115440786 | 0.167843587 | 0.131259025 | 0.400376242 | ns | TM | KS.1 |
| FIM1_1592 | 274.9599906 | 257.7004912 | 207.5354789 | -0.029869807 | 0.112131077 | 0.6321771 | 0.869527529 | ns | TM | KS.1 |
| FIM1_1593 | 3482.223387 | 2075.100825 | 3711.678997 | -0.213538095 | 0.277725314 | 0.035627031 | 0.188357575 | ns | TM | KS.1 |
| FIM1_1594 | 1782.497858 | 2486.741884 | 2701.158854 | 0.139401577 | 0.189606672 | 0.088105649 | 0.317685061 | ns | TM | KS.1 |
| FIM1_1595 | 1606.591597 | 1543.854524 | 1468.553781 | -0.020182678 | 0.10157454 | 0.736083297 | 0.912537853 | ns | TM | KS.1 |
| FIM1_1596 | 2242.375774 | 1618.077665 | 1618.955995 | -0.190426676 | 0.184128746 | 0.033514451 | 0.182822801 | ns | TM | KS.1 |
| FIM1_1597 | 822.4966779 | 1399.959332 | 1675.69388 | 0.258376463 | 0.294288822 | 0.020815421 | 0.140511155 | ns | TM | KS.1 |
| FIM1_1598 | 1767.981923 | 2522.112965 | 3069.22103 | 0.113920914 | 0.199184799 | 0.141128187 | 0.416559096 | ns | TM | KS.1 |
| FIM1_1599 | 2448.355978 | 2655.49748 | 2665.984911 | 0.066059514 | 0.077408438 | 0.205250515 | 0.512865935 | ns | TM | KS.1 |
| FIM1_1601 | 20748.21686 | 24225.97686 | 22763.71051 | 0.111496545 | 0.113177298 | 0.103643308 | 0.348223325 | ns | TM | KS.1 |
| FIM1_1602 | 3394.064107 | 2244.838389 | 2256.884439 | -0.549390919 | 0.109915819 | 1.30414263519 | 2.70337900420 | ns | TM | KS.1 |
| FIM1_1603 | 4107.247576 | 4332.53221 | 4657.555361 | 0.035816466 | 0.081326373 | 0.50119123 | 0.795351314 | ns | TM | KS.1 |
| FIM1_1604 | 2050.076903 | 2178.113342 | 2197.326057 | 0.034995499 | 0.095955512 | 0.550841119 | 0.822058746 | ns | TM | KS.1 |
| FIM1_1605 | 1309.411463 | 2300.940515 | 2906.898455 | 0.260208066 | 0.309440314 | 0.021771343 | 0.143841213 | ns | TM | KS.1 |
| FIM1_1606 | 6987.955087 | 7862.747761 | 7623.839318 | 0.077207543 | 0.10439044 | 0.222167333 | 0.536805479 | ns | TM | KS.1 |
| FIM1_1607 | 1405.393186 | 1409.100361 | 1263.574432 | 0.001422697 | 0.080177501 | 0.978418487 | 0.994408984 | ns | TM | KS.1 |
| FIM1_1608 | 2618.373989 | 2731.6816 | 2705.623237 | 0.02405983 | 0.093486416 | 0.67635007 | 0.887002016 | ns | TM | KS.1 |
| FIM1_1609 | 1231.131931 | 1352.165625 | 1307.238998 | 0.048918298 | 0.110328684 | 0.437803809 | 0.747832613 | ns | TM | KS.1 |
| FIM1_1610 | 1980.568778 | 2072.889202 | 2102.081972 | 0.018357776 | 0.119977489 | 0.770575003 | 0.925705893 | ns | TM | KS.1 |
| FIM1_1611 | 2126.835835 | 2086.300527 | 1829.231039 | -0.009501101 | 0.10261973 | 0.874089881 | 0.962759265 | ns | TM | KS.1 |
| FIM1_1612 | 1350.617264 | 1569.303795 | 1563.798557 | 0.064459151 | 0.132486642 | 0.338923085 | 0.667250632 | ns | TM | KS.1 |
| FIM1_1613 | 1061.970042 | 1209.717768 | 1328.538532 | 0.113190581 | 0.095038056 | 0.070150458 | 0.280239294 | ns | TM | KS.1 |
| FIM1_1614 | 728.6395534 | 666.4848209 | 538.7302173 | -0.054412786 | 0.098783997 | 0.366499278 | 0.690301888 | ns | TM | KS.1 |
| FIM1_1615 | 3143.468102 | 3087.127968 | 3138.809537 | -0.012222241 | 0.072389849 | 0.803780641 | 0.936908579 | ns | TM | KS.1 |
| FIM1_1616 | 775.5572485 | 857.6624111 | 848.8679576 | 0.063242729 | 0.100330589 | 0.301156802 | 0.630048398 | ns | TM | KS.1 |
| FIM1_1617 | 2463.37328 | 3036.517176 | 3551.417207 | 0.224350727 | 0.100336493 | 0.002763257 | 0.032809549 | ns | TM | KS.1 |
| FIM1_1618 | 147.400758 | 254.3220458 | 330.5806568 | 0.451062275 | 0.311190268 | 0.002157094 | 0.02746251 | ns | TM | KS.1 |
| FIM1_1619 | 1891.175573 | 1518.331307 | 1644.712945 | -0.206311688 | 0.118024388 | 0.009307058 | 0.079846584 | ns | TM | KS.1 |
| FIM1_1621 | 3333.043169 | 3313.172078 | 3449.992768 | -0.004077394 | 0.068390325 | 0.931324663 | 0.978943629 | ns | TM | KS.1 |
| FIM1_1622 | 685.4253995 | 482.3959031 | 889.5684964 | -0.327074912 | 0.17144722 | 0.001677152 | 0.022369518 | ns | TM | KS.1 |
| FIM1_1623 | 322.5251061 | 311.3309471 | 369.2683476 | -0.016994334 | 0.106017235 | 0.779967143 | 0.928976906 | ns | TM | KS.1 |
| FIM1_1624 | 972.8313236 | 702.5189591 | 674.7249446 | -0.274279713 | 0.167603462 | 0.005065618 | 0.0527227 | ns | TM | KS.1 |
| FIM1_1625 | 2038.937331 | 1831.356913 | 1770.134835 | -0.065088805 | 0.104887919 | 0.298777512 | 0.627569949 | ns | TM | KS.1 |
| FIM1_1627 | 2473.217366 | 2671.975048 | 2079.215773 | 0.0405441 | 0.105811359 | 0.510394613 | 0.802025647 | ns | TM | KS.1 |
| FIM1_1628 | 1650.969072 | 329.1605697 | 650.5022275 | -1.80265784 | 0.393938083 | 6.6885618899 | 7.5626353187 | Km<KS.1 | TM | KS.1 |
| FIM1_1629 | 379.7885445 | 395.5732327 | 370.7286386 | 0.023657829 | 0.093289213 | 0.681088189 | 0.887949093 | ns | TM | KS.1 |
| FIM1_1630 | 2708.506618 | 2547.683651 | 2888.50703 | -0.040942692 | 0.083586906 | 0.45059661 | 0.754323807 | ns | TM | KS.1 |
| FIM1_1631 | 764.0468921 | 699.4318312 | 737.8346959 | -0.040377067 | 0.116953927 | 0.526205675 | 0.808531286 | ns | TM | KS.1 |
| FIM1_1632 | 1431.786171 | 1308.766608 | 1564.129185 | -0.024702973 | 0.147669791 | 0.697632299 | 0.895901055 | ns | TM | KS.1 |
| FIM1_1633 | 1463.248953 | 1113.136259 | 1191.610824 | -0.0821517 | 0.175001459 | 0.24715227 | 0.57146079 | ns | TM | KS.1 |
| FIM1_1634 | 2079.845416 | 2187.54893 | 2116.876201 | 0.028245945 | 0.095715112 | 0.628762252 | 0.867709349 | ns | TM | KS.1 |
| FIM1_1635 | 5519.767338 | 6120.197903 | 6104.133034 | 0.099771489 | 0.076552454 | 0.061980781 | 0.261236705 | ns | TM | KS.1 |
| FIM1_1636 | 1125.99151 | 1022.634784 | 829.9480188 | -0.059826655 | 0.100298507 | 0.326584256 | 0.653825623 | ns | TM | KS.1 |

| | | | | | | | | | | |
|---|---|---|---|---|---|---|---|---|---|---|
| FIM1_1637 | 282.3438177 | 273.131494 | 238.0021329 | -0.014557153 | 0.107592661 | 0.811681517 | 0.939971031 | ns | TM | KS.1 |
| FIM1_1638 | 45.20680214 | 25.86150397 | 70.51401508 | -0.263233569 | 0.311571687 | 0.021155345 | 0.141841004 | ns | TM | KS.1 |
| FIM1_1640 | 2559.628625 | 2458.34608 | 2570.685489 | -0.026149028 | 0.081063558 | 0.622040275 | 0.86322186 | ns | TM | KS.1 |
| FIM1_1641 | 4708.644176 | 5071.1161 | 4617.052727 | 0.060405623 | 0.074252611 | 0.231588073 | 0.549231011 | ns | TM | KS.1 |
| FIM1_1642 | 2055.548181 | 1880.920963 | 1800.68227 | -0.070841141 | 0.082799588 | 0.195843713 | 0.50256523 | ns | TM | KS.1 |
| FIM1_1643 | 1522.026851 | 1521.092986 | 1577.718136 | -0.000271711 | 0.082012112 | 0.995935225 | 0.998142172 | ns | TM | KS.1 |
| FIM1_1644 | 599.0648544 | 663.1952002 | 640.2083275 | 0.072561808 | 0.095007839 | 0.225024009 | 0.541865655 | ns | TM | KS.1 |
| FIM1_1645 | 611.8611062 | 554.4118248 | 638.4829664 | -0.061642857 | 0.101295492 | 0.315290205 | 0.641836948 | ns | TM | KS.1 |
| FIM1_1646 | 2881.223732 | 2401.631215 | 3165.894573 | -0.110571085 | 0.128492554 | 0.124307998 | 0.387735747 | ns | TM | KS.1 |
| FIM1_1647 | 54.05793817 | 85.99841939 | 78.19150405 | 0.253166926 | 0.265033744 | 0.019185812 | 0.132384765 | ns | TM | KS.1 |
| FIM1_1648 | 4543.367272 | 3923.787808 | 3748.977745 | -0.08267936 | 0.120120993 | 0.221195059 | 0.535714496 | ns | TM | KS.1 |
| FIM1_1649 | 2735.165994 | 3243.820098 | 2915.235222 | 0.087662135 | 0.130637607 | 0.208949851 | 0.515893552 | ns | TM | KS.1 |
| FIM1_1650 | 1568.512707 | 997.3180154 | 1231.102463 | -0.270840421 | 0.25356039 | 0.013751279 | 0.104063892 | ns | TM | KS.1 |
| FIM1_1651 | 658.6797782 | 528.0795068 | 475.7090115 | -0.133405535 | 0.141602936 | 0.081991667 | 0.304774234 | ns | TM | KS.1 |
| FIM1_1652 | 3051.097903 | 3281.769952 | 2996.097767 | 0.043944575 | 0.095021509 | 0.453556762 | 0.757196272 | ns | TM | KS.1 |
| FIM1_1653 | 497.8239414 | 484.9246834 | 589.1797007 | -0.014978073 | 0.091590785 | 0.79233774 | 0.933303839 | ns | TM | KS.1 |
| FIM1_1654 | 591.1297719 | 463.2924102 | 472.4391557 | -0.119413781 | 0.154486991 | 0.117323809 | 0.375119503 | ns | TM | KS.1 |
| FIM1_1655 | 1427.585275 | 1291.740087 | 1447.568048 | -0.074261627 | 0.091004392 | 0.203195636 | 0.51148169 | ns | TM | KS.1 |
| FIM1_1656 | 1650.271504 | 1675.202486 | 1353.127567 | 0.008892293 | 0.085489691 | 0.870848658 | 0.96208467 | ns | TM | KS.1 |
| FIM1_1657 | 24805.18013 | 30313.35834 | 30073.46668 | 0.210001347 | 0.101667654 | 0.004639494 | 0.04900527 | ns | TM | KS.1 |
| FIM1_1658 | 1215.013338 | 1064.56724 | 1053.529141 | -0.08526978 | 0.109733827 | 0.192061012 | 0.496260756 | ns | TM | KS.1 |
| FIM1_1659 | 2638.931783 | 2808.320382 | 2925.916133 | 0.043674937 | 0.080040157 | 0.408171937 | 0.729140174 | ns | TM | KS.1 |
| FIM1_1660 | 2298.101203 | 2760.986892 | 2284.236219 | 0.179473011 | 0.104645337 | 0.0130654 | 0.101404627 | ns | TM | KS.1 |
| FIM1_1661 | 1502.576407 | 1794.45911 | 1659.451143 | 0.127372451 | 0.121217724 | 0.07746691 | 0.296004515 | ns | TM | KS.1 |
| FIM1_1664 | 2247.678552 | 2298.765126 | 2194.986819 | 0.017310829 | 0.061982874 | 0.694685465 | 0.893893954 | ns | TM | KS.1 |
| FIM1_1665 | 246.9983509 | 280.6381455 | 323.128888 | 0.052277574 | 0.12914809 | 0.427771783 | 0.742555695 | ns | TM | KS.1 |
| FIM1_1666 | 851.511237 | 884.2553878 | 919.0415446 | 0.027050047 | 0.064999748 | 0.553112377 | 0.822886984 | ns | TM | KS.1 |
| FIM1_1667 | 706.721787 | 1217.935839 | 1963.193442 | 0.341917738 | 0.312676347 | 0.008065402 | 0.072297973 | ns | TM | KS.1 |
| FIM1_1669 | 1725.969952 | 1781.843121 | 2080.923473 | 0.019694851 | 0.083832957 | 0.715919239 | 0.905084971 | ns | TM | KS.1 |
| FIM1_1670 | 1717.804293 | 1828.632613 | 1932.306517 | 0.026819111 | 0.117778604 | 0.670670947 | 0.882695228 | ns | TM | KS.1 |
| FIM1_1671 | 3179.111218 | 4883.038354 | 5657.092212 | 0.19293754 | 0.23468745 | 0.041160123 | 0.207258716 | ns | TM | KS.1 |
| FIM1_1672 | 4023.116452 | 4332.754727 | 4201.129198 | 0.041268574 | 0.101561357 | 0.495398 | 0.792222775 | ns | TM | KS.1 |
| FIM1_1673 | 2706.352066 | 3007.405091 | 3566.447103 | 0.095143443 | 0.08166281 | 0.08664169 | 0.313970477 | ns | TM | KS.1 |
| FIM1_1674 | 1905.370694 | 2340.737023 | 1673.238799 | 0.172413515 | 0.123005385 | 0.025666644 | 0.156676751 | ns | TM | KS.1 |
| FIM1_1675 | 15.29319008 | 20.65802725 | 18.27940201 | 0.067158172 | 0.19446712 | 0.324407837 | 0.651040335 | ns | TM | KS.1 |
| FIM1_1676 | 554.8476877 | 613.3333042 | 574.8902043 | 0.069843745 | 0.096371627 | 0.24568004 | 0.569551818 | ns | TM | KS.1 |
| FIM1_1677 | 892.2847603 | 514.057743 | 639.4036967 | -0.185340372 | 0.282601719 | 0.053368775 | 0.241250174 | ns | TM | KS.1 |
| FIM1_1678 | 188.8621426 | 170.8371098 | 212.1497853 | -0.036486353 | 0.132939692 | 0.572849096 | 0.832094673 | ns | TM | KS.1 |
| FIM1_1679 | 598.3741118 | 488.3096061 | 646.7479553 | -0.085194064 | 0.146767194 | 0.229193732 | 0.547402215 | ns | TM | KS.1 |
| FIM1_1680 | 449.7989217 | 468.6249035 | 512.5961361 | 0.019958348 | 0.104649396 | 0.741916785 | 0.914981657 | ns | TM | KS.1 |
| FIM1_1681 | 64.06327085 | 55.52898711 | 59.03730574 | -0.035582482 | 0.161246142 | 0.578470323 | 0.836595888 | ns | TM | KS.1 |
| FIM1_1682 | 696.1859544 | 798.8601793 | 870.1066741 | 0.068021178 | 0.123071019 | 0.308094619 | 0.636779799 | ns | TM | KS.1 |
| FIM1_1683 | 4265.053 | 3467.60493 | 4383.462853 | -0.117156329 | 0.138738776 | 0.114843244 | 0.37039344 | ns | TM | KS.1 |
| FIM1_1684 | 2949.313802 | 2986.474233 | 1764.440744 | 0.008342351 | 0.074941657 | 0.868403961 | 0.960282219 | ns | TM | KS.1 |
| FIM1_1685 | 3215.293709 | 2123.710787 | 2269.185769 | -0.551011757 | 0.111848596 | 1.93979197382[illegible] | 3.86018602791[illegible] | ns | TM | KS.1 |

| | | | | | | | | | | |
|---|---|---|---|---|---|---|---|---|---|---|
| FIM1_1686 | 1122.072399 | 1077.620309 | 1051.0058 | -0.023544597 | 0.090901984 | 0.678314304 | 0.887129504 | ns | TM | KS.1 |
| FIM1_1687 | 4406.604315 | 3806.944314 | 2583.761881 | -0.127545953 | 0.100539364 | 0.05262381 | 0.240005664 | ns | TM | KS.1 |
| FIM1_1688 | 509.9646607 | 155.1213728 | 317.1652885 | -1.533808473 | 0.25474086 | 1.5610309592 | 4.8538306387 | Km<KS.1 | TM | KS.1 |
| FIM1_1689 | 185.6000745 | 172.3818146 | 271.523357 | -0.030058205 | 0.120787907 | 0.636257694 | 0.869849417 | ns | TM | KS.1 |
| FIM1_1690 | 4485.309416 | 3870.710736 | 3733.747297 | -0.157909447 | 0.085105404 | 0.011349432 | 0.092109992 | ns | TM | KS.1 |
| FIM1_1691 | 166.5876816 | 369.8753078 | 2389.642711 | 0.359045578 | 0.431204889 | 0.011912973 | 0.095746435 | ns | TM | KS.1 |
| FIM1_1692 | 494.4294492 | 451.5407683 | 1205.142974 | -0.041317569 | 0.117100816 | 0.517139896 | 0.804981347 | ns | TM | KS.1 |
| FIM1_1693 | 1105.165738 | 442.8125603 | 768.7516542 | -1.07707753 | 0.351651522 | 2.7469708804 | 0.000923391 | Km<KS.1 | TM | KS.1 |
| FIM1_1694 | 44.13224663 | 54.4136587 | 53.63782133 | 0.056516395 | 0.159883569 | 0.397977273 | 0.721551361 | ns | TM | KS.1 |
| FIM1_1695 | 161.9325299 | 139.3387149 | 186.2138148 | -0.037271091 | 0.161289654 | 0.561770307 | 0.826391733 | ns | TM | KS.1 |
| FIM1_1697 | 178.1147785 | 123.5705213 | 205.3618799 | -0.156425376 | 0.204119343 | 0.067810174 | 0.274943454 | ns | TM | KS.1 |
| FIM1_1698 | 1399.355336 | 2094.46556 | 2099.503148 | 0.201645276 | 0.222997825 | 0.033972463 | 0.184512014 | ns | TM | KS.1 |
| FIM1_1701 | 431.1800076 | 409.4791942 | 548.8457104 | -0.020619012 | 0.119405453 | 0.743372675 | 0.91508234 | ns | TM | KS.1 |
| FIM1_1703 | 74.64234375 | 81.02406263 | 84.05414166 | 0.021979281 | 0.146059748 | 0.729055262 | 0.908876981 | ns | TM | KS.1 |
| FIM1_1705 | 18.49991091 | 17.56339468 | 22.09320857 | -0.007645077 | 0.163509851 | 0.90153159 | 0.968873116 | ns | TM | KS.1 |
| FIM1_1707 | 237.9453357 | 295.2124783 | 374.05025 | 0.120473246 | 0.141912986 | 0.108791169 | 0.357723772 | ns | TM | KS.1 |
| FIM1_1708 | 328.6619596 | 398.3672827 | 2951.506154 | 0.036402323 | 0.180845488 | 0.565600335 | 0.828581174 | ns | TM | KS.1 |
| FIM1_1709 | 766.3925782 | 803.8974149 | 1248.502807 | 0.015501541 | 0.134318167 | 0.806858514 | 0.937659684 | ns | TM | KS.1 |
| FIM1_1710 | 296.0376032 | 271.6008361 | 440.0704167 | -0.024352929 | 0.145033421 | 0.702044809 | 0.897397976 | ns | TM | KS.1 |
| FIM1_1711 | 1982.005194 | 1193.383592 | 1667.795688 | -0.172138469 | 0.262927204 | 0.061629845 | 0.260865744 | ns | TM | KS.1 |
| FIM1_1712 | 3433.629641 | 2061.104375 | 2816.95815 | -0.238700639 | 0.280150325 | 0.025400598 | 0.156140634 | ns | TM | KS.1 |
| FIM1_1713 | 62.84412938 | 43.81118949 | 91.60946069 | -0.15054616 | 0.197241153 | 0.073666919 | 0.288577104 | ns | TM | KS.1 |
| FIM1_1714 | 129.2475473 | 158.9474133 | 331.4930736 | 0.11007115 | 0.139437496 | 0.134315524 | 0.406212601 | ns | TM | KS.1 |
| FIM1_1715 | 276.2594736 | 250.8226399 | 205.5618188 | -0.037552081 | 0.126752295 | 0.560529593 | 0.826391733 | ns | TM | KS.1 |
| FIM1_1716 | 6704.859408 | 5705.727213 | 4759.1703 | -0.057947991 | 0.142699224 | 0.387890082 | 0.712272102 | ns | TM | KS.1 |
| FIM1_1717 | 129.9104578 | 169.5010463 | 180.8783533 | 0.200522033 | 0.148604065 | 0.019670921 | 0.134797287 | ns | TM | KS.1 |
| FIM1_1718 | 136.7484317 | 206.633938 | 178.6442181 | 0.424855735 | 0.183284616 | 0.00024694 | 0.005227779 | ns | TM | KS.1 |
| FIM1_1719 | 204.9637896 | 221.3857257 | 232.4189604 | 0.040392312 | 0.108545363 | 0.516137409 | 0.804981347 | ns | TM | KS.1 |
| FIM1_1721 | 18.8632283 | 23.12254471 | 37.01723315 | 0.045378091 | 0.183859123 | 0.482097743 | 0.783685372 | ns | TM | KS.1 |
| FIM1_1724 | 3268.983356 | 3167.263258 | 10659.98046 | -0.011595468 | 0.125689542 | 0.854250543 | 0.954818344 | ns | TM | KS.1 |
| FIM1_1725 | 15853.83762 | 17771.38025 | 18386.44368 | 0.047895183 | 0.126341146 | 0.463346192 | 0.765321151 | ns | TM | KS.1 |
| FIM1_1727 | 2696.699028 | 2425.738871 | 1983.948707 | -0.044799606 | 0.124481632 | 0.489970416 | 0.788868226 | ns | TM | KS.1 |
| FIM1_1728 | 6335.734995 | 5963.195154 | 6048.725339 | -0.035793623 | 0.093565824 | 0.536532771 | 0.813136987 | ns | TM | KS.1 |
| FIM1_1729 | 4183.650569 | 4196.553609 | 3532.289846 | 0.001863908 | 0.078685876 | 0.971390447 | 0.99362472 | ns | TM | KS.1 |
| FIM1_1730 | 4514.295116 | 4874.706368 | 4780.754088 | 0.071889607 | 0.065493506 | 0.122341002 | 0.38497564 | ns | TM | KS.1 |
| FIM1_1731 | 4690.684275 | 4179.442768 | 4615.337124 | -0.119144371 | 0.076215573 | 0.029511835 | 0.170326428 | ns | TM | KS.1 |
| FIM1_1732 | 1087.546045 | 1107.526588 | 1104.078117 | 0.009783671 | 0.097365067 | 0.867418142 | 0.960282219 | ns | TM | KS.1 |
| FIM1_1733 | 8889.793329 | 9191.350771 | 8127.148046 | 0.010048865 | 0.138463726 | 0.873566602 | 0.962759265 | ns | TM | KS.1 |
| FIM1_1734 | 542.9254512 | 582.3758946 | 542.6370371 | 0.027536809 | 0.123887336 | 0.665422224 | 0.880679852 | ns | TM | KS.1 |
| FIM1_1735 | 2618.623749 | 2144.470902 | 2353.005583 | -0.130016856 | 0.132015371 | 0.081916894 | 0.304774234 | ns | TM | KS.1 |
| FIM1_1736 | 686.9943826 | 730.3109959 | 664.8273505 | 0.03378596 | 0.099432777 | 0.571274436 | 0.832094673 | ns | TM | KS.1 |
| FIM1_1737 | 722.992202 | 542.3717354 | 484.8333214 | -0.200820099 | 0.161972975 | 0.022810262 | 0.147377994 | ns | TM | KS.1 |
| FIM1_1738 | 1775.077959 | 1163.200806 | 1392.724953 | -0.482397892 | 0.174756857 | 4.1807501836 | 0.001223484 | ns | TM | KS.1 |
| FIM1_1739 | 2233.696457 | 3025.282347 | 2986.10881 | 0.274056841 | 0.151078713 | 0.003681867 | 0.040978273 | ns | TM | KS.1 |
| FIM1_1740 | 1151.105359 | 1046.442952 | 944.6310137 | -0.067638173 | 0.092369406 | 0.248246099 | 0.573097143 | ns | TM | KS.1 |

| | | | | | | | | | | |
|---|---|---|---|---|---|---|---|---|---|---|
| FIM1_1741 | 2720.789391 | 3402.019424 | 4603.886683 | 0.142391494 | 0.141006242 | 0.066923216 | 0.272267737 | ns | TM | KS.1 |
| FIM1_1742 | 745.9147539 | 1038.842341 | 1223.137733 | 0.290749504 | 0.167054666 | 0.003477441 | 0.039377778 | ns | TM | KS.1 |
| FIM1_1743 | 3130.601106 | 3773.704475 | 3528.491098 | 0.07467217 | 0.143802065 | 0.281141769 | 0.606295641 | ns | TM | KS.1 |
| FIM1_1744 | 2089.998419 | 2379.472165 | 2700.92026 | 0.041148755 | 0.143597799 | 0.527932748 | 0.808531286 | ns | TM | KS.1 |
| FIM1_1745 | 1382.399904 | 1419.926 | 1396.955375 | 0.018704403 | 0.071502751 | 0.701376924 | 0.897235589 | ns | TM | KS.1 |
| FIM1_1746 | 477.3127892 | 506.8990417 | 512.8983152 | 0.029512652 | 0.10977083 | 0.633880654 | 0.869547581 | ns | TM | KS.1 |
| FIM1_1747 | 3109.918311 | 2988.355 | 3201.610679 | -0.019742956 | 0.104437329 | 0.744389594 | 0.91508234 | ns | TM | KS.1 |
| FIM1_1748 | 39.78421638 | 64.91332119 | 243.67275 | 0.235813587 | 0.270602191 | 0.025308388 | 0.156047257 | ns | TM | KS.1 |
| FIM1_1749 | 4002.939471 | 3881.448241 | 4469.165477 | -0.01651461 | 0.096672155 | 0.777671993 | 0.928020669 | ns | TM | KS.1 |
| FIM1_1750 | 4698.800058 | 5035.326984 | 4549.600814 | 0.022932143 | 0.134448543 | 0.719152219 | 0.905913973 | ns | TM | KS.1 |
| FIM1_1751 | 697.9220332 | 1120.894069 | 1262.995325 | 0.160489378 | 0.247661101 | 0.070974504 | 0.282027284 | ns | TM | KS.1 |
| FIM1_1752 | 965.8365883 | 964.4695144 | 1127.180991 | -0.000485191 | 0.140730697 | 0.993848293 | 0.99786059 | ns | TM | KS.1 |
| FIM1_1753 | 1019.6945 | 816.1135471 | 787.7175786 | -0.219006907 | 0.113999417 | 0.005755424 | 0.057381231 | ns | TM | KS.1 |
| FIM1_1754 | 307.2836017 | 279.7064574 | 560.3280302 | -0.03370812 | 0.13319913 | 0.601211874 | 0.851491516 | ns | TM | KS.1 |
| FIM1_1755 | 1755.845502 | 1762.474205 | 2207.912755 | 0.001007107 | 0.143572787 | 0.987207988 | 0.996167262 | ns | TM | KS.1 |
| FIM1_1756 | 1277.824258 | 1121.204619 | 1284.85607 | -0.052815029 | 0.131458959 | 0.424484808 | 0.740157476 | ns | TM | KS.1 |
| FIM1_1757 | 1420.620001 | 1418.737561 | 1650.215053 | -0.000586563 | 0.127427515 | 0.992575075 | 0.997449676 | ns | TM | KS.1 |
| FIM1_1758 | 343.358733 | 376.7757897 | 332.026574 | 0.033657821 | 0.131152418 | 0.601435415 | 0.851491516 | ns | TM | KS.1 |
| FIM1_1759 | 347.4040037 | 360.3903426 | 363.4795938 | 0.011771527 | 0.132577201 | 0.852420518 | 0.954011929 | ns | TM | KS.1 |
| FIM1_1760 | 3474.710186 | 3502.366123 | 3933.361456 | 0.002808682 | 0.126777991 | 0.964451559 | 0.993216478 | ns | TM | KS.1 |
| FIM1_1761 | 2345.390207 | 2443.480119 | 2837.073276 | 0.012966086 | 0.135428589 | 0.837702644 | 0.949337033 | ns | TM | KS.1 |
| FIM1_1762 | 2753.7388 | 4446.339656 | 5193.355965 | 0.262847953 | 0.267925863 | 0.016985274 | 0.121410542 | ns | TM | KS.1 |
| FIM1_1763 | 136.0683155 | 132.5710647 | 106.1058246 | -0.006938937 | 0.142210841 | 0.912220754 | 0.973674801 | ns | TM | KS.1 |
| FIM1_1764 | 3561.190236 | 3721.217975 | 3825.694864 | 0.030636248 | 0.075290245 | 0.544489719 | 0.817435092 | ns | TM | KS.1 |
| FIM1_1765 | 2234.627496 | 1714.193775 | 2218.508472 | -0.272089096 | 0.120315971 | 0.001533526 | 0.020788256 | ns | TM | KS.1 |
| FIM1_1766 | 1410.440156 | 1563.823256 | 1836.890909 | 0.031889764 | 0.141844424 | 0.620148175 | 0.861798092 | ns | TM | KS.1 |
| FIM1_1767 | 1916.300482 | 1810.176338 | 1753.236508 | -0.034168522 | 0.091856611 | 0.550908305 | 0.822058746 | ns | TM | KS.1 |
| FIM1_1769 | 18.58914958 | 17.68395728 | 21.2872472 | -0.005990844 | 0.188134784 | 0.920995877 | 0.975881018 | ns | TM | KS.1 |
| FIM1_1770 | 2789.375586 | 2959.651885 | 2963.922448 | 0.042308389 | 0.079156589 | 0.419383507 | 0.737776855 | ns | TM | KS.1 |
| FIM1_1771 | 1303.778758 | 1160.390449 | 1673.299133 | -0.071419004 | 0.106426352 | 0.260478733 | 0.588210523 | ns | TM | KS.1 |
| FIM1_1772 | 1644.153449 | 1727.920755 | 1718.984209 | 0.025899293 | 0.101724185 | 0.666304858 | 0.880910621 | ns | TM | KS.1 |
| FIM1_1774 | 1033.836288 | 949.2631547 | 925.4380031 | -0.054366689 | 0.095327536 | 0.357663488 | 0.682799636 | ns | TM | KS.1 |
| FIM1_1775 | 6972.615235 | 6811.279408 | 6632.013889 | -0.013223596 | 0.090902172 | 0.815415357 | 0.940791141 | ns | TM | KS.1 |
| FIM1_1776 | 2403.644545 | 2627.724453 | 2784.014127 | 0.048346004 | 0.106294533 | 0.436188575 | 0.747764597 | ns | TM | KS.1 |
| FIM1_1777 | 4641.236389 | 2200.091531 | 3183.634497 | -1.018843168 | 0.173843146 | 7.7693177413 | 1.8405883696 | Km<KS.1 | TM | KS.1 |
| FIM1_1778 | 2442.574418 | 2554.324772 | 2756.463574 | 0.023930599 | 0.098344496 | 0.685555187 | 0.890505759 | ns | TM | KS.1 |
| FIM1_1779 | 4224.501506 | 4704.662752 | 5361.145866 | 0.039797779 | 0.132251706 | 0.540129756 | 0.814399145 | ns | TM | KS.1 |
| FIM1_1780 | 791.3798412 | 874.4774572 | 982.3092643 | 0.066293578 | 0.097407849 | 0.271987609 | 0.598783418 | ns | TM | KS.1 |
| FIM1_1781 | 468.7812216 | 463.4987049 | 622.2993874 | -0.006045497 | 0.095208936 | 0.916923776 | 0.975634767 | ns | TM | KS.1 |
| FIM1_1782 | 1040.135633 | 964.2985975 | 1110.647824 | -0.063260173 | 0.072813382 | 0.2047514 | 0.512865935 | ns | TM | KS.1 |
| FIM1_1783 | 565.3948151 | 540.3075794 | 438.0035736 | -0.023433885 | 0.099860062 | 0.693740361 | 0.893671231 | ns | TM | KS.1 |
| FIM1_1784 | 1994.197973 | 2206.833109 | 2015.541673 | 0.065069552 | 0.099607936 | 0.28630935 | 0.612640436 | ns | TM | KS.1 |
| FIM1_1785 | 1037.840579 | 994.9986225 | 875.1746038 | -0.023085536 | 0.097490752 | 0.694811642 | 0.893893954 | ns | TM | KS.1 |
| FIM1_1786 | 50.37159789 | 47.04763114 | 53.71768876 | -0.015745423 | 0.133602943 | 0.803918045 | 0.936908579 | ns | TM | KS.1 |
| FIM1_1787 | 1632.940821 | 2186.827702 | 2637.023263 | 0.098890742 | 0.17752249 | 0.181497344 | 0.481472312 | ns | TM | KS.1 |

| | | | | | | | | | | |
|---|---|---|---|---|---|---|---|---|---|---|
| FIM1_1788 | 490.5518377 | 594.89472 | 541.1282185 | 0.133678871 | 0.127308877 | 0.071599554 | 0.283830902 | ns | TM | KS.1 |
| FIM1_1789 | 1302.84906 | 1519.703877 | 1310.35748 | 0.053627911 | 0.142598526 | 0.420629824 | 0.737778527 | ns | TM | KS.1 |
| FIM1_1790 | 1265.404348 | 774.8273424 | 819.0189478 | -0.54450924 | 0.225568873 | 0.000106932 | 0.002533262 | ns | TM | KS.1 |
| FIM1_1791 | 325.3522208 | 453.1213828 | 360.2382304 | 0.229278847 | 0.180857413 | 0.015408182 | 0.112895004 | ns | TM | KS.1 |
| FIM1_1792 | 6404.366035 | 6438.943305 | 6424.741287 | 0.00190646 | 0.126849423 | 0.975865346 | 0.993872484 | ns | TM | KS.1 |
| FIM1_1793 | 1745.615614 | 1622.934751 | 1819.898147 | -0.046842421 | 0.090921079 | 0.413463094 | 0.733587336 | ns | TM | KS.1 |
| FIM1_1794 | 889.6864513 | 778.8170072 | 681.1152267 | -0.031254436 | 0.161233591 | 0.62293637 | 0.863411908 | ns | TM | KS.1 |
| FIM1_1795 | 828.7162705 | 833.970065 | 820.1649257 | 0.002540203 | 0.053207941 | 0.948758714 | 0.986388143 | ns | TM | KS.1 |
| FIM1_1796 | 91.72099896 | 80.67194483 | 82.68238282 | -0.044183184 | 0.137582793 | 0.499942311 | 0.794415655 | ns | TM | KS.1 |
| FIM1_1797 | 1427.500453 | 1663.2439 | 1607.787515 | 0.058908655 | 0.137804327 | 0.379905699 | 0.703660035 | ns | TM | KS.1 |
| FIM1_1798 | 1412.839312 | 1366.934118 | 1934.363292 | -0.010190361 | 0.137242451 | 0.871868489 | 0.962615565 | ns | TM | KS.1 |
| FIM1_1799 | 1669.335434 | 1689.187159 | 1747.339532 | 0.00734713 | 0.082683657 | 0.890991286 | 0.967215407 | ns | TM | KS.1 |
| FIM1_1800 | 763.8112149 | 542.8856587 | 846.4389477 | -0.179690256 | 0.192446303 | 0.04285013 | 0.212118802 | ns | TM | KS.1 |
| FIM1_1801 | 1192.698363 | 1787.394654 | 2016.171127 | 0.147544743 | 0.21908997 | 0.081984819 | 0.304774234 | ns | TM | KS.1 |
| FIM1_1802 | 2320.223012 | 2230.903463 | 3150.560927 | -0.012416943 | 0.13585289 | 0.844427428 | 0.951966112 | ns | TM | KS.1 |
| FIM1_1803 | 2025.279214 | 2139.400387 | 4401.067168 | 0.012222039 | 0.158726378 | 0.844325484 | 0.951966112 | ns | TM | KS.1 |
| FIM1_1804 | 488.2799251 | 531.1075473 | 737.2140736 | 0.028870741 | 0.132963271 | 0.652607365 | 0.875599148 | ns | TM | KS.1 |
| FIM1_1805 | 38.11716587 | 26.74619851 | 87.50925761 | -0.096187713 | 0.199343011 | 0.193057276 | 0.496788132 | ns | TM | KS.1 |
| FIM1_1806 | 956.8217397 | 1053.618204 | 1470.141121 | 0.029501305 | 0.141406024 | 0.645610299 | 0.873039206 | ns | TM | KS.1 |
| FIM1_1807 | 74.80199791 | 84.142312 | 206.0439287 | 0.039335315 | 0.137628646 | 0.545268368 | 0.818313765 | ns | TM | KS.1 |
| FIM1_1808 | 2015.384247 | 2030.295165 | 2362.410885 | 0.002282575 | 0.133579397 | 0.971143762 | 0.99362472 | ns | TM | KS.1 |
| FIM1_1809 | 2854.530524 | 3292.927263 | 3503.93053 | 0.040652618 | 0.151262119 | 0.531582824 | 0.809001086 | ns | TM | KS.1 |
| FIM1_1810 | 1516.270384 | 1501.307912 | 1536.574471 | -0.005453062 | 0.093089622 | 0.924190281 | 0.976662213 | ns | TM | KS.1 |
| FIM1_1811 | 1421.062718 | 1475.061696 | 1497.327843 | 0.024086677 | 0.079964377 | 0.646817688 | 0.873595603 | ns | TM | KS.1 |
| FIM1_1812 | 742.3214371 | 664.1131465 | 669.3733824 | -0.050249925 | 0.121898743 | 0.439838498 | 0.747832613 | ns | TM | KS.1 |
| FIM1_1813 | 551.4734992 | 480.5902065 | 514.8656018 | -0.084003427 | 0.11353253 | 0.204848933 | 0.512865935 | ns | TM | KS.1 |
| FIM1_1814 | 965.6437865 | 950.9097818 | 879.8759774 | -0.008705592 | 0.076918091 | 0.865037414 | 0.959975149 | ns | TM | KS.1 |
| FIM1_1815 | 571.756571 | 448.7224893 | 661.3282019 | -0.213160204 | 0.129541134 | 0.010405531 | 0.087297666 | ns | TM | KS.1 |
| FIM1_1816 | 2185.179505 | 2090.635949 | 2257.114841 | -0.026292274 | 0.088965286 | 0.6394556 | 0.870968656 | ns | TM | KS.1 |
| FIM1_1817 | 1618.502178 | 1672.654704 | 1715.826269 | 0.018586562 | 0.091660876 | 0.74416882 | 0.91508234 | ns | TM | KS.1 |
| FIM1_1818 | 19932.78278 | 24291.20974 | 24018.39903 | 0.23466634 | 0.081239919 | 0.000486097 | 0.008793943 | ns | TM | KS.1 |
| FIM1_1819 | 3018.576373 | 3020.985839 | 2817.807409 | 0.000582644 | 0.081799376 | 0.991269922 | 0.996774593 | ns | TM | KS.1 |
| FIM1_1820 | 499.2483635 | 469.2889474 | 505.4351108 | -0.028754657 | 0.111271893 | 0.643911593 | 0.872978256 | ns | TM | KS.1 |
| FIM1_1821 | 1855.14253 | 1781.193727 | 1756.485633 | -0.023554522 | 0.090974299 | 0.678319726 | 0.887129504 | ns | TM | KS.1 |
| FIM1_1822 | 512.1554515 | 488.931795 | 470.8690079 | -0.026835758 | 0.091183043 | 0.637280814 | 0.870169358 | ns | TM | KS.1 |
| FIM1_1823 | 261.608135 | 254.2473565 | 249.6394333 | -0.006289326 | 0.154024301 | 0.919612484 | 0.975881018 | ns | TM | KS.1 |
| FIM1_1824 | 1077.788013 | 1081.9388 | 1074.180864 | 0.001767857 | 0.108017016 | 0.976891748 | 0.993872484 | ns | TM | KS.1 |
| FIM1_1825 | 4747.164053 | 4815.890381 | 4575.038093 | 0.007271248 | 0.101521031 | 0.903016343 | 0.970094214 | ns | TM | KS.1 |
| FIM1_1826 | 6848.73671 | 6786.394158 | 6200.367689 | -0.003537024 | 0.12145173 | 0.955047627 | 0.988678089 | ns | TM | KS.1 |
| FIM1_1827 | 5586.195527 | 5270.119112 | 5286.396663 | -0.039903435 | 0.080769087 | 0.452064584 | 0.755873008 | ns | TM | KS.1 |
| FIM1_1828 | 3599.710038 | 3594.898023 | 3775.938771 | -0.00074734 | 0.086364131 | 0.98915905 | 0.996167262 | ns | TM | KS.1 |
| FIM1_1829 | 309.371758 | 348.8909747 | 281.7080514 | 0.065363621 | 0.112199204 | 0.311053515 | 0.638961913 | ns | TM | KS.1 |
| FIM1_1830 | 1798.86556 | 1695.665311 | 1821.326502 | -0.040736106 | 0.080726486 | 0.442627344 | 0.748839657 | ns | TM | KS.1 |
| FIM1_1831 | 536.3182723 | 568.1201403 | 557.696713 | 0.03371 | 0.09570098 | 0.56480727 | 0.828150948 | ns | TM | KS.1 |
| FIM1_1832 | 432.7835939 | 438.7759472 | 410.2668753 | 0.007325351 | 0.092440488 | 0.897934973 | 0.967972723 | ns | TM | KS.1 |

| | | | | | | | | | | |
|---|---|---|---|---|---|---|---|---|---|---|
| FIM1_1833 | 1031.539493 | 817.4989745 | 903.3919552 | -0.279400813 | 0.082874015 | 7.3104334497$ | 0.001874712 | ns | TM | KS.1 |
| FIM1_1834 | 928.9588427 | 757.5297862 | 892.8793349 | -0.124098535 | 0.135840304 | 0.096705496 | 0.336440449 | ns | TM | KS.1 |
| FIM1_1835 | 849.5971712 | 780.1131471 | 680.737761 | -0.059210936 | 0.090836174 | 0.304380736 | 0.632801573 | ns | TM | KS.1 |
| FIM1_1836 | 2131.136097 | 2377.512192 | 2170.345231 | 0.053388116 | 0.117200284 | 0.408752428 | 0.729391439 | ns | TM | KS.1 |
| FIM1_1837 | 3647.849756 | 4776.463573 | 5368.555356 | 0.135490727 | 0.163011689 | 0.087586718 | 0.316214747 | ns | TM | KS.1 |
| FIM1_1838 | 158.7774698 | 157.2865578 | 170.4799956 | -0.004276263 | 0.125991864 | 0.94588649 | 0.985614597 | ns | TM | KS.1 |
| FIM1_1839 | 4303.362851 | 3590.168022 | 3181.795781 | -0.139221124 | 0.119274637 | 0.055978553 | 0.248654734 | ns | TM | KS.1 |
| FIM1_1840 | 5794.517282 | 6608.92879 | 7274.870358 | 0.067127963 | 0.120189 | 0.310678661 | 0.6389526 | ns | TM | KS.1 |
| FIM1_1841 | 1319.282451 | 1024.182406 | 1025.017145 | -0.298140355 | 0.09201052 | 8.5976373598 | 0.002149409 | ns | TM | KS.1 |
| FIM1_1842 | 1103.37383 | 1101.294895 | 1034.982679 | -0.000682874 | 0.13143661 | 0.991364962 | 0.996774593 | ns | TM | KS.1 |
| FIM1_1843 | 6740.201286 | 7786.271305 | 8375.507969 | 0.127740402 | 0.099057934 | 0.050375996 | 0.233221194 | ns | TM | KS.1 |
| FIM1_1844 | 55.70527345 | 66.99339046 | 62.51638684 | 0.059193762 | 0.150225509 | 0.379339368 | 0.70365897 | ns | TM | KS.1 |
| FIM1_1845 | 618.3664627 | 583.0060097 | 543.7782126 | -0.036419335 | 0.087079024 | 0.512351858 | 0.802566276 | ns | TM | KS.1 |
| FIM1_1846 | 3044.913213 | 3389.564812 | 3597.826933 | 0.071863101 | 0.099437541 | 0.241316729 | 0.563638838 | ns | TM | KS.1 |
| FIM1_1847 | 3252.900033 | 3143.182133 | 4023.427754 | -0.016512626 | 0.106649173 | 0.786460022 | 0.931802479 | ns | TM | KS.1 |
| FIM1_1848 | 6043.711924 | 7291.43628 | 7128.816569 | 0.097052363 | 0.135320885 | 0.174247358 | 0.470363867 | ns | TM | KS.1 |
| FIM1_1849 | 5041.763583 | 5350.464165 | 3645.990782 | 0.02480724 | 0.118726135 | 0.694144512 | 0.893729024 | ns | TM | KS.1 |
| FIM1_1850 | 762.1048708 | 760.1901481 | 680.5337264 | -0.00127334 | 0.108811358 | 0.983393507 | 0.996034691 | ns | TM | KS.1 |
| FIM1_1851 | 4859.215881 | 4039.622534 | 3843.198155 | -0.197095764 | 0.095401819 | 0.005196138 | 0.053410719 | ns | TM | KS.1 |
| FIM1_1853 | 41896.16543 | 52074.70197 | 51825.76471 | 0.170222443 | 0.130696337 | 0.031023961 | 0.176192016 | ns | TM | KS.1 |
| FIM1_1854 | 4406.168638 | 4771.291765 | 6421.667845 | 0.033160486 | 0.121278189 | 0.60313176 | 0.851547247 | ns | TM | KS.1 |
| FIM1_1855 | 772.3366115 | 612.5425038 | 833.8984346 | -0.096681304 | 0.154609659 | 0.184752372 | 0.487350505 | ns | TM | KS.1 |
| FIM1_1856 | 1786.858596 | 1637.313629 | 1627.702602 | -0.036442364 | 0.122469744 | 0.569692928 | 0.832094673 | ns | TM | KS.1 |
| FIM1_1857 | 5597.311003 | 5820.593225 | 6646.68315 | 0.022034917 | 0.093188592 | 0.701585482 | 0.897235589 | ns | TM | KS.1 |
| FIM1_1858 | 1530.895756 | 1140.579652 | 1277.778027 | -0.293452978 | 0.134237269 | 0.001381725 | 0.019255133 | ns | TM | KS.1 |
| FIM1_1859 | 2004.091664 | 2020.089998 | 1973.780227 | 0.004110529 | 0.099577275 | 0.944610638 | 0.984955766 | ns | TM | KS.1 |
| FIM1_1860 | 1352.811424 | 1359.928195 | 1463.585236 | 0.00199536 | 0.119809004 | 0.974576144 | 0.993872484 | ns | TM | KS.1 |
| FIM1_1861 | 614.4434598 | 548.7494744 | 621.8867677 | -0.070303327 | 0.104895042 | 0.264140511 | 0.590340989 | ns | TM | KS.1 |
| FIM1_1862 | 808.534476 | 816.5076993 | 889.5884481 | 0.005167059 | 0.08858337 | 0.926228331 | 0.976735299 | ns | TM | KS.1 |
| FIM1_1863 | 685.8278189 | 706.454941 | 657.76852 | 0.020529024 | 0.06816686 | 0.663490162 | 0.880619203 | ns | TM | KS.1 |
| FIM1_1864 | 1574.95102 | 1097.844131 | 1432.97111 | -0.420874127 | 0.133011866 | 1.7421319106 | 0.000637287 | ns | TM | KS.1 |
| FIM1_1865 | 805.1154548 | 947.264224 | 897.4371506 | 0.080996131 | 0.129314762 | 0.239590373 | 0.560395911 | ns | TM | KS.1 |
| FIM1_1866 | 234.4562275 | 207.5991416 | 472.7962783 | -0.03940923 | 0.142182354 | 0.544518572 | 0.817435092 | ns | TM | KS.1 |
| FIM1_1867 | 141.1398825 | 163.8477382 | 199.9969661 | 0.072569057 | 0.128698102 | 0.285232583 | 0.611828594 | ns | TM | KS.1 |
| FIM1_1868 | 1426.476393 | 1768.504655 | 1780.688788 | 0.172600252 | 0.127940598 | 0.027972844 | 0.165672497 | ns | TM | KS.1 |
| FIM1_1869 | 1615.624171 | 2152.916372 | 2045.223674 | 0.289956905 | 0.130240294 | 0.001324053 | 0.018660515 | ns | TM | KS.1 |
| FIM1_1870 | 1438.607448 | 1334.77501 | 1245.736782 | -0.045975718 | 0.094125491 | 0.431081071 | 0.743536592 | ns | TM | KS.1 |
| FIM1_1871 | 1522.344337 | 1328.833976 | 1710.552191 | -0.071750349 | 0.119983703 | 0.281196121 | 0.606295641 | ns | TM | KS.1 |
| FIM1_1872 | 672.6015208 | 671.5520207 | 908.4532218 | -0.000927331 | 0.097350025 | 0.987365072 | 0.996167262 | ns | TM | KS.1 |
| FIM1_1873 | 899.8280749 | 1062.702503 | 969.815206 | 0.132406255 | 0.112692533 | 0.059693534 | 0.256462934 | ns | TM | KS.1 |
| FIM1_1874 | 1870.479344 | 1683.268646 | 1626.170728 | -0.057192894 | 0.110861155 | 0.369048284 | 0.69186398 | ns | TM | KS.1 |
| FIM1_1875 | 64.6129773 | 60.07423133 | 58.7017236 | -0.025189153 | 0.13363923 | 0.693456749 | 0.893671231 | ns | TM | KS.1 |
| FIM1_1876 | 613.0841693 | 547.0075173 | 644.4371434 | -0.027270433 | 0.15832641 | 0.666649034 | 0.881131494 | ns | TM | KS.1 |
| FIM1_1877 | 595.4666007 | 946.2060929 | 1186.719207 | 0.157840505 | 0.243252738 | 0.073242869 | 0.288050017 | ns | TM | KS.1 |
| FIM1_1878 | 1993.0159 | 2363.67917 | 2956.918088 | 0.042248465 | 0.162582564 | 0.514467189 | 0.804256034 | ns | TM | KS.1 |

| | | | | | | | | | | |
|---|---|---|---|---|---|---|---|---|---|---|
| FIM1_1879 | 485.9773994 | 452.9895373 | 476.7881425 | -0.046172342 | 0.086224884 | 0.40502347 | 0.727171333 | ns | TM | KS.1 |
| FIM1_1880 | 3215.641178 | 3347.008195 | 3492.685695 | 0.024269381 | 0.087007277 | 0.661212192 | 0.879788889 | ns | TM | KS.1 |
| FIM1_1881 | 671.6257331 | 696.9842823 | 750.7455602 | 0.023271474 | 0.082405528 | 0.664019404 | 0.880619203 | ns | TM | KS.1 |
| FIM1_1882 | 3029.267621 | 3031.958686 | 3247.16289 | 0.000387927 | 0.112889709 | 0.994993705 | 0.99790453 | ns | TM | KS.1 |
| FIM1_1883 | 99.30835249 | 112.2359867 | 133.49796 | 0.035721966 | 0.145638325 | 0.580341474 | 0.838424819 | ns | TM | KS.1 |
| FIM1_1884 | 4680.210883 | 4602.860347 | 4422.520693 | -0.007720426 | 0.107797402 | 0.899336783 | 0.968452273 | ns | TM | KS.1 |
| FIM1_1885 | 1657.859558 | 1863.526402 | 1568.879703 | 0.096947949 | 0.091885264 | 0.106368187 | 0.353494809 | ns | TM | KS.1 |
| FIM1_1886 | 1010.310039 | 1000.498949 | 895.1978618 | -0.006043774 | 0.081807047 | 0.909648714 | 0.972251892 | ns | TM | KS.1 |
| FIM1_1887 | 5399.296898 | 6117.432608 | 5765.02779 | 0.066135877 | 0.116828665 | 0.312926233 | 0.639671112 | ns | TM | KS.1 |
| FIM1_1888 | 2762.793734 | 2811.900881 | 2288.547933 | 0.015119799 | 0.049015662 | 0.684458084 | 0.890505759 | ns | TM | KS.1 |
| FIM1_1889 | 1398.127881 | 1467.991028 | 1442.223128 | 0.031221244 | 0.08426785 | 0.566051663 | 0.828754274 | ns | TM | KS.1 |
| FIM1_1890 | 567.9964822 | 958.4629033 | 1155.395381 | 0.15044905 | 0.262215438 | 0.084692135 | 0.310267577 | ns | TM | KS.1 |
| FIM1_1891 | 1479.781495 | 1494.545854 | 1454.40925 | 0.002802919 | 0.141928791 | 0.96445389 | 0.993216478 | ns | TM | KS.1 |
| FIM1_1892 | 1121.731541 | 1015.652876 | 894.0127034 | -0.036864791 | 0.130689013 | 0.568624523 | 0.831053761 | ns | TM | KS.1 |
| FIM1_1893 | 1261.544902 | 1547.015113 | 1753.076194 | 0.177550825 | 0.120041397 | 0.021121592 | 0.141808262 | ns | TM | KS.1 |
| FIM1_1894 | 1623.264294 | 1792.325259 | 1675.682039 | 0.05703441 | 0.105325635 | 0.359968642 | 0.683664672 | ns | TM | KS.1 |
| FIM1_1895 | 1156.460956 | 1200.545456 | 1091.70638 | 0.023439298 | 0.081565943 | 0.659710563 | 0.878874681 | ns | TM | KS.1 |
| FIM1_1896 | 1135.871315 | 1244.070332 | 1174.592518 | 0.071942988 | 0.083787396 | 0.192840652 | 0.496788132 | ns | TM | KS.1 |
| FIM1_1897 | 1724.710488 | 1917.672856 | 1873.564261 | 0.085095148 | 0.089244474 | 0.143836683 | 0.420933821 | ns | TM | KS.1 |
| FIM1_1898 | 2328.384961 | 1737.802344 | 1951.388759 | -0.347894636 | 0.098490291 | 1.48047513668 | 0.000562242 | ns | TM | KS.1 |
| FIM1_1899 | 7621.689526 | 7800.195757 | 8322.433107 | 0.013289784 | 0.089773998 | 0.813268453 | 0.940696014 | ns | TM | KS.1 |
| FIM1_1900 | 433.8750951 | 447.7560356 | 403.9192299 | 0.019546234 | 0.085777725 | 0.721663267 | 0.906864045 | ns | TM | KS.1 |
| FIM1_1901 | 3783.618123 | 3942.077249 | 4115.123878 | 0.020599584 | 0.103079623 | 0.732461288 | 0.910998727 | ns | TM | KS.1 |
| FIM1_1902 | 759.4051363 | 789.7381478 | 880.7159253 | 0.020714482 | 0.098781405 | 0.726219919 | 0.907961357 | ns | TM | KS.1 |
| FIM1_1903 | 726.0193618 | 628.1384698 | 936.9536444 | -0.132167447 | 0.096857371 | 0.041518504 | 0.208364318 | ns | TM | KS.1 |
| FIM1_1904 | 2266.935494 | 2909.070894 | 5400.172419 | 0.214808266 | 0.135437884 | 0.011312435 | 0.091959747 | ns | TM | KS.1 |
| FIM1_1905 | 1081.519391 | 1060.756427 | 1071.12881 | -0.010973095 | 0.087206548 | 0.842816815 | 0.951443988 | ns | TM | KS.1 |
| FIM1_1907 | 1118.137166 | 1232.488552 | 1143.469546 | 0.048077679 | 0.113881983 | 0.450356212 | 0.754323807 | ns | TM | KS.1 |
| FIM1_1908 | 3852.767174 | 4509.62499 | 4344.102142 | 0.10105757 | 0.118570376 | 0.142716452 | 0.419382366 | ns | TM | KS.1 |
| FIM1_1909 | 4013.508284 | 3549.624619 | 4253.410281 | -0.106042838 | 0.0922177 | 0.081257093 | 0.304408161 | ns | TM | KS.1 |
| FIM1_1910 | 1900.698435 | 1824.267522 | 1422.179399 | -0.022391826 | 0.096944414 | 0.702710142 | 0.898017713 | ns | TM | KS.1 |
| FIM1_1911 | 5721.364077 | 7146.193138 | 7619.23478 | 0.271854198 | 0.077976321 | 5.49858540056 | 0.00148671 | ns | TM | KS.1 |
| FIM1_1912 | 2652.204066 | 2790.825792 | 3167.10355 | 0.032550711 | 0.084466655 | 0.550339661 | 0.822058746 | ns | TM | KS.1 |
| FIM1_1913 | 1634.766262 | 2540.083929 | 2600.344044 | 0.255479538 | 0.245820589 | 0.016454569 | 0.119173353 | ns | TM | KS.1 |
| FIM1_1914 | 627.6125581 | 614.8280114 | 540.0841743 | -0.010153406 | 0.102465081 | 0.865460776 | 0.959975149 | ns | TM | KS.1 |
| FIM1_1915 | 1909.923638 | 2059.314639 | 2001.753845 | 0.02603357 | 0.132360732 | 0.683905448 | 0.890494131 | ns | TM | KS.1 |
| FIM1_1916 | 1158.297165 | 1251.488589 | 1040.49571 | 0.033016616 | 0.119777871 | 0.603777154 | 0.851547247 | ns | TM | KS.1 |
| FIM1_1917 | 737.5466388 | 998.9425027 | 992.6185635 | 0.135076563 | 0.177681923 | 0.092501491 | 0.328710655 | ns | TM | KS.1 |
| FIM1_1918 | 868.5411379 | 939.2926432 | 883.3691243 | 0.041322424 | 0.105796732 | 0.502530606 | 0.796206932 | ns | TM | KS.1 |
| FIM1_1919 | 1281.882788 | 1037.009587 | 1482.241881 | -0.192665652 | 0.119442189 | 0.013969859 | 0.105143797 | ns | TM | KS.1 |
| FIM1_1920 | 1195.632789 | 1973.318201 | 2593.445431 | 0.21658366 | 0.271353104 | 0.033342557 | 0.182084765 | ns | TM | KS.1 |
| FIM1_1921 | 3283.443038 | 2954.296143 | 2806.918951 | -0.058887162 | 0.108941851 | 0.352478469 | 0.678105331 | ns | TM | KS.1 |
| FIM1_1922 | 2445.450369 | 2269.544484 | 2248.488215 | -0.072575498 | 0.060481521 | 0.098693795 | 0.339178757 | ns | TM | KS.1 |
| FIM1_1923 | 1929.952675 | 2443.972439 | 2652.797944 | 0.272330759 | 0.093097411 | 0.000300079 | 0.006019728 | ns | TM | KS.1 |
| FIM1_1924 | 1137.766694 | 1124.224822 | 1067.849095 | -0.005697275 | 0.101073571 | 0.923783274 | 0.976662213 | ns | TM | KS.1 |

| | | | | | | | | | | |
|---|---|---|---|---|---|---|---|---|---|---|
| FIM1_1925 | 2348.483302 | 2097.664656 | 2161.135795 | -0.100716649 | 0.086553173 | 0.082830956 | 0.305700302 | ns | TM | KS.1 |
| FIM1_1926 | 876.983987 | 555.7803485 | 614.4443236 | -0.441640956 | 0.231234975 | 0.000682235 | 0.011351568 | ns | TM | KS.1 |
| FIM1_1927 | 2052.151485 | 3209.261025 | 3871.751739 | 0.196423782 | 0.243417228 | 0.040327541 | 0.204589117 | ns | TM | KS.1 |
| FIM1_1928 | 1730.578567 | 1636.834916 | 1626.853045 | -0.036746419 | 0.083795677 | 0.498491406 | 0.793598318 | ns | TM | KS.1 |
| FIM1_1929 | 529.4918967 | 533.5193443 | 577.4460461 | 0.002492574 | 0.130748366 | 0.968490052 | 0.993374633 | ns | TM | KS.1 |
| FIM1_1930 | 770.6797221 | 696.361237 | 719.1019424 | -0.059128001 | 0.105718971 | 0.344330129 | 0.670129201 | ns | TM | KS.1 |
| FIM1_1931 | 18202.28428 | 15105.00816 | 12554.5319 | -0.225538624 | 0.075678672 | 0.000402254 | 0.007638225 | ns | TM | KS.1 |
| FIM1_1932 | 896.8476003 | 559.7629024 | 615.3969019 | -0.61727064 | 0.147229053 | 1.05917912468 | 1.1455252489 | ns | TM | KS.1 |
| FIM1_1933 | 4850.493622 | 4954.939408 | 5025.61618 | 0.012956318 | 0.083387286 | 0.810054664 | 0.939473859 | ns | TM | KS.1 |
| FIM1_1934 | 379.5753522 | 352.0812143 | 592.1345006 | -0.035441089 | 0.112634198 | 0.572211611 | 0.832094673 | ns | TM | KS.1 |
| FIM1_1935 | 892.3687634 | 918.1472962 | 2584.393175 | 0.014637017 | 0.101236589 | 0.80629475 | 0.937659684 | ns | TM | KS.1 |
| FIM1_1936 | 4251.722712 | 4787.502106 | 5334.781665 | 0.139877325 | 0.065082986 | 0.005971369 | 0.05906076 | ns | TM | KS.1 |
| FIM1_1937 | 2532.854916 | 2698.155552 | 2724.981043 | 0.045002536 | 0.079419349 | 0.391806411 | 0.715009037 | ns | TM | KS.1 |
| FIM1_1938 | 1342.007207 | 1393.816154 | 1245.841524 | 0.018907367 | 0.103837205 | 0.754240876 | 0.919693225 | ns | TM | KS.1 |
| FIM1_1939 | 229.684322 | 225.8527013 | 307.9868761 | -0.006112558 | 0.126042055 | 0.922740203 | 0.976369825 | ns | TM | KS.1 |
| FIM1_1940 | 3500.230171 | 3286.894429 | 3000.403003 | -0.02454287 | 0.123662051 | 0.699150837 | 0.896699645 | ns | TM | KS.1 |
| FIM1_1941 | 1583.924188 | 1324.083979 | 1548.488475 | -0.066145976 | 0.145219075 | 0.332144886 | 0.65912278 | ns | TM | KS.1 |
| FIM1_1942 | 1041.130504 | 813.975822 | 919.5417893 | -0.166541249 | 0.147265657 | 0.041413741 | 0.208324934 | ns | TM | KS.1 |
| FIM1_1943 | 1104.833134 | 1118.091046 | 1103.740139 | 0.006009103 | 0.097547512 | 0.918365092 | 0.975634767 | ns | TM | KS.1 |
| FIM1_1944 | 562.5599119 | 529.7526314 | 568.8789386 | -0.039792727 | 0.086776586 | 0.473645396 | 0.774362749 | ns | TM | KS.1 |
| FIM1_1945 | 796.4572758 | 766.1924164 | 776.3111489 | -0.020122866 | 0.099683239 | 0.734742273 | 0.91155681 | ns | TM | KS.1 |
| FIM1_1946 | 2309.389598 | 1832.006408 | 2199.170287 | -0.167654738 | 0.139225645 | 0.037058589 | 0.193255946 | ns | TM | KS.1 |
| FIM1_1947 | 1105.832269 | 1093.603483 | 995.8216166 | -0.005292075 | 0.103384853 | 0.929828867 | 0.97843743 | ns | TM | KS.1 |
| FIM1_1948 | 1319.435006 | 1738.981023 | 2939.991438 | 0.297474422 | 0.11725274 | 0.000591636 | 0.010184737 | ns | TM | KS.1 |
| FIM1_1949 | 500.0363114 | 470.0461435 | 448.1257848 | -0.030897501 | 0.107081013 | 0.615267318 | 0.858612877 | ns | TM | KS.1 |
| FIM1_1950 | 1337.903708 | 1186.241791 | 1066.766058 | -0.086833002 | 0.100630715 | 0.165740617 | 0.456381383 | ns | TM | KS.1 |
| FIM1_1951 | 2173.598293 | 1448.097855 | 1505.63966 | -0.456294457 | 0.168441347 | 5.8825284486 | 0.001573418 | ns | TM | KS.1 |
| FIM1_1952 | 1828.303803 | 1916.017005 | 2539.892516 | 0.032331448 | 0.075863763 | 0.524663291 | 0.808531286 | ns | TM | KS.1 |
| FIM1_1953 | 2138.943977 | 2128.009378 | 3001.487266 | -0.002099311 | 0.120067882 | 0.97326154 | 0.993872484 | ns | TM | KS.1 |
| FIM1_1954 | 1036.006565 | 1805.402806 | 2132.792289 | 0.153519584 | 0.273136496 | 0.082089944 | 0.304774234 | ns | TM | KS.1 |
| FIM1_1955 | 498.9442216 | 496.2527645 | 615.9159347 | -0.002952522 | 0.079786325 | 0.955092844 | 0.988678089 | ns | TM | KS.1 |
| FIM1_1956 | 807.8640805 | 631.7348215 | 737.4129827 | -0.128413798 | 0.154024182 | 0.097509974 | 0.3377271 | ns | TM | KS.1 |
| FIM1_1957 | 1142.91605 | 1226.511373 | 1281.058434 | 0.041247637 | 0.09663202 | 0.484995662 | 0.785178463 | ns | TM | KS.1 |
| FIM1_1958 | 1220.746527 | 1022.230668 | 1017.167306 | -0.110272763 | 0.126121264 | 0.12283172 | 0.385787757 | ns | TM | KS.1 |
| FIM1_1959 | 740.8971518 | 723.5579893 | 825.9505333 | -0.014140401 | 0.083225169 | 0.792854658 | 0.933303839 | ns | TM | KS.1 |
| FIM1_1960 | 2686.532549 | 2652.267232 | 2838.54034 | -0.009254658 | 0.057754893 | 0.825342411 | 0.945229857 | ns | TM | KS.1 |
| FIM1_1961 | 837.1170999 | 893.9009 | 3122.551629 | 0.029637234 | 0.115013834 | 0.637199257 | 0.870169358 | ns | TM | KS.1 |
| FIM1_1962 | 29339.40549 | 36327.37624 | 36346.13897 | 0.196987511 | 0.118482815 | 0.012137363 | 0.096768242 | ns | TM | KS.1 |
| FIM1_1963 | 541.0745211 | 574.7668992 | 534.5366959 | 0.031451906 | 0.104620798 | 0.605811102 | 0.851547247 | ns | TM | KS.1 |
| FIM1_1964 | 9628.314069 | 11194.17586 | 12314.88059 | 0.059325174 | 0.136433215 | 0.376468749 | 0.700423346 | ns | TM | KS.1 |
| FIM1_1965 | 1242.269235 | 1044.205688 | 1120.310801 | -0.125196695 | 0.119701802 | 0.080146865 | 0.301630805 | ns | TM | KS.1 |
| FIM1_1966 | 1695.791585 | 1286.652238 | 1545.494868 | -0.285514538 | 0.122733581 | 0.00112461 | 0.016554684 | ns | TM | KS.1 |
| FIM1_1967 | 2060.904317 | 1865.604666 | 2172.736869 | -0.067022257 | 0.096689954 | 0.265055475 | 0.591039896 | ns | TM | KS.1 |
| FIM1_1968 | 1591.06491 | 2083.609048 | 2275.329364 | 0.131514898 | 0.163670493 | 0.094979345 | 0.333005827 | ns | TM | KS.1 |
| FIM1_1970 | 674.2513656 | 605.9862868 | 570.7668803 | -0.055468987 | 0.113139729 | 0.386386782 | 0.710637426 | ns | TM | KS.1 |

| | | | | | | | | | | |
|---|---|---|---|---|---|---|---|---|---|---|
| FIM1_1971 | 476.8340184 | 459.0284787 | 411.3584587 | -0.019713501 | 0.102642227 | 0.74304107 | 0.91508234 | ns | TM | KS.1 |
| FIM1_1972 | 3130.809291 | 2204.141777 | 3532.771367 | -0.125940589 | 0.197139009 | 0.113836327 | 0.368468268 | ns | TM | KS.1 |
| FIM1_1973 | 4925.115457 | 6376.302141 | 7261.775774 | 0.105232351 | 0.163058242 | 0.15843275 | 0.445060944 | ns | TM | KS.1 |
| FIM1_1975 | 1557.781713 | 1444.277504 | 1218.097813 | -0.054838887 | 0.083095234 | 0.313085258 | 0.639671112 | ns | TM | KS.1 |
| FIM1_1976 | 2590.967816 | 2240.994241 | 2935.738703 | -0.094777787 | 0.113716483 | 0.158396633 | 0.445060944 | ns | TM | KS.1 |
| FIM1_1977 | 746.3395462 | 795.5253387 | 1034.614862 | 0.046762747 | 0.077104474 | 0.364238079 | 0.687959166 | ns | TM | KS.1 |
| FIM1_1978 | 2432.568125 | 2384.914049 | 2647.71356 | -0.007135453 | 0.126526931 | 0.909910515 | 0.972251892 | ns | TM | KS.1 |
| FIM1_1979 | 822.9746115 | 763.8984787 | 814.5126033 | -0.059942266 | 0.074110558 | 0.234445932 | 0.55356835 | ns | TM | KS.1 |
| FIM1_1980 | 10776.8042 | 9024.014192 | 8603.960782 | -0.095217155 | 0.131121685 | 0.17790113 | 0.476606421 | ns | TM | KS.1 |
| FIM1_1981 | 9846.902737 | 10666.50728 | 10239.13225 | 0.035076718 | 0.118126031 | 0.580945439 | 0.838424819 | ns | TM | KS.1 |
| FIM1_1982 | 23.36969219 | 13.49412252 | 64.93626001 | -0.11971321 | 0.261413988 | 0.133913783 | 0.40527648 | ns | TM | KS.1 |
| FIM1_1983 | 3742.627121 | 4141.597462 | 4897.795103 | 0.071743793 | 0.094389047 | 0.228184454 | 0.545762703 | ns | TM | KS.1 |
| FIM1_1984 | 529.9472223 | 532.431078 | 569.3680288 | 0.003005283 | 0.079414294 | 0.954156316 | 0.988678089 | ns | TM | KS.1 |
| FIM1_1985 | 2330.736613 | 2768.47537 | 2826.378344 | 0.145564253 | 0.111539698 | 0.041026884 | 0.206928663 | ns | TM | KS.1 |
| FIM1_1986 | 1872.618983 | 1936.604709 | 2153.313448 | 0.025075753 | 0.065354384 | 0.583932943 | 0.840829636 | ns | TM | KS.1 |
| FIM1_1987 | 761.5499972 | 806.105705 | 767.2735327 | 0.024860129 | 0.115991031 | 0.692034984 | 0.893131011 | ns | TM | KS.1 |
| FIM1_1988 | 2687.490387 | 2751.812408 | 2531.306992 | 0.011432026 | 0.104646342 | 0.849976749 | 0.953031594 | ns | TM | KS.1 |
| FIM1_1989 | 742.055692 | 693.1627878 | 1456.817408 | -0.030428106 | 0.116559941 | 0.629609492 | 0.868412743 | ns | TM | KS.1 |
| FIM1_1990 | 1517.532754 | 1799.047287 | 2173.972986 | 0.150285944 | 0.108612169 | 0.033696895 | 0.183415813 | ns | TM | KS.1 |
| FIM1_1991 | 2125.919645 | 2102.940188 | 2246.902047 | -0.004879851 | 0.110715815 | 0.936771933 | 0.982179213 | ns | TM | KS.1 |
| FIM1_1992 | 4530.974245 | 4393.692309 | 4555.279446 | -0.012115444 | 0.120629823 | 0.847070609 | 0.952348261 | ns | TM | KS.1 |
| FIM1_1993 | 2174.262419 | 1803.434763 | 1575.670143 | -0.118155034 | 0.128908893 | 0.104788835 | 0.350588067 | ns | TM | KS.1 |
| FIM1_1994 | 10759.50975 | 13290.24083 | 15695.74925 | 0.255578669 | 0.079011293 | 0.000146512 | 0.003343562 | ns | TM | KS.1 |
| FIM1_1995 | 2708.86079 | 3072.547967 | 3370.377933 | 0.049776068 | 0.131526065 | 0.44962557 | 0.754176403 | ns | TM | KS.1 |
| FIM1_1996 | 6128.351164 | 7584.922181 | 7937.596766 | 0.158108044 | 0.131886478 | 0.042269296 | 0.210079667 | ns | TM | KS.1 |
| FIM1_1997 | 1544.361913 | 1454.697185 | 1520.024576 | -0.043552554 | 0.075564639 | 0.391138606 | 0.71457503 | ns | TM | KS.1 |
| FIM1_1998 | 1437.271441 | 1457.272367 | 1350.747153 | 0.0102814 | 0.06466356 | 0.821144094 | 0.943898306 | ns | TM | KS.1 |
| FIM1_1999 | 262.0171337 | 217.6636122 | 411.8301405 | -0.084240822 | 0.13957957 | 0.230625178 | 0.548995638 | ns | TM | KS.1 |
| FIM1_2000 | 5493.878471 | 5676.165393 | 5751.658379 | 0.016000535 | 0.104546742 | 0.791401596 | 0.933303839 | ns | TM | KS.1 |
| FIM1_2001 | 1630.753265 | 2630.170598 | 3238.179242 | 0.257549568 | 0.266792058 | 0.018198999 | 0.127881388 | ns | TM | KS.1 |
| FIM1_2002 | 2735.218273 | 2927.140087 | 3293.617611 | 0.046052761 | 0.085281185 | 0.402972434 | 0.725058901 | ns | TM | KS.1 |
| FIM1_2003 | 4839.609839 | 5590.016706 | 4467.052273 | 0.089424983 | 0.11545805 | 0.18310111 | 0.48479405 | ns | TM | KS.1 |
| FIM1_2004 | 664.8815765 | 422.9190586 | 610.1034241 | -0.472555154 | 0.215137891 | 0.000262952 | 0.005450771 | ns | TM | KS.1 |
| FIM1_2005 | 28074.32149 | 32514.50907 | 32739.38233 | 0.079775682 | 0.121691864 | 0.237997684 | 0.558772287 | ns | TM | KS.1 |
| FIM1_2006 | 849.6112539 | 982.6690344 | 869.3747403 | 0.098960125 | 0.112200543 | 0.140524216 | 0.415393924 | ns | TM | KS.1 |
| FIM1_2007 | 548.285038 | 596.0499721 | 634.0629093 | 0.051712725 | 0.095459853 | 0.381164489 | 0.70494176 | ns | TM | KS.1 |
| FIM1_2008 | 588.1913306 | 559.0347861 | 575.9856178 | -0.026535297 | 0.100126894 | 0.656470784 | 0.877276998 | ns | TM | KS.1 |
| FIM1_2009 | 294.8793732 | 265.5312771 | 814.4961066 | -0.032242061 | 0.143329468 | 0.616290666 | 0.859446765 | ns | TM | KS.1 |
| FIM1_2010 | 39.78543691 | 28.70392744 | 78.59222337 | -0.099436852 | 0.183350053 | 0.180455624 | 0.480643537 | ns | TM | KS.1 |
| FIM1_2011 | 2058.884904 | 2337.501235 | 2751.075001 | 0.124983187 | 0.085307113 | 0.035429068 | 0.188204876 | ns | TM | KS.1 |
| FIM1_2012 | 2204.920949 | 2011.876517 | 3168.947475 | -0.062390442 | 0.092468295 | 0.285315043 | 0.611828594 | ns | TM | KS.1 |
| FIM1_2013 | 1310.276171 | 1021.384846 | 1113.463587 | -0.124339698 | 0.155999644 | 0.106749063 | 0.353815182 | ns | TM | KS.1 |
| FIM1_2014 | 1943.024652 | 1667.443699 | 1707.956662 | -0.137459893 | 0.101273352 | 0.040170684 | 0.204136011 | ns | TM | KS.1 |
| FIM1_2015 | 18.22583219 | 21.9355548 | 21.92099029 | 0.042436705 | 0.159852931 | 0.51335971 | 0.803386145 | ns | TM | KS.1 |
| FIM1_2016 | 2136.652146 | 2432.995264 | 2361.274511 | 0.115374354 | 0.093332194 | 0.062774439 | 0.262637093 | ns | TM | KS.1 |

| | | | | | | | | | | |
|---|---|---|---|---|---|---|---|---|---|---|
| FIM1_2017 | 1232.709302 | 905.6393599 | 1226.622931 | -0.198743453 | 0.173764483 | 0.026480942 | 0.159881904 | ns | TM | KS.1 |
| FIM1_2018 | 403.6866356 | 112.2947866 | 205.6604993 | -1.512971413 | 0.342185327 | 4.1738117693 | 3.9178704816 | Km<KS.1 | TM | KS.1 |
| FIM1_2019 | 1604.214515 | 1424.376165 | 1473.133084 | -0.047278411 | 0.130033394 | 0.470548173 | 0.771581134 | ns | TM | KS.1 |
| FIM1_2020 | 3506.079774 | 2694.12443 | 3206.016417 | -0.147849044 | 0.158624788 | 0.066931345 | 0.272267737 | ns | TM | KS.1 |
| FIM1_2021 | 4295.749324 | 3837.315977 | 4196.697368 | -0.053910755 | 0.119119235 | 0.406626682 | 0.728526463 | ns | TM | KS.1 |
| FIM1_2022 | 1671.767168 | 1698.43037 | 1584.664656 | 0.008662031 | 0.09528326 | 0.88127224 | 0.9656535 | ns | TM | KS.1 |
| FIM1_2023 | 2566.084458 | 1972.31337 | 2191.806579 | -0.188835663 | 0.151080743 | 0.026327924 | 0.159323047 | ns | TM | KS.1 |
| FIM1_2024 | 1003.651118 | 164.3651479 | 535.9085137 | -1.557679505 | 0.63674334 | 2.1056936279 | 0.000742966 | Km<KS.1 | TM | KS.1 |
| FIM1_2025 | 216.0464473 | 205.8610276 | 215.5552265 | -0.020268891 | 0.117575538 | 0.746785453 | 0.916442434 | ns | TM | KS.1 |
| FIM1_2026 | 1175.998928 | 1261.135323 | 1235.45582 | 0.051533732 | 0.07734178 | 0.319177634 | 0.647069572 | ns | TM | KS.1 |
| FIM1_2027 | 1097.289789 | 1039.152864 | 972.3894514 | -0.035015687 | 0.085142839 | 0.522884134 | 0.807872226 | ns | TM | KS.1 |
| FIM1_2028 | 1889.028668 | 1933.130661 | 2123.234624 | 0.010930427 | 0.106854501 | 0.857538545 | 0.955916258 | ns | TM | KS.1 |
| FIM1_2029 | 2723.651987 | 2676.39578 | 2108.789834 | -0.012329312 | 0.067805928 | 0.793160662 | 0.933303839 | ns | TM | KS.1 |
| FIM1_2030 | 448.2373552 | 409.7286805 | 1023.682246 | -0.040942515 | 0.118686364 | 0.522202024 | 0.807870838 | ns | TM | KS.1 |
| FIM1_2031 | 974.387691 | 983.6134418 | 1632.790141 | 0.002715985 | 0.138525593 | 0.965619848 | 0.993374633 | ns | TM | KS.1 |
| FIM1_2032 | 10.142242 | 9.04066601 | 27.57214771 | -0.015916715 | 0.192920166 | 0.79297425 | 0.933303839 | ns | TM | KS.1 |
| FIM1_2033 | 1472.526337 | 1391.428936 | 1731.269197 | -0.04018607 | 0.078165467 | 0.438993438 | 0.747832613 | ns | TM | KS.1 |
| FIM1_2034 | 1967.250169 | 1916.97088 | 2139.809632 | -0.018685497 | 0.067880882 | 0.691239016 | 0.893131011 | ns | TM | KS.1 |
| FIM1_2035 | 342.5992357 | 205.4369196 | 253.7108773 | -0.481927936 | 0.272797085 | 0.000848825 | 0.013521989 | ns | TM | KS.1 |
| FIM1_2037 | 506.6455046 | 439.639724 | 314.2585225 | -0.056008201 | 0.134543697 | 0.400546463 | 0.723750973 | ns | TM | KS.1 |
| FIM1_2038 | 1977.593701 | 2102.613419 | 1938.2021 | 0.029870903 | 0.108127844 | 0.628058938 | 0.867219877 | ns | TM | KS.1 |
| FIM1_2039 | 421.1715085 | 533.9663468 | 620.846494 | 0.13764547 | 0.148968406 | 0.078437851 | 0.298339685 | ns | TM | KS.1 |
| FIM1_2040 | 296.648853 | 348.7900148 | 447.3371595 | 0.086638604 | 0.126134404 | 0.209356005 | 0.516639943 | ns | TM | KS.1 |
| FIM1_2041 | 2038.301992 | 1630.625854 | 5070.218881 | -0.163393281 | 0.135531441 | 0.039104819 | 0.19994499 | ns | TM | KS.1 |
| FIM1_2042 | 3787.408022 | 3404.240003 | 3704.298972 | -0.0893807 | 0.086317961 | 0.118361383 | 0.377224778 | ns | TM | KS.1 |
| FIM1_2043 | 1475.028333 | 2096.994487 | 2435.732405 | 0.238992722 | 0.193368776 | 0.014419445 | 0.107712825 | ns | TM | KS.1 |
| FIM1_2044 | 757.635563 | 801.8286242 | 810.1224928 | 0.032036398 | 0.096921715 | 0.586414402 | 0.841613393 | ns | TM | KS.1 |
| FIM1_2045 | 422.533947 | 348.4580232 | 395.9114986 | -0.122162654 | 0.13166527 | 0.097810478 | 0.3377271 | ns | TM | KS.1 |
| FIM1_2046 | 237.2941603 | 298.8820703 | 265.7848338 | 0.098933488 | 0.15332909 | 0.17633129 | 0.473932019 | ns | TM | KS.1 |
| FIM1_2047 | 657.7242259 | 608.7736844 | 771.6282506 | -0.03838103 | 0.110256467 | 0.538891727 | 0.814399145 | ns | TM | KS.1 |
| FIM1_2048 | 2026.901258 | 1968.797315 | 1837.727007 | -0.020362875 | 0.071914413 | 0.677541352 | 0.887002016 | ns | TM | KS.1 |
| FIM1_2049 | 428.9859271 | 456.8392595 | 419.7850415 | 0.038522905 | 0.090370502 | 0.49809957 | 0.793454967 | ns | TM | KS.1 |
| FIM1_2050 | 3834.115801 | 3320.229093 | 3690.857324 | -0.133314819 | 0.095818743 | 0.038868122 | 0.198939204 | ns | TM | KS.1 |
| FIM1_2051 | 1949.421545 | 2075.658491 | 2042.719606 | 0.041871008 | 0.084251797 | 0.442831411 | 0.748839657 | ns | TM | KS.1 |
| FIM1_2052 | 3912.20657 | 4102.348116 | 3036.137461 | 0.022737368 | 0.107547563 | 0.710676606 | 0.901942886 | ns | TM | KS.1 |
| FIM1_2053 | 8692.297558 | 7279.965787 | 6991.329792 | -0.16081842 | 0.108740362 | 0.02521151 | 0.156001645 | ns | TM | KS.1 |
| FIM1_2054 | 2255.810431 | 2181.640861 | 1495.778769 | -0.0212342 | 0.080795475 | 0.68812921 | 0.891826565 | ns | TM | KS.1 |
| FIM1_2055 | 631.2310906 | 922.7038644 | 893.0585356 | 0.110951828 | 0.207992518 | 0.149927054 | 0.43114861 | ns | TM | KS.1 |
| FIM1_2056 | 753.3091254 | 960.1323795 | 844.1085238 | 0.127711582 | 0.152612241 | 0.098338574 | 0.338336381 | ns | TM | KS.1 |
| FIM1_2057 | 616.6741579 | 707.0935708 | 693.8576477 | 0.101013262 | 0.105504785 | 0.121919839 | 0.384126384 | ns | TM | KS.1 |
| FIM1_2058 | 269.5681982 | 438.3360146 | 517.6529092 | 0.266169831 | 0.272812789 | 0.01668795 | 0.120322541 | ns | TM | KS.1 |
| FIM1_2059 | 567.4795448 | 575.5167247 | 774.0261571 | 0.007903674 | 0.093673826 | 0.890673216 | 0.967215407 | ns | TM | KS.1 |
| FIM1_2060 | 6003.989208 | 4442.557869 | 3723.71952 | -0.093049686 | 0.182051266 | 0.202668142 | 0.510777105 | ns | TM | KS.1 |
| FIM1_2061 | 1644.872035 | 1870.28865 | 1823.673689 | 0.078631654 | 0.111102978 | 0.228023807 | 0.545655815 | ns | TM | KS.1 |
| FIM1_2062 | 762.29924 | 760.0915544 | 1165.197352 | -0.00086265 | 0.138149991 | 0.989077315 | 0.996167262 | ns | TM | KS.1 |

| | | | | | | | | | | |
|---|---|---|---|---|---|---|---|---|---|---|
| FIM1_2063 | 2805.149276 | 2610.63429 | 3667.742342 | -0.043690147 | 0.094505269 | 0.454778986 | 0.758218986 | ns | TM | KS.1 |
| FIM1_2064 | 1367.231728 | 524.3969645 | 1152.213554 | -1.215799884 | 0.273438373 | 5.9422076408 | 5.0969798298 | Km<KS.1 | TM | KS.1 |
| FIM1_2065 | 16.88395755 | 43.21195576 | 56.90405186 | 0.965722555 | 0.456746799 | 0.000216076 | 0.004673814 | ns | TM | KS.1 |
| FIM1_2066 | 973.4471818 | 421.7004649 | 3256.199129 | -0.464373721 | 0.471511575 | 0.005830454 | 0.058013014 | ns | TM | KS.1 |
| FIM1_2067 | 2255.567289 | 2636.449865 | 2073.776345 | 0.092331392 | 0.121017305 | 0.178695479 | 0.477526476 | ns | TM | KS.1 |
| FIM1_2068 | 962.6052885 | 1078.746105 | 1436.187791 | 0.085419304 | 0.096772737 | 0.162684349 | 0.45316609 | ns | TM | KS.1 |
| FIM1_2069 | 1440.724217 | 1211.860785 | 1555.198042 | -0.131240971 | 0.117000132 | 0.066180404 | 0.270807979 | ns | TM | KS.1 |
| FIM1_2070 | 76.79577331 | 78.5618506 | 80.28421507 | 0.007202676 | 0.13555266 | 0.909180262 | 0.972251892 | ns | TM | KS.1 |
| FIM1_2071 | 1798.890815 | 1992.342102 | 2107.247229 | 0.102290472 | 0.072767694 | 0.048281078 | 0.228978421 | ns | TM | KS.1 |
| FIM1_2072 | 1435.404482 | 1546.226791 | 1485.603666 | 0.043857912 | 0.097224375 | 0.459886991 | 0.763054265 | ns | TM | KS.1 |
| FIM1_2073 | 2558.78217 | 3352.647606 | 3368.971596 | 0.142671322 | 0.162496755 | 0.075743985 | 0.293478448 | ns | TM | KS.1 |
| FIM1_2074 | 693.2687774 | 795.0676502 | 831.1518814 | 0.113742773 | 0.101164033 | 0.079440446 | 0.300154562 | ns | TM | KS.1 |
| FIM1_2075 | 8026.080871 | 7335.782082 | 6104.843797 | -0.05427761 | 0.099906382 | 0.370432703 | 0.69308112 | ns | TM | KS.1 |
| FIM1_2076 | 1473.788212 | 1582.638811 | 1345.166444 | 0.047966252 | 0.087005271 | 0.390046379 | 0.713674416 | ns | TM | KS.1 |
| FIM1_2077 | 2300.22509 | 2298.918867 | 2543.597471 | -0.000269056 | 0.107685358 | 0.996478906 | 0.998284849 | ns | TM | KS.1 |
| FIM1_2078 | 1469.71962 | 1509.30416 | 1355.849322 | 0.012092377 | 0.109059536 | 0.843682921 | 0.951768052 | ns | TM | KS.1 |
| FIM1_2079 | 5503.333879 | 4302.471479 | 1965.955201 | -0.071355898 | 0.169378339 | 0.30184665 | 0.630695962 | ns | TM | KS.1 |
| FIM1_2080 | 25.40714817 | 23.87500016 | 36.46740299 | -0.008523188 | 0.160857202 | 0.890612427 | 0.967215407 | ns | TM | KS.1 |
| FIM1_2081 | 5637.069085 | 8949.686485 | 11195.6198 | 0.318143523 | 0.259373799 | 0.007147603 | 0.066581689 | ns | TM | KS.1 |
| FIM1_2083 | 712.074409 | 770.0636405 | 754.0544526 | 0.029365177 | 0.12791444 | 0.646451021 | 0.873463833 | ns | TM | KS.1 |
| FIM1_2084 | 6535.02701 | 7078.497719 | 6222.755823 | 0.077129487 | 0.065179409 | 0.097722655 | 0.3377271 | ns | TM | KS.1 |
| FIM1_2085 | 2844.965358 | 2825.863386 | 2721.631889 | -0.004786678 | 0.057663157 | 0.909044241 | 0.972251892 | ns | TM | KS.1 |
| FIM1_2086 | 4059.132579 | 3781.091756 | 3151.420719 | -0.054449207 | 0.07684782 | 0.291016967 | 0.618193599 | ns | TM | KS.1 |
| FIM1_2087 | 5187.07628 | 4557.940421 | 3960.918562 | -0.041458943 | 0.143029781 | 0.52506843 | 0.808531286 | ns | TM | KS.1 |
| FIM1_2088 | 1184.670051 | 1317.133545 | 1507.92372 | 0.064613301 | 0.104734433 | 0.301768377 | 0.630695962 | ns | TM | KS.1 |
| FIM1_2089 | 325.9318286 | 311.0356525 | 349.9156491 | -0.023435483 | 0.105343225 | 0.700144548 | 0.897138176 | ns | TM | KS.1 |
| FIM1_2090 | 687.67321 | 756.0545066 | 845.9294823 | 0.069580066 | 0.087433423 | 0.219743999 | 0.533281169 | ns | TM | KS.1 |
| FIM1_2091 | 1147.407749 | 1221.87717 | 1214.776412 | 0.036958866 | 0.094702797 | 0.526257186 | 0.808531286 | ns | TM | KS.1 |
| FIM1_2092 | 570.1567397 | 712.9505156 | 677.1625973 | 0.0757281 | 0.158825897 | 0.277912964 | 0.602049362 | ns | TM | KS.1 |
| FIM1_2093 | 1798.450899 | 2864.892335 | 3388.232517 | 0.196446794 | 0.251651896 | 0.041570401 | 0.208364318 | ns | TM | KS.1 |
| FIM1_2094 | 6264.835512 | 7723.067963 | 8107.293509 | 0.152610847 | 0.131238639 | 0.047788493 | 0.227509809 | ns | TM | KS.1 |
| FIM1_2095 | 1211.88645 | 811.0570375 | 1015.029635 | -0.362794968 | 0.203688355 | 0.001582453 | 0.021279312 | ns | TM | KS.1 |
| FIM1_2096 | 1605.25316 | 590.8811919 | 1200.383166 | -1.298099006 | 0.24377577 | 1.3388997447 | 1.8002773594 | Km<KS.1 | TM | KS.1 |
| FIM1_2097 | 1702.41664 | 1524.083955 | 1071.318606 | -0.101370588 | 0.082526737 | 0.071946043 | 0.284298304 | ns | TM | KS.1 |
| FIM1_2098 | 449.0004457 | 157.073416 | 1283.458349 | -0.876200712 | 0.55206692 | 0.000648251 | 0.010846109 | ns | TM | KS.1 |
| FIM1_2099 | 15.4217908 | 16.81186055 | 15.19172552 | 0.022338419 | 0.180623793 | 0.717632533 | 0.905458243 | ns | TM | KS.1 |
| FIM1_2100 | 1538.928455 | 1801.988803 | 1852.733756 | 0.090561858 | 0.123046718 | 0.188593486 | 0.491815863 | ns | TM | KS.1 |
| FIM1_2101 | 749.7754447 | 841.5388317 | 761.0685894 | 0.071840063 | 0.105690535 | 0.256306568 | 0.584919805 | ns | TM | KS.1 |
| FIM1_2102 | 1056.383812 | 1100.862536 | 1111.61358 | 0.025075201 | 0.086693655 | 0.65006118 | 0.874068749 | ns | TM | KS.1 |
| FIM1_2103 | 6538.902607 | 5265.095811 | 5729.274012 | -0.086397154 | 0.151474076 | 0.225179063 | 0.541907079 | ns | TM | KS.1 |
| FIM1_2104 | 565.2515181 | 689.6038143 | 766.9407679 | 0.090732943 | 0.142884591 | 0.203565454 | 0.511742362 | ns | TM | KS.1 |
| FIM1_2105 | 454.417109 | 558.0246585 | 459.8558034 | 0.1296168 | 0.135325748 | 0.085082503 | 0.31075147 | ns | TM | KS.1 |
| FIM1_2106 | 1028.026184 | 912.2856756 | 901.2688859 | -0.059691383 | 0.118387816 | 0.360678602 | 0.683750066 | ns | TM | KS.1 |
| FIM1_2107 | 628.2263173 | 741.9456957 | 817.3913675 | 0.126493301 | 0.115182056 | 0.072655766 | 0.286420314 | ns | TM | KS.1 |
| FIM1_2108 | 740.6507251 | 679.9506313 | 655.523916 | -0.048737801 | 0.102934036 | 0.426060621 | 0.741071977 | ns | TM | KS.1 |

| | | | | | | | | | | |
|---|---|---|---|---|---|---|---|---|---|---|
| FIM1_2109 | 195.7201602 | 215.6532926 | 233.2353216 | 0.042330256 | 0.119419843 | 0.509428877 | 0.801014116 | ns | TM | KS.1 |
| FIM1_2110 | 317.7879156 | 397.6917118 | 423.3050302 | 0.134013096 | 0.14418469 | 0.082326394 | 0.304891809 | ns | TM | KS.1 |
| FIM1_2111 | 40.73294071 | 46.72283922 | 44.30100201 | 0.03753491 | 0.153382028 | 0.561006821 | 0.826391733 | ns | TM | KS.1 |
| FIM1_2112 | 1307.988478 | 925.9560852 | 1126.871592 | -0.357871832 | 0.150713309 | 0.000406661 | 0.007658836 | ns | TM | KS.1 |
| FIM1_2113 | 1667.739182 | 1477.365292 | 1409.47737 | -0.08669672 | 0.10179869 | 0.169028446 | 0.463059758 | ns | TM | KS.1 |
| FIM1_2115 | 1183.795689 | 1216.5557 | 1230.235367 | 0.017439289 | 0.079001882 | 0.7379818 | 0.913453034 | ns | TM | KS.1 |
| FIM1_2116 | 1162.823933 | 1125.303653 | 1207.977676 | -0.026472007 | 0.059275485 | 0.535056962 | 0.812052589 | ns | TM | KS.1 |
| FIM1_2117 | 2180.101767 | 2201.207793 | 2470.402507 | 0.007354558 | 0.05546554 | 0.856706816 | 0.955916258 | ns | TM | KS.1 |
| FIM1_2118 | 1563.525789 | 1324.134658 | 1670.173672 | -0.207488839 | 0.064736817 | 0.000226892 | 0.004844585 | ns | TM | KS.1 |
| FIM1_2119 | 649.6288458 | 902.6369737 | 891.3692608 | 0.111566751 | 0.189670298 | 0.145630389 | 0.423690751 | ns | TM | KS.1 |
| FIM1_2120 | 2095.723096 | 1968.479764 | 1712.28105 | -0.036606721 | 0.094685008 | 0.530097562 | 0.808531286 | ns | TM | KS.1 |
| FIM1_2121 | 3569.815088 | 3227.498825 | 3712.965937 | -0.081371901 | 0.086433011 | 0.15221696 | 0.434578193 | ns | TM | KS.1 |
| FIM1_2122 | 1202.800777 | 1329.019094 | 1269.252022 | 0.072357666 | 0.093272083 | 0.221190098 | 0.535714496 | ns | TM | KS.1 |
| FIM1_2123 | 7261.242253 | 7550.035044 | 7668.136712 | 0.025634321 | 0.078915908 | 0.622913534 | 0.863411908 | ns | TM | KS.1 |
| FIM1_2124 | 3999.863627 | 1954.730044 | 2574.161137 | -0.93470641 | 0.228446648 | 1.2012742595[illegible] | 7.9684525881[illegible] | ns | TM | KS.1 |
| FIM1_2125 | 1090.052234 | 1080.242303 | 1307.919858 | -0.00483165 | 0.096969415 | 0.934132473 | 0.980651837 | ns | TM | KS.1 |
| FIM1_2126 | 1417.523968 | 956.030472 | 1140.724034 | -0.391837385 | 0.18534381 | 0.000558535 | 0.009784194 | ns | TM | KS.1 |
| FIM1_2127 | 932.917521 | 1003.054264 | 1172.311572 | 0.05374907 | 0.081105226 | 0.314796265 | 0.641585998 | ns | TM | KS.1 |
| FIM1_2128 | 757.9559001 | 781.0129026 | 763.6469738 | 0.015747994 | 0.101689842 | 0.792362929 | 0.933303839 | ns | TM | KS.1 |
| FIM1_2129 | 3038.741236 | 3027.439703 | 3207.282133 | -0.002129201 | 0.080584824 | 0.967808414 | 0.993374633 | ns | TM | KS.1 |
| FIM1_2130 | 2163.579827 | 1705.027107 | 4591.296402 | -0.111107196 | 0.153623471 | 0.138368646 | 0.412176572 | ns | TM | KS.1 |
| FIM1_2131 | 9573.722207 | 11197.93064 | 10641.61632 | 0.104494226 | 0.116840692 | 0.129072225 | 0.39662404 | ns | TM | KS.1 |
| FIM1_2132 | 1224.374764 | 1928.012495 | 2400.954067 | 0.141158963 | 0.236797551 | 0.094078189 | 0.331213943 | ns | TM | KS.1 |
| FIM1_2133 | 1067.045202 | 978.5567459 | 1195.454349 | -0.064676351 | 0.084995854 | 0.24372824 | 0.567141251 | ns | TM | KS.1 |
| FIM1_2134 | 975.0754474 | 1042.650228 | 923.6418429 | 0.039680898 | 0.09587732 | 0.49962206 | 0.79438151 | ns | TM | KS.1 |
| FIM1_2135 | 1878.20209 | 2279.387849 | 2432.676187 | 0.171203285 | 0.115912867 | 0.02272969 | 0.147048383 | ns | TM | KS.1 |
| FIM1_2136 | 1864.604426 | 2114.183023 | 1894.50463 | 0.111233108 | 0.091745002 | 0.068360306 | 0.275864055 | ns | TM | KS.1 |
| FIM1_2137 | 1140.930161 | 936.2579308 | 929.025873 | -0.145677374 | 0.12656608 | 0.053099311 | 0.24081046 | ns | TM | KS.1 |
| FIM1_2138 | 1002.238816 | 1072.642767 | 1077.161676 | 0.029098824 | 0.118215324 | 0.645375446 | 0.873039206 | ns | TM | KS.1 |
| FIM1_2139 | 3137.220324 | 3059.932859 | 2822.038722 | -0.013970667 | 0.091741908 | 0.806149948 | 0.937659684 | ns | TM | KS.1 |
| FIM1_2140 | 1049.82555 | 1000.089764 | 1049.093401 | -0.032697857 | 0.079190246 | 0.531847689 | 0.809156653 | ns | TM | KS.1 |
| FIM1_2141 | 1175.048297 | 1352.538449 | 1365.113948 | 0.088118631 | 0.113761814 | 0.186125979 | 0.488642081 | ns | TM | KS.1 |
| FIM1_2142 | 162.7933045 | 186.6692376 | 184.7391978 | 0.074376113 | 0.117859485 | 0.262767489 | 0.589200856 | ns | TM | KS.1 |
| FIM1_2143 | 898.6931716 | 933.9286153 | 887.3095082 | 0.023933659 | 0.088940719 | 0.669559365 | 0.882180579 | ns | TM | KS.1 |
| FIM1_2144 | 1272.544331 | 1379.375574 | 1324.008097 | 0.04703403 | 0.100126401 | 0.435805566 | 0.747374247 | ns | TM | KS.1 |
| FIM1_2145 | 1032.793443 | 783.6228454 | 768.7166833 | -0.123509662 | 0.167337248 | 0.11224617 | 0.364506981 | ns | TM | KS.1 |
| FIM1_2146 | 14108.6428 | 14465.76273 | 11469.12418 | 0.017594269 | 0.069251221 | 0.712296349 | 0.903537566 | ns | TM | KS.1 |
| FIM1_2147 | 479.1947728 | 664.134328 | 730.851883 | 0.253876036 | 0.174039489 | 0.008647118 | 0.076683448 | ns | TM | KS.1 |
| FIM1_2148 | 548.5415141 | 588.0137718 | 554.9139861 | 0.031066449 | 0.115400017 | 0.621738054 | 0.863043197 | ns | TM | KS.1 |
| FIM1_2149 | 934.2356904 | 979.3685288 | 947.8479098 | 0.031424772 | 0.078607491 | 0.545911511 | 0.818784976 | ns | TM | KS.1 |
| FIM1_2150 | 1544.13553 | 1483.880067 | 1219.572483 | -0.018080163 | 0.110876465 | 0.769761559 | 0.925705893 | ns | TM | KS.1 |
| FIM1_2151 | 546.3371242 | 725.1405762 | 1008.198779 | 0.240548678 | 0.149193113 | 0.007890274 | 0.070983926 | ns | TM | KS.1 |
| FIM1_2152 | 128.962954 | 178.8253954 | 256.6004841 | 0.288410939 | 0.165183164 | 0.0035335 | 0.03977186 | ns | TM | KS.1 |
| FIM1_2153 | 545.7106694 | 498.8787587 | 445.3975149 | -0.058952719 | 0.096315748 | 0.322871144 | 0.650844386 | ns | TM | KS.1 |
| FIM1_2154 | 739.5792127 | 1190.206502 | 1304.304847 | 0.147157724 | 0.245592152 | 0.086839586 | 0.314201411 | ns | TM | KS.1 |

| | | | | | | | | | | |
|---|---|---|---|---|---|---|---|---|---|---|
| FIM1_2155 | 752.5434571 | 785.0968749 | 789.1994816 | 0.028978986 | 0.076934332 | 0.572018979 | 0.832094673 | ns | TM | KS.1 |
| FIM1_2156 | 1217.421856 | 1378.375188 | 1257.674473 | 0.070080525 | 0.113775694 | 0.282597904 | 0.608625355 | ns | TM | KS.1 |
| FIM1_2157 | 5573.951776 | 5279.231981 | 3731.140676 | -0.041218174 | 0.070666559 | 0.394865729 | 0.718791439 | ns | TM | KS.1 |
| FIM1_2158 | 1996.733561 | 2597.440841 | 2815.198628 | 0.273110614 | 0.11840609 | 0.001371748 | 0.019169794 | ns | TM | KS.1 |
| FIM1_2159 | 50.24731571 | 65.76654177 | 86.34968685 | 0.075871736 | 0.177947754 | 0.277556161 | 0.602049362 | ns | TM | KS.1 |
| FIM1_2160 | 1731.37843 | 1735.961826 | 1771.477552 | 0.001551016 | 0.091017621 | 0.978148825 | 0.994338048 | ns | TM | KS.1 |
| FIM1_2161 | 1722.750275 | 1588.549463 | 1565.997904 | -0.082011785 | 0.060298956 | 0.063774335 | 0.265060417 | ns | TM | KS.1 |
| FIM1_2162 | 658.408469 | 665.3293091 | 767.1367039 | 0.006249465 | 0.080888238 | 0.905935549 | 0.971578849 | ns | TM | KS.1 |
| FIM1_2163 | 775.033855 | 793.0499362 | 761.4008034 | 0.010407078 | 0.1108201 | 0.86583337 | 0.959975149 | ns | TM | KS.1 |
| FIM1_2164 | 847.5238231 | 814.1839355 | 769.3607597 | -0.027018444 | 0.076457455 | 0.596705787 | 0.849348181 | ns | TM | KS.1 |
| FIM1_2165 | 2259.871354 | 2412.935043 | 2668.911091 | 0.043975135 | 0.084715069 | 0.422205914 | 0.73856344 | ns | TM | KS.1 |
| FIM1_2166 | 354.3645191 | 389.3551927 | 398.1753113 | 0.035513569 | 0.129739106 | 0.582047575 | 0.838843188 | ns | TM | KS.1 |
| FIM1_2167 | 5164.160969 | 5177.806122 | 4787.92655 | 0.001620841 | 0.080423096 | 0.975460698 | 0.993872484 | ns | TM | KS.1 |
| FIM1_2168 | 3859.683013 | 4446.000026 | 4647.44812 | 0.157585406 | 0.079572193 | 0.008413048 | 0.075008804 | ns | TM | KS.1 |
| FIM1_2169 | 2506.763372 | 2419.171761 | 2427.300803 | -0.02385033 | 0.07497118 | 0.635932438 | 0.869849417 | ns | TM | KS.1 |
| FIM1_2170 | 4098.994461 | 3746.552491 | 3230.549092 | -0.054863229 | 0.099191911 | 0.363731624 | 0.687524631 | ns | TM | KS.1 |
| FIM1_2171 | 1362.619196 | 1444.445361 | 1110.462969 | 0.027768228 | 0.109188639 | 0.653131616 | 0.875830132 | ns | TM | KS.1 |
| FIM1_2172 | 1793.249869 | 1502.920195 | 2057.017079 | -0.154679373 | 0.11059901 | 0.031325499 | 0.176980365 | ns | TM | KS.1 |
| FIM1_2173 | 953.9419239 | 1018.958234 | 774.6059075 | 0.030395168 | 0.112963783 | 0.627087306 | 0.866840608 | ns | TM | KS.1 |
| FIM1_2174 | 1296.207965 | 1031.058549 | 1263.218158 | -0.143298211 | 0.143468302 | 0.066892025 | 0.272267737 | ns | TM | KS.1 |
| FIM1_2175 | 3015.942786 | 3395.840929 | 3557.19596 | 0.083154202 | 0.101225972 | 0.184142872 | 0.486516617 | ns | TM | KS.1 |
| FIM1_2176 | 1557.222149 | 1193.400756 | 925.9356295 | -0.195542079 | 0.151098501 | 0.022696404 | 0.147024228 | ns | TM | KS.1 |
| FIM1_2177 | 48.42492959 | 45.49539794 | 79.54727981 | -0.018710433 | 0.13497454 | 0.768421803 | 0.925179691 | ns | TM | KS.1 |
| FIM1_2178 | 4836.247974 | 5426.132388 | 4144.390667 | 0.047626996 | 0.127204395 | 0.466174206 | 0.767952541 | ns | TM | KS.1 |
| FIM1_2179 | 10731.0576 | 9785.783583 | 9088.135023 | -0.062400281 | 0.093519203 | 0.288459245 | 0.61538797 | ns | TM | KS.1 |
| FIM1_2180 | 1671.514421 | 2584.011838 | 2978.44097 | 0.180994566 | 0.2360333 | 0.04996855 | 0.232979769 | ns | TM | KS.1 |
| FIM1_2181 | 2485.576409 | 2662.737049 | 2827.95635 | 0.025949802 | 0.126389277 | 0.683935292 | 0.890494131 | ns | TM | KS.1 |
| FIM1_2182 | 1717.260539 | 619.4070999 | 1562.092515 | -1.104938981 | 0.429357294 | 7.55908481122 | 0.001918696 | Km<KS.1 | TM | KS.1 |
| FIM1_2183 | 2565.902951 | 2095.11219 | 2582.084018 | -0.194661605 | 0.111675796 | 0.010634793 | 0.088198944 | ns | TM | KS.1 |
| FIM1_2184 | 3128.412275 | 2963.8692 | 2910.723947 | -0.031460567 | 0.093057958 | 0.585200526 | 0.841613393 | ns | TM | KS.1 |
| FIM1_2185 | 3081.230042 | 1966.658483 | 3145.59594 | -0.455880632 | 0.218406892 | 0.00039021 | 0.007466514 | ns | TM | KS.1 |
| FIM1_2186 | 1462.208896 | 1679.054436 | 1562.596842 | 0.106563416 | 0.104533448 | 0.103383347 | 0.348223325 | ns | TM | KS.1 |
| FIM1_2187 | 1292.303961 | 1592.383495 | 1825.116442 | 0.085974145 | 0.14843286 | 0.226204438 | 0.543918356 | ns | TM | KS.1 |
| FIM1_2188 | 4843.872007 | 4033.574991 | 4842.867628 | -0.190541409 | 0.098138691 | 0.007266437 | 0.06719428 | ns | TM | KS.1 |
| FIM1_2189 | 911.2444522 | 1008.048839 | 1001.661405 | 0.050609479 | 0.114169996 | 0.42843201 | 0.742665244 | ns | TM | KS.1 |
| FIM1_2190 | 1978.546484 | 1974.839321 | 2212.447532 | -0.001165393 | 0.078572925 | 0.982093327 | 0.995500061 | ns | TM | KS.1 |
| FIM1_2191 | 6819.900412 | 8123.924524 | 7904.87394 | 0.199914566 | 0.083693893 | 0.002346052 | 0.029179022 | ns | TM | KS.1 |
| FIM1_2192 | 681.2203812 | 916.3835691 | 907.4031976 | 0.214139228 | 0.164453316 | 0.017677512 | 0.124922762 | ns | TM | KS.1 |
| FIM1_2193 | 2146.994217 | 3541.502995 | 4372.129069 | 0.238566197 | 0.275420358 | 0.02490329 | 0.154938568 | ns | TM | KS.1 |
| FIM1_2194 | 514.796946 | 656.9666689 | 674.7807856 | 0.101424159 | 0.158568112 | 0.16939615 | 0.463542658 | ns | TM | KS.1 |
| FIM1_2195 | 698.2053126 | 701.8850658 | 827.4549321 | 0.003368959 | 0.089602057 | 0.952159343 | 0.987816044 | ns | TM | KS.1 |
| FIM1_2196 | 192.7982678 | 282.4999798 | 346.3742841 | 0.297322267 | 0.205461711 | 0.005583247 | 0.056456617 | ns | TM | KS.1 |
| FIM1_2197 | 1191.088523 | 1331.785031 | 1611.03317 | 0.08064109 | 0.096427025 | 0.184511737 | 0.487133708 | ns | TM | KS.1 |
| FIM1_2198 | 2869.026459 | 1820.856976 | 2316.489743 | -0.308948383 | 0.254486547 | 0.007830614 | 0.070772899 | ns | TM | KS.1 |
| FIM1_2199 | 65.14890449 | 80.31719286 | 102.5282884 | 0.072868663 | 0.157285446 | 0.293296507 | 0.620392168 | ns | TM | KS.1 |

| | | | | | | | | | | |
|---|---|---|---|---|---|---|---|---|---|---|
| FIM1_2200 | 1436.544405 | 1424.825404 | 1233.819232 | -0.004299692 | 0.097428671 | 0.941496745 | 0.98379092 | ns | TM | KS.1 |
| FIM1_2201 | 881.9389919 | 880.720295 | 1062.087388 | -0.000259408 | 0.067656568 | 0.995592573 | 0.997999809 | ns | TM | KS.1 |
| FIM1_2202 | 5141.646512 | 5172.500862 | 4287.758579 | 0.003403624 | 0.088701887 | 0.951398567 | 0.987490021 | ns | TM | KS.1 |
| FIM1_2203 | 1204.677674 | 1016.14658 | 970.0864016 | -0.134461729 | 0.114715768 | 0.058633559 | 0.254317312 | ns | TM | KS.1 |
| FIM1_2204 | 1287.53214 | 1022.186004 | 982.1773121 | -0.162263064 | 0.140321644 | 0.042473624 | 0.210464424 | ns | TM | KS.1 |
| FIM1_2205 | 965.2269309 | 925.0694831 | 972.1536766 | -0.023180604 | 0.095276272 | 0.690331715 | 0.892979792 | ns | TM | KS.1 |
| FIM1_2206 | 734.9022516 | 996.2869505 | 1280.748421 | 0.095705803 | 0.182763935 | 0.193123266 | 0.496788132 | ns | TM | KS.1 |
| FIM1_2207 | 467.6334811 | 426.9084834 | 462.4421567 | -0.022698052 | 0.153737802 | 0.719283961 | 0.905913973 | ns | TM | KS.1 |
| FIM1_2209 | 2074.258187 | 1615.708121 | 1758.944291 | -0.109283972 | 0.158926814 | 0.145305098 | 0.423239382 | ns | TM | KS.1 |
| FIM1_2210 | 2202.524669 | 3811.546112 | 4726.000485 | 0.435967703 | 0.313572962 | 0.002641887 | 0.031747309 | ns | TM | KS.1 |
| FIM1_2211 | 4859.735597 | 4390.289528 | 4800.855103 | -0.081497769 | 0.087011885 | 0.153403747 | 0.437104032 | ns | TM | KS.1 |
| FIM1_2212 | 1465.527092 | 1211.140924 | 1039.059064 | -0.154745239 | 0.120117308 | 0.038204807 | 0.196962604 | ns | TM | KS.1 |
| FIM1_2213 | 810.4301839 | 855.1106714 | 898.7242659 | 0.035081539 | 0.084768918 | 0.520975892 | 0.807566596 | ns | TM | KS.1 |
| FIM1_2214 | 692.1123314 | 685.6879997 | 624.8375672 | -0.005439091 | 0.077719533 | 0.915990626 | 0.975605516 | ns | TM | KS.1 |
| FIM1_2215 | 4820.529889 | 5917.874261 | 6323.786572 | 0.106966675 | 0.140455873 | 0.144157208 | 0.421623816 | ns | TM | KS.1 |
| FIM1_2216 | 923.0109561 | 1038.342553 | 919.6030042 | 0.086597212 | 0.099410433 | 0.163911956 | 0.45366779 | ns | TM | KS.1 |
| FIM1_2217 | 793.8283102 | 707.8733933 | 851.7880038 | -0.042139437 | 0.133556013 | 0.518154132 | 0.804981347 | ns | TM | KS.1 |
| FIM1_2218 | 9156.59755 | 9570.565858 | 8006.289705 | 0.032306548 | 0.070558915 | 0.503961722 | 0.796765282 | ns | TM | KS.1 |
| FIM1_2219 | 2530.192495 | 2567.358552 | 2623.452048 | 0.010227677 | 0.067878557 | 0.827947846 | 0.946035952 | ns | TM | KS.1 |
| FIM1_2220 | 1420.454238 | 1522.987304 | 1521.415233 | 0.066696222 | 0.06129566 | 0.131186539 | 0.400376242 | ns | TM | KS.1 |
| FIM1_2221 | 848.2932464 | 912.1203642 | 801.815568 | 0.055833498 | 0.07674106 | 0.278758159 | 0.602966017 | ns | TM | KS.1 |
| FIM1_2222 | 763.814313 | 608.6108278 | 622.0733073 | -0.146765381 | 0.142124896 | 0.061288072 | 0.260828192 | ns | TM | KS.1 |
| FIM1_2223 | 906.7778705 | 978.6276158 | 1068.373156 | 0.046053396 | 0.095904932 | 0.435020371 | 0.746542375 | ns | TM | KS.1 |
| FIM1_2224 | 3524.572555 | 3716.351551 | 3951.563321 | 0.030493194 | 0.094084414 | 0.598797698 | 0.849691542 | ns | TM | KS.1 |
| FIM1_2225 | 889.4106319 | 935.7573968 | 1245.852409 | 0.019144405 | 0.12466282 | 0.762405177 | 0.924230468 | ns | TM | KS.1 |
| FIM1_2226 | 9233.469563 | 9635.940627 | 10280.44126 | 0.03820664 | 0.052148705 | 0.325728915 | 0.653164592 | ns | TM | KS.1 |
| FIM1_2227 | 1234.425175 | 1489.342646 | 2047.478626 | 0.096625092 | 0.135449344 | 0.17593545 | 0.473568831 | ns | TM | KS.1 |
| FIM1_2228 | 959.6425101 | 956.0228787 | 941.2334906 | -0.001003007 | 0.146266461 | 0.987233823 | 0.996167262 | ns | TM | KS.1 |
| FIM1_2229 | 692.9426411 | 783.2944784 | 888.5419249 | 0.052922892 | 0.126302883 | 0.420837704 | 0.737778527 | ns | TM | KS.1 |
| FIM1_2230 | 288.4101873 | 274.5524744 | 429.8878394 | -0.01678107 | 0.133421648 | 0.791449046 | 0.933303839 | ns | TM | KS.1 |
| FIM1_2231 | 1086.943061 | 1118.125961 | 934.2259976 | 0.02002121 | 0.069052976 | 0.674114103 | 0.885586919 | ns | TM | KS.1 |
| FIM1_2232 | 313.0947341 | 284.6457198 | 872.8070102 | -0.032367447 | 0.13387423 | 0.615260815 | 0.858612877 | ns | TM | KS.1 |
| FIM1_2233 | 1847.905445 | 1732.824895 | 1744.585935 | -0.036955755 | 0.096184979 | 0.529648134 | 0.808531286 | ns | TM | KS.1 |
| FIM1_2234 | 505.0598793 | 531.4006014 | 525.7881748 | 0.033152321 | 0.080790179 | 0.53157217 | 0.809001086 | ns | TM | KS.1 |
| FIM1_2235 | 544.446832 | 622.2944104 | 542.9644863 | 0.063274105 | 0.124997499 | 0.341744378 | 0.668903606 | ns | TM | KS.1 |
| FIM1_2236 | 1330.644628 | 1510.336568 | 1267.879751 | 0.08590474 | 0.10573157 | 0.181182122 | 0.481249898 | ns | TM | KS.1 |
| FIM1_2237 | 24.25476062 | 21.91549349 | 40.526282 | -0.01813035 | 0.157877643 | 0.772154715 | 0.925733395 | ns | TM | KS.1 |
| FIM1_2238 | 1040.531882 | 762.9010829 | 875.0395528 | -0.288679506 | 0.150678442 | 0.002545602 | 0.030888701 | ns | TM | KS.1 |
| FIM1_2239 | 1084.169112 | 988.2759004 | 707.4400088 | -0.052817006 | 0.104456722 | 0.393099265 | 0.716470042 | ns | TM | KS.1 |
| FIM1_2240 | 839.2712846 | 836.8507231 | 670.9627868 | -0.001367369 | 0.098158497 | 0.981441024 | 0.995431301 | ns | TM | KS.1 |
| FIM1_2241 | 2933.224337 | 2762.036522 | 2657.322158 | -0.026999215 | 0.113824271 | 0.66599752 | 0.880910621 | ns | TM | KS.1 |
| FIM1_2243 | 1221.925619 | 1104.086575 | 1190.464514 | -0.063220322 | 0.100947241 | 0.302856726 | 0.631432423 | ns | TM | KS.1 |
| FIM1_2244 | 3995.527535 | 4400.966817 | 5009.643501 | 0.064871056 | 0.095667546 | 0.276978673 | 0.601532657 | ns | TM | KS.1 |
| FIM1_2245 | 1147.516758 | 1205.395962 | 1697.103408 | 0.023486437 | 0.107307326 | 0.701475681 | 0.897235589 | ns | TM | KS.1 |
| FIM1_2246 | 9320.255641 | 11979.2933 | 11412.85161 | 0.295034362 | 0.091741121 | 9.6740122142[illegible] | 0.002325034 | ns | TM | KS.1 |

| | | | | | | | | | | |
|---|---|---|---|---|---|---|---|---|---|---|
| FIM1_2247 | 1078.420459 | 1109.005049 | 1103.334595 | 0.017884282 | 0.076559912 | 0.726230331 | 0.907961357 | ns | TM | KS.1 |
| FIM1_2248 | 1484.514659 | 2365.338931 | 2883.141521 | 0.195682405 | 0.251658582 | 0.042052834 | 0.209871189 | ns | TM | KS.1 |
| FIM1_2249 | 2520.976716 | 3793.237939 | 4456.848732 | 0.18566903 | 0.224428152 | 0.044438175 | 0.217302694 | ns | TM | KS.1 |
| FIM1_2250 | 246.4501684 | 401.4698374 | 393.1173166 | 0.240573839 | 0.270685896 | 0.023667334 | 0.150569038 | ns | TM | KS.1 |
| FIM1_2251 | 450.8746977 | 457.6702474 | 559.9824934 | 0.011566069 | 0.071856585 | 0.813197596 | 0.940696014 | ns | TM | KS.1 |
| FIM1_2252 | 3710.073588 | 3592.50188 | 3579.467591 | -0.018066948 | 0.092688149 | 0.752472147 | 0.918662314 | ns | TM | KS.1 |
| FIM1_2253 | 10777.14047 | 13895.65396 | 15903.47299 | 0.322253073 | 0.073172399 | 7.2696677159[illegible] | 5.7407296645[illegible] | ns | TM | KS.1 |
| FIM1_2254 | 2706.007394 | 2897.464048 | 2607.848379 | 0.036292804 | 0.103538967 | 0.550964915 | 0.822058746 | ns | TM | KS.1 |
| FIM1_2255 | 13161.77074 | 13576.73886 | 13851.80021 | 0.026336632 | 0.052567153 | 0.500665916 | 0.795278714 | ns | TM | KS.1 |
| FIM1_2256 | 2368.990159 | 1552.205436 | 1342.415325 | -0.361015038 | 0.222914332 | 0.0023042 | 0.028802497 | ns | TM | KS.1 |
| FIM1_2257 | 1369.747661 | 900.0913308 | 1340.549922 | -0.116636325 | 0.221900014 | 0.137799131 | 0.411248158 | ns | TM | KS.1 |
| FIM1_2258 | 1733.052048 | 1572.069362 | 2130.826195 | -0.051632482 | 0.109674143 | 0.412936013 | 0.732913544 | ns | TM | KS.1 |
| FIM1_2259 | 523.7850462 | 438.7841331 | 421.2064794 | -0.122317733 | 0.122455493 | 0.089012411 | 0.319968746 | ns | TM | KS.1 |
| FIM1_2260 | 1825.446924 | 1779.314137 | 1913.190993 | -0.016765884 | 0.07580333 | 0.741079449 | 0.914629188 | ns | TM | KS.1 |
| FIM1_2261 | 2737.441579 | 3184.75185 | 3325.419316 | 0.125837169 | 0.105153451 | 0.061480317 | 0.260865744 | ns | TM | KS.1 |
| FIM1_2262 | 4435.049346 | 3073.893032 | 2968.684737 | -0.250644084 | 0.200885286 | 0.012475698 | 0.098206639 | ns | TM | KS.1 |
| FIM1_2263 | 5308.72395 | 5639.485672 | 4218.55243 | 0.033613634 | 0.098299062 | 0.571091287 | 0.832094673 | ns | TM | KS.1 |
| FIM1_2264 | 1263.710822 | 1284.197818 | 1340.425027 | 0.009877485 | 0.08078734 | 0.851749417 | 0.95392681 | ns | TM | KS.1 |
| FIM1_2265 | 931.181604 | 957.48925 | 992.9188256 | 0.018654473 | 0.074593686 | 0.710195344 | 0.901619224 | ns | TM | KS.1 |
| FIM1_2266 | 1425.857245 | 1558.124849 | 1334.817771 | 0.050620368 | 0.103394325 | 0.410056446 | 0.73093186 | ns | TM | KS.1 |
| FIM1_2267 | 1698.892587 | 1532.380073 | 1619.757706 | -0.06490945 | 0.101635912 | 0.29250984 | 0.619777024 | ns | TM | KS.1 |
| FIM1_2268 | 1330.723848 | 1385.206114 | 1336.216251 | 0.027427318 | 0.07537114 | 0.587598747 | 0.842693504 | ns | TM | KS.1 |
| FIM1_2270 | 724.1843485 | 869.7337713 | 840.3921184 | 0.089320933 | 0.135913909 | 0.205633096 | 0.512865935 | ns | TM | KS.1 |
| FIM1_2271 | 1437.273907 | 1271.053183 | 1494.176021 | -0.084643723 | 0.103954819 | 0.183249609 | 0.484929151 | ns | TM | KS.1 |
| FIM1_2272 | 1193.087906 | 1100.498673 | 737.2469833 | -0.041620449 | 0.107795338 | 0.502748363 | 0.796298346 | ns | TM | KS.1 |
| FIM1_2273 | 229.9599497 | 184.3753041 | 169.1592428 | -0.089694864 | 0.152128842 | 0.211228668 | 0.51871941 | ns | TM | KS.1 |
| FIM1_2274 | 3400.922211 | 2770.024297 | 2250.51838 | -0.08225509 | 0.148301683 | 0.243384833 | 0.566872447 | ns | TM | KS.1 |
| FIM1_2275 | 2631.058262 | 1781.187717 | 2090.895444 | -0.339139271 | 0.200247879 | 0.002343284 | 0.029179022 | ns | TM | KS.1 |
| FIM1_2276 | 815.9961351 | 757.2977033 | 871.5900199 | -0.0422169 | 0.099944779 | 0.48248564 | 0.783685372 | ns | TM | KS.1 |
| FIM1_2278 | 1624.714014 | 1678.826347 | 1870.660216 | 0.013682165 | 0.116964851 | 0.826731344 | 0.946035952 | ns | TM | KS.1 |
| FIM1_2279 | 335.1668459 | 217.3380288 | 338.5058139 | -0.525642301 | 0.170396349 | 1.0674943489[illegible] | 0.00043531 | ns | TM | KS.1 |
| FIM1_2280 | 2252.386925 | 2382.028101 | 1950.211584 | 0.038398567 | 0.079843423 | 0.465760437 | 0.767779382 | ns | TM | KS.1 |
| FIM1_2281 | 802.835042 | 606.18246 | 749.2942018 | -0.169540403 | 0.163955554 | 0.044372848 | 0.217278465 | ns | TM | KS.1 |
| FIM1_2282 | 404.7421762 | 318.0083193 | 305.8777883 | -0.20794999 | 0.133222757 | 0.01283096 | 0.100367965 | ns | TM | KS.1 |
| FIM1_2283 | 1905.281455 | 1807.851532 | 1312.087633 | -0.031594504 | 0.089532499 | 0.575462425 | 0.833942781 | ns | TM | KS.1 |
| FIM1_2284 | 2592.782713 | 2417.808261 | 2494.53559 | -0.027683989 | 0.123331456 | 0.663582363 | 0.880619203 | ns | TM | KS.1 |
| FIM1_2285 | 3418.106512 | 4520.570949 | 4091.590869 | 0.193180745 | 0.159186167 | 0.026084191 | 0.158641625 | ns | TM | KS.1 |
| FIM1_2286 | 991.0648074 | 959.5094878 | 1032.886163 | -0.01891944 | 0.088019071 | 0.734076015 | 0.911352378 | ns | TM | KS.1 |
| FIM1_2287 | 809.1501811 | 819.2913425 | 927.4687669 | 0.007008134 | 0.090957517 | 0.901498515 | 0.968873116 | ns | TM | KS.1 |
| FIM1_2288 | 1770.271877 | 1852.310827 | 1886.171791 | 0.028723479 | 0.08429977 | 0.597407859 | 0.849348181 | ns | TM | KS.1 |
| FIM1_2289 | 3179.1117 | 3456.675039 | 3305.0036 | 0.048048382 | 0.101658485 | 0.42969947 | 0.743536592 | ns | TM | KS.1 |
| FIM1_2290 | 1609.413407 | 1755.408894 | 2575.129183 | 0.041795545 | 0.113628597 | 0.508850838 | 0.800864574 | ns | TM | KS.1 |
| FIM1_2291 | 3116.850531 | 3392.075471 | 3516.64279 | 0.046583932 | 0.104529478 | 0.449195418 | 0.753709007 | ns | TM | KS.1 |
| FIM1_2292 | 847.63611 | 947.412733 | 872.7159229 | 0.057708703 | 0.114665406 | 0.370881588 | 0.693660112 | ns | TM | KS.1 |
| FIM1_2293 | 781.5867404 | 723.1110518 | 613.4994378 | -0.033330082 | 0.119077222 | 0.59999837 | 0.850484618 | ns | TM | KS.1 |

| | | | | | | | | | | |
|---|---|---|---|---|---|---|---|---|---|---|
| FIM1_2294 | 8085.427283 | 9073.132551 | 9396.152219 | 0.117800944 | 0.077648174 | 0.033196846 | 0.181488253 | ns | TM | KS.1 |
| FIM1_2295 | 873.8141622 | 782.8738156 | 746.0783644 | -0.060115922 | 0.110892369 | 0.346581214 | 0.671956953 | ns | TM | KS.1 |
| FIM1_2297 | 1171.991676 | 1152.770699 | 1216.926502 | -0.010635238 | 0.071640814 | 0.827640096 | 0.946035952 | ns | TM | KS.1 |
| FIM1_2298 | 1266.473487 | 1454.374325 | 1665.205528 | 0.073111029 | 0.120270462 | 0.273394811 | 0.599198132 | ns | TM | KS.1 |
| FIM1_2299 | 1587.484344 | 1677.913539 | 1448.848662 | 0.029534731 | 0.10027716 | 0.621272192 | 0.862872646 | ns | TM | KS.1 |
| FIM1_2300 | 586.1689428 | 540.8100251 | 538.300696 | -0.043676657 | 0.103833147 | 0.475742679 | 0.776238092 | ns | TM | KS.1 |
| FIM1_2301 | 2588.859933 | 2834.600836 | 2796.687825 | 0.040794359 | 0.11838646 | 0.523391833 | 0.807903931 | ns | TM | KS.1 |
| FIM1_2302 | 4483.569899 | 4572.795117 | 3328.172383 | 0.013080139 | 0.075237051 | 0.795563903 | 0.935238756 | ns | TM | KS.1 |
| FIM1_2303 | 183.266687 | 231.9918802 | 302.1341551 | 0.101114736 | 0.156451316 | 0.169830765 | 0.463725607 | ns | TM | KS.1 |
| FIM1_2304 | 475.4883204 | 448.5760694 | 326.5847066 | -0.029006799 | 0.106699061 | 0.636223046 | 0.869849417 | ns | TM | KS.1 |
| FIM1_2305 | 675.3218626 | 697.0933632 | 762.5268359 | 0.021876329 | 0.070079346 | 0.649265531 | 0.874068749 | ns | TM | KS.1 |
| FIM1_2306 | 359.8392857 | 211.0183672 | 275.4655839 | -0.687942771 | 0.189362302 | 1.71493050350 | 0.000109382 | ns | TM | KS.1 |
| FIM1_2307 | 2191.542055 | 2224.358743 | 2506.191909 | 0.009668893 | 0.078772634 | 0.852533394 | 0.954011929 | ns | TM | KS.1 |
| FIM1_2308 | 1454.629287 | 1842.895543 | 1907.014888 | 0.116831072 | 0.151976032 | 0.122610968 | 0.385581269 | ns | TM | KS.1 |
| FIM1_2309 | 411.9770304 | 254.9995481 | 295.1554919 | -0.282489523 | 0.269875645 | 0.013008924 | 0.101166593 | ns | TM | KS.1 |
| FIM1_2310 | 436.5188938 | 456.3436847 | 334.031678 | 0.01832314 | 0.120487439 | 0.771151761 | 0.925705893 | ns | TM | KS.1 |
| FIM1_2311 | 1602.228838 | 1673.461199 | 1672.056037 | 0.013485326 | 0.136809302 | 0.831294245 | 0.946802573 | ns | TM | KS.1 |
| FIM1_2312 | 4882.182992 | 4100.439194 | 4768.238598 | -0.168384262 | 0.103036666 | 0.017127814 | 0.122253765 | ns | TM | KS.1 |
| FIM1_2313 | 908.4314702 | 1174.191936 | 1339.623606 | 0.181525499 | 0.149179289 | 0.030311883 | 0.173534659 | ns | TM | KS.1 |
| FIM1_2314 | 1359.994785 | 1015.465083 | 1073.429431 | -0.342792705 | 0.102894521 | 3.21294366033 | 0.001024641 | ns | TM | KS.1 |
| FIM1_2315 | 849.2794553 | 656.8426526 | 1049.537091 | -0.128216866 | 0.158267781 | 0.099529305 | 0.340998887 | ns | TM | KS.1 |
| FIM1_2316 | 836.8124195 | 860.3128331 | 886.4830386 | 0.010184944 | 0.124088028 | 0.871589532 | 0.962521182 | ns | TM | KS.1 |
| FIM1_2317 | 1841.549556 | 1887.496871 | 2337.312787 | 0.014644175 | 0.086933313 | 0.791010487 | 0.933303839 | ns | TM | KS.1 |
| FIM1_2318 | 16501.8527 | 9630.729403 | 13391.12303 | -0.60577643 | 0.251534626 | 0.00010305 | 0.002452979 | ns | TM | KS.1 |
| FIM1_2319 | 174.4299872 | 157.4167553 | 164.3750103 | -0.039454877 | 0.130821113 | 0.543109519 | 0.816551785 | ns | TM | KS.1 |
| FIM1_2320 | 190.2929275 | 207.1722005 | 214.5370624 | 0.048332041 | 0.102645459 | 0.4291802 | 0.743181463 | ns | TM | KS.1 |
| FIM1_2321 | 1600.82889 | 1630.148697 | 1675.237695 | 0.014681564 | 0.053245719 | 0.71024035 | 0.901619224 | ns | TM | KS.1 |
| FIM1_2322 | 1063.188787 | 987.4662239 | 811.9243081 | -0.048169178 | 0.089944679 | 0.397662039 | 0.721505705 | ns | TM | KS.1 |
| FIM1_2323 | 2566.680932 | 2744.93739 | 3024.517997 | 0.044512614 | 0.086970309 | 0.424277713 | 0.740157476 | ns | TM | KS.1 |
| FIM1_2324 | 3802.175669 | 3902.46658 | 3890.555805 | 0.013019422 | 0.102149433 | 0.8279075 | 0.946035952 | ns | TM | KS.1 |
| FIM1_2325 | 41.71149794 | 41.00954614 | 40.96022622 | -0.013889834 | 0.127335996 | 0.826188861 | 0.94595497 | ns | TM | KS.1 |
| FIM1_2326 | 443.9450441 | 411.2868632 | 442.7879753 | -0.047358717 | 0.093130332 | 0.414928074 | 0.734631029 | ns | TM | KS.1 |
| FIM1_2327 | 3001.251855 | 2915.110711 | 3239.293127 | -0.020619384 | 0.068851435 | 0.664314346 | 0.880619203 | ns | TM | KS.1 |
| FIM1_2328 | 5927.646873 | 7077.198709 | 7241.459513 | 0.196296298 | 0.08913265 | 0.003801964 | 0.042032829 | ns | TM | KS.1 |
| FIM1_2329 | 306.4289982 | 311.0071414 | 371.0631883 | 0.005924061 | 0.117207734 | 0.924355752 | 0.976662213 | ns | TM | KS.1 |
| FIM1_2330 | 418.4846862 | 426.045563 | 471.7337333 | 0.007138498 | 0.1165198 | 0.908822212 | 0.972251892 | ns | TM | KS.1 |
| FIM1_2331 | 3061.685118 | 2756.60073 | 2684.208701 | -0.048554811 | 0.119263739 | 0.452156396 | 0.755873008 | ns | TM | KS.1 |
| FIM1_2332 | 9242.925255 | 10008.202 | 8886.548113 | 0.040109993 | 0.108935214 | 0.519537198 | 0.805989904 | ns | TM | KS.1 |
| FIM1_2333 | 387.4564313 | 393.5570327 | 387.8348301 | 0.006859301 | 0.104000467 | 0.909352295 | 0.972251892 | ns | TM | KS.1 |
| FIM1_2334 | 597.2128271 | 473.2015526 | 624.1776974 | -0.157330403 | 0.142743674 | 0.048682223 | 0.230174256 | ns | TM | KS.1 |
| FIM1_2335 | 6532.596796 | 6450.084857 | 6406.854433 | -0.006181711 | 0.103440398 | 0.918096621 | 0.975634767 | ns | TM | KS.1 |
| FIM1_2336 | 486.951378 | 1008.898569 | 1689.305228 | 0.895745641 | 0.286382469 | 1.96613511835 | 0.000708806 | ns | TM | KS.1 |
| FIM1_2337 | 125.3972343 | 155.7708917 | 216.0073424 | 0.083384606 | 0.151941598 | 0.23895577 | 0.559787835 | ns | TM | KS.1 |
| FIM1_2338 | 1831.59244 | 1378.012818 | 1538.303557 | -0.32975752 | 0.103472424 | 6.30023919439 | 0.001658396 | ns | TM | KS.1 |
| FIM1_2339 | 603.06743 | 579.3190792 | 640.8000692 | -0.024476012 | 0.090242302 | 0.665153847 | 0.880679852 | ns | TM | KS.1 |

| | | | | | | | | | | |
|---|---|---|---|---|---|---|---|---|---|---|
| FIM1_2340 | 1465.626158 | 1735.153854 | 1242.990799 | 0.078035151 | 0.133936867 | 0.258411327 | 0.58729847 | ns | TM | KS.1 |
| FIM1_2341 | 454.7424082 | 372.4735782 | 716.9177179 | -0.105219352 | 0.139135218 | 0.148772097 | 0.429441469 | ns | TM | KS.1 |
| FIM1_2342 | 1752.643775 | 1806.355042 | 1706.408292 | 0.016674697 | 0.093537946 | 0.771976785 | 0.925733395 | ns | TM | KS.1 |
| FIM1_2343 | 48.48980601 | 30.59025872 | 62.07038615 | -0.162059899 | 0.244448727 | 0.068785115 | 0.276450087 | ns | TM | KS.1 |
| FIM1_2344 | 531.9620568 | 378.9255438 | 771.224789 | -0.17391087 | 0.19175848 | 0.047452308 | 0.226777362 | ns | TM | KS.1 |
| FIM1_2345 | 1173.660659 | 540.2809801 | 917.6130562 | -1.071091191 | 0.156869121 | 2.2154416200 | 1.4841793695 | Km<KS.1 | TM | KS.1 |
| FIM1_2346 | 3470.161486 | 3714.273389 | 4009.822228 | 0.045143367 | 0.086618793 | 0.416696671 | 0.735551151 | ns | TM | KS.1 |
| FIM1_2347 | 621.1893272 | 662.8822146 | 620.2654514 | 0.031905306 | 0.108969346 | 0.60630273 | 0.851596861 | ns | TM | KS.1 |
| FIM1_2348 | 2568.880892 | 2823.460919 | 2895.085466 | 0.072502357 | 0.086810327 | 0.199925835 | 0.507723852 | ns | TM | KS.1 |
| FIM1_2349 | 2232.573256 | 2180.941085 | 2302.086355 | -0.01714938 | 0.065016283 | 0.707081625 | 0.89994363 | ns | TM | KS.1 |
| FIM1_2350 | 3479.57571 | 3245.275774 | 2886.681087 | -0.046115624 | 0.087127026 | 0.40861976 | 0.729391439 | ns | TM | KS.1 |
| FIM1_2351 | 1198.949597 | 1067.533797 | 1167.699938 | -0.05247296 | 0.122661094 | 0.421802803 | 0.738117813 | ns | TM | KS.1 |
| FIM1_2352 | 920.0464918 | 999.0094605 | 1125.829973 | 0.032269023 | 0.125403196 | 0.614469638 | 0.858462917 | ns | TM | KS.1 |
| FIM1_2353 | 1186.64374 | 945.7720687 | 972.6931714 | -0.208533632 | 0.122883403 | 0.010000119 | 0.084466202 | ns | TM | KS.1 |
| FIM1_2354 | 681.9530753 | 714.716918 | 595.0128552 | 0.021830702 | 0.109106841 | 0.722910541 | 0.907127048 | ns | TM | KS.1 |
| FIM1_2355 | 267.5596831 | 140.908704 | 214.1355932 | -0.798270973 | 0.252326926 | 1.3844398659 | 0.000532533 | ns | TM | KS.1 |
| FIM1_2356 | 496.5840559 | 366.8969361 | 428.712158 | -0.118060119 | 0.179009972 | 0.127451204 | 0.393587671 | ns | TM | KS.1 |
| FIM1_2357 | 1366.928677 | 1953.978583 | 2252.726222 | 0.237349087 | 0.196931644 | 0.015356736 | 0.112684017 | ns | TM | KS.1 |
| FIM1_2358 | 1534.032062 | 1152.641053 | 1063.905746 | -0.311571913 | 0.116042926 | 0.000339033 | 0.006614465 | ns | TM | KS.1 |
| FIM1_2359 | 4186.515293 | 4248.442224 | 4212.938516 | 0.010320024 | 0.068522922 | 0.827540565 | 0.946035952 | ns | TM | KS.1 |
| FIM1_2360 | 2248.215147 | 1518.986929 | 2317.011115 | -0.323454674 | 0.206231101 | 0.003482658 | 0.039377778 | ns | TM | KS.1 |
| FIM1_2361 | 3258.051919 | 3294.611195 | 3183.044409 | 0.008106647 | 0.065626059 | 0.859993144 | 0.957000476 | ns | TM | KS.1 |
| FIM1_2362 | 1324.07292 | 1403.332967 | 1599.294081 | 0.042491176 | 0.075225411 | 0.401210566 | 0.723750973 | ns | TM | KS.1 |
| FIM1_2363 | 1635.048184 | 1875.587117 | 2113.157998 | 0.082575518 | 0.114809324 | 0.213954414 | 0.521776083 | ns | TM | KS.1 |
| FIM1_2364 | 2348.349671 | 2385.48999 | 2512.202957 | 0.011847727 | 0.06030878 | 0.784186296 | 0.930812921 | ns | TM | KS.1 |
| FIM1_2365 | 18303.7264 | 20943.42621 | 19621.02589 | 0.095005859 | 0.106869939 | 0.14550858 | 0.423584075 | ns | TM | KS.1 |
| FIM1_2366 | 518.3265249 | 602.9629827 | 568.7596446 | 0.093012255 | 0.117756781 | 0.171507882 | 0.465692069 | ns | TM | KS.1 |
| FIM1_2367 | 422.0485968 | 492.1983956 | 488.8553682 | 0.068424416 | 0.132171468 | 0.312949954 | 0.639671112 | ns | TM | KS.1 |
| FIM1_2368 | 2040.639146 | 2430.660073 | 2404.573716 | 0.087504507 | 0.132641614 | 0.211270844 | 0.51871941 | ns | TM | KS.1 |
| FIM1_2369 | 1294.964995 | 1243.985128 | 1138.461136 | -0.023435702 | 0.089972594 | 0.677984437 | 0.887129504 | ns | TM | KS.1 |
| FIM1_2370 | 321.7708444 | 445.4023161 | 394.0827039 | 0.111232558 | 0.187908976 | 0.146220581 | 0.424167573 | ns | TM | KS.1 |
| FIM1_2371 | 2794.86438 | 2666.693464 | 4375.944249 | -0.013605491 | 0.141621885 | 0.829482006 | 0.946484629 | ns | TM | KS.1 |
| FIM1_2372 | 4141.268796 | 4604.920412 | 4975.428215 | 0.069415285 | 0.100488469 | 0.259370175 | 0.588210523 | ns | TM | KS.1 |
| FIM1_2373 | 2069.427449 | 807.7723802 | 648.7126715 | -1.067227755 | 0.385420828 | 5.2542347837 | 0.001437543 | Km<KS.1 | TM | KS.1 |
| FIM1_2374 | 1121.073636 | 938.9451558 | 1107.153194 | -0.151868525 | 0.112096109 | 0.035018872 | 0.187130923 | ns | TM | KS.1 |
| FIM1_2375 | 1670.793257 | 1865.405376 | 1708.348738 | 0.075268367 | 0.099913674 | 0.222392726 | 0.537089229 | ns | TM | KS.1 |
| FIM1_2376 | 1502.406566 | 1622.147039 | 1376.278378 | 0.04645313 | 0.096428029 | 0.432490261 | 0.74505982 | ns | TM | KS.1 |
| FIM1_2377 | 6474.804731 | 7474.761994 | 9042.424857 | 0.110425783 | 0.10611841 | 0.095660219 | 0.334441033 | ns | TM | KS.1 |
| FIM1_2378 | 811.0826049 | 701.9362074 | 689.8052412 | -0.08385134 | 0.118619391 | 0.213401366 | 0.521776083 | ns | TM | KS.1 |
| FIM1_2379 | 1372.714375 | 1274.6429 | 1764.115152 | -0.046494241 | 0.092153272 | 0.420492773 | 0.737778527 | ns | TM | KS.1 |
| FIM1_2380 | 778.347554 | 816.9335439 | 815.8202954 | 0.023863413 | 0.10682537 | 0.696507274 | 0.894950919 | ns | TM | KS.1 |
| FIM1_2381 | 1626.1825 | 1772.571618 | 1661.261604 | 0.072825549 | 0.076975249 | 0.16296393 | 0.45366779 | ns | TM | KS.1 |
| FIM1_2382 | 484.4986839 | 577.2729192 | 593.4255858 | 0.118677555 | 0.123204612 | 0.098001987 | 0.337879338 | ns | TM | KS.1 |
| FIM1_2383 | 702.9089402 | 637.8072742 | 893.42857 | -0.065587584 | 0.096297318 | 0.273764292 | 0.599198132 | ns | TM | KS.1 |
| FIM1_2384 | 8891.715107 | 10643.09177 | 9984.682313 | 0.167816745 | 0.107533862 | 0.02001944 | 0.136621004 | ns | TM | KS.1 |

| | | | | | | | | | | |
|---|---|---|---|---|---|---|---|---|---|---|
| FIM1_2385 | 966.5678451 | 1040.135032 | 919.9138713 | 0.059615317 | 0.073959631 | 0.23625892 | 0.556001952 | ns | TM | KS.1 |
| FIM1_2386 | 1200.694975 | 875.7138068 | 686.0131183 | -0.14928611 | 0.181747779 | 0.071786904 | 0.284128986 | ns | TM | KS.1 |
| FIM1_2387 | 466.8243672 | 458.6725416 | 480.6728324 | -0.011644518 | 0.076354634 | 0.819314469 | 0.94310261 | ns | TM | KS.1 |
| FIM1_2388 | 7871.319578 | 6465.595601 | 6074.465123 | -0.217359953 | 0.093275311 | 0.002364219 | 0.029258683 | ns | TM | KS.1 |
| FIM1_2389 | 1628.360527 | 1729.857518 | 1980.459844 | 0.031451362 | 0.1035102 | 0.604228015 | 0.851547247 | ns | TM | KS.1 |
| FIM1_2390 | 947.8808198 | 959.8598279 | 1057.863545 | 0.008604287 | 0.066549092 | 0.852930749 | 0.954011929 | ns | TM | KS.1 |
| FIM1_2391 | 890.1660593 | 1058.731208 | 1136.747479 | 0.099793304 | 0.128180416 | 0.158256584 | 0.445060944 | ns | TM | KS.1 |
| FIM1_2392 | 1064.495426 | 1202.470659 | 1043.287542 | 0.060450236 | 0.119355363 | 0.356202331 | 0.680951596 | ns | TM | KS.1 |
| FIM1_2393 | 486.8060917 | 565.5661643 | 366.9729904 | 0.063398414 | 0.132839578 | 0.346296813 | 0.671956953 | ns | TM | KS.1 |
| FIM1_2394 | 1117.285715 | 779.3907815 | 1032.077971 | -0.278403528 | 0.192075974 | 0.006707067 | 0.064540929 | ns | TM | KS.1 |
| FIM1_2395 | 12928.94877 | 17348.56939 | 17822.67553 | 0.339829742 | 0.106514622 | 5.3123244748[illegible] | 0.001444198 | ns | TM | KS.1 |
| FIM1_2396 | 2771.739486 | 2335.73905 | 2557.179755 | -0.178758368 | 0.094521491 | 0.008965754 | 0.078667767 | ns | TM | KS.1 |
| FIM1_2397 | 2741.664931 | 2869.163574 | 3026.433657 | 0.030570922 | 0.078571138 | 0.556705808 | 0.825232602 | ns | TM | KS.1 |
| FIM1_2398 | 3331.053755 | 3075.102042 | 2011.486547 | -0.046367832 | 0.099640932 | 0.441033344 | 0.747832613 | ns | TM | KS.1 |
| FIM1_2399 | 1674.515656 | 1363.288514 | 828.3418785 | -0.148522534 | 0.130319815 | 0.052074868 | 0.238705802 | ns | TM | KS.1 |
| FIM1_2400 | 602.9861715 | 528.8701082 | 510.3101681 | -0.077309084 | 0.113910303 | 0.240113029 | 0.561090803 | ns | TM | KS.1 |
| FIM1_2401 | 780.7641758 | 720.5616175 | 714.9424682 | -0.046916881 | 0.098996599 | 0.434318294 | 0.746009599 | ns | TM | KS.1 |
| FIM1_2402 | 1423.769912 | 1752.422972 | 1613.534404 | 0.174347988 | 0.123660226 | 0.024745361 | 0.154853047 | ns | TM | KS.1 |
| FIM1_2403 | 1815.361318 | 1944.870715 | 2146.319139 | 0.04697263 | 0.084937754 | 0.392609138 | 0.71599357 | ns | TM | KS.1 |
| FIM1_2404 | 5442.393631 | 4572.95847 | 5503.319899 | -0.136790125 | 0.115748413 | 0.056185323 | 0.249350562 | ns | TM | KS.1 |
| FIM1_2405 | 1863.649475 | 1829.871073 | 1827.205237 | -0.011477466 | 0.077906111 | 0.824143093 | 0.94482203 | ns | TM | KS.1 |
| FIM1_2406 | 6461.198696 | 6530.899158 | 8854.810501 | 0.003250533 | 0.137798144 | 0.958874783 | 0.989915345 | ns | TM | KS.1 |
| FIM1_2407 | 860.6717005 | 1264.40203 | 1434.130017 | 0.307693945 | 0.204265118 | 0.004535697 | 0.048319252 | ns | TM | KS.1 |
| FIM1_2408 | 313.9453476 | 291.1919083 | 317.70967 | -0.031718115 | 0.118467101 | 0.616860261 | 0.859871056 | ns | TM | KS.1 |
| FIM1_2409 | 634.156478 | 589.791152 | 558.8150676 | -0.040136769 | 0.10112125 | 0.506132824 | 0.798222804 | ns | TM | KS.1 |
| FIM1_2410 | 5337.755258 | 5712.058494 | 5763.511942 | 0.047327788 | 0.082655886 | 0.380781541 | 0.70494176 | ns | TM | KS.1 |
| FIM1_2411 | 593.0854742 | 649.0682635 | 686.7980707 | 0.054854918 | 0.099987484 | 0.365770642 | 0.689806272 | ns | TM | KS.1 |
| FIM1_2412 | 580.9483046 | 624.5591753 | 600.7828512 | 0.033820209 | 0.113367556 | 0.590156147 | 0.844171027 | ns | TM | KS.1 |
| FIM1_2413 | 1808.245511 | 2191.329292 | 2181.1896 | 0.111174037 | 0.133368022 | 0.126835724 | 0.39290643 | ns | TM | KS.1 |
| FIM1_2414 | 1397.436264 | 1393.905154 | 1461.678195 | -0.001110892 | 0.084062841 | 0.983628691 | 0.996034691 | ns | TM | KS.1 |
| FIM1_2415 | 679.7736112 | 831.8435536 | 948.249549 | 0.167277267 | 0.123131587 | 0.029330205 | 0.169869347 | ns | TM | KS.1 |
| FIM1_2416 | 3709.529626 | 4340.000447 | 4098.833739 | 0.163805971 | 0.090829824 | 0.012281603 | 0.097517084 | ns | TM | KS.1 |
| FIM1_2417 | 1381.046598 | 1162.328247 | 1140.405553 | -0.149140158 | 0.109441358 | 0.035484658 | 0.188204876 | ns | TM | KS.1 |
| FIM1_2418 | 11036.96939 | 13967.89368 | 15167.9165 | 0.294358678 | 0.074345921 | 7.4217414563[illegible] | 0.000329671 | ns | TM | KS.1 |
| FIM1_2419 | 701.5722077 | 541.0860499 | 811.3745993 | -0.168420447 | 0.154130562 | 0.042242178 | 0.210079667 | ns | TM | KS.1 |
| FIM1_2420 | 1456.267354 | 1491.665378 | 1741.522806 | 0.013907751 | 0.089192671 | 0.804085704 | 0.936908579 | ns | TM | KS.1 |
| FIM1_2422 | 3707.696682 | 4030.666458 | 4611.739681 | 0.076427497 | 0.070258326 | 0.119633796 | 0.379577892 | ns | TM | KS.1 |
| FIM1_2423 | 505.410638 | 570.4530112 | 573.2406725 | 0.051681591 | 0.126753492 | 0.431325951 | 0.743536592 | ns | TM | KS.1 |
| FIM1_2424 | 1311.19365 | 1113.151122 | 1761.714842 | -0.091663066 | 0.125636227 | 0.186883841 | 0.489513719 | ns | TM | KS.1 |
| FIM1_2425 | 636.2237669 | 658.7199047 | 549.0309316 | 0.014661229 | 0.116223329 | 0.81437398 | 0.940696014 | ns | TM | KS.1 |
| FIM1_2426 | 3157.868627 | 2896.114368 | 3547.211572 | -0.056232826 | 0.094342946 | 0.339172196 | 0.667476929 | ns | TM | KS.1 |
| FIM1_2427 | 237.1591035 | 244.7198292 | 261.1951881 | 0.017630871 | 0.107231685 | 0.772901414 | 0.925733395 | ns | TM | KS.1 |
| FIM1_2428 | 567.2336382 | 340.9682713 | 668.4622509 | -0.322567386 | 0.290381493 | 0.008731366 | 0.077292783 | ns | TM | KS.1 |
| FIM1_2429 | 1010.544428 | 1011.136013 | 1186.307702 | -0.000149372 | 0.06172664 | 0.997291852 | 0.998293524 | ns | TM | KS.1 |
| FIM1_2430 | 1530.162294 | 1587.597936 | 1665.928015 | 0.023937157 | 0.079430855 | 0.647441186 | 0.873595603 | ns | TM | KS.1 |

| | | | | | | | | | | |
|---|---|---|---|---|---|---|---|---|---|---|
| FIM1_2431 | 2300.68339 | 2552.447945 | 1976.13754 | 0.059659475 | 0.106924026 | 0.342697115 | 0.669908898 | ns | TM | KS.1 |
| FIM1_2432 | 1135.147641 | 1119.857005 | 1147.157163 | -0.007490525 | 0.094706328 | 0.896914131 | 0.967972723 | ns | TM | KS.1 |
| FIM1_2433 | 26200.66497 | 29942.08977 | 28152.16964 | 0.043793004 | 0.142419772 | 0.503638651 | 0.796765282 | ns | TM | KS.1 |
| FIM1_2434 | 4976.304255 | 5475.645778 | 5279.855016 | 0.036502157 | 0.128738024 | 0.57175333 | 0.832094673 | ns | TM | KS.1 |
| FIM1_2435 | 1726.568835 | 1609.574998 | 1608.339509 | -0.048300874 | 0.08499946 | 0.379845258 | 0.703660035 | ns | TM | KS.1 |
| FIM1_2436 | 318.9138283 | 275.2114984 | 317.0982153 | -0.068256316 | 0.128639982 | 0.311601568 | 0.638961913 | ns | TM | KS.1 |
| FIM1_2437 | 119.3021406 | 91.05078089 | 185.0976495 | -0.091843739 | 0.173305468 | 0.206313522 | 0.513461617 | ns | TM | KS.1 |
| FIM1_2438 | 1129.501324 | 1197.84603 | 1531.349559 | 0.034793533 | 0.09291323 | 0.546281129 | 0.818934867 | ns | TM | KS.1 |
| FIM1_2439 | 309.224966 | 299.5934818 | 443.2228972 | -0.009799865 | 0.135763658 | 0.876770048 | 0.964248917 | ns | TM | KS.1 |
| FIM1_2440 | 315.0418677 | 180.0874682 | 226.0552197 | -0.143107588 | 0.270176794 | 0.095092685 | 0.333005827 | ns | TM | KS.1 |
| FIM1_2444 | 9542.262665 | 20074.232 | 20677.52651 | 0.547447368 | 0.434642883 | 0.002604034 | 0.031368211 | ns | TM | KS.1 |
| FIM1_2445 | 339.7913025 | 719.4230052 | 805.4567149 | 0.74252174 | 0.399654262 | 0.000462551 | 0.008460261 | ns | TM | KS.1 |
| FIM1_2446 | 245.6392021 | 238.1687465 | 299.0757199 | -0.017853161 | 0.09705725 | 0.760706121 | 0.923262022 | ns | TM | KS.1 |
| FIM1_2447 | 479.1508321 | 531.6572225 | 473.4177694 | 0.067383926 | 0.101974916 | 0.276375331 | 0.601096377 | ns | TM | KS.1 |
| FIM1_2448 | 36.77374223 | 41.45608583 | 42.2117417 | 0.022038916 | 0.179695046 | 0.721331695 | 0.906676397 | ns | TM | KS.1 |
| FIM1_2449 | 31.46590952 | 37.07310565 | 43.40792016 | 0.032695133 | 0.177391197 | 0.603913 | 0.851547247 | ns | TM | KS.1 |
| FIM1_2450 | 271.4791911 | 387.3688836 | 632.294383 | 0.206481709 | 0.198181937 | 0.027229339 | 0.162624202 | ns | TM | KS.1 |
| FIM1_2451 | 219.3765583 | 554.9201879 | 931.4556968 | 0.273317759 | 0.426215584 | 0.023817258 | 0.151329319 | ns | TM | KS.1 |
| FIM1_2452 | 43.1902819 | 40.75674446 | 52.95915738 | -0.013138536 | 0.147857111 | 0.834502294 | 0.948141464 | ns | TM | KS.1 |
| FIM1_2453 | 172.1680006 | 190.2592461 | 209.9167577 | 0.044535877 | 0.115406204 | 0.484666621 | 0.785178463 | ns | TM | KS.1 |
| FIM1_2454 | 281.2248563 | 242.4931215 | 378.7276646 | -0.058217547 | 0.135270094 | 0.384179232 | 0.708394027 | ns | TM | KS.1 |
| FIM1_2455 | 630.0975262 | 777.5877233 | 1087.383669 | 0.047140991 | 0.173905971 | 0.469153041 | 0.77114073 | ns | TM | KS.1 |
| FIM1_2456 | 289.5077477 | 276.1248987 | 256.7192958 | -0.018659322 | 0.127124755 | 0.768596515 | 0.925179691 | ns | TM | KS.1 |
| FIM1_2457 | 330.2706582 | 349.3082118 | 402.8146455 | 0.025130571 | 0.110283149 | 0.684719986 | 0.890505759 | ns | TM | KS.1 |
| FIM1_2458 | 1688.45111 | 1610.178246 | 2445.15704 | -0.021748066 | 0.111107605 | 0.725421532 | 0.907961357 | ns | TM | KS.1 |
| FIM1_2459 | 1350.259858 | 1771.665199 | 2120.245717 | 0.119375357 | 0.166174662 | 0.121248802 | 0.382506526 | ns | TM | KS.1 |
| FIM1_2460 | 2485.27875 | 3462.038121 | 4269.400428 | 0.120568946 | 0.189820312 | 0.12384566 | 0.387735747 | ns | TM | KS.1 |
| FIM1_2461 | 2616.697973 | 2656.666098 | 2918.362486 | 0.011257006 | 0.065355994 | 0.805933284 | 0.937659684 | ns | TM | KS.1 |
| FIM1_2462 | 4049.446883 | 4733.727103 | 5382.854023 | 0.159887869 | 0.092170057 | 0.014787457 | 0.109802383 | ns | TM | KS.1 |
| FIM1_2463 | 837.9367582 | 783.8303014 | 944.635553 | -0.050523972 | 0.076022584 | 0.322851035 | 0.650844386 | ns | TM | KS.1 |
| FIM1_2464 | 2205.144966 | 2016.942532 | 1939.7883 | -0.0471958 | 0.108054678 | 0.449792848 | 0.754202703 | ns | TM | KS.1 |
| FIM1_2465 | 847.6271695 | 816.3046509 | 596.242639 | -0.023468878 | 0.083473421 | 0.663882526 | 0.880619203 | ns | TM | KS.1 |
| FIM1_2466 | 752.6778136 | 773.052114 | 842.4749687 | 0.008300111 | 0.134684997 | 0.895473503 | 0.967972723 | ns | TM | KS.1 |
| FIM1_2467 | 367.0351063 | 324.3026317 | 403.5880091 | -0.077900569 | 0.109773351 | 0.229506866 | 0.547690282 | ns | TM | KS.1 |
| FIM1_2468 | 2164.018051 | 2484.027737 | 2249.658896 | 0.099052309 | 0.107023967 | 0.13125511 | 0.400376242 | ns | TM | KS.1 |
| FIM1_2469 | 2251.858644 | 1984.344283 | 1894.960468 | -0.138251277 | 0.075559059 | 0.013684258 | 0.104063892 | ns | TM | KS.1 |
| FIM1_2470 | 6727.873227 | 7409.249219 | 6846.182636 | 0.077747142 | 0.084627251 | 0.164085094 | 0.45366779 | ns | TM | KS.1 |
| FIM1_2471 | 1371.277419 | 1384.078915 | 1109.614518 | 0.00464825 | 0.099417576 | 0.93733754 | 0.98242551 | ns | TM | KS.1 |
| FIM1_2472 | 1981.205568 | 1830.268887 | 2006.773113 | -0.057827539 | 0.084460672 | 0.293318477 | 0.620392168 | ns | TM | KS.1 |
| FIM1_2473 | 1013.160011 | 1368.13853 | 1554.022244 | 0.17901645 | 0.17229982 | 0.038756902 | 0.198583099 | ns | TM | KS.1 |
| FIM1_2474 | 2465.663687 | 2862.840163 | 3438.768156 | 0.148268204 | 0.093052574 | 0.022198679 | 0.145505177 | ns | TM | KS.1 |
| FIM1_2475 | 1616.301573 | 1450.430486 | 1268.99965 | -0.084355321 | 0.092218467 | 0.155428366 | 0.439849899 | ns | TM | KS.1 |
| FIM1_2476 | 9874.464416 | 10172.46728 | 9869.721191 | 0.020462277 | 0.07225889 | 0.677056658 | 0.887002016 | ns | TM | KS.1 |
| FIM1_2477 | 2137.026072 | 1955.990213 | 2092.768011 | -0.059960028 | 0.092278414 | 0.303102176 | 0.631432423 | ns | TM | KS.1 |
| FIM1_2478 | 1086.456818 | 1248.091823 | 1118.601481 | 0.093749574 | 0.110305425 | 0.156647657 | 0.441792571 | ns | TM | KS.1 |

| | | | | | | | | | | |
|---|---|---|---|---|---|---|---|---|---|---|
| FIM1_2479 | 770.7038243 | 809.8476851 | 825.2483291 | 0.033398757 | 0.082034773 | 0.532587927 | 0.809787573 | ns | TM | KS.1 |
| FIM1_2480 | 1741.797383 | 1871.045651 | 2053.053403 | 0.064154709 | 0.06551711 | 0.165765057 | 0.456381383 | ns | TM | KS.1 |
| FIM1_2481 | 1114.789384 | 1192.353648 | 1282.433671 | 0.050574455 | 0.07575605 | 0.321202971 | 0.649587309 | ns | TM | KS.1 |
| FIM1_2482 | 1335.802268 | 1424.615489 | 1389.641121 | 0.054678698 | 0.065736498 | 0.236378608 | 0.556020603 | ns | TM | KS.1 |
| FIM1_2483 | 1109.552295 | 1158.600272 | 1291.08699 | 0.030408798 | 0.073216926 | 0.539785425 | 0.814399145 | ns | TM | KS.1 |
| FIM1_2484 | 295.6050909 | 331.3502755 | 351.8027649 | 0.071575016 | 0.107427734 | 0.261601567 | 0.588461463 | ns | TM | KS.1 |
| FIM1_2485 | 1713.191054 | 1339.833293 | 1569.017362 | -0.230837699 | 0.126415996 | 0.006066712 | 0.059730875 | ns | TM | KS.1 |
| FIM1_2486 | 11114.77987 | 12664.00333 | 11891.10262 | 0.078382332 | 0.11258653 | 0.232007934 | 0.549507578 | ns | TM | KS.1 |
| FIM1_2487 | 2217.089991 | 2365.005644 | 2468.994696 | 0.051884695 | 0.070912229 | 0.286687553 | 0.612762705 | ns | TM | KS.1 |
| FIM1_2488 | 285.6145229 | 376.7938094 | 344.4512517 | 0.159606137 | 0.163537057 | 0.054148405 | 0.242809601 | ns | TM | KS.1 |
| FIM1_2489 | 778.1375829 | 786.3128386 | 585.8551589 | 0.004789318 | 0.104587125 | 0.936756188 | 0.982179213 | ns | TM | KS.1 |
| FIM1_2490 | 1143.955209 | 1282.138532 | 1138.043493 | 0.066366088 | 0.109522362 | 0.29943869 | 0.628303452 | ns | TM | KS.1 |
| FIM1_2491 | 329.6103804 | 100.1514319 | 811.3610947 | -1.292032177 | 0.436233777 | 2.70880053878 | 0.000916754 | Km<KS.1 | TM | KS.1 |
| FIM1_2492 | 2148.62548 | 2263.617395 | 2656.474913 | 0.020624639 | 0.12177677 | 0.744126113 | 0.91508234 | ns | TM | KS.1 |
| FIM1_2493 | 1565.046376 | 1646.234024 | 1708.561039 | 0.043715554 | 0.06133098 | 0.318007068 | 0.645485583 | ns | TM | KS.1 |
| FIM1_2494 | 824.4374102 | 874.0561739 | 863.5211927 | 0.035884188 | 0.091690069 | 0.530949088 | 0.808531286 | ns | TM | KS.1 |
| FIM1_2495 | 6185.962772 | 5492.117065 | 5237.240303 | -0.061045478 | 0.116826147 | 0.348684635 | 0.674195903 | ns | TM | KS.1 |
| FIM1_2496 | 1362.619525 | 1125.979109 | 1081.57116 | -0.140904712 | 0.124547059 | 0.058044973 | 0.253530772 | ns | TM | KS.1 |
| FIM1_2497 | 429.0415779 | 448.6612543 | 585.9993221 | 0.021463667 | 0.108679704 | 0.726993611 | 0.907961357 | ns | TM | KS.1 |
| FIM1_2498 | 106.7502964 | 112.1738303 | 134.5474941 | 0.015834718 | 0.133217942 | 0.802840151 | 0.936908579 | ns | TM | KS.1 |
| FIM1_2499 | 1061.280632 | 1078.054227 | 1237.132266 | 0.010855229 | 0.074795977 | 0.82908986 | 0.94647133 | ns | TM | KS.1 |
| FIM1_2500 | 1063.025663 | 781.1878294 | 1509.180097 | -0.167856156 | 0.177238056 | 0.049561263 | 0.232172585 | ns | TM | KS.1 |
| FIM1_2502 | 3944.598609 | 3452.456321 | 3124.580927 | -0.078446308 | 0.114327987 | 0.234592986 | 0.553652801 | ns | TM | KS.1 |
| FIM1_2503 | 560.7646576 | 649.4660082 | 574.3758891 | 0.061412057 | 0.13280209 | 0.360070859 | 0.683664672 | ns | TM | KS.1 |
| FIM1_2504 | 854.4881917 | 861.1201155 | 853.0960671 | 0.002622744 | 0.130407993 | 0.966844181 | 0.993374633 | ns | TM | KS.1 |
| FIM1_2505 | 735.9686902 | 668.7274871 | 596.8034414 | -0.046765735 | 0.114083514 | 0.462481652 | 0.764620503 | ns | TM | KS.1 |
| FIM1_2506 | 572.890092 | 390.7644966 | 537.1545035 | -0.28592971 | 0.206817981 | 0.006989952 | 0.066238115 | ns | TM | KS.1 |
| FIM1_2507 | 947.4111263 | 870.615706 | 1007.272581 | -0.053286698 | 0.095219021 | 0.366652618 | 0.690301888 | ns | TM | KS.1 |
| FIM1_2508 | 12.15998276 | 19.01400634 | 19.23299098 | 0.082372761 | 0.225223938 | 0.246103841 | 0.570003076 | ns | TM | KS.1 |
| FIM1_2510 | 246.5935339 | 314.0161068 | 247.8209449 | 0.126396929 | 0.153475911 | 0.101395474 | 0.343859906 | ns | TM | KS.1 |
| FIM1_2511 | 3095.300568 | 4393.577815 | 5768.840307 | 0.246522533 | 0.191175463 | 0.012241728 | 0.097444157 | ns | TM | KS.1 |
| FIM1_2512 | 1340.803159 | 1320.554683 | 1212.02316 | -0.009054454 | 0.084944692 | 0.868024492 | 0.960282219 | ns | TM | KS.1 |
| FIM1_2513 | 5731.05481 | 6362.708553 | 6384.272418 | 0.049658439 | 0.117720175 | 0.440928822 | 0.747832613 | ns | TM | KS.1 |
| FIM1_2514 | 449.0277143 | 414.0091178 | 481.8060497 | -0.046983149 | 0.101688045 | 0.439710512 | 0.747832613 | ns | TM | KS.1 |
| FIM1_2515 | 76.26970373 | 73.98519732 | 94.90197683 | -0.011872602 | 0.122324669 | 0.850383685 | 0.953065743 | ns | TM | KS.1 |
| FIM1_2516 | 3635.771885 | 3599.499158 | 4106.495846 | -0.006506751 | 0.075175016 | 0.8973853 | 0.967972723 | ns | TM | KS.1 |
| FIM1_2517 | 2318.700212 | 1335.178639 | 2074.916218 | -0.698525266 | 0.206157662 | 4.49512245367 | 0.000223632 | ns | TM | KS.1 |
| FIM1_2518 | 1930.799763 | 2048.839645 | 2038.925988 | 0.047355937 | 0.069038723 | 0.320953641 | 0.649347037 | ns | TM | KS.1 |
| FIM1_2519 | 1347.88479 | 2391.305761 | 2929.214204 | 0.199326015 | 0.295020375 | 0.045456079 | 0.220040594 | ns | TM | KS.1 |
| FIM1_2520 | 515.8464138 | 656.3604993 | 689.008901 | 0.156093943 | 0.147407656 | 0.052155419 | 0.238705802 | ns | TM | KS.1 |
| FIM1_2521 | 548.6240616 | 801.8065904 | 697.9728368 | 0.304536843 | 0.200601754 | 0.004569193 | 0.048468521 | ns | TM | KS.1 |
| FIM1_2522 | 489.8482512 | 460.1498628 | 490.9830642 | -0.032749013 | 0.10123432 | 0.586051115 | 0.841613393 | ns | TM | KS.1 |
| FIM1_2523 | 4192.741881 | 4278.504184 | 4516.161231 | 0.017605453 | 0.048604859 | 0.633919326 | 0.869547581 | ns | TM | KS.1 |
| FIM1_2524 | 1203.288687 | 1476.698216 | 1763.71944 | 0.07227341 | 0.152522223 | 0.296253028 | 0.623723579 | ns | TM | KS.1 |
| FIM1_2525 | 1100.339267 | 861.7641672 | 706.1662017 | -0.185085405 | 0.141471371 | 0.025473438 | 0.156140634 | ns | TM | KS.1 |

| | | | | | | | | | | |
|---|---|---|---|---|---|---|---|---|---|---|
| FIM1_2526 | 7947.87927 | 7318.26545 | 5219.96843 | -0.048884249 | 0.098883658 | 0.415636516 | 0.735011951 | ns | TM | KS.1 |
| FIM1_2527 | 135.554873 | 431.8224615 | 199.4878147 | 1.341685795 | 0.370619975 | 6.17513361253 | 0.000287115 | Km>KS.1 | TM | KS.1 |
| FIM1_2528 | 19739.54282 | 22277.89791 | 23970.31229 | 0.102137626 | 0.092595065 | 0.092382613 | 0.328522875 | ns | TM | KS.1 |
| FIM1_2529 | 7038.703875 | 6987.518258 | 8482.783516 | -0.001934129 | 0.146251101 | 0.975387512 | 0.993872484 | ns | TM | KS.1 |
| FIM1_2530 | 1054.876676 | 964.2126253 | 1352.405008 | -0.023831428 | 0.14979848 | 0.707111487 | 0.89994363 | ns | TM | KS.1 |
| FIM1_2531 | 2622.710599 | 2786.277143 | 3138.28937 | 0.028707983 | 0.109940454 | 0.64315735 | 0.872978256 | ns | TM | KS.1 |
| FIM1_2532 | 536.8635918 | 775.1892291 | 477.4910332 | 0.14905592 | 0.20396903 | 0.077036593 | 0.294813117 | ns | TM | KS.1 |
| FIM1_2533 | 3239.528079 | 3081.424262 | 2574.912595 | -0.029456782 | 0.09193091 | 0.606687908 | 0.851659878 | ns | TM | KS.1 |
| FIM1_2534 | 2017.821763 | 1833.565004 | 2106.468145 | -0.051608314 | 0.10856868 | 0.411277495 | 0.731735607 | ns | TM | KS.1 |
| FIM1_2535 | 389.2752926 | 415.1872643 | 461.4695118 | 0.030923116 | 0.110111096 | 0.618447291 | 0.860858199 | ns | TM | KS.1 |
| FIM1_2536 | 484.942231 | 343.2639867 | 460.6005252 | -0.116407356 | 0.195727644 | 0.134511452 | 0.406322117 | ns | TM | KS.1 |
| FIM1_2537 | 868.0677829 | 600.5833214 | 1077.293257 | -0.162527166 | 0.204776665 | 0.061120333 | 0.260560116 | ns | TM | KS.1 |
| FIM1_2538 | 981.9633528 | 789.3839646 | 815.3711556 | -0.11179507 | 0.145277048 | 0.132787921 | 0.403555229 | ns | TM | KS.1 |
| FIM1_2539 | 607.1231219 | 598.6227594 | 615.8128797 | -0.006337375 | 0.112734334 | 0.918371379 | 0.975634767 | ns | TM | KS.1 |
| FIM1_2540 | 2992.667833 | 3037.302615 | 3341.158667 | 0.006612426 | 0.11029031 | 0.914322713 | 0.974699214 | ns | TM | KS.1 |
| FIM1_2541 | 1233.29759 | 1217.249879 | 1315.902532 | -0.006703662 | 0.093743429 | 0.907214244 | 0.97188159 | ns | TM | KS.1 |
| FIM1_2542 | 452.1449364 | 447.980655 | 401.8720866 | -0.00412459 | 0.108399044 | 0.946190013 | 0.985614597 | ns | TM | KS.1 |
| FIM1_2543 | 1057.351458 | 1077.788231 | 1152.744984 | 0.009748781 | 0.100087155 | 0.869499181 | 0.96127965 | ns | TM | KS.1 |
| FIM1_2544 | 854.7408395 | 873.4895993 | 859.9406074 | 0.014106767 | 0.078102114 | 0.785117336 | 0.931320636 | ns | TM | KS.1 |
| FIM1_2546 | 1528.635015 | 1410.680447 | 1183.701345 | -0.050323679 | 0.094789 | 0.391833464 | 0.715009037 | ns | TM | KS.1 |
| FIM1_2547 | 1533.655212 | 1607.015974 | 2021.126718 | 0.014973827 | 0.134775952 | 0.813240694 | 0.940696014 | ns | TM | KS.1 |
| FIM1_2548 | 4880.298857 | 3656.902845 | 5162.908892 | -0.086960579 | 0.179008206 | 0.226374274 | 0.544063775 | ns | TM | KS.1 |
| FIM1_2549 | 5791.385207 | 5149.670941 | 5865.03692 | -0.045207015 | 0.131513947 | 0.489485816 | 0.788598425 | ns | TM | KS.1 |
| FIM1_2550 | 1965.264152 | 1802.046747 | 1720.231843 | -0.040932825 | 0.114489019 | 0.518262362 | 0.804981347 | ns | TM | KS.1 |
| FIM1_2552 | 1857.625103 | 1140.557618 | 2587.608678 | -0.213208726 | 0.264719475 | 0.03409973 | 0.184799735 | ns | TM | KS.1 |
| FIM1_2553 | 495.7688828 | 465.240833 | 530.5381407 | -0.021447795 | 0.132953965 | 0.736280657 | 0.912555125 | ns | TM | KS.1 |
| FIM1_2554 | 1415.499096 | 1416.588534 | 1249.548733 | 0.000194721 | 0.134898026 | 0.997537006 | 0.998293524 | ns | TM | KS.1 |
| FIM1_2555 | 5618.021013 | 1147.631679 | 14232.97528 | -1.827812374 | 0.369137058 | 1.18066905293 | 1.63161903843 | Km<KS.1 | TM | KS.1 |
| FIM1_2556 | 88.15959783 | 82.93460329 | 413.0204317 | -0.013202855 | 0.162502201 | 0.831555084 | 0.946802573 | ns | TM | KS.1 |
| FIM1_2557 | 10082.22443 | 9593.306401 | 12207.00935 | -0.020369571 | 0.119145958 | 0.746233338 | 0.916216895 | ns | TM | KS.1 |
| FIM1_2558 | 952.548432 | 1187.962964 | 1227.149074 | 0.086868608 | 0.153170451 | 0.223527236 | 0.539044111 | ns | TM | KS.1 |
| FIM1_2559 | 379.8424112 | 364.2177629 | 458.2771054 | -0.011325711 | 0.145008605 | 0.857217718 | 0.955916258 | ns | TM | KS.1 |
| FIM1_2560 | 2722.633564 | 3212.00806 | 3738.299824 | 0.115544847 | 0.117763618 | 0.099367207 | 0.340932312 | ns | TM | KS.1 |
| FIM1_2561 | 269.4927638 | 283.8374923 | 346.8324722 | 0.018440615 | 0.124788426 | 0.770816282 | 0.925705893 | ns | TM | KS.1 |
| FIM1_2562 | 2412.703501 | 2437.065791 | 2830.694731 | 0.005889684 | 0.085701739 | 0.914356027 | 0.974699214 | ns | TM | KS.1 |
| FIM1_2563 | 2348.318289 | 2508.931235 | 3001.103454 | 0.051073578 | 0.074494891 | 0.310899939 | 0.638961913 | ns | TM | KS.1 |
| FIM1_2564 | 1032.970811 | 1034.226589 | 1739.693656 | 0.00137534 | 0.094591771 | 0.981002523 | 0.995431301 | ns | TM | KS.1 |
| FIM1_2565 | 1219.506283 | 1137.244353 | 1138.975453 | -0.040741958 | 0.096466497 | 0.489858099 | 0.788868226 | ns | TM | KS.1 |
| FIM1_2566 | 2034.207612 | 2026.884881 | 2318.819173 | -0.001736417 | 0.106042781 | 0.977159578 | 0.993941709 | ns | TM | KS.1 |
| FIM1_2567 | 809.8601053 | 859.5364041 | 1056.417571 | 0.036846919 | 0.090926809 | 0.518172501 | 0.804981347 | ns | TM | KS.1 |
| FIM1_2568 | 1018.180878 | 1219.556702 | 1217.730461 | 0.070759412 | 0.143160043 | 0.303341405 | 0.631432423 | ns | TM | KS.1 |
| FIM1_2570 | 291.5120127 | 337.2409305 | 388.1636513 | 0.053989202 | 0.138002887 | 0.4171555 | 0.735551151 | ns | TM | KS.1 |
| FIM1_2571 | 246.3074599 | 349.4708737 | 421.0415144 | 0.144690082 | 0.197728903 | 0.081823009 | 0.304774234 | ns | TM | KS.1 |
| FIM1_2572 | 174.3648507 | 241.7198824 | 287.6389418 | 0.194843889 | 0.186165291 | 0.031235278 | 0.176980365 | ns | TM | KS.1 |
| FIM1_2573 | 566.126357 | 548.1207392 | 484.8169296 | -0.016673511 | 0.100441191 | 0.779391223 | 0.928513251 | ns | TM | KS.1 |

| | | | | | | | | | | |
|---|---|---|---|---|---|---|---|---|---|---|
| FIM1_2574 | 1746.658007 | 1732.444465 | 1794.752805 | -0.004723789 | 0.083618715 | 0.930253074 | 0.97843743 | ns | TM | KS.1 |
| FIM1_2575 | 922.7833275 | 929.8289894 | 933.2852499 | 0.005212253 | 0.08313042 | 0.922792144 | 0.976369825 | ns | TM | KS.1 |
| FIM1_2576 | 10991.19873 | 13732.54161 | 14153.82679 | 0.297050737 | 0.053712622 | 3.7958954940( | 6.98879304128 | ns | TM | KS.1 |
| FIM1_2577 | 2800.850563 | 4154.855576 | 5459.753975 | 0.387769205 | 0.187515604 | 0.000649676 | 0.010846109 | ns | TM | KS.1 |
| FIM1_2578 | 67.72099874 | 56.70667086 | 73.6616089 | -0.06822703 | 0.142181626 | 0.318413831 | 0.646047639 | ns | TM | KS.1 |
| FIM1_2579 | 222.6685277 | 148.8019751 | 269.9166729 | -0.357337794 | 0.208132323 | 0.001910307 | 0.024984234 | ns | TM | KS.1 |
| FIM1_2580 | 677.7555669 | 592.5517514 | 1231.593787 | -0.053242192 | 0.132996083 | 0.421656936 | 0.738117813 | ns | TM | KS.1 |
| FIM1_2581 | 16963.82689 | 18057.40618 | 20957.63236 | 0.03750119 | 0.092800149 | 0.515703563 | 0.804775792 | ns | TM | KS.1 |
| FIM1_2582 | 973.6376978 | 712.5603817 | 655.2949581 | -0.230772327 | 0.170285857 | 0.013334919 | 0.102378432 | ns | TM | KS.1 |
| FIM1_2583 | 824.3780295 | 659.0031941 | 481.033384 | -0.126466215 | 0.144624942 | 0.097096168 | 0.337101354 | ns | TM | KS.1 |
| FIM1_2584 | 22.50433899 | 64.60714158 | 103.7048874 | 0.565828205 | 0.564191959 | 0.003670742 | 0.040946059 | ns | TM | KS.1 |
| FIM1_2585 | 4738.93039 | 4201.666405 | 4837.266344 | -0.084087217 | 0.10203069 | 0.181556192 | 0.481472312 | ns | TM | KS.1 |
| FIM1_2586 | 1102.490546 | 1110.045563 | 1096.603075 | 0.003704923 | 0.077873781 | 0.942790717 | 0.984319543 | ns | TM | KS.1 |
| FIM1_2587 | 3954.88048 | 3452.194168 | 4513.054149 | -0.125691944 | 0.092933059 | 0.04503836 | 0.21945724 | ns | TM | KS.1 |
| FIM1_2588 | 7772.863562 | 10751.23705 | 5707.663931 | 0.183394097 | 0.183625892 | 0.038167217 | 0.196962604 | ns | TM | KS.1 |
| FIM1_2590 | 2099.15455 | 1295.781705 | 1955.598832 | -0.250634823 | 0.268277857 | 0.020259121 | 0.137690067 | ns | TM | KS.1 |
| FIM1_2591 | 705.0912172 | 826.6244275 | 888.9165065 | 0.056457737 | 0.142725845 | 0.398940231 | 0.722244413 | ns | TM | KS.1 |
| FIM1_2592 | 1379.645671 | 1624.484175 | 1822.145642 | 0.13128917 | 0.111092502 | 0.059695292 | 0.256462934 | ns | TM | KS.1 |
| FIM1_2593 | 1440.715016 | 1377.51823 | 1588.488274 | -0.032670191 | 0.070218156 | 0.497724121 | 0.793454967 | ns | TM | KS.1 |
| FIM1_2594 | 2623.922274 | 2233.007404 | 2182.106981 | -0.129322183 | 0.110410172 | 0.0621192 | 0.261236705 | ns | TM | KS.1 |
| FIM1_2595 | 1126.636818 | 1268.123157 | 1316.272804 | 0.098964771 | 0.091745628 | 0.099748073 | 0.341199031 | ns | TM | KS.1 |
| FIM1_2596 | 1926.871085 | 1815.042924 | 2024.028705 | -0.025158278 | 0.118335713 | 0.689914161 | 0.892671769 | ns | TM | KS.1 |
| FIM1_2597 | 934.8410476 | 1085.869684 | 977.7203132 | 0.070306755 | 0.128689106 | 0.298836826 | 0.627569949 | ns | TM | KS.1 |
| FIM1_2598 | 1256.332343 | 1221.130113 | 1013.907709 | -0.01693682 | 0.085654431 | 0.757276724 | 0.920753636 | ns | TM | KS.1 |
| FIM1_2599 | 786.5737842 | 1008.235707 | 964.505576 | 0.115102549 | 0.157242643 | 0.128983296 | 0.396596531 | ns | TM | KS.1 |
| FIM1_2600 | 1329.75974 | 1321.601466 | 1325.314155 | -0.002651963 | 0.106552725 | 0.965183155 | 0.993335994 | ns | TM | KS.1 |
| FIM1_2601 | 2056.363151 | 2043.433059 | 2074.916074 | -0.003728577 | 0.080946388 | 0.943816717 | 0.984585482 | ns | TM | KS.1 |
| FIM1_2602 | 1679.138368 | 1781.245795 | 1905.859368 | 0.04140041 | 0.080339737 | 0.433854796 | 0.745878978 | ns | TM | KS.1 |
| FIM1_2603 | 297.2428286 | 307.7424526 | 286.3863057 | 0.009053993 | 0.145965775 | 0.88542117 | 0.967028314 | ns | TM | KS.1 |
| FIM1_2604 | 625.1452909 | 612.5874558 | 543.7081307 | -0.008510352 | 0.117335157 | 0.891587911 | 0.967215407 | ns | TM | KS.1 |
| FIM1_2606 | 25.83567047 | 22.31102557 | 32.27504231 | -0.032720774 | 0.167981735 | 0.605925076 | 0.851547247 | ns | TM | KS.1 |
| FIM1_2607 | 2122.574108 | 2861.927965 | 3002.133013 | 0.17353852 | 0.171983654 | 0.043097092 | 0.212459177 | ns | TM | KS.1 |
| FIM1_2608 | 744.8903351 | 807.7430539 | 800.2735726 | 0.063209191 | 0.080207351 | 0.235970596 | 0.555586234 | ns | TM | KS.1 |
| FIM1_2609 | 531.6164104 | 476.9886556 | 502.6738705 | -0.062083968 | 0.109576738 | 0.329666053 | 0.657613719 | ns | TM | KS.1 |
| FIM1_2610 | 860.1616162 | 949.1114469 | 1233.092503 | 0.065888541 | 0.096888239 | 0.27333631 | 0.599198132 | ns | TM | KS.1 |
| FIM1_2611 | 1681.133363 | 2359.789801 | 1871.482649 | 0.204508907 | 0.189724367 | 0.026686996 | 0.160541482 | ns | TM | KS.1 |
| FIM1_2612 | 2495.354423 | 2730.299274 | 3394.826906 | 0.020782243 | 0.158432049 | 0.740565854 | 0.914311536 | ns | TM | KS.1 |
| FIM1_2613 | 2765.018129 | 2746.157391 | 3064.290298 | -0.00210497 | 0.137000838 | 0.973365921 | 0.993872484 | ns | TM | KS.1 |
| FIM1_2614 | 974.5460746 | 909.1508967 | 709.2168211 | -0.041035181 | 0.095827367 | 0.485289973 | 0.785236922 | ns | TM | KS.1 |
| FIM1_2615 | 75.15146764 | 82.09318902 | 64.47101098 | 0.024529041 | 0.142636694 | 0.70039893 | 0.897138176 | ns | TM | KS.1 |
| FIM1_2616 | 566.1552045 | 649.0765597 | 737.3979886 | 0.122455093 | 0.095482148 | 0.053607571 | 0.24174415 | ns | TM | KS.1 |
| FIM1_2617 | 1273.841001 | 1235.639914 | 1133.242136 | -0.022063054 | 0.065481158 | 0.630425186 | 0.868799252 | ns | TM | KS.1 |
| FIM1_2618 | 460.8249432 | 471.7397921 | 394.8937208 | 0.011628996 | 0.098631056 | 0.84367951 | 0.951768052 | ns | TM | KS.1 |
| FIM1_2619 | 330.0664064 | 322.529752 | 276.3989874 | -0.010803431 | 0.108900025 | 0.860050076 | 0.957000476 | ns | TM | KS.1 |
| FIM1_2620 | 872.7758023 | 714.0090858 | 670.0593794 | -0.143602729 | 0.129006972 | 0.057624732 | 0.253253569 | ns | TM | KS.1 |

| | | | | | | | | | | |
|---|---|---|---|---|---|---|---|---|---|---|
| FIM1_2621 | 6226.701416 | 7132.9854 | 8627.981714 | 0.072977644 | 0.119045817 | 0.272607652 | 0.599172403 | ns | TM | KS.1 |
| FIM1_2622 | 733.2826915 | 743.3527632 | 652.659282 | 0.007325975 | 0.089215331 | 0.895977822 | 0.967972723 | ns | TM | KS.1 |
| FIM1_2623 | 1536.370096 | 1756.079614 | 1779.418525 | 0.103534874 | 0.102146782 | 0.107779547 | 0.356044653 | ns | TM | KS.1 |
| FIM1_2624 | 6647.954724 | 6566.97734 | 7410.842857 | -0.007437349 | 0.082700336 | 0.889675129 | 0.967028314 | ns | TM | KS.1 |
| FIM1_2625 | 575.530626 | 586.7188171 | 576.4427825 | 0.009567559 | 0.102561706 | 0.87318957 | 0.962759265 | ns | TM | KS.1 |
| FIM1_2626 | 2054.88651 | 2083.118506 | 1584.582197 | 0.008294044 | 0.083541086 | 0.87781313 | 0.964470036 | ns | TM | KS.1 |
| FIM1_2627 | 1929.95314 | 1807.121235 | 1047.969986 | -0.033714889 | 0.105133786 | 0.58139998 | 0.838424819 | ns | TM | KS.1 |
| FIM1_2628 | 917.7225352 | 954.2813801 | 833.5519857 | 0.024219404 | 0.08591614 | 0.659487105 | 0.878874681 | ns | TM | KS.1 |
| FIM1_2629 | 3999.547839 | 5804.405473 | 7639.601927 | 0.170405181 | 0.206886244 | 0.053724446 | 0.24174415 | ns | TM | KS.1 |
| FIM1_2630 | 502.274401 | 630.3167349 | 544.7853279 | 0.135776299 | 0.144450968 | 0.07937985 | 0.300154562 | ns | TM | KS.1 |
| FIM1_2631 | 167.6092875 | 168.3543088 | 217.1279088 | 0.003166268 | 0.102952578 | 0.957904937 | 0.989353426 | ns | TM | KS.1 |
| FIM1_2632 | 645.5326011 | 769.0303588 | 622.236406 | 0.11957401 | 0.122577446 | 0.095257865 | 0.33326855 | ns | TM | KS.1 |
| FIM1_2633 | 865.1291114 | 692.3576726 | 808.6597244 | -0.148865915 | 0.138746952 | 0.056700754 | 0.251189893 | ns | TM | KS.1 |
| FIM1_2634 | 7544.642542 | 6695.586204 | 5047.092429 | -0.039040987 | 0.140598191 | 0.548131324 | 0.820629954 | ns | TM | KS.1 |
| FIM1_2635 | 32387.92418 | 32525.31462 | 31529.08503 | 0.002773273 | 0.075396946 | 0.956249443 | 0.989353426 | ns | TM | KS.1 |
| FIM1_2636 | 3550.055311 | 3719.245481 | 3879.739344 | 0.037785003 | 0.061559977 | 0.389009305 | 0.713088169 | ns | TM | KS.1 |
| FIM1_2637 | 6942.945855 | 5881.373558 | 4556.375614 | -0.092410613 | 0.1263692 | 0.184572269 | 0.487133708 | ns | TM | KS.1 |
| FIM1_2638 | 1393.525914 | 1422.199812 | 1535.1718 | 0.01260243 | 0.082157276 | 0.813485983 | 0.940696014 | ns | TM | KS.1 |
| FIM1_2639 | 5049.493758 | 3802.126553 | 3287.046383 | -0.169982781 | 0.165030413 | 0.044289626 | 0.217084621 | ns | TM | KS.1 |
| FIM1_2640 | 4106.910654 | 2824.090001 | 3174.526742 | -0.379425428 | 0.170305889 | 0.000467966 | 0.008527959 | ns | TM | KS.1 |
| FIM1_2641 | 1009.155539 | 1010.929385 | 974.5261912 | 0.000793128 | 0.075196776 | 0.987460006 | 0.996167262 | ns | TM | KS.1 |
| FIM1_2642 | 300.0453894 | 325.8019455 | 395.3580463 | 0.049511573 | 0.095401825 | 0.400961417 | 0.723750973 | ns | TM | KS.1 |
| FIM1_2643 | 2497.549767 | 2229.266609 | 2442.68225 | -0.069546898 | 0.106363483 | 0.272141849 | 0.598783418 | ns | TM | KS.1 |
| FIM1_2644 | 2142.73224 | 1960.824604 | 1769.190409 | -0.065904405 | 0.086017984 | 0.238910315 | 0.559787835 | ns | TM | KS.1 |
| FIM1_2645 | 704.9389796 | 660.5113966 | 693.2951935 | -0.039369773 | 0.093553338 | 0.497285903 | 0.793454967 | ns | TM | KS.1 |
| FIM1_2646 | 2800.640389 | 2752.357652 | 2678.289446 | -0.011629283 | 0.072036795 | 0.812509745 | 0.940492318 | ns | TM | KS.1 |
| FIM1_2647 | 3218.298947 | 3219.272526 | 3353.209645 | 0.000271592 | 0.065845643 | 0.99529694 | 0.99790453 | ns | TM | KS.1 |
| FIM1_2648 | 276.6486339 | 276.2039588 | 308.8158223 | -0.001242365 | 0.08271048 | 0.981511894 | 0.995431301 | ns | TM | KS.1 |
| FIM1_2649 | 760.0141801 | 789.696974 | 874.0597844 | 0.028203169 | 0.065764701 | 0.539711624 | 0.814399145 | ns | TM | KS.1 |
| FIM1_2650 | 12804.20752 | 15105.69771 | 13759.61847 | 0.122649481 | 0.115505235 | 0.080581073 | 0.302787644 | ns | TM | KS.1 |
| FIM1_2651 | 298.1159703 | 299.0005264 | 381.7676173 | 0.0020379 | 0.119170465 | 0.974011319 | 0.993872484 | ns | TM | KS.1 |
| FIM1_2652 | 5679.569457 | 7207.885853 | 7485.896065 | 0.284181127 | 0.08629125 | 8.8104561820 | 0.002159213 | ns | TM | KS.1 |
| FIM1_2653 | 1712.581367 | 1729.855764 | 1690.258184 | 0.006565303 | 0.078932706 | 0.899679216 | 0.968452273 | ns | TM | KS.1 |
| FIM1_2654 | 2164.003168 | 2107.88232 | 2235.239421 | -0.015766084 | 0.083915148 | 0.770831796 | 0.925705893 | ns | TM | KS.1 |
| FIM1_2655 | 1262.121492 | 1535.592389 | 1444.349023 | 0.043939042 | 0.171691259 | 0.497244115 | 0.793454967 | ns | TM | KS.1 |
| FIM1_2656 | 142.145243 | 168.4041313 | 282.8426353 | 0.04876691 | 0.155991971 | 0.459024731 | 0.762486824 | ns | TM | KS.1 |
| FIM1_2657 | 748.4122374 | 853.333555 | 919.771808 | 0.087121542 | 0.107490861 | 0.179222249 | 0.47808616 | ns | TM | KS.1 |
| FIM1_2659 | 1873.270747 | 1488.537336 | 1561.785365 | -0.194784322 | 0.130097453 | 0.016751921 | 0.120608985 | ns | TM | KS.1 |
| FIM1_2660 | 3946.063497 | 2274.284266 | 1832.770411 | -0.376955129 | 0.318052949 | 0.005519403 | 0.056268502 | ns | TM | KS.1 |
| FIM1_2661 | 1179.234261 | 1266.426351 | 1387.871653 | 0.01866307 | 0.149182412 | 0.767420593 | 0.924883104 | ns | TM | KS.1 |
| FIM1_2662 | 4815.942926 | 4720.597082 | 5391.751476 | -0.011575564 | 0.088834788 | 0.836017398 | 0.948719561 | ns | TM | KS.1 |
| FIM1_2663 | 403.651307 | 448.9128708 | 406.4431412 | 0.028383772 | 0.150757016 | 0.656175205 | 0.877276998 | ns | TM | KS.1 |
| FIM1_2664 | 3429.669282 | 2755.14742 | 2489.737755 | -0.248886217 | 0.092858952 | 0.000743168 | 0.012043191 | ns | TM | KS.1 |
| FIM1_2665 | 354.0435934 | 252.1394336 | 497.1695574 | -0.196239416 | 0.190644078 | 0.031308377 | 0.176980365 | ns | TM | KS.1 |
| FIM1_2666 | 582.5657386 | 531.0400988 | 1035.415032 | -0.042290473 | 0.117522522 | 0.508133787 | 0.800242353 | ns | TM | KS.1 |

| | | | | | | | | | | |
|---|---|---|---|---|---|---|---|---|---|---|
| FIM1_2667 | 2740.088693 | 2911.102609 | 3083.454991 | 0.050624051 | 0.065141129 | 0.269108963 | 0.596620808 | ns | TM | KS.1 |
| FIM1_2668 | 3201.350987 | 3142.841988 | 3118.802518 | -0.005866265 | 0.134655148 | 0.925963619 | 0.976735299 | ns | TM | KS.1 |
| FIM1_2669 | 48.85068235 | 41.53823864 | 47.98920621 | -0.0481877 | 0.15700159 | 0.463689038 | 0.765379219 | ns | TM | KS.1 |
| FIM1_2671 | 65.77677148 | 45.39311891 | 54.40823865 | -0.129060504 | 0.204191331 | 0.109047556 | 0.35800987 | ns | TM | KS.1 |
| FIM1_2672 | 793.1600957 | 890.0107349 | 872.2161792 | 0.090171867 | 0.094498942 | 0.136920515 | 0.409362716 | ns | TM | KS.1 |
| FIM1_2674 | 5758.863346 | 6347.372022 | 5072.522202 | 0.056719895 | 0.104222738 | 0.360167564 | 0.683664672 | ns | TM | KS.1 |
| FIM1_2675 | 1075.765005 | 119.7385313 | 10561.10044 | -1.708296829 | 0.777300504 | 1.1647803899[illegible] | 0.000467321 | Km<KS.1 | TM | KS.1 |
| FIM1_2676 | 12993.31779 | 10147.00408 | 23990.20896 | -0.079000216 | 0.166351339 | 0.261699333 | 0.588461463 | ns | TM | KS.1 |
| FIM1_2677 | 3643.463231 | 3679.161646 | 3814.504978 | 0.006456907 | 0.077143552 | 0.899868566 | 0.968452273 | ns | TM | KS.1 |
| FIM1_2678 | 568.2083172 | 583.043762 | 579.2869035 | 0.013715077 | 0.096275524 | 0.814128696 | 0.940696014 | ns | TM | KS.1 |
| FIM1_2679 | 1187.764348 | 1137.15862 | 1383.071188 | -0.022657223 | 0.100546144 | 0.704177875 | 0.898508574 | ns | TM | KS.1 |
| FIM1_2680 | 3093.912998 | 3678.45818 | 4303.406281 | 0.120732269 | 0.120812709 | 0.090746992 | 0.324562389 | ns | TM | KS.1 |
| FIM1_2681 | 2412.336497 | 2123.290498 | 4554.22403 | -0.064419734 | 0.119763862 | 0.3284067 | 0.655821244 | ns | TM | KS.1 |
| FIM1_2682 | 330.949789 | 446.1643415 | 945.3060971 | 0.101993559 | 0.178959011 | 0.171580616 | 0.465692069 | ns | TM | KS.1 |
| FIM1_2683 | 2729.172202 | 2233.937624 | 2297.684861 | -0.116377708 | 0.13570346 | 0.114706748 | 0.37039344 | ns | TM | KS.1 |
| FIM1_2684 | 2998.483626 | 4480.137082 | 5456.353642 | 0.228756464 | 0.222952979 | 0.021698688 | 0.143551826 | ns | TM | KS.1 |
| FIM1_2685 | 1353.740477 | 987.2283493 | 1047.779783 | -0.332155293 | 0.13376161 | 0.000426469 | 0.007916733 | ns | TM | KS.1 |
| FIM1_2686 | 186.1830404 | 142.6940064 | 211.4828374 | -0.139255117 | 0.161365617 | 0.08071583 | 0.303065098 | ns | TM | KS.1 |
| FIM1_2687 | 642.919459 | 679.4624551 | 777.3624947 | 0.01617119 | 0.141152581 | 0.798367659 | 0.936276987 | ns | TM | KS.1 |
| FIM1_2688 | 1493.771981 | 1620.353384 | 1954.134537 | 0.050835638 | 0.095080842 | 0.387991436 | 0.712272102 | ns | TM | KS.1 |
| FIM1_2689 | 1901.130726 | 1916.334644 | 1808.791337 | 0.005247118 | 0.07590902 | 0.917696425 | 0.975634767 | ns | TM | KS.1 |
| FIM1_2690 | 6090.735669 | 6826.411167 | 6279.00711 | 0.107509894 | 0.08282418 | 0.058866969 | 0.254885264 | ns | TM | KS.1 |
| FIM1_2691 | 1366.909345 | 1383.491334 | 1404.064091 | 0.008626267 | 0.059425973 | 0.840315113 | 0.949982862 | ns | TM | KS.1 |
| FIM1_2692 | 718.2425212 | 641.1416521 | 751.1524816 | -0.074690867 | 0.101977441 | 0.230799396 | 0.548995638 | ns | TM | KS.1 |
| FIM1_2693 | 2221.80648 | 2394.665313 | 2325.402404 | 0.058964823 | 0.076893473 | 0.254048226 | 0.581366111 | ns | TM | KS.1 |
| FIM1_2694 | 437.3629143 | 466.7390308 | 600.8518616 | 0.043840324 | 0.082444576 | 0.415512938 | 0.735011951 | ns | TM | KS.1 |
| FIM1_2695 | 2564.59578 | 2703.401984 | 2827.478016 | 0.034329208 | 0.084383491 | 0.52870417 | 0.808531286 | ns | TM | KS.1 |
| FIM1_2696 | 4787.226071 | 6165.731222 | 6745.90564 | 0.133209299 | 0.156158975 | 0.089207961 | 0.320208953 | ns | TM | KS.1 |
| FIM1_2697 | 423.9817459 | 475.3056323 | 428.8738886 | 0.039192746 | 0.137537468 | 0.546640551 | 0.819137573 | ns | TM | KS.1 |
| FIM1_2698 | 3513.109812 | 3546.058032 | 3296.22013 | 0.006116689 | 0.071607524 | 0.900319953 | 0.968452273 | ns | TM | KS.1 |
| FIM1_2699 | 1306.270976 | 1881.621383 | 2342.557362 | 0.190397417 | 0.203772743 | 0.037427764 | 0.194569622 | ns | TM | KS.1 |
| FIM1_2700 | 447.1323782 | 423.3795153 | 571.3223321 | -0.024700249 | 0.113037064 | 0.691928697 | 0.893131011 | ns | TM | KS.1 |
| FIM1_2701 | 2720.229813 | 2132.228686 | 2051.847764 | -0.136347401 | 0.151360455 | 0.081700481 | 0.304774234 | ns | TM | KS.1 |
| FIM1_2702 | 146.1428431 | 163.7082995 | 193.6273669 | 0.051153861 | 0.118424665 | 0.428828634 | 0.742835117 | ns | TM | KS.1 |
| FIM1_2703 | 606.0058077 | 96.25603536 | 206.2115144 | -2.076485722 | 0.415381725 | 1.0868322347[illegible] | 2.3508653773[illegible] | Km<KS.1 | TM | KS.1 |
| FIM1_2704 | 1441.246296 | 1364.563449 | 1576.395522 | -0.032345809 | 0.091822316 | 0.572013065 | 0.832094673 | ns | TM | KS.1 |
| FIM1_2705 | 2761.95483 | 2232.888712 | 3068.511263 | -0.107721614 | 0.143525776 | 0.143624011 | 0.420558832 | ns | TM | KS.1 |
| FIM1_2706 | 10.74027607 | 10.85652096 | 23.14872675 | 0.006966417 | 0.177982662 | 0.909022845 | 0.972251892 | ns | TM | KS.1 |
| FIM1_2708 | 3173.381004 | 4679.084413 | 5778.449195 | 0.170028789 | 0.214056251 | 0.055469439 | 0.246834043 | ns | TM | KS.1 |
| FIM1_2709 | 2850.065494 | 3288.155901 | 3182.780214 | 0.112988108 | 0.104530922 | 0.08671265 | 0.313970477 | ns | TM | KS.1 |
| FIM1_2710 | 7884.406216 | 9324.597116 | 8876.301857 | 0.094074148 | 0.126707439 | 0.178216753 | 0.476938324 | ns | TM | KS.1 |
| FIM1_2711 | 314.3981638 | 274.7764363 | 333.9744302 | -0.096720593 | 0.10681248 | 0.139073838 | 0.412823594 | ns | TM | KS.1 |
| FIM1_2712 | 2081.87768 | 3027.554117 | 2749.901888 | 0.369316697 | 0.175352157 | 0.000698423 | 0.011582185 | ns | TM | KS.1 |
| FIM1_2713 | 1944.99331 | 1975.446903 | 1923.715799 | 0.010234411 | 0.07382532 | 0.837331702 | 0.949337033 | ns | TM | KS.1 |
| FIM1_2714 | 1771.662104 | 1512.273648 | 1521.290893 | -0.183385869 | 0.078725663 | 0.003023101 | 0.035304992 | ns | TM | KS.1 |

| | | | | | | | | | | |
|---|---|---|---|---|---|---|---|---|---|---|
| FIM1_2715 | 2489.784377 | 3025.100048 | 2877.232405 | 0.210618654 | 0.096382185 | 0.003522035 | 0.039732712 | ns | TM | KS.1 |
| FIM1_2716 | 6246.895329 | 6398.28097 | 7226.098918 | 0.013337511 | 0.091753799 | 0.814765653 | 0.940696014 | ns | TM | KS.1 |
| FIM1_2718 | 4363.038394 | 4261.654303 | 5181.679644 | -0.016389473 | 0.070049172 | 0.733262752 | 0.911168134 | ns | TM | KS.1 |
| FIM1_2719 | 2274.665693 | 2341.946923 | 2146.318507 | 0.015167785 | 0.099885964 | 0.798304276 | 0.936276987 | ns | TM | KS.1 |
| FIM1_2720 | 4100.476411 | 4507.468093 | 3393.652226 | 0.090864074 | 0.072485574 | 0.074431058 | 0.290399848 | ns | TM | KS.1 |
| FIM1_2721 | 473.8518966 | 588.7707731 | 627.1013239 | 0.083554859 | 0.152692243 | 0.238328699 | 0.559285509 | ns | TM | KS.1 |
| FIM1_2722 | 2448.764798 | 2775.639586 | 2572.884247 | 0.082724922 | 0.106672249 | 0.198281082 | 0.505976981 | ns | TM | KS.1 |
| FIM1_2723 | 1002.194999 | 1049.784705 | 810.7871717 | 0.016778178 | 0.12699051 | 0.791097768 | 0.933303839 | ns | TM | KS.1 |
| FIM1_2724 | 2136.926157 | 2415.112387 | 1911.980482 | 0.083764495 | 0.103796715 | 0.187076829 | 0.489513719 | ns | TM | KS.1 |
| FIM1_2726 | 928.3649355 | 885.4945943 | 789.6774471 | -0.02586839 | 0.097089006 | 0.659896875 | 0.878874681 | ns | TM | KS.1 |
| FIM1_2727 | 129.2050873 | 86.0281156 | 134.5850001 | -0.237438027 | 0.225946412 | 0.019174164 | 0.132384765 | ns | TM | KS.1 |
| FIM1_2728 | 853.5247277 | 468.565243 | 592.1077316 | -0.541984285 | 0.336460652 | 0.001104956 | 0.016480869 | ns | TM | KS.1 |
| FIM1_2729 | 251.5749268 | 450.0585445 | 331.3030649 | 0.271041944 | 0.319028096 | 0.019842649 | 0.13560052 | ns | TM | KS.1 |
| FIM1_2730 | 124.3540608 | 62.07425429 | 150.8965523 | -0.620711673 | 0.395054897 | 0.001069378 | 0.016269592 | ns | TM | KS.1 |
| FIM1_2731 | 3606.783856 | 3775.427474 | 4238.867516 | 0.031893933 | 0.075367056 | 0.528435042 | 0.808531286 | ns | TM | KS.1 |
| FIM1_2732 | 7565.415911 | 8162.049629 | 8924.198354 | 0.058723276 | 0.078389823 | 0.26224309 | 0.588745204 | ns | TM | KS.1 |
| FIM1_2733 | 281.8146501 | 314.1348268 | 314.6935133 | 0.060323786 | 0.112718292 | 0.348037108 | 0.673729421 | ns | TM | KS.1 |
| FIM1_2734 | 278.3679336 | 259.3915057 | 239.54723 | -0.042174308 | 0.098462541 | 0.479689543 | 0.781163822 | ns | TM | KS.1 |
| FIM1_2735 | 600.0956609 | 563.6127175 | 651.349206 | -0.038702144 | 0.092399121 | 0.501547069 | 0.795417956 | ns | TM | KS.1 |
| FIM1_2736 | 651.0559006 | 703.4430497 | 595.3209154 | 0.044406963 | 0.101274434 | 0.463535697 | 0.765379219 | ns | TM | KS.1 |
| FIM1_2737 | 2470.628395 | 2330.226601 | 2475.593407 | -0.038411477 | 0.084754885 | 0.48257978 | 0.783685372 | ns | TM | KS.1 |
| FIM1_2738 | 662.7903223 | 599.8182681 | 482.4117522 | -0.035901238 | 0.13288937 | 0.578733784 | 0.836733675 | ns | TM | KS.1 |
| FIM1_2739 | 4371.924521 | 2569.148813 | 2229.439946 | -0.313579402 | 0.302534028 | 0.010693781 | 0.088521732 | ns | TM | KS.1 |
| FIM1_2740 | 6321.763321 | 6587.865422 | 5832.091995 | 0.032868716 | 0.061787836 | 0.454763508 | 0.758218986 | ns | TM | KS.1 |
| FIM1_2741 | 1060.659785 | 943.4293401 | 950.3022383 | -0.090231564 | 0.095275392 | 0.138613166 | 0.412292618 | ns | TM | KS.1 |
| FIM1_2742 | 6115.690855 | 7260.758759 | 6492.584788 | 0.066266555 | 0.141671335 | 0.330630073 | 0.657690769 | ns | TM | KS.1 |
| FIM1_2743 | 563.153299 | 805.4489257 | 767.070899 | 0.137826587 | 0.19972296 | 0.092778167 | 0.329235968 | ns | TM | KS.1 |
| FIM1_2744 | 8963.581757 | 9082.65927 | 6478.222438 | 0.010257141 | 0.057058361 | 0.805094513 | 0.937204068 | ns | TM | KS.1 |
| FIM1_2745 | 1393.057769 | 1705.02364 | 1426.642156 | 0.078041805 | 0.148800682 | 0.26444829 | 0.590354459 | ns | TM | KS.1 |
| FIM1_2746 | 1671.709664 | 1752.589538 | 1398.377701 | 0.028622946 | 0.088188773 | 0.60847103 | 0.852956713 | ns | TM | KS.1 |
| FIM1_2747 | 2486.620154 | 2800.094993 | 2989.429139 | 0.068309943 | 0.111184648 | 0.289549543 | 0.617085223 | ns | TM | KS.1 |
| FIM1_2748 | 482.0921113 | 496.7925409 | 558.3447268 | 0.018007697 | 0.091691377 | 0.751883135 | 0.918662314 | ns | TM | KS.1 |
| FIM1_2749 | 720.2686938 | 656.9868642 | 918.7957471 | -0.05733704 | 0.099933746 | 0.345263256 | 0.670970586 | ns | TM | KS.1 |
| FIM1_2750 | 78.76363744 | 70.85994721 | 94.23558957 | -0.042438521 | 0.12576414 | 0.51263601 | 0.802758625 | ns | TM | KS.1 |
| FIM1_2751 | 599.7039662 | 443.8930344 | 582.8865073 | -0.182065382 | 0.171650461 | 0.036326823 | 0.190639184 | ns | TM | KS.1 |
| FIM1_2752 | 1380.57989 | 1302.517154 | 1326.891207 | -0.044764959 | 0.07079221 | 0.356410808 | 0.680951596 | ns | TM | KS.1 |
| FIM1_2753 | 895.4922364 | 969.297914 | 1404.235879 | 0.041941178 | 0.104849858 | 0.494729422 | 0.792222775 | ns | TM | KS.1 |
| FIM1_2754 | 1753.299554 | 1599.81818 | 3605.0053 | -0.036497315 | 0.125831081 | 0.570769961 | 0.832094673 | ns | TM | KS.1 |
| FIM1_2755 | 1163.332728 | 1056.336116 | 1085.069687 | -0.066240923 | 0.094425087 | 0.263863056 | 0.590340989 | ns | TM | KS.1 |
| FIM1_2756 | 992.5123462 | 956.3658892 | 909.1344341 | -0.011257964 | 0.137635966 | 0.858631454 | 0.956277476 | ns | TM | KS.1 |
| FIM1_2757 | 3041.915231 | 4762.361633 | 5736.513145 | 0.191956557 | 0.243211879 | 0.043175121 | 0.212459177 | ns | TM | KS.1 |
| FIM1_2758 | 597.8491395 | 627.4523426 | 579.5831668 | 0.026100504 | 0.099333615 | 0.660532969 | 0.879309333 | ns | TM | KS.1 |
| FIM1_2759 | 318.9046961 | 171.2659585 | 236.8115605 | -0.793021312 | 0.226668499 | 4.67808305773 | 0.000225956 | ns | TM | KS.1 |
| FIM1_2760 | 6215.496703 | 6768.42738 | 8012.268097 | 0.035860464 | 0.121474604 | 0.575072657 | 0.833922256 | ns | TM | KS.1 |
| FIM1_2761 | 1358.996649 | 1444.071099 | 1486.793316 | 0.054006599 | 0.060198235 | 0.212459234 | 0.520438458 | ns | TM | KS.1 |

| | | | | | | | | | | |
|---|---|---|---|---|---|---|---|---|---|---|
| FIM1_2762 | 2269.004318 | 2478.01849 | 2559.5786 | 0.039431622 | 0.118126366 | 0.536588689 | 0.813136987 | ns | TM | KS.1 |
| FIM1_2763 | 9.269800719 | 17.15644995 | 643.8802253 | 0.076050625 | 0.264140555 | 0.261298825 | 0.588461463 | ns | TM | KS.1 |
| FIM1_2764 | 2588.373677 | 2057.443869 | 1403.205605 | -0.144262779 | 0.143390163 | 0.065447977 | 0.268649905 | ns | TM | KS.1 |
| FIM1_2765 | 1672.797317 | 1636.32659 | 1273.675358 | -0.013063306 | 0.083798426 | 0.809076267 | 0.938702059 | ns | TM | KS.1 |
| FIM1_2766 | 3244.518234 | 2988.969048 | 3595.941157 | -0.055444142 | 0.090111615 | 0.332721123 | 0.659583974 | ns | TM | KS.1 |
| FIM1_2767 | 2853.623455 | 2578.44724 | 2284.431879 | -0.044705297 | 0.121365634 | 0.48868156 | 0.788068318 | ns | TM | KS.1 |
| FIM1_2768 | 1481.456334 | 1148.287769 | 1224.0717 | -0.146402555 | 0.154820052 | 0.067516249 | 0.273974992 | ns | TM | KS.1 |
| FIM1_2769 | 2953.483802 | 2710.677511 | 2209.538832 | -0.069047566 | 0.079918907 | 0.196076203 | 0.50256523 | ns | TM | KS.1 |
| FIM1_2770 | 1555.674312 | 1321.932309 | 1412.254494 | -0.09538504 | 0.12343039 | 0.16948424 | 0.463542658 | ns | TM | KS.1 |
| FIM1_2771 | 545.2659952 | 610.6671915 | 445.9064314 | 0.049730208 | 0.123953247 | 0.445898118 | 0.751725902 | ns | TM | KS.1 |
| FIM1_2772 | 1269.939287 | 1118.101663 | 1609.891903 | -0.114588491 | 0.092059689 | 0.062107597 | 0.261236705 | ns | TM | KS.1 |
| FIM1_2773 | 464.401481 | 472.1322035 | 405.4531913 | 0.009084817 | 0.103463527 | 0.879941293 | 0.965258439 | ns | TM | KS.1 |
| FIM1_2774 | 705.9451089 | 636.0826144 | 928.7320496 | -0.035467193 | 0.135913446 | 0.583425309 | 0.840341897 | ns | TM | KS.1 |
| FIM1_2775 | 3379.12598 | 4431.513925 | 4011.411568 | 0.334462881 | 0.084243403 | 3.6261289648 | 0.000191915 | ns | TM | KS.1 |
| FIM1_2776 | 643.8732273 | 723.0688226 | 718.1820608 | 0.079449041 | 0.100983187 | 0.202182518 | 0.51032878 | ns | TM | KS.1 |
| FIM1_2777 | 1261.547658 | 774.951108 | 817.4033035 | -0.613622393 | 0.178178157 | 2.6242988490 | 0.000150068 | ns | TM | KS.1 |
| FIM1_2778 | 660.5562227 | 611.3198919 | 579.844977 | -0.053734256 | 0.088334042 | 0.341504648 | 0.668903606 | ns | TM | KS.1 |
| FIM1_2779 | 966.0445885 | 944.4850403 | 941.1714081 | -0.012102016 | 0.091886761 | 0.831749978 | 0.946802573 | ns | TM | KS.1 |
| FIM1_2780 | 546.4305446 | 164.0113215 | 312.6164471 | -1.541578588 | 0.261354292 | 3.1586845505 | 8.2707661257 | Km<KS.1 | TM | KS.1 |
| FIM1_2781 | 1505.839682 | 1119.921004 | 1642.397071 | -0.11333048 | 0.176690199 | 0.138667067 | 0.412292618 | ns | TM | KS.1 |
| FIM1_2782 | 1616.23622 | 1635.771805 | 1563.850147 | 0.006702804 | 0.094813808 | 0.907747715 | 0.972237864 | ns | TM | KS.1 |
| FIM1_2783 | 3460.790508 | 5302.937242 | 6384.891685 | 0.178585649 | 0.231840277 | 0.051194409 | 0.235826096 | ns | TM | KS.1 |
| FIM1_2784 | 894.2127672 | 920.5211462 | 907.651831 | 0.019882642 | 0.067011629 | 0.669747949 | 0.882180579 | ns | TM | KS.1 |
| FIM1_2785 | 3301.168892 | 3202.685203 | 2810.77649 | -0.019187468 | 0.079804546 | 0.71463642 | 0.905084971 | ns | TM | KS.1 |
| FIM1_2786 | 2470.684399 | 2876.278842 | 2411.717556 | 0.095742861 | 0.117324878 | 0.160170587 | 0.447416434 | ns | TM | KS.1 |
| FIM1_2787 | 4855.409997 | 5428.09923 | 5756.818924 | 0.03567957 | 0.140802282 | 0.581259703 | 0.838424819 | ns | TM | KS.1 |
| FIM1_2788 | 421.4367151 | 332.5587066 | 344.3151537 | -0.136571684 | 0.147849192 | 0.079718096 | 0.300680459 | ns | TM | KS.1 |
| FIM1_2789 | 332.7627828 | 357.6149049 | 274.6503493 | 0.031496226 | 0.117180885 | 0.618385877 | 0.860858199 | ns | TM | KS.1 |
| FIM1_2790 | 1433.871038 | 969.3531541 | 1088.731515 | -0.455821303 | 0.150219097 | 1.9206368767 | 0.000697458 | ns | TM | KS.1 |
| FIM1_2791 | 137.7990219 | 134.6805014 | 239.3295169 | -0.009846455 | 0.115804027 | 0.874398498 | 0.962844739 | ns | TM | KS.1 |
| FIM1_2792 | 937.9650543 | 931.8323278 | 699.8149305 | -0.003354675 | 0.09997002 | 0.954858515 | 0.988678089 | ns | TM | KS.1 |
| FIM1_2793 | 1528.148636 | 1270.418628 | 1821.019608 | -0.205858717 | 0.090197039 | 0.002916156 | 0.034216694 | ns | TM | KS.1 |
| FIM1_2794 | 1864.637083 | 2184.325679 | 2224.607093 | 0.114658382 | 0.113690336 | 0.09628613 | 0.336156841 | ns | TM | KS.1 |
| FIM1_2795 | 554.2954718 | 555.085484 | 653.7490727 | 0.000419577 | 0.135341873 | 0.994692348 | 0.99790453 | ns | TM | KS.1 |
| FIM1_2796 | 216.912129 | 157.0307932 | 654.7569723 | -0.096121112 | 0.189585679 | 0.192382632 | 0.496423026 | ns | TM | KS.1 |
| FIM1_2797 | 53.68443467 | 59.57668405 | 53.41088707 | 0.028468022 | 0.151847436 | 0.655024368 | 0.87699044 | ns | TM | KS.1 |
| FIM1_2798 | 2309.336991 | 2174.421999 | 1657.686551 | -0.038860667 | 0.086181976 | 0.482083383 | 0.783685372 | ns | TM | KS.1 |
| FIM1_2800 | 2084.108692 | 2545.440489 | 3329.2877 | 0.210440062 | 0.100827863 | 0.004405053 | 0.047230904 | ns | TM | KS.1 |
| FIM1_2801 | 4386.126746 | 4011.498508 | 4352.614611 | -0.059079788 | 0.093845734 | 0.314743961 | 0.641585998 | ns | TM | KS.1 |
| FIM1_2802 | 687.7688797 | 753.1118457 | 855.5880216 | 0.055889096 | 0.098187641 | 0.352621976 | 0.678119184 | ns | TM | KS.1 |
| FIM1_2803 | 983.1607783 | 1147.173033 | 1290.444468 | 0.142417236 | 0.099425572 | 0.032930143 | 0.181123861 | ns | TM | KS.1 |
| FIM1_2804 | 2454.453674 | 1961.551891 | 2299.026539 | -0.261649155 | 0.088310613 | 0.000305251 | 0.006098895 | ns | TM | KS.1 |
| FIM1_2805 | 2210.72492 | 1702.418147 | 1868.510163 | -0.258279267 | 0.124304195 | 0.002671248 | 0.032022791 | ns | TM | KS.1 |
| FIM1_2806 | 230.3165075 | 214.0101282 | 204.5735688 | -0.039360508 | 0.105233761 | 0.521526396 | 0.807711954 | ns | TM | KS.1 |
| FIM1_2807 | 1306.116024 | 1017.940883 | 1116.620185 | -0.201919559 | 0.13952739 | 0.016657668 | 0.120278515 | ns | TM | KS.1 |

| | | | | | | | | | | |
|---|---|---|---|---|---|---|---|---|---|---|
| FIM1_2808 | 1480.102749 | 1517.421311 | 1536.198656 | 0.01351464 | 0.094061499 | 0.814711604 | 0.940696014 | ns | TM | KS.1 |
| FIM1_2809 | 211.5430815 | 151.3702855 | 172.0515214 | -0.150831706 | 0.189527267 | 0.071592497 | 0.283830902 | ns | TM | KS.1 |
| FIM1_2810 | 139.5946731 | 219.4020096 | 237.534163 | 0.187099439 | 0.244403792 | 0.0467307 | 0.224840651 | ns | TM | KS.1 |
| FIM1_2811 | 381.8800589 | 464.2364972 | 460.8452583 | 0.146275397 | 0.124184602 | 0.050582063 | 0.23366998 | ns | TM | KS.1 |
| FIM1_2812 | 3031.615445 | 2467.696103 | 2959.470854 | -0.22326632 | 0.098003687 | 0.00253416 | 0.030888701 | ns | TM | KS.1 |
| FIM1_2813 | 1155.980048 | 911.3199248 | 1060.003627 | -0.193266499 | 0.135811237 | 0.019276569 | 0.132643057 | ns | TM | KS.1 |
| FIM1_2814 | 1925.480554 | 1577.403727 | 2193.081339 | -0.14814817 | 0.127062131 | 0.05033948 | 0.233221194 | ns | TM | KS.1 |
| FIM1_2815 | 453.3772685 | 392.0036143 | 337.3905196 | -0.086036693 | 0.117014705 | 0.20069857 | 0.508919034 | ns | TM | KS.1 |
| FIM1_2816 | 438.6737355 | 397.3120042 | 373.392304 | -0.055295597 | 0.107530303 | 0.37838885 | 0.702681796 | ns | TM | KS.1 |
| FIM1_2817 | 538.0871203 | 489.1483862 | 534.8311899 | -0.053008077 | 0.10648976 | 0.39550156 | 0.718896697 | ns | TM | KS.1 |
| FIM1_2818 | 414.6494415 | 385.0031588 | 428.9503715 | -0.046902268 | 0.09055278 | 0.411765867 | 0.732007978 | ns | TM | KS.1 |
| FIM1_2819 | 2318.919929 | 1918.724717 | 2042.335681 | -0.220722325 | 0.083771277 | 0.001075425 | 0.016290797 | ns | TM | KS.1 |
| FIM1_2820 | 1625.102499 | 1930.676975 | 2772.900424 | 0.062635103 | 0.144321269 | 0.355082087 | 0.679784721 | ns | TM | KS.1 |
| FIM1_2821 | 1816.576828 | 1838.669952 | 1743.736036 | 0.007171338 | 0.084201913 | 0.894737479 | 0.967972723 | ns | TM | KS.1 |
| FIM1_2822 | 422.1224388 | 1027.320935 | 639.5826493 | 1.231686601 | 0.152322887 | 1.41939099333 | 1.41229403836 | Km>KS.1 | TM | KS.1 |
| FIM1_2823 | 2896.274136 | 3319.223557 | 3390.920902 | 0.132273899 | 0.089941045 | 0.033049915 | 0.18140475 | ns | TM | KS.1 |
| FIM1_2824 | 961.0047569 | 1026.44375 | 1247.898865 | 0.050876367 | 0.073262774 | 0.3070999 | 0.635797754 | ns | TM | KS.1 |
| FIM1_2825 | 1115.724892 | 1112.039831 | 1050.128606 | -0.001861112 | 0.078410756 | 0.971369405 | 0.99362472 | ns | TM | KS.1 |
| FIM1_2826 | 1264.858618 | 896.5611364 | 888.972011 | -0.262784193 | 0.184249896 | 0.008242892 | 0.073623676 | ns | TM | KS.1 |
| FIM1_2827 | 2352.582237 | 3087.505063 | 3463.262847 | 0.120988702 | 0.165939891 | 0.117453839 | 0.375294059 | ns | TM | KS.1 |
| FIM1_2828 | 759.5943633 | 744.3739985 | 768.7387174 | -0.012146351 | 0.076827431 | 0.812418875 | 0.940492318 | ns | TM | KS.1 |
| FIM1_2829 | 2132.465853 | 2607.095108 | 2250.230989 | 0.166157455 | 0.122449039 | 0.029793598 | 0.171016808 | ns | TM | KS.1 |
| FIM1_2830 | 1818.282844 | 1564.257896 | 2405.026122 | -0.097887626 | 0.115974998 | 0.150218488 | 0.431365212 | ns | TM | KS.1 |
| FIM1_2831 | 260.7339645 | 93.8314416 | 339.8605135 | -0.796168187 | 0.554495903 | 0.001002327 | 0.015534505 | ns | TM | KS.1 |
| FIM1_2832 | 1169.806727 | 1203.398699 | 1539.108163 | 0.017917151 | 0.081182559 | 0.735526624 | 0.912302407 | ns | TM | KS.1 |
| FIM1_2833 | 1257.842949 | 1813.577215 | 1969.477907 | 0.129998975 | 0.202991569 | 0.107064413 | 0.354388193 | ns | TM | KS.1 |
| FIM1_2834 | 1128.175578 | 1340.015164 | 1728.387067 | 0.045434281 | 0.15929192 | 0.486782741 | 0.786535932 | ns | TM | KS.1 |
| FIM1_2835 | 1028.501417 | 1126.121446 | 1076.06063 | 0.026688118 | 0.143404716 | 0.676118898 | 0.887002016 | ns | TM | KS.1 |
| FIM1_2836 | 769.9776162 | 804.6443382 | 864.2023526 | 0.026329758 | 0.089565867 | 0.640263441 | 0.871255639 | ns | TM | KS.1 |
| FIM1_2837 | 1157.727563 | 1216.640024 | 858.4962492 | 0.015527567 | 0.13635965 | 0.806484119 | 0.937659684 | ns | TM | KS.1 |
| FIM1_2838 | 2320.090154 | 1973.761176 | 2749.047553 | -0.132997607 | 0.109292877 | 0.055029583 | 0.245563655 | ns | TM | KS.1 |
| FIM1_2839 | 662.4479406 | 618.5648983 | 734.8702793 | -0.049284633 | 0.080349454 | 0.352784115 | 0.678168846 | ns | TM | KS.1 |
| FIM1_2840 | 111.6268373 | 88.20954258 | 112.8839267 | -0.120683487 | 0.148879636 | 0.111952305 | 0.36402792 | ns | TM | KS.1 |
| FIM1_2841 | 854.1565847 | 974.6904529 | 1116.009165 | 0.095314963 | 0.104489784 | 0.139714992 | 0.413618606 | ns | TM | KS.1 |
| FIM1_2842 | 2977.937293 | 2981.300375 | 3122.825629 | 0.00096339 | 0.074126746 | 0.984622836 | 0.996034691 | ns | TM | KS.1 |
| FIM1_2843 | 213.8859981 | 260.9059866 | 223.4923183 | 0.113046398 | 0.136802808 | 0.124233056 | 0.387735747 | ns | TM | KS.1 |
| FIM1_2844 | 582.0298114 | 574.165226 | 525.3170241 | -0.006349496 | 0.107226261 | 0.916974674 | 0.975634767 | ns | TM | KS.1 |
| FIM1_2845 | 32745.38889 | 34428.80546 | 43296.34962 | 0.031664805 | 0.085431224 | 0.563884693 | 0.827926121 | ns | TM | KS.1 |
| FIM1_2846 | 700.1978674 | 1025.390952 | 1216.290288 | 0.156888117 | 0.210262342 | 0.068497996 | 0.275933222 | ns | TM | KS.1 |
| FIM1_2847 | 1534.277553 | 2175.121506 | 3194.810321 | 0.082774716 | 0.201064514 | 0.245033296 | 0.568582391 | ns | TM | KS.1 |
| FIM1_2848 | 740.1816202 | 527.4863937 | 706.3170098 | -0.322288091 | 0.161104843 | 0.001451253 | 0.019889761 | ns | TM | KS.1 |
| FIM1_2849 | 13958.5593 | 15342.45458 | 14759.95724 | 0.064918708 | 0.09365757 | 0.270740386 | 0.598244313 | ns | TM | KS.1 |
| FIM1_2850 | 2144.785985 | 2259.417969 | 2126.038905 | 0.028250697 | 0.098982614 | 0.63428908 | 0.869547581 | ns | TM | KS.1 |
| FIM1_2851 | 2437.213567 | 2334.708171 | 2482.05434 | -0.020476062 | 0.107915327 | 0.73836773 | 0.913453034 | ns | TM | KS.1 |
| FIM1_2852 | 1272.545483 | 905.4311124 | 869.8070042 | -0.31993575 | 0.162721219 | 0.001602135 | 0.021484157 | ns | TM | KS.1 |

| | | | | | | | | | | |
|---|---|---|---|---|---|---|---|---|---|---|
| FIM1_2853 | 1827.901484 | 3126.111201 | 4109.292691 | 0.310466416 | 0.3050861 | 0.011309308 | 0.091959747 | ns | TM | KS.1 |
| FIM1_2854 | 1927.175462 | 1816.929348 | 1743.33323 | -0.031188726 | 0.101805901 | 0.604627329 | 0.851547247 | ns | TM | KS.1 |
| FIM1_2855 | 139.5761784 | 123.0622189 | 121.4704813 | -0.050075586 | 0.131685936 | 0.447161419 | 0.752446492 | ns | TM | KS.1 |
| FIM1_2856 | 643.0982649 | 513.7997183 | 634.690912 | -0.131219379 | 0.143892008 | 0.087291654 | 0.315378344 | ns | TM | KS.1 |
| FIM1_2857 | 755.5784602 | 755.4487307 | 751.6658429 | -1.79E-05 | 0.10180579 | 0.999760607 | 0.999760607 | ns | TM | KS.1 |
| FIM1_2858 | 1132.422981 | 965.9309419 | 1021.325247 | -0.121009895 | 0.112423037 | 0.080171804 | 0.301630805 | ns | TM | KS.1 |
| FIM1_2859 | 4051.234479 | 4451.091766 | 3942.214549 | 0.049553974 | 0.1092141 | 0.43036436 | 0.743536592 | ns | TM | KS.1 |
| FIM1_2860 | 14448.35848 | 7559.152343 | 11384.25993 | -0.488884551 | 0.379024746 | 0.002840825 | 0.033570316 | ns | TM | KS.1 |
| FIM1_2862 | 235.6676924 | 261.8576019 | 207.0002085 | 0.0466539 | 0.122582016 | 0.471871751 | 0.772376515 | ns | TM | KS.1 |
| FIM1_2863 | 1020.824564 | 1385.081973 | 1263.715346 | 0.162106651 | 0.176085949 | 0.055035875 | 0.245563655 | ns | TM | KS.1 |
| FIM1_2864 | 2545.15961 | 3009.390035 | 3071.00938 | 0.119816469 | 0.118037921 | 0.08965747 | 0.321358726 | ns | TM | KS.1 |
| FIM1_2865 | 949.1856502 | 1162.241158 | 1040.611865 | 0.122562795 | 0.135679295 | 0.099965009 | 0.341335565 | ns | TM | KS.1 |
| FIM1_2866 | 652.8458759 | 596.6269082 | 682.8797352 | -0.051092822 | 0.104075521 | 0.407289938 | 0.728957685 | ns | TM | KS.1 |
| FIM1_2867 | 1370.718624 | 1558.370474 | 1333.668172 | 0.081323775 | 0.109718546 | 0.211345577 | 0.51871941 | ns | TM | KS.1 |
| FIM1_2868 | 42.25418473 | 38.83920307 | 53.04285322 | -0.01627943 | 0.146162314 | 0.796451077 | 0.935618443 | ns | TM | KS.1 |
| FIM1_2869 | 174.5368285 | 86.81085985 | 263.2725796 | -0.89204769 | 0.248611531 | 5.1607160803 | 0.00024452 | ns | TM | KS.1 |
| FIM1_2870 | 4250.056912 | 4672.609142 | 4619.041052 | 0.05093161 | 0.10836434 | 0.4168004 | 0.735551151 | ns | TM | KS.1 |
| FIM1_2871 | 883.4327756 | 929.4721959 | 1028.851863 | 0.033796301 | 0.08289436 | 0.530488812 | 0.808531286 | ns | TM | KS.1 |
| FIM1_2872 | 1417.740352 | 1381.066935 | 3341.988258 | -0.014370138 | 0.09443558 | 0.803632609 | 0.936908579 | ns | TM | KS.1 |
| FIM1_2873 | 383.5947801 | 266.62165 | 379.4106986 | -0.265576944 | 0.197047396 | 0.009094768 | 0.079482818 | ns | TM | KS.1 |
| FIM1_2874 | 6651.865625 | 3746.030391 | 5304.524092 | -0.748137131 | 0.195418556 | 1.1584067694 | 7.7879374023 | ns | TM | KS.1 |
| FIM1_2875 | 1782.495844 | 1820.310145 | 2510.602835 | 0.014972378 | 0.06719921 | 0.748614636 | 0.91742442 | ns | TM | KS.1 |
| FIM1_2876 | 1921.929296 | 2132.391566 | 2282.743622 | 0.07754781 | 0.091620066 | 0.186983912 | 0.489513719 | ns | TM | KS.1 |
| FIM1_2877 | 2079.589262 | 3093.829595 | 3588.738224 | 0.176703647 | 0.218531591 | 0.050389412 | 0.233221194 | ns | TM | KS.1 |
| FIM1_2878 | 1153.439967 | 1021.7342 | 1245.67938 | -0.082657824 | 0.103995308 | 0.192886841 | 0.496788132 | ns | TM | KS.1 |
| FIM1_2879 | 881.9105712 | 868.5355676 | 1558.259213 | -0.007842618 | 0.100009016 | 0.894782986 | 0.967972723 | ns | TM | KS.1 |
| FIM1_2880 | 957.497786 | 969.2279653 | 1078.416442 | 0.006623735 | 0.094006868 | 0.90844377 | 0.972251892 | ns | TM | KS.1 |
| FIM1_2881 | 1029.647156 | 1187.320909 | 1233.208042 | 0.07189491 | 0.123718288 | 0.284632526 | 0.611419177 | ns | TM | KS.1 |
| FIM1_2882 | 1062.121294 | 1102.690948 | 1469.445907 | 0.021993667 | 0.09040458 | 0.697448599 | 0.895896406 | ns | TM | KS.1 |
| FIM1_2883 | 2324.746179 | 2111.649807 | 2329.138013 | -0.079045836 | 0.08360723 | 0.154311122 | 0.437933732 | ns | TM | KS.1 |
| FIM1_2884 | 555.7934074 | 626.0948985 | 690.2018904 | 0.088424722 | 0.098531096 | 0.153912569 | 0.437933732 | ns | TM | KS.1 |
| FIM1_2885 | 947.5359172 | 1476.752183 | 1656.05392 | 0.156931147 | 0.235759079 | 0.073076573 | 0.287623378 | ns | TM | KS.1 |
| FIM1_2886 | 407.2972015 | 782.1585848 | 931.8734118 | 0.329087465 | 0.366093385 | 0.012674281 | 0.0992985 | ns | TM | KS.1 |
| FIM1_2887 | 2731.17459 | 3992.900774 | 4640.995791 | 0.162318454 | 0.20983721 | 0.062354568 | 0.261819279 | ns | TM | KS.1 |
| FIM1_2888 | 529.7130008 | 392.9983152 | 389.7551072 | -0.216865738 | 0.164933575 | 0.016778886 | 0.120628549 | ns | TM | KS.1 |
| FIM1_2889 | 1915.748216 | 2035.760852 | 2031.263372 | 0.044490317 | 0.07666727 | 0.385857044 | 0.709925589 | ns | TM | KS.1 |
| FIM1_2890 | 4838.129474 | 4743.393793 | 4347.024313 | -0.012774288 | 0.076168797 | 0.801825889 | 0.936908579 | ns | TM | KS.1 |
| FIM1_2891 | 4082.732521 | 2916.493152 | 4319.216258 | -0.423960196 | 0.097623309 | 1.2258486010 | 1.2975737851 | ns | TM | KS.1 |
| FIM1_2892 | 326.1677658 | 179.0956242 | 304.0822981 | -0.79964311 | 0.185077046 | 2.4913568037 | 2.4302941370 | ns | TM | KS.1 |
| FIM1_2893 | 716.2428098 | 503.5556906 | 642.7355356 | -0.339548947 | 0.16604274 | 0.001090189 | 0.016336421 | ns | TM | KS.1 |
| FIM1_2894 | 745.3042844 | 765.1224985 | 1040.38808 | 0.014752998 | 0.088930269 | 0.792134982 | 0.933303839 | ns | TM | KS.1 |
| FIM1_2895 | 3262.601543 | 3286.076943 | 3125.276002 | 0.004790072 | 0.076601047 | 0.92528661 | 0.976735299 | ns | TM | KS.1 |
| FIM1_2896 | 1187.616101 | 1224.321403 | 1441.378662 | 0.02156607 | 0.069220065 | 0.651120975 | 0.874784459 | ns | TM | KS.1 |
| FIM1_2897 | 5065.562497 | 3602.686491 | 5776.34956 | -0.250280435 | 0.184276017 | 0.010562913 | 0.088171963 | ns | TM | KS.1 |
| FIM1_2898 | 14180.03888 | 16917.01646 | 16191.39412 | 0.15170921 | 0.112017637 | 0.03510825 | 0.187206368 | ns | TM | KS.1 |

| | | | | | | | | | | |
|---|---|---|---|---|---|---|---|---|---|---|
| FIM1_2899 | 4150.909598 | 6520.386473 | 9340.909001 | 0.374109015 | 0.244901862 | 0.002589489 | 0.031270285 | ns | TM | KS.1 |
| FIM1_2900 | 3052.370503 | 3119.065067 | 6646.927741 | 0.011265653 | 0.098779868 | 0.848594943 | 0.952348261 | ns | TM | KS.1 |
| FIM1_2901 | 1435.511682 | 1553.778176 | 1677.89953 | 0.053666612 | 0.088158738 | 0.341502332 | 0.668903606 | ns | TM | KS.1 |
| FIM1_2903 | 2229.629233 | 2011.33075 | 2443.122967 | -0.066351126 | 0.100335761 | 0.279306407 | 0.603889341 | ns | TM | KS.1 |
| FIM1_2904 | 1507.720743 | 1308.587046 | 1819.270302 | -0.06996835 | 0.124502068 | 0.297222295 | 0.624970802 | ns | TM | KS.1 |
| FIM1_2905 | 660.4916064 | 752.7980036 | 846.2092377 | 0.035114502 | 0.152842939 | 0.585130484 | 0.841613393 | ns | TM | KS.1 |
| FIM1_2906 | 206.3800698 | 187.1250835 | 192.2999731 | -0.037691034 | 0.126290802 | 0.558955978 | 0.82614557 | ns | TM | KS.1 |
| FIM1_2907 | 1239.174238 | 1120.129937 | 990.5456322 | -0.045171737 | 0.120445689 | 0.48364836 | 0.784137657 | ns | TM | KS.1 |
| FIM1_2908 | 2415.817322 | 2561.213192 | 2481.130868 | 0.039366355 | 0.08165539 | 0.461304068 | 0.763468975 | ns | TM | KS.1 |
| FIM1_2909 | 1003.291779 | 1028.5824 | 1242.605634 | 0.016376204 | 0.077776329 | 0.750980404 | 0.918418758 | ns | TM | KS.1 |
| FIM1_2911 | 634.4357923 | 461.491811 | 424.6130829 | -0.127360163 | 0.184311244 | 0.108409906 | 0.357240387 | ns | TM | KS.1 |
| FIM1_2912 | 2075.748637 | 2144.015775 | 2982.181995 | 0.010073773 | 0.136150228 | 0.873356975 | 0.962759265 | ns | TM | KS.1 |
| FIM1_2913 | 12.8739904 | 9.435934258 | 14.35777334 | -0.054409729 | 0.203156687 | 0.406594646 | 0.728526463 | ns | TM | KS.1 |
| FIM1_2914 | 1391.852104 | 2043.056935 | 1880.422927 | 0.12645863 | 0.209626142 | 0.115026707 | 0.37039344 | ns | TM | KS.1 |
| FIM1_2915 | 3424.829509 | 2226.376035 | 1489.707832 | -0.441972691 | 0.203963179 | 0.000336865 | 0.006598054 | ns | TM | KS.1 |
| FIM1_2916 | 7241.86428 | 6728.410038 | 5132.835234 | -0.040091003 | 0.102559891 | 0.5093469 | 0.801014116 | ns | TM | KS.1 |
| FIM1_2917 | 1560.385028 | 1185.850694 | 1446.996132 | -0.310527273 | 0.105445888 | 0.000171514 | 0.003838806 | ns | TM | KS.1 |
| FIM1_2918 | 825.2107552 | 736.6924894 | 985.6773065 | -0.078918251 | 0.099609193 | 0.201589103 | 0.509840799 | ns | TM | KS.1 |
| FIM1_2919 | 2671.700887 | 2952.049475 | 3139.017352 | 0.073605352 | 0.09097296 | 0.206872172 | 0.513839538 | ns | TM | KS.1 |
| FIM1_2920 | 320.6138098 | 338.9905459 | 333.924461 | 0.019649099 | 0.129580487 | 0.757149562 | 0.920753636 | ns | TM | KS.1 |
| FIM1_2921 | 1932.309453 | 2609.158704 | 2861.91268 | 0.110352901 | 0.178754753 | 0.14697251 | 0.425671509 | ns | TM | KS.1 |
| FIM1_2922 | 1055.819768 | 991.1900503 | 940.8518398 | -0.031530049 | 0.106581009 | 0.607559958 | 0.852159794 | ns | TM | KS.1 |
| FIM1_2923 | 3280.773853 | 3775.790355 | 3964.760454 | 0.104744083 | 0.106327044 | 0.111795844 | 0.363946737 | ns | TM | KS.1 |
| FIM1_2924 | 5580.427126 | 6156.927377 | 6416.49765 | 0.064564691 | 0.097600863 | 0.284541464 | 0.611419177 | ns | TM | KS.1 |
| FIM1_2925 | 50.73369476 | 38.86949186 | 55.21287527 | -0.059954809 | 0.179971882 | 0.371223191 | 0.694038097 | ns | TM | KS.1 |
| FIM1_2926 | 2280.687049 | 2294.927122 | 2247.740019 | 0.002803059 | 0.108937189 | 0.963469605 | 0.99280474 | ns | TM | KS.1 |
| FIM1_2927 | 5835.518981 | 6169.765597 | 5522.650867 | 0.048519821 | 0.059946877 | 0.260328652 | 0.588210523 | ns | TM | KS.1 |
| FIM1_2928 | 834.1581062 | 481.0498717 | 1364.35551 | -0.333494481 | 0.315386525 | 0.009106574 | 0.079482818 | ns | TM | KS.1 |
| FIM1_2929 | 1551.212006 | 1444.845829 | 1704.738101 | -0.028398533 | 0.123026329 | 0.655533082 | 0.877039451 | ns | TM | KS.1 |
| FIM1_2930 | 243.8731237 | 333.728797 | 314.4429203 | 0.092306149 | 0.186813945 | 0.205867994 | 0.512865935 | ns | TM | KS.1 |
| FIM1_2931 | 2407.346834 | 2712.630138 | 3084.728023 | 0.072020522 | 0.109090394 | 0.262185026 | 0.588745204 | ns | TM | KS.1 |
| FIM1_2932 | 1049.080587 | 940.6021075 | 921.3139034 | -0.067677078 | 0.104661503 | 0.280655105 | 0.606278396 | ns | TM | KS.1 |
| FIM1_2933 | 2407.203152 | 2556.387812 | 2741.375958 | 0.045075945 | 0.074080506 | 0.368406356 | 0.691630801 | ns | TM | KS.1 |
| FIM1_2934 | 2060.547884 | 2828.646417 | 3436.724809 | 0.228311427 | 0.17366584 | 0.014575296 | 0.108551045 | ns | TM | KS.1 |
| FIM1_2935 | 11526.24895 | 13867.19558 | 14252.74012 | 0.116323038 | 0.12828629 | 0.108740704 | 0.357723772 | ns | TM | KS.1 |
| FIM1_2936 | 3229.058227 | 3850.351722 | 4841.660457 | 0.094527387 | 0.130692717 | 0.180216323 | 0.480480282 | ns | TM | KS.1 |
| FIM1_2937 | 1076.445976 | 817.9002377 | 929.7319652 | -0.162616291 | 0.161785895 | 0.050394529 | 0.233221194 | ns | TM | KS.1 |
| FIM1_2939 | 757.1975366 | 513.2089343 | 579.7654092 | -0.347747569 | 0.197459267 | 0.001879484 | 0.024671329 | ns | TM | KS.1 |
| FIM1_2941 | 1591.841613 | 1666.005843 | 1537.092147 | 0.026986713 | 0.090566595 | 0.634130115 | 0.869547581 | ns | TM | KS.1 |
| FIM1_2942 | 11908.07327 | 10573.06746 | 7572.433439 | -0.081494935 | 0.102445231 | 0.195155685 | 0.501497694 | ns | TM | KS.1 |
| FIM1_2943 | 1212.262791 | 1200.474192 | 1210.279347 | -0.006568265 | 0.074820701 | 0.896097816 | 0.967972723 | ns | TM | KS.1 |
| FIM1_2944 | 704.4738648 | 940.568393 | 950.1685742 | 0.247604919 | 0.149757445 | 0.006751392 | 0.064774438 | ns | TM | KS.1 |
| FIM1_2945 | 649.043934 | 822.9786708 | 787.5940644 | 0.133985485 | 0.149713922 | 0.085100318 | 0.31075147 | ns | TM | KS.1 |
| FIM1_2946 | 1891.197716 | 1845.474211 | 1978.107633 | -0.016774176 | 0.071140042 | 0.730027194 | 0.909157553 | ns | TM | KS.1 |
| FIM1_2947 | 3182.819434 | 3848.690926 | 3679.15989 | 0.16075482 | 0.117186352 | 0.030857042 | 0.175444324 | ns | TM | KS.1 |

| | | | | | | | | | | |
|---|---|---|---|---|---|---|---|---|---|---|
| FIM1_2948 | 1268.466302 | 1225.819601 | 1174.689626 | -0.021661186 | 0.084608352 | 0.690698274 | 0.892989583 | ns | TM | KS.1 |
| FIM1_2949 | 4048.533501 | 3669.442794 | 3327.628688 | -0.060137827 | 0.101576867 | 0.327320812 | 0.654510064 | ns | TM | KS.1 |
| FIM1_2950 | 74.13321987 | 73.94382462 | 91.11159101 | -0.000386802 | 0.126360555 | 0.995100976 | 0.99790453 | ns | TM | KS.1 |
| FIM1_2951 | 2268.617718 | 2322.894113 | 2696.25655 | 0.013714793 | 0.087227517 | 0.804345544 | 0.936927905 | ns | TM | KS.1 |
| FIM1_2952 | 982.6802668 | 857.7318969 | 991.1695776 | -0.099991865 | 0.105500007 | 0.125265199 | 0.389417837 | ns | TM | KS.1 |
| FIM1_2953 | 1709.388288 | 1648.125383 | 1696.349832 | -0.01703457 | 0.108606019 | 0.781255269 | 0.929216508 | ns | TM | KS.1 |
| FIM1_2954 | 550.7644671 | 611.0150315 | 669.8722657 | 0.088684027 | 0.083254396 | 0.112319565 | 0.364507394 | ns | TM | KS.1 |
| FIM1_2955 | 1084.973456 | 1193.563376 | 1135.911101 | 0.051914992 | 0.108046606 | 0.407718987 | 0.728957685 | ns | TM | KS.1 |
| FIM1_2956 | 284.5248058 | 346.2622207 | 356.0336572 | 0.107630922 | 0.136929655 | 0.139840502 | 0.413618606 | ns | TM | KS.1 |
| FIM1_2957 | 2008.288627 | 3200.024534 | 3906.709487 | 0.270576836 | 0.261122264 | 0.014547307 | 0.108505029 | ns | TM | KS.1 |
| FIM1_2958 | 1385.626272 | 1753.095446 | 2248.188588 | 0.215388817 | 0.125530502 | 0.008925217 | 0.078450448 | ns | TM | KS.1 |
| FIM1_2959 | 2248.306588 | 3622.770804 | 4410.205535 | 0.388329446 | 0.263755445 | 0.002703066 | 0.032248811 | ns | TM | KS.1 |
| FIM1_2960 | 1797.374466 | 2623.217554 | 3342.8738 | 0.201212461 | 0.210319787 | 0.032084615 | 0.179349393 | ns | TM | KS.1 |
| FIM1_2961 | 2496.720995 | 3556.493213 | 4279.333789 | 0.247361352 | 0.193288486 | 0.012309694 | 0.097517084 | ns | TM | KS.1 |
| FIM1_2962 | 325.774287 | 260.2246649 | 401.0509813 | -0.184869434 | 0.129005422 | 0.021014546 | 0.141433402 | ns | TM | KS.1 |
| FIM1_2963 | 434.3660528 | 412.7228117 | 393.8680425 | -0.03503105 | 0.081519939 | 0.511197441 | 0.802123385 | ns | TM | KS.1 |
| FIM1_2964 | 6894.706915 | 7529.83065 | 7234.558437 | 0.055254517 | 0.097048067 | 0.354887757 | 0.679784721 | ns | TM | KS.1 |
| FIM1_2965 | 1487.296035 | 1347.877026 | 1402.398664 | -0.067796664 | 0.094776272 | 0.254417432 | 0.581671847 | ns | TM | KS.1 |
| FIM1_2966 | 384.9995554 | 368.4539987 | 368.3976288 | -0.021017929 | 0.106147099 | 0.730249966 | 0.909157553 | ns | TM | KS.1 |
| FIM1_2967 | 1736.978699 | 1445.900867 | 1546.006294 | -0.204499298 | 0.088970331 | 0.002834301 | 0.033570316 | ns | TM | KS.1 |
| FIM1_2968 | 2401.139645 | 2110.899763 | 1890.686952 | -0.114905739 | 0.092363594 | 0.06201991 | 0.261236705 | ns | TM | KS.1 |
| FIM1_2969 | 5149.039043 | 6037.932871 | 5685.816099 | 0.105549923 | 0.117962486 | 0.127299786 | 0.393568488 | ns | TM | KS.1 |
| FIM1_2970 | 693.4887293 | 548.2049985 | 760.6778271 | -0.232278151 | 0.114299793 | 0.003965565 | 0.043455253 | ns | TM | KS.1 |
| FIM1_2971 | 460.9877638 | 599.7566352 | 637.3109641 | 0.113188204 | 0.163638829 | 0.135947086 | 0.407922054 | ns | TM | KS.1 |
| FIM1_2972 | 626.4801458 | 656.5605806 | 731.7172261 | 0.025361604 | 0.097129847 | 0.666156477 | 0.880910621 | ns | TM | KS.1 |
| FIM1_2973 | 286.5863206 | 315.3600275 | 901.5278038 | 0.041437369 | 0.120983931 | 0.519214665 | 0.805989904 | ns | TM | KS.1 |
| FIM1_2974 | 1692.050716 | 1853.814878 | 1991.706768 | 0.041376603 | 0.117874893 | 0.517277008 | 0.804981347 | ns | TM | KS.1 |
| FIM1_2975 | 1931.871401 | 2376.587517 | 2788.527927 | 0.123678296 | 0.137438071 | 0.098719767 | 0.339178757 | ns | TM | KS.1 |
| FIM1_2976 | 542.8156487 | 556.1382624 | 629.7327363 | 0.016046997 | 0.086619281 | 0.77109746 | 0.925705893 | ns | TM | KS.1 |
| FIM1_2977 | 1137.386059 | 856.7990928 | 839.6745868 | -0.234644009 | 0.150623278 | 0.009285968 | 0.079846584 | ns | TM | KS.1 |
| FIM1_2978 | 2670.919815 | 3040.260333 | 2827.370299 | 0.080716122 | 0.110650733 | 0.216207706 | 0.526496984 | ns | TM | KS.1 |
| FIM1_2979 | 178.7101114 | 179.9593361 | 219.4068092 | 0.004087427 | 0.108234351 | 0.946647729 | 0.985884959 | ns | TM | KS.1 |
| FIM1_2980 | 898.8743067 | 874.0324272 | 926.1346837 | -0.015000991 | 0.096200722 | 0.797046261 | 0.935875654 | ns | TM | KS.1 |
| FIM1_2981 | 2869.698976 | 4265.442187 | 5131.577162 | 0.218435091 | 0.220067779 | 0.02531259 | 0.156047257 | ns | TM | KS.1 |
| FIM1_2982 | 3029.221171 | 3405.822516 | 3129.856232 | 0.091434922 | 0.095275374 | 0.133924529 | 0.40527648 | ns | TM | KS.1 |
| FIM1_2983 | 5409.029269 | 6304.53994 | 6118.843466 | 0.081969289 | 0.124159442 | 0.229628722 | 0.547690282 | ns | TM | KS.1 |
| FIM1_2984 | 462.6970143 | 704.9390523 | 607.7647513 | 0.190354388 | 0.230893673 | 0.042252166 | 0.210079667 | ns | TM | KS.1 |
| FIM1_2985 | 18080.74208 | 19747.98661 | 15548.43375 | 0.024347837 | 0.147189654 | 0.701737423 | 0.897235589 | ns | TM | KS.1 |
| FIM1_2986 | 1134.987139 | 1008.928177 | 930.6546089 | -0.103737046 | 0.088054581 | 0.078322756 | 0.298177127 | ns | TM | KS.1 |
| FIM1_2987 | 1757.514065 | 1861.44348 | 1963.239033 | 0.044831682 | 0.071732895 | 0.360177911 | 0.683664672 | ns | TM | KS.1 |
| FIM1_2988 | 1115.646825 | 919.4483023 | 931.4687696 | -0.146490945 | 0.124002396 | 0.050176471 | 0.233221194 | ns | TM | KS.1 |
| FIM1_2989 | 1010.017603 | 990.9848389 | 992.0718388 | -0.008713925 | 0.108653696 | 0.886778374 | 0.967028314 | ns | TM | KS.1 |
| FIM1_2990 | 9001.897207 | 4418.264172 | 12583.20332 | -1.013611034 | 0.081177623 | 3.1705523881 | 1.5773498131 | Km<KS.1 | TM | KS.1 |
| FIM1_2991 | 436.6773274 | 622.2434439 | 416.6918034 | 0.216642289 | 0.197437995 | 0.022534344 | 0.146759546 | ns | TM | KS.1 |
| FIM1_2992 | 63.51722597 | 86.7298273 | 85.80505748 | 0.127889193 | 0.182663889 | 0.106999129 | 0.354388193 | ns | TM | KS.1 |

| | | | | | | | | | | |
|---|---|---|---|---|---|---|---|---|---|---|
| FIM1_2993 | 1084.116931 | 1040.166441 | 1039.204649 | -0.031350145 | 0.066068728 | 0.496867152 | 0.793297202 | ns | TM | KS.1 |
| FIM1_2994 | 1053.986043 | 983.0999485 | 988.7851955 | -0.043640368 | 0.090913961 | 0.445389708 | 0.751385662 | ns | TM | KS.1 |
| FIM1_2995 | 339.8849531 | 282.5996247 | 250.7650778 | -0.066813016 | 0.147491341 | 0.32825954 | 0.655821244 | ns | TM | KS.1 |
| FIM1_2996 | 602.6649173 | 459.5378972 | 702.7419252 | -0.073175818 | 0.177769634 | 0.291653377 | 0.618644347 | ns | TM | KS.1 |
| FIM1_2997 | 1530.81192 | 1467.954419 | 1624.884602 | -0.02750419 | 0.080099527 | 0.601389981 | 0.851491516 | ns | TM | KS.1 |
| FIM1_2998 | 5328.973872 | 4239.165896 | 3567.38224 | -0.216228145 | 0.120514957 | 0.007627172 | 0.069752169 | ns | TM | KS.1 |
| FIM1_2999 | 834.9459992 | 985.1184951 | 807.5264055 | 0.106753613 | 0.121092681 | 0.1275888 | 0.393768164 | ns | TM | KS.1 |
| FIM1_3000 | 1071.444442 | 1107.21288 | 1759.153242 | 0.018273176 | 0.096715319 | 0.754881103 | 0.920022902 | ns | TM | KS.1 |
| FIM1_3001 | 1109.417213 | 1253.23663 | 1124.216239 | 0.058405062 | 0.121265081 | 0.37355049 | 0.696295874 | ns | TM | KS.1 |
| FIM1_3002 | 489.7031701 | 450.962846 | 406.515444 | -0.047751824 | 0.097665261 | 0.423220493 | 0.739045122 | ns | TM | KS.1 |
| FIM1_3003 | 919.2242306 | 1150.295973 | 974.3824504 | 0.085108547 | 0.155071183 | 0.231725808 | 0.549231011 | ns | TM | KS.1 |
| FIM1_3004 | 5325.88757 | 5784.53949 | 5703.476767 | 0.066866137 | 0.078255837 | 0.203385021 | 0.511547259 | ns | TM | KS.1 |
| FIM1_3005 | 1554.966322 | 2151.180101 | 2134.188148 | 0.112668474 | 0.187826272 | 0.142455225 | 0.418862143 | ns | TM | KS.1 |
| FIM1_3006 | 3036.807238 | 4072.411074 | 4840.496114 | 0.262515228 | 0.147993601 | 0.004554818 | 0.048419271 | ns | TM | KS.1 |
| FIM1_3007 | 3304.090128 | 3692.178131 | 3697.877283 | 0.080231924 | 0.097096289 | 0.188342733 | 0.491608132 | ns | TM | KS.1 |
| FIM1_3008 | 1995.68527 | 2187.036294 | 2183.93518 | 0.080367054 | 0.077518672 | 0.127899029 | 0.394236475 | ns | TM | KS.1 |
| FIM1_3009 | 1440.590636 | 1694.530473 | 1567.872644 | 0.166038151 | 0.094189243 | 0.013231894 | 0.101775569 | ns | TM | KS.1 |
| FIM1_3010 | 691.6586232 | 716.4330827 | 653.7230729 | 0.022657153 | 0.082960622 | 0.673623923 | 0.885317825 | ns | TM | KS.1 |
| FIM1_3011 | 421.5590514 | 466.5544649 | 622.8877379 | 0.053272897 | 0.111564253 | 0.402035497 | 0.724158796 | ns | TM | KS.1 |
| FIM1_3012 | 2140.223542 | 2313.161894 | 2505.524487 | 0.058556514 | 0.081532497 | 0.276278968 | 0.601096377 | ns | TM | KS.1 |
| FIM1_3013 | 2173.0449 | 2456.671178 | 2299.118373 | 0.107984488 | 0.090709881 | 0.073580428 | 0.288556468 | ns | TM | KS.1 |
| FIM1_3014 | 2827.805816 | 4584.914247 | 6063.64154 | 0.251471731 | 0.26879164 | 0.020075327 | 0.136814727 | ns | TM | KS.1 |
| FIM1_3015 | 4605.103281 | 5278.157339 | 4634.54022 | 0.104708606 | 0.103669429 | 0.107151982 | 0.354442229 | ns | TM | KS.1 |
| FIM1_3016 | 616.0797853 | 686.5170758 | 759.6981876 | 0.06385667 | 0.107516281 | 0.312958362 | 0.639671112 | ns | TM | KS.1 |
| FIM1_3017 | 17763.51007 | 22180.77529 | 21330.23418 | 0.212612768 | 0.117376021 | 0.007679632 | 0.069974673 | ns | TM | KS.1 |
| FIM1_3018 | 526.7587361 | 590.478752 | 535.4983651 | 0.057945441 | 0.116932365 | 0.372102778 | 0.694598963 | ns | TM | KS.1 |
| FIM1_3019 | 5784.455865 | 5887.906422 | 4633.715476 | 0.011154437 | 0.079541521 | 0.831239577 | 0.946802573 | ns | TM | KS.1 |
| FIM1_3020 | 80.76637841 | 71.57043739 | 96.90294568 | -0.044920401 | 0.132981399 | 0.492484155 | 0.790867874 | ns | TM | KS.1 |
| FIM1_3021 | 3531.260516 | 3082.435209 | 3834.209133 | -0.112866502 | 0.099132504 | 0.077986931 | 0.297534494 | ns | TM | KS.1 |
| FIM1_3022 | 220.9701637 | 175.1298162 | 240.3611277 | -0.064237099 | 0.167781319 | 0.344447652 | 0.670129201 | ns | TM | KS.1 |
| FIM1_3023 | 1066.490997 | 537.1267562 | 752.7178282 | -0.806880293 | 0.302974222 | 5.95530866956 | 0.001575939 | ns | TM | KS.1 |
| FIM1_3024 | 19.57324589 | 33.60273847 | 74.4569741 | 0.108370907 | 0.255707188 | 0.159408377 | 0.44660975 | ns | TM | KS.1 |
| FIM1_3025 | 1032.211957 | 925.0116695 | 931.7120277 | -0.045423417 | 0.126323809 | 0.485347444 | 0.785236922 | ns | TM | KS.1 |
| FIM1_3026 | 2168.9894 | 2023.24723 | 2497.866158 | -0.033269473 | 0.110986999 | 0.593645069 | 0.846854399 | ns | TM | KS.1 |
| FIM1_3028 | 3579.472098 | 3926.899206 | 4185.317805 | 0.060148034 | 0.096230265 | 0.313339895 | 0.639928562 | ns | TM | KS.1 |
| FIM1_3029 | 1588.606992 | 1533.500226 | 1373.966389 | -0.022036571 | 0.084327387 | 0.685039281 | 0.890505759 | ns | TM | KS.1 |
| FIM1_3030 | 5426.832625 | 4554.440345 | 3644.827954 | -0.210765423 | 0.074877413 | 0.000708252 | 0.011628895 | ns | TM | KS.1 |
| FIM1_3031 | 2130.884576 | 1354.306426 | 1581.656361 | -0.576809078 | 0.153611557 | 6.50479390258 | 5.34667101033 | ns | TM | KS.1 |
| FIM1_3032 | 2741.290275 | 2913.554509 | 2951.006234 | 0.031322334 | 0.104329295 | 0.606861159 | 0.851659878 | ns | TM | KS.1 |
| FIM1_3033 | 1464.541651 | 1366.33018 | 1273.167036 | -0.043215347 | 0.091380884 | 0.451085655 | 0.754843974 | ns | TM | KS.1 |
| FIM1_3034 | 3553.727845 | 3096.674958 | 2931.69041 | -0.040988962 | 0.148202819 | 0.528926632 | 0.808531286 | ns | TM | KS.1 |
| FIM1_3035 | 13205.18068 | 12202.86178 | 11142.20316 | -0.041762045 | 0.105702869 | 0.49797844 | 0.793454967 | ns | TM | KS.1 |
| FIM1_3036 | 3264.701706 | 2714.315082 | 3404.866561 | -0.133585226 | 0.123201165 | 0.068226218 | 0.275864055 | ns | TM | KS.1 |
| FIM1_3037 | 1814.860695 | 1415.564729 | 1021.631874 | -0.068657963 | 0.171657717 | 0.317206345 | 0.644123088 | ns | TM | KS.1 |
| FIM1_3038 | 15522.28983 | 18312.15424 | 17148.25544 | 0.11582025 | 0.117843899 | 0.098788532 | 0.339180776 | ns | TM | KS.1 |

| | | | | | | | | | | |
|---|---|---|---|---|---|---|---|---|---|---|
| FIM1_3039 | 2782.55998 | 2882.711709 | 2869.090948 | 0.021193426 | 0.088025257 | 0.703677488 | 0.898252804 | ns | TM | KS.1 |
| FIM1_3040 | 1598.142262 | 2449.952319 | 2794.51789 | 0.155664273 | 0.229018971 | 0.073415768 | 0.288301464 | ns | TM | KS.1 |
| FIM1_3041 | 1610.219571 | 1711.823792 | 1740.792045 | 0.043524829 | 0.077932828 | 0.40137265 | 0.723750973 | ns | TM | KS.1 |
| FIM1_3043 | 141.5613167 | 164.2114676 | 207.6025605 | 0.061631153 | 0.132966648 | 0.358608167 | 0.683292083 | ns | TM | KS.1 |
| FIM1_3044 | 418.4902253 | 353.2133124 | 4677.982151 | -0.039394882 | 0.166089114 | 0.540171616 | 0.814399145 | ns | TM | KS.1 |
| FIM1_3045 | 571.7334543 | 474.0943032 | 549.2223036 | -0.091160597 | 0.137277842 | 0.19873751 | 0.506475009 | ns | TM | KS.1 |
| FIM1_3046 | 1587.271574 | 1551.301389 | 1642.342519 | -0.011521624 | 0.102926754 | 0.84785501 | 0.952348261 | ns | TM | KS.1 |
| FIM1_3047 | 856.945136 | 854.6830078 | 825.3510037 | -0.001705287 | 0.072444518 | 0.972362156 | 0.99362472 | ns | TM | KS.1 |
| FIM1_3048 | 2439.870942 | 3045.103835 | 3099.753428 | 0.137278195 | 0.141163861 | 0.075051803 | 0.291705249 | ns | TM | KS.1 |
| FIM1_3049 | 1053.32778 | 1138.61784 | 883.089918 | 0.050212204 | 0.092131262 | 0.385193443 | 0.70922923 | ns | TM | KS.1 |
| FIM1_3050 | 952.4345276 | 848.2000269 | 792.229839 | -0.087785811 | 0.095702555 | 0.149672491 | 0.430665498 | ns | TM | KS.1 |
| FIM1_3051 | 1468.146925 | 1062.489078 | 1014.141862 | -0.319622355 | 0.1479915 | 0.001060262 | 0.016180375 | ns | TM | KS.1 |
| FIM1_3052 | 819.317609 | 831.4090841 | 796.9924135 | 0.008036155 | 0.092126286 | 0.887903884 | 0.967028314 | ns | TM | KS.1 |
| FIM1_3053 | 3071.884715 | 2100.847631 | 2143.005932 | -0.392468703 | 0.170070649 | 0.000335307 | 0.006593486 | ns | TM | KS.1 |
| FIM1_3054 | 988.3712255 | 880.518168 | 1456.614948 | -0.034365307 | 0.145474895 | 0.594080451 | 0.847105258 | ns | TM | KS.1 |
| FIM1_3056 | 3219.763512 | 3904.173086 | 4038.32144 | 0.075423653 | 0.145989939 | 0.277628527 | 0.602049362 | ns | TM | KS.1 |
| FIM1_3057 | 618.1099616 | 544.5959429 | 483.5889983 | -0.044369965 | 0.137466405 | 0.498241873 | 0.793454967 | ns | TM | KS.1 |
| FIM1_3058 | 2884.454069 | 2702.667497 | 2914.185425 | -0.038816247 | 0.093634927 | 0.503445955 | 0.796765282 | ns | TM | KS.1 |
| FIM1_3059 | 873.7733445 | 888.4774427 | 849.149499 | 0.010563041 | 0.076428757 | 0.83594395 | 0.948719561 | ns | TM | KS.1 |
| FIM1_3060 | 724.6375365 | 716.4606075 | 717.4541639 | -0.00577957 | 0.085268571 | 0.915702132 | 0.975605516 | ns | TM | KS.1 |
| FIM1_3061 | 4423.09029 | 4343.715008 | 3894.558609 | -0.0120415 | 0.073773158 | 0.809033863 | 0.938702059 | ns | TM | KS.1 |
| FIM1_3062 | 1628.430064 | 1868.896068 | 1637.020043 | 0.067584149 | 0.123845024 | 0.311710666 | 0.638961913 | ns | TM | KS.1 |
| FIM1_3063 | 1276.650675 | 1160.455726 | 1105.968508 | -0.081390655 | 0.081949717 | 0.137996392 | 0.411379542 | ns | TM | KS.1 |
| FIM1_3064 | 4900.639674 | 5855.313037 | 6172.113327 | 0.171755297 | 0.104176822 | 0.016113144 | 0.117368798 | ns | TM | KS.1 |
| FIM1_3065 | 2399.831569 | 2530.218545 | 2380.210882 | 0.037181248 | 0.077752756 | 0.472119996 | 0.772376515 | ns | TM | KS.1 |
| FIM1_3066 | 6063.840035 | 5618.684886 | 4964.727518 | -0.055354676 | 0.082782703 | 0.307480429 | 0.636056189 | ns | TM | KS.1 |
| FIM1_3067 | 646.6108432 | 602.9091463 | 690.525503 | -0.042426759 | 0.09522854 | 0.469460998 | 0.77114073 | ns | TM | KS.1 |
| FIM1_3068 | 4741.755659 | 4898.263164 | 5347.153281 | 0.013847171 | 0.114828248 | 0.823971081 | 0.94482203 | ns | TM | KS.1 |
| FIM1_3069 | 1169.267647 | 1158.329978 | 1141.648876 | -0.004111323 | 0.106702513 | 0.946079849 | 0.985614597 | ns | TM | KS.1 |
| FIM1_3070 | 6263.128356 | 6936.225536 | 7452.029009 | 0.07781495 | 0.090520036 | 0.182291976 | 0.483166 | ns | TM | KS.1 |
| FIM1_3071 | 3828.635333 | 3746.806071 | 3315.501244 | -0.012570385 | 0.087485484 | 0.820654587 | 0.943851992 | ns | TM | KS.1 |
| FIM1_3072 | 2080.278001 | 2093.662082 | 1546.247319 | 0.003041757 | 0.107202706 | 0.960153742 | 0.990413616 | ns | TM | KS.1 |
| FIM1_3074 | 863.2664712 | 924.570422 | 871.4180666 | 0.046024274 | 0.086506122 | 0.40745439 | 0.728957685 | ns | TM | KS.1 |
| FIM1_3075 | 895.7147363 | 1080.649072 | 1321.472421 | 0.11486336 | 0.13056472 | 0.114512186 | 0.37017422 | ns | TM | KS.1 |
| FIM1_3076 | 833.1039663 | 922.2673812 | 1017.158479 | 0.08154963 | 0.087503634 | 0.154656792 | 0.438498334 | ns | TM | KS.1 |
| FIM1_3077 | 1963.879872 | 1819.143147 | 2215.349619 | -0.048042211 | 0.093591771 | 0.409710967 | 0.73083975 | ns | TM | KS.1 |
| FIM1_3078 | 744.5443901 | 819.1660845 | 790.6237241 | 0.071429952 | 0.088198116 | 0.21089752 | 0.51871941 | ns | TM | KS.1 |
| FIM1_3079 | 1693.928685 | 2186.614344 | 2045.88673 | 0.194559289 | 0.145205306 | 0.021532631 | 0.143215029 | ns | TM | KS.1 |
| FIM1_3080 | 1210.081475 | 1208.765495 | 1517.297272 | -0.000664076 | 0.076570868 | 0.989624295 | 0.996341203 | ns | TM | KS.1 |
| FIM1_3081 | 712.4750443 | 1243.499678 | 1776.031229 | 0.729925007 | 0.184948074 | 6.5557172187( | 5.3466710103: | ns | TM | KS.1 |
| FIM1_3082 | 1657.013783 | 1558.598232 | 1874.539366 | -0.04611198 | 0.074401489 | 0.359077462 | 0.683511403 | ns | TM | KS.1 |
| FIM1_3083 | 2670.96942 | 4086.185897 | 5074.020734 | 0.189571213 | 0.232472393 | 0.043049995 | 0.212459177 | ns | TM | KS.1 |
| FIM1_3085 | 21.52456122 | 24.03515771 | 43.4414597 | 0.030149447 | 0.160442479 | 0.634865278 | 0.869623553 | ns | TM | KS.1 |
| FIM1_3086 | 682.9560198 | 495.7447908 | 1403.542008 | -0.201094897 | 0.180565324 | 0.0266599 | 0.160541482 | ns | TM | KS.1 |
| FIM1_3087 | 5698.890783 | 5579.276005 | 7192.444108 | -0.008774271 | 0.115892508 | 0.887966562 | 0.967028314 | ns | TM | KS.1 |

| | | | | | | | | | | |
|---|---|---|---|---|---|---|---|---|---|---|
| FIM1_3088 | 500.9967024 | 496.2021506 | 879.435673 | -0.0035658 | 0.128798208 | 0.954920876 | 0.988678089 | ns | TM | KS.1 |
| FIM1_3089 | 506.5746672 | 497.8228876 | 526.3408947 | -0.007215713 | 0.11319925 | 0.907219719 | 0.97188159 | ns | TM | KS.1 |
| FIM1_3090 | 253.0155442 | 241.5740592 | 396.6058563 | -0.025793358 | 0.097740513 | 0.661838372 | 0.880170528 | ns | TM | KS.1 |
| FIM1_3092 | 273.8728764 | 278.1235418 | 500.4249974 | 0.006921798 | 0.110407098 | 0.910363411 | 0.972526942 | ns | TM | KS.1 |
| FIM1_3098 | 1116.770562 | 2027.776632 | 1200.264754 | 0.719004774 | 0.252703511 | 2.9828157599 | 0.000969903 | ns | TM | KS.1 |
| FIM1_3099 | 2293.933471 | 2014.92638 | 2350.021231 | -0.038065916 | 0.147954071 | 0.556846914 | 0.825232602 | ns | TM | KS.1 |
| FIM1_3100 | 875.6100051 | 897.4874263 | 982.5762384 | 0.012992967 | 0.096545766 | 0.823957333 | 0.94482203 | ns | TM | KS.1 |
| FIM1_3101 | 964.1435033 | 1038.141616 | 1052.947167 | 0.041959315 | 0.099830461 | 0.484849135 | 0.785178463 | ns | TM | KS.1 |
| FIM1_3102 | 14747.6073 | 17862.13669 | 16236.58602 | 0.141959395 | 0.124567386 | 0.056576153 | 0.250861283 | ns | TM | KS.1 |
| FIM1_3103 | 108.7606206 | 138.1283721 | 162.3837864 | 0.064421203 | 0.170728336 | 0.343037455 | 0.670129201 | ns | TM | KS.1 |
| FIM1_3104 | 1972.81682 | 1857.078779 | 1971.434572 | -0.038056701 | 0.088549223 | 0.498133893 | 0.793454967 | ns | TM | KS.1 |
| FIM1_3105 | 19.064122 | 12.66188458 | 14.7225111 | -0.080986703 | 0.215111599 | 0.252677162 | 0.579042268 | ns | TM | KS.1 |
| FIM1_3106 | 1479.126893 | 1457.178736 | 1352.376741 | -0.009687422 | 0.079809114 | 0.853463262 | 0.954152748 | ns | TM | KS.1 |
| FIM1_3107 | 2135.141529 | 574.4236485 | 1220.887682 | -1.400281804 | 0.44796555 | 1.5509862324 | 0.000580162 | Km<KS.1 | TM | KS.1 |
| FIM1_3108 | 2593.943067 | 1330.84331 | 2240.122344 | -0.922845292 | 0.149902423 | 1.0406056227 | 3.6978664093 | ns | TM | KS.1 |
| FIM1_3109 | 2620.666032 | 3350.721379 | 3375.384685 | 0.130380533 | 0.15351668 | 0.093446893 | 0.330183448 | ns | TM | KS.1 |
| FIM1_3110 | 1894.627745 | 599.9312981 | 4126.047925 | -0.846238331 | 0.60259086 | 0.000888377 | 0.013898345 | ns | TM | KS.1 |
| FIM1_3111 | 697.0718031 | 394.9951114 | 1294.786463 | -0.319234604 | 0.323441599 | 0.011370628 | 0.092131716 | ns | TM | KS.1 |
| FIM1_3112 | 792.9987558 | 882.7606353 | 703.4952814 | 0.049183983 | 0.119935343 | 0.447286073 | 0.752446492 | ns | TM | KS.1 |
| FIM1_3113 | 1058.678153 | 1770.482417 | 1621.420472 | 0.579485119 | 0.235014212 | 8.6781283957 | 0.002158684 | ns | TM | KS.1 |
| FIM1_3114 | 2074.311818 | 1513.007296 | 1719.038682 | -0.246308237 | 0.168736571 | 0.009433398 | 0.080637726 | ns | TM | KS.1 |
| FIM1_3115 | 336.9224781 | 259.0413603 | 260.7413305 | -0.177322198 | 0.153561539 | 0.03472254 | 0.186147237 | ns | TM | KS.1 |
| FIM1_3116 | 1459.631783 | 1462.940468 | 1724.004441 | 0.000923587 | 0.117954668 | 0.988197874 | 0.996167262 | ns | TM | KS.1 |
| FIM1_3117 | 21.15692528 | 18.86164103 | 19.37440892 | -0.030204363 | 0.158103487 | 0.634851613 | 0.869623553 | ns | TM | KS.1 |
| FIM1_3118 | 7750.339947 | 4980.911154 | 4526.624002 | -0.419638855 | 0.224264334 | 0.000852675 | 0.013521989 | ns | TM | KS.1 |
| FIM1_3119 | 916.4414785 | 946.276528 | 946.5580616 | 0.014233169 | 0.111339213 | 0.817717436 | 0.94235447 | ns | TM | KS.1 |
| FIM1_3120 | 2737.475329 | 2822.036822 | 2859.777166 | 0.021197398 | 0.07084492 | 0.661852853 | 0.880170528 | ns | TM | KS.1 |
| FIM1_3121 | 1004.895199 | 1025.463489 | 1170.818666 | 0.012985468 | 0.076108183 | 0.798517238 | 0.936276987 | ns | TM | KS.1 |
| FIM1_3122 | 2896.408066 | 2718.199702 | 1970.926694 | -0.033756391 | 0.10226766 | 0.576519778 | 0.834745603 | ns | TM | KS.1 |
| FIM1_3123 | 2965.470201 | 2780.365095 | 2384.973826 | -0.044994266 | 0.082212413 | 0.402665528 | 0.724768814 | ns | TM | KS.1 |
| FIM1_3124 | 3487.75875 | 3723.178886 | 4368.791076 | 0.065830756 | 0.05406204 | 0.10232558 | 0.345835436 | ns | TM | KS.1 |
| FIM1_3125 | 597.8690464 | 514.8116516 | 648.7975289 | -0.104259198 | 0.113519082 | 0.125006941 | 0.388936541 | ns | TM | KS.1 |
| FIM1_3126 | 2180.962528 | 2120.674934 | 2225.153781 | -0.012313942 | 0.113033331 | 0.842494709 | 0.951443988 | ns | TM | KS.1 |
| FIM1_3127 | 1858.62108 | 1945.719537 | 1641.556946 | 0.02003069 | 0.114047247 | 0.747895647 | 0.916900159 | ns | TM | KS.1 |
| FIM1_3128 | 9813.17007 | 12531.3658 | 9997.73968 | 0.289563689 | 0.088689409 | 8.9652347282 | 0.002175709 | ns | TM | KS.1 |
| FIM1_3129 | 3062.341356 | 3773.136901 | 3927.025455 | 0.236252649 | 0.09208566 | 0.001123896 | 0.016554684 | ns | TM | KS.1 |
| FIM1_3130 | 1024.470557 | 1276.103934 | 1143.616367 | 0.125537248 | 0.142701947 | 0.097981636 | 0.337879338 | ns | TM | KS.1 |
| FIM1_3131 | 1002.949614 | 1008.084068 | 1039.568712 | 0.002240531 | 0.093132367 | 0.968814818 | 0.993374633 | ns | TM | KS.1 |
| FIM1_3132 | 3224.689589 | 3253.306878 | 3367.068562 | 0.006657435 | 0.065237168 | 0.884352766 | 0.966744674 | ns | TM | KS.1 |
| FIM1_3133 | 386.7138796 | 408.5081792 | 594.2050343 | 0.026289629 | 0.108505843 | 0.669546128 | 0.882180579 | ns | TM | KS.1 |
| FIM1_3134 | 9605.767229 | 10464.10505 | 11415.25866 | 0.037824734 | 0.118503084 | 0.55299757 | 0.822886984 | ns | TM | KS.1 |
| FIM1_3135 | 1078.898572 | 1175.454102 | 1117.0395 | 0.060406235 | 0.089171401 | 0.289626934 | 0.617085223 | ns | TM | KS.1 |
| FIM1_3136 | 2677.891806 | 3892.081105 | 4630.98272 | 0.300721494 | 0.197163069 | 0.004677029 | 0.049202074 | ns | TM | KS.1 |
| FIM1_3137 | 17.15526674 | 25.08990854 | 24.44135902 | 0.077812825 | 0.216326093 | 0.266999685 | 0.593708608 | ns | TM | KS.1 |
| FIM1_3138 | 3543.15341 | 5399.397535 | 6167.117837 | 0.21454837 | 0.232899717 | 0.028947728 | 0.169342179 | ns | TM | KS.1 |

| | | | | | | | | | | |
|---|---|---|---|---|---|---|---|---|---|---|
| FIM1_3139 | 813.6658456 | 962.7721762 | 667.88201 | 0.08471168 | 0.130701713 | 0.222568244 | 0.537131242 | ns | TM | KS.1 |
| FIM1_3140 | 443.929579 | 458.3879372 | 459.0135077 | 0.017107619 | 0.093328128 | 0.765985439 | 0.924600573 | ns | TM | KS.1 |
| FIM1_3141 | 1121.427988 | 1272.954413 | 972.2596684 | 0.081443476 | 0.108472177 | 0.20832562 | 0.515579507 | ns | TM | KS.1 |
| FIM1_3142 | 203.6744927 | 177.9422796 | 264.7518978 | -0.056356234 | 0.130740609 | 0.396262917 | 0.719491976 | ns | TM | KS.1 |
| FIM1_3143 | 4850.794673 | 5525.666147 | 5562.129101 | 0.128801789 | 0.0865069 | 0.032612761 | 0.180382188 | ns | TM | KS.1 |
| FIM1_3144 | 486.6818779 | 427.608811 | 358.3171778 | -0.06483793 | 0.120357698 | 0.326235576 | 0.653613253 | ns | TM | KS.1 |
| FIM1_3145 | 1216.93715 | 1407.301738 | 1300.969336 | 0.109190512 | 0.10749421 | 0.101160772 | 0.343715475 | ns | TM | KS.1 |
| FIM1_3146 | 568.4291862 | 568.9100193 | 695.8146255 | -0.000180127 | 0.080952327 | 0.997283592 | 0.998293524 | ns | TM | KS.1 |
| FIM1_3147 | 1157.248192 | 962.3260951 | 1134.359183 | -0.135006587 | 0.122165239 | 0.064960601 | 0.267310992 | ns | TM | KS.1 |
| FIM1_3148 | 1068.141184 | 1098.964284 | 864.5807955 | 0.018400144 | 0.079833293 | 0.725907325 | 0.907961357 | ns | TM | KS.1 |
| FIM1_3149 | 1067.188705 | 1136.660444 | 1048.570728 | 0.03731449 | 0.094377297 | 0.521588808 | 0.807711954 | ns | TM | KS.1 |
| FIM1_3150 | 2429.147327 | 4354.070768 | 5060.087817 | 0.444066807 | 0.338403091 | 0.003102789 | 0.036150763 | ns | TM | KS.1 |
| FIM1_3151 | 904.2110118 | 553.0154475 | 787.651183 | -0.596737209 | 0.197342672 | 1.2743555534 | 0.000495306 | ns | TM | KS.1 |
| FIM1_3152 | 215.0033422 | 226.3883297 | 255.1933084 | 0.027632689 | 0.102742316 | 0.647128268 | 0.873595603 | ns | TM | KS.1 |
| FIM1_3153 | 832.0537046 | 735.1226301 | 769.2644039 | -0.091633801 | 0.100844168 | 0.14611787 | 0.424167573 | ns | TM | KS.1 |
| FIM1_3154 | 1985.091173 | 2318.991582 | 2146.446937 | 0.075542293 | 0.128609282 | 0.26806707 | 0.595106502 | ns | TM | KS.1 |
| FIM1_3155 | 1688.491403 | 2041.857963 | 2866.78671 | 0.143033384 | 0.123387748 | 0.054174604 | 0.242809601 | ns | TM | KS.1 |
| FIM1_3156 | 3730.066144 | 3757.068545 | 3973.487865 | 0.004423395 | 0.08398069 | 0.934846038 | 0.981103476 | ns | TM | KS.1 |
| FIM1_3157 | 2107.249937 | 2040.012453 | 1292.04226 | -0.010864083 | 0.131544093 | 0.863616325 | 0.959694581 | ns | TM | KS.1 |
| FIM1_3158 | 333.6481113 | 294.1960682 | 368.4361109 | -0.075772026 | 0.111512318 | 0.244587092 | 0.567815242 | ns | TM | KS.1 |
| FIM1_3159 | 533.7445607 | 379.9006461 | 404.0752316 | -0.326064204 | 0.160120087 | 0.001287945 | 0.01835967 | ns | TM | KS.1 |
| FIM1_3160 | 256.1724818 | 301.6231361 | 291.6823713 | 0.046206684 | 0.153903134 | 0.481068098 | 0.783152417 | ns | TM | KS.1 |
| FIM1_3161 | 962.171964 | 488.8790138 | 1401.267782 | -0.595919853 | 0.386207803 | 0.001193933 | 0.017266913 | ns | TM | KS.1 |
| FIM1_3162 | 2203.053652 | 1606.339737 | 2181.720415 | -0.118600572 | 0.18368519 | 0.127142524 | 0.393568488 | ns | TM | KS.1 |
| FIM1_3163 | 869.4655684 | 914.5650151 | 715.8975727 | 0.022485891 | 0.113722074 | 0.718402205 | 0.905913973 | ns | TM | KS.1 |
| FIM1_3164 | 2240.331479 | 2381.440016 | 2822.476065 | 0.060966773 | 0.052979941 | 0.123557339 | 0.387577402 | ns | TM | KS.1 |
| FIM1_3165 | 820.006294 | 814.1852505 | 793.2031306 | -0.004162891 | 0.096029406 | 0.942968229 | 0.984319543 | ns | TM | KS.1 |
| FIM1_3166 | 1227.622338 | 1148.198128 | 1168.677337 | -0.039293574 | 0.096040235 | 0.504110773 | 0.796765282 | ns | TM | KS.1 |
| FIM1_3167 | 1866.813544 | 2820.441452 | 3498.114776 | 0.158501893 | 0.223455839 | 0.069131694 | 0.27676212 | ns | TM | KS.1 |
| FIM1_3168 | 1285.237304 | 1249.220273 | 1110.963659 | -0.017708771 | 0.077816447 | 0.731574721 | 0.910578994 | ns | TM | KS.1 |
| FIM1_3169 | 1177.749624 | 1136.333781 | 1077.659817 | -0.018487944 | 0.099467536 | 0.755220038 | 0.920210554 | ns | TM | KS.1 |
| FIM1_3170 | 53.00335797 | 46.16537184 | 121.4952025 | -0.04600296 | 0.147086323 | 0.483722305 | 0.784137657 | ns | TM | KS.1 |
| FIM1_3171 | 1743.688278 | 1761.471999 | 1716.990805 | 0.005704115 | 0.092907585 | 0.920633497 | 0.975881018 | ns | TM | KS.1 |
| FIM1_3172 | 1670.8119 | 1825.113584 | 1513.119302 | 0.038245099 | 0.120094906 | 0.549956313 | 0.822058746 | ns | TM | KS.1 |
| FIM1_3173 | 1201.85201 | 1177.042176 | 1674.368048 | -0.011612198 | 0.092813882 | 0.839286196 | 0.949982862 | ns | TM | KS.1 |
| FIM1_3174 | 470.2718389 | 489.5383624 | 492.5185698 | 0.016054275 | 0.120145628 | 0.798507034 | 0.936276987 | ns | TM | KS.1 |
| FIM1_3175 | 1670.756925 | 1412.368739 | 1395.309944 | -0.125098059 | 0.116120503 | 0.076391939 | 0.294813117 | ns | TM | KS.1 |
| FIM1_3176 | 453.861535 | 355.7725804 | 464.5748983 | -0.204737417 | 0.134651593 | 0.014281349 | 0.107002579 | ns | TM | KS.1 |
| FIM1_3177 | 2067.905317 | 2166.420214 | 2248.049981 | 0.037939696 | 0.061943835 | 0.389213266 | 0.713199262 | ns | TM | KS.1 |
| FIM1_3178 | 2007.196289 | 2041.304904 | 2050.784747 | 0.009008464 | 0.095727251 | 0.876843176 | 0.964248917 | ns | TM | KS.1 |
| FIM1_3179 | 383.3127212 | 275.477949 | 479.2921944 | -0.158947607 | 0.188124673 | 0.061436285 | 0.260865744 | ns | TM | KS.1 |
| FIM1_3180 | 1441.328379 | 1189.132985 | 1588.096631 | -0.141454568 | 0.125150415 | 0.057742593 | 0.253323986 | ns | TM | KS.1 |
| FIM1_3181 | 1821.291189 | 2621.317485 | 3024.698306 | 0.150791865 | 0.202665128 | 0.074482454 | 0.290399848 | ns | TM | KS.1 |
| FIM1_3182 | 1804.582406 | 1841.418157 | 1881.758192 | 0.012202376 | 0.084671911 | 0.822514314 | 0.944618544 | ns | TM | KS.1 |
| FIM1_3183 | 6828.173184 | 5608.061802 | 6908.476124 | -0.136906537 | 0.128818617 | 0.06745293 | 0.273941491 | ns | TM | KS.1 |

| | | | | | | | | | | |
|---|---|---|---|---|---|---|---|---|---|---|
| FIM1_3184 | 773.0623293 | 718.6653875 | 690.2581567 | -0.028922345 | 0.123581812 | 0.649968963 | 0.874068749 | ns | TM | KS.1 |
| FIM1_3185 | 858.0416811 | 1060.442654 | 1068.012224 | 0.095730673 | 0.146515274 | 0.185445778 | 0.488144311 | ns | TM | KS.1 |
| FIM1_3186 | 4100.265605 | 4662.652081 | 3548.73503 | 0.092000497 | 0.103970205 | 0.151376806 | 0.433313929 | ns | TM | KS.1 |
| FIM1_3187 | 644.0446165 | 711.3611918 | 663.5778738 | 0.056002588 | 0.105963214 | 0.369583653 | 0.692013051 | ns | TM | KS.1 |
| FIM1_3188 | 660.6971905 | 694.0666419 | 690.8064976 | 0.029319075 | 0.090590936 | 0.605427977 | 0.851547247 | ns | TM | KS.1 |
| FIM1_3189 | 369.4715802 | 470.2231356 | 411.7370225 | 0.142524697 | 0.148984249 | 0.070711367 | 0.28143124 | ns | TM | KS.1 |
| FIM1_3190 | 682.1135232 | 663.7865768 | 680.2043265 | -0.014977386 | 0.093088932 | 0.794069342 | 0.934144 | ns | TM | KS.1 |
| FIM1_3191 | 900.4974165 | 947.3182832 | 883.2861558 | 0.023910179 | 0.109729015 | 0.698604309 | 0.896699645 | ns | TM | KS.1 |
| FIM1_3192 | 723.8149035 | 684.6613014 | 729.3971384 | -0.035297177 | 0.085135768 | 0.519559723 | 0.805989904 | ns | TM | KS.1 |
| FIM1_3193 | 31.37944038 | 29.6162117 | 43.75386283 | -0.010097442 | 0.147840985 | 0.872200419 | 0.962759265 | ns | TM | KS.1 |
| FIM1_3194 | 2095.55209 | 2202.391093 | 2395.124519 | 0.021152427 | 0.11639683 | 0.735720609 | 0.912315561 | ns | TM | KS.1 |
| FIM1_3195 | 1957.045382 | 2807.493018 | 3200.949999 | 0.188503233 | 0.201680248 | 0.03829454 | 0.197220846 | ns | TM | KS.1 |
| FIM1_3196 | 441.1376697 | 479.1988574 | 477.2816158 | 0.042857369 | 0.10789301 | 0.490717435 | 0.789815348 | ns | TM | KS.1 |
| FIM1_3197 | 511.1676252 | 528.8858679 | 1536.617768 | 0.0095686 | 0.142186427 | 0.879302919 | 0.965254197 | ns | TM | KS.1 |
| FIM1_3198 | 391.2550835 | 515.5635145 | 460.0031329 | 0.060024044 | 0.186737041 | 0.369916456 | 0.69237561 | ns | TM | KS.1 |
| FIM1_3199 | 2031.6768 | 2085.518014 | 2359.023598 | 0.014477508 | 0.093300855 | 0.801005489 | 0.936908579 | ns | TM | KS.1 |
| FIM1_3200 | 7298.988938 | 9223.961598 | 9302.489641 | 0.309235644 | 0.057888965 | 8.74893960668 | 1.28017572186 | ns | TM | KS.1 |
| FIM1_3201 | 547.7146993 | 606.8034225 | 501.8279529 | 0.055451946 | 0.110300879 | 0.38203865 | 0.706033538 | ns | TM | KS.1 |
| FIM1_3202 | 503.7703907 | 520.2316874 | 534.8867205 | 0.020187381 | 0.088048357 | 0.717111689 | 0.905260252 | ns | TM | KS.1 |
| FIM1_3203 | 1260.517812 | 1230.23425 | 1449.933299 | -0.014194476 | 0.088958177 | 0.799852272 | 0.936738478 | ns | TM | KS.1 |
| FIM1_3204 | 773.7139016 | 799.7086313 | 744.7715961 | 0.028726119 | 0.058194222 | 0.495078361 | 0.792222775 | ns | TM | KS.1 |
| FIM1_3205 | 4475.23803 | 4673.824434 | 4777.253445 | 0.029918905 | 0.075880826 | 0.556014528 | 0.825176412 | ns | TM | KS.1 |
| FIM1_3206 | 1721.21908 | 1782.38612 | 1815.968829 | 0.023845819 | 0.075542706 | 0.637716076 | 0.870169358 | ns | TM | KS.1 |
| FIM1_3207 | 957.5829662 | 986.1846586 | 891.1447567 | 0.012793702 | 0.113278627 | 0.836560433 | 0.949024751 | ns | TM | KS.1 |
| FIM1_3208 | 3342.70411 | 3895.997743 | 3247.202603 | 0.14302034 | 0.098701824 | 0.031641141 | 0.177624525 | ns | TM | KS.1 |
| FIM1_3209 | 7451.397581 | 7823.848061 | 7110.124517 | 0.029819544 | 0.087855626 | 0.592870993 | 0.846854399 | ns | TM | KS.1 |
| FIM1_3210 | 1056.742346 | 1163.608534 | 590.3318852 | 0.052558085 | 0.107836822 | 0.401832637 | 0.724158796 | ns | TM | KS.1 |
| FIM1_3211 | 446.9570673 | 507.2128146 | 437.1034821 | 0.062633354 | 0.120735764 | 0.342007163 | 0.669085975 | ns | TM | KS.1 |
| FIM1_3212 | 2781.152063 | 3225.844405 | 3159.226961 | 0.119476637 | 0.105689885 | 0.074144641 | 0.28953657 | ns | TM | KS.1 |
| FIM1_3213 | 3919.644884 | 3575.301245 | 3801.11775 | -0.070981983 | 0.085332445 | 0.203933833 | 0.512409504 | ns | TM | KS.1 |
| FIM1_3214 | 2008.179246 | 3177.954767 | 3474.386174 | 0.13325975 | 0.237077548 | 0.106745026 | 0.353815182 | ns | TM | KS.1 |
| FIM1_3215 | 2157.910801 | 1987.573399 | 2625.623474 | -0.068557689 | 0.076470405 | 0.185653453 | 0.48817438 | ns | TM | KS.1 |
| FIM1_3216 | 114.1875249 | 85.41059831 | 80.9259995 | -0.12043494 | 0.173829801 | 0.120770667 | 0.381584021 | ns | TM | KS.1 |
| FIM1_3217 | 2172.908276 | 1905.126602 | 2185.918002 | -0.046990035 | 0.137751534 | 0.475041446 | 0.775371126 | ns | TM | KS.1 |
| FIM1_3218 | 522.4746767 | 323.8991084 | 239.287081 | -0.233002498 | 0.264007099 | 0.025531128 | 0.156140634 | ns | TM | KS.1 |
| FIM1_3219 | 4212.141258 | 3485.619983 | 3158.916771 | -0.112954307 | 0.131444517 | 0.120336271 | 0.381077623 | ns | TM | KS.1 |
| FIM1_3220 | 833.2808012 | 847.3505607 | 2141.848212 | 0.006776172 | 0.120226893 | 0.913920131 | 0.974699214 | ns | TM | KS.1 |
| FIM1_3221 | 6573.263272 | 6592.839196 | 6394.062365 | 0.002037856 | 0.073237631 | 0.967216224 | 0.993374633 | ns | TM | KS.1 |
| FIM1_3222 | 4183.571696 | 3816.055859 | 3571.471653 | -0.097494995 | 0.063356371 | 0.036794746 | 0.192283467 | ns | TM | KS.1 |
| FIM1_3223 | 4157.194245 | 4793.568155 | 5186.784433 | 0.146395701 | 0.087618425 | 0.018918903 | 0.13145467 | ns | TM | KS.1 |
| FIM1_3224 | 1876.265799 | 1793.040842 | 1877.403515 | -0.014382266 | 0.13567691 | 0.820401211 | 0.943851992 | ns | TM | KS.1 |
| FIM1_3225 | 133.1809029 | 142.4916197 | 117.7906859 | 0.020335362 | 0.142646267 | 0.74869209 | 0.91742442 | ns | TM | KS.1 |
| FIM1_3226 | 883.7631871 | 877.9703579 | 829.6867632 | -0.003596561 | 0.089843509 | 0.949006787 | 0.986388143 | ns | TM | KS.1 |
| FIM1_3227 | 2035.876293 | 2232.376701 | 2360.123084 | 0.059122943 | 0.097089466 | 0.323671408 | 0.650941236 | ns | TM | KS.1 |
| FIM1_3228 | 2042.80203 | 2005.333145 | 1687.076866 | -0.010924556 | 0.083424332 | 0.839415906 | 0.949982862 | ns | TM | KS.1 |

| | | | | | | | | | | |
|---|---|---|---|---|---|---|---|---|---|---|
| FIM1_3229 | 3356.387458 | 3882.692655 | 3769.560011 | 0.140774869 | 0.093937535 | 0.029023108 | 0.169342179 | ns | TM | KS.1 |
| FIM1_3230 | 3982.007516 | 4538.266818 | 4397.562342 | 0.10669604 | 0.098347258 | 0.091655907 | 0.327107704 | ns | TM | KS.1 |
| FIM1_3231 | 4283.404706 | 4665.732627 | 4952.197279 | 0.080480159 | 0.069470869 | 0.099787857 | 0.341199031 | ns | TM | KS.1 |
| FIM1_3232 | 118.4082434 | 119.3593752 | 257.3779081 | 0.00338663 | 0.12924784 | 0.957188465 | 0.989353426 | ns | TM | KS.1 |
| FIM1_3233 | 4345.52378 | 2846.251124 | 1744.519258 | -0.534813795 | 0.140535868 | 5.18326698905 | 4.70464842814 | ns | TM | KS.1 |
| FIM1_3234 | 2043.765929 | 3353.669991 | 4010.201847 | 0.21643007 | 0.268705757 | 0.033072182 | 0.18140475 | ns | TM | KS.1 |
| FIM1_3235 | 1665.769578 | 1549.582387 | 1430.647497 | -0.049947258 | 0.085209645 | 0.364913489 | 0.688866363 | ns | TM | KS.1 |
| FIM1_3236 | 5952.882913 | 7999.741117 | 9277.944695 | 0.134216917 | 0.174216391 | 0.093116098 | 0.329798971 | ns | TM | KS.1 |
| FIM1_3237 | 5855.756023 | 5193.291786 | 5182.316218 | -0.080664179 | 0.103772754 | 0.20239579 | 0.510554281 | ns | TM | KS.1 |
| FIM1_3238 | 627.489168 | 487.7315498 | 470.235462 | -0.23734949 | 0.12730374 | 0.005222943 | 0.053465311 | ns | TM | KS.1 |
| FIM1_3239 | 3560.665926 | 2462.063864 | 3239.1185 | -0.447663811 | 0.12487765 | 2.48805218406 | 0.000143931 | ns | TM | KS.1 |
| FIM1_3240 | 2405.307086 | 2320.21393 | 2728.842769 | -0.020671536 | 0.091224198 | 0.716075358 | 0.905084971 | ns | TM | KS.1 |
| FIM1_3241 | 1550.999347 | 1186.784725 | 1207.718789 | -0.159504881 | 0.158852906 | 0.052728875 | 0.240005664 | ns | TM | KS.1 |
| FIM1_3242 | 652.2364353 | 675.2595639 | 739.1983989 | 0.019713281 | 0.090230708 | 0.727083284 | 0.907961357 | ns | TM | KS.1 |
| FIM1_3243 | 2021.48778 | 2338.249311 | 2177.611347 | 0.105501051 | 0.109502362 | 0.114899039 | 0.37039344 | ns | TM | KS.1 |
| FIM1_3244 | 3250.282257 | 1583.233171 | 1295.076085 | -0.9111724 | 0.258715226 | 6.67797032155 | 0.000302026 | ns | TM | KS.1 |
| FIM1_3245 | 1421.08335 | 1307.391602 | 1235.652559 | -0.048972024 | 0.100040602 | 0.417557179 | 0.73560445 | ns | TM | KS.1 |
| FIM1_3246 | 931.5917983 | 731.8857257 | 1352.307553 | -0.151711934 | 0.147540673 | 0.057436954 | 0.252651502 | ns | TM | KS.1 |
| FIM1_3247 | 4975.799722 | 5150.01178 | 5932.375847 | 0.010893213 | 0.135271401 | 0.863230839 | 0.959681212 | ns | TM | KS.1 |
| FIM1_3248 | 349.4804932 | 227.2927568 | 270.241428 | -0.477123808 | 0.187724147 | 8.90590000034 | 0.002171905 | ns | TM | KS.1 |
| FIM1_3249 | 4038.970278 | 3673.5282 | 3260.562503 | -0.059028188 | 0.09942324 | 0.330571854 | 0.657690769 | ns | TM | KS.1 |
| FIM1_3250 | 963.5983504 | 926.3142867 | 864.3547259 | -0.019693009 | 0.102952855 | 0.743603081 | 0.91508234 | ns | TM | KS.1 |
| FIM1_3251 | 592.8930807 | 633.3881485 | 786.1267761 | 0.042509336 | 0.085642349 | 0.440724088 | 0.747832613 | ns | TM | KS.1 |
| FIM1_3252 | 24.2166191 | 42.60348078 | 46.84830583 | 0.21842911 | 0.304140971 | 0.0362948 | 0.190639184 | ns | TM | KS.1 |
| FIM1_3253 | 4168.990974 | 3926.59584 | 3578.117444 | -0.018475966 | 0.138239923 | 0.771052974 | 0.925705893 | ns | TM | KS.1 |
| FIM1_3254 | 4523.443105 | 4623.230913 | 4440.083237 | 0.015400201 | 0.068205422 | 0.744264795 | 0.91508234 | ns | TM | KS.1 |
| FIM1_3255 | 647.8785474 | 705.932568 | 844.2191512 | 0.068297249 | 0.079508258 | 0.19916872 | 0.506836 | ns | TM | KS.1 |
| FIM1_3256 | 30.389805 | 33.68548386 | 40.43753559 | 0.028266732 | 0.154964599 | 0.656514464 | 0.877276998 | ns | TM | KS.1 |
| FIM1_3258 | 432.5024271 | 374.0011435 | 304.6439041 | -0.074245965 | 0.124289677 | 0.271370536 | 0.598713828 | ns | TM | KS.1 |
| FIM1_3259 | 619.2417717 | 616.4437851 | 420.9082459 | -0.00163335 | 0.123033338 | 0.97926386 | 0.99466856 | ns | TM | KS.1 |
| FIM1_3260 | 1182.723903 | 1191.22493 | 1532.399655 | 0.005102945 | 0.060095871 | 0.905897405 | 0.971578849 | ns | TM | KS.1 |
| FIM1_3261 | 672.4740424 | 615.0393517 | 583.1292843 | -0.049305892 | 0.105226007 | 0.425492088 | 0.741071977 | ns | TM | KS.1 |
| FIM1_3262 | 25743.35895 | 30461.67478 | 29873.94409 | 0.143004348 | 0.109911548 | 0.042446715 | 0.210464424 | ns | TM | KS.1 |
| FIM1_3263 | 1773.102908 | 1967.658935 | 1959.690367 | 0.080319935 | 0.090406956 | 0.169242895 | 0.463392076 | ns | TM | KS.1 |
| FIM1_3264 | 1597.441716 | 1708.361546 | 1677.565821 | 0.04348533 | 0.088315175 | 0.439144069 | 0.747832613 | ns | TM | KS.1 |
| FIM1_3265 | 1220.078214 | 1197.77391 | 1308.290788 | -0.012218606 | 0.069738963 | 0.798862391 | 0.936373441 | ns | TM | KS.1 |
| FIM1_3266 | 657.4999988 | 670.7782538 | 627.6223463 | 0.008042792 | 0.119757367 | 0.897877129 | 0.967972723 | ns | TM | KS.1 |
| FIM1_3267 | 3395.874219 | 3551.752664 | 3631.771014 | 0.029819462 | 0.079955199 | 0.570818382 | 0.832094673 | ns | TM | KS.1 |
| FIM1_3268 | 892.0796849 | 906.383848 | 861.409376 | 0.007792623 | 0.104569238 | 0.89730837 | 0.967972723 | ns | TM | KS.1 |
| FIM1_3269 | 644.1733839 | 514.3665609 | 594.8101771 | -0.141800474 | 0.142034752 | 0.068369926 | 0.275864055 | ns | TM | KS.1 |
| FIM1_3270 | 5707.335199 | 3269.936106 | 3849.519547 | -0.241357086 | 0.301596758 | 0.026803861 | 0.160661696 | ns | TM | KS.1 |
| FIM1_3271 | 4515.724539 | 3733.626684 | 3839.313539 | -0.155751323 | 0.119249102 | 0.036632181 | 0.191836946 | ns | TM | KS.1 |
| FIM1_3272 | 1283.973577 | 1216.793307 | 1169.173293 | -0.031307006 | 0.093639957 | 0.588260172 | 0.843155965 | ns | TM | KS.1 |
| FIM1_3273 | 1595.479738 | 1978.895342 | 2005.529265 | 0.076150579 | 0.155161981 | 0.275355183 | 0.600738415 | ns | TM | KS.1 |
| FIM1_3274 | 2765.348392 | 2821.037727 | 2580.125895 | 0.018465647 | 0.043261451 | 0.586676431 | 0.841613393 | ns | TM | KS.1 |

| | | | | | | | | | | |
|---|---|---|---|---|---|---|---|---|---|---|
| FIM1_3275 | 3135.515665 | 3384.882041 | 3593.207739 | 0.051012734 | 0.089382205 | 0.369282298 | 0.69186398 | ns | TM | KS.1 |
| FIM1_3276 | 479.209209 | 398.1828467 | 476.724671 | -0.085942447 | 0.138107267 | 0.221957197 | 0.536805479 | ns | TM | KS.1 |
| FIM1_3277 | 2204.65342 | 2336.178024 | 2368.234626 | 0.040036207 | 0.079590294 | 0.446179463 | 0.751945403 | ns | TM | KS.1 |
| FIM1_3278 | 6487.026829 | 6514.468519 | 6815.733062 | 0.002939142 | 0.06946859 | 0.951000551 | 0.987490021 | ns | TM | KS.1 |
| FIM1_3279 | 3372.30476 | 3441.182647 | 3833.314864 | 0.011965069 | 0.087316024 | 0.828956404 | 0.94647133 | ns | TM | KS.1 |
| FIM1_3280 | 2653.218955 | 2285.154321 | 2105.880803 | -0.112811012 | 0.10900642 | 0.094147954 | 0.331213943 | ns | TM | KS.1 |
| FIM1_3281 | 5894.058303 | 5868.451104 | 4292.32629 | -0.002324232 | 0.093361311 | 0.967690883 | 0.993374633 | ns | TM | KS.1 |
| FIM1_3282 | 5066.517739 | 5294.692628 | 5569.056616 | 0.011315925 | 0.149059333 | 0.85689031 | 0.955916258 | ns | TM | KS.1 |
| FIM1_3283 | 928.7946783 | 688.924729 | 807.0213661 | -0.240593412 | 0.158892144 | 0.009264643 | 0.079846584 | ns | TM | KS.1 |
| FIM1_3284 | 1559.234214 | 1413.772247 | 1342.972478 | -0.054952614 | 0.10680137 | 0.379820892 | 0.703660035 | ns | TM | KS.1 |
| FIM1_3286 | 1405.140906 | 1545.611315 | 1644.190823 | 0.067825122 | 0.091917087 | 0.245608121 | 0.569551818 | ns | TM | KS.1 |
| FIM1_3287 | 383.1675467 | 404.4612047 | 460.2756596 | 0.025811926 | 0.110729053 | 0.677161224 | 0.887002016 | ns | TM | KS.1 |
| FIM1_3288 | 26022.30775 | 26732.13385 | 24758.36259 | 0.011059492 | 0.117386055 | 0.859528781 | 0.956848441 | ns | TM | KS.1 |
| FIM1_3290 | 1719.12914 | 1755.80919 | 1587.834695 | 0.012495244 | 0.086489437 | 0.820548413 | 0.943851992 | ns | TM | KS.1 |
| FIM1_3291 | 14.26109464 | 9.242788839 | 18.39346275 | -0.078618884 | 0.230069218 | 0.261880136 | 0.588461463 | ns | TM | KS.1 |
| FIM1_3292 | 25859.19951 | 30016.41594 | 28547.75295 | 0.11561684 | 0.107754296 | 0.085548787 | 0.311772902 | ns | TM | KS.1 |
| FIM1_3293 | 356.2600218 | 391.1614776 | 402.3163413 | 0.043139108 | 0.117451938 | 0.499962898 | 0.794415655 | ns | TM | KS.1 |
| FIM1_3294 | 296.0720832 | 258.1370023 | 335.1507035 | -0.065097671 | 0.124891131 | 0.329158758 | 0.656865151 | ns | TM | KS.1 |
| FIM1_3295 | 920.8441557 | 1015.763516 | 1289.998448 | 0.041895493 | 0.121844673 | 0.515431182 | 0.804775792 | ns | TM | KS.1 |
| FIM1_3297 | 2392.935409 | 2350.074226 | 2133.083476 | -0.012367642 | 0.06617769 | 0.789041482 | 0.933303839 | ns | TM | KS.1 |
| FIM1_3298 | 2170.588558 | 3209.339711 | 3671.679717 | 0.201855006 | 0.216888831 | 0.032844053 | 0.180951453 | ns | TM | KS.1 |
| FIM1_3299 | 3839.617222 | 4487.477053 | 5009.030904 | 0.156185706 | 0.094439747 | 0.018233254 | 0.127941383 | ns | TM | KS.1 |
| FIM1_3300 | 11742.16946 | 13058.06907 | 12154.06664 | 0.061853561 | 0.106832509 | 0.326064124 | 0.653573335 | ns | TM | KS.1 |
| FIM1_3301 | 2176.984972 | 1710.899243 | 2184.731024 | -0.224078715 | 0.125934204 | 0.007169129 | 0.066581689 | ns | TM | KS.1 |
| FIM1_3302 | 443.9066425 | 669.3422404 | 695.5945618 | 0.12203135 | 0.219256061 | 0.125556933 | 0.389831665 | ns | TM | KS.1 |
| FIM1_3303 | 538.8884207 | 589.8546514 | 583.473781 | 0.046192175 | 0.110304402 | 0.462614616 | 0.764620503 | ns | TM | KS.1 |
| FIM1_3304 | 6349.436585 | 8860.273059 | 10610.29114 | 0.161379387 | 0.189057719 | 0.058970861 | 0.254891426 | ns | TM | KS.1 |
| FIM1_3305 | 12360.26944 | 11531.96232 | 11293.93362 | -0.043742031 | 0.090750142 | 0.443874017 | 0.74983811 | ns | TM | KS.1 |
| FIM1_3306 | 564.029372 | 420.678702 | 385.9812727 | -0.166182134 | 0.169707907 | 0.049177777 | 0.231028743 | ns | TM | KS.1 |
| FIM1_3307 | 1125.782803 | 867.9074941 | 1497.973183 | -0.200630903 | 0.146540438 | 0.019081962 | 0.132384765 | ns | TM | KS.1 |
| FIM1_3308 | 323.3212643 | 332.9720637 | 1459.473884 | 0.011496502 | 0.121457302 | 0.854907993 | 0.95512402 | ns | TM | KS.1 |
| FIM1_3309 | 3293.630832 | 4025.250653 | 4207.45595 | 0.145472181 | 0.128605584 | 0.054820691 | 0.245263433 | ns | TM | KS.1 |
| FIM1_3310 | 1659.762217 | 1623.973413 | 1586.122228 | -0.014032192 | 0.076725903 | 0.783797311 | 0.930812921 | ns | TM | KS.1 |
| FIM1_3311 | 5186.224245 | 6120.709056 | 5799.004745 | 0.209188008 | 0.0642355 | 0.000191441 | 0.004177267 | ns | TM | KS.1 |
| FIM1_3312 | 2021.491021 | 2003.806217 | 3030.972899 | -0.002689757 | 0.137807174 | 0.965962177 | 0.993374633 | ns | TM | KS.1 |
| FIM1_3313 | 363.193362 | 397.4749907 | 449.383105 | 0.018787794 | 0.164974756 | 0.762870723 | 0.924230468 | ns | TM | KS.1 |
| FIM1_3314 | 656.5793233 | 542.1788538 | 611.8314272 | -0.14744159 | 0.122852539 | 0.048149117 | 0.228570475 | ns | TM | KS.1 |
| FIM1_3315 | 647.5285141 | 714.5300509 | 625.3048471 | 0.049993489 | 0.11219263 | 0.431065905 | 0.743536592 | ns | TM | KS.1 |
| FIM1_3316 | 1535.011839 | 1550.495005 | 1921.22516 | 0.007231541 | 0.068065608 | 0.878114718 | 0.964588369 | ns | TM | KS.1 |
| FIM1_3317 | 248.8588785 | 192.7655299 | 156.9880462 | -0.169100065 | 0.151190145 | 0.040597574 | 0.205257043 | ns | TM | KS.1 |
| FIM1_3318 | 503.9187067 | 845.5328631 | 902.019129 | 0.259582027 | 0.287693026 | 0.01983043 | 0.13560052 | ns | TM | KS.1 |
| FIM1_3319 | 1382.45258 | 1485.647659 | 1424.631369 | 0.04076713 | 0.09923463 | 0.495774203 | 0.79247947 | ns | TM | KS.1 |
| FIM1_3320 | 1928.650992 | 1669.408076 | 3754.133926 | -0.044802812 | 0.146994708 | 0.494299349 | 0.792222775 | ns | TM | KS.1 |
| FIM1_3321 | 1301.137946 | 1380.183754 | 2821.384153 | 0.015658769 | 0.147649402 | 0.803744139 | 0.936908579 | ns | TM | KS.1 |
| FIM1_3322 | 453.8865841 | 350.2125091 | 576.8316103 | -0.095976322 | 0.165837608 | 0.189841896 | 0.49354318 | ns | TM | KS.1 |

| | | | | | | | | | | |
|---|---|---|---|---|---|---|---|---|---|---|
| FIM1_3323 | 1994.535144 | 1295.635151 | 1336.77278 | -0.521621129 | 0.162399549 | 6.5996755843E | 0.000301224 | ns | TM | KS.1 |
| FIM1_3324 | 1657.349008 | 1588.597577 | 2165.389234 | -0.02184829 | 0.1017624 | 0.715601724 | 0.905084971 | ns | TM | KS.1 |
| FIM1_3326 | 8022.132343 | 9892.005129 | 10761.74779 | 0.12485997 | 0.138125863 | 0.096624053 | 0.336392346 | ns | TM | KS.1 |
| FIM1_3327 | 1146.078762 | 1188.023542 | 1547.231501 | 0.018895776 | 0.098592146 | 0.749056767 | 0.917645263 | ns | TM | KS.1 |
| FIM1_3328 | 277.4786151 | 211.685642 | 250.7530417 | -0.15538209 | 0.160732831 | 0.058053602 | 0.253530772 | ns | TM | KS.1 |
| FIM1_3329 | 242.5767387 | 199.5617561 | 333.4106703 | -0.118099685 | 0.134274515 | 0.109358178 | 0.35800987 | ns | TM | KS.1 |
| FIM1_3331 | 565.4882355 | 445.0463789 | 427.7347597 | -0.222633813 | 0.126799121 | 0.007617851 | 0.069752169 | ns | TM | KS.1 |
| FIM1_3332 | 637.9501547 | 696.992732 | 563.365356 | 0.061613513 | 0.089881246 | 0.282824879 | 0.608850616 | ns | TM | KS.1 |
| FIM1_3333 | 2693.44428 | 3498.918772 | 3974.130456 | 0.149069389 | 0.157701065 | 0.06494581 | 0.267310992 | ns | TM | KS.1 |
| FIM1_3334 | 27766.91592 | 34223.08795 | 33685.66399 | 0.183674756 | 0.12090589 | 0.01834566 | 0.128548817 | ns | TM | KS.1 |
| FIM1_3335 | 1002.256654 | 1358.843724 | 1422.542747 | 0.189479028 | 0.173066251 | 0.031668935 | 0.177624525 | ns | TM | KS.1 |
| FIM1_3336 | 1144.291091 | 1166.131486 | 1146.475696 | 0.008630203 | 0.109454197 | 0.888114874 | 0.967028314 | ns | TM | KS.1 |
| FIM1_3337 | 3522.962116 | 2916.938109 | 2308.848679 | -0.219714115 | 0.082585309 | 0.001013252 | 0.015592815 | ns | TM | KS.1 |
| FIM1_3339 | 1479.354962 | 1339.170921 | 1165.07265 | -0.062604916 | 0.100188849 | 0.305443944 | 0.633687915 | ns | TM | KS.1 |
| FIM1_3340 | 11408.05679 | 9811.532753 | 5574.560663 | -0.068420925 | 0.130169166 | 0.311671182 | 0.638961913 | ns | TM | KS.1 |
| FIM1_3341 | 1197.483867 | 1266.990096 | 1121.771156 | 0.040302765 | 0.075285648 | 0.425937787 | 0.741071977 | ns | TM | KS.1 |
| FIM1_3342 | 208.3480273 | 152.9120274 | 158.9391856 | -0.146146763 | 0.180148564 | 0.075716041 | 0.293478448 | ns | TM | KS.1 |
| FIM1_3343 | 10076.5868 | 10181.61159 | 8696.620167 | 0.003314786 | 0.133898118 | 0.958108209 | 0.989353426 | ns | TM | KS.1 |
| FIM1_3344 | 20016.8576 | 23780.76097 | 18575.06962 | 0.046887273 | 0.157918789 | 0.474518808 | 0.775371126 | ns | TM | KS.1 |
| FIM1_3345 | 9307.016285 | 9653.457177 | 6481.108677 | 0.009406183 | 0.148704486 | 0.88075486 | 0.9656535 | ns | TM | KS.1 |
| FIM1_3346 | 203.9391107 | 215.5149742 | 215.6425215 | 0.019488006 | 0.130139917 | 0.759097949 | 0.922567322 | ns | TM | KS.1 |
| FIM1_3347 | 1483.876494 | 957.7722064 | 1658.200111 | -0.38908093 | 0.229052538 | 0.001580473 | 0.021279312 | ns | TM | KS.1 |
| FIM1_3348 | 794.1503197 | 591.6550928 | 957.1650824 | -0.167711097 | 0.170498091 | 0.047928143 | 0.227738787 | ns | TM | KS.1 |
| FIM1_3349 | 1926.954524 | 1631.341482 | 1789.716452 | -0.141341483 | 0.109427977 | 0.043949966 | 0.215632229 | ns | TM | KS.1 |
| FIM1_3350 | 8008.948328 | 5968.186806 | 9211.614553 | -0.176371847 | 0.169301699 | 0.040054411 | 0.203961818 | ns | TM | KS.1 |
| FIM1_3351 | 2036.577332 | 2003.02053 | 1694.361127 | -0.010610535 | 0.079015975 | 0.838680034 | 0.949982862 | ns | TM | KS.1 |
| FIM1_3353 | 7639.080516 | 9173.007105 | 8404.574634 | 0.129043983 | 0.123661127 | 0.076680736 | 0.294813117 | ns | TM | KS.1 |
| FIM1_3355 | 1860.043928 | 1361.408438 | 1440.681107 | -0.19463628 | 0.176201514 | 0.029270881 | 0.16972335 | ns | TM | KS.1 |
| FIM1_3356 | 1505.281872 | 1171.049019 | 1282.228097 | -0.128656369 | 0.156038671 | 0.09782206 | 0.3377271 | ns | TM | KS.1 |
| FIM1_3357 | 521.3896202 | 465.5311881 | 631.5619316 | -0.061190996 | 0.111702279 | 0.339941429 | 0.668193429 | ns | TM | KS.1 |
| FIM1_3358 | 10.43204589 | 12.23573496 | 16.18493289 | 0.029868501 | 0.201919003 | 0.627819594 | 0.867130061 | ns | TM | KS.1 |
| FIM1_3359 | 4505.284561 | 4971.408593 | 5452.627362 | 0.081614356 | 0.083991566 | 0.143496515 | 0.420432957 | ns | TM | KS.1 |
| FIM1_3360 | 4001.89926 | 4286.329527 | 3710.643881 | 0.046398937 | 0.085734773 | 0.401085859 | 0.723750973 | ns | TM | KS.1 |
| FIM1_3361 | 6335.286986 | 6348.17236 | 6897.756809 | 0.001563842 | 0.063395614 | 0.972187637 | 0.99362472 | ns | TM | KS.1 |
| FIM1_3362 | 2455.198295 | 2375.112274 | 3212.704601 | -0.015398957 | 0.108928476 | 0.801866514 | 0.936908579 | ns | TM | KS.1 |
| FIM1_3363 | 1461.951459 | 1466.460523 | 1497.899917 | 0.002167677 | 0.07839136 | 0.966650664 | 0.993374633 | ns | TM | KS.1 |
| FIM1_3364 | 1956.294464 | 1838.857083 | 1841.479941 | -0.047189188 | 0.07347239 | 0.343886034 | 0.670129201 | ns | TM | KS.1 |
| FIM1_3365 | 324.5624937 | 380.2933451 | 341.3008944 | 0.079336929 | 0.128185104 | 0.247147269 | 0.57146079 | ns | TM | KS.1 |
| FIM1_3366 | 332.1587443 | 373.9179115 | 426.0664148 | 0.024132477 | 0.168436534 | 0.699506692 | 0.896699645 | ns | TM | KS.1 |
| FIM1_3367 | 908.26935 | 755.3189506 | 713.2190129 | -0.105612023 | 0.130907484 | 0.141345975 | 0.416585441 | ns | TM | KS.1 |
| FIM1_3368 | 482.9782635 | 584.2546325 | 621.4381627 | 0.109414349 | 0.132860183 | 0.13146962 | 0.400772892 | ns | TM | KS.1 |
| FIM1_3369 | 1479.785665 | 923.1329358 | 1156.707981 | -0.422724042 | 0.251676938 | 0.001363585 | 0.019149579 | ns | TM | KS.1 |
| FIM1_3370 | 982.8261167 | 1105.542544 | 841.2917541 | 0.054589027 | 0.121927332 | 0.403934378 | 0.725739808 | ns | TM | KS.1 |
| FIM1_3371 | 1070.133974 | 973.0600161 | 821.7171968 | -0.048463569 | 0.111278183 | 0.443308553 | 0.749137246 | ns | TM | KS.1 |
| FIM1_3372 | 859.1577362 | 829.9711351 | 785.7021713 | -0.022318172 | 0.079364415 | 0.66960584 | 0.882180579 | ns | TM | KS.1 |

| | | | | | | | | | | |
|---|---|---|---|---|---|---|---|---|---|---|
| FIM1_3373 | 517.7486463 | 752.848025 | 864.5272455 | 0.12183999 | 0.206317842 | 0.1239713 | 0.387735747 | ns | TM | KS.1 |
| FIM1_3374 | 3716.920553 | 3596.293857 | 5453.42696 | -0.008761522 | 0.14663345 | 0.889025812 | 0.967028314 | ns | TM | KS.1 |
| FIM1_3375 | 5978.695093 | 4999.134851 | 8288.966713 | -0.141354122 | 0.117283334 | 0.051270568 | 0.235847526 | ns | TM | KS.1 |
| FIM1_3376 | 18.10554007 | 14.82325025 | 19.77240213 | -0.039029449 | 0.179319827 | 0.540531854 | 0.814399145 | ns | TM | KS.1 |
| FIM1_3377 | 1093.165466 | 1371.962732 | 1614.212457 | 0.079387314 | 0.158378093 | 0.259188384 | 0.588210523 | ns | TM | KS.1 |
| FIM1_3378 | 17.79177082 | 21.15432446 | 24.81555604 | 0.03821748 | 0.160757061 | 0.55270699 | 0.822775964 | ns | TM | KS.1 |
| FIM1_3379 | 655.6939948 | 806.8449051 | 602.1434 | 0.160249417 | 0.128127052 | 0.038041576 | 0.196732683 | ns | TM | KS.1 |
| FIM1_3380 | 380.9848728 | 488.6562329 | 384.328297 | 0.129845309 | 0.154816222 | 0.095008058 | 0.333005827 | ns | TM | KS.1 |
| FIM1_3381 | 16164.90093 | 19507.72638 | 18287.95914 | 0.147320125 | 0.120784055 | 0.046712207 | 0.224840651 | ns | TM | KS.1 |
| FIM1_3382 | 510.7948221 | 382.5760649 | 540.4338346 | -0.222610608 | 0.157179789 | 0.013466094 | 0.103226225 | ns | TM | KS.1 |
| FIM1_3383 | 2530.95017 | 2132.288305 | 2471.836453 | -0.137691261 | 0.11415989 | 0.053338681 | 0.241250174 | ns | TM | KS.1 |
| FIM1_3384 | 4891.099105 | 7890.714556 | 10102.61494 | 0.240850533 | 0.264785989 | 0.022883229 | 0.147657673 | ns | TM | KS.1 |
| FIM1_3385 | 62.40452894 | 81.71469042 | 83.71763438 | 0.091916557 | 0.170243833 | 0.205666912 | 0.512865935 | ns | TM | KS.1 |
| FIM1_3386 | 662.4700969 | 856.0479761 | 715.7074906 | 0.115159094 | 0.160609264 | 0.129938415 | 0.398547234 | ns | TM | KS.1 |
| FIM1_3387 | 1525.297184 | 1708.098294 | 1783.26767 | 0.041359467 | 0.133600006 | 0.525491732 | 0.808531286 | ns | TM | KS.1 |
| FIM1_3388 | 2814.476399 | 2846.185635 | 3075.192619 | 0.003741055 | 0.130657367 | 0.952729489 | 0.988081969 | ns | TM | KS.1 |
| FIM1_3389 | 1818.096411 | 1446.037083 | 1286.693949 | -0.071199307 | 0.162911674 | 0.302820159 | 0.631432423 | ns | TM | KS.1 |
| FIM1_3390 | 2942.460276 | 3091.686115 | 3612.230617 | 0.029727205 | 0.089333268 | 0.597565275 | 0.849348181 | ns | TM | KS.1 |
| FIM1_3391 | 1410.469469 | 1230.575792 | 1527.696093 | -0.061012962 | 0.127609137 | 0.35979721 | 0.683664672 | ns | TM | KS.1 |
| FIM1_3392 | 1352.971245 | 1031.987352 | 997.793671 | -0.23661554 | 0.141466928 | 0.007470994 | 0.068702762 | ns | TM | KS.1 |
| FIM1_3393 | 331.2095249 | 120.4030421 | 168.986438 | -1.037811207 | 0.465144382 | 0.00015717 | 0.003538111 | Km<KS.1 | TM | KS.1 |
| FIM1_3394 | 2569.819418 | 2338.784554 | 2718.688271 | -0.043263346 | 0.117435089 | 0.498767522 | 0.793783884 | ns | TM | KS.1 |
| FIM1_3395 | 1879.934562 | 2353.050901 | 2191.632266 | 0.11137788 | 0.147758624 | 0.135161295 | 0.407284943 | ns | TM | KS.1 |
| FIM1_3396 | 1060.856713 | 998.0289035 | 1060.624303 | -0.022553298 | 0.127190898 | 0.722891656 | 0.907127048 | ns | TM | KS.1 |
| FIM1_3397 | 7979.519134 | 9093.604208 | 7705.802668 | 0.038269415 | 0.148151287 | 0.554839643 | 0.824224313 | ns | TM | KS.1 |
| FIM1_3398 | 8.043899655 | 9.31385775 | 8.902902147 | 0.023250352 | 0.195829451 | 0.703739584 | 0.898252804 | ns | TM | KS.1 |
| FIM1_3399 | 1165.092047 | 1068.304845 | 1046.559827 | -0.039865406 | 0.116328767 | 0.530674479 | 0.808531286 | ns | TM | KS.1 |
| FIM1_3400 | 1503.017184 | 1410.549523 | 1585.761323 | -0.043954254 | 0.081853241 | 0.412131527 | 0.732007978 | ns | TM | KS.1 |
| FIM1_3401 | 1397.430118 | 1147.280706 | 1416.926159 | -0.138496023 | 0.128638275 | 0.064813423 | 0.267310992 | ns | TM | KS.1 |
| FIM1_3402 | 3625.029967 | 3863.996793 | 3669.48146 | 0.053038931 | 0.068160901 | 0.262801347 | 0.589200856 | ns | TM | KS.1 |
| FIM1_3403 | 2498.335233 | 3775.569519 | 4961.124225 | 0.20567538 | 0.228085319 | 0.032599048 | 0.180382188 | ns | TM | KS.1 |
| FIM1_3404 | 43.57789315 | 18.82086092 | 107.336423 | -0.396083058 | 0.457129757 | 0.009377607 | 0.08029879 | ns | TM | KS.1 |
| FIM1_3405 | 6464.729824 | 8761.931574 | 10136.20419 | 0.256746372 | 0.157588279 | 0.006300215 | 0.061337713 | ns | TM | KS.1 |
| FIM1_3407 | 1451.102465 | 1426.216164 | 1399.676544 | -0.00894336 | 0.097533576 | 0.878794732 | 0.965013767 | ns | TM | KS.1 |
| FIM1_3408 | 1035.61842 | 1000.806402 | 1106.331544 | -0.022768389 | 0.077730552 | 0.659059247 | 0.878874681 | ns | TM | KS.1 |
| FIM1_3409 | 8250.532697 | 10220.58281 | 9335.100371 | 0.123920509 | 0.140475793 | 0.100140229 | 0.341699342 | ns | TM | KS.1 |
| FIM1_3410 | 1149.073554 | 1409.833611 | 1501.152017 | 0.088569408 | 0.145782628 | 0.213918969 | 0.521776083 | ns | TM | KS.1 |
| FIM1_3411 | 1525.607869 | 1805.509631 | 1892.610748 | 0.067911326 | 0.139240768 | 0.319486554 | 0.647364498 | ns | TM | KS.1 |
| FIM1_3412 | 1532.36959 | 1469.297659 | 1428.368822 | -0.030640548 | 0.069849375 | 0.523229874 | 0.807903931 | ns | TM | KS.1 |
| FIM1_3413 | 2534.774145 | 2668.356398 | 2766.208354 | 0.02324856 | 0.112145651 | 0.708318309 | 0.90101856 | ns | TM | KS.1 |
| FIM1_3414 | 3323.674135 | 3230.973489 | 2553.054635 | -0.013127766 | 0.108381316 | 0.830207807 | 0.946661434 | ns | TM | KS.1 |
| FIM1_3415 | 763.3337331 | 984.6491992 | 1188.797963 | 0.07063484 | 0.172756926 | 0.305791491 | 0.634144504 | ns | TM | KS.1 |
| FIM1_3416 | 429.5878144 | 449.6254674 | 460.4770577 | 0.015355183 | 0.130285249 | 0.808563556 | 0.938680028 | ns | TM | KS.1 |
| FIM1_3417 | 13038.35769 | 15583.20073 | 14646.90247 | 0.151548337 | 0.113274554 | 0.036253525 | 0.190639184 | ns | TM | KS.1 |
| FIM1_3418 | 2429.996156 | 2523.479516 | 2580.809561 | 0.023461556 | 0.085383239 | 0.668296087 | 0.882180579 | ns | TM | KS.1 |

| | | | | | | | | | | |
|---|---|---|---|---|---|---|---|---|---|---|
| FIM1_3419 | 2503.349778 | 2599.499526 | 3343.998621 | 0.027968663 | 0.066396361 | 0.54583211 | 0.818784976 | ns | TM | KS.1 |
| FIM1_3420 | 158.499401 | 74.79763779 | 103.0118483 | -0.456164632 | 0.434347752 | 0.005371065 | 0.054868685 | ns | TM | KS.1 |
| FIM1_3421 | 140.0963804 | 120.6369676 | 127.9908184 | -0.038640757 | 0.157267746 | 0.549443474 | 0.821605435 | ns | TM | KS.1 |
| FIM1_3422 | 88.30108577 | 69.83792042 | 142.3187291 | -0.053974566 | 0.177496355 | 0.414169579 | 0.734555234 | ns | TM | KS.1 |
| FIM1_3423 | 381.3684256 | 339.3262919 | 604.0889833 | -0.033983798 | 0.147478025 | 0.597658704 | 0.849348181 | ns | TM | KS.1 |
| FIM1_3424 | 735.2568201 | 766.2715414 | 844.8276442 | 0.016993444 | 0.117645846 | 0.786278999 | 0.931802479 | ns | TM | KS.1 |
| FIM1_3425 | 1040.033793 | 810.5939145 | 1240.169645 | -0.113772851 | 0.157715022 | 0.132601233 | 0.40323419 | ns | TM | KS.1 |
| FIM1_3426 | 1742.890632 | 1489.797408 | 4545.849937 | -0.031597262 | 0.173177433 | 0.616383127 | 0.859446765 | ns | TM | KS.1 |
| FIM1_3427 | 935.6426581 | 1059.53037 | 1391.919756 | 0.054047864 | 0.125782174 | 0.411389848 | 0.731735607 | ns | TM | KS.1 |
| FIM1_3428 | 40.89137434 | 54.98049714 | 84.78845287 | 0.094137507 | 0.179719768 | 0.19842434 | 0.505976981 | ns | TM | KS.1 |
| FIM1_3429 | 2836.583587 | 3092.959077 | 3127.203162 | 0.060240607 | 0.08982602 | 0.29307657 | 0.620392168 | ns | TM | KS.1 |
| FIM1_3430 | 1922.680119 | 1430.658403 | 1770.705665 | -0.337073754 | 0.109008686 | 7.51315558712 | 0.001916818 | ns | TM | KS.1 |
| FIM1_3431 | 137.6532154 | 172.7195331 | 368.9373469 | 0.121236726 | 0.148052029 | 0.110280446 | 0.360239803 | ns | TM | KS.1 |
| FIM1_3432 | 1020.582103 | 798.5650794 | 922.3058643 | -0.162634039 | 0.147384079 | 0.045185479 | 0.219529062 | ns | TM | KS.1 |
| FIM1_3433 | 3387.714984 | 3652.822829 | 3764.933484 | 0.064621057 | 0.07060732 | 0.185888831 | 0.488277156 | ns | TM | KS.1 |
| FIM1_3434 | 809.4318431 | 845.1749256 | 717.5488816 | 0.03143058 | 0.071749595 | 0.52046186 | 0.807137704 | ns | TM | KS.1 |
| FIM1_3435 | 2842.71176 | 2448.024636 | 3127.965965 | -0.082805169 | 0.12173205 | 0.222626054 | 0.537131242 | ns | TM | KS.1 |
| FIM1_3437 | 20712.25627 | 19920.82879 | 16824.47536 | -0.018165333 | 0.108826269 | 0.767395109 | 0.924883104 | ns | TM | KS.1 |
| FIM1_3438 | 2650.016138 | 2943.763726 | 2603.795388 | 0.046833491 | 0.121245348 | 0.469216852 | 0.77114073 | ns | TM | KS.1 |
| FIM1_3439 | 1740.788425 | 1964.711882 | 1906.752553 | 0.041725042 | 0.13801197 | 0.522592296 | 0.807872226 | ns | TM | KS.1 |
| FIM1_3440 | 2426.783721 | 2682.884025 | 3000.803043 | 0.060292148 | 0.103357751 | 0.330293336 | 0.657690769 | ns | TM | KS.1 |
| FIM1_3441 | 913.4524352 | 789.0011839 | 987.7667858 | -0.091094637 | 0.116035369 | 0.176864844 | 0.474852995 | ns | TM | KS.1 |
| FIM1_3442 | 4476.489452 | 4625.412673 | 5052.080606 | 0.01776198 | 0.096377315 | 0.761104581 | 0.92331024 | ns | TM | KS.1 |
| FIM1_3444 | 4089.004235 | 3946.128982 | 4131.371059 | -0.01995524 | 0.092961946 | 0.728103252 | 0.908701505 | ns | TM | KS.1 |
| FIM1_3445 | 3319.455517 | 3535.041901 | 3482.366193 | 0.0291815 | 0.112121131 | 0.639877475 | 0.870968656 | ns | TM | KS.1 |
| FIM1_3446 | 1217.947789 | 1408.214221 | 1375.250604 | 0.162094619 | 0.079683904 | 0.007173424 | 0.066581689 | ns | TM | KS.1 |
| FIM1_3447 | 2549.294362 | 2929.044515 | 2612.233532 | 0.109713626 | 0.102987654 | 0.092231223 | 0.328219124 | ns | TM | KS.1 |
| FIM1_3448 | 651.958038 | 692.1432158 | 773.1970386 | 0.042576657 | 0.076717446 | 0.406598116 | 0.728526463 | ns | TM | KS.1 |
| FIM1_3449 | 1175.190348 | 836.3610134 | 936.3167794 | -0.389758605 | 0.12711597 | 3.68463192184 | 0.001112321 | ns | TM | KS.1 |
| FIM1_3450 | 2323.016751 | 3889.765174 | 4650.317217 | 0.293509791 | 0.291326261 | 0.012891899 | 0.100686336 | ns | TM | KS.1 |
| FIM1_3452 | 1703.993752 | 1679.259802 | 1780.430393 | -0.008991254 | 0.082669906 | 0.86680155 | 0.960282219 | ns | TM | KS.1 |
| FIM1_3454 | 1803.655781 | 1357.663393 | 12413.85407 | -0.132660228 | 0.169550974 | 0.094635761 | 0.332494994 | ns | TM | KS.1 |
| FIM1_3455 | 41.99421376 | 51.21773333 | 1212.410285 | 0.054159887 | 0.161687054 | 0.41529752 | 0.735007172 | ns | TM | KS.1 |
| FIM1_3456 | 16.51909114 | 24.19979208 | 220.5758679 | 0.087074845 | 0.210954896 | 0.227259357 | 0.54408821 | ns | TM | KS.1 |
| FIM1_3457 | 324.8133324 | 464.5798365 | 898.4169684 | 0.121740011 | 0.199997506 | 0.12311877 | 0.38644535 | ns | TM | KS.1 |
| FIM1_3458 | 2780.915453 | 3993.29859 | 8765.855031 | 0.239126797 | 0.199396028 | 0.015189648 | 0.112119433 | ns | TM | KS.1 |
| FIM1_3460 | 4982.241098 | 5131.951305 | 6233.295187 | 0.016969497 | 0.090108824 | 0.763552935 | 0.924230468 | ns | TM | KS.1 |
| FIM1_3461 | 1249.518948 | 1507.155347 | 1201.563314 | 0.048361924 | 0.162985625 | 0.461142169 | 0.763455006 | ns | TM | KS.1 |
| FIM1_3462 | 802.3182414 | 877.1293652 | 799.4370233 | 0.064518005 | 0.088832411 | 0.258207284 | 0.587268814 | ns | TM | KS.1 |
| FIM1_3463 | 2056.284959 | 2383.023111 | 3452.321499 | 0.046213077 | 0.146807956 | 0.481913946 | 0.783685372 | ns | TM | KS.1 |
| FIM1_3464 | 271.6927119 | 279.8615218 | 332.051609 | 0.010826947 | 0.126255104 | 0.863821574 | 0.959694581 | ns | TM | KS.1 |
| FIM1_3465 | 295.0583456 | 191.252498 | 228.7605552 | -0.171557904 | 0.234286385 | 0.05770824 | 0.253323986 | ns | TM | KS.1 |
| FIM1_3466 | 2589.529418 | 2832.531494 | 2895.335907 | 0.082652576 | 0.073199727 | 0.104003312 | 0.34904241 | ns | TM | KS.1 |
| FIM1_3467 | 613.5208634 | 777.5456931 | 819.1413989 | 0.052953392 | 0.178276781 | 0.42176046 | 0.738117813 | ns | TM | KS.1 |
| FIM1_3468 | 2295.078406 | 2588.957638 | 2493.683692 | 0.069048288 | 0.111810078 | 0.285863903 | 0.612477571 | ns | TM | KS.1 |

| | | | | | | | | | | |
|---|---|---|---|---|---|---|---|---|---|---|
| FIM1_3469 | 1943.141872 | 2057.350965 | 2023.438997 | 0.017333221 | 0.139013912 | 0.784609049 | 0.930939666 | ns | TM | KS.1 |
| FIM1_3470 | 663.2175556 | 731.3595375 | 741.3694133 | 0.033919691 | 0.134261652 | 0.599130821 | 0.84979238 | ns | TM | KS.1 |
| FIM1_3471 | 2216.874798 | 2635.19983 | 2447.582828 | 0.135554444 | 0.115827157 | 0.058095493 | 0.253530772 | ns | TM | KS.1 |
| FIM1_3472 | 602.394897 | 677.8481304 | 647.6004953 | 0.075544517 | 0.10534713 | 0.23361511 | 0.551868552 | ns | TM | KS.1 |
| FIM1_3473 | 1794.978203 | 2785.499951 | 3333.603423 | 0.157311446 | 0.234007865 | 0.072343283 | 0.285641137 | ns | TM | KS.1 |
| FIM1_3474 | 308.9921268 | 336.3749132 | 705.3237644 | 0.021944936 | 0.151182935 | 0.728513546 | 0.908701505 | ns | TM | KS.1 |
| FIM1_3475 | 3392.206527 | 3157.374193 | 3795.78794 | -0.052549071 | 0.080542768 | 0.323219369 | 0.650941236 | ns | TM | KS.1 |
| FIM1_3476 | 1429.437661 | 1424.840461 | 1593.52545 | -0.002142428 | 0.07119022 | 0.964867385 | 0.993216478 | ns | TM | KS.1 |
| FIM1_3477 | 1837.66897 | 1651.295034 | 1971.95525 | -0.062406636 | 0.107103948 | 0.322562172 | 0.650748908 | ns | TM | KS.1 |
| FIM1_3478 | 264.2920075 | 289.8557799 | 204.791761 | 0.03757011 | 0.123431266 | 0.558809804 | 0.82614557 | ns | TM | KS.1 |
| FIM1_3479 | 8019.775453 | 8687.683666 | 7511.9524 | 0.084783682 | 0.057434632 | 0.047751845 | 0.227509809 | ns | TM | KS.1 |
| FIM1_3480 | 1768.224317 | 2785.56576 | 3057.765141 | 0.153366134 | 0.239110669 | 0.07787509 | 0.29733582 | ns | TM | KS.1 |
| FIM1_3481 | 2941.659279 | 3167.00975 | 3386.072532 | 0.059102566 | 0.074869267 | 0.244216159 | 0.567522712 | ns | TM | KS.1 |
| FIM1_3482 | 5636.126607 | 5084.83618 | 3183.553039 | -0.049837647 | 0.116244275 | 0.437711099 | 0.747832613 | ns | TM | KS.1 |
| FIM1_3483 | 1046.100906 | 962.2524653 | 933.8487455 | -0.062401723 | 0.084195584 | 0.257079739 | 0.585794503 | ns | TM | KS.1 |
| FIM1_3484 | 1764.795121 | 1597.075377 | 1506.444146 | -0.089142355 | 0.08016438 | 0.101798277 | 0.34470873 | ns | TM | KS.1 |
| FIM1_3485 | 554.9433575 | 577.5841919 | 508.4485735 | 0.021002004 | 0.102734367 | 0.72712261 | 0.907961357 | ns | TM | KS.1 |
| FIM1_3486 | 4460.713159 | 4015.000913 | 3483.811659 | -0.061231595 | 0.106453603 | 0.329895705 | 0.657690769 | ns | TM | KS.1 |
| FIM1_3487 | 934.9245805 | 1014.767215 | 881.4953144 | 0.037667826 | 0.115535486 | 0.552027237 | 0.822689033 | ns | TM | KS.1 |
| FIM1_3488 | 414.9220646 | 393.0970388 | 422.2680436 | -0.036431601 | 0.086076677 | 0.509237852 | 0.801014116 | ns | TM | KS.1 |
| FIM1_3489 | 619.7571851 | 657.3771042 | 767.3091327 | 0.038314778 | 0.080830593 | 0.47031194 | 0.771448038 | ns | TM | KS.1 |
| FIM1_3490 | 1010.617681 | 1129.56508 | 1105.079606 | 0.084848758 | 0.094626108 | 0.15970226 | 0.44660975 | ns | TM | KS.1 |
| FIM1_3491 | 2043.642767 | 1878.729592 | 2077.042796 | -0.053073554 | 0.095451265 | 0.369153755 | 0.69186398 | ns | TM | KS.1 |
| FIM1_3492 | 1051.83881 | 856.6190687 | 782.9234932 | -0.11079729 | 0.139426701 | 0.132234042 | 0.402363522 | ns | TM | KS.1 |
| FIM1_3493 | 105.2755025 | 123.712724 | 111.3256679 | 0.049565549 | 0.150036677 | 0.453189517 | 0.756837143 | ns | TM | KS.1 |
| FIM1_3494 | 7425.905761 | 9258.120238 | 9553.046138 | 0.228508125 | 0.107653313 | 0.003409354 | 0.038902605 | ns | TM | KS.1 |
| FIM1_3495 | 186.0556602 | 131.6977179 | 257.5342315 | -0.254881861 | 0.187325424 | 0.009995273 | 0.084466202 | ns | TM | KS.1 |
| FIM1_3496 | 455.9168287 | 567.5110078 | 619.1826244 | 0.127394465 | 0.142835756 | 0.094204569 | 0.331213943 | ns | TM | KS.1 |
| FIM1_3497 | 1814.11995 | 972.2686881 | 1782.456726 | -0.33322623 | 0.353732455 | 0.011447654 | 0.092454675 | ns | TM | KS.1 |
| FIM1_3498 | 150.7437562 | 180.5381149 | 150.7250461 | 0.067151307 | 0.143778309 | 0.325506006 | 0.652980799 | ns | TM | KS.1 |
| FIM1_3499 | 2776.576459 | 2887.071794 | 2687.064173 | 0.029458531 | 0.06559883 | 0.52106307 | 0.807566596 | ns | TM | KS.1 |
| FIM1_3500 | 1996.141023 | 1897.687964 | 1777.103672 | -0.022424949 | 0.113660951 | 0.719084913 | 0.905913973 | ns | TM | KS.1 |
| FIM1_3501 | 2527.603206 | 2240.241716 | 2906.741389 | -0.042933719 | 0.136275651 | 0.511213484 | 0.802123385 | ns | TM | KS.1 |
| FIM1_3503 | 328.3431464 | 381.6287274 | 375.0021449 | 0.028469009 | 0.176259056 | 0.64919338 | 0.874068749 | ns | TM | KS.1 |
| FIM1_3504 | 401.580685 | 480.7892128 | 551.5915906 | 0.052229941 | 0.155869065 | 0.431027188 | 0.743536592 | ns | TM | KS.1 |
| FIM1_3505 | 4522.849613 | 3390.182686 | 3680.734522 | -0.33066894 | 0.10596077 | 7.61058833816 | 0.001921963 | ns | TM | KS.1 |
| FIM1_3506 | 674.2744389 | 756.5106163 | 702.3294603 | 0.055490158 | 0.118918024 | 0.393687704 | 0.717171852 | ns | TM | KS.1 |
| FIM1_3507 | 2644.21417 | 1959.497911 | 1893.918144 | -0.349375741 | 0.105810336 | 3.17701540886 | 0.001019719 | ns | TM | KS.1 |
| FIM1_3508 | 768.3498295 | 932.923488 | 926.6437892 | 0.160805738 | 0.12034865 | 0.032834957 | 0.180951453 | ns | TM | KS.1 |
| FIM1_3509 | 14.78129666 | 10.11547815 | 75.09163503 | -0.080436722 | 0.203054851 | 0.25542578 | 0.583177263 | ns | TM | KS.1 |
| FIM1_3511 | 6314.978694 | 5254.442209 | 4938.838614 | -0.140320344 | 0.120579799 | 0.055573546 | 0.247076311 | ns | TM | KS.1 |
| FIM1_3512 | 4806.494636 | 4304.652135 | 4645.559879 | -0.052222605 | 0.119015226 | 0.420446154 | 0.737778527 | ns | TM | KS.1 |
| FIM1_3513 | 881.5482712 | 687.9950825 | 986.2109913 | -0.053636975 | 0.18174095 | 0.416006714 | 0.735216129 | ns | TM | KS.1 |
| FIM1_3514 | 3151.923727 | 2235.146338 | 2916.016212 | -0.172873806 | 0.19359855 | 0.048754056 | 0.230174256 | ns | TM | KS.1 |
| FIM1_3516 | 5302.485429 | 7012.913629 | 7410.048664 | 0.089957055 | 0.174841994 | 0.213847818 | 0.521776083 | ns | TM | KS.1 |

| | | | | | | | | | | |
|---|---|---|---|---|---|---|---|---|---|---|
| FIM1_3517 | 744.8898698 | 521.7478812 | 512.655349 | -0.408035147 | 0.135052545 | 3.3512048868[illegible] | 0.001048569 | ns | TM | KS.1 |
| FIM1_3518 | 1626.370011 | 2625.84708 | 3047.35086 | 0.158482656 | 0.24918187 | 0.073422704 | 0.288301464 | ns | TM | KS.1 |
| FIM1_3519 | 665.1876942 | 858.4854357 | 757.2062454 | 0.103587907 | 0.161954376 | 0.163274386 | 0.45366779 | ns | TM | KS.1 |
| FIM1_3520 | 1230.348078 | 1093.857673 | 1141.632281 | -0.042348789 | 0.135357153 | 0.516508923 | 0.804981347 | ns | TM | KS.1 |
| FIM1_3522 | 1723.221011 | 2931.667361 | 3616.222836 | 0.20021333 | 0.279800459 | 0.043150301 | 0.212459177 | ns | TM | KS.1 |
| FIM1_3523 | 5161.038443 | 6742.636688 | 7519.610076 | 0.169493381 | 0.157059095 | 0.042267439 | 0.210079667 | ns | TM | KS.1 |
| FIM1_3524 | 1603.522349 | 1723.833967 | 1888.769054 | 0.036871374 | 0.107442552 | 0.550788811 | 0.822058746 | ns | TM | KS.1 |
| FIM1_3525 | 529.8380768 | 524.9282458 | 478.5321531 | -0.003016504 | 0.128407468 | 0.961851609 | 0.991548229 | ns | TM | KS.1 |
| FIM1_3526 | 3570.972352 | 2717.100665 | 3162.86158 | -0.321385703 | 0.096306375 | 4.4693038894[illegible] | 0.001277861 | ns | TM | KS.1 |
| FIM1_3527 | 2312.604183 | 1881.636022 | 1778.7921 | -0.085776958 | 0.147427895 | 0.22677227 | 0.54408821 | ns | TM | KS.1 |
| FIM1_3528 | 1671.891815 | 1727.611459 | 3188.141061 | 0.008299254 | 0.149375271 | 0.894585674 | 0.967972723 | ns | TM | KS.1 |
| FIM1_3529 | 698.4926754 | 658.4024227 | 616.9667557 | -0.025519909 | 0.116244367 | 0.684584594 | 0.890505759 | ns | TM | KS.1 |
| FIM1_3530 | 374.4752412 | 452.4952168 | 406.0561677 | 0.099590589 | 0.135920736 | 0.165464144 | 0.456057682 | ns | TM | KS.1 |
| FIM1_3531 | 1399.135217 | 1302.485486 | 1385.990734 | -0.057662385 | 0.073458614 | 0.249247841 | 0.574965989 | ns | TM | KS.1 |
| FIM1_3532 | 898.0796974 | 970.7529613 | 977.3404401 | 0.044794253 | 0.099321885 | 0.455514069 | 0.759118744 | ns | TM | KS.1 |
| FIM1_3533 | 401.7163738 | 398.59719 | 652.0218671 | -0.004226659 | 0.097477312 | 0.942501683 | 0.984319543 | ns | TM | KS.1 |
| FIM1_3534 | 2515.021411 | 2646.087171 | 2703.360744 | 0.039256602 | 0.067536049 | 0.402549453 | 0.724768814 | ns | TM | KS.1 |
| FIM1_3535 | 1561.101942 | 1617.048962 | 1527.632879 | 0.028582018 | 0.060862405 | 0.511107625 | 0.802123385 | ns | TM | KS.1 |
| FIM1_3536 | 481.653663 | 482.6860592 | 460.5971985 | 0.001360508 | 0.08962287 | 0.980671975 | 0.995431301 | ns | TM | KS.1 |
| FIM1_3537 | 4916.647387 | 7788.667917 | 8084.007475 | 0.24105618 | 0.255392892 | 0.02169602 | 0.143551826 | ns | TM | KS.1 |
| FIM1_3538 | 1015.653313 | 988.4108386 | 903.5957502 | -0.016634676 | 0.08236697 | 0.755932999 | 0.920354827 | ns | TM | KS.1 |
| FIM1_3539 | 630.3191206 | 620.9689031 | 579.1258978 | -0.008321286 | 0.09145274 | 0.883511909 | 0.966744674 | ns | TM | KS.1 |
| FIM1_3540 | 3316.757083 | 2848.251906 | 3361.474688 | -0.147709097 | 0.095411005 | 0.024617301 | 0.154245685 | ns | TM | KS.1 |
| FIM1_3541 | 234.2839462 | 240.8692959 | 306.9328249 | 0.009045221 | 0.126606784 | 0.886014978 | 0.967028314 | ns | TM | KS.1 |
| FIM1_3542 | 399.4029616 | 378.9571517 | 450.4154753 | -0.034112351 | 0.087481258 | 0.540252574 | 0.814399145 | ns | TM | KS.1 |
| FIM1_3544 | 2561.180295 | 2874.201017 | 2705.31606 | 0.06087351 | 0.114375108 | 0.346502028 | 0.671956953 | ns | TM | KS.1 |
| FIM1_3545 | 85.35426751 | 36.47512199 | 154.6222435 | -0.826871046 | 0.449495018 | 0.000445145 | 0.008202204 | ns | TM | KS.1 |
| FIM1_3546 | 1171.153553 | 1156.757554 | 1688.710571 | -0.00614303 | 0.099523494 | 0.917300297 | 0.975634767 | ns | TM | KS.1 |
| FIM1_3547 | 1248.985697 | 1287.120945 | 926.7482602 | 0.017634628 | 0.086734767 | 0.749420674 | 0.917865055 | ns | TM | KS.1 |
| FIM1_3548 | 222.3740768 | 444.9374237 | 437.2404573 | 0.510223327 | 0.406293577 | 0.002899803 | 0.03410525 | ns | TM | KS.1 |
| FIM1_3549 | 156.5466051 | 145.107845 | 171.1134703 | -0.035704016 | 0.114426584 | 0.571251816 | 0.832094673 | ns | TM | KS.1 |
| FIM1_3550 | 2874.681354 | 1767.085862 | 1459.170544 | -0.402740163 | 0.268770134 | 0.002353552 | 0.029199301 | ns | TM | KS.1 |
| FIM1_3551 | 1700.098909 | 1788.318264 | 1689.147242 | 0.022202571 | 0.114491465 | 0.722228229 | 0.907011644 | ns | TM | KS.1 |
| FIM1_3552 | 729.9815899 | 637.2925225 | 580.9714058 | -0.083558395 | 0.113576217 | 0.20708508 | 0.513839538 | ns | TM | KS.1 |
| FIM1_3553 | 443.6170303 | 470.9699744 | 438.2540512 | 0.021389064 | 0.128427871 | 0.736608668 | 0.912734277 | ns | TM | KS.1 |
| FIM1_3554 | 728.038323 | 780.8699084 | 602.7025135 | 0.037510513 | 0.10308724 | 0.537226183 | 0.81337601 | ns | TM | KS.1 |
| FIM1_3555 | 609.2600711 | 633.9784698 | 575.0273797 | 0.021430552 | 0.098245979 | 0.71653851 | 0.905084971 | ns | TM | KS.1 |
| FIM1_3556 | 1949.677624 | 1983.452852 | 2232.991527 | 0.011189202 | 0.075132877 | 0.824433629 | 0.94482203 | ns | TM | KS.1 |
| FIM1_3557 | 3114.064346 | 3942.824592 | 4774.948837 | 0.111825672 | 0.152699352 | 0.136028681 | 0.407922054 | ns | TM | KS.1 |
| FIM1_3558 | 1069.939443 | 1144.98579 | 1198.38517 | 0.038240192 | 0.099128755 | 0.522038842 | 0.807870838 | ns | TM | KS.1 |
| FIM1_3559 | 278.4537458 | 213.171348 | 243.8344568 | -0.182942475 | 0.153514999 | 0.030732209 | 0.174934485 | ns | TM | KS.1 |
| FIM1_3560 | 1370.3374 | 1120.281974 | 1344.499968 | -0.189981839 | 0.112983974 | 0.012610463 | 0.098954343 | ns | TM | KS.1 |
| FIM1_3561 | 794.1367071 | 600.5259252 | 872.0920139 | -0.140652788 | 0.166698901 | 0.080212574 | 0.301630805 | ns | TM | KS.1 |
| FIM1_3562 | 1196.862431 | 1172.730135 | 1735.131374 | -0.013229099 | 0.077930896 | 0.797900549 | 0.936276987 | ns | TM | KS.1 |
| FIM1_3563 | 1692.125097 | 1755.992368 | 1925.341098 | 0.023799561 | 0.081696024 | 0.655156492 | 0.87699044 | ns | TM | KS.1 |

| | | | | | | | | | | |
|---|---|---|---|---|---|---|---|---|---|---|
| FIM1_3565 | 73.52107804 | 76.82643562 | 110.8741976 | 0.011850133 | 0.153819569 | 0.849642134 | 0.95298643 | ns | TM | KS.1 |
| FIM1_3566 | 201.0926093 | 177.2312618 | 242.8917698 | -0.04844648 | 0.131622285 | 0.460998276 | 0.763455006 | ns | TM | KS.1 |
| FIM1_3567 | 5691.813774 | 7344.497437 | 7172.635201 | 0.178466471 | 0.14897253 | 0.032365485 | 0.18019422 | ns | TM | KS.1 |
| FIM1_3568 | 4560.909324 | 5314.572005 | 4147.484486 | 0.064881193 | 0.133538491 | 0.336652152 | 0.66488466 | ns | TM | KS.1 |
| FIM1_3569 | 1121.430919 | 1680.436833 | 2003.273747 | 0.102356992 | 0.216434221 | 0.17477065 | 0.471265031 | ns | TM | KS.1 |
| FIM1_3570 | 1278.453649 | 1475.143451 | 1301.032634 | 0.058878664 | 0.132884532 | 0.378351014 | 0.702681796 | ns | TM | KS.1 |
| FIM1_3571 | 4183.87973 | 3915.523867 | 4121.434755 | -0.048048166 | 0.079093722 | 0.359711909 | 0.683664672 | ns | TM | KS.1 |
| FIM1_3572 | 554.8578054 | 568.120469 | 519.58149 | 0.014030089 | 0.087887098 | 0.800790733 | 0.936908579 | ns | TM | KS.1 |
| FIM1_3573 | 3867.975916 | 4308.693225 | 4375.909811 | 0.087364726 | 0.089421134 | 0.134816038 | 0.406737289 | ns | TM | KS.1 |
| FIM1_3574 | 2161.700332 | 1689.084839 | 3146.18648 | -0.060298264 | 0.17622573 | 0.369364971 | 0.69186398 | ns | TM | KS.1 |
| FIM1_3575 | 2104.036457 | 957.4642253 | 1718.833637 | -0.99805744 | 0.268110784 | 4.60761212992 | 0.000225818 | ns | TM | KS.1 |
| FIM1_3576 | 4216.628702 | 3586.924247 | 3522.498511 | -0.126448794 | 0.111860659 | 0.068848574 | 0.276450087 | ns | TM | KS.1 |
| FIM1_3577 | 619.7558711 | 609.7025805 | 487.2915457 | -0.00888189 | 0.092659752 | 0.876576449 | 0.964248917 | ns | TM | KS.1 |
| FIM1_3579 | 2186.112294 | 2154.023117 | 2293.963276 | -0.006720929 | 0.109881005 | 0.912827055 | 0.973904053 | ns | TM | KS.1 |
| FIM1_3580 | 2796.526309 | 4503.56594 | 5248.873732 | 0.215429253 | 0.259944076 | 0.03238063 | 0.18019422 | ns | TM | KS.1 |
| FIM1_3581 | 4748.065146 | 6961.456618 | 4370.134313 | 0.228434433 | 0.212278121 | 0.020314905 | 0.137880836 | ns | TM | KS.1 |
| FIM1_3582 | 1918.935521 | 1613.094433 | 2298.735857 | -0.174574191 | 0.098533357 | 0.012095015 | 0.096768242 | ns | TM | KS.1 |
| FIM1_3583 | 2389.872369 | 1834.846068 | 1973.623488 | -0.108133456 | 0.164935169 | 0.150262495 | 0.431365212 | ns | TM | KS.1 |
| FIM1_3584 | 744.2115626 | 741.7097976 | 589.4158153 | -0.002111533 | 0.103123547 | 0.971936994 | 0.99362472 | ns | TM | KS.1 |
| FIM1_3585 | 1410.777699 | 1353.652369 | 1187.196258 | -0.01815918 | 0.113785605 | 0.770415253 | 0.925705893 | ns | TM | KS.1 |
| FIM1_3586 | 2179.201562 | 2718.80762 | 2909.673499 | 0.119986459 | 0.144476452 | 0.111377967 | 0.363109689 | ns | TM | KS.1 |
| FIM1_3587 | 63.3745175 | 62.84407198 | 85.38572007 | -0.000381881 | 0.149097187 | 0.995127468 | 0.99790453 | ns | TM | KS.1 |
| FIM1_3588 | 984.422243 | 1061.831503 | 1024.528335 | 0.023743693 | 0.138154557 | 0.709755297 | 0.901619224 | ns | TM | KS.1 |
| FIM1_3589 | 2667.903814 | 2478.226623 | 3873.387259 | -0.031836319 | 0.118353081 | 0.615508705 | 0.858708863 | ns | TM | KS.1 |
| FIM1_3591 | 1091.652271 | 556.7477885 | 629.3839924 | -0.701437847 | 0.348617659 | 0.00032035 | 0.006374966 | ns | TM | KS.1 |
| FIM1_3592 | 3655.782186 | 2928.510206 | 1972.23227 | -0.063109393 | 0.164783655 | 0.351944752 | 0.67786494 | ns | TM | KS.1 |
| FIM1_3594 | 2574.437054 | 2993.946919 | 3172.010947 | 0.121940957 | 0.106398724 | 0.070253897 | 0.280283191 | ns | TM | KS.1 |
| FIM1_3595 | 6156.813378 | 6373.863922 | 5378.255139 | 0.018068344 | 0.099399976 | 0.760502999 | 0.923255837 | ns | TM | KS.1 |
| FIM1_3596 | 34.99104663 | 33.55205962 | 37.01878554 | -0.008233402 | 0.165678077 | 0.893812732 | 0.967972723 | ns | TM | KS.1 |
| FIM1_3597 | 11535.83776 | 12032.49997 | 10687.94615 | 0.027919536 | 0.079366271 | 0.5937343 | 0.846854399 | ns | TM | KS.1 |
| FIM1_3598 | 1367.286152 | 1969.781149 | 2566.044627 | 0.214909355 | 0.203123803 | 0.024183522 | 0.152487988 | ns | TM | KS.1 |
| FIM1_3599 | 3943.949533 | 5227.69276 | 6212.493068 | 0.299121522 | 0.119701351 | 0.000634352 | 0.010704111 | ns | TM | KS.1 |
| FIM1_3600 | 421.6745982 | 479.8956296 | 386.4645009 | 0.062205741 | 0.123144603 | 0.347539556 | 0.673290223 | ns | TM | KS.1 |
| FIM1_3601 | 1979.654162 | 2094.272687 | 2242.782252 | 0.032638767 | 0.093586153 | 0.572573918 | 0.832094673 | ns | TM | KS.1 |
| FIM1_3603 | 693.4221921 | 655.141179 | 629.563039 | -0.035675263 | 0.089337157 | 0.527201292 | 0.808531286 | ns | TM | KS.1 |
| FIM1_3604 | 2491.440598 | 3482.14707 | 3903.096472 | 0.267310459 | 0.176523841 | 0.006770394 | 0.064774438 | ns | TM | KS.1 |
| FIM1_3605 | 1346.283298 | 1295.819851 | 1340.176429 | -0.031870564 | 0.058051263 | 0.448241332 | 0.753124157 | ns | TM | KS.1 |
| FIM1_3606 | 657.6463939 | 712.0898964 | 573.6389977 | 0.033143054 | 0.12096034 | 0.603137918 | 0.851547247 | ns | TM | KS.1 |
| FIM1_3607 | 1252.88689 | 1365.775241 | 1506.810348 | 0.053983168 | 0.097335808 | 0.366408217 | 0.690301888 | ns | TM | KS.1 |
| FIM1_3608 | 401.7655934 | 532.4198405 | 542.6673427 | 0.238664264 | 0.148297672 | 0.008115098 | 0.072612608 | ns | TM | KS.1 |
| FIM1_3609 | 2279.867777 | 1028.59249 | 1486.044669 | -0.963726877 | 0.311375653 | 2.38402522833 | 0.000829407 | ns | TM | KS.1 |
| FIM1_3610 | 1938.624863 | 2015.913223 | 1668.484706 | 0.024716182 | 0.082290312 | 0.644336916 | 0.872978256 | ns | TM | KS.1 |
| FIM1_3611 | 736.724129 | 860.3540718 | 868.238066 | 0.143572566 | 0.098685094 | 0.031105352 | 0.176452822 | ns | TM | KS.1 |
| FIM1_3612 | 1629.480529 | 1516.996152 | 1620.530005 | -0.053713173 | 0.080159721 | 0.311314798 | 0.638961913 | ns | TM | KS.1 |
| FIM1_3613 | 1272.114501 | 1422.307379 | 1052.78488 | 0.048058578 | 0.124197652 | 0.460565202 | 0.763054265 | ns | TM | KS.1 |

| | | | | | | | | | | |
|---|---|---|---|---|---|---|---|---|---|---|
| FIM1_3614 | 341.03187 | 433.1220466 | 312.8522437 | 0.117630885 | 0.152403108 | 0.120802981 | 0.381584021 | ns | TM | KS.1 |
| FIM1_3615 | 973.7151465 | 1293.061381 | 1455.642772 | 0.181545014 | 0.163576807 | 0.034599802 | 0.185689335 | ns | TM | KS.1 |
| FIM1_3616 | 2411.376585 | 3174.571752 | 3554.016383 | 0.115922518 | 0.167966288 | 0.130083188 | 0.398745447 | ns | TM | KS.1 |
| FIM1_3617 | 170.679991 | 223.517659 | 224.0788788 | 0.124540625 | 0.164501639 | 0.109198309 | 0.35800987 | ns | TM | KS.1 |
| FIM1_3618 | 3135.403007 | 3083.688528 | 2714.576116 | -0.01137973 | 0.070879411 | 0.814454868 | 0.940696014 | ns | TM | KS.1 |
| FIM1_3619 | 437.2104851 | 363.2592355 | 359.8464444 | -0.092404156 | 0.135020891 | 0.192119543 | 0.496260756 | ns | TM | KS.1 |
| FIM1_3620 | 1553.466781 | 1333.102344 | 1503.26998 | -0.10363593 | 0.114867338 | 0.128983555 | 0.396596531 | ns | TM | KS.1 |
| FIM1_3621 | 1703.13276 | 1639.006843 | 1545.844025 | -0.02147734 | 0.093112432 | 0.708625978 | 0.901179509 | ns | TM | KS.1 |
| FIM1_3622 | 2254.93545 | 2334.035169 | 2433.684599 | 0.025793479 | 0.065839603 | 0.575225587 | 0.833922256 | ns | TM | KS.1 |
| FIM1_3623 | 286.2234983 | 158.2390945 | 572.7992675 | -0.21707371 | 0.30822325 | 0.037337757 | 0.194304751 | ns | TM | KS.1 |
| FIM1_3624 | 700.1480457 | 413.5844821 | 462.092434 | -0.215567163 | 0.282051821 | 0.035169771 | 0.187333629 | ns | TM | KS.1 |
| FIM1_3625 | 13695.96561 | 18788.457 | 18717.45703 | 0.423746848 | 0.067700778 | 2.38662009176 | 1.48417936956 | ns | TM | KS.1 |
| FIM1_3626 | 332.7156073 | 250.2953964 | 201.9656237 | -0.247113406 | 0.148019449 | 0.006606428 | 0.063819379 | ns | TM | KS.1 |
| FIM1_3627 | 672.6826426 | 484.6194251 | 1302.001326 | -0.197269876 | 0.184874216 | 0.02958198 | 0.170502907 | ns | TM | KS.1 |
| FIM1_3628 | 8515.914953 | 11867.17137 | 11043.21588 | 0.102697219 | 0.191511167 | 0.171111885 | 0.465692069 | ns | TM | KS.1 |
| FIM1_3629 | 4426.506921 | 4883.129009 | 4731.292686 | 0.042493534 | 0.121768192 | 0.509681226 | 0.801157694 | ns | TM | KS.1 |
| FIM1_3630 | 200.1112141 | 156.4301105 | 197.6624048 | -0.11057488 | 0.15738581 | 0.141169575 | 0.416559096 | ns | TM | KS.1 |
| FIM1_3631 | 582.9355555 | 556.263983 | 557.9884718 | -0.026806735 | 0.088524496 | 0.631990497 | 0.869511262 | ns | TM | KS.1 |
| FIM1_3632 | 8044.421819 | 9068.694632 | 8567.738859 | 0.097702094 | 0.094037513 | 0.108927579 | 0.357935736 | ns | TM | KS.1 |
| FIM1_3633 | 3821.251502 | 1387.593507 | 2506.69561 | -1.231382807 | 0.32157734 | 4.06065976080 | 0.000208266 | Km<KS.1 | TM | KS.1 |
| FIM1_3634 | 5975.915299 | 3042.162249 | 3704.855705 | -0.685782372 | 0.357170335 | 0.000410203 | 0.007660186 | ns | TM | KS.1 |
| FIM1_3635 | 7682.765481 | 3405.339664 | 4861.765801 | -0.986557804 | 0.313456762 | 2.14315510213 | 0.000750859 | ns | TM | KS.1 |
| FIM1_3636 | 4804.540349 | 1801.164946 | 2761.731935 | -1.20956138 | 0.306207414 | 2.91212169540 | 0.000160976 | Km<KS.1 | TM | KS.1 |
| FIM1_3637 | 6032.147384 | 2319.385068 | 3180.97478 | -1.174307997 | 0.310217356 | 4.62983779042 | 0.000225818 | Km<KS.1 | TM | KS.1 |
| FIM1_3638 | 11881.46921 | 6048.785049 | 7351.407994 | -0.578951177 | 0.387555632 | 0.001391548 | 0.019268155 | ns | TM | KS.1 |
| FIM1_3639 | 643.2928507 | 277.685188 | 401.9788217 | -0.776644927 | 0.455922589 | 0.000634716 | 0.010704111 | ns | TM | KS.1 |
| FIM1_3640 | 1877.462684 | 768.8360324 | 1134.694664 | -1.039926897 | 0.358511648 | 3.96531451439 | 0.001167304 | Km<KS.1 | TM | KS.1 |
| FIM1_3641 | 1318.664056 | 658.5107162 | 956.6364595 | -0.704568853 | 0.367294578 | 0.000407958 | 0.007658836 | ns | TM | KS.1 |
| FIM1_3642 | 3839.584485 | 1936.743097 | 2394.135994 | -0.682964967 | 0.366551537 | 0.000476065 | 0.008643883 | ns | TM | KS.1 |
| FIM1_3643 | 2512.510276 | 953.4758796 | 1267.188752 | -1.147835616 | 0.348946504 | 1.61565048287 | 0.000595397 | Km<KS.1 | TM | KS.1 |
| FIM1_3644 | 5052.134856 | 1828.080807 | 2834.249379 | -1.298899233 | 0.263542337 | 8.51855159391 | 9.41773203993 | Km<KS.1 | TM | KS.1 |
| FIM1_3645 | 5498.207989 | 2068.259475 | 3118.831835 | -1.230290058 | 0.282498524 | 8.14952098599 | 6.11475692855 | Km<KS.1 | TM | KS.1 |
| FIM1_3646 | 2060.933951 | 916.0844593 | 1263.162388 | -0.931351067 | 0.35415068 | 7.21908384862 | 0.001860878 | ns | TM | KS.1 |
| FIM1_3647 | 31561.3314 | 24302.93819 | 25119.05881 | -0.12791466 | 0.160513615 | 0.100923925 | 0.343604292 | ns | TM | KS.1 |
| FIM1_3648 | 2404.083222 | 1599.930463 | 2447.029702 | -0.092105677 | 0.217793568 | 0.208408118 | 0.515579507 | ns | TM | KS.1 |
| FIM1_3649 | 2540.364152 | 1138.606204 | 1619.634937 | -0.885774889 | 0.375696136 | 0.000135084 | 0.003125782 | ns | TM | KS.1 |
| FIM1_3650 | 2593.81589 | 967.8321352 | 1797.310892 | -1.221069502 | 0.300885271 | 2.11337041598 | 0.000129026 | Km<KS.1 | TM | KS.1 |
| FIM1_3651 | 2227.774012 | 944.1302234 | 1422.069128 | -1.000971918 | 0.3528197 | 4.55935888577 | 0.001292076 | Km<KS.1 | TM | KS.1 |
| FIM1_3652 | 818.9402936 | 197.7959225 | 400.6113962 | -1.587449209 | 0.394017683 | 1.14480411107 | 7.78793740239 | Km<KS.1 | TM | KS.1 |
| FIM1_3653 | 7321.395751 | 3900.518491 | 5782.597739 | -0.509951728 | 0.365076969 | 0.002063369 | 0.026593944 | ns | TM | KS.1 |
| FIM1_3654 | 13310.68163 | 6342.996581 | 7828.351318 | -0.915976071 | 0.284379897 | 1.58775576634 | 0.000589484 | ns | TM | KS.1 |
| FIM1_3655 | 8275.642996 | 3608.199731 | 4492.996544 | -1.03542015 | 0.288981431 | 7.28516589119 | 0.00032652 | Km<KS.1 | TM | KS.1 |
| FIM1_3656 | 9237.994161 | 4506.854281 | 5621.544174 | -0.787256972 | 0.349486344 | 0.000172071 | 0.003838806 | ns | TM | KS.1 |
| FIM1_3657 | 1210.992589 | 436.1316695 | 773.3438998 | -1.226636641 | 0.3348049 | 6.30018740058 | 0.000290217 | Km<KS.1 | TM | KS.1 |
| FIM1_3658 | 2659.254171 | 1469.253717 | 2222.721408 | -0.270511383 | 0.325063147 | 0.02047876 | 0.138614734 | ns | TM | KS.1 |

| | | | | | | | | | | |
|---|---|---|---|---|---|---|---|---|---|---|
| FIM1_3659 | 3282.576127 | 1353.838815 | 1990.138433 | -1.047117308 | 0.344900073 | 2.91311272243 | 0.000953469 | Km<KS.1 | TM | KS.1 |
| FIM1_3660 | 2223.077144 | 970.1927949 | 1551.995642 | -0.900421217 | 0.392075953 | 0.000154953 | 0.003504058 | ns | TM | KS.1 |
| FIM1_3661 | 5726.622764 | 1668.039645 | 2798.114036 | -1.537285598 | 0.28904901 | 7.85080889169 | 1.18356891624 | Km<KS.1 | TM | KS.1 |
| FIM1_3662 | 2475.907006 | 877.5644232 | 1358.714328 | -1.28799696 | 0.296537283 | 8.23494902941 | 6.11475692855 | Km<KS.1 | TM | KS.1 |
| FIM1_3663 | 6363.198445 | 3090.159019 | 4056.103488 | -0.402961991 | 0.412672084 | 0.007653464 | 0.069864193 | ns | TM | KS.1 |
| FIM1_3664 | 2332.941849 | 977.5688314 | 1737.776234 | -0.974815009 | 0.383263009 | 8.80734930645 | 0.002159213 | ns | TM | KS.1 |
| FIM1_3665 | 2389.954234 | 931.7478005 | 1234.791245 | -1.011857969 | 0.424349212 | 0.000120594 | 0.002829986 | Km<KS.1 | TM | KS.1 |
| FIM1_3666 | 2435.570314 | 1069.464995 | 1461.323656 | -0.844066096 | 0.416462292 | 0.000291219 | 0.00591353 | ns | TM | KS.1 |
| FIM1_3667 | 3822.368346 | 1704.908915 | 2133.783649 | -0.951284942 | 0.335352244 | 4.46071155131 | 0.001277861 | ns | TM | KS.1 |
| FIM1_3668 | 2815.524925 | 1042.188039 | 1477.657812 | -1.230349361 | 0.301329068 | 1.95611570093 | 0.000121646 | Km<KS.1 | TM | KS.1 |
| FIM1_3669 | 2057.769323 | 770.1322783 | 1245.2853 | -1.138653332 | 0.370902889 | 2.65724378587 | 0.000913187 | Km<KS.1 | TM | KS.1 |
| FIM1_3670 | 1508.667013 | 560.4052454 | 785.9254473 | -1.184661419 | 0.339459303 | 9.91236781808 | 0.000414015 | Km<KS.1 | TM | KS.1 |
| FIM1_3671 | 1195.186354 | 409.7354865 | 649.025862 | -1.197931858 | 0.406376287 | 3.24882046175 | 0.00102915 | Km<KS.1 | TM | KS.1 |
| FIM1_3672 | 2976.529356 | 1269.882513 | 2288.424868 | -1.004156492 | 0.342854144 | 3.68919205459 | 0.001112321 | Km<KS.1 | TM | KS.1 |
| FIM1_3673 | 2706.93904 | 1521.187233 | 2448.216564 | -0.259321177 | 0.314759587 | 0.022503295 | 0.146759546 | ns | TM | KS.1 |
| FIM1_3674 | 109.1297372 | 32.77260702 | 71.71387351 | -1.366049661 | 0.390023263 | 7.77216622809 | 0.000342182 | Km<KS.1 | TM | KS.1 |
| FIM1_3675 | 7892.112591 | 3444.112206 | 4505.666315 | -0.992885252 | 0.326472475 | 2.80219699050 | 0.000929395 | ns | TM | KS.1 |
| FIM1_3676 | 211.4178388 | 70.50578783 | 187.0670644 | -1.317633825 | 0.335109023 | 2.84550004217 | 0.000160868 | Km<KS.1 | TM | KS.1 |
| FIM1_3677 | 2686.350793 | 1154.450495 | 1500.927836 | -0.996870099 | 0.340821414 | 3.71146279684 | 0.001112321 | ns | TM | KS.1 |
| FIM1_3678 | 3454.788807 | 1974.968208 | 2981.63063 | -0.150723431 | 0.273305948 | 0.08545362 | 0.311680174 | ns | TM | KS.1 |
| FIM1_3679 | 75571.47751 | 49057.13188 | 61159.05026 | -0.311626019 | 0.238629715 | 0.006430726 | 0.062364252 | ns | TM | KS.1 |
| FIM1_3680 | 164.5227904 | 106.8777095 | 178.89611 | -0.164533239 | 0.233039473 | 0.064314039 | 0.266080163 | ns | TM | KS.1 |
| FIM1_3681 | 2063.939381 | 816.9013845 | 1295.64304 | -1.164465894 | 0.28512382 | 1.94249069304 | 0.000121646 | Km<KS.1 | TM | KS.1 |
| FIM1_3682 | 3956.007194 | 1405.506148 | 3493.016974 | -1.258026748 | 0.320973636 | 3.11574966052 | 0.000168488 | Km<KS.1 | TM | KS.1 |
| FIM1_3683 | 2278.543911 | 1128.692873 | 1390.773785 | -0.756956392 | 0.350647505 | 0.000217621 | 0.004686868 | ns | TM | KS.1 |
| FIM1_3684 | 1039.524792 | 410.1661778 | 675.516524 | -1.135188847 | 0.317381962 | 8.00313477448 | 0.000346223 | Km<KS.1 | TM | KS.1 |
| FIM1_3685 | 1799.139632 | 676.4545408 | 967.9350028 | -1.131425559 | 0.372221558 | 2.84948061216 | 0.000938819 | Km<KS.1 | TM | KS.1 |
| FIM1_3686 | 1361.179551 | 816.7421071 | 1007.071036 | -0.142956715 | 0.256187585 | 0.094007317 | 0.331213943 | ns | TM | KS.1 |
| FIM1_3687 | 1399.523428 | 538.2258664 | 709.5175891 | -1.189239219 | 0.296155986 | 2.40536225057 | 0.000140799 | Km<KS.1 | TM | KS.1 |
| FIM1_3688 | 2870.913758 | 1028.074816 | 1342.19789 | -1.160419424 | 0.395715915 | 3.49521949174 | 0.001086795 | Km<KS.1 | TM | KS.1 |
| FIM1_3689 | 2791.254316 | 1286.44548 | 1805.398193 | -0.896619894 | 0.339978532 | 6.97032498475 | 0.001806113 | ns | TM | KS.1 |
| FIM1_3690 | 2074.120545 | 750.7330972 | 1028.481125 | -1.272430847 | 0.287612283 | 6.30415618469 | 5.31579271505 | Km<KS.1 | TM | KS.1 |
| FIM1_3691 | 6098.984683 | 2749.896117 | 3698.699931 | -0.970402322 | 0.306771544 | 2.02030802307 | 0.000723096 | ns | TM | KS.1 |
| FIM1_3692 | 3747.961729 | 1607.134311 | 2354.507026 | -1.007304327 | 0.334621654 | 3.04775871113 | 0.000984584 | Km<KS.1 | TM | KS.1 |
| FIM1_3693 | 3763.331109 | 1509.819117 | 2060.419245 | -1.112649262 | 0.318501423 | 9.83876124867 | 0.000414015 | Km<KS.1 | TM | KS.1 |
| FIM1_3694 | 1958.607806 | 775.971625 | 1198.114767 | -1.148687498 | 0.299575299 | 4.01973778404 | 0.000208266 | Km<KS.1 | TM | KS.1 |
| FIM1_3695 | 5861.064082 | 2838.625565 | 5682.653673 | -0.700698525 | 0.394383974 | 0.000583628 | 0.0101169 | ns | TM | KS.1 |
| FIM1_3696 | 8224.032419 | 2738.070687 | 4593.752869 | -1.38477952 | 0.279074454 | 6.46059531500 | 7.47475853305 | Km<KS.1 | TM | KS.1 |
| FIM1_3697 | 19565.92173 | 9318.687622 | 11978.24542 | -0.968528383 | 0.231789644 | 1.01166981612 | 7.08881314822 | ns | TM | KS.1 |
| FIM1_3698 | 4677.168162 | 2080.923665 | 3455.448409 | -0.934044742 | 0.351026066 | 6.71717501426 | 0.001749631 | ns | TM | KS.1 |
| FIM1_3699 | 535.1675137 | 115.9796105 | 1113.884538 | -1.663406889 | 0.417118274 | 9.48605031093 | 6.77808860802 | Km<KS.1 | TM | KS.1 |
| FIM1_3700 | 1455.133577 | 429.0089016 | 866.2872135 | -1.458937026 | 0.333249031 | 5.58687289823 | 4.96333797656 | Km<KS.1 | TM | KS.1 |
| FIM1_3701 | 1577.455244 | 732.3789302 | 989.2838633 | -0.686384844 | 0.430442138 | 0.000918092 | 0.014318212 | ns | TM | KS.1 |
| FIM1_3702 | 1324.793753 | 313.1985969 | 650.5353628 | -1.785194227 | 0.299304597 | 7.37491985567 | 2.88008403921 | Km<KS.1 | TM | KS.1 |
| FIM1_3703 | 4551.932391 | 1805.564302 | 2849.464825 | -1.151646606 | 0.294923689 | 3.29310546632 | 0.000176163 | Km<KS.1 | TM | KS.1 |

| | | | | | | | | | |
|---|---|---|---|---|---|---|---|---|---|
| FIM1_3704 | 2763.456492 | 988.4995573 | 1314.613492 | -1.242933499 | 0.327344649 | 4.40251712986 | 0.000221238 | Km<KS.1 | TM | KS.1 |
| FIM1_3705 | 4672.471128 | 2255.143855 | 2843.130502 | -0.635216835 | 0.41477517 | 0.001151871 | 0.016756014 | ns | TM | KS.1 |
| FIM1_3706 | 8506.105609 | 3704.280001 | 5158.641043 | -0.974149228 | 0.343988045 | 4.57096147617 | 0.001292076 | ns | TM | KS.1 |
| FIM1_3707 | 1644.165343 | 719.7039612 | 787.9552835 | -0.830826234 | 0.425001615 | 0.000345148 | 0.00668136 | ns | TM | KS.1 |
| FIM1_3708 | 9270.946237 | 4527.962671 | 5903.164291 | -0.916469407 | 0.249495628 | 4.30706333589 | 0.000218649 | ns | TM | KS.1 |
| FIM1_3709 | 3906.195565 | 1730.270104 | 2293.754854 | -0.892394417 | 0.382867929 | 0.000143627 | 0.003292826 | ns | TM | KS.1 |
| FIM1_3710 | 5595.874198 | 2365.677832 | 2787.408431 | -1.02683812 | 0.334323452 | 2.66155024692 | 0.000913187 | Km<KS.1 | TM | KS.1 |
| FIM1_3711 | 5381.395767 | 2254.27606 | 3620.269437 | -1.023943661 | 0.346859024 | 3.51839074927 | 0.001087205 | Km<KS.1 | TM | KS.1 |
| FIM1_3712 | 5387.221129 | 3041.658025 | 3898.949549 | -0.360644625 | 0.329961098 | 0.007312382 | 0.067493689 | ns | TM | KS.1 |
| FIM1_3713 | 5713.767665 | 2408.836261 | 3243.832749 | -1.055555248 | 0.312214426 | 1.26748503848 | 0.000495306 | Km<KS.1 | TM | KS.1 |
| FIM1_3714 | 2667.646332 | 441.5193833 | 1059.112359 | -2.101022861 | 0.381793515 | 6.92012881702 | 2.88008403921 | Km<KS.1 | TM | KS.1 |
| FIM1_3715 | 753.3678857 | 293.050585 | 396.9883538 | -0.919919299 | 0.474373147 | 0.000322273 | 0.00638768 | ns | TM | KS.1 |
| FIM1_3716 | 10963.14501 | 5086.858639 | 5631.398114 | -0.929243344 | 0.306777579 | 2.70072071039 | 0.000916754 | ns | TM | KS.1 |
| FIM1_3717 | 16096.20072 | 6919.190146 | 8897.841964 | -1.094158654 | 0.247027708 | 5.78616840917 | 5.05020839221 | Km<KS.1 | TM | KS.1 |
| FIM1_3718 | 3680.515124 | 1416.198558 | 1894.578112 | -1.089605269 | 0.3824419 | 4.38416554942 | 0.001268094 | Km<KS.1 | TM | KS.1 |
| FIM1_3719 | 2729.284633 | 951.7259635 | 1434.633716 | -1.155561363 | 0.422841723 | 5.22591281191 | 0.001437543 | Km<KS.1 | TM | KS.1 |
| FIM1_3720 | 2980.800168 | 1056.867795 | 1754.689686 | -1.253222753 | 0.327479198 | 4.03535862444 | 0.000208266 | Km<KS.1 | TM | KS.1 |
| FIM1_3722 | 213.1214819 | 88.64466962 | 144.6627501 | -0.885019485 | 0.442268878 | 0.000297239 | 0.005986897 | ns | TM | KS.1 |
| FIM1_3723 | 464.785814 | 216.0325827 | 374.7185674 | -0.382318159 | 0.426219205 | 0.009648627 | 0.082336056 | ns | TM | KS.1 |
| FIM1_3725 | 47.29927693 | 73.30499594 | 110.7036884 | 0.084322312 | 0.226657821 | 0.237929869 | 0.558772287 | ns | TM | KS.1 |
| FIM1_3727 | 805.4293039 | 423.1891576 | 551.350287 | -0.259934535 | 0.341965517 | 0.024573223 | 0.154163661 | ns | TM | KS.1 |
| FIM1_3731 | 70.50654542 | 140.2597128 | 113.1955211 | 0.34463055 | 0.382360581 | 0.011600587 | 0.093537962 | ns | TM | KS.1 |
| FIM1_3732 | 259.2174793 | 285.7036974 | 285.7986033 | 0.026461591 | 0.149521031 | 0.677574754 | 0.887002016 | ns | TM | KS.1 |
| FIM1_3733 | 1781.059997 | 1282.267387 | 1175.971791 | -0.304101677 | 0.159580478 | 0.002169106 | 0.027528828 | ns | TM | KS.1 |
| FIM1_3734 | 1096.340296 | 1479.268056 | 827.4544815 | 0.280131558 | 0.14426739 | 0.002682264 | 0.032077551 | ns | TM | KS.1 |
| FIM1_3735 | 1887.911707 | 1934.642702 | 1708.862647 | 0.01884911 | 0.0572362 | 0.650712397 | 0.874707964 | ns | TM | KS.1 |
| FIM1_3736 | 1016.645649 | 1215.682089 | 1412.888503 | 0.111049562 | 0.126640075 | 0.121161165 | 0.382472585 | ns | TM | KS.1 |
| FIM1_3737 | 424.5012476 | 219.8476691 | 793.9904804 | -0.648656571 | 0.355854803 | 0.000542991 | 0.009647782 | ns | TM | KS.1 |
| FIM1_3738 | 337.1537682 | 239.0217032 | 451.205716 | -0.325165253 | 0.164903489 | 0.001494191 | 0.020421975 | ns | TM | KS.1 |
| FIM1_3739 | 1466.356249 | 1105.742272 | 1405.266701 | -0.142868445 | 0.167689658 | 0.077095971 | 0.294813572 | ns | TM | KS.1 |
| FIM1_3740 | 2152.893267 | 1890.73891 | 2403.374038 | -0.077232538 | 0.112784976 | 0.238636754 | 0.55974439 | ns | TM | KS.1 |
| FIM1_3741 | 2842.110602 | 2430.929149 | 3060.206979 | -0.125278019 | 0.108664763 | 0.067017177 | 0.272394163 | ns | TM | KS.1 |
| FIM1_3742 | 2313.20104 | 2349.138195 | 2198.695666 | 0.009411222 | 0.0812359 | 0.859150909 | 0.956641847 | ns | TM | KS.1 |
| FIM1_3743 | 3075.949904 | 3173.484051 | 3417.466024 | 0.022437218 | 0.067661773 | 0.632603806 | 0.869547581 | ns | TM | KS.1 |
| FIM1_3744 | 1850.10561 | 1762.454472 | 1593.700646 | -0.030098289 | 0.086756978 | 0.58666607 | 0.841613393 | ns | TM | KS.1 |
| FIM1_3745 | 11926.96586 | 11692.30802 | 12179.15248 | -0.013487687 | 0.071380206 | 0.781914827 | 0.929216508 | ns | TM | KS.1 |
| FIM1_3746 | 2148.184262 | 1765.691197 | 1963.194845 | -0.167386386 | 0.118661398 | 0.026747897 | 0.160661696 | ns | TM | KS.1 |
| FIM1_3747 | 12954.41935 | 13728.55074 | 16972.88969 | 0.030988175 | 0.100942853 | 0.60554997 | 0.851547247 | ns | TM | KS.1 |
| FIM1_3748 | 7676.230265 | 9260.617414 | 7906.555294 | 0.135078966 | 0.124400164 | 0.066811678 | 0.272267737 | ns | TM | KS.1 |
| FIM1_3749 | 4354.715142 | 4652.851889 | 4416.571733 | 0.055523562 | 0.067717762 | 0.239363457 | 0.560128504 | ns | TM | KS.1 |
| FIM1_3750 | 830.1342799 | 1169.86045 | 1375.601999 | 0.117342545 | 0.194547276 | 0.13209437 | 0.402184512 | ns | TM | KS.1 |
| FIM1_3751 | 1943.078396 | 1958.791024 | 1850.963617 | 0.006316819 | 0.063029519 | 0.887532838 | 0.967028314 | ns | TM | KS.1 |
| FIM1_3752 | 2498.559706 | 2250.970632 | 2341.916908 | -0.082757294 | 0.089204016 | 0.154126093 | 0.437933732 | ns | TM | KS.1 |
| FIM1_3753 | 1412.546684 | 1934.715577 | 2476.245096 | 0.265866631 | 0.162120242 | 0.005550857 | 0.056358187 | ns | TM | KS.1 |
| FIM1_3754 | 1824.441849 | 1858.408897 | 2359.349226 | 0.006258069 | 0.129857764 | 0.921035943 | 0.975881018 | ns | TM | KS.1 |

| | | | | | | | | | | |
|---|---|---|---|---|---|---|---|---|---|---|
| FIM1_3755 | 231.2723884 | 235.5306172 | 213.6750868 | 0.008017483 | 0.119246056 | 0.898123033 | 0.967972723 | ns | TM | KS.1 |
| FIM1_3756 | 19945.39531 | 23509.62377 | 22678.61819 | 0.130160225 | 0.112262696 | 0.062857828 | 0.262637093 | ns | TM | KS.1 |
| FIM1_3757 | 696.0992252 | 687.9003318 | 660.2333306 | -0.006377678 | 0.082286649 | 0.905021077 | 0.971408815 | ns | TM | KS.1 |
| FIM1_3758 | 1756.526542 | 1781.606447 | 1542.398213 | 0.006694041 | 0.108352828 | 0.91279327 | 0.973904053 | ns | TM | KS.1 |
| FIM1_3759 | 8512.969583 | 9453.27427 | 9864.711821 | 0.038664684 | 0.131904239 | 0.551068526 | 0.822058746 | ns | TM | KS.1 |
| FIM1_3760 | 697.5762251 | 634.6300952 | 619.9495454 | -0.064668514 | 0.092854709 | 0.27006198 | 0.597663125 | ns | TM | KS.1 |
| FIM1_3761 | 739.1895188 | 683.0638112 | 735.6103109 | -0.061927917 | 0.077709249 | 0.235103632 | 0.554201572 | ns | TM | KS.1 |
| FIM1_3762 | 1792.975654 | 1820.030605 | 1837.459971 | 0.007969066 | 0.096302208 | 0.891255427 | 0.967215407 | ns | TM | KS.1 |
| FIM1_3763 | 3610.607181 | 3757.847444 | 3123.577917 | 0.016992897 | 0.115623421 | 0.785495389 | 0.931546975 | ns | TM | KS.1 |
| FIM1_3764 | 705.1010064 | 755.7117598 | 462.6643805 | 0.048482419 | 0.081058262 | 0.363334394 | 0.68729605 | ns | TM | KS.1 |
| FIM1_3765 | 10164.46209 | 10673.14351 | 9241.215501 | 0.026660484 | 0.097519077 | 0.650967072 | 0.874784459 | ns | TM | KS.1 |
| FIM1_3766 | 78.54431744 | 63.63546489 | 98.51668961 | -0.087183232 | 0.141241137 | 0.218011914 | 0.529594372 | ns | TM | KS.1 |
| FIM1_3767 | 2610.607184 | 2089.318656 | 2270.164745 | -0.179461337 | 0.130968328 | 0.024866494 | 0.154938568 | ns | TM | KS.1 |
| FIM1_3768 | 33501.99629 | 25789.53805 | 26488.78546 | -0.062470223 | 0.180186139 | 0.354581193 | 0.679784721 | ns | TM | KS.1 |
| FIM1_3769 | 1565.340455 | 1836.49702 | 2223.519758 | 0.167053239 | 0.091278163 | 0.01127608 | 0.091959747 | ns | TM | KS.1 |
| FIM1_3770 | 6667.63331 | 6838.060957 | 6474.110678 | 0.022799396 | 0.046959964 | 0.526956344 | 0.808531286 | ns | TM | KS.1 |
| FIM1_3771 | 1353.288813 | 1355.589485 | 1381.914983 | 0.000928314 | 0.097556433 | 0.987364032 | 0.996167262 | ns | TM | KS.1 |
| FIM1_3772 | 2288.669269 | 2158.43553 | 2235.125208 | -0.030123433 | 0.104064173 | 0.620093899 | 0.861798092 | ns | TM | KS.1 |
| FIM1_3773 | 9035.363081 | 9512.602587 | 9253.453081 | 0.02436551 | 0.109019707 | 0.692602902 | 0.893131011 | ns | TM | KS.1 |
| FIM1_3774 | 7366.489609 | 5338.139625 | 4738.193851 | -0.275437426 | 0.164560191 | 0.004677906 | 0.049202074 | ns | TM | KS.1 |
| FIM1_3775 | 254.9554529 | 266.3673993 | 237.9796987 | 0.02551778 | 0.092710212 | 0.656679305 | 0.877276998 | ns | TM | KS.1 |
| FIM1_3776 | 549.6632451 | 664.0923628 | 418.9431956 | 0.137679524 | 0.124740784 | 0.062980111 | 0.262637093 | ns | TM | KS.1 |
| FIM1_3777 | 9711.708377 | 11897.6173 | 9059.563685 | 0.050318072 | 0.167553477 | 0.44429833 | 0.750300134 | ns | TM | KS.1 |
| FIM1_3778 | 278.5980033 | 214.8857804 | 395.1968135 | -0.09926696 | 0.16499707 | 0.178140741 | 0.476938324 | ns | TM | KS.1 |
| FIM1_3779 | 1689.420152 | 1580.347179 | 1354.296373 | -0.038558547 | 0.096158211 | 0.512230414 | 0.802566276 | ns | TM | KS.1 |
| FIM1_3780 | 9439.468667 | 11607.78353 | 10131.22759 | 0.237373053 | 0.088910825 | 0.000853448 | 0.013521989 | ns | TM | KS.1 |
| FIM1_3782 | 3339.487787 | 3085.26529 | 3009.537331 | -0.050821568 | 0.09263883 | 0.381130479 | 0.70494176 | ns | TM | KS.1 |
| FIM1_3783 | 1245.348081 | 1450.651457 | 1425.301828 | 0.133011667 | 0.102868621 | 0.047614141 | 0.227332393 | ns | TM | KS.1 |
| FIM1_3784 | 4513.993105 | 4546.960654 | 3653.060382 | 0.004702859 | 0.079107398 | 0.928143958 | 0.977872975 | ns | TM | KS.1 |
| FIM1_3785 | 516.556388 | 441.9510847 | 374.948203 | -0.075256882 | 0.129375799 | 0.270299906 | 0.597663125 | ns | TM | KS.1 |
| FIM1_3786 | 5021.511906 | 4337.905645 | 4454.690071 | -0.170216093 | 0.07480646 | 0.003728009 | 0.041307001 | ns | TM | KS.1 |
| FIM1_3787 | 62.50761529 | 62.02173278 | 84.05890081 | 0.00572525 | 0.120876417 | 0.927274232 | 0.977370615 | ns | TM | KS.1 |
| FIM1_3788 | 6680.170483 | 6746.065066 | 6370.812734 | 0.006500077 | 0.074271656 | 0.896668275 | 0.967972723 | ns | TM | KS.1 |
| FIM1_3789 | 771.004266 | 652.3405219 | 690.6040231 | -0.130318086 | 0.114383558 | 0.064953503 | 0.267310992 | ns | TM | KS.1 |
| FIM1_3790 | 591.8708429 | 400.146988 | 421.9346332 | -0.382211195 | 0.187779187 | 0.000738369 | 0.012019452 | ns | TM | KS.1 |
| FIM1_3791 | 3198.668762 | 3042.762231 | 3342.107276 | -0.041848513 | 0.06153947 | 0.340158212 | 0.668193429 | ns | TM | KS.1 |
| FIM1_3792 | 2718.453405 | 2720.689293 | 3126.315086 | 0.0002269 | 0.121884529 | 0.997115578 | 0.998293524 | ns | TM | KS.1 |
| FIM1_3793 | 6731.428619 | 6212.719521 | 6434.595813 | -0.051336442 | 0.092960625 | 0.377419155 | 0.701404668 | ns | TM | KS.1 |
| FIM1_3794 | 8074.59924 | 10191.96153 | 9539.717368 | 0.218076191 | 0.12265186 | 0.007703364 | 0.07006259 | ns | TM | KS.1 |
| FIM1_3795 | 579.6886104 | 697.6703404 | 881.351855 | 0.083871597 | 0.138990202 | 0.232063401 | 0.549507578 | ns | TM | KS.1 |
| FIM1_3796 | 334.8536402 | 317.4608198 | 333.1994851 | -0.028030966 | 0.105116287 | 0.645549035 | 0.873039206 | ns | TM | KS.1 |
| FIM1_3797 | 4801.785005 | 3905.857754 | 4135.845504 | -0.167919176 | 0.124953676 | 0.029837706 | 0.171016808 | ns | TM | KS.1 |
| FIM1_3798 | 2431.792266 | 2215.856173 | 2050.789803 | -0.060036594 | 0.096973456 | 0.316260984 | 0.642990763 | ns | TM | KS.1 |
| FIM1_3799 | 16868.80331 | 15941.79466 | 14279.63957 | -0.028290909 | 0.105555198 | 0.643078112 | 0.872978256 | ns | TM | KS.1 |
| FIM1_3801 | 2110.968478 | 2203.132156 | 2398.246948 | 0.019017015 | 0.112675526 | 0.759379534 | 0.922567322 | ns | TM | KS.1 |

| | | | | | | | | | | |
|---|---|---|---|---|---|---|---|---|---|---|
| FIM1_3802 | 2834.835524 | 1861.328664 | 2210.137681 | -0.17948846 | 0.229255866 | 0.050014551 | 0.232979769 | ns | TM | KS.1 |
| FIM1_3803 | 326.4024824 | 308.797597 | 313.849329 | -0.024685793 | 0.113588499 | 0.692490243 | 0.893131011 | ns | TM | KS.1 |
| FIM1_3804 | 382.4178636 | 311.382742 | 257.7622274 | -0.150491893 | 0.130291297 | 0.049662733 | 0.232305651 | ns | TM | KS.1 |
| FIM1_3805 | 412.76089 | 327.3296406 | 471.9418112 | -0.161807394 | 0.141364291 | 0.043399769 | 0.213353609 | ns | TM | KS.1 |
| FIM1_3806 | 873.6033126 | 1017.818714 | 864.3943599 | 0.05246794 | 0.143353452 | 0.429845915 | 0.743536592 | ns | TM | KS.1 |
| FIM1_3807 | 7656.880486 | 9592.659838 | 10336.56045 | 0.290429937 | 0.064785816 | 8.71247509245 | 6.37420052720 | ns | TM | KS.1 |
| FIM1_3808 | 581.1647183 | 464.0706777 | 598.9609995 | -0.224357632 | 0.115377229 | 0.005178066 | 0.053400721 | ns | TM | KS.1 |
| FIM1_3809 | 1303.262218 | 1853.412678 | 2060.132763 | 0.11312488 | 0.198087443 | 0.142953869 | 0.419584366 | ns | TM | KS.1 |
| FIM1_3810 | 1990.746691 | 2047.98004 | 1956.994185 | 0.015757798 | 0.093435424 | 0.784036261 | 0.930812921 | ns | TM | KS.1 |
| FIM1_3811 | 3583.711491 | 2551.84428 | 5184.452121 | -0.307047498 | 0.167343025 | 0.002416995 | 0.02976374 | ns | TM | KS.1 |
| FIM1_3812 | 12.27872587 | 11.29256932 | 22.63589428 | -0.019347754 | 0.17704958 | 0.75380504 | 0.919612573 | ns | TM | KS.1 |
| FIM1_3813 | 3658.018861 | 3774.097925 | 4199.466434 | 0.024037521 | 0.062122864 | 0.586169541 | 0.841613393 | ns | TM | KS.1 |
| FIM1_3814 | 27.45472185 | 22.51406973 | 26.24877177 | -0.049652854 | 0.172058019 | 0.448806426 | 0.753357428 | ns | TM | KS.1 |
| FIM1_3815 | 19077.95546 | 16132.99136 | 13481.12748 | -0.062703032 | 0.142174116 | 0.354252888 | 0.679784721 | ns | TM | KS.1 |
| FIM1_3816 | 1738.088777 | 2398.223406 | 2349.079688 | 0.191305149 | 0.181927626 | 0.032496141 | 0.180382188 | ns | TM | KS.1 |
| FIM1_3817 | 3860.357559 | 5022.816602 | 3144.887224 | 0.252840004 | 0.128927575 | 0.003601619 | 0.040355983 | ns | TM | KS.1 |
| FIM1_3818 | 69.21266992 | 62.15548577 | 352.0551801 | -0.032018359 | 0.145978098 | 0.618254713 | 0.860858199 | ns | TM | KS.1 |
| FIM1_3819 | 2713.444826 | 1974.213114 | 2295.003418 | -0.223141679 | 0.175235697 | 0.016456702 | 0.119173353 | ns | TM | KS.1 |
| FIM1_3820 | 997.5194636 | 1321.712528 | 1699.424937 | 0.131107562 | 0.168738362 | 0.097281983 | 0.337501997 | ns | TM | KS.1 |
| FIM1_3821 | 731.4449771 | 929.5340236 | 934.0114174 | 0.147762578 | 0.147365009 | 0.062487597 | 0.262121245 | ns | TM | KS.1 |
| FIM1_3822 | 4607.034584 | 4197.442605 | 5347.315536 | -0.062443409 | 0.094458442 | 0.290946309 | 0.618193599 | ns | TM | KS.1 |
| FIM1_3823 | 66.40618748 | 79.12901779 | 134.5366798 | 0.077644889 | 0.1345862 | 0.260895437 | 0.588210523 | ns | TM | KS.1 |
| FIM1_3824 | 9835.919999 | 10149.36001 | 20601.80896 | 0.012196979 | 0.121529788 | 0.846228643 | 0.952348261 | ns | TM | KS.1 |
| FIM1_3825 | 402.8955646 | 335.2116288 | 253.0589104 | -0.051828369 | 0.157901445 | 0.433914176 | 0.745878978 | ns | TM | KS.1 |
| FIM1_3826 | 5038.000428 | 5561.151879 | 6236.632612 | 0.104571691 | 0.067365951 | 0.033693683 | 0.183415813 | ns | TM | KS.1 |
| FIM1_3827 | 582.5488613 | 503.7124225 | 477.8659757 | -0.093668714 | 0.113930101 | 0.163079138 | 0.45366779 | ns | TM | KS.1 |
| FIM1_3828 | 2911.032143 | 2911.978549 | 3076.068311 | 0.000615618 | 0.072181241 | 0.989996252 | 0.996341203 | ns | TM | KS.1 |
| FIM1_3829 | 2810.410398 | 2710.380666 | 2219.783123 | -0.024084833 | 0.077927057 | 0.641287598 | 0.872410664 | ns | TM | KS.1 |
| FIM1_3830 | 3642.420192 | 2885.995671 | 3249.216534 | -0.186968698 | 0.1344688 | 0.021951344 | 0.144454945 | ns | TM | KS.1 |
| FIM1_3831 | 875.5829667 | 637.8083296 | 740.3250707 | -0.240397596 | 0.171277284 | 0.011045451 | 0.090678411 | ns | TM | KS.1 |
| FIM1_3832 | 968.8290081 | 939.1421855 | 785.8791234 | -0.017586605 | 0.090510395 | 0.755828836 | 0.920354827 | ns | TM | KS.1 |
| FIM1_3833 | 66.40031993 | 60.76400271 | 255.9778003 | -0.022002036 | 0.154471254 | 0.727174533 | 0.907961357 | ns | TM | KS.1 |
| FIM1_3834 | 3367.330492 | 2815.456807 | 3235.127828 | -0.184436647 | 0.098142133 | 0.008786055 | 0.077638762 | ns | TM | KS.1 |
| FIM1_3835 | 327.0552068 | 291.0015311 | 381.0551814 | -0.07889026 | 0.104264107 | 0.212765072 | 0.520662191 | ns | TM | KS.1 |
| FIM1_3836 | 3625.888624 | 4032.515394 | 4396.370425 | 0.061827402 | 0.10687706 | 0.326346798 | 0.653613253 | ns | TM | KS.1 |
| FIM1_3837 | 2634.966854 | 2433.784929 | 2397.157586 | -0.056165561 | 0.085906931 | 0.312268203 | 0.63931779 | ns | TM | KS.1 |
| FIM1_3838 | 1748.181414 | 1775.018245 | 1857.047638 | 0.00947103 | 0.079910459 | 0.856813137 | 0.955916258 | ns | TM | KS.1 |
| FIM1_3839 | 2697.71126 | 2224.017014 | 2382.466326 | -0.077368911 | 0.145258139 | 0.267039379 | 0.593708608 | ns | TM | KS.1 |
| FIM1_3840 | 628.3831085 | 832.6865229 | 898.5308475 | 0.170463757 | 0.164384461 | 0.043671519 | 0.214477597 | ns | TM | KS.1 |
| FIM1_3841 | 1954.692482 | 2119.6375 | 2372.251608 | 0.035087428 | 0.118935163 | 0.581420226 | 0.838424819 | ns | TM | KS.1 |
| FIM1_3842 | 143.1828775 | 133.2395247 | 197.4601246 | -0.018555811 | 0.145959769 | 0.769219779 | 0.925705893 | ns | TM | KS.1 |
| FIM1_3843 | 9789.65696 | 11248.43718 | 11982.34472 | 0.157013862 | 0.076660805 | 0.007133089 | 0.066581689 | ns | TM | KS.1 |
| FIM1_3844 | 901.6744014 | 992.1116211 | 1232.990209 | 0.032815715 | 0.135139366 | 0.610651455 | 0.854808945 | ns | TM | KS.1 |
| FIM1_3845 | 677.8365404 | 805.879312 | 1007.305287 | 0.064827255 | 0.143439109 | 0.340342141 | 0.668193429 | ns | TM | KS.1 |
| FIM1_3846 | 790.0690884 | 716.9596729 | 975.6377112 | -0.064849962 | 0.095474753 | 0.276573448 | 0.601115292 | ns | TM | KS.1 |

| | | | | | | | | | | |
|---|---|---|---|---|---|---|---|---|---|---|
| FIM1_3847 | 25611.78242 | 32005.37484 | 31400.47508 | 0.262257167 | 0.086608348 | 0.000252165 | 0.005293345 | ns | TM | KS.1 |
| FIM1_3848 | 794.3869573 | 736.4463597 | 750.7664885 | -0.02667891 | 0.131069951 | 0.676582648 | 0.887002016 | ns | TM | KS.1 |
| FIM1_3849 | 26223.60713 | 31116.32816 | 30737.89021 | 0.115629833 | 0.121157899 | 0.103164303 | 0.347960953 | ns | TM | KS.1 |
| FIM1_3850 | 2362.202436 | 2425.528785 | 2505.267773 | 0.011694684 | 0.111443546 | 0.849715354 | 0.95298643 | ns | TM | KS.1 |
| FIM1_3851 | 785.0103152 | 772.4622513 | 1348.817359 | -0.008239501 | 0.091653955 | 0.884780819 | 0.967000126 | ns | TM | KS.1 |
| FIM1_3852 | 1470.71599 | 1129.734896 | 1300.114399 | -0.197899944 | 0.149207163 | 0.021034027 | 0.141433402 | ns | TM | KS.1 |
| FIM1_3853 | 1464.721281 | 1599.9985 | 1715.996936 | 0.059321564 | 0.092864272 | 0.309880875 | 0.638103209 | ns | TM | KS.1 |
| FIM1_3854 | 6051.254801 | 5953.749126 | 5777.754768 | -0.008757627 | 0.094841662 | 0.879701106 | 0.965258439 | ns | TM | KS.1 |
| FIM1_3855 | 4436.279544 | 3686.084099 | 3691.321226 | -0.155768652 | 0.116093383 | 0.034396906 | 0.185199792 | ns | TM | KS.1 |
| FIM1_3856 | 3029.652438 | 1980.900588 | 1475.589319 | -0.521758039 | 0.15270891 | 2.8843301662 | 0.000160976 | ns | TM | KS.1 |
| FIM1_3857 | 5852.034026 | 7175.765014 | 6700.010341 | 0.241171816 | 0.083024178 | 0.000439303 | 0.008124656 | ns | TM | KS.1 |
| FIM1_3858 | 1348.959797 | 1157.897773 | 1106.416192 | -0.125623265 | 0.106349373 | 0.063415488 | 0.264231199 | ns | TM | KS.1 |
| FIM1_3859 | 39.8675875 | 28.72372492 | 26.75156027 | -0.10706876 | 0.184008319 | 0.15705919 | 0.442201171 | ns | TM | KS.1 |
| FIM1_3860 | 2537.763197 | 2112.710192 | 2591.085474 | -0.077741621 | 0.140778109 | 0.263519728 | 0.590013793 | ns | TM | KS.1 |
| FIM1_3861 | 930.0047858 | 788.4768385 | 683.3851481 | -0.069090635 | 0.136963439 | 0.311276748 | 0.638961913 | ns | TM | KS.1 |
| FIM1_3862 | 6168.544268 | 4745.273913 | 4239.368737 | -0.079336482 | 0.171883375 | 0.260278789 | 0.588210523 | ns | TM | KS.1 |
| FIM1_3863 | 4842.729712 | 5305.452053 | 5834.400475 | 0.052150825 | 0.103671767 | 0.397207272 | 0.720943517 | ns | TM | KS.1 |
| FIM1_3864 | 236.5213355 | 219.7239447 | 239.0951616 | -0.020727379 | 0.146457076 | 0.743547534 | 0.91508234 | ns | TM | KS.1 |
| FIM1_3865 | 969.4434243 | 924.6065353 | 647.6237333 | -0.013280907 | 0.143287442 | 0.833292223 | 0.948005674 | ns | TM | KS.1 |
| FIM1_3866 | 1698.294071 | 1567.516216 | 1486.262765 | -0.045225699 | 0.100640573 | 0.454230136 | 0.757811846 | ns | TM | KS.1 |
| FIM1_3867 | 1450.734265 | 1444.603635 | 1536.73574 | -0.002269207 | 0.094540844 | 0.968653081 | 0.993374633 | ns | TM | KS.1 |
| FIM1_3868 | 457.8324002 | 341.435201 | 508.917387 | -0.181887259 | 0.168565574 | 0.035675587 | 0.188414062 | ns | TM | KS.1 |
| FIM1_3869 | 13275.81498 | 14468.7432 | 15450.81622 | 0.036573333 | 0.120975433 | 0.567463575 | 0.830088588 | ns | TM | KS.1 |
| FIM1_3870 | 2976.246549 | 1806.139744 | 1812.154242 | -0.632854785 | 0.18061763 | 2.2025113720 | 0.000132018 | ns | TM | KS.1 |
| FIM1_3871 | 1728.392646 | 2584.727287 | 2563.177529 | 0.255367971 | 0.222955354 | 0.013932755 | 0.105023419 | ns | TM | KS.1 |
| FIM1_3872 | 4060.727542 | 3488.832568 | 5529.200178 | -0.158093122 | 0.089168873 | 0.013738865 | 0.104063892 | ns | TM | KS.1 |
| FIM1_3873 | 1856.06525 | 2326.774514 | 1453.231919 | 0.152029349 | 0.140095868 | 0.05347146 | 0.241397925 | ns | TM | KS.1 |
| FIM1_3874 | 521.6513318 | 442.1932652 | 510.1569601 | -0.124971805 | 0.114832266 | 0.075148496 | 0.291853058 | ns | TM | KS.1 |
| FIM1_3875 | 308.712509 | 303.621844 | 283.7631909 | -0.010994018 | 0.090073158 | 0.845306952 | 0.952348261 | ns | TM | KS.1 |
| FIM1_3876 | 168.1891553 | 85.93751305 | 99.06440972 | -0.714084922 | 0.342930183 | 0.000265624 | 0.005483313 | ns | TM | KS.1 |
| FIM1_3877 | 124.2408567 | 72.62604463 | 174.706943 | -0.533190039 | 0.28725456 | 0.000577437 | 0.010044575 | ns | TM | KS.1 |
| FIM1_3879 | 12158.02837 | 12127.83238 | 20269.80526 | -0.000964195 | 0.12141874 | 0.987737141 | 0.996167262 | ns | TM | KS.1 |
| FIM1_3880 | 3676.045221 | 3443.204183 | 3181.816457 | -0.030361187 | 0.112234189 | 0.626800119 | 0.866840608 | ns | TM | KS.1 |
| FIM1_3881 | 8115.035787 | 8334.533699 | 6875.92404 | 0.023580772 | 0.048608365 | 0.523357825 | 0.807903931 | ns | TM | KS.1 |
| FIM1_3882 | 58.47516338 | 61.79954864 | 60.88332332 | 0.014455665 | 0.146748562 | 0.818506642 | 0.943045517 | ns | TM | KS.1 |
| FIM1_3883 | 587.5751303 | 454.6267522 | 422.477108 | -0.202609488 | 0.143504034 | 0.017446781 | 0.12351946 | ns | TM | KS.1 |
| FIM1_3884 | 489.1019012 | 412.3251732 | 508.2338864 | -0.083129447 | 0.13258491 | 0.231616634 | 0.549231011 | ns | TM | KS.1 |
| FIM1_3885 | 2650.618112 | 2726.114768 | 2432.3808 | 0.018494734 | 0.076139076 | 0.716299729 | 0.905084971 | ns | TM | KS.1 |
| FIM1_3886 | 4582.835091 | 4173.449002 | 3862.400914 | -0.062151072 | 0.095115312 | 0.295030141 | 0.622741988 | ns | TM | KS.1 |
| FIM1_3887 | 537.960397 | 489.3105574 | 388.6433825 | -0.070246973 | 0.088380718 | 0.218738996 | 0.531101271 | ns | TM | KS.1 |
| FIM1_3888 | 3252.816267 | 3227.677132 | 3161.601912 | -0.004597203 | 0.086267863 | 0.933351158 | 0.980245305 | ns | TM | KS.1 |
| FIM1_3889 | 1421.562764 | 1409.265981 | 1106.781719 | -0.004004658 | 0.11137635 | 0.948177003 | 0.986388143 | ns | TM | KS.1 |
| FIM1_3890 | 2629.471961 | 2211.019003 | 2590.63797 | -0.118617901 | 0.121552424 | 0.096376434 | 0.336236156 | ns | TM | KS.1 |
| FIM1_3891 | 312.9559039 | 256.4246116 | 393.9541856 | -0.083010619 | 0.146278836 | 0.239179697 | 0.55996188 | ns | TM | KS.1 |
| FIM1_3892 | 1346.481972 | 1416.036553 | 1681.82906 | 0.014594513 | 0.141381093 | 0.817438948 | 0.942251799 | ns | TM | KS.1 |

| | | | | | | | | | | |
|---|---|---|---|---|---|---|---|---|---|---|
| FIM1_3893 | 118.8973235 | 135.1489451 | 134.945167 | 0.063281722 | 0.120842359 | 0.337557622 | 0.665879925 | ns | TM | KS.1 |
| FIM1_3894 | 765.3014055 | 692.1518644 | 844.3097769 | -0.067251148 | 0.097875193 | 0.266770838 | 0.593708608 | ns | TM | KS.1 |
| FIM1_3895 | 138.2259951 | 154.991569 | 166.2505611 | 0.049360335 | 0.123620281 | 0.448834456 | 0.753357428 | ns | TM | KS.1 |
| FIM1_3896 | 1210.055495 | 1202.643086 | 1224.149328 | -0.004076057 | 0.079915396 | 0.938097651 | 0.982947729 | ns | TM | KS.1 |
| FIM1_3897 | 458.5976717 | 504.2605837 | 509.3109975 | 0.056499708 | 0.101755344 | 0.3564217 | 0.680951596 | ns | TM | KS.1 |
| FIM1_3898 | 2278.665042 | 4024.737062 | 4862.051882 | 0.249277596 | 0.308827692 | 0.024914745 | 0.154938568 | ns | TM | KS.1 |
| FIM1_3899 | 862.8215918 | 905.3709284 | 974.2576244 | 0.033419519 | 0.074470341 | 0.505508162 | 0.797875351 | ns | TM | KS.1 |
| FIM1_3900 | 312.0946092 | 324.9981537 | 381.9129481 | 0.02139931 | 0.097567199 | 0.716045157 | 0.905084971 | ns | TM | KS.1 |
| FIM1_3901 | 1301.2554 | 1367.895956 | 1554.936371 | 0.029128304 | 0.091646036 | 0.610058291 | 0.85421897 | ns | TM | KS.1 |
| FIM1_3902 | 640.3651043 | 467.0297557 | 1006.228109 | -0.197878902 | 0.177402219 | 0.027714798 | 0.164339833 | ns | TM | KS.1 |
| FIM1_3903 | 569.2011874 | 479.7891787 | 596.3936974 | -0.104512919 | 0.123713511 | 0.137644413 | 0.411032985 | ns | TM | KS.1 |
| FIM1_3904 | 7133.810059 | 8098.910712 | 7748.498376 | 0.096638014 | 0.10040856 | 0.126549954 | 0.392265433 | ns | TM | KS.1 |
| FIM1_3905 | 6266.820315 | 7060.216037 | 6693.299249 | 0.060337139 | 0.117581008 | 0.354879785 | 0.679784721 | ns | TM | KS.1 |
| FIM1_3906 | 1031.979559 | 1222.269845 | 747.9196432 | 0.105779189 | 0.12369724 | 0.133626025 | 0.405258563 | ns | TM | KS.1 |
| FIM1_3907 | 2582.746865 | 2407.152552 | 1759.155748 | -0.049570076 | 0.08264793 | 0.359020989 | 0.683511403 | ns | TM | KS.1 |
| FIM1_3908 | 1602.730025 | 1634.055376 | 1524.870191 | 0.014111146 | 0.058851353 | 0.739846136 | 0.914241064 | ns | TM | KS.1 |
| FIM1_3909 | 974.3678709 | 1258.228548 | 1174.845268 | 0.096792366 | 0.16404037 | 0.186580084 | 0.489513719 | ns | TM | KS.1 |
| FIM1_3910 | 5273.89956 | 6015.759797 | 6264.664992 | 0.130336258 | 0.086916785 | 0.031470622 | 0.17751286 | ns | TM | KS.1 |
| FIM1_3911 | 183.3583917 | 183.1437148 | 182.5380654 | -0.001754357 | 0.121303843 | 0.977687541 | 0.994223198 | ns | TM | KS.1 |
| FIM1_3912 | 1532.035747 | 1498.538173 | 4257.191824 | -0.008574033 | 0.12078787 | 0.891333881 | 0.967215407 | ns | TM | KS.1 |
| FIM1_3913 | 361.8743006 | 346.4785851 | 592.3381723 | -0.020670872 | 0.110114779 | 0.737676058 | 0.913453034 | ns | TM | KS.1 |
| FIM1_3914 | 386.4063064 | 403.2391615 | 465.4991353 | 0.02181126 | 0.102252129 | 0.716608985 | 0.905084971 | ns | TM | KS.1 |
| FIM1_3915 | 2350.248247 | 2311.094468 | 2899.055429 | -0.011082004 | 0.07285933 | 0.822530157 | 0.944618544 | ns | TM | KS.1 |
| FIM1_3916 | 824.6837936 | 772.4229893 | 852.7608378 | -0.03464894 | 0.10301648 | 0.567915758 | 0.830505848 | ns | TM | KS.1 |
| FIM1_3917 | 1604.359331 | 1739.984632 | 1846.392987 | 0.045820133 | 0.101612576 | 0.450585451 | 0.754323807 | ns | TM | KS.1 |
| FIM1_3918 | 1669.966732 | 1697.253121 | 1266.487463 | 0.006267625 | 0.121677119 | 0.920489602 | 0.975881018 | ns | TM | KS.1 |
| FIM1_3919 | 299.4458063 | 260.3889706 | 331.3315058 | -0.075897509 | 0.120693063 | 0.257827438 | 0.58677562 | ns | TM | KS.1 |
| FIM1_3920 | 294.5590794 | 229.2183543 | 231.2274604 | -0.181827406 | 0.147941267 | 0.029702821 | 0.170834141 | ns | TM | KS.1 |
| FIM1_3921 | 2519.427366 | 2591.187592 | 2798.983204 | 0.017141671 | 0.084434911 | 0.752401036 | 0.918662314 | ns | TM | KS.1 |
| FIM1_3922 | 516.4924036 | 559.2701845 | 500.5854168 | 0.041843228 | 0.105290573 | 0.496469027 | 0.792916022 | ns | TM | KS.1 |
| FIM1_3923 | 2664.396694 | 2775.94377 | 3006.232563 | 0.025523627 | 0.084793974 | 0.639840549 | 0.870968656 | ns | TM | KS.1 |
| FIM1_3924 | 290.1677404 | 539.9375759 | 705.9763253 | 0.232097875 | 0.324027412 | 0.032332211 | 0.18019422 | ns | TM | KS.1 |
| FIM1_3925 | 225.645439 | 141.173312 | 212.0267322 | -0.321442874 | 0.26562728 | 0.007216151 | 0.066853541 | ns | TM | KS.1 |
| FIM1_3926 | 1559.266322 | 1406.075153 | 1880.232738 | -0.065311467 | 0.101857493 | 0.290238294 | 0.618123079 | ns | TM | KS.1 |
| FIM1_3927 | 852.5853522 | 429.9232036 | 370.1716768 | -0.914023308 | 0.202548008 | 2.4150655491( | 2.4029902213 | ns | TM | KS.1 |
| FIM1_3928 | 1693.204158 | 1849.680731 | 2189.577524 | 0.049865411 | 0.10358867 | 0.41720067 | 0.735551151 | ns | TM | KS.1 |
| FIM1_3929 | 1467.181695 | 1655.070285 | 1378.074944 | 0.060088901 | 0.118796959 | 0.358213634 | 0.68306356 | ns | TM | KS.1 |
| FIM1_3930 | 979.0160828 | 1056.863424 | 1070.974216 | 0.057795549 | 0.080479704 | 0.278189141 | 0.602194469 | ns | TM | KS.1 |
| FIM1_3932 | 3355.682539 | 2388.395978 | 2228.120119 | -0.402600997 | 0.118042091 | 9.1586558627 | 0.000392796 | ns | TM | KS.1 |
| FIM1_3933 | 699.3816655 | 624.7455148 | 529.5501752 | -0.073081406 | 0.102873971 | 0.242411865 | 0.56513544 | ns | TM | KS.1 |
| FIM1_3934 | 731.1265358 | 782.2060369 | 1094.394944 | 0.046436717 | 0.085901132 | 0.401282827 | 0.723750973 | ns | TM | KS.1 |
| FIM1_3935 | 1466.269123 | 1326.646364 | 1625.570104 | -0.058662449 | 0.10476951 | 0.345943902 | 0.671768505 | ns | TM | KS.1 |
| FIM1_3936 | 1067.895909 | 789.2339243 | 891.8872015 | -0.3624734 | 0.101598468 | 1.0069522180 | 0.000414015 | ns | TM | KS.1 |
| FIM1_3937 | 52.83480664 | 35.01401901 | 34.38314948 | -0.100427319 | 0.216101041 | 0.180608934 | 0.480643537 | ns | TM | KS.1 |
| FIM1_3938 | 73.38229122 | 67.62232041 | 80.08171728 | -0.022890533 | 0.143001212 | 0.719017327 | 0.905913973 | ns | TM | KS.1 |

| | | | | | | | | | |
|---|---|---|---|---|---|---|---|---|---|
| FIM1_3939 | 546.4942255 | 253.4287026 | 588.5738225 | -0.91701219 | 0.318730029 | 3.9170767880 | 0.001159968 | ns | TM | KS.1 |
| FIM1_3940 | 1698.659376 | 2185.3756 | 2416.895088 | 0.080501118 | 0.167507568 | 0.254497525 | 0.581671847 | ns | TM | KS.1 |
| FIM1_3942 | 834.2187759 | 704.775668 | 686.7445039 | -0.142286463 | 0.109545123 | 0.0429385 | 0.212344969 | ns | TM | KS.1 |
| FIM1_3943 | 3805.367792 | 4159.72963 | 4597.163711 | 0.07698896 | 0.076900191 | 0.14131266 | 0.416585441 | ns | TM | KS.1 |
| FIM1_3944 | 1054.887122 | 1049.585147 | 950.884262 | -0.004507529 | 0.070974289 | 0.92601109 | 0.976735299 | ns | TM | KS.1 |
| FIM1_3945 | 935.8407708 | 974.9365373 | 915.1070056 | 0.015939801 | 0.122374992 | 0.800449714 | 0.936908579 | ns | TM | KS.1 |
| FIM1_3946 | 1248.560973 | 1408.065018 | 1684.428242 | 0.03314852 | 0.150329952 | 0.605707782 | 0.851547247 | ns | TM | KS.1 |
| FIM1_3947 | 1493.03568 | 1357.061871 | 1402.901195 | -0.061794296 | 0.097878523 | 0.305409281 | 0.633687915 | ns | TM | KS.1 |
| FIM1_3948 | 1569.599011 | 1601.248791 | 1624.501317 | 0.01384308 | 0.066802316 | 0.766071068 | 0.924600573 | ns | TM | KS.1 |
| FIM1_3949 | 1369.464042 | 1422.899302 | 1524.536322 | 0.028027777 | 0.071339006 | 0.564986855 | 0.828170184 | ns | TM | KS.1 |
| FIM1_3950 | 1413.471046 | 1364.467688 | 1387.801633 | -0.025289997 | 0.069204358 | 0.595838022 | 0.84839558 | ns | TM | KS.1 |
| FIM1_3951 | 1935.43901 | 1807.810126 | 2008.560255 | -0.062034823 | 0.063841922 | 0.171579967 | 0.465692069 | ns | TM | KS.1 |
| FIM1_3952 | 1370.981184 | 1472.665812 | 1720.543302 | 0.020439585 | 0.143756495 | 0.747324052 | 0.916738037 | ns | TM | KS.1 |
| FIM1_3953 | 2224.270675 | 2109.744974 | 1806.123057 | -0.034111216 | 0.084258924 | 0.530923807 | 0.808531286 | ns | TM | KS.1 |
| FIM1_3954 | 707.780834 | 546.1242068 | 635.473512 | -0.213891796 | 0.142254205 | 0.013113165 | 0.101458781 | ns | TM | KS.1 |
| FIM1_3955 | 794.9387329 | 667.0177263 | 717.4408983 | -0.09002926 | 0.131989821 | 0.199722675 | 0.50746696 | ns | TM | KS.1 |
| FIM1_3956 | 1802.333585 | 2416.699885 | 3147.383048 | 0.138425259 | 0.17293979 | 0.085604379 | 0.311772902 | ns | TM | KS.1 |
| FIM1_3957 | 2338.2309 | 2282.655542 | 2442.72559 | -0.019807217 | 0.052423647 | 0.612011254 | 0.855989876 | ns | TM | KS.1 |
| FIM1_3958 | 1499.907491 | 2557.490255 | 3999.83523 | 0.293084607 | 0.30113596 | 0.013763626 | 0.104063892 | ns | TM | KS.1 |
| FIM1_3959 | 918.1657288 | 653.6088731 | 783.6072442 | -0.386554041 | 0.127240039 | 4.2249070338 | 0.001229176 | ns | TM | KS.1 |
| FIM1_3960 | 176.9676015 | 175.4872024 | 230.82045 | -0.003049968 | 0.13156983 | 0.961454221 | 0.99134399 | ns | TM | KS.1 |
| FIM1_3961 | 925.4829936 | 902.3086025 | 941.5538678 | -0.00693426 | 0.144466705 | 0.912144438 | 0.973674801 | ns | TM | KS.1 |
| FIM1_3962 | 1958.870412 | 1187.842795 | 1344.310423 | -0.665697398 | 0.151274196 | 5.8259642756 | 6.900993397 | ns | TM | KS.1 |
| FIM1_3963 | 1457.615288 | 1395.041311 | 1670.37721 | -0.014920576 | 0.131260832 | 0.813873171 | 0.940696014 | ns | TM | KS.1 |
| FIM1_3964 | 1796.252845 | 2141.487777 | 2477.98014 | 0.05205364 | 0.154276881 | 0.432616573 | 0.74505982 | ns | TM | KS.1 |
| FIM1_3965 | 686.4911263 | 599.9881904 | 578.3806888 | -0.089382364 | 0.109185476 | 0.172542202 | 0.46779153 | ns | TM | KS.1 |
| FIM1_3966 | 289.4387878 | 410.8659778 | 404.0608576 | 0.071940472 | 0.204732584 | 0.297005771 | 0.624779582 | ns | TM | KS.1 |
| FIM1_3967 | 1459.850583 | 1583.793934 | 1412.758355 | 0.043375212 | 0.106143802 | 0.482813199 | 0.783685372 | ns | TM | KS.1 |
| FIM1_3968 | 1246.788983 | 1331.967769 | 1315.968476 | 0.04188629 | 0.088710887 | 0.457067298 | 0.761013991 | ns | TM | KS.1 |
| FIM1_3969 | 2324.511698 | 2267.664301 | 2221.040879 | -0.012349687 | 0.102245386 | 0.836667851 | 0.949024751 | ns | TM | KS.1 |
| FIM1_3970 | 692.9515382 | 759.1412171 | 906.8312738 | 0.037150096 | 0.123904544 | 0.56328471 | 0.827381586 | ns | TM | KS.1 |
| FIM1_3971 | 340.3774348 | 433.5826331 | 324.094424 | 0.079190121 | 0.164289278 | 0.260652502 | 0.588210523 | ns | TM | KS.1 |
| FIM1_3972 | 555.5383185 | 543.3154936 | 550.1518357 | -0.014400018 | 0.075978254 | 0.776883 | 0.9275241 | ns | TM | KS.1 |
| FIM1_3973 | 4863.171436 | 5726.929497 | 6210.686297 | 0.135025767 | 0.10989903 | 0.052728883 | 0.240005664 | ns | TM | KS.1 |
| FIM1_3974 | 1123.615854 | 1225.357371 | 1268.56398 | 0.051810402 | 0.098789503 | 0.389000924 | 0.713088169 | ns | TM | KS.1 |
| FIM1_3975 | 1212.861252 | 686.7355746 | 1007.307858 | -0.770602139 | 0.159556496 | 2.2976700910 | 2.9310022315 | ns | TM | KS.1 |
| FIM1_3976 | 6837.737225 | 6726.035746 | 8974.615497 | -0.008331829 | 0.100643462 | 0.888570334 | 0.967028314 | ns | TM | KS.1 |
| FIM1_3977 | 383.0441132 | 288.3131816 | 1321.053844 | -0.067369316 | 0.185422215 | 0.323878287 | 0.650941236 | ns | TM | KS.1 |
| FIM1_3978 | 918.5384086 | 1102.73976 | 1124.950898 | 0.112270102 | 0.128432397 | 0.119514048 | 0.379439943 | ns | TM | KS.1 |
| FIM1_3979 | 807.5320902 | 760.8009523 | 703.252064 | -0.040345764 | 0.080344298 | 0.445545267 | 0.751385662 | ns | TM | KS.1 |
| FIM1_3980 | 13896.34051 | 17140.00849 | 15468.15665 | 0.190084664 | 0.118827547 | 0.014735948 | 0.109583468 | ns | TM | KS.1 |
| FIM1_3981 | 314.3060872 | 146.2351786 | 506.7927928 | -0.666622702 | 0.433799761 | 0.001081881 | 0.016290797 | ns | TM | KS.1 |
| FIM1_3982 | 3707.805889 | 2985.494861 | 3579.231815 | -0.208484173 | 0.115090689 | 0.008029712 | 0.072107977 | ns | TM | KS.1 |
| FIM1_3983 | 1852.854155 | 1756.121445 | 1834.252194 | -0.032888161 | 0.087432538 | 0.554608797 | 0.824127468 | ns | TM | KS.1 |
| FIM1_3984 | 308.0280742 | 292.1439865 | 288.2195062 | -0.021481686 | 0.119797957 | 0.733330092 | 0.911168134 | ns | TM | KS.1 |

| | | | | | | | | | | |
|---|---|---|---|---|---|---|---|---|---|---|
| FIM1_3985 | 786.9939295 | 731.3524028 | 708.1430085 | -0.057726098 | 0.076377777 | 0.261647733 | 0.588461463 | ns | TM | KS.1 |
| FIM1_3986 | 1633.102321 | 1786.188126 | 1607.990702 | 0.040327149 | 0.117845548 | 0.527500122 | 0.808531286 | ns | TM | KS.1 |
| FIM1_3987 | 782.7941468 | 762.8363984 | 787.8348508 | -0.015907679 | 0.081885337 | 0.765441275 | 0.924600573 | ns | TM | KS.1 |
| FIM1_3988 | 1208.4436 | 1153.288859 | 1013.687429 | -0.029017518 | 0.086210574 | 0.598693351 | 0.849691542 | ns | TM | KS.1 |
| FIM1_3989 | 8996.010453 | 10493.41739 | 10098.21849 | 0.143427898 | 0.099189372 | 0.031725956 | 0.177743954 | ns | TM | KS.1 |
| FIM1_3990 | 2188.720596 | 1706.265961 | 1567.976856 | -0.097608415 | 0.161213306 | 0.183096276 | 0.48479405 | ns | TM | KS.1 |
| FIM1_3991 | 1058.776742 | 1345.879743 | 1565.42951 | 0.090891075 | 0.15944116 | 0.208028246 | 0.515497652 | ns | TM | KS.1 |
| FIM1_3992 | 738.1183898 | 731.1878333 | 725.4346143 | -0.006563709 | 0.068390867 | 0.889660784 | 0.967028314 | ns | TM | KS.1 |
| FIM1_3993 | 6317.985611 | 7497.588359 | 8151.152732 | 0.228222288 | 0.048946267 | 7.471988097[illegible] | 5.80830324764[illegible] | ns | TM | KS.1 |
| FIM1_3994 | 52.06017271 | 45.64565675 | 83.69121574 | -0.04702078 | 0.140201441 | 0.475001068 | 0.775371126 | ns | TM | KS.1 |
| FIM1_3995 | 173.3638087 | 127.9723776 | 185.7070297 | -0.109601504 | 0.179569861 | 0.149180984 | 0.429997333 | ns | TM | KS.1 |
| FIM1_3996 | 1024.576413 | 1420.676066 | 1615.367025 | 0.103921868 | 0.189567086 | 0.167130434 | 0.45887081 | ns | TM | KS.1 |
| FIM1_3997 | 1958.466884 | 2164.015459 | 1901.564487 | 0.065969428 | 0.097686644 | 0.274972591 | 0.600258288 | ns | TM | KS.1 |
| FIM1_3998 | 1528.012947 | 1323.743668 | 1173.428708 | -0.084490255 | 0.117542229 | 0.208779531 | 0.515893552 | ns | TM | KS.1 |
| FIM1_3999 | 336.4561427 | 289.9342474 | 239.0508598 | -0.059589565 | 0.136094887 | 0.374443376 | 0.69745885 | ns | TM | KS.1 |
| FIM1_4000 | 1018.868999 | 1060.207409 | 866.1570562 | 0.021075944 | 0.099080979 | 0.722042375 | 0.907011644 | ns | TM | KS.1 |
| FIM1_4001 | 95.57009142 | 116.0110845 | 118.8632034 | 0.076165756 | 0.146412603 | 0.273734889 | 0.599198132 | ns | TM | KS.1 |
| FIM1_4002 | 3820.864714 | 4731.458657 | 4722.895656 | 0.2550143 | 0.0823483 | 0.000222903 | 0.004779932 | ns | TM | KS.1 |
| FIM1_4003 | 4533.291957 | 4593.674098 | 4266.578467 | 0.008953234 | 0.074683244 | 0.858544958 | 0.956277476 | ns | TM | KS.1 |
| FIM1_4004 | 1460.060962 | 1400.540764 | 1181.977227 | -0.029511654 | 0.073326111 | 0.55219327 | 0.822689033 | ns | TM | KS.1 |
| FIM1_4005 | 3695.791608 | 3848.97655 | 2922.143101 | 0.021071444 | 0.100337406 | 0.723603316 | 0.907181536 | ns | TM | KS.1 |
| FIM1_4006 | 2805.242014 | 2366.404456 | 2646.572787 | -0.185488194 | 0.0896062 | 0.005637934 | 0.056841744 | ns | TM | KS.1 |
| FIM1_4007 | 1911.890046 | 1695.363896 | 1713.896372 | -0.120831062 | 0.081558294 | 0.035231257 | 0.187460433 | ns | TM | KS.1 |
| FIM1_4008 | 309.5337848 | 329.5346195 | 533.3020394 | 0.023142343 | 0.124356573 | 0.715452515 | 0.905084971 | ns | TM | KS.1 |
| FIM1_4009 | 2317.845497 | 2464.24361 | 3342.353643 | 0.033267718 | 0.099787882 | 0.57772797 | 0.836008333 | ns | TM | KS.1 |
| FIM1_4010 | 1723.522331 | 1830.187809 | 1854.404496 | 0.046949762 | 0.070565715 | 0.332589601 | 0.659583974 | ns | TM | KS.1 |
| FIM1_4011 | 1990.541671 | 1989.087113 | 1968.013867 | -0.00086608 | 0.050435272 | 0.981825406 | 0.995431301 | ns | TM | KS.1 |
| FIM1_4012 | 2057.757672 | 2222.629907 | 2395.085632 | 0.054024628 | 0.08613271 | 0.33133772 | 0.658571777 | ns | TM | KS.1 |
| FIM1_4013 | 1611.109355 | 1728.385879 | 1792.394935 | 0.061146351 | 0.066825949 | 0.19204178 | 0.496260756 | ns | TM | KS.1 |
| FIM1_4014 | 465.2941576 | 451.639427 | 366.1548634 | -0.01368264 | 0.11002908 | 0.824009359 | 0.94482203 | ns | TM | KS.1 |
| FIM1_4015 | 3817.852882 | 3559.605581 | 2846.782905 | -0.033641215 | 0.110556169 | 0.589151439 | 0.844030895 | ns | TM | KS.1 |
| FIM1_4016 | 2559.558723 | 2561.029214 | 2731.732684 | 0.000449325 | 0.090528781 | 0.99365073 | 0.99786059 | ns | TM | KS.1 |
| FIM1_4017 | 341.7862001 | 377.9501818 | 389.0880982 | 0.04600229 | 0.120243239 | 0.475884659 | 0.776238092 | ns | TM | KS.1 |
| FIM1_4018 | 2054.770358 | 3521.165815 | 4265.881815 | 0.227824796 | 0.29027367 | 0.030679515 | 0.174934485 | ns | TM | KS.1 |
| FIM1_4019 | 691.0420944 | 709.5617736 | 802.5732301 | 0.016742062 | 0.08079552 | 0.751538238 | 0.918662314 | ns | TM | KS.1 |
| FIM1_4020 | 3488.179831 | 2875.408527 | 3839.847189 | -0.220788345 | 0.087144738 | 0.001394279 | 0.019268155 | ns | TM | KS.1 |
| FIM1_4021 | 1153.032052 | 1157.959536 | 1069.033636 | 0.0024622 | 0.088431618 | 0.964770413 | 0.993216478 | ns | TM | KS.1 |
| FIM1_4022 | 30.63315877 | 28.97711918 | 31.43776702 | -0.011208337 | 0.159866545 | 0.856840763 | 0.955916258 | ns | TM | KS.1 |
| FIM1_4023 | 6465.852345 | 6015.120732 | 7102.690264 | -0.073233463 | 0.05633833 | 0.079493185 | 0.300154562 | ns | TM | KS.1 |
| FIM1_4024 | 3120.031346 | 2543.682253 | 2905.728058 | -0.054821726 | 0.16392025 | 0.40998136 | 0.73093186 | ns | TM | KS.1 |
| FIM1_4025 | 276.0918393 | 358.5420923 | 257.0080608 | 0.151760354 | 0.156585193 | 0.061034923 | 0.26041916 | ns | TM | KS.1 |
| FIM1_4026 | 108.4130284 | 159.9006724 | 370.4821554 | 0.11153862 | 0.211449894 | 0.148862607 | 0.429441469 | ns | TM | KS.1 |
| FIM1_4027 | 727.0870941 | 321.5931697 | 621.9376762 | -0.550501709 | 0.472061228 | 0.003135744 | 0.036399552 | ns | TM | KS.1 |
| FIM1_4028 | 1048.598173 | 1163.034134 | 1191.808455 | 0.088585913 | 0.085140245 | 0.118049469 | 0.376471865 | ns | TM | KS.1 |
| FIM1_4029 | 43.31611309 | 71.01112735 | 104.9791132 | 0.202825452 | 0.269970991 | 0.040342095 | 0.204589117 | ns | TM | KS.1 |

| | | | | | | | | | | |
|---|---|---|---|---|---|---|---|---|---|---|
| FIM1_4030 | 2845.919375 | 4543.550996 | 5795.456378 | 0.235621086 | 0.258803868 | 0.023944308 | 0.151692113 | ns | TM | KS.1 |
| FIM1_4031 | 3000.396001 | 3342.053197 | 3294.908715 | 0.073297406 | 0.09904321 | 0.231642836 | 0.549231011 | ns | TM | KS.1 |
| FIM1_4033 | 919.117948 | 859.3206973 | 824.2126454 | -0.040318488 | 0.093564741 | 0.487203883 | 0.786960818 | ns | TM | KS.1 |
| FIM1_4034 | 397.2520415 | 462.4922388 | 554.5582895 | 0.087451511 | 0.121398331 | 0.200220655 | 0.508213143 | ns | TM | KS.1 |
| FIM1_4035 | 329.5439984 | 381.5913318 | 436.1044789 | 0.072807359 | 0.125338364 | 0.280831686 | 0.606295641 | ns | TM | KS.1 |
| FIM1_4036 | 1601.309538 | 1744.006759 | 1895.044124 | 0.050541501 | 0.100064678 | 0.403368269 | 0.725246527 | ns | TM | KS.1 |
| FIM1_4037 | 1254.306788 | 1278.040416 | 1354.565124 | 0.012357251 | 0.077746941 | 0.810668853 | 0.939792751 | ns | TM | KS.1 |
| FIM1_4038 | 648.1422049 | 721.1678562 | 775.7880711 | 0.073253061 | 0.096290943 | 0.224517417 | 0.541169646 | ns | TM | KS.1 |
| FIM1_4039 | 721.8088044 | 743.770662 | 1193.073608 | 0.016809567 | 0.096138199 | 0.773235555 | 0.925733395 | ns | TM | KS.1 |
| FIM1_4041 | 578.9412451 | 954.8007057 | 1151.064925 | 0.218103307 | 0.271337898 | 0.032630439 | 0.180382188 | ns | TM | KS.1 |
| FIM1_4042 | 1159.22014 | 1328.914684 | 1666.45966 | 0.109084538 | 0.1017966 | 0.091764263 | 0.327259648 | ns | TM | KS.1 |
| FIM1_4043 | 1441.153793 | 1506.021832 | 1573.467723 | 0.023025825 | 0.10023878 | 0.699250282 | 0.896699645 | ns | TM | KS.1 |
| FIM1_4044 | 222.1544283 | 264.3985865 | 280.2021216 | 0.045877246 | 0.160133565 | 0.482769443 | 0.783685372 | ns | TM | KS.1 |
| FIM1_4045 | 3334.461673 | 4242.358757 | 4361.899764 | 0.165137189 | 0.144657 | 0.041650939 | 0.208464208 | ns | TM | KS.1 |
| FIM1_4046 | 1366.245159 | 1355.849441 | 1270.611571 | -0.003457133 | 0.105012244 | 0.954388254 | 0.988678089 | ns | TM | KS.1 |
| FIM1_4047 | 5935.756546 | 7188.744686 | 7888.412633 | 0.154840258 | 0.120440568 | 0.038336531 | 0.197232928 | ns | TM | KS.1 |
| FIM1_4048 | 410.6056513 | 437.6932779 | 405.4900098 | 0.027379809 | 0.117681401 | 0.664277121 | 0.880619203 | ns | TM | KS.1 |
| FIM1_4049 | 980.1824548 | 1503.055267 | 1563.26928 | 0.134954277 | 0.226379069 | 0.10231289 | 0.345835436 | ns | TM | KS.1 |
| FIM1_4050 | 3690.536303 | 3554.006849 | 3856.464842 | -0.029404591 | 0.062521096 | 0.507310189 | 0.799198286 | ns | TM | KS.1 |
| FIM1_4051 | 15032.89834 | 11559.19041 | 8292.48385 | -0.276012838 | 0.116072775 | 0.001128048 | 0.016554684 | ns | TM | KS.1 |
| FIM1_4052 | 41.32942577 | 49.58354625 | 75.04869013 | 0.070666413 | 0.150747212 | 0.305303322 | 0.633687915 | ns | TM | KS.1 |
| FIM1_4053 | 588.6149274 | 632.6555555 | 749.4921098 | 0.016388811 | 0.157905716 | 0.793167564 | 0.933303839 | ns | TM | KS.1 |
| FIM1_4054 | 2673.32788 | 2600.869868 | 2907.120408 | -0.013728388 | 0.102863515 | 0.81923057 | 0.94310261 | ns | TM | KS.1 |
| FIM1_4055 | 1389.711945 | 764.6395257 | 1475.51348 | -0.669269779 | 0.286331302 | 0.000129588 | 0.003026771 | ns | TM | KS.1 |
| FIM1_4056 | 1170.114463 | 1161.865664 | 1623.584067 | -0.004003241 | 0.090019171 | 0.943311006 | 0.98447079 | ns | TM | KS.1 |
| FIM1_4057 | 1170.301248 | 1084.456894 | 1189.612241 | -0.053623663 | 0.084856736 | 0.330232761 | 0.657690769 | ns | TM | KS.1 |
| FIM1_4058 | 460.5038258 | 422.915702 | 385.20337 | -0.044841286 | 0.10664006 | 0.469504577 | 0.77114073 | ns | TM | KS.1 |
| FIM1_4059 | 2783.796697 | 3214.527103 | 4790.05668 | 0.044931317 | 0.146494487 | 0.493204913 | 0.791514335 | ns | TM | KS.1 |
| FIM1_4060 | 1526.362409 | 2073.079231 | 2445.498945 | 0.128993628 | 0.179213682 | 0.10404621 | 0.34904241 | ns | TM | KS.1 |
| FIM1_4061 | 3317.265331 | 4864.010533 | 5649.129174 | 0.166251439 | 0.211366075 | 0.058600531 | 0.254317312 | ns | TM | KS.1 |
| FIM1_4062 | 915.0207999 | 1301.731553 | 1445.784866 | 0.115125965 | 0.19820846 | 0.138008534 | 0.411379542 | ns | TM | KS.1 |
| FIM1_4063 | 2075.798973 | 2095.732012 | 2330.080871 | 0.005274611 | 0.095949169 | 0.92775176 | 0.97766681 | ns | TM | KS.1 |
| FIM1_4064 | 5333.603593 | 5919.1903 | 5527.055317 | 0.049797871 | 0.117355471 | 0.439319978 | 0.747832613 | ns | TM | KS.1 |
| FIM1_4065 | 2368.381218 | 3692.037286 | 4374.277099 | 0.186808427 | 0.240516289 | 0.046317444 | 0.223284192 | ns | TM | KS.1 |
| FIM1_4066 | 2560.594599 | 2517.987551 | 3727.910755 | -0.010488481 | 0.079305079 | 0.840875731 | 0.950330932 | ns | TM | KS.1 |
| FIM1_4067 | 763.9571332 | 770.3106545 | 1068.184694 | 0.006646054 | 0.076325288 | 0.896237022 | 0.967972723 | ns | TM | KS.1 |
| FIM1_4068 | 337.7282339 | 350.3639531 | 406.8654862 | 0.017517768 | 0.109087926 | 0.775573234 | 0.926727011 | ns | TM | KS.1 |
| FIM1_4069 | 1954.266426 | 2240.766394 | 2644.230048 | 0.047740606 | 0.139669522 | 0.468750099 | 0.77114073 | ns | TM | KS.1 |
| FIM1_4070 | 2255.919862 | 2543.878001 | 2799.421974 | 0.040560407 | 0.138932765 | 0.533597069 | 0.810826335 | ns | TM | KS.1 |
| FIM1_4071 | 57.93653506 | 55.69243632 | 102.4320302 | -0.010566799 | 0.140446366 | 0.867018162 | 0.960282219 | ns | TM | KS.1 |
| FIM1_4072 | 1568.582219 | 1643.634698 | 1604.04108 | 0.031360381 | 0.079224821 | 0.548790205 | 0.82102634 | ns | TM | KS.1 |
| FIM1_4073 | 1050.651379 | 1060.217377 | 1025.178863 | 0.005428538 | 0.081918512 | 0.918878792 | 0.975757095 | ns | TM | KS.1 |
| FIM1_4074 | 1251.99599 | 1180.030621 | 1295.36021 | -0.016405087 | 0.145319222 | 0.795052755 | 0.93485877 | ns | TM | KS.1 |
| FIM1_4075 | 948.6548034 | 665.1475679 | 728.1756904 | -0.160877974 | 0.199090991 | 0.061716378 | 0.260865744 | ns | TM | KS.1 |
| FIM1_4076 | 15.0301895 | 12.01262949 | 18.61852671 | -0.0299676 | 0.183256426 | 0.631336763 | 0.86917233 | ns | TM | KS.1 |

| | | | | | | | | | | |
|---|---|---|---|---|---|---|---|---|---|---|
| FIM1_4077 | 22997.92297 | 26814.79744 | 36444.03178 | 0.047662245 | 0.148324928 | 0.469307587 | 0.77114073 | ns | TM | KS.1 |
| FIM1_4078 | 137.995597 | 167.045835 | 253.8581896 | 0.037311969 | 0.178749633 | 0.557177148 | 0.825232602 | ns | TM | KS.1 |
| FIM1_4080 | 2365.898775 | 1779.522524 | 3950.898547 | -0.094244128 | 0.175707946 | 0.197577853 | 0.505632622 | ns | TM | KS.1 |
| FIM1_4081 | 20.44568717 | 13.66714172 | 26.3944966 | -0.10939064 | 0.216621849 | 0.155048373 | 0.439274291 | ns | TM | KS.1 |
| FIM1_4082 | 13.16102476 | 14.83374157 | 53.98004836 | 0.024880828 | 0.180986817 | 0.688189162 | 0.891826565 | ns | TM | KS.1 |
| FIM1_4083 | 562.3345327 | 563.6197593 | 621.9285562 | 0.002160894 | 0.095488454 | 0.970293801 | 0.99362472 | ns | TM | KS.1 |
| FIM1_4084 | 3014.385761 | 3512.287775 | 3451.86931 | 0.101580754 | 0.115769313 | 0.137024259 | 0.409426841 | ns | TM | KS.1 |
| FIM1_4085 | 7490.463518 | 9500.559389 | 9779.713209 | 0.240860621 | 0.114906384 | 0.003153408 | 0.036399552 | ns | TM | KS.1 |
| FIM1_4086 | 6747.228927 | 9913.125006 | 11459.64232 | 0.237667578 | 0.213123773 | 0.017429357 | 0.12351946 | ns | TM | KS.1 |
| FIM1_4087 | 335.3576653 | 259.871163 | 342.0848434 | -0.082688631 | 0.167064464 | 0.244234397 | 0.567522712 | ns | TM | KS.1 |
| FIM1_4088 | 1761.963106 | 1953.643101 | 2166.353819 | 0.034875677 | 0.136648698 | 0.58946443 | 0.844122919 | ns | TM | KS.1 |
| FIM1_4089 | 192.9016142 | 145.9922107 | 243.3551311 | -0.195181707 | 0.159032636 | 0.024957408 | 0.155010117 | ns | TM | KS.1 |
| FIM1_4090 | 180.3583387 | 168.4757928 | 225.2102565 | -0.035182657 | 0.095173977 | 0.547047815 | 0.819501018 | ns | TM | KS.1 |
| FIM1_4091 | 1462.776974 | 2398.910723 | 2847.784754 | 0.192237569 | 0.263129773 | 0.045974815 | 0.221847435 | ns | TM | KS.1 |
| FIM1_4092 | 1658.379059 | 1248.535331 | 1310.075233 | -0.324848118 | 0.105864188 | 9.72846481867 | 0.00232688 | ns | TM | KS.1 |
| FIM1_4093 | 1180.335057 | 1291.032361 | 1490.982194 | 0.025642868 | 0.145041743 | 0.687516611 | 0.891826565 | ns | TM | KS.1 |
| FIM1_4094 | 3212.904164 | 2448.343381 | 6504.902137 | -0.107069022 | 0.168045566 | 0.154093075 | 0.437933732 | ns | TM | KS.1 |
| FIM1_4095 | 1805.859899 | 1824.169327 | 1717.111691 | 0.005119701 | 0.100445387 | 0.931305559 | 0.978943629 | ns | TM | KS.1 |
| FIM1_4096 | 2366.770326 | 2239.67767 | 1803.497877 | -0.028592177 | 0.102004276 | 0.634868893 | 0.869623553 | ns | TM | KS.1 |
| FIM1_4097 | 1067.796993 | 1600.435413 | 1921.574245 | 0.117184653 | 0.216771045 | 0.13587551 | 0.407922054 | ns | TM | KS.1 |
| FIM1_4098 | 2628.148303 | 2433.442288 | 1806.243203 | -0.030288003 | 0.124677636 | 0.635460588 | 0.869849417 | ns | TM | KS.1 |
| FIM1_4099 | 527.4163676 | 535.1064081 | 365.0170713 | 0.006397111 | 0.112114061 | 0.917484222 | 0.975634767 | ns | TM | KS.1 |
| FIM1_4100 | 306.8661576 | 314.9421329 | 354.4049164 | 0.014696516 | 0.084892984 | 0.787409578 | 0.932705393 | ns | TM | KS.1 |
| FIM1_4101 | 748.378978 | 749.3012319 | 806.4622174 | 0.000460918 | 0.112737164 | 0.994049666 | 0.99786059 | ns | TM | KS.1 |
| FIM1_4102 | 2257.911004 | 1981.963566 | 2062.510701 | -0.090365548 | 0.106138069 | 0.16247075 | 0.45282464 | ns | TM | KS.1 |
| FIM1_4103 | 9729.383105 | 9389.608339 | 4905.46933 | -0.015389976 | 0.114107893 | 0.804556287 | 0.936954009 | ns | TM | KS.1 |
| FIM1_4104 | 1246.329643 | 1545.854199 | 1512.017908 | 0.084975926 | 0.151451646 | 0.23153025 | 0.549231011 | ns | TM | KS.1 |
| FIM1_4105 | 295.4078838 | 259.8007558 | 221.2919914 | -0.046616195 | 0.137283873 | 0.478244078 | 0.779543161 | ns | TM | KS.1 |
| FIM1_4106 | 5326.20663 | 3697.502088 | 8180.424686 | -0.18040554 | 0.203658065 | 0.044567746 | 0.217376995 | ns | TM | KS.1 |
| FIM1_4107 | 1077.964613 | 1161.106954 | 1193.939476 | 0.05286623 | 0.083734416 | 0.33277503 | 0.659583974 | ns | TM | KS.1 |
| FIM1_4108 | 2656.311952 | 2543.575076 | 2811.311319 | -0.03324533 | 0.064903879 | 0.465569708 | 0.767719357 | ns | TM | KS.1 |
| FIM1_4109 | 328.7198847 | 379.6170929 | 326.6748471 | 0.063961519 | 0.130010527 | 0.340820617 | 0.668341572 | ns | TM | KS.1 |
| FIM1_4110 | 860.1864986 | 915.9690073 | 929.8709923 | 0.035811196 | 0.096086337 | 0.542034494 | 0.815921818 | ns | TM | KS.1 |
| FIM1_4111 | 2329.98815 | 2413.664174 | 2452.532085 | 0.03072331 | 0.051227472 | 0.423310759 | 0.739045122 | ns | TM | KS.1 |
| FIM1_4112 | 2012.923715 | 2057.100732 | 2371.194331 | 0.008116613 | 0.122759805 | 0.897312251 | 0.967972723 | ns | TM | KS.1 |
| FIM1_4113 | 877.730362 | 1373.626036 | 1778.14431 | 0.195412898 | 0.243436493 | 0.040964909 | 0.20690398 | ns | TM | KS.1 |
| FIM1_4114 | 855.6150899 | 1566.053292 | 2048.426331 | 0.212996427 | 0.310746777 | 0.039552749 | 0.202027643 | ns | TM | KS.1 |
| FIM1_4115 | 943.1955751 | 1139.2131 | 1095.168122 | 0.09827782 | 0.135290205 | 0.169719712 | 0.463676862 | ns | TM | KS.1 |
| FIM1_4116 | 254.153192 | 287.6629402 | 254.7718731 | 0.031049758 | 0.15565479 | 0.62645585 | 0.866840608 | ns | TM | KS.1 |
| FIM1_4117 | 181.3332344 | 167.9472992 | 180.9317383 | -0.018834343 | 0.15473449 | 0.764355613 | 0.924230468 | ns | TM | KS.1 |
| FIM1_4118 | 1949.975335 | 1962.953667 | 2095.015767 | 0.002179 | 0.131724564 | 0.972454023 | 0.99362472 | ns | TM | KS.1 |
| FIM1_4119 | 1609.346708 | 1603.226959 | 1782.112594 | -0.002187255 | 0.090807407 | 0.969151543 | 0.99351513 | ns | TM | KS.1 |
| FIM1_4120 | 1778.508865 | 2649.745165 | 2779.77814 | 0.117619274 | 0.214712789 | 0.134596754 | 0.406322117 | ns | TM | KS.1 |
| FIM1_4121 | 696.6767204 | 1028.094155 | 988.6484663 | 0.170739783 | 0.214508882 | 0.054893898 | 0.245370299 | ns | TM | KS.1 |
| FIM1_4122 | 745.0763273 | 668.9048729 | 644.6217503 | -0.064012969 | 0.106457011 | 0.30969251 | 0.637979394 | ns | TM | KS.1 |

| | | | | | | | | | | |
|---|---|---|---|---|---|---|---|---|---|---|
| FIM1_4123 | 1092.005192 | 995.1277451 | 1041.902857 | -0.051170327 | 0.106876542 | 0.412060494 | 0.732007978 | ns | TM | KS.1 |
| FIM1_4124 | 65.34393064 | 71.93449552 | 61.31069651 | 0.030692925 | 0.143707023 | 0.632578714 | 0.869547581 | ns | TM | KS.1 |
| FIM1_4125 | 5466.597461 | 6546.574097 | 7126.221652 | 0.095490502 | 0.132358123 | 0.177882053 | 0.476606421 | ns | TM | KS.1 |
| FIM1_4126 | 416.6369774 | 335.9501477 | 369.1405331 | -0.152597221 | 0.136128827 | 0.050661459 | 0.233804042 | ns | TM | KS.1 |
| FIM1_4127 | 63.50736835 | 57.97344326 | 56.92117253 | -0.026260876 | 0.147019305 | 0.680283269 | 0.887492141 | ns | TM | KS.1 |
| FIM1_4128 | 1101.004154 | 1276.101239 | 1315.209878 | 0.097847757 | 0.113958745 | 0.147295439 | 0.426043493 | ns | TM | KS.1 |
| FIM1_4129 | 2511.900516 | 2703.242796 | 2444.613147 | 0.039952472 | 0.102826473 | 0.511262965 | 0.802123385 | ns | TM | KS.1 |
| FIM1_4130 | 1649.91929 | 1641.130558 | 2108.407292 | -0.002453203 | 0.11074373 | 0.968184188 | 0.993374633 | ns | TM | KS.1 |
| FIM1_4132 | 246.7196251 | 236.0389846 | 198.7253864 | -0.025794523 | 0.092479279 | 0.652793363 | 0.875612558 | ns | TM | KS.1 |
| FIM1_4133 | 1045.243558 | 817.3483178 | 883.5089162 | -0.194202557 | 0.13999703 | 0.020150817 | 0.137141332 | ns | TM | KS.1 |
| FIM1_4134 | 2075.456244 | 1592.755475 | 1743.048971 | -0.273603892 | 0.119257517 | 0.001401032 | 0.019307857 | ns | TM | KS.1 |
| FIM1_4135 | 851.8634763 | 912.8553234 | 1187.428661 | 0.037912728 | 0.100311992 | 0.527876528 | 0.808531286 | ns | TM | KS.1 |
| FIM1_4136 | 2411.455978 | 2340.836335 | 2361.138264 | -0.019914206 | 0.074863067 | 0.692315559 | 0.893131011 | ns | TM | KS.1 |
| FIM1_4137 | 2536.227743 | 3045.076903 | 3197.217042 | 0.090156273 | 0.135430466 | 0.201739766 | 0.509840799 | ns | TM | KS.1 |
| FIM1_4138 | 1165.573847 | 1279.962702 | 1404.198415 | 0.072012999 | 0.086536755 | 0.201886708 | 0.509840799 | ns | TM | KS.1 |
| FIM1_4139 | 2052.205049 | 3070.750845 | 3528.830738 | 0.210863645 | 0.223354897 | 0.029236921 | 0.16972335 | ns | TM | KS.1 |
| FIM1_4140 | 1020.997998 | 1154.74717 | 1085.210523 | 0.047251119 | 0.132566544 | 0.47167524 | 0.772376515 | ns | TM | KS.1 |
| FIM1_4141 | 7735.755003 | 9059.252265 | 8667.610626 | 0.108154329 | 0.116227821 | 0.117158852 | 0.374832982 | ns | TM | KS.1 |
| FIM1_4142 | 771.0291918 | 844.0332813 | 850.6829663 | 0.064185765 | 0.090026886 | 0.264517672 | 0.590354459 | ns | TM | KS.1 |
| FIM1_4143 | 884.3858185 | 928.7646644 | 875.8348652 | 0.032875325 | 0.082527858 | 0.540531155 | 0.814399145 | ns | TM | KS.1 |
| FIM1_4144 | 504.4807368 | 448.7161008 | 395.2838946 | -0.055932842 | 0.119911884 | 0.39125676 | 0.71457503 | ns | TM | KS.1 |
| FIM1_4145 | 576.8496624 | 516.334777 | 511.0689433 | -0.054824965 | 0.11579496 | 0.395274862 | 0.718896697 | ns | TM | KS.1 |
| FIM1_4146 | 1065.957965 | 1568.072706 | 1621.817164 | 0.094211495 | 0.211027163 | 0.200703346 | 0.508919034 | ns | TM | KS.1 |
| FIM1_4147 | 1056.510796 | 871.7539896 | 825.2580876 | -0.102213006 | 0.135887223 | 0.15638859 | 0.441312101 | ns | TM | KS.1 |
| FIM1_4148 | 2724.806357 | 2775.828605 | 3486.418128 | 0.005623131 | 0.137259138 | 0.92896838 | 0.978304974 | ns | TM | KS.1 |
| FIM1_4149 | 13151.95877 | 13494.90618 | 12080.15282 | 0.016998617 | 0.076046146 | 0.738153646 | 0.913453034 | ns | TM | KS.1 |
| FIM1_4150 | 2150.129616 | 2259.595552 | 2153.571342 | 0.027108591 | 0.09799848 | 0.646337113 | 0.873463833 | ns | TM | KS.1 |
| FIM1_4151 | 2807.153801 | 2404.093302 | 3026.452796 | -0.11005791 | 0.113793975 | 0.108394509 | 0.357240387 | ns | TM | KS.1 |
| FIM1_4152 | 913.5123747 | 1464.696513 | 1951.519265 | 0.169641877 | 0.249081332 | 0.061907008 | 0.261227619 | ns | TM | KS.1 |
| FIM1_4153 | 1589.160454 | 1492.331638 | 1687.778482 | -0.032612949 | 0.10422817 | 0.592257066 | 0.846446108 | ns | TM | KS.1 |
| FIM1_4154 | 1188.048805 | 709.4466091 | 796.973279 | -0.502076218 | 0.27112381 | 0.000631432 | 0.010704111 | ns | TM | KS.1 |
| FIM1_4155 | 3652.304224 | 3930.545168 | 3767.699817 | 0.029994605 | 0.121878263 | 0.637489824 | 0.870169358 | ns | TM | KS.1 |
| FIM1_4156 | 2281.921781 | 3200.083749 | 3625.250613 | 0.129956204 | 0.192032177 | 0.105014566 | 0.351107167 | ns | TM | KS.1 |
| FIM1_4157 | 2045.390723 | 2447.792516 | 3012.997044 | 0.089965309 | 0.133986669 | 0.201530559 | 0.509840799 | ns | TM | KS.1 |
| FIM1_4158 | 4413.160775 | 3840.141038 | 4488.575145 | -0.06630889 | 0.125237045 | 0.321385083 | 0.649691502 | ns | TM | KS.1 |
| FIM1_4159 | 1486.248146 | 1146.400293 | 1271.519633 | -0.272830178 | 0.115377136 | 0.001208879 | 0.017381997 | ns | TM | KS.1 |
| FIM1_4160 | 443.047677 | 376.4290938 | 355.4111926 | -0.100608221 | 0.122208509 | 0.148901816 | 0.429441469 | ns | TM | KS.1 |
| FIM1_4161 | 13871.90451 | 15709.81339 | 15530.23963 | 0.11034185 | 0.091124516 | 0.069114019 | 0.27676212 | ns | TM | KS.1 |
| FIM1_4162 | 1511.41738 | 1820.311455 | 1415.571465 | 0.044195299 | 0.166961935 | 0.496021713 | 0.79247947 | ns | TM | KS.1 |
| FIM1_4163 | 9353.334201 | 7581.961301 | 7457.914414 | -0.183480378 | 0.121512496 | 0.018692601 | 0.130611925 | ns | TM | KS.1 |
| FIM1_4164 | 75.26711272 | 68.74001915 | 109.5779923 | -0.02027468 | 0.161853271 | 0.745819011 | 0.91614154 | ns | TM | KS.1 |
| FIM1_4165 | 545.540334 | 454.5759436 | 424.6697933 | -0.107164839 | 0.129929199 | 0.13566184 | 0.407805228 | ns | TM | KS.1 |
| FIM1_4166 | 1156.810289 | 1185.999874 | 2248.026681 | 0.008418016 | 0.129952333 | 0.893989845 | 0.967972723 | ns | TM | KS.1 |
| FIM1_4167 | 1167.008604 | 1172.134651 | 1085.647549 | 0.001076791 | 0.147800588 | 0.986277351 | 0.996167262 | ns | TM | KS.1 |
| FIM1_4169 | 624.151337 | 730.7644477 | 672.9160416 | 0.149440306 | 0.097696381 | 0.025242477 | 0.156001645 | ns | TM | KS.1 |

| | | | | | | | | | |
|---|---|---|---|---|---|---|---|---|---|
| FIM1_4170 | 934.4033246 | 856.7716329 | 738.4157999 | -0.04629769 | 0.106580669 | 0.455623833 | 0.759118744 | ns | TM | KS.1 |
| FIM1_4171 | 116.7457715 | 121.3243687 | 117.2484734 | 0.009321147 | 0.144259803 | 0.882226852 | 0.966130215 | ns | TM | KS.1 |
| FIM1_4172 | 1115.256636 | 931.7211125 | 1125.987572 | -0.071301974 | 0.142538309 | 0.299968652 | 0.628445786 | ns | TM | KS.1 |
| FIM1_4173 | 1268.043976 | 1022.792941 | 1128.570396 | -0.198959349 | 0.119184639 | 0.011726269 | 0.094398367 | ns | TM | KS.1 |
| FIM1_4174 | 2902.554593 | 4373.075866 | 5769.751469 | 0.20304522 | 0.226406351 | 0.033749378 | 0.183500714 | ns | TM | KS.1 |
| FIM1_4175 | 1664.271804 | 1850.895399 | 1627.710598 | 0.071915026 | 0.098247466 | 0.237880874 | 0.558772287 | ns | TM | KS.1 |
| FIM1_4176 | 1511.483013 | 1440.313141 | 1316.289622 | -0.033962091 | 0.075701194 | 0.50343691 | 0.796765282 | ns | TM | KS.1 |
| FIM1_4177 | 9013.642285 | 7825.822758 | 6783.080192 | -0.079853653 | 0.118445626 | 0.233337926 | 0.551737728 | ns | TM | KS.1 |
| FIM1_4178 | 8368.450372 | 7270.792772 | 6196.614027 | -0.088305485 | 0.113852805 | 0.185445706 | 0.488144311 | ns | TM | KS.1 |
| FIM1_4180 | 25.19794587 | 20.73840231 | 24.9312206 | -0.044596345 | 0.170495318 | 0.491718793 | 0.790857947 | ns | TM | KS.1 |
| FIM1_4181 | 2552.952888 | 2433.125553 | 3689.600522 | -0.026624733 | 0.095937004 | 0.648770885 | 0.874068749 | ns | TM | KS.1 |
| FIM1_4182 | 6872.028127 | 8198.088484 | 8869.187302 | 0.205203457 | 0.081722508 | 0.00165797 | 0.022173124 | ns | TM | KS.1 |
| FIM1_4183 | 1572.751767 | 1611.268577 | 1556.276746 | 0.015696437 | 0.075334105 | 0.756078427 | 0.920354827 | ns | TM | KS.1 |
| FIM1_4184 | 1960.403199 | 2014.776625 | 2060.641967 | 0.019359825 | 0.069243461 | 0.684867891 | 0.890505759 | ns | TM | KS.1 |
| FIM1_4185 | 915.658771 | 739.1030084 | 760.5632707 | -0.189256254 | 0.122231309 | 0.016341675 | 0.118859402 | ns | TM | KS.1 |
| FIM1_4186 | 2889.967164 | 3372.67715 | 2235.521867 | 0.062701005 | 0.13582122 | 0.352424008 | 0.678105331 | ns | TM | KS.1 |
| FIM1_4187 | 727.628884 | 673.4468017 | 693.7005976 | -0.04586491 | 0.097847189 | 0.441671763 | 0.747895515 | ns | TM | KS.1 |
| FIM1_4188 | 983.1447246 | 1144.272114 | 1297.856177 | 0.0878662 | 0.120559176 | 0.197276096 | 0.505120216 | ns | TM | KS.1 |
| FIM1_4189 | 3028.465174 | 3083.084382 | 3247.958278 | 0.00892969 | 0.101730589 | 0.881175882 | 0.9656535 | ns | TM | KS.1 |
| FIM1_4190 | 2458.597341 | 2309.613656 | 1947.08494 | -0.042040968 | 0.08402719 | 0.440234114 | 0.747832613 | ns | TM | KS.1 |
| FIM1_4191 | 1947.36251 | 2048.432408 | 2019.30619 | 0.029866645 | 0.092173966 | 0.602266672 | 0.851547247 | ns | TM | KS.1 |
| FIM1_4192 | 709.2172947 | 979.0835308 | 923.868796 | 0.163823897 | 0.184133979 | 0.05524614 | 0.246280956 | ns | TM | KS.1 |
| FIM1_4193 | 275.9783068 | 256.1354655 | 265.2306088 | -0.02960183 | 0.123894602 | 0.642637717 | 0.872815354 | ns | TM | KS.1 |
| FIM1_4194 | 1523.771528 | 1097.651948 | 1161.170013 | -0.2489524 | 0.176232427 | 0.009850017 | 0.083624294 | ns | TM | KS.1 |
| FIM1_4195 | 2136.651304 | 2290.181727 | 2259.970138 | 0.046023208 | 0.086267139 | 0.406656882 | 0.728526463 | ns | TM | KS.1 |
| FIM1_4196 | 3009.439528 | 2765.82899 | 3731.145569 | -0.040837958 | 0.112787618 | 0.517268207 | 0.804981347 | ns | TM | KS.1 |
| FIM1_4197 | 2429.879627 | 1449.319569 | 1354.845331 | -0.352321004 | 0.295434094 | 0.006176154 | 0.060484971 | ns | TM | KS.1 |
| FIM1_4198 | 1990.681664 | 1650.244821 | 1718.026555 | -0.080302278 | 0.141568418 | 0.250674397 | 0.576493979 | ns | TM | KS.1 |
| FIM1_4199 | 287.2811766 | 264.1643322 | 370.175666 | -0.034910562 | 0.122356496 | 0.585380918 | 0.841613393 | ns | TM | KS.1 |
| FIM1_4201 | 640.4585931 | 556.1705236 | 544.8274497 | -0.144034696 | 0.086600989 | 0.019587572 | 0.13459692 | ns | TM | KS.1 |
| FIM1_4203 | 270.0043972 | 316.1518832 | 324.5406144 | 0.063200693 | 0.136710883 | 0.349345611 | 0.675211505 | ns | TM | KS.1 |
| FIM1_4204 | 1212.845689 | 969.5583048 | 1148.355635 | -0.17963461 | 0.131689173 | 0.025060463 | 0.155262517 | ns | TM | KS.1 |
| FIM1_4205 | 1300.655185 | 1159.910175 | 3643.20249 | -0.028483767 | 0.15591131 | 0.653928105 | 0.876661903 | ns | TM | KS.1 |
| FIM1_4206 | 1227.69245 | 1415.163576 | 1485.64639 | 0.051421337 | 0.138678478 | 0.437823986 | 0.747832613 | ns | TM | KS.1 |
| FIM1_4207 | 2669.78011 | 2431.736492 | 2188.568411 | -0.046329737 | 0.11277615 | 0.464803307 | 0.766709699 | ns | TM | KS.1 |
| FIM1_4208 | 1025.673837 | 998.3084754 | 1001.676204 | -0.012161556 | 0.111466762 | 0.843822419 | 0.951768052 | ns | TM | KS.1 |
| FIM1_4209 | 700.0617818 | 606.2225135 | 664.3409912 | -0.094232298 | 0.113175878 | 0.159641587 | 0.44660975 | ns | TM | KS.1 |
| FIM1_4210 | 1928.913845 | 2065.521174 | 2048.586944 | 0.035500567 | 0.104780261 | 0.561457398 | 0.826391733 | ns | TM | KS.1 |
| FIM1_4211 | 680.3957724 | 700.8355833 | 725.4670799 | 0.011997238 | 0.11900765 | 0.848197947 | 0.952348261 | ns | TM | KS.1 |
| FIM1_4212 | 2424.682442 | 2980.464534 | 3234.851481 | 0.23075318 | 0.09356574 | 0.001511771 | 0.020549341 | ns | TM | KS.1 |
| FIM1_4213 | 398.5490834 | 405.9823576 | 423.953216 | 0.009913669 | 0.100756489 | 0.867691565 | 0.960282219 | ns | TM | KS.1 |
| FIM1_4214 | 1030.967562 | 1097.024949 | 1128.506859 | 0.038477205 | 0.089139268 | 0.495160142 | 0.792222775 | ns | TM | KS.1 |
| FIM1_4215 | 4225.81901 | 4416.033725 | 5631.325258 | 0.014215959 | 0.134461891 | 0.822457183 | 0.944618544 | ns | TM | KS.1 |
| FIM1_4216 | 768.1379125 | 629.5844748 | 743.7061317 | -0.09265619 | 0.142290415 | 0.195444464 | 0.50198049 | ns | TM | KS.1 |
| FIM1_4217 | 1002.458588 | 769.6533434 | 889.1683139 | -0.135589498 | 0.160843261 | 0.086684321 | 0.313970477 | ns | TM | KS.1 |

| | | | | | | | | | | |
|---|---|---|---|---|---|---|---|---|---|---|
| FIM1_4218 | 35.88468374 | 40.48322398 | 40.80047592 | 0.029655751 | 0.138972989 | 0.644142284 | 0.872978256 | ns | TM | KS.1 |
| FIM1_4219 | 468.4717709 | 471.9743751 | 472.1011329 | 0.003666477 | 0.104619505 | 0.951563248 | 0.987490021 | ns | TM | KS.1 |
| FIM1_4220 | 743.8079662 | 762.5403164 | 752.1887105 | 0.012467027 | 0.100957449 | 0.834282396 | 0.948141464 | ns | TM | KS.1 |
| FIM1_4221 | 307.4008642 | 289.7829332 | 215.5270907 | -0.026181444 | 0.114950795 | 0.676157953 | 0.887002016 | ns | TM | KS.1 |
| FIM1_4222 | 1540.484092 | 1629.568843 | 1521.313095 | 0.018069186 | 0.135324852 | 0.776029091 | 0.926727011 | ns | TM | KS.1 |
| FIM1_4223 | 2702.114181 | 2711.502688 | 2560.189644 | 0.001902429 | 0.094306947 | 0.973685176 | 0.993872484 | ns | TM | KS.1 |
| FIM1_4224 | 6123.8742 | 5999.128994 | 5080.741052 | -0.009520427 | 0.1079024 | 0.876122965 | 0.964103462 | ns | TM | KS.1 |
| FIM1_4225 | 439.5107364 | 626.4939254 | 581.9715583 | 0.093028003 | 0.200689895 | 0.204271982 | 0.512741227 | ns | TM | KS.1 |
| FIM1_4226 | 22931.02264 | 18676.3002 | 38582.63717 | -0.090710365 | 0.145214242 | 0.204681016 | 0.512865935 | ns | TM | KS.1 |
| FIM1_4227 | 1298.655097 | 1826.985211 | 2863.283207 | 0.087060496 | 0.197418405 | 0.226937225 | 0.54408821 | ns | TM | KS.1 |
| FIM1_4228 | 17.36047898 | 14.08280002 | 24.61517435 | -0.03971617 | 0.17918475 | 0.534074961 | 0.811304711 | ns | TM | KS.1 |
| FIM1_4229 | 354.42069 | 374.4026663 | 1314.53667 | 0.014731292 | 0.146431754 | 0.815187652 | 0.940791141 | ns | TM | KS.1 |
| FIM1_4230 | 2003.093421 | 2255.894295 | 2387.443186 | 0.04225136 | 0.135979701 | 0.517498265 | 0.804981347 | ns | TM | KS.1 |
| FIM1_4231 | 4402.115121 | 5056.938506 | 5772.270488 | 0.067794935 | 0.124028681 | 0.310529036 | 0.6389526 | ns | TM | KS.1 |
| FIM1_4232 | 652.1992792 | 466.3165016 | 487.0612049 | -0.242439239 | 0.182845896 | 0.012133868 | 0.096768242 | ns | TM | KS.1 |
| FIM1_4233 | 671.8813171 | 742.0736331 | 813.2059935 | 0.059642201 | 0.103053046 | 0.334569884 | 0.661823131 | ns | TM | KS.1 |
| FIM1_4234 | 1149.600938 | 1174.59647 | 1164.257849 | 0.011446828 | 0.097858901 | 0.845545773 | 0.952348261 | ns | TM | KS.1 |
| FIM1_4235 | 987.5461014 | 903.6159191 | 998.3452731 | -0.062455753 | 0.089250815 | 0.274475494 | 0.599699421 | ns | TM | KS.1 |
| FIM1_4236 | 2716.13632 | 2490.35462 | 2572.85615 | -0.060855968 | 0.089785676 | 0.288231512 | 0.61538797 | ns | TM | KS.1 |
| FIM1_4237 | 6260.93954 | 6839.041026 | 7971.332114 | 0.069107698 | 0.082977201 | 0.207082413 | 0.513839538 | ns | TM | KS.1 |
| FIM1_4238 | 2788.5684 | 2840.109112 | 2974.152138 | 0.011502948 | 0.081416881 | 0.828541275 | 0.94647133 | ns | TM | KS.1 |
| FIM1_4239 | 2129.637052 | 2237.608883 | 2512.177134 | 0.030969848 | 0.086497921 | 0.57528064 | 0.833922256 | ns | TM | KS.1 |
| FIM1_4240 | 959.3086422 | 1311.152787 | 1416.346129 | 0.129593781 | 0.181771342 | 0.103486341 | 0.348223325 | ns | TM | KS.1 |
| FIM1_4241 | 2152.45827 | 2233.134639 | 2297.8411 | 0.012464213 | 0.130841082 | 0.843869121 | 0.951768052 | ns | TM | KS.1 |
| FIM1_4242 | 203.0150087 | 171.6741534 | 256.0742388 | -0.072517846 | 0.137099657 | 0.290821464 | 0.618193599 | ns | TM | KS.1 |
| FIM1_4243 | 761.4988316 | 775.5777367 | 683.6635556 | 0.011759811 | 0.080285194 | 0.823260543 | 0.94482203 | ns | TM | KS.1 |
| FIM1_4244 | 970.5786609 | 840.4285447 | 691.9464541 | -0.065216968 | 0.128842806 | 0.331560198 | 0.658750792 | ns | TM | KS.1 |
| FIM1_4245 | 31.70217521 | 19.13516152 | 45.78680657 | -0.182646599 | 0.262923703 | 0.052848566 | 0.240330544 | ns | TM | KS.1 |
| FIM1_4246 | 585.4164902 | 543.6351512 | 598.2144239 | -0.049297486 | 0.090203324 | 0.38778438 | 0.712272102 | ns | TM | KS.1 |
| FIM1_4247 | 331.2562101 | 350.242534 | 496.5544882 | 0.022523437 | 0.120226131 | 0.721279673 | 0.906676397 | ns | TM | KS.1 |
| FIM1_4248 | 1609.422304 | 1183.655021 | 1281.443992 | -0.374205438 | 0.096374112 | 2.40561942687 | 0.000140799 | ns | TM | KS.1 |
| FIM1_4249 | 4476.726918 | 4603.022361 | 5696.017768 | 0.016589851 | 0.08625071 | 0.76302651 | 0.924230468 | ns | TM | KS.1 |
| FIM1_4250 | 987.2006467 | 784.4533747 | 1103.258929 | -0.288386231 | 0.072739644 | 7.91962816834 | 0.000345615 | ns | TM | KS.1 |
| FIM1_4251 | 3759.328989 | 3378.416636 | 3147.760199 | -0.097059194 | 0.081952163 | 0.081875758 | 0.304774234 | ns | TM | KS.1 |
| FIM1_4252 | 1142.517032 | 1157.681714 | 1133.022365 | 0.008153538 | 0.081205037 | 0.877779119 | 0.964470036 | ns | TM | KS.1 |
| FIM1_4253 | 2148.906152 | 3021.468697 | 2736.043885 | 0.222613798 | 0.188697426 | 0.018836826 | 0.131067425 | ns | TM | KS.1 |
| FIM1_4254 | 868.5991863 | 963.3015953 | 1081.144263 | 0.066836636 | 0.099439728 | 0.273709968 | 0.599198132 | ns | TM | KS.1 |
| FIM1_4255 | 1821.718329 | 1873.685026 | 1779.909196 | 0.011790254 | 0.116517845 | 0.850161651 | 0.953031594 | ns | TM | KS.1 |
| FIM1_4257 | 2237.795425 | 2433.926243 | 4226.826573 | 0.029552548 | 0.132111293 | 0.645122936 | 0.873039206 | ns | TM | KS.1 |
| FIM1_4258 | 31.62434316 | 39.65039346 | 42.59403129 | 0.06967281 | 0.164145782 | 0.311572869 | 0.638961913 | ns | TM | KS.1 |
| FIM1_4259 | 3116.67437 | 3289.656941 | 2675.666478 | 0.022788784 | 0.117450102 | 0.716942067 | 0.905260252 | ns | TM | KS.1 |
| FIM1_4260 | 789.8997683 | 836.8733948 | 767.4231294 | 0.021041079 | 0.127345469 | 0.740517614 | 0.914311536 | ns | TM | KS.1 |
| FIM1_4261 | 496.8939719 | 593.657352 | 920.3489585 | 0.07872107 | 0.137839295 | 0.25704522 | 0.585794503 | ns | TM | KS.1 |
| FIM1_4262 | 2425.653651 | 2652.295719 | 2964.984714 | 0.046285056 | 0.109388297 | 0.460364064 | 0.763054265 | ns | TM | KS.1 |
| FIM1_4263 | 785.3853243 | 524.2780694 | 569.9360588 | -0.500032307 | 0.13945443 | 1.56067729288 | 0.000102163 | ns | TM | KS.1 |

| | | | | | | | | | | |
|---|---|---|---|---|---|---|---|---|---|---|
| FIM1_4265 | 850.92158 | 937.4561344 | 891.4510669 | 0.062994007 | 0.097541735 | 0.295657603 | 0.623274449 | ns | TM | KS.1 |
| FIM1_4266 | 7346.668797 | 11156.67122 | 9953.457295 | 0.266379773 | 0.232011231 | 0.012508805 | 0.098311699 | ns | TM | KS.1 |
| FIM1_4267 | 1447.47527 | 1410.398741 | 1443.309574 | -0.013951503 | 0.093891957 | 0.80867992 | 0.938680028 | ns | TM | KS.1 |
| FIM1_4268 | 3186.331424 | 3265.0283 | 3576.938436 | 0.010341897 | 0.114963208 | 0.867936061 | 0.960282219 | ns | TM | KS.1 |
| FIM1_4269 | 317.7943033 | 367.1056011 | 394.092963 | 0.083143707 | 0.116293196 | 0.213462757 | 0.521776083 | ns | TM | KS.1 |
| FIM1_4270 | 2090.798022 | 1826.453662 | 2359.182223 | -0.105237091 | 0.102387536 | 0.103267985 | 0.348074679 | ns | TM | KS.1 |
| FIM1_4271 | 745.3055733 | 731.9562062 | 835.0517382 | -0.011873146 | 0.078531461 | 0.819105279 | 0.94310261 | ns | TM | KS.1 |
| FIM1_4272 | 663.6933517 | 584.2445349 | 712.7728461 | -0.053271885 | 0.129387078 | 0.419822875 | 0.737778527 | ns | TM | KS.1 |
| FIM1_4273 | 1098.893389 | 1676.317559 | 1940.035915 | 0.169545378 | 0.22881044 | 0.058632667 | 0.254317312 | ns | TM | KS.1 |
| FIM1_4274 | 187.3135751 | 208.7568939 | 458.2280169 | 0.065616173 | 0.111758646 | 0.30857104 | 0.636779799 | ns | TM | KS.1 |
| FIM1_4275 | 17.16113429 | 15.19812844 | 33.90945476 | -0.01953114 | 0.182204214 | 0.750626564 | 0.918418758 | ns | TM | KS.1 |
| FIM1_4276 | 2468.806846 | 2431.808344 | 2355.293366 | -0.009773819 | 0.074822168 | 0.845925117 | 0.952348261 | ns | TM | KS.1 |
| FIM1_4277 | 196.5329357 | 218.6686704 | 199.5390417 | 0.04861515 | 0.120920789 | 0.453144986 | 0.756837143 | ns | TM | KS.1 |
| FIM1_4278 | 3129.528472 | 3044.888762 | 2990.652528 | -0.020827301 | 0.059668729 | 0.627314655 | 0.866914003 | ns | TM | KS.1 |
| FIM1_4279 | 2145.479182 | 3154.912639 | 3477.358609 | 0.127011523 | 0.210371191 | 0.11406842 | 0.368979448 | ns | TM | KS.1 |
| FIM1_4280 | 3457.313201 | 3326.616484 | 2995.245579 | -0.022602347 | 0.089469066 | 0.687837129 | 0.891826565 | ns | TM | KS.1 |
| FIM1_4281 | 4673.462421 | 4984.490764 | 5184.495808 | 0.042474536 | 0.086138079 | 0.442730791 | 0.748839657 | ns | TM | KS.1 |
| FIM1_4282 | 1465.326415 | 537.8053096 | 2506.198991 | -1.179872392 | 0.355945148 | 1.50315768205 | 0.000566531 | Km<KS.1 | TM | KS.1 |
| FIM1_4283 | 47.05747217 | 29.86934213 | 72.17510083 | -0.171624828 | 0.243808327 | 0.059229054 | 0.255563352 | ns | TM | KS.1 |
| FIM1_4284 | 7459.214209 | 7087.793293 | 5208.636838 | -0.032517939 | 0.084571227 | 0.551042969 | 0.822058746 | ns | TM | KS.1 |
| FIM1_4285 | 2246.584144 | 2583.954636 | 2583.026233 | 0.101398375 | 0.107517485 | 0.12430925 | 0.387735747 | ns | TM | KS.1 |
| FIM1_4286 | 2545.545759 | 2539.416094 | 2558.176458 | -0.000734775 | 0.139142427 | 0.990692016 | 0.996500764 | ns | TM | KS.1 |
| FIM1_4287 | 10.78273614 | 11.44493462 | 12.20518445 | 0.017859004 | 0.180020962 | 0.771266518 | 0.925705893 | ns | TM | KS.1 |
| FIM1_4288 | 1318.081266 | 1204.903151 | 1283.172397 | -0.032687346 | 0.130943638 | 0.611530585 | 0.85579878 | ns | TM | KS.1 |
| FIM1_4289 | 2513.779451 | 2340.807232 | 2144.079177 | -0.039538187 | 0.100796586 | 0.511705193 | 0.802564103 | ns | TM | KS.1 |
| FIM1_4290 | 1285.386376 | 1316.541641 | 1397.806259 | 0.014695388 | 0.085601388 | 0.788444747 | 0.933067989 | ns | TM | KS.1 |
| FIM1_4291 | 775.7643132 | 833.0688646 | 822.1209317 | 0.058256484 | 0.074869334 | 0.250834085 | 0.576493979 | ns | TM | KS.1 |
| FIM1_4292 | 864.277209 | 991.9313611 | 926.2401936 | 0.105654033 | 0.104824585 | 0.106485776 | 0.353581815 | ns | TM | KS.1 |
| FIM1_4293 | 638.71622 | 658.512295 | 651.6130055 | 0.022680869 | 0.073215111 | 0.647316329 | 0.873595603 | ns | TM | KS.1 |
| FIM1_4294 | 1157.487022 | 1025.351904 | 1014.23371 | -0.083521127 | 0.103302245 | 0.187145458 | 0.489513719 | ns | TM | KS.1 |
| FIM1_4295 | 1013.773727 | 1075.221698 | 957.734905 | 0.045224127 | 0.070603153 | 0.350639327 | 0.676561755 | ns | TM | KS.1 |
| FIM1_4296 | 1425.70365 | 1118.787948 | 1234.676435 | -0.313327729 | 0.066640218 | 2.1619572908 | 2.1950484738 | ns | TM | KS.1 |
| FIM1_4297 | 686.3540551 | 677.3439111 | 795.4135375 | -0.006108047 | 0.105900622 | 0.919772864 | 0.975881018 | ns | TM | KS.1 |
| FIM1_4298 | 203.4597514 | 251.1655855 | 256.6677733 | 0.038875642 | 0.184486997 | 0.540813078 | 0.814576162 | ns | TM | KS.1 |
| FIM1_4299 | 221.3345648 | 279.984126 | 366.378426 | 0.152409341 | 0.143489817 | 0.054705291 | 0.244967437 | ns | TM | KS.1 |
| FIM1_4300 | 181.5691716 | 194.2775106 | 223.9361699 | 0.032071662 | 0.105549971 | 0.600155075 | 0.850484618 | ns | TM | KS.1 |
| FIM1_4303 | 45.81037528 | 58.61279961 | 82.48496482 | 0.091503387 | 0.164656285 | 0.206497303 | 0.513662041 | ns | TM | KS.1 |
| FIM1_4304 | 15.44885418 | 22.66702761 | 30.91326256 | 0.071259227 | 0.210425758 | 0.300138128 | 0.628445786 | ns | TM | KS.1 |
| FIM1_4307 | 351.9607161 | 384.5476417 | 476.7071757 | 0.041330836 | 0.117302163 | 0.517198893 | 0.804981347 | ns | TM | KS.1 |
| FIM1_4308 | 83.09878046 | 118.4657032 | 116.7686975 | 0.18815121 | 0.202858015 | 0.038758631 | 0.198583099 | ns | TM | KS.1 |
| FIM1_4314 | 344.1489986 | 591.5978997 | 646.7515623 | 0.538337023 | 0.287205951 | 0.000542874 | 0.009647782 | ns | TM | KS.1 |
| FIM1_4780 | 30.51009712 | 57.38502959 | 82.75928701 | 0.232154471 | 0.327943854 | 0.032632031 | 0.180382188 | ns | TM | KS.1 |
| FIM1_4781 | 41.60261245 | 28.99750924 | 87.57792087 | -0.151595356 | 0.204224156 | 0.073764873 | 0.288677781 | ns | TM | KS.1 |
| FIM1_4782 | 221.8068361 | 253.6514793 | 296.5047851 | 0.089299393 | 0.110884718 | 0.176005783 | 0.473568831 | ns | TM | KS.1 |
| FIM1_4783 | 1958.485844 | 2357.784445 | 1729.24346 | 0.146216815 | 0.119847273 | 0.047292139 | 0.226665116 | ns | TM | KS.1 |

| | | | | | | | | | | |
|---|---|---|---|---|---|---|---|---|---|---|
| FIM1_4785 | 725.936034 | 749.333919 | 1037.162625 | 0.019940111 | 0.088360588 | 0.720958913 | 0.906676397 | ns | TM | KS.1 |
| FIM1_4786 | 925.4289353 | 932.4957638 | 3276.771524 | 0.001435369 | 0.169963394 | 0.98132027 | 0.995431301 | ns | TM | KS.1 |
| FIM1_4787 | 118.8364372 | 120.4747036 | 235.4642185 | 0.005079723 | 0.134095865 | 0.935861333 | 0.981638231 | ns | TM | KS.1 |
| FIM1_4788 | 204.251331 | 208.4899156 | 744.3069426 | 0.005203647 | 0.15056937 | 0.933642742 | 0.980344585 | ns | TM | KS.1 |
| FIM1_4789 | 212.0556319 | 251.6895374 | 264.372092 | 0.034312624 | 0.174798876 | 0.587935498 | 0.842933459 | ns | TM | KS.1 |
| FIM1_4790 | 460.9033638 | 616.211966 | 614.4716867 | 0.177128682 | 0.166452203 | 0.038689276 | 0.198583099 | ns | TM | KS.1 |
| FIM1_4791 | 1025.193754 | 1489.74804 | 2073.260557 | 0.222465736 | 0.208060937 | 0.021885748 | 0.144214032 | ns | TM | KS.1 |
| FIM1_4792 | 449.1999272 | 444.005342 | 464.198878 | -0.003691989 | 0.136759993 | 0.953316998 | 0.988485216 | ns | TM | KS.1 |
| FIM1_4793 | 1716.568644 | 1579.730125 | 2070.580399 | -0.04542047 | 0.10452115 | 0.460187173 | 0.763054265 | ns | TM | KS.1 |
| FIM1_4794 | 2597.285271 | 1859.886803 | 2670.050722 | -0.180114323 | 0.188663675 | 0.041714414 | 0.208572069 | ns | TM | KS.1 |
| FIM1_4795 | 7009.975949 | 5271.743533 | 6331.766408 | -0.108069395 | 0.173101872 | 0.152255234 | 0.434578193 | ns | TM | KS.1 |
| FIM1_4796 | 3075.103851 | 1871.880426 | 2521.152864 | -0.452812412 | 0.266955809 | 0.001139612 | 0.016675198 | ns | TM | KS.1 |
| FIM1_4797 | 2751.513641 | 3682.21853 | 4622.760369 | 0.096699054 | 0.177663376 | 0.189028623 | 0.492365131 | ns | TM | KS.1 |
| FIM1_4798 | 1191.067093 | 1108.857206 | 907.3828427 | -0.027691725 | 0.125154447 | 0.664114478 | 0.880619203 | ns | TM | KS.1 |
| FIM1_4799 | 1002.501185 | 1042.697725 | 915.778069 | 0.015950358 | 0.118717387 | 0.799356738 | 0.936599099 | ns | TM | KS.1 |
| FIM1_4800 | 12.08923879 | 11.55619107 | 14.76985844 | -0.003331592 | 0.17448612 | 0.956495552 | 0.989353426 | ns | TM | KS.1 |
| FIM1_4801 | 351.4062942 | 286.2106157 | 369.4995752 | -0.077061549 | 0.150329111 | 0.269855393 | 0.597663125 | ns | TM | KS.1 |
| FIM1_4802 | 748.755251 | 758.047066 | 805.0057536 | 0.00811982 | 0.071218621 | 0.867449108 | 0.960282219 | ns | TM | KS.1 |
| FIM1_4803 | 1777.616682 | 1862.220275 | 1781.19925 | 0.017530305 | 0.124169039 | 0.781600659 | 0.929216508 | ns | TM | KS.1 |
| FIM1_4804 | 5042.020276 | 5860.543539 | 5526.854346 | 0.098260109 | 0.115575353 | 0.148277521 | 0.428385987 | ns | TM | KS.1 |
| FIM1_4805 | 2586.262598 | 4254.490781 | 4310.284211 | 0.617202137 | 0.190936885 | 6.03856979048 | 0.000283414 | ns | TM | KS.1 |
| FIM1_4806 | 1637.263677 | 1979.272392 | 2008.96905 | 0.229097284 | 0.076401196 | 0.00037173 | 0.007168052 | ns | TM | KS.1 |
| FIM1_4807 | 2245.247407 | 2699.472391 | 2654.061447 | 0.084906453 | 0.138095653 | 0.226724979 | 0.54408821 | ns | TM | KS.1 |
| FIM1_4808 | 2116.415921 | 2161.21744 | 2096.452981 | 0.008847797 | 0.115053478 | 0.886851674 | 0.967028314 | ns | TM | KS.1 |
| FIM1_4809 | 589.2943617 | 656.4030572 | 696.4791253 | 0.058753544 | 0.109841713 | 0.355127737 | 0.679784721 | ns | TM | KS.1 |
| FIM1_4810 | 10632.50589 | 12285.99945 | 12749.87612 | 0.140416452 | 0.092870555 | 0.02827523 | 0.166668193 | ns | TM | KS.1 |
| FIM1_4811 | 1902.55035 | 1596.375614 | 1936.960012 | -0.045193445 | 0.16127926 | 0.488478198 | 0.787995796 | ns | TM | KS.1 |
| FIM1_4812 | 242.680482 | 255.009183 | 213.5856373 | 0.022023239 | 0.109319322 | 0.720765391 | 0.906676397 | ns | TM | KS.1 |
| FIM1_4813 | 821.1846362 | 1335.421059 | 1787.613643 | 0.212464328 | 0.263871274 | 0.034354508 | 0.185199792 | ns | TM | KS.1 |
| FIM1_4814 | 1147.160597 | 1110.431168 | 1173.57424 | -0.024187969 | 0.068353674 | 0.60885662 | 0.853256813 | ns | TM | KS.1 |
| FIM1_4815 | 773.5779093 | 832.6333635 | 843.8898588 | 0.050479627 | 0.083861174 | 0.355016902 | 0.679784721 | ns | TM | KS.1 |
| FIM1_4816 | 1019.247179 | 1203.21572 | 1091.397534 | 0.154298723 | 0.102545872 | 0.025475091 | 0.156140634 | ns | TM | KS.1 |
| FIM1_4817 | 1448.719994 | 1581.574625 | 1649.15812 | 0.047858936 | 0.106119745 | 0.440324281 | 0.747832613 | ns | TM | KS.1 |
| FIM1_4818 | 1018.074816 | 974.1532458 | 1041.875757 | -0.031729259 | 0.073897206 | 0.525011495 | 0.808531286 | ns | TM | KS.1 |
| FIM1_4819 | 8167.88011 | 9625.849261 | 5748.894804 | 0.163698139 | 0.0972379 | 0.016064431 | 0.117185546 | ns | TM | KS.1 |
| FIM1_4820 | 2074.188212 | 1580.530912 | 1825.771121 | -0.305939969 | 0.105755569 | 0.000211376 | 0.00459213 | ns | TM | KS.1 |
| FIM1_4821 | 1371.45387 | 1087.362553 | 1098.858809 | -0.07645187 | 0.161491963 | 0.274298961 | 0.599699421 | ns | TM | KS.1 |
| FIM1_4822 | 2207.961895 | 1963.018875 | 1621.314464 | -0.083754897 | 0.100337055 | 0.179158978 | 0.47808616 | ns | TM | KS.1 |
| FIM1_4823 | 1075.301427 | 1380.262083 | 1166.873272 | 0.248862651 | 0.11981804 | 0.002992509 | 0.035029958 | ns | TM | KS.1 |
| FIM1_4824 | 2195.646305 | 2936.943699 | 3418.530375 | 0.16495541 | 0.169223562 | 0.050249963 | 0.233221194 | ns | TM | KS.1 |
| FIM1_4825 | 531.8219946 | 282.9637477 | 465.332088 | -0.699082414 | 0.305585685 | 0.000151488 | 0.003441335 | ns | TM | KS.1 |
| FIM1_4826 | 915.8772674 | 952.7870296 | 1047.673213 | 0.025678917 | 0.077819879 | 0.619109934 | 0.861076858 | ns | TM | KS.1 |
| FIM1_4827 | 1257.692904 | 1330.788812 | 1322.638848 | 0.037991234 | 0.081086919 | 0.474945457 | 0.775371126 | ns | TM | KS.1 |
| FIM1_4828 | 972.9176993 | 887.0342708 | 929.2585492 | -0.061523595 | 0.093894323 | 0.296113279 | 0.623693294 | ns | TM | KS.1 |
| FIM1_4829 | 2509.996467 | 1999.511997 | 2408.544241 | -0.068313014 | 0.163825991 | 0.319583358 | 0.647364498 | ns | TM | KS.1 |

| | | | | | | | | | | |
|---|---|---|---|---|---|---|---|---|---|---|
| FIM1_4830 | 2410.928929 | 2551.362707 | 2396.706495 | 0.018345422 | 0.134832644 | 0.77275552 | 0.925733395 | ns | TM | KS.1 |
| FIM1_4831 | 2184.759812 | 2088.587114 | 1966.919271 | -0.033388632 | 0.069126945 | 0.483524949 | 0.784137657 | ns | TM | KS.1 |
| FIM1_4832 | 4302.686581 | 3882.269408 | 3576.343722 | -0.109857125 | 0.068670547 | 0.028908417 | 0.169342179 | ns | TM | KS.1 |
| FIM1_4833 | 3455.86407 | 3853.716831 | 4511.192965 | 0.064899014 | 0.106241966 | 0.303008649 | 0.631432423 | ns | TM | KS.1 |
| FIM1_4834 | 893.5819255 | 1071.617189 | 1340.639535 | 0.043063635 | 0.166488213 | 0.506209637 | 0.798222804 | ns | TM | KS.1 |
| FIM1_4835 | 3397.77303 | 3847.177825 | 3751.275652 | 0.082510141 | 0.106019364 | 0.197982199 | 0.505976981 | ns | TM | KS.1 |
| FIM1_4836 | 761.8645033 | 819.8582262 | 802.9952928 | 0.02529997 | 0.131387339 | 0.692069886 | 0.893131011 | ns | TM | KS.1 |
| FIM1_4837 | 69.95928002 | 75.73057594 | 67.6932174 | 0.017374492 | 0.158210569 | 0.78118142 | 0.929216508 | ns | TM | KS.1 |
| FIM1_4839 | 776.6787492 | 629.9150585 | 576.3763114 | -0.071635013 | 0.154867021 | 0.300089594 | 0.628445786 | ns | TM | KS.1 |
| FIM1_4840 | 7446.96397 | 8454.917469 | 8620.223328 | 0.117034643 | 0.089826188 | 0.053838662 | 0.241957851 | ns | TM | KS.1 |
| FIM1_4841 | 212.4174002 | 149.9790663 | 248.0003937 | -0.301866877 | 0.17682429 | 0.003279472 | 0.037593029 | ns | TM | KS.1 |
| FIM1_4842 | 55678.10472 | 58878.07016 | 58807.41069 | 0.036648506 | 0.083763584 | 0.499521464 | 0.79438151 | ns | TM | KS.1 |
| FIM1_4844 | 629.7309875 | 277.1853115 | 1119.871416 | -0.358429332 | 0.437419223 | 0.012133912 | 0.096768242 | ns | TM | KS.1 |
| FIM1_4845 | 370.8007593 | 417.9652908 | 600.696593 | 0.031903882 | 0.151709758 | 0.618362026 | 0.860858199 | ns | TM | KS.1 |
| FIM1_4846 | 375.227582 | 617.4764092 | 542.9534828 | 0.187289892 | 0.263131943 | 0.049413806 | 0.231715567 | ns | TM | KS.1 |
| FIM1_4847 | 385.4110636 | 467.7463944 | 765.4735833 | 0.061159386 | 0.154364159 | 0.365586155 | 0.689719803 | ns | TM | KS.1 |
| FIM1_4848 | 669.5084009 | 2047.670781 | 2313.99835 | 1.426155132 | 0.264159388 | 7.1497851294[illegible] | 1.1115681568[illegible] | Km>KS.1 | TM | KS.1 |
| FIM1_4849 | 2571.822987 | 3122.617401 | 3562.945838 | 0.067033194 | 0.150955985 | 0.327231105 | 0.654510064 | ns | TM | KS.1 |
| FIM1_4850 | 421.6907067 | 348.2269915 | 797.3107309 | -0.042686142 | 0.171613963 | 0.508450914 | 0.800488385 | ns | TM | KS.1 |
| FIM1_4851 | 1042.376821 | 827.8917234 | 855.6509651 | -0.159112566 | 0.140369893 | 0.045659678 | 0.220755005 | ns | TM | KS.1 |
| FIM1_4852 | 604.3720552 | 562.7036558 | 468.2283045 | -0.035957721 | 0.108323763 | 0.561641497 | 0.826391733 | ns | TM | KS.1 |
| FIM1_4853 | 330.1928011 | 319.3883248 | 464.3356436 | -0.01311335 | 0.117667672 | 0.833991057 | 0.948141464 | ns | TM | KS.1 |
| FIM1_4854 | 2816.355212 | 2543.329787 | 3480.209368 | -0.027102575 | 0.150717767 | 0.670231558 | 0.882350357 | ns | TM | KS.1 |
| FIM1_4855 | 583.1187347 | 865.2844021 | 915.6515042 | 0.098148839 | 0.213364674 | 0.187622916 | 0.4902437 | ns | TM | KS.1 |
| FIM1_4856 | 122.0560452 | 112.4378457 | 90.94481753 | -0.028350535 | 0.134878423 | 0.658382282 | 0.87860833 | ns | TM | KS.1 |
| FIM1_4857 | 31.79817347 | 27.88218082 | 97.34138546 | -0.022007392 | 0.170827883 | 0.723546514 | 0.907181536 | ns | TM | KS.1 |
| FIM1_4858 | 1065.293505 | 1938.715928 | 1889.896656 | 0.373053182 | 0.346734761 | 0.00713976 | 0.066581689 | ns | TM | KS.1 |
| FIM1_4859 | 1355.183585 | 1451.162592 | 1667.293549 | 0.054448934 | 0.07266281 | 0.272250873 | 0.598783418 | ns | TM | KS.1 |
| FIM1_4860 | 715.4302694 | 621.0934148 | 551.1737234 | -0.049474595 | 0.140267582 | 0.454096133 | 0.757811846 | ns | TM | KS.1 |
| FIM1_4861 | 869.5009403 | 864.887287 | 825.2678814 | -0.003080263 | 0.075124571 | 0.951296091 | 0.987490021 | ns | TM | KS.1 |
| FIM1_4862 | 1538.816168 | 1571.430062 | 1646.755377 | 0.01096432 | 0.099255642 | 0.852902152 | 0.954011929 | ns | TM | KS.1 |
| FIM1_4863 | 3094.33151 | 4600.008713 | 5275.620652 | 0.168060143 | 0.2175133 | 0.057998399 | 0.253530772 | ns | TM | KS.1 |
| FIM1_4864 | 67.70715106 | 74.92836296 | 64.56806349 | 0.029314476 | 0.138958344 | 0.647825649 | 0.873595603 | ns | TM | KS.1 |
| FIM1_4865 | 709.9465622 | 692.2898304 | 766.4919362 | -0.009534012 | 0.122855014 | 0.879568423 | 0.965258439 | ns | TM | KS.1 |
| FIM1_4866 | 2245.968024 | 1872.060913 | 1468.118296 | -0.069815663 | 0.144345653 | 0.309262268 | 0.637621129 | ns | TM | KS.1 |
| FIM1_4867 | 1107.72784 | 1591.460613 | 1774.801931 | 0.257903289 | 0.197215351 | 0.010517482 | 0.088016463 | ns | TM | KS.1 |
| FIM1_4868 | 1295.171359 | 1348.609374 | 1300.526989 | 0.023645452 | 0.09028166 | 0.675866907 | 0.887002016 | ns | TM | KS.1 |
| FIM1_4869 | 2953.879466 | 2624.413813 | 2324.438527 | -0.066207894 | 0.112229473 | 0.305346834 | 0.633687915 | ns | TM | KS.1 |
| FIM1_4870 | 491.0855589 | 613.4528805 | 695.124892 | 0.074522313 | 0.158801257 | 0.284336398 | 0.611419177 | ns | TM | KS.1 |
| FIM1_4871 | 450.1677782 | 329.2287686 | 519.1620767 | -0.217067937 | 0.172982191 | 0.018191351 | 0.127881388 | ns | TM | KS.1 |
| FIM1_4872 | 300.382232 | 192.8374552 | 580.6762812 | -0.343421742 | 0.2448897 | 0.004180252 | 0.045081369 | ns | TM | KS.1 |
| FIM1_4873 | 1208.883666 | 1398.040402 | 1506.886808 | 0.07436732 | 0.123917053 | 0.270277823 | 0.597663125 | ns | TM | KS.1 |
| FIM1_4874 | 2011.560441 | 2707.591296 | 2687.585499 | 0.151849671 | 0.173353486 | 0.0662414 | 0.270807979 | ns | TM | KS.1 |
| FIM1_4875 | 1825.783301 | 1417.404355 | 1454.363538 | -0.101960802 | 0.161773171 | 0.168452516 | 0.461736235 | ns | TM | KS.1 |
| FIM1_4876 | 87.95870413 | 73.01049282 | 66.18190685 | -0.061900196 | 0.151451358 | 0.360580182 | 0.683750066 | ns | TM | KS.1 |

| | | | | | | | | | | |
|---|---|---|---|---|---|---|---|---|---|---|
| FIM1_4877 | 731.1387911 | 806.3335511 | 1003.914225 | 0.037945891 | 0.128007408 | 0.557071182 | 0.825232602 | ns | TM | KS.1 |
| FIM1_4878 | 16948.97945 | 20283.01002 | 19245.88169 | 0.140766016 | 0.117970296 | 0.052670342 | 0.240005664 | ns | TM | KS.1 |
| FIM1_4879 | 2349.885983 | 2552.836537 | 2685.028214 | 0.079401088 | 0.067500285 | 0.09709557 | 0.337101354 | ns | TM | KS.1 |
| FIM1_4880 | 3674.007726 | 4421.999726 | 4901.205305 | 0.232719527 | 0.067319954 | 8.7424817009 | 0.002159213 | ns | TM | KS.1 |
| FIM1_4881 | 1778.26113 | 1918.617183 | 2085.045205 | 0.047281088 | 0.094658091 | 0.419904657 | 0.737778527 | ns | TM | KS.1 |
| FIM1_4882 | 35.13807364 | 25.77935116 | 1008.366534 | -0.046009211 | 0.211889966 | 0.470103672 | 0.771448038 | ns | TM | KS.1 |
| FIM1_4883 | 3269.886094 | 3195.005971 | 3633.9843 | -0.011525876 | 0.102334456 | 0.847449785 | 0.952348261 | ns | TM | KS.1 |
| FIM1_4884 | 2244.610369 | 2384.934787 | 2324.54675 | 0.046914972 | 0.070872865 | 0.334429897 | 0.661809362 | ns | TM | KS.1 |
| FIM1_4885 | 1115.192392 | 955.4941862 | 961.0478088 | -0.128390346 | 0.106441529 | 0.058857518 | 0.254885264 | ns | TM | KS.1 |
| FIM1_4886 | 315.0269228 | 286.1989393 | 262.1975986 | -0.048856988 | 0.111295335 | 0.439811551 | 0.747832613 | ns | TM | KS.1 |
| FIM1_4887 | 466.3493948 | 493.4943365 | 366.9965181 | 0.035464913 | 0.086917558 | 0.522875373 | 0.807872226 | ns | TM | KS.1 |
| FIM1_4888 | 1062.518553 | 1015.537776 | 846.6949265 | -0.020389281 | 0.112522218 | 0.742686354 | 0.91508234 | ns | TM | KS.1 |
| FIM1_4889 | 849.77532 | 626.5378178 | 630.0680574 | -0.183894286 | 0.173850262 | 0.035563266 | 0.188357575 | ns | TM | KS.1 |
| FIM1_4890 | 587.2677238 | 638.6616388 | 592.0101904 | 0.039920406 | 0.113435792 | 0.527131604 | 0.808531286 | ns | TM | KS.1 |
| FIM1_4891 | 1078.595553 | 942.0559417 | 1119.446052 | -0.080874013 | 0.113869892 | 0.221037375 | 0.535714496 | ns | TM | KS.1 |
| FIM1_4892 | 3978.029036 | 4044.768091 | 4920.135313 | 0.005821551 | 0.127946038 | 0.92647515 | 0.976735299 | ns | TM | KS.1 |
| FIM1_4893 | 7043.625682 | 8572.426937 | 9611.487046 | 0.078691897 | 0.146062713 | 0.260407441 | 0.588210523 | ns | TM | KS.1 |
| FIM1_4894 | 1672.911835 | 1657.03519 | 1464.534456 | -0.00431404 | 0.108913597 | 0.943808304 | 0.984585482 | ns | TM | KS.1 |
| FIM1_4895 | 1550.909025 | 1674.722351 | 1695.610525 | 0.030092996 | 0.124802115 | 0.637621421 | 0.870169358 | ns | TM | KS.1 |
| FIM1_4896 | 981.7852039 | 1190.149716 | 1266.705427 | 0.080512799 | 0.143415063 | 0.25033986 | 0.576059577 | ns | TM | KS.1 |
| FIM1_4897 | 934.3805364 | 989.9148324 | 1069.82916 | 0.021123195 | 0.126807119 | 0.739468145 | 0.914051801 | ns | TM | KS.1 |
| FIM1_4898 | 1068.846349 | 971.5655274 | 920.0857228 | -0.031706952 | 0.1369169 | 0.62233391 | 0.863388512 | ns | TM | KS.1 |
| FIM1_4899 | 10626.15512 | 8967.129248 | 7998.009529 | -0.124318588 | 0.117519923 | 0.079518334 | 0.300154562 | ns | TM | KS.1 |
| FIM1_4900 | 5846.180235 | 5289.237636 | 4562.418037 | -0.057015276 | 0.106323619 | 0.362149814 | 0.685319085 | ns | TM | KS.1 |
| FIM1_4901 | 481.203573 | 419.6723246 | 385.1794082 | -0.056510323 | 0.132016303 | 0.395695261 | 0.718986094 | ns | TM | KS.1 |
| FIM1_4902 | 1053.522738 | 907.3258047 | 1002.088481 | -0.09371001 | 0.116820307 | 0.167354579 | 0.45923278 | ns | TM | KS.1 |
| FIM1_4903 | 1789.445541 | 957.2974844 | 1339.826362 | -0.861078097 | 0.151342703 | 8.6994678975 | 1.9672660359 | ns | TM | KS.1 |
| FIM1_4904 | 2104.714376 | 1804.84689 | 1271.668189 | -0.072660314 | 0.129057014 | 0.284982258 | 0.61164225 | ns | TM | KS.1 |
| FIM1_4905 | 472.6963676 | 482.0519021 | 445.754003 | 0.006647561 | 0.12443383 | 0.915916703 | 0.975605516 | ns | TM | KS.1 |
| FIM1_4906 | 1795.821759 | 1445.79011 | 1602.179037 | -0.203661413 | 0.117550665 | 0.009871506 | 0.08366396 | ns | TM | KS.1 |
| FIM1_4907 | 977.6328453 | 1689.125928 | 2107.583726 | 0.249398127 | 0.299301077 | 0.023936499 | 0.151692113 | ns | TM | KS.1 |
| FIM1_4908 | 633.1220099 | 646.2978952 | 559.5296818 | 0.009055628 | 0.112788543 | 0.883649801 | 0.966744674 | ns | TM | KS.1 |
| FIM1_4909 | 2604.433546 | 2719.51424 | 2808.153758 | 0.026367576 | 0.086943964 | 0.633995132 | 0.869547581 | ns | TM | KS.1 |
| FIM1_4910 | 487.8032804 | 681.1384085 | 642.7546195 | 0.247194616 | 0.181184837 | 0.010836293 | 0.08925589 | ns | TM | KS.1 |
| FIM1_4911 | 1103.951932 | 1346.349285 | 1207.163562 | 0.093454214 | 0.141509418 | 0.191874349 | 0.496260756 | ns | TM | KS.1 |
| FIM1_4912 | 949.1670437 | 1039.286093 | 1002.936123 | 0.041596376 | 0.116709211 | 0.514057984 | 0.804225935 | ns | TM | KS.1 |
| FIM1_4913 | 863.0121511 | 1415.994568 | 1779.518337 | 0.355585482 | 0.280999655 | 0.005181843 | 0.053400721 | ns | TM | KS.1 |
| FIM1_4914 | 1631.09802 | 2016.192933 | 2391.945927 | 0.174618792 | 0.126410832 | 0.025888755 | 0.157838916 | ns | TM | KS.1 |
| FIM1_4915 | 1761.451516 | 1861.029396 | 2017.395644 | 0.018278049 | 0.133122823 | 0.773562472 | 0.925733395 | ns | TM | KS.1 |
| FIM1_4916 | 758.54417 | 1386.64277 | 1357.458541 | 0.232440847 | 0.317327451 | 0.031610864 | 0.177624525 | ns | TM | KS.1 |
| FIM1_4917 | 1238.162118 | 1356.219665 | 1595.349315 | 0.059121514 | 0.095470337 | 0.31915978 | 0.647069572 | ns | TM | KS.1 |
| FIM1_4918 | 189.7133881 | 158.2788884 | 185.4433156 | -0.091404537 | 0.134667751 | 0.195972353 | 0.50256523 | ns | TM | KS.1 |
| FIM1_4919 | 1136.033547 | 1386.017679 | 1474.08409 | 0.094038581 | 0.14152027 | 0.189584636 | 0.493436406 | ns | TM | KS.1 |
| FIM1_4920 | 509.9513217 | 390.4391121 | 491.5382069 | -0.174011924 | 0.156002262 | 0.038087455 | 0.196765407 | ns | TM | KS.1 |
| FIM1_4921 | 8778.373758 | 10012.57328 | 10694.03251 | 0.082996398 | 0.110682938 | 0.20476459 | 0.512865935 | ns | TM | KS.1 |

| | | | | | | | | | |
|---|---|---|---|---|---|---|---|---|---|
| FIM1_4922 | 899.0542396 | 1075.525248 | 1043.283304 | 0.095181285 | 0.132133908 | 0.178890899 | 0.477714558 | ns | TM | KS.1 |
| FIM1_4923 | 822.7838355 | 604.9983919 | 779.2656105 | -0.354738767 | 0.113372757 | 5.08978512073 | 0.001414619 | ns | TM | KS.1 |
| FIM1_4924 | 5999.392536 | 3027.268563 | 2324.548194 | -0.675085365 | 0.368960219 | 0.000521939 | 0.009371875 | ns | TM | KS.1 |
| FIM1_4925 | 151.4968658 | 134.0587063 | 244.9278878 | -0.027496965 | 0.16196814 | 0.66327283 | 0.880619203 | ns | TM | KS.1 |
| FIM1_4926 | 1284.775603 | 1372.422986 | 1363.095632 | 0.050909178 | 0.072140345 | 0.301545484 | 0.630596378 | ns | TM | KS.1 |
| FIM1_4927 | 896.9550486 | 1280.954241 | 1324.343166 | 0.126986424 | 0.199358832 | 0.112166917 | 0.364487533 | ns | TM | KS.1 |
| FIM1_4928 | 682.4507377 | 1054.001796 | 1095.136634 | 0.16484882 | 0.233371549 | 0.064047907 | 0.265874687 | ns | TM | KS.1 |
| FIM1_4929 | 1044.70711 | 943.5043171 | 701.8857728 | -0.073844076 | 0.092885819 | 0.211277174 | 0.51871941 | ns | TM | KS.1 |
| FIM1_4931 | 4045.709224 | 3536.459274 | 3938.011143 | -0.071555178 | 0.119117731 | 0.28127243 | 0.606295641 | ns | TM | KS.1 |
| FIM1_4932 | 3811.726828 | 3235.437108 | 4932.710896 | -0.094049238 | 0.124710607 | 0.176175714 | 0.473769825 | ns | TM | KS.1 |
| FIM1_4933 | 2102.158253 | 1702.906388 | 1783.902986 | -0.131901221 | 0.137036587 | 0.08193676 | 0.304774234 | ns | TM | KS.1 |
| FIM1_4935 | 1116.072441 | 853.1914583 | 1921.499279 | -0.169573226 | 0.15783649 | 0.042449611 | 0.210464424 | ns | TM | KS.1 |
| FIM1_4936 | 1322.023617 | 1301.909207 | 1229.813806 | -0.004695621 | 0.1378551 | 0.940634434 | 0.983546982 | ns | TM | KS.1 |
| FIM1_4937 | 8033.791888 | 8785.049627 | 6854.844582 | 0.025637064 | 0.144955931 | 0.687595204 | 0.891826565 | ns | TM | KS.1 |
| FIM1_4938 | 900.0489189 | 1070.198579 | 1253.715977 | 0.096365089 | 0.12876095 | 0.171370115 | 0.465692069 | ns | TM | KS.1 |
| FIM1_4939 | 1199.598003 | 1214.704288 | 997.7523455 | 0.007175556 | 0.088224871 | 0.89746959 | 0.967972723 | ns | TM | KS.1 |
| FIM1_4940 | 1103.842924 | 1407.375177 | 1454.603785 | 0.096304079 | 0.159221639 | 0.187317751 | 0.489703527 | ns | TM | KS.1 |
| FIM1_4941 | 7217.038795 | 8156.047034 | 8851.194038 | 0.095349887 | 0.097399342 | 0.124492679 | 0.38782159 | ns | TM | KS.1 |
| FIM1_4942 | 1024.80393 | 1158.338696 | 1224.514551 | 0.030311498 | 0.156791651 | 0.634025077 | 0.869547581 | ns | TM | KS.1 |
| FIM1_4943 | 2481.026097 | 2718.375271 | 2318.517883 | 0.050594951 | 0.105672889 | 0.414849739 | 0.734631029 | ns | TM | KS.1 |
| FIM1_4944 | 776.8538251 | 885.7305614 | 1183.17116 | 0.038112155 | 0.148498194 | 0.556309886 | 0.825176412 | ns | TM | KS.1 |
| FIM1_4945 | 7583.857843 | 7485.169791 | 7429.713428 | -0.006005187 | 0.10866472 | 0.921800896 | 0.976151439 | ns | TM | KS.1 |
| FIM1_4946 | 2511.888954 | 2754.567525 | 3010.056622 | 0.049493996 | 0.107695676 | 0.428331465 | 0.742665244 | ns | TM | KS.1 |
| FIM1_4947 | 1816.54728 | 1885.275455 | 1835.211868 | 0.021932483 | 0.087175451 | 0.692265252 | 0.893131011 | ns | TM | KS.1 |
| FIM1_4948 | 517.2036417 | 507.6595532 | 502.0525132 | -0.011011822 | 0.093117385 | 0.847731001 | 0.952348261 | ns | TM | KS.1 |
| FIM1_4949 | 1901.138215 | 1779.09298 | 1482.360849 | -0.058774322 | 0.061302329 | 0.181460383 | 0.481472312 | ns | TM | KS.1 |
| FIM1_4950 | 4265.198448 | 3870.997354 | 3177.919928 | -0.08118927 | 0.082592178 | 0.140970337 | 0.416465218 | ns | TM | KS.1 |
| FIM1_4951 | 2283.151974 | 2239.649377 | 2430.954616 | -0.009335899 | 0.104463529 | 0.877106325 | 0.964248917 | ns | TM | KS.1 |
| FIM1_4952 | 3859.487716 | 6145.844355 | 6919.195298 | 0.239333563 | 0.257933966 | 0.022567046 | 0.146759546 | ns | TM | KS.1 |
| FIM1_4953 | 1054.614939 | 1818.772088 | 2320.49064 | 0.250747015 | 0.298753561 | 0.023466585 | 0.150446212 | ns | TM | KS.1 |
| FIM1_4954 | 198.8404803 | 311.2425809 | 227.721582 | 0.188638474 | 0.242902982 | 0.045397694 | 0.220040594 | ns | TM | KS.1 |
| FIM1_4955 | 7647.905992 | 9543.298234 | 10080.4091 | 0.145278408 | 0.13928847 | 0.061808842 | 0.261034796 | ns | TM | KS.1 |
| FIM1_4956 | 681.5811892 | 697.9733624 | 2186.763112 | 0.009869548 | 0.115888542 | 0.874127357 | 0.962759265 | ns | TM | KS.1 |
| FIM1_4957 | 277.7582329 | 347.2534073 | 753.7054491 | 0.103690038 | 0.149448239 | 0.158956946 | 0.446281495 | ns | TM | KS.1 |
| FIM1_4958 | 509.5472165 | 175.0392786 | 219.4088314 | -0.860601716 | 0.56460031 | 0.000739287 | 0.012019452 | ns | TM | KS.1 |
| FIM1_4960 | 3163.675459 | 3991.984962 | 4081.875028 | 0.258891227 | 0.099214379 | 0.000798426 | 0.012854914 | ns | TM | KS.1 |
| FIM1_4961 | 2664.202471 | 2665.761188 | 3685.306776 | 7.8453234371736e | 0.117905462 | 0.998997352 | 0.999198196 | ns | TM | KS.1 |
| FIM1_4962 | 527.8033219 | 647.1435874 | 811.1420474 | 0.130354574 | 0.134213868 | 0.082894003 | 0.305706201 | ns | TM | KS.1 |
| FIM1_4963 | 757.7561152 | 587.4873847 | 718.4865154 | -0.261551664 | 0.117909704 | 0.001913365 | 0.024984234 | ns | TM | KS.1 |
| FIM1_4964 | 45.24771319 | 59.30349231 | 90.85514937 | 0.136243798 | 0.162890265 | 0.086246549 | 0.312966141 | ns | TM | KS.1 |
| FIM1_4965 | 5381.49474 | 7095.724528 | 7135.825911 | 0.286142674 | 0.122806374 | 0.001106454 | 0.016480869 | ns | TM | KS.1 |
| FIM1_4966 | 5968.495489 | 4293.097458 | 4089.50433 | -0.405194929 | 0.102778266 | 1.05072321588 | 7.26020555422 | ns | TM | KS.1 |
| FIM1_4967 | 1771.885371 | 1573.062277 | 1829.677499 | -0.081100507 | 0.103282538 | 0.199064152 | 0.506829147 | ns | TM | KS.1 |
| FIM1_4968 | 1399.822367 | 1814.716112 | 1696.526117 | 0.077046261 | 0.17183925 | 0.271585577 | 0.598783418 | ns | TM | KS.1 |
| FIM1_4969 | 14425.15546 | 13086.51089 | 13025.51777 | -0.095487264 | 0.072254589 | 0.061666224 | 0.260865744 | ns | TM | KS.1 |

| | | | | | | | | | | |
|---|---|---|---|---|---|---|---|---|---|---|
| FIM1_4970 | 11229.11216 | 13363.84916 | 13733.02455 | 0.17660773 | 0.098414241 | 0.011310671 | 0.091959747 | ns | TM | KS.1 |
| FIM1_4971 | 5061.272449 | 4866.907625 | 5299.304992 | -0.01761442 | 0.111455283 | 0.775822873 | 0.926727011 | ns | TM | KS.1 |
| FIM1_4972 | 2219.92433 | 1702.357744 | 1907.458323 | -0.107877657 | 0.165410011 | 0.151120313 | 0.433078086 | ns | TM | KS.1 |
| FIM1_4973 | 2072.852998 | 2140.732173 | 1688.204938 | 0.013789513 | 0.11477429 | 0.824668875 | 0.94482203 | ns | TM | KS.1 |
| FIM1_4974 | 563.1609756 | 513.7165143 | 495.5369414 | -0.064388096 | 0.094503448 | 0.2770073 | 0.601532657 | ns | TM | KS.1 |
| FIM1_4976 | 6504.891854 | 7546.327572 | 7599.367736 | 0.119313733 | 0.105853754 | 0.074715191 | 0.290851389 | ns | TM | KS.1 |
| FIM1_4977 | 938.1701432 | 1391.733277 | 1707.165664 | 0.103746379 | 0.213169615 | 0.170376997 | 0.464961908 | ns | TM | KS.1 |
| FIM1_4978 | 939.1861416 | 1119.442179 | 947.8801932 | 0.12486706 | 0.120870731 | 0.082038636 | 0.304774234 | ns | TM | KS.1 |
| FIM1_4979 | 16826.31452 | 16065.35911 | 16927.46391 | -0.03377562 | 0.071181623 | 0.487495998 | 0.787177082 | ns | TM | KS.1 |
| FIM1_4980 | 564.8373636 | 610.2488342 | 556.3859116 | 0.038100322 | 0.110874362 | 0.542546563 | 0.815959873 | ns | TM | KS.1 |
| FIM1_4981 | 15195.36418 | 14608.92603 | 15859.94697 | -0.033362862 | 0.055210899 | 0.410295345 | 0.731095753 | ns | TM | KS.1 |
| FIM1_4982 | 917.9453549 | 985.7989841 | 1010.171553 | 0.064405068 | 0.067442605 | 0.173048445 | 0.468851247 | ns | TM | KS.1 |
| FIM1_4983 | 946.323517 | 939.1737935 | 976.9536087 | -0.004419309 | 0.073275155 | 0.929002971 | 0.978304974 | ns | TM | KS.1 |
| FIM1_4984 | 1368.077483 | 1133.35396 | 1549.734645 | -0.111531486 | 0.131441041 | 0.124246717 | 0.387735747 | ns | TM | KS.1 |
| FIM1_4985 | 2564.812906 | 2489.856013 | 2672.857618 | -0.021120683 | 0.068414759 | 0.655230848 | 0.87699044 | ns | TM | KS.1 |
| FIM1_4986 | 4328.582431 | 4252.472055 | 4455.986282 | -0.01281098 | 0.064617636 | 0.778039484 | 0.928236554 | ns | TM | KS.1 |
| FIM1_4987 | 4916.373327 | 3372.250862 | 2820.403304 | -0.308240548 | 0.197631189 | 0.004076022 | 0.044179107 | ns | TM | KS.1 |
| FIM1_4988 | 193.7795946 | 163.0463817 | 174.4108259 | -0.091295352 | 0.130356715 | 0.193004153 | 0.496788132 | ns | TM | KS.1 |
| FIM1_4989 | 6206.236071 | 4577.618686 | 7408.666665 | -0.240646301 | 0.16267163 | 0.009787232 | 0.083375823 | ns | TM | KS.1 |
| FIM1_4991 | 2970.913624 | 3555.282728 | 3383.67469 | 0.089080935 | 0.134278847 | 0.205543668 | 0.512865935 | ns | TM | KS.1 |
| FIM1_4992 | 1349.350247 | 1448.043685 | 1100.95135 | 0.032996774 | 0.112558902 | 0.598221911 | 0.849417898 | ns | TM | KS.1 |
| FIM1_4993 | 2037.592315 | 2284.311003 | 2479.73763 | 0.059117313 | 0.115158028 | 0.360772648 | 0.683750066 | ns | TM | KS.1 |
| FIM1_4995 | 1703.133584 | 1023.714665 | 1753.530347 | -0.501974303 | 0.264287505 | 0.000553269 | 0.009746554 | ns | TM | KS.1 |
| FIM1_4996 | 482.0390683 | 452.7957344 | 533.7981596 | -0.030608716 | 0.107551142 | 0.619078375 | 0.861076858 | ns | TM | KS.1 |
| FIM1_4997 | 1749.624032 | 1730.126188 | 1646.699176 | -0.006919327 | 0.083182679 | 0.897672882 | 0.967972723 | ns | TM | KS.1 |
| FIM1_4998 | 2457.293254 | 2199.854627 | 2531.997194 | -0.059222145 | 0.112493276 | 0.355985903 | 0.680903448 | ns | TM | KS.1 |
| FIM1_4999 | 2996.993901 | 3486.350054 | 3751.186881 | 0.094370663 | 0.117659476 | 0.165959105 | 0.456662914 | ns | TM | KS.1 |
| FIM1_5000 | 402.7881597 | 395.7674984 | 621.3457797 | -0.006021247 | 0.131094755 | 0.924026663 | 0.976662213 | ns | TM | KS.1 |
| FIM1_5001 | 606.1430841 | 488.0399657 | 1113.903104 | -0.078968446 | 0.154258833 | 0.260746881 | 0.588210523 | ns | TM | KS.1 |
| FIM1_5002 | 1652.307974 | 1618.920719 | 1846.561683 | -0.009979952 | 0.103480486 | 0.868245823 | 0.960282219 | ns | TM | KS.1 |
| FIM1_5003 | 1538.2268 | 1288.288308 | 1211.205547 | -0.124878045 | 0.121564892 | 0.082733536 | 0.305567439 | ns | TM | KS.1 |
| FIM1_5004 | 2288.220246 | 2739.151494 | 3222.634587 | 0.068568177 | 0.144001455 | 0.316754656 | 0.643468524 | ns | TM | KS.1 |
| FIM1_5005 | 18549.99565 | 21347.86138 | 20594.94691 | 0.113578987 | 0.102580125 | 0.082151489 | 0.304775287 | ns | TM | KS.1 |
| FIM1_5006 | 316.4088165 | 302.8896446 | 269.8336058 | -0.021390157 | 0.106853589 | 0.726341843 | 0.907961357 | ns | TM | KS.1 |
| FIM1_5007 | 1772.478194 | 2065.986464 | 2601.176588 | 0.041081636 | 0.155887159 | 0.526674681 | 0.808531286 | ns | TM | KS.1 |
| FIM1_5008 | 4716.530356 | 3751.635057 | 4296.317491 | -0.200525612 | 0.127063295 | 0.013606459 | 0.103981773 | ns | TM | KS.1 |
| FIM1_5009 | 144.9150645 | 44.8871868 | 119.1624388 | -1.277259697 | 0.431529364 | 2.7988168629[illegible] | 0.000929395 | Km<KS.1 | TM | KS.1 |
| FIM1_5010 | 1153.215108 | 1003.411126 | 1071.331298 | -0.063316316 | 0.12744736 | 0.343454846 | 0.670129201 | ns | TM | KS.1 |
| FIM1_5011 | 183.2980005 | 184.8632814 | 171.8248649 | 0.00313328 | 0.116018625 | 0.959848186 | 0.990303759 | ns | TM | KS.1 |
| FIM1_5012 | 1387.49522 | 1328.601787 | 962.7448218 | -0.020377983 | 0.107528586 | 0.739252206 | 0.914051801 | ns | TM | KS.1 |
| FIM1_5013 | 1392.670046 | 1286.205499 | 1207.073186 | -0.04842558 | 0.096662773 | 0.414371924 | 0.734555234 | ns | TM | KS.1 |
| FIM1_5014 | 2959.857169 | 3109.455439 | 2375.999414 | 0.034544392 | 0.077016948 | 0.501183179 | 0.795351314 | ns | TM | KS.1 |
| FIM1_5015 | 3150.471803 | 2916.88342 | 2910.350965 | -0.038739501 | 0.108684309 | 0.533121728 | 0.810351542 | ns | TM | KS.1 |
| FIM1_5016 | 869.0846483 | 483.189727 | 752.5604019 | -0.582733057 | 0.313069666 | 0.000523695 | 0.009371875 | ns | TM | KS.1 |
| FIM1_5017 | 2441.314434 | 2464.184473 | 2331.695015 | 0.003930869 | 0.114754071 | 0.949519696 | 0.986604111 | ns | TM | KS.1 |

| | | | | | | | | | | |
|---|---|---|---|---|---|---|---|---|---|---|
| FIM1_5018 | 494.5928583 | 408.3763987 | 382.6164024 | -0.115914103 | 0.131950045 | 0.112976055 | 0.366160178 | ns | TM | KS.1 |
| FIM1_5019 | 841.6492014 | 917.9213381 | 1121.73002 | 0.030861244 | 0.131490966 | 0.630963353 | 0.869059435 | ns | TM | KS.1 |
| FIM1_5020 | 724.4835013 | 860.8056809 | 1100.992639 | 0.050050845 | 0.154419967 | 0.448688284 | 0.753357428 | ns | TM | KS.1 |
| FIM1_5021 | 640.2025438 | 582.2854219 | 757.4105412 | -0.064565609 | 0.095264009 | 0.277971334 | 0.602049362 | ns | TM | KS.1 |
| FIM1_5022 | 2159.437417 | 2173.016312 | 2655.25825 | 0.004259533 | 0.069936838 | 0.929363105 | 0.978328703 | ns | TM | KS.1 |
| FIM1_5023 | 1687.746013 | 1810.074498 | 1964.965564 | 0.049642592 | 0.08285966 | 0.359135439 | 0.683511403 | ns | TM | KS.1 |
| FIM1_5024 | 2864.471478 | 3150.25162 | 3413.721675 | 0.065044179 | 0.093972107 | 0.270803255 | 0.598244313 | ns | TM | KS.1 |
| FIM1_5025 | 3309.976008 | 2962.406204 | 2924.38756 | -0.04805486 | 0.12408222 | 0.460518488 | 0.763054265 | ns | TM | KS.1 |
| FIM1_5026 | 1034.193571 | 1170.596136 | 2627.883575 | 0.079628696 | 0.107221276 | 0.215191094 | 0.524278007 | ns | TM | KS.1 |
| FIM1_5027 | 3051.284057 | 2910.404011 | 3523.227109 | -0.022836534 | 0.107263438 | 0.709225682 | 0.901619224 | ns | TM | KS.1 |
| FIM1_5028 | 2632.19192 | 3183.079948 | 3668.055305 | 0.191236196 | 0.103680995 | 0.008918808 | 0.078450448 | ns | TM | KS.1 |
| FIM1_5029 | 2710.244771 | 2867.506229 | 2781.189831 | 0.044374265 | 0.068200081 | 0.347824864 | 0.673580653 | ns | TM | KS.1 |
| FIM1_5030 | 5227.210943 | 4682.153407 | 3657.061808 | -0.112210278 | 0.075361622 | 0.036934713 | 0.192812378 | ns | TM | KS.1 |
| FIM1_5031 | 4055.929093 | 4184.863839 | 5372.95306 | 0.018792154 | 0.086030364 | 0.732397218 | 0.910998727 | ns | TM | KS.1 |
| FIM1_5032 | 1438.672995 | 1791.680578 | 2251.585274 | 0.069243214 | 0.160226444 | 0.314132561 | 0.640900344 | ns | TM | KS.1 |
| FIM1_5033 | 1934.096273 | 1945.922756 | 1764.733732 | 0.00411367 | 0.055604518 | 0.919709286 | 0.975881018 | ns | TM | KS.1 |
| FIM1_5034 | 6604.992386 | 6823.084757 | 4885.437389 | 0.019481169 | 0.08643275 | 0.7237394 | 0.907181536 | ns | TM | KS.1 |
| FIM1_5035 | 1057.808729 | 1053.072455 | 1070.618224 | -0.001738636 | 0.119481398 | 0.977836002 | 0.994223198 | ns | TM | KS.1 |
| FIM1_5036 | 1507.31294 | 1364.790175 | 1330.740806 | -0.051435297 | 0.111404877 | 0.417379075 | 0.735551151 | ns | TM | KS.1 |
| FIM1_5037 | 1221.365001 | 1322.531085 | 1385.984308 | 0.071444474 | 0.072109395 | 0.151535456 | 0.433367037 | ns | TM | KS.1 |
| FIM1_5038 | 3043.44628 | 3349.503159 | 3473.213098 | 0.07185657 | 0.088951342 | 0.210788635 | 0.51871941 | ns | TM | KS.1 |
| FIM1_5039 | 1461.797933 | 1414.477185 | 1431.105986 | -0.019502745 | 0.087664386 | 0.725620066 | 0.907961357 | ns | TM | KS.1 |
| FIM1_5040 | 13499.02005 | 16321.02543 | 15960.82951 | 0.147712453 | 0.121790527 | 0.047021544 | 0.225803262 | ns | TM | KS.1 |
| FIM1_5041 | 386.7344869 | 386.8013877 | 428.8023825 | 0.001836802 | 0.100499819 | 0.975328366 | 0.993872484 | ns | TM | KS.1 |
| FIM1_5042 | 13757.38501 | 16455.40954 | 18192.75871 | 0.125525967 | 0.122492295 | 0.082363029 | 0.304891809 | ns | TM | KS.1 |
| FIM1_5043 | 938.3100138 | 938.5075698 | 914.906838 | 0.000363157 | 0.106342421 | 0.995227076 | 0.99790453 | ns | TM | KS.1 |
| FIM1_5044 | 275.1910457 | 304.6137768 | 237.6434499 | 0.03927848 | 0.127885762 | 0.543902927 | 0.817250698 | ns | TM | KS.1 |
| FIM1_5045 | 3248.066096 | 3287.658293 | 2635.336793 | 0.005659243 | 0.107447364 | 0.926017142 | 0.976735299 | ns | TM | KS.1 |
| FIM1_5046 | 2907.895267 | 2015.163926 | 3462.930132 | -0.155440457 | 0.203890215 | 0.068933509 | 0.276567909 | ns | TM | KS.1 |
| FIM1_5047 | 1697.397539 | 1235.165659 | 1684.120007 | -0.34895229 | 0.127332085 | 0.000174054 | 0.003865702 | ns | TM | KS.1 |
| FIM1_5048 | 923.3592487 | 891.8251408 | 648.1284846 | -0.020449351 | 0.079688026 | 0.696532655 | 0.894950919 | ns | TM | KS.1 |
| FIM1_5049 | 2292.755136 | 2372.819591 | 2315.334913 | 0.023744781 | 0.073644126 | 0.633336877 | 0.869547581 | ns | TM | KS.1 |
| FIM1_5050 | 1888.986126 | 1747.576724 | 2158.68306 | -0.041840239 | 0.104604743 | 0.495307167 | 0.792222775 | ns | TM | KS.1 |
| FIM1_5051 | 18.42794641 | 16.69011279 | 26.75475964 | -0.017528513 | 0.160204301 | 0.778944245 | 0.928304554 | ns | TM | KS.1 |
| FIM1_5052 | 995.9854257 | 846.1148687 | 833.6461928 | -0.126983637 | 0.112089673 | 0.06815456 | 0.275864055 | ns | TM | KS.1 |
| FIM1_5053 | 1374.466179 | 1324.294422 | 1307.8111 | -0.021445755 | 0.090398259 | 0.704579896 | 0.898791021 | ns | TM | KS.1 |
| FIM1_5054 | 636.3083585 | 709.4955102 | 670.2714311 | 0.062194248 | 0.108953708 | 0.327706472 | 0.655017958 | ns | TM | KS.1 |
| FIM1_5055 | 3178.75078 | 3393.439777 | 3470.755075 | 0.043600468 | 0.085659332 | 0.429327342 | 0.743181463 | ns | TM | KS.1 |
| FIM1_5056 | 1308.032394 | 1412.871403 | 1420.427163 | 0.04870923 | 0.092836507 | 0.401253221 | 0.723750973 | ns | TM | KS.1 |
| FIM1_5057 | 3161.619301 | 5044.764427 | 6219.004623 | 0.173112442 | 0.247850739 | 0.058525553 | 0.254317312 | ns | TM | KS.1 |
| FIM1_5058 | 4911.924177 | 3291.712844 | 2622.191853 | -0.292898755 | 0.218048989 | 0.007025087 | 0.066444504 | ns | TM | KS.1 |
| FIM1_5059 | 694.4278811 | 705.8444656 | 686.9090204 | 0.009969246 | 0.065057091 | 0.827222156 | 0.946035952 | ns | TM | KS.1 |
| FIM1_5061 | 2482.094692 | 1916.146374 | 2153.880053 | -0.220487099 | 0.139239096 | 0.010598101 | 0.088198944 | ns | TM | KS.1 |
| FIM1_5062 | 726.4191349 | 658.7348728 | 633.3694877 | -0.068326475 | 0.094406806 | 0.249843415 | 0.575716068 | ns | TM | KS.1 |
| FIM1_5063 | 2006.091243 | 1920.317003 | 1990.11932 | -0.025242594 | 0.091677686 | 0.658193992 | 0.878592731 | ns | TM | KS.1 |

| | | | | | | | | | | |
|---|---|---|---|---|---|---|---|---|---|---|
| FIM1_5064 | 621.7091324 | 744.8973989 | 645.2072038 | 0.080254868 | 0.138610066 | 0.249566364 | 0.575344142 | ns | TM | KS.1 |
| FIM1_5065 | 3434.198256 | 3343.452688 | 3945.297633 | -0.011815728 | 0.112314443 | 0.848509478 | 0.952348261 | ns | TM | KS.1 |
| FIM1_5066 | 491.390196 | 449.2402278 | 475.1938222 | -0.061997795 | 0.088589967 | 0.275642102 | 0.600738415 | ns | TM | KS.1 |
| FIM1_5067 | 3278.460463 | 3148.286558 | 3488.00408 | -0.026023706 | 0.080723076 | 0.622771014 | 0.863411908 | ns | TM | KS.1 |
| FIM1_5068 | 1493.558788 | 1116.283705 | 1083.197022 | -0.199330049 | 0.164236786 | 0.024065067 | 0.152126693 | ns | TM | KS.1 |
| FIM1_5069 | 155.7873678 | 116.0807044 | 1090.397409 | -0.079504359 | 0.183461183 | 0.259700821 | 0.588210523 | ns | TM | KS.1 |
| FIM1_5070 | 95.9035759 | 90.11744908 | 66.93085334 | -0.021751081 | 0.119219736 | 0.729927851 | 0.909157553 | ns | TM | KS.1 |
| FIM1_5071 | 10812.39771 | 8727.196619 | 7131.434308 | -0.203984954 | 0.115657504 | 0.009258757 | 0.079846584 | ns | TM | KS.1 |
| FIM1_5072 | 187.4015249 | 188.2901694 | 128.7444275 | 0.001635484 | 0.125195153 | 0.97927369 | 0.99466856 | ns | TM | KS.1 |
| FIM1_5073 | 733.1928643 | 727.2003203 | 611.9560529 | -0.003893744 | 0.104896032 | 0.948616158 | 0.986388143 | ns | TM | KS.1 |
| FIM1_5074 | 631.748288 | 570.1194408 | 663.8664992 | -0.055967857 | 0.109740567 | 0.376911607 | 0.700723186 | ns | TM | KS.1 |
| FIM1_5075 | 864.2892976 | 862.5507308 | 1277.11883 | -0.001186104 | 0.102426559 | 0.984191119 | 0.996034691 | ns | TM | KS.1 |
| FIM1_5076 | 4188.289612 | 4337.232522 | 3651.790514 | 0.02204401 | 0.082259577 | 0.680351669 | 0.887492141 | ns | TM | KS.1 |
| FIM1_5077 | 2610.164225 | 2196.791955 | 2209.852189 | -0.107626846 | 0.124481026 | 0.128840201 | 0.396596531 | ns | TM | KS.1 |
| FIM1_5078 | 455.3227163 | 427.52982 | 454.6684508 | -0.034541716 | 0.09682357 | 0.557743987 | 0.825580582 | ns | TM | KS.1 |
| FIM1_5079 | 5200.327486 | 4843.652451 | 6011.899769 | -0.044375152 | 0.091657638 | 0.440080069 | 0.747832613 | ns | TM | KS.1 |
| FIM1_5080 | 1156.07454 | 1077.067185 | 1060.424088 | -0.064429627 | 0.061679033 | 0.146024592 | 0.424167573 | ns | TM | KS.1 |
| FIM1_5081 | 1586.091602 | 1584.635279 | 1610.532415 | -0.000635344 | 0.077507724 | 0.990151185 | 0.996341203 | ns | TM | KS.1 |
| FIM1_5082 | 498.0419474 | 618.6541858 | 557.9162857 | 0.172291529 | 0.130109219 | 0.029222939 | 0.16972335 | ns | TM | KS.1 |
| FIM1_5083 | 4696.457383 | 5499.905622 | 4870.532015 | 0.064306431 | 0.136423901 | 0.341779491 | 0.668903606 | ns | TM | KS.1 |
| FIM1_5084 | 697.4762117 | 875.3556745 | 802.1607 | 0.076850262 | 0.159610239 | 0.272116583 | 0.598783418 | ns | TM | KS.1 |
| FIM1_5085 | 1692.476921 | 1600.218359 | 1763.991479 | -0.021636185 | 0.123957834 | 0.732856694 | 0.911086503 | ns | TM | KS.1 |
| FIM1_5086 | 2526.195347 | 2674.686084 | 2732.781178 | 0.03474719 | 0.090307764 | 0.540397713 | 0.814399145 | ns | TM | KS.1 |
| FIM1_5087 | 2096.323874 | 2359.204888 | 1469.97806 | 0.057457739 | 0.119474931 | 0.378887717 | 0.703083325 | ns | TM | KS.1 |
| FIM1_5088 | 9181.01577 | 10424.21856 | 9428.848193 | 0.051012261 | 0.130912232 | 0.439025691 | 0.747832613 | ns | TM | KS.1 |
| FIM1_5089 | 3644.548212 | 3063.289946 | 2468.934404 | -0.06410317 | 0.144057745 | 0.345216196 | 0.670970586 | ns | TM | KS.1 |
| FIM1_5090 | 373.7342934 | 360.4412853 | 387.9351323 | -0.010665965 | 0.141652885 | 0.865699386 | 0.959975149 | ns | TM | KS.1 |
| FIM1_5091 | 668.3695576 | 592.6407514 | 492.4874872 | -0.041340332 | 0.138200768 | 0.526208183 | 0.808531286 | ns | TM | KS.1 |
| FIM1_5093 | 1822.77639 | 1426.629952 | 1273.576786 | -0.275014825 | 0.099775304 | 0.000456096 | 0.008372978 | ns | TM | KS.1 |
| FIM1_5094 | 86877.15499 | 93592.13466 | 89508.85285 | 0.054487754 | 0.081690173 | 0.310668406 | 0.6389526 | ns | TM | KS.1 |
| FIM1_5095 | 531.1069581 | 558.9510545 | 503.4258247 | 0.028390012 | 0.094294753 | 0.624377126 | 0.864597072 | ns | TM | KS.1 |
| FIM1_5096 | 1075.338392 | 1244.372299 | 1407.574219 | 0.077105729 | 0.12273197 | 0.253554853 | 0.580732538 | ns | TM | KS.1 |
| FIM1_5097 | 6852.427884 | 8499.352632 | 9492.802646 | 0.264634738 | 0.075894177 | 5.95349246582 | 0.001575939 | ns | TM | KS.1 |
| FIM1_5098 | 270.3221565 | 242.2404539 | 297.09687 | -0.043402752 | 0.128810159 | 0.505007379 | 0.797591019 | ns | TM | KS.1 |
| FIM1_5099 | 1846.946368 | 1924.469832 | 1756.661765 | 0.025429942 | 0.085740346 | 0.643293017 | 0.872978256 | ns | TM | KS.1 |
| FIM1_5100 | 1134.450648 | 1037.886091 | 1058.239174 | -0.091591162 | 0.065108369 | 0.052604508 | 0.240005664 | ns | TM | KS.1 |
| FIM1_5101 | 1187.198985 | 1035.728699 | 1189.433938 | -0.101551535 | 0.10496457 | 0.119187908 | 0.378888079 | ns | TM | KS.1 |
| FIM1_5102 | 2217.044483 | 1707.091828 | 3568.104286 | -0.106119422 | 0.164158588 | 0.156012095 | 0.441000098 | ns | TM | KS.1 |
| FIM1_5103 | 741.5121566 | 535.3376345 | 511.8376738 | -0.316808463 | 0.151618195 | 0.00127741 | 0.018261829 | ns | TM | KS.1 |
| FIM1_5104 | 774.0636314 | 681.5436446 | 743.3096505 | -0.11780814 | 0.090566035 | 0.053726121 | 0.24174415 | ns | TM | KS.1 |
| FIM1_5105 | 2258.393407 | 2335.314609 | 2556.219037 | 0.026987618 | 0.059296957 | 0.527230937 | 0.808531286 | ns | TM | KS.1 |
| FIM1_5106 | 960.7937021 | 773.7907864 | 873.3045665 | -0.163185634 | 0.132596089 | 0.037801718 | 0.196103802 | ns | TM | KS.1 |
| FIM1_5107 | 3655.100302 | 4019.698443 | 4318.641348 | 0.055913339 | 0.103070467 | 0.36419852 | 0.687959166 | ns | TM | KS.1 |
| FIM1_5108 | 1002.472969 | 965.8676414 | 919.5715219 | -0.019663571 | 0.097911441 | 0.73847399 | 0.913453034 | ns | TM | KS.1 |
| FIM1_5109 | 906.5299381 | 937.2491864 | 1055.894387 | 0.017469818 | 0.099346615 | 0.768110965 | 0.925179691 | ns | TM | KS.1 |

| | | | | | | | | | | |
|---|---|---|---|---|---|---|---|---|---|---|
| FIM1_5110 | 1129.252661 | 1301.53613 | 1771.777108 | 0.1114986 | 0.104678902 | 0.090580985 | 0.324201726 | ns | TM | KS.1 |
| FIM1_5111 | 1528.819433 | 1766.544292 | 1833.436691 | 0.124819986 | 0.100590497 | 0.057036322 | 0.252003289 | ns | TM | KS.1 |
| FIM1_5112 | 916.8340402 | 876.5725553 | 912.3996043 | -0.027831752 | 0.084723313 | 0.609869162 | 0.854194561 | ns | TM | KS.1 |
| FIM1_5113 | 127.5934524 | 149.9571623 | 185.8773579 | 0.071433862 | 0.133046679 | 0.295035752 | 0.622741988 | ns | TM | KS.1 |
| FIM1_5114 | 335.3065247 | 455.8022646 | 473.4386546 | 0.164094082 | 0.176354025 | 0.053041674 | 0.240768548 | ns | TM | KS.1 |
| FIM1_5115 | 1329.206662 | 1565.839025 | 1717.039464 | 0.127430123 | 0.112907493 | 0.068315475 | 0.275864055 | ns | TM | KS.1 |
| FIM1_5116 | 1467.454937 | 1502.67025 | 1592.914278 | 0.014170151 | 0.083470564 | 0.792792103 | 0.933303839 | ns | TM | KS.1 |
| FIM1_5117 | 634.9094758 | 589.9199696 | 553.3084487 | -0.026792864 | 0.129807237 | 0.675151748 | 0.886715931 | ns | TM | KS.1 |
| FIM1_5118 | 3267.009022 | 3269.185503 | 2917.278492 | 0.0005105 | 0.091385182 | 0.992822382 | 0.997449676 | ns | TM | KS.1 |
| FIM1_5119 | 1831.471069 | 1569.520026 | 1408.965236 | -0.128535415 | 0.1060667 | 0.058155801 | 0.253571527 | ns | TM | KS.1 |
| FIM1_5120 | 1216.628523 | 1243.161668 | 1218.891875 | 0.008456777 | 0.119977959 | 0.892695233 | 0.967822084 | ns | TM | KS.1 |
| FIM1_5121 | 3002.315047 | 3053.53473 | 2267.146921 | 0.009333252 | 0.0935602 | 0.871000875 | 0.96208467 | ns | TM | KS.1 |
| FIM1_5122 | 3136.736801 | 3308.680225 | 4430.879936 | 0.013924644 | 0.148420554 | 0.824774094 | 0.94482203 | ns | TM | KS.1 |
| FIM1_5123 | 4671.901297 | 5295.698036 | 4067.119462 | 0.031792976 | 0.155757529 | 0.618606645 | 0.860858199 | ns | TM | KS.1 |
| FIM1_5124 | 1030.683459 | 987.7920754 | 969.6212758 | -0.030472311 | 0.076657438 | 0.551485772 | 0.822434567 | ns | TM | KS.1 |
| FIM1_5125 | 2591.061894 | 3145.411095 | 2718.883313 | 0.063445377 | 0.153201334 | 0.350276281 | 0.676222158 | ns | TM | KS.1 |
| FIM1_5126 | 1439.799855 | 1522.374199 | 1546.701462 | 0.047987513 | 0.062497928 | 0.280033821 | 0.605199069 | ns | TM | KS.1 |
| FIM1_5127 | 4493.193759 | 4145.476629 | 4733.280554 | -0.045983116 | 0.100984061 | 0.447686757 | 0.752446492 | ns | TM | KS.1 |
| FIM1_5128 | 3141.826386 | 3409.134902 | 3394.131703 | 0.063336317 | 0.080126526 | 0.234746149 | 0.553751585 | ns | TM | KS.1 |
| FIM1_5129 | 1192.573734 | 1190.082799 | 1408.446859 | -0.001344004 | 0.089018163 | 0.980835328 | 0.995431301 | ns | TM | KS.1 |
| FIM1_5130 | 2923.634229 | 2673.824418 | 2394.966011 | -0.048101744 | 0.107008347 | 0.43969365 | 0.747832613 | ns | TM | KS.1 |
| FIM1_5131 | 4509.848079 | 2397.042284 | 2682.725242 | -0.223156073 | 0.324036108 | 0.035996762 | 0.189506763 | ns | TM | KS.1 |
| FIM1_5132 | 172.5992925 | 193.8415272 | 193.6028101 | 0.026092118 | 0.161178316 | 0.679050899 | 0.887492141 | ns | TM | KS.1 |
| FIM1_5133 | 1736.651039 | 1552.961166 | 1610.349592 | -0.066970047 | 0.107189545 | 0.290783307 | 0.618193599 | ns | TM | KS.1 |
| FIM1_5134 | 663.5237166 | 697.8779543 | 596.8297995 | 0.024765334 | 0.106588725 | 0.685533229 | 0.890505759 | ns | TM | KS.1 |
| FIM1_5135 | 1371.918942 | 1434.653384 | 1370.994416 | 0.02388084 | 0.099191876 | 0.687350628 | 0.891826565 | ns | TM | KS.1 |
| FIM1_5136 | 3320.200232 | 3051.117049 | 2722.172896 | -0.051612984 | 0.09750862 | 0.38750814 | 0.712272102 | ns | TM | KS.1 |
| FIM1_5137 | 1126.080395 | 1211.50564 | 955.2377765 | 0.036013992 | 0.109894182 | 0.563052401 | 0.827284612 | ns | TM | KS.1 |
| FIM1_5138 | 2138.142157 | 2266.013738 | 2129.186013 | 0.019066856 | 0.134155187 | 0.764212867 | 0.924230468 | ns | TM | KS.1 |
| FIM1_5139 | 3501.30404 | 3914.862782 | 3263.840028 | 0.036532005 | 0.13956624 | 0.572775664 | 0.832094673 | ns | TM | KS.1 |
| FIM1_5140 | 3782.56109 | 2978.810772 | 3280.874518 | -0.081820726 | 0.162015355 | 0.247830313 | 0.572402882 | ns | TM | KS.1 |
| FIM1_5141 | 1899.325272 | 1900.669696 | 2740.679032 | 0.000206371 | 0.097863709 | 0.997194847 | 0.998293524 | ns | TM | KS.1 |
| FIM1_5142 | 1942.447579 | 1690.863664 | 1638.967762 | -0.056004004 | 0.132912308 | 0.399959686 | 0.723667321 | ns | TM | KS.1 |
| FIM1_5143 | 2200.335526 | 2293.500062 | 2447.338046 | 0.031395114 | 0.067264083 | 0.501738212 | 0.795458128 | ns | TM | KS.1 |
| FIM1_5144 | 2334.859289 | 2080.51086 | 1888.397872 | -0.083763892 | 0.098095828 | 0.173629734 | 0.469348363 | ns | TM | KS.1 |
| FIM1_5145 | 3083.857663 | 2899.834018 | 2312.449822 | -0.037283866 | 0.09181329 | 0.515605144 | 0.804775792 | ns | TM | KS.1 |
| FIM1_5146 | 356.1093332 | 572.4610254 | 627.1426176 | 0.326806841 | 0.26732729 | 0.006787638 | 0.064814782 | ns | TM | KS.1 |
| FIM1_5147 | 26224.37159 | 30581.61356 | 29081.46093 | 0.122880265 | 0.10799449 | 0.070613596 | 0.281267124 | ns | TM | KS.1 |
| FIM1_5148 | 445.0764275 | 470.8022235 | 298.8760442 | 0.019167904 | 0.130827718 | 0.762928507 | 0.924230468 | ns | TM | KS.1 |
| FIM1_5149 | 9087.837689 | 3766.074719 | 4101.836441 | -0.644587117 | 0.50268415 | 0.001936117 | 0.02513155 | ns | TM | KS.1 |
| FIM1_5150 | 6343.869614 | 8909.768112 | 9268.607682 | 0.201626041 | 0.190112106 | 0.028233884 | 0.166623453 | ns | TM | KS.1 |
| FIM1_5151 | 216.3126143 | 221.6108728 | 223.4158694 | 0.012661897 | 0.101124615 | 0.83185408 | 0.946802573 | ns | TM | KS.1 |
| FIM1_5152 | 476.5478761 | 341.6592927 | 293.9752473 | -0.145506419 | 0.189308849 | 0.078811991 | 0.299076777 | ns | TM | KS.1 |
| FIM1_5153 | 4866.030396 | 5478.396597 | 6107.404169 | 0.102741044 | 0.089907846 | 0.08490944 | 0.31075147 | ns | TM | KS.1 |
| FIM1_5155 | 984.1664673 | 1120.515326 | 1027.34733 | 0.085934619 | 0.1078279 | 0.185296008 | 0.488144311 | ns | TM | KS.1 |

| | | | | | | | | | | |
|---|---|---|---|---|---|---|---|---|---|---|
| FIM1_5156 | 880.4747126 | 865.2210152 | 907.6906643 | -0.006577875 | 0.123198712 | 0.916704217 | 0.975634767 | ns | TM | KS.1 |
| FIM1_5157 | 2226.672669 | 2423.554375 | 2358.045634 | 0.079826334 | 0.06914437 | 0.101213767 | 0.343715475 | ns | TM | KS.1 |
| FIM1_5158 | 1464.288 | 1190.146826 | 1384.854333 | -0.099038176 | 0.143662134 | 0.172068532 | 0.466761693 | ns | TM | KS.1 |
| FIM1_5159 | 1310.282876 | 925.3215582 | 963.2899021 | -0.379881558 | 0.140935496 | 0.000131451 | 0.003055931 | ns | TM | KS.1 |
| FIM1_5160 | 6993.989014 | 10442.00919 | 11959.88436 | 0.229491856 | 0.222538981 | 0.021378528 | 0.14253455 | ns | TM | KS.1 |
| FIM1_5161 | 1402.209892 | 1558.141622 | 1637.126416 | 0.034057693 | 0.139399106 | 0.597811907 | 0.849348181 | ns | TM | KS.1 |
| FIM1_5162 | 2456.421328 | 2665.484631 | 2683.565014 | 0.060806601 | 0.083034919 | 0.264621194 | 0.590354459 | ns | TM | KS.1 |
| FIM1_5163 | 4526.133546 | 3780.06345 | 3379.838754 | -0.102565428 | 0.129982924 | 0.150412329 | 0.43154633 | ns | TM | KS.1 |
| FIM1_5164 | 267.8279627 | 175.3870947 | 390.2049425 | -0.462842286 | 0.180778292 | 9.11308580911 | 0.002200854 | ns | TM | KS.1 |
| FIM1_5165 | 1021.136951 | 930.1465126 | 1240.079075 | -0.056653596 | 0.100370349 | 0.35186369 | 0.67786494 | ns | TM | KS.1 |
| FIM1_5166 | 3131.736064 | 3086.465763 | 2961.30963 | -0.005852051 | 0.119042779 | 0.925487415 | 0.976735299 | ns | TM | KS.1 |
| FIM1_5167 | 732.4776361 | 703.6185041 | 637.241487 | -0.024275758 | 0.086465502 | 0.659957428 | 0.878874681 | ns | TM | KS.1 |
| FIM1_5168 | 664.0115079 | 719.1624071 | 781.3317534 | 0.043660816 | 0.103174801 | 0.474647961 | 0.775371126 | ns | TM | KS.1 |
| FIM1_5169 | 268.5490584 | 264.6903256 | 280.748438 | -0.00505109 | 0.117003055 | 0.93544449 | 0.981407916 | ns | TM | KS.1 |
| FIM1_5170 | 2073.230658 | 2008.298042 | 2146.695084 | -0.023088615 | 0.066497141 | 0.61845995 | 0.860858199 | ns | TM | KS.1 |
| FIM1_5171 | 739.503288 | 780.3200106 | 1014.350572 | 0.02711372 | 0.105050426 | 0.656152341 | 0.877276998 | ns | TM | KS.1 |
| FIM1_5172 | 5995.664002 | 6995.581241 | 7294.21892 | 0.123334609 | 0.10827803 | 0.070123208 | 0.280239294 | ns | TM | KS.1 |
| FIM1_5173 | 1987.448184 | 1448.276495 | 1704.099513 | -0.312946827 | 0.14483415 | 0.001146732 | 0.016730181 | ns | TM | KS.1 |
| FIM1_5174 | 4656.757133 | 4216.939264 | 4111.276212 | -0.089351349 | 0.078874719 | 0.097360879 | 0.337540328 | ns | TM | KS.1 |
| FIM1_5175 | 2600.544118 | 3121.521357 | 3329.456953 | 0.128703468 | 0.123284852 | 0.0769634 | 0.294813117 | ns | TM | KS.1 |
| FIM1_5176 | 449.3429391 | 459.376031 | 608.5618791 | 0.009043975 | 0.118570236 | 0.885080706 | 0.967028314 | ns | TM | KS.1 |
| FIM1_5177 | 940.9577341 | 1488.595945 | 1527.121631 | 0.150527608 | 0.240138729 | 0.081596691 | 0.304774234 | ns | TM | KS.1 |
| FIM1_5178 | 1052.393492 | 1059.769981 | 1130.801298 | 0.004418675 | 0.073477284 | 0.92914392 | 0.978304974 | ns | TM | KS.1 |
| FIM1_5179 | 2497.412748 | 4021.485482 | 4531.64748 | 0.236240997 | 0.263203995 | 0.024264999 | 0.152808065 | ns | TM | KS.1 |
| FIM1_5180 | 720.6586228 | 707.409913 | 802.0512042 | -0.010085248 | 0.089596012 | 0.857528363 | 0.955916258 | ns | TM | KS.1 |
| FIM1_5181 | 5343.099943 | 5379.704528 | 4942.464288 | 0.004049849 | 0.085872768 | 0.941113888 | 0.983672404 | ns | TM | KS.1 |
| FIM1_5182 | 1506.441156 | 1508.246142 | 1281.7074 | 0.001104811 | 0.070436183 | 0.981746058 | 0.995431301 | ns | TM | KS.1 |
| FIM1_5183 | 3987.101661 | 4852.160459 | 5023.530225 | 0.161514854 | 0.121038461 | 0.032668211 | 0.180382188 | ns | TM | KS.1 |
| FIM1_5184 | 572.9877077 | 700.3600287 | 724.6511252 | 0.147720002 | 0.128045014 | 0.051549048 | 0.236583501 | ns | TM | KS.1 |
| FIM1_5185 | 3369.498866 | 3409.118115 | 3451.371304 | 0.007874608 | 0.074247913 | 0.874955225 | 0.963031469 | ns | TM | KS.1 |
| FIM1_5186 | 1691.710614 | 1954.852546 | 1683.684437 | 0.122357613 | 0.101715079 | 0.062875344 | 0.262637093 | ns | TM | KS.1 |
| FIM1_5187 | 2117.480247 | 1692.749143 | 2096.002034 | -0.128165704 | 0.144119616 | 0.093355366 | 0.330094487 | ns | TM | KS.1 |
| FIM1_5188 | 7006.561956 | 8304.260368 | 8105.092407 | 0.097664851 | 0.126498837 | 0.164257228 | 0.45373665 | ns | TM | KS.1 |
| FIM1_5189 | 257.5673061 | 298.921994 | 255.0617251 | 0.053330579 | 0.140982017 | 0.422813638 | 0.739045122 | ns | TM | KS.1 |
| FIM1_5190 | 1100.380438 | 1213.103547 | 1079.103786 | 0.062772001 | 0.098534101 | 0.299943026 | 0.628445786 | ns | TM | KS.1 |
| FIM1_5191 | 1282.829678 | 1036.288337 | 1573.276048 | -0.134992261 | 0.137382184 | 0.076651158 | 0.294813117 | ns | TM | KS.1 |
| FIM1_5192 | 354.8259723 | 358.2576178 | 1933.053245 | 0.002163175 | 0.157488769 | 0.972189004 | 0.99362472 | ns | TM | KS.1 |
| FIM1_5193 | 13101.30335 | 16583.86842 | 15741.76877 | 0.248436988 | 0.10887995 | 0.001938741 | 0.02513155 | ns | TM | KS.1 |
| FIM1_5194 | 4088.145863 | 3375.202991 | 3900.873431 | -0.114762132 | 0.132079312 | 0.116056178 | 0.372502895 | ns | TM | KS.1 |
| FIM1_5195 | 5440.548414 | 3956.735138 | 5536.513644 | -0.335260452 | 0.134791552 | 0.000405515 | 0.007658836 | ns | TM | KS.1 |
| FIM1_5196 | 367.5119563 | 281.0062826 | 274.2174018 | -0.236668802 | 0.140802158 | 0.00735955 | 0.067803257 | ns | TM | KS.1 |
| FIM1_5197 | 356.7883658 | 368.9328687 | 384.7279617 | 0.021177789 | 0.085738805 | 0.699515844 | 0.896699645 | ns | TM | KS.1 |
| FIM1_5199 | 2258.689015 | 2762.13561 | 3164.931706 | 0.227305365 | 0.091062392 | 0.001449042 | 0.019889761 | ns | TM | KS.1 |
| FIM1_5200 | 200.6908219 | 165.8275701 | 329.1609456 | -0.053149797 | 0.159570564 | 0.423340195 | 0.739045122 | ns | TM | KS.1 |
| FIM1_5201 | 305.4510295 | 372.239259 | 502.8709471 | 0.086563912 | 0.142582452 | 0.221392765 | 0.535714496 | ns | TM | KS.1 |

| | | | | | | | | | | |
|---|---|---|---|---|---|---|---|---|---|---|
| FIM1_5202 | 890.8444464 | 648.6082458 | 778.6488792 | -0.362690018 | 0.118428213 | 5.5448566326 | 0.001491117 | ns | TM | KS.1 |
| FIM1_5203 | 641.294937 | 673.408327 | 745.3949667 | 0.025818231 | 0.099964349 | 0.664851898 | 0.880679852 | ns | TM | KS.1 |
| FIM1_5204 | 424.0723968 | 473.5174969 | 477.5816684 | 0.04956403 | 0.121265454 | 0.445189078 | 0.751385662 | ns | TM | KS.1 |
| FIM1_5205 | 1260.603159 | 1481.518888 | 1564.827119 | 0.163310129 | 0.093701368 | 0.014136353 | 0.10623619 | ns | TM | KS.1 |
| FIM1_5206 | 782.3304442 | 890.9283112 | 958.1753731 | 0.10258437 | 0.098175205 | 0.102795414 | 0.346951959 | ns | TM | KS.1 |
| FIM1_5207 | 272.5964667 | 282.1828461 | 269.4435916 | 0.014903498 | 0.108003102 | 0.807552925 | 0.937868053 | ns | TM | KS.1 |
| FIM1_5208 | 12.99982159 | 14.15386893 | 21.5559521 | 0.012087872 | 0.199855244 | 0.8403768 | 0.949982862 | ns | TM | KS.1 |
| FIM1_5209 | 6.613114832 | 36.95135788 | 37.8397702 | 1.560894047 | 0.551964958 | 1.3915430166 | 0.000532533 | Km>KS.1 | TM | KS.1 |
| FIM1_4316 | 2729.95145 | 4159.894537 | 4233.280858 | 0.336494281 | 0.22692942 | 0.003719256 | 0.041302007 | ns | TM | KS.1 |
| FIM1_4318 | 184.3064591 | 192.1464987 | 297.7479064 | 0.018983317 | 0.107060471 | 0.756009848 | 0.920354827 | ns | TM | KS.1 |
| FIM1_4319 | 642.599699 | 577.3793051 | 791.3821641 | -0.031119083 | 0.145615922 | 0.627778392 | 0.867130061 | ns | TM | KS.1 |
| FIM1_4320 | 3253.534897 | 3386.012251 | 3853.218499 | 0.025926204 | 0.080351982 | 0.623044975 | 0.863411908 | ns | TM | KS.1 |
| FIM1_4321 | 1715.126001 | 2026.554885 | 1574.930606 | 0.131092391 | 0.113722541 | 0.062936951 | 0.262637093 | ns | TM | KS.1 |
| FIM1_4322 | 1046.532855 | 1314.526034 | 1448.110775 | 0.194609089 | 0.128772277 | 0.016390338 | 0.119039318 | ns | TM | KS.1 |
| FIM1_4323 | 11041.1282 | 13819.691 | 16724.97566 | 0.280645662 | 0.072521027 | 1.2643678642 | 0.000495306 | ns | TM | KS.1 |
| FIM1_4324 | 30.34335488 | 34.97264631 | 38.88673509 | 0.033475154 | 0.153462289 | 0.601675012 | 0.851547247 | ns | TM | KS.1 |
| FIM1_4325 | 1821.904209 | 1913.629266 | 1983.336966 | 0.033511086 | 0.077739631 | 0.516632338 | 0.804981347 | ns | TM | KS.1 |
| FIM1_4326 | 1322.590449 | 1349.735633 | 1289.899005 | 0.010397725 | 0.100581244 | 0.861205574 | 0.958071943 | ns | TM | KS.1 |
| FIM1_4327 | 2168.577068 | 1810.747069 | 2328.645034 | -0.134482036 | 0.120567916 | 0.06434059 | 0.266080163 | ns | TM | KS.1 |
| FIM1_4328 | 722.7966557 | 816.3302033 | 872.2491588 | 0.056504252 | 0.122740967 | 0.389553029 | 0.713295089 | ns | TM | KS.1 |
| FIM1_4329 | 702.571509 | 902.9060543 | 704.8157214 | 0.181314826 | 0.146650916 | 0.029610957 | 0.170502907 | ns | TM | KS.1 |
| FIM1_4330 | 2375.199271 | 1917.017127 | 2374.448184 | -0.169557743 | 0.129074574 | 0.03073039 | 0.174934485 | ns | TM | KS.1 |
| FIM1_4331 | 4290.290566 | 2286.512399 | 5501.843841 | -0.784227555 | 0.246604109 | 1.2587978579 | 0.000495306 | ns | TM | KS.1 |
| FIM1_4332 | 1673.225041 | 1592.557654 | 1772.143234 | -0.039340529 | 0.064460477 | 0.385727221 | 0.709925589 | ns | TM | KS.1 |
| FIM1_4333 | 3008.142518 | 2859.759966 | 2923.975554 | -0.037364875 | 0.070884108 | 0.441267854 | 0.747837973 | ns | TM | KS.1 |
| FIM1_4334 | 903.5426057 | 911.4286162 | 898.2957246 | 0.004484797 | 0.101409423 | 0.940044551 | 0.983546982 | ns | TM | KS.1 |
| FIM1_4335 | 2707.2376 | 3035.06708 | 2760.392648 | 0.122736481 | 0.073103464 | 0.021831038 | 0.144044317 | ns | TM | KS.1 |
| FIM1_4336 | 1349.734593 | 1072.407087 | 963.1192203 | -0.167259114 | 0.138553542 | 0.037097373 | 0.193255946 | ns | TM | KS.1 |
| FIM1_4337 | 1418.39812 | 1544.670914 | 1516.12613 | 0.058456122 | 0.090202492 | 0.308301703 | 0.636779799 | ns | TM | KS.1 |
| FIM1_4338 | 3772.551948 | 4175.319508 | 4371.463693 | 0.107679819 | 0.069051539 | 0.032359648 | 0.18019422 | ns | TM | KS.1 |
| FIM1_4339 | 743.5087015 | 642.538488 | 495.8349819 | -0.083782075 | 0.119185483 | 0.214521473 | 0.522902659 | ns | TM | KS.1 |
| FIM1_4340 | 14648.55805 | 16633.15458 | 10345.93131 | 0.10083633 | 0.098352027 | 0.108428741 | 0.357240387 | ns | TM | KS.1 |
| FIM1_4341 | 387.3838782 | 445.6538677 | 351.1808717 | 0.050251349 | 0.139011155 | 0.447489899 | 0.752446492 | ns | TM | KS.1 |
| FIM1_4342 | 729.6852614 | 731.6893577 | 597.9498622 | 0.001066472 | 0.086299316 | 0.98452353 | 0.996034691 | ns | TM | KS.1 |
| FIM1_4343 | 23626.24603 | 26301.28176 | 25916.70843 | 0.082222462 | 0.09225663 | 0.165383136 | 0.456057682 | ns | TM | KS.1 |
| FIM1_4344 | 1739.570125 | 2318.281574 | 2485.044825 | 0.129605281 | 0.171262732 | 0.100850734 | 0.343604292 | ns | TM | KS.1 |
| FIM1_4345 | 807.7915275 | 737.5990879 | 544.9502246 | -0.049203926 | 0.107250346 | 0.430145719 | 0.743536592 | ns | TM | KS.1 |
| FIM1_4346 | 939.5638404 | 1408.349053 | 1382.390916 | 0.112101522 | 0.216639011 | 0.148075009 | 0.428049487 | ns | TM | KS.1 |
| FIM1_4347 | 1776.15316 | 2111.802501 | 1730.147006 | 0.148441497 | 0.111563049 | 0.037971401 | 0.196574108 | ns | TM | KS.1 |
| FIM1_4348 | 2223.37819 | 2229.62483 | 2477.390735 | 0.002166938 | 0.059742349 | 0.959796327 | 0.990303759 | ns | TM | KS.1 |
| FIM1_4349 | 116.7903441 | 71.68050867 | 484.6426657 | -0.185405088 | 0.25892363 | 0.050185841 | 0.233221194 | ns | TM | KS.1 |
| FIM1_4350 | 2394.97921 | 1892.193211 | 2577.991275 | -0.299035815 | 0.07038024 | 2.1266590078 | 0.000129026 | ns | TM | KS.1 |
| FIM1_4351 | 3972.903194 | 4421.28469 | 4666.112367 | 0.073301179 | 0.098095222 | 0.229129709 | 0.547402215 | ns | TM | KS.1 |
| FIM1_4352 | 995.221213 | 1278.760634 | 1263.868516 | 0.13163755 | 0.155524188 | 0.091865673 | 0.327386621 | ns | TM | KS.1 |
| FIM1_4353 | 1725.319272 | 2045.029059 | 2362.725098 | 0.125743836 | 0.117297167 | 0.076460458 | 0.294813117 | ns | TM | KS.1 |

| | | | | | | | | | | |
|---|---|---|---|---|---|---|---|---|---|---|
| FIM1_4354 | 2513.459746 | 2190.399669 | 1627.87739 | -0.082935795 | 0.114649139 | 0.211908532 | 0.519844649 | ns | TM | KS.1 |
| FIM1_4355 | 26.81976678 | 26.27926877 | 26.18476884 | -0.005985821 | 0.156024369 | 0.923325657 | 0.976662213 | ns | TM | KS.1 |
| FIM1_4356 | 22416.25101 | 26093.72442 | 23485.72274 | 0.181849494 | 0.071821677 | 0.001788912 | 0.023732899 | ns | TM | KS.1 |
| FIM1_4357 | 2686.086679 | 2396.993286 | 1923.814461 | -0.074912412 | 0.102611395 | 0.231027698 | 0.549146106 | ns | TM | KS.1 |
| FIM1_4358 | 4816.50163 | 4953.282715 | 4355.850489 | 0.016601469 | 0.087232434 | 0.764374174 | 0.924230468 | ns | TM | KS.1 |
| FIM1_4359 | 3239.647541 | 2315.373813 | 2335.321236 | -0.352982064 | 0.143166209 | 0.000342262 | 0.006651376 | ns | TM | KS.1 |
| FIM1_4360 | 7899.56767 | 7251.301195 | 8671.074157 | -0.068027804 | 0.080811454 | 0.205845049 | 0.512865935 | ns | TM | KS.1 |
| FIM1_4361 | 1712.397257 | 2352.843437 | 2926.548286 | 0.191214872 | 0.17965535 | 0.032063688 | 0.179349393 | ns | TM | KS.1 |
| FIM1_4363 | 531.5299847 | 822.5232186 | 979.5832279 | 0.286592011 | 0.243892108 | 0.010025525 | 0.084537265 | ns | TM | KS.1 |
| FIM1_4364 | 451.3410331 | 507.8254614 | 393.1670421 | 0.055982732 | 0.120945461 | 0.391925557 | 0.715009037 | ns | TM | KS.1 |
| FIM1_4365 | 1194.258337 | 1268.722678 | 1130.932375 | 0.036566122 | 0.093131434 | 0.526868838 | 0.808531286 | ns | TM | KS.1 |
| FIM1_4366 | 8626.606542 | 9222.885707 | 9010.826394 | 0.054093831 | 0.070851258 | 0.266769795 | 0.593708608 | ns | TM | KS.1 |
| FIM1_4367 | 1477.378157 | 1633.50814 | 1772.816264 | 0.062513407 | 0.101240565 | 0.308726206 | 0.636779799 | ns | TM | KS.1 |
| FIM1_4368 | 1534.877576 | 1619.935903 | 1621.621862 | 0.034183023 | 0.087305976 | 0.538936369 | 0.814399145 | ns | TM | KS.1 |
| FIM1_4369 | 1709.290358 | 2632.26053 | 3186.351165 | 0.202755798 | 0.237031331 | 0.035590719 | 0.188357575 | ns | TM | KS.1 |
| FIM1_4370 | 634.1875998 | 793.0098374 | 886.3623511 | 0.124661649 | 0.14474333 | 0.100974769 | 0.343604292 | ns | TM | KS.1 |
| FIM1_4371 | 3277.052193 | 3367.313466 | 3441.943324 | 0.013803848 | 0.101346751 | 0.817163238 | 0.942152285 | ns | TM | KS.1 |
| FIM1_4372 | 2411.133485 | 2618.852747 | 2499.143024 | 0.040638078 | 0.11180123 | 0.518052669 | 0.804981347 | ns | TM | KS.1 |
| FIM1_4373 | 4470.855056 | 6690.120154 | 8223.638193 | 0.274236849 | 0.222126039 | 0.01007512 | 0.084811716 | ns | TM | KS.1 |
| FIM1_4374 | 1210.830569 | 1186.765345 | 1187.237333 | -0.012194417 | 0.072346733 | 0.80414063 | 0.936908579 | ns | TM | KS.1 |
| FIM1_4376 | 258.9768267 | 244.9661578 | 275.9932316 | -0.026126362 | 0.109522868 | 0.672423007 | 0.884299354 | ns | TM | KS.1 |
| FIM1_4377 | 3740.783936 | 4769.156897 | 5376.730464 | 0.1201656 | 0.153946846 | 0.115339593 | 0.370766358 | ns | TM | KS.1 |
| FIM1_4378 | 245.2264049 | 241.1895461 | 200.5203965 | -0.00830219 | 0.106268011 | 0.891279875 | 0.967215407 | ns | TM | KS.1 |
| FIM1_4379 | 3896.738552 | 4766.925094 | 3467.010987 | 0.194512142 | 0.110807602 | 0.01039096 | 0.087297666 | ns | TM | KS.1 |
| FIM1_4380 | 1217.568418 | 1314.237322 | 1529.100328 | 0.061410132 | 0.076348509 | 0.233193279 | 0.5516579 | ns | TM | KS.1 |
| FIM1_4381 | 1401.415536 | 1431.308912 | 1501.693725 | 0.013298719 | 0.081576075 | 0.802548197 | 0.936908579 | ns | TM | KS.1 |
| FIM1_4382 | 861.4913588 | 928.470652 | 961.0533677 | 0.047148456 | 0.092426765 | 0.414937325 | 0.734631029 | ns | TM | KS.1 |
| FIM1_4383 | 4939.767399 | 5588.645505 | 5562.631831 | 0.140671169 | 0.070934122 | 0.009183634 | 0.07968336 | ns | TM | KS.1 |
| FIM1_4384 | 1019.128108 | 1063.088969 | 1504.303601 | 0.016078109 | 0.123363397 | 0.798975931 | 0.936373441 | ns | TM | KS.1 |
| FIM1_4385 | 1594.794751 | 1661.261763 | 1672.830195 | 0.026921232 | 0.078753405 | 0.605134525 | 0.851547247 | ns | TM | KS.1 |
| FIM1_4386 | 16434.04453 | 17676.27008 | 15426.11709 | 0.050572134 | 0.08492644 | 0.358058643 | 0.683043834 | ns | TM | KS.1 |
| FIM1_4387 | 27.78388778 | 29.53491531 | 32.08724951 | 0.02290362 | 0.150313517 | 0.717623963 | 0.905458243 | ns | TM | KS.1 |
| FIM1_4388 | 1146.920355 | 1393.998107 | 2828.08616 | 0.068737108 | 0.150573736 | 0.316752164 | 0.643468524 | ns | TM | KS.1 |
| FIM1_4389 | 299.7607277 | 231.2716902 | 261.3620366 | -0.158255456 | 0.156792943 | 0.05339024 | 0.241250174 | ns | TM | KS.1 |
| FIM1_4390 | 119.3554187 | 130.0342283 | 119.0014353 | 0.039390964 | 0.121083152 | 0.539243554 | 0.814399145 | ns | TM | KS.1 |
| FIM1_4391 | 6656.549968 | 9572.810596 | 11072.76012 | 0.267882496 | 0.196164449 | 0.00862204 | 0.076597584 | ns | TM | KS.1 |
| FIM1_4392 | 2289.499779 | 2062.324571 | 2446.787469 | -0.049040736 | 0.118374889 | 0.44700689 | 0.752446492 | ns | TM | KS.1 |
| FIM1_4393 | 109.9963793 | 111.0903695 | 102.1546705 | 0.004121632 | 0.132365418 | 0.947936516 | 0.986388143 | ns | TM | KS.1 |
| FIM1_4394 | 794.3242868 | 589.1341696 | 574.9786187 | -0.301744664 | 0.13307977 | 0.001043093 | 0.015967346 | ns | TM | KS.1 |
| FIM1_4396 | 4142.394317 | 3935.652962 | 3371.773329 | -0.027947784 | 0.097876058 | 0.636084933 | 0.869849417 | ns | TM | KS.1 |
| FIM1_4397 | 1515.480093 | 1311.768486 | 1309.553351 | -0.116259513 | 0.104255861 | 0.078800591 | 0.299076777 | ns | TM | KS.1 |
| FIM1_4398 | 127.333153 | 91.00967203 | 199.3745941 | -0.101716392 | 0.193716304 | 0.174419199 | 0.470572405 | ns | TM | KS.1 |
| FIM1_4399 | 11525.70022 | 13281.242 | 14052.33915 | 0.063146581 | 0.128939677 | 0.345700992 | 0.671558937 | ns | TM | KS.1 |
| FIM1_4400 | 1156.523366 | 1172.581913 | 1072.102269 | 0.009410822 | 0.071046483 | 0.846364304 | 0.952348261 | ns | TM | KS.1 |
| FIM1_4401 | 2165.709606 | 2468.740343 | 2429.574297 | 0.097454063 | 0.103054657 | 0.129164329 | 0.396662061 | ns | TM | KS.1 |

| | | | | | | | | | | |
|---|---|---|---|---|---|---|---|---|---|---|
| FIM1_4402 | 211.0016836 | 154.476002 | 172.6698895 | -0.213790232 | 0.17141366 | 0.019146349 | 0.132384765 | ns | TM | KS.1 |
| FIM1_4403 | 4095.440512 | 3839.723963 | 5429.59483 | -0.018975005 | 0.141567991 | 0.764835545 | 0.924230468 | ns | TM | KS.1 |
| FIM1_4404 | 465.9904643 | 460.9029346 | 460.0954742 | -0.005200977 | 0.100062716 | 0.930104289 | 0.97843743 | ns | TM | KS.1 |
| FIM1_4405 | 714.7695649 | 611.8988696 | 609.2161831 | -0.088851571 | 0.122242926 | 0.195024126 | 0.501418619 | ns | TM | KS.1 |
| FIM1_4406 | 1196.900009 | 1034.297268 | 1069.145318 | -0.110649503 | 0.107792673 | 0.097775241 | 0.3377271 | ns | TM | KS.1 |
| FIM1_4407 | 68.65831643 | 42.53300446 | 122.619116 | -0.229594557 | 0.263850942 | 0.026802059 | 0.160661696 | ns | TM | KS.1 |
| FIM1_4408 | 671.7988631 | 490.4319694 | 448.4685894 | -0.108925088 | 0.184554152 | 0.151913758 | 0.434101635 | ns | TM | KS.1 |
| FIM1_4409 | 4983.2207 | 3461.588374 | 6251.210809 | -0.135546113 | 0.202420433 | 0.097098742 | 0.337101354 | ns | TM | KS.1 |
| FIM1_4410 | 1023.775001 | 886.0433892 | 823.5708034 | -0.101922861 | 0.110984345 | 0.128500069 | 0.395843865 | ns | TM | KS.1 |
| FIM1_4412 | 191.4271488 | 154.8925167 | 240.9380461 | -0.063849461 | 0.160823003 | 0.347419371 | 0.673290223 | ns | TM | KS.1 |
| FIM1_4413 | 1448.731853 | 1531.228984 | 1434.163885 | 0.041614088 | 0.072461777 | 0.398709411 | 0.722244002 | ns | TM | KS.1 |
| FIM1_4414 | 3389.300904 | 3161.642228 | 3221.545165 | -0.044335721 | 0.090242202 | 0.436333794 | 0.747764597 | ns | TM | KS.1 |
| FIM1_4415 | 2877.637818 | 2982.385391 | 3335.323888 | 0.010763083 | 0.13848896 | 0.864707905 | 0.959975149 | ns | TM | KS.1 |
| FIM1_4416 | 1886.980424 | 1781.613802 | 1896.95476 | -0.024548055 | 0.117083528 | 0.696274376 | 0.894950919 | ns | TM | KS.1 |
| FIM1_4417 | 1109.829143 | 1169.735582 | 1298.851095 | 0.037606954 | 0.074935551 | 0.455876742 | 0.759285836 | ns | TM | KS.1 |
| FIM1_4418 | 5108.193105 | 5687.739747 | 5851.324223 | 0.091640023 | 0.086345547 | 0.11026335 | 0.360239803 | ns | TM | KS.1 |
| FIM1_4419 | 1233.221592 | 1246.988496 | 1039.942133 | 0.006995906 | 0.079171703 | 0.893342863 | 0.967972723 | ns | TM | KS.1 |
| FIM1_4420 | 1180.01612 | 1192.751107 | 1050.982712 | 0.007022694 | 0.074857339 | 0.888990412 | 0.967028314 | ns | TM | KS.1 |
| FIM1_4421 | 1039.197424 | 855.2579435 | 841.2736502 | -0.202047846 | 0.101746965 | 0.005951423 | 0.058980734 | ns | TM | KS.1 |
| FIM1_4422 | 15.37379167 | 11.72921026 | 14.57156587 | -0.043368907 | 0.199974226 | 0.496036396 | 0.79247947 | ns | TM | KS.1 |
| FIM1_4423 | 1919.506633 | 1852.613906 | 1754.733821 | -0.013437723 | 0.123071319 | 0.831180123 | 0.946802573 | ns | TM | KS.1 |
| FIM1_4424 | 763.8094743 | 890.400272 | 1053.413028 | 0.032035644 | 0.170481989 | 0.612441222 | 0.8563505 | ns | TM | KS.1 |
| FIM1_4425 | 645.4775823 | 603.9059537 | 643.1878699 | -0.051604773 | 0.07074628 | 0.288446863 | 0.61538797 | ns | TM | KS.1 |
| FIM1_4426 | 700.049223 | 650.9294403 | 728.6083259 | -0.044828576 | 0.093339852 | 0.440254499 | 0.747832613 | ns | TM | KS.1 |
| FIM1_4427 | 5068.047999 | 5094.185463 | 5440.023767 | 0.003245404 | 0.078216524 | 0.950017147 | 0.986914868 | ns | TM | KS.1 |
| FIM1_4428 | 5170.012102 | 6817.049015 | 6919.829657 | 0.128463047 | 0.166908428 | 0.101853635 | 0.34470873 | ns | TM | KS.1 |
| FIM1_4429 | 1836.533695 | 1828.466287 | 2044.53562 | -0.002286355 | 0.101116886 | 0.969372286 | 0.993536696 | ns | TM | KS.1 |
| FIM1_4430 | 369.7284531 | 379.0162803 | 343.7891576 | 0.012504602 | 0.103809126 | 0.835637455 | 0.948719561 | ns | TM | KS.1 |
| FIM1_4431 | 1519.033651 | 1639.233106 | 2195.196975 | 0.057093981 | 0.081186428 | 0.286717575 | 0.612762705 | ns | TM | KS.1 |
| FIM1_4432 | 4738.110853 | 5508.276092 | 5969.247125 | 0.15357838 | 0.091273596 | 0.017454107 | 0.12351946 | ns | TM | KS.1 |
| FIM1_4433 | 558.4393186 | 659.3397923 | 622.5379711 | 0.095577102 | 0.12589392 | 0.171492306 | 0.465692069 | ns | TM | KS.1 |
| FIM1_4434 | 1771.656305 | 1966.056619 | 1987.486864 | 0.115177183 | 0.06542408 | 0.018760394 | 0.130718431 | ns | TM | KS.1 |
| FIM1_4435 | 3677.208728 | 3722.800891 | 4237.227993 | 0.007937909 | 0.075141908 | 0.874949006 | 0.963031469 | ns | TM | KS.1 |
| FIM1_4436 | 1716.136764 | 1609.820308 | 1632.471748 | -0.053808063 | 0.065519744 | 0.242640508 | 0.565403527 | ns | TM | KS.1 |
| FIM1_4437 | 705.4210401 | 690.6099927 | 636.5513952 | -0.012037364 | 0.093330682 | 0.833965627 | 0.948141464 | ns | TM | KS.1 |
| FIM1_4438 | 1268.599593 | 1170.757585 | 1274.270211 | -0.043255594 | 0.104698721 | 0.481457409 | 0.783529804 | ns | TM | KS.1 |
| FIM1_4439 | 1982.18705 | 1550.296132 | 1607.302738 | -0.252987137 | 0.114322859 | 0.002141011 | 0.02738183 | ns | TM | KS.1 |
| FIM1_4440 | 2304.916611 | 1547.623279 | 1509.242747 | -0.262181398 | 0.220053737 | 0.012124466 | 0.096768242 | ns | TM | KS.1 |
| FIM1_4441 | 3308.671554 | 3546.198129 | 3396.226368 | 0.022760838 | 0.134996342 | 0.721119505 | 0.906676397 | ns | TM | KS.1 |
| FIM1_4443 | 473.7881039 | 297.9697211 | 510.6500014 | -0.146614092 | 0.241328718 | 0.086955591 | 0.314392489 | ns | TM | KS.1 |
| FIM1_4444 | 25287.01423 | 29523.33843 | 29229.62885 | 0.103411128 | 0.116149428 | 0.13155533 | 0.400788589 | ns | TM | KS.1 |
| FIM1_4445 | 1246.629701 | 1381.442406 | 1353.071427 | 0.077332808 | 0.091791137 | 0.188632007 | 0.491815863 | ns | TM | KS.1 |
| FIM1_4446 | 5774.705205 | 5456.084112 | 5081.720682 | -0.036584007 | 0.085361169 | 0.505305438 | 0.79780849 | ns | TM | KS.1 |
| FIM1_4447 | 1478.774722 | 1512.897947 | 1698.587544 | 0.017232254 | 0.060009381 | 0.689203616 | 0.892448722 | ns | TM | KS.1 |
| FIM1_4448 | 246.1424584 | 233.4904067 | 260.2087931 | -0.01607829 | 0.139673288 | 0.799623578 | 0.936691147 | ns | TM | KS.1 |

| | | | | | | | | | | |
|---|---|---|---|---|---|---|---|---|---|---|
| FIM1_4449 | 844.7696583 | 585.5122286 | 733.6709146 | -0.161834824 | 0.203949917 | 0.061690626 | 0.260865744 | ns | TM | KS.1 |
| FIM1_4450 | 3964.158874 | 4177.447538 | 3259.486394 | 0.03058614 | 0.092665599 | 0.594712025 | 0.847499664 | ns | TM | KS.1 |
| FIM1_4451 | 2956.340352 | 2705.565489 | 2267.005217 | -0.043509513 | 0.112531935 | 0.490966679 | 0.789960941 | ns | TM | KS.1 |
| FIM1_4452 | 9380.563897 | 10837.24994 | 10157.37843 | 0.087941559 | 0.116364093 | 0.191030091 | 0.495502973 | ns | TM | KS.1 |
| FIM1_4454 | 2999.49356 | 3423.073304 | 4077.397871 | 0.1323361 | 0.086143989 | 0.028537378 | 0.167619193 | ns | TM | KS.1 |
| FIM1_4455 | 1344.507009 | 1406.253793 | 1386.220544 | 0.022126276 | 0.105508095 | 0.716146275 | 0.905084971 | ns | TM | KS.1 |
| FIM1_4456 | 2378.919532 | 1588.983535 | 1521.687779 | -0.390104257 | 0.196036464 | 0.000774186 | 0.012505107 | ns | TM | KS.1 |
| FIM1_4457 | 2409.571788 | 2086.12793 | 1899.474495 | -0.123189771 | 0.100668818 | 0.059908863 | 0.257158405 | ns | TM | KS.1 |
| FIM1_4458 | 1601.344253 | 1714.086256 | 1523.401855 | 0.04882675 | 0.081341073 | 0.36110584 | 0.684120927 | ns | TM | KS.1 |
| FIM1_4460 | 567.4131194 | 676.736816 | 482.6566913 | 0.087715004 | 0.133762725 | 0.211168838 | 0.51871941 | ns | TM | KS.1 |
| FIM1_4461 | 291.9092215 | 310.3024668 | 235.8112276 | 0.026542514 | 0.117090027 | 0.673405895 | 0.885317825 | ns | TM | KS.1 |
| FIM1_4462 | 990.4962777 | 1052.022228 | 1086.853569 | 0.041372103 | 0.081842037 | 0.439727619 | 0.747832613 | ns | TM | KS.1 |
| FIM1_4463 | 801.7599229 | 839.2345088 | 1342.582568 | 0.026228479 | 0.090922492 | 0.644335638 | 0.872978256 | ns | TM | KS.1 |
| FIM1_4464 | 1043.530689 | 933.401108 | 1042.604224 | -0.066900558 | 0.10676184 | 0.290369194 | 0.618137245 | ns | TM | KS.1 |
| FIM1_4465 | 1618.756982 | 1483.617062 | 1372.504425 | -0.05619096 | 0.094503764 | 0.339986268 | 0.668193429 | ns | TM | KS.1 |
| FIM1_4466 | 1390.024742 | 1318.611089 | 1228.347793 | -0.036195384 | 0.078791322 | 0.487724795 | 0.787290997 | ns | TM | KS.1 |
| FIM1_4467 | 2573.976467 | 2744.147576 | 2880.810584 | 0.054634625 | 0.066718773 | 0.241763012 | 0.564151495 | ns | TM | KS.1 |
| FIM1_4468 | 320.6067217 | 300.6880265 | 290.415238 | -0.035220458 | 0.101879048 | 0.559768138 | 0.826391733 | ns | TM | KS.1 |
| FIM1_4469 | 1906.7854 | 3007.493197 | 3092.130005 | 0.214161442 | 0.249908468 | 0.031594657 | 0.177624525 | ns | TM | KS.1 |
| FIM1_4470 | 2090.27336 | 2883.858567 | 2972.149374 | 0.154980692 | 0.184113306 | 0.065117294 | 0.267734328 | ns | TM | KS.1 |
| FIM1_4471 | 847.0217872 | 927.7786813 | 801.3701326 | 0.0544301 | 0.101059508 | 0.371908842 | 0.694536971 | ns | TM | KS.1 |
| FIM1_4472 | 2250.835309 | 2319.78304 | 2324.054921 | 0.024250545 | 0.053477051 | 0.540334417 | 0.814399145 | ns | TM | KS.1 |
| FIM1_4473 | 919.3863509 | 720.6474556 | 825.0001088 | -0.196553004 | 0.137961116 | 0.018455757 | 0.129138383 | ns | TM | KS.1 |
| FIM1_4474 | 12606.29725 | 12036.5374 | 10933.34462 | -0.024307999 | 0.100298045 | 0.68361966 | 0.890494131 | ns | TM | KS.1 |
| FIM1_4475 | 1349.706544 | 1262.354379 | 812.4556507 | -0.032635514 | 0.108964413 | 0.59826338 | 0.849417898 | ns | TM | KS.1 |
| FIM1_4476 | 1521.879427 | 1727.545325 | 1386.57011 | 0.061805042 | 0.121250747 | 0.348472553 | 0.674195903 | ns | TM | KS.1 |
| FIM1_4477 | 2295.48085 | 2415.443879 | 2194.233526 | 0.036993515 | 0.073815152 | 0.458568324 | 0.762486824 | ns | TM | KS.1 |
| FIM1_4478 | 2190.621546 | 2011.814084 | 1911.228193 | -0.052011204 | 0.097517153 | 0.384006292 | 0.708354209 | ns | TM | KS.1 |
| FIM1_4479 | 1781.795789 | 2060.349374 | 2132.936426 | 0.060691252 | 0.132410343 | 0.364995323 | 0.688866363 | ns | TM | KS.1 |
| FIM1_4480 | 498.2736595 | 490.189724 | 436.876264 | -0.008333532 | 0.10363699 | 0.889860427 | 0.967028314 | ns | TM | KS.1 |
| FIM1_4481 | 2269.130473 | 2098.73571 | 1967.364171 | -0.066182151 | 0.073122287 | 0.186710391 | 0.489513719 | ns | TM | KS.1 |
| FIM1_4482 | 80.37348818 | 39.31636029 | 2864.161851 | -0.425167185 | 0.415839397 | 0.006388852 | 0.062079174 | ns | TM | KS.1 |
| FIM1_4484 | 9.037853576 | 19.69177248 | 72.63694843 | 0.137937486 | 0.321051528 | 0.098316685 | 0.338336381 | ns | TM | KS.1 |
| FIM1_4486 | 1163.877719 | 1133.634032 | 1769.97335 | -0.007201037 | 0.144783344 | 0.908766401 | 0.972251892 | ns | TM | KS.1 |
| FIM1_4487 | 279.7137299 | 193.0911782 | 435.3429884 | -0.131069034 | 0.205097679 | 0.10547849 | 0.351712793 | ns | TM | KS.1 |
| FIM1_4488 | 33.51625273 | 31.66566343 | 33.08812412 | -0.016638368 | 0.154446817 | 0.790765879 | 0.933303839 | ns | TM | KS.1 |
| FIM1_4489 | 1083.454144 | 1230.225999 | 1176.216034 | 0.040053447 | 0.143523392 | 0.53827457 | 0.814399145 | ns | TM | KS.1 |
| FIM1_4490 | 5821.48546 | 7728.039286 | 7550.038692 | 0.127495245 | 0.169769928 | 0.104599409 | 0.350588067 | ns | TM | KS.1 |
| FIM1_4491 | 833.1227211 | 759.422067 | 931.9821455 | -0.044158886 | 0.1151644 | 0.487940329 | 0.787383438 | ns | TM | KS.1 |
| FIM1_4492 | 800.6337251 | 722.2053097 | 821.0718333 | -0.068387205 | 0.097030026 | 0.25692183 | 0.585794503 | ns | TM | KS.1 |
| FIM1_4493 | 5026.261183 | 5056.115929 | 6962.732737 | 0.002479003 | 0.116894713 | 0.968275719 | 0.993374633 | ns | TM | KS.1 |
| FIM1_4494 | 24.65499901 | 29.63627301 | 44.75258721 | 0.054094018 | 0.16205577 | 0.415743443 | 0.735011951 | ns | TM | KS.1 |
| FIM1_4495 | 939.8157243 | 874.7273978 | 1200.271987 | -0.040895346 | 0.098480741 | 0.492825754 | 0.791161061 | ns | TM | KS.1 |
| FIM1_4496 | 593.8514026 | 610.03622 | 572.8180632 | 0.015794223 | 0.089793008 | 0.77902945 | 0.928304554 | ns | TM | KS.1 |
| FIM1_4497 | 2683.795385 | 2943.867915 | 4143.687139 | 0.063679479 | 0.092638275 | 0.276443922 | 0.601096377 | ns | TM | KS.1 |

| | | | | | | | | | | |
|---|---|---|---|---|---|---|---|---|---|---|
| FIM1_4498 | 17044.86207 | 20124.35728 | 18755.82401 | 0.132367983 | 0.112531418 | 0.059578601 | 0.256462934 | ns | TM | KS.1 |
| FIM1_4499 | 4827.373821 | 4271.720709 | 3375.850452 | -0.088131197 | 0.101223136 | 0.161433807 | 0.450439253 | ns | TM | KS.1 |
| FIM1_4500 | 2745.958736 | 2536.418213 | 2985.97264 | -0.067218379 | 0.073232129 | 0.180662298 | 0.480643537 | ns | TM | KS.1 |
| FIM1_4501 | 1726.141056 | 2030.678465 | 1516.750278 | 0.086981772 | 0.126955432 | 0.208624724 | 0.515858848 | ns | TM | KS.1 |
| FIM1_4502 | 230.5324694 | 241.1306123 | 211.4499236 | 0.017543417 | 0.124215794 | 0.781452296 | 0.929216508 | ns | TM | KS.1 |
| FIM1_4503 | 812.4773607 | 778.7869173 | 692.7756756 | -0.028437602 | 0.077449682 | 0.580881124 | 0.838424819 | ns | TM | KS.1 |
| FIM1_4504 | 1031.726799 | 959.3037208 | 1117.332412 | -0.035385355 | 0.110590779 | 0.570532848 | 0.832094673 | ns | TM | KS.1 |
| FIM1_4505 | 1556.166901 | 2365.729653 | 2753.53033 | 0.140652305 | 0.223873966 | 0.092781674 | 0.329235968 | ns | TM | KS.1 |
| FIM1_4506 | 1391.238878 | 1484.528669 | 1446.303108 | 0.031743683 | 0.108776304 | 0.607865802 | 0.852348468 | ns | TM | KS.1 |
| FIM1_4507 | 2807.453152 | 2595.977891 | 2743.829087 | -0.045574271 | 0.098887772 | 0.446936467 | 0.752446492 | ns | TM | KS.1 |
| FIM1_4508 | 1148.7223 | 1206.096485 | 1156.549221 | 0.034780686 | 0.076898875 | 0.497832246 | 0.793454967 | ns | TM | KS.1 |
| FIM1_4509 | 3859.71075 | 3158.055377 | 2958.750099 | -0.256336806 | 0.064041773 | 1.0022967805 | 0.000414015 | ns | TM | KS.1 |
| FIM1_4510 | 3995.542676 | 4761.580135 | 5040.076391 | 0.186537217 | 0.09383434 | 0.006765889 | 0.064774438 | ns | TM | KS.1 |
| FIM1_4512 | 1195.60759 | 1019.557842 | 894.7769578 | -0.140562629 | 0.104349942 | 0.039819552 | 0.202973639 | ns | TM | KS.1 |
| FIM1_4513 | 199.2354397 | 281.9691159 | 220.9323604 | 0.277693597 | 0.177669396 | 0.005548098 | 0.056358187 | ns | TM | KS.1 |
| FIM1_4514 | 382.4485885 | 375.3022187 | 316.521227 | -0.010816479 | 0.093764692 | 0.850917619 | 0.953390116 | ns | TM | KS.1 |
| FIM1_4515 | 1493.962688 | 1364.058767 | 1229.686415 | -0.071830668 | 0.08311781 | 0.191132888 | 0.495511264 | ns | TM | KS.1 |
| FIM1_4516 | 465.9242056 | 655.0264125 | 750.3798019 | 0.093871992 | 0.19601628 | 0.200890888 | 0.509135082 | ns | TM | KS.1 |
| FIM1_4517 | 1493.142992 | 1084.330974 | 1121.894738 | -0.190183639 | 0.180757702 | 0.03294816 | 0.181123861 | ns | TM | KS.1 |
| FIM1_4518 | 710.3071785 | 789.095171 | 733.9843654 | 0.058695438 | 0.109200899 | 0.354429601 | 0.679784721 | ns | TM | KS.1 |
| FIM1_4519 | 1101.985437 | 1034.889066 | 1030.65912 | -0.045658483 | 0.080789298 | 0.390438651 | 0.713987918 | ns | TM | KS.1 |
| FIM1_4520 | 1653.998333 | 1726.523631 | 2052.447947 | 0.020298966 | 0.108053482 | 0.740638289 | 0.914311536 | ns | TM | KS.1 |
| FIM1_4521 | 2292.049462 | 2416.227545 | 2375.04407 | 0.043065205 | 0.064075696 | 0.3405277 | 0.668294008 | ns | TM | KS.1 |
| FIM1_4522 | 574.1608942 | 308.7058742 | 643.0435648 | -0.57934245 | 0.344496003 | 0.000856521 | 0.013527595 | ns | TM | KS.1 |
| FIM1_4523 | 2628.064827 | 2907.339424 | 3284.485673 | 0.11179452 | 0.065113633 | 0.02140167 | 0.14253455 | ns | TM | KS.1 |
| FIM1_4524 | 2550.171425 | 1851.070983 | 2028.069343 | -0.348950215 | 0.128928906 | 0.000191021 | 0.004177267 | ns | TM | KS.1 |
| FIM1_4525 | 315.2598554 | 298.2519552 | 257.5943842 | -0.027055659 | 0.111022518 | 0.663049201 | 0.880619203 | ns | TM | KS.1 |
| FIM1_4526 | 6574.620432 | 6872.800536 | 6663.423493 | 0.039308698 | 0.053986937 | 0.324225922 | 0.651040335 | ns | TM | KS.1 |
| FIM1_4527 | 18328.84279 | 20558.12019 | 16071.36789 | 0.037187315 | 0.140540123 | 0.566239197 | 0.828784939 | ns | TM | KS.1 |
| FIM1_4528 | 169.9734 | 140.3914242 | 224.419495 | -0.054320546 | 0.15900381 | 0.414451566 | 0.734555234 | ns | TM | KS.1 |
| FIM1_4529 | 600.0344027 | 353.9756016 | 548.8525816 | -0.283100139 | 0.296876408 | 0.015274059 | 0.11240894 | ns | TM | KS.1 |
| FIM1_4530 | 2164.010938 | 2372.570429 | 2316.557567 | 0.0695565 | 0.086865609 | 0.217968445 | 0.529594372 | ns | TM | KS.1 |
| FIM1_4531 | 6350.514695 | 5477.392604 | 3611.620623 | -0.140591602 | 0.095322954 | 0.030589565 | 0.174722257 | ns | TM | KS.1 |
| FIM1_4532 | 1845.03522 | 1869.068519 | 1404.05236 | 0.006911331 | 0.092175338 | 0.903527747 | 0.970434055 | ns | TM | KS.1 |
| FIM1_4533 | 5034.642165 | 8190.647664 | 8237.061385 | 0.263568767 | 0.272041962 | 0.017230646 | 0.122811551 | ns | TM | KS.1 |
| FIM1_4534 | 1049.067371 | 907.7887162 | 795.2805864 | -0.094026688 | 0.113628466 | 0.161181916 | 0.449988794 | ns | TM | KS.1 |
| FIM1_4535 | 1438.324374 | 1812.666472 | 1795.554662 | 0.147216377 | 0.14380846 | 0.061530241 | 0.260865744 | ns | TM | KS.1 |
| FIM1_4536 | 2611.145372 | 2274.445141 | 2216.598367 | -0.05300236 | 0.135253296 | 0.424310741 | 0.740157476 | ns | TM | KS.1 |
| FIM1_4537 | 813.1797017 | 868.8223476 | 926.008992 | 0.044128702 | 0.086507742 | 0.426691244 | 0.741197254 | ns | TM | KS.1 |
| FIM1_4538 | 1603.366826 | 477.7943961 | 724.0558032 | -1.505668564 | 0.29120235 | 1.8141098412 | 2.3750517000 | Km<KS.1 | TM | KS.1 |
| FIM1_4539 | 1093.687874 | 1024.388941 | 1030.865874 | -0.04768583 | 0.078723346 | 0.361768471 | 0.685115394 | ns | TM | KS.1 |
| FIM1_4540 | 805.9041965 | 776.5566088 | 575.5351737 | -0.019051879 | 0.101313662 | 0.750001979 | 0.918124963 | ns | TM | KS.1 |
| FIM1_4542 | 4838.23439 | 4644.352992 | 4417.824229 | -0.021938617 | 0.097707468 | 0.709449598 | 0.901619224 | ns | TM | KS.1 |
| FIM1_4543 | 1834.937279 | 3277.04541 | 3931.707626 | 0.259336624 | 0.316294788 | 0.022637377 | 0.147024228 | ns | TM | KS.1 |
| FIM1_4544 | 800.0733922 | 598.724182 | 678.0470273 | -0.202962935 | 0.163030793 | 0.022037629 | 0.144831181 | ns | TM | KS.1 |

| | | | | | | | | | | |
|---|---|---|---|---|---|---|---|---|---|---|
| FIM1_4545 | 1342.531536 | 1938.83436 | 2193.460746 | 0.195390037 | 0.204997016 | 0.03450728 | 0.185392782 | ns | TM | KS.1 |
| FIM1_4546 | 2401.122193 | 3877.92104 | 5054.654355 | 0.248078581 | 0.266347084 | 0.02079392 | 0.140511155 | ns | TM | KS.1 |
| FIM1_4547 | 85.7880688 | 98.57121533 | 93.14022584 | 0.042684712 | 0.14924511 | 0.513103537 | 0.803237916 | ns | TM | KS.1 |
| FIM1_4548 | 1474.544735 | 1256.23109 | 1192.076338 | -0.110936826 | 0.116781075 | 0.110074625 | 0.360040276 | ns | TM | KS.1 |
| FIM1_4549 | 3268.302471 | 2686.731993 | 2090.131992 | -0.119191789 | 0.132890357 | 0.105649948 | 0.351822919 | ns | TM | KS.1 |
| FIM1_4550 | 63.70826206 | 48.32581618 | 58.11738184 | -0.083409583 | 0.176171957 | 0.24152329 | 0.563856579 | ns | TM | KS.1 |
| FIM1_4551 | 151.5140716 | 129.4433794 | 155.7175167 | -0.078511133 | 0.127493662 | 0.250876274 | 0.576493979 | ns | TM | KS.1 |
| FIM1_4552 | 1076.023744 | 1072.911986 | 991.3499352 | -0.001720476 | 0.096783708 | 0.976497422 | 0.993872484 | ns | TM | KS.1 |
| FIM1_4553 | 410.4262819 | 514.3781034 | 556.4867473 | 0.126765543 | 0.14488757 | 0.096616433 | 0.336392346 | ns | TM | KS.1 |
| FIM1_4554 | 2325.910482 | 2570.483705 | 2258.936361 | 0.053743649 | 0.109786248 | 0.395228515 | 0.718896697 | ns | TM | KS.1 |
| FIM1_4555 | 3790.965759 | 4770.022465 | 5241.801357 | 0.108634636 | 0.150668346 | 0.144270354 | 0.421706822 | ns | TM | KS.1 |
| FIM1_4556 | 2656.621223 | 2686.653355 | 2797.311843 | 0.004251045 | 0.122451813 | 0.94604924 | 0.985614597 | ns | TM | KS.1 |
| FIM1_4557 | 4845.300222 | 5022.255354 | 4748.96601 | 0.025455764 | 0.070781112 | 0.599210385 | 0.84979238 | ns | TM | KS.1 |
| FIM1_4558 | 2498.296872 | 2044.242472 | 1668.102854 | -0.137164769 | 0.130728381 | 0.068472435 | 0.275933222 | ns | TM | KS.1 |
| FIM1_4559 | 1564.410857 | 1403.26269 | 1020.512662 | -0.076781397 | 0.096907271 | 0.206075661 | 0.513126334 | ns | TM | KS.1 |
| FIM1_4560 | 246.1455564 | 199.6721561 | 161.0635889 | -0.086303878 | 0.148901033 | 0.224875957 | 0.541771374 | ns | TM | KS.1 |
| FIM1_4561 | 3429.924123 | 3602.548785 | 4062.928576 | 0.019350172 | 0.121612032 | 0.759198766 | 0.922567322 | ns | TM | KS.1 |
| FIM1_4562 | 315.441529 | 249.9189394 | 665.2961242 | -0.055375647 | 0.17510412 | 0.404156822 | 0.725877324 | ns | TM | KS.1 |
| FIM1_4563 | 1884.63829 | 2237.94658 | 2210.548623 | 0.070646038 | 0.139297002 | 0.302777666 | 0.631432423 | ns | TM | KS.1 |
| FIM1_4564 | 638.2889182 | 492.5074928 | 338.2719058 | -0.135378417 | 0.158138894 | 0.086088184 | 0.312966141 | ns | TM | KS.1 |
| FIM1_4565 | 5031.878263 | 4992.127747 | 4454.604135 | -0.002669728 | 0.130988741 | 0.966253472 | 0.993374633 | ns | TM | KS.1 |
| FIM1_4566 | 14626.44683 | 15020.30397 | 10289.83615 | 0.007878135 | 0.139267953 | 0.900593875 | 0.968537512 | ns | TM | KS.1 |
| FIM1_4567 | 5610.485071 | 5396.749267 | 6056.028043 | -0.025253411 | 0.079915375 | 0.630868024 | 0.869059435 | ns | TM | KS.1 |
| FIM1_4568 | 5007.507945 | 4743.779588 | 5307.079298 | -0.05035627 | 0.054719785 | 0.212089857 | 0.52003304 | ns | TM | KS.1 |
| FIM1_4569 | 2891.362595 | 2605.084183 | 2428.104162 | -0.061821218 | 0.105041387 | 0.322558763 | 0.650748908 | ns | TM | KS.1 |
| FIM1_4570 | 1375.559724 | 1298.106061 | 1569.40606 | -0.038883318 | 0.081424753 | 0.465945078 | 0.767829336 | ns | TM | KS.1 |
| FIM1_4571 | 1117.991526 | 1166.892501 | 1194.927305 | 0.026461712 | 0.086344742 | 0.631394734 | 0.86917233 | ns | TM | KS.1 |
| FIM1_4572 | 2952.540461 | 4174.462323 | 4823.600807 | 0.136643918 | 0.195095124 | 0.093783313 | 0.330902114 | ns | TM | KS.1 |
| FIM1_4573 | 1697.347868 | 1800.612637 | 1981.366376 | 0.028546574 | 0.10847135 | 0.643477226 | 0.872978256 | ns | TM | KS.1 |
| FIM1_4574 | 692.5445402 | 670.6848182 | 667.0720463 | -0.023437566 | 0.074062086 | 0.639025524 | 0.870968656 | ns | TM | KS.1 |
| FIM1_4575 | 7691.798364 | 8361.568534 | 8341.093234 | 0.061437391 | 0.084981358 | 0.267119788 | 0.593708608 | ns | TM | KS.1 |
| FIM1_4576 | 1738.835839 | 1767.711967 | 1594.398959 | 0.009886854 | 0.087161438 | 0.858146917 | 0.956277476 | ns | TM | KS.1 |
| FIM1_4577 | 543.3016558 | 518.288427 | 482.5281874 | -0.020020827 | 0.117265734 | 0.74961062 | 0.917871729 | ns | TM | KS.1 |
| FIM1_4578 | 449.1645552 | 396.1643455 | 446.1690164 | -0.076594487 | 0.111561382 | 0.240034713 | 0.561090803 | ns | TM | KS.1 |
| FIM1_4579 | 328.0181073 | 316.7456301 | 334.6155605 | -0.013057608 | 0.123827451 | 0.835977076 | 0.948719561 | ns | TM | KS.1 |
| FIM1_4580 | 2185.23717 | 2814.912007 | 3068.994516 | 0.176032732 | 0.148725843 | 0.034068146 | 0.184799735 | ns | TM | KS.1 |
| FIM1_4582 | 1300.312215 | 1471.546469 | 1719.339598 | 0.04730015 | 0.132801234 | 0.471316741 | 0.772376515 | ns | TM | KS.1 |
| FIM1_4583 | 1303.607586 | 1366.158041 | 1529.542087 | 0.036458284 | 0.066404543 | 0.43123498 | 0.743536592 | ns | TM | KS.1 |
| FIM1_4584 | 1666.925107 | 1769.864879 | 1894.691616 | 0.040014343 | 0.084222766 | 0.462952457 | 0.764924767 | ns | TM | KS.1 |
| FIM1_4585 | 1956.939772 | 2028.074046 | 2464.633392 | 0.027338265 | 0.063113722 | 0.54038633 | 0.814399145 | ns | TM | KS.1 |
| FIM1_4586 | 11011.56102 | 14966.56309 | 14634.50189 | 0.404812788 | 0.071426961 | 1.54325041699 | 4.85383063877 | ns | TM | KS.1 |
| FIM1_4587 | 1378.074892 | 1860.075411 | 2336.525261 | 0.086822358 | 0.183200602 | 0.22721108 | 0.54408821 | ns | TM | KS.1 |
| FIM1_4588 | 3743.384339 | 4166.336169 | 3198.471663 | 0.069908298 | 0.100866913 | 0.257160843 | 0.585794503 | ns | TM | KS.1 |
| FIM1_4589 | 1538.240593 | 2230.68327 | 2506.601279 | 0.20337038 | 0.207056659 | 0.030356469 | 0.173590153 | ns | TM | KS.1 |
| FIM1_4590 | 547.3744986 | 580.6323586 | 592.5620059 | 0.037817695 | 0.092690832 | 0.511949808 | 0.802566276 | ns | TM | KS.1 |

| | | | | | | | | | | |
|---|---|---|---|---|---|---|---|---|---|---|
| FIM1_4591 | 2347.577275 | 2580.192405 | 2635.107102 | 0.069368885 | 0.09041204 | 0.230854045 | 0.548995638 | ns | TM | KS.1 |
| FIM1_4592 | 1228.46802 | 1078.944936 | 1307.898843 | -0.121844027 | 0.089385515 | 0.045467685 | 0.220040594 | ns | TM | KS.1 |
| FIM1_4593 | 835.7327902 | 867.3445595 | 726.3664995 | 0.019691189 | 0.097890702 | 0.738097234 | 0.913453034 | ns | TM | KS.1 |
| FIM1_4594 | 411.6418303 | 554.6395369 | 742.0136083 | 0.140599851 | 0.174557996 | 0.082552135 | 0.305123977 | ns | TM | KS.1 |
| FIM1_4595 | 3910.16425 | 3842.901123 | 4025.083491 | -0.010529307 | 0.084594484 | 0.846419583 | 0.952348261 | ns | TM | KS.1 |
| FIM1_4596 | 2092.433747 | 3009.72396 | 3538.793639 | 0.205816087 | 0.202685198 | 0.028341968 | 0.166668193 | ns | TM | KS.1 |
| FIM1_4597 | 2101.407021 | 2108.693007 | 1866.817906 | 0.002071595 | 0.071734397 | 0.966203805 | 0.993374633 | ns | TM | KS.1 |
| FIM1_4598 | 501.6966707 | 512.8134022 | 478.6078264 | 0.010895299 | 0.102802103 | 0.855944249 | 0.955916258 | ns | TM | KS.1 |
| FIM1_4599 | 257.1387154 | 246.2966005 | 280.8019534 | -0.018989257 | 0.112885823 | 0.759836122 | 0.922896656 | ns | TM | KS.1 |
| FIM1_4600 | 2872.967183 | 3322.687511 | 3711.087624 | 0.127899704 | 0.099972255 | 0.051341281 | 0.235847526 | ns | TM | KS.1 |
| FIM1_4601 | 927.4180203 | 868.8906484 | 781.6834496 | -0.049388592 | 0.076578491 | 0.336075941 | 0.664010248 | ns | TM | KS.1 |
| FIM1_4602 | 515.3724269 | 554.2263376 | 450.4868022 | 0.024110291 | 0.135088844 | 0.705689918 | 0.899257553 | ns | TM | KS.1 |
| FIM1_4603 | 3874.968892 | 3980.422043 | 3864.028105 | 0.017475191 | 0.077317154 | 0.733904378 | 0.911352378 | ns | TM | KS.1 |
| FIM1_4604 | 166.7428804 | 169.8547429 | 140.3676843 | 0.007440436 | 0.128070724 | 0.906155952 | 0.971578849 | ns | TM | KS.1 |
| FIM1_4605 | 905.5107983 | 1043.352204 | 1105.088326 | 0.122094127 | 0.100107063 | 0.061020942 | 0.26041916 | ns | TM | KS.1 |
| FIM1_4606 | 752.7333411 | 744.7121797 | 706.6727291 | -0.00668581 | 0.077902466 | 0.896989405 | 0.967972723 | ns | TM | KS.1 |
| FIM1_4607 | 2466.014348 | 2259.243786 | 2344.433376 | -0.079804905 | 0.071430656 | 0.109310228 | 0.35800987 | ns | TM | KS.1 |
| FIM1_4608 | 480.0305532 | 359.4290232 | 390.933759 | -0.16650361 | 0.168231119 | 0.048450882 | 0.229564894 | ns | TM | KS.1 |
| FIM1_4609 | 2856.399746 | 3589.973311 | 3251.156676 | 0.115536035 | 0.148690183 | 0.124458797 | 0.38782159 | ns | TM | KS.1 |
| FIM1_4612 | 6852.007244 | 7317.644558 | 7107.770878 | 0.038280528 | 0.095941315 | 0.514723862 | 0.804256034 | ns | TM | KS.1 |
| FIM1_4613 | 23680.21354 | 13997.99126 | 15088.26707 | -0.405056688 | 0.299405052 | 0.003290789 | 0.037636031 | ns | TM | KS.1 |
| FIM1_4614 | 587.9941235 | 498.7236664 | 609.0939818 | -0.13988985 | 0.109441248 | 0.045744152 | 0.220948694 | ns | TM | KS.1 |
| FIM1_4615 | 1425.240716 | 1331.117029 | 1502.09062 | -0.051046176 | 0.077219194 | 0.323162611 | 0.650941236 | ns | TM | KS.1 |
| FIM1_4616 | 2939.498378 | 3039.263303 | 3201.066156 | 0.024491178 | 0.068021952 | 0.603131757 | 0.851547247 | ns | TM | KS.1 |
| FIM1_4617 | 3738.760989 | 3736.327214 | 3570.518226 | -0.000265024 | 0.103085215 | 0.996476352 | 0.998284849 | ns | TM | KS.1 |
| FIM1_4618 | 4690.977007 | 4481.18557 | 4057.374277 | -0.030105025 | 0.080424397 | 0.568620461 | 0.831053761 | ns | TM | KS.1 |
| FIM1_4619 | 957.0602297 | 975.2207892 | 857.5893191 | 0.012961032 | 0.072051281 | 0.791534319 | 0.933303839 | ns | TM | KS.1 |
| FIM1_4620 | 693.8984535 | 713.1694492 | 744.6249619 | 0.009624527 | 0.128238923 | 0.878889925 | 0.965013767 | ns | TM | KS.1 |
| FIM1_4621 | 79.50869852 | 134.6597177 | 125.0342392 | 0.127207953 | 0.256621522 | 0.119392068 | 0.379294726 | ns | TM | KS.1 |
| FIM1_4622 | 1108.869998 | 971.1516468 | 1327.016102 | -0.096396592 | 0.104947511 | 0.136668793 | 0.409101833 | ns | TM | KS.1 |
| FIM1_4623 | 1292.258761 | 1555.476362 | 1412.356005 | 0.161562599 | 0.114065515 | 0.028195275 | 0.16659322 | ns | TM | KS.1 |
| FIM1_4624 | 3112.569805 | 2849.976966 | 2292.836892 | -0.059216672 | 0.092484042 | 0.309533753 | 0.637916496 | ns | TM | KS.1 |
| FIM1_4625 | 887.3141671 | 906.7066395 | 926.2415632 | 0.015196493 | 0.063153973 | 0.733917102 | 0.911352378 | ns | TM | KS.1 |
| FIM1_4626 | 456.0898903 | 331.8796255 | 425.5798025 | -0.179629242 | 0.181143446 | 0.040412709 | 0.20473852 | ns | TM | KS.1 |
| FIM1_4627 | 2926.00263 | 2530.551369 | 2357.628351 | -0.083089235 | 0.119147823 | 0.217860711 | 0.529594372 | ns | TM | KS.1 |
| FIM1_4628 | 474.6058549 | 323.4009686 | 297.3288643 | -0.29169748 | 0.20674271 | 0.006288765 | 0.061337713 | ns | TM | KS.1 |
| FIM1_4629 | 831.5692715 | 736.3180697 | 744.8255768 | -0.061608126 | 0.118181148 | 0.346334574 | 0.671956953 | ns | TM | KS.1 |
| FIM1_4630 | 1274.327845 | 1170.68654 | 1317.615698 | -0.060642403 | 0.08640863 | 0.278280419 | 0.602194469 | ns | TM | KS.1 |
| FIM1_4631 | 2493.734664 | 2298.053163 | 2080.043648 | -0.046956278 | 0.100843363 | 0.438135572 | 0.747832613 | ns | TM | KS.1 |
| FIM1_4632 | 1907.075774 | 1906.899872 | 2347.234258 | -9.37E-05 | 0.119607665 | 0.99880566 | 0.999198196 | ns | TM | KS.1 |
| FIM1_4633 | 88.89081124 | 58.10601108 | 76.16514234 | -0.184108409 | 0.234973139 | 0.047393437 | 0.226713798 | ns | TM | KS.1 |
| FIM1_4634 | 2981.7848 | 3448.577052 | 2905.267577 | 0.040148047 | 0.153281285 | 0.535947402 | 0.812908026 | ns | TM | KS.1 |
| FIM1_4635 | 1226.538582 | 1140.692755 | 1175.173052 | -0.04609165 | 0.092588883 | 0.425678332 | 0.741071977 | ns | TM | KS.1 |
| FIM1_4636 | 624.496885 | 512.9550166 | 580.978079 | -0.208778118 | 0.101971804 | 0.004884521 | 0.051158935 | ns | TM | KS.1 |
| FIM1_4637 | 1105.349606 | 978.9420742 | 1091.158215 | -0.072183051 | 0.11077346 | 0.264370278 | 0.590354459 | ns | TM | KS.1 |

| | | | | | | | | | | |
|---|---|---|---|---|---|---|---|---|---|---|
| FIM1_4638 | 1315.646948 | 1187.077867 | 1228.748941 | -0.082388706 | 0.087271302 | 0.150078202 | 0.431333943 | ns | TM | KS.1 |
| FIM1_4639 | 2066.629235 | 2018.013748 | 2308.383022 | -0.014803605 | 0.082212956 | 0.78179771 | 0.929216508 | ns | TM | KS.1 |
| FIM1_4640 | 892.7319825 | 925.1682716 | 889.2184359 | 0.018087529 | 0.103586003 | 0.764287956 | 0.924230468 | ns | TM | KS.1 |
| FIM1_4641 | 6500.75611 | 6143.132169 | 6170.986705 | -0.038472146 | 0.080848031 | 0.468570692 | 0.771134367 | ns | TM | KS.1 |
| FIM1_4642 | 3133.46725 | 3305.175866 | 3031.349814 | 0.02932919 | 0.097789359 | 0.619551201 | 0.861449756 | ns | TM | KS.1 |
| FIM1_4643 | 521.2203685 | 519.4416786 | 499.9010469 | -0.001800724 | 0.096403523 | 0.9753559 | 0.993872484 | ns | TM | KS.1 |
| FIM1_4644 | 927.9380605 | 915.8460981 | 877.6145571 | -0.008199361 | 0.081340803 | 0.87722424 | 0.964248917 | ns | TM | KS.1 |
| FIM1_4645 | 1010.810595 | 1200.535626 | 1148.317039 | 0.119496627 | 0.120457029 | 0.093139207 | 0.329798971 | ns | TM | KS.1 |
| FIM1_4646 | 1644.47687 | 1660.02515 | 1583.751112 | 0.00627141 | 0.073165441 | 0.899282146 | 0.968452273 | ns | TM | KS.1 |
| FIM1_4647 | 2072.520052 | 2065.860698 | 1914.355097 | -0.001692858 | 0.085942471 | 0.97537709 | 0.993872484 | ns | TM | KS.1 |
| FIM1_4648 | 1710.695256 | 1689.201429 | 1698.51504 | -0.007370265 | 0.084909942 | 0.892369593 | 0.967822084 | ns | TM | KS.1 |
| FIM1_4649 | 2352.932115 | 2385.033353 | 2547.851914 | 0.00885539 | 0.073246037 | 0.858262393 | 0.956277476 | ns | TM | KS.1 |
| FIM1_4650 | 1427.93446 | 1572.684347 | 1549.062917 | 0.065610136 | 0.095513552 | 0.27137682 | 0.598713828 | ns | TM | KS.1 |
| FIM1_4651 | 3576.834717 | 2986.036078 | 4372.04566 | -0.063510893 | 0.147461792 | 0.349580935 | 0.675403942 | ns | TM | KS.1 |
| FIM1_4652 | 1535.002472 | 1587.576491 | 1412.47613 | 0.022644108 | 0.075853185 | 0.655620044 | 0.877039451 | ns | TM | KS.1 |
| FIM1_4653 | 827.2476477 | 1091.822593 | 1046.675546 | 0.156374396 | 0.163965594 | 0.0579386 | 0.253530772 | ns | TM | KS.1 |
| FIM1_4654 | 1388.318918 | 977.2833664 | 1079.974788 | -0.285381357 | 0.183564696 | 0.005181461 | 0.053400721 | ns | TM | KS.1 |
| FIM1_4655 | 1736.111847 | 1552.10564 | 1886.468117 | -0.041517759 | 0.132828488 | 0.523842073 | 0.808348112 | ns | TM | KS.1 |
| FIM1_4656 | 2225.311444 | 2534.703942 | 2415.83052 | 0.040874101 | 0.144189532 | 0.530464206 | 0.808531286 | ns | TM | KS.1 |
| FIM1_4657 | 969.032231 | 897.1964549 | 980.8930235 | -0.049334245 | 0.09372661 | 0.397962362 | 0.721551361 | ns | TM | KS.1 |
| FIM1_4658 | 18248.77016 | 20952.13942 | 20752.37853 | 0.074062913 | 0.11979674 | 0.267198708 | 0.593708608 | ns | TM | KS.1 |
| FIM1_4660 | 85.81790172 | 49.37218217 | 183.5974283 | -0.380611907 | 0.314148228 | 0.005112323 | 0.053097716 | ns | TM | KS.1 |
| FIM1_4661 | 191.1983681 | 178.3089719 | 320.9228849 | -0.024084833 | 0.131466314 | 0.705849396 | 0.899257553 | ns | TM | KS.1 |
| FIM1_4662 | 1842.843804 | 2624.279851 | 3115.470066 | 0.124109733 | 0.19816824 | 0.117770683 | 0.376064922 | ns | TM | KS.1 |
| FIM1_4664 | 60.69346935 | 41.39550824 | 39.5335887 | -0.163617587 | 0.211133644 | 0.061240053 | 0.260828192 | ns | TM | KS.1 |
| FIM1_4665 | 1995.331345 | 2873.238859 | 3582.114891 | 0.154241497 | 0.202922191 | 0.070186565 | 0.280239294 | ns | TM | KS.1 |
| FIM1_4666 | 953.4526635 | 1404.042142 | 786.747206 | 0.150303325 | 0.212010875 | 0.07697598 | 0.294813117 | ns | TM | KS.1 |
| FIM1_4667 | 2165.340925 | 4021.880108 | 5171.262894 | 0.339830865 | 0.352729298 | 0.010604768 | 0.088198944 | ns | TM | KS.1 |
| FIM1_4668 | 409.2730023 | 328.0867718 | 590.2764347 | -0.079665933 | 0.156135737 | 0.257540499 | 0.586390838 | ns | TM | KS.1 |
| FIM1_4669 | 30.90169842 | 34.18303115 | 39.49713505 | 0.022768204 | 0.163616736 | 0.716283688 | 0.905084971 | ns | TM | KS.1 |
| FIM1_4670 | 2384.601222 | 2357.329487 | 2421.800564 | -0.005047297 | 0.112888163 | 0.934957102 | 0.981103476 | ns | TM | KS.1 |
| FIM1_4671 | 1297.545658 | 1100.623935 | 1252.57698 | -0.125608272 | 0.114065147 | 0.073029859 | 0.287623378 | ns | TM | KS.1 |
| FIM1_4672 | 1182.628234 | 938.5852458 | 1166.076171 | -0.224669 | 0.118319727 | 0.005655868 | 0.056844331 | ns | TM | KS.1 |
| FIM1_4673 | 322.6595059 | 363.037579 | 398.5748486 | 0.072537586 | 0.106855852 | 0.254532585 | 0.581671847 | ns | TM | KS.1 |
| FIM1_4674 | 244.608724 | 225.6287395 | 279.8664622 | -0.048646691 | 0.092854206 | 0.401892997 | 0.724158796 | ns | TM | KS.1 |
| FIM1_4675 | 3463.724382 | 4031.669667 | 4876.118353 | 0.121447163 | 0.107423221 | 0.072628956 | 0.286420314 | ns | TM | KS.1 |
| FIM1_4676 | 1725.732986 | 1998.725146 | 2206.597425 | 0.164654163 | 0.079129604 | 0.006296004 | 0.061337713 | ns | TM | KS.1 |
| FIM1_4678 | 1462.932061 | 1929.044595 | 2206.929083 | 0.07857219 | 0.177442801 | 0.264104024 | 0.590340989 | ns | TM | KS.1 |
| FIM1_4679 | 842.2197318 | 1177.750934 | 1706.025796 | 0.147016016 | 0.189995317 | 0.076836492 | 0.294813117 | ns | TM | KS.1 |
| FIM1_4680 | 528.5536186 | 588.8690339 | 650.5430871 | 0.076201715 | 0.094449653 | 0.202477105 | 0.510554281 | ns | TM | KS.1 |
| FIM1_4681 | 2379.582977 | 1713.288423 | 3003.550954 | -0.264024352 | 0.172591246 | 0.006846368 | 0.065125582 | ns | TM | KS.1 |
| FIM1_4682 | 6782.349095 | 6069.189222 | 6334.836483 | -0.108400919 | 0.078741793 | 0.049138342 | 0.231028743 | ns | TM | KS.1 |
| FIM1_4683 | 193.3231168 | 218.6704481 | 149.174951 | 0.034701846 | 0.149753378 | 0.589932049 | 0.844166125 | ns | TM | KS.1 |
| FIM1_4684 | 1376.77837 | 1558.868392 | 1653.177765 | 0.063443591 | 0.118495827 | 0.333643727 | 0.660779276 | ns | TM | KS.1 |
| FIM1_4685 | 3285.383195 | 1331.724551 | 1895.683557 | -0.832498085 | 0.47959864 | 0.000546663 | 0.009678468 | ns | TM | KS.1 |

| | | | | | | | | | |
|---|---|---|---|---|---|---|---|---|---|
| FIM1_4686 | 229.8495152 | 81.52424408 | 374.4480036 | -1.115056471 | 0.436346774 | 7.72446807252 | 0.00194087 | Km<KS.1 | TM | KS.1 |
| FIM1_4688 | 904.8550991 | 1047.953858 | 1030.77713 | 0.077087937 | 0.12273373 | 0.253654634 | 0.580732538 | ns | TM | KS.1 |
| FIM1_4689 | 1162.107155 | 1137.163783 | 1260.925112 | -0.013894828 | 0.079098058 | 0.789944644 | 0.933303839 | ns | TM | KS.1 |
| FIM1_4690 | 2801.061917 | 3020.776711 | 3273.689111 | 0.044272654 | 0.097901496 | 0.457373662 | 0.761179252 | ns | TM | KS.1 |
| FIM1_4691 | 587.2251655 | 968.6340928 | 1392.037738 | 0.231564801 | 0.274001978 | 0.027291062 | 0.162797401 | ns | TM | KS.1 |
| FIM1_4692 | 1554.489101 | 1091.155208 | 1443.260619 | -0.283342809 | 0.185649455 | 0.005569179 | 0.056429059 | ns | TM | KS.1 |
| FIM1_4693 | 306.1936495 | 269.6548485 | 358.3793039 | -0.057582558 | 0.126692078 | 0.38445505 | 0.708394027 | ns | TM | KS.1 |
| FIM1_4694 | 976.6054768 | 772.7959557 | 868.552117 | -0.080362693 | 0.160811986 | 0.254666503 | 0.581710677 | ns | TM | KS.1 |
| FIM1_4695 | 608.2072316 | 628.0420105 | 611.148356 | 0.010018457 | 0.134360202 | 0.874088542 | 0.962759265 | ns | TM | KS.1 |
| FIM1_4697 | 675.285667 | 1104.46554 | 1508.15025 | 0.140899622 | 0.249538215 | 0.096208211 | 0.336120679 | ns | TM | KS.1 |
| FIM1_4698 | 1482.731014 | 1269.350464 | 1300.486597 | -0.096637351 | 0.119174739 | 0.159360275 | 0.44660975 | ns | TM | KS.1 |
| FIM1_4699 | 710.1188002 | 824.0614092 | 876.9958142 | 0.046601251 | 0.146886062 | 0.478537651 | 0.779543161 | ns | TM | KS.1 |
| FIM1_4700 | 39.06589019 | 27.68910031 | 31.51954424 | -0.105125977 | 0.193140427 | 0.164030993 | 0.45366779 | ns | TM | KS.1 |
| FIM1_4701 | 1824.283248 | 1228.285503 | 949.344502 | -0.245423541 | 0.219283569 | 0.015999278 | 0.116881654 | ns | TM | KS.1 |
| FIM1_4702 | 2460.014112 | 2292.25818 | 2349.804366 | -0.019249299 | 0.14673594 | 0.760879255 | 0.923262022 | ns | TM | KS.1 |
| FIM1_4703 | 9458.851456 | 9911.737762 | 8353.968002 | 0.01515862 | 0.134347497 | 0.811008753 | 0.939848252 | ns | TM | KS.1 |
| FIM1_4704 | 1469.288054 | 1412.745001 | 1487.561147 | -0.01284872 | 0.13387187 | 0.839173839 | 0.949982862 | ns | TM | KS.1 |
| FIM1_4705 | 1104.168179 | 1226.880805 | 1309.422391 | 0.064956831 | 0.103135668 | 0.295741165 | 0.623274449 | ns | TM | KS.1 |
| FIM1_4706 | 1395.328325 | 1271.642161 | 1323.038148 | -0.057242481 | 0.09911433 | 0.343960887 | 0.670129201 | ns | TM | KS.1 |
| FIM1_4707 | 692.5159278 | 713.1933948 | 651.8354446 | 0.011379072 | 0.120931708 | 0.856276571 | 0.955916258 | ns | TM | KS.1 |
| FIM1_4708 | 1003.318868 | 1135.55387 | 1232.431846 | 0.033353904 | 0.151984268 | 0.603245677 | 0.851547247 | ns | TM | KS.1 |
| FIM1_4709 | 1531.970182 | 1414.710036 | 1693.347632 | -0.027739521 | 0.132669046 | 0.664979835 | 0.880679852 | ns | TM | KS.1 |
| FIM1_4710 | 1285.483551 | 1361.275018 | 1338.096276 | 0.020181461 | 0.129625621 | 0.750883739 | 0.918418758 | ns | TM | KS.1 |
| FIM1_4711 | 2167.405627 | 2184.897193 | 2530.178698 | 0.003074535 | 0.12142468 | 0.960917212 | 0.990995674 | ns | TM | KS.1 |
| FIM1_4712 | 469.871694 | 510.1540307 | 488.8438043 | 0.032878552 | 0.123224912 | 0.607106867 | 0.851764429 | ns | TM | KS.1 |
| FIM1_4713 | 2374.442721 | 2249.306068 | 2362.926956 | -0.026255609 | 0.107938667 | 0.669374349 | 0.882180579 | ns | TM | KS.1 |
| FIM1_4714 | 1741.192918 | 1419.898244 | 1656.996978 | -0.101123575 | 0.141555688 | 0.163694561 | 0.45366779 | ns | TM | KS.1 |
| FIM1_4715 | 4384.582947 | 4115.035009 | 4836.093207 | -0.026607905 | 0.118997276 | 0.673731313 | 0.885317825 | ns | TM | KS.1 |
| FIM1_4716 | 2261.184367 | 2982.209454 | 3401.173216 | 0.066073303 | 0.183370905 | 0.331766875 | 0.658898284 | ns | TM | KS.1 |
| FIM1_4717 | 3734.507485 | 4975.436451 | 4812.049546 | 0.100027449 | 0.175285955 | 0.177383642 | 0.475989006 | ns | TM | KS.1 |
| FIM1_4718 | 1101.455694 | 1718.99176 | 1948.392689 | 0.119152724 | 0.230431936 | 0.133160928 | 0.404441767 | ns | TM | KS.1 |
| FIM1_4719 | 258.3941893 | 360.1345131 | 359.3621064 | 0.169479183 | 0.188721079 | 0.050762253 | 0.234052095 | ns | TM | KS.1 |
| FIM1_4720 | 1231.486091 | 2290.928172 | 2687.385742 | 0.463840221 | 0.362690169 | 0.003147107 | 0.036399552 | ns | TM | KS.1 |
| FIM1_4721 | 1489.13797 | 1537.372732 | 1004.264901 | 0.019345402 | 0.085523365 | 0.723931451 | 0.907193695 | ns | TM | KS.1 |
| FIM1_4722 | 839.1977962 | 1630.539213 | 1807.401918 | 0.199324951 | 0.323768667 | 0.047869742 | 0.227678742 | ns | TM | KS.1 |
| FIM1_4724 | 563.5822866 | 422.7242695 | 584.5339102 | -0.216712887 | 0.158204929 | 0.015535241 | 0.113658568 | ns | TM | KS.1 |
| FIM1_4725 | 223.0495711 | 188.588228 | 211.8274937 | -0.012298801 | 0.246166001 | 0.823549285 | 0.94482203 | ns | TM | KS.1 |
| FIM1_4726 | 67.49517923 | 59.13225086 | 61.86479955 | -0.012393403 | 0.228888489 | 0.829768185 | 0.946594066 | ns | TM | KS.1 |
| FIM1_4727 | 486.3084543 | 371.2580123 | 420.3880307 | -0.017833549 | 0.255111181 | 0.741535602 | 0.914738314 | ns | TM | KS.1 |
| FIM1_4728 | 151.1666711 | 114.2563997 | 140.8377942 | -0.021046968 | 0.248117029 | 0.706285201 | 0.899352156 | ns | TM | KS.1 |
| FIM1_4729 | 871.8563154 | 737.6460535 | 766.2970696 | -0.011225913 | 0.251552099 | 0.835486278 | 0.948719561 | ns | TM | KS.1 |
| FIM1_4730 | 456.9917127 | 356.931587 | 481.3915826 | -0.085469768 | 0.163685455 | 0.231429778 | 0.549231011 | ns | TM | KS.1 |
| FIM1_4731 | 1008.480374 | 1267.93528 | 1595.830275 | 0.084101848 | 0.15731562 | 0.236674286 | 0.556453012 | ns | TM | KS.1 |
| FIM1_4732 | 3855.964545 | 3641.888766 | 3415.38404 | -0.018383603 | 0.13549463 | 0.772286397 | 0.925733395 | ns | TM | KS.1 |
| FIM1_4733 | 1347.927962 | 1116.34119 | 1350.604209 | -0.100004858 | 0.134982841 | 0.16332235 | 0.45366779 | ns | TM | KS.1 |

| | | | | | | | | | | |
|---|---|---|---|---|---|---|---|---|---|---|
| FIM1_4734 | 362.3394155 | 273.0631242 | 494.3151773 | -0.142268424 | 0.168559792 | 0.07826886 | 0.298177127 | ns | TM | KS.1 |
| FIM1_4736 | 182.1275152 | 191.9236571 | 252.8171641 | 0.016921791 | 0.133845862 | 0.789760053 | 0.933303839 | ns | TM | KS.1 |
| FIM1_4737 | 542.6362359 | 455.1335408 | 807.1249009 | -0.073734734 | 0.139706455 | 0.284968891 | 0.61164225 | ns | TM | KS.1 |
| FIM1_4738 | 96.01523093 | 144.0519739 | 122.0593403 | 0.161017255 | 0.222520456 | 0.066165003 | 0.270807979 | ns | TM | KS.1 |
| FIM1_4739 | 343.6925459 | 395.3829853 | 449.3723635 | 0.077884599 | 0.118284706 | 0.243631948 | 0.567141251 | ns | TM | KS.1 |
| FIM1_4740 | 1300.072851 | 1457.171042 | 1494.907235 | 0.060249283 | 0.114126957 | 0.350764219 | 0.676561755 | ns | TM | KS.1 |
| FIM1_4741 | 575.8310427 | 559.588698 | 562.5953815 | -0.012438563 | 0.114170544 | 0.841329185 | 0.95052049 | ns | TM | KS.1 |
| FIM1_4742 | 232.4329477 | 229.5452527 | 286.3233181 | -0.004020631 | 0.139369681 | 0.949113576 | 0.986388143 | ns | TM | KS.1 |
| FIM1_4743 | 13.47202448 | 16.72040159 | 29.18473673 | 0.040369277 | 0.190402251 | 0.525434686 | 0.808531286 | ns | TM | KS.1 |
| FIM1_4744 | 2763.447845 | 2829.349096 | 2880.341186 | 0.010444407 | 0.111669652 | 0.865649671 | 0.959975149 | ns | TM | KS.1 |
| FIM1_4745 | 1334.058209 | 1035.218595 | 978.5767618 | -0.228743957 | 0.132690635 | 0.007509166 | 0.068926381 | ns | TM | KS.1 |
| FIM1_4746 | 2603.646294 | 2222.689386 | 2526.440432 | -0.061624268 | 0.138213625 | 0.360694119 | 0.683750066 | ns | TM | KS.1 |
| FIM1_4747 | 688.6356587 | 691.9660938 | 669.4574419 | 0.004061055 | 0.082629519 | 0.939584961 | 0.983546982 | ns | TM | KS.1 |
| FIM1_4748 | 4902.914386 | 4697.94249 | 4156.527977 | -0.030369725 | 0.072342951 | 0.536939315 | 0.81337601 | ns | TM | KS.1 |
| FIM1_4749 | 473.358101 | 420.4136312 | 434.7145254 | -0.083226714 | 0.10129052 | 0.183945232 | 0.486252673 | ns | TM | KS.1 |
| FIM1_4750 | 3970.101897 | 4204.686519 | 4683.332238 | 0.048207547 | 0.064004728 | 0.286160664 | 0.612585758 | ns | TM | KS.1 |
| FIM1_4751 | 199.2744733 | 176.2853322 | 204.7642249 | -0.054116496 | 0.126570245 | 0.411378716 | 0.731735607 | ns | TM | KS.1 |
| FIM1_4753 | 1258.374582 | 1877.119646 | 2190.71695 | 0.221014552 | 0.222037951 | 0.024533943 | 0.154111571 | ns | TM | KS.1 |
| FIM1_4754 | 1650.298474 | 2076.673934 | 1932.410155 | 0.218391481 | 0.120187325 | 0.007122378 | 0.066581689 | ns | TM | KS.1 |
| FIM1_4755 | 469.2112929 | 489.3827703 | 420.6909983 | 0.021268355 | 0.102588289 | 0.723674809 | 0.907181536 | ns | TM | KS.1 |
| FIM1_4756 | 1184.065493 | 1598.923887 | 1824.271366 | 0.251200003 | 0.157322998 | 0.007115638 | 0.066581689 | ns | TM | KS.1 |
| FIM1_4757 | 1300.621326 | 1856.06079 | 1744.657445 | 0.155391307 | 0.199214108 | 0.068020287 | 0.275570788 | ns | TM | KS.1 |
| FIM1_4758 | 690.0342928 | 504.9752219 | 689.9815069 | -0.337105464 | 0.1280579 | 0.000273437 | 0.005621273 | ns | TM | KS.1 |
| FIM1_4759 | 5887.022723 | 5139.527196 | 6178.264648 | -0.07602527 | 0.117263154 | 0.252477282 | 0.579042268 | ns | TM | KS.1 |
| FIM1_4760 | 4077.636854 | 3643.815209 | 4423.256503 | -0.03345614 | 0.145067364 | 0.603456845 | 0.851547247 | ns | TM | KS.1 |
| FIM1_4761 | 380.7957826 | 357.5909355 | 374.2602876 | -0.029813865 | 0.108955228 | 0.629621058 | 0.868412743 | ns | TM | KS.1 |
| FIM1_4762 | 112.0073605 | 107.8118977 | 127.7586392 | -0.018593393 | 0.114951453 | 0.765718033 | 0.924600573 | ns | TM | KS.1 |
| FIM1_4763 | 154.1230869 | 138.8009801 | 172.8569932 | -0.037760865 | 0.134360145 | 0.560359305 | 0.826391733 | ns | TM | KS.1 |
| FIM1_4764 | 345.5736124 | 309.6380251 | 328.2736609 | -0.036124998 | 0.139087275 | 0.576853845 | 0.834986291 | ns | TM | KS.1 |
| FIM1_4765 | 372.1482414 | 295.7467875 | 314.2749395 | -0.089554854 | 0.155896114 | 0.21273734 | 0.520662191 | ns | TM | KS.1 |
| FIM1_4766 | 164.6990617 | 177.8501632 | 195.8864339 | 0.031483474 | 0.122066908 | 0.621441747 | 0.862872646 | ns | TM | KS.1 |
| FIM1_4767 | 712.7161237 | 683.0611122 | 816.4770033 | -0.022435677 | 0.098933914 | 0.70486863 | 0.898928847 | ns | TM | KS.1 |
| FIM1_4769 | 59.92134488 | 82.97334182 | 118.1575199 | 0.093754243 | 0.193623148 | 0.201121233 | 0.509459336 | ns | TM | KS.1 |
| FIM1_4770 | 242.2749396 | 491.1132499 | 529.218885 | 0.280601172 | 0.37158543 | 0.02131992 | 0.14253455 | ns | TM | KS.1 |
| FIM1_4771 | 379.1226044 | 584.7756215 | 746.2717156 | 0.345546189 | 0.236412989 | 0.003633081 | 0.040617025 | ns | TM | KS.1 |
| FIM1_4772 | 46.23457903 | 47.18950512 | 75.1073442 | 0.006622093 | 0.133374522 | 0.916488035 | 0.975634767 | ns | TM | KS.1 |
| FIM1_4773 | 4071.828199 | 4205.70312 | 2745.278261 | 0.01612783 | 0.103039395 | 0.788616559 | 0.933067989 | ns | TM | KS.1 |
| FIM1_3 | 3075.343344 | 2973.550777 | 3037.529789 | -0.007984429 | 0.05702419 | 0.725908244 | 0.987157164 | ns | TM | KS.3 |
| FIM1_5 | 634.292505 | 625.0760985 | 633.8041188 | -0.001371295 | 0.079808378 | 0.946386464 | 0.994413583 | ns | TM | KS.3 |
| FIM1_6 | 444.7160679 | 405.6170494 | 405.4975614 | -0.014026889 | 0.073577572 | 0.541998389 | 0.95917717 | ns | TM | KS.3 |
| FIM1_7 | 20.23579022 | 39.42728799 | 60.8195808 | 0.185997302 | 0.316206845 | 0.00597202 | 0.16059892 | ns | TM | KS.3 |
| FIM1_8 | 288.162505 | 337.7417933 | 375.3172182 | 0.01958669 | 0.083000294 | 0.389363089 | 0.948637087 | ns | TM | KS.3 |
| FIM1_9 | 1124.181287 | 1078.663684 | 2901.971283 | -0.003132422 | 0.085030997 | 0.87219706 | 0.991141351 | ns | TM | KS.3 |
| FIM1_10 | 526.332748 | 953.0646011 | 2943.370395 | 0.141760399 | 0.266182134 | 0.009965199 | 0.221325297 | ns | TM | KS.3 |
| FIM1_11 | 253.3765851 | 212.377455 | 340.6940978 | -0.035428429 | 0.084178756 | 0.189329742 | 0.828217874 | ns | TM | KS.3 |

| | | | | | | | | | | |
|---|---|---|---|---|---|---|---|---|---|---|
| FIM1_12 | 357.8671807 | 571.9435466 | 1729.337393 | 0.104094493 | 0.195444748 | 0.017174322 | 0.302986707 | ns | TM | KS.3 |
| FIM1_13 | 37380.62821 | 34960.31201 | 35774.55391 | -0.003762773 | 0.09320922 | 0.837255566 | 0.991141351 | ns | TM | KS.3 |
| FIM1_14 | 10.08811561 | 14.37730315 | 16.92066841 | 0.018934069 | 0.114488281 | 0.337371798 | 0.940821016 | ns | TM | KS.3 |
| FIM1_15 | 15080.4237 | 15788.12038 | 14849.47076 | 0.001901164 | 0.102250374 | 0.912538127 | 0.991141351 | ns | TM | KS.3 |
| FIM1_16 | 15064.59561 | 14366.7213 | 13864.7932 | -0.004930831 | 0.078256937 | 0.81354069 | 0.988916276 | ns | TM | KS.3 |
| FIM1_17 | 21654.94777 | 21703.81208 | 21161.65642 | 0.000105396 | 0.098029887 | 0.995234423 | 0.999525119 | ns | TM | KS.3 |
| FIM1_18 | 1720.459557 | 1928.59211 | 2232.947735 | 0.012492007 | 0.08058039 | 0.562050473 | 0.961073589 | ns | TM | KS.3 |
| FIM1_19 | 1102.665297 | 1062.324111 | 965.1753006 | -0.011416503 | 0.048431065 | 0.589176892 | 0.963781856 | ns | TM | KS.3 |
| FIM1_20 | 4218.577684 | 3740.908132 | 3333.389345 | -0.060975649 | 0.076472803 | 0.066804006 | 0.566847784 | ns | TM | KS.3 |
| FIM1_21 | 640.3068026 | 612.0671481 | 585.7584931 | -0.010792585 | 0.059099418 | 0.641344368 | 0.975935248 | ns | TM | KS.3 |
| FIM1_22 | 1563.273417 | 1416.33382 | 1520.407378 | -0.058362008 | 0.069209675 | 0.064810389 | 0.561727671 | ns | TM | KS.3 |
| FIM1_23 | 240.1724688 | 210.5399599 | 230.7427362 | -0.025214191 | 0.076352153 | 0.315017445 | 0.927564309 | ns | TM | KS.3 |
| FIM1_24 | 935.1711412 | 981.6143765 | 1199.112786 | 0.0050221 | 0.078314449 | 0.810122431 | 0.988916276 | ns | TM | KS.3 |
| FIM1_25 | 478.5661284 | 497.8944151 | 454.9892691 | 0.004725527 | 0.074829275 | 0.826509116 | 0.991141351 | ns | TM | KS.3 |
| FIM1_26 | 314.6152736 | 284.9720257 | 284.9825584 | -0.014219131 | 0.07536617 | 0.53227265 | 0.959071703 | ns | TM | KS.3 |
| FIM1_27 | 683.867792 | 702.1323353 | 657.296323 | 0.005091437 | 0.062862568 | 0.824241777 | 0.991141351 | ns | TM | KS.3 |
| FIM1_28 | 12797.60721 | 12848.41434 | 11506.37819 | 0.00067062 | 0.066226069 | 0.976361694 | 0.998938406 | ns | TM | KS.3 |
| FIM1_29 | 3434.525736 | 3252.353971 | 2774.21492 | -0.009170635 | 0.068256172 | 0.689526194 | 0.980879562 | ns | TM | KS.3 |
| FIM1_30 | 2262.145374 | 2109.040388 | 2049.9388 | -0.018563351 | 0.060689766 | 0.440943814 | 0.950410534 | ns | TM | KS.3 |
| FIM1_31 | 3968.763651 | 4136.74939 | 4258.824352 | 0.013603917 | 0.046885828 | 0.510301544 | 0.953260948 | ns | TM | KS.3 |
| FIM1_32 | 3471.471457 | 3151.366704 | 2794.011073 | -0.026685397 | 0.066673516 | 0.300297005 | 0.926536521 | ns | TM | KS.3 |
| FIM1_33 | 736.7938059 | 803.145219 | 732.2157693 | 0.014666501 | 0.070803482 | 0.531953502 | 0.959071703 | ns | TM | KS.3 |
| FIM1_34 | 1629.559086 | 1566.308475 | 1643.975651 | -0.013395038 | 0.043614674 | 0.491477489 | 0.952591289 | ns | TM | KS.3 |
| FIM1_35 | 40.37412014 | 36.01776224 | 43.59138275 | -0.010952759 | 0.086626212 | 0.588401967 | 0.963781856 | ns | TM | KS.3 |
| FIM1_36 | 7087.992105 | 8764.592362 | 6124.361802 | 0.011960802 | 0.099929902 | 0.527141305 | 0.959071703 | ns | TM | KS.3 |
| FIM1_37 | 628.4519901 | 598.713297 | 700.1965119 | -0.010401767 | 0.061860491 | 0.655271953 | 0.976323032 | ns | TM | KS.3 |
| FIM1_38 | 981.3086511 | 1005.565519 | 1220.934355 | 0.004228818 | 0.065482069 | 0.852776889 | 0.991141351 | ns | TM | KS.3 |
| FIM1_39 | 1286.711828 | 1504.707474 | 1622.369305 | 0.021833973 | 0.081600266 | 0.353615487 | 0.948637087 | ns | TM | KS.3 |
| FIM1_40 | 2557.896741 | 2376.341907 | 1421.516646 | -0.019807557 | 0.061176809 | 0.414810165 | 0.950410534 | ns | TM | KS.3 |
| FIM1_41 | 808.0760119 | 895.0522114 | 1004.945188 | 0.026720057 | 0.068062792 | 0.300895959 | 0.926536521 | ns | TM | KS.3 |
| FIM1_42 | 1098.898578 | 1059.286967 | 1088.647994 | -0.007425237 | 0.062953999 | 0.747546613 | 0.988477371 | ns | TM | KS.3 |
| FIM1_43 | 2382.936713 | 2115.22232 | 2347.041188 | -0.076755611 | 0.076702673 | 0.035053646 | 0.42652037 | ns | TM | KS.3 |
| FIM1_44 | 315.6768078 | 281.623735 | 223.7839843 | -0.01768344 | 0.07579227 | 0.449932482 | 0.950410534 | ns | TM | KS.3 |
| FIM1_45 | 1446.513565 | 1259.362577 | 909.3249672 | -0.040545526 | 0.077839725 | 0.159992744 | 0.791498358 | ns | TM | KS.3 |
| FIM1_46 | 3147.699777 | 3130.415645 | 3895.846385 | -0.001285565 | 0.056336303 | 0.954411234 | 0.994413583 | ns | TM | KS.3 |
| FIM1_47 | 572.6947379 | 1232.982256 | 1453.094101 | 1.080566648 | 0.116491481 | 5.2774580057[illegible] | 1.3127676789[illegible] | Km>KS.3 | TM | KS.3 |
| FIM1_48 | 3163.231445 | 3367.076708 | 2574.536415 | 0.026094356 | 0.051155916 | 0.242810961 | 0.887413624 | ns | TM | KS.3 |
| FIM1_49 | 7148.727426 | 8055.8092 | 9518.429149 | 0.011727395 | 0.08278929 | 0.576782524 | 0.963382753 | ns | TM | KS.3 |
| FIM1_50 | 4983.192693 | 4484.855769 | 4848.123799 | -0.014910032 | 0.075744187 | 0.513897445 | 0.954240701 | ns | TM | KS.3 |
| FIM1_51 | 3073.743788 | 3065.936114 | 3374.321267 | -0.000563038 | 0.06005153 | 0.980266097 | 0.999333464 | ns | TM | KS.3 |
| FIM1_52 | 3629.347798 | 3524.569887 | 3261.822567 | -0.006775581 | 0.057503614 | 0.76599345 | 0.988477371 | ns | TM | KS.3 |
| FIM1_53 | 32740.89154 | 34161.16121 | 32534.50092 | 0.011602633 | 0.053654152 | 0.605076828 | 0.970937835 | ns | TM | KS.3 |
| FIM1_54 | 3139.654435 | 3606.686675 | 3260.43686 | 0.020295937 | 0.078930779 | 0.388673346 | 0.948637087 | ns | TM | KS.3 |
| FIM1_55 | 1323.686427 | 1168.64038 | 884.141548 | -0.009864826 | 0.087191138 | 0.621018709 | 0.973122675 | ns | TM | KS.3 |
| FIM1_56 | 32.6300034 | 15.8974698 | 110.1015889 | -0.113000448 | 0.269852986 | 0.016500566 | 0.298510235 | ns | TM | KS.3 |

| | | | | | | | | | | |
|---|---|---|---|---|---|---|---|---|---|---|
| FIM1_57 | 430.7858148 | 361.9690452 | 411.2940193 | -0.018496186 | 0.086299472 | 0.400417662 | 0.950410534 | ns | TM | KS.3 |
| FIM1_58 | 3406.402205 | 3687.691403 | 3663.533176 | 0.013307716 | 0.070462619 | 0.567578616 | 0.962265923 | ns | TM | KS.3 |
| FIM1_59 | 4454.268853 | 4572.198914 | 4886.402825 | 0.002550415 | 0.079180882 | 0.901287669 | 0.991141351 | ns | TM | KS.3 |
| FIM1_60 | 1881.509203 | 2011.939452 | 2097.26308 | 0.018570501 | 0.058995946 | 0.436657417 | 0.950410534 | ns | TM | KS.3 |
| FIM1_61 | 1480.029297 | 1329.293114 | 1514.364944 | -0.046019875 | 0.070082045 | 0.119371285 | 0.730090427 | ns | TM | KS.3 |
| FIM1_62 | 1303.41996 | 1236.183497 | 1248.976291 | -0.010581813 | 0.064346726 | 0.650158416 | 0.976323032 | ns | TM | KS.3 |
| FIM1_63 | 70.44288511 | 84.13967784 | 61.0388668 | 0.015255206 | 0.089320715 | 0.463011699 | 0.950410534 | ns | TM | KS.3 |
| FIM1_64 | 2330.889481 | 2177.572136 | 2027.713438 | -0.023411036 | 0.055963904 | 0.323148566 | 0.93251979 | ns | TM | KS.3 |
| FIM1_65 | 202.1962716 | 198.9679854 | 253.6940693 | -0.002958591 | 0.058426232 | 0.896332745 | 0.991141351 | ns | TM | KS.3 |
| FIM1_66 | 4794.123136 | 4537.989004 | 4634.515481 | -0.011405146 | 0.06338956 | 0.626324738 | 0.97344368 | ns | TM | KS.3 |
| FIM1_67 | 324.5857749 | 294.702463 | 516.1440968 | -0.016502765 | 0.072209151 | 0.485082151 | 0.952269291 | ns | TM | KS.3 |
| FIM1_68 | 15611.35878 | 16931.07465 | 15105.22591 | 0.015480822 | 0.068371076 | 0.515568478 | 0.955289825 | ns | TM | KS.3 |
| FIM1_69 | 14237.6434 | 15149.17464 | 14277.90827 | 0.015734602 | 0.05980248 | 0.506498475 | 0.953260948 | ns | TM | KS.3 |
| FIM1_70 | 868.2529692 | 1035.713776 | 1316.552668 | 0.0340572 | 0.084122864 | 0.200488437 | 0.845272486 | ns | TM | KS.3 |
| FIM1_71 | 688.0151063 | 726.2915504 | 738.570201 | 0.011519713 | 0.062437406 | 0.622827063 | 0.973167286 | ns | TM | KS.3 |
| FIM1_72 | 516.0228595 | 572.8221206 | 603.4281888 | 0.021666871 | 0.070413103 | 0.382476595 | 0.948637087 | ns | TM | KS.3 |
| FIM1_73 | 13890.28853 | 12936.36281 | 13298.10691 | -0.024908918 | 0.056817354 | 0.299845039 | 0.926536521 | ns | TM | KS.3 |
| FIM1_74 | 701.8788825 | 785.7038402 | 775.8522912 | 0.047253232 | 0.072015586 | 0.115692802 | 0.717670439 | ns | TM | KS.3 |
| FIM1_75 | 23572.13609 | 21876.02587 | 22825.45264 | -0.017731864 | 0.063538086 | 0.463780511 | 0.950410534 | ns | TM | KS.3 |
| FIM1_76 | 215.6869924 | 273.9535694 | 347.4029841 | 0.062481537 | 0.103802112 | 0.059789345 | 0.54403004 | ns | TM | KS.3 |
| FIM1_77 | 586.9761281 | 624.17644 | 859.8821874 | 0.017155602 | 0.05629936 | 0.461502723 | 0.950410534 | ns | TM | KS.3 |
| FIM1_78 | 3272.037285 | 3097.778633 | 3222.916234 | -0.013801983 | 0.058405684 | 0.554394586 | 0.95917717 | ns | TM | KS.3 |
| FIM1_79 | 919.5053149 | 1046.791296 | 3178.933275 | 0.018996994 | 0.077824369 | 0.416887706 | 0.950410534 | ns | TM | KS.3 |
| FIM1_80 | 84.96784741 | 74.04577491 | 115.301577 | -0.019255164 | 0.079523434 | 0.406631252 | 0.950410534 | ns | TM | KS.3 |
| FIM1_81 | 1201.235586 | 1105.738059 | 1455.405965 | -0.017347404 | 0.067019145 | 0.472964486 | 0.950410534 | ns | TM | KS.3 |
| FIM1_82 | 1887.057237 | 1957.057294 | 2001.056325 | 0.011142149 | 0.048822703 | 0.600045199 | 0.969543639 | ns | TM | KS.3 |
| FIM1_83 | 451.8410311 | 402.9913881 | 385.1962856 | -0.025850107 | 0.072463288 | 0.312200748 | 0.927416591 | ns | TM | KS.3 |
| FIM1_84 | 1022.453207 | 894.5121432 | 813.4542519 | -0.0345105 | 0.075919342 | 0.208512069 | 0.852551729 | ns | TM | KS.3 |
| FIM1_85 | 1226.70559 | 1173.645841 | 812.4313926 | -0.013089974 | 0.051105044 | 0.550304999 | 0.95917717 | ns | TM | KS.3 |
| FIM1_86 | 1454.128695 | 1438.877322 | 890.901293 | -0.002906108 | 0.051180127 | 0.893369453 | 0.991141351 | ns | TM | KS.3 |
| FIM1_87 | 450.0002389 | 549.810017 | 533.8379842 | 0.037714793 | 0.08911879 | 0.164457456 | 0.800584912 | ns | TM | KS.3 |
| FIM1_88 | 854.9715863 | 780.8128172 | 812.0837853 | -0.016819962 | 0.070094002 | 0.481733945 | 0.951416554 | ns | TM | KS.3 |
| FIM1_89 | 158.6289239 | 443.6438489 | 613.585864 | 0.9736534 | 0.542587787 | 9.13140074222 | 0.00797357 | ns | TM | KS.3 |
| FIM1_90 | 2327.727 | 2638.766652 | 2907.03043 | 0.083308576 | 0.079157272 | 0.028515567 | 0.390786333 | ns | TM | KS.3 |
| FIM1_91 | 1113.951693 | 1070.871715 | 2110.39745 | -0.004291818 | 0.077155239 | 0.838631741 | 0.991141351 | ns | TM | KS.3 |
| FIM1_92 | 363.2540623 | 385.1960404 | 320.0306234 | 0.012904546 | 0.062674019 | 0.584089505 | 0.963781856 | ns | TM | KS.3 |
| FIM1_93 | 286.6979951 | 319.6621741 | 258.6469668 | 0.036359731 | 0.071062665 | 0.191432164 | 0.829253069 | ns | TM | KS.3 |
| FIM1_94 | 436.0126698 | 406.8557693 | 357.0574842 | -0.015287319 | 0.064546999 | 0.522333909 | 0.958602567 | ns | TM | KS.3 |
| FIM1_95 | 5097.322106 | 4727.480924 | 5406.626742 | -0.027726939 | 0.058041727 | 0.259223036 | 0.895871714 | ns | TM | KS.3 |
| FIM1_96 | 497.4222369 | 313.983054 | 692.0664714 | -0.098807545 | 0.188145678 | 0.019148882 | 0.32184354 | ns | TM | KS.3 |
| FIM1_97 | 2649.351705 | 2035.626225 | 3895.884372 | -0.026908678 | 0.098504083 | 0.242358861 | 0.887413624 | ns | TM | KS.3 |
| FIM1_98 | 239.1122174 | 195.1583209 | 194.9117998 | -0.027028507 | 0.088945204 | 0.260007464 | 0.896422129 | ns | TM | KS.3 |
| FIM1_99 | 588.1745276 | 515.7010415 | 463.1981537 | -0.025670079 | 0.075762567 | 0.309697654 | 0.927416591 | ns | TM | KS.3 |
| FIM1_100 | 3421.690523 | 3425.045914 | 3763.677311 | 0.000265648 | 0.055684019 | 0.990540214 | 0.999525119 | ns | TM | KS.3 |
| FIM1_101 | 846.0646044 | 804.7445107 | 1500.413589 | -0.008521766 | 0.067492963 | 0.710475814 | 0.984652342 | ns | TM | KS.3 |

| | | | | | | | | | | |
|---|---|---|---|---|---|---|---|---|---|---|
| FIM1_102 | 220.194668 | 205.4882559 | 221.6806448 | -0.012294789 | 0.068807866 | 0.597654605 | 0.9678814 | ns | TM | KS.3 |
| FIM1_103 | 1974.948407 | 1878.887383 | 1767.344721 | -0.013787574 | 0.054957617 | 0.545556315 | 0.95917717 | ns | TM | KS.3 |
| FIM1_104 | 749.4503606 | 675.9740681 | 640.3642589 | -0.058559566 | 0.070405196 | 0.066178493 | 0.564730709 | ns | TM | KS.3 |
| FIM1_105 | 3097.991074 | 3581.472871 | 3825.164874 | 0.028043486 | 0.07815085 | 0.273643808 | 0.908468501 | ns | TM | KS.3 |
| FIM1_107 | 1086.291438 | 1020.278166 | 944.1502495 | -0.014363906 | 0.062392674 | 0.544944647 | 0.95917717 | ns | TM | KS.3 |
| FIM1_108 | 1928.25874 | 2166.353715 | 2134.243348 | 0.025185825 | 0.072385421 | 0.322089701 | 0.930003636 | ns | TM | KS.3 |
| FIM1_109 | 1112.220711 | 4978.030273 | 3009.743906 | 2.050742766 | 0.17266621 | 3.0330915469 | 1.5089630445 | Km>KS.3 | TM | KS.3 |
| FIM1_110 | 459.2811584 | 866.7297866 | 762.545038 | 0.880436609 | 0.145745795 | 4.6455270103 | 1.7778074520 | ns | TM | KS.3 |
| FIM1_111 | 8724.127283 | 7857.43515 | 5944.23296 | -0.043057888 | 0.069289757 | 0.136333968 | 0.768410055 | ns | TM | KS.3 |
| FIM1_112 | 859.2503121 | 941.9513615 | 919.1620486 | 0.017314471 | 0.070040427 | 0.470671791 | 0.950410534 | ns | TM | KS.3 |
| FIM1_113 | 1048.255885 | 953.7478862 | 1147.542679 | -0.026378441 | 0.065848148 | 0.303919595 | 0.926536521 | ns | TM | KS.3 |
| FIM1_114 | 1005.543158 | 1013.181312 | 985.3718464 | 0.00175649 | 0.055214186 | 0.937372494 | 0.994413583 | ns | TM | KS.3 |
| FIM1_115 | 861.3972255 | 873.4886779 | 1055.82097 | 0.001644956 | 0.075269598 | 0.938491744 | 0.994413583 | ns | TM | KS.3 |
| FIM1_116 | 474.2039034 | 392.4098367 | 432.800489 | -0.046697837 | 0.088949774 | 0.116942689 | 0.722720344 | ns | TM | KS.3 |
| FIM1_117 | 11565.17835 | 11218.60407 | 11163.9285 | -0.004826396 | 0.068357343 | 0.830671781 | 0.991141351 | ns | TM | KS.3 |
| FIM1_118 | 275.4155095 | 185.5820915 | 186.7466403 | -0.100854824 | 0.168471215 | 0.017503687 | 0.304361209 | ns | TM | KS.3 |
| FIM1_119 | 6410.363433 | 7062.973654 | 7368.886416 | 0.015767513 | 0.072790685 | 0.500946415 | 0.953260948 | ns | TM | KS.3 |
| FIM1_120 | 1315.091665 | 1351.89305 | 1362.100667 | 0.00752588 | 0.051411821 | 0.730144378 | 0.987157164 | ns | TM | KS.3 |
| FIM1_121 | 2703.416512 | 3133.694408 | 3572.133845 | 0.015827938 | 0.083606729 | 0.46676654 | 0.950410534 | ns | TM | KS.3 |
| FIM1_122 | 3427.953726 | 3198.817483 | 3375.700208 | -0.023934908 | 0.056248397 | 0.314702496 | 0.927564309 | ns | TM | KS.3 |
| FIM1_123 | 5318.050956 | 5022.621797 | 4792.686076 | -0.015525185 | 0.057167975 | 0.505557482 | 0.953260948 | ns | TM | KS.3 |
| FIM1_124 | 572.0958576 | 771.5669356 | 724.0275505 | 0.219586597 | 0.146356116 | 0.000888516 | 0.044072215 | ns | TM | KS.3 |
| FIM1_125 | 253.4957428 | 342.9734285 | 653.6319812 | 0.095613697 | 0.133466057 | 0.019839963 | 0.324535175 | ns | TM | KS.3 |
| FIM1_126 | 1296.257113 | 1346.988423 | 1309.752804 | 0.011539953 | 0.047843876 | 0.582037534 | 0.963781856 | ns | TM | KS.3 |
| FIM1_127 | 867.1644809 | 729.969962 | 779.7362617 | -0.031049736 | 0.083178405 | 0.229763458 | 0.879350777 | ns | TM | KS.3 |
| FIM1_128 | 18308.47246 | 17340.85182 | 15811.42649 | -0.015250932 | 0.055564164 | 0.507655831 | 0.953260948 | ns | TM | KS.3 |
| FIM1_129 | 813.9981456 | 968.1790301 | 1036.64876 | 0.029203025 | 0.0833283 | 0.248629041 | 0.888934767 | ns | TM | KS.3 |
| FIM1_130 | 12177.16834 | 11452.71749 | 9887.24238 | -0.017056895 | 0.057557906 | 0.468147646 | 0.950410534 | ns | TM | KS.3 |
| FIM1_131 | 1599.620269 | 2019.308942 | 2248.066324 | 0.071088963 | 0.105663688 | 0.045104298 | 0.487132212 | ns | TM | KS.3 |
| FIM1_132 | 22085.10545 | 21994.36638 | 21182.2828 | -0.001209786 | 0.041332856 | 0.948910011 | 0.994413583 | ns | TM | KS.3 |
| FIM1_133 | 3572.688213 | 4211.798991 | 4961.324517 | 0.0474484 | 0.084592305 | 0.117771923 | 0.724896022 | ns | TM | KS.3 |
| FIM1_134 | 4635.172044 | 4720.07677 | 4336.865324 | 0.004706793 | 0.051847811 | 0.829395755 | 0.991141351 | ns | TM | KS.3 |
| FIM1_135 | 1237.050311 | 1470.877417 | 1364.517315 | 0.119273374 | 0.093760689 | 0.009859796 | 0.219966293 | ns | TM | KS.3 |
| FIM1_136 | 71.91884616 | 84.25389719 | 244.4891628 | 0.015105815 | 0.086621766 | 0.474540379 | 0.950410534 | ns | TM | KS.3 |
| FIM1_137 | 736.5288591 | 452.2108762 | 852.1868397 | -0.231553864 | 0.276000857 | 0.002526971 | 0.093517802 | ns | TM | KS.3 |
| FIM1_138 | 11245.01444 | 12445.94793 | 11142.29983 | 0.022647779 | 0.069400572 | 0.365888849 | 0.948637087 | ns | TM | KS.3 |
| FIM1_140 | 17418.69705 | 18865.49261 | 17517.12602 | 0.020396319 | 0.06341011 | 0.406395644 | 0.950410534 | ns | TM | KS.3 |
| FIM1_141 | 604.5721702 | 780.1169253 | 718.8621058 | 0.107985792 | 0.12103977 | 0.014591504 | 0.273934835 | ns | TM | KS.3 |
| FIM1_142 | 12926.33447 | 12763.33607 | 12027.67333 | -0.004132712 | 0.032406534 | 0.785029563 | 0.988880259 | ns | TM | KS.3 |
| FIM1_143 | 116.6267177 | 133.6053606 | 247.9532581 | 0.014042401 | 0.08274575 | 0.51348601 | 0.954240701 | ns | TM | KS.3 |
| FIM1_144 | 1226.110983 | 822.2272707 | 911.1906525 | -0.035265012 | 0.12355307 | 0.146474133 | 0.780546912 | ns | TM | KS.3 |
| FIM1_145 | 11957.48678 | 12623.94068 | 13754.70225 | 0.008927246 | 0.068757706 | 0.696390125 | 0.981479944 | ns | TM | KS.3 |
| FIM1_146 | 2285.45936 | 2168.436172 | 2501.115526 | -0.017654916 | 0.050518862 | 0.419744524 | 0.950410534 | ns | TM | KS.3 |
| FIM1_147 | 2209.141032 | 1976.851442 | 2010.46277 | -0.060341785 | 0.073404757 | 0.06518525 | 0.563994123 | ns | TM | KS.3 |
| FIM1_148 | 714.8873772 | 808.6424116 | 787.1053614 | 0.062692221 | 0.077804645 | 0.063317217 | 0.553608357 | ns | TM | KS.3 |

| | | | | | | | | | | |
|---|---|---|---|---|---|---|---|---|---|---|
| FIM1_149 | 634.2398638 | 640.2786626 | 567.6685807 | 0.002331597 | 0.051390103 | 0.914504081 | 0.991141351 | ns | TM | KS.3 |
| FIM1_150 | 1122.803888 | 1203.162866 | 1020.816971 | 0.025430909 | 0.055620731 | 0.284252153 | 0.918450395 | ns | TM | KS.3 |
| FIM1_151 | 1000.122964 | 839.9831491 | 929.2554157 | -0.070347408 | 0.0924446 | 0.050326012 | 0.513412402 | ns | TM | KS.3 |
| FIM1_153 | 659.6326242 | 803.2441415 | 573.2743016 | 0.041945534 | 0.089261026 | 0.139251379 | 0.774219902 | ns | TM | KS.3 |
| FIM1_154 | 2011.222581 | 1908.786819 | 1999.653377 | -0.005416547 | 0.078457418 | 0.795595905 | 0.988916276 | ns | TM | KS.3 |
| FIM1_155 | 3951.664474 | 3612.844804 | 3525.482726 | -0.013601535 | 0.073176457 | 0.554016126 | 0.95917717 | ns | TM | KS.3 |
| FIM1_156 | 1061.704449 | 1064.471081 | 1443.315126 | 0.000522677 | 0.058175206 | 0.981593157 | 0.999333464 | ns | TM | KS.3 |
| FIM1_157 | 1516.453448 | 1349.395521 | 2207.756938 | -0.010080762 | 0.084981685 | 0.620178466 | 0.973122675 | ns | TM | KS.3 |
| FIM1_158 | 8067.892211 | 7605.860048 | 6922.7706 | -0.017027701 | 0.056027284 | 0.463664693 | 0.950410534 | ns | TM | KS.3 |
| FIM1_159 | 6874.960805 | 6735.447938 | 6425.530611 | -0.007215512 | 0.021017269 | 0.406744956 | 0.950410534 | ns | TM | KS.3 |
| FIM1_160 | 6084.979185 | 5852.079874 | 5656.953157 | -0.014035155 | 0.041760349 | 0.452498999 | 0.950410534 | ns | TM | KS.3 |
| FIM1_161 | 693.2183992 | 612.253163 | 678.5940738 | -0.021724698 | 0.075391719 | 0.372628799 | 0.948637087 | ns | TM | KS.3 |
| FIM1_162 | 1027.695589 | 862.3874805 | 1262.050445 | -0.044083943 | 0.085749854 | 0.132647318 | 0.761677364 | ns | TM | KS.3 |
| FIM1_163 | 450.4964022 | 297.5781382 | 516.5720716 | -0.102790697 | 0.17613806 | 0.01693204 | 0.302858265 | ns | TM | KS.3 |
| FIM1_164 | 803.8655098 | 825.6056512 | 893.1192894 | 0.007148243 | 0.046386857 | 0.727394222 | 0.987157164 | ns | TM | KS.3 |
| FIM1_165 | 564.2911979 | 603.099 | 587.4606674 | 0.014827821 | 0.062805276 | 0.533263864 | 0.959071703 | ns | TM | KS.3 |
| FIM1_166 | 678.1775202 | 875.0449489 | 737.7645358 | 0.057312253 | 0.105570469 | 0.069703431 | 0.581836521 | ns | TM | KS.3 |
| FIM1_167 | 447.1027606 | 363.9628548 | 515.6252124 | -0.046676257 | 0.092265281 | 0.113379959 | 0.713104042 | ns | TM | KS.3 |
| FIM1_168 | 1362.386785 | 1377.967691 | 1298.756978 | 0.002955485 | 0.052469607 | 0.8928134 | 0.991141351 | ns | TM | KS.3 |
| FIM1_169 | 1594.518522 | 1632.013647 | 2192.324479 | 0.005554303 | 0.054741536 | 0.803918303 | 0.988916276 | ns | TM | KS.3 |
| FIM1_170 | 12116.66828 | 11290.70089 | 11208.52652 | -0.050839337 | 0.05462012 | 0.054639977 | 0.531428718 | ns | TM | KS.3 |
| FIM1_171 | 1885.188365 | 1972.062597 | 1811.685558 | 0.01037331 | 0.059440127 | 0.654292186 | 0.976323032 | ns | TM | KS.3 |
| FIM1_172 | 795.7913317 | 808.8152968 | 709.2605182 | 0.003575985 | 0.057932381 | 0.874736423 | 0.991141351 | ns | TM | KS.3 |
| FIM1_173 | 910.8741874 | 867.3609117 | 894.9360852 | -0.012733706 | 0.05664498 | 0.580245907 | 0.963781856 | ns | TM | KS.3 |
| FIM1_174 | 383.0173307 | 391.7010551 | 327.707724 | 0.001462536 | 0.088583502 | 0.938033106 | 0.994413583 | ns | TM | KS.3 |
| FIM1_175 | 108.4844178 | 92.77669077 | 88.16067366 | -0.015520386 | 0.084815182 | 0.470160558 | 0.950410534 | ns | TM | KS.3 |
| FIM1_176 | 2027.077236 | 2010.843329 | 2120.157969 | -0.001808988 | 0.056133367 | 0.935833916 | 0.994413583 | ns | TM | KS.3 |
| FIM1_177 | 9213.103534 | 8916.825206 | 8182.819457 | -0.004161714 | 0.073507286 | 0.848354738 | 0.991141351 | ns | TM | KS.3 |
| FIM1_178 | 1041.318681 | 1008.453085 | 898.2572507 | -0.009864587 | 0.044943981 | 0.622423435 | 0.973122675 | ns | TM | KS.3 |
| FIM1_179 | 2440.999742 | 2189.205192 | 2568.32626 | -0.036502689 | 0.06981809 | 0.188756304 | 0.828217874 | ns | TM | KS.3 |
| FIM1_180 | 1509.995124 | 1565.233204 | 1767.514357 | 0.010485917 | 0.049673272 | 0.625300009 | 0.97344368 | ns | TM | KS.3 |
| FIM1_181 | 11932.59836 | 11389.90328 | 11963.46107 | -0.017769684 | 0.044094785 | 0.362552208 | 0.948637087 | ns | TM | KS.3 |
| FIM1_182 | 5537.681124 | 4834.003445 | 5639.517076 | -0.017833257 | 0.079810646 | 0.434563116 | 0.950410534 | ns | TM | KS.3 |
| FIM1_183 | 116.1283274 | 95.46490628 | 86.97005316 | -0.020459004 | 0.088580436 | 0.356842088 | 0.948637087 | ns | TM | KS.3 |
| FIM1_184 | 4942.192325 | 4383.969867 | 4253.522987 | -0.034104064 | 0.072620862 | 0.21341556 | 0.861104955 | ns | TM | KS.3 |
| FIM1_185 | 470.4132818 | 424.1783717 | 1297.052696 | -0.016518287 | 0.073820604 | 0.480981625 | 0.951416554 | ns | TM | KS.3 |
| FIM1_186 | 267.4639989 | 265.5884053 | 342.0256655 | -0.000695803 | 0.07625177 | 0.973698192 | 0.998607635 | ns | TM | KS.3 |
| FIM1_187 | 6560.957021 | 8242.911449 | 7421.616443 | 0.025793588 | 0.092797865 | 0.266497227 | 0.904370522 | ns | TM | KS.3 |
| FIM1_189 | 1168.133739 | 1217.801291 | 1349.127392 | 0.01159058 | 0.052460898 | 0.601363282 | 0.97002321 | ns | TM | KS.3 |
| FIM1_190 | 208.4069141 | 198.7729324 | 166.9618692 | -0.007610915 | 0.070306686 | 0.735804791 | 0.987655486 | ns | TM | KS.3 |
| FIM1_191 | 577.32139 | 519.5514212 | 673.3985639 | -0.031509167 | 0.068787614 | 0.238980433 | 0.886981413 | ns | TM | KS.3 |
| FIM1_192 | 3544.607641 | 3612.944116 | 3739.861374 | 0.001516971 | 0.083731916 | 0.938404596 | 0.994413583 | ns | TM | KS.3 |
| FIM1_193 | 31.28025926 | 16.84274193 | 567.2888931 | -0.067095124 | 0.195727006 | 0.043739304 | 0.479301841 | ns | TM | KS.3 |
| FIM1_194 | 2235.61634 | 2437.283669 | 2532.895346 | 0.013430262 | 0.072462581 | 0.560206254 | 0.961043487 | ns | TM | KS.3 |
| FIM1_195 | 1249.054141 | 1387.100409 | 1244.231713 | 0.021605228 | 0.070832824 | 0.383112303 | 0.948637087 | ns | TM | KS.3 |

| | | | | | | | | | |
|---|---|---|---|---|---|---|---|---|---|
| FIM1_196 | 2627.871338 | 2848.682913 | 2924.485626 | 0.02401197 | 0.061828597 | 0.334813038 | 0.938982061 | ns | TM | KS.3 |
| FIM1_197 | 1861.567574 | 1606.872402 | 2057.649556 | -0.022249858 | 0.079755411 | 0.352106028 | 0.948637087 | ns | TM | KS.3 |
| FIM1_198 | 3058.16247 | 3139.809456 | 4654.542393 | 0.00573851 | 0.059389717 | 0.801865891 | 0.988916276 | ns | TM | KS.3 |
| FIM1_199 | 995.3129439 | 1137.781231 | 1061.302746 | 0.026785808 | 0.0760916 | 0.293740108 | 0.923185001 | ns | TM | KS.3 |
| FIM1_200 | 2597.060632 | 2300.951095 | 2473.384623 | -0.02923142 | 0.072935949 | 0.26667569 | 0.904370522 | ns | TM | KS.3 |
| FIM1_201 | 1981.142666 | 2174.606386 | 2060.566516 | 0.029471013 | 0.065050887 | 0.258479466 | 0.895871714 | ns | TM | KS.3 |
| FIM1_203 | 1712.953269 | 1473.023084 | 1224.968132 | -0.06127899 | 0.084878723 | 0.070137937 | 0.582627046 | ns | TM | KS.3 |
| FIM1_204 | 140.5435965 | 145.240084 | 149.289268 | 0.003952948 | 0.073028367 | 0.856346391 | 0.991141351 | ns | TM | KS.3 |
| FIM1_205 | 1136.946311 | 1370.523793 | 1549.415675 | 0.020225638 | 0.08755698 | 0.363870819 | 0.948637087 | ns | TM | KS.3 |
| FIM1_206 | 2793.244647 | 3019.147645 | 2888.196518 | 0.025493058 | 0.059815275 | 0.302874712 | 0.926536521 | ns | TM | KS.3 |
| FIM1_207 | 1152.073138 | 946.2695922 | 798.6830117 | -0.05819823 | 0.093952611 | 0.074782706 | 0.599104608 | ns | TM | KS.3 |
| FIM1_208 | 3966.115458 | 3442.387064 | 3150.474754 | -0.049246268 | 0.080166091 | 0.111511265 | 0.71059097 | ns | TM | KS.3 |
| FIM1_209 | 3863.055348 | 3500.349752 | 4312.461468 | -0.028692205 | 0.066877892 | 0.272083383 | 0.908468501 | ns | TM | KS.3 |
| FIM1_210 | 3231.789783 | 3084.44636 | 2926.221732 | -0.010603146 | 0.060013878 | 0.648003924 | 0.976323032 | ns | TM | KS.3 |
| FIM1_211 | 1671.071322 | 1453.736149 | 1421.061327 | -0.049749493 | 0.079827637 | 0.10925445 | 0.705897255 | ns | TM | KS.3 |
| FIM1_212 | 1496.459302 | 1320.182369 | 1343.651911 | -0.031961948 | 0.073894366 | 0.234936088 | 0.883800031 | ns | TM | KS.3 |
| FIM1_213 | 1468.289187 | 1361.484535 | 1407.992985 | -0.029090577 | 0.058255059 | 0.239980407 | 0.887413624 | ns | TM | KS.3 |
| FIM1_214 | 456.9680017 | 359.0967915 | 339.0121516 | -0.05405698 | 0.10137352 | 0.080431938 | 0.61846815 | ns | TM | KS.3 |
| FIM1_215 | 811.5890942 | 720.5216289 | 763.6384068 | -0.064782334 | 0.076167677 | 0.056628245 | 0.535599847 | ns | TM | KS.3 |
| FIM1_216 | 3645.703069 | 3503.005224 | 3833.499387 | -0.006815671 | 0.066991889 | 0.765434178 | 0.988477371 | ns | TM | KS.3 |
| FIM1_217 | 2678.633992 | 2636.927716 | 2626.542711 | -0.003714758 | 0.055579163 | 0.868497454 | 0.991141351 | ns | TM | KS.3 |
| FIM1_218 | 1425.154799 | 860.0241218 | 1854.385565 | -0.122316046 | 0.221731685 | 0.012428967 | 0.254461363 | ns | TM | KS.3 |
| FIM1_219 | 1100.562785 | 971.1090283 | 1022.823757 | -0.061946375 | 0.078160119 | 0.065586597 | 0.564318013 | ns | TM | KS.3 |
| FIM1_220 | 382.7199363 | 437.2276632 | 417.3276213 | 0.032632586 | 0.075449295 | 0.226881252 | 0.876782875 | ns | TM | KS.3 |
| FIM1_221 | 1330.986864 | 1279.480893 | 1188.193571 | -0.008650057 | 0.060591014 | 0.70814705 | 0.983537569 | ns | TM | KS.3 |
| FIM1_222 | 1989.556474 | 1596.178937 | 1793.918807 | -0.244024121 | 0.071241024 | 8.55591430712 | 0.001064142 | ns | TM | KS.3 |
| FIM1_224 | 2082.895166 | 2174.501847 | 1990.069359 | 0.010257915 | 0.058086068 | 0.656044874 | 0.976323032 | ns | TM | KS.3 |
| FIM1_225 | 1334.295078 | 1367.478589 | 1133.049414 | 0.006721352 | 0.049635162 | 0.753340674 | 0.988477371 | ns | TM | KS.3 |
| FIM1_226 | 476.0946299 | 444.6020756 | 393.6260039 | -0.008987566 | 0.074373499 | 0.684059777 | 0.98018358 | ns | TM | KS.3 |
| FIM1_227 | 1739.286198 | 2121.166621 | 1808.330015 | 0.052919779 | 0.092682097 | 0.090525879 | 0.65689657 | ns | TM | KS.3 |
| FIM1_228 | 169.4231861 | 159.2512226 | 100.5731306 | -0.015035193 | 0.062459879 | 0.527733427 | 0.959071703 | ns | TM | KS.3 |
| FIM1_229 | 10679.78201 | 10839.21336 | 7860.361819 | 0.003534298 | 0.055312438 | 0.874614648 | 0.991141351 | ns | TM | KS.3 |
| FIM1_230 | 2293.760663 | 2143.30711 | 1941.403259 | -0.0228315 | 0.056377197 | 0.336456202 | 0.939848178 | ns | TM | KS.3 |
| FIM1_231 | 206.9415617 | 252.1306551 | 186.9610113 | 0.019552172 | 0.089506347 | 0.370803441 | 0.948637087 | ns | TM | KS.3 |
| FIM1_233 | 4858.450391 | 4969.296608 | 4732.029427 | 0.004769548 | 0.06010048 | 0.834748677 | 0.991141351 | ns | TM | KS.3 |
| FIM1_234 | 1861.003611 | 1812.547766 | 1866.763399 | -0.008974661 | 0.034963605 | 0.574172429 | 0.963192994 | ns | TM | KS.3 |
| FIM1_235 | 965.8757659 | 909.6309757 | 816.0938578 | -0.014542775 | 0.060301469 | 0.537960035 | 0.95917717 | ns | TM | KS.3 |
| FIM1_236 | 2427.218244 | 2389.351015 | 2878.122841 | -0.003689659 | 0.055908339 | 0.86961333 | 0.991141351 | ns | TM | KS.3 |
| FIM1_237 | 782.1451797 | 730.2500658 | 509.869959 | -0.006605585 | 0.080458328 | 0.748956319 | 0.988477371 | ns | TM | KS.3 |
| FIM1_238 | 533.0638009 | 530.1737527 | 439.7022536 | -0.000892114 | 0.057757599 | 0.96854474 | 0.996327965 | ns | TM | KS.3 |
| FIM1_239 | 94.0288204 | 97.0166118 | 93.79866522 | 0.004322956 | 0.072969001 | 0.843304018 | 0.991141351 | ns | TM | KS.3 |
| FIM1_240 | 658.6223467 | 543.718753 | 582.5653609 | -0.036534795 | 0.08717298 | 0.175810305 | 0.811391014 | ns | TM | KS.3 |
| FIM1_241 | 2565.363358 | 2214.943592 | 3621.35305 | -0.139729473 | 0.074330398 | 0.002976085 | 0.101288726 | ns | TM | KS.3 |
| FIM1_242 | 72.49303908 | 66.17263008 | 58.14148934 | -0.011790864 | 0.076049321 | 0.595648233 | 0.965889818 | ns | TM | KS.3 |
| FIM1_243 | 921.5925054 | 1137.039336 | 1207.709963 | 0.19194753 | 0.089368548 | 0.000647584 | 0.035018801 | ns | TM | KS.3 |

| | | | | | | | | | | |
|---|---|---|---|---|---|---|---|---|---|---|
| FIM1_245 | 4279.517273 | 4618.770728 | 4952.962543 | 0.009606458 | 0.075583359 | 0.661766096 | 0.976323032 | ns | TM | KS.3 |
| FIM1_246 | 2201.187491 | 2092.303652 | 2323.651007 | -0.016564335 | 0.050631103 | 0.449159304 | 0.950410534 | ns | TM | KS.3 |
| FIM1_247 | 1678.046089 | 1243.074093 | 1343.159356 | -0.061994629 | 0.117661304 | 0.055466229 | 0.532711374 | ns | TM | KS.3 |
| FIM1_248 | 849.0122364 | 848.9781407 | 1115.187354 | 0.000327993 | 0.051593578 | 0.987966917 | 0.999333464 | ns | TM | KS.3 |
| FIM1_249 | 651.4462458 | 884.5898796 | 995.5845502 | 0.02749994 | 0.105531556 | 0.225796881 | 0.876782875 | ns | TM | KS.3 |
| FIM1_250 | 905.4993387 | 738.4672766 | 804.4931944 | -0.089227066 | 0.102905146 | 0.026527569 | 0.382098641 | ns | TM | KS.3 |
| FIM1_251 | 633.5318749 | 649.0428452 | 522.5479514 | 0.006133403 | 0.055228927 | 0.784809604 | 0.988880259 | ns | TM | KS.3 |
| FIM1_252 | 2064.194155 | 2048.964485 | 2061.09628 | -0.002280562 | 0.03964834 | 0.900591658 | 0.991141351 | ns | TM | KS.3 |
| FIM1_253 | 4531.830135 | 4133.595082 | 4712.613198 | -0.033891352 | 0.064368843 | 0.203393416 | 0.847612283 | ns | TM | KS.3 |
| FIM1_254 | 629.6874088 | 667.5303304 | 630.1023059 | 0.013000578 | 0.062383198 | 0.581317827 | 0.963781856 | ns | TM | KS.3 |
| FIM1_255 | 4577.81768 | 4338.535522 | 3916.679253 | -0.011376363 | 0.062783858 | 0.627053771 | 0.97344368 | ns | TM | KS.3 |
| FIM1_256 | 2449.226237 | 2907.333028 | 2945.980669 | 0.036819729 | 0.083559456 | 0.179884648 | 0.813337273 | ns | TM | KS.3 |
| FIM1_257 | 829.9728779 | 950.623006 | 989.1991687 | 0.04975079 | 0.078749668 | 0.109053138 | 0.705512825 | ns | TM | KS.3 |
| FIM1_258 | 1460.981478 | 1305.273476 | 1326.343152 | -0.039511446 | 0.071155486 | 0.165076192 | 0.801223467 | ns | TM | KS.3 |
| FIM1_259 | 3310.258334 | 3208.557055 | 3182.099749 | -0.007039059 | 0.058725087 | 0.758435713 | 0.988477371 | ns | TM | KS.3 |
| FIM1_260 | 436.9922243 | 371.0793312 | 531.1506979 | -0.039884203 | 0.082973187 | 0.159811633 | 0.791498358 | ns | TM | KS.3 |
| FIM1_261 | 1702.872157 | 1609.417823 | 1417.32722 | -0.00823166 | 0.071417105 | 0.714085516 | 0.986826511 | ns | TM | KS.3 |
| FIM1_262 | 1013.630149 | 1093.035728 | 1256.727153 | 0.011806084 | 0.071293502 | 0.606983922 | 0.970937835 | ns | TM | KS.3 |
| FIM1_263 | 2270.898487 | 2562.493577 | 4356.538085 | 0.042779357 | 0.07399975 | 0.144556126 | 0.77783242 | ns | TM | KS.3 |
| FIM1_264 | 880.5722222 | 922.0570034 | 998.4399574 | 0.013872885 | 0.05215405 | 0.532160869 | 0.959071703 | ns | TM | KS.3 |
| FIM1_265 | 1452.009222 | 1520.328785 | 1462.899096 | 0.016440502 | 0.047413194 | 0.430099125 | 0.950410534 | ns | TM | KS.3 |
| FIM1_266 | 860.613852 | 857.9054438 | 972.3963985 | -0.000678382 | 0.057610199 | 0.976063492 | 0.998938406 | ns | TM | KS.3 |
| FIM1_267 | 4733.605585 | 4806.871193 | 5139.402025 | 0.0045494 | 0.043344276 | 0.815980163 | 0.988916276 | ns | TM | KS.3 |
| FIM1_268 | 7203.825349 | 6054.155049 | 5576.256847 | -0.053842359 | 0.088087497 | 0.090969954 | 0.657813255 | ns | TM | KS.3 |
| FIM1_269 | 1396.507569 | 1387.733751 | 1362.757812 | -0.001483584 | 0.052151519 | 0.945890385 | 0.994413583 | ns | TM | KS.3 |
| FIM1_270 | 150.3260155 | 185.3172846 | 135.2654486 | 0.037190808 | 0.089902978 | 0.166639771 | 0.80502665 | ns | TM | KS.3 |
| FIM1_271 | 2905.585807 | 2743.168176 | 2806.474049 | -0.013725787 | 0.060312546 | 0.559687407 | 0.960914203 | ns | TM | KS.3 |
| FIM1_272 | 677.6283532 | 664.3920651 | 690.4990834 | -0.004279344 | 0.058602631 | 0.850844702 | 0.991141351 | ns | TM | KS.3 |
| FIM1_273 | 808.7101485 | 539.9554574 | 446.7911323 | -0.140551536 | 0.193509903 | 0.007623927 | 0.188701684 | ns | TM | KS.3 |
| FIM1_274 | 1526.176766 | 1665.319891 | 1722.830084 | 0.012270607 | 0.074248338 | 0.587130058 | 0.963781856 | ns | TM | KS.3 |
| FIM1_275 | 1744.683048 | 1799.619941 | 1602.25596 | 0.005724724 | 0.064347216 | 0.802813592 | 0.988916276 | ns | TM | KS.3 |
| FIM1_276 | 947.1536568 | 978.7887596 | 906.7754852 | 0.007904725 | 0.056651232 | 0.727981284 | 0.987157164 | ns | TM | KS.3 |
| FIM1_277 | 312.6810903 | 315.948904 | 311.8922662 | 0.00329432 | 0.052566601 | 0.880735072 | 0.991141351 | ns | TM | KS.3 |
| FIM1_278 | 663.3783646 | 756.3340178 | 637.3066224 | 0.031801104 | 0.075235036 | 0.235612251 | 0.884880231 | ns | TM | KS.3 |
| FIM1_279 | 3415.822479 | 3447.992157 | 3531.57243 | 0.002570773 | 0.050181071 | 0.904655949 | 0.991141351 | ns | TM | KS.3 |
| FIM1_280 | 23.46180387 | 16.80196182 | 26.86602388 | -0.022140108 | 0.110015314 | 0.289067063 | 0.919993167 | ns | TM | KS.3 |
| FIM1_281 | 2342.734992 | 2289.612652 | 3055.708706 | -0.004217292 | 0.063989986 | 0.853623143 | 0.991141351 | ns | TM | KS.3 |
| FIM1_282 | 267.3244265 | 301.9677255 | 392.1436837 | 0.00928916 | 0.087761573 | 0.638396599 | 0.975246356 | ns | TM | KS.3 |
| FIM1_283 | 400.6069748 | 367.2576377 | 1789.244243 | -0.007341934 | 0.084072941 | 0.71443747 | 0.986942381 | ns | TM | KS.3 |
| FIM1_284 | 884.1641924 | 929.8788768 | 1082.482432 | 0.011073164 | 0.061698798 | 0.63541409 | 0.973922241 | ns | TM | KS.3 |
| FIM1_285 | 1224.368611 | 1528.15169 | 1445.590392 | 0.02127121 | 0.09282695 | 0.332078705 | 0.938982061 | ns | TM | KS.3 |
| FIM1_286 | 3332.414254 | 3080.723374 | 3409.995974 | -0.014462647 | 0.068670208 | 0.540472963 | 0.95917717 | ns | TM | KS.3 |
| FIM1_287 | 634.4186202 | 645.6871601 | 685.4018079 | 0.004269187 | 0.050742044 | 0.84335916 | 0.991141351 | ns | TM | KS.3 |
| FIM1_288 | 116.2298173 | 144.5182461 | 132.9791323 | 0.039380045 | 0.090416929 | 0.152001656 | 0.783447458 | ns | TM | KS.3 |
| FIM1_290 | 335.0630308 | 373.5515522 | 401.1793757 | 0.022261841 | 0.071412437 | 0.37069686 | 0.948637087 | ns | TM | KS.3 |

| | | | | | | | | | | |
|---|---|---|---|---|---|---|---|---|---|---|
| FIM1_291 | 1408.224951 | 1288.789305 | 1665.079634 | -0.01856983 | 0.067951099 | 0.445613063 | 0.950410534 | ns | TM | KS.3 |
| FIM1_292 | 1005.411085 | 306.8901533 | 1255.979238 | -1.477162479 | 0.290094168 | 2.96545955275 | 6.70598239769 | Km<KS.3 | TM | KS.3 |
| FIM1_293 | 2282.762082 | 2217.918888 | 3668.649036 | -0.006949701 | 0.056057583 | 0.758530723 | 0.988477371 | ns | TM | KS.3 |
| FIM1_294 | 139.2794212 | 158.6628089 | 2209.438311 | 0.008459094 | 0.091545758 | 0.657979212 | 0.976323032 | ns | TM | KS.3 |
| FIM1_295 | 20.02897815 | 26.10651341 | 54.48768277 | 0.017983519 | 0.101097308 | 0.375506339 | 0.948637087 | ns | TM | KS.3 |
| FIM1_296 | 57.79206709 | 64.28412741 | 99.09863344 | 0.007887257 | 0.086843739 | 0.688617471 | 0.980879562 | ns | TM | KS.3 |
| FIM1_297 | 20.70000985 | 18.66882436 | 31.93400973 | -0.005146297 | 0.092069856 | 0.781895908 | 0.988814136 | ns | TM | KS.3 |
| FIM1_298 | 28.02278199 | 33.34960804 | 43.81990865 | 0.017267827 | 0.088340441 | 0.419873451 | 0.950410534 | ns | TM | KS.3 |
| FIM1_299 | 639.6529984 | 644.6249458 | 625.4742574 | 0.000884283 | 0.076361813 | 0.966546248 | 0.995634078 | ns | TM | KS.3 |
| FIM1_300 | 1165.992105 | 1569.200965 | 1750.706662 | 0.024908814 | 0.104102801 | 0.258176862 | 0.895871714 | ns | TM | KS.3 |
| FIM1_301 | 214.7125307 | 238.9704967 | 194.6034244 | 0.012515512 | 0.079240955 | 0.565660708 | 0.962265923 | ns | TM | KS.3 |
| FIM1_302 | 788.1546227 | 789.5507942 | 737.3037931 | 0.000403173 | 0.062689145 | 0.985889469 | 0.999333464 | ns | TM | KS.3 |
| FIM1_303 | 27064.33492 | 26346.44688 | 31660.2633 | -0.005944018 | 0.058836432 | 0.794688602 | 0.988916276 | ns | TM | KS.3 |
| FIM1_304 | 516.5982175 | 516.1451871 | 807.8463057 | 4.56209872621263 | 0.070137303 | 0.998359399 | 0.999525119 | ns | TM | KS.3 |
| FIM1_305 | 450.0482847 | 200.6799825 | 843.7830533 | -0.418186873 | 0.486215886 | 0.001229319 | 0.055097839 | ns | TM | KS.3 |
| FIM1_306 | 217.195881 | 103.3387086 | 334.8470566 | -0.087990725 | 0.241686273 | 0.026144883 | 0.381439278 | ns | TM | KS.3 |
| FIM1_307 | 1184.183635 | 1284.814122 | 1116.511687 | 0.023087587 | 0.062548508 | 0.353096994 | 0.948637087 | ns | TM | KS.3 |
| FIM1_308 | 2259.393748 | 2264.504289 | 2450.767558 | 0.000484296 | 0.059183949 | 0.982994367 | 0.999333464 | ns | TM | KS.3 |
| FIM1_309 | 5701.994883 | 5732.402846 | 4752.360542 | 0.00162282 | 0.039501614 | 0.928982019 | 0.993697171 | ns | TM | KS.3 |
| FIM1_310 | 3271.414927 | 3648.471355 | 4156.181608 | 0.024414016 | 0.070853707 | 0.33550405 | 0.938982061 | ns | TM | KS.3 |
| FIM1_311 | 2293.293783 | 2651.270222 | 3860.09946 | 0.069191706 | 0.084832159 | 0.052261747 | 0.521046472 | ns | TM | KS.3 |
| FIM1_312 | 4585.082242 | 4052.403223 | 3656.8811 | -0.021573101 | 0.075252992 | 0.375612001 | 0.948637087 | ns | TM | KS.3 |
| FIM1_313 | 2761.089952 | 2750.071928 | 2693.26069 | -0.000699135 | 0.065598358 | 0.975407628 | 0.998880439 | ns | TM | KS.3 |
| FIM1_314 | 2645.208989 | 2993.177275 | 2126.457402 | 0.019377234 | 0.076382105 | 0.413522122 | 0.950410534 | ns | TM | KS.3 |
| FIM1_315 | 605.7656978 | 635.8770052 | 659.2698316 | 0.015742511 | 0.050607435 | 0.471388845 | 0.950410534 | ns | TM | KS.3 |
| FIM1_316 | 873.4184005 | 1028.114984 | 1101.901215 | 0.012118511 | 0.090827485 | 0.542301279 | 0.95917717 | ns | TM | KS.3 |
| FIM1_317 | 129.2340392 | 143.5367356 | 101.9596112 | 0.010842664 | 0.080277542 | 0.610669071 | 0.972185161 | ns | TM | KS.3 |
| FIM1_318 | 7492.058125 | 7227.978082 | 7843.167884 | -0.014008437 | 0.035824929 | 0.377556191 | 0.948637087 | ns | TM | KS.3 |
| FIM1_319 | 248.4544429 | 260.4189306 | 301.5180789 | 0.009166444 | 0.063628652 | 0.692823893 | 0.981188557 | ns | TM | KS.3 |
| FIM1_320 | 1292.308636 | 1390.14957 | 1810.685389 | 0.009539826 | 0.074702598 | 0.666042546 | 0.976323032 | ns | TM | KS.3 |
| FIM1_321 | 96.09034605 | 106.0907285 | 107.0083954 | 0.007256367 | 0.088969799 | 0.706505618 | 0.983265883 | ns | TM | KS.3 |
| FIM1_322 | 13936.9997 | 15261.79794 | 14723.54858 | 0.032430289 | 0.063993237 | 0.218786796 | 0.865538807 | ns | TM | KS.3 |
| FIM1_323 | 510.9391884 | 656.666813 | 571.029106 | 0.04099418 | 0.098904649 | 0.131873255 | 0.761283488 | ns | TM | KS.3 |
| FIM1_324 | 3704.080329 | 5331.450822 | 5869.511829 | 0.077639425 | 0.14343801 | 0.032149822 | 0.413606776 | ns | TM | KS.3 |
| FIM1_325 | 11.16088001 | 17.69504118 | 17.19711774 | 0.031261416 | 0.13266921 | 0.170125807 | 0.809599629 | ns | TM | KS.3 |
| FIM1_326 | 83.62099149 | 139.3628141 | 4974.569664 | 0.043634821 | 0.148743654 | 0.097752692 | 0.677062609 | ns | TM | KS.3 |
| FIM1_327 | 1238.91882 | 1236.688326 | 1256.923338 | -0.000479302 | 0.050108894 | 0.982160809 | 0.999333464 | ns | TM | KS.3 |
| FIM1_328 | 115.3170001 | 136.188598 | 197.3835281 | 0.013854658 | 0.088524323 | 0.500590014 | 0.953260948 | ns | TM | KS.3 |
| FIM1_329 | 702.9322783 | 685.3760932 | 674.5158635 | -0.002936972 | 0.074975883 | 0.890953275 | 0.991141351 | ns | TM | KS.3 |
| FIM1_330 | 11398.93544 | 9357.130501 | 10680.29304 | -0.055699727 | 0.093276829 | 0.0818464 | 0.621932675 | ns | TM | KS.3 |
| FIM1_331 | 370.3635624 | 319.5214853 | 526.2211276 | -0.017344685 | 0.082584609 | 0.436007871 | 0.950410534 | ns | TM | KS.3 |
| FIM1_332 | 1208.326537 | 1058.651884 | 1159.272756 | -0.027596343 | 0.07553926 | 0.284090833 | 0.918450395 | ns | TM | KS.3 |
| FIM1_333 | 1028.334512 | 913.0951985 | 1067.606057 | -0.027540018 | 0.072665713 | 0.288496255 | 0.919993167 | ns | TM | KS.3 |
| FIM1_334 | 1783.998082 | 1862.635994 | 1901.909592 | 0.009147515 | 0.061631102 | 0.693176986 | 0.981188557 | ns | TM | KS.3 |
| FIM1_335 | 993.6349409 | 1185.10772 | 1214.735243 | 0.01601827 | 0.088511296 | 0.447251479 | 0.950410534 | ns | TM | KS.3 |

| | | | | | | | | | | |
|---|---|---|---|---|---|---|---|---|---|---|
| FIM1_336 | 563.4367588 | 528.7872742 | 665.6980096 | -0.009845855 | 0.070400094 | 0.665821797 | 0.976323032 | ns | TM | KS.3 |
| FIM1_337 | 4401.595444 | 3938.279261 | 3669.238901 | -0.02909448 | 0.070474526 | 0.269210916 | 0.907401292 | ns | TM | KS.3 |
| FIM1_338 | 1286.848353 | 1333.315425 | 2051.336962 | 0.003292954 | 0.080361598 | 0.871539074 | 0.991141351 | ns | TM | KS.3 |
| FIM1_339 | 3200.706485 | 3275.224274 | 3976.175577 | 0.004481251 | 0.062689892 | 0.844835083 | 0.991141351 | ns | TM | KS.3 |
| FIM1_340 | 271.9245887 | 316.5980941 | 277.3983359 | 0.012364572 | 0.088284745 | 0.542043046 | 0.95917717 | ns | TM | KS.3 |
| FIM1_341 | 3811.159785 | 4029.140281 | 4523.907596 | 0.015420157 | 0.056314642 | 0.505696223 | 0.953260948 | ns | TM | KS.3 |
| FIM1_342 | 1152.210003 | 1112.767067 | 1112.983384 | -0.007618137 | 0.059852387 | 0.740462585 | 0.988051177 | ns | TM | KS.3 |
| FIM1_343 | 2532.0978 | 2330.789759 | 2331.160854 | -0.010339849 | 0.076129964 | 0.637905436 | 0.975246356 | ns | TM | KS.3 |
| FIM1_344 | 1857.874493 | 1823.788434 | 1776.822187 | -0.004703453 | 0.053101384 | 0.83131171 | 0.991141351 | ns | TM | KS.3 |
| FIM1_345 | 1413.422411 | 993.2536828 | 3734.663347 | -0.073891432 | 0.137865742 | 0.036100958 | 0.431736218 | ns | TM | KS.3 |
| FIM1_346 | 5127.130344 | 4636.305498 | 6352.248537 | -0.022168219 | 0.069377208 | 0.374364151 | 0.948637087 | ns | TM | KS.3 |
| FIM1_347 | 2073.04827 | 1949.29265 | 1962.428707 | -0.017294505 | 0.057802334 | 0.463118434 | 0.950410534 | ns | TM | KS.3 |
| FIM1_348 | 1426.975337 | 1756.292777 | 2060.195225 | 0.019183506 | 0.091531878 | 0.372437074 | 0.948637087 | ns | TM | KS.3 |
| FIM1_349 | 2658.84482 | 2564.178605 | 2501.27156 | -0.009577273 | 0.053386616 | 0.667481701 | 0.976323032 | ns | TM | KS.3 |
| FIM1_350 | 562.6920497 | 580.6811948 | 551.4005951 | 0.008705265 | 0.051624189 | 0.69097636 | 0.981188557 | ns | TM | KS.3 |
| FIM1_351 | 1728.001349 | 1651.366538 | 1800.110609 | -0.0172431 | 0.043879363 | 0.375158256 | 0.948637087 | ns | TM | KS.3 |
| FIM1_352 | 6072.197696 | 5923.381633 | 5972.361674 | -0.00212768 | 0.082032343 | 0.915172522 | 0.991141351 | ns | TM | KS.3 |
| FIM1_354 | 659.2917333 | 624.7641271 | 493.2727055 | -0.004810474 | 0.081721582 | 0.811878378 | 0.988916276 | ns | TM | KS.3 |
| FIM1_355 | 3151.916814 | 3008.625763 | 2709.148435 | -0.01217154 | 0.055860246 | 0.594524645 | 0.965889818 | ns | TM | KS.3 |
| FIM1_356 | 837.3391863 | 685.024702 | 2103.957352 | -0.030005507 | 0.088074415 | 0.229378114 | 0.879350777 | ns | TM | KS.3 |
| FIM1_357 | 5949.982978 | 4235.981531 | 3991.281274 | -0.041898564 | 0.116041638 | 0.114289743 | 0.714310891 | ns | TM | KS.3 |
| FIM1_358 | 4607.304402 | 5044.330479 | 6074.080717 | 0.008461828 | 0.082104122 | 0.680920302 | 0.97986479 | ns | TM | KS.3 |
| FIM1_359 | 7070.087783 | 7178.769763 | 7789.665494 | 0.005092212 | 0.025998265 | 0.668354965 | 0.976323032 | ns | TM | KS.3 |
| FIM1_360 | 1278.218752 | 1279.887749 | 1375.551515 | 0.000282999 | 0.045201538 | 0.988793638 | 0.999333464 | ns | TM | KS.3 |
| FIM1_361 | 562.3359387 | 547.2599944 | 560.8953301 | -0.004754067 | 0.065791094 | 0.834791585 | 0.991141351 | ns | TM | KS.3 |
| FIM1_362 | 2658.268237 | 2551.463513 | 2519.03007 | -0.006766069 | 0.067866305 | 0.766270078 | 0.988477371 | ns | TM | KS.3 |
| FIM1_363 | 1348.588824 | 1322.767778 | 1212.245779 | -0.004131419 | 0.059362904 | 0.856218308 | 0.991141351 | ns | TM | KS.3 |
| FIM1_364 | 3278.397807 | 2935.254093 | 2836.126551 | -0.088044566 | 0.071685053 | 0.019640221 | 0.324535175 | ns | TM | KS.3 |
| FIM1_365 | 1319.118296 | 1524.964043 | 1520.484683 | 0.028542439 | 0.078118023 | 0.267689945 | 0.90534159 | ns | TM | KS.3 |
| FIM1_366 | 2267.731354 | 1975.585194 | 1946.325552 | -0.031411562 | 0.076567848 | 0.238189663 | 0.88653074 | ns | TM | KS.3 |
| FIM1_367 | 1169.611848 | 2094.849405 | 4689.156278 | 0.160495554 | 0.277612976 | 0.007580594 | 0.188567287 | ns | TM | KS.3 |
| FIM1_368 | 420.7860148 | 415.9913199 | 489.1536025 | -0.001007161 | 0.081482503 | 0.959934967 | 0.994932595 | ns | TM | KS.3 |
| FIM1_369 | 35.41638303 | 42.29881508 | 35.23365655 | 0.010477762 | 0.097003059 | 0.578731367 | 0.963781856 | ns | TM | KS.3 |
| FIM1_370 | 961.7307183 | 973.5985144 | 767.2498702 | 0.003241279 | 0.053787574 | 0.883744117 | 0.991141351 | ns | TM | KS.3 |
| FIM1_371 | 78.68348782 | 93.3641182 | 98.26239274 | 0.016979453 | 0.08665659 | 0.431104042 | 0.950410534 | ns | TM | KS.3 |
| FIM1_372 | 15121.91311 | 15703.04195 | 15260.98212 | 0.005200282 | 0.071879348 | 0.814079728 | 0.988916276 | ns | TM | KS.3 |
| FIM1_373 | 1338.334444 | 1463.735129 | 1422.934025 | 0.015360089 | 0.071236002 | 0.514043232 | 0.954240701 | ns | TM | KS.3 |
| FIM1_374 | 1535.151091 | 1457.79129 | 1849.472422 | -0.012788665 | 0.058376014 | 0.582400083 | 0.963781856 | ns | TM | KS.3 |
| FIM1_375 | 1637.217132 | 1575.095467 | 1675.434603 | -0.005166402 | 0.072620513 | 0.814243223 | 0.988916276 | ns | TM | KS.3 |
| FIM1_376 | 1194.851054 | 906.9672096 | 754.4049239 | -0.032789955 | 0.101097313 | 0.182026412 | 0.818789693 | ns | TM | KS.3 |
| FIM1_377 | 1105.934141 | 1088.921106 | 809.5753069 | -0.00240362 | 0.068912498 | 0.914512906 | 0.991141351 | ns | TM | KS.3 |
| FIM1_378 | 40932.77024 | 42262.87679 | 46593.75577 | 0.007365788 | 0.057944072 | 0.74706459 | 0.988477371 | ns | TM | KS.3 |
| FIM1_379 | 484.5848403 | 498.3150779 | 566.4716419 | 0.002162693 | 0.084364215 | 0.911865322 | 0.991141351 | ns | TM | KS.3 |
| FIM1_380 | 2086.767411 | 2168.053067 | 2248.476043 | 0.004093785 | 0.077413429 | 0.845506405 | 0.991141351 | ns | TM | KS.3 |
| FIM1_381 | 2653.681713 | 2844.294576 | 2891.447937 | 0.006884598 | 0.079962217 | 0.740295247 | 0.988051177 | ns | TM | KS.3 |

| | | | | | | | | | | |
|---|---|---|---|---|---|---|---|---|---|---|
| FIM1_383 | 1210.10838 | 1773.676978 | 5227.277974 | 0.040470683 | 0.123422786 | 0.11769541 | 0.724896022 | ns | TM | KS.3 |
| FIM1_384 | 36222.10054 | 45479.24745 | 47056.6348 | 0.03990878 | 0.094237951 | 0.143450326 | 0.77783242 | ns | TM | KS.3 |
| FIM1_385 | 511.3084512 | 755.3687493 | 715.9322575 | 0.186570179 | 0.204834148 | 0.003230226 | 0.103770022 | ns | TM | KS.3 |
| FIM1_386 | 3621.597282 | 3071.260406 | 2795.587424 | -0.017767725 | 0.085172106 | 0.418789526 | 0.950410534 | ns | TM | KS.3 |
| FIM1_387 | 1093.356605 | 1059.028041 | 897.1630477 | -0.00657328 | 0.061097017 | 0.774935909 | 0.988814136 | ns | TM | KS.3 |
| FIM1_388 | 629.368538 | 640.6473965 | 624.7718793 | 0.003921617 | 0.059551056 | 0.863465374 | 0.991141351 | ns | TM | KS.3 |
| FIM1_389 | 3278.339831 | 3445.979569 | 3232.834531 | 0.010120094 | 0.06342926 | 0.663968276 | 0.976323032 | ns | TM | KS.3 |
| FIM1_390 | 467.5050375 | 467.0824308 | 386.9148257 | 1.46979494229778 | 0.070859298 | 0.999468862 | 0.999930384 | ns | TM | KS.3 |
| FIM1_391 | 1521.531083 | 1486.33631 | 1297.739817 | -0.005826939 | 0.054367502 | 0.794065428 | 0.988916276 | ns | TM | KS.3 |
| FIM1_392 | 4423.696373 | 4368.546749 | 4038.265816 | -0.002988466 | 0.055538586 | 0.893940764 | 0.991141351 | ns | TM | KS.3 |
| FIM1_393 | 1556.422348 | 1574.666902 | 1931.746616 | 0.003510179 | 0.042955445 | 0.856610837 | 0.991141351 | ns | TM | KS.3 |
| FIM1_394 | 153.4419764 | 173.2783155 | 215.3585894 | 0.015164311 | 0.079545314 | 0.495877475 | 0.953260948 | ns | TM | KS.3 |
| FIM1_395 | 1445.587235 | 1242.84309 | 1365.180795 | -0.040463245 | 0.080309565 | 0.158961204 | 0.790041949 | ns | TM | KS.3 |
| FIM1_396 | 2258.408695 | 2197.036376 | 2343.481647 | -0.00672708 | 0.05544957 | 0.765115107 | 0.988477371 | ns | TM | KS.3 |
| FIM1_397 | 34.66892718 | 37.33731997 | 44.24618426 | 0.005357619 | 0.08456503 | 0.78607858 | 0.988916276 | ns | TM | KS.3 |
| FIM1_398 | 2346.190355 | 2254.61457 | 2180.258315 | -0.01001856 | 0.05619347 | 0.660133566 | 0.976323032 | ns | TM | KS.3 |
| FIM1_399 | 3801.281045 | 3589.608717 | 2774.821104 | -0.01336009 | 0.060927387 | 0.570334618 | 0.962265923 | ns | TM | KS.3 |
| FIM1_401 | 1656.276101 | 1504.728716 | 1422.118783 | -0.026307869 | 0.066509275 | 0.305785917 | 0.926747341 | ns | TM | KS.3 |
| FIM1_402 | 265.3136378 | 221.0010137 | 416.4323931 | -0.031422603 | 0.084755598 | 0.222742176 | 0.871073186 | ns | TM | KS.3 |
| FIM1_403 | 734.0935784 | 635.2177215 | 967.7019184 | -0.029952687 | 0.077833531 | 0.251996424 | 0.888934767 | ns | TM | KS.3 |
| FIM1_404 | 502.5853618 | 453.1448006 | 546.760275 | -0.014352215 | 0.075970113 | 0.527154689 | 0.959071703 | ns | TM | KS.3 |
| FIM1_405 | 7049.679139 | 6831.822405 | 7626.635991 | -0.010014241 | 0.043584448 | 0.608519431 | 0.970937835 | ns | TM | KS.3 |
| FIM1_406 | 9746.627883 | 9143.275254 | 9920.880166 | -0.025383371 | 0.052511365 | 0.265312707 | 0.904062137 | ns | TM | KS.3 |
| FIM1_407 | 4013.324305 | 3008.424395 | 3459.852175 | -0.086587138 | 0.125435325 | 0.025875931 | 0.379742647 | ns | TM | KS.3 |
| FIM1_408 | 1691.009578 | 1459.752903 | 1724.87704 | -0.105154727 | 0.085530804 | 0.014354335 | 0.272568006 | ns | TM | KS.3 |
| FIM1_409 | 4867.574625 | 4708.35798 | 4668.864928 | -0.008576278 | 0.054072181 | 0.701516998 | 0.98151979 | ns | TM | KS.3 |
| FIM1_410 | 228.4745962 | 185.0743817 | 198.6579872 | -0.036163534 | 0.090240421 | 0.173130299 | 0.811391014 | ns | TM | KS.3 |
| FIM1_411 | 1405.13735 | 1277.679061 | 2144.197744 | -0.008172512 | 0.083994517 | 0.685743477 | 0.980861455 | ns | TM | KS.3 |
| FIM1_412 | 442.5741282 | 516.6622031 | 728.237725 | 0.053199988 | 0.083640081 | 0.095121346 | 0.673713046 | ns | TM | KS.3 |
| FIM1_413 | 3375.087227 | 3109.635095 | 4930.833769 | -0.031540219 | 0.060516576 | 0.217294118 | 0.865538807 | ns | TM | KS.3 |
| FIM1_414 | 518.388036 | 442.7160081 | 811.4516496 | -0.016524024 | 0.084795513 | 0.446979241 | 0.950410534 | ns | TM | KS.3 |
| FIM1_415 | 450.1927911 | 375.6496412 | 421.903523 | -0.043407699 | 0.086913064 | 0.134715614 | 0.766830871 | ns | TM | KS.3 |
| FIM1_416 | 1498.002463 | 1198.287083 | 1370.58872 | -0.125595156 | 0.112170738 | 0.008911175 | 0.204299984 | ns | TM | KS.3 |
| FIM1_417 | 5214.020234 | 5576.180398 | 6188.655595 | 0.014968062 | 0.06348404 | 0.530063136 | 0.959071703 | ns | TM | KS.3 |
| FIM1_418 | 1330.76421 | 1211.323972 | 901.0270708 | -0.017487804 | 0.07042808 | 0.466235055 | 0.950410534 | ns | TM | KS.3 |
| FIM1_419 | 905.6023322 | 900.6056066 | 844.6077708 | -0.001174398 | 0.058419374 | 0.958697383 | 0.994413583 | ns | TM | KS.3 |
| FIM1_420 | 755.515214 | 766.4134803 | 749.4564911 | 0.003224369 | 0.055748917 | 0.885782879 | 0.991141351 | ns | TM | KS.3 |
| FIM1_421 | 1128.399028 | 918.6580381 | 1164.997138 | -0.133079824 | 0.105017416 | 0.00690949 | 0.176553399 | ns | TM | KS.3 |
| FIM1_422 | 624.8473655 | 588.3949045 | 623.0930477 | -0.015400067 | 0.059047418 | 0.513482434 | 0.954240701 | ns | TM | KS.3 |
| FIM1_423 | 36.07760878 | 41.82198661 | 92.95811991 | 0.007915567 | 0.099123685 | 0.66385079 | 0.976323032 | ns | TM | KS.3 |
| FIM1_425 | 2440.385923 | 3073.185883 | 3568.612381 | 0.02474956 | 0.093327241 | 0.278923914 | 0.910529181 | ns | TM | KS.3 |
| FIM1_426 | 4281.732791 | 4005.652095 | 4407.258022 | -0.018591692 | 0.058831534 | 0.435709639 | 0.950410534 | ns | TM | KS.3 |
| FIM1_427 | 365.0987267 | 370.9317065 | 348.6366388 | 0.00299345 | 0.060428556 | 0.895612171 | 0.991141351 | ns | TM | KS.3 |
| FIM1_428 | 362.8019153 | 309.2417015 | 345.022864 | -0.028756874 | 0.080743171 | 0.259584195 | 0.895871714 | ns | TM | KS.3 |
| FIM1_429 | 1248.939995 | 1501.248519 | 1382.713337 | 0.093436896 | 0.097830621 | 0.02350128 | 0.359750361 | ns | TM | KS.3 |

| | | | | | | | | | | |
|---|---|---|---|---|---|---|---|---|---|---|
| FIM1_430 | 912.243464 | 950.3024508 | 1029.537073 | 0.011991091 | 0.050083139 | 0.579066586 | 0.963781856 | ns | TM | KS.3 |
| FIM1_431 | 545.4198918 | 570.558517 | 613.3133998 | 0.006899265 | 0.06976552 | 0.759733765 | 0.988477371 | ns | TM | KS.3 |
| FIM1_432 | 198.4829564 | 291.9956352 | 229.5730587 | 0.07218116 | 0.145789396 | 0.037654279 | 0.440708412 | ns | TM | KS.3 |
| FIM1_433 | 1970.666251 | 1942.11079 | 1934.95982 | -0.00360183 | 0.054763414 | 0.871814567 | 0.991141351 | ns | TM | KS.3 |
| FIM1_434 | 1023.352551 | 1010.389414 | 1025.508772 | -0.002880191 | 0.055885684 | 0.897949283 | 0.991141351 | ns | TM | KS.3 |
| FIM1_435 | 1054.554679 | 870.1599251 | 1007.262228 | -0.115092093 | 0.101471149 | 0.011929256 | 0.248318196 | ns | TM | KS.3 |
| FIM1_436 | 351.2926449 | 447.9321476 | 391.0584459 | 0.042180672 | 0.097871943 | 0.127088396 | 0.750017523 | ns | TM | KS.3 |
| FIM1_437 | 565.6332216 | 600.6774341 | 545.3349553 | 0.012147062 | 0.064418783 | 0.605265399 | 0.970937835 | ns | TM | KS.3 |
| FIM1_438 | 23.6413233 | 26.48231293 | 26.92615975 | 0.006647675 | 0.096590253 | 0.71588777 | 0.987123519 | ns | TM | KS.3 |
| FIM1_439 | 1273.432373 | 1744.05252 | 1895.423913 | 0.042484493 | 0.111400571 | 0.114222224 | 0.714310891 | ns | TM | KS.3 |
| FIM1_440 | 746.3612123 | 720.7893945 | 823.75812 | -0.00861007 | 0.055157237 | 0.702668083 | 0.98151979 | ns | TM | KS.3 |
| FIM1_441 | 710.0445871 | 713.4408632 | 966.3812468 | 0.001432099 | 0.050865451 | 0.947145207 | 0.994413583 | ns | TM | KS.3 |
| FIM1_442 | 1373.18281 | 1256.257032 | 1211.952098 | -0.015602591 | 0.070731441 | 0.509142101 | 0.953260948 | ns | TM | KS.3 |
| FIM1_443 | 182.6762527 | 180.956801 | 176.328668 | -0.000836685 | 0.081762143 | 0.966615597 | 0.995634078 | ns | TM | KS.3 |
| FIM1_444 | 1432.199366 | 1440.295673 | 1310.064377 | 0.001233812 | 0.05828444 | 0.956592208 | 0.994413583 | ns | TM | KS.3 |
| FIM1_445 | 2066.233581 | 1969.206533 | 1874.845332 | -0.012032813 | 0.057508169 | 0.602330484 | 0.970245665 | ns | TM | KS.3 |
| FIM1_446 | 478.4122803 | 452.831681 | 323.4614719 | -0.009628198 | 0.066048615 | 0.677745723 | 0.97986479 | ns | TM | KS.3 |
| FIM1_447 | 834.3242177 | 802.5185945 | 917.951041 | -0.012782222 | 0.045911936 | 0.529608108 | 0.959071703 | ns | TM | KS.3 |
| FIM1_448 | 2892.028727 | 2548.004254 | 2575.742673 | -0.038446231 | 0.074690506 | 0.175703525 | 0.811391014 | ns | TM | KS.3 |
| FIM1_449 | 4785.048524 | 4526.755583 | 4280.172542 | -0.008805423 | 0.069614909 | 0.698968562 | 0.98151979 | ns | TM | KS.3 |
| FIM1_450 | 193.1093285 | 123.6826991 | 132.0429841 | -0.112727378 | 0.194252051 | 0.0140266 | 0.269253438 | ns | TM | KS.3 |
| FIM1_451 | 1452.314337 | 1773.188389 | 2000.966981 | 0.028132323 | 0.087889226 | 0.249712991 | 0.888934767 | ns | TM | KS.3 |
| FIM1_452 | 685.5358252 | 815.5587139 | 780.8864883 | 0.040834769 | 0.084718112 | 0.151703943 | 0.783447458 | ns | TM | KS.3 |
| FIM1_453 | 3515.413297 | 3692.735283 | 2775.856282 | 0.00952757 | 0.064558108 | 0.681585051 | 0.97986479 | ns | TM | KS.3 |
| FIM1_454 | 240.0538929 | 268.3445969 | 268.9446485 | 0.007486007 | 0.089939857 | 0.696125209 | 0.981479944 | ns | TM | KS.3 |
| FIM1_455 | 1270.733586 | 1282.960806 | 1245.188269 | 0.001252295 | 0.072626105 | 0.954191312 | 0.994413583 | ns | TM | KS.3 |
| FIM1_456 | 1128.137246 | 1125.545171 | 1442.054858 | -0.000344956 | 0.071681234 | 0.987461081 | 0.999333464 | ns | TM | KS.3 |
| FIM1_457 | 14687.12414 | 15577.51038 | 14320.3747 | 0.014244878 | 0.060284962 | 0.545781469 | 0.95917717 | ns | TM | KS.3 |
| FIM1_458 | 20752.98366 | 21676.01053 | 20648.47836 | 0.011403723 | 0.055275405 | 0.615776822 | 0.973122675 | ns | TM | KS.3 |
| FIM1_459 | 1495.660754 | 1292.6779 | 1542.097326 | -0.025325666 | 0.078615284 | 0.308244079 | 0.927416591 | ns | TM | KS.3 |
| FIM1_460 | 397.0851723 | 439.1724655 | 413.5934671 | 0.018720927 | 0.071405195 | 0.438220417 | 0.950410534 | ns | TM | KS.3 |
| FIM1_461 | 890.5871404 | 845.8877861 | 842.2776394 | -0.012445224 | 0.059112537 | 0.593345578 | 0.965478443 | ns | TM | KS.3 |
| FIM1_462 | 762.10912 | 697.6475151 | 709.7308589 | -0.038350027 | 0.062378756 | 0.153153304 | 0.783447458 | ns | TM | KS.3 |
| FIM1_463 | 2603.383012 | 2747.767241 | 2168.265916 | 0.006547686 | 0.07523261 | 0.762277756 | 0.988477371 | ns | TM | KS.3 |
| FIM1_464 | 5639.369254 | 5525.073754 | 5385.754652 | -0.003992437 | 0.062465521 | 0.861463654 | 0.991141351 | ns | TM | KS.3 |
| FIM1_465 | 31.90713316 | 28.3368415 | 27.27999917 | -0.00785364 | 0.091354852 | 0.679917893 | 0.97986479 | ns | TM | KS.3 |
| FIM1_466 | 6463.277605 | 6759.306594 | 7328.141002 | 0.012255855 | 0.054187661 | 0.587207973 | 0.963781856 | ns | TM | KS.3 |
| FIM1_467 | 1764.535538 | 1949.177773 | 2370.570354 | 0.013887141 | 0.075469817 | 0.540487538 | 0.95917717 | ns | TM | KS.3 |
| FIM1_468 | 116.2516561 | 66.60986361 | 101.8914025 | -0.314206039 | 0.339115991 | 0.001407836 | 0.061438441 | ns | TM | KS.3 |
| FIM1_469 | 827.3201144 | 902.2424927 | 815.2713542 | 0.011945913 | 0.074665478 | 0.59509202 | 0.965889818 | ns | TM | KS.3 |
| FIM1_470 | 1352.479409 | 1299.476297 | 1474.85134 | -0.005460764 | 0.072212731 | 0.804656996 | 0.988916276 | ns | TM | KS.3 |
| FIM1_471 | 835.7482782 | 902.6871248 | 924.8653515 | 0.006621938 | 0.083214654 | 0.742128583 | 0.988477371 | ns | TM | KS.3 |
| FIM1_472 | 399.1218691 | 347.1484941 | 470.3324537 | -0.015426583 | 0.0825389 | 0.479744912 | 0.950972051 | ns | TM | KS.3 |
| FIM1_473 | 1608.71114 | 1420.085148 | 1561.354367 | -0.020822862 | 0.075838078 | 0.387619781 | 0.948637087 | ns | TM | KS.3 |
| FIM1_474 | 1711.197922 | 1727.655591 | 1664.142884 | 0.002326095 | 0.054338888 | 0.916693676 | 0.991141351 | ns | TM | KS.3 |

| | | | | | | | | | |
|---|---|---|---|---|---|---|---|---|---|
| FIM1_475 | 683.0212148 | 708.3496292 | 707.7649955 | 0.004621196 | 0.074090494 | 0.831279763 | 0.991141351 | ns | TM | KS.3 |
| FIM1_476 | 11008.48876 | 9494.535986 | 9708.260431 | -0.033688145 | 0.078754602 | 0.212915105 | 0.860185951 | ns | TM | KS.3 |
| FIM1_477 | 8700.462981 | 8634.796969 | 8762.515758 | -0.00212006 | 0.047079217 | 0.918378048 | 0.991141351 | ns | TM | KS.3 |
| FIM1_478 | 2427.40653 | 3119.32629 | 3891.826898 | 0.02563111 | 0.096528148 | 0.261142144 | 0.897845311 | ns | TM | KS.3 |
| FIM1_479 | 1460.85057 | 1435.868861 | 1402.401075 | -0.002156134 | 0.073411865 | 0.920792994 | 0.991141351 | ns | TM | KS.3 |
| FIM1_480 | 1470.184989 | 1296.04072 | 1130.206243 | -0.041001727 | 0.075045313 | 0.157024171 | 0.789034023 | ns | TM | KS.3 |
| FIM1_481 | 1099.740228 | 1000.443988 | 1343.131751 | -0.029131169 | 0.065505073 | 0.263950785 | 0.90372202 | ns | TM | KS.3 |
| FIM1_482 | 1186.171963 | 1080.900795 | 1188.457089 | -0.02095702 | 0.067726308 | 0.397521601 | 0.950345972 | ns | TM | KS.3 |
| FIM1_483 | 2055.022209 | 2036.327893 | 2054.337398 | -0.002620894 | 0.042565401 | 0.891984337 | 0.991141351 | ns | TM | KS.3 |
| FIM1_485 | 84.31723034 | 136.2300356 | 116.1817359 | 0.230063752 | 0.271887218 | 0.002508811 | 0.093517802 | ns | TM | KS.3 |
| FIM1_486 | 855.9731212 | 1103.839191 | 1003.7575 | 0.06930771 | 0.110054069 | 0.046156772 | 0.491712932 | ns | TM | KS.3 |
| FIM1_487 | 678.879111 | 781.8582619 | 668.4113705 | 0.0218717 | 0.078698495 | 0.361444286 | 0.948637087 | ns | TM | KS.3 |
| FIM1_488 | 6302.912569 | 6039.867952 | 5026.573853 | -0.007197424 | 0.067435864 | 0.752648229 | 0.988477371 | ns | TM | KS.3 |
| FIM1_489 | 2971.172647 | 3087.087761 | 3569.253078 | 0.004288415 | 0.076409097 | 0.839865207 | 0.991141351 | ns | TM | KS.3 |
| FIM1_490 | 1881.687179 | 1663.556488 | 1831.003238 | -0.038382455 | 0.073867574 | 0.176029515 | 0.811391014 | ns | TM | KS.3 |
| FIM1_491 | 11915.10037 | 12797.02562 | 12395.74351 | 0.026279332 | 0.056579921 | 0.27500892 | 0.908583046 | ns | TM | KS.3 |
| FIM1_492 | 2094.629436 | 1959.143974 | 2334.251937 | -0.02332802 | 0.055356337 | 0.321519425 | 0.929976243 | ns | TM | KS.3 |
| FIM1_493 | 1404.807448 | 1356.792518 | 1321.065216 | -0.008065459 | 0.05814716 | 0.724649445 | 0.987157164 | ns | TM | KS.3 |
| FIM1_494 | 1988.763233 | 2134.970474 | 1974.82989 | 0.020622077 | 0.059284492 | 0.392739677 | 0.948637087 | ns | TM | KS.3 |
| FIM1_495 | 4219.338786 | 4096.634279 | 3267.866095 | -0.008870223 | 0.04706353 | 0.668493071 | 0.976323032 | ns | TM | KS.3 |
| FIM1_496 | 5774.971493 | 5211.297983 | 5435.532296 | -0.080548184 | 0.068966594 | 0.024316108 | 0.368819013 | ns | TM | KS.3 |
| FIM1_497 | 1821.797851 | 1918.384754 | 1598.935476 | 0.00693227 | 0.073061328 | 0.753454985 | 0.988477371 | ns | TM | KS.3 |
| FIM1_498 | 717.2538142 | 701.3171762 | 630.0746192 | -0.004468556 | 0.061951195 | 0.845268188 | 0.991141351 | ns | TM | KS.3 |
| FIM1_499 | 3520.402846 | 3787.946388 | 3312.6156 | 0.021866236 | 0.059486085 | 0.368068369 | 0.948637087 | ns | TM | KS.3 |
| FIM1_500 | 343.5126105 | 310.5117006 | 352.4480144 | -0.021378978 | 0.07000197 | 0.388162594 | 0.948637087 | ns | TM | KS.3 |
| FIM1_501 | 632.7807679 | 592.5344583 | 642.575356 | -0.011699562 | 0.068176867 | 0.615102181 | 0.973122675 | ns | TM | KS.3 |
| FIM1_502 | 2158.886744 | 2090.829427 | 2460.250386 | -0.007460253 | 0.057800304 | 0.743838604 | 0.988477371 | ns | TM | KS.3 |
| FIM1_503 | 929.9536504 | 830.8989843 | 959.0798002 | -0.021835648 | 0.072770794 | 0.376106605 | 0.948637087 | ns | TM | KS.3 |
| FIM1_504 | 1111.367294 | 1113.614825 | 1291.733053 | 0.00053567 | 0.045081483 | 0.978751917 | 0.999157637 | ns | TM | KS.3 |
| FIM1_505 | 772.6857802 | 987.6703925 | 1053.730224 | 0.054534004 | 0.102444188 | 0.078428925 | 0.615432024 | ns | TM | KS.3 |
| FIM1_506 | 2300.064377 | 2703.063269 | 3207.087225 | 0.018728468 | 0.083971309 | 0.40295894 | 0.950410534 | ns | TM | KS.3 |
| FIM1_507 | 1584.03128 | 1786.272539 | 1545.803788 | 0.03364241 | 0.072621518 | 0.217976487 | 0.865538807 | ns | TM | KS.3 |
| FIM1_508 | 853.8946509 | 881.6238657 | 1038.605976 | 0.007068338 | 0.058921816 | 0.757632254 | 0.988477371 | ns | TM | KS.3 |
| FIM1_509 | 732.286933 | 562.5432993 | 964.2370488 | -0.046046676 | 0.102899153 | 0.105403512 | 0.698245636 | ns | TM | KS.3 |
| FIM1_510 | 825.9829056 | 888.7957616 | 854.0455006 | 0.017949565 | 0.062781746 | 0.458115327 | 0.950410534 | ns | TM | KS.3 |
| FIM1_511 | 1577.913226 | 1513.934016 | 1599.389843 | -0.006210127 | 0.070119442 | 0.782100502 | 0.988814136 | ns | TM | KS.3 |
| FIM1_512 | 325.0526619 | 201.3568922 | 324.3843601 | -0.227920615 | 0.27029993 | 0.00256075 | 0.093517802 | ns | TM | KS.3 |
| FIM1_513 | 3073.724296 | 3071.957775 | 3189.871274 | -0.00014283 | 0.053855241 | 0.994854746 | 0.999525119 | ns | TM | KS.3 |
| FIM1_514 | 2079.943447 | 1816.267643 | 1894.917338 | -0.027488807 | 0.076284719 | 0.284194115 | 0.918450395 | ns | TM | KS.3 |
| FIM1_515 | 1851.927577 | 1746.32347 | 1777.12397 | -0.023533872 | 0.049319577 | 0.275502095 | 0.908583046 | ns | TM | KS.3 |
| FIM1_516 | 1411.591763 | 1319.87994 | 1352.961096 | -0.020356314 | 0.057430713 | 0.391928469 | 0.948637087 | ns | TM | KS.3 |
| FIM1_517 | 1025.251942 | 1199.78699 | 1368.583645 | 0.016668568 | 0.084638963 | 0.44423522 | 0.950410534 | ns | TM | KS.3 |
| FIM1_518 | 23762.13269 | 24363.33976 | 24314.06549 | 0.008147258 | 0.039983797 | 0.654978834 | 0.976323032 | ns | TM | KS.3 |
| FIM1_519 | 177.1989477 | 206.4540479 | 587.3415799 | 0.010550459 | 0.091683068 | 0.587798132 | 0.963781856 | ns | TM | KS.3 |
| FIM1_522 | 2029.189594 | 2358.049191 | 2603.129516 | 0.018352474 | 0.082174821 | 0.416222896 | 0.950410534 | ns | TM | KS.3 |

| | | | | | | | | | | |
|---|---|---|---|---|---|---|---|---|---|---|
| FIM1_523 | 678.0770302 | 724.60455 | 636.3143509 | 0.011280063 | 0.068924943 | 0.626018577 | 0.97344368 | ns | TM | KS.3 |
| FIM1_524 | 2094.142424 | 1646.198045 | 1738.364328 | -0.143035089 | 0.118354551 | 0.005479897 | 0.152304407 | ns | TM | KS.3 |
| FIM1_525 | 1745.64255 | 1932.373413 | 1775.306378 | 0.061019812 | 0.070009379 | 0.058510753 | 0.540287513 | ns | TM | KS.3 |
| FIM1_526 | 76.90325438 | 130.9837371 | 136.6789371 | 0.067446126 | 0.177429241 | 0.043421563 | 0.476870369 | ns | TM | KS.3 |
| FIM1_527 | 5610.933337 | 6557.514136 | 14481.84218 | 0.01215659 | 0.089413233 | 0.544911201 | 0.95917717 | ns | TM | KS.3 |
| FIM1_528 | 2894.012718 | 2956.613382 | 3607.913838 | 0.001824328 | 0.082022143 | 0.927195442 | 0.993697171 | ns | TM | KS.3 |
| FIM1_529 | 15.03542523 | 15.96015389 | 26.20148965 | 0.005854095 | 0.087647925 | 0.761474819 | 0.988477371 | ns | TM | KS.3 |
| FIM1_530 | 3240.085271 | 3438.108145 | 3718.552015 | 0.011807719 | 0.064931441 | 0.614660777 | 0.973122675 | ns | TM | KS.3 |
| FIM1_531 | 10309.98813 | 8950.412551 | 10265.85102 | -0.014569183 | 0.083625752 | 0.497232955 | 0.953260948 | ns | TM | KS.3 |
| FIM1_532 | 2092.614624 | 2940.089478 | 4169.976391 | 0.025715487 | 0.110882009 | 0.240013859 | 0.887413624 | ns | TM | KS.3 |
| FIM1_533 | 953.1233563 | 949.9188591 | 1149.200053 | -0.000784282 | 0.049381505 | 0.970586174 | 0.997052226 | ns | TM | KS.3 |
| FIM1_534 | 1069.167455 | 985.6778051 | 1090.027217 | -0.021360639 | 0.063443494 | 0.387256103 | 0.948637087 | ns | TM | KS.3 |
| FIM1_535 | 1519.461057 | 1427.762318 | 1542.118228 | -0.004675582 | 0.085784398 | 0.810152296 | 0.988916276 | ns | TM | KS.3 |
| FIM1_536 | 406.0045271 | 862.0463126 | 1058.344059 | 0.609138569 | 0.47065385 | 0.000358401 | 0.022282532 | ns | TM | KS.3 |
| FIM1_537 | 823.8401062 | 860.4925655 | 995.1810894 | 0.00413505 | 0.080122458 | 0.839906002 | 0.991141351 | ns | TM | KS.3 |
| FIM1_538 | 1223.857447 | 1162.362184 | 1018.384598 | -0.010658833 | 0.063127236 | 0.647991958 | 0.976323032 | ns | TM | KS.3 |
| FIM1_539 | 450.2558995 | 457.7991299 | 440.214433 | 0.003466252 | 0.055500426 | 0.877136147 | 0.991141351 | ns | TM | KS.3 |
| FIM1_540 | 2548.92188 | 2332.64501 | 2288.724654 | -0.033439281 | 0.063037125 | 0.204213148 | 0.847612283 | ns | TM | KS.3 |
| FIM1_541 | 651.4516202 | 568.251458 | 552.7711887 | -0.097794953 | 0.082572285 | 0.017747698 | 0.304361209 | ns | TM | KS.3 |
| FIM1_542 | 850.2717766 | 895.699332 | 827.8097266 | 0.013407301 | 0.057057292 | 0.562242852 | 0.961073589 | ns | TM | KS.3 |
| FIM1_543 | 461.306207 | 466.5903508 | 548.75533 | 0.001683118 | 0.071352744 | 0.939079415 | 0.994413583 | ns | TM | KS.3 |
| FIM1_544 | 1484.215592 | 1714.968816 | 1915.488907 | 0.039925696 | 0.078960571 | 0.163576554 | 0.800496208 | ns | TM | KS.3 |
| FIM1_545 | 3680.703641 | 3535.640454 | 1877.281489 | -0.007532736 | 0.064656653 | 0.74372955 | 0.988477371 | ns | TM | KS.3 |
| FIM1_546 | 3365.037294 | 3128.122656 | 4994.01369 | -0.024843777 | 0.057946808 | 0.306369979 | 0.927241586 | ns | TM | KS.3 |
| FIM1_547 | 2077.859523 | 2001.988909 | 2150.754405 | -0.006993289 | 0.064346673 | 0.7611977 | 0.988477371 | ns | TM | KS.3 |
| FIM1_549 | 745.6242236 | 869.0121365 | 1034.980657 | 0.031753541 | 0.079572836 | 0.229995998 | 0.879350777 | ns | TM | KS.3 |
| FIM1_550 | 10390.44708 | 11322.84086 | 11820.81753 | 0.024208546 | 0.063729146 | 0.335624341 | 0.938982061 | ns | TM | KS.3 |
| FIM1_551 | 678.1682961 | 694.9960008 | 566.140344 | 0.004380671 | 0.06530243 | 0.847674135 | 0.991141351 | ns | TM | KS.3 |
| FIM1_552 | 1203.144382 | 1213.996562 | 1160.952802 | 0.002549585 | 0.050051649 | 0.905309606 | 0.991141351 | ns | TM | KS.3 |
| FIM1_553 | 1729.32288 | 1727.439357 | 1591.648014 | -0.000110643 | 0.043987255 | 0.995535522 | 0.999525119 | ns | TM | KS.3 |
| FIM1_554 | 2041.291984 | 2071.435895 | 1887.284174 | 0.003932469 | 0.046925082 | 0.848978677 | 0.991141351 | ns | TM | KS.3 |
| FIM1_555 | 630.8435151 | 685.0866595 | 575.5339994 | 0.019170025 | 0.065223551 | 0.43330738 | 0.950410534 | ns | TM | KS.3 |
| FIM1_556 | 1134.257408 | 1066.545026 | 1202.338386 | -0.015982319 | 0.059592087 | 0.499871087 | 0.953260948 | ns | TM | KS.3 |
| FIM1_557 | 818.2602653 | 839.830654 | 994.4919767 | 0.007096007 | 0.049216959 | 0.738919101 | 0.988051177 | ns | TM | KS.3 |
| FIM1_558 | 1372.726069 | 1281.545289 | 1298.567273 | -0.01478278 | 0.06438622 | 0.534996258 | 0.959071703 | ns | TM | KS.3 |
| FIM1_559 | 908.9148986 | 792.3225424 | 786.9371466 | -0.049138994 | 0.07873679 | 0.111837735 | 0.71059097 | ns | TM | KS.3 |
| FIM1_560 | 3413.597958 | 3166.543076 | 2200.518531 | -0.020222459 | 0.061504926 | 0.40677419 | 0.950410534 | ns | TM | KS.3 |
| FIM1_561 | 717.9086643 | 767.48188 | 945.6953352 | 0.015556182 | 0.062797448 | 0.514907897 | 0.954689943 | ns | TM | KS.3 |
| FIM1_562 | 1682.442926 | 1703.547522 | 1953.332671 | 0.003517566 | 0.046127316 | 0.86325147 | 0.991141351 | ns | TM | KS.3 |
| FIM1_563 | 201.3211746 | 160.1569647 | 220.2215981 | -0.029741606 | 0.092822863 | 0.222235478 | 0.870568114 | ns | TM | KS.3 |
| FIM1_564 | 1692.901328 | 1690.997486 | 1556.202246 | -0.000201754 | 0.066101381 | 0.992889868 | 0.999525119 | ns | TM | KS.3 |
| FIM1_565 | 1164.748824 | 1394.350156 | 1520.594745 | 0.033114008 | 0.084663024 | 0.207489067 | 0.852551729 | ns | TM | KS.3 |
| FIM1_566 | 799.09022 | 866.8383028 | 826.6143196 | 0.018117457 | 0.065653415 | 0.456097205 | 0.950410534 | ns | TM | KS.3 |
| FIM1_567 | 1669.649948 | 1740.389336 | 1648.299936 | 0.011160738 | 0.053938191 | 0.619421575 | 0.973122675 | ns | TM | KS.3 |
| FIM1_568 | 2008.386026 | 2751.667434 | 3256.95093 | 0.023139546 | 0.107109814 | 0.278058439 | 0.909494236 | ns | TM | KS.3 |

| | | | | | | | | | | |
|---|---|---|---|---|---|---|---|---|---|---|
| FIM1_569 | 2155.159956 | 2310.629326 | 2490.001199 | 0.006543419 | 0.081092319 | 0.749726437 | 0.988477371 | ns | TM | KS.3 |
| FIM1_570 | 889.8655753 | 1070.166026 | 1243.278647 | 0.036619253 | 0.08587371 | 0.177459421 | 0.811391014 | ns | TM | KS.3 |
| FIM1_571 | 1703.740952 | 2705.281135 | 2026.460833 | 0.062229395 | 0.157035988 | 0.050911674 | 0.514808085 | ns | TM | KS.3 |
| FIM1_572 | 532.7416794 | 998.9973296 | 510.8466022 | 0.074183177 | 0.204430344 | 0.036253425 | 0.432519877 | ns | TM | KS.3 |
| FIM1_573 | 3249.53496 | 3596.544202 | 4434.991654 | 0.027143983 | 0.068030607 | 0.294744268 | 0.92509331 | ns | TM | KS.3 |
| FIM1_574 | 2536.242786 | 3027.964001 | 3334.420775 | 0.015212371 | 0.089454967 | 0.463674168 | 0.950410534 | ns | TM | KS.3 |
| FIM1_575 | 2559.005457 | 2505.752149 | 2703.430966 | -0.005553883 | 0.051531546 | 0.798872184 | 0.988916276 | ns | TM | KS.3 |
| FIM1_576 | 5367.309442 | 5180.728694 | 5823.24603 | -0.011065051 | 0.04638723 | 0.589462454 | 0.963781856 | ns | TM | KS.3 |
| FIM1_577 | 885.6895838 | 839.7674422 | 905.1260683 | -0.012856668 | 0.05967222 | 0.582617168 | 0.963781856 | ns | TM | KS.3 |
| FIM1_578 | 5526.868347 | 5189.849545 | 4422.84059 | -0.016335785 | 0.059449198 | 0.49079061 | 0.952591289 | ns | TM | KS.3 |
| FIM1_579 | 917.2608618 | 871.3923904 | 850.9124711 | -0.013057953 | 0.05773281 | 0.573554559 | 0.963192994 | ns | TM | KS.3 |
| FIM1_580 | 249.0671984 | 279.3798878 | 260.9150765 | 0.029465311 | 0.070605326 | 0.264552249 | 0.90372202 | ns | TM | KS.3 |
| FIM1_581 | 719.141438 | 734.2146886 | 835.7897162 | 0.004249724 | 0.062680533 | 0.852693777 | 0.991141351 | ns | TM | KS.3 |
| FIM1_582 | 2269.136757 | 2902.153078 | 3226.125734 | 0.021665975 | 0.096623508 | 0.31737963 | 0.92825612 | ns | TM | KS.3 |
| FIM1_583 | 216.0951627 | 294.9654273 | 256.7458051 | 0.07057416 | 0.124979527 | 0.041161775 | 0.462993409 | ns | TM | KS.3 |
| FIM1_584 | 1022.079672 | 1054.709124 | 916.6185616 | 0.007966876 | 0.054303547 | 0.721978358 | 0.987157164 | ns | TM | KS.3 |
| FIM1_585 | 10076.51384 | 9955.921277 | 10000.39067 | -0.00199695 | 0.066724927 | 0.929628253 | 0.993993515 | ns | TM | KS.3 |
| FIM1_586 | 1354.009679 | 1568.633428 | 2018.87692 | 0.024680109 | 0.079078021 | 0.316308047 | 0.927564309 | ns | TM | KS.3 |
| FIM1_587 | 677.2741466 | 735.7842192 | 743.9692453 | 0.013189013 | 0.071600736 | 0.56849188 | 0.962265923 | ns | TM | KS.3 |
| FIM1_588 | 629.2441459 | 701.5148222 | 685.0246758 | 0.022779827 | 0.071158587 | 0.362061992 | 0.948637087 | ns | TM | KS.3 |
| FIM1_589 | 917.9124405 | 878.723235 | 965.9544655 | -0.012361284 | 0.052545021 | 0.578272752 | 0.963781856 | ns | TM | KS.3 |
| FIM1_590 | 793.6773286 | 735.9854856 | 896.1242943 | -0.025413281 | 0.059061401 | 0.301274032 | 0.926536521 | ns | TM | KS.3 |
| FIM1_591 | 617.6290334 | 611.4874892 | 799.1978164 | -0.002292535 | 0.054494174 | 0.917974377 | 0.991141351 | ns | TM | KS.3 |
| FIM1_592 | 6005.546424 | 6053.004126 | 5793.999017 | 0.001806486 | 0.05689716 | 0.936157282 | 0.994413583 | ns | TM | KS.3 |
| FIM1_593 | 1150.822939 | 1241.56885 | 1160.875021 | 0.015263069 | 0.0666885 | 0.522374809 | 0.958602567 | ns | TM | KS.3 |
| FIM1_594 | 1842.272861 | 1781.422503 | 2091.221904 | -0.005804592 | 0.066277289 | 0.799257501 | 0.988916276 | ns | TM | KS.3 |
| FIM1_595 | 2284.35243 | 2097.26828 | 2258.61927 | -0.018360288 | 0.067078194 | 0.450573031 | 0.950410534 | ns | TM | KS.3 |
| FIM1_596 | 4635.367802 | 4211.036424 | 3810.382358 | -0.020548275 | 0.06887534 | 0.404809772 | 0.950410534 | ns | TM | KS.3 |
| FIM1_597 | 1205.099721 | 1338.100979 | 1847.660826 | 0.034157327 | 0.068496864 | 0.209637486 | 0.852551729 | ns | TM | KS.3 |
| FIM1_598 | 992.1827851 | 997.09616 | 942.4573859 | 0.001065623 | 0.058690821 | 0.962549047 | 0.995175461 | ns | TM | KS.3 |
| FIM1_599 | 642.6031339 | 620.7332647 | 612.5166413 | -0.009720934 | 0.051983941 | 0.658737822 | 0.976323032 | ns | TM | KS.3 |
| FIM1_600 | 2346.785529 | 1753.310232 | 1731.684023 | -0.087338163 | 0.126700248 | 0.025241614 | 0.375979132 | ns | TM | KS.3 |
| FIM1_601 | 2030.141579 | 2434.626181 | 2786.680377 | 0.016782901 | 0.088829336 | 0.429042299 | 0.950410534 | ns | TM | KS.3 |
| FIM1_602 | 2021.32265 | 1882.289158 | 2163.783996 | -0.028361969 | 0.055539801 | 0.236051354 | 0.884880231 | ns | TM | KS.3 |
| FIM1_603 | 8099.77067 | 5519.133923 | 6877.477411 | -0.413314411 | 0.184327948 | 2.73974708006 | 0.003097782 | ns | TM | KS.3 |
| FIM1_604 | 2505.119399 | 2142.859995 | 2260.077787 | -0.057307328 | 0.085228562 | 0.081253576 | 0.619994692 | ns | TM | KS.3 |
| FIM1_605 | 720.6104749 | 613.1049995 | 511.9090931 | -0.034921019 | 0.081715276 | 0.197640701 | 0.838676184 | ns | TM | KS.3 |
| FIM1_606 | 1880.316739 | 1747.203444 | 1345.168045 | -0.01606274 | 0.064753266 | 0.503371115 | 0.953260948 | ns | TM | KS.3 |
| FIM1_607 | 510.3100876 | 572.8269261 | 715.6332677 | 0.012411012 | 0.081012189 | 0.56292833 | 0.961073589 | ns | TM | KS.3 |
| FIM1_608 | 545.1407248 | 636.8570018 | 591.9919784 | 0.047253149 | 0.082321051 | 0.12010338 | 0.730457601 | ns | TM | KS.3 |
| FIM1_609 | 5731.479464 | 6039.913856 | 7419.754934 | 0.007373911 | 0.072074983 | 0.740629965 | 0.988051177 | ns | TM | KS.3 |
| FIM1_610 | 641.1296207 | 591.7950592 | 549.4681379 | -0.022838228 | 0.061329695 | 0.354794855 | 0.948637087 | ns | TM | KS.3 |
| FIM1_611 | 164.4598141 | 150.4649371 | 193.9310292 | -0.014138902 | 0.0718395 | 0.543205025 | 0.95917717 | ns | TM | KS.3 |
| FIM1_612 | 1588.863546 | 1491.225575 | 1807.422097 | -0.025020123 | 0.052544059 | 0.272094319 | 0.908468501 | ns | TM | KS.3 |
| FIM1_613 | 1363.426339 | 1343.526307 | 1297.621499 | -0.004446661 | 0.044514639 | 0.823459383 | 0.991141351 | ns | TM | KS.3 |

| | | | | | | | | | | |
|---|---|---|---|---|---|---|---|---|---|---|
| FIM1_615 | 872.0919195 | 802.3354128 | 988.5036954 | -0.022758899 | 0.062935814 | 0.359905986 | 0.948637087 | ns | TM | KS.3 |
| FIM1_616 | 2810.502222 | 2406.033932 | 2442.610828 | -0.068241066 | 0.087327006 | 0.054519426 | 0.531428718 | ns | TM | KS.3 |
| FIM1_617 | 409.0627316 | 435.4591305 | 457.5838486 | 0.004971647 | 0.084335448 | 0.801172885 | 0.988916276 | ns | TM | KS.3 |
| FIM1_618 | 975.5920864 | 1070.764362 | 1419.095574 | 0.012570013 | 0.075384742 | 0.575792015 | 0.963192994 | ns | TM | KS.3 |
| FIM1_619 | 10102.86766 | 10998.2416 | 13755.79097 | 0.017759192 | 0.067386592 | 0.463561476 | 0.950410534 | ns | TM | KS.3 |
| FIM1_620 | 1077.128794 | 1029.241286 | 905.0081186 | -0.004523421 | 0.079176649 | 0.826924164 | 0.991141351 | ns | TM | KS.3 |
| FIM1_621 | 1011.426927 | 1155.893513 | 628.3279031 | 0.014259352 | 0.082454701 | 0.508868922 | 0.953260948 | ns | TM | KS.3 |
| FIM1_622 | 437.5932298 | 438.0049444 | 308.1685685 | 4.0949672193582є | 0.053458799 | 0.998520571 | 0.999525119 | ns | TM | KS.3 |
| FIM1_623 | 3042.735841 | 2961.161892 | 3212.491914 | -0.007241968 | 0.051642095 | 0.740430348 | 0.988051177 | ns | TM | KS.3 |
| FIM1_625 | 215.3947529 | 218.4612143 | 1665.134305 | 0.001379969 | 0.080754405 | 0.945540249 | 0.994413583 | ns | TM | KS.3 |
| FIM1_626 | 3028.820382 | 3348.244661 | 3460.275943 | 0.013763471 | 0.075741952 | 0.542909345 | 0.95917717 | ns | TM | KS.3 |
| FIM1_627 | 950.3299809 | 872.0727021 | 1097.322672 | -0.021483339 | 0.064274093 | 0.38596015 | 0.948637087 | ns | TM | KS.3 |
| FIM1_628 | 3932.657642 | 3307.52408 | 2869.052745 | -0.036246817 | 0.08380781 | 0.183719108 | 0.821657502 | ns | TM | KS.3 |
| FIM1_629 | 2241.219964 | 2155.556788 | 2629.388905 | -0.00645968 | 0.067693062 | 0.776489806 | 0.988814136 | ns | TM | KS.3 |
| FIM1_630 | 846.8733424 | 869.6461316 | 786.9538102 | 0.005941044 | 0.057879241 | 0.794100978 | 0.988916276 | ns | TM | KS.3 |
| FIM1_631 | 1756.751817 | 1627.773471 | 2014.126863 | -0.008038629 | 0.079007313 | 0.70323565 | 0.98151979 | ns | TM | KS.3 |
| FIM1_632 | 340.0178797 | 264.1335806 | 338.091691 | -0.054038345 | 0.103887134 | 0.078820591 | 0.616697988 | ns | TM | KS.3 |
| FIM1_633 | 5950.408901 | 5546.892572 | 5665.138552 | -0.022641311 | 0.057470679 | 0.345202067 | 0.946736649 | ns | TM | KS.3 |
| FIM1_634 | 477.6259789 | 468.5489361 | 504.229274 | -0.005202203 | 0.051533874 | 0.811302018 | 0.988916276 | ns | TM | KS.3 |
| FIM1_635 | 3916.94156 | 3801.728471 | 3749.836985 | -0.008329897 | 0.050137882 | 0.699012232 | 0.98151979 | ns | TM | KS.3 |
| FIM1_636 | 413.1806677 | 497.0751686 | 505.1129773 | 0.115102085 | 0.098431376 | 0.011753439 | 0.245686378 | ns | TM | KS.3 |
| FIM1_637 | 8978.539753 | 8586.201668 | 8546.879484 | -0.011681748 | 0.055573256 | 0.608232567 | 0.970937835 | ns | TM | KS.3 |
| FIM1_638 | 1475.251536 | 1556.363691 | 1509.720033 | 0.015614405 | 0.053904368 | 0.491548433 | 0.952591289 | ns | TM | KS.3 |
| FIM1_639 | 46.81136776 | 74.13874779 | 79.66084673 | 0.079199611 | 0.168784471 | 0.030892689 | 0.403388782 | ns | TM | KS.3 |
| FIM1_640 | 1155.343473 | 1104.477496 | 1173.532448 | -0.015058231 | 0.046004055 | 0.459090558 | 0.950410534 | ns | TM | KS.3 |
| FIM1_641 | 491.2491731 | 550.5897984 | 514.1290684 | 0.026525785 | 0.071432971 | 0.303414356 | 0.926536521 | ns | TM | KS.3 |
| FIM1_642 | 3145.67477 | 3214.909867 | 3393.308776 | 0.002933719 | 0.072013404 | 0.893710253 | 0.991141351 | ns | TM | KS.3 |
| FIM1_644 | 1578.904575 | 1784.178779 | 1767.328646 | 0.01832699 | 0.076698423 | 0.433792566 | 0.950410534 | ns | TM | KS.3 |
| FIM1_645 | 655.7246092 | 970.1028909 | 1058.851261 | 0.132394977 | 0.18446673 | 0.008766651 | 0.202237047 | ns | TM | KS.3 |
| FIM1_646 | 1362.656542 | 2508.004839 | 2889.920807 | 0.846294319 | 0.142387544 | 7.5321910665 | 2.6766178968 | ns | TM | KS.3 |
| FIM1_647 | 1255.60044 | 1446.006924 | 1610.557098 | 0.02455602 | 0.077917255 | 0.321033786 | 0.929652552 | ns | TM | KS.3 |
| FIM1_648 | 189.683562 | 158.3587358 | 180.6262444 | -0.056588876 | 0.090635711 | 0.081115146 | 0.619889174 | ns | TM | KS.3 |
| FIM1_650 | 686.8842866 | 553.06652 | 655.9854887 | -0.089984972 | 0.106383449 | 0.025557019 | 0.378411211 | ns | TM | KS.3 |
| FIM1_651 | 1030.143022 | 871.1214392 | 1317.559511 | -0.021302282 | 0.083787142 | 0.356058152 | 0.948637087 | ns | TM | KS.3 |
| FIM1_652 | 2569.198098 | 2311.96738 | 2526.977574 | -0.043893537 | 0.069657304 | 0.131550876 | 0.761283488 | ns | TM | KS.3 |
| FIM1_653 | 2660.534513 | 3173.826408 | 3738.84694 | 0.016494201 | 0.088152706 | 0.437429188 | 0.950410534 | ns | TM | KS.3 |
| FIM1_654 | 648.950602 | 635.85543 | 618.8363942 | -0.005362531 | 0.0533254 | 0.808580022 | 0.988916276 | ns | TM | KS.3 |
| FIM1_655 | 501.5222619 | 457.8357619 | 491.6011962 | -0.019173232 | 0.068915324 | 0.432212015 | 0.950410534 | ns | TM | KS.3 |
| FIM1_656 | 1807.137826 | 1779.777228 | 1796.04403 | -0.003797298 | 0.053962655 | 0.864215713 | 0.991141351 | ns | TM | KS.3 |
| FIM1_657 | 1035.456429 | 859.2209875 | 840.9049317 | -0.047009138 | 0.088649742 | 0.115931854 | 0.718257753 | ns | TM | KS.3 |
| FIM1_658 | 802.2146644 | 675.6721015 | 678.8625973 | -0.036875373 | 0.083562935 | 0.179472683 | 0.813196759 | ns | TM | KS.3 |
| FIM1_659 | 3000.881624 | 2801.0304 | 2457.652158 | -0.020005939 | 0.058656408 | 0.403829097 | 0.950410534 | ns | TM | KS.3 |
| FIM1_660 | 890.1678813 | 866.3344081 | 935.7586285 | -0.008033006 | 0.047006051 | 0.697873429 | 0.98151979 | ns | TM | KS.3 |
| FIM1_661 | 770.9353493 | 711.8837965 | 869.1390861 | -0.012957499 | 0.070672648 | 0.576533645 | 0.963382753 | ns | TM | KS.3 |
| FIM1_662 | 11354.01102 | 12475.58426 | 13329.56743 | 0.018887776 | 0.069416102 | 0.437607896 | 0.950410534 | ns | TM | KS.3 |

| | | | | | | | | | | |
|---|---|---|---|---|---|---|---|---|---|---|
| FIM1_663 | 609.818095 | 542.7692442 | 728.0410642 | -0.009062696 | 0.087123732 | 0.647568877 | 0.976323032 | ns | TM | KS.3 |
| FIM1_664 | 2748.378251 | 2937.285529 | 3029.072366 | 0.015827608 | 0.061868198 | 0.507303177 | 0.953260948 | ns | TM | KS.3 |
| FIM1_665 | 836.9009368 | 659.9767545 | 624.5284246 | -0.181538699 | 0.111456766 | 0.001601477 | 0.067706867 | ns | TM | KS.3 |
| FIM1_666 | 225.7815456 | 228.2770962 | 221.6060011 | 0.001373393 | 0.07614546 | 0.948193363 | 0.994413583 | ns | TM | KS.3 |
| FIM1_667 | 268.9826538 | 235.1009121 | 255.920656 | -0.022373505 | 0.077324695 | 0.356748633 | 0.948637087 | ns | TM | KS.3 |
| FIM1_668 | 3055.531266 | 3502.834465 | 3917.240784 | 0.042882644 | 0.077726083 | 0.145036411 | 0.778377716 | ns | TM | KS.3 |
| FIM1_669 | 1100.863548 | 1815.206045 | 1242.978932 | 0.162065023 | 0.247887037 | 0.006486584 | 0.170744739 | ns | TM | KS.3 |
| FIM1_670 | 1495.480089 | 2299.356337 | 2971.716088 | 0.035909194 | 0.129054691 | 0.140030035 | 0.775302706 | ns | TM | KS.3 |
| FIM1_671 | 571.600214 | 606.1276331 | 622.0532644 | 0.014829413 | 0.059472161 | 0.529082421 | 0.959071703 | ns | TM | KS.3 |
| FIM1_672 | 696.3226499 | 625.4730384 | 662.1761727 | -0.024004201 | 0.070166084 | 0.342575568 | 0.944808611 | ns | TM | KS.3 |
| FIM1_673 | 184.6051635 | 162.9054939 | 349.8445058 | -0.011297776 | 0.084497568 | 0.584193904 | 0.963781856 | ns | TM | KS.3 |
| FIM1_674 | 5755.640703 | 4832.931834 | 5215.570752 | -0.058287425 | 0.089436644 | 0.077080249 | 0.609656977 | ns | TM | KS.3 |
| FIM1_675 | 1749.265142 | 1554.691751 | 1473.001452 | -0.039283496 | 0.072697973 | 0.168361549 | 0.805573568 | ns | TM | KS.3 |
| FIM1_676 | 4736.820756 | 4455.136606 | 3264.916752 | -0.007710085 | 0.074800389 | 0.724219818 | 0.987157164 | ns | TM | KS.3 |
| FIM1_677 | 1431.125657 | 1268.052611 | 1919.82398 | -0.027380947 | 0.073130006 | 0.290168131 | 0.919993167 | ns | TM | KS.3 |
| FIM1_678 | 2346.951856 | 2226.041191 | 2766.754028 | -0.011037689 | 0.062965735 | 0.636891376 | 0.974658588 | ns | TM | KS.3 |
| FIM1_679 | 3282.788443 | 2414.680932 | 2876.776366 | -0.047107258 | 0.112004883 | 0.095425084 | 0.673713046 | ns | TM | KS.3 |
| FIM1_680 | 4851.442261 | 6386.762088 | 5497.70073 | 0.03505751 | 0.101457954 | 0.164622787 | 0.800584912 | ns | TM | KS.3 |
| FIM1_681 | 1347.076386 | 1418.387295 | 2011.579786 | 0.004096769 | 0.084212671 | 0.835150966 | 0.991141351 | ns | TM | KS.3 |
| FIM1_682 | 1030.663872 | 1144.558756 | 1325.94449 | 0.009924052 | 0.082565455 | 0.632085543 | 0.97344368 | ns | TM | KS.3 |
| FIM1_683 | 2464.089994 | 2966.576986 | 3487.012945 | 0.077410063 | 0.096292206 | 0.039354196 | 0.450085339 | ns | TM | KS.3 |
| FIM1_684 | 1436.503361 | 1513.976858 | 1704.721099 | 0.016694173 | 0.052762483 | 0.45772153 | 0.950410534 | ns | TM | KS.3 |
| FIM1_685 | 623.7387455 | 492.9389107 | 571.6491387 | -0.085145627 | 0.110424787 | 0.028838473 | 0.390786333 | ns | TM | KS.3 |
| FIM1_686 | 3904.072761 | 3637.952261 | 3809.941275 | -0.033252598 | 0.055082438 | 0.169077506 | 0.806481872 | ns | TM | KS.3 |
| FIM1_687 | 1066.791172 | 928.0518081 | 1055.214549 | -0.049226444 | 0.079732517 | 0.111587478 | 0.71059097 | ns | TM | KS.3 |
| FIM1_688 | 1387.229135 | 1295.162169 | 1036.037415 | -0.021833218 | 0.057035798 | 0.359416425 | 0.948637087 | ns | TM | KS.3 |
| FIM1_689 | 119.648666 | 121.4994295 | 143.7030848 | 0.001215842 | 0.077898323 | 0.953338663 | 0.994413583 | ns | TM | KS.3 |
| FIM1_690 | 1150.366141 | 1264.625816 | 1524.061999 | 0.030330733 | 0.065563298 | 0.248365132 | 0.888934767 | ns | TM | KS.3 |
| FIM1_691 | 1600.685652 | 1626.807675 | 1611.399486 | 0.003819643 | 0.055449101 | 0.864733314 | 0.991141351 | ns | TM | KS.3 |
| FIM1_692 | 1206.220876 | 1327.973954 | 1227.666889 | 0.024419586 | 0.067194155 | 0.336080345 | 0.939325685 | ns | TM | KS.3 |
| FIM1_693 | 1285.163716 | 1155.657745 | 1255.218501 | -0.041225377 | 0.069604396 | 0.149774998 | 0.783447458 | ns | TM | KS.3 |
| FIM1_694 | 628.4695389 | 560.2094658 | 543.4684132 | -0.032930576 | 0.071193855 | 0.224956162 | 0.875024944 | ns | TM | KS.3 |
| FIM1_695 | 971.3120619 | 918.4673475 | 1000.654697 | -0.018639519 | 0.050227635 | 0.392790337 | 0.948637087 | ns | TM | KS.3 |
| FIM1_696 | 1523.129993 | 1411.308021 | 1557.699312 | -0.022553219 | 0.060809667 | 0.358749182 | 0.948637087 | ns | TM | KS.3 |
| FIM1_697 | 827.8765203 | 824.2466283 | 898.8205216 | -0.001264062 | 0.051754896 | 0.953723571 | 0.994413583 | ns | TM | KS.3 |
| FIM1_698 | 531.8880827 | 716.5867672 | 812.8636411 | 0.163020467 | 0.148283173 | 0.003736751 | 0.115467924 | ns | TM | KS.3 |
| FIM1_699 | 2733.164478 | 2788.879166 | 2938.740619 | 0.002632444 | 0.072779069 | 0.903947426 | 0.991141351 | ns | TM | KS.3 |
| FIM1_700 | 6796.965531 | 6380.797816 | 6849.827476 | -0.024394056 | 0.052483123 | 0.283252856 | 0.918450395 | ns | TM | KS.3 |
| FIM1_701 | 1502.665253 | 1812.924802 | 1751.349444 | 0.030024768 | 0.085800967 | 0.234246954 | 0.883800031 | ns | TM | KS.3 |
| FIM1_702 | 1205.444923 | 1124.236235 | 1186.573425 | -0.019357552 | 0.060041841 | 0.421878301 | 0.950410534 | ns | TM | KS.3 |
| FIM1_703 | 2649.826439 | 2581.06457 | 2874.929683 | -0.00561468 | 0.059951701 | 0.806262296 | 0.988916276 | ns | TM | KS.3 |
| FIM1_704 | 3877.591219 | 5235.165198 | 6127.943018 | 0.046913514 | 0.110513571 | 0.096930998 | 0.676341814 | ns | TM | KS.3 |
| FIM1_705 | 1039.837905 | 1079.598932 | 1247.382072 | 0.003904438 | 0.077989623 | 0.851669582 | 0.991141351 | ns | TM | KS.3 |
| FIM1_706 | 2673.68013 | 2084.393139 | 2196.0927 | -0.163948519 | 0.121084809 | 0.003053931 | 0.101288726 | ns | TM | KS.3 |
| FIM1_707 | 908.7649768 | 881.4309841 | 994.1467093 | -0.009595781 | 0.046723711 | 0.641559973 | 0.975935248 | ns | TM | KS.3 |

| | | | | | | | | | | |
|---|---|---|---|---|---|---|---|---|---|---|
| FIM1_708 | 685.5681536 | 666.4133583 | 661.6962084 | -0.006760662 | 0.055934625 | 0.764658986 | 0.988477371 | ns | TM | KS.3 |
| FIM1_709 | 1404.561191 | 1275.720041 | 1294.26645 | -0.032540544 | 0.06602975 | 0.222695337 | 0.871073186 | ns | TM | KS.3 |
| FIM1_710 | 762.4964339 | 696.6961224 | 649.6720697 | -0.023412274 | 0.065938826 | 0.35228168 | 0.948637087 | ns | TM | KS.3 |
| FIM1_711 | 489.7334064 | 550.1664407 | 444.1594199 | 0.021808502 | 0.073468045 | 0.375303851 | 0.948637087 | ns | TM | KS.3 |
| FIM1_712 | 9.501228528 | 33.39130948 | 21.06410889 | 0.480035684 | 0.631312316 | 0.000903591 | 0.044072215 | ns | TM | KS.3 |
| FIM1_713 | 1511.456629 | 1434.175069 | 1449.725859 | -0.004836657 | 0.080950514 | 0.812256787 | 0.988916276 | ns | TM | KS.3 |
| FIM1_714 | 716.1489631 | 650.2822959 | 955.3241447 | -0.021176097 | 0.069186611 | 0.392636647 | 0.948637087 | ns | TM | KS.3 |
| FIM1_715 | 2462.73502 | 2289.242145 | 1366.728225 | -0.009642956 | 0.074469836 | 0.663430996 | 0.976323032 | ns | TM | KS.3 |
| FIM1_718 | 5554.487271 | 5051.657643 | 4056.178042 | -0.018977142 | 0.069556283 | 0.435594581 | 0.950410534 | ns | TM | KS.3 |
| FIM1_719 | 7716.968101 | 7672.373175 | 7126.365091 | -0.001292269 | 0.057399316 | 0.954392055 | 0.994413583 | ns | TM | KS.3 |
| FIM1_720 | 488.6544252 | 511.912137 | 588.1164666 | 0.010407282 | 0.060769996 | 0.654484565 | 0.976323032 | ns | TM | KS.3 |
| FIM1_721 | 1997.320762 | 2133.985484 | 2180.455325 | 0.016348894 | 0.06114723 | 0.493559097 | 0.952949531 | ns | TM | KS.3 |
| FIM1_722 | 4269.779376 | 4636.412325 | 4671.506458 | 0.033891923 | 0.060571708 | 0.189797799 | 0.828284256 | ns | TM | KS.3 |
| FIM1_723 | 640.2986021 | 732.0988026 | 877.597854 | 0.042971818 | 0.077204979 | 0.144518991 | 0.77783242 | ns | TM | KS.3 |
| FIM1_724 | 330.0175997 | 375.3980484 | 388.5643602 | 0.020494463 | 0.076841606 | 0.390995979 | 0.948637087 | ns | TM | KS.3 |
| FIM1_725 | 4491.453713 | 4847.582603 | 5829.750812 | 0.011342515 | 0.072426374 | 0.617740739 | 0.973122675 | ns | TM | KS.3 |
| FIM1_726 | 801.2083767 | 856.4708341 | 898.0738828 | 0.017337966 | 0.060129748 | 0.468005537 | 0.950410534 | ns | TM | KS.3 |
| FIM1_727 | 1463.447883 | 1357.631586 | 1229.378744 | -0.018208978 | 0.063265961 | 0.452805892 | 0.950410534 | ns | TM | KS.3 |
| FIM1_728 | 1989.125742 | 2005.200118 | 2356.280893 | 0.001452906 | 0.064220143 | 0.949111306 | 0.994413583 | ns | TM | KS.3 |
| FIM1_729 | 2262.134184 | 2425.658567 | 2429.537675 | 0.019094998 | 0.059893676 | 0.427275998 | 0.950410534 | ns | TM | KS.3 |
| FIM1_730 | 1884.551787 | 1925.265174 | 1867.201485 | 0.006242266 | 0.046748868 | 0.761988397 | 0.988477371 | ns | TM | KS.3 |
| FIM1_731 | 385.2069775 | 447.3731663 | 460.353051 | 0.064385768 | 0.085315941 | 0.062544016 | 0.551491823 | ns | TM | KS.3 |
| FIM1_732 | 9301.51769 | 9089.056098 | 8995.403339 | -0.005300815 | 0.057249602 | 0.815177512 | 0.988916276 | ns | TM | KS.3 |
| FIM1_733 | 1738.520396 | 1890.220298 | 2077.4532 | 0.010573535 | 0.075970222 | 0.631329184 | 0.97344368 | ns | TM | KS.3 |
| FIM1_734 | 2584.89641 | 2833.295279 | 2175.091979 | 0.068316578 | 0.064752195 | 0.035422963 | 0.42652037 | ns | TM | KS.3 |
| FIM1_735 | 557.4543031 | 721.167852 | 672.6776497 | 0.218717564 | 0.112947559 | 0.000480078 | 0.027140746 | ns | TM | KS.3 |
| FIM1_736 | 3499.559295 | 2588.82802 | 2714.657046 | -0.243642079 | 0.144255599 | 0.000451244 | 0.02580389 | ns | TM | KS.3 |
| FIM1_737 | 2260.258575 | 2357.977536 | 2322.058371 | 0.013905183 | 0.046316836 | 0.496846659 | 0.953260948 | ns | TM | KS.3 |
| FIM1_738 | 2247.054782 | 2126.791554 | 1834.221427 | -0.010627596 | 0.065167304 | 0.648519564 | 0.976323032 | ns | TM | KS.3 |
| FIM1_739 | 505.5888395 | 542.5369861 | 482.0774618 | 0.010450692 | 0.071641881 | 0.645414051 | 0.976323032 | ns | TM | KS.3 |
| FIM1_740 | 825.2428475 | 873.7828761 | 827.4917073 | 0.01349326 | 0.059975837 | 0.565518159 | 0.962265923 | ns | TM | KS.3 |
| FIM1_741 | 1922.01487 | 1740.069661 | 1896.080384 | -0.037421372 | 0.067303261 | 0.176013887 | 0.811391014 | ns | TM | KS.3 |
| FIM1_742 | 1827.081739 | 2053.821688 | 1995.452123 | 0.036016127 | 0.072094323 | 0.195276362 | 0.837467988 | ns | TM | KS.3 |
| FIM1_743 | 2146.135702 | 2108.434437 | 2245.619871 | -0.005305835 | 0.044795715 | 0.79096027 | 0.988916276 | ns | TM | KS.3 |
| FIM1_744 | 612.8730329 | 655.1586062 | 472.0112613 | 0.012988497 | 0.065960921 | 0.581583159 | 0.963781856 | ns | TM | KS.3 |
| FIM1_745 | 467.4313998 | 466.3834224 | 570.5548511 | -0.000315606 | 0.049815023 | 0.988216384 | 0.999333464 | ns | TM | KS.3 |
| FIM1_746 | 1197.940185 | 1031.706948 | 1985.127197 | -0.037800677 | 0.079584431 | 0.178098289 | 0.811391014 | ns | TM | KS.3 |
| FIM1_747 | 1598.815921 | 1514.321341 | 1178.845108 | -0.011351178 | 0.063058839 | 0.627836128 | 0.97344368 | ns | TM | KS.3 |
| FIM1_748 | 854.0337432 | 826.9285006 | 794.7275218 | -0.003809738 | 0.075052871 | 0.859062422 | 0.991141351 | ns | TM | KS.3 |
| FIM1_749 | 1221.045838 | 1170.002121 | 1109.475622 | -0.009764549 | 0.059242952 | 0.67255046 | 0.976861203 | ns | TM | KS.3 |
| FIM1_750 | 1526.305537 | 1345.989922 | 1422.150315 | -0.049994372 | 0.076205059 | 0.106631083 | 0.700394017 | ns | TM | KS.3 |
| FIM1_751 | 4144.000704 | 3880.982442 | 3565.154167 | -0.02710275 | 0.052824496 | 0.238212006 | 0.88653074 | ns | TM | KS.3 |
| FIM1_752 | 3286.973079 | 3918.551841 | 5517.589437 | 0.118904274 | 0.094807866 | 0.01008975 | 0.223094117 | ns | TM | KS.3 |
| FIM1_753 | 1323.623383 | 1493.554668 | 2091.90095 | 0.02927529 | 0.072932632 | 0.266141741 | 0.904370522 | ns | TM | KS.3 |
| FIM1_754 | 2023.901415 | 2013.828284 | 2116.637094 | -0.000833008 | 0.066216973 | 0.970642916 | 0.997052226 | ns | TM | KS.3 |

| | | | | | | | | | | |
|---|---|---|---|---|---|---|---|---|---|---|
| FIM1_755 | 3224.853601 | 2804.095578 | 2519.886863 | -0.136309761 | 0.071148114 | 0.002988405 | 0.101288726 | ns | TM | KS.3 |
| FIM1_756 | 893.8927653 | 880.0004575 | 822.4926349 | -0.004544206 | 0.052025282 | 0.835440439 | 0.991141351 | ns | TM | KS.3 |
| FIM1_757 | 2079.652513 | 2370.458678 | 2005.582828 | 0.04463882 | 0.076749121 | 0.134555187 | 0.766830871 | ns | TM | KS.3 |
| FIM1_758 | 299.7166343 | 363.3405968 | 378.9891539 | 0.085578365 | 0.098598258 | 0.030122382 | 0.400453344 | ns | TM | KS.3 |
| FIM1_759 | 705.4028327 | 724.4634913 | 738.316749 | 0.006907609 | 0.051845743 | 0.752381877 | 0.988477371 | ns | TM | KS.3 |
| FIM1_760 | 2512.387733 | 2417.785423 | 2381.124377 | -0.007896921 | 0.061917019 | 0.732410355 | 0.987157164 | ns | TM | KS.3 |
| FIM1_761 | 3544.363923 | 3406.430353 | 2800.818831 | -0.007366517 | 0.064870233 | 0.748965602 | 0.988477371 | ns | TM | KS.3 |
| FIM1_762 | 1570.096454 | 1330.024022 | 1122.594086 | -0.044156874 | 0.084126966 | 0.13398345 | 0.766169728 | ns | TM | KS.3 |
| FIM1_763 | 643.6092404 | 709.9218588 | 775.0109219 | 0.027404969 | 0.066664312 | 0.289767981 | 0.919993167 | ns | TM | KS.3 |
| FIM1_764 | 2343.689858 | 2159.202778 | 2454.664853 | -0.029081519 | 0.060844517 | 0.251340941 | 0.888934767 | ns | TM | KS.3 |
| FIM1_765 | 353.7545187 | 371.9951709 | 354.7709332 | 0.005916004 | 0.076325601 | 0.781961034 | 0.988814136 | ns | TM | KS.3 |
| FIM1_767 | 569.9136308 | 468.5648494 | 1201.766306 | -0.018749045 | 0.089869434 | 0.385185891 | 0.948637087 | ns | TM | KS.3 |
| FIM1_768 | 390.7468489 | 385.5271755 | 370.6882992 | -0.002554604 | 0.060484802 | 0.91081087 | 0.991141351 | ns | TM | KS.3 |
| FIM1_769 | 2656.990744 | 2343.602579 | 2602.665713 | -0.101292432 | 0.076695147 | 0.013672584 | 0.265707448 | ns | TM | KS.3 |
| FIM1_770 | 1126.178641 | 3023.787599 | 2444.516323 | 1.299443198 | 0.227481331 | 1.85483533182 | 6.15187051722 | Km>KS.3 | TM | KS.3 |
| FIM1_772 | 27.75911794 | 122.108497 | 53.38291118 | 1.683477479 | 0.370164122 | 1.34836301415 | 2.58004076746 | Km>KS.3 | TM | KS.3 |
| FIM1_773 | 1803.157417 | 2249.681387 | 2254.322242 | 0.04209144 | 0.093671476 | 0.132584207 | 0.761677364 | ns | TM | KS.3 |
| FIM1_774 | 829.383527 | 732.8729869 | 784.9324631 | -0.036534292 | 0.07357988 | 0.191224844 | 0.829253069 | ns | TM | KS.3 |
| FIM1_776 | 1810.99605 | 2272.76805 | 2078.601108 | 0.097810906 | 0.11086441 | 0.01997419 | 0.324535175 | ns | TM | KS.3 |
| FIM1_777 | 2461.085113 | 2597.734263 | 2381.458233 | 0.016006694 | 0.054147479 | 0.482204837 | 0.951594234 | ns | TM | KS.3 |
| FIM1_778 | 617.6732706 | 651.6828171 | 665.8322517 | 0.011212348 | 0.063230921 | 0.631863939 | 0.97344368 | ns | TM | KS.3 |
| FIM1_779 | 10131.83177 | 18282.99214 | 14674.89046 | 0.836136224 | 0.09788125 | 6.05815287478 | 1.00464368506 | ns | TM | KS.3 |
| FIM1_780 | 220.9317361 | 333.1220586 | 453.4928227 | 0.086778997 | 0.163836203 | 0.025009618 | 0.373642194 | ns | TM | KS.3 |
| FIM1_781 | 254.5522635 | 274.5021243 | 472.8245672 | 0.008950315 | 0.07641762 | 0.680215186 | 0.97986479 | ns | TM | KS.3 |
| FIM1_782 | 1862.10156 | 1279.538289 | 1671.123472 | -0.079573499 | 0.147894388 | 0.030340267 | 0.40080992 | ns | TM | KS.3 |
| FIM1_783 | 838.7227004 | 918.3876341 | 935.7181848 | 0.017670734 | 0.06926278 | 0.46382848 | 0.950410534 | ns | TM | KS.3 |
| FIM1_784 | 365.5672191 | 316.5373176 | 488.4529459 | -0.020503228 | 0.07971159 | 0.382531557 | 0.948637087 | ns | TM | KS.3 |
| FIM1_785 | 3231.049109 | 4201.351547 | 3305.208004 | 0.11242509 | 0.124825596 | 0.01286145 | 0.257037736 | ns | TM | KS.3 |
| FIM1_786 | 2214.784896 | 2370.151216 | 2337.434031 | 0.019681356 | 0.058269639 | 0.409612528 | 0.950410534 | ns | TM | KS.3 |
| FIM1_787 | 2572.711191 | 3467.702687 | 4819.528209 | 0.095928861 | 0.132179918 | 0.019717329 | 0.324535175 | ns | TM | KS.3 |
| FIM1_788 | 161.8263225 | 153.3453118 | 200.3153645 | -0.005583993 | 0.078625931 | 0.78925115 | 0.988916276 | ns | TM | KS.3 |
| FIM1_789 | 1280.507221 | 1473.028774 | 1008.009036 | 0.02360517 | 0.07792636 | 0.335393494 | 0.938982061 | ns | TM | KS.3 |
| FIM1_790 | 208.1259445 | 181.5026568 | 368.2901065 | -0.018677327 | 0.07905369 | 0.419549674 | 0.950410534 | ns | TM | KS.3 |
| FIM1_791 | 1231.440933 | 2095.533534 | 3356.780222 | 0.470384358 | 0.325423288 | 0.000270387 | 0.018427072 | ns | TM | KS.3 |
| FIM1_792 | 1146.942239 | 1110.078088 | 1088.768169 | -0.008840565 | 0.052239786 | 0.688266871 | 0.980879562 | ns | TM | KS.3 |
| FIM1_793 | 3145.678059 | 3173.513623 | 3888.038793 | 0.002233406 | 0.052219937 | 0.918692716 | 0.991141351 | ns | TM | KS.3 |
| FIM1_794 | 777.9051356 | 865.7541154 | 1027.485245 | 0.039675812 | 0.069719263 | 0.161731418 | 0.795859349 | ns | TM | KS.3 |
| FIM1_795 | 2637.383395 | 2815.84286 | 2824.487688 | 0.014800857 | 0.062946059 | 0.534039977 | 0.959071703 | ns | TM | KS.3 |
| FIM1_796 | 1419.74461 | 1574.842899 | 1281.943094 | 0.04418832 | 0.06906269 | 0.128493567 | 0.753101833 | ns | TM | KS.3 |
| FIM1_797 | 371.072416 | 450.6503469 | 336.0054251 | 0.033119919 | 0.086873394 | 0.203006922 | 0.847612283 | ns | TM | KS.3 |
| FIM1_798 | 717.540859 | 767.9894838 | 648.7373743 | 0.011966188 | 0.068003242 | 0.607828776 | 0.970937835 | ns | TM | KS.3 |
| FIM1_799 | 1555.773228 | 1139.581854 | 1734.643283 | -0.039300977 | 0.109469415 | 0.131124944 | 0.761283488 | ns | TM | KS.3 |
| FIM1_800 | 2864.539428 | 3171.880224 | 2792.052274 | 0.022661268 | 0.069506317 | 0.365588636 | 0.948637087 | ns | TM | KS.3 |
| FIM1_801 | 348.3288313 | 740.4917274 | 2056.706458 | 0.685937111 | 0.455118634 | 0.000216967 | 0.015734323 | ns | TM | KS.3 |
| FIM1_802 | 556.6200974 | 785.2213455 | 852.4081707 | 0.110576128 | 0.154543713 | 0.013484569 | 0.263081302 | ns | TM | KS.3 |

| | | | | | | | | | | |
|---|---|---|---|---|---|---|---|---|---|---|
| FIM1_803 | 1120.014762 | 1116.973695 | 1145.42991 | -0.000757678 | 0.043064278 | 0.968965067 | 0.996327965 | ns | TM | KS.3 |
| FIM1_804 | 3495.12552 | 4602.963636 | 5144.829841 | 0.059383813 | 0.110782771 | 0.062759806 | 0.551491823 | ns | TM | KS.3 |
| FIM1_805 | 821.6490747 | 784.4758259 | 1049.39095 | -0.010752328 | 0.05905043 | 0.642481784 | 0.976237372 | ns | TM | KS.3 |
| FIM1_806 | 973.9466551 | 779.2363097 | 1397.046938 | -0.028984931 | 0.091759674 | 0.232039348 | 0.880810476 | ns | TM | KS.3 |
| FIM1_807 | 225.7463514 | 244.6117346 | 604.1760257 | 0.014130005 | 0.06927158 | 0.54811835 | 0.95917717 | ns | TM | KS.3 |
| FIM1_808 | 1312.686243 | 1467.764849 | 1277.977105 | 0.026497982 | 0.071078858 | 0.304028143 | 0.926536521 | ns | TM | KS.3 |
| FIM1_809 | 36952.08719 | 33559.46611 | 20737.53121 | -0.028597909 | 0.066176301 | 0.2723577 | 0.908468501 | ns | TM | KS.3 |
| FIM1_810 | 1315.823798 | 1248.273405 | 1437.83775 | -0.016862377 | 0.05259065 | 0.452460464 | 0.950410534 | ns | TM | KS.3 |
| FIM1_811 | 2211.543227 | 2101.880716 | 2368.765038 | -0.009434957 | 0.065679418 | 0.683849891 | 0.98018358 | ns | TM | KS.3 |
| FIM1_812 | 2609.637428 | 2455.086699 | 2025.255696 | -0.014463444 | 0.061235147 | 0.541266622 | 0.95917717 | ns | TM | KS.3 |
| FIM1_813 | 560.5346691 | 610.1835808 | 733.4134806 | 0.026848861 | 0.062911502 | 0.29091892 | 0.919993167 | ns | TM | KS.3 |
| FIM1_814 | 40.02360445 | 27.95239331 | 59.6130047 | -0.034219724 | 0.116093901 | 0.157625995 | 0.789048157 | ns | TM | KS.3 |
| FIM1_815 | 56.17031693 | 66.57713958 | 102.8411246 | 0.015620019 | 0.087183193 | 0.460481973 | 0.950410534 | ns | TM | KS.3 |
| FIM1_816 | 273.5982569 | 329.2153793 | 391.056969 | 0.026522992 | 0.085320981 | 0.275075498 | 0.908583046 | ns | TM | KS.3 |
| FIM1_817 | 1077.683799 | 1207.583534 | 1487.348653 | 0.022483081 | 0.072751918 | 0.364886136 | 0.948637087 | ns | TM | KS.3 |
| FIM1_818 | 374.2054232 | 409.4956113 | 327.2956797 | 0.015592784 | 0.071332427 | 0.508242532 | 0.953260948 | ns | TM | KS.3 |
| FIM1_819 | 1064.220026 | 1343.751737 | 1264.18978 | 0.108176633 | 0.114154668 | 0.014719121 | 0.275291831 | ns | TM | KS.3 |
| FIM1_820 | 4998.105908 | 5089.963251 | 6602.112733 | 0.00392097 | 0.059170618 | 0.863359928 | 0.991141351 | ns | TM | KS.3 |
| FIM1_821 | 8637.483097 | 8179.501741 | 8074.419783 | -0.010742536 | 0.064594577 | 0.645376889 | 0.976323032 | ns | TM | KS.3 |
| FIM1_822 | 1108.793 | 1100.591016 | 963.4226905 | -0.001322561 | 0.06444282 | 0.953647287 | 0.994413583 | ns | TM | KS.3 |
| FIM1_823 | 1316.837365 | 1354.981951 | 1053.613319 | 0.008212727 | 0.048942533 | 0.698970018 | 0.98151979 | ns | TM | KS.3 |
| FIM1_824 | 3040.686371 | 3890.42595 | 4683.444448 | 0.021050538 | 0.096920254 | 0.32688265 | 0.934739762 | ns | TM | KS.3 |
| FIM1_825 | 4124.836219 | 3316.205227 | 3341.939333 | -0.126242781 | 0.110029594 | 0.008700092 | 0.202237047 | ns | TM | KS.3 |
| FIM1_826 | 95.52018265 | 92.82710579 | 94.20694577 | -0.002084291 | 0.083139425 | 0.916001435 | 0.991141351 | ns | TM | KS.3 |
| FIM1_828 | 1531.45105 | 1942.594265 | 1423.450385 | 0.0951625 | 0.113244008 | 0.021349721 | 0.338445603 | ns | TM | KS.3 |
| FIM1_829 | 747.6842059 | 701.8299792 | 492.427216 | -0.013956139 | 0.063215375 | 0.556078408 | 0.959760553 | ns | TM | KS.3 |
| FIM1_830 | 3224.457073 | 2822.606051 | 2874.1411 | -0.011626412 | 0.085769611 | 0.570675447 | 0.962265923 | ns | TM | KS.3 |
| FIM1_831 | 4710.529794 | 4660.043152 | 4234.203468 | -0.002400832 | 0.057486743 | 0.915467241 | 0.991141351 | ns | TM | KS.3 |
| FIM1_832 | 682.9886655 | 784.6304966 | 754.9707349 | 0.026975121 | 0.077025861 | 0.289434774 | 0.919993167 | ns | TM | KS.3 |
| FIM1_833 | 3357.417151 | 3677.387832 | 3921.404092 | 0.012212819 | 0.075404427 | 0.585605033 | 0.963781856 | ns | TM | KS.3 |
| FIM1_834 | 1172.406631 | 1363.951816 | 1221.750856 | 0.062460566 | 0.085255116 | 0.067149782 | 0.567220699 | ns | TM | KS.3 |
| FIM1_835 | 691.1486333 | 842.8261198 | 953.2250199 | 0.07145435 | 0.098032882 | 0.047135337 | 0.496748336 | ns | TM | KS.3 |
| FIM1_836 | 1057.387064 | 316.1250396 | 1089.792055 | -1.542228128 | 0.262603327 | 3.7592430947[illegible] | 1.5585195330[illegible] | Km<KS.3 | TM | KS.3 |
| FIM1_840 | 630.5424139 | 565.7904549 | 485.8406089 | -0.0232542 | 0.070692446 | 0.354526858 | 0.948637087 | ns | TM | KS.3 |
| FIM1_841 | 45.48488675 | 44.47093591 | 116.3551879 | -0.001102583 | 0.08529792 | 0.954532099 | 0.994413583 | ns | TM | KS.3 |
| FIM1_842 | 136.0332139 | 134.4360197 | 255.2486176 | -0.001246842 | 0.075499115 | 0.953238795 | 0.994413583 | ns | TM | KS.3 |
| FIM1_843 | 259.7487008 | 195.975701 | 1017.35679 | -0.017802161 | 0.104126191 | 0.374045359 | 0.948637087 | ns | TM | KS.3 |
| FIM1_844 | 79.44843037 | 81.33109866 | 112.3709184 | 0.002666101 | 0.07380232 | 0.90190901 | 0.991141351 | ns | TM | KS.3 |
| FIM1_846 | 52.76833331 | 47.60364952 | 84.39783745 | -0.010391055 | 0.083056697 | 0.616047027 | 0.973122675 | ns | TM | KS.3 |
| FIM1_847 | 9.414998418 | 10.09508809 | 15.97427327 | 0.002567169 | 0.09297095 | 0.888228787 | 0.991141351 | ns | TM | KS.3 |
| FIM1_850 | 510.7461721 | 605.1253389 | 698.1875307 | 0.01641618 | 0.087039193 | 0.442482977 | 0.950410534 | ns | TM | KS.3 |
| FIM1_851 | 316.9260063 | 319.0074971 | 305.0748288 | 0.000941545 | 0.076251432 | 0.964420203 | 0.995175461 | ns | TM | KS.3 |
| FIM1_852 | 77.64653924 | 67.70447322 | 63.46054071 | -0.018562506 | 0.078392062 | 0.423913868 | 0.950410534 | ns | TM | KS.3 |
| FIM1_853 | 69.47947247 | 57.72123416 | 101.2122781 | -0.024777665 | 0.087095296 | 0.293373019 | 0.923185001 | ns | TM | KS.3 |
| FIM1_854 | 1251.44531 | 970.5857297 | 1559.299267 | -0.032728219 | 0.097313371 | 0.187714047 | 0.828217874 | ns | TM | KS.3 |

| | | | | | | | | | | |
|---|---|---|---|---|---|---|---|---|---|---|
| FIM1_855 | 274.3421633 | 254.7286618 | 276.8693972 | -0.011667712 | 0.07178401 | 0.609874118 | 0.971747391 | ns | TM | KS.3 |
| FIM1_856 | 248.2328909 | 283.5729493 | 299.5086226 | 0.025523596 | 0.075851396 | 0.311605425 | 0.927416591 | ns | TM | KS.3 |
| FIM1_857 | 136.8100082 | 124.1496288 | 194.7328761 | -0.020030742 | 0.069454975 | 0.414415741 | 0.950410534 | ns | TM | KS.3 |
| FIM1_858 | 957.8755308 | 403.5727166 | 643.7882858 | -1.020872024 | 0.361516897 | 1.8783362174 | 0.002224934 | Km<KS.3 | TM | KS.3 |
| FIM1_859 | 577.6423861 | 452.595917 | 460.6221006 | -0.200939857 | 0.112574738 | 0.000868399 | 0.044072215 | ns | TM | KS.3 |
| FIM1_860 | 508.5388176 | 507.7808382 | 530.8493055 | -0.000531156 | 0.047771954 | 0.979695596 | 0.999333464 | ns | TM | KS.3 |
| FIM1_861 | 800.1373593 | 757.4479579 | 689.1671746 | -0.01141504 | 0.062422531 | 0.625830177 | 0.97344368 | ns | TM | KS.3 |
| FIM1_862 | 134.1871252 | 129.8593664 | 158.4303416 | -0.004096962 | 0.069740306 | 0.854604265 | 0.991141351 | ns | TM | KS.3 |
| FIM1_863 | 52.65192235 | 56.93010508 | 32.18866186 | 0.004671341 | 0.089828375 | 0.804153902 | 0.988916276 | ns | TM | KS.3 |
| FIM1_864 | 749.2273446 | 722.5471762 | 574.4896196 | -0.003878878 | 0.077429546 | 0.853398868 | 0.991141351 | ns | TM | KS.3 |
| FIM1_865 | 1121.53163 | 990.1465823 | 947.753543 | -0.019352347 | 0.07663729 | 0.413300286 | 0.950410534 | ns | TM | KS.3 |
| FIM1_866 | 596.9255138 | 473.7854656 | 514.4551748 | -0.056525809 | 0.100250108 | 0.074767743 | 0.599104608 | ns | TM | KS.3 |
| FIM1_867 | 1103.357569 | 1079.492436 | 1299.862932 | -0.003356643 | 0.068706036 | 0.881157951 | 0.991141351 | ns | TM | KS.3 |
| FIM1_868 | 938.3477627 | 832.6668791 | 901.8214115 | -0.026793591 | 0.072952375 | 0.298326678 | 0.926536521 | ns | TM | KS.3 |
| FIM1_869 | 722.792724 | 775.4575028 | 674.8613506 | 0.019746436 | 0.059580335 | 0.41221079 | 0.950410534 | ns | TM | KS.3 |
| FIM1_870 | 971.5008055 | 1297.847634 | 1337.424702 | 0.062524355 | 0.115579688 | 0.055119039 | 0.531428718 | ns | TM | KS.3 |
| FIM1_871 | 1159.061417 | 1323.563672 | 1085.450442 | 0.069629651 | 0.081027896 | 0.049836623 | 0.511210716 | ns | TM | KS.3 |
| FIM1_872 | 2923.899904 | 2686.148478 | 2799.267011 | -0.029780805 | 0.061789656 | 0.245120293 | 0.888934767 | ns | TM | KS.3 |
| FIM1_873 | 883.1183186 | 786.72051 | 998.9041111 | -0.028134747 | 0.07191332 | 0.281199446 | 0.916154055 | ns | TM | KS.3 |
| FIM1_874 | 3124.94667 | 2803.267862 | 4884.448618 | -0.015813891 | 0.075627434 | 0.492462202 | 0.952591289 | ns | TM | KS.3 |
| FIM1_875 | 325.3107307 | 325.2564185 | 368.4667161 | -0.000126324 | 0.070460481 | 0.995447547 | 0.999525119 | ns | TM | KS.3 |
| FIM1_876 | 2461.149922 | 2611.842735 | 2803.00508 | 0.006430485 | 0.077799133 | 0.760997265 | 0.988477371 | ns | TM | KS.3 |
| FIM1_877 | 1485.415573 | 1371.714075 | 1254.645693 | -0.018996186 | 0.064472472 | 0.436608135 | 0.950410534 | ns | TM | KS.3 |
| FIM1_878 | 4070.746786 | 3668.062996 | 3660.528863 | -0.034957841 | 0.068382472 | 0.201355445 | 0.84606701 | ns | TM | KS.3 |
| FIM1_879 | 17.90240002 | 11.63715871 | 27.8276178 | -0.030736113 | 0.126721877 | 0.177635133 | 0.811391014 | ns | TM | KS.3 |
| FIM1_880 | 1412.274087 | 1240.28202 | 1216.289069 | -0.056709581 | 0.078655359 | 0.081795841 | 0.621932675 | ns | TM | KS.3 |
| FIM1_881 | 1644.472184 | 1389.362938 | 1551.362047 | -0.105888328 | 0.093164931 | 0.015341555 | 0.284118339 | ns | TM | KS.3 |
| FIM1_882 | 515.2466073 | 428.2495103 | 510.0564056 | -0.06353125 | 0.093402006 | 0.06285344 | 0.551491823 | ns | TM | KS.3 |
| FIM1_883 | 1774.416659 | 1613.033749 | 1562.433082 | -0.015755522 | 0.072374868 | 0.502173791 | 0.953260948 | ns | TM | KS.3 |
| FIM1_884 | 456.3439763 | 321.4163312 | 330.5532116 | -0.059997512 | 0.128700766 | 0.056781796 | 0.535878428 | ns | TM | KS.3 |
| FIM1_885 | 1788.66442 | 1877.925715 | 2050.872708 | 0.016557796 | 0.047639547 | 0.428582218 | 0.950410534 | ns | TM | KS.3 |
| FIM1_886 | 4869.540204 | 4191.53619 | 4045.572158 | -0.022361597 | 0.080216032 | 0.348943557 | 0.948637087 | ns | TM | KS.3 |
| FIM1_887 | 879.2701232 | 846.1430468 | 602.4879608 | -0.00925573 | 0.056882307 | 0.685164925 | 0.980355335 | ns | TM | KS.3 |
| FIM1_888 | 1643.015574 | 1665.479979 | 1495.16734 | 0.003123914 | 0.057003146 | 0.889997384 | 0.991141351 | ns | TM | KS.3 |
| FIM1_889 | 1010.624805 | 964.6300614 | 968.179931 | -0.014377309 | 0.051492022 | 0.514546335 | 0.954689943 | ns | TM | KS.3 |
| FIM1_890 | 321.6935136 | 329.8592774 | 335.6733714 | 0.004936894 | 0.061960351 | 0.829444786 | 0.991141351 | ns | TM | KS.3 |
| FIM1_891 | 4066.792752 | 3225.672429 | 4113.37712 | -0.082940157 | 0.108765616 | 0.031042698 | 0.403710444 | ns | TM | KS.3 |
| FIM1_892 | 236.8519228 | 172.0540598 | 170.0596042 | -0.03526043 | 0.10958453 | 0.155256752 | 0.786819827 | ns | TM | KS.3 |
| FIM1_893 | 12.84782413 | 13.16992318 | 28.89626081 | 0.001119899 | 0.093637944 | 0.950739288 | 0.994413583 | ns | TM | KS.3 |
| FIM1_894 | 2358.121978 | 1700.612766 | 3726.986747 | -0.043335659 | 0.114203245 | 0.108915329 | 0.705512825 | ns | TM | KS.3 |
| FIM1_895 | 708.6907117 | 676.4678652 | 707.5695422 | -0.011403593 | 0.05747952 | 0.620694603 | 0.973122675 | ns | TM | KS.3 |
| FIM1_896 | 133.8196029 | 158.4925537 | 140.4085589 | 0.037235938 | 0.082855061 | 0.177959273 | 0.811391014 | ns | TM | KS.3 |
| FIM1_897 | 611.9091402 | 643.6089984 | 566.0714161 | 0.007030387 | 0.072259202 | 0.751617485 | 0.988477371 | ns | TM | KS.3 |
| FIM1_898 | 6316.84383 | 5396.514697 | 5129.727209 | -0.030361862 | 0.080433691 | 0.242590848 | 0.887413624 | ns | TM | KS.3 |
| FIM1_899 | 707.5865854 | 694.4139952 | 688.9489661 | -0.003053555 | 0.066148162 | 0.892954231 | 0.991141351 | ns | TM | KS.3 |

| | | | | | | | | | | |
|---|---|---|---|---|---|---|---|---|---|---|
| FIM1_900 | 2181.621331 | 1822.27149 | 1909.136672 | -0.039514647 | 0.085659974 | 0.158418525 | 0.789048157 | ns | TM | KS.3 |
| FIM1_901 | 1939.909031 | 3159.423007 | 3456.279029 | 0.675672721 | 0.13032248 | 7.65222906123 | 2.00367576734 | ns | TM | KS.3 |
| FIM1_902 | 546.4043803 | 547.6247058 | 486.7253884 | 0.000686904 | 0.052801238 | 0.975062454 | 0.998751433 | ns | TM | KS.3 |
| FIM1_903 | 1193.921953 | 1420.546351 | 1651.287398 | 0.014936571 | 0.089130483 | 0.471333768 | 0.950410534 | ns | TM | KS.3 |
| FIM1_904 | 952.976685 | 1168.070513 | 1256.333312 | 0.021053698 | 0.089944223 | 0.34286916 | 0.944808611 | ns | TM | KS.3 |
| FIM1_905 | 678.1373125 | 627.7884131 | 879.7972504 | -0.018596755 | 0.063675599 | 0.444629587 | 0.950410534 | ns | TM | KS.3 |
| FIM1_906 | 12189.89102 | 13251.15864 | 12090.97668 | 0.020699355 | 0.064633073 | 0.401636859 | 0.950410534 | ns | TM | KS.3 |
| FIM1_907 | 16356.09979 | 16970.57943 | 16591.70845 | 0.009704014 | 0.053965611 | 0.664957102 | 0.976323032 | ns | TM | KS.3 |
| FIM1_909 | 1041.955841 | 1196.082299 | 1077.98299 | 0.035750209 | 0.076896007 | 0.196854384 | 0.83848507 | ns | TM | KS.3 |
| FIM1_910 | 214.9963885 | 189.5797022 | 267.2848716 | -0.018528748 | 0.077367261 | 0.427687773 | 0.950410534 | ns | TM | KS.3 |
| FIM1_912 | 83.55020229 | 53.59230573 | 111.3884135 | -0.076915456 | 0.163613609 | 0.032812908 | 0.417169416 | ns | TM | KS.3 |
| FIM1_913 | 1765.686693 | 1857.875914 | 2001.170019 | 0.006169895 | 0.075212404 | 0.775273241 | 0.988814136 | ns | TM | KS.3 |
| FIM1_914 | 493.5639353 | 590.1005542 | 592.0757581 | 0.073700321 | 0.094524553 | 0.044690204 | 0.486507146 | ns | TM | KS.3 |
| FIM1_915 | 3966.763753 | 3697.242498 | 4187.559889 | -0.020940941 | 0.058654163 | 0.384043815 | 0.948637087 | ns | TM | KS.3 |
| FIM1_916 | 1052.61767 | 1020.59668 | 1234.286868 | -0.006212883 | 0.061919005 | 0.786931798 | 0.988916276 | ns | TM | KS.3 |
| FIM1_917 | 5796.069411 | 5408.883683 | 10820.5135 | -0.006486843 | 0.081099635 | 0.751725186 | 0.988477371 | ns | TM | KS.3 |
| FIM1_918 | 207.8893919 | 295.971277 | 235.7526735 | 0.063333319 | 0.130755462 | 0.05067045 | 0.513412402 | ns | TM | KS.3 |
| FIM1_919 | 227.1564969 | 271.3974632 | 300.1039835 | 0.019583037 | 0.08645927 | 0.378867466 | 0.948637087 | ns | TM | KS.3 |
| FIM1_920 | 1476.729703 | 1798.997505 | 1881.743709 | 0.043283332 | 0.089728085 | 0.131763604 | 0.761283488 | ns | TM | KS.3 |
| FIM1_921 | 3141.078712 | 2917.020571 | 2693.04147 | -0.017549356 | 0.063449896 | 0.467866445 | 0.950410534 | ns | TM | KS.3 |
| FIM1_922 | 533.7803128 | 590.9291068 | 488.257024 | 0.018914518 | 0.07161572 | 0.433856483 | 0.950410534 | ns | TM | KS.3 |
| FIM1_923 | 403.4247784 | 379.0186507 | 409.4636735 | -0.010198839 | 0.070036902 | 0.655860232 | 0.976323032 | ns | TM | KS.3 |
| FIM1_924 | 695.0160176 | 695.616282 | 926.2492243 | 5.56109911754634 | 0.057234662 | 0.998034395 | 0.999525119 | ns | TM | KS.3 |
| FIM1_925 | 166.5189713 | 199.788283 | 227.6234873 | 0.03217925 | 0.08461886 | 0.215946644 | 0.865538807 | ns | TM | KS.3 |
| FIM1_926 | 1811.74992 | 1627.57108 | 4723.754127 | -0.008607046 | 0.08596009 | 0.665898212 | 0.976323032 | ns | TM | KS.3 |
| FIM1_927 | 6917.777308 | 6618.878663 | 6918.70648 | -0.015199439 | 0.045847706 | 0.453560474 | 0.950410534 | ns | TM | KS.3 |
| FIM1_928 | 1056.63732 | 1125.594179 | 996.4788868 | 0.013358492 | 0.064084722 | 0.572137206 | 0.962265923 | ns | TM | KS.3 |
| FIM1_929 | 1333.472742 | 1370.033487 | 1444.466614 | 0.003503431 | 0.072976687 | 0.872439242 | 0.991141351 | ns | TM | KS.3 |
| FIM1_930 | 1560.185677 | 1629.620309 | 1751.919322 | 0.007621702 | 0.066748886 | 0.739610039 | 0.988051177 | ns | TM | KS.3 |
| FIM1_931 | 1702.330948 | 1683.981117 | 1672.699267 | -0.00276156 | 0.052733721 | 0.900014432 | 0.991141351 | ns | TM | KS.3 |
| FIM1_932 | 534.0317851 | 492.2285504 | 497.2585964 | -0.025200129 | 0.061797112 | 0.314316577 | 0.927564309 | ns | TM | KS.3 |
| FIM1_933 | 2152.197532 | 1942.430837 | 2148.831949 | -0.054752059 | 0.069665417 | 0.077742536 | 0.612946304 | ns | TM | KS.3 |
| FIM1_934 | 1360.543381 | 1467.366964 | 1323.273902 | 0.020919933 | 0.061292044 | 0.391996542 | 0.948637087 | ns | TM | KS.3 |
| FIM1_935 | 850.0806089 | 777.4232878 | 1205.089316 | -0.006692865 | 0.08667261 | 0.731991152 | 0.987157164 | ns | TM | KS.3 |
| FIM1_936 | 9269.911164 | 8966.507202 | 8369.136041 | -0.003971109 | 0.074928382 | 0.853387771 | 0.991141351 | ns | TM | KS.3 |
| FIM1_937 | 2820.829236 | 2631.066924 | 2593.518284 | -0.020399548 | 0.058790758 | 0.395853386 | 0.948637087 | ns | TM | KS.3 |
| FIM1_938 | 3253.157551 | 2918.885655 | 3427.445661 | -0.017565179 | 0.074136965 | 0.456879028 | 0.950410534 | ns | TM | KS.3 |
| FIM1_939 | 1951.315907 | 1983.917651 | 1869.916936 | 0.005390517 | 0.039155155 | 0.764615271 | 0.988477371 | ns | TM | KS.3 |
| FIM1_940 | 3059.063621 | 2407.583641 | 2972.43512 | -0.063199247 | 0.104361697 | 0.058180735 | 0.540287513 | ns | TM | KS.3 |
| FIM1_941 | 453.9563235 | 390.6375258 | 506.1071318 | -0.020039319 | 0.080939872 | 0.387303333 | 0.948637087 | ns | TM | KS.3 |
| FIM1_942 | 491.257035 | 794.4379936 | 1787.929814 | 0.246262582 | 0.276776675 | 0.002056535 | 0.081850075 | ns | TM | KS.3 |
| FIM1_943 | 8863.725291 | 8937.827657 | 8646.69721 | 0.001830394 | 0.05829344 | 0.935663928 | 0.994413583 | ns | TM | KS.3 |
| FIM1_944 | 1243.65942 | 1430.196348 | 1283.016317 | 0.01621311 | 0.081789197 | 0.463762268 | 0.950410534 | ns | TM | KS.3 |
| FIM1_945 | 1754.383109 | 1785.685005 | 2338.598098 | 0.005039213 | 0.047400792 | 0.808469101 | 0.988916276 | ns | TM | KS.3 |
| FIM1_946 | 44.8103453 | 47.10820873 | 45.99085676 | 0.006149516 | 0.073459399 | 0.779134723 | 0.988814136 | ns | TM | KS.3 |

| | | | | | | | | | | |
|---|---|---|---|---|---|---|---|---|---|---|
| FIM1_947 | 5108.425595 | 4823.708472 | 5270.519313 | -0.014575227 | 0.058795915 | 0.534439172 | 0.959071703 | ns | TM | KS.3 |
| FIM1_948 | 4586.54291 | 4769.789856 | 5590.866524 | 0.011616346 | 0.050067793 | 0.590757781 | 0.963781856 | ns | TM | KS.3 |
| FIM1_949 | 1471.506158 | 1211.098565 | 1384.197803 | -0.027489446 | 0.087071784 | 0.259019911 | 0.895871714 | ns | TM | KS.3 |
| FIM1_950 | 420.0211517 | 467.393684 | 465.2174882 | 0.013621022 | 0.077487386 | 0.541389519 | 0.95917717 | ns | TM | KS.3 |
| FIM1_951 | 14376.46583 | 14368.93313 | 20186.18416 | -0.000103605 | 0.058032546 | 0.996349133 | 0.999525119 | ns | TM | KS.3 |
| FIM1_952 | 30.09397212 | 19.16663548 | 41.24879713 | -0.036346053 | 0.131742314 | 0.136383131 | 0.768410055 | ns | TM | KS.3 |
| FIM1_953 | 3550.135151 | 3447.977096 | 3895.405561 | -0.006876637 | 0.057065108 | 0.762187816 | 0.988477371 | ns | TM | KS.3 |
| FIM1_954 | 1854.21344 | 1909.408335 | 1865.356409 | 0.003927396 | 0.072213955 | 0.858150524 | 0.991141351 | ns | TM | KS.3 |
| FIM1_955 | 819.5871708 | 1076.375149 | 1946.284371 | 0.029790486 | 0.100175726 | 0.209524185 | 0.852551729 | ns | TM | KS.3 |
| FIM1_957 | 14604.43452 | 12824.12218 | 11679.93381 | -0.012391557 | 0.08401596 | 0.553961653 | 0.95917717 | ns | TM | KS.3 |
| FIM1_958 | 3972.187727 | 3687.169095 | 4172.932193 | -0.013816017 | 0.068081934 | 0.557802944 | 0.959899566 | ns | TM | KS.3 |
| FIM1_959 | 870.2929563 | 978.8175799 | 1630.320037 | 0.014925472 | 0.078542896 | 0.50494513 | 0.953260948 | ns | TM | KS.3 |
| FIM1_960 | 2090.023842 | 2262.435494 | 2383.03718 | 0.02214924 | 0.062102153 | 0.369633465 | 0.948637087 | ns | TM | KS.3 |
| FIM1_961 | 5614.676962 | 5432.482615 | 4809.67305 | -0.010226255 | 0.045955124 | 0.615607445 | 0.973122675 | ns | TM | KS.3 |
| FIM1_962 | 3064.338263 | 2743.209678 | 2784.4084 | -0.047395227 | 0.071715323 | 0.114473314 | 0.714461279 | ns | TM | KS.3 |
| FIM1_963 | 721.5867231 | 693.8586343 | 705.9682752 | -0.007990142 | 0.062508052 | 0.729533948 | 0.987157164 | ns | TM | KS.3 |
| FIM1_964 | 248.6009329 | 287.086069 | 229.22586 | 0.038948572 | 0.078143811 | 0.170950889 | 0.809734625 | ns | TM | KS.3 |
| FIM1_965 | 1129.619729 | 899.5683282 | 1032.537501 | -0.193265565 | 0.101467365 | 0.000884279 | 0.044072215 | ns | TM | KS.3 |
| FIM1_966 | 5072.773762 | 4761.675234 | 3950.198171 | -0.012725007 | 0.065136234 | 0.589116352 | 0.963781856 | ns | TM | KS.3 |
| FIM1_967 | 1178.067005 | 1117.471617 | 1015.043269 | -0.014225313 | 0.056130087 | 0.537136722 | 0.95917717 | ns | TM | KS.3 |
| FIM1_968 | 657.2718793 | 634.4372785 | 752.0076583 | -0.007252181 | 0.062117864 | 0.753079724 | 0.988477371 | ns | TM | KS.3 |
| FIM1_969 | 305.8412675 | 312.7012486 | 440.0148115 | 0.00386343 | 0.064506258 | 0.865576218 | 0.991141351 | ns | TM | KS.3 |
| FIM1_970 | 1000.076484 | 913.9131258 | 965.1903519 | -0.028039484 | 0.063912532 | 0.275723519 | 0.908583046 | ns | TM | KS.3 |
| FIM1_971 | 97.39707369 | 124.0965796 | 131.9005992 | 0.04538033 | 0.097849663 | 0.112716302 | 0.711904568 | ns | TM | KS.3 |
| FIM1_972 | 187.339083 | 168.6166805 | 157.1682748 | -0.015177257 | 0.074572306 | 0.510641292 | 0.953260948 | ns | TM | KS.3 |
| FIM1_973 | 945.6887845 | 1022.80622 | 1188.232828 | 0.02905155 | 0.058847844 | 0.243353737 | 0.888250069 | ns | TM | KS.3 |
| FIM1_974 | 790.8736832 | 1032.778222 | 1062.195313 | 0.081681411 | 0.117714454 | 0.030656749 | 0.402578636 | ns | TM | KS.3 |
| FIM1_975 | 3136.026955 | 3472.714923 | 2546.823218 | 0.017650428 | 0.072486018 | 0.458854172 | 0.950410534 | ns | TM | KS.3 |
| FIM1_976 | 41.55912451 | 37.67293196 | 101.3528942 | -0.007222881 | 0.086234015 | 0.713662637 | 0.986826511 | ns | TM | KS.3 |
| FIM1_977 | 2987.76286 | 3067.107874 | 3415.972487 | 0.002168706 | 0.082845951 | 0.912871909 | 0.991141351 | ns | TM | KS.3 |
| FIM1_978 | 2832.863677 | 3661.484263 | 4313.41991 | 0.016510556 | 0.101313759 | 0.406491428 | 0.950410534 | ns | TM | KS.3 |
| FIM1_979 | 1521.598624 | 1835.282709 | 2201.409289 | 0.017595024 | 0.089175524 | 0.410543081 | 0.950410534 | ns | TM | KS.3 |
| FIM1_980 | 1978.128252 | 1965.917509 | 2473.290594 | -0.000787714 | 0.073334087 | 0.9709986 | 0.997052226 | ns | TM | KS.3 |
| FIM1_981 | 826.9152552 | 573.8107379 | 1418.350743 | -0.03526752 | 0.117638382 | 0.149617916 | 0.783447458 | ns | TM | KS.3 |
| FIM1_982 | 1356.449267 | 1205.20209 | 1544.672182 | -0.021688408 | 0.074067585 | 0.376215465 | 0.948637087 | ns | TM | KS.3 |
| FIM1_983 | 14580.10514 | 12281.83772 | 12436.09678 | -0.097495115 | 0.094551408 | 0.020450741 | 0.327146095 | ns | TM | KS.3 |
| FIM1_984 | 31.49350902 | 14.58001856 | 91.15760918 | -0.039891284 | 0.174821756 | 0.102109781 | 0.687410227 | ns | TM | KS.3 |
| FIM1_988 | 2420.510708 | 2266.786324 | 2634.034784 | -0.016796603 | 0.060457898 | 0.481565696 | 0.951416554 | ns | TM | KS.3 |
| FIM1_989 | 26.93106693 | 25.52714206 | 71.54174204 | -0.005112019 | 0.081754348 | 0.800536184 | 0.988916276 | ns | TM | KS.3 |
| FIM1_991 | 1573.028063 | 1315.991155 | 1301.236003 | -0.02580588 | 0.084479412 | 0.286451829 | 0.919993167 | ns | TM | KS.3 |
| FIM1_992 | 2338.577272 | 2141.104458 | 2183.789848 | -0.013124272 | 0.073377258 | 0.566131993 | 0.962265923 | ns | TM | KS.3 |
| FIM1_993 | 1525.021365 | 1508.545061 | 1362.119294 | -0.001270243 | 0.075102164 | 0.95253676 | 0.994413583 | ns | TM | KS.3 |
| FIM1_994 | 2009.756247 | 2052.848015 | 2129.332173 | 0.003518523 | 0.067096811 | 0.876443395 | 0.991141351 | ns | TM | KS.3 |
| FIM1_995 | 1693.684662 | 1897.319081 | 2185.273744 | 0.012812994 | 0.079996559 | 0.555069562 | 0.95917717 | ns | TM | KS.3 |
| FIM1_996 | 2637.449494 | 2846.806104 | 2898.353279 | 0.005276107 | 0.088146387 | 0.782865441 | 0.988814136 | ns | TM | KS.3 |

| | | | | | | | | | | |
|---|---|---|---|---|---|---|---|---|---|---|
| FIM1_997 | 166.7188656 | 117.9294823 | 234.4595379 | -0.052202063 | 0.123120423 | 0.075451722 | 0.603492469 | ns | TM | KS.3 |
| FIM1_998 | 1047.840294 | 1047.087443 | 1120.239119 | 4.62814117799333 | 0.053048245 | 0.998322707 | 0.999525119 | ns | TM | KS.3 |
| FIM1_999 | 1316.230804 | 1389.805768 | 1425.907567 | 0.013177085 | 0.059345637 | 0.573237052 | 0.963192994 | ns | TM | KS.3 |
| FIM1_1000 | 556.4274037 | 552.4861624 | 961.7356256 | -0.000588933 | 0.081952512 | 0.976449742 | 0.998938406 | ns | TM | KS.3 |
| FIM1_1001 | 307.5893761 | 339.7412927 | 323.4595592 | 0.032155796 | 0.067733442 | 0.230213227 | 0.879350777 | ns | TM | KS.3 |
| FIM1_1002 | 283.9800362 | 334.8308701 | 328.5117821 | 0.043974616 | 0.082817091 | 0.136145918 | 0.768410055 | ns | TM | KS.3 |
| FIM1_1003 | 2639.952937 | 3166.890016 | 3583.500302 | 0.022156669 | 0.085947946 | 0.335543825 | 0.938982061 | ns | TM | KS.3 |
| FIM1_1004 | 4061.766533 | 3632.334097 | 3041.230843 | -0.016479805 | 0.075766598 | 0.476629861 | 0.950410534 | ns | TM | KS.3 |
| FIM1_1005 | 1063.575988 | 1134.505791 | 1178.634649 | 0.013507836 | 0.0645298 | 0.568139731 | 0.962265923 | ns | TM | KS.3 |
| FIM1_1006 | 3711.954436 | 3427.144243 | 3500.720483 | -0.015619789 | 0.067768249 | 0.512819775 | 0.954240701 | ns | TM | KS.3 |
| FIM1_1007 | 754.7938474 | 856.0793852 | 680.5163647 | 0.05965 | 0.077926526 | 0.071932008 | 0.590530926 | ns | TM | KS.3 |
| FIM1_1008 | 328.7554081 | 290.1293003 | 350.3893167 | -0.025223665 | 0.074831893 | 0.317865949 | 0.928601572 | ns | TM | KS.3 |
| FIM1_1009 | 726.1247608 | 685.7242381 | 736.7478592 | -0.013732899 | 0.059819261 | 0.558786439 | 0.960263397 | ns | TM | KS.3 |
| FIM1_1010 | 2008.215395 | 1797.103994 | 2615.082822 | -0.01155709 | 0.081196936 | 0.586707979 | 0.963781856 | ns | TM | KS.3 |
| FIM1_1011 | 16993.71409 | 13244.13004 | 10635.097 | -0.044989039 | 0.099754547 | 0.112462911 | 0.711904568 | ns | TM | KS.3 |
| FIM1_1012 | 2468.118889 | 1844.524243 | 2599.211892 | -0.023912127 | 0.103285292 | 0.272538011 | 0.908468501 | ns | TM | KS.3 |
| FIM1_1013 | 3729.363725 | 3558.618417 | 2768.382107 | -0.006265776 | 0.072893544 | 0.776139337 | 0.988814136 | ns | TM | KS.3 |
| FIM1_1014 | 1550.144744 | 755.6273036 | 2115.903474 | -0.166834596 | 0.322385461 | 0.007866559 | 0.191538592 | ns | TM | KS.3 |
| FIM1_1015 | 1512.761147 | 2039.893442 | 2155.16098 | 0.035988189 | 0.106011549 | 0.153568023 | 0.783447458 | ns | TM | KS.3 |
| FIM1_1016 | 213.1235269 | 200.2337067 | 196.0072282 | -0.007194012 | 0.077458903 | 0.735472027 | 0.987655486 | ns | TM | KS.3 |
| FIM1_1017 | 3685.025032 | 3726.957617 | 4189.521793 | 0.002616611 | 0.055865887 | 0.907214611 | 0.991141351 | ns | TM | KS.3 |
| FIM1_1018 | 1521.362133 | 1372.858641 | 1304.181682 | -0.04509835 | 0.06902969 | 0.122770817 | 0.737662821 | ns | TM | KS.3 |
| FIM1_1019 | 702.5662596 | 789.7121607 | 854.6122633 | 0.024324592 | 0.072922911 | 0.33455222 | 0.938982061 | ns | TM | KS.3 |
| FIM1_1020 | 175.521951 | 144.1835555 | 134.9193748 | -0.016860294 | 0.091017421 | 0.421330974 | 0.950410534 | ns | TM | KS.3 |
| FIM1_1021 | 244.9262423 | 261.261396 | 651.045516 | 0.005140278 | 0.084424614 | 0.79458505 | 0.988916276 | ns | TM | KS.3 |
| FIM1_1022 | 6827.54268 | 6770.390994 | 7715.193246 | -0.002278325 | 0.048502232 | 0.913857877 | 0.991141351 | ns | TM | KS.3 |
| FIM1_1024 | 1538.642333 | 1232.945559 | 1785.986354 | -0.030426243 | 0.091572049 | 0.21797231 | 0.865538807 | ns | TM | KS.3 |
| FIM1_1025 | 4189.11102 | 5232.142144 | 5296.423224 | 0.073383203 | 0.103849218 | 0.04262482 | 0.473090953 | ns | TM | KS.3 |
| FIM1_1026 | 6666.599295 | 7148.932591 | 6893.761453 | 0.014989939 | 0.064675841 | 0.529779636 | 0.959071703 | ns | TM | KS.3 |
| FIM1_1027 | 1310.358068 | 1104.168399 | 1025.936303 | -0.032454731 | 0.083007994 | 0.216715813 | 0.865538807 | ns | TM | KS.3 |
| FIM1_1028 | 132.63474 | 125.5091103 | 216.8554548 | -0.007079687 | 0.072469888 | 0.749619429 | 0.988477371 | ns | TM | KS.3 |
| FIM1_1029 | 15174.0733 | 12492.49297 | 12682.87503 | -0.02758197 | 0.087009063 | 0.258090213 | 0.895871714 | ns | TM | KS.3 |
| FIM1_1030 | 2817.423174 | 3548.00551 | 4336.793453 | 0.018796632 | 0.095496736 | 0.370492037 | 0.948637087 | ns | TM | KS.3 |
| FIM1_1031 | 94.55193778 | 66.24396283 | 81.79274124 | -0.052734447 | 0.124262003 | 0.073751345 | 0.597271765 | ns | TM | KS.3 |
| FIM1_1032 | 662.0426197 | 680.5177336 | 743.9069697 | 0.008128931 | 0.049017676 | 0.702140523 | 0.98151979 | ns | TM | KS.3 |
| FIM1_1033 | 232.6482987 | 209.1581117 | 215.5946391 | -0.015802935 | 0.075950584 | 0.491788674 | 0.952591289 | ns | TM | KS.3 |
| FIM1_1034 | 643.9355329 | 666.7477851 | 648.1549468 | 0.010622876 | 0.049678844 | 0.620860776 | 0.973122675 | ns | TM | KS.3 |
| FIM1_1035 | 162.8478301 | 192.8764576 | 164.1932233 | 0.035748865 | 0.083009511 | 0.188860294 | 0.828217874 | ns | TM | KS.3 |
| FIM1_1036 | 837.9876384 | 712.9414084 | 581.8946502 | -0.058766212 | 0.086970563 | 0.076670156 | 0.609319531 | ns | TM | KS.3 |
| FIM1_1037 | 1352.054577 | 1337.914501 | 1268.132034 | -0.003200548 | 0.034843894 | 0.844203438 | 0.991141351 | ns | TM | KS.3 |
| FIM1_1038 | 3498.683855 | 3669.128474 | 3371.647619 | 0.017150404 | 0.045995503 | 0.398278111 | 0.950410534 | ns | TM | KS.3 |
| FIM1_1039 | 2128.513752 | 2203.339397 | 3203.341312 | 0.009288715 | 0.052443443 | 0.674166821 | 0.976966447 | ns | TM | KS.3 |
| FIM1_1040 | 1526.082696 | 1514.498554 | 1696.149232 | -0.001128734 | 0.069984525 | 0.959482415 | 0.994849847 | ns | TM | KS.3 |
| FIM1_1041 | 579.6346262 | 1565.229013 | 2064.977982 | 1.215139544 | 0.322598086 | 3.06796259503 | 0.000401661 | Km>KS.3 | TM | KS.3 |
| FIM1_1042 | 56.19347829 | 63.24078923 | 77.10730827 | 0.015337214 | 0.076774277 | 0.500440554 | 0.953260948 | ns | TM | KS.3 |

| | | | | | | | | | | |
|---|---|---|---|---|---|---|---|---|---|---|
| FIM1_1043 | 136.3374863 | 202.7504169 | 188.1524893 | 0.071977144 | 0.148777313 | 0.037819651 | 0.440708412 | ns | TM | KS.3 |
| FIM1_1044 | 1682.451449 | 1767.129191 | 2014.807761 | 0.006387064 | 0.073576946 | 0.770854731 | 0.988814136 | ns | TM | KS.3 |
| FIM1_1045 | 495.90699 | 534.1998096 | 478.7153123 | 0.017456187 | 0.063582859 | 0.4701044 | 0.950410534 | ns | TM | KS.3 |
| FIM1_1046 | 884.8198849 | 897.2765208 | 813.5084914 | 0.003727345 | 0.054632009 | 0.867291526 | 0.991141351 | ns | TM | KS.3 |
| FIM1_1047 | 1532.439654 | 1466.34716 | 1493.593789 | -0.015930062 | 0.043481185 | 0.409747298 | 0.950410534 | ns | TM | KS.3 |
| FIM1_1048 | 1785.987896 | 1773.723513 | 1727.147317 | -0.001841488 | 0.04851831 | 0.930334301 | 0.993993515 | ns | TM | KS.3 |
| FIM1_1049 | 1782.579939 | 2307.856707 | 2601.580577 | 0.02342963 | 0.098100515 | 0.287766675 | 0.919993167 | ns | TM | KS.3 |
| FIM1_1050 | 140.8197735 | 151.2358101 | 107.7630703 | 0.00644501 | 0.080548234 | 0.754447059 | 0.988477371 | ns | TM | KS.3 |
| FIM1_1051 | 658.8152216 | 638.3946367 | 591.553105 | -0.005859529 | 0.064125538 | 0.798406411 | 0.988916276 | ns | TM | KS.3 |
| FIM1_1052 | 1786.269363 | 1894.825637 | 1052.997585 | 0.008766846 | 0.071184631 | 0.697598883 | 0.98151979 | ns | TM | KS.3 |
| FIM1_1053 | 107.0905158 | 104.4125994 | 89.13311434 | -0.003385746 | 0.070505465 | 0.878857972 | 0.991141351 | ns | TM | KS.3 |
| FIM1_1054 | 1998.937358 | 2039.134695 | 2238.829363 | 0.005647983 | 0.046215072 | 0.782427085 | 0.988814136 | ns | TM | KS.3 |
| FIM1_1055 | 240.3023767 | 252.1016813 | 204.8296976 | 0.009218417 | 0.06652822 | 0.68983495 | 0.980879562 | ns | TM | KS.3 |
| FIM1_1056 | 551.3840978 | 549.5133425 | 564.8640782 | -0.000434504 | 0.060215467 | 0.984774177 | 0.999333464 | ns | TM | KS.3 |
| FIM1_1057 | 1294.617898 | 1284.954421 | 1415.285574 | -0.00215573 | 0.04767616 | 0.917645976 | 0.991141351 | ns | TM | KS.3 |
| FIM1_1058 | 3295.984524 | 3215.964315 | 2535.405671 | -0.00688133 | 0.048936373 | 0.745677388 | 0.988477371 | ns | TM | KS.3 |
| FIM1_1060 | 3189.27066 | 3765.23557 | 4530.229662 | 0.028147946 | 0.082070557 | 0.263541416 | 0.90372202 | ns | TM | KS.3 |
| FIM1_1061 | 704.0818848 | 683.2209821 | 795.8899212 | -0.006714283 | 0.05901344 | 0.769297945 | 0.988814136 | ns | TM | KS.3 |
| FIM1_1062 | 748.5773491 | 745.775908 | 854.6353268 | -0.001310556 | 0.048253016 | 0.950227135 | 0.994413583 | ns | TM | KS.3 |
| FIM1_1063 | 761.5212093 | 714.2705968 | 789.7390678 | -0.014828835 | 0.062330172 | 0.532890881 | 0.959071703 | ns | TM | KS.3 |
| FIM1_1064 | 1003.750891 | 1154.405872 | 1260.751839 | 0.03276155 | 0.077006373 | 0.223966319 | 0.873580587 | ns | TM | KS.3 |
| FIM1_1065 | 1085.464331 | 1183.572386 | 1242.392912 | 0.03132088 | 0.062522686 | 0.227589919 | 0.877720812 | ns | TM | KS.3 |
| FIM1_1066 | 465.9350241 | 496.6631307 | 442.4206693 | 0.017040692 | 0.059260916 | 0.473170919 | 0.950410534 | ns | TM | KS.3 |
| FIM1_1067 | 383.7500069 | 438.1885198 | 409.8270158 | 0.036504901 | 0.075823561 | 0.19115987 | 0.829253069 | ns | TM | KS.3 |
| FIM1_1068 | 101.5465132 | 153.7640628 | 234.1648906 | 0.07595843 | 0.156857073 | 0.033628514 | 0.424624004 | ns | TM | KS.3 |
| FIM1_1069 | 437.3002496 | 417.1080973 | 417.8969508 | -0.004210277 | 0.081458731 | 0.834990204 | 0.991141351 | ns | TM | KS.3 |
| FIM1_1070 | 1353.391769 | 1274.863371 | 1189.605498 | -0.017898744 | 0.055256625 | 0.43921273 | 0.950410534 | ns | TM | KS.3 |
| FIM1_1071 | 1499.711286 | 1439.629782 | 1522.70044 | -0.015476814 | 0.041826513 | 0.406225266 | 0.950410534 | ns | TM | KS.3 |
| FIM1_1072 | 787.9639351 | 965.5635963 | 1457.758092 | 0.016103265 | 0.09305222 | 0.433163109 | 0.950410534 | ns | TM | KS.3 |
| FIM1_1074 | 1200.717239 | 1213.800693 | 1221.662544 | 0.002565103 | 0.053454444 | 0.907604773 | 0.991141351 | ns | TM | KS.3 |
| FIM1_1075 | 2660.562861 | 2489.160606 | 3065.067165 | -0.016742393 | 0.060943185 | 0.48371489 | 0.952269291 | ns | TM | KS.3 |
| FIM1_1076 | 498.639185 | 493.0969618 | 428.2304838 | -0.001431385 | 0.074100243 | 0.947008621 | 0.994413583 | ns | TM | KS.3 |
| FIM1_1077 | 109.0305153 | 104.1885077 | 125.6029179 | -0.003835615 | 0.084154833 | 0.845445447 | 0.991141351 | ns | TM | KS.3 |
| FIM1_1078 | 239.8983773 | 291.3888731 | 294.9642808 | 0.044375211 | 0.089395091 | 0.126895286 | 0.749767277 | ns | TM | KS.3 |
| FIM1_1079 | 246.6510878 | 365.3808115 | 387.3355343 | 0.138950928 | 0.187732695 | 0.007701876 | 0.18968728 | ns | TM | KS.3 |
| FIM1_1080 | 972.8775817 | 1105.618292 | 1084.702492 | 0.016010257 | 0.079632739 | 0.475496245 | 0.950410534 | ns | TM | KS.3 |
| FIM1_1081 | 832.6369748 | 1462.323519 | 1316.689336 | 0.150464641 | 0.263448987 | 0.008492485 | 0.200237497 | ns | TM | KS.3 |
| FIM1_1082 | 776.972848 | 760.6991772 | 1101.869711 | -0.002825804 | 0.071798177 | 0.89774138 | 0.991141351 | ns | TM | KS.3 |
| FIM1_1083 | 1329.282592 | 1408.564756 | 1554.803504 | 0.017584444 | 0.054281596 | 0.442548801 | 0.950410534 | ns | TM | KS.3 |
| FIM1_1084 | 267.9877775 | 697.1605241 | 682.8854909 | 1.297242814 | 0.185337392 | 4.6931715462[illegible] | 3.3355040632[illegible] | Km>KS.3 | TM | KS.3 |
| FIM1_1085 | 608.009569 | 618.412334 | 580.9697454 | 0.005144822 | 0.050458104 | 0.81136151 | 0.988916276 | ns | TM | KS.3 |
| FIM1_1086 | 811.0139174 | 832.9009993 | 635.0051541 | 0.005719544 | 0.059911265 | 0.802743459 | 0.988916276 | ns | TM | KS.3 |
| FIM1_1087 | 700.7524771 | 691.4407537 | 755.4464923 | -0.00260282 | 0.056793771 | 0.908127519 | 0.991141351 | ns | TM | KS.3 |
| FIM1_1088 | 2199.275587 | 2178.681121 | 2300.985618 | -0.002967779 | 0.040523649 | 0.873058071 | 0.991141351 | ns | TM | KS.3 |
| FIM1_1089 | 127.1503361 | 143.6173745 | 127.3354343 | 0.017925416 | 0.076932738 | 0.441505121 | 0.950410534 | ns | TM | KS.3 |

| | | | | | | | | | | |
|---|---|---|---|---|---|---|---|---|---|---|
| FIM1_1090 | 5096.695913 | 4358.876269 | 4998.7929 | -0.066750508 | 0.087187614 | 0.05750769 | 0.536774409 | ns | TM | KS.3 |
| FIM1_1091 | 1719.035673 | 2058.33429 | 2368.850307 | 0.024567124 | 0.08499213 | 0.301901819 | 0.926536521 | ns | TM | KS.3 |
| FIM1_1092 | 938.9982621 | 1036.330039 | 980.3221085 | 0.044923281 | 0.068213624 | 0.122080713 | 0.736051706 | ns | TM | KS.3 |
| FIM1_1093 | 1212.098388 | 1006.806735 | 1159.130995 | -0.057492883 | 0.091456903 | 0.078144901 | 0.614172013 | ns | TM | KS.3 |
| FIM1_1094 | 1688.141383 | 1459.180667 | 1676.028457 | -0.154000048 | 0.064753647 | 0.000998346 | 0.04634941 | ns | TM | KS.3 |
| FIM1_1095 | 1157.095832 | 1072.11651 | 1197.186712 | -0.024694924 | 0.058976742 | 0.313364279 | 0.927416591 | ns | TM | KS.3 |
| FIM1_1096 | 389.7271041 | 417.2129003 | 436.598392 | 0.01212191 | 0.067272132 | 0.604291351 | 0.970729568 | ns | TM | KS.3 |
| FIM1_1097 | 5259.029237 | 5526.157987 | 8368.502534 | 0.00882028 | 0.066661444 | 0.701973917 | 0.98151979 | ns | TM | KS.3 |
| FIM1_1098 | 4031.820673 | 3395.988068 | 3522.134248 | -0.037743317 | 0.083773557 | 0.17295337 | 0.811391014 | ns | TM | KS.3 |
| FIM1_1099 | 799.9551725 | 769.8373124 | 822.1322244 | -0.00755973 | 0.062081431 | 0.743194377 | 0.988477371 | ns | TM | KS.3 |
| FIM1_1100 | 7305.356617 | 7295.592749 | 7850.830426 | -0.000391663 | 0.041897787 | 0.983622936 | 0.999333464 | ns | TM | KS.3 |
| FIM1_1101 | 50.97767239 | 72.80718482 | 78.11312331 | 0.017181603 | 0.115456802 | 0.37072469 | 0.948637087 | ns | TM | KS.3 |
| FIM1_1102 | 1360.324056 | 1434.222076 | 1220.438134 | 0.013068903 | 0.058705545 | 0.575154636 | 0.963192994 | ns | TM | KS.3 |
| FIM1_1103 | 985.8857313 | 1066.378395 | 973.7573959 | 0.009622568 | 0.076181793 | 0.659705558 | 0.976323032 | ns | TM | KS.3 |
| FIM1_1104 | 1652.316847 | 1701.876415 | 1469.220352 | 0.005166735 | 0.065843964 | 0.820839864 | 0.990886949 | ns | TM | KS.3 |
| FIM1_1105 | 4279.057687 | 4379.741756 | 4789.31649 | 0.007752138 | 0.037542664 | 0.652839807 | 0.976323032 | ns | TM | KS.3 |
| FIM1_1106 | 7282.731826 | 6877.60752 | 6673.423361 | -0.011744074 | 0.063970903 | 0.616658139 | 0.973122675 | ns | TM | KS.3 |
| FIM1_1107 | 1045.169461 | 1132.358009 | 984.9406452 | 0.017004431 | 0.066468623 | 0.481011617 | 0.951416554 | ns | TM | KS.3 |
| FIM1_1108 | 1088.035132 | 1547.902266 | 1788.027479 | 0.044082782 | 0.119668143 | 0.103315045 | 0.689041746 | ns | TM | KS.3 |
| FIM1_1109 | 4409.394868 | 4127.585698 | 4698.985873 | -0.014100166 | 0.064209468 | 0.552540821 | 0.95917717 | ns | TM | KS.3 |
| FIM1_1110 | 1404.196596 | 1444.675098 | 1348.385957 | 0.005905643 | 0.060612625 | 0.796810112 | 0.988916276 | ns | TM | KS.3 |
| FIM1_1111 | 1207.220963 | 1242.166046 | 1127.910016 | 0.007189215 | 0.053022326 | 0.745338203 | 0.988477371 | ns | TM | KS.3 |
| FIM1_1112 | 2916.321047 | 2936.108633 | 3978.287789 | 0.0018385 | 0.049628964 | 0.931317486 | 0.994058033 | ns | TM | KS.3 |
| FIM1_1113 | 1134.527894 | 1341.695622 | 1419.349751 | 0.018996001 | 0.084914121 | 0.394763715 | 0.948637087 | ns | TM | KS.3 |
| FIM1_1114 | 382.6836007 | 252.3323152 | 2368.019001 | -0.035180258 | 0.126297361 | 0.145726635 | 0.779559149 | ns | TM | KS.3 |
| FIM1_1115 | 191.2394569 | 204.658717 | 361.7384784 | 0.007871788 | 0.076965207 | 0.713889612 | 0.986826511 | ns | TM | KS.3 |
| FIM1_1116 | 7730.838582 | 7688.946828 | 6624.83979 | -0.001110725 | 0.060227023 | 0.961099895 | 0.995175461 | ns | TM | KS.3 |
| FIM1_1117 | 2771.085926 | 2513.576797 | 1887.62211 | -0.020998021 | 0.069014965 | 0.396112017 | 0.948781719 | ns | TM | KS.3 |
| FIM1_1118 | 13.94371018 | 16.77167302 | 20.87135697 | 0.012541038 | 0.091650896 | 0.528072883 | 0.959071703 | ns | TM | KS.3 |
| FIM1_1119 | 2516.928714 | 2501.273603 | 2937.395474 | -0.001416438 | 0.056918061 | 0.94991596 | 0.994413583 | ns | TM | KS.3 |
| FIM1_1120 | 33.69779409 | 26.04475066 | 43.48615303 | -0.033956453 | 0.102967392 | 0.170818323 | 0.809734625 | ns | TM | KS.3 |
| FIM1_1121 | 14730.0869 | 12652.31126 | 12996.19495 | -0.042730751 | 0.08099453 | 0.144511483 | 0.77783242 | ns | TM | KS.3 |
| FIM1_1122 | 4769.774784 | 5195.334511 | 5340.92505 | 0.042305342 | 0.062305321 | 0.122206776 | 0.736051706 | ns | TM | KS.3 |
| FIM1_1123 | 384.6994823 | 446.6773804 | 448.2279881 | 0.041757944 | 0.080607662 | 0.150583235 | 0.783447458 | ns | TM | KS.3 |
| FIM1_1124 | 2874.04072 | 2902.18902 | 3235.534849 | 0.002302792 | 0.056338346 | 0.918482506 | 0.991141351 | ns | TM | KS.3 |
| FIM1_1125 | 21029.29822 | 23133.37669 | 21740.66123 | 0.025870481 | 0.066446911 | 0.31241472 | 0.927416591 | ns | TM | KS.3 |
| FIM1_1126 | 543.72352 | 544.5726311 | 764.9335658 | 9.19676276225745 | 0.079503803 | 0.996409497 | 0.999525119 | ns | TM | KS.3 |
| FIM1_1127 | 448.2219495 | 426.1860529 | 411.6442297 | -0.005930759 | 0.075745965 | 0.782548439 | 0.988814136 | ns | TM | KS.3 |
| FIM1_1128 | 247.980457 | 169.3888685 | 192.2800568 | -0.054339112 | 0.132334374 | 0.068160282 | 0.572799666 | ns | TM | KS.3 |
| FIM1_1129 | 3542.916401 | 3079.230566 | 3127.035772 | -0.055469378 | 0.081163225 | 0.087106732 | 0.645836051 | ns | TM | KS.3 |
| FIM1_1130 | 376.2231297 | 490.354752 | 445.2423528 | 0.087686947 | 0.118927352 | 0.025717713 | 0.379660596 | ns | TM | KS.3 |
| FIM1_1131 | 1402.99874 | 1359.329988 | 1327.843283 | -0.008589496 | 0.050955127 | 0.69282387 | 0.981188557 | ns | TM | KS.3 |
| FIM1_1132 | 726.2165862 | 440.9737222 | 990.3518956 | -0.304619095 | 0.301491422 | 0.001166202 | 0.052744145 | ns | TM | KS.3 |
| FIM1_1133 | 540.064531 | 555.3175669 | 653.9017944 | 0.003229148 | 0.075239048 | 0.879982061 | 0.991141351 | ns | TM | KS.3 |
| FIM1_1134 | 276.6098795 | 336.8305643 | 307.6470545 | 0.023633421 | 0.088172785 | 0.306781839 | 0.927241586 | ns | TM | KS.3 |

| | | | | | | | | | | |
|---|---|---|---|---|---|---|---|---|---|---|
| FIM1_1135 | 15191.56432 | 12985.56765 | 14001.82884 | -0.021057057 | 0.08198416 | 0.365750168 | 0.948637087 | ns | TM | KS.3 |
| FIM1_1136 | 1149.538208 | 1165.852876 | 1073.687292 | 0.002993716 | 0.05711095 | 0.894591305 | 0.991141351 | ns | TM | KS.3 |
| FIM1_1137 | 3639.940363 | 4425.379962 | 3821.963831 | 0.023325451 | 0.087836188 | 0.312216571 | 0.927416591 | ns | TM | KS.3 |
| FIM1_1138 | 670.4263442 | 607.6655556 | 592.9896694 | -0.027727372 | 0.066656251 | 0.285156333 | 0.919509407 | ns | TM | KS.3 |
| FIM1_1139 | 10582.46288 | 8606.233105 | 9084.169046 | -0.172067763 | 0.096422836 | 0.001681834 | 0.069149779 | ns | TM | KS.3 |
| FIM1_1140 | 4971.920765 | 3397.012278 | 3299.452156 | -0.030702877 | 0.118393351 | 0.183149581 | 0.82087312 | ns | TM | KS.3 |
| FIM1_1141 | 5293.645777 | 3802.289531 | 3765.705268 | -0.034471106 | 0.111246315 | 0.159324731 | 0.791058419 | ns | TM | KS.3 |
| FIM1_1142 | 824.3745108 | 817.6858416 | 869.6513995 | -0.002023688 | 0.048342811 | 0.923304124 | 0.992531983 | ns | TM | KS.3 |
| FIM1_1143 | 1642.389944 | 1669.242414 | 1700.876473 | 0.004794462 | 0.04410322 | 0.808645237 | 0.988916276 | ns | TM | KS.3 |
| FIM1_1144 | 939.1198041 | 976.0083628 | 867.3689057 | 0.005214232 | 0.072222777 | 0.813137127 | 0.988916276 | ns | TM | KS.3 |
| FIM1_1145 | 7845.387106 | 8048.359456 | 9504.782481 | 0.007760868 | 0.04502283 | 0.699093293 | 0.98151979 | ns | TM | KS.3 |
| FIM1_1146 | 748.7169214 | 665.7261562 | 629.5973332 | -0.030841599 | 0.072145976 | 0.247982203 | 0.888934767 | ns | TM | KS.3 |
| FIM1_1147 | 3241.445719 | 3042.691445 | 3609.849336 | -0.019880291 | 0.055649075 | 0.394830719 | 0.948637087 | ns | TM | KS.3 |
| FIM1_1148 | 744.9868825 | 680.61942 | 672.4674812 | -0.019816037 | 0.067611953 | 0.419973561 | 0.950410534 | ns | TM | KS.3 |
| FIM1_1149 | 255.2934629 | 278.9470661 | 286.7529395 | 0.016526593 | 0.069456579 | 0.489399505 | 0.952591289 | ns | TM | KS.3 |
| FIM1_1150 | 929.8864107 | 976.257743 | 929.2402497 | 0.009896926 | 0.062965172 | 0.670647394 | 0.976861203 | ns | TM | KS.3 |
| FIM1_1151 | 5889.737401 | 4972.809732 | 6165.423123 | -0.049332248 | 0.08598067 | 0.108742322 | 0.705336441 | ns | TM | KS.3 |
| FIM1_1152 | 751.7137643 | 725.982648 | 803.2191854 | -0.004444266 | 0.073534776 | 0.838296979 | 0.991141351 | ns | TM | KS.3 |
| FIM1_1153 | 118.0802184 | 76.9682447 | 75.4474694 | -0.141032316 | 0.206239021 | 0.007969545 | 0.191538592 | ns | TM | KS.3 |
| FIM1_1154 | 1628.303271 | 1523.791494 | 1354.118306 | -0.01953672 | 0.057743512 | 0.410988097 | 0.950410534 | ns | TM | KS.3 |
| FIM1_1155 | 88.34253838 | 77.69008261 | 96.14274037 | -0.008824612 | 0.089874729 | 0.648979588 | 0.976323032 | ns | TM | KS.3 |
| FIM1_1156 | 9907.465431 | 9044.600881 | 9766.387008 | -0.026394187 | 0.064852994 | 0.302144198 | 0.926536521 | ns | TM | KS.3 |
| FIM1_1157 | 2895.014678 | 2689.694355 | 2947.350756 | -0.025841223 | 0.057675075 | 0.287886724 | 0.919993167 | ns | TM | KS.3 |
| FIM1_1158 | 1565.599692 | 1270.412082 | 1034.973562 | -0.023693736 | 0.089972346 | 0.3013397 | 0.926536521 | ns | TM | KS.3 |
| FIM1_1159 | 1542.630842 | 1941.4519 | 1818.015142 | 0.230330857 | 0.087308101 | 0.000103742 | 0.00889853 | ns | TM | KS.3 |
| FIM1_1160 | 10295.9451 | 8741.625416 | 13109.57996 | -0.1362807 | 0.085509629 | 0.004762708 | 0.138564175 | ns | TM | KS.3 |
| FIM1_1161 | 80.3206789 | 49.91915823 | 118.611153 | -0.047943573 | 0.146470132 | 0.08358992 | 0.629137448 | ns | TM | KS.3 |
| FIM1_1162 | 6044.829846 | 7173.980331 | 6707.310668 | 0.01559964 | 0.088045812 | 0.458411254 | 0.950410534 | ns | TM | KS.3 |
| FIM1_1163 | 996.6630886 | 907.8688316 | 1155.211628 | -0.036712905 | 0.065104042 | 0.176949557 | 0.811391014 | ns | TM | KS.3 |
| FIM1_1164 | 38.80922194 | 41.85135407 | 38.21550992 | 0.006362983 | 0.082518313 | 0.753023424 | 0.988477371 | ns | TM | KS.3 |
| FIM1_1165 | 959.8810182 | 975.7102392 | 916.3283181 | 0.003107505 | 0.063115417 | 0.891770118 | 0.991141351 | ns | TM | KS.3 |
| FIM1_1166 | 3452.808591 | 3212.519982 | 6055.591906 | -0.016302973 | 0.063959113 | 0.497373127 | 0.953260948 | ns | TM | KS.3 |
| FIM1_1167 | 712.0190576 | 828.2955104 | 888.8186405 | 0.042752996 | 0.080896454 | 0.144453014 | 0.77783242 | ns | TM | KS.3 |
| FIM1_1168 | 1393.325894 | 1553.588294 | 1564.529 | 0.036756362 | 0.070086691 | 0.186810395 | 0.827669994 | ns | TM | KS.3 |
| FIM1_1169 | 1207.213344 | 963.8963522 | 944.2420511 | -0.069557214 | 0.103407418 | 0.048077029 | 0.499686002 | ns | TM | KS.3 |
| FIM1_1170 | 1115.73008 | 1098.299384 | 1068.3569 | -0.003493318 | 0.058230858 | 0.87772603 | 0.991141351 | ns | TM | KS.3 |
| FIM1_1171 | 27.93863737 | 38.12844905 | 54.07909024 | 0.041672 | 0.111692599 | 0.117764374 | 0.724896022 | ns | TM | KS.3 |
| FIM1_1172 | 2437.993956 | 2701.491526 | 2917.458834 | 0.060547785 | 0.070221746 | 0.060117666 | 0.544665909 | ns | TM | KS.3 |
| FIM1_1173 | 556.6220811 | 702.8618161 | 611.6997115 | 0.049636935 | 0.098157576 | 0.096418832 | 0.673713046 | ns | TM | KS.3 |
| FIM1_1174 | 352.606392 | 311.2403445 | 552.607912 | -0.016044499 | 0.079143908 | 0.476281482 | 0.950410534 | ns | TM | KS.3 |
| FIM1_1175 | 113.5381909 | 156.574091 | 154.079058 | 0.048020937 | 0.114909527 | 0.090866207 | 0.657813255 | ns | TM | KS.3 |
| FIM1_1177 | 2265.314655 | 2292.539688 | 2264.533237 | 0.00357423 | 0.041357563 | 0.849769083 | 0.991141351 | ns | TM | KS.3 |
| FIM1_1178 | 11224.27066 | 12736.0968 | 14048.49393 | 0.032031462 | 0.074109861 | 0.234073525 | 0.883800031 | ns | TM | KS.3 |
| FIM1_1179 | 990.0848841 | 910.404284 | 940.2419267 | -0.028907667 | 0.061786627 | 0.257186343 | 0.895871714 | ns | TM | KS.3 |
| FIM1_1180 | 1097.746422 | 954.5925272 | 1300.328035 | -0.036508727 | 0.077596826 | 0.189992107 | 0.828405552 | ns | TM | KS.3 |

| | | | | | | | | | | |
|---|---|---|---|---|---|---|---|---|---|---|
| FIM1_1181 | 2568.404416 | 2175.592812 | 2233.461848 | -0.03321196 | 0.082137888 | 0.211545707 | 0.856547038 | ns | TM | KS.3 |
| FIM1_1182 | 62.58027191 | 55.42703682 | 106.0153218 | -0.014708208 | 0.077973885 | 0.512121063 | 0.953967865 | ns | TM | KS.3 |
| FIM1_1183 | 470.1898255 | 550.9966133 | 1349.595068 | 0.032589836 | 0.080963315 | 0.219415821 | 0.865538807 | ns | TM | KS.3 |
| FIM1_1184 | 600.7790626 | 654.9967769 | 592.4635588 | 0.018533085 | 0.067364907 | 0.446734008 | 0.950410534 | ns | TM | KS.3 |
| FIM1_1186 | 621.3648487 | 671.5014717 | 717.6328512 | 0.033008617 | 0.059721783 | 0.196114273 | 0.837834055 | ns | TM | KS.3 |
| FIM1_1187 | 5799.200694 | 5472.425358 | 6179.320756 | -0.028841232 | 0.047627019 | 0.17106951 | 0.809734625 | ns | TM | KS.3 |
| FIM1_1188 | 388.5633791 | 366.8636844 | 1370.245573 | -0.007769502 | 0.073113914 | 0.725781354 | 0.987157164 | ns | TM | KS.3 |
| FIM1_1189 | 972.6452795 | 789.7637371 | 1684.525144 | -0.025790391 | 0.089501009 | 0.273964202 | 0.908468501 | ns | TM | KS.3 |
| FIM1_1190 | 291.9425179 | 236.9946831 | 919.1149451 | -0.025047196 | 0.089567595 | 0.283511534 | 0.918450395 | ns | TM | KS.3 |
| FIM1_1191 | 80.92559478 | 32.89257704 | 87.82668476 | -0.10697047 | 0.290264659 | 0.017501756 | 0.304361209 | ns | TM | KS.3 |
| FIM1_1192 | 951.1141889 | 1036.074278 | 1024.594112 | 0.015332176 | 0.069782498 | 0.517280964 | 0.956681337 | ns | TM | KS.3 |
| FIM1_1193 | 4774.214372 | 4796.553199 | 4984.269302 | 0.001515594 | 0.032268541 | 0.920529024 | 0.991141351 | ns | TM | KS.3 |
| FIM1_1194 | 2701.2952 | 2840.089708 | 3125.589537 | 0.013825179 | 0.05490332 | 0.544319899 | 0.95917717 | ns | TM | KS.3 |
| FIM1_1195 | 350.1649725 | 415.8351353 | 301.4078431 | 0.040657798 | 0.084094437 | 0.153577681 | 0.783447458 | ns | TM | KS.3 |
| FIM1_1196 | 16057.05587 | 17318.69138 | 16415.82363 | 0.020906952 | 0.061287893 | 0.392249998 | 0.948637087 | ns | TM | KS.3 |
| FIM1_1197 | 5137.537574 | 4846.04499 | 4171.618958 | -0.015830729 | 0.05744788 | 0.49848997 | 0.953260948 | ns | TM | KS.3 |
| FIM1_1198 | 1166.470662 | 1264.372621 | 1483.304579 | 0.012941059 | 0.071380889 | 0.575581726 | 0.963192994 | ns | TM | KS.3 |
| FIM1_1199 | 1340.513622 | 1341.160253 | 1383.38834 | 7.73652490606455 | 0.048521465 | 0.997069052 | 0.999525119 | ns | TM | KS.3 |
| FIM1_1200 | 3533.807782 | 3551.825158 | 4476.768721 | 0.000717473 | 0.070662963 | 0.974116769 | 0.998607635 | ns | TM | KS.3 |
| FIM1_1201 | 228.3476163 | 225.7603211 | 205.2568071 | -0.002368805 | 0.063946096 | 0.917232458 | 0.991141351 | ns | TM | KS.3 |
| FIM1_1202 | 303.7686531 | 351.2363785 | 329.0836406 | 0.019473811 | 0.080459356 | 0.399485985 | 0.950410534 | ns | TM | KS.3 |
| FIM1_1203 | 1344.264653 | 1120.833686 | 1165.203984 | -0.019569235 | 0.087018431 | 0.377485369 | 0.948637087 | ns | TM | KS.3 |
| FIM1_1204 | 3824.349121 | 3330.371614 | 4190.507741 | -0.032167525 | 0.076692564 | 0.230249046 | 0.879350777 | ns | TM | KS.3 |
| FIM1_1205 | 704.259217 | 825.2221893 | 847.5103943 | 0.053024452 | 0.08451977 | 0.095501675 | 0.673713046 | ns | TM | KS.3 |
| FIM1_1206 | 780.8422383 | 780.7989086 | 779.5023305 | -0.000182889 | 0.051652702 | 0.993293721 | 0.999525119 | ns | TM | KS.3 |
| FIM1_1207 | 1927.432159 | 1808.574587 | 1871.703589 | -0.018802001 | 0.056632375 | 0.423477229 | 0.950410534 | ns | TM | KS.3 |
| FIM1_1208 | 1773.916245 | 1578.027726 | 1630.311595 | -0.041479402 | 0.072712218 | 0.15222262 | 0.783447458 | ns | TM | KS.3 |
| FIM1_1209 | 1013.954819 | 1075.386532 | 1031.406901 | 0.019678839 | 0.052932256 | 0.385218099 | 0.948637087 | ns | TM | KS.3 |
| FIM1_1210 | 508.0850255 | 468.6612026 | 392.2183859 | -0.012293947 | 0.072605363 | 0.59054092 | 0.963781856 | ns | TM | KS.3 |
| FIM1_1211 | 2202.478163 | 1883.372975 | 1425.858515 | -0.042961817 | 0.081927217 | 0.142456527 | 0.777709768 | ns | TM | KS.3 |
| FIM1_1212 | 8344.100208 | 8939.514156 | 9933.849973 | 0.021406258 | 0.057664833 | 0.370765784 | 0.948637087 | ns | TM | KS.3 |
| FIM1_1213 | 8371.819592 | 9034.632634 | 8591.439274 | 0.019070256 | 0.063003532 | 0.433633756 | 0.950410534 | ns | TM | KS.3 |
| FIM1_1214 | 18334.49374 | 18675.26191 | 18090.46764 | 0.005491183 | 0.044643499 | 0.783316439 | 0.988814136 | ns | TM | KS.3 |
| FIM1_1215 | 869.4723651 | 892.8846972 | 930.5813569 | 0.008183235 | 0.047183865 | 0.693256458 | 0.981188557 | ns | TM | KS.3 |
| FIM1_1216 | 806.8158885 | 811.6559467 | 932.4318863 | 0.001008494 | 0.0614844 | 0.964721854 | 0.995175461 | ns | TM | KS.3 |
| FIM1_1217 | 2775.28231 | 3431.068663 | 3883.21751 | 0.021325642 | 0.091249579 | 0.334972472 | 0.938982061 | ns | TM | KS.3 |
| FIM1_1218 | 576.8230394 | 583.3605627 | 430.1686544 | 0.002434064 | 0.060593925 | 0.915005373 | 0.991141351 | ns | TM | KS.3 |
| FIM1_1219 | 1095.647098 | 1129.009215 | 1348.642288 | 0.008422354 | 0.050274523 | 0.696310143 | 0.981479944 | ns | TM | KS.3 |
| FIM1_1220 | 589.3994394 | 475.9737635 | 623.5037668 | -0.038886121 | 0.091526747 | 0.153288146 | 0.783447458 | ns | TM | KS.3 |
| FIM1_1221 | 879.8917231 | 806.0217095 | 911.9478004 | -0.0358495 | 0.063300687 | 0.17965681 | 0.813278098 | ns | TM | KS.3 |
| FIM1_1222 | 5847.04263 | 5506.943646 | 5569.113365 | -0.031796362 | 0.048666819 | 0.142704139 | 0.777709768 | ns | TM | KS.3 |
| FIM1_1223 | 342.7698678 | 304.3300288 | 358.2853896 | -0.025086856 | 0.073571473 | 0.321977302 | 0.930003636 | ns | TM | KS.3 |
| FIM1_1224 | 297.1924944 | 292.4685795 | 283.1675351 | -0.004025381 | 0.059854094 | 0.860002927 | 0.991141351 | ns | TM | KS.3 |
| FIM1_1225 | 1807.955614 | 1840.364327 | 2384.065711 | 0.003726227 | 0.060273062 | 0.87035487 | 0.991141351 | ns | TM | KS.3 |
| FIM1_1227 | 3338.29261 | 3422.133213 | 4298.800342 | 0.003934417 | 0.068268217 | 0.861370883 | 0.991141351 | ns | TM | KS.3 |

| | | | | | | | | | | |
|---|---|---|---|---|---|---|---|---|---|---|
| FIM1_1228 | 1148.811693 | 1275.909209 | 1288.517643 | 0.023870431 | 0.069996166 | 0.344863843 | 0.946330732 | ns | TM | KS.3 |
| FIM1_1230 | 1960.425451 | 1880.001863 | 2123.311934 | -0.008751321 | 0.062008127 | 0.705559654 | 0.982962554 | ns | TM | KS.3 |
| FIM1_1231 | 3437.035742 | 3330.487241 | 3724.980557 | -0.006025615 | 0.063441097 | 0.79306357 | 0.988916276 | ns | TM | KS.3 |
| FIM1_1232 | 2615.745887 | 2566.217152 | 3261.024632 | -0.003867799 | 0.060849272 | 0.865613199 | 0.991141351 | ns | TM | KS.3 |
| FIM1_1233 | 1341.941854 | 1428.386179 | 1200.732231 | 0.009265224 | 0.071454623 | 0.681617249 | 0.97986479 | ns | TM | KS.3 |
| FIM1_1234 | 8381.015761 | 9032.325348 | 8850.688557 | 0.026104122 | 0.058262205 | 0.286214831 | 0.919993167 | ns | TM | KS.3 |
| FIM1_1235 | 1805.051213 | 1816.232812 | 2019.227918 | 0.001468771 | 0.053927318 | 0.947168778 | 0.994413583 | ns | TM | KS.3 |
| FIM1_1236 | 447.4624607 | 355.9339232 | 389.0121877 | -0.058267825 | 0.100685585 | 0.070321851 | 0.582627046 | ns | TM | KS.3 |
| FIM1_1239 | 2854.047896 | 2890.48451 | 2726.193429 | 0.002038733 | 0.067633827 | 0.927886152 | 0.993697171 | ns | TM | KS.3 |
| FIM1_1240 | 258.5425982 | 309.1406725 | 277.3851919 | 0.042143214 | 0.086034249 | 0.142518905 | 0.777709768 | ns | TM | KS.3 |
| FIM1_1242 | 717.1689686 | 803.469812 | 774.0457167 | 0.030475164 | 0.071223999 | 0.252333591 | 0.889064882 | ns | TM | KS.3 |
| FIM1_1243 | 3423.613396 | 3328.681257 | 3450.124753 | -0.009853841 | 0.036499892 | 0.553177426 | 0.95917717 | ns | TM | KS.3 |
| FIM1_1244 | 1018.724011 | 939.4366886 | 884.1417876 | -0.024036904 | 0.061876856 | 0.334499579 | 0.938982061 | ns | TM | KS.3 |
| FIM1_1245 | 1181.742939 | 1103.824885 | 1256.399244 | -0.016616031 | 0.061825688 | 0.4880387 | 0.952591289 | ns | TM | KS.3 |
| FIM1_1246 | 634.7265438 | 593.2537269 | 571.5752836 | -0.021277999 | 0.056917768 | 0.370324144 | 0.948637087 | ns | TM | KS.3 |
| FIM1_1247 | 927.8479669 | 894.4679416 | 887.4453676 | -0.008820048 | 0.05789513 | 0.700340625 | 0.98151979 | ns | TM | KS.3 |
| FIM1_1248 | 1547.579215 | 1745.418807 | 1548.077761 | 0.030965572 | 0.072744767 | 0.246391406 | 0.888934767 | ns | TM | KS.3 |
| FIM1_1249 | 2595.754424 | 3140.359959 | 4249.76047 | 0.019347172 | 0.088581988 | 0.377203657 | 0.948637087 | ns | TM | KS.3 |
| FIM1_1250 | 1340.137499 | 1499.233171 | 2503.527131 | 0.022888131 | 0.072196804 | 0.358874578 | 0.948637087 | ns | TM | KS.3 |
| FIM1_1251 | 3559.75482 | 3412.41412 | 3902.077065 | -0.011817606 | 0.052733146 | 0.595409136 | 0.965889818 | ns | TM | KS.3 |
| FIM1_1252 | 1547.591846 | 1699.570366 | 1817.874988 | 0.021525878 | 0.067544333 | 0.386805773 | 0.948637087 | ns | TM | KS.3 |
| FIM1_1253 | 403.6969259 | 418.3146858 | 403.9005518 | 0.009852573 | 0.05071084 | 0.650108054 | 0.976323032 | ns | TM | KS.3 |
| FIM1_1254 | 1222.892509 | 1249.544894 | 1261.360215 | 0.003493106 | 0.067352353 | 0.877192779 | 0.991141351 | ns | TM | KS.3 |
| FIM1_1255 | 1708.43615 | 1979.624311 | 2162.44669 | 0.014137488 | 0.085214232 | 0.503155608 | 0.953260948 | ns | TM | KS.3 |
| FIM1_1256 | 1356.523091 | 1195.937726 | 1269.138431 | -0.042063194 | 0.07499459 | 0.14983203 | 0.783447458 | ns | TM | KS.3 |
| FIM1_1257 | 1884.655882 | 1663.042788 | 1885.744787 | -0.035481894 | 0.074029544 | 0.200507054 | 0.845272486 | ns | TM | KS.3 |
| FIM1_1258 | 1354.397751 | 1424.414445 | 1152.889444 | 0.012812966 | 0.057264509 | 0.579440288 | 0.963781856 | ns | TM | KS.3 |
| FIM1_1259 | 349.2562866 | 360.821979 | 347.9151025 | 0.004810971 | 0.069427936 | 0.830249165 | 0.991141351 | ns | TM | KS.3 |
| FIM1_1261 | 2239.966375 | 2184.057909 | 2347.620967 | -0.004382909 | 0.065929791 | 0.847332606 | 0.991141351 | ns | TM | KS.3 |
| FIM1_1262 | 3683.628421 | 3682.671461 | 2501.382779 | -5.25E-05 | 0.069006771 | 0.998124645 | 0.999525119 | ns | TM | KS.3 |
| FIM1_1263 | 2136.953645 | 2073.760751 | 2066.89919 | -0.005821332 | 0.063150202 | 0.799862428 | 0.988916276 | ns | TM | KS.3 |
| FIM1_1264 | 2702.366263 | 2853.510632 | 2953.260636 | 0.009018277 | 0.068624998 | 0.693750658 | 0.981346978 | ns | TM | KS.3 |
| FIM1_1265 | 5376.024855 | 4924.421375 | 5046.128415 | -0.017237968 | 0.068747025 | 0.474047559 | 0.950410534 | ns | TM | KS.3 |
| FIM1_1266 | 786.6374318 | 839.4411775 | 805.668196 | 0.011283702 | 0.068613228 | 0.62635137 | 0.97344368 | ns | TM | KS.3 |
| FIM1_1267 | 3066.629983 | 3371.772297 | 4172.094383 | 0.008892987 | 0.082235296 | 0.666256177 | 0.976323032 | ns | TM | KS.3 |
| FIM1_1268 | 871.9556137 | 839.8602811 | 779.0367007 | -0.008252225 | 0.059899539 | 0.720245418 | 0.987157164 | ns | TM | KS.3 |
| FIM1_1269 | 2408.233879 | 2820.462045 | 3666.06524 | 0.014563749 | 0.086722232 | 0.487667805 | 0.952591289 | ns | TM | KS.3 |
| FIM1_1270 | 2879.176212 | 2730.870127 | 2826.111165 | -0.013546301 | 0.057633847 | 0.559744595 | 0.960914203 | ns | TM | KS.3 |
| FIM1_1271 | 3578.401444 | 3186.656567 | 3309.465527 | -0.038434069 | 0.072004779 | 0.174525087 | 0.811391014 | ns | TM | KS.3 |
| FIM1_1272 | 6798.732795 | 6932.762856 | 7662.356285 | 0.005624869 | 0.046957268 | 0.785500339 | 0.988916276 | ns | TM | KS.3 |
| FIM1_1273 | 577.6351695 | 617.5555991 | 549.5565453 | 0.012286974 | 0.068013938 | 0.598904815 | 0.968980748 | ns | TM | KS.3 |
| FIM1_1274 | 72.6080258 | 84.32470227 | 79.67623041 | 0.015606892 | 0.083697748 | 0.471681148 | 0.950410534 | ns | TM | KS.3 |
| FIM1_1275 | 3017.673379 | 2979.988869 | 2704.232452 | -0.003056921 | 0.054572289 | 0.890888944 | 0.991141351 | ns | TM | KS.3 |
| FIM1_1276 | 3530.34832 | 3300.124601 | 3207.325652 | -0.052624987 | 0.052576233 | 0.044229555 | 0.48254832 | ns | TM | KS.3 |
| FIM1_1277 | 2653.745961 | 2269.995788 | 2437.463365 | -0.033992647 | 0.080401581 | 0.207684864 | 0.852551729 | ns | TM | KS.3 |

| | | | | | | | | | | |
|---|---|---|---|---|---|---|---|---|---|---|
| FIM1_1279 | 1159.140548 | 1179.424869 | 1120.380077 | 0.00415347 | 0.053822531 | 0.851522309 | 0.991141351 | ns | TM | KS.3 |
| FIM1_1280 | 12096.65579 | 10384.19372 | 7070.403574 | -0.066357833 | 0.086244934 | 0.058272679 | 0.540287513 | ns | TM | KS.3 |
| FIM1_1281 | 1944.357508 | 2257.898241 | 2186.258833 | 0.065201965 | 0.085273337 | 0.060681432 | 0.544665909 | ns | TM | KS.3 |
| FIM1_1282 | 626.0209187 | 616.4258302 | 725.1315443 | -0.001500104 | 0.079419581 | 0.941594594 | 0.994413583 | ns | TM | KS.3 |
| FIM1_1283 | 4686.335748 | 3332.141434 | 3946.865104 | -0.046025131 | 0.118422239 | 0.096369702 | 0.673713046 | ns | TM | KS.3 |
| FIM1_1284 | 648.7403199 | 472.5861418 | 937.7489032 | -0.061690326 | 0.121596487 | 0.055036767 | 0.531428718 | ns | TM | KS.3 |
| FIM1_1285 | 4381.045137 | 3833.479912 | 4577.267726 | -0.062384001 | 0.080642651 | 0.06603915 | 0.564509918 | ns | TM | KS.3 |
| FIM1_1286 | 475.4873282 | 601.5977095 | 438.3162663 | 0.06233303 | 0.102898891 | 0.06051722 | 0.544665909 | ns | TM | KS.3 |
| FIM1_1287 | 1675.517094 | 1557.910984 | 1377.703421 | -0.009912741 | 0.07385558 | 0.656658429 | 0.976323032 | ns | TM | KS.3 |
| FIM1_1288 | 514.377948 | 547.2253586 | 608.8673932 | 0.014096931 | 0.06151683 | 0.551194489 | 0.95917717 | ns | TM | KS.3 |
| FIM1_1289 | 2186.88238 | 2145.945127 | 2209.892379 | -0.005009405 | 0.050840548 | 0.816924849 | 0.989579041 | ns | TM | KS.3 |
| FIM1_1290 | 4670.234109 | 4400.506953 | 4253.796382 | -0.014773574 | 0.059683611 | 0.530920831 | 0.959071703 | ns | TM | KS.3 |
| FIM1_1291 | 1132.70374 | 1134.354546 | 1057.93371 | 0.000185388 | 0.064931173 | 0.993489078 | 0.999525119 | ns | TM | KS.3 |
| FIM1_1292 | 233.4442248 | 239.1002356 | 242.678051 | 0.003745013 | 0.065940652 | 0.869158893 | 0.991141351 | ns | TM | KS.3 |
| FIM1_1293 | 2166.968452 | 2315.896073 | 2514.956573 | 0.009021763 | 0.073592189 | 0.684768373 | 0.980234354 | ns | TM | KS.3 |
| FIM1_1294 | 1022.213649 | 1089.341597 | 1168.931978 | 0.005998509 | 0.080775884 | 0.769949373 | 0.988814136 | ns | TM | KS.3 |
| FIM1_1295 | 3391.855137 | 3072.410593 | 3331.000546 | -0.027383668 | 0.067266731 | 0.290715985 | 0.919993167 | ns | TM | KS.3 |
| FIM1_1296 | 807.3721147 | 790.9812796 | 921.9107713 | -0.001443435 | 0.086649823 | 0.939813735 | 0.994413583 | ns | TM | KS.3 |
| FIM1_1297 | 1484.564004 | 603.6479486 | 1960.959462 | -1.041333286 | 0.384916723 | 2.23837416265 | 0.002589747 | Km<KS.3 | TM | KS.3 |
| FIM1_1298 | 3874.070258 | 3385.457219 | 2757.110115 | -0.020020936 | 0.078307257 | 0.395645895 | 0.948637087 | ns | TM | KS.3 |
| FIM1_1299 | 992.7582211 | 1028.472414 | 985.6162665 | 0.010092825 | 0.048929464 | 0.635033777 | 0.973888113 | ns | TM | KS.3 |
| FIM1_1300 | 1258.023373 | 1207.336329 | 1455.409244 | -0.008477025 | 0.061798779 | 0.714072829 | 0.986826511 | ns | TM | KS.3 |
| FIM1_1301 | 330.9850418 | 336.0100678 | 342.8589182 | 0.002723546 | 0.059650961 | 0.904815851 | 0.991141351 | ns | TM | KS.3 |
| FIM1_1302 | 723.0158814 | 656.8969192 | 736.6684656 | -0.02940308 | 0.065679006 | 0.260647927 | 0.897386461 | ns | TM | KS.3 |
| FIM1_1303 | 1635.605447 | 1825.286011 | 2040.5456 | 0.016725827 | 0.075080599 | 0.472969421 | 0.950410534 | ns | TM | KS.3 |
| FIM1_1304 | 1436.277156 | 1409.555577 | 1576.531809 | -0.0019502 | 0.077731283 | 0.925417935 | 0.99348841 | ns | TM | KS.3 |
| FIM1_1305 | 4081.960572 | 3602.381585 | 4427.895953 | -0.029831638 | 0.073804488 | 0.258791953 | 0.895871714 | ns | TM | KS.3 |
| FIM1_1306 | 113.6385394 | 110.5217533 | 117.1812301 | -0.003415648 | 0.072560197 | 0.875990533 | 0.991141351 | ns | TM | KS.3 |
| FIM1_1307 | 1135.657849 | 1164.613425 | 1081.113627 | 0.006036251 | 0.055647267 | 0.788634649 | 0.988916276 | ns | TM | KS.3 |
| FIM1_1308 | 768.4550844 | 789.4272038 | 883.5704345 | 0.006905436 | 0.053778213 | 0.756408614 | 0.988477371 | ns | TM | KS.3 |
| FIM1_1309 | 1117.960996 | 1094.082487 | 1078.806146 | -0.004311001 | 0.062108475 | 0.850620315 | 0.991141351 | ns | TM | KS.3 |
| FIM1_1310 | 1965.204953 | 2079.566489 | 2115.586315 | 0.017105228 | 0.054434041 | 0.455210118 | 0.950410534 | ns | TM | KS.3 |
| FIM1_1311 | 766.4602787 | 471.8352651 | 2063.53469 | -0.144238943 | 0.230382095 | 0.008305649 | 0.197706228 | ns | TM | KS.3 |
| FIM1_1312 | 1601.872741 | 1589.612529 | 1879.694692 | -0.001014067 | 0.07215998 | 0.963027682 | 0.995175461 | ns | TM | KS.3 |
| FIM1_1313 | 41277.59324 | 40081.23572 | 32699.65459 | -0.006323163 | 0.059940375 | 0.782721685 | 0.988814136 | ns | TM | KS.3 |
| FIM1_1314 | 732.537321 | 751.906836 | 895.8018927 | 0.003718337 | 0.070572074 | 0.867103519 | 0.991141351 | ns | TM | KS.3 |
| FIM1_1315 | 3483.716717 | 3456.452244 | 3495.315727 | -0.001425259 | 0.064021562 | 0.95009484 | 0.994413583 | ns | TM | KS.3 |
| FIM1_1316 | 769.8938848 | 917.2807881 | 918.0455375 | 0.029356776 | 0.083684889 | 0.246128325 | 0.888934767 | ns | TM | KS.3 |
| FIM1_1317 | 615.0977947 | 635.6234162 | 631.2855543 | 0.005454737 | 0.066937037 | 0.81050721 | 0.988916276 | ns | TM | KS.3 |
| FIM1_1318 | 1585.300451 | 1585.346362 | 1423.226802 | 1.96226927952269 | 0.056410549 | 0.999930384 | 0.999930384 | ns | TM | KS.3 |
| FIM1_1319 | 3325.440138 | 3317.284585 | 3635.495891 | -0.000317067 | 0.072113677 | 0.988436666 | 0.999333464 | ns | TM | KS.3 |
| FIM1_1320 | 13260.34227 | 12960.62696 | 13061.52587 | -0.00346197 | 0.069271444 | 0.877103026 | 0.991141351 | ns | TM | KS.3 |
| FIM1_1321 | 795.9784921 | 729.9815736 | 788.6044429 | -0.024687172 | 0.06391971 | 0.327863494 | 0.934739762 | ns | TM | KS.3 |
| FIM1_1322 | 9880.434983 | 8417.265241 | 7929.863146 | -0.023538932 | 0.081746205 | 0.325988544 | 0.934739762 | ns | TM | KS.3 |
| FIM1_1323 | 445.6942204 | 352.8211368 | 817.2490221 | -0.055572929 | 0.100437146 | 0.07705413 | 0.609656977 | ns | TM | KS.3 |

| | | | | | | | | | | |
|---|---|---|---|---|---|---|---|---|---|---|
| FIM1_1324 | 961.5831823 | 1076.820795 | 1164.683031 | 0.012294783 | 0.080578753 | 0.567565479 | 0.962265923 | ns | TM | KS.3 |
| FIM1_1325 | 1450.232798 | 1352.128619 | 1963.284815 | -0.014966539 | 0.064705269 | 0.5303688 | 0.959071703 | ns | TM | KS.3 |
| FIM1_1326 | 955.7707231 | 972.6061427 | 1051.740856 | 0.004884925 | 0.047589231 | 0.814667356 | 0.988916276 | ns | TM | KS.3 |
| FIM1_1327 | 2149.329936 | 1824.272527 | 2187.925136 | -0.082982123 | 0.091423451 | 0.032870434 | 0.417169416 | ns | TM | KS.3 |
| FIM1_1328 | 1573.77618 | 1543.644215 | 1573.607562 | -0.006622523 | 0.037434519 | 0.701023699 | 0.98151979 | ns | TM | KS.3 |
| FIM1_1329 | 718.0030552 | 588.8555148 | 1818.801545 | -0.028498374 | 0.087652247 | 0.246193826 | 0.888934767 | ns | TM | KS.3 |
| FIM1_1331 | 181.5195249 | 177.0506493 | 153.9760707 | -0.002155184 | 0.079921581 | 0.915842927 | 0.991141351 | ns | TM | KS.3 |
| FIM1_1332 | 758.7778461 | 684.3550806 | 565.8234931 | -0.028459681 | 0.068358261 | 0.276707405 | 0.908583046 | ns | TM | KS.3 |
| FIM1_1333 | 5703.546589 | 6572.617591 | 7441.272709 | 0.020437095 | 0.079479283 | 0.384437438 | 0.948637087 | ns | TM | KS.3 |
| FIM1_1334 | 4299.83136 | 3462.077242 | 3113.798748 | -0.040771216 | 0.092517076 | 0.141042223 | 0.776200289 | ns | TM | KS.3 |
| FIM1_1335 | 1158.387859 | 1047.196856 | 988.3723608 | -0.019652531 | 0.070789884 | 0.420273543 | 0.950410534 | ns | TM | KS.3 |
| FIM1_1336 | 655.2578241 | 913.0924108 | 1587.422816 | 0.044608518 | 0.11588089 | 0.102861126 | 0.689041746 | ns | TM | KS.3 |
| FIM1_1337 | 2165.17033 | 3032.495151 | 3774.680074 | 0.038602148 | 0.113962338 | 0.131904797 | 0.761283488 | ns | TM | KS.3 |
| FIM1_1338 | 665.9583725 | 575.7720223 | 857.7782986 | -0.024698984 | 0.078769459 | 0.316820042 | 0.927798616 | ns | TM | KS.3 |
| FIM1_1339 | 4463.318206 | 4346.720621 | 5129.626516 | -0.004515684 | 0.066391116 | 0.842582065 | 0.991141351 | ns | TM | KS.3 |
| FIM1_1340 | 2066.373714 | 2218.701365 | 2228.020992 | 0.019800057 | 0.060127138 | 0.412517518 | 0.950410534 | ns | TM | KS.3 |
| FIM1_1341 | 763.4538507 | 859.6536102 | 703.1858564 | 0.01626519 | 0.077517367 | 0.476325739 | 0.950410534 | ns | TM | KS.3 |
| FIM1_1342 | 2493.119265 | 2380.712866 | 2388.158002 | -0.00978324 | 0.061875411 | 0.673841843 | 0.976861203 | ns | TM | KS.3 |
| FIM1_1344 | 309.2978761 | 303.0833177 | 431.1494487 | -0.005674606 | 0.05528564 | 0.800482373 | 0.988916276 | ns | TM | KS.3 |
| FIM1_1345 | 804.6554003 | 1075.901571 | 1529.858814 | 0.191335892 | 0.143236848 | 0.001772965 | 0.071711409 | ns | TM | KS.3 |
| FIM1_1346 | 7930.837237 | 7580.333901 | 8008.702261 | -0.012175365 | 0.054661033 | 0.591059647 | 0.963781856 | ns | TM | KS.3 |
| FIM1_1347 | 17200.83514 | 17768.75977 | 16702.93723 | 0.00762887 | 0.057369866 | 0.737870581 | 0.988051177 | ns | TM | KS.3 |
| FIM1_1348 | 9631.685584 | 10074.08091 | 10458.08256 | 0.015330107 | 0.046494044 | 0.454896651 | 0.950410534 | ns | TM | KS.3 |
| FIM1_1349 | 1559.852184 | 1356.42001 | 1253.590956 | -0.079656895 | 0.084188011 | 0.035285508 | 0.42652037 | ns | TM | KS.3 |
| FIM1_1350 | 9031.719042 | 9025.011838 | 8388.563378 | -0.000135679 | 0.062464027 | 0.995251517 | 0.999525119 | ns | TM | KS.3 |
| FIM1_1352 | 2875.780282 | 2638.805797 | 2375.419536 | -0.032614437 | 0.062164253 | 0.210648173 | 0.854791728 | ns | TM | KS.3 |
| FIM1_1353 | 11331.26218 | 11003.33574 | 8281.703894 | -0.008888339 | 0.046107073 | 0.663367177 | 0.976323032 | ns | TM | KS.3 |
| FIM1_1354 | 1171.529787 | 1106.151282 | 1056.633872 | -0.012281049 | 0.063261892 | 0.60151188 | 0.97002321 | ns | TM | KS.3 |
| FIM1_1355 | 1513.166812 | 1419.197162 | 1650.334581 | -0.030949759 | 0.052087961 | 0.177416321 | 0.811391014 | ns | TM | KS.3 |
| FIM1_1356 | 764.2881185 | 734.2407645 | 718.3923749 | -0.010993581 | 0.052736243 | 0.620781128 | 0.973122675 | ns | TM | KS.3 |
| FIM1_1357 | 1105.253987 | 1035.854659 | 1025.871029 | -0.013778267 | 0.064262614 | 0.560983656 | 0.961073589 | ns | TM | KS.3 |
| FIM1_1358 | 1296.714256 | 1329.526581 | 1315.886861 | 0.006201361 | 0.054199406 | 0.781019028 | 0.988814136 | ns | TM | KS.3 |
| FIM1_1359 | 463.4783052 | 490.3108792 | 495.7212686 | 0.012716014 | 0.061906644 | 0.588850019 | 0.963781856 | ns | TM | KS.3 |
| FIM1_1360 | 1774.247489 | 1620.18187 | 2675.62504 | -0.011728824 | 0.076028561 | 0.597471172 | 0.9678814 | ns | TM | KS.3 |
| FIM1_1361 | 284.0713815 | 227.596631 | 279.5368154 | -0.05623856 | 0.098067134 | 0.076974671 | 0.609656977 | ns | TM | KS.3 |
| FIM1_1362 | 529.3713009 | 456.7026776 | 484.6343487 | -0.036670682 | 0.079276242 | 0.187039423 | 0.82775868 | ns | TM | KS.3 |
| FIM1_1363 | 2782.220055 | 3141.458927 | 3829.713585 | 0.012251945 | 0.082447431 | 0.562830422 | 0.961073589 | ns | TM | KS.3 |
| FIM1_1364 | 330.0630402 | 380.8358206 | 299.4874911 | 0.018674027 | 0.080524764 | 0.41499213 | 0.950410534 | ns | TM | KS.3 |
| FIM1_1365 | 2496.899593 | 2519.821624 | 2515.753909 | 0.001768307 | 0.061565091 | 0.938214365 | 0.994413583 | ns | TM | KS.3 |
| FIM1_1366 | 1263.846819 | 1307.438948 | 1390.123103 | 0.007441559 | 0.05978412 | 0.746110712 | 0.988477371 | ns | TM | KS.3 |
| FIM1_1367 | 959.740926 | 904.1802294 | 982.0969972 | -0.019413918 | 0.052788354 | 0.390551781 | 0.948637087 | ns | TM | KS.3 |
| FIM1_1368 | 6879.128534 | 5987.557917 | 5774.293505 | -0.077281908 | 0.08369422 | 0.038331495 | 0.444235187 | ns | TM | KS.3 |
| FIM1_1369 | 1549.068994 | 1737.688568 | 3750.884313 | 0.019890066 | 0.074177493 | 0.409165195 | 0.950410534 | ns | TM | KS.3 |
| FIM1_1370 | 13.14792553 | 14.37638182 | 34.13613517 | 0.004862221 | 0.092710182 | 0.792465871 | 0.988916276 | ns | TM | KS.3 |
| FIM1_1371 | 1253.254875 | 1607.150886 | 1244.944897 | 0.195355946 | 0.114289622 | 0.00107673 | 0.049144341 | ns | TM | KS.3 |

| | | | | | | | | | | |
|---|---|---|---|---|---|---|---|---|---|---|
| FIM1_1372 | 326.7093854 | 422.4007276 | 292.5230999 | 0.068543866 | 0.109701079 | 0.047358548 | 0.497064927 | ns | TM | KS.3 |
| FIM1_1373 | 1533.618119 | 2860.380094 | 2413.523477 | 0.876850092 | 0.114827684 | 9.48347507598 | 1.17777544215 | ns | TM | KS.3 |
| FIM1_1374 | 2325.465098 | 2224.169916 | 2208.060344 | -0.013495426 | 0.050976485 | 0.537638568 | 0.95917717 | ns | TM | KS.3 |
| FIM1_1375 | 1653.477571 | 1393.903852 | 1570.463931 | -0.016193989 | 0.087400035 | 0.446461375 | 0.950410534 | ns | TM | KS.3 |
| FIM1_1376 | 106.7118651 | 59.3433511 | 96.17648643 | -0.628718294 | 0.327472641 | 8.86232655611 | 0.00797357 | ns | TM | KS.3 |
| FIM1_1377 | 2930.205102 | 3087.735096 | 3826.541247 | 0.01551007 | 0.053779148 | 0.493801932 | 0.952949531 | ns | TM | KS.3 |
| FIM1_1378 | 347.8259076 | 301.4451815 | 321.4348399 | -0.030570987 | 0.078203009 | 0.24460522 | 0.88883001 | ns | TM | KS.3 |
| FIM1_1379 | 258.027303 | 237.7663435 | 272.0880493 | -0.011053726 | 0.074271283 | 0.621665051 | 0.973122675 | ns | TM | KS.3 |
| FIM1_1380 | 1418.086004 | 1533.322767 | 2315.659662 | 0.008976992 | 0.07746562 | 0.676624391 | 0.979149565 | ns | TM | KS.3 |
| FIM1_1382 | 3905.441439 | 3813.951868 | 4035.366892 | -0.007930174 | 0.034761387 | 0.619470228 | 0.973122675 | ns | TM | KS.3 |
| FIM1_1383 | 2830.073182 | 2634.768851 | 2587.870545 | -0.020300928 | 0.059803888 | 0.401006674 | 0.950410534 | ns | TM | KS.3 |
| FIM1_1384 | 1030.947889 | 1091.300179 | 1200.455473 | 0.011827082 | 0.063573252 | 0.614308986 | 0.973122675 | ns | TM | KS.3 |
| FIM1_1385 | 4850.532043 | 4082.89287 | 5001.549149 | -0.037437495 | 0.08385683 | 0.17497048 | 0.811391014 | ns | TM | KS.3 |
| FIM1_1386 | 474.9182901 | 631.4022373 | 571.1931895 | 0.160857557 | 0.14052451 | 0.003774734 | 0.115921601 | ns | TM | KS.3 |
| FIM1_1387 | 2866.293493 | 3004.752452 | 2965.730192 | 0.024164524 | 0.04019947 | 0.169660022 | 0.808485258 | ns | TM | KS.3 |
| FIM1_1388 | 411.2167041 | 373.424735 | 383.3514424 | -0.033791483 | 0.066561934 | 0.209998035 | 0.852849163 | ns | TM | KS.3 |
| FIM1_1389 | 759.0972368 | 704.137034 | 608.2360905 | -0.021877036 | 0.060935601 | 0.372101074 | 0.948637087 | ns | TM | KS.3 |
| FIM1_1390 | 2728.915187 | 2599.14076 | 2724.645374 | -0.007646072 | 0.069463957 | 0.735871153 | 0.987655486 | ns | TM | KS.3 |
| FIM1_1391 | 5620.524174 | 5000.060479 | 3696.141313 | -0.009057031 | 0.08719733 | 0.647571876 | 0.976323032 | ns | TM | KS.3 |
| FIM1_1392 | 1201.09264 | 1528.933578 | 1101.82239 | 0.06296686 | 0.104692715 | 0.058456771 | 0.540287513 | ns | TM | KS.3 |
| FIM1_1393 | 1285.137785 | 1322.140626 | 1062.710524 | 0.004445997 | 0.068566069 | 0.843522129 | 0.991141351 | ns | TM | KS.3 |
| FIM1_1394 | 556.1501808 | 405.3251762 | 587.753791 | -0.045818041 | 0.113290205 | 0.099464351 | 0.677856366 | ns | TM | KS.3 |
| FIM1_1395 | 869.1505266 | 905.347024 | 753.8510585 | 0.004663538 | 0.076206386 | 0.826616035 | 0.991141351 | ns | TM | KS.3 |
| FIM1_1396 | 1470.524615 | 1310.908963 | 1338.63058 | -0.023123547 | 0.072716031 | 0.354162062 | 0.948637087 | ns | TM | KS.3 |
| FIM1_1397 | 5692.445397 | 5007.190515 | 4469.632152 | -0.034809113 | 0.074677221 | 0.206464305 | 0.850925432 | ns | TM | KS.3 |
| FIM1_1399 | 2982.426026 | 2509.683821 | 2805.973654 | -0.028661951 | 0.083196099 | 0.254944497 | 0.892600401 | ns | TM | KS.3 |
| FIM1_1400 | 1484.640509 | 1704.413933 | 1902.072314 | 0.019857637 | 0.079056854 | 0.396486796 | 0.948783939 | ns | TM | KS.3 |
| FIM1_1401 | 2158.8283 | 3138.185931 | 3542.249151 | 0.055425855 | 0.131353686 | 0.065791541 | 0.564318013 | ns | TM | KS.3 |
| FIM1_1402 | 733.7981901 | 773.679664 | 872.8792913 | 0.016984727 | 0.052825152 | 0.450524489 | 0.950410534 | ns | TM | KS.3 |
| FIM1_1403 | 689.959237 | 608.5101924 | 747.890367 | -0.029316023 | 0.074059438 | 0.264680966 | 0.90372202 | ns | TM | KS.3 |
| FIM1_1404 | 986.5177175 | 520.5324969 | 894.6827909 | -0.172947667 | 0.305545151 | 0.006920183 | 0.176553399 | ns | TM | KS.3 |
| FIM1_1405 | 818.7503931 | 1182.977296 | 849.0081097 | 0.09993903 | 0.157886483 | 0.017652549 | 0.304361209 | ns | TM | KS.3 |
| FIM1_1406 | 6636.439211 | 6307.328519 | 6026.309982 | -0.006431842 | 0.07423978 | 0.76813136 | 0.988477371 | ns | TM | KS.3 |
| FIM1_1407 | 1390.951115 | 1361.919106 | 1369.341169 | -0.005404281 | 0.050434068 | 0.802031537 | 0.988916276 | ns | TM | KS.3 |
| FIM1_1408 | 5003.444488 | 4809.395168 | 7942.031075 | -0.010032029 | 0.05563246 | 0.658562408 | 0.976323032 | ns | TM | KS.3 |
| FIM1_1409 | 1922.15212 | 2155.996874 | 2107.260258 | 0.00904595 | 0.086718882 | 0.649163121 | 0.976323032 | ns | TM | KS.3 |
| FIM1_1410 | 3796.94647 | 4325.137147 | 3987.427979 | 0.031385467 | 0.074974054 | 0.240269335 | 0.887413624 | ns | TM | KS.3 |
| FIM1_1411 | 1382.045454 | 1357.678402 | 1205.529616 | -0.00160902 | 0.080778845 | 0.936523538 | 0.994413583 | ns | TM | KS.3 |
| FIM1_1412 | 3772.328156 | 3454.658378 | 4924.602873 | -0.012994546 | 0.073500669 | 0.569297467 | 0.962265923 | ns | TM | KS.3 |
| FIM1_1413 | 541.5617317 | 576.5225053 | 623.2557519 | 0.005877012 | 0.080971973 | 0.773935553 | 0.988814136 | ns | TM | KS.3 |
| FIM1_1414 | 334.6570874 | 326.7744137 | 305.0673911 | -0.003923724 | 0.067322147 | 0.86233335 | 0.991141351 | ns | TM | KS.3 |
| FIM1_1415 | 1627.900695 | 1852.281456 | 1607.211688 | 0.074280474 | 0.080361806 | 0.041227353 | 0.462993409 | ns | TM | KS.3 |
| FIM1_1416 | 1421.325373 | 1457.708804 | 1016.078387 | 0.00737786 | 0.048424986 | 0.726443751 | 0.987157164 | ns | TM | KS.3 |
| FIM1_1418 | 1066.286084 | 1116.917002 | 1019.213384 | 0.014283548 | 0.04922695 | 0.504839473 | 0.953260948 | ns | TM | KS.3 |
| FIM1_1420 | 1307.558768 | 1490.230744 | 1437.044234 | 0.047769301 | 0.077117975 | 0.117877564 | 0.724896022 | ns | TM | KS.3 |

| | | | | | | | | | | |
|---|---|---|---|---|---|---|---|---|---|---|
| FIM1_1421 | 574.4455932 | 593.8257235 | 553.3328384 | 0.004118107 | 0.074487825 | 0.848670131 | 0.991141351 | ns | TM | KS.3 |
| FIM1_1422 | 477.8230248 | 619.5016516 | 486.6103039 | 0.057203871 | 0.106462795 | 0.06948202 | 0.581836521 | ns | TM | KS.3 |
| FIM1_1423 | 709.0228983 | 786.2568265 | 793.3963286 | 0.048131537 | 0.0695332 | 0.106713299 | 0.700394017 | ns | TM | KS.3 |
| FIM1_1424 | 1372.17646 | 1317.580085 | 1449.15332 | -0.014325865 | 0.04555177 | 0.47786131 | 0.950410534 | ns | TM | KS.3 |
| FIM1_1425 | 1117.879717 | 1097.553025 | 1331.415899 | -0.003633366 | 0.062117121 | 0.873742437 | 0.991141351 | ns | TM | KS.3 |
| FIM1_1426 | 3177.773899 | 2930.988883 | 3731.823932 | -0.006041905 | 0.086445883 | 0.756659651 | 0.988477371 | ns | TM | KS.3 |
| FIM1_1427 | 28.8816751 | 38.94174594 | 38.6833852 | 0.027635322 | 0.104160675 | 0.226043781 | 0.876782875 | ns | TM | KS.3 |
| FIM1_1428 | 2421.070394 | 2338.028457 | 2119.551811 | -0.007092485 | 0.061883957 | 0.758204961 | 0.988477371 | ns | TM | KS.3 |
| FIM1_1429 | 2028.685827 | 2039.599953 | 2233.881656 | 0.000999159 | 0.063109685 | 0.965035846 | 0.995175461 | ns | TM | KS.3 |
| FIM1_1430 | 1231.125651 | 1187.869699 | 1121.095545 | -0.006386716 | 0.065542564 | 0.780448089 | 0.988814136 | ns | TM | KS.3 |
| FIM1_1431 | 2109.367569 | 2083.783996 | 2212.056116 | -0.002081326 | 0.066091019 | 0.926837779 | 0.993697171 | ns | TM | KS.3 |
| FIM1_1432 | 14797.70775 | 12732.15517 | 13282.27271 | -0.054042766 | 0.083187523 | 0.092190803 | 0.665673792 | ns | TM | KS.3 |
| FIM1_1433 | 1911.357236 | 1850.834335 | 1941.906229 | -0.009585798 | 0.047743095 | 0.646802447 | 0.976323032 | ns | TM | KS.3 |
| FIM1_1434 | 1881.185918 | 1785.891776 | 1918.794881 | -0.006021053 | 0.076134598 | 0.778663334 | 0.988814136 | ns | TM | KS.3 |
| FIM1_1435 | 5741.18369 | 4902.349331 | 5628.031343 | -0.035857958 | 0.080941918 | 0.191273758 | 0.829253069 | ns | TM | KS.3 |
| FIM1_1436 | 548.9415132 | 494.5953305 | 560.7691604 | -0.023310549 | 0.070265388 | 0.353951089 | 0.948637087 | ns | TM | KS.3 |
| FIM1_1437 | 373.2911024 | 362.9967989 | 339.2170924 | -0.006828993 | 0.049748562 | 0.749856158 | 0.988477371 | ns | TM | KS.3 |
| FIM1_1438 | 56986.20218 | 59973.43748 | 60172.18324 | 0.017073962 | 0.049865571 | 0.43062947 | 0.950410534 | ns | TM | KS.3 |
| FIM1_1439 | 856.0418249 | 906.6128751 | 1220.359772 | 0.014046722 | 0.060154191 | 0.55085819 | 0.95917717 | ns | TM | KS.3 |
| FIM1_1440 | 351.9297253 | 356.3825693 | 401.8067455 | 0.002473821 | 0.054038141 | 0.911249573 | 0.991141351 | ns | TM | KS.3 |
| FIM1_1441 | 113.6546019 | 134.8519418 | 322.5394479 | 0.015306944 | 0.087961918 | 0.465659018 | 0.950410534 | ns | TM | KS.3 |
| FIM1_1442 | 56.3154845 | 48.274216 | 578.1003268 | -0.012255401 | 0.088903945 | 0.543448667 | 0.95917717 | ns | TM | KS.3 |
| FIM1_1443 | 453.2656178 | 474.0875621 | 494.6692785 | 0.009961122 | 0.062638368 | 0.668676209 | 0.976323032 | ns | TM | KS.3 |
| FIM1_1444 | 12747.88942 | 13398.22951 | 12306.94573 | 0.011254624 | 0.060614027 | 0.629356891 | 0.97344368 | ns | TM | KS.3 |
| FIM1_1445 | 759.1222865 | 912.5741447 | 1243.352178 | 0.019224573 | 0.087616594 | 0.382278829 | 0.948637087 | ns | TM | KS.3 |
| FIM1_1446 | 643.5383097 | 675.3665145 | 640.2136342 | 0.009559735 | 0.063319543 | 0.680846867 | 0.97986479 | ns | TM | KS.3 |
| FIM1_1447 | 839.7446882 | 945.6162777 | 788.7167144 | 0.016660999 | 0.07709765 | 0.468602351 | 0.950410534 | ns | TM | KS.3 |
| FIM1_1448 | 495.7279109 | 581.2771369 | 525.5591259 | 0.027280237 | 0.080649486 | 0.277431837 | 0.908583046 | ns | TM | KS.3 |
| FIM1_1449 | 2519.68233 | 2407.019697 | 2542.263048 | -0.012378208 | 0.05448565 | 0.584539398 | 0.963781856 | ns | TM | KS.3 |
| FIM1_1450 | 820.35105 | 791.2716389 | 920.6746159 | -0.008028793 | 0.059846061 | 0.727310425 | 0.987157164 | ns | TM | KS.3 |
| FIM1_1451 | 11534.66962 | 10806.83993 | 8214.938459 | -0.007688071 | 0.076285187 | 0.721598305 | 0.987157164 | ns | TM | KS.3 |
| FIM1_1453 | 3255.531591 | 3763.594417 | 3848.212792 | 0.015971836 | 0.082977727 | 0.465449924 | 0.950410534 | ns | TM | KS.3 |
| FIM1_1454 | 825.2498273 | 869.6586234 | 1125.145479 | 0.011794677 | 0.060541087 | 0.613674549 | 0.973122675 | ns | TM | KS.3 |
| FIM1_1455 | 1283.915846 | 1571.333385 | 1192.667858 | 0.036620149 | 0.089013592 | 0.172028899 | 0.811391014 | ns | TM | KS.3 |
| FIM1_1456 | 430.9210747 | 533.6486205 | 560.0456439 | 0.060432672 | 0.097871514 | 0.06715437 | 0.567220699 | ns | TM | KS.3 |
| FIM1_1457 | 3484.116305 | 3204.968403 | 2947.262511 | -0.018332336 | 0.066393247 | 0.451369427 | 0.950410534 | ns | TM | KS.3 |
| FIM1_1458 | 1393.620179 | 1434.102718 | 1399.980782 | 0.006866091 | 0.056520001 | 0.761899175 | 0.988477371 | ns | TM | KS.3 |
| FIM1_1460 | 1210.538966 | 1267.914187 | 1103.395768 | 0.015046843 | 0.048454518 | 0.477559848 | 0.950410534 | ns | TM | KS.3 |
| FIM1_1461 | 1610.954326 | 1558.62835 | 986.9451075 | -0.003372577 | 0.078379509 | 0.870996615 | 0.991141351 | ns | TM | KS.3 |
| FIM1_1462 | 1525.121612 | 1496.405744 | 1477.699355 | -0.005271281 | 0.049317501 | 0.804447531 | 0.988916276 | ns | TM | KS.3 |
| FIM1_1463 | 224.2037721 | 242.7512153 | 239.1789131 | 0.012873744 | 0.070206715 | 0.57963816 | 0.963781856 | ns | TM | KS.3 |
| FIM1_1464 | 26.72634036 | 15.30007873 | 24.69833755 | -0.030066586 | 0.142542916 | 0.173477833 | 0.811391014 | ns | TM | KS.3 |
| FIM1_1465 | 1168.439679 | 1014.100578 | 1340.776731 | -0.057363453 | 0.081770292 | 0.080989436 | 0.619880684 | ns | TM | KS.3 |
| FIM1_1466 | 1329.823676 | 1314.542405 | 1195.574456 | -0.003266973 | 0.050105519 | 0.878954156 | 0.991141351 | ns | TM | KS.3 |
| FIM1_1467 | 2629.1538 | 2347.226497 | 2349.248935 | -0.026969519 | 0.071375228 | 0.297216436 | 0.926536521 | ns | TM | KS.3 |

| | | | | | | | | | | |
|---|---|---|---|---|---|---|---|---|---|---|
| FIM1_1468 | 3368.964441 | 3182.294788 | 3250.144257 | -0.01805079 | 0.053418568 | 0.42678999 | 0.950410534 | ns | TM | KS.3 |
| FIM1_1469 | 441.9381717 | 323.053745 | 198.5580107 | -0.079984976 | 0.12992374 | 0.030749725 | 0.402578636 | ns | TM | KS.3 |
| FIM1_1470 | 1766.944548 | 2000.393526 | 1860.483098 | 0.083024664 | 0.078437267 | 0.028420933 | 0.390786333 | ns | TM | KS.3 |
| FIM1_1471 | 5651.933274 | 5032.888146 | 4424.68561 | -0.036724298 | 0.071857689 | 0.188827452 | 0.828217874 | ns | TM | KS.3 |
| FIM1_1472 | 1637.832696 | 1666.938517 | 1066.927055 | 0.003606526 | 0.06094849 | 0.874578572 | 0.991141351 | ns | TM | KS.3 |
| FIM1_1473 | 969.5448612 | 933.9471216 | 1273.656244 | -0.009018902 | 0.05742499 | 0.693438988 | 0.981188557 | ns | TM | KS.3 |
| FIM1_1474 | 2655.735415 | 2581.393924 | 2614.068586 | -0.005863209 | 0.061191732 | 0.79837406 | 0.988916276 | ns | TM | KS.3 |
| FIM1_1475 | 1349.247721 | 1125.089261 | 1427.26863 | -0.116975553 | 0.097548889 | 0.010999117 | 0.234852386 | ns | TM | KS.3 |
| FIM1_1476 | 104.7479809 | 58.95890292 | 1083.108515 | -0.185208378 | 0.291726509 | 0.005476799 | 0.152304407 | ns | TM | KS.3 |
| FIM1_1477 | 1724.72079 | 1990.652333 | 1843.773248 | 0.016724564 | 0.082050069 | 0.451337492 | 0.950410534 | ns | TM | KS.3 |
| FIM1_1478 | 1211.091162 | 1439.39144 | 1710.692423 | 0.021936524 | 0.084453898 | 0.343446326 | 0.944808611 | ns | TM | KS.3 |
| FIM1_1479 | 1107.558926 | 863.0066415 | 809.3494458 | -0.228097592 | 0.103716204 | 0.000255003 | 0.017944075 | ns | TM | KS.3 |
| FIM1_1480 | 840.9210899 | 972.1835074 | 905.3788425 | 0.020679845 | 0.079901761 | 0.37876096 | 0.948637087 | ns | TM | KS.3 |
| FIM1_1481 | 540.4836312 | 505.5947397 | 507.7294437 | -0.011929642 | 0.068482543 | 0.608244827 | 0.970937835 | ns | TM | KS.3 |
| FIM1_1482 | 2439.965036 | 5099.32711 | 6288.323436 | 1.024261205 | 0.141766796 | 1.67074966841 | 1.38532993339 | Km>KS.3 | TM | KS.3 |
| FIM1_1483 | 349.3439807 | 419.5881302 | 398.8084871 | 0.029584632 | 0.084963743 | 0.240730973 | 0.887413624 | ns | TM | KS.3 |
| FIM1_1484 | 886.6606772 | 856.7801518 | 742.8195055 | -0.008052602 | 0.057909384 | 0.724799676 | 0.987157164 | ns | TM | KS.3 |
| FIM1_1485 | 3581.827374 | 3505.658573 | 3134.913706 | -0.006773076 | 0.039879545 | 0.710440107 | 0.984652342 | ns | TM | KS.3 |
| FIM1_1486 | 3322.518934 | 3230.937307 | 2021.714419 | -0.004355335 | 0.06869909 | 0.846534054 | 0.991141351 | ns | TM | KS.3 |
| FIM1_1487 | 1487.018758 | 1749.355512 | 1944.335533 | 0.017357663 | 0.085051179 | 0.427856959 | 0.950410534 | ns | TM | KS.3 |
| FIM1_1488 | 5131.181594 | 5130.658749 | 5118.618575 | 6.36761832201824 | 0.036221332 | 0.999699705 | 0.999930384 | ns | TM | KS.3 |
| FIM1_1489 | 528.794881 | 516.2123717 | 496.6185416 | -0.005203701 | 0.060915619 | 0.820330835 | 0.990886949 | ns | TM | KS.3 |
| FIM1_1490 | 646.1219068 | 629.6448535 | 542.9241269 | -0.005480699 | 0.062376617 | 0.811231048 | 0.988916276 | ns | TM | KS.3 |
| FIM1_1491 | 1120.275599 | 1082.230738 | 1008.296104 | -0.01081848 | 0.044893868 | 0.588627296 | 0.963781856 | ns | TM | KS.3 |
| FIM1_1492 | 1871.915883 | 1897.719721 | 1549.742596 | 0.003328602 | 0.053399208 | 0.880299782 | 0.991141351 | ns | TM | KS.3 |
| FIM1_1493 | 871.5895156 | 725.2436425 | 639.7551531 | -0.049339291 | 0.088852465 | 0.106312374 | 0.700394017 | ns | TM | KS.3 |
| FIM1_1494 | 266.6837966 | 254.5856675 | 226.628268 | -0.008023295 | 0.064906846 | 0.728032366 | 0.987157164 | ns | TM | KS.3 |
| FIM1_1495 | 67.55344992 | 72.08804598 | 72.26210851 | 0.005702553 | 0.081278788 | 0.77965858 | 0.988814136 | ns | TM | KS.3 |
| FIM1_1496 | 731.2609937 | 781.7969619 | 888.9900235 | 0.018778743 | 0.057927404 | 0.42865986 | 0.950410534 | ns | TM | KS.3 |
| FIM1_1497 | 472.3314822 | 581.248362 | 541.8235652 | 0.103489057 | 0.105251176 | 0.017137538 | 0.302986707 | ns | TM | KS.3 |
| FIM1_1498 | 604.0365002 | 525.3506452 | 589.1756182 | -0.03480235 | 0.077327981 | 0.204667951 | 0.847812702 | ns | TM | KS.3 |
| FIM1_1499 | 2537.523629 | 3017.918625 | 1990.863723 | 0.031059014 | 0.083361056 | 0.229278293 | 0.879350777 | ns | TM | KS.3 |
| FIM1_1500 | 1647.734295 | 1631.036419 | 1542.250736 | -0.002295809 | 0.056272552 | 0.918701671 | 0.991141351 | ns | TM | KS.3 |
| FIM1_1501 | 1474.278901 | 1585.209622 | 1400.018085 | 0.017605367 | 0.062868182 | 0.466043784 | 0.950410534 | ns | TM | KS.3 |
| FIM1_1503 | 2034.867233 | 2516.415784 | 2153.953742 | 0.17799536 | 0.09766036 | 0.001398751 | 0.061438441 | ns | TM | KS.3 |
| FIM1_1504 | 922.9332463 | 867.3264099 | 912.8558115 | -0.021844424 | 0.053326946 | 0.33969194 | 0.943910292 | ns | TM | KS.3 |
| FIM1_1505 | 588.1800434 | 514.6748017 | 530.282422 | -0.039573306 | 0.076559921 | 0.167154277 | 0.80502665 | ns | TM | KS.3 |
| FIM1_1506 | 133.7327116 | 134.7282174 | 147.5683477 | 0.000945669 | 0.076549974 | 0.96416268 | 0.995175461 | ns | TM | KS.3 |
| FIM1_1507 | 6464.116776 | 6112.255272 | 4989.294237 | -0.013726931 | 0.059451391 | 0.558363642 | 0.960200179 | ns | TM | KS.3 |
| FIM1_1508 | 1804.753298 | 1842.734492 | 1713.87884 | 0.00527505 | 0.052638981 | 0.810580615 | 0.988916276 | ns | TM | KS.3 |
| FIM1_1509 | 1635.114879 | 1625.716644 | 1673.109911 | -0.001325574 | 0.056975473 | 0.953135017 | 0.994413583 | ns | TM | KS.3 |
| FIM1_1510 | 1410.998359 | 1337.432806 | 1336.410245 | -0.014438067 | 0.056834671 | 0.533382381 | 0.959071703 | ns | TM | KS.3 |
| FIM1_1511 | 220.2669211 | 223.4048886 | 223.070927 | 0.002270112 | 0.066021219 | 0.920258459 | 0.991141351 | ns | TM | KS.3 |
| FIM1_1512 | 638.6051582 | 615.1757626 | 639.4952849 | -0.008655182 | 0.059395886 | 0.707129496 | 0.983265883 | ns | TM | KS.3 |
| FIM1_1513 | 5823.453377 | 5373.060334 | 4753.957716 | -0.033760707 | 0.059797729 | 0.187888042 | 0.828217874 | ns | TM | KS.3 |

| | | | | | | | | | | |
|---|---|---|---|---|---|---|---|---|---|---|
| FIM1_1514 | 14475.85675 | 13651.04367 | 11778.31794 | -0.015684256 | 0.057834554 | 0.503303744 | 0.953260948 | ns | TM | KS.3 |
| FIM1_1515 | 349.5894174 | 275.3342973 | 365.2917633 | -0.067842341 | 0.105735486 | 0.049792078 | 0.511210716 | ns | TM | KS.3 |
| FIM1_1516 | 71.41255423 | 49.34097205 | 209.8896237 | -0.025114055 | 0.115331482 | 0.242810831 | 0.887413624 | ns | TM | KS.3 |
| FIM1_1518 | 1446.336572 | 1505.897053 | 1559.237096 | 0.009298709 | 0.05884241 | 0.686477416 | 0.980879562 | ns | TM | KS.3 |
| FIM1_1519 | 667.3530772 | 683.5164066 | 582.3236775 | 0.005410195 | 0.058123423 | 0.812128834 | 0.988916276 | ns | TM | KS.3 |
| FIM1_1520 | 5086.805535 | 4613.476859 | 4137.183329 | -0.022596995 | 0.068328339 | 0.367225882 | 0.948637087 | ns | TM | KS.3 |
| FIM1_1521 | 2727.287525 | 3310.53979 | 4002.233212 | 0.015797479 | 0.091735415 | 0.443582982 | 0.950410534 | ns | TM | KS.3 |
| FIM1_1523 | 436.2552977 | 367.2373774 | 369.144513 | -0.059231837 | 0.089473714 | 0.074563601 | 0.599104608 | ns | TM | KS.3 |
| FIM1_1524 | 1264.997731 | 1247.736438 | 1103.512142 | -0.003322033 | 0.056221755 | 0.882654995 | 0.991141351 | ns | TM | KS.3 |
| FIM1_1525 | 2153.604624 | 1853.304627 | 1670.902005 | -0.123689704 | 0.083075564 | 0.007076115 | 0.178698832 | ns | TM | KS.3 |
| FIM1_1526 | 16188.63432 | 11450.31636 | 7536.909485 | -0.331481765 | 0.168186347 | 8.7055080600 | 0.00797357 | ns | TM | KS.3 |
| FIM1_1527 | 521.9181806 | 518.7733527 | 531.9941973 | -0.001830374 | 0.049065351 | 0.931191296 | 0.994058033 | ns | TM | KS.3 |
| FIM1_1528 | 619.7477893 | 729.2409245 | 668.6270175 | 0.013599089 | 0.088658385 | 0.506965153 | 0.953260948 | ns | TM | KS.3 |
| FIM1_1529 | 1402.650129 | 1193.962765 | 1461.92079 | -0.052793658 | 0.085161722 | 0.096080756 | 0.673713046 | ns | TM | KS.3 |
| FIM1_1530 | 1965.240587 | 1916.873009 | 2471.08405 | -0.005385734 | 0.059223711 | 0.813605221 | 0.988916276 | ns | TM | KS.3 |
| FIM1_1531 | 984.5895503 | 869.9647101 | 741.0917151 | -0.02056259 | 0.075717135 | 0.39270776 | 0.948637087 | ns | TM | KS.3 |
| FIM1_1532 | 1043.617476 | 880.9732307 | 768.8674388 | -0.02237752 | 0.083741117 | 0.338367177 | 0.942012706 | ns | TM | KS.3 |
| FIM1_1533 | 754.8926525 | 719.3655853 | 731.9239523 | -0.012832241 | 0.055849265 | 0.57536066 | 0.963192994 | ns | TM | KS.3 |
| FIM1_1534 | 427.6532717 | 426.024974 | 392.2872536 | -0.000117489 | 0.052015434 | 0.995705642 | 0.999525119 | ns | TM | KS.3 |
| FIM1_1535 | 859.344223 | 810.6634821 | 781.0826287 | -0.020532463 | 0.050449018 | 0.3490569 | 0.948637087 | ns | TM | KS.3 |
| FIM1_1536 | 568.7623568 | 612.9652969 | 720.7602547 | 0.019424684 | 0.061455442 | 0.423556247 | 0.950410534 | ns | TM | KS.3 |
| FIM1_1537 | 1796.785787 | 1815.926544 | 1969.124807 | 0.002915516 | 0.050036155 | 0.891795566 | 0.991141351 | ns | TM | KS.3 |
| FIM1_1538 | 2111.493508 | 1819.974593 | 1902.472634 | -0.0554805 | 0.083097045 | 0.087261191 | 0.646018488 | ns | TM | KS.3 |
| FIM1_1540 | 1017.644391 | 939.1884524 | 957.5712073 | -0.017309554 | 0.065990933 | 0.474160081 | 0.950410534 | ns | TM | KS.3 |
| FIM1_1541 | 1825.047727 | 1745.230926 | 2516.562676 | -0.00703009 | 0.069152677 | 0.756230047 | 0.988477371 | ns | TM | KS.3 |
| FIM1_1542 | 2881.459392 | 3400.561762 | 4261.464166 | 0.018056744 | 0.085126426 | 0.412930206 | 0.950410534 | ns | TM | KS.3 |
| FIM1_1543 | 2443.667647 | 2977.811047 | 3473.59313 | 0.016880995 | 0.091444082 | 0.419757828 | 0.950410534 | ns | TM | KS.3 |
| FIM1_1544 | 278.5982078 | 257.5767019 | 191.583174 | -0.013591972 | 0.069424188 | 0.56184008 | 0.961073589 | ns | TM | KS.3 |
| FIM1_1545 | 1317.112802 | 1104.164649 | 1128.202135 | -0.057394292 | 0.089571283 | 0.079457972 | 0.616697988 | ns | TM | KS.3 |
| FIM1_1546 | 351.9907483 | 358.6138837 | 298.4181561 | 0.003620182 | 0.061015839 | 0.874118963 | 0.991141351 | ns | TM | KS.3 |
| FIM1_1547 | 929.0981256 | 983.6370907 | 968.3615341 | 0.012870282 | 0.061186816 | 0.584015261 | 0.963781856 | ns | TM | KS.3 |
| FIM1_1548 | 1472.494937 | 1594.224528 | 1539.925275 | 0.012599314 | 0.071459306 | 0.584673223 | 0.963781856 | ns | TM | KS.3 |
| FIM1_1549 | 2913.281531 | 2833.089383 | 1949.095125 | -0.006641556 | 0.056513131 | 0.769355098 | 0.988814136 | ns | TM | KS.3 |
| FIM1_1550 | 2101.202327 | 2157.213973 | 1478.150675 | 0.004343484 | 0.067049096 | 0.848086056 | 0.991141351 | ns | TM | KS.3 |
| FIM1_1551 | 43302.14685 | 39016.55492 | 21078.24352 | -0.013036213 | 0.077597409 | 0.556584499 | 0.959760553 | ns | TM | KS.3 |
| FIM1_1552 | 286.8017119 | 265.1158826 | 398.9322626 | -0.011332686 | 0.073016751 | 0.61666921 | 0.973122675 | ns | TM | KS.3 |
| FIM1_1553 | 2499.118225 | 3063.996098 | 3566.150704 | 0.017382751 | 0.092072578 | 0.407304495 | 0.950410534 | ns | TM | KS.3 |
| FIM1_1554 | 883.3183941 | 907.0363947 | 1029.671585 | 0.005728993 | 0.059420682 | 0.802199682 | 0.988916276 | ns | TM | KS.3 |
| FIM1_1556 | 479.1879492 | 471.1224046 | 412.6890585 | -0.002851852 | 0.067907023 | 0.899240131 | 0.991141351 | ns | TM | KS.3 |
| FIM1_1557 | 460.0777458 | 391.4613378 | 370.8202114 | -0.033980985 | 0.081344819 | 0.206208983 | 0.85082987 | ns | TM | KS.3 |
| FIM1_1558 | 2203.005892 | 2062.932737 | 2132.708685 | -0.01860109 | 0.058245246 | 0.433739172 | 0.950410534 | ns | TM | KS.3 |
| FIM1_1559 | 5919.768391 | 6120.959459 | 5562.228255 | 0.005660439 | 0.06690015 | 0.803646849 | 0.988916276 | ns | TM | KS.3 |
| FIM1_1560 | 1903.230399 | 1990.832353 | 1399.809743 | 0.012878646 | 0.052978686 | 0.564562839 | 0.962265923 | ns | TM | KS.3 |
| FIM1_1561 | 1947.306164 | 1654.756952 | 1903.390235 | -0.043693964 | 0.083311433 | 0.137199353 | 0.769522863 | ns | TM | KS.3 |
| FIM1_1562 | 513.5869956 | 582.3992713 | 554.8368495 | 0.018693777 | 0.077056293 | 0.425240554 | 0.950410534 | ns | TM | KS.3 |

| | | | | | | | | | | |
|---|---|---|---|---|---|---|---|---|---|---|
| FIM1_1563 | 773.9109129 | 845.5779861 | 1104.718313 | 0.032814009 | 0.063912829 | 0.214096701 | 0.862454321 | ns | TM | KS.3 |
| FIM1_1564 | 52.55654753 | 40.35065615 | 328.3333144 | -0.023813577 | 0.10091194 | 0.277539918 | 0.908583046 | ns | TM | KS.3 |
| FIM1_1566 | 1902.294619 | 1974.171982 | 2607.517748 | 0.006304671 | 0.067053181 | 0.782143554 | 0.988814136 | ns | TM | KS.3 |
| FIM1_1567 | 617.7300829 | 684.2614223 | 1116.362786 | 0.013256201 | 0.076915249 | 0.552776588 | 0.95917717 | ns | TM | KS.3 |
| FIM1_1568 | 7739.505675 | 7345.035613 | 7098.304888 | -0.018432495 | 0.04899756 | 0.389004733 | 0.948637087 | ns | TM | KS.3 |
| FIM1_1569 | 196.9411401 | 187.3401329 | 191.7269476 | -0.005825563 | 0.075714909 | 0.786272575 | 0.988916276 | ns | TM | KS.3 |
| FIM1_1570 | 1015.877734 | 1080.380552 | 1038.993896 | 0.017525637 | 0.057863618 | 0.457744422 | 0.950410534 | ns | TM | KS.3 |
| FIM1_1571 | 391.0537266 | 371.2946444 | 365.0476731 | -0.011394333 | 0.063000043 | 0.626584072 | 0.97344368 | ns | TM | KS.3 |
| FIM1_1572 | 448.4653404 | 502.2321146 | 549.9838849 | 0.029119821 | 0.071333466 | 0.2688158 | 0.906922649 | ns | TM | KS.3 |
| FIM1_1573 | 1066.934254 | 1013.40512 | 1173.957086 | -0.017894364 | 0.049426683 | 0.406030031 | 0.950410534 | ns | TM | KS.3 |
| FIM1_1574 | 2909.196264 | 2931.592996 | 3203.394133 | 0.002434371 | 0.034869894 | 0.881423165 | 0.991141351 | ns | TM | KS.3 |
| FIM1_1575 | 953.6061642 | 899.4536461 | 851.4544938 | -0.016874889 | 0.056288232 | 0.468417031 | 0.950410534 | ns | TM | KS.3 |
| FIM1_1576 | 45.60612995 | 43.67921425 | 38.68437699 | -0.005176435 | 0.077633702 | 0.805759694 | 0.988916276 | ns | TM | KS.3 |
| FIM1_1577 | 791.3944321 | 897.0342146 | 674.6265987 | 0.032282168 | 0.073765444 | 0.231630832 | 0.880810476 | ns | TM | KS.3 |
| FIM1_1578 | 8550.159241 | 9143.04587 | 9826.523273 | 0.024219256 | 0.054912881 | 0.302089974 | 0.926536521 | ns | TM | KS.3 |
| FIM1_1579 | 1366.044029 | 1600.854956 | 1877.861067 | 0.015924473 | 0.085499699 | 0.458488741 | 0.950410534 | ns | TM | KS.3 |
| FIM1_1580 | 1456.271484 | 1383.241718 | 1777.236402 | -0.008017404 | 0.069701357 | 0.723656387 | 0.987157164 | ns | TM | KS.3 |
| FIM1_1581 | 4111.174874 | 4052.449384 | 4862.540858 | -0.004470305 | 0.040749025 | 0.81041442 | 0.988916276 | ns | TM | KS.3 |
| FIM1_1582 | 114.0359597 | 74.16870784 | 111.7929124 | -0.064921693 | 0.152745541 | 0.046837904 | 0.495784193 | ns | TM | KS.3 |
| FIM1_1583 | 759.7215054 | 756.9559708 | 887.9140318 | -0.000941792 | 0.053778582 | 0.966068251 | 0.995634078 | ns | TM | KS.3 |
| FIM1_1584 | 4771.322013 | 4720.524411 | 5326.866002 | -0.002548809 | 0.055285066 | 0.909306734 | 0.991141351 | ns | TM | KS.3 |
| FIM1_1585 | 893.4228327 | 950.3745102 | 919.5679531 | 0.014408277 | 0.061720778 | 0.54321964 | 0.95917717 | ns | TM | KS.3 |
| FIM1_1586 | 1158.806678 | 1075.794597 | 2013.747848 | -0.009229046 | 0.075694506 | 0.673248468 | 0.976861203 | ns | TM | KS.3 |
| FIM1_1587 | 7491.807521 | 7361.253434 | 7941.355191 | -0.004701175 | 0.049970041 | 0.826421972 | 0.991141351 | ns | TM | KS.3 |
| FIM1_1588 | 6119.046917 | 5854.990648 | 6573.148533 | -0.007785226 | 0.06640559 | 0.734667642 | 0.987655486 | ns | TM | KS.3 |
| FIM1_1589 | 175.3408261 | 180.6106328 | 174.21158 | 0.004872915 | 0.058927596 | 0.830775821 | 0.991141351 | ns | TM | KS.3 |
| FIM1_1590 | 901.2025841 | 1004.021005 | 907.6081619 | 0.01656059 | 0.074826778 | 0.477400606 | 0.950410534 | ns | TM | KS.3 |
| FIM1_1591 | 214.1192059 | 173.9116811 | 193.798865 | -0.027818502 | 0.089425862 | 0.249730014 | 0.888934767 | ns | TM | KS.3 |
| FIM1_1592 | 280.8860159 | 257.7004912 | 207.5354789 | -0.015276062 | 0.070236213 | 0.517910677 | 0.95713433 | ns | TM | KS.3 |
| FIM1_1593 | 2698.83153 | 2075.100825 | 3711.678997 | -0.0234468 | 0.098811668 | 0.28624564 | 0.919993167 | ns | TM | KS.3 |
| FIM1_1594 | 1874.622754 | 2486.741884 | 2701.158854 | 0.037326369 | 0.103453527 | 0.147604893 | 0.780546912 | ns | TM | KS.3 |
| FIM1_1595 | 1514.769313 | 1543.854524 | 1468.553781 | 0.003735789 | 0.062307511 | 0.870235214 | 0.991141351 | ns | TM | KS.3 |
| FIM1_1596 | 1632.524443 | 1618.077665 | 1618.955995 | -0.00127192 | 0.07017029 | 0.954298238 | 0.994413583 | ns | TM | KS.3 |
| FIM1_1597 | 1010.046041 | 1399.959332 | 1675.69388 | 0.03510638 | 0.110666796 | 0.155426248 | 0.786819827 | ns | TM | KS.3 |
| FIM1_1598 | 2034.808636 | 2522.112965 | 3069.22103 | 0.018949642 | 0.092821836 | 0.373712391 | 0.948637087 | ns | TM | KS.3 |
| FIM1_1599 | 2673.452379 | 2655.49748 | 2665.984911 | -0.001952899 | 0.042076397 | 0.918717993 | 0.991141351 | ns | TM | KS.3 |
| FIM1_1601 | 22147.54164 | 24225.97686 | 22763.71051 | 0.023731615 | 0.065269828 | 0.346107048 | 0.947377991 | ns | TM | KS.3 |
| FIM1_1602 | 2028.003583 | 2244.838389 | 2256.884439 | 0.042282092 | 0.06815065 | 0.139281771 | 0.774219902 | ns | TM | KS.3 |
| FIM1_1603 | 4077.188792 | 4332.53221 | 4657.555361 | 0.017858574 | 0.056229489 | 0.444135823 | 0.950410534 | ns | TM | KS.3 |
| FIM1_1604 | 2217.395306 | 2178.113342 | 2197.326057 | -0.004020568 | 0.058049302 | 0.859455225 | 0.991141351 | ns | TM | KS.3 |
| FIM1_1605 | 1807.663391 | 2300.940515 | 2906.898455 | 0.021240144 | 0.096021172 | 0.325557337 | 0.934591893 | ns | TM | KS.3 |
| FIM1_1606 | 8440.060728 | 7862.747761 | 7623.839318 | -0.017690819 | 0.061880833 | 0.462854874 | 0.950410534 | ns | TM | KS.3 |
| FIM1_1607 | 1650.304395 | 1409.100361 | 1263.574432 | -0.065410285 | 0.087381212 | 0.060340036 | 0.544665909 | ns | TM | KS.3 |
| FIM1_1608 | 2892.157333 | 2731.6816 | 2705.623237 | -0.013213241 | 0.061054599 | 0.574469887 | 0.963192994 | ns | TM | KS.3 |
| FIM1_1609 | 1239.591915 | 1352.165625 | 1307.238998 | 0.017255383 | 0.068455294 | 0.473959408 | 0.950410534 | ns | TM | KS.3 |

| | | | | | | | | | | |
|---|---|---|---|---|---|---|---|---|---|---|
| FIM1_1610 | 2327.07658 | 2072.889202 | 2102.081972 | -0.017257458 | 0.076048092 | 0.458444186 | 0.950410534 | ns | TM | KS.3 |
| FIM1_1611 | 1907.720588 | 2086.300527 | 1829.231039 | 0.017898122 | 0.06879215 | 0.459371812 | 0.950410534 | ns | TM | KS.3 |
| FIM1_1612 | 1368.228498 | 1569.303795 | 1563.798557 | 0.020697514 | 0.078403145 | 0.382878467 | 0.948637087 | ns | TM | KS.3 |
| FIM1_1613 | 1240.641722 | 1209.717768 | 1328.538532 | -0.007080187 | 0.047116555 | 0.732579749 | 0.987157164 | ns | TM | KS.3 |
| FIM1_1614 | 664.0463889 | 666.4848209 | 538.7302173 | 0.000870237 | 0.056507961 | 0.969152089 | 0.996327965 | ns | TM | KS.3 |
| FIM1_1615 | 2387.265508 | 3087.127968 | 3138.809537 | 0.221177594 | 0.11304012 | 0.00044188 | 0.02580389 | ns | TM | KS.3 |
| FIM1_1616 | 783.6938516 | 857.6624111 | 848.8679576 | 0.023585888 | 0.065510535 | 0.348881976 | 0.948637087 | ns | TM | KS.3 |
| FIM1_1617 | 2997.742449 | 3036.517176 | 3551.417207 | 0.003760094 | 0.04522997 | 0.852011606 | 0.991141351 | ns | TM | KS.3 |
| FIM1_1618 | 228.0110776 | 254.3220458 | 330.5806568 | 0.013871318 | 0.077527145 | 0.534748632 | 0.959071703 | ns | TM | KS.3 |
| FIM1_1619 | 1449.112823 | 1518.331307 | 1644.712945 | 0.012459885 | 0.054919689 | 0.583472047 | 0.963781856 | ns | TM | KS.3 |
| FIM1_1621 | 3524.540937 | 3313.172078 | 3449.992768 | -0.020843844 | 0.053859882 | 0.364339416 | 0.948637087 | ns | TM | KS.3 |
| FIM1_1622 | 501.3562791 | 482.3959031 | 889.5684964 | -0.00815622 | 0.062239353 | 0.724253556 | 0.987157164 | ns | TM | KS.3 |
| FIM1_1623 | 379.3712552 | 311.3309471 | 369.2683476 | -0.047859691 | 0.091356448 | 0.109590305 | 0.707148855 | ns | TM | KS.3 |
| FIM1_1624 | 648.1450511 | 702.5189591 | 674.7249446 | 0.016500887 | 0.066929057 | 0.492475336 | 0.952591289 | ns | TM | KS.3 |
| FIM1_1625 | 2017.730855 | 1831.356913 | 1770.134835 | -0.023819416 | 0.067661011 | 0.346197626 | 0.947377991 | ns | TM | KS.3 |
| FIM1_1627 | 2564.770575 | 2671.975048 | 2079.215773 | 0.008078723 | 0.063346969 | 0.726718983 | 0.987157164 | ns | TM | KS.3 |
| FIM1_1628 | 250.6812281 | 329.1605697 | 650.5022275 | 0.018361476 | 0.102360963 | 0.365732088 | 0.948637087 | ns | TM | KS.3 |
| FIM1_1629 | 442.2713819 | 395.5732327 | 370.7286386 | -0.029312372 | 0.0704325 | 0.266452648 | 0.904370522 | ns | TM | KS.3 |
| FIM1_1630 | 2496.479979 | 2547.683651 | 2888.50703 | 0.00549998 | 0.050775929 | 0.799312939 | 0.988916276 | ns | TM | KS.3 |
| FIM1_1631 | 705.980513 | 699.4318312 | 737.8346959 | -0.001580237 | 0.067356251 | 0.944117451 | 0.994413583 | ns | TM | KS.3 |
| FIM1_1632 | 1192.093374 | 1308.766608 | 1564.129185 | 0.008172427 | 0.083520275 | 0.686967585 | 0.980879562 | ns | TM | KS.3 |
| FIM1_1633 | 1099.173897 | 1113.136259 | 1191.610824 | 0.001067433 | 0.082365101 | 0.957174345 | 0.994413583 | ns | TM | KS.3 |
| FIM1_1634 | 2011.896462 | 2187.54893 | 2116.876201 | 0.019748174 | 0.065385406 | 0.421330231 | 0.950410534 | ns | TM | KS.3 |
| FIM1_1635 | 6446.481071 | 6120.197903 | 6104.133034 | -0.019529483 | 0.047358237 | 0.34784758 | 0.948637087 | ns | TM | KS.3 |
| FIM1_1636 | 937.29894 | 1022.634784 | 829.9480188 | 0.021788632 | 0.064991048 | 0.380879825 | 0.948637087 | ns | TM | KS.3 |
| FIM1_1637 | 258.9695778 | 273.131494 | 238.0021329 | 0.009237486 | 0.066782396 | 0.689033815 | 0.980879562 | ns | TM | KS.3 |
| FIM1_1638 | 38.33311087 | 25.86150397 | 70.51401508 | -0.043801338 | 0.129569061 | 0.101189166 | 0.684438124 | ns | TM | KS.3 |
| FIM1_1640 | 3033.905123 | 2458.34608 | 2570.685489 | -0.120870393 | 0.107376319 | 0.010156367 | 0.223094117 | ns | TM | KS.3 |
| FIM1_1641 | 5335.580324 | 5071.1161 | 4617.052727 | -0.017511342 | 0.048915385 | 0.412329063 | 0.950410534 | ns | TM | KS.3 |
| FIM1_1642 | 2044.453849 | 1880.920963 | 1800.68227 | -0.030981691 | 0.061143542 | 0.2268949 | 0.876782875 | ns | TM | KS.3 |
| FIM1_1643 | 1630.203244 | 1521.092986 | 1577.718136 | -0.019176965 | 0.059722179 | 0.425025947 | 0.950410534 | ns | TM | KS.3 |
| FIM1_1644 | 609.9621833 | 663.1952002 | 640.2083275 | 0.025730394 | 0.06299302 | 0.308737054 | 0.927416591 | ns | TM | KS.3 |
| FIM1_1645 | 603.0681535 | 554.4118248 | 638.4829664 | -0.021485387 | 0.0648394 | 0.386502309 | 0.948637087 | ns | TM | KS.3 |
| FIM1_1646 | 2207.307299 | 2401.631215 | 3165.894573 | 0.017223422 | 0.067756341 | 0.475295816 | 0.950410534 | ns | TM | KS.3 |
| FIM1_1647 | 57.07483181 | 85.99841939 | 78.19150405 | 0.071189182 | 0.15192257 | 0.038678788 | 0.444462459 | ns | TM | KS.3 |
| FIM1_1648 | 3879.521147 | 3923.787808 | 3748.977745 | 0.002159948 | 0.062874174 | 0.924581116 | 0.993271864 | ns | TM | KS.3 |
| FIM1_1649 | 2745.937918 | 3243.820098 | 2915.235222 | 0.032309845 | 0.082196933 | 0.219674527 | 0.865538807 | ns | TM | KS.3 |
| FIM1_1650 | 1259.419336 | 997.3180154 | 1231.102463 | -0.031461852 | 0.093657829 | 0.204476702 | 0.847726326 | ns | TM | KS.3 |
| FIM1_1651 | 541.1718904 | 528.0795068 | 475.7090115 | -0.004239059 | 0.064941951 | 0.852623651 | 0.991141351 | ns | TM | KS.3 |
| FIM1_1652 | 3201.915029 | 3281.769952 | 2996.097767 | 0.00573471 | 0.056764336 | 0.800054624 | 0.988916276 | ns | TM | KS.3 |
| FIM1_1653 | 494.3874927 | 484.9246834 | 589.1797007 | -0.004731792 | 0.057612244 | 0.83481773 | 0.991141351 | ns | TM | KS.3 |
| FIM1_1654 | 685.4869766 | 463.2924102 | 472.4391557 | -0.119029756 | 0.177351723 | 0.011564412 | 0.24513135 | ns | TM | KS.3 |
| FIM1_1655 | 1241.792571 | 1291.740087 | 1447.568048 | 0.01091704 | 0.051897992 | 0.620258975 | 0.973122675 | ns | TM | KS.3 |
| FIM1_1656 | 2211.931938 | 1675.202486 | 1353.127567 | -0.177039117 | 0.13598298 | 0.002405211 | 0.091342955 | ns | TM | KS.3 |
| FIM1_1657 | 28310.75114 | 30313.35834 | 30073.46668 | 0.022938601 | 0.056334645 | 0.334154464 | 0.938982061 | ns | TM | KS.3 |

| | | | | | | | | | | |
|---|---|---|---|---|---|---|---|---|---|---|
| FIM1_1658 | 1036.279082 | 1064.56724 | 1053.529141 | 0.005854624 | 0.058150582 | 0.797205086 | 0.988916276 | ns | TM | KS.3 |
| FIM1_1659 | 2901.47762 | 2808.320382 | 2925.916133 | -0.009287293 | 0.049601316 | 0.66471957 | 0.976323032 | ns | TM | KS.3 |
| FIM1_1660 | 2780.221638 | 2760.986892 | 2284.236219 | -0.001921089 | 0.046853055 | 0.92579141 | 0.99348841 | ns | TM | KS.3 |
| FIM1_1661 | 1831.340845 | 1794.45911 | 1659.451143 | -0.004736427 | 0.057602301 | 0.834652819 | 0.991141351 | ns | TM | KS.3 |
| FIM1_1664 | 2261.967619 | 2298.765126 | 2194.986819 | 0.005040206 | 0.039976293 | 0.783359481 | 0.988814136 | ns | TM | KS.3 |
| FIM1_1665 | 310.5775544 | 280.6381455 | 323.128888 | -0.014875045 | 0.074891555 | 0.517172378 | 0.956681337 | ns | TM | KS.3 |
| FIM1_1666 | 995.7909195 | 884.2553878 | 919.0415446 | -0.070559354 | 0.076764766 | 0.045062614 | 0.487132212 | ns | TM | KS.3 |
| FIM1_1667 | 1172.12079 | 1217.935839 | 1963.193442 | 0.003896702 | 0.078567862 | 0.851135687 | 0.991141351 | ns | TM | KS.3 |
| FIM1_1669 | 1837.689501 | 1781.843121 | 2080.923473 | -0.007955654 | 0.054113154 | 0.721965859 | 0.987157164 | ns | TM | KS.3 |
| FIM1_1670 | 1699.811901 | 1828.632613 | 1932.306517 | 0.011287244 | 0.071432897 | 0.621445667 | 0.973122675 | ns | TM | KS.3 |
| FIM1_1671 | 4190.36552 | 4883.038354 | 5657.092212 | 0.015692008 | 0.08469165 | 0.466483716 | 0.950410534 | ns | TM | KS.3 |
| FIM1_1672 | 4237.885859 | 4332.754727 | 4201.129198 | 0.004637418 | 0.060304944 | 0.83928483 | 0.991141351 | ns | TM | KS.3 |
| FIM1_1673 | 3155.913512 | 3007.405091 | 3566.447103 | -0.016504906 | 0.047802403 | 0.431314082 | 0.950410534 | ns | TM | KS.3 |
| FIM1_1674 | 3006.241781 | 2340.737023 | 1673.238799 | -0.138890309 | 0.122883936 | 0.006259175 | 0.166520831 | ns | TM | KS.3 |
| FIM1_1675 | 23.3895507 | 20.65802725 | 18.27940201 | -0.009092591 | 0.091825315 | 0.635447557 | 0.973922241 | ns | TM | KS.3 |
| FIM1_1676 | 631.4020859 | 613.3333042 | 574.8902043 | -0.008012075 | 0.053772628 | 0.719391912 | 0.987157164 | ns | TM | KS.3 |
| FIM1_1677 | 662.4421255 | 514.057743 | 639.4036967 | -0.018168713 | 0.099562746 | 0.374592166 | 0.948637087 | ns | TM | KS.3 |
| FIM1_1678 | 192.6963258 | 170.8371098 | 212.1497853 | -0.015330026 | 0.079122257 | 0.493240704 | 0.952949531 | ns | TM | KS.3 |
| FIM1_1679 | 588.4276847 | 488.3096061 | 646.7479553 | -0.02674092 | 0.08574064 | 0.271301881 | 0.908468501 | ns | TM | KS.3 |
| FIM1_1680 | 381.5432739 | 468.6249035 | 512.5961361 | 0.050097312 | 0.092533257 | 0.1000548 | 0.680017255 | ns | TM | KS.3 |
| FIM1_1681 | 89.95371959 | 55.52898711 | 59.03730574 | -0.05582161 | 0.155646849 | 0.062785761 | 0.551491823 | ns | TM | KS.3 |
| FIM1_1682 | 808.5262149 | 798.8601793 | 870.1066741 | -0.002112418 | 0.066244102 | 0.925711617 | 0.99348841 | ns | TM | KS.3 |
| FIM1_1683 | 3885.255498 | 3467.60493 | 4383.462853 | -0.021227693 | 0.073190065 | 0.386221713 | 0.948637087 | ns | TM | KS.3 |
| FIM1_1684 | 3694.582393 | 2986.474233 | 1764.440744 | -0.141566953 | 0.106971525 | 0.005373338 | 0.151888395 | ns | TM | KS.3 |
| FIM1_1685 | 2353.017822 | 2123.710787 | 2269.185769 | -0.04288928 | 0.068828787 | 0.136544688 | 0.768450024 | ns | TM | KS.3 |
| FIM1_1686 | 1191.210517 | 1077.620309 | 1051.0058 | -0.026643267 | 0.067857368 | 0.301904015 | 0.926536521 | ns | TM | KS.3 |
| FIM1_1687 | 4929.523068 | 3806.944314 | 2583.761881 | -0.215173066 | 0.11628564 | 0.00058706 | 0.032094739 | ns | TM | KS.3 |
| FIM1_1688 | 201.5083747 | 155.1213728 | 317.1652885 | -0.035326362 | 0.099359866 | 0.165303942 | 0.801546891 | ns | TM | KS.3 |
| FIM1_1689 | 183.802399 | 172.3818146 | 271.523357 | -0.009545277 | 0.071623491 | 0.672695714 | 0.976861203 | ns | TM | KS.3 |
| FIM1_1690 | 4257.347879 | 3870.710736 | 3733.747297 | -0.047798139 | 0.066561481 | 0.101804913 | 0.686685387 | ns | TM | KS.3 |
| FIM1_1691 | 412.2159723 | 369.8753078 | 2389.642711 | -0.006392139 | 0.093108259 | 0.73125085 | 0.987157164 | ns | TM | KS.3 |
| FIM1_1692 | 611.9482449 | 451.5407683 | 1205.142974 | -0.083679445 | 0.128927457 | 0.027772406 | 0.388845019 | ns | TM | KS.3 |
| FIM1_1693 | 466.4811216 | 442.8125603 | 768.7516542 | -0.004872179 | 0.080749984 | 0.811289171 | 0.988916276 | ns | TM | KS.3 |
| FIM1_1694 | 45.45053489 | 54.4136587 | 53.63782133 | 0.01727099 | 0.089351136 | 0.416939083 | 0.950410534 | ns | TM | KS.3 |
| FIM1_1695 | 116.661912 | 139.3387149 | 186.2138148 | 0.013576644 | 0.091617746 | 0.499587194 | 0.953260948 | ns | TM | KS.3 |
| FIM1_1697 | 133.3953701 | 123.5705213 | 205.3618799 | -0.008248438 | 0.07939842 | 0.695237011 | 0.981479944 | ns | TM | KS.3 |
| FIM1_1698 | 2039.586097 | 2094.46556 | 2099.503148 | 0.002971118 | 0.076271642 | 0.888388136 | 0.991141351 | ns | TM | KS.3 |
| FIM1_1701 | 460.650656 | 409.4791942 | 548.8457104 | -0.017855594 | 0.075882093 | 0.446004391 | 0.950410534 | ns | TM | KS.3 |
| FIM1_1703 | 92.72191157 | 81.02406263 | 84.05414166 | -0.012233929 | 0.084552763 | 0.556762837 | 0.959760553 | ns | TM | KS.3 |
| FIM1_1705 | 18.71713533 | 17.56339468 | 22.09320857 | -0.004261921 | 0.08948215 | 0.821117364 | 0.990886949 | ns | TM | KS.3 |
| FIM1_1707 | 279.0513387 | 295.2124783 | 374.05025 | 0.01031035 | 0.067792484 | 0.655722539 | 0.976323032 | ns | TM | KS.3 |
| FIM1_1708 | 401.2312037 | 398.3672827 | 2951.506154 | -0.000379696 | 0.093423852 | 0.98330108 | 0.999333464 | ns | TM | KS.3 |
| FIM1_1709 | 829.98022 | 803.8974149 | 1248.502807 | -0.003426292 | 0.077087745 | 0.870589093 | 0.991141351 | ns | TM | KS.3 |
| FIM1_1710 | 274.6401792 | 271.6008361 | 440.0704167 | -0.001053636 | 0.080678537 | 0.958421128 | 0.994413583 | ns | TM | KS.3 |
| FIM1_1711 | 1094.003609 | 1193.383592 | 1667.795688 | 0.006163912 | 0.087825676 | 0.749386736 | 0.988477371 | ns | TM | KS.3 |

| | | | | | | | | | | |
|---|---|---|---|---|---|---|---|---|---|---|
| FIM1_1712 | 1033.027207 | 2061.104375 | 2816.95815 | 0.285078079 | 0.39176238 | 0.002548153 | 0.093517802 | ns | TM | KS.3 |
| FIM1_1713 | 62.50384775 | 43.81118949 | 91.60946069 | -0.052708183 | 0.123296993 | 0.074064905 | 0.597271765 | ns | TM | KS.3 |
| FIM1_1714 | 169.9960963 | 158.9474133 | 331.4930736 | -0.011648477 | 0.068199384 | 0.616516499 | 0.973122675 | ns | TM | KS.3 |
| FIM1_1715 | 246.7856467 | 250.8226399 | 205.5618188 | 0.002058468 | 0.071949992 | 0.925234692 | 0.99348841 | ns | TM | KS.3 |
| FIM1_1716 | 6185.453136 | 5705.727213 | 4759.1703 | -0.009320065 | 0.077489151 | 0.665662425 | 0.976323032 | ns | TM | KS.3 |
| FIM1_1717 | 181.9835633 | 169.5010463 | 180.8783533 | -0.016028994 | 0.063908669 | 0.503955971 | 0.953260948 | ns | TM | KS.3 |
| FIM1_1718 | 202.4332645 | 206.633938 | 178.6442181 | 0.005006449 | 0.058748886 | 0.826158374 | 0.991141351 | ns | TM | KS.3 |
| FIM1_1719 | 181.0559665 | 221.3857257 | 232.4189604 | 0.050377621 | 0.09247979 | 0.09912043 | 0.677856366 | ns | TM | KS.3 |
| FIM1_1721 | 33.87105703 | 23.12254471 | 37.01723315 | -0.027888474 | 0.117780383 | 0.20975343 | 0.852551729 | ns | TM | KS.3 |
| FIM1_1724 | 3153.808849 | 3167.263258 | 10659.98046 | 0.000546076 | 0.073404326 | 0.979878524 | 0.999333464 | ns | TM | KS.3 |
| FIM1_1725 | 20419.38151 | 17771.38025 | 18386.44368 | -0.021270032 | 0.078536186 | 0.372287155 | 0.948637087 | ns | TM | KS.3 |
| FIM1_1727 | 2624.955332 | 2425.738871 | 1983.948707 | -0.011692929 | 0.072685785 | 0.607165228 | 0.970937835 | ns | TM | KS.3 |
| FIM1_1728 | 6913.957933 | 5963.195154 | 6048.725339 | -0.045182211 | 0.080658592 | 0.131166844 | 0.761283488 | ns | TM | KS.3 |
| FIM1_1729 | 4590.431525 | 4196.553609 | 3532.289846 | -0.027966185 | 0.06397614 | 0.276931207 | 0.908583046 | ns | TM | KS.3 |
| FIM1_1730 | 5210.112796 | 4874.706368 | 4780.754088 | -0.031289529 | 0.05292578 | 0.178844226 | 0.811815714 | ns | TM | KS.3 |
| FIM1_1731 | 4986.136572 | 4179.442768 | 4615.337124 | -0.173770277 | 0.076392594 | 0.000804558 | 0.04169454 | ns | TM | KS.3 |
| FIM1_1732 | 1051.091854 | 1107.526588 | 1104.078117 | 0.010952128 | 0.062829452 | 0.63935859 | 0.975246356 | ns | TM | KS.3 |
| FIM1_1733 | 10466.96848 | 9191.350771 | 8127.148046 | -0.013198816 | 0.08294825 | 0.535085621 | 0.959071703 | ns | TM | KS.3 |
| FIM1_1734 | 603.3446135 | 582.3758946 | 542.6370371 | -0.004815808 | 0.072000112 | 0.827237373 | 0.991141351 | ns | TM | KS.3 |
| FIM1_1735 | 2265.820529 | 2144.470902 | 2353.005583 | -0.011225804 | 0.0638492 | 0.631486096 | 0.97344368 | ns | TM | KS.3 |
| FIM1_1736 | 642.0739015 | 730.3109959 | 664.8273505 | 0.031983519 | 0.074696256 | 0.234166417 | 0.883800031 | ns | TM | KS.3 |
| FIM1_1737 | 617.1865509 | 542.3717354 | 484.8333214 | -0.025974695 | 0.075190693 | 0.306468839 | 0.927241586 | ns | TM | KS.3 |
| FIM1_1738 | 1197.100203 | 1163.200806 | 1392.724953 | -0.006348383 | 0.059024569 | 0.781373589 | 0.988814136 | ns | TM | KS.3 |
| FIM1_1739 | 2908.968495 | 3025.282347 | 2986.10881 | 0.008676218 | 0.059937035 | 0.706900639 | 0.983265883 | ns | TM | KS.3 |
| FIM1_1740 | 1060.13157 | 1046.442952 | 944.6310137 | -0.003590914 | 0.051108385 | 0.868410395 | 0.991141351 | ns | TM | KS.3 |
| FIM1_1741 | 3139.09777 | 3402.019424 | 4603.886683 | 0.015775452 | 0.067812968 | 0.50902195 | 0.953260948 | ns | TM | KS.3 |
| FIM1_1742 | 1044.780863 | 1038.842341 | 1223.137733 | -0.001191057 | 0.060833624 | 0.958323648 | 0.994413583 | ns | TM | KS.3 |
| FIM1_1743 | 3005.13358 | 3773.704475 | 3528.491098 | 0.033456738 | 0.092919685 | 0.188868647 | 0.828217874 | ns | TM | KS.3 |
| FIM1_1744 | 2094.205226 | 2379.472165 | 2700.92026 | 0.01331688 | 0.082343267 | 0.533860302 | 0.959071703 | ns | TM | KS.3 |
| FIM1_1745 | 1381.091164 | 1419.926 | 1396.955375 | 0.008116799 | 0.047117091 | 0.695343045 | 0.981479944 | ns | TM | KS.3 |
| FIM1_1746 | 522.217581 | 506.8990417 | 512.8983152 | -0.005264282 | 0.065533613 | 0.81771632 | 0.98978971 | ns | TM | KS.3 |
| FIM1_1747 | 3340.101802 | 2988.355 | 3201.610679 | -0.022615089 | 0.071979725 | 0.363818185 | 0.948637087 | ns | TM | KS.3 |
| FIM1_1748 | 25.67252662 | 64.91332119 | 243.67275 | 0.994822818 | 0.451151275 | 5.0123190100[illegible] | 0.004987257 | ns | TM | KS.3 |
| FIM1_1749 | 4097.907928 | 3881.448241 | 4469.165477 | -0.011640168 | 0.062493904 | 0.619398325 | 0.973122675 | ns | TM | KS.3 |
| FIM1_1750 | 5197.217304 | 5035.326984 | 4549.600814 | -0.003473262 | 0.076708291 | 0.869315583 | 0.991141351 | ns | TM | KS.3 |
| FIM1_1751 | 920.2346402 | 1120.894069 | 1262.995325 | 0.015364421 | 0.092737092 | 0.451332238 | 0.950410534 | ns | TM | KS.3 |
| FIM1_1752 | 1029.396566 | 964.4695144 | 1127.180991 | -0.006151023 | 0.080914011 | 0.764159473 | 0.988477371 | ns | TM | KS.3 |
| FIM1_1753 | 1038.50172 | 816.1135471 | 787.7175786 | -0.166268308 | 0.115788253 | 0.002718476 | 0.096373426 | ns | TM | KS.3 |
| FIM1_1754 | 256.2891022 | 279.7064574 | 560.3280302 | 0.010465435 | 0.077283855 | 0.630896238 | 0.97344368 | ns | TM | KS.3 |
| FIM1_1755 | 1694.478582 | 1762.474205 | 2207.912755 | 0.00348955 | 0.081514475 | 0.862504082 | 0.991141351 | ns | TM | KS.3 |
| FIM1_1756 | 1050.728927 | 1121.204619 | 1284.85607 | 0.008803749 | 0.073519556 | 0.691784077 | 0.981188557 | ns | TM | KS.3 |
| FIM1_1757 | 1417.416697 | 1418.737561 | 1650.215053 | 0.000130859 | 0.074348045 | 0.995137683 | 0.999525119 | ns | TM | KS.3 |
| FIM1_1758 | 364.7763461 | 376.7757897 | 332.026574 | 0.003982861 | 0.074344916 | 0.853704065 | 0.991141351 | ns | TM | KS.3 |
| FIM1_1759 | 282.8086702 | 360.3903426 | 363.4795938 | 0.03108151 | 0.095214772 | 0.205208114 | 0.849343067 | ns | TM | KS.3 |
| FIM1_1760 | 3376.952649 | 3502.366123 | 3933.361456 | 0.004457076 | 0.074483878 | 0.836559618 | 0.991141351 | ns | TM | KS.3 |

| | | | | | | | | | | |
|---|---|---|---|---|---|---|---|---|---|---|
| FIM1_1761 | 2687.6234 | 2443.480119 | 2837.073276 | -0.010162607 | 0.07977072 | 0.6328767 | 0.97344368 | ns | TM | KS.3 |
| FIM1_1762 | 3728.857075 | 4446.339656 | 5193.355965 | 0.019582044 | 0.086021439 | 0.380186434 | 0.948637087 | ns | TM | KS.3 |
| FIM1_1763 | 106.5261288 | 132.5710647 | 106.1058246 | 0.022034799 | 0.092559504 | 0.320281182 | 0.929108338 | ns | TM | KS.3 |
| FIM1_1764 | 3638.142943 | 3721.217975 | 3825.694864 | 0.006533009 | 0.047568986 | 0.753975486 | 0.988477371 | ns | TM | KS.3 |
| FIM1_1765 | 1579.723883 | 1714.193775 | 2218.508472 | 0.024472422 | 0.06212225 | 0.327549769 | 0.934739762 | ns | TM | KS.3 |
| FIM1_1766 | 1511.267201 | 1563.823256 | 1836.890909 | 0.003405131 | 0.078913284 | 0.869100553 | 0.991141351 | ns | TM | KS.3 |
| FIM1_1767 | 2025.785421 | 1810.176338 | 1753.236508 | -0.031708049 | 0.070787742 | 0.238080339 | 0.88653074 | ns | TM | KS.3 |
| FIM1_1769 | 17.35341425 | 17.68395728 | 21.2872472 | 0.000922581 | 0.100598931 | 0.957760357 | 0.994413583 | ns | TM | KS.3 |
| FIM1_1770 | 2910.441874 | 2959.651885 | 2963.922448 | 0.004845553 | 0.04769608 | 0.816382589 | 0.989163025 | ns | TM | KS.3 |
| FIM1_1771 | 1136.728128 | 1160.390449 | 1673.299133 | 0.004640207 | 0.058340206 | 0.838383783 | 0.991141351 | ns | TM | KS.3 |
| FIM1_1772 | 1648.741985 | 1727.920755 | 1718.984209 | 0.009461499 | 0.063371058 | 0.683831032 | 0.98018358 | ns | TM | KS.3 |
| FIM1_1774 | 1084.974 | 949.2631547 | 925.4380031 | -0.042557617 | 0.077136526 | 0.147082896 | 0.780546912 | ns | TM | KS.3 |
| FIM1_1775 | 7739.033752 | 6811.279408 | 6632.013889 | -0.03533477 | 0.074595264 | 0.201721499 | 0.846889839 | ns | TM | KS.3 |
| FIM1_1776 | 2654.709957 | 2627.724453 | 2784.014127 | -0.002034282 | 0.061345891 | 0.928950719 | 0.993697171 | ns | TM | KS.3 |
| FIM1_1777 | 2456.667472 | 2200.091531 | 3183.634497 | -0.0225587 | 0.071746612 | 0.365108818 | 0.948637087 | ns | TM | KS.3 |
| FIM1_1778 | 2526.972899 | 2554.324772 | 2756.463574 | 0.002295331 | 0.060225705 | 0.919779487 | 0.991141351 | ns | TM | KS.3 |
| FIM1_1779 | 4285.583512 | 4704.662752 | 5361.145866 | 0.011740535 | 0.076706103 | 0.595183914 | 0.965889818 | ns | TM | KS.3 |
| FIM1_1780 | 1073.856017 | 874.4774572 | 982.3092643 | -0.09561948 | 0.104379487 | 0.021760067 | 0.33950953 | ns | TM | KS.3 |
| FIM1_1781 | 566.0177506 | 463.4987049 | 622.2993874 | -0.063106788 | 0.095740085 | 0.062652871 | 0.551491823 | ns | TM | KS.3 |
| FIM1_1782 | 1042.420242 | 964.2985975 | 1110.647824 | -0.032385347 | 0.058036086 | 0.195437253 | 0.837467988 | ns | TM | KS.3 |
| FIM1_1783 | 649.2946179 | 540.3075794 | 438.0035736 | -0.051378745 | 0.089674079 | 0.09812635 | 0.677062609 | ns | TM | KS.3 |
| FIM1_1784 | 2496.26665 | 2206.833109 | 2015.541673 | -0.036381886 | 0.07359031 | 0.192545691 | 0.832245711 | ns | TM | KS.3 |
| FIM1_1785 | 1132.460513 | 994.9986225 | 875.1746038 | -0.032169726 | 0.074845548 | 0.232109052 | 0.880810476 | ns | TM | KS.3 |
| FIM1_1786 | 69.16330864 | 47.04763114 | 53.71768876 | -0.074993875 | 0.14800597 | 0.03461149 | 0.42621823 | ns | TM | KS.3 |
| FIM1_1787 | 2019.025173 | 2186.827702 | 2637.023263 | 0.007514639 | 0.081461873 | 0.7147898 | 0.986942381 | ns | TM | KS.3 |
| FIM1_1788 | 458.1164669 | 594.89472 | 541.1282185 | 0.107893978 | 0.122904811 | 0.014575047 | 0.273934835 | ns | TM | KS.3 |
| FIM1_1789 | 1626.741102 | 1519.703877 | 1310.35748 | -0.007630503 | 0.077441324 | 0.720782765 | 0.987157164 | ns | TM | KS.3 |
| FIM1_1790 | 623.6602756 | 774.8273424 | 819.0189478 | 0.052361593 | 0.096036297 | 0.089443802 | 0.653903013 | ns | TM | KS.3 |
| FIM1_1791 | 404.6274109 | 453.1213828 | 360.2382304 | 0.020497311 | 0.073401732 | 0.399294312 | 0.950410534 | ns | TM | KS.3 |
| FIM1_1792 | 7331.957723 | 6438.943305 | 6424.741287 | -0.016577775 | 0.079554056 | 0.462760248 | 0.950410534 | ns | TM | KS.3 |
| FIM1_1793 | 2122.61803 | 1622.934751 | 1819.898147 | -0.172863656 | 0.130778113 | 0.002575264 | 0.093517802 | ns | TM | KS.3 |
| FIM1_1794 | 814.7145384 | 778.8170072 | 681.1152267 | -0.003190507 | 0.086801081 | 0.867859581 | 0.991141351 | ns | TM | KS.3 |
| FIM1_1795 | 748.7646287 | 833.970065 | 820.1649257 | 0.075610074 | 0.071173905 | 0.031977791 | 0.413219513 | ns | TM | KS.3 |
| FIM1_1796 | 104.5901192 | 80.67194483 | 82.68238282 | -0.037957607 | 0.100387769 | 0.147064398 | 0.780546912 | ns | TM | KS.3 |
| FIM1_1797 | 1685.924979 | 1663.2439 | 1607.787515 | -0.001710275 | 0.0736628 | 0.936961568 | 0.994413583 | ns | TM | KS.3 |
| FIM1_1798 | 1481.551179 | 1366.934118 | 1934.363292 | -0.00820299 | 0.079887769 | 0.695464362 | 0.981479944 | ns | TM | KS.3 |
| FIM1_1799 | 1664.032427 | 1689.187159 | 1747.339532 | 0.003776169 | 0.05310617 | 0.864093696 | 0.991141351 | ns | TM | KS.3 |
| FIM1_1800 | 642.2925653 | 542.8856587 | 846.4389477 | -0.023748151 | 0.08310205 | 0.319000203 | 0.929077912 | ns | TM | KS.3 |
| FIM1_1801 | 1740.050298 | 1787.394654 | 2016.171127 | 0.002309922 | 0.081964913 | 0.908033633 | 0.991141351 | ns | TM | KS.3 |
| FIM1_1802 | 2372.488236 | 2230.903463 | 3150.560927 | -0.006429414 | 0.078599778 | 0.75930272 | 0.988477371 | ns | TM | KS.3 |
| FIM1_1803 | 2621.907675 | 2139.400387 | 4401.067168 | -0.014627649 | 0.094565532 | 0.465089768 | 0.950410534 | ns | TM | KS.3 |
| FIM1_1804 | 525.5768087 | 531.1075473 | 737.2140736 | 0.001199924 | 0.075291148 | 0.955080769 | 0.994413583 | ns | TM | KS.3 |
| FIM1_1805 | 50.62071903 | 26.74619851 | 87.50925761 | -0.121202051 | 0.260071048 | 0.014125658 | 0.269253438 | ns | TM | KS.3 |
| FIM1_1806 | 1186.29032 | 1053.618204 | 1470.141121 | -0.012067139 | 0.082086798 | 0.569215248 | 0.962265923 | ns | TM | KS.3 |
| FIM1_1807 | 92.46326096 | 84.142312 | 206.0439287 | -0.010114879 | 0.077532137 | 0.640836423 | 0.975868137 | ns | TM | KS.3 |

| | | | | | | | | | | |
|---|---|---|---|---|---|---|---|---|---|---|
| FIM1_1808 | 2058.155652 | 2030.295165 | 2362.410885 | -0.001457975 | 0.077003166 | 0.944560599 | 0.994413583 | ns | TM | KS.3 |
| FIM1_1809 | 3372.485344 | 3292.927263 | 3503.93053 | -0.002106255 | 0.081402093 | 0.916533149 | 0.991141351 | ns | TM | KS.3 |
| FIM1_1810 | 1616.745795 | 1501.307912 | 1536.574471 | -0.01707063 | 0.063862281 | 0.479196186 | 0.950972051 | ns | TM | KS.3 |
| FIM1_1811 | 1590.482889 | 1475.061696 | 1497.327843 | -0.022765417 | 0.059853493 | 0.351829699 | 0.948637087 | ns | TM | KS.3 |
| FIM1_1812 | 706.185381 | 664.1131465 | 669.3733824 | -0.009743887 | 0.070059015 | 0.669476399 | 0.976323032 | ns | TM | KS.3 |
| FIM1_1813 | 456.2115202 | 480.5902065 | 514.8656018 | 0.011145045 | 0.061984689 | 0.633462825 | 0.97344368 | ns | TM | KS.3 |
| FIM1_1814 | 976.9245251 | 950.9097818 | 879.8759774 | -0.007245986 | 0.050581696 | 0.737570339 | 0.988051177 | ns | TM | KS.3 |
| FIM1_1815 | 572.7773168 | 448.7224893 | 661.3282019 | -0.127861876 | 0.120179775 | 0.008449856 | 0.200181112 | ns | TM | KS.3 |
| FIM1_1816 | 2435.940332 | 2090.635949 | 2257.114841 | -0.05123141 | 0.083094778 | 0.102693917 | 0.689041746 | ns | TM | KS.3 |
| FIM1_1817 | 1877.934862 | 1672.654704 | 1715.826269 | -0.030796282 | 0.071425524 | 0.248582097 | 0.888934767 | ns | TM | KS.3 |
| FIM1_1818 | 22685.27581 | 24291.20974 | 24018.39903 | 0.02847888 | 0.054356569 | 0.22699424 | 0.876782875 | ns | TM | KS.3 |
| FIM1_1819 | 3023.632176 | 3020.985839 | 2817.807409 | -0.000188369 | 0.05242473 | 0.993138949 | 0.999525119 | ns | TM | KS.3 |
| FIM1_1820 | 374.8909119 | 469.2889474 | 505.4351108 | 0.050531173 | 0.096690856 | 0.09472083 | 0.673663379 | ns | TM | KS.3 |
| FIM1_1821 | 1628.395837 | 1781.193727 | 1756.485633 | 0.022977954 | 0.065586669 | 0.359623372 | 0.948637087 | ns | TM | KS.3 |
| FIM1_1822 | 427.4889102 | 488.931795 | 470.8690079 | 0.038812541 | 0.076194901 | 0.172811413 | 0.811391014 | ns | TM | KS.3 |
| FIM1_1823 | 253.6309629 | 254.2473565 | 249.6394333 | 0.000114098 | 0.08553438 | 0.995279274 | 0.999525119 | ns | TM | KS.3 |
| FIM1_1824 | 978.8658073 | 1081.9388 | 1074.180864 | 0.018136999 | 0.07157568 | 0.450191009 | 0.950410534 | ns | TM | KS.3 |
| FIM1_1825 | 5336.967834 | 4815.890381 | 4575.038093 | -0.021325101 | 0.070378118 | 0.388761105 | 0.948637087 | ns | TM | KS.3 |
| FIM1_1826 | 6424.748086 | 6786.394158 | 6200.367689 | 0.007454502 | 0.072877946 | 0.7365282 | 0.987928766 | ns | TM | KS.3 |
| FIM1_1827 | 6052.293909 | 5270.119112 | 5286.396663 | -0.059809571 | 0.081494566 | 0.07350599 | 0.597271765 | ns | TM | KS.3 |
| FIM1_1828 | 4034.864607 | 3594.898023 | 3775.938771 | -0.033490678 | 0.071435205 | 0.219241608 | 0.865538807 | ns | TM | KS.3 |
| FIM1_1829 | 311.7674307 | 348.8909747 | 281.7080514 | 0.025051023 | 0.071638268 | 0.324923379 | 0.934066145 | ns | TM | KS.3 |
| FIM1_1830 | 1881.917313 | 1695.665311 | 1821.326502 | -0.037530157 | 0.068387375 | 0.177261123 | 0.811391014 | ns | TM | KS.3 |
| FIM1_1831 | 506.5571241 | 568.1201403 | 557.696713 | 0.028816945 | 0.07066408 | 0.272756058 | 0.908468501 | ns | TM | KS.3 |
| FIM1_1832 | 477.6582453 | 438.7759472 | 410.2668753 | -0.020559094 | 0.065306927 | 0.404907344 | 0.950410534 | ns | TM | KS.3 |
| FIM1_1833 | 873.1243347 | 817.4989745 | 903.3919552 | -0.026543955 | 0.053856626 | 0.254420676 | 0.892600401 | ns | TM | KS.3 |
| FIM1_1834 | 853.774832 | 757.5297862 | 892.8793349 | -0.024513796 | 0.073340115 | 0.331002977 | 0.938313283 | ns | TM | KS.3 |
| FIM1_1835 | 741.0058738 | 780.1131471 | 680.737761 | 0.014189082 | 0.055775992 | 0.537049045 | 0.95917717 | ns | TM | KS.3 |
| FIM1_1836 | 2143.676378 | 2377.512192 | 2170.345231 | 0.018898177 | 0.071994545 | 0.433499678 | 0.950410534 | ns | TM | KS.3 |
| FIM1_1837 | 4534.163775 | 4776.463573 | 5368.555356 | 0.007366585 | 0.071929289 | 0.74112346 | 0.988051177 | ns | TM | KS.3 |
| FIM1_1838 | 160.5388443 | 157.2865578 | 170.4799956 | -0.003056796 | 0.073701473 | 0.887809058 | 0.991141351 | ns | TM | KS.3 |
| FIM1_1839 | 3539.559978 | 3590.168022 | 3181.795781 | 0.003370779 | 0.055526503 | 0.880501471 | 0.991141351 | ns | TM | KS.3 |
| FIM1_1840 | 7015.714283 | 6608.92879 | 7274.870358 | -0.010701302 | 0.067298419 | 0.64476614 | 0.976323032 | ns | TM | KS.3 |
| FIM1_1841 | 1010.764599 | 1024.182406 | 1025.017145 | 0.003835848 | 0.042982844 | 0.843525843 | 0.991141351 | ns | TM | KS.3 |
| FIM1_1842 | 1074.959805 | 1101.294895 | 1034.982679 | 0.002716431 | 0.076212093 | 0.897899276 | 0.991141351 | ns | TM | KS.3 |
| FIM1_1843 | 7570.503277 | 7786.271305 | 8375.507969 | 0.008090073 | 0.048578491 | 0.701927956 | 0.98151979 | ns | TM | KS.3 |
| FIM1_1844 | 59.1649726 | 66.99339046 | 62.51638684 | 0.013438794 | 0.081691572 | 0.532718494 | 0.959071703 | ns | TM | KS.3 |
| FIM1_1845 | 492.5758353 | 583.0060097 | 543.7782126 | 0.067117857 | 0.090041521 | 0.056587847 | 0.535599847 | ns | TM | KS.3 |
| FIM1_1846 | 3064.032786 | 3389.564812 | 3597.826933 | 0.028996111 | 0.067604358 | 0.268886614 | 0.906922649 | ns | TM | KS.3 |
| FIM1_1847 | 3972.026681 | 3143.182133 | 4023.427754 | -0.059601176 | 0.101929126 | 0.066613149 | 0.566847784 | ns | TM | KS.3 |
| FIM1_1848 | 7890.17586 | 7291.43628 | 7128.816569 | -0.013329948 | 0.070288026 | 0.567304684 | 0.962265923 | ns | TM | KS.3 |
| FIM1_1849 | 5755.207155 | 5350.464165 | 3645.990782 | -0.010991794 | 0.071910191 | 0.628987832 | 0.97344368 | ns | TM | KS.3 |
| FIM1_1850 | 683.4877964 | 760.1901481 | 680.5337264 | 0.019087333 | 0.072505494 | 0.428654177 | 0.950410534 | ns | TM | KS.3 |
| FIM1_1851 | 4161.596339 | 4039.622534 | 3843.198155 | -0.009111858 | 0.045552574 | 0.652603331 | 0.976323032 | ns | TM | KS.3 |
| FIM1_1853 | 48399.41286 | 52074.70197 | 51825.76471 | 0.0175394 | 0.063104359 | 0.467790833 | 0.950410534 | ns | TM | KS.3 |

| | | | | | | | | | | |
|---|---|---|---|---|---|---|---|---|---|---|
| FIM1_1854 | 4981.277042 | 4771.291765 | 6421.667845 | -0.006295996 | 0.070769811 | 0.77836633 | 0.988814136 | ns | TM | KS.3 |
| FIM1_1855 | 572.2363344 | 612.5425038 | 833.8984346 | 0.008582078 | 0.075366154 | 0.694592661 | 0.981425301 | ns | TM | KS.3 |
| FIM1_1856 | 1755.811905 | 1637.313629 | 1627.702602 | -0.010254637 | 0.072187123 | 0.650114959 | 0.976323032 | ns | TM | KS.3 |
| FIM1_1857 | 6113.418543 | 5820.593225 | 6646.68315 | -0.011215842 | 0.060198082 | 0.630098695 | 0.97344368 | ns | TM | KS.3 |
| FIM1_1858 | 1141.949387 | 1140.579652 | 1277.778027 | -0.000305245 | 0.054379234 | 0.989043835 | 0.999333464 | ns | TM | KS.3 |
| FIM1_1859 | 1884.088805 | 2020.089998 | 1973.780227 | 0.014250129 | 0.06554871 | 0.548476003 | 0.95917717 | ns | TM | KS.3 |
| FIM1_1860 | 1570.943942 | 1359.928195 | 1463.585236 | -0.02164165 | 0.079397949 | 0.36336385 | 0.948637087 | ns | TM | KS.3 |
| FIM1_1861 | 548.1948601 | 548.7494744 | 621.8867677 | -0.000192552 | 0.057226167 | 0.993194013 | 0.999525119 | ns | TM | KS.3 |
| FIM1_1862 | 843.6861346 | 816.5076993 | 889.5884481 | -0.007677996 | 0.057246971 | 0.736125938 | 0.987655486 | ns | TM | KS.3 |
| FIM1_1863 | 838.5024932 | 706.454941 | 657.76852 | -0.126956672 | 0.092118878 | 0.007452965 | 0.186324131 | ns | TM | KS.3 |
| FIM1_1864 | 1128.96909 | 1097.844131 | 1432.97111 | -0.007318062 | 0.052742025 | 0.740380607 | 0.988051177 | ns | TM | KS.3 |
| FIM1_1865 | 850.7237264 | 947.264224 | 897.4371506 | 0.018841173 | 0.073144448 | 0.432332058 | 0.950410534 | ns | TM | KS.3 |
| FIM1_1866 | 226.8499579 | 207.5991416 | 472.7962783 | -0.009362375 | 0.079867954 | 0.657693359 | 0.976323032 | ns | TM | KS.3 |
| FIM1_1867 | 153.4265752 | 163.8477382 | 199.9969661 | 0.011193538 | 0.069836274 | 0.627086379 | 0.97344368 | ns | TM | KS.3 |
| FIM1_1868 | 1954.169661 | 1768.504655 | 1780.688788 | -0.026776676 | 0.067749005 | 0.299880054 | 0.926536521 | ns | TM | KS.3 |
| FIM1_1869 | 2071.743114 | 2152.916372 | 2045.223674 | 0.009836145 | 0.055358336 | 0.664082841 | 0.976323032 | ns | TM | KS.3 |
| FIM1_1870 | 1395.089962 | 1334.77501 | 1245.736782 | -0.010870142 | 0.057753947 | 0.637050869 | 0.974658588 | ns | TM | KS.3 |
| FIM1_1871 | 1569.583087 | 1328.833976 | 1710.552191 | -0.036066847 | 0.082535334 | 0.187181611 | 0.82775868 | ns | TM | KS.3 |
| FIM1_1872 | 764.1453193 | 671.5520207 | 908.4532218 | -0.030536474 | 0.074497774 | 0.250054497 | 0.888934767 | ns | TM | KS.3 |
| FIM1_1873 | 1027.095764 | 1062.702503 | 969.815206 | 0.008715911 | 0.054425196 | 0.697750091 | 0.98151979 | ns | TM | KS.3 |
| FIM1_1874 | 1657.336212 | 1683.268646 | 1626.170728 | 0.00308775 | 0.062470444 | 0.892488599 | 0.991141351 | ns | TM | KS.3 |
| FIM1_1875 | 48.8300581 | 60.07423133 | 58.7017236 | 0.026354383 | 0.090357886 | 0.264899124 | 0.90372202 | ns | TM | KS.3 |
| FIM1_1876 | 386.2994333 | 547.0075173 | 644.4371434 | 0.030324202 | 0.112881133 | 0.190638744 | 0.829253069 | ns | TM | KS.3 |
| FIM1_1877 | 703.4890242 | 946.2060929 | 1186.719207 | 0.02551485 | 0.103989001 | 0.250718965 | 0.888934767 | ns | TM | KS.3 |
| FIM1_1878 | 2606.905837 | 2363.67917 | 2956.918088 | -0.007337371 | 0.087032758 | 0.707751625 | 0.983273233 | ns | TM | KS.3 |
| FIM1_1879 | 450.8323591 | 452.9895373 | 476.7881425 | 0.001271427 | 0.050646461 | 0.952965035 | 0.994413583 | ns | TM | KS.3 |
| FIM1_1880 | 3922.047288 | 3347.008195 | 3492.685695 | -0.057049283 | 0.085729432 | 0.081929854 | 0.621932675 | ns | TM | KS.3 |
| FIM1_1881 | 869.6300457 | 696.9842823 | 750.7455602 | -0.129014034 | 0.111146349 | 0.008036655 | 0.192222881 | ns | TM | KS.3 |
| FIM1_1882 | 2861.126955 | 3031.958686 | 3247.16289 | 0.009262811 | 0.069652294 | 0.684777421 | 0.980234354 | ns | TM | KS.3 |
| FIM1_1883 | 105.3635849 | 112.2359867 | 133.49796 | 0.006199727 | 0.080395136 | 0.763531548 | 0.988477371 | ns | TM | KS.3 |
| FIM1_1884 | 4899.816893 | 4602.860347 | 4422.520693 | -0.011037787 | 0.067883388 | 0.634349241 | 0.973738807 | ns | TM | KS.3 |
| FIM1_1885 | 1997.145595 | 1863.526402 | 1568.879703 | -0.02299401 | 0.056761668 | 0.335133686 | 0.938982061 | ns | TM | KS.3 |
| FIM1_1886 | 977.3226065 | 1000.498949 | 895.1978618 | 0.00586687 | 0.053097668 | 0.790728589 | 0.988916276 | ns | TM | KS.3 |
| FIM1_1887 | 6350.62362 | 6117.432608 | 5765.02779 | -0.006957583 | 0.06469876 | 0.762225825 | 0.988477371 | ns | TM | KS.3 |
| FIM1_1888 | 2166.741982 | 2811.900881 | 2288.547933 | 0.324233015 | 0.070382688 | 3.1689487030[illegible] | 8.8674837302[illegible] | ns | TM | KS.3 |
| FIM1_1889 | 1287.784332 | 1467.991028 | 1442.223128 | 0.046265048 | 0.076951403 | 0.125596839 | 0.744748835 | ns | TM | KS.3 |
| FIM1_1890 | 649.0822965 | 958.4629033 | 1155.395381 | 0.02894437 | 0.119225058 | 0.198313768 | 0.840384151 | ns | TM | KS.3 |
| FIM1_1891 | 1229.989518 | 1494.545854 | 1454.40925 | 0.019064966 | 0.089467032 | 0.380144725 | 0.948637087 | ns | TM | KS.3 |
| FIM1_1892 | 1001.289843 | 1015.652876 | 894.0127034 | 0.001758136 | 0.073670043 | 0.935201209 | 0.994413583 | ns | TM | KS.3 |
| FIM1_1893 | 1620.747875 | 1547.015113 | 1753.076194 | -0.012170885 | 0.055999354 | 0.594899897 | 0.965889818 | ns | TM | KS.3 |
| FIM1_1894 | 2187.127574 | 1792.325259 | 1675.682039 | -0.062287738 | 0.095555921 | 0.064447877 | 0.560538788 | ns | TM | KS.3 |
| FIM1_1895 | 1066.956896 | 1200.545456 | 1091.70638 | 0.041078945 | 0.073061705 | 0.155357698 | 0.786819827 | ns | TM | KS.3 |
| FIM1_1896 | 1309.416833 | 1244.070332 | 1174.592518 | -0.016505574 | 0.051353444 | 0.455025314 | 0.950410534 | ns | TM | KS.3 |
| FIM1_1897 | 1862.974375 | 1917.672856 | 1873.564261 | 0.008639017 | 0.048174004 | 0.681281116 | 0.97986479 | ns | TM | KS.3 |
| FIM1_1898 | 1704.907684 | 1737.802344 | 1951.388759 | 0.005529392 | 0.044864864 | 0.782592373 | 0.988814136 | ns | TM | KS.3 |

| | | | | | | | | | | |
|---|---|---|---|---|---|---|---|---|---|---|
| FIM1_1899 | 6452.260094 | 7800.195757 | 8322.433107 | 0.068042239 | 0.095116875 | 0.05358423 | 0.528931635 | ns | TM | KS.3 |
| FIM1_1900 | 393.1756139 | 447.7560356 | 403.9192299 | 0.042390767 | 0.076326176 | 0.148086329 | 0.780546912 | ns | TM | KS.3 |
| FIM1_1901 | 4434.583349 | 3942.077249 | 4115.123878 | -0.024945058 | 0.072810041 | 0.325186445 | 0.934066145 | ns | TM | KS.3 |
| FIM1_1902 | 805.4060432 | 789.7381478 | 880.7159253 | -0.004174612 | 0.060839436 | 0.855135431 | 0.991141351 | ns | TM | KS.3 |
| FIM1_1903 | 873.6670243 | 628.1384698 | 936.9536444 | -0.383887219 | 0.128180133 | 1.95049501835 | 0.00026368 | ns | TM | KS.3 |
| FIM1_1904 | 2991.490373 | 2909.070894 | 5400.172419 | -0.006447883 | 0.057354254 | 0.776712929 | 0.988814136 | ns | TM | KS.3 |
| FIM1_1905 | 1046.185294 | 1060.756427 | 1071.12881 | 0.003325009 | 0.055370237 | 0.88200025 | 0.991141351 | ns | TM | KS.3 |
| FIM1_1907 | 1153.64818 | 1232.488552 | 1143.469546 | 0.011866824 | 0.067549975 | 0.611126998 | 0.972453674 | ns | TM | KS.3 |
| FIM1_1908 | 4798.533848 | 4509.62499 | 4344.102142 | -0.013698453 | 0.062930475 | 0.562775671 | 0.961073589 | ns | TM | KS.3 |
| FIM1_1909 | 4373.990106 | 3549.624619 | 4253.410281 | -0.15758444 | 0.10179694 | 0.003041897 | 0.101288726 | ns | TM | KS.3 |
| FIM1_1910 | 2338.138423 | 1824.267522 | 1422.179399 | -0.094574964 | 0.116186373 | 0.021450061 | 0.338445603 | ns | TM | KS.3 |
| FIM1_1911 | 7133.316706 | 7146.193138 | 7619.23478 | 0.000563116 | 0.036928978 | 0.973893563 | 0.998607635 | ns | TM | KS.3 |
| FIM1_1912 | 2758.58653 | 2790.825792 | 3167.10355 | 0.002953134 | 0.051874753 | 0.892346893 | 0.991141351 | ns | TM | KS.3 |
| FIM1_1913 | 2234.255191 | 2540.083929 | 2600.344044 | 0.015574829 | 0.080099398 | 0.484208766 | 0.952269291 | ns | TM | KS.3 |
| FIM1_1914 | 625.835324 | 614.8280114 | 540.0841743 | -0.003333905 | 0.063136002 | 0.883974953 | 0.991141351 | ns | TM | KS.3 |
| FIM1_1915 | 1800.909436 | 2059.314639 | 2001.753845 | 0.016169796 | 0.080705972 | 0.468306834 | 0.950410534 | ns | TM | KS.3 |
| FIM1_1916 | 1018.87235 | 1251.488589 | 1040.49571 | 0.036961343 | 0.089773445 | 0.168401309 | 0.805573568 | ns | TM | KS.3 |
| FIM1_1917 | 1000.682038 | 998.9425027 | 992.6185635 | -0.000215228 | 0.07459187 | 0.991985571 | 0.999525119 | ns | TM | KS.3 |
| FIM1_1918 | 914.5207826 | 939.2926432 | 883.3691243 | 0.005176949 | 0.062416608 | 0.821390508 | 0.990886949 | ns | TM | KS.3 |
| FIM1_1919 | 1059.517673 | 1037.009587 | 1482.241881 | -0.005443521 | 0.052954727 | 0.805193431 | 0.988916276 | ns | TM | KS.3 |
| FIM1_1920 | 1655.457573 | 1973.318201 | 2593.445431 | 0.015636693 | 0.08875278 | 0.455488801 | 0.950410534 | ns | TM | KS.3 |
| FIM1_1921 | 3288.720062 | 2954.296143 | 2806.918951 | -0.023884486 | 0.070515619 | 0.344270925 | 0.945225083 | ns | TM | KS.3 |
| FIM1_1922 | 2298.295598 | 2269.544484 | 2248.488215 | -0.003863897 | 0.031230575 | 0.791510854 | 0.988916276 | ns | TM | KS.3 |
| FIM1_1923 | 2261.973577 | 2443.972439 | 2652.797944 | 0.029971459 | 0.058820451 | 0.230336176 | 0.879350777 | ns | TM | KS.3 |
| FIM1_1924 | 1075.824191 | 1124.224822 | 1067.849095 | 0.008601533 | 0.063884655 | 0.710240065 | 0.984652342 | ns | TM | KS.3 |
| FIM1_1925 | 2479.330049 | 2097.664656 | 2161.135795 | -0.116591771 | 0.091650835 | 0.010534831 | 0.229871871 | ns | TM | KS.3 |
| FIM1_1926 | 486.5969912 | 555.7803485 | 614.4443236 | 0.024236581 | 0.076158642 | 0.329918444 | 0.936305909 | ns | TM | KS.3 |
| FIM1_1927 | 2746.226569 | 3209.261025 | 3871.751739 | 0.01522401 | 0.085643042 | 0.474679028 | 0.950410534 | ns | TM | KS.3 |
| FIM1_1928 | 1511.966249 | 1636.834916 | 1626.853045 | 0.023924765 | 0.061371057 | 0.335118667 | 0.938982061 | ns | TM | KS.3 |
| FIM1_1929 | 496.9629513 | 533.5193443 | 577.4460461 | 0.008150635 | 0.077164222 | 0.704196598 | 0.982408703 | ns | TM | KS.3 |
| FIM1_1930 | 717.8004538 | 696.361237 | 719.1019424 | -0.006714196 | 0.060307575 | 0.770037429 | 0.988814136 | ns | TM | KS.3 |
| FIM1_1931 | 17344.60985 | 15105.00816 | 12554.5319 | -0.115362579 | 0.079016345 | 0.00865515 | 0.202156667 | ns | TM | KS.3 |
| FIM1_1932 | 608.3834655 | 559.7629024 | 615.3969019 | -0.023836334 | 0.06351557 | 0.341670643 | 0.944808611 | ns | TM | KS.3 |
| FIM1_1933 | 5040.888644 | 4954.939408 | 5025.61618 | -0.004300762 | 0.053394955 | 0.845854254 | 0.991141351 | ns | TM | KS.3 |
| FIM1_1934 | 350.4302023 | 352.0812143 | 592.1345006 | 0.001024229 | 0.06587519 | 0.963952975 | 0.995175461 | ns | TM | KS.3 |
| FIM1_1935 | 898.4156847 | 918.1472962 | 2584.393175 | 0.004292683 | 0.062514559 | 0.85124062 | 0.991141351 | ns | TM | KS.3 |
| FIM1_1936 | 4328.672417 | 4787.502106 | 5334.781665 | 0.083965423 | 0.066921748 | 0.019655055 | 0.324535175 | ns | TM | KS.3 |
| FIM1_1937 | 2778.471587 | 2698.155552 | 2724.981043 | -0.00832252 | 0.048608503 | 0.693945461 | 0.981346978 | ns | TM | KS.3 |
| FIM1_1938 | 1221.827725 | 1393.816154 | 1245.841524 | 0.028033582 | 0.075429143 | 0.278699712 | 0.910394659 | ns | TM | KS.3 |
| FIM1_1939 | 149.8540754 | 225.8527013 | 307.9868761 | 0.093421694 | 0.167058516 | 0.021083484 | 0.336186958 | ns | TM | KS.3 |
| FIM1_1940 | 3064.352777 | 3286.894429 | 3000.403003 | 0.009598255 | 0.073597479 | 0.666870694 | 0.976323032 | ns | TM | KS.3 |
| FIM1_1941 | 1513.763664 | 1324.083979 | 1548.488475 | -0.016155942 | 0.080700086 | 0.468645134 | 0.950410534 | ns | TM | KS.3 |
| FIM1_1942 | 831.6573359 | 813.975822 | 919.5417893 | -0.004024157 | 0.063419428 | 0.860302923 | 0.991141351 | ns | TM | KS.3 |
| FIM1_1943 | 1296.77893 | 1118.091046 | 1103.740139 | -0.037029019 | 0.079225989 | 0.184309164 | 0.822366001 | ns | TM | KS.3 |
| FIM1_1944 | 582.9404323 | 529.7526314 | 568.8789386 | -0.029686371 | 0.06608655 | 0.257667153 | 0.895871714 | ns | TM | KS.3 |

| | | | | | | | | | | |
|---|---|---|---|---|---|---|---|---|---|---|
| FIM1_1945 | 811.4744858 | 766.1924164 | 776.3111489 | -0.01205507 | 0.063805641 | 0.607890567 | 0.970937835 | ns | TM | KS.3 |
| FIM1_1946 | 1769.982101 | 1832.006408 | 2199.170287 | 0.007142315 | 0.061482705 | 0.756512288 | 0.988477371 | ns | TM | KS.3 |
| FIM1_1947 | 1083.77161 | 1093.603483 | 995.8216166 | 0.001665801 | 0.063602431 | 0.941732903 | 0.994413583 | ns | TM | KS.3 |
| FIM1_1948 | 1471.183014 | 1738.981023 | 2939.991438 | 0.078952843 | 0.091852051 | 0.037707271 | 0.440708412 | ns | TM | KS.3 |
| FIM1_1949 | 546.6301828 | 470.0461435 | 448.1257848 | -0.032787186 | 0.079243923 | 0.220537889 | 0.867332804 | ns | TM | KS.3 |
| FIM1_1950 | 1404.41135 | 1186.241791 | 1066.766058 | -0.070049242 | 0.090857717 | 0.051068484 | 0.5150643 | ns | TM | KS.3 |
| FIM1_1951 | 1836.942301 | 1448.097855 | 1505.63966 | -0.099857409 | 0.114373063 | 0.018614409 | 0.314988715 | ns | TM | KS.3 |
| FIM1_1952 | 1848.944129 | 1916.017005 | 2539.892516 | 0.010502553 | 0.049350852 | 0.623357648 | 0.973384902 | ns | TM | KS.3 |
| FIM1_1953 | 2005.104222 | 2128.009378 | 3001.487266 | 0.008319156 | 0.072537359 | 0.709203454 | 0.984180526 | ns | TM | KS.3 |
| FIM1_1954 | 1337.949135 | 1805.402806 | 2132.792289 | 0.018972607 | 0.106185267 | 0.347846584 | 0.948637087 | ns | TM | KS.3 |
| FIM1_1955 | 588.7093552 | 496.2527645 | 615.9159347 | -0.080032045 | 0.093457836 | 0.036366137 | 0.432826634 | ns | TM | KS.3 |
| FIM1_1956 | 659.9622849 | 631.7348215 | 737.4129827 | -0.006591136 | 0.070056609 | 0.769508074 | 0.988814136 | ns | TM | KS.3 |
| FIM1_1957 | 1223.489682 | 1226.511373 | 1281.058434 | 0.000399281 | 0.057199141 | 0.985886141 | 0.999333464 | ns | TM | KS.3 |
| FIM1_1958 | 1269.853836 | 1022.230668 | 1017.167306 | -0.064433371 | 0.100002182 | 0.057985918 | 0.54022461 | ns | TM | KS.3 |
| FIM1_1959 | 671.4007041 | 723.5579893 | 825.9505333 | 0.02043465 | 0.06078066 | 0.400749687 | 0.950410534 | ns | TM | KS.3 |
| FIM1_1960 | 2891.368023 | 2652.267232 | 2838.54034 | -0.043630537 | 0.062499338 | 0.114081535 | 0.714310891 | ns | TM | KS.3 |
| FIM1_1961 | 1042.606279 | 893.9009 | 3122.551629 | -0.027742608 | 0.079739164 | 0.273871227 | 0.908468501 | ns | TM | KS.3 |
| FIM1_1962 | 33644.74738 | 36327.37624 | 36346.13897 | 0.022116961 | 0.06098122 | 0.367564787 | 0.948637087 | ns | TM | KS.3 |
| FIM1_1963 | 570.4622557 | 574.7668992 | 534.5366959 | 0.001503893 | 0.062416177 | 0.947431776 | 0.994413583 | ns | TM | KS.3 |
| FIM1_1964 | 12266.94956 | 11194.17586 | 12314.88059 | -0.011969244 | 0.075885161 | 0.591071099 | 0.963781856 | ns | TM | KS.3 |
| FIM1_1965 | 589.0741705 | 1044.205688 | 1120.310801 | 0.790267501 | 0.149476277 | 1.44020951554 | 3.41192492372 | ns | TM | KS.3 |
| FIM1_1966 | 1377.630666 | 1286.652238 | 1545.494868 | -0.019447763 | 0.059166015 | 0.4175349 | 0.950410534 | ns | TM | KS.3 |
| FIM1_1967 | 2093.674071 | 1865.604666 | 2172.736869 | -0.035738984 | 0.071523574 | 0.197456069 | 0.838676184 | ns | TM | KS.3 |
| FIM1_1968 | 1731.595767 | 2083.609048 | 2275.329364 | 0.028495842 | 0.085373186 | 0.25159408 | 0.888934767 | ns | TM | KS.3 |
| FIM1_1970 | 658.1475184 | 605.9862868 | 570.7668803 | -0.016216253 | 0.068275424 | 0.498072948 | 0.953260948 | ns | TM | KS.3 |
| FIM1_1971 | 464.1170705 | 459.0284787 | 411.3584587 | -0.002011228 | 0.062561486 | 0.929762146 | 0.993993515 | ns | TM | KS.3 |
| FIM1_1972 | 2307.108687 | 2204.141777 | 3532.771367 | -0.004175411 | 0.080953367 | 0.837099972 | 0.991141351 | ns | TM | KS.3 |
| FIM1_1973 | 5266.677194 | 6376.302141 | 7261.775774 | 0.024623719 | 0.08683732 | 0.296172372 | 0.926536521 | ns | TM | KS.3 |
| FIM1_1975 | 1301.15936 | 1444.277504 | 1218.097813 | 0.03823282 | 0.068281993 | 0.170963174 | 0.809734625 | ns | TM | KS.3 |
| FIM1_1976 | 2280.341156 | 2240.994241 | 2935.738703 | -0.00384219 | 0.058274987 | 0.865704083 | 0.991141351 | ns | TM | KS.3 |
| FIM1_1977 | 776.8035749 | 795.5253387 | 1034.614862 | 0.00702792 | 0.046176628 | 0.731001642 | 0.987157164 | ns | TM | KS.3 |
| FIM1_1978 | 2453.318314 | 2384.914049 | 2647.71356 | -0.003484469 | 0.074138237 | 0.871895309 | 0.991141351 | ns | TM | KS.3 |
| FIM1_1979 | 760.8257731 | 763.8984787 | 814.5126033 | 0.001235271 | 0.040908007 | 0.947413525 | 0.994413583 | ns | TM | KS.3 |
| FIM1_1980 | 7873.268031 | 9024.014192 | 8603.960782 | 0.026194525 | 0.076650775 | 0.300629349 | 0.926536521 | ns | TM | KS.3 |
| FIM1_1981 | 11634.82413 | 10666.50728 | 10239.13225 | -0.013913228 | 0.071998696 | 0.548655217 | 0.95917717 | ns | TM | KS.3 |
| FIM1_1982 | 17.65977213 | 13.49412252 | 64.93626001 | -0.014272966 | 0.107485432 | 0.450057556 | 0.950410534 | ns | TM | KS.3 |
| FIM1_1983 | 4379.143114 | 4141.597462 | 4897.795103 | -0.015416181 | 0.056382159 | 0.506016068 | 0.953260948 | ns | TM | KS.3 |
| FIM1_1984 | 562.5160797 | 532.431078 | 569.3680288 | -0.015132642 | 0.055683534 | 0.511213134 | 0.953818112 | ns | TM | KS.3 |
| FIM1_1985 | 2702.315528 | 2768.47537 | 2826.378344 | 0.006396476 | 0.052249937 | 0.770863312 | 0.988814136 | ns | TM | KS.3 |
| FIM1_1986 | 2137.976559 | 1936.604709 | 2153.313448 | -0.047762069 | 0.067900783 | 0.105183091 | 0.697714507 | ns | TM | KS.3 |
| FIM1_1987 | 628.7871046 | 806.105705 | 767.2735327 | 0.051766019 | 0.102006178 | 0.086534444 | 0.643510999 | ns | TM | KS.3 |
| FIM1_1988 | 2352.529724 | 2751.812408 | 2531.306992 | 0.033815736 | 0.080477994 | 0.209118197 | 0.852551729 | ns | TM | KS.3 |
| FIM1_1989 | 689.8550402 | 693.1627878 | 1456.817408 | 0.000743684 | 0.068179531 | 0.97356668 | 0.998607635 | ns | TM | KS.3 |
| FIM1_1990 | 1909.607686 | 1799.047287 | 2173.972986 | -0.017476904 | 0.05608654 | 0.452830349 | 0.950410534 | ns | TM | KS.3 |
| FIM1_1991 | 1933.839147 | 2102.940188 | 2246.902047 | 0.014244372 | 0.070710678 | 0.542782741 | 0.95917717 | ns | TM | KS.3 |

| | | | | | | | | | | |
|---|---|---|---|---|---|---|---|---|---|---|
| FIM1_1992 | 4436.048261 | 4393.692309 | 4555.279446 | -0.001320191 | 0.071236454 | 0.952220657 | 0.994413583 | ns | TM | KS.3 |
| FIM1_1993 | 1867.689021 | 1803.434763 | 1575.670143 | -0.006937395 | 0.062560576 | 0.763288415 | 0.988477371 | ns | TM | KS.3 |
| FIM1_1994 | 12785.59895 | 13290.24083 | 15695.74925 | 0.013475024 | 0.043114454 | 0.484365539 | 0.952269291 | ns | TM | KS.3 |
| FIM1_1995 | 2726.556194 | 3072.547967 | 3370.377933 | 0.016423991 | 0.0775386 | 0.472625885 | 0.950410534 | ns | TM | KS.3 |
| FIM1_1996 | 6882.910008 | 7584.922181 | 7937.596766 | 0.023302479 | 0.067838823 | 0.354992355 | 0.948637087 | ns | TM | KS.3 |
| FIM1_1997 | 1501.416519 | 1454.697185 | 1520.024576 | -0.009550512 | 0.047128647 | 0.64511811 | 0.976323032 | ns | TM | KS.3 |
| FIM1_1998 | 1313.192557 | 1457.272367 | 1350.747153 | 0.051296166 | 0.070678846 | 0.093612481 | 0.671406965 | ns | TM | KS.3 |
| FIM1_1999 | 241.1127201 | 217.6636122 | 411.8301405 | -0.015389328 | 0.074879583 | 0.504672232 | 0.953260948 | ns | TM | KS.3 |
| FIM1_2000 | 6138.194136 | 5676.165393 | 5751.658379 | -0.015034977 | 0.067853304 | 0.527067982 | 0.959071703 | ns | TM | KS.3 |
| FIM1_2001 | 2272.899249 | 2630.170598 | 3238.179242 | 0.01571116 | 0.083386956 | 0.470221681 | 0.950410534 | ns | TM | KS.3 |
| FIM1_2002 | 2685.200184 | 2927.140087 | 3293.617611 | 0.026889228 | 0.062901902 | 0.290274209 | 0.919993167 | ns | TM | KS.3 |
| FIM1_2003 | 5002.231552 | 5590.016706 | 4467.052273 | 0.026494099 | 0.070834318 | 0.304211064 | 0.926536521 | ns | TM | KS.3 |
| FIM1_2004 | 433.017472 | 422.9190586 | 610.1034241 | -0.004435708 | 0.064433005 | 0.846091435 | 0.991141351 | ns | TM | KS.3 |
| FIM1_2005 | 34267.17987 | 32514.50907 | 32739.38233 | -0.009791379 | 0.06575158 | 0.673027704 | 0.976861203 | ns | TM | KS.3 |
| FIM1_2006 | 695.791332 | 982.6690344 | 869.3747403 | 0.273176419 | 0.181183265 | 0.000441194 | 0.02580389 | ns | TM | KS.3 |
| FIM1_2007 | 536.7544745 | 596.0499721 | 634.0629093 | 0.029364213 | 0.068633497 | 0.265031242 | 0.90372202 | ns | TM | KS.3 |
| FIM1_2008 | 551.3966902 | 559.0347861 | 575.9856178 | 0.003005839 | 0.060922709 | 0.895249634 | 0.991141351 | ns | TM | KS.3 |
| FIM1_2009 | 293.7044619 | 265.5312771 | 814.4961066 | -0.009967188 | 0.081528342 | 0.633765644 | 0.97344368 | ns | TM | KS.3 |
| FIM1_2010 | 38.85835346 | 28.70392744 | 78.59222337 | -0.029628629 | 0.102289982 | 0.208189706 | 0.852551729 | ns | TM | KS.3 |
| FIM1_2011 | 2531.901234 | 2337.501235 | 2751.075001 | -0.033978183 | 0.059372084 | 0.18354173 | 0.821657502 | ns | TM | KS.3 |
| FIM1_2012 | 2195.476608 | 2011.876517 | 3168.947475 | -0.02593394 | 0.063512788 | 0.306680144 | 0.927241586 | ns | TM | KS.3 |
| FIM1_2013 | 1447.097807 | 1021.384846 | 1113.463587 | -0.088088624 | 0.144768533 | 0.023995714 | 0.365072411 | ns | TM | KS.3 |
| FIM1_2014 | 1779.501415 | 1667.443699 | 1707.956662 | -0.02056422 | 0.056123958 | 0.381965154 | 0.948637087 | ns | TM | KS.3 |
| FIM1_2015 | 37.45673106 | 21.9355548 | 21.92099029 | -0.112274581 | 0.222476515 | 0.01538283 | 0.284118339 | ns | TM | KS.3 |
| FIM1_2016 | 2828.259501 | 2432.995264 | 2361.274511 | -0.085008965 | 0.087653363 | 0.030038597 | 0.400453344 | ns | TM | KS.3 |
| FIM1_2017 | 970.6183886 | 905.6393599 | 1226.622931 | -0.011496133 | 0.069736241 | 0.618601676 | 0.973122675 | ns | TM | KS.3 |
| FIM1_2018 | 189.7543512 | 112.2947866 | 205.6604993 | -0.0706654 | 0.179431736 | 0.039599264 | 0.4518494 | ns | TM | KS.3 |
| FIM1_2019 | 1290.218551 | 1424.376165 | 1473.133084 | 0.013526236 | 0.075664349 | 0.549331292 | 0.95917717 | ns | TM | KS.3 |
| FIM1_2020 | 2881.087334 | 2694.12443 | 3206.016417 | -0.010695707 | 0.070385831 | 0.640586312 | 0.975785947 | ns | TM | KS.3 |
| FIM1_2021 | 3719.720692 | 3837.315977 | 4196.697368 | 0.005195023 | 0.067110453 | 0.819121604 | 0.990886949 | ns | TM | KS.3 |
| FIM1_2022 | 1674.075242 | 1698.43037 | 1584.664656 | 0.002992221 | 0.059671835 | 0.895512263 | 0.991141351 | ns | TM | KS.3 |
| FIM1_2023 | 1807.05639 | 1972.31337 | 2191.806579 | 0.017914642 | 0.068178894 | 0.459581682 | 0.950410534 | ns | TM | KS.3 |
| FIM1_2024 | 135.1699289 | 164.3651479 | 535.9085137 | 0.008019816 | 0.106872484 | 0.64889705 | 0.976323032 | ns | TM | KS.3 |
| FIM1_2025 | 384.62412 | 205.8610276 | 215.5552265 | -0.769915689 | 0.277420841 | 1.74892973740 | 0.002122177 | ns | TM | KS.3 |
| FIM1_2026 | 1270.946496 | 1261.135323 | 1235.45582 | -0.002267755 | 0.044344005 | 0.9091986 | 0.991141351 | ns | TM | KS.3 |
| FIM1_2027 | 1236.39767 | 1039.152864 | 972.3894514 | -0.076457152 | 0.093047725 | 0.040973996 | 0.462234994 | ns | TM | KS.3 |
| FIM1_2028 | 1796.332572 | 1933.130661 | 2123.234624 | 0.013320338 | 0.068322794 | 0.570544133 | 0.962265923 | ns | TM | KS.3 |
| FIM1_2029 | 3027.36616 | 2676.39578 | 2108.789834 | -0.062306041 | 0.077640953 | 0.064201101 | 0.559370367 | ns | TM | KS.3 |
| FIM1_2030 | 483.5022699 | 409.7286805 | 1023.682246 | -0.029896561 | 0.081901941 | 0.24439406 | 0.88883001 | ns | TM | KS.3 |
| FIM1_2031 | 938.6839469 | 983.6134418 | 1632.790141 | 0.004638008 | 0.079556111 | 0.822050395 | 0.991141351 | ns | TM | KS.3 |
| FIM1_2032 | 16.86398747 | 9.04066601 | 27.57214771 | -0.039360385 | 0.159250432 | 0.111786624 | 0.71059097 | ns | TM | KS.3 |
| FIM1_2033 | 1450.178031 | 1391.428936 | 1731.269197 | -0.012204894 | 0.050501446 | 0.574380194 | 0.963192994 | ns | TM | KS.3 |
| FIM1_2034 | 1998.405 | 1916.97088 | 2139.809632 | -0.01304166 | 0.047617154 | 0.532702979 | 0.959071703 | ns | TM | KS.3 |
| FIM1_2035 | 229.528127 | 205.4369196 | 253.7108773 | -0.017663793 | 0.075978401 | 0.449830825 | 0.950410534 | ns | TM | KS.3 |
| FIM1_2037 | 346.8968468 | 439.639724 | 314.2585225 | 0.03643905 | 0.094894284 | 0.163825521 | 0.800496208 | ns | TM | KS.3 |

| | | | | | | | | | | |
|---|---|---|---|---|---|---|---|---|---|---|
| FIM1_2038 | 1704.067945 | 2102.613419 | 1938.2021 | 0.050745072 | 0.094234848 | 0.096232109 | 0.673713046 | ns | TM | KS.3 |
| FIM1_2039 | 557.6717239 | 533.9663468 | 620.846494 | -0.007524773 | 0.067252614 | 0.742274698 | 0.988477371 | ns | TM | KS.3 |
| FIM1_2040 | 350.7119745 | 348.7900148 | 447.3371595 | -0.001051615 | 0.064435949 | 0.963130419 | 0.995175461 | ns | TM | KS.3 |
| FIM1_2041 | 1471.606185 | 1630.625854 | 5070.218881 | 0.024099596 | 0.06924685 | 0.341470762 | 0.944808611 | ns | TM | KS.3 |
| FIM1_2042 | 3420.10527 | 3404.240003 | 3704.298972 | -0.001415123 | 0.044082855 | 0.943031802 | 0.994413583 | ns | TM | KS.3 |
| FIM1_2043 | 1888.724907 | 2096.994487 | 2435.732405 | 0.017206207 | 0.073474328 | 0.466362685 | 0.950410534 | ns | TM | KS.3 |
| FIM1_2044 | 931.5603617 | 801.8286242 | 810.1224928 | -0.041708594 | 0.080249409 | 0.151151282 | 0.783447458 | ns | TM | KS.3 |
| FIM1_2045 | 452.864184 | 348.4580232 | 395.9114986 | -0.090125153 | 0.119427208 | 0.023972298 | 0.365072411 | ns | TM | KS.3 |
| FIM1_2046 | 285.7526907 | 298.8820703 | 265.7848338 | 0.005792359 | 0.073453515 | 0.791370602 | 0.988916276 | ns | TM | KS.3 |
| FIM1_2047 | 630.1618747 | 608.7736844 | 771.6282506 | -0.006388159 | 0.065215452 | 0.780566526 | 0.988814136 | ns | TM | KS.3 |
| FIM1_2048 | 1383.812695 | 1968.797315 | 1837.727007 | 0.431186316 | 0.133873982 | 7.54512874232 | 0.000131451 | ns | TM | KS.3 |
| FIM1_2049 | 441.7455575 | 456.8392595 | 419.7850415 | 0.008650754 | 0.055601624 | 0.702191516 | 0.98151979 | ns | TM | KS.3 |
| FIM1_2050 | 3597.306603 | 3320.229093 | 3690.857324 | -0.029053911 | 0.060049076 | 0.248577023 | 0.888934767 | ns | TM | KS.3 |
| FIM1_2051 | 2112.144527 | 2075.658491 | 2042.719606 | -0.004556815 | 0.05068788 | 0.832899436 | 0.991141351 | ns | TM | KS.3 |
| FIM1_2052 | 3287.854604 | 4102.348116 | 3036.137461 | 0.054170458 | 0.097554073 | 0.08290573 | 0.625881647 | ns | TM | KS.3 |
| FIM1_2053 | 5408.913199 | 7279.965787 | 6991.329792 | 0.266260726 | 0.134327442 | 0.000181924 | 0.013924187 | ns | TM | KS.3 |
| FIM1_2054 | 2130.739114 | 2181.640861 | 1495.778769 | 0.006293536 | 0.05180158 | 0.773515112 | 0.988814136 | ns | TM | KS.3 |
| FIM1_2055 | 593.3709648 | 922.7038644 | 893.0585356 | 0.04526596 | 0.137923747 | 0.093794541 | 0.671406965 | ns | TM | KS.3 |
| FIM1_2056 | 568.1828292 | 960.1323795 | 844.1085238 | 0.435675263 | 0.32496314 | 0.000365558 | 0.022282532 | ns | TM | KS.3 |
| FIM1_2057 | 668.3006468 | 707.0935708 | 693.8576477 | 0.014357353 | 0.057038499 | 0.536116263 | 0.959071703 | ns | TM | KS.3 |
| FIM1_2058 | 326.9142535 | 438.3360146 | 517.6529092 | 0.037042626 | 0.105272112 | 0.147578532 | 0.780546912 | ns | TM | KS.3 |
| FIM1_2059 | 714.3241315 | 575.5167247 | 774.0261571 | -0.07840236 | 0.104112496 | 0.036552778 | 0.434009714 | ns | TM | KS.3 |
| FIM1_2060 | 3463.050163 | 4442.557869 | 3723.71952 | 0.023310069 | 0.096636411 | 0.292237694 | 0.921929314 | ns | TM | KS.3 |
| FIM1_2061 | 1834.710146 | 1870.28865 | 1823.673689 | 0.004142155 | 0.059378957 | 0.855856783 | 0.991141351 | ns | TM | KS.3 |
| FIM1_2062 | 754.3470971 | 760.0915544 | 1165.197352 | 0.00075096 | 0.078894854 | 0.970872627 | 0.997052226 | ns | TM | KS.3 |
| FIM1_2063 | 2524.500591 | 2610.63429 | 3667.742342 | 0.00794095 | 0.057203664 | 0.727547816 | 0.987157164 | ns | TM | KS.3 |
| FIM1_2064 | 350.2687911 | 524.3969645 | 1152.213554 | 0.071949928 | 0.150498783 | 0.037825626 | 0.440708412 | ns | TM | KS.3 |
| FIM1_2065 | 28.57176776 | 43.21195576 | 56.90405186 | 0.053778447 | 0.138736708 | 0.068646221 | 0.575910535 | ns | TM | KS.3 |
| FIM1_2066 | 737.2657683 | 421.7004649 | 3256.199129 | -0.054118647 | 0.167805817 | 0.065728628 | 0.564318013 | ns | TM | KS.3 |
| FIM1_2067 | 2865.718419 | 2636.449865 | 2073.776345 | -0.017413388 | 0.067111931 | 0.471429643 | 0.950410534 | ns | TM | KS.3 |
| FIM1_2068 | 1131.109747 | 1078.746105 | 1436.187791 | -0.01310306 | 0.054125973 | 0.562216307 | 0.961073589 | ns | TM | KS.3 |
| FIM1_2069 | 1446.991383 | 1211.860785 | 1555.198042 | -0.065777691 | 0.091804941 | 0.058843158 | 0.541117769 | ns | TM | KS.3 |
| FIM1_2070 | 88.70640948 | 78.5618506 | 80.28421507 | -0.013362126 | 0.081075176 | 0.536760662 | 0.95917717 | ns | TM | KS.3 |
| FIM1_2071 | 2117.680509 | 1992.342102 | 2107.247229 | -0.026435553 | 0.050328056 | 0.230487118 | 0.879350777 | ns | TM | KS.3 |
| FIM1_2072 | 1440.776507 | 1546.226791 | 1485.603666 | 0.01692077 | 0.062655221 | 0.481739829 | 0.951416554 | ns | TM | KS.3 |
| FIM1_2073 | 2934.373326 | 3352.647606 | 3368.971596 | 0.020912826 | 0.077567411 | 0.381370485 | 0.948637087 | ns | TM | KS.3 |
| FIM1_2074 | 816.2752433 | 795.0676502 | 831.1518814 | -0.007668279 | 0.050208564 | 0.721918441 | 0.987157164 | ns | TM | KS.3 |
| FIM1_2075 | 8155.660805 | 7335.782082 | 6104.843797 | -0.027441024 | 0.069269248 | 0.291059283 | 0.919993167 | ns | TM | KS.3 |
| FIM1_2076 | 1579.348114 | 1582.638811 | 1345.166444 | 0.000378294 | 0.050805165 | 0.98601819 | 0.999333464 | ns | TM | KS.3 |
| FIM1_2077 | 2107.815025 | 2298.918867 | 2543.597471 | 0.015598128 | 0.070041126 | 0.510431939 | 0.953260948 | ns | TM | KS.3 |
| FIM1_2078 | 1356.024295 | 1509.30416 | 1355.849322 | 0.019456041 | 0.072596273 | 0.421070626 | 0.950410534 | ns | TM | KS.3 |
| FIM1_2079 | 3546.494956 | 4302.471479 | 1965.955201 | 0.017084798 | 0.090539889 | 0.417701689 | 0.950410534 | ns | TM | KS.3 |
| FIM1_2080 | 26.58472224 | 23.87500016 | 36.46740299 | -0.006889556 | 0.089362861 | 0.719208323 | 0.987157164 | ns | TM | KS.3 |
| FIM1_2081 | 6880.658179 | 8949.686485 | 11195.6198 | 0.04169151 | 0.101209829 | 0.125959129 | 0.745120886 | ns | TM | KS.3 |
| FIM1_2083 | 557.2667644 | 770.0636405 | 754.0544526 | 0.059669601 | 0.122223719 | 0.058644273 | 0.540287513 | ns | TM | KS.3 |

| | | | | | | | | | | |
|---|---|---|---|---|---|---|---|---|---|---|
| FIM1_2084 | 6807.448687 | 7078.497719 | 6222.755823 | 0.014598756 | 0.040347284 | 0.417947283 | 0.950410534 | ns | TM | KS.3 |
| FIM1_2085 | 3209.835459 | 2825.863386 | 2721.631889 | -0.093744985 | 0.078866153 | 0.019109426 | 0.32184354 | ns | TM | KS.3 |
| FIM1_2086 | 3306.668224 | 3781.091756 | 3151.420719 | 0.071260495 | 0.081670419 | 0.047117243 | 0.496748336 | ns | TM | KS.3 |
| FIM1_2087 | 4097.987058 | 4557.940421 | 3960.918562 | 0.011103711 | 0.080848145 | 0.601127421 | 0.97002321 | ns | TM | KS.3 |
| FIM1_2088 | 1128.925372 | 1317.133545 | 1507.92372 | 0.044220686 | 0.081670806 | 0.135713083 | 0.768410055 | ns | TM | KS.3 |
| FIM1_2089 | 340.8923949 | 311.0356525 | 349.9156491 | -0.01829164 | 0.06925397 | 0.450366459 | 0.950410534 | ns | TM | KS.3 |
| FIM1_2090 | 749.5771991 | 756.0545066 | 845.9294823 | 0.002126072 | 0.047327734 | 0.918412411 | 0.991141351 | ns | TM | KS.3 |
| FIM1_2091 | 1190.214035 | 1221.87717 | 1214.776412 | 0.005912021 | 0.057504086 | 0.79476116 | 0.988916276 | ns | TM | KS.3 |
| FIM1_2092 | 555.9267468 | 712.9505156 | 677.1625973 | 0.028606345 | 0.096104836 | 0.227803385 | 0.877863547 | ns | TM | KS.3 |
| FIM1_2093 | 2430.324761 | 2864.892335 | 3388.232517 | 0.015165866 | 0.087267823 | 0.471130271 | 0.950410534 | ns | TM | KS.3 |
| FIM1_2094 | 7384.801383 | 7723.067963 | 8107.293509 | 0.009839794 | 0.06078657 | 0.671579341 | 0.976861203 | ns | TM | KS.3 |
| FIM1_2095 | 1040.890576 | 811.0570375 | 1015.029635 | -0.067390147 | 0.108179788 | 0.049589952 | 0.510786778 | ns | TM | KS.3 |
| FIM1_2096 | 640.5158416 | 590.8811919 | 1200.383166 | -0.009968422 | 0.076144386 | 0.649150179 | 0.976323032 | ns | TM | KS.3 |
| FIM1_2097 | 1485.550312 | 1524.083955 | 1071.318606 | 0.007455749 | 0.042309143 | 0.696958177 | 0.98151979 | ns | TM | KS.3 |
| FIM1_2098 | 384.1831238 | 157.073416 | 1283.458349 | -0.252934681 | 0.433676353 | 0.003693957 | 0.115135887 | ns | TM | KS.3 |
| FIM1_2099 | 12.55686745 | 16.81186055 | 15.19172552 | 0.016901346 | 0.109101704 | 0.385412771 | 0.948637087 | ns | TM | KS.3 |
| FIM1_2100 | 1592.250349 | 1801.988803 | 1852.733756 | 0.02638756 | 0.073917421 | 0.302739123 | 0.926536521 | ns | TM | KS.3 |
| FIM1_2101 | 650.4947867 | 841.5388317 | 761.0685894 | 0.119015397 | 0.124262139 | 0.010778327 | 0.231130071 | ns | TM | KS.3 |
| FIM1_2102 | 1033.570874 | 1100.862536 | 1111.61358 | 0.016507021 | 0.059266713 | 0.486169824 | 0.952269291 | ns | TM | KS.3 |
| FIM1_2103 | 5645.157535 | 5265.095811 | 5729.274012 | -0.008487991 | 0.075867832 | 0.696406875 | 0.981479944 | ns | TM | KS.3 |
| FIM1_2104 | 914.8405803 | 689.6038143 | 766.9407679 | -0.056199904 | 0.111016304 | 0.069636814 | 0.581836521 | ns | TM | KS.3 |
| FIM1_2105 | 512.1224062 | 558.0246585 | 459.8558034 | 0.017428441 | 0.067880272 | 0.470586512 | 0.950410534 | ns | TM | KS.3 |
| FIM1_2106 | 1080.34005 | 912.2856756 | 901.2688859 | -0.03498548 | 0.082906837 | 0.195084205 | 0.837467988 | ns | TM | KS.3 |
| FIM1_2107 | 679.1956993 | 741.9456957 | 817.3913675 | 0.024752165 | 0.065006159 | 0.328547125 | 0.935147309 | ns | TM | KS.3 |
| FIM1_2108 | 741.6601421 | 679.9506313 | 655.523916 | -0.019607276 | 0.066295027 | 0.424442644 | 0.950410534 | ns | TM | KS.3 |
| FIM1_2109 | 215.5207886 | 215.6532926 | 233.2353216 | -5.66E-05 | 0.067817064 | 0.997991954 | 0.999525119 | ns | TM | KS.3 |
| FIM1_2110 | 410.2301301 | 397.6917118 | 423.3050302 | -0.005716322 | 0.065530774 | 0.802613585 | 0.988916276 | ns | TM | KS.3 |
| FIM1_2111 | 43.23483843 | 46.72283922 | 44.30100201 | 0.00598415 | 0.082869841 | 0.766071249 | 0.988477371 | ns | TM | KS.3 |
| FIM1_2112 | 1027.324376 | 925.9560852 | 1126.871592 | -0.027477743 | 0.068582428 | 0.290313308 | 0.919993167 | ns | TM | KS.3 |
| FIM1_2113 | 1681.816257 | 1477.365292 | 1409.47737 | -0.042926967 | 0.075975586 | 0.144622108 | 0.77783242 | ns | TM | KS.3 |
| FIM1_2115 | 1317.427038 | 1216.5557 | 1230.235367 | -0.025084417 | 0.061430526 | 0.315196569 | 0.927564309 | ns | TM | KS.3 |
| FIM1_2116 | 1316.10812 | 1125.303653 | 1207.977676 | -0.143662456 | 0.078829976 | 0.003007528 | 0.101288726 | ns | TM | KS.3 |
| FIM1_2117 | 2224.07326 | 2201.207793 | 2470.402507 | -0.003359715 | 0.035493481 | 0.839236123 | 0.991141351 | ns | TM | KS.3 |
| FIM1_2118 | 1463.556111 | 1324.134658 | 1670.173672 | -0.078366313 | 0.067551118 | 0.025351761 | 0.376492575 | ns | TM | KS.3 |
| FIM1_2119 | 790.7390225 | 902.6369737 | 891.3692608 | 0.012169588 | 0.084815638 | 0.557791096 | 0.959899566 | ns | TM | KS.3 |
| FIM1_2120 | 2207.178147 | 1968.479764 | 1712.28105 | -0.030801415 | 0.071269913 | 0.248517862 | 0.888934767 | ns | TM | KS.3 |
| FIM1_2121 | 3488.659113 | 3227.498825 | 3712.965937 | -0.027524691 | 0.05927296 | 0.267950518 | 0.905607219 | ns | TM | KS.3 |
| FIM1_2122 | 1283.561018 | 1329.019094 | 1269.252022 | 0.00948096 | 0.052565404 | 0.668279597 | 0.976323032 | ns | TM | KS.3 |
| FIM1_2123 | 7493.644517 | 7550.035044 | 7668.136712 | 0.002020865 | 0.049373878 | 0.924316466 | 0.993271864 | ns | TM | KS.3 |
| FIM1_2124 | 1719.318868 | 1954.730044 | 2574.161137 | 0.020828669 | 0.07660184 | 0.38552541 | 0.948637087 | ns | TM | KS.3 |
| FIM1_2125 | 1123.211656 | 1080.242303 | 1307.919858 | -0.008202934 | 0.061740539 | 0.7226784 | 0.987157164 | ns | TM | KS.3 |
| FIM1_2126 | 1354.983456 | 956.030472 | 1140.724034 | -0.197515608 | 0.181846859 | 0.002183836 | 0.084221572 | ns | TM | KS.3 |
| FIM1_2127 | 809.6264134 | 1003.054264 | 1172.311572 | 0.156983588 | 0.104985044 | 0.003233036 | 0.103770022 | ns | TM | KS.3 |
| FIM1_2128 | 878.8742673 | 781.0129026 | 763.6469738 | -0.025781241 | 0.07287007 | 0.312729892 | 0.927416591 | ns | TM | KS.3 |
| FIM1_2129 | 2992.98464 | 3027.439703 | 3207.282133 | 0.002889249 | 0.05192949 | 0.89470929 | 0.991141351 | ns | TM | KS.3 |

| | | | | | | | | | | |
|---|---|---|---|---|---|---|---|---|---|---|
| FIM1_2130 | 1825.042997 | 1705.027107 | 4591.296402 | -0.009447571 | 0.073133288 | 0.672552752 | 0.976861203 | ns | TM | KS.3 |
| FIM1_2131 | 11864.10124 | 11197.93064 | 10641.61632 | -0.013186429 | 0.061500395 | 0.575620819 | 0.963192994 | ns | TM | KS.3 |
| FIM1_2132 | 1556.543394 | 1928.012495 | 2400.954067 | 0.01606856 | 0.09486928 | 0.429645734 | 0.950410534 | ns | TM | KS.3 |
| FIM1_2133 | 1025.518511 | 978.5567459 | 1195.454349 | -0.014639311 | 0.052025976 | 0.509728838 | 0.953260948 | ns | TM | KS.3 |
| FIM1_2134 | 1098.696983 | 1042.650228 | 923.6418429 | -0.012259371 | 0.059998692 | 0.599831806 | 0.969513722 | ns | TM | KS.3 |
| FIM1_2135 | 2329.668125 | 2279.387849 | 2432.676187 | -0.005564886 | 0.052290261 | 0.799856272 | 0.988916276 | ns | TM | KS.3 |
| FIM1_2136 | 2288.365785 | 2114.183023 | 1894.50463 | -0.029002106 | 0.059569666 | 0.247290308 | 0.888934767 | ns | TM | KS.3 |
| FIM1_2137 | 836.9557033 | 936.2579308 | 929.025873 | 0.028703742 | 0.070705417 | 0.274210413 | 0.908468501 | ns | TM | KS.3 |
| FIM1_2138 | 1099.380527 | 1072.642767 | 1077.161676 | -0.003743925 | 0.069258467 | 0.867272626 | 0.991141351 | ns | TM | KS.3 |
| FIM1_2139 | 3276.304196 | 3059.932859 | 2822.038722 | -0.01618741 | 0.062457373 | 0.49910211 | 0.953260948 | ns | TM | KS.3 |
| FIM1_2140 | 977.9847368 | 1000.089764 | 1049.093401 | 0.00640362 | 0.049557582 | 0.764388739 | 0.988477371 | ns | TM | KS.3 |
| FIM1_2141 | 1397.035275 | 1352.538449 | 1365.113948 | -0.007116772 | 0.059849932 | 0.756673781 | 0.988477371 | ns | TM | KS.3 |
| FIM1_2142 | 220.6576048 | 186.6692376 | 184.7391978 | -0.039368191 | 0.082203108 | 0.164121836 | 0.800496208 | ns | TM | KS.3 |
| FIM1_2143 | 936.0557835 | 933.9286153 | 887.3095082 | -0.000677499 | 0.055055002 | 0.975790741 | 0.998880439 | ns | TM | KS.3 |
| FIM1_2144 | 1464.429563 | 1379.375574 | 1324.008097 | -0.013642249 | 0.06186333 | 0.56360239 | 0.961234792 | ns | TM | KS.3 |
| FIM1_2145 | 667.7274378 | 783.6228454 | 768.7166833 | 0.021401376 | 0.082418189 | 0.358483784 | 0.948637087 | ns | TM | KS.3 |
| FIM1_2146 | 14765.6346 | 14465.76273 | 11469.12418 | -0.006060555 | 0.045161247 | 0.763358597 | 0.988477371 | ns | TM | KS.3 |
| FIM1_2147 | 665.4326101 | 664.134328 | 730.851883 | -0.000370252 | 0.063526868 | 0.98703146 | 0.999333464 | ns | TM | KS.3 |
| FIM1_2148 | 614.3797169 | 588.0137718 | 554.9139861 | -0.007182235 | 0.068456038 | 0.752095492 | 0.988477371 | ns | TM | KS.3 |
| FIM1_2149 | 984.211882 | 979.3685288 | 947.8479098 | -0.001696101 | 0.048193861 | 0.935570382 | 0.994413583 | ns | TM | KS.3 |
| FIM1_2150 | 1402.178528 | 1483.880067 | 1219.572483 | 0.009543238 | 0.068291364 | 0.678112409 | 0.97986479 | ns | TM | KS.3 |
| FIM1_2151 | 721.7439236 | 725.1405762 | 1008.198779 | 0.001010438 | 0.058786822 | 0.964496223 | 0.995175461 | ns | TM | KS.3 |
| FIM1_2152 | 187.2428658 | 178.8253954 | 256.6004841 | -0.010314989 | 0.060911075 | 0.657342264 | 0.976323032 | ns | TM | KS.3 |
| FIM1_2153 | 523.2691678 | 498.8787587 | 445.3975149 | -0.012624865 | 0.057693486 | 0.585702786 | 0.963781856 | ns | TM | KS.3 |
| FIM1_2154 | 925.3165311 | 1190.206502 | 1304.304847 | 0.018746813 | 0.098910438 | 0.364441629 | 0.948637087 | ns | TM | KS.3 |
| FIM1_2155 | 741.2603534 | 785.0968749 | 789.1994816 | 0.017824248 | 0.054528412 | 0.437787083 | 0.950410534 | ns | TM | KS.3 |
| FIM1_2156 | 1462.786715 | 1378.375188 | 1257.674473 | -0.012125817 | 0.064413976 | 0.605859818 | 0.970937835 | ns | TM | KS.3 |
| FIM1_2157 | 5217.072804 | 5279.231981 | 3731.140676 | 0.003583948 | 0.042296482 | 0.851930611 | 0.991141351 | ns | TM | KS.3 |
| FIM1_2158 | 2619.21755 | 2597.440841 | 2815.198628 | -0.002234365 | 0.050478156 | 0.917325679 | 0.991141351 | ns | TM | KS.3 |
| FIM1_2159 | 61.37214588 | 65.76654177 | 86.34968685 | 0.00545249 | 0.084536777 | 0.782544956 | 0.988814136 | ns | TM | KS.3 |
| FIM1_2160 | 1877.00094 | 1735.961826 | 1771.477552 | -0.018892457 | 0.06397756 | 0.438469297 | 0.950410534 | ns | TM | KS.3 |
| FIM1_2161 | 1683.764756 | 1588.549463 | 1565.997904 | -0.028555315 | 0.047814357 | 0.176632001 | 0.811391014 | ns | TM | KS.3 |
| FIM1_2162 | 833.6629299 | 665.3293091 | 767.1367039 | -0.134489221 | 0.113390104 | 0.006886785 | 0.176553399 | ns | TM | KS.3 |
| FIM1_2163 | 1023.729659 | 793.0499362 | 761.4008034 | -0.060609068 | 0.106821541 | 0.06200352 | 0.549852965 | ns | TM | KS.3 |
| FIM1_2164 | 784.0365054 | 814.1839355 | 769.3607597 | 0.01054495 | 0.050354438 | 0.626213676 | 0.97344368 | ns | TM | KS.3 |
| FIM1_2165 | 2222.36905 | 2412.935043 | 2668.911091 | 0.025432686 | 0.06189423 | 0.310722676 | 0.927416591 | ns | TM | KS.3 |
| FIM1_2166 | 393.3676066 | 389.3551927 | 398.1753113 | -0.001334768 | 0.073250052 | 0.950932637 | 0.994413583 | ns | TM | KS.3 |
| FIM1_2167 | 5744.556848 | 5177.806122 | 4787.92655 | -0.033006467 | 0.068246065 | 0.221473403 | 0.870323997 | ns | TM | KS.3 |
| FIM1_2168 | 4777.233415 | 4446.000026 | 4647.44812 | -0.032593961 | 0.055675384 | 0.180123696 | 0.813337273 | ns | TM | KS.3 |
| FIM1_2169 | 2393.992248 | 2419.171761 | 2427.300803 | 0.00293634 | 0.047265997 | 0.887421211 | 0.991141351 | ns | TM | KS.3 |
| FIM1_2170 | 3839.646417 | 3746.552491 | 3230.549092 | -0.005635487 | 0.057376144 | 0.803957735 | 0.988916276 | ns | TM | KS.3 |
| FIM1_2171 | 1069.614933 | 1444.445361 | 1110.462969 | 0.095022915 | 0.132544785 | 0.020188065 | 0.324535175 | ns | TM | KS.3 |
| FIM1_2172 | 1269.209515 | 1502.920195 | 2057.017079 | 0.076753176 | 0.091888906 | 0.040616833 | 0.459247141 | ns | TM | KS.3 |
| FIM1_2173 | 761.3775454 | 1018.958234 | 774.6059075 | 0.078824398 | 0.123107467 | 0.032502302 | 0.415678547 | ns | TM | KS.3 |
| FIM1_2174 | 1201.409125 | 1031.058549 | 1263.218158 | -0.032791854 | 0.079760344 | 0.219644953 | 0.865538807 | ns | TM | KS.3 |

| | | | | | | | | | | |
|---|---|---|---|---|---|---|---|---|---|---|
| FIM1_2175 | 3357.958423 | 3395.840929 | 3557.19596 | 0.002852634 | 0.0534216 | 0.897295723 | 0.991141351 | ns | TM | KS.3 |
| FIM1_2176 | 1430.619828 | 1193.400756 | 925.9356295 | -0.046383344 | 0.087582434 | 0.119771343 | 0.730090427 | ns | TM | KS.3 |
| FIM1_2177 | 59.86472119 | 45.49539794 | 79.54727981 | -0.036450842 | 0.100246281 | 0.156641441 | 0.789034023 | ns | TM | KS.3 |
| FIM1_2178 | 4365.125518 | 5426.132388 | 4144.390667 | 0.036548807 | 0.091710786 | 0.168014183 | 0.805573568 | ns | TM | KS.3 |
| FIM1_2179 | 10992.88416 | 9785.783583 | 9088.135023 | -0.03775188 | 0.072032988 | 0.180160337 | 0.813337273 | ns | TM | KS.3 |
| FIM1_2180 | 2230.332695 | 2584.011838 | 2978.44097 | 0.013954312 | 0.085394463 | 0.507331787 | 0.953260948 | ns | TM | KS.3 |
| FIM1_2181 | 2657.326261 | 2662.737049 | 2827.95635 | 0.000256593 | 0.072828489 | 0.99058866 | 0.999525119 | ns | TM | KS.3 |
| FIM1_2182 | 460.939408 | 619.4070999 | 1562.092515 | 0.025322475 | 0.103858798 | 0.253283572 | 0.889891082 | ns | TM | KS.3 |
| FIM1_2183 | 2032.494157 | 2095.11219 | 2582.084018 | 0.008386793 | 0.051145925 | 0.700217924 | 0.98151979 | ns | TM | KS.3 |
| FIM1_2184 | 3002.317061 | 2963.8692 | 2910.723947 | -0.002918351 | 0.056818351 | 0.897088402 | 0.991141351 | ns | TM | KS.3 |
| FIM1_2185 | 1889.207054 | 1966.658483 | 3145.59594 | 0.00726818 | 0.065534839 | 0.751792333 | 0.988477371 | ns | TM | KS.3 |
| FIM1_2186 | 1558.044076 | 1679.054436 | 1562.596842 | 0.021465752 | 0.060474912 | 0.378982353 | 0.948637087 | ns | TM | KS.3 |
| FIM1_2187 | 1344.066771 | 1592.383495 | 1825.116442 | 0.023268078 | 0.083481303 | 0.325163534 | 0.934066145 | ns | TM | KS.3 |
| FIM1_2188 | 4448.118544 | 4033.574991 | 4842.867628 | -0.040273536 | 0.066795815 | 0.150944565 | 0.783447458 | ns | TM | KS.3 |
| FIM1_2189 | 905.7272266 | 1008.048839 | 1001.661405 | 0.020540043 | 0.071934632 | 0.401242326 | 0.950410534 | ns | TM | KS.3 |
| FIM1_2190 | 2024.56623 | 1974.839321 | 2212.447532 | -0.006520306 | 0.051568551 | 0.765141135 | 0.988477371 | ns | TM | KS.3 |
| FIM1_2191 | 8416.65468 | 8123.924524 | 7904.87394 | -0.011913698 | 0.042894099 | 0.535743002 | 0.959071703 | ns | TM | KS.3 |
| FIM1_2192 | 866.1816996 | 916.3835691 | 907.4031976 | 0.010592503 | 0.065993017 | 0.649070484 | 0.976323032 | ns | TM | KS.3 |
| FIM1_2193 | 3080.266493 | 3541.502995 | 4372.129069 | 0.013100279 | 0.084968992 | 0.531371966 | 0.959071703 | ns | TM | KS.3 |
| FIM1_2194 | 445.045107 | 656.9666689 | 674.7807856 | 0.081375328 | 0.153349171 | 0.028822626 | 0.390786333 | ns | TM | KS.3 |
| FIM1_2195 | 672.7305534 | 701.8850658 | 827.4549321 | 0.009527858 | 0.058512328 | 0.678996618 | 0.97986479 | ns | TM | KS.3 |
| FIM1_2196 | 272.0213878 | 282.4999798 | 346.3742841 | 0.00634139 | 0.067119573 | 0.780879822 | 0.988814136 | ns | TM | KS.3 |
| FIM1_2197 | 1247.066094 | 1331.785031 | 1611.03317 | 0.018693821 | 0.058049594 | 0.430997102 | 0.950410534 | ns | TM | KS.3 |
| FIM1_2198 | 1406.785935 | 1820.856976 | 2316.489743 | 0.04069132 | 0.099958848 | 0.13228803 | 0.761677364 | ns | TM | KS.3 |
| FIM1_2199 | 57.99551089 | 80.31719286 | 102.5282884 | 0.045658125 | 0.118231097 | 0.097801353 | 0.677062609 | ns | TM | KS.3 |
| FIM1_2200 | 936.5399377 | 1424.825404 | 1233.819232 | 0.37867294 | 0.236628185 | 0.000193848 | 0.014393914 | ns | TM | KS.3 |
| FIM1_2201 | 897.4507684 | 880.720295 | 1062.087388 | -0.005518598 | 0.044469421 | 0.781664569 | 0.988814136 | ns | TM | KS.3 |
| FIM1_2202 | 5986.080261 | 5172.500862 | 4287.758579 | -0.044754169 | 0.080202736 | 0.133637465 | 0.765070643 | ns | TM | KS.3 |
| FIM1_2203 | 1173.092137 | 1016.14658 | 970.0864016 | -0.04963639 | 0.080795055 | 0.109756128 | 0.707301473 | ns | TM | KS.3 |
| FIM1_2204 | 1177.393747 | 1022.186004 | 982.1773121 | -0.033679602 | 0.077416797 | 0.214731611 | 0.864312109 | ns | TM | KS.3 |
| FIM1_2205 | 1036.79422 | 925.0694831 | 972.1536766 | -0.028475024 | 0.071201972 | 0.277076067 | 0.908583046 | ns | TM | KS.3 |
| FIM1_2206 | 872.7311649 | 996.2869505 | 1280.748421 | 0.011625412 | 0.085629717 | 0.571117873 | 0.962265923 | ns | TM | KS.3 |
| FIM1_2207 | 383.7226745 | 426.9084834 | 462.4421567 | 0.008333165 | 0.086512907 | 0.673886965 | 0.976861203 | ns | TM | KS.3 |
| FIM1_2209 | 1703.952555 | 1615.708121 | 1758.944291 | -0.006806763 | 0.074209239 | 0.755503867 | 0.988477371 | ns | TM | KS.3 |
| FIM1_2210 | 2938.846533 | 3811.546112 | 4726.000485 | 0.037094894 | 0.09931332 | 0.153631002 | 0.783447458 | ns | TM | KS.3 |
| FIM1_2211 | 5158.695016 | 4390.289528 | 4800.855103 | -0.09345727 | 0.091240303 | 0.023000471 | 0.356471474 | ns | TM | KS.3 |
| FIM1_2212 | 1276.111746 | 1211.140924 | 1039.059064 | -0.01323462 | 0.058038895 | 0.569260616 | 0.962265923 | ns | TM | KS.3 |
| FIM1_2213 | 864.6034237 | 855.1106714 | 898.7242659 | -0.002865563 | 0.05149896 | 0.895159477 | 0.991141351 | ns | TM | KS.3 |
| FIM1_2214 | 705.8585688 | 685.6879997 | 624.8375672 | -0.007890959 | 0.05130389 | 0.71742842 | 0.987157164 | ns | TM | KS.3 |
| FIM1_2215 | 5825.697595 | 5917.874261 | 6323.786572 | 0.002523419 | 0.067650005 | 0.910855064 | 0.991141351 | ns | TM | KS.3 |
| FIM1_2216 | 951.1430473 | 1038.342553 | 919.6030042 | 0.026039039 | 0.06349428 | 0.304970629 | 0.926747341 | ns | TM | KS.3 |
| FIM1_2217 | 751.8813129 | 707.8733933 | 851.7880038 | -0.007370825 | 0.07528187 | 0.734408009 | 0.987655486 | ns | TM | KS.3 |
| FIM1_2218 | 10912.59804 | 9570.565858 | 8006.289705 | -0.070973958 | 0.080910645 | 0.047228535 | 0.496748336 | ns | TM | KS.3 |
| FIM1_2219 | 2343.604272 | 2567.358552 | 2623.452048 | 0.036600777 | 0.064112806 | 0.175024612 | 0.811391014 | ns | TM | KS.3 |
| FIM1_2220 | 1535.77528 | 1522.987304 | 1521.415233 | -0.002082743 | 0.031517133 | 0.888332168 | 0.991141351 | ns | TM | KS.3 |

| | | | | | | | | | | |
|---|---|---|---|---|---|---|---|---|---|---|
| FIM1_2221 | 889.3536261 | 912.1203642 | 801.815568 | 0.006835013 | 0.044427912 | 0.731109925 | 0.987157164 | ns | TM | KS.3 |
| FIM1_2222 | 652.2343721 | 608.6108278 | 622.0733073 | -0.013242358 | 0.066543566 | 0.574385444 | 0.963192994 | ns | TM | KS.3 |
| FIM1_2223 | 1010.660283 | 978.6276158 | 1068.373156 | -0.007469361 | 0.057260229 | 0.742937707 | 0.988477371 | ns | TM | KS.3 |
| FIM1_2224 | 3712.651683 | 3716.351551 | 3951.563321 | 0.0002295 | 0.057274062 | 0.991889687 | 0.999525119 | ns | TM | KS.3 |
| FIM1_2225 | 927.3141629 | 935.7573968 | 1245.852409 | 0.001185354 | 0.072572981 | 0.956653453 | 0.994413583 | ns | TM | KS.3 |
| FIM1_2226 | 10040.7464 | 9635.940627 | 10280.44126 | -0.016574047 | 0.039357246 | 0.341295849 | 0.944808611 | ns | TM | KS.3 |
| FIM1_2227 | 1369.544857 | 1489.342646 | 2047.478626 | 0.014196634 | 0.070765654 | 0.543896911 | 0.95917717 | ns | TM | KS.3 |
| FIM1_2228 | 850.6526543 | 956.0228787 | 941.2334906 | 0.010025813 | 0.085097957 | 0.621592314 | 0.973122675 | ns | TM | KS.3 |
| FIM1_2229 | 743.3278923 | 783.2944784 | 888.5419249 | 0.00771657 | 0.07089875 | 0.731474824 | 0.987157164 | ns | TM | KS.3 |
| FIM1_2230 | 310.4857687 | 274.5524744 | 429.8878394 | -0.014012491 | 0.080499513 | 0.521586243 | 0.958602567 | ns | TM | KS.3 |
| FIM1_2231 | 970.6940895 | 1118.125961 | 934.2259976 | 0.07997934 | 0.08470437 | 0.035057505 | 0.42652037 | ns | TM | KS.3 |
| FIM1_2232 | 290.1271299 | 284.6457198 | 872.8070102 | -0.002281026 | 0.075450182 | 0.914726718 | 0.991141351 | ns | TM | KS.3 |
| FIM1_2233 | 1535.462681 | 1732.824895 | 1744.585935 | 0.031855015 | 0.072759308 | 0.236550634 | 0.88502866 | ns | TM | KS.3 |
| FIM1_2234 | 512.1378471 | 531.4006014 | 525.7881748 | 0.01021715 | 0.051660731 | 0.641665165 | 0.975935248 | ns | TM | KS.3 |
| FIM1_2235 | 511.7942094 | 622.2944104 | 542.9644863 | 0.037372946 | 0.08789059 | 0.168771981 | 0.806481872 | ns | TM | KS.3 |
| FIM1_2236 | 1508.140072 | 1510.336568 | 1267.879751 | 0.00042778 | 0.055108229 | 0.984716921 | 0.999333464 | ns | TM | KS.3 |
| FIM1_2237 | 24.97628777 | 21.91549349 | 40.526282 | -0.009011091 | 0.088353794 | 0.646239123 | 0.976323032 | ns | TM | KS.3 |
| FIM1_2238 | 836.3687765 | 762.9010829 | 875.0395528 | -0.022401379 | 0.066743311 | 0.370646532 | 0.948637087 | ns | TM | KS.3 |
| FIM1_2239 | 992.0368148 | 988.2759004 | 707.4400088 | -0.000859014 | 0.059734977 | 0.969881265 | 0.996727803 | ns | TM | KS.3 |
| FIM1_2240 | 811.3267129 | 836.8507231 | 670.9627868 | 0.006283964 | 0.061834188 | 0.784581994 | 0.988880259 | ns | TM | KS.3 |
| FIM1_2241 | 3085.580697 | 2762.036522 | 2657.322158 | -0.019094876 | 0.073630671 | 0.426102193 | 0.950410534 | ns | TM | KS.3 |
| FIM1_2243 | 1032.237515 | 1104.086575 | 1190.464514 | 0.016608946 | 0.061676461 | 0.488013729 | 0.952591289 | ns | TM | KS.3 |
| FIM1_2244 | 4270.707409 | 4400.966817 | 5009.643501 | 0.007579903 | 0.054447471 | 0.735052684 | 0.987655486 | ns | TM | KS.3 |
| FIM1_2245 | 1369.810843 | 1205.395962 | 1697.103408 | -0.025670545 | 0.075029288 | 0.311057907 | 0.927416591 | ns | TM | KS.3 |
| FIM1_2246 | 11480.92231 | 11979.2933 | 11412.85161 | 0.013787047 | 0.04743275 | 0.508367324 | 0.953260948 | ns | TM | KS.3 |
| FIM1_2247 | 1139.739308 | 1109.005049 | 1103.334595 | -0.007492721 | 0.049866384 | 0.726977645 | 0.987157164 | ns | TM | KS.3 |
| FIM1_2248 | 2146.739619 | 2365.338931 | 2883.141521 | 0.008637481 | 0.083357349 | 0.671682584 | 0.976861203 | ns | TM | KS.3 |
| FIM1_2249 | 3078.459968 | 3793.237939 | 4456.848732 | 0.023500411 | 0.090021841 | 0.304031119 | 0.926536521 | ns | TM | KS.3 |
| FIM1_2250 | 288.1113277 | 401.4698374 | 393.1173166 | 0.040163042 | 0.113615428 | 0.123922956 | 0.739893493 | ns | TM | KS.3 |
| FIM1_2251 | 499.0853981 | 457.6702474 | 559.9824934 | -0.030809513 | 0.061675601 | 0.231111818 | 0.880810476 | ns | TM | KS.3 |
| FIM1_2252 | 3949.259716 | 3592.50188 | 3579.467591 | -0.023397384 | 0.067049189 | 0.353191472 | 0.948637087 | ns | TM | KS.3 |
| FIM1_2253 | 14221.55996 | 13895.65396 | 15903.47299 | -0.007705694 | 0.037079001 | 0.651023262 | 0.976323032 | ns | TM | KS.3 |
| FIM1_2254 | 3218.421383 | 2897.464048 | 2607.848379 | -0.022988228 | 0.070278713 | 0.359387495 | 0.948637087 | ns | TM | KS.3 |
| FIM1_2255 | 13484.65016 | 13576.73886 | 13851.80021 | 0.002218569 | 0.031643365 | 0.881528378 | 0.991141351 | ns | TM | KS.3 |
| FIM1_2256 | 2563.644864 | 1552.205436 | 1342.415325 | -0.434613657 | 0.305185794 | 0.000291764 | 0.019353666 | ns | TM | KS.3 |
| FIM1_2257 | 915.2946713 | 900.0913308 | 1340.549922 | -0.001120631 | 0.087912646 | 0.952734357 | 0.994413583 | ns | TM | KS.3 |
| FIM1_2258 | 1866.474315 | 1572.069362 | 2130.826195 | -0.041683856 | 0.084648851 | 0.146844148 | 0.780546912 | ns | TM | KS.3 |
| FIM1_2259 | 438.1863336 | 438.7841331 | 421.2064794 | 0.00011718 | 0.058420048 | 0.995876047 | 0.999525119 | ns | TM | KS.3 |
| FIM1_2260 | 2028.260595 | 1779.314137 | 1913.190993 | -0.054966638 | 0.078622759 | 0.087874706 | 0.64839159 | ns | TM | KS.3 |
| FIM1_2261 | 3590.794944 | 3184.75185 | 3325.419316 | -0.043987496 | 0.073865498 | 0.13680251 | 0.769031059 | ns | TM | KS.3 |
| FIM1_2262 | 3892.96921 | 3073.893032 | 2968.684737 | -0.046729889 | 0.097892385 | 0.107244255 | 0.702423202 | ns | TM | KS.3 |
| FIM1_2263 | 5814.648793 | 5639.485672 | 4218.55243 | -0.006586766 | 0.060029542 | 0.774085863 | 0.988814136 | ns | TM | KS.3 |
| FIM1_2264 | 1482.472999 | 1284.197818 | 1340.425027 | -0.05496155 | 0.081833424 | 0.088978422 | 0.651940574 | ns | TM | KS.3 |
| FIM1_2265 | 882.0573438 | 957.48925 | 992.9188256 | 0.026591994 | 0.060268138 | 0.286556447 | 0.919993167 | ns | TM | KS.3 |
| FIM1_2266 | 1088.205507 | 1558.124849 | 1334.817771 | 0.244717963 | 0.192006374 | 0.000928452 | 0.044845145 | ns | TM | KS.3 |

| | | | | | | | | | | |
|---|---|---|---|---|---|---|---|---|---|---|
| FIM1_2267 | 1496.442578 | 1532.380073 | 1619.757706 | 0.005436789 | 0.057054818 | 0.810402934 | 0.988916276 | ns | TM | KS.3 |
| FIM1_2268 | 1199.648566 | 1385.206114 | 1336.216251 | 0.068717289 | 0.084056641 | 0.052967178 | 0.527023426 | ns | TM | KS.3 |
| FIM1_2270 | 792.8086692 | 869.7337713 | 840.3921184 | 0.015000689 | 0.072410758 | 0.520338106 | 0.958509403 | ns | TM | KS.3 |
| FIM1_2271 | 1636.285765 | 1271.053183 | 1494.176021 | -0.140072079 | 0.123998396 | 0.006079041 | 0.162598018 | ns | TM | KS.3 |
| FIM1_2272 | 1083.222624 | 1100.498673 | 737.2469833 | 0.003000786 | 0.063092053 | 0.895451932 | 0.991141351 | ns | TM | KS.3 |
| FIM1_2273 | 230.7820163 | 184.3753041 | 169.1592428 | -0.032155735 | 0.092290981 | 0.200826548 | 0.845272486 | ns | TM | KS.3 |
| FIM1_2274 | 2618.735765 | 2770.024297 | 2250.51838 | 0.006940728 | 0.074951881 | 0.749516302 | 0.988477371 | ns | TM | KS.3 |
| FIM1_2275 | 2005.611819 | 1781.187717 | 2090.895444 | -0.02299165 | 0.073630582 | 0.354801215 | 0.948637087 | ns | TM | KS.3 |
| FIM1_2276 | 698.7745142 | 757.2977033 | 871.5900199 | 0.018719678 | 0.065300907 | 0.442928174 | 0.950410534 | ns | TM | KS.3 |
| FIM1_2278 | 1690.558736 | 1678.826347 | 1870.660216 | -0.00103751 | 0.069528516 | 0.962857551 | 0.995175461 | ns | TM | KS.3 |
| FIM1_2279 | 232.8748641 | 217.3380288 | 338.5058139 | -0.017612422 | 0.063341534 | 0.466334375 | 0.950410534 | ns | TM | KS.3 |
| FIM1_2280 | 2140.937965 | 2382.028101 | 1950.211584 | 0.039267736 | 0.069414111 | 0.164464953 | 0.800584912 | ns | TM | KS.3 |
| FIM1_2281 | 868.5927744 | 606.18246 | 749.2942018 | -0.12477675 | 0.16718403 | 0.009848056 | 0.219966293 | ns | TM | KS.3 |
| FIM1_2282 | 351.3905852 | 318.0083193 | 305.8777883 | -0.026813401 | 0.067964824 | 0.299479495 | 0.926536521 | ns | TM | KS.3 |
| FIM1_2283 | 1828.797095 | 1807.851532 | 1312.087633 | -0.002729164 | 0.054843093 | 0.902670736 | 0.991141351 | ns | TM | KS.3 |
| FIM1_2284 | 2432.988411 | 2417.808261 | 2494.53559 | -0.000846861 | 0.071407108 | 0.96929193 | 0.996327965 | ns | TM | KS.3 |
| FIM1_2285 | 4530.318137 | 4520.570949 | 4091.590869 | -0.000364286 | 0.064201604 | 0.987226946 | 0.999333464 | ns | TM | KS.3 |
| FIM1_2286 | 1109.631377 | 959.5094878 | 1032.886163 | -0.047262348 | 0.08060492 | 0.120699274 | 0.730509595 | ns | TM | KS.3 |
| FIM1_2287 | 815.7775763 | 819.2913425 | 927.4687669 | 0.000979469 | 0.057242615 | 0.965397423 | 0.995203518 | ns | TM | KS.3 |
| FIM1_2288 | 1795.661003 | 1852.310827 | 1886.171791 | 0.008052263 | 0.053541826 | 0.717561133 | 0.987157164 | ns | TM | KS.3 |
| FIM1_2289 | 3148.132088 | 3456.675039 | 3305.0036 | 0.021956415 | 0.067322297 | 0.37884866 | 0.948637087 | ns | TM | KS.3 |
| FIM1_2290 | 1674.797994 | 1755.408894 | 2575.129183 | 0.008218068 | 0.066907012 | 0.720557617 | 0.987157164 | ns | TM | KS.3 |
| FIM1_2291 | 3404.39482 | 3392.075471 | 3516.64279 | -0.00073915 | 0.06064844 | 0.974119167 | 0.998607635 | ns | TM | KS.3 |
| FIM1_2292 | 837.1021536 | 947.412733 | 872.7159229 | 0.025028999 | 0.074083811 | 0.322028522 | 0.930003636 | ns | TM | KS.3 |
| FIM1_2293 | 709.6334268 | 723.1110518 | 613.4994378 | 0.002782277 | 0.069168108 | 0.901025937 | 0.991141351 | ns | TM | KS.3 |
| FIM1_2294 | 8652.924477 | 9073.132551 | 9396.152219 | 0.017629461 | 0.04542726 | 0.379734504 | 0.948637087 | ns | TM | KS.3 |
| FIM1_2295 | 873.0731574 | 782.8738156 | 746.0783644 | -0.023767409 | 0.071113095 | 0.345645015 | 0.94690746 | ns | TM | KS.3 |
| FIM1_2297 | 1282.858421 | 1152.770699 | 1216.926502 | -0.043267893 | 0.069971885 | 0.13621451 | 0.768410055 | ns | TM | KS.3 |
| FIM1_2298 | 1348.998357 | 1454.374325 | 1665.205528 | 0.01414912 | 0.067902193 | 0.549386297 | 0.95917717 | ns | TM | KS.3 |
| FIM1_2299 | 1878.965807 | 1677.913539 | 1448.848662 | -0.026199061 | 0.071511599 | 0.308031424 | 0.927416591 | ns | TM | KS.3 |
| FIM1_2300 | 536.8538391 | 540.8100251 | 538.300696 | 0.001176241 | 0.060685779 | 0.958834564 | 0.994413583 | ns | TM | KS.3 |
| FIM1_2301 | 3000.78489 | 2834.600836 | 2796.687825 | -0.009147767 | 0.06948307 | 0.688568188 | 0.980879562 | ns | TM | KS.3 |
| FIM1_2302 | 5558.005044 | 4572.795117 | 3328.172383 | -0.118407074 | 0.102048722 | 0.010773068 | 0.231130071 | ns | TM | KS.3 |
| FIM1_2303 | 204.3888464 | 231.9918802 | 302.1341551 | 0.016969329 | 0.079025433 | 0.455715021 | 0.950410534 | ns | TM | KS.3 |
| FIM1_2304 | 459.1247607 | 448.5760694 | 326.5847066 | -0.004276287 | 0.064044632 | 0.851604347 | 0.991141351 | ns | TM | KS.3 |
| FIM1_2305 | 817.5150762 | 697.0933632 | 762.5268359 | -0.101513532 | 0.090201027 | 0.017322281 | 0.304361209 | ns | TM | KS.3 |
| FIM1_2306 | 285.7652434 | 211.0183672 | 275.4655839 | -0.139181742 | 0.147301835 | 0.006638207 | 0.173816208 | ns | TM | KS.3 |
| FIM1_2307 | 2124.565772 | 2224.358743 | 2506.191909 | 0.012627506 | 0.0541125 | 0.576011124 | 0.963192994 | ns | TM | KS.3 |
| FIM1_2308 | 1668.581638 | 1842.895543 | 1907.014888 | 0.015000006 | 0.074200753 | 0.515962933 | 0.955340251 | ns | TM | KS.3 |
| FIM1_2309 | 269.6955766 | 254.9995481 | 295.1554919 | -0.006267688 | 0.077581254 | 0.76714193 | 0.988477371 | ns | TM | KS.3 |
| FIM1_2310 | 329.5500118 | 456.3436847 | 334.031678 | 0.072148268 | 0.129684753 | 0.038683868 | 0.444462459 | ns | TM | KS.3 |
| FIM1_2311 | 1374.947885 | 1673.461199 | 1672.056037 | 0.021818466 | 0.088493245 | 0.333974647 | 0.938982061 | ns | TM | KS.3 |
| FIM1_2312 | 4363.146846 | 4100.439194 | 4768.238598 | -0.019751537 | 0.055055961 | 0.394925114 | 0.948637087 | ns | TM | KS.3 |
| FIM1_2313 | 927.8792335 | 1174.191936 | 1339.623606 | 0.070346405 | 0.105973916 | 0.046030799 | 0.491423234 | ns | TM | KS.3 |
| FIM1_2314 | 984.7217631 | 1015.465083 | 1073.429431 | 0.008860694 | 0.047679499 | 0.671532027 | 0.976861203 | ns | TM | KS.3 |

| | | | | | | | | | | |
|---|---|---|---|---|---|---|---|---|---|---|
| FIM1_2315 | 536.2962125 | 656.8426526 | 1049.537091 | 0.034067908 | 0.088969639 | 0.191117217 | 0.829253069 | ns | TM | KS.3 |
| FIM1_2316 | 893.1066292 | 860.3128331 | 886.4830386 | -0.004850184 | 0.073188735 | 0.824472233 | 0.991141351 | ns | TM | KS.3 |
| FIM1_2317 | 1602.875638 | 1887.496871 | 2337.312787 | 0.057527377 | 0.086910452 | 0.080193872 | 0.618428374 | ns | TM | KS.3 |
| FIM1_2318 | 9056.827567 | 9630.729403 | 13391.12303 | 0.010189799 | 0.069132838 | 0.657546112 | 0.976323032 | ns | TM | KS.3 |
| FIM1_2319 | 150.6500015 | 157.4167553 | 164.3750103 | 0.005646819 | 0.074032652 | 0.795440556 | 0.988916276 | ns | TM | KS.3 |
| FIM1_2320 | 234.5036735 | 207.1722005 | 214.5370624 | -0.032682749 | 0.073971179 | 0.227385031 | 0.87761096 | ns | TM | KS.3 |
| FIM1_2321 | 1729.487998 | 1630.148697 | 1675.237695 | -0.026438605 | 0.049024449 | 0.219645657 | 0.865538807 | ns | TM | KS.3 |
| FIM1_2322 | 929.0935936 | 987.4662239 | 811.9243081 | 0.016409303 | 0.058187255 | 0.485927545 | 0.952269291 | ns | TM | KS.3 |
| FIM1_2323 | 2433.831806 | 2744.93739 | 3024.517997 | 0.041375489 | 0.07360571 | 0.153697431 | 0.783447458 | ns | TM | KS.3 |
| FIM1_2324 | 3677.906095 | 3902.46658 | 3890.555805 | 0.011621332 | 0.065304047 | 0.61978514 | 0.973122675 | ns | TM | KS.3 |
| FIM1_2325 | 45.11044669 | 41.00954614 | 40.96022622 | -0.012133031 | 0.075924992 | 0.586360589 | 0.963781856 | ns | TM | KS.3 |
| FIM1_2326 | 416.5749309 | 411.2868632 | 442.7879753 | -0.003321246 | 0.054623748 | 0.881586942 | 0.991141351 | ns | TM | KS.3 |
| FIM1_2327 | 2978.084402 | 2915.110711 | 3239.293127 | -0.00630915 | 0.044660308 | 0.75204238 | 0.988477371 | ns | TM | KS.3 |
| FIM1_2328 | 7099.10378 | 7077.198709 | 7241.459513 | -0.000916513 | 0.040498463 | 0.960662411 | 0.995175461 | ns | TM | KS.3 |
| FIM1_2329 | 278.1935868 | 311.0071414 | 371.0631883 | 0.016939735 | 0.075072759 | 0.468192403 | 0.950410534 | ns | TM | KS.3 |
| FIM1_2330 | 365.5270908 | 426.045563 | 471.7337333 | 0.024806473 | 0.079977224 | 0.312150393 | 0.927416591 | ns | TM | KS.3 |
| FIM1_2331 | 2577.538399 | 2756.60073 | 2684.208701 | 0.01106427 | 0.069663299 | 0.631094208 | 0.97344368 | ns | TM | KS.3 |
| FIM1_2332 | 11097.78662 | 10008.202 | 8886.548113 | -0.02057964 | 0.070903521 | 0.402066732 | 0.950410534 | ns | TM | KS.3 |
| FIM1_2333 | 352.8621941 | 393.5570327 | 387.8348301 | 0.021228965 | 0.07144447 | 0.389160453 | 0.948637087 | ns | TM | KS.3 |
| FIM1_2334 | 452.4204187 | 473.2015526 | 624.1776974 | 0.008617595 | 0.064023418 | 0.709708237 | 0.98460638 | ns | TM | KS.3 |
| FIM1_2335 | 5844.760136 | 6450.084857 | 6406.854433 | 0.019537661 | 0.070228104 | 0.423343116 | 0.950410534 | ns | TM | KS.3 |
| FIM1_2336 | 698.7520364 | 1008.898569 | 1689.305228 | 0.079834494 | 0.145839041 | 0.03014519 | 0.400453344 | ns | TM | KS.3 |
| FIM1_2337 | 151.0650896 | 155.7708917 | 216.0073424 | 0.003652698 | 0.07480157 | 0.865055097 | 0.991141351 | ns | TM | KS.3 |
| FIM1_2338 | 1030.506751 | 1378.012818 | 1538.303557 | 0.284295784 | 0.121424989 | 5.9396498590[illegible] | 0.00579407 | ns | TM | KS.3 |
| FIM1_2339 | 602.5844635 | 579.3190792 | 640.8000692 | -0.009493663 | 0.057613211 | 0.678851036 | 0.97986479 | ns | TM | KS.3 |
| FIM1_2340 | 1713.204918 | 1735.153854 | 1242.990799 | 0.001890307 | 0.069338905 | 0.932511923 | 0.994413583 | ns | TM | KS.3 |
| FIM1_2341 | 491.8579388 | 372.4735782 | 716.9177179 | -0.068957128 | 0.115420542 | 0.045139286 | 0.487132212 | ns | TM | KS.3 |
| FIM1_2342 | 1830.075648 | 1806.355042 | 1706.408292 | -0.002943923 | 0.058363935 | 0.896821514 | 0.991141351 | ns | TM | KS.3 |
| FIM1_2343 | 30.04981431 | 30.59025872 | 62.07038615 | 0.001984702 | 0.08567475 | 0.918097144 | 0.991141351 | ns | TM | KS.3 |
| FIM1_2344 | 408.9800112 | 378.9255438 | 771.224789 | -0.009849365 | 0.075240748 | 0.655167369 | 0.976323032 | ns | TM | KS.3 |
| FIM1_2345 | 594.1279437 | 540.2809801 | 917.6130562 | -0.021120785 | 0.068359711 | 0.394176966 | 0.948637087 | ns | TM | KS.3 |
| FIM1_2346 | 4092.306378 | 3714.273389 | 4009.822228 | -0.030567231 | 0.066141794 | 0.246520148 | 0.888934767 | ns | TM | KS.3 |
| FIM1_2347 | 561.4735756 | 662.8822146 | 620.2654514 | 0.035773666 | 0.082570083 | 0.189396611 | 0.828217874 | ns | TM | KS.3 |
| FIM1_2348 | 2662.869744 | 2823.460919 | 2895.085466 | 0.018253556 | 0.054192451 | 0.425747686 | 0.950410534 | ns | TM | KS.3 |
| FIM1_2349 | 2675.918988 | 2180.941085 | 2302.086355 | -0.182166517 | 0.091005016 | 0.001006178 | 0.04634941 | ns | TM | KS.3 |
| FIM1_2350 | 3481.995578 | 3245.275774 | 2886.681087 | -0.02016792 | 0.059330272 | 0.402455099 | 0.950410534 | ns | TM | KS.3 |
| FIM1_2351 | 1091.349523 | 1067.533797 | 1167.699938 | -0.003440072 | 0.068557616 | 0.878341228 | 0.991141351 | ns | TM | KS.3 |
| FIM1_2352 | 934.8046648 | 999.0094605 | 1125.829973 | 0.009120567 | 0.073459889 | 0.681972579 | 0.97986479 | ns | TM | KS.3 |
| FIM1_2353 | 967.5691874 | 945.7720687 | 972.6931714 | -0.005656036 | 0.053711423 | 0.798985603 | 0.988916276 | ns | TM | KS.3 |
| FIM1_2354 | 727.081216 | 714.716918 | 595.0128552 | -0.002859034 | 0.06556085 | 0.899889447 | 0.991141351 | ns | TM | KS.3 |
| FIM1_2355 | 155.0946084 | 140.908704 | 214.1355932 | -0.01452544 | 0.073480688 | 0.529590466 | 0.959071703 | ns | TM | KS.3 |
| FIM1_2356 | 437.5680226 | 366.8969361 | 428.712158 | -0.019813375 | 0.085920117 | 0.376171649 | 0.948637087 | ns | TM | KS.3 |
| FIM1_2357 | 1656.613121 | 1953.978583 | 2252.726222 | 0.028337365 | 0.081895704 | 0.261751103 | 0.899317499 | ns | TM | KS.3 |
| FIM1_2358 | 1218.764105 | 1152.641053 | 1063.905746 | -0.016294655 | 0.055399331 | 0.479800321 | 0.950972051 | ns | TM | KS.3 |
| FIM1_2359 | 4067.829978 | 4248.442224 | 4212.938516 | 0.013587451 | 0.048961167 | 0.524061195 | 0.958883577 | ns | TM | KS.3 |

| | | | | | | | | | | |
|---|---|---|---|---|---|---|---|---|---|---|
| FIM1_2360 | 1738.340655 | 1518.986929 | 2317.011115 | -0.025159632 | 0.076535234 | 0.315413922 | 0.927564309 | ns | TM | KS.3 |
| FIM1_2361 | 3368.423374 | 3294.611195 | 3183.044409 | -0.006797607 | 0.043692854 | 0.729309662 | 0.987157164 | ns | TM | KS.3 |
| FIM1_2362 | 1520.26276 | 1403.332967 | 1599.294081 | -0.029400718 | 0.059872847 | 0.242946202 | 0.887413624 | ns | TM | KS.3 |
| FIM1_2363 | 1909.014622 | 1875.587117 | 2113.157998 | -0.003663571 | 0.06048785 | 0.872547465 | 0.991141351 | ns | TM | KS.3 |
| FIM1_2364 | 2319.384286 | 2385.48999 | 2512.202957 | 0.009376828 | 0.041361757 | 0.616648436 | 0.973122675 | ns | TM | KS.3 |
| FIM1_2365 | 19322.80864 | 20943.42621 | 19621.02589 | 0.021480014 | 0.062987038 | 0.384218362 | 0.948637087 | ns | TM | KS.3 |
| FIM1_2366 | 604.2497896 | 602.9629827 | 568.7596446 | -0.000355049 | 0.059816425 | 0.987549339 | 0.999333464 | ns | TM | KS.3 |
| FIM1_2367 | 422.3469232 | 492.1983956 | 488.8553682 | 0.024604336 | 0.08029552 | 0.314213453 | 0.927564309 | ns | TM | KS.3 |
| FIM1_2368 | 2348.710387 | 2430.660073 | 2404.573716 | 0.005579587 | 0.067949236 | 0.805529283 | 0.988916276 | ns | TM | KS.3 |
| FIM1_2369 | 1408.136178 | 1243.985128 | 1138.461136 | -0.03579199 | 0.073664241 | 0.197742144 | 0.838676184 | ns | TM | KS.3 |
| FIM1_2370 | 364.3329417 | 445.4023161 | 394.0827039 | 0.019578682 | 0.090058617 | 0.368827218 | 0.948637087 | ns | TM | KS.3 |
| FIM1_2371 | 2614.945889 | 2666.693464 | 4375.944249 | 0.001824544 | 0.080106523 | 0.928543424 | 0.993697171 | ns | TM | KS.3 |
| FIM1_2372 | 3719.261131 | 4604.920412 | 4975.428215 | 0.093295232 | 0.106287949 | 0.02318017 | 0.35748181 | ns | TM | KS.3 |
| FIM1_2373 | 955.0516043 | 807.7723802 | 648.7126715 | -0.015055204 | 0.087943665 | 0.471831026 | 0.950410534 | ns | TM | KS.3 |
| FIM1_2374 | 967.4560032 | 938.9451558 | 1107.153194 | -0.008004181 | 0.052740885 | 0.717315462 | 0.987157164 | ns | TM | KS.3 |
| FIM1_2375 | 1676.620362 | 1865.405376 | 1708.348738 | 0.031750282 | 0.069196873 | 0.236600627 | 0.88502866 | ns | TM | KS.3 |
| FIM1_2376 | 1712.219429 | 1622.147039 | 1376.278378 | -0.012893257 | 0.059736896 | 0.581690312 | 0.963781856 | ns | TM | KS.3 |
| FIM1_2377 | 7490.541754 | 7474.761994 | 9042.424857 | -0.000527828 | 0.052081969 | 0.980720682 | 0.999333464 | ns | TM | KS.3 |
| FIM1_2378 | 775.0408281 | 701.9362074 | 689.8052412 | -0.021145706 | 0.069501385 | 0.392954439 | 0.948637087 | ns | TM | KS.3 |
| FIM1_2379 | 1286.095782 | 1274.6429 | 1764.115152 | -0.002354987 | 0.054209227 | 0.915590163 | 0.991141351 | ns | TM | KS.3 |
| FIM1_2380 | 878.6706437 | 816.9335439 | 815.8202954 | -0.013716538 | 0.067622734 | 0.560900349 | 0.961073589 | ns | TM | KS.3 |
| FIM1_2381 | 1643.361015 | 1772.571618 | 1661.261604 | 0.030425662 | 0.057694677 | 0.218622147 | 0.865538807 | ns | TM | KS.3 |
| FIM1_2382 | 502.8505295 | 577.2729192 | 593.4255858 | 0.036373904 | 0.077055274 | 0.191519955 | 0.829253069 | ns | TM | KS.3 |
| FIM1_2383 | 555.0806272 | 637.8072742 | 893.42857 | 0.048429045 | 0.07999214 | 0.115251359 | 0.717000304 | ns | TM | KS.3 |
| FIM1_2384 | 10012.90697 | 10643.09177 | 9984.682313 | 0.018384031 | 0.055740129 | 0.429642814 | 0.950410534 | ns | TM | KS.3 |
| FIM1_2385 | 968.7372248 | 1040.135032 | 919.9138713 | 0.028007719 | 0.056107926 | 0.244736947 | 0.88883001 | ns | TM | KS.3 |
| FIM1_2386 | 782.345492 | 875.7138068 | 686.0131183 | 0.01434293 | 0.078022974 | 0.52112664 | 0.958602567 | ns | TM | KS.3 |
| FIM1_2387 | 529.0380907 | 458.6725416 | 480.6728324 | -0.066641522 | 0.084613086 | 0.057403836 | 0.536774409 | ns | TM | KS.3 |
| FIM1_2388 | 6569.832523 | 6465.595601 | 6074.465123 | -0.004810123 | 0.043133894 | 0.804923527 | 0.988916276 | ns | TM | KS.3 |
| FIM1_2389 | 1568.885864 | 1729.857518 | 1980.459844 | 0.020618807 | 0.069303095 | 0.403111919 | 0.950410534 | ns | TM | KS.3 |
| FIM1_2390 | 1008.266531 | 959.8598279 | 1057.863545 | -0.016137616 | 0.049278211 | 0.452111074 | 0.950410534 | ns | TM | KS.3 |
| FIM1_2391 | 1026.630419 | 1058.731208 | 1136.747479 | 0.0055895 | 0.064244161 | 0.807341992 | 0.988916276 | ns | TM | KS.3 |
| FIM1_2392 | 1201.844289 | 1202.470659 | 1043.287542 | 9.69807724207602 | 0.065678223 | 0.996586878 | 0.999525119 | ns | TM | KS.3 |
| FIM1_2393 | 538.2366523 | 565.5661643 | 366.9729904 | 0.006951051 | 0.071875676 | 0.754890168 | 0.988477371 | ns | TM | KS.3 |
| FIM1_2394 | 857.9270182 | 779.3907815 | 1032.077971 | -0.017187268 | 0.07121426 | 0.471565131 | 0.950410534 | ns | TM | KS.3 |
| FIM1_2395 | 16943.21697 | 17348.56939 | 17822.67553 | 0.006800376 | 0.047639483 | 0.744509032 | 0.988477371 | ns | TM | KS.3 |
| FIM1_2396 | 2392.503118 | 2335.73905 | 2557.179755 | -0.007255115 | 0.044412022 | 0.715161263 | 0.986942381 | ns | TM | KS.3 |
| FIM1_2397 | 2939.909294 | 2869.163574 | 3026.433657 | -0.006658581 | 0.049549612 | 0.75532595 | 0.988477371 | ns | TM | KS.3 |
| FIM1_2398 | 3155.722603 | 3075.102042 | 2011.486547 | -0.005703527 | 0.058872542 | 0.802748718 | 0.988916276 | ns | TM | KS.3 |
| FIM1_2399 | 1293.343012 | 1363.288514 | 828.3418785 | 0.011524412 | 0.061562712 | 0.622257428 | 0.973122675 | ns | TM | KS.3 |
| FIM1_2400 | 636.3245111 | 528.8701082 | 510.3101681 | -0.051548155 | 0.089565508 | 0.097625704 | 0.677062609 | ns | TM | KS.3 |
| FIM1_2401 | 748.7784244 | 720.5616175 | 714.9424682 | -0.008976308 | 0.059361019 | 0.697034869 | 0.98151979 | ns | TM | KS.3 |
| FIM1_2402 | 1737.494054 | 1752.422972 | 1613.534404 | 0.001996533 | 0.05441336 | 0.928484678 | 0.993697171 | ns | TM | KS.3 |
| FIM1_2403 | 2077.551244 | 1944.870715 | 2146.319139 | -0.019445803 | 0.057529021 | 0.41224006 | 0.950410534 | ns | TM | KS.3 |
| FIM1_2404 | 4736.33613 | 4572.95847 | 5503.319899 | -0.008698827 | 0.055638113 | 0.700713387 | 0.98151979 | ns | TM | KS.3 |

| | | | | | | | | | | |
|---|---|---|---|---|---|---|---|---|---|---|
| FIM1_2405 | 1855.029278 | 1829.871073 | 1827.205237 | -0.003628764 | 0.050150095 | 0.86575706 | 0.991141351 | ns | TM | KS.3 |
| FIM1_2406 | 6562.422439 | 6530.899158 | 8854.810501 | -0.000472406 | 0.078743429 | 0.981699191 | 0.999333464 | ns | TM | KS.3 |
| FIM1_2407 | 1065.072386 | 1264.40203 | 1434.130017 | 0.03394873 | 0.083304594 | 0.202945583 | 0.847612283 | ns | TM | KS.3 |
| FIM1_2408 | 316.0913762 | 291.1919083 | 317.70967 | -0.012856888 | 0.071846712 | 0.576867783 | 0.963382753 | ns | TM | KS.3 |
| FIM1_2409 | 601.2186966 | 589.791152 | 558.8150676 | -0.00402654 | 0.060053135 | 0.860017779 | 0.991141351 | ns | TM | KS.3 |
| FIM1_2410 | 6561.22452 | 5712.058494 | 5763.511942 | -0.06018828 | 0.08163602 | 0.072472937 | 0.593991535 | ns | TM | KS.3 |
| FIM1_2411 | 608.4467552 | 649.0682635 | 686.7980707 | 0.015286409 | 0.061476402 | 0.520388723 | 0.958509403 | ns | TM | KS.3 |
| FIM1_2412 | 635.9585942 | 624.5591753 | 600.7828512 | -0.002997895 | 0.066458896 | 0.894771522 | 0.991141351 | ns | TM | KS.3 |
| FIM1_2413 | 2031.890245 | 2191.329292 | 2181.1896 | 0.014213626 | 0.067939836 | 0.547689457 | 0.95917717 | ns | TM | KS.3 |
| FIM1_2414 | 1412.329311 | 1393.905154 | 1461.678195 | -0.003296421 | 0.053892454 | 0.88187497 | 0.991141351 | ns | TM | KS.3 |
| FIM1_2415 | 800.7237822 | 831.8435536 | 948.249549 | 0.009229291 | 0.056226693 | 0.68488334 | 0.980234354 | ns | TM | KS.3 |
| FIM1_2416 | 4338.586767 | 4340.000447 | 4098.833739 | 8.5819268444839 | 0.04121546 | 0.996365805 | 0.999525119 | ns | TM | KS.3 |
| FIM1_2417 | 1195.836169 | 1162.328247 | 1140.405553 | -0.007487978 | 0.051603215 | 0.731927271 | 0.987157164 | ns | TM | KS.3 |
| FIM1_2418 | 14686.319 | 13967.89368 | 15167.9165 | -0.019316823 | 0.045902549 | 0.339633999 | 0.943910292 | ns | TM | KS.3 |
| FIM1_2419 | 650.0697512 | 541.0860499 | 811.3745993 | -0.040418521 | 0.086571655 | 0.151600451 | 0.783447458 | ns | TM | KS.3 |
| FIM1_2420 | 1680.797134 | 1491.665378 | 1741.522806 | -0.033582909 | 0.072412807 | 0.21855556 | 0.865538807 | ns | TM | KS.3 |
| FIM1_2422 | 4336.354862 | 4030.666458 | 4611.739681 | -0.03332784 | 0.056016782 | 0.173604897 | 0.811391014 | ns | TM | KS.3 |
| FIM1_2423 | 455.4557447 | 570.4530112 | 573.2406725 | 0.040116934 | 0.093916729 | 0.142723421 | 0.777709768 | ns | TM | KS.3 |
| FIM1_2424 | 1088.679775 | 1113.151122 | 1761.714842 | 0.004093815 | 0.064220149 | 0.857777708 | 0.991141351 | ns | TM | KS.3 |
| FIM1_2425 | 535.0071289 | 658.7199047 | 549.0309316 | 0.037795682 | 0.090241182 | 0.162108293 | 0.796138951 | ns | TM | KS.3 |
| FIM1_2426 | 2941.577383 | 2896.114368 | 3547.211572 | -0.003806293 | 0.054151441 | 0.864078729 | 0.991141351 | ns | TM | KS.3 |
| FIM1_2427 | 299.5897958 | 244.7198292 | 261.1951881 | -0.048219023 | 0.092061101 | 0.107453747 | 0.702423202 | ns | TM | KS.3 |
| FIM1_2428 | 372.6374159 | 340.9682713 | 668.4622509 | -0.009763531 | 0.078967061 | 0.647603396 | 0.976323032 | ns | TM | KS.3 |
| FIM1_2429 | 1220.839026 | 1011.136013 | 1186.307702 | -0.171528224 | 0.086040074 | 0.001300646 | 0.057774225 | ns | TM | KS.3 |
| FIM1_2430 | 1764.678437 | 1587.597936 | 1665.928015 | -0.036776392 | 0.069031101 | 0.185121743 | 0.825251496 | ns | TM | KS.3 |
| FIM1_2431 | 2734.031508 | 2552.447945 | 1976.13754 | -0.01502508 | 0.064150687 | 0.528835401 | 0.959071703 | ns | TM | KS.3 |
| FIM1_2432 | 1416.843447 | 1119.857005 | 1147.157163 | -0.086823659 | 0.110641272 | 0.027439722 | 0.38781993 | ns | TM | KS.3 |
| FIM1_2433 | 33988.8552 | 29942.08977 | 28152.16964 | -0.013719839 | 0.081680209 | 0.525338546 | 0.959071703 | ns | TM | KS.3 |
| FIM1_2434 | 5494.651005 | 5475.645778 | 5279.855016 | -0.000442181 | 0.072837329 | 0.983781941 | 0.999333464 | ns | TM | KS.3 |
| FIM1_2435 | 1438.722346 | 1609.574998 | 1608.339509 | 0.040142786 | 0.071360473 | 0.160477079 | 0.792037169 | ns | TM | KS.3 |
| FIM1_2436 | 287.7089338 | 275.2114984 | 317.0982153 | -0.006997965 | 0.069535958 | 0.75679828 | 0.988477371 | ns | TM | KS.3 |
| FIM1_2437 | 107.1759432 | 91.05078089 | 185.0976495 | -0.016263941 | 0.086541416 | 0.447448001 | 0.950410534 | ns | TM | KS.3 |
| FIM1_2438 | 1217.067977 | 1197.84603 | 1531.349559 | -0.003825479 | 0.056351905 | 0.865192124 | 0.991141351 | ns | TM | KS.3 |
| FIM1_2439 | 308.0372106 | 299.5934818 | 443.2228972 | -0.002928911 | 0.077851881 | 0.88828603 | 0.991141351 | ns | TM | KS.3 |
| FIM1_2440 | 131.4307453 | 180.0874682 | 226.0552197 | 0.018692242 | 0.108567454 | 0.349733123 | 0.948637087 | ns | TM | KS.3 |
| FIM1_2444 | 23251.25233 | 20074.232 | 20677.52651 | -0.012066683 | 0.087828852 | 0.551878843 | 0.95917717 | ns | TM | KS.3 |
| FIM1_2445 | 812.2153863 | 719.4230052 | 805.4567149 | -0.012061213 | 0.082664471 | 0.567563308 | 0.962265923 | ns | TM | KS.3 |
| FIM1_2446 | 295.2118866 | 238.1687465 | 299.0757199 | -0.073303786 | 0.102425609 | 0.043165916 | 0.476165039 | ns | TM | KS.3 |
| FIM1_2447 | 519.7253848 | 531.6572225 | 473.4177694 | 0.004646104 | 0.056302159 | 0.836770554 | 0.991141351 | ns | TM | KS.3 |
| FIM1_2448 | 58.82484468 | 41.45608583 | 42.2117417 | -0.020845766 | 0.113063491 | 0.305754323 | 0.926747341 | ns | TM | KS.3 |
| FIM1_2449 | 35.53905049 | 37.07310565 | 43.40792016 | 0.002151571 | 0.092321208 | 0.90653128 | 0.991141351 | ns | TM | KS.3 |
| FIM1_2450 | 360.9775241 | 387.3688836 | 632.294383 | 0.010109862 | 0.073064561 | 0.652532015 | 0.976323032 | ns | TM | KS.3 |
| FIM1_2451 | 628.6568185 | 554.9201879 | 931.4556968 | -0.005351312 | 0.102781296 | 0.76035215 | 0.988477371 | ns | TM | KS.3 |
| FIM1_2452 | 37.9165985 | 40.75674446 | 52.95915738 | 0.004844069 | 0.08319591 | 0.808026668 | 0.988916276 | ns | TM | KS.3 |
| FIM1_2453 | 206.082047 | 190.2592461 | 209.9167577 | -0.014219083 | 0.069165692 | 0.545992366 | 0.95917717 | ns | TM | KS.3 |

| | | | | | | | | | | |
|---|---|---|---|---|---|---|---|---|---|---|
| FIM1_2454 | 289.6407725 | 242.4931215 | 378.7276646 | -0.025148261 | 0.083995776 | 0.296548451 | 0.926536521 | ns | TM | KS.3 |
| FIM1_2455 | 865.4166171 | 777.5877233 | 1087.383669 | -0.00700408 | 0.090447716 | 0.712827259 | 0.986183429 | ns | TM | KS.3 |
| FIM1_2456 | 325.2279706 | 276.1248987 | 256.7192958 | -0.022217308 | 0.082773941 | 0.343801686 | 0.944808611 | ns | TM | KS.3 |
| FIM1_2457 | 323.8202729 | 349.3082118 | 402.8146455 | 0.013913128 | 0.069208089 | 0.553800556 | 0.95917717 | ns | TM | KS.3 |
| FIM1_2458 | 1846.041258 | 1610.178246 | 2445.15704 | -0.025412222 | 0.076893515 | 0.311017524 | 0.927416591 | ns | TM | KS.3 |
| FIM1_2459 | 1529.570029 | 1771.665199 | 2120.245717 | 0.019265659 | 0.08103451 | 0.401700132 | 0.950410534 | ns | TM | KS.3 |
| FIM1_2460 | 2996.776307 | 3462.038121 | 4269.400428 | 0.014239264 | 0.084533929 | 0.502686465 | 0.953260948 | ns | TM | KS.3 |
| FIM1_2461 | 2474.036649 | 2656.666098 | 2918.362486 | 0.026803398 | 0.056226266 | 0.264447957 | 0.90372202 | ns | TM | KS.3 |
| FIM1_2462 | 5169.766906 | 4733.727103 | 5382.854023 | -0.037192431 | 0.063060886 | 0.165969991 | 0.803992898 | ns | TM | KS.3 |
| FIM1_2463 | 845.3646349 | 783.8303014 | 944.635553 | -0.027724153 | 0.058096717 | 0.259531133 | 0.895871714 | ns | TM | KS.3 |
| FIM1_2464 | 1940.846971 | 2016.942532 | 1939.7883 | 0.007510508 | 0.063440836 | 0.74476871 | 0.988477371 | ns | TM | KS.3 |
| FIM1_2465 | 736.5856093 | 816.3046509 | 596.242639 | 0.031466764 | 0.067679208 | 0.238124479 | 0.88653074 | ns | TM | KS.3 |
| FIM1_2466 | 822.1805341 | 773.052114 | 842.4749687 | -0.006537458 | 0.078239444 | 0.756313411 | 0.988477371 | ns | TM | KS.3 |
| FIM1_2467 | 381.7210305 | 324.3026317 | 403.5880091 | -0.047256598 | 0.083961789 | 0.11909522 | 0.729678224 | ns | TM | KS.3 |
| FIM1_2468 | 2382.569947 | 2484.027737 | 2249.658896 | 0.010369154 | 0.056299344 | 0.649539891 | 0.976323032 | ns | TM | KS.3 |
| FIM1_2469 | 1926.026342 | 1984.344283 | 1894.960468 | 0.010057232 | 0.038558489 | 0.566203508 | 0.962265923 | ns | TM | KS.3 |
| FIM1_2470 | 7269.065007 | 7409.249219 | 6846.182636 | 0.005641135 | 0.04559536 | 0.78073332 | 0.988814136 | ns | TM | KS.3 |
| FIM1_2471 | 1353.446394 | 1384.078915 | 1109.614518 | 0.00442778 | 0.062041575 | 0.846654386 | 0.991141351 | ns | TM | KS.3 |
| FIM1_2472 | 2160.237374 | 1830.268887 | 2006.773113 | -0.088055704 | 0.092584354 | 0.027788089 | 0.388845019 | ns | TM | KS.3 |
| FIM1_2473 | 1119.317539 | 1368.13853 | 1554.022244 | 0.037386959 | 0.0888887 | 0.167009586 | 0.80502665 | ns | TM | KS.3 |
| FIM1_2474 | 2647.446863 | 2862.840163 | 3438.768156 | 0.030096336 | 0.059035399 | 0.229636185 | 0.879350777 | ns | TM | KS.3 |
| FIM1_2475 | 1592.216721 | 1450.430486 | 1268.99965 | -0.032194904 | 0.064720695 | 0.223565366 | 0.87302802 | ns | TM | KS.3 |
| FIM1_2476 | 10500.72136 | 10172.46728 | 9869.721191 | -0.009318114 | 0.048082605 | 0.657485003 | 0.976323032 | ns | TM | KS.3 |
| FIM1_2477 | 2069.347514 | 1955.990213 | 2092.768011 | -0.015279172 | 0.056984432 | 0.511437447 | 0.953818112 | ns | TM | KS.3 |
| FIM1_2478 | 1325.604414 | 1248.091823 | 1118.601481 | -0.014645851 | 0.060484833 | 0.535535668 | 0.959071703 | ns | TM | KS.3 |
| FIM1_2479 | 901.680803 | 809.8476851 | 825.2483291 | -0.037236095 | 0.069480309 | 0.181702957 | 0.818789693 | ns | TM | KS.3 |
| FIM1_2480 | 2153.300619 | 1871.045651 | 2053.053403 | -0.124242334 | 0.076720136 | 0.005836913 | 0.159552978 | ns | TM | KS.3 |
| FIM1_2481 | 1368.898332 | 1192.353648 | 1282.433671 | -0.078275012 | 0.083524039 | 0.036871452 | 0.436751124 | ns | TM | KS.3 |
| FIM1_2482 | 1461.871071 | 1424.615489 | 1389.641121 | -0.008509531 | 0.038768862 | 0.630968325 | 0.97344368 | ns | TM | KS.3 |
| FIM1_2483 | 1224.69691 | 1158.600272 | 1291.08699 | -0.017459022 | 0.052080631 | 0.434324731 | 0.950410534 | ns | TM | KS.3 |
| FIM1_2484 | 258.6631802 | 331.3502755 | 351.8027649 | 0.101388614 | 0.117369205 | 0.017662423 | 0.304361209 | ns | TM | KS.3 |
| FIM1_2485 | 1442.83745 | 1339.833293 | 1569.017362 | -0.020146516 | 0.061227709 | 0.40777462 | 0.950410534 | ns | TM | KS.3 |
| FIM1_2486 | 12681.29705 | 12664.00333 | 11891.10262 | -0.000273739 | 0.059871875 | 0.990401432 | 0.999525119 | ns | TM | KS.3 |
| FIM1_2487 | 2307.61265 | 2365.005644 | 2468.994696 | 0.007555251 | 0.042049134 | 0.691626313 | 0.981188557 | ns | TM | KS.3 |
| FIM1_2488 | 354.5247338 | 376.7938094 | 344.4512517 | 0.009673766 | 0.069689173 | 0.672192047 | 0.976861203 | ns | TM | KS.3 |
| FIM1_2489 | 818.1537398 | 786.3128386 | 585.8551589 | -0.007342101 | 0.065112585 | 0.749637781 | 0.988477371 | ns | TM | KS.3 |
| FIM1_2490 | 1104.472502 | 1282.138532 | 1138.043493 | 0.038108004 | 0.079572716 | 0.175852069 | 0.811391014 | ns | TM | KS.3 |
| FIM1_2491 | 308.8862978 | 100.1514319 | 811.3610947 | -1.05073085 | 0.572869544 | 7.27588376223 | 0.006961062 | Km<KS.3 | TM | KS.3 |
| FIM1_2492 | 2197.973433 | 2263.617395 | 2656.474913 | 0.00407535 | 0.071569237 | 0.853614889 | 0.991141351 | ns | TM | KS.3 |
| FIM1_2493 | 1870.384355 | 1646.234024 | 1708.561039 | -0.103839038 | 0.0769015 | 0.012505103 | 0.25497086 | ns | TM | KS.3 |
| FIM1_2494 | 824.2925535 | 874.0561739 | 863.5211927 | 0.014905067 | 0.059712444 | 0.527548917 | 0.959071703 | ns | TM | KS.3 |
| FIM1_2495 | 6434.307348 | 5492.117065 | 5237.240303 | -0.03341628 | 0.080736756 | 0.212243231 | 0.858463475 | ns | TM | KS.3 |
| FIM1_2496 | 1267.663489 | 1125.979109 | 1081.57116 | -0.031707555 | 0.072282912 | 0.238249568 | 0.88653074 | ns | TM | KS.3 |
| FIM1_2497 | 541.131581 | 448.6612543 | 585.9993221 | -0.041498968 | 0.087642837 | 0.143909747 | 0.77783242 | ns | TM | KS.3 |
| FIM1_2498 | 131.4132983 | 112.1738303 | 134.5474941 | -0.019264658 | 0.083491569 | 0.39397737 | 0.948637087 | ns | TM | KS.3 |

| | | | | | | | | | | |
|---|---|---|---|---|---|---|---|---|---|---|
| FIM1_2499 | 1202.230723 | 1078.054227 | 1237.132266 | -0.041198177 | 0.070317792 | 0.151178513 | 0.783447458 | ns | TM | KS.3 |
| FIM1_2500 | 1105.925041 | 781.1878294 | 1509.180097 | -0.084290045 | 0.142430025 | 0.026670727 | 0.382098641 | ns | TM | KS.3 |
| FIM1_2502 | 4377.891826 | 3452.456321 | 3124.580927 | -0.079724439 | 0.109357034 | 0.033988537 | 0.42621823 | ns | TM | KS.3 |
| FIM1_2503 | 637.4857102 | 649.4660082 | 574.3758891 | 0.002619522 | 0.071345989 | 0.90544646 | 0.991141351 | ns | TM | KS.3 |
| FIM1_2504 | 1105.003763 | 861.1201155 | 853.0960671 | -0.033748606 | 0.096711286 | 0.180510122 | 0.814177566 | ns | TM | KS.3 |
| FIM1_2505 | 827.5784647 | 668.7274871 | 596.8034414 | -0.049883057 | 0.094613428 | 0.098828866 | 0.677856366 | ns | TM | KS.3 |
| FIM1_2506 | 473.4414469 | 390.7644966 | 537.1545035 | -0.03640777 | 0.087213764 | 0.176645234 | 0.811391014 | ns | TM | KS.3 |
| FIM1_2507 | 874.3976944 | 870.615706 | 1007.272581 | -0.001152851 | 0.054755184 | 0.958745248 | 0.994413583 | ns | TM | KS.3 |
| FIM1_2508 | 15.972828 | 19.01400634 | 19.23299098 | 0.008805164 | 0.099034101 | 0.631722781 | 0.97344368 | ns | TM | KS.3 |
| FIM1_2510 | 272.1651708 | 314.0161068 | 247.8209449 | 0.023709891 | 0.078536173 | 0.332200247 | 0.938982061 | ns | TM | KS.3 |
| FIM1_2511 | 4055.369396 | 4393.577815 | 5768.840307 | 0.013348232 | 0.070659332 | 0.566142212 | 0.962265923 | ns | TM | KS.3 |
| FIM1_2512 | 1591.070861 | 1320.554683 | 1212.02316 | -0.07693909 | 0.096356833 | 0.039943526 | 0.454734648 | ns | TM | KS.3 |
| FIM1_2513 | 6956.909412 | 6362.708553 | 6384.272418 | -0.015539922 | 0.070870661 | 0.510394147 | 0.953260948 | ns | TM | KS.3 |
| FIM1_2514 | 401.4238974 | 414.0091178 | 481.8060497 | 0.006816561 | 0.060085577 | 0.766585387 | 0.988477371 | ns | TM | KS.3 |
| FIM1_2515 | 66.48774473 | 73.98519732 | 94.90197683 | 0.015335649 | 0.076389098 | 0.501634807 | 0.953260948 | ns | TM | KS.3 |
| FIM1_2516 | 4299.150784 | 3599.499158 | 4106.495846 | -0.096219524 | 0.096332312 | 0.021447758 | 0.338445603 | ns | TM | KS.3 |
| FIM1_2517 | 1633.871536 | 1335.178639 | 2074.916218 | -0.051852465 | 0.09302663 | 0.09364723 | 0.671406965 | ns | TM | KS.3 |
| FIM1_2518 | 1989.299863 | 2048.839645 | 2038.925988 | 0.009212568 | 0.042436154 | 0.630431739 | 0.97344368 | ns | TM | KS.3 |
| FIM1_2519 | 2083.832416 | 2391.305761 | 2929.214204 | 0.009268927 | 0.09128905 | 0.630649394 | 0.97344368 | ns | TM | KS.3 |
| FIM1_2520 | 560.1004889 | 656.3604993 | 689.008901 | 0.034477518 | 0.080803136 | 0.202845225 | 0.847612283 | ns | TM | KS.3 |
| FIM1_2521 | 683.7545522 | 801.8065904 | 697.9728368 | 0.031676243 | 0.08094791 | 0.228190868 | 0.877996572 | ns | TM | KS.3 |
| FIM1_2522 | 478.5696382 | 460.1498628 | 490.9830642 | -0.008233216 | 0.061949045 | 0.72177285 | 0.987157164 | ns | TM | KS.3 |
| FIM1_2523 | 4512.6607 | 4278.504184 | 4516.161231 | -0.024469627 | 0.045045485 | 0.217066167 | 0.865538807 | ns | TM | KS.3 |
| FIM1_2524 | 1359.801023 | 1476.698216 | 1763.71944 | 0.008983598 | 0.078701097 | 0.673078769 | 0.976861203 | ns | TM | KS.3 |
| FIM1_2525 | 700.7613785 | 861.7641672 | 706.1662017 | 0.063892488 | 0.097635934 | 0.06015703 | 0.544665909 | ns | TM | KS.3 |
| FIM1_2526 | 9625.345352 | 7318.26545 | 5219.96843 | -0.136991305 | 0.132933937 | 0.006756998 | 0.175083682 | ns | TM | KS.3 |
| FIM1_2527 | 316.0191627 | 431.8224615 | 199.4878147 | 0.028359064 | 0.106602093 | 0.215505638 | 0.865538807 | ns | TM | KS.3 |
| FIM1_2528 | 22166.95556 | 22277.89791 | 23970.31229 | 0.00143053 | 0.046026117 | 0.9440714 | 0.994413583 | ns | TM | KS.3 |
| FIM1_2529 | 6828.318797 | 6987.518258 | 8482.783516 | 0.001940202 | 0.08243737 | 0.922290285 | 0.992085225 | ns | TM | KS.3 |
| FIM1_2530 | 947.1248223 | 964.2126253 | 1352.405008 | 0.001505142 | 0.082740327 | 0.93945658 | 0.994413583 | ns | TM | KS.3 |
| FIM1_2531 | 2437.626255 | 2786.277143 | 3138.28937 | 0.025976523 | 0.076164459 | 0.304611269 | 0.926536521 | ns | TM | KS.3 |
| FIM1_2532 | 520.0631284 | 775.1892291 | 477.4910332 | 0.060613012 | 0.140961391 | 0.054298058 | 0.531428718 | ns | TM | KS.3 |
| FIM1_2533 | 3207.459093 | 3081.424262 | 2574.912595 | -0.009443906 | 0.058136691 | 0.681121741 | 0.97986479 | ns | TM | KS.3 |
| FIM1_2534 | 2054.850529 | 1833.565004 | 2106.468145 | -0.024740044 | 0.071906936 | 0.32940869 | 0.936229083 | ns | TM | KS.3 |
| FIM1_2535 | 428.6178257 | 415.1872643 | 461.4695118 | -0.005764038 | 0.065613641 | 0.800984481 | 0.988916276 | ns | TM | KS.3 |
| FIM1_2536 | 484.8118698 | 343.2639867 | 460.6005252 | -0.037735024 | 0.114994445 | 0.136128988 | 0.768410055 | ns | TM | KS.3 |
| FIM1_2537 | 760.0309104 | 600.5833214 | 1077.293257 | -0.028669724 | 0.09384139 | 0.231254877 | 0.880810476 | ns | TM | KS.3 |
| FIM1_2538 | 733.0849223 | 789.3839646 | 815.3711556 | 0.011604105 | 0.07115227 | 0.61296967 | 0.973122675 | ns | TM | KS.3 |
| FIM1_2539 | 615.0677727 | 598.6227594 | 615.8128797 | -0.004338326 | 0.068174843 | 0.847517454 | 0.991141351 | ns | TM | KS.3 |
| FIM1_2540 | 3379.59475 | 3037.302615 | 3341.158667 | -0.018727352 | 0.072897082 | 0.435199663 | 0.950410534 | ns | TM | KS.3 |
| FIM1_2541 | 1287.509156 | 1217.249879 | 1315.902532 | -0.012506663 | 0.061796896 | 0.594587132 | 0.965889818 | ns | TM | KS.3 |
| FIM1_2542 | 428.0505505 | 447.980655 | 401.8720866 | 0.00773502 | 0.067027139 | 0.735754899 | 0.987655486 | ns | TM | KS.3 |
| FIM1_2543 | 1100.808522 | 1077.788231 | 1152.744984 | -0.004038612 | 0.062127262 | 0.859891154 | 0.991141351 | ns | TM | KS.3 |
| FIM1_2544 | 720.3585449 | 873.4895993 | 859.9406074 | 0.105164518 | 0.101687184 | 0.016284199 | 0.29567114 | ns | TM | KS.3 |
| FIM1_2546 | 1419.687057 | 1410.680447 | 1183.701345 | -0.001584611 | 0.055029395 | 0.943422665 | 0.994413583 | ns | TM | KS.3 |

| | | | | | | | | | | |
|---|---|---|---|---|---|---|---|---|---|---|
| FIM1_2547 | 1965.281178 | 1607.015974 | 2021.126718 | -0.023594521 | 0.088744167 | 0.305873193 | 0.926747341 | ns | TM | KS.3 |
| FIM1_2548 | 3472.606353 | 3656.902845 | 5162.908892 | 0.004315082 | 0.083120579 | 0.828369697 | 0.991141351 | ns | TM | KS.3 |
| FIM1_2549 | 5539.509743 | 5149.670941 | 5865.03692 | -0.009481765 | 0.074778941 | 0.667652084 | 0.976323032 | ns | TM | KS.3 |
| FIM1_2550 | 1987.625529 | 1802.046747 | 1720.231843 | -0.017538705 | 0.07157083 | 0.463130132 | 0.950410534 | ns | TM | KS.3 |
| FIM1_2552 | 1515.745339 | 1140.557618 | 2587.608678 | -0.028788706 | 0.102052771 | 0.216777142 | 0.865538807 | ns | TM | KS.3 |
| FIM1_2553 | 545.1591557 | 465.240833 | 530.5381407 | -0.018984582 | 0.083279178 | 0.400085587 | 0.950410534 | ns | TM | KS.3 |
| FIM1_2554 | 1457.263189 | 1416.588534 | 1249.548733 | -0.002957987 | 0.077715723 | 0.887340422 | 0.991141351 | ns | TM | KS.3 |
| FIM1_2555 | 4548.104625 | 1147.631679 | 14232.97528 | -1.562557881 | 0.384518743 | 7.66248282742 | 0.000131451 | Km<KS.3 | TM | KS.3 |
| FIM1_2556 | 123.6165006 | 82.93460329 | 413.0204317 | -0.032675686 | 0.122423091 | 0.164919645 | 0.801223467 | ns | TM | KS.3 |
| FIM1_2557 | 8982.447394 | 9593.306401 | 12207.00935 | 0.009664148 | 0.071923848 | 0.668472866 | 0.976323032 | ns | TM | KS.3 |
| FIM1_2558 | 1062.061102 | 1187.962964 | 1227.149074 | 0.013632835 | 0.078618783 | 0.537528415 | 0.95917717 | ns | TM | KS.3 |
| FIM1_2559 | 410.8274447 | 364.2177629 | 458.2771054 | -0.010789644 | 0.084552444 | 0.599367461 | 0.969393081 | ns | TM | KS.3 |
| FIM1_2560 | 3856.349791 | 3212.00806 | 3738.299824 | -0.06196692 | 0.092107877 | 0.066806428 | 0.566847784 | ns | TM | KS.3 |
| FIM1_2561 | 392.665151 | 283.8374923 | 346.8324722 | -0.063985165 | 0.124637199 | 0.050546009 | 0.513412402 | ns | TM | KS.3 |
| FIM1_2562 | 2800.808187 | 2437.065791 | 2830.694731 | -0.044973062 | 0.078675799 | 0.132830503 | 0.761677364 | ns | TM | KS.3 |
| FIM1_2563 | 2369.854744 | 2508.931235 | 3001.103454 | 0.019626896 | 0.051763007 | 0.379219739 | 0.948637087 | ns | TM | KS.3 |
| FIM1_2564 | 1083.361637 | 1034.226589 | 1739.693656 | -0.009926897 | 0.061193172 | 0.669186715 | 0.976323032 | ns | TM | KS.3 |
| FIM1_2565 | 1205.130744 | 1137.244353 | 1138.975453 | -0.013486304 | 0.060852736 | 0.566826776 | 0.962265923 | ns | TM | KS.3 |
| FIM1_2566 | 2060.210902 | 2026.884881 | 2318.819173 | -0.00288129 | 0.065028226 | 0.899260108 | 0.991141351 | ns | TM | KS.3 |
| FIM1_2567 | 837.4011137 | 859.5364041 | 1056.417571 | 0.006526322 | 0.055594121 | 0.772046031 | 0.988814136 | ns | TM | KS.3 |
| FIM1_2568 | 1230.184376 | 1219.556702 | 1217.730461 | -0.001091352 | 0.07386223 | 0.959655159 | 0.994849847 | ns | TM | KS.3 |
| FIM1_2570 | 342.5948817 | 337.2409305 | 388.1636513 | -0.001779205 | 0.074280216 | 0.934081944 | 0.994413583 | ns | TM | KS.3 |
| FIM1_2571 | 328.4807968 | 349.4708737 | 421.0415144 | 0.006538108 | 0.078642478 | 0.75540343 | 0.988477371 | ns | TM | KS.3 |
| FIM1_2572 | 212.0796606 | 241.7198824 | 287.6389418 | 0.020791859 | 0.077227011 | 0.384498289 | 0.948637087 | ns | TM | KS.3 |
| FIM1_2573 | 504.91869 | 548.1207392 | 484.8169296 | 0.01682325 | 0.066822957 | 0.485034209 | 0.952269291 | ns | TM | KS.3 |
| FIM1_2574 | 1477.816144 | 1732.444465 | 1794.752805 | 0.059014653 | 0.086256596 | 0.076139665 | 0.607043 | ns | TM | KS.3 |
| FIM1_2575 | 840.5584237 | 929.8289894 | 933.2852499 | 0.030252723 | 0.067627315 | 0.252775969 | 0.889363822 | ns | TM | KS.3 |
| FIM1_2576 | 13682.14706 | 13732.54161 | 14153.82679 | 0.001206793 | 0.025636694 | 0.920085867 | 0.991141351 | ns | TM | KS.3 |
| FIM1_2577 | 3576.786669 | 4154.855576 | 5459.753975 | 0.035784666 | 0.079355395 | 0.194030859 | 0.836484854 | ns | TM | KS.3 |
| FIM1_2578 | 45.9272675 | 56.70667086 | 73.6616089 | 0.027676404 | 0.089193808 | 0.251873812 | 0.888934767 | ns | TM | KS.3 |
| FIM1_2579 | 122.4639438 | 148.8019751 | 269.9166729 | 0.042388799 | 0.090270134 | 0.135563617 | 0.768410055 | ns | TM | KS.3 |
| FIM1_2580 | 564.8747565 | 592.5517514 | 1231.593787 | 0.006290601 | 0.073350818 | 0.774522491 | 0.988814136 | ns | TM | KS.3 |
| FIM1_2581 | 18919.54953 | 18057.40618 | 20957.63236 | -0.011219253 | 0.058404792 | 0.627719652 | 0.97344368 | ns | TM | KS.3 |
| FIM1_2582 | 585.6468383 | 712.5603817 | 655.2949581 | 0.044963613 | 0.089850147 | 0.123604327 | 0.739893493 | ns | TM | KS.3 |
| FIM1_2583 | 417.7941853 | 659.0031941 | 481.033384 | 0.303730159 | 0.271118469 | 0.000882071 | 0.044072215 | ns | TM | KS.3 |
| FIM1_2584 | 63.37748081 | 64.60714158 | 103.7048874 | 0.00113733 | 0.094308483 | 0.949759806 | 0.994413583 | ns | TM | KS.3 |
| FIM1_2585 | 4897.5619 | 4201.666405 | 4837.266344 | -0.055097162 | 0.08407273 | 0.088448362 | 0.649971345 | ns | TM | KS.3 |
| FIM1_2586 | 1189.143458 | 1110.045563 | 1096.603075 | -0.020651619 | 0.058493978 | 0.389536281 | 0.948637087 | ns | TM | KS.3 |
| FIM1_2587 | 3412.964567 | 3452.194168 | 4513.054149 | 0.003355596 | 0.044296875 | 0.865867846 | 0.991141351 | ns | TM | KS.3 |
| FIM1_2588 | 13274.80198 | 10751.23705 | 5707.663931 | -0.035537896 | 0.090341225 | 0.177439004 | 0.811391014 | ns | TM | KS.3 |
| FIM1_2590 | 1359.405366 | 1295.781705 | 1955.598832 | -0.004767378 | 0.079193422 | 0.817894572 | 0.98978971 | ns | TM | KS.3 |
| FIM1_2591 | 727.870981 | 826.6244275 | 888.9165065 | 0.014877079 | 0.080464803 | 0.499902906 | 0.953260948 | ns | TM | KS.3 |
| FIM1_2592 | 1749.266188 | 1624.484175 | 1822.145642 | -0.020869064 | 0.060468507 | 0.391014824 | 0.948637087 | ns | TM | KS.3 |
| FIM1_2593 | 1605.195397 | 1377.51823 | 1588.488274 | -0.096319787 | 0.087857954 | 0.020222292 | 0.324535175 | ns | TM | KS.3 |
| FIM1_2594 | 2468.31155 | 2233.007404 | 2182.106981 | -0.030651706 | 0.067101366 | 0.247098447 | 0.888934767 | ns | TM | KS.3 |

| | | | | | | | | | | |
|---|---|---|---|---|---|---|---|---|---|---|
| FIM1_2595 | 1276.26612 | 1268.123157 | 1316.272804 | -0.001776031 | 0.045635557 | 0.930206117 | 0.993993515 | ns | TM | KS.3 |
| FIM1_2596 | 1486.18978 | 1815.042924 | 2024.028705 | 0.035260608 | 0.088387628 | 0.182960542 | 0.820765281 | ns | TM | KS.3 |
| FIM1_2597 | 888.5553776 | 1085.869684 | 977.7203132 | 0.03780907 | 0.089020408 | 0.164000923 | 0.800496208 | ns | TM | KS.3 |
| FIM1_2598 | 1217.93068 | 1221.130113 | 1013.907709 | 0.000618955 | 0.053923512 | 0.977717478 | 0.999036302 | ns | TM | KS.3 |
| FIM1_2599 | 863.3546099 | 1008.235707 | 964.505576 | 0.022622361 | 0.081088117 | 0.342192415 | 0.944808611 | ns | TM | KS.3 |
| FIM1_2600 | 1179.324059 | 1321.601466 | 1325.314155 | 0.021637522 | 0.07302964 | 0.379156005 | 0.948637087 | ns | TM | KS.3 |
| FIM1_2601 | 1756.303818 | 2043.433059 | 2074.916074 | 0.058462955 | 0.084353335 | 0.077908723 | 0.613284649 | ns | TM | KS.3 |
| FIM1_2602 | 1717.15838 | 1781.245795 | 1905.859368 | 0.010503177 | 0.050665945 | 0.628800807 | 0.97344368 | ns | TM | KS.3 |
| FIM1_2603 | 322.3343247 | 307.7424526 | 286.3863057 | -0.004000568 | 0.082396805 | 0.841577711 | 0.991141351 | ns | TM | KS.3 |
| FIM1_2604 | 597.6681772 | 612.5874558 | 543.7081307 | 0.00365523 | 0.070103758 | 0.869710386 | 0.991141351 | ns | TM | KS.3 |
| FIM1_2606 | 19.95255399 | 22.31102557 | 32.27504231 | 0.006544403 | 0.090962931 | 0.729126967 | 0.987157164 | ns | TM | KS.3 |
| FIM1_2607 | 2586.937781 | 2861.927965 | 3002.133013 | 0.016004167 | 0.073617762 | 0.493381464 | 0.952949531 | ns | TM | KS.3 |
| FIM1_2608 | 757.8695783 | 807.7430539 | 800.2735726 | 0.020811771 | 0.053644995 | 0.36380959 | 0.948637087 | ns | TM | KS.3 |
| FIM1_2609 | 520.7485377 | 476.9886556 | 502.6738705 | -0.019309425 | 0.067380406 | 0.430404248 | 0.950410534 | ns | TM | KS.3 |
| FIM1_2610 | 918.6156194 | 949.1114469 | 1233.092503 | 0.008204228 | 0.0549865 | 0.715515184 | 0.987012589 | ns | TM | KS.3 |
| FIM1_2611 | 2408.813265 | 2359.789801 | 1871.482649 | -0.002984032 | 0.070137554 | 0.893307516 | 0.991141351 | ns | TM | KS.3 |
| FIM1_2612 | 2653.889045 | 2730.299274 | 3394.826906 | 0.002000724 | 0.086666516 | 0.916724196 | 0.991141351 | ns | TM | KS.3 |
| FIM1_2613 | 3101.741431 | 2746.157391 | 3064.290298 | -0.012577707 | 0.082064649 | 0.554919516 | 0.95917717 | ns | TM | KS.3 |
| FIM1_2614 | 977.1005968 | 909.1508967 | 709.2168211 | -0.017325335 | 0.062523597 | 0.472129352 | 0.950410534 | ns | TM | KS.3 |
| FIM1_2615 | 76.59832075 | 82.09318902 | 64.47101098 | 0.007244194 | 0.080671002 | 0.726093956 | 0.987157164 | ns | TM | KS.3 |
| FIM1_2616 | 847.6854327 | 649.0765597 | 737.3979886 | -0.257773418 | 0.108123165 | 9.13554699343 | 0.00797357 | ns | TM | KS.3 |
| FIM1_2617 | 1363.419483 | 1235.639914 | 1133.242136 | -0.046118996 | 0.0673669 | 0.113001204 | 0.711904568 | ns | TM | KS.3 |
| FIM1_2618 | 436.3161791 | 471.7397921 | 394.8937208 | 0.015869496 | 0.065624002 | 0.508002743 | 0.953260948 | ns | TM | KS.3 |
| FIM1_2619 | 326.9184642 | 322.529752 | 276.3989874 | -0.002480833 | 0.06598197 | 0.912918546 | 0.991141351 | ns | TM | KS.3 |
| FIM1_2620 | 749.3344297 | 714.0090858 | 670.0593794 | -0.010932127 | 0.061221002 | 0.639256032 | 0.975246356 | ns | TM | KS.3 |
| FIM1_2621 | 7706.158222 | 7132.9854 | 8627.981714 | -0.014836841 | 0.067762306 | 0.532093424 | 0.959071703 | ns | TM | KS.3 |
| FIM1_2622 | 826.1323062 | 743.3527632 | 652.659282 | -0.028114313 | 0.068835841 | 0.281652951 | 0.917031043 | ns | TM | KS.3 |
| FIM1_2623 | 1771.231253 | 1756.079614 | 1779.418525 | -0.002320998 | 0.050863082 | 0.914462535 | 0.991141351 | ns | TM | KS.3 |
| FIM1_2624 | 9168.773436 | 6566.97734 | 7410.842857 | -0.302552272 | 0.162169102 | 0.00014758 | 0.011842135 | ns | TM | KS.3 |
| FIM1_2625 | 539.3305323 | 586.7188171 | 576.4427825 | 0.016808692 | 0.068094211 | 0.484463937 | 0.952269291 | ns | TM | KS.3 |
| FIM1_2626 | 1963.730235 | 2083.118506 | 1584.582197 | 0.015546435 | 0.058261961 | 0.507930857 | 0.953260948 | ns | TM | KS.3 |
| FIM1_2627 | 1881.194938 | 1807.121235 | 1047.969986 | -0.007846222 | 0.063738887 | 0.734018255 | 0.987655486 | ns | TM | KS.3 |
| FIM1_2628 | 870.6732507 | 954.2813801 | 833.5519857 | 0.026340184 | 0.06524391 | 0.303622458 | 0.926536521 | ns | TM | KS.3 |
| FIM1_2629 | 5540.248922 | 5804.405473 | 7639.601927 | 0.005107618 | 0.077114185 | 0.809152936 | 0.988916276 | ns | TM | KS.3 |
| FIM1_2630 | 502.254458 | 630.3167349 | 544.7853279 | 0.055009073 | 0.098906252 | 0.079603906 | 0.616765353 | ns | TM | KS.3 |
| FIM1_2631 | 196.6116209 | 168.3543088 | 217.1279088 | -0.035120097 | 0.079920099 | 0.198763658 | 0.841573784 | ns | TM | KS.3 |
| FIM1_2632 | 698.9279046 | 769.0303588 | 622.236406 | 0.023150547 | 0.067664508 | 0.357596168 | 0.948637087 | ns | TM | KS.3 |
| FIM1_2633 | 548.0969197 | 692.3576726 | 808.6597244 | 0.070142152 | 0.105477733 | 0.046489169 | 0.493142038 | ns | TM | KS.3 |
| FIM1_2634 | 5073.872398 | 6695.586204 | 5047.092429 | 0.035227495 | 0.101934683 | 0.162889171 | 0.799185036 | ns | TM | KS.3 |
| FIM1_2635 | 31123.57394 | 32525.31462 | 31529.08503 | 0.012591826 | 0.052502674 | 0.571207151 | 0.962265923 | ns | TM | KS.3 |
| FIM1_2636 | 4006.556796 | 3719.245481 | 3879.739344 | -0.034760458 | 0.057027242 | 0.16356277 | 0.800496208 | ns | TM | KS.3 |
| FIM1_2637 | 6168.328577 | 5881.373558 | 4556.375614 | -0.008801846 | 0.065564082 | 0.703340312 | 0.98151979 | ns | TM | KS.3 |
| FIM1_2638 | 1521.749951 | 1422.199812 | 1535.1718 | -0.018748829 | 0.059183126 | 0.433135393 | 0.950410534 | ns | TM | KS.3 |
| FIM1_2639 | 3366.308357 | 3802.126553 | 3287.046383 | 0.020867828 | 0.075180535 | 0.388429134 | 0.948637087 | ns | TM | KS.3 |
| FIM1_2640 | 2819.845777 | 2824.090001 | 3174.526742 | 0.000332307 | 0.059264787 | 0.988332892 | 0.999333464 | ns | TM | KS.3 |

| | | | | | | | | | | |
|---|---|---|---|---|---|---|---|---|---|---|
| FIM1_2641 | 1144.207778 | 1010.929385 | 974.5261912 | -0.049699083 | 0.075855901 | 0.107728423 | 0.702423202 | ns | TM | KS.3 |
| FIM1_2642 | 307.3726166 | 325.8019455 | 395.3580463 | 0.014711071 | 0.059401576 | 0.532045627 | 0.959071703 | ns | TM | KS.3 |
| FIM1_2643 | 2649.682669 | 2229.266609 | 2442.68225 | -0.052895875 | 0.087618032 | 0.094411295 | 0.673663379 | ns | TM | KS.3 |
| FIM1_2644 | 1985.461513 | 1960.824604 | 1769.190409 | -0.003508104 | 0.047709953 | 0.866468728 | 0.991141351 | ns | TM | KS.3 |
| FIM1_2645 | 661.6023642 | 660.5113966 | 693.2951935 | -0.000159559 | 0.05583298 | 0.994322551 | 0.999525119 | ns | TM | KS.3 |
| FIM1_2646 | 2900.609679 | 2752.357652 | 2678.289446 | -0.016168979 | 0.052415336 | 0.46976618 | 0.950410534 | ns | TM | KS.3 |
| FIM1_2647 | 3617.115799 | 3219.272526 | 3353.209645 | -0.058351295 | 0.075015676 | 0.073165012 | 0.595738026 | ns | TM | KS.3 |
| FIM1_2648 | 266.6542373 | 276.2039588 | 308.8158223 | 0.009621216 | 0.054696706 | 0.669363111 | 0.976323032 | ns | TM | KS.3 |
| FIM1_2649 | 826.0408591 | 789.696974 | 874.0597844 | -0.0163216 | 0.047270041 | 0.432325266 | 0.950410534 | ns | TM | KS.3 |
| FIM1_2650 | 14817.08426 | 15105.69771 | 13759.61847 | 0.004599391 | 0.055697987 | 0.83785102 | 0.991141351 | ns | TM | KS.3 |
| FIM1_2651 | 269.739294 | 299.0005264 | 381.7676173 | 0.015133029 | 0.075038139 | 0.510441167 | 0.953260948 | ns | TM | KS.3 |
| FIM1_2652 | 7227.88054 | 7207.885853 | 7485.896065 | -0.000845109 | 0.04020626 | 0.963515587 | 0.995175461 | ns | TM | KS.3 |
| FIM1_2653 | 1778.297862 | 1729.855764 | 1690.258184 | -0.007320898 | 0.051916562 | 0.738395105 | 0.988051177 | ns | TM | KS.3 |
| FIM1_2654 | 2242.871912 | 2107.88232 | 2235.239421 | -0.016531568 | 0.058627331 | 0.484038771 | 0.952269291 | ns | TM | KS.3 |
| FIM1_2655 | 1316.454442 | 1535.592389 | 1444.349023 | 0.01031927 | 0.09231204 | 0.593715514 | 0.965588324 | ns | TM | KS.3 |
| FIM1_2656 | 266.2566358 | 168.4041313 | 282.8426353 | -0.065752952 | 0.159073168 | 0.045580091 | 0.489573998 | ns | TM | KS.3 |
| FIM1_2657 | 774.6566218 | 853.333555 | 919.771808 | 0.025637387 | 0.066845388 | 0.316397313 | 0.927564309 | ns | TM | KS.3 |
| FIM1_2659 | 1528.830955 | 1488.537336 | 1561.785365 | -0.006249372 | 0.056645764 | 0.78261129 | 0.988814136 | ns | TM | KS.3 |
| FIM1_2660 | 2392.031223 | 2274.284266 | 1832.770411 | -0.005328091 | 0.078006311 | 0.799625118 | 0.988916276 | ns | TM | KS.3 |
| FIM1_2661 | 1189.034146 | 1266.426351 | 1387.871653 | 0.005211657 | 0.083614698 | 0.793356817 | 0.988916276 | ns | TM | KS.3 |
| FIM1_2662 | 4640.470256 | 4720.597082 | 5391.751476 | 0.003998882 | 0.056399473 | 0.859206019 | 0.991141351 | ns | TM | KS.3 |
| FIM1_2663 | 348.9906786 | 448.9128708 | 406.4431412 | 0.023307108 | 0.097074989 | 0.291439042 | 0.919993167 | ns | TM | KS.3 |
| FIM1_2664 | 2968.675456 | 2755.14742 | 2489.737755 | -0.028141806 | 0.057617728 | 0.25079768 | 0.888934767 | ns | TM | KS.3 |
| FIM1_2665 | 329.3241076 | 252.1394336 | 497.1695574 | -0.050079169 | 0.105205108 | 0.089509136 | 0.653903013 | ns | TM | KS.3 |
| FIM1_2666 | 502.5208291 | 531.0400988 | 1035.415032 | 0.008980496 | 0.068899899 | 0.694553919 | 0.981425301 | ns | TM | KS.3 |
| FIM1_2667 | 2893.46023 | 2911.102609 | 3083.454991 | 0.001909582 | 0.036662301 | 0.911006556 | 0.991141351 | ns | TM | KS.3 |
| FIM1_2668 | 2130.086631 | 3142.841988 | 3118.802518 | 0.062866373 | 0.14007049 | 0.050529994 | 0.513412402 | ns | TM | KS.3 |
| FIM1_2669 | 22.24950685 | 41.53823864 | 47.98920621 | 0.112594563 | 0.244205989 | 0.016108956 | 0.294639904 | ns | TM | KS.3 |
| FIM1_2671 | 55.35349612 | 45.39311891 | 54.40823865 | -0.018417447 | 0.090439151 | 0.390219871 | 0.948637087 | ns | TM | KS.3 |
| FIM1_2672 | 932.2002907 | 890.0107349 | 872.2161792 | -0.013531423 | 0.052481041 | 0.543468173 | 0.95917717 | ns | TM | KS.3 |
| FIM1_2674 | 6901.740058 | 6347.372022 | 5072.522202 | -0.019278644 | 0.065701248 | 0.431172906 | 0.950410534 | ns | TM | KS.3 |
| FIM1_2675 | 331.1116832 | 119.7385313 | 10561.10044 | -0.061449036 | 0.23058883 | 0.042697053 | 0.473090953 | ns | TM | KS.3 |
| FIM1_2676 | 8541.793921 | 10147.00408 | 23990.20896 | 0.016755835 | 0.087209026 | 0.43436858 | 0.950410534 | ns | TM | KS.3 |
| FIM1_2677 | 3454.525653 | 3679.161646 | 3814.504978 | 0.018476391 | 0.056829086 | 0.431778083 | 0.950410534 | ns | TM | KS.3 |
| FIM1_2678 | 621.3058095 | 583.043762 | 579.2869035 | -0.014031256 | 0.063343987 | 0.554153041 | 0.95917717 | ns | TM | KS.3 |
| FIM1_2679 | 1150.516723 | 1137.15862 | 1383.071188 | -0.002300665 | 0.061417933 | 0.919699525 | 0.991141351 | ns | TM | KS.3 |
| FIM1_2680 | 3511.679757 | 3678.45818 | 4303.406281 | 0.010525772 | 0.060049344 | 0.650345604 | 0.976323032 | ns | TM | KS.3 |
| FIM1_2681 | 2219.336227 | 2123.290498 | 4554.22403 | -0.007798871 | 0.066530505 | 0.734136874 | 0.987655486 | ns | TM | KS.3 |
| FIM1_2682 | 389.1129244 | 446.1643415 | 945.3060971 | 0.01322794 | 0.084096043 | 0.530645702 | 0.959071703 | ns | TM | KS.3 |
| FIM1_2683 | 2381.42696 | 2233.937624 | 2297.684861 | -0.01178008 | 0.06710459 | 0.614018371 | 0.973122675 | ns | TM | KS.3 |
| FIM1_2684 | 3754.79922 | 4480.137082 | 5456.353642 | 0.023939712 | 0.084513788 | 0.312210886 | 0.927416591 | ns | TM | KS.3 |
| FIM1_2685 | 1019.680285 | 987.2283493 | 1047.779783 | -0.008181089 | 0.055043083 | 0.716375633 | 0.987157164 | ns | TM | KS.3 |
| FIM1_2686 | 196.7168414 | 142.6940064 | 211.4828374 | -0.076041119 | 0.129808013 | 0.03446779 | 0.42621823 | ns | TM | KS.3 |
| FIM1_2687 | 683.5994372 | 679.4624551 | 777.3624947 | -0.000558935 | 0.079652558 | 0.978151925 | 0.999036302 | ns | TM | KS.3 |
| FIM1_2688 | 1591.952994 | 1620.353384 | 1954.134537 | 0.004196703 | 0.055377583 | 0.851527983 | 0.991141351 | ns | TM | KS.3 |

| | | | | | | | | | | |
|---|---|---|---|---|---|---|---|---|---|---|
| FIM1_2689 | 1963.344022 | 1916.334644 | 1808.791337 | -0.006633626 | 0.049986161 | 0.757373795 | 0.988477371 | ns | TM | KS.3 |
| FIM1_2690 | 7203.834658 | 6826.411167 | 6279.00711 | -0.019228229 | 0.049343472 | 0.371724855 | 0.948637087 | ns | TM | KS.3 |
| FIM1_2691 | 1632.171841 | 1383.491334 | 1404.064091 | -0.147678469 | 0.082714972 | 0.002917271 | 0.100787663 | ns | TM | KS.3 |
| FIM1_2692 | 797.3699303 | 641.1416521 | 751.1524816 | -0.09799645 | 0.107853012 | 0.020068528 | 0.324535175 | ns | TM | KS.3 |
| FIM1_2693 | 2464.417629 | 2394.665313 | 2325.402404 | -0.008821446 | 0.045113977 | 0.660748426 | 0.976323032 | ns | TM | KS.3 |
| FIM1_2694 | 573.4804336 | 466.7390308 | 600.8518616 | -0.138015483 | 0.105513528 | 0.005940574 | 0.16059892 | ns | TM | KS.3 |
| FIM1_2695 | 2656.205869 | 2703.401984 | 2827.478016 | 0.004524361 | 0.051879658 | 0.835933586 | 0.991141351 | ns | TM | KS.3 |
| FIM1_2696 | 5585.743192 | 6165.731222 | 6745.90564 | 0.015451527 | 0.073588752 | 0.506569264 | 0.953260948 | ns | TM | KS.3 |
| FIM1_2697 | 398.3395019 | 475.3056323 | 428.8738886 | 0.021256411 | 0.085366571 | 0.352152332 | 0.948637087 | ns | TM | KS.3 |
| FIM1_2698 | 4365.5332 | 3546.058032 | 3296.22013 | -0.14974205 | 0.103276566 | 0.004006751 | 0.120809617 | ns | TM | KS.3 |
| FIM1_2699 | 1742.055074 | 1881.621383 | 2342.557362 | 0.009665925 | 0.07572756 | 0.659553972 | 0.976323032 | ns | TM | KS.3 |
| FIM1_2700 | 432.7287423 | 423.3795153 | 571.3223321 | -0.003558407 | 0.067261096 | 0.874979403 | 0.991141351 | ns | TM | KS.3 |
| FIM1_2701 | 1883.906743 | 2132.228686 | 2051.847764 | 0.021521331 | 0.075354282 | 0.376284479 | 0.948637087 | ns | TM | KS.3 |
| FIM1_2702 | 160.0965033 | 163.7082995 | 193.6273669 | 0.003447771 | 0.066305348 | 0.879250689 | 0.991141351 | ns | TM | KS.3 |
| FIM1_2703 | 205.8637441 | 96.25603536 | 206.2115144 | -0.13261225 | 0.298515705 | 0.01227708 | 0.253437653 | ns | TM | KS.3 |
| FIM1_2704 | 1312.397213 | 1364.563449 | 1576.395522 | 0.00938165 | 0.057738522 | 0.682517422 | 0.97986479 | ns | TM | KS.3 |
| FIM1_2705 | 2134.315783 | 2232.888712 | 3068.511263 | 0.007009432 | 0.069493416 | 0.756477914 | 0.988477371 | ns | TM | KS.3 |
| FIM1_2706 | 10.10351681 | 10.85652096 | 23.14872675 | 0.002398686 | 0.095465789 | 0.893819939 | 0.991141351 | ns | TM | KS.3 |
| FIM1_2708 | 4178.022147 | 4679.084413 | 5778.449195 | 0.011928131 | 0.08115845 | 0.576117884 | 0.963192994 | ns | TM | KS.3 |
| FIM1_2709 | 3509.995396 | 3288.155901 | 3182.780214 | -0.01870552 | 0.058064871 | 0.430779149 | 0.950410534 | ns | TM | KS.3 |
| FIM1_2710 | 9349.422336 | 9324.597116 | 8876.301857 | -0.000486477 | 0.063967749 | 0.982950515 | 0.999333464 | ns | TM | KS.3 |
| FIM1_2711 | 320.64746 | 274.7764363 | 333.9744302 | -0.055897659 | 0.084754578 | 0.08571169 | 0.641226554 | ns | TM | KS.3 |
| FIM1_2712 | 2685.933647 | 3027.554117 | 2749.901888 | 0.028415774 | 0.072705233 | 0.277040711 | 0.908583046 | ns | TM | KS.3 |
| FIM1_2713 | 2274.522541 | 1975.446903 | 1923.715799 | -0.065935587 | 0.083275675 | 0.058567136 | 0.540287513 | ns | TM | KS.3 |
| FIM1_2714 | 1559.593234 | 1512.273648 | 1521.290893 | -0.010065547 | 0.039767433 | 0.577111112 | 0.963465698 | ns | TM | KS.3 |
| FIM1_2715 | 2607.611742 | 3025.100048 | 2877.232405 | 0.089151202 | 0.087115569 | 0.02581669 | 0.379742647 | ns | TM | KS.3 |
| FIM1_2716 | 8178.184214 | 6398.28097 | 7226.098918 | -0.106230597 | 0.117874311 | 0.015419488 | 0.284118339 | ns | TM | KS.3 |
| FIM1_2718 | 4121.773563 | 4261.654303 | 5181.679644 | 0.010260608 | 0.047586509 | 0.623120234 | 0.973319675 | ns | TM | KS.3 |
| FIM1_2719 | 1713.252811 | 2341.946923 | 2146.318507 | 0.139141023 | 0.150771861 | 0.0067136 | 0.17486996 | ns | TM | KS.3 |
| FIM1_2720 | 4261.810844 | 4507.468093 | 3393.652226 | 0.022815128 | 0.048654879 | 0.284448642 | 0.918450395 | ns | TM | KS.3 |
| FIM1_2721 | 339.7790842 | 588.7707731 | 627.1013239 | 0.238604843 | 0.310630017 | 0.002870775 | 0.100578206 | ns | TM | KS.3 |
| FIM1_2722 | 2600.423369 | 2775.639586 | 2572.884247 | 0.015951651 | 0.061161958 | 0.503333884 | 0.953260948 | ns | TM | KS.3 |
| FIM1_2723 | 1153.894742 | 1049.784705 | 810.7871717 | -0.012058678 | 0.076579082 | 0.5865517 | 0.963781856 | ns | TM | KS.3 |
| FIM1_2724 | 2788.224885 | 2415.112387 | 1911.980482 | -0.047470797 | 0.080240879 | 0.119766064 | 0.730090427 | ns | TM | KS.3 |
| FIM1_2726 | 913.2629656 | 885.4945943 | 789.6774471 | -0.006701597 | 0.060181943 | 0.770392783 | 0.988814136 | ns | TM | KS.3 |
| FIM1_2727 | 183.6123727 | 86.0281156 | 134.5850001 | -0.947818409 | 0.287202244 | 8.07116844597 | 0.001029591 | ns | TM | KS.3 |
| FIM1_2728 | 541.683455 | 468.565243 | 592.1077316 | -0.017710223 | 0.081653848 | 0.43122383 | 0.950410534 | ns | TM | KS.3 |
| FIM1_2729 | 244.1412476 | 450.0585445 | 331.3030649 | 0.123178925 | 0.254883662 | 0.013471536 | 0.263081302 | ns | TM | KS.3 |
| FIM1_2730 | 63.01076121 | 62.07425429 | 150.8965523 | -0.00093167 | 0.080203581 | 0.963399388 | 0.995175461 | ns | TM | KS.3 |
| FIM1_2731 | 4139.527278 | 3775.427474 | 4238.867516 | -0.033811734 | 0.064340516 | 0.204178285 | 0.847612283 | ns | TM | KS.3 |
| FIM1_2732 | 8819.950913 | 8162.049629 | 8924.198354 | -0.028470665 | 0.058905939 | 0.252068786 | 0.888934767 | ns | TM | KS.3 |
| FIM1_2733 | 242.6259212 | 314.1348268 | 314.6935133 | 0.084024448 | 0.116949291 | 0.028780274 | 0.390786333 | ns | TM | KS.3 |
| FIM1_2734 | 207.1386075 | 259.3915057 | 239.54723 | 0.080224977 | 0.105642517 | 0.034276595 | 0.42621823 | ns | TM | KS.3 |
| FIM1_2735 | 716.1309964 | 563.6127175 | 651.349206 | -0.121138511 | 0.118359539 | 0.01017937 | 0.223094117 | ns | TM | KS.3 |
| FIM1_2736 | 645.8551177 | 703.4430497 | 595.3209154 | 0.019626458 | 0.066065579 | 0.424024977 | 0.950410534 | ns | TM | KS.3 |

| | | | | | | | | | | |
|---|---|---|---|---|---|---|---|---|---|---|
| FIM1_2737 | 2274.536038 | 2330.226601 | 2475.593407 | 0.006312577 | 0.052047798 | 0.773358864 | 0.988814136 | ns | TM | KS.3 |
| FIM1_2738 | 615.4753041 | 599.8182681 | 482.4117522 | -0.00303821 | 0.074790448 | 0.887430208 | 0.991141351 | ns | TM | KS.3 |
| FIM1_2739 | 2765.224192 | 2569.148813 | 2229.439946 | -0.007352768 | 0.079980742 | 0.7240273 | 0.987157164 | ns | TM | KS.3 |
| FIM1_2740 | 6608.56107 | 6587.865422 | 5832.091995 | -0.000948251 | 0.037136258 | 0.95626031 | 0.994413583 | ns | TM | KS.3 |
| FIM1_2741 | 1147.391357 | 943.4293401 | 950.3022383 | -0.115906025 | 0.10265769 | 0.011677614 | 0.24513135 | ns | TM | KS.3 |
| FIM1_2742 | 6862.338991 | 7260.758759 | 6492.584788 | 0.006953435 | 0.075019134 | 0.748950533 | 0.988477371 | ns | TM | KS.3 |
| FIM1_2743 | 547.8381673 | 805.4489257 | 767.070899 | 0.053584599 | 0.132564337 | 0.069946143 | 0.582627046 | ns | TM | KS.3 |
| FIM1_2744 | 9338.120335 | 9082.65927 | 6478.222438 | -0.009249064 | 0.039368695 | 0.605915346 | 0.970937835 | ns | TM | KS.3 |
| FIM1_2745 | 1580.641933 | 1705.02364 | 1426.642156 | 0.009091 | 0.076459387 | 0.67564034 | 0.978262715 | ns | TM | KS.3 |
| FIM1_2746 | 1830.755988 | 1752.589538 | 1398.377701 | -0.010931784 | 0.056794363 | 0.633440179 | 0.97344368 | ns | TM | KS.3 |
| FIM1_2747 | 2688.037042 | 2800.094993 | 2989.429139 | 0.0084304 | 0.062125929 | 0.715608926 | 0.987012589 | ns | TM | KS.3 |
| FIM1_2748 | 451.5831435 | 496.7925409 | 558.3447268 | 0.023721092 | 0.06670071 | 0.347490239 | 0.948637087 | ns | TM | KS.3 |
| FIM1_2749 | 740.5625091 | 656.9868642 | 918.7957471 | -0.033463847 | 0.072823068 | 0.219765954 | 0.865538807 | ns | TM | KS.3 |
| FIM1_2750 | 60.91147572 | 70.85994721 | 94.23558957 | 0.02121072 | 0.079389509 | 0.370852885 | 0.948637087 | ns | TM | KS.3 |
| FIM1_2751 | 569.8926344 | 443.8930344 | 582.8865073 | -0.053708831 | 0.103161234 | 0.080178201 | 0.618428374 | ns | TM | KS.3 |
| FIM1_2752 | 1494.814012 | 1302.517154 | 1326.891207 | -0.085684048 | 0.083192642 | 0.02790305 | 0.388845019 | ns | TM | KS.3 |
| FIM1_2753 | 1206.508261 | 969.297914 | 1404.235879 | -0.06614893 | 0.100961188 | 0.054496035 | 0.531428718 | ns | TM | KS.3 |
| FIM1_2754 | 1819.98096 | 1599.81818 | 3605.0053 | -0.018548671 | 0.077961889 | 0.425503308 | 0.950410534 | ns | TM | KS.3 |
| FIM1_2755 | 1013.452454 | 1056.336116 | 1085.069687 | 0.010473167 | 0.054739372 | 0.642794387 | 0.976237372 | ns | TM | KS.3 |
| FIM1_2756 | 921.2370557 | 956.3658892 | 909.1344341 | 0.003734515 | 0.078780397 | 0.856896947 | 0.991141351 | ns | TM | KS.3 |
| FIM1_2757 | 4084.854863 | 4762.361633 | 5736.513145 | 0.014640219 | 0.085805893 | 0.488552784 | 0.952591289 | ns | TM | KS.3 |
| FIM1_2758 | 480.6262298 | 627.4523426 | 579.5831668 | 0.102442557 | 0.123844625 | 0.016855981 | 0.302738295 | ns | TM | KS.3 |
| FIM1_2759 | 147.2178371 | 171.2659585 | 236.8115605 | 0.026854286 | 0.079189999 | 0.286287353 | 0.919993167 | ns | TM | KS.3 |
| FIM1_2760 | 7324.822168 | 6768.42738 | 8012.268097 | -0.011848321 | 0.072465164 | 0.603284816 | 0.970245665 | ns | TM | KS.3 |
| FIM1_2761 | 1371.794543 | 1444.071099 | 1486.793316 | 0.020958272 | 0.044936371 | 0.289663744 | 0.919993167 | ns | TM | KS.3 |
| FIM1_2762 | 2303.185632 | 2478.01849 | 2559.5786 | 0.011827402 | 0.070557489 | 0.60780645 | 0.970937835 | ns | TM | KS.3 |
| FIM1_2763 | 20.28283624 | 17.15644995 | 643.8802253 | -0.005084274 | 0.113045496 | 0.761374171 | 0.988477371 | ns | TM | KS.3 |
| FIM1_2764 | 1982.794517 | 2057.443869 | 1403.205605 | 0.00681956 | 0.064886073 | 0.766627214 | 0.988477371 | ns | TM | KS.3 |
| FIM1_2765 | 1649.76351 | 1636.32659 | 1273.675358 | -0.001976652 | 0.053232074 | 0.928593384 | 0.993697171 | ns | TM | KS.3 |
| FIM1_2766 | 3222.237688 | 2988.969048 | 3595.941157 | -0.021739362 | 0.060553679 | 0.373775715 | 0.948637087 | ns | TM | KS.3 |
| FIM1_2767 | 2717.81828 | 2578.44724 | 2284.431879 | -0.008127907 | 0.070029251 | 0.719658886 | 0.987157164 | ns | TM | KS.3 |
| FIM1_2768 | 1143.557019 | 1148.287769 | 1224.0717 | 0.000637712 | 0.06733778 | 0.977421574 | 0.999036302 | ns | TM | KS.3 |
| FIM1_2769 | 2849.255198 | 2710.677511 | 2209.538832 | -0.016316454 | 0.049903201 | 0.451293904 | 0.950410534 | ns | TM | KS.3 |
| FIM1_2770 | 1528.60046 | 1321.932309 | 1412.254494 | -0.033165701 | 0.078198993 | 0.21853419 | 0.865538807 | ns | TM | KS.3 |
| FIM1_2771 | 490.8780616 | 610.6671915 | 445.9064314 | 0.040102898 | 0.09241596 | 0.144885859 | 0.778377716 | ns | TM | KS.3 |
| FIM1_2772 | 1270.785887 | 1118.101663 | 1609.891903 | -0.063939926 | 0.079183508 | 0.061183024 | 0.545493804 | ns | TM | KS.3 |
| FIM1_2773 | 433.6870794 | 472.1322035 | 405.4531913 | 0.016457355 | 0.068324907 | 0.492371066 | 0.952591289 | ns | TM | KS.3 |
| FIM1_2774 | 452.0902382 | 636.0826144 | 928.7320496 | 0.053960656 | 0.123049684 | 0.070941732 | 0.584329662 | ns | TM | KS.3 |
| FIM1_2775 | 4474.782184 | 4431.513925 | 4011.411568 | -0.002908935 | 0.039128699 | 0.871971018 | 0.991141351 | ns | TM | KS.3 |
| FIM1_2776 | 652.968331 | 723.0688226 | 718.1820608 | 0.030261982 | 0.067984943 | 0.253088403 | 0.889833785 | ns | TM | KS.3 |
| FIM1_2777 | 824.9807492 | 774.951108 | 817.4033035 | -0.014212746 | 0.062367177 | 0.548873374 | 0.95917717 | ns | TM | KS.3 |
| FIM1_2778 | 605.3625011 | 611.3198919 | 579.844977 | 0.002771189 | 0.050896356 | 0.89800749 | 0.991141351 | ns | TM | KS.3 |
| FIM1_2779 | 912.8290286 | 944.4850403 | 941.1714081 | 0.007620324 | 0.058803862 | 0.73961154 | 0.988051177 | ns | TM | KS.3 |
| FIM1_2780 | 170.6086308 | 164.0113215 | 312.6164471 | -0.004851652 | 0.07745769 | 0.817787679 | 0.98978971 | ns | TM | KS.3 |
| FIM1_2781 | 897.0072861 | 1119.921004 | 1642.397071 | 0.025905094 | 0.091804457 | 0.267242031 | 0.905057253 | ns | TM | KS.3 |

| | | | | | | | | | | |
|---|---|---|---|---|---|---|---|---|---|---|
| FIM1_2782 | 1523.00436 | 1635.771805 | 1563.850147 | 0.015992393 | 0.064119064 | 0.504923512 | 0.953260948 | ns | TM | KS.3 |
| FIM1_2783 | 4520.585643 | 5302.937242 | 6384.891685 | 0.015553304 | 0.086023092 | 0.465596156 | 0.950410534 | ns | TM | KS.3 |
| FIM1_2784 | 953.8689078 | 920.5211462 | 907.651831 | -0.010773102 | 0.045405882 | 0.593453885 | 0.965478443 | ns | TM | KS.3 |
| FIM1_2785 | 3490.193408 | 3202.685203 | 2810.77649 | -0.026647834 | 0.06290987 | 0.294016301 | 0.923441351 | ns | TM | KS.3 |
| FIM1_2786 | 2956.339462 | 2876.278842 | 2411.717556 | -0.005823628 | 0.060166575 | 0.799378418 | 0.988916276 | ns | TM | KS.3 |
| FIM1_2787 | 5407.452787 | 5428.09923 | 5756.818924 | 0.000392651 | 0.077827049 | 0.984925302 | 0.999333464 | ns | TM | KS.3 |
| FIM1_2788 | 381.1341434 | 332.5587066 | 344.3151537 | -0.025111484 | 0.076501386 | 0.316183327 | 0.927564309 | ns | TM | KS.3 |
| FIM1_2789 | 318.2480334 | 357.6149049 | 274.6503493 | 0.019128471 | 0.074714644 | 0.422844906 | 0.950410534 | ns | TM | KS.3 |
| FIM1_2790 | 984.1764286 | 969.3531541 | 1088.731515 | -0.003621595 | 0.054647657 | 0.871023392 | 0.991141351 | ns | TM | KS.3 |
| FIM1_2791 | 188.1255418 | 134.6805014 | 239.3295169 | -0.095508547 | 0.144513665 | 0.019687576 | 0.324535175 | ns | TM | KS.3 |
| FIM1_2792 | 921.0578193 | 931.8323278 | 699.8149305 | 0.00228807 | 0.062021007 | 0.920152458 | 0.991141351 | ns | TM | KS.3 |
| FIM1_2793 | 1173.939082 | 1270.418628 | 1821.019608 | 0.031643071 | 0.05900721 | 0.209310638 | 0.852551729 | ns | TM | KS.3 |
| FIM1_2794 | 2204.601848 | 2184.325679 | 2224.607093 | -0.002212487 | 0.055288169 | 0.921215855 | 0.991358183 | ns | TM | KS.3 |
| FIM1_2795 | 584.9471518 | 555.085484 | 653.7490727 | -0.005444909 | 0.078344982 | 0.794789078 | 0.988916276 | ns | TM | KS.3 |
| FIM1_2796 | 193.9227413 | 157.0307932 | 654.7569723 | -0.017841069 | 0.092890949 | 0.39566032 | 0.948637087 | ns | TM | KS.3 |
| FIM1_2797 | 50.50156132 | 59.57668405 | 53.41088707 | 0.014274174 | 0.088489029 | 0.489812768 | 0.952591289 | ns | TM | KS.3 |
| FIM1_2798 | 2240.288055 | 2174.421999 | 1657.686551 | -0.00776173 | 0.053243549 | 0.726565838 | 0.987157164 | ns | TM | KS.3 |
| FIM1_2800 | 2449.36682 | 2545.440489 | 3329.2877 | 0.011665681 | 0.048597214 | 0.581999917 | 0.963781856 | ns | TM | KS.3 |
| FIM1_2801 | 4267.995729 | 4011.498508 | 4352.614611 | -0.016533596 | 0.058664662 | 0.484082488 | 0.952269291 | ns | TM | KS.3 |
| FIM1_2802 | 897.0784535 | 753.1118457 | 855.5880216 | -0.062208805 | 0.090515926 | 0.066882342 | 0.566847784 | ns | TM | KS.3 |
| FIM1_2803 | 1236.775439 | 1147.173033 | 1290.444468 | -0.025922675 | 0.058445474 | 0.290060014 | 0.919993167 | ns | TM | KS.3 |
| FIM1_2804 | 2112.897437 | 1961.551891 | 2299.026539 | -0.029272622 | 0.057204319 | 0.232027129 | 0.880810476 | ns | TM | KS.3 |
| FIM1_2805 | 1636.879948 | 1702.418147 | 1868.510163 | 0.010027224 | 0.054747897 | 0.656701831 | 0.976323032 | ns | TM | KS.3 |
| FIM1_2806 | 177.9653542 | 214.0101282 | 204.5735688 | 0.047682494 | 0.088921937 | 0.112833707 | 0.711904568 | ns | TM | KS.3 |
| FIM1_2807 | 1024.061783 | 1017.940883 | 1116.620185 | -0.001368337 | 0.058440924 | 0.951891844 | 0.994413583 | ns | TM | KS.3 |
| FIM1_2808 | 1408.295582 | 1517.421311 | 1536.198656 | 0.017078925 | 0.064095238 | 0.479130215 | 0.950972051 | ns | TM | KS.3 |
| FIM1_2809 | 181.3467817 | 151.3702855 | 172.0515214 | -0.023084764 | 0.085950591 | 0.320974837 | 0.929652552 | ns | TM | KS.3 |
| FIM1_2810 | 256.3506673 | 219.4020096 | 237.534163 | -0.014715025 | 0.086159363 | 0.485593646 | 0.952269291 | ns | TM | KS.3 |
| FIM1_2811 | 474.1144227 | 464.2364972 | 460.8452583 | -0.005393125 | 0.056637577 | 0.811511794 | 0.988916276 | ns | TM | KS.3 |
| FIM1_2812 | 2839.764136 | 2467.696103 | 2959.470854 | -0.076970028 | 0.084230192 | 0.038987608 | 0.446920158 | ns | TM | KS.3 |
| FIM1_2813 | 887.4784819 | 911.3199248 | 1060.003627 | 0.006025004 | 0.058513448 | 0.791771988 | 0.988916276 | ns | TM | KS.3 |
| FIM1_2814 | 1749.442016 | 1577.403727 | 2193.081339 | -0.02612796 | 0.068814452 | 0.309890329 | 0.927416591 | ns | TM | KS.3 |
| FIM1_2815 | 307.4341593 | 392.0036143 | 337.3905196 | 0.078340915 | 0.109446964 | 0.035379553 | 0.42652037 | ns | TM | KS.3 |
| FIM1_2816 | 410.1115319 | 397.3120042 | 373.392304 | -0.007091313 | 0.061829723 | 0.758233972 | 0.988477371 | ns | TM | KS.3 |
| FIM1_2817 | 542.710677 | 489.1483862 | 534.8311899 | -0.023758416 | 0.069967102 | 0.346720436 | 0.948287064 | ns | TM | KS.3 |
| FIM1_2818 | 413.7901567 | 385.0031588 | 428.9503715 | -0.018786708 | 0.05980764 | 0.433920951 | 0.950410534 | ns | TM | KS.3 |
| FIM1_2819 | 2095.022069 | 1918.724717 | 2042.335681 | -0.042043622 | 0.063387726 | 0.127721273 | 0.75107959 | ns | TM | KS.3 |
| FIM1_2820 | 1277.809062 | 1930.676975 | 2772.900424 | 0.084338273 | 0.16246938 | 0.026679859 | 0.382098641 | ns | TM | KS.3 |
| FIM1_2821 | 1776.504652 | 1838.669952 | 1743.736036 | 0.008565755 | 0.055300547 | 0.70437223 | 0.982408703 | ns | TM | KS.3 |
| FIM1_2822 | 618.0532662 | 1027.320935 | 639.5826493 | 0.669390007 | 0.184268193 | 1.4374803305[illegible] | 0.000204328 | ns | TM | KS.3 |
| FIM1_2823 | 3528.59019 | 3319.223557 | 3390.920902 | -0.021940689 | 0.052733759 | 0.333906981 | 0.938982061 | ns | TM | KS.3 |
| FIM1_2824 | 1094.03708 | 1026.44375 | 1247.898865 | -0.023542778 | 0.053601751 | 0.307009048 | 0.927364916 | ns | TM | KS.3 |
| FIM1_2825 | 1204.533129 | 1112.039831 | 1050.128606 | -0.024279884 | 0.061205459 | 0.32837396 | 0.935147309 | ns | TM | KS.3 |
| FIM1_2826 | 991.4516523 | 896.5611364 | 888.972011 | -0.018560472 | 0.071408783 | 0.441554717 | 0.950410534 | ns | TM | KS.3 |
| FIM1_2827 | 2606.372413 | 3087.505063 | 3463.262847 | 0.023013704 | 0.083509485 | 0.328997781 | 0.93582845 | ns | TM | KS.3 |

| | | | | | | | | | | |
|---|---|---|---|---|---|---|---|---|---|---|
| FIM1_2828 | 558.3617062 | 744.3739985 | 768.7387174 | 0.245079302 | 0.131012939 | 0.000320851 | 0.021003105 | ns | TM | KS.3 |
| FIM1_2829 | 1975.917094 | 2607.095108 | 2250.230989 | 0.172255701 | 0.13588846 | 0.002727192 | 0.096373426 | ns | TM | KS.3 |
| FIM1_2830 | 2073.103725 | 1564.257896 | 2405.026122 | -0.13598628 | 0.136403098 | 0.006988917 | 0.177397248 | ns | TM | KS.3 |
| FIM1_2831 | 88.46023213 | 93.8314416 | 339.8605135 | 0.003409669 | 0.09195188 | 0.853354593 | 0.991141351 | ns | TM | KS.3 |
| FIM1_2832 | 1056.613639 | 1203.398699 | 1539.108163 | 0.047382196 | 0.07727908 | 0.119896257 | 0.730090427 | ns | TM | KS.3 |
| FIM1_2833 | 1397.376591 | 1813.577215 | 1969.477907 | 0.026266366 | 0.098114176 | 0.250549224 | 0.888934767 | ns | TM | KS.3 |
| FIM1_2834 | 1310.425025 | 1340.015164 | 1728.387067 | 0.001772791 | 0.083686578 | 0.92810471 | 0.993697171 | ns | TM | KS.3 |
| FIM1_2835 | 1036.439974 | 1126.121446 | 1076.06063 | 0.007911832 | 0.081332137 | 0.7015065 | 0.98151979 | ns | TM | KS.3 |
| FIM1_2836 | 740.8474922 | 804.6443382 | 864.2023526 | 0.021958855 | 0.06390755 | 0.376432421 | 0.948637087 | ns | TM | KS.3 |
| FIM1_2837 | 1275.752928 | 1216.640024 | 858.4962492 | -0.004933476 | 0.078252756 | 0.813451795 | 0.988916276 | ns | TM | KS.3 |
| FIM1_2838 | 2109.037066 | 1973.761176 | 2749.047553 | -0.018857352 | 0.058531684 | 0.428794901 | 0.950410534 | ns | TM | KS.3 |
| FIM1_2839 | 691.9486287 | 618.5648983 | 734.8702793 | -0.046106057 | 0.072078337 | 0.122151833 | 0.736051706 | ns | TM | KS.3 |
| FIM1_2840 | 107.8140473 | 88.20954258 | 112.8839267 | -0.038718768 | 0.089667883 | 0.157172405 | 0.789034023 | ns | TM | KS.3 |
| FIM1_2841 | 1067.954298 | 974.6904529 | 1116.009165 | -0.026753918 | 0.065048066 | 0.297007609 | 0.926536521 | ns | TM | KS.3 |
| FIM1_2842 | 2889.375962 | 2981.300375 | 3122.825629 | 0.008818723 | 0.049912104 | 0.68165343 | 0.97986479 | ns | TM | KS.3 |
| FIM1_2843 | 270.2748847 | 260.9059866 | 223.4923183 | -0.006278878 | 0.065460343 | 0.784035363 | 0.988880259 | ns | TM | KS.3 |
| FIM1_2844 | 590.964417 | 574.165226 | 525.3170241 | -0.005020893 | 0.065760316 | 0.825792008 | 0.991141351 | ns | TM | KS.3 |
| FIM1_2845 | 31739.07823 | 34428.80546 | 43296.34962 | 0.023280035 | 0.062327597 | 0.349112796 | 0.948637087 | ns | TM | KS.3 |
| FIM1_2846 | 756.5591041 | 1025.390952 | 1216.290288 | 0.037095244 | 0.107370636 | 0.145380488 | 0.77938354 | ns | TM | KS.3 |
| FIM1_2847 | 1817.72973 | 2175.121506 | 3194.810321 | 0.011551161 | 0.094822545 | 0.549667238 | 0.95917717 | ns | TM | KS.3 |
| FIM1_2848 | 508.0444171 | 527.4863937 | 706.3170098 | 0.007883966 | 0.060593074 | 0.732364472 | 0.987157164 | ns | TM | KS.3 |
| FIM1_2849 | 14207.84705 | 15342.45458 | 14759.95724 | 0.021962368 | 0.061134468 | 0.370953277 | 0.948637087 | ns | TM | KS.3 |
| FIM1_2850 | 1848.414535 | 2259.417969 | 2126.038905 | 0.060742753 | 0.095454194 | 0.067859682 | 0.571238443 | ns | TM | KS.3 |
| FIM1_2851 | 2374.115079 | 2334.708171 | 2482.05434 | -0.002930304 | 0.065210315 | 0.897516218 | 0.991141351 | ns | TM | KS.3 |
| FIM1_2852 | 934.4479085 | 905.4311124 | 869.8070042 | -0.006901992 | 0.060509815 | 0.763991713 | 0.988477371 | ns | TM | KS.3 |
| FIM1_2853 | 2754.052886 | 3126.111201 | 4109.292691 | 0.01267468 | 0.082964409 | 0.549361146 | 0.95917717 | ns | TM | KS.3 |
| FIM1_2854 | 1514.992448 | 1816.929348 | 1743.33323 | 0.048394184 | 0.088184851 | 0.110655747 | 0.71059097 | ns | TM | KS.3 |
| FIM1_2855 | 145.9389218 | 123.0622189 | 121.4704813 | -0.025446002 | 0.083589514 | 0.293600781 | 0.923185001 | ns | TM | KS.3 |
| FIM1_2856 | 609.051129 | 513.7997183 | 634.690912 | -0.034986511 | 0.08310645 | 0.194725892 | 0.837304507 | ns | TM | KS.3 |
| FIM1_2857 | 632.9651195 | 755.4487307 | 751.6658429 | 0.042136911 | 0.085540278 | 0.143182383 | 0.77783242 | ns | TM | KS.3 |
| FIM1_2858 | 840.8199004 | 965.9309419 | 1021.325247 | 0.045785951 | 0.078872998 | 0.128431815 | 0.753101833 | ns | TM | KS.3 |
| FIM1_2859 | 4245.680544 | 4451.091766 | 3942.214549 | 0.009208052 | 0.064197891 | 0.691434501 | 0.981188557 | ns | TM | KS.3 |
| FIM1_2860 | 7872.678888 | 7559.152343 | 11384.25993 | -0.003818344 | 0.080228315 | 0.851704512 | 0.991141351 | ns | TM | KS.3 |
| FIM1_2862 | 222.7938474 | 261.8576019 | 207.0002085 | 0.027318909 | 0.081147259 | 0.275745269 | 0.908583046 | ns | TM | KS.3 |
| FIM1_2863 | 990.9004222 | 1385.081973 | 1263.715346 | 0.075215575 | 0.133606576 | 0.03499012 | 0.42652037 | ns | TM | KS.3 |
| FIM1_2864 | 2703.788661 | 3009.390035 | 3071.00938 | 0.028376554 | 0.069449884 | 0.27836704 | 0.909905404 | ns | TM | KS.3 |
| FIM1_2865 | 1024.86491 | 1162.241158 | 1040.611865 | 0.026119461 | 0.074487103 | 0.305626339 | 0.926747341 | ns | TM | KS.3 |
| FIM1_2866 | 645.4138386 | 596.6269082 | 682.8797352 | -0.017897499 | 0.065511945 | 0.460951768 | 0.950410534 | ns | TM | KS.3 |
| FIM1_2867 | 1387.606944 | 1558.370474 | 1333.668172 | 0.029942032 | 0.071672179 | 0.258639925 | 0.895871714 | ns | TM | KS.3 |
| FIM1_2868 | 35.2501793 | 38.83920307 | 53.04285322 | 0.008124739 | 0.083548381 | 0.688523901 | 0.980879562 | ns | TM | KS.3 |
| FIM1_2869 | 110.9194393 | 86.81085985 | 263.2725796 | -0.043326768 | 0.097966009 | 0.121566167 | 0.734862308 | ns | TM | KS.3 |
| FIM1_2870 | 4807.641265 | 4672.609142 | 4619.041052 | -0.005616149 | 0.06257106 | 0.806716346 | 0.988916276 | ns | TM | KS.3 |
| FIM1_2871 | 961.0180944 | 929.4721959 | 1028.851863 | -0.009138682 | 0.05237886 | 0.678875813 | 0.97986479 | ns | TM | KS.3 |
| FIM1_2872 | 1218.42382 | 1381.066935 | 3341.988258 | 0.031999027 | 0.073958941 | 0.234506337 | 0.883800031 | ns | TM | KS.3 |
| FIM1_2873 | 262.3848374 | 266.62165 | 379.4106986 | 0.002450347 | 0.067723336 | 0.913388194 | 0.991141351 | ns | TM | KS.3 |

| | | | | | | | | | | |
|---|---|---|---|---|---|---|---|---|---|---|
| FIM1_2874 | 3792.377455 | 3746.030391 | 5304.524092 | -0.002354199 | 0.062475304 | 0.917853628 | 0.991141351 | ns | TM | KS.3 |
| FIM1_2875 | 1892.296542 | 1820.310145 | 2510.602835 | -0.012374655 | 0.047049618 | 0.5503385 | 0.95917717 | ns | TM | KS.3 |
| FIM1_2876 | 2058.517595 | 2132.391566 | 2282.743622 | 0.009808181 | 0.051096879 | 0.652939261 | 0.976323032 | ns | TM | KS.3 |
| FIM1_2877 | 2580.406346 | 3093.829595 | 3588.738224 | 0.019956759 | 0.086762989 | 0.371042984 | 0.948637087 | ns | TM | KS.3 |
| FIM1_2878 | 1217.857087 | 1021.7342 | 1245.67938 | -0.065467639 | 0.091358131 | 0.05958986 | 0.543962484 | ns | TM | KS.3 |
| FIM1_2879 | 1019.368332 | 868.5355676 | 1558.259213 | -0.039009623 | 0.081876803 | 0.166918611 | 0.80502665 | ns | TM | KS.3 |
| FIM1_2880 | 1014.301683 | 969.2279653 | 1078.416442 | -0.010144347 | 0.060889424 | 0.662447016 | 0.976323032 | ns | TM | KS.3 |
| FIM1_2881 | 1255.121582 | 1187.320909 | 1233.208042 | -0.009661483 | 0.06765851 | 0.67527559 | 0.978019231 | ns | TM | KS.3 |
| FIM1_2882 | 1311.139396 | 1102.690948 | 1469.445907 | -0.059462685 | 0.089318442 | 0.074027884 | 0.597271765 | ns | TM | KS.3 |
| FIM1_2883 | 2276.145998 | 2111.649807 | 2329.138013 | -0.027288558 | 0.058034277 | 0.266006792 | 0.904370522 | ns | TM | KS.3 |
| FIM1_2884 | 630.8202743 | 626.0948985 | 690.2018904 | -0.002036831 | 0.050630397 | 0.924721758 | 0.993271864 | ns | TM | KS.3 |
| FIM1_2885 | 1290.394686 | 1476.752183 | 1656.05392 | 0.011002531 | 0.087192498 | 0.585295452 | 0.963781856 | ns | TM | KS.3 |
| FIM1_2886 | 620.5493504 | 782.1585848 | 931.8734118 | 0.018530036 | 0.09579406 | 0.37505149 | 0.948637087 | ns | TM | KS.3 |
| FIM1_2887 | 3435.071033 | 3992.900774 | 4640.995791 | 0.016081393 | 0.083893557 | 0.459947984 | 0.950410534 | ns | TM | KS.3 |
| FIM1_2888 | 465.593952 | 392.9983152 | 389.7551072 | -0.038391258 | 0.083964416 | 0.168224247 | 0.805573568 | ns | TM | KS.3 |
| FIM1_2889 | 2102.429105 | 2035.760852 | 2031.263372 | -0.009638507 | 0.047620067 | 0.644427925 | 0.976323032 | ns | TM | KS.3 |
| FIM1_2890 | 5231.223338 | 4743.393793 | 4347.024313 | -0.033847516 | 0.066398747 | 0.209046441 | 0.852551729 | ns | TM | KS.3 |
| FIM1_2891 | 3195.988729 | 2916.493152 | 4319.216258 | -0.039559744 | 0.06447339 | 0.150099948 | 0.783447458 | ns | TM | KS.3 |
| FIM1_2892 | 211.4955203 | 179.0956242 | 304.0822981 | -0.041067713 | 0.08416326 | 0.151036998 | 0.783447458 | ns | TM | KS.3 |
| FIM1_2893 | 403.4464758 | 503.5556906 | 642.7355356 | 0.07134855 | 0.102469762 | 0.045821473 | 0.490240488 | ns | TM | KS.3 |
| FIM1_2894 | 799.3020679 | 765.1224985 | 1040.38808 | -0.01040993 | 0.058009056 | 0.651318021 | 0.976323032 | ns | TM | KS.3 |
| FIM1_2895 | 3338.418541 | 3286.076943 | 3125.276002 | -0.00431895 | 0.049813084 | 0.840030287 | 0.991141351 | ns | TM | KS.3 |
| FIM1_2896 | 1160.291822 | 1224.321403 | 1441.378662 | 0.017425426 | 0.050933186 | 0.428332919 | 0.950410534 | ns | TM | KS.3 |
| FIM1_2897 | 4426.351629 | 3602.686491 | 5776.34956 | -0.044758852 | 0.091643689 | 0.122377208 | 0.736186951 | ns | TM | KS.3 |
| FIM1_2898 | 16590.91494 | 16917.01646 | 16191.39412 | 0.005091284 | 0.051797458 | 0.815680242 | 0.988916276 | ns | TM | KS.3 |
| FIM1_2899 | 6091.635342 | 6520.386473 | 9340.909001 | 0.010323077 | 0.071423261 | 0.64963767 | 0.976323032 | ns | TM | KS.3 |
| FIM1_2900 | 2774.06037 | 3119.065067 | 6646.927741 | 0.026522542 | 0.072381186 | 0.302684121 | 0.926536521 | ns | TM | KS.3 |
| FIM1_2901 | 1631.610981 | 1553.778176 | 1677.89953 | -0.013695785 | 0.054745387 | 0.547423881 | 0.95917717 | ns | TM | KS.3 |
| FIM1_2903 | 2094.88478 | 2011.33075 | 2443.122967 | -0.009787778 | 0.057516033 | 0.669590791 | 0.976323032 | ns | TM | KS.3 |
| FIM1_2904 | 1332.567267 | 1308.587046 | 1819.270302 | -0.002979478 | 0.066807129 | 0.895275232 | 0.991141351 | ns | TM | KS.3 |
| FIM1_2905 | 677.5911529 | 752.7980036 | 846.2092377 | 0.008887723 | 0.084805299 | 0.659453542 | 0.976323032 | ns | TM | KS.3 |
| FIM1_2906 | 161.252271 | 187.1250835 | 192.2999731 | 0.023013414 | 0.081340822 | 0.335257126 | 0.938982061 | ns | TM | KS.3 |
| FIM1_2907 | 903.9728616 | 1120.129937 | 990.5456322 | 0.041405398 | 0.092261031 | 0.137990582 | 0.770139466 | ns | TM | KS.3 |
| FIM1_2908 | 2296.46954 | 2561.213192 | 2481.130868 | 0.039114006 | 0.070120543 | 0.166827681 | 0.80502665 | ns | TM | KS.3 |
| FIM1_2909 | 1080.482448 | 1028.5824 | 1242.605634 | -0.01428706 | 0.053926569 | 0.52786432 | 0.959071703 | ns | TM | KS.3 |
| FIM1_2911 | 463.8617485 | 461.491811 | 424.6130829 | -0.000547617 | 0.077392814 | 0.979067604 | 0.999157637 | ns | TM | KS.3 |
| FIM1_2912 | 1912.146906 | 2144.015775 | 2982.181995 | 0.012079833 | 0.081196847 | 0.57167147 | 0.962265923 | ns | TM | KS.3 |
| FIM1_2913 | 7.930695373 | 9.435934258 | 14.35777334 | 0.009949361 | 0.105463893 | 0.581889752 | 0.963781856 | ns | TM | KS.3 |
| FIM1_2914 | 1563.887802 | 2043.056935 | 1880.422927 | 0.024405783 | 0.099372202 | 0.272004934 | 0.908468501 | ns | TM | KS.3 |
| FIM1_2915 | 2354.020664 | 2226.376035 | 1489.707832 | -0.010815771 | 0.065117496 | 0.64304118 | 0.976237372 | ns | TM | KS.3 |
| FIM1_2916 | 6269.337087 | 6728.410038 | 5132.835234 | 0.015116118 | 0.064815981 | 0.52661614 | 0.959071703 | ns | TM | KS.3 |
| FIM1_2917 | 1198.043099 | 1185.850694 | 1446.996132 | -0.003075138 | 0.046914248 | 0.881592942 | 0.991141351 | ns | TM | KS.3 |
| FIM1_2918 | 664.6778742 | 736.6924894 | 985.6773065 | 0.029838472 | 0.06740645 | 0.257718077 | 0.895871714 | ns | TM | KS.3 |
| FIM1_2919 | 3203.182741 | 2952.049475 | 3139.017352 | -0.02610118 | 0.061324853 | 0.298083766 | 0.926536521 | ns | TM | KS.3 |
| FIM1_2920 | 262.9865041 | 338.9905459 | 333.924461 | 0.036538814 | 0.098198396 | 0.158602645 | 0.789048157 | ns | TM | KS.3 |

| | | | | | | | | | | |
|---|---|---|---|---|---|---|---|---|---|---|
| FIM1_2921 | 2409.757265 | 2609.158704 | 2861.91268 | 0.008032861 | 0.080022239 | 0.700807074 | 0.98151979 | ns | TM | KS.3 |
| FIM1_2922 | 1105.36625 | 991.1900503 | 940.8518398 | -0.022109363 | 0.071625182 | 0.373082586 | 0.948637087 | ns | TM | KS.3 |
| FIM1_2923 | 3941.454557 | 3775.790355 | 3964.760454 | -0.011055821 | 0.05556266 | 0.626988777 | 0.97344368 | ns | TM | KS.3 |
| FIM1_2924 | 6506.644998 | 6156.927377 | 6416.49765 | -0.014021647 | 0.058342596 | 0.548302311 | 0.95917717 | ns | TM | KS.3 |
| FIM1_2925 | 37.68249379 | 38.86949186 | 55.21287527 | 0.002234281 | 0.088737926 | 0.905496815 | 0.991141351 | ns | TM | KS.3 |
| FIM1_2926 | 2340.867041 | 2294.927122 | 2247.740019 | -0.003352952 | 0.06641961 | 0.882480572 | 0.991141351 | ns | TM | KS.3 |
| FIM1_2927 | 6130.187912 | 6169.765597 | 5522.650867 | 0.002052083 | 0.033618871 | 0.896553915 | 0.991141351 | ns | TM | KS.3 |
| FIM1_2928 | 576.7866641 | 481.0498717 | 1364.35551 | -0.018772016 | 0.087333942 | 0.391874639 | 0.948637087 | ns | TM | KS.3 |
| FIM1_2929 | 1637.353657 | 1444.845829 | 1704.738101 | -0.018284111 | 0.077323936 | 0.432847949 | 0.950410534 | ns | TM | KS.3 |
| FIM1_2930 | 258.3694767 | 333.728797 | 314.4429203 | 0.022621878 | 0.098002739 | 0.299820428 | 0.926536521 | ns | TM | KS.3 |
| FIM1_2931 | 2701.307816 | 2712.630138 | 3084.728023 | 0.000902719 | 0.059094913 | 0.968304077 | 0.996327965 | ns | TM | KS.3 |
| FIM1_2932 | 981.8239462 | 940.6021075 | 921.3139034 | -0.009897273 | 0.059532372 | 0.668796518 | 0.976323032 | ns | TM | KS.3 |
| FIM1_2933 | 2725.228204 | 2556.387812 | 2741.375958 | -0.022133289 | 0.054047013 | 0.337999685 | 0.942012706 | ns | TM | KS.3 |
| FIM1_2934 | 2502.461831 | 2828.646417 | 3436.724809 | 0.023585954 | 0.074332509 | 0.343828919 | 0.944808611 | ns | TM | KS.3 |
| FIM1_2935 | 15729.75009 | 13867.19558 | 14252.74012 | -0.028541593 | 0.074136324 | 0.274077346 | 0.908468501 | ns | TM | KS.3 |
| FIM1_2936 | 3453.118158 | 3850.351722 | 4841.660457 | 0.02019544 | 0.072555252 | 0.406703434 | 0.950410534 | ns | TM | KS.3 |
| FIM1_2937 | 829.8565687 | 817.9002377 | 929.7319652 | -0.00236354 | 0.067685887 | 0.916449754 | 0.991141351 | ns | TM | KS.3 |
| FIM1_2939 | 544.664574 | 513.2089343 | 579.7654092 | -0.011199028 | 0.066643694 | 0.630961614 | 0.97344368 | ns | TM | KS.3 |
| FIM1_2941 | 1453.948656 | 1666.005843 | 1537.092147 | 0.040872413 | 0.077188277 | 0.157993062 | 0.789048157 | ns | TM | KS.3 |
| FIM1_2942 | 10870.32988 | 10573.06746 | 7572.433439 | -0.006792782 | 0.055257559 | 0.762637033 | 0.988477371 | ns | TM | KS.3 |
| FIM1_2943 | 1212.509926 | 1200.474192 | 1210.279347 | -0.002858558 | 0.048323517 | 0.891825422 | 0.991141351 | ns | TM | KS.3 |
| FIM1_2944 | 833.5738737 | 940.568393 | 950.1685742 | 0.030203498 | 0.072667952 | 0.255192677 | 0.892600401 | ns | TM | KS.3 |
| FIM1_2945 | 745.9180288 | 822.9786708 | 787.5940644 | 0.016883008 | 0.072111716 | 0.476633981 | 0.950410534 | ns | TM | KS.3 |
| FIM1_2946 | 2121.270677 | 1845.474211 | 1978.107633 | -0.071090497 | 0.083186902 | 0.048130084 | 0.499686002 | ns | TM | KS.3 |
| FIM1_2947 | 3838.749442 | 3848.690926 | 3679.15989 | 0.000611434 | 0.052696048 | 0.977782497 | 0.999036302 | ns | TM | KS.3 |
| FIM1_2948 | 1361.985555 | 1225.819601 | 1174.689626 | -0.032101632 | 0.068676909 | 0.232074205 | 0.880810476 | ns | TM | KS.3 |
| FIM1_2949 | 3733.639093 | 3669.442794 | 3327.628688 | -0.003928895 | 0.05746794 | 0.862304132 | 0.991141351 | ns | TM | KS.3 |
| FIM1_2950 | 52.38906102 | 73.94382462 | 91.11159101 | 0.062890039 | 0.130446268 | 0.051436928 | 0.515921463 | ns | TM | KS.3 |
| FIM1_2951 | 2543.895557 | 2322.894113 | 2696.25655 | -0.02451055 | 0.065380957 | 0.333042996 | 0.938982061 | ns | TM | KS.3 |
| FIM1_2952 | 976.5104988 | 857.7318969 | 991.1695776 | -0.043583533 | 0.076396785 | 0.14069026 | 0.775632767 | ns | TM | KS.3 |
| FIM1_2953 | 1867.292182 | 1648.125383 | 1696.349832 | -0.023675022 | 0.074801385 | 0.341495571 | 0.944808611 | ns | TM | KS.3 |
| FIM1_2954 | 694.6987124 | 611.0150315 | 669.8722657 | -0.07298743 | 0.079468801 | 0.04279668 | 0.473141074 | ns | TM | KS.3 |
| FIM1_2955 | 1216.291764 | 1193.563376 | 1135.911101 | -0.003823978 | 0.061849412 | 0.867211538 | 0.991141351 | ns | TM | KS.3 |
| FIM1_2956 | 376.0190643 | 346.2622207 | 356.0336572 | -0.015117159 | 0.069771954 | 0.522558827 | 0.958602567 | ns | TM | KS.3 |
| FIM1_2957 | 2903.032028 | 3200.024534 | 3906.709487 | 0.011049915 | 0.078800844 | 0.608983003 | 0.971365963 | ns | TM | KS.3 |
| FIM1_2958 | 2047.380746 | 1753.095446 | 2248.188588 | -0.057074704 | 0.084916715 | 0.082007605 | 0.621932675 | ns | TM | KS.3 |
| FIM1_2959 | 3562.259451 | 3622.770804 | 4410.205535 | 0.002345107 | 0.071189631 | 0.915368935 | 0.991141351 | ns | TM | KS.3 |
| FIM1_2960 | 2263.356034 | 2623.217554 | 3342.8738 | 0.019226502 | 0.081193264 | 0.401959865 | 0.950410534 | ns | TM | KS.3 |
| FIM1_2961 | 3457.146236 | 3556.493213 | 4279.333789 | 0.00453509 | 0.068077627 | 0.84084247 | 0.991141351 | ns | TM | KS.3 |
| FIM1_2962 | 269.2617587 | 260.2246649 | 401.0509813 | -0.007975089 | 0.057348993 | 0.726628649 | 0.987157164 | ns | TM | KS.3 |
| FIM1_2963 | 368.3086382 | 412.7228117 | 393.8680425 | 0.040819376 | 0.071756372 | 0.155923145 | 0.78780435 | ns | TM | KS.3 |
| FIM1_2964 | 7704.255751 | 7529.83065 | 7234.558437 | -0.005423708 | 0.056110693 | 0.809958586 | 0.988916276 | ns | TM | KS.3 |
| FIM1_2965 | 1326.445617 | 1347.877026 | 1402.398664 | 0.003906036 | 0.052538786 | 0.858835409 | 0.991141351 | ns | TM | KS.3 |
| FIM1_2966 | 310.5522455 | 368.4539987 | 368.3976288 | 0.037134368 | 0.083297785 | 0.17801224 | 0.811391014 | ns | TM | KS.3 |
| FIM1_2967 | 1550.464518 | 1445.900867 | 1546.006294 | -0.02727618 | 0.055498049 | 0.25266034 | 0.889363822 | ns | TM | KS.3 |

| | | | | | | | | | | |
|---|---|---|---|---|---|---|---|---|---|---|
| FIM1_2968 | 2418.671464 | 2110.899763 | 1890.686952 | -0.071021891 | 0.082355879 | 0.047890415 | 0.499686002 | ns | TM | KS.3 |
| FIM1_2969 | 6361.259871 | 6037.932871 | 5685.816099 | -0.011677713 | 0.061151522 | 0.617563383 | 0.973122675 | ns | TM | KS.3 |
| FIM1_2970 | 557.3029585 | 548.2049985 | 760.6778271 | -0.004558472 | 0.050374265 | 0.832306773 | 0.991141351 | ns | TM | KS.3 |
| FIM1_2971 | 525.1566054 | 599.7566352 | 637.3109641 | 0.017019685 | 0.079643447 | 0.452624694 | 0.950410534 | ns | TM | KS.3 |
| FIM1_2972 | 571.5032958 | 656.5605806 | 731.7172261 | 0.035735265 | 0.077061462 | 0.196848026 | 0.83848507 | ns | TM | KS.3 |
| FIM1_2973 | 174.8999093 | 315.3600275 | 901.5278038 | 0.636365585 | 0.32592622 | 8.2860641457 | 0.007777956 | ns | TM | KS.3 |
| FIM1_2974 | 2111.245568 | 1853.814878 | 1991.706768 | -0.022619839 | 0.076218739 | 0.355439634 | 0.948637087 | ns | TM | KS.3 |
| FIM1_2975 | 2174.612236 | 2376.587517 | 2788.527927 | 0.01697884 | 0.069310909 | 0.479230186 | 0.950972051 | ns | TM | KS.3 |
| FIM1_2976 | 647.9040684 | 556.1382624 | 629.7327363 | -0.052964147 | 0.083340491 | 0.096045081 | 0.673713046 | ns | TM | KS.3 |
| FIM1_2977 | 1004.334952 | 856.7990928 | 839.6745868 | -0.042622459 | 0.082383339 | 0.143976353 | 0.77783242 | ns | TM | KS.3 |
| FIM1_2978 | 2924.856612 | 3040.260333 | 2827.370299 | 0.008491884 | 0.059993934 | 0.712736922 | 0.986183429 | ns | TM | KS.3 |
| FIM1_2979 | 209.1285015 | 179.9593361 | 219.4068092 | -0.030014131 | 0.078991097 | 0.249191956 | 0.888934767 | ns | TM | KS.3 |
| FIM1_2980 | 965.9317754 | 874.0324272 | 926.1346837 | -0.023255514 | 0.068694773 | 0.355710228 | 0.948637087 | ns | TM | KS.3 |
| FIM1_2981 | 3859.022502 | 4265.442187 | 5131.577162 | 0.012516044 | 0.077310686 | 0.571637829 | 0.962265923 | ns | TM | KS.3 |
| FIM1_2982 | 2923.062289 | 3405.822516 | 3129.856232 | 0.070591315 | 0.087029133 | 0.050079207 | 0.512642091 | ns | TM | KS.3 |
| FIM1_2983 | 5667.989546 | 6304.53994 | 6118.843466 | 0.020407773 | 0.071806497 | 0.403970109 | 0.950410534 | ns | TM | KS.3 |
| FIM1_2984 | 581.4169452 | 704.9390523 | 607.7647513 | 0.020819051 | 0.088243891 | 0.35146067 | 0.948637087 | ns | TM | KS.3 |
| FIM1_2985 | 16711.47998 | 19747.98661 | 15548.43375 | 0.015228738 | 0.087637031 | 0.468506411 | 0.950410534 | ns | TM | KS.3 |
| FIM1_2986 | 933.8166665 | 1008.928177 | 930.6546089 | 0.0294926 | 0.059219162 | 0.238801551 | 0.886981413 | ns | TM | KS.3 |
| FIM1_2987 | 1906.913432 | 1861.44348 | 1963.239033 | -0.007469848 | 0.043543786 | 0.702883141 | 0.98151979 | ns | TM | KS.3 |
| FIM1_2988 | 420.8576862 | 919.4483023 | 931.4687696 | 1.087316411 | 0.143805909 | 1.1836939117 | 1.1777754421 | Km>KS.3 | TM | KS.3 |
| FIM1_2989 | 831.862684 | 990.9848389 | 992.0718388 | 0.035432939 | 0.084013864 | 0.189586548 | 0.828217874 | ns | TM | KS.3 |
| FIM1_2990 | 3023.562647 | 4418.264172 | 12583.20332 | 0.525285151 | 0.092940711 | 3.7953948475 | 1.8882089366 | ns | TM | KS.3 |
| FIM1_2991 | 1554.50773 | 622.2434439 | 416.6918034 | -1.217991074 | 0.21284983 | 2.0970390369 | 6.5204807556 | Km<KS.3 | TM | KS.3 |
| FIM1_2992 | 144.4635823 | 86.7298273 | 85.80505748 | -0.189818621 | 0.271081169 | 0.004605316 | 0.134937815 | ns | TM | KS.3 |
| FIM1_2993 | 1238.507061 | 1040.166441 | 1039.204649 | -0.146492541 | 0.088003393 | 0.00349316 | 0.110690892 | ns | TM | KS.3 |
| FIM1_2994 | 1108.016284 | 983.0999485 | 988.7851955 | -0.037535215 | 0.073063167 | 0.182590993 | 0.820587346 | ns | TM | KS.3 |
| FIM1_2995 | 340.3336032 | 282.5996247 | 250.7650778 | -0.022642401 | 0.086379443 | 0.32661438 | 0.934739762 | ns | TM | KS.3 |
| FIM1_2996 | 541.2857518 | 459.5378972 | 702.7419252 | -0.012853623 | 0.08988362 | 0.523876785 | 0.958883577 | ns | TM | KS.3 |
| FIM1_2997 | 1536.128361 | 1467.954419 | 1624.884602 | -0.012705582 | 0.053445091 | 0.571398019 | 0.962265923 | ns | TM | KS.3 |
| FIM1_2998 | 4943.252011 | 4239.165896 | 3567.38224 | -0.060223457 | 0.085298222 | 0.072939206 | 0.594873028 | ns | TM | KS.3 |
| FIM1_2999 | 1113.481297 | 985.1184951 | 807.5264055 | -0.030165758 | 0.073238759 | 0.255310627 | 0.892600401 | ns | TM | KS.3 |
| FIM1_3000 | 1070.132873 | 1107.21288 | 1759.153242 | 0.007167409 | 0.060735284 | 0.755460967 | 0.988477371 | ns | TM | KS.3 |
| FIM1_3001 | 1152.861343 | 1253.23663 | 1124.216239 | 0.014247404 | 0.070560899 | 0.542982077 | 0.95917717 | ns | TM | KS.3 |
| FIM1_3002 | 543.3593501 | 450.962846 | 406.515444 | -0.065144607 | 0.093532957 | 0.059523861 | 0.543962484 | ns | TM | KS.3 |
| FIM1_3003 | 934.017103 | 1150.295973 | 974.3824504 | 0.026437851 | 0.089450413 | 0.2659699 | 0.904370522 | ns | TM | KS.3 |
| FIM1_3004 | 5654.178517 | 5784.53949 | 5703.476767 | 0.006968187 | 0.043977358 | 0.724064781 | 0.987157164 | ns | TM | KS.3 |
| FIM1_3005 | 1866.960074 | 2151.180101 | 2134.188148 | 0.013458033 | 0.084937662 | 0.521828516 | 0.958602567 | ns | TM | KS.3 |
| FIM1_3006 | 3923.473865 | 4072.411074 | 4840.496114 | 0.008309727 | 0.059526679 | 0.718147124 | 0.987157164 | ns | TM | KS.3 |
| FIM1_3007 | 3364.609377 | 3692.178131 | 3697.877283 | 0.028544216 | 0.065067724 | 0.271121509 | 0.908468501 | ns | TM | KS.3 |
| FIM1_3008 | 2301.582392 | 2187.036294 | 2183.93518 | -0.017996514 | 0.048309039 | 0.394925598 | 0.948637087 | ns | TM | KS.3 |
| FIM1_3009 | 1825.936061 | 1694.530473 | 1567.872644 | -0.028023452 | 0.057422967 | 0.251601628 | 0.888934767 | ns | TM | KS.3 |
| FIM1_3010 | 624.9077272 | 716.4330827 | 653.7230729 | 0.049867903 | 0.079320761 | 0.108660837 | 0.705336441 | ns | TM | KS.3 |
| FIM1_3011 | 441.9465534 | 466.5544649 | 622.8877379 | 0.010282816 | 0.064991243 | 0.658764312 | 0.976323032 | ns | TM | KS.3 |
| FIM1_3012 | 2321.382097 | 2313.161894 | 2505.524487 | -0.001005283 | 0.045793189 | 0.960549827 | 0.995175461 | ns | TM | KS.3 |

| | | | | | | | | | |
|---|---|---|---|---|---|---|---|---|---|
| FIM1_3013 | 2657.893701 | 2456.671178 | 2299.118373 | -0.028746814 | 0.059244521 | 0.249546806 | 0.888934767 ns | TM | KS.3 |
| FIM1_3014 | 4135.617318 | 4584.914247 | 6063.64154 | 0.010460712 | 0.081175274 | 0.619526415 | 0.973122675 ns | TM | KS.3 |
| FIM1_3015 | 5474.550879 | 5278.157339 | 4634.54022 | -0.009648717 | 0.053681684 | 0.665982271 | 0.976323032 ns | TM | KS.3 |
| FIM1_3016 | 685.6137754 | 686.5170758 | 759.6981876 | 0.00045109 | 0.05938339 | 0.984167647 | 0.999333464 ns | TM | KS.3 |
| FIM1_3017 | 20250.27977 | 22180.77529 | 21330.23418 | 0.028244992 | 0.064467022 | 0.274060381 | 0.908468501 ns | TM | KS.3 |
| FIM1_3018 | 499.719851 | 590.478752 | 535.4983651 | 0.035060924 | 0.082482416 | 0.195207424 | 0.837467988 ns | TM | KS.3 |
| FIM1_3019 | 5832.696085 | 5887.906422 | 4633.715476 | 0.002480267 | 0.051008811 | 0.908752727 | 0.991141351 ns | TM | KS.3 |
| FIM1_3020 | 34.10099075 | 71.57043739 | 96.90294568 | 0.735180594 | 0.417147061 | 0.0001282 | 0.010455632 ns | TM | KS.3 |
| FIM1_3021 | 3252.710376 | 3082.435209 | 3834.209133 | -0.016007322 | 0.054081148 | 0.481903014 | 0.951416554 ns | TM | KS.3 |
| FIM1_3022 | 185.5047046 | 175.1298162 | 240.3611277 | -0.004672807 | 0.084006413 | 0.813182543 | 0.988916276 ns | TM | KS.3 |
| FIM1_3023 | 680.1473382 | 537.1267562 | 752.7178282 | -0.035847205 | 0.094847983 | 0.167933068 | 0.805573568 ns | TM | KS.3 |
| FIM1_3024 | 49.75761521 | 33.60273847 | 74.4569741 | -0.022116147 | 0.118404253 | 0.280056799 | 0.913627918 ns | TM | KS.3 |
| FIM1_3025 | 883.7988573 | 925.0116695 | 931.7120277 | 0.006478806 | 0.071634767 | 0.770988026 | 0.988814136 ns | TM | KS.3 |
| FIM1_3026 | 2261.153149 | 2023.24723 | 2497.866158 | -0.020812872 | 0.072748853 | 0.394679408 | 0.948637087 ns | TM | KS.3 |
| FIM1_3028 | 4088.234788 | 3926.899206 | 4185.317805 | -0.009968284 | 0.056371221 | 0.662035408 | 0.976323032 ns | TM | KS.3 |
| FIM1_3029 | 1583.225672 | 1533.500226 | 1373.966389 | -0.008181405 | 0.054290404 | 0.71489637 | 0.986942381 ns | TM | KS.3 |
| FIM1_3030 | 4571.03482 | 4554.440345 | 3644.827954 | -0.001104999 | 0.035020593 | 0.946309534 | 0.994413583 ns | TM | KS.3 |
| FIM1_3031 | 1501.128789 | 1354.306426 | 1581.656361 | -0.029860405 | 0.068187037 | 0.258327945 | 0.895871714 ns | TM | KS.3 |
| FIM1_3032 | 2969.449527 | 2913.554509 | 2951.006234 | -0.003643138 | 0.062652261 | 0.873397254 | 0.991141351 ns | TM | KS.3 |
| FIM1_3033 | 1493.254145 | 1366.33018 | 1273.167036 | -0.024412172 | 0.064535063 | 0.333525261 | 0.938982061 ns | TM | KS.3 |
| FIM1_3034 | 3826.158313 | 3096.674958 | 2931.69041 | -0.021439769 | 0.091104541 | 0.333436604 | 0.938982061 ns | TM | KS.3 |
| FIM1_3035 | 12724.85911 | 12202.86178 | 11142.20316 | -0.00831767 | 0.063252539 | 0.719206007 | 0.987157164 ns | TM | KS.3 |
| FIM1_3036 | 2660.312072 | 2714.315082 | 3404.866561 | 0.004504384 | 0.057917139 | 0.842782624 | 0.991141351 ns | TM | KS.3 |
| FIM1_3037 | 1164.233947 | 1415.564729 | 1021.631874 | 0.016330747 | 0.091522606 | 0.431833339 | 0.950410534 ns | TM | KS.3 |
| FIM1_3038 | 17470.89922 | 18312.15424 | 17148.25544 | 0.010910472 | 0.0595386 | 0.638374269 | 0.975246356 ns | TM | KS.3 |
| FIM1_3039 | 2481.844738 | 2882.711709 | 2869.090948 | 0.049399218 | 0.081993952 | 0.110571685 | 0.71059097 ns | TM | KS.3 |
| FIM1_3040 | 2078.258131 | 2449.952319 | 2794.51789 | 0.014290819 | 0.088200443 | 0.490217807 | 0.952591289 ns | TM | KS.3 |
| FIM1_3041 | 1739.226644 | 1711.823792 | 1740.792045 | -0.004329079 | 0.046521928 | 0.833059865 | 0.991141351 ns | TM | KS.3 |
| FIM1_3043 | 130.8755427 | 164.2114676 | 207.6025605 | 0.038016898 | 0.093643782 | 0.155466109 | 0.786819827 ns | TM | KS.3 |
| FIM1_3044 | 423.7561643 | 353.2133124 | 4677.982151 | -0.013064968 | 0.092871141 | 0.5103795 | 0.953260948 ns | TM | KS.3 |
| FIM1_3045 | 653.6812936 | 474.0943032 | 549.2223036 | -0.084478689 | 0.134510998 | 0.026812964 | 0.382098641 ns | TM | KS.3 |
| FIM1_3046 | 1582.249691 | 1551.301389 | 1642.342519 | -0.003650158 | 0.063417474 | 0.873088481 | 0.991141351 ns | TM | KS.3 |
| FIM1_3047 | 774.6370099 | 854.6830078 | 825.3510037 | 0.036351236 | 0.066487332 | 0.184000534 | 0.821725903 ns | TM | KS.3 |
| FIM1_3048 | 2896.212858 | 3045.103835 | 3099.753428 | 0.009348765 | 0.065578777 | 0.686534809 | 0.980879562 ns | TM | KS.3 |
| FIM1_3049 | 1112.81985 | 1138.61784 | 883.089918 | 0.005537185 | 0.054087351 | 0.803630109 | 0.988916276 ns | TM | KS.3 |
| FIM1_3050 | 723.124289 | 848.2000269 | 792.229839 | 0.071889117 | 0.088759508 | 0.047974841 | 0.499686002 ns | TM | KS.3 |
| FIM1_3051 | 1425.040292 | 1062.489078 | 1014.141862 | -0.172619682 | 0.145314995 | 0.002895876 | 0.100748141 ns | TM | KS.3 |
| FIM1_3052 | 910.8174372 | 831.4090841 | 796.9924135 | -0.022316473 | 0.066403558 | 0.372054851 | 0.948637087 ns | TM | KS.3 |
| FIM1_3053 | 2406.847793 | 2100.847631 | 2143.005932 | -0.035516632 | 0.076473546 | 0.199173189 | 0.842123734 ns | TM | KS.3 |
| FIM1_3054 | 643.3647876 | 880.518168 | 1456.614948 | 0.036397548 | 0.108863287 | 0.148458273 | 0.781566041 ns | TM | KS.3 |
| FIM1_3056 | 3562.823312 | 3904.173086 | 4038.32144 | 0.011353841 | 0.076813078 | 0.605988484 | 0.970937835 ns | TM | KS.3 |
| FIM1_3057 | 566.6814401 | 544.5959429 | 483.5889983 | -0.004608605 | 0.075910899 | 0.829049192 | 0.991141351 ns | TM | KS.3 |
| FIM1_3058 | 3203.963328 | 2702.667497 | 2914.185425 | -0.057741514 | 0.088304877 | 0.079074594 | 0.616697988 ns | TM | KS.3 |
| FIM1_3059 | 892.6975767 | 888.4774427 | 849.149499 | -0.001368704 | 0.048872631 | 0.948400533 | 0.994413583 ns | TM | KS.3 |
| FIM1_3060 | 700.8453643 | 716.4606075 | 717.4541639 | 0.005352155 | 0.054847899 | 0.810995185 | 0.988916276 ns | TM | KS.3 |

| | | | | | | | | | | |
|---|---|---|---|---|---|---|---|---|---|---|
| FIM1_3061 | 4938.723984 | 4343.715008 | 3894.558609 | -0.055369614 | 0.078067138 | 0.086074865 | 0.642805774 | ns | TM | KS.3 |
| FIM1_3062 | 1782.199119 | 1868.896068 | 1637.020043 | 0.007978557 | 0.067873056 | 0.72718156 | 0.987157164 | ns | TM | KS.3 |
| FIM1_3063 | 1251.18447 | 1160.455726 | 1105.968508 | -0.02901259 | 0.058110756 | 0.24039647 | 0.887413624 | ns | TM | KS.3 |
| FIM1_3064 | 5313.462287 | 5855.313037 | 6172.113327 | 0.035908759 | 0.066252094 | 0.187680782 | 0.828217874 | ns | TM | KS.3 |
| FIM1_3065 | 2557.217932 | 2530.218545 | 2380.210882 | -0.002955721 | 0.047179991 | 0.886556247 | 0.991141351 | ns | TM | KS.3 |
| FIM1_3066 | 5656.620428 | 5618.684886 | 4964.727518 | -0.001900864 | 0.047308723 | 0.927009075 | 0.993697171 | ns | TM | KS.3 |
| FIM1_3067 | 624.0429559 | 602.9091463 | 690.525503 | -0.008489262 | 0.057819592 | 0.710730967 | 0.984652342 | ns | TM | KS.3 |
| FIM1_3068 | 5142.489004 | 4898.263164 | 5347.153281 | -0.007563673 | 0.069667951 | 0.738246648 | 0.988051177 | ns | TM | KS.3 |
| FIM1_3069 | 1029.521964 | 1158.329978 | 1141.648876 | 0.022562005 | 0.073681841 | 0.361895791 | 0.948637087 | ns | TM | KS.3 |
| FIM1_3070 | 6989.505363 | 6936.225536 | 7452.029009 | -0.002086439 | 0.048217648 | 0.920815141 | 0.991141351 | ns | TM | KS.3 |
| FIM1_3071 | 4039.305599 | 3746.806071 | 3315.501244 | -0.019346903 | 0.062297062 | 0.426697451 | 0.950410534 | ns | TM | KS.3 |
| FIM1_3072 | 2394.663562 | 2093.662082 | 1546.247319 | -0.026058155 | 0.076261331 | 0.303288269 | 0.926536521 | ns | TM | KS.3 |
| FIM1_3074 | 1005.463907 | 924.570422 | 871.4180666 | -0.025681317 | 0.062424718 | 0.308095973 | 0.927416591 | ns | TM | KS.3 |
| FIM1_3075 | 928.561818 | 1080.649072 | 1321.472421 | 0.034951513 | 0.079786812 | 0.200368803 | 0.845272486 | ns | TM | KS.3 |
| FIM1_3076 | 918.6567475 | 922.2673812 | 1017.158479 | 0.00067529 | 0.045400913 | 0.973342312 | 0.998607635 | ns | TM | KS.3 |
| FIM1_3077 | 2003.4013 | 1819.143147 | 2215.349619 | -0.026704909 | 0.066536714 | 0.299857495 | 0.926536521 | ns | TM | KS.3 |
| FIM1_3078 | 859.1386937 | 819.1660845 | 790.6237241 | -0.014016646 | 0.05204993 | 0.527550695 | 0.959071703 | ns | TM | KS.3 |
| FIM1_3079 | 1942.344232 | 2186.614344 | 2045.88673 | 0.027804972 | 0.072421698 | 0.285186536 | 0.919509407 | ns | TM | KS.3 |
| FIM1_3080 | 1200.505233 | 1208.765495 | 1517.297272 | 0.001939561 | 0.049382491 | 0.927358211 | 0.993697171 | ns | TM | KS.3 |
| FIM1_3081 | 1252.716419 | 1243.499678 | 1776.031229 | -0.001627938 | 0.060270291 | 0.943031548 | 0.994413583 | ns | TM | KS.3 |
| FIM1_3082 | 1659.097053 | 1558.598232 | 1874.539366 | -0.021421657 | 0.053601599 | 0.350333447 | 0.948637087 | ns | TM | KS.3 |
| FIM1_3083 | 3361.108493 | 4086.185897 | 5074.020734 | 0.020689572 | 0.088748206 | 0.352330139 | 0.948637087 | ns | TM | KS.3 |
| FIM1_3085 | 28.18335076 | 24.03515771 | 43.4414597 | -0.011758154 | 0.088639009 | 0.558637537 | 0.960263397 | ns | TM | KS.3 |
| FIM1_3086 | 609.0856223 | 495.7447908 | 1403.542008 | -0.03875286 | 0.090065163 | 0.156341443 | 0.788842474 | ns | TM | KS.3 |
| FIM1_3087 | 5442.128039 | 5579.276005 | 7192.444108 | 0.003746382 | 0.069514656 | 0.866991872 | 0.991141351 | ns | TM | KS.3 |
| FIM1_3088 | 514.9494339 | 496.2021506 | 879.435673 | -0.004391159 | 0.075270753 | 0.837806811 | 0.991141351 | ns | TM | KS.3 |
| FIM1_3089 | 518.4570783 | 497.8228876 | 526.3408947 | -0.006689907 | 0.068834852 | 0.767748141 | 0.988477371 | ns | TM | KS.3 |
| FIM1_3090 | 320.1573323 | 241.5740592 | 396.6058563 | -0.133931505 | 0.136503499 | 0.007366406 | 0.185090242 | ns | TM | KS.3 |
| FIM1_3092 | 282.5604073 | 278.1235418 | 500.4249974 | -0.002656517 | 0.066733258 | 0.906552768 | 0.991141351 | ns | TM | KS.3 |
| FIM1_3098 | 1936.09604 | 2027.776632 | 1200.264754 | 0.007450424 | 0.068919597 | 0.742874598 | 0.988477371 | ns | TM | KS.3 |
| FIM1_3099 | 2249.688912 | 2014.92638 | 2350.021231 | -0.010339229 | 0.08304089 | 0.617699921 | 0.973122675 | ns | TM | KS.3 |
| FIM1_3100 | 930.953823 | 897.4874263 | 982.5762384 | -0.007685532 | 0.061053562 | 0.738902962 | 0.988051177 | ns | TM | KS.3 |
| FIM1_3101 | 1014.851991 | 1038.141616 | 1052.947167 | 0.004754379 | 0.059252263 | 0.834947115 | 0.991141351 | ns | TM | KS.3 |
| FIM1_3102 | 16827.97439 | 17862.13669 | 16236.58602 | 0.014024038 | 0.061096764 | 0.552680317 | 0.95917717 | ns | TM | KS.3 |
| FIM1_3103 | 142.8259509 | 138.1283721 | 162.3837864 | -0.002543913 | 0.08411204 | 0.896745315 | 0.991141351 | ns | TM | KS.3 |
| FIM1_3104 | 2064.004569 | 1857.078779 | 1971.434572 | -0.031857062 | 0.068837209 | 0.235028631 | 0.883800031 | ns | TM | KS.3 |
| FIM1_3105 | 15.8803414 | 12.66188458 | 14.7225111 | -0.010529584 | 0.09975959 | 0.571947588 | 0.962265923 | ns | TM | KS.3 |
| FIM1_3106 | 1604.492477 | 1457.178736 | 1352.376741 | -0.030432721 | 0.065930129 | 0.247806569 | 0.888934767 | ns | TM | KS.3 |
| FIM1_3107 | 1164.51179 | 574.4236485 | 1220.887682 | -0.101626785 | 0.252614629 | 0.020094467 | 0.324535175 | ns | TM | KS.3 |
| FIM1_3108 | 2274.107194 | 1330.84331 | 2240.122344 | -0.727215884 | 0.164355112 | 2.1582505543[illegible] | 3.9767764844[illegible] | ns | TM | KS.3 |
| FIM1_3109 | 2738.081319 | 3350.721379 | 3375.384685 | 0.037151225 | 0.08907727 | 0.168279993 | 0.805573568 | ns | TM | KS.3 |
| FIM1_3110 | 774.6554011 | 599.9312981 | 4126.047925 | -0.014091017 | 0.103754799 | 0.460791721 | 0.950410534 | ns | TM | KS.3 |
| FIM1_3111 | 566.6814401 | 394.9951114 | 1294.786463 | -0.042835423 | 0.120696129 | 0.108069194 | 0.703722825 | ns | TM | KS.3 |
| FIM1_3112 | 831.2823363 | 882.7606353 | 703.4952814 | 0.009908629 | 0.069353907 | 0.665622503 | 0.976323032 | ns | TM | KS.3 |
| FIM1_3113 | 1452.688896 | 1770.482417 | 1621.420472 | 0.044085683 | 0.09006257 | 0.12744226 | 0.75107959 | ns | TM | KS.3 |

| | | | | | | | | | | |
|---|---|---|---|---|---|---|---|---|---|---|
| FIM1_3114 | 1728.712712 | 1513.007296 | 1719.038682 | -0.028377087 | 0.075706928 | 0.273987111 | 0.908468501 | ns | TM | KS.3 |
| FIM1_3115 | 313.206875 | 259.0413603 | 260.7413305 | -0.043937177 | 0.088113162 | 0.1305844 | 0.761283488 | ns | TM | KS.3 |
| FIM1_3116 | 1825.241481 | 1462.940468 | 1724.004441 | -0.038986184 | 0.092924124 | 0.150621238 | 0.783447458 | ns | TM | KS.3 |
| FIM1_3117 | 31.02928939 | 18.86164103 | 19.37440892 | -0.069848201 | 0.17242956 | 0.040416464 | 0.458022575 | ns | TM | KS.3 |
| FIM1_3118 | 3860.983314 | 4980.911154 | 4526.624002 | 0.064096414 | 0.108069872 | 0.054985244 | 0.531428718 | ns | TM | KS.3 |
| FIM1_3119 | 1041.298786 | 946.276528 | 946.5580616 | -0.016527258 | 0.071784408 | 0.485404022 | 0.952269291 | ns | TM | KS.3 |
| FIM1_3120 | 3009.450829 | 2822.036822 | 2859.777166 | -0.021628382 | 0.054749849 | 0.352475307 | 0.948637087 | ns | TM | KS.3 |
| FIM1_3121 | 1219.689278 | 1025.463489 | 1170.818666 | -0.091648865 | 0.094745862 | 0.024808379 | 0.373642194 | ns | TM | KS.3 |
| FIM1_3122 | 2548.019427 | 2718.199702 | 1970.926694 | 0.013440186 | 0.064566341 | 0.569938473 | 0.962265923 | ns | TM | KS.3 |
| FIM1_3123 | 3027.5511 | 2780.365095 | 2384.973826 | -0.027965767 | 0.062251035 | 0.272255969 | 0.908468501 | ns | TM | KS.3 |
| FIM1_3124 | 3871.493114 | 3723.178886 | 4368.791076 | -0.016118014 | 0.03717413 | 0.324641489 | 0.934066145 | ns | TM | KS.3 |
| FIM1_3125 | 546.2450546 | 514.8116516 | 648.7975289 | -0.014680868 | 0.060746325 | 0.534990368 | 0.959071703 | ns | TM | KS.3 |
| FIM1_3126 | 2446.044132 | 2120.674934 | 2225.153781 | -0.024997013 | 0.078134778 | 0.314085238 | 0.927564309 | ns | TM | KS.3 |
| FIM1_3127 | 2064.370928 | 1945.719537 | 1641.556946 | -0.00952082 | 0.069570116 | 0.677019036 | 0.97940381 | ns | TM | KS.3 |
| FIM1_3128 | 13040.1316 | 12531.3658 | 9997.73968 | -0.01300541 | 0.046011517 | 0.523108339 | 0.958732165 | ns | TM | KS.3 |
| FIM1_3129 | 4144.178854 | 3773.136901 | 3927.025455 | -0.041127655 | 0.065448655 | 0.140783194 | 0.775632767 | ns | TM | KS.3 |
| FIM1_3130 | 928.8959883 | 1276.103934 | 1143.616367 | 0.102085916 | 0.141170354 | 0.016607084 | 0.299348715 | ns | TM | KS.3 |
| FIM1_3131 | 1063.722155 | 1008.084068 | 1039.568712 | -0.012209257 | 0.061405261 | 0.602639411 | 0.970245665 | ns | TM | KS.3 |
| FIM1_3132 | 3002.121233 | 3253.306878 | 3367.068562 | 0.031564003 | 0.059596101 | 0.213014893 | 0.860185951 | ns | TM | KS.3 |
| FIM1_3133 | 429.1232354 | 408.5081792 | 594.2050343 | -0.009251762 | 0.066257262 | 0.689031065 | 0.980879562 | ns | TM | KS.3 |
| FIM1_3134 | 9193.05849 | 10464.10505 | 11415.25866 | 0.021768477 | 0.076449289 | 0.369271476 | 0.948637087 | ns | TM | KS.3 |
| FIM1_3135 | 1219.548722 | 1175.454102 | 1117.0395 | -0.010192312 | 0.05242267 | 0.644992744 | 0.976323032 | ns | TM | KS.3 |
| FIM1_3136 | 3675.014353 | 3892.081105 | 4630.98272 | 0.010059474 | 0.067609891 | 0.663381492 | 0.976323032 | ns | TM | KS.3 |
| FIM1_3137 | 11.36560659 | 25.08990854 | 24.44135902 | 0.080668916 | 0.239446279 | 0.0301484 | 0.400453344 | ns | TM | KS.3 |
| FIM1_3138 | 4148.509907 | 5399.397535 | 6167.117837 | 0.034453663 | 0.099320615 | 0.171584954 | 0.810669655 | ns | TM | KS.3 |
| FIM1_3139 | 421.7621216 | 962.7721762 | 667.88201 | 1.111817654 | 0.195759266 | 3.20833582198 | 8.86748373021 | Km>KS.3 | TM | KS.3 |
| FIM1_3140 | 511.7433151 | 458.3879372 | 459.0135077 | -0.027728625 | 0.069940913 | 0.287172759 | 0.919993167 | ns | TM | KS.3 |
| FIM1_3141 | 1050.239058 | 1272.954413 | 972.2596684 | 0.066209475 | 0.095139323 | 0.056873128 | 0.535878428 | ns | TM | KS.3 |
| FIM1_3142 | 186.7716266 | 177.9422796 | 264.7518978 | -0.007031508 | 0.072093021 | 0.751864752 | 0.988477371 | ns | TM | KS.3 |
| FIM1_3143 | 5030.667632 | 5525.666147 | 5562.129101 | 0.04364018 | 0.065685066 | 0.123860051 | 0.739893493 | ns | TM | KS.3 |
| FIM1_3144 | 389.9690707 | 427.608811 | 358.3171778 | 0.016434136 | 0.070474642 | 0.48998241 | 0.952591289 | ns | TM | KS.3 |
| FIM1_3145 | 1254.3123 | 1407.301738 | 1300.969336 | 0.036289941 | 0.071458538 | 0.192392119 | 0.832245711 | ns | TM | KS.3 |
| FIM1_3146 | 614.1612343 | 568.9100193 | 695.8146255 | -0.02190425 | 0.060723348 | 0.371005203 | 0.948637087 | ns | TM | KS.3 |
| FIM1_3147 | 967.677595 | 962.3260951 | 1134.359183 | -0.001261573 | 0.056943193 | 0.955387785 | 0.994413583 | ns | TM | KS.3 |
| FIM1_3148 | 1111.64303 | 1098.964284 | 864.5807955 | -0.002882652 | 0.050569098 | 0.89359061 | 0.991141351 | ns | TM | KS.3 |
| FIM1_3149 | 1258.081625 | 1136.660444 | 1048.570728 | -0.026862962 | 0.068233783 | 0.29890575 | 0.926536521 | ns | TM | KS.3 |
| FIM1_3150 | 3823.013342 | 4354.070768 | 5060.087817 | 0.014346769 | 0.081659904 | 0.509243315 | 0.953260948 | ns | TM | KS.3 |
| FIM1_3151 | 546.9742608 | 553.0154475 | 787.651183 | 0.001983951 | 0.061268304 | 0.930697609 | 0.993993515 | ns | TM | KS.3 |
| FIM1_3152 | 248.8491561 | 226.3883297 | 255.1933084 | -0.020839425 | 0.068809453 | 0.399264941 | 0.950410534 | ns | TM | KS.3 |
| FIM1_3153 | 844.7291584 | 735.1226301 | 769.2644039 | -0.052160533 | 0.080414387 | 0.099212638 | 0.677856366 | ns | TM | KS.3 |
| FIM1_3154 | 2041.970939 | 2318.991582 | 2146.446937 | 0.022106551 | 0.075818693 | 0.365026038 | 0.948637087 | ns | TM | KS.3 |
| FIM1_3155 | 2215.661141 | 2041.857963 | 2866.78671 | -0.020474098 | 0.064095435 | 0.40562598 | 0.950410534 | ns | TM | KS.3 |
| FIM1_3156 | 4596.813947 | 3757.068545 | 3973.487865 | -0.091210247 | 0.102692747 | 0.024993412 | 0.373642194 | ns | TM | KS.3 |
| FIM1_3157 | 2514.166558 | 2040.012453 | 1292.04226 | -0.026286611 | 0.089564672 | 0.267569393 | 0.90534159 | ns | TM | KS.3 |
| FIM1_3158 | 289.5774828 | 294.1960682 | 368.4361109 | 0.003624104 | 0.060023868 | 0.873804344 | 0.991141351 | ns | TM | KS.3 |

| | | | | | | | | | | |
|---|---|---|---|---|---|---|---|---|---|---|
| FIM1_3159 | 450.1234261 | 379.9006461 | 404.0752316 | -0.04979734 | 0.086365449 | 0.106606553 | 0.700394017 | ns | TM | KS.3 |
| FIM1_3160 | 308.0198654 | 301.6231361 | 291.6823713 | -0.001798816 | 0.081438465 | 0.928615056 | 0.993697171 | ns | TM | KS.3 |
| FIM1_3161 | 429.2796109 | 488.8790138 | 1401.267782 | 0.013052414 | 0.082984735 | 0.538931997 | 0.95917717 | ns | TM | KS.3 |
| FIM1_3162 | 2207.386825 | 1606.339737 | 2181.720415 | -0.040418667 | 0.111138034 | 0.124183129 | 0.739893493 | ns | TM | KS.3 |
| FIM1_3163 | 876.8038178 | 914.5650151 | 715.8975727 | 0.006749265 | 0.068396722 | 0.766280462 | 0.988477371 | ns | TM | KS.3 |
| FIM1_3164 | 2608.930485 | 2381.440016 | 2822.476065 | -0.079357581 | 0.062393723 | 0.019836375 | 0.324535175 | ns | TM | KS.3 |
| FIM1_3165 | 771.122063 | 814.1852505 | 793.2031306 | 0.011529064 | 0.062564076 | 0.622602909 | 0.973122675 | ns | TM | KS.3 |
| FIM1_3166 | 1220.547147 | 1148.198128 | 1168.677337 | -0.014441528 | 0.061299458 | 0.541911986 | 0.95917717 | ns | TM | KS.3 |
| FIM1_3167 | 2240.18847 | 2820.441452 | 3498.114776 | 0.02251151 | 0.09382065 | 0.309974914 | 0.927416591 | ns | TM | KS.3 |
| FIM1_3168 | 1094.635717 | 1249.220273 | 1110.963659 | 0.052233937 | 0.078366062 | 0.098259136 | 0.677062609 | ns | TM | KS.3 |
| FIM1_3169 | 943.5770034 | 1136.333781 | 1077.659817 | 0.050022924 | 0.089306319 | 0.103321637 | 0.689041746 | ns | TM | KS.3 |
| FIM1_3170 | 60.92270592 | 46.16537184 | 121.4952025 | -0.033198423 | 0.101194133 | 0.178681252 | 0.811815714 | ns | TM | KS.3 |
| FIM1_3171 | 1608.566311 | 1761.471999 | 1716.990805 | 0.021605973 | 0.066547116 | 0.385228185 | 0.948637087 | ns | TM | KS.3 |
| FIM1_3172 | 1844.523107 | 1825.113584 | 1513.119302 | -0.001604361 | 0.06897865 | 0.942803134 | 0.994413583 | ns | TM | KS.3 |
| FIM1_3173 | 1260.561308 | 1177.042176 | 1674.368048 | -0.015830183 | 0.062832244 | 0.508162388 | 0.953260948 | ns | TM | KS.3 |
| FIM1_3174 | 452.7631742 | 489.5383624 | 492.5185698 | 0.011147297 | 0.073135496 | 0.62173638 | 0.973122675 | ns | TM | KS.3 |
| FIM1_3175 | 1440.456613 | 1412.368739 | 1395.309944 | -0.004711352 | 0.055754776 | 0.834037894 | 0.991141351 | ns | TM | KS.3 |
| FIM1_3176 | 300.698376 | 355.7725804 | 464.5748983 | 0.052644252 | 0.086536695 | 0.095926873 | 0.673713046 | ns | TM | KS.3 |
| FIM1_3177 | 2052.072592 | 2166.420214 | 2248.049981 | 0.021071044 | 0.047895468 | 0.315872687 | 0.927564309 | ns | TM | KS.3 |
| FIM1_3178 | 2113.283846 | 2041.304904 | 2050.784747 | -0.00739772 | 0.060807581 | 0.748049419 | 0.988477371 | ns | TM | KS.3 |
| FIM1_3179 | 377.9487778 | 275.477949 | 479.2921944 | -0.053614591 | 0.117067775 | 0.073598821 | 0.597271765 | ns | TM | KS.3 |
| FIM1_3180 | 1388.507248 | 1189.132985 | 1588.096631 | -0.046066473 | 0.082326154 | 0.12579772 | 0.745051976 | ns | TM | KS.3 |
| FIM1_3181 | 2230.120611 | 2621.317485 | 3024.698306 | 0.01737337 | 0.084891301 | 0.428021475 | 0.950410534 | ns | TM | KS.3 |
| FIM1_3182 | 1772.018437 | 1841.418157 | 1881.758192 | 0.009693608 | 0.055781145 | 0.669432648 | 0.976323032 | ns | TM | KS.3 |
| FIM1_3183 | 5939.219603 | 5608.061802 | 6908.476124 | -0.012536781 | 0.062482712 | 0.594140605 | 0.965889818 | ns | TM | KS.3 |
| FIM1_3184 | 738.3043 | 718.6653875 | 690.2581567 | -0.003702606 | 0.071639422 | 0.866680335 | 0.991141351 | ns | TM | KS.3 |
| FIM1_3185 | 801.691547 | 1060.442654 | 1068.012224 | 0.052463715 | 0.108728376 | 0.080302458 | 0.618428374 | ns | TM | KS.3 |
| FIM1_3186 | 4463.33012 | 4662.652081 | 3548.73503 | 0.011148146 | 0.056133016 | 0.625532457 | 0.97344368 | ns | TM | KS.3 |
| FIM1_3187 | 829.4955858 | 711.3611918 | 663.5778738 | -0.040663452 | 0.080907621 | 0.157139576 | 0.789034023 | ns | TM | KS.3 |
| FIM1_3188 | 736.4812536 | 694.0666419 | 690.8064976 | -0.014830799 | 0.059701776 | 0.529461202 | 0.959071703 | ns | TM | KS.3 |
| FIM1_3189 | 433.7417665 | 470.2231356 | 411.7370225 | 0.014335653 | 0.06940298 | 0.542669597 | 0.95917717 | ns | TM | KS.3 |
| FIM1_3190 | 618.4431472 | 663.7865768 | 680.2043265 | 0.016609557 | 0.063280678 | 0.489655892 | 0.952591289 | ns | TM | KS.3 |
| FIM1_3191 | 879.9366439 | 947.3182832 | 883.2861558 | 0.013110355 | 0.068767921 | 0.57555309 | 0.963192994 | ns | TM | KS.3 |
| FIM1_3192 | 742.5741005 | 684.6613014 | 729.3971384 | -0.024225097 | 0.062190866 | 0.332005599 | 0.938982061 | ns | TM | KS.3 |
| FIM1_3193 | 33.32415675 | 29.6162117 | 43.75386283 | -0.009918751 | 0.084612818 | 0.626365703 | 0.97344368 | ns | TM | KS.3 |
| FIM1_3194 | 2343.632831 | 2202.391093 | 2395.124519 | -0.009608136 | 0.070589267 | 0.672692508 | 0.976861203 | ns | TM | KS.3 |
| FIM1_3195 | 2427.44451 | 2807.493018 | 3200.949999 | 0.01923059 | 0.080781054 | 0.403179812 | 0.950410534 | ns | TM | KS.3 |
| FIM1_3196 | 516.1281023 | 479.1988574 | 477.2816158 | -0.014864731 | 0.066571514 | 0.532355122 | 0.959071703 | ns | TM | KS.3 |
| FIM1_3197 | 447.4822141 | 528.8858679 | 1536.617768 | 0.016072862 | 0.086861054 | 0.450863327 | 0.950410534 | ns | TM | KS.3 |
| FIM1_3198 | 449.6214069 | 515.5635145 | 460.0031329 | 0.008430048 | 0.093284163 | 0.655449189 | 0.976323032 | ns | TM | KS.3 |
| FIM1_3199 | 2354.496718 | 2085.518014 | 2359.023598 | -0.031412438 | 0.072910582 | 0.241341108 | 0.887413624 | ns | TM | KS.3 |
| FIM1_3200 | 10041.90028 | 9223.961598 | 9302.489641 | -0.065461269 | 0.061485825 | 0.03545647 | 0.42652037 | ns | TM | KS.3 |
| FIM1_3201 | 445.3880437 | 606.8034225 | 501.8279529 | 0.128616714 | 0.147191532 | 0.00858258 | 0.201407251 | ns | TM | KS.3 |
| FIM1_3202 | 474.2340445 | 520.2316874 | 534.8867205 | 0.026326162 | 0.066532188 | 0.305533364 | 0.926747341 | ns | TM | KS.3 |
| FIM1_3203 | 1221.815155 | 1230.23425 | 1449.933299 | 0.001720215 | 0.055979238 | 0.93892462 | 0.994413583 | ns | TM | KS.3 |

| | | | | | | | | | | |
|---|---|---|---|---|---|---|---|---|---|---|
| FIM1_3204 | 811.7347418 | 799.7086313 | 744.7715961 | -0.004793976 | 0.035526152 | 0.771646825 | 0.988814136 | ns | TM | KS.3 |
| FIM1_3205 | 5174.065934 | 4673.824434 | 4777.253445 | -0.038439644 | 0.067831574 | 0.16828327 | 0.805573568 | ns | TM | KS.3 |
| FIM1_3206 | 1983.771788 | 1782.38612 | 1815.968829 | -0.041041019 | 0.0697881 | 0.151451546 | 0.783447458 | ns | TM | KS.3 |
| FIM1_3207 | 973.4110645 | 986.1846586 | 891.1447567 | 0.002080958 | 0.067966392 | 0.926284858 | 0.993697171 | ns | TM | KS.3 |
| FIM1_3208 | 4381.859152 | 3895.997743 | 3247.202603 | -0.05153168 | 0.074388046 | 0.097984602 | 0.677062609 | ns | TM | KS.3 |
| FIM1_3209 | 7974.687498 | 7823.848061 | 7110.124517 | -0.004688569 | 0.054482723 | 0.833524155 | 0.991141351 | ns | TM | KS.3 |
| FIM1_3210 | 921.0838291 | 1163.608534 | 590.3318852 | 0.073649476 | 0.106739956 | 0.041421052 | 0.464121025 | ns | TM | KS.3 |
| FIM1_3211 | 556.5042062 | 507.2128146 | 437.1034821 | -0.016837863 | 0.070831711 | 0.480113299 | 0.950972051 | ns | TM | KS.3 |
| FIM1_3212 | 3284.284142 | 3225.844405 | 3159.226961 | -0.004745961 | 0.051346038 | 0.827206199 | 0.991141351 | ns | TM | KS.3 |
| FIM1_3213 | 3701.812702 | 3575.301245 | 3801.11775 | -0.010234784 | 0.048934621 | 0.630341572 | 0.97344368 | ns | TM | KS.3 |
| FIM1_3214 | 2341.91119 | 3177.954767 | 3474.386174 | 0.022912704 | 0.105660505 | 0.283259158 | 0.918450395 | ns | TM | KS.3 |
| FIM1_3215 | 2012.474746 | 1987.573399 | 2625.623474 | -0.003747195 | 0.041550389 | 0.843140248 | 0.991141351 | ns | TM | KS.3 |
| FIM1_3216 | 74.45762419 | 85.41059831 | 80.9259995 | 0.015045559 | 0.081103644 | 0.493671496 | 0.952949531 | ns | TM | KS.3 |
| FIM1_3217 | 1986.044692 | 1905.126602 | 2185.918002 | -0.004838139 | 0.075747745 | 0.821082435 | 0.990886949 | ns | TM | KS.3 |
| FIM1_3218 | 376.0898535 | 323.8991084 | 239.287081 | -0.014851505 | 0.084891333 | 0.486156533 | 0.952269291 | ns | TM | KS.3 |
| FIM1_3219 | 4272.141596 | 3485.619983 | 3158.916771 | -0.05147927 | 0.093200526 | 0.09471917 | 0.673663379 | ns | TM | KS.3 |
| FIM1_3220 | 717.3879551 | 847.3505607 | 2141.848212 | 0.025559464 | 0.082526221 | 0.294914169 | 0.92509331 | ns | TM | KS.3 |
| FIM1_3221 | 6818.393938 | 6592.839196 | 6394.062365 | -0.009624963 | 0.049766284 | 0.654061217 | 0.976323032 | ns | TM | KS.3 |
| FIM1_3222 | 3966.070601 | 3816.055859 | 3571.471653 | -0.014696667 | 0.038146439 | 0.386605369 | 0.948637087 | ns | TM | KS.3 |
| FIM1_3223 | 4638.926007 | 4793.568155 | 5186.784433 | 0.010573402 | 0.0436952 | 0.589184249 | 0.963781856 | ns | TM | KS.3 |
| FIM1_3224 | 2052.939488 | 1793.040842 | 1877.403515 | -0.014719897 | 0.082294791 | 0.497792956 | 0.953260948 | ns | TM | KS.3 |
| FIM1_3225 | 146.0715367 | 142.4916197 | 117.7906859 | -0.002185404 | 0.079906835 | 0.914684804 | 0.991141351 | ns | TM | KS.3 |
| FIM1_3226 | 943.6965458 | 877.9703579 | 829.6867632 | -0.017432349 | 0.062455047 | 0.469569213 | 0.950410534 | ns | TM | KS.3 |
| FIM1_3227 | 2174.585503 | 2232.376701 | 2360.123084 | 0.006284003 | 0.055689897 | 0.780321513 | 0.988814136 | ns | TM | KS.3 |
| FIM1_3228 | 1913.373038 | 2005.333145 | 1687.076866 | 0.01211931 | 0.056296582 | 0.597142517 | 0.967682091 | ns | TM | KS.3 |
| FIM1_3229 | 3977.544795 | 3882.692655 | 3769.560011 | -0.007268199 | 0.044977843 | 0.717191159 | 0.987157164 | ns | TM | KS.3 |
| FIM1_3230 | 4883.523174 | 4538.266818 | 4397.562342 | -0.023007833 | 0.058817369 | 0.34358894 | 0.944808611 | ns | TM | KS.3 |
| FIM1_3231 | 4852.67104 | 4665.732627 | 4952.197279 | -0.014079149 | 0.041657212 | 0.449951427 | 0.950410534 | ns | TM | KS.3 |
| FIM1_3232 | 88.48898873 | 119.3593752 | 257.3779081 | 0.043689369 | 0.108063612 | 0.111118011 | 0.71059097 | ns | TM | KS.3 |
| FIM1_3233 | 3064.172012 | 2846.251124 | 1744.519258 | -0.021393696 | 0.060266593 | 0.37984531 | 0.948637087 | ns | TM | KS.3 |
| FIM1_3234 | 2653.10028 | 3353.669991 | 4010.201847 | 0.022211316 | 0.09455324 | 0.313032434 | 0.927416591 | ns | TM | KS.3 |
| FIM1_3235 | 1650.281636 | 1549.582387 | 1430.647497 | -0.018403681 | 0.05665063 | 0.432808432 | 0.950410534 | ns | TM | KS.3 |
| FIM1_3236 | 7660.300479 | 7999.741117 | 9277.944695 | 0.005340484 | 0.07454079 | 0.805269907 | 0.988916276 | ns | TM | KS.3 |
| FIM1_3237 | 5633.44683 | 5193.291786 | 5182.316218 | -0.021361648 | 0.063372829 | 0.387132381 | 0.948637087 | ns | TM | KS.3 |
| FIM1_3238 | 514.1136624 | 487.7315498 | 470.235462 | -0.013722753 | 0.057643395 | 0.554906292 | 0.95917717 | ns | TM | KS.3 |
| FIM1_3239 | 2699.964284 | 2462.063864 | 3239.1185 | -0.030826779 | 0.064523615 | 0.239698434 | 0.887413624 | ns | TM | KS.3 |
| FIM1_3240 | 2368.993518 | 2320.21393 | 2728.842769 | -0.004790571 | 0.057202883 | 0.832523185 | 0.991141351 | ns | TM | KS.3 |
| FIM1_3241 | 1477.505914 | 1186.784725 | 1207.718789 | -0.046283228 | 0.094527879 | 0.112503851 | 0.711904568 | ns | TM | KS.3 |
| FIM1_3242 | 658.0342166 | 675.2595639 | 739.1983989 | 0.006129111 | 0.056922642 | 0.786933125 | 0.988916276 | ns | TM | KS.3 |
| FIM1_3243 | 2267.326025 | 2338.249311 | 2177.611347 | 0.007477373 | 0.055567397 | 0.740355138 | 0.988051177 | ns | TM | KS.3 |
| FIM1_3244 | 1806.906751 | 1583.233171 | 1295.076085 | -0.019344343 | 0.078121179 | 0.409158111 | 0.950410534 | ns | TM | KS.3 |
| FIM1_3245 | 1364.704876 | 1307.391602 | 1235.652559 | -0.009557104 | 0.059862853 | 0.679504133 | 0.97986479 | ns | TM | KS.3 |
| FIM1_3246 | 760.0012716 | 731.8857257 | 1352.307553 | -0.00681244 | 0.065458258 | 0.766581584 | 0.988477371 | ns | TM | KS.3 |
| FIM1_3247 | 5553.782094 | 5150.01178 | 5932.375847 | -0.007986334 | 0.07892146 | 0.705201732 | 0.982962554 | ns | TM | KS.3 |
| FIM1_3248 | 248.8814224 | 227.2927568 | 270.241428 | -0.02057131 | 0.067839415 | 0.404927289 | 0.950410534 | ns | TM | KS.3 |

| | | | | | | | | | | |
|---|---|---|---|---|---|---|---|---|---|---|
| FIM1_3249 | 3938.007927 | 3673.5282 | 3260.562503 | -0.017169305 | 0.06182378 | 0.474897392 | 0.950410534 | ns | TM | KS.3 |
| FIM1_3250 | 1014.859492 | 926.3142867 | 864.3547259 | -0.0185847 | 0.068743569 | 0.444659057 | 0.950410534 | ns | TM | KS.3 |
| FIM1_3251 | 687.2711999 | 633.3881485 | 786.1267761 | -0.025048771 | 0.061894344 | 0.317128669 | 0.928067724 | ns | TM | KS.3 |
| FIM1_3252 | 44.69682255 | 42.60348078 | 46.84830583 | -0.002844642 | 0.085022139 | 0.883757526 | 0.991141351 | ns | TM | KS.3 |
| FIM1_3253 | 3382.56509 | 3926.59584 | 3578.117444 | 0.01570034 | 0.083972688 | 0.468585513 | 0.950410534 | ns | TM | KS.3 |
| FIM1_3254 | 4204.239542 | 4623.230913 | 4440.083237 | 0.039384622 | 0.065756591 | 0.155309506 | 0.786819827 | ns | TM | KS.3 |
| FIM1_3255 | 689.6872707 | 705.932568 | 844.2191512 | 0.006524056 | 0.043833508 | 0.740445128 | 0.988051177 | ns | TM | KS.3 |
| FIM1_3256 | 39.3644642 | 33.68548386 | 40.43753559 | -0.014900304 | 0.08907209 | 0.472394132 | 0.950410534 | ns | TM | KS.3 |
| FIM1_3258 | 414.2891065 | 374.0011435 | 304.6439041 | -0.018870764 | 0.072020117 | 0.434013923 | 0.950410534 | ns | TM | KS.3 |
| FIM1_3259 | 608.185617 | 616.4437851 | 420.9082459 | 0.001658583 | 0.072488277 | 0.939445683 | 0.994413583 | ns | TM | KS.3 |
| FIM1_3260 | 1397.240024 | 1191.22493 | 1532.399655 | -0.134366132 | 0.084162044 | 0.004943324 | 0.140896498 | ns | TM | KS.3 |
| FIM1_3261 | 561.2232272 | 615.0393517 | 583.1292843 | 0.020256334 | 0.067847193 | 0.411095876 | 0.950410534 | ns | TM | KS.3 |
| FIM1_3262 | 29775.90851 | 30461.67478 | 29873.94409 | 0.006002354 | 0.051720625 | 0.783498557 | 0.988814136 | ns | TM | KS.3 |
| FIM1_3263 | 1988.385787 | 1967.658935 | 1959.690367 | -0.002770799 | 0.04771948 | 0.894341265 | 0.991141351 | ns | TM | KS.3 |
| FIM1_3264 | 1829.515178 | 1708.361546 | 1677.565821 | -0.018920971 | 0.059378337 | 0.429800691 | 0.950410534 | ns | TM | KS.3 |
| FIM1_3265 | 1278.008412 | 1197.77391 | 1308.290788 | -0.022113727 | 0.055029943 | 0.343929326 | 0.944808611 | ns | TM | KS.3 |
| FIM1_3266 | 480.3591975 | 670.7782538 | 627.6223463 | 0.075150777 | 0.133404116 | 0.035071784 | 0.42652037 | ns | TM | KS.3 |
| FIM1_3267 | 3270.644286 | 3551.752664 | 3631.771014 | 0.026107819 | 0.061694682 | 0.299126182 | 0.926536521 | ns | TM | KS.3 |
| FIM1_3268 | 804.9006955 | 906.383848 | 861.409376 | 0.023776386 | 0.073378663 | 0.342486934 | 0.944808611 | ns | TM | KS.3 |
| FIM1_3269 | 858.0199451 | 514.3665609 | 594.8101771 | -0.597982384 | 0.253565131 | 2.90295301984 | 0.003209376 | ns | TM | KS.3 |
| FIM1_3270 | 1760.031871 | 3269.936106 | 3849.519547 | 0.121400742 | 0.254693363 | 0.013917783 | 0.268375845 | ns | TM | KS.3 |
| FIM1_3271 | 3740.7655 | 3733.626684 | 3839.313539 | -0.000461337 | 0.053955861 | 0.983394206 | 0.999333464 | ns | TM | KS.3 |
| FIM1_3272 | 1361.141182 | 1216.793307 | 1169.173293 | -0.029564596 | 0.070453112 | 0.263300639 | 0.90372202 | ns | TM | KS.3 |
| FIM1_3273 | 1921.748421 | 1978.895342 | 2005.529265 | 0.003110421 | 0.077394252 | 0.881986069 | 0.991141351 | ns | TM | KS.3 |
| FIM1_3274 | 3188.755436 | 2821.037727 | 2580.125895 | -0.141135367 | 0.053478088 | 0.000745094 | 0.039019373 | ns | TM | KS.3 |
| FIM1_3275 | 3180.070636 | 3384.882041 | 3593.207739 | 0.017384258 | 0.058043569 | 0.461675244 | 0.950410534 | ns | TM | KS.3 |
| FIM1_3276 | 442.6570124 | 398.1828467 | 476.724671 | -0.016108057 | 0.074452838 | 0.488811905 | 0.952591289 | ns | TM | KS.3 |
| FIM1_3277 | 2691.654644 | 2336.178024 | 2368.234626 | -0.065151041 | 0.083193877 | 0.060315239 | 0.544665909 | ns | TM | KS.3 |
| FIM1_3278 | 6668.314937 | 6514.468519 | 6815.733062 | -0.006821852 | 0.04631457 | 0.739105007 | 0.988051177 | ns | TM | KS.3 |
| FIM1_3279 | 3388.586453 | 3441.182647 | 3833.314864 | 0.003653227 | 0.055487142 | 0.870579752 | 0.991141351 | ns | TM | KS.3 |
| FIM1_3280 | 2411.9655 | 2285.154321 | 2105.880803 | -0.014065473 | 0.057448362 | 0.545077049 | 0.95917717 | ns | TM | KS.3 |
| FIM1_3281 | 5598.695766 | 5868.451104 | 4292.32629 | 0.010407259 | 0.060871122 | 0.654557069 | 0.976323032 | ns | TM | KS.3 |
| FIM1_3282 | 5665.031852 | 5294.692628 | 5569.056616 | -0.005495586 | 0.084129145 | 0.781696123 | 0.988814136 | ns | TM | KS.3 |
| FIM1_3283 | 720.2532226 | 688.924729 | 807.0213661 | -0.008990545 | 0.06309616 | 0.698272946 | 0.98151979 | ns | TM | KS.3 |
| FIM1_3284 | 1562.385146 | 1413.772247 | 1342.972478 | -0.022456892 | 0.069112443 | 0.369399859 | 0.948637087 | ns | TM | KS.3 |
| FIM1_3286 | 1305.395154 | 1545.611315 | 1644.190823 | 0.077123454 | 0.091998186 | 0.040110844 | 0.455596911 | ns | TM | KS.3 |
| FIM1_3287 | 388.174482 | 404.4612047 | 460.2756596 | 0.007396113 | 0.066849736 | 0.746757896 | 0.988477371 | ns | TM | KS.3 |
| FIM1_3288 | 27764.6382 | 26732.13385 | 24758.36259 | -0.005572815 | 0.070433539 | 0.803174344 | 0.988916276 | ns | TM | KS.3 |
| FIM1_3290 | 1627.948743 | 1755.80919 | 1587.834695 | 0.019804976 | 0.062092543 | 0.416627408 | 0.950410534 | ns | TM | KS.3 |
| FIM1_3291 | 10.34823019 | 9.242788839 | 18.39346275 | -0.005182946 | 0.102403504 | 0.767818146 | 0.988477371 | ns | TM | KS.3 |
| FIM1_3292 | 27539.75708 | 30016.41594 | 28547.75295 | 0.025077435 | 0.063463696 | 0.320468018 | 0.929108338 | ns | TM | KS.3 |
| FIM1_3293 | 319.8572706 | 391.1614776 | 402.3163413 | 0.040357758 | 0.089623775 | 0.147482152 | 0.780546912 | ns | TM | KS.3 |
| FIM1_3294 | 268.535339 | 258.1370023 | 335.1507035 | -0.006530056 | 0.068282072 | 0.773602624 | 0.988814136 | ns | TM | KS.3 |
| FIM1_3295 | 894.2221272 | 1015.763516 | 1289.998448 | 0.020451473 | 0.076630265 | 0.392365135 | 0.948637087 | ns | TM | KS.3 |
| FIM1_3297 | 2454.83554 | 2350.074226 | 2133.083476 | -0.0142418 | 0.047856493 | 0.497376508 | 0.953260948 | ns | TM | KS.3 |

| | | | | | | | | | | |
|---|---|---|---|---|---|---|---|---|---|---|
| FIM1_3298 | 3022.463857 | 3209.339711 | 3671.679717 | 0.007063117 | 0.076115789 | 0.742926607 | 0.988477371 | ns | TM | KS.3 |
| FIM1_3299 | 4185.909981 | 4487.477053 | 5009.030904 | 0.025138544 | 0.055955921 | 0.2912599 | 0.919993167 | ns | TM | KS.3 |
| FIM1_3300 | 12752.36317 | 13058.06907 | 12154.06664 | 0.005013243 | 0.060101807 | 0.826513956 | 0.991141351 | ns | TM | KS.3 |
| FIM1_3301 | 1898.407378 | 1710.899243 | 2184.731024 | -0.030995843 | 0.068443481 | 0.244660849 | 0.88883001 | ns | TM | KS.3 |
| FIM1_3302 | 539.1586142 | 669.3422404 | 695.5945618 | 0.016521313 | 0.094803054 | 0.419509937 | 0.950410534 | ns | TM | KS.3 |
| FIM1_3303 | 670.5521591 | 589.8546514 | 583.473781 | -0.027314121 | 0.074969672 | 0.288660117 | 0.919993167 | ns | TM | KS.3 |
| FIM1_3304 | 7841.935516 | 8860.273059 | 10610.29114 | 0.015552436 | 0.078839297 | 0.48882659 | 0.952591289 | ns | TM | KS.3 |
| FIM1_3305 | 9177.013072 | 11531.96232 | 11293.93362 | 0.116437439 | 0.113517846 | 0.011635609 | 0.24513135 | ns | TM | KS.3 |
| FIM1_3306 | 377.911498 | 420.678702 | 385.9812727 | 0.016711551 | 0.074414277 | 0.475072097 | 0.950410534 | ns | TM | KS.3 |
| FIM1_3307 | 1092.694696 | 867.9074941 | 1497.973183 | -0.076582629 | 0.106611396 | 0.03796989 | 0.44135561 | ns | TM | KS.3 |
| FIM1_3308 | 268.862733 | 332.9720637 | 1459.473884 | 0.034007912 | 0.090425452 | 0.188901196 | 0.828217874 | ns | TM | KS.3 |
| FIM1_3309 | 3861.43843 | 4025.250653 | 4207.45595 | 0.009209649 | 0.060173503 | 0.690457293 | 0.981155393 | ns | TM | KS.3 |
| FIM1_3310 | 1616.942519 | 1623.973413 | 1586.122228 | 0.001170179 | 0.048981117 | 0.95593009 | 0.994413583 | ns | TM | KS.3 |
| FIM1_3311 | 6266.225201 | 6120.709056 | 5799.004745 | -0.008060254 | 0.033281662 | 0.597109626 | 0.967682091 | ns | TM | KS.3 |
| FIM1_3312 | 1946.427133 | 2003.806217 | 3030.972899 | 0.00286824 | 0.078939725 | 0.889405989 | 0.991141351 | ns | TM | KS.3 |
| FIM1_3313 | 356.1355367 | 397.4749907 | 449.383105 | 0.00702066 | 0.091325407 | 0.71053494 | 0.984652342 | ns | TM | KS.3 |
| FIM1_3314 | 535.696388 | 542.1788538 | 611.8314272 | 0.002623748 | 0.05626346 | 0.907156375 | 0.991141351 | ns | TM | KS.3 |
| FIM1_3315 | 528.8841805 | 714.5300509 | 625.3048471 | 0.104349502 | 0.136123125 | 0.01570935 | 0.2883912 | ns | TM | KS.3 |
| FIM1_3316 | 1752.758329 | 1550.495005 | 1921.22516 | -0.061026538 | 0.077365465 | 0.067472769 | 0.568944115 | ns | TM | KS.3 |
| FIM1_3317 | 202.0798606 | 192.7655299 | 156.9880462 | -0.008659463 | 0.065557992 | 0.707759085 | 0.983273233 | ns | TM | KS.3 |
| FIM1_3318 | 762.1980174 | 845.5328631 | 902.019129 | 0.009675311 | 0.082788862 | 0.639333312 | 0.975246356 | ns | TM | KS.3 |
| FIM1_3319 | 1435.092413 | 1485.647659 | 1424.631369 | 0.007621996 | 0.059873388 | 0.740351631 | 0.988051177 | ns | TM | KS.3 |
| FIM1_3320 | 2520.275543 | 1669.408076 | 3754.133926 | -0.062305409 | 0.145258841 | 0.051144073 | 0.5150643 | ns | TM | KS.3 |
| FIM1_3321 | 1105.437881 | 1380.183754 | 2821.384153 | 0.020192765 | 0.093349104 | 0.349427815 | 0.948637087 | ns | TM | KS.3 |
| FIM1_3322 | 432.0822962 | 350.2125091 | 576.8316103 | -0.024565843 | 0.089975469 | 0.28904198 | 0.919993167 | ns | TM | KS.3 |
| FIM1_3323 | 1388.260387 | 1295.635151 | 1336.77278 | -0.017211346 | 0.061470554 | 0.473398229 | 0.950410534 | ns | TM | KS.3 |
| FIM1_3324 | 1480.227224 | 1588.597577 | 2165.389234 | 0.014364322 | 0.065847535 | 0.545424418 | 0.95917717 | ns | TM | KS.3 |
| FIM1_3326 | 10850.4471 | 9892.005129 | 10761.74779 | -0.017668049 | 0.069836309 | 0.463144502 | 0.950410534 | ns | TM | KS.3 |
| FIM1_3327 | 1608.334756 | 1188.023542 | 1547.231501 | -0.140050916 | 0.146446295 | 0.006482899 | 0.170744739 | ns | TM | KS.3 |
| FIM1_3328 | 204.5542474 | 211.685642 | 250.7530417 | 0.005462785 | 0.068784627 | 0.808696802 | 0.988916276 | ns | TM | KS.3 |
| FIM1_3329 | 196.1235166 | 199.5617561 | 333.4106703 | 0.002520667 | 0.06399424 | 0.911955963 | 0.991141351 | ns | TM | KS.3 |
| FIM1_3331 | 391.7436295 | 445.0463789 | 427.7347597 | 0.038297804 | 0.074013154 | 0.176742109 | 0.811391014 | ns | TM | KS.3 |
| FIM1_3332 | 568.5150158 | 696.992732 | 563.365356 | 0.112292937 | 0.104750738 | 0.01311847 | 0.261057546 | ns | TM | KS.3 |
| FIM1_3333 | 2885.732776 | 3498.918772 | 3974.130456 | 0.0367449 | 0.087393554 | 0.173943876 | 0.811391014 | ns | TM | KS.3 |
| FIM1_3334 | 31776.07992 | 34223.08795 | 33685.66399 | 0.020384328 | 0.061012155 | 0.402338146 | 0.950410534 | ns | TM | KS.3 |
| FIM1_3335 | 1196.395 | 1358.843724 | 1422.542747 | 0.021948023 | 0.075940364 | 0.367445257 | 0.948637087 | ns | TM | KS.3 |
| FIM1_3336 | 1089.557319 | 1166.131486 | 1146.475696 | 0.011667191 | 0.068855814 | 0.615107493 | 0.973122675 | ns | TM | KS.3 |
| FIM1_3337 | 2712.829701 | 2916.938109 | 2308.848679 | 0.0310821 | 0.056411364 | 0.203033482 | 0.847612283 | ns | TM | KS.3 |
| FIM1_3339 | 1550.693531 | 1339.170921 | 1165.07265 | -0.045567245 | 0.080450344 | 0.129237698 | 0.756420644 | ns | TM | KS.3 |
| FIM1_3340 | 11070.57568 | 9811.532753 | 5574.560663 | -0.019244485 | 0.0758196 | 0.417671035 | 0.950410534 | ns | TM | KS.3 |
| FIM1_3341 | 1385.617191 | 1266.990096 | 1121.771156 | -0.034281254 | 0.063406361 | 0.196196316 | 0.837834055 | ns | TM | KS.3 |
| FIM1_3342 | 137.2811455 | 152.9120274 | 158.9391856 | 0.01318115 | 0.077711799 | 0.552340286 | 0.95917717 | ns | TM | KS.3 |
| FIM1_3343 | 8313.745792 | 10181.61159 | 8696.620167 | 0.023744575 | 0.088942002 | 0.303183648 | 0.926536521 | ns | TM | KS.3 |
| FIM1_3344 | 22664.19357 | 23780.76097 | 18575.06962 | 0.003960026 | 0.083328849 | 0.841772094 | 0.991141351 | ns | TM | KS.3 |
| FIM1_3345 | 9795.953251 | 9653.457177 | 6481.108677 | -0.001186208 | 0.083239292 | 0.952019004 | 0.994413583 | ns | TM | KS.3 |

| | | | | | | | | | | |
|---|---|---|---|---|---|---|---|---|---|---|
| FIM1_3346 | 192.2756028 | 215.5149742 | 215.6425215 | 0.013696912 | 0.078642248 | 0.535776673 | 0.959071703 | ns | TM | KS.3 |
| FIM1_3347 | 1118.33911 | 957.7722064 | 1658.200111 | -0.028487432 | 0.079973259 | 0.264429946 | 0.90372202 | ns | TM | KS.3 |
| FIM1_3348 | 784.8357204 | 591.6550928 | 957.1650824 | -0.061792318 | 0.11350013 | 0.05709014 | 0.535913008 | ns | TM | KS.3 |
| FIM1_3349 | 1614.70027 | 1631.341482 | 1789.716452 | 0.002651098 | 0.050940083 | 0.902440866 | 0.991141351 | ns | TM | KS.3 |
| FIM1_3350 | 5059.039269 | 5968.186806 | 9211.614553 | 0.029505872 | 0.081876698 | 0.248659585 | 0.888934767 | ns | TM | KS.3 |
| FIM1_3351 | 2155.75601 | 2003.02053 | 1694.361127 | -0.021668877 | 0.059634877 | 0.372464532 | 0.948637087 | ns | TM | KS.3 |
| FIM1_3353 | 8487.133396 | 9173.007105 | 8404.574634 | 0.018288316 | 0.064281227 | 0.451860191 | 0.950410534 | ns | TM | KS.3 |
| FIM1_3355 | 1310.965827 | 1361.408438 | 1440.681107 | 0.005934792 | 0.068815175 | 0.792808022 | 0.988916276 | ns | TM | KS.3 |
| FIM1_3356 | 1299.872691 | 1171.049019 | 1282.228097 | -0.015961271 | 0.074442362 | 0.492267645 | 0.952591289 | ns | TM | KS.3 |
| FIM1_3357 | 460.3491303 | 465.5311881 | 631.5619316 | 0.002392338 | 0.062188539 | 0.916535561 | 0.991141351 | ns | TM | KS.3 |
| FIM1_3358 | 21.62552117 | 12.23573496 | 16.18493289 | -0.032264532 | 0.145247839 | 0.155977344 | 0.78780435 | ns | TM | KS.3 |
| FIM1_3359 | 4805.079849 | 4971.408593 | 5452.627362 | 0.010491979 | 0.046807346 | 0.611228671 | 0.972453674 | ns | TM | KS.3 |
| FIM1_3360 | 4821.405713 | 4286.329527 | 3710.643881 | -0.041740854 | 0.072911655 | 0.150595161 | 0.783447458 | ns | TM | KS.3 |
| FIM1_3361 | 5955.255719 | 6348.17236 | 6897.756809 | 0.023607832 | 0.053258494 | 0.303536801 | 0.926536521 | ns | TM | KS.3 |
| FIM1_3362 | 2459.060827 | 2375.112274 | 3212.704601 | -0.005982075 | 0.066504328 | 0.793215145 | 0.988916276 | ns | TM | KS.3 |
| FIM1_3363 | 1547.865299 | 1466.460523 | 1497.899917 | -0.015259728 | 0.055474977 | 0.507110471 | 0.953260948 | ns | TM | KS.3 |
| FIM1_3364 | 1776.281579 | 1838.857083 | 1841.479941 | 0.010749444 | 0.046064612 | 0.598257753 | 0.968542897 | ns | TM | KS.3 |
| FIM1_3365 | 349.1266617 | 380.2933451 | 341.3008944 | 0.014865408 | 0.070569004 | 0.527429944 | 0.959071703 | ns | TM | KS.3 |
| FIM1_3366 | 341.8012843 | 373.9179115 | 426.0664148 | 0.005548799 | 0.09152302 | 0.76671124 | 0.988477371 | ns | TM | KS.3 |
| FIM1_3367 | 725.6142755 | 755.3189506 | 713.2190129 | 0.007345141 | 0.065163348 | 0.749515229 | 0.988477371 | ns | TM | KS.3 |
| FIM1_3368 | 697.0890564 | 584.2546325 | 621.4381627 | -0.039723171 | 0.084896801 | 0.158226865 | 0.789048157 | ns | TM | KS.3 |
| FIM1_3369 | 931.6064859 | 923.1329358 | 1156.707981 | -0.00137582 | 0.069035681 | 0.950913885 | 0.994413583 | ns | TM | KS.3 |
| FIM1_3370 | 1153.174398 | 1105.542544 | 841.2917541 | -0.00684198 | 0.068535851 | 0.763093699 | 0.988477371 | ns | TM | KS.3 |
| FIM1_3371 | 866.1626646 | 973.0600161 | 821.7171968 | 0.023466575 | 0.072839087 | 0.348338634 | 0.948637087 | ns | TM | KS.3 |
| FIM1_3372 | 697.0343058 | 829.9711351 | 785.7021713 | 0.082283817 | 0.093562675 | 0.033747362 | 0.425045894 | ns | TM | KS.3 |
| FIM1_3373 | 534.4628736 | 752.848025 | 864.5272455 | 0.034165799 | 0.113121913 | 0.160049718 | 0.791498358 | ns | TM | KS.3 |
| FIM1_3374 | 3609.587799 | 3596.293857 | 5453.42696 | -0.000307554 | 0.082346451 | 0.987651303 | 0.999333464 | ns | TM | KS.3 |
| FIM1_3375 | 4549.699011 | 4999.134851 | 8288.966713 | 0.0263804 | 0.065980569 | 0.304063828 | 0.926536521 | ns | TM | KS.3 |
| FIM1_3376 | 15.49112168 | 14.82325025 | 19.77240213 | -0.004163927 | 0.092983885 | 0.820774811 | 0.990886949 | ns | TM | KS.3 |
| FIM1_3377 | 1292.837224 | 1371.962732 | 1614.212457 | 0.006197106 | 0.078561695 | 0.767556238 | 0.988477371 | ns | TM | KS.3 |
| FIM1_3378 | 23.78294141 | 21.15432446 | 24.81555604 | -0.009774209 | 0.085741026 | 0.627852112 | 0.97344368 | ns | TM | KS.3 |
| FIM1_3379 | 778.7802326 | 806.8449051 | 602.1434 | 0.008257448 | 0.05819181 | 0.718553267 | 0.987157164 | ns | TM | KS.3 |
| FIM1_3380 | 510.177433 | 488.6562329 | 384.328297 | -0.006565709 | 0.069778787 | 0.770709446 | 0.988814136 | ns | TM | KS.3 |
| FIM1_3381 | 18336.86603 | 19507.72638 | 18287.95914 | 0.015525599 | 0.060016386 | 0.512170486 | 0.953967865 | ns | TM | KS.3 |
| FIM1_3382 | 350.233015 | 382.5760649 | 540.4338346 | 0.017984777 | 0.067961703 | 0.458219668 | 0.950410534 | ns | TM | KS.3 |
| FIM1_3383 | 2121.746182 | 2132.288305 | 2471.836453 | 0.001170361 | 0.053205384 | 0.957662819 | 0.994413583 | ns | TM | KS.3 |
| FIM1_3384 | 7004.052058 | 7890.714556 | 10102.61494 | 0.01199576 | 0.082338709 | 0.570458929 | 0.962265923 | ns | TM | KS.3 |
| FIM1_3385 | 69.63023529 | 81.71469042 | 83.71763438 | 0.016257538 | 0.084748402 | 0.453171611 | 0.950410534 | ns | TM | KS.3 |
| FIM1_3386 | 702.544381 | 856.0479761 | 715.7074906 | 0.028885912 | 0.087518034 | 0.242282954 | 0.887413624 | ns | TM | KS.3 |
| FIM1_3387 | 1514.4039 | 1708.098294 | 1783.26767 | 0.015165326 | 0.078888984 | 0.497975164 | 0.953260948 | ns | TM | KS.3 |
| FIM1_3388 | 3205.798965 | 2846.185635 | 3075.192619 | -0.013965275 | 0.079795993 | 0.525094178 | 0.959071703 | ns | TM | KS.3 |
| FIM1_3389 | 1600.127419 | 1446.037083 | 1286.693949 | -0.009387597 | 0.08278248 | 0.648594443 | 0.976323032 | ns | TM | KS.3 |
| FIM1_3390 | 3405.515336 | 3091.686115 | 3612.230617 | -0.026686174 | 0.066628984 | 0.30023767 | 0.926536521 | ns | TM | KS.3 |
| FIM1_3391 | 1236.134486 | 1230.575792 | 1527.696093 | -0.000678658 | 0.069271685 | 0.975731948 | 0.998880439 | ns | TM | KS.3 |
| FIM1_3392 | 1122.595052 | 1031.987352 | 997.793671 | -0.020303175 | 0.064899478 | 0.409765404 | 0.950410534 | ns | TM | KS.3 |

| | | | | | | | | | | |
|---|---|---|---|---|---|---|---|---|---|---|
| FIM1_3393 | 138.2031471 | 120.4030421 | 168.986438 | -0.010075346 | 0.089809667 | 0.607516659 | 0.970937835 | ns | TM | KS.3 |
| FIM1_3394 | 2366.742813 | 2338.784554 | 2718.688271 | -0.00192889 | 0.067278487 | 0.931861405 | 0.994413583 | ns | TM | KS.3 |
| FIM1_3395 | 2147.874327 | 2353.050901 | 2191.632266 | 0.014043652 | 0.073082992 | 0.542794193 | 0.95917717 | ns | TM | KS.3 |
| FIM1_3396 | 975.2228807 | 998.0289035 | 1060.624303 | 0.002899617 | 0.073601435 | 0.89359623 | 0.991141351 | ns | TM | KS.3 |
| FIM1_3397 | 9484.904452 | 9093.604208 | 7705.802668 | -0.003868725 | 0.080786883 | 0.848988055 | 0.991141351 | ns | TM | KS.3 |
| FIM1_3398 | 12.07724663 | 9.31385775 | 8.902902147 | -0.013842469 | 0.108724229 | 0.459446392 | 0.950410534 | ns | TM | KS.3 |
| FIM1_3399 | 1080.241426 | 1068.304845 | 1046.559827 | -0.001791972 | 0.067158422 | 0.936711102 | 0.994413583 | ns | TM | KS.3 |
| FIM1_3400 | 1407.015503 | 1410.549523 | 1585.761323 | 0.000699691 | 0.048469214 | 0.973481607 | 0.998607635 | ns | TM | KS.3 |
| FIM1_3401 | 678.4473787 | 1147.280706 | 1416.926159 | 0.696357268 | 0.184572861 | 1.18531996572 | 0.00018428 | ns | TM | KS.3 |
| FIM1_3402 | 3863.665713 | 3863.996793 | 3669.48146 | 9.13811547554588 | 0.038175681 | 0.999588204 | 0.999930384 | ns | TM | KS.3 |
| FIM1_3403 | 3233.857124 | 3775.569519 | 4961.124225 | 0.017872322 | 0.083336364 | 0.422401057 | 0.950410534 | ns | TM | KS.3 |
| FIM1_3404 | 42.09270909 | 18.82086092 | 107.336423 | -0.129447289 | 0.302975796 | 0.012819189 | 0.257037736 | ns | TM | KS.3 |
| FIM1_3405 | 8276.457444 | 8761.931574 | 10136.20419 | 0.01205487 | 0.063260275 | 0.607847304 | 0.970937835 | ns | TM | KS.3 |
| FIM1_3407 | 1449.348173 | 1426.216164 | 1399.676544 | -0.003274859 | 0.060822358 | 0.885961978 | 0.991141351 | ns | TM | KS.3 |
| FIM1_3408 | 952.7860545 | 1000.806402 | 1106.331544 | 0.014184628 | 0.053488427 | 0.528994876 | 0.959071703 | ns | TM | KS.3 |
| FIM1_3409 | 10836.99458 | 10220.58281 | 9335.100371 | -0.010422896 | 0.067332074 | 0.652901001 | 0.976323032 | ns | TM | KS.3 |
| FIM1_3410 | 1179.251326 | 1409.833611 | 1501.152017 | 0.0265058 | 0.084422671 | 0.277597232 | 0.908583046 | ns | TM | KS.3 |
| FIM1_3411 | 1755.846314 | 1805.509631 | 1892.610748 | 0.003593017 | 0.073100438 | 0.869105561 | 0.991141351 | ns | TM | KS.3 |
| FIM1_3412 | 1414.278095 | 1469.297659 | 1428.368822 | 0.012061244 | 0.046973624 | 0.560013488 | 0.961043487 | ns | TM | KS.3 |
| FIM1_3413 | 2496.326826 | 2668.356398 | 2766.208354 | 0.011247056 | 0.069159753 | 0.626619309 | 0.97344368 | ns | TM | KS.3 |
| FIM1_3414 | 3219.165206 | 3230.973489 | 2553.054635 | 0.000622663 | 0.065687729 | 0.978090313 | 0.999036302 | ns | TM | KS.3 |
| FIM1_3415 | 682.3413769 | 984.6491992 | 1188.797963 | 0.036788374 | 0.118571784 | 0.139619234 | 0.775229562 | ns | TM | KS.3 |
| FIM1_3416 | 415.2063776 | 449.6254674 | 460.4770577 | 0.009429631 | 0.077005479 | 0.66352353 | 0.976323032 | ns | TM | KS.3 |
| FIM1_3417 | 15143.28194 | 15583.20073 | 14646.90247 | 0.007468784 | 0.053038906 | 0.735940272 | 0.987655486 | ns | TM | KS.3 |
| FIM1_3418 | 2285.304489 | 2523.479516 | 2580.809561 | 0.029134451 | 0.067000348 | 0.266330192 | 0.904370522 | ns | TM | KS.3 |
| FIM1_3419 | 2681.380649 | 2599.499526 | 3343.998621 | -0.009869963 | 0.043906415 | 0.61583883 | 0.973122675 | ns | TM | KS.3 |
| FIM1_3420 | 73.89883247 | 74.79763779 | 103.0118483 | 0.0008052 | 0.08694489 | 0.966292347 | 0.995634078 | ns | TM | KS.3 |
| FIM1_3421 | 107.0167987 | 120.6369676 | 127.9908184 | 0.009418371 | 0.086607472 | 0.637103463 | 0.974658588 | ns | TM | KS.3 |
| FIM1_3422 | 144.9294074 | 69.83792042 | 142.3187291 | -0.14205595 | 0.301421127 | 0.010738533 | 0.231130071 | ns | TM | KS.3 |
| FIM1_3423 | 343.4985542 | 339.3262919 | 604.0889833 | -0.001107531 | 0.080686463 | 0.95629531 | 0.994413583 | ns | TM | KS.3 |
| FIM1_3424 | 791.7290665 | 766.2715414 | 844.8276442 | -0.004837896 | 0.070047404 | 0.828709834 | 0.991141351 | ns | TM | KS.3 |
| FIM1_3425 | 702.804513 | 810.5939145 | 1240.169645 | 0.020144722 | 0.079860733 | 0.388667196 | 0.948637087 | ns | TM | KS.3 |
| FIM1_3426 | 2188.709435 | 1489.797408 | 4545.849937 | -0.026950207 | 0.117860911 | 0.21948832 | 0.865538807 | ns | TM | KS.3 |
| FIM1_3427 | 1010.145768 | 1059.53037 | 1391.919756 | 0.007201533 | 0.070398764 | 0.748960699 | 0.988477371 | ns | TM | KS.3 |
| FIM1_3428 | 26.12613757 | 54.98049714 | 84.78845287 | 0.313019659 | 0.419125015 | 0.002183582 | 0.084221572 | ns | TM | KS.3 |
| FIM1_3429 | 2975.97418 | 3092.959077 | 3127.203162 | 0.010412843 | 0.053100523 | 0.640120219 | 0.975673434 | ns | TM | KS.3 |
| FIM1_3430 | 1496.63547 | 1430.658403 | 1770.705665 | -0.013359398 | 0.05161721 | 0.544683917 | 0.95917717 | ns | TM | KS.3 |
| FIM1_3431 | 233.557889 | 172.7195331 | 368.9373469 | -0.082652954 | 0.127527406 | 0.028701599 | 0.390786333 | ns | TM | KS.3 |
| FIM1_3432 | 782.4373571 | 798.5650794 | 922.3058643 | 0.00380769 | 0.063787812 | 0.867638624 | 0.991141351 | ns | TM | KS.3 |
| FIM1_3433 | 3327.876579 | 3652.822829 | 3764.933484 | 0.048063007 | 0.065952379 | 0.098894264 | 0.677856366 | ns | TM | KS.3 |
| FIM1_3434 | 885.2823577 | 845.1749256 | 717.5488816 | -0.014623908 | 0.049189904 | 0.494623351 | 0.953260948 | ns | TM | KS.3 |
| FIM1_3435 | 2732.403487 | 2448.024636 | 3127.965965 | -0.022230427 | 0.071762052 | 0.370770839 | 0.948637087 | ns | TM | KS.3 |
| FIM1_3437 | 21697.94449 | 19920.82879 | 16824.47536 | -0.015362498 | 0.069895125 | 0.516365568 | 0.955698921 | ns | TM | KS.3 |
| FIM1_3438 | 3117.292486 | 2943.763726 | 2603.795388 | -0.008998065 | 0.06996521 | 0.692452728 | 0.981188557 | ns | TM | KS.3 |
| FIM1_3439 | 1520.688916 | 1964.711882 | 1906.752553 | 0.034125245 | 0.097998291 | 0.175798197 | 0.811391014 | ns | TM | KS.3 |

| | | | | | | | | | | |
|---|---|---|---|---|---|---|---|---|---|---|
| FIM1_3440 | 2815.359318 | 2682.884025 | 3000.803043 | -0.010866589 | 0.060370076 | 0.640522797 | 0.975785947 | ns | TM | KS.3 |
| FIM1_3441 | 995.3687722 | 789.0011839 | 987.7667858 | -0.082785715 | 0.108888857 | 0.031160766 | 0.403710444 | ns | TM | KS.3 |
| FIM1_3442 | 5251.744947 | 4625.412673 | 5052.080606 | -0.031382864 | 0.074229498 | 0.240926806 | 0.887413624 | ns | TM | KS.3 |
| FIM1_3444 | 3760.567336 | 3946.128982 | 4131.371059 | 0.010988676 | 0.06016567 | 0.636732026 | 0.974658588 | ns | TM | KS.3 |
| FIM1_3445 | 3622.188659 | 3535.041901 | 3482.366193 | -0.004107741 | 0.066613553 | 0.856398968 | 0.991141351 | ns | TM | KS.3 |
| FIM1_3446 | 1464.208672 | 1408.214221 | 1375.250604 | -0.014137022 | 0.042280682 | 0.454425704 | 0.950410534 | ns | TM | KS.3 |
| FIM1_3447 | 3081.129369 | 2929.044515 | 2612.233532 | -0.014148702 | 0.054964144 | 0.535506719 | 0.959071703 | ns | TM | KS.3 |
| FIM1_3448 | 743.9424567 | 692.1432158 | 773.1970386 | -0.023761853 | 0.056487819 | 0.319154603 | 0.929077912 | ns | TM | KS.3 |
| FIM1_3449 | 940.9946717 | 836.3610134 | 936.3167794 | -0.041684289 | 0.07319177 | 0.151235579 | 0.783447458 | ns | TM | KS.3 |
| FIM1_3450 | 3286.688385 | 3889.765174 | 4650.317217 | 0.017812902 | 0.085778033 | 0.415972038 | 0.950410534 | ns | TM | KS.3 |
| FIM1_3452 | 1944.092884 | 1679.259802 | 1780.430393 | -0.053254817 | 0.082035144 | 0.09511962 | 0.673713046 | ns | TM | KS.3 |
| FIM1_3454 | 4039.200201 | 1357.663393 | 12413.85407 | -1.42737422 | 0.233651021 | 1.2584955627[illegible] | 5.6918322041[illegible] | Km<KS.3 | TM | KS.3 |
| FIM1_3455 | 128.6058825 | 51.21773333 | 1212.410285 | -1.004591502 | 0.434482403 | 4.2270299321[illegible] | 0.00438114 | Km<KS.3 | TM | KS.3 |
| FIM1_3456 | 26.47954147 | 24.19979208 | 220.5758679 | -0.005402259 | 0.09027089 | 0.774455308 | 0.988814136 | ns | TM | KS.3 |
| FIM1_3457 | 320.7678608 | 464.5798365 | 898.4169684 | 0.042025302 | 0.121877025 | 0.111128706 | 0.71059097 | ns | TM | KS.3 |
| FIM1_3458 | 3624.434497 | 3993.29859 | 8765.855031 | 0.015119433 | 0.073433193 | 0.515003859 | 0.954689943 | ns | TM | KS.3 |
| FIM1_3460 | 5763.631232 | 5131.951305 | 6233.295187 | -0.031894417 | 0.071617256 | 0.236205215 | 0.884880231 | ns | TM | KS.3 |
| FIM1_3461 | 1226.730616 | 1507.155347 | 1201.563314 | 0.016797796 | 0.092868943 | 0.417975018 | 0.950410534 | ns | TM | KS.3 |
| FIM1_3462 | 940.3209727 | 877.1293652 | 799.4370233 | -0.021355963 | 0.05805573 | 0.373318231 | 0.948637087 | ns | TM | KS.3 |
| FIM1_3463 | 2405.651082 | 2383.023111 | 3452.321499 | -0.00093389 | 0.078843572 | 0.963805107 | 0.995175461 | ns | TM | KS.3 |
| FIM1_3464 | 329.3171898 | 279.8615218 | 332.051609 | -0.022006717 | 0.082539257 | 0.347950079 | 0.948637087 | ns | TM | KS.3 |
| FIM1_3465 | 234.3614338 | 191.252498 | 228.7605552 | -0.019106372 | 0.090843832 | 0.375677985 | 0.948637087 | ns | TM | KS.3 |
| FIM1_3466 | 2953.743827 | 2832.531494 | 2895.335907 | -0.014806998 | 0.043722213 | 0.446849754 | 0.950410534 | ns | TM | KS.3 |
| FIM1_3467 | 815.0386262 | 777.5456931 | 819.1413989 | -0.003001613 | 0.089420017 | 0.872893887 | 0.991141351 | ns | TM | KS.3 |
| FIM1_3468 | 2556.623281 | 2588.957638 | 2493.683692 | 0.002560378 | 0.06102668 | 0.910667321 | 0.991141351 | ns | TM | KS.3 |
| FIM1_3469 | 2278.245025 | 2057.350965 | 2023.438997 | -0.010288224 | 0.081200956 | 0.624751643 | 0.97344368 | ns | TM | KS.3 |
| FIM1_3470 | 716.7591531 | 731.3595375 | 741.3694133 | 0.002282662 | 0.075436563 | 0.914676661 | 0.991141351 | ns | TM | KS.3 |
| FIM1_3471 | 2474.853554 | 2635.19983 | 2447.582828 | 0.016328106 | 0.059373829 | 0.490817224 | 0.952591289 | ns | TM | KS.3 |
| FIM1_3472 | 684.371439 | 677.8481304 | 647.6004953 | -0.001952663 | 0.056426697 | 0.930857445 | 0.993993515 | ns | TM | KS.3 |
| FIM1_3473 | 2133.030178 | 2785.499951 | 3333.603423 | 0.024208745 | 0.099347407 | 0.274705998 | 0.908583046 | ns | TM | KS.3 |
| FIM1_3474 | 281.5740145 | 336.3749132 | 705.3237644 | 0.015087194 | 0.089869143 | 0.465575126 | 0.950410534 | ns | TM | KS.3 |
| FIM1_3475 | 3714.653705 | 3157.374193 | 3795.78794 | -0.091357099 | 0.091652488 | 0.024760478 | 0.373642194 | ns | TM | KS.3 |
| FIM1_3476 | 1533.233691 | 1424.840461 | 1593.52545 | -0.024974523 | 0.058084396 | 0.304686181 | 0.926536521 | ns | TM | KS.3 |
| FIM1_3477 | 2006.777771 | 1651.295034 | 1971.95525 | -0.059622422 | 0.093972427 | 0.071289238 | 0.586221418 | ns | TM | KS.3 |
| FIM1_3478 | 284.9289918 | 289.8557799 | 204.791761 | 0.00265385 | 0.070406351 | 0.904833821 | 0.991141351 | ns | TM | KS.3 |
| FIM1_3479 | 10936.65844 | 8687.683666 | 7511.9524 | -0.283434116 | 0.060206538 | 9.4377471260[illegible] | 2.3476395976[illegible] | ns | TM | KS.3 |
| FIM1_3480 | 2508.974196 | 2785.56576 | 3057.765141 | 0.007994238 | 0.086870821 | 0.684798506 | 0.980234354 | ns | TM | KS.3 |
| FIM1_3481 | 3254.103964 | 3167.00975 | 3386.072532 | -0.008386536 | 0.043726983 | 0.669265892 | 0.976323032 | ns | TM | KS.3 |
| FIM1_3482 | 4937.820362 | 5084.83618 | 3183.553039 | 0.005033904 | 0.066331726 | 0.825050066 | 0.991141351 | ns | TM | KS.3 |
| FIM1_3483 | 1200.407291 | 962.2524653 | 933.8487455 | -0.167446256 | 0.106446747 | 0.002346839 | 0.089811734 | ns | TM | KS.3 |
| FIM1_3484 | 1979.424037 | 1597.075377 | 1506.444146 | -0.202894152 | 0.088576436 | 0.000397649 | 0.023835005 | ns | TM | KS.3 |
| FIM1_3485 | 644.3048577 | 577.5841919 | 508.4485735 | -0.023311031 | 0.071324769 | 0.35292583 | 0.948637087 | ns | TM | KS.3 |
| FIM1_3486 | 4257.149934 | 4015.000913 | 3483.811659 | -0.012786248 | 0.06266081 | 0.587329692 | 0.963781856 | ns | TM | KS.3 |
| FIM1_3487 | 952.0083781 | 1014.767215 | 881.4953144 | 0.010670663 | 0.069080865 | 0.643434573 | 0.976323032 | ns | TM | KS.3 |
| FIM1_3488 | 446.2122623 | 393.0970388 | 422.2680436 | -0.046487057 | 0.076966445 | 0.124420384 | 0.739983875 | ns | TM | KS.3 |

| | | | | | | | | | | |
|---|---|---|---|---|---|---|---|---|---|---|
| FIM1_3489 | 850.9581697 | 657.3771042 | 767.3091327 | -0.212244772 | 0.117496367 | 0.000663836 | 0.035511648 | ns | TM | KS.3 |
| FIM1_3490 | 1091.880259 | 1129.56508 | 1105.079606 | 0.009198842 | 0.051262723 | 0.673501397 | 0.976861203 | ns | TM | KS.3 |
| FIM1_3491 | 2012.219802 | 1878.729592 | 2077.042796 | -0.017610841 | 0.060941926 | 0.463181954 | 0.950410534 | ns | TM | KS.3 |
| FIM1_3492 | 978.8820112 | 856.6190687 | 782.9234932 | -0.024370568 | 0.07639717 | 0.327360535 | 0.934739762 | ns | TM | KS.3 |
| FIM1_3493 | 86.61769294 | 123.712724 | 111.3256679 | 0.045901046 | 0.12208297 | 0.095473212 | 0.673713046 | ns | TM | KS.3 |
| FIM1_3494 | 9025.670769 | 9258.120238 | 9553.046138 | 0.007183809 | 0.048877531 | 0.734777962 | 0.987655486 | ns | TM | KS.3 |
| FIM1_3495 | 149.2560996 | 131.6977179 | 257.5342315 | -0.022592084 | 0.075015017 | 0.358671499 | 0.948637087 | ns | TM | KS.3 |
| FIM1_3496 | 575.7838242 | 567.5110078 | 619.1826244 | -0.002542196 | 0.065523935 | 0.910905413 | 0.991141351 | ns | TM | KS.3 |
| FIM1_3497 | 1471.564655 | 972.2686881 | 1782.456726 | -0.045735519 | 0.133038232 | 0.093104404 | 0.671036206 | ns | TM | KS.3 |
| FIM1_3498 | 153.5736868 | 180.5381149 | 150.7250461 | 0.020367905 | 0.082570313 | 0.376221068 | 0.948637087 | ns | TM | KS.3 |
| FIM1_3499 | 3079.425465 | 2887.071794 | 2687.064173 | -0.024342054 | 0.053103586 | 0.288478585 | 0.919993167 | ns | TM | KS.3 |
| FIM1_3500 | 1949.158825 | 1897.687964 | 1777.103672 | -0.004313461 | 0.067900185 | 0.848566363 | 0.991141351 | ns | TM | KS.3 |
| FIM1_3501 | 2543.399213 | 2240.241716 | 2906.741389 | -0.01535142 | 0.080009496 | 0.489843489 | 0.952591289 | ns | TM | KS.3 |
| FIM1_3503 | 429.0666441 | 381.6287274 | 375.0021449 | -0.006564427 | 0.094800481 | 0.721866965 | 0.987157164 | ns | TM | KS.3 |
| FIM1_3504 | 413.6093545 | 480.7892128 | 551.5915906 | 0.013882255 | 0.086238165 | 0.506613947 | 0.953260948 | ns | TM | KS.3 |
| FIM1_3505 | 3131.473291 | 3390.182686 | 3680.734522 | 0.027234585 | 0.060113716 | 0.27586527 | 0.908583046 | ns | TM | KS.3 |
| FIM1_3506 | 734.2873339 | 756.5106163 | 702.3294603 | 0.005007166 | 0.066639087 | 0.825771112 | 0.991141351 | ns | TM | KS.3 |
| FIM1_3507 | 1770.652466 | 1959.497911 | 1893.918144 | 0.039362845 | 0.067886902 | 0.160743225 | 0.792564466 | ns | TM | KS.3 |
| FIM1_3508 | 933.2198941 | 932.923488 | 926.6437892 | 5.50198125331872 | 0.053667641 | 0.998015296 | 0.999525119 | ns | TM | KS.3 |
| FIM1_3509 | 23.45971837 | 10.11547815 | 75.09163503 | -0.272037762 | 0.431846871 | 0.00318079 | 0.103770022 | ns | TM | KS.3 |
| FIM1_3511 | 5590.088221 | 5254.442209 | 4938.838614 | -0.015296768 | 0.060373699 | 0.518587939 | 0.957674461 | ns | TM | KS.3 |
| FIM1_3512 | 4988.978843 | 4304.652135 | 4645.559879 | -0.027371714 | 0.078693735 | 0.280821612 | 0.915522622 | ns | TM | KS.3 |
| FIM1_3513 | 969.1646845 | 687.9950825 | 986.2109913 | -0.024120599 | 0.111271563 | 0.259607366 | 0.895871714 | ns | TM | KS.3 |
| FIM1_3514 | 2234.260441 | 2235.146338 | 2916.016212 | 5.33263935581525 | 0.073730773 | 0.99802918 | 0.999525119 | ns | TM | KS.3 |
| FIM1_3516 | 6200.176153 | 7012.913629 | 7410.048664 | 0.011344575 | 0.084170151 | 0.583786379 | 0.963781856 | ns | TM | KS.3 |
| FIM1_3517 | 496.4831666 | 521.7478812 | 512.655349 | 0.012455179 | 0.055886544 | 0.586333338 | 0.963781856 | ns | TM | KS.3 |
| FIM1_3518 | 2203.195896 | 2625.84708 | 3047.35086 | 0.013134178 | 0.091596165 | 0.511639917 | 0.953818112 | ns | TM | KS.3 |
| FIM1_3519 | 840.1867303 | 858.4854357 | 757.2062454 | 0.002524971 | 0.075030101 | 0.906048728 | 0.991141351 | ns | TM | KS.3 |
| FIM1_3520 | 1018.036216 | 1093.857673 | 1141.632281 | 0.008541949 | 0.076420025 | 0.693302898 | 0.981188557 | ns | TM | KS.3 |
| FIM1_3522 | 2536.691186 | 2931.667361 | 3616.222836 | 0.010944546 | 0.089262115 | 0.581494168 | 0.963781856 | ns | TM | KS.3 |
| FIM1_3523 | 6619.218914 | 6742.636688 | 7519.610076 | 0.003197158 | 0.065789333 | 0.888106339 | 0.991141351 | ns | TM | KS.3 |
| FIM1_3524 | 1735.757414 | 1723.833967 | 1888.769054 | -0.001334605 | 0.063301356 | 0.953308672 | 0.994413583 | ns | TM | KS.3 |
| FIM1_3525 | 444.616042 | 524.9282458 | 478.5321531 | 0.021189154 | 0.083612559 | 0.358524453 | 0.948637087 | ns | TM | KS.3 |
| FIM1_3526 | 2739.143271 | 2717.100665 | 3162.86158 | -0.002470028 | 0.043795061 | 0.900264446 | 0.991141351 | ns | TM | KS.3 |
| FIM1_3527 | 1787.240016 | 1881.636022 | 1778.7921 | 0.006585405 | 0.074003329 | 0.763363336 | 0.988477371 | ns | TM | KS.3 |
| FIM1_3528 | 1862.616838 | 1727.611459 | 3188.141061 | -0.006062138 | 0.084576961 | 0.759659656 | 0.988477371 | ns | TM | KS.3 |
| FIM1_3529 | 690.985216 | 658.4024227 | 616.9667557 | -0.007575682 | 0.069520053 | 0.738061249 | 0.988051177 | ns | TM | KS.3 |
| FIM1_3530 | 464.0193893 | 452.4952168 | 406.0561677 | -0.004314237 | 0.066942564 | 0.849148887 | 0.991141351 | ns | TM | KS.3 |
| FIM1_3531 | 1595.921112 | 1302.485486 | 1385.990734 | -0.181814875 | 0.089790257 | 0.000982315 | 0.04634941 | ns | TM | KS.3 |
| FIM1_3532 | 1090.290656 | 970.7529613 | 977.3404401 | -0.030082916 | 0.071853027 | 0.256912798 | 0.895871714 | ns | TM | KS.3 |
| FIM1_3533 | 375.6497395 | 398.59719 | 652.0218671 | 0.01275027 | 0.063746937 | 0.588601666 | 0.963781856 | ns | TM | KS.3 |
| FIM1_3534 | 3068.647547 | 2646.087171 | 2703.360744 | -0.10712367 | 0.085649616 | 0.013416665 | 0.263081302 | ns | TM | KS.3 |
| FIM1_3535 | 1664.109455 | 1617.048962 | 1527.632879 | -0.009483193 | 0.040055029 | 0.602350069 | 0.970245665 | ns | TM | KS.3 |
| FIM1_3536 | 494.3195917 | 482.6860592 | 460.5971985 | -0.005632231 | 0.057171674 | 0.803891792 | 0.988916276 | ns | TM | KS.3 |
| FIM1_3537 | 7003.095937 | 7788.667917 | 8084.007475 | 0.011330235 | 0.080482427 | 0.595567359 | 0.965889818 | ns | TM | KS.3 |

| | | | | | | | | | | |
|---|---|---|---|---|---|---|---|---|---|---|
| FIM1_3538 | 1032.797306 | 988.4108386 | 903.5957502 | -0.011773942 | 0.055119833 | 0.604286878 | 0.970729568 | ns | TM | KS.3 |
| FIM1_3539 | 654.044362 | 620.9689031 | 579.1258978 | -0.01228394 | 0.060413568 | 0.599614899 | 0.969478103 | ns | TM | KS.3 |
| FIM1_3540 | 3145.526636 | 2848.251906 | 3361.474688 | -0.041417719 | 0.067241961 | 0.143380665 | 0.77783242 | ns | TM | KS.3 |
| FIM1_3541 | 300.3610346 | 240.8692959 | 306.9328249 | -0.03226572 | 0.091785835 | 0.200802003 | 0.845272486 | ns | TM | KS.3 |
| FIM1_3542 | 442.9594202 | 378.9571517 | 450.4154753 | -0.057017193 | 0.084281278 | 0.082257627 | 0.62282947 | ns | TM | KS.3 |
| FIM1_3544 | 2776.847546 | 2874.201017 | 2705.31606 | 0.006464451 | 0.064197622 | 0.778461358 | 0.988814136 | ns | TM | KS.3 |
| FIM1_3545 | 67.28352938 | 36.47512199 | 154.6222435 | -0.128392402 | 0.259654463 | 0.01238942 | 0.254461363 | ns | TM | KS.3 |
| FIM1_3546 | 1150.698051 | 1156.757554 | 1688.710571 | 0.001183033 | 0.061690241 | 0.958628115 | 0.994413583 | ns | TM | KS.3 |
| FIM1_3547 | 819.0923856 | 1287.120945 | 926.7482602 | 0.582987605 | 0.171285547 | 1.36940238101 | 0.000200376 | ns | TM | KS.3 |
| FIM1_3548 | 487.2246678 | 444.9374237 | 437.2404573 | -0.008024967 | 0.083201878 | 0.692828723 | 0.981188557 | ns | TM | KS.3 |
| FIM1_3549 | 130.7422666 | 145.107845 | 171.1134703 | 0.017630479 | 0.072291934 | 0.459694955 | 0.950410534 | ns | TM | KS.3 |
| FIM1_3550 | 1773.213488 | 1767.085862 | 1459.170544 | -0.000495569 | 0.071313592 | 0.98203667 | 0.999333464 | ns | TM | KS.3 |
| FIM1_3551 | 1552.270682 | 1788.318264 | 1689.147242 | 0.024583358 | 0.077982156 | 0.320472343 | 0.929108338 | ns | TM | KS.3 |
| FIM1_3552 | 682.1870026 | 637.2925225 | 580.9714058 | -0.015362904 | 0.063946552 | 0.520327588 | 0.958509403 | ns | TM | KS.3 |
| FIM1_3553 | 455.4857841 | 470.9699744 | 438.2540512 | 0.004063834 | 0.07430645 | 0.850848198 | 0.991141351 | ns | TM | KS.3 |
| FIM1_3554 | 785.7606101 | 780.8699084 | 602.7025135 | -0.001368001 | 0.060997503 | 0.952151245 | 0.994413583 | ns | TM | KS.3 |
| FIM1_3555 | 701.0424498 | 633.9784698 | 575.0273797 | -0.022983781 | 0.068965193 | 0.360280496 | 0.948637087 | ns | TM | KS.3 |
| FIM1_3556 | 2215.478632 | 1983.452852 | 2232.991527 | -0.04176226 | 0.070954617 | 0.148107796 | 0.780546912 | ns | TM | KS.3 |
| FIM1_3557 | 3982.561628 | 3942.824592 | 4774.948837 | -0.00139925 | 0.071011027 | 0.949447774 | 0.994413583 | ns | TM | KS.3 |
| FIM1_3558 | 1106.83848 | 1144.98579 | 1198.38517 | 0.007245632 | 0.060004059 | 0.752578546 | 0.988477371 | ns | TM | KS.3 |
| FIM1_3559 | 251.5192265 | 213.171348 | 243.8344568 | -0.036625421 | 0.081959839 | 0.18382006 | 0.821657502 | ns | TM | KS.3 |
| FIM1_3560 | 1145.27922 | 1120.281974 | 1344.499968 | -0.005865711 | 0.050833387 | 0.786471384 | 0.988916276 | ns | TM | KS.3 |
| FIM1_3561 | 758.493626 | 600.5259252 | 872.0920139 | -0.039746662 | 0.095307005 | 0.142932247 | 0.77783242 | ns | TM | KS.3 |
| FIM1_3562 | 1573.235559 | 1172.730135 | 1735.131374 | -0.25968978 | 0.133534852 | 0.000218225 | 0.015734323 | ns | TM | KS.3 |
| FIM1_3563 | 2079.055639 | 1755.992368 | 1925.341098 | -0.075479033 | 0.091804243 | 0.042405565 | 0.472993959 | ns | TM | KS.3 |
| FIM1_3565 | 49.46944497 | 76.82643562 | 110.8741976 | 0.04418782 | 0.136156661 | 0.09812274 | 0.677062609 | ns | TM | KS.3 |
| FIM1_3566 | 165.3272303 | 177.2312618 | 242.8917698 | 0.009692205 | 0.074548351 | 0.661725029 | 0.976323032 | ns | TM | KS.3 |
| FIM1_3567 | 7051.438929 | 7344.497437 | 7172.635201 | 0.007901085 | 0.063783508 | 0.732269 | 0.987157164 | ns | TM | KS.3 |
| FIM1_3568 | 4312.55527 | 5314.572005 | 4147.484486 | 0.034149311 | 0.08979625 | 0.188936045 | 0.828217874 | ns | TM | KS.3 |
| FIM1_3569 | 1350.831634 | 1680.436833 | 2003.273747 | 0.014458843 | 0.097282706 | 0.463392243 | 0.950410534 | ns | TM | KS.3 |
| FIM1_3570 | 1322.768299 | 1475.143451 | 1301.032634 | 0.015353265 | 0.076149546 | 0.501919934 | 0.953260948 | ns | TM | KS.3 |
| FIM1_3571 | 3964.10338 | 3915.523867 | 4121.434755 | -0.003555757 | 0.046502221 | 0.862501421 | 0.991141351 | ns | TM | KS.3 |
| FIM1_3572 | 577.2054988 | 568.120469 | 519.58149 | -0.004235246 | 0.055539566 | 0.850328725 | 0.991141351 | ns | TM | KS.3 |
| FIM1_3573 | 4235.049184 | 4308.693225 | 4375.909811 | 0.004950216 | 0.046715885 | 0.810060181 | 0.988916276 | ns | TM | KS.3 |
| FIM1_3574 | 1716.666748 | 1689.084839 | 3146.18648 | -0.001121363 | 0.086973686 | 0.953074239 | 0.994413583 | ns | TM | KS.3 |
| FIM1_3575 | 871.876001 | 957.4642253 | 1718.833637 | 0.012013785 | 0.0762915 | 0.588651563 | 0.963781856 | ns | TM | KS.3 |
| FIM1_3576 | 3836.784278 | 3586.924247 | 3522.498511 | -0.01817265 | 0.059707383 | 0.447604707 | 0.950410534 | ns | TM | KS.3 |
| FIM1_3577 | 648.6507835 | 609.7025805 | 487.2915457 | -0.013882147 | 0.061754535 | 0.557108291 | 0.959760553 | ns | TM | KS.3 |
| FIM1_3579 | 2002.644779 | 2154.023117 | 2293.963276 | 0.012470939 | 0.069467306 | 0.591825188 | 0.964028601 | ns | TM | KS.3 |
| FIM1_3580 | 3904.19708 | 4503.56594 | 5248.873732 | 0.013462553 | 0.085146588 | 0.52105749 | 0.958602567 | ns | TM | KS.3 |
| FIM1_3581 | 5762.473024 | 6961.456618 | 4370.134313 | 0.028357574 | 0.086035361 | 0.251559556 | 0.888934767 | ns | TM | KS.3 |
| FIM1_3582 | 1439.209342 | 1613.094433 | 2298.735857 | 0.051122841 | 0.073412642 | 0.098594003 | 0.677856366 | ns | TM | KS.3 |
| FIM1_3583 | 1032.154133 | 1834.846068 | 1973.623488 | 0.2952285 | 0.344245893 | 0.001793517 | 0.071957633 | ns | TM | KS.3 |
| FIM1_3584 | 548.5260245 | 741.7097976 | 589.4158153 | 0.107445584 | 0.137236612 | 0.014488041 | 0.273934835 | ns | TM | KS.3 |
| FIM1_3585 | 1588.669851 | 1353.652369 | 1187.196258 | -0.028776081 | 0.080971299 | 0.25884813 | 0.895871714 | ns | TM | KS.3 |

| | | | | | | | | | | |
|---|---|---|---|---|---|---|---|---|---|---|
| FIM1_3586 | 2733.252787 | 2718.80762 | 2909.673499 | -0.000878487 | 0.06716326 | 0.968927283 | 0.996327965 | ns | TM | KS.3 |
| FIM1_3587 | 63.71951304 | 62.84407198 | 85.38572007 | -0.001724445 | 0.083337658 | 0.930284268 | 0.993993515 | ns | TM | KS.3 |
| FIM1_3588 | 1004.792955 | 1061.831503 | 1024.528335 | 0.005697867 | 0.078640002 | 0.78514111 | 0.988880259 | ns | TM | KS.3 |
| FIM1_3589 | 2670.017229 | 2478.226623 | 3873.387259 | -0.011618831 | 0.071380996 | 0.612094094 | 0.973122675 | ns | TM | KS.3 |
| FIM1_3591 | 615.5589846 | 556.7477885 | 629.3839924 | -0.011672982 | 0.078659488 | 0.591248689 | 0.963781856 | ns | TM | KS.3 |
| FIM1_3592 | 1837.27724 | 2928.510206 | 1972.23227 | 0.062391323 | 0.158070085 | 0.050634818 | 0.513412402 | ns | TM | KS.3 |
| FIM1_3594 | 2653.130093 | 2993.946919 | 3172.010947 | 0.042431236 | 0.073822729 | 0.146703353 | 0.780546912 | ns | TM | KS.3 |
| FIM1_3595 | 6578.172428 | 6373.863922 | 5378.255139 | -0.006385996 | 0.061932128 | 0.781240724 | 0.988814136 | ns | TM | KS.3 |
| FIM1_3596 | 30.02735391 | 33.55205962 | 37.01878554 | 0.006701111 | 0.092355477 | 0.720697307 | 0.987157164 | ns | TM | KS.3 |
| FIM1_3597 | 12076.07417 | 12032.49997 | 10687.94615 | -0.00096788 | 0.04930409 | 0.963674423 | 0.995175461 | ns | TM | KS.3 |
| FIM1_3598 | 1741.851528 | 1969.781149 | 2566.044627 | 0.017617491 | 0.077351186 | 0.446810509 | 0.950410534 | ns | TM | KS.3 |
| FIM1_3599 | 6008.158088 | 5227.69276 | 6212.493068 | -0.054604731 | 0.080699844 | 0.090035795 | 0.655824418 | ns | TM | KS.3 |
| FIM1_3600 | 429.2172654 | 479.8956296 | 386.4645009 | 0.019314134 | 0.073555226 | 0.421877472 | 0.950410534 | ns | TM | KS.3 |
| FIM1_3601 | 2320.840153 | 2094.272687 | 2242.782252 | -0.026832181 | 0.068440912 | 0.29944566 | 0.926536521 | ns | TM | KS.3 |
| FIM1_3603 | 732.4593535 | 655.141179 | 629.563039 | -0.033755415 | 0.070637545 | 0.216095848 | 0.865538807 | ns | TM | KS.3 |
| FIM1_3604 | 3358.734588 | 3482.14707 | 3903.096472 | 0.006705003 | 0.064565185 | 0.770483089 | 0.988814136 | ns | TM | KS.3 |
| FIM1_3605 | 1446.501713 | 1295.819851 | 1340.176429 | -0.079981961 | 0.073068477 | 0.028202466 | 0.390786333 | ns | TM | KS.3 |
| FIM1_3606 | 649.5540713 | 712.0898964 | 573.6389977 | 0.013771405 | 0.07335636 | 0.549146718 | 0.95917717 | ns | TM | KS.3 |
| FIM1_3607 | 1347.285905 | 1365.775241 | 1506.810348 | 0.003164372 | 0.055923825 | 0.887994665 | 0.991141351 | ns | TM | KS.3 |
| FIM1_3608 | 526.911174 | 532.4198405 | 542.6673427 | 0.002498333 | 0.058596097 | 0.912396772 | 0.991141351 | ns | TM | KS.3 |
| FIM1_3609 | 951.9610715 | 1028.59249 | 1486.044669 | 0.008777007 | 0.077605529 | 0.682673808 | 0.97986479 | ns | TM | KS.3 |
| FIM1_3610 | 1959.270707 | 2015.913223 | 1668.484706 | 0.007443615 | 0.052641477 | 0.735912293 | 0.987655486 | ns | TM | KS.3 |
| FIM1_3611 | 896.3999605 | 860.3540718 | 868.238066 | -0.013458828 | 0.049655191 | 0.531878146 | 0.959071703 | ns | TM | KS.3 |
| FIM1_3612 | 1737.315479 | 1516.996152 | 1620.530005 | -0.066581945 | 0.081877189 | 0.056575872 | 0.535599847 | ns | TM | KS.3 |
| FIM1_3613 | 2086.569278 | 1422.307379 | 1052.78488 | -0.122429953 | 0.175862833 | 0.010651297 | 0.231130071 | ns | TM | KS.3 |
| FIM1_3614 | 450.8041843 | 433.1220466 | 312.8522437 | -0.005952818 | 0.070485239 | 0.790263623 | 0.988916276 | ns | TM | KS.3 |
| FIM1_3615 | 1250.047151 | 1293.061381 | 1455.642772 | 0.005713832 | 0.066750661 | 0.801969146 | 0.988916276 | ns | TM | KS.3 |
| FIM1_3616 | 2957.888795 | 3174.571752 | 3554.016383 | 0.008433365 | 0.076323941 | 0.697054107 | 0.98151979 | ns | TM | KS.3 |
| FIM1_3617 | 217.8254619 | 223.517659 | 224.0788788 | 0.003207523 | 0.072445374 | 0.88354038 | 0.991141351 | ns | TM | KS.3 |
| FIM1_3618 | 3544.344016 | 3083.688528 | 2714.576116 | -0.070533228 | 0.083296706 | 0.049200534 | 0.508882862 | ns | TM | KS.3 |
| FIM1_3619 | 365.6231666 | 363.2592355 | 359.8464444 | -0.001194565 | 0.067655846 | 0.957671322 | 0.994413583 | ns | TM | KS.3 |
| FIM1_3620 | 1590.267316 | 1333.102344 | 1503.26998 | -0.057454561 | 0.089538613 | 0.079307311 | 0.616697988 | ns | TM | KS.3 |
| FIM1_3621 | 1706.294085 | 1639.006843 | 1545.844025 | -0.009257019 | 0.059390243 | 0.688330515 | 0.980879562 | ns | TM | KS.3 |
| FIM1_3622 | 2422.861833 | 2334.035169 | 2433.684599 | -0.01219797 | 0.045180165 | 0.543625403 | 0.95917717 | ns | TM | KS.3 |
| FIM1_3623 | 230.7644899 | 158.2390945 | 572.7992675 | -0.030471791 | 0.1177353 | 0.185592751 | 0.825871141 | ns | TM | KS.3 |
| FIM1_3624 | 553.9100004 | 413.5844821 | 462.092434 | -0.026254289 | 0.103252137 | 0.242817367 | 0.887413624 | ns | TM | KS.3 |
| FIM1_3625 | 16442.78909 | 18788.457 | 18717.45703 | 0.133060945 | 0.068459657 | 0.003015126 | 0.101288726 | ns | TM | KS.3 |
| FIM1_3626 | 288.2298242 | 250.2953964 | 201.9656237 | -0.038138571 | 0.078093355 | 0.176920912 | 0.811391014 | ns | TM | KS.3 |
| FIM1_3627 | 614.182129 | 484.6194251 | 1302.001326 | -0.044633259 | 0.097379832 | 0.116370477 | 0.720078512 | ns | TM | KS.3 |
| FIM1_3628 | 10819.35697 | 11867.17137 | 11043.21588 | 0.007623642 | 0.084718958 | 0.702961089 | 0.98151979 | ns | TM | KS.3 |
| FIM1_3629 | 5151.029406 | 4883.129009 | 4731.292686 | -0.008111783 | 0.070459867 | 0.719507255 | 0.987157164 | ns | TM | KS.3 |
| FIM1_3630 | 156.8346738 | 156.4301105 | 197.6624048 | -0.000950749 | 0.072752838 | 0.96517017 | 0.995175461 | ns | TM | KS.3 |
| FIM1_3631 | 572.8825993 | 556.263983 | 557.9884718 | -0.006996267 | 0.055602287 | 0.756347811 | 0.988477371 | ns | TM | KS.3 |
| FIM1_3632 | 8712.768068 | 9068.694632 | 8567.738859 | 0.011686224 | 0.05072411 | 0.59148671 | 0.96385404 | ns | TM | KS.3 |
| FIM1_3633 | 2417.030003 | 1387.593507 | 2506.69561 | -0.129337367 | 0.24368205 | 0.011620684 | 0.24513135 | ns | TM | KS.3 |

| | | | | | | | | | | |
|---|---|---|---|---|---|---|---|---|---|---|
| FIM1_3634 | 2503.890865 | 3042.162249 | 3704.855705 | 0.023594559 | 0.087664552 | 0.308685742 | 0.927416591 | ns | TM | KS.3 |
| FIM1_3635 | 3701.270367 | 3405.339664 | 4861.765801 | -0.009326407 | 0.078235394 | 0.663416286 | 0.976323032 | ns | TM | KS.3 |
| FIM1_3636 | 1983.736916 | 1801.164946 | 2761.731935 | -0.009616412 | 0.081127858 | 0.645999071 | 0.976323032 | ns | TM | KS.3 |
| FIM1_3637 | 2452.038738 | 2319.385068 | 3180.97478 | -0.005514265 | 0.079491684 | 0.790073274 | 0.988916276 | ns | TM | KS.3 |
| FIM1_3638 | 6992.818697 | 6048.785049 | 7351.407994 | -0.014691175 | 0.084187111 | 0.492377898 | 0.952591289 | ns | TM | KS.3 |
| FIM1_3639 | 268.2876977 | 277.685188 | 401.9788217 | 0.002671947 | 0.08393728 | 0.891791238 | 0.991141351 | ns | TM | KS.3 |
| FIM1_3640 | 853.3008858 | 768.8360324 | 1134.694664 | -0.009933327 | 0.082410241 | 0.632245785 | 0.97344368 | ns | TM | KS.3 |
| FIM1_3641 | 745.2538354 | 658.5107162 | 956.6364595 | -0.013624538 | 0.081218296 | 0.529345283 | 0.959071703 | ns | TM | KS.3 |
| FIM1_3642 | 2265.15369 | 1936.743097 | 2394.135994 | -0.017710127 | 0.083748346 | 0.424486133 | 0.950410534 | ns | TM | KS.3 |
| FIM1_3643 | 930.8056258 | 953.4758796 | 1267.188752 | 0.002147028 | 0.081051752 | 0.915221466 | 0.991141351 | ns | TM | KS.3 |
| FIM1_3644 | 1919.430932 | 1828.080807 | 2834.249379 | -0.005458445 | 0.076777324 | 0.797234418 | 0.988916276 | ns | TM | KS.3 |
| FIM1_3645 | 2324.408702 | 2068.259475 | 3118.831835 | -0.012658673 | 0.08089844 | 0.556414889 | 0.959760553 | ns | TM | KS.3 |
| FIM1_3646 | 793.4592863 | 916.0844593 | 1263.162388 | 0.015130509 | 0.083477632 | 0.483837111 | 0.952269291 | ns | TM | KS.3 |
| FIM1_3647 | 22182.69748 | 24302.93819 | 25119.05881 | 0.013007519 | 0.074387309 | 0.566654747 | 0.962265923 | ns | TM | KS.3 |
| FIM1_3648 | 1401.644887 | 1599.930463 | 2447.029702 | 0.00759384 | 0.094831731 | 0.682852508 | 0.97986479 | ns | TM | KS.3 |
| FIM1_3649 | 898.2591455 | 1138.606204 | 1619.634937 | 0.025990294 | 0.094235485 | 0.261077899 | 0.897845311 | ns | TM | KS.3 |
| FIM1_3650 | 1051.720694 | 967.8321352 | 1797.310892 | -0.008334217 | 0.0803512 | 0.689867137 | 0.980879562 | ns | TM | KS.3 |
| FIM1_3651 | 1161.069238 | 944.1302234 | 1422.069128 | -0.022016087 | 0.090165198 | 0.326410321 | 0.934739762 | ns | TM | KS.3 |
| FIM1_3652 | 247.8354706 | 197.7959225 | 400.6113962 | -0.015218216 | 0.097533049 | 0.443960037 | 0.950410534 | ns | TM | KS.3 |
| FIM1_3653 | 3446.089727 | 3900.518491 | 5782.597739 | 0.012947789 | 0.082017177 | 0.544885335 | 0.95917717 | ns | TM | KS.3 |
| FIM1_3654 | 6695.092734 | 6342.996581 | 7828.351318 | -0.006874414 | 0.074250537 | 0.753148353 | 0.988477371 | ns | TM | KS.3 |
| FIM1_3655 | 3726.653164 | 3608.199731 | 4492.996544 | -0.003722155 | 0.075693115 | 0.861443777 | 0.991141351 | ns | TM | KS.3 |
| FIM1_3656 | 4782.548622 | 4506.854281 | 5621.544174 | -0.006610749 | 0.077229372 | 0.755968769 | 0.988477371 | ns | TM | KS.3 |
| FIM1_3657 | 413.4560023 | 436.1316695 | 773.3438998 | 0.004702576 | 0.081800743 | 0.815799496 | 0.988916276 | ns | TM | KS.3 |
| FIM1_3658 | 1444.447529 | 1469.253717 | 2222.721408 | 0.001290402 | 0.084794011 | 0.947053144 | 0.994413583 | ns | TM | KS.3 |
| FIM1_3659 | 1357.248465 | 1353.838815 | 1990.138433 | -0.000245613 | 0.079229389 | 0.990437656 | 0.999525119 | ns | TM | KS.3 |
| FIM1_3660 | 1037.196732 | 970.1927949 | 1551.995642 | -0.006246275 | 0.081182847 | 0.760152719 | 0.988477371 | ns | TM | KS.3 |
| FIM1_3661 | 2146.165424 | 1668.039645 | 2798.114036 | -0.027505442 | 0.096621005 | 0.238799308 | 0.886981413 | ns | TM | KS.3 |
| FIM1_3662 | 931.2410491 | 877.5644232 | 1358.714328 | -0.00581222 | 0.079873084 | 0.778545577 | 0.988814136 | ns | TM | KS.3 |
| FIM1_3663 | 2520.61741 | 3090.159019 | 4056.103488 | 0.015205055 | 0.094001991 | 0.452116048 | 0.950410534 | ns | TM | KS.3 |
| FIM1_3664 | 1224.401979 | 977.5688314 | 1737.776234 | -0.022592425 | 0.092963684 | 0.310648672 | 0.927416591 | ns | TM | KS.3 |
| FIM1_3665 | 929.1960262 | 931.7478005 | 1234.791245 | 0.000201698 | 0.083455601 | 0.991814218 | 0.999525119 | ns | TM | KS.3 |
| FIM1_3666 | 1036.727923 | 1069.464995 | 1461.323656 | 0.002739968 | 0.08143165 | 0.891683033 | 0.991141351 | ns | TM | KS.3 |
| FIM1_3667 | 1850.079282 | 1704.908915 | 2133.783649 | -0.008754808 | 0.078990599 | 0.679689833 | 0.97986479 | ns | TM | KS.3 |
| FIM1_3668 | 1178.981224 | 1042.188039 | 1477.657812 | -0.012492226 | 0.082507287 | 0.55588955 | 0.959760553 | ns | TM | KS.3 |
| FIM1_3669 | 881.0283987 | 770.1322783 | 1245.2853 | -0.011773245 | 0.085839655 | 0.566162334 | 0.962265923 | ns | TM | KS.3 |
| FIM1_3670 | 620.3009461 | 560.4052454 | 785.9254473 | -0.009286052 | 0.083109293 | 0.650967158 | 0.976323032 | ns | TM | KS.3 |
| FIM1_3671 | 461.2590593 | 409.7354865 | 649.025862 | -0.009053158 | 0.087572168 | 0.646756964 | 0.976323032 | ns | TM | KS.3 |
| FIM1_3672 | 1266.902757 | 1269.882513 | 2288.424868 | 0.000228791 | 0.078497823 | 0.991157062 | 0.999525119 | ns | TM | KS.3 |
| FIM1_3673 | 1380.569595 | 1521.187233 | 2448.216564 | 0.007581238 | 0.086066246 | 0.701249187 | 0.98151979 | ns | TM | KS.3 |
| FIM1_3674 | 47.06939684 | 32.77260702 | 71.71387351 | -0.030573062 | 0.116144849 | 0.185877939 | 0.825903583 | ns | TM | KS.3 |
| FIM1_3675 | 3622.850929 | 3444.112206 | 4505.666315 | -0.005366402 | 0.07792554 | 0.798398817 | 0.988916276 | ns | TM | KS.3 |
| FIM1_3676 | 62.75832737 | 70.50578783 | 187.0670644 | 0.009721391 | 0.086009893 | 0.628823047 | 0.97344368 | ns | TM | KS.3 |
| FIM1_3677 | 1090.750044 | 1154.450495 | 1500.927836 | 0.005752679 | 0.07905572 | 0.782346243 | 0.988814136 | ns | TM | KS.3 |
| FIM1_3678 | 1618.587572 | 1974.968208 | 2981.63063 | 0.011613886 | 0.098230257 | 0.540778572 | 0.95917717 | ns | TM | KS.3 |

| | | | | | | | | | | |
|---|---|---|---|---|---|---|---|---|---|---|
| FIM1_3679 | 48922.22115 | 49057.13188 | 61159.05026 | 0.000383907 | 0.071032929 | 0.986111245 | 0.999333464 | ns | TM | KS.3 |
| FIM1_3680 | 90.33233064 | 106.8777095 | 178.89611 | 0.014461873 | 0.088311459 | 0.485541051 | 0.952269291 | ns | TM | KS.3 |
| FIM1_3681 | 860.660898 | 816.9013845 | 1295.64304 | -0.005620717 | 0.077603998 | 0.789917936 | 0.988916276 | ns | TM | KS.3 |
| FIM1_3682 | 1539.347495 | 1405.506148 | 3493.016974 | -0.008410418 | 0.082341854 | 0.682007886 | 0.97986479 | ns | TM | KS.3 |
| FIM1_3683 | 1269.058854 | 1128.692873 | 1390.773785 | -0.013449008 | 0.080015767 | 0.537886613 | 0.95917717 | ns | TM | KS.3 |
| FIM1_3684 | 456.4877593 | 410.1661778 | 675.516524 | -0.010759552 | 0.081588288 | 0.609180347 | 0.971367716 | ns | TM | KS.3 |
| FIM1_3685 | 673.5782784 | 676.4545408 | 967.9350028 | 0.000345061 | 0.082103266 | 0.986178475 | 0.999333464 | ns | TM | KS.3 |
| FIM1_3686 | 715.9103488 | 816.7421071 | 1007.071036 | 0.008146046 | 0.092970012 | 0.666186132 | 0.976323032 | ns | TM | KS.3 |
| FIM1_3687 | 643.0583106 | 538.2258664 | 709.5175891 | -0.019423418 | 0.086425006 | 0.381978696 | 0.948637087 | ns | TM | KS.3 |
| FIM1_3688 | 989.3464315 | 1028.074816 | 1342.19789 | 0.00302594 | 0.084105594 | 0.877495856 | 0.991141351 | ns | TM | KS.3 |
| FIM1_3689 | 1348.419111 | 1286.44548 | 1805.398193 | -0.005078271 | 0.07743173 | 0.809651303 | 0.988916276 | ns | TM | KS.3 |
| FIM1_3690 | 911.9720398 | 750.7330972 | 1028.481125 | -0.0213782 | 0.088326643 | 0.341702847 | 0.944808611 | ns | TM | KS.3 |
| FIM1_3691 | 2760.500599 | 2749.896117 | 3698.699931 | -0.00044049 | 0.075677742 | 0.983435404 | 0.999333464 | ns | TM | KS.3 |
| FIM1_3692 | 1438.133531 | 1607.134311 | 2354.507026 | 0.011617289 | 0.081120768 | 0.585195773 | 0.963781856 | ns | TM | KS.3 |
| FIM1_3693 | 1692.832647 | 1509.819117 | 2060.419245 | -0.0117237 | 0.081702076 | 0.580289042 | 0.963781856 | ns | TM | KS.3 |
| FIM1_3694 | 1047.441205 | 775.971625 | 1198.114767 | -0.038778502 | 0.10719547 | 0.135733528 | 0.768410055 | ns | TM | KS.3 |
| FIM1_3695 | 2884.571735 | 2838.625565 | 5682.653673 | -0.00156287 | 0.07910968 | 0.939347235 | 0.994413583 | ns | TM | KS.3 |
| FIM1_3696 | 2890.617377 | 2738.070687 | 4593.752869 | -0.005556073 | 0.078772365 | 0.789968894 | 0.988916276 | ns | TM | KS.3 |
| FIM1_3697 | 8710.402723 | 9318.687622 | 11978.24542 | 0.010073021 | 0.071718962 | 0.65651954 | 0.976323032 | ns | TM | KS.3 |
| FIM1_3698 | 2100.056632 | 2080.923665 | 3455.448409 | -0.00093025 | 0.078043011 | 0.964230402 | 0.995175461 | ns | TM | KS.3 |
| FIM1_3699 | 110.5905813 | 115.9796105 | 1113.884538 | 0.002761335 | 0.091926618 | 0.880751364 | 0.991141351 | ns | TM | KS.3 |
| FIM1_3700 | 461.6446898 | 429.0089016 | 866.2872135 | -0.00593009 | 0.084445413 | 0.764829439 | 0.988477371 | ns | TM | KS.3 |
| FIM1_3701 | 743.0922442 | 732.3789302 | 989.2838633 | -0.001262242 | 0.08184067 | 0.949637286 | 0.994413583 | ns | TM | KS.3 |
| FIM1_3702 | 480.2648846 | 313.1985969 | 650.5353628 | -0.060173198 | 0.147300903 | 0.054708644 | 0.531428718 | ns | TM | KS.3 |
| FIM1_3703 | 2076.982646 | 1805.564302 | 2849.464825 | -0.015327046 | 0.082603549 | 0.481923561 | 0.951416554 | ns | TM | KS.3 |
| FIM1_3704 | 1376.258705 | 988.4995573 | 1314.613492 | -0.037366415 | 0.112338151 | 0.139967697 | 0.775302706 | ns | TM | KS.3 |
| FIM1_3705 | 2511.816946 | 2255.143855 | 2843.130502 | -0.009898942 | 0.083310155 | 0.630719346 | 0.97344368 | ns | TM | KS.3 |
| FIM1_3706 | 3604.481195 | 3704.280001 | 5158.641043 | 0.002774453 | 0.078290277 | 0.89365145 | 0.991141351 | ns | TM | KS.3 |
| FIM1_3707 | 710.6024346 | 719.7039612 | 787.9552835 | 0.001088633 | 0.081755466 | 0.956584468 | 0.994413583 | ns | TM | KS.3 |
| FIM1_3708 | 5132.991385 | 4527.962671 | 5903.164291 | -0.018879905 | 0.077031937 | 0.421554871 | 0.950410534 | ns | TM | KS.3 |
| FIM1_3709 | 1555.548573 | 1730.270104 | 2293.754854 | 0.010356849 | 0.082136512 | 0.619840547 | 0.973122675 | ns | TM | KS.3 |
| FIM1_3710 | 2145.090275 | 2365.677832 | 2787.408431 | 0.010062839 | 0.080621122 | 0.633453828 | 0.97344368 | ns | TM | KS.3 |
| FIM1_3711 | 2143.235075 | 2254.27606 | 3620.269437 | 0.004941138 | 0.079586536 | 0.810812959 | 0.988916276 | ns | TM | KS.3 |
| FIM1_3712 | 2974.807974 | 3041.658025 | 3898.949549 | 0.002137654 | 0.079466747 | 0.916895853 | 0.991141351 | ns | TM | KS.3 |
| FIM1_3713 | 2523.908271 | 2408.836261 | 3243.832749 | -0.004949228 | 0.077825947 | 0.813623379 | 0.988916276 | ns | TM | KS.3 |
| FIM1_3714 | 465.1594729 | 441.5193833 | 1059.112359 | -0.003599379 | 0.087623472 | 0.850319191 | 0.991141351 | ns | TM | KS.3 |
| FIM1_3715 | 308.3485422 | 293.050585 | 396.9883538 | -0.003740611 | 0.086110216 | 0.84655104 | 0.991141351 | ns | TM | KS.3 |
| FIM1_3716 | 5613.873636 | 5086.858639 | 5631.398114 | -0.011787068 | 0.077951203 | 0.590144524 | 0.963781856 | ns | TM | KS.3 |
| FIM1_3717 | 7604.717977 | 6919.190146 | 8897.841964 | -0.0125028 | 0.075900204 | 0.576173337 | 0.963192994 | ns | TM | KS.3 |
| FIM1_3718 | 1538.872463 | 1416.198558 | 1894.578112 | -0.007167714 | 0.083434219 | 0.72212072 | 0.987157164 | ns | TM | KS.3 |
| FIM1_3719 | 1312.588984 | 951.7259635 | 1434.633716 | -0.027997241 | 0.108113731 | 0.217559538 | 0.865538807 | ns | TM | KS.3 |
| FIM1_3720 | 1180.616408 | 1056.867795 | 1754.689686 | -0.010130675 | 0.083590952 | 0.622596551 | 0.973122675 | ns | TM | KS.3 |
| FIM1_3722 | 102.526791 | 88.64466962 | 144.6627501 | -0.011734625 | 0.087604674 | 0.5621989 | 0.961073589 | ns | TM | KS.3 |
| FIM1_3723 | 230.9467561 | 216.0325827 | 374.7185674 | -0.004225251 | 0.090174038 | 0.821695123 | 0.991014118 | ns | TM | KS.3 |
| FIM1_3725 | 77.59612495 | 73.30499594 | 110.7036884 | -0.00266583 | 0.096840482 | 0.881232929 | 0.991141351 | ns | TM | KS.3 |

| | | | | | | | | | | |
|---|---|---|---|---|---|---|---|---|---|---|
| FIM1_3727 | 342.0381755 | 423.1891576 | 551.350287 | 0.014748027 | 0.096071335 | 0.458676299 | 0.950410534 | ns | TM | KS.3 |
| FIM1_3731 | 127.155971 | 140.2597128 | 113.1955211 | 0.007293168 | 0.087260842 | 0.708839052 | 0.983990871 | ns | TM | KS.3 |
| FIM1_3732 | 262.9879284 | 285.7036974 | 285.7986033 | 0.007099606 | 0.083615762 | 0.724109028 | 0.987157164 | ns | TM | KS.3 |
| FIM1_3733 | 1020.73476 | 1282.267387 | 1175.971791 | 0.079475755 | 0.106894024 | 0.034773604 | 0.42652037 | ns | TM | KS.3 |
| FIM1_3734 | 1056.990464 | 1479.268056 | 827.4544815 | 0.245775446 | 0.173398053 | 0.000706046 | 0.03736785 | ns | TM | KS.3 |
| FIM1_3735 | 2161.289867 | 1934.642702 | 1708.862647 | -0.075270925 | 0.073334925 | 0.034413023 | 0.42621823 | ns | TM | KS.3 |
| FIM1_3736 | 1163.302721 | 1215.682089 | 1412.888503 | 0.008847454 | 0.062972374 | 0.702696124 | 0.98151979 | ns | TM | KS.3 |
| FIM1_3737 | 289.1238719 | 219.8476691 | 793.9904804 | -0.036736031 | 0.101538831 | 0.153318198 | 0.783447458 | ns | TM | KS.3 |
| FIM1_3738 | 256.1958526 | 239.0217032 | 451.205716 | -0.01508316 | 0.064227618 | 0.527390198 | 0.959071703 | ns | TM | KS.3 |
| FIM1_3739 | 1343.546456 | 1105.742272 | 1405.266701 | -0.03098217 | 0.087095849 | 0.221905074 | 0.870513238 | ns | TM | KS.3 |
| FIM1_3740 | 2295.133308 | 1890.73891 | 2403.374038 | -0.057274053 | 0.093080457 | 0.077715613 | 0.612946304 | ns | TM | KS.3 |
| FIM1_3741 | 2816.75374 | 2430.929149 | 3060.206979 | -0.056453901 | 0.082966111 | 0.084064191 | 0.631751287 | ns | TM | KS.3 |
| FIM1_3742 | 2664.075689 | 2349.138195 | 2198.695666 | -0.043287242 | 0.075163463 | 0.142048701 | 0.777709768 | ns | TM | KS.3 |
| FIM1_3743 | 2864.665928 | 3173.484051 | 3417.466024 | 0.046633684 | 0.069097364 | 0.113937803 | 0.714310891 | ns | TM | KS.3 |
| FIM1_3744 | 2051.633117 | 1762.454472 | 1593.700646 | -0.055233962 | 0.083806656 | 0.088030612 | 0.64839159 | ns | TM | KS.3 |
| FIM1_3745 | 10353.48071 | 11692.30802 | 12179.15248 | 0.054176404 | 0.076027154 | 0.088861446 | 0.651940574 | ns | TM | KS.3 |
| FIM1_3746 | 1945.895453 | 1765.691197 | 1963.194845 | -0.028761934 | 0.06650244 | 0.270619461 | 0.908468501 | ns | TM | KS.3 |
| FIM1_3747 | 16242.10054 | 13728.55074 | 16972.88969 | -0.043983435 | 0.084443503 | 0.134563089 | 0.766830871 | ns | TM | KS.3 |
| FIM1_3748 | 8137.371981 | 9260.617414 | 7906.555294 | 0.034821225 | 0.074930123 | 0.206251422 | 0.85082987 | ns | TM | KS.3 |
| FIM1_3749 | 4595.274544 | 4652.851889 | 4416.571733 | 0.004003392 | 0.038134825 | 0.820432478 | 0.990886949 | ns | TM | KS.3 |
| FIM1_3750 | 1073.68554 | 1169.86045 | 1375.601999 | 0.007661673 | 0.082762582 | 0.706455853 | 0.983265883 | ns | TM | KS.3 |
| FIM1_3751 | 1901.373262 | 1958.791024 | 1850.963617 | 0.009752863 | 0.043532807 | 0.617672007 | 0.973122675 | ns | TM | KS.3 |
| FIM1_3752 | 2702.343426 | 2250.970632 | 2341.916908 | -0.112542635 | 0.097725079 | 0.01271438 | 0.257037736 | ns | TM | KS.3 |
| FIM1_3753 | 1328.483643 | 1934.715577 | 2476.245096 | 0.255513829 | 0.206386253 | 0.000901759 | 0.044072215 | ns | TM | KS.3 |
| FIM1_3754 | 1814.700961 | 1858.408897 | 2359.349226 | 0.002751736 | 0.075483474 | 0.897281247 | 0.991141351 | ns | TM | KS.3 |
| FIM1_3755 | 176.8455438 | 235.5306172 | 213.6750868 | 0.055744426 | 0.111387643 | 0.07055137 | 0.582627046 | ns | TM | KS.3 |
| FIM1_3756 | 22483.85473 | 23509.62377 | 22678.61819 | 0.011532053 | 0.055909836 | 0.613525018 | 0.973122675 | ns | TM | KS.3 |
| FIM1_3757 | 682.9562576 | 687.9003318 | 660.2333306 | 0.001594941 | 0.052524834 | 0.942020926 | 0.994413583 | ns | TM | KS.3 |
| FIM1_3758 | 1581.612682 | 1781.606447 | 1542.398213 | 0.021988856 | 0.074038191 | 0.371018613 | 0.948637087 | ns | TM | KS.3 |
| FIM1_3759 | 10798.62334 | 9453.27427 | 9864.711821 | -0.017188154 | 0.079710133 | 0.448717402 | 0.950410534 | ns | TM | KS.3 |
| FIM1_3760 | 530.2780428 | 634.6300952 | 619.9495454 | 0.081510259 | 0.095726148 | 0.034538853 | 0.42621823 | ns | TM | KS.3 |
| FIM1_3761 | 655.3156982 | 683.0638112 | 735.6103109 | 0.013167062 | 0.047677139 | 0.529159521 | 0.959071703 | ns | TM | KS.3 |
| FIM1_3762 | 1553.265615 | 1820.030605 | 1837.459971 | 0.041694724 | 0.082032278 | 0.14971405 | 0.783447458 | ns | TM | KS.3 |
| FIM1_3763 | 3667.295392 | 3757.847444 | 3123.577917 | 0.003729054 | 0.069039278 | 0.867951825 | 0.991141351 | ns | TM | KS.3 |
| FIM1_3764 | 638.831684 | 755.7117598 | 462.6643805 | 0.091333665 | 0.09303602 | 0.024938142 | 0.373642194 | ns | TM | KS.3 |
| FIM1_3765 | 11477.71707 | 10673.14351 | 9241.215501 | -0.016179091 | 0.064364082 | 0.500492359 | 0.953260948 | ns | TM | KS.3 |
| FIM1_3766 | 76.97195808 | 63.63546489 | 98.51668961 | -0.032956287 | 0.087378174 | 0.203399071 | 0.847612283 | ns | TM | KS.3 |
| FIM1_3767 | 2357.362325 | 2089.318656 | 2270.164745 | -0.033023614 | 0.072787366 | 0.22423365 | 0.873580587 | ns | TM | KS.3 |
| FIM1_3768 | 24784.22115 | 25789.53805 | 26488.78546 | 0.002665406 | 0.088057555 | 0.888114572 | 0.991141351 | ns | TM | KS.3 |
| FIM1_3769 | 1906.254552 | 1836.49702 | 2223.519758 | -0.011922005 | 0.045350551 | 0.554042251 | 0.95917717 | ns | TM | KS.3 |
| FIM1_3770 | 7007.640285 | 6838.060957 | 6474.110678 | -0.008605911 | 0.031459138 | 0.546552351 | 0.95917717 | ns | TM | KS.3 |
| FIM1_3771 | 1475.931051 | 1355.589485 | 1381.914983 | -0.018508847 | 0.066850298 | 0.447452911 | 0.950410534 | ns | TM | KS.3 |
| FIM1_3772 | 2373.52679 | 2158.43553 | 2235.125208 | -0.019640322 | 0.069095629 | 0.422537221 | 0.950410534 | ns | TM | KS.3 |
| FIM1_3773 | 9426.842791 | 9512.602587 | 9253.453081 | 0.001582482 | 0.065331705 | 0.944443243 | 0.994413583 | ns | TM | KS.3 |
| FIM1_3774 | 6966.010993 | 5338.139625 | 4738.193851 | -0.097481503 | 0.122417605 | 0.019363331 | 0.32435209 | ns | TM | KS.3 |

| | | | | | | | | | | |
|---|---|---|---|---|---|---|---|---|---|---|
| FIM1_3775 | 259.4218662 | 266.3673993 | 237.9796987 | 0.006451453 | 0.057511721 | 0.776746396 | 0.988814136 | ns | TM | KS.3 |
| FIM1_3776 | 686.5727738 | 664.0923628 | 418.9431956 | -0.007783105 | 0.058531017 | 0.734114631 | 0.987655486 | ns | TM | KS.3 |
| FIM1_3777 | 12583.61347 | 11897.6173 | 9059.563685 | -0.004081174 | 0.086347886 | 0.83267084 | 0.991141351 | ns | TM | KS.3 |
| FIM1_3778 | 264.2394095 | 214.8857804 | 395.1968135 | -0.025103945 | 0.089317118 | 0.283361971 | 0.918450395 | ns | TM | KS.3 |
| FIM1_3779 | 1594.051596 | 1580.347179 | 1354.296373 | -0.001915786 | 0.057477457 | 0.932473837 | 0.994413583 | ns | TM | KS.3 |
| FIM1_3780 | 12853.87913 | 11607.78353 | 10131.22759 | -0.049439627 | 0.069194003 | 0.099446445 | 0.677856366 | ns | TM | KS.3 |
| FIM1_3782 | 3174.559535 | 3085.26529 | 3009.537331 | -0.007079129 | 0.05486134 | 0.752445217 | 0.988477371 | ns | TM | KS.3 |
| FIM1_3783 | 1456.884516 | 1450.651457 | 1425.301828 | -0.001137785 | 0.048041502 | 0.956669592 | 0.994413583 | ns | TM | KS.3 |
| FIM1_3784 | 6153.646449 | 4546.960654 | 3653.060382 | -0.260973837 | 0.141114447 | 0.0002579 | 0.017944075 | ns | TM | KS.3 |
| FIM1_3785 | 550.0294593 | 441.9510847 | 374.948203 | -0.043964702 | 0.09363259 | 0.123600131 | 0.739893493 | ns | TM | KS.3 |
| FIM1_3786 | 4879.070021 | 4337.905645 | 4454.690071 | -0.089883409 | 0.074621581 | 0.019918375 | 0.324535175 | ns | TM | KS.3 |
| FIM1_3787 | 41.82487405 | 62.02173278 | 84.05890081 | 0.100232896 | 0.168348781 | 0.017773487 | 0.304361209 | ns | TM | KS.3 |
| FIM1_3788 | 7454.003293 | 6746.065066 | 6370.812734 | -0.035921667 | 0.067075325 | 0.189445656 | 0.828217874 | ns | TM | KS.3 |
| FIM1_3789 | 637.5598677 | 652.3405219 | 690.6040231 | 0.005587353 | 0.054707887 | 0.802741019 | 0.988916276 | ns | TM | KS.3 |
| FIM1_3790 | 421.0959619 | 400.146988 | 421.9346332 | -0.010213217 | 0.064057533 | 0.661123218 | 0.976323032 | ns | TM | KS.3 |
| FIM1_3791 | 3589.361812 | 3042.762231 | 3342.107276 | -0.158174926 | 0.07724038 | 0.001605912 | 0.067706867 | ns | TM | KS.3 |
| FIM1_3792 | 3242.274026 | 2720.689293 | 3126.315086 | -0.026071456 | 0.083908214 | 0.284488856 | 0.918450395 | ns | TM | KS.3 |
| FIM1_3793 | 7123.680319 | 6212.719521 | 6434.595813 | -0.045267063 | 0.078245404 | 0.131238198 | 0.761283488 | ns | TM | KS.3 |
| FIM1_3794 | 9621.765692 | 10191.96153 | 9539.717368 | 0.015435922 | 0.057569944 | 0.508995219 | 0.953260948 | ns | TM | KS.3 |
| FIM1_3795 | 600.8541643 | 697.6703404 | 881.351855 | 0.023463477 | 0.079856109 | 0.332442749 | 0.938982061 | ns | TM | KS.3 |
| FIM1_3796 | 330.1506946 | 317.4608198 | 333.1994851 | -0.007898513 | 0.064148575 | 0.732260037 | 0.987157164 | ns | TM | KS.3 |
| FIM1_3797 | 3655.270746 | 3905.857754 | 4135.845504 | 0.01662135 | 0.060825904 | 0.486438984 | 0.952394312 | ns | TM | KS.3 |
| FIM1_3798 | 1863.790189 | 2215.856173 | 2050.789803 | 0.063903839 | 0.090498828 | 0.063104723 | 0.552721826 | ns | TM | KS.3 |
| FIM1_3799 | 20176.66403 | 15941.79466 | 14279.63957 | -0.065925842 | 0.104481149 | 0.053353698 | 0.528754276 | ns | TM | KS.3 |
| FIM1_3801 | 2178.532003 | 2203.132156 | 2398.246948 | 0.001830126 | 0.067379923 | 0.935307157 | 0.994413583 | ns | TM | KS.3 |
| FIM1_3802 | 2288.517168 | 1861.328664 | 2210.137681 | -0.021419059 | 0.09031559 | 0.335768661 | 0.938982061 | ns | TM | KS.3 |
| FIM1_3803 | 341.2175223 | 308.797597 | 313.849329 | -0.017426272 | 0.072547999 | 0.463617379 | 0.950410534 | ns | TM | KS.3 |
| FIM1_3804 | 348.8505244 | 311.382742 | 257.7622274 | -0.027493789 | 0.070607463 | 0.290304645 | 0.919993167 | ns | TM | KS.3 |
| FIM1_3805 | 350.0507885 | 327.3296406 | 471.9418112 | -0.013764275 | 0.065181168 | 0.561240667 | 0.961073589 | ns | TM | KS.3 |
| FIM1_3806 | 849.0848679 | 1017.818714 | 864.3943599 | 0.021287176 | 0.086129922 | 0.349407171 | 0.948637087 | ns | TM | KS.3 |
| FIM1_3807 | 9638.862899 | 9592.659838 | 10336.56045 | -0.001495562 | 0.031200906 | 0.918981014 | 0.991141351 | ns | TM | KS.3 |
| FIM1_3808 | 503.2220417 | 464.0706777 | 598.9609995 | -0.02506904 | 0.061617961 | 0.316005567 | 0.927564309 | ns | TM | KS.3 |
| FIM1_3809 | 1529.474677 | 1853.412678 | 2060.132763 | 0.016793307 | 0.090579873 | 0.423971452 | 0.950410534 | ns | TM | KS.3 |
| FIM1_3810 | 1963.37015 | 2047.98004 | 1956.994185 | 0.009352607 | 0.060004974 | 0.685912965 | 0.980861455 | ns | TM | KS.3 |
| FIM1_3811 | 2409.407246 | 2551.84428 | 5184.452121 | 0.01203766 | 0.06349015 | 0.608362519 | 0.970937835 | ns | TM | KS.3 |
| FIM1_3812 | 21.43477154 | 11.29256932 | 22.63589428 | -0.07178083 | 0.205649606 | 0.038493395 | 0.444326306 | ns | TM | KS.3 |
| FIM1_3813 | 3771.360817 | 3774.097925 | 4199.466434 | 0.00028913 | 0.038476944 | 0.987057035 | 0.999333464 | ns | TM | KS.3 |
| FIM1_3814 | 22.20179959 | 22.51406973 | 26.24877177 | -0.00035573 | 0.087774028 | 0.984995932 | 0.999333464 | ns | TM | KS.3 |
| FIM1_3815 | 13430.60882 | 16132.99136 | 13481.12748 | 0.023791766 | 0.08569239 | 0.311114923 | 0.927416591 | ns | TM | KS.3 |
| FIM1_3816 | 2233.810176 | 2398.223406 | 2349.079688 | 0.010733797 | 0.071707631 | 0.636938492 | 0.974658588 | ns | TM | KS.3 |
| FIM1_3817 | 5529.415591 | 5022.816602 | 3144.887224 | -0.027599843 | 0.066265603 | 0.286461862 | 0.919993167 | ns | TM | KS.3 |
| FIM1_3818 | 69.80478101 | 62.15548577 | 352.0551801 | -0.011854536 | 0.083446315 | 0.571058441 | 0.962265923 | ns | TM | KS.3 |
| FIM1_3819 | 1929.941237 | 1974.213114 | 2295.003418 | 0.003888047 | 0.06617376 | 0.864146208 | 0.991141351 | ns | TM | KS.3 |
| FIM1_3820 | 1147.715784 | 1321.712528 | 1699.424937 | 0.019268001 | 0.079942849 | 0.405078294 | 0.950410534 | ns | TM | KS.3 |
| FIM1_3821 | 941.9630978 | 929.5340236 | 934.0114174 | -0.00236491 | 0.064785542 | 0.917234561 | 0.991141351 | ns | TM | KS.3 |

| | | | | | | | | | | |
|---|---|---|---|---|---|---|---|---|---|---|
| FIM1_3822 | 4483.196185 | 4197.442605 | 5347.315536 | -0.017785084 | 0.059226981 | 0.455406137 | 0.950410534 | ns | TM | KS.3 |
| FIM1_3823 | 35.5832083 | 79.12901779 | 134.5366798 | 1.012230502 | 0.253124188 | 1.96103588911 | 0.00026368 | Km>KS.3 | TM | KS.3 |
| FIM1_3824 | 8801.647606 | 10149.36001 | 20601.80896 | 0.020776321 | 0.079443846 | 0.378397722 | 0.948637087 | ns | TM | KS.3 |
| FIM1_3825 | 335.1218268 | 335.2116288 | 253.0589104 | -4.67E-06 | 0.082108461 | 0.999812975 | 0.999930384 | ns | TM | KS.3 |
| FIM1_3826 | 5755.673791 | 5561.151879 | 6236.632612 | -0.012568444 | 0.037789229 | 0.459399267 | 0.950410534 | ns | TM | KS.3 |
| FIM1_3827 | 444.0605168 | 503.7124225 | 477.8659757 | 0.032168963 | 0.073641172 | 0.232878632 | 0.883057311 | ns | TM | KS.3 |
| FIM1_3828 | 2625.146659 | 2911.978549 | 3076.068311 | 0.039832955 | 0.068626495 | 0.158513439 | 0.789048157 | ns | TM | KS.3 |
| FIM1_3829 | 2173.586434 | 2710.380666 | 2219.783123 | 0.144964151 | 0.109707407 | 0.004933873 | 0.140896498 | ns | TM | KS.3 |
| FIM1_3830 | 3120.159845 | 2885.995671 | 3249.216534 | -0.018692635 | 0.064019844 | 0.442827254 | 0.950410534 | ns | TM | KS.3 |
| FIM1_3831 | 820.4702648 | 637.8083296 | 740.3250707 | -0.071386871 | 0.110198347 | 0.04329755 | 0.476560428 | ns | TM | KS.3 |
| FIM1_3832 | 888.5389702 | 939.1421855 | 785.8791234 | 0.013008615 | 0.059976877 | 0.578820875 | 0.963781856 | ns | TM | KS.3 |
| FIM1_3833 | 76.74410986 | 60.76400271 | 255.9778003 | -0.019470418 | 0.095406901 | 0.357908771 | 0.948637087 | ns | TM | KS.3 |
| FIM1_3834 | 2743.194775 | 2815.456807 | 3235.127828 | 0.007771909 | 0.046029908 | 0.703184892 | 0.98151979 | ns | TM | KS.3 |
| FIM1_3835 | 305.9379648 | 291.0015311 | 381.0551814 | -0.013751086 | 0.05923149 | 0.557336329 | 0.959760553 | ns | TM | KS.3 |
| FIM1_3836 | 3869.410813 | 4032.515394 | 4396.370425 | 0.008820021 | 0.061153318 | 0.703127515 | 0.98151979 | ns | TM | KS.3 |
| FIM1_3837 | 2761.548355 | 2433.784929 | 2397.157586 | -0.050205521 | 0.076705188 | 0.10602484 | 0.700394017 | ns | TM | KS.3 |
| FIM1_3838 | 1590.650257 | 1775.018245 | 1857.047638 | 0.036452572 | 0.070138177 | 0.189616112 | 0.828217874 | ns | TM | KS.3 |
| FIM1_3839 | 2466.576096 | 2224.017014 | 2382.466326 | -0.013264338 | 0.077148061 | 0.551852833 | 0.95917717 | ns | TM | KS.3 |
| FIM1_3840 | 820.6301947 | 832.6865229 | 898.5308475 | 0.0023759 | 0.067355085 | 0.916140122 | 0.991141351 | ns | TM | KS.3 |
| FIM1_3841 | 2079.315171 | 2119.6375 | 2372.251608 | 0.002941614 | 0.068934708 | 0.895553952 | 0.991141351 | ns | TM | KS.3 |
| FIM1_3842 | 125.5341811 | 133.2395247 | 197.4601246 | 0.005303138 | 0.082261416 | 0.792464044 | 0.988916276 | ns | TM | KS.3 |
| FIM1_3843 | 10829.14196 | 11248.43718 | 11982.34472 | 0.013630624 | 0.041272689 | 0.461107285 | 0.950410534 | ns | TM | KS.3 |
| FIM1_3844 | 865.5935695 | 992.1116211 | 1232.990209 | 0.016106677 | 0.081254177 | 0.467965789 | 0.950410534 | ns | TM | KS.3 |
| FIM1_3845 | 794.3106401 | 805.879312 | 1007.305287 | 0.00170573 | 0.074876353 | 0.93645635 | 0.994413583 | ns | TM | KS.3 |
| FIM1_3846 | 704.5716343 | 716.9596729 | 975.6377112 | 0.004646068 | 0.053534215 | 0.833885475 | 0.991141351 | ns | TM | KS.3 |
| FIM1_3847 | 29921.38602 | 32005.37484 | 31400.47508 | 0.025935424 | 0.054406348 | 0.268262329 | 0.906045545 | ns | TM | KS.3 |
| FIM1_3848 | 685.9143568 | 736.4463597 | 750.7664885 | 0.008476858 | 0.076249089 | 0.695831164 | 0.981479944 | ns | TM | KS.3 |
| FIM1_3849 | 33205.25253 | 31116.32816 | 30737.89021 | -0.014722418 | 0.062772459 | 0.535940251 | 0.959071703 | ns | TM | KS.3 |
| FIM1_3850 | 2345.192072 | 2425.528785 | 2505.267773 | 0.005523443 | 0.067722495 | 0.807605101 | 0.988916276 | ns | TM | KS.3 |
| FIM1_3851 | 772.1383602 | 772.4622513 | 1348.817359 | 0.000330198 | 0.057569806 | 0.988345245 | 0.999333464 | ns | TM | KS.3 |
| FIM1_3852 | 1077.646898 | 1129.734896 | 1300.114399 | 0.009511226 | 0.063156507 | 0.682323349 | 0.97986479 | ns | TM | KS.3 |
| FIM1_3853 | 1567.071861 | 1599.9985 | 1715.996936 | 0.005270686 | 0.053021401 | 0.81132477 | 0.988916276 | ns | TM | KS.3 |
| FIM1_3854 | 7183.993557 | 5953.749126 | 5777.754768 | -0.056853528 | 0.091719224 | 0.079714597 | 0.616765353 | ns | TM | KS.3 |
| FIM1_3855 | 4196.260132 | 3686.084099 | 3691.321226 | -0.04401587 | 0.076345517 | 0.138083297 | 0.770139466 | ns | TM | KS.3 |
| FIM1_3856 | 2102.97071 | 1980.900588 | 1475.589319 | -0.015451778 | 0.058839611 | 0.511706642 | 0.953818112 | ns | TM | KS.3 |
| FIM1_3857 | 7354.157878 | 7175.765014 | 6700.010341 | -0.007867196 | 0.040823507 | 0.67189999 | 0.976861203 | ns | TM | KS.3 |
| FIM1_3858 | 1236.646232 | 1157.897773 | 1106.416192 | -0.019072662 | 0.058008715 | 0.422268805 | 0.950410534 | ns | TM | KS.3 |
| FIM1_3859 | 29.10886208 | 28.72372492 | 26.75156027 | 0.000275228 | 0.081048758 | 0.98908904 | 0.999333464 | ns | TM | KS.3 |
| FIM1_3860 | 1930.349124 | 2112.710192 | 2591.085474 | 0.012382791 | 0.074957594 | 0.582127284 | 0.963781856 | ns | TM | KS.3 |
| FIM1_3861 | 720.1534162 | 788.4768385 | 683.3851481 | 0.012451888 | 0.074866662 | 0.580470368 | 0.963781856 | ns | TM | KS.3 |
| FIM1_3862 | 4347.917553 | 4745.273913 | 4239.368737 | 0.007584872 | 0.083461481 | 0.707390246 | 0.983273233 | ns | TM | KS.3 |
| FIM1_3863 | 5440.617335 | 5305.452053 | 5834.400475 | -0.005349448 | 0.060012925 | 0.815169898 | 0.988916276 | ns | TM | KS.3 |
| FIM1_3864 | 142.5778692 | 219.7239447 | 239.0951616 | 0.057479237 | 0.147228338 | 0.059815966 | 0.54403004 | ns | TM | KS.3 |
| FIM1_3865 | 695.3967935 | 924.6065353 | 647.6237333 | 0.030126495 | 0.102265048 | 0.203516319 | 0.847612283 | ns | TM | KS.3 |
| FIM1_3866 | 1208.732543 | 1567.516216 | 1486.262765 | 0.109060082 | 0.123391079 | 0.014110296 | 0.269253438 | ns | TM | KS.3 |

| | | | | | | | | | | |
|---|---|---|---|---|---|---|---|---|---|---|
| FIM1_3867 | 1334.238023 | 1444.603635 | 1536.73574 | 0.017866464 | 0.065110645 | 0.461575033 | 0.950410534 | ns | TM | KS.3 |
| FIM1_3868 | 340.9977172 | 341.435201 | 508.917387 | 0.000433194 | 0.067572938 | 0.984644279 | 0.999333464 | ns | TM | KS.3 |
| FIM1_3869 | 15940.46659 | 14468.7432 | 15450.81622 | -0.014909449 | 0.073629338 | 0.519665166 | 0.958509403 | ns | TM | KS.3 |
| FIM1_3870 | 1797.313713 | 1806.139744 | 1812.154242 | 0.000993176 | 0.05890593 | 0.965113666 | 0.995175461 | ns | TM | KS.3 |
| FIM1_3871 | 2197.260673 | 2584.727287 | 2563.177529 | 0.023883962 | 0.082060298 | 0.319917055 | 0.929108338 | ns | TM | KS.3 |
| FIM1_3872 | 3619.899268 | 3488.832568 | 5529.200178 | -0.012033464 | 0.045054312 | 0.548232007 | 0.95917717 | ns | TM | KS.3 |
| FIM1_3873 | 2663.47891 | 2326.774514 | 1453.231919 | -0.03068526 | 0.076025381 | 0.24668469 | 0.888934767 | ns | TM | KS.3 |
| FIM1_3874 | 545.0827315 | 442.1932652 | 510.1569601 | -0.093390206 | 0.105034117 | 0.023219876 | 0.35748181 | ns | TM | KS.3 |
| FIM1_3875 | 353.4002326 | 303.621844 | 283.7631909 | -0.047370355 | 0.081809955 | 0.119787257 | 0.730090427 | ns | TM | KS.3 |
| FIM1_3876 | 114.5016036 | 85.93751305 | 99.06440972 | -0.04142788 | 0.105831026 | 0.123090354 | 0.738690602 | ns | TM | KS.3 |
| FIM1_3877 | 106.657941 | 72.62604463 | 174.706943 | -0.103608577 | 0.164031496 | 0.016229233 | 0.29567114 | ns | TM | KS.3 |
| FIM1_3879 | 14178.9614 | 12127.83238 | 20269.80526 | -0.022905727 | 0.081202284 | 0.337350176 | 0.940821016 | ns | TM | KS.3 |
| FIM1_3880 | 3761.738408 | 3443.204183 | 3181.816457 | -0.01555556 | 0.07060038 | 0.510507544 | 0.953260948 | ns | TM | KS.3 |
| FIM1_3881 | 9250.057864 | 8334.533699 | 6875.92404 | -0.093106633 | 0.066408796 | 0.013332774 | 0.263081302 | ns | TM | KS.3 |
| FIM1_3882 | 58.83047964 | 61.79954864 | 60.88332332 | 0.004282075 | 0.082298922 | 0.830933154 | 0.991141351 | ns | TM | KS.3 |
| FIM1_3883 | 474.6242641 | 454.6267522 | 422.477108 | -0.009346881 | 0.061287908 | 0.686902153 | 0.980879562 | ns | TM | KS.3 |
| FIM1_3884 | 391.5403272 | 412.3251732 | 508.2338864 | 0.008066743 | 0.069323688 | 0.722614707 | 0.987157164 | ns | TM | KS.3 |
| FIM1_3885 | 2698.40035 | 2726.114768 | 2432.3808 | 0.002791182 | 0.048465343 | 0.894537398 | 0.991141351 | ns | TM | KS.3 |
| FIM1_3886 | 4920.588757 | 4173.449002 | 3862.400914 | -0.063353194 | 0.088477572 | 0.064802695 | 0.561727671 | ns | TM | KS.3 |
| FIM1_3887 | 504.6074602 | 489.3105574 | 388.6433825 | -0.008893236 | 0.04993329 | 0.679210233 | 0.97986479 | ns | TM | KS.3 |
| FIM1_3888 | 3599.12778 | 3227.677132 | 3161.601912 | -0.031152962 | 0.069751018 | 0.243915211 | 0.88883001 | ns | TM | KS.3 |
| FIM1_3889 | 1754.921904 | 1409.265981 | 1106.781719 | -0.045801446 | 0.094435278 | 0.114601025 | 0.714461279 | ns | TM | KS.3 |
| FIM1_3890 | 2691.114774 | 2211.019003 | 2590.63797 | -0.064845558 | 0.095771667 | 0.059187999 | 0.543284675 | ns | TM | KS.3 |
| FIM1_3891 | 296.0468951 | 256.4246116 | 393.9541856 | -0.019692083 | 0.080415475 | 0.395453729 | 0.948637087 | ns | TM | KS.3 |
| FIM1_3892 | 1478.014032 | 1416.036553 | 1681.82906 | -0.004029232 | 0.080254245 | 0.843692575 | 0.991141351 | ns | TM | KS.3 |
| FIM1_3893 | 127.4104507 | 135.1489451 | 134.945167 | 0.010874622 | 0.067553762 | 0.639447962 | 0.975246356 | ns | TM | KS.3 |
| FIM1_3894 | 738.0175143 | 692.1518644 | 844.3097769 | -0.016831455 | 0.059558546 | 0.478905232 | 0.950972051 | ns | TM | KS.3 |
| FIM1_3895 | 161.4125171 | 154.991569 | 166.2505611 | -0.006676235 | 0.069395271 | 0.767538158 | 0.988477371 | ns | TM | KS.3 |
| FIM1_3896 | 1326.258439 | 1202.643086 | 1224.149328 | -0.031105201 | 0.066553801 | 0.240644858 | 0.887413624 | ns | TM | KS.3 |
| FIM1_3897 | 487.6769959 | 504.2605837 | 509.3109975 | 0.007646782 | 0.058718914 | 0.738683184 | 0.988051177 | ns | TM | KS.3 |
| FIM1_3898 | 3446.547778 | 4024.737062 | 4862.051882 | 0.012348416 | 0.088972949 | 0.540595782 | 0.95917717 | ns | TM | KS.3 |
| FIM1_3899 | 859.4934201 | 905.3709284 | 974.2576244 | 0.01641925 | 0.051783632 | 0.459746405 | 0.950410534 | ns | TM | KS.3 |
| FIM1_3900 | 341.7974916 | 324.9981537 | 381.9129481 | -0.011325446 | 0.062048792 | 0.628261634 | 0.97344368 | ns | TM | KS.3 |
| FIM1_3901 | 1316.320708 | 1367.895956 | 1554.936371 | 0.009169575 | 0.057857909 | 0.68930476 | 0.980879562 | ns | TM | KS.3 |
| FIM1_3902 | 505.6217115 | 467.0297557 | 1006.228109 | -0.012505651 | 0.071047005 | 0.588064516 | 0.963781856 | ns | TM | KS.3 |
| FIM1_3903 | 549.0083523 | 479.7891787 | 596.3936974 | -0.031854815 | 0.076003787 | 0.234256627 | 0.883800031 | ns | TM | KS.3 |
| FIM1_3904 | 8236.810995 | 8098.910712 | 7748.498376 | -0.004418456 | 0.051357345 | 0.838936262 | 0.991141351 | ns | TM | KS.3 |
| FIM1_3905 | 6314.705969 | 7060.216037 | 6693.299249 | 0.021281449 | 0.072657467 | 0.386234947 | 0.948637087 | ns | TM | KS.3 |
| FIM1_3906 | 985.6957447 | 1222.269845 | 747.9196432 | 0.064992605 | 0.099731264 | 0.05709224 | 0.535913008 | ns | TM | KS.3 |
| FIM1_3907 | 2412.23299 | 2407.152552 | 1759.155748 | -0.000661404 | 0.048074627 | 0.974812718 | 0.99870125 | ns | TM | KS.3 |
| FIM1_3908 | 1554.619112 | 1634.055376 | 1524.870191 | 0.018729171 | 0.046733994 | 0.362510325 | 0.948637087 | ns | TM | KS.3 |
| FIM1_3909 | 1032.652862 | 1258.228548 | 1174.845268 | 0.023590758 | 0.088109945 | 0.307570809 | 0.927416591 | ns | TM | KS.3 |
| FIM1_3910 | 6118.772428 | 6015.759797 | 6264.664992 | -0.005265183 | 0.041455351 | 0.780332137 | 0.988814136 | ns | TM | KS.3 |
| FIM1_3911 | 206.1679545 | 183.1437148 | 182.5380654 | -0.018000373 | 0.077383062 | 0.438589212 | 0.950410534 | ns | TM | KS.3 |
| FIM1_3912 | 1395.545383 | 1498.538173 | 4257.191824 | 0.009989318 | 0.073325485 | 0.655574542 | 0.976323032 | ns | TM | KS.3 |

| | | | | | | | | | | |
|---|---|---|---|---|---|---|---|---|---|---|
| FIM1_3913 | 316.895684 | 346.4785851 | 592.3381723 | 0.016089915 | 0.070697826 | 0.497626344 | 0.953260948 | ns | TM | KS.3 |
| FIM1_3914 | 430.6504532 | 403.2391615 | 465.4991353 | -0.013374184 | 0.065297665 | 0.571536318 | 0.962265923 | ns | TM | KS.3 |
| FIM1_3915 | 2620.017207 | 2311.094468 | 2899.055429 | -0.054645283 | 0.077137045 | 0.088047685 | 0.64839159 | ns | TM | KS.3 |
| FIM1_3916 | 946.409609 | 772.4229893 | 852.7608378 | -0.058100841 | 0.095252984 | 0.074193759 | 0.597271765 | ns | TM | KS.3 |
| FIM1_3917 | 1744.947899 | 1739.984632 | 1846.392987 | -0.000563927 | 0.059222287 | 0.980200901 | 0.999333464 | ns | TM | KS.3 |
| FIM1_3918 | 2097.723202 | 1697.253121 | 1266.487463 | -0.033483401 | 0.090188488 | 0.193549858 | 0.835859849 | ns | TM | KS.3 |
| FIM1_3919 | 306.3797861 | 260.3889706 | 331.3315058 | -0.035799593 | 0.081751505 | 0.1905067 | 0.829253069 | ns | TM | KS.3 |
| FIM1_3920 | 290.9782628 | 229.2183543 | 231.2274604 | -0.076404965 | 0.109328914 | 0.037489058 | 0.440708412 | ns | TM | KS.3 |
| FIM1_3921 | 2801.600044 | 2591.187592 | 2798.983204 | -0.021483055 | 0.061852307 | 0.381997383 | 0.948637087 | ns | TM | KS.3 |
| FIM1_3922 | 606.7387352 | 559.2701845 | 500.5854168 | -0.017254407 | 0.06676063 | 0.475189674 | 0.950410534 | ns | TM | KS.3 |
| FIM1_3923 | 2965.436766 | 2775.94377 | 3006.232563 | -0.018000864 | 0.059173874 | 0.450236523 | 0.950410534 | ns | TM | KS.3 |
| FIM1_3924 | 407.2484293 | 539.9375759 | 705.9763253 | 0.020254633 | 0.102900249 | 0.329703187 | 0.936229083 | ns | TM | KS.3 |
| FIM1_3925 | 177.3489078 | 141.173312 | 212.0267322 | -0.032746869 | 0.092968976 | 0.194589152 | 0.837304507 | ns | TM | KS.3 |
| FIM1_3926 | 1339.5085 | 1406.075153 | 1880.232738 | 0.011546157 | 0.059229041 | 0.619329371 | 0.973122675 | ns | TM | KS.3 |
| FIM1_3927 | 632.4847977 | 429.9232036 | 370.1716768 | -0.177993503 | 0.200377258 | 0.003702863 | 0.115135887 | ns | TM | KS.3 |
| FIM1_3928 | 1854.449269 | 1849.680731 | 2189.577524 | -0.000555758 | 0.059665233 | 0.980506982 | 0.999333464 | ns | TM | KS.3 |
| FIM1_3929 | 1325.124211 | 1655.070285 | 1378.074944 | 0.051763459 | 0.096891926 | 0.090579105 | 0.65689657 | ns | TM | KS.3 |
| FIM1_3930 | 1098.226325 | 1056.863424 | 1070.974216 | -0.011888009 | 0.048766664 | 0.575772864 | 0.963192994 | ns | TM | KS.3 |
| FIM1_3932 | 2432.66252 | 2388.395978 | 2228.120119 | -0.00509972 | 0.04891828 | 0.809803313 | 0.988916276 | ns | TM | KS.3 |
| FIM1_3933 | 560.2646071 | 624.7455148 | 529.5501752 | 0.029772172 | 0.069699055 | 0.260506596 | 0.897386461 | ns | TM | KS.3 |
| FIM1_3934 | 725.2009329 | 782.2060369 | 1094.394944 | 0.021958031 | 0.059688374 | 0.366906171 | 0.948637087 | ns | TM | KS.3 |
| FIM1_3935 | 1478.952316 | 1326.646364 | 1625.570104 | -0.026478485 | 0.070266009 | 0.304655627 | 0.926536521 | ns | TM | KS.3 |
| FIM1_3936 | 779.3723526 | 789.2339243 | 891.8872015 | 0.003242885 | 0.045073984 | 0.871882248 | 0.991141351 | ns | TM | KS.3 |
| FIM1_3937 | 33.83107022 | 35.01401901 | 34.38314948 | 0.001971234 | 0.090622258 | 0.915319766 | 0.991141351 | ns | TM | KS.3 |
| FIM1_3938 | 57.18769332 | 67.62232041 | 80.08171728 | 0.01628018 | 0.086695822 | 0.446608161 | 0.950410534 | ns | TM | KS.3 |
| FIM1_3939 | 233.904895 | 253.4287026 | 588.5738225 | 0.009019925 | 0.077801989 | 0.674352317 | 0.976966447 | ns | TM | KS.3 |
| FIM1_3940 | 1493.66665 | 2185.3756 | 2416.895088 | 0.048487187 | 0.128122817 | 0.084999845 | 0.636858776 | ns | TM | KS.3 |
| FIM1_3942 | 576.3953986 | 704.775668 | 686.7445039 | 0.107121962 | 0.104505137 | 0.015356878 | 0.284118339 | ns | TM | KS.3 |
| FIM1_3943 | 4162.362129 | 4159.72963 | 4597.163711 | -0.000160156 | 0.040022678 | 0.993058682 | 0.999525119 | ns | TM | KS.3 |
| FIM1_3944 | 1180.346471 | 1049.585147 | 950.884262 | -0.051049272 | 0.074192469 | 0.09988975 | 0.679824224 | ns | TM | KS.3 |
| FIM1_3945 | 909.4124465 | 974.9365373 | 915.1070056 | 0.009551467 | 0.073544199 | 0.66843085 | 0.976323032 | ns | TM | KS.3 |
| FIM1_3946 | 1346.179711 | 1408.065018 | 1684.428242 | 0.003865437 | 0.082279344 | 0.846954029 | 0.991141351 | ns | TM | KS.3 |
| FIM1_3947 | 1490.728563 | 1357.061871 | 1402.901195 | -0.025615044 | 0.065961531 | 0.315867647 | 0.927564309 | ns | TM | KS.3 |
| FIM1_3948 | 1614.962612 | 1601.248791 | 1624.501317 | -0.002440496 | 0.042764707 | 0.899731985 | 0.991141351 | ns | TM | KS.3 |
| FIM1_3949 | 1499.077414 | 1422.899302 | 1524.536322 | -0.016897108 | 0.051007396 | 0.442601001 | 0.950410534 | ns | TM | KS.3 |
| FIM1_3950 | 1493.120387 | 1364.467688 | 1387.801633 | -0.037181282 | 0.063928256 | 0.168994324 | 0.806481872 | ns | TM | KS.3 |
| FIM1_3951 | 2022.311405 | 1807.810126 | 2008.560255 | -0.085022239 | 0.073404643 | 0.023281227 | 0.35748181 | ns | TM | KS.3 |
| FIM1_3952 | 1364.497482 | 1472.665812 | 1720.543302 | 0.00704457 | 0.08172932 | 0.730519689 | 0.987157164 | ns | TM | KS.3 |
| FIM1_3953 | 2108.511044 | 2109.744974 | 1806.123057 | 0.000165327 | 0.051360763 | 0.993921434 | 0.999525119 | ns | TM | KS.3 |
| FIM1_3954 | 644.2137333 | 546.1242068 | 635.473512 | -0.048453486 | 0.085067786 | 0.113046153 | 0.711904568 | ns | TM | KS.3 |
| FIM1_3955 | 695.4071019 | 667.0177263 | 717.4408983 | -0.007025265 | 0.067697518 | 0.757981908 | 0.988477371 | ns | TM | KS.3 |
| FIM1_3956 | 2786.476059 | 2416.699885 | 3147.383048 | -0.019341115 | 0.080145153 | 0.403027373 | 0.950410534 | ns | TM | KS.3 |
| FIM1_3957 | 2156.686173 | 2282.655542 | 2442.72559 | 0.02558501 | 0.047875855 | 0.22420412 | 0.873580587 | ns | TM | KS.3 |
| FIM1_3958 | 2265.702611 | 2557.490255 | 3999.83523 | 0.011740732 | 0.083133562 | 0.575325373 | 0.963192994 | ns | TM | KS.3 |
| FIM1_3959 | 586.1336201 | 653.6088731 | 783.6072442 | 0.036912092 | 0.070511143 | 0.185932063 | 0.825903583 | ns | TM | KS.3 |

| | | | | | | | | | | |
|---|---|---|---|---|---|---|---|---|---|---|
| FIM1_3960 | 167.3738349 | 175.4872024 | 230.82045 | 0.005425453 | 0.076656804 | 0.798624773 | 0.988916276 | ns | TM | KS.3 |
| FIM1_3961 | 915.1372116 | 902.3086025 | 941.5538678 | -0.001229698 | 0.081509488 | 0.951091178 | 0.994413583 | ns | TM | KS.3 |
| FIM1_3962 | 1111.331818 | 1187.842795 | 1344.310423 | 0.017151319 | 0.06046173 | 0.473089503 | 0.950410534 | ns | TM | KS.3 |
| FIM1_3963 | 1217.798427 | 1395.041311 | 1670.37721 | 0.016190156 | 0.08106257 | 0.46667068 | 0.950410534 | ns | TM | KS.3 |
| FIM1_3964 | 1845.932671 | 2141.487777 | 2477.98014 | 0.013938181 | 0.085666321 | 0.506909165 | 0.953260948 | ns | TM | KS.3 |
| FIM1_3965 | 604.443987 | 599.9881904 | 578.3806888 | -0.001778476 | 0.05655898 | 0.937045744 | 0.994413583 | ns | TM | KS.3 |
| FIM1_3966 | 354.2080899 | 410.8659778 | 404.0608576 | 0.008230963 | 0.096530988 | 0.656558666 | 0.976323032 | ns | TM | KS.3 |
| FIM1_3967 | 1537.164044 | 1583.793934 | 1412.758355 | 0.005903458 | 0.062532162 | 0.797179847 | 0.988916276 | ns | TM | KS.3 |
| FIM1_3968 | 1273.304834 | 1331.967769 | 1315.968476 | 0.011883025 | 0.055871244 | 0.603026898 | 0.970245665 | ns | TM | KS.3 |
| FIM1_3969 | 1854.281183 | 2267.664301 | 2221.040879 | 0.051869478 | 0.092899405 | 0.093702724 | 0.671406965 | ns | TM | KS.3 |
| FIM1_3970 | 899.7652477 | 759.1412171 | 906.8312738 | -0.026927183 | 0.082861653 | 0.276320916 | 0.908583046 | ns | TM | KS.3 |
| FIM1_3971 | 369.0308472 | 433.5826331 | 324.094424 | 0.016171377 | 0.085778822 | 0.451911768 | 0.950410534 | ns | TM | KS.3 |
| FIM1_3972 | 538.0979224 | 543.3154936 | 550.1518357 | 0.003079611 | 0.048540069 | 0.883826145 | 0.991141351 | ns | TM | KS.3 |
| FIM1_3973 | 5453.829747 | 5726.929497 | 6210.686297 | 0.013203824 | 0.055293178 | 0.563129171 | 0.961086664 | ns | TM | KS.3 |
| FIM1_3974 | 1295.464718 | 1225.357371 | 1268.56398 | -0.013281502 | 0.060321227 | 0.571764075 | 0.962265923 | ns | TM | KS.3 |
| FIM1_3975 | 749.5998009 | 686.7355746 | 1007.307858 | -0.021008898 | 0.065707364 | 0.396295158 | 0.948781719 | ns | TM | KS.3 |
| FIM1_3976 | 8382.827645 | 6726.035746 | 8974.615497 | -0.062775855 | 0.100119391 | 0.061072556 | 0.545486476 | ns | TM | KS.3 |
| FIM1_3977 | 454.4782137 | 288.3131816 | 1321.053844 | -0.040405545 | 0.136440727 | 0.113862217 | 0.714310891 | ns | TM | KS.3 |
| FIM1_3978 | 1045.757613 | 1102.73976 | 1124.950898 | 0.010602967 | 0.064180415 | 0.649574229 | 0.976323032 | ns | TM | KS.3 |
| FIM1_3979 | 809.4208444 | 760.8009523 | 703.252064 | -0.019253457 | 0.055867105 | 0.409965619 | 0.950410534 | ns | TM | KS.3 |
| FIM1_3980 | 14738.02329 | 17140.00849 | 15468.15665 | 0.05806257 | 0.084228204 | 0.07909171 | 0.616697988 | ns | TM | KS.3 |
| FIM1_3981 | 220.2149014 | 146.2351786 | 506.7927928 | -0.049420551 | 0.134997844 | 0.08078241 | 0.619248827 | ns | TM | KS.3 |
| FIM1_3982 | 3819.217916 | 2985.494861 | 3579.231815 | -0.169973373 | 0.118825104 | 0.002508248 | 0.093517802 | ns | TM | KS.3 |
| FIM1_3983 | 2001.127542 | 1756.121445 | 1834.252194 | -0.043372986 | 0.076527303 | 0.141997765 | 0.777709768 | ns | TM | KS.3 |
| FIM1_3984 | 282.5461474 | 292.1439865 | 288.2195062 | 0.00434052 | 0.070652721 | 0.845286284 | 0.991141351 | ns | TM | KS.3 |
| FIM1_3985 | 858.8986711 | 731.3524028 | 708.1430085 | -0.112275635 | 0.090430477 | 0.012020825 | 0.249181684 | ns | TM | KS.3 |
| FIM1_3986 | 1660.967201 | 1786.188126 | 1607.990702 | 0.011834466 | 0.070321235 | 0.608037754 | 0.970937835 | ns | TM | KS.3 |
| FIM1_3987 | 755.8905759 | 762.8363984 | 787.8348508 | 0.002458298 | 0.05193766 | 0.910328212 | 0.991141351 | ns | TM | KS.3 |
| FIM1_3988 | 1108.435622 | 1153.288859 | 1013.687429 | 0.00986842 | 0.05530693 | 0.662960058 | 0.976323032 | ns | TM | KS.3 |
| FIM1_3989 | 10270.69435 | 10493.41739 | 10098.21849 | 0.006193486 | 0.046931321 | 0.76437552 | 0.988477371 | ns | TM | KS.3 |
| FIM1_3990 | 1524.309223 | 1706.265961 | 1567.976856 | 0.013177918 | 0.079373381 | 0.547183216 | 0.95917717 | ns | TM | KS.3 |
| FIM1_3991 | 1059.621894 | 1345.879743 | 1565.42951 | 0.030798298 | 0.09462051 | 0.208820873 | 0.852551729 | ns | TM | KS.3 |
| FIM1_3992 | 659.6746344 | 731.1878333 | 725.4346143 | 0.042488408 | 0.0680967 | 0.137722086 | 0.770139466 | ns | TM | KS.3 |
| FIM1_3993 | 7730.521473 | 7497.588359 | 8151.152732 | -0.011877284 | 0.030691163 | 0.375193378 | 0.948637087 | ns | TM | KS.3 |
| FIM1_3994 | 59.0085351 | 45.64565675 | 83.69121574 | -0.036807272 | 0.0996939 | 0.154999987 | 0.786819827 | ns | TM | KS.3 |
| FIM1_3995 | 128.0414955 | 127.9723776 | 185.7070297 | -7.54E-05 | 0.079012332 | 0.997068888 | 0.999525119 | ns | TM | KS.3 |
| FIM1_3996 | 1145.426614 | 1420.676066 | 1615.367025 | 0.019594045 | 0.092579624 | 0.362130345 | 0.948637087 | ns | TM | KS.3 |
| FIM1_3997 | 1569.786491 | 2164.015459 | 1901.564487 | 0.255504956 | 0.159694452 | 0.000448187 | 0.02580389 | ns | TM | KS.3 |
| FIM1_3998 | 1312.786691 | 1323.743668 | 1173.428708 | 0.001659784 | 0.061374115 | 0.941989053 | 0.994413583 | ns | TM | KS.3 |
| FIM1_3999 | 278.1774847 | 289.9342474 | 239.0508598 | 0.00522951 | 0.073472024 | 0.810816794 | 0.988916276 | ns | TM | KS.3 |
| FIM1_4000 | 578.4515262 | 1060.207409 | 866.1570562 | 0.82466519 | 0.170016544 | 9.71690282217[illegible] | 1.93366366161[illegible] | ns | TM | KS.3 |
| FIM1_4001 | 117.5824496 | 116.0110845 | 118.8632034 | -0.00164636 | 0.0735849 | 0.939348138 | 0.994413583 | ns | TM | KS.3 |
| FIM1_4002 | 4733.027523 | 4731.458657 | 4722.895656 | -0.000111548 | 0.038479399 | 0.995006674 | 0.999525119 | ns | TM | KS.3 |
| FIM1_4003 | 4938.066721 | 4593.674098 | 4266.578467 | -0.022933472 | 0.05841807 | 0.343466581 | 0.944808611 | ns | TM | KS.3 |
| FIM1_4004 | 1483.690949 | 1400.540764 | 1181.977227 | -0.018733554 | 0.053227936 | 0.409193176 | 0.950410534 | ns | TM | KS.3 |

| | | | | | | | | | | |
|---|---|---|---|---|---|---|---|---|---|---|
| FIM1_4005 | 3920.027086 | 3848.97655 | 2922.143101 | -0.00362679 | 0.061621527 | 0.873949968 | 0.991141351 | ns | TM | KS.3 |
| FIM1_4006 | 2996.798304 | 2366.404456 | 2646.572787 | -0.233051305 | 0.091261255 | 0.000115409 | 0.009731481 | ns | TM | KS.3 |
| FIM1_4007 | 1729.963227 | 1695.363896 | 1713.896372 | -0.00651224 | 0.039881296 | 0.721255285 | 0.987157164 | ns | TM | KS.3 |
| FIM1_4008 | 431.8246915 | 329.5346195 | 533.3020394 | -0.047045094 | 0.104164432 | 0.100626223 | 0.682037412 | ns | TM | KS.3 |
| FIM1_4009 | 2629.450292 | 2464.24361 | 3342.353643 | -0.014107289 | 0.063683946 | 0.552267242 | 0.95917717 | ns | TM | KS.3 |
| FIM1_4010 | 1820.613146 | 1830.187809 | 1854.404496 | 0.001510631 | 0.04074122 | 0.935502596 | 0.994413583 | ns | TM | KS.3 |
| FIM1_4011 | 2145.324782 | 1989.087113 | 1968.013867 | -0.042629513 | 0.057405766 | 0.10125557 | 0.684438124 | ns | TM | KS.3 |
| FIM1_4012 | 2112.096559 | 2222.629907 | 2395.085632 | 0.014616361 | 0.054230623 | 0.519921085 | 0.958509403 | ns | TM | KS.3 |
| FIM1_4013 | 1701.709488 | 1728.385879 | 1792.394935 | 0.004714201 | 0.036751079 | 0.78211555 | 0.988814136 | ns | TM | KS.3 |
| FIM1_4014 | 379.8725337 | 451.639427 | 366.1548634 | 0.033796576 | 0.083284468 | 0.20427882 | 0.847612283 | ns | TM | KS.3 |
| FIM1_4015 | 3132.376465 | 3559.605581 | 2846.782905 | 0.024782534 | 0.075154392 | 0.323777713 | 0.933252679 | ns | TM | KS.3 |
| FIM1_4016 | 2605.355035 | 2561.029214 | 2731.732684 | -0.003903511 | 0.05752302 | 0.863211212 | 0.991141351 | ns | TM | KS.3 |
| FIM1_4017 | 332.9932252 | 377.9501818 | 389.0880982 | 0.021529749 | 0.075957411 | 0.374668795 | 0.948637087 | ns | TM | KS.3 |
| FIM1_4018 | 3031.107396 | 3521.165815 | 4265.881815 | 0.012185932 | 0.088219538 | 0.547340859 | 0.95917717 | ns | TM | KS.3 |
| FIM1_4019 | 830.5230908 | 709.5617736 | 802.5732301 | -0.065285708 | 0.086698621 | 0.060611691 | 0.544665909 | ns | TM | KS.3 |
| FIM1_4020 | 2959.657058 | 2875.408527 | 3839.847189 | -0.009277017 | 0.042889471 | 0.631011911 | 0.97344368 | ns | TM | KS.3 |
| FIM1_4021 | 1217.863942 | 1157.959536 | 1069.033636 | -0.012159686 | 0.059036984 | 0.601356303 | 0.97002321 | ns | TM | KS.3 |
| FIM1_4022 | 36.66014369 | 28.97711918 | 31.43776702 | -0.018851144 | 0.098553595 | 0.363119727 | 0.948637087 | ns | TM | KS.3 |
| FIM1_4023 | 7033.047627 | 6015.120732 | 7102.690264 | -0.178065141 | 0.057057287 | 0.000127795 | 0.010455632 | ns | TM | KS.3 |
| FIM1_4024 | 2389.911675 | 2543.682253 | 2905.728058 | 0.004957751 | 0.084455823 | 0.801493565 | 0.988916276 | ns | TM | KS.3 |
| FIM1_4025 | 333.3583394 | 358.5420923 | 257.0080608 | 0.011991194 | 0.069501669 | 0.605037232 | 0.970937835 | ns | TM | KS.3 |
| FIM1_4026 | 166.7060523 | 159.9006724 | 370.4821554 | -0.003060198 | 0.086213068 | 0.873788682 | 0.991141351 | ns | TM | KS.3 |
| FIM1_4027 | 482.4343204 | 321.5931697 | 621.9376762 | -0.035801281 | 0.124850961 | 0.142501358 | 0.777709768 | ns | TM | KS.3 |
| FIM1_4028 | 1402.060314 | 1163.034134 | 1191.808455 | -0.144875402 | 0.09477076 | 0.004208118 | 0.125361591 | ns | TM | KS.3 |
| FIM1_4029 | 53.64617709 | 71.01112735 | 104.9791132 | 0.026104209 | 0.100575363 | 0.248505817 | 0.888934767 | ns | TM | KS.3 |
| FIM1_4030 | 4175.060559 | 4543.550996 | 5795.456378 | 0.008557301 | 0.080236991 | 0.68278283 | 0.97986479 | ns | TM | KS.3 |
| FIM1_4031 | 3318.443369 | 3342.053197 | 3294.908715 | 0.001729471 | 0.053667214 | 0.937699472 | 0.994413583 | ns | TM | KS.3 |
| FIM1_4033 | 773.0776449 | 859.3206973 | 824.2126454 | 0.028668512 | 0.068919953 | 0.274275314 | 0.908468501 | ns | TM | KS.3 |
| FIM1_4034 | 490.162392 | 462.4922388 | 554.5582895 | -0.011718996 | 0.064834003 | 0.617200893 | 0.973122675 | ns | TM | KS.3 |
| FIM1_4035 | 409.0390902 | 381.5913318 | 436.1044789 | -0.012097389 | 0.068918294 | 0.602964088 | 0.970245665 | ns | TM | KS.3 |
| FIM1_4036 | 1861.178779 | 1744.006759 | 1895.044124 | -0.015196997 | 0.062165504 | 0.523379845 | 0.958732165 | ns | TM | KS.3 |
| FIM1_4037 | 1423.100474 | 1278.040416 | 1354.565124 | -0.037481611 | 0.06943407 | 0.179475385 | 0.813196759 | ns | TM | KS.3 |
| FIM1_4038 | 776.9668109 | 721.1678562 | 775.7880711 | -0.021719197 | 0.060669399 | 0.374488151 | 0.948637087 | ns | TM | KS.3 |
| FIM1_4039 | 773.3029896 | 743.770662 | 1193.073608 | -0.008223867 | 0.060767178 | 0.721653942 | 0.987157164 | ns | TM | KS.3 |
| FIM1_4041 | 808.8896059 | 954.8007057 | 1151.064925 | 0.014834503 | 0.087843117 | 0.477563704 | 0.950410534 | ns | TM | KS.3 |
| FIM1_4042 | 1427.353009 | 1328.914684 | 1666.45966 | -0.021392082 | 0.058815463 | 0.375333557 | 0.948637087 | ns | TM | KS.3 |
| FIM1_4043 | 1480.275073 | 1506.021832 | 1573.467723 | 0.00349645 | 0.061364802 | 0.878402117 | 0.991141351 | ns | TM | KS.3 |
| FIM1_4044 | 237.4066056 | 264.3985865 | 280.2021216 | 0.008736751 | 0.085989754 | 0.661428456 | 0.976323032 | ns | TM | KS.3 |
| FIM1_4045 | 3924.680009 | 4242.358757 | 4361.899764 | 0.015788284 | 0.066875544 | 0.509444682 | 0.953260948 | ns | TM | KS.3 |
| FIM1_4046 | 1331.356591 | 1355.849441 | 1270.611571 | 0.003270733 | 0.064570345 | 0.885914903 | 0.991141351 | ns | TM | KS.3 |
| FIM1_4047 | 7142.602986 | 7188.744686 | 7888.412633 | 0.001544449 | 0.054497082 | 0.944667412 | 0.994413583 | ns | TM | KS.3 |
| FIM1_4048 | 307.3270345 | 437.6932779 | 405.4900098 | 0.101038697 | 0.153300734 | 0.017097704 | 0.302986707 | ns | TM | KS.3 |
| FIM1_4049 | 1409.21257 | 1503.055267 | 1563.26928 | 0.004787324 | 0.086069492 | 0.805339925 | 0.988916276 | ns | TM | KS.3 |
| FIM1_4050 | 3543.721354 | 3554.006849 | 3856.464842 | 0.000991129 | 0.038140203 | 0.955316358 | 0.994413583 | ns | TM | KS.3 |
| FIM1_4051 | 11487.30911 | 11559.19041 | 8292.48385 | 0.001649183 | 0.049970009 | 0.938596476 | 0.994413583 | ns | TM | KS.3 |

| | | | | | | | | | | |
|---|---|---|---|---|---|---|---|---|---|---|
| FIM1_4052 | 54.81848729 | 49.58354625 | 75.04869013 | -0.013527785 | 0.078570871 | 0.540441297 | 0.95917717 | ns | TM | KS.3 |
| FIM1_4053 | 586.9475527 | 632.6555555 | 749.4921098 | 0.005288313 | 0.087612871 | 0.783292725 | 0.988814136 | ns | TM | KS.3 |
| FIM1_4054 | 2332.843961 | 2600.869868 | 2907.120408 | 0.022391737 | 0.071469931 | 0.368359961 | 0.948637087 | ns | TM | KS.3 |
| FIM1_4055 | 962.7883407 | 764.6395257 | 1475.51348 | -0.040300024 | 0.094907799 | 0.14038997 | 0.775632767 | ns | TM | KS.3 |
| FIM1_4056 | 1300.857021 | 1161.865664 | 1623.584067 | -0.030082836 | 0.070730452 | 0.256983132 | 0.895871714 | ns | TM | KS.3 |
| FIM1_4057 | 1149.317883 | 1084.456894 | 1189.612241 | -0.017062216 | 0.05506066 | 0.458991729 | 0.950410534 | ns | TM | KS.3 |
| FIM1_4058 | 369.3878403 | 422.915702 | 385.20337 | 0.031043031 | 0.076388845 | 0.242333023 | 0.887413624 | ns | TM | KS.3 |
| FIM1_4059 | 2694.600772 | 3214.527103 | 4790.05668 | 0.018433502 | 0.086751638 | 0.400320097 | 0.950410534 | ns | TM | KS.3 |
| FIM1_4060 | 1822.980465 | 2073.079231 | 2445.498945 | 0.015063995 | 0.080617734 | 0.494816253 | 0.953260948 | ns | TM | KS.3 |
| FIM1_4061 | 3739.55612 | 4864.010533 | 5649.129174 | 0.031751029 | 0.098729137 | 0.193819695 | 0.836299203 | ns | TM | KS.3 |
| FIM1_4062 | 879.7660274 | 1301.731553 | 1445.784866 | 0.044397532 | 0.127700735 | 0.099374187 | 0.677856366 | ns | TM | KS.3 |
| FIM1_4063 | 2238.0725 | 2095.732012 | 2330.080871 | -0.014269418 | 0.0637756 | 0.548069001 | 0.95917717 | ns | TM | KS.3 |
| FIM1_4064 | 5664.630026 | 5919.1903 | 5527.055317 | 0.007451601 | 0.067399555 | 0.744487219 | 0.988477371 | ns | TM | KS.3 |
| FIM1_4065 | 3047.900449 | 3692.037286 | 4374.277099 | 0.018637742 | 0.089213218 | 0.389171514 | 0.948637087 | ns | TM | KS.3 |
| FIM1_4066 | 2347.13925 | 2517.987551 | 3727.910755 | 0.020519569 | 0.059103342 | 0.394322789 | 0.948637087 | ns | TM | KS.3 |
| FIM1_4067 | 756.2011048 | 770.3106545 | 1068.184694 | 0.004992686 | 0.04959248 | 0.815125911 | 0.988916276 | ns | TM | KS.3 |
| FIM1_4068 | 338.1949921 | 350.3639531 | 406.8654862 | 0.005865089 | 0.066280183 | 0.797244504 | 0.988916276 | ns | TM | KS.3 |
| FIM1_4069 | 2363.249807 | 2240.766394 | 2644.230048 | -0.006019286 | 0.076659741 | 0.777699435 | 0.988814136 | ns | TM | KS.3 |
| FIM1_4070 | 2949.386509 | 2543.878001 | 2799.421974 | -0.017024692 | 0.082655182 | 0.442731593 | 0.950410534 | ns | TM | KS.3 |
| FIM1_4071 | 71.11955176 | 55.69243632 | 102.4320302 | -0.026850306 | 0.094876908 | 0.249479343 | 0.888934767 | ns | TM | KS.3 |
| FIM1_4072 | 1864.939462 | 1643.634698 | 1604.04108 | -0.050533826 | 0.076575868 | 0.104454289 | 0.69380519 | ns | TM | KS.3 |
| FIM1_4073 | 1245.429383 | 1060.217377 | 1025.178863 | -0.064931824 | 0.088105822 | 0.061333789 | 0.545665039 | ns | TM | KS.3 |
| FIM1_4074 | 1187.228603 | 1180.030621 | 1295.36021 | -0.000539051 | 0.081410171 | 0.978557872 | 0.999157637 | ns | TM | KS.3 |
| FIM1_4075 | 534.5157358 | 665.1475679 | 728.1756904 | 0.027285848 | 0.09110303 | 0.252117981 | 0.888934767 | ns | TM | KS.3 |
| FIM1_4076 | 9.583287645 | 12.01262949 | 18.61852671 | 0.011172514 | 0.103178984 | 0.545607614 | 0.95917717 | ns | TM | KS.3 |
| FIM1_4077 | 24176.93869 | 26814.79744 | 36444.03178 | 0.010214114 | 0.081718095 | 0.625504781 | 0.97344368 | ns | TM | KS.3 |
| FIM1_4078 | 140.4685966 | 167.045835 | 253.8581896 | 0.0100693 | 0.09664902 | 0.592744357 | 0.964955229 | ns | TM | KS.3 |
| FIM1_4080 | 1995.57751 | 1779.522524 | 3950.898547 | -0.010744425 | 0.083280877 | 0.604524471 | 0.970790588 | ns | TM | KS.3 |
| FIM1_4081 | 14.53268276 | 13.66714172 | 26.3944966 | -0.004468493 | 0.089743717 | 0.812424758 | 0.988916276 | ns | TM | KS.3 |
| FIM1_4082 | 9.701784111 | 14.83374157 | 53.98004836 | 0.025056012 | 0.123633248 | 0.236107671 | 0.884880231 | ns | TM | KS.3 |
| FIM1_4083 | 455.3981695 | 563.6197593 | 621.9285562 | 0.068137148 | 0.100118233 | 0.051540295 | 0.515921463 | ns | TM | KS.3 |
| FIM1_4084 | 3268.274155 | 3512.287775 | 3451.86931 | 0.016887956 | 0.063309362 | 0.483094447 | 0.952269291 | ns | TM | KS.3 |
| FIM1_4085 | 9051.65942 | 9500.559389 | 9779.713209 | 0.013634915 | 0.054051852 | 0.546705533 | 0.95917717 | ns | TM | KS.3 |
| FIM1_4086 | 8708.612379 | 9913.125006 | 11459.64232 | 0.018655208 | 0.078022563 | 0.42315586 | 0.950410534 | ns | TM | KS.3 |
| FIM1_4087 | 435.3769738 | 259.871163 | 342.0848434 | -0.106415938 | 0.211710986 | 0.016984413 | 0.302858265 | ns | TM | KS.3 |
| FIM1_4088 | 1690.664861 | 1953.643101 | 2166.353819 | 0.016813769 | 0.082209077 | 0.44883434 | 0.950410534 | ns | TM | KS.3 |
| FIM1_4089 | 165.5605323 | 145.9922107 | 243.3551311 | -0.026656418 | 0.075323449 | 0.296848945 | 0.926536521 | ns | TM | KS.3 |
| FIM1_4090 | 209.9524213 | 168.4757928 | 225.2102565 | -0.087219624 | 0.105214431 | 0.027896386 | 0.388845019 | ns | TM | KS.3 |
| FIM1_4091 | 1964.688414 | 2398.910723 | 2847.784754 | 0.016818986 | 0.091828422 | 0.420127853 | 0.950410534 | ns | TM | KS.3 |
| FIM1_4092 | 1364.087644 | 1248.535331 | 1310.075233 | -0.031709401 | 0.063171287 | 0.224907332 | 0.875024944 | ns | TM | KS.3 |
| FIM1_4093 | 1161.502396 | 1291.032361 | 1490.982194 | 0.00981335 | 0.083000851 | 0.634327337 | 0.973738807 | ns | TM | KS.3 |
| FIM1_4094 | 1398.939545 | 2448.343381 | 6504.902137 | 0.225549863 | 0.308862032 | 0.003377173 | 0.107701496 | ns | TM | KS.3 |
| FIM1_4095 | 1487.021561 | 1824.169327 | 1717.111691 | 0.055189345 | 0.094498373 | 0.082376239 | 0.62282947 | ns | TM | KS.3 |
| FIM1_4096 | 2110.094453 | 2239.67767 | 1803.497877 | 0.012187474 | 0.064385798 | 0.60414272 | 0.970729568 | ns | TM | KS.3 |
| FIM1_4097 | 1309.707751 | 1600.435413 | 1921.574245 | 0.014924443 | 0.093753014 | 0.459591285 | 0.950410534 | ns | TM | KS.3 |

| | | | | | | | | | | |
|---|---|---|---|---|---|---|---|---|---|---|
| FIM1_4098 | 2181.326057 | 2433.442288 | 1806.243203 | 0.015349789 | 0.076218032 | 0.501801598 | 0.953260948 | ns | TM | KS.3 |
| FIM1_4099 | 447.9219275 | 535.1064081 | 365.0170713 | 0.033297169 | 0.084299813 | 0.206616668 | 0.850925432 | ns | TM | KS.3 |
| FIM1_4100 | 285.1502609 | 314.9421329 | 354.4049164 | 0.026689921 | 0.065528815 | 0.298751523 | 0.926536521 | ns | TM | KS.3 |
| FIM1_4101 | 798.3176191 | 749.3012319 | 806.4622174 | -0.010189732 | 0.06983214 | 0.656468456 | 0.976323032 | ns | TM | KS.3 |
| FIM1_4102 | 2300.003499 | 1981.963566 | 2062.510701 | -0.049348616 | 0.081750044 | 0.110855135 | 0.71059097 | ns | TM | KS.3 |
| FIM1_4103 | 10884.96977 | 9389.608339 | 4905.46933 | -0.025604975 | 0.078990911 | 0.303432349 | 0.926536521 | ns | TM | KS.3 |
| FIM1_4104 | 1292.417808 | 1545.854199 | 1512.017908 | 0.023399727 | 0.085110439 | 0.318550114 | 0.929077912 | ns | TM | KS.3 |
| FIM1_4105 | 196.3873618 | 259.8007558 | 221.2919914 | 0.040614427 | 0.104206841 | 0.128519288 | 0.753101833 | ns | TM | KS.3 |
| FIM1_4106 | 3091.81878 | 3697.502088 | 8180.424686 | 0.022954989 | 0.085168766 | 0.325161679 | 0.934066145 | ns | TM | KS.3 |
| FIM1_4107 | 1144.647892 | 1161.106954 | 1193.939476 | 0.003770949 | 0.048353048 | 0.857712986 | 0.991141351 | ns | TM | KS.3 |
| FIM1_4108 | 2814.203951 | 2543.575076 | 2811.311319 | -0.052846951 | 0.069280919 | 0.084424004 | 0.63349837 | ns | TM | KS.3 |
| FIM1_4109 | 321.643139 | 379.6170929 | 326.6748471 | 0.027403337 | 0.082157835 | 0.27222308 | 0.908468501 | ns | TM | KS.3 |
| FIM1_4110 | 812.2872771 | 915.9690073 | 929.8709923 | 0.031626225 | 0.072598071 | 0.239083834 | 0.886981413 | ns | TM | KS.3 |
| FIM1_4111 | 2392.998638 | 2413.664174 | 2452.532085 | 0.002737677 | 0.030062081 | 0.846316081 | 0.991141351 | ns | TM | KS.3 |
| FIM1_4112 | 2139.016572 | 2057.100732 | 2371.194331 | -0.005187651 | 0.072776027 | 0.813287437 | 0.988916276 | ns | TM | KS.3 |
| FIM1_4113 | 1204.333834 | 1373.626036 | 1778.14431 | 0.012627283 | 0.083992645 | 0.547433137 | 0.95917717 | ns | TM | KS.3 |
| FIM1_4114 | 1322.026942 | 1566.053292 | 2048.426331 | 0.01119128 | 0.093593776 | 0.563371707 | 0.961170864 | ns | TM | KS.3 |
| FIM1_4115 | 1019.122997 | 1139.2131 | 1095.168122 | 0.019644347 | 0.073461427 | 0.41555318 | 0.950410534 | ns | TM | KS.3 |
| FIM1_4116 | 220.5768285 | 287.6629402 | 254.7718731 | 0.022929921 | 0.099352386 | 0.292792629 | 0.923094632 | ns | TM | KS.3 |
| FIM1_4117 | 144.5472469 | 167.9472992 | 180.9317383 | 0.011560893 | 0.089373198 | 0.562511547 | 0.961073589 | ns | TM | KS.3 |
| FIM1_4118 | 1895.002548 | 1962.953667 | 2095.015767 | 0.003919691 | 0.076511015 | 0.85316015 | 0.991141351 | ns | TM | KS.3 |
| FIM1_4119 | 1550.244798 | 1603.226959 | 1782.112594 | 0.007844575 | 0.058624512 | 0.73222362 | 0.987157164 | ns | TM | KS.3 |
| FIM1_4120 | 1886.738021 | 2649.745165 | 2779.77814 | 0.029201116 | 0.111263007 | 0.202402631 | 0.847612283 | ns | TM | KS.3 |
| FIM1_4121 | 897.0813021 | 1028.094155 | 988.6484663 | 0.014582037 | 0.082645292 | 0.500090543 | 0.953260948 | ns | TM | KS.3 |
| FIM1_4122 | 703.4543894 | 668.9048729 | 644.6217503 | -0.01113292 | 0.06145465 | 0.63353116 | 0.97344368 | ns | TM | KS.3 |
| FIM1_4123 | 1083.198079 | 995.1277451 | 1041.902857 | -0.018171295 | 0.067043374 | 0.45468529 | 0.950410534 | ns | TM | KS.3 |
| FIM1_4124 | 60.06092459 | 71.93449552 | 61.31069651 | 0.01722046 | 0.086361044 | 0.426762555 | 0.950410534 | ns | TM | KS.3 |
| FIM1_4125 | 7073.698163 | 6546.574097 | 7126.221652 | -0.013567297 | 0.069456377 | 0.562437345 | 0.961073589 | ns | TM | KS.3 |
| FIM1_4126 | 223.3464844 | 335.9501477 | 369.1405331 | 0.339493652 | 0.235542574 | 0.000335738 | 0.021692164 | ns | TM | KS.3 |
| FIM1_4127 | 45.925182 | 57.97344326 | 56.92117253 | 0.02225004 | 0.094469089 | 0.312609264 | 0.927416591 | ns | TM | KS.3 |
| FIM1_4128 | 1099.42585 | 1276.101239 | 1315.209878 | 0.043450699 | 0.080653116 | 0.140614816 | 0.775632767 | ns | TM | KS.3 |
| FIM1_4129 | 2566.493313 | 2703.242796 | 2444.613147 | 0.010809201 | 0.062941238 | 0.643556036 | 0.976323032 | ns | TM | KS.3 |
| FIM1_4130 | 1606.069308 | 1641.130558 | 2108.407292 | 0.003610402 | 0.06728359 | 0.873168861 | 0.991141351 | ns | TM | KS.3 |
| FIM1_4132 | 220.6030195 | 236.0389846 | 198.7253864 | 0.019467825 | 0.063839217 | 0.426087344 | 0.950410534 | ns | TM | KS.3 |
| FIM1_4133 | 967.1833757 | 817.3483178 | 883.5089162 | -0.048670882 | 0.08560502 | 0.111734639 | 0.71059097 | ns | TM | KS.3 |
| FIM1_4134 | 1759.39481 | 1592.755475 | 1743.048971 | -0.032058475 | 0.066914921 | 0.229989726 | 0.879350777 | ns | TM | KS.3 |
| FIM1_4135 | 860.6532173 | 912.8553234 | 1187.428661 | 0.012896315 | 0.062789326 | 0.584364023 | 0.963781856 | ns | TM | KS.3 |
| FIM1_4136 | 2372.977036 | 2340.836335 | 2361.138264 | -0.003786635 | 0.047742919 | 0.856048938 | 0.991141351 | ns | TM | KS.3 |
| FIM1_4137 | 2856.666653 | 3045.076903 | 3197.217042 | 0.010233926 | 0.070037946 | 0.654814278 | 0.976323032 | ns | TM | KS.3 |
| FIM1_4138 | 1201.534822 | 1279.962702 | 1404.198415 | 0.019939657 | 0.05514278 | 0.391113778 | 0.948637087 | ns | TM | KS.3 |
| FIM1_4139 | 2743.062065 | 3070.750845 | 3528.830738 | 0.013446515 | 0.079070263 | 0.540995896 | 0.95917717 | ns | TM | KS.3 |
| FIM1_4140 | 1143.748673 | 1154.74717 | 1085.210523 | 0.001235663 | 0.073133911 | 0.954611364 | 0.994413583 | ns | TM | KS.3 |
| FIM1_4141 | 10001.25792 | 9059.252265 | 8667.610626 | -0.025049431 | 0.067779928 | 0.326181246 | 0.934739762 | ns | TM | KS.3 |
| FIM1_4142 | 748.9903517 | 844.0332813 | 850.6829663 | 0.043412152 | 0.073528003 | 0.140099926 | 0.775302706 | ns | TM | KS.3 |
| FIM1_4143 | 839.984892 | 928.7646644 | 875.8348652 | 0.033504895 | 0.067570637 | 0.215014924 | 0.864752826 | ns | TM | KS.3 |

| | | | | | | | | | |
|---|---|---|---|---|---|---|---|---|---|
| FIM1_4144 | 455.4808501 | 448.7161008 | 395.2838946 | -0.002576533 | 0.066756024 | 0.909335899 | 0.991141351 | ns | TM | KS.3 |
| FIM1_4145 | 546.8158569 | 516.334777 | 511.0689433 | -0.010464956 | 0.066948017 | 0.652040698 | 0.976323032 | ns | TM | KS.3 |
| FIM1_4146 | 1259.259271 | 1568.072706 | 1621.817164 | 0.014290816 | 0.097648021 | 0.466942309 | 0.950410534 | ns | TM | KS.3 |
| FIM1_4147 | 898.6484049 | 871.7539896 | 825.2580876 | -0.005076078 | 0.067019255 | 0.823198928 | 0.991141351 | ns | TM | KS.3 |
| FIM1_4148 | 2671.31686 | 2775.828605 | 3486.418128 | 0.003847645 | 0.078818296 | 0.852610706 | 0.991141351 | ns | TM | KS.3 |
| FIM1_4149 | 16489.97216 | 13494.90618 | 12080.15282 | -0.123035038 | 0.103688604 | 0.009386859 | 0.211310513 | ns | TM | KS.3 |
| FIM1_4150 | 2307.367916 | 2259.595552 | 2153.571342 | -0.004400575 | 0.060035696 | 0.847253277 | 0.991141351 | ns | TM | KS.3 |
| FIM1_4151 | 2956.711858 | 2404.093302 | 3026.452796 | -0.082935106 | 0.102488684 | 0.032174035 | 0.413606776 | ns | TM | KS.3 |
| FIM1_4152 | 1411.628264 | 1464.696513 | 1951.519265 | 0.002864988 | 0.084538606 | 0.883448754 | 0.991141351 | ns | TM | KS.3 |
| FIM1_4153 | 1676.017333 | 1492.331638 | 1687.778482 | -0.025025304 | 0.072334751 | 0.32456757 | 0.934066145 | ns | TM | KS.3 |
| FIM1_4154 | 844.2637355 | 709.4466091 | 796.973279 | -0.029075605 | 0.083433176 | 0.249779851 | 0.888934767 | ns | TM | KS.3 |
| FIM1_4155 | 4065.482493 | 3930.545168 | 3767.699817 | -0.004808674 | 0.071009643 | 0.828670174 | 0.991141351 | ns | TM | KS.3 |
| FIM1_4156 | 2746.916314 | 3200.083749 | 3625.250613 | 0.01582429 | 0.084527204 | 0.463897405 | 0.950410534 | ns | TM | KS.3 |
| FIM1_4157 | 2553.263832 | 2447.792516 | 3012.997044 | -0.006828392 | 0.068399729 | 0.763685887 | 0.988477371 | ns | TM | KS.3 |
| FIM1_4158 | 3898.518753 | 3840.141038 | 4488.575145 | -0.002427169 | 0.067610657 | 0.914245908 | 0.991141351 | ns | TM | KS.3 |
| FIM1_4159 | 1215.243744 | 1146.400293 | 1271.519633 | -0.016981481 | 0.056005358 | 0.464731691 | 0.950410534 | ns | TM | KS.3 |
| FIM1_4160 | 370.339921 | 376.4290938 | 355.4111926 | 0.0032352 | 0.06133577 | 0.887381867 | 0.991141351 | ns | TM | KS.3 |
| FIM1_4161 | 16797.72824 | 15709.81339 | 15530.23963 | -0.02302968 | 0.055443937 | 0.327801716 | 0.934739762 | ns | TM | KS.3 |
| FIM1_4162 | 2017.724994 | 1820.311455 | 1415.571465 | -0.007315447 | 0.088438972 | 0.705474779 | 0.982962554 | ns | TM | KS.3 |
| FIM1_4163 | 7117.39484 | 7581.961301 | 7457.914414 | 0.016821396 | 0.059049444 | 0.477975308 | 0.950410534 | ns | TM | KS.3 |
| FIM1_4164 | 95.10575579 | 68.74001915 | 109.5779923 | -0.02531213 | 0.108474903 | 0.247570958 | 0.888934767 | ns | TM | KS.3 |
| FIM1_4165 | 415.0290231 | 454.5759436 | 424.6697933 | 0.017990813 | 0.069305751 | 0.456785562 | 0.950410534 | ns | TM | KS.3 |
| FIM1_4166 | 1116.516222 | 1185.999874 | 2248.026681 | 0.007048146 | 0.076359803 | 0.742899607 | 0.988477371 | ns | TM | KS.3 |
| FIM1_4167 | 1107.03994 | 1172.134651 | 1085.647549 | 0.004692021 | 0.083615905 | 0.813102007 | 0.988916276 | ns | TM | KS.3 |
| FIM1_4169 | 698.6632567 | 730.7644477 | 672.9160416 | 0.01360634 | 0.049112573 | 0.52437317 | 0.959071703 | ns | TM | KS.3 |
| FIM1_4170 | 1220.340494 | 856.7716329 | 738.4157999 | -0.200158602 | 0.185110729 | 0.002125248 | 0.083913551 | ns | TM | KS.3 |
| FIM1_4171 | 525.6241536 | 121.3243687 | 117.2484734 | -1.814071397 | 0.295766589 | 3.07358696029 | 1.82115719802 | Km<KS.3 | TM | KS.3 |
| FIM1_4172 | 935.9817055 | 931.7211125 | 1125.987572 | -0.00056695 | 0.073605613 | 0.979074066 | 0.999157637 | ns | TM | KS.3 |
| FIM1_4173 | 1030.407325 | 1022.792941 | 1128.570396 | -0.001904032 | 0.05206755 | 0.930549869 | 0.993993515 | ns | TM | KS.3 |
| FIM1_4174 | 4673.581972 | 4373.075866 | 5769.751469 | -0.007319501 | 0.07770939 | 0.730628559 | 0.987157164 | ns | TM | KS.3 |
| FIM1_4175 | 1849.981636 | 1850.895399 | 1627.710598 | -5.40E-05 | 0.053317314 | 0.998048111 | 0.999525119 | ns | TM | KS.3 |
| FIM1_4176 | 1571.391754 | 1440.313141 | 1316.289622 | -0.031443514 | 0.062641587 | 0.226460664 | 0.876782875 | ns | TM | KS.3 |
| FIM1_4177 | 6940.358154 | 7825.822758 | 6783.080192 | 0.025863379 | 0.073163105 | 0.311176278 | 0.927416591 | ns | TM | KS.3 |
| FIM1_4178 | 6143.495864 | 7270.792772 | 6196.614027 | 0.048407375 | 0.085594554 | 0.112847253 | 0.711904568 | ns | TM | KS.3 |
| FIM1_4180 | 18.52363895 | 20.73840231 | 24.9312206 | 0.008669044 | 0.091393258 | 0.650970381 | 0.976323032 | ns | TM | KS.3 |
| FIM1_4181 | 2887.73264 | 2433.125553 | 3689.600522 | -0.050637907 | 0.086694983 | 0.103128648 | 0.689041746 | ns | TM | KS.3 |
| FIM1_4182 | 9701.457496 | 8198.088484 | 8869.187302 | -0.15063919 | 0.082678108 | 0.002607368 | 0.093997493 | ns | TM | KS.3 |
| FIM1_4183 | 1558.908927 | 1611.268577 | 1556.276746 | 0.009486983 | 0.050264975 | 0.660528379 | 0.976323032 | ns | TM | KS.3 |
| FIM1_4184 | 2021.29967 | 2014.776625 | 2060.641967 | -0.000762133 | 0.043768044 | 0.969145541 | 0.996327965 | ns | TM | KS.3 |
| FIM1_4185 | 729.7319113 | 739.1030084 | 760.5632707 | 0.003046327 | 0.053627834 | 0.890547683 | 0.991141351 | ns | TM | KS.3 |
| FIM1_4186 | 3781.723436 | 3372.67715 | 2235.521867 | -0.01574544 | 0.077039922 | 0.489897202 | 0.952591289 | ns | TM | KS.3 |
| FIM1_4187 | 508.2249205 | 673.4468017 | 693.7005976 | 0.146165798 | 0.137914721 | 0.005416179 | 0.152234402 | ns | TM | KS.3 |
| FIM1_4188 | 1150.363937 | 1144.272114 | 1297.856177 | -0.001007701 | 0.06206643 | 0.96475436 | 0.995175461 | ns | TM | KS.3 |
| FIM1_4189 | 3905.617329 | 3083.084382 | 3247.958278 | -0.0693197 | 0.105845278 | 0.047543085 | 0.497951255 | ns | TM | KS.3 |
| FIM1_4190 | 2392.560948 | 2309.613656 | 1947.08494 | -0.00964675 | 0.05233009 | 0.662215063 | 0.976323032 | ns | TM | KS.3 |

| | | | | | | | | | | |
|---|---|---|---|---|---|---|---|---|---|---|
| FIM1_4191 | 2231.03922 | 2048.432408 | 2019.30619 | -0.021693646 | 0.064672802 | 0.382371124 | 0.948637087 | ns | TM | KS.3 |
| FIM1_4192 | 722.2669393 | 979.0835308 | 923.868796 | 0.055189284 | 0.115261722 | 0.070336036 | 0.582627046 | ns | TM | KS.3 |
| FIM1_4193 | 307.1524888 | 256.1354655 | 265.2306088 | -0.028322629 | 0.084843225 | 0.25481873 | 0.892600401 | ns | TM | KS.3 |
| FIM1_4194 | 1215.792171 | 1097.651948 | 1161.170013 | -0.019537617 | 0.071210521 | 0.421923928 | 0.950410534 | ns | TM | KS.3 |
| FIM1_4195 | 2197.478205 | 2290.181727 | 2259.970138 | 0.011077301 | 0.053616379 | 0.621010413 | 0.973122675 | ns | TM | KS.3 |
| FIM1_4196 | 3396.934729 | 2765.82899 | 3731.145569 | -0.046286964 | 0.09199923 | 0.115296531 | 0.717000304 | ns | TM | KS.3 |
| FIM1_4197 | 1884.521383 | 1449.319569 | 1354.845331 | -0.035491636 | 0.099393906 | 0.16411519 | 0.800496208 | ns | TM | KS.3 |
| FIM1_4198 | 1530.2708 | 1650.244821 | 1718.026555 | 0.010311211 | 0.073969444 | 0.64435934 | 0.976323032 | ns | TM | KS.3 |
| FIM1_4199 | 274.1088613 | 264.1643322 | 370.175666 | -0.005203471 | 0.07073644 | 0.815403784 | 0.988916276 | ns | TM | KS.3 |
| FIM1_4201 | 560.8922043 | 556.1705236 | 544.8274497 | -0.001894767 | 0.039946038 | 0.917859399 | 0.991141351 | ns | TM | KS.3 |
| FIM1_4203 | 330.5622332 | 316.1518832 | 324.5406144 | -0.006027557 | 0.073056779 | 0.783965524 | 0.988880259 | ns | TM | KS.3 |
| FIM1_4204 | 1101.509253 | 969.5583048 | 1148.355635 | -0.035191519 | 0.074481323 | 0.203042523 | 0.847612283 | ns | TM | KS.3 |
| FIM1_4205 | 1078.191232 | 1159.910175 | 3643.20249 | 0.005594228 | 0.085639265 | 0.775105429 | 0.988814136 | ns | TM | KS.3 |
| FIM1_4206 | 1209.386088 | 1415.163576 | 1485.64639 | 0.019640425 | 0.082715812 | 0.38923629 | 0.948637087 | ns | TM | KS.3 |
| FIM1_4207 | 2582.661046 | 2431.736492 | 2188.568411 | -0.010942848 | 0.066937155 | 0.638092534 | 0.975246356 | ns | TM | KS.3 |
| FIM1_4208 | 1028.398383 | 998.3084754 | 1001.676204 | -0.004847881 | 0.067549531 | 0.830562045 | 0.991141351 | ns | TM | KS.3 |
| FIM1_4209 | 793.7544934 | 606.2225135 | 664.3409912 | -0.13080146 | 0.130308487 | 0.007903791 | 0.191538592 | ns | TM | KS.3 |
| FIM1_4210 | 1910.268873 | 2065.521174 | 2048.586944 | 0.015996181 | 0.066822241 | 0.504482503 | 0.953260948 | ns | TM | KS.3 |
| FIM1_4211 | 622.3582387 | 700.8355833 | 725.4670799 | 0.017796755 | 0.076264049 | 0.44617544 | 0.950410534 | ns | TM | KS.3 |
| FIM1_4212 | 3004.46646 | 2980.464534 | 3234.851481 | -0.002321406 | 0.042535149 | 0.904221839 | 0.991141351 | ns | TM | KS.3 |
| FIM1_4213 | 424.7251722 | 405.9823576 | 423.953216 | -0.008896596 | 0.063487769 | 0.701150609 | 0.98151979 | ns | TM | KS.3 |
| FIM1_4214 | 1120.059061 | 1097.024949 | 1128.506859 | -0.005028594 | 0.053997794 | 0.821119011 | 0.990886949 | ns | TM | KS.3 |
| FIM1_4215 | 4357.139887 | 4416.033725 | 5631.325258 | 0.001437535 | 0.077039384 | 0.94531617 | 0.994413583 | ns | TM | KS.3 |
| FIM1_4216 | 768.1689599 | 629.5844748 | 743.7061317 | -0.03362548 | 0.088028439 | 0.196494901 | 0.838389481 | ns | TM | KS.3 |
| FIM1_4217 | 964.1073395 | 769.6533434 | 889.1683139 | -0.039921611 | 0.093870089 | 0.143879492 | 0.77783242 | ns | TM | KS.3 |
| FIM1_4218 | 51.75242089 | 40.48322398 | 40.80047592 | -0.03208289 | 0.095398707 | 0.196084217 | 0.837834055 | ns | TM | KS.3 |
| FIM1_4219 | 452.7801188 | 471.9743751 | 472.1011329 | 0.007530767 | 0.06519703 | 0.743548851 | 0.988477371 | ns | TM | KS.3 |
| FIM1_4220 | 777.7746459 | 762.5403164 | 752.1887105 | -0.003938078 | 0.062384634 | 0.863320341 | 0.991141351 | ns | TM | KS.3 |
| FIM1_4221 | 320.7095448 | 289.7829332 | 215.5270907 | -0.017685122 | 0.073199398 | 0.456528615 | 0.950410534 | ns | TM | KS.3 |
| FIM1_4222 | 1602.679475 | 1629.568843 | 1521.313095 | 0.00177416 | 0.077199231 | 0.932464444 | 0.994413583 | ns | TM | KS.3 |
| FIM1_4223 | 2896.954973 | 2711.502688 | 2560.189644 | -0.014783138 | 0.063303637 | 0.534682193 | 0.959071703 | ns | TM | KS.3 |
| FIM1_4224 | 5846.473541 | 5999.128994 | 5080.741052 | 0.004457349 | 0.06599091 | 0.844769243 | 0.991141351 | ns | TM | KS.3 |
| FIM1_4225 | 516.7074898 | 626.4939254 | 581.9715583 | 0.013791323 | 0.093783269 | 0.488537536 | 0.952591289 | ns | TM | KS.3 |
| FIM1_4226 | 17643.47909 | 18676.3002 | 38582.63717 | 0.007808671 | 0.072802177 | 0.725116599 | 0.987157164 | ns | TM | KS.3 |
| FIM1_4227 | 1175.868632 | 1826.985211 | 2863.283207 | 0.039603019 | 0.133339856 | 0.118532152 | 0.727123865 | ns | TM | KS.3 |
| FIM1_4228 | 14.69325153 | 14.08280002 | 24.61517435 | -0.002455152 | 0.092461958 | 0.89340309 | 0.991141351 | ns | TM | KS.3 |
| FIM1_4229 | 434.5727454 | 374.4026663 | 1314.53667 | -0.013273575 | 0.086563884 | 0.521806952 | 0.958602567 | ns | TM | KS.3 |
| FIM1_4230 | 1918.614493 | 2255.894295 | 2387.443186 | 0.020177556 | 0.08327983 | 0.377497729 | 0.948637087 | ns | TM | KS.3 |
| FIM1_4231 | 4610.860454 | 5056.938506 | 5772.270488 | 0.01597023 | 0.071328627 | 0.499272366 | 0.953260948 | ns | TM | KS.3 |
| FIM1_4232 | 489.9559026 | 466.3165016 | 487.0612049 | -0.008204499 | 0.067543365 | 0.720380209 | 0.987157164 | ns | TM | KS.3 |
| FIM1_4233 | 592.375467 | 742.0736331 | 813.2059935 | 0.087732209 | 0.108908904 | 0.026969522 | 0.382260313 | ns | TM | KS.3 |
| FIM1_4234 | 1091.315132 | 1174.59647 | 1164.257849 | 0.015651667 | 0.065218976 | 0.513355743 | 0.954240701 | ns | TM | KS.3 |
| FIM1_4235 | 1135.737942 | 903.6159191 | 998.3452731 | -0.151565985 | 0.112832519 | 0.004128801 | 0.123739677 | ns | TM | KS.3 |
| FIM1_4236 | 2716.366865 | 2490.35462 | 2572.85615 | -0.026975338 | 0.063142982 | 0.289583395 | 0.919993167 | ns | TM | KS.3 |
| FIM1_4237 | 8368.659986 | 6839.041026 | 7971.332114 | -0.150064545 | 0.100326714 | 0.003826384 | 0.116725767 | ns | TM | KS.3 |

| | | | | | | | | | | |
|---|---|---|---|---|---|---|---|---|---|---|
| FIM1_4238 | 2797.498967 | 2840.109112 | 2974.152138 | 0.003826787 | 0.052251992 | 0.861324978 | 0.991141351 | ns | TM | KS.3 |
| FIM1_4239 | 2062.328054 | 2237.608883 | 2512.177134 | 0.022930949 | 0.062650257 | 0.356174745 | 0.948637087 | ns | TM | KS.3 |
| FIM1_4240 | 1124.295973 | 1311.152787 | 1416.346129 | 0.017897216 | 0.08311793 | 0.422580716 | 0.950410534 | ns | TM | KS.3 |
| FIM1_4241 | 1900.895003 | 2233.134639 | 2297.8411 | 0.019641529 | 0.083399785 | 0.387080537 | 0.948637087 | ns | TM | KS.3 |
| FIM1_4242 | 179.620032 | 171.6741534 | 256.0742388 | -0.006113458 | 0.072084466 | 0.782588578 | 0.988814136 | ns | TM | KS.3 |
| FIM1_4243 | 677.3672945 | 775.5777367 | 683.6635556 | 0.049149996 | 0.078248036 | 0.111649803 | 0.71059097 | ns | TM | KS.3 |
| FIM1_4244 | 876.4509575 | 840.4285447 | 691.9464541 | -0.006287739 | 0.069921555 | 0.77976037 | 0.988814136 | ns | TM | KS.3 |
| FIM1_4245 | 24.71617319 | 19.13516152 | 45.78680657 | -0.019527405 | 0.097045884 | 0.353392154 | 0.948637087 | ns | TM | KS.3 |
| FIM1_4246 | 606.4759136 | 543.6351512 | 598.2144239 | -0.034506836 | 0.069813293 | 0.207776627 | 0.852551729 | ns | TM | KS.3 |
| FIM1_4247 | 397.5576942 | 350.242534 | 496.5544882 | -0.019454305 | 0.076997794 | 0.410288856 | 0.950410534 | ns | TM | KS.3 |
| FIM1_4248 | 1272.129092 | 1183.655021 | 1281.443992 | -0.027066298 | 0.056968856 | 0.264195815 | 0.90372202 | ns | TM | KS.3 |
| FIM1_4249 | 4507.592948 | 4603.022361 | 5696.017768 | 0.005111223 | 0.054925594 | 0.819384716 | 0.990886949 | ns | TM | KS.3 |
| FIM1_4250 | 853.7545762 | 784.4533747 | 1103.258929 | -0.046479574 | 0.062452388 | 0.097235666 | 0.676927258 | ns | TM | KS.3 |
| FIM1_4251 | 3429.145609 | 3378.416636 | 3147.760199 | -0.004548316 | 0.041288651 | 0.809143593 | 0.988916276 | ns | TM | KS.3 |
| FIM1_4252 | 1281.325066 | 1157.681714 | 1133.022365 | -0.031767704 | 0.06750847 | 0.23434724 | 0.883800031 | ns | TM | KS.3 |
| FIM1_4253 | 2480.359 | 3021.468697 | 2736.043885 | 0.036693259 | 0.088226797 | 0.172869627 | 0.811391014 | ns | TM | KS.3 |
| FIM1_4254 | 837.547084 | 963.3015953 | 1081.144263 | 0.045173612 | 0.079132643 | 0.131676728 | 0.761283488 | ns | TM | KS.3 |
| FIM1_4255 | 1788.167817 | 1873.685026 | 1779.909196 | 0.007022372 | 0.070351776 | 0.754891492 | 0.988477371 | ns | TM | KS.3 |
| FIM1_4257 | 2101.11043 | 2433.926243 | 4226.826573 | 0.018302234 | 0.081649887 | 0.418931252 | 0.950410534 | ns | TM | KS.3 |
| FIM1_4258 | 40.87766533 | 39.65039346 | 42.59403129 | -0.001539219 | 0.079765427 | 0.939871837 | 0.994413583 | ns | TM | KS.3 |
| FIM1_4259 | 3268.071077 | 3289.656941 | 2675.666478 | 0.000991745 | 0.069225997 | 0.96455758 | 0.995175461 | ns | TM | KS.3 |
| FIM1_4260 | 863.0521638 | 836.8733948 | 767.4231294 | -0.003799185 | 0.073846904 | 0.860894838 | 0.991141351 | ns | TM | KS.3 |
| FIM1_4261 | 442.8126869 | 593.657352 | 920.3489585 | 0.059542645 | 0.115061497 | 0.060871205 | 0.544665909 | ns | TM | KS.3 |
| FIM1_4262 | 2381.902953 | 2652.295719 | 2964.984714 | 0.022106945 | 0.071299289 | 0.373561587 | 0.948637087 | ns | TM | KS.3 |
| FIM1_4263 | 625.1103683 | 524.2780694 | 569.9360588 | -0.07837626 | 0.093874936 | 0.038396207 | 0.444235187 | ns | TM | KS.3 |
| FIM1_4265 | 832.8954442 | 937.4561344 | 891.4510669 | 0.03577623 | 0.072482705 | 0.197628393 | 0.838676184 | ns | TM | KS.3 |
| FIM1_4266 | 8524.852277 | 11156.67122 | 9953.457295 | 0.045330891 | 0.103672651 | 0.107605524 | 0.702423202 | ns | TM | KS.3 |
| FIM1_4267 | 1428.945161 | 1410.398741 | 1443.309574 | -0.002691835 | 0.05868874 | 0.9056852 | 0.991141351 | ns | TM | KS.3 |
| FIM1_4268 | 3041.17457 | 3265.0283 | 3576.938436 | 0.011082476 | 0.071086544 | 0.628031438 | 0.97344368 | ns | TM | KS.3 |
| FIM1_4269 | 321.2884126 | 367.1056011 | 394.092963 | 0.032426226 | 0.075986966 | 0.228441421 | 0.87828135 | ns | TM | KS.3 |
| FIM1_4270 | 1715.089733 | 1826.453662 | 2359.182223 | 0.017814782 | 0.05748651 | 0.44965358 | 0.950410534 | ns | TM | KS.3 |
| FIM1_4271 | 818.3827516 | 731.9562062 | 835.0517382 | -0.040489225 | 0.071687869 | 0.15827158 | 0.789048157 | ns | TM | KS.3 |
| FIM1_4272 | 732.0884232 | 584.2445349 | 712.7728461 | -0.038333015 | 0.093540798 | 0.153665618 | 0.783447458 | ns | TM | KS.3 |
| FIM1_4273 | 1430.805667 | 1676.317559 | 1940.035915 | 0.015020914 | 0.086318534 | 0.477541995 | 0.950410534 | ns | TM | KS.3 |
| FIM1_4274 | 192.3844745 | 208.7568939 | 458.2280169 | 0.018898635 | 0.067269698 | 0.439016538 | 0.950410534 | ns | TM | KS.3 |
| FIM1_4275 | 19.78426477 | 15.19812844 | 33.90945476 | -0.014355144 | 0.105391209 | 0.451242811 | 0.950410534 | ns | TM | KS.3 |
| FIM1_4276 | 2462.859549 | 2431.808344 | 2355.293366 | -0.003410521 | 0.048371646 | 0.871201115 | 0.991141351 | ns | TM | KS.3 |
| FIM1_4277 | 242.7918023 | 218.6686704 | 199.5390417 | -0.018470298 | 0.073540456 | 0.439091794 | 0.950410534 | ns | TM | KS.3 |
| FIM1_4278 | 3298.586908 | 3044.888762 | 2990.652528 | -0.03799561 | 0.059737772 | 0.145653567 | 0.779559149 | ns | TM | KS.3 |
| FIM1_4279 | 2788.691933 | 3154.912639 | 3477.358609 | 0.010198481 | 0.086226189 | 0.612970389 | 0.973122675 | ns | TM | KS.3 |
| FIM1_4280 | 3603.704945 | 3326.616484 | 2995.245579 | -0.020563385 | 0.063365466 | 0.4029543 | 0.950410534 | ns | TM | KS.3 |
| FIM1_4281 | 5157.792981 | 4984.490764 | 5184.495808 | -0.009051188 | 0.053198885 | 0.683964323 | 0.98018358 | ns | TM | KS.3 |
| FIM1_4282 | 663.2752693 | 537.8053096 | 2506.198991 | -0.019449162 | 0.091687465 | 0.367031472 | 0.948637087 | ns | TM | KS.3 |
| FIM1_4283 | 43.57492663 | 29.86934213 | 72.17510083 | -0.040011944 | 0.121711501 | 0.120621489 | 0.730509595 | ns | TM | KS.3 |
| FIM1_4284 | 7102.224727 | 7087.793293 | 5208.636838 | -0.000510556 | 0.051790923 | 0.981303681 | 0.999333464 | ns | TM | KS.3 |

| | | | | | | | | | | |
|---|---|---|---|---|---|---|---|---|---|---|
| FIM1_4285 | 2194.58777 | 2583.954636 | 2583.026233 | 0.060192826 | 0.087431756 | 0.072728853 | 0.594131438 | ns | TM | KS.3 |
| FIM1_4286 | 2605.211592 | 2539.416094 | 2558.176458 | -0.002461794 | 0.07946642 | 0.904414589 | 0.991141351 | ns | TM | KS.3 |
| FIM1_4287 | 10.25076988 | 11.44493462 | 12.20518445 | 0.005448961 | 0.096559828 | 0.763579107 | 0.988477371 | ns | TM | KS.3 |
| FIM1_4288 | 1005.400053 | 1204.903151 | 1283.172397 | 0.024122361 | 0.085194161 | 0.307679262 | 0.927416591 | ns | TM | KS.3 |
| FIM1_4289 | 2120.645945 | 2340.807232 | 2144.079177 | 0.022830876 | 0.068569676 | 0.3630636 | 0.948637087 | ns | TM | KS.3 |
| FIM1_4290 | 1415.574298 | 1316.541641 | 1397.806259 | -0.019302996 | 0.061125497 | 0.42552971 | 0.950410534 | ns | TM | KS.3 |
| FIM1_4291 | 893.8259897 | 833.0688646 | 822.1209317 | -0.026522859 | 0.055412355 | 0.264427926 | 0.90372202 | ns | TM | KS.3 |
| FIM1_4292 | 910.2346813 | 991.9313611 | 926.2401936 | 0.025253732 | 0.063139408 | 0.316850221 | 0.927798616 | ns | TM | KS.3 |
| FIM1_4293 | 653.7780926 | 658.512295 | 651.6130055 | 0.002130093 | 0.04583069 | 0.916574469 | 0.991141351 | ns | TM | KS.3 |
| FIM1_4294 | 1223.844069 | 1025.351904 | 1014.23371 | -0.06851539 | 0.092420815 | 0.053516298 | 0.528931635 | ns | TM | KS.3 |
| FIM1_4295 | 1079.171329 | 1075.221698 | 957.734905 | -0.001066988 | 0.040838311 | 0.954510045 | 0.994413583 | ns | TM | KS.3 |
| FIM1_4296 | 1171.883474 | 1118.787948 | 1234.676435 | -0.018569432 | 0.042779605 | 0.326090466 | 0.934739762 | ns | TM | KS.3 |
| FIM1_4297 | 758.0923177 | 677.3439111 | 795.4135375 | -0.021641101 | 0.072533696 | 0.379971473 | 0.948637087 | ns | TM | KS.3 |
| FIM1_4298 | 239.6052954 | 251.1655855 | 256.6677733 | 0.002483476 | 0.094629813 | 0.890672568 | 0.991141351 | ns | TM | KS.3 |
| FIM1_4299 | 272.5872784 | 279.984126 | 366.378426 | 0.005209233 | 0.062986748 | 0.820280819 | 0.990886949 | ns | TM | KS.3 |
| FIM1_4300 | 156.0466096 | 194.2775106 | 223.9361699 | 0.055855447 | 0.096076906 | 0.079394529 | 0.616697988 | ns | TM | KS.3 |
| FIM1_4303 | 59.87240193 | 58.61279961 | 82.48496482 | -0.002327057 | 0.076426682 | 0.912225481 | 0.991141351 | ns | TM | KS.3 |
| FIM1_4304 | 27.32654314 | 22.66702761 | 30.91326256 | -0.010035624 | 0.097639197 | 0.591981382 | 0.964028601 | ns | TM | KS.3 |
| FIM1_4307 | 380.3450953 | 384.5476417 | 476.7071757 | 0.001621102 | 0.067216964 | 0.942711504 | 0.994413583 | ns | TM | KS.3 |
| FIM1_4308 | 106.4651059 | 118.4657032 | 116.7686975 | 0.014412794 | 0.077060338 | 0.522355817 | 0.958602567 | ns | TM | KS.3 |
| FIM1_4314 | 529.4632454 | 591.5978997 | 646.7515623 | 0.016526243 | 0.075468303 | 0.476416778 | 0.950410534 | ns | TM | KS.3 |
| FIM1_4780 | 43.64012059 | 57.38502959 | 82.75928701 | 0.019631571 | 0.101316946 | 0.343442935 | 0.944808611 | ns | TM | KS.3 |
| FIM1_4781 | 36.95815958 | 28.99750924 | 87.57792087 | -0.030304099 | 0.097597552 | 0.208505161 | 0.852551729 | ns | TM | KS.3 |
| FIM1_4782 | 236.1991408 | 253.6514793 | 296.5047851 | 0.017487129 | 0.062489236 | 0.468343201 | 0.950410534 | ns | TM | KS.3 |
| FIM1_4783 | 2877.254054 | 2357.784445 | 1729.24346 | -0.083044688 | 0.100667297 | 0.032351766 | 0.414819679 | ns | TM | KS.3 |
| FIM1_4785 | 666.4756737 | 749.333919 | 1037.162625 | 0.033135319 | 0.07139192 | 0.222889681 | 0.871073186 | ns | TM | KS.3 |
| FIM1_4786 | 729.130669 | 932.4957638 | 3276.771524 | 0.014607102 | 0.101616578 | 0.451234939 | 0.950410534 | ns | TM | KS.3 |
| FIM1_4787 | 118.9082694 | 120.4747036 | 235.4642185 | 0.001133487 | 0.076962134 | 0.956887697 | 0.994413583 | ns | TM | KS.3 |
| FIM1_4788 | 323.8642495 | 208.4899156 | 744.3069426 | -0.050529922 | 0.142054875 | 0.076618676 | 0.609319531 | ns | TM | KS.3 |
| FIM1_4789 | 262.5599028 | 251.6895374 | 264.372092 | -0.002551703 | 0.091582391 | 0.889936032 | 0.991141351 | ns | TM | KS.3 |
| FIM1_4790 | 611.6132893 | 616.211966 | 614.4716867 | 0.0012226 | 0.067093596 | 0.956787381 | 0.994413583 | ns | TM | KS.3 |
| FIM1_4791 | 1333.329598 | 1489.74804 | 2073.260557 | 0.015534994 | 0.076376888 | 0.496889678 | 0.953260948 | ns | TM | KS.3 |
| FIM1_4792 | 399.1654674 | 444.005342 | 464.198878 | 0.011005765 | 0.081091312 | 0.603297065 | 0.970245665 | ns | TM | KS.3 |
| FIM1_4793 | 1900.114518 | 1579.730125 | 2070.580399 | -0.050856236 | 0.089382515 | 0.100218011 | 0.680197281 | ns | TM | KS.3 |
| FIM1_4794 | 2074.851457 | 1859.886803 | 2670.050722 | -0.015164572 | 0.076381741 | 0.505797615 | 0.953260948 | ns | TM | KS.3 |
| FIM1_4795 | 4607.421875 | 5271.743533 | 6331.766408 | 0.014694043 | 0.082207887 | 0.498720789 | 0.953260948 | ns | TM | KS.3 |
| FIM1_4796 | 1931.35561 | 1871.880426 | 2521.152864 | -0.004556337 | 0.070377024 | 0.838079396 | 0.991141351 | ns | TM | KS.3 |
| FIM1_4797 | 3172.614322 | 3682.21853 | 4622.760369 | 0.014157915 | 0.085498992 | 0.501742892 | 0.953260948 | ns | TM | KS.3 |
| FIM1_4798 | 1039.159295 | 1108.857206 | 907.3828427 | 0.008740525 | 0.073686404 | 0.693415855 | 0.981188557 | ns | TM | KS.3 |
| FIM1_4799 | 1320.322832 | 1042.697725 | 915.778069 | -0.04310034 | 0.096673859 | 0.124045699 | 0.739893493 | ns | TM | KS.3 |
| FIM1_4800 | 13.89746688 | 11.55619107 | 14.76985844 | -0.008932982 | 0.096480981 | 0.631667341 | 0.97344368 | ns | TM | KS.3 |
| FIM1_4801 | 423.336306 | 286.2106157 | 369.4995752 | -0.07524692 | 0.149325171 | 0.0343394 | 0.42621823 | ns | TM | KS.3 |
| FIM1_4802 | 817.1015538 | 758.047066 | 805.0057536 | -0.025070835 | 0.057846304 | 0.301926622 | 0.926536521 | ns | TM | KS.3 |
| FIM1_4803 | 1978.108085 | 1862.220275 | 1781.19925 | -0.007990036 | 0.07371553 | 0.717396127 | 0.987157164 | ns | TM | KS.3 |
| FIM1_4804 | 6120.490985 | 5860.543539 | 5526.854346 | -0.009660064 | 0.060221427 | 0.676646474 | 0.979149565 | ns | TM | KS.3 |

| | | | | | | | | | | |
|---|---|---|---|---|---|---|---|---|---|---|
| FIM1_4805 | 3884.455768 | 4254.490781 | 4310.284211 | 0.020482098 | 0.067368467 | 0.406802922 | 0.950410534 | ns | TM | KS.3 |
| FIM1_4806 | 1812.182014 | 1979.272392 | 2008.96905 | 0.047725112 | 0.063676923 | 0.094414549 | 0.673663379 | ns | TM | KS.3 |
| FIM1_4807 | 2616.693492 | 2699.472391 | 2654.061447 | 0.004574492 | 0.070268419 | 0.837561792 | 0.991141351 | ns | TM | KS.3 |
| FIM1_4808 | 2121.854253 | 2161.21744 | 2096.452981 | 0.00281811 | 0.069005139 | 0.899855612 | 0.991141351 | ns | TM | KS.3 |
| FIM1_4809 | 635.1324251 | 656.4030572 | 696.4791253 | 0.006463356 | 0.062195829 | 0.778727602 | 0.988814136 | ns | TM | KS.3 |
| FIM1_4810 | 14269.08705 | 12285.99945 | 12749.87612 | -0.092566397 | 0.087476013 | 0.022970818 | 0.356471474 | ns | TM | KS.3 |
| FIM1_4811 | 1833.826446 | 1596.375614 | 1936.960012 | -0.010967714 | 0.087997043 | 0.584148376 | 0.963781856 | ns | TM | KS.3 |
| FIM1_4812 | 256.7653945 | 255.009183 | 213.5856373 | -0.000394392 | 0.06528938 | 0.986136779 | 0.999333464 | ns | TM | KS.3 |
| FIM1_4813 | 1095.0628 | 1335.421059 | 1787.613643 | 0.01860604 | 0.090373619 | 0.386646842 | 0.948637087 | ns | TM | KS.3 |
| FIM1_4814 | 1304.654938 | 1110.431168 | 1173.57424 | -0.110215451 | 0.090220884 | 0.012864803 | 0.257037736 | ns | TM | KS.3 |
| FIM1_4815 | 841.2869289 | 832.6333635 | 843.8898588 | -0.002480635 | 0.048483255 | 0.906225928 | 0.991141351 | ns | TM | KS.3 |
| FIM1_4816 | 1392.298011 | 1203.21572 | 1091.397534 | -0.073006242 | 0.085250138 | 0.045501409 | 0.489573998 | ns | TM | KS.3 |
| FIM1_4817 | 1661.451977 | 1581.574625 | 1649.15812 | -0.010117959 | 0.063192205 | 0.664031611 | 0.976323032 | ns | TM | KS.3 |
| FIM1_4818 | 1263.857269 | 974.1532458 | 1041.875757 | -0.240766033 | 0.108706916 | 0.000186891 | 0.014087652 | ns | TM | KS.3 |
| FIM1_4819 | 9272.326028 | 9625.849261 | 5748.894804 | 0.011530072 | 0.04780488 | 0.582145288 | 0.963781856 | ns | TM | KS.3 |
| FIM1_4820 | 1781.232557 | 1580.530912 | 1825.771121 | -0.050618537 | 0.074933655 | 0.102642834 | 0.689041746 | ns | TM | KS.3 |
| FIM1_4821 | 1289.76881 | 1087.362553 | 1098.858809 | -0.017519026 | 0.086401413 | 0.420251077 | 0.950410534 | ns | TM | KS.3 |
| FIM1_4822 | 1931.894348 | 1963.018875 | 1621.314464 | 0.003959235 | 0.053103206 | 0.857592827 | 0.991141351 | ns | TM | KS.3 |
| FIM1_4823 | 1365.57193 | 1380.262083 | 1166.873272 | 0.002858593 | 0.051073178 | 0.89499305 | 0.991141351 | ns | TM | KS.3 |
| FIM1_4824 | 2666.055794 | 2936.943699 | 3418.530375 | 0.015169144 | 0.073363654 | 0.513963133 | 0.954240701 | ns | TM | KS.3 |
| FIM1_4825 | 252.3877841 | 282.9637477 | 465.332088 | 0.015219012 | 0.077222649 | 0.501927892 | 0.953260948 | ns | TM | KS.3 |
| FIM1_4826 | 930.1304231 | 952.7870296 | 1047.673213 | 0.006502132 | 0.04950827 | 0.760759296 | 0.988477371 | ns | TM | KS.3 |
| FIM1_4827 | 1247.645362 | 1330.788812 | 1322.638848 | 0.01957929 | 0.056857337 | 0.406715477 | 0.950410534 | ns | TM | KS.3 |
| FIM1_4828 | 1016.697199 | 887.0342708 | 929.2585492 | -0.047738139 | 0.078597147 | 0.118473904 | 0.727123865 | ns | TM | KS.3 |
| FIM1_4829 | 2647.324296 | 1999.511997 | 2408.544241 | -0.028632949 | 0.101403888 | 0.219251754 | 0.865538807 | ns | TM | KS.3 |
| FIM1_4830 | 2898.868172 | 2551.362707 | 2396.706495 | -0.014211671 | 0.08130996 | 0.513826775 | 0.954240701 | ns | TM | KS.3 |
| FIM1_4831 | 2071.675291 | 2088.587114 | 1966.919271 | 0.002432975 | 0.042198482 | 0.899039448 | 0.991141351 | ns | TM | KS.3 |
| FIM1_4832 | 4460.859588 | 3882.269408 | 3576.343722 | -0.134821235 | 0.071270035 | 0.003186725 | 0.103770022 | ns | TM | KS.3 |
| FIM1_4833 | 4175.615762 | 3853.716831 | 4511.192965 | -0.018544222 | 0.065027997 | 0.446628116 | 0.950410534 | ns | TM | KS.3 |
| FIM1_4834 | 1039.834157 | 1071.617189 | 1340.639535 | 0.002108355 | 0.08681423 | 0.912172172 | 0.991141351 | ns | TM | KS.3 |
| FIM1_4835 | 3924.270312 | 3847.177825 | 3751.275652 | -0.004616643 | 0.056345976 | 0.837821082 | 0.991141351 | ns | TM | KS.3 |
| FIM1_4836 | 1620.580371 | 819.8582262 | 802.9952928 | -0.656805078 | 0.405397473 | 0.000174634 | 0.013620889 | ns | TM | KS.3 |
| FIM1_4837 | 170.5534637 | 75.73057594 | 67.6932174 | -0.325272757 | 0.453457631 | 0.002176969 | 0.084221572 | ns | TM | KS.3 |
| FIM1_4839 | 717.8834968 | 629.9150585 | 576.3763114 | -0.013930693 | 0.082200046 | 0.518149301 | 0.957219745 | ns | TM | KS.3 |
| FIM1_4840 | 8635.991521 | 8454.917469 | 8620.223328 | -0.006360812 | 0.044205485 | 0.748264268 | 0.988477371 | ns | TM | KS.3 |
| FIM1_4841 | 146.7136703 | 149.9790663 | 248.0003937 | 0.003032726 | 0.063237984 | 0.8943369 | 0.991141351 | ns | TM | KS.3 |
| FIM1_4842 | 57697.17695 | 58878.07016 | 58807.41069 | 0.005335554 | 0.051476065 | 0.806478815 | 0.988916276 | ns | TM | KS.3 |
| FIM1_4844 | 133.5420637 | 277.1853115 | 1119.871416 | 0.086747377 | 0.238337977 | 0.026881311 | 0.382098641 | ns | TM | KS.3 |
| FIM1_4845 | 438.4323298 | 417.9652908 | 600.696593 | -0.004043418 | 0.08295162 | 0.83909926 | 0.991141351 | ns | TM | KS.3 |
| FIM1_4846 | 641.4366622 | 617.4764092 | 542.9534828 | -0.002934221 | 0.084493452 | 0.880726412 | 0.991141351 | ns | TM | KS.3 |
| FIM1_4847 | 408.5736275 | 467.7463944 | 765.4735833 | 0.013543882 | 0.0836558 | 0.523593024 | 0.958732165 | ns | TM | KS.3 |
| FIM1_4848 | 1374.082753 | 2047.670781 | 2313.99835 | 0.072042897 | 0.149373543 | 0.037736163 | 0.440708412 | ns | TM | KS.3 |
| FIM1_4849 | 2938.769627 | 3122.617401 | 3562.945838 | 0.006462224 | 0.078161866 | 0.759107394 | 0.988477371 | ns | TM | KS.3 |
| FIM1_4850 | 478.4140431 | 348.2269915 | 797.3107309 | -0.023431457 | 0.10748623 | 0.273456204 | 0.908468501 | ns | TM | KS.3 |
| FIM1_4851 | 903.6782934 | 827.8917234 | 855.6509651 | -0.01857007 | 0.067626701 | 0.445808288 | 0.950410534 | ns | TM | KS.3 |

| | | | | | | | | | | |
|---|---|---|---|---|---|---|---|---|---|---|
| FIM1_4852 | 596.906722 | 562.7036558 | 468.2283045 | -0.011127851 | 0.066091312 | 0.633437561 | 0.97344368 | ns | TM | KS.3 |
| FIM1_4853 | 345.4429059 | 319.3883248 | 464.3356436 | -0.011745456 | 0.07231217 | 0.606527684 | 0.970937835 | ns | TM | KS.3 |
| FIM1_4854 | 2205.247876 | 2543.329787 | 3480.209368 | 0.012167899 | 0.086847647 | 0.551774836 | 0.95917717 | ns | TM | KS.3 |
| FIM1_4855 | 715.0886735 | 865.2844021 | 915.6515042 | 0.012391894 | 0.095404039 | 0.523557026 | 0.958732165 | ns | TM | KS.3 |
| FIM1_4856 | 91.29574327 | 112.4378457 | 90.94481753 | 0.025690418 | 0.089787298 | 0.274565829 | 0.908583046 | ns | TM | KS.3 |
| FIM1_4857 | 19.50102853 | 27.88218082 | 97.34138546 | 0.023148258 | 0.113893892 | 0.269970666 | 0.908468501 | ns | TM | KS.3 |
| FIM1_4858 | 1627.789236 | 1938.715928 | 1889.896656 | 0.016831026 | 0.087584427 | 0.431590487 | 0.950410534 | ns | TM | KS.3 |
| FIM1_4859 | 1306.273902 | 1451.162592 | 1667.293549 | 0.052965037 | 0.070924545 | 0.086918312 | 0.645400896 | ns | TM | KS.3 |
| FIM1_4860 | 624.3543495 | 621.0934148 | 551.1737234 | -0.000599657 | 0.075909213 | 0.977403752 | 0.999036302 | ns | TM | KS.3 |
| FIM1_4861 | 914.0535174 | 864.887287 | 825.2678814 | -0.016079317 | 0.054117107 | 0.480169204 | 0.950972051 | ns | TM | KS.3 |
| FIM1_4862 | 1517.513677 | 1571.430062 | 1646.755377 | 0.007028749 | 0.062374342 | 0.760320903 | 0.988477371 | ns | TM | KS.3 |
| FIM1_4863 | 3771.007409 | 4600.008713 | 5275.620652 | 0.021245254 | 0.089069855 | 0.341932291 | 0.944808611 | ns | TM | KS.3 |
| FIM1_4864 | 55.43058152 | 74.92836296 | 64.56806349 | 0.037162466 | 0.105455736 | 0.146671928 | 0.780546912 | ns | TM | KS.3 |
| FIM1_4865 | 757.2339444 | 692.2898304 | 766.4919362 | -0.012151142 | 0.075065638 | 0.588261333 | 0.963781856 | ns | TM | KS.3 |
| FIM1_4866 | 1890.717469 | 1872.060913 | 1468.118296 | -0.001187215 | 0.074506066 | 0.955870089 | 0.994413583 | ns | TM | KS.3 |
| FIM1_4867 | 1410.253289 | 1591.460613 | 1774.801931 | 0.020673407 | 0.075065412 | 0.392276621 | 0.948637087 | ns | TM | KS.3 |
| FIM1_4868 | 1297.127196 | 1348.609374 | 1300.526989 | 0.009161433 | 0.05768448 | 0.689324265 | 0.980879562 | ns | TM | KS.3 |
| FIM1_4869 | 2760.027251 | 2624.413813 | 2324.438527 | -0.010254664 | 0.063620037 | 0.659948149 | 0.976323032 | ns | TM | KS.3 |
| FIM1_4870 | 445.9455527 | 613.4528805 | 695.124892 | 0.040320726 | 0.11141208 | 0.12449578 | 0.739983875 | ns | TM | KS.3 |
| FIM1_4871 | 252.1026436 | 329.2287686 | 519.1620767 | 0.068490774 | 0.112857437 | 0.046476096 | 0.493142038 | ns | TM | KS.3 |
| FIM1_4872 | 158.8456437 | 192.8374552 | 580.6762812 | 0.030650431 | 0.086603301 | 0.226193782 | 0.876782875 | ns | TM | KS.3 |
| FIM1_4873 | 1121.439506 | 1398.040402 | 1506.886808 | 0.050649368 | 0.096178595 | 0.094786807 | 0.673663379 | ns | TM | KS.3 |
| FIM1_4874 | 2403.177297 | 2707.591296 | 2687.585499 | 0.017055944 | 0.076992094 | 0.460109334 | 0.950410534 | ns | TM | KS.3 |
| FIM1_4875 | 1815.822415 | 1417.404355 | 1454.363538 | -0.034159357 | 0.096489236 | 0.177703168 | 0.811391014 | ns | TM | KS.3 |
| FIM1_4876 | 117.0663914 | 73.01049282 | 66.18190685 | -0.103337259 | 0.196570637 | 0.017531402 | 0.304361209 | ns | TM | KS.3 |
| FIM1_4877 | 715.2812877 | 806.3335511 | 1003.914225 | 0.016488395 | 0.077516165 | 0.471229472 | 0.950410534 | ns | TM | KS.3 |
| FIM1_4878 | 19204.80271 | 20283.01002 | 19245.88169 | 0.01376999 | 0.058491178 | 0.555438919 | 0.95948216 | ns | TM | KS.3 |
| FIM1_4879 | 2741.279644 | 2552.836537 | 2685.028214 | -0.034236954 | 0.054866452 | 0.157110905 | 0.789034023 | ns | TM | KS.3 |
| FIM1_4880 | 5161.210729 | 4421.999726 | 4901.205305 | -0.159834022 | 0.068155338 | 0.000950196 | 0.04545411 | ns | TM | KS.3 |
| FIM1_4881 | 1840.738704 | 1918.617183 | 2085.045205 | 0.010102159 | 0.057509373 | 0.659911251 | 0.976323032 | ns | TM | KS.3 |
| FIM1_4882 | 38.20272295 | 25.77935116 | 1008.366534 | -0.017305604 | 0.119888503 | 0.362291839 | 0.948637087 | ns | TM | KS.3 |
| FIM1_4883 | 3310.675792 | 3195.005971 | 3633.9843 | -0.006796158 | 0.063794837 | 0.767745261 | 0.988477371 | ns | TM | KS.3 |
| FIM1_4884 | 2645.385139 | 2384.934787 | 2324.54675 | -0.051117484 | 0.070012817 | 0.09320322 | 0.671036206 | ns | TM | KS.3 |
| FIM1_4885 | 1079.151695 | 955.4941862 | 961.0478088 | -0.044449622 | 0.074526977 | 0.134535203 | 0.766830871 | ns | TM | KS.3 |
| FIM1_4886 | 300.9914182 | 286.1989393 | 262.1975986 | -0.009651312 | 0.065296198 | 0.677515677 | 0.979837352 | ns | TM | KS.3 |
| FIM1_4887 | 374.0377953 | 493.4943365 | 366.9965181 | 0.191314894 | 0.134966129 | 0.001623795 | 0.067857575 | ns | TM | KS.3 |
| FIM1_4888 | 979.4938622 | 1015.537776 | 846.6949265 | 0.005834909 | 0.067819331 | 0.797093954 | 0.988916276 | ns | TM | KS.3 |
| FIM1_4889 | 824.7547432 | 626.5378178 | 630.0680574 | -0.061974794 | 0.111907751 | 0.057307626 | 0.536774409 | ns | TM | KS.3 |
| FIM1_4890 | 600.1318535 | 638.6616388 | 592.0101904 | 0.010881876 | 0.06790389 | 0.638839523 | 0.975246356 | ns | TM | KS.3 |
| FIM1_4891 | 898.7015039 | 942.0559417 | 1119.446052 | 0.009847471 | 0.062067211 | 0.671963489 | 0.976861203 | ns | TM | KS.3 |
| FIM1_4892 | 4455.086949 | 4044.768091 | 4920.135313 | -0.011840688 | 0.077394399 | 0.590296346 | 0.963781856 | ns | TM | KS.3 |
| FIM1_4893 | 9036.050582 | 8572.426937 | 9611.487046 | -0.006570584 | 0.07460004 | 0.762735003 | 0.988477371 | ns | TM | KS.3 |
| FIM1_4894 | 1595.683734 | 1657.03519 | 1464.534456 | 0.006408832 | 0.066967114 | 0.778780039 | 0.988814136 | ns | TM | KS.3 |
| FIM1_4895 | 1706.073563 | 1674.722351 | 1695.610525 | -0.002483504 | 0.071904756 | 0.909946149 | 0.991141351 | ns | TM | KS.3 |
| FIM1_4896 | 1011.538025 | 1190.149716 | 1266.705427 | 0.023208274 | 0.082269753 | 0.329556817 | 0.936229083 | ns | TM | KS.3 |

| | | | | | | | | | | |
|---|---|---|---|---|---|---|---|---|---|---|
| FIM1_4897 | 1024.44226 | 989.9148324 | 1069.82916 | -0.004313176 | 0.073705124 | 0.842722619 | 0.991141351 | ns | TM | KS.3 |
| FIM1_4898 | 939.7112246 | 971.5655274 | 920.0857228 | 0.003635995 | 0.076982393 | 0.862962495 | 0.991141351 | ns | TM | KS.3 |
| FIM1_4899 | 7945.805315 | 8967.129248 | 7998.009529 | 0.034293891 | 0.072904887 | 0.211597248 | 0.856547038 | ns | TM | KS.3 |
| FIM1_4900 | 4361.59296 | 5289.237636 | 4562.418037 | 0.056746174 | 0.092701221 | 0.079371684 | 0.616697988 | ns | TM | KS.3 |
| FIM1_4901 | 388.2948208 | 419.6723246 | 385.1794082 | 0.010636415 | 0.073944325 | 0.634737737 | 0.973888113 | ns | TM | KS.3 |
| FIM1_4902 | 987.0974068 | 907.3258047 | 1002.088481 | -0.019084713 | 0.066082656 | 0.435299165 | 0.950410534 | ns | TM | KS.3 |
| FIM1_4903 | 1037.879276 | 957.2974844 | 1339.826362 | -0.019319691 | 0.064685929 | 0.429880554 | 0.950410534 | ns | TM | KS.3 |
| FIM1_4904 | 1910.466738 | 1804.84689 | 1271.668189 | -0.009023879 | 0.069701209 | 0.692062433 | 0.981188557 | ns | TM | KS.3 |
| FIM1_4905 | 436.4679259 | 482.0519021 | 445.754003 | 0.01311715 | 0.076366101 | 0.558133539 | 0.960136361 | ns | TM | KS.3 |
| FIM1_4906 | 1370.431264 | 1445.79011 | 1602.179037 | 0.01489236 | 0.056147618 | 0.519145146 | 0.957992249 | ns | TM | KS.3 |
| FIM1_4907 | 1404.293136 | 1689.125928 | 2107.583726 | 0.015927282 | 0.090063237 | 0.445000417 | 0.950410534 | ns | TM | KS.3 |
| FIM1_4908 | 586.5561854 | 646.2978952 | 559.5296818 | 0.015996935 | 0.072431984 | 0.496343953 | 0.953260948 | ns | TM | KS.3 |
| FIM1_4909 | 2759.316477 | 2719.51424 | 2808.153758 | -0.003606059 | 0.054106161 | 0.871100596 | 0.991141351 | ns | TM | KS.3 |
| FIM1_4910 | 451.909436 | 681.1384085 | 642.7546195 | 0.23264504 | 0.22968595 | 0.001719025 | 0.070099605 | ns | TM | KS.3 |
| FIM1_4911 | 1209.393587 | 1346.349285 | 1207.163562 | 0.016456768 | 0.074771566 | 0.479917108 | 0.950972051 | ns | TM | KS.3 |
| FIM1_4912 | 1163.110808 | 1039.286093 | 1002.936123 | -0.019719689 | 0.073715435 | 0.413520364 | 0.950410534 | ns | TM | KS.3 |
| FIM1_4913 | 1205.014214 | 1415.994568 | 1779.518337 | 0.021187165 | 0.082727296 | 0.361232962 | 0.948637087 | ns | TM | KS.3 |
| FIM1_4914 | 1984.788803 | 2016.192933 | 2391.945927 | 0.003765515 | 0.055668369 | 0.866791777 | 0.991141351 | ns | TM | KS.3 |
| FIM1_4915 | 1933.433723 | 1861.029396 | 2017.395644 | -0.004237569 | 0.076586126 | 0.841454216 | 0.991141351 | ns | TM | KS.3 |
| FIM1_4916 | 1076.502605 | 1386.64277 | 1357.458541 | 0.018628427 | 0.099217892 | 0.366152371 | 0.948637087 | ns | TM | KS.3 |
| FIM1_4917 | 1289.345161 | 1356.219665 | 1595.349315 | 0.01284587 | 0.057380195 | 0.578765313 | 0.963781856 | ns | TM | KS.3 |
| FIM1_4918 | 148.2471049 | 158.2788884 | 185.4433156 | 0.009825228 | 0.069624328 | 0.667717743 | 0.976323032 | ns | TM | KS.3 |
| FIM1_4919 | 1305.306216 | 1386.017679 | 1474.08409 | 0.008926302 | 0.071255696 | 0.692491001 | 0.981188557 | ns | TM | KS.3 |
| FIM1_4920 | 555.1272712 | 390.4391121 | 491.5382069 | -0.148256477 | 0.172355406 | 0.005901204 | 0.16042891 | ns | TM | KS.3 |
| FIM1_4921 | 12316.55432 | 10012.57328 | 10694.03251 | -0.072037518 | 0.100013911 | 0.045660771 | 0.489573998 | ns | TM | KS.3 |
| FIM1_4922 | 951.7637441 | 1075.525248 | 1043.283304 | 0.022612587 | 0.074533728 | 0.35935443 | 0.948637087 | ns | TM | KS.3 |
| FIM1_4923 | 720.744514 | 604.9983919 | 779.2656105 | -0.098548443 | 0.096457251 | 0.019893956 | 0.324535175 | ns | TM | KS.3 |
| FIM1_4924 | 2228.250855 | 3027.268563 | 2324.548194 | 0.041377112 | 0.109345489 | 0.12069614 | 0.730509595 | ns | TM | KS.3 |
| FIM1_4925 | 124.2215356 | 134.0587063 | 244.9278878 | 0.005058093 | 0.088169311 | 0.79134623 | 0.988916276 | ns | TM | KS.3 |
| FIM1_4926 | 1403.260098 | 1372.422986 | 1363.095632 | -0.006871103 | 0.042478638 | 0.72069897 | 0.987157164 | ns | TM | KS.3 |
| FIM1_4927 | 1065.460662 | 1280.954241 | 1324.343166 | 0.017640353 | 0.088609974 | 0.411187453 | 0.950410534 | ns | TM | KS.3 |
| FIM1_4928 | 832.1292983 | 1054.001796 | 1095.136634 | 0.022160071 | 0.094921466 | 0.313047125 | 0.927416591 | ns | TM | KS.3 |
| FIM1_4929 | 884.0223931 | 943.5043171 | 701.8857728 | 0.01893202 | 0.057572841 | 0.42396563 | 0.950410534 | ns | TM | KS.3 |
| FIM1_4931 | 3179.155951 | 3536.459274 | 3938.011143 | 0.021001368 | 0.071535411 | 0.393229043 | 0.948637087 | ns | TM | KS.3 |
| FIM1_4932 | 2973.122408 | 3235.437108 | 4932.710896 | 0.016743222 | 0.068172029 | 0.485897504 | 0.952269291 | ns | TM | KS.3 |
| FIM1_4933 | 1765.128623 | 1702.906388 | 1783.902986 | -0.00680814 | 0.064013838 | 0.767302164 | 0.988477371 | ns | TM | KS.3 |
| FIM1_4935 | 965.7721682 | 853.1914583 | 1921.499279 | -0.023104993 | 0.07470249 | 0.350861029 | 0.948637087 | ns | TM | KS.3 |
| FIM1_4936 | 1343.321841 | 1301.909207 | 1229.813806 | -0.003093961 | 0.078954584 | 0.880817381 | 0.991141351 | ns | TM | KS.3 |
| FIM1_4937 | 8751.352868 | 8785.049627 | 6854.844582 | 0.000350233 | 0.080536343 | 0.986186566 | 0.999333464 | ns | TM | KS.3 |
| FIM1_4938 | 1038.62142 | 1070.198579 | 1253.715977 | 0.005444835 | 0.064955107 | 0.811923103 | 0.988916276 | ns | TM | KS.3 |
| FIM1_4939 | 1255.552834 | 1214.704288 | 997.7523455 | -0.007819941 | 0.057109781 | 0.731346119 | 0.987157164 | ns | TM | KS.3 |
| FIM1_4940 | 1172.420971 | 1407.375177 | 1454.603785 | 0.023096035 | 0.085791863 | 0.321245204 | 0.929723612 | ns | TM | KS.3 |
| FIM1_4941 | 8726.245487 | 8156.047034 | 8851.194038 | -0.020234698 | 0.057775148 | 0.395847595 | 0.948637087 | ns | TM | KS.3 |
| FIM1_4942 | 1104.26535 | 1158.338696 | 1224.514551 | 0.003623904 | 0.085194189 | 0.852395728 | 0.991141351 | ns | TM | KS.3 |
| FIM1_4943 | 2880.788247 | 2718.375271 | 2318.517883 | -0.012230512 | 0.063507801 | 0.602960457 | 0.970245665 | ns | TM | KS.3 |

| | | | | | | | | | | |
|---|---|---|---|---|---|---|---|---|---|---|
| FIM1_4944 | 737.2736699 | 885.7305614 | 1183.17116 | 0.017767813 | 0.088393985 | 0.409137388 | 0.950410534 | ns | TM | KS.3 |
| FIM1_4945 | 7480.400937 | 7485.169791 | 7429.713428 | 0.000112962 | 0.066049988 | 0.996019656 | 0.999525119 | ns | TM | KS.3 |
| FIM1_4946 | 2632.57923 | 2754.567525 | 3010.056622 | 0.009067656 | 0.063414305 | 0.695882185 | 0.981479944 | ns | TM | KS.3 |
| FIM1_4947 | 2249.848758 | 1885.275455 | 1835.211868 | -0.068095273 | 0.092284654 | 0.054307239 | 0.531428718 | ns | TM | KS.3 |
| FIM1_4948 | 520.1645785 | 507.6595532 | 502.0525132 | -0.005501336 | 0.058777903 | 0.809476049 | 0.988916276 | ns | TM | KS.3 |
| FIM1_4949 | 1788.841957 | 1779.09298 | 1482.360849 | -0.001202065 | 0.032813416 | 0.937954201 | 0.994413583 | ns | TM | KS.3 |
| FIM1_4950 | 4454.026295 | 3870.997354 | 3177.919928 | -0.081686974 | 0.084335663 | 0.032798106 | 0.417169416 | ns | TM | KS.3 |
| FIM1_4951 | 2365.035737 | 2239.649377 | 2430.954616 | -0.010139525 | 0.065928343 | 0.662496492 | 0.976323032 | ns | TM | KS.3 |
| FIM1_4952 | 5677.534917 | 6145.844355 | 6919.195298 | 0.008165022 | 0.079627494 | 0.697412453 | 0.98151979 | ns | TM | KS.3 |
| FIM1_4953 | 1503.03021 | 1818.772088 | 2320.49064 | 0.01666508 | 0.090447257 | 0.427169717 | 0.950410534 | ns | TM | KS.3 |
| FIM1_4954 | 277.873252 | 311.2425809 | 227.721582 | 0.010578947 | 0.08335247 | 0.609374602 | 0.971367716 | ns | TM | KS.3 |
| FIM1_4955 | 10717.17011 | 9543.298234 | 10080.4091 | -0.02482217 | 0.072438896 | 0.327529088 | 0.934739762 | ns | TM | KS.3 |
| FIM1_4956 | 664.7105029 | 697.9733624 | 2186.763112 | 0.007448359 | 0.070222903 | 0.741187336 | 0.988051177 | ns | TM | KS.3 |
| FIM1_4957 | 251.5101215 | 347.2534073 | 753.7054491 | 0.069681846 | 0.127336224 | 0.041961081 | 0.469115458 | ns | TM | KS.3 |
| FIM1_4958 | 201.0941291 | 175.0392786 | 219.4088314 | -0.008120061 | 0.094536709 | 0.664139875 | 0.976323032 | ns | TM | KS.3 |
| FIM1_4960 | 4774.565328 | 3991.984962 | 4081.875028 | -0.121545953 | 0.095824422 | 0.009334836 | 0.211094585 | ns | TM | KS.3 |
| FIM1_4961 | 3912.196796 | 2665.761188 | 3685.306776 | -0.114025019 | 0.171499955 | 0.012800434 | 0.257037736 | ns | TM | KS.3 |
| FIM1_4962 | 646.8810173 | 647.1435874 | 811.1420474 | 0.000165439 | 0.061908677 | 0.994210459 | 0.999525119 | ns | TM | KS.3 |
| FIM1_4963 | 578.6683254 | 587.4873847 | 718.4865154 | 0.003771717 | 0.051107032 | 0.861863763 | 0.991141351 | ns | TM | KS.3 |
| FIM1_4964 | 49.77163186 | 59.30349231 | 90.85514937 | 0.027180427 | 0.08143346 | 0.27675741 | 0.908583046 | ns | TM | KS.3 |
| FIM1_4965 | 6242.229309 | 7095.724528 | 7135.825911 | 0.045353443 | 0.076204075 | 0.13029479 | 0.76081758 | ns | TM | KS.3 |
| FIM1_4966 | 5172.746205 | 4293.097458 | 4089.50433 | -0.136471523 | 0.096294736 | 0.005714058 | 0.157057677 | ns | TM | KS.3 |
| FIM1_4967 | 1656.055188 | 1573.062277 | 1829.677499 | -0.012911947 | 0.058178649 | 0.578559275 | 0.963781856 | ns | TM | KS.3 |
| FIM1_4968 | 1517.044105 | 1814.716112 | 1696.526117 | 0.01592232 | 0.089125067 | 0.447726502 | 0.950410534 | ns | TM | KS.3 |
| FIM1_4969 | 14799.27045 | 13086.51089 | 13025.51777 | -0.093860113 | 0.076664002 | 0.018056168 | 0.307635054 | ns | TM | KS.3 |
| FIM1_4970 | 12992.72592 | 13363.84916 | 13733.02455 | 0.008434474 | 0.04645154 | 0.6811562 | 0.97986479 | ns | TM | KS.3 |
| FIM1_4971 | 4644.370809 | 4866.907625 | 5299.304992 | 0.007788175 | 0.068027438 | 0.733072678 | 0.987553905 | ns | TM | KS.3 |
| FIM1_4972 | 1517.473593 | 1702.357744 | 1907.458323 | 0.013679404 | 0.07925895 | 0.534260908 | 0.959071703 | ns | TM | KS.3 |
| FIM1_4973 | 1870.666189 | 2140.732173 | 1688.204938 | 0.022549981 | 0.077243089 | 0.354035506 | 0.948637087 | ns | TM | KS.3 |
| FIM1_4974 | 515.1020613 | 513.7165143 | 495.5369414 | -0.001445697 | 0.052536162 | 0.947440325 | 0.994413583 | ns | TM | KS.3 |
| FIM1_4976 | 7366.736735 | 7546.327572 | 7599.367736 | 0.00634638 | 0.051945544 | 0.771978006 | 0.988814136 | ns | TM | KS.3 |
| FIM1_4977 | 988.8012955 | 1391.733277 | 1707.165664 | 0.026146114 | 0.111176942 | 0.234646216 | 0.883800031 | ns | TM | KS.3 |
| FIM1_4978 | 1814.399857 | 1119.442179 | 947.8801932 | -0.633711868 | 0.176903632 | 1.24573221591 | 0.000187804 | ns | TM | KS.3 |
| FIM1_4979 | 17206.02486 | 16065.35911 | 16927.46391 | -0.024399387 | 0.055680466 | 0.303086941 | 0.926536521 | ns | TM | KS.3 |
| FIM1_4980 | 571.3624804 | 610.2488342 | 556.3859116 | 0.012147393 | 0.067446955 | 0.603405042 | 0.970245665 | ns | TM | KS.3 |
| FIM1_4981 | 14992.6904 | 14608.92603 | 15859.94697 | -0.008860869 | 0.035824905 | 0.588686383 | 0.963781856 | ns | TM | KS.3 |
| FIM1_4982 | 998.1214215 | 985.7989841 | 1010.171553 | -0.004326818 | 0.036201139 | 0.797088273 | 0.988916276 | ns | TM | KS.3 |
| FIM1_4983 | 941.3048 | 939.1737935 | 976.9536087 | -0.000578453 | 0.047169704 | 0.977717334 | 0.999036302 | ns | TM | KS.3 |
| FIM1_4984 | 1354.353056 | 1133.35396 | 1549.734645 | -0.041552055 | 0.085715152 | 0.146229213 | 0.780546912 | ns | TM | KS.3 |
| FIM1_4985 | 2728.67547 | 2489.856013 | 2672.857618 | -0.037591654 | 0.064236955 | 0.16623504 | 0.804493506 | ns | TM | KS.3 |
| FIM1_4986 | 5069.575782 | 4252.472055 | 4455.986282 | -0.138277551 | 0.090951302 | 0.004956158 | 0.140896498 | ns | TM | KS.3 |
| FIM1_4987 | 3599.998733 | 3372.250862 | 2820.403304 | -0.011573844 | 0.068049291 | 0.618814259 | 0.973122675 | ns | TM | KS.3 |
| FIM1_4988 | 182.4792466 | 163.0463817 | 174.4108259 | -0.021231898 | 0.073380933 | 0.385772649 | 0.948637087 | ns | TM | KS.3 |
| FIM1_4989 | 5263.726432 | 4577.618686 | 7408.666665 | -0.031578757 | 0.076982524 | 0.235890745 | 0.884880231 | ns | TM | KS.3 |
| FIM1_4991 | 3245.047082 | 3555.282728 | 3383.67469 | 0.015101782 | 0.071928446 | 0.518918579 | 0.957929472 | ns | TM | KS.3 |

| | | | | | | | | | | |
|---|---|---|---|---|---|---|---|---|---|---|
| FIM1_4992 | 1516.564965 | 1448.043685 | 1100.95135 | -0.007915749 | 0.067271582 | 0.729781083 | 0.987157164 | ns | TM | KS.3 |
| FIM1_4993 | 2063.747044 | 2284.311003 | 2479.73763 | 0.01989248 | 0.070836705 | 0.415457211 | 0.950410534 | ns | TM | KS.3 |
| FIM1_4995 | 1414.295745 | 1023.714665 | 1753.530347 | -0.082452145 | 0.133967762 | 0.028406827 | 0.390786333 | ns | TM | KS.3 |
| FIM1_4996 | 509.0617935 | 452.7957344 | 533.7981596 | -0.023886457 | 0.073229929 | 0.340976263 | 0.944808611 | ns | TM | KS.3 |
| FIM1_4997 | 1757.976322 | 1730.126188 | 1646.699176 | -0.003986813 | 0.053439025 | 0.856986212 | 0.991141351 | ns | TM | KS.3 |
| FIM1_4998 | 2475.526828 | 2199.854627 | 2531.997194 | -0.025042557 | 0.072894811 | 0.323603768 | 0.933252679 | ns | TM | KS.3 |
| FIM1_4999 | 3326.490674 | 3486.350054 | 3751.186881 | 0.00999652 | 0.061837292 | 0.66738923 | 0.976323032 | ns | TM | KS.3 |
| FIM1_5000 | 460.0559069 | 395.7674984 | 621.3457797 | -0.018043263 | 0.08246217 | 0.421650165 | 0.950410534 | ns | TM | KS.3 |
| FIM1_5001 | 413.4205489 | 488.0399657 | 1113.903104 | 0.019381454 | 0.084452494 | 0.388750468 | 0.948637087 | ns | TM | KS.3 |
| FIM1_5002 | 1677.622077 | 1618.920719 | 1846.561683 | -0.006648216 | 0.064337286 | 0.772413124 | 0.988814136 | ns | TM | KS.3 |
| FIM1_5003 | 1131.639618 | 1288.288308 | 1211.205547 | 0.034869539 | 0.074954652 | 0.205799863 | 0.85037734 | ns | TM | KS.3 |
| FIM1_5004 | 2726.679117 | 2739.151494 | 3222.634587 | 0.000549951 | 0.074519045 | 0.979538742 | 0.999333464 | ns | TM | KS.3 |
| FIM1_5005 | 20604.55797 | 21347.86138 | 20594.94691 | 0.009687651 | 0.052235961 | 0.66060437 | 0.976323032 | ns | TM | KS.3 |
| FIM1_5006 | 344.5978482 | 302.8896446 | 269.8336058 | -0.026483885 | 0.075330829 | 0.299179708 | 0.926536521 | ns | TM | KS.3 |
| FIM1_5007 | 1980.254999 | 2065.986464 | 2601.176588 | 0.003473088 | 0.083328493 | 0.860772157 | 0.991141351 | ns | TM | KS.3 |
| FIM1_5008 | 3596.424623 | 3751.635057 | 4296.317491 | 0.010423075 | 0.056917266 | 0.649084403 | 0.976323032 | ns | TM | KS.3 |
| FIM1_5009 | 108.5606607 | 44.8871868 | 119.1624388 | -0.382579456 | 0.502224334 | 0.001598923 | 0.067706867 | ns | TM | KS.3 |
| FIM1_5010 | 896.3079158 | 1003.411126 | 1071.331298 | 0.018124864 | 0.074767635 | 0.443313025 | 0.950410534 | ns | TM | KS.3 |
| FIM1_5011 | 187.1741621 | 184.8632814 | 171.8248649 | -0.001760321 | 0.069201981 | 0.937183901 | 0.994413583 | ns | TM | KS.3 |
| FIM1_5012 | 1382.829427 | 1328.601787 | 962.7448218 | -0.007148705 | 0.065829972 | 0.755468657 | 0.988477371 | ns | TM | KS.3 |
| FIM1_5013 | 1295.514812 | 1286.205499 | 1207.073186 | -0.001665612 | 0.056341543 | 0.94096748 | 0.994413583 | ns | TM | KS.3 |
| FIM1_5014 | 3507.765306 | 3109.455439 | 2375.999414 | -0.050542459 | 0.075049569 | 0.103105469 | 0.689041746 | ns | TM | KS.3 |
| FIM1_5015 | 2919.087634 | 2916.88342 | 2910.350965 | -0.000136879 | 0.063698847 | 0.995204211 | 0.999525119 | ns | TM | KS.3 |
| FIM1_5016 | 526.2168792 | 483.189727 | 752.5604019 | -0.011136363 | 0.075431859 | 0.616197398 | 0.973122675 | ns | TM | KS.3 |
| FIM1_5017 | 1843.222014 | 2464.184473 | 2331.695015 | 0.066465144 | 0.117461137 | 0.048210911 | 0.499686002 | ns | TM | KS.3 |
| FIM1_5018 | 385.6397732 | 408.3763987 | 382.6164024 | 0.011233209 | 0.065616217 | 0.630707205 | 0.97344368 | ns | TM | KS.3 |
| FIM1_5019 | 803.803967 | 917.9213381 | 1121.73002 | 0.016620922 | 0.080062289 | 0.460143215 | 0.950410534 | ns | TM | KS.3 |
| FIM1_5020 | 807.2577892 | 860.8056809 | 1100.992639 | 0.005741532 | 0.081931218 | 0.776906909 | 0.988814136 | ns | TM | KS.3 |
| FIM1_5021 | 648.6406549 | 582.2854219 | 757.4105412 | -0.033604721 | 0.069596338 | 0.216749731 | 0.865538807 | ns | TM | KS.3 |
| FIM1_5022 | 1896.474619 | 2173.016312 | 2655.25825 | 0.068202162 | 0.082118022 | 0.053249885 | 0.528754276 | ns | TM | KS.3 |
| FIM1_5023 | 1796.274318 | 1810.074498 | 1964.965564 | 0.002026967 | 0.048171797 | 0.923021409 | 0.992531983 | ns | TM | KS.3 |
| FIM1_5024 | 2972.460322 | 3150.25162 | 3413.721675 | 0.015695003 | 0.057328933 | 0.501633419 | 0.953260948 | ns | TM | KS.3 |
| FIM1_5025 | 2620.511183 | 2962.406204 | 2924.38756 | 0.019316417 | 0.076183228 | 0.415264525 | 0.950410534 | ns | TM | KS.3 |
| FIM1_5026 | 1166.05419 | 1170.596136 | 2627.883575 | 0.000908138 | 0.056836633 | 0.967858794 | 0.996327965 | ns | TM | KS.3 |
| FIM1_5027 | 2126.883761 | 2910.404011 | 3523.227109 | 0.109324316 | 0.142378052 | 0.01379494 | 0.267042128 | ns | TM | KS.3 |
| FIM1_5028 | 3041.246903 | 3183.079948 | 3668.055305 | 0.013775837 | 0.050935595 | 0.529179892 | 0.959071703 | ns | TM | KS.3 |
| FIM1_5029 | 3111.787074 | 2867.506229 | 2781.189831 | -0.03547199 | 0.060148565 | 0.171224287 | 0.809734625 | ns | TM | KS.3 |
| FIM1_5030 | 4798.311471 | 4682.153407 | 3657.061808 | -0.008105216 | 0.038166546 | 0.642900352 | 0.976237372 | ns | TM | KS.3 |
| FIM1_5031 | 4163.455584 | 4184.863839 | 5372.95306 | 0.001229349 | 0.054067337 | 0.955810714 | 0.994413583 | ns | TM | KS.3 |
| FIM1_5032 | 1496.522394 | 1791.680578 | 2251.585274 | 0.017876668 | 0.087737461 | 0.408773131 | 0.950410534 | ns | TM | KS.3 |
| FIM1_5033 | 1973.195948 | 1945.922756 | 1764.733732 | -0.004249049 | 0.035961838 | 0.799504289 | 0.988916276 | ns | TM | KS.3 |
| FIM1_5034 | 8134.267864 | 6823.084757 | 4885.437389 | -0.067612807 | 0.091918982 | 0.055281703 | 0.5319661 | ns | TM | KS.3 |
| FIM1_5035 | 1042.850731 | 1053.072455 | 1070.618224 | 0.001386602 | 0.070948342 | 0.949925541 | 0.994413583 | ns | TM | KS.3 |
| FIM1_5036 | 1288.209849 | 1364.790175 | 1330.740806 | 0.011087249 | 0.065703562 | 0.634860195 | 0.973888113 | ns | TM | KS.3 |
| FIM1_5037 | 1358.605529 | 1322.531085 | 1385.984308 | -0.008498075 | 0.040120071 | 0.641877538 | 0.975959887 | ns | TM | KS.3 |

| | | | | | | | | | | |
|---|---|---|---|---|---|---|---|---|---|---|
| FIM1_5038 | 3455.146733 | 3349.503159 | 3473.213098 | -0.008783691 | 0.050060931 | 0.683355653 | 0.98018358 | ns | TM | KS.3 |
| FIM1_5039 | 1449.145333 | 1414.477185 | 1431.105986 | -0.005805502 | 0.055592098 | 0.796393623 | 0.988916276 | ns | TM | KS.3 |
| FIM1_5040 | 16169.47391 | 16321.02543 | 15960.82951 | 0.002194498 | 0.055672273 | 0.922024991 | 0.992014345 | ns | TM | KS.3 |
| FIM1_5041 | 374.5711986 | 386.8013877 | 428.8023825 | 0.006131011 | 0.062842234 | 0.789652442 | 0.988916276 | ns | TM | KS.3 |
| FIM1_5042 | 18380.85763 | 16455.40954 | 18192.75871 | -0.028050277 | 0.070452942 | 0.282811288 | 0.918450395 | ns | TM | KS.3 |
| FIM1_5043 | 853.1679895 | 938.5075698 | 914.906838 | 0.017844888 | 0.070631943 | 0.458097225 | 0.950410534 | ns | TM | KS.3 |
| FIM1_5044 | 205.5672716 | 304.6137768 | 237.6434499 | 0.098567336 | 0.165225876 | 0.018443046 | 0.313154106 | ns | TM | KS.3 |
| FIM1_5045 | 2133.725081 | 3287.658293 | 2635.336793 | 0.280703417 | 0.251765289 | 0.000996476 | 0.04634941 | ns | TM | KS.3 |
| FIM1_5046 | 1783.665555 | 2015.163926 | 3462.930132 | 0.012976629 | 0.081604765 | 0.545428625 | 0.95917717 | ns | TM | KS.3 |
| FIM1_5047 | 1349.854928 | 1235.165659 | 1684.120007 | -0.026298667 | 0.063736173 | 0.301425679 | 0.926536521 | ns | TM | KS.3 |
| FIM1_5048 | 879.82675 | 891.8251408 | 648.1284846 | 0.003549913 | 0.050478649 | 0.869085272 | 0.991141351 | ns | TM | KS.3 |
| FIM1_5049 | 2526.715649 | 2372.819591 | 2315.334913 | -0.020123757 | 0.054937212 | 0.385989661 | 0.948637087 | ns | TM | KS.3 |
| FIM1_5050 | 1861.960406 | 1747.576724 | 2158.68306 | -0.013050303 | 0.06453643 | 0.580414928 | 0.963781856 | ns | TM | KS.3 |
| FIM1_5051 | 16.07800877 | 16.69011279 | 26.75475964 | 0.002164636 | 0.087525021 | 0.909309237 | 0.991141351 | ns | TM | KS.3 |
| FIM1_5052 | 656.8246663 | 846.1148687 | 833.6461928 | 0.148276812 | 0.124143428 | 0.004850267 | 0.140291152 | ns | TM | KS.3 |
| FIM1_5053 | 1285.987934 | 1324.294422 | 1307.8111 | 0.006705834 | 0.057168174 | 0.767954925 | 0.988477371 | ns | TM | KS.3 |
| FIM1_5054 | 718.6803607 | 709.4955102 | 670.2714311 | -0.002834519 | 0.060531342 | 0.901121695 | 0.991141351 | ns | TM | KS.3 |
| FIM1_5055 | 3289.434234 | 3393.439777 | 3470.755075 | 0.008287729 | 0.052457539 | 0.70716087 | 0.983265883 | ns | TM | KS.3 |
| FIM1_5056 | 1598.615304 | 1412.871403 | 1420.427163 | -0.038545852 | 0.073970776 | 0.1747761 | 0.811391014 | ns | TM | KS.3 |
| FIM1_5057 | 4504.880933 | 5044.764427 | 6219.004623 | 0.0091751 | 0.086062562 | 0.646568246 | 0.976323032 | ns | TM | KS.3 |
| FIM1_5058 | 3191.804237 | 3291.712844 | 2622.191853 | 0.004647286 | 0.069591686 | 0.835713812 | 0.991141351 | ns | TM | KS.3 |
| FIM1_5059 | 669.4587209 | 705.8444656 | 686.9090204 | 0.018934938 | 0.050402034 | 0.386699849 | 0.948637087 | ns | TM | KS.3 |
| FIM1_5061 | 1932.316558 | 1916.146374 | 2153.880053 | -0.001904974 | 0.057553645 | 0.932873755 | 0.994413583 | ns | TM | KS.3 |
| FIM1_5062 | 700.1100208 | 658.7348728 | 633.3694877 | -0.016545058 | 0.057476531 | 0.480533794 | 0.951315411 | ns | TM | KS.3 |
| FIM1_5063 | 2354.802912 | 1920.317003 | 1990.11932 | -0.078001008 | 0.100786751 | 0.037771506 | 0.440708412 | ns | TM | KS.3 |
| FIM1_5064 | 673.0837602 | 744.8973989 | 645.2072038 | 0.014962539 | 0.074879123 | 0.515052825 | 0.954689943 | ns | TM | KS.3 |
| FIM1_5065 | 3454.813101 | 3343.452688 | 3945.297633 | -0.00527477 | 0.06806629 | 0.815695256 | 0.988916276 | ns | TM | KS.3 |
| FIM1_5066 | 412.9316816 | 449.2402278 | 475.1938222 | 0.02540353 | 0.061431883 | 0.309852096 | 0.927416591 | ns | TM | KS.3 |
| FIM1_5067 | 3009.705545 | 3148.286558 | 3488.00408 | 0.0123693 | 0.053853719 | 0.5827525 | 0.963781856 | ns | TM | KS.3 |
| FIM1_5068 | 1093.599713 | 1116.283705 | 1083.197022 | 0.003617944 | 0.065344814 | 0.8737467 | 0.991141351 | ns | TM | KS.3 |
| FIM1_5069 | 190.9822703 | 116.0807044 | 1090.397409 | -0.059290907 | 0.161490547 | 0.05583317 | 0.535202356 | ns | TM | KS.3 |
| FIM1_5070 | 87.58886575 | 90.11744908 | 66.93085334 | 0.005530602 | 0.070484636 | 0.804545865 | 0.988916276 | ns | TM | KS.3 |
| FIM1_5071 | 8372.490185 | 8727.196619 | 7131.434308 | 0.011306471 | 0.053575765 | 0.613850675 | 0.973122675 | ns | TM | KS.3 |
| FIM1_5072 | 165.1769079 | 188.2901694 | 128.7444275 | 0.016739666 | 0.078824852 | 0.461461507 | 0.950410534 | ns | TM | KS.3 |
| FIM1_5073 | 751.9675207 | 727.2003203 | 611.9560529 | -0.006149095 | 0.064971604 | 0.788539883 | 0.988916276 | ns | TM | KS.3 |
| FIM1_5074 | 677.2659858 | 570.1194408 | 663.8664992 | -0.043222578 | 0.084985187 | 0.13796704 | 0.770139466 | ns | TM | KS.3 |
| FIM1_5075 | 1065.447465 | 862.5507308 | 1277.11883 | -0.055452558 | 0.095881399 | 0.080622576 | 0.618977341 | ns | TM | KS.3 |
| FIM1_5076 | 4606.068774 | 4337.232522 | 3651.790514 | -0.016623353 | 0.05739204 | 0.478329221 | 0.950734269 | ns | TM | KS.3 |
| FIM1_5077 | 2222.668468 | 2196.791955 | 2209.852189 | -0.002344078 | 0.061225147 | 0.918182376 | 0.991141351 | ns | TM | KS.3 |
| FIM1_5078 | 424.513307 | 427.52982 | 454.6684508 | 0.001524043 | 0.058210323 | 0.946389652 | 0.994413583 | ns | TM | KS.3 |
| FIM1_5079 | 5831.589343 | 4843.652451 | 6011.899769 | -0.075084677 | 0.095806678 | 0.042498151 | 0.472993959 | ns | TM | KS.3 |
| FIM1_5080 | 1069.008474 | 1077.067185 | 1060.424088 | 0.003042059 | 0.032602576 | 0.842345359 | 0.991141351 | ns | TM | KS.3 |
| FIM1_5081 | 1613.542259 | 1584.635279 | 1610.532415 | -0.00474195 | 0.050438448 | 0.825800122 | 0.991141351 | ns | TM | KS.3 |
| FIM1_5082 | 626.4075332 | 618.6541858 | 557.9162857 | -0.002741696 | 0.056615348 | 0.903179468 | 0.991141351 | ns | TM | KS.3 |
| FIM1_5083 | 5265.418208 | 5499.905622 | 4870.532015 | 0.00577178 | 0.072988757 | 0.792821111 | 0.988916276 | ns | TM | KS.3 |

| | | | | | | | | | | |
|---|---|---|---|---|---|---|---|---|---|---|
| FIM1_5084 | 745.3061158 | 875.3556745 | 802.1607 | 0.016957501 | 0.085071326 | 0.436475777 | 0.950410534 | ns | TM | KS.3 |
| FIM1_5085 | 1685.315254 | 1600.218359 | 1763.991479 | -0.006933643 | 0.07306617 | 0.753400434 | 0.988477371 | ns | TM | KS.3 |
| FIM1_5086 | 2640.985753 | 2674.686084 | 2732.781178 | 0.003077984 | 0.054912933 | 0.890382764 | 0.991141351 | ns | TM | KS.3 |
| FIM1_5087 | 2381.264887 | 2359.204888 | 1469.97806 | -0.001578849 | 0.06625573 | 0.944409242 | 0.994413583 | ns | TM | KS.3 |
| FIM1_5088 | 9566.435949 | 10424.21856 | 9428.848193 | 0.011744803 | 0.074676189 | 0.600742657 | 0.97002321 | ns | TM | KS.3 |
| FIM1_5089 | 2913.189544 | 3063.289946 | 2468.934404 | 0.005847973 | 0.075964327 | 0.785025092 | 0.988880259 | ns | TM | KS.3 |
| FIM1_5090 | 412.8736486 | 360.4412853 | 387.9351323 | -0.013048704 | 0.084295532 | 0.534877496 | 0.959071703 | ns | TM | KS.3 |
| FIM1_5091 | 632.5348511 | 592.6407514 | 492.4874872 | -0.007301353 | 0.077271316 | 0.732266605 | 0.987157164 | ns | TM | KS.3 |
| FIM1_5093 | 1326.214723 | 1426.629952 | 1273.576786 | 0.02556202 | 0.057902769 | 0.293698435 | 0.923185001 | ns | TM | KS.3 |
| FIM1_5094 | 94921.09439 | 93592.13466 | 89508.85285 | -0.003971212 | 0.047146077 | 0.847956562 | 0.991141351 | ns | TM | KS.3 |
| FIM1_5095 | 531.0588666 | 558.9510545 | 503.4258247 | 0.011491413 | 0.060131909 | 0.622005095 | 0.973122675 | ns | TM | KS.3 |
| FIM1_5096 | 1213.43602 | 1244.372299 | 1407.574219 | 0.004456543 | 0.065081289 | 0.845177341 | 0.991141351 | ns | TM | KS.3 |
| FIM1_5097 | 9104.150486 | 8499.352632 | 9492.802646 | -0.030515016 | 0.054269705 | 0.197630786 | 0.838676184 | ns | TM | KS.3 |
| FIM1_5098 | 243.9405108 | 242.2404539 | 297.09687 | -0.000856 | 0.072065904 | 0.968807042 | 0.996327965 | ns | TM | KS.3 |
| FIM1_5099 | 1692.563947 | 1924.469832 | 1756.661765 | 0.041930609 | 0.075580895 | 0.150912576 | 0.783447458 | ns | TM | KS.3 |
| FIM1_5100 | 1020.444685 | 1037.886091 | 1058.239174 | 0.005623095 | 0.032852582 | 0.712507603 | 0.986183429 | ns | TM | KS.3 |
| FIM1_5101 | 1108.783674 | 1035.728699 | 1189.433938 | -0.018687617 | 0.059521133 | 0.435419051 | 0.950410534 | ns | TM | KS.3 |
| FIM1_5102 | 1708.195528 | 1707.091828 | 3568.104286 | -6.88E-05 | 0.075260712 | 0.997421713 | 0.999525119 | ns | TM | KS.3 |
| FIM1_5103 | 622.6639354 | 535.3376345 | 511.8376738 | -0.04438119 | 0.081216549 | 0.135140492 | 0.768370223 | ns | TM | KS.3 |
| FIM1_5104 | 743.6344314 | 681.5436446 | 743.3096505 | -0.034907137 | 0.062567762 | 0.186683388 | 0.827669994 | ns | TM | KS.3 |
| FIM1_5105 | 2568.799932 | 2335.314609 | 2556.219037 | -0.054325647 | 0.067000411 | 0.074107378 | 0.597271765 | ns | TM | KS.3 |
| FIM1_5106 | 812.1460993 | 773.7907864 | 873.3045665 | -0.010918715 | 0.060906064 | 0.639428485 | 0.975246356 | ns | TM | KS.3 |
| FIM1_5107 | 4509.267101 | 4019.698443 | 4318.641348 | -0.028792966 | 0.071429262 | 0.272922762 | 0.908468501 | ns | TM | KS.3 |
| FIM1_5108 | 755.7266388 | 965.8676414 | 919.5715219 | 0.087443932 | 0.113791362 | 0.026521804 | 0.382098641 | ns | TM | KS.3 |
| FIM1_5109 | 930.748955 | 937.2491864 | 1055.894387 | 0.001361634 | 0.061074749 | 0.95237676 | 0.994413583 | ns | TM | KS.3 |
| FIM1_5110 | 1274.784422 | 1301.53613 | 1771.777108 | 0.005360135 | 0.051728061 | 0.806062507 | 0.988916276 | ns | TM | KS.3 |
| FIM1_5111 | 1718.878614 | 1766.544292 | 1833.436691 | 0.00786283 | 0.049277344 | 0.712262673 | 0.986183429 | ns | TM | KS.3 |
| FIM1_5112 | 791.0697848 | 876.5725553 | 912.3996043 | 0.031344527 | 0.067816065 | 0.239757654 | 0.887413624 | ns | TM | KS.3 |
| FIM1_5113 | 138.6421596 | 149.9571623 | 185.8773579 | 0.011951407 | 0.072147522 | 0.601102409 | 0.97002321 | ns | TM | KS.3 |
| FIM1_5114 | 338.9271486 | 455.8022646 | 473.4386546 | 0.06013995 | 0.116044698 | 0.059383279 | 0.543962484 | ns | TM | KS.3 |
| FIM1_5115 | 1614.910853 | 1565.839025 | 1717.039464 | -0.007741994 | 0.054690937 | 0.730136443 | 0.987157164 | ns | TM | KS.3 |
| FIM1_5116 | 1769.497586 | 1502.67025 | 1592.914278 | -0.064341943 | 0.088440399 | 0.062587269 | 0.551491823 | ns | TM | KS.3 |
| FIM1_5117 | 425.0084084 | 589.9199696 | 553.3084487 | 0.05632854 | 0.12123191 | 0.065838627 | 0.564318013 | ns | TM | KS.3 |
| FIM1_5118 | 2879.77689 | 3269.185503 | 2917.278492 | 0.034033647 | 0.074277943 | 0.213880324 | 0.862280884 | ns | TM | KS.3 |
| FIM1_5119 | 1458.809193 | 1569.520026 | 1408.965236 | 0.021731475 | 0.059491451 | 0.370761184 | 0.948637087 | ns | TM | KS.3 |
| FIM1_5120 | 1230.07577 | 1243.161668 | 1218.891875 | 0.001471124 | 0.071056918 | 0.946841205 | 0.994413583 | ns | TM | KS.3 |
| FIM1_5121 | 2840.874779 | 3053.53473 | 2267.146921 | 0.016508042 | 0.063734895 | 0.492362847 | 0.952591289 | ns | TM | KS.3 |
| FIM1_5122 | 3123.143575 | 3308.680225 | 4430.879936 | 0.004768219 | 0.083497216 | 0.810392807 | 0.988916276 | ns | TM | KS.3 |
| FIM1_5123 | 4043.621937 | 5295.698036 | 4067.119462 | 0.023460744 | 0.099936263 | 0.284103848 | 0.918450395 | ns | TM | KS.3 |
| FIM1_5124 | 1092.643541 | 987.7920754 | 969.6212758 | -0.037476743 | 0.067400602 | 0.175732911 | 0.811391014 | ns | TM | KS.3 |
| FIM1_5125 | 2625.132722 | 3145.411095 | 2718.883313 | 0.019255762 | 0.087010571 | 0.383443484 | 0.948637087 | ns | TM | KS.3 |
| FIM1_5126 | 1330.41877 | 1522.374199 | 1546.701462 | 0.113773561 | 0.078306696 | 0.009001856 | 0.205432275 | ns | TM | KS.3 |
| FIM1_5127 | 4389.912569 | 4145.476629 | 4733.280554 | -0.012720619 | 0.062111142 | 0.588852799 | 0.963781856 | ns | TM | KS.3 |
| FIM1_5128 | 3568.420436 | 3409.134902 | 3394.131703 | -0.014399658 | 0.049496272 | 0.503034265 | 0.953260948 | ns | TM | KS.3 |
| FIM1_5129 | 1078.12792 | 1190.082799 | 1408.446859 | 0.025695496 | 0.067422931 | 0.315910737 | 0.927564309 | ns | TM | KS.3 |

| | | | | | | | | | | |
|---|---|---|---|---|---|---|---|---|---|---|
| FIM1_5130 | 2438.280837 | 2673.824418 | 2394.966011 | 0.019451897 | 0.068339824 | 0.426965403 | 0.950410534 | ns | TM | KS.3 |
| FIM1_5131 | 3085.7299 | 2397.042284 | 2682.725242 | -0.016774099 | 0.100444214 | 0.402242621 | 0.950410534 | ns | TM | KS.3 |
| FIM1_5132 | 156.9279632 | 193.8415272 | 193.6028101 | 0.015336662 | 0.094934933 | 0.446754961 | 0.950410534 | ns | TM | KS.3 |
| FIM1_5133 | 1808.763189 | 1552.961166 | 1610.349592 | -0.041632103 | 0.080772226 | 0.151220921 | 0.783447458 | ns | TM | KS.3 |
| FIM1_5134 | 634.2026634 | 697.8779543 | 596.8297995 | 0.01838849 | 0.069981113 | 0.447397704 | 0.950410534 | ns | TM | KS.3 |
| FIM1_5135 | 1424.672544 | 1434.653384 | 1370.994416 | 0.001282223 | 0.060546159 | 0.955123766 | 0.994413583 | ns | TM | KS.3 |
| FIM1_5136 | 3241.091094 | 3051.117049 | 2722.172896 | -0.014611878 | 0.060526083 | 0.536480943 | 0.95917717 | ns | TM | KS.3 |
| FIM1_5137 | 904.4579362 | 1211.50564 | 955.2377765 | 0.09251579 | 0.128879024 | 0.021769556 | 0.33950953 | ns | TM | KS.3 |
| FIM1_5138 | 1991.746156 | 2266.013738 | 2129.186013 | 0.014595972 | 0.081182725 | 0.504529899 | 0.953260948 | ns | TM | KS.3 |
| FIM1_5139 | 4710.874634 | 3914.862782 | 3263.840028 | -0.021176758 | 0.086821792 | 0.349300867 | 0.948637087 | ns | TM | KS.3 |
| FIM1_5140 | 3602.91576 | 2978.810772 | 3280.874518 | -0.020528414 | 0.087960873 | 0.357332674 | 0.948637087 | ns | TM | KS.3 |
| FIM1_5141 | 1936.316218 | 1900.669696 | 2740.679032 | -0.003748884 | 0.06121335 | 0.869731948 | 0.991141351 | ns | TM | KS.3 |
| FIM1_5142 | 1733.44298 | 1690.863664 | 1638.967762 | -0.003304657 | 0.072348202 | 0.880155155 | 0.991141351 | ns | TM | KS.3 |
| FIM1_5143 | 2058.713962 | 2293.500062 | 2447.338046 | 0.054253835 | 0.07173225 | 0.083124882 | 0.626585283 | ns | TM | KS.3 |
| FIM1_5144 | 2162.567917 | 2080.51086 | 1888.397872 | -0.010266456 | 0.053927299 | 0.647101173 | 0.976323032 | ns | TM | KS.3 |
| FIM1_5145 | 3066.412097 | 2899.834018 | 2312.449822 | -0.013880167 | 0.059171811 | 0.55373776 | 0.95917717 | ns | TM | KS.3 |
| FIM1_5146 | 452.4747831 | 572.4610254 | 627.1426176 | 0.034487528 | 0.094150704 | 0.178827915 | 0.811815714 | ns | TM | KS.3 |
| FIM1_5147 | 29735.20007 | 30581.61356 | 29081.46093 | 0.007310815 | 0.052947935 | 0.741070632 | 0.988051177 | ns | TM | KS.3 |
| FIM1_5148 | 447.935345 | 470.8022235 | 298.8760442 | 0.005857979 | 0.075861864 | 0.78486773 | 0.988880259 | ns | TM | KS.3 |
| FIM1_5149 | 5542.341147 | 3766.074719 | 4101.836441 | -0.031714674 | 0.119674289 | 0.173973255 | 0.811391014 | ns | TM | KS.3 |
| FIM1_5150 | 8545.973181 | 8909.768112 | 9268.607682 | 0.006012564 | 0.071032855 | 0.787523283 | 0.988916276 | ns | TM | KS.3 |
| FIM1_5151 | 188.5210577 | 221.6108728 | 223.4158694 | 0.037529504 | 0.081310276 | 0.178030807 | 0.811391014 | ns | TM | KS.3 |
| FIM1_5152 | 389.3479335 | 341.6592927 | 293.9752473 | -0.015208034 | 0.080908251 | 0.490359756 | 0.952591289 | ns | TM | KS.3 |
| FIM1_5153 | 5926.773839 | 5478.396597 | 6107.404169 | -0.0286587 | 0.059388457 | 0.251461106 | 0.888934767 | ns | TM | KS.3 |
| FIM1_5155 | 1049.542339 | 1120.515326 | 1027.34733 | 0.01611107 | 0.061252034 | 0.499530987 | 0.953260948 | ns | TM | KS.3 |
| FIM1_5156 | 772.492883 | 865.2210152 | 907.6906643 | 0.015399156 | 0.077038329 | 0.498147451 | 0.953260948 | ns | TM | KS.3 |
| FIM1_5157 | 2438.609561 | 2423.554375 | 2358.045634 | -0.001802877 | 0.034889782 | 0.912156443 | 0.991141351 | ns | TM | KS.3 |
| FIM1_5158 | 1397.987338 | 1190.146826 | 1384.854333 | -0.02623645 | 0.081350576 | 0.288999748 | 0.919993167 | ns | TM | KS.3 |
| FIM1_5159 | 797.43071 | 925.3215582 | 963.2899021 | 0.050943624 | 0.082208949 | 0.104059682 | 0.693034698 | ns | TM | KS.3 |
| FIM1_5160 | 9340.738212 | 10442.00919 | 11959.88436 | 0.01433966 | 0.077741861 | 0.522092324 | 0.958602567 | ns | TM | KS.3 |
| FIM1_5161 | 1393.481912 | 1558.141622 | 1637.126416 | 0.012021832 | 0.080667576 | 0.574996473 | 0.963192994 | ns | TM | KS.3 |
| FIM1_5162 | 2629.920966 | 2665.484631 | 2683.565014 | 0.003816817 | 0.046755414 | 0.853031169 | 0.991141351 | ns | TM | KS.3 |
| FIM1_5163 | 3719.577758 | 3780.06345 | 3379.838754 | 0.002948835 | 0.064303882 | 0.897080413 | 0.991141351 | ns | TM | KS.3 |
| FIM1_5164 | 202.325337 | 175.3870947 | 390.2049425 | -0.035451446 | 0.077104181 | 0.199232087 | 0.842123734 | ns | TM | KS.3 |
| FIM1_5165 | 1032.881955 | 930.1465126 | 1240.079075 | -0.027031912 | 0.068791997 | 0.296703242 | 0.926536521 | ns | TM | KS.3 |
| FIM1_5166 | 3305.449077 | 3086.465763 | 2961.30963 | -0.009843407 | 0.072551207 | 0.661716446 | 0.976323032 | ns | TM | KS.3 |
| FIM1_5167 | 693.4006654 | 703.6185041 | 637.241487 | 0.003609154 | 0.054002597 | 0.870897494 | 0.991141351 | ns | TM | KS.3 |
| FIM1_5168 | 798.2826457 | 719.1624071 | 781.3317534 | -0.02384973 | 0.069593477 | 0.345423938 | 0.946823191 | ns | TM | KS.3 |
| FIM1_5169 | 286.8197804 | 264.6903256 | 280.748438 | -0.012294109 | 0.072433873 | 0.590929731 | 0.963781856 | ns | TM | KS.3 |
| FIM1_5170 | 2208.543576 | 2008.298042 | 2146.695084 | -0.042762867 | 0.06608751 | 0.130845757 | 0.761283488 | ns | TM | KS.3 |
| FIM1_5171 | 755.7564573 | 780.3200106 | 1014.350572 | 0.006025295 | 0.063784931 | 0.793008232 | 0.988916276 | ns | TM | KS.3 |
| FIM1_5172 | 6799.662375 | 6995.581241 | 7294.21892 | 0.007392529 | 0.053028969 | 0.738486815 | 0.988051177 | ns | TM | KS.3 |
| FIM1_5173 | 1579.3554 | 1448.276495 | 1704.099513 | -0.022138409 | 0.064923693 | 0.374268845 | 0.948637087 | ns | TM | KS.3 |
| FIM1_5174 | 4195.655675 | 4216.939264 | 4111.276212 | 0.001513933 | 0.039769705 | 0.93410257 | 0.994413583 | ns | TM | KS.3 |
| FIM1_5175 | 4174.100978 | 3121.521357 | 3329.456953 | -0.153285463 | 0.142465776 | 0.00461094 | 0.134937815 | ns | TM | KS.3 |

| | | | | | | | | | | |
|---|---|---|---|---|---|---|---|---|---|---|
| FIM1_5176 | 400.8658066 | 459.376031 | 608.5618791 | 0.020938007 | 0.07818572 | 0.379173444 | 0.948637087 | ns | TM | KS.3 |
| FIM1_5177 | 1272.004363 | 1488.595945 | 1527.121631 | 0.011930515 | 0.090033607 | 0.549842022 | 0.95917717 | ns | TM | KS.3 |
| FIM1_5178 | 1107.300654 | 1059.769981 | 1130.801298 | -0.013067998 | 0.051448936 | 0.552547819 | 0.95917717 | ns | TM | KS.3 |
| FIM1_5179 | 3345.846117 | 4021.485482 | 4531.64748 | 0.019132041 | 0.087607826 | 0.384078538 | 0.948637087 | ns | TM | KS.3 |
| FIM1_5180 | 754.1612988 | 707.409913 | 802.0512042 | -0.015744969 | 0.061188067 | 0.508502372 | 0.953260948 | ns | TM | KS.3 |
| FIM1_5181 | 6026.698956 | 5379.704528 | 4942.464288 | -0.033199239 | 0.070985591 | 0.222046492 | 0.870513238 | ns | TM | KS.3 |
| FIM1_5182 | 1618.159687 | 1508.246142 | 1281.7074 | -0.023846119 | 0.056932541 | 0.319753493 | 0.929108338 | ns | TM | KS.3 |
| FIM1_5183 | 5247.266363 | 4852.160459 | 5023.530225 | -0.020959992 | 0.062373228 | 0.39340915 | 0.948637087 | ns | TM | KS.3 |
| FIM1_5184 | 621.6837593 | 700.3600287 | 724.6511252 | 0.031120387 | 0.072369595 | 0.244763239 | 0.88883001 | ns | TM | KS.3 |
| FIM1_5185 | 3417.437205 | 3409.118115 | 3451.371304 | -0.000724974 | 0.047815892 | 0.972304627 | 0.998187272 | ns | TM | KS.3 |
| FIM1_5186 | 1849.194657 | 1954.852546 | 1683.684437 | 0.016302401 | 0.054455477 | 0.475804508 | 0.950410534 | ns | TM | KS.3 |
| FIM1_5187 | 2040.130408 | 1692.749143 | 2096.002034 | -0.038689405 | 0.086662697 | 0.162104805 | 0.796138951 | ns | TM | KS.3 |
| FIM1_5188 | 8801.746959 | 8304.260368 | 8105.092407 | -0.011129218 | 0.065694142 | 0.633655017 | 0.97344368 | ns | TM | KS.3 |
| FIM1_5189 | 290.0835936 | 298.921994 | 255.0617251 | 0.003333647 | 0.075703748 | 0.875639342 | 0.991141351 | ns | TM | KS.3 |
| FIM1_5190 | 1133.737982 | 1213.103547 | 1079.103786 | 0.017131312 | 0.060714489 | 0.47403305 | 0.950410534 | ns | TM | KS.3 |
| FIM1_5191 | 1147.312407 | 1036.288337 | 1573.276048 | -0.020892028 | 0.070323423 | 0.39688875 | 0.949289197 | ns | TM | KS.3 |
| FIM1_5192 | 446.1220424 | 358.2576178 | 1933.053245 | -0.016089558 | 0.095700821 | 0.427284262 | 0.950410534 | ns | TM | KS.3 |
| FIM1_5193 | 15306.60492 | 16583.86842 | 15741.76877 | 0.026388624 | 0.060655322 | 0.291211789 | 0.919993167 | ns | TM | KS.3 |
| FIM1_5194 | 3440.155267 | 3375.202991 | 3900.873431 | -0.003582196 | 0.063571475 | 0.875397003 | 0.991141351 | ns | TM | KS.3 |
| FIM1_5195 | 4171.283292 | 3956.735138 | 5536.513644 | -0.013559748 | 0.05750094 | 0.559068275 | 0.960415976 | ns | TM | KS.3 |
| FIM1_5196 | 312.1166636 | 281.0062826 | 274.2174018 | -0.026043604 | 0.068467187 | 0.311075444 | 0.927416591 | ns | TM | KS.3 |
| FIM1_5197 | 353.2165819 | 368.9328687 | 384.7279617 | 0.011677801 | 0.056322104 | 0.610173244 | 0.971747391 | ns | TM | KS.3 |
| FIM1_5199 | 2874.716943 | 2762.13561 | 3164.931706 | -0.013134634 | 0.046303639 | 0.521033804 | 0.958602567 | ns | TM | KS.3 |
| FIM1_5200 | 154.8148819 | 165.8275701 | 329.1609456 | 0.005828718 | 0.083498815 | 0.770508106 | 0.988814136 | ns | TM | KS.3 |
| FIM1_5201 | 284.6243571 | 372.239259 | 502.8709471 | 0.048415401 | 0.104261181 | 0.095669825 | 0.673713046 | ns | TM | KS.3 |
| FIM1_5202 | 622.7915368 | 648.6082458 | 778.6488792 | 0.011205786 | 0.052532976 | 0.613494967 | 0.973122675 | ns | TM | KS.3 |
| FIM1_5203 | 671.9973177 | 673.408327 | 745.3949667 | 0.000324503 | 0.060636447 | 0.988634945 | 0.999333464 | ns | TM | KS.3 |
| FIM1_5204 | 232.2082243 | 473.5174969 | 477.5816684 | 0.941236037 | 0.222078924 | 9.77845980153 | 0.000156929 | ns | TM | KS.3 |
| FIM1_5205 | 1552.456457 | 1481.518888 | 1564.827119 | -0.015623758 | 0.048637156 | 0.462199003 | 0.950410534 | ns | TM | KS.3 |
| FIM1_5206 | 885.1708807 | 890.9283112 | 958.1753731 | 0.001703572 | 0.0486069 | 0.935605695 | 0.994413583 | ns | TM | KS.3 |
| FIM1_5207 | 230.419428 | 282.1828461 | 269.4435916 | 0.044396909 | 0.090595983 | 0.125315993 | 0.743970245 | ns | TM | KS.3 |
| FIM1_5208 | 21.02323289 | 14.15386893 | 21.5559521 | -0.01696733 | 0.119954967 | 0.369226924 | 0.948637087 | ns | TM | KS.3 |
| FIM1_5209 | 26.88482363 | 36.95135788 | 37.8397702 | 0.022734194 | 0.1086281 | 0.281964911 | 0.917446325 | ns | TM | KS.3 |
| FIM1_4316 | 4090.394936 | 4159.894537 | 4233.280858 | 0.002625983 | 0.068587796 | 0.906832912 | 0.991141351 | ns | TM | KS.3 |
| FIM1_4318 | 231.4348603 | 192.1464987 | 297.7479064 | -0.042544832 | 0.08713467 | 0.138928694 | 0.773986848 | ns | TM | KS.3 |
| FIM1_4319 | 871.3116552 | 577.3793051 | 791.3821641 | -0.0571925 | 0.141137153 | 0.060826851 | 0.544665909 | ns | TM | KS.3 |
| FIM1_4320 | 3794.160135 | 3386.012251 | 3853.218499 | -0.040190987 | 0.071617011 | 0.160416674 | 0.792037169 | ns | TM | KS.3 |
| FIM1_4321 | 2411.541464 | 2026.554885 | 1574.930606 | -0.069777729 | 0.091999543 | 0.051377282 | 0.515921463 | ns | TM | KS.3 |
| FIM1_4322 | 1263.82801 | 1314.526034 | 1448.110775 | 0.009643332 | 0.057094671 | 0.673396593 | 0.976861203 | ns | TM | KS.3 |
| FIM1_4323 | 14641.94869 | 13819.691 | 16724.97566 | -0.02415331 | 0.048891937 | 0.259667741 | 0.895871714 | ns | TM | KS.3 |
| FIM1_4324 | 40.54174812 | 34.97264631 | 38.88673509 | -0.013444185 | 0.08640572 | 0.517692802 | 0.957087213 | ns | TM | KS.3 |
| FIM1_4325 | 2209.834745 | 1913.629266 | 1983.336966 | -0.067735328 | 0.084141678 | 0.054981191 | 0.531428718 | ns | TM | KS.3 |
| FIM1_4326 | 1511.041658 | 1349.735633 | 1289.899005 | -0.024481731 | 0.071802075 | 0.333531114 | 0.938982061 | ns | TM | KS.3 |
| FIM1_4327 | 1264.014525 | 1810.747069 | 2328.645034 | 0.304280847 | 0.187462797 | 0.000259693 | 0.017944075 | ns | TM | KS.3 |
| FIM1_4328 | 763.7967874 | 816.3302033 | 872.2491588 | 0.010859636 | 0.06994731 | 0.636557636 | 0.974658588 | ns | TM | KS.3 |

| | | | | | | | | | | |
|---|---|---|---|---|---|---|---|---|---|---|
| FIM1_4329 | 1159.086544 | 902.9060543 | 704.8157214 | -0.084423651 | 0.113877265 | 0.028899523 | 0.390786333 | ns | TM | KS.3 |
| FIM1_4330 | 2028.982662 | 1917.017127 | 2374.448184 | -0.013419572 | 0.060402295 | 0.56809923 | 0.962265923 | ns | TM | KS.3 |
| FIM1_4331 | 3351.525237 | 2286.512399 | 5501.843841 | -0.13637472 | 0.182208877 | 0.007968624 | 0.191538592 | ns | TM | KS.3 |
| FIM1_4332 | 1646.861702 | 1592.557654 | 1772.143234 | -0.011465293 | 0.04222113 | 0.546008131 | 0.95917717 | ns | TM | KS.3 |
| FIM1_4333 | 3462.054857 | 2859.759966 | 2923.975554 | -0.1526964 | 0.094403024 | 0.003193593 | 0.103770022 | ns | TM | KS.3 |
| FIM1_4334 | 974.9933873 | 911.4286162 | 898.2957246 | -0.013478089 | 0.065939836 | 0.568493306 | 0.962265923 | ns | TM | KS.3 |
| FIM1_4335 | 4205.330895 | 3035.06708 | 2760.392648 | -0.437951652 | 0.083272762 | 3.2945557351[illegible] | 1.8211571980[illegible] | ns | TM | KS.3 |
| FIM1_4336 | 837.8915403 | 1072.407087 | 963.1192203 | 0.087097182 | 0.113656945 | 0.026803431 | 0.382098641 | ns | TM | KS.3 |
| FIM1_4337 | 1564.554678 | 1544.670914 | 1516.12613 | -0.003417795 | 0.051225955 | 0.874827931 | 0.991141351 | ns | TM | KS.3 |
| FIM1_4338 | 4624.811063 | 4175.319508 | 4371.463693 | -0.075379906 | 0.069508293 | 0.030691029 | 0.402578636 | ns | TM | KS.3 |
| FIM1_4339 | 695.7291836 | 642.538488 | 495.8349819 | -0.016196395 | 0.066895656 | 0.499669961 | 0.953260948 | ns | TM | KS.3 |
| FIM1_4340 | 17071.86209 | 16633.15458 | 10345.93131 | -0.007106792 | 0.05052916 | 0.742256266 | 0.988477371 | ns | TM | KS.3 |
| FIM1_4341 | 467.3420605 | 445.6538677 | 351.1808717 | -0.00564609 | 0.075839506 | 0.792323263 | 0.988916276 | ns | TM | KS.3 |
| FIM1_4342 | 766.7050318 | 731.6893577 | 597.9498622 | -0.011602196 | 0.05761491 | 0.615099248 | 0.973122675 | ns | TM | KS.3 |
| FIM1_4343 | 26107.28218 | 26301.28176 | 25916.70843 | 0.002030776 | 0.048572356 | 0.923246375 | 0.992531983 | ns | TM | KS.3 |
| FIM1_4344 | 1831.182025 | 2318.281574 | 2485.044825 | 0.034565832 | 0.094513089 | 0.177667016 | 0.811391014 | ns | TM | KS.3 |
| FIM1_4345 | 750.2066385 | 737.5990879 | 544.9502246 | -0.003596882 | 0.061877783 | 0.874987583 | 0.991141351 | ns | TM | KS.3 |
| FIM1_4346 | 1010.16157 | 1408.349053 | 1382.390916 | 0.026254613 | 0.10968509 | 0.234967168 | 0.883800031 | ns | TM | KS.3 |
| FIM1_4347 | 1943.590361 | 2111.802501 | 1730.147006 | 0.024944994 | 0.062296107 | 0.319948725 | 0.929108338 | ns | TM | KS.3 |
| FIM1_4348 | 2346.843766 | 2229.62483 | 2477.390735 | -0.018143724 | 0.04721185 | 0.382020757 | 0.948637087 | ns | TM | KS.3 |
| FIM1_4349 | 107.4557714 | 71.68050867 | 484.6426657 | -0.043026811 | 0.129129436 | 0.104434981 | 0.69380519 | ns | TM | KS.3 |
| FIM1_4350 | 2172.593789 | 1892.193211 | 2577.991275 | -0.124641963 | 0.075441478 | 0.005517862 | 0.152507586 | ns | TM | KS.3 |
| FIM1_4351 | 4920.553614 | 4421.28469 | 4666.112367 | -0.032502508 | 0.069126691 | 0.228088336 | 0.877996572 | ns | TM | KS.3 |
| FIM1_4352 | 1211.212903 | 1278.760634 | 1263.868516 | 0.008253294 | 0.070450448 | 0.715024634 | 0.986942381 | ns | TM | KS.3 |
| FIM1_4353 | 2065.886739 | 2045.029059 | 2362.725098 | -0.002363528 | 0.055699103 | 0.916062227 | 0.991141351 | ns | TM | KS.3 |
| FIM1_4354 | 2266.731418 | 2190.399669 | 1627.87739 | -0.007189898 | 0.061217956 | 0.754901513 | 0.988477371 | ns | TM | KS.3 |
| FIM1_4355 | 27.71287464 | 26.27926877 | 26.18476884 | -0.003009979 | 0.086141392 | 0.875899694 | 0.991141351 | ns | TM | KS.3 |
| FIM1_4356 | 27013.96755 | 26093.72442 | 23485.72274 | -0.012460807 | 0.038874681 | 0.476511255 | 0.950410534 | ns | TM | KS.3 |
| FIM1_4357 | 3137.989804 | 2396.993286 | 1923.814461 | -0.153206145 | 0.131960767 | 0.004401767 | 0.130349945 | ns | TM | KS.3 |
| FIM1_4358 | 5300.813112 | 4953.282715 | 4355.850489 | -0.017378791 | 0.060769126 | 0.468290346 | 0.950410534 | ns | TM | KS.3 |
| FIM1_4359 | 2565.495548 | 2315.373813 | 2335.321236 | -0.028673444 | 0.068164195 | 0.273699501 | 0.908468501 | ns | TM | KS.3 |
| FIM1_4360 | 8678.458779 | 7251.301195 | 8671.074157 | -0.126557444 | 0.095269065 | 0.007859248 | 0.191538592 | ns | TM | KS.3 |
| FIM1_4361 | 2042.354407 | 2352.843437 | 2926.548286 | 0.023319466 | 0.078337931 | 0.338786452 | 0.942652461 | ns | TM | KS.3 |
| FIM1_4363 | 626.4168591 | 822.5232186 | 979.5832279 | 0.045482944 | 0.104521155 | 0.106327517 | 0.700394017 | ns | TM | KS.3 |
| FIM1_4364 | 540.0112905 | 507.8254614 | 393.1670421 | -0.010331518 | 0.068881236 | 0.653702789 | 0.976323032 | ns | TM | KS.3 |
| FIM1_4365 | 1307.474606 | 1268.722678 | 1130.932375 | -0.007173558 | 0.057039678 | 0.752368873 | 0.988477371 | ns | TM | KS.3 |
| FIM1_4366 | 9507.960245 | 9222.885707 | 9010.826394 | -0.009858751 | 0.042994245 | 0.610198764 | 0.971747391 | ns | TM | KS.3 |
| FIM1_4367 | 1769.514893 | 1633.50814 | 1772.816264 | -0.019918138 | 0.063719794 | 0.416567141 | 0.950410534 | ns | TM | KS.3 |
| FIM1_4368 | 1748.266236 | 1619.935903 | 1621.621862 | -0.021114701 | 0.061415146 | 0.388350764 | 0.948637087 | ns | TM | KS.3 |
| FIM1_4369 | 2451.435089 | 2632.26053 | 3186.351165 | 0.007255338 | 0.079493656 | 0.728560703 | 0.987157164 | ns | TM | KS.3 |
| FIM1_4370 | 727.3648305 | 793.0098374 | 886.3623511 | 0.014906211 | 0.070568715 | 0.526419097 | 0.959071703 | ns | TM | KS.3 |
| FIM1_4371 | 3187.888082 | 3367.313466 | 3441.943324 | 0.010823274 | 0.064558763 | 0.643029843 | 0.976237372 | ns | TM | KS.3 |
| FIM1_4372 | 2434.360417 | 2618.852747 | 2499.143024 | 0.0133599 | 0.068216217 | 0.569626209 | 0.962265923 | ns | TM | KS.3 |
| FIM1_4373 | 6108.692763 | 6690.120154 | 8223.638193 | 0.013247363 | 0.073978722 | 0.561324436 | 0.961073589 | ns | TM | KS.3 |
| FIM1_4374 | 1032.502461 | 1186.765345 | 1187.237333 | 0.067654518 | 0.083179678 | 0.054828374 | 0.531428718 | ns | TM | KS.3 |

| | | | | | | | | | | |
|---|---|---|---|---|---|---|---|---|---|---|
| FIM1_4376 | 276.9575069 | 244.9661578 | 275.9932316 | -0.023989057 | 0.074498336 | 0.337146235 | 0.940821016 | ns | TM | KS.3 |
| FIM1_4377 | 4523.482956 | 4769.156897 | 5376.730464 | 0.007694515 | 0.071369908 | 0.731409688 | 0.987157164 | ns | TM | KS.3 |
| FIM1_4378 | 267.5306568 | 241.1895461 | 200.5203965 | -0.020106033 | 0.071446412 | 0.410360392 | 0.950410534 | ns | TM | KS.3 |
| FIM1_4379 | 4945.592111 | 4766.925094 | 3467.010987 | -0.010248943 | 0.051504242 | 0.640097817 | 0.975673434 | ns | TM | KS.3 |
| FIM1_4380 | 1483.032995 | 1314.237322 | 1529.100328 | -0.065202253 | 0.077170197 | 0.056557534 | 0.535599847 | ns | TM | KS.3 |
| FIM1_4381 | 1474.201814 | 1431.308912 | 1501.693725 | -0.007831515 | 0.053249404 | 0.724249939 | 0.987157164 | ns | TM | KS.3 |
| FIM1_4382 | 976.5020774 | 928.470652 | 961.0533677 | -0.013137018 | 0.057577729 | 0.571003222 | 0.962265923 | ns | TM | KS.3 |
| FIM1_4383 | 5628.241638 | 5588.645505 | 5562.631831 | -0.002262835 | 0.032440596 | 0.882037636 | 0.991141351 | ns | TM | KS.3 |
| FIM1_4384 | 1085.008634 | 1063.088969 | 1504.303601 | -0.002693922 | 0.072306831 | 0.902089786 | 0.991141351 | ns | TM | KS.3 |
| FIM1_4385 | 1654.476335 | 1661.261763 | 1672.830195 | 0.001131001 | 0.048937991 | 0.957382925 | 0.994413583 | ns | TM | KS.3 |
| FIM1_4386 | 17294.02624 | 17676.27008 | 15426.11709 | 0.005943731 | 0.050211593 | 0.782333845 | 0.988814136 | ns | TM | KS.3 |
| FIM1_4387 | 29.15865484 | 29.53491531 | 32.08724951 | 0.00180754 | 0.082141878 | 0.927777602 | 0.993697171 | ns | TM | KS.3 |
| FIM1_4388 | 943.2880989 | 1393.998107 | 2828.08616 | 0.066335995 | 0.143070089 | 0.045107751 | 0.487132212 | ns | TM | KS.3 |
| FIM1_4389 | 309.8503319 | 231.2716902 | 261.3620366 | -0.082343015 | 0.125636563 | 0.029128433 | 0.392720739 | ns | TM | KS.3 |
| FIM1_4390 | 131.5717816 | 130.0342283 | 119.0014353 | -0.001417706 | 0.068703843 | 0.949516417 | 0.994413583 | ns | TM | KS.3 |
| FIM1_4391 | 10084.33561 | 9572.810596 | 11072.76012 | -0.008608958 | 0.068478663 | 0.706639246 | 0.983265883 | ns | TM | KS.3 |
| FIM1_4392 | 1731.349086 | 2062.324571 | 2446.787469 | 0.033714134 | 0.083775403 | 0.204048838 | 0.847612283 | ns | TM | KS.3 |
| FIM1_4393 | 98.80561382 | 111.0903695 | 102.1546705 | 0.012366738 | 0.079644304 | 0.568519934 | 0.962265923 | ns | TM | KS.3 |
| FIM1_4394 | 476.0433731 | 589.1341696 | 574.9786187 | 0.095844555 | 0.106208932 | 0.021497248 | 0.338445603 | ns | TM | KS.3 |
| FIM1_4396 | 4382.281575 | 3935.652962 | 3371.773329 | -0.025477908 | 0.07011073 | 0.319492663 | 0.929108338 | ns | TM | KS.3 |
| FIM1_4397 | 1251.009422 | 1311.768486 | 1309.553351 | 0.013043646 | 0.054441746 | 0.565005168 | 0.962265923 | ns | TM | KS.3 |
| FIM1_4398 | 69.01188457 | 91.00967203 | 199.3745941 | 0.024281496 | 0.101265785 | 0.270588886 | 0.908468501 | ns | TM | KS.3 |
| FIM1_4399 | 13941.54965 | 13281.242 | 14052.33915 | -0.007259861 | 0.070521245 | 0.74687672 | 0.988477371 | ns | TM | KS.3 |
| FIM1_4400 | 1104.589495 | 1172.581913 | 1072.102269 | 0.019293621 | 0.054005304 | 0.400126888 | 0.950410534 | ns | TM | KS.3 |
| FIM1_4401 | 2264.999192 | 2468.740343 | 2429.574297 | 0.025211422 | 0.063370008 | 0.318057704 | 0.928601572 | ns | TM | KS.3 |
| FIM1_4402 | 94.00006379 | 154.476002 | 172.6698895 | 0.411043052 | 0.304918505 | 0.00036727 | 0.022282532 | ns | TM | KS.3 |
| FIM1_4403 | 4161.759038 | 3839.723963 | 5429.59483 | -0.007710482 | 0.081139037 | 0.708869001 | 0.983990871 | ns | TM | KS.3 |
| FIM1_4404 | 459.6577237 | 460.9029346 | 460.0954742 | 0.000630219 | 0.061858546 | 0.977949668 | 0.999036302 | ns | TM | KS.3 |
| FIM1_4405 | 675.2211044 | 611.8988696 | 609.2161831 | -0.020398058 | 0.069903314 | 0.406780233 | 0.950410534 | ns | TM | KS.3 |
| FIM1_4406 | 1250.844123 | 1034.297268 | 1069.145318 | -0.0845882 | 0.098452973 | 0.031089914 | 0.403710444 | ns | TM | KS.3 |
| FIM1_4407 | 59.52463299 | 42.53300446 | 122.619116 | -0.040704906 | 0.114779025 | 0.120575475 | 0.730509595 | ns | TM | KS.3 |
| FIM1_4408 | 533.4871118 | 490.4319694 | 448.4685894 | -0.007857484 | 0.081806913 | 0.702139238 | 0.98151979 | ns | TM | KS.3 |
| FIM1_4409 | 3648.050433 | 3461.588374 | 6251.210809 | -0.004863116 | 0.080802158 | 0.811533709 | 0.988916276 | ns | TM | KS.3 |
| FIM1_4410 | 932.6490868 | 886.0433892 | 823.5708034 | -0.012526895 | 0.058836173 | 0.590593277 | 0.963781856 | ns | TM | KS.3 |
| FIM1_4412 | 150.0666639 | 154.8925167 | 240.9380461 | 0.00286263 | 0.081404888 | 0.886931587 | 0.991141351 | ns | TM | KS.3 |
| FIM1_4413 | 1428.500649 | 1531.228984 | 1434.163885 | 0.025079964 | 0.055743115 | 0.291129194 | 0.919993167 | ns | TM | KS.3 |
| FIM1_4414 | 2619.507288 | 3161.642228 | 3221.545165 | 0.080500396 | 0.097503714 | 0.035493353 | 0.42652037 | ns | TM | KS.3 |
| FIM1_4415 | 3091.428672 | 2982.385391 | 3335.323888 | -0.003538083 | 0.079129521 | 0.863807535 | 0.991141351 | ns | TM | KS.3 |
| FIM1_4416 | 2063.112189 | 1781.613802 | 1896.95476 | -0.024518588 | 0.079007184 | 0.318845148 | 0.929077912 | ns | TM | KS.3 |
| FIM1_4417 | 1279.384365 | 1169.735582 | 1298.851095 | -0.033746148 | 0.063193675 | 0.20131993 | 0.84606701 | ns | TM | KS.3 |
| FIM1_4418 | 5316.646221 | 5687.739747 | 5851.324223 | 0.023610187 | 0.055397178 | 0.316318711 | 0.927564309 | ns | TM | KS.3 |
| FIM1_4419 | 1174.831345 | 1246.988496 | 1039.942133 | 0.017050751 | 0.056912707 | 0.46622633 | 0.950410534 | ns | TM | KS.3 |
| FIM1_4420 | 1158.510447 | 1192.751107 | 1050.982712 | 0.008196101 | 0.049869893 | 0.70271154 | 0.98151979 | ns | TM | KS.3 |
| FIM1_4421 | 795.5761379 | 855.2579435 | 841.2736502 | 0.024538258 | 0.057656316 | 0.310489118 | 0.927416591 | ns | TM | KS.3 |
| FIM1_4422 | 15.28430961 | 11.72921026 | 14.57156587 | -0.013204204 | 0.107954023 | 0.477950535 | 0.950410534 | ns | TM | KS.3 |

| | | | | | | | | | | |
|---|---|---|---|---|---|---|---|---|---|---|
| FIM1_4423 | 1818.111625 | 1852.613906 | 1754.733821 | 0.002489949 | 0.072321775 | 0.909427014 | 0.991141351 | ns | TM | KS.3 |
| FIM1_4424 | 904.5726067 | 890.400272 | 1053.413028 | -0.000961291 | 0.090080433 | 0.958741359 | 0.994413583 | ns | TM | KS.3 |
| FIM1_4425 | 588.4569214 | 603.9059537 | 643.1878699 | 0.007014065 | 0.041703998 | 0.711038262 | 0.98480383 | ns | TM | KS.3 |
| FIM1_4426 | 705.6944506 | 650.9294403 | 728.6083259 | -0.021501089 | 0.063300226 | 0.38431146 | 0.948637087 | ns | TM | KS.3 |
| FIM1_4427 | 5662.110732 | 5094.185463 | 5440.023767 | -0.035615447 | 0.068891792 | 0.195732882 | 0.837834055 | ns | TM | KS.3 |
| FIM1_4428 | 6438.086752 | 6817.049015 | 6919.829657 | 0.007444324 | 0.073943653 | 0.734778236 | 0.987655486 | ns | TM | KS.3 |
| FIM1_4429 | 1924.980872 | 1828.466287 | 2044.53562 | -0.010057146 | 0.064451944 | 0.665678074 | 0.976323032 | ns | TM | KS.3 |
| FIM1_4430 | 288.9177211 | 379.0162803 | 343.7891576 | 0.084331065 | 0.119730261 | 0.028169335 | 0.390786333 | ns | TM | KS.3 |
| FIM1_4431 | 1407.560718 | 1639.233106 | 2195.196975 | 0.081386004 | 0.087913615 | 0.034209311 | 0.42621823 | ns | TM | KS.3 |
| FIM1_4432 | 5662.922897 | 5508.276092 | 5969.247125 | -0.008625287 | 0.043997077 | 0.661890665 | 0.976323032 | ns | TM | KS.3 |
| FIM1_4433 | 558.9612081 | 659.3397923 | 622.5379711 | 0.037739525 | 0.082625513 | 0.174694083 | 0.811391014 | ns | TM | KS.3 |
| FIM1_4434 | 2281.168285 | 1966.056619 | 1987.486864 | -0.162248197 | 0.062203719 | 0.000554134 | 0.030975465 | ns | TM | KS.3 |
| FIM1_4435 | 4312.647196 | 3722.800891 | 4237.227993 | -0.067221023 | 0.084955767 | 0.056256494 | 0.535599847 | ns | TM | KS.3 |
| FIM1_4436 | 1678.23828 | 1609.820308 | 1632.471748 | -0.014757316 | 0.043164784 | 0.443069533 | 0.950410534 | ns | TM | KS.3 |
| FIM1_4437 | 808.9451354 | 690.6099927 | 636.5513952 | -0.045633995 | 0.082761629 | 0.12765234 | 0.75107959 | ns | TM | KS.3 |
| FIM1_4438 | 1362.308315 | 1170.757585 | 1274.270211 | -0.037130365 | 0.079829761 | 0.182854705 | 0.820765281 | ns | TM | KS.3 |
| FIM1_4439 | 1733.744744 | 1550.296132 | 1607.302738 | -0.04001087 | 0.071103093 | 0.161158169 | 0.793823652 | ns | TM | KS.3 |
| FIM1_4440 | 2166.529133 | 1547.623279 | 1509.242747 | -0.076026076 | 0.134491996 | 0.034108583 | 0.42621823 | ns | TM | KS.3 |
| FIM1_4441 | 3515.9126 | 3546.198129 | 3396.226368 | 0.000928332 | 0.076699189 | 0.964768398 | 0.995175461 | ns | TM | KS.3 |
| FIM1_4443 | 407.94573 | 297.9697211 | 510.6500014 | -0.025595796 | 0.106869218 | 0.246011076 | 0.888934767 | ns | TM | KS.3 |
| FIM1_4444 | 31485.63271 | 29523.33843 | 29229.62885 | -0.014901312 | 0.062205891 | 0.530932262 | 0.959071703 | ns | TM | KS.3 |
| FIM1_4445 | 1326.794211 | 1381.442406 | 1353.071427 | 0.011374586 | 0.051908739 | 0.605991655 | 0.970937835 | ns | TM | KS.3 |
| FIM1_4446 | 6589.22152 | 5456.084112 | 5081.720682 | -0.089618813 | 0.098943401 | 0.026503323 | 0.382098641 | ns | TM | KS.3 |
| FIM1_4447 | 1846.89402 | 1512.897947 | 1698.587544 | -0.196784474 | 0.079936315 | 0.000343046 | 0.021880147 | ns | TM | KS.3 |
| FIM1_4448 | 197.2251791 | 233.4904067 | 260.2087931 | 0.017410264 | 0.086233909 | 0.423075163 | 0.950410534 | ns | TM | KS.3 |
| FIM1_4449 | 715.0339467 | 585.5122286 | 733.6709146 | -0.023459738 | 0.088539398 | 0.308381575 | 0.927416591 | ns | TM | KS.3 |
| FIM1_4450 | 4548.081694 | 4177.447538 | 3259.486394 | -0.021425582 | 0.064717707 | 0.387535125 | 0.948637087 | ns | TM | KS.3 |
| FIM1_4451 | 3068.095873 | 2705.565489 | 2267.005217 | -0.024518085 | 0.0747266 | 0.328570351 | 0.935147309 | ns | TM | KS.3 |
| FIM1_4452 | 9947.876025 | 10837.24994 | 10157.37843 | 0.018952137 | 0.066640146 | 0.43806428 | 0.950410534 | ns | TM | KS.3 |
| FIM1_4454 | 3375.400887 | 3423.073304 | 4077.397871 | 0.004373024 | 0.040619792 | 0.814012145 | 0.988916276 | ns | TM | KS.3 |
| FIM1_4455 | 1281.791252 | 1406.253793 | 1386.220544 | 0.018097934 | 0.069549949 | 0.454172448 | 0.950410534 | ns | TM | KS.3 |
| FIM1_4456 | 1518.386908 | 1588.983535 | 1521.687779 | 0.008711245 | 0.064376991 | 0.706697023 | 0.983265883 | ns | TM | KS.3 |
| FIM1_4457 | 1990.833739 | 2086.12793 | 1899.474495 | 0.013607917 | 0.052503047 | 0.541345585 | 0.95917717 | ns | TM | KS.3 |
| FIM1_4458 | 2009.153138 | 1714.086256 | 1523.401855 | -0.082515867 | 0.089901342 | 0.033227265 | 0.420625047 | ns | TM | KS.3 |
| FIM1_4460 | 603.5809849 | 676.736816 | 482.6566913 | 0.019572683 | 0.074370954 | 0.414885115 | 0.950410534 | ns | TM | KS.3 |
| FIM1_4461 | 294.5737825 | 310.3024668 | 235.8112276 | 0.007900227 | 0.069800644 | 0.727253183 | 0.987157164 | ns | TM | KS.3 |
| FIM1_4462 | 1115.724388 | 1052.022228 | 1086.853569 | -0.017438791 | 0.055274195 | 0.450525908 | 0.950410534 | ns | TM | KS.3 |
| FIM1_4463 | 712.3367872 | 839.2345088 | 1342.582568 | 0.054963371 | 0.086645961 | 0.088103058 | 0.64839159 | ns | TM | KS.3 |
| FIM1_4464 | 890.927308 | 933.401108 | 1042.604224 | 0.010211245 | 0.060880638 | 0.660431209 | 0.976323032 | ns | TM | KS.3 |
| FIM1_4465 | 1623.605444 | 1483.617062 | 1372.504425 | -0.02533069 | 0.06493574 | 0.319008159 | 0.929077912 | ns | TM | KS.3 |
| FIM1_4466 | 1450.449716 | 1318.611089 | 1228.347793 | -0.033703525 | 0.065432103 | 0.208325889 | 0.852551729 | ns | TM | KS.3 |
| FIM1_4467 | 2740.205651 | 2744.147576 | 2880.810584 | 0.000248132 | 0.036901741 | 0.988488041 | 0.999333464 | ns | TM | KS.3 |
| FIM1_4468 | 290.7294181 | 300.6880265 | 290.415238 | 0.006565996 | 0.061565149 | 0.775284645 | 0.988814136 | ns | TM | KS.3 |
| FIM1_4469 | 2574.435466 | 3007.493197 | 3092.130005 | 0.015801778 | 0.085053496 | 0.462756371 | 0.950410534 | ns | TM | KS.3 |
| FIM1_4470 | 2600.893264 | 2883.858567 | 2972.149374 | 0.013206698 | 0.077156803 | 0.553363239 | 0.95917717 | ns | TM | KS.3 |

| | | | | | | | | | | |
|---|---|---|---|---|---|---|---|---|---|---|
| FIM1_4471 | 1003.63408 | 927.7786813 | 801.3701326 | -0.018897977 | 0.06408101 | 0.438433779 | 0.950410534 | ns | TM | KS.3 |
| FIM1_4472 | 2503.51436 | 2319.78304 | 2324.054921 | -0.042171989 | 0.05787101 | 0.10606045 | 0.700394017 | ns | TM | KS.3 |
| FIM1_4473 | 772.3519088 | 720.6474556 | 825.0001088 | -0.016040881 | 0.063062679 | 0.503181294 | 0.953260948 | ns | TM | KS.3 |
| FIM1_4474 | 9357.292212 | 12036.5374 | 10933.34462 | 0.087038373 | 0.115228635 | 0.026640186 | 0.382098641 | ns | TM | KS.3 |
| FIM1_4475 | 1288.776061 | 1262.354379 | 812.4556507 | -0.003724682 | 0.0647508 | 0.870263277 | 0.991141351 | ns | TM | KS.3 |
| FIM1_4476 | 1571.55507 | 1727.545325 | 1386.57011 | 0.016634598 | 0.071317971 | 0.483859256 | 0.952269291 | ns | TM | KS.3 |
| FIM1_4477 | 2519.793405 | 2415.443879 | 2194.233526 | -0.013150545 | 0.049003458 | 0.537607881 | 0.95917717 | ns | TM | KS.3 |
| FIM1_4478 | 1983.416072 | 2011.814084 | 1911.228193 | 0.003217664 | 0.056453981 | 0.886437116 | 0.991141351 | ns | TM | KS.3 |
| FIM1_4479 | 2203.739687 | 2060.349374 | 2132.936426 | -0.009441168 | 0.072960112 | 0.673129645 | 0.976861203 | ns | TM | KS.3 |
| FIM1_4480 | 477.2566541 | 490.189724 | 436.876264 | 0.005174121 | 0.064075046 | 0.821269926 | 0.990886949 | ns | TM | KS.3 |
| FIM1_4481 | 1952.325625 | 2098.73571 | 1967.364171 | 0.030051079 | 0.056532215 | 0.217459155 | 0.865538807 | ns | TM | KS.3 |
| FIM1_4482 | 70.41559246 | 39.31636029 | 2864.161851 | -0.083247195 | 0.204721451 | 0.028906406 | 0.390786333 | ns | TM | KS.3 |
| FIM1_4484 | 14.79488284 | 19.69177248 | 72.63694843 | 0.010431315 | 0.112953869 | 0.554730426 | 0.95917717 | ns | TM | KS.3 |
| FIM1_4486 | 1242.771029 | 1133.634032 | 1769.97335 | -0.008095677 | 0.083335315 | 0.690066023 | 0.980879562 | ns | TM | KS.3 |
| FIM1_4487 | 211.4477733 | 193.0911782 | 435.3429884 | -0.008102822 | 0.083273028 | 0.689982372 | 0.980879562 | ns | TM | KS.3 |
| FIM1_4488 | 49.36009321 | 31.66566343 | 33.08812412 | -0.052685185 | 0.145635865 | 0.070617911 | 0.582627046 | ns | TM | KS.3 |
| FIM1_4489 | 1192.022545 | 1230.225999 | 1176.216034 | 0.003174745 | 0.078590738 | 0.878187624 | 0.991141351 | ns | TM | KS.3 |
| FIM1_4490 | 5538.201645 | 7728.039286 | 7550.038692 | 0.060669841 | 0.124937522 | 0.056190793 | 0.535599847 | ns | TM | KS.3 |
| FIM1_4491 | 778.8384691 | 759.422067 | 931.9821455 | -0.00424294 | 0.066389867 | 0.85188613 | 0.991141351 | ns | TM | KS.3 |
| FIM1_4492 | 642.2287336 | 722.2053097 | 821.0718333 | 0.036118088 | 0.071895834 | 0.194242227 | 0.836671065 | ns | TM | KS.3 |
| FIM1_4493 | 2236.932622 | 5056.115929 | 6962.732737 | 1.082896757 | 0.215966778 | 8.3193837415 | 1.7995188745 | Km>KS.3 | TM | KS.3 |
| FIM1_4494 | 20.50153976 | 29.63627301 | 44.75258721 | 0.035073876 | 0.114483183 | 0.152878038 | 0.783447458 | ns | TM | KS.3 |
| FIM1_4495 | 821.1327336 | 874.7273978 | 1200.271987 | 0.014294338 | 0.062304788 | 0.546694101 | 0.95917717 | ns | TM | KS.3 |
| FIM1_4496 | 459.1591919 | 610.03622 | 572.8180632 | 0.159901249 | 0.139602691 | 0.003847844 | 0.116725767 | ns | TM | KS.3 |
| FIM1_4497 | 4623.17834 | 2943.867915 | 4143.687139 | -0.61027033 | 0.141931175 | 9.0687418420 | 1.8798746110 | ns | TM | KS.3 |
| FIM1_4498 | 18469.43271 | 20124.35728 | 18755.82401 | 0.024509573 | 0.063561105 | 0.330173384 | 0.936495203 | ns | TM | KS.3 |
| FIM1_4499 | 4652.376419 | 4271.720709 | 3375.850452 | -0.024647899 | 0.063350818 | 0.327409301 | 0.934739762 | ns | TM | KS.3 |
| FIM1_4500 | 2801.902463 | 2536.418213 | 2985.97264 | -0.051088428 | 0.068520684 | 0.090390195 | 0.65689657 | ns | TM | KS.3 |
| FIM1_4501 | 2002.243809 | 2030.678465 | 1516.750278 | 0.002499092 | 0.065190582 | 0.912484848 | 0.991141351 | ns | TM | KS.3 |
| FIM1_4502 | 226.6425243 | 241.1306123 | 211.4499236 | 0.008369378 | 0.073750265 | 0.705095267 | 0.982962554 | ns | TM | KS.3 |
| FIM1_4503 | 814.4873117 | 778.7869173 | 692.7756756 | -0.01329505 | 0.052094641 | 0.548703613 | 0.95917717 | ns | TM | KS.3 |
| FIM1_4504 | 1073.648442 | 959.3037208 | 1117.332412 | -0.021571125 | 0.072776468 | 0.380799269 | 0.948637087 | ns | TM | KS.3 |
| FIM1_4505 | 1776.675915 | 2365.729653 | 2753.53033 | 0.025742727 | 0.102328421 | 0.250259634 | 0.888934767 | ns | TM | KS.3 |
| FIM1_4506 | 1444.932564 | 1484.528669 | 1446.303108 | 0.004840869 | 0.064914409 | 0.83222348 | 0.991141351 | ns | TM | KS.3 |
| FIM1_4507 | 2777.927731 | 2595.977891 | 2743.829087 | -0.015812096 | 0.062674096 | 0.508479523 | 0.953260948 | ns | TM | KS.3 |
| FIM1_4508 | 1261.448958 | 1206.096485 | 1156.549221 | -0.013179406 | 0.050923011 | 0.546769161 | 0.95917717 | ns | TM | KS.3 |
| FIM1_4509 | 3295.022655 | 3158.055377 | 2958.750099 | -0.016672269 | 0.040484001 | 0.352983551 | 0.948637087 | ns | TM | KS.3 |
| FIM1_4510 | 4930.028324 | 4761.580135 | 5040.076391 | -0.010926813 | 0.045813236 | 0.590696048 | 0.963781856 | ns | TM | KS.3 |
| FIM1_4512 | 1182.653969 | 1019.557842 | 894.7769578 | -0.0696314 | 0.0860638 | 0.051696094 | 0.516441907 | ns | TM | KS.3 |
| FIM1_4513 | 218.6970492 | 281.9691159 | 220.9323604 | 0.077060372 | 0.112358877 | 0.036080811 | 0.431736218 | ns | TM | KS.3 |
| FIM1_4514 | 340.6182637 | 375.3022187 | 316.521227 | 0.023549593 | 0.068014007 | 0.350788931 | 0.948637087 | ns | TM | KS.3 |
| FIM1_4515 | 1343.330121 | 1364.058767 | 1229.686415 | 0.004404464 | 0.045245493 | 0.82705592 | 0.991141351 | ns | TM | KS.3 |
| FIM1_4516 | 516.0922642 | 655.0264125 | 750.3798019 | 0.018641538 | 0.096805993 | 0.370706694 | 0.948637087 | ns | TM | KS.3 |
| FIM1_4517 | 1202.717764 | 1084.330974 | 1121.894738 | -0.016080514 | 0.074174399 | 0.490183193 | 0.952591289 | ns | TM | KS.3 |
| FIM1_4518 | 943.2788922 | 789.095171 | 733.9843654 | -0.047439935 | 0.087157842 | 0.115595057 | 0.717670439 | ns | TM | KS.3 |

| | | | | | | | | | | |
|---|---|---|---|---|---|---|---|---|---|---|
| FIM1_4519 | 1126.479342 | 1034.889066 | 1030.65912 | -0.028968467 | 0.062035488 | 0.257162842 | 0.895871714 | ns | TM | KS.3 |
| FIM1_4520 | 2018.203638 | 1726.523631 | 2052.447947 | -0.031981338 | 0.080249162 | 0.226533596 | 0.876782875 | ns | TM | KS.3 |
| FIM1_4521 | 2510.30447 | 2416.227545 | 2375.04407 | -0.013136612 | 0.042578588 | 0.490823653 | 0.952591289 | ns | TM | KS.3 |
| FIM1_4522 | 480.0695236 | 308.7058742 | 643.0435648 | -0.101443867 | 0.184137833 | 0.017802837 | 0.304361209 | ns | TM | KS.3 |
| FIM1_4523 | 2941.297356 | 2907.339424 | 3284.485673 | -0.003793883 | 0.030588438 | 0.791001085 | 0.988916276 | ns | TM | KS.3 |
| FIM1_4524 | 2026.641766 | 1851.070983 | 2028.069343 | -0.027180755 | 0.064625359 | 0.289838035 | 0.919993167 | ns | TM | KS.3 |
| FIM1_4525 | 222.615792 | 298.2519552 | 257.5943842 | 0.081052968 | 0.124146404 | 0.030372933 | 0.40080992 | ns | TM | KS.3 |
| FIM1_4526 | 7156.139159 | 6872.800536 | 6663.423493 | -0.01571441 | 0.039517186 | 0.370752767 | 0.948637087 | ns | TM | KS.3 |
| FIM1_4527 | 17319.41572 | 20558.12019 | 16071.36789 | 0.019065889 | 0.085510615 | 0.391578864 | 0.948637087 | ns | TM | KS.3 |
| FIM1_4528 | 65.00969817 | 140.3914242 | 224.419495 | 0.377158353 | 0.458333289 | 0.001498206 | 0.064813696 | ns | TM | KS.3 |
| FIM1_4529 | 366.3832769 | 353.9756016 | 548.8525816 | -0.003227755 | 0.080324229 | 0.874082751 | 0.991141351 | ns | TM | KS.3 |
| FIM1_4530 | 2406.309556 | 2372.570429 | 2316.557567 | -0.004016546 | 0.047614551 | 0.847174294 | 0.991141351 | ns | TM | KS.3 |
| FIM1_4531 | 5497.656572 | 5477.392604 | 3611.620623 | -0.001053352 | 0.044236817 | 0.957684077 | 0.994413583 | ns | TM | KS.3 |
| FIM1_4532 | 1604.319293 | 1869.068519 | 1404.05236 | 0.043867105 | 0.081397214 | 0.137819078 | 0.770139466 | ns | TM | KS.3 |
| FIM1_4533 | 6319.611671 | 8190.647664 | 8237.061385 | 0.030705573 | 0.097984324 | 0.204185953 | 0.847612283 | ns | TM | KS.3 |
| FIM1_4534 | 959.6261364 | 907.7887162 | 795.2805864 | -0.012813327 | 0.061248139 | 0.585644636 | 0.963781856 | ns | TM | KS.3 |
| FIM1_4535 | 1777.682219 | 1812.666472 | 1795.554662 | 0.003603709 | 0.063940455 | 0.874598459 | 0.991141351 | ns | TM | KS.3 |
| FIM1_4536 | 2781.573647 | 2274.445141 | 2216.598367 | -0.028586573 | 0.088123287 | 0.244145509 | 0.88883001 | ns | TM | KS.3 |
| FIM1_4537 | 981.0442861 | 868.8223476 | 926.008992 | -0.04323924 | 0.074269283 | 0.141803242 | 0.777709768 | ns | TM | KS.3 |
| FIM1_4538 | 710.5917863 | 477.7943961 | 724.0558032 | -0.055077693 | 0.13624395 | 0.065903269 | 0.564318013 | ns | TM | KS.3 |
| FIM1_4539 | 1095.715581 | 1024.388941 | 1030.865874 | -0.022500238 | 0.056298771 | 0.34261403 | 0.944808611 | ns | TM | KS.3 |
| FIM1_4540 | 741.2153531 | 776.5566088 | 575.5351737 | 0.009288435 | 0.063645133 | 0.689085612 | 0.980879562 | ns | TM | KS.3 |
| FIM1_4542 | 5274.268918 | 4644.352992 | 4417.824229 | -0.030941534 | 0.074281976 | 0.245722829 | 0.888934767 | ns | TM | KS.3 |
| FIM1_4543 | 2750.936286 | 3277.04541 | 3931.707626 | 0.014017925 | 0.090378529 | 0.491184184 | 0.952591289 | ns | TM | KS.3 |
| FIM1_4544 | 723.1658812 | 598.724182 | 678.0470273 | -0.042715518 | 0.087904466 | 0.137026967 | 0.769423431 | ns | TM | KS.3 |
| FIM1_4545 | 1732.272717 | 1938.83436 | 2193.460746 | 0.014577393 | 0.077783845 | 0.515979749 | 0.955340251 | ns | TM | KS.3 |
| FIM1_4546 | 3459.237678 | 3877.92104 | 5054.654355 | 0.011687137 | 0.081725514 | 0.581274606 | 0.963781856 | ns | TM | KS.3 |
| FIM1_4547 | 77.10092174 | 98.57121533 | 93.14022584 | 0.024753892 | 0.09470273 | 0.276005605 | 0.908583046 | ns | TM | KS.3 |
| FIM1_4548 | 1188.500358 | 1256.23109 | 1192.076338 | 0.01276447 | 0.060529938 | 0.586274558 | 0.963781856 | ns | TM | KS.3 |
| FIM1_4549 | 2419.699047 | 2686.731993 | 2090.131992 | 0.021372548 | 0.070847385 | 0.38733407 | 0.948637087 | ns | TM | KS.3 |
| FIM1_4550 | 42.67813221 | 48.32581618 | 58.11738184 | 0.01016364 | 0.085908946 | 0.614952757 | 0.973122675 | ns | TM | KS.3 |
| FIM1_4551 | 220.6097958 | 129.4433794 | 155.7175167 | -0.605630057 | 0.27724674 | 4.56465640458 | 0.004634524 | ns | TM | KS.3 |
| FIM1_4552 | 954.5755344 | 1072.911986 | 991.3499352 | 0.027362324 | 0.072225088 | 0.291250102 | 0.919993167 | ns | TM | KS.3 |
| FIM1_4553 | 446.9333698 | 514.3781034 | 556.4867473 | 0.026561073 | 0.077575454 | 0.293749117 | 0.923185001 | ns | TM | KS.3 |
| FIM1_4554 | 2698.093813 | 2570.483705 | 2258.936361 | -0.009541369 | 0.063970795 | 0.681327633 | 0.97986479 | ns | TM | KS.3 |
| FIM1_4555 | 4867.79243 | 4770.022465 | 5241.801357 | -0.002854477 | 0.070940778 | 0.89734579 | 0.991141351 | ns | TM | KS.3 |
| FIM1_4556 | 3291.784285 | 2686.653355 | 2797.311843 | -0.031025904 | 0.088493508 | 0.218516473 | 0.865538807 | ns | TM | KS.3 |
| FIM1_4557 | 5442.845177 | 5022.255354 | 4748.96601 | -0.029844034 | 0.060050808 | 0.237562074 | 0.88653074 | ns | TM | KS.3 |
| FIM1_4558 | 2596.081558 | 2044.242472 | 1668.102854 | -0.083998764 | 0.110956817 | 0.029724408 | 0.399672788 | ns | TM | KS.3 |
| FIM1_4559 | 1931.401033 | 1403.26269 | 1020.512662 | -0.284884639 | 0.15177008 | 0.000175224 | 0.013620889 | ns | TM | KS.3 |
| FIM1_4560 | 214.7144747 | 199.6721561 | 161.0635889 | -0.009391631 | 0.075304881 | 0.66914601 | 0.976323032 | ns | TM | KS.3 |
| FIM1_4561 | 3898.834432 | 3602.548785 | 4062.928576 | -0.011134733 | 0.073640591 | 0.620885985 | 0.973122675 | ns | TM | KS.3 |
| FIM1_4562 | 295.2720671 | 249.9189394 | 665.2961242 | -0.011735257 | 0.092235878 | 0.550071712 | 0.95917717 | ns | TM | KS.3 |
| FIM1_4563 | 1740.590642 | 2237.94658 | 2210.548623 | 0.040941438 | 0.09882339 | 0.132235142 | 0.761677364 | ns | TM | KS.3 |
| FIM1_4564 | 406.9530186 | 492.5074928 | 338.2719058 | 0.032521816 | 0.086364312 | 0.209243686 | 0.852551729 | ns | TM | KS.3 |

| | | | | | | | | | | |
|---|---|---|---|---|---|---|---|---|---|---|
| FIM1_4565 | 5140.444586 | 4992.127747 | 4454.604135 | -0.003296149 | 0.076099873 | 0.876567037 | 0.991141351 | ns | TM | KS.3 |
| FIM1_4566 | 10608.01606 | 15020.30397 | 10289.83615 | 0.044516559 | 0.118990381 | 0.101864084 | 0.686685387 | ns | TM | KS.3 |
| FIM1_4567 | 6494.47334 | 5396.749267 | 6056.028043 | -0.09588601 | 0.098552277 | 0.021744463 | 0.33950953 | ns | TM | KS.3 |
| FIM1_4568 | 5436.24893 | 4743.779588 | 5307.079298 | -0.142971877 | 0.064755793 | 0.001636766 | 0.067857575 | ns | TM | KS.3 |
| FIM1_4569 | 2618.251471 | 2605.084183 | 2428.104162 | -0.001093798 | 0.058700014 | 0.961561068 | 0.995175461 | ns | TM | KS.3 |
| FIM1_4570 | 1699.494657 | 1298.106061 | 1569.40606 | -0.220623629 | 0.12254048 | 0.000565435 | 0.031255978 | ns | TM | KS.3 |
| FIM1_4571 | 1255.260555 | 1166.892501 | 1194.927305 | -0.019715912 | 0.060813438 | 0.415969258 | 0.950410534 | ns | TM | KS.3 |
| FIM1_4572 | 3626.511026 | 4174.462323 | 4823.600807 | 0.014664859 | 0.083394424 | 0.495585203 | 0.953260948 | ns | TM | KS.3 |
| FIM1_4573 | 2004.833364 | 1800.612637 | 1981.366376 | -0.020636434 | 0.071880942 | 0.399510393 | 0.950410534 | ns | TM | KS.3 |
| FIM1_4574 | 763.6359754 | 670.6848182 | 667.0720463 | -0.060504473 | 0.079463397 | 0.070512152 | 0.582627046 | ns | TM | KS.3 |
| FIM1_4575 | 9660.492998 | 8361.568534 | 8341.093234 | -0.067924517 | 0.084375449 | 0.05465984 | 0.531428718 | ns | TM | KS.3 |
| FIM1_4576 | 1656.158086 | 1767.711967 | 1594.398959 | 0.016514469 | 0.060569765 | 0.488593701 | 0.952591289 | ns | TM | KS.3 |
| FIM1_4577 | 616.5072566 | 518.288427 | 482.5281874 | -0.029765807 | 0.083383533 | 0.242449063 | 0.887413624 | ns | TM | KS.3 |
| FIM1_4578 | 446.0770819 | 396.1643455 | 446.1690164 | -0.029207416 | 0.07258869 | 0.267198964 | 0.905057253 | ns | TM | KS.3 |
| FIM1_4579 | 312.5940177 | 316.7456301 | 334.6155605 | 0.001689516 | 0.072534161 | 0.938298387 | 0.994413583 | ns | TM | KS.3 |
| FIM1_4580 | 2540.554879 | 2814.912007 | 3068.994516 | 0.021265757 | 0.070348314 | 0.389889552 | 0.948637087 | ns | TM | KS.3 |
| FIM1_4582 | 1538.576722 | 1471.546469 | 1719.339598 | -0.005646652 | 0.073843606 | 0.79575758 | 0.988916276 | ns | TM | KS.3 |
| FIM1_4583 | 1377.997907 | 1366.158041 | 1529.542087 | -0.002464452 | 0.039697428 | 0.89271809 | 0.991141351 | ns | TM | KS.3 |
| FIM1_4584 | 1629.270012 | 1769.864879 | 1894.691616 | 0.025277653 | 0.062031166 | 0.313676335 | 0.927564309 | ns | TM | KS.3 |
| FIM1_4585 | 2255.086266 | 2028.074046 | 2464.633392 | -0.05936928 | 0.071618721 | 0.06559 | 0.564318013 | ns | TM | KS.3 |
| FIM1_4586 | 16037.46348 | 14966.56309 | 14634.50189 | -0.034712299 | 0.054156106 | 0.148079921 | 0.780546912 | ns | TM | KS.3 |
| FIM1_4587 | 1564.829517 | 1860.075411 | 2336.525261 | 0.014283193 | 0.089697194 | 0.486183718 | 0.952269291 | ns | TM | KS.3 |
| FIM1_4588 | 4405.788163 | 4166.336169 | 3198.471663 | -0.013876167 | 0.059093526 | 0.553705404 | 0.95917717 | ns | TM | KS.3 |
| FIM1_4589 | 1963.324332 | 2230.68327 | 2506.601279 | 0.016986446 | 0.07881372 | 0.45600991 | 0.950410534 | ns | TM | KS.3 |
| FIM1_4590 | 619.9022034 | 580.6323586 | 592.5620059 | -0.016273943 | 0.060510883 | 0.494368963 | 0.953260948 | ns | TM | KS.3 |
| FIM1_4591 | 2888.407757 | 2580.192405 | 2635.107102 | -0.041100896 | 0.071553979 | 0.153636218 | 0.783447458 | ns | TM | KS.3 |
| FIM1_4592 | 1227.051714 | 1078.944936 | 1307.898843 | -0.07249754 | 0.080424015 | 0.0442294 | 0.48254832 | ns | TM | KS.3 |
| FIM1_4593 | 828.7417717 | 867.3445595 | 726.3664995 | 0.009617403 | 0.0620042 | 0.678916188 | 0.97986479 | ns | TM | KS.3 |
| FIM1_4594 | 545.3654654 | 554.6395369 | 742.0136083 | 0.002202811 | 0.073128029 | 0.919279568 | 0.991141351 | ns | TM | KS.3 |
| FIM1_4595 | 3271.900997 | 3842.901123 | 4025.083491 | 0.059048747 | 0.086649846 | 0.075966816 | 0.606637094 | ns | TM | KS.3 |
| FIM1_4596 | 2581.73068 | 3009.72396 | 3538.793639 | 0.021886798 | 0.081031808 | 0.354418882 | 0.948637087 | ns | TM | KS.3 |
| FIM1_4597 | 1694.56389 | 2108.693007 | 1866.817906 | 0.16251141 | 0.105812941 | 0.002731388 | 0.096373426 | ns | TM | KS.3 |
| FIM1_4598 | 497.7790488 | 512.8134022 | 478.6078264 | 0.005667 | 0.063658479 | 0.804906288 | 0.988916276 | ns | TM | KS.3 |
| FIM1_4599 | 244.7938087 | 246.2966005 | 280.8019534 | 0.001087964 | 0.067325825 | 0.961500422 | 0.995175461 | ns | TM | KS.3 |
| FIM1_4600 | 3202.299822 | 3322.687511 | 3711.087624 | 0.010687175 | 0.050120336 | 0.620655142 | 0.973122675 | ns | TM | KS.3 |
| FIM1_4601 | 783.4526877 | 868.8906484 | 781.6834496 | 0.04407439 | 0.06926843 | 0.129634943 | 0.757854102 | ns | TM | KS.3 |
| FIM1_4602 | 470.2881045 | 554.2263376 | 450.4868022 | 0.018902784 | 0.084162582 | 0.398918597 | 0.950410534 | ns | TM | KS.3 |
| FIM1_4603 | 3871.668893 | 3980.422043 | 3864.028105 | 0.00754242 | 0.050541737 | 0.727221786 | 0.987157164 | ns | TM | KS.3 |
| FIM1_4604 | 87.98353924 | 169.8547429 | 140.3676843 | 0.54802153 | 0.415280431 | 0.000359119 | 0.022282532 | ns | TM | KS.3 |
| FIM1_4605 | 1088.957014 | 1043.352204 | 1105.088326 | -0.012921792 | 0.051507607 | 0.557173156 | 0.959760553 | ns | TM | KS.3 |
| FIM1_4606 | 712.9368882 | 744.7121797 | 706.6727291 | 0.012270556 | 0.053480906 | 0.584413367 | 0.963781856 | ns | TM | KS.3 |
| FIM1_4607 | 2245.678983 | 2259.243786 | 2344.433376 | 0.001716639 | 0.036489458 | 0.919646691 | 0.991141351 | ns | TM | KS.3 |
| FIM1_4608 | 526.5806485 | 359.4290232 | 390.933759 | -0.131584393 | 0.179909187 | 0.008780543 | 0.202237047 | ns | TM | KS.3 |
| FIM1_4609 | 3157.604125 | 3589.973311 | 3251.156676 | 0.020696234 | 0.076679158 | 0.387730701 | 0.948637087 | ns | TM | KS.3 |
| FIM1_4612 | 7437.010364 | 7317.644558 | 7107.770878 | -0.003631081 | 0.057621026 | 0.872680611 | 0.991141351 | ns | TM | KS.3 |

| | | | | | | | | | | |
|---|---|---|---|---|---|---|---|---|---|---|
| FIM1_4613 | 11333.13935 | 13997.99126 | 15088.26707 | 0.028829038 | 0.089783017 | 0.237835248 | 0.88653074 | ns | TM | KS.3 |
| FIM1_4614 | 538.7833714 | 498.7236664 | 609.0939818 | -0.023627154 | 0.061143112 | 0.339807705 | 0.943910292 | ns | TM | KS.3 |
| FIM1_4615 | 1361.607362 | 1331.117029 | 1502.09062 | -0.007031499 | 0.045830679 | 0.729523219 | 0.987157164 | ns | TM | KS.3 |
| FIM1_4616 | 2822.024414 | 3039.263303 | 3201.066156 | 0.028121175 | 0.057569987 | 0.250867991 | 0.888934767 | ns | TM | KS.3 |
| FIM1_4617 | 3652.210141 | 3736.327214 | 3570.518226 | 0.004256681 | 0.063851403 | 0.85231154 | 0.991141351 | ns | TM | KS.3 |
| FIM1_4618 | 4776.862207 | 4481.18557 | 4057.374277 | -0.018805923 | 0.0570278 | 0.424887209 | 0.950410534 | ns | TM | KS.3 |
| FIM1_4619 | 968.4504392 | 975.2207892 | 857.5893191 | 0.002052646 | 0.046101769 | 0.919905765 | 0.991141351 | ns | TM | KS.3 |
| FIM1_4620 | 685.352378 | 713.1694492 | 744.6249619 | 0.004808913 | 0.075004488 | 0.823281881 | 0.991141351 | ns | TM | KS.3 |
| FIM1_4621 | 107.3287915 | 134.6597177 | 125.0342392 | 0.012791715 | 0.100696442 | 0.501573731 | 0.953260948 | ns | TM | KS.3 |
| FIM1_4622 | 972.8095392 | 971.1516468 | 1327.016102 | -0.000257549 | 0.053319531 | 0.990685815 | 0.999525119 | ns | TM | KS.3 |
| FIM1_4623 | 1652.128386 | 1555.476362 | 1412.356005 | -0.016807503 | 0.057128335 | 0.472935413 | 0.950410534 | ns | TM | KS.3 |
| FIM1_4624 | 3109.912901 | 2849.976966 | 2292.836892 | -0.025614918 | 0.063764662 | 0.31232536 | 0.927416591 | ns | TM | KS.3 |
| FIM1_4625 | 669.1674018 | 906.7066395 | 926.2415632 | 0.351102742 | 0.10726027 | 8.37088757515 | 0.000138817 | ns | TM | KS.3 |
| FIM1_4626 | 413.0879999 | 331.8796255 | 425.5798025 | -0.038011641 | 0.092363399 | 0.157409839 | 0.789048157 | ns | TM | KS.3 |
| FIM1_4627 | 2722.645385 | 2530.551369 | 2357.628351 | -0.014806517 | 0.066303544 | 0.53397176 | 0.959071703 | ns | TM | KS.3 |
| FIM1_4628 | 329.7685738 | 323.4009686 | 297.3288643 | -0.003245436 | 0.067969666 | 0.885467597 | 0.991141351 | ns | TM | KS.3 |
| FIM1_4629 | 727.2226703 | 736.3180697 | 744.8255768 | 0.002145851 | 0.065156399 | 0.92479101 | 0.993271864 | ns | TM | KS.3 |
| FIM1_4630 | 1251.465961 | 1170.68654 | 1317.615698 | -0.020395881 | 0.057018343 | 0.38944015 | 0.948637087 | ns | TM | KS.3 |
| FIM1_4631 | 2371.073372 | 2298.053163 | 2080.043648 | -0.006814496 | 0.059685799 | 0.766438822 | 0.988477371 | ns | TM | KS.3 |
| FIM1_4632 | 2008.148235 | 1906.899872 | 2347.234258 | -0.007266608 | 0.072009248 | 0.744262718 | 0.988477371 | ns | TM | KS.3 |
| FIM1_4633 | 53.92670629 | 58.10601108 | 76.16514234 | 0.007219626 | 0.082031881 | 0.723650962 | 0.987157164 | ns | TM | KS.3 |
| FIM1_4634 | 3305.337697 | 3448.577052 | 2905.267577 | 0.00360795 | 0.082486482 | 0.856624891 | 0.991141351 | ns | TM | KS.3 |
| FIM1_4635 | 1304.351586 | 1140.692755 | 1175.173052 | -0.043341836 | 0.07744316 | 0.142272937 | 0.777709768 | ns | TM | KS.3 |
| FIM1_4636 | 682.8379026 | 512.9550166 | 580.978079 | -0.288251958 | 0.116821319 | 3.99471334939 | 0.004228447 | ns | TM | KS.3 |
| FIM1_4637 | 1103.426029 | 978.9420742 | 1091.158215 | -0.028856416 | 0.072737565 | 0.271452603 | 0.908468501 | ns | TM | KS.3 |
| FIM1_4638 | 1188.509785 | 1187.077867 | 1228.748941 | -0.000281409 | 0.045513663 | 0.988907726 | 0.999333464 | ns | TM | KS.3 |
| FIM1_4639 | 1980.586502 | 2018.013748 | 2308.383022 | 0.004791887 | 0.052710661 | 0.827661157 | 0.991141351 | ns | TM | KS.3 |
| FIM1_4640 | 788.78651 | 925.1682716 | 889.2184359 | 0.036335045 | 0.081425173 | 0.186828825 | 0.827669994 | ns | TM | KS.3 |
| FIM1_4641 | 5530.113473 | 6143.132169 | 6170.986705 | 0.03761406 | 0.068904608 | 0.177454686 | 0.811391014 | ns | TM | KS.3 |
| FIM1_4642 | 2497.634838 | 3305.175866 | 3031.349814 | 0.125220527 | 0.133801046 | 0.009173506 | 0.208393578 | ns | TM | KS.3 |
| FIM1_4643 | 439.6153281 | 519.4416786 | 499.9010469 | 0.044363991 | 0.08415438 | 0.132891649 | 0.761677364 | ns | TM | KS.3 |
| FIM1_4644 | 1101.608878 | 915.8460981 | 877.6145571 | -0.085604208 | 0.097346683 | 0.030184925 | 0.400453344 | ns | TM | KS.3 |
| FIM1_4645 | 1142.868704 | 1200.535626 | 1148.317039 | 0.011314364 | 0.060251641 | 0.627276005 | 0.97344368 | ns | TM | KS.3 |
| FIM1_4646 | 1581.941522 | 1660.02515 | 1583.751112 | 0.014175367 | 0.052018537 | 0.522857292 | 0.958732165 | ns | TM | KS.3 |
| FIM1_4647 | 1925.652643 | 2065.860698 | 1914.355097 | 0.018065527 | 0.061075696 | 0.452956673 | 0.950410534 | ns | TM | KS.3 |
| FIM1_4648 | 1586.607982 | 1689.201429 | 1698.51504 | 0.016206607 | 0.059365685 | 0.493809828 | 0.952949531 | ns | TM | KS.3 |
| FIM1_4649 | 2601.506782 | 2385.033353 | 2547.851914 | -0.030184578 | 0.062592951 | 0.242419578 | 0.887413624 | ns | TM | KS.3 |
| FIM1_4650 | 1506.577819 | 1572.684347 | 1549.062917 | 0.011114569 | 0.055583841 | 0.625263081 | 0.97344368 | ns | TM | KS.3 |
| FIM1_4651 | 3224.975911 | 2986.036078 | 4372.04566 | -0.008509537 | 0.078139685 | 0.689945892 | 0.980879562 | ns | TM | KS.3 |
| FIM1_4652 | 1155.634367 | 1587.576491 | 1412.47613 | 0.325294306 | 0.13725089 | 2.97712411153 | 0.003219824 | ns | TM | KS.3 |
| FIM1_4653 | 1133.303084 | 1091.822593 | 1046.675546 | -0.005723846 | 0.069228294 | 0.799447902 | 0.988916276 | ns | TM | KS.3 |
| FIM1_4654 | 1139.888523 | 977.2833664 | 1079.974788 | -0.032736935 | 0.079910063 | 0.219907749 | 0.865538807 | ns | TM | KS.3 |
| FIM1_4655 | 1383.408996 | 1552.10564 | 1886.468117 | 0.014646188 | 0.078255557 | 0.512774011 | 0.954240701 | ns | TM | KS.3 |
| FIM1_4656 | 2114.005457 | 2534.703942 | 2415.83052 | 0.019378347 | 0.08706496 | 0.380950335 | 0.948637087 | ns | TM | KS.3 |
| FIM1_4657 | 835.6129166 | 897.1964549 | 980.8930235 | 0.018292016 | 0.06106454 | 0.44779424 | 0.950410534 | ns | TM | KS.3 |

| | | | | | | | | | | |
|---|---|---|---|---|---|---|---|---|---|---|
| FIM1_4658 | 21767.45567 | 20952.13942 | 20752.37853 | -0.007049794 | 0.064900285 | 0.759156923 | 0.988477371 | ns | TM | KS.3 |
| FIM1_4660 | 58.98753866 | 49.37218217 | 183.5974283 | -0.019852558 | 0.08581093 | 0.375771783 | 0.948637087 | ns | TM | KS.3 |
| FIM1_4661 | 141.4992887 | 178.3089719 | 320.9228849 | 0.030850344 | 0.093182267 | 0.210909093 | 0.855153004 | ns | TM | KS.3 |
| FIM1_4662 | 2234.253078 | 2624.279851 | 3115.470066 | 0.015021767 | 0.086766795 | 0.476156034 | 0.950410534 | ns | TM | KS.3 |
| FIM1_4664 | 46.12284937 | 41.39550824 | 39.5335887 | -0.012511633 | 0.082111781 | 0.556607845 | 0.959760553 | ns | TM | KS.3 |
| FIM1_4665 | 2369.739537 | 2873.238859 | 3582.114891 | 0.021654635 | 0.087908028 | 0.338250493 | 0.942012706 | ns | TM | KS.3 |
| FIM1_4666 | 1095.002477 | 1404.042142 | 786.747206 | 0.026190561 | 0.096095643 | 0.25504456 | 0.892600401 | ns | TM | KS.3 |
| FIM1_4667 | 3158.172304 | 4021.880108 | 5171.262894 | 0.021716365 | 0.095921584 | 0.317994863 | 0.928601572 | ns | TM | KS.3 |
| FIM1_4668 | 303.1875028 | 328.0867718 | 590.2764347 | 0.008571932 | 0.07867159 | 0.686504461 | 0.980879562 | ns | TM | KS.3 |
| FIM1_4669 | 27.86429871 | 34.18303115 | 39.49713505 | 0.014075247 | 0.095362244 | 0.477438521 | 0.950410534 | ns | TM | KS.3 |
| FIM1_4670 | 2456.5482 | 2357.329487 | 2421.800564 | -0.006542731 | 0.068790681 | 0.772643844 | 0.988814136 | ns | TM | KS.3 |
| FIM1_4671 | 1232.014284 | 1100.623935 | 1252.57698 | -0.033737066 | 0.070934465 | 0.2164763 | 0.865538807 | ns | TM | KS.3 |
| FIM1_4672 | 975.8755436 | 938.5852458 | 1166.076171 | -0.010425014 | 0.053678624 | 0.641441001 | 0.975935248 | ns | TM | KS.3 |
| FIM1_4673 | 381.8458232 | 363.037579 | 398.5748486 | -0.011300802 | 0.05989009 | 0.627292069 | 0.97344368 | ns | TM | KS.3 |
| FIM1_4674 | 233.720583 | 225.6287395 | 279.8664622 | -0.007908759 | 0.055214032 | 0.725588165 | 0.987157164 | ns | TM | KS.3 |
| FIM1_4675 | 4334.565269 | 4031.669667 | 4876.118353 | -0.020782402 | 0.059866301 | 0.391142946 | 0.948637087 | ns | TM | KS.3 |
| FIM1_4676 | 2007.870751 | 1998.725146 | 2206.597425 | -0.001526646 | 0.03578564 | 0.927279116 | 0.993697171 | ns | TM | KS.3 |
| FIM1_4678 | 1770.964725 | 1929.044595 | 2206.929083 | 0.00685406 | 0.085060171 | 0.729608459 | 0.987157164 | ns | TM | KS.3 |
| FIM1_4679 | 1608.94897 | 1177.750934 | 1706.025796 | -0.046485571 | 0.112719515 | 0.097287033 | 0.676927258 | ns | TM | KS.3 |
| FIM1_4680 | 515.363759 | 588.8690339 | 650.5430871 | 0.049985955 | 0.077955058 | 0.107712113 | 0.702423202 | ns | TM | KS.3 |
| FIM1_4681 | 2244.643961 | 1713.288423 | 3003.550954 | -0.087093709 | 0.120388142 | 0.02598973 | 0.380290896 | ns | TM | KS.3 |
| FIM1_4682 | 6546.472469 | 6069.189222 | 6334.836483 | -0.033624138 | 0.057555514 | 0.178584964 | 0.811815714 | ns | TM | KS.3 |
| FIM1_4683 | 166.286952 | 218.6704481 | 149.174951 | 0.027400897 | 0.100114547 | 0.234252905 | 0.883800031 | ns | TM | KS.3 |
| FIM1_4684 | 1309.788985 | 1558.868392 | 1653.177765 | 0.037322233 | 0.084197833 | 0.175244762 | 0.811391014 | ns | TM | KS.3 |
| FIM1_4685 | 1276.765387 | 1331.724551 | 1895.683557 | 0.003141988 | 0.0856845 | 0.871071647 | 0.991141351 | ns | TM | KS.3 |
| FIM1_4686 | 108.099369 | 81.52424408 | 374.4480036 | -0.023169901 | 0.102313906 | 0.284421502 | 0.918450395 | ns | TM | KS.3 |
| FIM1_4688 | 826.9731308 | 1047.953858 | 1030.77713 | 0.060733686 | 0.103037045 | 0.063625849 | 0.555330875 | ns | TM | KS.3 |
| FIM1_4689 | 1274.477222 | 1137.163783 | 1260.925112 | -0.040299299 | 0.071866117 | 0.15988368 | 0.791498358 | ns | TM | KS.3 |
| FIM1_4690 | 3252.333315 | 3020.776711 | 3273.689111 | -0.017568691 | 0.063308565 | 0.467305139 | 0.950410534 | ns | TM | KS.3 |
| FIM1_4691 | 704.0632171 | 968.6340928 | 1392.037738 | 0.034854097 | 0.109273309 | 0.158269422 | 0.789048157 | ns | TM | KS.3 |
| FIM1_4692 | 1444.85999 | 1091.155208 | 1443.260619 | -0.084737411 | 0.122556501 | 0.02752901 | 0.387979669 | ns | TM | KS.3 |
| FIM1_4693 | 315.5300586 | 269.6548485 | 358.3793039 | -0.025581821 | 0.080138251 | 0.300873568 | 0.926536521 | ns | TM | KS.3 |
| FIM1_4694 | 799.6113934 | 772.7959557 | 868.552117 | -0.003419382 | 0.078619545 | 0.868936316 | 0.991141351 | ns | TM | KS.3 |
| FIM1_4695 | 447.8203582 | 628.0420105 | 611.148356 | 0.048820261 | 0.119432461 | 0.086459667 | 0.643510999 | ns | TM | KS.3 |
| FIM1_4697 | 932.3093039 | 1104.46554 | 1508.15025 | 0.011097079 | 0.093793343 | 0.565831742 | 0.962265923 | ns | TM | KS.3 |
| FIM1_4698 | 1155.065527 | 1269.350464 | 1300.486597 | 0.021042274 | 0.067991005 | 0.395814747 | 0.948637087 | ns | TM | KS.3 |
| FIM1_4699 | 645.2443063 | 824.0614092 | 876.9958142 | 0.027360866 | 0.095345403 | 0.242722733 | 0.887413624 | ns | TM | KS.3 |
| FIM1_4700 | 22.85179513 | 27.68910031 | 31.51954424 | 0.015715241 | 0.091810599 | 0.445321951 | 0.950410534 | ns | TM | KS.3 |
| FIM1_4701 | 987.2424106 | 1228.285503 | 949.344502 | 0.034220674 | 0.091391111 | 0.185504561 | 0.825871141 | ns | TM | KS.3 |
| FIM1_4702 | 2383.280707 | 2292.25818 | 2349.804366 | -0.003361154 | 0.082103391 | 0.866712518 | 0.991141351 | ns | TM | KS.3 |
| FIM1_4703 | 10239.17594 | 9911.737762 | 8353.968002 | -0.003494147 | 0.077174152 | 0.867966543 | 0.991141351 | ns | TM | KS.3 |
| FIM1_4704 | 1477.745756 | 1412.745001 | 1487.561147 | -0.004875802 | 0.077327795 | 0.817133819 | 0.989591224 | ns | TM | KS.3 |
| FIM1_4705 | 1413.252014 | 1226.880805 | 1309.422391 | -0.041194581 | 0.078539619 | 0.155407687 | 0.786819827 | ns | TM | KS.3 |
| FIM1_4706 | 1290.36494 | 1271.642161 | 1323.038148 | -0.003386032 | 0.05650183 | 0.880586394 | 0.991141351 | ns | TM | KS.3 |
| FIM1_4707 | 650.5605401 | 713.1933948 | 651.8354446 | 0.012981331 | 0.074554616 | 0.566910095 | 0.962265923 | ns | TM | KS.3 |

| | | | | | | | | | | |
|---|---|---|---|---|---|---|---|---|---|---|
| FIM1_4708 | 963.122433 | 1135.55387 | 1232.431846 | 0.01435535 | 0.088150332 | 0.488714757 | 0.952591289 | ns | TM | KS.3 |
| FIM1_4709 | 1238.612564 | 1414.710036 | 1693.347632 | 0.016036573 | 0.080622103 | 0.471659073 | 0.950410534 | ns | TM | KS.3 |
| FIM1_4710 | 1490.213465 | 1361.275018 | 1338.096276 | -0.011030617 | 0.07709385 | 0.614613398 | 0.973122675 | ns | TM | KS.3 |
| FIM1_4711 | 2417.627417 | 2184.897193 | 2530.178698 | -0.014138913 | 0.075567285 | 0.533742385 | 0.959071703 | ns | TM | KS.3 |
| FIM1_4712 | 410.7195569 | 510.1540307 | 488.8438043 | 0.036267494 | 0.091400894 | 0.170457542 | 0.809734625 | ns | TM | KS.3 |
| FIM1_4713 | 2557.117958 | 2249.306068 | 2362.926956 | -0.025513154 | 0.07506973 | 0.313238886 | 0.927416591 | ns | TM | KS.3 |
| FIM1_4714 | 1250.676354 | 1419.898244 | 1656.996978 | 0.020759393 | 0.076323403 | 0.387523123 | 0.948637087 | ns | TM | KS.3 |
| FIM1_4715 | 4858.723239 | 4115.035009 | 4836.093207 | -0.027536908 | 0.082109889 | 0.270719157 | 0.908468501 | ns | TM | KS.3 |
| FIM1_4716 | 2955.03361 | 2982.209454 | 3401.173216 | 0.000605696 | 0.088087172 | 0.974393123 | 0.998682692 | ns | TM | KS.3 |
| FIM1_4717 | 4539.509037 | 4975.436451 | 4812.049546 | 0.008976951 | 0.081219764 | 0.666320301 | 0.976323032 | ns | TM | KS.3 |
| FIM1_4718 | 1297.676845 | 1718.99176 | 1948.392689 | 0.019518903 | 0.10308555 | 0.342526621 | 0.944808611 | ns | TM | KS.3 |
| FIM1_4719 | 312.5800805 | 360.1345131 | 359.3621064 | 0.019435954 | 0.079932395 | 0.401856037 | 0.950410534 | ns | TM | KS.3 |
| FIM1_4720 | 1938.464257 | 2290.928172 | 2687.385742 | 0.017334931 | 0.085881281 | 0.425767659 | 0.950410534 | ns | TM | KS.3 |
| FIM1_4721 | 1497.907829 | 1537.372732 | 1004.264901 | 0.006354875 | 0.054576449 | 0.776376042 | 0.988814136 | ns | TM | KS.3 |
| FIM1_4722 | 1259.526326 | 1630.539213 | 1807.401918 | 0.014894304 | 0.103190341 | 0.441276935 | 0.950410534 | ns | TM | KS.3 |
| FIM1_4724 | 497.0813857 | 422.7242695 | 584.5339102 | -0.038454571 | 0.082252231 | 0.170219339 | 0.809599629 | ns | TM | KS.3 |
| FIM1_4725 | 200.1342262 | 188.588228 | 211.8274937 | -0.001185574 | 0.128155064 | 0.937290859 | 0.994413583 | ns | TM | KS.3 |
| FIM1_4726 | 60.3618287 | 59.13225086 | 61.86479955 | -0.00050529 | 0.119614192 | 0.974641757 | 0.99870125 | ns | TM | KS.3 |
| FIM1_4727 | 390.8328978 | 371.2580123 | 420.3880307 | -0.000919573 | 0.131604767 | 0.949945114 | 0.994413583 | ns | TM | KS.3 |
| FIM1_4728 | 135.3032447 | 114.2563997 | 140.8377942 | -0.003454239 | 0.128403658 | 0.82027173 | 0.990886949 | ns | TM | KS.3 |
| FIM1_4729 | 814.7608214 | 737.6460535 | 766.2970696 | -0.001821557 | 0.131100511 | 0.901731155 | 0.991141351 | ns | TM | KS.3 |
| FIM1_4730 | 376.0866665 | 356.931587 | 481.3915826 | -0.005215917 | 0.079033757 | 0.801772923 | 0.988916276 | ns | TM | KS.3 |
| FIM1_4731 | 1200.928584 | 1267.93528 | 1595.830275 | 0.005950283 | 0.077357318 | 0.778736224 | 0.988814136 | ns | TM | KS.3 |
| FIM1_4732 | 3670.067093 | 3641.888766 | 3415.38404 | -0.000819093 | 0.077210274 | 0.96875546 | 0.996327965 | ns | TM | KS.3 |
| FIM1_4733 | 1020.694616 | 1116.34119 | 1350.604209 | 0.015642968 | 0.07093535 | 0.507799688 | 0.953260948 | ns | TM | KS.3 |
| FIM1_4734 | 302.245929 | 273.0631242 | 494.3151773 | -0.014140001 | 0.07536376 | 0.534295984 | 0.959071703 | ns | TM | KS.3 |
| FIM1_4736 | 108.9219663 | 191.9236571 | 252.8171641 | 0.219296961 | 0.310252942 | 0.00370012 | 0.115135887 | ns | TM | KS.3 |
| FIM1_4737 | 824.7109701 | 455.1335408 | 807.1249009 | -0.521438939 | 0.369123912 | 0.000290186 | 0.019353666 | ns | TM | KS.3 |
| FIM1_4738 | 94.46281945 | 144.0519739 | 122.0593403 | 0.05928184 | 0.1453944 | 0.056437098 | 0.535599847 | ns | TM | KS.3 |
| FIM1_4739 | 319.283038 | 395.3829853 | 449.3723635 | 0.05819049 | 0.097273297 | 0.072662883 | 0.594131438 | ns | TM | KS.3 |
| FIM1_4740 | 1249.523655 | 1457.171042 | 1494.907235 | 0.034086627 | 0.079961669 | 0.207556663 | 0.852551729 | ns | TM | KS.3 |
| FIM1_4741 | 595.455148 | 559.588698 | 562.5953815 | -0.009872112 | 0.070095611 | 0.66554654 | 0.976323032 | ns | TM | KS.3 |
| FIM1_4742 | 219.8378958 | 229.5452527 | 286.3233181 | 0.004211178 | 0.07976882 | 0.837595899 | 0.991141351 | ns | TM | KS.3 |
| FIM1_4743 | 29.63121645 | 16.72040159 | 29.18473673 | -0.046468994 | 0.160246608 | 0.086181196 | 0.642805774 | ns | TM | KS.3 |
| FIM1_4744 | 3195.985276 | 2829.349096 | 2880.341186 | -0.021292797 | 0.075023198 | 0.381123434 | 0.948637087 | ns | TM | KS.3 |
| FIM1_4745 | 1308.45775 | 1035.218595 | 978.5767618 | -0.111766638 | 0.115070992 | 0.01327539 | 0.263081302 | ns | TM | KS.3 |
| FIM1_4746 | 1583.103892 | 2222.689386 | 2526.440432 | 0.064252761 | 0.128546618 | 0.049497738 | 0.510786778 | ns | TM | KS.3 |
| FIM1_4747 | 631.2053628 | 691.9660938 | 669.4574419 | 0.026775628 | 0.065020153 | 0.296631971 | 0.926536521 | ns | TM | KS.3 |
| FIM1_4748 | 4986.703966 | 4697.94249 | 4156.527977 | -0.019700242 | 0.053372384 | 0.387260939 | 0.948637087 | ns | TM | KS.3 |
| FIM1_4749 | 477.3987522 | 420.4136312 | 434.7145254 | -0.042376453 | 0.075637577 | 0.148023379 | 0.780546912 | ns | TM | KS.3 |
| FIM1_4750 | 4399.57867 | 4204.686519 | 4683.332238 | -0.016414336 | 0.044311541 | 0.403555655 | 0.950410534 | ns | TM | KS.3 |
| FIM1_4751 | 155.9926459 | 176.2853322 | 204.7642249 | 0.01831554 | 0.075994822 | 0.436026509 | 0.950410534 | ns | TM | KS.3 |
| FIM1_4753 | 1580.540286 | 1877.119646 | 2190.71695 | 0.02269317 | 0.084074946 | 0.332374766 | 0.938982061 | ns | TM | KS.3 |
| FIM1_4754 | 1884.812138 | 2076.673934 | 1932.410155 | 0.029875293 | 0.066122707 | 0.255280089 | 0.892600401 | ns | TM | KS.3 |
| FIM1_4755 | 413.0236087 | 489.3827703 | 420.6909983 | 0.040743201 | 0.083940639 | 0.153258645 | 0.783447458 | ns | TM | KS.3 |

| | | | | | | | | | | |
|---|---|---|---|---|---|---|---|---|---|---|
| FIM1_4756 | 1561.034882 | 1598.923887 | 1824.271366 | 0.004911022 | 0.060949583 | 0.830186932 | 0.991141351 | ns | TM | KS.3 |
| FIM1_4757 | 1758.461499 | 1856.06079 | 1744.657445 | 0.005862685 | 0.077553877 | 0.781435176 | 0.988814136 | ns | TM | KS.3 |
| FIM1_4758 | 584.0019268 | 504.9752219 | 689.9815069 | -0.054747064 | 0.082335207 | 0.089760512 | 0.654777926 | ns | TM | KS.3 |
| FIM1_4759 | 5627.712326 | 5139.527196 | 6178.264648 | -0.01860016 | 0.068569334 | 0.444489738 | 0.950410534 | ns | TM | KS.3 |
| FIM1_4760 | 2797.170872 | 3643.815209 | 4423.256503 | 0.028753267 | 0.098688062 | 0.222024844 | 0.870513238 | ns | TM | KS.3 |
| FIM1_4761 | 303.4271407 | 357.5909355 | 374.2602876 | 0.033984273 | 0.081568232 | 0.20579307 | 0.85037734 | ns | TM | KS.3 |
| FIM1_4762 | 85.12073544 | 107.8118977 | 127.7586392 | 0.046439126 | 0.09760054 | 0.10867311 | 0.705336441 | ns | TM | KS.3 |
| FIM1_4763 | 126.0467296 | 138.8009801 | 172.8569932 | 0.011493194 | 0.077841095 | 0.598917749 | 0.968980748 | ns | TM | KS.3 |
| FIM1_4764 | 301.8854659 | 309.6380251 | 328.2736609 | 0.002693029 | 0.077254315 | 0.897764427 | 0.991141351 | ns | TM | KS.3 |
| FIM1_4765 | 356.5032561 | 295.7467875 | 314.2749395 | -0.023799483 | 0.086394133 | 0.309086975 | 0.927416591 | ns | TM | KS.3 |
| FIM1_4766 | 218.0176755 | 177.8501632 | 195.8864339 | -0.035251805 | 0.088989074 | 0.181939903 | 0.818789693 | ns | TM | KS.3 |
| FIM1_4767 | 752.1006777 | 683.0611122 | 816.4770033 | -0.021602062 | 0.068432671 | 0.385154632 | 0.948637087 | ns | TM | KS.3 |
| FIM1_4769 | 74.86082086 | 82.97334182 | 118.1575199 | 0.007743951 | 0.086552535 | 0.694352636 | 0.981425301 | ns | TM | KS.3 |
| FIM1_4770 | 559.1417511 | 491.1132499 | 529.218885 | -0.008028849 | 0.092807162 | 0.670714403 | 0.976861203 | ns | TM | KS.3 |
| FIM1_4771 | 770.2994102 | 584.7756215 | 746.2717156 | -0.060006799 | 0.110979178 | 0.061421592 | 0.545665039 | ns | TM | KS.3 |
| FIM1_4772 | 56.41791853 | 47.18950512 | 75.1073442 | -0.019403349 | 0.083360503 | 0.391725552 | 0.948637087 | ns | TM | KS.3 |
| FIM1_4773 | 4867.857103 | 4205.70312 | 2745.278261 | -0.032403301 | 0.078332851 | 0.225649465 | 0.876782875 | ns | TM | KS.3 |

**TF_targets**

| TF | Targets | w303_id | w303_annotation | Ambiguous gene model | km.gene | Orthogroup | Ortholog_gene_length_ratio | DEG.KSR1 | DEG.KSR3 |
|---|---|---|---|---|---|---|---|---|---|
| Oaf1 | YBR126C | W303.asm02.HP0_( | YBR126C | n | FIM1_5102 | OG0000677 | 0.98019802 | Km<KS.1 | Km>KS.3 |
| Oaf1 | YBR135W | W303.asm02.HP0_( | YBR135W | n | FIM1_4519 | OG0001275 | 1.136363636 | ns | ns |
| Oaf1 | YBR159W | W303.asm02.HP0_( | YBR159W | n | FIM1_2050 | OG0001770 | 1.002890173 | ns | ns |
| Oaf1 | YCR010C | W303.asm02.HP0_( | YCR010C | n | FIM1_859 | OG0000037 | 1.025362319 | Km<KS.1 | ns |
| Oaf1 | YDL110C | W303.asm02.HP0_( | YDL110C | n | FIM1_4871 | OG0004239 | 1 | Km<KS.1 | ns |
| Oaf1 | YDL070W | W303.asm02.HP0_( | YDL070W | n | NA | NA | NA | NA | NA |
| Oaf1 | YDL037C | NA | NA | NA | NA | NA | NA | NA | NA |
| Oaf1 | YDR072C | W303.asm02.HP0_( | YDR072C | n | FIM1_876 | OG0003617 | 0.966972477 | ns | ns |
| Oaf1 | YDR077W | W303.asm02.HP0_( | YDR077W | n | FIM1_1283 | OG0000029 | 1.475982533 | Km<KS.1 | ns |
| Oaf1 | YDR207C | W303.asm02.HP0_( | YDR207C | n | FIM1_2661 | OG0000014 | 0.896995708 | ns | Km>KS.3 |
| Oaf1 | YDR244W | W303.asm02.HP0_( | YDR244W | n | FIM1_2212 | OG0001744 | 1.071803853 | ns | ns |
| Oaf1 | YDR256C | W303.asm02.HP0_( | YDR256C | n | FIM1_3892 | OG0000841 | 1.007827789 | ns | Km>KS.3 |
| Oaf1 | YDR384C | W303.asm02.HP0_( | YDR384C | n | FIM1_1442 | OG0000037 | 1.018518519 | Km<KS.1 | Km>KS.3 |
| Oaf1 | YEL060C | W303.asm02.HP0_( | YEL060C | n | FIM1_4811 | OG0000749 | 1.148282098 | ns | Km>KS.3 |
| Oaf1 | YEL011W | W303.asm02.HP0_( | YEL011W | n | FIM1_292 | OG0002853 | 0.981868898 | Km<KS.1 | Km>KS.3 |
| Oaf1 | YER015W | W303.asm02.HP0_( | YER015W | n | FIM1_4620 | OG0000315 | 0.989361702 | ns | ns |
| Oaf1 | YER065C | W303.asm02.HP0_( | YER065C | n | FIM1_4484 | OG0001550 | 1.005415162 | Km<KS.1 | ns |
| Oaf1 | YGL205W | W303.asm02.HP0_( | YGL205W | n | FIM1_1120 | OG0000496 | 1.017687075 | Km<KS.1 | ns |
| Oaf1 | YGL096W | W303.asm02.HP0_( | YGL096W | n | NA | NA | NA | NA | NA |
| Oaf1 | YGL045W | W303.asm02.HP0_( | YGL045W | n | FIM1_2083 | OG0002817 | 0.87987013 | ns | ns |
| Oaf1 | YGR035C | W303.asm02.HP0_( | YGR035C | n | NA | NA | NA | NA | NA |
| Oaf1 | YGR052W | W303.asm02.HP0_( | YGR052W | n | FIM1_2330 | OG0004098 | 0.884892086 | ns | ns |
| Oaf1 | YGR117C | W303.asm02.HP0_( | YGR117C | n | FIM1_4605 | OG0002778 | 1.00845666 | ns | ns |
| Oaf1 | YGR161C | W303.asm02.HP0_( | YGR161C | n | FIM1_2079 | OG0002752 | 0.859477124 | ns | ns |
| Oaf1 | YHL027W | W303.asm02.HP0_( | YHL027W | n | FIM1_1295 | OG0000187 | 1.560081466 | ns | Km>KS.3 |
| Oaf1 | YHR096C | W303.asm02.HP0_( | YHR096C | n | FIM1_3458 | OG0000000 | 1.038596491 | ns | Km>KS.3 |
| Oaf1 | YHR160C | W303.asm02.HP0_( | YHR160C | n | NA | NA | NA | NA | NA |
| Oaf1 | YIL160C | W303.asm02.HP0_( | YIL160C | n | FIM1_1077 | OG0002755 | 1.01459854 | ns | Km>KS.3 |
| Oaf1 | YIL120W | W303.asm02.HP0_( | YIL120W | n | NA | NA | NA | NA | NA |
| Oaf1 | YIL118W | W303.asm02.HP0_( | YIL118W | n | FIM1_4950 | OG0001023 | 1.017621145 | ns | ns |
| Oaf1 | YJL218W | W303.asm02.HP0_( | YJL218W | n | FIM1_1582 | OG0002280 | 1.037037037 | Km<KS.1 | ns |
| Oaf1 | YJL196C | W303.asm02.HP0_( | YJL196C | n | NA | NA | NA | NA | NA |
| Oaf1 | YJL153C | W303.asm02.HP0_( | YJL153C | n | FIM1_3581 | OG0003867 | 1.005660377 | Km>KS.1 | Km<KS.3 |
| Oaf1 | YJR019C | W303.asm02.HP0_( | YJR019C | n | FIM1_4216 | OG0000250 | 1.357771261 | ns | ns |
| Oaf1 | YJR071W | NA | NA | NA | NA | NA | NA | NA | NA |
| Oaf1 | YJR113C | W303.asm02.HP0_( | YJR113C | n | FIM1_4401 | OG0003400 | 0.875886525 | Km>KS.1 | ns |
| Oaf1 | YKL201C | W303.asm02.HP0_( | YKL201C | n | NA | NA | NA | NA | NA |
| Oaf1 | YKL188C | W303.asm02.HP0_( | YKL188C | n | FIM1_2992 | OG0002336 | 1.044063647 | ns | ns |
| Oaf1 | YKL096W-A | NA | NA | NA | NA | NA | NA | NA | NA |
| Oaf1 | YKL087C | W303.asm02.HP0_( | YKL087C | n | FIM1_4128 | OG0003507 | 0.832713755 | ns | ns |
| Oaf1 | YKL085W | W303.asm02.HP0_( | YKL085W | n | FIM1_4125 | OG0000053 | 0.98816568 | ns | Km>KS.3 |
| Oaf1 | YKL052C | W303.asm02.HP0_( | YKL052C | n | FIM1_5044 | OG0003542 | 1.20661157 | ns | Km>KS.3 |
| Oaf1 | YKR009C | W303.asm02.HP0_( | YKR009C | n | FIM1_4734 | OG0001189 | 0.969827586 | Km<KS.1 | Km>KS.3 |
| Oaf1 | YKR080W | W303.asm02.HP0_( | YKR080W | n | NA | NA | NA | NA | NA |

| | | | | | | | | | | |
|---|---|---|---|---|---|---|---|---|---|---|
| Oaf1 | YKR093W | W303.asm02.HP0_( | YKR093W | n | FIM1_4685 | OG0000627 | | 0.949447077 | ns | ns |
| Oaf1 | YKR097W | W303.asm02.HP0_( | YKR097W | n | FIM1_2428 | OG0002754 | | 1.011049724 | Km<KS.1 | ns |
| Oaf1 | YLR284C | W303.asm02.HP0_( | YLR284C | n | FIM1_648 | OG0000262 | | 1 | ns | ns |
| Oaf1 | YLR285W | W303.asm02.HP0_( | YLR285W | n | FIM1_647 | OG0002090 | | 0.966666667 | ns | ns |
| Oaf1 | YML128C | W303.asm02.HP0_( | YML128C | n | FIM1_1516 | OG0002351 | | 1.059917355 | Km<KS.1 | ns |
| Oaf1 | YML063W | W303.asm02.HP0_( | YLR441C/YML063W | y | FIM1_3262 | OG0000552 | | 1 | ns | ns |
| Oaf1 | YML007W | W303.asm02.HP0_( | YML007W | n | FIM1_2371 | OG0000555 | | 1.063829787 | ns | Km>KS.3 |
| Oaf1 | YML004C | W303.asm02.HP0_( | YML004C | n | FIM1_3822 | OG0002805 | | 0.964497041 | ns | ns |
| Oaf1 | YMR009W | W303.asm02.HP0_( | YMR009W | n | FIM1_3506 | OG0001851 | | 1.005617978 | ns | ns |
| Oaf1 | YMR017W | W303.asm02.HP0_( | YMR017W | n | NA | NA | NA | | NA | NA |
| Oaf1 | YMR018W | W303.asm02.HP0_( | YMR018W | n | NA | NA | NA | | NA | NA |
| Oaf1 | YMR102C | W303.asm02.HP0_( | YMR102C | n | FIM1_1798 | OG0000449 | | 0.966396292 | ns | Km>KS.3 |
| Oaf1 | YMR135C | W303.asm02.HP0_( | YMR135C | n | FIM1_2506 | OG0002491 | | 1.261111111 | ns | ns |
| Oaf1 | YMR210W | W303.asm02.HP0_( | YMR210W | n | FIM1_4326 | OG0001716 | | 1.025114155 | Km>KS.1 | ns |
| Oaf1 | YMR272C | W303.asm02.HP0_( | YMR272C | n | FIM1_263 | OG0002495 | | 1.007874016 | Km<KS.1 | ns |
| Oaf1 | YMR303C | W303.asm02.HP0_( | YMR303C | n | NA | NA | NA | | NA | NA |
| Oaf1 | YMR316W | NA | NA | NA | NA | NA | NA | | NA | NA |
| Oaf1 | YNL202W | W303.asm02.HP0_( | YNL202W | n | FIM1_3674 | OG0000700 | | 1 | Km<KS.1 | ns |
| Oaf1 | YNL117W | W303.asm02.HP0_( | YNL117W | n | FIM1_4053 | OG0000576 | | 0.998198198 | ns | Km>KS.3 |
| Oaf1 | YNL065W | W303.asm02.HP0_( | YNL065W | n | FIM1_4956 | OG0000101 | | 1.069343066 | Km<KS.1 | Km>KS.3 |
| Oaf1 | YNL009W | W303.asm02.HP0_( | YNL009W | n | NA | NA | NA | | NA | NA |
| Oaf1 | YNR018W | W303.asm02.HP0_( | YNR018W | n | FIM1_3811 | OG0000686 | | 1.013574661 | Km<KS.1 | ns |
| Oaf1 | YOL147C | W303.asm02.HP0_( | YOL147C | n | FIM1_1490 | OG0000688 | | 1.008547009 | Km>KS.1 | ns |
| Oaf1 | YOL126C | W303.asm02.HP0_( | YOL126C | n | FIM1_2847 | OG0000053 | | 1.00802139 | ns | ns |
| Oaf1 | YOL111C | W303.asm02.HP0_( | YOL111C | n | FIM1_444 | OG0003967 | | 0.986046512 | ns | ns |
| Oaf1 | YOL101C | W303.asm02.HP0_( | YOL101C | n | NA | NA | NA | | NA | NA |
| Oaf1 | YOL084W | W303.asm02.HP0_( | YOL084W | n | FIM1_2096 | OG0000193 | | 1.026943005 | ns | Km>KS.3 |
| Oaf1 | YOL083W | W303.asm02.HP0_( | YOL083W | n | FIM1_2095 | OG0000507 | | 0.710344828 | ns | ns |
| Oaf1 | YOL002C | W303.asm02.HP0_( | YOL002C | n | FIM1_4825 | OG0000223 | | 1.068627451 | Km<KS.1 | ns |
| Oaf1 | YOR049C | W303.asm02.HP0_( | YOR049C | n | FIM1_1945 | OG0000299 | | 0.894613583 | ns | ns |
| Oaf1 | YOR084W | W303.asm02.HP0_( | YOR084W | n | FIM1_4693 | OG0001202 | | 0.943902439 | ns | ns |
| Oaf1 | YOR100C | W303.asm02.HP0_( | YOR100C | n | FIM1_5009 | OG0002705 | | 0.947826087 | Km<KS.1 | ns |
| Oaf1 | YOR101W | W303.asm02.HP0_( | YOR101W | n | NA | NA | NA | | NA | NA |
| Oaf1 | YOR153W | W303.asm02.HP0_( | YOR153W | n | FIM1_2701 | OG0000009 | | 0.997359736 | ns | ns |
| Oaf1 | YOR180C | W303.asm02.HP0_( | YOR180C | n | NA | NA | NA | | NA | NA |
| Oaf1 | YOR220W | W303.asm02.HP0_( | YOR220W | n | FIM1_1656 | OG0003242 | | 1.039215686 | ns | ns |
| Oaf1 | YOR273C | W303.asm02.HP0_( | YOR273C | n | FIM1_2879 | OG0002506 | | 1.083881579 | ns | ns |
| Oaf1 | YOR302W | NA | NA | NA | NA | NA | NA | | NA | NA |
| Oaf1 | YOR317W | W303.asm02.HP0_( | YOR317W | n | FIM1_1792 | OG0000522 | | 0.998573466 | ns | ns |
| Oaf1 | YOR342C | W303.asm02.HP0_( | YOR342C | n | FIM1_4083 | OG0000575 | | 1.015923567 | Km<KS.1 | ns |
| Oaf1 | YOR348C | W303.asm02.HP0_( | YOR348C | n | FIM1_4075 | OG0000002 | | 1.094240838 | Km>KS.1 | Km>KS.3 |
| Oaf1 | YOR363C | W303.asm02.HP0_( | YOR363C | n | NA | NA | NA | | NA | NA |
| Oaf1 | YPL177C | W303.asm02.HP0_( | YPL177C | n | FIM1_2023 | OG0000548 | | 0.690744921 | ns | ns |
| Oaf1 | YPL112C | W303.asm02.HP0_( | YPL112C | n | FIM1_3119 | OG0000851 | | 0.992443325 | Km>KS.1 | ns |
| Oaf1 | YPL095C | W303.asm02.HP0_( | YPL095C | n | NA | NA | NA | | NA | NA |
| Oaf1 | YPL016W | W303.asm02.HP0_( | YPL016W | n | FIM1_3917 | OG0001751 | | 1.086848635 | ns | ns |

| | | | | | | | | | | |
|---|---|---|---|---|---|---|---|---|---|---|
| Oaf1 | YPR128C | W303.asm02.HP0_( | YPR128C | n | FIM1_4608 | OG0002777 | | 0.913649025 | ns | ns |
| Oaf1 | YAR035W | W303.asm02.HP0_( | YAR035W | n | FIM1_673 | OG0003793 | | 0.984240688 | Km<KS.1 | Km>KS.3 |
| Oaf1 | YAR071W | NA | NA | NA | NA | NA | NA | | NA | NA |
| Oaf1 | YBR053C | W303.asm02.HP0_( | YBR053C | n | FIM1_3739 | OG0003117 | | 1.06547619 | ns | ns |
| Oaf1 | YBR072W | W303.asm02.HP0_( | YBR072W | n | FIM1_2584 | OG0004415 | | 1.10880829 | Km<KS.1 | ns |
| Oaf1 | YJL076W | W303.asm02.HP0_( | YJL076W | n | NA | NA | NA | | NA | NA |
| Oaf1 | YJL044C | W303.asm02.HP0_( | YJL044C | n | FIM1_950 | OG0001452 | | 1.491856678 | ns | Km>KS.3 |
| Kar4 | YCL055W | W303.asm02.HP0_( | YCL055W | n | FIM1_826 | OG0001765 | | 1.037151703 | ns | ns |
| Kar4 | YCL027W | W303.asm02.HP0_( | YCL027W | n | FIM1_790 | OG0001691 | | 0.975238095 | ns | Km>KS.3 |
| Kar4 | YCR089W | W303.asm02.HP0_( | YCR089W | n | NA | NA | NA | | NA | NA |
| Kar4 | YDL127W | W303.asm02.HP0_( | YDL127W | n | FIM1_3798 | OG0000683 | | 0.922155689 | ns | ns |
| Kar4 | YDR309C | W303.asm02.HP0_( | YDR309C | n | FIM1_2241 | OG0003681 | | 1.026809651 | Km>KS.1 | ns |
| Kar4 | YER164W | W303.asm02.HP0_( | YER164W | n | FIM1_1995 | OG0003297 | | 0.938019169 | ns | Km>KS.3 |
| Kar4 | YFL026W | W303.asm02.HP0_( | YFL026W | n | FIM1_3999 | OG0001477 | | 0.995381062 | ns | ns |
| Kar4 | YGL028C | W303.asm02.HP0_( | YGL028C | n | FIM1_901 | OG0003345 | | 1.054474708 | ns | ns |
| Kar4 | YIL117C | W303.asm02.HP0_( | YIL117C | n | NA | NA | NA | | NA | NA |
| Kar4 | YIL102C | W303.asm02.HP0_( | YIL102C | n | NA | NA | NA | | NA | NA |
| Kar4 | YIL101C | W303.asm02.HP0_( | YIL101C | n | FIM1_4925 | OG0000649 | | 2.246527778 | Km<KS.1 | ns |
| Kar4 | YIL015W | W303.asm02.HP0_( | YIL015W | n | FIM1_3559 | OG0000015 | | 1.171656687 | ns | ns |
| Kar4 | YJL162C | W303.asm02.HP0_( | YJL162C | n | FIM1_3341 | OG0004168 | | 0.898305085 | Km>KS.1 | ns |
| Kar4 | YJL108C | NA | NA | NA | NA | NA | NA | | NA | NA |
| Kar4 | YJL107C | W303.asm02.HP0_( | YJL107C | n | FIM1_2274 | OG0003900 | | 0.399380805 | ns | ns |
| Kar4 | YKL127W | W303.asm02.HP0_( | YKL127W | n | NA | NA | NA | | NA | NA |
| Kar4 | YKR041W | W303.asm02.HP0_( | YKR041W | n | NA | NA | NA | | NA | NA |
| Kar4 | YKR042W | W303.asm02.HP0_( | YKR042W | n | FIM1_4286 | OG0000063 | | 0.921717172 | ns | ns |
| Kar4 | YML056C | W303.asm02.HP0_( | YML056C | n | FIM1_3275 | OG0000352 | | 3.402597403 | ns | ns |
| Kar4 | YMR198W | W303.asm02.HP0_( | YMR198W | n | NA | NA | NA | | NA | NA |
| Kar4 | YNL279W | W303.asm02.HP0_( | YNL279W | n | FIM1_772 | OG0001826 | | 1.024806202 | ns | Km>KS.3 |
| Kar4 | YNL146W | W303.asm02.HP0_( | YNL146W | n | NA | NA | NA | | NA | NA |
| Kar4 | YNR044W | W303.asm02.HP0_( | YNR044W | n | FIM1_1682 | OG0000026 | | 0.60720268 | ns | ns |
| Kar4 | YOL109W | W303.asm02.HP0_( | YOL109W | n | NA | NA | NA | | NA | NA |
| Kar4 | TOS7 | NA | NA | NA | NA | NA | NA | | NA | NA |
| Kar4 | YPL192C | W303.asm02.HP0_( | YPL192C | n | NA | NA | NA | | NA | NA |
| Kar4 | YPL156C | W303.asm02.HP0_( | YPL156C | n | FIM1_4329 | OG0004264 | | 0.975945017 | ns | ns |
| Kar4 | YPR141C | W303.asm02.HP0_( | YPR141C | n | FIM1_4590 | OG0001893 | | 1.059593023 | ns | Km<KS.3 |
| Kar4 | YPR201W | W303.asm02.HP0_( | YPR201W | n | FIM1_5202 | OG0003827 | | 0.955082742 | ns | ns |
| Kar4 | YBL016W | W303.asm02.HP0_( | YBL016W | n | FIM1_892 | OG0000190 | | 0.964480874 | ns | ns |
| Kar4 | YBR040W | W303.asm02.HP0_( | YBR040W | n | FIM1_3325 | OG0001920 | | 1 | NA | NA |
| Kar4 | YBR083W | W303.asm02.HP0_( | YBR083W | n | FIM1_4666 | OG0000926 | | 0.722139673 | ns | Km>KS.3 |
| Kar4 | YBR088C | W303.asm02.HP0_( | YBR088C | n | FIM1_4642 | OG0003523 | | 1 | Km>KS.1 | ns |
| Hcm1 | YEL061C | W303.asm02.HP0_( | YEL061C | n | FIM1_4814 | OG0000253 | | 1.004016064 | ns | ns |
| Hcm1 | YGL033W | W303.asm02.HP0_( | YGL033W | n | NA | NA | NA | | NA | NA |
| Hcm1 | YIL131C | W303.asm02.HP0_( | YIL131C | n | NA | NA | NA | | NA | NA |
| Hcm1 | YLR004C | W303.asm02.HP0_( | YLR004C | n | FIM1_4857 | OG0001430 | | 1.015533981 | Km<KS.1 | Km>KS.3 |
| Hcm1 | YML085C | W303.asm02.HP0_( | YML085C | n | FIM1_4567 | OG0000364 | | 1.839506173 | ns | ns |
| Hcm1 | YMR202W | W303.asm02.HP0_( | YMR202W | n | FIM1_3828 | OG0002793 | | 0.995515695 | ns | ns |

| | | | | | | | | | | |
|---|---|---|---|---|---|---|---|---|---|---|
| Hcm1 | YMR250W | W303.asm02.HP0_( | YMR250W | n | FIM1_2257 | OG0001690 | | 1.031746032 | Km<KS.1 | Km>KS.3 |
| Hcm1 | YNL068C | W303.asm02.HP0_( | YNL068C | n | FIM1_3754 | OG0000287 | | 1.062885327 | ns | ns |
| Hcm1 | YNL013C | NA | NA | NA | NA | NA | NA | | NA | NA |
| Hcm1 | YNR076W | W303.asm02.HP0_( | YAL068C/YBR301W/ | y | NA | NA | NA | | NA | NA |
| Hcm1 | YOL164W | W303.asm02.HP0_( | YOL164W | n | NA | NA | NA | | NA | NA |
| Hcm1 | YOR070C | W303.asm02.HP0_( | YOR070C | n | FIM1_422 | OG0000805 | | 1.329853862 | ns | ns |
| Hcm1 | YOR077W | W303.asm02.HP0_( | YOR077W | n | FIM1_436 | OG0002494 | | 0.970711297 | ns | ns |
| Hcm1 | YOR372C | W303.asm02.HP0_( | YOR372C | n | FIM1_3104 | OG0004157 | | 0.994614004 | ns | ns |
| Hcm1 | YPL171C | W303.asm02.HP0_( | YPL171C | n | NA | NA | NA | | NA | NA |
| Hcm1 | YBL032W | W303.asm02.HP0_( | YBL032W | n | FIM1_4656 | OG0000854 | | 1.026954178 | ns | Km>KS.3 |
| Tup1 | YAL064W | W303.asm02.HP0_( | YAL064W | n | NA | NA | NA | | NA | NA |
| Tup1 | YAL063C | W303.asm02.HP0_( | YAL063C | n | FIM1_4786 | OG0000010 | | 1.156605424 | ns | Km>KS.3 |
| Tup1 | YBR092C | W303.asm02.HP0_( | YBR092C | n | NA | NA | NA | | NA | NA |
| Tup1 | YBR093C | W303.asm02.HP0_( | YBR093C | n | NA | NA | NA | | NA | NA |
| Tup1 | YBR118W | NA | NA | NA | NA | NA | NA | | NA | NA |
| Tup1 | YBR158W | W303.asm02.HP0_( | YBR158W | n | FIM1_2048 | OG0003918 | | 1.27972028 | ns | ns |
| Tup1 | YBR181C | NA | NA | NA | NA | NA | NA | | NA | NA |
| Tup1 | YBR182C | W303.asm02.HP0_( | YBR182C | n | NA | NA | NA | | NA | NA |
| Tup1 | YBR238C | W303.asm02.HP0_( | YBR238C | n | FIM1_1097 | OG0000465 | | 1.071847507 | ns | ns |
| Tup1 | YBR240C | W303.asm02.HP0_( | YBR240C | n | FIM1_1101 | OG0004016 | | 1.229508197 | ns | ns |
| Tup1 | YCL025C | W303.asm02.HP0_( | YCL025C | n | FIM1_787 | OG0000001 | | 1.024271845 | ns | ns |
| Tup1 | YDL228C | NA | NA | NA | NA | NA | NA | | NA | NA |
| Tup1 | YDL227C | W303.asm02.HP0_( | YDL227C | n | NA | NA | NA | | NA | NA |
| Tup1 | YDL185W | W303.asm02.HP0_( | YDL185W | n | FIM1_3788 | OG0001045 | | 1.735818476 | ns | ns |
| Tup1 | YDL037C | NA | NA | NA | NA | NA | NA | | NA | NA |
| Tup1 | YDL022W | W303.asm02.HP0_( | YDL022W | n | FIM1_680 | OG0000313 | | 0.900921659 | Km<KS.1 | ns |
| Tup1 | YDR009W | W303.asm02.HP0_( | YDR009W | n | FIM1_1190 | OG0000311 | | 1.027667984 | Km<KS.1 | Km>KS.3 |
| Tup1 | YDR041W | W303.asm02.HP0_( | YDR041W | n | FIM1_2574 | OG0002975 | | 0.995098039 | ns | ns |
| Tup1 | YDR043C | W303.asm02.HP0_( | YDR043C | n | FIM1_2570 | OG0000451 | | 0.620967742 | Km<KS.1 | ns |
| Tup1 | YDR072C | W303.asm02.HP0_( | YDR072C | n | FIM1_876 | OG0003617 | | 0.966972477 | ns | ns |
| Tup1 | CCW22 | NA | NA | NA | NA | NA | NA | | NA | NA |
| Tup1 | YDR186C | W303.asm02.HP0_( | YDR186C | n | FIM1_3651 | OG0004194 | | 1.274709302 | Km<KS.1 | ns |
| Tup1 | YDR215C | W303.asm02.HP0_( | W303.asm02.HP0_G( | y | NA | NA | NA | | NA | NA |
| Tup1 | YDR258C | W303.asm02.HP0_( | YDR258C | n | FIM1_4197 | OG0003204 | | 0.970095694 | Km>KS.1 | ns |
| Tup1 | YDR259C | W303.asm02.HP0_( | YDR259C | n | NA | NA | NA | | NA | NA |
| Tup1 | YDR311W | W303.asm02.HP0_( | YDR311W | n | FIM1_2247 | OG0002458 | | 1.038834951 | ns | ns |
| Tup1 | YDR342C | NA | NA | NA | NA | NA | NA | | NA | NA |
| Tup1 | YDR343C | NA | NA | NA | NA | NA | NA | | NA | NA |
| Tup1 | YDR441C | W303.asm02.HP0_( | YDR441C | n | NA | NA | NA | | NA | NA |
| Tup1 | YDR449C | W303.asm02.HP0_( | YDR449C | n | FIM1_3040 | OG0001862 | | 0.977777778 | ns | Km<KS.3 |
| Tup1 | YDR452W | W303.asm02.HP0_( | YDR452W | n | FIM1_3036 | OG0003045 | | 0.972582973 | ns | ns |
| Tup1 | YDR508C | W303.asm02.HP0_( | YDR508C | n | NA | NA | NA | | NA | NA |
| Tup1 | YDR525W | NA | NA | NA | NA | NA | NA | | NA | NA |
| Tup1 | YEL060C | W303.asm02.HP0_( | YEL060C | n | FIM1_4811 | OG0000749 | | 1.148282098 | ns | Km>KS.3 |
| Tup1 | YEL046C | W303.asm02.HP0_( | YEL046C | n | FIM1_1973 | OG0000654 | | 0.989769821 | ns | Km>KS.3 |
| Tup1 | YEL045C | NA | NA | NA | NA | NA | NA | | NA | NA |

| | | | | | | | | | | |
|---|---|---|---|---|---|---|---|---|---|---|
| Tup1 | YEL040W | W303.asm02.HP0_( | YEL040W | n | FIM1_1980 | OG0000241 | | 0.993617021 | Km>KS.1 | ns |
| Tup1 | YEL034W | NA | NA | NA | NA | NA | NA | | NA | NA |
| Tup1 | YEL007W | W303.asm02.HP0_( | YEL007W | n | NA | NA | NA | | NA | NA |
| Tup1 | YER011W | W303.asm02.HP0_( | W303.asm02.HP0_G( | y | NA | NA | NA | | NA | NA |
| Tup1 | YER036C | W303.asm02.HP0_( | YER036C | n | FIM1_2384 | OG0002251 | | 1.004942339 | ns | ns |
| Tup1 | YER055C | W303.asm02.HP0_( | YER055C | n | FIM1_2794 | OG0003405 | | 3.030612245 | ns | ns |
| Tup1 | YER056C | W303.asm02.HP0_( | YER056C | n | FIM1_4493 | OG0000068 | | 1.026974952 | ns | Km>KS.3 |
| Tup1 | YER145C | NA | NA | NA | NA | NA | NA | | NA | NA |
| Tup1 | YER158C | W303.asm02.HP0_( | YER158C | n | NA | NA | NA | | NA | NA |
| Tup1 | YFL054C | NA | NA | NA | NA | NA | NA | | NA | NA |
| Tup1 | ZNF1 | NA | NA | NA | NA | NA | NA | | NA | NA |
| Tup1 | YFL051C | NA | NA | NA | NA | NA | NA | | NA | NA |
| Tup1 | YFL026W | W303.asm02.HP0_( | YFL026W | n | FIM1_3999 | OG0001477 | | 0.995381062 | ns | ns |
| Tup1 | YFL022C | W303.asm02.HP0_( | YFL022C | n | FIM1_3993 | OG0003902 | | 1.008016032 | ns | ns |
| Tup1 | YFL021W | W303.asm02.HP0_( | YFL021W | n | FIM1_3990 | OG0000035 | | 0.672823219 | ns | ns |
| Tup1 | YFL016C | W303.asm02.HP0_( | YFL016C | n | FIM1_1627 | OG0001409 | | 1.096566524 | Km>KS.1 | ns |
| Tup1 | YFL015C | W303.asm02.HP0_( | W303.asm02.HP0_G( | y | NA | NA | NA | | NA | NA |
| Tup1 | YFL014W | W303.asm02.HP0_( | YFL014W | n | FIM1_1628 | OG0001024 | | 1.048076923 | Km<KS.1 | ns |
| Tup1 | YFR032C | W303.asm02.HP0_( | YFR032C | n | FIM1_3565 | OG0000118 | | 0.598343685 | ns | Km>KS.3 |
| Tup1 | YGL253W | W303.asm02.HP0_( | YGL253W | n | FIM1_3879 | OG0000486 | | 1.002061856 | ns | Km>KS.3 |
| Tup1 | YGL158W | W303.asm02.HP0_( | YGL158W | n | NA | NA | NA | | NA | NA |
| Tup1 | YGL123W | W303.asm02.HP0_( | YGL123W | n | FIM1_1125 | OG0002666 | | 0.988326848 | ns | ns |
| Tup1 | YGL103W | NA | NA | NA | NA | NA | NA | | NA | NA |
| Tup1 | YGL097W | W303.asm02.HP0_( | YGL097W | n | FIM1_1270 | OG0002453 | | 1.006263048 | ns | ns |
| Tup1 | YGL055W | W303.asm02.HP0_( | YGL055W | n | FIM1_619 | OG0000750 | | 1.06918239 | Km<KS.1 | ns |
| Tup1 | YGL030W | W303.asm02.HP0_( | YGL030W | n | FIM1_906 | OG0001085 | | 0.898305085 | ns | ns |
| Tup1 | YGL008C | W303.asm02.HP0_( | YGL008C | n | FIM1_384 | OG0000490 | | 1.021134594 | ns | ns |
| Tup1 | YGR013W | W303.asm02.HP0_( | YGR013W | n | FIM1_2494 | OG0003469 | | 1.083916084 | ns | ns |
| Tup1 | YGR033C | W303.asm02.HP0_( | YGR033C | n | FIM1_3286 | OG0001821 | | 1.004201681 | ns | ns |
| Tup1 | YGR036C | W303.asm02.HP0_( | YGR036C | n | FIM1_3283 | OG0001815 | | 1.034632035 | ns | ns |
| Tup1 | YGR040W | W303.asm02.HP0_( | YGR040W | n | FIM1_2340 | OG0000190 | | 1.016574586 | ns | ns |
| Tup1 | YGR041W | W303.asm02.HP0_( | YGR041W | n | FIM1_2338 | OG0000947 | | 1.035984848 | ns | ns |
| Tup1 | YGR052W | W303.asm02.HP0_( | YGR052W | n | FIM1_2330 | OG0004098 | | 0.884892086 | ns | ns |
| Tup1 | YGR060W | W303.asm02.HP0_( | YGR060W | n | FIM1_3375 | OG0000410 | | 0.974763407 | ns | ns |
| Tup1 | YGR085C | W303.asm02.HP0_( | YGR085C/YPR102C | y | FIM1_1196 | OG0000607 | | 1 | ns | ns |
| Tup1 | YGR086C | W303.asm02.HP0_( | YGR086C | n | FIM1_1200 | OG0000121 | | 1.100649351 | ns | Km>KS.3 |
| Tup1 | YGR108W | W303.asm02.HP0_( | YGR108W | n | NA | NA | NA | | NA | NA |
| Tup1 | YGR139W | NA | NA | NA | NA | NA | NA | | NA | NA |
| Tup1 | YGR143W | W303.asm02.HP0_( | YGR143W | n | NA | NA | NA | | NA | NA |
| Tup1 | YGR180C | W303.asm02.HP0_( | YGR180C | n | NA | NA | NA | | NA | NA |
| Tup1 | YGR210C | W303.asm02.HP0_( | YGR210C | n | FIM1_4230 | OG0003644 | | 1.004889976 | ns | Km>KS.3 |
| Tup1 | YGR211W | W303.asm02.HP0_( | YGR211W | n | FIM1_4231 | OG0002655 | | 0.997946612 | ns | ns |
| Tup1 | YGR213C | W303.asm02.HP0_( | YGR213C | n | NA | NA | NA | | NA | NA |
| Tup1 | YGR243W | W303.asm02.HP0_( | YGR243W | n | NA | NA | NA | | NA | NA |
| Tup1 | YGR248W | W303.asm02.HP0_( | YGR248W | n | NA | NA | NA | | NA | NA |
| Tup1 | YGR287C | W303.asm02.HP0_( | YGR287C | n | NA | NA | NA | | NA | NA |

| | | | | | | | | | | |
|---|---|---|---|---|---|---|---|---|---|---|
| Tup1 | YHL033C | W303.asm02.HP0_( | YHL033C/YLL045C | y | NA | NA | NA | | NA | NA |
| Tup1 | YHL029C | W303.asm02.HP0_( | YHL029C | n | FIM1_169 | OG0002978 | | 1.290874525 | Km<KS.1 | ns |
| Tup1 | YHL028W | W303.asm02.HP0_( | YHL028W | n | FIM1_1296 | OG0000050 | | 1.198019802 | ns | Km>KS.3 |
| Tup1 | YHL027W | W303.asm02.HP0_( | YHL027W | n | FIM1_1295 | OG0000187 | | 1.560081466 | ns | Km>KS.3 |
| Tup1 | YHL016C | W303.asm02.HP0_( | YHL016C | n | FIM1_1999 | OG0000309 | | 0.977931034 | ns | Km>KS.3 |
| Tup1 | YHL001W | W303.asm02.HP0_( | YHL001W/YKL006W | y | NA | NA | NA | | NA | NA |
| Tup1 | YHR007C | W303.asm02.HP0_( | YHR007C | n | FIM1_1408 | OG0002417 | | 1.007604563 | Km<KS.1 | ns |
| Tup1 | YHR010W | W303.asm02.HP0_( | YDR471W/YHR010W | y | NA | NA | NA | | NA | NA |
| Tup1 | YHR032W | W303.asm02.HP0_( | YHR032W | n | FIM1_2705 | OG0000065 | | 0.952459016 | ns | ns |
| Tup1 | YHR033W | W303.asm02.HP0_( | YHR033W | n | NA | NA | NA | | NA | NA |
| Tup1 | YHR049W | W303.asm02.HP0_( | YHR049W | n | FIM1_2104 | OG0002961 | | 1.008298755 | ns | Km<KS.3 |
| Tup1 | YHR070W | W303.asm02.HP0_( | YHR070W | n | FIM1_4678 | OG0001174 | | 0.994023904 | ns | ns |
| Tup1 | YHR071W | W303.asm02.HP0_( | YHR071W | n | FIM1_4679 | OG0000561 | | 1.031531532 | ns | ns |
| Tup1 | YHR092C | W303.asm02.HP0_( | YHR092C | n | FIM1_3457 | OG0000000 | | 1.019469027 | ns | Km>KS.3 |
| Tup1 | YHR095W | NA | NA | NA | NA | NA | NA | | NA | NA |
| Tup1 | YHR137W | W303.asm02.HP0_( | YHR137W | n | FIM1_3866 | OG0002367 | | 0.967924528 | ns | ns |
| Tup1 | YHR139C | W303.asm02.HP0_( | YHR139C | n | NA | NA | NA | | NA | NA |
| Tup1 | YHR203C | NA | NA | NA | NA | NA | NA | | NA | NA |
| Tup1 | YHR210C | W303.asm02.HP0_( | YHR210C | n | NA | NA | NA | | NA | NA |
| Tup1 | YHR211W | W303.asm02.HP0_( | YHR211W | n | NA | NA | NA | | NA | NA |
| Tup1 | YHR215W | W303.asm02.HP0_( | YAR071W/YHR215W | y | NA | NA | NA | | NA | NA |
| Tup1 | YIL172C | NA | NA | NA | NA | NA | NA | | NA | NA |
| Tup1 | YIL162W | W303.asm02.HP0_( | YIL162W | n | FIM1_326 | OG0003957 | | 0.956834532 | Km<KS.1 | Km>KS.3 |
| Tup1 | YIL123W | W303.asm02.HP0_( | YIL123W | n | NA | NA | NA | | NA | NA |
| Tup1 | YIL117C | W303.asm02.HP0_( | YIL117C | n | NA | NA | NA | | NA | NA |
| Tup1 | YIL099W | W303.asm02.HP0_( | YIL099W | n | FIM1_4923 | OG0003342 | | 0.922689076 | ns | ns |
| Tup1 | YIL055C | W303.asm02.HP0_( | YIL055C | n | FIM1_4487 | OG0004274 | | 1.208092486 | Km<KS.1 | Km>KS.3 |
| Tup1 | YIL047C | W303.asm02.HP0_( | YIL047C | n | FIM1_2793 | OG0000723 | | 1.066193853 | ns | ns |
| Tup1 | YIR013C | W303.asm02.HP0_( | YIR013C | n | FIM1_1128 | OG0000062 | | 0.47826087 | Km<KS.1 | ns |
| Tup1 | YJL225C | W303.asm02.HP0_( | YPR204W | y | NA | NA | NA | | NA | NA |
| Tup1 | YJL218W | W303.asm02.HP0_( | YJL218W | n | FIM1_1582 | OG0002280 | | 1.037037037 | Km<KS.1 | ns |
| Tup1 | YJL217W | W303.asm02.HP0_( | YJL217W | n | NA | NA | NA | | NA | NA |
| Tup1 | YJL216C | W303.asm02.HP0_( | YJL216C | n | NA | NA | NA | | NA | NA |
| Tup1 | YJR094C | W303.asm02.HP0_( | YJR094C | n | FIM1_4412 | OG0004270 | | 1.049562682 | ns | ns |
| Tup1 | YJR145C | W303.asm02.HP0_( | YHR203C/YJR145C | y | NA | NA | NA | | NA | NA |
| Tup1 | YJR146W | NA | NA | NA | NA | NA | NA | | NA | NA |
| Tup1 | YJR147W | NA | NA | NA | NA | NA | NA | | NA | NA |
| Tup1 | YJR157W | NA | NA | NA | NA | NA | NA | | NA | NA |
| Tup1 | YJR158W | NA | NA | NA | NA | NA | NA | | NA | NA |
| Tup1 | YJR159W | W303.asm02.HP0_( | YDL246C/YJR159W | y | NA | NA | NA | | NA | NA |
| Tup1 | YKL186C | W303.asm02.HP0_( | YKL186C | n | FIM1_2989 | OG0001041 | | 1.051428571 | ns | ns |
| Tup1 | YKL182W | W303.asm02.HP0_( | YKL182W | n | FIM1_2985 | OG0002436 | | 1.001464844 | ns | ns |
| Tup1 | YKL097C | NA | NA | NA | NA | NA | NA | | NA | NA |
| Tup1 | YKL096W | W303.asm02.HP0_( | YKL096W | n | FIM1_3305 | OG0000583 | | 0.952191235 | Km<KS.1 | ns |
| Tup1 | YKL062W | W303.asm02.HP0_( | YKL062W | n | NA | NA | NA | | NA | NA |
| Tup1 | YKL060C | W303.asm02.HP0_( | YKL060C | n | FIM1_2845 | OG0003525 | | 0.994459834 | ns | ns |

| | | | | | | | | | | |
|---|---|---|---|---|---|---|---|---|---|---|
| Tup1 | YKL044W | W303.asm02.HP0_( | YKL044W | n | NA | NA | NA | | NA | NA |
| Tup1 | YKL043W | W303.asm02.HP0_( | YKL043W | n | NA | NA | NA | | NA | NA |
| Tup1 | YKR075C | W303.asm02.HP0_( | YKR075C | n | FIM1_1961 | OG0000438 | | 1.180769231 | Km<KS.1 | Km>KS.3 |
| Tup1 | YKR080W | W303.asm02.HP0_( | YKR080W | n | NA | NA | NA | | NA | NA |
| Tup1 | YKR093W | W303.asm02.HP0_( | YKR093W | n | FIM1_4685 | OG0000627 | | 0.949447077 | ns | ns |
| Tup1 | YKR096W | W303.asm02.HP0_( | YKR096W | n | FIM1_2426 | OG0000924 | | 0.967611336 | Km<KS.1 | ns |
| Tup1 | YKR101W | W303.asm02.HP0_( | YKR101W | n | NA | NA | NA | | NA | NA |
| Tup1 | YKR102W | W303.asm02.HP0_( | YKR102W | n | NA | NA | NA | | NA | NA |
| Tup1 | YLL045C | W303.asm02.HP0_( | YHL033C/YLL045C | y | FIM1_5193 | OG0000283 | | 1.003921569 | ns | ns |
| Tup1 | YLL028W | W303.asm02.HP0_( | YLL028W | n | FIM1_2548 | OG0000764 | | 1.017361111 | ns | Km>KS.3 |
| Tup1 | YLR044C | W303.asm02.HP0_( | YLR044C | n | FIM1_4226 | OG0000024 | | 0.99822695 | ns | Km>KS.3 |
| Tup1 | YLR045C | W303.asm02.HP0_( | YLR045C | n | FIM1_4225 | OG0001387 | | 1.002257336 | ns | Km>KS.3 |
| Tup1 | YLR076C | NA | NA | NA | NA | NA | NA | | NA | NA |
| Tup1 | YLR109W | W303.asm02.HP0_( | YLR109W | n | FIM1_2634 | OG0000322 | | 1.041420118 | ns | ns |
| Tup1 | YLR110C | W303.asm02.HP0_( | W303.asm02.HP0_G( | y | FIM1_2635 | OG0000026 | | 0.528 | ns | ns |
| Tup1 | YLR113W | W303.asm02.HP0_( | YLR113W | n | FIM1_2636 | OG0003206 | | 0.935483871 | ns | ns |
| Tup1 | YLR141W | W303.asm02.HP0_( | YLR141W | n | FIM1_3608 | OG0001600 | | 0.67472119 | ns | ns |
| Tup1 | YLR236C | W303.asm02.HP0_( | W303.asm02.HP0_G( | y | NA | NA | NA | | NA | NA |
| Tup1 | YLR237W | W303.asm02.HP0_( | YLR237W | n | NA | NA | NA | | NA | NA |
| Tup1 | YLR264W | W303.asm02.HP0_( | YLR264W/YOR167C | y | FIM1_2650 | OG0000624 | | 1 | ns | ns |
| Tup1 | YLR267W | W303.asm02.HP0_( | YLR267W | n | NA | NA | NA | | NA | NA |
| Tup1 | YLR279W | NA | NA | NA | NA | NA | NA | | NA | NA |
| Tup1 | RSO55 | NA | NA | NA | NA | NA | NA | | NA | NA |
| Tup1 | YLR287C | W303.asm02.HP0_( | YLR287C | n | FIM1_644 | OG0001383 | | 1.011396011 | ns | ns |
| Tup1 | YLR296W | NA | NA | NA | NA | NA | NA | | NA | NA |
| Tup1 | YLR299W | W303.asm02.HP0_( | YLR299W | n | FIM1_632 | OG0000630 | | 1.092715232 | ns | ns |
| Tup1 | YLR300W | W303.asm02.HP0_( | YLR300W | n | FIM1_629 | OG0000551 | | 1.039443155 | ns | Km>KS.3 |
| Tup1 | YLR302C | NA | NA | NA | NA | NA | NA | | NA | NA |
| Tup1 | YLR304C | W303.asm02.HP0_( | YLR304C | n | FIM1_2005 | OG0000169 | | 0.997435897 | Km>KS.1 | ns |
| Tup1 | ATG39 | NA | NA | NA | NA | NA | NA | | NA | NA |
| Tup1 | YLR332W | W303.asm02.HP0_( | YLR332W | n | FIM1_2311 | OG0000945 | | 1.035812672 | ns | Km>KS.3 |
| Tup1 | YLR342W | W303.asm02.HP0_( | YLR342W | n | FIM1_3288 | OG0000071 | | 0.99154334 | ns | ns |
| Tup1 | YLR402W | NA | NA | NA | NA | NA | NA | | NA | NA |
| Tup1 | YLR413W | W303.asm02.HP0_( | YLR413W | n | NA | NA | NA | | NA | NA |
| Tup1 | YLR441C | W303.asm02.HP0_( | YLR441C/YML063W | y | NA | NA | NA | | NA | NA |
| Tup1 | YLR461W | W303.asm02.HP0_( | YAL068C/YBR301W/ | y | NA | NA | NA | | NA | NA |
| Tup1 | YML131W | W303.asm02.HP0_( | YML131W | n | NA | NA | NA | | NA | NA |
| Tup1 | YML122C | W303.asm02.HP0_( | W303.asm02.HP0_G( | y | NA | NA | NA | | NA | NA |
| Tup1 | YML117W | W303.asm02.HP0_( | YML117W | n | FIM1_4830 | OG0004237 | | 0.979274611 | ns | Km>KS.3 |
| Tup1 | YML116W | W303.asm02.HP0_( | YML116W | n | NA | NA | NA | | NA | NA |
| Tup1 | YML091C | W303.asm02.HP0_( | YML091C | n | FIM1_3839 | OG0001779 | | 0.991749175 | ns | ns |
| Tup1 | YMR006C | W303.asm02.HP0_( | YMR006C | n | NA | NA | NA | | NA | NA |
| Tup1 | YMR007W | NA | NA | NA | NA | NA | NA | | NA | NA |
| Tup1 | YMR011W | W303.asm02.HP0_( | YMR011W | n | NA | NA | NA | | NA | NA |
| Tup1 | YMR016C | W303.asm02.HP0_( | YMR016C | n | FIM1_3763 | OG0000422 | | 1.382716049 | ns | ns |
| Tup1 | YMR017W | W303.asm02.HP0_( | YMR017W | n | NA | NA | NA | | NA | NA |

| | | | | | | | | | | |
|---|---|---|---|---|---|---|---|---|---|---|
| Tup1 | YMR057C | NA | NA | NA | NA | NA | NA | | NA | NA |
| Tup1 | YMR136W | W303.asm02.HP0_( | YMR136W | n | NA | NA | NA | | NA | NA |
| Tup1 | YMR142C | W303.asm02.HP0_( | YDL082W/YMR142C | y | NA | NA | NA | | NA | NA |
| Tup1 | YMR188C | W303.asm02.HP0_( | YMR188C | n | FIM1_3429 | OG0001627 | | 1.021551724 | ns | ns |
| Tup1 | YMR195W | W303.asm02.HP0_( | YMR195W | n | NA | NA | NA | | NA | NA |
| Tup1 | YMR242C | NA | NA | NA | NA | NA | NA | | NA | NA |
| Tup1 | YMR243C | W303.asm02.HP0_( | YMR243C | n | FIM1_1172 | OG0000566 | | 1 | ns | ns |
| Tup1 | YMR265C | W303.asm02.HP0_( | YMR265C | n | FIM1_919 | OG0003673 | | 0.91468254 | ns | Km>KS.3 |
| Tup1 | YMR278W | W303.asm02.HP0_( | YMR278W | n | FIM1_4834 | OG0002969 | | 1.014681892 | ns | Km>KS.3 |
| Tup1 | YMR319C | W303.asm02.HP0_( | YMR319C | n | FIM1_3712 | OG0004202 | | 1.009140768 | Km<KS.1 | ns |
| Tup1 | YNL069C | W303.asm02.HP0_( | YNL069C | n | NA | NA | NA | | NA | NA |
| Tup1 | YNL067W | W303.asm02.HP0_( | YGL147C/YNL067W | y | FIM1_4878 | OG0000240 | | 1 | ns | ns |
| Tup1 | YNL066W | W303.asm02.HP0_( | YNL066W | n | FIM1_3745 | OG0000063 | | 1.042183623 | ns | ns |
| Tup1 | YNL036W | W303.asm02.HP0_( | YNL036W | n | FIM1_4924 | OG0003343 | | 1.01843318 | ns | ns |
| Tup1 | YNR059W | NA | NA | NA | NA | NA | NA | | NA | NA |
| Tup1 | YNR067C | W303.asm02.HP0_( | YNR067C | n | FIM1_1041 | OG0000236 | | 1.104846686 | ns | Km>KS.3 |
| Tup1 | YNR071C | NA | NA | NA | NA | NA | NA | | NA | NA |
| Tup1 | YNR072W | NA | NA | NA | NA | NA | NA | | NA | NA |
| Tup1 | YNR073C | W303.asm02.HP0_( | YEL070W/YNR073C | y | NA | NA | NA | | NA | NA |
| Tup1 | YOL158C | W303.asm02.HP0_( | YOL158C | n | NA | NA | NA | | NA | NA |
| Tup1 | YOL157C | W303.asm02.HP0_( | YIL172C/YJL221C/YC | y | NA | NA | NA | | NA | NA |
| Tup1 | YOL156W | W303.asm02.HP0_( | W303.asm02.HP0_G( | y | NA | NA | NA | | NA | NA |
| Tup1 | YOL151W | W303.asm02.HP0_( | YOL151W | n | NA | NA | NA | | NA | NA |
| Tup1 | YOL125W | W303.asm02.HP0_( | YOL125W | n | FIM1_2846 | OG0002735 | | 1.112149533 | ns | ns |
| Tup1 | YOL052C-A | W303.asm02.HP0_( | YOL052C-A | n | NA | NA | NA | | NA | NA |
| Tup1 | YOL040C | W303.asm02.HP0_( | YOL040C | n | FIM1_1214 | OG0001062 | | 1 | ns | ns |
| Tup1 | YOR010C | W303.asm02.HP0_( | YOR010C | n | NA | NA | NA | | NA | NA |
| Tup1 | YOR027W | W303.asm02.HP0_( | YOR027W | n | FIM1_464 | OG0001989 | | 1.019031142 | ns | ns |
| Tup1 | YOR032C | W303.asm02.HP0_( | YOR032C | n | FIM1_4205 | OG0000096 | | 0.4375 | ns | Km>KS.3 |
| Tup1 | YOR050C | NA | NA | NA | NA | NA | NA | | NA | NA |
| Tup1 | YOR062C | W303.asm02.HP0_( | YOR062C | n | NA | NA | NA | | NA | NA |
| Tup1 | YOR063W | W303.asm02.HP0_( | YOR063W | n | FIM1_1962 | OG0001279 | | 1 | ns | ns |
| Tup1 | YOR134W | W303.asm02.HP0_( | YOR134W | n | NA | NA | NA | | NA | NA |
| Tup1 | YOR229W | W303.asm02.HP0_( | YOR229W | n | NA | NA | NA | | NA | NA |
| Tup1 | YOR230W | W303.asm02.HP0_( | YOR230W | n | NA | NA | NA | | NA | NA |
| Tup1 | YOR268C | W303.asm02.HP0_( | YOR268C | n | NA | NA | NA | | NA | NA |
| Tup1 | YOR317W | W303.asm02.HP0_( | YOR317W | n | FIM1_1792 | OG0000522 | | 0.998573466 | ns | ns |
| Tup1 | YOR344C | W303.asm02.HP0_( | YOR344C | n | FIM1_4080 | OG0000096 | | 0.954098361 | ns | Km>KS.3 |
| Tup1 | YOR348C | W303.asm02.HP0_( | YOR348C | n | FIM1_4075 | OG0000002 | | 1.094240838 | Km>KS.1 | Km>KS.3 |
| Tup1 | YOR374W | W303.asm02.HP0_( | YOR374W | n | FIM1_3107 | OG0000069 | | 0.996161228 | ns | ns |
| Tup1 | YOR375C | W303.asm02.HP0_( | YOR375C | n | NA | NA | NA | | NA | NA |
| Tup1 | YPL277C | NA | NA | NA | NA | NA | NA | | NA | NA |
| Tup1 | YPL265W | W303.asm02.HP0_( | YPL265W | n | FIM1_4227 | OG0000002 | | 1.057391304 | ns | Km>KS.3 |
| Tup1 | YPL231W | W303.asm02.HP0_( | YPL231W | n | FIM1_4527 | OG0000568 | | 0.996304118 | ns | Km>KS.3 |
| Tup1 | YPL190C | W303.asm02.HP0_( | YPL190C | n | FIM1_3928 | OG0002830 | | 1.04973822 | ns | ns |
| Tup1 | YPL189W | W303.asm02.HP0_( | YPL189W | n | NA | NA | NA | | NA | NA |

| | | | | | | | | | | |
|---|---|---|---|---|---|---|---|---|---|---|
| Tup1 | YPL177C | W303.asm02.HP0_( | YPL177C | n | FIM1_2023 | OG0000548 | | 0.690744921 | ns | ns |
| Tup1 | YPL137C | W303.asm02.HP0_( | YPL137C | n | FIM1_1646 | OG0000042 | | 1.01754386 | ns | Km>KS.3 |
| Tup1 | YPL135W | W303.asm02.HP0_( | YPL135W | n | FIM1_1648 | OG0000549 | | 0.926966292 | ns | ns |
| Tup1 | YPL111W | W303.asm02.HP0_( | YPL111W | n | FIM1_3118 | OG0003354 | | 1.027777778 | ns | ns |
| Tup1 | YPL024W | W303.asm02.HP0_( | YPL024W | n | FIM1_3914 | OG0002461 | | 1.061674009 | ns | ns |
| Tup1 | YPL016W | W303.asm02.HP0_( | YPL016W | n | FIM1_3917 | OG0001751 | | 1.086848635 | ns | ns |
| Tup1 | YPR005C | W303.asm02.HP0_( | YPR005C | n | NA | NA | NA | | NA | NA |
| Tup1 | YPR013C | W303.asm02.HP0_( | YPR013C | n | FIM1_3232 | OG0000006 | | 0.788557214 | ns | Km>KS.3 |
| Tup1 | YPR015C | W303.asm02.HP0_( | YPR015C | n | NA | NA | NA | | NA | NA |
| Tup1 | YPR035W | W303.asm02.HP0_( | YPR035W | n | FIM1_4428 | OG0001518 | | 1.002710027 | ns | ns |
| Tup1 | YPR064W | NA | NA | NA | NA | NA | NA | | NA | NA |
| Tup1 | YPR065W | W303.asm02.HP0_( | YPR065W | n | FIM1_4617 | OG0000459 | | 0.924623116 | ns | ns |
| Tup1 | YAL061W | W303.asm02.HP0_( | YAL061W | n | FIM1_3111 | OG0000248 | | 1.091623037 | Km<KS.1 | Km>KS.3 |
| Tup1 | YAL039C | W303.asm02.HP0_( | YAL039C | n | FIM1_5023 | OG0002200 | | 0.93728223 | ns | ns |
| Tup1 | YAL003W | W303.asm02.HP0_( | YAL003W | n | FIM1_2476 | OG0001084 | | 1.775862069 | ns | ns |
| Tup1 | YAR028W | W303.asm02.HP0_( | YAR028W | n | NA | NA | NA | | NA | NA |
| Tup1 | YAR035W | W303.asm02.HP0_( | YAR035W | n | FIM1_673 | OG0003793 | | 0.984240688 | Km<KS.1 | Km>KS.3 |
| Tup1 | YAR047C | NA | NA | NA | NA | NA | NA | | NA | NA |
| Tup1 | YAR060C | NA | NA | NA | NA | NA | NA | | NA | NA |
| Tup1 | YBL072C | NA | NA | NA | NA | NA | NA | | NA | NA |
| Tup1 | YBL064C | W303.asm02.HP0_( | YBL064C | n | FIM1_5140 | OG0000325 | | 1.048192771 | ns | ns |
| Tup1 | YBL043W | W303.asm02.HP0_( | YBL043W | n | FIM1_4398 | OG0000957 | | 1.048979592 | ns | ns |
| Tup1 | YBL041W | W303.asm02.HP0_( | YBL041W | n | FIM1_4396 | OG0003126 | | 1.004166667 | ns | ns |
| Tup1 | YBL030C | W303.asm02.HP0_( | YBL030C | n | FIM1_4658 | OG0000264 | | 1.042622951 | Km>KS.1 | ns |
| Tup1 | YBL016W | W303.asm02.HP0_( | YBL016W | n | FIM1_892 | OG0000190 | | 0.964480874 | ns | ns |
| Tup1 | YBR066C | W303.asm02.HP0_( | YBR066C | n | NA | NA | NA | | NA | NA |
| Tup1 | YBR067C | W303.asm02.HP0_( | YBR067C | n | NA | NA | NA | | NA | NA |
| Tup1 | YBR069C | W303.asm02.HP0_( | YBR069C | n | NA | NA | NA | | NA | NA |
| Tup1 | YBR078W | W303.asm02.HP0_( | YBR078W | n | FIM1_2255 | OG0000056 | | 1.198324022 | ns | ns |
| Tup1 | YBR082C | W303.asm02.HP0_( | YBR082C | n | NA | NA | NA | | NA | NA |
| Tup1 | YBR084C-A | W303.asm02.HP0_( | YBL027W/YBR084C- | y | NA | NA | NA | | NA | NA |
| Tup1 | YBR085W | W303.asm02.HP0_( | YBR085W | n | NA | NA | NA | | NA | NA |
| Tup1 | YJL079C | W303.asm02.HP0_( | YJL079C | n | NA | NA | NA | | NA | NA |
| Tup1 | YJL078C | W303.asm02.HP0_( | YJL078C | n | NA | NA | NA | | NA | NA |
| Tup1 | YDL182W | W303.asm02.HP0_( | YDL182W | n | FIM1_67 | OG0000178 | | 1.089058524 | ns | ns |
| Tup1 | YIL170W | NA | NA | NA | NA | NA | NA | | NA | NA |
| Rds1 | YCR106W | W303.asm02.HP0_( | YCR106W | n | FIM1_4973 | OG0000043 | | 0.796934866 | ns | ns |
| Rds1 | YCR107W | W303.asm02.HP0_( | YCR107W | n | NA | NA | NA | | NA | NA |
| Rds1 | YMR103C | NA | NA | NA | NA | NA | NA | | NA | NA |
