## Supplementary material for "Intergeneric chromosomal transfer in yeast results in improved phenotypes and widespread transcriptional responses": TableS5

Table S5. Orthologous genes in Km and Sc (W303).

| Kmar_cds | Scer_W303 | Orthogroup | chr.w303 | start.w303 | end.w303 | chr.km | start.km | end.km | aa.similarity | sc.length | km.length | length.ratio | reciprocal.hit | single.copy.ortholog | one.to.one.ortholog | FIM1 | W303_annotation | Scer_systematic_name | Scer_standard_name | Ambiguous_gene_model |
|---|---|---|---|---|---|---|---|---|---|---|---|---|---|---|---|---|---|---|---|---|
| Kmar_QGN17960.1 | W303.asm02.HP0_G0043810 | OG0000000 | chrXIV | 39139 | 40761 | CP015060.1 | 831859 | 833448 | 44.38305709 | 540 | 529 | 1.020793951 | 1 | 0 | 1 | FIM1_5169 | YNL318C | YNL318C | HXT14 | n |
| Kmar_QGN16737.1 | W303.asm02.HP0_G0024260 | OG0000000 | chrVIII | 300642 | 302420 | CP015058.1 | 782298 | 784010 | 71.16357504 | 592 | 570 | 1.038596491 | 1 | 0 | 1 | FIM1_3458 | YHR096C | YHR096C | HXT5 | n |
| Kmar_QGN16736.1 | W303.asm02.HP0_G0024230 | OG0000000 | chrVIII | 293054 | 294784 | CP015058.1 | 778532 | 780229 | 73.13691508 | 576 | 565 | 1.019469027 | 1 | 0 | 0 | FIM1_3457 | YHR092C | YHR092C | HXT4 | n |
| Kmar_QGN16734.1 | W303.asm02.HP0_G0025580 | OG0000000 | chrIX | 8379 | 8708 | CP015058.1 | 772166 | 772456 | 42.70833333 | 109 | 96 | 1.135416667 | 1 | 0 | 0 | FIM1_3455 | W303.asm02.HP0_G0025580 | YOL156W | HXT11 | y |
| Kmar_QGN15879.1 | W303.asm02.HP0_G0009150 | OG0000001 | chrIV | 552308 | 554122 | CP015057.1 | 284057 | 285919 | 57.09677419 | 604 | 620 | 0.974193548 | 1 | 0 | 0 | FIM1_2577 | YDR046C | YDR046C | BAP3 | n |
| Kmar_QGN14134.1 | W303.asm02.HP0_G0005340 | OG0000001 | chrIII | 75669 | 77570 | CP015054.1 | 1610634 | 1612490 | 58.59375 | 633 | 618 | 1.024271845 | 1 | 0 | 0 | FIM1_787 | YCL025C | YCL025C | AGP1 | n |
| Kmar_QGN13825.1 | W303.asm02.HP0_G0033640 | OG0000001 | chrXI | 524724 | 526532 | CP015054.1 | 946442 | 948214 | 68.49087894 | 602 | 590 | 1.020338983 | 1 | 1 | 1 | FIM1_471 | YKR039W | YKR039W | GAP1 | n |
| Kmar_QGN16605.1 | W303.asm02.HP0_G0052760 | OG0000001 | chrXVI | 23490 | 25253 | CP015058.1 | 483226 | 484977 | 67.17687075 | 587 | 583 | 1.006861063 | 1 | 0 | 0 | FIM1_3321 | YPL274W | YPL274W | SAM3 | n |
| Kmar_QGN18198.1 | W303.asm02.HP0_G0003260 | OG0000002 | chrII | 515938 | 517728 | CP015061.1 | 427170 | 428885 | 62 | 596 | 571 | 1.043782837 | 1 | 1 | 1 | FIM1_4522 | YBR132C | YBR132C | AGP2 | n |
| Kmar_QGN17343.1 | W303.asm02.HP0_G0052250 | OG0000002 | chrXV | 989180 | 991063 | CP015059.1 | 734845 | 736566 | 58.98251192 | 627 | 573 | 1.094240838 | 1 | 0 | 1 | FIM1_4075 | YOR348C | YOR348C | PUT4 | n |
| Kmar_QGN17491.1 | W303.asm02.HP0_G0052850 | OG0000002 | chrXVI | 41595 | 43421 | CP015059.1 | 1020708 | 1022435 | 61.18421053 | 608 | 575 | 1.057391304 | 1 | 1 | 1 | FIM1_4227 | YPL265W | YPL265W | DIP5 | n |
| Kmar_QGN14101.1 | W303.asm02.HP0_G0044280 | OG0000002 | chrXIV | 139104 | 140939 | CP015054.1 | 1522307 | 1524040 | 71.61500816 | 611 | 577 | 1.058925477 | 1 | 0 | 1 | FIM1_752 | YNL268W | YNL268W | LYP1 | n |
| Kmar_QGN15034.1 | W303.asm02.HP0_G0013750 | OG0000004 | chrIV | 1525339 | 1526373 | CP015056.1 | 61375 | 62400 | 48.10495627 | 344 | 341 | 1.008797654 | 1 | 0 | 0 | FIM1_1718 | YDR541C | YDR541C | | n |
| Kmar_QGN15616.1 | W303.asm02.HP0_G0006950 | OG0000005 | chrIV | 101364 | 103427 | CP015056.1 | 1312188 | 1314215 | 26.79083095 | 687 | 675 | 1.017777778 | 1 | 0 | 1 | FIM1_2309 | YDL199C | YDL199C | | n |
| Kmar_QGN18039.1 | W303.asm02.HP0_G0016510 | OG0000005 | chrVI | 19797 | 21419 | CP015061.1 | 79437 | 80936 | 26.2962963 | 540 | 499 | 1.082164329 | 1 | 0 | 1 | FIM1_4355 | YFL040W | YFL040W | | n |
| Kmar_QGN17381.1 | W303.asm02.HP0_G0013710 | OG0000005 | chrIV | 1513680 | 1515389 | CP015059.1 | 807069 | 808910 | 63.37115073 | 569 | 613 | 0.92822186 | 1 | 0 | 1 | FIM1_4113 | YDR536W | YDR536W | STL1 | n |
| Kmar_QGN16374.1 | W303.asm02.HP0_G0018010 | OG0000006 | chrVII | 91371 | 92519 | CP015057.1 | 1387116 | 1387850 | 25.94752187 | 382 | 244 | 1.56557377 | 1 | 0 | 0 | FIM1_3081 | YGL209W | YGL209W | MIG2 | n |
| Kmar_QGN14256.1 | W303.asm02.HP0_G0019570 | OG0000006 | chrVII | 428634 | 430148 | CP015055.1 | 123625 | 125274 | 24.59605027 | 504 | 549 | 0.918032787 | 1 | 0 | 1 | FIM1_913 | YGL035C | YGL035C | MIG1 | n |
| Kmar_QGN16519.1 | W303.asm02.HP0_G0055520 | OG0000006 | chrXVI | 591128 | 592081 | CP015058.1 | 288496 | 289704 | 24.35897436 | 317 | 402 | 0.788557214 | 1 | 0 | 0 | FIM1_3232 | YPR013C | YPR013C | CMR3 | n |
| Kmar_QGN16401.1 | W303.asm02.HP0_G0052560 | OG0000008 | chrXV | 1057826 | 1059961 | CP015058.1 | 34966 | 36882 | 26.11111111 | 711 | 638 | 1.114420063 | 1 | 0 | 0 | FIM1_3114 | YOR381W | YOR381W | FRE3 | n |
| Kmar_QGN13547.1 | W303.asm02.HP0_G0008800 | OG0000009 | chrIV | 465578 | 470083 | CP015054.1 | 387023 | 391408 | 62.88180611 | 1501 | 1461 | 1.027378508 | 1 | 0 | 1 | FIM1_187 | YDR011W | YDR011W | SNQ2 | n |
| Kmar_QGN15365.1 | W303.asm02.HP0_G0054830 | OG0000009 | chrXVI | 452201 | 456736 | CP015056.1 | 805659 | 808136 | 47.85276074 | 1511 | 825 | 1.831515152 | 1 | 0 | 0 | FIM1_2055 | YPL058C | YPL058C | PDR12 | n |
| Kmar_QGN16001.1 | W303.asm02.HP0_G0050470 | OG0000009 | chrXV | 620715 | 625250 | CP015057.1 | 586003 | 590550 | 64.00791035 | 1511 | 1515 | 0.997359736 | 1 | 0 | 0 | FIM1_2701 | YOR153W | YOR153W | PDR5 | n |
| Kmar_QGN17579.1 | W303.asm02.HP0_G0000100 | OG0000010 | chrI | 24635 | 28603 | CP015060.1 | 23568 | 26999 | 33.632287 | 1322 | 1143 | 1.156605424 | 1 | 0 | 0 | FIM1_4786 | YAL063C | YAL063C | FLO9 | n |
| Kmar_QGN16952.1 | W303.asm02.HP0_G0006670 | OG0000013 | chrIV | 44138 | 45979 | CP015058.1 | 1246499 | 1248340 | 91.68026101 | 613 | 613 | 1 | 1 | 0 | 0 | FIM1_3679 | YDL229W/YNL209W | YDL229W | SSB1 | y |
| Kmar_QGN14878.1 | W303.asm02.HP0_G0029360 | OG0000013 | chrX | 376976 | 379024 | CP015055.1 | 1408067 | 1410106 | 77.01317716 | 682 | 679 | 1.004418262 | 1 | 1 | 1 | FIM1_1551 | YJL034W | YJL034W | KAR2 | n |
| Kmar_QGN17940.1 | W303.asm02.HP0_G0001190 | OG0000013 | chrII | 85652 | 87601 | CP015060.1 | 780229 | 782181 | 86.76923077 | 649 | 650 | 0.998461538 | 1 | 0 | 0 | FIM1_5149 | YBL075C | YBL075C | SSA3 | n |
| Kmar_QGN14649.1 | W303.asm02.HP0_G0034690 | OG0000013 | chrXII | 95575 | 97494 | CP015055.1 | 945687 | 947609 | 90.15625 | 639 | 640 | 0.9984375 | 1 | 0 | 0 | FIM1_1313 | YAL005C/YLL024C | YLL024C | SSA2 | y |
| Kmar_QGN15963.1 | W303.asm02.HP0_G0010660 | OG0000014 | chrIV | 868728 | 871238 | CP015057.1 | 488081 | 490879 | 24.23280423 | 836 | 932 | 0.896995708 | 1 | 0 | 1 | FIM1_2661 | YDR207C | YDR207C | UME6 | n |
| Kmar_QGN16835.1 | W303.asm02.HP0_G0027050 | OG0000015 | chrIX | 312971 | 314734 | CP015058.1 | 991707 | 993212 | 28.28947368 | 587 | 501 | 1.171656687 | 1 | 0 | 1 | FIM1_3559 | YIL015W | YIL015W | BAR1 | n |
| Kmar_QGN14723.1 | W303.asm02.HP0_G0036080 | OG0000015 | chrXII | 396697 | 398223 | CP015055.1 | 1099091 | 1100893 | 41.65289256 | 508 | 600 | 0.846666667 | 1 | 0 | 1 | FIM1_1391 | YLR121C | YLR121C | YPS3 | n |
| Kmar_QGN13381.1 | W303.asm02.HP0_G0013840 | OG0000016 | chrV | 28021 | 29907 | CP015054.1 | 40329 | 42233 | 24.88335925 | 628 | 634 | 0.990536278 | 1 | 0 | 0 | FIM1_18 | YEL065W | YEL065W | SIT1 | n |
| Kmar_QGN14190.1 | W303.asm02.HP0_G0022660 | OG0000016 | chrVIII | 5974 | 7836 | CP015054.1 | 1721497 | 1723356 | 59.51612903 | 620 | 619 | 1.001615509 | 1 | 0 | 0 | FIM1_843 | YHL047C | YHL047C | ARN2 | n |
| Kmar_QGN13681.1 | W303.asm02.HP0_G0022710 | OG0000016 | chrVIII | 16712 | 18595 | CP015054.1 | 652835 | 654652 | 57.6433121 | 627 | 605 | 1.036363636 | 1 | 0 | 1 | FIM1_324 | YHL040C | YHL040C | ARN1 | n |
| Kmar_QGN16383.1 | W303.asm02.HP0_G0004930 | OG0000016 | chrIII | 6132 | 7979 | CP015057.1 | 1408245 | 1410113 | 41.28 | 615 | 622 | 0.988745981 | 1 | 0 | 0 | FIM1_3092 | YCL073C/YKR106W | YCL073C | GEX1 | y |
| Kmar_QGN18403.1 | W303.asm02.HP0_G0019430 | OG0000017 | chrVII | 400772 | 401476 | CP015061.1 | 853232 | 853810 | 22.64957265 | 234 | 192 | 1.21875 | 1 | 0 | 0 | FIM1_4730 | YAR033W/YGL051W | YGL051W | MST27 | y |
| Kmar_QGN14267.1 | W303.asm02.HP0_G0019420 | OG0000017 | chrVII | 399674 | 400387 | CP015055.1 | 146340 | 146951 | 24.38016529 | 237 | 203 | 1.167487685 | 1 | 0 | 0 | FIM1_924 | YGL053W | YGL053W | PRM8 | n |
| Kmar_QGN15018.1 | W303.asm02.HP0_G0000810 | OG0000017 | chrI | 184970 | 185677 | CP015056.1 | 4963 | 5544 | 23.1884058 | 235 | 193 | 1.21761658 | 1 | 0 | 0 | FIM1_1701 | YAR027W | YAR027W | UIP3 | n |
| Kmar_QGN18398.1 | W303.asm02.HP0_G0005580 | OG0000017 | chrIII | 131251 | 131970 | CP015061.1 | 848407 | 849006 | 20.49180328 | 239 | 199 | 1.201005025 | 1 | 0 | 0 | FIM1_4725 | YCR007C | YCR007C | DFP3 | n |
| Kmar_QGN13870.1 | W303.asm02.HP0_G0028400 | OG0000018 | chrX | 141241 | 142527 | CP015054.1 | 1027369 | 1028637 | 59.38967136 | 428 | 422 | 1.014218009 | 1 | 0 | 0 | FIM1_516 | YJL139C | YJL139C | YUR1 | n |
| Kmar_QGN15352.1 | W303.asm02.HP0_G0013220 | OG0000018 | chrIV | 1426851 | 1428179 | CP015056.1 | 773782 | 775155 | 48.14814815 | 442 | 457 | 0.967177243 | 1 | 1 | 1 | FIM1_2042 | YDR483W | YDR483W | KRE2 | n |
| Kmar_QGN13463.1 | W303.asm02.HP0_G0003990 | OG0000018 | chrII | 650401 | 651615 | CP015054.1 | 191487 | 192776 | 59.06976744 | 404 | 429 | 0.941724942 | 1 | 0 | 1 | FIM1_100 | YBR205W | YBR205W | KTR3 | n |
| Kmar_QGN17799.1 | W303.asm02.HP0_G0049990 | OG0000018 | chrXV | 512560 | 513741 | CP015060.1 | 511218 | 512447 | 61.55717762 | 393 | 409 | 0.960880196 | 1 | 0 | 1 | FIM1_5008 | YOR099W | YOR099W | KTR1 | n |
| Kmar_QGN13739.1 | W303.asm02.HP0_G0025080 | OG0000019 | chrVIII | 468608 | 469810 | CP015054.1 | 778638 | 779849 | 71.25 | 400 | 403 | 0.992555831 | 1 | 0 | 0 | FIM1_383 | YHR179W | YHR179W | OYE2 | n |
| Kmar_QGN17090.1 | W303.asm02.HP0_G0046820 | OG0000020 | chrXIV | 656473 | 659103 | CP015059.1 | 214626 | 217178 | 53.79310345 | 876 | 850 | 1.030588235 | 1 | 1 | 1 | FIM1_3821 | YNR011C | YNR011C | PRP2 | n |
| Kmar_QGN16376.1 | W303.asm02.HP0_G0018830 | OG0000020 | chrVII | 278589 | 280892 | CP015057.1 | 1390005 | 1392311 | 88.42652796 | 767 | 768 | 0.998697917 | 1 | 1 | 1 | FIM1_3083 | YGL120C | YGL120C | PRP43 | n |
| Kmar_QGN18297.1 | W303.asm02.HP0_G0014600 | OG0000020 | chrV | 179988 | 183425 | CP015061.1 | 643203 | 646541 | 60.70811744 | 1145 | 1112 | 1.029676259 | 1 | 1 | 1 | FIM1_4623 | YER013W | YER013W | PRP22 | n |
| Kmar_QGN13788.1 | W303.asm02.HP0_G0034100 | OG0000020 | chrXI | 618180 | 621395 | CP015054.1 | 880783 | 884046 | 49.18330309 | 1071 | 1087 | 0.985280589 | 1 | 1 | 1 | FIM1_434 | YKR086W | YKR086W | PRP16 | n |
| Kmar_QGN18186.1 | W303.asm02.HP0_G0006120 | OG0000021 | chrIII | 262036 | 262992 | CP015061.1 | 396024 | 396950 | 38.19875776 | 318 | 308 | 1.032467532 | 1 | 0 | 1 | FIM1_4509 | YCR069W | YCR069W | CPR4 | n |
| Kmar_QGN15448.1 | W303.asm02.HP0_G0011560 | OG0000021 | chrIV | 1076379 | 1077056 | CP015056.1 | 980029 | 980631 | 60.44444444 | 225 | 200 | 1.125 | 1 | 0 | 1 | FIM1_2139 | YDR304C | YDR304C | CPR5 | n |
| Kmar_QGN16273.1 | W303.asm02.HP0_G0039760 | OG0000021 | chrXIII | 105542 | 106090 | CP015057.1 | 1172228 | 1172806 | 73.15789474 | 182 | 192 | 0.947916667 | 1 | 1 | 1 | FIM1_2978 | YML078W | YML078W | CPR3 | n |
| Kmar_QGN16871.1 | W303.asm02.HP0_G0010150 | OG0000021 | chrIV | 772226 | 772714 | CP015058.1 | 1074196 | 1074684 | 87.65432099 | 162 | 162 | 1 | 1 | 1 | 1 | FIM1_3597 | YDR155C | YDR155C | CPR1 | n |
| Kmar_QGN14781.1 | W303.asm02.HP0_G0012310 | OG0000024 | chrIV | 1239811 | 1241718 | CP015055.1 | 1215011 | 1216903 | 48.05598756 | 635 | 630 | 1.007936508 | 1 | 1 | 1 | FIM1_1451 | YDR380W | YDR380W | ARO10 | n |
| Kmar_QGN15803.1 | W303.asm02.HP0_G0008040 | OG0000024 | chrIV | 310972 | 312801 | CP015057.1 | 124663 | 126399 | 34.09090909 | 609 | 578 | 1.053633218 | 1 | 0 | 1 | FIM1_2499 | YDL080C | YDL080C | THI3 | n |
| Kmar_QGN17490.1 | W303.asm02.HP0_G0035350 | OG0000024 | chrXII | 240203 | 241894 | CP015059.1 | 1018663 | 1020357 | 86.14564831 | 563 | 564 | 0.99822695 | 1 | 0 | 0 | FIM1_4226 | YLR044C | YLR044C | PDC1 | n |
| Kmar_QGN16781.1 | W303.asm02.HP0_G0027400 | OG0000025 | chrIX | 375777 | 376514 | CP015058.1 | 878404 | 879885 | 18.60940695 | 245 | 493 | 0.496957404 | 1 | 0 | 0 | FIM1_3504 | YIR018W | YIR018W | YAP5 | n |
| Kmar_QGN14340.1 | W303.asm02.HP0_G0032180 | OG0000025 | chrXI | 240086 | 241750 | CP015055.1 | 310260 | 312116 | 17.3566879 | 554 | 618 | 0.896440129 | 1 | 1 | 1 | FIM1_1000 | YKL109W | YKL109W | HAP4 | n |
| Kmar_QGN15072.1 | W303.asm02.HP0_G0023020 | OG0000025 | chrVIII | 82773 | 83765 | CP015056.1 | 143603 | 144772 | 26.99228792 | 330 | 389 | 0.848329049 | 1 | 0 | 1 | FIM1_1757 | YHL009C | YHL009C | YAP3 | n |
| Kmar_QGN15937.1 | W303.asm02.HP0_G0009970 | OG0000026 | chrIV | 726190 | 726390 | CP015057.1 | 424339 | 424716 | 78.78787879 | 66 | 125 | 0.528 | 1 | 0 | 0 | FIM1_2635 | W303.asm02.HP0_G0009970 | YLR110C | CCW12 | y |
| Kmar_QGN15001.1 | W303.asm02.HP0_G0047160 | OG0000026 | chrXIV | 716281 | 718458 | CP015055.1 | 1658728 | 1662312 | 25.27196653 | 725 | 1194 | 0.60720268 | 1 | 0 | 0 | FIM1_1682 | YNR044W | YNR044W | AGA1 | n |
| Kmar_QGN16740.1 | W303.asm02.HP0_G0027070 | OG0000027 | chrIX | 316732 | 318624 | CP015058.1 | 793757 | 795622 | 34.99222395 | 630 | 621 | 1.014492754 | 1 | 0 | 1 | FIM1_3462 | YIL014W | YIL014W | MNT3 | n |
| Kmar_QGN17639.1 | W303.asm02.HP0_G0014470 | OG0000027 | chrV | 154703 | 156991 | CP015060.1 | 161004 | 163295 | 32.27040816 | 762 | 763 | 0.998689384 | 1 | 0 | 0 | FIM1_4846 | YER001W | YER001W | MNN1 | n |
| Kmar_QGN14619.1 | W303.asm02.HP0_G0009440 | OG0000029 | chrIV | 605069 | 606085 | CP015055.1 | 870422 | 871111 | 35.44668588 | 338 | 229 | 1.475982533 | 1 | 0 | 0 | FIM1_1283 | YDR077W | YDR077W | SED1 | n |
| Kmar_QGN14110.1 | W303.asm02.HP0_G0044220 | OG0000030 | chrXIV | 121672 | 122724 | CP015054.1 | 1554660 | 1555709 | 62.85714286 | 350 | 349 | 1.00286533 | 1 | 0 | 1 | FIM1_761 | YNL274C | YNL274C | GOR1 | n |
| Kmar_QGN16409.1 | W303.asm02.HP0_G0054310 | OG0000030 | chrXVI | 342311 | 343501 | CP015058.1 | 53924 | 55099 | 48.50746269 | 396 | 391 | 1.012787724 | 1 | 0 | 1 | FIM1_3122 | YPL113C | YPL113C | | n |
| Kmar_QGN14491.1 | W303.asm02.HP0_G0018240 | OG0000030 | chrVII | 148680 | 149819 | CP015055.1 | 621579 | 622847 | 35.6979405 | 379 | 422 | 0.898104265 | 1 | 0 | 1 | FIM1_1152 | YGL185C | YGL185C | | n |
| Kmar_QGN16628.1 | W303.asm02.HP0_G0028220 | OG0000031 | chrX | 107489 | 108352 | CP015058.1 | 533564 | 534592 | 44.15204678 | 287 | 342 | 0.839181287 | 1 | 0 | 0 | FIM1_3345 | YJL160C | YJL160C | PIR5 | n |
| Kmar_QGN16627.1 | W303.asm02.HP0_G0031680 | OG0000031 | chrXI | 142961 | 143986 | CP015058.1 | 530759 | 531934 | 65.56962025 | 341 | 391 | 0.872122762 | 1 | 0 | 0 | FIM1_3344 | YKL164C | YKL164C | PIR1 | n |
| Kmar_QGN15976.1 | W303.asm02.HP0_G0048250 | OG0000032 | chrXV | 160003 | 161049 | CP015057.1 | 529296 | 530342 | 86.20689655 | 348 | 348 | 1 | 1 | 0 | 0 | FIM1_2676 | YOL086C | YOL086C | ADH1 | n |
| Kmar_QGN17882.1 | W303.asm02.HP0_G0041250 | OG0000032 | chrXIII | 435293 | 436420 | CP015060.1 | 672847 | 673974 | 77.86666667 | 375 | 375 | 1 | 1 | 0 | 1 | FIM1_5091 | YMR083W | YMR083W | ADH3 | n |
| Kmar_QGN14922.1 | W303.asm02.HP0_G0045030 | OG0000034 | chrXIV | 293833 | 296205 | CP015055.1 | 1499254 | 1501641 | 57.52427184 | 790 | 795 | 0.993710692 | 1 | 0 | 0 | FIM1_1596 | YNL183C | YNL183C | NPR1 | n |
| Kmar_QGN14031.1 | W303.asm02.HP0_G0008510 | OG0000034 | chrIV | 405386 | 407248 | CP015054.1 | 1373118 | 1374980 | 42.65625 | 620 | 620 | 1 | 1 | 1 | 1 | FIM1_682 | YDL025C | YDL025C | RTK1 | n |
| Kmar_QGN16170.1 | W303.asm02.HP0_G0051500 | OG0000034 | chrXV | 824310 | 826586 | CP015057.1 | 949393 | 951486 | 44.50194049 | 758 | 697 | 1.087517934 | 1 | 1 | 1 | FIM1_2872 | YOR267C | YOR267C | HRK1 | n |
| Kmar_QGN15696.1 | W303.asm02.HP0_G0014890 | OG0000035 | chrV | 230934 | 233126 | CP015056.1 | 1486074 | 1488467 | 27.63320942 | 730 | 797 | 0.915934755 | 1 | 1 | 1 | FIM1_2389 | YER040W | YER040W | GLN3 | n |
| Kmar_QGN17259.1 | W303.asm02.HP0_G0016700 | OG0000035 | chrVI | 64446 | 65978 | CP015059.1 | 565175 | 567451 | 21.76781003 | 510 | 758 | 0.672823219 | 1 | 1 | 1 | FIM1_3990 | YFL021W | YFL021W | GAT1 | n |
| Kmar_QGN15988.1 | W303.asm02.HP0_G0033590 | OG0000035 | chrXI | 516587 | 517396 | CP015057.1 | 556848 | 558425 | 22.20113852 | 269 | 525 | 0.512380952 | 1 | 0 | 1 | FIM1_2688 | YKR034W | YKR034W | DAL80 | n |
| Kmar_QGN14718.1 | W303.asm02.HP0_G0017300 | OG0000036 | chrVI | 198284 | 200845 | CP015055.1 | 1089205 | 1091700 | 58.77598152 | 853 | 831 | 1.026474128 | 1 | 1 | 1 | FIM1_1386 | YFR038W | YFR038W | IRC5 | n |
| Kmar_QGN17280.1 | W303.asm02.HP0_G0051840 | OG0000036 | chrXV | 886233 | 889595 | CP015059.1 | 607617 | 610844 | 63.64444444 | 1120 | 1075 | 1.041860465 | 1 | 1 | 1 | FIM1_4011 | YOR304W | YOR304W | ISW2 | n |
| Kmar_QGN14602.1 | W303.asm02.HP0_G0004350 | OG0000036 | chrII | 731505 | 734894 | CP015055.1 | 836399 | 839680 | 69.03114187 | 1129 | 1093 | 1.032936871 | 1 | 0 | 1 | FIM1_1265 | YBR245C | YBR245C | ISW1 | n |
| Kmar_QGN14772.1 | W303.asm02.HP0_G0012360 | OG0000037 | chrIV | 1246798 | 1247625 | CP015055.1 | 1202668 | 1203480 | 58.33333333 | 275 | 270 | 1.018518519 | 1 | 0 | 0 | FIM1_1442 | YDR384C | YDR384C | ATO3 | n |
| Kmar_QGN14203.1 | W303.asm02.HP0_G0005610 | OG0000037 | chrIII | 137513 | 138364 | CP015055.1 | 29706 | 30536 | 64.66431095 | 283 | 276 | 1.025362319 | 1 | 0 | 0 | FIM1_859 | YCR010C | YCR010C | ADY2 | n |
| Kmar_QGN17426.1 | W303.asm02.HP0_G0002030 | OG0000038 | chrII | 262796 | 264589 | CP015059.1 | 897199 | 899061 | 50.31746032 | 597 | 620 | 0.962903226 | 1 | 0 | 1 | FIM1_4158 | YBR015C | YBR015C | MNN2 | n |
| Kmar_QGN18224.1 | W303.asm02.HP0_G0027990 | OG0000038 | chrX | 69685 | 71445 | CP015061.1 | 490058 | 491872 | 34.76112026 | 586 | 604 | 0.970198675 | 1 | 0 | 1 | FIM1_4549 | YJL186W | YJL186W | MNN5 | n |
| Kmar_QGN14162.1 | W303.asm02.HP0_G0013590 | OG0000039 | chrIV | 1491259 | 1492731 | CP015054.1 | 1660100 | 1661590 | 31.81818182 | 490 | 496 | 0.987903226 | 1 | 1 | 1 | FIM1_815 | YDR523C | YDR523C | SPS1 | n |
| Kmar_QGN14021.1 | W303.asm02.HP0_G0000780 | OG0000039 | chrI | 173415 | 176339 | CP015054.1 | 1345091 | 1347598 | 30.48289738 | 974 | 835 | 1.166467066 | 1 | 1 | 1 | FIM1_672 | YAR019C | YAR019C | CDC15 | n |
| Kmar_QGN13514.1 | W303.asm02.HP0_G0024320 | OG0000039 | chrVIII | 322545 | 325787 | CP015054.1 | 306791 | 310000 | 40.10647737 | 1080 | 1069 | 1.010289991 | 1 | 0 | 1 | FIM1_154 | YHR102W | YHR102W | KIC1 | n |
| Kmar_QGN14235.1 | W303.asm02.HP0_G0001740 | OG0000040 | chrII | 189131 | 193870 | CP015055.1 | 81396 | 86099 | 42.48284467 | 1579 | 1567 | 1.007657945 | 1 | 0 | 0 | FIM1_891 | YBL017C | YBL017C | PEP1 | n |
| Kmar_QGN17839.1 | W303.asm02.HP0_G0032770 | OG0000041 | chrXI | 354679 | 355368 | CP015060.1 | 595369 | 595947 | 52.40174672 | 229 | 192 | 1.192708333 | 1 | 1 | 1 | FIM1_5048 | YKL049C | YKL049C | CSE4 | n |
| Kmar_QGN17693.1 | W303.asm02.HP0_G0002000 | OG0000041 | chrII | 258670 | 259080 | CP015060.1 | 267112 | 267588 | 86.07594937 | 136 | 158 | 0.860759494 | 1 | 0 | 0 | FIM1_4900 | YBR010W/YNL031C | YNL031C | HHT2 | y |
| Kmar_QGN14291.1 | W303.asm02.HP0_G0029300 | OG0000042 | chrX | 357183 | 364022 | CP015055.1 | 201146 | 203488 | 17.86553832 | 2123 | 780 | 2.721794872 | 1 | 1 | 1 | FIM1_948 | YJL042W | YJL042W | MHP1 | n |
| Kmar_QGN14968.1 | W303.asm02.HP0_G0054080 | OG0000042 | chrXVI | 298750 | 302580 | CP015055.1 | 1593366 | 1597130 | 42.52265861 | 1276 | 1254 | 1.01754386 | 1 | 0 | 0 | FIM1_1646 | YPL137C | YPL137C | GIP3 | n |
| Kmar_QGN15605.1 | W303.asm02.HP0_G0000200 | OG0000043 | chrI | 49199 | 52342 | CP015056.1 | 1285674 | 1288751 | 29.76958525 | 1047 | 1025 | 1.021463415 | 1 | 0 | 0 | FIM1_2298 | YAL051W | YAL051W | OAF1 | n |
| Kmar_QGN17764.1 | W303.asm02.HP0_G0006450 | OG0000043 | chrIII | 334987 | 337485 | CP015060.1 | 422688 | 425822 | 20 | 832 | 1044 | 0.796934866 | 1 | 0 | 0 | FIM1_4973 | YCR106W | YCR106W | RDS1 | n |
| Kmar_QGN13955.1 | W303.asm02.HP0_G0048970 | OG0000044 | chrXV | 305804 | 307864 | CP015054.1 | 1212217 | 1214169 | 58.22416303 | 686 | 650 | 1.055384615 | 1 | 0 | 1 | FIM1_603 | YOL011W | YOL011W | PLB3 | n |
| Kmar_QGN17646.1 | W303.asm02.HP0_G0029650 | OG0000045 | chrX | 438558 | 440510 | CP015060.1 | 177845 | 180244 | 30.08739076 | 650 | 799 | 0.813516896 | 1 | 0 | 1 | FIM1_4853 | YJR004C | YJR004C | SAG1 | n |
| Kmar_QGN15332.1 | W303.asm02.HP0_G0053670 | OG0000047 | chrXVI | 213708 | 215357 | CP015056.1 | 727231 | 728760 | 56.45756458 | 549 | 509 | 1.078585462 | 1 | 1 | 1 | FIM1_2021 | YPL179W | YPL179W | PPQ1 | n |
| Kmar_QGN16352.1 | W303.asm02.HP0_G0040350 | OG0000047 | chrXIII | 239964 | 242042 | CP015057.1 | 1344943 | 1346967 | 59.25414365 | 692 | 674 | 1.026706231 | 1 | 0 | 0 | FIM1_3058 | YML016C | YML016C | PPZ1 | n |

| | | | | | | | | | | | | | | | | | | | | |
|---|---|---|---|---|---|---|---|---|---|---|---|---|---|---|---|---|---|---|---|---|
| Kmar_QGN17296.1 | W303.asm02.HP0_G0045480 | OG0000048 | chrXIV | 373284 | 374414 | CP015059.1 | 640872 | 641987 | 55.2 | 376 | 371 | 1.013477089 | 1 | 0 | 0 | FIM1_4027 | YNL134C | YNL134C | | n |
| Kmar_QGN15821.1 | W303.asm02.HP0_G0035260 | OG0000049 | chrXII | 218325 | 219746 | CP015057.1 | 157166 | 158617 | 69.15113872 | 473 | 483 | 0.979296066 | 1 | 1 | 1 | FIM1_2517 | YLR034C | YLR034C | SMF3 | n |
| Kmar_QGN15593.1 | W303.asm02.HP0_G0047900 | OG0000049 | chrXV | 90032 | 91759 | CP015056.1 | 1262937 | 1264739 | 56.27062706 | 575 | 600 | 0.958333333 | 1 | 0 | 1 | FIM1_2285 | YOL122C | YOL122C | SMF1 | n |
| Kmar_QGN14877.1 | W303.asm02.HP0_G0023650 | OG0000049 | chrVIII | 201498 | 203147 | CP015055.1 | 1404773 | 1406551 | 63.85135135 | 549 | 592 | 0.927364865 | 1 | 1 | 1 | FIM1_1550 | YHR050W | YHR050W | SMF2 | n |
| Kmar_QGN14632.1 | W303.asm02.HP0_G0022830 | OG0000050 | chrVIII | 46386 | 48203 | CP015055.1 | 904615 | 906132 | 35.09060956 | 605 | 505 | 1.198019802 | 1 | 1 | 1 | FIM1_1296 | YHL028W | YHL028W | WSC4 | n |
| Kmar_QGN17856.1 | W303.asm02.HP0_G0044130 | OG0000050 | chrXIV | 105742 | 107250 | CP015060.1 | 624165 | 625460 | 35.78528827 | 502 | 431 | 1.164733179 | 1 | 0 | 0 | FIM1_5065 | YNL283C | YNL283C | WSC2 | n |
| Kmar_QGN15069.1 | W303.asm02.HP0_G0030150 | OG0000051 | chrX | 559166 | 561973 | CP015056.1 | 135445 | 138249 | 34.75103734 | 935 | 934 | 1.001070664 | 1 | 0 | 0 | FIM1_1754 | YJR061W | YJR061W | MNN14 | n |
| Kmar_QGN17746.1 | W303.asm02.HP0_G0046120 | OG0000052 | chrXIV | 508055 | 509284 | CP015060.1 | 377052 | 378284 | 74.57627119 | 409 | 410 | 0.997560976 | 1 | 1 | 1 | FIM1_4955 | YNL064C | YNL064C | YDJ1 | n |
| Kmar_QGN15899.1 | W303.asm02.HP0_G0035790 | OG0000052 | chrXII | 328650 | 330029 | CP015057.1 | 338820 | 340160 | 38.54389722 | 459 | 446 | 1.029147982 | 1 | 1 | 1 | FIM1_2597 | YLR090W | YLR090W | XDJ1 | n |
| Kmar_QGN16129.1 | W303.asm02.HP0_G0045990 | OG0000052 | chrXIV | 483579 | 485165 | CP015057.1 | 871429 | 872871 | 42.85714286 | 528 | 480 | 1.1 | 1 | 1 | 1 | FIM1_2830 | YNL077W | YNL077W | APJ1 | n |
| Kmar_QGN15806.1 | W303.asm02.HP0_G0008070 | OG0000053 | chrIV | 315687 | 316718 | CP015057.1 | 130410 | 131426 | 53.46820809 | 343 | 338 | 1.014792899 | 1 | 1 | 1 | FIM1_2502 | YDL078C | YDL078C | MDH3 | n |
| Kmar_QGN16146.1 | W303.asm02.HP0_G0047860 | OG0000053 | chrXV | 82127 | 83260 | CP015057.1 | 904076 | 905200 | 55.4973822 | 377 | 374 | 1.00802139 | 1 | 1 | 1 | FIM1_2847 | YOL126C | YOL126C | MDH2 | n |
| Kmar_QGN17393.1 | W303.asm02.HP0_G0032410 | OG0000053 | chrXI | 287599 | 288603 | CP015059.1 | 830458 | 831474 | 78.40236686 | 334 | 338 | 0.98816568 | 1 | 1 | 1 | FIM1_4125 | YKL085W | YKL085W | MDH1 | n |
| Kmar_QGN18237.1 | W303.asm02.HP0_G0027410 | OG0000054 | chrIX | 380873 | 384976 | CP015061.1 | 516019 | 519609 | 31.58650395 | 1367 | 1196 | 1.142976589 | 1 | 0 | 0 | FIM1_4562 | YIR019C | YIR019C | FLO11 | n |
| Kmar_QGN15564.1 | W303.asm02.HP0_G0002740 | OG0000056 | chrII | 409410 | 411028 | CP015056.1 | 1193025 | 1194101 | 47.78554779 | 429 | 358 | 1.198324022 | 1 | 0 | 0 | FIM1_2255 | YBR078W | YBR078W | ECM33 | n |
| Kmar_QGN14161.1 | W303.asm02.HP0_G0005130 | OG0000056 | chrIII | 41817 | 43208 | CP015054.1 | 1658100 | 1659392 | 39.74082073 | 463 | 430 | 1.076744186 | 1 | 0 | 0 | FIM1_814 | YCL048W | YCL048W | SPS22 | n |
| Kmar_QGN18003.1 | W303.asm02.HP0_G0004800 | OG0000058 | chrII | 810375 | 811799 | CP015061.1 | 13517 | 15214 | 45.64831261 | 474 | 565 | 0.838938053 | 1 | 0 | 0 | FIM1_4319 | YBR293W | YBR293W | VBA2 | n |
| Kmar_QGN14347.1 | W303.asm02.HP0_G0041290 | OG0000058 | chrXIII | 443919 | 445607 | CP015055.1 | 324314 | 326011 | 51.74216028 | 562 | 565 | 0.994690265 | 1 | 0 | 1 | FIM1_1007 | YMR088C | YMR088C | VBA1 | n |
| Kmar_QGN18259.1 | W303.asm02.HP0_G0056960 | OG0000060 | chrXVI | 892356 | 893669 | CP015061.1 | 568848 | 570146 | 84.21052632 | 437 | 432 | 1.011574074 | 1 | 1 | 1 | FIM1_4585 | YPR173C | YPR173C | VPS4 | n |
| Kmar_QGN13443.1 | W303.asm02.HP0_G0054690 | OG0000060 | chrXVI | 422121 | 424385 | CP015054.1 | 156711 | 158681 | 42.85714286 | 754 | 656 | 1.149390244 | 1 | 0 | 1 | FIM1_80 | YPL074W | YPL074W | YTA6 | n |
| Kmar_QGN14708.1 | W303.asm02.HP0_G0014970 | OG0000060 | chrV | 245081 | 247774 | CP015055.1 | 1068707 | 1071040 | 45.37346711 | 897 | 777 | 1.154440154 | 1 | 0 | 1 | FIM1_1375 | YER047C | YER047C | SAP1 | n |
| Kmar_QGN14186.1 | W303.asm02.HP0_G0004970 | OG0000061 | chrIII | 12935 | 13772 | CP015054.1 | 1710095 | 1710874 | 22.47191011 | 271 | 259 | 1.046332046 | 1 | 0 | 1 | FIM1_839 | YCL066W/YCR040W | YCR040W | MATALPHA1 | y |
| Kmar_QGN14066.1 | W303.asm02.HP0_G0005910 | OG0000061 | chrIII | 223288 | 223950 | CP015054.1 | 1453171 | 1453812 | 14.08450704 | 165 | 213 | 0.774647887 | 1 | 0 | 0 | FIM1_717 | YCR097W | YCR097W | HMRA1 | n |
| Kmar_QGN15809.1 | W303.asm02.HP0_G0055190 | OG0000062 | chrXVI | 517458 | 518021 | CP015057.1 | 134518 | 135753 | 16.30170316 | 187 | 411 | 0.454987835 | 1 | 0 | 0 | FIM1_2505 | YPL021W | YPL021W | ECM23 | n |
| Kmar_QGN14467.1 | W303.asm02.HP0_G0027350 | OG0000062 | chrIX | 371187 | 371552 | CP015055.1 | 577507 | 578268 | 17.64705882 | 121 | 253 | 0.47826087 | 1 | 0 | 0 | FIM1_1128 | YIR013C | YIR013C | GAT4 | n |
| Kmar_QGN17550.1 | W303.asm02.HP0_G0033660 | OG0000063 | chrXI | 529215 | 530312 | CP015059.1 | 1146524 | 1147714 | 62.97229219 | 365 | 396 | 0.921717172 | 1 | 0 | 0 | FIM1_4286 | YKR042W | YKR042W | UTH1 | n |
| Kmar_QGN17016.1 | W303.asm02.HP0_G0046100 | OG0000063 | chrXIV | 503702 | 504964 | CP015059.1 | 38082 | 39293 | 64.13301663 | 420 | 403 | 1.042183623 | 1 | 0 | 0 | FIM1_3745 | YNL066W | YNL066W | SUN4 | n |
| Kmar_QGN15704.1 | W303.asm02.HP0_G0017320 | OG0000064 | chrVI | 203146 | 206154 | CP015056.1 | 1501548 | 1504484 | 47.15370019 | 1002 | 978 | 1.024539877 | 1 | 0 | 0 | FIM1_2397 | YFR040W | YFR040W | SAP155 | n |
| Kmar_QGN15169.1 | W303.asm02.HP0_G0033540 | OG0000064 | chrXI | 503937 | 507041 | CP015056.1 | 349639 | 352968 | 44.72984942 | 1034 | 1109 | 0.932371506 | 1 | 0 | 0 | FIM1_1857 | YKR028W | YKR028W | SAP190 | n |
| Kmar_QGN13824.1 | W303.asm02.HP0_G0011910 | OG0000065 | chrIV | 1157979 | 1160066 | CP015054.1 | 942951 | 945065 | 58.70786517 | 695 | 704 | 0.987215909 | 1 | 1 | 1 | FIM1_470 | YDR338C | YDR338C | | n |
| Kmar_QGN16005.1 | W303.asm02.HP0_G0023450 | OG0000065 | chrVIII | 167188 | 168933 | CP015057.1 | 601775 | 603607 | 58.43071786 | 581 | 610 | 0.952459016 | 1 | 0 | 1 | FIM1_2705 | YHR032W | YHR032W | ERC1 | n |
| Kmar_QGN17936.1 | W303.asm02.HP0_G0015570 | OG0000066 | chrV | 362172 | 363464 | CP015060.1 | 772985 | 774271 | 48.14814815 | 430 | 428 | 1.004672897 | 1 | 0 | 0 | FIM1_5145 | YER101C | YER101C | AST2 | n |
| Kmar_QGN18031.1 | W303.asm02.HP0_G0016470 | OG0000067 | chrVI | 14167 | 16191 | CP015061.1 | 60862 | 63114 | 30.1061008 | 674 | 750 | 0.898666667 | 1 | 0 | 0 | FIM1_4347 | YFL042C | YFL042C | LAM5 | n |
| Kmar_QGN17462.1 | W303.asm02.HP0_G0011800 | OG0000067 | chrIV | 1131209 | 1135525 | CP015059.1 | 967236 | 971423 | 32.45614035 | 1438 | 1395 | 1.030824373 | 1 | 0 | 0 | FIM1_4196 | YDR326C | YDR326C | YSP2 | n |
| Kmar_QGN18170.1 | W303.asm02.HP0_G0015070 | OG0000068 | chrV | 268183 | 269784 | CP015061.1 | 365396 | 366955 | 64.23220974 | 533 | 519 | 1.026974952 | 1 | 0 | 0 | FIM1_4493 | YER056C | YER056C | FCY2 | n |
| Kmar_QGN13460.1 | W303.asm02.HP0_G0054800 | OG0000069 | chrXVI | 438946 | 440448 | CP015054.1 | 187053 | 188576 | 72.38658777 | 500 | 507 | 0.986193294 | 1 | 0 | 1 | FIM1_97 | YPL061W | YPL061W | ALD6 | n |
| Kmar_QGN16394.1 | W303.asm02.HP0_G0052490 | OG0000069 | chrXV | 1042121 | 1043680 | CP015058.1 | 19940 | 21505 | 75.52581262 | 519 | 521 | 0.996161228 | 1 | 0 | 1 | FIM1_3107 | YOR374W | YOR374W | ALD4 | n |
| Kmar_QGN16159.1 | W303.asm02.HP0_G0015250 | OG0000069 | chrV | 305700 | 307262 | CP015057.1 | 925048 | 926592 | 77.11538462 | 520 | 514 | 1.011673152 | 1 | 0 | 1 | FIM1_2860 | YER073W | YER073W | ALD5 | n |
| Kmar_QGN17771.1 | W303.asm02.HP0_G0043460 | OG0000071 | chrXIII | 888079 | 893436 | CP015060.1 | 441665 | 446992 | 67.43144936 | 1785 | 1775 | 1.005633803 | 1 | 1 | 1 | FIM1_4980 | YMR306W | YMR306W | FKS3 | n |
| Kmar_QGN16573.1 | W303.asm02.HP0_G0038090 | OG0000071 | chrXII | 806175 | 811805 | CP015058.1 | 407759 | 413437 | 80.79957917 | 1876 | 1892 | 0.99154334 | 1 | 0 | 0 | FIM1_3288 | YLR342W | YLR342W | FKS1 | n |
| Kmar_QGN13600.1 | W303.asm02.HP0_G0048620 | OG0000074 | chrXV | 240659 | 241765 | CP015054.1 | 496312 | 497010 | 22.80701754 | 342 | 232 | 1.474137931 | 1 | 0 | 1 | FIM1_242 | YOL048C | YOL048C | RRT8 | n |
| Kmar_QGN14564.1 | W303.asm02.HP0_G0000530 | OG0000074 | chrI | 119328 | 120305 | CP015055.1 | 765502 | 766431 | 34.08360129 | 325 | 309 | 1.051779935 | 1 | 0 | 1 | FIM1_1226 | YAL018C | YAL018C | LDS1 | n |
| Kmar_QGN15045.1 | W303.asm02.HP0_G0030300 | OG0000075 | chrX | 584641 | 585888 | CP015056.1 | 83552 | 84820 | 72 | 415 | 422 | 0.983412322 | 1 | 1 | 1 | FIM1_1730 | YJR076C | YJR076C | CDC11 | n |
| Kmar_QGN14315.1 | W303.asm02.HP0_G0010770 | OG0000075 | chrIV | 908282 | 909553 | CP015055.1 | 260193 | 261098 | 30.96926714 | 423 | 301 | 1.405315615 | 1 | 0 | 1 | FIM1_972 | YDR218C | YDR218C | SPR28 | n |
| Kmar_QGN15969.1 | W303.asm02.HP0_G0006700 | OG0000075 | chrIV | 52518 | 54173 | CP015057.1 | 504116 | 505783 | 54.75352113 | 551 | 555 | 0.992792793 | 1 | 0 | 1 | FIM1_2667 | YDL225W | YDL225W | SHS1 | n |
| Kmar_QGN15819.1 | W303.asm02.HP0_G0035270 | OG0000076 | chrXII | 220182 | 222269 | CP015057.1 | 150416 | 152215 | 21 | 695 | 599 | 1.160267112 | 1 | 0 | 0 | FIM1_2515 | YLR035C | YLR035C | MLH2 | n |
| Kmar_QGN15398.1 | W303.asm02.HP0_G0045940 | OG0000076 | chrXIV | 473005 | 477639 | CP015056.1 | 879328 | 880929 | 24.0981241 | 1390 | 533 | 2.607879925 | 1 | 0 | 1 | FIM1_2088 | YNL082W | YNL082W | PMS1 | n |
| Kmar_QGN17274.1 | W303.asm02.HP0_G0051810 | OG0000077 | chrXV | 880154 | 882394 | CP015059.1 | 594822 | 596873 | 40.50131926 | 746 | 647 | 1.15301391 | 1 | 0 | 0 | FIM1_4005 | YOR299W | YOR299W | BUD7 | n |
| Kmar_QGN15168.1 | W303.asm02.HP0_G0028760 | OG0000077 | chrX | 229430 | 231670 | CP015056.1 | 346798 | 348951 | 38.60589812 | 746 | 717 | 1.040446304 | 1 | 0 | 1 | FIM1_1856 | YJL099W | YJL099W | CHS6 | n |
| Kmar_QGN13585.1 | W303.asm02.HP0_G0033390 | OG0000078 | chrXI | 473650 | 474639 | CP015054.1 | 468279 | 469196 | 50.45592705 | 329 | 305 | 1.078688525 | 1 | 0 | 1 | FIM1_227 | YKR013W | YKR013W | PRY2 | n |
| Kmar_QGN15327.1 | W303.asm02.HP0_G0015850 | OG0000079 | chrV | 423094 | 424425 | CP015056.1 | 711926 | 713299 | 27.21382289 | 443 | 457 | 0.969365427 | 1 | 0 | 1 | FIM1_2016 | YER130C | YER130C | COM2 | n |
| Kmar_QGN16070.1 | W303.asm02.HP0_G0040870 | OG0000079 | chrXIII | 344906 | 347020 | CP015057.1 | 743791 | 745920 | 24.50980392 | 704 | 709 | 0.992947814 | 1 | 0 | 0 | FIM1_2770 | YMR037C | YMR037C | MSN2 | n |
| Kmar_QGN13563.1 | W303.asm02.HP0_G0031020 | OG0000080 | chrX | 729489 | 731120 | CP015054.1 | 423615 | 425270 | 55.97826087 | 543 | 551 | 0.985480944 | 1 | 0 | 1 | FIM1_204 | YJR152W | YJR152W | DAL5 | n |
| Kmar_QGN13723.1 | W303.asm02.HP0_G0022290 | OG0000081 | chrVII | 1025399 | 1027003 | CP015054.1 | 740795 | 742321 | 64.23220974 | 534 | 508 | 1.051181102 | 1 | 0 | 1 | FIM1_367 | YGR260W | YGR260W | TNA1 | n |
| Kmar_QGN14521.1 | W303.asm02.HP0_G0051870 | OG0000082 | chrXV | 891587 | 893152 | CP015055.1 | 686617 | 688089 | 49.90439771 | 521 | 490 | 1.063265306 | 1 | 0 | 0 | FIM1_1183 | YOR306C | YOR306C | MCH5 | n |
| Kmar_QGN14805.1 | W303.asm02.HP0_G0012170 | OG0000083 | chrIV | 1219497 | 1220435 | CP015055.1 | 1262979 | 1263905 | 60.25641026 | 312 | 308 | 1.012987013 | 1 | 0 | 0 | FIM1_1476 | YDR368W | YDR368W | YPR1 | n |
| Kmar_QGN13803.1 | W303.asm02.HP0_G0044000 | OG0000084 | chrXIV | 69362 | 71890 | CP015054.1 | 908819 | 911476 | 56.9213732 | 842 | 885 | 0.951412429 | 1 | 0 | 0 | FIM1_449 | YNL298W | YNL298W | CLA4 | n |
| Kmar_QGN15075.1 | W303.asm02.HP0_G0023040 | OG0000084 | chrVIII | 87480 | 90299 | CP015056.1 | 148487 | 151882 | 42.6975945 | 939 | 1131 | 0.830238727 | 1 | 0 | 1 | FIM1_1760 | YHL007C | YHL007C | STE20 | n |
| Kmar_QGN18439.1 | W303.asm02.HP0_G0031140 | OG0000085 | chrXI | 4387 | 6504 | CP015061.1 | 932761 | 934764 | 26.75070028 | 705 | 667 | 1.056971514 | 1 | 0 | 1 | FIM1_4767 | YKL222C | YKL222C | | n |
| Kmar_QGN14476.1 | W303.asm02.HP0_G0050560 | OG0000085 | chrXV | 640435 | 642867 | CP015055.1 | 597025 | 599418 | 24.41860465 | 810 | 797 | 1.016311167 | 1 | 0 | 0 | FIM1_1137 | YOR162C | YOR162C | YRR1 | n |
| Kmar_QGN16108.1 | W303.asm02.HP0_G0026960 | OG0000086 | chrIX | 299231 | 299800 | CP015057.1 | 830364 | 831074 | 23.72881356 | 189 | 236 | 0.800847458 | 1 | 0 | 1 | FIM1_2809 | YIL024C | YIL024C | | n |
| Kmar_QGN14931.1 | W303.asm02.HP0_G0024750 | OG0000086 | chrVIII | 396405 | 397802 | CP015055.1 | 1517468 | 1518853 | 30.94736842 | 465 | 461 | 1.00867679 | 1 | 0 | 0 | FIM1_1606 | YHR146W | YHR146W | CRP1 | n |
| Kmar_QGN16128.1 | W303.asm02.HP0_G0025910 | OG0000087 | chrIX | 78979 | 81819 | CP015057.1 | 866843 | 869554 | 43.78947368 | 946 | 903 | 1.047619048 | 1 | 1 | 1 | FIM1_2829 | YIL137C | YIL137C | TMA108 | n |
| Kmar_QGN15413.1 | W303.asm02.HP0_G0031760 | OG0000087 | chrXI | 156926 | 159511 | CP015056.1 | 906652 | 909243 | 71.41203704 | 861 | 863 | 0.997682503 | 1 | 0 | 0 | FIM1_2103 | YKL157W | YKL157W | APE2 | n |
| Kmar_QGN13833.1 | W303.asm02.HP0_G0031600 | OG0000088 | chrXI | 123687 | 126713 | CP015054.1 | 962670 | 965495 | 43.79344588 | 1008 | 941 | 1.07120085 | 1 | 1 | 1 | FIM1_479 | YKL173W | YKL173W | SNU114 | n |
| Kmar_QGN14768.1 | W303.asm02.HP0_G0012370 | OG0000088 | chrIV | 1248824 | 1251352 | CP015055.1 | 1192064 | 1194592 | 94.06175772 | 842 | 842 | 1 | 1 | 0 | 0 | FIM1_1438 | YDR385W/YOR133W | YOR133W | EFT1 | y |
| Kmar_QGN14939.1 | W303.asm02.HP0_G0045180 | OG0000089 | chrXIV | 324524 | 325744 | CP015055.1 | 1534521 | 1535609 | 32.93556086 | 406 | 362 | 1.121546961 | 1 | 0 | 0 | FIM1_1614 | YNL165W | YNL165W | | n |
| Kmar_QGN16054.1 | W303.asm02.HP0_G0037340 | OG0000090 | chrXII | 641663 | 646114 | CP015057.1 | 711568 | 715410 | 42.45660881 | 1483 | 1280 | 1.15859375 | 1 | 1 | 1 | FIM1_2754 | YLR256W | YLR256W | HAP1 | n |
| Kmar_QGN17212.1 | W303.asm02.HP0_G0019070 | OG0000091 | chrVII | 334858 | 338811 | CP015059.1 | 456879 | 460871 | 34.63485177 | 1317 | 1330 | 0.990225564 | 1 | 1 | 1 | FIM1_3943 | YGL092W | YGL092W | NUP145 | n |
| Kmar_QGN16060.1 | W303.asm02.HP0_G0040940 | OG0000091 | chrXIII | 363867 | 367208 | CP015057.1 | 724757 | 728167 | 40.2173913 | 1113 | 1136 | 0.979753521 | 1 | 0 | 0 | FIM1_2760 | YMR047C | YMR047C | NUP116 | n |
| Kmar_QGN14283.1 | W303.asm02.HP0_G0023360 | OG0000092 | chrVIII | 145507 | 151293 | CP015055.1 | 179027 | 184612 | 45.76446281 | 1928 | 1861 | 1.036002149 | 1 | 1 | 1 | FIM1_940 | YHR023W | YHR023W | MYO1 | n |
| Kmar_QGN16496.1 | W303.asm02.HP0_G0052050 | OG0000092 | chrXV | 927441 | 932165 | CP015058.1 | 231329 | 235987 | 63.33122229 | 1574 | 1552 | 1.014175258 | 1 | 0 | 0 | FIM1_3209 | YOR326W | YOR326W | MYO2 | n |
| Kmar_QGN18034.1 | W303.asm02.HP0_G0016500 | OG0000093 | chrVI | 17586 | 19454 | CP015061.1 | 69300 | 71144 | 58.12101911 | 622 | 614 | 1.013029316 | 1 | 0 | 1 | FIM1_4350 | YFL041W | YFL041W | FET5 | n |
| Kmar_QGN17374.1 | W303.asm02.HP0_G0041020 | OG0000093 | chrXIII | 389325 | 391235 | CP015059.1 | 795427 | 797304 | 68.31210191 | 636 | 625 | 1.0176 | 1 | 0 | 1 | FIM1_4106 | YMR058W | YMR058W | FET3 | n |
| Kmar_QGN15189.1 | W303.asm02.HP0_G0021420 | OG0000094 | chrVII | 846470 | 847684 | CP015056.1 | 396068 | 397189 | 45.43209877 | 404 | 373 | 1.08310992 | 1 | 1 | 1 | FIM1_1877 | YGR169C | YGR169C | PUS6 | n |
| Kmar_QGN17282.1 | W303.asm02.HP0_G0048440 | OG0000094 | chrXV | 203151 | 204926 | CP015059.1 | 612670 | 614454 | 64.32160804 | 591 | 594 | 0.994949495 | 1 | 0 | 0 | FIM1_4013 | YOL066C | YOL066C | RIB2 | n |
| Kmar_QGN14151.1 | W303.asm02.HP0_G0005210 | OG0000095 | chrIII | 57025 | 58425 | CP015054.1 | 1638733 | 1640184 | 25.94059406 | 466 | 483 | 0.964803313 | 1 | 0 | 1 | FIM1_804 | YCL037C | YCL037C | SRO9 | n |
| Kmar_QGN16377.1 | W303.asm02.HP0_G0013520 | OG0000095 | chrIV | 1479123 | 1480466 | CP015057.1 | 1394559 | 1395170 | 13.12335958 | 447 | 203 | 2.201970443 | 1 | 0 | 1 | FIM1_3085 | YDR515W | YDR515W | SLF1 | n |
| Kmar_QGN17105.1 | W303.asm02.HP0_G0008310 | OG0000096 | chrIV | 364287 | 365114 | CP015059.1 | 239452 | 240240 | 47.68683274 | 275 | 262 | 1.049618321 | 1 | 1 | 1 | FIM1_3836 | YDL051W | YDL051W | LHP1 | n |
| Kmar_QGN17348.1 | W303.asm02.HP0_G0052220 | OG0000096 | chrXV | 979475 | 980350 | CP015059.1 | 744327 | 745244 | 32.38993711 | 291 | 305 | 0.954098361 | 1 | 0 | 1 | FIM1_4080 | YOR344C | YOR344C | TYE7 | n |
| Kmar_QGN17469.1 | W303.asm02.HP0_G0049380 | OG0000096 | chrXV | 390227 | 391531 | CP015059.1 | 984433 | 987411 | 12.90322581 | 434 | 992 | 0.4375 | 1 | 0 | 0 | FIM1_4205 | YOR032C | YOR032C | HMS1 | n |
| Kmar_QGN18229.1 | W303.asm02.HP0_G0007290 | OG0000097 | chrIV | 172693 | 174240 | CP015061.1 | 500401 | 501885 | 48.96421846 | 515 | 494 | 1.042510121 | 1 | 1 | 1 | FIM1_4554 | YDL159W | YDL159W | STE7 | n |
| Kmar_QGN14964.1 | W303.asm02.HP0_G0051160 | OG0000097 | chrXV | 774325 | 775851 | CP015055.1 | 1586929 | 1588446 | 52.60115607 | 508 | 505 | 1.005940594 | 1 | 0 | 0 | FIM1_1642 | YOR231W | YOR231W | MKK1 | n |
| Kmar_QGN15010.1 | W303.asm02.HP0_G0008520 | OG0000098 | chrIV | 408490 | 409896 | CP015055.1 | 1682469 | 1683884 | 53.81355932 | 468 | 471 | 0.993630573 | 1 | 0 | 0 | FIM1_1691 | YDL024C | YDL024C | DIA3 | n |
| Kmar_QGN17088.1 | W303.asm02.HP0_G0046840 | OG0000099 | chrXIV | 661181 | 663865 | CP015059.1 | 210014 | 212590 | 57.20620843 | 894 | 858 | 1.041958042 | 1 | 1 | 1 | FIM1_3819 | YNR013C | YNR013C | PHO91 | n |
| Kmar_QGN14053.1 | W303.asm02.HP0_G0005890 | OG0000099 | chrIII | 217500 | 220271 | CP015054.1 | 1419303 | 1422059 | 57.72532189 | 923 | 918 | 1.005446623 | 1 | 0 | 0 | FIM1_704 | YCR037C | YCR037C | PHO87 | n |
| Kmar_QGN16762.1 | W303.asm02.HP0_G0027210 | OG0000100 | chrIX | 341646 | 344486 | CP015058.1 | 839503 | 842289 | 52.26554268 | 946 | 928 | 1.019396552 | 1 | 1 | 1 | FIM1_3484 | YIL002C | YIL002C | INP51 | n |
| Kmar_QGN17337.1 | W303.asm02.HP0_G0050070 | OG0000100 | chrXV | 526153 | 529476 | CP015059.1 | 717928 | 721368 | 56.35888502 | 1107 | 1146 | 0.965968586 | 1 | 0 | 0 | FIM1_4069 | YOR109W | YOR109W | INP53 | n |
| Kmar_QGN17747.1 | W303.asm02.HP0_G0046110 | OG0000101 | chrXIV | 505911 | 507671 | CP015060.1 | 379102 | 380748 | 55.97269625 | 586 | 548 | 1.069343066 | 1 | 0 | 1 | FIM1_4956 | YNL065W | YNL065W | AQR1 | n |
| Kmar_QGN17748.1 | W303.asm02.HP0_G0026080 | OG0000101 | chrIX | 121964 | 123592 | CP015060.1 | 381478 | 383397 | 44.09836066 | 542 | 639 | 0.848200313 | 1 | 0 | 0 | FIM1_4957 | YIL121W | YIL121W | QDR2 | n |
| Kmar_QGN18386.1 | W303.asm02.HP0_G0015780 | OG0000102 | chrV | 406836 | 408410 | CP015061.1 | 823862 | 825295 | 59.54198473 | 524 | 477 | 1.098532495 | 1 | 1 | 1 | FIM1_4713 | YER123W | YER123W | YCK3 | n |
| Kmar_QGN17132.1 | W303.asm02.HP0_G0045300 | OG0000102 | chrXIV | 344328 | 345968 | CP015059.1 | 303035 | 304669 | 65.64748201 | 546 | 544 | 1.003676471 | 1 | 0 | 0 | FIM1_3863 | YNL154C | YNL154C | YCK2 | n |
| Kmar_QGN15625.1 | W303.asm02.HP0_G0015930 | OG0000103 | chrV | 465371 | 469305 | CP015056.1 | 1338198 | 1339397 | 23.57320099 | 1269 | 399 | 3.180451128 | 1 | 0 | 1 | FIM1_2318 | YER144C | YER144C | UBP5 | n |
| Kmar_QGN14224.1 | W303.asm02.HP0_G0009360 | OG0000103 | chrIV | 589088 | 591868 | CP015055.1 | 63633 | 66032 | 41.53354633 | 926 | 799 | 1.158948686 | 1 | 0 | 1 | FIM1_880 | YDR069C | YDR069C | DOA4 | n |
| Kmar_QGN13876.1 | W303.asm02.HP0_G0033780 | OG0000104 | chrXI | 557882 | 558757 | CP015054.1 | 1038725 | 1039381 | 46.39175258 | 291 | 218 | 1.334862385 | 1 | 1 | 1 | FIM1_523 | YKR055W | YKR055W | RHO4 | n |
| Kmar_QGN18250.1 | W303.asm02.HP0_G0056880 | OG0000104 | chrXVI | 881200 | 881829 | CP015061.1 | 556540 | 557166 | 80.56872038 | 209 | 208 | 1.004807692 | 1 | 0 | 1 | FIM1_4575 | YPR165W | YPR165W | RHO1 | n |
| Kmar_QGN13768.1 | W303.asm02.HP0_G0040040 | OG0000105 | chrXIII | 165076 | 166085 | CP015054.1 | 842361 | 843311 | 29.24528302 | 274 | 316 | 0.867088608 | 1 | 1 | 1 | FIM1_413 | YML052W | YML052W | SUR7 | n |
| Kmar_QGN15972.1 | W303.asm02.HP0_G0006730 | OG0000105 | chrIV | 60945 | 61874 | CP015057.1 | 515086 | 515901 | 45.27687296 | 309 | 271 | 1.140221402 | 1 | 0 | 0 | FIM1_2671 | YDL222C | YDL222C | FMP45 | n |
| Kmar_QGN15008.1 | W303.asm02.HP0_G0010980 | OG0000106 | chrIV | 951308 | 952957 | CP015055.1 | 1677254 | 1678927 | 29.8951049 | 549 | 557 | 0.985637343 | 1 | 0 | 0 | FIM1_1689 | YDR242W | YDR242W | AMD2 | n |

| | | | | | | | | | | | | | | | | | | | | |
|---|---|---|---|---|---|---|---|---|---|---|---|---|---|---|---|---|---|---|---|---|
| Kmar_QGN15547.1 | W303.asm02.HP0_G0011590 | OG0000107 | chrIV | 1080364 | 1082352 | CP015056.1 | 1162956 | 1164893 | 25.71001495 | 662 | 645 | 1.026356589 | 1 | 1 | 1 | FIM1_2238 | YDR307W | YDR307W | PMT7 | n |
| Kmar_QGN14843.1 | W303.asm02.HP0_G0007890 | OG0000107 | chrIV | 287389 | 289842 | CP015055.1 | 1331791 | 1334292 | 58.87290168 | 817 | 833 | 0.980792317 | 1 | 0 | 0 | FIM1_1514 | YDL095W | YDL095W | PMT1 | n |
| Kmar_QGN15869.1 | W303.asm02.HP0_G0002540 | OG0000108 | chrII | 373148 | 376474 | CP015057.1 | 265145 | 267721 | 35.66308244 | 1108 | 858 | 1.291375291 | 1 | 1 | 1 | FIM1_2566 | YBR059C | YBR059C | AKL1 | n |
| Kmar_QGN17708.1 | W303.asm02.HP0_G0026330 | OG0000108 | chrIX | 174295 | 176727 | CP015060.1 | 290794 | 293295 | 42.89017341 | 810 | 833 | 0.972388956 | 1 | 0 | 0 | FIM1_4915 | YIL095W | YIL095W | PRK1 | n |
| Kmar_QGN16014.1 | W303.asm02.HP0_G0021710 | OG0000109 | chrVII | 909917 | 912196 | CP015057.1 | 614572 | 616848 | 54.85564304 | 759 | 758 | 1.001319261 | 1 | 1 | 1 | FIM1_2714 | YGR199W | YGR199W | PMT6 | n |
| Kmar_QGN15242.1 | W303.asm02.HP0_G0000480 | OG0000109 | chrI | 107036 | 109315 | CP015056.1 | 510003 | 512240 | 72.45033113 | 759 | 745 | 1.018791946 | 1 | 0 | 0 | FIM1_1931 | YAL023C | YAL023C | PMT2 | n |
| Kmar_QGN16572.1 | W303.asm02.HP0_G0038110 | OG0000110 | chrXII | 812272 | 813939 | CP015058.1 | 405592 | 407172 | 54.23423423 | 555 | 526 | 1.05513308 | 1 | 1 | 1 | FIM1_3287 | YLR343W | YLR343W | GAS2 | n |
| Kmar_QGN17770.1 | W303.asm02.HP0_G0043470 | OG0000110 | chrXIII | 893922 | 895601 | CP015060.1 | 438658 | 440364 | 64.49912127 | 559 | 568 | 0.98415493 | 1 | 0 | 0 | FIM1_4979 | YMR307W | YMR307W | GAS1 | n |
| Kmar_QGN13937.1 | W303.asm02.HP0_G0039360 | OG0000111 | chrXIII | 22817 | 24358 | CP015054.1 | 1176531 | 1178087 | 73.16602317 | 513 | 518 | 0.99034749 | 1 | 0 | 1 | FIM1_585 | YML120C | YML120C | NDI1 | n |
| Kmar_QGN13506.1 | W303.asm02.HP0_G0041840 | OG0000111 | chrXIII | 555416 | 557098 | CP015054.1 | 290384 | 292018 | 70.46263345 | 560 | 544 | 1.029411765 | 1 | 0 | 1 | FIM1_145 | YMR145C | YMR145C | NDE1 | n |
| Kmar_QGN16137.1 | W303.asm02.HP0_G0032800 | OG0000112 | chrXI | 359879 | 361228 | CP015057.1 | 890470 | 891828 | 65.78366446 | 449 | 452 | 0.993362832 | 1 | 0 | 1 | FIM1_2838 | YKL046C | YKL046C | DCW1 | n |
| Kmar_QGN17275.1 | W303.asm02.HP0_G0042790 | OG0000112 | chrXIII | 746931 | 748307 | CP015059.1 | 597517 | 598881 | 66.59436009 | 458 | 454 | 1.008810573 | 1 | 0 | 1 | FIM1_4006 | YMR238W | YMR238W | DFG5 | n |
| Kmar_QGN17635.1 | W303.asm02.HP0_G0021640 | OG0000113 | chrVII | 895227 | 896225 | CP015060.1 | 154965 | 155960 | 85.24096386 | 332 | 331 | 1.003021148 | 1 | 0 | 0 | FIM1_4842 | YGR192C/YJR009C | YGR192C | TDH3 | y |
| Kmar_QGN18070.1 | W303.asm02.HP0_G0001470 | OG0000114 | chrII | 139980 | 141899 | CP015061.1 | 149176 | 151080 | 52.05479452 | 639 | 634 | 1.007886435 | 1 | 0 | 1 | FIM1_4388 | YBL042C | YBL042C | FUI1 | n |
| Kmar_QGN13546.1 | W303.asm02.HP0_G0002090 | OG0000114 | chrII | 276540 | 278441 | CP015054.1 | 384294 | 386168 | 60.47244094 | 633 | 624 | 1.014423077 | 1 | 0 | 0 | FIM1_186 | YBR021W | YBR021W | FUR4 | n |
| Kmar_QGN16135.1 | W303.asm02.HP0_G0040690 | OG0000115 | chrXIII | 317668 | 318921 | CP015057.1 | 887451 | 888950 | 27.70137525 | 417 | 499 | 0.835671343 | 1 | 1 | 1 | FIM1_2836 | YMR021C | YMR021C | MAC1 | n |
| Kmar_QGN15624.1 | W303.asm02.HP0_G0055470 | OG0000115 | chrXVI | 579375 | 581459 | CP015056.1 | 1334598 | 1336400 | 29.32761087 | 694 | 600 | 1.156666667 | 1 | 0 | 0 | FIM1_2317 | YPR008W | YPR008W | HAA1 | n |
| Kmar_QGN16647.1 | W303.asm02.HP0_G0020490 | OG0000116 | chrVII | 623080 | 624840 | CP015058.1 | 583678 | 585840 | 29.9744898 | 586 | 720 | 0.813888889 | 1 | 1 | 1 | FIM1_3366 | YGR068C | YGR068C | ART5 | n |
| Kmar_QGN14665.1 | W303.asm02.HP0_G0049250 | OG0000116 | chrXV | 364825 | 367338 | CP015055.1 | 982236 | 984566 | 37.79342723 | 837 | 776 | 1.078608247 | 1 | 0 | 0 | FIM1_1329 | YOR018W | YOR018W | ROD1 | n |
| Kmar_QGN17985.1 | W303.asm02.HP0_G0022770 | OG0000118 | chrVIII | 30817 | 31701 | CP015060.1 | 890986 | 891714 | 39.79591837 | 294 | 242 | 1.214876033 | 1 | 0 | 0 | FIM1_5194 | YHL034C | YHL034C | SBP1 | n |
| Kmar_QGN16840.1 | W303.asm02.HP0_G0017220 | OG0000118 | chrVI | 190995 | 191864 | CP015058.1 | 1007625 | 1009076 | 22.42063492 | 289 | 483 | 0.598343685 | 1 | 1 | 1 | FIM1_3565 | YFR032C | YFR032C | RRT5 | n |
| Kmar_QGN15875.1 | W303.asm02.HP0_G0002590 | OG0000119 | chrII | 382263 | 382887 | CP015057.1 | 279420 | 279761 | 33.33333333 | 180 | 113 | 1.592920354 | 1 | 0 | 1 | FIM1_2573 | YBR062C | YBR062C | | n |
| Kmar_QGN14722.1 | W303.asm02.HP0_G0010050 | OG0000119 | chrIV | 746951 | 748782 | CP015055.1 | 1096335 | 1098545 | 21.77419355 | 588 | 736 | 0.798913043 | 1 | 1 | 1 | FIM1_1390 | YDR143C | YDR143C | SAN1 | n |
| Kmar_QGN14551.1 | W303.asm02.HP0_G0048700 | OG0000120 | chrXV | 254752 | 255072 | CP015055.1 | 742411 | 742734 | 72.89719626 | 106 | 107 | 0.990654206 | 1 | 1 | 1 | FIM1_1213 | YOL039W | YOL039W | RPP2A | n |
| Kmar_QGN14774.1 | W303.asm02.HP0_G0012340 | OG0000120 | chrIV | 1245085 | 1245417 | CP015055.1 | 1204793 | 1205122 | 67.27272727 | 110 | 109 | 1.009174312 | 1 | 0 | 1 | FIM1_1444 | YDR382W | YDR382W | RPP2B | n |
| Kmar_QGN18431.1 | W303.asm02.HP0_G0055360 | OG0000121 | chrXVI | 556989 | 558014 | CP015061.1 | 915667 | 916623 | 76.83284457 | 341 | 318 | 1.072327044 | 1 | 0 | 1 | FIM1_4759 | YPL004C | YPL004C | LSP1 | n |
| Kmar_QGN14538.1 | W303.asm02.HP0_G0020630 | OG0000121 | chrVII | 647635 | 648654 | CP015055.1 | 719585 | 720511 | 80.2359882 | 339 | 308 | 1.100649351 | 1 | 0 | 1 | FIM1_1200 | YGR086C | YGR086C | PIL1 | n |
| Kmar_QGN17696.1 | W303.asm02.HP0_G0026400 | OG0000122 | chrIX | 188426 | 189898 | CP015060.1 | 269617 | 271053 | 49.79674797 | 490 | 478 | 1.025104603 | 1 | 1 | 1 | FIM1_4903 | YIL088C | YIL088C | AVT7 | n |
| Kmar_QGN18134.1 | W303.asm02.HP0_G0015740 | OG0000122 | chrV | 401518 | 402864 | CP015061.1 | 295421 | 296821 | 56.92640693 | 448 | 466 | 0.961373391 | 1 | 0 | 0 | FIM1_4456 | YER119C | YER119C | AVT6 | n |
| Kmar_QGN16222.1 | W303.asm02.HP0_G0008170 | OG0000123 | chrIV | 335165 | 336451 | CP015057.1 | 1064748 | 1066037 | 80.7424594 | 428 | 429 | 0.997668998 | 1 | 1 | 1 | FIM1_2926 | YDL066W | YDL066W | IDP1 | n |
| Kmar_QGN15161.1 | W303.asm02.HP0_G0036610 | OG0000123 | chrXII | 504852 | 506090 | CP015056.1 | 329017 | 330264 | 80 | 412 | 415 | 0.992771084 | 1 | 0 | 0 | FIM1_1848 | YLR174W | YLR174W | IDP2 | n |
| Kmar_QGN15178.1 | W303.asm02.HP0_G0028860 | OG0000124 | chrX | 255942 | 258431 | CP015056.1 | 373810 | 376059 | 26.36363636 | 829 | 749 | 1.106809079 | 1 | 1 | 1 | FIM1_1866 | YJL089W | YJL089W | SIP4 | n |
| Kmar_QGN17622.1 | W303.asm02.HP0_G0043190 | OG0000124 | chrXIII | 833950 | 838251 | CP015060.1 | 124511 | 128890 | 33.12262958 | 1433 | 1459 | 0.982179575 | 1 | 0 | 1 | FIM1_4829 | YMR280C | YMR280C | CAT8 | n |
| Kmar_QGN14049.1 | W303.asm02.HP0_G0005860 | OG0000125 | chrIII | 213678 | 214721 | CP015054.1 | 1410958 | 1412019 | 74.78753541 | 347 | 353 | 0.983002833 | 1 | 1 | 1 | FIM1_700 | YCR034W | YCR034W | ELO2 | n |
| Kmar_QGN16642.1 | W303.asm02.HP0_G0038380 | OG0000125 | chrXII | 864020 | 865057 | CP015058.1 | 571600 | 572634 | 66.66666667 | 345 | 344 | 1.002906977 | 1 | 0 | 1 | FIM1_3361 | YLR372W | YLR372W | ELO3 | n |
| Kmar_QGN17269.1 | W303.asm02.HP0_G0051780 | OG0000127 | chrXV | 876462 | 877040 | CP015059.1 | 588975 | 589520 | 37.18592965 | 192 | 181 | 1.060773481 | 1 | 1 | 1 | FIM1_4000 | YOR297C | YOR297C | TIM18 | n |
| Kmar_QGN16911.1 | W303.asm02.HP0_G0036530 | OG0000127 | chrXII | 494144 | 494650 | CP015058.1 | 1176891 | 1177385 | 50.89820359 | 168 | 164 | 1.024390244 | 1 | 0 | 0 | FIM1_3638 | YLR164W | YLR164W | SHH4 | n |
| Kmar_QGN14742.1 | W303.asm02.HP0_G0013070 | OG0000128 | chrIV | 1400814 | 1403510 | CP015055.1 | 1141349 | 1143418 | 37.20930233 | 898 | 689 | 1.303338171 | 1 | 1 | 1 | FIM1_1411 | YDR466W | YDR466W | PKH3 | n |
| Kmar_QGN17841.1 | W303.asm02.HP0_G0048100 | OG0000128 | chrXV | 129692 | 132937 | CP015060.1 | 597742 | 600660 | 35.86459286 | 1081 | 972 | 1.112139918 | 1 | 0 | 0 | FIM1_5050 | YOL100W | YOL100W | PKH2 | n |
| Kmar_QGN17248.1 | W303.asm02.HP0_G0022970 | OG0000129 | chrVIII | 74435 | 75652 | CP015059.1 | 546045 | 547274 | 62.28710462 | 405 | 409 | 0.990220049 | 1 | 1 | 1 | FIM1_3979 | YHL014C | YHL014C | YLF2 | n |
| Kmar_QGN13541.1 | W303.asm02.HP0_G0002130 | OG0000129 | chrII | 285778 | 286962 | CP015054.1 | 365100 | 366281 | 89.34010152 | 394 | 393 | 1.002544529 | 1 | 0 | 1 | FIM1_181 | YBR025C | YBR025C | OLA1 | n |
| Kmar_QGN16117.1 | W303.asm02.HP0_G0054810 | OG0000130 | chrXVI | 440881 | 442122 | CP015057.1 | 841794 | 842990 | 53.47721823 | 413 | 398 | 1.037688442 | 1 | 1 | 1 | FIM1_2818 | YPL060W | YPL060W | MFM1 | n |
| Kmar_QGN16491.1 | W303.asm02.HP0_G0052120 | OG0000130 | chrXV | 946743 | 948155 | CP015058.1 | 220207 | 221547 | 51.98329854 | 470 | 446 | 1.053811659 | 1 | 0 | 0 | FIM1_3204 | YOR334W | YOR334W | MRS2 | n |
| Kmar_QGN15537.1 | W303.asm02.HP0_G0006910 | OG0000131 | chrIV | 96156 | 98027 | CP015056.1 | 1145327 | 1146934 | 38.17619784 | 623 | 535 | 1.164485981 | 1 | 1 | 1 | FIM1_2228 | YDL203C | YDL203C | ACK1 | n |
| Kmar_QGN17930.1 | W303.asm02.HP0_G0001290 | OG0000131 | chrII | 106464 | 108554 | CP015060.1 | 758216 | 760324 | 44.69496021 | 696 | 702 | 0.991452991 | 1 | 0 | 0 | FIM1_5139 | YBL061C | YBL061C | SKT5 | n |
| Kmar_QGN13806.1 | W303.asm02.HP0_G0047950 | OG0000132 | chrXV | 100264 | 101412 | CP015054.1 | 915328 | 916620 | 32.50564334 | 382 | 430 | 0.888372093 | 1 | 0 | 1 | FIM1_452 | YOL116W | YOL116W | MSN1 | n |
| Kmar_QGN17142.1 | W303.asm02.HP0_G0042100 | OG0000132 | chrXIII | 606565 | 608724 | CP015059.1 | 324534 | 326822 | 22.94654498 | 719 | 762 | 0.943569554 | 1 | 0 | 1 | FIM1_3873 | YMR172W | YMR172W | HOT1 | n |
| Kmar_QGN14353.1 | W303.asm02.HP0_G0032240 | OG0000133 | chrXI | 257396 | 261952 | CP015055.1 | 337869 | 341780 | 33.55176933 | 1518 | 1303 | 1.165003837 | 1 | 1 | 1 | FIM1_1013 | YKL101W | YKL101W | HSL1 | n |
| Kmar_QGN14132.1 | W303.asm02.HP0_G0013460 | OG0000133 | chrIV | 1468052 | 1471480 | CP015054.1 | 1603819 | 1607160 | 52.62252794 | 1142 | 1113 | 1.026055705 | 1 | 0 | 0 | FIM1_785 | YDR507C | YDR507C | GIN4 | n |
| Kmar_QGN14191.1 | W303.asm02.HP0_G0052540 | OG0000134 | chrXV | 1051792 | 1053339 | CP015054.1 | 1724682 | 1726214 | 63.49809886 | 515 | 510 | 1.009803922 | 1 | 0 | 1 | FIM1_844 | YOR378W | YOR378W | AMF1 | n |
| Kmar_QGN15800.1 | W303.asm02.HP0_G0020020 | OG0000135 | chrVII | 518624 | 519610 | CP015057.1 | 121040 | 121963 | 41.21212121 | 328 | 307 | 1.068403909 | 1 | 0 | 0 | FIM1_2496 | YGR015C | YGR015C | EAT1 | n |
| Kmar_QGN16512.1 | W303.asm02.HP0_G0041070 | OG0000136 | chrXIII | 397580 | 398299 | CP015058.1 | 277980 | 278609 | 26.97095436 | 239 | 209 | 1.14354067 | 1 | 1 | 1 | FIM1_3225 | YMR063W | YMR063W | RIM9 | n |
| Kmar_QGN13671.1 | W303.asm02.HP0_G0048890 | OG0000136 | chrXV | 289353 | 291008 | CP015054.1 | 636351 | 638378 | 25.76866764 | 551 | 675 | 0.816296296 | 1 | 0 | 0 | FIM1_314 | YOL019W | YOL019W | TOS7 | n |
| Kmar_QGN14015.1 | W303.asm02.HP0_G0008570 | OG0000137 | chrIV | 422870 | 423547 | CP015054.1 | 1332885 | 1333514 | 37.33333333 | 225 | 209 | 1.076555024 | 1 | 0 | 1 | FIM1_666 | YDL018C | YDL018C | ERP3 | n |
| Kmar_QGN16788.1 | W303.asm02.HP0_G0049230 | OG0000137 | chrXV | 360917 | 361540 | CP015058.1 | 890995 | 891624 | 67.14285714 | 207 | 209 | 0.990430622 | 1 | 0 | 1 | FIM1_3511 | YOR016C | YOR016C | ERP4 | n |
| Kmar_QGN17064.1 | W303.asm02.HP0_G0007560 | OG0000138 | chrIV | 230112 | 230282 | CP015059.1 | 156115 | 156429 | 34.73684211 | 56 | 104 | 0.538461538 | 1 | 1 | 1 | FIM1_3794 | W303.asm02.HP0_G0007560 | YDL130W | RPP1B | y |
| Kmar_QGN13500.1 | W303.asm02.HP0_G0008030 | OG0000138 | chrIV | 310132 | 310452 | CP015054.1 | 273723 | 274046 | 76.63551402 | 106 | 107 | 0.990654206 | 1 | 0 | 1 | FIM1_138 | YDL081C | YDL081C | RPP1A | n |
| Kmar_QGN13542.1 | W303.asm02.HP0_G0002240 | OG0000139 | chrII | 306999 | 309890 | CP015054.1 | 367249 | 369996 | 64.90134995 | 963 | 915 | 1.052459016 | 1 | 1 | 1 | FIM1_182 | YBR038W | YBR038W | CHS2 | n |
| Kmar_QGN15974.1 | W303.asm02.HP0_G0044950 | OG0000139 | chrXIV | 277202 | 280597 | CP015057.1 | 522235 | 525288 | 50.70422535 | 1131 | 1017 | 1.112094395 | 1 | 0 | 1 | FIM1_2674 | YNL192W | YNL192W | CHS1 | n |
| Kmar_QGN18307.1 | W303.asm02.HP0_G0027390 | OG0000140 | chrIX | 374724 | 375287 | CP015061.1 | 661527 | 662384 | 31.35888502 | 187 | 285 | 0.656140351 | 1 | 0 | 1 | FIM1_4633 | YIR017C | YIR017C | MET28 | n |
| Kmar_QGN17810.1 | W303.asm02.HP0_G0045750 | OG0000140 | chrXIV | 429390 | 431408 | CP015060.1 | 530986 | 533109 | 26.8 | 672 | 707 | 0.95049505 | 1 | 0 | 1 | FIM1_5019 | YNL103W | YNL103W | MET4 | n |
| Kmar_QGN17049.1 | W303.asm02.HP0_G0006990 | OG0000141 | chrIV | 111654 | 114308 | CP015059.1 | 122311 | 123741 | 48.49445325 | 884 | 476 | 1.857142857 | 1 | 0 | 0 | FIM1_3778 | YDL194W | YDL194W | SNF3 | n |
| Kmar_QGN16900.1 | W303.asm02.HP0_G0036460 | OG0000142 | chrXII | 461546 | 463276 | CP015058.1 | 1130710 | 1132632 | 47.65625 | 576 | 640 | 0.9 | 1 | 0 | 1 | FIM1_3627 | YLR152C | YLR152C | | n |
| Kmar_QGN17792.1 | W303.asm02.HP0_G0049920 | OG0000142 | chrXV | 495862 | 497703 | CP015060.1 | 493167 | 494807 | 56.60685155 | 613 | 546 | 1.122710623 | 1 | 0 | 0 | FIM1_5001 | YOR092W | YOR092W | ECM3 | n |
| Kmar_QGN15611.1 | W303.asm02.HP0_G0000180 | OG0000143 | chrI | 43516 | 45657 | CP015056.1 | 1304876 | 1306999 | 76.75070028 | 713 | 707 | 1.008486563 | 1 | 0 | 1 | FIM1_2304 | YAL054C | YAL054C | ACS1 | n |
| Kmar_QGN16901.1 | W303.asm02.HP0_G0036470 | OG0000143 | chrXII | 464112 | 466163 | CP015058.1 | 1133969 | 1136023 | 83.47953216 | 683 | 684 | 0.998538012 | 1 | 0 | 1 | FIM1_3628 | YLR153C | YLR153C | ACS2 | n |
| Kmar_QGN13748.1 | W303.asm02.HP0_G0000710 | OG0000144 | chrI | 155269 | 155928 | CP015054.1 | 803667 | 804317 | 56.16438356 | 219 | 216 | 1.013888889 | 1 | 0 | 0 | FIM1_392 | YAR002C-A | YAR002C-A | ERP1 | n |
| Kmar_QGN14637.1 | W303.asm02.HP0_G0024400 | OG0000144 | chrVIII | 338255 | 338893 | CP015055.1 | 919247 | 919888 | 39.43661972 | 212 | 213 | 0.995305164 | 1 | 0 | 1 | FIM1_1301 | YHR110W | YHR110W | ERP5 | n |
| Kmar_QGN15109.1 | W303.asm02.HP0_G0041420 | OG0000145 | chrXIII | 468950 | 469981 | CP015056.1 | 222195 | 223352 | 41.4507772 | 343 | 385 | 0.890909091 | 1 | 1 | 1 | FIM1_1796 | YMR101C | YMR101C | SRT1 | n |
| Kmar_QGN17449.1 | W303.asm02.HP0_G0001920 | OG0000145 | chrII | 244047 | 244907 | CP015059.1 | 943089 | 943919 | 54.48275862 | 286 | 276 | 1.036231884 | 1 | 1 | 1 | FIM1_4183 | YBR002C | YBR002C | RER2 | n |
| Kmar_QGN17423.1 | W303.asm02.HP0_G0048330 | OG0000146 | chrXV | 185893 | 186099 | CP015059.1 | 892031 | 892240 | 44.92753623 | 68 | 69 | 0.985507246 | 1 | 1 | 1 | FIM1_4155 | YOL077W-A | YOL077W-A | ATP19 | n |
| Kmar_QGN17661.1 | W303.asm02.HP0_G0049280 | OG0000146 | chrXV | 372141 | 372413 | CP015060.1 | 208308 | 208535 | 39.50617284 | 90 | 75 | 1.2 | 1 | 1 | 1 | FIM1_4868 | YOR020W-A | YOR020W-A | MCO10 | n |
| Kmar_QGN14159.1 | W303.asm02.HP0_G0013560 | OG0000147 | chrIV | 1486118 | 1486525 | CP015054.1 | 1655206 | 1655628 | 56.33802817 | 135 | 140 | 0.964285714 | 1 | 1 | 1 | FIM1_812 | YDR519W | YDR519W | FPR2 | n |
| Kmar_QGN17628.1 | W303.asm02.HP0_G0045470 | OG0000147 | chrXIV | 372715 | 373059 | CP015060.1 | 141725 | 142069 | 90.35087719 | 114 | 114 | 1 | 1 | 1 | 1 | FIM1_4835 | YNL135C | YNL135C | FPR1 | n |
| Kmar_QGN13467.1 | W303.asm02.HP0_G0024580 | OG0000148 | chrVIII | 368972 | 370126 | CP015054.1 | 199114 | 200304 | 55.38847118 | 384 | 396 | 0.96969697 | 1 | 1 | 1 | FIM1_104 | YHR129C | YHR129C | ARP1 | n |
| Kmar_QGN18040.1 | W303.asm02.HP0_G0016520 | OG0000148 | chrVI | 21707 | 22789 | CP015061.1 | 81567 | 82703 | 92.59259259 | 360 | 378 | 0.952380952 | 1 | 1 | 1 | FIM1_4356 | YFL039C | YFL039C | ACT1 | n |
| Kmar_QGN17877.1 | W303.asm02.HP0_G0056050 | OG0000149 | chrXVI | 696605 | 698287 | CP015060.1 | 661645 | 663441 | 51.20772947 | 560 | 598 | 0.936454849 | 1 | 1 | 1 | FIM1_5086 | YPR072W | YPR072W | NOT5 | n |
| Kmar_QGN14710.1 | W303.asm02.HP0_G0026830 | OG0000149 | chrIX | 270771 | 273281 | CP015055.1 | 1072894 | 1075185 | 42.97619048 | 836 | 763 | 1.095674967 | 1 | 1 | 1 | FIM1_1377 | YIL038C | YIL038C | NOT3 | n |
| Kmar_QGN18353.1 | W303.asm02.HP0_G0054900 | OG0000150 | chrXVI | 466322 | 466918 | CP015061.1 | 754840 | 755439 | 37.31343284 | 198 | 199 | 0.994974874 | 1 | 1 | 1 | FIM1_4680 | YPL051W | YPL051W | ARL3 | n |
| Kmar_QGN17794.1 | W303.asm02.HP0_G0049940 | OG0000150 | chrXV | 503530 | 504081 | CP015060.1 | 501640 | 502188 | 82.5136612 | 183 | 182 | 1.005494505 | 1 | 1 | 1 | FIM1_5003 | YOR094W | YOR094W | ARF3 | n |
| Kmar_QGN17159.1 | W303.asm02.HP0_G0054930 | OG0000151 | chrXVI | 470761 | 472008 | CP015059.1 | 360354 | 361025 | 50.22222222 | 415 | 223 | 1.860986547 | 1 | 0 | 0 | FIM1_3890 | YPL048W | YPL048W | CAM1 | n |
| Kmar_QGN18314.1 | W303.asm02.HP0_G0002880 | OG0000152 | chrII | 443441 | 443770 | CP015061.1 | 677006 | 677329 | 61.46788991 | 109 | 107 | 1.018691589 | 1 | 1 | 1 | FIM1_4640 | YBR091C | YBR091C | TIM12 | n |
| Kmar_QGN17397.1 | W303.asm02.HP0_G0023160 | OG0000152 | chrVIII | 108604 | 108885 | CP015059.1 | 836298 | 836567 | 77.41935484 | 93 | 89 | 1.04494382 | 1 | 1 | 1 | FIM1_4129 | YHR005C-A | YHR005C-A | TIM10 | n |
| Kmar_QGN16228.1 | W303.asm02.HP0_G0057100 | OG0000153 | chrXVI | 917820 | 918311 | CP015057.1 | 1078196 | 1078660 | 33.12883436 | 163 | 154 | 1.058441558 | 1 | 0 | 1 | FIM1_2932 | YPR188C | YPR188C | MLC2 | n |
| Kmar_QGN17867.1 | W303.asm02.HP0_G0003040 | OG0000153 | chrII | 474205 | 474648 | CP015060.1 | 639667 | 640110 | 93.87755102 | 147 | 147 | 1 | 1 | 0 | 1 | FIM1_5076 | YBR109C | YBR109C | CMD1 | n |
| Kmar_QGN17329.1 | W303.asm02.HP0_G0045680 | OG0000154 | chrXIV | 414846 | 417936 | CP015059.1 | 706163 | 707542 | 65.63071298 | 546 | 459 | 1.189542484 | 1 | 1 | 1 | FIM1_4061 | YNL112W | YNL112W | DBP2 | n |
| Kmar_QGN14434.1 | W303.asm02.HP0_G0004270 | OG0000154 | chrII | 714812 | 717361 | CP015055.1 | 509996 | 512449 | 47.18969555 | 849 | 817 | 1.039167687 | 1 | 1 | 1 | FIM1_1095 | YBR237W | YBR237W | PRP5 | n |
| Kmar_QGN14598.1 | W303.asm02.HP0_G0004320 | OG0000155 | chrII | 728025 | 728741 | CP015055.1 | 830860 | 831636 | 55.42635659 | 238 | 258 | 0.92248062 | 1 | 1 | 1 | FIM1_1261 | YBR242W | YBR242W | | n |
| Kmar_QGN14724.1 | W303.asm02.HP0_G0021080 | OG0000156 | chrVII | 768623 | 769174 | CP015055.1 | 1101781 | 1102311 | 31.55080214 | 183 | 176 | 1.039772727 | 1 | 1 | 1 | FIM1_1392 | YGR133W | YGR133W | PEX4 | n |
| Kmar_QGN16366.1 | W303.asm02.HP0_G0012720 | OG0000157 | chrIV | 1325620 | 1327497 | CP015057.1 | 1370356 | 1372095 | 35.02377179 | 625 | 579 | 1.079447323 | 1 | 1 | 1 | FIM1_3072 | YDR425W | YDR425W | SNX41 | n |
| Kmar_QGN17668.1 | W303.asm02.HP0_G0007720 | OG0000157 | chrIV | 256839 | 258761 | CP015060.1 | 219655 | 221457 | 44.89164087 | 640 | 600 | 1.066666667 | 1 | 1 | 1 | FIM1_4875 | YDL113C | YDL113C | ATG20 | n |
| Kmar_QGN15257.1 | W303.asm02.HP0_G0033900 | OG0000158 | chrXI | 577585 | 579816 | CP015056.1 | 544031 | 546316 | 47.94344473 | 743 | 761 | 0.976346912 | 1 | 1 | 1 | FIM1_1946 | YKR067W | YKR067W | GPT2 | n |
| Kmar_QGN15331.1 | W303.asm02.HP0_G0001790 | OG0000158 | chrII | 205823 | 208102 | CP015056.1 | 723581 | 725734 | 58.20895522 | 759 | 717 | 1.058577406 | 1 | 1 | 1 | FIM1_2020 | YBL011W | YBL011W | SCT1 | n |
| Kmar_QGN18151.1 | W303.asm02.HP0_G0015220 | OG0000159 | chrV | 300620 | 303286 | CP015061.1 | 333573 | 336212 | 78.69955157 | 888 | 879 | 1.010238908 | 1 | 0 | 0 | FIM1_4474 | YER070W | YER070W | RNR1 | n |
| Kmar_QGN14980.1 | W303.asm02.HP0_G0051050 | OG0000160 | chrXV | 753509 | 756304 | CP015055.1 | 1613450 | 1616065 | 39.09774436 | 931 | 871 | 1.068886338 | 1 | 1 | 1 | FIM1_1658 | YOR219C | YOR219C | STE13 | n |
| Kmar_QGN14278.1 | W303.asm02.HP0_G0023410 | OG0000160 | chrVIII | 158821 | 161277 | CP015055.1 | 168337 | 170802 | 56.74698795 | 818 | 821 | 0.99634592 | 1 | 1 | 1 | FIM1_935 | YHR028C | YHR028C | DAP2 | n |
| Kmar_QGN13488.1 | W303.asm02.HP0_G0014190 | OG0000161 | chrV | 97651 | 98589 | CP015054.1 | 245480 | 246433 | 52.17391304 | 312 | 317 | 0.984227129 | 1 | 1 | 1 | FIM1_126 | YEL029C | YEL029C | BUD16 | n |
| Kmar_QGN15512.1 | W303.asm02.HP0_G0046980 | OG0000161 | chrXIV | 687505 | 688458 | CP015056.1 | 1096928 | 1097836 | 46.37223975 | 317 | 302 | 1.049668874 | 1 | 1 | 1 | FIM1_2203 | YNR027W | YNR027W | BUD17 | n |
| Kmar_QGN14807.1 | W303.asm02.HP0_G0021690 | OG0000162 | chrVII | 904912 | 906555 | CP015055.1 | 1267737 | 1269356 | 39.61937716 | 547 | 539 | 1.014842301 | 1 | 0 | 0 | FIM1_1478 | YGR197C | YGR197C | SNG1 | n |

| | | | | | | | | | | | | | | | | | | | | |
|---|---|---|---|---|---|---|---|---|---|---|---|---|---|---|---|---|---|---|---|---|
| Kmar_QGN17692.1 | W303.asm02.HP0_G0001990 | OG0000163 | chrII | 257711 | 258022 | CP015060.1 | 266036 | 266347 | 97.08737864 | 103 | 103 | 1 | 1 | 0 | 0 | FIM1_4899 | YBR009C/YNL030W | YNL030W | HHF2 | y |
| Kmar_QGN13857.1 | W303.asm02.HP0_G0028310 | OG0000164 | chrX | 126907 | 128898 | CP015054.1 | 1006803 | 1008401 | 46.72897196 | 663 | 532 | 1.246240602 | 1 | 1 | 1 | FIM1_503 | YJL149W | YJL149W | DAS1 | n |
| Kmar_QGN15934.1 | W303.asm02.HP0_G0009950 | OG0000164 | chrIV | 721699 | 723369 | CP015057.1 | 417318 | 418907 | 31.79396092 | 556 | 529 | 1.051039698 | 1 | 1 | 1 | FIM1_2632 | YDR131C | YDR131C | | n |
| Kmar_QGN14349.1 | W303.asm02.HP0_G0032210 | OG0000165 | chrXI | 245395 | 246750 | CP015055.1 | 327296 | 328582 | 56.98447894 | 451 | 428 | 1.053738318 | 1 | 1 | 1 | FIM1_1009 | YKL106W | YKL106W | AAT1 | n |
| Kmar_QGN15835.1 | W303.asm02.HP0_G0035200 | OG0000165 | chrXII | 204214 | 205512 | CP015057.1 | 188448 | 189737 | 66.13995485 | 432 | 429 | 1.006993007 | 1 | 1 | 1 | FIM1_2531 | YLR027C | YLR027C | AAT2 | n |
| Kmar_QGN14706.1 | W303.asm02.HP0_G0014980 | OG0000166 | chrV | 248253 | 249428 | CP015055.1 | 1065310 | 1066581 | 43.97163121 | 391 | 423 | 0.924349882 | 1 | 1 | 1 | FIM1_1373 | YER048C | YER048C | CAJ1 | n |
| Kmar_QGN16744.1 | W303.asm02.HP0_G0027260 | OG0000166 | chrIX | 354275 | 355573 | CP015058.1 | 802198 | 803469 | 64.84018265 | 432 | 423 | 1.021276596 | 1 | 1 | 1 | FIM1_3466 | YIR004W | YIR004W | DJP1 | n |
| Kmar_QGN17422.1 | W303.asm02.HP0_G0003320 | OG0000167 | chrII | 531950 | 533476 | CP015059.1 | 889872 | 891350 | 52.84872299 | 508 | 492 | 1.032520325 | 1 | 0 | 1 | FIM1_4154 | YBR139W | YBR139W | ATG42 | n |
| Kmar_QGN13702.1 | W303.asm02.HP0_G0043370 | OG0000167 | chrXIII | 868842 | 870440 | CP015054.1 | 689459 | 691081 | 65.56169429 | 532 | 540 | 0.985185185 | 1 | 1 | 1 | FIM1_345 | YMR297W | YMR297W | PRC1 | n |
| Kmar_QGN14501.1 | W303.asm02.HP0_G0035490 | OG0000168 | chrXII | 265804 | 267213 | CP015055.1 | 644648 | 646057 | 85.71428571 | 469 | 469 | 1 | 1 | 1 | 1 | FIM1_1162 | YLR058C | YLR058C | SHM2 | n |
| Kmar_QGN16430.1 | W303.asm02.HP0_G0004530 | OG0000168 | chrII | 759620 | 761092 | CP015058.1 | 102803 | 104296 | 69.41649899 | 490 | 497 | 0.985915493 | 1 | 1 | 1 | FIM1_3143 | YBR263W | YBR263W | SHM1 | n |
| Kmar_QGN15316.1 | W303.asm02.HP0_G0037750 | OG0000169 | chrXII | 730469 | 732805 | CP015056.1 | 684998 | 687340 | 86.6837388 | 778 | 780 | 0.997435897 | 1 | 1 | 1 | FIM1_2005 | YLR304C | YLR304C | ACO1 | n |
| Kmar_QGN14054.1 | W303.asm02.HP0_G0027870 | OG0000169 | chrX | 45975 | 48344 | CP015054.1 | 1423584 | 1425977 | 80.02512563 | 789 | 797 | 0.989962359 | 1 | 1 | 1 | FIM1_705 | YJL200C | YJL200C | ACO2 | n |
| Kmar_QGN15922.1 | W303.asm02.HP0_G0009900 | OG0000170 | chrIV | 708143 | 709153 | CP015057.1 | 380937 | 381902 | 38.3480826 | 336 | 321 | 1.046728972 | 1 | 1 | 1 | FIM1_2620 | YDR126W | YDR126W | SWF1 | n |
| Kmar_QGN16923.1 | W303.asm02.HP0_G0037260 | OG0000170 | chrXII | 622356 | 623435 | CP015058.1 | 1196548 | 1197627 | 44.23076923 | 359 | 359 | 1 | 1 | 1 | 1 | FIM1_3650 | YLR246W | YLR246W | ERF2 | n |
| Kmar_QGN16172.1 | W303.asm02.HP0_G0051530 | OG0000171 | chrXV | 829771 | 832293 | CP015057.1 | 955058 | 957577 | 64.15981199 | 840 | 839 | 1.001191895 | 1 | 1 | 1 | FIM1_2874 | YOR270C | YOR270C | VPH1 | n |
| Kmar_QGN17370.1 | W303.asm02.HP0_G0040990 | OG0000171 | chrXIII | 383806 | 386478 | CP015059.1 | 785481 | 788093 | 61.03183315 | 890 | 870 | 1.022988506 | 1 | 1 | 1 | FIM1_4102 | YMR054W | YMR054W | STV1 | n |
| Kmar_QGN16739.1 | W303.asm02.HP0_G0023440 | OG0000172 | chrVIII | 164642 | 166813 | CP015058.1 | 791343 | 793454 | 42.46947083 | 723 | 703 | 1.028449502 | 1 | 1 | 1 | FIM1_3461 | YHR031C | YHR031C | RRM3 | n |
| Kmar_QGN16551.1 | W303.asm02.HP0_G0039930 | OG0000172 | chrXIII | 143493 | 146072 | CP015058.1 | 354419 | 356977 | 41.76334107 | 859 | 852 | 1.008215962 | 1 | 1 | 1 | FIM1_3265 | YML061C | YML061C | PIF1 | n |
| Kmar_QGN14368.1 | W303.asm02.HP0_G0009680 | OG0000173 | chrIV | 657180 | 659933 | CP015055.1 | 371167 | 373458 | 25.2972973 | 917 | 763 | 1.201834862 | 1 | 1 | 1 | FIM1_1028 | YDR103W | YDR103W | STE5 | n |
| Kmar_QGN16629.1 | W303.asm02.HP0_G0028250 | OG0000173 | chrX | 113364 | 115856 | CP015058.1 | 535698 | 538463 | 20.61965812 | 830 | 921 | 0.901194354 | 1 | 1 | 1 | FIM1_3346 | YJL157C | YJL157C | FAR1 | n |
| Kmar_QGN17752.1 | W303.asm02.HP0_G0030610 | OG0000174 | chrX | 639358 | 642714 | CP015060.1 | 391542 | 394895 | 81.6399287 | 1118 | 1117 | 1.000895255 | 1 | 1 | 1 | FIM1_4961 | YJR109C | YJR109C | CPA2 | n |
| Kmar_QGN17530.1 | W303.asm02.HP0_G0028500 | OG0000174 | chrX | 155741 | 162385 | CP015059.1 | 1101662 | 1108348 | 81.35365307 | 2214 | 2228 | 0.993716338 | 1 | 1 | 1 | FIM1_4266 | YJL130C | YJL130C | URA2 | n |
| Kmar_QGN18338.1 | W303.asm02.HP0_G0002800 | OG0000175 | chrII | 427340 | 430267 | CP015061.1 | 721074 | 723995 | 73.51738241 | 975 | 973 | 1.002055498 | 1 | 0 | 0 | FIM1_4665 | YBR084W | YBR084W | MIS1 | n |
| Kmar_QGN14365.1 | W303.asm02.HP0_G0001880 | OG0000176 | chrII | 237733 | 238131 | CP015055.1 | 365977 | 366369 | 96.96969697 | 132 | 130 | 1.015384615 | 1 | 0 | 0 | FIM1_1025 | YBL003C/YDR225W | YBL003C | HTA2 | y |
| Kmar_QGN15919.1 | W303.asm02.HP0_G0009890 | OG0000177 | chrIV | 706306 | 707667 | CP015057.1 | 378137 | 379366 | 41.37168142 | 453 | 409 | 1.107579462 | 1 | 0 | 0 | FIM1_2617 | YDR125C | YDR125C | ECM18 | n |
| Kmar_QGN13430.1 | W303.asm02.HP0_G0007090 | OG0000178 | chrIV | 133657 | 134943 | CP015054.1 | 133982 | 135163 | 89.08629442 | 428 | 393 | 1.089058524 | 1 | 0 | 0 | FIM1_67 | YDL182W | YDL182W | LYS20 | n |
| Kmar_QGN17062.1 | W303.asm02.HP0_G0007540 | OG0000178 | chrIV | 227599 | 228921 | CP015059.1 | 152278 | 153606 | 90.51918736 | 440 | 442 | 0.995475113 | 1 | 0 | 0 | FIM1_3792 | YDL131W | YDL131W | LYS21 | n |
| Kmar_QGN18406.1 | W303.asm02.HP0_G0031850 | OG0000179 | chrXI | 173051 | 175129 | CP015061.1 | 856590 | 858293 | 45.16594517 | 692 | 567 | 1.220458554 | 1 | 1 | 1 | FIM1_4733 | YKL146W | YKL146W | AVT3 | n |
| Kmar_QGN17807.1 | W303.asm02.HP0_G0045770 | OG0000179 | chrXIV | 436654 | 438795 | CP015060.1 | 522381 | 524555 | 52.67489712 | 713 | 724 | 0.98480663 | 1 | 1 | 1 | FIM1_5016 | YNL101W | YNL101W | AVT4 | n |
| Kmar_QGN16645.1 | W303.asm02.HP0_G0038350 | OG0000180 | chrXII | 855732 | 857705 | CP015058.1 | 578621 | 580564 | 68.23708207 | 657 | 647 | 1.015455951 | 1 | 1 | 1 | FIM1_3364 | YLR369W | YLR369W | SSQ1 | n |
| Kmar_QGN13490.1 | W303.asm02.HP0_G0030000 | OG0000180 | chrX | 528169 | 530133 | CP015054.1 | 248213 | 250138 | 89.29663609 | 654 | 641 | 1.020280811 | 1 | 1 | 1 | FIM1_128 | YJR045C | YJR045C | SSC1 | n |
| Kmar_QGN17110.1 | W303.asm02.HP0_G0056840 | OG0000181 | chrXVI | 870281 | 872254 | CP015059.1 | 254038 | 256047 | 48.45814978 | 657 | 669 | 0.98206278 | 1 | 1 | 1 | FIM1_3841 | YPR161C | YPR161C | SGV1 | n |
| Kmar_QGN15361.1 | W303.asm02.HP0_G0003530 | OG0000181 | chrII | 576855 | 577751 | CP015056.1 | 794401 | 795288 | 86.24161074 | 298 | 295 | 1.010169492 | 1 | 1 | 1 | FIM1_2051 | YBR160W | YBR160W | CDC28 | n |
| Kmar_QGN15841.1 | W303.asm02.HP0_G0002990 | OG0000182 | chrII | 465953 | 466942 | CP015057.1 | 197930 | 198871 | 47.40061162 | 329 | 313 | 1.051118211 | 1 | 0 | 1 | FIM1_2537 | YBR104W | YBR104W | YMC2 | n |
| Kmar_QGN17906.1 | W303.asm02.HP0_G0055930 | OG0000182 | chrXVI | 680249 | 681172 | CP015060.1 | 713324 | 714241 | 71.66123779 | 307 | 305 | 1.006557377 | 1 | 0 | 1 | FIM1_5115 | YPR058W | YPR058W | YMC1 | n |
| Kmar_QGN14259.1 | W303.asm02.HP0_G0043060 | OG0000183 | chrXIII | 808272 | 809204 | CP015055.1 | 132182 | 133060 | 54.48717949 | 310 | 292 | 1.061643836 | 1 | 1 | 1 | FIM1_916 | YMR267W | YMR267W | PPA2 | n |
| Kmar_QGN17429.1 | W303.asm02.HP0_G0002010 | OG0000183 | chrII | 259452 | 260315 | CP015059.1 | 901525 | 902388 | 85.36585366 | 287 | 287 | 1 | 1 | 1 | 1 | FIM1_4161 | YBR011C | YBR011C | IPP1 | n |
| Kmar_QGN15586.1 | W303.asm02.HP0_G0048210 | OG0000184 | chrXV | 150853 | 153945 | CP015056.1 | 1245103 | 1248525 | 20.81526453 | 1030 | 1140 | 0.903508772 | 1 | 0 | 0 | FIM1_2278 | YOL089C | YOL089C | HAL9 | n |
| Kmar_QGN16888.1 | W303.asm02.HP0_G0036300 | OG0000185 | chrXII | 440868 | 441770 | CP015058.1 | 1109000 | 1109884 | 62.66666667 | 300 | 294 | 1.020408163 | 1 | 0 | 0 | FIM1_3615 | YLR146C | YLR146C | SPE4 | n |
| Kmar_QGN17870.1 | W303.asm02.HP0_G0056020 | OG0000185 | chrXVI | 691054 | 691935 | CP015060.1 | 643847 | 644728 | 86.00682594 | 293 | 293 | 1 | 1 | 0 | 1 | FIM1_5079 | YPR069C | YPR069C | SPE3 | n |
| Kmar_QGN16620.1 | W303.asm02.HP0_G0028180 | OG0000186 | chrX | 99494 | 100687 | CP015058.1 | 513610 | 515061 | 62.52587992 | 397 | 483 | 0.82194617 | 1 | 0 | 1 | FIM1_3337 | YJL164C | YJL164C | TPK1 | n |
| Kmar_QGN14631.1 | W303.asm02.HP0_G0022840 | OG0000187 | chrVIII | 48734 | 51192 | CP015055.1 | 901696 | 903171 | 26.0756193 | 766 | 491 | 1.560081466 | 1 | 1 | 1 | FIM1_1295 | YHL027W | YHL027W | RIM101 | n |
| Kmar_QGN16442.1 | W303.asm02.HP0_G0029160 | OG0000187 | chrX | 320450 | 323092 | CP015058.1 | 127698 | 129755 | 29.42528736 | 880 | 685 | 1.284671533 | 1 | 1 | 1 | FIM1_3155 | YJL056C | YJL056C | ZAP1 | n |
| Kmar_QGN17580.1 | W303.asm02.HP0_G0006840 | OG0000188 | chrIV | 84343 | 86058 | CP015060.1 | 31177 | 32886 | 54.02097902 | 571 | 569 | 1.003514938 | 1 | 0 | 0 | FIM1_4787 | YDL210W | YDL210W | UGA4 | n |
| Kmar_QGN15296.1 | W303.asm02.HP0_G0016190 | OG0000189 | chrV | 537232 | 543723 | CP015056.1 | 622146 | 628577 | 47.83808648 | 2163 | 2143 | 1.009332711 | 1 | 1 | 1 | FIM1_1985 | YER172C | YER172C | BRR2 | n |
| Kmar_QGN13706.1 | W303.asm02.HP0_G0022380 | OG0000189 | chrVII | 1045198 | 1051101 | CP015054.1 | 696227 | 702103 | 70.42110604 | 1967 | 1958 | 1.004596527 | 1 | 1 | 1 | FIM1_349 | YGR271W | YGR271W | SLH1 | n |
| Kmar_QGN14236.1 | W303.asm02.HP0_G0001750 | OG0000190 | chrII | 194738 | 195799 | CP015055.1 | 86766 | 87866 | 68.57923497 | 353 | 366 | 0.964480874 | 1 | 1 | 1 | FIM1_892 | YBL016W | YBL016W | FUS3 | n |
| Kmar_QGN15647.1 | W303.asm02.HP0_G0020240 | OG0000190 | chrVII | 573156 | 574262 | CP015056.1 | 1385427 | 1386515 | 74.45652174 | 368 | 362 | 1.016574586 | 1 | 1 | 1 | FIM1_2340 | YGR040W | YGR040W | KSS1 | n |
| Kmar_QGN13469.1 | W303.asm02.HP0_G0045350 | OG0000191 | chrXIV | 350599 | 351363 | CP015054.1 | 202582 | 203334 | 29.58801498 | 254 | 250 | 1.016 | 1 | 1 | 1 | FIM1_107 | YNL148C | YNL148C | ALF1 | n |
| Kmar_QGN15472.1 | W303.asm02.HP0_G0053720 | OG0000191 | chrXVI | 226100 | 228706 | CP015056.1 | 1017357 | 1019846 | 22.75784753 | 868 | 829 | 1.047044632 | 1 | 1 | 1 | FIM1_2163 | YPL174C | YPL174C | NIP100 | n |
| Kmar_QGN13889.1 | W303.asm02.HP0_G0022180 | OG0000192 | chrVII | 1000490 | 1001860 | CP015054.1 | 1075892 | 1077349 | 27.66393443 | 456 | 485 | 0.940206186 | 1 | 1 | 1 | FIM1_536 | YGR249W | YGR249W | MGA1 | n |
| Kmar_QGN14759.1 | W303.asm02.HP0_G0050350 | OG0000192 | chrXV | 587856 | 590156 | CP015055.1 | 1168860 | 1171187 | 22.02970297 | 766 | 775 | 0.988387097 | 1 | 1 | 1 | FIM1_1429 | YOR140W | YOR140W | SFL1 | n |
| Kmar_QGN14260.1 | W303.asm02.HP0_G0043050 | OG0000193 | chrXIII | 805018 | 807879 | CP015055.1 | 133417 | 136140 | 60.66945607 | 953 | 907 | 1.050716648 | 1 | 1 | 1 | FIM1_917 | YMR266W | YMR266W | RSN1 | n |
| Kmar_QGN15406.1 | W303.asm02.HP0_G0048270 | OG0000193 | chrXV | 162811 | 165786 | CP015056.1 | 894365 | 897262 | 51.20240481 | 991 | 965 | 1.026943005 | 1 | 1 | 1 | FIM1_2096 | YOL084W | YOL084W | PHM7 | n |
| Kmar_QGN13522.1 | W303.asm02.HP0_G0034780 | OG0000194 | chrXII | 117192 | 121871 | CP015054.1 | 327119 | 331849 | 54.13533835 | 1559 | 1576 | 0.989213198 | 1 | 0 | 1 | FIM1_162 | YLL015W | YLL015W | BPT1 | n |
| Kmar_QGN15939.1 | W303.asm02.HP0_G0009980 | OG0000194 | chrIV | 727913 | 732460 | CP015057.1 | 429946 | 434493 | 68.90920967 | 1515 | 1515 | 1 | 1 | 0 | 1 | FIM1_2637 | YDR135C | YDR135C | YCF1 | n |
| Kmar_QGN16809.1 | W303.asm02.HP0_G0056450 | OG0000195 | chrXVI | 766954 | 774423 | CP015058.1 | 938013 | 945194 | 35.4324432 | 2489 | 2393 | 1.040117008 | 1 | 1 | 1 | FIM1_3533 | YPR117W | YPR117W | HOB2 | n |
| Kmar_QGN18248.1 | W303.asm02.HP0_G0039150 | OG0000195 | chrXII | 1040665 | 1048551 | CP015061.1 | 543927 | 551765 | 34.60682493 | 2628 | 2612 | 1.006125574 | 1 | 1 | 1 | FIM1_4573 | YLR454W | YLR454W | FMP27 | n |
| Kmar_QGN14359.1 | W303.asm02.HP0_G0010850 | OG0000196 | chrIV | 922072 | 926148 | CP015055.1 | 350528 | 354061 | 17.03216374 | 1358 | 1177 | 1.153780799 | 1 | 0 | 1 | FIM1_1019 | YDR227W | YDR227W | SIR4 | n |
| Kmar_QGN14384.1 | W303.asm02.HP0_G0009200 | OG0000196 | chrIV | 562660 | 564774 | CP015055.1 | 412067 | 414028 | 32.53333333 | 704 | 653 | 1.078101072 | 1 | 0 | 1 | FIM1_1044 | YDR052C | YDR052C | DBF4 | n |
| Kmar_QGN14402.1 | W303.asm02.HP0_G0051280 | OG0000197 | chrXV | 794254 | 795591 | CP015055.1 | 444626 | 445942 | 77.97752809 | 445 | 438 | 1.015981735 | 1 | 1 | 1 | FIM1_1062 | YOR244W | YOR244W | ESA1 | n |
| Kmar_QGN17915.1 | W303.asm02.HP0_G0001380 | OG0000197 | chrII | 120527 | 123022 | CP015060.1 | 728108 | 730252 | 34.46601942 | 831 | 714 | 1.163865546 | 1 | 1 | 1 | FIM1_5124 | YBL052C | YBL052C | SAS3 | n |
| Kmar_QGN17596.1 | W303.asm02.HP0_G0052800 | OG0000198 | chrXVI | 31034 | 33355 | CP015060.1 | 67646 | 70003 | 55.54156171 | 773 | 785 | 0.984713376 | 1 | 1 | 1 | FIM1_4803 | YPL270W | YPL270W | MDL2 | n |
| Kmar_QGN14117.1 | W303.asm02.HP0_G0036740 | OG0000198 | chrXII | 528428 | 530515 | CP015054.1 | 1569708 | 1571780 | 66.42651297 | 695 | 690 | 1.007246377 | 1 | 1 | 1 | FIM1_769 | YLR188W | YLR188W | MDL1 | n |
| Kmar_QGN15293.1 | W303.asm02.HP0_G0016220 | OG0000199 | chrV | 547166 | 548065 | CP015056.1 | 616781 | 617638 | 38.53820598 | 299 | 285 | 1.049122807 | 1 | 0 | 0 | FIM1_1982 | YER175C | YER175C | TMT1 | n |
| Kmar_QGN14415.1 | W303.asm02.HP0_G0025700 | OG0000200 | chrIX | 30856 | 34983 | CP015055.1 | 469904 | 473851 | 31.64285714 | 1375 | 1315 | 1.045627376 | 1 | 1 | 1 | FIM1_1076 | YIL159W | YIL159W | BNR1 | n |
| Kmar_QGN14103.1 | W303.asm02.HP0_G0044250 | OG0000200 | chrXIV | 130076 | 135937 | CP015054.1 | 1530075 | 1535660 | 43.83767535 | 1953 | 1861 | 1.049435787 | 1 | 1 | 1 | FIM1_754 | YNL271C | YNL271C | BNI1 | n |
| Kmar_QGN14319.1 | W303.asm02.HP0_G0010800 | OG0000201 | chrIV | 914555 | 915802 | CP015055.1 | 268115 | 269398 | 29.25170068 | 415 | 427 | 0.971896956 | 1 | 0 | 0 | FIM1_976 | YDR222W | YDR222W | | n |
| Kmar_QGN16738.1 | W303.asm02.HP0_G0011960 | OG0000201 | chrIV | 1178250 | 1179695 | CP015058.1 | 789619 | 790857 | 49.07216495 | 481 | 412 | 1.167475728 | 1 | 0 | 1 | FIM1_3460 | YDR346C | YDR346C | SVF1 | n |
| Kmar_QGN14441.1 | W303.asm02.HP0_G0006240 | OG0000202 | chrIII | 282558 | 282941 | CP015055.1 | 522271 | 522720 | 46.76258993 | 127 | 149 | 0.852348993 | 1 | 0 | 1 | FIM1_1102 | YCR083W | YCR083W | TRX3 | n |
| Kmar_QGN17488.1 | W303.asm02.HP0_G0021810 | OG0000202 | chrVII | 925328 | 925642 | CP015059.1 | 1013885 | 1014199 | 71.42857143 | 104 | 104 | 1 | 1 | 0 | 0 | FIM1_4224 | YGR209C | YGR209C | TRX2 | n |
| Kmar_QGN16669.1 | W303.asm02.HP0_G0051700 | OG0000203 | chrXV | 856866 | 861977 | CP015058.1 | 632125 | 637005 | 53.27868852 | 1703 | 1626 | 1.047355474 | 1 | 1 | 1 | FIM1_3388 | YOR290C | YOR290C | SNF2 | n |
| Kmar_QGN17020.1 | W303.asm02.HP0_G0026030 | OG0000203 | chrIX | 107026 | 111105 | CP015059.1 | 49174 | 53154 | 61.65951359 | 1359 | 1326 | 1.024886878 | 1 | 1 | 1 | FIM1_3749 | YIL126W | YIL126W | STH1 | n |
| Kmar_QGN13840.1 | W303.asm02.HP0_G0028110 | OG0000204 | chrX | 87267 | 88997 | CP015054.1 | 978035 | 979774 | 49.65397924 | 576 | 579 | 0.994818653 | 1 | 0 | 1 | FIM1_486 | YJL172W | YJL172W | CPS1 | n |
| Kmar_QGN15221.1 | W303.asm02.HP0_G0035170 | OG0000205 | chrXII | 195054 | 200672 | CP015056.1 | 462393 | 467531 | 36.06819393 | 1872 | 1712 | 1.093457944 | 1 | 1 | 1 | FIM1_1910 | YLR024C | YLR024C | UBR2 | n |
| Kmar_QGN15220.1 | W303.asm02.HP0_G0021570 | OG0000205 | chrVII | 872316 | 878168 | CP015056.1 | 455339 | 461227 | 46.88279302 | 1950 | 1962 | 0.993883792 | 1 | 1 | 1 | FIM1_1909 | YGR184C | YGR184C | UBR1 | n |
| Kmar_QGN16044.1 | W303.asm02.HP0_G0021620 | OG0000206 | chrVII | 889084 | 890607 | CP015057.1 | 692416 | 693804 | 44.37869822 | 507 | 462 | 1.097402597 | 1 | 0 | 1 | FIM1_2744 | YGR189C | YGR189C | CRH1 | n |
| Kmar_QGN14075.1 | W303.asm02.HP0_G0044450 | OG0000207 | chrXIV | 179949 | 181577 | CP015054.1 | 1466627 | 1468177 | 34.55882353 | 542 | 516 | 1.050387597 | 1 | 1 | 1 | FIM1_726 | YNL249C | YNL249C | MPA43 | n |
| Kmar_QGN15892.1 | W303.asm02.HP0_G0009740 | OG0000207 | chrIV | 678271 | 680418 | CP015057.1 | 313631 | 315970 | 55.58375635 | 715 | 779 | 0.917843389 | 1 | 1 | 1 | FIM1_2590 | YDR109C | YDR109C | | n |
| Kmar_QGN13512.1 | W303.asm02.HP0_G0049440 | OG0000208 | chrXV | 400804 | 401670 | CP015054.1 | 301460 | 302314 | 44.02730375 | 288 | 284 | 1.014084507 | 1 | 1 | 1 | FIM1_151 | YOR036W | YOR036W | PEP12 | n |
| Kmar_QGN17809.1 | W303.asm02.HP0_G0050040 | OG0000208 | chrXV | 519856 | 520707 | CP015060.1 | 530111 | 530896 | 32.15547703 | 283 | 261 | 1.084291188 | 1 | 1 | 1 | FIM1_5018 | YOR106W | YOR106W | VAM3 | n |
| Kmar_QGN13880.1 | W303.asm02.HP0_G0022090 | OG0000209 | chrVII | 983186 | 986149 | CP015054.1 | 1054965 | 1057901 | 73.04964539 | 987 | 978 | 1.009202454 | 1 | 1 | 1 | FIM1_527 | YGR240C | YGR240C | PFK1 | n |
| Kmar_QGN17093.1 | W303.asm02.HP0_G0042450 | OG0000209 | chrXIII | 672468 | 675347 | CP015059.1 | 219773 | 222589 | 73.02083333 | 959 | 938 | 1.02238806 | 1 | 1 | 1 | FIM1_3824 | YMR205C | YMR205C | PFK2 | n |
| Kmar_QGN17106.1 | W303.asm02.HP0_G0028090 | OG0000210 | chrX | 84622 | 85452 | CP015059.1 | 240619 | 241446 | 43.47826087 | 276 | 275 | 1.003636364 | 1 | 0 | 0 | FIM1_3837 | YJL174W | YJL174W | KRE9 | n |
| Kmar_QGN13736.1 | W303.asm02.HP0_G0019770 | OG0000211 | chrVII | 466626 | 469832 | CP015054.1 | 770147 | 773758 | 19.81981982 | 1068 | 1203 | 0.887780549 | 1 | 0 | 0 | FIM1_380 | YGL013C | YGL013C | PDR1 | n |
| Kmar_QGN16635.1 | W303.asm02.HP0_G0027960 | OG0000212 | chrX | 64441 | 64833 | CP015058.1 | 554636 | 555028 | 96.92307692 | 130 | 130 | 1 | 1 | 0 | 0 | FIM1_3353 | YJL190C/YLR367W | YJL190C | RPS22A | y |
| Kmar_QGN16121.1 | W303.asm02.HP0_G0027590 | OG0000213 | chrIX | 412327 | 413091 | CP015057.1 | 853931 | 854692 | 53.35968379 | 254 | 253 | 1.003952569 | 1 | 0 | 0 | FIM1_2822 | YIR035C | YIR035C | NRE1 | n |
| Kmar_QGN17412.1 | W303.asm02.HP0_G0033080 | OG0000214 | chrXI | 412672 | 412971 | CP015059.1 | 864337 | 864674 | 33.70786517 | 99 | 84 | 1.178571429 | 1 | 1 | 1 | FIM1_4144 | YKL018C-A | YKL018C-A | MCO12 | n |
| Kmar_QGN14984.1 | W303.asm02.HP0_G0054150 | OG0000215 | chrXVI | 311231 | 312070 | CP015055.1 | 1622098 | 1622775 | 62.45059289 | 244 | 225 | 1.084444444 | 1 | 0 | 1 | FIM1_1664 | YPL129W | YPL129W | TAF14 | n |
| Kmar_QGN15846.1 | W303.asm02.HP0_G0010010 | OG0000216 | chrIV | 738527 | 738833 | CP015057.1 | 208368 | 208538 | 53.24675325 | 77 | 56 | 1.375 | 1 | 0 | 1 | FIM1_2542 | YDR139C | YDR139C | RUB1 | n |
| Kmar_QGN16386.1 | W303.asm02.HP0_G0027620 | OG0000217 | chrIX | 415110 | 415814 | CP015058.1 | 4941 | 5645 | 62.82051282 | 234 | 234 | 1 | 1 | 0 | 0 | FIM1_3099 | YIR038C | YIR038C | GTT1 | n |
| Kmar_QGN14285.1 | W303.asm02.HP0_G0023350 | OG0000218 | chrVIII | 143134 | 144190 | CP015055.1 | 188902 | 190083 | 22.13930348 | 315 | 393 | 0.801526718 | 1 | 0 | 1 | FIM1_942 | YHR022C | YHR022C | | n |
| Kmar_QGN17366.1 | W303.asm02.HP0_G0021250 | OG0000218 | chrVII | 806401 | 807219 | CP015059.1 | 779518 | 780363 | 64.76868327 | 272 | 281 | 0.96797153 | 1 | 0 | 1 | FIM1_4098 | YGR152C | YGR152C | RSR1 | n |
| Kmar_QGN14337.1 | W303.asm02.HP0_G0028720 | OG0000219 | chrX | 219041 | 220897 | CP015055.1 | 304453 | 306249 | 33.44 | 618 | 598 | 1.033444816 | 1 | 0 | 1 | FIM1_997 | YJL103C | YJL103C | GSM1 | n |
| Kmar_QGN14438.1 | W303.asm02.HP0_G0004290 | OG0000219 | chrII | 721194 | 722783 | CP015055.1 | 517061 | 518563 | 45.21575985 | 529 | 500 | 1.058 | 1 | 1 | 1 | FIM1_1099 | YBR239C | YBR239C | ERT1 | n |
| Kmar_QGN18424.1 | W303.asm02.HP0_G0055400 | OG0000220 | chrXVI | 562734 | 564194 | CP015061.1 | 903161 | 904630 | 63.46938776 | 486 | 489 | 0.993865031 | 1 | 0 | 1 | FIM1_4752 | YPR001W | YPR001W | CIT3 | n |
| Kmar_QGN14300.1 | W303.asm02.HP0_G0046730 | OG0000220 | chrXIV | 641275 | 642714 | CP015055.1 | 226657 | 228108 | 85.06224066 | 479 | 483 | 0.991718427 | 1 | 0 | 0 | FIM1_957 | YNR001C | YNR001C | CIT1 | n |
| Kmar_QGN14301.1 | W303.asm02.HP0_G0005550 | OG0000221 | chrIII | 124815 | 125558 | CP015055.1 | 228928 | 229671 | 62.09677419 | 247 | 247 | 1 | 1 | 0 | 1 | FIM1_958 | YCR004C | YCR004C | YCP4 | n |

| | | | | | | | | | | | | | | | | | | | | |
|---|---|---|---|---|---|---|---|---|---|---|---|---|---|---|---|---|---|---|---|---|
| Kmar_QGN17011.1 | W303.asm02.HP0_G0008990 | OG0000221 | chrIV | 503758 | 504354 | CP015059.1 | 25290 | 25886 | 81.31313131 | 198 | 198 | 1 | 1 | 0 | 1 | FIM1_3740 | YDR032C | YDR032C | PST2 | n |
| Kmar_QGN13462.1 | W303.asm02.HP0_G0022320 | OG0000222 | chrVII | 1030879 | 1032153 | CP015054.1 | 190155 | 191225 | 20.74592075 | 424 | 356 | 1.191011236 | 1 | 0 | 1 | FIM1_99 | YGR263C | YGR263C | SAY1 | n |
| Kmar_QGN17843.1 | W303.asm02.HP0_G0013310 | OG0000223 | chrIV | 1440618 | 1441568 | CP015060.1 | 602796 | 603743 | 55.69620253 | 316 | 315 | 1.003174603 | 1 | 0 | 1 | FIM1_5052 | YDR492W | YDR492W | IZH1 | n |
| Kmar_QGN17618.1 | W303.asm02.HP0_G0049060 | OG0000223 | chrXV | 323866 | 324849 | CP015060.1 | 117299 | 118219 | 53.65853659 | 327 | 306 | 1.068627451 | 1 | 0 | 1 | FIM1_4825 | YOL002C | YOL002C | IZH2 | n |
| Kmar_QGN16098.1 | W303.asm02.HP0_G0026780 | OG0000224 | chrIX | 263582 | 265555 | CP015057.1 | 808466 | 809314 | 20.4248366 | 604 | 282 | 2.141843972 | 1 | 0 | 1 | FIM1_2798 | W303.asm02.HP0_G0026780 | YIL044C | AGE2 | y |
| Kmar_QGN16509.1 | W303.asm02.HP0_G0037120 | OG0000226 | chrXII | 598989 | 599564 | CP015058.1 | 269679 | 270254 | 97.38219895 | 191 | 191 | 1 | 1 | 1 | 1 | FIM1_3222 | YLR229C | YLR229C | CDC42 | n |
| Kmar_QGN13915.1 | W303.asm02.HP0_G0051650 | OG0000227 | chrXV | 851354 | 851773 | CP015054.1 | 1140579 | 1141016 | 47.5862069 | 139 | 145 | 0.95862069 | 1 | 0 | 1 | FIM1_563 | YOR285W | YOR285W | RDL1 | n |
| Kmar_QGN13916.1 | W303.asm02.HP0_G0051660 | OG0000227 | chrXV | 851999 | 852448 | CP015054.1 | 1141432 | 1141860 | 48.32214765 | 149 | 142 | 1.049295775 | 1 | 0 | 1 | FIM1_564 | YOR286W | YOR286W | RDL2 | n |
| Kmar_QGN14366.1 | W303.asm02.HP0_G0010820 | OG0000228 | chrIV | 918818 | 919213 | CP015055.1 | 367077 | 367475 | 93.18181818 | 131 | 132 | 0.992424242 | 1 | 0 | 0 | FIM1_1026 | YBL002W/YDR224C | YDR224C | HTB1 | y |
| Kmar_QGN13972.1 | W303.asm02.HP0_G0004040 | OG0000229 | chrII | 670171 | 672183 | CP015054.1 | 1243690 | 1245582 | 32.54086181 | 670 | 630 | 1.063492063 | 1 | 0 | 1 | FIM1_621 | YBR212W | YBR212W | NGR1 | n |
| Kmar_QGN15203.1 | W303.asm02.HP0_G0024160 | OG0000229 | chrVIII | 284126 | 285697 | CP015056.1 | 421698 | 423458 | 41.26455907 | 523 | 586 | 0.892491468 | 1 | 1 | 1 | FIM1_1891 | YHR086W | YHR086W | NAM8 | n |
| Kmar_QGN16431.1 | W303.asm02.HP0_G0004540 | OG0000230 | chrII | 761126 | 761782 | CP015058.1 | 104397 | 105029 | 37.93103448 | 218 | 210 | 1.038095238 | 1 | 0 | 1 | FIM1_3144 | YBR264C | YBR264C | YPT10 | n |
| Kmar_QGN17788.1 | W303.asm02.HP0_G0049890 | OG0000230 | chrXV | 490931 | 491563 | CP015060.1 | 484870 | 485496 | 75.23809524 | 210 | 208 | 1.009615385 | 1 | 0 | 1 | FIM1_4997 | YOR089C | YOR089C | VPS21 | n |
| Kmar_QGN17816.1 | W303.asm02.HP0_G0000300 | OG0000231 | chrI | 66413 | 68155 | CP015060.1 | 549723 | 550976 | 25.9057971 | 580 | 417 | 1.39088729 | 1 | 0 | 1 | FIM1_5025 | YAL040C | YAL040C | CLN3 | n |
| Kmar_QGN15918.1 | W303.asm02.HP0_G0035870 | OG0000232 | chrXII | 345489 | 347435 | CP015057.1 | 375953 | 377767 | 55.28330781 | 648 | 604 | 1.072847682 | 1 | 0 | 1 | FIM1_2616 | YLR098C | YLR098C | CHA4 | n |
| Kmar_QGN16485.1 | W303.asm02.HP0_G0052150 | OG0000232 | chrXV | 956626 | 958905 | CP015058.1 | 204894 | 207038 | 46.44670051 | 759 | 714 | 1.06302521 | 1 | 0 | 1 | FIM1_3198 | YOR337W | YOR337W | TEA1 | n |
| Kmar_QGN16296.1 | W303.asm02.HP0_G0053400 | OG0000233 | chrXVI | 163396 | 165147 | CP015057.1 | 1224776 | 1226545 | 37.78877888 | 583 | 589 | 0.989813243 | 1 | 0 | 1 | FIM1_3001 | YPL208W | YPL208W | RKM1 | n |
| Kmar_QGN17989.1 | W303.asm02.HP0_G0022720 | OG0000233 | chrVIII | 19407 | 21164 | CP015060.1 | 902181 | 903845 | 35.59322034 | 585 | 554 | 1.055956679 | 1 | 0 | 1 | FIM1_5199 | YHL039W | YHL039W | EFM1 | n |
| Kmar_QGN15437.1 | W303.asm02.HP0_G0023480 | OG0000234 | chrVIII | 172063 | 173955 | CP015056.1 | 960032 | 962023 | 24.96473907 | 630 | 663 | 0.950226244 | 1 | 0 | 1 | FIM1_2128 | YHR035W | YHR035W | NEL1 | n |
| Kmar_QGN14686.1 | W303.asm02.HP0_G0057030 | OG0000234 | chrXVI | 903193 | 905499 | CP015055.1 | 1031766 | 1034048 | 79.71391417 | 768 | 760 | 1.010526316 | 1 | 1 | 1 | FIM1_1353 | YPR181C | YPR181C | SEC23 | n |
| Kmar_QGN14737.1 | W303.asm02.HP0_G0056820 | OG0000235 | chrXVI | 863415 | 865577 | CP015055.1 | 1128426 | 1130702 | 60.34255599 | 720 | 758 | 0.949868074 | 1 | 0 | 0 | FIM1_1406 | YPR159W | YPR159W | KRE6 | n |
| Kmar_QGN14381.1 | W303.asm02.HP0_G0047380 | OG0000236 | chrXIV | 768324 | 771677 | CP015055.1 | 399918 | 402953 | 47.41992883 | 1117 | 1011 | 1.104846686 | 1 | 0 | 1 | FIM1_1041 | YNR067C | YNR067C | DSE4 | n |
| Kmar_QGN16886.1 | W303.asm02.HP0_G0036280 | OG0000236 | chrXII | 437723 | 440062 | CP015058.1 | 1105363 | 1107756 | 49.62871287 | 779 | 797 | 0.977415307 | 1 | 1 | 1 | FIM1_3613 | YLR144C | YLR144C | ACF2 | n |
| Kmar_QGN15962.1 | W303.asm02.HP0_G0009090 | OG0000237 | chrIV | 538737 | 542012 | CP015057.1 | 483361 | 486612 | 63.51228389 | 1091 | 1083 | 1.007386888 | 1 | 0 | 0 | FIM1_2660 | YDR038C/YDR039C/YDR040C | YDR040C | ENA1 | y |
| Kmar_QGN18292.1 | W303.asm02.HP0_G0014640 | OG0000238 | chrV | 190650 | 192935 | CP015061.1 | 634115 | 636451 | 66.24203822 | 761 | 778 | 0.9781491 | 1 | 0 | 1 | FIM1_4618 | YER017C | YER017C | AFG3 | n |
| Kmar_QGN14346.1 | W303.asm02.HP0_G0041300 | OG0000238 | chrXIII | 446114 | 448591 | CP015055.1 | 321291 | 323816 | 65.96736597 | 825 | 841 | 0.98097503 | 1 | 1 | 1 | FIM1_1006 | YMR089C | YMR089C | YTA12 | n |
| Kmar_QGN13552.1 | W303.asm02.HP0_G0053460 | OG0000239 | chrXVI | 175322 | 176707 | CP015054.1 | 403155 | 404330 | 17.84946237 | 461 | 391 | 1.179028133 | 1 | 0 | 1 | FIM1_193 | YPL201C | YPL201C | YIG1 | n |
| Kmar_QGN13528.1 | W303.asm02.HP0_G0034580 | OG0000239 | chrXII | 66754 | 68265 | CP015054.1 | 347411 | 348880 | 29.24901186 | 503 | 489 | 1.028629857 | 1 | 0 | 1 | FIM1_168 | YLL036C | YLL036C | PRP19 | n |
| Kmar_QGN17671.1 | W303.asm02.HP0_G0046090 | OG0000240 | chrXIV | 501868 | 502443 | CP015060.1 | 224409 | 224984 | 83.2460733 | 191 | 191 | 1 | 1 | 0 | 0 | FIM1_4878 | YGL147C/YNL067W | YNL067W | RPL9B | y |
| Kmar_QGN17037.1 | W303.asm02.HP0_G0036970 | OG0000241 | chrXII | 566778 | 568046 | CP015059.1 | 94406 | 95968 | 38.21656051 | 422 | 520 | 0.811538462 | 1 | 0 | 1 | FIM1_3766 | YLR213C | YLR213C | CRR1 | n |
| Kmar_QGN15291.1 | W303.asm02.HP0_G0014090 | OG0000241 | chrV | 78414 | 79817 | CP015056.1 | 613131 | 614543 | 46.92144374 | 467 | 470 | 0.993617021 | 1 | 0 | 1 | FIM1_1980 | YEL040W | YEL040W | UTR2 | n |
| Kmar_QGN13577.1 | W303.asm02.HP0_G0040010 | OG0000242 | chrXIII | 160207 | 161982 | CP015054.1 | 449817 | 451595 | 64.52702703 | 591 | 592 | 0.998310811 | 1 | 0 | 1 | FIM1_218 | YML054C | YML054C | CYB2 | n |
| Kmar_QGN13579.1 | W303.asm02.HP0_G0038890 | OG0000244 | chrXII | 983731 | 984782 | CP015054.1 | 456957 | 457955 | 35.62874251 | 326 | 332 | 0.981927711 | 1 | 1 | 1 | FIM1_220 | YLR426W | YLR426W | TDA5 | n |
| Kmar_QGN14439.1 | W303.asm02.HP0_G0018960 | OG0000245 | chrVII | 304378 | 305508 | CP015055.1 | 519259 | 520392 | 66.84073107 | 376 | 377 | 0.99734748 | 1 | 1 | 1 | FIM1_1100 | YGL105W | YGL105W | ARC1 | n |
| Kmar_QGN16893.1 | W303.asm02.HP0_G0036330 | OG0000246 | chrXII | 442686 | 445442 | CP015058.1 | 1116073 | 1118838 | 35.17169615 | 918 | 921 | 0.996742671 | 1 | 0 | 0 | FIM1_3620 | YLR148W | YLR148W | PEP3 | n |
| Kmar_QGN17440.1 | W303.asm02.HP0_G0001950 | OG0000247 | chrII | 248245 | 248886 | CP015059.1 | 920705 | 921364 | 31.98198198 | 213 | 219 | 0.97260274 | 1 | 0 | 0 | FIM1_4173 | YBR005W | YBR005W | RCR1 | n |
| Kmar_QGN16398.1 | W303.asm02.HP0_G0000120 | OG0000248 | chrI | 34083 | 35336 | CP015058.1 | 27307 | 28455 | 55.82524272 | 417 | 382 | 1.091623037 | 1 | 0 | 1 | FIM1_3111 | YAL061W | YAL061W | BDH2 | n |
| Kmar_QGN16397.1 | W303.asm02.HP0_G0000130 | OG0000248 | chrI | 35790 | 36938 | CP015058.1 | 25200 | 26564 | 53.31858407 | 382 | 454 | 0.841409692 | 1 | 0 | 1 | FIM1_3110 | YAL060W | YAL060W | BDH1 | n |
| Kmar_QGN14765.1 | W303.asm02.HP0_G0012400 | OG0000249 | chrIV | 1255780 | 1257359 | CP015055.1 | 1185098 | 1186498 | 61.02783726 | 379 | 466 | 0.813304721 | 1 | 0 | 1 | FIM1_1435 | YDR388W | YDR388W | RVS167 | n |
| Kmar_QGN17480.1 | W303.asm02.HP0_G0029820 | OG0000250 | chrX | 463995 | 465620 | CP015059.1 | 1006928 | 1007953 | 35.99137931 | 463 | 341 | 1.357771261 | 1 | 0 | 1 | FIM1_4216 | YJR019C | YJR019C | TES1 | n |
| Kmar_QGN17437.1 | W303.asm02.HP0_G0001960 | OG0000252 | chrII | 249349 | 250842 | CP015059.1 | 916654 | 918150 | 70.88353414 | 497 | 498 | 0.997991968 | 1 | 1 | 1 | FIM1_4170 | YBR006W | YBR006W | UGA2 | n |
| Kmar_QGN17607.1 | W303.asm02.HP0_G0013880 | OG0000253 | chrV | 36898 | 39900 | CP015060.1 | 89733 | 92723 | 47.2195122 | 1000 | 996 | 1.004016064 | 1 | 1 | 1 | FIM1_4814 | YEL061C | YEL061C | CIN8 | n |
| Kmar_QGN14198.1 | W303.asm02.HP0_G0027770 | OG0000254 | chrX | 23381 | 25780 | CP015055.1 | 13722 | 16103 | 80.72590738 | 799 | 793 | 1.007566204 | 1 | 0 | 1 | FIM1_854 | YJL212C | YJL212C | OPT1 | n |
| Kmar_QGN16263.1 | W303.asm02.HP0_G0030810 | OG0000255 | chrX | 674991 | 676910 | CP015057.1 | 1152526 | 1154604 | 56.34005764 | 639 | 692 | 0.923410405 | 1 | 0 | 0 | FIM1_2968 | YJR130C | YJR130C | STR2 | n |
| Kmar_QGN15469.1 | W303.asm02.HP0_G0053760 | OG0000256 | chrXVI | 234247 | 234705 | CP015056.1 | 1012211 | 1012669 | 71.71052632 | 152 | 152 | 1 | 1 | 1 | 1 | FIM1_2160 | YPL170W | YPL170W | DAP1 | n |
| Kmar_QGN17772.1 | W303.asm02.HP0_G0022460 | OG0000257 | chrVII | 1062205 | 1063365 | CP015060.1 | 448606 | 449739 | 57.14285714 | 386 | 377 | 1.023872679 | 1 | 0 | 0 | FIM1_4981 | YGR279C | YGR279C | SCW4 | n |
| Kmar_QGN14230.1 | W303.asm02.HP0_G0001690 | OG0000258 | chrII | 180161 | 183562 | CP015055.1 | 72343 | 75717 | 67.93760832 | 1133 | 1124 | 1.008007117 | 1 | 1 | 1 | FIM1_886 | YBL022C | YBL022C | PIM1 | n |
| Kmar_QGN18191.1 | W303.asm02.HP0_G0020570 | OG0000259 | chrVII | 638758 | 641046 | CP015061.1 | 405568 | 406515 | 17.76223776 | 715 | 315 | 2.26984127 | 1 | 0 | 0 | FIM1_4515 | YGR079W | YGR079W | | n |
| Kmar_QGN17659.1 | W303.asm02.HP0_G0049300 | OG0000260 | chrXV | 374166 | 378165 | CP015060.1 | 205304 | 207262 | 23.9046887 | 1286 | 652 | 1.972392638 | 1 | 0 | 0 | FIM1_4866 | YOR022C | YOR022C | DDL1 | n |
| Kmar_QGN14822.1 | W303.asm02.HP0_G0025620 | OG0000261 | chrIX | 18446 | 19078 | CP015055.1 | 1295195 | 1296172 | 34.73053892 | 210 | 325 | 0.646153846 | 1 | 0 | 1 | FIM1_1493 | W303.asm02.HP0_G0025620 | YCL064C | CHA1 | y |
| Kmar_QGN14183.1 | W303.asm02.HP0_G0004980 | OG0000261 | chrIII | 15451 | 16533 | CP015054.1 | 1702402 | 1703478 | 50.40871935 | 360 | 358 | 1.005586592 | 1 | 0 | 1 | FIM1_836 | YCL064C | YCL064C | CHA1 | n |
| Kmar_QGN13998.1 | W303.asm02.HP0_G0037570 | OG0000262 | chrXII | 700967 | 701809 | CP015054.1 | 1296051 | 1296893 | 47.55244755 | 280 | 280 | 1 | 1 | 0 | 0 | FIM1_648 | YLR284C | YLR284C | ECI1 | n |
| Kmar_QGN17051.1 | W303.asm02.HP0_G0007480 | OG0000263 | chrIV | 216736 | 217281 | CP015059.1 | 127976 | 128521 | 87.84530387 | 181 | 181 | 1 | 1 | 0 | 0 | FIM1_3780 | YDL137W/YDL192W | YDL137W | ARF2 | y |
| Kmar_QGN18332.1 | W303.asm02.HP0_G0001600 | OG0000264 | chrII | 165343 | 166299 | CP015061.1 | 712379 | 713296 | 84.90566038 | 318 | 305 | 1.042622951 | 1 | 0 | 0 | FIM1_4658 | YBL030C | YBL030C | PET9 | n |
| Kmar_QGN15811.1 | W303.asm02.HP0_G0041740 | OG0000265 | chrXIII | 536824 | 539169 | CP015057.1 | 137804 | 138526 | 19.1904048 | 664 | 240 | 2.766666667 | 1 | 0 | 1 | FIM1_2507 | YMR133W | YMR133W | REC114 | n |
| Kmar_QGN18167.1 | W303.asm02.HP0_G0026690 | OG0000267 | chrIX | 245743 | 246495 | CP015061.1 | 361425 | 362171 | 77.6 | 250 | 248 | 1.008064516 | 1 | 0 | 0 | FIM1_4490 | YIL053W | YIL053W | GPP1 | n |
| Kmar_QGN17038.1 | W303.asm02.HP0_G0007410 | OG0000268 | chrIV | 198868 | 199938 | CP015059.1 | 96527 | 97615 | 55.61643836 | 356 | 362 | 0.983425414 | 1 | 0 | 1 | FIM1_3767 | YDL144C | YDL144C | | n |
| Kmar_QGN17914.1 | W303.asm02.HP0_G0043580 | OG0000269 | chrXIII | 919089 | 920171 | CP015060.1 | 726362 | 727462 | 62.98342541 | 360 | 366 | 0.983606557 | 1 | 0 | 1 | FIM1_5123 | YMR318C | YMR318C | ADH6 | n |
| Kmar_QGN13851.1 | W303.asm02.HP0_G0005800 | OG0000270 | chrIII | 193972 | 195510 | CP015054.1 | 999154 | 1000470 | 52.66393443 | 512 | 438 | 1.168949772 | 1 | 0 | 1 | FIM1_497 | YCR028C | YCR028C | FEN2 | n |
| Kmar_QGN17070.1 | W303.asm02.HP0_G0007130 | OG0000271 | chrIV | 139742 | 141334 | CP015059.1 | 169569 | 171215 | 69.89051095 | 530 | 548 | 0.967153285 | 1 | 0 | 0 | FIM1_3801 | YDL178W | YDL178W | DLD2 | n |
| Kmar_QGN14890.1 | W303.asm02.HP0_G0037490 | OG0000272 | chrXII | 683733 | 685679 | CP015055.1 | 1439030 | 1441336 | 15.68627451 | 648 | 768 | 0.84375 | 1 | 0 | 0 | FIM1_1564 | YLR273C | YLR273C | PIG1 | n |
| Kmar_QGN17682.1 | W303.asm02.HP0_G0046590 | OG0000273 | chrXIV | 616567 | 616794 | CP015060.1 | 250039 | 250302 | 33.33333333 | 75 | 87 | 0.862068966 | 1 | 0 | 1 | FIM1_4889 | YNL015W | YNL015W | PBI2 | n |
| Kmar_QGN17137.1 | W303.asm02.HP0_G0024670 | OG0000273 | chrVIII | 383328 | 383672 | CP015059.1 | 313016 | 313312 | 33.04347826 | 114 | 98 | 1.163265306 | 1 | 0 | 1 | FIM1_3868 | YHR138C | YHR138C | | n |
| Kmar_QGN13910.1 | W303.asm02.HP0_G0042620 | OG0000274 | chrXIII | 716217 | 716888 | CP015054.1 | 1133411 | 1134070 | 56.05381166 | 223 | 219 | 1.01826484 | 1 | 0 | 1 | FIM1_558 | YMR222C | YMR222C | FSH2 | n |
| Kmar_QGN13909.1 | W303.asm02.HP0_G0051610 | OG0000274 | chrXV | 845417 | 847511 | CP015054.1 | 1132367 | 1133146 | 45.52238806 | 586 | 259 | 2.262548263 | 1 | 0 | 1 | FIM1_557 | YOR279C | YOR279C | RFM1 | n |
| Kmar_QGN14451.1 | W303.asm02.HP0_G0013040 | OG0000275 | chrIV | 1392509 | 1394068 | CP015055.1 | 537067 | 538779 | 33.50340136 | 519 | 570 | 0.910526316 | 1 | 0 | 0 | FIM1_1112 | YDR463W | YDR463W | STP1 | n |
| Kmar_QGN15799.1 | W303.asm02.HP0_G0020010 | OG0000276 | chrVII | 514477 | 518397 | CP015057.1 | 118189 | 120774 | 26.40239342 | 1306 | 861 | 1.516840883 | 1 | 0 | 1 | FIM1_2495 | YGR014W | YGR014W | MSB2 | n |
| Kmar_QGN15675.1 | W303.asm02.HP0_G0012670 | OG0000276 | chrIV | 1311991 | 1317231 | CP015056.1 | 1436471 | 1440661 | 19.7021764 | 1746 | 1396 | 1.250716332 | 1 | 0 | 1 | FIM1_2368 | YDR420W | YDR420W | HKR1 | n |
| Kmar_QGN16856.1 | W303.asm02.HP0_G0036150 | OG0000277 | chrXII | 410747 | 412015 | CP015058.1 | 1041965 | 1043161 | 53.9503386 | 422 | 398 | 1.060301508 | 1 | 0 | 0 | FIM1_3582 | YLR130C | YLR130C | ZRT2 | n |
| Kmar_QGN13616.1 | W303.asm02.HP0_G0039450 | OG0000278 | chrXIII | 41494 | 44256 | CP015054.1 | 524484 | 527240 | 45 | 920 | 918 | 1.002178649 | 1 | 0 | 0 | FIM1_258 | YML111W | YML111W | BUL2 | n |
| Kmar_QGN16659.1 | W303.asm02.HP0_G0020410 | OG0000282 | chrVII | 605322 | 606860 | CP015058.1 | 608134 | 609786 | 36.5248227 | 512 | 550 | 0.930909091 | 1 | 0 | 1 | FIM1_3378 | YGR059W | YGR059W | SPR3 | n |
| Kmar_QGN13519.1 | W303.asm02.HP0_G0024370 | OG0000282 | chrVIII | 332786 | 334009 | CP015054.1 | 319028 | 320209 | 72.39709443 | 407 | 393 | 1.03562341 | 1 | 0 | 1 | FIM1_159 | YHR107C | YHR107C | CDC12 | n |
| Kmar_QGN17984.1 | W303.asm02.HP0_G0034520 | OG0000283 | chrXII | 47729 | 48499 | CP015060.1 | 888580 | 889347 | 85.15625 | 256 | 255 | 1.003921569 | 1 | 0 | 0 | FIM1_5193 | YHL033C/YLL045C | YLL045C | RPL8B | y |
| Kmar_QGN14025.1 | W303.asm02.HP0_G0008550 | OG0000284 | chrIV | 415150 | 416745 | CP015054.1 | 1357030 | 1358886 | 25.12077295 | 531 | 618 | 0.859223301 | 1 | 0 | 1 | FIM1_676 | YDL020C | YDL020C | RPN4 | n |
| Kmar_QGN15804.1 | W303.asm02.HP0_G0008050 | OG0000285 | chrIV | 313281 | 315078 | CP015057.1 | 127020 | 128180 | 40.6 | 501 | 386 | 1.297927461 | 1 | 0 | 1 | FIM1_2500 | YDL079C | YDL079C | MRK1 | n |
| Kmar_QGN17025.1 | W303.asm02.HP0_G0046080 | OG0000287 | chrXIV | 497887 | 500475 | CP015059.1 | 69536 | 71971 | 31.66109253 | 862 | 811 | 1.062885327 | 1 | 0 | 0 | FIM1_3754 | YNL068C | YNL068C | FKH2 | n |
| Kmar_QGN16021.1 | W303.asm02.HP0_G0011980 | OG0000288 | chrIV | 1181362 | 1182861 | CP015057.1 | 642438 | 642767 | 16.44736842 | 499 | 109 | 4.577981651 | 1 | 0 | 0 | FIM1_2721 | YDR348C | YDR348C | PAL1 | n |
| Kmar_QGN16330.1 | W303.asm02.HP0_G0040230 | OG0000289 | chrXIII | 220646 | 221236 | CP015057.1 | 1304200 | 1304793 | 89.79591837 | 196 | 197 | 0.994923858 | 1 | 0 | 0 | FIM1_3035 | YML028W | YML028W | TSA1 | n |
| Kmar_QGN14163.1 | W303.asm02.HP0_G0013620 | OG0000290 | chrIV | 1496289 | 1496528 | CP015054.1 | 1666132 | 1666359 | 72.15189873 | 79 | 75 | 1.053333333 | 1 | 0 | 0 | FIM1_816 | YDR525W-A | YDR525W-A | SNA2 | n |
| Kmar_QGN13679.1 | W303.asm02.HP0_G0002370 | OG0000291 | chrII | 340044 | 341024 | CP015054.1 | 650064 | 650950 | 89.1025641 | 156 | 156 | 1 | 1 | 0 | 0 | FIM1_322 | YBR048W/YDR025W | YDR025W | RPS11A | y |
| Kmar_QGN14324.1 | W303.asm02.HP0_G0033990 | OG0000292 | chrXI | 600104 | 601216 | CP015055.1 | 279531 | 280631 | 62.26415094 | 370 | 366 | 1.010928962 | 1 | 0 | 0 | FIM1_981 | YKR076W | YKR076W | ECM4 | n |
| Kmar_QGN17885.1 | W303.asm02.HP0_G0003120 | OG0000294 | chrII | 493957 | 495333 | CP015060.1 | 678563 | 679939 | 95.41484716 | 458 | 458 | 1 | 1 | 0 | 0 | FIM1_5094 | YBR118W/YPR080W | YPR080W | TEF1 | y |
| Kmar_QGN14154.1 | W303.asm02.HP0_G0005180 | OG0000295 | chrIII | 50490 | 51992 | CP015054.1 | 1644828 | 1646273 | 60.83499006 | 500 | 481 | 1.03950104 | 1 | 0 | 0 | FIM1_807 | YCL040W | YCL040W | GLK1 | n |
| Kmar_QGN15609.1 | W303.asm02.HP0_G0000190 | OG0000296 | chrI | 46534 | 48885 | CP015056.1 | 1297674 | 1300025 | 57.94044665 | 783 | 783 | 1 | 1 | 0 | 0 | FIM1_2302 | YAL053W | YAL053W | FLC2 | n |
| Kmar_QGN14063.1 | W303.asm02.HP0_G0027820 | OG0000297 | chrX | 37190 | 39466 | CP015054.1 | 1445124 | 1447100 | 36.71052632 | 758 | 658 | 1.151975684 | 1 | 0 | 1 | FIM1_714 | YJL206C | YJL206C | | n |
| Kmar_QGN17024.1 | W303.asm02.HP0_G0025990 | OG0000297 | chrIX | 91813 | 94710 | CP015059.1 | 64998 | 67586 | 42.95918367 | 965 | 862 | 1.119489559 | 1 | 0 | 1 | FIM1_3753 | YIL130W | YIL130W | ASG1 | n |
| Kmar_QGN14859.1 | W303.asm02.HP0_G0042080 | OG0000298 | chrXIII | 602146 | 603666 | CP015055.1 | 1363754 | 1365259 | 66.79841897 | 506 | 501 | 1.00998004 | 1 | 0 | 0 | FIM1_1532 | YMR170C | YMR170C | ALD2 | n |
| Kmar_QGN15256.1 | W303.asm02.HP0_G0049560 | OG0000299 | chrXV | 423124 | 424272 | CP015056.1 | 540073 | 541356 | 40.65420561 | 382 | 427 | 0.894613583 | 1 | 0 | 1 | FIM1_1945 | YOR049C | YOR049C | RSB1 | n |
| Kmar_QGN13464.1 | W303.asm02.HP0_G0024590 | OG0000300 | chrVIII | 371313 | 373865 | CP015054.1 | 193650 | 196247 | 29.42502818 | 850 | 865 | 0.98265896 | 1 | 0 | 1 | FIM1_101 | YHR131C | YHR131C | | n |
| Kmar_QGN14332.1 | W303.asm02.HP0_G0037280 | OG0000301 | chrXII | 629486 | 631318 | CP015055.1 | 294957 | 296588 | 52.0661157 | 610 | 543 | 1.123388582 | 1 | 0 | 1 | FIM1_992 | YLR248W | YLR248W | RCK2 | n |
| Kmar_QGN17626.1 | W303.asm02.HP0_G0052770 | OG0000302 | chrXVI | 25639 | 26616 | CP015060.1 | 138296 | 139294 | 67.27828746 | 325 | 332 | 0.978915663 | 1 | 0 | 1 | FIM1_4833 | YPL273W | YPL273W | SAM4 | n |
| Kmar_QGN16984.1 | W303.asm02.HP0_G0044580 | OG0000303 | chrXIV | 210929 | 212209 | CP015058.1 | 1308199 | 1309482 | 28.31460674 | 426 | 427 | 0.99765808 | 1 | 0 | 1 | FIM1_3711 | YNL234W | YNL234W | | n |
| Kmar_QGN14379.1 | W303.asm02.HP0_G0002710 | OG0000304 | chrII | 402575 | 405505 | CP015055.1 | 392118 | 394883 | 40.06085193 | 976 | 921 | 1.059717698 | 1 | 0 | 1 | FIM1_1039 | YBR074W | YBR074W | PFF1 | n |
| Kmar_QGN14887.1 | W303.asm02.HP0_G0037820 | OG0000305 | chrXII | 747481 | 752250 | CP015055.1 | 1427528 | 1432090 | 32.86413709 | 1589 | 1520 | 1.045394737 | 1 | 0 | 0 | FIM1_1561 | YLR310C | YLR310C | CDC25 | n |
| Kmar_QGN14597.1 | W303.asm02.HP0_G0004310 | OG0000307 | chrII | 725445 | 726911 | CP015055.1 | 826995 | 828587 | 46.19771863 | 488 | 530 | 0.920754717 | 1 | 0 | 0 | FIM1_1259 | YBR241C | YBR241C | VVS1 | n |
| Kmar_QGN16557.1 | W303.asm02.HP0_G0038980 | OG0000308 | chrXII | 1008282 | 1008680 | CP015058.1 | 368182 | 368649 | 19.87179487 | 132 | 155 | 0.851612903 | 1 | 0 | 0 | FIM1_3271 | YLR437C | YLR437C | DIF1 | n |
| Kmar_QGN15310.1 | W303.asm02.HP0_G0022950 | OG0000309 | chrVIII | 70237 | 72584 | CP015056.1 | 670888 | 673065 | 61.37281292 | 709 | 725 | 0.977931034 | 1 | 0 | 1 | FIM1_1999 | YHL016C | YHL016C | DUR3 | n |
| Kmar_QGN15685.1 | W303.asm02.HP0_G0017950 | OG0000310 | chrVII | 80397 | 82814 | CP015056.1 | 1466844 | 1469351 | 43.74269006 | 805 | 835 | 0.964071856 | 1 | 1 | 1 | FIM1_2378 | YGL216W | YGL216W | KIP3 | n |
| Kmar_QGN14528.1 | W303.asm02.HP0_G0008790 | OG0000311 | chrIV | 463093 | 464655 | CP015055.1 | 700643 | 702163 | 57.9047619 | 520 | 506 | 1.027667984 | 1 | 0 | 0 | FIM1_1190 | YDR009W | YDR009W | GAL3 | n |

| | | | | | | | | | | | | | | | | | | | | |
|---|---|---|---|---|---|---|---|---|---|---|---|---|---|---|---|---|---|---|---|---|
| Kmar_QGN14527.1 | W303.asm02.HP0_G0002070 | OG0000312 | chrII | 271350 | 273449 | CP015055.1 | 696976 | 699051 | 57.04125178 | 699 | 691 | 1.011577424 | 1 | 0 | 1 | FIM1_1189 | YBR019C | YBR019C | GAL10 | n |
| Kmar_QGN14029.1 | W303.asm02.HP0_G0008530 | OG0000313 | chrIV | 411862 | 413037 | CP015054.1 | 1366439 | 1367743 | 63.82488479 | 391 | 434 | 0.900921659 | 1 | 0 | 0 | FIM1_680 | YDL022W | YDL022W | GPD1 | n |
| Kmar_QGN15789.1 | W303.asm02.HP0_G0037950 | OG0000314 | chrXII | 774900 | 776471 | CP015057.1 | 103405 | 105042 | 49.00181488 | 523 | 545 | 0.959633028 | 1 | 0 | 0 | FIM1_2485 | YLR324W | YLR324W | PEX30 | n |
| Kmar_QGN18294.1 | W303.asm02.HP0_G0014620 | OG0000315 | chrV | 185688 | 187922 | CP015061.1 | 637876 | 640134 | 51.97368421 | 744 | 752 | 0.989361702 | 1 | 0 | 1 | FIM1_4620 | YER015W | YER015W | FAA2 | n |
| Kmar_QGN17338.1 | W303.asm02.HP0_G0045740 | OG0000316 | chrXIV | 426550 | 428409 | CP015059.1 | 721939 | 723768 | 77.54442649 | 619 | 609 | 1.016420361 | 1 | 0 | 0 | FIM1_4070 | YNL104C | YNL104C | LEU4 | n |
| Kmar_QGN15494.1 | W303.asm02.HP0_G0046900 | OG0000317 | chrXIV | 677493 | 679421 | CP015056.1 | 1059162 | 1060967 | 42.92379471 | 642 | 601 | 1.068219634 | 1 | 0 | 0 | FIM1_2185 | YNR019W | YNR019W | ARE2 | n |
| Kmar_QGN14118.1 | W303.asm02.HP0_G0036730 | OG0000319 | chrXII | 524994 | 528074 | CP015054.1 | 1572197 | 1575571 | 37.20316623 | 1026 | 1124 | 0.912811388 | 1 | 0 | 0 | FIM1_770 | YLR187W | YLR187W | SKG3 | n |
| Kmar_QGN15759.1 | W303.asm02.HP0_G0031210 | OG0000320 | chrXI | 27849 | 31709 | CP015057.1 | 30813 | 34673 | 73.374613 | 1286 | 1286 | 1 | 1 | 0 | 0 | FIM1_2455 | YKL215C | YKL215C | OXP1 | n |
| Kmar_QGN15116.1 | W303.asm02.HP0_G0041450 | OG0000321 | chrXIII | 476403 | 478112 | CP015056.1 | 236708 | 238408 | 74.38596491 | 569 | 566 | 1.005300353 | 1 | 0 | 0 | FIM1_1803 | YMR105C | YMR105C | PGM2 | n |
| Kmar_QGN15936.1 | W303.asm02.HP0_G0035970 | OG0000322 | chrXII | 376735 | 377265 | CP015057.1 | 422576 | 423085 | 50 | 176 | 169 | 1.041420118 | 1 | 0 | 1 | FIM1_2634 | YLR109W | YLR109W | AHP1 | n |
| Kmar_QGN14039.1 | W303.asm02.HP0_G0048180 | OG0000323 | chrXV | 144659 | 145585 | CP015054.1 | 1389955 | 1390875 | 63.17460317 | 308 | 306 | 1.006535948 | 1 | 0 | 0 | FIM1_690 | YOL092W | YOL092W | YPQ1 | n |
| Kmar_QGN15303.1 | W303.asm02.HP0_G0016130 | OG0000324 | chrV | 520446 | 525161 | CP015056.1 | 645652 | 650409 | 64.22864167 | 1571 | 1585 | 0.991167192 | 1 | 0 | 0 | FIM1_1992 | YER166W | YER166W | DNF1 | n |
| Kmar_QGN17931.1 | W303.asm02.HP0_G0001270 | OG0000325 | chrII | 101519 | 102304 | CP015060.1 | 761053 | 761802 | 63.84615385 | 261 | 249 | 1.048192771 | 1 | 0 | 0 | FIM1_5140 | YBL064C | YBL064C | PRX1 | n |
| Kmar_QGN15119.1 | W303.asm02.HP0_G0032020 | OG0000326 | chrXI | 202189 | 203076 | CP015056.1 | 241601 | 242629 | 37.00564972 | 295 | 342 | 0.862573099 | 1 | 0 | 1 | FIM1_1806 | YKL128C | YKL128C | PMU1 | n |
| Kmar_QGN13796.1 | W303.asm02.HP0_G0034130 | OG0000327 | chrXI | 623956 | 626688 | CP015054.1 | 893334 | 895655 | 48.30601093 | 910 | 773 | 1.177231565 | 1 | 0 | 0 | FIM1_442 | YKR089C | YKR089C | TGL4 | n |
| Kmar_QGN17014.1 | W303.asm02.HP0_G0002380 | OG0000328 | chrII | 341597 | 344029 | CP015059.1 | 33141 | 34955 | 36.69501823 | 810 | 604 | 1.341059603 | 1 | 0 | 0 | FIM1_3743 | YBR049C | YBR049C | REB1 | n |
| Kmar_QGN15012.1 | W303.asm02.HP0_G0047300 | OG0000329 | chrXIV | 746858 | 748300 | CP015055.1 | 1688099 | 1689565 | 50.70993915 | 480 | 488 | 0.983606557 | 1 | 0 | 0 | FIM1_1693 | YNR058W | YNR058W | BIO3 | n |
| Kmar_QGN17007.1 | W303.asm02.HP0_G0009010 | OG0000331 | chrIV | 509396 | 511768 | CP015059.1 | 17277 | 19526 | 56.98113208 | 790 | 749 | 1.054739653 | 1 | 0 | 1 | FIM1_3736 | YDR034C | YDR034C | LYS14 | n |
| Kmar_QGN15104.1 | W303.asm02.HP0_G0042750 | OG0000332 | chrXIII | 740844 | 741890 | CP015056.1 | 212101 | 213201 | 40.48913043 | 348 | 366 | 0.950819672 | 1 | 0 | 1 | FIM1_1791 | YMR234W | YMR234W | RNH1 | n |
| Kmar_QGN16675.1 | W303.asm02.HP0_G0007160 | OG0000333 | chrIV | 146046 | 147809 | CP015058.1 | 646762 | 648558 | 57.33333333 | 587 | 598 | 0.981605351 | 1 | 0 | 0 | FIM1_3394 | YDL174C | YDL174C | DLD1 | n |
| Kmar_QGN16062.1 | W303.asm02.HP0_G0040920 | OG0000334 | chrXIII | 353106 | 353639 | CP015057.1 | 731579 | 732253 | 14.67889908 | 177 | 224 | 0.790178571 | 1 | 0 | 1 | FIM1_2762 | YMR042W | YMR042W | ARG80 | n |
| Kmar_QGN14149.1 | W303.asm02.HP0_G0013510 | OG0000335 | chrIV | 1477242 | 1478693 | CP015054.1 | 1635668 | 1636834 | 28.721174 | 483 | 388 | 1.244845361 | 1 | 0 | 0 | FIM1_802 | YDR514C | YDR514C | | n |
| Kmar_QGN14996.1 | W303.asm02.HP0_G0054210 | OG0000336 | chrXVI | 323572 | 324876 | CP015055.1 | 1648947 | 1650224 | 47.72727273 | 434 | 425 | 1.021176471 | 1 | 0 | 0 | FIM1_1677 | YPL123C | YPL123C | RNY1 | n |
| Kmar_QGN17307.1 | W303.asm02.HP0_G0045570 | OG0000337 | chrXIV | 390979 | 393375 | CP015059.1 | 661939 | 663903 | 41.41791045 | 798 | 654 | 1.220183486 | 1 | 0 | 0 | FIM1_4039 | YNL125C | YNL125C | ESBP6 | n |
| Kmar_QGN13676.1 | W303.asm02.HP0_G0002320 | OG0000339 | chrII | 329771 | 331117 | CP015054.1 | 644190 | 645905 | 16.45796064 | 448 | 571 | 0.784588441 | 1 | 0 | 0 | FIM1_319 | W303.asm02.HP0_G0002320 | YBR045C | GIP1 | y |
| Kmar_QGN14361.1 | W303.asm02.HP0_G0026100 | OG0000340 | chrIX | 126915 | 128138 | CP015055.1 | 357509 | 358477 | 14.4638404 | 407 | 322 | 1.263975155 | 1 | 0 | 1 | FIM1_1021 | YIL119C | YIL119C | RPI1 | n |
| Kmar_QGN14965.1 | W303.asm02.HP0_G0054060 | OG0000341 | chrXVI | 295602 | 296984 | CP015055.1 | 1589356 | 1590723 | 32.7198364 | 460 | 455 | 1.010989011 | 1 | 0 | 0 | FIM1_1643 | YPL139C | YPL139C | UME1 | n |
| Kmar_QGN17986.1 | W303.asm02.HP0_G0034500 | OG0000342 | chrXII | 41151 | 46136 | CP015060.1 | 893353 | 898299 | 63.69617225 | 1661 | 1648 | 1.00788835 | 1 | 0 | 1 | FIM1_5195 | YLL048C | YLL048C | YBT1 | n |
| Kmar_QGN17063.1 | W303.asm02.HP0_G0007100 | OG0000344 | chrIV | 135399 | 135656 | CP015059.1 | 154330 | 154596 | 53.40909091 | 85 | 88 | 0.965909091 | 1 | 0 | 0 | FIM1_3793 | YDL181W | YDL181W | INH1 | n |
| Kmar_QGN18175.1 | W303.asm02.HP0_G0026700 | OG0000345 | chrIX | 246856 | 247694 | CP015061.1 | 373806 | 374632 | 81.6 | 121 | 125 | 0.968 | 1 | 0 | 0 | FIM1_4498 | YER056C-A/YIL052C | YIL052C | RPL34B | y |
| Kmar_QGN18174.1 | W303.asm02.HP0_G0026710 | OG0000346 | chrIX | 248475 | 248912 | CP015061.1 | 372553 | 372975 | 71.03448276 | 145 | 140 | 1.035714286 | 1 | 0 | 0 | FIM1_4497 | YIL051C | YIL051C | MMF1 | n |
| Kmar_QGN17780.1 | W303.asm02.HP0_G0049870 | OG0000347 | chrXV | 483954 | 487514 | CP015060.1 | 464737 | 468258 | 58.36837679 | 1186 | 1173 | 1.011082694 | 1 | 0 | 0 | FIM1_4989 | YOR086C | YOR086C | TCB1 | n |
| Kmar_QGN15084.1 | W303.asm02.HP0_G0023090 | OG0000348 | chrVIII | 96639 | 97453 | CP015056.1 | 168702 | 168842 | 82.60869565 | 138 | 46 | 3 | 1 | 0 | 0 | FIM1_1770 | YHL001W/YKL006W | YHL001W | RPL14B | y |
| Kmar_QGN15643.1 | W303.asm02.HP0_G0057180 | OG0000349 | chrXVI | 937208 | 938620 | CP015056.1 | 1377909 | 1379111 | 16.91648822 | 470 | 400 | 1.175 | 1 | 0 | 0 | FIM1_2336 | YPR196W | YPR196W | | n |
| Kmar_QGN13431.1 | W303.asm02.HP0_G0003820 | OG0000350 | chrII | 621283 | 622283 | CP015054.1 | 135654 | 136786 | 91.79487179 | 195 | 189 | 1.031746032 | 1 | 0 | 0 | FIM1_68 | YBR189W/YPL081W | YBR189W | RPS9B | y |
| Kmar_QGN16697.1 | W303.asm02.HP0_G0052990 | OG0000351 | chrXVI | 81266 | 81806 | CP015058.1 | 686278 | 686977 | 80 | 100 | 100 | 1 | 1 | 0 | 0 | FIM1_3417 | YMR194W/YPL249C-A | YPL249C-A | RPL36B | y |
| Kmar_QGN16561.1 | W303.asm02.HP0_G0039990 | OG0000352 | chrXIII | 156734 | 158716 | CP015058.1 | 376931 | 377395 | 28.60759494 | 524 | 154 | 3.402597403 | 1 | 0 | 0 | FIM1_3275 | YML056C | YML056C | IMD4 | n |
| Kmar_QGN16662.1 | W303.asm02.HP0_G0051730 | OG0000353 | chrXV | 868817 | 869571 | CP015058.1 | 612290 | 612783 | 36.19047619 | 105 | 55 | 1.909090909 | 1 | 0 | 0 | FIM1_3381 | YMR230W/YOR293W | YOR293W | RPS10A | y |
| Kmar_QGN18142.1 | W303.asm02.HP0_G0047790 | OG0000354 | chrXV | 70666 | 71031 | CP015061.1 | 317525 | 317851 | 58.67768595 | 121 | 108 | 1.12037037 | 1 | 1 | 1 | FIM1_4465 | YOL133W | YOL133W | HRT1 | n |
| Kmar_QGN13812.1 | W303.asm02.HP0_G0047910 | OG0000355 | chrXV | 92366 | 93190 | CP015054.1 | 925939 | 926893 | 88.88888889 | 144 | 144 | 1 | 1 | 0 | 0 | FIM1_458 | YNL302C/YOL121C | YOL121C | RPS19A | y |
| Kmar_QGN17497.1 | W303.asm02.HP0_G0021860 | OG0000356 | chrVII | 932990 | 934203 | CP015059.1 | 1033916 | 1034014 | 17.28395062 | 252 | 32 | 7.875 | 1 | 0 | 0 | FIM1_4233 | YGR214W | YGR214W | RPS0A | n |
| Kmar_QGN14684.1 | W303.asm02.HP0_G0035520 | OG0000357 | chrXII | 271006 | 271760 | CP015055.1 | 1028090 | 1028807 | 89.25619835 | 121 | 120 | 1.008333333 | 1 | 0 | 0 | FIM1_1350 | YLR061W | YLR061W | RPL22A | n |
| Kmar_QGN16333.1 | W303.asm02.HP0_G0012920 | OG0000358 | chrIV | 1365616 | 1366491 | CP015057.1 | 1310486 | 1310878 | 82.19178082 | 146 | 130 | 1.123076923 | 1 | 0 | 0 | FIM1_3038 | YDR450W/YML026C | YML026C | RPS18B | y |
| Kmar_QGN17334.1 | W303.asm02.HP0_G0050080 | OG0000359 | chrXV | 529816 | 531123 | CP015059.1 | 713666 | 716070 | 46.32034632 | 435 | 456 | 0.953947368 | 1 | 0 | 0 | FIM1_4066 | YOR110W | YOR110W | TFC7 | n |
| Kmar_QGN17912.1 | W303.asm02.HP0_G0014670 | OG0000360 | chrV | 195420 | 195686 | CP015060.1 | 721068 | 721334 | 68.53932584 | 88 | 88 | 1 | 1 | 0 | 0 | FIM1_5121 | YER019C-A | YER019C-A | SBH2 | n |
| Kmar_QGN14286.1 | W303.asm02.HP0_G0031770 | OG0000361 | chrXI | 159892 | 160491 | CP015055.1 | 190717 | 191548 | 88.76404494 | 89 | 82 | 1.085365854 | 1 | 0 | 0 | FIM1_943 | YHR021C/YKL156W | YKL156W | RPS27A | y |
| Kmar_QGN17417.1 | W303.asm02.HP0_G0025900 | OG0000362 | chrIX | 78261 | 78746 | CP015059.1 | 872892 | 873457 | 63.35403727 | 161 | 161 | 1 | 1 | 0 | 0 | FIM1_4149 | YIL138C | YIL138C | TPM2 | n |
| Kmar_QGN18055.1 | W303.asm02.HP0_G0038520 | OG0000363 | chrXII | 895321 | 895491 | CP015061.1 | 109733 | 109903 | 80.35714286 | 56 | 56 | 1 | 1 | 0 | 0 | FIM1_4372 | YLR388W | YLR388W | RPS29A | n |
| Kmar_QGN18242.1 | W303.asm02.HP0_G0039690 | OG0000364 | chrXIII | 92481 | 93940 | CP015061.1 | 532355 | 533086 | 49.44071588 | 447 | 243 | 1.839506173 | 1 | 0 | 0 | FIM1_4567 | YML085C | YML085C | TUB1 | n |
| Kmar_QGN15004.1 | W303.asm02.HP0_G0004750 | OG0000365 | chrII | 799940 | 801223 | CP015055.1 | 1667970 | 1669262 | 65.27777778 | 427 | 430 | 0.993023256 | 1 | 0 | 0 | FIM1_1685 | YBR287W | YBR287W | | n |
| Kmar_QGN18159.1 | W303.asm02.HP0_G0015190 | OG0000366 | chrV | 293736 | 294221 | CP015061.1 | 348130 | 348615 | 67.28395062 | 161 | 161 | 1 | 1 | 0 | 0 | FIM1_4482 | YER067W | YER067W | RGI1 | n |
| Kmar_QGN16797.1 | W303.asm02.HP0_G0020690 | OG0000368 | chrVII | 666657 | 668375 | CP015058.1 | 909714 | 911477 | 68.18950931 | 572 | 587 | 0.974446337 | 1 | 0 | 0 | FIM1_3520 | YGR092W | YGR092W | DBF2 | n |
| Kmar_QGN15470.1 | W303.asm02.HP0_G0051390 | OG0000369 | chrXV | 809977 | 812406 | CP015056.1 | 1013208 | 1015580 | 42.66337855 | 809 | 790 | 1.024050633 | 1 | 0 | 0 | FIM1_2161 | YOR256C | YOR256C | TRE2 | n |
| Kmar_QGN14169.1 | W303.asm02.HP0_G0013640 | OG0000370 | chrIV | 1500279 | 1501550 | CP015054.1 | 1674976 | 1676382 | 25.26096033 | 423 | 468 | 0.903846154 | 1 | 0 | 0 | FIM1_822 | YDR528W | YDR528W | HLR1 | n |
| Kmar_QGN13365.1 | W303.asm02.HP0_G0028120 | OG0000371 | chrX | 89232 | 90422 | CP015054.1 | 915 | 2102 | 57.43073048 | 396 | 395 | 1.002531646 | 1 | 0 | 0 | FIM1_1 | YJL171C | YJL171C | TOH1 | n |
| Kmar_QGN14946.1 | W303.asm02.HP0_G0024690 | OG0000372 | chrVIII | 386546 | 387265 | CP015055.1 | 1548991 | 1549659 | 42.67782427 | 239 | 222 | 1.076576577 | 1 | 0 | 1 | FIM1_1622 | YHR140W | YHR140W | | n |
| Kmar_QGN17837.1 | W303.asm02.HP0_G0040790 | OG0000373 | chrXIII | 332715 | 335246 | CP015060.1 | 589410 | 591893 | 32.43559719 | 843 | 827 | 1.019347037 | 1 | 0 | 0 | FIM1_5046 | YMR031C | YMR031C | EIS1 | n |
| Kmar_QGN17017.1 | W303.asm02.HP0_G0026050 | OG0000374 | chrIX | 115238 | 116131 | CP015059.1 | 41678 | 42571 | 57.75577558 | 297 | 297 | 1 | 1 | 0 | 1 | FIM1_3746 | YIL124W | YIL124W | AYR1 | n |
| Kmar_QGN17315.1 | W303.asm02.HP0_G0045610 | OG0000375 | chrXIV | 399892 | 401745 | CP015059.1 | 678517 | 680358 | 57.21077655 | 617 | 613 | 1.006525285 | 1 | 0 | 0 | FIM1_4047 | YNL121C | YNL121C | TOM70 | n |
| Kmar_QGN18212.1 | W303.asm02.HP0_G0042160 | OG0000376 | chrXIII | 617150 | 618682 | CP015061.1 | 458779 | 460278 | 52.58126195 | 510 | 499 | 1.022044088 | 1 | 0 | 0 | FIM1_4536 | YMR177W | YMR177W | MMT1 | n |
| Kmar_QGN15246.1 | W303.asm02.HP0_G0011300 | OG0000377 | chrIV | 1018901 | 1020202 | CP015056.1 | 521907 | 523340 | 52.35173824 | 433 | 477 | 0.907756813 | 1 | 0 | 1 | FIM1_1935 | YDR277C | YDR277C | MTH1 | n |
| Kmar_QGN15589.1 | W303.asm02.HP0_G0013140 | OG0000379 | chrIV | 1413157 | 1415787 | CP015056.1 | 1253569 | 1255908 | 27.13963964 | 876 | 779 | 1.124518614 | 1 | 0 | 0 | FIM1_2281 | YDR475C | YDR475C | JIP4 | n |
| Kmar_QGN15040.1 | W303.asm02.HP0_G0022030 | OG0000380 | chrVII | 972319 | 973518 | CP015056.1 | 70800 | 71987 | 63.7279597 | 399 | 395 | 1.010126582 | 1 | 0 | 1 | FIM1_1724 | YGR234W | YGR234W | YHB1 | n |
| Kmar_QGN14468.1 | W303.asm02.HP0_G0004180 | OG0000381 | chrII | 699200 | 702064 | CP015055.1 | 578911 | 581574 | 48.32985386 | 954 | 887 | 1.075535513 | 1 | 0 | 1 | FIM1_1129 | YBR229C | YBR229C | ROT2 | n |
| Kmar_QGN16899.1 | W303.asm02.HP0_G0036450 | OG0000382 | chrXII | 460303 | 461325 | CP015058.1 | 1128718 | 1129842 | 42.51968504 | 340 | 374 | 0.909090909 | 1 | 0 | 0 | FIM1_3626 | YLR151C | YLR151C | PCD1 | n |
| Kmar_QGN15145.1 | W303.asm02.HP0_G0015880 | OG0000383 | chrV | 458128 | 458808 | CP015056.1 | 292274 | 292906 | 49.55752212 | 226 | 210 | 1.076190476 | 1 | 0 | 0 | FIM1_1832 | YER139C | YER139C | RTR1 | n |
| Kmar_QGN15165.1 | W303.asm02.HP0_G0037290 | OG0000384 | chrXII | 632012 | 635146 | CP015056.1 | 337431 | 340565 | 86.01532567 | 1044 | 1044 | 1 | 1 | 0 | 0 | FIM1_1853 | YLR249W | YLR249W | YEF3 | n |
| Kmar_QGN14264.1 | W303.asm02.HP0_G0043030 | OG0000385 | chrXIII | 802307 | 802918 | CP015055.1 | 143314 | 143919 | 44.76190476 | 203 | 201 | 1.009950249 | 1 | 0 | 0 | FIM1_921 | YMR264W | YMR264W | CUE1 | n |
| Kmar_QGN14734.1 | W303.asm02.HP0_G0021140 | OG0000386 | chrVII | 782296 | 783699 | CP015055.1 | 1121918 | 1123204 | 54.25531915 | 467 | 428 | 1.091121495 | 1 | 0 | 1 | FIM1_1403 | YGR141W | YGR141W | VPS62 | n |
| Kmar_QGN13988.1 | W303.asm02.HP0_G0037670 | OG0000387 | chrXII | 715540 | 716199 | CP015054.1 | 1277261 | 1277905 | 96.34703196 | 219 | 214 | 1.023364486 | 1 | 0 | 0 | FIM1_637 | YLR293C/YOR185C | YLR293C | GSP1 | y |
| Kmar_QGN17505.1 | W303.asm02.HP0_G0024780 | OG0000388 | chrVIII | 400560 | 402764 | CP015059.1 | 1052282 | 1054390 | 31.67539267 | 734 | 702 | 1.045584046 | 1 | 0 | 0 | FIM1_4241 | YHR149C | YHR149C | SKG6 | n |
| Kmar_QGN16507.1 | W303.asm02.HP0_G0010720 | OG0000389 | chrIV | 894103 | 896844 | CP015058.1 | 263614 | 266001 | 43.58695652 | 913 | 795 | 1.148427673 | 1 | 0 | 0 | FIM1_3220 | YDR213W | YDR213W | UPC2 | n |
| Kmar_QGN15115.1 | W303.asm02.HP0_G0032040 | OG0000390 | chrXI | 212941 | 214983 | CP015056.1 | 233963 | 236263 | 60.10430248 | 680 | 766 | 0.88772846 | 1 | 0 | 0 | FIM1_1802 | YKL126W | YKL126W | YPK1 | n |
| Kmar_QGN16942.1 | W303.asm02.HP0_G0037180 | OG0000391 | chrXII | 609985 | 611421 | CP015058.1 | 1230764 | 1232194 | 40.90909091 | 478 | 476 | 1.004201681 | 1 | 0 | 0 | FIM1_3669 | YLR238W | YLR238W | FAR10 | n |
| Kmar_QGN15170.1 | W303.asm02.HP0_G0028780 | OG0000392 | chrX | 235607 | 236260 | CP015056.1 | 353510 | 354163 | 61.18721461 | 217 | 217 | 1 | 1 | 0 | 1 | FIM1_1858 | YJL097W | YJL097W | PHS1 | n |
| Kmar_QGN15703.1 | W303.asm02.HP0_G0017830 | OG0000393 | chrVII | 62968 | 64701 | CP015056.1 | 1497920 | 1499881 | 24.02597403 | 577 | 653 | 0.883614089 | 1 | 0 | 0 | FIM1_2396 | YGL228W | YGL228W | SHE10 | n |
| Kmar_QGN17322.1 | W303.asm02.HP0_G0024450 | OG0000394 | chrVIII | 346082 | 347332 | CP015059.1 | 692668 | 694302 | 40.58500914 | 416 | 544 | 0.764705882 | 1 | 0 | 0 | FIM1_4054 | YHR115C | YHR115C | DMA1 | n |
| Kmar_QGN16857.1 | W303.asm02.HP0_G0036160 | OG0000395 | chrXII | 412463 | 414775 | CP015058.1 | 1044131 | 1046734 | 26.16609784 | 770 | 867 | 0.888119954 | 1 | 0 | 0 | FIM1_3583 | YLR131C | YLR131C | ACE2 | n |
| Kmar_QGN15945.1 | W303.asm02.HP0_G0037460 | OG0000396 | chrXII | 675836 | 676888 | CP015057.1 | 445432 | 446487 | 65.52706553 | 350 | 351 | 0.997150997 | 1 | 0 | 0 | FIM1_2643 | YLR270W | YLR270W | DCS1 | n |
| Kmar_QGN15776.1 | W303.asm02.HP0_G0047190 | OG0000397 | chrXIV | 721105 | 723786 | CP015057.1 | 73380 | 75722 | 50.33557047 | 893 | 780 | 1.144871795 | 1 | 0 | 1 | FIM1_2472 | YNR047W | YNR047W | FPK1 | n |
| Kmar_QGN13881.1 | W303.asm02.HP0_G0024890 | OG0000398 | chrVIII | 426479 | 428392 | CP015054.1 | 1058897 | 1060624 | 39.96865204 | 637 | 575 | 1.107826087 | 1 | 0 | 0 | FIM1_528 | YHR161C | YHR161C | YAP1801 | n |
| Kmar_QGN13595.1 | W303.asm02.HP0_G0028910 | OG0000399 | chrX | 268858 | 270672 | CP015054.1 | 488339 | 490069 | 29.08496732 | 604 | 576 | 1.048611111 | 1 | 0 | 0 | FIM1_237 | YJL083W | YJL083W | TAX4 | n |
| Kmar_QGN17938.1 | W303.asm02.HP0_G0001210 | OG0000400 | chrII | 89671 | 90273 | CP015060.1 | 776316 | 776921 | 89.55223881 | 200 | 201 | 0.995024876 | 1 | 0 | 0 | FIM1_5147 | YBL072C/YER102W | YER102W | RPS8B | y |
| Kmar_QGN15283.1 | W303.asm02.HP0_G0014020 | OG0000401 | chrV | 65748 | 67160 | CP015056.1 | 596503 | 597966 | 58.3501006 | 470 | 487 | 0.965092402 | 1 | 0 | 0 | FIM1_1972 | YEL047C | YEL047C | FRD1 | n |
| Kmar_QGN14958.1 | W303.asm02.HP0_G0054010 | OG0000402 | chrXVI | 284328 | 285632 | CP015055.1 | 1572915 | 1574162 | 67.43119266 | 434 | 415 | 1.045783133 | 1 | 0 | 0 | FIM1_1635 | YPL145C | YPL145C | KES1 | n |
| Kmar_QGN13845.1 | W303.asm02.HP0_G0028070 | OG0000403 | chrX | 80316 | 81187 | CP015054.1 | 990187 | 990504 | 53.26086957 | 184 | 105 | 1.752380952 | 1 | 0 | 0 | FIM1_491 | YJL177W/YKL180W | YJL177W | RPL17B | y |
| Kmar_QGN17174.1 | W303.asm02.HP0_G0055080 | OG0000404 | chrXVI | 495246 | 497723 | CP015059.1 | 384872 | 387550 | 35.91160221 | 825 | 892 | 0.924887892 | 1 | 0 | 0 | FIM1_3905 | YPL032C | YPL032C | SVL3 | n |
| Kmar_QGN17173.1 | W303.asm02.HP0_G0055040 | OG0000405 | chrXVI | 487787 | 488260 | CP015059.1 | 383862 | 384335 | 75 | 157 | 157 | 1 | 1 | 0 | 0 | FIM1_3904 | YPL037C | YPL037C | EGD1 | n |
| Kmar_QGN13414.1 | W303.asm02.HP0_G0054520 | OG0000406 | chrXVI | 380151 | 381527 | CP015054.1 | 91875 | 93227 | 40.63829787 | 458 | 450 | 1.017777778 | 1 | 0 | 1 | FIM1_51 | YPL092W | YPL092W | SSU1 | n |
| Kmar_QGN15230.1 | W303.asm02.HP0_G0035130 | OG0000407 | chrXII | 189179 | 190795 | CP015056.1 | 482751 | 484370 | 49.81818182 | 538 | 539 | 0.998144712 | 1 | 0 | 0 | FIM1_1919 | YLR020C | YLR020C | YEH2 | n |
| Kmar_QGN16556.1 | W303.asm02.HP0_G0038990 | OG0000408 | chrXII | 1009165 | 1010439 | CP015058.1 | 365631 | 366944 | 70.30162413 | 424 | 437 | 0.970251716 | 1 | 0 | 1 | FIM1_3270 | YLR438W | YLR438W | CAR2 | n |
| Kmar_QGN16064.1 | W303.asm02.HP0_G0032610 | OG0000409 | chrXI | 324913 | 325533 | CP015057.1 | 734704 | 735360 | 39.7260274 | 206 | 218 | 0.944954128 | 1 | 0 | 1 | FIM1_2764 | YKL065C | YKL065C | YET1 | n |
| Kmar_QGN16656.1 | W303.asm02.HP0_G0020420 | OG0000410 | chrVII | 608322 | 609251 | CP015058.1 | 603718 | 604671 | 81.13207547 | 309 | 317 | 0.974763407 | 1 | 0 | 1 | FIM1_3375 | YGR060W | YGR060W | ERG25 | n |
| Kmar_QGN14962.1 | W303.asm02.HP0_G0054040 | OG0000411 | chrXVI | 289397 | 291994 | CP015055.1 | 1582373 | 1585075 | 45.65217391 | 865 | 900 | 0.961111111 | 1 | 0 | 0 | FIM1_1640 | YPL141C | YPL141C | FRK1 | n |
| Kmar_QGN16981.1 | W303.asm02.HP0_G0044610 | OG0000413 | chrXIV | 216683 | 217738 | CP015058.1 | 1302676 | 1303710 | 67.91907514 | 351 | 344 | 1.020348837 | 1 | 0 | 0 | FIM1_3708 | YNL231C | YNL231C | PDR16 | n |
| Kmar_QGN16197.1 | W303.asm02.HP0_G0039800 | OG0000414 | chrXIII | 114629 | 115864 | CP015057.1 | 1007868 | 1009112 | 64.31924883 | 411 | 414 | 0.992753623 | 1 | 0 | 0 | FIM1_2899 | YML074C | YML074C | FPR3 | n |
| Kmar_QGN17181.1 | W303.asm02.HP0_G0055140 | OG0000415 | chrXVI | 507034 | 508542 | CP015059.1 | 399943 | 401379 | 46.82539683 | 502 | 478 | 1.050209205 | 1 | 0 | 1 | FIM1_3912 | YPL026C | YPL026C | SKS1 | n |
| Kmar_QGN14126.1 | W303.asm02.HP0_G0036670 | OG0000416 | chrXII | 515391 | 516749 | CP015054.1 | 1591167 | 1592321 | 53.37423313 | 200 | 384 | 0.520833333 | 1 | 0 | 0 | FIM1_779 | W303.asm02.HP0_G0036670 | YLR180W | SAM1 | y |
| Kmar_QGN14123.1 | W303.asm02.HP0_G0013400 | OG0000417 | chrIV | 1457047 | 1458612 | CP015054.1 | 1585404 | 1586996 | 38.01498127 | 521 | 530 | 0.983018868 | 1 | 0 | 1 | FIM1_776 | YDR501W | YDR501W | PLM2 | n |

| | | | | | | | | | | | | | | | | | | | | |
|---|---|---|---|---|---|---|---|---|---|---|---|---|---|---|---|---|---|---|---|---|
| Kmar_QGN14129.1 | W303.asm02.HP0_G0036640 | OG0000418 | chrXII | 511315 | 513069 | CP015054.1 | 1596420 | 1598921 | 38.38862559 | 584 | 833 | 0.701080432 | 1 | 0 | 0 | FIM1_782 | YLR177W | YLR177W | | n |
| Kmar_QGN18268.1 | W303.asm02.HP0_G0056630 | OG0000419 | chrXVI | 822617 | 824086 | CP015061.1 | 585537 | 586892 | 73.79454927 | 489 | 451 | 1.084257206 | 1 | 0 | 0 | FIM1_4594 | YPR138C | YPR138C | MEP3 | n |
| Kmar_QGN13799.1 | W303.asm02.HP0_G0044040 | OG0000420 | chrXIV | 81087 | 82988 | CP015054.1 | 897581 | 899410 | 49.00763359 | 633 | 609 | 1.039408867 | 1 | 0 | 1 | FIM1_445 | YNL293W | YNL293W | MSB3 | n |
| Kmar_QGN17531.1 | W303.asm02.HP0_G0033730 | OG0000421 | chrXI | 537503 | 540172 | CP015059.1 | 1110431 | 1113520 | 39.9047619 | 889 | 1029 | 0.863945578 | 1 | 0 | 0 | FIM1_4267 | YKR050W | YKR050W | TRK2 | n |
| Kmar_QGN17034.1 | W303.asm02.HP0_G0040640 | OG0000422 | chrXIII | 303742 | 306096 | CP015059.1 | 86985 | 88688 | 26.02040816 | 784 | 567 | 1.382716049 | 1 | 0 | 0 | FIM1_3763 | YMR016C | YMR016C | SOK2 | n |
| Kmar_QGN15362.1 | W303.asm02.HP0_G0054840 | OG0000423 | chrXVI | 458268 | 459416 | CP015056.1 | 798763 | 799836 | 57.55208333 | 382 | 357 | 1.070028011 | 1 | 0 | 0 | FIM1_2052 | YPL057C | YPL057C | SUR1 | n |
| Kmar_QGN16375.1 | W303.asm02.HP0_G0014750 | OG0000424 | chrV | 210118 | 211371 | CP015057.1 | 1388272 | 1389765 | 47.70459082 | 417 | 497 | 0.839034205 | 1 | 0 | 0 | FIM1_3082 | YER027C | YER027C | GAL83 | n |
| Kmar_QGN14601.1 | W303.asm02.HP0_G0027610 | OG0000425 | chrIX | 414429 | 414920 | CP015055.1 | 835529 | 836113 | 66.49484536 | 163 | 194 | 0.840206186 | 1 | 0 | 1 | FIM1_1264 | YIR037W | YIR037W | HYR1 | n |
| Kmar_QGN13604.1 | W303.asm02.HP0_G0019970 | OG0000426 | chrVII | 506581 | 508536 | CP015054.1 | 503296 | 504972 | 45.09202454 | 651 | 558 | 1.166666667 | 1 | 0 | 0 | FIM1_246 | YGR009C | YGR009C | SEC9 | n |
| Kmar_QGN16607.1 | W303.asm02.HP0_G0002280 | OG0000427 | chrII | 315517 | 316710 | CP015058.1 | 488221 | 489432 | 51.13350126 | 397 | 403 | 0.985111663 | 1 | 0 | 0 | FIM1_3323 | YBR042C | YBR042C | CST26 | n |
| Kmar_QGN15377.1 | W303.asm02.HP0_G0048920 | OG0000428 | chrXV | 295232 | 296575 | CP015056.1 | 836325 | 837602 | 60.98654709 | 447 | 425 | 1.051764706 | 1 | 0 | 0 | FIM1_2067 | YOL016C | YOL016C | CMK2 | n |
| Kmar_QGN17172.1 | W303.asm02.HP0_G0011100 | OG0000429 | chrIV | 968491 | 969066 | CP015059.1 | 382748 | 383590 | 31.69014085 | 191 | 280 | 0.682142857 | 1 | 0 | 0 | FIM1_3903 | YDR253C | YDR253C | MET32 | n |
| Kmar_QGN16523.1 | W303.asm02.HP0_G0021360 | OG0000430 | chrVII | 835834 | 838692 | CP015058.1 | 299620 | 302763 | 40.90480679 | 952 | 1047 | 0.909264565 | 1 | 0 | 0 | FIM1_3236 | YGR162W | YGR162W | TIF4631 | n |
| Kmar_QGN17195.1 | W303.asm02.HP0_G0019160 | OG0000431 | chrVII | 352779 | 353924 | CP015059.1 | 428112 | 429146 | 45.14435696 | 381 | 344 | 1.10755814 | 1 | 0 | 0 | FIM1_3926 | YGL082W | YGL082W | | n |
| Kmar_QGN18177.1 | W303.asm02.HP0_G0047060 | OG0000432 | chrXIV | 702903 | 703868 | CP015061.1 | 379208 | 380179 | 76.54320988 | 321 | 323 | 0.99380805 | 1 | 0 | 0 | FIM1_4500 | YNR034W | YNR034W | SOL1 | n |
| Kmar_QGN16484.1 | W303.asm02.HP0_G0052160 | OG0000433 | chrXV | 959180 | 960271 | CP015058.1 | 202141 | 203484 | 37.30684327 | 363 | 447 | 0.812080537 | 1 | 0 | 0 | FIM1_3197 | YOR338W | YOR338W | | n |
| Kmar_QGN18202.1 | W303.asm02.HP0_G0053160 | OG0000435 | chrXVI | 112840 | 113712 | CP015061.1 | 433123 | 433980 | 67.5862069 | 290 | 285 | 1.01754386 | 1 | 0 | 0 | FIM1_4526 | YPL232W | YPL232W | SSO1 | n |
| Kmar_QGN15634.1 | W303.asm02.HP0_G0023100 | OG0000436 | chrVIII | 98417 | 99730 | CP015056.1 | 1354302 | 1355660 | 73.45132743 | 437 | 452 | 0.966814159 | 1 | 0 | 0 | FIM1_2327 | YHR001W | YHR001W | OSH7 | n |
| Kmar_QGN16859.1 | W303.asm02.HP0_G0036180 | OG0000437 | chrXII | 416398 | 418146 | CP015058.1 | 1049296 | 1050915 | 45.81196581 | 582 | 539 | 1.079777365 | 1 | 0 | 0 | FIM1_3585 | YLR133W | YLR133W | CKI1 | n |
| Kmar_QGN15272.1 | W303.asm02.HP0_G0033980 | OG0000438 | chrXI | 597649 | 598572 | CP015056.1 | 570404 | 571186 | 34.08360129 | 307 | 260 | 1.180769231 | 1 | 0 | 0 | FIM1_1961 | YKR075C | YKR075C | | n |
| Kmar_QGN13687.1 | W303.asm02.HP0_G0025380 | OG0000439 | chrVIII | 523635 | 524816 | CP015054.1 | 663976 | 665175 | 77.94486216 | 393 | 399 | 0.984962406 | 1 | 0 | 0 | FIM1_330 | YHR208W | YHR208W | BAT1 | n |
| Kmar_QGN13677.1 | W303.asm02.HP0_G0002330 | OG0000440 | chrII | 331533 | 332537 | CP015054.1 | 646527 | 647585 | 62.78409091 | 334 | 352 | 0.948863636 | 1 | 0 | 0 | FIM1_320 | YBR046C | YBR046C | ZTA1 | n |
| Kmar_QGN17356.1 | W303.asm02.HP0_G0017020 | OG0000441 | chrVI | 138405 | 140768 | CP015059.1 | 762955 | 765957 | 31.77387914 | 787 | 1000 | 0.787 | 1 | 0 | 0 | FIM1_4088 | YFR013W | YFR013W | IOC3 | n |
| Kmar_QGN13553.1 | W303.asm02.HP0_G0019250 | OG0000442 | chrVII | 368965 | 371037 | CP015054.1 | 405323 | 407632 | 27.2611465 | 690 | 769 | 0.897269181 | 1 | 0 | 0 | FIM1_194 | YGL071W | YGL071W | AFT1 | n |
| Kmar_QGN15062.1 | W303.asm02.HP0_G0031330 | OG0000443 | chrXI | 56960 | 64384 | CP015056.1 | 114753 | 122126 | 70.89220832 | 2474 | 2457 | 1.006919007 | 1 | 0 | 0 | FIM1_1747 | YKL203C | YKL203C | TOR2 | n |
| Kmar_QGN17849.1 | W303.asm02.HP0_G0013360 | OG0000444 | chrIV | 1449407 | 1451161 | CP015060.1 | 612350 | 614068 | 67.17171717 | 584 | 572 | 1.020979021 | 1 | 0 | 0 | FIM1_5058 | YDR497C | YDR497C | ITR1 | n |
| Kmar_QGN17740.1 | W303.asm02.HP0_G0046170 | OG0000445 | chrXIV | 517950 | 518900 | CP015060.1 | 366579 | 367742 | 23.45360825 | 316 | 387 | 0.816537468 | 1 | 0 | 0 | FIM1_4949 | YNL058C | YNL058C | | n |
| Kmar_QGN15569.1 | W303.asm02.HP0_G0052000 | OG0000446 | chrXV | 915339 | 916814 | CP015056.1 | 1206035 | 1207390 | 40.12219959 | 491 | 451 | 1.088691796 | 1 | 0 | 1 | FIM1_2260 | YOR320C | YOR320C | GNT1 | n |
| Kmar_QGN15579.1 | W303.asm02.HP0_G0028700 | OG0000448 | chrX | 215071 | 216753 | CP015056.1 | 1226729 | 1229092 | 23.52201258 | 560 | 787 | 0.711562897 | 1 | 0 | 1 | FIM1_2271 | YJL105W | YJL105W | SET4 | n |
| Kmar_QGN15111.1 | W303.asm02.HP0_G0041430 | OG0000449 | chrXIII | 470353 | 472857 | CP015056.1 | 225656 | 228247 | 28.05139186 | 834 | 863 | 0.966396292 | 1 | 0 | 0 | FIM1_1798 | YMR102C | YMR102C | LAF1 | n |
| Kmar_QGN15852.1 | W303.asm02.HP0_G0034640 | OG0000450 | chrXII | 81470 | 83719 | CP015057.1 | 222752 | 224932 | 59.31307794 | 749 | 726 | 1.031680441 | 1 | 0 | 1 | FIM1_2549 | YLL029W | YLL029W | FRA1 | n |
| Kmar_QGN15872.1 | W303.asm02.HP0_G0009120 | OG0000451 | chrIV | 546219 | 546914 | CP015057.1 | 275157 | 276275 | 24.66487936 | 231 | 372 | 0.620967742 | 1 | 0 | 0 | FIM1_2570 | YDR043C | YDR043C | NRG1 | n |
| Kmar_QGN14480.1 | W303.asm02.HP0_G0025630 | OG0000452 | chrIX | 19968 | 21596 | CP015055.1 | 606385 | 608103 | 43.55400697 | 542 | 572 | 0.947552448 | 1 | 0 | 1 | FIM1_1141 | YIL166C | YIL166C | SOA1 | n |
| Kmar_QGN18223.1 | W303.asm02.HP0_G0007320 | OG0000453 | chrIV | 176985 | 178268 | CP015061.1 | 487348 | 488706 | 44.02654867 | 427 | 452 | 0.944690265 | 1 | 0 | 0 | FIM1_4548 | YDL155W | YDL155W | CLB3 | n |
| Kmar_QGN13419.1 | W303.asm02.HP0_G0054570 | OG0000454 | chrXVI | 389813 | 390766 | CP015054.1 | 103476 | 104429 | 51.875 | 317 | 317 | 1 | 1 | 0 | 0 | FIM1_56 | YPL087W | YPL087W | YDC1 | n |
| Kmar_QGN13394.1 | W303.asm02.HP0_G0054390 | OG0000455 | chrXVI | 359221 | 361770 | CP015054.1 | 59856 | 62099 | 26.70588235 | 849 | 747 | 1.136546185 | 1 | 0 | 0 | FIM1_31 | YPL105C | YPL105C | SYH1 | n |
| Kmar_QGN14656.1 | W303.asm02.HP0_G0031840 | OG0000456 | chrXI | 170474 | 172396 | CP015055.1 | 959146 | 961089 | 86.55332303 | 640 | 647 | 0.989180835 | 1 | 0 | 0 | FIM1_1320 | YKL148C | YKL148C | SDH1 | n |
| Kmar_QGN13515.1 | W303.asm02.HP0_G0024330 | OG0000457 | chrVIII | 326387 | 328945 | CP015054.1 | 311315 | 313957 | 30.19079686 | 852 | 880 | 0.968181818 | 1 | 0 | 0 | FIM1_155 | YHR103W | YHR103W | SBE22 | n |
| Kmar_QGN14128.1 | W303.asm02.HP0_G0036650 | OG0000458 | chrXII | 513291 | 513950 | CP015054.1 | 1595097 | 1595702 | 42.00913242 | 219 | 201 | 1.089552239 | 1 | 0 | 0 | FIM1_781 | YLR178C | YLR178C | TFS1 | n |
| Kmar_QGN18291.1 | W303.asm02.HP0_G0055980 | OG0000459 | chrXVI | 686191 | 687297 | CP015061.1 | 628668 | 629864 | 16.00985222 | 368 | 398 | 0.924623116 | 1 | 0 | 1 | FIM1_4617 | YPR065W | YPR065W | ROX1 | n |
| Kmar_QGN13805.1 | W303.asm02.HP0_G0043990 | OG0000460 | chrXIV | 66964 | 68892 | CP015054.1 | 912583 | 914793 | 47.49661705 | 642 | 736 | 0.872282609 | 1 | 0 | 0 | FIM1_451 | YNL299W | YNL299W | TRF5 | n |
| Kmar_QGN17526.1 | W303.asm02.HP0_G0033750 | OG0000461 | chrXI | 542216 | 543130 | CP015059.1 | 1094861 | 1095901 | 61.83431953 | 304 | 346 | 0.878612717 | 1 | 0 | 0 | FIM1_4262 | YKR052C | YKR052C | MRS4 | n |
| Kmar_QGN13873.1 | W303.asm02.HP0_G0028420 | OG0000462 | chrX | 144513 | 145655 | CP015054.1 | 1032123 | 1033484 | 32.33404711 | 380 | 453 | 0.838852097 | 1 | 0 | 0 | FIM1_519 | YJL137C | YJL137C | GLG2 | n |
| Kmar_QGN17860.1 | W303.asm02.HP0_G0003000 | OG0000463 | chrII | 467167 | 468255 | CP015060.1 | 630901 | 631962 | 37.8238342 | 362 | 353 | 1.025495751 | 1 | 0 | 0 | FIM1_5069 | YBR105C | YBR105C | VID24 | n |
| Kmar_QGN14563.1 | W303.asm02.HP0_G0000540 | OG0000464 | chrI | 120989 | 125059 | CP015055.1 | 759603 | 763223 | 47.16157205 | 1356 | 1206 | 1.124378109 | 1 | 0 | 0 | FIM1_1225 | YAL017W | YAL017W | PSK1 | n |
| Kmar_QGN14436.1 | W303.asm02.HP0_G0004280 | OG0000465 | chrII | 717950 | 720145 | CP015055.1 | 513350 | 515398 | 43.75 | 731 | 682 | 1.071847507 | 1 | 0 | 0 | FIM1_1097 | YBR238C | YBR238C | | n |
| Kmar_QGN13883.1 | W303.asm02.HP0_G0024900 | OG0000466 | chrVIII | 429178 | 429567 | CP015054.1 | 1062292 | 1062708 | 67.44186047 | 129 | 138 | 0.934782609 | 1 | 0 | 0 | FIM1_530 | YHR162W | YHR162W | MPC2 | n |
| Kmar_QGN15834.1 | W303.asm02.HP0_G0041600 | OG0000467 | chrXIII | 505958 | 507832 | CP015057.1 | 185583 | 187565 | 29.65722802 | 624 | 660 | 0.945454545 | 1 | 0 | 0 | FIM1_2530 | YMR119W | YMR119W | ASI1 | n |
| Kmar_QGN17419.1 | W303.asm02.HP0_G0003330 | OG0000468 | chrII | 533636 | 542914 | CP015059.1 | 875927 | 884638 | 34.42307692 | 3092 | 2903 | 1.065105064 | 1 | 0 | 0 | FIM1_4151 | YBR140C | YBR140C | IRA1 | n |
| Kmar_QGN15956.1 | W303.asm02.HP0_G0037380 | OG0000469 | chrXII | 660528 | 662564 | CP015057.1 | 464505 | 466268 | 48.08259587 | 678 | 587 | 1.155025554 | 1 | 0 | 0 | FIM1_2654 | YLR260W | YLR260W | LCB5 | n |
| Kmar_QGN18086.1 | W303.asm02.HP0_G0030630 | OG0000470 | chrX | 645628 | 646479 | CP015061.1 | 178598 | 179470 | 43.52159468 | 283 | 290 | 0.975862069 | 1 | 0 | 1 | FIM1_4405 | YJR111C | YJR111C | PXP2 | n |
| Kmar_QGN15695.1 | W303.asm02.HP0_G0014870 | OG0000471 | chrV | 229595 | 230344 | CP015056.1 | 1483895 | 1484908 | 54.22740525 | 249 | 337 | 0.738872404 | 1 | 0 | 1 | FIM1_2388 | YER039C | YER039C | HVG1 | n |
| Kmar_QGN15264.1 | W303.asm02.HP0_G0033950 | OG0000472 | chrXI | 586097 | 587773 | CP015056.1 | 555484 | 557589 | 42.85714286 | 558 | 701 | 0.796005706 | 1 | 0 | 0 | FIM1_1953 | YKR072C | YKR072C | SIS2 | n |
| Kmar_QGN16421.1 | W303.asm02.HP0_G0050910 | OG0000473 | chrXV | 724206 | 726020 | CP015058.1 | 81717 | 83600 | 66.45468998 | 604 | 627 | 0.963317384 | 1 | 0 | 0 | FIM1_3134 | YOR204W | YOR204W | DED1 | n |
| Kmar_QGN15290.1 | W303.asm02.HP0_G0030030 | OG0000474 | chrX | 535539 | 537131 | CP015056.1 | 610680 | 612224 | 50.58365759 | 530 | 514 | 1.031128405 | 1 | 0 | 0 | FIM1_1979 | YJR049C | YJR049C | UTR1 | n |
| Kmar_QGN15511.1 | W303.asm02.HP0_G0006100 | OG0000475 | chrIII | 256106 | 259303 | CP015056.1 | 1093359 | 1096442 | 28.10457516 | 1065 | 1027 | 1.037000974 | 1 | 0 | 0 | FIM1_2202 | YCR067C | YCR067C | SED4 | n |
| Kmar_QGN16596.1 | W303.asm02.HP0_G0031350 | OG0000476 | chrXI | 69398 | 71389 | CP015058.1 | 465505 | 468123 | 29.84356197 | 663 | 872 | 0.760321101 | 1 | 0 | 0 | FIM1_3312 | YKL198C | YKL198C | PTK1 | n |
| Kmar_QGN17470.1 | W303.asm02.HP0_G0049400 | OG0000478 | chrXV | 392872 | 394980 | CP015059.1 | 988681 | 990579 | 40.42553191 | 702 | 632 | 1.110759494 | 1 | 0 | 0 | FIM1_4206 | YOR033C | YOR033C | EXO1 | n |
| Kmar_QGN16866.1 | W303.asm02.HP0_G0036210 | OG0000479 | chrXII | 423754 | 424611 | CP015058.1 | 1068929 | 1069864 | 35.90504451 | 285 | 311 | 0.916398714 | 1 | 0 | 1 | FIM1_3592 | YLR136C | YLR136C | TIS11 | n |
| Kmar_QGN18181.1 | W303.asm02.HP0_G0047020 | OG0000480 | chrXIV | 693276 | 698015 | CP015061.1 | 385103 | 389764 | 47.53894081 | 1579 | 1553 | 1.01674179 | 1 | 0 | 0 | FIM1_4504 | YNR031C | YNR031C | SSK2 | n |
| Kmar_QGN14251.1 | W303.asm02.HP0_G0019600 | OG0000481 | chrVII | 434553 | 435020 | CP015055.1 | 115004 | 115471 | 87.74193548 | 155 | 155 | 1 | 1 | 0 | 0 | FIM1_907 | YGL031C/YGR148C | YGL031C | RPL24A | y |
| Kmar_QGN14414.1 | W303.asm02.HP0_G0034240 | OG0000482 | chrXI | 657167 | 658234 | CP015055.1 | 468249 | 469313 | 30.02754821 | 355 | 354 | 1.002824859 | 1 | 0 | 0 | FIM1_1075 | YKR100C | YKR100C | SKG1 | n |
| Kmar_QGN14886.1 | W303.asm02.HP0_G0015840 | OG0000483 | chrV | 419302 | 422730 | CP015055.1 | 1424290 | 1427433 | 37.98785776 | 1142 | 1047 | 1.090735435 | 1 | 0 | 0 | FIM1_1560 | YER129W | YER129W | SAK1 | n |
| Kmar_QGN15833.1 | W303.asm02.HP0_G0041610 | OG0000484 | chrXIII | 508127 | 509905 | CP015057.1 | 182737 | 184509 | 87.66891892 | 592 | 590 | 1.003389831 | 1 | 0 | 0 | FIM1_2529 | YMR120C | YMR120C | ADE17 | n |
| Kmar_QGN13439.1 | W303.asm02.HP0_G0003900 | OG0000485 | chrII | 631978 | 632631 | CP015054.1 | 147793 | 148206 | 19.3236715 | 217 | 137 | 1.583941606 | 1 | 0 | 0 | FIM1_76 | YBR197C | YBR197C | | n |
| Kmar_QGN17148.1 | W303.asm02.HP0_G0017600 | OG0000486 | chrVII | 19305 | 20765 | CP015059.1 | 339971 | 341428 | 72.83950617 | 486 | 485 | 1.002061856 | 1 | 0 | 0 | FIM1_3879 | YGL253W | YGL253W | HXK2 | n |
| Kmar_QGN14023.1 | W303.asm02.HP0_G0000850 | OG0000487 | chrI | 193819 | 197385 | CP015054.1 | 1350532 | 1354221 | 54.14673046 | 1188 | 1229 | 0.966639544 | 1 | 0 | 0 | FIM1_674 | YAR042W | YAR042W | SWH1 | n |
| Kmar_QGN17879.1 | W303.asm02.HP0_G0056070 | OG0000488 | chrXVI | 699294 | 701336 | CP015060.1 | 664462 | 666501 | 77.94117647 | 680 | 679 | 1.001472754 | 1 | 0 | 0 | FIM1_5088 | YPR074C | YPR074C | TKL1 | n |
| Kmar_QGN13969.1 | W303.asm02.HP0_G0004060 | OG0000489 | chrII | 673696 | 675279 | CP015054.1 | 1236874 | 1238406 | 50.51194539 | 527 | 510 | 1.033333333 | 1 | 0 | 0 | FIM1_618 | YBR214W | YBR214W | SDS24 | n |
| Kmar_QGN13740.1 | W303.asm02.HP0_G0019820 | OG0000490 | chrVII | 477444 | 480200 | CP015054.1 | 781604 | 784303 | 86.49237473 | 918 | 899 | 1.021134594 | 1 | 0 | 0 | FIM1_384 | YGL008C | YGL008C | PMA1 | n |
| Kmar_QGN15232.1 | W303.asm02.HP0_G0034840 | OG0000491 | chrXII | 130516 | 131787 | CP015056.1 | 486678 | 487964 | 47.6635514 | 423 | 428 | 0.988317757 | 1 | 0 | 0 | FIM1_1921 | YLL010C | YLL010C | PSR1 | n |
| Kmar_QGN14016.1 | W303.asm02.HP0_G0027510 | OG0000492 | chrIX | 397561 | 398943 | CP015054.1 | 1333594 | 1335192 | 54.5112782 | 460 | 532 | 0.864661654 | 1 | 0 | 1 | FIM1_667 | YIR027C | YIR027C | DAL1 | n |
| Kmar_QGN17957.1 | W303.asm02.HP0_G0015690 | OG0000493 | chrV | 392612 | 395734 | CP015060.1 | 824348 | 827482 | 41.06324473 | 1040 | 1044 | 0.996168582 | 1 | 0 | 0 | FIM1_5166 | YER114C | YER114C | BOI2 | n |
| Kmar_QGN15080.1 | W303.asm02.HP0_G0023070 | OG0000494 | chrVIII | 93010 | 94245 | CP015056.1 | 161231 | 162460 | 63.94230769 | 411 | 409 | 1.004889976 | 1 | 0 | 0 | FIM1_1765 | YHL003C | YHL003C | LAG1 | n |
| Kmar_QGN15376.1 | W303.asm02.HP0_G0037360 | OG0000495 | chrXII | 655400 | 657517 | CP015056.1 | 833249 | 835354 | 77.29196051 | 705 | 701 | 1.005706134 | 1 | 0 | 0 | FIM1_2066 | YLR258W | YLR258W | GSY2 | n |
| Kmar_QGN14459.1 | W303.asm02.HP0_G0018050 | OG0000496 | chrVII | 103671 | 105917 | CP015055.1 | 557321 | 559528 | 63.5026738 | 748 | 735 | 1.017687075 | 1 | 0 | 1 | FIM1_1120 | YGL205W | YGL205W | POX1 | n |
| Kmar_QGN15773.1 | W303.asm02.HP0_G0006340 | OG0000497 | chrIII | 310354 | 311529 | CP015057.1 | 61751 | 62911 | 58.97435897 | 391 | 386 | 1.012953368 | 1 | 0 | 0 | FIM1_2469 | YCR094W | YCR094W | CDC50 | n |
| Kmar_QGN16096.1 | W303.asm02.HP0_G0015050 | OG0000498 | chrV | 263723 | 265369 | CP015057.1 | 803993 | 805609 | 22.29965157 | 548 | 538 | 1.018587361 | 1 | 0 | 0 | FIM1_2796 | YER054C | YER054C | GIP2 | n |
| Kmar_QGN13732.1 | W303.asm02.HP0_G0025050 | OG0000499 | chrVIII | 460335 | 461633 | CP015054.1 | 762348 | 763616 | 37.1559633 | 432 | 422 | 1.023696682 | 1 | 0 | 1 | FIM1_376 | YHR176W | YHR176W | FMO1 | n |
| Kmar_QGN13878.1 | W303.asm02.HP0_G0028450 | OG0000500 | chrX | 147716 | 148945 | CP015054.1 | 1052228 | 1053448 | 46.39423077 | 409 | 406 | 1.007389163 | 1 | 0 | 0 | FIM1_525 | YJL134W | YJL134W | LCB3 | n |
| Kmar_QGN15659.1 | W303.asm02.HP0_G0012590 | OG0000501 | chrIV | 1295131 | 1297845 | CP015056.1 | 1406743 | 1409160 | 30.08849558 | 904 | 805 | 1.122981366 | 1 | 0 | 0 | FIM1_2352 | YDR409W | YDR409W | SIZ1 | n |
| Kmar_QGN17918.1 | W303.asm02.HP0_G0001350 | OG0000502 | chrII | 114910 | 116316 | CP015060.1 | 736429 | 737742 | 56.62393162 | 468 | 437 | 1.070938215 | 1 | 0 | 0 | FIM1_5127 | YBL056W | YBL056W | PTC3 | n |
| Kmar_QGN17916.1 | W303.asm02.HP0_G0015430 | OG0000503 | chrV | 334846 | 336858 | CP015060.1 | 730857 | 732887 | 25.88401697 | 670 | 676 | 0.99112426 | 1 | 0 | 0 | FIM1_5125 | YER088C | YER088C | DOT6 | n |
| Kmar_QGN15518.1 | W303.asm02.HP0_G0019210 | OG0000504 | chrVII | 359177 | 360868 | CP015056.1 | 1107384 | 1109045 | 65.90909091 | 563 | 553 | 1.018083183 | 1 | 0 | 1 | FIM1_2209 | YGL077C | YGL077C | HNM1 | n |
| Kmar_QGN13391.1 | W303.asm02.HP0_G0054380 | OG0000505 | chrXVI | 356552 | 358633 | CP015054.1 | 54532 | 56616 | 81.26801153 | 693 | 694 | 0.998559078 | 1 | 0 | 0 | FIM1_28 | YPL106C | YPL106C | SSE1 | n |
| Kmar_QGN16316.1 | W303.asm02.HP0_G0053270 | OG0000506 | chrXVI | 138607 | 140988 | CP015057.1 | 1266820 | 1269195 | 57.17852684 | 793 | 791 | 1.002528445 | 1 | 0 | 0 | FIM1_3021 | YPL221W | YPL221W | FLC1 | n |
| Kmar_QGN15405.1 | W303.asm02.HP0_G0048280 | OG0000507 | chrXV | 166169 | 167407 | CP015056.1 | 891992 | 893734 | 16.55172414 | 412 | 580 | 0.710344828 | 1 | 0 | 0 | FIM1_2095 | YOL083W | YOL083W | ATG34 | n |
| Kmar_QGN18163.1 | W303.asm02.HP0_G0026660 | OG0000508 | chrIX | 240620 | 242542 | CP015061.1 | 354288 | 356351 | 33.86880857 | 640 | 687 | 0.931586608 | 1 | 0 | 0 | FIM1_4486 | YIL056W | YIL056W | VHR1 | n |
| Kmar_QGN15915.1 | W303.asm02.HP0_G0009860 | OG0000509 | chrIV | 699609 | 702803 | CP015057.1 | 369045 | 372557 | 47.16049383 | 1064 | 1170 | 0.909401709 | 1 | 0 | 0 | FIM1_2613 | YDR122W | YDR122W | KIN1 | n |
| Kmar_QGN13996.1 | W303.asm02.HP0_G0037600 | OG0000510 | chrXII | 703217 | 704905 | CP015054.1 | 1292543 | 1294201 | 53.98230088 | 562 | 552 | 1.018115942 | 1 | 0 | 1 | FIM1_646 | YLR286C | YLR286C | CTS1 | n |
| Kmar_QGN18438.1 | W303.asm02.HP0_G0031150 | OG0000511 | chrXI | 6991 | 8412 | CP015061.1 | 930720 | 932156 | 55.48654244 | 473 | 478 | 0.989539749 | 1 | 0 | 0 | FIM1_4766 | YKL221W | YKL221W | MCH2 | n |
| Kmar_QGN18172.1 | W303.asm02.HP0_G0015110 | OG0000512 | chrV | 274294 | 275556 | CP015061.1 | 369542 | 370840 | 36.78646934 | 420 | 432 | 0.972222222 | 1 | 0 | 1 | FIM1_4495 | YER059W | YER059W | PCL6 | n |
| Kmar_QGN18278.1 | W303.asm02.HP0_G0020920 | OG0000513 | chrVII | 726748 | 727502 | CP015061.1 | 604920 | 605015 | 17.93103448 | 145 | 31 | 4.677419355 | 1 | 0 | 0 | FIM1_4604 | YGR118W/YPR132W | YPR132W | RPS23B | y |
| Kmar_QGN14612.1 | W303.asm02.HP0_G0052930 | OG0000514 | chrXVI | 70545 | 72182 | CP015055.1 | 856052 | 857647 | 48.56115108 | 545 | 531 | 1.026365348 | 1 | 0 | 1 | FIM1_1275 | YPL256C | YPL256C | CLN2 | n |
| Kmar_QGN16560.1 | W303.asm02.HP0_G0038950 | OG0000515 | chrXII | 1001004 | 1002665 | CP015058.1 | 373480 | 375117 | 63.3273703 | 553 | 545 | 1.014678899 | 1 | 0 | 0 | FIM1_3274 | YLR433C | YLR433C | CNA1 | n |
| Kmar_QGN14147.1 | W303.asm02.HP0_G0005230 | OG0000516 | chrIII | 60492 | 60824 | CP015054.1 | 1632585 | 1632908 | 59.81308411 | 110 | 107 | 1.028037383 | 1 | 0 | 1 | FIM1_800 | YCL035C | YCL035C | GRX1 | n |
| Kmar_QGN17027.1 | W303.asm02.HP0_G0025960 | OG0000517 | chrIX | 87558 | 88447 | CP015059.1 | 73708 | 74781 | 84 | 199 | 199 | 1 | 1 | 0 | 0 | FIM1_3756 | YIL133C | YIL133C | RPL16A | n |
| Kmar_QGN14713.1 | W303.asm02.HP0_G0026850 | OG0000518 | chrIX | 276295 | 278058 | CP015055.1 | 1079789 | 1081324 | 27.75041051 | 587 | 511 | 1.148727984 | 1 | 0 | 0 | FIM1_1380 | YIL036W | YIL036W | CST6 | n |
| Kmar_QGN16308.1 | W303.asm02.HP0_G0018710 | OG0000519 | chrVII | 254175 | 257969 | CP015057.1 | 1248724 | 1252377 | 37.94162826 | 1264 | 1217 | 1.038619556 | 1 | 0 | 0 | FIM1_3013 | YGL133W | YGL133W | ITC1 | n |

| | | | | | | | | | | | | | | | | | | | | |
|---|---|---|---|---|---|---|---|---|---|---|---|---|---|---|---|---|---|---|---|---|
| Kmar_QGN15183.1 | W303.asm02.HP0_G0033470 | OG0000520 | chrXI | 488927 | 491674 | CP015056.1 | 384124 | 386886 | 39.87274655 | 915 | 920 | 0.994565217 | 1 | 0 | 0 | FIM1_1871 | YKR021W | YKR021W | ALY1 | n |
| Kmar_QGN14167.1 | W303.asm02.HP0_G0005100 | OG0000521 | chrIII | 37490 | 38455 | CP015054.1 | 1672378 | 1673361 | 51.20481928 | 321 | 327 | 0.981651376 | 1 | 0 | 0 | FIM1_820 | YCL050C | YCL050C | APA1 | n |
| Kmar_QGN15105.1 | W303.asm02.HP0_G0051970 | OG0000522 | chrXV | 911063 | 913165 | CP015056.1 | 214171 | 216276 | 64.15362731 | 700 | 701 | 0.998573466 | 1 | 0 | 0 | FIM1_1792 | YOR317W | YOR317W | FAA1 | n |
| Kmar_QGN15444.1 | W303.asm02.HP0_G0011520 | OG0000523 | chrIV | 1066004 | 1067290 | CP015056.1 | 972200 | 973477 | 75.70093458 | 428 | 425 | 1.007058824 | 1 | 0 | 0 | FIM1_2135 | YDR300C | YDR300C | PRO1 | n |
| Kmar_QGN17933.1 | W303.asm02.HP0_G0015540 | OG0000524 | chrV | 357136 | 359400 | CP015060.1 | 767861 | 770251 | 38.04347826 | 754 | 796 | 0.947236181 | 1 | 0 | 1 | FIM1_5142 | YER098W | YER098W | UBP9 | n |
| Kmar_QGN14455.1 | W303.asm02.HP0_G0018080 | OG0000525 | chrVII | 111572 | 113074 | CP015055.1 | 549333 | 550826 | 61.6 | 500 | 497 | 1.006036217 | 1 | 0 | 1 | FIM1_1116 | YGL202W | YGL202W | ARO8 | n |
| Kmar_QGN15155.1 | W303.asm02.HP0_G0018120 | OG0000526 | chrVII | 120211 | 124674 | CP015056.1 | 310713 | 315185 | 35.15775918 | 1487 | 1490 | 0.997986577 | 1 | 0 | 0 | FIM1_1842 | YGL197W | YGL197W | MDS3 | n |
| Kmar_QGN15484.1 | W303.asm02.HP0_G0033370 | OG0000527 | chrXI | 468486 | 470801 | CP015056.1 | 1035928 | 1039569 | 19.49013158 | 771 | 1213 | 0.63561418 | 1 | 0 | 1 | FIM1_2175 | YKR010C | YKR010C | TOF2 | n |
| Kmar_QGN16111.1 | W303.asm02.HP0_G0045280 | OG0000528 | chrXIV | 341761 | 342660 | CP015057.1 | 834546 | 835322 | 45.36423841 | 299 | 258 | 1.158914729 | 1 | 0 | 0 | FIM1_2812 | YNL156C | YNL156C | NSG2 | n |
| Kmar_QGN17369.1 | W303.asm02.HP0_G0032520 | OG0000529 | chrXI | 308058 | 310358 | CP015059.1 | 782876 | 785077 | 33.03341902 | 766 | 733 | 1.045020464 | 1 | 0 | 0 | FIM1_4101 | YKL072W | YKL072W | STB6 | n |
| Kmar_QGN15898.1 | W303.asm02.HP0_G0035780 | OG0000530 | chrXII | 326186 | 327964 | CP015057.1 | 335427 | 337055 | 65.03378378 | 592 | 542 | 1.092250923 | 1 | 0 | 0 | FIM1_2596 | YLR089C | YLR089C | ALT1 | n |
| Kmar_QGN17332.1 | W303.asm02.HP0_G0045690 | OG0000531 | chrXIV | 418595 | 418957 | CP015059.1 | 711909 | 712277 | 49.18032787 | 120 | 122 | 0.983606557 | 1 | 0 | 1 | FIM1_4064 | YNL111C | YNL111C | CYB5 | n |
| Kmar_QGN16272.1 | W303.asm02.HP0_G0030780 | OG0000532 | chrX | 666028 | 668463 | CP015057.1 | 1169568 | 1171892 | 43.03329223 | 811 | 774 | 1.047803618 | 1 | 0 | 0 | FIM1_2977 | YJR126C | YJR126C | VPS70 | n |
| Kmar_QGN14764.1 | W303.asm02.HP0_G0012410 | OG0000533 | chrIV | 1258262 | 1260226 | CP015055.1 | 1181784 | 1183703 | 39.82430454 | 654 | 639 | 1.023474178 | 1 | 0 | 0 | FIM1_1434 | YDR389W | YDR389W | SAC7 | n |
| Kmar_QGN15887.1 | W303.asm02.HP0_G0035720 | OG0000534 | chrXII | 302044 | 304047 | CP015057.1 | 295927 | 297897 | 67.6161919 | 667 | 656 | 1.016768293 | 1 | 0 | 0 | FIM1_2585 | YLR083C | YLR083C | EMP70 | n |
| Kmar_QGN17246.1 | W303.asm02.HP0_G0032880 | OG0000535 | chrXI | 374208 | 377720 | CP015059.1 | 539192 | 542461 | 29.42097027 | 1170 | 1089 | 1.074380165 | 1 | 0 | 0 | FIM1_3977 | YKL038W | YKL038W | RGT1 | n |
| Kmar_QGN15935.1 | W303.asm02.HP0_G0009960 | OG0000536 | chrIV | 723725 | 725212 | CP015057.1 | 419678 | 421036 | 35.41247485 | 495 | 452 | 1.095132743 | 1 | 0 | 0 | FIM1_2633 | YDR132C | YDR132C | MRX16 | n |
| Kmar_QGN18258.1 | W303.asm02.HP0_G0056950 | OG0000537 | chrXVI | 891628 | 892230 | CP015061.1 | 568114 | 568698 | 52.04081633 | 200 | 194 | 1.030927835 | 1 | 0 | 1 | FIM1_4584 | YPR172W | YPR172W | | n |
| Kmar_QGN18095.1 | W303.asm02.HP0_G0030410 | OG0000538 | chrX | 604835 | 608110 | CP015061.1 | 205797 | 209204 | 50.08474576 | 1091 | 1135 | 0.96123348 | 1 | 0 | 0 | FIM1_4415 | YJR091C | YJR091C | JSN1 | n |
| Kmar_QGN14320.1 | W303.asm02.HP0_G0037060 | OG0000539 | chrXII | 582360 | 585617 | CP015055.1 | 270008 | 273223 | 40 | 1085 | 1071 | 1.013071895 | 1 | 0 | 0 | FIM1_977 | YLR223C | YLR223C | IFH1 | n |
| Kmar_QGN13518.1 | W303.asm02.HP0_G0012040 | OG0000540 | chrIV | 1194815 | 1195774 | CP015054.1 | 317596 | 318588 | 85.45454545 | 319 | 330 | 0.966666667 | 1 | 0 | 0 | FIM1_158 | YDR353W | YDR353W | TRR1 | n |
| Kmar_QGN15622.1 | W303.asm02.HP0_G0018440 | OG0000541 | chrVII | 194606 | 195505 | CP015056.1 | 1327472 | 1328665 | 24.38423645 | 299 | 397 | 0.753148615 | 1 | 0 | 1 | FIM1_2315 | YGL162W | YGL162W | SUT1 | n |
| Kmar_QGN13961.1 | W303.asm02.HP0_G0019330 | OG0000542 | chrVII | 382149 | 385685 | CP015054.1 | 1223197 | 1226724 | 85.41136556 | 1178 | 1175 | 1.002553191 | 1 | 0 | 0 | FIM1_609 | YGL062W | YGL062W | PYC1 | n |
| Kmar_QGN16876.1 | W303.asm02.HP0_G0010190 | OG0000543 | chrIV | 779877 | 782435 | CP015058.1 | 1086512 | 1089004 | 44.80746791 | 852 | 830 | 1.026506024 | 1 | 0 | 1 | FIM1_3603 | YDR160W | YDR160W | SSY1 | n |
| Kmar_QGN17464.1 | W303.asm02.HP0_G0049340 | OG0000544 | chrXV | 383989 | 384876 | CP015059.1 | 974903 | 976681 | 17.56756757 | 295 | 592 | 0.498310811 | 1 | 0 | 0 | FIM1_4198 | YOR028C | YOR028C | CIN5 | n |
| Kmar_QGN15063.1 | W303.asm02.HP0_G0048550 | OG0000545 | chrXV | 224875 | 226530 | CP015056.1 | 122662 | 124224 | 36.23188406 | 551 | 520 | 1.059615385 | 1 | 0 | 0 | FIM1_1748 | YOL055C | YOL055C | THI20 | n |
| Kmar_QGN14588.1 | W303.asm02.HP0_G0038400 | OG0000546 | chrXII | 868363 | 869394 | CP015055.1 | 807690 | 809162 | 33.33333333 | 343 | 490 | 0.7 | 1 | 0 | 0 | FIM1_1250 | YLR375W | YLR375W | STP3 | n |
| Kmar_QGN15970.1 | W303.asm02.HP0_G0006710 | OG0000547 | chrIV | 54470 | 56419 | CP015057.1 | 506494 | 508542 | 35.38681948 | 649 | 682 | 0.951612903 | 1 | 0 | 0 | FIM1_2668 | YDL224C | YDL224C | WHI4 | n |
| Kmar_QGN15334.1 | W303.asm02.HP0_G0053690 | OG0000548 | chrXVI | 218592 | 219512 | CP015056.1 | 731471 | 732802 | 26.28062361 | 306 | 443 | 0.690744921 | 1 | 0 | 0 | FIM1_2023 | YPL177C | YPL177C | CUP9 | n |
| Kmar_QGN14970.1 | W303.asm02.HP0_G0054090 | OG0000549 | chrXVI | 303486 | 303983 | CP015055.1 | 1598902 | 1599438 | 72.06703911 | 165 | 178 | 0.926966292 | 1 | 0 | 0 | FIM1_1648 | YPL135W | YPL135W | ISU1 | n |
| Kmar_QGN13773.1 | W303.asm02.HP0_G0030080 | OG0000550 | chrX | 544711 | 546204 | CP015054.1 | 854735 | 856306 | 41.87380497 | 497 | 523 | 0.950286807 | 1 | 0 | 0 | FIM1_418 | YJR054W | YJR054W | KCH1 | n |
| Kmar_QGN13980.1 | W303.asm02.HP0_G0037720 | OG0000551 | chrXII | 723724 | 725070 | CP015054.1 | 1261348 | 1262643 | 60 | 448 | 431 | 1.039443155 | 1 | 0 | 0 | FIM1_629 | YLR300W | YLR300W | EXG1 | n |
| Kmar_QGN16548.1 | W303.asm02.HP0_G0039910 | OG0000552 | chrXIII | 141022 | 141789 | CP015058.1 | 349903 | 350670 | 93.33333333 | 255 | 255 | 1 | 1 | 0 | 0 | FIM1_3262 | YLR441C/YML063W | YML063W | RPS1B | y |
| Kmar_QGN16461.1 | W303.asm02.HP0_G0029070 | OG0000553 | chrX | 305073 | 305576 | CP015058.1 | 161400 | 161888 | 33.33333333 | 167 | 162 | 1.030864198 | 1 | 0 | 1 | FIM1_3174 | YJL065C | YJL065C | DLS1 | n |
| Kmar_QGN18147.1 | W303.asm02.HP0_G0047820 | OG0000554 | chrXV | 74740 | 77319 | CP015061.1 | 324563 | 326944 | 49.48096886 | 859 | 793 | 1.083228247 | 1 | 0 | 0 | FIM1_4470 | YOL130W | YOL130W | ALR1 | n |
| Kmar_QGN15678.1 | W303.asm02.HP0_G0040440 | OG0000555 | chrXIII | 254354 | 256306 | CP015056.1 | 1447304 | 1449139 | 32.31707317 | 650 | 611 | 1.063829787 | 1 | 0 | 0 | FIM1_2371 | YML007W | YML007W | YAP1 | n |
| Kmar_QGN17056.1 | W303.asm02.HP0_G0007510 | OG0000556 | chrIV | 219868 | 220977 | CP015059.1 | 138593 | 139675 | 84.01084011 | 369 | 360 | 1.025 | 1 | 0 | 0 | FIM1_3786 | YDL134C | YDL134C | PPH21 | n |
| Kmar_QGN13473.1 | W303.asm02.HP0_G0014260 | OG0000557 | chrV | 112604 | 116983 | CP015054.1 | 207735 | 211883 | 53.58885017 | 1459 | 1382 | 1.055716353 | 1 | 0 | 0 | FIM1_111 | YEL022W | YEL022W | GEA2 | n |
| Kmar_QGN15693.1 | W303.asm02.HP0_G0017890 | OG0000558 | chrVII | 73387 | 74229 | CP015056.1 | 1481329 | 1482204 | 59.02777778 | 280 | 291 | 0.962199313 | 1 | 0 | 0 | FIM1_2386 | YGL224C | YGL224C | SDT1 | n |
| Kmar_QGN13708.1 | W303.asm02.HP0_G0043330 | OG0000559 | chrXIII | 862319 | 863713 | CP015054.1 | 703765 | 705162 | 58.06451613 | 464 | 465 | 0.997849462 | 1 | 0 | 1 | FIM1_351 | YMR293C | YMR293C | HER2 | n |
| Kmar_QGN17555.1 | W303.asm02.HP0_G0018600 | OG0000560 | chrVII | 228918 | 230975 | CP015059.1 | 1155577 | 1157625 | 47.15672677 | 685 | 682 | 1.004398827 | 1 | 0 | 0 | FIM1_4291 | YGL144C | YGL144C | ROG1 | n |
| Kmar_QGN18352.1 | W303.asm02.HP0_G0023990 | OG0000561 | chrVIII | 242978 | 243667 | CP015061.1 | 753859 | 754527 | 32.36514523 | 229 | 222 | 1.031531532 | 1 | 0 | 1 | FIM1_4679 | YHR071W | YHR071W | PCL5 | n |
| Kmar_QGN15964.1 | W303.asm02.HP0_G0010650 | OG0000562 | chrIV | 865770 | 868424 | CP015057.1 | 491648 | 494521 | 25.38540596 | 884 | 957 | 0.923719958 | 1 | 0 | 0 | FIM1_2662 | YDR206W | YDR206W | EBS1 | n |
| Kmar_QGN16269.1 | W303.asm02.HP0_G0039730 | OG0000563 | chrXIII | 99317 | 103072 | CP015057.1 | 1162558 | 1166610 | 39.82365907 | 1251 | 1350 | 0.926666667 | 1 | 0 | 0 | FIM1_2974 | YML081W | YML081W | TDA9 | n |
| Kmar_QGN13964.1 | W303.asm02.HP0_G0019350 | OG0000564 | chrVII | 386925 | 388943 | CP015054.1 | 1228636 | 1230750 | 31.83098592 | 672 | 704 | 0.954545455 | 1 | 0 | 0 | FIM1_612 | YGL060W | YGL060W | YBP2 | n |
| Kmar_QGN16822.1 | W303.asm02.HP0_G0056470 | OG0000565 | chrXVI | 778582 | 780057 | CP015058.1 | 967592 | 969223 | 55.04587156 | 491 | 543 | 0.904235727 | 1 | 0 | 1 | FIM1_3546 | YPR119W | YPR119W | CLB2 | n |
| Kmar_QGN14511.1 | W303.asm02.HP0_G0042840 | OG0000566 | chrXIII | 761344 | 762672 | CP015055.1 | 661239 | 662567 | 56.74518201 | 442 | 442 | 1 | 1 | 0 | 0 | FIM1_1172 | YMR243C | YMR243C | ZRC1 | n |
| Kmar_QGN13417.1 | W303.asm02.HP0_G0054550 | OG0000567 | chrXVI | 385478 | 387508 | CP015054.1 | 97473 | 99704 | 29.18848168 | 676 | 743 | 0.909825034 | 1 | 0 | 0 | FIM1_54 | YPL089C | YPL089C | RLM1 | n |
| Kmar_QGN18203.1 | W303.asm02.HP0_G0053170 | OG0000568 | chrXVI | 114217 | 119880 | CP015061.1 | 435843 | 441527 | 81.23352662 | 1887 | 1894 | 0.996304118 | 1 | 0 | 1 | FIM1_4527 | YPL231W | YPL231W | FAS2 | n |
| Kmar_QGN16850.1 | W303.asm02.HP0_G0056730 | OG0000569 | chrXVI | 841551 | 842072 | CP015058.1 | 1027609 | 1028118 | 54.9132948 | 173 | 169 | 1.023668639 | 1 | 0 | 0 | FIM1_3575 | YPR149W | YPR149W | NCE102 | n |
| Kmar_QGN17728.1 | W303.asm02.HP0_G0026200 | OG0000570 | chrIX | 147742 | 150522 | CP015060.1 | 338555 | 341308 | 60.39915966 | 926 | 917 | 1.009814613 | 1 | 0 | 0 | FIM1_4937 | YIL109C | YIL109C | SEC24 | n |
| Kmar_QGN14732.1 | W303.asm02.HP0_G0056790 | OG0000571 | chrXVI | 849542 | 851410 | CP015055.1 | 1115631 | 1117466 | 74.83974359 | 622 | 611 | 1.018003273 | 1 | 0 | 0 | FIM1_1401 | YPR156C | YPR156C | TPO3 | n |
| Kmar_QGN16990.1 | W303.asm02.HP0_G0044520 | OG0000572 | chrXIV | 196980 | 198497 | CP015058.1 | 1324483 | 1325976 | 70.0990099 | 505 | 497 | 1.016096579 | 1 | 0 | 1 | FIM1_3717 | YNL241C | YNL241C | ZWF1 | n |
| Kmar_QGN15194.1 | W303.asm02.HP0_G0008400 | OG0000573 | chrIV | 377644 | 379332 | CP015056.1 | 404525 | 406519 | 48.27067669 | 562 | 664 | 0.846385542 | 1 | 0 | 0 | FIM1_1882 | YDL042C | YDL042C | SIR2 | n |
| Kmar_QGN16540.1 | W303.asm02.HP0_G0039840 | OG0000574 | chrXIII | 128015 | 129769 | CP015058.1 | 334751 | 336517 | 66.60988075 | 584 | 588 | 0.993197279 | 1 | 0 | 1 | FIM1_3253 | YML070W | YML070W | DAK1 | n |
| Kmar_QGN17351.1 | W303.asm02.HP0_G0052200 | OG0000575 | chrXV | 968948 | 969907 | CP015059.1 | 748502 | 749446 | 28.44827586 | 319 | 314 | 1.015923567 | 1 | 0 | 0 | FIM1_4083 | YOR342C | YOR342C | | n |
| Kmar_QGN17321.1 | W303.asm02.HP0_G0045640 | OG0000576 | chrXIV | 407566 | 409230 | CP015059.1 | 690207 | 691874 | 79.60288809 | 554 | 555 | 0.998198198 | 1 | 0 | 0 | FIM1_4053 | YNL117W | YNL117W | MLS1 | n |
| Kmar_QGN16771.1 | W303.asm02.HP0_G0025060 | OG0000577 | chrVIII | 462697 | 464058 | CP015058.1 | 852589 | 853953 | 26.17021277 | 453 | 454 | 0.997797357 | 1 | 0 | 0 | FIM1_3493 | YHR177W | YHR177W | ROF1 | n |
| Kmar_QGN18120.1 | W303.asm02.HP0_G0000970 | OG0000578 | chrII | 25250 | 29108 | CP015061.1 | 255666 | 259067 | 31.70914543 | 1244 | 1133 | 1.097969991 | 1 | 0 | 0 | FIM1_4440 | YBL101C | YBL101C | ECM21 | n |
| Kmar_QGN15771.1 | W303.asm02.HP0_G0034440 | OG0000579 | chrXII | 29980 | 31575 | CP015057.1 | 58848 | 60533 | 49.73913043 | 531 | 561 | 0.946524064 | 1 | 0 | 1 | FIM1_2467 | YLL055W | YLL055W | YCT1 | n |
| Kmar_QGN14136.1 | W303.asm02.HP0_G0005320 | OG0000580 | chrIII | 73056 | 73637 | CP015054.1 | 1614593 | 1615183 | 67.34693878 | 193 | 196 | 0.984693878 | 1 | 0 | 0 | FIM1_789 | YCL026C-B | YCL026C-B | HBN1 | n |
| Kmar_QGN17895.1 | W303.asm02.HP0_G0042920 | OG0000581 | chrXIII | 781818 | 782222 | CP015060.1 | 694942 | 695310 | 42.25352113 | 134 | 122 | 1.098360656 | 1 | 0 | 1 | FIM1_5104 | YMR252C | YMR252C | MLO1 | n |
| Kmar_QGN16443.1 | W303.asm02.HP0_G0041150 | OG0000582 | chrXIII | 412073 | 412624 | CP015058.1 | 129936 | 130436 | 29.94652406 | 183 | 166 | 1.102409639 | 1 | 0 | 1 | FIM1_3156 | YMR072W | YMR072W | ABF2 | n |
| Kmar_QGN17233.1 | W303.asm02.HP0_G0032940 | OG0000582 | chrXI | 389012 | 389689 | CP015059.1 | 509024 | 510379 | 10.13513514 | 225 | 451 | 0.498891353 | 1 | 0 | 1 | FIM1_3964 | W303.asm02.HP0_G0032940 | YKL032C | IXR1 | y |
| Kmar_QGN16589.1 | W303.asm02.HP0_G0032300 | OG0000583 | chrXI | 269608 | 270327 | CP015058.1 | 449524 | 450279 | 41.66666667 | 239 | 251 | 0.952191235 | 1 | 0 | 1 | FIM1_3305 | YKL096W | YKL096W | CWP1 | n |
| Kmar_QGN17207.1 | W303.asm02.HP0_G0053600 | OG0000584 | chrXVI | 199199 | 199696 | CP015059.1 | 448348 | 448899 | 47.2826087 | 165 | 183 | 0.901639344 | 1 | 0 | 0 | FIM1_3938 | YPL187W | YPL187W | MF(ALPHA)1 | n |
| Kmar_QGN17982.1 | W303.asm02.HP0_G0034530 | OG0000585 | chrXII | 49945 | 51954 | CP015060.1 | 882858 | 884576 | 37.9671151 | 669 | 572 | 1.16958042 | 1 | 0 | 1 | FIM1_5191 | YLL043W | YLL043W | FPS1 | n |
| Kmar_QGN18322.1 | W303.asm02.HP0_G0002850 | OG0000588 | chrII | 440051 | 441115 | CP015061.1 | 689877 | 690944 | 71.18644068 | 354 | 355 | 0.997183099 | 1 | 0 | 1 | FIM1_4648 | YBR087W | YBR087W | RFC5 | n |
| Kmar_QGN13501.1 | W303.asm02.HP0_G0041820 | OG0000591 | chrXIII | 552550 | 553526 | CP015054.1 | 276815 | 277722 | 96.5034965 | 143 | 143 | 1 | 1 | 0 | 1 | FIM1_140 | YDL083C/YMR143W | YMR143W | RPS16A | y |
| Kmar_QGN16547.1 | W303.asm02.HP0_G0039900 | OG0000592 | chrXIII | 139679 | 140416 | CP015058.1 | 348231 | 348899 | 60 | 245 | 222 | 1.103603604 | 1 | 0 | 1 | FIM1_3261 | YML064C | YML064C | TEM1 | n |
| Kmar_QGN16196.1 | W303.asm02.HP0_G0039810 | OG0000593 | chrXIII | 117767 | 118712 | CP015057.1 | 1006237 | 1007351 | 80.79096045 | 176 | 175 | 1.005714286 | 1 | 0 | 1 | FIM1_2898 | YML073C | YML073C | RPL6A | n |
| Kmar_QGN13728.1 | W303.asm02.HP0_G0025120 | OG0000600 | chrVIII | 477064 | 478533 | CP015054.1 | 751838 | 753316 | 85.56910569 | 489 | 492 | 0.993902439 | 1 | 0 | 0 | FIM1_372 | YHR183W | YHR183W | GND1 | n |
| Kmar_QGN15795.1 | W303.asm02.HP0_G0019960 | OG0000601 | chrVII | 505644 | 505898 | CP015057.1 | 110451 | 110678 | 70.66666667 | 84 | 75 | 1.12 | 1 | 0 | 0 | FIM1_2491 | YGR008C | YGR008C | STF2 | n |
| Kmar_QGN16449.1 | W303.asm02.HP0_G0029120 | OG0000603 | chrX | 313405 | 314739 | CP015058.1 | 139624 | 140964 | 70.91722595 | 444 | 446 | 0.995515695 | 1 | 0 | 1 | FIM1_3162 | YJL060W | YJL060W | BNA3 | n |
| Kmar_QGN13432.1 | W303.asm02.HP0_G0054640 | OG0000604 | chrXVI | 412994 | 413897 | CP015054.1 | 137703 | 138605 | 94.375 | 160 | 160 | 1 | 1 | 0 | 1 | FIM1_69 | YBR191W/YPL079W | YPL079W | RPL21B | y |
| Kmar_QGN16517.1 | W303.asm02.HP0_G0037680 | OG0000606 | chrXII | 716768 | 717142 | CP015058.1 | 284235 | 284567 | 46.77419355 | 124 | 110 | 1.127272727 | 1 | 0 | 0 | FIM1_3230 | YLR295C | YLR295C | ATP14 | n |
| Kmar_QGN14534.1 | W303.asm02.HP0_G0020620 | OG0000607 | chrVII | 646419 | 646943 | CP015055.1 | 710966 | 711490 | 90.22988506 | 174 | 174 | 1 | 1 | 0 | 0 | FIM1_1196 | YGR085C/YPR102C | YGR085C | RPL11B | y |
| Kmar_QGN14269.1 | W303.asm02.HP0_G0043000 | OG0000608 | chrXIII | 796707 | 799871 | CP015055.1 | 148794 | 152531 | 46.19085806 | 1054 | 1245 | 0.846586345 | 1 | 0 | 0 | FIM1_926 | YMR261C | YMR261C | TPS3 | n |
| Kmar_QGN15248.1 | W303.asm02.HP0_G0011290 | OG0000609 | chrIV | 1017976 | 1018143 | CP015056.1 | 526186 | 526528 | 74.54545455 | 55 | 55 | 1 | 1 | 0 | 1 | FIM1_1937 | YDR276C | YDR276C | PMP3 | n |
| Kmar_QGN14911.1 | W303.asm02.HP0_G0006750 | OG0000610 | chrIV | 65315 | 65838 | CP015055.1 | 1478446 | 1478898 | 69.33333333 | 150 | 150 | 1 | 1 | 0 | 1 | FIM1_1585 | YDL219W | YDL219W | DTD1 | n |
| Kmar_QGN16898.1 | W303.asm02.HP0_G0036350 | OG0000611 | chrXII | 448511 | 449332 | CP015058.1 | 1127088 | 1128218 | 53.47985348 | 273 | 248 | 1.100806452 | 1 | 0 | 0 | FIM1_3625 | YLR150W | YLR150W | STM1 | n |
| Kmar_QGN13811.1 | W303.asm02.HP0_G0043970 | OG0000612 | chrXIV | 64005 | 64997 | CP015054.1 | 923838 | 924927 | 91.93548387 | 186 | 186 | 1 | 1 | 0 | 0 | FIM1_457 | YNL301C/YOL120C | YOL120C | RPL18A | y |
| Kmar_QGN15449.1 | W303.asm02.HP0_G0023860 | OG0000613 | chrVIII | 224827 | 225714 | CP015056.1 | 980868 | 981725 | 43.79084967 | 295 | 285 | 1.035087719 | 1 | 0 | 1 | FIM1_2140 | YHR058C | YHR058C | MED6 | n |
| Kmar_QGN13416.1 | W303.asm02.HP0_G0003730 | OG0000614 | chrII | 608487 | 609549 | CP015054.1 | 94633 | 96130 | 88.55932203 | 236 | 239 | 0.987447699 | 1 | 0 | 0 | FIM1_53 | YBR181C/YPL090C | YPL090C | RPS6A | y |
| Kmar_QGN18315.1 | W303.asm02.HP0_G0055880 | OG0000615 | chrXVI | 672191 | 672472 | CP015061.1 | 678728 | 679009 | 80.64516129 | 93 | 93 | 1 | 1 | 0 | 1 | FIM1_4641 | YPR052C | YPR052C | NHP6A | n |
| Kmar_QGN18094.1 | W303.asm02.HP0_G0030420 | OG0000616 | chrX | 608509 | 612851 | CP015061.1 | 200689 | 204303 | 27.65363128 | 1420 | 1204 | 1.179401993 | 1 | 0 | 1 | FIM1_4414 | YJR092W | YJR092W | BUD4 | n |
| Kmar_QGN16600.1 | W303.asm02.HP0_G0017170 | OG0000618 | chrVI | 177318 | 178973 | CP015058.1 | 474682 | 476223 | 66.60714286 | 551 | 513 | 1.074074074 | 1 | 0 | 1 | FIM1_3316 | YFR028C | YFR028C | CDC14 | n |
| Kmar_QGN18131.1 | W303.asm02.HP0_G0001030 | OG0000619 | chrII | 47131 | 47523 | CP015061.1 | 289815 | 290207 | 80.76923077 | 130 | 130 | 1 | 1 | 0 | 1 | FIM1_4452 | YBL092W | YBL092W | RPL32 | n |
| Kmar_QGN16301.1 | W303.asm02.HP0_G0053360 | OG0000621 | chrXVI | 157070 | 158704 | CP015057.1 | 1234591 | 1236210 | 61.64874552 | 544 | 539 | 1.009276438 | 1 | 0 | 1 | FIM1_3006 | YPL212C | YPL212C | PUS1 | n |
| Kmar_QGN16321.1 | W303.asm02.HP0_G0040180 | OG0000622 | chrXIII | 210033 | 212663 | CP015057.1 | 1277686 | 1279638 | 25.88757396 | 834 | 650 | 1.283076923 | 1 | 0 | 0 | FIM1_3026 | YML034W | YML034W | SRC1 | n |
| Kmar_QGN14741.1 | W303.asm02.HP0_G0023210 | OG0000623 | chrVIII | 116362 | 117063 | CP015055.1 | 1139591 | 1140277 | 73.27586207 | 233 | 228 | 1.021929825 | 1 | 0 | 1 | FIM1_1410 | YHR008C | YHR008C | SOD2 | n |
| Kmar_QGN15952.1 | W303.asm02.HP0_G0037410 | OG0000624 | chrXII | 667782 | 667985 | CP015057.1 | 456493 | 456696 | 97.01492537 | 67 | 67 | 1 | 1 | 0 | 1 | FIM1_2650 | YLR264W/YOR167C | YLR264W | RPS28B | y |
| Kmar_QGN15131.1 | W303.asm02.HP0_G0041570 | OG0000625 | chrXIII | 500081 | 501313 | CP015056.1 | 264673 | 266038 | 66.87116564 | 319 | 326 | 0.978527607 | 1 | 0 | 1 | FIM1_1818 | YMR116C | YMR116C | ASC1 | n |
| Kmar_QGN14370.1 | W303.asm02.HP0_G0009670 | OG0000626 | chrIV | 654516 | 656297 | CP015055.1 | 376800 | 378515 | 52.47524752 | 593 | 571 | 1.038528897 | 1 | 0 | 1 | FIM1_1030 | YDR101C | YDR101C | ARX1 | n |
| Kmar_QGN18358.1 | W303.asm02.HP0_G0034170 | OG0000627 | chrXI | 633997 | 635802 | CP015061.1 | 761477 | 763378 | 57.66192733 | 601 | 633 | 0.949447077 | 1 | 0 | 1 | FIM1_4685 | YKR093W | YKR093W | PTR2 | n |
| Kmar_QGN18368.1 | W303.asm02.HP0_G0034140 | OG0000628 | chrXI | 627269 | 629389 | CP015061.1 | 789980 | 792073 | 25.73018081 | 706 | 697 | 1.012912482 | 1 | 0 | 1 | FIM1_4695 | YKR090W | YKR090W | PXL1 | n |
| Kmar_QGN13724.1 | W303.asm02.HP0_G0022270 | OG0000629 | chrVII | 1020111 | 1023206 | CP015054.1 | 743561 | 746698 | 45.74275362 | 1031 | 1045 | 0.986602871 | 1 | 0 | 0 | FIM1_368 | YGR258C | YGR258C | RAD2 | n |
| Kmar_QGN13983.1 | W303.asm02.HP0_G0037710 | OG0000630 | chrXII | 720838 | 722820 | CP015054.1 | 1270442 | 1272256 | 54.4494721 | 660 | 604 | 1.092715232 | 1 | 0 | 1 | FIM1_632 | YLR299W | YLR299W | ECM38 | n |
| Kmar_QGN14974.1 | W303.asm02.HP0_G0054100 | OG0000631 | chrXVI | 304504 | 305436 | CP015055.1 | 1603562 | 1604458 | 62.37942122 | 310 | 298 | 1.040268456 | 1 | 0 | 0 | FIM1_1652 | YPL134C | YPL134C | ODC1 | n |

| | | | | | | | | | | | | | | | | | | | | |
|---|---|---|---|---|---|---|---|---|---|---|---|---|---|---|---|---|---|---|---|---|
| Kmar_QGN16753.1 | W303.asm02.HP0_G0056180 | OG0000632 | chrXVI | 716600 | 717637 | CP015058.1 | 824852 | 825925 | 64.70588235 | 345 | 357 | 0.966386555 | 1 | 0 | 1 | FIM1_3475 | YPR086W | YPR086W | SUA7 | n |
| Kmar_QGN14916.1 | W303.asm02.HP0_G0006770 | OG0000634 | chrIV | 68056 | 68679 | CP015055.1 | 1486396 | 1486986 | 70.53140097 | 207 | 196 | 1.056122449 | 1 | 0 | 1 | FIM1_1590 | YDL217C | YDL217C | TIM22 | n |
| Kmar_QGN18111.1 | W303.asm02.HP0_G0030330 | OG0000635 | chrX | 589719 | 590903 | CP015061.1 | 235489 | 236556 | 27.5 | 394 | 355 | 1.109859155 | 1 | 0 | 1 | FIM1_4431 | YJR080C | YJR080C | AIM24 | n |
| Kmar_QGN16400.1 | W303.asm02.HP0_G0000110 | OG0000636 | chrI | 32202 | 33575 | CP015058.1 | 31742 | 33127 | 75.05422993 | 457 | 461 | 0.99132321 | 1 | 0 | 0 | FIM1_3113 | YAL062W | YAL062W | GDH3 | n |
| Kmar_QGN18019.1 | W303.asm02.HP0_G0016430 | OG0000637 | chrVI | 8868 | 11012 | CP015061.1 | 43898 | 45961 | 50.55555556 | 714 | 687 | 1.03930131 | 1 | 0 | 1 | FIM1_4335 | YFL047W | YFL047W | RGD2 | n |
| Kmar_QGN16511.1 | W303.asm02.HP0_G0010670 | OG0000638 | chrIV | 871940 | 874279 | CP015058.1 | 274495 | 276948 | 43.46289753 | 779 | 817 | 0.953488372 | 1 | 0 | 1 | FIM1_3224 | YDR208W | YDR208W | MSS4 | n |
| Kmar_QGN14914.1 | W303.asm02.HP0_G0044980 | OG0000639 | chrXIV | 284956 | 286584 | CP015055.1 | 1483539 | 1485155 | 82.47232472 | 542 | 538 | 1.007434944 | 1 | 0 | 1 | FIM1_1588 | YNL189W | YNL189W | SRP1 | n |
| Kmar_QGN17077.1 | W303.asm02.HP0_G0026470 | OG0000640 | chrIX | 205486 | 206448 | CP015059.1 | 181104 | 182057 | 29.51289398 | 320 | 317 | 1.009463722 | 1 | 0 | 1 | FIM1_3808 | YIL077C | YIL077C | RCI37 | n |
| Kmar_QGN14581.1 | W303.asm02.HP0_G0007950 | OG0000641 | chrIV | 298747 | 300333 | CP015055.1 | 795441 | 796709 | 35.72778828 | 528 | 422 | 1.251184834 | 1 | 0 | 0 | FIM1_1243 | YDL088C | YDL088C | ASM4 | n |
| Kmar_QGN13868.1 | W303.asm02.HP0_G0028380 | OG0000642 | chrX | 137495 | 139918 | CP015054.1 | 1024052 | 1026214 | 48.65853659 | 807 | 720 | 1.120833333 | 1 | 0 | 1 | FIM1_514 | YJL141C | YJL141C | YAK1 | n |
| Kmar_QGN14545.1 | W303.asm02.HP0_G0000600 | OG0000643 | chrI | 132965 | 134842 | CP015055.1 | 729625 | 731973 | 30.82524272 | 625 | 782 | 0.799232737 | 1 | 0 | 1 | FIM1_1207 | YAL011W | YAL011W | SWC3 | n |
| Kmar_QGN16818.1 | W303.asm02.HP0_G0057160 | OG0000644 | chrXVI | 930136 | 932769 | CP015058.1 | 957877 | 960486 | 43.06651635 | 877 | 869 | 1.009205984 | 1 | 0 | 1 | FIM1_3542 | YPR194C | YPR194C | OPT2 | n |
| Kmar_QGN16934.1 | W303.asm02.HP0_G0037210 | OG0000646 | chrXII | 615703 | 618051 | CP015058.1 | 1215057 | 1217393 | 67.93409379 | 782 | 778 | 1.005141388 | 1 | 0 | 1 | FIM1_3661 | YLR241W | YLR241W | CSC1 | n |
| Kmar_QGN17718.1 | W303.asm02.HP0_G0026280 | OG0000649 | chrIX | 165665 | 167608 | CP015060.1 | 312619 | 313485 | 19.16537867 | 647 | 288 | 2.246527778 | 1 | 0 | 1 | FIM1_4925 | YIL101C | YIL101C | XBP1 | n |
| Kmar_QGN14358.1 | W303.asm02.HP0_G0010860 | OG0000650 | chrIV | 926427 | 928307 | CP015055.1 | 348406 | 350268 | 45.31722054 | 626 | 620 | 1.009677419 | 1 | 0 | 1 | FIM1_1018 | YDR228C | YDR228C | PCF11 | n |
| Kmar_QGN15122.1 | W303.asm02.HP0_G0041500 | OG0000651 | chrXIII | 487212 | 490871 | CP015056.1 | 248893 | 252684 | 71.80094787 | 1219 | 1263 | 0.965162312 | 1 | 0 | 0 | FIM1_1809 | YMR109W | YMR109W | MYO5 | n |
| Kmar_QGN14055.1 | W303.asm02.HP0_G0005900 | OG0000652 | chrIII | 220707 | 222635 | CP015054.1 | 1426260 | 1430357 | 15.73938507 | 642 | 1365 | 0.47032967 | 1 | 0 | 0 | FIM1_706 | YCR038C | YCR038C | BUD5 | n |
| Kmar_QGN16705.1 | W303.asm02.HP0_G0000040 | OG0000653 | chrI | 7481 | 9262 | CP015058.1 | 707311 | 709086 | 69.73244147 | 593 | 591 | 1.003384095 | 1 | 0 | 1 | FIM1_3425 | YAL067C | YAL067C | SEO1 | n |
| Kmar_QGN15284.1 | W303.asm02.HP0_G0014030 | OG0000654 | chrV | 67992 | 69155 | CP015056.1 | 599296 | 600471 | 76.72634271 | 387 | 391 | 0.989769821 | 1 | 0 | 1 | FIM1_1973 | YEL046C | YEL046C | GLY1 | n |
| Kmar_QGN14826.1 | W303.asm02.HP0_G0042020 | OG0000655 | chrXIII | 587882 | 590134 | CP015055.1 | 1301417 | 1302760 | 22.20689655 | 750 | 447 | 1.677852349 | 1 | 0 | 1 | FIM1_1497 | YMR164C | YMR164C | MSS11 | n |
| Kmar_QGN13908.1 | W303.asm02.HP0_G0042610 | OG0000656 | chrXIII | 714510 | 716024 | CP015054.1 | 1130298 | 1131836 | 58.05825243 | 504 | 512 | 0.984375 | 1 | 0 | 1 | FIM1_556 | YMR221C | YMR221C | FMP42 | n |
| Kmar_QGN18354.1 | W303.asm02.HP0_G0013200 | OG0000657 | chrIV | 1424244 | 1425944 | CP015061.1 | 755914 | 757554 | 56.18374558 | 566 | 546 | 1.036630037 | 1 | 0 | 1 | FIM1_4681 | YDR481C | YDR481C | PHO8 | n |
| Kmar_QGN15558.1 | W303.asm02.HP0_G0011640 | OG0000658 | chrIV | 1092211 | 1093572 | CP015056.1 | 1181508 | 1182929 | 67.94055202 | 453 | 473 | 0.957716702 | 1 | 0 | 0 | FIM1_2249 | YDR312W | YDR312W | SSF2 | n |
| Kmar_QGN16663.1 | W303.asm02.HP0_G0051720 | OG0000659 | chrXV | 867372 | 868301 | CP015058.1 | 613360 | 614487 | 42.89544236 | 309 | 375 | 0.824 | 1 | 0 | 1 | FIM1_3382 | YOR292C | YOR292C | | n |
| Kmar_QGN13660.1 | W303.asm02.HP0_G0005630 | OG0000660 | chrIII | 142986 | 144236 | CP015054.1 | 612722 | 613972 | 82.69230769 | 416 | 416 | 1 | 1 | 0 | 1 | FIM1_303 | YCR012W | YCR012W | PGK1 | n |
| Kmar_QGN17932.1 | W303.asm02.HP0_G0001260 | OG0000661 | chrII | 97817 | 101263 | CP015060.1 | 762806 | 766300 | 49.63971177 | 1148 | 1164 | 0.986254296 | 1 | 0 | 1 | FIM1_5141 | YBL066C | YBL066C | SEF1 | n |
| Kmar_QGN16052.1 | W303.asm02.HP0_G0037320 | OG0000662 | chrXII | 637862 | 639571 | CP015057.1 | 707498 | 709201 | 61.52512998 | 569 | 567 | 1.003527337 | 1 | 0 | 1 | FIM1_2752 | YLR253W | YLR253W | CQD2 | n |
| Kmar_QGN16892.1 | W303.asm02.HP0_G0036320 | OG0000663 | chrXII | 442203 | 442508 | CP015058.1 | 1115498 | 1115800 | 66.66666667 | 101 | 100 | 1.01 | 1 | 0 | 0 | FIM1_3619 | YLR147C | YLR147C | SMD3 | n |
| Kmar_QGN16584.1 | W303.asm02.HP0_G0038030 | OG0000664 | chrXII | 791259 | 791585 | CP015058.1 | 437538 | 437864 | 87.96296296 | 108 | 108 | 1 | 1 | 0 | 0 | FIM1_3300 | YGR027C/YLR333C | YLR333C | RPS25B | y |
| Kmar_QGN14352.1 | W303.asm02.HP0_G0032230 | OG0000665 | chrXI | 254614 | 256158 | CP015055.1 | 335261 | 336832 | 59.88593156 | 514 | 523 | 0.982791587 | 1 | 0 | 1 | FIM1_1012 | YKL103C | YKL103C | APE1 | n |
| Kmar_QGN16047.1 | W303.asm02.HP0_G0036590 | OG0000666 | chrXII | 501521 | 502423 | CP015057.1 | 696880 | 697779 | 84.66666667 | 300 | 299 | 1.003344482 | 1 | 0 | 1 | FIM1_2747 | YLR172C | YLR172C | DPH5 | n |
| Kmar_QGN15226.1 | W303.asm02.HP0_G0034810 | OG0000668 | chrXII | 123259 | 125895 | CP015056.1 | 474100 | 476964 | 39.81576254 | 878 | 954 | 0.92033543 | 1 | 0 | 0 | FIM1_1915 | YLL013C | YLL013C | PUF3 | n |
| Kmar_QGN14001.1 | W303.asm02.HP0_G0037530 | OG0000669 | chrXII | 691806 | 694145 | CP015054.1 | 1303440 | 1305743 | 69.29936306 | 779 | 767 | 1.015645372 | 1 | 0 | 0 | FIM1_652 | YLR277C | YLR277C | YSH1 | n |
| Kmar_QGN17209.1 | W303.asm02.HP0_G0053620 | OG0000670 | chrXVI | 201501 | 203339 | CP015059.1 | 451616 | 453664 | 50.58479532 | 612 | 682 | 0.897360704 | 1 | 0 | 1 | FIM1_3940 | YPL184C | YPL184C | MRN1 | n |
| Kmar_QGN17735.1 | W303.asm02.HP0_G0046210 | OG0000671 | chrXIV | 537269 | 540766 | CP015060.1 | 352738 | 356121 | 27.96610169 | 1165 | 1127 | 1.033717835 | 1 | 0 | 0 | FIM1_4944 | YNL054W | YNL054W | VAC7 | n |
| Kmar_QGN17683.1 | W303.asm02.HP0_G0007660 | OG0000672 | chrIV | 246129 | 246653 | CP015060.1 | 250669 | 251295 | 38.38862559 | 174 | 208 | 0.836538462 | 1 | 0 | 1 | FIM1_4890 | YDL120W | YDL120W | YFH1 | n |
| Kmar_QGN14844.1 | W303.asm02.HP0_G0041930 | OG0000673 | chrXIII | 569166 | 570809 | CP015055.1 | 1334481 | 1336328 | 37.32057416 | 547 | 615 | 0.889430894 | 1 | 0 | 1 | FIM1_1515 | YMR155W | YMR155W | | n |
| Kmar_QGN15314.1 | W303.asm02.HP0_G0034720 | OG0000674 | chrXII | 100956 | 105356 | CP015056.1 | 677892 | 681644 | 26.85248131 | 1466 | 1250 | 1.1728 | 1 | 0 | 1 | FIM1_2003 | YLL021W | YLL021W | SPA2 | n |
| Kmar_QGN18088.1 | W303.asm02.HP0_G0001410 | OG0000675 | chrII | 128469 | 128885 | CP015061.1 | 182525 | 182899 | 50.72463768 | 138 | 124 | 1.112903226 | 1 | 0 | 1 | FIM1_4407 | YBL049W | YBL049W | MOH1 | n |
| Kmar_QGN17071.1 | W303.asm02.HP0_G0007620 | OG0000676 | chrIV | 240465 | 241403 | CP015059.1 | 171723 | 172655 | 66.66666667 | 312 | 310 | 1.006451613 | 1 | 0 | 1 | FIM1_3802 | YDL124W | YDL124W | | n |
| Kmar_QGN17893.1 | W303.asm02.HP0_G0003190 | OG0000677 | chrII | 505190 | 506677 | CP015060.1 | 690333 | 691850 | 77.1484375 | 495 | 505 | 0.98019802 | 1 | 0 | 1 | FIM1_5102 | YBR126C | YBR126C | TPS1 | n |
| Kmar_QGN18325.1 | W303.asm02.HP0_G0002840 | OG0000678 | chrII | 436487 | 439327 | CP015061.1 | 695984 | 698692 | 29.36842105 | 946 | 902 | 1.048780488 | 1 | 0 | 1 | FIM1_4651 | YBR086C | YBR086C | IST2 | n |
| Kmar_QGN17454.1 | W303.asm02.HP0_G0011850 | OG0000679 | chrIV | 1141601 | 1143670 | CP015059.1 | 952794 | 954800 | 47.47474747 | 689 | 668 | 1.031437126 | 1 | 0 | 1 | FIM1_4188 | YDR332W | YDR332W | IRC3 | n |
| Kmar_QGN13888.1 | W303.asm02.HP0_G0024910 | OG0000680 | chrVIII | 429830 | 430579 | CP015054.1 | 1071612 | 1072412 | 48.12030075 | 249 | 266 | 0.936090226 | 1 | 0 | 0 | FIM1_535 | YHR163W | YHR163W | SOL3 | n |
| Kmar_QGN15827.1 | W303.asm02.HP0_G0041660 | OG0000681 | chrXIII | 518155 | 521062 | CP015057.1 | 171004 | 173580 | 53.58796296 | 861 | 858 | 1.003496503 | 1 | 0 | 1 | FIM1_2523 | YMR125W | YMR125W | STO1 | n |
| Kmar_QGN13734.1 | W303.asm02.HP0_G0025020 | OG0000682 | chrVIII | 457433 | 458746 | CP015054.1 | 764652 | 765965 | 86.04118993 | 437 | 437 | 1 | 1 | 0 | 0 | FIM1_378 | YHR174W | YHR174W | ENO2 | n |
| Kmar_QGN17068.1 | W303.asm02.HP0_G0007590 | OG0000683 | chrIV | 235133 | 236059 | CP015059.1 | 163581 | 164585 | 42.26190476 | 308 | 334 | 0.922155689 | 1 | 0 | 0 | FIM1_3798 | YDL127W | YDL127W | PCL2 | n |
| Kmar_QGN17067.1 | W303.asm02.HP0_G0007580 | OG0000684 | chrIV | 232858 | 234093 | CP015059.1 | 161360 | 162580 | 70.34313725 | 411 | 406 | 1.012315271 | 1 | 0 | 0 | FIM1_3797 | YDL128W | YDL128W | VCX1 | n |
| Kmar_QGN16510.1 | W303.asm02.HP0_G0010700 | OG0000685 | chrIV | 889079 | 891217 | CP015058.1 | 270648 | 272852 | 63.59945873 | 712 | 734 | 0.970027248 | 1 | 0 | 0 | FIM1_3223 | YDR211W | YDR211W | GCD6 | n |
| Kmar_QGN17080.1 | W303.asm02.HP0_G0046890 | OG0000686 | chrXIV | 676424 | 677098 | CP015059.1 | 184853 | 185518 | 64.44444444 | 224 | 221 | 1.013574661 | 1 | 0 | 1 | FIM1_3811 | YNR018W | YNR018W | RCF2 | n |
| Kmar_QGN14113.1 | W303.asm02.HP0_G0005940 | OG0000687 | chrIII | 229858 | 230931 | CP015054.1 | 1558705 | 1559748 | 53.86740331 | 357 | 347 | 1.028818444 | 1 | 0 | 1 | FIM1_764 | YCR044C | YCR044C | PER1 | n |
| Kmar_QGN14819.1 | W303.asm02.HP0_G0047670 | OG0000688 | chrXV | 48141 | 48851 | CP015055.1 | 1291218 | 1291922 | 54.66101695 | 236 | 234 | 1.008547009 | 1 | 0 | 1 | FIM1_1490 | YOL147C | YOL147C | PEX11 | n |
| Kmar_QGN14623.1 | W303.asm02.HP0_G0016000 | OG0000689 | chrV | 480222 | 481553 | CP015055.1 | 881204 | 882559 | 57.52212389 | 443 | 451 | 0.982261641 | 1 | 0 | 1 | FIM1_1287 | YER152C | YER152C | | n |
| Kmar_QGN16355.1 | W303.asm02.HP0_G0012780 | OG0000690 | chrIV | 1336793 | 1338397 | CP015057.1 | 1350301 | 1351902 | 48.38129496 | 534 | 533 | 1.001876173 | 1 | 0 | 1 | FIM1_3061 | YDR434W | YDR434W | GPI17 | n |
| Kmar_QGN18114.1 | W303.asm02.HP0_G0000920 | OG0000691 | chrII | 11151 | 14183 | CP015061.1 | 239629 | 242595 | 48.03536346 | 1010 | 988 | 1.022267206 | 1 | 0 | 0 | FIM1_4434 | YBL106C | YBL106C | SRO77 | n |
| Kmar_QGN15127.1 | W303.asm02.HP0_G0041540 | OG0000692 | chrXIII | 495624 | 496907 | CP015056.1 | 257622 | 258932 | 52.59593679 | 427 | 436 | 0.979357798 | 1 | 0 | 0 | FIM1_1814 | YMR113W | YMR113W | FOL3 | n |
| Kmar_QGN16312.1 | W303.asm02.HP0_G0018690 | OG0000693 | chrVII | 251109 | 251762 | CP015057.1 | 1259861 | 1260514 | 92.16589862 | 217 | 217 | 1 | 1 | 0 | 0 | FIM1_3017 | YGL135W/YPL220W | YPL220W | RPL1A | y |
| Kmar_QGN13557.1 | W303.asm02.HP0_G0053440 | OG0000694 | chrXVI | 169828 | 171312 | CP015054.1 | 411142 | 412617 | 68.30708661 | 494 | 491 | 1.00610998 | 1 | 0 | 1 | FIM1_198 | YPL204W | YPL204W | HRR25 | n |
| Kmar_QGN13558.1 | W303.asm02.HP0_G0019280 | OG0000695 | chrVII | 373054 | 374208 | CP015054.1 | 413786 | 414964 | 46.34760705 | 384 | 392 | 0.979591837 | 1 | 0 | 0 | FIM1_199 | YGL067W | YGL067W | NPY1 | n |
| Kmar_QGN18260.1 | W303.asm02.HP0_G0020990 | OG0000696 | chrVII | 751616 | 753334 | CP015061.1 | 570512 | 572233 | 86.03839442 | 572 | 573 | 0.998254799 | 1 | 0 | 0 | FIM1_4586 | YGR124W | YGR124W | ASN2 | n |
| Kmar_QGN16897.1 | W303.asm02.HP0_G0010280 | OG0000697 | chrIV | 796062 | 797603 | CP015058.1 | 1124296 | 1125936 | 28.73194222 | 513 | 546 | 0.93956044 | 1 | 0 | 1 | FIM1_3624 | YDR169C | YDR169C | STB3 | n |
| Kmar_QGN13455.1 | W303.asm02.HP0_G0054760 | OG0000698 | chrXVI | 432591 | 434030 | CP015054.1 | 177851 | 178485 | 18.37160752 | 479 | 159 | 3.012578616 | 1 | 0 | 0 | FIM1_92 | YPL066W | YPL066W | RGL1 | n |
| Kmar_QGN17391.1 | W303.asm02.HP0_G0041120 | OG0000699 | chrXIII | 406808 | 408088 | CP015059.1 | 828290 | 829723 | 31.97424893 | 426 | 477 | 0.893081761 | 1 | 0 | 1 | FIM1_4123 | YMR068W | YMR068W | AVO2 | n |
| Kmar_QGN16947.1 | W303.asm02.HP0_G0044860 | OG0000700 | chrXIV | 260280 | 261158 | CP015058.1 | 1238381 | 1239259 | 64.96598639 | 292 | 292 | 1 | 1 | 0 | 1 | FIM1_3674 | YNL202W | YNL202W | SPS19 | n |
| Kmar_QGN13564.1 | W303.asm02.HP0_G0031420 | OG0000701 | chrXI | 82204 | 83808 | CP015054.1 | 426309 | 428084 | 46.32107023 | 534 | 591 | 0.903553299 | 1 | 0 | 1 | FIM1_205 | YKL191W | YKL191W | DPH2 | n |
| Kmar_QGN17689.1 | W303.asm02.HP0_G0046470 | OG0000702 | chrXIV | 590763 | 592799 | CP015060.1 | 260267 | 262060 | 27.36535662 | 678 | 597 | 1.135678392 | 1 | 0 | 1 | FIM1_4896 | YNL027W | YNL027W | CRZ1 | n |
| Kmar_QGN16851.1 | W303.asm02.HP0_G0021070 | OG0000703 | chrVII | 767316 | 768179 | CP015058.1 | 1029273 | 1030118 | 79.09407666 | 287 | 281 | 1.021352313 | 1 | 0 | 1 | FIM1_3576 | YGR132C | YGR132C | PHB1 | n |
| Kmar_QGN14323.1 | W303.asm02.HP0_G0037030 | OG0000704 | chrXII | 576949 | 577917 | CP015055.1 | 277160 | 278032 | 62.42236025 | 322 | 290 | 1.110344828 | 1 | 0 | 1 | FIM1_980 | YLR220W | YLR220W | CCC1 | n |
| Kmar_QGN15828.1 | W303.asm02.HP0_G0041650 | OG0000705 | chrXIII | 515084 | 517903 | CP015057.1 | 174395 | 177415 | 28.87391723 | 939 | 1006 | 0.933399602 | 1 | 0 | 0 | FIM1_2524 | YMR124W | YMR124W | EPO1 | n |
| Kmar_QGN16632.1 | W303.asm02.HP0_G0028280 | OG0000706 | chrX | 120629 | 123463 | CP015058.1 | 544631 | 547279 | 51.01832994 | 944 | 882 | 1.070294785 | 1 | 0 | 1 | FIM1_3349 | YJL154C | YJL154C | VPS35 | n |
| Kmar_QGN16434.1 | W303.asm02.HP0_G0004620 | OG0000707 | chrII | 771410 | 772720 | CP015058.1 | 111735 | 112967 | 33.18872017 | 436 | 410 | 1.063414634 | 1 | 0 | 0 | FIM1_3147 | YBR273C | YBR273C | UBX7 | n |
| Kmar_QGN17519.1 | W303.asm02.HP0_G0022040 | OG0000708 | chrVII | 973775 | 974476 | CP015059.1 | 1080149 | 1080841 | 40.94827586 | 233 | 230 | 1.013043478 | 1 | 0 | 1 | FIM1_4255 | YGR235C | YGR235C | MIC26 | n |
| Kmar_QGN15914.1 | W303.asm02.HP0_G0035840 | OG0000709 | chrXII | 337634 | 340078 | CP015057.1 | 365214 | 367463 | 32.19512195 | 814 | 749 | 1.086782377 | 1 | 0 | 1 | FIM1_2612 | YLR095C | YLR095C | IOC2 | n |
| Kmar_QGN17293.1 | W303.asm02.HP0_G0002140 | OG0000710 | chrII | 287974 | 289116 | CP015059.1 | 635896 | 637038 | 63.28125 | 380 | 380 | 1 | 1 | 0 | 1 | FIM1_4024 | YBR026C | YBR026C | ETR1 | n |
| Kmar_QGN18241.1 | W303.asm02.HP0_G0039330 | OG0000711 | chrXIII | 18589 | 20352 | CP015061.1 | 528667 | 530385 | 69.33560477 | 587 | 572 | 1.026223776 | 1 | 0 | 1 | FIM1_4566 | YML123C | YML123C | PHO84 | n |
| Kmar_QGN18017.1 | W303.asm02.HP0_G0016420 | OG0000712 | chrVI | 7290 | 8627 | CP015061.1 | 41456 | 42757 | 46.34146341 | 445 | 433 | 1.027713626 | 1 | 0 | 0 | FIM1_4333 | YFL048C | YFL048C | EMP47 | n |
| Kmar_QGN15211.1 | W303.asm02.HP0_G0029430 | OG0000713 | chrX | 388053 | 389252 | CP015056.1 | 439011 | 440228 | 79.06403941 | 399 | 405 | 0.985185185 | 1 | 0 | 1 | FIM1_1899 | YJL026W | YJL026W | RNR2 | n |
| Kmar_QGN15941.1 | W303.asm02.HP0_G0050670 | OG0000714 | chrXV | 663276 | 664457 | CP015057.1 | 438869 | 440026 | 70.37974684 | 393 | 385 | 1.020779221 | 1 | 0 | 1 | FIM1_2639 | YOR176W | YOR176W | HEM15 | n |
| Kmar_QGN17549.1 | W303.asm02.HP0_G0033670 | OG0000715 | chrXI | 530585 | 531400 | CP015059.1 | 1144788 | 1145603 | 73.43173432 | 271 | 271 | 1 | 1 | 0 | 1 | FIM1_4285 | YKR043C | YKR043C | SHB17 | n |
| Kmar_QGN15475.1 | W303.asm02.HP0_G0051410 | OG0000716 | chrXV | 813394 | 814047 | CP015056.1 | 1022388 | 1022954 | 14.28571429 | 217 | 188 | 1.154255319 | 1 | 0 | 1 | FIM1_2166 | YOR258W | YOR258W | HNT3 | n |
| Kmar_QGN13754.1 | W303.asm02.HP0_G0046720 | OG0000717 | chrXIV | 639109 | 640269 | CP015054.1 | 813619 | 814794 | 59.07928389 | 386 | 391 | 0.987212276 | 1 | 0 | 1 | FIM1_398 | YNL001W | YNL001W | DOM34 | n |
| Kmar_QGN13459.1 | W303.asm02.HP0_G0003970 | OG0000718 | chrII | 645947 | 648721 | CP015054.1 | 182548 | 184614 | 25.5091104 | 924 | 688 | 1.343023256 | 1 | 0 | 1 | FIM1_96 | YBR203W | YBR203W | COS111 | n |
| Kmar_QGN16074.1 | W303.asm02.HP0_G0040840 | OG0000719 | chrXIII | 339921 | 341225 | CP015057.1 | 751996 | 753408 | 55.41666667 | 434 | 470 | 0.923404255 | 1 | 0 | 0 | FIM1_2774 | YMR034C | YMR034C | RCH1 | n |
| Kmar_QGN14669.1 | W303.asm02.HP0_G0046660 | OG0000720 | chrXIV | 629690 | 631219 | CP015055.1 | 993232 | 994299 | 58.38926174 | 440 | 355 | 1.23943662 | 1 | 0 | 1 | FIM1_1334 | YNL007C | YNL007C | SIS1 | n |
| Kmar_QGN14671.1 | W303.asm02.HP0_G0036630 | OG0000721 | chrXII | 508058 | 510493 | CP015055.1 | 996334 | 998832 | 26.6589057 | 811 | 832 | 0.974759615 | 1 | 0 | 1 | FIM1_1336 | YLR176C | YLR176C | RFX1 | n |
| Kmar_QGN17884.1 | W303.asm02.HP0_G0056110 | OG0000722 | chrXVI | 705368 | 706513 | CP015060.1 | 675822 | 676871 | 39.48717949 | 381 | 349 | 1.091690544 | 1 | 0 | 0 | FIM1_5093 | YPR079W | YPR079W | MRL1 | n |
| Kmar_QGN16093.1 | W303.asm02.HP0_G0026750 | OG0000723 | chrIX | 255748 | 258456 | CP015057.1 | 795139 | 797679 | 39.12568306 | 902 | 846 | 1.066193853 | 1 | 0 | 1 | FIM1_2793 | YIL047C | YIL047C | SYG1 | n |
| Kmar_QGN16198.1 | W303.asm02.HP0_G0039790 | OG0000724 | chrXIII | 110274 | 113438 | CP015057.1 | 1010528 | 1013749 | 59.74025974 | 1054 | 1073 | 0.982292637 | 1 | 0 | 0 | FIM1_2900 | YML075C | YML075C | HMG1 | n |
| Kmar_QGN17318.1 | W303.asm02.HP0_G0045630 | OG0000725 | chrXIV | 403860 | 406772 | CP015059.1 | 683203 | 685695 | 29.51991828 | 970 | 830 | 1.168674699 | 1 | 0 | 1 | FIM1_4050 | YNL118C | YNL118C | DCP2 | n |
| Kmar_QGN14526.1 | W303.asm02.HP0_G0002060 | OG0000726 | chrII | 269524 | 270624 | CP015055.1 | 694352 | 695470 | 66.93989071 | 366 | 372 | 0.983870968 | 1 | 0 | 1 | FIM1_1188 | YBR018C | YBR018C | GAL7 | n |
| Kmar_QGN16570.1 | W303.asm02.HP0_G0038130 | OG0000727 | chrXII | 816689 | 818218 | CP015058.1 | 400123 | 401784 | 56.13718412 | 509 | 553 | 0.920433996 | 1 | 0 | 1 | FIM1_3284 | YLR345W | YLR345W | | n |
| Kmar_QGN14458.1 | W303.asm02.HP0_G0014740 | OG0000728 | chrV | 208783 | 209613 | CP015055.1 | 555848 | 556672 | 70.28985507 | 276 | 274 | 1.00729927 | 1 | 0 | 1 | FIM1_1119 | YER026C | YER026C | CHO1 | n |
| Kmar_QGN17452.1 | W303.asm02.HP0_G0024140 | OG0000730 | chrVIII | 280148 | 282214 | CP015059.1 | 947856 | 949973 | 32.73231623 | 688 | 705 | 0.975886525 | 1 | 0 | 1 | FIM1_4186 | YHR084W | YHR084W | STE12 | n |
| Kmar_QGN16966.1 | W303.asm02.HP0_G0006620 | OG0000731 | chrIV | 33488 | 33991 | CP015058.1 | 1272150 | 1272557 | 46.47058824 | 167 | 135 | 1.237037037 | 1 | 0 | 1 | FIM1_3693 | YDL235C | YDL235C | YPD1 | n |
| Kmar_QGN16348.1 | W303.asm02.HP0_G0040330 | OG0000732 | chrXIII | 235277 | 236458 | CP015057.1 | 1339140 | 1340345 | 58.70646766 | 393 | 401 | 0.980049875 | 1 | 0 | 1 | FIM1_3054 | YML018C | YML018C | | n |
| Kmar_QGN17069.1 | W303.asm02.HP0_G0007600 | OG0000733 | chrIV | 236363 | 238870 | CP015059.1 | 164889 | 167387 | 88.55780691 | 835 | 832 | 1.003605769 | 1 | 0 | 1 | FIM1_3799 | YDL126C | YDL126C | CDC48 | n |
| Kmar_QGN17821.1 | W303.asm02.HP0_G0000280 | OG0000734 | chrI | 61951 | 63198 | CP015060.1 | 561948 | 563171 | 62.01923077 | 415 | 407 | 1.01965602 | 1 | 0 | 1 | FIM1_5030 | YAL042W | YAL042W | ERV46 | n |
| Kmar_QGN15541.1 | W303.asm02.HP0_G0047760 | OG0000735 | chrXV | 65962 | 67455 | CP015056.1 | 1153526 | 1155292 | 20.30201342 | 497 | 588 | 0.845238095 | 1 | 0 | 1 | FIM1_2232 | YOL137W | YOL137W | BSC6 | n |
| Kmar_QGN17285.1 | W303.asm02.HP0_G0048460 | OG0000736 | chrXV | 206557 | 207630 | CP015059.1 | 619462 | 620553 | 64.93150685 | 357 | 363 | 0.983471074 | 1 | 0 | 1 | FIM1_4016 | YOL064C | YOL064C | MET22 | n |

| | | | | | | | | | | | | | | | | | | | | |
|---|---|---|---|---|---|---|---|---|---|---|---|---|---|---|---|---|---|---|---|---|
| Kmar_QGN17416.1 | W303.asm02.HP0_G0027570 | OG0000738 | chrIX | 407425 | 410766 | CP015059.1 | 868820 | 872521 | 35.05976096 | 1113 | 1233 | 0.902676399 | 1 | 0 | 0 | FIM1_4148 | YIR033W | YIR033W | MGA2 | n |
| Kmar_QGN15292.1 | W303.asm02.HP0_G0030020 | OG0000739 | chrX | 534990 | 535319 | CP015056.1 | 615192 | 615524 | 85.45454545 | 109 | 110 | 0.990909091 | 1 | 0 | 1 | FIM1_1981 | YJR048W | YJR048W | CYC1 | n |
| Kmar_QGN16760.1 | W303.asm02.HP0_G0014460 | OG0000740 | chrV | 151484 | 152161 | CP015058.1 | 836774 | 837499 | 33.88429752 | 225 | 241 | 0.933609959 | 1 | 0 | 1 | FIM1_3482 | YEL001C | YEL001C | IRC22 | n |
| Kmar_QGN16961.1 | W303.asm02.HP0_G0006640 | OG0000741 | chrIV | 36870 | 38246 | CP015058.1 | 1260641 | 1262602 | 28.0245023 | 458 | 653 | 0.701378254 | 1 | 0 | 1 | FIM1_3688 | YDL233W | YDL233W | MFG1 | n |
| Kmar_QGN13938.1 | W303.asm02.HP0_G0043260 | OG0000742 | chrXIII | 849358 | 850815 | CP015054.1 | 1179721 | 1182699 | 21.92192192 | 485 | 992 | 0.48891129 | 1 | 0 | 0 | FIM1_586 | W303.asm02.HP0_G0043260 | YMR287C | DSS1 | y |
| Kmar_QGN16759.1 | W303.asm02.HP0_G0027230 | OG0000744 | chrIX | 347197 | 347949 | CP015058.1 | 833823 | 834881 | 28.40909091 | 250 | 352 | 0.710227273 | 1 | 0 | 1 | FIM1_3481 | YIR001C | YIR001C | SGN1 | n |
| Kmar_QGN16770.1 | W303.asm02.HP0_G0027160 | OG0000745 | chrIX | 335117 | 336238 | CP015058.1 | 850713 | 851822 | 51.39240506 | 373 | 369 | 1.010840108 | 1 | 0 | 0 | FIM1_3492 | YIL006W | YIL006W | YIA6 | n |
| Kmar_QGN16622.1 | W303.asm02.HP0_G0028190 | OG0000746 | chrX | 101194 | 102861 | CP015058.1 | 517041 | 518885 | 32.65306122 | 555 | 614 | 0.903908795 | 1 | 0 | 0 | FIM1_3339 | YJL163C | YJL163C | | n |
| Kmar_QGN16810.1 | W303.asm02.HP0_G0020770 | OG0000747 | chrVII | 688715 | 691567 | CP015058.1 | 945658 | 948522 | 58.87265136 | 950 | 954 | 0.995807128 | 1 | 0 | 1 | FIM1_3534 | YGR100W | YGR100W | MDR1 | n |
| Kmar_QGN15560.1 | W303.asm02.HP0_G0011650 | OG0000748 | chrIV | 1093849 | 1094709 | CP015056.1 | 1184166 | 1185152 | 36.49851632 | 286 | 328 | 0.87195122 | 1 | 0 | 1 | FIM1_2251 | YDR313C | YDR313C | PIB1 | n |
| Kmar_QGN17604.1 | W303.asm02.HP0_G0013890 | OG0000749 | chrV | 40409 | 42316 | CP015060.1 | 84651 | 86312 | 54.48916409 | 635 | 553 | 1.148282098 | 1 | 0 | 1 | FIM1_4811 | YEL060C | YEL060C | PRB1 | n |
| Kmar_QGN13970.1 | W303.asm02.HP0_G0019400 | OG0000750 | chrVII | 395581 | 397113 | CP015054.1 | 1240356 | 1241789 | 65.87771203 | 510 | 477 | 1.06918239 | 1 | 0 | 1 | FIM1_619 | YGL055W | YGL055W | OLE1 | n |
| Kmar_QGN15499.1 | W303.asm02.HP0_G0046950 | OG0000751 | chrXIV | 683000 | 684700 | CP015056.1 | 1066032 | 1067423 | 35.33568905 | 566 | 463 | 1.222462203 | 1 | 0 | 0 | FIM1_2190 | YNR023W | YNR023W | SNF12 | n |
| Kmar_QGN17583.1 | W303.asm02.HP0_G0027780 | OG0000752 | chrX | 26450 | 27265 | CP015060.1 | 40984 | 41688 | 27.57352941 | 271 | 234 | 1.158119658 | 1 | 0 | 1 | FIM1_4790 | YJL210W | YJL210W | PEX2 | n |
| Kmar_QGN13588.1 | W303.asm02.HP0_G0033400 | OG0000753 | chrXI | 475060 | 475764 | CP015054.1 | 475373 | 475996 | 60.68376068 | 234 | 207 | 1.130434783 | 1 | 0 | 1 | FIM1_230 | YKR014C | YKR014C | YPT52 | n |
| Kmar_QGN18421.1 | W303.asm02.HP0_G0016920 | OG0000754 | chrVI | 121140 | 121607 | CP015061.1 | 896890 | 897345 | 50.63291139 | 155 | 151 | 1.026490066 | 1 | 0 | 1 | FIM1_4749 | YFR003C | YFR003C | YPI1 | n |
| Kmar_QGN13594.1 | W303.asm02.HP0_G0033440 | OG0000755 | chrXI | 483418 | 485595 | CP015054.1 | 485724 | 487904 | 61.8852459 | 725 | 726 | 0.99862259 | 1 | 0 | 0 | FIM1_236 | YKR018C | YKR018C | | n |
| Kmar_QGN14935.1 | W303.asm02.HP0_G0045150 | OG0000756 | chrXIV | 318723 | 319502 | CP015055.1 | 1528870 | 1529661 | 69.31818182 | 259 | 263 | 0.984790875 | 1 | 0 | 1 | FIM1_1610 | YNL168C | YNL168C | FMP41 | n |
| Kmar_QGN14222.1 | W303.asm02.HP0_G0009380 | OG0000757 | chrIV | 593103 | 593678 | CP015055.1 | 61495 | 62079 | 58.79396985 | 191 | 194 | 0.984536082 | 1 | 0 | 1 | FIM1_878 | YDR071C | YDR071C | PAA1 | n |
| Kmar_QGN17776.1 | W303.asm02.HP0_G0043420 | OG0000758 | chrXIII | 876993 | 879545 | CP015060.1 | 457146 | 459590 | 65.29411765 | 850 | 814 | 1.044226044 | 1 | 0 | 1 | FIM1_4985 | YMR302C | YMR302C | YME2 | n |
| Kmar_QGN15129.1 | W303.asm02.HP0_G0041560 | OG0000759 | chrXIII | 498329 | 499834 | CP015056.1 | 260164 | 261672 | 45.91439689 | 501 | 502 | 0.998007968 | 1 | 0 | 0 | FIM1_1816 | YMR115W | YMR115W | MGR3 | n |
| Kmar_QGN13875.1 | W303.asm02.HP0_G0033790 | OG0000760 | chrXI | 559224 | 561032 | CP015054.1 | 1036669 | 1038405 | 59.86733002 | 602 | 578 | 1.041522491 | 1 | 0 | 1 | FIM1_522 | YKR056W | YKR056W | TRM2 | n |
| Kmar_QGN15531.1 | W303.asm02.HP0_G0006880 | OG0000761 | chrIV | 90249 | 92537 | CP015056.1 | 1131394 | 1133496 | 34.36692506 | 762 | 700 | 1.088571429 | 1 | 0 | 1 | FIM1_2222 | YDL206W | YDL206W | YCX1 | n |
| Kmar_QGN17965.1 | W303.asm02.HP0_G0043780 | OG0000762 | chrXIV | 33726 | 34667 | CP015060.1 | 845024 | 845908 | 35.66878981 | 313 | 294 | 1.06462585 | 1 | 0 | 1 | FIM1_5174 | YNL322C | YNL322C | KRE1 | n |
| Kmar_QGN15139.1 | W303.asm02.HP0_G0016860 | OG0000763 | chrVI | 100293 | 102779 | CP015056.1 | 273929 | 276532 | 56.10599078 | 828 | 867 | 0.955017301 | 1 | 0 | 0 | FIM1_1826 | YFL004W | YFL004W | VTC2 | n |
| Kmar_QGN15851.1 | W303.asm02.HP0_G0034650 | OG0000764 | chrXII | 84813 | 86573 | CP015057.1 | 220341 | 222071 | 65.36502547 | 586 | 576 | 1.017361111 | 1 | 0 | 1 | FIM1_2548 | YLL028W | YLL028W | TPO1 | n |
| Kmar_QGN14067.1 | W303.asm02.HP0_G0044500 | OG0000765 | chrXIV | 188606 | 191512 | CP015054.1 | 1454235 | 1455965 | 27.13797035 | 968 | 576 | 1.680555556 | 1 | 0 | 1 | FIM1_718 | YNL243W | YNL243W | SLA2 | n |
| Kmar_QGN17612.1 | W303.asm02.HP0_G0049140 | OG0000766 | chrXV | 339393 | 340433 | CP015060.1 | 104062 | 105060 | 57.10227273 | 346 | 332 | 1.042168675 | 1 | 0 | 1 | FIM1_4819 | YOR007C | YOR007C | SGT2 | n |
| Kmar_QGN16917.1 | W303.asm02.HP0_G0010430 | OG0000767 | chrIV | 831297 | 832772 | CP015058.1 | 1187263 | 1188708 | 54.03225806 | 491 | 481 | 1.020790021 | 1 | 0 | 1 | FIM1_3644 | YDR182W | YDR182W | CDC1 | n |
| Kmar_QGN16922.1 | W303.asm02.HP0_G0036570 | OG0000768 | chrXII | 499839 | 500531 | CP015058.1 | 1195158 | 1195856 | 57.93991416 | 230 | 232 | 0.99137931 | 1 | 0 | 0 | FIM1_3649 | YLR168C | YLR168C | UPS2 | n |
| Kmar_QGN15278.1 | W303.asm02.HP0_G0034010 | OG0000770 | chrXI | 602773 | 604530 | CP015056.1 | 582712 | 584715 | 18.96296296 | 585 | 667 | 0.877061469 | 1 | 0 | 1 | FIM1_1967 | YKR078W | YKR078W | VPS501 | n |
| Kmar_QGN15670.1 | W303.asm02.HP0_G0013970 | OG0000771 | chrV | 56934 | 58463 | CP015056.1 | 1425717 | 1427171 | 64.50980392 | 509 | 484 | 1.051652893 | 1 | 0 | 1 | FIM1_2363 | YEL052W | YEL052W | AFG1 | n |
| Kmar_QGN15085.1 | W303.asm02.HP0_G0033240 | OG0000773 | chrXI | 444655 | 445860 | CP015056.1 | 170478 | 171728 | 66.83046683 | 401 | 416 | 0.963942308 | 1 | 0 | 1 | FIM1_1771 | YKL004W | YKL004W | AUR1 | n |
| Kmar_QGN14814.1 | W303.asm02.HP0_G0043700 | OG0000774 | chrXIV | 18433 | 19734 | CP015055.1 | 1281898 | 1283196 | 89.58333333 | 433 | 432 | 1.002314815 | 1 | 0 | 1 | FIM1_1485 | YNL330C | YNL330C | RPD3 | n |
| Kmar_QGN15252.1 | W303.asm02.HP0_G0049490 | OG0000775 | chrXV | 408881 | 410116 | CP015056.1 | 531947 | 533158 | 39.95485327 | 411 | 403 | 1.019851117 | 1 | 0 | 0 | FIM1_1941 | YOR042W | YOR042W | CUE5 | n |
| Kmar_QGN17434.1 | W303.asm02.HP0_G0008770 | OG0000776 | chrIV | 458201 | 460906 | CP015059.1 | 905960 | 909715 | 38.24701195 | 901 | 1251 | 0.720223821 | 1 | 0 | 1 | FIM1_4166 | YDR006C | YDR006C | SOK1 | n |
| Kmar_QGN16360.1 | W303.asm02.HP0_G0040390 | OG0000777 | chrXIII | 246622 | 247257 | CP015057.1 | 1361864 | 1362502 | 74.05660377 | 211 | 212 | 0.995283019 | 1 | 0 | 1 | FIM1_3066 | YML012W | YML012W | ERV25 | n |
| Kmar_QGN18390.1 | W303.asm02.HP0_G0015810 | OG0000778 | chrV | 415416 | 416201 | CP015061.1 | 837519 | 838304 | 93.8697318 | 261 | 261 | 1 | 1 | 0 | 1 | FIM1_4717 | YER126C | YER126C | NSA2 | n |
| Kmar_QGN17493.1 | W303.asm02.HP0_G0031190 | OG0000779 | chrXI | 23256 | 25106 | CP015059.1 | 1024993 | 1026801 | 63.40288925 | 616 | 602 | 1.023255814 | 1 | 0 | 1 | FIM1_4229 | YKL217W | YKL217W | JEN1 | n |
| Kmar_QGN14048.1 | W303.asm02.HP0_G0005850 | OG0000780 | chrIII | 209575 | 213255 | CP015054.1 | 1405879 | 1410354 | 27.6298269 | 1226 | 1491 | 0.822266935 | 1 | 0 | 1 | FIM1_699 | YCR033W | YCR033W | SNT1 | n |
| Kmar_QGN15487.1 | W303.asm02.HP0_G0015350 | OG0000781 | chrV | 324356 | 325765 | CP015056.1 | 1045554 | 1046966 | 83.82978723 | 469 | 470 | 0.99787234 | 1 | 0 | 0 | FIM1_2178 | YER081W | YER081W | SER3 | n |
| Kmar_QGN15090.1 | W303.asm02.HP0_G0020370 | OG0000782 | chrVII | 597176 | 598900 | CP015056.1 | 183749 | 185488 | 71.97231834 | 574 | 579 | 0.991364421 | 1 | 0 | 0 | FIM1_1777 | YGR055W | YGR055W | MUP1 | n |
| Kmar_QGN14700.1 | W303.asm02.HP0_G0026790 | OG0000783 | chrIX | 265737 | 266921 | CP015055.1 | 1057269 | 1058483 | 44.17475728 | 394 | 404 | 0.975247525 | 1 | 0 | 0 | FIM1_1367 | YIL042C | YIL042C | PKP1 | n |
| Kmar_QGN14036.1 | W303.asm02.HP0_G0008490 | OG0000784 | chrIV | 401040 | 403334 | CP015054.1 | 1382189 | 1384606 | 37.74509804 | 764 | 805 | 0.949068323 | 1 | 0 | 1 | FIM1_687 | YDL028C | YDL028C | MPS1 | n |
| Kmar_QGN17977.1 | W303.asm02.HP0_G0024420 | OG0000785 | chrVIII | 340500 | 341636 | CP015060.1 | 867591 | 868739 | 70.52631579 | 378 | 382 | 0.989528796 | 1 | 0 | 1 | FIM1_5186 | YHR112C | YHR112C | | n |
| Kmar_QGN15370.1 | W303.asm02.HP0_G0056530 | OG0000786 | chrXVI | 797012 | 798049 | CP015056.1 | 818809 | 819840 | 50.28735632 | 345 | 343 | 1.005830904 | 1 | 0 | 1 | FIM1_2060 | YPR127W | YPR127W | | n |
| Kmar_QGN18244.1 | W303.asm02.HP0_G0039310 | OG0000787 | chrXIII | 15313 | 16251 | CP015061.1 | 535891 | 536823 | 59.93589744 | 312 | 310 | 1.006451613 | 1 | 0 | 0 | FIM1_4569 | YML125C | YML125C | PGA3 | n |
| Kmar_QGN16680.1 | W303.asm02.HP0_G0007190 | OG0000788 | chrIV | 156538 | 158124 | CP015058.1 | 659275 | 661212 | 32.37951807 | 528 | 645 | 0.818604651 | 1 | 0 | 1 | FIM1_3399 | YDL170W | YDL170W | UGA3 | n |
| Kmar_QGN16134.1 | W303.asm02.HP0_G0040680 | OG0000789 | chrXIII | 315880 | 317406 | CP015057.1 | 885888 | 887393 | 44.27184466 | 508 | 501 | 1.013972056 | 1 | 0 | 1 | FIM1_2835 | YMR020W | YMR020W | FMS1 | n |
| Kmar_QGN15756.1 | W303.asm02.HP0_G0006530 | OG0000790 | chrIV | 16287 | 17309 | CP015057.1 | 25748 | 26770 | 82.05882353 | 340 | 340 | 1 | 1 | 0 | 0 | FIM1_2452 | YDL244W/YFL058W/YJR156C/YI | YDL244W | THI13 | y |
| Kmar_QGN13990.1 | W303.asm02.HP0_G0005700 | OG0000791 | chrIII | 165926 | 166573 | CP015054.1 | 1280538 | 1281179 | 46.04651163 | 215 | 213 | 1.009389671 | 1 | 0 | 1 | FIM1_639 | YCR020C | YCR020C | PET18 | n |
| Kmar_QGN18054.1 | W303.asm02.HP0_G0038530 | OG0000792 | chrXII | 896247 | 899330 | CP015061.1 | 105953 | 108973 | 58.51272016 | 1027 | 1006 | 1.020874751 | 1 | 0 | 1 | FIM1_4371 | YLR389C | YLR389C | STE23 | n |
| Kmar_QGN17935.1 | W303.asm02.HP0_G0015560 | OG0000793 | chrV | 361232 | 361984 | CP015060.1 | 772096 | 772854 | 59.52380952 | 250 | 252 | 0.992063492 | 1 | 0 | 1 | FIM1_5144 | YER100W | YER100W | UBC6 | n |
| Kmar_QGN17688.1 | W303.asm02.HP0_G0046480 | OG0000794 | chrXIV | 593103 | 594557 | CP015060.1 | 258518 | 260020 | 34.98098859 | 484 | 500 | 0.968 | 1 | 0 | 1 | FIM1_4895 | YNL026W | YNL026W | SAM50 | n |
| Kmar_QGN15267.1 | W303.asm02.HP0_G0049620 | OG0000795 | chrXV | 434284 | 436941 | CP015056.1 | 562376 | 564217 | 26.8403171 | 885 | 613 | 1.443719413 | 1 | 0 | 1 | FIM1_1956 | YOR058C | YOR058C | ASE1 | n |
| Kmar_QGN15046.1 | W303.asm02.HP0_G0030290 | OG0000796 | chrX | 583268 | 584458 | CP015056.1 | 85612 | 86823 | 48.7804878 | 396 | 403 | 0.982630273 | 1 | 0 | 1 | FIM1_1731 | YJR075W | YJR075W | HOC1 | n |
| Kmar_QGN17474.1 | W303.asm02.HP0_G0011230 | OG0000797 | chrIV | 1004605 | 1006524 | CP015059.1 | 997013 | 998947 | 39.7881997 | 639 | 644 | 0.992236025 | 1 | 0 | 1 | FIM1_4210 | YDR266C | YDR266C | HEL2 | n |
| Kmar_QGN16896.1 | W303.asm02.HP0_G0036340 | OG0000798 | chrXII | 445676 | 447868 | CP015058.1 | 1121677 | 1123737 | 41.18438762 | 730 | 686 | 1.064139942 | 1 | 0 | 0 | FIM1_3623 | YLR149C | YLR149C | GID11 | n |
| Kmar_QGN14156.1 | W303.asm02.HP0_G0005170 | OG0000799 | chrIII | 48305 | 49873 | CP015054.1 | 1649243 | 1650805 | 51.60680529 | 522 | 520 | 1.003846154 | 1 | 0 | 1 | FIM1_809 | YCL043C | YCL043C | PDI1 | n |
| Kmar_QGN16103.1 | W303.asm02.HP0_G0026910 | OG0000800 | chrIX | 286679 | 290638 | CP015057.1 | 820203 | 822974 | 25.74404762 | 1319 | 923 | 1.429035753 | 1 | 0 | 0 | FIM1_2804 | YIL030C | YIL030C | SSM4 | n |
| Kmar_QGN16704.1 | W303.asm02.HP0_G0042290 | OG0000801 | chrXIII | 641498 | 645841 | CP015058.1 | 702971 | 707113 | 41.10885734 | 1447 | 1380 | 1.048550725 | 1 | 0 | 1 | FIM1_3424 | YMR190C | YMR190C | SGS1 | n |
| Kmar_QGN16703.1 | W303.asm02.HP0_G0053010 | OG0000802 | chrXVI | 85276 | 87921 | CP015058.1 | 698974 | 701661 | 13.14917127 | 881 | 895 | 0.984357542 | 1 | 0 | 1 | FIM1_3423 | YPL248C | YPL248C | GAL4 | n |
| Kmar_QGN16699.1 | W303.asm02.HP0_G0053000 | OG0000803 | chrXVI | 82234 | 84918 | CP015058.1 | 688978 | 691164 | 37.36017897 | 894 | 728 | 1.228021978 | 1 | 0 | 1 | FIM1_3419 | YPL249C | YPL249C | GYP5 | n |
| Kmar_QGN15535.1 | W303.asm02.HP0_G0010900 | OG0000804 | chrIV | 933971 | 934858 | CP015056.1 | 1141033 | 1141896 | 44.44444444 | 295 | 287 | 1.027874564 | 1 | 0 | 1 | FIM1_2226 | YDR233C | YDR233C | RTN1 | n |
| Kmar_QGN13777.1 | W303.asm02.HP0_G0049730 | OG0000805 | chrXV | 456642 | 458555 | CP015054.1 | 863079 | 864518 | 48.75 | 637 | 479 | 1.329853862 | 1 | 0 | 0 | FIM1_422 | YOR070C | YOR070C | GYP1 | n |
| Kmar_QGN15738.1 | W303.asm02.HP0_G0025740 | OG0000806 | chrIX | 37122 | 40337 | CP015056.1 | 1575216 | 1578272 | 41.91780822 | 1071 | 1018 | 1.052062868 | 1 | 0 | 0 | FIM1_2432 | YIL156W | YIL156W | UBP7 | n |
| Kmar_QGN13619.1 | W303.asm02.HP0_G0043120 | OG0000807 | chrXIII | 817732 | 820479 | CP015054.1 | 531295 | 534168 | 29.79591837 | 915 | 957 | 0.956112853 | 1 | 0 | 0 | FIM1_261 | YMR273C | YMR273C | ZDS1 | n |
| Kmar_QGN15465.1 | W303.asm02.HP0_G0051360 | OG0000808 | chrXV | 806100 | 806630 | CP015056.1 | 1006093 | 1006584 | 35.22727273 | 176 | 163 | 1.079754601 | 1 | 0 | 1 | FIM1_2156 | YOR253W | YOR253W | NAT5 | n |
| Kmar_QGN15832.1 | W303.asm02.HP0_G0035220 | OG0000809 | chrXII | 209361 | 209975 | CP015057.1 | 181329 | 181943 | 95.58823529 | 204 | 204 | 1 | 1 | 0 | 0 | FIM1_2528 | YLR029C/YMR121C | YLR029C | RPL15A | y |
| Kmar_QGN17298.1 | W303.asm02.HP0_G0024540 | OG0000810 | chrVIII | 362534 | 364417 | CP015059.1 | 643524 | 645284 | 34.87261146 | 627 | 586 | 1.06996587 | 1 | 0 | 1 | FIM1_4029 | YHR124W | YHR124W | NDT80 | n |
| Kmar_QGN14232.1 | W303.asm02.HP0_G0001710 | OG0000811 | chrII | 184688 | 186412 | CP015055.1 | 77177 | 78868 | 55.99315068 | 574 | 563 | 1.019538188 | 1 | 0 | 1 | FIM1_888 | YBL020W | YBL020W | RFT1 | n |
| Kmar_QGN17603.1 | W303.asm02.HP0_G0052880 | OG0000812 | chrXVI | 47885 | 49351 | CP015060.1 | 81653 | 83104 | 79.09836066 | 488 | 483 | 1.010351967 | 1 | 0 | 1 | FIM1_4810 | YPL262W | YPL262W | FUM1 | n |
| Kmar_QGN14152.1 | W303.asm02.HP0_G0005200 | OG0000813 | chrIII | 54593 | 56179 | CP015054.1 | 1640587 | 1642077 | 55.06692161 | 528 | 496 | 1.064516129 | 1 | 0 | 0 | FIM1_805 | YCL038C | YCL038C | ATG22 | n |
| Kmar_QGN15783.1 | W303.asm02.HP0_G0037900 | OG0000814 | chrXII | 762227 | 764881 | CP015057.1 | 88587 | 91166 | 34.30821147 | 884 | 859 | 1.029103609 | 1 | 0 | 0 | FIM1_2479 | YLR318W | YLR318W | EST2 | n |
| Kmar_QGN14559.1 | W303.asm02.HP0_G0048660 | OG0000815 | chrXV | 248847 | 249989 | CP015055.1 | 752704 | 753804 | 48.03149606 | 380 | 366 | 1.038251366 | 1 | 0 | 0 | FIM1_1221 | YOL043C | YOL043C | NTG2 | n |
| Kmar_QGN14565.1 | W303.asm02.HP0_G0048600 | OG0000816 | chrXV | 235395 | 238640 | CP015055.1 | 767923 | 771105 | 34.15929204 | 1081 | 1060 | 1.019811321 | 1 | 0 | 1 | FIM1_1227 | YOL051W | YOL051W | GAL11 | n |
| Kmar_QGN14064.1 | W303.asm02.HP0_G0027810 | OG0000818 | chrX | 30920 | 36964 | CP015054.1 | 1447371 | 1451489 | 23.73259053 | 2014 | 1372 | 1.467930029 | 1 | 0 | 0 | FIM1_715 | YJL207C | YJL207C | LAA1 | n |
| Kmar_QGN17400.1 | W303.asm02.HP0_G0033160 | OG0000819 | chrXI | 429249 | 430310 | CP015059.1 | 839902 | 840975 | 27.52043597 | 353 | 357 | 0.988795518 | 1 | 0 | 0 | FIM1_4132 | YKL011C | YKL011C | CCE1 | n |
| Kmar_QGN16566.1 | W303.asm02.HP0_G0020230 | OG0000820 | chrVII | 558440 | 559108 | CP015058.1 | 393203 | 393814 | 72.19730942 | 222 | 203 | 1.093596059 | 1 | 0 | 0 | FIM1_3280 | YGR038W | YGR038W | ORM1 | n |
| Kmar_QGN16616.1 | W303.asm02.HP0_G0002200 | OG0000821 | chrII | 300032 | 301078 | CP015058.1 | 507364 | 508410 | 79.31034483 | 348 | 348 | 1 | 1 | 0 | 1 | FIM1_3333 | YBR034C | YBR034C | HMT1 | n |
| Kmar_QGN13872.1 | W303.asm02.HP0_G0028410 | OG0000822 | chrX | 143032 | 144219 | CP015054.1 | 1030083 | 1031273 | 83.83838384 | 395 | 396 | 0.997474747 | 1 | 0 | 0 | FIM1_518 | YJL138C/YKR059W | YKR059W | TIF1 | y |
| Kmar_QGN13820.1 | W303.asm02.HP0_G0011940 | OG0000823 | chrIV | 1162839 | 1164662 | CP015054.1 | 937916 | 939874 | 78.06748466 | 607 | 652 | 0.930981595 | 1 | 0 | 0 | FIM1_466 | YDR341C | YDR341C | | n |
| Kmar_QGN15619.1 | W303.asm02.HP0_G0018400 | OG0000824 | chrVII | 184084 | 186936 | CP015056.1 | 1320903 | 1323716 | 71.42857143 | 950 | 937 | 1.013874066 | 1 | 0 | 1 | FIM1_2312 | YGL167C | YGL167C | PMR1 | n |
| Kmar_QGN13611.1 | W303.asm02.HP0_G0039500 | OG0000825 | chrXIII | 51332 | 52012 | CP015054.1 | 516898 | 517575 | 78.76106195 | 226 | 225 | 1.004444444 | 1 | 0 | 1 | FIM1_253 | YML106W | YML106W | URA5 | n |
| Kmar_QGN14955.1 | W303.asm02.HP0_G0016820 | OG0000826 | chrVI | 84628 | 86967 | CP015055.1 | 1564937 | 1566901 | 46.55612245 | 779 | 654 | 1.191131498 | 1 | 0 | 0 | FIM1_1632 | YFL009W | YFL009W | CDC4 | n |
| Kmar_QGN15140.1 | W303.asm02.HP0_G0016850 | OG0000827 | chrVI | 98817 | 99464 | CP015056.1 | 277328 | 277972 | 75.81395349 | 215 | 214 | 1.004672897 | 1 | 0 | 1 | FIM1_1827 | YFL005W | YFL005W | SEC4 | n |
| Kmar_QGN14193.1 | W303.asm02.HP0_G0047290 | OG0000828 | chrXIV | 745923 | 746636 | CP015054.1 | 1732752 | 1733117 | 51.53846154 | 237 | 121 | 1.958677686 | 1 | 0 | 0 | FIM1_847 | YNR057C | YNR057C | BIO4 | n |
| Kmar_QGN18206.1 | W303.asm02.HP0_G0053200 | OG0000829 | chrXVI | 123947 | 125596 | CP015061.1 | 447090 | 448778 | 46.31578947 | 549 | 562 | 0.976868327 | 1 | 0 | 0 | FIM1_4530 | YPL228W | YPL228W | CET1 | n |
| Kmar_QGN17061.1 | W303.asm02.HP0_G0007530 | OG0000830 | chrIV | 224510 | 226957 | CP015059.1 | 149266 | 151695 | 56.76004872 | 815 | 809 | 1.007416564 | 1 | 0 | 1 | FIM1_3791 | YDL132W | YDL132W | CDC53 | n |
| Kmar_QGN17944.1 | W303.asm02.HP0_G0001160 | OG0000831 | chrII | 76412 | 80920 | CP015060.1 | 789255 | 793622 | 56.58504302 | 1502 | 1455 | 1.032302405 | 1 | 0 | 1 | FIM1_5153 | YBL079W | YBL079W | NUP170 | n |
| Kmar_QGN17081.1 | W303.asm02.HP0_G0005950 | OG0000832 | chrIII | 231116 | 232815 | CP015059.1 | 187634 | 189133 | 36.75373134 | 518 | 499 | 1.038076152 | 1 | 0 | 1 | FIM1_3812 | YCR045C | YCR045C | RRT12 | n |
| Kmar_QGN17447.1 | W303.asm02.HP0_G0008720 | OG0000833 | chrIV | 449879 | 452134 | CP015059.1 | 938441 | 940702 | 70.5026455 | 751 | 753 | 0.997343958 | 1 | 0 | 0 | FIM1_4181 | YDR001C | YDR001C | NTH1 | n |
| Kmar_QGN17501.1 | W303.asm02.HP0_G0021900 | OG0000834 | chrVII | 944956 | 948210 | CP015059.1 | 1045204 | 1048467 | 76.90892364 | 1084 | 1087 | 0.99724011 | 1 | 0 | 1 | FIM1_4237 | YGR218W | YGR218W | CRM1 | n |
| Kmar_QGN16533.1 | W303.asm02.HP0_G0019810 | OG0000835 | chrVII | 473847 | 476186 | CP015058.1 | 320456 | 322810 | 76.55259823 | 779 | 784 | 0.993622449 | 1 | 0 | 1 | FIM1_3246 | YGL009C | YGL009C | LEU1 | n |
| Kmar_QGN15796.1 | W303.asm02.HP0_G0037990 | OG0000836 | chrXII | 780599 | 781804 | CP015057.1 | 112345 | 113772 | 61.00628931 | 401 | 475 | 0.844210526 | 1 | 0 | 0 | FIM1_2492 | YLR328W | YLR328W | NMA1 | n |
| Kmar_QGN15321.1 | W303.asm02.HP0_G0037800 | OG0000837 | chrXII | 743194 | 744132 | CP015056.1 | 699270 | 700250 | 55.52147239 | 312 | 326 | 0.957055215 | 1 | 0 | 0 | FIM1_2010 | YLR308W | YLR308W | CDA2 | n |
| Kmar_QGN18308.1 | W303.asm02.HP0_G0023430 | OG0000838 | chrVIII | 162733 | 164187 | CP015061.1 | 663633 | 665186 | 73.55212355 | 484 | 517 | 0.936170213 | 1 | 0 | 1 | FIM1_4634 | YHR030C | YHR030C | SLT2 | n |
| Kmar_QGN17266.1 | W303.asm02.HP0_G0016670 | OG0000839 | chrVI | 57649 | 58822 | CP015059.1 | 581143 | 583578 | 21.98830409 | 375 | 811 | 0.462392109 | 1 | 0 | 0 | FIM1_3997 | W303.asm02.HP0_G0016670 | YFL024C | EPL1 | y |
| Kmar_QGN17633.1 | W303.asm02.HP0_G0029690 | OG0000840 | chrX | 446669 | 447583 | CP015060.1 | 152582 | 153490 | 79.60526316 | 304 | 302 | 1.006622517 | 1 | 0 | 1 | FIM1_4840 | YJR007W | YJR007W | SUI2 | n |

| | | | | | | | | | | | | | | | | | | | | |
|---|---|---|---|---|---|---|---|---|---|---|---|---|---|---|---|---|---|---|---|---|
| Kmar_QGN17161.1 | W303.asm02.HP0_G0011130 | OG0000841 | chrIV | 972634 | 974181 | CP015059.1 | 363795 | 365330 | 77.56286267 | 515 | 511 | 1.007827789 | 1 | 0 | 1 | FIM1_3892 | YDR256C | YDR256C | CTA1 | n |
| Kmar_QGN15604.1 | W303.asm02.HP0_G0000220 | OG0000842 | chrI | 53436 | 55424 | CP015056.1 | 1283654 | 1285639 | 67.0694864 | 662 | 661 | 1.001512859 | 1 | 0 | 1 | FIM1_2297 | YAL048C | YAL048C | GEM1 | n |
| Kmar_QGN18045.1 | W303.asm02.HP0_G0016540 | OG0000843 | chrVI | 27230 | 31285 | CP015061.1 | 90387 | 94175 | 53.77218935 | 1351 | 1262 | 1.070522979 | 1 | 0 | 1 | FIM1_4361 | YFL036W | YFL036W | RPO41 | n |
| Kmar_QGN13984.1 | W303.asm02.HP0_G0050770 | OG0000844 | chrXV | 685325 | 686638 | CP015054.1 | 1272712 | 1274004 | 86.49885584 | 437 | 430 | 1.01627907 | 1 | 0 | 0 | FIM1_633 | YOR187W | YOR187W | TUF1 | n |
| Kmar_QGN14664.1 | W303.asm02.HP0_G0017100 | OG0000845 | chrVI | 163715 | 165217 | CP015055.1 | 979354 | 980895 | 59.49612403 | 500 | 513 | 0.974658869 | 1 | 0 | 1 | FIM1_1328 | YFR021W | YFR021W | ATG18 | n |
| Kmar_QGN15123.1 | W303.asm02.HP0_G0041510 | OG0000846 | chrXIII | 491019 | 492617 | CP015056.1 | 252949 | 254514 | 47.83427495 | 532 | 521 | 1.021113244 | 1 | 0 | 1 | FIM1_1810 | YMR110C | YMR110C | HFD1 | n |
| Kmar_QGN17523.1 | W303.asm02.HP0_G0024860 | OG0000847 | chrVIII | 419791 | 423285 | CP015059.1 | 1087202 | 1090084 | 38.2832618 | 1164 | 960 | 1.2125 | 1 | 0 | 0 | FIM1_4259 | YHR158C | YHR158C | KEL1 | n |
| Kmar_QGN14445.1 | W303.asm02.HP0_G0006250 | OG0000848 | chrIII | 283292 | 285433 | CP015055.1 | 527500 | 529581 | 60.50420168 | 713 | 693 | 1.028860029 | 1 | 0 | 1 | FIM1_1106 | YCR084C | YCR084C | TUP1 | n |
| Kmar_QGN13941.1 | W303.asm02.HP0_G0043240 | OG0000849 | chrXIII | 847066 | 848613 | CP015054.1 | 1184872 | 1186455 | 58.38041431 | 515 | 527 | 0.977229602 | 1 | 0 | 1 | FIM1_589 | YMR285C | YMR285C | NGL2 | n |
| Kmar_QGN15074.1 | W303.asm02.HP0_G0032370 | OG0000850 | chrXI | 281870 | 283519 | CP015056.1 | 146627 | 148282 | 38.0952381 | 549 | 551 | 0.996370236 | 1 | 0 | 1 | FIM1_1759 | YKL089W | YKL089W | MIF2 | n |
| Kmar_QGN16406.1 | W303.asm02.HP0_G0054320 | OG0000851 | chrXVI | 343794 | 344978 | CP015058.1 | 48424 | 49617 | 32.51231527 | 394 | 397 | 0.992443325 | 1 | 0 | 0 | FIM1_3119 | YPL112C | YPL112C | PEX25 | n |
| Kmar_QGN14899.1 | W303.asm02.HP0_G0003240 | OG0000852 | chrII | 511878 | 513155 | CP015055.1 | 1456388 | 1457455 | 28.21782178 | 425 | 355 | 1.197183099 | 1 | 0 | 1 | FIM1_1573 | YBR130C | YBR130C | SHE3 | n |
| Kmar_QGN17219.1 | W303.asm02.HP0_G0053650 | OG0000853 | chrXVI | 208972 | 210492 | CP015059.1 | 476231 | 477565 | 35.77075099 | 506 | 444 | 1.13963964 | 1 | 0 | 1 | FIM1_3950 | YPL181W | YPL181W | CTI6 | n |
| Kmar_QGN18330.1 | W303.asm02.HP0_G0001580 | OG0000854 | chrII | 162487 | 163632 | CP015061.1 | 708237 | 709352 | 38.25 | 381 | 371 | 1.026954178 | 1 | 0 | 1 | FIM1_4656 | YBL032W | YBL032W | HEK2 | n |
| Kmar_QGN16693.1 | W303.asm02.HP0_G0036870 | OG0000855 | chrXII | 550777 | 552087 | CP015058.1 | 678854 | 680137 | 70.41284404 | 436 | 427 | 1.021077283 | 1 | 0 | 1 | FIM1_3413 | YLR203C | YLR203C | MSS51 | n |
| Kmar_QGN16161.1 | W303.asm02.HP0_G0015280 | OG0000856 | chrV | 310083 | 312869 | CP015057.1 | 929869 | 933015 | 30.69216758 | 928 | 1048 | 0.885496183 | 1 | 0 | 1 | FIM1_2863 | YER075C | YER075C | PTP3 | n |
| Kmar_QGN16258.1 | W303.asm02.HP0_G0025230 | OG0000857 | chrVIII | 494759 | 496498 | CP015057.1 | 1145062 | 1146906 | 59.03225806 | 579 | 614 | 0.942996743 | 1 | 0 | 1 | FIM1_2963 | YHR194W | YHR194W | MDM31 | n |
| Kmar_QGN17963.1 | W303.asm02.HP0_G0047740 | OG0000858 | chrXV | 60727 | 61368 | CP015060.1 | 840151 | 840783 | 64.78873239 | 213 | 210 | 1.014285714 | 1 | 0 | 1 | FIM1_5172 | YOL139C | YOL139C | CDC33 | n |
| Kmar_QGN18033.1 | W303.asm02.HP0_G0006500 | OG0000859 | chrIV | 8769 | 9842 | CP015061.1 | 67838 | 68902 | 66.38655462 | 357 | 354 | 1.008474576 | 1 | 0 | 0 | FIM1_4349 | YDL246C/YJR159W | YJR159W | SOR1 | y |
| Kmar_QGN17251.1 | W303.asm02.HP0_G0032870 | OG0000860 | chrXI | 371225 | 372796 | CP015059.1 | 550934 | 552439 | 51.24282983 | 523 | 501 | 1.043912176 | 1 | 0 | 0 | FIM1_3982 | YKL039W | YKL039W | PTM1 | n |
| Kmar_QGN17012.1 | W303.asm02.HP0_G0008960 | OG0000861 | chrIV | 497494 | 500538 | CP015059.1 | 27176 | 29713 | 31.11919927 | 1014 | 845 | 1.2 | 1 | 0 | 1 | FIM1_3741 | YDR028C | YDR028C | REG1 | n |
| Kmar_QGN13584.1 | W303.asm02.HP0_G0051690 | OG0000862 | chrXV | 855076 | 855831 | CP015054.1 | 466125 | 466826 | 50 | 251 | 233 | 1.077253219 | 1 | 0 | 1 | FIM1_226 | YOR289W | YOR289W | | n |
| Kmar_QGN17864.1 | W303.asm02.HP0_G0056000 | OG0000864 | chrXVI | 688718 | 689275 | CP015060.1 | 635304 | 635891 | 46.875 | 185 | 195 | 0.948717949 | 1 | 0 | 1 | FIM1_5073 | YPR067W | YPR067W | ISA2 | n |
| Kmar_QGN17749.1 | W303.asm02.HP0_G0030470 | OG0000865 | chrX | 620968 | 621816 | CP015060.1 | 385050 | 385922 | 64.48275862 | 282 | 290 | 0.972413793 | 1 | 0 | 1 | FIM1_4958 | YJR096W | YJR096W | | n |
| Kmar_QGN16633.1 | W303.asm02.HP0_G0033890 | OG0000866 | chrXI | 575780 | 576865 | CP015058.1 | 548325 | 549362 | 61.21883657 | 361 | 345 | 1.046376812 | 1 | 0 | 1 | FIM1_3350 | YKR066C | YKR066C | CCP1 | n |
| Kmar_QGN16112.1 | W303.asm02.HP0_G0024610 | OG0000867 | chrVIII | 376027 | 376422 | CP015057.1 | 835647 | 836144 | 51.80722892 | 131 | 165 | 0.793939394 | 1 | 0 | 0 | FIM1_2813 | YHR132W-A | YHR132W-A | IGO2 | n |
| Kmar_QGN15341.1 | W303.asm02.HP0_G0042360 | OG0000868 | chrXIII | 655659 | 658925 | CP015056.1 | 746868 | 750089 | 66 | 1088 | 1073 | 1.013979497 | 1 | 0 | 1 | FIM1_2030 | YMR196W | YMR196W | | n |
| Kmar_QGN16332.1 | W303.asm02.HP0_G0040240 | OG0000869 | chrXIII | 221914 | 223071 | CP015057.1 | 1308652 | 1310061 | 31.78947368 | 385 | 469 | 0.820895522 | 1 | 0 | 0 | FIM1_3037 | YML027W | YML027W | YOX1 | n |
| Kmar_QGN15251.1 | W303.asm02.HP0_G0049500 | OG0000870 | chrXV | 411326 | 412786 | CP015056.1 | 529243 | 530679 | 49.50298211 | 486 | 478 | 1.016736402 | 1 | 0 | 1 | FIM1_1940 | YOR043W | YOR043W | WHI2 | n |
| Kmar_QGN14910.1 | W303.asm02.HP0_G0023560 | OG0000871 | chrVIII | 184387 | 186462 | CP015055.1 | 1475946 | 1478003 | 59.85611511 | 691 | 685 | 1.008759124 | 1 | 0 | 1 | FIM1_1584 | YHR042W | YHR042W | NCP1 | n |
| Kmar_QGN17836.1 | W303.asm02.HP0_G0032750 | OG0000872 | chrXI | 349916 | 350977 | CP015060.1 | 588215 | 589225 | 49.47643979 | 353 | 336 | 1.050595238 | 1 | 0 | 1 | FIM1_5045 | YKL051W | YKL051W | SFK1 | n |
| Kmar_QGN15294.1 | W303.asm02.HP0_G0016210 | OG0000873 | chrV | 546136 | 546870 | CP015056.1 | 618680 | 619426 | 65.99190283 | 244 | 248 | 0.983870968 | 1 | 0 | 0 | FIM1_1983 | YER174C | YER174C | GRX4 | n |
| Kmar_QGN15300.1 | W303.asm02.HP0_G0016160 | OG0000874 | chrV | 531071 | 533461 | CP015056.1 | 636499 | 639006 | 46.9273743 | 796 | 835 | 0.953293413 | 1 | 0 | 1 | FIM1_1989 | YER169W | YER169W | RPH1 | n |
| Kmar_QGN15906.1 | W303.asm02.HP0_G0004810 | OG0000875 | chrII | 812604 | 815183 | CP015057.1 | 349582 | 352329 | 57.08154506 | 859 | 915 | 0.938797814 | 1 | 0 | 0 | FIM1_2604 | YBR294W | YBR294W | SUL1 | n |
| Kmar_QGN14855.1 | W303.asm02.HP0_G0039570 | OG0000876 | chrXIII | 68948 | 71590 | CP015055.1 | 1353456 | 1356236 | 47.49478079 | 880 | 926 | 0.950323974 | 1 | 0 | 1 | FIM1_1528 | YML099C | YML099C | ARG81 | n |
| Kmar_QGN14312.1 | W303.asm02.HP0_G0010750 | OG0000877 | chrIV | 899536 | 903507 | CP015055.1 | 250394 | 254590 | 26.52777778 | 1323 | 1398 | 0.946351931 | 1 | 0 | 1 | FIM1_969 | YDR216W | YDR216W | ADR1 | n |
| Kmar_QGN17113.1 | W303.asm02.HP0_G0039120 | OG0000878 | chrXII | 1032760 | 1035420 | CP015059.1 | 261631 | 264435 | 48.08510638 | 886 | 934 | 0.948608137 | 1 | 0 | 1 | FIM1_3844 | YLR451W | YLR451W | LEU3 | n |
| Kmar_QGN15500.1 | W303.asm02.HP0_G0006010 | OG0000879 | chrIII | 239678 | 241222 | CP015056.1 | 1067815 | 1069347 | 79.37743191 | 514 | 510 | 1.007843137 | 1 | 0 | 1 | FIM1_2191 | YCR053W | YCR053W | THR4 | n |
| Kmar_QGN17116.1 | W303.asm02.HP0_G0030750 | OG0000880 | chrX | 661675 | 662352 | CP015059.1 | 270982 | 271665 | 88.98678414 | 225 | 227 | 0.991189427 | 1 | 0 | 1 | FIM1_3847 | YJR123W | YJR123W | RPS5 | n |
| Kmar_QGN15958.1 | W303.asm02.HP0_G0020050 | OG0000881 | chrVII | 522763 | 524178 | CP015057.1 | 469789 | 471207 | 77.96610169 | 471 | 472 | 0.997881356 | 1 | 0 | 1 | FIM1_2656 | YGR019W | YGR019W | UGA1 | n |
| Kmar_QGN13568.1 | W303.asm02.HP0_G0038850 | OG0000882 | chrXII | 962566 | 968364 | CP015054.1 | 431422 | 437304 | 36.3086233 | 1932 | 1960 | 0.985714286 | 1 | 0 | 1 | FIM1_209 | YLR422W | YLR422W | DCK1 | n |
| Kmar_QGN13989.1 | W303.asm02.HP0_G0037650 | OG0000883 | chrXII | 713086 | 714231 | CP015054.1 | 1278914 | 1279984 | 64.15584416 | 381 | 356 | 1.070224719 | 1 | 0 | 0 | FIM1_638 | YLR291C | YLR291C | GCD7 | n |
| Kmar_QGN16741.1 | W303.asm02.HP0_G0027280 | OG0000884 | chrIX | 356520 | 360962 | CP015058.1 | 796031 | 800416 | 41.0044501 | 1480 | 1461 | 1.013004791 | 1 | 0 | 1 | FIM1_3463 | YIR006C | YIR006C | PAN1 | n |
| Kmar_QGN16643.1 | W303.asm02.HP0_G0038370 | OG0000885 | chrXII | 859381 | 863451 | CP015058.1 | 573426 | 577121 | 58.87096774 | 1356 | 1231 | 1.101543461 | 1 | 0 | 0 | FIM1_3362 | YLR371W | YLR371W | ROM2 | n |
| Kmar_QGN16716.1 | W303.asm02.HP0_G0042250 | OG0000886 | chrXIII | 632938 | 635055 | CP015058.1 | 732362 | 734503 | 89.34081346 | 705 | 713 | 0.988779804 | 1 | 0 | 0 | FIM1_3437 | YMR186W/YPL240C | YMR186W | HSC82 | y |
| Kmar_QGN14892.1 | W303.asm02.HP0_G0009210 | OG0000887 | chrIV | 565589 | 566476 | CP015055.1 | 1444016 | 1444918 | 63.12292359 | 295 | 300 | 0.983333333 | 1 | 0 | 1 | FIM1_1566 | YDR054C | YDR054C | CDC34 | n |
| Kmar_QGN15739.1 | W303.asm02.HP0_G0043670 | OG0000888 | chrXIV | 13698 | 14594 | CP015056.1 | 1578980 | 1579873 | 76.58862876 | 298 | 297 | 1.003367003 | 1 | 0 | 0 | FIM1_2433 | YFL059W/YNL333W | YNL333W | SNZ2 | y |
| Kmar_QGN15740.1 | W303.asm02.HP0_G0043660 | OG0000889 | chrXIV | 12637 | 13305 | CP015056.1 | 1580910 | 1581548 | 53.4562212 | 222 | 212 | 1.047169811 | 1 | 0 | 0 | FIM1_2434 | YFL060C/YNL334C | YNL334C | SNO2 | y |
| Kmar_QGN18187.1 | W303.asm02.HP0_G0020590 | OG0000890 | chrVII | 642082 | 642633 | CP015061.1 | 397446 | 398000 | 54.34782609 | 183 | 184 | 0.994565217 | 1 | 0 | 1 | FIM1_4510 | YGR082W | YGR082W | TOM20 | n |
| Kmar_QGN15091.1 | W303.asm02.HP0_G0020380 | OG0000891 | chrVII | 599420 | 602206 | CP015056.1 | 185883 | 188621 | 40.0203666 | 928 | 912 | 1.01754386 | 1 | 0 | 1 | FIM1_1778 | YGR056W | YGR056W | RSC1 | n |
| Kmar_QGN17875.1 | W303.asm02.HP0_G0003090 | OG0000892 | chrII | 483534 | 485906 | CP015060.1 | 654208 | 656457 | 72.79596977 | 790 | 749 | 1.054739653 | 1 | 0 | 1 | FIM1_5084 | YBR114W | YBR114W | RAD16 | n |
| Kmar_QGN16546.1 | W303.asm02.HP0_G0039890 | OG0000894 | chrXIII | 136750 | 139494 | CP015058.1 | 345422 | 348052 | 44.82758621 | 914 | 876 | 1.043378995 | 1 | 0 | 1 | FIM1_3260 | YML065W | YML065W | ORC1 | n |
| Kmar_QGN14641.1 | W303.asm02.HP0_G0024380 | OG0000895 | chrVIII | 334276 | 336033 | CP015055.1 | 926595 | 928343 | 62.62626263 | 585 | 582 | 1.005154639 | 1 | 0 | 0 | FIM1_1305 | YHR108W | YHR108W | GGA2 | n |
| Kmar_QGN14326.1 | W303.asm02.HP0_G0009650 | OG0000896 | chrIV | 652437 | 653258 | CP015055.1 | 281836 | 282588 | 86.1423221 | 273 | 250 | 1.092 | 1 | 0 | 1 | FIM1_983 | YDR099W | YDR099W | BMH2 | n |
| Kmar_QGN13409.1 | W303.asm02.HP0_G0003700 | OG0000897 | chrII | 601582 | 602937 | CP015054.1 | 80672 | 82018 | 53.40659341 | 451 | 448 | 1.006696429 | 1 | 0 | 0 | FIM1_46 | YBR177C | YBR177C | EHT1 | n |
| Kmar_QGN14085.1 | W303.asm02.HP0_G0044380 | OG0000898 | chrXIV | 161186 | 164875 | CP015054.1 | 1486464 | 1490021 | 28.77930477 | 1229 | 1185 | 1.037130802 | 1 | 0 | 0 | FIM1_736 | YNL257C | YNL257C | SIP3 | n |
| Kmar_QGN15121.1 | W303.asm02.HP0_G0041490 | OG0000899 | chrXIII | 484709 | 486772 | CP015056.1 | 246314 | 248416 | 72 | 687 | 700 | 0.981428571 | 1 | 0 | 1 | FIM1_1808 | YMR108W | YMR108W | ILV2 | n |
| Kmar_QGN15672.1 | W303.asm02.HP0_G0013950 | OG0000900 | chrV | 53084 | 53581 | CP015056.1 | 1431149 | 1431646 | 92.72727273 | 165 | 165 | 1 | 1 | 0 | 1 | FIM1_2365 | YDR418W/YEL054C | YEL054C | RPL12A | y |
| Kmar_QGN17796.1 | W303.asm02.HP0_G0049960 | OG0000901 | chrXV | 506529 | 507502 | CP015060.1 | 504981 | 505970 | 91.57894737 | 190 | 188 | 1.010638298 | 1 | 0 | 0 | FIM1_5005 | YOR096W | YOR096W | RPS7A | n |
| Kmar_QGN18073.1 | W303.asm02.HP0_G0001510 | OG0000902 | chrII | 146155 | 147894 | CP015061.1 | 154725 | 156461 | 79.00172117 | 579 | 578 | 1.001730104 | 1 | 0 | 0 | FIM1_4391 | YBL039C | YBL039C | URA7 | n |
| Kmar_QGN15864.1 | W303.asm02.HP0_G0002510 | OG0000903 | chrII | 368478 | 369578 | CP015057.1 | 256260 | 257171 | 35.82887701 | 366 | 303 | 1.207920792 | 1 | 0 | 1 | FIM1_2561 | YBR057C | YBR057C | MUM2 | n |
| Kmar_QGN17471.1 | W303.asm02.HP0_G0011210 | OG0000904 | chrIV | 1000530 | 1002824 | CP015059.1 | 991146 | 993401 | 44.50194049 | 764 | 751 | 1.017310253 | 1 | 0 | 0 | FIM1_4207 | YDR264C | YDR264C | AKR1 | n |
| Kmar_QGN17201.1 | W303.asm02.HP0_G0019140 | OG0000905 | chrVII | 347568 | 349250 | CP015059.1 | 439092 | 440861 | 57.55517827 | 560 | 589 | 0.950764007 | 1 | 0 | 0 | FIM1_3932 | YGL084C | YGL084C | GUP1 | n |
| Kmar_QGN18389.1 | W303.asm02.HP0_G0018310 | OG0000906 | chrVII | 164126 | 167039 | CP015061.1 | 833439 | 836168 | 40.41501976 | 932 | 909 | 1.02530253 | 1 | 0 | 0 | FIM1_4716 | YGL178W | YGL178W | MPT5 | n |
| Kmar_QGN18236.1 | W303.asm02.HP0_G0007260 | OG0000907 | chrIV | 167934 | 169313 | CP015061.1 | 513232 | 514788 | 54.89443378 | 459 | 518 | 0.886100386 | 1 | 0 | 0 | FIM1_4561 | YDL161W | YDL161W | ENT1 | n |
| Kmar_QGN17518.1 | W303.asm02.HP0_G0024830 | OG0000908 | chrVIII | 409072 | 412284 | CP015059.1 | 1077380 | 1080070 | 31.22087605 | 1070 | 896 | 1.194196429 | 1 | 0 | 1 | FIM1_4254 | YHR154W | YHR154W | RTT107 | n |
| Kmar_QGN14496.1 | W303.asm02.HP0_G0018200 | OG0000909 | chrVII | 141322 | 142902 | CP015055.1 | 630863 | 632221 | 73.38403042 | 526 | 452 | 1.163716814 | 1 | 0 | 1 | FIM1_1157 | YGL190C | YGL190C | CDC55 | n |
| Kmar_QGN14681.1 | W303.asm02.HP0_G0015860 | OG0000910 | chrV | 425929 | 426288 | CP015055.1 | 1022073 | 1022432 | 92.43697479 | 119 | 119 | 1 | 1 | 0 | 0 | FIM1_1347 | YER131W/YGL189C | YER131W | RPS26B | y |
| Kmar_QGN15138.1 | W303.asm02.HP0_G0031900 | OG0000911 | chrXI | 180946 | 181542 | CP015056.1 | 272995 | 273540 | 48.48484848 | 198 | 181 | 1.093922652 | 1 | 0 | 0 | FIM1_1825 | YKL141W | YKL141W | SDH3 | n |
| Kmar_QGN17376.1 | W303.asm02.HP0_G0032480 | OG0000912 | chrXI | 299929 | 301107 | CP015059.1 | 799541 | 800716 | 57.50636132 | 392 | 391 | 1.002557545 | 1 | 0 | 1 | FIM1_4108 | YKL077W | YKL077W | PSG1 | n |
| Kmar_QGN17241.1 | W303.asm02.HP0_G0023010 | OG0000913 | chrVIII | 80669 | 82426 | CP015059.1 | 529326 | 530924 | 38.99159664 | 585 | 532 | 1.09962406 | 1 | 0 | 1 | FIM1_3972 | YHL010C | YHL010C | ETP1 | n |
| Kmar_QGN17759.1 | W303.asm02.HP0_G0043500 | OG0000914 | chrXIII | 902632 | 903585 | CP015060.1 | 405012 | 406265 | 38.424821 | 317 | 417 | 0.760191847 | 1 | 0 | 0 | FIM1_4968 | YMR310C | YMR310C | UPA2 | n |
| Kmar_QGN17760.1 | W303.asm02.HP0_G0022490 | OG0000915 | chrVII | 1071189 | 1072130 | CP015060.1 | 406732 | 407670 | 74.12140575 | 313 | 312 | 1.003205128 | 1 | 0 | 1 | FIM1_4969 | YGR282C | YGR282C | BGL2 | n |
| Kmar_QGN17757.1 | W303.asm02.HP0_G0022510 | OG0000917 | chrVII | 1074064 | 1074996 | CP015060.1 | 402105 | 403022 | 66.77419355 | 310 | 305 | 1.016393443 | 1 | 0 | 1 | FIM1_4966 | YGR284C | YGR284C | ERV29 | n |
| Kmar_QGN14787.1 | W303.asm02.HP0_G0050240 | OG0000918 | chrXV | 562046 | 565069 | CP015055.1 | 1230486 | 1233599 | 33.30161751 | 1007 | 1037 | 0.971070395 | 1 | 0 | 0 | FIM1_1457 | YOR127W | YOR127W | RGA1 | n |
| Kmar_QGN15608.1 | W303.asm02.HP0_G0000210 | OG0000919 | chrI | 52490 | 53230 | CP015056.1 | 1296620 | 1297381 | 54.72440945 | 246 | 253 | 0.972332016 | 1 | 0 | 1 | FIM1_2301 | YAL049C | YAL049C | AIM2 | n |
| Kmar_QGN17934.1 | W303.asm02.HP0_G0015550 | OG0000920 | chrV | 359775 | 360731 | CP015060.1 | 770430 | 771398 | 85.71428571 | 318 | 322 | 0.98757764 | 1 | 0 | 0 | FIM1_5143 | YER099C | YER099C | PRS2 | n |
| Kmar_QGN17833.1 | W303.asm02.HP0_G0032720 | OG0000921 | chrXI | 345141 | 347357 | CP015060.1 | 582814 | 584823 | 32.79569892 | 738 | 669 | 1.103139013 | 1 | 0 | 1 | FIM1_5042 | YKL054C | YKL054C | DEF1 | n |
| Kmar_QGN15728.1 | W303.asm02.HP0_G0034180 | OG0000923 | chrXI | 638072 | 643699 | CP015056.1 | 1549780 | 1555026 | 30.41622199 | 1875 | 1748 | 1.072654462 | 1 | 0 | 0 | FIM1_2422 | YKR095W | YKR095W | MLP1 | n |
| Kmar_QGN15732.1 | W303.asm02.HP0_G0034200 | OG0000924 | chrXI | 645060 | 648647 | CP015056.1 | 1560267 | 1563974 | 42.11783439 | 1195 | 1235 | 0.967611336 | 1 | 0 | 0 | FIM1_2426 | YKR096W | YKR096W | ESL2 | n |
| Kmar_QGN15923.1 | W303.asm02.HP0_G0009910 | OG0000925 | chrIV | 709393 | 714159 | CP015057.1 | 382386 | 387128 | 71.23977344 | 1588 | 1580 | 1.005063291 | 1 | 0 | 1 | FIM1_2621 | YDR127W | YDR127W | ARO1 | n |
| Kmar_QGN18339.1 | W303.asm02.HP0_G0002790 | OG0000926 | chrII | 425455 | 426915 | CP015061.1 | 724434 | 726455 | 22.3042836 | 486 | 673 | 0.722139673 | 1 | 0 | 1 | FIM1_4666 | YBR083W | YBR083W | TEC1 | n |
| Kmar_QGN14775.1 | W303.asm02.HP0_G0050270 | OG0000927 | chrXV | 570805 | 571683 | CP015055.1 | 1205758 | 1206612 | 60.75085324 | 292 | 284 | 1.028169014 | 1 | 0 | 1 | FIM1_1445 | YOR130C | YOR130C | ORT1 | n |
| Kmar_QGN16208.1 | W303.asm02.HP0_G0038650 | OG0000929 | chrXII | 922238 | 924289 | CP015057.1 | 1037094 | 1039172 | 34.91620112 | 683 | 692 | 0.98699422 | 1 | 0 | 1 | FIM1_2912 | YLR403W | YLR403W | SFP1 | n |
| Kmar_QGN14730.1 | W303.asm02.HP0_G0056770 | OG0000930 | chrXVI | 846198 | 846845 | CP015055.1 | 1111949 | 1112623 | 47.36842105 | 215 | 224 | 0.959821429 | 1 | 0 | 0 | FIM1_1399 | YPR154W | YPR154W | PIN3 | n |
| Kmar_QGN15174.1 | W303.asm02.HP0_G0028830 | OG0000931 | chrX | 247439 | 250963 | CP015056.1 | 361944 | 365159 | 33.38898164 | 1174 | 1071 | 1.096171802 | 1 | 0 | 0 | FIM1_1862 | YJL092W | YJL092W | SRS2 | n |
| Kmar_QGN17822.1 | W303.asm02.HP0_G0047890 | OG0000932 | chrXV | 88184 | 89788 | CP015060.1 | 563751 | 565496 | 48.98305085 | 534 | 581 | 0.919104991 | 1 | 0 | 1 | FIM1_5031 | YOL123W | YOL123W | HRP1 | n |
| Kmar_QGN16414.1 | W303.asm02.HP0_G0050870 | OG0000933 | chrXV | 718318 | 719679 | CP015058.1 | 69830 | 71110 | 62.09150327 | 453 | 426 | 1.063380282 | 1 | 0 | 1 | FIM1_3127 | YOR197W | YOR197W | MCA1 | n |
| Kmar_QGN17732.1 | W303.asm02.HP0_G0046230 | OG0000934 | chrXIV | 542908 | 543369 | CP015060.1 | 346149 | 346601 | 67.32026144 | 153 | 150 | 1.02 | 1 | 0 | 0 | FIM1_4941 | YNL052W | YNL052W | COX5A | n |
| Kmar_QGN15043.1 | W303.asm02.HP0_G0031240 | OG0000935 | chrXI | 35565 | 37436 | CP015056.1 | 79267 | 81138 | 60.8974359 | 623 | 623 | 1 | 1 | 0 | 1 | FIM1_1728 | YKL212W | YKL212W | SAC1 | n |
| Kmar_QGN17734.1 | W303.asm02.HP0_G0046220 | OG0000936 | chrXIV | 541125 | 542594 | CP015060.1 | 350687 | 351859 | 29.35222672 | 489 | 390 | 1.253846154 | 1 | 0 | 0 | FIM1_4943 | YNL053W | YNL053W | MSG5 | n |
| Kmar_QGN14853.1 | W303.asm02.HP0_G0039260 | OG0000937 | chrXIII | 6034 | 7725 | CP015055.1 | 1350580 | 1352235 | 52.82685512 | 563 | 551 | 1.021778584 | 1 | 0 | 1 | FIM1_1526 | YML130C | YML130C | ERO1 | n |
| Kmar_QGN16755.1 | W303.asm02.HP0_G0014500 | OG0000938 | chrV | 160763 | 161458 | CP015058.1 | 828738 | 829415 | 57.01754386 | 231 | 225 | 1.026666667 | 1 | 0 | 1 | FIM1_3477 | YER004W | YER004W | FMP52 | n |
| Kmar_QGN14929.1 | W303.asm02.HP0_G0045100 | OG0000939 | chrXIV | 305766 | 307676 | CP015055.1 | 1513847 | 1515136 | 23.74213836 | 636 | 429 | 1.482517483 | 1 | 0 | 1 | FIM1_1604 | YNL176C | YNL176C | TDA7 | n |
| Kmar_QGN13765.1 | W303.asm02.HP0_G0017120 | OG0000940 | chrVI | 168777 | 170612 | CP015054.1 | 834646 | 837420 | 28.54054054 | 611 | 924 | 0.661255411 | 1 | 0 | 0 | FIM1_410 | YFR023W | YFR023W | PES4 | n |
| Kmar_QGN13827.1 | W303.asm02.HP0_G0031640 | OG0000941 | chrXI | 132458 | 134659 | CP015054.1 | 952202 | 954529 | 35.88761175 | 733 | 775 | 0.945806452 | 1 | 0 | 0 | FIM1_473 | YKL168C | YKL168C | KKQ8 | n |
| Kmar_QGN14692.1 | W303.asm02.HP0_G0039180 | OG0000942 | chrXII | 1052479 | 1053438 | CP015055.1 | 1042878 | 1043900 | 24.63343109 | 319 | 340 | 0.938235294 | 1 | 0 | 0 | FIM1_1359 | YLR457C | YLR457C | NBP1 | n |
| Kmar_QGN15276.1 | W303.asm02.HP0_G0049700 | OG0000943 | chrXV | 450171 | 452060 | CP015056.1 | 578965 | 580560 | 23.31288344 | 629 | 531 | 1.184557439 | 1 | 0 | 0 | FIM1_1965 | YOR066W | YOR066W | MSA1 | n |
| Kmar_QGN15338.1 | W303.asm02.HP0_G0052960 | OG0000944 | chrXVI | 76631 | 78574 | CP015056.1 | 742497 | 744365 | 22.96072508 | 647 | 622 | 1.040192926 | 1 | 0 | 0 | FIM1_2027 | YPL253C | YPL253C | VIK1 | n |

| | | | | | | | | | | | | | | | | | | | | |
|---|---|---|---|---|---|---|---|---|---|---|---|---|---|---|---|---|---|---|---|---|
| Kmar_QGN15618.1 | W303.asm02.HP0_G0038020 | OG0000945 | chrXII | 786362 | 787492 | CP015056.1 | 1318514 | 1319605 | 38.93129771 | 376 | 363 | 1.035812672 | 1 | 0 | 0 | FIM1_2311 | YLR332W | YLR332W | MID2 | n |
| Kmar_QGN15638.1 | W303.asm02.HP0_G0038240 | OG0000946 | chrXII | 836501 | 837094 | CP015056.1 | 1363762 | 1364442 | 24.7826087 | 197 | 226 | 0.871681416 | 1 | 0 | 0 | FIM1_2331 | YLR356W | YLR356W | ATG33 | n |
| Kmar_QGN15645.1 | W303.asm02.HP0_G0020250 | OG0000947 | chrVII | 575245 | 576888 | CP015056.1 | 1380454 | 1382040 | 21.90305206 | 547 | 528 | 1.035984848 | 1 | 0 | 0 | FIM1_2338 | YGR041W | YGR041W | BUD9 | n |
| Kmar_QGN16089.1 | W303.asm02.HP0_G0006180 | OG0000948 | chrIII | 271956 | 272183 | CP015057.1 | 789297 | 789542 | 40.74074074 | 75 | 81 | 0.925925926 | 1 | 0 | 0 | FIM1_2789 | YCR075W-A | YCR075W-A | EGO2 | n |
| Kmar_QGN16285.1 | W303.asm02.HP0_G0031460 | OG0000949 | chrXI | 90453 | 92705 | CP015057.1 | 1204745 | 1206802 | 36.51006711 | 750 | 685 | 1.094890511 | 1 | 0 | 0 | FIM1_2990 | YKL187C | YKL187C | FAT3 | n |
| Kmar_QGN16311.1 | W303.asm02.HP0_G0018700 | OG0000950 | chrVII | 252131 | 253432 | CP015057.1 | 1257639 | 1259117 | 24.7464503 | 433 | 492 | 0.880081301 | 1 | 0 | 0 | FIM1_3016 | YGL134W | YGL134W | PCL10 | n |
| Kmar_QGN16445.1 | W303.asm02.HP0_G0029140 | OG0000951 | chrX | 316261 | 317892 | CP015058.1 | 132931 | 133650 | 19.52117864 | 543 | 239 | 2.271966527 | 1 | 0 | 1 | FIM1_3158 | YJL058C | YJL058C | BIT61 | n |
| Kmar_QGN16497.1 | W303.asm02.HP0_G0052040 | OG0000952 | chrXV | 924952 | 926760 | CP015058.1 | 237165 | 238568 | 19.07131012 | 602 | 467 | 1.289079229 | 1 | 0 | 0 | FIM1_3210 | YOR324C | YOR324C | FRT1 | n |
| Kmar_QGN16823.1 | W303.asm02.HP0_G0056480 | OG0000953 | chrXVI | 780804 | 782111 | CP015058.1 | 970029 | 971378 | 36.97104677 | 435 | 449 | 0.968819599 | 1 | 0 | 0 | FIM1_3547 | YPR120C | YPR120C | CLB5 | n |
| Kmar_QGN17087.1 | W303.asm02.HP0_G0042460 | OG0000954 | chrXIII | 676475 | 677416 | CP015059.1 | 207038 | 208078 | 27.66570605 | 313 | 346 | 0.904624277 | 1 | 0 | 0 | FIM1_3818 | YMR206W | YMR206W | | n |
| Kmar_QGN17319.1 | W303.asm02.HP0_G0016320 | OG0000955 | chrV | 567156 | 568067 | CP015059.1 | 686431 | 687399 | 38.80126183 | 303 | 322 | 0.940993789 | 1 | 0 | 0 | FIM1_4051 | YER185W | YER185W | PUG1 | n |
| Kmar_QGN17691.1 | W303.asm02.HP0_G0046460 | OG0000956 | chrXIV | 588388 | 589956 | CP015060.1 | 264298 | 265866 | 45.84837545 | 522 | 522 | 1 | 1 | 0 | 0 | FIM1_4898 | YNL029C | YNL029C | KTR5 | n |
| Kmar_QGN18079.1 | W303.asm02.HP0_G0001460 | OG0000957 | chrII | 138329 | 139102 | CP015061.1 | 164769 | 165506 | 20.62256809 | 257 | 245 | 1.048979592 | 1 | 0 | 0 | FIM1_4398 | YBL043W | YBL043W | ECM13 | n |
| Kmar_QGN18306.1 | W303.asm02.HP0_G0048720 | OG0000958 | chrXV | 257200 | 259485 | CP015061.1 | 658669 | 661146 | 19.87878788 | 761 | 825 | 0.922424242 | 1 | 0 | 0 | FIM1_4632 | YOL036W | YOL036W | | n |
| Kmar_QGN14444.1 | W303.asm02.HP0_G0047150 | OG0000961 | chrXIV | 714477 | 715667 | CP015055.1 | 525731 | 526787 | 46.55647383 | 396 | 242 | 1.636363636 | 1 | 1 | 1 | FIM1_1105 | YNR043W | YNR043W | MVD1 | n |
| Kmar_QGN13669.1 | W303.asm02.HP0_G0016990 | OG0000962 | chrVI | 133550 | 135049 | CP015054.1 | 632494 | 634088 | 58.31663327 | 499 | 484 | 1.030991736 | 1 | 1 | 1 | FIM1_312 | YFR010W | YFR010W | UBP6 | n |
| Kmar_QGN13670.1 | W303.asm02.HP0_G0017000 | OG0000963 | chrVI | 135229 | 135741 | CP015054.1 | 634420 | 634887 | 38.95348837 | 170 | 155 | 1.096774194 | 1 | 1 | 1 | FIM1_313 | YFR011C | YFR011C | MIC19 | n |
| Kmar_QGN15101.1 | W303.asm02.HP0_G0042740 | OG0000964 | chrXIII | 739763 | 740443 | CP015056.1 | 205329 | 206157 | 41.84100418 | 226 | 230 | 0.982608696 | 1 | 1 | 1 | FIM1_1788 | YMR233W | YMR233W | TRI1 | n |
| Kmar_QGN13944.1 | W303.asm02.HP0_G0049040 | OG0000965 | chrXV | 317393 | 322003 | CP015054.1 | 1189978 | 1194447 | 49.59145192 | 1536 | 1489 | 1.031564809 | 1 | 1 | 1 | FIM1_592 | YOL004W | YOL004W | SIN3 | n |
| Kmar_QGN13945.1 | W303.asm02.HP0_G0049030 | OG0000966 | chrXV | 316268 | 316630 | CP015054.1 | 1195076 | 1195441 | 80.99173554 | 120 | 121 | 0.991735537 | 1 | 1 | 1 | FIM1_593 | YOL005C | YOL005C | RPB11 | n |
| Kmar_QGN17314.1 | W303.asm02.HP0_G0045600 | OG0000967 | chrXIV | 399231 | 399578 | CP015059.1 | 677437 | 677724 | 45.2173913 | 115 | 95 | 1.210526316 | 1 | 1 | 1 | FIM1_4046 | YNL122C | YNL122C | MRP35 | n |
| Kmar_QGN17952.1 | W303.asm02.HP0_G0015660 | OG0000968 | chrV | 384617 | 387898 | CP015060.1 | 811554 | 814430 | 33.24764353 | 1093 | 958 | 1.14091858 | 1 | 1 | 1 | FIM1_5161 | YER111C | YER111C | SWI4 | n |
| Kmar_QGN14571.1 | W303.asm02.HP0_G0007980 | OG0000969 | chrIV | 302794 | 303000 | CP015055.1 | 785104 | 785307 | 77.94117647 | 68 | 67 | 1.014925373 | 1 | 1 | 1 | FIM1_1233 | YDL085C-A | YDL085C-A | | n |
| Kmar_QGN16195.1 | W303.asm02.HP0_G0039820 | OG0000970 | chrXIII | 119270 | 123907 | CP015057.1 | 1000912 | 1005381 | 47.68041237 | 1545 | 1489 | 1.037609134 | 1 | 1 | 1 | FIM1_2897 | YML072C | YML072C | TCB3 | n |
| Kmar_QGN15128.1 | W303.asm02.HP0_G0041550 | OG0000971 | chrXIII | 496968 | 498074 | CP015056.1 | 258955 | 259854 | 31.93717277 | 368 | 299 | 1.230769231 | 1 | 1 | 1 | FIM1_1815 | YMR114C | YMR114C | | n |
| Kmar_QGN17122.1 | W303.asm02.HP0_G0014700 | OG0000972 | chrV | 199951 | 202014 | CP015059.1 | 282781 | 284790 | 39.40256046 | 687 | 669 | 1.02690583 | 1 | 1 | 1 | FIM1_3853 | YER022W | YER022W | SRB4 | n |
| Kmar_QGN14936.1 | W303.asm02.HP0_G0045160 | OG0000973 | chrXIV | 320109 | 322052 | CP015055.1 | 1530182 | 1531879 | 27.59197324 | 647 | 565 | 1.145132743 | 1 | 1 | 1 | FIM1_1611 | YNL167C | YNL167C | SKO1 | n |
| Kmar_QGN13520.1 | W303.asm02.HP0_G0037870 | OG0000974 | chrXII | 757832 | 759394 | CP015054.1 | 321062 | 322546 | 54.89443378 | 520 | 494 | 1.052631579 | 1 | 1 | 1 | FIM1_160 | YLR314C | YLR314C | CDC3 | n |
| Kmar_QGN13567.1 | W303.asm02.HP0_G0038840 | OG0000975 | chrXII | 961759 | 962229 | CP015054.1 | 430356 | 430883 | 50.31847134 | 156 | 175 | 0.891428571 | 1 | 1 | 1 | FIM1_208 | YLR421C | YLR421C | RPN13 | n |
| Kmar_QGN17176.1 | W303.asm02.HP0_G0055090 | OG0000976 | chrXVI | 498377 | 499396 | CP015059.1 | 390889 | 391827 | 80.51118211 | 305 | 312 | 0.977564103 | 1 | 1 | 1 | FIM1_3907 | YPL031C | YPL031C | PHO85 | n |
| Kmar_QGN18123.1 | W303.asm02.HP0_G0000980 | OG0000977 | chrII | 37862 | 39499 | CP015061.1 | 269217 | 270863 | 88.16029144 | 545 | 548 | 0.994525547 | 1 | 1 | 1 | FIM1_4444 | YBL099W | YBL099W | ATP1 | n |
| Kmar_QGN14999.1 | W303.asm02.HP0_G0029010 | OG0000978 | chrX | 295240 | 295881 | CP015055.1 | 1655194 | 1655799 | 59.33014354 | 213 | 201 | 1.059701493 | 1 | 1 | 1 | FIM1_1680 | YJL072C | YJL072C | PSF2 | n |
| Kmar_QGN16470.1 | W303.asm02.HP0_G0004710 | OG0000979 | chrII | 792312 | 793784 | CP015058.1 | 176872 | 178311 | 56.85071575 | 490 | 479 | 1.022964509 | 1 | 1 | 1 | FIM1_3183 | YBR283C | YBR283C | SSH1 | n |
| Kmar_QGN14815.1 | W303.asm02.HP0_G0047710 | OG0000980 | chrXV | 54795 | 55304 | CP015055.1 | 1283627 | 1284124 | 74.54545455 | 169 | 165 | 1.024242424 | 1 | 1 | 1 | FIM1_1486 | YOL143C | YOL143C | RIB4 | n |
| Kmar_QGN17943.1 | W303.asm02.HP0_G0001170 | OG0000981 | chrII | 81531 | 81884 | CP015060.1 | 787903 | 788277 | 79.83870968 | 117 | 124 | 0.943548387 | 1 | 1 | 1 | FIM1_5152 | YBL078C | YBL078C | ATG8 | n |
| Kmar_QGN14499.1 | W303.asm02.HP0_G0016590 | OG0000983 | chrVI | 43633 | 44319 | CP015055.1 | 640858 | 641769 | 31.84713376 | 228 | 303 | 0.752475248 | 1 | 1 | 1 | FIM1_1160 | YFL031W | YFL031W | HAC1 | n |
| Kmar_QGN18276.1 | W303.asm02.HP0_G0056590 | OG0000984 | chrXVI | 804486 | 804638 | CP015061.1 | 602207 | 602356 | 66 | 50 | 49 | 1.020408163 | 1 | 1 | 1 | FIM1_4602 | YPR133W-A | YPR133W-A | TOM5 | n |
| Kmar_QGN18173.1 | W303.asm02.HP0_G0015100 | OG0000985 | chrV | 273439 | 273762 | CP015061.1 | 371697 | 372014 | 67.28971963 | 107 | 105 | 1.019047619 | 1 | 1 | 1 | FIM1_4496 | YER058W | YER058W | PET117 | n |
| Kmar_QGN14943.1 | W303.asm02.HP0_G0045210 | OG0000986 | chrXIV | 331021 | 331239 | CP015055.1 | 1542487 | 1542708 | 45.20547945 | 72 | 73 | 0.98630137 | 1 | 1 | 1 | FIM1_1618 | YNL162W-A | YNL162W-A | | n |
| Kmar_QGN17878.1 | W303.asm02.HP0_G0056060 | OG0000987 | chrXVI | 698431 | 698916 | CP015060.1 | 663618 | 664094 | 62.11180124 | 161 | 158 | 1.018987342 | 1 | 1 | 1 | FIM1_5087 | YPR073C | YPR073C | LTP1 | n |
| Kmar_QGN18380.1 | W303.asm02.HP0_G0008370 | OG0000988 | chrIV | 373573 | 374493 | CP015061.1 | 809302 | 810378 | 44.65753425 | 306 | 358 | 0.854748603 | 1 | 1 | 1 | FIM1_4707 | YDL045C | YDL045C | FAD1 | n |
| Kmar_QGN17729.1 | W303.asm02.HP0_G0046250 | OG0000989 | chrXIV | 545262 | 546165 | CP015060.1 | 341748 | 342566 | 40.54982818 | 270 | 272 | 0.992647059 | 1 | 1 | 1 | FIM1_4938 | YNL050C | YNL050C | | n |
| Kmar_QGN16543.1 | W303.asm02.HP0_G0039060 | OG0000990 | chrXII | 1020856 | 1021504 | CP015058.1 | 340241 | 340750 | 27.12765957 | 188 | 169 | 1.112426036 | 1 | 1 | 1 | FIM1_3256 | YLR445W | YLR445W | GMC2 | n |
| Kmar_QGN17714.1 | W303.asm02.HP0_G0046380 | OG0000991 | chrXIV | 569103 | 570185 | CP015060.1 | 301509 | 302594 | 78.17679558 | 360 | 361 | 0.997229917 | 1 | 1 | 1 | FIM1_4921 | YNL037C | YNL037C | IDH1 | n |
| Kmar_QGN15382.1 | W303.asm02.HP0_G0009460 | OG0000992 | chrIV | 607340 | 607675 | CP015056.1 | 846230 | 846604 | 72.13114754 | 111 | 124 | 0.89516129 | 1 | 1 | 1 | FIM1_2072 | YDR079W | YDR079W | PET100 | n |
| Kmar_QGN16746.1 | W303.asm02.HP0_G0014560 | OG0000993 | chrV | 173298 | 173675 | CP015058.1 | 806073 | 806450 | 87.2 | 125 | 125 | 1 | 1 | 1 | 1 | FIM1_3468 | YER009W | YER009W | NTF2 | n |
| Kmar_QGN18084.1 | W303.asm02.HP0_G0001420 | OG0000995 | chrII | 129536 | 133681 | CP015061.1 | 173845 | 177639 | 38.66197183 | 1381 | 1264 | 1.092563291 | 1 | 1 | 1 | FIM1_4403 | YBL047C | YBL047C | EDE1 | n |
| Kmar_QGN18083.1 | W303.asm02.HP0_G0030650 | OG0000996 | chrX | 647702 | 647941 | CP015061.1 | 172869 | 173324 | 23.17880795 | 79 | 151 | 0.523178808 | 1 | 1 | 1 | FIM1_4402 | YJR112W-A | YJR112W-A | | n |
| Kmar_QGN14234.1 | W303.asm02.HP0_G0001730 | OG0000997 | chrII | 188285 | 188761 | CP015055.1 | 80619 | 81014 | 45.86466165 | 133 | 131 | 1.015267176 | 1 | 1 | 1 | FIM1_890 | YBL018C | YBL018C | POP8 | n |
| Kmar_QGN14371.1 | W303.asm02.HP0_G0016260 | OG0000998 | chrV | 556123 | 557219 | CP015055.1 | 379025 | 380014 | 82.03592814 | 334 | 329 | 1.015197568 | 1 | 1 | 1 | FIM1_1031 | YER179W | YER179W | DMC1 | n |
| Kmar_QGN17204.1 | W303.asm02.HP0_G0019110 | OG0000999 | chrVII | 343358 | 343856 | CP015059.1 | 444647 | 444952 | 57.97101449 | 137 | 101 | 1.356435644 | 1 | 1 | 1 | FIM1_3935 | YGL087C | YGL087C | MMS2 | n |
| Kmar_QGN15980.1 | W303.asm02.HP0_G0003470 | OG0001000 | chrII | 565139 | 565786 | CP015057.1 | 535840 | 536487 | 84.65116279 | 215 | 215 | 1 | 1 | 1 | 1 | FIM1_2680 | YBR154C | YBR154C | RPB5 | n |
| Kmar_QGN15621.1 | W303.asm02.HP0_G0018450 | OG0001001 | chrVII | 195678 | 196610 | CP015056.1 | 1326402 | 1327211 | 24.84076433 | 310 | 269 | 1.152416357 | 1 | 1 | 1 | FIM1_2314 | YGL161C | YGL161C | YIP5 | n |
| Kmar_QGN13525.1 | W303.asm02.HP0_G0034740 | OG0001002 | chrXI | 108687 | 108983 | CP015054.1 | 342938 | 343234 | 67.34693878 | 98 | 98 | 1 | 1 | 1 | 1 | FIM1_165 | YLL018C-A | YLL018C-A | COX19 | n |
| Kmar_QGN16032.1 | W303.asm02.HP0_G0018350 | OG0001003 | chrVII | 171995 | 176581 | CP015057.1 | 667276 | 671697 | 56.76028739 | 1528 | 1473 | 1.037338764 | 1 | 1 | 1 | FIM1_2732 | YGL173C | YGL173C | XRN1 | n |
| Kmar_QGN16495.1 | W303.asm02.HP0_G0052060 | OG0001004 | chrXV | 932454 | 932801 | CP015058.1 | 230160 | 230634 | 80.86956522 | 115 | 110 | 1.045454545 | 1 | 1 | 1 | FIM1_3208 | YOR327C | YOR327C | SNC2 | n |
| Kmar_QGN16612.1 | W303.asm02.HP0_G0008850 | OG0001005 | chrIV | 478133 | 478417 | CP015058.1 | 499755 | 500072 | 46.36363636 | 94 | 105 | 0.895238095 | 1 | 1 | 1 | FIM1_3328 | YDR016C | YDR016C | DAD1 | n |
| Kmar_QGN18310.1 | W303.asm02.HP0_G0055890 | OG0001006 | chrXVI | 672778 | 673944 | CP015061.1 | 670293 | 671612 | 52.38095238 | 388 | 439 | 0.883826879 | 1 | 1 | 1 | FIM1_4636 | YPR054W | YPR054W | SMK1 | n |
| Kmar_QGN17662.1 | W303.asm02.HP0_G0049270 | OG0001007 | chrXV | 370980 | 371300 | CP015060.1 | 209348 | 209665 | 84.90566038 | 106 | 105 | 1.00952381 | 1 | 1 | 1 | FIM1_4869 | YOR020C | YOR020C | HSP10 | n |
| Kmar_QGN15790.1 | W303.asm02.HP0_G0037960 | OG0001008 | chrXII | 776828 | 777064 | CP015057.1 | 105592 | 105828 | 75.64102564 | 78 | 78 | 1 | 1 | 1 | 1 | FIM1_2486 | YLR325C | YLR325C | RPL38 | n |
| Kmar_QGN15794.1 | W303.asm02.HP0_G0019950 | OG0001009 | chrVII | 504503 | 505474 | CP015057.1 | 109410 | 110363 | 46.38554217 | 323 | 317 | 1.018927445 | 1 | 1 | 1 | FIM1_2490 | YGR007W | YGR007W | ECT1 | n |
| Kmar_QGN16532.1 | W303.asm02.HP0_G0001830 | OG0001010 | chrII | 211938 | 214460 | CP015058.1 | 317282 | 319870 | 49.60272418 | 840 | 862 | 0.974477958 | 1 | 1 | 1 | FIM1_3245 | YBL008W | YBL008W | HIR1 | n |
| Kmar_QGN15822.1 | W303.asm02.HP0_G0035250 | OG0001011 | chrXI | 216145 | 217653 | CP015057.1 | 158842 | 160323 | 44.22700587 | 502 | 493 | 1.018255578 | 1 | 1 | 1 | FIM1_2518 | YLR033W | YLR033W | RSC58 | n |
| Kmar_QGN16264.1 | W303.asm02.HP0_G0039720 | OG0001012 | chrXIII | 98523 | 98702 | CP015057.1 | 1154937 | 1155113 | 72.4137931 | 59 | 58 | 1.017241379 | 1 | 1 | 1 | FIM1_2969 | YML081C-A | YML081C-A | ATP18 | n |
| Kmar_QGN17257.1 | W303.asm02.HP0_G0032850 | OG0001013 | chrXI | 368746 | 369420 | CP015059.1 | 559219 | 559911 | 62.06896552 | 224 | 230 | 0.973913043 | 1 | 1 | 1 | FIM1_3988 | YKL041W | YKL041W | VPS24 | n |
| Kmar_QGN13496.1 | W303.asm02.HP0_G0014120 | OG0001014 | chrV | 81765 | 82961 | CP015054.1 | 265481 | 266650 | 48.38709677 | 398 | 389 | 1.023136247 | 1 | 1 | 1 | FIM1_134 | YEL037C | YEL037C | RAD23 | n |
| Kmar_QGN15863.1 | W303.asm02.HP0_G0009030 | OG0001015 | chrIV | 521475 | 522587 | CP015057.1 | 254653 | 255771 | 79.56989247 | 370 | 372 | 0.994623656 | 1 | 1 | 1 | FIM1_2560 | YDR035W | YDR035W | ARO3 | n |
| Kmar_QGN16200.1 | W303.asm02.HP0_G0030770 | OG0001016 | chrX | 664511 | 665737 | CP015057.1 | 1020490 | 1021716 | 52.95508274 | 408 | 408 | 1 | 1 | 1 | 1 | FIM1_2903 | YJR125C | YJR125C | ENT3 | n |
| Kmar_QGN16160.1 | W303.asm02.HP0_G0015270 | OG0001017 | chrV | 309323 | 309793 | CP015057.1 | 929106 | 929404 | 71.23287671 | 85 | 70 | 1.214285714 | 1 | 1 | 1 | FIM1_2862 | YER074W-A | YER074W-A | YOS1 | n |
| Kmar_QGN15258.1 | W303.asm02.HP0_G0033910 | OG0001018 | chrXI | 579993 | 580574 | CP015056.1 | 546589 | 547215 | 70.67307692 | 193 | 208 | 0.927884615 | 1 | 1 | 1 | FIM1_1947 | YKR068C | YKR068C | BET3 | n |
| Kmar_QGN16010.1 | W303.asm02.HP0_G0023670 | OG0001019 | chrVIII | 203549 | 203995 | CP015057.1 | 608610 | 609056 | 72.18543046 | 148 | 148 | 1 | 1 | 1 | 1 | FIM1_2710 | YHR051W | YHR051W | COX6 | n |
| Kmar_QGN17294.1 | W303.asm02.HP0_G0048030 | OG0001020 | chrXV | 111886 | 112341 | CP015059.1 | 637687 | 638042 | 21.19205298 | 151 | 96 | 1.572916667 | 1 | 1 | 1 | FIM1_4025 | YOL108C | YOL108C | INO4 | n |
| Kmar_QGN14651.1 | W303.asm02.HP0_G0055500 | OG0001021 | chrXVI | 588869 | 589230 | CP015055.1 | 950340 | 950558 | 54.79452055 | 72 | 72 | 1 | 1 | 1 | 1 | FIM1_1315 | YPR010C-A | YPR010C-A | MRA1 | n |
| Kmar_QGN13830.1 | W303.asm02.HP0_G0028160 | OG0001022 | chrX | 95962 | 96246 | CP015054.1 | 958889 | 959173 | 71.27659574 | 94 | 94 | 1 | 1 | 1 | 1 | FIM1_476 | YJL166W | YJL166W | QCR8 | n |
| Kmar_QGN17741.1 | W303.asm02.HP0_G0026110 | OG0001023 | chrIX | 130108 | 130803 | CP015060.1 | 368102 | 368785 | 75.84745763 | 231 | 227 | 1.017621145 | 1 | 1 | 1 | FIM1_4950 | YIL118W | YIL118W | RHO3 | n |
| Kmar_QGN14951.1 | W303.asm02.HP0_G0016770 | OG0001024 | chrVI | 75740 | 76069 | CP015055.1 | 1558582 | 1558896 | 61.46788991 | 109 | 104 | 1.048076923 | 1 | 1 | 1 | FIM1_1628 | YFL014W | YFL014W | HSP12 | n |
| Kmar_QGN16037.1 | W303.asm02.HP0_G0021380 | OG0001025 | chrVII | 840891 | 841928 | CP015057.1 | 677571 | 678194 | 28.9017341 | 345 | 207 | 1.666666667 | 1 | 1 | 1 | FIM1_2737 | YGR165W | YGR165W | MRPS35 | n |
| Kmar_QGN18153.1 | W303.asm02.HP0_G0026600 | OG0001027 | chrIX | 232573 | 233346 | CP015061.1 | 338467 | 339156 | 53.307393 | 257 | 229 | 1.122270742 | 1 | 1 | 1 | FIM1_4476 | YIL064W | YIL064W | EFM4 | n |
| Kmar_QGN18157.1 | W303.asm02.HP0_G0026630 | OG0001028 | chrIX | 235287 | 236189 | CP015061.1 | 344578 | 345489 | 40.06309148 | 300 | 303 | 0.99009901 | 1 | 1 | 1 | FIM1_4480 | YIL061C | YIL061C | SNP1 | n |
| Kmar_QGN17937.1 | W303.asm02.HP0_G0001220 | OG0001029 | chrII | 91126 | 91374 | CP015060.1 | 774747 | 774995 | 75.6097561 | 82 | 82 | 1 | 1 | 1 | 1 | FIM1_5146 | YBL071W-A | YBL071W-A | KTI11 | n |
| Kmar_QGN14981.1 | W303.asm02.HP0_G0051040 | OG0001030 | chrXV | 750596 | 753181 | CP015055.1 | 1616094 | 1618583 | 56.49651972 | 861 | 829 | 1.038600724 | 1 | 1 | 1 | FIM1_1659 | YOR217W | YOR217W | RFC1 | n |
| Kmar_QGN16220.1 | W303.asm02.HP0_G0038590 | OG0001031 | chrXII | 906399 | 906635 | CP015057.1 | 1062233 | 1062451 | 53.84615385 | 78 | 72 | 1.083333333 | 1 | 1 | 1 | FIM1_2924 | YLR395C | YLR395C | COX8 | n |
| Kmar_QGN16219.1 | W303.asm02.HP0_G0008160 | OG0001032 | chrIV | 334547 | 334726 | CP015057.1 | 1061010 | 1061192 | 63.33333333 | 59 | 60 | 0.983333333 | 1 | 1 | 1 | FIM1_2923 | YDL067C | YDL067C | COX9 | n |
| Kmar_QGN14888.1 | W303.asm02.HP0_G0037510 | OG0001033 | chrXII | 689028 | 689450 | CP015055.1 | 1432551 | 1432883 | 69.09090909 | 110 | 110 | 1 | 1 | 1 | 1 | FIM1_1562 | YLR275W | YLR275W | SMD2 | n |
| Kmar_QGN15515.1 | W303.asm02.HP0_G0020580 | OG0001034 | chrVII | 641186 | 641818 | CP015056.1 | 1102886 | 1103470 | 41.9047619 | 210 | 194 | 1.082474227 | 1 | 1 | 1 | FIM1_2206 | YGR081C | YGR081C | SLX9 | n |
| Kmar_QGN15497.1 | W303.asm02.HP0_G0046930 | OG0001035 | chrXIV | 680959 | 682173 | CP015056.1 | 1063570 | 1064796 | 43.27628362 | 404 | 408 | 0.990196078 | 1 | 1 | 1 | FIM1_2188 | YNR021W | YNR021W | | n |
| Kmar_QGN17339.1 | W303.asm02.HP0_G0052280 | OG0001036 | chrXV | 997299 | 998792 | CP015059.1 | 724767 | 726239 | 50.68226121 | 497 | 490 | 1.014285714 | 1 | 1 | 1 | FIM1_4071 | YOR351C | YOR351C | MEK1 | n |
| Kmar_QGN17783.1 | W303.asm02.HP0_G0045880 | OG0001037 | chrXIV | 458192 | 458770 | CP015060.1 | 474474 | 475037 | 78.64583333 | 192 | 187 | 1.026737968 | 1 | 1 | 1 | FIM1_4992 | YNL090W | YNL090W | RHO2 | n |
| Kmar_QGN17388.1 | W303.asm02.HP0_G0032430 | OG0001038 | chrXI | 289501 | 290805 | CP015059.1 | 822979 | 824298 | 57.33634312 | 434 | 439 | 0.988610478 | 1 | 1 | 1 | FIM1_4120 | YKL082C | YKL082C | RRP14 | n |
| Kmar_QGN16815.1 | W303.asm02.HP0_G0020820 | OG0001039 | chrVII | 697069 | 697302 | CP015058.1 | 954146 | 954520 | 63.41463415 | 77 | 82 | 0.93902439 | 1 | 1 | 1 | FIM1_3539 | YGR105W | YGR105W | VMA21 | n |
| Kmar_QGN16284.1 | W303.asm02.HP0_G0031470 | OG0001041 | chrXI | 93913 | 94467 | CP015057.1 | 1202221 | 1202748 | 56.98924731 | 184 | 175 | 1.051428571 | 1 | 1 | 1 | FIM1_2989 | YKL186C | YKL186C | MTR2 | n |
| Kmar_QGN13476.1 | W303.asm02.HP0_G0029910 | OG0001042 | chrX | 505214 | 505540 | CP015054.1 | 217915 | 218241 | 76.85185185 | 108 | 108 | 1 | 1 | 1 | 1 | FIM1_114 | YJR034W | YJR034W | PET191 | n |
| Kmar_QGN13940.1 | W303.asm02.HP0_G0043250 | OG0001043 | chrXIII | 848864 | 849124 | CP015054.1 | 1184091 | 1184345 | 58.33333333 | 86 | 84 | 1.023809524 | 1 | 1 | 1 | FIM1_588 | YMR286W | YMR286W | MRPL33 | n |
| Kmar_QGN17442.1 | W303.asm02.HP0_G0008700 | OG0001044 | chrIV | 446626 | 447237 | CP015059.1 | 930011 | 930811 | 40.66390041 | 203 | 266 | 0.763157895 | 1 | 1 | 1 | FIM1_4175 | YDL002C | YDL002C | NHP10 | n |
| Kmar_QGN17058.1 | W303.asm02.HP0_G0007070 | OG0001045 | chrIV | 127007 | 130222 | CP015059.1 | 142232 | 144085 | 52.38095238 | 1071 | 617 | 1.735818476 | 1 | 1 | 1 | FIM1_3788 | YDL185W | YDL185W | VMA1 | n |
| Kmar_QGN17047.1 | W303.asm02.HP0_G0037010 | OG0001046 | chrXII | 573590 | 574042 | CP015059.1 | 116938 | 117198 | 36.66666667 | 150 | 86 | 1.744186047 | 1 | 1 | 1 | FIM1_3776 | YLR218C | YLR218C | COA4 | n |
| Kmar_QGN17270.1 | W303.asm02.HP0_G0051790 | OG0001047 | chrXV | 877318 | 878757 | CP015059.1 | 590050 | 591066 | 22.5 | 479 | 338 | 1.417159763 | 1 | 1 | 1 | FIM1_4001 | YOR298W | YOR298W | MUM3 | n |
| Kmar_QGN16350.1 | W303.asm02.HP0_G0040340 | OG0001048 | chrXIII | 237501 | 239237 | CP015057.1 | 1341444 | 1343162 | 31.74342105 | 578 | 572 | 1.01048951 | 1 | 1 | 1 | FIM1_3056 | YML017W | YML017W | PSP2 | n |
| Kmar_QGN15671.1 | W303.asm02.HP0_G0013960 | OG0001049 | chrV | 54264 | 56465 | CP015056.1 | 1427988 | 1430339 | 39.00889454 | 733 | 783 | 0.93614304 | 1 | 1 | 1 | FIM1_2364 | YEL053C | YEL053C | MAK10 | n |

| | | | | | | | | | | | | | | | | | | | | |
|---|---|---|---|---|---|---|---|---|---|---|---|---|---|---|---|---|---|---|---|---|
| Kmar_QGN17899.1 | W303.asm02.HP0_G0048560 | OG0001050 | chrXV | 229069 | 230289 | CP015060.1 | 699052 | 700113 | 35.22458629 | 406 | 353 | 1.150141643 | 1 | 1 | 1 | FIM1_5108 | YOL054W | YOL054W | PSH1 | n |
| Kmar_QGN18143.1 | W303.asm02.HP0_G0043830 | OG0001051 | chrXIV | 42503 | 43507 | CP015061.1 | 318078 | 319013 | 47.80058651 | 334 | 311 | 1.073954984 | 1 | 1 | 1 | FIM1_4466 | YNL316C | YNL316C | PHA2 | n |
| Kmar_QGN15364.1 | W303.asm02.HP0_G0003560 | OG0001052 | chrII | 582007 | 582204 | CP015056.1 | 803920 | 804114 | 60.60606061 | 65 | 64 | 1.015625 | 1 | 1 | 1 | FIM1_2054 | YBR162W-A | YBR162W-A | YSY6 | n |
| Kmar_QGN15340.1 | W303.asm02.HP0_G0042370 | OG0001053 | chrXIII | 659128 | 659781 | CP015056.1 | 745985 | 746650 | 61.43497758 | 217 | 221 | 0.981900452 | 1 | 1 | 1 | FIM1_2029 | YMR197C | YMR197C | VTI1 | n |
| Kmar_QGN16842.1 | W303.asm02.HP0_G0017250 | OG0001054 | chrVI | 193231 | 193674 | CP015058.1 | 1011736 | 1012113 | 56.46258503 | 147 | 125 | 1.176 | 1 | 1 | 1 | FIM1_3567 | YFR033C | YFR033C | QCR6 | n |
| Kmar_QGN17222.1 | W303.asm02.HP0_G0004780 | OG0001055 | chrII | 805960 | 806925 | CP015059.1 | 482859 | 483779 | 49.22600619 | 321 | 306 | 1.049019608 | 1 | 1 | 1 | FIM1_3953 | YBR290W | YBR290W | BSD2 | n |
| Kmar_QGN17223.1 | W303.asm02.HP0_G0004790 | OG0001056 | chrII | 807042 | 807941 | CP015059.1 | 484164 | 485060 | 85.28428094 | 299 | 298 | 1.003355705 | 1 | 1 | 1 | FIM1_3954 | YBR291C | YBR291C | CTP1 | n |
| Kmar_QGN13485.1 | W303.asm02.HP0_G0014210 | OG0001057 | chrV | 101952 | 102434 | CP015054.1 | 239989 | 240471 | 95.625 | 160 | 160 | 1 | 1 | 1 | 1 | FIM1_123 | YEL027W | YEL027W | VMA3 | n |
| Kmar_QGN13479.1 | W303.asm02.HP0_G0014240 | OG0001058 | chrV | 108443 | 109090 | CP015054.1 | 225657 | 226298 | 79.53488372 | 215 | 213 | 1.009389671 | 1 | 1 | 1 | FIM1_117 | YEL024W | YEL024W | RIP1 | n |
| Kmar_QGN13478.1 | W303.asm02.HP0_G0014250 | OG0001059 | chrV | 109687 | 111735 | CP015054.1 | 222800 | 224818 | 42.2005571 | 682 | 672 | 1.014880952 | 1 | 1 | 1 | FIM1_116 | YEL023C | YEL023C | | n |
| Kmar_QGN14779.1 | W303.asm02.HP0_G0012320 | OG0001061 | chrIV | 1242151 | 1243597 | CP015055.1 | 1211003 | 1211290 | 25.56818182 | 226 | 95 | 2.378947368 | 1 | 1 | 1 | FIM1_1449 | YDR381W | YDR381W | YRA1 | n |
| Kmar_QGN14552.1 | W303.asm02.HP0_G0048690 | OG0001062 | chrXV | 253604 | 254032 | CP015055.1 | 743606 | 744034 | 83.8028169 | 142 | 142 | 1 | 1 | 1 | 1 | FIM1_1214 | YOL040C | YOL040C | RPS15 | n |
| Kmar_QGN17509.1 | W303.asm02.HP0_G0021960 | OG0001063 | chrVII | 957560 | 959434 | CP015059.1 | 1058668 | 1060323 | 36.82487725 | 593 | 551 | 1.076225045 | 1 | 1 | 1 | FIM1_4245 | YGR225W | YGR225W | AMA1 | n |
| Kmar_QGN14005.1 | W303.asm02.HP0_G0020560 | OG0001065 | chrVII | 637675 | 638274 | CP015054.1 | 1310983 | 1311585 | 64.87804878 | 199 | 200 | 0.995 | 1 | 1 | 1 | FIM1_656 | YGR078C | YGR078C | PAC10 | n |
| Kmar_QGN17162.1 | W303.asm02.HP0_G0054940 | OG0001066 | chrXVI | 472322 | 472621 | CP015059.1 | 365748 | 366041 | 53.53535354 | 99 | 97 | 1.020618557 | 1 | 1 | 1 | FIM1_3893 | YPL047W | YPL047W | SGF11 | n |
| Kmar_QGN16363.1 | W303.asm02.HP0_G0040420 | OG0001067 | chrXIII | 251810 | 252022 | CP015057.1 | 1366766 | 1366981 | 74.28571429 | 70 | 71 | 0.985915493 | 1 | 1 | 1 | FIM1_3069 | YML009C | YML009C | MRPL39 | n |
| Kmar_QGN15574.1 | W303.asm02.HP0_G0032140 | OG0001068 | chrXI | 231211 | 232314 | CP015056.1 | 1217191 | 1218264 | 66.21253406 | 367 | 357 | 1.028011204 | 1 | 1 | 1 | FIM1_2265 | YKL114C | YKL114C | APN1 | n |
| Kmar_QGN17364.1 | W303.asm02.HP0_G0009540 | OG0001069 | chrIV | 621205 | 621447 | CP015059.1 | 778008 | 778220 | 67.5 | 80 | 70 | 1.142857143 | 1 | 1 | 1 | FIM1_4096 | YDR086C | YDR086C | SSS1 | n |
| Kmar_QGN13974.1 | W303.asm02.HP0_G0019410 | OG0001070 | chrVII | 397956 | 398372 | CP015054.1 | 1249459 | 1249866 | 82.96296296 | 138 | 135 | 1.022222222 | 1 | 1 | 1 | FIM1_623 | YGL054C | YGL054C | ERV14 | n |
| Kmar_QGN15882.1 | W303.asm02.HP0_G0009190 | OG0001071 | chrIV | 561205 | 562209 | CP015057.1 | 288505 | 289554 | 49.86595174 | 334 | 349 | 0.957020057 | 1 | 1 | 1 | FIM1_2580 | YDR051C | YDR051C | DET1 | n |
| Kmar_QGN17625.1 | W303.asm02.HP0_G0039410 | OG0001072 | chrXIII | 34738 | 36345 | CP015060.1 | 136325 | 137833 | 60.18691589 | 535 | 502 | 1.065737052 | 1 | 1 | 1 | FIM1_4832 | YML115C | YML115C | VAN1 | n |
| Kmar_QGN16241.1 | W303.asm02.HP0_G0030920 | OG0001073 | chrX | 700520 | 705466 | CP015057.1 | 1107440 | 1112398 | 36.2396204 | 1648 | 1652 | 0.997578692 | 1 | 1 | 1 | FIM1_2946 | YJR140C | YJR140C | HIR3 | n |
| Kmar_QGN16154.1 | W303.asm02.HP0_G0047800 | OG0001074 | chrXV | 71640 | 73055 | CP015057.1 | 916504 | 917919 | 55.85774059 | 471 | 471 | 1 | 1 | 1 | 1 | FIM1_2855 | YOL132W | YOL132W | GAS4 | n |
| Kmar_QGN17118.1 | W303.asm02.HP0_G0030730 | OG0001075 | chrX | 657381 | 658916 | CP015059.1 | 274957 | 276474 | 91.5851272 | 511 | 505 | 1.011881188 | 1 | 1 | 1 | FIM1_3849 | YJR121W | YJR121W | ATP2 | n |
| Kmar_QGN17124.1 | W303.asm02.HP0_G0030690 | OG0001076 | chrX | 651781 | 653142 | CP015059.1 | 287136 | 288503 | 66.88596491 | 453 | 455 | 0.995604396 | 1 | 1 | 1 | FIM1_3855 | YJR117W | YJR117W | STE24 | n |
| Kmar_QGN17473.1 | W303.asm02.HP0_G0011220 | OG0001077 | chrIV | 1003365 | 1004378 | CP015059.1 | 996009 | 996926 | 43.1547619 | 337 | 305 | 1.104918033 | 1 | 1 | 1 | FIM1_4209 | YDR265W | YDR265W | PEX10 | n |
| Kmar_QGN16724.1 | W303.asm02.HP0_G0011760 | OG0001078 | chrIV | 1122603 | 1122893 | CP015058.1 | 746954 | 747235 | 56.98924731 | 96 | 93 | 1.032258065 | 1 | 1 | 1 | FIM1_3445 | YDR322C-A | YDR322C-A | TIM11 | n |
| Kmar_QGN14786.1 | W303.asm02.HP0_G0012300 | OG0001079 | chrIV | 1238871 | 1239110 | CP015055.1 | 1230016 | 1230255 | 64.55696203 | 79 | 79 | 1 | 1 | 1 | 1 | FIM1_1456 | YDR379C-A | YDR379C-A | SDH6 | n |
| Kmar_QGN13382.1 | W303.asm02.HP0_G0054350 | OG0001080 | chrXVI | 351622 | 353747 | CP015054.1 | 43519 | 45324 | 55.62805873 | 657 | 601 | 1.093178037 | 1 | 1 | 1 | FIM1_19 | YPL109C | YPL109C | CQD1 | n |
| Kmar_QGN18074.1 | W303.asm02.HP0_G0030550 | OG0001081 | chrX | 632323 | 632787 | CP015061.1 | 156918 | 157382 | 70.77922078 | 154 | 154 | 1 | 1 | 1 | 1 | FIM1_4392 | YJR104C | YJR104C | SOD1 | n |
| Kmar_QGN15219.1 | W303.asm02.HP0_G0021560 | OG0001082 | chrVII | 871478 | 871891 | CP015056.1 | 454326 | 454613 | 50.52631579 | 66 | 95 | 0.694736842 | 1 | 1 | 1 | FIM1_1908 | YGR183C | YGR183C | QCR9 | n |
| Kmar_QGN14989.1 | W303.asm02.HP0_G0050970 | OG0001083 | chrXV | 739615 | 739827 | CP015055.1 | 1633689 | 1633901 | 92.85714286 | 70 | 70 | 1 | 1 | 1 | 1 | FIM1_1670 | YOR210W | YOR210W | RPB10 | n |
| Kmar_QGN15780.1 | W303.asm02.HP0_G0000660 | OG0001084 | chrI | 142940 | 143926 | CP015057.1 | 83938 | 84677 | 19.90291262 | 206 | 116 | 1.775862069 | 1 | 1 | 1 | FIM1_2476 | YAL003W | YAL003W | EFB1 | n |
| Kmar_QGN14250.1 | W303.asm02.HP0_G0019610 | OG0001085 | chrVII | 436319 | 436724 | CP015055.1 | 113154 | 113510 | 77.96610169 | 106 | 118 | 0.898305085 | 1 | 1 | 1 | FIM1_906 | YGL030W | YGL030W | RPL30 | n |
| Kmar_QGN14178.1 | W303.asm02.HP0_G0005030 | OG0001086 | chrIII | 23685 | 23978 | CP015054.1 | 1694304 | 1694552 | 60.82474227 | 97 | 82 | 1.182926829 | 1 | 1 | 1 | FIM1_831 | YCL057C-A | YCL057C-A | MIC10 | n |
| Kmar_QGN18179.1 | W303.asm02.HP0_G0047040 | OG0001087 | chrXIV | 699825 | 700046 | CP015061.1 | 383382 | 383606 | 77.02702703 | 73 | 74 | 0.986486486 | 1 | 1 | 1 | FIM1_4502 | YNR032C-A | YNR032C-A | HUB1 | n |
| Kmar_QGN17644.1 | W303.asm02.HP0_G0029730 | OG0001088 | chrX | 454314 | 454598 | CP015060.1 | 175538 | 175816 | 36.17021277 | 94 | 92 | 1.02173913 | 1 | 1 | 1 | FIM1_4851 | YJR010C-A | YJR010C-A | SPC1 | n |
| Kmar_QGN18215.1 | W303.asm02.HP0_G0036960 | OG0001089 | chrXII | 564987 | 566408 | CP015061.1 | 468851 | 470263 | 52.32067511 | 473 | 470 | 1.006382979 | 1 | 1 | 1 | FIM1_4539 | YLR212C | YLR212C | TUB4 | n |
| Kmar_QGN14704.1 | W303.asm02.HP0_G0014990 | OG0001090 | chrV | 251989 | 252273 | CP015055.1 | 1063692 | 1063967 | 59.57446809 | 94 | 91 | 1.032967033 | 1 | 1 | 1 | FIM1_1371 | YER048W-A | YER048W-A | ISD11 | n |
| Kmar_QGN15688.1 | W303.asm02.HP0_G0017920 | OG0001091 | chrVII | 76938 | 77804 | CP015056.1 | 1472562 | 1473623 | 47.07520891 | 288 | 353 | 0.815864023 | 1 | 1 | 1 | FIM1_2381 | YGL221C | YGL221C | NIF3 | n |
| Kmar_QGN15731.1 | W303.asm02.HP0_G0034190 | OG0001092 | chrXI | 644131 | 644472 | CP015056.1 | 1558565 | 1558825 | 51.13636364 | 88 | 86 | 1.023255814 | 1 | 1 | 1 | FIM1_2425 | YKR095W-A | YKR095W-A | PCC1 | n |
| Kmar_QGN15070.1 | W303.asm02.HP0_G0030140 | OG0001093 | chrX | 557414 | 558469 | CP015056.1 | 138935 | 140050 | 33.87096774 | 351 | 371 | 0.946091644 | 1 | 1 | 1 | FIM1_1755 | YJR060W | YJR060W | CBF1 | n |
| Kmar_QGN13395.1 | W303.asm02.HP0_G0003670 | OG0001094 | chrII | 598501 | 598947 | CP015054.1 | 62507 | 62947 | 64.18918919 | 148 | 146 | 1.01369863 | 1 | 1 | 1 | FIM1_32 | YBR173C | YBR173C | UMP1 | n |
| Kmar_QGN16925.1 | W303.asm02.HP0_G0037250 | OG0001095 | chrXII | 621740 | 622168 | CP015058.1 | 1201059 | 1201505 | 44.21768707 | 142 | 148 | 0.959459459 | 1 | 1 | 1 | FIM1_3652 | YLR245C | YLR245C | CDD1 | n |
| Kmar_QGN15545.1 | W303.asm02.HP0_G0013010 | OG0001096 | chrIV | 1389504 | 1390469 | CP015056.1 | 1159281 | 1160237 | 72.89719626 | 321 | 318 | 1.009433962 | 1 | 1 | 1 | FIM1_2236 | YDR460W | YDR460W | TFB3 | n |
| Kmar_QGN17079.1 | W303.asm02.HP0_G0005960 | OG0001097 | chrIII | 232895 | 233404 | CP015059.1 | 183805 | 184326 | 57.30994152 | 169 | 173 | 0.976878613 | 1 | 1 | 1 | FIM1_3810 | YCR046C | YCR046C | IMG1 | n |
| Kmar_QGN13797.1 | W303.asm02.HP0_G0048010 | OG0001098 | chrXV | 109631 | 110344 | CP015054.1 | 895910 | 896704 | 33.21167883 | 237 | 264 | 0.897727273 | 1 | 1 | 1 | FIM1_443 | YOL110W | YOL110W | SHR5 | n |
| Kmar_QGN17483.1 | W303.asm02.HP0_G0029840 | OG0001099 | chrX | 466489 | 466875 | CP015059.1 | 1010131 | 1010421 | 27.34375 | 128 | 96 | 1.333333333 | 1 | 1 | 1 | FIM1_4219 | YJR022W | YJR022W | LSM8 | n |
| Kmar_QGN18379.1 | W303.asm02.HP0_G0008360 | OG0001100 | chrIV | 373133 | 373420 | CP015061.1 | 808862 | 809149 | 74.73684211 | 95 | 95 | 1 | 1 | 1 | 1 | FIM1_4706 | YDL045W-A | YDL045W-A | MRP10 | n |
| Kmar_QGN16328.1 | W303.asm02.HP0_G0040210 | OG0001101 | chrXIII | 216943 | 217422 | CP015057.1 | 1300035 | 1300514 | 60.37735849 | 159 | 159 | 1 | 1 | 1 | 1 | FIM1_3033 | YML030W | YML030W | RCF1 | n |
| Kmar_QGN18372.1 | W303.asm02.HP0_G0044070 | OG0001102 | chrXIV | 86665 | 87687 | CP015061.1 | 796096 | 797100 | 73.62318841 | 340 | 334 | 1.017964072 | 1 | 1 | 1 | FIM1_4699 | YNL290W | YNL290W | RFC3 | n |
| Kmar_QGN14833.1 | W303.asm02.HP0_G0007840 | OG0001103 | chrIV | 280636 | 282177 | CP015055.1 | 1318290 | 1319540 | 56.42023346 | 513 | 416 | 1.233173077 | 1 | 1 | 1 | FIM1_1504 | YDL101C | YDL101C | DUN1 | n |
| Kmar_QGN16957.1 | W303.asm02.HP0_G0044790 | OG0001104 | chrXIV | 250754 | 251014 | CP015058.1 | 1256104 | 1256364 | 56.81818182 | 86 | 86 | 1 | 1 | 1 | 1 | FIM1_3684 | YNL211C | YNL211C | MRX7 | n |
| Kmar_QGN15654.1 | W303.asm02.HP0_G0050530 | OG0001105 | chrXV | 634157 | 634441 | CP015056.1 | 1399083 | 1399364 | 63.82978723 | 94 | 93 | 1.010752688 | 1 | 1 | 1 | FIM1_2347 | YOR159C | YOR159C | SME1 | n |
| Kmar_QGN17909.1 | W303.asm02.HP0_G0055960 | OG0001106 | chrXVI | 683663 | 684139 | CP015060.1 | 717181 | 717639 | 72.15189873 | 158 | 152 | 1.039473684 | 1 | 1 | 1 | FIM1_5118 | YPR062W | YPR062W | FCY1 | n |
| Kmar_QGN15849.1 | W303.asm02.HP0_G0009300 | OG0001107 | chrIV | 582812 | 583261 | CP015057.1 | 214370 | 214813 | 63.08724832 | 149 | 147 | 1.013605442 | 1 | 1 | 1 | FIM1_2546 | YDR063W | YDR063W | AIM7 | n |
| Kmar_QGN17910.1 | W303.asm02.HP0_G0055970 | OG0001108 | chrXVI | 684310 | 684818 | CP015060.1 | 717819 | 718211 | 36.87943262 | 140 | 130 | 1.076923077 | 1 | 1 | 1 | FIM1_5119 | YPR063C | YPR063C | | n |
| Kmar_QGN17475.1 | W303.asm02.HP0_G0024300 | OG0001109 | chrVIII | 320089 | 320646 | CP015059.1 | 999207 | 999776 | 51.85185185 | 185 | 189 | 0.978835979 | 1 | 1 | 1 | FIM1_4211 | YHR100C | YHR100C | GEP4 | n |
| Kmar_QGN18130.1 | W303.asm02.HP0_G0001020 | OG0001110 | chrII | 45409 | 46071 | CP015061.1 | 287074 | 287691 | 58.55855856 | 220 | 205 | 1.073170732 | 1 | 1 | 1 | FIM1_4451 | YBL093C | YBL093C | ROX3 | n |
| Kmar_QGN17854.1 | W303.asm02.HP0_G0044150 | OG0001111 | chrXIV | 109021 | 109482 | CP015060.1 | 621779 | 622207 | 47.05882353 | 153 | 142 | 1.077464789 | 1 | 1 | 1 | FIM1_5063 | YNL281W | YNL281W | HCH1 | n |
| Kmar_QGN16148.1 | W303.asm02.HP0_G0047850 | OG0001112 | chrXV | 80688 | 81530 | CP015057.1 | 906813 | 907250 | 92.4137931 | 142 | 145 | 0.979310345 | 1 | 1 | 1 | FIM1_2849 | YOL127W | YOL127W | RPL25 | n |
| Kmar_QGN16310.1 | W303.asm02.HP0_G0053300 | OG0001113 | chrXVI | 144262 | 144973 | CP015057.1 | 1256605 | 1257177 | 87.36842105 | 190 | 190 | 1 | 1 | 1 | 1 | FIM1_3015 | YPL218W | YPL218W | SAR1 | n |
| Kmar_QGN15764.1 | W303.asm02.HP0_G0047250 | OG0001114 | chrXIV | 733687 | 735679 | CP015057.1 | 43604 | 44985 | 42.97029703 | 486 | 272 | 1.786764706 | 1 | 1 | 1 | FIM1_2460 | YNR053C | YNR053C | NOG2 | n |
| Kmar_QGN15817.1 | W303.asm02.HP0_G0035300 | OG0001115 | chrXII | 232734 | 232985 | CP015057.1 | 148135 | 148383 | 82.92682927 | 83 | 82 | 1.012195122 | 1 | 1 | 1 | FIM1_2513 | YLR038C | YLR038C | COX12 | n |
| Kmar_QGN16567.1 | W303.asm02.HP0_G0020220 | OG0001117 | chrVII | 557489 | 557752 | CP015058.1 | 394814 | 395178 | 78.16091954 | 87 | 87 | 1 | 1 | 1 | 1 | FIM1_3281 | YGR037C | YGR037C | ACB1 | n |
| Kmar_QGN13952.1 | W303.asm02.HP0_G0040540 | OG0001118 | chrXIII | 274523 | 276058 | CP015054.1 | 1207212 | 1208756 | 49.32301741 | 511 | 514 | 0.994163424 | 1 | 1 | 1 | FIM1_600 | YMR004W | YMR004W | MVP1 | n |
| Kmar_QGN16456.1 | W303.asm02.HP0_G0029090 | OG0001119 | chrX | 306742 | 306999 | CP015058.1 | 150777 | 151043 | 68.53932584 | 85 | 88 | 0.965909091 | 1 | 1 | 1 | FIM1_3169 | YJL062W-A | YJL062W-A | COA3 | n |
| Kmar_QGN18102.1 | W303.asm02.HP0_G0030370 | OG0001120 | chrX | 595843 | 596175 | CP015061.1 | 221777 | 222049 | 46.36363636 | 110 | 90 | 1.222222222 | 1 | 1 | 1 | FIM1_4422 | YJR086W | YJR086W | STE18 | n |
| Kmar_QGN15053.1 | W303.asm02.HP0_G0031290 | OG0001122 | chrXI | 49132 | 49980 | CP015056.1 | 100590 | 101402 | 44.56140351 | 282 | 270 | 1.044444444 | 1 | 1 | 1 | FIM1_1738 | YKL207W | YKL207W | EMC3 | n |
| Kmar_QGN17953.1 | W303.asm02.HP0_G0015670 | OG0001123 | chrV | 389254 | 389817 | CP015060.1 | 815073 | 815597 | 38.29787234 | 187 | 174 | 1.074712644 | 1 | 1 | 1 | FIM1_5162 | YER112W | YER112W | LSM4 | n |
| Kmar_QGN15239.1 | W303.asm02.HP0_G0034870 | OG0001124 | chrXII | 135496 | 137493 | CP015056.1 | 502759 | 504738 | 24.77876106 | 665 | 659 | 1.009104704 | 1 | 1 | 1 | FIM1_1928 | YLL007C | YLL007C | LMO1 | n |
| Kmar_QGN17926.1 | W303.asm02.HP0_G0001310 | OG0001125 | chrII | 111273 | 111687 | CP015060.1 | 752151 | 752459 | 63.30275229 | 109 | 102 | 1.068627451 | 1 | 1 | 1 | FIM1_5135 | YBL059C-A | YBL059C-A | CMC2 | n |
| Kmar_QGN13470.1 | W303.asm02.HP0_G0014280 | OG0001126 | chrV | 118393 | 118656 | CP015054.1 | 204294 | 204557 | 86.20689655 | 87 | 87 | 1 | 1 | 1 | 1 | FIM1_108 | YEL020W-A | YEL020W-A | TIM9 | n |
| Kmar_QGN13967.1 | W303.asm02.HP0_G0019370 | OG0001127 | chrVII | 390939 | 391457 | CP015054.1 | 1235035 | 1235532 | 95.15151515 | 172 | 165 | 1.042424242 | 1 | 1 | 1 | FIM1_616 | YGL058W | YGL058W | RAD6 | n |
| Kmar_QGN14788.1 | W303.asm02.HP0_G0012280 | OG0001129 | chrIV | 1234942 | 1235313 | CP015055.1 | 1234893 | 1235135 | 39.66942149 | 123 | 80 | 1.5375 | 1 | 1 | 1 | FIM1_1458 | YDR378C | YDR378C | LSM6 | n |
| Kmar_QGN15687.1 | W303.asm02.HP0_G0017930 | OG0001130 | chrVII | 77886 | 78248 | CP015056.1 | 1472138 | 1472395 | 36.66666667 | 120 | 85 | 1.411764706 | 1 | 1 | 1 | FIM1_2380 | YGL220W | YGL220W | BOL2 | n |
| Kmar_QGN15504.1 | W303.asm02.HP0_G0006040 | OG0001131 | chrIII | 246435 | 247211 | CP015056.1 | 1075408 | 1076238 | 59.02255639 | 258 | 276 | 0.934782609 | 1 | 1 | 1 | FIM1_2195 | YCR059C | YCR059C | YIH1 | n |
| Kmar_QGN15920.1 | W303.asm02.HP0_G0035890 | OG0001132 | chrXII | 349281 | 349544 | CP015057.1 | 379772 | 380089 | 30.47619048 | 87 | 105 | 0.828571429 | 1 | 1 | 1 | FIM1_2618 | YLR099W-A | YLR099W-A | MIM2 | n |
| Kmar_QGN17404.1 | W303.asm02.HP0_G0033140 | OG0001133 | chrXI | 426602 | 427117 | CP015059.1 | 844925 | 845440 | 87.71929825 | 171 | 171 | 1 | 1 | 1 | 1 | FIM1_4136 | YKL013C | YKL013C | ARC19 | n |
| Kmar_QGN16610.1 | W303.asm02.HP0_G0002250 | OG0001134 | chrII | 310676 | 311611 | CP015058.1 | 493757 | 494626 | 69.45337621 | 311 | 289 | 1.076124567 | 1 | 1 | 1 | FIM1_3326 | YBR039W | YBR039W | ATP3 | n |
| Kmar_QGN17919.1 | W303.asm02.HP0_G0015450 | OG0001135 | chrV | 339619 | 341142 | CP015060.1 | 738518 | 740041 | 75.63850688 | 507 | 507 | 1 | 1 | 1 | 1 | FIM1_5128 | YER090W | YER090W | TRP2 | n |
| Kmar_QGN16492.1 | W303.asm02.HP0_G0052100 | OG0001136 | chrXV | 945376 | 946077 | CP015058.1 | 221932 | 222621 | 65.2360515 | 233 | 229 | 1.017467249 | 1 | 1 | 1 | FIM1_3205 | YOR332W | YOR332W | VMA4 | n |
| Kmar_QGN14437.1 | W303.asm02.HP0_G0018950 | OG0001137 | chrVII | 303507 | 303956 | CP015055.1 | 516584 | 517012 | 63.75838926 | 149 | 142 | 1.049295775 | 1 | 1 | 1 | FIM1_1098 | YGL106W | YGL106W | MLC1 | n |
| Kmar_QGN14695.1 | W303.asm02.HP0_G0021020 | OG0001138 | chrVII | 758459 | 759397 | CP015055.1 | 1051119 | 1052042 | 45.79439252 | 312 | 307 | 1.016286645 | 1 | 1 | 1 | FIM1_1362 | YGR127W | YGR127W | | n |
| Kmar_QGN18158.1 | W303.asm02.HP0_G0015200 | OG0001139 | chrV | 294720 | 296483 | CP015061.1 | 345791 | 347503 | 50.67567568 | 587 | 570 | 1.029824561 | 1 | 1 | 1 | FIM1_4481 | YER068W | YER068W | MOT2 | n |
| Kmar_QGN14214.1 | W303.asm02.HP0_G0009420 | OG0001140 | chrIV | 601432 | 602358 | CP015055.1 | 47098 | 48024 | 83.44155844 | 308 | 308 | 1 | 1 | 1 | 1 | FIM1_870 | YDR075W | YDR075W | PPH3 | n |
| Kmar_QGN15946.1 | W303.asm02.HP0_G0037450 | OG0001141 | chrXII | 674850 | 675494 | CP015057.1 | 447560 | 448204 | 69.62616822 | 214 | 214 | 1 | 1 | 1 | 1 | FIM1_2644 | YLR268W | YLR268W | SEC22 | n |
| Kmar_QGN15473.1 | W303.asm02.HP0_G0051400 | OG0001142 | chrXV | 812731 | 813216 | CP015056.1 | 1020160 | 1020663 | 73.05389222 | 161 | 167 | 0.964071856 | 1 | 1 | 1 | FIM1_2164 | YOR257W | YOR257W | CDC31 | n |
| Kmar_QGN14489.1 | W303.asm02.HP0_G0001650 | OG0001143 | chrII | 172918 | 173333 | CP015055.1 | 618693 | 618980 | 82.10526316 | 95 | 95 | 1 | 1 | 1 | 1 | FIM1_1150 | YBL026W | YBL026W | LSM2 | n |
| Kmar_QGN18304.1 | W303.asm02.HP0_G0048810 | OG0001144 | chrXV | 274467 | 274808 | CP015061.1 | 656284 | 656727 | 38.0952381 | 113 | 147 | 0.768707483 | 1 | 1 | 1 | FIM1_4630 | YOL026C | YOL026C | MIM1 | n |
| Kmar_QGN15482.1 | W303.asm02.HP0_G0051480 | OG0001145 | chrXV | 822176 | 822496 | CP015056.1 | 1033471 | 1033788 | 65.09433962 | 106 | 105 | 1.00952381 | 1 | 1 | 1 | FIM1_2173 | YOR265W | YOR265W | RBL2 | n |
| Kmar_QGN17245.1 | W303.asm02.HP0_G0032900 | OG0001146 | chrXI | 378494 | 379993 | CP015059.1 | 536611 | 538119 | 86.48111332 | 499 | 502 | 0.994023904 | 1 | 1 | 1 | FIM1_3976 | YKL035W | YKL035W | UGP1 | n |
| Kmar_QGN16654.1 | W303.asm02.HP0_G0038310 | OG0001147 | chrXII | 849641 | 849898 | CP015058.1 | 598087 | 598359 | 75.55555556 | 85 | 90 | 0.944444444 | 1 | 1 | 1 | FIM1_3373 | YLR363W-A | YLR363W-A | | n |
| Kmar_QGN18107.1 | W303.asm02.HP0_G0055750 | OG0001148 | chrXVI | 650333 | 651769 | CP015061.1 | 227695 | 229077 | 44.12371134 | 478 | 460 | 1.039130435 | 1 | 1 | 1 | FIM1_4427 | YPR036W | YPR036W | VMA13 | n |
| Kmar_QGN13918.1 | W303.asm02.HP0_G0042650 | OG0001149 | chrXIII | 721539 | 721982 | CP015054.1 | 1143406 | 1143783 | 54.62962963 | 108 | 98 | 1.102040816 | 1 | 1 | 1 | FIM1_566 | YMR225C | YMR225C | MRPL44 | n |
| Kmar_QGN16424.1 | W303.asm02.HP0_G0054230 | OG0001150 | chrXVI | 326893 | 327561 | CP015058.1 | 90240 | 90896 | 33.48214286 | 222 | 218 | 1.018348624 | 1 | 1 | 1 | FIM1_3137 | YPL121C | YPL121C | MEI5 | n |
| Kmar_QGN13887.1 | W303.asm02.HP0_G0022160 | OG0001151 | chrVII | 997389 | 998108 | CP015054.1 | 1070565 | 1071209 | 43.33333333 | 239 | 214 | 1.11682243 | 1 | 1 | 1 | FIM1_534 | YGR247W | YGR247W | CPD1 | n |
| Kmar_QGN13884.1 | W303.asm02.HP0_G0022130 | OG0001152 | chrVII | 990451 | 991734 | CP015054.1 | 1063113 | 1064387 | 68.37209302 | 427 | 424 | 1.007075472 | 1 | 1 | 1 | FIM1_531 | YGR244C | YGR244C | LSC2 | n |
| Kmar_QGN17690.1 | W303.asm02.HP0_G0026440 | OG0001153 | chrIX | 192771 | 193754 | CP015060.1 | 263086 | 264099 | 49.04632153 | 327 | 337 | 0.970326409 | 1 | 1 | 1 | FIM1_4897 | YIL084C | YIL084C | SDS3 | n |

| | | | | | | | | | | | | | | | | | | | | |
|---|---|---|---|---|---|---|---|---|---|---|---|---|---|---|---|---|---|---|---|---|
| Kmar_QGN17701.1 | W303.asm02.HP0_G0046500 | OG0001154 | chrXIV | 597785 | 598003 | CP015060.1 | 279333 | 279551 | 80.55555556 | 72 | 72 | 1 | 1 | 1 | 1 | FIM1_4908 | YNL024C-A | YNL024C-A | KSH1 | n |
| Kmar_QGN14087.1 | W303.asm02.HP0_G0044360 | OG0001155 | chrXIV | 158199 | 158420 | CP015054.1 | 1493233 | 1493451 | 71.23287671 | 73 | 72 | 1.013888889 | 1 | 1 | 1 | FIM1_738 | YNL259C | YNL259C | ATX1 | n |
| Kmar_QGN16554.1 | W303.asm02.HP0_G0039000 | OG0001156 | chrXII | 1010573 | 1010842 | CP015058.1 | 360086 | 360541 | 37.74834437 | 89 | 151 | 0.589403974 | 1 | 1 | 1 | FIM1_3268 | YLR438C-A | YLR438C-A | LSM3 | n |
| Kmar_QGN16373.1 | W303.asm02.HP0_G0014770 | OG0001157 | chrV | 213726 | 214316 | CP015057.1 | 1385252 | 1385830 | 43.93939394 | 196 | 192 | 1.020833333 | 1 | 1 | 1 | FIM1_3080 | YER029C | YER029C | SMB1 | n |
| Kmar_QGN14592.1 | W303.asm02.HP0_G0047100 | OG0001158 | chrXIV | 707634 | 707909 | CP015055.1 | 816316 | 816591 | 84.61538462 | 91 | 91 | 1 | 1 | 1 | 1 | FIM1_1254 | YNR037C | YNR037C | RSM19 | n |
| Kmar_QGN18239.1 | W303.asm02.HP0_G0011730 | OG0001159 | chrIV | 1118346 | 1118564 | CP015061.1 | 523227 | 523445 | 66.66666667 | 72 | 72 | 1 | 1 | 1 | 1 | FIM1_4564 | YDR320C-A | YDR320C-A | DAD4 | n |
| Kmar_QGN18238.1 | W303.asm02.HP0_G0024010 | OG0001160 | chrVIII | 247638 | 247814 | CP015061.1 | 522697 | 522870 | 84.21052632 | 58 | 57 | 1.01754386 | 1 | 1 | 1 | FIM1_4563 | YHR072W-A | YHR072W-A | NOP10 | n |
| Kmar_QGN18053.1 | W303.asm02.HP0_G0038540 | OG0001161 | chrXII | 899736 | 900074 | CP015061.1 | 104752 | 105024 | 33.89830508 | 112 | 90 | 1.244444444 | 1 | 1 | 1 | FIM1_4370 | YLR390W | YLR390W | ECM19 | n |
| Kmar_QGN14035.1 | W303.asm02.HP0_G0008480 | OG0001162 | chrIV | 399383 | 400681 | CP015054.1 | 1380835 | 1382010 | 91.81585678 | 391 | 391 | 1 | 1 | 1 | 1 | FIM1_686 | YDL029W | YDL029W | ARP2 | n |
| Kmar_QGN14848.1 | W303.asm02.HP0_G0039630 | OG0001163 | chrXIII | 80537 | 81289 | CP015055.1 | 1341435 | 1342187 | 88 | 250 | 250 | 1 | 1 | 1 | 1 | FIM1_1520 | YML092C | YML092C | PRE8 | n |
| Kmar_QGN16687.1 | W303.asm02.HP0_G0036850 | OG0001164 | chrXII | 549140 | 549484 | CP015058.1 | 671570 | 671905 | 64.65517241 | 114 | 111 | 1.027027027 | 1 | 1 | 1 | FIM1_3407 | YLR200W | YLR200W | YKE2 | n |
| Kmar_QGN16689.1 | W303.asm02.HP0_G0007210 | OG0001165 | chrIV | 159824 | 160984 | CP015058.1 | 673774 | 674928 | 80.56994819 | 386 | 384 | 1.005208333 | 1 | 1 | 1 | FIM1_3409 | YDL168W | YDL168W | SFA1 | n |
| Kmar_QGN14670.1 | W303.asm02.HP0_G0046670 | OG0001166 | chrXIV | 631720 | 632631 | CP015055.1 | 994834 | 995745 | 85.80858086 | 303 | 303 | 1 | 1 | 1 | 1 | FIM1_1335 | YNL006W | YNL006W | LST8 | n |
| Kmar_QGN14672.1 | W303.asm02.HP0_G0046710 | OG0001167 | chrXIV | 637827 | 638795 | CP015055.1 | 1000566 | 1001531 | 65.12345679 | 322 | 321 | 1.003115265 | 1 | 1 | 1 | FIM1_1337 | YNL002C | YNL002C | RLP7 | n |
| Kmar_QGN16125.1 | W303.asm02.HP0_G0027580 | OG0001168 | chrIX | 410916 | 412037 | CP015057.1 | 861883 | 863001 | 77.21179625 | 373 | 372 | 1.002688172 | 1 | 1 | 1 | FIM1_2826 | YIR034C | YIR034C | LYS1 | n |
| Kmar_QGN16681.1 | W303.asm02.HP0_G0036790 | OG0001169 | chrXII | 540137 | 540664 | CP015058.1 | 661584 | 662111 | 60 | 175 | 175 | 1 | 1 | 1 | 1 | FIM1_3400 | YLR193C | YLR193C | UPS1 | n |
| Kmar_QGN17672.1 | W303.asm02.HP0_G0007690 | OG0001170 | chrIV | 251772 | 253952 | CP015060.1 | 225220 | 227388 | 44.15405777 | 726 | 722 | 1.005540166 | 1 | 1 | 1 | FIM1_4879 | YDL116W | YDL116W | NUP84 | n |
| Kmar_QGN14842.1 | W303.asm02.HP0_G0007880 | OG0001171 | chrIV | 285720 | 287024 | CP015055.1 | 1329716 | 1330990 | 72.11981567 | 434 | 424 | 1.023584906 | 1 | 1 | 1 | FIM1_1513 | YDL097C | YDL097C | RPN6 | n |
| Kmar_QGN18349.1 | W303.asm02.HP0_G0011680 | OG0001172 | chrIV | 1098393 | 1099808 | CP015061.1 | 748720 | 750021 | 60.43478261 | 471 | 433 | 1.087759815 | 1 | 1 | 1 | FIM1_4676 | YDR316W | YDR316W | OMS1 | n |
| Kmar_QGN17556.1 | W303.asm02.HP0_G0018610 | OG0001173 | chrVII | 231185 | 232426 | CP015059.1 | 1157822 | 1159081 | 60.33254157 | 413 | 419 | 0.985680191 | 1 | 1 | 1 | FIM1_4292 | YGL143C | YGL143C | MRF1 | n |
| Kmar_QGN18351.1 | W303.asm02.HP0_G0023980 | OG0001174 | chrVIII | 240712 | 242211 | CP015061.1 | 751476 | 752984 | 50.19455253 | 499 | 502 | 0.994023904 | 1 | 1 | 1 | FIM1_4678 | YHR070W | YHR070W | TRM5 | n |
| Kmar_QGN15241.1 | W303.asm02.HP0_G0034890 | OG0001175 | chrXII | 139132 | 141738 | CP015056.1 | 506738 | 509527 | 33.72340426 | 868 | 929 | 0.934337998 | 1 | 1 | 1 | FIM1_1930 | YLL005C | YLL005C | SPO75 | n |
| Kmar_QGN18240.1 | W303.asm02.HP0_G0024020 | OG0001176 | chrVIII | 248556 | 251546 | CP015061.1 | 524080 | 527154 | 37.67705382 | 996 | 1024 | 0.97265625 | 1 | 1 | 1 | FIM1_4565 | YHR073W | YHR073W | OSH3 | n |
| Kmar_QGN13718.1 | W303.asm02.HP0_G0022300 | OG0001177 | chrVII | 1027236 | 1029665 | CP015054.1 | 726649 | 729141 | 40.35294118 | 809 | 830 | 0.974698795 | 1 | 1 | 1 | FIM1_362 | YGR261C | YGR261C | APL6 | n |
| Kmar_QGN13733.1 | W303.asm02.HP0_G0025030 | OG0001178 | chrVIII | 458978 | 459547 | CP015054.1 | 763780 | 764244 | 34.39153439 | 189 | 154 | 1.227272727 | 1 | 1 | 1 | FIM1_377 | YHR175W | YHR175W | CTR2 | n |
| Kmar_QGN14155.1 | W303.asm02.HP0_G0013540 | OG0001179 | chrIV | 1482932 | 1484050 | CP015054.1 | 1647626 | 1648798 | 44.36090226 | 372 | 390 | 0.953846154 | 1 | 1 | 1 | FIM1_808 | YDR517W | YDR517W | GRH1 | n |
| Kmar_QGN16800.1 | W303.asm02.HP0_G0020720 | OG0001180 | chrVII | 674141 | 674812 | CP015058.1 | 920617 | 921294 | 51.7699115 | 223 | 225 | 0.991111111 | 1 | 1 | 1 | FIM1_3524 | YGR095C | YGR095C | RRP46 | n |
| Kmar_QGN16796.1 | W303.asm02.HP0_G0020680 | OG0001181 | chrVII | 664809 | 666293 | CP015058.1 | 908130 | 909350 | 29.27835052 | 494 | 406 | 1.216748768 | 1 | 1 | 1 | FIM1_3519 | YGR091W | YGR091W | PRP31 | n |
| Kmar_QGN15584.1 | W303.asm02.HP0_G0003440 | OG0001182 | chrII | 561805 | 562755 | CP015056.1 | 1242406 | 1243329 | 51.41955836 | 316 | 307 | 1.029315961 | 1 | 1 | 1 | FIM1_2276 | YBR151W | YBR151W | APD1 | n |
| Kmar_QGN18263.1 | W303.asm02.HP0_G0056670 | OG0001183 | chrXVI | 829956 | 830708 | CP015061.1 | 577051 | 577824 | 56.42023346 | 250 | 257 | 0.972762646 | 1 | 1 | 1 | FIM1_4589 | YPR143W | YPR143W | RRP15 | n |
| Kmar_QGN17561.1 | W303.asm02.HP0_G0018630 | OG0001184 | chrVII | 234821 | 237553 | CP015059.1 | 1165048 | 1167765 | 34.77789816 | 910 | 905 | 1.005524862 | 1 | 1 | 1 | FIM1_4297 | YGL141W | YGL141W | HUL5 | n |
| Kmar_QGN15202.1 | W303.asm02.HP0_G0024150 | OG0001185 | chrVIII | 282737 | 283741 | CP015056.1 | 418688 | 419833 | 36.97916667 | 334 | 381 | 0.87664042 | 1 | 1 | 1 | FIM1_1890 | YHR085W | YHR085W | IPI1 | n |
| Kmar_QGN16147.1 | W303.asm02.HP0_G0043940 | OG0001186 | chrXIV | 59329 | 60225 | CP015057.1 | 905806 | 906567 | 40.93959732 | 298 | 253 | 1.177865613 | 1 | 1 | 1 | FIM1_2848 | YNL305C | YNL305C | BXI1 | n |
| Kmar_QGN16773.1 | W303.asm02.HP0_G0027150 | OG0001187 | chrIX | 334049 | 334711 | CP015058.1 | 859731 | 860345 | 29.59641256 | 220 | 204 | 1.078431373 | 1 | 1 | 1 | FIM1_3495 | YIL007C | YIL007C | NAS2 | n |
| Kmar_QGN16799.1 | W303.asm02.HP0_G0020710 | OG0001188 | chrVII | 670654 | 673968 | CP015058.1 | 917176 | 920481 | 79.74683544 | 1104 | 1101 | 1.002724796 | 1 | 1 | 1 | FIM1_3523 | YGR094W | YGR094W | VAS1 | n |
| Kmar_QGN18407.1 | W303.asm02.HP0_G0033360 | OG0001189 | chrXI | 463914 | 466616 | CP015061.1 | 861816 | 864602 | 63.38329764 | 900 | 928 | 0.969827586 | 1 | 1 | 1 | FIM1_4734 | YKR009C | YKR009C | FOX2 | n |
| Kmar_QGN18385.1 | W303.asm02.HP0_G0055620 | OG0001190 | chrXVI | 615324 | 616529 | CP015061.1 | 821792 | 822919 | 36.31840796 | 401 | 375 | 1.069333333 | 1 | 1 | 1 | FIM1_4712 | YPR023C | YPR023C | EAF3 | n |
| Kmar_QGN14585.1 | W303.asm02.HP0_G0007920 | OG0001191 | chrIV | 293722 | 295089 | CP015055.1 | 802570 | 803829 | 36.60130719 | 455 | 419 | 1.085918854 | 1 | 1 | 1 | FIM1_1247 | YDL091C | YDL091C | UBX3 | n |
| Kmar_QGN17643.1 | W303.asm02.HP0_G0021660 | OG0001192 | chrVII | 898488 | 900290 | CP015060.1 | 173464 | 175272 | 54.58937198 | 600 | 602 | 0.996677741 | 1 | 1 | 1 | FIM1_4850 | YGR194C | YGR194C | XKS1 | n |
| Kmar_QGN18178.1 | W303.asm02.HP0_G0047050 | OG0001193 | chrXIV | 700217 | 702580 | CP015061.1 | 380858 | 383113 | 53.33333333 | 787 | 751 | 1.047936085 | 1 | 1 | 1 | FIM1_4501 | YNR033W | YNR033W | ABZ1 | n |
| Kmar_QGN17221.1 | W303.asm02.HP0_G0004770 | OG0001194 | chrII | 803036 | 805753 | CP015059.1 | 479801 | 482458 | 39.34588702 | 905 | 885 | 1.02259887 | 1 | 1 | 1 | FIM1_3952 | YBR289W | YBR289W | SNF5 | n |
| Kmar_QGN15928.1 | W303.asm02.HP0_G0035920 | OG0001195 | chrXII | 351944 | 353896 | CP015057.1 | 395347 | 397356 | 62.81481481 | 650 | 669 | 0.971599402 | 1 | 1 | 1 | FIM1_2626 | YLR103C | YLR103C | CDC45 | n |
| Kmar_QGN18061.1 | W303.asm02.HP0_G0008240 | OG0001196 | chrIV | 345997 | 351369 | CP015061.1 | 124517 | 129790 | 38.26955075 | 1790 | 1757 | 1.018782015 | 1 | 1 | 1 | FIM1_4379 | YDL058W | YDL058W | USO1 | n |
| Kmar_QGN18060.1 | W303.asm02.HP0_G0008230 | OG0001197 | chrIV | 344569 | 345285 | CP015061.1 | 123147 | 123776 | 47.05882353 | 238 | 209 | 1.138755981 | 1 | 1 | 1 | FIM1_4378 | YDL059C | YDL059C | RAD59 | n |
| Kmar_QGN18059.1 | W303.asm02.HP0_G0038480 | OG0001198 | chrXII | 885521 | 889570 | CP015061.1 | 119071 | 123048 | 67.7037037 | 1349 | 1325 | 1.018113208 | 1 | 1 | 1 | FIM1_4377 | YLR384C | YLR384C | IKI3 | n |
| Kmar_QGN17179.1 | W303.asm02.HP0_G0055120 | OG0001199 | chrXVI | 504455 | 505651 | CP015059.1 | 397380 | 398576 | 76.94235589 | 398 | 398 | 1 | 1 | 1 | 1 | FIM1_3910 | YPL028W | YPL028W | ERG10 | n |
| Kmar_QGN17169.1 | W303.asm02.HP0_G0054990 | OG0001200 | chrXVI | 481478 | 482101 | CP015059.1 | 376440 | 377042 | 40.29126214 | 207 | 200 | 1.035 | 1 | 1 | 1 | FIM1_3900 | YPL041C | YPL041C | MRX11 | n |
| Kmar_QGN15304.1 | W303.asm02.HP0_G0009600 | OG0001201 | chrIV | 634148 | 634877 | CP015056.1 | 651292 | 651774 | 83.125 | 153 | 160 | 0.95625 | 1 | 1 | 1 | FIM1_1993 | YDR092W | YDR092W | UBC13 | n |
| Kmar_QGN18366.1 | W303.asm02.HP0_G0049850 | OG0001202 | chrXV | 481322 | 482485 | CP015061.1 | 786555 | 787787 | 35.25179856 | 387 | 410 | 0.943902439 | 1 | 0 | 1 | FIM1_4693 | YOR084W | YOR084W | LPX1 | n |
| Kmar_QGN15177.1 | W303.asm02.HP0_G0028850 | OG0001203 | chrX | 252773 | 255067 | CP015056.1 | 370156 | 372240 | 29.98729352 | 764 | 694 | 1.100864553 | 1 | 1 | 1 | FIM1_1865 | YJL090C | YJL090C | DPB11 | n |
| Kmar_QGN14973.1 | W303.asm02.HP0_G0051090 | OG0001204 | chrXV | 761077 | 761955 | CP015055.1 | 1601831 | 1602571 | 47.78156997 | 292 | 246 | 1.18699187 | 1 | 1 | 1 | FIM1_1651 | YOR223W | YOR223W | DSC3 | n |
| Kmar_QGN13982.1 | W303.asm02.HP0_G0050780 | OG0001205 | chrXV | 687062 | 690475 | CP015054.1 | 1266192 | 1270043 | 32.56880734 | 1137 | 1283 | 0.886204209 | 1 | 1 | 1 | FIM1_631 | YOR188W | YOR188W | MSB1 | n |
| Kmar_QGN17681.1 | W303.asm02.HP0_G0018510 | OG0001206 | chrVII | 210549 | 211679 | CP015060.1 | 248527 | 249666 | 44.76439791 | 376 | 379 | 0.992084433 | 1 | 1 | 1 | FIM1_4888 | YGL155W | YGL155W | CDC43 | n |
| Kmar_QGN17680.1 | W303.asm02.HP0_G0007670 | OG0001207 | chrIV | 246895 | 247818 | CP015060.1 | 247573 | 248457 | 55.70032573 | 307 | 294 | 1.044217687 | 1 | 1 | 1 | FIM1_4887 | YDL119C | YDL119C | HEM25 | n |
| Kmar_QGN17678.1 | W303.asm02.HP0_G0018530 | OG0001208 | chrVII | 212741 | 213766 | CP015060.1 | 245345 | 246397 | 41.24293785 | 341 | 350 | 0.974285714 | 1 | 1 | 1 | FIM1_4885 | YGL153W | YGL153W | PEX14 | n |
| Kmar_QGN13942.1 | W303.asm02.HP0_G0043230 | OG0001209 | chrXIII | 845109 | 846917 | CP015054.1 | 1186484 | 1188259 | 44.81054366 | 602 | 591 | 1.018612521 | 1 | 1 | 1 | FIM1_590 | YMR284W | YMR284W | YKU70 | n |
| Kmar_QGN17673.1 | W303.asm02.HP0_G0018560 | OG0001210 | chrVII | 222867 | 223997 | CP015060.1 | 228030 | 229160 | 86.17021277 | 376 | 376 | 1 | 1 | 1 | 1 | FIM1_4880 | YGL148W | YGL148W | ARO2 | n |
| Kmar_QGN16072.1 | W303.asm02.HP0_G0040860 | OG0001211 | chrXIII | 342359 | 344023 | CP015057.1 | 748387 | 749934 | 27.35346359 | 554 | 515 | 1.075728155 | 1 | 1 | 1 | FIM1_2772 | YMR036C | YMR036C | MIH1 | n |
| Kmar_QGN16891.1 | W303.asm02.HP0_G0010250 | OG0001212 | chrIV | 790020 | 792935 | CP015058.1 | 1112460 | 1115225 | 44.35401831 | 971 | 921 | 1.054288817 | 1 | 1 | 1 | FIM1_3618 | YDR166C | YDR166C | SEC5 | n |
| Kmar_QGN14171.1 | W303.asm02.HP0_G0005070 | OG0001213 | chrIII | 31103 | 33628 | CP015054.1 | 1679666 | 1682173 | 77.8173191 | 841 | 835 | 1.007185629 | 1 | 1 | 1 | FIM1_824 | YCL054W | YCL054W | SPB1 | n |
| Kmar_QGN17676.1 | W303.asm02.HP0_G0018550 | OG0001214 | chrVII | 217572 | 222041 | CP015060.1 | 237242 | 241684 | 53.33333333 | 1489 | 1480 | 1.006081081 | 1 | 1 | 1 | FIM1_4883 | YGL150C | YGL150C | INO80 | n |
| Kmar_QGN13920.1 | W303.asm02.HP0_G0012530 | OG0001215 | chrIV | 1280327 | 1281937 | CP015054.1 | 1145434 | 1147023 | 54.88029466 | 536 | 529 | 1.013232514 | 1 | 1 | 1 | FIM1_568 | YDR403W | YDR403W | DIT1 | n |
| Kmar_QGN16611.1 | W303.asm02.HP0_G0008860 | OG0001216 | chrIV | 478774 | 481926 | CP015058.1 | 496249 | 498969 | 36.94207028 | 1050 | 906 | 1.158940397 | 1 | 1 | 1 | FIM1_3327 | YDR017C | YDR017C | KCS1 | n |
| Kmar_QGN16046.1 | W303.asm02.HP0_G0036580 | OG0001217 | chrXII | 500840 | 501310 | CP015057.1 | 696142 | 696612 | 68.58974359 | 156 | 156 | 1 | 1 | 1 | 1 | FIM1_2746 | YLR170C | YLR170C | APS1 | n |
| Kmar_QGN13629.1 | W303.asm02.HP0_G0045420 | OG0001218 | chrXIV | 361757 | 366550 | CP015054.1 | 548111 | 552784 | 32.50466128 | 1597 | 1557 | 1.02569043 | 1 | 1 | 1 | FIM1_271 | YNL139C | YNL139C | THO2 | n |
| Kmar_QGN16226.1 | W303.asm02.HP0_G0057080 | OG0001219 | chrXVI | 915565 | 916854 | CP015057.1 | 1075712 | 1076992 | 51.34529148 | 429 | 426 | 1.007042254 | 1 | 1 | 1 | FIM1_2930 | YPR186C | YPR186C | PZF1 | n |
| Kmar_QGN18347.1 | W303.asm02.HP0_G0011670 | OG0001220 | chrIV | 1097371 | 1098216 | CP015061.1 | 745935 | 746774 | 31.75675676 | 281 | 279 | 1.007168459 | 1 | 1 | 1 | FIM1_4674 | YDR315C | YDR315C | IPK1 | n |
| Kmar_QGN16863.1 | W303.asm02.HP0_G0010110 | OG0001221 | chrIV | 760536 | 767588 | CP015058.1 | 1056857 | 1065277 | 24.49336129 | 2350 | 2806 | 0.837491091 | 1 | 1 | 1 | FIM1_3589 | YDR150W | YDR150W | NUM1 | n |
| Kmar_QGN14242.1 | W303.asm02.HP0_G0019650 | OG0001222 | chrVII | 443498 | 445621 | CP015055.1 | 95606 | 97735 | 78.3592645 | 707 | 709 | 0.997179126 | 1 | 1 | 1 | FIM1_898 | YGL026C | YGL026C | TRP5 | n |
| Kmar_QGN17983.1 | W303.asm02.HP0_G0022790 | OG0001223 | chrVIII | 34003 | 36132 | CP015060.1 | 885848 | 888049 | 59.73333333 | 709 | 733 | 0.967257844 | 1 | 1 | 1 | FIM1_5192 | YHL032C | YHL032C | GUT1 | n |
| Kmar_QGN16193.1 | W303.asm02.HP0_G0039080 | OG0001224 | chrXII | 1023523 | 1024560 | CP015057.1 | 997364 | 998407 | 77.80979827 | 345 | 347 | 0.994236311 | 1 | 1 | 1 | FIM1_2895 | YLR447C | YLR447C | VMA6 | n |
| Kmar_QGN16388.1 | W303.asm02.HP0_G0052430 | OG0001225 | chrXV | 1029124 | 1030329 | CP015058.1 | 6716 | 7837 | 44.41687345 | 401 | 373 | 1.075067024 | 1 | 1 | 1 | FIM1_3101 | YOR368W | YOR368W | RAD17 | n |
| Kmar_QGN16122.1 | W303.asm02.HP0_G0033010 | OG0001227 | chrXI | 401618 | 402232 | CP015057.1 | 855999 | 856910 | 48.50498339 | 204 | 303 | 0.673267327 | 1 | 1 | 1 | FIM1_2823 | YKL024C | YKL024C | URA6 | n |
| Kmar_QGN17478.1 | W303.asm02.HP0_G0029810 | OG0001228 | chrX | 463284 | 463796 | CP015059.1 | 1002354 | 1002845 | 68.23529412 | 170 | 163 | 1.042944785 | 1 | 1 | 1 | FIM1_4214 | YJR017C | YJR017C | ESS1 | n |
| Kmar_QGN17167.1 | W303.asm02.HP0_G0054970 | OG0001229 | chrXVI | 476299 | 478356 | CP015059.1 | 372156 | 374204 | 63.63636364 | 685 | 682 | 1.004398827 | 1 | 1 | 1 | FIM1_3898 | YPL043W | YPL043W | NOP4 | n |
| Kmar_QGN13539.1 | W303.asm02.HP0_G0008340 | OG0001230 | chrIV | 370656 | 371591 | CP015054.1 | 362238 | 363167 | 93.56913183 | 311 | 309 | 1.006472492 | 1 | 1 | 1 | FIM1_179 | YDL047W | YDL047W | SIT4 | n |
| Kmar_QGN14699.1 | W303.asm02.HP0_G0015010 | OG0001231 | chrV | 255242 | 255694 | CP015055.1 | 1056674 | 1057144 | 41.33333333 | 150 | 156 | 0.961538462 | 1 | 1 | 1 | FIM1_1366 | YER050C | YER050C | RSM18 | n |
| Kmar_QGN17213.1 | W303.asm02.HP0_G0019060 | OG0001232 | chrVII | 331838 | 334591 | CP015059.1 | 461202 | 463742 | 22.6927253 | 917 | 846 | 1.08392435 | 1 | 1 | 1 | FIM1_3944 | YGL093W | YGL093W | SPC105 | n |
| Kmar_QGN15317.1 | W303.asm02.HP0_G0037760 | OG0001233 | chrXII | 733418 | 739120 | CP015056.1 | 688549 | 694272 | 48.49428868 | 1900 | 1907 | 0.996329313 | 1 | 1 | 1 | FIM1_2006 | YLR305C | YLR305C | STT4 | n |
| Kmar_QGN13603.1 | W303.asm02.HP0_G0055540 | OG0001234 | chrXVI | 598830 | 599567 | CP015054.1 | 502159 | 502896 | 93.87755102 | 245 | 245 | 1 | 1 | 1 | 1 | FIM1_245 | YPR016C | YPR016C | TIF6 | n |
| Kmar_QGN17443.1 | W303.asm02.HP0_G0008710 | OG0001235 | chrIV | 447643 | 448935 | CP015059.1 | 931277 | 932644 | 72.80701754 | 430 | 455 | 0.945054945 | 1 | 1 | 1 | FIM1_4176 | YDL001W | YDL001W | RMD1 | n |
| Kmar_QGN17441.1 | W303.asm02.HP0_G0001870 | OG0001236 | chrII | 229975 | 237456 | CP015059.1 | 922279 | 929769 | 50.09972078 | 2493 | 2496 | 0.998798077 | 1 | 1 | 1 | FIM1_4174 | YBL004W | YBL004W | UTP20 | n |
| Kmar_QGN16791.1 | W303.asm02.HP0_G0020660 | OG0001237 | chrVII | 655429 | 658239 | CP015058.1 | 896133 | 899105 | 34.36893204 | 936 | 990 | 0.945454545 | 1 | 1 | 1 | FIM1_3514 | YGR089W | YGR089W | NNF2 | n |
| Kmar_QGN14869.1 | W303.asm02.HP0_G0029630 | OG0001238 | chrX | 434514 | 436295 | CP015055.1 | 1391716 | 1393575 | 50 | 593 | 619 | 0.957996769 | 1 | 1 | 1 | FIM1_1542 | YJR002W | YJR002W | MPP10 | n |
| Kmar_QGN15114.1 | W303.asm02.HP0_G0032050 | OG0001239 | chrXI | 215481 | 217364 | CP015056.1 | 231431 | 233329 | 60.12658228 | 627 | 632 | 0.992088608 | 1 | 1 | 1 | FIM1_1801 | YKL125W | YKL125W | RRN3 | n |
| Kmar_QGN14621.1 | W303.asm02.HP0_G0015990 | OG0001240 | chrV | 477251 | 479989 | CP015055.1 | 877094 | 879457 | 38.46994536 | 912 | 787 | 1.158831004 | 1 | 1 | 1 | FIM1_1285 | YER151C | YER151C | UBP3 | n |
| Kmar_QGN16958.1 | W303.asm02.HP0_G0044780 | OG0001241 | chrXIV | 248160 | 250508 | CP015058.1 | 1256793 | 1257752 | 14.65053763 | 782 | 319 | 2.451410658 | 1 | 1 | 1 | FIM1_3685 | YNL212W | YNL212W | VID27 | n |
| Kmar_QGN14512.1 | W303.asm02.HP0_G0051940 | OG0001242 | chrXV | 906480 | 907520 | CP015055.1 | 663494 | 664549 | 26.96335079 | 346 | 351 | 0.985754986 | 1 | 1 | 1 | FIM1_1173 | YOR315W | YOR315W | SFG1 | n |
| Kmar_QGN14141.1 | W303.asm02.HP0_G0013490 | OG0001243 | chrIV | 1475711 | 1476112 | CP015054.1 | 1626284 | 1626670 | 60.15625 | 133 | 128 | 1.0390625 | 1 | 1 | 1 | FIM1_794 | YDR511W | YDR511W | SDH7 | n |
| Kmar_QGN13828.1 | W303.asm02.HP0_G0031630 | OG0001244 | chrXI | 131804 | 132220 | CP015054.1 | 954790 | 955206 | 76.8115942 | 138 | 138 | 1 | 1 | 1 | 1 | FIM1_474 | YKL170W | YKL170W | MRPL38 | n |
| Kmar_QGN17076.1 | W303.asm02.HP0_G0026460 | OG0001245 | chrIX | 202994 | 205198 | CP015059.1 | 178628 | 180859 | 81.74496644 | 734 | 743 | 0.987886945 | 1 | 1 | 1 | FIM1_3807 | YIL078W | YIL078W | THS1 | n |
| Kmar_QGN15896.1 | W303.asm02.HP0_G0035770 | OG0001246 | chrXII | 324056 | 325900 | CP015057.1 | 331459 | 333201 | 52.8363047 | 614 | 580 | 1.05862069 | 1 | 1 | 1 | FIM1_2594 | YLR088W | YLR088W | GAA1 | n |
| Kmar_QGN15903.1 | W303.asm02.HP0_G0009790 | OG0001247 | chrIV | 687631 | 688488 | CP015057.1 | 343634 | 344434 | 52.63157895 | 285 | 266 | 1.071428571 | 1 | 1 | 1 | FIM1_2601 | YDR116C | YDR116C | MRPL1 | n |
| Kmar_QGN17335.1 | W303.asm02.HP0_G0045720 | OG0001248 | chrXIV | 421753 | 422433 | CP015059.1 | 716385 | 717059 | 60.35242291 | 226 | 224 | 1.008928571 | 1 | 1 | 1 | FIM1_4067 | YNL107W | YNL107W | YAF9 | n |
| Kmar_QGN14608.1 | W303.asm02.HP0_G0004380 | OG0001249 | chrII | 738162 | 739820 | CP015055.1 | 848796 | 850454 | 74.14104882 | 552 | 552 | 1 | 1 | 1 | 1 | FIM1_1271 | YBR248C | YBR248C | HIS7 | n |
| Kmar_QGN17592.1 | W303.asm02.HP0_G0052820 | OG0001250 | chrXVI | 35788 | 38397 | CP015060.1 | 60542 | 63319 | 40.02145923 | 869 | 925 | 0.939459459 | 1 | 1 | 1 | FIM1_4799 | YPL268W | YPL268W | PLC1 | n |
| Kmar_QGN14606.1 | W303.asm02.HP0_G0004370 | OG0001251 | chrII | 736359 | 737810 | CP015055.1 | 845201 | 846622 | 67.56198347 | 483 | 473 | 1.021141649 | 1 | 1 | 1 | FIM1_1269 | YBR247C | YBR247C | ENP1 | n |
| Kmar_QGN17667.1 | W303.asm02.HP0_G0007730 | OG0001253 | chrIV | 259121 | 263431 | CP015060.1 | 214959 | 219179 | 38.53083853 | 1436 | 1406 | 1.021337127 | 1 | 1 | 1 | FIM1_4874 | YDL112W | YDL112W | TRM3 | n |
| Kmar_QGN17666.1 | W303.asm02.HP0_G0007740 | OG0001254 | chrIV | 263519 | 264316 | CP015060.1 | 214077 | 214871 | 61.88679245 | 265 | 264 | 1.003787879 | 1 | 1 | 1 | FIM1_4873 | YDL111C | YDL111C | RRP42 | n |

| | | | | | | | | | | | | | | | | | | | | |
|---|---|---|---|---|---|---|---|---|---|---|---|---|---|---|---|---|---|---|---|---|
| Kmar_QGN14582.1 | W303.asm02.HP0_G0007940 | OG0001255 | chrIV | 297150 | 298604 | CP015055.1 | 796831 | 798462 | 29.52898551 | 484 | 543 | 0.891344383 | 1 | 1 | 1 | FIM1_1244 | YDL089W | YDL089W | NUR1 | n |
| Kmar_QGN14993.1 | W303.asm02.HP0_G0050950 | OG0001256 | chrXV | 735220 | 737472 | CP015055.1 | 1642667 | 1645354 | 33.81057269 | 750 | 895 | 0.837988827 | 1 | 1 | 1 | FIM1_1674 | YOR208W | YOR208W | PTP2 | n |
| Kmar_QGN14584.1 | W303.asm02.HP0_G0007930 | OG0001257 | chrIV | 295364 | 296659 | CP015055.1 | 800952 | 802241 | 48.96073903 | 431 | 429 | 1.004662005 | 1 | 1 | 1 | FIM1_1246 | YDL090C | YDL090C | RAM1 | n |
| Kmar_QGN18132.1 | W303.asm02.HP0_G0001050 | OG0001258 | chrII | 48516 | 49781 | CP015061.1 | 292829 | 294112 | 70.62937063 | 421 | 427 | 0.985948478 | 1 | 1 | 1 | FIM1_4454 | YBL091C | YBL091C | MAP2 | n |
| Kmar_QGN15570.1 | W303.asm02.HP0_G0042880 | OG0001259 | chrXIII | 769856 | 774544 | CP015056.1 | 1207679 | 1212241 | 33.0994898 | 1562 | 1520 | 1.027631579 | 1 | 1 | 1 | FIM1_2261 | YMR247C | YMR247C | RKR1 | n |
| Kmar_QGN16293.1 | W303.asm02.HP0_G0019300 | OG0001260 | chrVII | 376713 | 378224 | CP015057.1 | 1220550 | 1222079 | 45.98825832 | 503 | 509 | 0.988212181 | 1 | 1 | 1 | FIM1_2998 | YGL065C | YGL065C | ALG2 | n |
| Kmar_QGN13869.1 | W303.asm02.HP0_G0028390 | OG0001261 | chrX | 140489 | 141154 | CP015054.1 | 1026668 | 1027270 | 66.06334842 | 221 | 200 | 1.105 | 1 | 1 | 1 | FIM1_515 | YJL140W | YJL140W | RPB4 | n |
| Kmar_QGN13867.1 | W303.asm02.HP0_G0028370 | OG0001262 | chrX | 136629 | 137105 | CP015054.1 | 1023258 | 1023722 | 91.13924051 | 158 | 154 | 1.025974026 | 1 | 1 | 1 | FIM1_513 | YJL143W | YJL143W | TIM17 | n |
| Kmar_QGN13834.1 | W303.asm02.HP0_G0031590 | OG0001263 | chrXI | 121550 | 123406 | CP015054.1 | 965647 | 967308 | 54.87364621 | 618 | 553 | 1.117540687 | 1 | 1 | 1 | FIM1_480 | YKL174C | YKL174C | TPO5 | n |
| Kmar_QGN13871.1 | W303.asm02.HP0_G0033830 | OG0001264 | chrXI | 566181 | 567005 | CP015054.1 | 1028917 | 1029729 | 56.36363636 | 274 | 270 | 1.014814815 | 1 | 1 | 1 | FIM1_517 | YKR060W | YKR060W | UTP30 | n |
| Kmar_QGN16979.1 | W303.asm02.HP0_G0044630 | OG0001265 | chrXIV | 219833 | 220897 | CP015058.1 | 1299360 | 1300565 | 67.24565757 | 354 | 401 | 0.882793017 | 1 | 1 | 1 | FIM1_3706 | YNL229C | YNL229C | URE2 | n |
| Kmar_QGN16417.1 | W303.asm02.HP0_G0050890 | OG0001266 | chrXV | 721765 | 723003 | CP015058.1 | 74966 | 76165 | 43.85542169 | 412 | 399 | 1.032581454 | 1 | 1 | 1 | FIM1_3130 | YOR201C | YOR201C | MRM1 | n |
| Kmar_QGN16418.1 | W303.asm02.HP0_G0050900 | OG0001267 | chrXV | 723241 | 723903 | CP015058.1 | 76510 | 77175 | 83.33333333 | 220 | 221 | 0.995475113 | 1 | 1 | 1 | FIM1_3131 | YOR202W | YOR202W | HIS3 | n |
| Kmar_QGN14544.1 | W303.asm02.HP0_G0000610 | OG0001268 | chrI | 134950 | 136431 | CP015055.1 | 728128 | 729525 | 44.12955466 | 493 | 465 | 1.060215054 | 1 | 1 | 1 | FIM1_1206 | YAL010C | YAL010C | MDM10 | n |
| Kmar_QGN16419.1 | W303.asm02.HP0_G0054270 | OG0001269 | chrXVI | 332986 | 334020 | CP015058.1 | 77380 | 78441 | 39.83516484 | 344 | 353 | 0.974504249 | 1 | 1 | 1 | FIM1_3132 | YPL118W | YPL118W | MRP51 | n |
| Kmar_QGN16695.1 | W303.asm02.HP0_G0007250 | OG0001270 | chrIV | 165208 | 167626 | CP015058.1 | 682431 | 684569 | 57.69230769 | 798 | 712 | 1.120786517 | 1 | 1 | 1 | FIM1_3415 | YDL164C | YDL164C | CDC9 | n |
| Kmar_QGN17925.1 | W303.asm02.HP0_G0015500 | OG0001271 | chrV | 349582 | 350070 | CP015060.1 | 751338 | 751860 | 40.78947368 | 137 | 152 | 0.901315789 | 1 | 1 | 1 | FIM1_5134 | YER093C-A | YER093C-A | AIM11 | n |
| Kmar_QGN17018.1 | W303.asm02.HP0_G0026040 | OG0001272 | chrIX | 111723 | 114767 | CP015059.1 | 43630 | 46692 | 78.84427032 | 1014 | 1020 | 0.994117647 | 1 | 1 | 1 | FIM1_3747 | YIL125W | YIL125W | KGD1 | n |
| Kmar_QGN17289.1 | W303.asm02.HP0_G0002160 | OG0001273 | chrII | 291465 | 292838 | CP015059.1 | 627913 | 629256 | 67.46724891 | 457 | 447 | 1.022371365 | 1 | 1 | 1 | FIM1_4020 | YBR029C | YBR029C | CDS1 | n |
| Kmar_QGN14068.1 | W303.asm02.HP0_G0044490 | OG0001274 | chrXIV | 187725 | 188051 | CP015054.1 | 1456931 | 1457257 | 92.59259259 | 108 | 108 | 1 | 1 | 1 | 1 | FIM1_719 | YNL244C | YNL244C | SUI1 | n |
| Kmar_QGN18195.1 | W303.asm02.HP0_G0003280 | OG0001275 | chrII | 521140 | 521592 | CP015061.1 | 420183 | 420581 | 83.45864662 | 150 | 132 | 1.136363636 | 1 | 1 | 1 | FIM1_4519 | YBR135W | YBR135W | CKS1 | n |
| Kmar_QGN18194.1 | W303.asm02.HP0_G0003290 | OG0001276 | chrII | 521954 | 529060 | CP015061.1 | 412619 | 419530 | 45.2909167 | 2368 | 2303 | 1.028224056 | 1 | 1 | 1 | FIM1_4518 | YBR136W | YBR136W | MEC1 | n |
| Kmar_QGN18193.1 | W303.asm02.HP0_G0056210 | OG0001277 | chrXVI | 722655 | 724967 | CP015061.1 | 410187 | 412574 | 34.44976077 | 770 | 795 | 0.968553459 | 1 | 1 | 1 | FIM1_4517 | YPR091C | YPR091C | NVJ2 | n |
| Kmar_QGN17852.1 | W303.asm02.HP0_G0044160 | OG0001278 | chrXIV | 109649 | 110965 | CP015060.1 | 618532 | 619842 | 73.05936073 | 438 | 436 | 1.004587156 | 1 | 1 | 1 | FIM1_5061 | YNL280C | YNL280C | ERG24 | n |
| Kmar_QGN15273.1 | W303.asm02.HP0_G0049670 | OG0001279 | chrXV | 445282 | 446445 | CP015056.1 | 573391 | 574554 | 93.28165375 | 387 | 387 | 1 | 1 | 1 | 1 | FIM1_1962 | YOR063W | YOR063W | RPL3 | n |
| Kmar_QGN18361.1 | W303.asm02.HP0_G0025840 | OG0001280 | chrIX | 65385 | 66314 | CP015061.1 | 777092 | 778027 | 59.42492013 | 309 | 311 | 0.993569132 | 1 | 1 | 1 | FIM1_4688 | YIL145C | YIL145C | PAN6 | n |
| Kmar_QGN15861.1 | W303.asm02.HP0_G0021310 | OG0001281 | chrVII | 816998 | 817750 | CP015057.1 | 249826 | 250524 | 39.91935484 | 250 | 232 | 1.077586207 | 1 | 1 | 1 | FIM1_2558 | YGR158C | YGR158C | MTR3 | n |
| Kmar_QGN15860.1 | W303.asm02.HP0_G0021300 | OG0001282 | chrVII | 814170 | 816779 | CP015057.1 | 246910 | 249603 | 41.21475054 | 869 | 897 | 0.968784838 | 1 | 1 | 1 | FIM1_2557 | YGR157W | YGR157W | CHO2 | n |
| Kmar_QGN17328.1 | W303.asm02.HP0_G0045670 | OG0001283 | chrXIV | 413979 | 414407 | CP015059.1 | 705208 | 705591 | 65.49295775 | 142 | 127 | 1.118110236 | 1 | 1 | 1 | FIM1_4060 | YNL113W | YNL113W | RPC19 | n |
| Kmar_QGN17327.1 | W303.asm02.HP0_G0050110 | OG0001284 | chrXV | 534950 | 537694 | CP015059.1 | 701750 | 704248 | 33.04628633 | 914 | 832 | 1.098557692 | 1 | 1 | 1 | FIM1_4059 | YOR113W | YOR113W | AZF1 | n |
| Kmar_QGN14679.1 | W303.asm02.HP0_G0018230 | OG0001285 | chrVII | 146550 | 148289 | CP015055.1 | 1017690 | 1019369 | 58.17555938 | 579 | 559 | 1.035778175 | 1 | 1 | 1 | FIM1_1345 | YGL186C | YGL186C | TPN1 | n |
| Kmar_QGN14668.1 | W303.asm02.HP0_G0036620 | OG0001286 | chrXII | 506395 | 507846 | CP015055.1 | 990921 | 992357 | 84.47204969 | 483 | 478 | 1.010460251 | 1 | 1 | 1 | FIM1_1333 | YLR175W | YLR175W | CBF5 | n |
| Kmar_QGN14215.1 | W303.asm02.HP0_G0015970 | OG0001287 | chrV | 473773 | 475035 | CP015055.1 | 48744 | 49826 | 26.66666667 | 420 | 360 | 1.166666667 | 1 | 1 | 1 | FIM1_871 | YER149C | YER149C | PEA2 | n |
| Kmar_QGN14233.1 | W303.asm02.HP0_G0001720 | OG0001288 | chrII | 186640 | 188202 | CP015055.1 | 79079 | 80569 | 50.47984645 | 520 | 496 | 1.048387097 | 1 | 1 | 1 | FIM1_889 | YBL019W | YBL019W | APN2 | n |
| Kmar_QGN14216.1 | W303.asm02.HP0_G0015960 | OG0001289 | chrV | 472868 | 473590 | CP015055.1 | 50164 | 50865 | 82.91666667 | 240 | 233 | 1.030042918 | 1 | 1 | 1 | FIM1_872 | YER148W | YER148W | SPT15 | n |
| Kmar_QGN17354.1 | W303.asm02.HP0_G0052190 | OG0001290 | chrXV | 963269 | 968263 | CP015059.1 | 754696 | 759678 | 80.37103531 | 1664 | 1660 | 1.002409639 | 1 | 1 | 1 | FIM1_4086 | YOR341W | YOR341W | RPA190 | n |
| Kmar_QGN16446.1 | W303.asm02.HP0_G0029130 | OG0001291 | chrX | 314983 | 316209 | CP015058.1 | 133682 | 134866 | 54.98783455 | 408 | 394 | 1.035532995 | 1 | 1 | 1 | FIM1_3159 | YJL059W | YJL059W | YHC3 | n |
| Kmar_QGN14609.1 | W303.asm02.HP0_G0004390 | OG0001292 | chrII | 740237 | 741349 | CP015055.1 | 850842 | 851936 | 82.16216216 | 370 | 364 | 1.016483516 | 1 | 1 | 1 | FIM1_1272 | YBR249C | YBR249C | ARO4 | n |
| Kmar_QGN16453.1 | W303.asm02.HP0_G0029100 | OG0001293 | chrX | 307303 | 309795 | CP015058.1 | 145132 | 147501 | 42.43146603 | 830 | 789 | 1.051964512 | 1 | 1 | 1 | FIM1_3166 | YJL062W | YJL062W | LAS21 | n |
| Kmar_QGN17300.1 | W303.asm02.HP0_G0045510 | OG0001294 | chrXIV | 379600 | 380058 | CP015059.1 | 649474 | 649932 | 53.84615385 | 152 | 152 | 1 | 1 | 1 | 1 | FIM1_4031 | YNL131W | YNL131W | TOM22 | n |
| Kmar_QGN17305.1 | W303.asm02.HP0_G0045550 | OG0001295 | chrXIV | 384820 | 387681 | CP015059.1 | 656269 | 659088 | 41.96891192 | 953 | 939 | 1.014909478 | 1 | 1 | 1 | FIM1_4037 | YNL127W | YNL127W | FAR11 | n |
| Kmar_QGN17304.1 | W303.asm02.HP0_G0024510 | OG0001296 | chrVIII | 358729 | 359292 | CP015059.1 | 655279 | 655836 | 52.65957447 | 187 | 185 | 1.010810811 | 1 | 1 | 1 | FIM1_4036 | YHR121W | YHR121W | LSM12 | n |
| Kmar_QGN17303.1 | W303.asm02.HP0_G0045540 | OG0001297 | chrXIV | 383192 | 384496 | CP015059.1 | 653813 | 655117 | 33.99558499 | 434 | 434 | 1 | 1 | 1 | 1 | FIM1_4035 | YNL128W | YNL128W | TEP1 | n |
| Kmar_QGN15313.1 | W303.asm02.HP0_G0037840 | OG0001299 | chrXII | 754737 | 755498 | CP015056.1 | 676551 | 677303 | 35.20599251 | 253 | 250 | 1.012 | 1 | 1 | 1 | FIM1_2002 | YLR312W-A | YLR312W-A | MRPL15 | n |
| Kmar_QGN15312.1 | W303.asm02.HP0_G0018370 | OG0001300 | chrVII | 178858 | 180552 | CP015056.1 | 674145 | 675908 | 69.21768707 | 564 | 587 | 0.960817717 | 1 | 1 | 1 | FIM1_2001 | YGL171W | YGL171W | ROK1 | n |
| Kmar_QGN15311.1 | W303.asm02.HP0_G0055590 | OG0001301 | chrXVI | 606368 | 606715 | CP015056.1 | 673551 | 673913 | 51.66666667 | 115 | 120 | 0.958333333 | 1 | 1 | 1 | FIM1_2000 | YPR020W | YPR020W | ATP20 | n |
| Kmar_QGN14372.1 | W303.asm02.HP0_G0009700 | OG0001302 | chrIV | 664179 | 665600 | CP015055.1 | 380082 | 381491 | 65.18987342 | 473 | 469 | 1.008528785 | 1 | 1 | 1 | FIM1_1032 | YDR105C | YDR105C | TMS1 | n |
| Kmar_QGN13402.1 | W303.asm02.HP0_G0054420 | OG0001303 | chrXVI | 366567 | 367937 | CP015054.1 | 72988 | 74406 | 52.41090147 | 456 | 472 | 0.966101695 | 1 | 1 | 1 | FIM1_39 | YPL101W | YPL101W | ELP4 | n |
| Kmar_QGN15309.1 | W303.asm02.HP0_G0055580 | OG0001304 | chrXVI | 603248 | 606049 | CP015056.1 | 667370 | 670033 | 70.20364416 | 933 | 887 | 1.051860203 | 1 | 1 | 1 | FIM1_1998 | YPR019W | YPR019W | MCM4 | n |
| Kmar_QGN15888.1 | W303.asm02.HP0_G0009730 | OG0001305 | chrIV | 676020 | 678116 | CP015057.1 | 298762 | 300720 | 38.77266388 | 698 | 652 | 1.070552147 | 1 | 1 | 1 | FIM1_2586 | YDR108W | YDR108W | TRS85 | n |
| Kmar_QGN15889.1 | W303.asm02.HP0_G0035730 | OG0001306 | chrXII | 304542 | 308204 | CP015057.1 | 300837 | 304451 | 36.85897436 | 1220 | 1204 | 1.013289037 | 1 | 1 | 1 | FIM1_2587 | YLR084C | YLR084C | RAX2 | n |
| Kmar_QGN16422.1 | W303.asm02.HP0_G0054220 | OG0001307 | chrXVI | 325157 | 326698 | CP015058.1 | 84631 | 86121 | 64.0625 | 513 | 496 | 1.034274194 | 1 | 1 | 1 | FIM1_3135 | YPL122C | YPL122C | TFB2 | n |
| Kmar_QGN16423.1 | W303.asm02.HP0_G0050940 | OG0001308 | chrXV | 731303 | 734752 | CP015058.1 | 86671 | 90099 | 86.94516971 | 1149 | 1142 | 1.006129597 | 1 | 1 | 1 | FIM1_3136 | YOR207C | YOR207C | RET1 | n |
| Kmar_QGN14976.1 | W303.asm02.HP0_G0054110 | OG0001309 | chrXVI | 305821 | 307161 | CP015055.1 | 1606706 | 1608145 | 56.34095634 | 446 | 479 | 0.931106472 | 1 | 1 | 1 | FIM1_1654 | YPL133C | YPL133C | RDS2 | n |
| Kmar_QGN16425.1 | W303.asm02.HP0_G0050930 | OG0001310 | chrXV | 728807 | 730939 | CP015058.1 | 91157 | 93286 | 65.54621849 | 710 | 709 | 1.001410437 | 1 | 1 | 1 | FIM1_3138 | YOR206W | YOR206W | NOC2 | n |
| Kmar_QGN14460.1 | W303.asm02.HP0_G0018040 | OG0001311 | chrVII | 98056 | 103017 | CP015055.1 | 560869 | 565833 | 79.00786449 | 1653 | 1654 | 0.999395405 | 1 | 1 | 1 | FIM1_1121 | YGL206C | YGL206C | CHC1 | n |
| Kmar_QGN17724.1 | W303.asm02.HP0_G0046280 | OG0001312 | chrXIV | 551094 | 553064 | CP015060.1 | 326740 | 328845 | 43.92655367 | 656 | 701 | 0.935805991 | 1 | 1 | 1 | FIM1_4932 | YNL047C | YNL047C | SLM2 | n |
| Kmar_QGN14657.1 | W303.asm02.HP0_G0031830 | OG0001313 | chrXI | 168871 | 170088 | CP015055.1 | 961895 | 963049 | 42.50614251 | 405 | 384 | 1.0546875 | 1 | 1 | 1 | FIM1_1321 | YKL149C | YKL149C | DBR1 | n |
| Kmar_QGN14658.1 | W303.asm02.HP0_G0031820 | OG0001314 | chrXI | 167803 | 168711 | CP015055.1 | 963227 | 964114 | 64.2384106 | 302 | 295 | 1.023728814 | 1 | 1 | 1 | FIM1_1322 | YKL150W | YKL150W | MCR1 | n |
| Kmar_QGN14659.1 | W303.asm02.HP0_G0031810 | OG0001315 | chrXI | 166176 | 167189 | CP015055.1 | 964853 | 965851 | 45.32163743 | 337 | 332 | 1.015060241 | 1 | 1 | 1 | FIM1_1323 | YKL151C | YKL151C | NNR2 | n |
| Kmar_QGN18245.1 | W303.asm02.HP0_G0039660 | OG0001316 | chrXIII | 86775 | 88781 | CP015061.1 | 536925 | 538850 | 39.22734027 | 668 | 641 | 1.042121685 | 1 | 1 | 1 | FIM1_4570 | YML088W | YML088W | UFO1 | n |
| Kmar_QGN17258.1 | W303.asm02.HP0_G0040600 | OG0001317 | chrXIII | 291640 | 295473 | CP015059.1 | 560833 | 564666 | 39.80509745 | 1277 | 1277 | 1 | 1 | 1 | 1 | FIM1_3989 | YMR012W | YMR012W | CLU1 | n |
| Kmar_QGN14998.1 | W303.asm02.HP0_G0029020 | OG0001318 | chrX | 296151 | 297875 | CP015055.1 | 1653185 | 1654876 | 45.37671233 | 574 | 563 | 1.019538188 | 1 | 1 | 1 | FIM1_1679 | YJL071W | YJL071W | ARG2 | n |
| Kmar_QGN17617.1 | W303.asm02.HP0_G0049080 | OG0001319 | chrXV | 327287 | 329488 | CP015060.1 | 114574 | 116814 | 50.9383378 | 733 | 746 | 0.982573727 | 1 | 1 | 1 | FIM1_4824 | YOR001W | YOR001W | RRP6 | n |
| Kmar_QGN16454.1 | W303.asm02.HP0_G0004560 | OG0001320 | chrII | 763197 | 764378 | CP015058.1 | 147923 | 149173 | 55.21327014 | 393 | 416 | 0.944711538 | 1 | 1 | 1 | FIM1_3167 | YBR267W | YBR267W | REI1 | n |
| Kmar_QGN17722.1 | W303.asm02.HP0_G0046330 | OG0001321 | chrXIV | 559283 | 560473 | CP015060.1 | 319769 | 320857 | 28.39805825 | 396 | 362 | 1.093922652 | 1 | 1 | 1 | FIM1_4929 | YNL042W | YNL042W | BOP3 | n |
| Kmar_QGN17721.1 | W303.asm02.HP0_G0026240 | OG0001322 | chrIX | 160340 | 161863 | CP015060.1 | 318256 | 319677 | 54.296875 | 507 | 473 | 1.071881607 | 1 | 1 | 1 | FIM1_4928 | YIL104C | YIL104C | SHQ1 | n |
| Kmar_QGN17720.1 | W303.asm02.HP0_G0026250 | OG0001323 | chrIX | 162109 | 163386 | CP015060.1 | 316605 | 317885 | 72.3255814 | 425 | 426 | 0.997652582 | 1 | 1 | 1 | FIM1_4927 | YIL103W | YIL103W | DPH1 | n |
| Kmar_QGN17719.1 | W303.asm02.HP0_G0046340 | OG0001324 | chrXIV | 560651 | 563170 | CP015060.1 | 314217 | 316565 | 35.39192399 | 839 | 782 | 1.072890026 | 1 | 1 | 1 | FIM1_4926 | YNL041C | YNL041C | COG6 | n |
| Kmar_QGN16218.1 | W303.asm02.HP0_G0038600 | OG0001325 | chrXII | 906905 | 908980 | CP015057.1 | 1058583 | 1060454 | 30.96590909 | 691 | 623 | 1.109149278 | 1 | 1 | 1 | FIM1_2922 | YLR396C | YLR396C | VPS33 | n |
| Kmar_QGN14139.1 | W303.asm02.HP0_G0005290 | OG0001326 | chrIII | 68250 | 69572 | CP015054.1 | 1622010 | 1623242 | 33.95833333 | 440 | 410 | 1.073170732 | 1 | 1 | 1 | FIM1_792 | YCL029C | YCL029C | BIK1 | n |
| Kmar_QGN17737.1 | W303.asm02.HP0_G0026140 | OG0001327 | chrIX | 134683 | 139065 | CP015060.1 | 359668 | 364179 | 24.23051735 | 1460 | 1503 | 0.971390552 | 1 | 1 | 1 | FIM1_4946 | YIL115C | YIL115C | NUP159 | n |
| Kmar_QGN17260.1 | W303.asm02.HP0_G0035420 | OG0001328 | chrXII | 254137 | 254790 | CP015059.1 | 568866 | 569555 | 48.93617021 | 217 | 229 | 0.947598253 | 1 | 1 | 1 | FIM1_3991 | YLR051C | YLR051C | FCF2 | n |
| Kmar_QGN14616.1 | W303.asm02.HP0_G0038430 | OG0001329 | chrXII | 872402 | 873844 | CP015055.1 | 864760 | 866202 | 82.12058212 | 480 | 480 | 1 | 1 | 1 | 1 | FIM1_1280 | YLR378C | YLR378C | SEC61 | n |
| Kmar_QGN17102.1 | W303.asm02.HP0_G0038420 | OG0001330 | chrXII | 870412 | 871458 | CP015059.1 | 235884 | 236939 | 73.29545455 | 348 | 351 | 0.991452991 | 1 | 1 | 1 | FIM1_3833 | YLR377C | YLR377C | FBP1 | n |
| Kmar_QGN14615.1 | W303.asm02.HP0_G0008280 | OG0001331 | chrIV | 360160 | 361620 | CP015055.1 | 862919 | 864367 | 37.5249501 | 486 | 482 | 1.008298755 | 1 | 1 | 1 | FIM1_1279 | YDL054C | YDL054C | MCH1 | n |
| Kmar_QGN16250.1 | W303.asm02.HP0_G0030870 | OG0001332 | chrX | 686743 | 687006 | CP015057.1 | 1130915 | 1131178 | 74.71264368 | 87 | 87 | 1 | 1 | 1 | 1 | FIM1_2955 | YJR135W-A | YJR135W-A | TIM8 | n |
| Kmar_QGN16247.1 | W303.asm02.HP0_G0025270 | OG0001333 | chrVIII | 502417 | 503382 | CP015057.1 | 1125972 | 1126922 | 40.37267081 | 321 | 316 | 1.015822785 | 1 | 1 | 1 | FIM1_2952 | YHR198C | YHR198C | AIM18 | n |
| Kmar_QGN18371.1 | W303.asm02.HP0_G0044060 | OG0001334 | chrXIV | 84616 | 86262 | CP015061.1 | 794132 | 795778 | 43.40949033 | 548 | 548 | 1 | 1 | 1 | 1 | FIM1_4698 | YNL291C | YNL291C | MID1 | n |
| Kmar_QGN14008.1 | W303.asm02.HP0_G0002040 | OG0001335 | chrII | 265333 | 265731 | CP015054.1 | 1317824 | 1318177 | 43.18181818 | 132 | 117 | 1.128205128 | 1 | 1 | 1 | FIM1_659 | YBR016W | YBR016W | CPP1 | n |
| Kmar_QGN17114.1 | W303.asm02.HP0_G0030760 | OG0001336 | chrX | 662666 | 664012 | CP015059.1 | 265369 | 266721 | 57.52212389 | 448 | 450 | 0.995555556 | 1 | 1 | 1 | FIM1_3845 | YJR124C | YJR124C | | n |
| Kmar_QGN14761.1 | W303.asm02.HP0_G0050330 | OG0001337 | chrXV | 582688 | 584556 | CP015055.1 | 1174836 | 1176575 | 26.40625 | 622 | 579 | 1.074265976 | 1 | 1 | 1 | FIM1_1431 | YOR137C | YOR137C | SIA1 | n |
| Kmar_QGN13821.1 | W303.asm02.HP0_G0011920 | OG0001338 | chrIV | 1160551 | 1161120 | CP015054.1 | 940233 | 940802 | 87.3015873 | 189 | 189 | 1 | 1 | 1 | 1 | FIM1_467 | YDR339C | YDR339C | FCF1 | n |
| Kmar_QGN16224.1 | W303.asm02.HP0_G0057060 | OG0001339 | chrXVI | 907876 | 912486 | CP015057.1 | 1068100 | 1072668 | 67.23127036 | 1536 | 1522 | 1.009198423 | 1 | 1 | 1 | FIM1_2928 | YPR184W | YPR184W | GDB1 | n |
| Kmar_QGN13644.1 | W303.asm02.HP0_G0014350 | OG0001340 | chrV | 130008 | 131744 | CP015054.1 | 572723 | 574465 | 73.10344828 | 578 | 580 | 0.996551724 | 1 | 1 | 1 | FIM1_286 | YEL013W | YEL013W | VAC8 | n |
| Kmar_QGN16435.1 | W303.asm02.HP0_G0029230 | OG0001341 | chrX | 335992 | 337344 | CP015058.1 | 113152 | 114495 | 26.79738562 | 450 | 447 | 1.006711409 | 1 | 1 | 1 | FIM1_3148 | YJL049W | YJL049W | CHM7 | n |
| Kmar_QGN14003.1 | W303.asm02.HP0_G0020550 | OG0001342 | chrVII | 635646 | 637415 | CP015054.1 | 1308108 | 1309805 | 24.32885906 | 589 | 565 | 1.042477876 | 1 | 1 | 1 | FIM1_654 | YGR077C | YGR077C | PEX8 | n |
| Kmar_QGN14593.1 | W303.asm02.HP0_G0047110 | OG0001343 | chrXIV | 708177 | 710066 | CP015055.1 | 816879 | 818810 | 49.15514593 | 629 | 643 | 0.978227061 | 1 | 1 | 1 | FIM1_1255 | YNR038W | YNR038W | DBP6 | n |
| Kmar_QGN14591.1 | W303.asm02.HP0_G0006210 | OG0001344 | chrIII | 275826 | 277154 | CP015055.1 | 814550 | 816043 | 41.24513619 | 442 | 497 | 0.889336016 | 1 | 1 | 1 | FIM1_1253 | YCR079W | YCR079W | PTC6 | n |
| Kmar_QGN17553.1 | W303.asm02.HP0_G0028640 | OG0001345 | chrX | 186303 | 187142 | CP015059.1 | 1151633 | 1152550 | 63.07189542 | 279 | 305 | 0.914754098 | 1 | 1 | 1 | FIM1_4289 | YJL115W | YJL115W | ASF1 | n |
| Kmar_QGN17613.1 | W303.asm02.HP0_G0049090 | OG0001346 | chrXV | 329872 | 331506 | CP015060.1 | 105987 | 107711 | 64.29809359 | 544 | 574 | 0.947735192 | 1 | 1 | 1 | FIM1_4820 | YOR002W | YOR002W | ALG6 | n |
| Kmar_QGN17700.1 | W303.asm02.HP0_G0026370 | OG0001347 | chrIX | 181525 | 183690 | CP015060.1 | 276920 | 279034 | 64.21636616 | 721 | 704 | 1.024147727 | 1 | 1 | 1 | FIM1_4907 | YIL091C | YIL091C | UTP25 | n |
| Kmar_QGN17695.1 | W303.asm02.HP0_G0046430 | OG0001348 | chrXIV | 585689 | 586534 | CP015060.1 | 268392 | 269252 | 55.94405594 | 281 | 286 | 0.982517483 | 1 | 1 | 1 | FIM1_4902 | YNL032W | YNL032W | SIW14 | n |
| Kmar_QGN17694.1 | W303.asm02.HP0_G0026410 | OG0001349 | chrIX | 190141 | 190614 | CP015060.1 | 267924 | 268337 | 42.67515924 | 157 | 137 | 1.145985401 | 1 | 1 | 1 | FIM1_4901 | YIL087C | YIL087C | AIM19 | n |
| Kmar_QGN17679.1 | W303.asm02.HP0_G0018520 | OG0001350 | chrVII | 211746 | 212564 | CP015060.1 | 246613 | 247380 | 29.5620438 | 272 | 255 | 1.066666667 | 1 | 1 | 1 | FIM1_4886 | YGL154C | YGL154C | LYS5 | n |
| Kmar_QGN14926.1 | W303.asm02.HP0_G0045080 | OG0001351 | chrXIV | 303376 | 304098 | CP015055.1 | 1510498 | 1511211 | 87.08333333 | 240 | 237 | 1.012658228 | 1 | 1 | 1 | FIM1_1601 | YNL178W | YNL178W | RPS3 | n |
| Kmar_QGN14971.1 | W303.asm02.HP0_G0051100 | OG0001352 | chrXV | 762120 | 762560 | CP015055.1 | 1600086 | 1600523 | 71.62162162 | 146 | 145 | 1.006896552 | 1 | 1 | 1 | FIM1_1649 | YOR224C | YOR224C | RPB8 | n |
| Kmar_QGN16655.1 | W303.asm02.HP0_G0020430 | OG0001353 | chrVII | 609645 | 613721 | CP015058.1 | 599237 | 603286 | 77.24594993 | 1358 | 1349 | 1.006671609 | 1 | 1 | 1 | FIM1_3374 | YGR061C | YGR061C | ADE6 | n |
| Kmar_QGN18103.1 | W303.asm02.HP0_G0030360 | OG0001354 | chrX | 594895 | 595212 | CP015061.1 | 223056 | 223373 | 70.47619048 | 105 | 105 | 1 | 1 | 1 | 1 | FIM1_4423 | YJR085C | YJR085C | TMH11 | n |

| | | | | | | | | | | | | | | | | | | | | |
|---|---|---|---|---|---|---|---|---|---|---|---|---|---|---|---|---|---|---|---|---|
| Kmar_QGN14288.1 | W303.asm02.HP0_G0031790 | OG0001355 | chrXI | 162880 | 163614 | CP015055.1 | 194376 | 195125 | 41.6988417 | 244 | 249 | 0.979919679 | 1 | 1 | 1 | FIM1_945 | YKL154W | YKL154W | SRP102 | n |
| Kmar_QGN18392.1 | W303.asm02.HP0_G0015830 | OG0001356 | chrV | 417880 | 418491 | CP015061.1 | 839895 | 840524 | 40.88669951 | 203 | 209 | 0.971291866 | 1 | 1 | 1 | FIM1_4719 | YER128W | YER128W | VFA1 | n |
| Kmar_QGN14282.1 | W303.asm02.HP0_G0023370 | OG0001357 | chrVIII | 151585 | 153033 | CP015055.1 | 177345 | 178811 | 71.51639344 | 482 | 488 | 0.987704918 | 1 | 1 | 1 | FIM1_939 | YHR024C | YHR024C | MAS2 | n |
| Kmar_QGN18427.1 | W303.asm02.HP0_G0055380 | OG0001358 | chrXVI | 559984 | 560685 | CP015061.1 | 908994 | 909692 | 58.72340426 | 233 | 232 | 1.004310345 | 1 | 1 | 1 | FIM1_4755 | YPL002C | YPL002C | SNF8 | n |
| Kmar_QGN18428.1 | W303.asm02.HP0_G0016890 | OG0001359 | chrVI | 115614 | 116942 | CP015061.1 | 909788 | 911140 | 57.9064588 | 442 | 450 | 0.982222222 | 1 | 1 | 1 | FIM1_4756 | YFL001W | YFL001W | DEG1 | n |
| Kmar_QGN16677.1 | W303.asm02.HP0_G0007170 | OG0001360 | chrIV | 148411 | 149298 | CP015058.1 | 650664 | 651149 | 24.24242424 | 295 | 161 | 1.832298137 | 1 | 1 | 1 | FIM1_3396 | YDL173W | YDL173W | PAR32 | n |
| Kmar_QGN15057.1 | W303.asm02.HP0_G0030240 | OG0001361 | chrX | 578093 | 578686 | CP015056.1 | 106055 | 106639 | 63.63636364 | 197 | 194 | 1.015463918 | 1 | 1 | 1 | FIM1_1742 | YJR069C | YJR069C | HAM1 | n |
| Kmar_QGN14755.1 | W303.asm02.HP0_G0050380 | OG0001362 | chrXV | 602258 | 603217 | CP015055.1 | 1162510 | 1163448 | 40.3030303 | 319 | 312 | 1.022435897 | 1 | 1 | 1 | FIM1_1425 | YOR143C | YOR143C | THI80 | n |
| Kmar_QGN17273.1 | W303.asm02.HP0_G0042770 | OG0001363 | chrXIII | 743549 | 744022 | CP015059.1 | 594123 | 594527 | 66.0130719 | 157 | 134 | 1.171641791 | 1 | 1 | 1 | FIM1_4004 | YMR236W | YMR236W | TAF9 | n |
| Kmar_QGN16327.1 | W303.asm02.HP0_G0012960 | OG0001364 | chrIV | 1371957 | 1372520 | CP015057.1 | 1299106 | 1299684 | 76.68393782 | 187 | 192 | 0.973958333 | 1 | 1 | 1 | FIM1_3032 | YDR454C | YDR454C | GUK1 | n |
| Kmar_QGN15653.1 | W303.asm02.HP0_G0050540 | OG0001365 | chrXV | 634714 | 637632 | CP015056.1 | 1395929 | 1398814 | 47.44897959 | 972 | 961 | 1.01144641 | 1 | 1 | 1 | FIM1_2346 | YOR160W | YOR160W | MTR10 | n |
| Kmar_QGN16504.1 | W303.asm02.HP0_G0052020 | OG0001366 | chrXV | 920326 | 922782 | CP015058.1 | 255303 | 257708 | 43.41637011 | 818 | 801 | 1.021223471 | 1 | 1 | 1 | FIM1_3217 | YOR322C | YOR322C | LDB19 | n |
| Kmar_QGN15287.1 | W303.asm02.HP0_G0014060 | OG0001367 | chrV | 70839 | 73709 | CP015056.1 | 605104 | 607467 | 30.8411215 | 956 | 787 | 1.214739517 | 1 | 1 | 1 | FIM1_1976 | YEL043W | YEL043W | GTA1 | n |
| Kmar_QGN14249.1 | W303.asm02.HP0_G0021230 | OG0001368 | chrVII | 800758 | 802056 | CP015055.1 | 111455 | 112684 | 52.65588915 | 432 | 409 | 1.056234719 | 1 | 1 | 1 | FIM1_905 | YGR149W | YGR149W | GPC1 | n |
| Kmar_QGN18290.1 | W303.asm02.HP0_G0004410 | OG0001369 | chrII | 744745 | 745668 | CP015061.1 | 625598 | 626482 | 63.51791531 | 307 | 294 | 1.044217687 | 1 | 1 | 1 | FIM1_4616 | YBR251W | YBR251W | MRPS5 | n |
| Kmar_QGN16836.1 | W303.asm02.HP0_G0027040 | OG0001370 | chrIX | 312083 | 312562 | CP015058.1 | 994009 | 994527 | 33.13953488 | 159 | 172 | 0.924418605 | 1 | 1 | 1 | FIM1_3560 | YIL016W | YIL016W | SNL1 | n |
| Kmar_QGN17903.1 | W303.asm02.HP0_G0055910 | OG0001371 | chrXVI | 677624 | 678640 | CP015060.1 | 710342 | 711337 | 62.57309942 | 338 | 331 | 1.021148036 | 1 | 1 | 1 | FIM1_5112 | YPR056W | YPR056W | TFB4 | n |
| Kmar_QGN14519.1 | W303.asm02.HP0_G0042820 | OG0001372 | chrXIII | 758467 | 759411 | CP015055.1 | 684458 | 685384 | 78.98089172 | 314 | 308 | 1.019480519 | 1 | 1 | 1 | FIM1_1181 | YMR241W | YMR241W | YHM2 | n |
| Kmar_QGN14518.1 | W303.asm02.HP0_G0051880 | OG0001373 | chrXV | 894451 | 895812 | CP015055.1 | 682642 | 683973 | 51.53508772 | 453 | 443 | 1.022573363 | 1 | 1 | 1 | FIM1_1180 | YOR307C | YOR307C | SLY41 | n |
| Kmar_QGN16326.1 | W303.asm02.HP0_G0012970 | OG0001374 | chrIV | 1373178 | 1375079 | CP015057.1 | 1296673 | 1298496 | 70.21943574 | 633 | 607 | 1.042833608 | 1 | 1 | 1 | FIM1_3031 | YDR456W | YDR456W | NHX1 | n |
| Kmar_QGN16325.1 | W303.asm02.HP0_G0040200 | OG0001375 | chrXIII | 214697 | 216664 | CP015057.1 | 1294533 | 1296362 | 32.88490284 | 655 | 609 | 1.075533662 | 1 | 1 | 1 | FIM1_3030 | YML031W | YML031W | NDC1 | n |
| Kmar_QGN15655.1 | W303.asm02.HP0_G0050520 | OG0001376 | chrXV | 633039 | 633995 | CP015056.1 | 1399495 | 1400409 | 45.28301887 | 318 | 304 | 1.046052632 | 1 | 1 | 1 | FIM1_2348 | YOR158W | YOR158W | PET123 | n |
| Kmar_QGN14767.1 | W303.asm02.HP0_G0012380 | OG0001377 | chrIV | 1251678 | 1253576 | CP015055.1 | 1189793 | 1191646 | 41.84952978 | 632 | 617 | 1.024311183 | 1 | 1 | 1 | FIM1_1437 | YDR386W | YDR386W | MUS81 | n |
| Kmar_QGN16324.1 | W303.asm02.HP0_G0040190 | OG0001378 | chrXIII | 213023 | 214537 | CP015057.1 | 1292218 | 1293555 | 47.38878143 | 504 | 445 | 1.13258427 | 1 | 1 | 1 | FIM1_3029 | YML032C | YML032C | RAD52 | n |
| Kmar_QGN17863.1 | W303.asm02.HP0_G0055990 | OG0001379 | chrXVI | 687711 | 688610 | CP015060.1 | 634156 | 635088 | 49.51768489 | 299 | 310 | 0.964516129 | 1 | 1 | 1 | FIM1_5072 | YPR066W | YPR066W | UBA3 | n |
| Kmar_QGN15106.1 | W303.asm02.HP0_G0020860 | OG0001380 | chrVII | 713496 | 714833 | CP015056.1 | 217290 | 218591 | 48.0349345 | 445 | 433 | 1.027713626 | 1 | 1 | 1 | FIM1_1793 | YGR110W | YGR110W | CLD1 | n |
| Kmar_QGN15549.1 | W303.asm02.HP0_G0023880 | OG0001381 | chrVIII | 226555 | 227100 | CP015056.1 | 1165665 | 1166171 | 41.75824176 | 181 | 168 | 1.077380952 | 1 | 1 | 1 | FIM1_2240 | YHR060W | YHR060W | VMA22 | n |
| Kmar_QGN16465.1 | W303.asm02.HP0_G0029050 | OG0001382 | chrX | 303035 | 303934 | CP015058.1 | 167265 | 168152 | 64.90066225 | 299 | 295 | 1.013559322 | 1 | 1 | 1 | FIM1_3178 | YJL068C | YJL068C | | n |
| Kmar_QGN13994.1 | W303.asm02.HP0_G0037610 | OG0001383 | chrXII | 705760 | 706827 | CP015054.1 | 1287771 | 1288826 | 30.08130081 | 355 | 351 | 1.011396011 | 1 | 1 | 1 | FIM1_644 | YLR287C | YLR287C | | n |
| Kmar_QGN13995.1 | W303.asm02.HP0_G0050720 | OG0001384 | chrXV | 677234 | 679135 | CP015054.1 | 1288937 | 1291165 | 43.47240915 | 633 | 742 | 0.85309973 | 1 | 1 | 1 | FIM1_645 | YOR181W | YOR181W | LAS17 | n |
| Kmar_QGN16685.1 | W303.asm02.HP0_G0007200 | OG0001385 | chrIV | 158284 | 158955 | CP015058.1 | 667360 | 667872 | 27.96610169 | 223 | 170 | 1.311764706 | 1 | 1 | 1 | FIM1_3404 | YDL169C | YDL169C | UGX2 | n |
| Kmar_QGN16688.1 | W303.asm02.HP0_G0036860 | OG0001386 | chrXII | 549639 | 550421 | CP015058.1 | 672014 | 672769 | 53.84615385 | 260 | 251 | 1.035856574 | 1 | 1 | 1 | FIM1_3408 | YLR201C | YLR201C | COQ9 | n |
| Kmar_QGN17489.1 | W303.asm02.HP0_G0035360 | OG0001387 | chrXII | 242850 | 245516 | CP015059.1 | 1014892 | 1017552 | 32.7173913 | 888 | 886 | 1.002257336 | 1 | 1 | 1 | FIM1_4225 | YLR045C | YLR045C | STU2 | n |
| Kmar_QGN17486.1 | W303.asm02.HP0_G0021790 | OG0001388 | chrVII | 923258 | 924043 | CP015059.1 | 1011666 | 1012442 | 67.17557252 | 261 | 258 | 1.011627907 | 1 | 1 | 1 | FIM1_4222 | YGR207C | YGR207C | CIR1 | n |
| Kmar_QGN18287.1 | W303.asm02.HP0_G0056510 | OG0001389 | chrXVI | 793137 | 794357 | CP015061.1 | 620971 | 622020 | 39.75308642 | 406 | 349 | 1.163323782 | 1 | 1 | 1 | FIM1_4613 | YPR124W | YPR124W | CTR1 | n |
| Kmar_QGN14276.1 | W303.asm02.HP0_G0055900 | OG0001390 | chrXVI | 674174 | 677371 | CP015055.1 | 163236 | 166310 | 35.41472507 | 1065 | 1024 | 1.040039063 | 1 | 1 | 1 | FIM1_933 | YPR055W | YPR055W | SEC8 | n |
| Kmar_QGN14274.1 | W303.asm02.HP0_G0002960 | OG0001391 | chrII | 460107 | 460979 | CP015055.1 | 159770 | 160642 | 60.48109966 | 290 | 290 | 1 | 1 | 1 | 1 | FIM1_931 | YBR101C | YBR101C | FES1 | n |
| Kmar_QGN16608.1 | W303.asm02.HP0_G0002270 | OG0001392 | chrII | 313367 | 315376 | CP015058.1 | 489581 | 491617 | 56.63716814 | 669 | 678 | 0.986725664 | 1 | 1 | 1 | FIM1_3324 | YBR041W | YBR041W | FAT1 | n |
| Kmar_QGN14229.1 | W303.asm02.HP0_G0015900 | OG0001393 | chrV | 461024 | 462484 | CP015055.1 | 70525 | 72003 | 63.43434343 | 486 | 492 | 0.987804878 | 1 | 1 | 1 | FIM1_885 | YER141W | YER141W | COX15 | n |
| Kmar_QGN15146.1 | W303.asm02.HP0_G0015890 | OG0001394 | chrV | 459130 | 460800 | CP015056.1 | 293178 | 294803 | 35.13986014 | 556 | 541 | 1.027726433 | 1 | 1 | 1 | FIM1_1833 | YER140W | YER140W | EMP65 | n |
| Kmar_QGN14158.1 | W303.asm02.HP0_G0005150 | OG0001395 | chrIII | 44275 | 46557 | CP015054.1 | 1652755 | 1654917 | 34.29319372 | 760 | 720 | 1.055555556 | 1 | 1 | 1 | FIM1_811 | YCL045C | YCL045C | EMC1 | n |
| Kmar_QGN14164.1 | W303.asm02.HP0_G0013630 | OG0001396 | chrIV | 1496787 | 1498106 | CP015054.1 | 1666786 | 1668222 | 48.04928131 | 439 | 478 | 0.918410042 | 1 | 1 | 1 | FIM1_817 | YDR527W | YDR527W | RBA50 | n |
| Kmar_QGN18005.1 | W303.asm02.HP0_G0013000 | OG0001397 | chrIV | 1388012 | 1389136 | CP015061.1 | 18442 | 19515 | 40.16172507 | 374 | 357 | 1.047619048 | 1 | 1 | 1 | FIM1_4321 | YDR459C | YDR459C | PFA5 | n |
| Kmar_QGN14106.1 | W303.asm02.HP0_G0044230 | OG0001398 | chrXIV | 123437 | 127153 | CP015054.1 | 1539707 | 1543177 | 39.36 | 1238 | 1156 | 1.070934256 | 1 | 1 | 1 | FIM1_757 | YNL273W | YNL273W | TOF1 | n |
| Kmar_QGN14105.1 | W303.asm02.HP0_G0013850 | OG0001399 | chrV | 30161 | 31603 | CP015054.1 | 1538261 | 1539658 | 51.58562368 | 480 | 465 | 1.032258065 | 1 | 1 | 1 | FIM1_756 | YEL064C | YEL064C | AVT2 | n |
| Kmar_QGN18036.1 | W303.asm02.HP0_G0035580 | OG0001400 | chrXII | 278959 | 279414 | CP015061.1 | 74023 | 74457 | 51.65562914 | 151 | 144 | 1.048611111 | 1 | 1 | 1 | FIM1_4352 | YLR068W | YLR068W | FYV7 | n |
| Kmar_QGN13415.1 | W303.asm02.HP0_G0054530 | OG0001401 | chrXVI | 381860 | 383311 | CP015054.1 | 93583 | 95037 | 73.40206186 | 483 | 484 | 0.997933884 | 1 | 1 | 1 | FIM1_52 | YPL091W | YPL091W | GLR1 | n |
| Kmar_QGN17427.1 | W303.asm02.HP0_G0002020 | OG0001402 | chrII | 261810 | 262421 | CP015059.1 | 899510 | 900196 | 37.06896552 | 203 | 228 | 0.890350877 | 1 | 1 | 1 | FIM1_4159 | YBR014C | YBR014C | GRX7 | n |
| Kmar_QGN18430.1 | W303.asm02.HP0_G0055370 | OG0001403 | chrXVI | 558377 | 559765 | CP015061.1 | 913322 | 914689 | 25.20661157 | 462 | 455 | 1.015384615 | 1 | 1 | 1 | FIM1_4758 | YPL003W | YPL003W | ULA1 | n |
| Kmar_QGN18429.1 | W303.asm02.HP0_G0016880 | OG0001404 | chrVI | 113597 | 115417 | CP015061.1 | 911494 | 913287 | 65.34653465 | 606 | 597 | 1.015075377 | 1 | 1 | 1 | FIM1_4757 | YFL002C | YFL002C | SPB4 | n |
| Kmar_QGN13950.1 | W303.asm02.HP0_G0049000 | OG0001405 | chrXV | 310767 | 311390 | CP015054.1 | 1205218 | 1205805 | 48.07692308 | 207 | 195 | 1.061538462 | 1 | 1 | 1 | FIM1_598 | YOL008W | YOL008W | COQ10 | n |
| Kmar_QGN16583.1 | W303.asm02.HP0_G0038040 | OG0001406 | chrXII | 793117 | 795279 | CP015058.1 | 433812 | 435980 | 35.43307087 | 720 | 722 | 0.997229917 | 1 | 1 | 1 | FIM1_3299 | YLR335W | YLR335W | NUP2 | n |
| Kmar_QGN14265.1 | W303.asm02.HP0_G0043020 | OG0001407 | chrXIII | 801421 | 802026 | CP015055.1 | 144206 | 144718 | 45.27363184 | 201 | 170 | 1.182352941 | 1 | 1 | 1 | FIM1_922 | YMR263W | YMR263W | SAP30 | n |
| Kmar_QGN14263.1 | W303.asm02.HP0_G0039540 | OG0001408 | chrXIII | 62843 | 64249 | CP015055.1 | 140745 | 142295 | 44.33962264 | 468 | 516 | 0.906976744 | 1 | 1 | 1 | FIM1_920 | YML102W | YML102W | CAC2 | n |
| Kmar_QGN14950.1 | W303.asm02.HP0_G0016750 | OG0001409 | chrVI | 73184 | 74719 | CP015055.1 | 1555676 | 1557076 | 50.68226121 | 511 | 466 | 1.096566524 | 1 | 1 | 1 | FIM1_1627 | YFL016C | YFL016C | MDJ1 | n |
| Kmar_QGN13448.1 | W303.asm02.HP0_G0054730 | OG0001410 | chrXVI | 429243 | 430250 | CP015054.1 | 166601 | 167536 | 38.87240356 | 335 | 311 | 1.077170418 | 1 | 1 | 1 | FIM1_85 | YPL069C | YPL069C | BTS1 | n |
| Kmar_QGN14872.1 | W303.asm02.HP0_G0029640 | OG0001411 | chrX | 436643 | 438202 | CP015055.1 | 1396792 | 1398351 | 32.69961977 | 519 | 519 | 1 | 1 | 1 | 1 | FIM1_1545 | YJR003C | YJR003C | MRX12 | n |
| Kmar_QGN16126.1 | W303.asm02.HP0_G0033050 | OG0001412 | chrXI | 406436 | 407842 | CP015057.1 | 863177 | 864409 | 52.68595041 | 468 | 410 | 1.141463415 | 1 | 1 | 1 | FIM1_2827 | YKL021C | YKL021C | MAK11 | n |
| Kmar_QGN16124.1 | W303.asm02.HP0_G0033040 | OG0001413 | chrXI | 403818 | 406340 | CP015057.1 | 858951 | 861131 | 41.63701068 | 840 | 726 | 1.157024793 | 1 | 1 | 1 | FIM1_2825 | YKL022C | YKL022C | CDC16 | n |
| Kmar_QGN17508.1 | W303.asm02.HP0_G0021940 | OG0001414 | chrVII | 953284 | 954630 | CP015059.1 | 1056501 | 1057556 | 30.80357143 | 448 | 351 | 1.276353276 | 1 | 1 | 1 | FIM1_4244 | YGR223C | YGR223C | HSV2 | n |
| Kmar_QGN14702.1 | W303.asm02.HP0_G0015000 | OG0001415 | chrV | 252999 | 254933 | CP015055.1 | 1060951 | 1062795 | 60.03062787 | 644 | 614 | 1.048859935 | 1 | 1 | 1 | FIM1_1369 | YER049W | YER049W | TPA1 | n |
| Kmar_QGN16671.1 | W303.asm02.HP0_G0015400 | OG0001416 | chrV | 330147 | 331877 | CP015058.1 | 639483 | 641210 | 76.04166667 | 576 | 575 | 1.00173913 | 1 | 1 | 1 | FIM1_3390 | YER086W | YER086W | ILV1 | n |
| Kmar_QGN13991.1 | W303.asm02.HP0_G0037630 | OG0001417 | chrXII | 709858 | 711795 | CP015054.1 | 1281472 | 1283367 | 73.17829457 | 645 | 631 | 1.022187005 | 1 | 1 | 1 | FIM1_640 | YLR289W | YLR289W | GUF1 | n |
| Kmar_QGN13993.1 | W303.asm02.HP0_G0050740 | OG0001418 | chrXV | 680652 | 681839 | CP015054.1 | 1284812 | 1285990 | 66.16161616 | 395 | 392 | 1.007653061 | 1 | 1 | 1 | FIM1_642 | YOR184W | YOR184W | SER1 | n |
| Kmar_QGN13992.1 | W303.asm02.HP0_G0037620 | OG0001419 | chrXII | 708249 | 709673 | CP015054.1 | 1283601 | 1284803 | 36.70886076 | 474 | 400 | 1.185 | 1 | 1 | 1 | FIM1_641 | YLR288C | YLR288C | MEC3 | n |
| Kmar_QGN14889.1 | W303.asm02.HP0_G0037500 | OG0001420 | chrXII | 686205 | 688532 | CP015055.1 | 1433177 | 1435417 | 71.90721649 | 775 | 746 | 1.038873995 | 1 | 1 | 1 | FIM1_1563 | YLR274W | YLR274W | MCM5 | n |
| Kmar_QGN13999.1 | W303.asm02.HP0_G0037550 | OG0001421 | chrXII | 699262 | 699729 | CP015054.1 | 1298001 | 1298537 | 47.19101124 | 155 | 178 | 0.870786517 | 1 | 1 | 1 | FIM1_650 | YLR281C | YLR281C | RSO55 | n |
| Kmar_QGN13985.1 | W303.asm02.HP0_G0037700 | OG0001422 | chrXII | 719490 | 720185 | CP015054.1 | 1274552 | 1275151 | 35.8974359 | 231 | 199 | 1.16080402 | 1 | 1 | 1 | FIM1_634 | YLR298C | YLR298C | YHC1 | n |
| Kmar_QGN16501.1 | W303.asm02.HP0_G0000460 | OG0001423 | chrI | 100988 | 101908 | CP015058.1 | 245984 | 246889 | 71.33550489 | 306 | 301 | 1.016611296 | 1 | 1 | 1 | FIM1_3214 | YAL025C | YAL025C | MAK16 | n |
| Kmar_QGN17684.1 | W303.asm02.HP0_G0007650 | OG0001424 | chrIV | 245339 | 245788 | CP015060.1 | 251603 | 252019 | 36.91275168 | 149 | 138 | 1.079710145 | 1 | 1 | 1 | FIM1_4891 | YDL121C | YDL121C | EXP1 | n |
| Kmar_QGN14977.1 | W303.asm02.HP0_G0054120 | OG0001425 | chrXVI | 307649 | 308551 | CP015055.1 | 1608419 | 1609198 | 62.83783784 | 300 | 259 | 1.158301158 | 1 | 1 | 1 | FIM1_1655 | YPL132W | YPL132W | COX11 | n |
| Kmar_QGN15710.1 | W303.asm02.HP0_G0017780 | OG0001426 | chrVII | 54492 | 57224 | CP015056.1 | 1510124 | 1512706 | 47.14912281 | 910 | 860 | 1.058139535 | 1 | 1 | 1 | FIM1_2403 | YGL233W | YGL233W | SEC15 | n |
| Kmar_QGN15707.1 | W303.asm02.HP0_G0017350 | OG0001427 | chrVI | 208018 | 208731 | CP015056.1 | 1506381 | 1507127 | 28.17460317 | 237 | 248 | 0.955645161 | 1 | 1 | 1 | FIM1_2400 | YFR043C | YFR043C | IRC6 | n |
| Kmar_QGN13386.1 | W303.asm02.HP0_G0027560 | OG0001428 | chrIX | 406331 | 406918 | CP015054.1 | 50028 | 50660 | 48.05825243 | 195 | 210 | 0.928571429 | 1 | 1 | 1 | FIM1_23 | YIR032C | YIR032C | DAL3 | n |
| Kmar_QGN13410.1 | W303.asm02.HP0_G0003710 | OG0001429 | chrII | 603322 | 605889 | CP015054.1 | 82396 | 84888 | 48.02325581 | 855 | 830 | 1.030120482 | 1 | 1 | 1 | FIM1_47 | YBR179C | YBR179C | FZO1 | n |
| Kmar_QGN17650.1 | W303.asm02.HP0_G0034970 | OG0001430 | chrXII | 159358 | 160929 | CP015060.1 | 190934 | 192481 | 69.33333333 | 523 | 515 | 1.015533981 | 1 | 1 | 1 | FIM1_4857 | YLR004C | YLR004C | THI73 | n |
| Kmar_QGN13960.1 | W303.asm02.HP0_G0018760 | OG0001431 | chrVII | 266860 | 267243 | CP015054.1 | 1221293 | 1221673 | 62.99212598 | 127 | 126 | 1.007936508 | 1 | 1 | 1 | FIM1_608 | YGL127C | YGL127C | SOH1 | n |
| Kmar_QGN15981.1 | W303.asm02.HP0_G0003460 | OG0001432 | chrII | 564237 | 564971 | CP015057.1 | 536925 | 537674 | 52.4 | 244 | 249 | 0.979919679 | 1 | 1 | 1 | FIM1_2681 | YBR153W | YBR153W | RIB7 | n |
| Kmar_QGN14051.1 | W303.asm02.HP0_G0005880 | OG0001433 | chrIII | 216383 | 217384 | CP015054.1 | 1414033 | 1415040 | 51.32743363 | 333 | 335 | 0.994029851 | 1 | 1 | 1 | FIM1_702 | YCR036W | YCR036W | RBK1 | n |
| Kmar_QGN15562.1 | W303.asm02.HP0_G0002750 | OG0001434 | chrII | 411669 | 414563 | CP015056.1 | 1188129 | 1190909 | 49.69008264 | 964 | 926 | 1.041036717 | 1 | 1 | 1 | FIM1_2253 | YBR079C | YBR079C | RPG1 | n |
| Kmar_QGN14639.1 | W303.asm02.HP0_G0012100 | OG0001435 | chrIV | 1206928 | 1207779 | CP015055.1 | 922051 | 922863 | 66.07773852 | 283 | 270 | 1.048148148 | 1 | 1 | 1 | FIM1_1303 | YDR361C | YDR361C | BCP1 | n |
| Kmar_QGN17971.1 | W303.asm02.HP0_G0012140 | OG0001436 | chrIV | 1214359 | 1215726 | CP015060.1 | 855440 | 856729 | 52.63157895 | 455 | 429 | 1.060606061 | 1 | 1 | 1 | FIM1_5180 | YDR364C | YDR364C | CDC40 | n |
| Kmar_QGN15815.1 | W303.asm02.HP0_G0041720 | OG0001437 | chrXIII | 533779 | 535314 | CP015057.1 | 142647 | 144218 | 74.0952381 | 511 | 523 | 0.977055449 | 1 | 1 | 1 | FIM1_2511 | YMR131C | YMR131C | RRB1 | n |
| Kmar_QGN17006.1 | W303.asm02.HP0_G0002480 | OG0001438 | chrII | 360888 | 363587 | CP015059.1 | 14282 | 16912 | 34.27947598 | 899 | 876 | 1.026255708 | 1 | 1 | 1 | FIM1_3735 | YBR055C | YBR055C | PRP6 | n |
| Kmar_QGN17203.1 | W303.asm02.HP0_G0019120 | OG0001439 | chrVII | 344071 | 346320 | CP015059.1 | 441984 | 443948 | 39.36170213 | 749 | 654 | 1.145259939 | 1 | 1 | 1 | FIM1_3934 | YGL086W | YGL086W | MAD1 | n |
| Kmar_QGN16042.1 | W303.asm02.HP0_G0029520 | OG0001440 | chrX | 406883 | 409048 | CP015057.1 | 689029 | 691215 | 76.51098901 | 721 | 728 | 0.990384615 | 1 | 1 | 1 | FIM1_2742 | YJL012C | YJL012C | VTC4 | n |
| Kmar_QGN14927.1 | W303.asm02.HP0_G0006820 | OG0001441 | chrIV | 78499 | 79131 | CP015055.1 | 1511676 | 1512290 | 55.45023697 | 210 | 204 | 1.029411765 | 1 | 1 | 1 | FIM1_1602 | YDL212W | YDL212W | SHR3 | n |
| Kmar_QGN14928.1 | W303.asm02.HP0_G0045090 | OG0001442 | chrXIV | 304382 | 305311 | CP015055.1 | 1512468 | 1513364 | 50.48231511 | 309 | 298 | 1.036912752 | 1 | 1 | 1 | FIM1_1603 | YNL177C | YNL177C | MRPL22 | n |
| Kmar_QGN14920.1 | W303.asm02.HP0_G0045010 | OG0001443 | chrXIV | 290194 | 292572 | CP015055.1 | 1495699 | 1498014 | 45.63345633 | 792 | 771 | 1.027237354 | 1 | 1 | 1 | FIM1_1594 | YNL186W | YNL186W | UBP10 | n |
| Kmar_QGN14918.1 | W303.asm02.HP0_G0006780 | OG0001444 | chrIV | 69070 | 70437 | CP015055.1 | 1488508 | 1489650 | 30.71895425 | 455 | 380 | 1.197368421 | 1 | 1 | 1 | FIM1_1592 | YDL216C | YDL216C | RRI1 | n |
| Kmar_QGN15856.1 | W303.asm02.HP0_G0022870 | OG0001445 | chrVIII | 54853 | 56994 | CP015057.1 | 238228 | 240357 | 29.79274611 | 713 | 709 | 1.005641749 | 1 | 1 | 1 | FIM1_2553 | YHL024W | YHL024W | RIM4 | n |
| Kmar_QGN16436.1 | W303.asm02.HP0_G0004610 | OG0001446 | chrII | 769704 | 771152 | CP015058.1 | 114761 | 116173 | 32.71221532 | 482 | 470 | 1.025531915 | 1 | 1 | 1 | FIM1_3149 | YBR272C | YBR272C | HSM3 | n |
| Kmar_QGN16437.1 | W303.asm02.HP0_G0029220 | OG0001447 | chrX | 332541 | 335762 | CP015058.1 | 116232 | 119441 | 80.72625698 | 1073 | 1069 | 1.003741815 | 1 | 1 | 1 | FIM1_3150 | YJL050W | YJL050W | MTR4 | n |
| Kmar_QGN17927.1 | W303.asm02.HP0_G0015510 | OG0001448 | chrV | 350399 | 351016 | CP015060.1 | 752721 | 753338 | 91.70731707 | 205 | 205 | 1 | 1 | 1 | 1 | FIM1_5136 | YER094C | YER094C | PUP3 | n |
| Kmar_QGN17542.1 | W303.asm02.HP0_G0028580 | OG0001449 | chrX | 178021 | 179457 | CP015059.1 | 1132794 | 1133990 | 41.2244898 | 478 | 398 | 1.201005025 | 1 | 1 | 1 | FIM1_4278 | YJL123C | YJL123C | MTC1 | n |
| Kmar_QGN16986.1 | W303.asm02.HP0_G0044570 | OG0001450 | chrXIV | 207626 | 210550 | CP015058.1 | 1313680 | 1316544 | 36.09172483 | 974 | 954 | 1.020964361 | 1 | 1 | 1 | FIM1_3713 | YNL236W | YNL236W | SIN4 | n |
| Kmar_QGN16009.1 | W303.asm02.HP0_G0031730 | OG0001451 | chrXI | 154548 | 154985 | CP015057.1 | 607614 | 608138 | 57.38636364 | 145 | 174 | 0.833333333 | 1 | 1 | 1 | FIM1_2709 | YKL160W | YKL160W | ELF1 | n |
| Kmar_QGN14293.1 | W303.asm02.HP0_G0029280 | OG0001452 | chrX | 354235 | 355611 | CP015055.1 | 209109 | 210032 | 16.15720524 | 458 | 307 | 1.491856678 | 1 | 1 | 1 | FIM1_950 | YJL044C | YJL044C | GYP6 | n |
| Kmar_QGN14070.1 | W303.asm02.HP0_G0044480 | OG0001453 | chrXIV | 186015 | 186904 | CP015054.1 | 1458191 | 1458958 | 48.74551971 | 264 | 255 | 1.035294118 | 1 | 1 | 1 | FIM1_721 | YNL246W | YNL246W | VPS75 | n |

| | | | | | | | | | | | | | | | | | | | | |
|---|---|---|---|---|---|---|---|---|---|---|---|---|---|---|---|---|---|---|---|---|
| Kmar_QGN15591.1 | W303.asm02.HP0_G0013150 | OG0001454 | chrIV | 1416146 | 1416820 | CP015056.1 | 1257690 | 1258331 | 31.73913043 | 224 | 213 | 1.051643192 | 1 | 1 | 1 | FIM1_2283 | YDR476C | YDR476C | | n |
| Kmar_QGN14114.1 | W303.asm02.HP0_G0014650 | OG0001455 | chrV | 193106 | 193771 | CP015056.1 | 1560176 | 1560835 | 29.27927928 | 221 | 219 | 1.00913242 | 1 | 1 | 1 | FIM1_765 | YER018C | YER018C | SPC25 | n |
| Kmar_QGN15929.1 | W303.asm02.HP0_G0035930 | OG0001456 | chrXII | 354540 | 354935 | CP015057.1 | 397458 | 398039 | 38.02083333 | 131 | 193 | 0.678756477 | 1 | 1 | 1 | FIM1_2627 | YLR104W | YLR104W | LCL2 | n |
| Kmar_QGN14894.1 | W303.asm02.HP0_G0003210 | OG0001457 | chrII | 507554 | 509107 | CP015055.1 | 1448846 | 1450396 | 93.05019305 | 517 | 516 | 1.001937984 | 1 | 1 | 1 | FIM1_1568 | YBR127C | YBR127C | VMA2 | n |
| Kmar_QGN14896.1 | W303.asm02.HP0_G0056160 | OG0001458 | chrXVI | 713471 | 714841 | CP015055.1 | 1452165 | 1453523 | 44.28571429 | 456 | 452 | 1.008849558 | 1 | 1 | 1 | FIM1_1570 | YPR084W | YPR084W | | n |
| Kmar_QGN17892.1 | W303.asm02.HP0_G0003180 | OG0001459 | chrII | 503484 | 504665 | CP015060.1 | 688604 | 689818 | 58.43520782 | 393 | 404 | 0.972772277 | 1 | 1 | 1 | FIM1_5101 | YBR125C | YBR125C | PTC4 | n |
| Kmar_QGN14893.1 | W303.asm02.HP0_G0056150 | OG0001460 | chrXVI | 711353 | 713092 | CP015055.1 | 1446845 | 1448548 | 29.41176471 | 579 | 567 | 1.021164021 | 1 | 1 | 1 | FIM1_1567 | YPR083W | YPR083W | MDM36 | n |
| Kmar_QGN14092.1 | W303.asm02.HP0_G0044320 | OG0001461 | chrXIV | 147450 | 148394 | CP015054.1 | 1504166 | 1505056 | 54.74683544 | 314 | 296 | 1.060810811 | 1 | 1 | 1 | FIM1_743 | YNL263C | YNL263C | YIF1 | n |
| Kmar_QGN14975.1 | W303.asm02.HP0_G0051070 | OG0001462 | chrXV | 757771 | 758853 | CP015055.1 | 1605675 | 1606631 | 34.7826087 | 360 | 318 | 1.132075472 | 1 | 1 | 1 | FIM1_1653 | YOR221C | YOR221C | MCT1 | n |
| Kmar_QGN16907.1 | W303.asm02.HP0_G0010340 | OG0001463 | chrIV | 815824 | 816564 | CP015058.1 | 1169907 | 1170825 | 69.47791165 | 246 | 245 | 1.004081633 | 1 | 1 | 1 | FIM1_3634 | YDR174W | YDR174W | HMO1 | n |
| Kmar_QGN13885.1 | W303.asm02.HP0_G0022140 | OG0001464 | chrVII | 992185 | 994488 | CP015054.1 | 1065523 | 1067841 | 65.71072319 | 767 | 772 | 0.993523316 | 1 | 1 | 1 | FIM1_532 | YGR245C | YGR245C | SDA1 | n |
| Kmar_QGN13886.1 | W303.asm02.HP0_G0022150 | OG0001465 | chrVII | 994905 | 996695 | CP015054.1 | 1068470 | 1070149 | 63.88888889 | 596 | 559 | 1.066189624 | 1 | 1 | 1 | FIM1_533 | YGR246C | YGR246C | BRF1 | n |
| Kmar_QGN16183.1 | W303.asm02.HP0_G0024970 | OG0001466 | chrVIII | 448287 | 449582 | CP015057.1 | 979059 | 980366 | 76.32183908 | 431 | 435 | 0.990804598 | 1 | 1 | 1 | FIM1_2885 | YHR169W | YHR169W | DBP8 | n |
| Kmar_QGN13896.1 | W303.asm02.HP0_G0024960 | OG0001467 | chrVIII | 446483 | 448039 | CP015054.1 | 1102940 | 1104586 | 53.97111913 | 518 | 548 | 0.945255474 | 1 | 1 | 1 | FIM1_543 | YHR168W | YHR168W | MTG2 | n |
| Kmar_QGN17407.1 | W303.asm02.HP0_G0033130 | OG0001468 | chrXI | 420714 | 426008 | CP015059.1 | 851244 | 856310 | 34.23423423 | 1764 | 1688 | 1.045023697 | 1 | 1 | 1 | FIM1_4139 | YKL014C | YKL014C | URB1 | n |
| Kmar_QGN15371.1 | W303.asm02.HP0_G0054820 | OG0001469 | chrXVI | 450938 | 451390 | CP015056.1 | 821030 | 821470 | 69.33333333 | 150 | 146 | 1.02739726 | 1 | 1 | 1 | FIM1_2061 | YPL059W | YPL059W | GRX5 | n |
| Kmar_QGN17409.1 | W303.asm02.HP0_G0033110 | OG0001470 | chrXI | 416557 | 417081 | CP015059.1 | 860071 | 860595 | 76.43678161 | 174 | 174 | 1 | 1 | 1 | 1 | FIM1_4141 | YKL016C | YKL016C | ATP7 | n |
| Kmar_QGN18323.1 | W303.asm02.HP0_G0001530 | OG0001471 | chrII | 149512 | 152340 | CP015061.1 | 691228 | 694107 | 37.72102161 | 942 | 959 | 0.982273201 | 1 | 1 | 1 | FIM1_4649 | YBL037W | YBL037W | APL3 | n |
| Kmar_QGN17411.1 | W303.asm02.HP0_G0033090 | OG0001472 | chrXI | 413194 | 414183 | CP015059.1 | 863102 | 864178 | 63.88888889 | 329 | 358 | 0.918994413 | 1 | 1 | 1 | FIM1_4143 | YKL018W | YKL018W | SWD2 | n |
| Kmar_QGN15190.1 | W303.asm02.HP0_G0029470 | OG0001473 | chrX | 394586 | 398059 | CP015056.1 | 397369 | 400290 | 27.26488352 | 1157 | 973 | 1.189105858 | 1 | 1 | 1 | FIM1_1878 | YJL020C | YJL020C | BBC1 | n |
| Kmar_QGN18324.1 | W303.asm02.HP0_G0001540 | OG0001474 | chrII | 152750 | 153523 | CP015061.1 | 694177 | 694974 | 59.62962963 | 257 | 265 | 0.969811321 | 1 | 1 | 1 | FIM1_4650 | YBL036C | YBL036C | | n |
| Kmar_QGN18326.1 | W303.asm02.HP0_G0001550 | OG0001475 | chrII | 153796 | 155913 | CP015061.1 | 698811 | 700739 | 47.95486601 | 705 | 642 | 1.098130841 | 1 | 1 | 1 | FIM1_4652 | YBL035C | YBL035C | POL12 | n |
| Kmar_QGN17456.1 | W303.asm02.HP0_G0011840 | OG0001476 | chrIV | 1140188 | 1141423 | CP015059.1 | 959031 | 960260 | 61.48325359 | 411 | 409 | 1.004889976 | 1 | 1 | 1 | FIM1_4190 | YDR331W | YDR331W | GPI8 | n |
| Kmar_QGN17268.1 | W303.asm02.HP0_G0016640 | OG0001477 | chrVI | 51028 | 52323 | CP015059.1 | 586984 | 588285 | 47.1395881 | 431 | 433 | 0.995381062 | 1 | 1 | 1 | FIM1_3999 | YFL026W | YFL026W | STE2 | n |
| Kmar_QGN17699.1 | W303.asm02.HP0_G0026380 | OG0001478 | chrIX | 184090 | 185565 | CP015060.1 | 275004 | 276359 | 51.52749491 | 491 | 451 | 1.088691796 | 1 | 1 | 1 | FIM1_4906 | YIL090W | YIL090W | ICE2 | n |
| Kmar_QGN16187.1 | W303.asm02.HP0_G0022210 | OG0001479 | chrVII | 1009309 | 1010628 | CP015057.1 | 985538 | 987076 | 69.140625 | 439 | 512 | 0.857421875 | 1 | 1 | 1 | FIM1_2889 | YGR252W | YGR252W | GCN5 | n |
| Kmar_QGN16185.1 | W303.asm02.HP0_G0024980 | OG0001480 | chrVIII | 449934 | 451490 | CP015057.1 | 981459 | 983006 | 80.15414258 | 518 | 515 | 1.005825243 | 1 | 1 | 1 | FIM1_2887 | YHR170W | YHR170W | NMD3 | n |
| Kmar_QGN16186.1 | W303.asm02.HP0_G0024990 | OG0001481 | chrVIII | 451819 | 453711 | CP015057.1 | 983324 | 985144 | 48.96988906 | 630 | 606 | 1.03960396 | 1 | 1 | 1 | FIM1_2888 | YHR171W | YHR171W | ATG7 | n |
| Kmar_QGN13735.1 | W303.asm02.HP0_G0025000 | OG0001482 | chrVIII | 454441 | 456912 | CP015054.1 | 767166 | 769502 | 27.20763723 | 823 | 778 | 1.057840617 | 1 | 1 | 1 | FIM1_379 | YHR172W | YHR172W | SPC97 | n |
| Kmar_QGN16188.1 | W303.asm02.HP0_G0022220 | OG0001483 | chrVII | 1010798 | 1011580 | CP015057.1 | 987437 | 988240 | 83.89513109 | 260 | 267 | 0.973782772 | 1 | 1 | 1 | FIM1_2890 | YGR253C | YGR253C | PUP2 | n |
| Kmar_QGN15005.1 | W303.asm02.HP0_G0052420 | OG0001484 | chrXV | 1028288 | 1028890 | CP015055.1 | 1670361 | 1670975 | 68.4729064 | 200 | 204 | 0.980392157 | 1 | 1 | 1 | FIM1_1686 | YOR367W | YOR367W | SCP1 | n |
| Kmar_QGN14542.1 | W303.asm02.HP0_G0049220 | OG0001485 | chrXV | 358130 | 360403 | CP015055.1 | 723232 | 724752 | 60.30428769 | 757 | 506 | 1.496047431 | 1 | 1 | 1 | FIM1_1204 | YOR014W | YOR014W | RTS1 | n |
| Kmar_QGN16051.1 | W303.asm02.HP0_G0037310 | OG0001486 | chrXII | 636699 | 637292 | CP015057.1 | 706560 | 707150 | 56.28140704 | 197 | 196 | 1.005102041 | 1 | 1 | 1 | FIM1_2751 | YLR251W | YLR251W | SYM1 | n |
| Kmar_QGN14040.1 | W303.asm02.HP0_G0048170 | OG0001487 | chrXV | 143270 | 144151 | CP015054.1 | 1391344 | 1392300 | 52.40963855 | 293 | 318 | 0.921383648 | 1 | 1 | 1 | FIM1_691 | YOL093W | YOL093W | TRM10 | n |
| Kmar_QGN13408.1 | W303.asm02.HP0_G0054480 | OG0001488 | chrXVI | 373285 | 374376 | CP015054.1 | 79478 | 80542 | 59.34065934 | 363 | 354 | 1.025423729 | 1 | 1 | 1 | FIM1_45 | YPL096W | YPL096W | PNG1 | n |
| Kmar_QGN16801.1 | W303.asm02.HP0_G0020730 | OG0001489 | chrVII | 675091 | 676035 | CP015058.1 | 921648 | 922595 | 43.4375 | 314 | 315 | 0.996825397 | 1 | 1 | 1 | FIM1_3525 | YGR096W | YGR096W | TPC1 | n |
| Kmar_QGN16194.1 | W303.asm02.HP0_G0039830 | OG0001490 | chrXIII | 124289 | 126112 | CP015057.1 | 999043 | 1000299 | 30.79470199 | 607 | 418 | 1.45215311 | 1 | 1 | 1 | FIM1_2896 | YML071C | YML071C | COG8 | n |
| Kmar_QGN15098.1 | W303.asm02.HP0_G0042720 | OG0001491 | chrXIII | 734123 | 737212 | CP015056.1 | 198668 | 201742 | 44.52975048 | 1029 | 1024 | 1.004882813 | 1 | 1 | 1 | FIM1_1785 | YMR231W | YMR231W | PEP5 | n |
| Kmar_QGN13640.1 | W303.asm02.HP0_G0005780 | OG0001492 | chrIII | 176025 | 178253 | CP015054.1 | 565197 | 567122 | 25.59681698 | 742 | 641 | 1.157566303 | 1 | 0 | 0 | FIM1_282 | YCR026C | YCR026C | NPP1 | n |
| Kmar_QGN17868.1 | W303.asm02.HP0_G0003050 | OG0001493 | chrII | 475158 | 476507 | CP015060.1 | 640565 | 642115 | 53.7428023 | 449 | 516 | 0.870155039 | 1 | 1 | 1 | FIM1_5077 | YBR110W | YBR110W | ALG1 | n |
| Kmar_QGN17869.1 | W303.asm02.HP0_G0056010 | OG0001494 | chrXVI | 689439 | 690851 | CP015060.1 | 642106 | 643419 | 47.65957447 | 470 | 437 | 1.075514874 | 1 | 1 | 1 | FIM1_5078 | YPR068C | YPR068C | HOS1 | n |
| Kmar_QGN17346.1 | W303.asm02.HP0_G0052230 | OG0001495 | chrXV | 984109 | 987066 | CP015059.1 | 739688 | 742144 | 31.56843157 | 985 | 818 | 1.204156479 | 1 | 1 | 1 | FIM1_4078 | YOR346W | YOR346W | REV1 | n |
| Kmar_QGN17978.1 | W303.asm02.HP0_G0034560 | OG0001496 | chrXII | 54220 | 63654 | CP015060.1 | 869087 | 878470 | 43.08560677 | 3144 | 3127 | 1.005436521 | 1 | 1 | 1 | FIM1_5187 | YLL040C | YLL040C | VPS13 | n |
| Kmar_QGN14239.1 | W303.asm02.HP0_G0001780 | OG0001497 | chrII | 204341 | 205546 | CP015055.1 | 91847 | 92938 | 40.94292804 | 401 | 363 | 1.104683196 | 1 | 1 | 1 | FIM1_895 | YBL013W | YBL013W | FMT1 | n |
| Kmar_QGN14243.1 | W303.asm02.HP0_G0001770 | OG0001498 | chrII | 201349 | 204033 | CP015055.1 | 98153 | 100792 | 24.55947137 | 894 | 879 | 1.017064846 | 1 | 1 | 1 | FIM1_899 | YBL014C | YBL014C | RRN6 | n |
| Kmar_QGN14244.1 | W303.asm02.HP0_G0019640 | OG0001499 | chrVII | 440728 | 443229 | CP015055.1 | 101195 | 103654 | 49.46996466 | 833 | 819 | 1.017094017 | 1 | 1 | 1 | FIM1_900 | YGL027C | YGL027C | CWH41 | n |
| Kmar_QGN16707.1 | W303.asm02.HP0_G0053020 | OG0001500 | chrXVI | 88190 | 89761 | CP015058.1 | 713548 | 715050 | 38.48739496 | 523 | 500 | 1.046 | 1 | 1 | 1 | FIM1_3427 | YPL247C | YPL247C | | n |
| Kmar_QGN16706.1 | W303.asm02.HP0_G0042280 | OG0001501 | chrXIII | 638083 | 641187 | CP015058.1 | 709790 | 712882 | 69.46564885 | 1034 | 1030 | 1.003883495 | 1 | 1 | 1 | FIM1_3426 | YMR189W | YMR189W | GCV2 | n |
| Kmar_QGN14506.1 | W303.asm02.HP0_G0016620 | OG0001502 | chrVI | 47792 | 48661 | CP015055.1 | 652549 | 653394 | 56.41891892 | 289 | 281 | 1.028469751 | 1 | 1 | 1 | FIM1_1167 | YFL028C | YFL028C | CAF16 | n |
| Kmar_QGN15217.1 | W303.asm02.HP0_G0021480 | OG0001503 | chrVII | 857674 | 858186 | CP015056.1 | 451327 | 451761 | 47.64705882 | 170 | 144 | 1.180555556 | 1 | 1 | 1 | FIM1_1905 | YGR174C | YGR174C | CBP4 | n |
| Kmar_QGN15542.1 | W303.asm02.HP0_G0013030 | OG0001504 | chrIV | 1391766 | 1392209 | CP015056.1 | 1155741 | 1156184 | 55.10204082 | 147 | 147 | 1 | 1 | 1 | 1 | FIM1_2233 | YDR462W | YDR462W | MRPL28 | n |
| Kmar_QGN16508.1 | W303.asm02.HP0_G0010710 | OG0001505 | chrIV | 891584 | 893263 | CP015058.1 | 267048 | 268727 | 88.01431127 | 559 | 559 | 1 | 1 | 1 | 1 | FIM1_3221 | YDR212W | YDR212W | TCP1 | n |
| Kmar_QGN15216.1 | W303.asm02.HP0_G0021490 | OG0001506 | chrVII | 858714 | 860204 | CP015056.1 | 449310 | 450812 | 74.2 | 496 | 500 | 0.992 | 1 | 1 | 1 | FIM1_1904 | YGR175C | YGR175C | ERG1 | n |
| Kmar_QGN16983.1 | W303.asm02.HP0_G0044590 | OG0001507 | chrXIV | 212618 | 215296 | CP015058.1 | 1305137 | 1307590 | 32.07343413 | 892 | 817 | 1.091799266 | 1 | 1 | 1 | FIM1_3710 | YNL233W | YNL233W | BNI4 | n |
| Kmar_QGN17377.1 | W303.asm02.HP0_G0041030 | OG0001508 | chrXIII | 391602 | 391988 | CP015059.1 | 800996 | 801349 | 43.75 | 128 | 117 | 1.094017094 | 1 | 1 | 1 | FIM1_4109 | YMR059W | YMR059W | SEN15 | n |
| Kmar_QGN15826.1 | W303.asm02.HP0_G0041670 | OG0001509 | chrXIII | 521377 | 522405 | CP015057.1 | 169770 | 170747 | 31.48688047 | 342 | 325 | 1.052307692 | 1 | 1 | 1 | FIM1_2522 | YMR126C | YMR126C | DLT1 | n |
| Kmar_QGN16943.1 | W303.asm02.HP0_G0010610 | OG0001510 | chrIV | 860033 | 860530 | CP015058.1 | 1232590 | 1233057 | 41.21212121 | 165 | 155 | 1.064516129 | 1 | 1 | 1 | FIM1_3670 | YDR201W | YDR201W | SPC19 | n |
| Kmar_QGN17130.1 | W303.asm02.HP0_G0045290 | OG0001511 | chrXIV | 343208 | 344032 | CP015059.1 | 299362 | 300177 | 41.99288256 | 274 | 271 | 1.011070111 | 1 | 1 | 1 | FIM1_3861 | YNL155W | YNL155W | CUZ1 | n |
| Kmar_QGN13397.1 | W303.asm02.HP0_G0054400 | OG0001512 | chrXVI | 362058 | 364034 | CP015054.1 | 64400 | 66592 | 53.58592693 | 658 | 730 | 0.901369863 | 1 | 1 | 1 | FIM1_34 | YPL104W | YPL104W | MSD1 | n |
| Kmar_QGN18393.1 | W303.asm02.HP0_G0029540 | OG0001513 | chrX | 413211 | 415211 | CP015061.1 | 840779 | 842800 | 64.6971935 | 666 | 673 | 0.989598811 | 1 | 1 | 1 | FIM1_4720 | YJL010C | YJL010C | NOP9 | n |
| Kmar_QGN17979.1 | W303.asm02.HP0_G0034550 | OG0001514 | chrXII | 53140 | 53940 | CP015060.1 | 879853 | 880617 | 78.57142857 | 266 | 254 | 1.047244094 | 1 | 1 | 1 | FIM1_5188 | YLL041C | YLL041C | SDH2 | n |
| Kmar_QGN14802.1 | W303.asm02.HP0_G0012190 | OG0001515 | chrIV | 1223375 | 1224703 | CP015055.1 | 1257089 | 1258336 | 26.23318386 | 442 | 415 | 1.065060241 | 1 | 1 | 1 | FIM1_1473 | YDR370C | YDR370C | DXO1 | n |
| Kmar_QGN15705.1 | W303.asm02.HP0_G0017330 | OG0001516 | chrVI | 206272 | 207159 | CP015056.1 | 1504637 | 1505458 | 43.2885906 | 295 | 273 | 1.080586081 | 1 | 1 | 1 | FIM1_2398 | YFR041C | YFR041C | ERJ5 | n |
| Kmar_QGN18097.1 | W303.asm02.HP0_G0030390 | OG0001517 | chrX | 597492 | 600356 | CP015061.1 | 213924 | 216167 | 18.72384937 | 954 | 747 | 1.277108434 | 1 | 1 | 1 | FIM1_4417 | YJR089W | YJR089W | BIR1 | n |
| Kmar_QGN18108.1 | W303.asm02.HP0_G0055740 | OG0001518 | chrXVI | 648705 | 649817 | CP015061.1 | 230147 | 231256 | 86.48648649 | 370 | 369 | 1.002710027 | 1 | 1 | 1 | FIM1_4428 | YPR035W | YPR035W | GLN1 | n |
| Kmar_QGN15697.1 | W303.asm02.HP0_G0014900 | OG0001519 | chrV | 233600 | 235879 | CP015056.1 | 1488703 | 1490481 | 24.47368421 | 759 | 592 | 1.282094595 | 1 | 1 | 1 | FIM1_2390 | YER041W | YER041W | YEN1 | n |
| Kmar_QGN13787.1 | W303.asm02.HP0_G0034090 | OG0001520 | chrXI | 617417 | 618004 | CP015054.1 | 879729 | 880322 | 42.42424242 | 195 | 197 | 0.989847716 | 1 | 1 | 1 | FIM1_433 | YKR085C | YKR085C | MRPL20 | n |
| Kmar_QGN13468.1 | W303.asm02.HP0_G0024570 | OG0001521 | chrVIII | 368088 | 368738 | CP015054.1 | 200568 | 201221 | 83.87096774 | 216 | 217 | 0.995391705 | 1 | 1 | 1 | FIM1_105 | YHR128W | YHR128W | FUR1 | n |
| Kmar_QGN16227.1 | W303.asm02.HP0_G0057090 | OG0001522 | chrXVI | 917089 | 917632 | CP015057.1 | 1077543 | 1078019 | 80.37974684 | 155 | 158 | 0.981012658 | 1 | 1 | 1 | FIM1_2931 | YPR187W | YPR187W | RPO26 | n |
| Kmar_QGN16231.1 | W303.asm02.HP0_G0057130 | OG0001523 | chrXVI | 925213 | 926319 | CP015057.1 | 1086215 | 1087297 | 63.04347826 | 368 | 360 | 1.022222222 | 1 | 1 | 1 | FIM1_2935 | YPR191W | YPR191W | QCR2 | n |
| Kmar_QGN18374.1 | W303.asm02.HP0_G0044080 | OG0001524 | chrXIV | 88343 | 89182 | CP015061.1 | 797724 | 798569 | 43.72881356 | 279 | 281 | 0.992882562 | 1 | 1 | 1 | FIM1_4701 | YNL289W | YNL289W | PCL1 | n |
| Kmar_QGN13817.1 | W303.asm02.HP0_G0011140 | OG0001525 | chrIV | 974491 | 975975 | CP015054.1 | 932778 | 934325 | 41.16504854 | 494 | 515 | 0.959223301 | 1 | 1 | 1 | FIM1_463 | YDR257C | YDR257C | RKM4 | n |
| Kmar_QGN13816.1 | W303.asm02.HP0_G0049320 | OG0001526 | chrXV | 380237 | 381262 | CP015054.1 | 931756 | 932697 | 37.79069767 | 341 | 313 | 1.089456869 | 1 | 1 | 1 | FIM1_462 | YOR026W | YOR026W | BUB3 | n |
| Kmar_QGN13695.1 | W303.asm02.HP0_G0043410 | OG0001527 | chrXIII | 874475 | 876547 | CP015054.1 | 675927 | 678053 | 72.24606581 | 690 | 708 | 0.974576271 | 1 | 1 | 1 | FIM1_338 | YMR301C | YMR301C | ATM1 | n |
| Kmar_QGN15814.1 | W303.asm02.HP0_G0041730 | OG0001528 | chrXIII | 535560 | 536186 | CP015057.1 | 141728 | 142357 | 61.53846154 | 208 | 209 | 0.995215311 | 1 | 1 | 1 | FIM1_2510 | YMR132C | YMR132C | JLP2 | n |
| Kmar_QGN15774.1 | W303.asm02.HP0_G0006330 | OG0001529 | chrIII | 303709 | 310035 | CP015057.1 | 63215 | 69568 | 47.84222738 | 2108 | 2117 | 0.995748701 | 1 | 1 | 1 | FIM1_2470 | YCR093W | YCR093W | CDC39 | n |
| Kmar_QGN15772.1 | W303.asm02.HP0_G0010370 | OG0001530 | chrIV | 820592 | 821239 | CP015057.1 | 60729 | 61406 | 68.75 | 215 | 225 | 0.955555556 | 1 | 1 | 1 | FIM1_2468 | YDR177W | YDR177W | UBC1 | n |
| Kmar_QGN15766.1 | W303.asm02.HP0_G0047230 | OG0001531 | chrXIV | 729363 | 730910 | CP015057.1 | 47466 | 48986 | 30.70017953 | 515 | 506 | 1.017786561 | 1 | 1 | 1 | FIM1_2462 | YNR051C | YNR051C | BRE5 | n |
| Kmar_QGN14816.1 | W303.asm02.HP0_G0047700 | OG0001532 | chrXV | 53301 | 54752 | CP015055.1 | 1284218 | 1285708 | 42.58188825 | 483 | 496 | 0.973790323 | 1 | 1 | 1 | FIM1_1487 | YOL144W | YOL144W | NOP8 | n |
| Kmar_QGN17950.1 | W303.asm02.HP0_G0001130 | OG0001533 | chrII | 70904 | 72280 | CP015060.1 | 805797 | 807206 | 46.23430962 | 458 | 469 | 0.976545842 | 1 | 1 | 1 | FIM1_5159 | YBL082C | YBL082C | ALG3 | n |
| Kmar_QGN17948.1 | W303.asm02.HP0_G0015630 | OG0001535 | chrV | 375118 | 376215 | CP015060.1 | 799492 | 800604 | 68.73315364 | 365 | 370 | 0.986486486 | 1 | 1 | 1 | FIM1_5157 | YER107C | YER107C | GLE2 | n |
| Kmar_QGN16026.1 | W303.asm02.HP0_G0019550 | OG0001536 | chrVII | 424383 | 425033 | CP015057.1 | 653160 | 653810 | 46.36363636 | 216 | 216 | 1 | 1 | 1 | 1 | FIM1_2726 | YGL037C | YGL037C | PNC1 | n |
| Kmar_QGN15288.1 | W303.asm02.HP0_G0030040 | OG0001537 | chrX | 537352 | 538059 | CP015056.1 | 607588 | 608304 | 41.8699187 | 235 | 238 | 0.987394958 | 1 | 1 | 1 | FIM1_1977 | YJR050W | YJR050W | ISY1 | n |
| Kmar_QGN16521.1 | W303.asm02.HP0_G0021180 | OG0001538 | chrVII | 793494 | 795617 | CP015058.1 | 295196 | 297295 | 75.38896747 | 707 | 699 | 1.011444921 | 1 | 1 | 1 | FIM1_3234 | YGR145W | YGR145W | ENP2 | n |
| Kmar_QGN13762.1 | W303.asm02.HP0_G0023310 | OG0001539 | chrVIII | 133842 | 135233 | CP015054.1 | 829264 | 830652 | 82.46753247 | 463 | 462 | 1.002164502 | 1 | 1 | 1 | FIM1_407 | YHR018C | YHR018C | ARG4 | n |
| Kmar_QGN16075.1 | W303.asm02.HP0_G0048110 | OG0001540 | chrXV | 133180 | 136293 | CP015057.1 | 753801 | 756932 | 53.14285714 | 1037 | 1043 | 0.994247363 | 1 | 1 | 1 | FIM1_2775 | YOL098C | YOL098C | SDD3 | n |
| Kmar_QGN13634.1 | W303.asm02.HP0_G0005760 | OG0001541 | chrIII | 172658 | 174136 | CP015054.1 | 557620 | 558951 | 44.24242424 | 492 | 443 | 1.110609481 | 1 | 1 | 1 | FIM1_276 | YCR024C | YCR024C | SLM5 | n |
| Kmar_QGN13759.1 | W303.asm02.HP0_G0017140 | OG0001542 | chrVI | 172648 | 173655 | CP015054.1 | 823477 | 824499 | 44.6685879 | 335 | 340 | 0.985294118 | 1 | 1 | 1 | FIM1_404 | YFR025C | YFR025C | HIS2 | n |
| Kmar_QGN13760.1 | W303.asm02.HP0_G0023330 | OG0001543 | chrVIII | 137828 | 139906 | CP015054.1 | 824743 | 826776 | 76.04617605 | 692 | 677 | 1.022156573 | 1 | 1 | 1 | FIM1_405 | YHR020W | YHR020W | | n |
| Kmar_QGN13761.1 | W303.asm02.HP0_G0023320 | OG0001544 | chrVIII | 135725 | 137389 | CP015054.1 | 827115 | 828773 | 75.27075812 | 554 | 552 | 1.003623188 | 1 | 1 | 1 | FIM1_406 | YHR019C | YHR019C | DED81 | n |
| Kmar_QGN17928.1 | W303.asm02.HP0_G0001300 | OG0001545 | chrII | 109080 | 111143 | CP015060.1 | 753686 | 755650 | 27.57660167 | 687 | 654 | 1.050458716 | 1 | 1 | 1 | FIM1_5137 | YBL060W | YBL060W | YEL1 | n |
| Kmar_QGN16963.1 | W303.asm02.HP0_G0006630 | OG0001546 | chrIV | 34310 | 36550 | CP015058.1 | 1264111 | 1266351 | 46.25322997 | 746 | 746 | 1 | 1 | 1 | 1 | FIM1_3690 | YDL234C | YDL234C | GYP7 | n |
| Kmar_QGN16965.1 | W303.asm02.HP0_G0044730 | OG0001547 | chrXIV | 241030 | 242010 | CP015058.1 | 1269193 | 1270128 | 33.92857143 | 326 | 311 | 1.048231511 | 1 | 1 | 1 | FIM1_3692 | YNL217W | YNL217W | PPN2 | n |
| Kmar_QGN13578.1 | W303.asm02.HP0_G0038880 | OG0001548 | chrXII | 979563 | 983486 | CP015054.1 | 452474 | 456367 | 43.62670713 | 1307 | 1297 | 1.0077101 | 1 | 1 | 1 | FIM1_219 | YLR425W | YLR425W | TUS1 | n |
| Kmar_QGN14646.1 | W303.asm02.HP0_G0034710 | OG0001549 | chrXII | 99053 | 100210 | CP015055.1 | 935060 | 936145 | 21.6957606 | 385 | 361 | 1.066481994 | 1 | 1 | 1 | FIM1_1310 | YLL022C | YLL022C | HIF1 | n |
| Kmar_QGN18161.1 | W303.asm02.HP0_G0015170 | OG0001550 | chrV | 286911 | 288584 | CP015061.1 | 350798 | 352462 | 69.87522282 | 557 | 554 | 1.005415162 | 1 | 1 | 1 | FIM1_4484 | YER065C | YER065C | ICL1 | n |
| Kmar_QGN17177.1 | W303.asm02.HP0_G0055100 | OG0001551 | chrXVI | 499902 | 501605 | CP015059.1 | 392602 | 394212 | 59.40246046 | 567 | 536 | 1.057835821 | 1 | 1 | 1 | FIM1_3908 | YPL030W | YPL030W | TRM44 | n |
| Kmar_QGN17165.1 | W303.asm02.HP0_G0054960 | OG0001552 | chrXVI | 473620 | 476016 | CP015059.1 | 368167 | 370548 | 35.93556382 | 798 | 793 | 1.00630517 | 1 | 1 | 1 | FIM1_3896 | YPL045W | YPL045W | VPS16 | n |
| Kmar_QGN17164.1 | W303.asm02.HP0_G0054950 | OG0001553 | chrXVI | 473004 | 473303 | CP015059.1 | 367633 | 367923 | 50.50505051 | 99 | 96 | 1.03125 | 1 | 1 | 1 | FIM1_3895 | YPL046C | YPL046C | ELC1 | n |

| | | | | | | | | | | | | | | | | | | | | |
|---|---|---|---|---|---|---|---|---|---|---|---|---|---|---|---|---|---|---|---|---|
| Kmar_QGN17163.1 | W303.asm02.HP0_G0011120 | OG0001554 | chrIV | 971058 | 972323 | CP015059.1 | 366372 | 367562 | 41.84397163 | 421 | 396 | 1.063131313 | 1 | 1 | 1 | FIM1_3894 | YDR255C | YDR255C | RMD5 | n |
| Kmar_QGN13643.1 | W303.asm02.HP0_G0014340 | OG0001555 | chrV | 127812 | 129467 | CP015054.1 | 570621 | 572123 | 28.57142857 | 551 | 500 | 1.102 | 1 | 1 | 1 | FIM1_285 | YEL015W | YEL015W | EDC3 | n |
| Kmar_QGN16802.1 | W303.asm02.HP0_G0056410 | OG0001556 | chrXVI | 759186 | 759848 | CP015058.1 | 923042 | 923749 | 56.3559322 | 220 | 235 | 0.936170213 | 1 | 1 | 1 | FIM1_3526 | YPR113W | YPR113W | PIS1 | n |
| Kmar_QGN16252.1 | W303.asm02.HP0_G0025260 | OG0001557 | chrVIII | 499998 | 502289 | CP015057.1 | 1132249 | 1134570 | 40.72164948 | 763 | 773 | 0.987063389 | 1 | 1 | 1 | FIM1_2957 | YHR197W | YHR197W | RIX1 | n |
| Kmar_QGN16950.1 | W303.asm02.HP0_G0044840 | OG0001558 | chrXIV | 257487 | 258854 | CP015058.1 | 1243203 | 1244615 | 42.33128834 | 455 | 470 | 0.968085106 | 1 | 1 | 1 | FIM1_3677 | YNL206C | YNL206C | RTT106 | n |
| Kmar_QGN15681.1 | W303.asm02.HP0_G0017980 | OG0001559 | chrVII | 86945 | 87895 | CP015056.1 | 1456078 | 1457133 | 25.35211268 | 316 | 351 | 0.9002849 | 1 | 1 | 1 | FIM1_2374 | YGL212W | YGL212W | VAM7 | n |
| Kmar_QGN17323.1 | W303.asm02.HP0_G0045660 | OG0001560 | chrXIV | 411327 | 413261 | CP015059.1 | 694629 | 696602 | 50.56980057 | 644 | 657 | 0.98021309 | 1 | 1 | 1 | FIM1_4055 | YNL115C | YNL115C | | n |
| Kmar_QGN18233.1 | W303.asm02.HP0_G0036910 | OG0001561 | chrXII | 556915 | 559416 | CP015061.1 | 508315 | 510759 | 34.07494145 | 833 | 814 | 1.023341523 | 1 | 1 | 1 | FIM1_4558 | YLR207W | YLR207W | HRD3 | n |
| Kmar_QGN13891.1 | W303.asm02.HP0_G0024920 | OG0001562 | chrVIII | 430716 | 435284 | CP015054.1 | 1085706 | 1090142 | 35.77806122 | 1522 | 1478 | 1.029769959 | 1 | 1 | 1 | FIM1_538 | YHR164C | YHR164C | DNA2 | n |
| Kmar_QGN15130.1 | W303.asm02.HP0_G0031970 | OG0001563 | chrXI | 190429 | 192747 | CP015056.1 | 262038 | 264386 | 52.67175573 | 772 | 782 | 0.987212276 | 1 | 1 | 1 | FIM1_1817 | YKL134C | YKL134C | OCT1 | n |
| Kmar_QGN18113.1 | W303.asm02.HP0_G0000910 | OG0001564 | chrII | 10265 | 10855 | CP015061.1 | 238811 | 239407 | 37 | 196 | 198 | 0.98989899 | 1 | 1 | 1 | FIM1_4433 | YBL107C | YBL107C | MIX23 | n |
| Kmar_QGN14516.1 | W303.asm02.HP0_G0051900 | OG0001565 | chrXV | 898545 | 900080 | CP015055.1 | 678558 | 680099 | 89.66861598 | 511 | 513 | 0.996101365 | 1 | 1 | 1 | FIM1_1178 | YOR310C | YOR310C | NOP58 | n |
| Kmar_QGN14748.1 | W303.asm02.HP0_G0023240 | OG0001566 | chrVIII | 121246 | 122586 | CP015055.1 | 1152945 | 1154279 | 57.71812081 | 446 | 444 | 1.004504505 | 1 | 1 | 1 | FIM1_1418 | YHR011W | YHR011W | DIA4 | n |
| Kmar_QGN17220.1 | W303.asm02.HP0_G0004760 | OG0001567 | chrII | 801381 | 802832 | CP015059.1 | 477884 | 479452 | 28.67513612 | 483 | 522 | 0.925287356 | 1 | 1 | 1 | FIM1_3951 | YBR288C | YBR288C | APM3 | n |
| Kmar_QGN16860.1 | W303.asm02.HP0_G0010100 | OG0001568 | chrIV | 758583 | 759974 | CP015058.1 | 1051664 | 1053031 | 68.25053996 | 463 | 455 | 1.017582418 | 1 | 1 | 1 | FIM1_3586 | YDR148C | YDR148C | KGD2 | n |
| Kmar_QGN17138.1 | W303.asm02.HP0_G0016720 | OG0001569 | chrVI | 70111 | 71610 | CP015059.1 | 314520 | 316001 | 80.76152305 | 499 | 493 | 1.012170385 | 1 | 1 | 1 | FIM1_3869 | YFL018C | YFL018C | LPD1 | n |
| Kmar_QGN16458.1 | W303.asm02.HP0_G0029080 | OG0001570 | chrX | 305778 | 306494 | CP015058.1 | 155896 | 156615 | 40.08264463 | 238 | 239 | 0.9958159 | 1 | 1 | 1 | FIM1_3171 | YJL063C | YJL063C | MRPL8 | n |
| Kmar_QGN16455.1 | W303.asm02.HP0_G0004550 | OG0001571 | chrII | 761938 | 762900 | CP015058.1 | 149500 | 150444 | 44.89164087 | 320 | 314 | 1.01910828 | 1 | 1 | 1 | FIM1_3168 | YBR265W | YBR265W | TSC10 | n |
| Kmar_QGN16471.1 | W303.asm02.HP0_G0004520 | OG0001572 | chrII | 759076 | 759396 | CP015058.1 | 179162 | 179446 | 29.52380952 | 106 | 94 | 1.127659574 | 1 | 1 | 1 | FIM1_3184 | YBR262C | YBR262C | MIC12 | n |
| Kmar_QGN16550.1 | W303.asm02.HP0_G0039020 | OG0001573 | chrXII | 1012234 | 1014363 | CP015058.1 | 352294 | 354327 | 29.37931034 | 709 | 677 | 1.047267356 | 1 | 1 | 1 | FIM1_3264 | YLR440C | YLR440C | SEC39 | n |
| Kmar_QGN16553.1 | W303.asm02.HP0_G0039010 | OG0001574 | chrXII | 1011155 | 1012114 | CP015058.1 | 358818 | 359816 | 49.38271605 | 319 | 332 | 0.960843373 | 1 | 1 | 1 | FIM1_3267 | YLR439W | YLR439W | MRPL4 | n |
| Kmar_QGN15326.1 | W303.asm02.HP0_G0018260 | OG0001575 | chrVII | 152056 | 153109 | CP015056.1 | 710797 | 711474 | 28.39506173 | 324 | 225 | 1.44 | 1 | 1 | 1 | FIM1_2015 | YGL183C | YGL183C | MND1 | n |
| Kmar_QGN15684.1 | W303.asm02.HP0_G0017960 | OG0001576 | chrVII | 83493 | 84851 | CP015056.1 | 1463273 | 1464535 | 30.85106383 | 452 | 420 | 1.076190476 | 1 | 1 | 1 | FIM1_2377 | YGL215W | YGL215W | CLG1 | n |
| Kmar_QGN16372.1 | W303.asm02.HP0_G0014780 | OG0001577 | chrV | 214555 | 215037 | CP015057.1 | 1384316 | 1384810 | 47.97687861 | 160 | 164 | 0.975609756 | 1 | 1 | 1 | FIM1_3079 | YER030W | YER030W | CHZ1 | n |
| Kmar_QGN18155.1 | W303.asm02.HP0_G0026620 | OG0001578 | chrIX | 234628 | 235092 | CP015061.1 | 340550 | 341011 | 70.77922078 | 154 | 153 | 1.006535948 | 1 | 1 | 1 | FIM1_4478 | YIL062C | YIL062C | ARC15 | n |
| Kmar_QGN14367.1 | W303.asm02.HP0_G0009690 | OG0001579 | chrIV | 660003 | 663740 | CP015055.1 | 367810 | 371172 | 30.18124507 | 1245 | 1120 | 1.111607143 | 1 | 1 | 1 | FIM1_1027 | YDR104C | YDR104C | SPO71 | n |
| Kmar_QGN17244.1 | W303.asm02.HP0_G0032910 | OG0001580 | chrXI | 380432 | 382708 | CP015059.1 | 533516 | 535789 | 56.62337662 | 758 | 757 | 1.001321004 | 1 | 1 | 1 | FIM1_3975 | YKL034W | YKL034W | TUL1 | n |
| Kmar_QGN14109.1 | W303.asm02.HP0_G0005920 | OG0001581 | chrIII | 224155 | 228378 | CP015054.1 | 1549611 | 1553792 | 47.44827586 | 1407 | 1393 | 1.010050251 | 1 | 1 | 1 | FIM1_760 | YCR042C | YCR042C | TAF2 | n |
| Kmar_QGN17941.1 | W303.asm02.HP0_G0001180 | OG0001582 | chrII | 82196 | 85414 | CP015060.1 | 783102 | 786323 | 81.36067102 | 1072 | 1073 | 0.999068034 | 1 | 1 | 1 | FIM1_5150 | YBL076C | YBL076C | ILS1 | n |
| Kmar_QGN17215.1 | W303.asm02.HP0_G0053640 | OG0001583 | chrXVI | 205046 | 208087 | CP015059.1 | 464769 | 467744 | 43.52941176 | 1013 | 991 | 1.022199798 | 1 | 1 | 1 | FIM1_3946 | YPL183C | YPL183C | RTT10 | n |
| Kmar_QGN17599.1 | W303.asm02.HP0_G0013930 | OG0001584 | chrV | 47531 | 48736 | CP015060.1 | 74616 | 75845 | 50.84337349 | 401 | 409 | 0.980440098 | 1 | 1 | 1 | FIM1_4806 | YEL056W | YEL056W | HAT2 | n |
| Kmar_QGN18417.1 | W303.asm02.HP0_G0055420 | OG0001585 | chrXVI | 567861 | 570125 | CP015061.1 | 888773 | 890893 | 50.39787798 | 754 | 706 | 1.067988669 | 1 | 1 | 1 | FIM1_4745 | YPR003C | YPR003C | | n |
| Kmar_QGN18416.1 | W303.asm02.HP0_G0055430 | OG0001586 | chrXVI | 570364 | 571398 | CP015061.1 | 887279 | 888313 | 54.44126074 | 344 | 344 | 1 | 1 | 1 | 1 | FIM1_4744 | YPR004C | YPR004C | AIM45 | n |
| Kmar_QGN15949.1 | W303.asm02.HP0_G0050590 | OG0001587 | chrXV | 645441 | 647771 | CP015057.1 | 450678 | 453053 | 45.9697733 | 776 | 791 | 0.981036662 | 1 | 1 | 1 | FIM1_2647 | YOR165W | YOR165W | SEY1 | n |
| Kmar_QGN13951.1 | W303.asm02.HP0_G0048990 | OG0001588 | chrXV | 309779 | 310594 | CP015054.1 | 1206036 | 1206938 | 49.07749077 | 271 | 300 | 0.903333333 | 1 | 1 | 1 | FIM1_599 | YOL009C | YOL009C | MDM12 | n |
| Kmar_QGN14336.1 | W303.asm02.HP0_G0033500 | OG0001589 | chrXI | 494836 | 497064 | CP015055.1 | 301906 | 304116 | 59.94659546 | 742 | 736 | 1.008152174 | 1 | 1 | 1 | FIM1_996 | YKR024C | YKR024C | DBP7 | n |
| Kmar_QGN14335.1 | W303.asm02.HP0_G0033510 | OG0001590 | chrXI | 497462 | 498310 | CP015055.1 | 300595 | 301359 | 41.46341463 | 282 | 254 | 1.11023622 | 1 | 1 | 1 | FIM1_995 | YKR025W | YKR025W | RPC37 | n |
| Kmar_QGN14334.1 | W303.asm02.HP0_G0033520 | OG0001591 | chrXI | 498430 | 499347 | CP015055.1 | 299599 | 300528 | 77.66990291 | 305 | 309 | 0.987055016 | 1 | 1 | 1 | FIM1_994 | YKR026C | YKR026C | GCN3 | n |
| Kmar_QGN15163.1 | W303.asm02.HP0_G0046620 | OG0001592 | chrXIV | 623019 | 624353 | CP015056.1 | 332177 | 333556 | 27.27272727 | 444 | 459 | 0.967320261 | 1 | 1 | 1 | FIM1_1850 | YNL011C | YNL011C | | n |
| Kmar_QGN18227.1 | W303.asm02.HP0_G0007310 | OG0001593 | chrIV | 175130 | 176698 | CP015061.1 | 497975 | 499531 | 67.80952381 | 522 | 518 | 1.007722008 | 1 | 1 | 1 | FIM1_4552 | YDL156W | YDL156W | CMR1 | n |
| Kmar_QGN15126.1 | W303.asm02.HP0_G0041530 | OG0001594 | chrXIII | 494725 | 495120 | CP015056.1 | 257070 | 257423 | 44.6969697 | 131 | 117 | 1.11965812 | 1 | 1 | 1 | FIM1_1813 | YMR112C | YMR112C | MED11 | n |
| Kmar_QGN16615.1 | W303.asm02.HP0_G0008820 | OG0001595 | chrIV | 472816 | 473442 | CP015058.1 | 506381 | 507016 | 64.1509434 | 208 | 211 | 0.985781991 | 1 | 1 | 1 | FIM1_3332 | YDR013W | YDR013W | PSF1 | n |
| Kmar_QGN15423.1 | W303.asm02.HP0_G0011350 | OG0001596 | chrIV | 1029569 | 1034548 | CP015056.1 | 925359 | 930380 | 61.50178784 | 1659 | 1673 | 0.991631799 | 1 | 1 | 1 | FIM1_2113 | YDR283C | YDR283C | GCN2 | n |
| Kmar_QGN13913.1 | W303.asm02.HP0_G0042640 | OG0001597 | chrXIII | 719154 | 721232 | CP015054.1 | 1137365 | 1139425 | 54.46808511 | 692 | 686 | 1.008746356 | 1 | 1 | 1 | FIM1_561 | YMR224C | YMR224C | MRE11 | n |
| Kmar_QGN18144.1 | W303.asm02.HP0_G0043840 | OG0001598 | chrXIV | 43756 | 44712 | CP015061.1 | 319358 | 320329 | 56.70731707 | 318 | 323 | 0.984520124 | 1 | 1 | 1 | FIM1_4467 | YNL315C | YNL315C | ATP11 | n |
| Kmar_QGN16156.1 | W303.asm02.HP0_G0043890 | OG0001599 | chrXIV | 52339 | 52956 | CP015057.1 | 920320 | 920889 | 44.85981308 | 205 | 189 | 1.084656085 | 1 | 1 | 1 | FIM1_2857 | YNL310C | YNL310C | ZIM17 | n |
| Kmar_QGN16881.1 | W303.asm02.HP0_G0036250 | OG0001600 | chrXII | 431729 | 432820 | CP015058.1 | 1095085 | 1096701 | 22.61208577 | 363 | 538 | 0.67472119 | 1 | 1 | 1 | FIM1_3608 | YLR141W | YLR141W | RRN5 | n |
| Kmar_QGN13729.1 | W303.asm02.HP0_G0025110 | OG0001601 | chrVIII | 474323 | 476680 | CP015054.1 | 756036 | 758240 | 16.05562579 | 785 | 734 | 1.069482289 | 1 | 1 | 1 | FIM1_373 | YHR182W | YHR182W | RGD3 | n |
| Kmar_QGN13396.1 | W303.asm02.HP0_G0003680 | OG0001602 | chrII | 599183 | 600130 | CP015054.1 | 63430 | 64371 | 52.53164557 | 315 | 313 | 1.006389776 | 1 | 1 | 1 | FIM1_33 | YBR175W | YBR175W | SWD3 | n |
| Kmar_QGN16151.1 | W303.asm02.HP0_G0047830 | OG0001603 | chrXV | 77900 | 78454 | CP015057.1 | 912010 | 912540 | 63.58695652 | 184 | 176 | 1.045454545 | 1 | 1 | 1 | FIM1_2852 | YOL129W | YOL129W | VPS68 | n |
| Kmar_QGN16150.1 | W303.asm02.HP0_G0043920 | OG0001604 | chrXIV | 56878 | 58005 | CP015057.1 | 910320 | 911417 | 73.33333333 | 375 | 365 | 1.02739726 | 1 | 1 | 1 | FIM1_2851 | YNL307C | YNL307C | MCK1 | n |
| Kmar_QGN14604.1 | W303.asm02.HP0_G0019000 | OG0001605 | chrVII | 311574 | 313496 | CP015055.1 | 841365 | 843314 | 73.77300613 | 640 | 649 | 0.986132512 | 1 | 1 | 1 | FIM1_1267 | YGL099W | YGL099W | LSG1 | n |
| Kmar_QGN16314.1 | W303.asm02.HP0_G0018670 | OG0001606 | chrVII | 246337 | 249206 | CP015057.1 | 1262197 | 1264431 | 53.97536394 | 889 | 744 | 1.194892473 | 1 | 1 | 1 | FIM1_3019 | YGL137W | YGL137W | SEC27 | n |
| Kmar_QGN17725.1 | W303.asm02.HP0_G0046270 | OG0001607 | chrXIV | 549356 | 551002 | CP015060.1 | 329096 | 330808 | 43.2885906 | 548 | 570 | 0.961403509 | 1 | 1 | 1 | FIM1_4933 | YNL048W | YNL048W | ALG11 | n |
| Kmar_QGN16292.1 | W303.asm02.HP0_G0019290 | OG0001608 | chrVII | 374562 | 376535 | CP015057.1 | 1218803 | 1220398 | 36.83409437 | 657 | 531 | 1.237288136 | 1 | 1 | 1 | FIM1_2997 | YGL066W | YGL066W | SGF73 | n |
| Kmar_QGN16297.1 | W303.asm02.HP0_G0053390 | OG0001609 | chrXVI | 162044 | 163147 | CP015057.1 | 1227867 | 1229024 | 50.78125 | 367 | 385 | 0.953246753 | 1 | 1 | 1 | FIM1_3002 | YPL209C | YPL209C | IPL1 | n |
| Kmar_QGN16300.1 | W303.asm02.HP0_G0053370 | OG0001610 | chrXVI | 159049 | 159594 | CP015057.1 | 1233500 | 1234045 | 89.50276243 | 181 | 181 | 1 | 1 | 1 | 1 | FIM1_3005 | YPL211W | YPL211W | NIP7 | n |
| Kmar_QGN16302.1 | W303.asm02.HP0_G0018740 | OG0001611 | chrVII | 264191 | 265543 | CP015057.1 | 1236583 | 1237890 | 46.88888889 | 450 | 435 | 1.034482759 | 1 | 1 | 1 | FIM1_3007 | YGL129C | YGL129C | RSM23 | n |
| Kmar_QGN16298.1 | W303.asm02.HP0_G0019310 | OG0001612 | chrVII | 378476 | 380161 | CP015057.1 | 1229100 | 1230794 | 47.9020979 | 561 | 564 | 0.994680851 | 1 | 1 | 1 | FIM1_3003 | YGL064C | YGL064C | MRH4 | n |
| Kmar_QGN16291.1 | W303.asm02.HP0_G0053420 | OG0001613 | chrXVI | 168184 | 169149 | CP015057.1 | 1217266 | 1218234 | 43.42507645 | 321 | 322 | 0.99689441 | 1 | 1 | 1 | FIM1_2996 | YPL206C | YPL206C | PGC1 | n |
| Kmar_QGN13453.1 | W303.asm02.HP0_G0003960 | OG0001614 | chrII | 642548 | 645085 | CP015054.1 | 173322 | 175787 | 66.90140845 | 845 | 821 | 1.029232643 | 1 | 1 | 1 | FIM1_90 | YBR202W | YBR202W | MCM7 | n |
| Kmar_QGN18169.1 | W303.asm02.HP0_G0026730 | OG0001615 | chrIX | 250791 | 251552 | CP015061.1 | 364059 | 364760 | 34.63035019 | 253 | 233 | 1.08583691 | 1 | 1 | 1 | FIM1_4492 | YIL049W | YIL049W | DFG10 | n |
| Kmar_QGN13457.1 | W303.asm02.HP0_G0054770 | OG0001616 | chrXVI | 434285 | 435013 | CP015054.1 | 179381 | 180133 | 49.40711462 | 242 | 250 | 0.968 | 1 | 1 | 1 | FIM1_94 | YPL065W | YPL065W | VPS28 | n |
| Kmar_QGN17149.1 | W303.asm02.HP0_G0017440 | OG0001617 | chrVI | 221410 | 222234 | CP015059.1 | 342053 | 342850 | 53.62318841 | 274 | 265 | 1.033962264 | 1 | 1 | 1 | FIM1_3880 | YFR052W | YFR052W | RPN12 | n |
| Kmar_QGN14864.1 | W303.asm02.HP0_G0042040 | OG0001618 | chrXIII | 593951 | 595057 | CP015055.1 | 1372945 | 1374045 | 64.78494624 | 368 | 366 | 1.005464481 | 1 | 1 | 1 | FIM1_1537 | YMR166C | YMR166C | MME1 | n |
| Kmar_QGN14863.1 | W303.asm02.HP0_G0007810 | OG0001619 | chrIV | 274063 | 275205 | CP015055.1 | 1371162 | 1372343 | 49.61038961 | 380 | 393 | 0.96692112 | 1 | 1 | 1 | FIM1_1536 | YDL104C | YDL104C | QRI7 | n |
| Kmar_QGN14862.1 | W303.asm02.HP0_G0007800 | OG0001620 | chrIV | 272595 | 273803 | CP015055.1 | 1369923 | 1371074 | 39.10891089 | 402 | 383 | 1.049608355 | 1 | 1 | 1 | FIM1_1535 | YDL105W | YDL105W | NSE4 | n |
| Kmar_QGN14861.1 | W303.asm02.HP0_G0042050 | OG0001621 | chrXIII | 595470 | 597779 | CP015055.1 | 1367488 | 1369650 | 53.94566624 | 769 | 720 | 1.068055556 | 1 | 1 | 1 | FIM1_1534 | YMR167W | YMR167W | MLH1 | n |
| Kmar_QGN16013.1 | W303.asm02.HP0_G0021700 | OG0001622 | chrVII | 907108 | 909561 | CP015057.1 | 612063 | 614249 | 33.37393423 | 817 | 728 | 1.122252747 | 1 | 1 | 1 | FIM1_2713 | YGR198W | YGR198W | YPP1 | n |
| Kmar_QGN16012.1 | W303.asm02.HP0_G0029760 | OG0001623 | chrX | 458066 | 458662 | CP015057.1 | 611057 | 611659 | 71.2195122 | 198 | 200 | 0.99 | 1 | 1 | 1 | FIM1_2712 | YJR014W | YJR014W | TMA22 | n |
| Kmar_QGN13388.1 | W303.asm02.HP0_G0003610 | OG0001624 | chrII | 588243 | 588665 | CP015054.1 | 51470 | 51880 | 42.55319149 | 140 | 136 | 1.029411765 | 1 | 1 | 1 | FIM1_25 | YBR167C | YBR167C | POP7 | n |
| Kmar_QGN15243.1 | W303.asm02.HP0_G0000490 | OG0001625 | chrI | 109641 | 111194 | CP015056.1 | 512621 | 513910 | 42.7184466 | 517 | 429 | 1.205128205 | 1 | 1 | 1 | FIM1_1932 | YAL022C | YAL022C | FUN26 | n |
| Kmar_QGN15173.1 | W303.asm02.HP0_G0028820 | OG0001626 | chrX | 244753 | 246828 | CP015056.1 | 359573 | 361492 | 35.45586107 | 691 | 639 | 1.081377152 | 1 | 1 | 1 | FIM1_1861 | YJL093C | YJL093C | TOK1 | n |
| Kmar_QGN16709.1 | W303.asm02.HP0_G0042270 | OG0001627 | chrXIII | 636874 | 637587 | CP015058.1 | 717211 | 717909 | 59.24369748 | 237 | 232 | 1.021551724 | 1 | 1 | 1 | FIM1_3429 | YMR188C | YMR188C | MRPS17 | n |
| Kmar_QGN17738.1 | W303.asm02.HP0_G0026130 | OG0001628 | chrIX | 133284 | 134441 | CP015060.1 | 364268 | 365437 | 61.06870229 | 385 | 389 | 0.989717224 | 1 | 1 | 1 | FIM1_4947 | YIL116W | YIL116W | HIS5 | n |
| Kmar_QGN16515.1 | W303.asm02.HP0_G0037660 | OG0001629 | chrXII | 714558 | 715139 | CP015058.1 | 281517 | 282098 | 46.11398964 | 193 | 193 | 1 | 1 | 1 | 1 | FIM1_3228 | YLR292C | YLR292C | SEC72 | n |
| Kmar_QGN13580.1 | W303.asm02.HP0_G0040000 | OG0001630 | chrXIII | 159331 | 159867 | CP015054.1 | 458115 | 458624 | 39.3258427 | 178 | 169 | 1.053254438 | 1 | 1 | 1 | FIM1_221 | YML055W | YML055W | SPC2 | n |
| Kmar_QGN17477.1 | W303.asm02.HP0_G0021750 | OG0001631 | chrVII | 917652 | 918098 | CP015059.1 | 1001875 | 1002318 | 56.08108108 | 148 | 147 | 1.006802721 | 1 | 1 | 1 | FIM1_4213 | YGR203W | YGR203W | YCH1 | n |
| Kmar_QGN17476.1 | W303.asm02.HP0_G0021740 | OG0001632 | chrVII | 915889 | 917163 | CP015059.1 | 1000334 | 1001638 | 55.09761388 | 424 | 434 | 0.976958525 | 1 | 1 | 1 | FIM1_4212 | YGR202C | YGR202C | PCT1 | n |
| Kmar_QGN15240.1 | W303.asm02.HP0_G0034880 | OG0001633 | chrXII | 137779 | 139059 | CP015056.1 | 505428 | 506741 | 51.12612613 | 426 | 437 | 0.974828375 | 1 | 1 | 1 | FIM1_1929 | YLL006W | YLL006W | MMM1 | n |
| Kmar_QGN16256.1 | W303.asm02.HP0_G0030830 | OG0001634 | chrX | 679295 | 682441 | CP015057.1 | 1140263 | 1143421 | 51.89393939 | 1048 | 1052 | 0.996197719 | 1 | 1 | 1 | FIM1_2961 | YJR132W | YJR132W | NMD5 | n |
| Kmar_QGN17655.1 | W303.asm02.HP0_G0035010 | OG0001635 | chrXII | 167002 | 167508 | CP015060.1 | 200648 | 201160 | 65.14285714 | 168 | 170 | 0.988235294 | 1 | 1 | 1 | FIM1_4862 | YLR008C | YLR008C | PAM18 | n |
| Kmar_QGN17656.1 | W303.asm02.HP0_G0035020 | OG0001636 | chrXII | 167962 | 168561 | CP015060.1 | 201674 | 202282 | 80.69306931 | 199 | 202 | 0.985148515 | 1 | 1 | 1 | FIM1_4863 | YLR009W | YLR009W | RLP24 | n |
| Kmar_QGN17535.1 | W303.asm02.HP0_G0028540 | OG0001637 | chrX | 172315 | 174237 | CP015059.1 | 1120897 | 1122615 | 44.34108527 | 640 | 572 | 1.118881119 | 1 | 1 | 1 | FIM1_4271 | YJL127C | YJL127C | SPT10 | n |
| Kmar_QGN13917.1 | W303.asm02.HP0_G0051670 | OG0001638 | chrXV | 852656 | 853558 | CP015054.1 | 1141977 | 1143014 | 46.72364672 | 300 | 345 | 0.869565217 | 1 | 1 | 1 | FIM1_565 | YOR287C | YOR287C | RRP36 | n |
| Kmar_QGN14091.1 | W303.asm02.HP0_G0044330 | OG0001639 | chrXIV | 148766 | 155434 | CP015054.1 | 1497138 | 1503695 | 64.34352518 | 2222 | 2185 | 1.016933638 | 1 | 1 | 1 | FIM1_742 | YNL262W | YNL262W | POL2 | n |
| Kmar_QGN14088.1 | W303.asm02.HP0_G0044350 | OG0001640 | chrXIV | 157414 | 158010 | CP015054.1 | 1493735 | 1494232 | 36.13861386 | 198 | 165 | 1.2 | 1 | 1 | 1 | FIM1_739 | YNL260C | YNL260C | LTO1 | n |
| Kmar_QGN14090.1 | W303.asm02.HP0_G0044340 | OG0001641 | chrXIV | 155655 | 157094 | CP015054.1 | 1495498 | 1496922 | 34.3035343 | 479 | 474 | 1.010548523 | 1 | 1 | 1 | FIM1_741 | YNL261W | YNL261W | ORC5 | n |
| Kmar_QGN14086.1 | W303.asm02.HP0_G0044370 | OG0001642 | chrXIV | 158664 | 160928 | CP015054.1 | 1490788 | 1492830 | 29.56636005 | 754 | 680 | 1.108823529 | 1 | 1 | 1 | FIM1_737 | YNL258C | YNL258C | DSL1 | n |
| Kmar_QGN16354.1 | W303.asm02.HP0_G0040360 | OG0001643 | chrXIII | 242495 | 243535 | CP015057.1 | 1348809 | 1349861 | 41.45658263 | 346 | 350 | 0.988571429 | 1 | 1 | 1 | FIM1_3060 | YML015C | YML015C | TAF11 | n |
| Kmar_QGN16440.1 | W303.asm02.HP0_G0029190 | OG0001644 | chrX | 325917 | 327056 | CP015058.1 | 124456 | 125370 | 55.93667546 | 379 | 304 | 1.246710526 | 1 | 1 | 1 | FIM1_3153 | YJL053W | YJL053W | PEP8 | n |
| Kmar_QGN15458.1 | W303.asm02.HP0_G0051330 | OG0001645 | chrXV | 802693 | 804030 | CP015056.1 | 994128 | 995441 | 46.46017699 | 445 | 437 | 1.018306636 | 1 | 1 | 1 | FIM1_2149 | YOR250C | YOR250C | CLP1 | n |
| Kmar_QGN14403.1 | W303.asm02.HP0_G0053930 | OG0001646 | chrXVI | 270962 | 272038 | CP015055.1 | 445992 | 447083 | 62.91208791 | 358 | 363 | 0.986225895 | 1 | 1 | 1 | FIM1_1063 | YPL152W | YPL152W | RRD2 | n |
| Kmar_QGN16169.1 | W303.asm02.HP0_G0051490 | OG0001647 | chrXV | 822745 | 824016 | CP015057.1 | 947723 | 948913 | 29.46635731 | 423 | 396 | 1.068181818 | 1 | 1 | 1 | FIM1_2871 | YOR266W | YOR266W | PNT1 | n |
| Kmar_QGN14405.1 | W303.asm02.HP0_G0053920 | OG0001648 | chrXVI | 267661 | 270126 | CP015055.1 | 449598 | 452021 | 56.68257757 | 821 | 807 | 1.017348203 | 1 | 1 | 1 | FIM1_1065 | YPL153C | YPL153C | RAD53 | n |
| Kmar_QGN15234.1 | W303.asm02.HP0_G0035100 | OG0001649 | chrXII | 184998 | 186011 | CP015056.1 | 490292 | 491314 | 73.23529412 | 337 | 340 | 0.991176471 | 1 | 1 | 1 | FIM1_1923 | YLR017W | YLR017W | MEU1 | n |
| Kmar_QGN14404.1 | W303.asm02.HP0_G0051290 | OG0001650 | chrXV | 795799 | 797055 | CP015055.1 | 447429 | 448691 | 46.44549763 | 418 | 420 | 0.995238095 | 1 | 1 | 1 | FIM1_1064 | YOR245C | YOR245C | DGA1 | n |
| Kmar_QGN14310.1 | W303.asm02.HP0_G0034900 | OG0001651 | chrXII | 142265 | 144115 | CP015055.1 | 246267 | 248198 | 41.11282844 | 616 | 643 | 0.958009331 | 1 | 1 | 1 | FIM1_967 | YLL004W | YLL004W | ORC3 | n |
| Kmar_QGN13516.1 | W303.asm02.HP0_G0012030 | OG0001652 | chrIV | 1193317 | 1194270 | CP015054.1 | 314365 | 315270 | 47.94952681 | 317 | 301 | 1.053156146 | 1 | 1 | 1 | FIM1_156 | YDR352W | YDR352W | YPQ2 | n |

| | | | | | | | | | | | | | | | | | | | | |
|---|---|---|---|---|---|---|---|---|---|---|---|---|---|---|---|---|---|---|---|---|
| Kmar_QGN15435.1 | W303.asm02.HP0_G0011460 | OG0001653 | chrIV | 1054958 | 1056727 | CP015056.1 | 955464 | 957200 | 60.44142615 | 589 | 578 | 1.019031142 | 1 | 1 | 1 | FIM1_2126 | YDR294C | YDR294C | DPL1 | n |
| Kmar_QGN15436.1 | W303.asm02.HP0_G0011470 | OG0001654 | chrIV | 1057122 | 1059146 | CP015056.1 | 957784 | 959973 | 31.2987013 | 674 | 729 | 0.924554184 | 1 | 1 | 1 | FIM1_2127 | YDR295C | YDR295C | HDA2 | n |
| Kmar_QGN15265.1 | W303.asm02.HP0_G0049600 | OG0001655 | chrXV | 430843 | 432222 | CP015056.1 | 558350 | 559720 | 68.04347826 | 459 | 456 | 1.006578947 | 1 | 1 | 1 | FIM1_1954 | YOR056C | YOR056C | NOB1 | n |
| Kmar_QGN13741.1 | W303.asm02.HP0_G0000740 | OG0001657 | chrI | 160095 | 160997 | CP015054.1 | 788542 | 789423 | 37.13355049 | 300 | 293 | 1.023890785 | 1 | 1 | 1 | FIM1_385 | YAR008W | YAR008W | SEN34 | n |
| Kmar_QGN14537.1 | W303.asm02.HP0_G0056330 | OG0001658 | chrXVI | 743483 | 746068 | CP015055.1 | 716287 | 718629 | 34.60648148 | 861 | 780 | 1.103846154 | 1 | 1 | 1 | FIM1_1199 | YPR105C | YPR105C | COG4 | n |
| Kmar_QGN14540.1 | W303.asm02.HP0_G0056350 | OG0001659 | chrXVI | 747937 | 748563 | CP015055.1 | 721331 | 721963 | 77.88461538 | 208 | 210 | 0.99047619 | 1 | 1 | 1 | FIM1_1202 | YPR107C | YPR107C | YTH1 | n |
| Kmar_QGN15176.1 | W303.asm02.HP0_G0028840 | OG0001660 | chrX | 251096 | 252592 | CP015056.1 | 368514 | 369842 | 41.16465863 | 498 | 442 | 1.126696833 | 1 | 1 | 1 | FIM1_1864 | YJL091C | YJL091C | GWT1 | n |
| Kmar_QGN14543.1 | W303.asm02.HP0_G0000620 | OG0001661 | chrI | 136620 | 137399 | CP015055.1 | 726651 | 727520 | 53.76712329 | 259 | 289 | 0.896193772 | 1 | 1 | 1 | FIM1_1205 | YAL009W | YAL009W | SPO7 | n |
| Kmar_QGN14389.1 | W303.asm02.HP0_G0054000 | OG0001662 | chrXVI | 282095 | 283462 | CP015055.1 | 422253 | 423587 | 58.42450766 | 455 | 444 | 1.024774775 | 1 | 1 | 1 | FIM1_1049 | YPL146C | YPL146C | NOP53 | n |
| Kmar_QGN14982.1 | W303.asm02.HP0_G0051030 | OG0001663 | chrXV | 748820 | 750274 | CP015055.1 | 1618994 | 1620445 | 40.57377049 | 484 | 483 | 1.002070393 | 1 | 1 | 1 | FIM1_1660 | YOR216C | YOR216C | RUD3 | n |
| Kmar_QGN13843.1 | W303.asm02.HP0_G0028080 | OG0001664 | chrX | 81583 | 84060 | CP015054.1 | 984103 | 987045 | 30.28515241 | 825 | 980 | 0.841836735 | 1 | 1 | 1 | FIM1_489 | YJL176C | YJL176C | SWI3 | n |
| Kmar_QGN13831.1 | W303.asm02.HP0_G0028150 | OG0001665 | chrX | 94542 | 95600 | CP015054.1 | 959667 | 960716 | 77.05382436 | 352 | 349 | 1.008595989 | 1 | 1 | 1 | FIM1_477 | YJL167W | YJL167W | ERG20 | n |
| Kmar_QGN13832.1 | W303.asm02.HP0_G0031610 | OG0001666 | chrXI | 126929 | 128212 | CP015054.1 | 961139 | 962385 | 63.44827586 | 427 | 415 | 1.028915663 | 1 | 1 | 1 | FIM1_478 | YKL172W | YKL172W | EBP2 | n |
| Kmar_QGN13427.1 | W303.asm02.HP0_G0003800 | OG0001667 | chrII | 619409 | 620251 | CP015054.1 | 125202 | 126041 | 51.06382979 | 280 | 279 | 1.003584229 | 1 | 1 | 1 | FIM1_64 | YBR187W | YBR187W | GDT1 | n |
| Kmar_QGN13434.1 | W303.asm02.HP0_G0003850 | OG0001668 | chrII | 625857 | 626528 | CP015054.1 | 140408 | 141052 | 49.77578475 | 223 | 214 | 1.042056075 | 1 | 1 | 1 | FIM1_71 | YBR193C | YBR193C | MED8 | n |
| Kmar_QGN13433.1 | W303.asm02.HP0_G0003840 | OG0001669 | chrII | 624427 | 625560 | CP015054.1 | 139193 | 140299 | 73.74005305 | 377 | 368 | 1.024456522 | 1 | 1 | 1 | FIM1_70 | YBR192W | YBR192W | RIM2 | n |
| Kmar_QGN13429.1 | W303.asm02.HP0_G0054620 | OG0001670 | chrXVI | 404838 | 410441 | CP015054.1 | 126739 | 132525 | 64.11311054 | 1867 | 1928 | 0.968360996 | 1 | 1 | 1 | FIM1_66 | YPL082C | YPL082C | MOT1 | n |
| Kmar_QGN14398.1 | W303.asm02.HP0_G0051250 | OG0001671 | chrXV | 788718 | 790364 | CP015055.1 | 437423 | 438913 | 59.70961887 | 548 | 496 | 1.10483871 | 1 | 1 | 1 | FIM1_1058 | YOR241W | YOR241W | MET7 | n |
| Kmar_QGN17178.1 | W303.asm02.HP0_G0055110 | OG0001672 | chrXVI | 501865 | 504078 | CP015059.1 | 394605 | 396851 | 48.80952381 | 737 | 748 | 0.985294118 | 1 | 1 | 1 | FIM1_3909 | YPL029W | YPL029W | SUV3 | n |
| Kmar_QGN17180.1 | W303.asm02.HP0_G0011050 | OG0001673 | chrIV | 962259 | 962840 | CP015059.1 | 399079 | 399627 | 45.54973822 | 193 | 182 | 1.06043956 | 1 | 1 | 1 | FIM1_3911 | YDR248C | YDR248C | | n |
| Kmar_QGN18006.1 | W303.asm02.HP0_G0051300 | OG0001674 | chrXV | 797524 | 798516 | CP015061.1 | 20195 | 21172 | 59.15915916 | 330 | 325 | 1.015384615 | 1 | 1 | 1 | FIM1_4322 | YOR246C | YOR246C | ENV9 | n |
| Kmar_QGN17234.1 | W303.asm02.HP0_G0041160 | OG0001675 | chrXIII | 412772 | 413377 | CP015059.1 | 511481 | 511915 | 29.85074627 | 201 | 144 | 1.395833333 | 1 | 1 | 1 | FIM1_3965 | YMR073C | YMR073C | IRC21 | n |
| Kmar_QGN13688.1 | W303.asm02.HP0_G0016300 | OG0001676 | chrV | 561036 | 561671 | CP015054.1 | 665771 | 666493 | 42.32365145 | 211 | 240 | 0.879166667 | 1 | 1 | 1 | FIM1_331 | YER183C | YER183C | FAU1 | n |
| Kmar_QGN15112.1 | W303.asm02.HP0_G0032070 | OG0001677 | chrXI | 219729 | 220232 | CP015056.1 | 228648 | 229109 | 32.74853801 | 167 | 153 | 1.091503268 | 1 | 1 | 1 | FIM1_1799 | YKL122C | YKL122C | SRP21 | n |
| Kmar_QGN14006.1 | W303.asm02.HP0_G0048360 | OG0001678 | chrXV | 190113 | 193997 | CP015054.1 | 1311846 | 1315616 | 50.42471042 | 1294 | 1256 | 1.030254777 | 1 | 0 | 1 | FIM1_657 | YOL075C | YOL075C | | n |
| Kmar_QGN15383.1 | W303.asm02.HP0_G0016040 | OG0001679 | chrV | 491027 | 492043 | CP015056.1 | 847149 | 848150 | 67.75147929 | 338 | 333 | 1.015015015 | 1 | 1 | 1 | FIM1_2073 | YER156C | YER156C | MYG1 | n |
| Kmar_QGN15904.1 | W303.asm02.HP0_G0009800 | OG0001680 | chrIV | 688853 | 690550 | CP015057.1 | 344789 | 346534 | 48.02065404 | 565 | 581 | 0.972461274 | 1 | 1 | 1 | FIM1_2602 | YDR117C | YDR117C | TMA64 | n |
| Kmar_QGN16479.1 | W303.asm02.HP0_G0017180 | OG0001681 | chrVI | 179842 | 181878 | CP015058.1 | 190960 | 192897 | 46.72364672 | 678 | 645 | 1.051162791 | 1 | 1 | 1 | FIM1_3192 | YFR029W | YFR029W | PTR3 | n |
| Kmar_QGN14484.1 | W303.asm02.HP0_G0021280 | OG0001682 | chrVII | 810270 | 811793 | CP015055.1 | 611955 | 613436 | 76.13412229 | 507 | 493 | 1.028397566 | 1 | 1 | 1 | FIM1_1145 | YGR155W | YGR155W | CYS4 | n |
| Kmar_QGN14482.1 | W303.asm02.HP0_G0019500 | OG0001683 | chrVII | 414569 | 415498 | CP015055.1 | 610124 | 611017 | 60.96774194 | 309 | 297 | 1.04040404 | 1 | 1 | 1 | FIM1_1143 | YGL043W | YGL043W | DST1 | n |
| Kmar_QGN14481.1 | W303.asm02.HP0_G0019490 | OG0001684 | chrVII | 413232 | 414122 | CP015055.1 | 608971 | 609759 | 51.68918919 | 296 | 262 | 1.129770992 | 1 | 1 | 1 | FIM1_1142 | YGL044C | YGL044C | RNA15 | n |
| Kmar_QGN13648.1 | W303.asm02.HP0_G0014360 | OG0001685 | chrV | 132955 | 133978 | CP015054.1 | 578634 | 579227 | 64.62264151 | 212 | 197 | 1.076142132 | 1 | 1 | 1 | FIM1_291 | YEL012W | YEL012W | UBC8 | n |
| Kmar_QGN18124.1 | W303.asm02.HP0_G0055640 | OG0001686 | chrXVI | 619875 | 621056 | CP015061.1 | 271663 | 272877 | 49.64370546 | 393 | 404 | 0.972772277 | 1 | 1 | 1 | FIM1_4445 | YPR025C | YPR025C | CCL1 | n |
| Kmar_QGN13645.1 | W303.asm02.HP0_G0005690 | OG0001687 | chrIII | 164751 | 165842 | CP015054.1 | 575932 | 576966 | 45.32967033 | 363 | 344 | 1.055232558 | 1 | 1 | 1 | FIM1_287 | YCR019W | YCR019W | MAK32 | n |
| Kmar_QGN16598.1 | W303.asm02.HP0_G0030120 | OG0001688 | chrX | 553387 | 553830 | CP015058.1 | 472860 | 473306 | 63.51351351 | 147 | 148 | 0.993243243 | 1 | 1 | 1 | FIM1_3314 | YJR058C | YJR058C | APS2 | n |
| Kmar_QGN16599.1 | W303.asm02.HP0_G0030110 | OG0001689 | chrX | 552717 | 553367 | CP015058.1 | 473354 | 473986 | 56.01851852 | 216 | 210 | 1.028571429 | 1 | 1 | 1 | FIM1_3315 | YJR057W | YJR057W | CDC8 | n |
| Kmar_QGN15566.1 | W303.asm02.HP0_G0042890 | OG0001690 | chrXIII | 777305 | 779062 | CP015056.1 | 1200119 | 1201822 | 58.4040747 | 585 | 567 | 1.031746032 | 1 | 1 | 1 | FIM1_2257 | YMR250W | YMR250W | GAD1 | n |
| Kmar_QGN14137.1 | W303.asm02.HP0_G0005310 | OG0001691 | chrIII | 71454 | 72992 | CP015054.1 | 1615474 | 1617051 | 24.90909091 | 512 | 525 | 0.975238095 | 1 | 1 | 1 | FIM1_790 | YCL027W | YCL027W | FUS1 | n |
| Kmar_QGN15346.1 | W303.asm02.HP0_G0011710 | OG0001692 | chrIV | 1115069 | 1115893 | CP015056.1 | 758192 | 758995 | 37.80918728 | 274 | 267 | 1.026217228 | 1 | 1 | 1 | FIM1_2035 | YDR319C | YDR319C | YFT2 | n |
| Kmar_QGN13730.1 | W303.asm02.HP0_G0025100 | OG0001693 | chrVIII | 473332 | 474018 | CP015054.1 | 758379 | 759080 | 60.51502146 | 228 | 233 | 0.978540773 | 1 | 1 | 1 | FIM1_374 | YHR181W | YHR181W | SVP26 | n |
| Kmar_QGN15987.1 | W303.asm02.HP0_G0033570 | OG0001694 | chrXI | 511027 | 516078 | CP015057.1 | 550026 | 554969 | 47.39819005 | 1683 | 1647 | 1.021857923 | 1 | 1 | 1 | FIM1_2687 | YKR031C | YKR031C | SPO14 | n |
| Kmar_QGN15997.1 | W303.asm02.HP0_G0011890 | OG0001695 | chrIV | 1155692 | 1156636 | CP015057.1 | 572965 | 574056 | 37.98882682 | 314 | 363 | 0.865013774 | 1 | 1 | 1 | FIM1_2697 | YDR336W | YDR336W | MRX8 | n |
| Kmar_QGN15996.1 | W303.asm02.HP0_G0024200 | OG0001696 | chrVIII | 288654 | 289271 | CP015057.1 | 572078 | 572728 | 77.16894977 | 205 | 216 | 0.949074074 | 1 | 1 | 1 | FIM1_2696 | YHR089C | YHR089C | GAR1 | n |
| Kmar_QGN17908.1 | W303.asm02.HP0_G0055950 | OG0001697 | chrXVI | 682475 | 683380 | CP015060.1 | 715718 | 716599 | 37.89808917 | 301 | 293 | 1.027303754 | 1 | 1 | 1 | FIM1_5117 | YPR061C | YPR061C | JID1 | n |
| Kmar_QGN17907.1 | W303.asm02.HP0_G0055940 | OG0001698 | chrXVI | 681359 | 682129 | CP015060.1 | 714401 | 715180 | 76.171875 | 256 | 259 | 0.988416988 | 1 | 1 | 1 | FIM1_5116 | YPR060C | YPR060C | ARO7 | n |
| Kmar_QGN17236.1 | W303.asm02.HP0_G0041170 | OG0001699 | chrXIII | 413540 | 414109 | CP015059.1 | 515639 | 516040 | 29.1005291 | 189 | 133 | 1.421052632 | 1 | 1 | 1 | FIM1_3967 | YMR074C | YMR074C | SDD2 | n |
| Kmar_QGN14478.1 | W303.asm02.HP0_G0037370 | OG0001700 | chrXII | 657968 | 659686 | CP015055.1 | 601642 | 603366 | 87.17504333 | 572 | 574 | 0.996515679 | 1 | 1 | 1 | FIM1_1139 | YLR259C | YLR259C | HSP60 | n |
| Kmar_QGN13959.1 | W303.asm02.HP0_G0040110 | OG0001701 | chrXIII | 190601 | 192124 | CP015054.1 | 1218990 | 1220435 | 32.82149712 | 507 | 481 | 1.054054054 | 1 | 1 | 1 | FIM1_607 | YML043C | YML043C | RRN11 | n |
| Kmar_QGN13958.1 | W303.asm02.HP0_G0048950 | OG0001702 | chrXV | 301835 | 303490 | CP015054.1 | 1217228 | 1218862 | 27.33686067 | 551 | 544 | 1.012867647 | 1 | 1 | 1 | FIM1_606 | YOL013C | YOL013C | HRD1 | n |
| Kmar_QGN14630.1 | W303.asm02.HP0_G0034590 | OG0001703 | chrXII | 68589 | 70487 | CP015055.1 | 899382 | 901274 | 38.33075734 | 632 | 630 | 1.003174603 | 1 | 1 | 1 | FIM1_1294 | YLL035W | YLL035W | GRC3 | n |
| Kmar_QGN17924.1 | W303.asm02.HP0_G0015490 | OG0001704 | chrV | 344990 | 349282 | CP015060.1 | 747076 | 751089 | 36.06102635 | 1430 | 1337 | 1.069558714 | 1 | 1 | 1 | FIM1_5133 | YER093C | YER093C | TSC11 | n |
| Kmar_QGN16003.1 | W303.asm02.HP0_G0024180 | OG0001705 | chrVIII | 286793 | 287128 | CP015057.1 | 594763 | 595101 | 56.25 | 111 | 112 | 0.991071429 | 1 | 1 | 1 | FIM1_2703 | YHR087W | YHR087W | RTC3 | n |
| Kmar_QGN15201.1 | W303.asm02.HP0_G0011860 | OG0001706 | chrIV | 1143860 | 1146031 | CP015056.1 | 416047 | 418260 | 45.36912752 | 723 | 737 | 0.981004071 | 1 | 1 | 1 | FIM1_1889 | YDR333C | YDR333C | RQC1 | n |
| Kmar_QGN15430.1 | W303.asm02.HP0_G0023520 | OG0001707 | chrVIII | 178719 | 180653 | CP015056.1 | 943272 | 945194 | 68.82716049 | 644 | 640 | 1.00625 | 1 | 1 | 1 | FIM1_2121 | YHR039C | YHR039C | MSC7 | n |
| Kmar_QGN15429.1 | W303.asm02.HP0_G0023530 | OG0001708 | chrVIII | 181018 | 181524 | CP015056.1 | 942463 | 942804 | 65.78947368 | 114 | 113 | 1.008849558 | 1 | 1 | 1 | FIM1_2120 | YHR039C-A | YHR039C-A | VMA10 | n |
| Kmar_QGN16004.1 | W303.asm02.HP0_G0011870 | OG0001709 | chrIV | 1146532 | 1151076 | CP015057.1 | 595819 | 600435 | 54.24588086 | 1514 | 1538 | 0.984395319 | 1 | 1 | 1 | FIM1_2704 | YDR334W | YDR334W | SWR1 | n |
| Kmar_QGN15943.1 | W303.asm02.HP0_G0050650 | OG0001710 | chrXV | 659622 | 660476 | CP015057.1 | 443015 | 443833 | 41.1971831 | 284 | 272 | 1.044117647 | 1 | 1 | 1 | FIM1_2641 | YOR174W | YOR174W | MED4 | n |
| Kmar_QGN15942.1 | W303.asm02.HP0_G0050660 | OG0001711 | chrXV | 660690 | 662549 | CP015057.1 | 441010 | 442869 | 59.48963317 | 619 | 619 | 1 | 1 | 1 | 1 | FIM1_2640 | YOR175C | YOR175C | ALE1 | n |
| Kmar_QGN15940.1 | W303.asm02.HP0_G0037480 | OG0001712 | chrXII | 678322 | 681852 | CP015057.1 | 435381 | 438641 | 52.79187817 | 1176 | 1086 | 1.082872928 | 1 | 1 | 1 | FIM1_2638 | YLR272C | YLR272C | YCS4 | n |
| Kmar_QGN13954.1 | W303.asm02.HP0_G0040550 | OG0001713 | chrXIII | 276551 | 277717 | CP015054.1 | 1210633 | 1211715 | 46.15384615 | 388 | 360 | 1.077777778 | 1 | 1 | 1 | FIM1_602 | YMR005W | YMR005W | TAF4 | n |
| Kmar_QGN13953.1 | W303.asm02.HP0_G0048980 | OG0001714 | chrXV | 308393 | 309496 | CP015054.1 | 1208885 | 1209982 | 81.74386921 | 367 | 365 | 1.005479452 | 1 | 1 | 1 | FIM1_601 | YOL010W | YOL010W | RCL1 | n |
| Kmar_QGN13922.1 | W303.asm02.HP0_G0012510 | OG0001715 | chrIV | 1276788 | 1277810 | CP015054.1 | 1149524 | 1150540 | 57.771261 | 340 | 338 | 1.00591716 | 1 | 1 | 1 | FIM1_570 | YDR400W | YDR400W | URH1 | n |
| Kmar_QGN18010.1 | W303.asm02.HP0_G0042500 | OG0001716 | chrXIII | 688095 | 689444 | CP015061.1 | 27340 | 28656 | 43.48785872 | 449 | 438 | 1.025114155 | 1 | 1 | 1 | FIM1_4326 | YMR210W | YMR210W | MGL2 | n |
| Kmar_QGN18015.1 | W303.asm02.HP0_G0053910 | OG0001717 | chrXVI | 265648 | 266865 | CP015061.1 | 37828 | 39054 | 74.08312958 | 405 | 408 | 0.992647059 | 1 | 1 | 1 | FIM1_4331 | YPL154C | YPL154C | PEP4 | n |
| Kmar_QGN18012.1 | W303.asm02.HP0_G0053880 | OG0001718 | chrXVI | 260748 | 261695 | CP015061.1 | 32026 | 32922 | 43.49206349 | 315 | 298 | 1.05704698 | 1 | 1 | 1 | FIM1_4328 | YPL157W | YPL157W | TGS1 | n |
| Kmar_QGN13493.1 | W303.asm02.HP0_G0014160 | OG0001719 | chrV | 87295 | 90210 | CP015054.1 | 256889 | 259717 | 64.60446247 | 971 | 942 | 1.030785563 | 1 | 1 | 1 | FIM1_131 | YEL032W | YEL032W | MCM3 | n |
| Kmar_QGN18261.1 | W303.asm02.HP0_G0056680 | OG0001720 | chrXVI | 831398 | 833056 | CP015061.1 | 573471 | 575039 | 52.3465704 | 552 | 522 | 1.057471264 | 1 | 1 | 1 | FIM1_4587 | YPR144C | YPR144C | NOC4 | n |
| Kmar_QGN15829.1 | W303.asm02.HP0_G0041640 | OG0001721 | chrXIII | 514220 | 514588 | CP015057.1 | 177948 | 178325 | 47.61904762 | 122 | 125 | 0.976 | 1 | 1 | 1 | FIM1_2525 | YMR123W | YMR123W | PKR1 | n |
| Kmar_QGN17170.1 | W303.asm02.HP0_G0055000 | OG0001722 | chrXVI | 482353 | 485361 | CP015059.1 | 377276 | 380254 | 56.14552606 | 1002 | 992 | 1.010080645 | 1 | 1 | 1 | FIM1_3901 | YPL040C | YPL040C | ISM1 | n |
| Kmar_QGN15117.1 | W303.asm02.HP0_G0041470 | OG0001723 | chrXIII | 478807 | 480696 | CP015056.1 | 238800 | 240713 | 25.18518519 | 629 | 637 | 0.98744113 | 1 | 1 | 1 | FIM1_1804 | YMR106C | YMR106C | YKU80 | n |
| Kmar_QGN15578.1 | W303.asm02.HP0_G0041380 | OG0001724 | chrXIII | 459929 | 461032 | CP015056.1 | 1225273 | 1226421 | 42.82115869 | 367 | 382 | 0.960732984 | 1 | 1 | 1 | FIM1_2270 | YMR097C | YMR097C | MTG1 | n |
| Kmar_QGN14355.1 | W303.asm02.HP0_G0032260 | OG0001725 | chrXI | 264196 | 264948 | CP015055.1 | 344745 | 345497 | 56.2992126 | 250 | 250 | 1 | 1 | 1 | 1 | FIM1_1015 | YKL099C | YKL099C | UTP11 | n |
| Kmar_QGN15577.1 | W303.asm02.HP0_G0041390 | OG0001726 | chrXIII | 461276 | 463114 | CP015056.1 | 1222237 | 1223940 | 25.59241706 | 612 | 567 | 1.079365079 | 1 | 1 | 1 | FIM1_2268 | YMR098C | YMR098C | ATP25 | n |
| Kmar_QGN14342.1 | W303.asm02.HP0_G0032190 | OG0001727 | chrXI | 242285 | 243646 | CP015055.1 | 313765 | 315066 | 20.82429501 | 453 | 433 | 1.046189376 | 1 | 1 | 1 | FIM1_1002 | YKL108W | YKL108W | SLD2 | n |
| Kmar_QGN14338.1 | W303.asm02.HP0_G0033490 | OG0001728 | chrXI | 493112 | 494704 | CP015055.1 | 306395 | 307891 | 44.34137291 | 530 | 498 | 1.064257028 | 1 | 1 | 1 | FIM1_998 | YKR023W | YKR023W | RQT4 | n |
| Kmar_QGN15581.1 | W303.asm02.HP0_G0028690 | OG0001729 | chrX | 211406 | 213343 | CP015056.1 | 1231864 | 1234221 | 35.75909661 | 645 | 785 | 0.821656051 | 1 | 1 | 1 | FIM1_2273 | YJL106W | YJL106W | IME2 | n |
| Kmar_QGN16978.1 | W303.asm02.HP0_G0044640 | OG0001730 | chrXIV | 221355 | 223127 | CP015058.1 | 1297101 | 1298945 | 47.10610932 | 590 | 614 | 0.960912052 | 1 | 1 | 1 | FIM1_3705 | YNL227C | YNL227C | JJJ1 | n |
| Kmar_QGN16976.1 | W303.asm02.HP0_G0044660 | OG0001731 | chrXIV | 225492 | 227795 | CP015058.1 | 1291383 | 1293662 | 39.375 | 767 | 759 | 1.010540184 | 1 | 1 | 1 | FIM1_3703 | YNL224C | YNL224C | SQS1 | n |
| Kmar_QGN16975.1 | W303.asm02.HP0_G0044670 | OG0001732 | chrXIV | 228066 | 229550 | CP015058.1 | 1289746 | 1291101 | 38.90020367 | 494 | 451 | 1.095343681 | 1 | 1 | 1 | FIM1_3702 | YNL223W | YNL223W | ATG4 | n |
| Kmar_QGN16974.1 | W303.asm02.HP0_G0044680 | OG0001733 | chrXIV | 229790 | 230410 | CP015058.1 | 1288672 | 1289307 | 76.30331754 | 206 | 211 | 0.976303318 | 1 | 1 | 1 | FIM1_3701 | YNL222W | YNL222W | SSU72 | n |
| Kmar_QGN16971.1 | W303.asm02.HP0_G0044690 | OG0001734 | chrXIV | 231764 | 234391 | CP015058.1 | 1280590 | 1283049 | 43.78563284 | 875 | 819 | 1.068376068 | 1 | 1 | 1 | FIM1_3698 | YNL221C | YNL221C | POP1 | n |
| Kmar_QGN16522.1 | W303.asm02.HP0_G0021370 | OG0001735 | chrVII | 839327 | 840352 | CP015058.1 | 298034 | 299080 | 70.0280112 | 341 | 348 | 0.979885057 | 1 | 1 | 1 | FIM1_3235 | YGR163W | YGR163W | GTR2 | n |
| Kmar_QGN13530.1 | W303.asm02.HP0_G0034750 | OG0001736 | chrXII | 109914 | 111587 | CP015054.1 | 351297 | 352964 | 82.76481149 | 557 | 555 | 1.003603604 | 1 | 1 | 1 | FIM1_170 | YLL018C | YLL018C | DPS1 | n |
| Kmar_QGN15322.1 | W303.asm02.HP0_G0037810 | OG0001737 | chrXII | 744291 | 747026 | CP015056.1 | 700376 | 702715 | 37.39035088 | 911 | 779 | 1.16944801 | 1 | 1 | 1 | FIM1_2011 | YLR309C | YLR309C | IMH1 | n |
| Kmar_QGN14556.1 | W303.asm02.HP0_G0000570 | OG0001738 | chrI | 129018 | 129785 | CP015055.1 | 749227 | 750006 | 35.47169811 | 255 | 259 | 0.984555985 | 1 | 1 | 1 | FIM1_1218 | YAL014C | YAL014C | SYN8 | n |
| Kmar_QGN13489.1 | W303.asm02.HP0_G0029990 | OG0001739 | chrX | 527294 | 527716 | CP015054.1 | 247032 | 247460 | 58.45070423 | 140 | 142 | 0.985915493 | 1 | 1 | 1 | FIM1_127 | YJR044C | YJR044C | VPS55 | n |
| Kmar_QGN16462.1 | W303.asm02.HP0_G0029060 | OG0001740 | chrX | 304133 | 304891 | CP015058.1 | 162219 | 162923 | 33.2046332 | 252 | 234 | 1.076923077 | 1 | 1 | 1 | FIM1_3175 | YJL066C | YJL066C | MPM1 | n |
| Kmar_QGN14547.1 | W303.asm02.HP0_G0048740 | OG0001741 | chrXV | 263930 | 265540 | CP015055.1 | 735858 | 737465 | 54.71349353 | 536 | 535 | 1.001869159 | 1 | 1 | 1 | FIM1_1209 | YOL033W | YOL033W | MSE1 | n |
| Kmar_QGN14546.1 | W303.asm02.HP0_G0048730 | OG0001742 | chrXV | 260378 | 263659 | CP015055.1 | 732233 | 735571 | 40.73741007 | 1093 | 1112 | 0.982913669 | 1 | 1 | 1 | FIM1_1208 | YOL034W | YOL034W | SMC5 | n |
| Kmar_QGN14140.1 | W303.asm02.HP0_G0005280 | OG0001743 | chrIII | 65585 | 67984 | CP015054.1 | 1623589 | 1625997 | 69.39281289 | 799 | 802 | 0.996259352 | 1 | 1 | 1 | FIM1_793 | YCL030C | YCL030C | HIS4 | n |
| Kmar_QGN15521.1 | W303.asm02.HP0_G0011000 | OG0001744 | chrIV | 955064 | 956902 | CP015056.1 | 1113040 | 1114755 | 44.77124183 | 612 | 571 | 1.071803853 | 1 | 1 | 1 | FIM1_2212 | YDR244W | YDR244W | PEX5 | n |
| Kmar_QGN16545.1 | W303.asm02.HP0_G0039050 | OG0001745 | chrXII | 1019292 | 1020638 | CP015058.1 | 343781 | 344965 | 30.48245614 | 448 | 394 | 1.137055838 | 1 | 1 | 1 | FIM1_3259 | YLR443W | YLR443W | ECM7 | n |
| Kmar_QGN16544.1 | W303.asm02.HP0_G0039880 | OG0001746 | chrXIII | 134964 | 136073 | CP015058.1 | 342418 | 343572 | 21.93877551 | 369 | 384 | 0.9609375 | 1 | 1 | 1 | FIM1_3258 | YML066C | YML066C | SMA2 | n |
| Kmar_QGN13523.1 | W303.asm02.HP0_G0037740 | OG0001747 | chrXII | 727311 | 728645 | CP015054.1 | 332305 | 333639 | 82.20720721 | 444 | 444 | 1 | 1 | 1 | 1 | FIM1_163 | YLR303W | YLR303W | MET17 | n |
| Kmar_QGN14574.1 | W303.asm02.HP0_G0055060 | OG0001748 | chrXVI | 493071 | 493568 | CP015055.1 | 788141 | 788938 | 19.70260223 | 165 | 265 | 0.622641509 | 1 | 1 | 1 | FIM1_1236 | YPL034W | YPL034W | | n |
| Kmar_QGN17119.1 | W303.asm02.HP0_G0014710 | OG0001749 | chrV | 202215 | 203075 | CP015059.1 | 277421 | 278281 | 66.55052265 | 286 | 286 | 1 | 1 | 1 | 1 | FIM1_3850 | YER023W | YER023W | PRO3 | n |
| Kmar_QGN17120.1 | W303.asm02.HP0_G0030710 | OG0001750 | chrX | 654078 | 656264 | CP015059.1 | 279336 | 281822 | 40.33613445 | 728 | 828 | 0.879227053 | 1 | 1 | 1 | FIM1_3851 | YJR119C | YJR119C | JHD2 | n |
| Kmar_QGN17186.1 | W303.asm02.HP0_G0055240 | OG0001751 | chrXVI | 527371 | 531315 | CP015059.1 | 410589 | 414218 | 31.09118087 | 1314 | 1209 | 1.086848635 | 1 | 1 | 1 | FIM1_3917 | YPL016W | YPL016W | SWI1 | n |
| Kmar_QGN15520.1 | W303.asm02.HP0_G0011010 | OG0001752 | chrIV | 957301 | 958482 | CP015056.1 | 1111513 | 1112508 | 57.39795918 | 393 | 331 | 1.187311178 | 1 | 1 | 1 | FIM1_2211 | YDR245W | YDR245W | MNN10 | n |

| | | | | | | | | | | | | | | | | | | | | |
|---|---|---|---|---|---|---|---|---|---|---|---|---|---|---|---|---|---|---|---|---|
| Kmar_QGN16725.1 | W303.asm02.HP0_G0024050 | OG0001753 | chrVIII | 257075 | 258199 | CP015058.1 | 747982 | 749028 | 54.27807487 | 374 | 348 | 1.074712644 | 1 | 1 | 1 | FIM1_3446 | YHR076W | YHR076W | PTC7 | n |
| Kmar_QGN16727.1 | W303.asm02.HP0_G0011770 | OG0001754 | chrIV | 1123080 | 1124627 | CP015058.1 | 752739 | 754346 | 38.54545455 | 515 | 535 | 0.962616822 | 1 | 1 | 1 | FIM1_3448 | YDR323C | YDR323C | PEP7 | n |
| Kmar_QGN16065.1 | W303.asm02.HP0_G0040910 | OG0001755 | chrXIII | 351469 | 352476 | CP015057.1 | 735361 | 736371 | 53.70919881 | 335 | 336 | 0.99702381 | 1 | 1 | 1 | FIM1_2765 | YMR041C | YMR041C | ARA2 | n |
| Kmar_QGN16066.1 | W303.asm02.HP0_G0040890 | OG0001756 | chrXIII | 349147 | 350025 | CP015057.1 | 736971 | 737732 | 28.08219178 | 292 | 253 | 1.154150198 | 1 | 1 | 1 | FIM1_2766 | YMR039C | YMR039C | SUB1 | n |
| Kmar_QGN16067.1 | W303.asm02.HP0_G0032620 | OG0001757 | chrXI | 326240 | 329149 | CP015057.1 | 738435 | 741338 | 52.62135922 | 969 | 967 | 1.002068252 | 1 | 1 | 1 | FIM1_2767 | YKL064W | YKL064W | MNR2 | n |
| Kmar_QGN14947.1 | W303.asm02.HP0_G0045400 | OG0001758 | chrXIV | 358286 | 359785 | CP015055.1 | 1551628 | 1553109 | 72.2 | 499 | 493 | 1.012170385 | 1 | 1 | 1 | FIM1_1623 | YNL142W | YNL142W | MEP2 | n |
| Kmar_QGN13978.1 | W303.asm02.HP0_G0050810 | OG0001759 | chrXV | 693770 | 698629 | CP015054.1 | 1254221 | 1259113 | 32.72727273 | 1619 | 1630 | 0.993251534 | 1 | 1 | 1 | FIM1_627 | YOR191W | YOR191W | ULS1 | n |
| Kmar_QGN13581.1 | W303.asm02.HP0_G0038900 | OG0001760 | chrXII | 985097 | 987109 | CP015054.1 | 459340 | 461208 | 38.3480826 | 670 | 622 | 1.077170418 | 1 | 1 | 1 | FIM1_222 | YLR427W | YLR427W | MAG2 | n |
| Kmar_QGN17704.1 | W303.asm02.HP0_G0046520 | OG0001761 | chrXIV | 599446 | 602343 | CP015060.1 | 282124 | 284820 | 49.38016529 | 965 | 898 | 1.074610245 | 1 | 1 | 1 | FIM1_4911 | YNL023C | YNL023C | FAP1 | n |
| Kmar_QGN17606.1 | W303.asm02.HP0_G0049110 | OG0001762 | chrXV | 334047 | 334811 | CP015060.1 | 88542 | 89390 | 60.80586081 | 254 | 282 | 0.90070922 | 1 | 1 | 1 | FIM1_4813 | YOR004W | YOR004W | UTP23 | n |
| Kmar_QGN16490.1 | W303.asm02.HP0_G0000400 | OG0001763 | chrI | 85512 | 87794 | CP015058.1 | 217556 | 219835 | 30.32581454 | 760 | 759 | 1.001317523 | 1 | 1 | 1 | FIM1_3203 | YAL031C | YAL031C | GIP4 | n |
| Kmar_QGN16106.1 | W303.asm02.HP0_G0026940 | OG0001764 | chrIX | 294308 | 294733 | CP015057.1 | 825348 | 825758 | 41.84397163 | 141 | 136 | 1.036764706 | 1 | 1 | 1 | FIM1_2807 | YIL027C | YIL027C | EMC5 | n |
| Kmar_QGN14173.1 | W303.asm02.HP0_G0005060 | OG0001765 | chrIII | 27583 | 28590 | CP015054.1 | 1684706 | 1685677 | 64.88095238 | 335 | 323 | 1.037151703 | 1 | 1 | 1 | FIM1_826 | YCL055W | YCL055W | KAR4 | n |
| Kmar_QGN13447.1 | W303.asm02.HP0_G0054720 | OG0001766 | chrXVI | 427306 | 429144 | CP015054.1 | 164913 | 166556 | 29.18660287 | 612 | 547 | 1.118829982 | 1 | 1 | 1 | FIM1_84 | YPL070W | YPL070W | MUK1 | n |
| Kmar_QGN17331.1 | W303.asm02.HP0_G0050100 | OG0001767 | chrXV | 532383 | 534668 | CP015059.1 | 709275 | 711470 | 36.67953668 | 761 | 731 | 1.041039672 | 1 | 1 | 1 | FIM1_4063 | YOR112W | YOR112W | CEX1 | n |
| Kmar_QGN17929.1 | W303.asm02.HP0_G0015520 | OG0001768 | chrV | 351650 | 352852 | CP015060.1 | 756514 | 757644 | 77.75 | 400 | 376 | 1.063829787 | 1 | 1 | 1 | FIM1_5138 | YER095W | YER095W | RAD51 | n |
| Kmar_QGN14298.1 | W303.asm02.HP0_G0005590 | OG0001769 | chrIII | 133710 | 135521 | CP015055.1 | 220841 | 222493 | 54.21487603 | 603 | 550 | 1.096363636 | 1 | 1 | 1 | FIM1_955 | YCR008W | YCR008W | SAT4 | n |
| Kmar_QGN15360.1 | W303.asm02.HP0_G0003520 | OG0001770 | chrII | 575462 | 576505 | CP015056.1 | 792697 | 793737 | 64.28571429 | 347 | 346 | 1.002890173 | 1 | 1 | 1 | FIM1_2050 | YBR159W | YBR159W | IFA38 | n |
| Kmar_QGN15092.1 | W303.asm02.HP0_G0038260 | OG0001771 | chrXII | 840462 | 841910 | CP015056.1 | 189136 | 190584 | 90.66390041 | 482 | 482 | 1 | 1 | 1 | 1 | FIM1_1779 | YLR359W | YLR359W | ADE13 | n |
| Kmar_QGN14428.1 | W303.asm02.HP0_G0004230 | OG0001772 | chrII | 707820 | 708104 | CP015055.1 | 499264 | 499512 | 43.61702128 | 94 | 82 | 1.146341463 | 1 | 1 | 1 | FIM1_1089 | YBR233W-A | YBR233W-A | DAD3 | n |
| Kmar_QGN16585.1 | W303.asm02.HP0_G0020100 | OG0001773 | chrVII | 530606 | 531442 | CP015058.1 | 439183 | 439998 | 40.35714286 | 278 | 271 | 1.025830258 | 1 | 1 | 1 | FIM1_3301 | YGR026W | YGR026W | | n |
| Kmar_QGN14429.1 | W303.asm02.HP0_G0004240 | OG0001774 | chrII | 708280 | 709434 | CP015055.1 | 499604 | 500725 | 66.66666667 | 384 | 373 | 1.029490617 | 1 | 1 | 1 | FIM1_1090 | YBR234C | YBR234C | ARC40 | n |
| Kmar_QGN13620.1 | W303.asm02.HP0_G0039480 | OG0001775 | chrXIII | 49352 | 49669 | CP015054.1 | 534422 | 534742 | 61.81818182 | 105 | 106 | 0.990566038 | 1 | 1 | 1 | FIM1_262 | YML108W | YML108W | | n |
| Kmar_QGN13438.1 | W303.asm02.HP0_G0003880 | OG0001776 | chrII | 629011 | 630675 | CP015054.1 | 145163 | 146830 | 87.38738739 | 554 | 555 | 0.998198198 | 1 | 1 | 1 | FIM1_75 | YBR196C | YBR196C | PGI1 | n |
| Kmar_QGN15152.1 | W303.asm02.HP0_G0018100 | OG0001777 | chrVII | 118207 | 118818 | CP015056.1 | 305122 | 305721 | 74.38423645 | 203 | 199 | 1.020100503 | 1 | 1 | 1 | FIM1_1839 | YGL200C | YGL200C | EMP24 | n |
| Kmar_QGN14568.1 | W303.asm02.HP0_G0000520 | OG0001778 | chrI | 115683 | 119078 | CP015055.1 | 773897 | 777022 | 49.82547993 | 1131 | 1041 | 1.086455331 | 1 | 1 | 1 | FIM1_1230 | YAL019W | YAL019W | FUN30 | n |
| Kmar_QGN17108.1 | W303.asm02.HP0_G0039640 | OG0001779 | chrXIII | 81664 | 85272 | CP015059.1 | 244552 | 248190 | 40.11299435 | 1202 | 1212 | 0.991749175 | 1 | 1 | 1 | FIM1_3839 | YML091C | YML091C | RPM2 | n |
| Kmar_QGN16452.1 | W303.asm02.HP0_G0004570 | OG0001780 | chrII | 764655 | 764972 | CP015058.1 | 144674 | 144970 | 49.52380952 | 105 | 98 | 1.071428571 | 1 | 1 | 1 | FIM1_3165 | YBR268W | YBR268W | MRPL37 | n |
| Kmar_QGN18197.1 | W303.asm02.HP0_G0056200 | OG0001781 | chrXVI | 719774 | 722440 | CP015061.1 | 423799 | 426057 | 35.08379888 | 888 | 752 | 1.180851064 | 1 | 1 | 1 | FIM1_4521 | YPR089W | YPR089W | | n |
| Kmar_QGN18376.1 | W303.asm02.HP0_G0044100 | OG0001782 | chrXIV | 92438 | 95245 | CP015061.1 | 803890 | 806760 | 60.72916667 | 935 | 956 | 0.978033473 | 1 | 1 | 1 | FIM1_4703 | YNL287W | YNL287W | SEC21 | n |
| Kmar_QGN17214.1 | W303.asm02.HP0_G0053630 | OG0001783 | chrXVI | 204646 | 204927 | CP015059.1 | 464332 | 464565 | 46.23655914 | 93 | 77 | 1.207792208 | 1 | 1 | 1 | FIM1_3945 | YPL183W-A | YPL183W-A | RTC6 | n |
| Kmar_QGN17217.1 | W303.asm02.HP0_G0053660 | OG0001784 | chrXVI | 210799 | 213198 | CP015059.1 | 471800 | 474091 | 25.91687042 | 799 | 763 | 1.047182176 | 1 | 1 | 1 | FIM1_3948 | YPL180W | YPL180W | TCO89 | n |
| Kmar_QGN15333.1 | W303.asm02.HP0_G0053680 | OG0001785 | chrXVI | 217709 | 218335 | CP015056.1 | 730019 | 730615 | 65.23809524 | 208 | 198 | 1.050505051 | 1 | 1 | 1 | FIM1_2022 | YPL178W | YPL178W | CBC2 | n |
| Kmar_QGN17199.1 | W303.asm02.HP0_G0019150 | OG0001786 | chrVII | 350010 | 352424 | CP015059.1 | 435871 | 438252 | 47.75561097 | 804 | 793 | 1.013871375 | 1 | 1 | 1 | FIM1_3930 | YGL083W | YGL083W | SCY1 | n |
| Kmar_QGN14322.1 | W303.asm02.HP0_G0037040 | OG0001787 | chrXII | 578486 | 579148 | CP015055.1 | 276374 | 276955 | 37.96296296 | 220 | 193 | 1.139896373 | 1 | 1 | 1 | FIM1_979 | YLR221C | YLR221C | RSA3 | n |
| Kmar_QGN15627.1 | W303.asm02.HP0_G0055460 | OG0001788 | chrXVI | 575693 | 577735 | CP015056.1 | 1341542 | 1343437 | 27.87356322 | 680 | 631 | 1.077654517 | 1 | 1 | 1 | FIM1_2320 | YPR007C | YPR007C | REC8 | n |
| Kmar_QGN18426.1 | W303.asm02.HP0_G0055390 | OG0001789 | chrXVI | 560962 | 562086 | CP015061.1 | 907553 | 908731 | 53.58090186 | 374 | 392 | 0.954081633 | 1 | 1 | 1 | FIM1_4754 | YPL001W | YPL001W | HAT1 | n |
| Kmar_QGN16119.1 | W303.asm02.HP0_G0004010 | OG0001790 | chrII | 653482 | 658989 | CP015057.1 | 844806 | 850301 | 71.91743618 | 1835 | 1831 | 1.002184599 | 1 | 1 | 1 | FIM1_2820 | YBR208C | YBR208C | DUR12 | n |
| Kmar_QGN13461.1 | W303.asm02.HP0_G0003980 | OG0001791 | chrII | 649033 | 650160 | CP015054.1 | 188846 | 189838 | 29.49061662 | 375 | 330 | 1.136363636 | 1 | 1 | 1 | FIM1_98 | YBR204C | YBR204C | LDH1 | n |
| Kmar_QGN18176.1 | W303.asm02.HP0_G0026740 | OG0001792 | chrIX | 252070 | 255525 | CP015061.1 | 375492 | 378938 | 70.36395147 | 1151 | 1148 | 1.00261324 | 1 | 1 | 1 | FIM1_4499 | YIL048W | YIL048W | NEO1 | n |
| Kmar_QGN16982.1 | W303.asm02.HP0_G0044600 | OG0001793 | chrXIV | 215619 | 216497 | CP015058.1 | 1303909 | 1304790 | 57.09459459 | 292 | 293 | 0.996587031 | 1 | 1 | 1 | FIM1_3709 | YNL232W | YNL232W | CSL4 | n |
| Kmar_QGN16980.1 | W303.asm02.HP0_G0044620 | OG0001794 | chrXIV | 218219 | 219358 | CP015058.1 | 1300963 | 1301952 | 27.76349614 | 379 | 329 | 1.151975684 | 1 | 1 | 1 | FIM1_3707 | YNL230C | YNL230C | ELA1 | n |
| Kmar_QGN15552.1 | W303.asm02.HP0_G0023900 | OG0001795 | chrVIII | 228707 | 229588 | CP015056.1 | 1169256 | 1170113 | 57.67918089 | 293 | 285 | 1.028070175 | 1 | 1 | 1 | FIM1_2243 | YHR062C | YHR062C | RPP1 | n |
| Kmar_QGN16199.1 | W303.asm02.HP0_G0039780 | OG0001796 | chrXIII | 107053 | 109887 | CP015057.1 | 1016570 | 1019614 | 27.08883026 | 944 | 1014 | 0.930966469 | 1 | 1 | 1 | FIM1_2901 | YML076C | YML076C | WAR1 | n |
| Kmar_QGN16262.1 | W303.asm02.HP0_G0030820 | OG0001797 | chrX | 677417 | 679066 | CP015057.1 | 1150465 | 1152111 | 60.83032491 | 549 | 548 | 1.001824818 | 1 | 1 | 1 | FIM1_2967 | YJR131W | YJR131W | MNS1 | n |
| Kmar_QGN14882.1 | W303.asm02.HP0_G0029390 | OG0001798 | chrX | 383306 | 383896 | CP015055.1 | 1416939 | 1417532 | 68.68686869 | 196 | 197 | 0.994923858 | 1 | 1 | 1 | FIM1_1556 | YJL030W | YJL030W | MAD2 | n |
| Kmar_QGN14881.1 | W303.asm02.HP0_G0029560 | OG0001799 | chrX | 418787 | 419758 | CP015055.1 | 1414631 | 1415602 | 55.52147239 | 323 | 323 | 1 | 1 | 1 | 1 | FIM1_1554 | YJL006C | YJL006C | CTK2 | n |
| Kmar_QGN17736.1 | W303.asm02.HP0_G0046190 | OG0001800 | chrXIV | 520181 | 521032 | CP015060.1 | 358241 | 359089 | 66.90140845 | 283 | 282 | 1.003546099 | 1 | 1 | 1 | FIM1_4945 | YNL055C | YNL055C | POR1 | n |
| Kmar_QGN18266.1 | W303.asm02.HP0_G0056640 | OG0001801 | chrXVI | 824789 | 825691 | CP015061.1 | 582186 | 583043 | 34.11371237 | 300 | 285 | 1.052631579 | 1 | 1 | 1 | FIM1_4592 | YPR139C | YPR139C | LOA1 | n |
| Kmar_QGN16742.1 | W303.asm02.HP0_G0027270 | OG0001802 | chrIX | 355943 | 356389 | CP015058.1 | 800525 | 801043 | 53.64238411 | 148 | 172 | 0.860465116 | 1 | 1 | 1 | FIM1_3464 | YIR005W | YIR005W | IST3 | n |
| Kmar_QGN16057.1 | W303.asm02.HP0_G0040960 | OG0001803 | chrXIII | 368597 | 371020 | CP015057.1 | 719541 | 721946 | 77.94117647 | 807 | 801 | 1.007490637 | 1 | 1 | 1 | FIM1_2757 | YMR049C | YMR049C | ERB1 | n |
| Kmar_QGN14348.1 | W303.asm02.HP0_G0041280 | OG0001804 | chrXIII | 443032 | 443886 | CP015055.1 | 326085 | 326837 | 37.41258741 | 284 | 250 | 1.136 | 1 | 1 | 1 | FIM1_1008 | YMR087W | YMR087W | PDL32 | n |
| Kmar_QGN17267.1 | W303.asm02.HP0_G0016650 | OG0001805 | chrVI | 52593 | 55682 | CP015059.1 | 583904 | 586831 | 43.780135 | 1029 | 975 | 1.055384615 | 1 | 1 | 1 | FIM1_3998 | YFL025C | YFL025C | BST1 | n |
| Kmar_QGN14648.1 | W303.asm02.HP0_G0034700 | OG0001806 | chrXII | 98006 | 98845 | CP015055.1 | 943985 | 944809 | 31.22807018 | 279 | 274 | 1.018248175 | 1 | 1 | 1 | FIM1_1312 | YLL023C | YLL023C | POM33 | n |
| Kmar_QGN17451.1 | W303.asm02.HP0_G0001940 | OG0001807 | chrII | 246705 | 248006 | CP015059.1 | 946269 | 947522 | 43.59550562 | 433 | 417 | 1.038369305 | 1 | 1 | 1 | FIM1_4185 | YBR004C | YBR004C | GPI18 | n |
| Kmar_QGN13451.1 | W303.asm02.HP0_G0054750 | OG0001808 | chrXVI | 431609 | 432205 | CP015054.1 | 169960 | 170571 | 29.95391705 | 198 | 203 | 0.975369458 | 1 | 1 | 1 | FIM1_88 | YPL067C | YPL067C | HTC1 | n |
| Kmar_QGN16784.1 | W303.asm02.HP0_G0040580 | OG0001809 | chrXIII | 285606 | 286823 | CP015058.1 | 883792 | 884940 | 55.18072289 | 405 | 382 | 1.060209424 | 1 | 1 | 1 | FIM1_3507 | YMR010W | YMR010W | ANY1 | n |
| Kmar_QGN13444.1 | W303.asm02.HP0_G0054700 | OG0001810 | chrXVI | 424870 | 426369 | CP015054.1 | 159197 | 161596 | 24.875 | 499 | 799 | 0.624530663 | 1 | 1 | 1 | FIM1_81 | YPL072W | YPL072W | UBP16 | n |
| Kmar_QGN17182.1 | W303.asm02.HP0_G0011020 | OG0001811 | chrIV | 958789 | 959448 | CP015059.1 | 403167 | 403739 | 50.45454545 | 219 | 190 | 1.152631579 | 1 | 1 | 1 | FIM1_3913 | YDR246W | YDR246W | TRS23 | n |
| Kmar_QGN14395.1 | W303.asm02.HP0_G0053960 | OG0001812 | chrXVI | 277244 | 278128 | CP015055.1 | 431520 | 432335 | 31.09540636 | 294 | 271 | 1.084870849 | 1 | 1 | 1 | FIM1_1055 | YPL149W | YPL149W | ATG5 | n |
| Kmar_QGN15967.1 | W303.asm02.HP0_G0044880 | OG0001813 | chrXIV | 264413 | 265153 | CP015057.1 | 501125 | 501847 | 64.91935484 | 246 | 240 | 1.025 | 1 | 1 | 1 | FIM1_2665 | YNL200C | YNL200C | NNR1 | n |
| Kmar_QGN16568.1 | W303.asm02.HP0_G0038150 | OG0001814 | chrXII | 820008 | 822593 | CP015058.1 | 395959 | 398544 | 70.96399535 | 861 | 861 | 1 | 1 | 1 | 1 | FIM1_3282 | YLR347C | YLR347C | KAP95 | n |
| Kmar_QGN16569.1 | W303.asm02.HP0_G0020210 | OG0001815 | chrVII | 555907 | 556626 | CP015058.1 | 399215 | 399910 | 48.53556485 | 239 | 231 | 1.034632035 | 1 | 1 | 1 | FIM1_3283 | YGR036C | YGR036C | CAX4 | n |
| Kmar_QGN16948.1 | W303.asm02.HP0_G0006690 | OG0001816 | chrIV | 51188 | 52246 | CP015058.1 | 1240232 | 1241305 | 55.52560647 | 352 | 357 | 0.985994398 | 1 | 1 | 1 | FIM1_3675 | YDL226C | YDL226C | GCS1 | n |
| Kmar_QGN16946.1 | W303.asm02.HP0_G0044870 | OG0001817 | chrXIV | 261329 | 263905 | CP015058.1 | 1235964 | 1238225 | 42.59681093 | 858 | 753 | 1.139442231 | 1 | 1 | 1 | FIM1_3673 | YNL201C | YNL201C | PSY2 | n |
| Kmar_QGN13848.1 | W303.asm02.HP0_G0028040 | OG0001818 | chrX | 77113 | 78090 | CP015054.1 | 993052 | 994041 | 57.70392749 | 325 | 329 | 0.987841945 | 1 | 1 | 1 | FIM1_494 | YJL180C | YJL180C | ATP12 | n |
| Kmar_QGN13847.1 | W303.asm02.HP0_G0028050 | OG0001819 | chrX | 78317 | 78646 | CP015054.1 | 992424 | 992765 | 51.32743363 | 109 | 113 | 0.96460177 | 1 | 1 | 1 | FIM1_493 | YJL179W | YJL179W | PFD1 | n |
| Kmar_QGN13852.1 | W303.asm02.HP0_G0028010 | OG0001820 | chrX | 72978 | 73349 | CP015054.1 | 1000695 | 1001048 | 45.96774194 | 123 | 117 | 1.051282051 | 1 | 1 | 1 | FIM1_498 | YJL184W | YJL184W | GON7 | n |
| Kmar_QGN16571.1 | W303.asm02.HP0_G0020170 | OG0001821 | chrVII | 551870 | 552589 | CP015058.1 | 404587 | 405303 | 54.35684647 | 239 | 238 | 1.004201681 | 1 | 1 | 1 | FIM1_3286 | YGR033C | YGR033C | TIM21 | n |
| Kmar_QGN14181.1 | W303.asm02.HP0_G0005000 | OG0001822 | chrIII | 18469 | 21759 | CP015054.1 | 1697188 | 1700028 | 31.8018018 | 1096 | 946 | 1.158562368 | 1 | 1 | 1 | FIM1_834 | YCL061C | YCL061C | MRC1 | n |
| Kmar_QGN14180.1 | W303.asm02.HP0_G0005010 | OG0001823 | chrIII | 22082 | 23032 | CP015054.1 | 1695850 | 1696809 | 84.11214953 | 316 | 319 | 0.990595611 | 1 | 1 | 1 | FIM1_833 | YCL059C | YCL059C | KRR1 | n |
| Kmar_QGN14121.1 | W303.asm02.HP0_G0036720 | OG0001824 | chrXII | 523761 | 524519 | CP015054.1 | 1580204 | 1580962 | 91.26984127 | 252 | 252 | 1 | 1 | 1 | 1 | FIM1_773 | YLR186W | YLR186W | EMG1 | n |
| Kmar_QGN14122.1 | W303.asm02.HP0_G0013380 | OG0001825 | chrIV | 1453524 | 1455767 | CP015054.1 | 1581726 | 1583735 | 27.77777778 | 747 | 669 | 1.116591928 | 1 | 1 | 1 | FIM1_774 | YDR499W | YDR499W | LCD1 | n |
| Kmar_QGN14120.1 | W303.asm02.HP0_G0044170 | OG0001826 | chrXIV | 111471 | 113456 | CP015054.1 | 1577799 | 1579736 | 41.43070045 | 661 | 645 | 1.024806202 | 1 | 1 | 1 | FIM1_772 | YNL279W | YNL279W | PRM1 | n |
| Kmar_QGN14133.1 | W303.asm02.HP0_G0013480 | OG0001827 | chrIV | 1475094 | 1475399 | CP015054.1 | 1608219 | 1608500 | 76 | 101 | 93 | 1.086021505 | 1 | 1 | 1 | FIM1_786 | YDR510W | YDR510W | SMT3 | n |
| Kmar_QGN15522.1 | W303.asm02.HP0_G0010990 | OG0001828 | chrIV | 953019 | 954785 | CP015056.1 | 1115212 | 1116834 | 50.6779661 | 588 | 540 | 1.088888889 | 1 | 1 | 1 | FIM1_2213 | YDR243C | YDR243C | PRP28 | n |
| Kmar_QGN15523.1 | W303.asm02.HP0_G0006850 | OG0001829 | chrIV | 86280 | 87414 | CP015056.1 | 1117126 | 1118154 | 50.27777778 | 356 | 342 | 1.040935673 | 1 | 1 | 1 | FIM1_2214 | YDL209C | YDL209C | CWC2 | n |
| Kmar_QGN18293.1 | W303.asm02.HP0_G0014630 | OG0001830 | chrV | 189424 | 190458 | CP015061.1 | 636679 | 637521 | 45.34883721 | 344 | 280 | 1.228571429 | 1 | 1 | 1 | FIM1_4619 | YER016W | YER016W | BIM1 | n |
| Kmar_QGN13458.1 | W303.asm02.HP0_G0054790 | OG0001831 | chrXVI | 436297 | 437727 | CP015054.1 | 180527 | 181984 | 64.28571429 | 476 | 485 | 0.981443299 | 1 | 1 | 1 | FIM1_95 | YPL063W | YPL063W | TIM50 | n |
| Kmar_QGN14357.1 | W303.asm02.HP0_G0006960 | OG0001832 | chrIV | 103723 | 104625 | CP015055.1 | 346929 | 347831 | 86.04651163 | 300 | 300 | 1 | 1 | 1 | 1 | FIM1_1017 | YDL198C | YDL198C | GGC1 | n |
| Kmar_QGN16149.1 | W303.asm02.HP0_G0043930 | OG0001833 | chrXIV | 58587 | 59240 | CP015057.1 | 908526 | 909143 | 48.62385321 | 217 | 205 | 1.058536585 | 1 | 1 | 1 | FIM1_2850 | YNL306W | YNL306W | MRPS18 | n |
| Kmar_QGN17460.1 | W303.asm02.HP0_G0011820 | OG0001834 | chrIV | 1136870 | 1138195 | CP015059.1 | 964548 | 966032 | 41.66666667 | 441 | 494 | 0.892712551 | 1 | 1 | 1 | FIM1_4194 | YDR329C | YDR329C | PEX3 | n |
| Kmar_QGN17461.1 | W303.asm02.HP0_G0011810 | OG0001835 | chrIV | 1136034 | 1136618 | CP015059.1 | 966314 | 966859 | 78.57142857 | 194 | 181 | 1.071823204 | 1 | 1 | 1 | FIM1_4195 | YDR328C | YDR328C | SKP1 | n |
| Kmar_QGN17457.1 | W303.asm02.HP0_G0011830 | OG0001836 | chrIV | 1138472 | 1139974 | CP015059.1 | 960570 | 962135 | 54.28571429 | 500 | 521 | 0.959692898 | 1 | 1 | 1 | FIM1_4191 | YDR330W | YDR330W | UBX5 | n |
| Kmar_QGN17455.1 | W303.asm02.HP0_G0024120 | OG0001837 | chrVIII | 274433 | 277522 | CP015059.1 | 956084 | 958774 | 43.36118849 | 1029 | 896 | 1.1484375 | 1 | 1 | 1 | FIM1_4189 | YHR082C | YHR082C | KSP1 | n |
| Kmar_QGN18327.1 | W303.asm02.HP0_G0001560 | OG0001838 | chrII | 156151 | 160692 | CP015061.1 | 700949 | 705379 | 22.62443439 | 1513 | 1476 | 1.025067751 | 1 | 1 | 1 | FIM1_4653 | YBL034C | YBL034C | STU1 | n |
| Kmar_QGN15641.1 | W303.asm02.HP0_G0020280 | OG0001839 | chrVII | 580749 | 581651 | CP015056.1 | 1371374 | 1372999 | 20.88724584 | 300 | 541 | 0.554528651 | 1 | 1 | 1 | FIM1_2334 | YGR044C | YGR044C | RME1 | n |
| Kmar_QGN15381.1 | W303.asm02.HP0_G0016050 | OG0001840 | chrV | 492490 | 494895 | CP015056.1 | 843778 | 845892 | 38.35616438 | 801 | 704 | 1.137784091 | 1 | 1 | 1 | FIM1_2071 | YER157W | YER157W | COG3 | n |
| Kmar_QGN15729.1 | W303.asm02.HP0_G0025800 | OG0001841 | chrIX | 50045 | 51760 | CP015056.1 | 1555252 | 1557087 | 36.9047619 | 571 | 582 | 0.981099656 | 1 | 1 | 1 | FIM1_2423 | YIL150C | YIL150C | MCM10 | n |
| Kmar_QGN16969.1 | W303.asm02.HP0_G0044710 | OG0001842 | chrXIV | 236695 | 238362 | CP015058.1 | 1276443 | 1278095 | 52.85714286 | 555 | 550 | 1.009090909 | 1 | 1 | 1 | FIM1_3696 | YNL219C | YNL219C | ALG9 | n |
| Kmar_QGN16968.1 | W303.asm02.HP0_G0006610 | OG0001843 | chrIV | 32371 | 33309 | CP015058.1 | 1274830 | 1275765 | 70.73954984 | 312 | 311 | 1.003215434 | 1 | 1 | 1 | FIM1_3695 | YDL236W | YDL236W | PHO13 | n |
| Kmar_QGN17813.1 | W303.asm02.HP0_G0052300 | OG0001844 | chrXV | 1000734 | 1003109 | CP015060.1 | 537659 | 540232 | 33.25892857 | 791 | 857 | 0.922987165 | 1 | 1 | 1 | FIM1_5022 | YOR353C | YOR353C | SOG2 | n |
| Kmar_QGN16068.1 | W303.asm02.HP0_G0040880 | OG0001845 | chrXIII | 348014 | 348763 | CP015057.1 | 741693 | 742427 | 50.20080321 | 249 | 244 | 1.020491803 | 1 | 1 | 1 | FIM1_2768 | YMR038C | YMR038C | CCS1 | n |
| Kmar_QGN14942.1 | W303.asm02.HP0_G0045200 | OG0001846 | chrXIV | 327433 | 330765 | CP015055.1 | 1538776 | 1542204 | 63.31877729 | 1110 | 1142 | 0.971978984 | 1 | 1 | 1 | FIM1_1617 | YNL163C | YNL163C | RIA1 | n |
| Kmar_QGN16764.1 | W303.asm02.HP0_G0014450 | OG0001847 | chrV | 149905 | 151197 | CP015058.1 | 842901 | 844205 | 46.55963303 | 430 | 434 | 0.99078341 | 1 | 1 | 1 | FIM1_3486 | YEL002C | YEL002C | WBP1 | n |
| Kmar_QGN16774.1 | W303.asm02.HP0_G0027140 | OG0001848 | chrIX | 333591 | 333890 | CP015058.1 | 860389 | 860694 | 67.32673267 | 99 | 101 | 0.98019802 | 1 | 1 | 1 | FIM1_3496 | YIL008W | YIL008W | URM1 | n |
| Kmar_QGN13532.1 | W303.asm02.HP0_G0020530 | OG0001849 | chrVII | 634049 | 634777 | CP015054.1 | 354222 | 354959 | 42.62948207 | 242 | 245 | 0.987755102 | 1 | 1 | 1 | FIM1_172 | YGR075C | YGR075C | PRP38 | n |
| Kmar_QGN16765.1 | W303.asm02.HP0_G0014440 | OG0001850 | chrV | 149359 | 149782 | CP015058.1 | 844308 | 844634 | 62.16216216 | 111 | 108 | 1.027777778 | 1 | 1 | 1 | FIM1_3487 | YEL003W | YEL003W | GIM4 | n |
| Kmar_QGN16783.1 | W303.asm02.HP0_G0040570 | OG0001851 | chrXIII | 284608 | 285147 | CP015058.1 | 882894 | 883430 | 58.01104972 | 179 | 178 | 1.005617978 | 1 | 1 | 1 | FIM1_3506 | YMR009W | YMR009W | ADI1 | n |

| | | | | | | | | | | | | | | | | | | | | |
|---|---|---|---|---|---|---|---|---|---|---|---|---|---|---|---|---|---|---|---|---|
| Kmar_QGN18139.1 | W303.asm02.HP0_G0043820 | OG0001852 | chrXIV | 41051 | 42448 | CP015061.1 | 311073 | 312431 | 60.63829787 | 465 | 452 | 1.028761062 | 1 | 1 | 1 | FIM1_4462 | YNL317W | YNL317W | PFS2 | n |
| Kmar_QGN18140.1 | W303.asm02.HP0_G0047770 | OG0001853 | chrXV | 67902 | 69095 | CP015061.1 | 313194 | 314525 | 36.16071429 | 397 | 443 | 0.896162528 | 1 | 1 | 1 | FIM1_4463 | YOL136C | YOL136C | PFK27 | n |
| Kmar_QGN16785.1 | W303.asm02.HP0_G0049240 | OG0001854 | chrXV | 361868 | 364270 | CP015058.1 | 885046 | 887169 | 30.51869723 | 800 | 707 | 1.131541726 | 1 | 1 | 1 | FIM1_3508 | YOR017W | YOR017W | PET127 | n |
| Kmar_QGN14247.1 | W303.asm02.HP0_G0019620 | OG0001855 | chrVII | 437149 | 437511 | CP015055.1 | 107932 | 108300 | 72.13114754 | 120 | 122 | 0.983606557 | 1 | 1 | 1 | FIM1_903 | YGL029W | YGL029W | CGR1 | n |
| Kmar_QGN18118.1 | W303.asm02.HP0_G0000950 | OG0001856 | chrII | 22380 | 23840 | CP015061.1 | 252917 | 254356 | 32.99798793 | 486 | 479 | 1.014613779 | 1 | 1 | 1 | FIM1_4438 | YBL103C | YBL103C | RTG3 | n |
| Kmar_QGN17970.1 | W303.asm02.HP0_G0012150 | OG0001857 | chrIV | 1216013 | 1217899 | CP015060.1 | 853275 | 855191 | 60.79027356 | 628 | 638 | 0.984326019 | 1 | 1 | 1 | FIM1_5179 | YDR365C | YDR365C | ESF1 | n |
| Kmar_QGN17968.1 | W303.asm02.HP0_G0050170 | OG0001858 | chrXV | 549667 | 551121 | CP015060.1 | 849042 | 850670 | 65.44117647 | 484 | 542 | 0.89298893 | 1 | 1 | 1 | FIM1_5177 | YOR119C | YOR119C | RIO1 | n |
| Kmar_QGN14636.1 | W303.asm02.HP0_G0024410 | OG0001859 | chrVIII | 339045 | 340367 | CP015055.1 | 917571 | 918905 | 63.73873874 | 440 | 444 | 0.990990991 | 1 | 1 | 1 | FIM1_1300 | YHR111W | YHR111W | UBA4 | n |
| Kmar_QGN14635.1 | W303.asm02.HP0_G0012110 | OG0001860 | chrIV | 1208195 | 1210213 | CP015055.1 | 915667 | 917505 | 34.27299703 | 672 | 612 | 1.098039216 | 1 | 1 | 1 | FIM1_1299 | YDR362C | YDR362C | TFC6 | n |
| Kmar_QGN17975.1 | W303.asm02.HP0_G0012120 | OG0001861 | chrIV | 1210699 | 1212069 | CP015060.1 | 863695 | 864873 | 27.51091703 | 456 | 392 | 1.163265306 | 1 | 1 | 1 | FIM1_5184 | YDR363W | YDR363W | ESC2 | n |
| Kmar_QGN16335.1 | W303.asm02.HP0_G0012910 | OG0001862 | chrIV | 1363273 | 1364595 | CP015057.1 | 1313559 | 1314911 | 64.30155211 | 440 | 450 | 0.977777778 | 1 | 1 | 1 | FIM1_3040 | YDR449C | YDR449C | UTP6 | n |
| Kmar_QGN16336.1 | W303.asm02.HP0_G0012900 | OG0001863 | chrIV | 1361620 | 1362924 | CP015057.1 | 1314993 | 1316300 | 78.85057471 | 434 | 435 | 0.997701149 | 1 | 1 | 1 | FIM1_3041 | YDR448W | YDR448W | ADA2 | n |
| Kmar_QGN16337.1 | W303.asm02.HP0_G0012880 | OG0001864 | chrIV | 1359280 | 1360188 | CP015057.1 | 1318145 | 1319098 | 23.6196319 | 302 | 317 | 0.952681388 | 1 | 1 | 1 | FIM1_3043 | YDR446W | YDR446W | ECM11 | n |
| Kmar_QGN14743.1 | W303.asm02.HP0_G0023220 | OG0001865 | chrVIII | 117575 | 119146 | CP015055.1 | 1144142 | 1145734 | 55.7168784 | 523 | 530 | 0.986792453 | 1 | 1 | 1 | FIM1_1412 | YHR009C | YHR009C | TDA3 | n |
| Kmar_QGN17976.1 | W303.asm02.HP0_G0024430 | OG0001866 | chrVIII | 342310 | 343782 | CP015060.1 | 865174 | 866700 | 64.56692913 | 490 | 508 | 0.964566929 | 1 | 1 | 1 | FIM1_5185 | YHR113W | YHR113W | APE4 | n |
| Kmar_QGN18119.1 | W303.asm02.HP0_G0000960 | OG0001867 | chrII | 24402 | 25049 | CP015061.1 | 254901 | 255542 | 72.09302326 | 215 | 213 | 1.009389671 | 1 | 1 | 1 | FIM1_4439 | YBL102W | YBL102W | SFT2 | n |
| Kmar_QGN14666.1 | W303.asm02.HP0_G0005450 | OG0001868 | chrIII | 110936 | 112093 | CP015055.1 | 986110 | 986536 | 13.75661376 | 385 | 94 | 4.095744681 | 1 | 1 | 1 | FIM1_1331 | YCL008C | YCL008C | STP22 | n |
| Kmar_QGN16320.1 | W303.asm02.HP0_G0053260 | OG0001869 | chrXVI | 135727 | 137793 | CP015057.1 | 1274835 | 1276847 | 47.1942446 | 688 | 670 | 1.026865672 | 1 | 1 | 1 | FIM1_3025 | YPL222W | YPL222W | FMP40 | n |
| Kmar_QGN15649.1 | W303.asm02.HP0_G0038190 | OG0001870 | chrXII | 825669 | 826544 | CP015056.1 | 1388963 | 1389835 | 72.60273973 | 291 | 290 | 1.003448276 | 1 | 1 | 1 | FIM1_2342 | YLR351C | YLR351C | NIT3 | n |
| Kmar_QGN15567.1 | W303.asm02.HP0_G0048590 | OG0001871 | chrXV | 232901 | 234091 | CP015056.1 | 1202514 | 1203605 | 45.84382872 | 396 | 363 | 1.090909091 | 1 | 1 | 1 | FIM1_2258 | YOL052C | YOL052C | SPE2 | n |
| Kmar_QGN16261.1 | W303.asm02.HP0_G0025200 | OG0001872 | chrVIII | 492332 | 492733 | CP015057.1 | 1150021 | 1150422 | 44.85294118 | 133 | 133 | 1 | 1 | 1 | 1 | FIM1_2966 | YHR191C | YHR191C | CTF8 | n |
| Kmar_QGN14082.1 | W303.asm02.HP0_G0044390 | OG0001873 | chrXIV | 165058 | 167652 | CP015054.1 | 1480008 | 1482401 | 46.59746251 | 864 | 797 | 1.084065245 | 1 | 1 | 1 | FIM1_733 | YNL256W | YNL256W | FOL1 | n |
| Kmar_QGN14080.1 | W303.asm02.HP0_G0044400 | OG0001874 | chrXIV | 168598 | 169803 | CP015054.1 | 1477883 | 1478989 | 26.43391521 | 401 | 368 | 1.089673913 | 1 | 1 | 1 | FIM1_731 | YNL254C | YNL254C | RTC4 | n |
| Kmar_QGN14077.1 | W303.asm02.HP0_G0044430 | OG0001875 | chrXIV | 173143 | 174870 | CP015054.1 | 1472748 | 1474571 | 44.57050243 | 575 | 607 | 0.947281713 | 1 | 1 | 1 | FIM1_728 | YNL251C | YNL251C | NRD1 | n |
| Kmar_QGN14078.1 | W303.asm02.HP0_G0044420 | OG0001876 | chrXIV | 171996 | 172841 | CP015054.1 | 1474877 | 1475668 | 51.57894737 | 281 | 263 | 1.068441065 | 1 | 1 | 1 | FIM1_729 | YNL252C | YNL252C | MRPL17 | n |
| Kmar_QGN14079.1 | W303.asm02.HP0_G0044410 | OG0001877 | chrXIV | 170573 | 171841 | CP015054.1 | 1475867 | 1477186 | 22.34513274 | 422 | 439 | 0.961275626 | 1 | 1 | 1 | FIM1_730 | YNL253W | YNL253W | TEX1 | n |
| Kmar_QGN15344.1 | W303.asm02.HP0_G0011720 | OG0001878 | chrIV | 1116161 | 1118167 | CP015056.1 | 753749 | 755677 | 32.73542601 | 668 | 642 | 1.040498442 | 1 | 1 | 1 | FIM1_2033 | YDR320C | YDR320C | SWA2 | n |
| Kmar_QGN16132.1 | W303.asm02.HP0_G0046010 | OG0001879 | chrXIV | 487794 | 488666 | CP015057.1 | 881373 | 882224 | 85.51724138 | 290 | 283 | 1.024734982 | 1 | 1 | 1 | FIM1_2833 | YNL075W | YNL075W | IMP4 | n |
| Kmar_QGN15646.1 | W303.asm02.HP0_G0038200 | OG0001880 | chrXII | 827296 | 829719 | CP015056.1 | 1382716 | 1385064 | 52.15252153 | 807 | 782 | 1.031969309 | 1 | 1 | 1 | FIM1_2339 | YLR352W | YLR352W | LUG1 | n |
| Kmar_QGN13477.1 | W303.asm02.HP0_G0029920 | OG0001881 | chrX | 505886 | 509143 | CP015054.1 | 219380 | 222493 | 56.26715462 | 1085 | 1037 | 1.046287367 | 1 | 1 | 1 | FIM1_115 | YJR035W | YJR035W | RAD26 | n |
| Kmar_QGN13770.1 | W303.asm02.HP0_G0040060 | OG0001882 | chrXIII | 167914 | 168849 | CP015054.1 | 846911 | 847837 | 33.22981366 | 311 | 308 | 1.00974026 | 1 | 1 | 1 | FIM1_415 | YML050W | YML050W | AIM32 | n |
| Kmar_QGN16838.1 | W303.asm02.HP0_G0017190 | OG0001883 | chrVI | 182217 | 185324 | CP015058.1 | 998455 | 1001553 | 60.6903164 | 1035 | 1032 | 1.002906977 | 1 | 1 | 1 | FIM1_3562 | YFR030W | YFR030W | MET10 | n |
| Kmar_QGN15410.1 | W303.asm02.HP0_G0011320 | OG0001884 | chrIV | 1025246 | 1026163 | CP015056.1 | 902069 | 902986 | 78.68852459 | 305 | 305 | 1 | 1 | 1 | 1 | FIM1_2100 | YDR280W | YDR280W | RRP45 | n |
| Kmar_QGN17739.1 | W303.asm02.HP0_G0046180 | OG0001885 | chrXIV | 519436 | 520029 | CP015060.1 | 365765 | 366301 | 63.02083333 | 197 | 178 | 1.106741573 | 1 | 1 | 1 | FIM1_4948 | YNL056W | YNL056W | OCA2 | n |
| Kmar_QGN15253.1 | W303.asm02.HP0_G0049480 | OG0001886 | chrXV | 407520 | 408377 | CP015056.1 | 533999 | 534868 | 59.31034483 | 285 | 289 | 0.98615917 | 1 | 1 | 1 | FIM1_1942 | YOR040W | YOR040W | GLO4 | n |
| Kmar_QGN15266.1 | W303.asm02.HP0_G0049610 | OG0001887 | chrXV | 432782 | 433969 | CP015056.1 | 560382 | 561533 | 49.74747475 | 395 | 383 | 1.031331593 | 1 | 1 | 1 | FIM1_1955 | YOR057W | YOR057W | SGT1 | n |
| Kmar_QGN16814.1 | W303.asm02.HP0_G0020810 | OG0001888 | chrVII | 695915 | 696838 | CP015058.1 | 953106 | 953873 | 37.62057878 | 307 | 255 | 1.203921569 | 1 | 1 | 1 | FIM1_3538 | YGR104C | YGR104C | SRB5 | n |
| Kmar_QGN16812.1 | W303.asm02.HP0_G0020790 | OG0001889 | chrVII | 693054 | 693605 | CP015058.1 | 950022 | 950543 | 35.51912568 | 183 | 173 | 1.057803468 | 1 | 1 | 1 | FIM1_3536 | YGR102C | YGR102C | GTF1 | n |
| Kmar_QGN16813.1 | W303.asm02.HP0_G0020800 | OG0001890 | chrVII | 693887 | 695704 | CP015058.1 | 951201 | 953015 | 75.36704731 | 605 | 604 | 1.001655629 | 1 | 1 | 1 | FIM1_3537 | YGR103W | YGR103W | NOP7 | n |
| Kmar_QGN18265.1 | W303.asm02.HP0_G0056650 | OG0001891 | chrXVI | 826024 | 827169 | CP015061.1 | 580708 | 581916 | 65.67164179 | 381 | 402 | 0.947761194 | 1 | 1 | 1 | FIM1_4591 | YPR140W | YPR140W | TAZ1 | n |
| Kmar_QGN16246.1 | W303.asm02.HP0_G0030890 | OG0001892 | chrX | 688727 | 693055 | CP015057.1 | 1121379 | 1125665 | 63.09606082 | 1442 | 1428 | 1.009803922 | 1 | 1 | 1 | FIM1_2951 | YJR137C | YJR137C | MET5 | n |
| Kmar_QGN18264.1 | W303.asm02.HP0_G0056660 | OG0001893 | chrXVI | 827367 | 829556 | CP015061.1 | 578319 | 580385 | 44.71879287 | 729 | 688 | 1.059593023 | 1 | 1 | 1 | FIM1_4590 | YPR141C | YPR141C | KAR3 | n |
| Kmar_QGN16244.1 | W303.asm02.HP0_G0025300 | OG0001894 | chrVIII | 505181 | 505987 | CP015057.1 | 1118948 | 1119694 | 69.68503937 | 268 | 248 | 1.080645161 | 1 | 1 | 1 | FIM1_2949 | YHR200W | YHR200W | RPN10 | n |
| Kmar_QGN16058.1 | W303.asm02.HP0_G0040950 | OG0001895 | chrXIII | 367484 | 368437 | CP015057.1 | 722055 | 722867 | 32.60869565 | 317 | 270 | 1.174074074 | 1 | 1 | 1 | FIM1_2758 | YMR048W | YMR048W | CSM3 | n |
| Kmar_QGN16240.1 | W303.asm02.HP0_G0030930 | OG0001896 | chrX | 705676 | 706719 | CP015057.1 | 1106193 | 1107263 | 23.39832869 | 347 | 356 | 0.974719101 | 1 | 1 | 1 | FIM1_2945 | YJR141W | YJR141W | IPA1 | n |
| Kmar_QGN16239.1 | W303.asm02.HP0_G0025310 | OG0001897 | chrVIII | 506052 | 507245 | CP015057.1 | 1104977 | 1106161 | 35.3221957 | 397 | 394 | 1.007614213 | 1 | 1 | 1 | FIM1_2944 | YHR201C | YHR201C | PPX1 | n |
| Kmar_QGN16237.1 | W303.asm02.HP0_G0030950 | OG0001898 | chrX | 708095 | 710383 | CP015057.1 | 1101341 | 1103647 | 60.12987013 | 762 | 768 | 0.9921875 | 1 | 1 | 1 | FIM1_2942 | YJR143C | YJR143C | PMT4 | n |
| Kmar_QGN16238.1 | W303.asm02.HP0_G0030940 | OG0001899 | chrX | 706910 | 707938 | CP015057.1 | 1103811 | 1104839 | 51.75438596 | 342 | 342 | 1 | 1 | 1 | 1 | FIM1_2943 | YJR142W | YJR142W | | n |
| Kmar_QGN16992.1 | W303.asm02.HP0_G0044510 | OG0001900 | chrXIV | 191879 | 196657 | CP015058.1 | 1329782 | 1334281 | 31.09452736 | 1592 | 1499 | 1.062041361 | 1 | 1 | 1 | FIM1_3719 | YNL242W | YNL242W | ATG2 | n |
| Kmar_QGN13859.1 | W303.asm02.HP0_G0033860 | OG0001901 | chrXI | 570450 | 571961 | CP015054.1 | 1011051 | 1012421 | 32.01581028 | 503 | 456 | 1.103070175 | 1 | 1 | 1 | FIM1_505 | YKR063C | YKR063C | LAS1 | n |
| Kmar_QGN15639.1 | W303.asm02.HP0_G0038230 | OG0001902 | chrXII | 834246 | 835433 | CP015056.1 | 1365904 | 1367109 | 86.28428928 | 395 | 401 | 0.985037406 | 1 | 1 | 1 | FIM1_2332 | YLR355C | YLR355C | ILV5 | n |
| Kmar_QGN14721.1 | W303.asm02.HP0_G0010040 | OG0001903 | chrIV | 745382 | 746509 | CP015055.1 | 1094723 | 1095841 | 60.20942408 | 375 | 372 | 1.008064516 | 1 | 1 | 1 | FIM1_1389 | YDR142C | YDR142C | PEX7 | n |
| Kmar_QGN15636.1 | W303.asm02.HP0_G0020350 | OG0001904 | chrVII | 592745 | 593596 | CP015056.1 | 1359349 | 1359993 | 17.20430108 | 283 | 214 | 1.322429907 | 1 | 1 | 1 | FIM1_2329 | YGR053C | YGR053C | MCO32 | n |
| Kmar_QGN15373.1 | W303.asm02.HP0_G0017050 | OG0001905 | chrVI | 145949 | 149650 | CP015056.1 | 823792 | 825552 | 19.04376013 | 1233 | 586 | 2.104095563 | 1 | 1 | 1 | FIM1_2063 | YFR016C | YFR016C | AIP5 | n |
| Kmar_QGN13948.1 | W303.asm02.HP0_G0040520 | OG0001906 | chrXIII | 272699 | 273169 | CP015054.1 | 1202924 | 1203385 | 67.29559748 | 156 | 153 | 1.019607843 | 1 | 1 | 1 | FIM1_596 | YMR002W | YMR002W | MIX17 | n |
| Kmar_QGN13947.1 | W303.asm02.HP0_G0049020 | OG0001907 | chrXV | 313533 | 315842 | CP015054.1 | 1200000 | 1202303 | 67.74611399 | 769 | 767 | 1.002607562 | 1 | 1 | 1 | FIM1_595 | YOL006C | YOL006C | TOP1 | n |
| Kmar_QGN17128.1 | W303.asm02.HP0_G0030580 | OG0001908 | chrX | 637068 | 638099 | CP015059.1 | 294416 | 295585 | 46.01542416 | 343 | 389 | 0.881748072 | 1 | 0 | 1 | FIM1_3859 | YJR107W | YJR107W | LIH1 | n |
| Kmar_QGN17384.1 | W303.asm02.HP0_G0041090 | OG0001909 | chrXIII | 400205 | 401719 | CP015059.1 | 814892 | 816295 | 25.7751938 | 504 | 467 | 1.079229122 | 1 | 1 | 1 | FIM1_4116 | YMR065W | YMR065W | KAR5 | n |
| Kmar_QGN17382.1 | W303.asm02.HP0_G0032470 | OG0001910 | chrXI | 297321 | 299528 | CP015059.1 | 810085 | 812295 | 59.3625498 | 735 | 736 | 0.998641304 | 1 | 1 | 1 | FIM1_4114 | YKL078W | YKL078W | DHR2 | n |
| Kmar_QGN13406.1 | W303.asm02.HP0_G0054460 | OG0001911 | chrXVI | 371310 | 372788 | CP015054.1 | 77423 | 78847 | 54.01606426 | 492 | 474 | 1.037974684 | 1 | 1 | 1 | FIM1_43 | YPL097W | YPL097W | MSY1 | n |
| Kmar_QGN15660.1 | W303.asm02.HP0_G0012600 | OG0001912 | chrIV | 1298097 | 1298816 | CP015056.1 | 1409254 | 1409997 | 52.63157895 | 239 | 247 | 0.967611336 | 1 | 1 | 1 | FIM1_2353 | YDR410C | YDR410C | STE14 | n |
| Kmar_QGN16644.1 | W303.asm02.HP0_G0038360 | OG0001913 | chrXII | 857898 | 858434 | CP015058.1 | 577988 | 578518 | 79.7752809 | 178 | 176 | 1.011363636 | 1 | 1 | 1 | FIM1_3363 | YLR370C | YLR370C | ARC18 | n |
| Kmar_QGN14421.1 | W303.asm02.HP0_G0018860 | OG0001914 | chrVII | 286762 | 288594 | CP015055.1 | 486050 | 487693 | 53.84615385 | 610 | 547 | 1.115173675 | 1 | 1 | 1 | FIM1_1082 | YGL116W | YGL116W | CDC20 | n |
| Kmar_QGN16011.1 | W303.asm02.HP0_G0031740 | OG0001915 | chrXI | 155095 | 155730 | CP015057.1 | 610094 | 610765 | 31.55555556 | 211 | 223 | 0.946188341 | 1 | 1 | 1 | FIM1_2711 | YKL159C | YKL159C | RCN1 | n |
| Kmar_QGN16008.1 | W303.asm02.HP0_G0023680 | OG0001916 | chrVIII | 204691 | 205821 | CP015057.1 | 605745 | 606908 | 49.74489796 | 376 | 387 | 0.971576227 | 1 | 1 | 1 | FIM1_2708 | YHR052W | YHR052W | CIC1 | n |
| Kmar_QGN18309.1 | W303.asm02.HP0_G0002940 | OG0001917 | chrII | 453237 | 457601 | CP015061.1 | 665434 | 669672 | 42.79863481 | 1454 | 1412 | 1.029745042 | 1 | 1 | 1 | FIM1_4635 | YBR097W | YBR097W | VPS15 | n |
| Kmar_QGN17905.1 | W303.asm02.HP0_G0055920 | OG0001918 | chrXVI | 678969 | 679994 | CP015060.1 | 712015 | 712989 | 27.98833819 | 341 | 324 | 1.052469136 | 1 | 1 | 1 | FIM1_5114 | YPR057W | YPR057W | BRR1 | n |
| Kmar_QGN14273.1 | W303.asm02.HP0_G0002970 | OG0001919 | chrII | 461348 | 463609 | CP015055.1 | 157218 | 159380 | 38.60103627 | 753 | 720 | 1.045833333 | 1 | 1 | 1 | FIM1_930 | YBR102C | YBR102C | EXO84 | n |
| Kmar_QGN16609.1 | W303.asm02.HP0_G0002260 | OG0001920 | chrII | 312069 | 312965 | CP015058.1 | 491974 | 492870 | 46.84385382 | 298 | 298 | 1 | 1 | 1 | 1 | FIM1_3325 | YBR040W | YBR040W | FIG1 | n |
| Kmar_QGN15533.1 | W303.asm02.HP0_G0006890 | OG0001921 | chrIV | 92836 | 93819 | CP015056.1 | 1135595 | 1136578 | 67.07317073 | 327 | 327 | 1 | 1 | 1 | 1 | FIM1_2224 | YDL205C | YDL205C | HEM3 | n |
| Kmar_QGN13903.1 | W303.asm02.HP0_G0051600 | OG0001922 | chrXV | 844536 | 845363 | CP015054.1 | 1118324 | 1119109 | 48.02867384 | 275 | 261 | 1.053639847 | 1 | 1 | 1 | FIM1_551 | YOR278W | YOR278W | HEM4 | n |
| Kmar_QGN14399.1 | W303.asm02.HP0_G0051260 | OG0001923 | chrXV | 790465 | 791580 | CP015055.1 | 438994 | 440133 | 28.98172324 | 371 | 379 | 0.978891821 | 1 | 1 | 1 | FIM1_1059 | YOR242C | YOR242C | SSP2 | n |
| Kmar_QGN18014.1 | W303.asm02.HP0_G0053900 | OG0001924 | chrXVI | 263150 | 265270 | CP015061.1 | 34799 | 36865 | 40 | 706 | 688 | 1.026162791 | 1 | 1 | 1 | FIM1_4330 | YPL155C | YPL155C | KIP2 | n |
| Kmar_QGN14400.1 | W303.asm02.HP0_G0051270 | OG0001925 | chrXV | 791934 | 794200 | CP015055.1 | 440686 | 442716 | 55.54054054 | 737 | 676 | 1.090236686 | 1 | 1 | 1 | FIM1_1060 | YOR243C | YOR243C | PUS7 | n |
| Kmar_QGN13837.1 | W303.asm02.HP0_G0028140 | OG0001926 | chrX | 91755 | 93956 | CP015054.1 | 972323 | 974461 | 52.31175694 | 733 | 712 | 1.029494382 | 1 | 1 | 1 | FIM1_483 | YJL168C | YJL168C | SET2 | n |
| Kmar_QGN17801.1 | W303.asm02.HP0_G0045810 | OG0001927 | chrXIV | 443000 | 443992 | CP015060.1 | 514765 | 515994 | 45.49878345 | 330 | 409 | 0.806845966 | 1 | 1 | 1 | FIM1_5010 | YNL097C | YNL097C | PHO23 | n |
| Kmar_QGN17803.1 | W303.asm02.HP0_G0050020 | OG0001929 | chrXV | 517184 | 517576 | CP015060.1 | 518332 | 518706 | 56.92307692 | 130 | 124 | 1.048387097 | 1 | 1 | 1 | FIM1_5012 | YOR103C | YOR103C | OST2 | n |
| Kmar_QGN17804.1 | W303.asm02.HP0_G0045790 | OG0001930 | chrXIV | 440222 | 440938 | CP015060.1 | 519049 | 519702 | 71.42857143 | 238 | 217 | 1.096774194 | 1 | 1 | 1 | FIM1_5013 | YNL099C | YNL099C | OCA1 | n |
| Kmar_QGN13911.1 | W303.asm02.HP0_G0042630 | OG0001931 | chrXIII | 717294 | 718709 | CP015054.1 | 1134739 | 1136082 | 49.57264957 | 471 | 447 | 1.053691275 | 1 | 1 | 1 | FIM1_559 | YMR223W | YMR223W | UBP8 | n |
| Kmar_QGN13912.1 | W303.asm02.HP0_G0051620 | OG0001932 | chrXV | 847988 | 848848 | CP015054.1 | 1136149 | 1136991 | 68.88111888 | 286 | 280 | 1.021428571 | 1 | 1 | 1 | FIM1_560 | YOR281C | YOR281C | PLP2 | n |
| Kmar_QGN17773.1 | W303.asm02.HP0_G0022450 | OG0001933 | chrVII | 1060138 | 1061871 | CP015060.1 | 450249 | 451838 | 33.2761578 | 577 | 529 | 1.09073724 | 1 | 1 | 1 | FIM1_4982 | YGR278W | YGR278W | CWC22 | n |
| Kmar_QGN16163.1 | W303.asm02.HP0_G0015300 | OG0001934 | chrV | 316204 | 318270 | CP015057.1 | 935953 | 937896 | 29.08309456 | 688 | 647 | 1.063369397 | 1 | 1 | 1 | FIM1_2865 | YER077C | YER077C | MRX1 | n |
| Kmar_QGN16964.1 | W303.asm02.HP0_G0044740 | OG0001935 | chrXIV | 242388 | 244871 | CP015058.1 | 1266517 | 1268574 | 38.68263473 | 827 | 685 | 1.20729927 | 1 | 1 | 1 | FIM1_3691 | YNL216W | YNL216W | RAP1 | n |
| Kmar_QGN17774.1 | W303.asm02.HP0_G0022440 | OG0001936 | chrVII | 1059051 | 1059968 | CP015060.1 | 452058 | 452930 | 43.77104377 | 305 | 290 | 1.051724138 | 1 | 1 | 1 | FIM1_4983 | YGR277C | YGR277C | CAB4 | n |
| Kmar_QGN17775.1 | W303.asm02.HP0_G0043440 | OG0001937 | chrXIII | 881907 | 885599 | CP015060.1 | 453000 | 456512 | 49.26948052 | 1230 | 1170 | 1.051282051 | 1 | 1 | 1 | FIM1_4984 | YMR304W | YMR304W | UBP15 | n |
| Kmar_QGN17763.1 | W303.asm02.HP0_G0022480 | OG0001938 | chrVII | 1066230 | 1070663 | CP015060.1 | 416067 | 420428 | 58.72483221 | 1477 | 1453 | 1.01651755 | 1 | 1 | 1 | FIM1_4972 | YGR281W | YGR281W | YOR1 | n |
| Kmar_QGN16088.1 | W303.asm02.HP0_G0006190 | OG0001939 | chrIII | 272274 | 273026 | CP015057.1 | 788205 | 789212 | 21.72619048 | 250 | 335 | 0.746268657 | 1 | 1 | 1 | FIM1_2788 | YCR076C | YCR076C | FUB1 | n |
| Kmar_QGN15777.1 | W303.asm02.HP0_G0047180 | OG0001940 | chrXIV | 720370 | 720777 | CP015057.1 | 76406 | 76804 | 74.81481481 | 135 | 132 | 1.022727273 | 1 | 1 | 1 | FIM1_2473 | YNR046W | YNR046W | TRM112 | n |
| Kmar_QGN16787.1 | W303.asm02.HP0_G0022890 | OG0001941 | chrVIII | 61165 | 62361 | CP015058.1 | 889247 | 890380 | 29.22705314 | 398 | 377 | 1.055702918 | 1 | 1 | 1 | FIM1_3510 | YHL022C | YHL022C | SPO11 | n |
| Kmar_QGN15002.1 | W303.asm02.HP0_G0004740 | OG0001942 | chrII | 798069 | 799682 | CP015055.1 | 1662814 | 1664361 | 64.54033771 | 537 | 515 | 1.042718447 | 1 | 1 | 1 | FIM1_1683 | YBR286W | YBR286W | APE3 | n |
| Kmar_QGN15003.1 | W303.asm02.HP0_G0029000 | OG0001943 | chrX | 293200 | 295137 | CP015055.1 | 1664600 | 1666498 | 28.20512821 | 645 | 632 | 1.02056962 | 1 | 1 | 1 | FIM1_1684 | YJL073W | YJL073W | JEM1 | n |
| Kmar_QGN16524.1 | W303.asm02.HP0_G0019460 | OG0001944 | chrVII | 407155 | 408372 | CP015058.1 | 303495 | 304709 | 92.09876543 | 405 | 404 | 1.002475248 | 1 | 1 | 1 | FIM1_3237 | YGL048C | YGL048C | RPT6 | n |
| Kmar_QGN15775.1 | W303.asm02.HP0_G0006320 | OG0001945 | chrIII | 299746 | 303500 | CP015057.1 | 70250 | 73369 | 42.21840069 | 1158 | 1039 | 1.114533205 | 1 | 1 | 1 | FIM1_2471 | YCR092C | YCR092C | MSH3 | n |
| Kmar_QGN14343.1 | W303.asm02.HP0_G0041340 | OG0001946 | chrXIII | 454520 | 456061 | CP015055.1 | 315184 | 316746 | 73.84615385 | 513 | 520 | 0.986538462 | 1 | 1 | 1 | FIM1_1003 | YMR093W | YMR093W | UTP15 | n |
| Kmar_QGN16464.1 | W303.asm02.HP0_G0004670 | OG0001947 | chrII | 784609 | 785964 | CP015058.1 | 165663 | 166886 | 45.15418502 | 451 | 407 | 1.108108108 | 1 | 1 | 1 | FIM1_3177 | YBR279W | YBR279W | PAF1 | n |
| Kmar_QGN14907.1 | W303.asm02.HP0_G0025190 | OG0001948 | chrVIII | 490947 | 492281 | CP015055.1 | 1470299 | 1471624 | 68.70748299 | 444 | 441 | 1.006802721 | 1 | 1 | 1 | FIM1_1581 | YHR190W | YHR190W | ERG9 | n |
| Kmar_QGN13936.1 | W303.asm02.HP0_G0039350 | OG0001949 | chrXIII | 21481 | 22413 | CP015054.1 | 1175125 | 1176024 | 77.09677419 | 310 | 299 | 1.036789298 | 1 | 1 | 1 | FIM1_584 | YML121W | YML121W | GTR1 | n |
| Kmar_QGN13495.1 | W303.asm02.HP0_G0014130 | OG0001950 | chrV | 83408 | 84910 | CP015054.1 | 263490 | 264842 | 65.66866267 | 500 | 450 | 1.111111111 | 1 | 1 | 1 | FIM1_133 | YEL036C | YEL036C | ANP1 | n |
| Kmar_QGN16839.1 | W303.asm02.HP0_G0017200 | OG0001951 | chrVI | 185499 | 189011 | CP015058.1 | 1001623 | 1005135 | 57.43589744 | 1170 | 1170 | 1 | 1 | 1 | 1 | FIM1_3563 | YFR031C | YFR031C | SMC2 | n |

| | | | | | | | | | | | | | | | | | | | | |
|---|---|---|---|---|---|---|---|---|---|---|---|---|---|---|---|---|---|---|---|---|
| Kmar_QGN18388.1 | W303.asm02.HP0_G0015800 | OG0001952 | chrV | 412213 | 414642 | CP015061.1 | 829687 | 832200 | 79.97630332 | 809 | 837 | 0.966547192 | 1 | 1 | 1 | FIM1_4715 | YER125W | YER125W | RSP5 | n |
| Kmar_QGN18387.1 | W303.asm02.HP0_G0015790 | OG0001953 | chrV | 409368 | 411089 | CP015061.1 | 826447 | 828159 | 28.30508475 | 573 | 570 | 1.005263158 | 1 | 1 | 1 | FIM1_4714 | YER124C | YER124C | DSE1 | n |
| Kmar_QGN17359.1 | W303.asm02.HP0_G0009510 | OG0001954 | chrIV | 616350 | 617528 | CP015059.1 | 768803 | 769975 | 71.28463476 | 392 | 390 | 1.005128205 | 1 | 1 | 1 | FIM1_4091 | YDR083W | YDR083W | RRP8 | n |
| Kmar_QGN14228.1 | W303.asm02.HP0_G0015910 | OG0001955 | chrV | 462711 | 463601 | CP015055.1 | 69526 | 70365 | 45.97315436 | 296 | 279 | 1.0609319 | 1 | 1 | 1 | FIM1_884 | YER142C | YER142C | MAG1 | n |
| Kmar_QGN15459.1 | W303.asm02.HP0_G0051340 | OG0001956 | chrXV | 804273 | 805187 | CP015056.1 | 995792 | 996691 | 53.15531353 | 304 | 299 | 1.016722408 | 1 | 1 | 1 | FIM1_2150 | YOR251C | YOR251C | TUM1 | n |
| Kmar_QGN16772.1 | W303.asm02.HP0_G0014380 | OG0001957 | chrV | 140101 | 140946 | CP015058.1 | 858039 | 859142 | 38.69209809 | 281 | 367 | 0.765667575 | 1 | 1 | 1 | FIM1_3494 | YEL009C | YEL009C | GCN4 | n |
| Kmar_QGN17390.1 | W303.asm02.HP0_G0041110 | OG0001958 | chrXIII | 404826 | 406076 | CP015059.1 | 827367 | 828008 | 16.9451074 | 416 | 213 | 1.953051643 | 1 | 1 | 1 | FIM1_4122 | YMR067C | YMR067C | UBX4 | n |
| Kmar_QGN16959.1 | W303.asm02.HP0_G0044770 | OG0001959 | chrXIV | 247159 | 247803 | CP015058.1 | 1258736 | 1259446 | 35.16949153 | 214 | 236 | 0.906779661 | 1 | 1 | 1 | FIM1_3686 | YNL213C | YNL213C | RRG9 | n |
| Kmar_QGN17082.1 | W303.asm02.HP0_G0046880 | OG0001960 | chrXIV | 675067 | 675735 | CP015059.1 | 189715 | 190374 | 68.75 | 222 | 219 | 1.01369863 | 1 | 1 | 1 | FIM1_3813 | YNR017W | YNR017W | TIM23 | n |
| Kmar_QGN17085.1 | W303.asm02.HP0_G0046860 | OG0001961 | chrXIV | 665540 | 666694 | CP015059.1 | 200712 | 201848 | 53.64583333 | 384 | 378 | 1.015873016 | 1 | 1 | 1 | FIM1_3816 | YNR015W | YNR015W | SMM1 | n |
| Kmar_QGN16483.1 | W303.asm02.HP0_G0000360 | OG0001962 | chrI | 80481 | 81350 | CP015058.1 | 200748 | 201617 | 40.75342466 | 289 | 289 | 1 | 1 | 1 | 1 | FIM1_3196 | YAL034W-A | YAL034W-A | MTW1 | n |
| Kmar_QGN16904.1 | W303.asm02.HP0_G0010330 | OG0001963 | chrIV | 814279 | 815346 | CP015058.1 | 1145963 | 1147021 | 48.45070423 | 355 | 352 | 1.008522727 | 1 | 1 | 1 | FIM1_3631 | YDR173C | YDR173C | ARG82 | n |
| Kmar_QGN16905.1 | W303.asm02.HP0_G0010320 | OG0001964 | chrIV | 812038 | 814095 | CP015058.1 | 1147079 | 1149214 | 69.69273743 | 685 | 711 | 0.963431786 | 1 | 1 | 1 | FIM1_3632 | YDR172W | YDR172W | SUP35 | n |
| Kmar_QGN14146.1 | W303.asm02.HP0_G0005240 | OG0001965 | chrIII | 61309 | 62373 | CP015054.1 | 1630294 | 1631292 | 41.24293785 | 354 | 332 | 1.06626506 | 1 | 1 | 1 | FIM1_799 | YCL034W | YCL034W | LSB5 | n |
| Kmar_QGN16768.1 | W303.asm02.HP0_G0027180 | OG0001966 | chrIX | 339001 | 339560 | CP015058.1 | 847491 | 847934 | 58.50340136 | 142 | 147 | 0.965986395 | 1 | 1 | 1 | FIM1_3490 | YIL004C | YIL004C | BET1 | n |
| Kmar_QGN16767.1 | W303.asm02.HP0_G0027190 | OG0001967 | chrIX | 340177 | 341058 | CP015058.1 | 846256 | 847092 | 64.86486486 | 293 | 278 | 1.053956835 | 1 | 1 | 1 | FIM1_3489 | YIL003W | YIL003W | CFD1 | n |
| Kmar_QGN16758.1 | W303.asm02.HP0_G0014480 | OG0001968 | chrV | 157986 | 158681 | CP015058.1 | 832322 | 833053 | 55.51020408 | 231 | 243 | 0.950617284 | 1 | 1 | 1 | FIM1_3480 | YER002W | YER002W | NOP16 | n |
| Kmar_QGN16757.1 | W303.asm02.HP0_G0014490 | OG0001969 | chrV | 158919 | 160301 | CP015058.1 | 830839 | 832131 | 76.97674419 | 429 | 430 | 0.997674419 | 1 | 1 | 1 | FIM1_3479 | YER003C | YER003C | PMI40 | n |
| Kmar_QGN16769.1 | W303.asm02.HP0_G0027170 | OG0001970 | chrIX | 336747 | 338852 | CP015058.1 | 848141 | 850249 | 38.23529412 | 701 | 702 | 0.998575499 | 1 | 1 | 1 | FIM1_3491 | YIL005W | YIL005W | EPS1 | n |
| Kmar_QGN14449.1 | W303.asm02.HP0_G0006300 | OG0001971 | chrIII | 295298 | 295846 | CP015055.1 | 534038 | 534598 | 47.59358289 | 182 | 186 | 0.978494624 | 1 | 1 | 1 | FIM1_1110 | YCR090C | YCR090C | | n |
| Kmar_QGN15394.1 | W303.asm02.HP0_G0025870 | OG0001972 | chrIX | 72333 | 73916 | CP015056.1 | 868516 | 870105 | 85.82230624 | 527 | 529 | 0.996219282 | 1 | 1 | 1 | FIM1_2084 | YIL142W | YIL142W | CCT2 | n |
| Kmar_QGN16270.1 | W303.asm02.HP0_G0039740 | OG0001973 | chrXIII | 103346 | 104617 | CP015057.1 | 1167043 | 1168293 | 71.72897196 | 423 | 416 | 1.016826923 | 1 | 1 | 1 | FIM1_2975 | YML080W | YML080W | DUS1 | n |
| Kmar_QGN15399.1 | W303.asm02.HP0_G0025890 | OG0001974 | chrIX | 77010 | 77747 | CP015056.1 | 881151 | 881849 | 37.09677419 | 245 | 232 | 1.056034483 | 1 | 1 | 1 | FIM1_2089 | YIL139C | YIL139C | REV7 | n |
| Kmar_QGN15763.1 | W303.asm02.HP0_G0047260 | OG0001975 | chrXIV | 735923 | 736873 | CP015057.1 | 41590 | 42501 | 59.74842767 | 316 | 303 | 1.04290429 | 1 | 1 | 1 | FIM1_2459 | YNR054C | YNR054C | ESF2 | n |
| Kmar_QGN15844.1 | W303.asm02.HP0_G0010030 | OG0001976 | chrIV | 739810 | 744906 | CP015057.1 | 202312 | 207408 | 43.96999423 | 1698 | 1698 | 1 | 1 | 1 | 1 | FIM1_2540 | YDR141C | YDR141C | DOP1 | n |
| Kmar_QGN15478.1 | W303.asm02.HP0_G0053740 | OG0001977 | chrXVI | 230286 | 231674 | CP015056.1 | 1026983 | 1028335 | 62.17391304 | 462 | 450 | 1.026666667 | 1 | 1 | 1 | FIM1_2169 | YPL172C | YPL172C | COX10 | n |
| Kmar_QGN15479.1 | W303.asm02.HP0_G0051440 | OG0001978 | chrXV | 817638 | 818654 | CP015056.1 | 1028765 | 1029772 | 70.29411765 | 338 | 335 | 1.008955224 | 1 | 1 | 1 | FIM1_2170 | YOR261C | YOR261C | RPN8 | n |
| Kmar_QGN15480.1 | W303.asm02.HP0_G0051450 | OG0001979 | chrXV | 819015 | 820058 | CP015056.1 | 1030037 | 1031080 | 82.42074928 | 347 | 347 | 1 | 1 | 1 | 1 | FIM1_2171 | YOR262W | YOR262W | GPN2 | n |
| Kmar_QGN17265.1 | W303.asm02.HP0_G0016680 | OG0001980 | chrVI | 59463 | 61856 | CP015059.1 | 577827 | 580058 | 31.8016287 | 797 | 743 | 1.072678331 | 1 | 1 | 1 | FIM1_3996 | YFL023W | YFL023W | BUD27 | n |
| Kmar_QGN14238.1 | W303.asm02.HP0_G0001760 | OG0001981 | chrII | 196407 | 197987 | CP015055.1 | 89622 | 91193 | 83.65019011 | 526 | 523 | 1.005736138 | 1 | 1 | 1 | FIM1_894 | YBL015W | YBL015W | ACH1 | n |
| Kmar_QGN15164.1 | W303.asm02.HP0_G0037300 | OG0001982 | chrXII | 635552 | 636256 | CP015056.1 | 333694 | 334368 | 55.60344828 | 234 | 224 | 1.044642857 | 1 | 1 | 1 | FIM1_1851 | YLR250W | YLR250W | SSP120 | n |
| Kmar_QGN17820.1 | W303.asm02.HP0_G0000290 | OG0001983 | chrI | 63475 | 66039 | CP015060.1 | 559139 | 561430 | 53.65005794 | 854 | 763 | 1.119266055 | 1 | 1 | 1 | FIM1_5029 | YAL041W | YAL041W | CDC24 | n |
| Kmar_QGN17819.1 | W303.asm02.HP0_G0052330 | OG0001984 | chrXV | 1009502 | 1011397 | CP015060.1 | 556884 | 558842 | 64.32926829 | 631 | 652 | 0.967791411 | 1 | 1 | 1 | FIM1_5028 | YOR356W | YOR356W | CIR2 | n |
| Kmar_QGN13800.1 | W303.asm02.HP0_G0044030 | OG0001985 | chrXIV | 79106 | 80707 | CP015054.1 | 900040 | 901845 | 32.08469055 | 533 | 601 | 0.886855241 | 1 | 1 | 1 | FIM1_446 | YNL294C | YNL294C | RIM21 | n |
| Kmar_QGN17288.1 | W303.asm02.HP0_G0008470 | OG0001986 | chrIV | 397580 | 399172 | CP015059.1 | 626067 | 627644 | 36.44859813 | 530 | 525 | 1.00952381 | 1 | 1 | 1 | FIM1_4019 | YDL030W | YDL030W | PRP9 | n |
| Kmar_QGN17283.1 | W303.asm02.HP0_G0048450 | OG0001987 | chrXV | 205185 | 206339 | CP015059.1 | 614637 | 615794 | 32.67813268 | 384 | 385 | 0.997402597 | 1 | 1 | 1 | FIM1_4014 | YOL065C | YOL065C | INP54 | n |
| Kmar_QGN14525.1 | W303.asm02.HP0_G0008410 | OG0001988 | chrIV | 379761 | 382325 | CP015055.1 | 691458 | 693980 | 54.25904317 | 854 | 840 | 1.016666667 | 1 | 1 | 1 | FIM1_1187 | YDL040C | YDL040C | NAT1 | n |
| Kmar_QGN13818.1 | W303.asm02.HP0_G0049330 | OG0001989 | chrXV | 381509 | 383278 | CP015054.1 | 935034 | 936770 | 73.85398981 | 589 | 578 | 1.019031142 | 1 | 1 | 1 | FIM1_464 | YOR027W | YOR027W | STI1 | n |
| Kmar_QGN15089.1 | W303.asm02.HP0_G0033290 | OG0001990 | chrXI | 452306 | 454012 | CP015056.1 | 180589 | 182445 | 65.10500808 | 568 | 618 | 0.919093851 | 1 | 1 | 1 | FIM1_1776 | YKR002W | YKR002W | PAP1 | n |
| Kmar_QGN14100.1 | W303.asm02.HP0_G0044290 | OG0001991 | chrXIV | 141432 | 144632 | CP015054.1 | 1518646 | 1521543 | 59.4972067 | 1066 | 965 | 1.104663212 | 1 | 1 | 1 | FIM1_751 | YNL267W | YNL267W | PIK1 | n |
| Kmar_QGN14058.1 | W303.asm02.HP0_G0027840 | OG0001992 | chrX | 40160 | 42682 | CP015054.1 | 1434592 | 1436943 | 44.69339623 | 840 | 783 | 1.072796935 | 1 | 1 | 1 | FIM1_709 | YJL204C | YJL204C | RCY1 | n |
| Kmar_QGN17890.1 | W303.asm02.HP0_G0003160 | OG0001993 | chrII | 500255 | 500845 | CP015060.1 | 685569 | 686075 | 48.73096447 | 196 | 168 | 1.166666667 | 1 | 1 | 1 | FIM1_5099 | YBR122C | YBR122C | MRPL36 | n |
| Kmar_QGN15990.1 | W303.asm02.HP0_G0028660 | OG0001994 | chrX | 197893 | 199545 | CP015057.1 | 559682 | 561334 | 86.72727273 | 550 | 550 | 1 | 1 | 1 | 1 | FIM1_2690 | YJL111W | YJL111W | CCT7 | n |
| Kmar_QGN17512.1 | W303.asm02.HP0_G0021970 | OG0001995 | chrVII | 959835 | 961412 | CP015059.1 | 1064222 | 1065751 | 54.66666667 | 525 | 509 | 1.031434185 | 1 | 1 | 1 | FIM1_4248 | YGR227W | YGR227W | DIE2 | n |
| Kmar_QGN17513.1 | W303.asm02.HP0_G0021980 | OG0001996 | chrVII | 961797 | 963314 | CP015059.1 | 1066062 | 1067771 | 40.91680815 | 505 | 569 | 0.887521968 | 1 | 1 | 1 | FIM1_4249 | YGR229C | YGR229C | SMI1 | n |
| Kmar_QGN17499.1 | W303.asm02.HP0_G0021880 | OG0001997 | chrVII | 935048 | 936877 | CP015059.1 | 1036421 | 1038280 | 36.16352201 | 609 | 619 | 0.983844911 | 1 | 1 | 1 | FIM1_4235 | YGR216C | YGR216C | GPI1 | n |
| Kmar_QGN14030.1 | W303.asm02.HP0_G0048500 | OG0001998 | chrXV | 214471 | 216591 | CP015054.1 | 1370285 | 1372510 | 52.75590551 | 706 | 741 | 0.952766532 | 1 | 1 | 1 | FIM1_681 | YOL060C | YOL060C | MAM3 | n |
| Kmar_QGN17527.1 | W303.asm02.HP0_G0033740 | OG0001999 | chrXI | 540633 | 541889 | CP015059.1 | 1096129 | 1097385 | 47.05882353 | 418 | 418 | 1 | 1 | 1 | 1 | FIM1_4263 | YKR051W | YKR051W | HFL1 | n |
| Kmar_QGN17529.1 | W303.asm02.HP0_G0028490 | OG0002000 | chrX | 154296 | 155366 | CP015059.1 | 1099840 | 1100874 | 32.68156425 | 356 | 344 | 1.034883721 | 1 | 1 | 1 | FIM1_4265 | YJL131C | YJL131C | AIM23 | n |
| Kmar_QGN14305.1 | W303.asm02.HP0_G0034930 | OG0002001 | chrXII | 149082 | 151355 | CP015055.1 | 235802 | 238132 | 70.3989704 | 757 | 776 | 0.975515464 | 1 | 1 | 1 | FIM1_962 | YLL001W | YLL001W | DNM1 | n |
| Kmar_QGN14309.1 | W303.asm02.HP0_G0029600 | OG0002002 | chrX | 428870 | 430300 | CP015055.1 | 244726 | 246108 | 38.65546218 | 476 | 460 | 1.034782609 | 1 | 1 | 1 | FIM1_966 | YJL002C | YJL002C | OST1 | n |
| Kmar_QGN14307.1 | W303.asm02.HP0_G0029610 | OG0002003 | chrX | 430812 | 431575 | CP015055.1 | 240895 | 240969 | 8.837209302 | 215 | 24 | 8.958333333 | 1 | 1 | 1 | FIM1_964 | YJL001W | YJL001W | PRE3 | n |
| Kmar_QGN18189.1 | W303.asm02.HP0_G0056230 | OG0002004 | chrXVI | 727136 | 727459 | CP015061.1 | 404003 | 404344 | 50.87719298 | 107 | 113 | 0.946902655 | 1 | 1 | 1 | FIM1_4513 | YPR094W | YPR094W | RDS3 | n |
| Kmar_QGN14535.1 | W303.asm02.HP0_G0056310 | OG0002005 | chrXVI | 738848 | 739711 | CP015055.1 | 712262 | 713116 | 85.0174216 | 287 | 284 | 1.01056338 | 1 | 1 | 1 | FIM1_1197 | YPR103W | YPR103W | PRE2 | n |
| Kmar_QGN14536.1 | W303.asm02.HP0_G0056320 | OG0002006 | chrXVI | 740122 | 742932 | CP015055.1 | 713433 | 716054 | 43.44608879 | 936 | 873 | 1.072164948 | 1 | 1 | 1 | FIM1_1198 | YPR104C | YPR104C | FHL1 | n |
| Kmar_QGN16502.1 | W303.asm02.HP0_G0000470 | OG0002007 | chrI | 102329 | 106636 | CP015058.1 | 247326 | 251474 | 36.54618474 | 1435 | 1382 | 1.038350217 | 1 | 1 | 1 | FIM1_3215 | YAL024C | YAL024C | LTE1 | n |
| Kmar_QGN13548.1 | W303.asm02.HP0_G0053480 | OG0002008 | chrXVI | 177585 | 178307 | CP015054.1 | 395443 | 396219 | 44.01544402 | 240 | 258 | 0.930232558 | 1 | 1 | 1 | FIM1_189 | YPL199C | YPL199C | | n |
| Kmar_QGN15495.1 | W303.asm02.HP0_G0046920 | OG0002009 | chrXIV | 679991 | 680674 | CP015056.1 | 1061223 | 1061969 | 67.74193548 | 227 | 248 | 0.915322581 | 1 | 1 | 1 | FIM1_2186 | YNR020C | YNR020C | ATP23 | n |
| Kmar_QGN15496.1 | W303.asm02.HP0_G0005990 | OG0002010 | chrIII | 237052 | 237720 | CP015056.1 | 1062516 | 1063148 | 47.53363229 | 222 | 210 | 1.057142857 | 1 | 1 | 1 | FIM1_2187 | YCR051W | YCR051W | | n |
| Kmar_QGN15498.1 | W303.asm02.HP0_G0046940 | OG0002011 | chrXIV | 682356 | 682775 | CP015056.1 | 1065059 | 1065463 | 48.20143885 | 139 | 134 | 1.037313433 | 1 | 1 | 1 | FIM1_2189 | YNR022C | YNR022C | MRPL50 | n |
| Kmar_QGN15428.1 | W303.asm02.HP0_G0023540 | OG0002012 | chrVIII | 181769 | 182869 | CP015056.1 | 940750 | 941943 | 39.26829268 | 366 | 397 | 0.921914358 | 1 | 1 | 1 | FIM1_2119 | YHR040W | YHR040W | BCD1 | n |
| Kmar_QGN15568.1 | W303.asm02.HP0_G0051990 | OG0002013 | chrXV | 914542 | 915183 | CP015056.1 | 1205406 | 1206029 | 44.29223744 | 213 | 207 | 1.028985507 | 1 | 1 | 1 | FIM1_2259 | YOR319W | YOR319W | HSH49 | n |
| Kmar_QGN16153.1 | W303.asm02.HP0_G0043870 | OG0002014 | chrXIV | 48719 | 49648 | CP015057.1 | 915223 | 916145 | 52.17391304 | 273 | 273 | 1 | 1 | 1 | 1 | FIM1_2854 | YNL312W | YNL312W | RFA2 | n |
| Kmar_QGN15162.1 | W303.asm02.HP0_G0046630 | OG0002015 | chrXIV | 624818 | 625543 | CP015056.1 | 331087 | 331764 | 52.45901639 | 241 | 225 | 1.071111111 | 1 | 1 | 1 | FIM1_1849 | YNL010W | YNL010W | PYP1 | n |
| Kmar_QGN13555.1 | W303.asm02.HP0_G0019270 | OG0002016 | chrVII | 372040 | 372624 | CP015054.1 | 408626 | 409159 | 53.09278351 | 194 | 177 | 1.096045198 | 1 | 1 | 1 | FIM1_196 | YGL068W | YGL068W | MNP1 | n |
| Kmar_QGN13554.1 | W303.asm02.HP0_G0019260 | OG0002017 | chrVII | 371412 | 371780 | CP015054.1 | 407815 | 408183 | 90.98360656 | 122 | 122 | 1 | 1 | 1 | 1 | FIM1_195 | YGL070C | YGL070C | RPB9 | n |
| Kmar_QGN15055.1 | W303.asm02.HP0_G0031300 | OG0002018 | chrXI | 50031 | 50834 | CP015056.1 | 103021 | 103845 | 34.17266187 | 267 | 274 | 0.974452555 | 1 | 1 | 1 | FIM1_1740 | YKL206C | YKL206C | ADD66 | n |
| Kmar_QGN15056.1 | W303.asm02.HP0_G0030250 | OG0002019 | chrX | 578908 | 579885 | CP015056.1 | 104836 | 105804 | 72.13622291 | 325 | 322 | 1.00931677 | 1 | 1 | 1 | FIM1_1741 | YJR070C | YJR070C | LIA1 | n |
| Kmar_QGN15066.1 | W303.asm02.HP0_G0030180 | OG0002020 | chrX | 564482 | 564859 | CP015056.1 | 128711 | 129097 | 67.1875 | 125 | 128 | 0.9765625 | 1 | 0 | 0 | FIM1_1751 | YJR063W | YJR063W | RPA12 | n |
| Kmar_QGN15065.1 | W303.asm02.HP0_G0030190 | OG0002021 | chrX | 565201 | 566889 | CP015056.1 | 126699 | 128351 | 81.49466192 | 562 | 550 | 1.021818182 | 1 | 1 | 1 | FIM1_1750 | YJR064W | YJR064W | CCT5 | n |
| Kmar_QGN15064.1 | W303.asm02.HP0_G0030200 | OG0002022 | chrX | 567096 | 568445 | CP015056.1 | 124814 | 126181 | 87.69230769 | 449 | 455 | 0.986813187 | 1 | 1 | 1 | FIM1_1749 | YJR065C | YJR065C | ARP3 | n |
| Kmar_QGN14789.1 | W303.asm02.HP0_G0012260 | OG0002023 | chrIV | 1232415 | 1233896 | CP015055.1 | 1236893 | 1238329 | 53.03643725 | 493 | 478 | 1.031380753 | 1 | 1 | 1 | FIM1_1460 | YDR376W | YDR376W | ARH1 | n |
| Kmar_QGN15275.1 | W303.asm02.HP0_G0049690 | OG0002024 | chrXV | 448035 | 448964 | CP015056.1 | 576451 | 577329 | 72.81553398 | 309 | 265 | 1.166037736 | 1 | 1 | 1 | FIM1_1964 | YOR065W | YOR065W | CYT1 | n |
| Kmar_QGN15268.1 | W303.asm02.HP0_G0033970 | OG0002025 | chrXI | 596650 | 597117 | CP015056.1 | 565321 | 565688 | 47.12643678 | 155 | 61 | 2.540983607 | 1 | 1 | 1 | FIM1_1957 | YKR074W | YKR074W | AIM29 | n |
| Kmar_QGN15271.1 | W303.asm02.HP0_G0049650 | OG0002026 | chrXV | 442130 | 443149 | CP015056.1 | 568841 | 569863 | 80.88235294 | 339 | 340 | 0.997058824 | 1 | 1 | 1 | FIM1_1960 | YOR061W | YOR061W | CKA2 | n |
| Kmar_QGN15270.1 | W303.asm02.HP0_G0049640 | OG0002027 | chrXV | 440986 | 441759 | CP015056.1 | 567635 | 568441 | 31.89964158 | 257 | 268 | 0.958955224 | 1 | 1 | 1 | FIM1_1959 | YOR060C | YOR060C | SLD7 | n |
| Kmar_QGN17858.1 | W303.asm02.HP0_G0044120 | OG0002028 | chrXIV | 103693 | 104661 | CP015060.1 | 627906 | 628748 | 56.21118012 | 322 | 280 | 1.15 | 1 | 1 | 1 | FIM1_5067 | YNL284C | YNL284C | MRPL10 | n |
| Kmar_QGN14144.1 | W303.asm02.HP0_G0013500 | OG0002029 | chrIV | 1476194 | 1476757 | CP015054.1 | 1629258 | 1629518 | 30.21978022 | 187 | 86 | 2.174418605 | 1 | 1 | 1 | FIM1_797 | YDR512C | YDR512C | EMI1 | n |
| Kmar_QGN17394.1 | W303.asm02.HP0_G0032400 | OG0002030 | chrXI | 286757 | 287140 | CP015059.1 | 832408 | 832800 | 49.23076923 | 127 | 130 | 0.976923077 | 1 | 1 | 1 | FIM1_4126 | YKL086W | YKL086W | SRX1 | n |
| Kmar_QGN15402.1 | W303.asm02.HP0_G0003340 | OG0002031 | chrII | 543311 | 544324 | CP015056.1 | 885606 | 886616 | 40.34582133 | 337 | 336 | 1.00297619 | 1 | 1 | 1 | FIM1_2092 | YBR141C | YBR141C | BMT2 | n |
| Kmar_QGN15403.1 | W303.asm02.HP0_G0003350 | OG0002032 | chrII | 544603 | 546924 | CP015056.1 | 886993 | 889347 | 58.39506173 | 773 | 784 | 0.985969388 | 1 | 1 | 1 | FIM1_2093 | YBR142W | YBR142W | MAK5 | n |
| Kmar_QGN15404.1 | W303.asm02.HP0_G0003360 | OG0002033 | chrII | 547155 | 548468 | CP015056.1 | 889597 | 890913 | 85.84474886 | 437 | 438 | 0.997716895 | 1 | 1 | 1 | FIM1_2094 | YBR143C | YBR143C | SUP45 | n |
| Kmar_QGN14806.1 | W303.asm02.HP0_G0043770 | OG0002034 | chrXIV | 32376 | 33620 | CP015055.1 | 1264907 | 1266226 | 52.04545455 | 414 | 439 | 0.943052392 | 1 | 1 | 1 | FIM1_1477 | YNL323W | YNL323W | LEM3 | n |
| Kmar_QGN17848.1 | W303.asm02.HP0_G0013350 | OG0002035 | chrIV | 1447127 | 1449097 | CP015060.1 | 609844 | 611817 | 65.80547112 | 656 | 657 | 0.99847793 | 1 | 1 | 1 | FIM1_5057 | YDR496C | YDR496C | PUF6 | n |
| Kmar_QGN18128.1 | W303.asm02.HP0_G0001010 | OG0002036 | chrII | 44430 | 45242 | CP015061.1 | 283301 | 284050 | 46.66666667 | 270 | 249 | 1.084337349 | 1 | 1 | 1 | FIM1_4449 | YBL095W | YBL095W | MRX3 | n |
| Kmar_QGN13783.1 | W303.asm02.HP0_G0049760 | OG0002037 | chrXV | 467410 | 468324 | CP015054.1 | 873362 | 874276 | 81.37254902 | 304 | 304 | 1 | 1 | 1 | 1 | FIM1_429 | YOR074C | YOR074C | CDC21 | n |
| Kmar_QGN13781.1 | W303.asm02.HP0_G0034060 | OG0002038 | chrXI | 614243 | 614644 | CP015054.1 | 870436 | 870804 | 41.04477612 | 133 | 122 | 1.090163934 | 1 | 1 | 1 | FIM1_427 | YKR083C | YKR083C | DAD2 | n |
| Kmar_QGN13780.1 | W303.asm02.HP0_G0034050 | OG0002039 | chrXI | 610647 | 614120 | CP015054.1 | 867027 | 870380 | 30.6122449 | 1157 | 1117 | 1.035810206 | 1 | 1 | 1 | FIM1_426 | YKR082W | YKR082W | NUP133 | n |
| Kmar_QGN13779.1 | W303.asm02.HP0_G0034040 | OG0002040 | chrXI | 609328 | 610362 | CP015054.1 | 865485 | 866513 | 71.88405797 | 344 | 342 | 1.005847953 | 1 | 1 | 1 | FIM1_425 | YKR081C | YKR081C | RPF2 | n |
| Kmar_QGN14018.1 | W303.asm02.HP0_G0000750 | OG0002041 | chrI | 167946 | 170075 | CP015054.1 | 1338833 | 1340992 | 31.05756359 | 709 | 719 | 0.986091794 | 1 | 1 | 1 | FIM1_669 | YAR014C | YAR014C | BUD14 | n |
| Kmar_QGN14017.1 | W303.asm02.HP0_G0002050 | OG0002042 | chrII | 266041 | 268800 | CP015054.1 | 1335825 | 1338491 | 52.80172414 | 919 | 888 | 1.03490991 | 1 | 1 | 1 | FIM1_668 | YBR017C | YBR017C | KAP104 | n |
| Kmar_QGN16351.1 | W303.asm02.HP0_G0012810 | OG0002043 | chrIV | 1342908 | 1343330 | CP015057.1 | 1343810 | 1344235 | 37.93103448 | 140 | 141 | 0.992907801 | 1 | 1 | 1 | FIM1_3057 | YDR437W | YDR437W | GPI19 | n |
| Kmar_QGN14014.1 | W303.asm02.HP0_G0008580 | OG0002044 | chrIV | 424246 | 425769 | CP015054.1 | 1331150 | 1332589 | 49.21259843 | 507 | 479 | 1.058455115 | 1 | 1 | 1 | FIM1_665 | YDL017W | YDL017W | CDC7 | n |
| Kmar_QGN16795.1 | W303.asm02.HP0_G0056380 | OG0002045 | chrXVI | 752327 | 753334 | CP015058.1 | 905136 | 906143 | 78.27380952 | 335 | 335 | 1 | 1 | 1 | 1 | FIM1_3518 | YPR110C | YPR110C | RPC40 | n |
| Kmar_QGN16202.1 | W303.asm02.HP0_G0008090 | OG0002046 | chrIV | 320998 | 321882 | CP015057.1 | 1023473 | 1024339 | 38.48684211 | 294 | 288 | 1.020833333 | 1 | 1 | 1 | FIM1_2905 | YDL076C | YDL076C | RXT3 | n |
| Kmar_QGN16201.1 | W303.asm02.HP0_G0038700 | OG0002047 | chrXII | 929636 | 930322 | CP015057.1 | 1022198 | 1022899 | 22.78481013 | 228 | 233 | 0.978540773 | 1 | 1 | 1 | FIM1_2904 | YLR407W | YLR407W | GAG1 | n |
| Kmar_QGN14813.1 | W303.asm02.HP0_G0043710 | OG0002048 | chrXIV | 19973 | 23065 | CP015055.1 | 1278567 | 1281566 | 47.18446602 | 1030 | 999 | 1.031031031 | 1 | 1 | 1 | FIM1_1484 | YNL329C | YNL329C | PEX6 | n |
| Kmar_QGN14812.1 | W303.asm02.HP0_G0043720 | OG0002049 | chrXIV | 23266 | 23706 | CP015055.1 | 1277794 | 1278240 | 49.33333333 | 146 | 148 | 0.986486486 | 1 | 1 | 1 | FIM1_1483 | YNL328C | YNL328C | MDJ2 | n |
| Kmar_QGN17345.1 | W303.asm02.HP0_G0000320 | OG0002050 | chrI | 72549 | 74051 | CP015059.1 | 737842 | 739347 | 85.42914172 | 500 | 501 | 0.998003992 | 1 | 1 | 1 | FIM1_4077 | YAL038W | YAL038W | CDC19 | n |

| | | | | | | | | | | | | | | | | | | | | |
|---|---|---|---|---|---|---|---|---|---|---|---|---|---|---|---|---|---|---|---|---|
| Kmar_QGN13617.1 | W303.asm02.HP0_G0039460 | OG0002051 | chrXIII | 44586 | 45509 | CP015054.1 | 527512 | 528435 | 72.31270358 | 307 | 307 | 1 | 1 | 1 | 1 | FIM1_259 | YML110C | YML110C | COQ5 | n |
| Kmar_QGN13436.1 | W303.asm02.HP0_G0054650 | OG0002052 | chrXVI | 414368 | 415102 | CP015054.1 | 142048 | 142758 | 59.83606557 | 244 | 236 | 1.033898305 | 1 | 1 | 1 | FIM1_73 | YPL078C | YPL078C | ATP4 | n |
| Kmar_QGN13437.1 | W303.asm02.HP0_G0003870 | OG0002053 | chrII | 627389 | 628657 | CP015054.1 | 143499 | 144791 | 48.05491991 | 422 | 430 | 0.981395349 | 1 | 1 | 1 | FIM1_74 | YBR195C | YBR195C | MSI1 | n |
| Kmar_QGN18154.1 | W303.asm02.HP0_G0026610 | OG0002054 | chrIX | 233391 | 234374 | CP015061.1 | 339255 | 340310 | 36.90140845 | 327 | 351 | 0.931623932 | 1 | 1 | 1 | FIM1_4477 | YIL063C | YIL063C | YRB2 | n |
| Kmar_QGN14226.1 | W303.asm02.HP0_G0009340 | OG0002055 | chrIV | 586940 | 587614 | CP015055.1 | 67293 | 67985 | 53.21888412 | 224 | 230 | 0.973913043 | 1 | 1 | 1 | FIM1_882 | YDR067C | YDR067C | OCA6 | n |
| Kmar_QGN17634.1 | W303.asm02.HP0_G0029700 | OG0002056 | chrX | 448079 | 449095 | CP015060.1 | 153868 | 154797 | 43.31395349 | 338 | 309 | 1.093851133 | 1 | 1 | 1 | FIM1_4841 | YJR008W | YJR008W | MHO1 | n |
| Kmar_QGN17652.1 | W303.asm02.HP0_G0029660 | OG0002057 | chrX | 441572 | 443674 | CP015060.1 | 193869 | 195962 | 48.50640114 | 700 | 697 | 1.004304161 | 1 | 1 | 1 | FIM1_4859 | YJR005W | YJR005W | APL1 | n |
| Kmar_QGN15425.1 | W303.asm02.HP0_G0011420 | OG0002058 | chrIV | 1044227 | 1047460 | CP015056.1 | 933172 | 936441 | 67.43970315 | 1077 | 1089 | 0.988980716 | 1 | 1 | 1 | FIM1_2116 | YDR291W | YDR291W | HRQ1 | n |
| Kmar_QGN15862.1 | W303.asm02.HP0_G0002490 | OG0002059 | chrII | 364165 | 365670 | CP015057.1 | 251424 | 252893 | 47.81746032 | 501 | 489 | 1.024539877 | 1 | 1 | 1 | FIM1_2559 | YBR056W | YBR056W | MRX18 | n |
| Kmar_QGN15984.1 | W303.asm02.HP0_G0028670 | OG0002060 | chrX | 199938 | 207325 | CP015057.1 | 541010 | 546349 | 51.18811881 | 2299 | 1779 | 1.292299044 | 1 | 1 | 1 | FIM1_2684 | YJL109C | YJL109C | UTP10 | n |
| Kmar_QGN16617.1 | W303.asm02.HP0_G0008810 | OG0002061 | chrIV | 471512 | 472600 | CP015058.1 | 508986 | 510077 | 85.12396694 | 362 | 363 | 0.997245179 | 1 | 1 | 1 | FIM1_3334 | YBR031W/YDR012W | YDR012W | RPL4B | y |
| Kmar_QGN14339.1 | W303.asm02.HP0_G0028710 | OG0002062 | chrX | 217343 | 217792 | CP015055.1 | 309045 | 309458 | 63.75838926 | 149 | 137 | 1.087591241 | 1 | 1 | 1 | FIM1_999 | YJL104W | YJL104W | PAM16 | n |
| Kmar_QGN16618.1 | W303.asm02.HP0_G0002170 | OG0002063 | chrII | 293388 | 295058 | CP015058.1 | 510441 | 512099 | 42.38178634 | 556 | 552 | 1.007246377 | 1 | 1 | 1 | FIM1_3335 | YBR030W | YBR030W | RKM3 | n |
| Kmar_QGN13593.1 | W303.asm02.HP0_G0033430 | OG0002064 | chrXI | 481388 | 483043 | CP015054.1 | 483926 | 485428 | 39.24050633 | 551 | 500 | 1.102 | 1 | 1 | 1 | FIM1_235 | YKR017C | YKR017C | HEL1 | n |
| Kmar_QGN15985.1 | W303.asm02.HP0_G0033560 | OG0002065 | chrXI | 509965 | 510786 | CP015057.1 | 546716 | 547453 | 54.94505495 | 273 | 245 | 1.114285714 | 1 | 1 | 1 | FIM1_2685 | YKR030W | YKR030W | GMH1 | n |
| Kmar_QGN16107.1 | W303.asm02.HP0_G0026950 | OG0002066 | chrIX | 295106 | 298558 | CP015057.1 | 826230 | 829442 | 36.07878245 | 1150 | 1070 | 1.074766355 | 1 | 1 | 1 | FIM1_2808 | YIL026C | YIL026C | IRR1 | n |
| Kmar_QGN14719.1 | W303.asm02.HP0_G0017290 | OG0002067 | chrVI | 196417 | 198090 | CP015055.1 | 1092101 | 1093762 | 51.73010381 | 557 | 553 | 1.007233273 | 1 | 1 | 1 | FIM1_1387 | YFR037C | YFR037C | RSC8 | n |
| Kmar_QGN16364.1 | W303.asm02.HP0_G0012750 | OG0002068 | chrIV | 1330033 | 1330857 | CP015057.1 | 1367586 | 1368398 | 54.74452555 | 274 | 270 | 1.014814815 | 1 | 1 | 1 | FIM1_3070 | YDR429C | YDR429C | TIF35 | n |
| Kmar_QGN16358.1 | W303.asm02.HP0_G0012770 | OG0002069 | chrIV | 1334339 | 1335583 | CP015057.1 | 1356696 | 1357706 | 50.46728972 | 414 | 336 | 1.232142857 | 1 | 1 | 1 | FIM1_3064 | YDR432W | YDR432W | NPL3 | n |
| Kmar_QGN14934.1 | W303.asm02.HP0_G0024740 | OG0002070 | chrVIII | 393893 | 394831 | CP015055.1 | 1527814 | 1528776 | 53.01587302 | 312 | 320 | 0.975 | 1 | 1 | 1 | FIM1_1609 | YHR144C | YHR144C | DCD1 | n |
| Kmar_QGN17286.1 | W303.asm02.HP0_G0008450 | OG0002071 | chrIV | 392702 | 393955 | CP015059.1 | 620881 | 622128 | 57.75656325 | 417 | 415 | 1.004819277 | 1 | 1 | 1 | FIM1_4017 | YDL033C | YDL033C | SLM3 | n |
| Kmar_QGN16076.1 | W303.asm02.HP0_G0055350 | OG0002072 | chrXVI | 554843 | 556663 | CP015057.1 | 758092 | 759933 | 31.35048232 | 606 | 613 | 0.98858075 | 1 | 1 | 1 | FIM1_2776 | YPL005W | YPL005W | AEP3 | n |
| Kmar_QGN14937.1 | W303.asm02.HP0_G0024730 | OG0002073 | chrVIII | 393338 | 393550 | CP015055.1 | 1532461 | 1532673 | 88.57142857 | 70 | 70 | 1 | 1 | 1 | 1 | FIM1_1612 | YHR143W-A | YHR143W-A | RPC10 | n |
| Kmar_QGN14221.1 | W303.asm02.HP0_G0015940 | OG0002074 | chrV | 470150 | 470431 | CP015055.1 | 60337 | 60606 | 55.20833333 | 93 | 89 | 1.04494382 | 1 | 1 | 1 | FIM1_877 | YER146W | YER146W | LSM5 | n |
| Kmar_QGN15456.1 | W303.asm02.HP0_G0053810 | OG0002075 | chrXVI | 245283 | 247430 | CP015056.1 | 989977 | 992103 | 36.23789765 | 715 | 708 | 1.009887006 | 1 | 1 | 1 | FIM1_2147 | YPL164C | YPL164C | MLH3 | n |
| Kmar_QGN14227.1 | W303.asm02.HP0_G0015920 | OG0002076 | chrV | 463884 | 465170 | CP015055.1 | 68067 | 69326 | 48.13953488 | 428 | 419 | 1.021479714 | 1 | 1 | 1 | FIM1_883 | YER143W | YER143W | DDI1 | n |
| Kmar_QGN18418.1 | W303.asm02.HP0_G0016950 | OG0002077 | chrVI | 124628 | 126235 | CP015061.1 | 891822 | 893375 | 56.52985075 | 535 | 517 | 1.034816248 | 1 | 1 | 1 | FIM1_4746 | YFR006W | YFR006W | | n |
| Kmar_QGN18420.1 | W303.asm02.HP0_G0016930 | OG0002078 | chrVI | 121876 | 122796 | CP015061.1 | 895528 | 896463 | 82.31511254 | 306 | 311 | 0.98392283 | 1 | 1 | 1 | FIM1_4748 | YFR004W | YFR004W | RPN11 | n |
| Kmar_QGN18419.1 | W303.asm02.HP0_G0016940 | OG0002079 | chrVI | 123010 | 124356 | CP015061.1 | 894045 | 895487 | 35.12396694 | 448 | 480 | 0.933333333 | 1 | 1 | 1 | FIM1_4747 | YFR005C | YFR005C | SAD1 | n |
| Kmar_QGN17453.1 | W303.asm02.HP0_G0024130 | OG0002080 | chrVIII | 278601 | 279590 | CP015059.1 | 951544 | 952548 | 30.97345133 | 329 | 334 | 0.98502994 | 1 | 1 | 1 | FIM1_4187 | YHR083W | YHR083W | SAM35 | n |
| Kmar_QGN14411.1 | W303.asm02.HP0_G0055200 | OG0002081 | chrXVI | 518670 | 520535 | CP015055.1 | 462016 | 463683 | 40.063593 | 621 | 555 | 1.118918919 | 1 | 1 | 1 | FIM1_1071 | YPL020C | YPL020C | ULP1 | n |
| Kmar_QGN15149.1 | W303.asm02.HP0_G0001680 | OG0002082 | chrII | 177207 | 179813 | CP015056.1 | 298558 | 301218 | 78.37837838 | 868 | 886 | 0.979683973 | 1 | 1 | 1 | FIM1_1836 | YBL023C | YBL023C | MCM2 | n |
| Kmar_QGN18228.1 | W303.asm02.HP0_G0007300 | OG0002083 | chrIV | 174444 | 174800 | CP015061.1 | 499838 | 500164 | 66.66666667 | 118 | 108 | 1.092592593 | 1 | 1 | 1 | FIM1_4553 | YDL157C | YDL157C | DMO2 | n |
| Kmar_QGN16581.1 | W303.asm02.HP0_G0020120 | OG0002084 | chrVII | 539825 | 540913 | CP015058.1 | 429103 | 430185 | 67.40331492 | 362 | 360 | 1.005555556 | 1 | 1 | 1 | FIM1_3297 | YGR028W | YGR028W | MSP1 | n |
| Kmar_QGN16582.1 | W303.asm02.HP0_G0038050 | OG0002085 | chrXII | 795384 | 798083 | CP015058.1 | 430499 | 433714 | 40.98513011 | 899 | 1071 | 0.839402428 | 1 | 1 | 1 | FIM1_3298 | YLR336C | YLR336C | SGD1 | n |
| Kmar_QGN14957.1 | W303.asm02.HP0_G0016830 | OG0002086 | chrVI | 87912 | 91589 | CP015055.1 | 1568910 | 1572638 | 52.65700483 | 1225 | 1242 | 0.986312399 | 1 | 1 | 1 | FIM1_1634 | YFL008W | YFL008W | SMC1 | n |
| Kmar_QGN15642.1 | W303.asm02.HP0_G0038220 | OG0002087 | chrXII | 832530 | 833537 | CP015056.1 | 1374884 | 1375888 | 77.91044776 | 335 | 334 | 1.002994012 | 1 | 1 | 1 | FIM1_2335 | YLR354C | YLR354C | TAL1 | n |
| Kmar_QGN15530.1 | W303.asm02.HP0_G0010930 | OG0002088 | chrIV | 939737 | 940393 | CP015056.1 | 1130477 | 1131040 | 51.37614679 | 218 | 187 | 1.165775401 | 1 | 1 | 1 | FIM1_2221 | YDR236C | YDR236C | FMN1 | n |
| Kmar_QGN15532.1 | W303.asm02.HP0_G0010920 | OG0002089 | chrIV | 938005 | 939639 | CP015056.1 | 1133570 | 1135210 | 44.24131627 | 544 | 546 | 0.996336996 | 1 | 1 | 1 | FIM1_2223 | YDR235W | YDR235W | PRP42 | n |
| Kmar_QGN13997.1 | W303.asm02.HP0_G0037580 | OG0002090 | chrXII | 702129 | 702914 | CP015054.1 | 1294698 | 1295510 | 59.05797101 | 261 | 270 | 0.966666667 | 1 | 1 | 1 | FIM1_647 | YLR285W | YLR285W | NNT1 | n |
| Kmar_QGN16085.1 | W303.asm02.HP0_G0055300 | OG0002091 | chrXVI | 541375 | 541944 | CP015057.1 | 780942 | 781511 | 56.31578947 | 189 | 189 | 1 | 1 | 1 | 1 | FIM1_2785 | YPL010W | YPL010W | RET3 | n |
| Kmar_QGN16084.1 | W303.asm02.HP0_G0055290 | OG0002092 | chrXVI | 539999 | 541060 | CP015057.1 | 779358 | 780407 | 38.99721448 | 353 | 349 | 1.011461318 | 1 | 1 | 1 | FIM1_2784 | YPL011C | YPL011C | TAF3 | n |
| Kmar_QGN16081.1 | W303.asm02.HP0_G0055260 | OG0002093 | chrXVI | 533904 | 535049 | CP015057.1 | 772501 | 773925 | 29.32489451 | 381 | 474 | 0.803797468 | 1 | 1 | 1 | FIM1_2781 | YPL014W | YPL014W | CIP1 | n |
| Kmar_QGN15369.1 | W303.asm02.HP0_G0003570 | OG0002094 | chrII | 582499 | 584256 | CP015056.1 | 816626 | 818257 | 31.93277311 | 585 | 543 | 1.077348066 | 1 | 1 | 1 | FIM1_2059 | YBR163W | YBR163W | EXO5 | n |
| Kmar_QGN14344.1 | W303.asm02.HP0_G0041330 | OG0002095 | chrXIII | 452137 | 453984 | CP015055.1 | 317366 | 319234 | 57.00787402 | 615 | 622 | 0.988745981 | 1 | 1 | 1 | FIM1_1004 | YMR092C | YMR092C | AIP1 | n |
| Kmar_QGN13591.1 | W303.asm02.HP0_G0033420 | OG0002096 | chrXI | 479414 | 481033 | CP015054.1 | 480165 | 481757 | 36.55536028 | 539 | 530 | 1.016981132 | 1 | 1 | 1 | FIM1_233 | YKR016W | YKR016W | MIC60 | n |
| Kmar_QGN17317.1 | W303.asm02.HP0_G0045620 | OG0002097 | chrXIV | 402248 | 403729 | CP015059.1 | 681656 | 683044 | 36.07214429 | 493 | 462 | 1.067099567 | 1 | 1 | 1 | FIM1_4049 | YNL119W | YNL119W | NCS2 | n |
| Kmar_QGN17414.1 | W303.asm02.HP0_G0027500 | OG0002098 | chrIX | 396174 | 397268 | CP015059.1 | 865806 | 866876 | 66.66666667 | 364 | 356 | 1.02247191 | 1 | 1 | 1 | FIM1_4146 | YIR026C | YIR026C | YVH1 | n |
| Kmar_QGN15237.1 | W303.asm02.HP0_G0029580 | OG0002099 | chrX | 427238 | 427849 | CP015056.1 | 499307 | 499900 | 65.02463054 | 203 | 197 | 1.030456853 | 1 | 1 | 1 | FIM1_1926 | YJL004C | YJL004C | SYS1 | n |
| Kmar_QGN18286.1 | W303.asm02.HP0_G0020900 | OG0002100 | chrVII | 720200 | 724555 | CP015061.1 | 616075 | 620457 | 69.36691627 | 1451 | 1460 | 0.993835616 | 1 | 1 | 1 | FIM1_4612 | YGR116W | YGR116W | SPT6 | n |
| Kmar_QGN14697.1 | W303.asm02.HP0_G0021040 | OG0002101 | chrVII | 762061 | 762708 | CP015055.1 | 1054346 | 1054987 | 21.00456621 | 215 | 213 | 1.009389671 | 1 | 1 | 1 | FIM1_1364 | YGR129W | YGR129W | SYF2 | n |
| Kmar_QGN13784.1 | W303.asm02.HP0_G0049770 | OG0002102 | chrXV | 468947 | 469987 | CP015054.1 | 874735 | 875724 | 40.34582133 | 346 | 329 | 1.051671733 | 1 | 1 | 1 | FIM1_430 | YOR075W | YOR075W | UFE1 | n |
| Kmar_QGN15058.1 | W303.asm02.HP0_G0031310 | OG0002103 | chrXI | 51076 | 54378 | CP015056.1 | 107042 | 110329 | 42.02401372 | 1100 | 1095 | 1.00456621 | 1 | 1 | 1 | FIM1_1743 | YKL205W | YKL205W | LOS1 | n |
| Kmar_QGN16468.1 | W303.asm02.HP0_G0029040 | OG0002104 | chrX | 300941 | 302725 | CP015058.1 | 173925 | 175640 | 62.37458194 | 594 | 571 | 1.04028021 | 1 | 1 | 1 | FIM1_3181 | YJL069C | YJL069C | UTP18 | n |
| Kmar_QGN14906.1 | W303.asm02.HP0_G0042240 | OG0002105 | chrXIII | 629608 | 632553 | CP015055.1 | 1466968 | 1469883 | 35.84720862 | 981 | 971 | 1.010298661 | 1 | 1 | 1 | FIM1_1580 | YMR185W | YMR185W | RTP1 | n |
| Kmar_QGN14905.1 | W303.asm02.HP0_G0053100 | OG0002106 | chrXVI | 105049 | 105651 | CP015055.1 | 1466014 | 1466619 | 65.85365854 | 200 | 201 | 0.995024876 | 1 | 1 | 1 | FIM1_1579 | YPL239W | YPL239W | YAR1 | n |
| Kmar_QGN14904.1 | W303.asm02.HP0_G0053110 | OG0002107 | chrXVI | 106061 | 106918 | CP015055.1 | 1464684 | 1465523 | 74.21602787 | 285 | 279 | 1.021505376 | 1 | 1 | 1 | FIM1_1578 | YPL237W | YPL237W | SUI3 | n |
| Kmar_QGN14830.1 | W303.asm02.HP0_G0041990 | OG0002108 | chrXIII | 578326 | 579000 | CP015055.1 | 1312793 | 1313527 | 47.38955823 | 224 | 244 | 0.918032787 | 1 | 1 | 1 | FIM1_1501 | YMR161W | YMR161W | HLJ1 | n |
| Kmar_QGN14991.1 | W303.asm02.HP0_G0050960 | OG0002109 | chrXV | 737732 | 739021 | CP015055.1 | 1637672 | 1638955 | 67.83216783 | 429 | 427 | 1.004683841 | 1 | 1 | 1 | FIM1_1672 | YOR209C | YOR209C | NPT1 | n |
| Kmar_QGN14992.1 | W303.asm02.HP0_G0054190 | OG0002110 | chrXVI | 319319 | 322417 | CP015055.1 | 1639506 | 1642577 | 55.15564202 | 1032 | 1023 | 1.008797654 | 1 | 1 | 1 | FIM1_1673 | YPL125W | YPL125W | KAP120 | n |
| Kmar_QGN13694.1 | W303.asm02.HP0_G0043540 | OG0002111 | chrXIII | 908623 | 909327 | CP015054.1 | 674770 | 675471 | 82.05128205 | 234 | 233 | 1.004291845 | 1 | 1 | 1 | FIM1_337 | YMR314W | YMR314W | PRE5 | n |
| Kmar_QGN15709.1 | W303.asm02.HP0_G0017790 | OG0002112 | chrVII | 57445 | 58372 | CP015056.1 | 1508941 | 1509849 | 46.3022508 | 289 | 302 | 0.956953642 | 1 | 1 | 1 | FIM1_2402 | YGL232W | YGL232W | TAN1 | n |
| Kmar_QGN15708.1 | W303.asm02.HP0_G0017800 | OG0002113 | chrVII | 58418 | 58990 | CP015056.1 | 1508379 | 1508921 | 40.83769634 | 190 | 180 | 1.055555556 | 1 | 1 | 1 | FIM1_2401 | YGL231C | YGL231C | EMC4 | n |
| Kmar_QGN16162.1 | W303.asm02.HP0_G0026540 | OG0002114 | chrIX | 220767 | 221567 | CP015057.1 | 934905 | 935690 | 51.86567164 | 266 | 261 | 1.019157088 | 1 | 1 | 1 | FIM1_2864 | YIL070C | YIL070C | MAM33 | n |
| Kmar_QGN16552.1 | W303.asm02.HP0_G0039940 | OG0002115 | chrXIII | 146411 | 147541 | CP015058.1 | 357436 | 358599 | 42.67676768 | 376 | 387 | 0.971576227 | 1 | 1 | 1 | FIM1_3266 | YML060W | YML060W | OGG1 | n |
| Kmar_QGN15113.1 | W303.asm02.HP0_G0032060 | OG0002116 | chrXI | 217827 | 219566 | CP015056.1 | 229197 | 230906 | 52.85234899 | 579 | 569 | 1.017574692 | 1 | 1 | 1 | FIM1_1800 | YKL124W | YKL124W | SSH4 | n |
| Kmar_QGN14870.1 | W303.asm02.HP0_G0034950 | OG0002117 | chrXII | 155767 | 157758 | CP015055.1 | 1393690 | 1395669 | 64.30722892 | 663 | 659 | 1.006069803 | 1 | 1 | 1 | FIM1_1543 | YLR002C | YLR002C | NOC3 | n |
| Kmar_QGN18375.1 | W303.asm02.HP0_G0044090 | OG0002118 | chrXIV | 90747 | 91868 | CP015061.1 | 802173 | 803411 | 59.46601942 | 373 | 412 | 0.905339806 | 1 | 1 | 1 | FIM1_4702 | YNL288W | YNL288W | CAF40 | n |
| Kmar_QGN13755.1 | W303.asm02.HP0_G0005500 | OG0002119 | chrIII | 117156 | 117722 | CP015054.1 | 815174 | 815722 | 65.59139785 | 188 | 182 | 1.032967033 | 1 | 1 | 1 | FIM1_399 | YCL001W | YCL001W | RER1 | n |
| Kmar_QGN15159.1 | W303.asm02.HP0_G0046680 | OG0002120 | chrXIV | 632966 | 634081 | CP015056.1 | 325978 | 327069 | 52.01072386 | 371 | 363 | 1.022038567 | 1 | 1 | 1 | FIM1_1846 | YNL005C | YNL005C | MRP7 | n |
| Kmar_QGN16323.1 | W303.asm02.HP0_G0012980 | OG0002121 | chrIV | 1375483 | 1385289 | CP015057.1 | 1281922 | 1291650 | 49.36325045 | 3268 | 3242 | 1.008019741 | 1 | 1 | 1 | FIM1_3028 | YDR457W | YDR457W | TOM1 | n |
| Kmar_QGN18311.1 | W303.asm02.HP0_G0002930 | OG0002122 | chrII | 452307 | 452999 | CP015061.1 | 671989 | 672681 | 55.84415584 | 230 | 230 | 1 | 1 | 1 | 1 | FIM1_4637 | YBR096W | YBR096W | | n |
| Kmar_QGN18150.1 | W303.asm02.HP0_G0026570 | OG0002123 | chrIX | 226219 | 228255 | CP015061.1 | 331132 | 333102 | 51.31964809 | 678 | 656 | 1.033536585 | 1 | 1 | 1 | FIM1_4473 | YIL067C | YIL067C | | n |
| Kmar_QGN17660.1 | W303.asm02.HP0_G0049290 | OG0002124 | chrXV | 373254 | 373895 | CP015060.1 | 207513 | 208151 | 74.05660377 | 213 | 212 | 1.004716981 | 1 | 1 | 1 | FIM1_4867 | YOR021C | YOR021C | SFM1 | n |
| Kmar_QGN15658.1 | W303.asm02.HP0_G0012580 | OG0002125 | chrIV | 1293940 | 1294584 | CP015056.1 | 1405484 | 1406131 | 71.1627907 | 214 | 215 | 0.995348837 | 1 | 1 | 1 | FIM1_2351 | YDR408C | YDR408C | ADE8 | n |
| Kmar_QGN14762.1 | W303.asm02.HP0_G0050320 | OG0002126 | chrXV | 581126 | 582235 | CP015055.1 | 1177293 | 1178399 | 84.82384824 | 369 | 368 | 1.002717391 | 1 | 1 | 1 | FIM1_1432 | YOR136W | YOR136W | IDH2 | n |
| Kmar_QGN13929.1 | W303.asm02.HP0_G0050430 | OG0002127 | chrXV | 610713 | 612263 | CP015054.1 | 1165217 | 1166734 | 40 | 516 | 505 | 1.021782178 | 1 | 1 | 1 | FIM1_577 | YOR149C | YOR149C | SMP3 | n |
| Kmar_QGN13930.1 | W303.asm02.HP0_G0012460 | OG0002128 | chrIV | 1267406 | 1268692 | CP015054.1 | 1166846 | 1168132 | 88.86310905 | 428 | 428 | 1 | 1 | 1 | 1 | FIM1_578 | YDR394W | YDR394W | RPT3 | n |
| Kmar_QGN13931.1 | W303.asm02.HP0_G0012450 | OG0002129 | chrIV | 1265626 | 1266996 | CP015054.1 | 1168527 | 1169768 | 43.20175439 | 456 | 413 | 1.104116223 | 1 | 1 | 1 | FIM1_579 | YDR393W | YDR393W | SHE9 | n |
| Kmar_QGN17609.1 | W303.asm02.HP0_G0049120 | OG0002130 | chrXV | 334964 | 337798 | CP015060.1 | 96643 | 99363 | 42.75210084 | 944 | 906 | 1.041942605 | 1 | 1 | 1 | FIM1_4816 | YOR005C | YOR005C | DNL4 | n |
| Kmar_QGN17610.1 | W303.asm02.HP0_G0049130 | OG0002131 | chrXV | 338135 | 339076 | CP015060.1 | 99526 | 100458 | 68.03797468 | 313 | 310 | 1.009677419 | 1 | 1 | 1 | FIM1_4817 | YOR006C | YOR006C | TSR3 | n |
| Kmar_QGN14644.1 | W303.asm02.HP0_G0004600 | OG0002132 | chrII | 768208 | 769467 | CP015055.1 | 931980 | 933179 | 44.60093897 | 419 | 399 | 1.050125313 | 1 | 1 | 1 | FIM1_1308 | YBR271W | YBR271W | EFM2 | n |
| Kmar_QGN15260.1 | W303.asm02.HP0_G0033920 | OG0002133 | chrXI | 581276 | 583051 | CP015056.1 | 549439 | 551286 | 55.71658615 | 591 | 615 | 0.96097561 | 1 | 1 | 1 | FIM1_1949 | YKR069W | YKR069W | MET1 | n |
| Kmar_QGN15259.1 | W303.asm02.HP0_G0049570 | OG0002134 | chrXV | 425302 | 426540 | CP015056.1 | 547591 | 548856 | 47.55244755 | 412 | 421 | 0.978622328 | 1 | 1 | 1 | FIM1_1948 | YOR051C | YOR051C | ETT1 | n |
| Kmar_QGN15262.1 | W303.asm02.HP0_G0049580 | OG0002135 | chrXV | 427228 | 427680 | CP015056.1 | 553087 | 553422 | 43.33333333 | 150 | 111 | 1.351351351 | 1 | 1 | 1 | FIM1_1951 | YOR052C | YOR052C | TMC1 | n |
| Kmar_QGN15439.1 | W303.asm02.HP0_G0011490 | OG0002136 | chrIV | 1061050 | 1062099 | CP015056.1 | 964285 | 965304 | 61.03151862 | 349 | 339 | 1.029498525 | 1 | 1 | 1 | FIM1_2130 | YDR297W | YDR297W | SUR2 | n |
| Kmar_QGN17611.1 | W303.asm02.HP0_G0040480 | OG0002137 | chrXIII | 263989 | 267260 | CP015060.1 | 100749 | 103958 | 31.54545455 | 1077 | 1069 | 1.00748363 | 1 | 1 | 1 | FIM1_4818 | YML002W | YML002W | VRL1 | n |
| Kmar_QGN13927.1 | W303.asm02.HP0_G0012480 | OG0002138 | chrIV | 1272091 | 1272623 | CP015054.1 | 1161116 | 1161580 | 67.34693878 | 146 | 154 | 0.948051948 | 1 | 1 | 1 | FIM1_575 | YDR397C | YDR397C | NCB2 | n |
| Kmar_QGN17373.1 | W303.asm02.HP0_G0041000 | OG0002139 | chrXIII | 386604 | 387524 | CP015059.1 | 792993 | 793901 | 62.41830065 | 306 | 302 | 1.013245033 | 1 | 1 | 1 | FIM1_4105 | YMR055C | YMR055C | BUB2 | n |
| Kmar_QGN17372.1 | W303.asm02.HP0_G0032500 | OG0002140 | chrXI | 303086 | 304669 | CP015059.1 | 791327 | 792928 | 23.09090909 | 527 | 533 | 0.988742964 | 1 | 1 | 1 | FIM1_4104 | YKL074C | YKL074C | MUD2 | n |
| Kmar_QGN17371.1 | W303.asm02.HP0_G0032510 | OG0002141 | chrXI | 304906 | 307551 | CP015059.1 | 788193 | 790844 | 35.61190739 | 881 | 883 | 0.997734994 | 1 | 1 | 1 | FIM1_4103 | YKL073W | YKL073W | LHS1 | n |
| Kmar_QGN13925.1 | W303.asm02.HP0_G0012500 | OG0002142 | chrIV | 1275793 | 1276458 | CP015054.1 | 1156636 | 1157283 | 83.25581395 | 221 | 215 | 1.027906977 | 1 | 1 | 1 | FIM1_573 | YDR399W | YDR399W | HPT1 | n |
| Kmar_QGN14096.1 | W303.asm02.HP0_G0020310 | OG0002143 | chrVII | 587585 | 588670 | CP015054.1 | 1511383 | 1512429 | 67.86703601 | 361 | 348 | 1.037356322 | 1 | 1 | 1 | FIM1_747 | YGR048W | YGR048W | UFD1 | n |
| Kmar_QGN13928.1 | W303.asm02.HP0_G0012470 | OG0002144 | chrIV | 1269049 | 1271883 | CP015054.1 | 1161896 | 1164754 | 50.89192025 | 944 | 952 | 0.991596639 | 1 | 1 | 1 | FIM1_576 | YDR395W | YDR395W | SXM1 | n |
| Kmar_QGN14142.1 | W303.asm02.HP0_G0005270 | OG0002145 | chrIII | 64326 | 65219 | CP015054.1 | 1626843 | 1627820 | 59.45121951 | 297 | 325 | 0.913846154 | 1 | 1 | 1 | FIM1_795 | YCL031C | YCL031C | RRP7 | n |
| Kmar_QGN13717.1 | W303.asm02.HP0_G0043290 | OG0002146 | chrXIII | 855606 | 856730 | CP015054.1 | 724387 | 725634 | 32.62411348 | 374 | 415 | 0.901204819 | 1 | 1 | 1 | FIM1_361 | YMR289W | YMR289W | ABZ2 | n |
| Kmar_QGN17487.1 | W303.asm02.HP0_G0021800 | OG0002147 | chrVII | 924298 | 925227 | CP015059.1 | 1012809 | 1013732 | 55.97484277 | 309 | 307 | 1.006514658 | 1 | 1 | 1 | FIM1_4223 | YGR208W | YGR208W | SER2 | n |
| Kmar_QGN18148.1 | W303.asm02.HP0_G0015230 | OG0002148 | chrV | 303617 | 303997 | CP015061.1 | 328044 | 328385 | 33.07086614 | 126 | 113 | 1.115044248 | 1 | 1 | 1 | FIM1_4471 | YER071C | YER071C | TDA2 | n |
| Kmar_QGN14325.1 | W303.asm02.HP0_G0009660 | OG0002149 | chrIV | 653843 | 654274 | CP015055.1 | 280987 | 281406 | 31.46853147 | 143 | 139 | 1.028776978 | 1 | 1 | 1 | FIM1_982 | YDR100W | YDR100W | TVP15 | n |

| | | | | | | | | | | | | | | | | | | | |
|---|---|---|---|---|---|---|---|---|---|---|---|---|---|---|---|---|---|---|---|
| Kmar_QGN14640.1 | W303.asm02.HP0_G0012090 | OG0002150 | chrIV | 1203452 | 1206400 | CP015055.1 | 923426 | 926407 | 39.6353167 | 982 | 993 | 0.988922457 | 1 | 1 | 1 | FIM1_1304 | YDR359C | YDR359C | EAF1 | n |
| Kmar_QGN16915.1 | W303.asm02.HP0_G0010410 | OG0002151 | chrIV | 825010 | 829491 | CP015058.1 | 1181161 | 1185582 | 32.07171315 | 1493 | 1473 | 1.013577733 | 1 | 1 | 1 | FIM1_3642 | YDR180W | YDR180W | SCC2 | n |
| Kmar_QGN16916.1 | W303.asm02.HP0_G0010420 | OG0002152 | chrIV | 829625 | 831070 | CP015058.1 | 1185655 | 1186734 | 25.91006424 | 481 | 359 | 1.339832869 | 1 | 1 | 1 | FIM1_3643 | YDR181C | YDR181C | SAS4 | n |
| Kmar_QGN16918.1 | W303.asm02.HP0_G0036550 | OG0002153 | chrXII | 495690 | 498305 | CP015058.1 | 1188727 | 1191180 | 42.25028703 | 871 | 817 | 1.066095471 | 1 | 1 | 1 | FIM1_3645 | YLR166C | YLR166C | SEC10 | n |
| Kmar_QGN16920.1 | W303.asm02.HP0_G0036560 | OG0002154 | chrXII | 499208 | 499666 | CP015058.1 | 1192624 | 1193079 | 89.54248366 | 152 | 151 | 1.006622517 | 1 | 1 | 1 | FIM1_3647 | YLR167W | YLR167W | RPS31 | n |
| Kmar_QGN15588.1 | W303.asm02.HP0_G0048200 | OG0002155 | chrXV | 147837 | 150731 | CP015056.1 | 1250181 | 1253051 | 61.6255144 | 964 | 956 | 1.008368201 | 1 | 1 | 1 | FIM1_2280 | YOL090W | YOL090W | MSH2 | n |
| Kmar_QGN14369.1 | W303.asm02.HP0_G0016250 | OG0002156 | chrV | 553313 | 555781 | CP015055.1 | 374690 | 375931 | 51.64179104 | 670 | 413 | 1.622276029 | 1 | 1 | 1 | FIM1_1029 | YER178W | YER178W | PDA1 | n |
| Kmar_QGN14645.1 | W303.asm02.HP0_G0012050 | OG0002157 | chrIV | 1196263 | 1197405 | CP015055.1 | 933420 | 934619 | 56.96821516 | 380 | 399 | 0.952380952 | 1 | 1 | 1 | FIM1_1309 | YDR354W | YDR354W | TRP4 | n |
| Kmar_QGN17974.1 | W303.asm02.HP0_G0050140 | OG0002158 | chrXV | 540638 | 545020 | CP015060.1 | 859213 | 863586 | 83.49315068 | 1460 | 1457 | 1.002059025 | 1 | 1 | 1 | FIM1_5183 | YOR116C | YOR116C | RPO31 | n |
| Kmar_QGN18253.1 | W303.asm02.HP0_G0056900 | OG0002159 | chrXVI | 882679 | 883464 | CP015061.1 | 559643 | 560434 | 70.45454545 | 261 | 263 | 0.992395437 | 1 | 1 | 1 | FIM1_4578 | YPR167C | YPR167C | MET16 | n |
| Kmar_QGN16914.1 | W303.asm02.HP0_G0036540 | OG0002160 | chrXII | 494755 | 495519 | CP015058.1 | 1179838 | 1180764 | 30.53691275 | 254 | 308 | 0.824675325 | 1 | 1 | 1 | FIM1_3641 | YLR165C | YLR165C | PUS5 | n |
| Kmar_QGN14580.1 | W303.asm02.HP0_G0007960 | OG0002161 | chrIV | 300544 | 301329 | CP015055.1 | 794348 | 795148 | 45.03546099 | 261 | 266 | 0.981203008 | 1 | 1 | 1 | FIM1_1242 | YDL087C | YDL087C | LUC7 | n |
| Kmar_QGN17972.1 | W303.asm02.HP0_G0050150 | OG0002162 | chrXV | 545904 | 547208 | CP015060.1 | 856824 | 858122 | 88.70967742 | 434 | 432 | 1.00462963 | 1 | 1 | 1 | FIM1_5181 | YOR117W | YOR117W | RPT5 | n |
| Kmar_QGN16926.1 | W303.asm02.HP0_G0037240 | OG0002163 | chrXII | 620402 | 621565 | CP015058.1 | 1201873 | 1202988 | 74.67700258 | 387 | 371 | 1.043126685 | 1 | 1 | 1 | FIM1_3653 | YLR244C | YLR244C | MAP1 | n |
| Kmar_QGN16927.1 | W303.asm02.HP0_G0010490 | OG0002164 | chrIV | 840136 | 841776 | CP015058.1 | 1203553 | 1205187 | 85.40145985 | 546 | 544 | 1.003676471 | 1 | 1 | 1 | FIM1_3654 | YDR188W | YDR188W | CCT6 | n |
| Kmar_QGN16928.1 | W303.asm02.HP0_G0010500 | OG0002165 | chrIV | 842107 | 844107 | CP015058.1 | 1205505 | 1207433 | 60.83086053 | 666 | 642 | 1.037383178 | 1 | 1 | 1 | FIM1_3655 | YDR189W | YDR189W | SLY1 | n |
| Kmar_QGN16929.1 | W303.asm02.HP0_G0010510 | OG0002166 | chrIV | 844319 | 845710 | CP015058.1 | 1207546 | 1208919 | 87.04103672 | 463 | 457 | 1.013129103 | 1 | 1 | 1 | FIM1_3656 | YDR190C | YDR190C | RVB1 | n |
| Kmar_QGN16931.1 | W303.asm02.HP0_G0037230 | OG0002167 | chrXII | 619433 | 620251 | CP015058.1 | 1210987 | 1211802 | 81.25 | 272 | 271 | 1.003690037 | 1 | 1 | 1 | FIM1_3658 | YLR243W | YLR243W | GPN3 | n |
| Kmar_QGN15966.1 | W303.asm02.HP0_G0010640 | OG0002168 | chrIV | 863062 | 865236 | CP015057.1 | 498365 | 500500 | 41.76394293 | 724 | 711 | 1.018284107 | 1 | 1 | 1 | FIM1_2664 | YDR205W | YDR205W | MSC2 | n |
| Kmar_QGN15737.1 | W303.asm02.HP0_G0025750 | OG0002169 | chrIX | 40790 | 42739 | CP015056.1 | 1573062 | 1574990 | 64.26380368 | 649 | 642 | 1.010903427 | 1 | 1 | 1 | FIM1_2431 | YIL155C | YIL155C | GUT2 | n |
| Kmar_QGN16357.1 | W303.asm02.HP0_G0040380 | OG0002170 | chrXIII | 244655 | 246409 | CP015057.1 | 1353462 | 1355330 | 34.80314961 | 584 | 622 | 0.938906752 | 1 | 1 | 1 | FIM1_3063 | YML013W | YML013W | UBX2 | n |
| Kmar_QGN15736.1 | W303.asm02.HP0_G0025760 | OG0002171 | chrIX | 43012 | 44052 | CP015056.1 | 1570030 | 1571124 | 38.27493261 | 346 | 364 | 0.950549451 | 1 | 1 | 1 | FIM1_2430 | YIL154C | YIL154C | IMP21 | n |
| Kmar_QGN14011.1 | W303.asm02.HP0_G0008600 | OG0002172 | chrIV | 427400 | 428383 | CP015054.1 | 1322290 | 1323273 | 90.82568807 | 327 | 327 | 1 | 1 | 1 | 1 | FIM1_662 | YDL014W | YDL014W | NOP1 | n |
| Kmar_QGN17782.1 | W303.asm02.HP0_G0045890 | OG0002173 | chrXIV | 459332 | 463618 | CP015060.1 | 469448 | 473797 | 68.08219178 | 1428 | 1449 | 0.985507246 | 1 | 1 | 1 | FIM1_4991 | YNL088W | YNL088W | TOP2 | n |
| Kmar_QGN14461.1 | W303.asm02.HP0_G0018030 | OG0002174 | chrVII | 94482 | 97589 | CP015055.1 | 566046 | 569141 | 71.73076923 | 1035 | 1031 | 1.003879728 | 1 | 1 | 1 | FIM1_1122 | YGL207W | YGL207W | SPT16 | n |
| Kmar_QGN17033.1 | W303.asm02.HP0_G0032810 | OG0002175 | chrXI | 362097 | 363683 | CP015059.1 | 82883 | 84355 | 59.3385214 | 528 | 490 | 1.07755102 | 1 | 1 | 1 | FIM1_3762 | YKL045W | YKL045W | PRI2 | n |
| Kmar_QGN13704.1 | W303.asm02.HP0_G0043350 | OG0002176 | chrXIII | 865217 | 865810 | CP015054.1 | 694291 | 694857 | 53.8071066 | 197 | 188 | 1.04787234 | 1 | 1 | 1 | FIM1_347 | YMR295C | YMR295C | GSR1 | n |
| Kmar_QGN15282.1 | W303.asm02.HP0_G0014010 | OG0002177 | chrV | 65072 | 65530 | CP015056.1 | 595770 | 596189 | 31.21019108 | 152 | 139 | 1.09352518 | 1 | 1 | 1 | FIM1_1971 | YEL048C | YEL048C | TCA17 | n |
| Kmar_QGN15286.1 | W303.asm02.HP0_G0014050 | OG0002178 | chrV | 70120 | 70620 | CP015056.1 | 604284 | 604712 | 41.42011834 | 166 | 142 | 1.169014085 | 1 | 1 | 1 | FIM1_1975 | YEL044W | YEL044W | IES6 | n |
| Kmar_QGN15669.1 | W303.asm02.HP0_G0012650 | OG0002179 | chrIV | 1304157 | 1306736 | CP015056.1 | 1423147 | 1425648 | 28.45804989 | 859 | 833 | 1.031212485 | 1 | 1 | 1 | FIM1_2362 | YDR416W | YDR416W | SYF1 | n |
| Kmar_QGN15668.1 | W303.asm02.HP0_G0013980 | OG0002180 | chrV | 58741 | 59511 | CP015056.1 | 1421936 | 1422829 | 73.4006734 | 256 | 297 | 0.861952862 | 1 | 1 | 1 | FIM1_2361 | YEL051W | YEL051W | VMA8 | n |
| Kmar_QGN14112.1 | W303.asm02.HP0_G0044210 | OG0002181 | chrXIV | 119822 | 121552 | CP015054.1 | 1556556 | 1558244 | 63.04728546 | 576 | 562 | 1.024911032 | 1 | 1 | 1 | FIM1_763 | YNL275W | YNL275W | BOR1 | n |
| Kmar_QGN13749.1 | W303.asm02.HP0_G0000700 | OG0002182 | chrI | 153461 | 155080 | CP015054.1 | 804412 | 805566 | 17.98165138 | 539 | 384 | 1.403645833 | 1 | 0 | 0 | FIM1_393 | YAR002W | YAR002W | NUP60 | n |
| Kmar_QGN15778.1 | W303.asm02.HP0_G0019880 | OG0002183 | chrVII | 492986 | 494035 | CP015057.1 | 78385 | 79440 | 77.20797721 | 349 | 351 | 0.994301994 | 1 | 1 | 1 | FIM1_2474 | YGL001C | YGL001C | ERG26 | n |
| Kmar_QGN16747.1 | W303.asm02.HP0_G0014550 | OG0002184 | chrV | 168991 | 173001 | CP015058.1 | 807276 | 811262 | 32.62927895 | 1336 | 1328 | 1.006024096 | 1 | 1 | 1 | FIM1_3469 | YER008C | YER008C | SEC3 | n |
| Kmar_QGN16754.1 | W303.asm02.HP0_G0014510 | OG0002185 | chrV | 161733 | 163625 | CP015058.1 | 826483 | 828459 | 50.2276176 | 630 | 658 | 0.957446809 | 1 | 1 | 1 | FIM1_3476 | YER005W | YER005W | YND1 | n |
| Kmar_QGN16751.1 | W303.asm02.HP0_G0014520 | OG0002186 | chrV | 163906 | 165468 | CP015058.1 | 817920 | 819509 | 69.75425331 | 520 | 529 | 0.982986767 | 1 | 1 | 1 | FIM1_3473 | YER006W | YER006W | NUG1 | n |
| Kmar_QGN16748.1 | W303.asm02.HP0_G0027240 | OG0002187 | chrIX | 348469 | 351450 | CP015058.1 | 811530 | 814649 | 53.21805956 | 993 | 1039 | 0.95572666 | 1 | 1 | 1 | FIM1_3470 | YIR002C | YIR002C | MPH1 | n |
| Kmar_QGN16749.1 | W303.asm02.HP0_G0014540 | OG0002188 | chrV | 167420 | 168068 | CP015058.1 | 815450 | 815995 | 77.3480663 | 181 | 181 | 1 | 1 | 1 | 1 | FIM1_3471 | YER007C-A | YER007C-A | TMA20 | n |
| Kmar_QGN16750.1 | W303.asm02.HP0_G0014530 | OG0002189 | chrV | 165710 | 167266 | CP015058.1 | 816021 | 817556 | 28.62453532 | 518 | 511 | 1.01369863 | 1 | 1 | 1 | FIM1_3472 | YER007W | YER007W | PAC2 | n |
| Kmar_QGN13815.1 | W303.asm02.HP0_G0013180 | OG0002190 | chrIV | 1420904 | 1422568 | CP015054.1 | 929846 | 931480 | 38.02345059 | 554 | 544 | 1.018382353 | 1 | 1 | 1 | FIM1_461 | YDR479C | YDR479C | PEX29 | n |
| Kmar_QGN13813.1 | W303.asm02.HP0_G0049310 | OG0002191 | chrXV | 378675 | 380018 | CP015054.1 | 927380 | 928789 | 51.25523013 | 447 | 469 | 0.953091684 | 1 | 1 | 1 | FIM1_459 | YOR025W | YOR025W | HST3 | n |
| Kmar_QGN15592.1 | W303.asm02.HP0_G0013160 | OG0002192 | chrIV | 1418067 | 1419968 | CP015056.1 | 1259671 | 1261491 | 72.57053292 | 633 | 606 | 1.044554455 | 1 | 1 | 1 | FIM1_2284 | YDR477W | YDR477W | SNF1 | n |
| Kmar_QGN14793.1 | W303.asm02.HP0_G0012230 | OG0002193 | chrIV | 1229067 | 1229987 | CP015055.1 | 1241889 | 1242689 | 25.94936709 | 306 | 266 | 1.15037594 | 1 | 1 | 1 | FIM1_1464 | YDR374C | YDR374C | PHO92 | n |
| Kmar_QGN14794.1 | W303.asm02.HP0_G0050220 | OG0002194 | chrXV | 559905 | 560606 | CP015055.1 | 1243144 | 1243992 | 57.80141844 | 233 | 282 | 0.826241135 | 1 | 1 | 1 | FIM1_1465 | YOR125C | YOR125C | CAT5 | n |
| Kmar_QGN16035.1 | W303.asm02.HP0_G0048410 | OG0002195 | chrXV | 199397 | 200752 | CP015057.1 | 674359 | 675708 | 49.00221729 | 451 | 449 | 1.004454343 | 1 | 1 | 1 | FIM1_2735 | YOL069W | YOL069W | NUF2 | n |
| Kmar_QGN17284.1 | W303.asm02.HP0_G0008440 | OG0002196 | chrIV | 389221 | 392106 | CP015059.1 | 616176 | 618743 | 35.4338843 | 961 | 855 | 1.123976608 | 1 | 1 | 1 | FIM1_4015 | YDL035C | YDL035C | GPR1 | n |
| Kmar_QGN15572.1 | W303.asm02.HP0_G0041400 | OG0002197 | chrXIII | 464439 | 465332 | CP015056.1 | 1213577 | 1214467 | 75 | 297 | 296 | 1.003378378 | 1 | 1 | 1 | FIM1_2263 | YMR099C | YMR099C | | n |
| Kmar_QGN15088.1 | W303.asm02.HP0_G0033280 | OG0002198 | chrXI | 449682 | 451796 | CP015056.1 | 176989 | 179043 | 79.26136364 | 704 | 684 | 1.029239766 | 1 | 1 | 1 | FIM1_1775 | YKR001C | YKR001C | VPS1 | n |
| Kmar_QGN18370.1 | W303.asm02.HP0_G0044050 | OG0002199 | chrXIV | 83253 | 84464 | CP015061.1 | 792838 | 794061 | 58.96805897 | 403 | 407 | 0.99017199 | 1 | 1 | 1 | FIM1_4697 | YNL292W | YNL292W | PUS4 | n |
| Kmar_QGN17814.1 | W303.asm02.HP0_G0000310 | OG0002200 | chrI | 69351 | 70160 | CP015060.1 | 541164 | 542027 | 55.40069686 | 269 | 287 | 0.93728223 | 1 | 1 | 1 | FIM1_5023 | YAL039C | YAL039C | CYC3 | n |
| Kmar_QGN17247.1 | W303.asm02.HP0_G0022980 | OG0002201 | chrVIII | 75768 | 76691 | CP015059.1 | 544789 | 545787 | 55 | 307 | 332 | 0.924698795 | 1 | 1 | 1 | FIM1_3978 | YHL013C | YHL013C | OTU2 | n |
| Kmar_QGN17818.1 | W303.asm02.HP0_G0052320 | OG0002202 | chrXV | 1007418 | 1008986 | CP015060.1 | 554820 | 556202 | 37.80718336 | 522 | 460 | 1.134782609 | 1 | 1 | 1 | FIM1_5027 | YOR355W | YOR355W | GDS1 | n |
| Kmar_QGN17078.1 | W303.asm02.HP0_G0005970 | OG0002203 | chrIII | 233699 | 234526 | CP015059.1 | 182715 | 183518 | 76.36363636 | 275 | 267 | 1.029962547 | 1 | 1 | 1 | FIM1_3809 | YCR047C | YCR047C | BUD23 | n |
| Kmar_QGN13690.1 | W303.asm02.HP0_G0025370 | OG0002204 | chrVIII | 521008 | 522588 | CP015054.1 | 667657 | 669126 | 36.96060038 | 526 | 489 | 1.075664622 | 1 | 1 | 1 | FIM1_333 | YHR207C | YHR207C | SET5 | n |
| Kmar_QGN15595.1 | W303.asm02.HP0_G0052350 | OG0002205 | chrXV | 1012442 | 1013170 | CP015056.1 | 1265690 | 1266166 | 50 | 242 | 158 | 1.53164557 | 1 | 1 | 1 | FIM1_2287 | YOR358W | YOR358W | HAP5 | n |
| Kmar_QGN17889.1 | W303.asm02.HP0_G0056140 | OG0002206 | chrXVI | 710724 | 711155 | CP015060.1 | 684845 | 685276 | 81.81818182 | 143 | 143 | 1 | 1 | 1 | 1 | FIM1_5098 | YPR082C | YPR082C | DIB1 | n |
| Kmar_QGN13565.1 | W303.asm02.HP0_G0038830 | OG0002207 | chrXII | 960454 | 961548 | CP015054.1 | 428636 | 429721 | 69.06077348 | 364 | 361 | 1.008310249 | 1 | 1 | 1 | FIM1_206 | YLR420W | YLR420W | URA4 | n |
| Kmar_QGN16073.1 | W303.asm02.HP0_G0040850 | OG0002208 | chrXIII | 341645 | 342178 | CP015057.1 | 750029 | 750565 | 51.66666667 | 177 | 178 | 0.994382022 | 1 | 1 | 1 | FIM1_2773 | YMR035W | YMR035W | IMP2 | n |
| Kmar_QGN18274.1 | W303.asm02.HP0_G0020930 | OG0002209 | chrVII | 727818 | 729446 | CP015061.1 | 599523 | 601193 | 49.03339192 | 542 | 556 | 0.974820144 | 1 | 1 | 1 | FIM1_4600 | YGR119C | YGR119C | NUP57 | n |
| Kmar_QGN18272.1 | W303.asm02.HP0_G0020940 | OG0002210 | chrVII | 729807 | 730595 | CP015061.1 | 595506 | 596237 | 29.27756654 | 262 | 243 | 1.0781893 | 1 | 1 | 1 | FIM1_4598 | YGR120C | YGR120C | COG2 | n |
| Kmar_QGN15263.1 | W303.asm02.HP0_G0033940 | OG0002211 | chrXI | 584597 | 585643 | CP015056.1 | 554001 | 555026 | 48.71794872 | 348 | 341 | 1.020527859 | 1 | 1 | 1 | FIM1_1952 | YKR071C | YKR071C | DRE2 | n |
| Kmar_QGN18362.1 | W303.asm02.HP0_G0025850 | OG0002212 | chrIX | 67105 | 69180 | CP015061.1 | 778491 | 780521 | 41.66666667 | 691 | 676 | 1.022189349 | 1 | 1 | 1 | FIM1_4689 | YIL144W | YIL144W | NDC80 | n |
| Kmar_QGN13633.1 | W303.asm02.HP0_G0045340 | OG0002213 | chrXIV | 350059 | 350448 | CP015054.1 | 556906 | 557355 | 36.91275168 | 129 | 149 | 0.865771812 | 1 | 1 | 1 | FIM1_275 | YNL149C | YNL149C | PGA2 | n |
| Kmar_QGN18270.1 | W303.asm02.HP0_G0056620 | OG0002214 | chrXVI | 809288 | 811009 | CP015061.1 | 590149 | 591945 | 66.33499171 | 573 | 598 | 0.95819398 | 1 | 1 | 1 | FIM1_4596 | YPR137W | YPR137W | RRP9 | n |
| Kmar_QGN13632.1 | W303.asm02.HP0_G0045330 | OG0002215 | chrXIV | 348458 | 349213 | CP015054.1 | 555890 | 556639 | 50.38461538 | 251 | 249 | 1.008032129 | 1 | 1 | 1 | FIM1_274 | YNL151C | YNL151C | RPC31 | n |
| Kmar_QGN13630.1 | W303.asm02.HP0_G0045320 | OG0002216 | chrXIV | 347005 | 348234 | CP015054.1 | 553108 | 554283 | 35.90361446 | 409 | 391 | 1.046035806 | 1 | 1 | 1 | FIM1_272 | YNL152W | YNL152W | INN1 | n |
| Kmar_QGN14687.1 | W303.asm02.HP0_G0057020 | OG0002217 | chrXVI | 901793 | 902836 | CP015055.1 | 1034229 | 1035242 | 53.0259366 | 347 | 337 | 1.029673591 | 1 | 1 | 1 | FIM1_1354 | YPR180W | YPR180W | AOS1 | n |
| Kmar_QGN18207.1 | W303.asm02.HP0_G0053210 | OG0002218 | chrXVI | 125728 | 126732 | CP015061.1 | 448927 | 449874 | 58.6102719 | 334 | 315 | 1.06031746 | 1 | 1 | 1 | FIM1_4531 | YPL227C | YPL227C | ALG5 | n |
| Kmar_QGN18121.1 | W303.asm02.HP0_G0055680 | OG0002219 | chrXVI | 630966 | 633464 | CP015061.1 | 259684 | 262107 | 47.68133175 | 832 | 807 | 1.030978934 | 1 | 1 | 1 | FIM1_4441 | YPR029C | YPR029C | APL4 | n |
| Kmar_QGN16230.1 | W303.asm02.HP0_G0057120 | OG0002220 | chrXVI | 922909 | 924873 | CP015057.1 | 1083257 | 1085272 | 47.19764012 | 654 | 671 | 0.97466468 | 1 | 1 | 1 | FIM1_2934 | YPR190C | YPR190C | RPC82 | n |
| Kmar_QGN16225.1 | W303.asm02.HP0_G0057070 | OG0002221 | chrXVI | 913050 | 915266 | CP015057.1 | 1073271 | 1075409 | 37.10526316 | 738 | 712 | 1.036516854 | 1 | 1 | 1 | FIM1_2929 | YPR185W | YPR185W | ATG13 | n |
| Kmar_QGN15351.1 | W303.asm02.HP0_G0057140 | OG0002222 | chrXVI | 927692 | 928609 | CP015056.1 | 771709 | 772638 | 49.20127796 | 305 | 309 | 0.987055016 | 1 | 0 | 0 | FIM1_2041 | YPR192W | YPR192W | AQY1 | n |
| Kmar_QGN18133.1 | W303.asm02.HP0_G0001060 | OG0002223 | chrII | 49978 | 50511 | CP015061.1 | 294597 | 295046 | 35.59322034 | 177 | 149 | 1.187919463 | 1 | 1 | 1 | FIM1_4455 | YBL090W | YBL090W | MRP21 | n |
| Kmar_QGN18363.1 | W303.asm02.HP0_G0025860 | OG0002224 | chrIX | 69541 | 72072 | CP015061.1 | 780879 | 783479 | 79.86191024 | 843 | 866 | 0.973441109 | 1 | 1 | 1 | FIM1_4690 | YIL143C | YIL143C | SSL2 | n |
| Kmar_QGN18364.1 | W303.asm02.HP0_G0034160 | OG0002225 | chrXI | 630988 | 632148 | CP015061.1 | 784340 | 785485 | 60.51282051 | 386 | 381 | 1.01312336 | 1 | 1 | 1 | FIM1_4691 | W303.asm02.HP0_G0034160 | YKR092C | SRP40 | y |
| Kmar_QGN18329.1 | W303.asm02.HP0_G0001570 | OG0002226 | chrII | 160959 | 161996 | CP015061.1 | 706456 | 707448 | 66.38176638 | 345 | 330 | 1.045454545 | 1 | 1 | 1 | FIM1_4655 | YBL033C | YBL033C | RIB1 | n |
| Kmar_QGN14287.1 | W303.asm02.HP0_G0031780 | OG0002227 | chrXI | 160735 | 162621 | CP015055.1 | 191822 | 194020 | 47.73662551 | 628 | 732 | 0.857923497 | 1 | 1 | 1 | FIM1_944 | YKL155C | YKL155C | RSM22 | n |
| Kmar_QGN13722.1 | W303.asm02.HP0_G0025150 | OG0002228 | chrVIII | 482101 | 486774 | CP015054.1 | 736079 | 740590 | 59.52685422 | 1557 | 1502 | 1.036617843 | 1 | 1 | 1 | FIM1_366 | YHR186C | YHR186C | KOG1 | n |
| Kmar_QGN14820.1 | W303.asm02.HP0_G0047660 | OG0002229 | chrXV | 45971 | 47782 | CP015055.1 | 1292456 | 1294186 | 45.17133956 | 603 | 576 | 1.046875 | 1 | 1 | 1 | FIM1_1491 | YOL148C | YOL148C | SPT20 | n |
| Kmar_QGN14821.1 | W303.asm02.HP0_G0047650 | OG0002230 | chrXV | 45165 | 45860 | CP015055.1 | 1294278 | 1294844 | 54.54545455 | 231 | 188 | 1.228723404 | 1 | 1 | 1 | FIM1_1492 | YOL149W | YOL149W | DCP1 | n |
| Kmar_QGN13804.1 | W303.asm02.HP0_G0047970 | OG0002231 | chrXV | 103772 | 104380 | CP015054.1 | 912036 | 912530 | 43 | 202 | 164 | 1.231707317 | 1 | 1 | 1 | FIM1_450 | YOL114C | YOL114C | PTH4 | n |
| Kmar_QGN16432.1 | W303.asm02.HP0_G0004640 | OG0002232 | chrII | 774705 | 780455 | CP015058.1 | 105414 | 109778 | 19.61598339 | 1916 | 1454 | 1.317744154 | 1 | 1 | 1 | FIM1_3145 | YBR275C | YBR275C | RIF1 | n |
| Kmar_QGN16444.1 | W303.asm02.HP0_G0029150 | OG0002233 | chrX | 318137 | 320140 | CP015058.1 | 131303 | 132748 | 29.35916542 | 667 | 481 | 1.386694387 | 1 | 1 | 1 | FIM1_3157 | YJL057C | YJL057C | IKS1 | n |
| Kmar_QGN15452.1 | W303.asm02.HP0_G0011580 | OG0002234 | chrIV | 1078234 | 1079670 | CP015056.1 | 983353 | 984633 | 51.98329854 | 478 | 426 | 1.122065728 | 1 | 1 | 1 | FIM1_2143 | YDR306C | YDR306C | PFU1 | n |
| Kmar_QGN15193.1 | W303.asm02.HP0_G0021390 | OG0002235 | chrVII | 842290 | 843972 | CP015056.1 | 402768 | 404054 | 26.19469027 | 560 | 428 | 1.308411215 | 1 | 1 | 1 | FIM1_1881 | YGR166W | YGR166W | TRS65 | n |
| Kmar_QGN17888.1 | W303.asm02.HP0_G0003150 | OG0002236 | chrII | 497649 | 499652 | CP015060.1 | 682368 | 684368 | 80.78078078 | 667 | 666 | 1.001501502 | 1 | 1 | 1 | FIM1_5097 | YBR121C | YBR121C | GRS1 | n |
| Kmar_QGN16728.1 | W303.asm02.HP0_G0024070 | OG0002237 | chrVIII | 262334 | 263992 | CP015058.1 | 754547 | 756175 | 38.98601399 | 552 | 542 | 1.018450185 | 1 | 1 | 1 | FIM1_3449 | YHR078W | YHR078W | | n |
| Kmar_QGN15438.1 | W303.asm02.HP0_G0011480 | OG0002238 | chrIV | 1059711 | 1060391 | CP015056.1 | 962502 | 963161 | 53.64806867 | 226 | 219 | 1.03196347 | 1 | 1 | 1 | FIM1_2129 | YDR296W | YDR296W | MHR1 | n |
| Kmar_QGN18405.1 | W303.asm02.HP0_G0031860 | OG0002239 | chrXI | 175481 | 176884 | CP015061.1 | 854817 | 856235 | 87.07627119 | 467 | 472 | 0.98940678 | 1 | 1 | 1 | FIM1_4732 | YKL145W | YKL145W | RPT1 | n |
| Kmar_QGN18404.1 | W303.asm02.HP0_G0031870 | OG0002240 | chrXI | 177111 | 177749 | CP015061.1 | 854061 | 854693 | 75 | 212 | 210 | 1.00952381 | 1 | 1 | 1 | FIM1_4731 | YKL144C | YKL144C | RPC25 | n |
| Kmar_QGN14442.1 | W303.asm02.HP0_G0047130 | OG0002241 | chrXIV | 712272 | 713042 | CP015055.1 | 522997 | 523815 | 29.31034483 | 256 | 272 | 0.941176471 | 1 | 1 | 1 | FIM1_1103 | YNR040W | YNR040W | MRX15 | n |
| Kmar_QGN15261.1 | W303.asm02.HP0_G0033930 | OG0002242 | chrXI | 583238 | 584296 | CP015056.1 | 551656 | 552714 | 60.05665722 | 352 | 352 | 1 | 1 | 1 | 1 | FIM1_1950 | YKR070W | YKR070W | | n |
| Kmar_QGN14796.1 | W303.asm02.HP0_G0050210 | OG0002243 | chrXV | 555699 | 559517 | CP015055.1 | 1245898 | 1249569 | 46.35865309 | 1272 | 1223 | 1.040065413 | 1 | 1 | 1 | FIM1_1467 | YOR124C | YOR124C | UBP2 | n |
| Kmar_QGN14791.1 | W303.asm02.HP0_G0012250 | OG0002244 | chrIV | 1230759 | 1232129 | CP015055.1 | 1239913 | 1241253 | 66.44736842 | 456 | 446 | 1.022421525 | 1 | 1 | 1 | FIM1_1462 | YDR375C | YDR375C | BCS1 | n |
| Kmar_QGN14661.1 | W303.asm02.HP0_G0005620 | OG0002245 | chrIII | 138964 | 142113 | CP015055.1 | 967317 | 970385 | 64.95238095 | 1049 | 1022 | 1.026418787 | 1 | 1 | 1 | FIM1_1325 | YCR011C | YCR011C | ADP1 | n |
| Kmar_QGN17386.1 | W303.asm02.HP0_G0032450 | OG0002246 | chrXI | 293506 | 294684 | CP015059.1 | 818338 | 819513 | 67.09183673 | 392 | 391 | 1.002557545 | 1 | 1 | 1 | FIM1_4118 | YKL080W | YKL080W | VMA5 | n |
| Kmar_QGN15587.1 | W303.asm02.HP0_G0003420 | OG0002247 | chrII | 556764 | 557798 | CP015056.1 | 1248706 | 1249722 | 56.10465116 | 344 | 338 | 1.017751479 | 1 | 1 | 1 | FIM1_2279 | YBR149W | YBR149W | ARA1 | n |
| Kmar_QGN17436.1 | W303.asm02.HP0_G0008750 | OG0002248 | chrIV | 454441 | 456242 | CP015059.1 | 914709 | 916145 | 42.5 | 485 | 478 | 1.014644351 | 1 | 1 | 1 | FIM1_4169 | YDR004W | YDR004W | RAD57 | n |

| | | | | | | | | | | | | | | | | | | | | |
|---|---|---|---|---|---|---|---|---|---|---|---|---|---|---|---|---|---|---|---|---|
| Kmar_QGN16660.1 | W303.asm02.HP0_G0020400 | OG0002249 | chrVII | 603895 | 604902 | CP015058.1 | 610282 | 611088 | 44.7761194 | 335 | 268 | 1.25 | 1 | 1 | 1 | FIM1_3379 | YGR058W | YGR058W | PEF1 | n |
| Kmar_QGN15141.1 | W303.asm02.HP0_G0016840 | OG0002250 | chrVI | 91962 | 98393 | CP015056.1 | 278398 | 284850 | 44.4598338 | 2143 | 2150 | 0.996744186 | 1 | 1 | 1 | FIM1_1828 | YFL007W | YFL007W | BLM10 | n |
| Kmar_QGN15691.1 | W303.asm02.HP0_G0014840 | OG0002251 | chrV | 224506 | 226338 | CP015056.1 | 1476704 | 1478527 | 88.36065574 | 610 | 607 | 1.004942339 | 1 | 1 | 1 | FIM1_2384 | YER036C | YER036C | ARB1 | n |
| Kmar_QGN14430.1 | W303.asm02.HP0_G0018910 | OG0002252 | chrVII | 296926 | 298317 | CP015055.1 | 501302 | 502600 | 42.43070362 | 463 | 432 | 1.071759259 | 1 | 1 | 1 | FIM1_1091 | YGL111W | YGL111W | NSA1 | n |
| Kmar_QGN14600.1 | W303.asm02.HP0_G0018990 | OG0002253 | chrVII | 310177 | 311226 | CP015055.1 | 833450 | 834457 | 68.48137536 | 349 | 335 | 1.041791045 | 1 | 1 | 1 | FIM1_1263 | YGL100W | YGL100W | SEH1 | n |
| Kmar_QGN14599.1 | W303.asm02.HP0_G0004330 | OG0002254 | chrII | 728802 | 730148 | CP015055.1 | 831773 | 833116 | 67.92035398 | 448 | 447 | 1.002237136 | 1 | 1 | 1 | FIM1_1262 | YBR243C | YBR243C | ALG7 | n |
| Kmar_QGN16580.1 | W303.asm02.HP0_G0038060 | OG0002255 | chrXII | 798340 | 800793 | CP015058.1 | 425477 | 427933 | 35.15715949 | 817 | 818 | 0.998777506 | 1 | 1 | 1 | FIM1_3295 | YLR337C | YLR337C | VRP1 | n |
| Kmar_QGN14603.1 | W303.asm02.HP0_G0004360 | OG0002256 | chrII | 734946 | 736109 | CP015055.1 | 840011 | 841141 | 54.43037975 | 387 | 376 | 1.029255319 | 1 | 1 | 1 | FIM1_1266 | YBR246W | YBR246W | RRT2 | n |
| Kmar_QGN14919.1 | W303.asm02.HP0_G0006790 | OG0002257 | chrIV | 70713 | 73991 | CP015055.1 | 1490052 | 1493156 | 61.88747731 | 1092 | 1034 | 1.056092843 | 1 | 1 | 1 | FIM1_1593 | YDL215C | YDL215C | GDH2 | n |
| Kmar_QGN14032.1 | W303.asm02.HP0_G0048490 | OG0002258 | chrXV | 212698 | 214188 | CP015054.1 | 1375229 | 1376710 | 62.69230769 | 496 | 493 | 1.006085193 | 1 | 1 | 1 | FIM1_683 | YOL061W | YOL061W | PRS5 | n |
| Kmar_QGN14033.1 | W303.asm02.HP0_G0048480 | OG0002259 | chrXV | 210974 | 212449 | CP015054.1 | 1377405 | 1378850 | 45.58232932 | 491 | 481 | 1.020790021 | 1 | 1 | 1 | FIM1_684 | YOL062C | YOL062C | APM4 | n |
| Kmar_QGN14431.1 | W303.asm02.HP0_G0018920 | OG0002260 | chrVII | 298485 | 300359 | CP015055.1 | 502673 | 504604 | 45.49689441 | 624 | 643 | 0.970451011 | 1 | 1 | 1 | FIM1_1092 | YGL110C | YGL110C | CUE3 | n |
| Kmar_QGN17548.1 | W303.asm02.HP0_G0028620 | OG0002262 | chrX | 182548 | 183483 | CP015059.1 | 1143275 | 1144162 | 45.54140127 | 311 | 295 | 1.054237288 | 1 | 1 | 1 | FIM1_4284 | YJL117W | YJL117W | PHO86 | n |
| Kmar_QGN17887.1 | W303.asm02.HP0_G0003140 | OG0002263 | chrII | 496720 | 497208 | CP015060.1 | 681585 | 682055 | 58.02469136 | 162 | 156 | 1.038461538 | 1 | 1 | 1 | FIM1_5096 | YBR120C | YBR120C | CBP6 | n |
| Kmar_QGN17784.1 | W303.asm02.HP0_G0045870 | OG0002265 | chrXIV | 454035 | 457757 | CP015060.1 | 475651 | 479457 | 25.48262548 | 1240 | 1268 | 0.977917981 | 1 | 1 | 1 | FIM1_4993 | YNL091W | YNL091W | NST1 | n |
| Kmar_QGN17786.1 | W303.asm02.HP0_G0049880 | OG0002266 | chrXV | 488442 | 490469 | CP015060.1 | 481220 | 483250 | 54.70588235 | 675 | 676 | 0.99852071 | 1 | 1 | 1 | FIM1_4995 | YOR087W | YOR087W | YVC1 | n |
| Kmar_QGN15644.1 | W303.asm02.HP0_G0020260 | OG0002267 | chrVII | 577234 | 578049 | CP015056.1 | 1379204 | 1379965 | 27.30627306 | 271 | 253 | 1.071146245 | 1 | 1 | 1 | FIM1_2337 | YGR042W | YGR042W | MTE1 | n |
| Kmar_QGN15528.1 | W303.asm02.HP0_G0010940 | OG0002268 | chrIV | 941116 | 941994 | CP015056.1 | 1127334 | 1128206 | 71.91780822 | 292 | 290 | 1.006896552 | 1 | 1 | 1 | FIM1_2219 | YDR237W | YDR237W | MRPL7 | n |
| Kmar_QGN15529.1 | W303.asm02.HP0_G0006870 | OG0002269 | chrIV | 88321 | 89937 | CP015056.1 | 1128662 | 1130278 | 45.93128391 | 538 | 538 | 1 | 1 | 1 | 1 | FIM1_2220 | YDL207W | YDL207W | GLE1 | n |
| Kmar_QGN17238.1 | W303.asm02.HP0_G0041190 | OG0002270 | chrXIII | 416701 | 420534 | CP015059.1 | 518576 | 522316 | 41.36607828 | 1277 | 1246 | 1.024879615 | 1 | 1 | 1 | FIM1_3969 | YMR076C | YMR076C | PDS5 | n |
| Kmar_QGN16343.1 | W303.asm02.HP0_G0012840 | OG0002271 | chrIV | 1348048 | 1349796 | CP015057.1 | 1331371 | 1333104 | 38.81355932 | 582 | 577 | 1.008665511 | 1 | 1 | 1 | FIM1_3049 | YDR440W | YDR440W | DOT1 | n |
| Kmar_QGN16345.1 | W303.asm02.HP0_G0040310 | OG0002272 | chrXIII | 231657 | 233651 | CP015057.1 | 1334830 | 1336755 | 46.54919236 | 664 | 641 | 1.035881435 | 1 | 1 | 1 | FIM1_3051 | YML020W | YML020W | | n |
| Kmar_QGN16359.1 | W303.asm02.HP0_G0012760 | OG0002273 | chrIV | 1331057 | 1334026 | CP015057.1 | 1358469 | 1361423 | 63.370333 | 989 | 984 | 1.005081301 | 1 | 1 | 1 | FIM1_3065 | YDR430C | YDR430C | CYM1 | n |
| Kmar_QGN15110.1 | W303.asm02.HP0_G0032090 | OG0002274 | chrXI | 225203 | 226177 | CP015056.1 | 223827 | 224786 | 72.89719626 | 324 | 319 | 1.015673981 | 1 | 1 | 1 | FIM1_1797 | YKL120W | YKL120W | OAC1 | n |
| Kmar_QGN15107.1 | W303.asm02.HP0_G0041410 | OG0002275 | chrXIII | 466805 | 468667 | CP015056.1 | 219003 | 220925 | 34.10138249 | 620 | 640 | 0.96875 | 1 | 1 | 1 | FIM1_1794 | YMR100W | YMR100W | MUB1 | n |
| Kmar_QGN13635.1 | W303.asm02.HP0_G0014300 | OG0002276 | chrV | 121680 | 122483 | CP015054.1 | 559146 | 559907 | 27.51937984 | 267 | 253 | 1.055335968 | 1 | 1 | 1 | FIM1_277 | YEL019C | YEL019C | MMS21 | n |
| Kmar_QGN18391.1 | W303.asm02.HP0_G0015820 | OG0002277 | chrV | 416502 | 417575 | CP015061.1 | 838715 | 839746 | 50.96418733 | 357 | 343 | 1.040816327 | 1 | 1 | 1 | FIM1_4718 | YER127W | YER127W | LCP5 | n |
| Kmar_QGN16344.1 | W303.asm02.HP0_G0040300 | OG0002278 | chrXIII | 230242 | 231321 | CP015057.1 | 1333336 | 1334382 | 52.7027027 | 359 | 348 | 1.031609195 | 1 | 1 | 1 | FIM1_3050 | YML021C | YML021C | UNG1 | n |
| Kmar_QGN16347.1 | W303.asm02.HP0_G0040320 | OG0002279 | chrXIII | 233964 | 234962 | CP015057.1 | 1337922 | 1338917 | 36.68639053 | 332 | 331 | 1.003021148 | 1 | 1 | 1 | FIM1_3053 | YML019W | YML019W | OST6 | n |
| Kmar_QGN14908.1 | W303.asm02.HP0_G0027720 | OG0002280 | chrX | 11504 | 12094 | CP015055.1 | 1472572 | 1473141 | 47.47474747 | 196 | 189 | 1.037037037 | 1 | 1 | 1 | FIM1_1582 | YJL218W | YJL218W | | n |
| Kmar_QGN15048.1 | W303.asm02.HP0_G0031260 | OG0002281 | chrXI | 40188 | 43262 | CP015056.1 | 88268 | 91330 | 76.85546875 | 1024 | 1020 | 1.003921569 | 1 | 1 | 1 | FIM1_1733 | YKL210W | YKL210W | UBA1 | n |
| Kmar_QGN15049.1 | W303.asm02.HP0_G0031270 | OG0002282 | chrXI | 43448 | 47320 | CP015056.1 | 91562 | 95452 | 47.07668945 | 1290 | 1296 | 0.99537037 | 1 | 1 | 1 | FIM1_1734 | YKL209C | YKL209C | STE6 | n |
| Kmar_QGN18356.1 | W303.asm02.HP0_G0017660 | OG0002283 | chrVII | 32303 | 32896 | CP015061.1 | 759436 | 759807 | 21.22905028 | 197 | 123 | 1.601626016 | 1 | 1 | 1 | FIM1_4683 | YGL247W | YGL247W | BRR6 | n |
| Kmar_QGN17812.1 | W303.asm02.HP0_G0052290 | OG0002284 | chrXV | 999494 | 1000525 | CP015060.1 | 536492 | 537541 | 36.82795699 | 343 | 349 | 0.982808023 | 1 | 1 | 1 | FIM1_5021 | YOR352W | YOR352W | TFB6 | n |
| Kmar_QGN15050.1 | W303.asm02.HP0_G0030270 | OG0002285 | chrX | 581602 | 582222 | CP015056.1 | 96264 | 96917 | 56.68202765 | 206 | 217 | 0.949308756 | 1 | 1 | 1 | FIM1_1735 | YJR073C | YJR073C | OPI3 | n |
| Kmar_QGN15768.1 | W303.asm02.HP0_G0047220 | OG0002286 | chrXIV | 726630 | 727970 | CP015057.1 | 51719 | 53056 | 85.87443946 | 446 | 445 | 1.002247191 | 1 | 1 | 1 | FIM1_2464 | YNR050C | YNR050C | LYS9 | n |
| Kmar_QGN16178.1 | W303.asm02.HP0_G0042510 | OG0002287 | chrXIII | 689662 | 691089 | CP015057.1 | 969224 | 970636 | 45.28688525 | 475 | 470 | 1.010638298 | 1 | 1 | 1 | FIM1_2880 | YMR211W | YMR211W | DML1 | n |
| Kmar_QGN14938.1 | W303.asm02.HP0_G0045170 | OG0002288 | chrXIV | 322911 | 324257 | CP015055.1 | 1533083 | 1534210 | 29.3986637 | 448 | 375 | 1.194666667 | 1 | 1 | 1 | FIM1_1613 | YNL166C | YNL166C | BNI5 | n |
| Kmar_QGN14426.1 | W303.asm02.HP0_G0018890 | OG0002289 | chrVII | 292884 | 294890 | CP015055.1 | 495414 | 497303 | 30.04418262 | 668 | 629 | 1.06200318 | 1 | 1 | 1 | FIM1_1087 | YGL113W | YGL113W | SLD3 | n |
| Kmar_QGN14427.1 | W303.asm02.HP0_G0018900 | OG0002290 | chrVII | 295126 | 296676 | CP015055.1 | 497404 | 498966 | 62.83018868 | 516 | 520 | 0.992307692 | 1 | 1 | 1 | FIM1_1088 | YGL112C | YGL112C | TAF6 | n |
| Kmar_QGN16954.1 | W303.asm02.HP0_G0006660 | OG0002291 | chrIV | 42773 | 43780 | CP015058.1 | 1250224 | 1251222 | 46.64723032 | 335 | 332 | 1.009036145 | 1 | 1 | 1 | FIM1_3681 | YDL230W | YDL230W | PTP1 | n |
| Kmar_QGN16951.1 | W303.asm02.HP0_G0044830 | OG0002292 | chrXIV | 256052 | 257329 | CP015058.1 | 1244669 | 1245919 | 75.29411765 | 425 | 416 | 1.021634615 | 1 | 1 | 1 | FIM1_3678 | YNL207W | YNL207W | RIO2 | n |
| Kmar_QGN18048.1 | W303.asm02.HP0_G0038570 | OG0002293 | chrXII | 903749 | 904588 | CP015061.1 | 98027 | 98881 | 43.85964912 | 279 | 284 | 0.982394366 | 1 | 1 | 1 | FIM1_4365 | YLR393W | YLR393W | ATP10 | n |
| Kmar_QGN17249.1 | W303.asm02.HP0_G0022960 | OG0002294 | chrVIII | 73754 | 74119 | CP015059.1 | 548339 | 548692 | 79.16666667 | 121 | 117 | 1.034188034 | 1 | 1 | 1 | FIM1_3980 | YHL015W | YHL015W | RPS20 | n |
| Kmar_QGN17789.1 | W303.asm02.HP0_G0049900 | OG0002295 | chrXV | 491858 | 493576 | CP015060.1 | 485914 | 487566 | 53.72616984 | 572 | 550 | 1.04 | 1 | 1 | 1 | FIM1_4998 | YOR090C | YOR090C | PTC5 | n |
| Kmar_QGN16353.1 | W303.asm02.HP0_G0012790 | OG0002296 | chrIV | 1338539 | 1339525 | CP015057.1 | 1347692 | 1348690 | 47.14714715 | 328 | 332 | 0.987951807 | 1 | 1 | 1 | FIM1_3059 | YDR435C | YDR435C | PPM1 | n |
| Kmar_QGN16356.1 | W303.asm02.HP0_G0040370 | OG0002297 | chrXIII | 243731 | 244570 | CP015057.1 | 1352256 | 1353080 | 64.41281139 | 279 | 274 | 1.018248175 | 1 | 1 | 1 | FIM1_3062 | YML014W | YML014W | TRM9 | n |
| Kmar_QGN15855.1 | W303.asm02.HP0_G0034670 | OG0002298 | chrXII | 88632 | 91358 | CP015057.1 | 229095 | 231791 | 76.31578947 | 908 | 898 | 1.011135857 | 1 | 1 | 1 | FIM1_2552 | YLL026W | YLL026W | HSP104 | n |
| Kmar_QGN15325.1 | W303.asm02.HP0_G0018270 | OG0002299 | chrVII | 153419 | 154609 | CP015056.1 | 708978 | 710069 | 35.1010101 | 396 | 363 | 1.090909091 | 1 | 1 | 1 | FIM1_2014 | YGL181W | YGL181W | GTS1 | n |
| Kmar_QGN16555.1 | W303.asm02.HP0_G0039950 | OG0002300 | chrXIII | 147759 | 152798 | CP015058.1 | 360757 | 365196 | 44.7752481 | 1679 | 1479 | 1.135226504 | 1 | 1 | 1 | FIM1_3269 | YML059C | YML059C | NTE1 | n |
| Kmar_QGN15245.1 | W303.asm02.HP0_G0000510 | OG0002301 | chrI | 114378 | 115379 | CP015056.1 | 518297 | 519412 | 35.71428571 | 333 | 371 | 0.897574124 | 1 | 1 | 1 | FIM1_1934 | YAL020C | YAL020C | ATS1 | n |
| Kmar_QGN14548.1 | W303.asm02.HP0_G0048750 | OG0002302 | chrXV | 265884 | 266624 | CP015055.1 | 737743 | 738495 | 64.96062992 | 246 | 250 | 0.984 | 1 | 1 | 1 | FIM1_1210 | YOL032W | YOL032W | OPI10 | n |
| Kmar_QGN14550.1 | W303.asm02.HP0_G0000590 | OG0002303 | chrI | 131565 | 132749 | CP015055.1 | 740269 | 741441 | 83.50253807 | 394 | 390 | 1.01025641 | 1 | 1 | 1 | FIM1_1212 | YAL012W | YAL012W | CYS3 | n |
| Kmar_QGN14549.1 | W303.asm02.HP0_G0048760 | OG0002304 | chrXV | 266719 | 267984 | CP015055.1 | 738531 | 739781 | 34.72222222 | 421 | 416 | 1.012019231 | 1 | 1 | 1 | FIM1_1211 | YOL031C | YOL031C | SIL1 | n |
| Kmar_QGN14566.1 | W303.asm02.HP0_G0048610 | OG0002305 | chrXV | 239074 | 240549 | CP015055.1 | 771488 | 772951 | 51.90380762 | 491 | 487 | 1.008213552 | 1 | 1 | 1 | FIM1_1228 | YOL049W | YOL049W | GSH2 | n |
| Kmar_QGN15401.1 | W303.asm02.HP0_G0045950 | OG0002306 | chrXIV | 477815 | 478246 | CP015056.1 | 884952 | 885365 | 74.82517483 | 143 | 137 | 1.04379562 | 1 | 1 | 1 | FIM1_2091 | YNL081C | YNL081C | SWS2 | n |
| Kmar_QGN18057.1 | W303.asm02.HP0_G0038510 | OG0002307 | chrXII | 893044 | 894342 | CP015061.1 | 113665 | 114879 | 47.46543779 | 432 | 404 | 1.069306931 | 1 | 1 | 1 | FIM1_4374 | YLR387C | YLR387C | REH1 | n |
| Kmar_QGN17418.1 | W303.asm02.HP0_G0045960 | OG0002308 | chrXIV | 478558 | 479658 | CP015059.1 | 874264 | 875274 | 43.25068871 | 366 | 336 | 1.089285714 | 1 | 1 | 1 | FIM1_4150 | YNL080C | YNL080C | EOS1 | n |
| Kmar_QGN18056.1 | W303.asm02.HP0_G0008220 | OG0002309 | chrIV | 341949 | 344315 | CP015061.1 | 110993 | 113374 | 64.16978777 | 788 | 793 | 0.99369483 | 1 | 1 | 1 | FIM1_4373 | YDL060W | YDL060W | TSR1 | n |
| Kmar_QGN18052.1 | W303.asm02.HP0_G0008200 | OG0002310 | chrIV | 338602 | 340464 | CP015061.1 | 102472 | 104319 | 46.14173228 | 620 | 615 | 1.008130081 | 1 | 1 | 1 | FIM1_4369 | YDL063C | YDL063C | SYO1 | n |
| Kmar_QGN16630.1 | W303.asm02.HP0_G0028260 | OG0002311 | chrX | 116381 | 118480 | CP015058.1 | 539388 | 541535 | 48.63387978 | 699 | 715 | 0.977622378 | 1 | 1 | 1 | FIM1_3347 | YJL156C | YJL156C | SSY5 | n |
| Kmar_QGN16631.1 | W303.asm02.HP0_G0028270 | OG0002312 | chrX | 118813 | 120171 | CP015058.1 | 542828 | 544198 | 69.07894737 | 452 | 456 | 0.99122807 | 1 | 1 | 1 | FIM1_3348 | YJL155C | YJL155C | FBP26 | n |
| Kmar_QGN18051.1 | W303.asm02.HP0_G0008190 | OG0002313 | chrIV | 337817 | 338400 | CP015061.1 | 101828 | 102301 | 83.43949045 | 157 | 157 | 1 | 1 | 1 | 1 | FIM1_4368 | YDL064W | YDL064W | UBC9 | n |
| Kmar_QGN16215.1 | W303.asm02.HP0_G0038620 | OG0002314 | chrXII | 911826 | 915689 | CP015057.1 | 1051301 | 1055098 | 56.90184049 | 1287 | 1265 | 1.017391304 | 1 | 1 | 1 | FIM1_2919 | YLR398C | YLR398C | SKI2 | n |
| Kmar_QGN16217.1 | W303.asm02.HP0_G0038610 | OG0002315 | chrXII | 909220 | 911562 | CP015057.1 | 1056019 | 1058355 | 69.13265306 | 780 | 778 | 1.002570694 | 1 | 1 | 1 | FIM1_2921 | YLR397C | YLR397C | AFG2 | n |
| Kmar_QGN18050.1 | W303.asm02.HP0_G0008180 | OG0002316 | chrIV | 336579 | 337631 | CP015061.1 | 100623 | 101633 | 37.74647887 | 350 | 336 | 1.041666667 | 1 | 1 | 1 | FIM1_4367 | YDL065C | YDL065C | PEX19 | n |
| Kmar_QGN13894.1 | W303.asm02.HP0_G0024940 | OG0002317 | chrVIII | 443276 | 445156 | CP015054.1 | 1099730 | 1101628 | 46.8412943 | 626 | 632 | 0.990506329 | 1 | 1 | 1 | FIM1_541 | YHR166C | YHR166C | CDC23 | n |
| Kmar_QGN13703.1 | W303.asm02.HP0_G0043360 | OG0002318 | chrXIII | 866135 | 867811 | CP015054.1 | 692182 | 693843 | 60.502693 | 558 | 553 | 1.009041591 | 1 | 1 | 1 | FIM1_346 | YMR296C | YMR296C | LCB1 | n |
| Kmar_QGN13893.1 | W303.asm02.HP0_G0024930 | OG0002319 | chrVIII | 435813 | 443054 | CP015054.1 | 1091998 | 1099248 | 71.95222405 | 2413 | 2416 | 0.998758278 | 1 | 1 | 1 | FIM1_540 | YHR165C | YHR165C | PRP8 | n |
| Kmar_QGN15857.1 | W303.asm02.HP0_G0022880 | OG0002320 | chrVIII | 57327 | 60767 | CP015057.1 | 240962 | 243562 | 37.17277487 | 1146 | 866 | 1.323325635 | 1 | 1 | 1 | FIM1_2554 | YHL023C | YHL023C | NPR3 | n |
| Kmar_QGN17420.1 | W303.asm02.HP0_G0048310 | OG0002321 | chrXV | 181013 | 181882 | CP015059.1 | 884951 | 885820 | 56.80272109 | 289 | 289 | 1 | 1 | 1 | 1 | FIM1_4152 | YOL080C | YOL080C | REX4 | n |
| Kmar_QGN17421.1 | W303.asm02.HP0_G0048320 | OG0002322 | chrXV | 182137 | 185667 | CP015059.1 | 886448 | 889723 | 34.63166808 | 1176 | 1091 | 1.077910174 | 1 | 1 | 1 | FIM1_4153 | YOL078W | YOL078W | AVO1 | n |
| Kmar_QGN16640.1 | W303.asm02.HP0_G0048130 | OG0002323 | chrXV | 136982 | 138280 | CP015058.1 | 568511 | 569809 | 82.15102975 | 432 | 432 | 1 | 1 | 1 | 1 | FIM1_3359 | YOL097C | YOL097C | WRS1 | n |
| Kmar_QGN15229.1 | W303.asm02.HP0_G0035140 | OG0002324 | chrXII | 191013 | 191552 | CP015056.1 | 481991 | 482476 | 42.45810056 | 179 | 161 | 1.111801242 | 1 | 1 | 1 | FIM1_1918 | YLR021W | YLR021W | IRC25 | n |
| Kmar_QGN16641.1 | W303.asm02.HP0_G0013260 | OG0002325 | chrIV | 1434048 | 1434674 | CP015058.1 | 569990 | 570622 | 67.61904762 | 208 | 210 | 0.99047619 | 1 | 1 | 1 | FIM1_3360 | YDR487C | YDR487C | RIB3 | n |
| Kmar_QGN13667.1 | W303.asm02.HP0_G0048860 | OG0002326 | chrXV | 282876 | 285881 | CP015054.1 | 626306 | 629338 | 76.77165354 | 1001 | 1010 | 0.991089109 | 1 | 1 | 1 | FIM1_310 | YOL021C | YOL021C | DIS3 | n |
| Kmar_QGN15977.1 | W303.asm02.HP0_G0003380 | OG0002327 | chrII | 551546 | 552382 | CP015057.1 | 530878 | 531756 | 60.47297297 | 278 | 292 | 0.952054795 | 1 | 1 | 1 | FIM1_2677 | YBR146W | YBR146W | MRPS9 | n |
| Kmar_QGN17424.1 | W303.asm02.HP0_G0048340 | OG0002328 | chrXV | 186303 | 187178 | CP015059.1 | 892541 | 893419 | 83.56164384 | 291 | 292 | 0.996575342 | 1 | 1 | 1 | FIM1_4156 | YOL077C | YOL077C | BRX1 | n |
| Kmar_QGN15630.1 | W303.asm02.HP0_G0033330 | OG0002329 | chrXI | 459991 | 460818 | CP015056.1 | 1346703 | 1347473 | 33.57142857 | 275 | 256 | 1.07421875 | 1 | 1 | 1 | FIM1_2323 | YKR006C | YKR006C | MRPL13 | n |
| Kmar_QGN15108.1 | W303.asm02.HP0_G0032100 | OG0002330 | chrXI | 226429 | 227076 | CP015056.1 | 221271 | 221888 | 37.03703704 | 215 | 205 | 1.048780488 | 1 | 1 | 1 | FIM1_1795 | YKL119C | YKL119C | VPH2 | n |
| Kmar_QGN14047.1 | W303.asm02.HP0_G0027920 | OG0002331 | chrX | 58869 | 60410 | CP015054.1 | 1402802 | 1404385 | 36.93693694 | 513 | 527 | 0.973434535 | 1 | 1 | 1 | FIM1_698 | YJL194W | YJL194W | CDC6 | n |
| Kmar_QGN15044.1 | W303.asm02.HP0_G0031250 | OG0002332 | chrXI | 37724 | 39178 | CP015056.1 | 81315 | 82793 | 69.73947896 | 484 | 492 | 0.983739837 | 1 | 1 | 1 | FIM1_1729 | YKL211C | YKL211C | TRP3 | n |
| Kmar_QGN16623.1 | W303.asm02.HP0_G0031670 | OG0002333 | chrXI | 139100 | 141859 | CP015058.1 | 519354 | 522119 | 69.6969697 | 919 | 921 | 0.997828447 | 1 | 1 | 1 | FIM1_3340 | YKL165C | YKL165C | MCD4 | n |
| Kmar_QGN15767.1 | W303.asm02.HP0_G0006350 | OG0002334 | chrIII | 311762 | 312850 | CP015057.1 | 49498 | 50580 | 44.38642298 | 362 | 360 | 1.005555556 | 1 | 1 | 1 | FIM1_2463 | YCR095C | YCR095C | OCA4 | n |
| Kmar_QGN15769.1 | W303.asm02.HP0_G0047210 | OG0002335 | chrXIV | 725605 | 726237 | CP015057.1 | 53457 | 54080 | 30.04694836 | 210 | 207 | 1.014492754 | 1 | 1 | 1 | FIM1_2465 | YNR049C | YNR049C | MSO1 | n |
| Kmar_QGN16287.1 | W303.asm02.HP0_G0031450 | OG0002336 | chrXI | 87394 | 89955 | CP015057.1 | 1209635 | 1212088 | 48.00884956 | 853 | 817 | 1.044063647 | 1 | 1 | 1 | FIM1_2992 | YKL188C | YKL188C | PXA2 | n |
| Kmar_QGN15488.1 | W303.asm02.HP0_G0026490 | OG0002337 | chrIX | 208358 | 211195 | CP015056.1 | 1047301 | 1050105 | 64.75583864 | 945 | 934 | 1.011777302 | 1 | 1 | 1 | FIM1_2179 | YIL075C | YIL075C | RPN2 | n |
| Kmar_QGN18254.1 | W303.asm02.HP0_G0056910 | OG0002338 | chrXVI | 883908 | 884381 | CP015061.1 | 561052 | 561504 | 62.5 | 157 | 150 | 1.046666667 | 1 | 1 | 1 | FIM1_4579 | YPR168W | YPR168W | NUT2 | n |
| Kmar_QGN14125.1 | W303.asm02.HP0_G0036680 | OG0002339 | chrXII | 516807 | 517799 | CP015054.1 | 1589825 | 1590922 | 32.43967828 | 330 | 365 | 0.904109589 | 1 | 1 | 1 | FIM1_778 | YLR181C | YLR181C | VTA1 | n |
| Kmar_QGN13538.1 | W303.asm02.HP0_G0048380 | OG0002340 | chrXV | 195425 | 196792 | CP015054.1 | 360125 | 361504 | 40.30172414 | 455 | 459 | 0.991285403 | 1 | 1 | 1 | FIM1_178 | YOL072W | YOL072W | THP1 | n |
| Kmar_QGN13536.1 | W303.asm02.HP0_G0020520 | OG0002341 | chrVII | 632206 | 633369 | CP015054.1 | 357307 | 358536 | 16.78832117 | 387 | 409 | 0.946210269 | 1 | 1 | 1 | FIM1_176 | YGR072W | YGR072W | UPF3 | n |
| Kmar_QGN14530.1 | W303.asm02.HP0_G0056280 | OG0002342 | chrXVI | 736290 | 736712 | CP015055.1 | 706193 | 706621 | 67.6056338 | 140 | 142 | 0.985915493 | 1 | 1 | 1 | FIM1_1192 | YPR100W | YPR100W | MRPL51 | n |
| Kmar_QGN13587.1 | W303.asm02.HP0_G0028940 | OG0002343 | chrX | 275576 | 279244 | CP015054.1 | 471244 | 474927 | 46.88 | 1222 | 1227 | 0.99592502 | 1 | 1 | 1 | FIM1_229 | YJL080C | YJL080C | SCP160 | n |
| Kmar_QGN17536.1 | W303.asm02.HP0_G0028550 | OG0002344 | chrX | 174518 | 175441 | CP015059.1 | 1123071 | 1123973 | 61.56351792 | 307 | 300 | 1.023333333 | 1 | 1 | 1 | FIM1_4272 | YJL126W | YJL126W | NIT2 | n |
| Kmar_QGN16673.1 | W303.asm02.HP0_G0036770 | OG0002345 | chrXII | 537400 | 538560 | CP015058.1 | 644079 | 645299 | 48.62155388 | 386 | 406 | 0.950738916 | 1 | 1 | 1 | FIM1_3392 | YLR191W | YLR191W | PEX13 | n |
| Kmar_QGN15957.1 | W303.asm02.HP0_G0021290 | OG0002346 | chrVII | 812273 | 813550 | CP015057.1 | 467680 | 469011 | 35.97560976 | 425 | 443 | 0.959367946 | 1 | 1 | 1 | FIM1_2655 | YGR156W | YGR156W | PTI1 | n |
| Kmar_QGN16789.1 | W303.asm02.HP0_G0056360 | OG0002347 | chrXVI | 748953 | 750242 | CP015058.1 | 891979 | 893205 | 69.46386946 | 429 | 408 | 1.051470588 | 1 | 1 | 1 | FIM1_3512 | YPR108W | YPR108W | RPN7 | n |
| Kmar_QGN17540.1 | W303.asm02.HP0_G0028570 | OG0002348 | chrX | 177143 | 177661 | CP015059.1 | 1131075 | 1131611 | 66.29213483 | 172 | 178 | 0.966292135 | 1 | 1 | 1 | FIM1_4276 | YJL124C | YJL124C | LSM1 | n |
| Kmar_QGN17537.1 | W303.asm02.HP0_G0028560 | OG0002349 | chrX | 175545 | 176696 | CP015059.1 | 1124012 | 1125238 | 64.76683938 | 383 | 408 | 0.93872549 | 1 | 1 | 1 | FIM1_4273 | YJL125C | YJL125C | GCD14 | n |

| | | | | | | | | | | | | | | | | | | | | |
|---|---|---|---|---|---|---|---|---|---|---|---|---|---|---|---|---|---|---|---|---|
| Kmar_QGN16494.1 | W303.asm02.HP0_G0052080 | OG0002350 | chrXV | 938451 | 941069 | CP015058.1 | 227486 | 229639 | 34.64566929 | 872 | 717 | 1.216178522 | 1 | 1 | 1 | FIM1_3207 | YOR329C | YOR329C | SCD5 | n |
| Kmar_QGN14845.1 | W303.asm02.HP0_G0039280 | OG0002351 | chrXIII | 9687 | 11228 | CP015055.1 | 1337574 | 1339028 | 39.84526112 | 513 | 484 | 1.059917355 | 1 | 1 | 1 | FIM1_1516 | YML128C | YML128C | MSC1 | n |
| Kmar_QGN17109.1 | W303.asm02.HP0_G0039300 | OG0002352 | chrXIII | 13612 | 15087 | CP015059.1 | 252051 | 252161 | 12.29946524 | 491 | 36 | 13.63888889 | 1 | 1 | 1 | FIM1_3840 | YML126C | YML126C | ERG13 | n |
| Kmar_QGN17830.1 | W303.asm02.HP0_G0040770 | OG0002353 | chrXIII | 329163 | 330734 | CP015060.1 | 578448 | 580181 | 29.54159593 | 523 | 577 | 0.906412478 | 1 | 1 | 1 | FIM1_5039 | YMR029C | YMR029C | FAR8 | n |
| Kmar_QGN18211.1 | W303.asm02.HP0_G0053230 | OG0002354 | chrXVI | 131571 | 132011 | CP015061.1 | 458194 | 458631 | 79.31034483 | 146 | 145 | 1.006896552 | 1 | 1 | 1 | FIM1_4535 | YPL225W | YPL225W | CHP1 | n |
| Kmar_QGN13540.1 | W303.asm02.HP0_G0002230 | OG0002355 | chrII | 305666 | 306553 | CP015054.1 | 363937 | 364878 | 60.50955414 | 295 | 313 | 0.942492013 | 1 | 1 | 1 | FIM1_180 | YBR037C | YBR037C | SCO1 | n |
| Kmar_QGN17862.1 | W303.asm02.HP0_G0003010 | OG0002356 | chrII | 468944 | 469510 | CP015060.1 | 633330 | 633902 | 72.10526316 | 188 | 190 | 0.989473684 | 1 | 1 | 1 | FIM1_5071 | YBR106W | YBR106W | SND3 | n |
| Kmar_QGN16136.1 | W303.asm02.HP0_G0040700 | OG0002357 | chrXIII | 319183 | 319680 | CP015057.1 | 889524 | 890021 | 83.63636364 | 165 | 165 | 1 | 1 | 1 | 1 | FIM1_2837 | YMR022W | YMR022W | UBC7 | n |
| Kmar_QGN18209.1 | W303.asm02.HP0_G0053220 | OG0002358 | chrXVI | 127332 | 130922 | CP015061.1 | 453344 | 456895 | 72.66666667 | 1196 | 1183 | 1.010989011 | 1 | 1 | 1 | FIM1_4533 | YPL226W | YPL226W | NEW1 | n |
| Kmar_QGN14689.1 | W303.asm02.HP0_G0057000 | OG0002359 | chrXVI | 898164 | 899561 | CP015055.1 | 1037709 | 1039127 | 35.74380165 | 465 | 472 | 0.985169492 | 1 | 1 | 1 | FIM1_1356 | YPR178W | YPR178W | PRP4 | n |
| Kmar_QGN14688.1 | W303.asm02.HP0_G0057010 | OG0002360 | chrXVI | 899629 | 901596 | CP015055.1 | 1035722 | 1037632 | 46.93251534 | 655 | 636 | 1.029874214 | 1 | 1 | 1 | FIM1_1355 | YPR179C | YPR179C | HDA3 | n |
| Kmar_QGN17481.1 | W303.asm02.HP0_G0021770 | OG0002361 | chrVII | 921628 | 922500 | CP015059.1 | 1008364 | 1009194 | 43.15068493 | 290 | 276 | 1.050724638 | 1 | 1 | 1 | FIM1_4217 | YGR205W | YGR205W | TDA10 | n |
| Kmar_QGN15716.1 | W303.asm02.HP0_G0017750 | OG0002362 | chrVII | 48101 | 48898 | CP015056.1 | 1523411 | 1524160 | 32.83018868 | 265 | 249 | 1.064257028 | 1 | 1 | 1 | FIM1_2409 | YGL237C | YGL237C | HAP2 | n |
| Kmar_QGN15717.1 | W303.asm02.HP0_G0017740 | OG0002363 | chrVII | 44922 | 47804 | CP015056.1 | 1524275 | 1527157 | 63.55140187 | 960 | 960 | 1 | 1 | 1 | 1 | FIM1_2410 | YGL238W | YGL238W | CSE1 | n |
| Kmar_QGN13696.1 | W303.asm02.HP0_G0043400 | OG0002364 | chrXIII | 872479 | 874011 | CP015054.1 | 678528 | 680057 | 81.56862745 | 510 | 509 | 1.001964637 | 1 | 1 | 1 | FIM1_339 | YMR300C | YMR300C | ADE4 | n |
| Kmar_QGN16099.1 | W303.asm02.HP0_G0015030 | OG0002365 | chrV | 257648 | 259232 | CP015057.1 | 812321 | 813889 | 66.9793621 | 515 | 522 | 0.986590038 | 1 | 1 | 1 | FIM1_2800 | YER052C | YER052C | HOM3 | n |
| Kmar_QGN14701.1 | W303.asm02.HP0_G0026800 | OG0002366 | chrIX | 267154 | 268134 | CP015055.1 | 1059678 | 1060583 | 51.61290323 | 326 | 301 | 1.083056478 | 1 | 1 | 1 | FIM1_1368 | YIL041W | YIL041W | GVP36 | n |
| Kmar_QGN17135.1 | W303.asm02.HP0_G0024660 | OG0002367 | chrVIII | 381682 | 383223 | CP015059.1 | 310404 | 311996 | 42.04753199 | 513 | 530 | 0.967924528 | 1 | 1 | 1 | FIM1_3866 | YHR137W | YHR137W | ARO9 | n |
| Kmar_QGN14350.1 | W303.asm02.HP0_G0041270 | OG0002368 | chrXIII | 439714 | 442596 | CP015055.1 | 328693 | 331674 | 19.31034483 | 960 | 993 | 0.966767372 | 1 | 1 | 1 | FIM1_1010 | YMR086W | YMR086W | SEG1 | n |
| Kmar_QGN16846.1 | W303.asm02.HP0_G0026990 | OG0002369 | chrIX | 303534 | 304490 | CP015058.1 | 1018732 | 1019655 | 74.52830189 | 318 | 307 | 1.035830619 | 1 | 1 | 1 | FIM1_3571 | YIL021W | YIL021W | RPB3 | n |
| Kmar_QGN16845.1 | W303.asm02.HP0_G0027000 | OG0002370 | chrIX | 304664 | 305449 | CP015058.1 | 1017670 | 1018521 | 60 | 261 | 283 | 0.922261484 | 1 | 1 | 1 | FIM1_3570 | YIL020C | YIL020C | HIS6 | n |
| Kmar_QGN16844.1 | W303.asm02.HP0_G0027010 | OG0002371 | chrIX | 305722 | 306762 | CP015058.1 | 1016590 | 1017564 | 40.82191781 | 346 | 324 | 1.067901235 | 1 | 1 | 1 | FIM1_3569 | YIL019W | YIL019W | FAF1 | n |
| Kmar_QGN18126.1 | W303.asm02.HP0_G0001000 | OG0002372 | chrII | 41640 | 43904 | CP015061.1 | 275973 | 278309 | 44.15422886 | 754 | 778 | 0.969151671 | 1 | 1 | 1 | FIM1_4447 | YBL097W | YBL097W | BRN1 | n |
| Kmar_QGN18125.1 | W303.asm02.HP0_G0055630 | OG0002373 | chrXVI | 616979 | 619222 | CP015061.1 | 273353 | 275611 | 71.42857143 | 747 | 752 | 0.993351064 | 1 | 1 | 1 | FIM1_4446 | YPR024W | YPR024W | YME1 | n |
| Kmar_QGN16843.1 | W303.asm02.HP0_G0017260 | OG0002374 | chrVI | 193925 | 194863 | CP015058.1 | 1013386 | 1015488 | 16.21621622 | 312 | 700 | 0.445714286 | 1 | 1 | 1 | FIM1_3568 | YFR034C | YFR034C | PHO4 | n |
| Kmar_QGN16304.1 | W303.asm02.HP0_G0018720 | OG0002375 | chrVII | 258116 | 262327 | CP015057.1 | 1239901 | 1244442 | 32.89986996 | 1403 | 1513 | 0.927296761 | 1 | 1 | 1 | FIM1_3009 | YGL131C | YGL131C | SNT2 | n |
| Kmar_QGN16305.1 | W303.asm02.HP0_G0053350 | OG0002376 | chrXVI | 156169 | 156885 | CP015057.1 | 1244504 | 1245226 | 46.47302905 | 238 | 240 | 0.991666667 | 1 | 1 | 1 | FIM1_3010 | YPL213W | YPL213W | LEA1 | n |
| Kmar_QGN16303.1 | W303.asm02.HP0_G0018730 | OG0002377 | chrVII | 262613 | 263992 | CP015057.1 | 1238040 | 1239461 | 68.6695279 | 459 | 473 | 0.970401691 | 1 | 1 | 1 | FIM1_3008 | YGL130W | YGL130W | CEG1 | n |
| Kmar_QGN13904.1 | W303.asm02.HP0_G0042580 | OG0002378 | chrXIII | 704158 | 707466 | CP015054.1 | 1119138 | 1122341 | 26.46264626 | 1102 | 1067 | 1.032802249 | 1 | 1 | 1 | FIM1_552 | YMR218C | YMR218C | TRS130 | n |
| Kmar_QGN16826.1 | W303.asm02.HP0_G0020870 | OG0002379 | chrVII | 715611 | 716813 | CP015058.1 | 978232 | 979434 | 44.03892944 | 400 | 400 | 1 | 1 | 1 | 1 | FIM1_3550 | YGR111W | YGR111W | | n |
| Kmar_QGN17136.1 | W303.asm02.HP0_G0045310 | OG0002380 | chrXIV | 346361 | 346750 | CP015059.1 | 312347 | 312736 | 69.76744186 | 129 | 129 | 1 | 1 | 1 | 1 | FIM1_3867 | YNL153C | YNL153C | GIM3 | n |
| Kmar_QGN17698.1 | W303.asm02.HP0_G0046490 | OG0002381 | chrXIV | 595503 | 596474 | CP015060.1 | 272884 | 273795 | 66.35802469 | 323 | 303 | 1.066006601 | 1 | 1 | 1 | FIM1_4905 | YNL025C | YNL025C | SSN8 | n |
| Kmar_QGN17687.1 | W303.asm02.HP0_G0026450 | OG0002382 | chrIX | 194051 | 195148 | CP015060.1 | 257304 | 258440 | 62.20472441 | 365 | 378 | 0.965608466 | 1 | 1 | 1 | FIM1_4894 | YIL083C | YIL083C | CAB2 | n |
| Kmar_QGN13425.1 | W303.asm02.HP0_G0054610 | OG0002383 | chrXVI | 403060 | 404463 | CP015054.1 | 120247 | 121575 | 33.68200837 | 467 | 442 | 1.056561086 | 1 | 1 | 1 | FIM1_62 | YPL083C | YPL083C | SEN54 | n |
| Kmar_QGN17713.1 | W303.asm02.HP0_G0026310 | OG0002384 | chrIX | 170785 | 172335 | CP015060.1 | 299520 | 300929 | 39.26499033 | 516 | 469 | 1.10021322 | 1 | 1 | 1 | FIM1_4920 | YIL097W | YIL097W | FYV10 | n |
| Kmar_QGN17712.1 | W303.asm02.HP0_G0026320 | OG0002385 | chrIX | 172475 | 173485 | CP015060.1 | 298510 | 299484 | 49.10714286 | 336 | 324 | 1.037037037 | 1 | 1 | 1 | FIM1_4919 | YIL096C | YIL096C | BMT5 | n |
| Kmar_QGN17685.1 | W303.asm02.HP0_G0007640 | OG0002386 | chrIV | 242758 | 245187 | CP015060.1 | 252118 | 254484 | 52.1945433 | 809 | 788 | 1.026649746 | 1 | 1 | 1 | FIM1_4892 | YDL122W | YDL122W | UBP1 | n |
| Kmar_QGN17686.1 | W303.asm02.HP0_G0046580 | OG0002387 | chrXIV | 614090 | 615451 | CP015060.1 | 255089 | 256459 | 52.72331155 | 453 | 456 | 0.993421053 | 1 | 1 | 1 | FIM1_4893 | YNL016W | YNL016W | PUB1 | n |
| Kmar_QGN18355.1 | W303.asm02.HP0_G0054910 | OG0002388 | chrXVI | 467138 | 468325 | CP015061.1 | 757686 | 758864 | 71.50127226 | 395 | 392 | 1.007653061 | 1 | 1 | 1 | FIM1_4682 | YPL050C | YPL050C | MNN9 | n |
| Kmar_QGN13412.1 | W303.asm02.HP0_G0054510 | OG0002389 | chrXVI | 377336 | 379279 | CP015054.1 | 86736 | 88667 | 83.64197531 | 647 | 643 | 1.00622084 | 1 | 1 | 1 | FIM1_49 | YPL093W | YPL093W | NOG1 | n |
| Kmar_QGN15357.1 | W303.asm02.HP0_G0054870 | OG0002390 | chrXVI | 462118 | 463023 | CP015056.1 | 786525 | 787985 | 24.52025586 | 301 | 486 | 0.619341564 | 1 | 1 | 1 | FIM1_2047 | YPL054W | YPL054W | LEE1 | n |
| Kmar_QGN13423.1 | W303.asm02.HP0_G0003780 | OG0002391 | chrII | 615898 | 616734 | CP015054.1 | 116532 | 117350 | 54.96453901 | 278 | 272 | 1.022058824 | 1 | 1 | 1 | FIM1_60 | YBR185C | YBR185C | MBA1 | n |
| Kmar_QGN13421.1 | W303.asm02.HP0_G0054580 | OG0002392 | chrXVI | 391131 | 392804 | CP015054.1 | 106150 | 107835 | 93.89587074 | 557 | 561 | 0.992869875 | 1 | 1 | 1 | FIM1_58 | YPL086C | YPL086C | ELP3 | n |
| Kmar_QGN17532.1 | W303.asm02.HP0_G0028520 | OG0002393 | chrX | 168114 | 170120 | CP015059.1 | 1114314 | 1116785 | 50.59952038 | 668 | 823 | 0.811664642 | 1 | 1 | 1 | FIM1_4268 | YJL128C | YJL128C | PBS2 | n |
| Kmar_QGN15462.1 | W303.asm02.HP0_G0053790 | OG0002394 | chrXVI | 238526 | 243253 | CP015056.1 | 999309 | 1003685 | 42.62820513 | 1552 | 1458 | 1.064471879 | 1 | 1 | 1 | FIM1_2153 | YPL167C | YPL167C | REV3 | n |
| Kmar_QGN16024.1 | W303.asm02.HP0_G0019540 | OG0002395 | chrVII | 422448 | 423890 | CP015057.1 | 650793 | 652220 | 48.4 | 480 | 475 | 1.010526316 | 1 | 1 | 1 | FIM1_2724 | YGL038C | YGL038C | OCH1 | n |
| Kmar_QGN15289.1 | W303.asm02.HP0_G0014070 | OG0002396 | chrV | 74132 | 75688 | CP015056.1 | 608676 | 610256 | 66.15969582 | 518 | 526 | 0.984790875 | 1 | 1 | 1 | FIM1_1978 | YEL042W | YEL042W | GDA1 | n |
| Kmar_QGN18384.1 | W303.asm02.HP0_G0055610 | OG0002397 | chrXVI | 610409 | 613810 | CP015061.1 | 817771 | 820941 | 37.68366465 | 1133 | 1056 | 1.072916667 | 1 | 1 | 1 | FIM1_4711 | YPR022C | YPR022C | SDD4 | n |
| Kmar_QGN14362.1 | W303.asm02.HP0_G0010840 | OG0002398 | chrIV | 920987 | 921655 | CP015055.1 | 361624 | 362307 | 80.17621145 | 222 | 227 | 0.977973568 | 1 | 1 | 1 | FIM1_1022 | YDR226W | YDR226W | ADK1 | n |
| Kmar_QGN15408.1 | W303.asm02.HP0_G0023570 | OG0002399 | chrVIII | 186649 | 187389 | CP015056.1 | 899848 | 900594 | 52.82258065 | 246 | 248 | 0.991935484 | 1 | 0 | 0 | FIM1_2098 | YHR043C | YHR043C | DOG2 | n |
| Kmar_QGN14002.1 | W303.asm02.HP0_G0037520 | OG0002400 | chrXII | 689696 | 691480 | CP015054.1 | 1306266 | 1308044 | 73.36683417 | 594 | 592 | 1.003378378 | 1 | 1 | 1 | FIM1_653 | YLR276C | YLR276C | DBP9 | n |
| Kmar_QGN14044.1 | W303.asm02.HP0_G0013240 | OG0002401 | chrIV | 1430514 | 1432946 | CP015054.1 | 1398108 | 1400255 | 35.4679803 | 810 | 715 | 1.132867133 | 1 | 1 | 1 | FIM1_695 | YDR485C | YDR485C | VPS72 | n |
| Kmar_QGN14045.1 | W303.asm02.HP0_G0048140 | OG0002402 | chrXV | 138562 | 139500 | CP015054.1 | 1400519 | 1401502 | 53.49544073 | 312 | 327 | 0.95412844 | 1 | 1 | 1 | FIM1_696 | YOL096C | YOL096C | COQ3 | n |
| Kmar_QGN14046.1 | W303.asm02.HP0_G0013250 | OG0002403 | chrIV | 1433125 | 1433814 | CP015054.1 | 1401663 | 1402340 | 60.51502146 | 229 | 225 | 1.017777778 | 1 | 1 | 1 | FIM1_697 | YDR486C | YDR486C | VPS60 | n |
| Kmar_QGN17239.1 | W303.asm02.HP0_G0023030 | OG0002404 | chrVIII | 84763 | 86661 | CP015059.1 | 522893 | 524938 | 43.67283951 | 632 | 681 | 0.92804699 | 1 | 1 | 1 | FIM1_3970 | YHL008C | YHL008C | | n |
| Kmar_QGN15510.1 | W303.asm02.HP0_G0006090 | OG0002405 | chrIII | 254481 | 255944 | CP015056.1 | 1092060 | 1093238 | 33.26572008 | 487 | 392 | 1.242346939 | 1 | 1 | 1 | FIM1_2201 | YCR066W | YCR066W | RAD18 | n |
| Kmar_QGN15513.1 | W303.asm02.HP0_G0006110 | OG0002406 | chrIII | 260195 | 261757 | CP015056.1 | 1098195 | 1099829 | 52.8440367 | 520 | 544 | 0.955882353 | 1 | 1 | 1 | FIM1_2204 | YCR068W | YCR068W | ATG15 | n |
| Kmar_QGN17227.1 | W303.asm02.HP0_G0032960 | OG0002407 | chrXI | 391660 | 393669 | CP015059.1 | 491886 | 493814 | 71.53502235 | 669 | 642 | 1.042056075 | 1 | 1 | 1 | FIM1_3958 | YKL029C | YKL029C | MAE1 | n |
| Kmar_QGN18247.1 | W303.asm02.HP0_G0056860 | OG0002408 | chrXVI | 874477 | 875787 | CP015061.1 | 541455 | 542786 | 47.21030043 | 436 | 443 | 0.984198646 | 1 | 1 | 1 | FIM1_4572 | YPR163C | YPR163C | TIF3 | n |
| Kmar_QGN18257.1 | W303.asm02.HP0_G0039160 | OG0002409 | chrXII | 1050296 | 1051210 | CP015061.1 | 566605 | 567672 | 37.35955056 | 304 | 355 | 0.856338028 | 1 | 1 | 1 | FIM1_4583 | YLR455W | YLR455W | PDP3 | n |
| Kmar_QGN13639.1 | W303.asm02.HP0_G0014320 | OG0002410 | chrV | 124840 | 125853 | CP015054.1 | 563574 | 564527 | 28.8184438 | 337 | 317 | 1.063091483 | 1 | 1 | 1 | FIM1_281 | YEL017W | YEL017W | GTT3 | n |
| Kmar_QGN16493.1 | W303.asm02.HP0_G0052090 | OG0002411 | chrXV | 941341 | 945183 | CP015058.1 | 223499 | 227098 | 52.28859581 | 1280 | 1199 | 1.067556297 | 1 | 1 | 1 | FIM1_3206 | YOR330C | YOR330C | MIP1 | n |
| Kmar_QGN13898.1 | W303.asm02.HP0_G0042540 | OG0002412 | chrXIII | 695929 | 697062 | CP015054.1 | 1105750 | 1106859 | 57.25593668 | 377 | 369 | 1.021680217 | 1 | 1 | 1 | FIM1_545 | YMR214W | YMR214W | SCJ1 | n |
| Kmar_QGN15047.1 | W303.asm02.HP0_G0030280 | OG0002413 | chrX | 582382 | 583038 | CP015056.1 | 87255 | 87848 | 41.22807018 | 218 | 197 | 1.106598985 | 1 | 1 | 1 | FIM1_1732 | YJR074W | YJR074W | MOG1 | n |
| Kmar_QGN14170.1 | W303.asm02.HP0_G0005080 | OG0002414 | chrIII | 33797 | 35047 | CP015054.1 | 1678312 | 1679586 | 35 | 416 | 424 | 0.981132075 | 1 | 1 | 1 | FIM1_823 | YCL052C | YCL052C | PBN1 | n |
| Kmar_QGN16018.1 | W303.asm02.HP0_G0024290 | OG0002415 | chrVIII | 308734 | 319968 | CP015057.1 | 622867 | 634218 | 61.81627846 | 3744 | 3783 | 0.989690722 | 1 | 1 | 1 | FIM1_2718 | YHR099W | YHR099W | TRA1 | n |
| Kmar_QGN14740.1 | W303.asm02.HP0_G0013060 | OG0002416 | chrIV | 1399029 | 1400267 | CP015055.1 | 1138014 | 1139267 | 60.46511628 | 412 | 417 | 0.988009592 | 1 | 1 | 1 | FIM1_1409 | YDR465C | YDR465C | RMT2 | n |
| Kmar_QGN14739.1 | W303.asm02.HP0_G0023180 | OG0002417 | chrVIII | 113564 | 115156 | CP015055.1 | 1135802 | 1137382 | 78.3269962 | 530 | 526 | 1.007604563 | 1 | 1 | 1 | FIM1_1408 | YHR007C | YHR007C | ERG11 | n |
| Kmar_QGN14281.1 | W303.asm02.HP0_G0023380 | OG0002418 | chrVIII | 153279 | 154352 | CP015055.1 | 175659 | 176732 | 82.91316527 | 357 | 357 | 1 | 1 | 1 | 1 | FIM1_938 | YHR025W | YHR025W | THR1 | n |
| Kmar_QGN15544.1 | W303.asm02.HP0_G0040160 | OG0002419 | chrXIII | 206149 | 206800 | CP015056.1 | 1158689 | 1159144 | 39.44444444 | 181 | 151 | 1.198675497 | 1 | 1 | 1 | FIM1_2235 | YML036W | YML036W | CGI121 | n |
| Kmar_QGN16593.1 | W303.asm02.HP0_G0041230 | OG0002420 | chrXIII | 427217 | 430132 | CP015058.1 | 456932 | 459805 | 69.56967213 | 971 | 957 | 1.014629049 | 1 | 1 | 1 | FIM1_3309 | YMR080C | YMR080C | NAM7 | n |
| Kmar_QGN16594.1 | W303.asm02.HP0_G0032340 | OG0002421 | chrXI | 274621 | 277935 | CP015058.1 | 459970 | 463113 | 37.36933798 | 1104 | 1047 | 1.054441261 | 1 | 1 | 1 | FIM1_3310 | YKL092C | YKL092C | BUD2 | n |
| Kmar_QGN16595.1 | W303.asm02.HP0_G0041220 | OG0002422 | chrXIII | 425494 | 426564 | CP015058.1 | 463865 | 464770 | 79.5379538 | 304 | 301 | 1.009966777 | 1 | 1 | 1 | FIM1_3311 | YMR079W | YMR079W | SEC14 | n |
| Kmar_QGN15087.1 | W303.asm02.HP0_G0033270 | OG0002423 | chrXI | 447851 | 448459 | CP015056.1 | 174132 | 174734 | 86 | 202 | 200 | 1.01 | 1 | 1 | 1 | FIM1_1774 | YKL001C | YKL001C | MET14 | n |
| Kmar_QGN15086.1 | W303.asm02.HP0_G0033250 | OG0002424 | chrXI | 446172 | 446567 | CP015056.1 | 172253 | 172642 | 61.83206107 | 131 | 129 | 1.015503876 | 1 | 1 | 1 | FIM1_1772 | YKL003C | YKL003C | MRP17 | n |
| Kmar_QGN15895.1 | W303.asm02.HP0_G0035760 | OG0002425 | chrXII | 314804 | 323680 | CP015057.1 | 322390 | 331233 | 30.62913907 | 2958 | 2947 | 1.003732609 | 1 | 1 | 1 | FIM1_2593 | YLR087C | YLR087C | CSF1 | n |
| Kmar_QGN16872.1 | W303.asm02.HP0_G0010160 | OG0002426 | chrIV | 773239 | 774046 | CP015058.1 | 1075552 | 1076055 | 30.83333333 | 236 | 167 | 1.413173653 | 1 | 1 | 1 | FIM1_3598 | YDR156W | YDR156W | RPA14 | n |
| Kmar_QGN15894.1 | W303.asm02.HP0_G0035750 | OG0002427 | chrXII | 310195 | 314451 | CP015057.1 | 318259 | 322404 | 49.75369458 | 1418 | 1381 | 1.02679218 | 1 | 1 | 1 | FIM1_2592 | YLR086W | YLR086W | SMC4 | n |
| Kmar_QGN16204.1 | W303.asm02.HP0_G0038670 | OG0002428 | chrXII | 926459 | 927562 | CP015057.1 | 1027167 | 1028228 | 64.03269755 | 367 | 353 | 1.039660057 | 1 | 1 | 1 | FIM1_2907 | YLR405W | YLR405W | DUS4 | n |
| Kmar_QGN17584.1 | W303.asm02.HP0_G0027800 | OG0002429 | chrX | 29725 | 30714 | CP015060.1 | 41736 | 42764 | 63.11239193 | 329 | 342 | 0.961988304 | 1 | 1 | 1 | FIM1_4791 | YJL208C | YJL208C | NUC1 | n |
| Kmar_QGN16278.1 | W303.asm02.HP0_G0031530 | OG0002430 | chrXI | 108485 | 109768 | CP015057.1 | 1184193 | 1185446 | 83.60655738 | 427 | 417 | 1.023980815 | 1 | 1 | 1 | FIM1_2983 | YKL181W | YKL181W | PRS1 | n |
| Kmar_QGN16279.1 | W303.asm02.HP0_G0038750 | OG0002431 | chrXII | 945037 | 945861 | CP015057.1 | 1186137 | 1186940 | 40.433213 | 274 | 267 | 1.026217228 | 1 | 1 | 1 | FIM1_2984 | YLR412W | YLR412W | BER1 | n |
| Kmar_QGN13691.1 | W303.asm02.HP0_G0025360 | OG0002432 | chrVIII | 518835 | 520703 | CP015054.1 | 669539 | 671020 | 43.95604396 | 622 | 493 | 1.261663286 | 1 | 1 | 1 | FIM1_334 | YHR206W | YHR206W | SKN7 | n |
| Kmar_QGN14469.1 | W303.asm02.HP0_G0004170 | OG0002433 | chrII | 698156 | 699070 | CP015055.1 | 581622 | 582485 | 40.32786885 | 304 | 287 | 1.059233449 | 1 | 1 | 1 | FIM1_1130 | YBR228W | YBR228W | SLX1 | n |
| Kmar_QGN16205.1 | W303.asm02.HP0_G0008110 | OG0002434 | chrIV | 324377 | 326479 | CP015057.1 | 1028748 | 1030772 | 42.57425743 | 700 | 674 | 1.038575668 | 1 | 1 | 1 | FIM1_2908 | YDL074C | YDL074C | BRE1 | n |
| Kmar_QGN16212.1 | W303.asm02.HP0_G0038630 | OG0002435 | chrXII | 916206 | 918266 | CP015057.1 | 1045312 | 1047417 | 56.76056338 | 686 | 701 | 0.978601997 | 1 | 1 | 1 | FIM1_2916 | YLR399C | YLR399C | BDF1 | n |
| Kmar_QGN16280.1 | W303.asm02.HP0_G0031520 | OG0002436 | chrXI | 101840 | 107995 | CP015057.1 | 1187460 | 1193606 | 74.33024842 | 2051 | 2048 | 1.001464844 | 1 | 1 | 1 | FIM1_2985 | YKL182W | YKL182W | FAS1 | n |
| Kmar_QGN16277.1 | W303.asm02.HP0_G0038730 | OG0002437 | chrXII | 934210 | 937650 | CP015057.1 | 1179867 | 1183289 | 73.53448276 | 1146 | 1140 | 1.005263158 | 1 | 1 | 1 | FIM1_2982 | YLR410W | YLR410W | VIP1 | n |
| Kmar_QGN14153.1 | W303.asm02.HP0_G0005190 | OG0002438 | chrIII | 52297 | 54534 | CP015054.1 | 1642070 | 1644220 | 34.0232859 | 745 | 716 | 1.040502793 | 1 | 1 | 1 | FIM1_806 | YCL039W | YCL039W | GID7 | n |
| Kmar_QGN14959.1 | W303.asm02.HP0_G0051200 | OG0002439 | chrXV | 782630 | 783265 | CP015055.1 | 1574872 | 1575552 | 45.65217391 | 211 | 226 | 0.933628319 | 1 | 1 | 1 | FIM1_1636 | YOR236W | YOR236W | DFR1 | n |
| Kmar_QGN16441.1 | W303.asm02.HP0_G0029180 | OG0002440 | chrX | 324284 | 325720 | CP015058.1 | 125638 | 127038 | 36.98630137 | 478 | 466 | 1.025751073 | 1 | 1 | 1 | FIM1_3154 | YJL054W | YJL054W | TIM54 | n |
| Kmar_QGN15236.1 | W303.asm02.HP0_G0029570 | OG0002441 | chrX | 420806 | 426886 | CP015056.1 | 493252 | 499029 | 48.41075795 | 2026 | 1925 | 1.052467532 | 1 | 1 | 1 | FIM1_1925 | YJL005W | YJL005W | CYR1 | n |
| Kmar_QGN14912.1 | W303.asm02.HP0_G0044960 | OG0002442 | chrXIV | 281132 | 282205 | CP015055.1 | 1479332 | 1480540 | 66.74816626 | 357 | 402 | 0.888059701 | 1 | 1 | 1 | FIM1_1586 | YNL191W | YNL191W | DUG3 | n |
| Kmar_QGN15171.1 | W303.asm02.HP0_G0028790 | OG0002443 | chrX | 236506 | 236991 | CP015056.1 | 354527 | 355015 | 62.34567901 | 161 | 162 | 0.99382716 | 1 | 1 | 1 | FIM1_1859 | YJL096W | YJL096W | MRPL49 | n |
| Kmar_QGN15172.1 | W303.asm02.HP0_G0028800 | OG0002444 | chrX | 237271 | 241707 | CP015056.1 | 355359 | 359393 | 46.05781866 | 1478 | 1344 | 1.099702381 | 1 | 1 | 1 | FIM1_1860 | YJL095W | YJL095W | BCK1 | n |
| Kmar_QGN14308.1 | W303.asm02.HP0_G0034910 | OG0002445 | chrXII | 144393 | 147233 | CP015055.1 | 241350 | 244382 | 23.46534653 | 946 | 1010 | 0.936633663 | 1 | 1 | 1 | FIM1_965 | YLL003W | YLL003W | SFI1 | n |
| Kmar_QGN18184.1 | W303.asm02.HP0_G0047010 | OG0002446 | chrXIV | 691381 | 693036 | CP015061.1 | 392769 | 394385 | 44.86486486 | 551 | 538 | 1.024163569 | 1 | 1 | 1 | FIM1_4507 | YNR030W | YNR030W | ALG12 | n |
| Kmar_QGN18185.1 | W303.asm02.HP0_G0047000 | OG0002447 | chrXIV | 689781 | 691070 | CP015061.1 | 394573 | 395835 | 57.53424658 | 429 | 420 | 1.021428571 | 1 | 1 | 1 | FIM1_4508 | YNR029C | YNR029C | ZNG1 | n |
| Kmar_QGN13664.1 | W303.asm02.HP0_G0048840 | OG0002448 | chrXV | 278511 | 280541 | CP015054.1 | 619534 | 621555 | 59.61820852 | 676 | 673 | 1.004457652 | 1 | 1 | 1 | FIM1_307 | YOL023W | YOL023W | IFM1 | n |

| | | | | | | | | | | | | | | | | | | | | |
|---|---|---|---|---|---|---|---|---|---|---|---|---|---|---|---|---|---|---|---|---|
| Kmar_QGN13665.1 | W303.asm02.HP0_G0048850 | OG0002449 | chrXV | 280727 | 281953 | CP015054.1 | 621623 | 622927 | 53.56321839 | 408 | 434 | 0.940092166 | 1 | 1 | 1 | FIM1_308 | YOL022C | YOL022C | TSR4 | n |
| Kmar_QGN14854.1 | W303.asm02.HP0_G0039580 | OG0002450 | chrXIII | 71817 | 72320 | CP015055.1 | 1352421 | 1352939 | 45.66473988 | 167 | 172 | 0.970930233 | 1 | 1 | 1 | FIM1_1527 | YML098W | YML098W | TAF13 | n |
| Kmar_QGN17141.1 | W303.asm02.HP0_G0040630 | OG0002451 | chrXIII | 301375 | 302991 | CP015059.1 | 322572 | 324179 | 77.50929368 | 538 | 535 | 1.005607477 | 1 | 1 | 1 | FIM1_3872 | YMR015C | YMR015C | ERG5 | n |
| Kmar_QGN14832.1 | W303.asm02.HP0_G0007830 | OG0002452 | chrIV | 277201 | 280494 | CP015055.1 | 1314934 | 1318239 | 75.22603978 | 1097 | 1101 | 0.996366939 | 1 | 1 | 1 | FIM1_1503 | YDL102W | YDL102W | POL3 | n |
| Kmar_QGN14607.1 | W303.asm02.HP0_G0019020 | OG0002453 | chrVII | 318731 | 320179 | CP015055.1 | 847225 | 848664 | 73.70600414 | 482 | 479 | 1.006263048 | 1 | 1 | 1 | FIM1_1270 | YGL097W | YGL097W | SRM1 | n |
| Kmar_QGN14477.1 | W303.asm02.HP0_G0018770 | OG0002454 | chrVII | 267465 | 268607 | CP015055.1 | 599588 | 600592 | 31.68831169 | 380 | 334 | 1.137724551 | 1 | 1 | 1 | FIM1_1138 | YGL126W | YGL126W | SCS3 | n |
| Kmar_QGN15396.1 | W303.asm02.HP0_G0025880 | OG0002455 | chrIX | 74397 | 76868 | CP015056.1 | 873115 | 875472 | 44.8856799 | 823 | 785 | 1.048407643 | 1 | 1 | 1 | FIM1_2086 | YIL140W | YIL140W | AXL2 | n |
| Kmar_QGN14474.1 | W303.asm02.HP0_G0004120 | OG0002456 | chrII | 687437 | 688537 | CP015055.1 | 592132 | 593193 | 82.78688525 | 366 | 353 | 1.036827195 | 1 | 1 | 1 | FIM1_1135 | YBR221C | YBR221C | PDB1 | n |
| Kmar_QGN14413.1 | W303.asm02.HP0_G0025720 | OG0002457 | chrIX | 35980 | 36573 | CP015055.1 | 467471 | 468013 | 49.49494949 | 197 | 180 | 1.094444444 | 1 | 1 | 1 | FIM1_1074 | YIL157C | YIL157C | COA1 | n |
| Kmar_QGN15556.1 | W303.asm02.HP0_G0011630 | OG0002458 | chrIV | 1089695 | 1091623 | CP015056.1 | 1177664 | 1179520 | 55.57275542 | 642 | 618 | 1.038834951 | 1 | 1 | 1 | FIM1_2247 | YDR311W | YDR311W | TFB1 | n |
| Kmar_QGN16396.1 | W303.asm02.HP0_G0000140 | OG0002459 | chrI | 37144 | 37782 | CP015058.1 | 24190 | 24798 | 41.48471616 | 212 | 202 | 1.04950495 | 1 | 1 | 1 | FIM1_3109 | YAL059W | YAL059W | ECM1 | n |
| Kmar_QGN17185.1 | W303.asm02.HP0_G0055180 | OG0002460 | chrXVI | 513054 | 516356 | CP015059.1 | 407232 | 410405 | 48.55595668 | 1100 | 1057 | 1.040681173 | 1 | 1 | 1 | FIM1_3916 | YPL022W | YPL022W | RAD1 | n |
| Kmar_QGN17183.1 | W303.asm02.HP0_G0055160 | OG0002461 | chrXVI | 509874 | 510599 | CP015059.1 | 404129 | 404812 | 35.56485356 | 241 | 227 | 1.061674009 | 1 | 1 | 1 | FIM1_3914 | YPL024W | YPL024W | RMI1 | n |
| Kmar_QGN17184.1 | W303.asm02.HP0_G0055170 | OG0002462 | chrXVI | 510696 | 512669 | CP015059.1 | 404876 | 406804 | 65.29680365 | 657 | 642 | 1.023364486 | 1 | 1 | 1 | FIM1_3915 | YPL023C | YPL023C | MET12 | n |
| Kmar_QGN13846.1 | W303.asm02.HP0_G0028060 | OG0002463 | chrX | 78812 | 79627 | CP015054.1 | 991420 | 992259 | 41.07142857 | 271 | 279 | 0.971326165 | 1 | 1 | 1 | FIM1_492 | YJL178C | YJL178C | ATG27 | n |
| Kmar_QGN14020.1 | W303.asm02.HP0_G0000770 | OG0002464 | chrI | 171600 | 172907 | CP015054.1 | 1343676 | 1344887 | 52.14446953 | 435 | 403 | 1.079404467 | 1 | 1 | 1 | FIM1_671 | YAR018C | YAR018C | KIN3 | n |
| Kmar_QGN15156.1 | W303.asm02.HP0_G0018140 | OG0002465 | chrVII | 127038 | 135056 | CP015056.1 | 315941 | 323947 | 58.71593878 | 2672 | 2668 | 1.00149925 | 1 | 1 | 1 | FIM1_1843 | YGL195W | YGL195W | GCN1 | n |
| Kmar_QGN16234.1 | W303.asm02.HP0_G0025320 | OG0002466 | chrVIII | 508490 | 510298 | CP015057.1 | 1096068 | 1097870 | 47.38562092 | 602 | 600 | 1.003333333 | 1 | 1 | 1 | FIM1_2939 | YHR202W | YHR202W | SMN1 | n |
| Kmar_QGN16283.1 | W303.asm02.HP0_G0031480 | OG0002467 | chrXI | 95668 | 97434 | CP015057.1 | 1199383 | 1200744 | 30.32367973 | 588 | 453 | 1.298013245 | 1 | 1 | 1 | FIM1_2988 | YKL185W | YKL185W | ASH1 | n |
| Kmar_QGN13613.1 | W303.asm02.HP0_G0043150 | OG0002468 | chrXIII | 825326 | 826447 | CP015054.1 | 519553 | 520713 | 57.69230769 | 373 | 386 | 0.966321244 | 1 | 1 | 1 | FIM1_255 | YMR276W | YMR276W | DSK2 | n |
| Kmar_QGN13622.1 | W303.asm02.HP0_G0039420 | OG0002469 | chrXIII | 36595 | 38127 | CP015054.1 | 537075 | 538649 | 22.77039848 | 510 | 524 | 0.973282443 | 1 | 1 | 1 | FIM1_264 | YML114C | YML114C | TAF8 | n |
| Kmar_QGN14270.1 | W303.asm02.HP0_G0042990 | OG0002470 | chrXIII | 795880 | 796341 | CP015055.1 | 153241 | 153711 | 89.74358974 | 153 | 156 | 0.980769231 | 1 | 1 | 1 | FIM1_927 | YMR260C | YMR260C | TIF11 | n |
| Kmar_QGN18146.1 | W303.asm02.HP0_G0043860 | OG0002471 | chrXIV | 45740 | 48454 | CP015061.1 | 321524 | 324214 | 60.63947078 | 904 | 896 | 1.008928571 | 1 | 1 | 1 | FIM1_4469 | YNL313C | YNL313C | EMW1 | n |
| Kmar_QGN15870.1 | W303.asm02.HP0_G0002550 | OG0002472 | chrII | 376939 | 378801 | CP015057.1 | 268187 | 269929 | 35.45310016 | 620 | 580 | 1.068965517 | 1 | 1 | 1 | FIM1_2567 | YBR060C | YBR060C | ORC2 | n |
| Kmar_QGN15871.1 | W303.asm02.HP0_G0009140 | OG0002473 | chrIV | 551523 | 551855 | CP015057.1 | 270321 | 270659 | 79.46428571 | 110 | 112 | 0.982142857 | 1 | 1 | 1 | FIM1_2568 | YDR045C | YDR045C | RPC11 | n |
| Kmar_QGN15884.1 | W303.asm02.HP0_G0002660 | OG0002474 | chrII | 395508 | 396221 | CP015057.1 | 291602 | 292291 | 45.26748971 | 237 | 229 | 1.034934498 | 1 | 1 | 1 | FIM1_2582 | YBR070C | YBR070C | ALG14 | n |
| Kmar_QGN15873.1 | W303.asm02.HP0_G0002610 | OG0002475 | chrII | 384871 | 385965 | CP015057.1 | 277288 | 278190 | 23.7704918 | 364 | 300 | 1.213333333 | 1 | 1 | 1 | FIM1_2571 | YBR065C | YBR065C | ECM2 | n |
| Kmar_QGN17902.1 | W303.asm02.HP0_G0042980 | OG0002476 | chrXIII | 791123 | 795385 | CP015060.1 | 705646 | 709875 | 36.25 | 1420 | 1409 | 1.007806955 | 1 | 1 | 1 | FIM1_5111 | YMR259C | YMR259C | TRM732 | n |
| Kmar_QGN14376.1 | W303.asm02.HP0_G0002700 | OG0002477 | chrII | 399502 | 402276 | CP015055.1 | 384680 | 387502 | 51.86335404 | 924 | 940 | 0.982978723 | 1 | 1 | 1 | FIM1_1036 | YBR073W | YBR073W | RDH54 | n |
| Kmar_QGN14377.1 | W303.asm02.HP0_G0009170 | OG0002478 | chrIV | 557403 | 559301 | CP015055.1 | 387557 | 389455 | 47.13178295 | 632 | 632 | 1 | 1 | 1 | 1 | FIM1_1037 | YDR049W | YDR049W | VMS1 | n |
| Kmar_QGN14378.1 | W303.asm02.HP0_G0009160 | OG0002479 | chrIV | 556009 | 557097 | CP015055.1 | 389805 | 390914 | 79.67479675 | 362 | 369 | 0.98102981 | 1 | 1 | 1 | FIM1_1038 | YDR047W | YDR047W | HEM12 | n |
| Kmar_QGN14594.1 | W303.asm02.HP0_G0006220 | OG0002480 | chrIII | 277352 | 281635 | CP015055.1 | 819092 | 823060 | 37.53481894 | 1427 | 1322 | 1.079425113 | 1 | 1 | 1 | FIM1_1256 | YCR081W | YCR081W | SRB8 | n |
| Kmar_QGN14595.1 | W303.asm02.HP0_G0047120 | OG0002481 | chrXIV | 710196 | 712013 | CP015055.1 | 823148 | 825046 | 40.88820827 | 605 | 632 | 0.957278481 | 1 | 1 | 1 | FIM1_1257 | YNR039C | YNR039C | ZRG17 | n |
| Kmar_QGN18180.1 | W303.asm02.HP0_G0047030 | OG0002482 | chrXIV | 698592 | 699698 | CP015061.1 | 383692 | 384726 | 68.47826087 | 368 | 344 | 1.069767442 | 1 | 1 | 1 | FIM1_4503 | YNR032W | YNR032W | PPG1 | n |
| Kmar_QGN18183.1 | W303.asm02.HP0_G0006130 | OG0002483 | chrIII | 263084 | 263524 | CP015061.1 | 392110 | 392649 | 40.55555556 | 146 | 179 | 0.815642458 | 1 | 1 | 1 | FIM1_4506 | YCR071C | YCR071C | IMG2 | n |
| Kmar_QGN18182.1 | W303.asm02.HP0_G0006140 | OG0002484 | chrIII | 263786 | 265333 | CP015061.1 | 390262 | 391809 | 80.77669903 | 515 | 515 | 1 | 1 | 1 | 1 | FIM1_4505 | YCR072C | YCR072C | RSA4 | n |
| Kmar_QGN14475.1 | W303.asm02.HP0_G0004110 | OG0002485 | chrII | 684784 | 686962 | CP015055.1 | 594630 | 596183 | 52.60416667 | 575 | 517 | 1.112185687 | 1 | 1 | 1 | FIM1_1136 | YBR220C | YBR220C | | n |
| Kmar_QGN15993.1 | W303.asm02.HP0_G0033630 | OG0002486 | chrXI | 521695 | 522855 | CP015057.1 | 565853 | 567010 | 86.52849741 | 386 | 385 | 1.002597403 | 1 | 1 | 1 | FIM1_2693 | YKR038C | YKR038C | KAE1 | n |
| Kmar_QGN16087.1 | W303.asm02.HP0_G0006200 | OG0002487 | chrIII | 273219 | 275609 | CP015057.1 | 785254 | 787908 | 47.49455338 | 796 | 884 | 0.900452489 | 1 | 1 | 1 | FIM1_2787 | YCR077C | YCR077C | PAT1 | n |
| Kmar_QGN17281.1 | W303.asm02.HP0_G0042800 | OG0002488 | chrXIII | 754769 | 756184 | CP015059.1 | 610884 | 612278 | 46.70912951 | 471 | 464 | 1.015086207 | 1 | 1 | 1 | FIM1_4012 | YMR239C | YMR239C | RNT1 | n |
| Kmar_QGN15614.1 | W303.asm02.HP0_G0006920 | OG0002489 | chrIV | 98549 | 99298 | CP015056.1 | 1310087 | 1310815 | 48.79032258 | 249 | 242 | 1.02892562 | 1 | 1 | 1 | FIM1_2307 | YDL202W | YDL202W | MRPL11 | n |
| Kmar_QGN15615.1 | W303.asm02.HP0_G0006930 | OG0002490 | chrIV | 99635 | 100495 | CP015056.1 | 1311104 | 1311946 | 76.04166667 | 286 | 280 | 1.021428571 | 1 | 1 | 1 | FIM1_2308 | YDL201W | YDL201W | TRM8 | n |
| Kmar_QGN15810.1 | W303.asm02.HP0_G0041750 | OG0002491 | chrXIII | 539289 | 540653 | CP015057.1 | 136630 | 137712 | 38.52813853 | 454 | 360 | 1.261111111 | 1 | 1 | 1 | FIM1_2506 | YMR135C | YMR135C | GID8 | n |
| Kmar_QGN14412.1 | W303.asm02.HP0_G0034230 | OG0002492 | chrXI | 654117 | 656552 | CP015055.1 | 464298 | 466673 | 34.69626168 | 811 | 791 | 1.02528445 | 1 | 1 | 1 | FIM1_1072 | YKR099W | YKR099W | BAS1 | n |
| Kmar_QGN15555.1 | W303.asm02.HP0_G0023920 | OG0002493 | chrVIII | 231355 | 233064 | CP015056.1 | 1175474 | 1177108 | 68.34782609 | 569 | 544 | 1.045955882 | 1 | 1 | 1 | FIM1_2246 | YHR064C | YHR064C | SSZ1 | n |
| Kmar_QGN13790.1 | W303.asm02.HP0_G0049790 | OG0002494 | chrXV | 472634 | 473332 | CP015054.1 | 885382 | 886101 | 40.33613445 | 232 | 239 | 0.970711297 | 1 | 1 | 1 | FIM1_436 | YOR077W | YOR077W | RTS2 | n |
| Kmar_QGN13621.1 | W303.asm02.HP0_G0043110 | OG0002495 | chrXIII | 816122 | 817276 | CP015054.1 | 535283 | 536428 | 73.4375 | 384 | 381 | 1.007874016 | 1 | 1 | 1 | FIM1_263 | YMR272C | YMR272C | SCS7 | n |
| Kmar_QGN13618.1 | W303.asm02.HP0_G0043130 | OG0002496 | chrXIII | 820863 | 821810 | CP015054.1 | 529622 | 530596 | 30.76923077 | 315 | 324 | 0.972222222 | 1 | 1 | 1 | FIM1_260 | YMR274C | YMR274C | RCE1 | n |
| Kmar_QGN13789.1 | W303.asm02.HP0_G0034110 | OG0002497 | chrXI | 621518 | 622555 | CP015054.1 | 884087 | 885127 | 53.2967033 | 345 | 346 | 0.997109827 | 1 | 1 | 1 | FIM1_435 | YKR087C | YKR087C | OMA1 | n |
| Kmar_QGN17768.1 | W303.asm02.HP0_G0022470 | OG0002498 | chrVII | 1064317 | 1065132 | CP015060.1 | 434977 | 435846 | 63.79310345 | 271 | 289 | 0.937716263 | 1 | 1 | 1 | FIM1_4977 | YGR280C | YGR280C | PXR1 | n |
| Kmar_QGN13751.1 | W303.asm02.HP0_G0000680 | OG0002499 | chrI | 148360 | 151932 | CP015054.1 | 806485 | 809964 | 41.78601516 | 1160 | 1159 | 1.000862813 | 1 | 1 | 1 | FIM1_395 | YAL001C | YAL001C | TFC3 | n |
| Kmar_QGN13752.1 | W303.asm02.HP0_G0019890 | OG0002500 | chrVII | 494667 | 495568 | CP015054.1 | 810503 | 811112 | 20 | 248 | 61 | 4.06557377 | 1 | 1 | 1 | FIM1_396 | YGR001C | YGR001C | EFM5 | n |
| Kmar_QGN15324.1 | W303.asm02.HP0_G0018280 | OG0002501 | chrVII | 156005 | 158698 | CP015056.1 | 705898 | 708408 | 51.56599553 | 897 | 836 | 1.072966507 | 1 | 1 | 1 | FIM1_2013 | YGL180W | YGL180W | ATG1 | n |
| Kmar_QGN13393.1 | W303.asm02.HP0_G0003650 | OG0002502 | chrII | 595139 | 595759 | CP015054.1 | 59110 | 59718 | 60.67961165 | 206 | 202 | 1.01980198 | 1 | 1 | 1 | FIM1_30 | YBR171W | YBR171W | SEC66 | n |
| Kmar_QGN18394.1 | W303.asm02.HP0_G0031890 | OG0002503 | chrXI | 179783 | 180442 | CP015061.1 | 842910 | 843584 | 43.11111111 | 219 | 224 | 0.977678571 | 1 | 1 | 1 | FIM1_4721 | YKL142W | YKL142W | MRP8 | n |
| Kmar_QGN15534.1 | W303.asm02.HP0_G0010910 | OG0002504 | chrIV | 935630 | 937711 | CP015056.1 | 1137345 | 1139462 | 80.05657709 | 693 | 705 | 0.982978723 | 1 | 1 | 1 | FIM1_2225 | YDR234W | YDR234W | LYS4 | n |
| Kmar_QGN16853.1 | W303.asm02.HP0_G0010070 | OG0002505 | chrIV | 751646 | 753265 | CP015058.1 | 1034178 | 1035800 | 42.63157895 | 539 | 540 | 0.998148148 | 1 | 1 | 1 | FIM1_3579 | YDR145W | YDR145W | TAF12 | n |
| Kmar_QGN16177.1 | W303.asm02.HP0_G0051560 | OG0002506 | chrXV | 836171 | 838150 | CP015057.1 | 965886 | 967712 | 55.75757576 | 659 | 608 | 1.083881579 | 1 | 1 | 1 | FIM1_2879 | YOR273C | YOR273C | TPO4 | n |
| Kmar_QGN15434.1 | W303.asm02.HP0_G0023490 | OG0002507 | chrVIII | 174189 | 175604 | CP015056.1 | 953714 | 955117 | 33.13131313 | 471 | 467 | 1.00856531 | 1 | 1 | 1 | FIM1_2125 | YHR036W | YHR036W | BRL1 | n |
| Kmar_QGN15431.1 | W303.asm02.HP0_G0023510 | OG0002508 | chrVIII | 177910 | 178602 | CP015056.1 | 945474 | 946196 | 47.91666667 | 230 | 240 | 0.958333333 | 1 | 1 | 1 | FIM1_2122 | YHR038W | YHR038W | RRF1 | n |
| Kmar_QGN14466.1 | W303.asm02.HP0_G0018790 | OG0002509 | chrVII | 271250 | 273184 | CP015055.1 | 575930 | 577564 | 29.42989214 | 644 | 544 | 1.183823529 | 1 | 1 | 1 | FIM1_1127 | YGL124C | YGL124C | MON1 | n |
| Kmar_QGN15477.1 | W303.asm02.HP0_G0051430 | OG0002510 | chrXV | 815707 | 817443 | CP015056.1 | 1025169 | 1026701 | 47.40484429 | 578 | 510 | 1.133333333 | 1 | 1 | 1 | FIM1_2168 | YOR260W | YOR260W | GCD1 | n |
| Kmar_QGN15476.1 | W303.asm02.HP0_G0051420 | OG0002511 | chrXV | 814118 | 815431 | CP015056.1 | 1023446 | 1024759 | 85.81235698 | 437 | 437 | 1 | 1 | 1 | 1 | FIM1_2167 | YOR259C | YOR259C | RPT4 | n |
| Kmar_QGN15474.1 | W303.asm02.HP0_G0053730 | OG0002512 | chrXVI | 229076 | 229969 | CP015056.1 | 1021120 | 1022010 | 52.01342282 | 297 | 296 | 1.003378378 | 1 | 1 | 1 | FIM1_2165 | YPL173W | YPL173W | MRPL40 | n |
| Kmar_QGN15471.1 | W303.asm02.HP0_G0053710 | OG0002513 | chrXVI | 224563 | 226021 | CP015056.1 | 1015959 | 1017308 | 64.01766004 | 452 | 449 | 1.006681514 | 1 | 1 | 1 | FIM1_2162 | YPL175W | YPL175W | SPT14 | n |
| Kmar_QGN15467.1 | W303.asm02.HP0_G0053770 | OG0002514 | chrXVI | 234972 | 236771 | CP015056.1 | 1009322 | 1011094 | 56.07321131 | 599 | 590 | 1.015254237 | 1 | 1 | 1 | FIM1_2158 | YPL169C | YPL169C | MEX67 | n |
| Kmar_QGN15466.1 | W303.asm02.HP0_G0051370 | OG0002515 | chrXV | 806755 | 808746 | CP015056.1 | 1006707 | 1008728 | 47.33044733 | 663 | 673 | 0.985141159 | 1 | 1 | 1 | FIM1_2157 | YOR254C | YOR254C | SEC63 | n |
| Kmar_QGN17309.1 | W303.asm02.HP0_G0045580 | OG0002516 | chrXIV | 394100 | 395578 | CP015059.1 | 664866 | 666371 | 40.42145594 | 492 | 501 | 0.982035928 | 1 | 1 | 1 | FIM1_4041 | YNL124W | YNL124W | NAF1 | n |
| Kmar_QGN17310.1 | W303.asm02.HP0_G0024500 | OG0002517 | chrVIII | 355547 | 358426 | CP015059.1 | 666485 | 669316 | 52.02492212 | 959 | 943 | 1.016967126 | 1 | 1 | 1 | FIM1_4042 | YHR120W | YHR120W | MSH1 | n |
| Kmar_QGN13624.1 | W303.asm02.HP0_G0045460 | OG0002518 | chrXIV | 371201 | 372478 | CP015054.1 | 541375 | 542844 | 28.3095723 | 425 | 489 | 0.869120654 | 1 | 1 | 1 | FIM1_266 | YNL136W | YNL136W | EAF7 | n |
| Kmar_QGN13625.1 | W303.asm02.HP0_G0045450 | OG0002519 | chrXIV | 369428 | 370888 | CP015054.1 | 543232 | 544680 | 56.14754098 | 486 | 482 | 1.008298755 | 1 | 1 | 1 | FIM1_267 | YNL137C | YNL137C | NAM9 | n |
| Kmar_QGN17745.1 | W303.asm02.HP0_G0046130 | OG0002520 | chrXIV | 509945 | 510889 | CP015060.1 | 375453 | 376373 | 41.06583072 | 314 | 306 | 1.026143791 | 1 | 1 | 1 | FIM1_4954 | YNL063W | YNL063W | MTQ1 | n |
| Kmar_QGN17744.1 | W303.asm02.HP0_G0046140 | OG0002521 | chrXIV | 510963 | 512399 | CP015060.1 | 374008 | 375432 | 54.87077535 | 478 | 474 | 1.008438819 | 1 | 1 | 1 | FIM1_4953 | YNL062C | YNL062C | GCD10 | n |
| Kmar_QGN17743.1 | W303.asm02.HP0_G0046150 | OG0002522 | chrXIV | 512727 | 514583 | CP015060.1 | 371794 | 373677 | 80 | 618 | 627 | 0.985645933 | 1 | 1 | 1 | FIM1_4952 | YNL061W | YNL061W | NOP2 | n |
| Kmar_QGN17742.1 | W303.asm02.HP0_G0046160 | OG0002523 | chrXIV | 514855 | 517122 | CP015060.1 | 369313 | 371562 | 61.24338624 | 755 | 749 | 1.008010681 | 1 | 1 | 1 | FIM1_4951 | YNL059C | YNL059C | ARP5 | n |
| Kmar_QGN13626.1 | W303.asm02.HP0_G0045440 | OG0002524 | chrXIV | 367574 | 369154 | CP015054.1 | 544899 | 546500 | 58.17843866 | 526 | 533 | 0.986866792 | 1 | 1 | 1 | FIM1_268 | YNL138W | YNL138W | SRV2 | n |
| Kmar_QGN17302.1 | W303.asm02.HP0_G0045530 | OG0002525 | chrXIV | 382313 | 383035 | CP015059.1 | 652779 | 653495 | 56.84647303 | 240 | 238 | 1.008403361 | 1 | 1 | 1 | FIM1_4034 | YNL129W | YNL129W | NRK1 | n |
| Kmar_QGN17299.1 | W303.asm02.HP0_G0045500 | OG0002526 | chrXIV | 376154 | 379324 | CP015059.1 | 645974 | 649147 | 81.62111216 | 1056 | 1057 | 0.999053926 | 1 | 1 | 1 | FIM1_4030 | YNL132W | YNL132W | KRE33 | n |
| Kmar_QGN14231.1 | W303.asm02.HP0_G0001700 | OG0002527 | chrII | 183947 | 184381 | CP015055.1 | 76244 | 76900 | 47.39336493 | 144 | 218 | 0.660550459 | 1 | 1 | 1 | FIM1_887 | YBL021C | YBL021C | HAP3 | n |
| Kmar_QGN15150.1 | W303.asm02.HP0_G0001670 | OG0002528 | chrII | 174828 | 176876 | CP015056.1 | 301452 | 303569 | 69.32952924 | 682 | 705 | 0.967375887 | 1 | 1 | 1 | FIM1_1837 | YBL024W | YBL024W | NCL1 | n |
| Kmar_QGN15151.1 | W303.asm02.HP0_G0001660 | OG0002529 | chrII | 173776 | 174213 | CP015056.1 | 303855 | 304232 | 39.31034483 | 145 | 125 | 1.16 | 1 | 1 | 1 | FIM1_1838 | YBL025W | YBL025W | RRN10 | n |
| Kmar_QGN15524.1 | W303.asm02.HP0_G0006860 | OG0002530 | chrIV | 87534 | 88055 | CP015056.1 | 1118953 | 1119402 | 73.41040462 | 173 | 149 | 1.161073826 | 1 | 1 | 1 | FIM1_2215 | YDL208W | YDL208W | NHP2 | n |
| Kmar_QGN15527.1 | W303.asm02.HP0_G0010950 | OG0002531 | chrIV | 942396 | 945317 | CP015056.1 | 1124130 | 1127048 | 65.12820513 | 973 | 972 | 1.001028807 | 1 | 1 | 1 | FIM1_2218 | YDR238C | YDR238C | SEC26 | n |
| Kmar_QGN13441.1 | W303.asm02.HP0_G0003910 | OG0002532 | chrII | 632902 | 635298 | CP015054.1 | 149447 | 151882 | 58.88754534 | 798 | 811 | 0.983970407 | 1 | 1 | 1 | FIM1_78 | YBR198C | YBR198C | TAF5 | n |
| Kmar_QGN13440.1 | W303.asm02.HP0_G0054670 | OG0002533 | chrXVI | 416801 | 417643 | CP015054.1 | 148509 | 149327 | 43.25259516 | 280 | 272 | 1.029411765 | 1 | 1 | 1 | FIM1_77 | YPL076W | YPL076W | GPI2 | n |
| Kmar_QGN18303.1 | W303.asm02.HP0_G0027370 | OG0002535 | chrIX | 373116 | 373550 | CP015061.1 | 655672 | 656118 | 38.85350318 | 144 | 148 | 0.972972973 | 1 | 1 | 1 | FIM1_4629 | YIR015W | YIR015W | RPR2 | n |
| Kmar_QGN15575.1 | W303.asm02.HP0_G0032150 | OG0002536 | chrXI | 232586 | 233734 | CP015056.1 | 1218579 | 1219724 | 69.10994764 | 382 | 381 | 1.002624672 | 1 | 1 | 1 | FIM1_2266 | YKL113C | YKL113C | RAD27 | n |
| Kmar_QGN14304.1 | W303.asm02.HP0_G0005530 | OG0002537 | chrIII | 122620 | 123588 | CP015055.1 | 233266 | 234282 | 73.07692308 | 322 | 338 | 0.952662722 | 1 | 1 | 1 | FIM1_961 | YCR002C | YCR002C | CDC10 | n |
| Kmar_QGN14303.1 | W303.asm02.HP0_G0005540 | OG0002538 | chrIII | 123860 | 124411 | CP015055.1 | 232142 | 232699 | 52.65957447 | 183 | 185 | 0.989189189 | 1 | 1 | 1 | FIM1_960 | YCR003W | YCR003W | MRPL32 | n |
| Kmar_QGN13413.1 | W303.asm02.HP0_G0003720 | OG0002539 | chrII | 606516 | 608234 | CP015054.1 | 89549 | 91300 | 44.93243243 | 572 | 583 | 0.981132075 | 1 | 1 | 1 | FIM1_50 | YBR180W | YBR180W | DTR1 | n |
| Kmar_QGN13921.1 | W303.asm02.HP0_G0012520 | OG0002540 | chrIV | 1277959 | 1279428 | CP015054.1 | 1147814 | 1149358 | 47.96116505 | 489 | 514 | 0.951361868 | 1 | 1 | 1 | FIM1_569 | YDR402C | YDR402C | DIT2 | n |
| Kmar_QGN17651.1 | W303.asm02.HP0_G0034960 | OG0002541 | chrXII | 158279 | 159154 | CP015060.1 | 192825 | 193775 | 51.875 | 291 | 316 | 0.920886076 | 1 | 1 | 1 | FIM1_4858 | YLR003C | YLR003C | CMS1 | n |
| Kmar_QGN15714.1 | W303.asm02.HP0_G0017760 | OG0002542 | chrVII | 49157 | 51166 | CP015056.1 | 1520136 | 1522178 | 57.32946299 | 669 | 680 | 0.983823529 | 1 | 1 | 1 | FIM1_2407 | YGL236C | YGL236C | MTO1 | n |
| Kmar_QGN15712.1 | W303.asm02.HP0_G0017370 | OG0002543 | chrVI | 211047 | 211904 | CP015056.1 | 1515385 | 1516416 | 37.75216138 | 285 | 343 | 0.83090379 | 1 | 1 | 1 | FIM1_2405 | YFR045W | YFR045W | MRX20 | n |
| Kmar_QGN16601.1 | W303.asm02.HP0_G0017160 | OG0002544 | chrVI | 176357 | 177202 | CP015058.1 | 476426 | 477178 | 42.25352113 | 281 | 250 | 1.124 | 1 | 1 | 1 | FIM1_3317 | YFR027W | YFR027W | ECO1 | n |
| Kmar_QGN15718.1 | W303.asm02.HP0_G0017730 | OG0002545 | chrVII | 43884 | 44735 | CP015056.1 | 1527481 | 1528245 | 35.78947368 | 283 | 254 | 1.114173228 | 1 | 1 | 1 | FIM1_2411 | YGL240W | YGL240W | DOC1 | n |
| Kmar_QGN15166.1 | W303.asm02.HP0_G0028740 | OG0002546 | chrX | 224336 | 226372 | CP015056.1 | 341940 | 343946 | 64.36950147 | 678 | 668 | 1.01497006 | 1 | 1 | 1 | FIM1_1854 | YJL101C | YJL101C | GSH1 | n |
| Kmar_QGN15713.1 | W303.asm02.HP0_G0017770 | OG0002547 | chrVII | 51852 | 54260 | CP015056.1 | 1516665 | 1519046 | 68.28358209 | 802 | 793 | 1.011349306 | 1 | 1 | 1 | FIM1_2406 | YGL234W | YGL234W | ADE57 | n |
| Kmar_QGN16141.1 | W303.asm02.HP0_G0040720 | OG0002548 | chrXIII | 322378 | 323550 | CP015057.1 | 895642 | 896598 | 45.2685422 | 390 | 318 | 1.226415094 | 1 | 1 | 1 | FIM1_2842 | YMR024W | YMR024W | MRPL3 | n |

| | | | | | | | | | | | | | | | | | | | | |
|---|---|---|---|---|---|---|---|---|---|---|---|---|---|---|---|---|---|---|---|---|
| Kmar_QGN15665.1 | W303.asm02.HP0_G0012630 | OG0002549 | chrIV | 1301323 | 1302411 | CP015056.1 | 1416597 | 1417730 | 30.69053708 | 362 | 377 | 0.960212202 | 1 | 1 | 1 | FIM1_2358 | YDR414C | YDR414C | ERD1 | n |
| Kmar_QGN15664.1 | W303.asm02.HP0_G0012620 | OG0002550 | chrIV | 1300418 | 1301125 | CP015056.1 | 1415723 | 1416490 | 57.19844358 | 235 | 255 | 0.921568627 | 1 | 1 | 1 | FIM1_2357 | YDR412W | YDR412W | RRP17 | n |
| Kmar_QGN17458.1 | W303.asm02.HP0_G0024110 | OG0002551 | chrVIII | 273512 | 274066 | CP015059.1 | 962426 | 963073 | 38.70967742 | 184 | 215 | 0.855813953 | 1 | 1 | 1 | FIM1_4192 | YHR081W | YHR081W | LRP1 | n |
| Kmar_QGN16731.1 | W303.asm02.HP0_G0024080 | OG0002552 | chrVIII | 264218 | 267565 | CP015058.1 | 761988 | 765404 | 45.66724437 | 1115 | 1138 | 0.979789104 | 1 | 1 | 1 | FIM1_3452 | YHR079C | YHR079C | IRE1 | n |
| Kmar_QGN16690.1 | W303.asm02.HP0_G0007220 | OG0002553 | chrIV | 161215 | 163374 | CP015058.1 | 675165 | 676832 | 37.41307371 | 719 | 555 | 1.295495495 | 1 | 1 | 1 | FIM1_3410 | YDL167C | YDL167C | NRP1 | n |
| Kmar_QGN16223.1 | W303.asm02.HP0_G0057050 | OG0002554 | chrXVI | 906587 | 907390 | CP015057.1 | 1066586 | 1067383 | 78.27715356 | 267 | 265 | 1.00754717 | 1 | 1 | 1 | FIM1_2927 | YPR183W | YPR183W | DPM1 | n |
| Kmar_QGN14763.1 | W303.asm02.HP0_G0012420 | OG0002555 | chrIV | 1260662 | 1262572 | CP015055.1 | 1179393 | 1181288 | 56.69781931 | 636 | 631 | 1.00792393 | 1 | 1 | 1 | FIM1_1433 | YDR390C | YDR390C | UBA2 | n |
| Kmar_QGN14800.1 | W303.asm02.HP0_G0050190 | OG0002556 | chrXV | 553173 | 553762 | CP015055.1 | 1254760 | 1255376 | 80.95238095 | 126 | 126 | 1 | 1 | 1 | 1 | FIM1_1471 | YOR122C | YOR122C | PFY1 | n |
| Kmar_QGN17353.1 | W303.asm02.HP0_G0000350 | OG0002557 | chrI | 77190 | 80198 | CP015059.1 | 751545 | 754517 | 73.93034826 | 1002 | 990 | 1.012121212 | 1 | 1 | 1 | FIM1_4085 | YAL035W | YAL035W | FUN12 | n |
| Kmar_QGN17352.1 | W303.asm02.HP0_G0000340 | OG0002558 | chrI | 75806 | 76915 | CP015059.1 | 749789 | 750898 | 95.12195122 | 369 | 369 | 1 | 1 | 1 | 1 | FIM1_4084 | YAL036C | YAL036C | RBG1 | n |
| Kmar_QGN17767.1 | W303.asm02.HP0_G0043480 | OG0002559 | chrXIII | 895871 | 899140 | CP015060.1 | 431244 | 434516 | 66.84981685 | 1089 | 1090 | 0.999082569 | 1 | 1 | 1 | FIM1_4976 | YMR308C | YMR308C | PSE1 | n |
| Kmar_QGN15784.1 | W303.asm02.HP0_G0037910 | OG0002560 | chrXII | 765003 | 767369 | CP015057.1 | 91202 | 93451 | 44.48669202 | 788 | 749 | 1.052069426 | 1 | 1 | 1 | FIM1_2480 | YLR319C | YLR319C | BUD6 | n |
| Kmar_QGN15782.1 | W303.asm02.HP0_G0037880 | OG0002561 | chrXII | 760523 | 761615 | CP015057.1 | 86888 | 88275 | 44.79768786 | 332 | 333 | 0.996996997 | 1 | 1 | 1 | FIM1_2478 | YLR316C | YLR316C | TAD3 | n |
| Kmar_QGN15781.1 | W303.asm02.HP0_G0037930 | OG0002562 | chrXII | 772269 | 773549 | CP015057.1 | 85303 | 86616 | 43.82022472 | 426 | 437 | 0.974828375 | 1 | 1 | 1 | FIM1_2477 | YLR321C | YLR321C | SFH1 | n |
| Kmar_QGN15244.1 | W303.asm02.HP0_G0000500 | OG0002563 | chrI | 111610 | 114123 | CP015056.1 | 514512 | 516962 | 56.97399527 | 837 | 816 | 1.025735294 | 1 | 1 | 1 | FIM1_1933 | YAL021C | YAL021C | CCR4 | n |
| Kmar_QGN16854.1 | W303.asm02.HP0_G0036140 | OG0002564 | chrXII | 407610 | 410441 | CP015058.1 | 1036210 | 1039020 | 65.89229145 | 943 | 936 | 1.007478632 | 1 | 1 | 1 | FIM1_3580 | YLR129W | YLR129W | DIP2 | n |
| Kmar_QGN17880.1 | W303.asm02.HP0_G0056090 | OG0002565 | chrXVI | 702236 | 703318 | CP015060.1 | 667343 | 668638 | 32.66219239 | 360 | 431 | 0.835266821 | 1 | 1 | 1 | FIM1_5089 | YPR075C | YPR075C | OPY2 | n |
| Kmar_QGN15594.1 | W303.asm02.HP0_G0052340 | OG0002566 | chrXV | 1011505 | 1011993 | CP015056.1 | 1264875 | 1265369 | 68.48484848 | 162 | 164 | 0.987804878 | 1 | 1 | 1 | FIM1_2286 | YOR357C | YOR357C | SNX3 | n |
| Kmar_QGN14493.1 | W303.asm02.HP0_G0018160 | OG0002567 | chrVII | 135881 | 137239 | CP015055.1 | 625508 | 626866 | 79.34065934 | 452 | 452 | 1 | 1 | 1 | 1 | FIM1_1154 | YGL194C | YGL194C | HOS2 | n |
| Kmar_QGN16536.1 | W303.asm02.HP0_G0019790 | OG0002568 | chrVII | 472023 | 472781 | CP015058.1 | 328705 | 329445 | 76.98412698 | 252 | 246 | 1.024390244 | 1 | 1 | 1 | FIM1_3249 | YGL011C | YGL011C | SCL1 | n |
| Kmar_QGN16189.1 | W303.asm02.HP0_G0019780 | OG0002569 | chrVII | 470389 | 471810 | CP015057.1 | 989078 | 990457 | 74.41860465 | 473 | 459 | 1.030501089 | 1 | 1 | 1 | FIM1_2891 | YGL012W | YGL012W | ERG4 | n |
| Kmar_QGN13737.1 | W303.asm02.HP0_G0019760 | OG0002570 | chrVII | 463675 | 466341 | CP015054.1 | 774252 | 776702 | 35.28761062 | 888 | 816 | 1.088235294 | 1 | 1 | 1 | FIM1_381 | YGL014W | YGL014W | PUF4 | n |
| Kmar_QGN15779.1 | W303.asm02.HP0_G0000670 | OG0002571 | chrI | 144473 | 148297 | CP015057.1 | 79478 | 83077 | 29.140625 | 1274 | 1199 | 1.062552127 | 1 | 1 | 1 | FIM1_2475 | YAL002W | YAL002W | VPS8 | n |
| Kmar_QGN13747.1 | W303.asm02.HP0_G0000720 | OG0002572 | chrI | 156209 | 157489 | CP015054.1 | 801651 | 803021 | 50.43668122 | 426 | 456 | 0.934210526 | 1 | 1 | 1 | FIM1_391 | YAR003W | YAR003W | SWD1 | n |
| Kmar_QGN13771.1 | W303.asm02.HP0_G0040070 | OG0002573 | chrXIII | 168995 | 173080 | CP015054.1 | 847828 | 851709 | 32.0318149 | 1361 | 1293 | 1.052590874 | 1 | 1 | 1 | FIM1_416 | YML049C | YML049C | RSE1 | n |
| Kmar_QGN13475.1 | W303.asm02.HP0_G0029900 | OG0002574 | chrX | 500909 | 504982 | CP015054.1 | 213462 | 217592 | 47.84482759 | 1357 | 1376 | 0.98619186 | 1 | 1 | 1 | FIM1_113 | YJR033C | YJR033C | RAV1 | n |
| Kmar_QGN13743.1 | W303.asm02.HP0_G0019840 | OG0002575 | chrVII | 487241 | 488080 | CP015054.1 | 794607 | 795377 | 32.26950355 | 279 | 256 | 1.08984375 | 1 | 1 | 1 | FIM1_387 | YGL005C | YGL005C | COG7 | n |
| Kmar_QGN15231.1 | W303.asm02.HP0_G0034830 | OG0002576 | chrXII | 128706 | 130175 | CP015056.1 | 484814 | 486247 | 76.89161554 | 489 | 477 | 1.025157233 | 1 | 1 | 1 | FIM1_1920 | YLL011W | YLL011W | SOF1 | n |
| Kmar_QGN14098.1 | W303.asm02.HP0_G0044300 | OG0002577 | chrXIV | 144834 | 145835 | CP015054.1 | 1516197 | 1517048 | 53.46534653 | 298 | 283 | 1.053003534 | 1 | 1 | 1 | FIM1_749 | YNL265C | YNL265C | IST1 | n |
| Kmar_QGN15319.1 | W303.asm02.HP0_G0037770 | OG0002578 | chrXII | 739408 | 740108 | CP015056.1 | 696326 | 696898 | 46.66666667 | 188 | 190 | 0.989473684 | 1 | 1 | 1 | FIM1_2008 | YLR306W | YLR306W | UBC12 | n |
| Kmar_QGN15633.1 | W303.asm02.HP0_G0033310 | OG0002579 | chrXI | 456367 | 457604 | CP015056.1 | 1353297 | 1354208 | 23.01587302 | 377 | 303 | 1.244224422 | 1 | 1 | 1 | FIM1_2326 | YKR004C | YKR004C | ECM9 | n |
| Kmar_QGN14401.1 | W303.asm02.HP0_G0053940 | OG0002580 | chrXVI | 272114 | 273469 | CP015055.1 | 442999 | 444282 | 56.76274945 | 451 | 427 | 1.056206089 | 1 | 1 | 1 | FIM1_1061 | YPL151C | YPL151C | PRP46 | n |
| Kmar_QGN16778.1 | W303.asm02.HP0_G0027110 | OG0002581 | chrIX | 325511 | 326158 | CP015058.1 | 866524 | 867159 | 53.95348837 | 215 | 211 | 1.018957346 | 1 | 1 | 1 | FIM1_3500 | YIL010W | YIL010W | DOT5 | n |
| Kmar_QGN14707.1 | W303.asm02.HP0_G0026820 | OG0002582 | chrIX | 269055 | 270476 | CP015055.1 | 1066999 | 1068381 | 58.40336134 | 473 | 460 | 1.02826087 | 1 | 1 | 1 | FIM1_1374 | YIL039W | YIL039W | TED1 | n |
| Kmar_QGN15440.1 | W303.asm02.HP0_G0011500 | OG0002583 | chrIV | 1062675 | 1063313 | CP015056.1 | 966000 | 966620 | 57.54716981 | 212 | 206 | 1.029126214 | 1 | 1 | 1 | FIM1_2131 | YDR298C | YDR298C | ATP5 | n |
| Kmar_QGN15666.1 | W303.asm02.HP0_G0013990 | OG0002584 | chrV | 60033 | 61214 | CP015056.1 | 1419151 | 1420347 | 71.85929648 | 393 | 398 | 0.987437186 | 1 | 1 | 1 | FIM1_2359 | YEL050C | YEL050C | RML2 | n |
| Kmar_QGN15279.1 | W303.asm02.HP0_G0034020 | OG0002585 | chrXI | 604610 | 607126 | CP015056.1 | 584771 | 587221 | 51.92307692 | 838 | 816 | 1.026960784 | 1 | 1 | 1 | FIM1_1968 | YKR079C | YKR079C | TRZ1 | n |
| Kmar_QGN13794.1 | W303.asm02.HP0_G0049810 | OG0002586 | chrXV | 474210 | 475151 | CP015054.1 | 889885 | 890787 | 37.0030581 | 313 | 300 | 1.043333333 | 1 | 1 | 1 | FIM1_440 | YOR079C | YOR079C | ATX2 | n |
| Kmar_QGN13795.1 | W303.asm02.HP0_G0049820 | OG0002587 | chrXV | 475287 | 477527 | CP015054.1 | 891107 | 893215 | 23.94736842 | 746 | 702 | 1.062678063 | 1 | 1 | 1 | FIM1_441 | YOR080W | YOR080W | DIA2 | n |
| Kmar_QGN13793.1 | W303.asm02.HP0_G0049800 | OG0002588 | chrXV | 473460 | 474104 | CP015054.1 | 889040 | 889813 | 31.12840467 | 214 | 257 | 0.832684825 | 1 | 1 | 1 | FIM1_439 | YOR078W | YOR078W | BUD21 | n |
| Kmar_QGN17845.1 | W303.asm02.HP0_G0048080 | OG0002589 | chrXV | 126452 | 127144 | CP015060.1 | 604880 | 605593 | 48.34710744 | 230 | 237 | 0.970464135 | 1 | 1 | 1 | FIM1_5054 | YOL102C | YOL102C | TPT1 | n |
| Kmar_QGN17844.1 | W303.asm02.HP0_G0013320 | OG0002590 | chrIV | 1441911 | 1442282 | CP015060.1 | 604171 | 604518 | 50.40650407 | 123 | 115 | 1.069565217 | 1 | 1 | 1 | FIM1_5053 | YDR493W | YDR493W | MZM1 | n |
| Kmar_QGN14696.1 | W303.asm02.HP0_G0021030 | OG0002591 | chrVII | 759611 | 761752 | CP015055.1 | 1052113 | 1054050 | 42.53835425 | 713 | 645 | 1.105426357 | 1 | 1 | 1 | FIM1_1363 | YGR128C | YGR128C | UTP8 | n |
| Kmar_QGN18137.1 | W303.asm02.HP0_G0015710 | OG0002592 | chrV | 397370 | 398194 | CP015061.1 | 308746 | 309567 | 23.48754448 | 274 | 273 | 1.003663004 | 1 | 1 | 1 | FIM1_4460 | YER116C | YER116C | SLX8 | n |
| Kmar_QGN15196.1 | W303.asm02.HP0_G0008650 | OG0002593 | chrIV | 437706 | 439019 | CP015056.1 | 408018 | 409322 | 91.53318078 | 437 | 434 | 1.006912442 | 1 | 1 | 1 | FIM1_1884 | YDL007W | YDL007W | RPT2 | n |
| Kmar_QGN16102.1 | W303.asm02.HP0_G0026900 | OG0002594 | chrIX | 283262 | 286366 | CP015057.1 | 816912 | 820058 | 26.38762511 | 1034 | 1048 | 0.986641221 | 1 | 1 | 1 | FIM1_2803 | YIL031W | YIL031W | ULP2 | n |
| Kmar_QGN16848.1 | W303.asm02.HP0_G0026980 | OG0002595 | chrIX | 301794 | 303089 | CP015058.1 | 1020751 | 1022007 | 74.47795824 | 431 | 418 | 1.031100478 | 1 | 1 | 1 | FIM1_3573 | YIL022W | YIL022W | TIM44 | n |
| Kmar_QGN18129.1 | W303.asm02.HP0_G0015770 | OG0002596 | chrV | 404892 | 406373 | CP015061.1 | 284857 | 286410 | 47.91666667 | 493 | 517 | 0.953578337 | 1 | 1 | 1 | FIM1_4450 | YER122C | YER122C | GLO3 | n |
| Kmar_QGN16158.1 | W303.asm02.HP0_G0015240 | OG0002597 | chrV | 304476 | 304865 | CP015057.1 | 923269 | 923649 | 83.72093023 | 129 | 126 | 1.023809524 | 1 | 1 | 1 | FIM1_2859 | YER072W | YER072W | VTC1 | n |
| Kmar_QGN16138.1 | W303.asm02.HP0_G0032790 | OG0002598 | chrXI | 358067 | 359617 | CP015057.1 | 892087 | 893235 | 24.6124031 | 516 | 382 | 1.35078534 | 1 | 1 | 1 | FIM1_2839 | YKL047W | YKL047W | ANR2 | n |
| Kmar_QGN15213.1 | W303.asm02.HP0_G0021510 | OG0002599 | chrVII | 862825 | 864993 | CP015056.1 | 442283 | 444523 | 40.85603113 | 722 | 746 | 0.967828418 | 1 | 1 | 1 | FIM1_1901 | YGR178C | YGR178C | PBP1 | n |
| Kmar_QGN18149.1 | W303.asm02.HP0_G0026560 | OG0002600 | chrIX | 223552 | 225969 | CP015061.1 | 328441 | 330825 | 50.7518797 | 805 | 794 | 1.013853904 | 1 | 1 | 1 | FIM1_4472 | YIL068C | YIL068C | SEC6 | n |
| Kmar_QGN16590.1 | W303.asm02.HP0_G0032310 | OG0002601 | chrXI | 270753 | 271589 | CP015058.1 | 451128 | 451994 | 42.06896552 | 278 | 288 | 0.965277778 | 1 | 1 | 1 | FIM1_3306 | YKL095W | YKL095W | YJU2 | n |
| Kmar_QGN14693.1 | W303.asm02.HP0_G0021000 | OG0002602 | chrVII | 753995 | 757105 | CP015055.1 | 1047040 | 1050150 | 51.95530726 | 1036 | 1036 | 1 | 1 | 1 | 1 | FIM1_1360 | YGR125W | YGR125W | VSB1 | n |
| Kmar_QGN13650.1 | W303.asm02.HP0_G0005670 | OG0002603 | chrIII | 150013 | 152874 | CP015054.1 | 582999 | 585938 | 57.33063701 | 953 | 979 | 0.973442288 | 1 | 1 | 1 | FIM1_293 | YCR017C | YCR017C | CWH43 | n |
| Kmar_QGN18377.1 | W303.asm02.HP0_G0008350 | OG0002604 | chrIV | 372125 | 372646 | CP015061.1 | 807125 | 807640 | 53.17919075 | 173 | 171 | 1.011695906 | 1 | 1 | 1 | FIM1_4704 | YDL046W | YDL046W | NPC2 | n |
| Kmar_QGN16118.1 | W303.asm02.HP0_G0004000 | OG0002605 | chrII | 651925 | 653322 | CP015057.1 | 843269 | 844594 | 59.41422594 | 465 | 441 | 1.054421769 | 1 | 0 | 1 | FIM1_2819 | YBR207W | YBR207W | FTH1 | n |
| Kmar_QGN17954.1 | W303.asm02.HP0_G0015680 | OG0002606 | chrV | 389954 | 392074 | CP015060.1 | 816153 | 818180 | 39.88995873 | 706 | 675 | 1.045925926 | 1 | 1 | 1 | FIM1_5163 | YER113C | YER113C | TMN3 | n |
| Kmar_QGN17956.1 | W303.asm02.HP0_G0001120 | OG0002607 | chrII | 68322 | 70598 | CP015060.1 | 821697 | 824084 | 36.76646707 | 758 | 795 | 0.953459119 | 1 | 1 | 1 | FIM1_5165 | YBL084C | YBL084C | CDC27 | n |
| Kmar_QGN15081.1 | W303.asm02.HP0_G0023080 | OG0002608 | chrVIII | 94974 | 96332 | CP015056.1 | 164075 | 165640 | 47.22792608 | 452 | 521 | 0.86756238 | 1 | 1 | 1 | FIM1_1766 | YHL002W | YHL002W | HSE1 | n |
| Kmar_QGN15082.1 | W303.asm02.HP0_G0033200 | OG0002609 | chrXI | 438394 | 439200 | CP015056.1 | 166028 | 166828 | 39.77695167 | 268 | 266 | 1.007518797 | 1 | 1 | 1 | FIM1_1767 | YKL007W | YKL007W | CAP1 | n |
| Kmar_QGN15076.1 | W303.asm02.HP0_G0033170 | OG0002610 | chrXI | 430518 | 434969 | CP015056.1 | 152710 | 157143 | 39.56975228 | 1483 | 1477 | 1.004062288 | 1 | 1 | 1 | FIM1_1761 | YKL010C | YKL010C | UFD4 | n |
| Kmar_QGN15071.1 | W303.asm02.HP0_G0041210 | OG0002611 | chrXIII | 423008 | 425233 | CP015056.1 | 140672 | 142990 | 40.27603513 | 741 | 772 | 0.95984456 | 1 | 1 | 1 | FIM1_1756 | YMR078C | YMR078C | CTF18 | n |
| Kmar_QGN15925.1 | W303.asm02.HP0_G0035900 | OG0002612 | chrXII | 349766 | 350809 | CP015057.1 | 390997 | 392037 | 68.3908046 | 347 | 346 | 1.002890173 | 1 | 1 | 1 | FIM1_2623 | YLR100W | YLR100W | ERG27 | n |
| Kmar_QGN15924.1 | W303.asm02.HP0_G0009920 | OG0002613 | chrIV | 714458 | 717904 | CP015057.1 | 387357 | 390764 | 46.51762683 | 1148 | 1135 | 1.011453744 | 1 | 1 | 1 | FIM1_2622 | YDR128W | YDR128W | MTC5 | n |
| Kmar_QGN16894.1 | W303.asm02.HP0_G0010260 | OG0002614 | chrIV | 793163 | 793783 | CP015058.1 | 1119118 | 1119696 | 58.25242718 | 206 | 192 | 1.072916667 | 1 | 1 | 1 | FIM1_3621 | YDR167W | YDR167W | TAF10 | n |
| Kmar_QGN14900.1 | W303.asm02.HP0_G0056190 | OG0002615 | chrXVI | 717902 | 719527 | CP015055.1 | 1457862 | 1459463 | 75.8302583 | 541 | 533 | 1.015009381 | 1 | 1 | 1 | FIM1_1574 | YPR088C | YPR088C | SRP54 | n |
| Kmar_QGN17316.1 | W303.asm02.HP0_G0024460 | OG0002616 | chrVIII | 347638 | 348093 | CP015059.1 | 680955 | 681362 | 43.70860927 | 151 | 135 | 1.118518519 | 1 | 1 | 1 | FIM1_4048 | YHR116W | YHR116W | COX23 | n |
| Kmar_QGN17403.1 | W303.asm02.HP0_G0033150 | OG0002617 | chrXI | 427404 | 429155 | CP015059.1 | 842623 | 844323 | 39.04109589 | 583 | 566 | 1.030035336 | 1 | 1 | 1 | FIM1_4135 | YKL012W | YKL012W | PRP40 | n |
| Kmar_QGN17405.1 | W303.asm02.HP0_G0027470 | OG0002618 | chrIX | 391078 | 393990 | CP015059.1 | 846374 | 849880 | 40.73763621 | 970 | 1168 | 0.830479452 | 1 | 1 | 1 | FIM1_4137 | YIR023W | YIR023W | DAL81 | n |
| Kmar_QGN15727.1 | W303.asm02.HP0_G0025820 | OG0002619 | chrIX | 58822 | 62484 | CP015056.1 | 1544457 | 1548056 | 52.61923378 | 1220 | 1199 | 1.017514595 | 1 | 1 | 1 | FIM1_2420 | YIL147C | YIL147C | SLN1 | n |
| Kmar_QGN17402.1 | W303.asm02.HP0_G0027460 | OG0002620 | chrIX | 390034 | 390537 | CP015059.1 | 842034 | 842543 | 67.26190476 | 167 | 169 | 0.98816568 | 1 | 1 | 1 | FIM1_4134 | YIR022W | YIR022W | SEC11 | n |
| Kmar_QGN17398.1 | W303.asm02.HP0_G0032380 | OG0002621 | chrXI | 283759 | 285474 | CP015059.1 | 836984 | 838585 | 35.2530541 | 571 | 533 | 1.071294559 | 1 | 1 | 1 | FIM1_4130 | YKL088W | YKL088W | CAB3 | n |
| Kmar_QGN15909.1 | W303.asm02.HP0_G0009840 | OG0002622 | chrIV | 696458 | 698170 | CP015057.1 | 356630 | 358324 | 63.69863014 | 570 | 564 | 1.010638298 | 1 | 1 | 1 | FIM1_2607 | YDR120C | YDR120C | TRM1 | n |
| Kmar_QGN17831.1 | W303.asm02.HP0_G0032700 | OG0002623 | chrXI | 343016 | 343519 | CP015060.1 | 580682 | 581185 | 80.83832335 | 167 | 167 | 1 | 1 | 1 | 1 | FIM1_5040 | YKL056C | YKL056C | TMA19 | n |
| Kmar_QGN15503.1 | W303.asm02.HP0_G0046960 | OG0002624 | chrXIV | 684991 | 685551 | CP015056.1 | 1074814 | 1075320 | 28.94736842 | 186 | 168 | 1.107142857 | 1 | 1 | 1 | FIM1_2194 | YNR024W | YNR024W | MPP6 | n |
| Kmar_QGN15502.1 | W303.asm02.HP0_G0006030 | OG0002625 | chrIII | 243438 | 246209 | CP015056.1 | 1071403 | 1074144 | 70.53087757 | 923 | 913 | 1.010952903 | 1 | 1 | 1 | FIM1_2193 | YCR057C | YCR057C | PWP2 | n |
| Kmar_QGN17832.1 | W303.asm02.HP0_G0032710 | OG0002626 | chrXI | 343925 | 344761 | CP015060.1 | 581596 | 582438 | 32.87197232 | 278 | 280 | 0.992857143 | 1 | 1 | 1 | FIM1_5041 | YKL055C | YKL055C | OAR1 | n |
| Kmar_QGN14318.1 | W303.asm02.HP0_G0010790 | OG0002627 | chrIV | 911831 | 913939 | CP015055.1 | 265050 | 267077 | 31.49717514 | 702 | 675 | 1.04 | 1 | 1 | 1 | FIM1_975 | YDR221W | YDR221W | GTB1 | n |
| Kmar_QGN17140.1 | W303.asm02.HP0_G0040620 | OG0002628 | chrXIII | 299374 | 300933 | CP015059.1 | 320824 | 322254 | 33.33333333 | 519 | 476 | 1.090336134 | 1 | 1 | 1 | FIM1_3871 | YMR014W | YMR014W | BUD22 | n |
| Kmar_QGN17793.1 | W303.asm02.HP0_G0049930 | OG0002629 | chrXV | 498241 | 503187 | CP015060.1 | 495041 | 499939 | 51.33372851 | 1648 | 1632 | 1.009803922 | 1 | 1 | 1 | FIM1_5002 | YOR093C | YOR093C | CMR2 | n |
| Kmar_QGN14248.1 | W303.asm02.HP0_G0021240 | OG0002630 | chrVII | 802186 | 804780 | CP015055.1 | 108570 | 111167 | 32.47480403 | 864 | 865 | 0.998843931 | 1 | 1 | 1 | FIM1_904 | YGR150C | YGR150C | CCM1 | n |
| Kmar_QGN18063.1 | W303.asm02.HP0_G0038460 | OG0002631 | chrXII | 878737 | 881421 | CP015061.1 | 133660 | 136308 | 56.63716814 | 894 | 882 | 1.013605442 | 1 | 1 | 1 | FIM1_4381 | YLR382C | YLR382C | NAM2 | n |
| Kmar_QGN18065.1 | W303.asm02.HP0_G0038440 | OG0002632 | chrXII | 874948 | 876174 | CP015061.1 | 138583 | 139929 | 46.54788419 | 408 | 448 | 0.910714286 | 1 | 1 | 1 | FIM1_4383 | YLR380W | YLR380W | CSR1 | n |
| Kmar_QGN15765.1 | W303.asm02.HP0_G0047240 | OG0002633 | chrXIV | 731928 | 733214 | CP015057.1 | 45339 | 46700 | 50.32537961 | 428 | 453 | 0.944812362 | 1 | 1 | 1 | FIM1_2461 | YNR052C | YNR052C | POP2 | n |
| Kmar_QGN16487.1 | W303.asm02.HP0_G0052130 | OG0002634 | chrXV | 948380 | 951256 | CP015058.1 | 212154 | 215024 | 80.27139875 | 958 | 956 | 1.00209205 | 1 | 1 | 1 | FIM1_3200 | YOR335C | YOR335C | ALA1 | n |
| Kmar_QGN15079.1 | W303.asm02.HP0_G0023060 | OG0002635 | chrVIII | 91580 | 92764 | CP015056.1 | 159409 | 160626 | 53.58024691 | 394 | 405 | 0.972839506 | 1 | 1 | 1 | FIM1_1764 | YHL004W | YHL004W | MRP4 | n |
| Kmar_QGN15571.1 | W303.asm02.HP0_G0032110 | OG0002636 | chrXI | 228183 | 228833 | CP015056.1 | 1212818 | 1213366 | 46.75925926 | 216 | 182 | 1.186813187 | 1 | 1 | 1 | FIM1_2262 | YKL117W | YKL117W | SBA1 | n |
| Kmar_QGN16466.1 | W303.asm02.HP0_G0004680 | OG0002637 | chrII | 786154 | 788082 | CP015058.1 | 168291 | 170102 | 44.90740741 | 642 | 603 | 1.064676617 | 1 | 1 | 1 | FIM1_3179 | YBR280C | YBR280C | SAF1 | n |
| Kmar_QGN15227.1 | W303.asm02.HP0_G0035160 | OG0002638 | chrXII | 192889 | 194520 | CP015056.1 | 478617 | 480626 | 35.45051699 | 543 | 669 | 0.811659193 | 1 | 1 | 1 | FIM1_1916 | YLR023C | YLR023C | IZH3 | n |
| Kmar_QGN17616.1 | W303.asm02.HP0_G0049070 | OG0002639 | chrXV | 325704 | 326585 | CP015060.1 | 112038 | 113105 | 42 | 293 | 355 | 0.825352113 | 1 | 1 | 1 | FIM1_4823 | YOL001W | YOL001W | PHO80 | n |
| Kmar_QGN15683.1 | W303.asm02.HP0_G0017970 | OG0002640 | chrVII | 85567 | 86760 | CP015056.1 | 1461192 | 1461798 | 24.78632479 | 397 | 187 | 2.122994652 | 1 | 1 | 1 | FIM1_2376 | YGL213C | YGL213C | SKI8 | n |
| Kmar_QGN16451.1 | W303.asm02.HP0_G0029110 | OG0002641 | chrX | 310035 | 312176 | CP015058.1 | 142335 | 144395 | 26.53061224 | 713 | 686 | 1.039358601 | 1 | 1 | 1 | FIM1_3164 | YJL061W | YJL061W | NUP82 | n |
| Kmar_QGN14422.1 | W303.asm02.HP0_G0018870 | OG0002642 | chrVII | 288985 | 289953 | CP015055.1 | 488647 | 489636 | 75.68389058 | 322 | 329 | 0.978723404 | 1 | 1 | 1 | FIM1_1083 | YGL115W | YGL115W | SNF4 | n |
| Kmar_QGN14423.1 | W303.asm02.HP0_G0018880 | OG0002643 | chrVII | 290412 | 292589 | CP015055.1 | 490240 | 492495 | 64.61336828 | 725 | 751 | 0.965379494 | 1 | 1 | 1 | FIM1_1084 | YGL114W | YGL114W | | n |
| Kmar_QGN14425.1 | W303.asm02.HP0_G0004220 | OG0002644 | chrII | 706270 | 707511 | CP015055.1 | 493808 | 495055 | 41.35514019 | 413 | 415 | 0.995180723 | 1 | 1 | 1 | FIM1_1086 | YBR233W | YBR233W | PBP2 | n |
| Kmar_QGN14424.1 | W303.asm02.HP0_G0004210 | OG0002645 | chrII | 705021 | 705932 | CP015055.1 | 492568 | 493515 | 46.15384615 | 303 | 315 | 0.961904762 | 1 | 1 | 1 | FIM1_1085 | YBR231C | YBR231C | SWC5 | n |
| Kmar_QGN14027.1 | W303.asm02.HP0_G0048530 | OG0002646 | chrXV | 221222 | 223357 | CP015054.1 | 1360483 | 1362495 | 58.96358543 | 711 | 670 | 1.06119403 | 1 | 1 | 1 | FIM1_678 | YOL057W | YOL057W | | n |
| Kmar_QGN17313.1 | W303.asm02.HP0_G0045590 | OG0002647 | chrXIV | 395893 | 398886 | CP015059.1 | 674270 | 677209 | 65.43086172 | 997 | 979 | 1.018386108 | 1 | 1 | 1 | FIM1_4045 | YNL123W | YNL123W | NMA111 | n |

| | | | | | | | | | | | | | | | | | | | | |
|---|---|---|---|---|---|---|---|---|---|---|---|---|---|---|---|---|---|---|---|---|
| Kmar_QGN17312.1 | W303.asm02.HP0_G0024480 | OG0002648 | chrVIII | 350294 | 351601 | CP015059.1 | 672737 | 674023 | 37.24604966 | 435 | 428 | 1.01635514 | 1 | 1 | 1 | FIM1_4044 | YHR118C | YHR118C | ORC6 | n |
| Kmar_QGN14145.1 | W303.asm02.HP0_G0005250 | OG0002649 | chrIII | 62427 | 62933 | CP015055.1 | 1629793 | 1630224 | 50 | 168 | 143 | 1.174825175 | 1 | 1 | 1 | FIM1_798 | YCL033C | YCL033C | MXR2 | n |
| Kmar_QGN14013.1 | W303.asm02.HP0_G0008590 | OG0002650 | chrIV | 426039 | 426971 | CP015054.1 | 1329941 | 1330975 | 53.33333333 | 310 | 344 | 0.901162791 | 1 | 1 | 1 | FIM1_664 | YDL015C | YDL015C | TSC13 | n |
| Kmar_QGN17272.1 | W303.asm02.HP0_G0051800 | OG0002651 | chrXV | 878949 | 879404 | CP015059.1 | 593301 | 593753 | 71.33333333 | 151 | 150 | 1.006666667 | 1 | 1 | 1 | FIM1_4003 | YOR298C-A | YOR298C-A | MBF1 | n |
| Kmar_QGN16829.1 | W303.asm02.HP0_G0027320 | OG0002652 | chrIX | 366599 | 368329 | CP015058.1 | 981653 | 983296 | 22.35494881 | 576 | 547 | 1.053016453 | 1 | 1 | 1 | FIM1_3553 | YIR010W | YIR010W | DSN1 | n |
| Kmar_QGN17498.1 | W303.asm02.HP0_G0021870 | OG0002653 | chrVII | 934590 | 934922 | CP015059.1 | 1036006 | 1036332 | 70.90909091 | 110 | 108 | 1.018518519 | 1 | 1 | 1 | FIM1_4234 | YGR215W | YGR215W | RSM27 | n |
| Kmar_QGN17500.1 | W303.asm02.HP0_G0021890 | OG0002654 | chrVII | 937111 | 943230 | CP015059.1 | 1038672 | 1044659 | 43.1884058 | 2039 | 1995 | 1.022055138 | 1 | 1 | 1 | FIM1_4236 | YGR217W | YGR217W | CCH1 | n |
| Kmar_QGN17495.1 | W303.asm02.HP0_G0021830 | OG0002655 | chrVII | 927656 | 929116 | CP015059.1 | 1028806 | 1030269 | 73.06122449 | 486 | 487 | 0.997946612 | 1 | 1 | 1 | FIM1_4231 | YGR211W | YGR211W | ZPR1 | n |
| Kmar_QGN16095.1 | W303.asm02.HP0_G0026760 | OG0002656 | chrIX | 259284 | 261206 | CP015057.1 | 800161 | 802185 | 55.53956835 | 640 | 674 | 0.949554896 | 1 | 1 | 1 | FIM1_2795 | YIL046W | YIL046W | MET30 | n |
| Kmar_QGN14809.1 | W303.asm02.HP0_G0047720 | OG0002657 | chrXV | 55758 | 56480 | CP015055.1 | 1272360 | 1273076 | 53.30578512 | 240 | 238 | 1.008403361 | 1 | 1 | 1 | FIM1_1480 | YOL142W | YOL142W | RRP40 | n |
| Kmar_QGN16166.1 | W303.asm02.HP0_G0026520 | OG0002658 | chrIX | 217097 | 218914 | CP015057.1 | 941277 | 943019 | 41.62601626 | 605 | 580 | 1.043103448 | 1 | 1 | 1 | FIM1_2868 | YIL072W | YIL072W | HOP1 | n |
| Kmar_QGN13731.1 | W303.asm02.HP0_G0025070 | OG0002659 | chrVIII | 465405 | 467636 | CP015054.1 | 760118 | 762103 | 47.80292943 | 743 | 661 | 1.124054463 | 1 | 1 | 1 | FIM1_375 | YHR178W | YHR178W | STB5 | n |
| Kmar_QGN16167.1 | W303.asm02.HP0_G0015330 | OG0002660 | chrV | 320590 | 321222 | CP015057.1 | 943951 | 944628 | 27.92792793 | 210 | 225 | 0.933333333 | 1 | 1 | 1 | FIM1_2869 | YER079W | YER079W | | n |
| Kmar_QGN16168.1 | W303.asm02.HP0_G0015340 | OG0002661 | chrV | 321633 | 323516 | CP015057.1 | 945512 | 947380 | 58.7955626 | 627 | 622 | 1.008038585 | 1 | 1 | 1 | FIM1_2870 | YER080W | YER080W | AIM9 | n |
| Kmar_QGN16830.1 | W303.asm02.HP0_G0004430 | OG0002662 | chrII | 747092 | 747619 | CP015058.1 | 983453 | 983929 | 52.57142857 | 175 | 158 | 1.107594937 | 1 | 1 | 1 | FIM1_3554 | YBR254C | YBR254C | TRS20 | n |
| Kmar_QGN16828.1 | W303.asm02.HP0_G0004420 | OG0002663 | chrII | 746625 | 746990 | CP015058.1 | 981001 | 981366 | 34.95934959 | 121 | 121 | 1 | 1 | 1 | 1 | FIM1_3552 | YBR253W | YBR253W | SRB6 | n |
| Kmar_QGN18283.1 | W303.asm02.HP0_G0056520 | OG0002664 | chrXVI | 794890 | 796254 | CP015061.1 | 611817 | 613193 | 59.73741794 | 454 | 458 | 0.991266376 | 1 | 1 | 1 | FIM1_4609 | YPR125W | YPR125W | YLH47 | n |
| Kmar_QGN14463.1 | W303.asm02.HP0_G0018810 | OG0002665 | chrVII | 275901 | 277478 | CP015055.1 | 570856 | 572448 | 40.56939502 | 525 | 530 | 0.990566038 | 1 | 1 | 1 | FIM1_1124 | YGL122C | YGL122C | NAB2 | n |
| Kmar_QGN14464.1 | W303.asm02.HP0_G0018800 | OG0002666 | chrVII | 274087 | 274851 | CP015055.1 | 572934 | 573707 | 85.21400778 | 254 | 257 | 0.988326848 | 1 | 1 | 1 | FIM1_1125 | YGL123W | YGL123W | RPS2 | n |
| Kmar_QGN14465.1 | W303.asm02.HP0_G0004190 | OG0002667 | chrII | 702392 | 702808 | CP015055.1 | 574991 | 575512 | 26.43678161 | 138 | 173 | 0.797687861 | 1 | 1 | 1 | FIM1_1126 | YBR230C | YBR230C | OM14 | n |
| Kmar_QGN14760.1 | W303.asm02.HP0_G0050340 | OG0002668 | chrXV | 585184 | 587199 | CP015055.1 | 1172267 | 1174006 | 27.0863836 | 671 | 579 | 1.158894646 | 1 | 1 | 1 | FIM1_1430 | YOR138C | YOR138C | RUP1 | n |
| Kmar_QGN14769.1 | W303.asm02.HP0_G0050290 | OG0002669 | chrXV | 574051 | 575706 | CP015055.1 | 1195262 | 1196944 | 50.08818342 | 551 | 560 | 0.983928571 | 1 | 1 | 1 | FIM1_1439 | YOR132W | YOR132W | VPS17 | n |
| Kmar_QGN13919.1 | W303.asm02.HP0_G0012540 | OG0002670 | chrIV | 1282379 | 1282894 | CP015054.1 | 1144287 | 1144802 | 78.3625731 | 171 | 171 | 1 | 1 | 1 | 1 | FIM1_567 | YDR404C | YDR404C | RPB7 | n |
| Kmar_QGN13582.1 | W303.asm02.HP0_G0050440 | OG0002671 | chrXV | 612874 | 613365 | CP015054.1 | 464029 | 464514 | 70.55214724 | 163 | 161 | 1.01242236 | 1 | 1 | 1 | FIM1_224 | YOR150W | YOR150W | MRPL23 | n |
| Kmar_QGN16564.1 | W303.asm02.HP0_G0050450 | OG0002672 | chrXV | 613872 | 617546 | CP015058.1 | 387255 | 390920 | 89.0790546 | 1224 | 1221 | 1.002457002 | 1 | 1 | 1 | FIM1_3278 | YOR151C | YOR151C | RPB2 | n |
| Kmar_QGN16565.1 | W303.asm02.HP0_G0012550 | OG0002673 | chrIV | 1283371 | 1284162 | CP015058.1 | 391680 | 392399 | 48.67924528 | 263 | 239 | 1.10041841 | 1 | 1 | 1 | FIM1_3279 | YDR405W | YDR405W | MRP20 | n |
| Kmar_QGN15651.1 | W303.asm02.HP0_G0050490 | OG0002674 | chrXV | 627503 | 628855 | CP015056.1 | 1391776 | 1393044 | 55.55555556 | 450 | 422 | 1.066350711 | 1 | 1 | 1 | FIM1_2344 | YOR155C | YOR155C | ISN1 | n |
| Kmar_QGN16807.1 | W303.asm02.HP0_G0020750 | OG0002675 | chrVII | 681037 | 685929 | CP015058.1 | 930659 | 935431 | 33.47457627 | 1630 | 1590 | 1.025157233 | 1 | 1 | 1 | FIM1_3531 | YGR098C | YGR098C | ESP1 | n |
| Kmar_QGN15450.1 | W303.asm02.HP0_G0023870 | OG0002676 | chrVIII | 225938 | 226330 | CP015056.1 | 982026 | 982403 | 52.30769231 | 130 | 125 | 1.04 | 1 | 1 | 1 | FIM1_2141 | YHR059W | YHR059W | FYV4 | n |
| Kmar_QGN14756.1 | W303.asm02.HP0_G0050370 | OG0002677 | chrXV | 593932 | 594921 | CP015055.1 | 1163998 | 1164954 | 80.92307692 | 329 | 318 | 1.034591195 | 1 | 1 | 1 | FIM1_1426 | YOR142W | YOR142W | LSC1 | n |
| Kmar_QGN14758.1 | W303.asm02.HP0_G0050360 | OG0002678 | chrXV | 590817 | 593462 | CP015055.1 | 1166308 | 1168686 | 40.85903084 | 881 | 792 | 1.112373737 | 1 | 1 | 1 | FIM1_1428 | YOR141C | YOR141C | ARP8 | n |
| Kmar_QGN18085.1 | W303.asm02.HP0_G0030640 | OG0002679 | chrX | 646803 | 647408 | CP015061.1 | 177735 | 178346 | 36.40776699 | 201 | 203 | 0.990147783 | 1 | 1 | 1 | FIM1_4404 | YJR112W | YJR112W | NNF1 | n |
| Kmar_QGN15995.1 | W303.asm02.HP0_G0011900 | OG0002680 | chrIV | 1156919 | 1157779 | CP015057.1 | 570467 | 571315 | 59.09090909 | 286 | 282 | 1.014184397 | 1 | 1 | 1 | FIM1_2695 | YDR337W | YDR337W | MRPS28 | n |
| Kmar_QGN14963.1 | W303.asm02.HP0_G0051170 | OG0002681 | chrXV | 776297 | 776983 | CP015055.1 | 1585853 | 1586623 | 64.453125 | 228 | 256 | 0.890625 | 1 | 1 | 1 | FIM1_1641 | YOR232W | YOR232W | MGE1 | n |
| Kmar_QGN17705.1 | W303.asm02.HP0_G0026340 | OG0002682 | chrIX | 176875 | 177990 | CP015060.1 | 285450 | 286559 | 79.5148248 | 371 | 369 | 1.005420054 | 1 | 1 | 1 | FIM1_4912 | YIL094C | YIL094C | LYS12 | n |
| Kmar_QGN17709.1 | W303.asm02.HP0_G0046400 | OG0002683 | chrXIV | 579705 | 580874 | CP015060.1 | 294098 | 295381 | 51.1682243 | 389 | 427 | 0.911007026 | 1 | 1 | 1 | FIM1_4916 | YNL035C | YNL035C | | n |
| Kmar_QGN17710.1 | W303.asm02.HP0_G0046360 | OG0002684 | chrXIV | 566231 | 568015 | CP015060.1 | 295673 | 297484 | 42.02898551 | 594 | 603 | 0.985074627 | 1 | 1 | 1 | FIM1_4917 | YNL039W | YNL039W | BDP1 | n |
| Kmar_QGN17711.1 | W303.asm02.HP0_G0046370 | OG0002685 | chrXIV | 568203 | 568966 | CP015060.1 | 297745 | 298227 | 22.01834862 | 229 | 160 | 1.43125 | 1 | 1 | 1 | FIM1_4918 | YNL038W | YNL038W | GPI15 | n |
| Kmar_QGN17911.1 | W303.asm02.HP0_G0015410 | OG0002686 | chrV | 332246 | 333976 | CP015060.1 | 719183 | 720838 | 52.43055556 | 576 | 551 | 1.045372051 | 1 | 1 | 1 | FIM1_5120 | YER087W | YER087W | AIM10 | n |
| Kmar_QGN17913.1 | W303.asm02.HP0_G0001390 | OG0002687 | chrII | 124394 | 126400 | CP015060.1 | 722717 | 724621 | 44.50867052 | 668 | 634 | 1.05362776 | 1 | 1 | 1 | FIM1_5122 | YBL051C | YBL051C | PIN4 | n |
| Kmar_QGN17917.1 | W303.asm02.HP0_G0001360 | OG0002688 | chrII | 116721 | 117977 | CP015060.1 | 734300 | 735568 | 43.59550562 | 418 | 422 | 0.990521327 | 1 | 1 | 1 | FIM1_5126 | YBL055C | YBL055C | | n |
| Kmar_QGN17920.1 | W303.asm02.HP0_G0001340 | OG0002689 | chrII | 113949 | 114575 | CP015060.1 | 740463 | 741083 | 63.33333333 | 208 | 206 | 1.009708738 | 1 | 1 | 1 | FIM1_5129 | YBL057C | YBL057C | PTH2 | n |
| Kmar_QGN17921.1 | W303.asm02.HP0_G0001330 | OG0002690 | chrII | 112585 | 113856 | CP015060.1 | 741218 | 742435 | 53.95348837 | 423 | 405 | 1.044444444 | 1 | 1 | 1 | FIM1_5130 | YBL058W | YBL058W | SHP1 | n |
| Kmar_QGN17922.1 | W303.asm02.HP0_G0015460 | OG0002691 | chrV | 341534 | 343837 | CP015060.1 | 743296 | 745602 | 78.25520833 | 767 | 768 | 0.998697917 | 1 | 1 | 1 | FIM1_5131 | YER091C | YER091C | MET6 | n |
| Kmar_QGN13610.1 | W303.asm02.HP0_G0039510 | OG0002692 | chrXIII | 52425 | 53246 | CP015054.1 | 515858 | 516685 | 55.4770318 | 273 | 275 | 0.992727273 | 1 | 1 | 1 | FIM1_252 | YML105C | YML105C | SEC65 | n |
| Kmar_QGN18007.1 | W303.asm02.HP0_G0053850 | OG0002693 | chrXVI | 252924 | 256196 | CP015061.1 | 21573 | 24854 | 77.6459854 | 1090 | 1093 | 0.997255261 | 1 | 1 | 1 | FIM1_4323 | YPL160W | YPL160W | CDC60 | n |
| Kmar_QGN14257.1 | W303.asm02.HP0_G0043070 | OG0002694 | chrXIII | 809388 | 810722 | CP015055.1 | 125656 | 126978 | 39.29359823 | 444 | 440 | 1.009090909 | 1 | 1 | 1 | FIM1_914 | YMR268C | YMR268C | PRP24 | n |
| Kmar_QGN13607.1 | W303.asm02.HP0_G0043080 | OG0002695 | chrXIII | 810956 | 811591 | CP015054.1 | 510137 | 510712 | 53.08056872 | 211 | 191 | 1.104712042 | 1 | 1 | 1 | FIM1_249 | YMR269W | YMR269W | TMA23 | n |
| Kmar_QGN13608.1 | W303.asm02.HP0_G0039520 | OG0002696 | chrXIII | 53497 | 56880 | CP015054.1 | 510952 | 514359 | 34.27350427 | 1127 | 1135 | 0.992951542 | 1 | 1 | 1 | FIM1_250 | YML104C | YML104C | MDM1 | n |
| Kmar_QGN13979.1 | W303.asm02.HP0_G0016810 | OG0002697 | chrVI | 83591 | 84226 | CP015054.1 | 1259575 | 1260186 | 47.98206278 | 211 | 203 | 1.039408867 | 1 | 1 | 1 | FIM1_628 | YFL010C | YFL010C | WWM1 | n |
| Kmar_QGN14682.1 | W303.asm02.HP0_G0035510 | OG0002698 | chrXII | 268791 | 270578 | CP015055.1 | 1022960 | 1024747 | 74.62184874 | 595 | 595 | 1 | 1 | 1 | 1 | FIM1_1348 | YLR060W | YLR060W | FRS1 | n |
| Kmar_QGN14683.1 | W303.asm02.HP0_G0016570 | OG0002699 | chrVI | 33925 | 37146 | CP015055.1 | 1024865 | 1027582 | 43.8547486 | 1073 | 905 | 1.185635359 | 1 | 1 | 1 | FIM1_1349 | YFL034W | YFL034W | MIL1 | n |
| Kmar_QGN18046.1 | W303.asm02.HP0_G0035530 | OG0002700 | chrXII | 272108 | 273205 | CP015061.1 | 95718 | 96755 | 47.0437018 | 365 | 345 | 1.057971014 | 1 | 1 | 1 | FIM1_4363 | YLR063W | YLR063W | BMT6 | n |
| Kmar_QGN14409.1 | W303.asm02.HP0_G0055250 | OG0002701 | chrXVI | 532167 | 533240 | CP015055.1 | 458231 | 459301 | 46.0704607 | 357 | 356 | 1.002808989 | 1 | 1 | 1 | FIM1_1069 | YPL015C | YPL015C | HST2 | n |
| Kmar_QGN16086.1 | W303.asm02.HP0_G0055310 | OG0002702 | chrXVI | 542177 | 545293 | CP015057.1 | 781591 | 784668 | 49.85994398 | 1038 | 1025 | 1.012682927 | 1 | 1 | 1 | FIM1_2786 | YPL009C | YPL009C | RQC2 | n |
| Kmar_QGN13865.1 | W303.asm02.HP0_G0022530 | OG0002703 | chrVII | 1077219 | 1078346 | CP015054.1 | 1020190 | 1021344 | 75.2 | 375 | 384 | 0.9765625 | 1 | 1 | 1 | FIM1_511 | YGR286C | YGR286C | BIO2 | n |
| Kmar_QGN16482.1 | W303.asm02.HP0_G0052180 | OG0002704 | chrXV | 961484 | 962464 | CP015058.1 | 199419 | 200285 | 55.21472393 | 326 | 288 | 1.131944444 | 1 | 1 | 1 | FIM1_3195 | YOR340C | YOR340C | RPA43 | n |
| Kmar_QGN17800.1 | W303.asm02.HP0_G0050000 | OG0002705 | chrXV | 514030 | 515013 | CP015060.1 | 512654 | 513691 | 59.14285714 | 327 | 345 | 0.947826087 | 1 | 1 | 1 | FIM1_5009 | YOR100C | YOR100C | CRC1 | n |
| Kmar_QGN17306.1 | W303.asm02.HP0_G0045560 | OG0002706 | chrXIV | 388060 | 390600 | CP015059.1 | 659336 | 661861 | 34.30232558 | 846 | 841 | 1.005945303 | 1 | 1 | 1 | FIM1_4038 | YNL126W | YNL126W | SPC98 | n |
| Kmar_QGN15536.1 | W303.asm02.HP0_G0010890 | OG0002707 | chrIV | 931953 | 933599 | CP015056.1 | 1143011 | 1144726 | 69.32409012 | 548 | 571 | 0.95971979 | 1 | 1 | 1 | FIM1_2227 | YDR232W | YDR232W | HEM1 | n |
| Kmar_QGN17706.1 | W303.asm02.HP0_G0046530 | OG0002708 | chrXIV | 602610 | 604082 | CP015060.1 | 286617 | 288101 | 64.12825651 | 490 | 494 | 0.991902834 | 1 | 1 | 1 | FIM1_4913 | YNL022C | YNL022C | RCM1 | n |
| Kmar_QGN17707.1 | W303.asm02.HP0_G0046540 | OG0002709 | chrXIV | 604410 | 606530 | CP015060.1 | 288617 | 290737 | 69.49152542 | 706 | 706 | 1 | 1 | 1 | 1 | FIM1_4914 | YNL021W | YNL021W | HDA1 | n |
| Kmar_QGN14452.1 | W303.asm02.HP0_G0018090 | OG0002710 | chrVII | 113367 | 116420 | CP015055.1 | 539576 | 542710 | 63.31753555 | 1017 | 1044 | 0.974137931 | 1 | 1 | 1 | FIM1_1113 | YGL201C | YGL201C | MCM6 | n |
| Kmar_QGN14115.1 | W303.asm02.HP0_G0036750 | OG0002711 | chrXII | 530925 | 534521 | CP015054.1 | 1563585 | 1567280 | 45.1561249 | 1198 | 1231 | 0.973192526 | 1 | 1 | 1 | FIM1_767 | YLR189C | YLR189C | ATG26 | n |
| Kmar_QGN14116.1 | W303.asm02.HP0_G0044200 | OG0002712 | chrXIV | 117903 | 119363 | CP015054.1 | 1567643 | 1569121 | 68.07228916 | 486 | 492 | 0.987804878 | 1 | 1 | 1 | FIM1_768 | YNL277W | YNL277W | MET2 | n |
| Kmar_QGN13392.1 | W303.asm02.HP0_G0003640 | OG0002713 | chrII | 593119 | 594861 | CP015054.1 | 56965 | 58755 | 63.71237458 | 580 | 596 | 0.973154362 | 1 | 1 | 1 | FIM1_29 | YBR170C | YBR170C | NPL4 | n |
| Kmar_QGN13900.1 | W303.asm02.HP0_G0042560 | OG0002714 | chrXIII | 699390 | 701618 | CP015054.1 | 1111183 | 1113327 | 51.98938992 | 742 | 714 | 1.039215686 | 1 | 1 | 1 | FIM1_547 | YMR216C | YMR216C | SKY1 | n |
| Kmar_QGN14589.1 | W303.asm02.HP0_G0047080 | OG0002715 | chrXIV | 705601 | 706629 | CP015055.1 | 812119 | 813099 | 57.92507205 | 342 | 326 | 1.049079755 | 1 | 1 | 1 | FIM1_1251 | YNR035C | YNR035C | ARC35 | n |
| Kmar_QGN14590.1 | W303.asm02.HP0_G0047090 | OG0002716 | chrXIV | 706943 | 707404 | CP015055.1 | 813426 | 813929 | 69.46107784 | 153 | 167 | 0.916167665 | 1 | 1 | 1 | FIM1_1252 | YNR036C | YNR036C | MRPS12 | n |
| Kmar_QGN16476.1 | W303.asm02.HP0_G0004480 | OG0002717 | chrII | 753090 | 753518 | CP015058.1 | 188261 | 188644 | 31.54362416 | 142 | 127 | 1.118110236 | 1 | 1 | 1 | FIM1_3189 | YBR258C | YBR258C | SHG1 | n |
| Kmar_QGN16473.1 | W303.asm02.HP0_G0004500 | OG0002718 | chrII | 755995 | 757995 | CP015058.1 | 181581 | 183734 | 57.98898072 | 666 | 717 | 0.928870293 | 1 | 1 | 1 | FIM1_3186 | YBR260C | YBR260C | RGD1 | n |
| Kmar_QGN14944.1 | W303.asm02.HP0_G0024710 | OG0002719 | chrVIII | 389511 | 390461 | CP015055.1 | 1542909 | 1543862 | 56.46687697 | 316 | 317 | 0.996845426 | 1 | 1 | 1 | FIM1_1619 | YHR142W | YHR142W | CHS7 | n |
| Kmar_QGN15167.1 | W303.asm02.HP0_G0028750 | OG0002720 | chrX | 227279 | 229102 | CP015056.1 | 344724 | 346400 | 46.53784219 | 607 | 558 | 1.08781362 | 1 | 1 | 1 | FIM1_1855 | YJL100W | YJL100W | LSB6 | n |
| Kmar_QGN16467.1 | W303.asm02.HP0_G0004690 | OG0002721 | chrII | 788340 | 790976 | CP015058.1 | 170482 | 173091 | 50.16797312 | 878 | 869 | 1.010356732 | 1 | 1 | 1 | FIM1_3180 | YBR281C | YBR281C | DUG2 | n |
| Kmar_QGN13499.1 | W303.asm02.HP0_G0041800 | OG0002722 | chrXIII | 548335 | 549804 | CP015054.1 | 271626 | 273074 | 53.87673956 | 489 | 482 | 1.014522822 | 1 | 1 | 1 | FIM1_137 | YMR140W | YMR140W | SIP5 | n |
| Kmar_QGN14850.1 | W303.asm02.HP0_G0039600 | OG0002723 | chrXIII | 76031 | 76663 | CP015055.1 | 1346309 | 1346980 | 40.17857143 | 210 | 223 | 0.941704036 | 1 | 1 | 1 | FIM1_1523 | YML095C | YML095C | RAD10 | n |
| Kmar_QGN17107.1 | W303.asm02.HP0_G0039290 | OG0002724 | chrXIII | 11616 | 13361 | CP015059.1 | 242511 | 244205 | 46.64371773 | 581 | 564 | 1.030141844 | 1 | 1 | 1 | FIM1_3838 | YML127W | YML127W | RSC9 | n |
| Kmar_QGN16602.1 | W303.asm02.HP0_G0008890 | OG0002725 | chrIV | 485405 | 486103 | CP015058.1 | 477658 | 478287 | 38.79310345 | 232 | 209 | 1.110047847 | 1 | 1 | 1 | FIM1_3318 | YDR020C | YDR020C | DAS2 | n |
| Kmar_QGN13673.1 | W303.asm02.HP0_G0008900 | OG0002726 | chrIV | 486463 | 487662 | CP015054.1 | 640032 | 641228 | 80.20050125 | 399 | 398 | 1.002512563 | 1 | 1 | 1 | FIM1_316 | YDR021W | YDR021W | FAL1 | n |
| Kmar_QGN17951.1 | W303.asm02.HP0_G0015650 | OG0002727 | chrV | 380432 | 383773 | CP015060.1 | 807636 | 810977 | 69.12028725 | 1113 | 1113 | 1 | 1 | 1 | 1 | FIM1_5160 | YER110C | YER110C | KAP123 | n |
| Kmar_QGN13839.1 | W303.asm02.HP0_G0031560 | OG0002728 | chrXI | 114385 | 115797 | CP015054.1 | 976534 | 977853 | 47.77070064 | 470 | 439 | 1.070615034 | 1 | 1 | 1 | FIM1_485 | YKL178C | YKL178C | STE3 | n |
| Kmar_QGN17946.1 | W303.asm02.HP0_G0001150 | OG0002729 | chrII | 74223 | 75848 | CP015060.1 | 795585 | 797204 | 60.36697248 | 541 | 539 | 1.003710575 | 1 | 1 | 1 | FIM1_5155 | YBL080C | YBL080C | PET112 | n |
| Kmar_QGN13474.1 | W303.asm02.HP0_G0029890 | OG0002730 | chrX | 499612 | 500793 | CP015054.1 | 212266 | 213435 | 55.18987342 | 393 | 389 | 1.010282776 | 1 | 1 | 1 | FIM1_112 | YJR032W | YJR032W | CPR7 | n |
| Kmar_QGN16315.1 | W303.asm02.HP0_G0018660 | OG0002731 | chrVII | 244962 | 245999 | CP015057.1 | 1265474 | 1266709 | 19.80676329 | 345 | 411 | 0.839416058 | 1 | 1 | 1 | FIM1_3020 | YGL138C | YGL138C | | n |
| Kmar_QGN16309.1 | W303.asm02.HP0_G0053310 | OG0002732 | chrXVI | 145184 | 148735 | CP015057.1 | 1252953 | 1256471 | 70.2020202 | 1183 | 1172 | 1.009385666 | 1 | 1 | 1 | FIM1_3014 | YPL217C | YPL217C | BMS1 | n |
| Kmar_QGN17840.1 | W303.asm02.HP0_G0040830 | OG0002733 | chrXIII | 338291 | 339780 | CP015060.1 | 596213 | 597577 | 49.46466809 | 467 | 454 | 1.028634361 | 1 | 1 | 1 | FIM1_5049 | YMR033W | YMR033W | ARP9 | n |
| Kmar_QGN17341.1 | W303.asm02.HP0_G0052260 | OG0002734 | chrXV | 992070 | 995114 | CP015059.1 | 728527 | 731568 | 24.56978967 | 1014 | 1013 | 1.000987167 | 1 | 1 | 1 | FIM1_4073 | YOR349W | YOR349W | CIN1 | n |
| Kmar_QGN16145.1 | W303.asm02.HP0_G0047870 | OG0002735 | chrXV | 84174 | 85604 | CP015057.1 | 901226 | 902512 | 38.03680982 | 476 | 428 | 1.112149533 | 1 | 1 | 1 | FIM1_2846 | YOL125W | YOL125W | TRM13 | n |
| Kmar_QGN17084.1 | W303.asm02.HP0_G0046870 | OG0002736 | chrXIV | 666826 | 673527 | CP015059.1 | 193714 | 200412 | 80.80536913 | 2233 | 2232 | 1.000448029 | 1 | 1 | 1 | FIM1_3815 | YNR016C | YNR016C | ACC1 | n |
| Kmar_QGN14028.1 | W303.asm02.HP0_G0048520 | OG0002737 | chrXV | 219666 | 220928 | CP015054.1 | 1363585 | 1364847 | 83.57142857 | 420 | 420 | 1 | 1 | 1 | 1 | FIM1_679 | YOL058W | YOL058W | ARG1 | n |
| Kmar_QGN18216.1 | W303.asm02.HP0_G0007390 | OG0002738 | chrIV | 192956 | 194431 | CP015061.1 | 470463 | 472010 | 40.29850746 | 491 | 515 | 0.953398058 | 1 | 1 | 1 | FIM1_4540 | YDL146W | YDL146W | LDB17 | n |
| Kmar_QGN17517.1 | W303.asm02.HP0_G0022020 | OG0002739 | chrVII | 967089 | 970625 | CP015059.1 | 1071758 | 1075171 | 43.86989158 | 1178 | 1137 | 1.036059807 | 1 | 1 | 1 | FIM1_4253 | YGR233C | YGR233C | PHO81 | n |
| Kmar_QGN17448.1 | W303.asm02.HP0_G0008730 | OG0002740 | chrIV | 452704 | 453309 | CP015059.1 | 942094 | 942702 | 73.55769231 | 201 | 202 | 0.995049505 | 1 | 1 | 1 | FIM1_4182 | YDR002W | YDR002W | YRB1 | n |
| Kmar_QGN17450.1 | W303.asm02.HP0_G0001930 | OG0002741 | chrII | 245148 | 246569 | CP015059.1 | 944620 | 946053 | 73.22175732 | 473 | 477 | 0.991614256 | 1 | 1 | 1 | FIM1_4184 | YBR003W | YBR003W | COQ1 | n |
| Kmar_QGN14019.1 | W303.asm02.HP0_G0000760 | OG0002742 | chrI | 170579 | 171499 | CP015054.1 | 1342673 | 1343590 | 79.41176471 | 306 | 305 | 1.003278689 | 1 | 1 | 1 | FIM1_670 | YAR015W | YAR015W | ADE1 | n |
| Kmar_QGN17502.1 | W303.asm02.HP0_G0021910 | OG0002743 | chrVII | 948489 | 949298 | CP015059.1 | 1049011 | 1049835 | 62.18181818 | 269 | 274 | 0.981751825 | 1 | 1 | 1 | FIM1_4238 | YGR220C | YGR220C | MRPL9 | n |
| Kmar_QGN17503.1 | W303.asm02.HP0_G0024760 | OG0002744 | chrVIII | 398744 | 399388 | CP015059.1 | 1050236 | 1050880 | 62.61682243 | 214 | 214 | 1 | 1 | 1 | 1 | FIM1_4239 | YHR147C | YHR147C | MRPL6 | n |
| Kmar_QGN17504.1 | W303.asm02.HP0_G0024770 | OG0002745 | chrVIII | 399639 | 400190 | CP015059.1 | 1051465 | 1052016 | 85.79234973 | 183 | 183 | 1 | 1 | 1 | 1 | FIM1_4240 | YHR148W | YHR148W | IMP3 | n |
| Kmar_QGN16991.1 | W303.asm02.HP0_G0006570 | OG0002746 | chrIV | 22897 | 25950 | CP015058.1 | 1326232 | 1329207 | 46.52777778 | 1017 | 991 | 1.026236125 | 1 | 1 | 1 | FIM1_3718 | YDL240W | YDL240W | LRG1 | n |

| | | | | | | | | | | | | | | | | | | | | |
|---|---|---|---|---|---|---|---|---|---|---|---|---|---|---|---|---|---|---|---|---|
| Kmar_QGN14306.1 | W303.asm02.HP0_G0034920 | OG0002747 | chrXII | 147483 | 148793 | CP015055.1 | 238638 | 239933 | 44.94382022 | 436 | 431 | 1.011600928 | 1 | 1 | 1 | FIM1_963 | YLL002W | YLL002W | RTT109 | n |
| Kmar_QGN15786.1 | W303.asm02.HP0_G0019900 | OG0002748 | chrVII | 496010 | 497440 | CP015057.1 | 98226 | 99764 | 51.49812734 | 476 | 512 | 0.9296875 | 1 | 1 | 1 | FIM1_2482 | YGR002C | YGR002C | SWC4 | n |
| Kmar_QGN15787.1 | W303.asm02.HP0_G0019910 | OG0002749 | chrVII | 497666 | 499900 | CP015057.1 | 100070 | 102277 | 29.51251647 | 744 | 735 | 1.012244898 | 1 | 1 | 1 | FIM1_2483 | YGR003W | YGR003W | CUL3 | n |
| Kmar_QGN14390.1 | W303.asm02.HP0_G0053990 | OG0002750 | chrXVI | 279758 | 281800 | CP015055.1 | 423786 | 426320 | 53.06603774 | 680 | 844 | 0.805687204 | 1 | 1 | 1 | FIM1_1050 | YPL147W | YPL147W | PXA1 | n |
| Kmar_QGN17545.1 | W303.asm02.HP0_G0033710 | OG0002751 | chrXI | 535075 | 536328 | CP015059.1 | 1136864 | 1138123 | 53.02325581 | 417 | 419 | 0.99522673 | 1 | 1 | 1 | FIM1_4281 | YKR048C | YKR048C | NAP1 | n |
| Kmar_QGN15389.1 | W303.asm02.HP0_G0021330 | OG0002752 | chrVII | 820402 | 821193 | CP015056.1 | 858590 | 859510 | 24.50980392 | 263 | 306 | 0.859477124 | 1 | 1 | 1 | FIM1_2079 | YGR161C | YGR161C | RTS3 | n |
| Kmar_QGN14290.1 | W303.asm02.HP0_G0029310 | OG0002753 | chrX | 364397 | 369448 | CP015055.1 | 196050 | 200996 | 46.96523276 | 1683 | 1648 | 1.021237864 | 1 | 1 | 1 | FIM1_947 | YJL039C | YJL039C | NUP192 | n |
| Kmar_QGN15734.1 | W303.asm02.HP0_G0034210 | OG0002754 | chrXI | 649418 | 651067 | CP015056.1 | 1566215 | 1567846 | 83.06010929 | 549 | 543 | 1.011049724 | 1 | 1 | 1 | FIM1_2428 | YKR097W | YKR097W | PCK1 | n |
| Kmar_QGN14416.1 | W303.asm02.HP0_G0025690 | OG0002755 | chrIX | 29222 | 30475 | CP015055.1 | 474757 | 475992 | 65.94724221 | 417 | 411 | 1.01459854 | 1 | 1 | 1 | FIM1_1077 | YIL160C | YIL160C | POT1 | n |
| Kmar_QGN17401.1 | W303.asm02.HP0_G0041310 | OG0002756 | chrXIII | 449750 | 450433 | CP015059.1 | 841034 | 841711 | 54.58515284 | 227 | 225 | 1.008888889 | 1 | 0 | 1 | FIM1_4133 | YMR090W | YMR090W | | n |
| Kmar_QGN13700.1 | W303.asm02.HP0_G0022410 | OG0002757 | chrVII | 1053302 | 1056502 | CP015054.1 | 685439 | 688606 | 48.11151079 | 1066 | 1055 | 1.01042654 | 1 | 1 | 1 | FIM1_343 | YGR274C | YGR274C | TAF1 | n |
| Kmar_QGN13698.1 | W303.asm02.HP0_G0022430 | OG0002758 | chrVII | 1057226 | 1058887 | CP015054.1 | 682071 | 684209 | 36.61360347 | 553 | 712 | 0.776685393 | 1 | 1 | 1 | FIM1_341 | YGR276C | YGR276C | RNH70 | n |
| Kmar_QGN18230.1 | W303.asm02.HP0_G0007280 | OG0002759 | chrIV | 170622 | 172142 | CP015061.1 | 503087 | 504685 | 74.86238532 | 506 | 532 | 0.95112782 | 1 | 1 | 1 | FIM1_4555 | YDL160C | YDL160C | DHH1 | n |
| Kmar_QGN17410.1 | W303.asm02.HP0_G0033100 | OG0002760 | chrXI | 414282 | 416333 | CP015059.1 | 861045 | 863120 | 50.97493036 | 683 | 691 | 0.988422576 | 1 | 1 | 1 | FIM1_4142 | YKL017C | YKL017C | HCS1 | n |
| Kmar_QGN17408.1 | W303.asm02.HP0_G0033120 | OG0002761 | chrXI | 417637 | 420576 | CP015059.1 | 856423 | 858987 | 45.64777328 | 979 | 854 | 1.146370023 | 1 | 1 | 1 | FIM1_4140 | YKL015W | YKL015W | PUT3 | n |
| Kmar_QGN14333.1 | W303.asm02.HP0_G0028730 | OG0002762 | chrX | 221317 | 223776 | CP015055.1 | 296874 | 299270 | 52.77777778 | 819 | 798 | 1.026315789 | 1 | 1 | 1 | FIM1_993 | YJL102W | YJL102W | MEF2 | n |
| Kmar_QGN18122.1 | W303.asm02.HP0_G0055650 | OG0002763 | chrXVI | 621877 | 625512 | CP015061.1 | 264047 | 267520 | 49.47453517 | 1211 | 1157 | 1.046672429 | 1 | 1 | 1 | FIM1_4443 | YPR026W | YPR026W | ATH1 | n |
| Kmar_QGN17048.1 | W303.asm02.HP0_G0006980 | OG0002764 | chrIV | 107282 | 111103 | CP015059.1 | 117854 | 121624 | 48.25716499 | 1273 | 1256 | 1.013535032 | 1 | 1 | 1 | FIM1_3777 | YDL195W | YDL195W | SEC31 | n |
| Kmar_QGN14727.1 | W303.asm02.HP0_G0021090 | OG0002765 | chrVII | 769497 | 772865 | CP015055.1 | 1104665 | 1108306 | 22.88477035 | 1122 | 1213 | 0.92497939 | 1 | 1 | 1 | FIM1_1395 | YGR134W | YGR134W | CAF130 | n |
| Kmar_QGN18357.1 | W303.asm02.HP0_G0017670 | OG0002766 | chrVII | 32987 | 34150 | CP015061.1 | 759852 | 761018 | 62.53229974 | 387 | 388 | 0.99742268 | 1 | 1 | 1 | FIM1_4684 | YGL246C | YGL246C | RAI1 | n |
| Kmar_QGN14921.1 | W303.asm02.HP0_G0045020 | OG0002767 | chrXIV | 292889 | 293365 | CP015055.1 | 1498315 | 1498764 | 77.21518987 | 158 | 149 | 1.060402685 | 1 | 1 | 1 | FIM1_1595 | YNL185C | YNL185C | MRPL19 | n |
| Kmar_QGN14925.1 | W303.asm02.HP0_G0045050 | OG0002768 | chrXIV | 299031 | 300254 | CP015055.1 | 1505861 | 1507114 | 60.8490566 | 407 | 417 | 0.976019185 | 1 | 1 | 1 | FIM1_1599 | YNL181W | YNL181W | PBR1 | n |
| Kmar_QGN14923.1 | W303.asm02.HP0_G0006810 | OG0002769 | chrIV | 77362 | 78039 | CP015055.1 | 1502997 | 1503656 | 60.96491228 | 225 | 219 | 1.02739726 | 1 | 1 | 1 | FIM1_1597 | YDL213C | YDL213C | NOP6 | n |
| Kmar_QGN17557.1 | W303.asm02.HP0_G0007770 | OG0002770 | chrIV | 267904 | 268905 | CP015059.1 | 1159326 | 1160258 | 75.80645161 | 306 | 310 | 0.987096774 | 1 | 1 | 1 | FIM1_4293 | YDL108W | YDL108W | KIN28 | n |
| Kmar_QGN17674.1 | W303.asm02.HP0_G0007680 | OG0002771 | chrIV | 248787 | 251444 | CP015060.1 | 229768 | 232284 | 43.06651635 | 885 | 838 | 1.056085919 | 1 | 1 | 1 | FIM1_4881 | YDL117W | YDL117W | CYK3 | n |
| Kmar_QGN16935.1 | W303.asm02.HP0_G0037200 | OG0002772 | chrXII | 612763 | 615390 | CP015058.1 | 1217698 | 1220298 | 60.61643836 | 875 | 866 | 1.01039261 | 1 | 1 | 1 | FIM1_3662 | YLR240W | YLR240W | VPS34 | n |
| Kmar_QGN18280.1 | W303.asm02.HP0_G0056560 | OG0002773 | chrXVI | 800840 | 801595 | CP015061.1 | 607727 | 608320 | 52.56916996 | 251 | 197 | 1.274111675 | 1 | 1 | 1 | FIM1_4606 | YPR131C | YPR131C | NAT3 | n |
| Kmar_QGN18271.1 | W303.asm02.HP0_G0056610 | OG0002774 | chrXVI | 806163 | 808946 | CP015061.1 | 592243 | 595092 | 42.11618257 | 927 | 949 | 0.976817703 | 1 | 1 | 1 | FIM1_4597 | YPR135W | YPR135W | CTF4 | n |
| Kmar_QGN18281.1 | W303.asm02.HP0_G0056550 | OG0002775 | chrXVI | 799616 | 800665 | CP015061.1 | 608424 | 609398 | 44.85714286 | 349 | 324 | 1.077160494 | 1 | 1 | 1 | FIM1_4607 | YPR129W | YPR129W | SCD6 | n |
| Kmar_QGN18277.1 | W303.asm02.HP0_G0056580 | OG0002776 | chrXVI | 802907 | 804139 | CP015061.1 | 602761 | 603858 | 56.73076923 | 410 | 365 | 1.123287671 | 1 | 1 | 1 | FIM1_4603 | YPR133C | YPR133C | SPN1 | n |
| Kmar_QGN18282.1 | W303.asm02.HP0_G0056540 | OG0002777 | chrXVI | 798147 | 799133 | CP015061.1 | 610333 | 611412 | 43.5483871 | 328 | 359 | 0.913649025 | 1 | 1 | 1 | FIM1_4608 | YPR128C | YPR128C | ANT1 | n |
| Kmar_QGN18279.1 | W303.asm02.HP0_G0020910 | OG0002778 | chrVII | 724816 | 726249 | CP015061.1 | 605814 | 607235 | 28.43137255 | 477 | 473 | 1.00845666 | 1 | 1 | 1 | FIM1_4605 | YGR117C | YGR117C | | n |
| Kmar_QGN16678.1 | W303.asm02.HP0_G0007180 | OG0002779 | chrIV | 149423 | 155860 | CP015058.1 | 651333 | 657752 | 80.65862709 | 2145 | 2139 | 1.002805049 | 1 | 1 | 1 | FIM1_3397 | YDL171C | YDL171C | GLT1 | n |
| Kmar_QGN14050.1 | W303.asm02.HP0_G0005870 | OG0002780 | chrIII | 214920 | 216104 | CP015054.1 | 1412312 | 1413604 | 47.82608696 | 394 | 430 | 0.91627907 | 1 | 1 | 1 | FIM1_701 | YCR035C | YCR035C | RRP43 | n |
| Kmar_QGN14052.1 | W303.asm02.HP0_G0027900 | OG0002781 | chrX | 53336 | 57100 | CP015054.1 | 1415164 | 1418895 | 39.16030534 | 1254 | 1243 | 1.008849558 | 1 | 1 | 1 | FIM1_703 | YJL197W | YJL197W | UBP12 | n |
| Kmar_QGN16637.1 | W303.asm02.HP0_G0005840 | OG0002782 | chrIII | 202606 | 209109 | CP015058.1 | 558377 | 564358 | 27.26021111 | 2167 | 1993 | 1.087305569 | 1 | 1 | 1 | FIM1_3356 | YCR032W | YCR032W | BPH1 | n |
| Kmar_QGN17435.1 | W303.asm02.HP0_G0001970 | OG0002783 | chrII | 251145 | 253355 | CP015059.1 | 910419 | 912524 | 27.89115646 | 736 | 701 | 1.049928673 | 1 | 1 | 1 | FIM1_4167 | YBR007C | YBR007C | DSF2 | n |
| Kmar_QGN14210.1 | W303.asm02.HP0_G0030500 | OG0002784 | chrX | 625349 | 626059 | CP015055.1 | 41439 | 42194 | 42.29249012 | 236 | 251 | 0.940239044 | 1 | 1 | 1 | FIM1_866 | YJR099W | YJR099W | YUH1 | n |
| Kmar_QGN17290.1 | W303.asm02.HP0_G0002150 | OG0002785 | chrII | 289522 | 291099 | CP015059.1 | 629804 | 631477 | 53.31010453 | 525 | 557 | 0.942549372 | 1 | 1 | 1 | FIM1_4021 | YBR028C | YBR028C | YPK3 | n |
| Kmar_QGN17311.1 | W303.asm02.HP0_G0024490 | OG0002786 | chrVIII | 352016 | 355258 | CP015059.1 | 669514 | 672477 | 40.97605893 | 1080 | 987 | 1.094224924 | 1 | 1 | 1 | FIM1_4043 | YHR119W | YHR119W | SET1 | n |
| Kmar_QGN13946.1 | W303.asm02.HP0_G0040500 | OG0002787 | chrXIII | 269525 | 271642 | CP015054.1 | 1196233 | 1198347 | 66.1971831 | 705 | 704 | 1.001420455 | 1 | 1 | 1 | FIM1_594 | YMR001C | YMR001C | CDC5 | n |
| Kmar_QGN15102.1 | W303.asm02.HP0_G0051770 | OG0002788 | chrXV | 871921 | 875790 | CP015056.1 | 206459 | 210565 | 24.38162544 | 1289 | 1368 | 0.942251462 | 1 | 1 | 1 | FIM1_1789 | YOR296W | YOR296W | | n |
| Kmar_QGN15100.1 | W303.asm02.HP0_G0051750 | OG0002789 | chrXV | 870059 | 870670 | CP015056.1 | 204360 | 204980 | 79.51219512 | 203 | 206 | 0.985436893 | 1 | 1 | 1 | FIM1_1787 | YOR294W | YOR294W | RRS1 | n |
| Kmar_QGN17099.1 | W303.asm02.HP0_G0046790 | OG0002790 | chrXIV | 652549 | 654534 | CP015059.1 | 230220 | 232142 | 62.33183857 | 661 | 640 | 1.0328125 | 1 | 1 | 1 | FIM1_3830 | YNR008W | YNR008W | LRO1 | n |
| Kmar_QGN17100.1 | W303.asm02.HP0_G0046780 | OG0002791 | chrXIV | 651335 | 652267 | CP015059.1 | 233018 | 233944 | 56.54952077 | 310 | 308 | 1.006493506 | 1 | 1 | 1 | FIM1_3831 | YNR007C | YNR007C | ATG3 | n |
| Kmar_QGN16525.1 | W303.asm02.HP0_G0019470 | OG0002792 | chrVII | 408638 | 409246 | CP015058.1 | 305164 | 305745 | 41.08910891 | 202 | 193 | 1.046632124 | 1 | 1 | 1 | FIM1_3238 | YGL047W | YGL047W | ALG13 | n |
| Kmar_QGN17097.1 | W303.asm02.HP0_G0042420 | OG0002793 | chrXIII | 668118 | 668786 | CP015059.1 | 227239 | 227910 | 70.40358744 | 222 | 223 | 0.995515695 | 1 | 1 | 1 | FIM1_3828 | YMR202W | YMR202W | ERG2 | n |
| Kmar_QGN14613.1 | W303.asm02.HP0_G0042400 | OG0002794 | chrXIII | 665335 | 666105 | CP015055.1 | 858332 | 859057 | 56.640625 | 256 | 241 | 1.062240664 | 1 | 1 | 1 | FIM1_1276 | YMR200W | YMR200W | ROT1 | n |
| Kmar_QGN14614.1 | W303.asm02.HP0_G0046770 | OG0002795 | chrXIV | 649139 | 651007 | CP015055.1 | 859487 | 861301 | 46.31410256 | 622 | 604 | 1.029801325 | 1 | 1 | 1 | FIM1_1277 | YNR006W | YNR006W | VPS27 | n |
| Kmar_QGN16282.1 | W303.asm02.HP0_G0031490 | OG0002796 | chrXI | 97926 | 99326 | CP015057.1 | 1197491 | 1198852 | 59.52890792 | 466 | 453 | 1.028697572 | 1 | 1 | 1 | FIM1_2987 | YKL184W | YKL184W | SPE1 | n |
| Kmar_QGN15816.1 | W303.asm02.HP0_G0035310 | OG0002797 | chrXII | 233239 | 236409 | CP015057.1 | 144569 | 147592 | 33.57933579 | 1056 | 1007 | 1.048659384 | 1 | 1 | 1 | FIM1_2512 | YLR039C | YLR039C | RIC1 | n |
| Kmar_QGN15837.1 | W303.asm02.HP0_G0035190 | OG0002798 | chrXII | 202838 | 203860 | CP015057.1 | 192241 | 193215 | 51.84135977 | 340 | 324 | 1.049382716 | 1 | 1 | 1 | FIM1_2533 | YLR026C | YLR026C | SED5 | n |
| Kmar_QGN15838.1 | W303.asm02.HP0_G0035180 | OG0002799 | chrXII | 201843 | 202565 | CP015057.1 | 193281 | 194021 | 54.4 | 240 | 246 | 0.975609756 | 1 | 1 | 1 | FIM1_2534 | YLR025W | YLR025W | SNF7 | n |
| Kmar_QGN14933.1 | W303.asm02.HP0_G0045140 | OG0002800 | chrXIV | 316862 | 318364 | CP015055.1 | 1525800 | 1527353 | 52.89719626 | 500 | 517 | 0.967117988 | 1 | 1 | 1 | FIM1_1608 | YNL169C | YNL169C | PSD1 | n |
| Kmar_QGN14932.1 | W303.asm02.HP0_G0045130 | OG0002801 | chrXIV | 311332 | 316575 | CP015055.1 | 1520570 | 1525567 | 29.47368421 | 1747 | 1665 | 1.049249249 | 1 | 1 | 1 | FIM1_1607 | YNL172W | YNL172W | APC1 | n |
| Kmar_QGN15353.1 | W303.asm02.HP0_G0003480 | OG0002802 | chrII | 566548 | 567705 | CP015056.1 | 775624 | 776736 | 51.94805195 | 385 | 370 | 1.040540541 | 1 | 1 | 1 | FIM1_2043 | YBR155W | YBR155W | CNS1 | n |
| Kmar_QGN13914.1 | W303.asm02.HP0_G0051630 | OG0002803 | chrXV | 849172 | 849864 | CP015054.1 | 1139641 | 1140333 | 59.05172414 | 230 | 230 | 1 | 1 | 1 | 1 | FIM1_562 | YOR283W | YOR283W | | n |
| Kmar_QGN13400.1 | W303.asm02.HP0_G0003690 | OG0002804 | chrII | 600495 | 601433 | CP015054.1 | 70202 | 71143 | 50.95541401 | 312 | 313 | 0.996805112 | 1 | 1 | 1 | FIM1_37 | YBR176W | YBR176W | ECM31 | n |
| Kmar_QGN17091.1 | W303.asm02.HP0_G0040470 | OG0002805 | chrXIII | 262211 | 263191 | CP015059.1 | 217694 | 218710 | 61.62790698 | 326 | 338 | 0.964497041 | 1 | 1 | 1 | FIM1_3822 | YML004C | YML004C | GLO1 | n |
| Kmar_QGN13666.1 | W303.asm02.HP0_G0016980 | OG0002806 | chrVI | 130971 | 133229 | CP015054.1 | 624002 | 626260 | 82.84574468 | 752 | 752 | 1 | 1 | 1 | 1 | FIM1_309 | YFR009W | YFR009W | GCN20 | n |
| Kmar_QGN13401.1 | W303.asm02.HP0_G0054410 | OG0002807 | chrXVI | 364358 | 365764 | CP015054.1 | 71174 | 72520 | 51.77453027 | 468 | 448 | 1.044642857 | 1 | 1 | 1 | FIM1_38 | YPL103C | YPL103C | FMP30 | n |
| Kmar_QGN13405.1 | W303.asm02.HP0_G0054450 | OG0002808 | chrXVI | 370744 | 371085 | CP015054.1 | 76777 | 77139 | 69.16666667 | 113 | 120 | 0.941666667 | 1 | 1 | 1 | FIM1_42 | YPL098C | YPL098C | MGR2 | n |
| Kmar_QGN17425.1 | W303.asm02.HP0_G0048350 | OG0002809 | chrXV | 187479 | 189869 | CP015059.1 | 894076 | 896436 | 34.28217822 | 796 | 786 | 1.012722646 | 1 | 1 | 1 | FIM1_4157 | YOL076W | YOL076W | MDM20 | n |
| Kmar_QGN16694.1 | W303.asm02.HP0_G0036890 | OG0002810 | chrXII | 552851 | 553804 | CP015058.1 | 681129 | 682175 | 50.85227273 | 317 | 348 | 0.91091954 | 1 | 1 | 1 | FIM1_3414 | YLR205C | YLR205C | HMX1 | n |
| Kmar_QGN16691.1 | W303.asm02.HP0_G0007230 | OG0002811 | chrIV | 163669 | 164262 | CP015058.1 | 677191 | 677829 | 66.03773585 | 197 | 212 | 0.929245283 | 1 | 1 | 1 | FIM1_3411 | YDL166C | YDL166C | FAP7 | n |
| Kmar_QGN16692.1 | W303.asm02.HP0_G0007240 | OG0002812 | chrIV | 164510 | 165085 | CP015058.1 | 678156 | 678710 | 67.01570681 | 191 | 184 | 1.038043478 | 1 | 1 | 1 | FIM1_3412 | YDL165W | YDL165W | CDC36 | n |
| Kmar_QGN15652.1 | W303.asm02.HP0_G0050550 | OG0002813 | chrXV | 637814 | 639433 | CP015056.1 | 1394093 | 1395712 | 63.83763838 | 539 | 539 | 1 | 1 | 1 | 1 | FIM1_2345 | YOR161C | YOR161C | PNS1 | n |
| Kmar_QGN13774.1 | W303.asm02.HP0_G0030070 | OG0002814 | chrX | 542682 | 544406 | CP015054.1 | 856908 | 858776 | 28.72841444 | 574 | 622 | 0.922829582 | 1 | 1 | 1 | FIM1_419 | YJR053W | YJR053W | BFA1 | n |
| Kmar_QGN13772.1 | W303.asm02.HP0_G0040080 | OG0002815 | chrXIII | 173201 | 174412 | CP015054.1 | 852994 | 854151 | 48.26732673 | 403 | 385 | 1.046753247 | 1 | 1 | 1 | FIM1_417 | YML048W | YML048W | GSF2 | n |
| Kmar_QGN15391.1 | W303.asm02.HP0_G0021320 | OG0002816 | chrVII | 818141 | 819427 | CP015056.1 | 862995 | 864374 | 45.43478261 | 275 | 459 | 0.59912854 | 1 | 1 | 1 | FIM1_2081 | YGR159C | YGR159C | NSR1 | n |
| Kmar_QGN15393.1 | W303.asm02.HP0_G0019480 | OG0002817 | chrVII | 411188 | 412816 | CP015056.1 | 866073 | 867923 | 34.41358025 | 542 | 616 | 0.87987013 | 1 | 1 | 1 | FIM1_2083 | YGL045W | YGL045W | RIM8 | n |
| Kmar_QGN14884.1 | W303.asm02.HP0_G0029400 | OG0002818 | chrX | 384034 | 386502 | CP015055.1 | 1418621 | 1421032 | 45.92326139 | 822 | 803 | 1.02366127 | 1 | 1 | 1 | FIM1_1558 | YJL029C | YJL029C | VPS53 | n |
| Kmar_QGN17430.1 | W303.asm02.HP0_G0008780 | OG0002819 | chrIV | 461501 | 462175 | CP015059.1 | 902463 | 903128 | 45.88744589 | 216 | 221 | 0.977375566 | 1 | 1 | 1 | FIM1_4162 | YDR007W | YDR007W | TRP1 | n |
| Kmar_QGN14885.1 | W303.asm02.HP0_G0029550 | OG0002820 | chrX | 415606 | 417312 | CP015055.1 | 1421731 | 1423446 | 80.55555556 | 568 | 571 | 0.99474606 | 1 | 1 | 1 | FIM1_1559 | YJL008C | YJL008C | CCT8 | n |
| Kmar_QGN17042.1 | W303.asm02.HP0_G0007430 | OG0002821 | chrIV | 201926 | 202777 | CP015059.1 | 104513 | 105382 | 44.17808219 | 283 | 289 | 0.979238754 | 1 | 1 | 1 | FIM1_3771 | YDL142C | YDL142C | CRD1 | n |
| Kmar_QGN14712.1 | W303.asm02.HP0_G0014960 | OG0002822 | chrV | 244319 | 244750 | CP015055.1 | 1077723 | 1078574 | 28.64864865 | 143 | 283 | 0.505300353 | 1 | 1 | 1 | FIM1_1379 | YER046W | YER046W | SPO73 | n |
| Kmar_QGN16489.1 | W303.asm02.HP0_G0000390 | OG0002823 | chrI | 84098 | 85237 | CP015058.1 | 216008 | 217333 | 32.11009174 | 379 | 441 | 0.859410431 | 1 | 1 | 1 | FIM1_3202 | YAL032C | YAL032C | PRP45 | n |
| Kmar_QGN16307.1 | W303.asm02.HP0_G0053330 | OG0002824 | chrXVI | 152971 | 153978 | CP015057.1 | 1247370 | 1248338 | 60.59701493 | 335 | 322 | 1.040372671 | 1 | 1 | 1 | FIM1_3012 | YPL215W | YPL215W | CBP3 | n |
| Kmar_QGN16299.1 | W303.asm02.HP0_G0053380 | OG0002825 | chrXVI | 159844 | 161766 | CP015057.1 | 1231362 | 1233275 | 28.74617737 | 640 | 637 | 1.004709576 | 1 | 1 | 1 | FIM1_3004 | YPL210C | YPL210C | SRP72 | n |
| Kmar_QGN16604.1 | W303.asm02.HP0_G0008880 | OG0002826 | chrIV | 483822 | 485024 | CP015058.1 | 480868 | 482049 | 64.25 | 400 | 393 | 1.017811705 | 1 | 1 | 1 | FIM1_3320 | YDR019C | YDR019C | GCV1 | n |
| Kmar_QGN15853.1 | W303.asm02.HP0_G0034660 | OG0002827 | chrXII | 87412 | 88164 | CP015057.1 | 225823 | 226512 | 47.2 | 250 | 229 | 1.091703057 | 1 | 1 | 1 | FIM1_2550 | YLL027W | YLL027W | ISA1 | n |
| Kmar_QGN15762.1 | W303.asm02.HP0_G0043550 | OG0002828 | chrXIII | 909714 | 910763 | CP015057.1 | 39882 | 40940 | 57.67045455 | 349 | 352 | 0.991477273 | 1 | 1 | 1 | FIM1_2458 | YMR315W | YMR315W | | n |
| Kmar_QGN17850.1 | W303.asm02.HP0_G0013370 | OG0002829 | chrIV | 1451537 | 1452688 | CP015060.1 | 615474 | 616523 | 27.39018088 | 383 | 349 | 1.097421203 | 1 | 1 | 1 | FIM1_5059 | YDR498C | YDR498C | SEC20 | n |
| Kmar_QGN17197.1 | W303.asm02.HP0_G0053560 | OG0002830 | chrXVI | 190869 | 193277 | CP015059.1 | 431900 | 434194 | 42.82238443 | 802 | 764 | 1.04973822 | 1 | 1 | 1 | FIM1_3928 | YPL190C | YPL190C | NAB3 | n |
| Kmar_QGN16895.1 | W303.asm02.HP0_G0010270 | OG0002831 | chrIV | 794042 | 795562 | CP015058.1 | 1119992 | 1121485 | 54.94071146 | 506 | 497 | 1.018108652 | 1 | 1 | 1 | FIM1_3622 | YDR168W | YDR168W | CDC37 | n |
| Kmar_QGN14852.1 | W303.asm02.HP0_G0039590 | OG0002832 | chrXIII | 72885 | 74240 | CP015055.1 | 1348927 | 1350237 | 43.37606838 | 451 | 436 | 1.03440367 | 1 | 1 | 1 | FIM1_1525 | YML097C | YML097C | VPS9 | n |
| Kmar_QGN13791.1 | W303.asm02.HP0_G0034120 | OG0002833 | chrXI | 622728 | 623741 | CP015054.1 | 886245 | 887162 | 43.32344214 | 337 | 305 | 1.104918033 | 1 | 1 | 1 | FIM1_437 | YKR088C | YKR088C | TVP38 | n |
| Kmar_QGN15926.1 | W303.asm02.HP0_G0009930 | OG0002834 | chrIV | 718249 | 720288 | CP015057.1 | 392188 | 394137 | 80.70987654 | 642 | 649 | 0.989214176 | 1 | 1 | 1 | FIM1_2624 | YDR129C | YDR129C | SAC6 | n |
| Kmar_QGN15077.1 | W303.asm02.HP0_G0033180 | OG0002835 | chrXI | 435336 | 436046 | CP015056.1 | 157711 | 158421 | 81.77966102 | 236 | 236 | 1 | 1 | 1 | 1 | FIM1_1762 | YKL009W | YKL009W | MRT4 | n |
| Kmar_QGN16056.1 | W303.asm02.HP0_G0032550 | OG0002836 | chrXI | 315761 | 316303 | CP015057.1 | 718220 | 718750 | 68.15642458 | 180 | 176 | 1.022727273 | 1 | 1 | 1 | FIM1_2756 | YKL069W | YKL069W | | n |
| Kmar_QGN16944.1 | W303.asm02.HP0_G0010620 | OG0002837 | chrIV | 860614 | 861669 | CP015058.1 | 1233100 | 1234224 | 42.89473684 | 351 | 374 | 0.938502674 | 1 | 1 | 1 | FIM1_3671 | YDR202C | YDR202C | RAV2 | n |
| Kmar_QGN16478.1 | W303.asm02.HP0_G0029260 | OG0002838 | chrX | 342411 | 343634 | CP015058.1 | 189744 | 190937 | 42.18009479 | 407 | 397 | 1.025188917 | 1 | 1 | 1 | FIM1_3191 | YJL046W | YJL046W | AIM22 | n |
| Kmar_QGN16477.1 | W303.asm02.HP0_G0004470 | OG0002839 | chrII | 752241 | 753080 | CP015058.1 | 188646 | 189476 | 51.05633803 | 279 | 276 | 1.010869565 | 1 | 1 | 1 | FIM1_3190 | YBR257W | YBR257W | POP4 | n |
| Kmar_QGN13596.1 | W303.asm02.HP0_G0033460 | OG0002840 | chrXI | 488031 | 488525 | CP015054.1 | 490620 | 491108 | 36.74698795 | 164 | 162 | 1.012345679 | 1 | 1 | 1 | FIM1_238 | YKR020W | YKR020W | VPS51 | n |
| Kmar_QGN15797.1 | W303.asm02.HP0_G0019990 | OG0002841 | chrVII | 510693 | 511874 | CP015057.1 | 113985 | 115235 | 60 | 393 | 416 | 0.944711538 | 1 | 1 | 1 | FIM1_2493 | YGR012W | YGR012W | MCY1 | n |
| Kmar_QGN14166.1 | W303.asm02.HP0_G0013670 | OG0002842 | chrIV | 1503925 | 1505028 | CP015054.1 | 1670089 | 1671183 | 60.8 | 367 | 364 | 1.008241758 | 1 | 1 | 1 | FIM1_819 | YDR531W | YDR531W | CAB1 | n |
| Kmar_QGN18136.1 | W303.asm02.HP0_G0001080 | OG0002843 | chrII | 52172 | 60535 | CP015061.1 | 298434 | 306746 | 31.26989742 | 2787 | 2770 | 1.006137184 | 1 | 1 | 1 | FIM1_4458 | YBL088C | YBL088C | TEL1 | n |
| Kmar_QGN14808.1 | W303.asm02.HP0_G0043750 | OG0002844 | chrXIV | 29171 | 31810 | CP015055.1 | 1269727 | 1272330 | 60.76662909 | 879 | 867 | 1.01384083 | 1 | 1 | 1 | FIM1_1479 | YNL325C | YNL325C | FIG4 | n |
| Kmar_QGN17126.1 | W303.asm02.HP0_G0030560 | OG0002845 | chrX | 633354 | 634376 | CP015059.1 | 290404 | 291726 | 53.28798186 | 340 | 440 | 0.772727273 | 1 | 1 | 1 | FIM1_3857 | YJR105W | YJR105W | ADO1 | n |

| | | | | | | | | | | | | | | | | | | | | |
|---|---|---|---|---|---|---|---|---|---|---|---|---|---|---|---|---|---|---|---|---|
| Kmar_QGN14897.1 | W303.asm02.HP0_G0056170 | OG0002846 | chrXVI | 714996 | 716342 | CP015055.1 | 1453548 | 1454813 | 28.73051225 | 448 | 421 | 1.064133017 | 1 | 1 | 1 | FIM1_1571 | YPR085C | YPR085C | ASA1 | n |
| Kmar_QGN17598.1 | W303.asm02.HP0_G0013940 | OG0002847 | chrV | 48834 | 51902 | CP015060.1 | 71465 | 74533 | 46.70433145 | 1022 | 1022 | 1 | 1 | 1 | 1 | FIM1_4805 | YEL055C | YEL055C | POL5 | n |
| Kmar_QGN14660.1 | W303.asm02.HP0_G0046750 | OG0002849 | chrXIV | 646497 | 647450 | CP015055.1 | 966006 | 966942 | 65.29968454 | 317 | 312 | 1.016025641 | 1 | 1 | 1 | FIM1_1324 | YNR003C | YNR003C | RPC34 | n |
| Kmar_QGN15180.1 | W303.asm02.HP0_G0028870 | OG0002850 | chrX | 258816 | 259832 | CP015056.1 | 378207 | 379229 | 71.26436782 | 338 | 340 | 0.994117647 | 1 | 1 | 1 | FIM1_1868 | YJL088W | YJL088W | ARG3 | n |
| Kmar_QGN17218.1 | W303.asm02.HP0_G0019040 | OG0002851 | chrVII | 325826 | 327559 | CP015059.1 | 474418 | 476151 | 52.4137931 | 577 | 577 | 1 | 1 | 1 | 1 | FIM1_3949 | YGL095C | YGL095C | VPS45 | n |
| Kmar_QGN15516.1 | W303.asm02.HP0_G0056260 | OG0002852 | chrXVI | 731893 | 735114 | CP015056.1 | 1103518 | 1106499 | 40.38112523 | 1073 | 993 | 1.080563948 | 1 | 1 | 1 | FIM1_2207 | YPR097W | YPR097W | LEC1 | n |
| Kmar_QGN13649.1 | W303.asm02.HP0_G0014370 | OG0002853 | chrV | 134303 | 136417 | CP015054.1 | 579843 | 581996 | 75.97222222 | 704 | 717 | 0.981868898 | 1 | 1 | 1 | FIM1_292 | YEL011W | YEL011W | GLC3 | n |
| Kmar_QGN13592.1 | W303.asm02.HP0_G0028930 | OG0002854 | chrX | 273820 | 275289 | CP015054.1 | 482215 | 483789 | 56.46387833 | 489 | 524 | 0.933206107 | 1 | 1 | 1 | FIM1_234 | YJL081C | YJL081C | ARP4 | n |
| Kmar_QGN14168.1 | W303.asm02.HP0_G0013650 | OG0002855 | chrIV | 1501858 | 1502241 | CP015054.1 | 1673858 | 1674241 | 71.65354331 | 127 | 127 | 1 | 1 | 1 | 1 | FIM1_821 | YDR529C | YDR529C | QCR7 | n |
| Kmar_QGN18064.1 | W303.asm02.HP0_G0038450 | OG0002856 | chrXII | 876389 | 878590 | CP015061.1 | 136363 | 138228 | 21.26200274 | 733 | 621 | 1.180354267 | 1 | 1 | 1 | FIM1_4382 | YLR381W | YLR381W | CTF3 | n |
| Kmar_QGN15557.1 | W303.asm02.HP0_G0023930 | OG0002857 | chrVIII | 233362 | 234993 | CP015056.1 | 1179647 | 1181110 | 75.8747698 | 543 | 487 | 1.114989733 | 1 | 1 | 1 | FIM1_2248 | YHR065C | YHR065C | RRP3 | n |
| Kmar_QGN15548.1 | W303.asm02.HP0_G0011600 | OG0002858 | chrIV | 1082526 | 1082948 | CP015056.1 | 1164989 | 1165423 | 65.97222222 | 140 | 144 | 0.972222222 | 1 | 1 | 1 | FIM1_2239 | YDR308C | YDR308C | SRB7 | n |
| Kmar_QGN18243.1 | W303.asm02.HP0_G0039680 | OG0002859 | chrXIII | 90331 | 91911 | CP015061.1 | 533882 | 535462 | 70.37735849 | 526 | 526 | 1 | 1 | 1 | 1 | FIM1_4568 | YML086C | YML086C | ALO1 | n |
| Kmar_QGN14849.1 | W303.asm02.HP0_G0039620 | OG0002860 | chrXIII | 77640 | 80339 | CP015055.1 | 1342467 | 1345103 | 53.81263617 | 899 | 878 | 1.023917995 | 1 | 1 | 1 | FIM1_1521 | YML093W | YML093W | UTP14 | n |
| Kmar_QGN14690.1 | W303.asm02.HP0_G0056990 | OG0002861 | chrXVI | 896933 | 897910 | CP015055.1 | 1039288 | 1040262 | 69.53846154 | 325 | 324 | 1.00308642 | 1 | 1 | 1 | FIM1_1357 | YPR176C | YPR176C | BET2 | n |
| Kmar_QGN14691.1 | W303.asm02.HP0_G0056980 | OG0002862 | chrXVI | 894806 | 896875 | CP015055.1 | 1040323 | 1042440 | 51.27118644 | 689 | 705 | 0.977304965 | 1 | 1 | 1 | FIM1_1358 | YPR175W | YPR175W | DPB2 | n |
| Kmar_QGN14514.1 | W303.asm02.HP0_G0051930 | OG0002863 | chrXV | 903578 | 904594 | CP015055.1 | 673232 | 674287 | 33.87096774 | 338 | 351 | 0.962962963 | 1 | 1 | 1 | FIM1_1175 | YOR313C | YOR313C | SPS4 | n |
| Kmar_QGN16686.1 | W303.asm02.HP0_G0036830 | OG0002864 | chrXII | 546225 | 547739 | CP015058.1 | 668516 | 670066 | 84.94208494 | 504 | 516 | 0.976744186 | 1 | 1 | 1 | FIM1_3405 | YLR197W | YLR197W | NOP56 | n |
| Kmar_QGN18343.1 | W303.asm02.HP0_G0002770 | OG0002865 | chrII | 417539 | 421537 | CP015061.1 | 738634 | 742635 | 43.19009468 | 1332 | 1333 | 0.999249812 | 1 | 1 | 1 | FIM1_4670 | YBR081C | YBR081C | SPT7 | n |
| Kmar_QGN18344.1 | W303.asm02.HP0_G0009240 | OG0002866 | chrIV | 570076 | 571704 | CP015061.1 | 742743 | 744044 | 21.24542125 | 542 | 433 | 1.251732102 | 1 | 1 | 1 | FIM1_4671 | YDR057W | YDR057W | YOS9 | n |
| Kmar_QGN14524.1 | W303.asm02.HP0_G0042810 | OG0002867 | chrXIII | 756437 | 757747 | CP015055.1 | 689682 | 691145 | 39.59731544 | 436 | 487 | 0.895277207 | 1 | 1 | 1 | FIM1_1186 | YMR240C | YMR240C | CUS1 | n |
| Kmar_QGN14522.1 | W303.asm02.HP0_G0051860 | OG0002868 | chrXV | 890742 | 891470 | CP015055.1 | 688328 | 689035 | 42.56198347 | 242 | 235 | 1.029787234 | 1 | 1 | 1 | FIM1_1184 | YOR305W | YOR305W | RRG7 | n |
| Kmar_QGN18345.1 | W303.asm02.HP0_G0009230 | OG0002869 | chrIV | 569172 | 569789 | CP015061.1 | 744301 | 744891 | 27.48815166 | 205 | 196 | 1.045918367 | 1 | 1 | 1 | FIM1_4672 | YDR056C | YDR056C | EMC10 | n |
| Kmar_QGN16717.1 | W303.asm02.HP0_G0011740 | OG0002870 | chrIV | 1118768 | 1119913 | CP015058.1 | 734945 | 736075 | 70.15706806 | 381 | 376 | 1.013297872 | 1 | 1 | 1 | FIM1_3438 | YDR321W | YDR321W | ASP1 | n |
| Kmar_QGN13926.1 | W303.asm02.HP0_G0012490 | OG0002871 | chrIV | 1273196 | 1275127 | CP015054.1 | 1158216 | 1160096 | 45.24886878 | 643 | 626 | 1.02715655 | 1 | 1 | 1 | FIM1_574 | YDR398W | YDR398W | UTP5 | n |
| Kmar_QGN16313.1 | W303.asm02.HP0_G0018680 | OG0002872 | chrVII | 249365 | 250327 | CP015057.1 | 1261126 | 1261998 | 55.72755418 | 320 | 290 | 1.103448276 | 1 | 1 | 1 | FIM1_3018 | YGL136C | YGL136C | MRM2 | n |
| Kmar_QGN16506.1 | W303.asm02.HP0_G0010730 | OG0002873 | chrIV | 897376 | 898428 | CP015058.1 | 262223 | 263311 | 54.76839237 | 350 | 362 | 0.966850829 | 1 | 1 | 1 | FIM1_3219 | YDR214W | YDR214W | AHA1 | n |
| Kmar_QGN14509.1 | W303.asm02.HP0_G0042860 | OG0002874 | chrXIII | 765022 | 765336 | CP015055.1 | 657474 | 657782 | 51.92307692 | 104 | 102 | 1.019607843 | 1 | 1 | 1 | FIM1_1170 | YMR244C-A | YMR244C-A | COA6 | n |
| Kmar_QGN15656.1 | W303.asm02.HP0_G0012570 | OG0002875 | chrIV | 1289794 | 1293663 | CP015056.1 | 1400693 | 1404364 | 35.17665131 | 1289 | 1223 | 1.053965658 | 1 | 1 | 1 | FIM1_2349 | YDR407C | YDR407C | TRS120 | n |
| Kmar_QGN15657.1 | W303.asm02.HP0_G0050510 | OG0002876 | chrXV | 631841 | 632626 | CP015056.1 | 1404544 | 1405401 | 79.29824561 | 261 | 285 | 0.915789474 | 1 | 1 | 1 | FIM1_2350 | YOR157C | YOR157C | PUP1 | n |
| Kmar_QGN14638.1 | W303.asm02.HP0_G0024390 | OG0002877 | chrVIII | 336283 | 338040 | CP015055.1 | 920182 | 921717 | 25.58528428 | 585 | 511 | 1.14481409 | 1 | 1 | 1 | FIM1_1302 | YHR109W | YHR109W | CTM1 | n |
| Kmar_QGN16676.1 | W303.asm02.HP0_G0036780 | OG0002878 | chrXII | 538920 | 539717 | CP015058.1 | 649090 | 649905 | 64.94464945 | 265 | 271 | 0.977859779 | 1 | 1 | 1 | FIM1_3395 | YLR192C | YLR192C | HCR1 | n |
| Kmar_QGN14751.1 | W303.asm02.HP0_G0013120 | OG0002879 | chrIV | 1409015 | 1409866 | CP015055.1 | 1157000 | 1157737 | 42.56055363 | 283 | 245 | 1.155102041 | 1 | 1 | 1 | FIM1_1421 | YDR472W | YDR472W | TRS31 | n |
| Kmar_QGN14750.1 | W303.asm02.HP0_G0023260 | OG0002880 | chrVIII | 124196 | 124912 | CP015055.1 | 1156054 | 1156743 | 66.39004149 | 238 | 229 | 1.03930131 | 1 | 1 | 1 | FIM1_1420 | YHR013C | YHR013C | ARD1 | n |
| Kmar_QGN13934.1 | W303.asm02.HP0_G0050400 | OG0002881 | chrXV | 606222 | 607046 | CP015054.1 | 1172963 | 1173784 | 77.45454545 | 274 | 273 | 1.003663004 | 1 | 1 | 1 | FIM1_582 | YOR145C | YOR145C | PNO1 | n |
| Kmar_QGN13932.1 | W303.asm02.HP0_G0050420 | OG0002882 | chrXV | 609515 | 610072 | CP015054.1 | 1169959 | 1170492 | 29.3814433 | 185 | 177 | 1.04519774 | 1 | 1 | 1 | FIM1_580 | YOR148C | YOR148C | SPP2 | n |
| Kmar_QGN14753.1 | W303.asm02.HP0_G0050390 | OG0002883 | chrXV | 603592 | 605967 | CP015055.1 | 1158850 | 1161189 | 26.32226322 | 791 | 779 | 1.015404365 | 1 | 1 | 1 | FIM1_1423 | YOR144C | YOR144C | ELG1 | n |
| Kmar_QGN14754.1 | W303.asm02.HP0_G0012440 | OG0002884 | chrIV | 1264421 | 1265434 | CP015055.1 | 1161283 | 1162272 | 72.40356083 | 337 | 329 | 1.024316109 | 1 | 1 | 1 | FIM1_1424 | YDR392W | YDR392W | SPT3 | n |
| Kmar_QGN15441.1 | W303.asm02.HP0_G0011510 | OG0002885 | chrIV | 1064126 | 1065730 | CP015056.1 | 967427 | 969028 | 54.02930403 | 534 | 533 | 1.001876173 | 1 | 1 | 1 | FIM1_2132 | YDR299W | YDR299W | BFR2 | n |
| Kmar_QGN17962.1 | W303.asm02.HP0_G0043800 | OG0002886 | chrXIV | 38132 | 38986 | CP015060.1 | 837887 | 838747 | 61.53846154 | 284 | 286 | 0.993006993 | 1 | 1 | 1 | FIM1_5171 | YNL320W | YNL320W | | n |
| Kmar_QGN17961.1 | W303.asm02.HP0_G0047750 | OG0002887 | chrXV | 61669 | 65691 | CP015060.1 | 833910 | 837863 | 36.92528736 | 1340 | 1317 | 1.017463933 | 1 | 1 | 1 | FIM1_5170 | YOL138C | YOL138C | RTC1 | n |
| Kmar_QGN14406.1 | W303.asm02.HP0_G0013280 | OG0002888 | chrIV | 1436706 | 1437590 | CP015055.1 | 452211 | 453089 | 51.67785235 | 294 | 292 | 1.006849315 | 1 | 1 | 1 | FIM1_1066 | YDR489W | YDR489W | SLD5 | n |
| Kmar_QGN15200.1 | W303.asm02.HP0_G0008690 | OG0002889 | chrIV | 444342 | 446042 | CP015056.1 | 414129 | 415934 | 35.31353135 | 566 | 601 | 0.941763727 | 1 | 1 | 1 | FIM1_1888 | YDL003W | YDL003W | MCD1 | n |
| Kmar_QGN15238.1 | W303.asm02.HP0_G0034860 | OG0002890 | chrXII | 132914 | 135181 | CP015056.1 | 500368 | 502647 | 70.03891051 | 755 | 759 | 0.994729908 | 1 | 1 | 1 | FIM1_1927 | YLL008W | YLL008W | DRS1 | n |
| Kmar_QGN17847.1 | W303.asm02.HP0_G0013340 | OG0002891 | chrIV | 1443809 | 1446844 | CP015060.1 | 607001 | 609631 | 27.21893491 | 1011 | 876 | 1.154109589 | 1 | 1 | 1 | FIM1_5056 | YDR495C | YDR495C | VPS3 | n |
| Kmar_QGN14443.1 | W303.asm02.HP0_G0047140 | OG0002892 | chrXIV | 713123 | 714241 | CP015055.1 | 523917 | 525041 | 65.24064171 | 372 | 374 | 0.994652406 | 1 | 1 | 1 | FIM1_1104 | YNR041C | YNR041C | COQ2 | n |
| Kmar_QGN17428.1 | W303.asm02.HP0_G0008630 | OG0002893 | chrIV | 433533 | 434030 | CP015059.1 | 900411 | 900911 | 37.43315508 | 165 | 166 | 0.993975904 | 1 | 1 | 1 | FIM1_4160 | YDL008W | YDL008W | APC11 | n |
| Kmar_QGN17043.1 | W303.asm02.HP0_G0007440 | OG0002894 | chrIV | 203246 | 205318 | CP015059.1 | 106118 | 108187 | 54.11255411 | 690 | 689 | 1.001451379 | 1 | 1 | 1 | FIM1_3772 | YDL141W | YDL141W | BPL1 | n |
| Kmar_QGN15443.1 | W303.asm02.HP0_G0023470 | OG0002895 | chrVIII | 170809 | 171843 | CP015056.1 | 971090 | 972097 | 38.13559322 | 344 | 335 | 1.026865672 | 1 | 1 | 1 | FIM1_2134 | YHR034C | YHR034C | PIH1 | n |
| Kmar_QGN18395.1 | W303.asm02.HP0_G0031880 | OG0002896 | chrXI | 178049 | 179440 | CP015061.1 | 843912 | 845396 | 56 | 463 | 494 | 0.937246964 | 1 | 1 | 1 | FIM1_4722 | YKL143W | YKL143W | LTV1 | n |
| Kmar_QGN14714.1 | W303.asm02.HP0_G0026860 | OG0002897 | chrIX | 278419 | 279537 | CP015055.1 | 1083392 | 1084513 | 74.79892761 | 372 | 373 | 0.997319035 | 1 | 1 | 1 | FIM1_1382 | YIL035C | YIL035C | CKA1 | n |
| Kmar_QGN17045.1 | W303.asm02.HP0_G0037000 | OG0002898 | chrXII | 572220 | 573335 | CP015059.1 | 114659 | 115777 | 67.02412869 | 371 | 372 | 0.997311828 | 1 | 1 | 1 | FIM1_3774 | YLR216C | YLR216C | CPR6 | n |
| Kmar_QGN17044.1 | W303.asm02.HP0_G0007450 | OG0002899 | chrIV | 205567 | 210768 | CP015059.1 | 108487 | 113670 | 85.92848904 | 1733 | 1727 | 1.003474233 | 1 | 1 | 1 | FIM1_3773 | YDL140C | YDL140C | RPO21 | n |
| Kmar_QGN17855.1 | W303.asm02.HP0_G0044140 | OG0002900 | chrXIV | 108241 | 108828 | CP015060.1 | 622488 | 623105 | 37.32057416 | 195 | 205 | 0.951219512 | 1 | 1 | 1 | FIM1_5064 | YNL282W | YNL282W | POP3 | n |
| Kmar_QGN14041.1 | W303.asm02.HP0_G0048160 | OG0002901 | chrXV | 142039 | 143010 | CP015054.1 | 1392700 | 1393665 | 80.74534161 | 323 | 321 | 1.00623053 | 1 | 1 | 1 | FIM1_692 | YOL094C | YOL094C | RFC4 | n |
| Kmar_QGN14042.1 | W303.asm02.HP0_G0013230 | OG0002902 | chrIV | 1428457 | 1430382 | CP015054.1 | 1393962 | 1395743 | 49.6875 | 641 | 593 | 1.080944351 | 1 | 1 | 1 | FIM1_693 | YDR484W | YDR484W | VPS52 | n |
| Kmar_QGN14043.1 | W303.asm02.HP0_G0048150 | OG0002903 | chrXV | 139682 | 141802 | CP015054.1 | 1395975 | 1398095 | 41.35546335 | 706 | 706 | 1 | 1 | 1 | 1 | FIM1_694 | YOL095C | YOL095C | HMI1 | n |
| Kmar_QGN15424.1 | W303.asm02.HP0_G0011410 | OG0002904 | chrIV | 1042779 | 1044008 | CP015056.1 | 931530 | 932657 | 33.48946136 | 409 | 375 | 1.090666667 | 1 | 1 | 1 | FIM1_2115 | YDR289C | YDR289C | RTT103 | n |
| Kmar_QGN14446.1 | W303.asm02.HP0_G0006260 | OG0002905 | chrIII | 286375 | 286947 | CP015055.1 | 530177 | 530707 | 50 | 190 | 176 | 1.079545455 | 1 | 1 | 1 | FIM1_1107 | YCR086W | YCR086W | CSM1 | n |
| Kmar_QGN14447.1 | W303.asm02.HP0_G0006270 | OG0002906 | chrIII | 286989 | 287435 | CP015055.1 | 530778 | 531239 | 58.70967742 | 148 | 153 | 0.967320261 | 1 | 1 | 1 | FIM1_1108 | YCR087C-A | YCR087C-A | | n |
| Kmar_QGN14311.1 | W303.asm02.HP0_G0029590 | OG0002907 | chrX | 428293 | 428649 | CP015055.1 | 248592 | 248948 | 72.88135593 | 118 | 118 | 1 | 1 | 1 | 1 | FIM1_968 | YJL003W | YJL003W | COX16 | n |
| Kmar_QGN14790.1 | W303.asm02.HP0_G0050230 | OG0002908 | chrXV | 560836 | 561552 | CP015055.1 | 1238891 | 1239616 | 50.20576132 | 238 | 241 | 0.987551867 | 1 | 1 | 1 | FIM1_1461 | YOR126C | YOR126C | IAH1 | n |
| Kmar_QGN17964.1 | W303.asm02.HP0_G0043790 | OG0002909 | chrXIV | 35128 | 37854 | CP015060.1 | 841190 | 844228 | 50.04878049 | 908 | 1012 | 0.897233202 | 1 | 1 | 1 | FIM1_5173 | YNL321W | YNL321W | VNX1 | n |
| Kmar_QGN15663.1 | W303.asm02.HP0_G0012610 | OG0002910 | chrIV | 1299094 | 1300119 | CP015056.1 | 1414145 | 1415002 | 32.18390805 | 341 | 285 | 1.196491228 | 1 | 1 | 1 | FIM1_2356 | YDR411C | YDR411C | DFM1 | n |
| Kmar_QGN13531.1 | W303.asm02.HP0_G0020540 | OG0002911 | chrVII | 635006 | 635479 | CP015054.1 | 353471 | 353944 | 56.875 | 157 | 157 | 1 | 1 | 1 | 1 | FIM1_171 | YGR076C | YGR076C | MRPL25 | n |
| Kmar_QGN15199.1 | W303.asm02.HP0_G0008680 | OG0002912 | chrIV | 442688 | 443170 | CP015056.1 | 412666 | 413139 | 72.5 | 160 | 157 | 1.01910828 | 1 | 1 | 1 | FIM1_1887 | YDL004W | YDL004W | ATP16 | n |
| Kmar_QGN15416.1 | W303.asm02.HP0_G0029340 | OG0002913 | chrX | 374474 | 375745 | CP015056.1 | 913278 | 914471 | 49.88179669 | 423 | 397 | 1.065491184 | 1 | 1 | 1 | FIM1_2106 | YJL036W | YJL036W | SNX4 | n |
| Kmar_QGN16365.1 | W303.asm02.HP0_G0012730 | OG0002914 | chrIV | 1327761 | 1328942 | CP015057.1 | 1368829 | 1370013 | 62.62626263 | 393 | 394 | 0.997461929 | 1 | 1 | 1 | FIM1_3071 | YDR427W | YDR427W | RPN9 | n |
| Kmar_QGN16079.1 | W303.asm02.HP0_G0055320 | OG0002915 | chrXVI | 545742 | 548327 | CP015057.1 | 765866 | 768343 | 47.17416378 | 861 | 825 | 1.043636364 | 1 | 1 | 1 | FIM1_2779 | YPL008W | YPL008W | CHL1 | n |
| Kmar_QGN18422.1 | W303.asm02.HP0_G0016910 | OG0002916 | chrVI | 118499 | 121018 | CP015061.1 | 897526 | 900084 | 55.73770492 | 839 | 852 | 0.984741784 | 1 | 1 | 1 | FIM1_4750 | YFR002W | YFR002W | NIC96 | n |
| Kmar_QGN15454.1 | W303.asm02.HP0_G0053830 | OG0002917 | chrXVI | 249139 | 249960 | CP015056.1 | 987146 | 987952 | 50.90252708 | 273 | 268 | 1.018656716 | 1 | 1 | 1 | FIM1_2145 | YPL162C | YPL162C | | n |
| Kmar_QGN17013.1 | W303.asm02.HP0_G0008950 | OG0002918 | chrIV | 494308 | 496977 | CP015059.1 | 30130 | 32664 | 41.68539326 | 889 | 844 | 1.053317536 | 1 | 1 | 1 | FIM1_3742 | YDR027C | YDR027C | VPS54 | n |
| Kmar_QGN16820.1 | W303.asm02.HP0_G0056460 | OG0002919 | chrXVI | 774681 | 775916 | CP015058.1 | 963289 | 964527 | 61.72248804 | 411 | 412 | 0.997572816 | 1 | 1 | 1 | FIM1_3544 | YPR118W | YPR118W | MRI1 | n |
| Kmar_QGN16143.1 | W303.asm02.HP0_G0040740 | OG0002920 | chrXIII | 324739 | 325938 | CP015057.1 | 897525 | 898532 | 23.94014963 | 399 | 335 | 1.191044776 | 1 | 1 | 1 | FIM1_2844 | YMR026C | YMR026C | PEX12 | n |
| Kmar_QGN14717.1 | W303.asm02.HP0_G0026880 | OG0002921 | chrIX | 281048 | 282298 | CP015055.1 | 1087271 | 1088671 | 58.17409766 | 416 | 466 | 0.892703863 | 1 | 1 | 1 | FIM1_1385 | YIL033C | YIL033C | BCY1 | n |
| Kmar_QGN14716.1 | W303.asm02.HP0_G0014930 | OG0002922 | chrV | 238709 | 239155 | CP015055.1 | 1086511 | 1086960 | 66.21621622 | 148 | 149 | 0.993288591 | 1 | 1 | 1 | FIM1_1384 | YER044C | YER044C | ERG28 | n |
| Kmar_QGN14074.1 | W303.asm02.HP0_G0044460 | OG0002923 | chrXIV | 181916 | 183163 | CP015054.1 | 1465182 | 1466420 | 71.80722892 | 415 | 412 | 1.007281553 | 1 | 1 | 1 | FIM1_725 | YNL248C | YNL248C | RPA49 | n |
| Kmar_QGN14073.1 | W303.asm02.HP0_G0030480 | OG0002924 | chrX | 622186 | 622704 | CP015054.1 | 1464210 | 1464692 | 45.56213018 | 172 | 160 | 1.075 | 1 | 1 | 1 | FIM1_724 | YJR097W | YJR097W | JJJ3 | n |
| Kmar_QGN18321.1 | W303.asm02.HP0_G0001520 | OG0002925 | chrII | 148490 | 149188 | CP015061.1 | 689130 | 689795 | 67.67241379 | 232 | 221 | 1.049773756 | 1 | 1 | 1 | FIM1_4647 | YBL038W | YBL038W | MRPL16 | n |
| Kmar_QGN15698.1 | W303.asm02.HP0_G0014910 | OG0002926 | chrV | 236076 | 236630 | CP015056.1 | 1490952 | 1491506 | 67.93478261 | 184 | 184 | 1 | 1 | 1 | 1 | FIM1_2391 | YER042W | YER042W | MXR1 | n |
| Kmar_QGN18320.1 | W303.asm02.HP0_G0055840 | OG0002927 | chrXVI | 664027 | 665436 | CP015061.1 | 687302 | 688717 | 62.4742268 | 469 | 471 | 0.995753715 | 1 | 1 | 1 | FIM1_4646 | YPR047W | YPR047W | MSF1 | n |
| Kmar_QGN18319.1 | W303.asm02.HP0_G0055850 | OG0002928 | chrXVI | 665680 | 667551 | CP015061.1 | 685040 | 686914 | 59.80861244 | 623 | 624 | 0.998397436 | 1 | 1 | 1 | FIM1_4645 | YPR048W | YPR048W | TAH18 | n |
| Kmar_QGN15628.1 | W303.asm02.HP0_G0033350 | OG0002929 | chrXI | 461763 | 463640 | CP015056.1 | 1343594 | 1345267 | 40 | 625 | 557 | 1.122082585 | 1 | 1 | 1 | FIM1_2321 | YKR008W | YKR008W | RSC4 | n |
| Kmar_QGN16140.1 | W303.asm02.HP0_G0040710 | OG0002930 | chrXIII | 319940 | 321520 | CP015057.1 | 893758 | 895239 | 46.3878327 | 526 | 493 | 1.06693712 | 1 | 1 | 1 | FIM1_2841 | YMR023C | YMR023C | MSS1 | n |
| Kmar_QGN14515.1 | W303.asm02.HP0_G0051910 | OG0002931 | chrXV | 900776 | 901648 | CP015055.1 | 677096 | 677941 | 67.93103448 | 290 | 281 | 1.03202847 | 1 | 1 | 1 | FIM1_1177 | YOR311C | YOR311C | DGK1 | n |
| Kmar_QGN14345.1 | W303.asm02.HP0_G0041320 | OG0002932 | chrXIII | 450563 | 451870 | CP015055.1 | 319617 | 320903 | 45.72072072 | 435 | 428 | 1.01635514 | 1 | 1 | 1 | FIM1_1005 | YMR091C | YMR091C | NPL6 | n |
| Kmar_QGN16265.1 | W303.asm02.HP0_G0027530 | OG0002933 | chrIX | 402108 | 403139 | CP015057.1 | 1156317 | 1157360 | 59.1954023 | 343 | 347 | 0.988472622 | 1 | 1 | 1 | FIM1_2970 | YIR029W | YIR029W | DAL2 | n |
| Kmar_QGN16266.1 | W303.asm02.HP0_G0030800 | OG0002934 | chrX | 673773 | 674792 | CP015057.1 | 1157643 | 1158584 | 34.09090909 | 339 | 313 | 1.083067093 | 1 | 1 | 1 | FIM1_2971 | YJR129C | YJR129C | EFM3 | n |
| Kmar_QGN17252.1 | W303.asm02.HP0_G0032860 | OG0002935 | chrXI | 369661 | 370431 | CP015059.1 | 552819 | 553586 | 59.16030534 | 256 | 255 | 1.003921569 | 1 | 1 | 1 | FIM1_3983 | YKL040C | YKL040C | NFU1 | n |
| Kmar_QGN15818.1 | W303.asm02.HP0_G0041710 | OG0002936 | chrXIII | 532735 | 533643 | CP015057.1 | 149195 | 150124 | 50 | 302 | 309 | 0.977346278 | 1 | 1 | 1 | FIM1_2514 | YMR130W | YMR130W | DPI35 | n |
| Kmar_QGN17254.1 | W303.asm02.HP0_G0022920 | OG0002937 | chrVIII | 65935 | 67752 | CP015059.1 | 554322 | 555839 | 33.92568659 | 605 | 505 | 1.198019802 | 1 | 1 | 1 | FIM1_3985 | YHL019C | YHL019C | APM2 | n |
| Kmar_QGN17253.1 | W303.asm02.HP0_G0022930 | OG0002938 | chrVIII | 67912 | 68274 | CP015059.1 | 553700 | 554017 | 38.65546218 | 120 | 105 | 1.142857143 | 1 | 1 | 1 | FIM1_3984 | YHL018W | YHL018W | MCO14 | n |
| Kmar_QGN15820.1 | W303.asm02.HP0_G0041700 | OG0002939 | chrXIII | 528420 | 532433 | CP015057.1 | 152307 | 156248 | 50.07462687 | 1337 | 1313 | 1.018278751 | 1 | 1 | 1 | FIM1_2516 | YMR129W | YMR129W | POM152 | n |
| Kmar_QGN13746.1 | W303.asm02.HP0_G0019860 | OG0002940 | chrVII | 490007 | 491707 | CP015054.1 | 799574 | 801316 | 57.70547945 | 566 | 580 | 0.975862069 | 1 | 1 | 1 | FIM1_390 | YGL003C | YGL003C | CDH1 | n |
| Kmar_QGN15824.1 | W303.asm02.HP0_G0041680 | OG0002941 | chrXIII | 522945 | 523961 | CP015057.1 | 164816 | 165790 | 50.85714286 | 338 | 324 | 1.043209877 | 1 | 1 | 1 | FIM1_2520 | YMR127C | YMR127C | SAS2 | n |
| Kmar_QGN15910.1 | W303.asm02.HP0_G0009850 | OG0002942 | chrIV | 698494 | 699084 | CP015057.1 | 358860 | 359441 | 45.04950495 | 196 | 193 | 1.015544041 | 1 | 1 | 1 | FIM1_2608 | YDR121W | YDR121W | DPB4 | n |
| Kmar_QGN15823.1 | W303.asm02.HP0_G0041690 | OG0002943 | chrXIII | 524312 | 528115 | CP015057.1 | 160680 | 164483 | 65.63706564 | 1267 | 1267 | 1 | 1 | 1 | 1 | FIM1_2519 | YMR128W | YMR128W | ECM16 | n |
| Kmar_QGN13745.1 | W303.asm02.HP0_G0000730 | OG0002944 | chrI | 157958 | 159823 | CP015054.1 | 797483 | 799348 | 59.14149444 | 621 | 621 | 1 | 1 | 1 | 1 | FIM1_389 | YAR007C | YAR007C | RFA1 | n |
| Kmar_QGN16530.1 | W303.asm02.HP0_G0019740 | OG0002945 | chrVII | 459200 | 462445 | CP015058.1 | 311143 | 314325 | 44.55626715 | 1081 | 1060 | 1.019811321 | 1 | 1 | 1 | FIM1_3243 | YGL016W | YGL016W | KAP122 | n |

| | | | | | | | | | | | | | | | | | | | | |
|---|---|---|---|---|---|---|---|---|---|---|---|---|---|---|---|---|---|---|---|---|
| Kmar_QGN15825.1 | W303.asm02.HP0_G0035240 | OG0002946 | chrXII | 212375 | 215884 | CP015057.1 | 165852 | 169250 | 41.37931034 | 1169 | 1132 | 1.032685512 | 1 | 1 | 1 | FIM1_2521 | YLR032W | YLR032W | RAD5 | n |
| Kmar_QGN16526.1 | W303.asm02.HP0_G0019700 | OG0002947 | chrVII | 454697 | 455404 | CP015058.1 | 305793 | 306452 | 31.14754098 | 235 | 219 | 1.073059361 | 1 | 1 | 1 | FIM1_3239 | YGL020C | YGL020C | GET1 | n |
| Kmar_QGN16527.1 | W303.asm02.HP0_G0019710 | OG0002948 | chrVII | 455690 | 456526 | CP015058.1 | 307176 | 308006 | 69.78417266 | 278 | 276 | 1.007246377 | 1 | 1 | 1 | FIM1_3240 | YGL019W | YGL019W | CKB1 | n |
| Kmar_QGN16528.1 | W303.asm02.HP0_G0019720 | OG0002949 | chrVII | 456644 | 457198 | CP015058.1 | 308255 | 308818 | 44.14893617 | 184 | 187 | 0.983957219 | 1 | 1 | 1 | FIM1_3241 | YGL018C | YGL018C | JAC1 | n |
| Kmar_QGN15788.1 | W303.asm02.HP0_G0037940 | OG0002950 | chrXII | 773858 | 774637 | CP015057.1 | 102306 | 103007 | 38.996139 | 259 | 233 | 1.111587983 | 1 | 1 | 1 | FIM1_2484 | YLR323C | YLR323C | CWC24 | n |
| Kmar_QGN15793.1 | W303.asm02.HP0_G0037970 | OG0002951 | chrXII | 777859 | 778581 | CP015057.1 | 108535 | 109191 | 43.49593496 | 240 | 218 | 1.100917431 | 1 | 1 | 1 | FIM1_2489 | YLR326W | YLR326W | | n |
| Kmar_QGN16529.1 | W303.asm02.HP0_G0019730 | OG0002952 | chrVII | 457388 | 458899 | CP015058.1 | 309367 | 310866 | 45.02923977 | 503 | 499 | 1.008016032 | 1 | 1 | 1 | FIM1_3242 | YGL017W | YGL017W | ATE1 | n |
| Kmar_QGN15791.1 | W303.asm02.HP0_G0019930 | OG0002953 | chrVII | 502191 | 503393 | CP015057.1 | 106552 | 107709 | 66.91176471 | 400 | 385 | 1.038961039 | 1 | 1 | 1 | FIM1_2487 | YGR005C | YGR005C | TFG2 | n |
| Kmar_QGN15792.1 | W303.asm02.HP0_G0019940 | OG0002954 | chrVII | 503604 | 504359 | CP015057.1 | 107993 | 108523 | 27.09163347 | 251 | 176 | 1.426136364 | 1 | 1 | 1 | FIM1_2488 | YGR006W | YGR006W | PRP18 | n |
| Kmar_QGN17225.1 | W303.asm02.HP0_G0032980 | OG0002955 | chrXI | 396654 | 397997 | CP015059.1 | 488157 | 489656 | 57.11422846 | 447 | 499 | 0.895791583 | 1 | 1 | 1 | FIM1_3956 | YKL027W | YKL027W | TCD2 | n |
| Kmar_QGN14253.1 | W303.asm02.HP0_G0021210 | OG0002956 | chrVII | 797781 | 798647 | CP015055.1 | 116996 | 117649 | 38.54166667 | 288 | 217 | 1.32718894 | 1 | 1 | 1 | FIM1_909 | YGR147C | YGR147C | NAT2 | n |
| Kmar_QGN17224.1 | W303.asm02.HP0_G0023120 | OG0002957 | chrVIII | 101174 | 102247 | CP015059.1 | 486445 | 487569 | 65.69148936 | 357 | 374 | 0.954545455 | 1 | 1 | 1 | FIM1_3955 | YHR002W | YHR002W | LEU5 | n |
| Kmar_QGN16255.1 | W303.asm02.HP0_G0025250 | OG0002958 | chrVIII | 498033 | 499760 | CP015057.1 | 1138317 | 1139954 | 37.39279588 | 575 | 545 | 1.055045872 | 1 | 1 | 1 | FIM1_2960 | YHR196W | YHR196W | UTP9 | n |
| Kmar_QGN16253.1 | W303.asm02.HP0_G0030850 | OG0002959 | chrX | 683501 | 685624 | CP015057.1 | 1135253 | 1137205 | 28.33333333 | 707 | 650 | 1.087692308 | 1 | 1 | 1 | FIM1_2958 | YJR134C | YJR134C | SGM1 | n |
| Kmar_QGN14433.1 | W303.asm02.HP0_G0004260 | OG0002960 | chrII | 713226 | 714536 | CP015055.1 | 508251 | 509513 | 70.62937063 | 436 | 420 | 1.038095238 | 1 | 1 | 1 | FIM1_1094 | YBR236C | YBR236C | ABD1 | n |
| Kmar_QGN15414.1 | W303.asm02.HP0_G0023640 | OG0002961 | chrVIII | 200306 | 201037 | CP015056.1 | 911422 | 912147 | 74.18032787 | 243 | 241 | 1.008298755 | 1 | 1 | 1 | FIM1_2104 | YHR049W | YHR049W | FSH1 | n |
| Kmar_QGN13597.1 | W303.asm02.HP0_G0009580 | OG0002962 | chrIV | 629338 | 630270 | CP015054.1 | 491119 | 491928 | 53.37620579 | 310 | 269 | 1.152416357 | 1 | 1 | 1 | FIM1_239 | YDR090C | YDR090C | ILT1 | n |
| Kmar_QGN15184.1 | W303.asm02.HP0_G0016090 | OG0002963 | chrV | 508332 | 510596 | CP015056.1 | 387174 | 389525 | 39.89962359 | 754 | 783 | 0.962962963 | 1 | 1 | 1 | FIM1_1872 | YER162C | YER162C | RAD4 | n |
| Kmar_QGN15185.1 | W303.asm02.HP0_G0016080 | OG0002964 | chrV | 507049 | 508050 | CP015056.1 | 389739 | 390704 | 43.56725146 | 333 | 321 | 1.037383178 | 1 | 1 | 1 | FIM1_1873 | YER161C | YER161C | SPT2 | n |
| Kmar_QGN17619.1 | W303.asm02.HP0_G0043220 | OG0002965 | chrXIII | 842248 | 843789 | CP015060.1 | 118817 | 120283 | 49.90366089 | 513 | 488 | 1.051229508 | 1 | 1 | 1 | FIM1_4826 | YMR283C | YMR283C | RIT1 | n |
| Kmar_QGN13657.1 | W303.asm02.HP0_G0005660 | OG0002966 | chrIII | 148874 | 149746 | CP015054.1 | 608642 | 609463 | 41.05263158 | 290 | 273 | 1.062271062 | 1 | 1 | 1 | FIM1_300 | YCR016W | YCR016W | RBP95 | n |
| Kmar_QGN13658.1 | W303.asm02.HP0_G0005650 | OG0002967 | chrIII | 146457 | 147410 | CP015054.1 | 610276 | 610965 | 26.41509434 | 317 | 229 | 1.384279476 | 1 | 1 | 1 | FIM1_301 | YCR015C | YCR015C | CTO1 | n |
| Kmar_QGN16426.1 | W303.asm02.HP0_G0050920 | OG0002968 | chrXV | 726859 | 728529 | CP015058.1 | 93624 | 95291 | 37.65541741 | 556 | 555 | 1.001801802 | 1 | 1 | 1 | FIM1_3139 | YOR205C | YOR205C | GEP3 | n |
| Kmar_QGN17627.1 | W303.asm02.HP0_G0043170 | OG0002969 | chrXIII | 829685 | 831553 | CP015060.1 | 139619 | 141460 | 56.46687697 | 622 | 613 | 1.014681892 | 1 | 1 | 1 | FIM1_4834 | YMR278W | YMR278W | PRM15 | n |
| Kmar_QGN13623.1 | W303.asm02.HP0_G0043160 | OG0002970 | chrXIII | 826755 | 828953 | CP015054.1 | 539016 | 541205 | 48.41688654 | 732 | 729 | 1.004115226 | 1 | 1 | 1 | FIM1_265 | YMR277W | YMR277W | FCP1 | n |
| Kmar_QGN14266.1 | W303.asm02.HP0_G0043010 | OG0002971 | chrXIII | 800228 | 801169 | CP015055.1 | 144972 | 146207 | 28.26603325 | 313 | 411 | 0.761557178 | 1 | 1 | 1 | FIM1_923 | YMR262W | YMR262W | | n |
| Kmar_QGN15883.1 | W303.asm02.HP0_G0009180 | OG0002972 | chrIV | 559875 | 560621 | CP015057.1 | 290519 | 291265 | 82.25806452 | 248 | 248 | 1 | 1 | 1 | 1 | FIM1_2581 | YDR050C | YDR050C | TPI1 | n |
| Kmar_QGN15878.1 | W303.asm02.HP0_G0009050 | OG0002973 | chrIV | 525100 | 526875 | CP015057.1 | 281714 | 283666 | 80.36809816 | 591 | 650 | 0.909230769 | 1 | 1 | 1 | FIM1_2576 | YDR037W | YDR037W | KRS1 | n |
| Kmar_QGN15877.1 | W303.asm02.HP0_G0002580 | OG0002974 | chrII | 381074 | 382006 | CP015057.1 | 280731 | 281645 | 75.39936102 | 310 | 304 | 1.019736842 | 1 | 1 | 1 | FIM1_2575 | YBR061C | YBR061C | TRM7 | n |
| Kmar_QGN15876.1 | W303.asm02.HP0_G0009100 | OG0002975 | chrIV | 543348 | 543959 | CP015057.1 | 279820 | 280434 | 55.55555556 | 203 | 204 | 0.995098039 | 1 | 1 | 1 | FIM1_2574 | YDR041W | YDR041W | RSM10 | n |
| Kmar_QGN15154.1 | W303.asm02.HP0_G0018110 | OG0002976 | chrVII | 119104 | 119811 | CP015056.1 | 309637 | 310305 | 45.76271186 | 235 | 222 | 1.058558559 | 1 | 1 | 1 | FIM1_1841 | YGL198W | YGL198W | YIP4 | n |
| Kmar_QGN14930.1 | W303.asm02.HP0_G0045110 | OG0002977 | chrXIV | 308097 | 309308 | CP015055.1 | 1515667 | 1516839 | 60.14851485 | 403 | 390 | 1.033333333 | 1 | 1 | 1 | FIM1_1605 | YNL175C | YNL175C | NOP13 | n |
| Kmar_QGN13529.1 | W303.asm02.HP0_G0022820 | OG0002978 | chrVIII | 43552 | 45591 | CP015054.1 | 349303 | 350883 | 37.64534884 | 679 | 526 | 1.290874525 | 1 | 1 | 1 | FIM1_169 | YHL029C | YHL029C | OCA5 | n |
| Kmar_QGN13526.1 | W303.asm02.HP0_G0034570 | OG0002979 | chrXII | 65784 | 66527 | CP015054.1 | 343721 | 344398 | 40.16064257 | 247 | 225 | 1.097777778 | 1 | 1 | 1 | FIM1_166 | YLL038C | YLL038C | ENT4 | n |
| Kmar_QGN14634.1 | W303.asm02.HP0_G0022810 | OG0002980 | chrVIII | 37707 | 43313 | CP015055.1 | 909798 | 915350 | 39.67161017 | 1868 | 1850 | 1.00972973 | 1 | 1 | 1 | FIM1_1298 | YHL030W | YHL030W | ECM29 | n |
| Kmar_QGN14531.1 | W303.asm02.HP0_G0020600 | OG0002981 | chrVII | 642896 | 644851 | CP015055.1 | 706830 | 708671 | 57.16463415 | 651 | 613 | 1.061990212 | 1 | 1 | 1 | FIM1_1193 | YGR083C | YGR083C | GCD2 | n |
| Kmar_QGN16500.1 | W303.asm02.HP0_G0000450 | OG0002982 | chrI | 96393 | 100460 | CP015058.1 | 241633 | 245613 | 70.38123167 | 1355 | 1326 | 1.021870287 | 1 | 1 | 1 | FIM1_3213 | YAL026C | YAL026C | DRS2 | n |
| Kmar_QGN16499.1 | W303.asm02.HP0_G0052030 | OG0002983 | chrXV | 923255 | 924625 | CP015058.1 | 240214 | 241557 | 72.5877193 | 456 | 447 | 1.020134228 | 1 | 1 | 1 | FIM1_3212 | YOR323C | YOR323C | PRO2 | n |
| Kmar_QGN13971.1 | W303.asm02.HP0_G0004050 | OG0002984 | chrII | 672649 | 673473 | CP015054.1 | 1242355 | 1243185 | 36.51877133 | 274 | 276 | 0.992753623 | 1 | 1 | 1 | FIM1_620 | YBR213W | YBR213W | MET8 | n |
| Kmar_QGN15009.1 | W303.asm02.HP0_G0052450 | OG0002985 | chrXV | 1031464 | 1033275 | CP015055.1 | 1680126 | 1681961 | 66.61237785 | 603 | 611 | 0.98690671 | 1 | 1 | 1 | FIM1_1690 | YOR370C | YOR370C | MRS6 | n |
| Kmar_QGN16389.1 | W303.asm02.HP0_G0052440 | OG0002986 | chrXV | 1030475 | 1030906 | CP015058.1 | 8037 | 8441 | 76.22377622 | 143 | 134 | 1.067164179 | 1 | 1 | 1 | FIM1_3102 | YOR369C | YOR369C | RPS12 | n |
| Kmar_QGN16393.1 | W303.asm02.HP0_G0052480 | OG0002987 | chrXV | 1039115 | 1041670 | CP015058.1 | 16058 | 18868 | 23.82934443 | 851 | 936 | 0.909188034 | 1 | 1 | 1 | FIM1_3106 | YOR373W | YOR373W | NUD1 | n |
| Kmar_QGN15735.1 | W303.asm02.HP0_G0025770 | OG0002988 | chrIX | 44229 | 45410 | CP015056.1 | 1568295 | 1569398 | 45.03816794 | 393 | 367 | 1.070844687 | 1 | 1 | 1 | FIM1_2429 | YIL153W | YIL153W | RRD1 | n |
| Kmar_QGN17872.1 | W303.asm02.HP0_G0003060 | OG0002989 | chrII | 477464 | 478159 | CP015060.1 | 647354 | 648028 | 69.56521739 | 231 | 224 | 1.03125 | 1 | 1 | 1 | FIM1_5081 | YBR111C | YBR111C | YSA1 | n |
| Kmar_QGN17874.1 | W303.asm02.HP0_G0003080 | OG0002990 | chrII | 479156 | 482056 | CP015060.1 | 649036 | 652491 | 48.82710686 | 966 | 1151 | 0.8392702 | 1 | 1 | 1 | FIM1_5083 | YBR112C | YBR112C | CYC8 | n |
| Kmar_QGN17205.1 | W303.asm02.HP0_G0053590 | OG0002991 | chrXVI | 196958 | 198202 | CP015059.1 | 445561 | 446796 | 67.07021792 | 414 | 411 | 1.00729927 | 1 | 1 | 1 | FIM1_3936 | YPL188W | YPL188W | POS5 | n |
| Kmar_QGN14810.1 | W303.asm02.HP0_G0043740 | OG0002992 | chrXIV | 27769 | 28779 | CP015055.1 | 1273389 | 1274366 | 41.59021407 | 336 | 325 | 1.033846154 | 1 | 1 | 1 | FIM1_1481 | YNL326C | YNL326C | PFA3 | n |
| Kmar_QGN16712.1 | W303.asm02.HP0_G0053050 | OG0002993 | chrXVI | 92579 | 93598 | CP015058.1 | 721053 | 722054 | 53.33333333 | 339 | 333 | 1.018018018 | 1 | 1 | 1 | FIM1_3432 | YPL244C | YPL244C | HUT1 | n |
| Kmar_QGN16713.1 | W303.asm02.HP0_G0053060 | OG0002994 | chrXVI | 94082 | 95881 | CP015058.1 | 722592 | 724358 | 36.7816092 | 599 | 588 | 1.018707483 | 1 | 1 | 1 | FIM1_3433 | YPL243W | YPL243W | SRP68 | n |
| Kmar_QGN15186.1 | W303.asm02.HP0_G0009570 | OG0002995 | chrIV | 626516 | 628993 | CP015056.1 | 390862 | 393396 | 30.9336333 | 825 | 844 | 0.977488152 | 1 | 1 | 1 | FIM1_1874 | YDR089W | YDR089W | VTC5 | n |
| Kmar_QGN14560.1 | W303.asm02.HP0_G0000550 | OG0002996 | chrI | 125643 | 127550 | CP015055.1 | 753969 | 755813 | 67.45283019 | 635 | 614 | 1.034201954 | 1 | 1 | 1 | FIM1_1222 | YAL016W | YAL016W | TPD3 | n |
| Kmar_QGN13559.1 | W303.asm02.HP0_G0038820 | OG0002997 | chrXII | 955097 | 959404 | CP015054.1 | 415562 | 419764 | 54.58970793 | 1435 | 1400 | 1.025 | 1 | 1 | 1 | FIM1_200 | YLR419W | YLR419W | | n |
| Kmar_QGN16852.1 | W303.asm02.HP0_G0036120 | OG0002998 | chrXII | 403711 | 406272 | CP015058.1 | 1030523 | 1032613 | 21.38728324 | 853 | 696 | 1.225574713 | 1 | 1 | 1 | FIM1_3577 | YLR127C | YLR127C | APC2 | n |
| Kmar_QGN15188.1 | W303.asm02.HP0_G0009560 | OG0002999 | chrIV | 622774 | 623922 | CP015056.1 | 394692 | 395813 | 32.11678832 | 382 | 373 | 1.024128686 | 1 | 1 | 1 | FIM1_1876 | YDR088C | YDR088C | SLU7 | n |
| Kmar_QGN17365.1 | W303.asm02.HP0_G0009550 | OG0003000 | chrIV | 621747 | 622583 | CP015059.1 | 778452 | 779444 | 42.27405248 | 278 | 330 | 0.842424242 | 1 | 1 | 1 | FIM1_4097 | YDR087C | YDR087C | RRP1 | n |
| Kmar_QGN17620.1 | W303.asm02.HP0_G0043210 | OG0003001 | chrXIII | 840278 | 842020 | CP015060.1 | 120918 | 122504 | 23.19587629 | 580 | 528 | 1.098484848 | 1 | 1 | 1 | FIM1_4827 | YMR282C | YMR282C | AEP2 | n |
| Kmar_QGN14432.1 | W303.asm02.HP0_G0004250 | OG0003002 | chrII | 709744 | 713106 | CP015055.1 | 504838 | 508116 | 60.0177305 | 1120 | 1092 | 1.025641026 | 1 | 1 | 1 | FIM1_1093 | YBR235W | YBR235W | VHC1 | n |
| Kmar_QGN17898.1 | W303.asm02.HP0_G0042950 | OG0003003 | chrXIII | 785448 | 785630 | CP015060.1 | 698307 | 698495 | 72.58064516 | 60 | 62 | 0.967741935 | 1 | 1 | 1 | FIM1_5107 | YMR256C | YMR256C | COX7 | n |
| Kmar_QGN15132.1 | W303.asm02.HP0_G0031960 | OG0003004 | chrXI | 187764 | 189944 | CP015056.1 | 266824 | 268947 | 54.12719892 | 726 | 707 | 1.026874116 | 1 | 1 | 1 | FIM1_1819 | YKL135C | YKL135C | APL2 | n |
| Kmar_QGN18141.1 | W303.asm02.HP0_G0047780 | OG0003005 | chrXV | 69717 | 70385 | CP015061.1 | 316385 | 317026 | 48.44444444 | 222 | 213 | 1.042253521 | 1 | 1 | 1 | FIM1_4464 | YOL135C | YOL135C | MED7 | n |
| Kmar_QGN13856.1 | W303.asm02.HP0_G0033880 | OG0003006 | chrXI | 574966 | 575559 | CP015054.1 | 1004937 | 1005497 | 47.20812183 | 197 | 186 | 1.059139785 | 1 | 1 | 1 | FIM1_502 | YKR065C | YKR065C | PAM17 | n |
| Kmar_QGN17188.1 | W303.asm02.HP0_G0053500 | OG0003007 | chrXVI | 180595 | 181416 | CP015059.1 | 416948 | 417748 | 53.84615385 | 273 | 266 | 1.026315789 | 1 | 1 | 1 | FIM1_3919 | YPL196W | YPL196W | OXR1 | n |
| Kmar_QGN18252.1 | W303.asm02.HP0_G0027540 | OG0003008 | chrIX | 403337 | 404071 | CP015061.1 | 558447 | 559274 | 19.09722222 | 244 | 275 | 0.887272727 | 1 | 1 | 1 | FIM1_4577 | YIR030C | YIR030C | DCG1 | n |
| Kmar_QGN17191.1 | W303.asm02.HP0_G0053520 | OG0003009 | chrXVI | 184829 | 186667 | CP015059.1 | 421626 | 423317 | 34.73344103 | 612 | 563 | 1.087033748 | 1 | 1 | 1 | FIM1_3922 | YPL194W | YPL194W | DDC1 | n |
| Kmar_QGN17727.1 | W303.asm02.HP0_G0026210 | OG0003010 | chrIX | 151244 | 153334 | CP015060.1 | 335522 | 337576 | 56.38450502 | 696 | 684 | 1.01754386 | 1 | 1 | 1 | FIM1_4936 | YIL108W | YIL108W | | n |
| Kmar_QGN17189.1 | W303.asm02.HP0_G0019190 | OG0003011 | chrVII | 356396 | 357052 | CP015059.1 | 417847 | 418380 | 36.27906977 | 218 | 177 | 1.231638418 | 1 | 1 | 1 | FIM1_3920 | YGL079W | YGL079W | KXD1 | n |
| Kmar_QGN17190.1 | W303.asm02.HP0_G0053510 | OG0003012 | chrXVI | 181775 | 184573 | CP015059.1 | 418689 | 421397 | 50.74786325 | 932 | 902 | 1.033259424 | 1 | 1 | 1 | FIM1_3921 | YPL195W | YPL195W | APL5 | n |
| Kmar_QGN17103.1 | W303.asm02.HP0_G0008300 | OG0003013 | chrIV | 363007 | 363918 | CP015059.1 | 237112 | 238074 | 49.0625 | 303 | 320 | 0.946875 | 1 | 1 | 1 | FIM1_3834 | YDL052C | YDL052C | SLC1 | n |
| Kmar_QGN17139.1 | W303.asm02.HP0_G0040610 | OG0003014 | chrXIII | 295685 | 297244 | CP015059.1 | 316892 | 318475 | 32.83302064 | 519 | 527 | 0.984819734 | 1 | 1 | 1 | FIM1_3870 | YMR013C | YMR013C | SEC59 | n |
| Kmar_QGN15741.1 | W303.asm02.HP0_G0028990 | OG0003015 | chrX | 289176 | 292868 | CP015056.1 | 1582179 | 1585853 | 48.33468725 | 1230 | 1224 | 1.004901961 | 1 | 1 | 1 | FIM1_2435 | YJL074C | YJL074C | SMC3 | n |
| Kmar_QGN13675.1 | W303.asm02.HP0_G0008920 | OG0003016 | chrIV | 489167 | 490555 | CP015054.1 | 642403 | 643782 | 84.63203463 | 462 | 459 | 1.006535948 | 1 | 1 | 1 | FIM1_318 | YDR023W | YDR023W | SES1 | n |
| Kmar_QGN14097.1 | W303.asm02.HP0_G0020300 | OG0003017 | chrVII | 584151 | 587228 | CP015054.1 | 1512854 | 1514563 | 37.00340522 | 1025 | 569 | 1.801405975 | 1 | 1 | 1 | FIM1_748 | YGR047C | YGR047C | TFC4 | n |
| Kmar_QGN16488.1 | W303.asm02.HP0_G0000380 | OG0003018 | chrI | 83469 | 83990 | CP015058.1 | 215479 | 215967 | 50.28571429 | 173 | 162 | 1.067901235 | 1 | 1 | 1 | FIM1_3201 | YAL033W | YAL033W | POP5 | n |
| Kmar_QGN16306.1 | W303.asm02.HP0_G0053340 | OG0003019 | chrXVI | 154124 | 155746 | CP015057.1 | 1245607 | 1247163 | 55.63770795 | 540 | 518 | 1.042471042 | 1 | 1 | 1 | FIM1_3011 | YPL214C | YPL214C | THI6 | n |
| Kmar_QGN14472.1 | W303.asm02.HP0_G0018780 | OG0003020 | chrVII | 268990 | 270792 | CP015055.1 | 587382 | 589235 | 62.58064516 | 600 | 617 | 0.972447326 | 1 | 1 | 1 | FIM1_1133 | YGL125W | YGL125W | MET13 | n |
| Kmar_QGN16486.1 | W303.asm02.HP0_G0052140 | OG0003021 | chrXV | 951920 | 956017 | CP015058.1 | 207840 | 211592 | 29.36449963 | 1365 | 1250 | 1.092 | 1 | 1 | 1 | FIM1_3199 | YOR336W | YOR336W | KRE5 | n |
| Kmar_QGN16110.1 | W303.asm02.HP0_G0024630 | OG0003022 | chrVIII | 377722 | 378531 | CP015057.1 | 833240 | 833992 | 48.89705882 | 269 | 250 | 1.076 | 1 | 1 | 1 | FIM1_2811 | YHR134W | YHR134W | WSS1 | n |
| Kmar_QGN16113.1 | W303.asm02.HP0_G0024600 | OG0003023 | chrVIII | 374475 | 375767 | CP015057.1 | 836875 | 838233 | 56.3876652 | 430 | 452 | 0.951327434 | 1 | 1 | 1 | FIM1_2814 | YHR132C | YHR132C | ECM14 | n |
| Kmar_QGN15337.1 | W303.asm02.HP0_G0052950 | OG0003024 | chrXVI | 75053 | 76519 | CP015056.1 | 741079 | 742374 | 49.3852459 | 488 | 431 | 1.13225058 | 1 | 1 | 1 | FIM1_2026 | YPL254W | YPL254W | HFI1 | n |
| Kmar_QGN17368.1 | W303.asm02.HP0_G0040970 | OG0003025 | chrXIII | 380089 | 380703 | CP015059.1 | 782216 | 782845 | 34.43396226 | 204 | 209 | 0.976076555 | 1 | 1 | 1 | FIM1_4100 | YMR052W | YMR052W | FAR3 | n |
| Kmar_QGN18232.1 | W303.asm02.HP0_G0036920 | OG0003026 | chrXII | 559678 | 560571 | CP015061.1 | 507011 | 507916 | 82.72425249 | 297 | 301 | 0.986710963 | 1 | 1 | 1 | FIM1_4557 | YLR208W | YLR208W | SEC13 | n |
| Kmar_QGN18231.1 | W303.asm02.HP0_G0036930 | OG0003027 | chrXII | 560924 | 561859 | CP015061.1 | 505814 | 506734 | 66.55948553 | 311 | 306 | 1.016339869 | 1 | 1 | 1 | FIM1_4556 | YLR209C | YLR209C | PNP1 | n |
| Kmar_QGN17117.1 | W303.asm02.HP0_G0030740 | OG0003028 | chrX | 659550 | 661043 | CP015059.1 | 272641 | 274005 | 35.01006036 | 497 | 454 | 1.094713656 | 1 | 1 | 1 | FIM1_3848 | YJR122W | YJR122W | IBA57 | n |
| Kmar_QGN15419.1 | W303.asm02.HP0_G0011380 | OG0003029 | chrIV | 1039730 | 1040074 | CP015056.1 | 918948 | 919280 | 37.16814159 | 114 | 110 | 1.036363636 | 1 | 1 | 1 | FIM1_2109 | YDR286C | YDR286C | MGP12 | n |
| Kmar_QGN15143.1 | W303.asm02.HP0_G0036020 | OG0003030 | chrXII | 385938 | 388517 | CP015056.1 | 287439 | 289898 | 41.89814815 | 859 | 819 | 1.048840049 | 1 | 1 | 1 | FIM1_1830 | YLR115W | YLR115W | CFT2 | n |
| Kmar_QGN13956.1 | W303.asm02.HP0_G0048960 | OG0003031 | chrXV | 304034 | 304438 | CP015054.1 | 1215646 | 1216044 | 92.53731343 | 134 | 132 | 1.015151515 | 1 | 1 | 1 | FIM1_604 | YOL012C | YOL012C | HTZ1 | n |
| Kmar_QGN17608.1 | W303.asm02.HP0_G0013870 | OG0003032 | chrV | 34743 | 36617 | CP015060.1 | 92822 | 94795 | 44.86803519 | 624 | 657 | 0.949771689 | 1 | 1 | 1 | FIM1_4815 | YEL062W | YEL062W | NPR2 | n |
| Kmar_QGN15848.1 | W303.asm02.HP0_G0036030 | OG0003033 | chrXII | 388775 | 390205 | CP015057.1 | 211030 | 212385 | 42.17118998 | 476 | 451 | 1.055432373 | 1 | 1 | 1 | FIM1_2544 | YLR116W | YLR116W | MSL5 | n |
| Kmar_QGN15845.1 | W303.asm02.HP0_G0010020 | OG0003034 | chrIV | 739047 | 739712 | CP015057.1 | 207464 | 208132 | 55.35714286 | 221 | 222 | 0.995495495 | 1 | 1 | 1 | FIM1_2541 | YDR140W | YDR140W | MTQ2 | n |
| Kmar_QGN15847.1 | W303.asm02.HP0_G0036040 | OG0003035 | chrXII | 390424 | 392487 | CP015057.1 | 208896 | 210965 | 47.98270893 | 687 | 689 | 0.997097242 | 1 | 1 | 1 | FIM1_2543 | YLR117C | YLR117C | CLF1 | n |
| Kmar_QGN15339.1 | W303.asm02.HP0_G0052970 | OG0003036 | chrXVI | 78931 | 79449 | CP015056.1 | 744893 | 745396 | 64.53488372 | 172 | 167 | 1.02994012 | 1 | 1 | 1 | FIM1_2028 | YPL252C | YPL252C | YAH1 | n |
| Kmar_QGN14720.1 | W303.asm02.HP0_G0036060 | OG0003037 | chrXII | 393487 | 394128 | CP015055.1 | 1094025 | 1094585 | 29.30232558 | 213 | 186 | 1.14516129 | 1 | 1 | 1 | FIM1_1388 | YLR119W | YLR119W | SRN2 | n |
| Kmar_QGN14731.1 | W303.asm02.HP0_G0056780 | OG0003038 | chrXVI | 847196 | 849046 | CP015055.1 | 1112796 | 1114604 | 26.16372392 | 616 | 602 | 1.023255814 | 1 | 1 | 1 | FIM1_1400 | YPR155C | YPR155C | NCA2 | n |
| Kmar_QGN14729.1 | W303.asm02.HP0_G0021100 | OG0003039 | chrVII | 773119 | 773895 | CP015055.1 | 1109896 | 1110651 | 84.52380952 | 258 | 251 | 1.027888446 | 1 | 1 | 1 | FIM1_1397 | YGR135W | YGR135W | PRE9 | n |
| Kmar_QGN16165.1 | W303.asm02.HP0_G0015310 | OG0003040 | chrV | 318477 | 320012 | CP015057.1 | 939487 | 941013 | 64.27184466 | 511 | 508 | 1.005905512 | 1 | 1 | 1 | FIM1_2867 | YER078C | YER078C | ICP55 | n |
| Kmar_QGN16761.1 | W303.asm02.HP0_G0027220 | OG0003041 | chrIX | 344995 | 346536 | CP015058.1 | 837596 | 839140 | 46.3878327 | 513 | 514 | 0.998054475 | 1 | 1 | 1 | FIM1_3483 | YIL001W | YIL001W | | n |
| Kmar_QGN16083.1 | W303.asm02.HP0_G0055280 | OG0003042 | chrXVI | 536080 | 539766 | CP015057.1 | 775490 | 779137 | 61.12012987 | 1228 | 1215 | 1.010699588 | 1 | 1 | 1 | FIM1_2783 | YPL012W | YPL012W | RRP12 | n |
| Kmar_QGN16082.1 | W303.asm02.HP0_G0055270 | OG0003043 | chrXVI | 535342 | 535707 | CP015057.1 | 774477 | 774839 | 65.2892562 | 121 | 120 | 1.008333333 | 1 | 1 | 1 | FIM1_2782 | YPL013C | YPL013C | MRPS16 | n |
| Kmar_QGN16577.1 | W303.asm02.HP0_G0038070 | OG0003044 | chrXII | 801574 | 802512 | CP015058.1 | 420211 | 421146 | 86.53846154 | 312 | 311 | 1.003215434 | 1 | 1 | 1 | FIM1_3292 | YLR340W | YLR340W | RPP0 | n |

| | | | | | | | | | | | | | | | | | | | | |
|---|---|---|---|---|---|---|---|---|---|---|---|---|---|---|---|---|---|---|---|---|
| Kmar_QGN16331.1 | W303.asm02.HP0_G0012940 | OG0003045 | chrIV | 1368571 | 1370595 | CP015057.1 | 1305135 | 1307216 | 46.10123119 | 674 | 693 | 0.972582973 | 1 | 1 | 1 | FIM1_3036 | YDR452W | YDR452W | PPN1 | n |
| Kmar_QGN16334.1 | W303.asm02.HP0_G0040260 | OG0003046 | chrXIII | 224914 | 225873 | CP015057.1 | 1311736 | 1312941 | 56.64335664 | 286 | 272 | 1.051470588 | 1 | 1 | 1 | FIM1_3039 | YML025C | YML025C | YML6 | n |
| Kmar_QGN14456.1 | W303.asm02.HP0_G0018070 | OG0003047 | chrVII | 107988 | 110177 | CP015055.1 | 552028 | 553914 | 42.85714286 | 729 | 628 | 1.160828025 | 1 | 1 | 1 | FIM1_1117 | YGL203C | YGL203C | KEX1 | n |
| Kmar_QGN14453.1 | W303.asm02.HP0_G0056830 | OG0003048 | chrXVI | 867138 | 869846 | CP015055.1 | 543503 | 546214 | 76.4254386 | 902 | 903 | 0.99889258 | 1 | 1 | 1 | FIM1_1114 | YPR160W | YPR160W | GPH1 | n |
| Kmar_QGN16061.1 | W303.asm02.HP0_G0040930 | OG0003049 | chrXIII | 355887 | 357314 | CP015057.1 | 728790 | 730172 | 35.8974359 | 475 | 460 | 1.032608696 | 1 | 1 | 1 | FIM1_2761 | YMR044W | YMR044W | IOC4 | n |
| Kmar_QGN16973.1 | W303.asm02.HP0_G0006590 | OG0003050 | chrIV | 29060 | 30529 | CP015058.1 | 1287063 | 1288565 | 48.6328125 | 489 | 500 | 0.978 | 1 | 1 | 1 | FIM1_3700 | YDL238C | YDL238C | GUD1 | n |
| Kmar_QGN18049.1 | W303.asm02.HP0_G0038550 | OG0003051 | chrXII | 900394 | 901110 | CP015061.1 | 99401 | 100087 | 49.79079498 | 238 | 228 | 1.043859649 | 1 | 1 | 1 | FIM1_4366 | YLR390W-A | YLR390W-A | CCW14 | n |
| Kmar_QGN16460.1 | W303.asm02.HP0_G0004650 | OG0003052 | chrII | 780969 | 783392 | CP015058.1 | 158197 | 160524 | 30.35495716 | 807 | 775 | 1.041290323 | 1 | 1 | 1 | FIM1_3173 | YBR276C | YBR276C | PPS1 | n |
| Kmar_QGN15060.1 | W303.asm02.HP0_G0030230 | OG0003053 | chrX | 576930 | 577991 | CP015056.1 | 112921 | 113976 | 69.68838527 | 353 | 351 | 1.005698006 | 1 | 1 | 1 | FIM1_1745 | YJR068W | YJR068W | RFC2 | n |
| Kmar_QGN15061.1 | W303.asm02.HP0_G0030220 | OG0003054 | chrX | 576306 | 576731 | CP015056.1 | 114215 | 114631 | 44.20289855 | 141 | 138 | 1.02173913 | 1 | 1 | 1 | FIM1_1746 | YJR067C | YJR067C | YAE1 | n |
| Kmar_QGN15596.1 | W303.asm02.HP0_G0000270 | OG0003055 | chrI | 59330 | 61687 | CP015056.1 | 1266704 | 1269193 | 36.71782763 | 785 | 829 | 0.946924005 | 1 | 1 | 1 | FIM1_2288 | YAL043C | YAL043C | PTA1 | n |
| Kmar_QGN14497.1 | W303.asm02.HP0_G0016580 | OG0003056 | chrVI | 37563 | 42875 | CP015055.1 | 633638 | 638428 | 40.31728665 | 1770 | 1596 | 1.109022556 | 1 | 1 | 1 | FIM1_1158 | YFL033C | YFL033C | RIM15 | n |
| Kmar_QGN15723.1 | W303.asm02.HP0_G0017690 | OG0003057 | chrVII | 36868 | 38544 | CP015056.1 | 1535026 | 1536735 | 57.06806283 | 558 | 569 | 0.980667838 | 1 | 1 | 1 | FIM1_2416 | YGL244W | YGL244W | RTF1 | n |
| Kmar_QGN17333.1 | W303.asm02.HP0_G0045700 | OG0003058 | chrXIV | 419481 | 420143 | CP015059.1 | 712859 | 713566 | 50.63829787 | 220 | 235 | 0.936170213 | 1 | 1 | 1 | FIM1_4065 | YNL110C | YNL110C | NOP15 | n |
| Kmar_QGN15725.1 | W303.asm02.HP0_G0017680 | OG0003059 | chrVII | 34345 | 36519 | CP015056.1 | 1539009 | 1541129 | 78.17679558 | 724 | 706 | 1.025495751 | 1 | 1 | 1 | FIM1_2418 | YGL245W | YGL245W | GUS1 | n |
| Kmar_QGN15724.1 | W303.asm02.HP0_G0017400 | OG0003060 | chrVI | 215051 | 217039 | CP015056.1 | 1537155 | 1538954 | 57.14285714 | 662 | 599 | 1.105175292 | 1 | 1 | 1 | FIM1_2417 | YFR048W | YFR048W | RMD8 | n |
| Kmar_QGN15951.1 | W303.asm02.HP0_G0050600 | OG0003061 | chrXV | 648001 | 649377 | CP015057.1 | 454330 | 455652 | 32.67973856 | 458 | 440 | 1.040909091 | 1 | 1 | 1 | FIM1_2649 | YOR166C | YOR166C | SWT1 | n |
| Kmar_QGN18413.1 | W303.asm02.HP0_G0016960 | OG0003062 | chrVI | 127782 | 128843 | CP015061.1 | 881869 | 882939 | 47.81420765 | 353 | 356 | 0.991573034 | 1 | 1 | 1 | FIM1_4741 | YFR007W | YFR007W | YFH7 | n |
| Kmar_QGN16588.1 | W303.asm02.HP0_G0055490 | OG0003063 | chrXVI | 583942 | 587553 | CP015058.1 | 442932 | 446543 | 86.26964433 | 1203 | 1203 | 1 | 1 | 1 | 1 | FIM1_3304 | YPR010C | YPR010C | RPA135 | n |
| Kmar_QGN15954.1 | W303.asm02.HP0_G0050610 | OG0003064 | chrXV | 650178 | 652607 | CP015057.1 | 460222 | 462636 | 77.13226205 | 809 | 804 | 1.006218905 | 1 | 1 | 1 | FIM1_2652 | YOR168W | YOR168W | GLN4 | n |
| Kmar_QGN14818.1 | W303.asm02.HP0_G0047680 | OG0003065 | chrXV | 49067 | 49651 | CP015055.1 | 1289693 | 1290259 | 50 | 194 | 188 | 1.031914894 | 1 | 1 | 1 | FIM1_1489 | YOL146W | YOL146W | PSF3 | n |
| Kmar_QGN14817.1 | W303.asm02.HP0_G0047690 | OG0003066 | chrXV | 49758 | 52991 | CP015055.1 | 1286099 | 1289422 | 51.78890877 | 1077 | 1107 | 0.972899729 | 1 | 1 | 1 | FIM1_1488 | YOL145C | YOL145C | CTR9 | n |
| Kmar_QGN16472.1 | W303.asm02.HP0_G0004510 | OG0003067 | chrII | 758188 | 758886 | CP015058.1 | 179802 | 180491 | 65.94827586 | 232 | 229 | 1.013100437 | 1 | 1 | 1 | FIM1_3185 | YBR261C | YBR261C | TAE1 | n |
| Kmar_QGN13756.1 | W303.asm02.HP0_G0005480 | OG0003068 | chrIII | 114345 | 115910 | CP015054.1 | 817323 | 818948 | 51.18829982 | 521 | 541 | 0.963031423 | 1 | 1 | 1 | FIM1_401 | YCL004W | YCL004W | PGS1 | n |
| Kmar_QGN15158.1 | W303.asm02.HP0_G0005430 | OG0003069 | chrIII | 108811 | 109590 | CP015056.1 | 324544 | 325341 | 48.72727273 | 259 | 265 | 0.977358491 | 1 | 1 | 1 | FIM1_1845 | YCL010C | YCL010C | SGF29 | n |
| Kmar_QGN14677.1 | W303.asm02.HP0_G0005400 | OG0003070 | chrIII | 101521 | 106431 | CP015055.1 | 1008432 | 1012799 | 35.31553398 | 1636 | 1455 | 1.124398625 | 1 | 1 | 1 | FIM1_1342 | YCL014W | YCL014W | BUD3 | n |
| Kmar_QGN14673.1 | W303.asm02.HP0_G0005370 | OG0003071 | chrIII | 96562 | 97658 | CP015055.1 | 1002117 | 1003205 | 74.72527473 | 313 | 362 | 0.864640884 | 1 | 1 | 1 | FIM1_1338 | YCL018W | YCL018W | LEU2 | n |
| Kmar_QGN14674.1 | W303.asm02.HP0_G0005380 | OG0003072 | chrIII | 98017 | 99510 | CP015055.1 | 1003729 | 1005207 | 77.86720322 | 497 | 492 | 1.010162602 | 1 | 1 | 1 | FIM1_1339 | YCL017C | YCL017C | NFS1 | n |
| Kmar_QGN14675.1 | W303.asm02.HP0_G0046700 | OG0003073 | chrXIV | 636628 | 637482 | CP015055.1 | 1005743 | 1006567 | 72.62773723 | 284 | 274 | 1.03649635 | 1 | 1 | 1 | FIM1_1340 | YNL003C | YNL003C | PET8 | n |
| Kmar_QGN14676.1 | W303.asm02.HP0_G0005390 | OG0003074 | chrIII | 99861 | 101003 | CP015055.1 | 1006780 | 1007892 | 42.1875 | 380 | 370 | 1.027027027 | 1 | 1 | 1 | FIM1_1341 | YCL016C | YCL016C | DCC1 | n |
| Kmar_QGN15720.1 | W303.asm02.HP0_G0017720 | OG0003075 | chrVII | 40815 | 43829 | CP015056.1 | 1529772 | 1532612 | 26.28968254 | 1004 | 946 | 1.061310782 | 1 | 1 | 1 | FIM1_2413 | YGL241W | YGL241W | KAP114 | n |
| Kmar_QGN15620.1 | W303.asm02.HP0_G0018390 | OG0003076 | chrVII | 182527 | 183807 | CP015056.1 | 1323868 | 1325076 | 60.56338028 | 426 | 402 | 1.059701493 | 1 | 1 | 1 | FIM1_2313 | YGL169W | YGL169W | SUA5 | n |
| Kmar_QGN16513.1 | W303.asm02.HP0_G0037640 | OG0003077 | chrXII | 711914 | 712747 | CP015058.1 | 278652 | 279488 | 45.54794521 | 277 | 278 | 0.996402878 | 1 | 1 | 1 | FIM1_3226 | YLR290C | YLR290C | COQ11 | n |
| Kmar_QGN16613.1 | W303.asm02.HP0_G0052730 | OG0003078 | chrXVI | 13783 | 14910 | CP015058.1 | 502258 | 503256 | 29.36507937 | 375 | 332 | 1.129518072 | 1 | 1 | 1 | FIM1_3329 | YOR390W/YPL279C | YPL279C | FEX2 | y |
| Kmar_QGN14867.1 | W303.asm02.HP0_G0034940 | OG0003079 | chrXII | 152813 | 155401 | CP015055.1 | 1385860 | 1388475 | 37.13318284 | 862 | 871 | 0.989667049 | 1 | 1 | 1 | FIM1_1540 | YLR001C | YLR001C | | n |
| Kmar_QGN14868.1 | W303.asm02.HP0_G0029620 | OG0003080 | chrX | 432451 | 434259 | CP015055.1 | 1389408 | 1391294 | 56.59777424 | 602 | 628 | 0.958598726 | 1 | 1 | 1 | FIM1_1541 | YJR001W | YJR001W | AVT1 | n |
| Kmar_QGN16808.1 | W303.asm02.HP0_G0020760 | OG0003081 | chrVII | 686370 | 688436 | CP015058.1 | 935686 | 937717 | 37.46397695 | 688 | 677 | 1.016248154 | 1 | 1 | 1 | FIM1_3532 | YGR099W | YGR099W | TEL2 | n |
| Kmar_QGN17543.1 | W303.asm02.HP0_G0028590 | OG0003082 | chrX | 179733 | 180260 | CP015059.1 | 1134775 | 1135260 | 50.28248588 | 175 | 161 | 1.086956522 | 1 | 1 | 1 | FIM1_4279 | YJL122W | YJL122W | ALB1 | n |
| Kmar_QGN17544.1 | W303.asm02.HP0_G0028600 | OG0003083 | chrX | 180391 | 181107 | CP015059.1 | 1135446 | 1136156 | 71.42857143 | 238 | 236 | 1.008474576 | 1 | 1 | 1 | FIM1_4280 | YJL121C | YJL121C | RPE1 | n |
| Kmar_QGN15721.1 | W303.asm02.HP0_G0017710 | OG0003084 | chrVII | 40022 | 40567 | CP015056.1 | 1533021 | 1533536 | 61.49425287 | 181 | 171 | 1.058479532 | 1 | 1 | 1 | FIM1_2414 | YGL242C | YGL242C | ANK1 | n |
| Kmar_QGN15722.1 | W303.asm02.HP0_G0017700 | OG0003085 | chrVII | 38677 | 39879 | CP015056.1 | 1533637 | 1534788 | 41.83168317 | 400 | 383 | 1.044386423 | 1 | 1 | 1 | FIM1_2415 | YGL243W | YGL243W | TAD1 | n |
| Kmar_QGN18152.1 | W303.asm02.HP0_G0026590 | OG0003086 | chrIX | 231938 | 232405 | CP015061.1 | 337638 | 338105 | 57.05128205 | 155 | 155 | 1 | 1 | 1 | 1 | FIM1_4475 | YIL065C | YIL065C | FIS1 | n |
| Kmar_QGN14799.1 | W303.asm02.HP0_G0050200 | OG0003087 | chrXV | 554051 | 555445 | CP015055.1 | 1252929 | 1254170 | 45.80645161 | 464 | 413 | 1.123486683 | 1 | 1 | 1 | FIM1_1470 | YOR123C | YOR123C | LEO1 | n |
| Kmar_QGN15298.1 | W303.asm02.HP0_G0016170 | OG0003088 | chrV | 533676 | 534353 | CP015056.1 | 631322 | 632005 | 64 | 225 | 227 | 0.991189427 | 1 | 1 | 1 | FIM1_1987 | YER170W | YER170W | ADK2 | n |
| Kmar_QGN15297.1 | W303.asm02.HP0_G0016180 | OG0003089 | chrV | 534784 | 537120 | CP015056.1 | 628664 | 631000 | 85.3470437 | 778 | 778 | 1 | 1 | 1 | 1 | FIM1_1986 | YER171W | YER171W | RAD3 | n |
| Kmar_QGN14296.1 | W303.asm02.HP0_G0005600 | OG0003090 | chrIII | 135985 | 136782 | CP015055.1 | 214408 | 215205 | 83.77358491 | 265 | 265 | 1 | 1 | 1 | 1 | FIM1_953 | YCR009C | YCR009C | RVS161 | n |
| Kmar_QGN17777.1 | W303.asm02.HP0_G0045920 | OG0003091 | chrXIV | 468759 | 471251 | CP015060.1 | 460004 | 462358 | 50.72289157 | 830 | 784 | 1.058673469 | 1 | 1 | 1 | FIM1_4986 | YNL085W | YNL085W | MKT1 | n |
| Kmar_QGN15998.1 | W303.asm02.HP0_G0011880 | OG0003092 | chrIV | 1151768 | 1155442 | CP015057.1 | 574440 | 578105 | 66.50326797 | 1224 | 1221 | 1.002457002 | 1 | 1 | 1 | FIM1_2698 | YDR335W | YDR335W | MSN5 | n |
| Kmar_QGN15999.1 | W303.asm02.HP0_G0024190 | OG0003093 | chrVIII | 287468 | 288355 | CP015057.1 | 578657 | 579538 | 78.98305085 | 295 | 293 | 1.006825939 | 1 | 1 | 1 | FIM1_2699 | YHR088W | YHR088W | RPF1 | n |
| Kmar_QGN18346.1 | W303.asm02.HP0_G0023950 | OG0003094 | chrVIII | 236801 | 237643 | CP015061.1 | 745120 | 745938 | 30.38869258 | 280 | 272 | 1.029411765 | 1 | 1 | 1 | FIM1_4673 | YHR067W | YHR067W | HTD2 | n |
| Kmar_QGN14909.1 | W303.asm02.HP0_G0011400 | OG0003095 | chrIV | 1041694 | 1042605 | CP015055.1 | 1474173 | 1475198 | 25.21489971 | 303 | 341 | 0.88856305 | 1 | 1 | 1 | FIM1_1583 | YDR288W | YDR288W | NSE3 | n |
| Kmar_QGN13719.1 | W303.asm02.HP0_G0043280 | OG0003096 | chrXIII | 852492 | 855407 | CP015054.1 | 729218 | 732127 | 56.30081301 | 971 | 969 | 1.002063983 | 1 | 1 | 1 | FIM1_363 | YMR288W | YMR288W | HSH155 | n |
| Kmar_QGN13721.1 | W303.asm02.HP0_G0025160 | OG0003097 | chrVIII | 487092 | 488021 | CP015054.1 | 734421 | 735308 | 49.51456311 | 309 | 295 | 1.047457627 | 1 | 1 | 1 | FIM1_365 | YHR187W | YHR187W | IKI1 | n |
| Kmar_QGN13720.1 | W303.asm02.HP0_G0025170 | OG0003098 | chrVIII | 488107 | 489939 | CP015054.1 | 732399 | 734300 | 51.78294574 | 610 | 633 | 0.963665087 | 1 | 1 | 1 | FIM1_364 | YHR188C | YHR188C | GPI16 | n |
| Kmar_QGN16041.1 | W303.asm02.HP0_G0021610 | OG0003099 | chrVII | 884459 | 887524 | CP015057.1 | 685775 | 688675 | 42.35517569 | 1021 | 966 | 1.056935818 | 1 | 1 | 1 | FIM1_2741 | YGR188C | YGR188C | BUB1 | n |
| Kmar_QGN14829.1 | W303.asm02.HP0_G0042000 | OG0003100 | chrXIII | 579559 | 584529 | CP015055.1 | 1307814 | 1312517 | 55.6422569 | 1656 | 1567 | 1.056796426 | 1 | 1 | 1 | FIM1_1500 | YMR162C | YMR162C | DNF3 | n |
| Kmar_QGN16407.1 | W303.asm02.HP0_G0050840 | OG0003101 | chrXV | 712978 | 713838 | CP015058.1 | 49981 | 50682 | 41.75438596 | 286 | 233 | 1.227467811 | 1 | 1 | 1 | FIM1_3120 | YOR194C | YOR194C | TOA1 | n |
| Kmar_QGN14990.1 | W303.asm02.HP0_G0054180 | OG0003102 | chrXVI | 316143 | 318833 | CP015055.1 | 1634290 | 1636857 | 49.33628319 | 896 | 855 | 1.047953216 | 1 | 1 | 1 | FIM1_1671 | YPL126W | YPL126W | NAN1 | n |
| Kmar_QGN14988.1 | W303.asm02.HP0_G0050980 | OG0003103 | chrXV | 740219 | 742927 | CP015055.1 | 1630786 | 1633419 | 66.04189636 | 902 | 877 | 1.028506271 | 1 | 1 | 1 | FIM1_1669 | YOR211C | YOR211C | MGM1 | n |
| Kmar_QGN15222.1 | W303.asm02.HP0_G0021580 | OG0003104 | chrVII | 878751 | 879935 | CP015056.1 | 467843 | 469036 | 79.24050633 | 394 | 397 | 0.992443325 | 1 | 1 | 1 | FIM1_1911 | YGR185C | YGR185C | TYS1 | n |
| Kmar_QGN15225.1 | W303.asm02.HP0_G0034790 | OG0003105 | chrXII | 122082 | 122408 | CP015056.1 | 473534 | 473851 | 37.03703704 | 108 | 105 | 1.028571429 | 1 | 1 | 1 | FIM1_1914 | YLL014W | YLL014W | EMC6 | n |
| Kmar_QGN14448.1 | W303.asm02.HP0_G0006280 | OG0003106 | chrIII | 288036 | 289814 | CP015055.1 | 532055 | 533908 | 52.22929936 | 592 | 617 | 0.959481361 | 1 | 1 | 1 | FIM1_1109 | YCR088W | YCR088W | ABP1 | n |
| Kmar_QGN14856.1 | W303.asm02.HP0_G0042090 | OG0003107 | chrXIII | 604452 | 606104 | CP015055.1 | 1356813 | 1358234 | 45.81818182 | 550 | 473 | 1.162790698 | 1 | 1 | 1 | FIM1_1529 | YMR171C | YMR171C | EAR1 | n |
| Kmar_QGN15228.1 | W303.asm02.HP0_G0035150 | OG0003108 | chrXII | 191564 | 192316 | CP015056.1 | 481196 | 481939 | 72.61904762 | 250 | 247 | 1.012145749 | 1 | 1 | 1 | FIM1_1917 | YLR022C | YLR022C | SDO1 | n |
| Kmar_QGN17756.1 | W303.asm02.HP0_G0022520 | OG0003109 | chrVII | 1075258 | 1076559 | CP015060.1 | 400132 | 401433 | 80.13856813 | 433 | 433 | 1 | 1 | 1 | 1 | FIM1_4965 | YGR285C | YGR285C | ZUO1 | n |
| Kmar_QGN17758.1 | W303.asm02.HP0_G0043510 | OG0003110 | chrXIII | 903830 | 904519 | CP015060.1 | 403798 | 404403 | 41.12554113 | 229 | 201 | 1.139303483 | 1 | 1 | 1 | FIM1_4967 | YMR311C | YMR311C | GLC8 | n |
| Kmar_QGN16015.1 | W303.asm02.HP0_G0021720 | OG0003111 | chrVII | 912319 | 914685 | CP015057.1 | 616890 | 619262 | 58.64661654 | 788 | 790 | 0.997468354 | 1 | 1 | 1 | FIM1_2715 | YGR200C | YGR200C | ELP2 | n |
| Kmar_QGN14880.1 | W303.asm02.HP0_G0029370 | OG0003112 | chrX | 379486 | 381798 | CP015055.1 | 1411904 | 1414234 | 73.91304348 | 770 | 776 | 0.992268041 | 1 | 1 | 1 | FIM1_1553 | YJL033W | YJL033W | HCA4 | n |
| Kmar_QGN18220.1 | W303.asm02.HP0_G0007350 | OG0003113 | chrIV | 183549 | 184817 | CP015061.1 | 480628 | 481869 | 44.74885845 | 422 | 413 | 1.021791768 | 1 | 1 | 1 | FIM1_4545 | YDL150W | YDL150W | RPC53 | n |
| Kmar_QGN18221.1 | W303.asm02.HP0_G0007340 | OG0003114 | chrIV | 181398 | 183224 | CP015061.1 | 482280 | 484109 | 58.6038961 | 608 | 609 | 0.998357964 | 1 | 1 | 1 | FIM1_4546 | YDL153C | YDL153C | SAS10 | n |
| Kmar_QGN17041.1 | W303.asm02.HP0_G0007420 | OG0003115 | chrIV | 200202 | 201788 | CP015059.1 | 102753 | 104345 | 84.33962264 | 528 | 530 | 0.996226415 | 1 | 1 | 1 | FIM1_3770 | YDL143W | YDL143W | CCT4 | n |
| Kmar_QGN17040.1 | W303.asm02.HP0_G0036990 | OG0003116 | chrXII | 570900 | 571982 | CP015059.1 | 100733 | 101839 | 51.89189189 | 360 | 368 | 0.97826087 | 1 | 1 | 1 | FIM1_3769 | YLR215C | YLR215C | CDC123 | n |
| Kmar_QGN17010.1 | W303.asm02.HP0_G0002460 | OG0003117 | chrII | 355961 | 357037 | CP015059.1 | 23603 | 24613 | 39.89071038 | 358 | 336 | 1.06547619 | 1 | 1 | 1 | FIM1_3739 | YBR053C | YBR053C | | n |
| Kmar_QGN15559.1 | W303.asm02.HP0_G0030090 | OG0003118 | chrX | 547427 | 547921 | CP015056.1 | 1183044 | 1183508 | 43.03030303 | 164 | 154 | 1.064935065 | 1 | 1 | 1 | FIM1_2250 | YJR055W | YJR055W | HIT1 | n |
| Kmar_QGN15295.1 | W303.asm02.HP0_G0016200 | OG0003119 | chrV | 544002 | 545981 | CP015056.1 | 619822 | 621816 | 38.94582723 | 659 | 664 | 0.99246988 | 1 | 1 | 1 | FIM1_1984 | YER173W | YER173W | RAD24 | n |
| Kmar_QGN16427.1 | W303.asm02.HP0_G0054240 | OG0003120 | chrXVI | 328429 | 330102 | CP015058.1 | 95591 | 96991 | 36.47912886 | 557 | 466 | 1.19527897 | 1 | 1 | 1 | FIM1_3140 | YPL120W | YPL120W | VPS30 | n |
| Kmar_QGN15930.1 | W303.asm02.HP0_G0035940 | OG0003121 | chrXII | 355002 | 356135 | CP015057.1 | 398120 | 399175 | 48.54881266 | 377 | 351 | 1.074074074 | 1 | 1 | 1 | FIM1_2628 | YLR105C | YLR105C | SEN2 | n |
| Kmar_QGN17363.1 | W303.asm02.HP0_G0016070 | OG0003122 | chrV | 499232 | 499660 | CP015059.1 | 777216 | 777641 | 50.69444444 | 142 | 141 | 1.007092199 | 1 | 1 | 1 | FIM1_4095 | YER159C | YER159C | BUR6 | n |
| Kmar_QGN13601.1 | W303.asm02.HP0_G0055570 | OG0003123 | chrXVI | 600974 | 602794 | CP015054.1 | 497707 | 499521 | 40.09661836 | 606 | 604 | 1.003311258 | 1 | 1 | 1 | FIM1_243 | YPR018W | YPR018W | RLF2 | n |
| Kmar_QGN16862.1 | W303.asm02.HP0_G0036200 | OG0003124 | chrXII | 421234 | 423480 | CP015058.1 | 1054195 | 1056408 | 20 | 748 | 737 | 1.014925373 | 1 | 1 | 1 | FIM1_3588 | YLR135W | YLR135W | SLX4 | n |
| Kmar_QGN15218.1 | W303.asm02.HP0_G0021550 | OG0003125 | chrVII | 870702 | 871019 | CP015056.1 | 453354 | 453659 | 65.71428571 | 105 | 101 | 1.03960396 | 1 | 1 | 1 | FIM1_1907 | YGR181W | YGR181W | TIM13 | n |
| Kmar_QGN18077.1 | W303.asm02.HP0_G0001480 | OG0003126 | chrII | 142886 | 143611 | CP015061.1 | 161157 | 161879 | 86.30705394 | 241 | 240 | 1.004166667 | 1 | 1 | 1 | FIM1_4396 | YBL041W | YBL041W | PRE7 | n |
| Kmar_QGN15445.1 | W303.asm02.HP0_G0011530 | OG0003127 | chrIV | 1067851 | 1071924 | CP015056.1 | 973842 | 977756 | 48.4915379 | 1357 | 1304 | 1.040644172 | 1 | 1 | 1 | FIM1_2136 | YDR301W | YDR301W | CFT1 | n |
| Kmar_QGN15446.1 | W303.asm02.HP0_G0011540 | OG0003128 | chrIV | 1072230 | 1072889 | CP015056.1 | 978058 | 978711 | 56.56108597 | 219 | 217 | 1.00921659 | 1 | 1 | 1 | FIM1_2137 | YDR302W | YDR302W | GPI11 | n |
| Kmar_QGN16410.1 | W303.asm02.HP0_G0054300 | OG0003129 | chrXVI | 338458 | 341844 | CP015058.1 | 55700 | 58930 | 37.42384682 | 1128 | 1076 | 1.048327138 | 1 | 1 | 1 | FIM1_3123 | YPL115C | YPL115C | BEM3 | n |
| Kmar_QGN16413.1 | W303.asm02.HP0_G0054290 | OG0003130 | chrXVI | 335964 | 338057 | CP015058.1 | 66824 | 68989 | 58.85486019 | 697 | 721 | 0.966712899 | 1 | 1 | 1 | FIM1_3126 | YPL116W | YPL116W | HOS3 | n |
| Kmar_QGN16411.1 | W303.asm02.HP0_G0050860 | OG0003131 | chrXV | 716888 | 718132 | CP015058.1 | 59246 | 60364 | 69.56521739 | 414 | 372 | 1.112903226 | 1 | 1 | 1 | FIM1_3124 | YOR196C | YOR196C | LIP5 | n |
| Kmar_QGN16416.1 | W303.asm02.HP0_G0054280 | OG0003132 | chrXVI | 334222 | 335088 | CP015058.1 | 73678 | 74514 | 67.01388889 | 288 | 278 | 1.035971223 | 1 | 1 | 1 | FIM1_3129 | YPL117C | YPL117C | IDI1 | n |
| Kmar_QGN14629.1 | W303.asm02.HP0_G0034600 | OG0003133 | chrXII | 70642 | 73155 | CP015055.1 | 896902 | 899346 | 70.90261283 | 837 | 814 | 1.028255528 | 1 | 1 | 1 | FIM1_1293 | YLL034C | YLL034C | RIX7 | n |
| Kmar_QGN18246.1 | W303.asm02.HP0_G0056850 | OG0003134 | chrXVI | 872547 | 874136 | CP015061.1 | 539293 | 540927 | 38.29787234 | 529 | 544 | 0.972426471 | 1 | 1 | 1 | FIM1_4571 | YPR162C | YPR162C | ORC4 | n |
| Kmar_QGN17111.1 | W303.asm02.HP0_G0039130 | OG0003135 | chrXII | 1035937 | 1038033 | CP015059.1 | 257367 | 259472 | 40.23746702 | 698 | 701 | 0.995720399 | 1 | 1 | 1 | FIM1_3842 | YLR452C | YLR452C | SST2 | n |
| Kmar_QGN17112.1 | W303.asm02.HP0_G0014730 | OG0003136 | chrV | 206390 | 207973 | CP015059.1 | 259617 | 261191 | 88.23529412 | 527 | 524 | 1.005725191 | 1 | 1 | 1 | FIM1_3843 | YER025W | YER025W | GCD11 | n |
| Kmar_QGN15948.1 | W303.asm02.HP0_G0050580 | OG0003137 | chrXV | 644269 | 645207 | CP015057.1 | 449410 | 450318 | 59.29487179 | 312 | 302 | 1.033112583 | 1 | 1 | 1 | FIM1_2646 | YOR164C | YOR164C | GET4 | n |
| Kmar_QGN15947.1 | W303.asm02.HP0_G0050570 | OG0003138 | chrXV | 643616 | 644182 | CP015057.1 | 448822 | 449358 | 59.67741935 | 188 | 178 | 1.056179775 | 1 | 1 | 1 | FIM1_2645 | YOR163W | YOR163W | DDP1 | n |
| Kmar_QGN17155.1 | W303.asm02.HP0_G0017420 | OG0003139 | chrVI | 217971 | 218771 | CP015059.1 | 354883 | 355680 | 71.05263158 | 266 | 265 | 1.003773585 | 1 | 1 | 1 | FIM1_3886 | YFR050C | YFR050C | PRE4 | n |
| Kmar_QGN14099.1 | W303.asm02.HP0_G0020290 | OG0003140 | chrVII | 582654 | 583811 | CP015054.1 | 1517158 | 1518342 | 60.40609137 | 385 | 394 | 0.97715736 | 1 | 1 | 1 | FIM1_750 | YGR046W | YGR046W | TAM41 | n |
| Kmar_QGN17156.1 | W303.asm02.HP0_G0017650 | OG0003141 | chrVII | 31023 | 32132 | CP015059.1 | 356178 | 357296 | 36.48293963 | 369 | 372 | 0.991935484 | 1 | 1 | 1 | FIM1_3887 | YGL248W | YGL248W | PDE1 | n |
| Kmar_QGN17150.1 | W303.asm02.HP0_G0017430 | OG0003142 | chrVI | 219068 | 220708 | CP015059.1 | 343280 | 344908 | 52.42369838 | 546 | 542 | 1.007380074 | 1 | 1 | 1 | FIM1_3881 | YFR051C | YFR051C | RET2 | n |
| Kmar_QGN15073.1 | W303.asm02.HP0_G0032360 | OG0003143 | chrXI | 280354 | 281685 | CP015056.1 | 145123 | 146385 | 23.00884956 | 443 | 420 | 1.054761905 | 1 | 1 | 1 | FIM1_1758 | YKL090W | YKL090W | CUE2 | n |

| | | | | | | | | | | | | | | | | | | | | |
|---|---|---|---|---|---|---|---|---|---|---|---|---|---|---|---|---|---|---|---|---|
| Kmar_QGN15843.1 | W303.asm02.HP0_G0036050 | OG0003144 | chrXII | 392678 | 393361 | CP015057.1 | 201358 | 202044 | 49.56521739 | 227 | 228 | 0.995614035 | 1 | 1 | 1 | FIM1_2539 | YLR118C | YLR118C | TML25 | n |
| Kmar_QGN13716.1 | W303.asm02.HP0_G0022310 | OG0003145 | chrVII | 1029889 | 1030674 | CP015054.1 | 723527 | 724321 | 70.07575758 | 261 | 264 | 0.988636364 | 1 | 1 | 1 | FIM1_360 | YGR262C | YGR262C | BUD32 | n |
| Kmar_QGN18093.1 | W303.asm02.HP0_G0030430 | OG0003146 | chrX | 612993 | 613976 | CP015061.1 | 199630 | 200481 | 37.6899696 | 327 | 283 | 1.155477032 | 1 | 1 | 1 | FIM1_4413 | YJR093C | YJR093C | FIP1 | n |
| Kmar_QGN18098.1 | W303.asm02.HP0_G0055790 | OG0003147 | chrXVI | 655202 | 656419 | CP015061.1 | 216671 | 217876 | 76.34146341 | 405 | 401 | 1.009975062 | 1 | 1 | 1 | FIM1_4418 | YPR041W | YPR041W | TIF5 | n |
| Kmar_QGN14828.1 | W303.asm02.HP0_G0007820 | OG0003148 | chrIV | 275477 | 276910 | CP015055.1 | 1305486 | 1306898 | 64.57023061 | 477 | 470 | 1.014893617 | 1 | 1 | 1 | FIM1_1499 | YDL103C | YDL103C | QRI1 | n |
| Kmar_QGN14321.1 | W303.asm02.HP0_G0037050 | OG0003149 | chrXII | 579445 | 581898 | CP015055.1 | 273617 | 276094 | 62.23021583 | 817 | 825 | 0.99030303 | 1 | 1 | 1 | FIM1_978 | YLR222C | YLR222C | UTP13 | n |
| Kmar_QGN16882.1 | W303.asm02.HP0_G0036260 | OG0003150 | chrXII | 433232 | 434662 | CP015058.1 | 1097418 | 1098818 | 58.15899582 | 476 | 466 | 1.021459227 | 1 | 1 | 1 | FIM1_3609 | YLR142W | YLR142W | PUT1 | n |
| Kmar_QGN14313.1 | W303.asm02.HP0_G0010760 | OG0003151 | chrIV | 904052 | 907981 | CP015055.1 | 254940 | 257831 | 23.74323279 | 1309 | 963 | 1.359293873 | 1 | 1 | 1 | FIM1_970 | YDR217C | YDR217C | RAD9 | n |
| Kmar_QGN14316.1 | W303.asm02.HP0_G0037090 | OG0003152 | chrXII | 590042 | 591229 | CP015055.1 | 261468 | 262673 | 48.51485149 | 395 | 401 | 0.985037406 | 1 | 1 | 1 | FIM1_973 | YLR226W | YLR226W | BUR2 | n |
| Kmar_QGN13511.1 | W303.asm02.HP0_G0011240 | OG0003153 | chrIV | 1007011 | 1008003 | CP015054.1 | 300311 | 301315 | 60.11904762 | 330 | 334 | 0.988023952 | 1 | 1 | 1 | FIM1_150 | YDR267C | YDR267C | CIA1 | n |
| Kmar_QGN13510.1 | W303.asm02.HP0_G0049450 | OG0003154 | chrXV | 401897 | 403111 | CP015054.1 | 298959 | 300038 | 43.10344828 | 404 | 359 | 1.125348189 | 1 | 1 | 1 | FIM1_149 | YOR037W | YOR037W | CYC2 | n |
| Kmar_QGN13508.1 | W303.asm02.HP0_G0049460 | OG0003155 | chrXV | 403217 | 405844 | CP015054.1 | 294978 | 297527 | 44.05436014 | 875 | 849 | 1.030624264 | 1 | 1 | 1 | FIM1_147 | YOR038C | YOR038C | HIR2 | n |
| Kmar_QGN13507.1 | W303.asm02.HP0_G0049470 | OG0003156 | chrXV | 406224 | 407000 | CP015054.1 | 293721 | 294509 | 70.22900763 | 258 | 262 | 0.984732824 | 1 | 1 | 1 | FIM1_146 | YOR039W | YOR039W | CKB2 | n |
| Kmar_QGN13509.1 | W303.asm02.HP0_G0011250 | OG0003157 | chrIV | 1008505 | 1009644 | CP015054.1 | 297582 | 298721 | 59.11458333 | 379 | 379 | 1 | 1 | 1 | 1 | FIM1_148 | YDR268W | YDR268W | MSW1 | n |
| Kmar_QGN18208.1 | W303.asm02.HP0_G0042180 | OG0003158 | chrXIII | 620442 | 622718 | CP015061.1 | 450279 | 452210 | 29.12873862 | 758 | 643 | 1.178849145 | 1 | 1 | 1 | FIM1_4532 | YMR179W | YMR179W | SPT21 | n |
| Kmar_QGN17484.1 | W303.asm02.HP0_G0029850 | OG0003159 | chrX | 466989 | 467723 | CP015059.1 | 1010464 | 1011087 | 43.85964912 | 244 | 207 | 1.178743961 | 1 | 1 | 1 | FIM1_4220 | YJR024C | YJR024C | MDE1 | n |
| Kmar_QGN15989.1 | W303.asm02.HP0_G0033600 | OG0003160 | chrXI | 517628 | 518242 | CP015057.1 | 558938 | 559552 | 76.09756098 | 204 | 204 | 1 | 1 | 1 | 1 | FIM1_2689 | YKR035W-A | YKR035W-A | DID2 | n |
| Kmar_QGN13537.1 | W303.asm02.HP0_G0032580 | OG0003161 | chrXI | 323288 | 323749 | CP015054.1 | 358966 | 359424 | 73.85620915 | 153 | 152 | 1.006578947 | 1 | 1 | 1 | FIM1_177 | YKL067W | YKL067W | YNK1 | n |
| Kmar_QGN14507.1 | W303.asm02.HP0_G0035460 | OG0003162 | chrXII | 259085 | 260893 | CP015055.1 | 653736 | 655652 | 48.2388974 | 602 | 638 | 0.943573668 | 1 | 1 | 1 | FIM1_1168 | YLR055C | YLR055C | SPT8 | n |
| Kmar_QGN14508.1 | W303.asm02.HP0_G0016630 | OG0003163 | chrVI | 48867 | 50360 | CP015055.1 | 655828 | 657033 | 30.32128514 | 497 | 401 | 1.239401496 | 1 | 1 | 1 | FIM1_1169 | YFL027C | YFL027C | GYP8 | n |
| Kmar_QGN13517.1 | W303.asm02.HP0_G0024340 | OG0003164 | chrVIII | 329382 | 330365 | CP015054.1 | 316117 | 317106 | 67.27272727 | 327 | 329 | 0.993920973 | 1 | 1 | 1 | FIM1_157 | YHR104W | YHR104W | GRE3 | n |
| Kmar_QGN14517.1 | W303.asm02.HP0_G0051890 | OG0003165 | chrXV | 896344 | 898107 | CP015055.1 | 680457 | 682100 | 23.4006734 | 587 | 547 | 1.073126143 | 1 | 1 | 1 | FIM1_1179 | YOR308C | YOR308C | SNU66 | n |
| Kmar_QGN14570.1 | W303.asm02.HP0_G0011260 | OG0003166 | chrIV | 1010176 | 1014335 | CP015055.1 | 781145 | 784072 | 50 | 1313 | 975 | 1.346666667 | 1 | 1 | 1 | FIM1_1232 | YDR270W | YDR270W | CCC2 | n |
| Kmar_QGN14685.1 | W303.asm02.HP0_G0039190 | OG0003167 | chrXII | 1054001 | 1055185 | CP015055.1 | 1030147 | 1031325 | 58.08080808 | 394 | 392 | 1.005102041 | 1 | 1 | 1 | FIM1_1352 | YLR459W | YLR459W | GAB1 | n |
| Kmar_QGN13422.1 | W303.asm02.HP0_G0054590 | OG0003168 | chrXVI | 393425 | 400012 | CP015054.1 | 109490 | 116374 | 26.42398287 | 2195 | 2294 | 0.956843941 | 1 | 1 | 1 | FIM1_59 | YPL085W | YPL085W | SEC16 | n |
| Kmar_QGN15573.1 | W303.asm02.HP0_G0032120 | OG0003169 | chrXI | 229203 | 230759 | CP015056.1 | 1214897 | 1216558 | 41.60714286 | 518 | 553 | 0.936708861 | 1 | 1 | 1 | FIM1_2264 | YKL116C | YKL116C | PRR1 | n |
| Kmar_QGN18032.1 | W303.asm02.HP0_G0035610 | OG0003170 | chrXII | 283329 | 286577 | CP015061.1 | 63902 | 66850 | 42.39423942 | 1082 | 982 | 1.101832994 | 1 | 1 | 1 | FIM1_4348 | YLR071C | YLR071C | RGR1 | n |
| Kmar_QGN18382.1 | W303.asm02.HP0_G0048400 | OG0003171 | chrXV | 197675 | 199180 | CP015061.1 | 812065 | 813810 | 34.80475382 | 501 | 581 | 0.862306368 | 1 | 1 | 1 | FIM1_4709 | YOL070C | YOL070C | NBA1 | n |
| Kmar_QGN14504.1 | W303.asm02.HP0_G0016610 | OG0003172 | chrVI | 46503 | 47609 | CP015055.1 | 649075 | 650028 | 28.5326087 | 368 | 317 | 1.160883281 | 1 | 1 | 1 | FIM1_1165 | YFL029C | YFL029C | CAK1 | n |
| Kmar_QGN17168.1 | W303.asm02.HP0_G0054980 | OG0003173 | chrXVI | 479400 | 481067 | CP015059.1 | 374254 | 375996 | 58.41924399 | 555 | 580 | 0.956896552 | 1 | 1 | 1 | FIM1_3899 | YPL042C | YPL042C | SSN3 | n |
| Kmar_QGN16288.1 | W303.asm02.HP0_G0038800 | OG0003174 | chrXII | 951679 | 953379 | CP015057.1 | 1212360 | 1213943 | 44.03508772 | 566 | 527 | 1.074003795 | 1 | 1 | 1 | FIM1_2993 | YLR417W | YLR417W | VPS36 | n |
| Kmar_QGN16289.1 | W303.asm02.HP0_G0031440 | OG0003175 | chrXI | 85873 | 87072 | CP015057.1 | 1214047 | 1215141 | 56.53333333 | 399 | 364 | 1.096153846 | 1 | 1 | 1 | FIM1_2994 | YKL189W | YKL189W | HYM1 | n |
| Kmar_QGN14986.1 | W303.asm02.HP0_G0050990 | OG0003176 | chrXV | 744205 | 745476 | CP015055.1 | 1624011 | 1625291 | 50.46082949 | 423 | 426 | 0.992957746 | 1 | 1 | 1 | FIM1_1666 | YOR212W | YOR212W | STE4 | n |
| Kmar_QGN14987.1 | W303.asm02.HP0_G0054160 | OG0003177 | chrXVI | 312466 | 314154 | CP015055.1 | 1626044 | 1627549 | 30.01745201 | 562 | 501 | 1.121756487 | 1 | 1 | 1 | FIM1_1667 | YPL128C | YPL128C | TBF1 | n |
| Kmar_QGN14653.1 | W303.asm02.HP0_G0020070 | OG0003178 | chrVII | 525166 | 526038 | CP015055.1 | 953773 | 954642 | 49.35064935 | 290 | 289 | 1.003460208 | 1 | 1 | 1 | FIM1_1317 | YGR021W | YGR021W | DPC29 | n |
| Kmar_QGN14654.1 | W303.asm02.HP0_G0020060 | OG0003179 | chrVII | 524507 | 524863 | CP015055.1 | 955161 | 955517 | 77.96610169 | 118 | 118 | 1 | 1 | 1 | 1 | FIM1_1318 | YGR020C | YGR020C | VMA7 | n |
| Kmar_QGN14655.1 | W303.asm02.HP0_G0038010 | OG0003180 | chrXII | 783350 | 785365 | CP015055.1 | 955696 | 957456 | 39.27536232 | 671 | 586 | 1.145051195 | 1 | 1 | 1 | FIM1_1319 | YLR330W | YLR330W | CHS5 | n |
| Kmar_QGN14553.1 | W303.asm02.HP0_G0000580 | OG0003181 | chrI | 130036 | 131253 | CP015055.1 | 744790 | 745590 | 40.39408867 | 405 | 266 | 1.522556391 | 1 | 1 | 1 | FIM1_1215 | YAL013W | YAL013W | DEP1 | n |
| Kmar_QGN13569.1 | W303.asm02.HP0_G0031410 | OG0003182 | chrXI | 81329 | 81646 | CP015054.1 | 437564 | 437929 | 59.67741935 | 105 | 121 | 0.867768595 | 1 | 1 | 1 | FIM1_210 | YKL192C | YKL192C | ACP1 | n |
| Kmar_QGN13570.1 | W303.asm02.HP0_G0031400 | OG0003183 | chrXI | 80035 | 81051 | CP015054.1 | 438288 | 439376 | 64.36464088 | 338 | 362 | 0.933701657 | 1 | 1 | 1 | FIM1_211 | YKL193C | YKL193C | SDS22 | n |
| Kmar_QGN16698.1 | W303.asm02.HP0_G0042320 | OG0003184 | chrXIII | 650619 | 651395 | CP015058.1 | 687793 | 688560 | 60.46511628 | 258 | 255 | 1.011764706 | 1 | 1 | 1 | FIM1_3418 | YMR193W | YMR193W | MRPL24 | n |
| Kmar_QGN16232.1 | W303.asm02.HP0_G0025350 | OG0003185 | chrVIII | 515466 | 517940 | CP015057.1 | 1088376 | 1090652 | 61.23188406 | 824 | 758 | 1.08707124 | 1 | 1 | 1 | FIM1_2936 | YHR205W | YHR205W | SCH9 | n |
| Kmar_QGN15308.1 | W303.asm02.HP0_G0034630 | OG0003186 | chrXII | 77161 | 80214 | CP015056.1 | 663662 | 666676 | 59.51456311 | 1017 | 1004 | 1.012948207 | 1 | 1 | 1 | FIM1_1997 | YLL031C | YLL031C | GPI13 | n |
| Kmar_QGN16233.1 | W303.asm02.HP0_G0025340 | OG0003187 | chrVIII | 512421 | 514811 | CP015057.1 | 1091406 | 1093748 | 40.24691358 | 796 | 780 | 1.020512821 | 1 | 1 | 1 | FIM1_2937 | YHR204W | YHR204W | MNL1 | n |
| Kmar_QGN14953.1 | W303.asm02.HP0_G0016780 | OG0003188 | chrVI | 76335 | 78413 | CP015055.1 | 1560084 | 1562168 | 31.43631436 | 692 | 694 | 0.997118156 | 1 | 1 | 1 | FIM1_1630 | YFL013C | YFL013C | IES1 | n |
| Kmar_QGN14949.1 | W303.asm02.HP0_G0016740 | OG0003189 | chrVI | 72466 | 72945 | CP015055.1 | 1554471 | 1554938 | 49.68944099 | 159 | 155 | 1.025806452 | 1 | 1 | 1 | FIM1_1625 | YFL017C | YFL017C | GNA1 | n |
| Kmar_QGN13844.1 | W303.asm02.HP0_G0031550 | OG0003190 | chrXI | 111634 | 113673 | CP015054.1 | 987401 | 989431 | 45.88068182 | 679 | 676 | 1.00443787 | 1 | 1 | 1 | FIM1_490 | YKL179C | YKL179C | COY1 | n |
| Kmar_QGN15095.1 | W303.asm02.HP0_G0038270 | OG0003191 | chrXII | 843568 | 845304 | CP015056.1 | 193607 | 195352 | 46.8907563 | 578 | 581 | 0.994836489 | 1 | 1 | 1 | FIM1_1782 | YLR361C | YLR361C | DCR2 | n |
| Kmar_QGN15097.1 | W303.asm02.HP0_G0038290 | OG0003192 | chrXII | 846016 | 848199 | CP015056.1 | 196238 | 198445 | 56.25806452 | 727 | 735 | 0.989115646 | 1 | 1 | 1 | FIM1_1784 | YLR362W | YLR362W | STE11 | n |
| Kmar_QGN13829.1 | W303.asm02.HP0_G0031620 | OG0003193 | chrXI | 128645 | 131431 | CP015054.1 | 955641 | 958061 | 26.63157895 | 928 | 806 | 1.151364764 | 1 | 1 | 1 | FIM1_475 | YKL171W | YKL171W | NNK1 | n |
| Kmar_QGN15068.1 | W303.asm02.HP0_G0030160 | OG0003194 | chrX | 562131 | 563504 | CP015056.1 | 134058 | 135284 | 42.76457883 | 457 | 408 | 1.120098039 | 1 | 1 | 1 | FIM1_1753 | YJR062C | YJR062C | NTA1 | n |
| Kmar_QGN16726.1 | W303.asm02.HP0_G0024060 | OG0003195 | chrVIII | 258348 | 261623 | CP015058.1 | 749176 | 752385 | 44.78976234 | 1091 | 1069 | 1.020579981 | 1 | 1 | 1 | FIM1_3447 | YHR077C | YHR077C | NMD2 | n |
| Kmar_QGN15674.1 | W303.asm02.HP0_G0012660 | OG0003196 | chrIV | 1308899 | 1310797 | CP015056.1 | 1434009 | 1435910 | 47.53846154 | 632 | 633 | 0.998420221 | 1 | 1 | 1 | FIM1_2367 | YDR419W | YDR419W | RAD30 | n |
| Kmar_QGN16055.1 | W303.asm02.HP0_G0018460 | OG0003197 | chrVII | 197029 | 198741 | CP015057.1 | 715839 | 717377 | 35.7267951 | 570 | 512 | 1.11328125 | 1 | 1 | 1 | FIM1_2755 | YGL160W | YGL160W | AIM14 | n |
| Kmar_QGN13836.1 | W303.asm02.HP0_G0031570 | OG0003198 | chrXI | 116670 | 119156 | CP015054.1 | 969754 | 971940 | 31.13983549 | 828 | 728 | 1.137362637 | 1 | 1 | 1 | FIM1_482 | YKL176C | YKL176C | LST4 | n |
| Kmar_QGN14157.1 | W303.asm02.HP0_G0005160 | OG0003199 | chrIII | 46763 | 48016 | CP015054.1 | 1651258 | 1652418 | 37.88235294 | 417 | 386 | 1.080310881 | 1 | 1 | 1 | FIM1_810 | YCL044C | YCL044C | MGR1 | n |
| Kmar_QGN13823.1 | W303.asm02.HP0_G0024210 | OG0003200 | chrVIII | 289750 | 290598 | CP015054.1 | 941499 | 942800 | 30.78703704 | 282 | 433 | 0.651270208 | 1 | 1 | 1 | FIM1_469 | YHR090C | YHR090C | YNG2 | n |
| Kmar_QGN17472.1 | W303.asm02.HP0_G0049430 | OG0003201 | chrXV | 398191 | 400560 | CP015059.1 | 993752 | 995773 | 33.04130163 | 789 | 673 | 1.172362556 | 1 | 1 | 1 | FIM1_4208 | YOR035C | YOR035C | SHE4 | n |
| Kmar_QGN16020.1 | W303.asm02.HP0_G0024280 | OG0003202 | chrVIII | 305118 | 307907 | CP015057.1 | 638962 | 641679 | 52.94748124 | 929 | 905 | 1.026519337 | 1 | 1 | 1 | FIM1_2720 | YHR098C | YHR098C | SFB3 | n |
| Kmar_QGN17466.1 | W303.asm02.HP0_G0049360 | OG0003203 | chrXV | 387281 | 389140 | CP015059.1 | 978678 | 980117 | 22.75641026 | 619 | 479 | 1.292275574 | 1 | 1 | 1 | FIM1_4201 | YOR030W | YOR030W | DFG16 | n |
| Kmar_QGN17463.1 | W303.asm02.HP0_G0011150 | OG0003204 | chrIV | 976309 | 978744 | CP015059.1 | 971767 | 974277 | 73.57142857 | 811 | 836 | 0.970095694 | 1 | 1 | 1 | FIM1_4197 | YDR258C | YDR258C | HSP78 | n |
| Kmar_QGN18004.1 | W303.asm02.HP0_G0040170 | OG0003205 | chrXIII | 206935 | 209367 | CP015061.1 | 15876 | 18293 | 65.02463054 | 810 | 805 | 1.00621118 | 1 | 1 | 1 | FIM1_4320 | YML035C | YML035C | AMD1 | n |
| Kmar_QGN15938.1 | W303.asm02.HP0_G0036000 | OG0003206 | chrXII | 379574 | 380881 | CP015057.1 | 428230 | 429627 | 77.77777778 | 435 | 465 | 0.935483871 | 1 | 1 | 1 | FIM1_2636 | YLR113W | YLR113W | HOG1 | n |
| Kmar_QGN14485.1 | W303.asm02.HP0_G0041200 | OG0003207 | chrXIII | 421989 | 422654 | CP015055.1 | 614626 | 615294 | 42.22222222 | 221 | 222 | 0.995495495 | 1 | 1 | 1 | FIM1_1146 | YMR077C | YMR077C | VPS20 | n |
| Kmar_QGN14486.1 | W303.asm02.HP0_G0019520 | OG0003208 | chrVII | 416613 | 417641 | CP015055.1 | 615610 | 616641 | 79.30029155 | 342 | 343 | 0.997084548 | 1 | 1 | 1 | FIM1_1147 | YGL040C | YGL040C | HEM2 | n |
| Kmar_QGN14294.1 | W303.asm02.HP0_G0031800 | OG0003209 | chrXI | 164921 | 165664 | CP015055.1 | 210292 | 211035 | 86.63967611 | 247 | 247 | 1 | 1 | 1 | 1 | FIM1_951 | YKL152C | YKL152C | GPM1 | n |
| Kmar_QGN17663.1 | W303.asm02.HP0_G0013130 | OG0003210 | chrIV | 1410138 | 1411547 | CP015060.1 | 209783 | 211075 | 38.15513627 | 469 | 430 | 1.090697674 | 1 | 1 | 1 | FIM1_4870 | YDR473C | YDR473C | PRP3 | n |
| Kmar_QGN17301.1 | W303.asm02.HP0_G0024520 | OG0003211 | chrVIII | 359598 | 360293 | CP015059.1 | 651764 | 652447 | 64.93506494 | 231 | 227 | 1.017621145 | 1 | 1 | 1 | FIM1_4033 | YHR122W | YHR122W | CIA2 | n |
| Kmar_QGN17959.1 | W303.asm02.HP0_G0001100 | OG0003212 | chrII | 62355 | 63755 | CP015060.1 | 829864 | 831258 | 38.35341365 | 466 | 464 | 1.004310345 | 1 | 1 | 1 | FIM1_5168 | YBL086C | YBL086C | | n |
| Kmar_QGN15422.1 | W303.asm02.HP0_G0011360 | OG0003213 | chrIV | 1035049 | 1035918 | CP015056.1 | 923981 | 924868 | 56.80272109 | 289 | 295 | 0.979661017 | 1 | 1 | 1 | FIM1_2112 | YDR284C | YDR284C | DPP1 | n |
| Kmar_QGN16597.1 | W303.asm02.HP0_G0031360 | OG0003214 | chrXI | 71903 | 75034 | CP015058.1 | 469338 | 472505 | 37.1559633 | 1043 | 1055 | 0.988625592 | 1 | 1 | 1 | FIM1_3313 | YKL197C | YKL197C | PEX1 | n |
| Kmar_QGN15538.1 | W303.asm02.HP0_G0010880 | OG0003215 | chrIV | 930794 | 931411 | CP015056.1 | 1147594 | 1148178 | 48.58490566 | 205 | 194 | 1.056701031 | 1 | 1 | 1 | FIM1_2229 | YDR231C | YDR231C | COX20 | n |
| Kmar_QGN15054.1 | W303.asm02.HP0_G0030260 | OG0003216 | chrX | 580254 | 581411 | CP015056.1 | 101758 | 102939 | 75.82697201 | 385 | 393 | 0.979643766 | 1 | 1 | 1 | FIM1_1739 | YJR072C | YJR072C | NPA3 | n |
| Kmar_QGN14387.1 | W303.asm02.HP0_G0036010 | OG0003217 | chrXII | 382897 | 385191 | CP015055.1 | 417513 | 419225 | 39.91825613 | 764 | 570 | 1.340350877 | 1 | 1 | 1 | FIM1_1047 | YLR114C | YLR114C | AVL9 | n |
| Kmar_QGN16889.1 | W303.asm02.HP0_G0010240 | OG0003218 | chrIV | 788585 | 789919 | CP015058.1 | 1110465 | 1111820 | 47.50542299 | 444 | 451 | 0.984478936 | 1 | 1 | 1 | FIM1_3616 | YDR165W | YDR165W | TRM82 | n |
| Kmar_QGN16885.1 | W303.asm02.HP0_G0010230 | OG0003219 | chrIV | 785755 | 787929 | CP015058.1 | 1102814 | 1104910 | 52.99026426 | 724 | 698 | 1.037249284 | 1 | 1 | 1 | FIM1_3612 | YDR164C | YDR164C | SEC1 | n |
| Kmar_QGN14450.1 | W303.asm02.HP0_G0040140 | OG0003220 | chrXIII | 203282 | 204610 | CP015055.1 | 534997 | 536292 | 40.34334764 | 442 | 431 | 1.025522042 | 1 | 1 | 1 | FIM1_1111 | YML038C | YML038C | YMD8 | n |
| Kmar_QGN16019.1 | W303.asm02.HP0_G0012000 | OG0003221 | chrIV | 1183902 | 1185692 | CP015057.1 | 636683 | 638371 | 35.6187291 | 596 | 562 | 1.060498221 | 1 | 1 | 1 | FIM1_2719 | YDR349C | YDR349C | YPS7 | n |
| Kmar_QGN17325.1 | W303.asm02.HP0_G0050130 | OG0003222 | chrXV | 539534 | 540340 | CP015059.1 | 699319 | 700131 | 43.33333333 | 268 | 270 | 0.992592593 | 1 | 1 | 1 | FIM1_4057 | YOR115C | YOR115C | TRS33 | n |
| Kmar_QGN17670.1 | W303.asm02.HP0_G0007700 | OG0003223 | chrIV | 254201 | 255332 | CP015060.1 | 222804 | 223928 | 36.59793814 | 353 | 374 | 0.943850267 | 1 | 1 | 1 | FIM1_4877 | YDL115C | YDL115C | IWR1 | n |
| Kmar_QGN17380.1 | W303.asm02.HP0_G0041060 | OG0003224 | chrXIII | 395557 | 396882 | CP015059.1 | 804973 | 806307 | 76.12612613 | 441 | 444 | 0.993243243 | 1 | 1 | 1 | FIM1_4112 | YMR062C | YMR062C | ARG7 | n |
| Kmar_QGN16619.1 | W303.asm02.HP0_G0031650 | OG0003225 | chrXI | 134891 | 135304 | CP015058.1 | 512416 | 512808 | 53.07692308 | 137 | 130 | 1.053846154 | 1 | 1 | 1 | FIM1_3336 | YKL167C | YKL167C | MRP49 | n |
| Kmar_QGN17237.1 | W303.asm02.HP0_G0041180 | OG0003226 | chrXIII | 414487 | 416541 | CP015059.1 | 516410 | 518374 | 31.6970547 | 684 | 654 | 1.04587156 | 1 | 1 | 1 | FIM1_3968 | YMR075W | YMR075W | RCO1 | n |
| Kmar_QGN15093.1 | W303.asm02.HP0_G0020390 | OG0003227 | chrVII | 602532 | 603260 | CP015056.1 | 190779 | 191534 | 46 | 242 | 251 | 0.964143426 | 1 | 1 | 1 | FIM1_1780 | YGR057C | YGR057C | LST7 | n |
| Kmar_QGN15181.1 | W303.asm02.HP0_G0028880 | OG0003228 | chrX | 260019 | 262502 | CP015056.1 | 379362 | 381803 | 47.1291866 | 827 | 813 | 1.017220172 | 1 | 1 | 1 | FIM1_1869 | YJL087C | YJL087C | TRL1 | n |
| Kmar_QGN15182.1 | W303.asm02.HP0_G0028890 | OG0003229 | chrX | 262844 | 264715 | CP015056.1 | 382256 | 384115 | 50.64102564 | 623 | 619 | 1.006462036 | 1 | 1 | 1 | FIM1_1870 | YJL085W | YJL085W | EXO70 | n |
| Kmar_QGN17095.1 | W303.asm02.HP0_G0042430 | OG0003230 | chrXIII | 669073 | 670236 | CP015059.1 | 224808 | 225971 | 71.64948454 | 387 | 387 | 1 | 1 | 1 | 1 | FIM1_3826 | YMR203W | YMR203W | TOM40 | n |
| Kmar_QGN17094.1 | W303.asm02.HP0_G0042440 | OG0003231 | chrXIII | 670644 | 671906 | CP015059.1 | 223094 | 224152 | 15.71428571 | 420 | 352 | 1.193181818 | 1 | 1 | 1 | FIM1_3825 | YMR204C | YMR204C | INP1 | n |
| Kmar_QGN16710.1 | W303.asm02.HP0_G0053030 | OG0003232 | chrXVI | 90074 | 90862 | CP015058.1 | 718073 | 718900 | 42.96028881 | 262 | 275 | 0.952727273 | 1 | 1 | 1 | FIM1_3430 | YPL246C | YPL246C | RBD2 | n |
| Kmar_QGN17028.1 | W303.asm02.HP0_G0025950 | OG0003233 | chrIX | 86426 | 87361 | CP015059.1 | 75041 | 75964 | 49.52380952 | 311 | 307 | 1.013029316 | 1 | 1 | 1 | FIM1_3757 | YIL134W | YIL134W | FLX1 | n |
| Kmar_QGN17032.1 | W303.asm02.HP0_G0046030 | OG0003234 | chrXIV | 490573 | 492303 | CP015059.1 | 80188 | 81756 | 45.76856649 | 576 | 522 | 1.103448276 | 1 | 1 | 1 | FIM1_3761 | YNL073W | YNL073W | MSK1 | n |
| Kmar_QGN16715.1 | W303.asm02.HP0_G0053070 | OG0003235 | chrXVI | 96187 | 100674 | CP015058.1 | 726184 | 730656 | 31.00724161 | 1495 | 1490 | 1.003355705 | 1 | 1 | 1 | FIM1_3435 | YPL242C | YPL242C | IQG1 | n |
| Kmar_QGN18201.1 | W303.asm02.HP0_G0053150 | OG0003236 | chrXVI | 111737 | 112387 | CP015061.1 | 431981 | 432649 | 40.54054054 | 216 | 222 | 0.972972973 | 1 | 1 | 1 | FIM1_4525 | YPL233W | YPL233W | NSL1 | n |
| Kmar_QGN18200.1 | W303.asm02.HP0_G0053140 | OG0003237 | chrXVI | 110511 | 111005 | CP015061.1 | 431069 | 431566 | 89.63414634 | 164 | 165 | 0.993939394 | 1 | 1 | 1 | FIM1_4524 | YPL234C | YPL234C | VMA11 | n |
| Kmar_QGN14979.1 | W303.asm02.HP0_G0054130 | OG0003238 | chrXVI | 309054 | 309947 | CP015055.1 | 1612022 | 1612915 | 87.87878788 | 297 | 297 | 1 | 1 | 1 | 1 | FIM1_1657 | YPL131W | YPL131W | RPL5 | n |
| Kmar_QGN14967.1 | W303.asm02.HP0_G0051130 | OG0003239 | chrXV | 768164 | 769072 | CP015055.1 | 1592376 | 1593239 | 31.04575163 | 302 | 287 | 1.052264808 | 1 | 1 | 1 | FIM1_1645 | YOR228C | YOR228C | MCP1 | n |
| Kmar_QGN17981.1 | W303.asm02.HP0_G0022800 | OG0003240 | chrVII | 36440 | 37111 | CP015060.1 | 882127 | 882774 | 58.74439462 | 223 | 215 | 1.037209302 | 1 | 1 | 1 | FIM1_5190 | YHL031C | YHL031C | GOS1 | n |
| Kmar_QGN18101.1 | W303.asm02.HP0_G0055770 | OG0003241 | chrXVI | 652945 | 653535 | CP015061.1 | 221102 | 221704 | 52.63157895 | 196 | 200 | 0.98 | 1 | 1 | 1 | FIM1_4421 | YPR037C | YPR037C | ERV2 | n |
| Kmar_QGN14978.1 | W303.asm02.HP0_G0051060 | OG0003242 | chrXV | 756623 | 757420 | CP015055.1 | 1609522 | 1610289 | 29.05660377 | 265 | 255 | 1.039215686 | 1 | 1 | 1 | FIM1_1656 | YOR220W | YOR220W | RCN2 | n |

| | | | | | | | | | | | | | | | | | | | | |
|---|---|---|---|---|---|---|---|---|---|---|---|---|---|---|---|---|---|---|---|---|
| Kmar_QGN17392.1 | W303.asm02.HP0_G0032420 | OG0003243 | chrXI | 288985 | 289335 | CP015059.1 | 829762 | 830103 | 53.84615385 | 116 | 113 | 1.026548673 | 1 | 1 | 1 | FIM1_4124 | YKL084W | YKL084W | HOT13 | n |
| Kmar_QGN17030.1 | W303.asm02.HP0_G0046050 | OG0003244 | chrXIV | 493710 | 495158 | CP015059.1 | 76967 | 78370 | 69.56521739 | 482 | 467 | 1.032119914 | 1 | 1 | 1 | FIM1_3759 | YNL071W | YNL071W | LAT1 | n |
| Kmar_QGN17031.1 | W303.asm02.HP0_G0046040 | OG0003245 | chrXIV | 492504 | 493427 | CP015059.1 | 79010 | 79885 | 58.11688312 | 307 | 291 | 1.054982818 | 1 | 1 | 1 | FIM1_3760 | YNL072W | YNL072W | RNH201 | n |
| Kmar_QGN14625.1 | W303.asm02.HP0_G0016020 | OG0003246 | chrV | 482722 | 483930 | CP015055.1 | 884288 | 885403 | 49.75124378 | 402 | 371 | 1.083557951 | 1 | 1 | 1 | FIM1_1289 | YER154W | YER154W | OXA1 | n |
| Kmar_QGN18062.1 | W303.asm02.HP0_G0038470 | OG0003247 | chrXII | 881958 | 885302 | CP015061.1 | 129857 | 133180 | 39.82222222 | 1114 | 1107 | 1.006323397 | 1 | 1 | 1 | FIM1_4380 | YLR383W | YLR383W | SMC6 | n |
| Kmar_QGN14626.1 | W303.asm02.HP0_G0016030 | OG0003248 | chrV | 484047 | 490550 | CP015055.1 | 885547 | 891675 | 44.01101423 | 2167 | 2042 | 1.061214496 | 1 | 1 | 1 | FIM1_1290 | YER155C | YER155C | BEM2 | n |
| Kmar_QGN13943.1 | W303.asm02.HP0_G0049050 | OG0003249 | chrXV | 322313 | 323449 | CP015054.1 | 1188767 | 1189894 | 52.38095238 | 378 | 375 | 1.008 | 1 | 1 | 1 | FIM1_591 | YOL003C | YOL003C | PFA4 | n |
| Kmar_QGN15603.1 | W303.asm02.HP0_G0000230 | OG0003250 | chrI | 55624 | 57492 | CP015056.1 | 1279837 | 1282026 | 21.81571816 | 622 | 729 | 0.853223594 | 1 | 1 | 1 | FIM1_2295 | YAL047C | YAL047C | SPC72 | n |
| Kmar_QGN15602.1 | W303.asm02.HP0_G0052380 | OG0003251 | chrXV | 1017642 | 1019933 | CP015056.1 | 1277126 | 1279336 | 50.4516129 | 763 | 736 | 1.036684783 | 1 | 1 | 1 | FIM1_2294 | YOR361C | YOR361C | PRT1 | n |
| Kmar_QGN15601.1 | W303.asm02.HP0_G0000240 | OG0003252 | chrI | 57664 | 58020 | CP015056.1 | 1276530 | 1276805 | 50.90909091 | 118 | 91 | 1.296703297 | 1 | 1 | 1 | FIM1_2293 | YAL046C | YAL046C | BOL3 | n |
| Kmar_QGN16868.1 | W303.asm02.HP0_G0010130 | OG0003253 | chrIV | 769420 | 770217 | CP015058.1 | 1070679 | 1071479 | 50 | 265 | 266 | 0.996240602 | 1 | 1 | 1 | FIM1_3594 | YDR152W | YDR152W | GIR2 | n |
| Kmar_QGN16878.1 | W303.asm02.HP0_G0010210 | OG0003254 | chrIV | 784104 | 784814 | CP015058.1 | 1090528 | 1091331 | 33.5483871 | 236 | 267 | 0.883895131 | 1 | 1 | 1 | FIM1_3605 | YDR162C | YDR162C | NBP2 | n |
| Kmar_QGN15807.1 | W303.asm02.HP0_G0041780 | OG0003255 | chrXIII | 545776 | 546351 | CP015057.1 | 131874 | 132473 | 38.27751196 | 191 | 199 | 0.959798995 | 1 | 1 | 1 | FIM1_2503 | YMR138W | YMR138W | CIN4 | n |
| Kmar_QGN15808.1 | W303.asm02.HP0_G0041770 | OG0003256 | chrXIII | 543593 | 545578 | CP015057.1 | 132769 | 134250 | 28.61491629 | 661 | 493 | 1.340770791 | 1 | 1 | 1 | FIM1_2504 | YMR137C | YMR137C | PSO2 | n |
| Kmar_QGN13850.1 | W303.asm02.HP0_G0028020 | OG0003257 | chrX | 73598 | 74866 | CP015054.1 | 997291 | 998520 | 45.75471698 | 422 | 409 | 1.031784841 | 1 | 1 | 1 | FIM1_496 | YJL183W | YJL183W | MNN11 | n |
| Kmar_QGN17859.1 | W303.asm02.HP0_G0048040 | OG0003258 | chrXV | 112557 | 113585 | CP015060.1 | 628938 | 629984 | 49.12790698 | 342 | 348 | 0.982758621 | 1 | 1 | 1 | FIM1_5068 | YOL107W | YOL107W | | n |
| Kmar_QGN15635.1 | W303.asm02.HP0_G0020360 | OG0003259 | chrVII | 594452 | 596380 | CP015056.1 | 1356805 | 1358583 | 54.93171472 | 642 | 592 | 1.084459459 | 1 | 1 | 1 | FIM1_2328 | YGR054W | YGR054W | | n |
| Kmar_QGN18029.1 | W303.asm02.HP0_G0016460 | OG0003260 | chrVI | 13102 | 14007 | CP015061.1 | 58616 | 59539 | 38.78205128 | 301 | 307 | 0.980456026 | 1 | 1 | 1 | FIM1_4345 | YFL044C | YFL044C | OTU1 | n |
| Kmar_QGN18028.1 | W303.asm02.HP0_G0035640 | OG0003261 | chrXII | 289907 | 290407 | CP015061.1 | 57944 | 58447 | 64.67065868 | 166 | 167 | 0.994011976 | 1 | 1 | 1 | FIM1_4344 | YLR074C | YLR074C | BUD20 | n |
| Kmar_QGN18027.1 | W303.asm02.HP0_G0035650 | OG0003262 | chrXII | 290878 | 291543 | CP015061.1 | 56645 | 57307 | 88.23529412 | 221 | 220 | 1.004545455 | 1 | 1 | 1 | FIM1_4343 | YLR075W | YLR075W | RPL10 | n |
| Kmar_QGN18024.1 | W303.asm02.HP0_G0016450 | OG0003263 | chrVI | 12075 | 12839 | CP015061.1 | 51467 | 52231 | 84.2519685 | 254 | 254 | 1 | 1 | 1 | 1 | FIM1_4340 | YFL045C | YFL045C | SEC53 | n |
| Kmar_QGN18022.1 | W303.asm02.HP0_G0035660 | OG0003264 | chrXII | 291823 | 293574 | CP015061.1 | 48478 | 50298 | 40.64 | 583 | 606 | 0.962046205 | 1 | 1 | 1 | FIM1_4338 | YLR077W | YLR077W | FMP25 | n |
| Kmar_QGN18023.1 | W303.asm02.HP0_G0016440 | OG0003265 | chrVI | 11262 | 11885 | CP015061.1 | 50667 | 51350 | 67.40088106 | 207 | 227 | 0.911894273 | 1 | 1 | 1 | FIM1_4339 | YFL046W | YFL046W | FMP32 | n |
| Kmar_QGN18021.1 | W303.asm02.HP0_G0035670 | OG0003266 | chrXII | 293687 | 294510 | CP015061.1 | 47639 | 48394 | 47.80876494 | 244 | 251 | 0.972111554 | 1 | 1 | 1 | FIM1_4337 | YLR078C | YLR078C | BOS1 | n |
| Kmar_QGN16243.1 | W303.asm02.HP0_G0030900 | OG0003267 | chrX | 694337 | 699091 | CP015057.1 | 1113916 | 1118571 | 46.02484472 | 1584 | 1551 | 1.021276596 | 1 | 1 | 1 | FIM1_2948 | YJR138W | YJR138W | IML1 | n |
| Kmar_QGN16242.1 | W303.asm02.HP0_G0030910 | OG0003268 | chrX | 699215 | 700294 | CP015057.1 | 1112730 | 1113886 | 73.05555556 | 359 | 360 | 0.997222222 | 1 | 1 | 1 | FIM1_2947 | YJR139C | YJR139C | HOM6 | n |
| Kmar_QGN16236.1 | W303.asm02.HP0_G0030960 | OG0003269 | chrX | 710651 | 711460 | CP015057.1 | 1099894 | 1100691 | 58.45588235 | 269 | 265 | 1.01509434 | 1 | 1 | 1 | FIM1_2941 | YJR144W | YJR144W | MGM101 | n |
| Kmar_QGN14836.1 | W303.asm02.HP0_G0007850 | OG0003270 | chrIV | 282441 | 283505 | CP015055.1 | 1323592 | 1324641 | 83.61581921 | 354 | 349 | 1.014326648 | 1 | 1 | 1 | FIM1_1507 | YDL100C | YDL100C | GET3 | n |
| Kmar_QGN14572.1 | W303.asm02.HP0_G0041850 | OG0003271 | chrXIII | 558106 | 559149 | CP015055.1 | 785521 | 786564 | 63.03724928 | 347 | 347 | 1 | 1 | 1 | 1 | FIM1_1234 | YMR146C | YMR146C | TIF34 | n |
| Kmar_QGN17876.1 | W303.asm02.HP0_G0003100 | OG0003272 | chrII | 486034 | 490212 | CP015060.1 | 656590 | 660750 | 73.5483871 | 1392 | 1386 | 1.004329004 | 1 | 1 | 1 | FIM1_5085 | YBR115C | YBR115C | LYS2 | n |
| Kmar_QGN15598.1 | W303.asm02.HP0_G0052370 | OG0003273 | chrXV | 1015522 | 1017102 | CP015056.1 | 1272213 | 1273823 | 34.96376812 | 526 | 536 | 0.981343284 | 1 | 1 | 1 | FIM1_2290 | YOR360C | YOR360C | PDE2 | n |
| Kmar_QGN15597.1 | W303.asm02.HP0_G0052360 | OG0003274 | chrXV | 1013470 | 1015041 | CP015056.1 | 1270021 | 1271430 | 38.67924528 | 523 | 469 | 1.115138593 | 1 | 1 | 1 | FIM1_2289 | YOR359W | YOR359W | VTS1 | n |
| Kmar_QGN13576.1 | W303.asm02.HP0_G0031370 | OG0003275 | chrXI | 76101 | 76703 | CP015054.1 | 448089 | 448691 | 84 | 200 | 200 | 1 | 1 | 1 | 1 | FIM1_217 | YKL196C | YKL196C | YKT6 | n |
| Kmar_QGN13575.1 | W303.asm02.HP0_G0031380 | OG0003276 | chrXI | 76918 | 78201 | CP015054.1 | 446521 | 447684 | 29.66292135 | 427 | 387 | 1.103359173 | 1 | 1 | 1 | FIM1_216 | YKL195W | YKL195W | MIA40 | n |
| Kmar_QGN13574.1 | W303.asm02.HP0_G0038870 | OG0003277 | chrXII | 970064 | 972190 | CP015054.1 | 444171 | 446177 | 27.93296089 | 708 | 668 | 1.05988024 | 1 | 1 | 1 | FIM1_215 | YLR424W | YLR424W | SPP382 | n |
| Kmar_QGN13560.1 | W303.asm02.HP0_G0038810 | OG0003278 | chrXII | 953583 | 954764 | CP015054.1 | 420250 | 421347 | 54.06091371 | 393 | 365 | 1.076712329 | 1 | 1 | 1 | FIM1_201 | YLR418C | YLR418C | CDC73 | n |
| Kmar_QGN13572.1 | W303.asm02.HP0_G0031390 | OG0003279 | chrXI | 78427 | 79815 | CP015054.1 | 441229 | 442614 | 58.49462366 | 462 | 461 | 1.002169197 | 1 | 1 | 1 | FIM1_213 | YKL194C | YKL194C | MST1 | n |
| Kmar_QGN16281.1 | W303.asm02.HP0_G0031510 | OG0003280 | chrXI | 99890 | 100810 | CP015057.1 | 1196162 | 1197070 | 38.43648208 | 306 | 302 | 1.013245033 | 1 | 1 | 1 | FIM1_2986 | YKL183W | YKL183W | LOT5 | n |
| Kmar_QGN16558.1 | W303.asm02.HP0_G0038970 | OG0003281 | chrXII | 1004082 | 1007906 | CP015058.1 | 369394 | 372477 | 19.78193146 | 1274 | 1027 | 1.240506329 | 1 | 1 | 1 | FIM1_3272 | YLR436C | YLR436C | ECM30 | n |
| Kmar_QGN16559.1 | W303.asm02.HP0_G0038960 | OG0003282 | chrXII | 1003039 | 1003656 | CP015058.1 | 372498 | 373133 | 57.34597156 | 205 | 211 | 0.971563981 | 1 | 1 | 1 | FIM1_3273 | YLR435W | YLR435W | TSR2 | n |
| Kmar_QGN17468.1 | W303.asm02.HP0_G0011190 | OG0003283 | chrIV | 997635 | 998453 | CP015059.1 | 982508 | 983323 | 30.03533569 | 272 | 271 | 1.003690037 | 1 | 1 | 1 | FIM1_4204 | YDR262W | YDR262W | | n |
| Kmar_QGN14642.1 | W303.asm02.HP0_G0012070 | OG0003284 | chrIV | 1200715 | 1201083 | CP015055.1 | 928699 | 929031 | 44.26229508 | 122 | 110 | 1.109090909 | 1 | 1 | 1 | FIM1_1306 | YDR357C | YDR357C | CNL1 | n |
| Kmar_QGN16945.1 | W303.asm02.HP0_G0010630 | OG0003285 | chrIV | 861853 | 862860 | CP015058.1 | 1234499 | 1235512 | 60.65088757 | 335 | 337 | 0.994065282 | 1 | 1 | 1 | FIM1_3672 | YDR204W | YDR204W | COQ4 | n |
| Kmar_QGN16941.1 | W303.asm02.HP0_G0037190 | OG0003286 | chrXII | 611562 | 612548 | CP015058.1 | 1229542 | 1230705 | 46.52956298 | 328 | 387 | 0.84754522 | 1 | 1 | 1 | FIM1_3668 | YLR239C | YLR239C | LIP2 | n |
| Kmar_QGN15676.1 | W303.asm02.HP0_G0012680 | OG0003287 | chrIV | 1317596 | 1320448 | CP015056.1 | 1441251 | 1444025 | 42.78187566 | 950 | 924 | 1.028138528 | 1 | 1 | 1 | FIM1_2369 | YDR421W | YDR421W | ARO80 | n |
| Kmar_QGN14966.1 | W303.asm02.HP0_G0054070 | OG0003288 | chrXVI | 297299 | 298537 | CP015055.1 | 1591020 | 1592111 | 33.41013825 | 402 | 363 | 1.107438017 | 1 | 1 | 1 | FIM1_1644 | YPL138C | YPL138C | SPP1 | n |
| Kmar_QGN16967.1 | W303.asm02.HP0_G0044720 | OG0003289 | chrXIV | 238937 | 240700 | CP015058.1 | 1272911 | 1274590 | 58.43270869 | 587 | 559 | 1.050089445 | 1 | 1 | 1 | FIM1_3694 | YNL218W | YNL218W | MGS1 | n |
| Kmar_QGN16908.1 | W303.asm02.HP0_G0010350 | OG0003290 | chrIV | 816907 | 817866 | CP015058.1 | 1171345 | 1172253 | 54.51713396 | 319 | 302 | 1.056291391 | 1 | 1 | 1 | FIM1_3635 | YDR175C | YDR175C | RSM24 | n |
| Kmar_QGN16909.1 | W303.asm02.HP0_G0010360 | OG0003291 | chrIV | 818166 | 820274 | CP015058.1 | 1172591 | 1174519 | 51.83615819 | 702 | 642 | 1.093457944 | 1 | 1 | 1 | FIM1_3636 | YDR176W | YDR176W | NGG1 | n |
| Kmar_QGN16910.1 | W303.asm02.HP0_G0036520 | OG0003292 | chrXII | 492127 | 493515 | CP015058.1 | 1174701 | 1176107 | 78.41880342 | 462 | 468 | 0.987179487 | 1 | 1 | 1 | FIM1_3637 | YLR163C | YLR163C | MAS1 | n |
| Kmar_QGN18109.1 | W303.asm02.HP0_G0055730 | OG0003293 | chrXVI | 646022 | 647455 | CP015061.1 | 233011 | 234390 | 27.67676768 | 477 | 459 | 1.039215686 | 1 | 1 | 1 | FIM1_4429 | YPR034W | YPR034W | ARP7 | n |
| Kmar_QGN15866.1 | W303.asm02.HP0_G0009040 | OG0003294 | chrIV | 522870 | 524372 | CP015057.1 | 260455 | 261963 | 45.65217391 | 500 | 502 | 0.996015936 | 1 | 1 | 1 | FIM1_2563 | YDR036C | YDR036C | EHD3 | n |
| Kmar_QGN13411.1 | W303.asm02.HP0_G0054500 | OG0003295 | chrXVI | 376197 | 377021 | CP015054.1 | 85383 | 86123 | 56.22641509 | 274 | 246 | 1.113821138 | 1 | 1 | 1 | FIM1_48 | YPL094C | YPL094C | SEC62 | n |
| Kmar_QGN15307.1 | W303.asm02.HP0_G0009590 | OG0003296 | chrIV | 630980 | 632806 | CP015056.1 | 661413 | 663236 | 90.13157895 | 608 | 607 | 1.001647446 | 1 | 1 | 1 | FIM1_1996 | YDR091C | YDR091C | RLI1 | n |
| Kmar_QGN15306.1 | W303.asm02.HP0_G0016110 | OG0003297 | chrV | 513094 | 517500 | CP015056.1 | 655324 | 660021 | 61.65605096 | 1468 | 1565 | 0.938019169 | 1 | 1 | 1 | FIM1_1995 | YER164W | YER164W | CHD1 | n |
| Kmar_QGN15305.1 | W303.asm02.HP0_G0016120 | OG0003298 | chrV | 518075 | 519808 | CP015056.1 | 652565 | 654337 | 68.95973154 | 577 | 590 | 0.977966102 | 1 | 1 | 1 | FIM1_1994 | YER165W | YER165W | PAB1 | n |
| Kmar_QGN14218.1 | W303.asm02.HP0_G0009410 | OG0003299 | chrIV | 598169 | 600859 | CP015055.1 | 53580 | 56378 | 60.26970954 | 896 | 932 | 0.961373391 | 1 | 1 | 1 | FIM1_874 | YDR074W | YDR074W | TPS2 | n |
| Kmar_QGN14084.1 | W303.asm02.HP0_G0009250 | OG0003300 | chrIV | 571872 | 572852 | CP015054.1 | 1484973 | 1486157 | 45.56962025 | 326 | 394 | 0.827411168 | 1 | 1 | 1 | FIM1_735 | YDR058C | YDR058C | TGL2 | n |
| Kmar_QGN17125.1 | W303.asm02.HP0_G0030680 | OG0003301 | chrX | 650599 | 651438 | CP015059.1 | 288955 | 289806 | 47.53521127 | 279 | 283 | 0.985865724 | 1 | 1 | 1 | FIM1_3856 | YJR116W | YJR116W | TDA4 | n |
| Kmar_QGN18042.1 | W303.asm02.HP0_G0016530 | OG0003302 | chrVI | 23814 | 24434 | CP015061.1 | 85178 | 85792 | 77.18446602 | 206 | 204 | 1.009803922 | 1 | 1 | 1 | FIM1_4358 | YFL038C | YFL038C | YPT1 | n |
| Kmar_QGN17123.1 | W303.asm02.HP0_G0014690 | OG0003303 | chrV | 198092 | 199663 | CP015059.1 | 285172 | 286695 | 55.61904762 | 523 | 507 | 1.031558185 | 1 | 1 | 1 | FIM1_3854 | YER021W | YER021W | RPN3 | n |
| Kmar_QGN17121.1 | W303.asm02.HP0_G0030700 | OG0003304 | chrX | 653264 | 653875 | CP015059.1 | 282197 | 282715 | 44.44444444 | 203 | 172 | 1.180232558 | 1 | 1 | 1 | FIM1_3852 | YJR118C | YJR118C | ILM1 | n |
| Kmar_QGN14083.1 | W303.asm02.HP0_G0022240 | OG0003305 | chrVII | 1014963 | 1016402 | CP015054.1 | 1483307 | 1484728 | 60.33402923 | 479 | 473 | 1.012684989 | 1 | 1 | 1 | FIM1_734 | YGR255C | YGR255C | COQ6 | n |
| Kmar_QGN14577.1 | W303.asm02.HP0_G0041880 | OG0003306 | chrXIII | 561617 | 562477 | CP015055.1 | 791933 | 792730 | 31.01045296 | 286 | 265 | 1.079245283 | 1 | 1 | 1 | FIM1_1239 | YMR149W | YMR149W | SWP1 | n |
| Kmar_QGN14578.1 | W303.asm02.HP0_G0041890 | OG0003307 | chrXIII | 562571 | 563143 | CP015055.1 | 792788 | 793297 | 59.47368421 | 190 | 169 | 1.124260355 | 1 | 1 | 1 | FIM1_1240 | YMR150C | YMR150C | IMP1 | n |
| Kmar_QGN17240.1 | W303.asm02.HP0_G0032930 | OG0003308 | chrXI | 384055 | 387171 | CP015059.1 | 525921 | 529010 | 26.78405931 | 1038 | 1029 | 1.008746356 | 1 | 1 | 1 | FIM1_3971 | YKL033W | YKL033W | TTI1 | n |
| Kmar_QGN16469.1 | W303.asm02.HP0_G0004700 | OG0003309 | chrII | 791609 | 792049 | CP015058.1 | 176019 | 176456 | 59.5890411 | 146 | 145 | 1.006896552 | 1 | 1 | 1 | FIM1_3182 | YBR282W | YBR282W | MRPL27 | n |
| Kmar_QGN16152.1 | W303.asm02.HP0_G0043910 | OG0003310 | chrXIV | 54554 | 56329 | CP015057.1 | 913148 | 914887 | 57.85953177 | 591 | 579 | 1.020725389 | 1 | 1 | 1 | FIM1_2853 | YNL308C | YNL308C | KRI1 | n |
| Kmar_QGN14351.1 | W303.asm02.HP0_G0032220 | OG0003311 | chrXI | 251697 | 253850 | CP015055.1 | 332390 | 334510 | 81.08484006 | 717 | 706 | 1.015580737 | 1 | 1 | 1 | FIM1_1011 | YKL104C | YKL104C | GFA1 | n |
| Kmar_QGN15137.1 | W303.asm02.HP0_G0031920 | OG0003312 | chrXI | 184238 | 185848 | CP015056.1 | 271257 | 272621 | 41.87866928 | 536 | 454 | 1.18061674 | 1 | 1 | 1 | FIM1_1824 | YKL139W | YKL139W | CTK1 | n |
| Kmar_QGN17129.1 | W303.asm02.HP0_G0014680 | OG0003313 | chrV | 196313 | 197662 | CP015059.1 | 296200 | 297705 | 51.89620758 | 449 | 501 | 0.896207585 | 1 | 1 | 1 | FIM1_3860 | YER020W | YER020W | GPA2 | n |
| Kmar_QGN18348.1 | W303.asm02.HP0_G0023960 | OG0003314 | chrVIII | 237964 | 239127 | CP015061.1 | 747084 | 748223 | 80.36175711 | 387 | 379 | 1.021108179 | 1 | 1 | 1 | FIM1_4675 | YHR068W | YHR068W | DYS1 | n |
| Kmar_QGN15679.1 | W303.asm02.HP0_G0040430 | OG0003315 | chrXIII | 252345 | 253496 | CP015056.1 | 1451041 | 1452168 | 79.31034483 | 383 | 375 | 1.021333333 | 1 | 1 | 1 | FIM1_2372 | YML008C | YML008C | ERG6 | n |
| Kmar_QGN16729.1 | W303.asm02.HP0_G0011780 | OG0003316 | chrIV | 1125033 | 1127363 | CP015058.1 | 756257 | 758467 | 54.95495495 | 776 | 736 | 1.054347826 | 1 | 1 | 1 | FIM1_3450 | YDR324C | YDR324C | UTP4 | n |
| Kmar_QGN14143.1 | W303.asm02.HP0_G0005260 | OG0003317 | chrIII | 63092 | 64132 | CP015054.1 | 1627991 | 1628890 | 39.48126801 | 346 | 299 | 1.157190635 | 1 | 1 | 1 | FIM1_796 | YCL032W | YCL032W | STE50 | n |
| Kmar_QGN18425.1 | W303.asm02.HP0_G0016900 | OG0003318 | chrVI | 117593 | 118207 | CP015061.1 | 905708 | 906331 | 64.42307692 | 204 | 207 | 0.985507246 | 1 | 1 | 1 | FIM1_4753 | YFR001W | YFR001W | LOC1 | n |
| Kmar_QGN16811.1 | W303.asm02.HP0_G0020780 | OG0003319 | chrVII | 691833 | 692873 | CP015058.1 | 948700 | 949890 | 47.2361809 | 346 | 396 | 0.873737374 | 1 | 1 | 1 | FIM1_3535 | YGR101W | YGR101W | PCP1 | n |
| Kmar_QGN16798.1 | W303.asm02.HP0_G0056400 | OG0003320 | chrXVI | 756185 | 758848 | CP015058.1 | 913408 | 916101 | 62.36203091 | 887 | 897 | 0.988851728 | 1 | 1 | 1 | FIM1_3522 | YPR112C | YPR112C | MRD1 | n |
| Kmar_QGN14076.1 | W303.asm02.HP0_G0044440 | OG0003321 | chrXIV | 175965 | 179903 | CP015054.1 | 1468208 | 1472098 | 50.57121097 | 1312 | 1296 | 1.012345679 | 1 | 1 | 1 | FIM1_727 | YNL250W | YNL250W | RAD50 | n |
| Kmar_QGN17271.1 | W303.asm02.HP0_G0042760 | OG0003322 | chrXIII | 742090 | 743313 | CP015059.1 | 591297 | 592466 | 55.25672372 | 407 | 389 | 1.046272494 | 1 | 1 | 1 | FIM1_4002 | YMR235C | YMR235C | RNA1 | n |
| Kmar_QGN15249.1 | W303.asm02.HP0_G0049520 | OG0003323 | chrXV | 414308 | 414493 | CP015056.1 | 527271 | 527444 | 50.81967213 | 61 | 57 | 1.070175439 | 1 | 1 | 1 | FIM1_1938 | YOR045W | YOR045W | TOM6 | n |
| Kmar_QGN15277.1 | W303.asm02.HP0_G0049710 | OG0003324 | chrXV | 452464 | 454197 | CP015056.1 | 580844 | 582529 | 64.81802426 | 577 | 561 | 1.028520499 | 1 | 1 | 1 | FIM1_1966 | YOR067C | YOR067C | ALG8 | n |
| Kmar_QGN15247.1 | W303.asm02.HP0_G0049530 | OG0003325 | chrXV | 414915 | 416363 | CP015056.1 | 524196 | 525620 | 72.82157676 | 482 | 474 | 1.016877637 | 1 | 1 | 1 | FIM1_1936 | YOR046C | YOR046C | DBP5 | n |
| Kmar_QGN16022.1 | W303.asm02.HP0_G0011970 | OG0003326 | chrIV | 1180214 | 1181179 | CP015057.1 | 646285 | 647172 | 52.32198142 | 321 | 295 | 1.088135593 | 1 | 1 | 1 | FIM1_2722 | YDR347W | YDR347W | MRP1 | n |
| Kmar_QGN17046.1 | W303.asm02.HP0_G0007460 | OG0003327 | chrIV | 211582 | 212253 | CP015059.1 | 116099 | 116710 | 40.69264069 | 223 | 203 | 1.098522167 | 1 | 1 | 1 | FIM1_3775 | YDL139C | YDL139C | SCM3 | n |
| Kmar_QGN13424.1 | W303.asm02.HP0_G0054600 | OG0003328 | chrXVI | 400396 | 402930 | CP015054.1 | 117772 | 120207 | 44.99411072 | 844 | 811 | 1.040690506 | 1 | 1 | 1 | FIM1_61 | YPL084W | YPL084W | BRO1 | n |
| Kmar_QGN17036.1 | W303.asm02.HP0_G0007400 | OG0003329 | chrIV | 194778 | 198383 | CP015059.1 | 90221 | 93856 | 70.33057851 | 1201 | 1211 | 0.991742362 | 1 | 1 | 1 | FIM1_3765 | YDL145C | YDL145C | COP1 | n |
| Kmar_QGN18262.1 | W303.asm02.HP0_G0020980 | OG0003330 | chrVII | 748334 | 749875 | CP015061.1 | 575309 | 576847 | 64.20233463 | 513 | 512 | 1.001953125 | 1 | 1 | 1 | FIM1_4588 | YGR123C | YGR123C | PPT1 | n |
| Kmar_QGN17060.1 | W303.asm02.HP0_G0007520 | OG0003331 | chrIV | 222633 | 223946 | CP015059.1 | 147261 | 148700 | 35.6846473 | 437 | 479 | 0.912317328 | 1 | 1 | 1 | FIM1_3790 | YDL133W | YDL133W | SRF1 | n |
| Kmar_QGN17059.1 | W303.asm02.HP0_G0007080 | OG0003332 | chrIV | 131091 | 132053 | CP015059.1 | 145754 | 146680 | 39.7515528 | 320 | 308 | 1.038961039 | 1 | 1 | 1 | FIM1_3789 | YDL183C | YDL183C | MRX19 | n |
| Kmar_QGN15363.1 | W303.asm02.HP0_G0003550 | OG0003333 | chrII | 579979 | 581346 | CP015056.1 | 800771 | 802159 | 59.02335456 | 455 | 462 | 0.984848485 | 1 | 1 | 1 | FIM1_2053 | YBR162C | YBR162C | TOS1 | n |
| Kmar_QGN17023.1 | W303.asm02.HP0_G0026000 | OG0003334 | chrIX | 95141 | 102271 | CP015059.1 | 58067 | 64858 | 54.64090718 | 2376 | 2263 | 1.049933716 | 1 | 1 | 1 | FIM1_3752 | YIL129C | YIL129C | TAO3 | n |
| Kmar_QGN17790.1 | W303.asm02.HP0_G0049910 | OG0003335 | chrXV | 494168 | 495205 | CP015060.1 | 488446 | 489492 | 74.14772727 | 345 | 348 | 0.99137931 | 1 | 1 | 1 | FIM1_4999 | YOR091W | YOR091W | TMA46 | n |
| Kmar_QGN17791.1 | W303.asm02.HP0_G0045840 | OG0003336 | chrXIV | 449238 | 451001 | CP015060.1 | 489770 | 491470 | 37.41610738 | 587 | 566 | 1.037102473 | 1 | 1 | 1 | FIM1_5000 | YNL094W | YNL094W | APP1 | n |
| Kmar_QGN17324.1 | W303.asm02.HP0_G0024440 | OG0003337 | chrVIII | 344056 | 345957 | CP015059.1 | 697139 | 699031 | 46.86520376 | 633 | 630 | 1.004761905 | 1 | 1 | 1 | FIM1_4056 | YHR114W | YHR114W | BZZ1 | n |
| Kmar_QGN17795.1 | W303.asm02.HP0_G0049950 | OG0003338 | chrXV | 504287 | 505063 | CP015060.1 | 502748 | 503566 | 66.54676259 | 258 | 272 | 0.948529412 | 1 | 1 | 1 | FIM1_5004 | YOR095C | YOR095C | RKI1 | n |
| Kmar_QGN17022.1 | W303.asm02.HP0_G0026010 | OG0003339 | chrIX | 102840 | 105938 | CP015059.1 | 54538 | 57627 | 45.62078922 | 1032 | 1029 | 1.002915452 | 1 | 1 | 1 | FIM1_3751 | YIL128W | YIL128W | MET18 | n |
| Kmar_QGN15395.1 | W303.asm02.HP0_G0045930 | OG0003340 | chrXIV | 471681 | 472730 | CP015056.1 | 870338 | 871489 | 47.90575916 | 349 | 383 | 0.911227154 | 1 | 1 | 1 | FIM1_2085 | YNL084C | YNL084C | END3 | n |
| Kmar_QGN13471.1 | W303.asm02.HP0_G0014270 | OG0003341 | chrV | 117349 | 118152 | CP015054.1 | 204969 | 205772 | 80.52434457 | 267 | 267 | 1 | 1 | 1 | 1 | FIM1_109 | YEL021W | YEL021W | URA3 | n |

| | | | | | | | | | | | | | | | | | | | | |
|---|---|---|---|---|---|---|---|---|---|---|---|---|---|---|---|---|---|---|---|---|
| Kmar_QGN17716.1 | W303.asm02.HP0_G0026290 | OG0003342 | chrIX | 168362 | 170011 | CP015060.1 | 304353 | 306140 | 33.9314845 | 549 | 595 | 0.922689076 | 1 | 1 | 1 | FIM1_4923 | YIL099W | YIL099W | SGA1 | n |
| Kmar_QGN17717.1 | W303.asm02.HP0_G0046390 | OG0003343 | chrXIV | 570997 | 571662 | CP015060.1 | 308607 | 309260 | 53.66972477 | 221 | 217 | 1.01843318 | 1 | 1 | 1 | FIM1_4924 | YNL036W | YNL036W | NCE103 | n |
| Kmar_QGN13742.1 | W303.asm02.HP0_G0019830 | OG0003344 | chrVII | 483454 | 486975 | CP015054.1 | 790482 | 794306 | 56.70682731 | 1173 | 1274 | 0.920722135 | 1 | 1 | 1 | FIM1_386 | YGL006W | YGL006W | PMC1 | n |
| Kmar_QGN14245.1 | W303.asm02.HP0_G0019630 | OG0003345 | chrVII | 438367 | 439995 | CP015055.1 | 104589 | 106133 | 50.09107468 | 542 | 514 | 1.054474708 | 1 | 1 | 1 | FIM1_901 | YGL028C | YGL028C | SCW11 | n |
| Kmar_QGN14744.1 | W303.asm02.HP0_G0013080 | OG0003346 | chrIV | 1403719 | 1404393 | CP015055.1 | 1146430 | 1147239 | 37.5464684 | 224 | 269 | 0.832713755 | 1 | 1 | 1 | FIM1_1413 | YDR468C | YDR468C | TLG1 | n |
| Kmar_QGN14745.1 | W303.asm02.HP0_G0013090 | OG0003347 | chrIV | 1404708 | 1405235 | CP015055.1 | 1147412 | 1147810 | 30.85714286 | 175 | 132 | 1.325757576 | 1 | 1 | 1 | FIM1_1414 | YDR469W | YDR469W | SDC1 | n |
| Kmar_QGN14746.1 | W303.asm02.HP0_G0013100 | OG0003348 | chrIV | 1405399 | 1406907 | CP015055.1 | 1147924 | 1149381 | 40.15748031 | 502 | 485 | 1.035051546 | 1 | 1 | 1 | FIM1_1415 | YDR470C | YDR470C | UGO1 | n |
| Kmar_QGN15702.1 | W303.asm02.HP0_G0014920 | OG0003349 | chrV | 236909 | 238258 | CP015056.1 | 1496113 | 1497462 | 90.64587973 | 449 | 449 | 1 | 1 | 1 | 1 | FIM1_2395 | YER043C | YER043C | SAH1 | n |
| Kmar_QGN15711.1 | W303.asm02.HP0_G0017360 | OG0003350 | chrVI | 208896 | 210341 | CP015056.1 | 1513144 | 1514700 | 65.83011583 | 481 | 518 | 0.928571429 | 1 | 1 | 1 | FIM1_2404 | YFR044C | YFR044C | DUG1 | n |
| Kmar_QGN13583.1 | W303.asm02.HP0_G0051680 | OG0003351 | chrXV | 853840 | 854796 | CP015054.1 | 464759 | 465871 | 38.60589812 | 318 | 370 | 0.859459459 | 1 | 1 | 1 | FIM1_225 | YOR288C | YOR288C | MPD1 | n |
| Kmar_QGN14213.1 | W303.asm02.HP0_G0030510 | OG0003352 | chrX | 626124 | 627107 | CP015055.1 | 45671 | 46585 | 59.02140673 | 327 | 304 | 1.075657895 | 1 | 1 | 1 | FIM1_869 | YJR100C | YJR100C | AIM25 | n |
| Kmar_QGN17751.1 | W303.asm02.HP0_G0030460 | OG0003353 | chrX | 619546 | 620514 | CP015060.1 | 388511 | 389425 | 78.70967742 | 322 | 304 | 1.059210526 | 1 | 1 | 1 | FIM1_4960 | YJR095W | YJR095W | SFC1 | n |
| Kmar_QGN16405.1 | W303.asm02.HP0_G0054330 | OG0003354 | chrXVI | 346302 | 347303 | CP015058.1 | 46259 | 47233 | 63.60946746 | 333 | 324 | 1.027777778 | 1 | 1 | 1 | FIM1_3118 | YPL111W | YPL111W | CAR1 | n |
| Kmar_QGN16403.1 | W303.asm02.HP0_G0054340 | OG0003355 | chrXVI | 347426 | 351097 | CP015058.1 | 40074 | 44186 | 48.90350877 | 1223 | 1370 | 0.89270073 | 1 | 1 | 1 | FIM1_3116 | YPL110C | YPL110C | GDE1 | n |
| Kmar_QGN18100.1 | W303.asm02.HP0_G0055780 | OG0003356 | chrXVI | 653802 | 654872 | CP015061.1 | 219445 | 220641 | 44.98777506 | 356 | 398 | 0.894472362 | 1 | 1 | 1 | FIM1_4420 | YPR040W | YPR040W | TIP41 | n |
| Kmar_QGN14617.1 | W303.asm02.HP0_G0030520 | OG0003357 | chrX | 628007 | 628807 | CP015055.1 | 866907 | 867665 | 54.68164794 | 266 | 252 | 1.055555556 | 1 | 1 | 1 | FIM1_1281 | YJR101W | YJR101W | RSM26 | n |
| Kmar_QGN18099.1 | W303.asm02.HP0_G0030380 | OG0003358 | chrX | 596269 | 597147 | CP015061.1 | 218519 | 219376 | 39.79591837 | 292 | 285 | 1.024561404 | 1 | 1 | 1 | FIM1_4419 | YJR088C | YJR088C | EMC2 | n |
| Kmar_QGN18071.1 | W303.asm02.HP0_G0030530 | OG0003359 | chrX | 628930 | 629538 | CP015061.1 | 152882 | 153445 | 43.90243902 | 202 | 187 | 1.080213904 | 1 | 1 | 1 | FIM1_4389 | YJR102C | YJR102C | VPS25 | n |
| Kmar_QGN18096.1 | W303.asm02.HP0_G0030400 | OG0003360 | chrX | 600646 | 604101 | CP015061.1 | 209946 | 213731 | 37.89883268 | 1151 | 1261 | 0.912767645 | 1 | 1 | 1 | FIM1_4416 | YJR090C | YJR090C | GRR1 | n |
| Kmar_QGN16653.1 | W303.asm02.HP0_G0020440 | OG0003361 | chrVII | 614085 | 615035 | CP015058.1 | 597045 | 597989 | 44.87951807 | 316 | 314 | 1.006369427 | 1 | 1 | 1 | FIM1_3372 | YGR062C | YGR062C | COX18 | n |
| Kmar_QGN16652.1 | W303.asm02.HP0_G0020450 | OG0003362 | chrVII | 615272 | 615580 | CP015058.1 | 596351 | 596659 | 77.45098039 | 102 | 102 | 1 | 1 | 1 | 1 | FIM1_3371 | YGR063C | YGR063C | SPT4 | n |
| Kmar_QGN13709.1 | W303.asm02.HP0_G0022370 | OG0003363 | chrVII | 1040777 | 1044916 | CP015054.1 | 705329 | 709474 | 47.72413793 | 1379 | 1381 | 0.998551774 | 1 | 1 | 1 | FIM1_352 | YGR270W | YGR270W | YTA7 | n |
| Kmar_QGN15208.1 | W303.asm02.HP0_G0021450 | OG0003364 | chrVII | 852605 | 854332 | CP015056.1 | 433054 | 434793 | 50.42158516 | 575 | 579 | 0.993091537 | 1 | 1 | 1 | FIM1_1896 | YGR171C | YGR171C | MSM1 | n |
| Kmar_QGN15210.1 | W303.asm02.HP0_G0021460 | OG0003365 | chrVII | 854626 | 855372 | CP015056.1 | 436793 | 437617 | 55.27272727 | 248 | 274 | 0.905109489 | 1 | 1 | 1 | FIM1_1898 | YGR172C | YGR172C | YIP1 | n |
| Kmar_QGN15426.1 | W303.asm02.HP0_G0011430 | OG0003366 | chrIV | 1047645 | 1049510 | CP015056.1 | 936526 | 938355 | 59.71563981 | 621 | 609 | 1.019704433 | 1 | 1 | 1 | FIM1_2117 | YDR292C | YDR292C | SRP101 | n |
| Kmar_QGN15206.1 | W303.asm02.HP0_G0021440 | OG0003367 | chrVII | 848923 | 852339 | CP015056.1 | 428147 | 431296 | 40.31141869 | 1138 | 1049 | 1.084842707 | 1 | 1 | 1 | FIM1_1894 | YGR170W | YGR170W | PSD2 | n |
| Kmar_QGN13715.1 | W303.asm02.HP0_G0022330 | OG0003368 | chrVII | 1032510 | 1034765 | CP015054.1 | 720891 | 723152 | 71.44754316 | 751 | 753 | 0.997343958 | 1 | 1 | 1 | FIM1_359 | YGR264C | YGR264C | MES1 | n |
| Kmar_QGN13711.1 | W303.asm02.HP0_G0022350 | OG0003369 | chrVII | 1037916 | 1038647 | CP015054.1 | 712194 | 712931 | 81.63265306 | 243 | 245 | 0.991836735 | 1 | 1 | 1 | FIM1_355 | YGR267C | YGR267C | FOL2 | n |
| Kmar_QGN15432.1 | W303.asm02.HP0_G0011440 | OG0003370 | chrIV | 1050139 | 1051626 | CP015056.1 | 946823 | 950362 | 24.7257384 | 495 | 1179 | 0.419847328 | 1 | 1 | 1 | FIM1_2123 | W303.asm02.HP0_G0011440 | YDR293C | SSD1 | y |
| Kmar_QGN15207.1 | W303.asm02.HP0_G0029680 | OG0003371 | chrX | 444851 | 446314 | CP015056.1 | 431546 | 433009 | 52.66393443 | 487 | 462 | 1.054112554 | 1 | 1 | 1 | FIM1_1895 | YJR006W | YJR006W | POL31 | n |
| Kmar_QGN13714.1 | W303.asm02.HP0_G0043300 | OG0003372 | chrXIII | 856995 | 858512 | CP015054.1 | 718700 | 720235 | 86.6927593 | 505 | 511 | 0.988258317 | 1 | 1 | 1 | FIM1_358 | YMR290C | YMR290C | HAS1 | n |
| Kmar_QGN13689.1 | W303.asm02.HP0_G0016290 | OG0003373 | chrV | 560227 | 560961 | CP015054.1 | 666532 | 667371 | 41.17647059 | 244 | 279 | 0.874551971 | 1 | 1 | 1 | FIM1_332 | YER182W | YER182W | FMP10 | n |
| Kmar_QGN15417.1 | W303.asm02.HP0_G0035080 | OG0003374 | chrXII | 182617 | 184134 | CP015056.1 | 914785 | 916023 | 43.19526627 | 505 | 412 | 1.225728155 | 1 | 1 | 1 | FIM1_2107 | YLR015W | YLR015W | BRE2 | n |
| Kmar_QGN16184.1 | W303.asm02.HP0_G0022200 | OG0003375 | chrVII | 1008079 | 1008669 | CP015057.1 | 980637 | 981221 | 39.71291866 | 196 | 194 | 1.010309278 | 1 | 1 | 1 | FIM1_2886 | YGR251W | YGR251W | NOP19 | n |
| Kmar_QGN13710.1 | W303.asm02.HP0_G0022360 | OG0003376 | chrVII | 1038969 | 1040060 | CP015054.1 | 711163 | 711792 | 31.59609121 | 304 | 209 | 1.454545455 | 1 | 1 | 1 | FIM1_354 | YGR268C | YGR268C | HUA1 | n |
| Kmar_QGN15433.1 | W303.asm02.HP0_G0023500 | OG0003377 | chrVIII | 175821 | 177548 | CP015056.1 | 951410 | 953137 | 74.39446367 | 575 | 575 | 1 | 1 | 1 | 1 | FIM1_2124 | YHR037W | YHR037W | PUT2 | n |
| Kmar_QGN13707.1 | W303.asm02.HP0_G0043340 | OG0003378 | chrXIII | 863886 | 865007 | CP015054.1 | 702379 | 703482 | 22.22222222 | 373 | 367 | 1.016348774 | 1 | 1 | 1 | FIM1_350 | YMR294W | YMR294W | JNM1 | n |
| Kmar_QGN15209.1 | W303.asm02.HP0_G0021470 | OG0003379 | chrVII | 855635 | 856741 | CP015056.1 | 435056 | 436165 | 91.05691057 | 368 | 369 | 0.997289973 | 1 | 1 | 1 | FIM1_1897 | YGR173W | YGR173W | RBG2 | n |
| Kmar_QGN13705.1 | W303.asm02.HP0_G0022390 | OG0003380 | chrVII | 1051207 | 1051908 | CP015054.1 | 695485 | 696180 | 46.44351464 | 233 | 231 | 1.008658009 | 1 | 1 | 1 | FIM1_348 | YGR271C-A | YGR271C-A | EFG1 | n |
| Kmar_QGN13713.1 | W303.asm02.HP0_G0022340 | OG0003381 | chrVII | 1035568 | 1037673 | CP015054.1 | 715674 | 717827 | 52.47933884 | 701 | 717 | 0.977684798 | 1 | 1 | 1 | FIM1_357 | YGR266W | YGR266W | | n |
| Kmar_QGN17521.1 | W303.asm02.HP0_G0022060 | OG0003382 | chrVII | 975713 | 978070 | CP015059.1 | 1082503 | 1084857 | 29.05569007 | 785 | 784 | 1.00127551 | 1 | 1 | 1 | FIM1_4257 | YGR237C | YGR237C | | n |
| Kmar_QGN15418.1 | W303.asm02.HP0_G0035070 | OG0003383 | chrXII | 179658 | 182372 | CP015056.1 | 916327 | 918657 | 38.29787234 | 904 | 776 | 1.164948454 | 1 | 1 | 1 | FIM1_2108 | YLR014C | YLR014C | PPR1 | n |
| Kmar_QGN16043.1 | W303.asm02.HP0_G0029530 | OG0003384 | chrX | 409934 | 410419 | CP015057.1 | 691550 | 692041 | 47.59036145 | 161 | 163 | 0.987730061 | 1 | 1 | 1 | FIM1_2743 | YJL011C | YJL011C | RPC17 | n |
| Kmar_QGN17779.1 | W303.asm02.HP0_G0045910 | OG0003385 | chrXIV | 467962 | 468270 | CP015060.1 | 464158 | 464457 | 41.17647059 | 102 | 99 | 1.03030303 | 1 | 1 | 1 | FIM1_4988 | YNL086W | YNL086W | SNN1 | n |
| Kmar_QGN17778.1 | W303.asm02.HP0_G0049860 | OG0003386 | chrXV | 482768 | 483820 | CP015060.1 | 463007 | 464044 | 30.44692737 | 350 | 345 | 1.014492754 | 1 | 1 | 1 | FIM1_4987 | YOR085W | YOR085W | OST3 | n |
| Kmar_QGN14583.1 | W303.asm02.HP0_G0041920 | OG0003387 | chrXIII | 566614 | 568797 | CP015055.1 | 798671 | 800764 | 24.09152086 | 727 | 697 | 1.043041607 | 1 | 1 | 1 | FIM1_1245 | YMR154C | YMR154C | RIM13 | n |
| Kmar_QGN17560.1 | W303.asm02.HP0_G0018620 | OG0003388 | chrVII | 232737 | 234587 | CP015059.1 | 1163021 | 1164838 | 48.16 | 616 | 605 | 1.018181818 | 1 | 1 | 1 | FIM1_4296 | YGL142C | YGL142C | GPI10 | n |
| Kmar_QGN14857.1 | W303.asm02.HP0_G0007790 | OG0003389 | chrIV | 270428 | 272107 | CP015055.1 | 1358684 | 1360225 | 36.10648918 | 559 | 513 | 1.089668616 | 1 | 1 | 1 | FIM1_1530 | YDL106C | YDL106C | PHO2 | n |
| Kmar_QGN17559.1 | W303.asm02.HP0_G0007780 | OG0003390 | chrIV | 269127 | 270182 | CP015059.1 | 1161764 | 1162906 | 36.08247423 | 351 | 380 | 0.923684211 | 1 | 1 | 1 | FIM1_4295 | YDL107W | YDL107W | MSS2 | n |
| Kmar_QGN16040.1 | W303.asm02.HP0_G0029500 | OG0003391 | chrX | 403207 | 404811 | CP015057.1 | 684056 | 685663 | 89.88764045 | 534 | 535 | 0.998130841 | 1 | 1 | 1 | FIM1_2740 | YJL014W | YJL014W | CCT3 | n |
| Kmar_QGN16938.1 | W303.asm02.HP0_G0010570 | OG0003392 | chrIV | 853989 | 854714 | CP015058.1 | 1225343 | 1226314 | 41.17647059 | 241 | 323 | 0.746130031 | 1 | 1 | 1 | FIM1_3665 | YDR196C | YDR196C | CAB5 | n |
| Kmar_QGN15554.1 | W303.asm02.HP0_G0023910 | OG0003393 | chrVIII | 229860 | 230999 | CP015056.1 | 1173997 | 1175154 | 43.03797468 | 379 | 385 | 0.984415584 | 1 | 1 | 1 | FIM1_2245 | YHR063C | YHR063C | PAN5 | n |
| Kmar_QGN16940.1 | W303.asm02.HP0_G0010590 | OG0003394 | chrIV | 856249 | 857688 | CP015058.1 | 1228053 | 1229414 | 42.37995825 | 479 | 453 | 1.057395143 | 1 | 1 | 1 | FIM1_3667 | YDR198C | YDR198C | RKM2 | n |
| Kmar_QGN13562.1 | W303.asm02.HP0_G0031430 | OG0003395 | chrXI | 84116 | 84719 | CP015054.1 | 422809 | 423336 | 83.42857143 | 175 | 175 | 1 | 1 | 1 | 1 | FIM1_203 | YKL190W | YKL190W | CNB1 | n |
| Kmar_QGN16120.1 | W303.asm02.HP0_G0033000 | OG0003396 | chrXI | 399332 | 401371 | CP015057.1 | 851545 | 853587 | 39.26553672 | 679 | 680 | 0.998529412 | 1 | 1 | 1 | FIM1_2821 | YKL025C | YKL025C | PAN3 | n |
| Kmar_QGN15412.1 | W303.asm02.HP0_G0023590 | OG0003397 | chrVIII | 189395 | 191077 | CP015056.1 | 904652 | 906310 | 43.18584071 | 560 | 552 | 1.014492754 | 1 | 1 | 1 | FIM1_2102 | YHR045W | YHR045W | DDE1 | n |
| Kmar_QGN17395.1 | W303.asm02.HP0_G0023150 | OG0003398 | chrVIII | 106483 | 107901 | CP015059.1 | 833120 | 834442 | 60.59322034 | 472 | 440 | 1.072727273 | 1 | 1 | 1 | FIM1_4127 | YHR005C | YHR005C | GPA1 | n |
| Kmar_QGN18080.1 | W303.asm02.HP0_G0001440 | OG0003399 | chrII | 135784 | 137157 | CP015061.1 | 167431 | 168768 | 61.26914661 | 457 | 445 | 1.026966292 | 1 | 1 | 1 | FIM1_4399 | YBL045C | YBL045C | COR1 | n |
| Kmar_QGN18082.1 | W303.asm02.HP0_G0030660 | OG0003400 | chrX | 648005 | 648748 | CP015061.1 | 171861 | 172709 | 54.60992908 | 247 | 282 | 0.875886525 | 1 | 1 | 1 | FIM1_4401 | YJR113C | YJR113C | RSM7 | n |
| Kmar_QGN18081.1 | W303.asm02.HP0_G0001430 | OG0003401 | chrII | 134065 | 135390 | CP015061.1 | 169752 | 170912 | 27.99097065 | 441 | 386 | 1.142487047 | 1 | 1 | 1 | FIM1_4400 | YBL046W | YBL046W | PSY4 | n |
| Kmar_QGN18251.1 | W303.asm02.HP0_G0056890 | OG0003402 | chrXVI | 882114 | 882461 | CP015061.1 | 557845 | 558192 | 69.56521739 | 115 | 115 | 1 | 1 | 1 | 1 | FIM1_4576 | YPR166C | YPR166C | MRP2 | n |
| Kmar_QGN16834.1 | W303.asm02.HP0_G0004460 | OG0003403 | chrII | 750742 | 751458 | CP015058.1 | 990628 | 991341 | 65.96638655 | 238 | 237 | 1.004219409 | 1 | 1 | 1 | FIM1_3558 | YBR256C | YBR256C | RIB5 | n |
| Kmar_QGN18298.1 | W303.asm02.HP0_G0014590 | OG0003404 | chrV | 178982 | 179578 | CP015061.1 | 646864 | 647451 | 83.58974359 | 198 | 195 | 1.015384615 | 1 | 1 | 1 | FIM1_4624 | YER012W | YER012W | PRE1 | n |
| Kmar_QGN16094.1 | W303.asm02.HP0_G0015060 | OG0003405 | chrV | 266563 | 267456 | CP015057.1 | 798474 | 799367 | 26.26262626 | 297 | 98 | 3.030612245 | 1 | 1 | 1 | FIM1_2794 | YER055C | YER055C | HIS1 | n |
| Kmar_QGN16833.1 | W303.asm02.HP0_G0027340 | OG0003406 | chrIX | 369654 | 370949 | CP015058.1 | 989198 | 990490 | 64.1723356 | 431 | 430 | 1.002325581 | 1 | 1 | 1 | FIM1_3557 | YIR012W | YIR012W | SQT1 | n |
| Kmar_QGN18299.1 | W303.asm02.HP0_G0027300 | OG0003407 | chrIX | 364245 | 365474 | CP015061.1 | 647807 | 649123 | 64.00911162 | 409 | 438 | 0.933789954 | 1 | 1 | 1 | FIM1_4625 | YIR008C | YIR008C | PRI1 | n |
| Kmar_QGN18300.1 | W303.asm02.HP0_G0027290 | OG0003408 | chrIX | 361872 | 364166 | CP015061.1 | 649482 | 651698 | 50.19556714 | 764 | 738 | 1.035230352 | 1 | 1 | 1 | FIM1_4626 | YIR007W | YIR007W | EGH1 | n |
| Kmar_QGN18217.1 | W303.asm02.HP0_G0007380 | OG0003409 | chrIV | 191131 | 192468 | CP015061.1 | 472969 | 474294 | 77.52808989 | 445 | 441 | 1.009070295 | 1 | 1 | 1 | FIM1_4542 | YDL147W | YDL147W | RPN5 | n |
| Kmar_QGN16782.1 | W303.asm02.HP0_G0048770 | OG0003410 | chrXV | 268642 | 270096 | CP015058.1 | 880010 | 881473 | 63.56435644 | 484 | 487 | 0.993839836 | 1 | 1 | 1 | FIM1_3505 | YOL030W | YOL030W | GAS5 | n |
| Kmar_QGN16777.1 | W303.asm02.HP0_G0052900 | OG0003411 | chrXVI | 51793 | 53220 | CP015058.1 | 865106 | 866464 | 68.77637131 | 475 | 452 | 1.050884956 | 1 | 1 | 1 | FIM1_3499 | YPL259C | YPL259C | APM1 | n |
| Kmar_QGN16016.1 | W303.asm02.HP0_G0029800 | OG0003412 | chrX | 461214 | 462971 | CP015057.1 | 619782 | 621533 | 82.73504274 | 585 | 583 | 1.003430532 | 1 | 1 | 1 | FIM1_2716 | YJR016C | YJR016C | ILV3 | n |
| Kmar_QGN18112.1 | W303.asm02.HP0_G0055720 | OG0003413 | chrXVI | 643876 | 645516 | CP015061.1 | 237083 | 238687 | 77.09090909 | 546 | 534 | 1.02247191 | 1 | 1 | 1 | FIM1_4432 | YPR033C | YPR033C | HTS1 | n |
| Kmar_QGN15979.1 | W303.asm02.HP0_G0048230 | OG0003414 | chrXV | 155742 | 159092 | CP015057.1 | 532666 | 535317 | 32.24092117 | 1116 | 883 | 1.26387316 | 1 | 1 | 1 | FIM1_2679 | YOL087C | YOL087C | DUF1 | n |
| Kmar_QGN15978.1 | W303.asm02.HP0_G0048240 | OG0003415 | chrXV | 159628 | 159900 | CP015057.1 | 531944 | 532261 | 37.73584906 | 90 | 105 | 0.857142857 | 1 | 1 | 1 | FIM1_2678 | YOL086W-A | YOL086W-A | MHF1 | n |
| Kmar_QGN14104.1 | W303.asm02.HP0_G0044240 | OG0003416 | chrXIV | 127359 | 129807 | CP015054.1 | 1535904 | 1537982 | 40.58524173 | 779 | 692 | 1.125722543 | 1 | 1 | 1 | FIM1_755 | YNL272C | YNL272C | SEC2 | n |
| Kmar_QGN18037.1 | W303.asm02.HP0_G0035570 | OG0003417 | chrXII | 275764 | 278661 | CP015061.1 | 75009 | 77864 | 31.50826446 | 965 | 951 | 1.014721346 | 1 | 1 | 1 | FIM1_4353 | YLR067C | YLR067C | PET309 | n |
| Kmar_QGN18038.1 | W303.asm02.HP0_G0035560 | OG0003418 | chrXII | 275120 | 275674 | CP015061.1 | 78050 | 78625 | 43.22916667 | 184 | 191 | 0.963350785 | 1 | 1 | 1 | FIM1_4354 | YLR066W | YLR066W | SPC3 | n |
| Kmar_QGN14071.1 | W303.asm02.HP0_G0044470 | OG0003419 | chrXIV | 183430 | 185733 | CP015054.1 | 1459430 | 1461730 | 64.81241915 | 767 | 766 | 1.001305483 | 1 | 1 | 1 | FIM1_722 | YNL247W | YNL247W | CRS1 | n |
| Kmar_QGN18035.1 | W303.asm02.HP0_G0035590 | OG0003420 | chrXII | 279581 | 281866 | CP015061.1 | 71450 | 73714 | 78.44940867 | 761 | 754 | 1.00928382 | 1 | 1 | 1 | FIM1_4351 | YLR069C | YLR069C | MEF1 | n |
| Kmar_QGN13544.1 | W303.asm02.HP0_G0002110 | OG0003421 | chrII | 279525 | 283022 | CP015054.1 | 372586 | 376098 | 59.2654424 | 1165 | 1170 | 0.995726496 | 1 | 1 | 1 | FIM1_184 | YBR023C | YBR023C | CHS3 | n |
| Kmar_QGN13802.1 | W303.asm02.HP0_G0044010 | OG0003422 | chrXIV | 72120 | 77030 | CP015054.1 | 903840 | 908747 | 35.46056592 | 1636 | 1635 | 1.000611621 | 1 | 1 | 1 | FIM1_448 | YNL297C | YNL297C | MON2 | n |
| Kmar_QGN15299.1 | W303.asm02.HP0_G0009630 | OG0003423 | chrIV | 644379 | 648107 | CP015056.1 | 632575 | 636156 | 58.16572808 | 1242 | 1193 | 1.041072925 | 1 | 1 | 1 | FIM1_1988 | YDR097C | YDR097C | MSH6 | n |
| Kmar_QGN15932.1 | W303.asm02.HP0_G0035950 | OG0003424 | chrXII | 372070 | 373284 | CP015057.1 | 415175 | 416350 | 45.83333333 | 404 | 391 | 1.033248082 | 1 | 1 | 1 | FIM1_2630 | YLR107W | YLR107W | REX3 | n |
| Kmar_QGN14280.1 | W303.asm02.HP0_G0023390 | OG0003425 | chrVIII | 154685 | 155326 | CP015055.1 | 174508 | 175143 | 79.342723 | 213 | 211 | 1.009478673 | 1 | 1 | 1 | FIM1_937 | YHR026W | YHR026W | VMA16 | n |
| Kmar_QGN14279.1 | W303.asm02.HP0_G0023400 | OG0003426 | chrVIII | 155573 | 158554 | CP015055.1 | 171248 | 174265 | 64.76377953 | 993 | 1005 | 0.988059701 | 1 | 1 | 1 | FIM1_936 | YHR027C | YHR027C | RPN1 | n |
| Kmar_QGN14056.1 | W303.asm02.HP0_G0027860 | OG0003427 | chrX | 43910 | 45709 | CP015054.1 | 1430673 | 1432400 | 47.94069193 | 599 | 575 | 1.04173913 | 1 | 1 | 1 | FIM1_707 | YJL201W | YJL201W | ECM25 | n |
| Kmar_QGN16887.1 | W303.asm02.HP0_G0036290 | OG0003428 | chrXII | 440213 | 440818 | CP015058.1 | 1108234 | 1108881 | 36.86635945 | 201 | 215 | 0.934883721 | 1 | 1 | 1 | FIM1_3614 | YLR145W | YLR145W | RMP1 | n |
| Kmar_QGN16877.1 | W303.asm02.HP0_G0010200 | OG0003429 | chrIV | 782757 | 783920 | CP015058.1 | 1089290 | 1090423 | 55.55555556 | 387 | 377 | 1.026525199 | 1 | 1 | 1 | FIM1_3604 | YDR161W | YDR161W | ACL4 | n |
| Kmar_QGN16033.1 | W303.asm02.HP0_G0018340 | OG0003430 | chrVII | 170831 | 171813 | CP015057.1 | 671846 | 672673 | 30.35714286 | 272 | 275 | 0.989090909 | 1 | 1 | 1 | FIM1_2733 | YGL174W | YGL174W | BUD13 | n |
| Kmar_QGN18383.1 | W303.asm02.HP0_G0055600 | OG0003431 | chrXVI | 607147 | 609855 | CP015061.1 | 814229 | 816964 | 57.21812434 | 902 | 911 | 0.990120746 | 1 | 1 | 1 | FIM1_4710 | YPR021C | YPR021C | AGC1 | n |
| Kmar_QGN16028.1 | W303.asm02.HP0_G0034730 | OG0003432 | chrXII | 105697 | 107910 | CP015057.1 | 657870 | 660014 | 48.36601307 | 737 | 714 | 1.032212885 | 1 | 1 | 1 | FIM1_2728 | YLL019C | YLL019C | KNS1 | n |
| Kmar_QGN16031.1 | W303.asm02.HP0_G0018360 | OG0003433 | chrVII | 177168 | 178586 | CP015057.1 | 665082 | 666725 | 41.56305506 | 472 | 547 | 0.862888483 | 1 | 1 | 1 | FIM1_2731 | YGL172W | YGL172W | NUP49 | n |
| Kmar_QGN15561.1 | W303.asm02.HP0_G0002760 | OG0003434 | chrII | 414900 | 417176 | CP015056.1 | 1185587 | 1187881 | 82.32984293 | 758 | 764 | 0.992146597 | 1 | 1 | 1 | FIM1_2252 | YBR080C | YBR080C | SEC18 | n |
| Kmar_QGN18335.1 | W303.asm02.HP0_G0001620 | OG0003435 | chrII | 168446 | 169576 | CP015061.1 | 716450 | 717376 | 31.91489362 | 376 | 308 | 1.220779221 | 1 | 1 | 1 | FIM1_4661 | YBL029W | YBL029W | | n |
| Kmar_QGN15623.1 | W303.asm02.HP0_G0018430 | OG0003436 | chrVII | 190175 | 192871 | CP015056.1 | 1331538 | 1334288 | 64.97354497 | 898 | 916 | 0.980349345 | 1 | 1 | 1 | FIM1_2316 | YGL163C | YGL163C | RAD54 | n |
| Kmar_QGN15599.1 | W303.asm02.HP0_G0000260 | OG0003437 | chrI | 58585 | 59097 | CP015056.1 | 1275168 | 1275695 | 54.54545455 | 170 | 175 | 0.971428571 | 1 | 1 | 1 | FIM1_2291 | YAL044C | YAL044C | GCV3 | n |
| Kmar_QGN17600.1 | W303.asm02.HP0_G0052870 | OG0003438 | chrXVI | 45102 | 47057 | CP015060.1 | 76210 | 78171 | 52.82738095 | 651 | 653 | 0.996937213 | 1 | 1 | 1 | FIM1_4807 | YPL263C | YPL263C | KEL3 | n |
| Kmar_QGN17590.1 | W303.asm02.HP0_G0052840 | OG0003439 | chrXVI | 39673 | 40629 | CP015060.1 | 58146 | 59138 | 82.72727273 | 318 | 330 | 0.963636364 | 1 | 1 | 1 | FIM1_4797 | YPL266W | YPL266W | DIM1 | n |
| Kmar_QGN17595.1 | W303.asm02.HP0_G0052810 | OG0003440 | chrXVI | 33565 | 35499 | CP015060.1 | 65384 | 67228 | 28.52713178 | 644 | 614 | 1.048859935 | 1 | 1 | 1 | FIM1_4802 | YPL269W | YPL269W | KAR9 | n |

| | | | | | | | | | | | | | | | | | | | | |
|---|---|---|---|---|---|---|---|---|---|---|---|---|---|---|---|---|---|---|---|---|
| Kmar_QGN17601.1 | W303.asm02.HP0_G0013910 | OG0003441 | chrV | 43615 | 45288 | CP015060.1 | 78401 | 80029 | 54.2039356 | 557 | 542 | 1.027675277 | 1 | 1 | 1 | FIM1_4808 | YEL058W | YEL058W | PCM1 | n |
| Kmar_QGN17202.1 | W303.asm02.HP0_G0019130 | OG0003442 | chrVII | 346568 | 347392 | CP015059.1 | 440949 | 441761 | 52.3297491 | 274 | 270 | 1.014814815 | 1 | 1 | 1 | FIM1_3933 | YGL085W | YGL085W | LCL3 | n |
| Kmar_QGN15600.1 | W303.asm02.HP0_G0000250 | OG0003443 | chrI | 58153 | 58485 | CP015056.1 | 1275826 | 1276185 | 42.27642276 | 110 | 119 | 0.924369748 | 1 | 1 | 1 | FIM1_2292 | YAL044W-A | YAL044W-A | BOL1 | n |
| Kmar_QGN17642.1 | W303.asm02.HP0_G0021650 | OG0003444 | chrVII | 896924 | 898156 | CP015060.1 | 171857 | 173107 | 43.42723005 | 410 | 416 | 0.985576923 | 1 | 1 | 1 | FIM1_4849 | YGR193C | YGR193C | PDX1 | n |
| Kmar_QGN17638.1 | W303.asm02.HP0_G0029720 | OG0003445 | chrX | 452476 | 454011 | CP015060.1 | 159232 | 160740 | 74.609375 | 511 | 502 | 1.017928287 | 1 | 1 | 1 | FIM1_4845 | YJR010W | YJR010W | MET3 | n |
| Kmar_QGN13599.1 | W303.asm02.HP0_G0009290 | OG0003446 | chrIV | 580623 | 582308 | CP015054.1 | 494329 | 496017 | 77.97513321 | 561 | 562 | 0.998220641 | 1 | 1 | 1 | FIM1_241 | YDR062W | YDR062W | LCB2 | n |
| Kmar_QGN13598.1 | W303.asm02.HP0_G0009280 | OG0003447 | chrIV | 578313 | 579932 | CP015054.1 | 492260 | 493837 | 57.09023941 | 539 | 525 | 1.026666667 | 1 | 1 | 1 | FIM1_240 | YDR061W | YDR061W | | n |
| Kmar_QGN14225.1 | W303.asm02.HP0_G0009350 | OG0003448 | chrIV | 587862 | 588794 | CP015055.1 | 66097 | 66990 | 34.79623824 | 310 | 297 | 1.043771044 | 1 | 1 | 1 | FIM1_881 | YDR068W | YDR068W | DOS2 | n |
| Kmar_QGN17653.1 | W303.asm02.HP0_G0034990 | OG0003449 | chrXII | 163180 | 165318 | CP015060.1 | 196641 | 198833 | 25.65789474 | 712 | 730 | 0.975342466 | 1 | 1 | 1 | FIM1_4860 | YLR006C | YLR006C | SSK1 | n |
| Kmar_QGN13963.1 | W303.asm02.HP0_G0004090 | OG0003450 | chrII | 680117 | 680677 | CP015054.1 | 1227777 | 1228364 | 34.87179487 | 186 | 195 | 0.953846154 | 1 | 1 | 1 | FIM1_611 | YBR217W | YBR217W | ATG12 | n |
| Kmar_QGN13965.1 | W303.asm02.HP0_G0004070 | OG0003451 | chrII | 675638 | 677599 | CP015054.1 | 1230852 | 1232303 | 27.82193959 | 625 | 483 | 1.293995859 | 1 | 1 | 1 | FIM1_613 | YBR215W | YBR215W | HPC2 | n |
| Kmar_QGN16342.1 | W303.asm02.HP0_G0040290 | OG0003452 | chrXIII | 229445 | 230008 | CP015057.1 | 1330594 | 1331157 | 74.86631016 | 187 | 187 | 1 | 1 | 1 | 1 | FIM1_3048 | YML022W | YML022W | APT1 | n |
| Kmar_QGN16340.1 | W303.asm02.HP0_G0012860 | OG0003453 | chrIV | 1351231 | 1355493 | CP015057.1 | 1324521 | 1328711 | 40.74074074 | 1420 | 1396 | 1.017191977 | 1 | 1 | 1 | FIM1_3046 | YDR443C | YDR443C | SSN2 | n |
| Kmar_QGN16362.1 | W303.asm02.HP0_G0040410 | OG0003454 | chrXIII | 248183 | 251374 | CP015057.1 | 1363460 | 1366552 | 63.79468378 | 1063 | 1030 | 1.032038835 | 1 | 1 | 1 | FIM1_3068 | YML010W | YML010W | SPT5 | n |
| Kmar_QGN17648.1 | W303.asm02.HP0_G0034980 | OG0003455 | chrXII | 161473 | 162858 | CP015060.1 | 185958 | 187601 | 60.32906764 | 461 | 547 | 0.842778793 | 1 | 1 | 1 | FIM1_4855 | YLR005W | YLR005W | SSL1 | n |
| Kmar_QGN15415.1 | W303.asm02.HP0_G0029350 | OG0003456 | chrX | 375898 | 376650 | CP015056.1 | 912355 | 913101 | 58.984375 | 250 | 248 | 1.008064516 | 1 | 1 | 1 | FIM1_2105 | YJL035C | YJL035C | TAD2 | n |
| Kmar_QGN16719.1 | W303.asm02.HP0_G0024030 | OG0003457 | chrVIII | 252167 | 254311 | CP015058.1 | 737978 | 740119 | 82.77310924 | 714 | 713 | 1.001402525 | 1 | 1 | 1 | FIM1_3440 | YHR074W | YHR074W | QNS1 | n |
| Kmar_QGN14865.1 | W303.asm02.HP0_G0042030 | OG0003458 | chrXIII | 590624 | 593212 | CP015055.1 | 1374622 | 1377147 | 48.25708061 | 862 | 841 | 1.024970273 | 1 | 1 | 1 | FIM1_1538 | YMR165C | YMR165C | PAH1 | n |
| Kmar_QGN14498.1 | W303.asm02.HP0_G0035500 | OG0003459 | chrXII | 267550 | 268359 | CP015055.1 | 639069 | 640070 | 43.65781711 | 269 | 333 | 0.807807808 | 1 | 1 | 1 | FIM1_1159 | YLR059C | YLR059C | REX2 | n |
| Kmar_QGN14495.1 | W303.asm02.HP0_G0018190 | OG0003460 | chrVII | 140321 | 140710 | CP015055.1 | 629904 | 630287 | 73.4375 | 129 | 127 | 1.015748031 | 1 | 1 | 1 | FIM1_1156 | YGL191W | YGL191W | COX13 | n |
| Kmar_QGN14680.1 | W303.asm02.HP0_G0018220 | OG0003461 | chrVII | 145217 | 145684 | CP015055.1 | 1020078 | 1020542 | 72.90322581 | 155 | 154 | 1.006493506 | 1 | 1 | 1 | FIM1_1346 | YGL187C | YGL187C | COX4 | n |
| Kmar_QGN16534.1 | W303.asm02.HP0_G0001840 | OG0003462 | chrII | 214917 | 218651 | CP015058.1 | 323091 | 326810 | 44.09020218 | 1244 | 1239 | 1.004035513 | 1 | 1 | 1 | FIM1_3247 | YBL007C | YBL007C | SLA1 | n |
| Kmar_QGN14494.1 | W303.asm02.HP0_G0018180 | OG0003463 | chrVII | 137759 | 139561 | CP015055.1 | 627680 | 629287 | 48.69281046 | 600 | 535 | 1.121495327 | 1 | 1 | 1 | FIM1_1155 | YGL192W | YGL192W | IME4 | n |
| Kmar_QGN16535.1 | W303.asm02.HP0_G0019800 | OG0003464 | chrVII | 473079 | 473603 | CP015058.1 | 327864 | 328379 | 50.56818182 | 174 | 171 | 1.01754386 | 1 | 1 | 1 | FIM1_3248 | YGL010W | YGL010W | MPO1 | n |
| Kmar_QGN15385.1 | W303.asm02.HP0_G0019680 | OG0003465 | chrVII | 449490 | 452094 | CP015056.1 | 849427 | 851724 | 63.64692218 | 813 | 765 | 1.062745098 | 1 | 1 | 1 | FIM1_2075 | YGL022W | YGL022W | STT3 | n |
| Kmar_QGN15330.1 | W303.asm02.HP0_G0019670 | OG0003466 | chrVII | 447283 | 449190 | CP015056.1 | 720189 | 722201 | 29.62447844 | 635 | 670 | 0.947761194 | 1 | 1 | 1 | FIM1_2019 | YGL023C | YGL023C | PIB2 | n |
| Kmar_QGN15133.1 | W303.asm02.HP0_G0031950 | OG0003467 | chrXI | 187261 | 187596 | CP015056.1 | 269065 | 269394 | 45.2173913 | 111 | 109 | 1.018348624 | 1 | 1 | 1 | FIM1_1820 | YKL137W | YKL137W | CMC1 | n |
| Kmar_QGN15134.1 | W303.asm02.HP0_G0031940 | OG0003468 | chrXI | 186590 | 186985 | CP015056.1 | 269745 | 270134 | 54.96183206 | 131 | 129 | 1.015503876 | 1 | 1 | 1 | FIM1_1821 | YKL138C | YKL138C | MRPL31 | n |
| Kmar_QGN15798.1 | W303.asm02.HP0_G0020000 | OG0003469 | chrVII | 512088 | 513950 | CP015057.1 | 115462 | 117180 | 31.76100629 | 620 | 572 | 1.083916084 | 1 | 1 | 1 | FIM1_2494 | YGR013W | YGR013W | SNU71 | n |
| Kmar_QGN13775.1 | W303.asm02.HP0_G0040100 | OG0003470 | chrXIII | 176249 | 178138 | CP015054.1 | 859185 | 861044 | 39.6875 | 629 | 619 | 1.016155089 | 1 | 1 | 1 | FIM1_420 | YML046W | YML046W | PRP39 | n |
| Kmar_QGN13776.1 | W303.asm02.HP0_G0030060 | OG0003471 | chrX | 540717 | 542414 | CP015054.1 | 861113 | 862813 | 50.52264808 | 565 | 566 | 0.998233216 | 1 | 1 | 1 | FIM1_421 | YJR052W | YJR052W | RAD7 | n |
| Kmar_QGN13497.1 | W303.asm02.HP0_G0014110 | OG0003472 | chrV | 80823 | 81506 | CP015054.1 | 267099 | 267764 | 55.60344828 | 227 | 221 | 1.027149321 | 1 | 1 | 1 | FIM1_135 | YEL038W | YEL038W | UTR4 | n |
| Kmar_QGN14663.1 | W303.asm02.HP0_G0017080 | OG0003473 | chrVI | 153409 | 160242 | CP015055.1 | 972143 | 978313 | 46.03036876 | 2277 | 2056 | 1.107490272 | 1 | 1 | 1 | FIM1_1327 | YFR019W | YFR019W | FAB1 | n |
| Kmar_QGN14797.1 | W303.asm02.HP0_G0012210 | OG0003474 | chrIV | 1226705 | 1227742 | CP015055.1 | 1249947 | 1250927 | 71.30434783 | 345 | 326 | 1.058282209 | 1 | 1 | 1 | FIM1_1468 | YDR372C | YDR372C | VPS74 | n |
| Kmar_QGN15865.1 | W303.asm02.HP0_G0002520 | OG0003475 | chrII | 369959 | 372304 | CP015057.1 | 257761 | 260091 | 47.03656999 | 781 | 776 | 1.006443299 | 1 | 1 | 1 | FIM1_2562 | YBR058C | YBR058C | UBP14 | n |
| Kmar_QGN16369.1 | W303.asm02.HP0_G0014810 | OG0003476 | chrV | 219196 | 222426 | CP015057.1 | 1376404 | 1379004 | 25.09259259 | 1076 | 866 | 1.242494226 | 1 | 1 | 1 | FIM1_3076 | YER033C | YER033C | ZRG8 | n |
| Kmar_QGN16658.1 | W303.asm02.HP0_G0038300 | OG0003477 | chrXII | 848675 | 849331 | CP015058.1 | 606315 | 607022 | 38.72340426 | 218 | 235 | 0.927659574 | 1 | 1 | 1 | FIM1_3377 | YLR363C | YLR363C | NMD4 | n |
| Kmar_QGN16667.1 | W303.asm02.HP0_G0042680 | OG0003478 | chrXIII | 725205 | 726230 | CP015058.1 | 626685 | 627692 | 37.10144928 | 341 | 335 | 1.017910448 | 1 | 1 | 1 | FIM1_3386 | YMR228W | YMR228W | MTF1 | n |
| Kmar_QGN17379.1 | W303.asm02.HP0_G0041050 | OG0003479 | chrXIII | 393258 | 395291 | CP015059.1 | 802832 | 804862 | 44.63519313 | 677 | 676 | 1.00147929 | 1 | 1 | 1 | FIM1_4111 | YMR061W | YMR061W | RNA14 | n |
| Kmar_QGN17378.1 | W303.asm02.HP0_G0041040 | OG0003480 | chrXIII | 392035 | 393018 | CP015059.1 | 801383 | 802312 | 35.92814371 | 327 | 309 | 1.058252427 | 1 | 1 | 1 | FIM1_4110 | YMR060C | YMR060C | SAM37 | n |
| Kmar_QGN13835.1 | W303.asm02.HP0_G0031580 | OG0003481 | chrXI | 119963 | 121474 | CP015054.1 | 967486 | 969066 | 39.1221374 | 503 | 526 | 0.956273764 | 1 | 1 | 1 | FIM1_481 | YKL175W | YKL175W | ZRT3 | n |
| Kmar_QGN13672.1 | W303.asm02.HP0_G0048900 | OG0003482 | chrXV | 291336 | 292529 | CP015054.1 | 638587 | 639681 | 48.86649874 | 397 | 364 | 1.090659341 | 1 | 1 | 1 | FIM1_315 | YOL018C | YOL018C | TLG2 | n |
| Kmar_QGN14010.1 | W303.asm02.HP0_G0008610 | OG0003483 | chrIV | 429103 | 430962 | CP015054.1 | 1319809 | 1321647 | 36.49068323 | 619 | 612 | 1.011437908 | 1 | 1 | 1 | FIM1_661 | YDL013W | YDL013W | SLX5 | n |
| Kmar_QGN17256.1 | W303.asm02.HP0_G0022910 | OG0003484 | chrVIII | 64446 | 65660 | CP015059.1 | 557727 | 558992 | 24.83370288 | 404 | 421 | 0.959619952 | 1 | 1 | 1 | FIM1_3987 | YHL020C | YHL020C | OPI1 | n |
| Kmar_QGN14838.1 | W303.asm02.HP0_G0041960 | OG0003485 | chrXIII | 572859 | 573326 | CP015055.1 | 1326363 | 1326830 | 71.61290323 | 155 | 155 | 1 | 1 | 1 | 1 | FIM1_1509 | YMR158W | YMR158W | MRPS8 | n |
| Kmar_QGN14839.1 | W303.asm02.HP0_G0041950 | OG0003486 | chrXIII | 571888 | 572655 | CP015055.1 | 1327369 | 1328022 | 38.82352941 | 255 | 217 | 1.175115207 | 1 | 1 | 1 | FIM1_1510 | YMR157C | YMR157C | AIM36 | n |
| Kmar_QGN14841.1 | W303.asm02.HP0_G0041940 | OG0003487 | chrXIII | 570915 | 571631 | CP015055.1 | 1328949 | 1329665 | 36.94779116 | 238 | 238 | 1 | 1 | 1 | 1 | FIM1_1512 | YMR156C | YMR156C | TPP1 | n |
| Kmar_QGN14840.1 | W303.asm02.HP0_G0007870 | OG0003488 | chrIV | 284910 | 285494 | CP015055.1 | 1328158 | 1328721 | 43.29896907 | 194 | 187 | 1.037433155 | 1 | 1 | 1 | FIM1_1511 | YDL098C | YDL098C | SNU23 | n |
| Kmar_QGN16875.1 | W303.asm02.HP0_G0010180 | OG0003489 | chrIV | 775591 | 779496 | CP015058.1 | 1078927 | 1082919 | 39.13043478 | 1301 | 1330 | 0.978195489 | 1 | 1 | 1 | FIM1_3601 | YDR159W | YDR159W | SAC3 | n |
| Kmar_QGN16874.1 | W303.asm02.HP0_G0036220 | OG0003490 | chrXII | 424959 | 426062 | CP015058.1 | 1077748 | 1078710 | 37.05722071 | 367 | 320 | 1.146875 | 1 | 1 | 1 | FIM1_3600 | YLR137W | YLR137W | RKM5 | n |
| Kmar_QGN16873.1 | W303.asm02.HP0_G0010170 | OG0003491 | chrIV | 774071 | 775168 | CP015058.1 | 1076407 | 1077501 | 81.64383562 | 365 | 364 | 1.002747253 | 1 | 1 | 1 | FIM1_3599 | YDR158W | YDR158W | HOM2 | n |
| Kmar_QGN16541.1 | W303.asm02.HP0_G0039850 | OG0003492 | chrXIII | 130040 | 131698 | CP015058.1 | 336867 | 338540 | 72.62969589 | 552 | 557 | 0.991023339 | 1 | 1 | 1 | FIM1_3254 | YML069W | YML069W | POB3 | n |
| Kmar_QGN16211.1 | W303.asm02.HP0_G0008130 | OG0003493 | chrIV | 330166 | 330777 | CP015057.1 | 1042807 | 1043436 | 41.20171674 | 203 | 209 | 0.971291866 | 1 | 1 | 1 | FIM1_2915 | YDL072C | YDL072C | YET3 | n |
| Kmar_QGN16210.1 | W303.asm02.HP0_G0038640 | OG0003494 | chrXII | 919112 | 921118 | CP015057.1 | 1040597 | 1042654 | 63.05916306 | 668 | 685 | 0.975182482 | 1 | 1 | 1 | FIM1_2914 | YLR401C | YLR401C | DUS3 | n |
| Kmar_QGN16276.1 | W303.asm02.HP0_G0038720 | OG0003495 | chrXII | 931083 | 933902 | CP015057.1 | 1176480 | 1179305 | 67.1957672 | 939 | 941 | 0.997874601 | 1 | 1 | 1 | FIM1_2981 | YLR409C | YLR409C | UTP21 | n |
| Kmar_QGN16275.1 | W303.asm02.HP0_G0008080 | OG0003496 | chrIV | 317301 | 320450 | CP015057.1 | 1173461 | 1176448 | 30.85004776 | 1049 | 995 | 1.054271357 | 1 | 1 | 1 | FIM1_2980 | YDL077C | YDL077C | VAM6 | n |
| Kmar_QGN16793.1 | W303.asm02.HP0_G0020670 | OG0003497 | chrVII | 660826 | 664539 | CP015058.1 | 900393 | 904004 | 63.14516129 | 1237 | 1203 | 1.028262677 | 1 | 1 | 1 | FIM1_3516 | YGR090W | YGR090W | UTP22 | n |
| Kmar_QGN15345.1 | W303.asm02.HP0_G0024000 | OG0003498 | chrVIII | 245072 | 247267 | CP015056.1 | 755741 | 757936 | 70.04103967 | 731 | 731 | 1 | 1 | 1 | 1 | FIM1_2034 | YHR072W | YHR072W | ERG7 | n |
| Kmar_QGN16664.1 | W303.asm02.HP0_G0051710 | OG0003499 | chrXV | 862894 | 867312 | CP015058.1 | 614568 | 618884 | 60.3646185 | 1472 | 1438 | 1.02364395 | 1 | 1 | 1 | FIM1_3383 | YOR291W | YOR291W | YPK9 | n |
| Kmar_QGN16668.1 | W303.asm02.HP0_G0042670 | OG0003500 | chrXIII | 723192 | 724964 | CP015058.1 | 628176 | 630323 | 41.72461752 | 590 | 715 | 0.825174825 | 1 | 1 | 1 | FIM1_3387 | YMR227C | YMR227C | TAF7 | n |
| Kmar_QGN17242.1 | W303.asm02.HP0_G0023000 | OG0003501 | chrVIII | 78992 | 79954 | CP015059.1 | 531340 | 532302 | 92.1875 | 320 | 320 | 1 | 1 | 1 | 1 | FIM1_3973 | YHL011C | YHL011C | PRS3 | n |
| Kmar_QGN16665.1 | W303.asm02.HP0_G0042690 | OG0003502 | chrXIII | 726513 | 731702 | CP015058.1 | 620061 | 625265 | 57.44438106 | 1729 | 1734 | 0.997116494 | 1 | 1 | 1 | FIM1_3384 | YMR229C | YMR229C | RRP5 | n |
| Kmar_QGN16450.1 | W303.asm02.HP0_G0004580 | OG0003503 | chrII | 765516 | 765932 | CP015058.1 | 141857 | 142255 | 61.5942029 | 138 | 132 | 1.045454545 | 1 | 1 | 1 | FIM1_3163 | YBR269C | YBR269C | SDH8 | n |
| Kmar_QGN15214.1 | W303.asm02.HP0_G0029440 | OG0003504 | chrX | 389616 | 391160 | CP015056.1 | 444972 | 446558 | 36.82310469 | 514 | 528 | 0.973484848 | 1 | 1 | 1 | FIM1_1902 | YJL025W | YJL025W | RRN7 | n |
| Kmar_QGN16415.1 | W303.asm02.HP0_G0050880 | OG0003505 | chrXV | 719948 | 721360 | CP015058.1 | 71473 | 72852 | 52.85412262 | 470 | 459 | 1.023965142 | 1 | 1 | 1 | FIM1_3128 | YOR198C | YOR198C | BFR1 | n |
| Kmar_QGN15269.1 | W303.asm02.HP0_G0049630 | OG0003506 | chrXV | 439502 | 440854 | CP015056.1 | 565873 | 567291 | 39.07563025 | 450 | 472 | 0.953389831 | 1 | 1 | 1 | FIM1_1958 | YOR059C | YOR059C | LPL1 | n |
| Kmar_QGN17396.1 | W303.asm02.HP0_G0032390 | OG0003507 | chrXI | 285665 | 286339 | CP015059.1 | 835271 | 836080 | 52.91970803 | 224 | 269 | 0.832713755 | 1 | 1 | 1 | FIM1_4128 | YKL087C | YKL087C | CYT2 | n |
| Kmar_QGN14874.1 | W303.asm02.HP0_G0029750 | OG0003508 | chrX | 456619 | 457830 | CP015055.1 | 1399591 | 1400799 | 55.66502463 | 403 | 402 | 1.002487562 | 1 | 1 | 1 | FIM1_1547 | YJR013W | YJR013W | GPI14 | n |
| Kmar_QGN14875.1 | W303.asm02.HP0_G0021670 | OG0003509 | chrVII | 901297 | 902037 | CP015055.1 | 1401134 | 1401871 | 74.3902439 | 246 | 245 | 1.004081633 | 1 | 1 | 1 | FIM1_1548 | YGR195W | YGR195W | SKI6 | n |
| Kmar_QGN15223.1 | W303.asm02.HP0_G0021590 | OG0003510 | chrVII | 880189 | 882396 | CP015056.1 | 469742 | 471778 | 53.10810811 | 735 | 678 | 1.084070796 | 1 | 1 | 1 | FIM1_1912 | YGR186W | YGR186W | TFG1 | n |
| Kmar_QGN15224.1 | W303.asm02.HP0_G0021600 | OG0003511 | chrVII | 882647 | 883831 | CP015056.1 | 471937 | 473091 | 67.42424242 | 394 | 384 | 1.026041667 | 1 | 1 | 1 | FIM1_1913 | YGR187C | YGR187C | HGH1 | n |
| Kmar_QGN15662.1 | W303.asm02.HP0_G0040120 | OG0003512 | chrXIII | 193146 | 195158 | CP015056.1 | 1411423 | 1413423 | 56.21301775 | 670 | 666 | 1.006006006 | 1 | 1 | 1 | FIM1_2355 | YML042W | YML042W | CAT2 | n |
| Kmar_QGN15192.1 | W303.asm02.HP0_G0021400 | OG0003513 | chrVII | 844231 | 844938 | CP015056.1 | 401761 | 402486 | 44.91525424 | 235 | 241 | 0.975103734 | 1 | 1 | 1 | FIM1_1880 | YGR167W | YGR167W | CLC1 | n |
| Kmar_QGN14715.1 | W303.asm02.HP0_G0026870 | OG0003514 | chrIX | 279855 | 280718 | CP015055.1 | 1084955 | 1085794 | 52.77777778 | 287 | 279 | 1.028673835 | 1 | 1 | 1 | FIM1_1383 | YIL034C | YIL034C | CAP2 | n |
| Kmar_QGN17005.1 | W303.asm02.HP0_G0004830 | OG0003515 | chrII | 820167 | 821891 | CP015059.1 | 12166 | 13923 | 70.5982906 | 574 | 585 | 0.981196581 | 1 | 0 | 1 | FIM1_3734 | YBR296C | YBR296C | PHO89 | n |
| Kmar_QGN13390.1 | W303.asm02.HP0_G0054370 | OG0003516 | chrXVI | 355477 | 356223 | CP015054.1 | 53586 | 54311 | 47.22222222 | 248 | 241 | 1.029045643 | 1 | 1 | 1 | FIM1_27 | YPL107W | YPL107W | DPC25 | n |
| Kmar_QGN17891.1 | W303.asm02.HP0_G0003170 | OG0003517 | chrII | 501027 | 502976 | CP015060.1 | 686403 | 688259 | 52.38828968 | 649 | 618 | 1.050161812 | 1 | 1 | 1 | FIM1_5100 | YBR123C | YBR123C | TFC1 | n |
| Kmar_QGN13383.1 | W303.asm02.HP0_G0003580 | OG0003518 | chrII | 584651 | 585202 | CP015054.1 | 45931 | 46482 | 93.44262295 | 183 | 183 | 1 | 1 | 1 | 1 | FIM1_20 | YBR164C | YBR164C | ARL1 | n |
| Kmar_QGN13385.1 | W303.asm02.HP0_G0003600 | OG0003519 | chrII | 586618 | 587976 | CP015054.1 | 48164 | 49492 | 72.74774775 | 452 | 442 | 1.022624434 | 1 | 1 | 1 | FIM1_22 | YBR166C | YBR166C | TYR1 | n |
| Kmar_QGN13906.1 | W303.asm02.HP0_G0042600 | OG0003520 | chrXIII | 712895 | 714250 | CP015054.1 | 1127476 | 1128825 | 48.01762115 | 451 | 449 | 1.004454343 | 1 | 1 | 1 | FIM1_554 | YMR220W | YMR220W | ERG8 | n |
| Kmar_QGN18318.1 | W303.asm02.HP0_G0055860 | OG0003521 | chrXVI | 667635 | 671171 | CP015061.1 | 681768 | 685022 | 27.25806452 | 1178 | 1084 | 1.086715867 | 1 | 1 | 1 | FIM1_4644 | YPR049C | YPR049C | ATG11 | n |
| Kmar_QGN18312.1 | W303.asm02.HP0_G0002920 | OG0003522 | chrII | 450692 | 451984 | CP015061.1 | 673290 | 674405 | 41.95804196 | 430 | 371 | 1.15902965 | 1 | 1 | 1 | FIM1_4638 | YBR095C | YBR095C | RXT2 | n |
| Kmar_QGN18316.1 | W303.asm02.HP0_G0002860 | OG0003523 | chrII | 441276 | 442052 | CP015061.1 | 679830 | 680606 | 69.37984496 | 258 | 258 | 1 | 1 | 1 | 1 | FIM1_4642 | YBR088C | YBR088C | POL30 | n |
| Kmar_QGN18317.1 | W303.asm02.HP0_G0055870 | OG0003524 | chrXVI | 671458 | 671988 | CP015061.1 | 680703 | 681341 | 56.74418605 | 176 | 212 | 0.830188679 | 1 | 1 | 1 | FIM1_4643 | YPR051W | YPR051W | MAK3 | n |
| Kmar_QGN16144.1 | W303.asm02.HP0_G0032660 | OG0003525 | chrXI | 335012 | 336091 | CP015057.1 | 898929 | 900014 | 77.22222222 | 359 | 361 | 0.994459834 | 1 | 1 | 1 | FIM1_2845 | YKL060C | YKL060C | FBA1 | n |
| Kmar_QGN17825.1 | W303.asm02.HP0_G0040750 | OG0003526 | chrXIII | 326380 | 327792 | CP015060.1 | 570038 | 571396 | 50.31578947 | 470 | 452 | 1.039823009 | 1 | 1 | 1 | FIM1_5034 | YMR027W | YMR027W | | n |
| Kmar_QGN17827.1 | W303.asm02.HP0_G0032680 | OG0003527 | chrXI | 338770 | 339138 | CP015060.1 | 573092 | 573433 | 62.60162602 | 122 | 113 | 1.079646018 | 1 | 1 | 1 | FIM1_5036 | YKL058W | YKL058W | TOA2 | n |
| Kmar_QGN17828.1 | W303.asm02.HP0_G0040760 | OG0003528 | chrXIII | 327985 | 329085 | CP015060.1 | 573688 | 574857 | 44.89795918 | 366 | 389 | 0.940874036 | 1 | 1 | 1 | FIM1_5037 | YMR028W | YMR028W | TAP42 | n |
| Kmar_QGN17829.1 | W303.asm02.HP0_G0032690 | OG0003529 | chrXI | 339460 | 342573 | CP015060.1 | 574867 | 577905 | 32.41965974 | 1037 | 1012 | 1.024703557 | 1 | 1 | 1 | FIM1_5038 | YKL057C | YKL057C | NUP120 | n |
| Kmar_QGN17826.1 | W303.asm02.HP0_G0032670 | OG0003530 | chrXI | 336722 | 338047 | CP015060.1 | 571498 | 572844 | 52.51641138 | 441 | 448 | 0.984375 | 1 | 1 | 1 | FIM1_5035 | YKL059C | YKL059C | MPE1 | n |
| Kmar_QGN17278.1 | W303.asm02.HP0_G0051830 | OG0003531 | chrXV | 884618 | 885853 | CP015059.1 | 601763 | 602992 | 64.55399061 | 411 | 409 | 1.004889976 | 1 | 1 | 1 | FIM1_4009 | YOR303W | YOR303W | CPA1 | n |
| Kmar_QGN16670.1 | W303.asm02.HP0_G0042660 | OG0003532 | chrXIII | 722172 | 722975 | CP015058.1 | 637630 | 638436 | 74.15730337 | 267 | 268 | 0.996268657 | 1 | 1 | 1 | FIM1_3389 | YMR226C | YMR226C | ORA1 | n |
| Kmar_QGN16368.1 | W303.asm02.HP0_G0017990 | OG0003533 | chrVII | 88025 | 89104 | CP015057.1 | 1374926 | 1376122 | 72.18045113 | 359 | 398 | 0.90201005 | 1 | 1 | 1 | FIM1_3075 | YGL211W | YGL211W | NCS6 | n |
| Kmar_QGN17761.1 | W303.asm02.HP0_G0043490 | OG0003534 | chrXIII | 899904 | 902342 | CP015060.1 | 408409 | 410868 | 56.07701564 | 812 | 819 | 0.991452991 | 1 | 1 | 1 | FIM1_4970 | YMR309C | YMR309C | NIP1 | n |
| Kmar_QGN15917.1 | W303.asm02.HP0_G0035860 | OG0003535 | chrXII | 344193 | 345227 | CP015057.1 | 374495 | 375514 | 48.99135447 | 344 | 339 | 1.014749263 | 1 | 1 | 1 | FIM1_2615 | YLR097C | YLR097C | HRT3 | n |
| Kmar_QGN16684.1 | W303.asm02.HP0_G0036820 | OG0003536 | chrXII | 544096 | 545826 | CP015058.1 | 665449 | 667179 | 69.16524702 | 576 | 576 | 1 | 1 | 1 | 1 | FIM1_3403 | YLR196W | YLR196W | PWP1 | n |
| Kmar_QGN16683.1 | W303.asm02.HP0_G0036810 | OG0003537 | chrXII | 542065 | 543432 | CP015058.1 | 663558 | 664907 | 62.80087527 | 455 | 449 | 1.013363029 | 1 | 1 | 1 | FIM1_3402 | YLR195C | YLR195C | NMT1 | n |
| Kmar_QGN13445.1 | W303.asm02.HP0_G0003930 | OG0003538 | chrII | 637647 | 639302 | CP015054.1 | 161842 | 163857 | 53.34420881 | 551 | 604 | 0.912251656 | 1 | 1 | 1 | FIM1_82 | YBR200W | YBR200W | BEM1 | n |
| Kmar_QGN15160.1 | W303.asm02.HP0_G0005440 | OG0003539 | chrIII | 109859 | 110788 | CP015056.1 | 327266 | 328153 | 71.84466019 | 309 | 295 | 1.047457627 | 1 | 1 | 1 | FIM1_1847 | YCL009C | YCL009C | ILV6 | n |

| | | | | | | | | | | | | | | | | | | | | |
|---|---|---|---|---|---|---|---|---|---|---|---|---|---|---|---|---|---|---|---|---|
| Kmar_QGN14057.1 | W303.asm02.HP0_G0027850 | OG0003540 | chrX | 42872 | 43714 | CP015054.1 | 1433250 | 1434359 | 22.43243243 | 280 | 369 | 0.758807588 | 1 | 1 | 1 | FIM1_708 | YJL203W | YJL203W | PRP21 | n |
| Kmar_QGN17229.1 | W303.asm02.HP0_G0041130 | OG0003541 | chrXIII | 408213 | 409070 | CP015059.1 | 497112 | 497762 | 22.34042553 | 285 | 216 | 1.319444444 | 1 | 1 | 1 | FIM1_3960 | YMR069W | YMR069W | NAT4 | n |
| Kmar_QGN17835.1 | W303.asm02.HP0_G0032740 | OG0003542 | chrXI | 348272 | 349150 | CP015060.1 | 586326 | 587054 | 30.84745763 | 292 | 242 | 1.20661157 | 1 | 1 | 1 | FIM1_5044 | YKL052C | YKL052C | ASK1 | n |
| Kmar_QGN17834.1 | W303.asm02.HP0_G0032730 | OG0003543 | chrXI | 347786 | 348046 | CP015060.1 | 585854 | 586108 | 75.58139535 | 86 | 84 | 1.023809524 | 1 | 1 | 1 | FIM1_5043 | YKL053C-A | YKL053C-A | MDM35 | n |
| Kmar_QGN15661.1 | W303.asm02.HP0_G0040130 | OG0003544 | chrXIII | 195271 | 196113 | CP015056.1 | 1410288 | 1411040 | 46.42857143 | 280 | 250 | 1.12 | 1 | 1 | 1 | FIM1_2354 | YML041C | YML041C | VPS71 | n |
| Kmar_QGN14569.1 | W303.asm02.HP0_G0049550 | OG0003545 | chrXV | 419086 | 422106 | CP015055.1 | 777976 | 780699 | 60.1980198 | 1006 | 907 | 1.109151047 | 1 | 1 | 1 | FIM1_1231 | YOR048C | YOR048C | RAT1 | n |
| Kmar_QGN13693.1 | W303.asm02.HP0_G0043530 | OG0003546 | chrXIII | 906295 | 908223 | CP015054.1 | 672492 | 674273 | 54.4049459 | 642 | 593 | 1.082630691 | 1 | 1 | 1 | FIM1_336 | YMR313C | YMR313C | TGL3 | n |
| Kmar_QGN13692.1 | W303.asm02.HP0_G0043520 | OG0003547 | chrXIII | 905321 | 906142 | CP015054.1 | 671641 | 672447 | 37.09090909 | 273 | 268 | 1.018656716 | 1 | 1 | 1 | FIM1_335 | YMR312W | YMR312W | ELP6 | n |
| Kmar_QGN15301.1 | W303.asm02.HP0_G0016150 | OG0003548 | chrV | 528731 | 530371 | CP015056.1 | 640124 | 641599 | 42.49084249 | 546 | 491 | 1.112016293 | 1 | 1 | 1 | FIM1_1990 | YER168C | YER168C | CCA1 | n |
| Kmar_QGN17805.1 | W303.asm02.HP0_G0050030 | OG0003549 | chrXV | 518377 | 519225 | CP015060.1 | 520199 | 521077 | 32.1192053 | 282 | 292 | 0.965753425 | 1 | 1 | 1 | FIM1_5014 | YOR104W | YOR104W | PIN2 | n |
| Kmar_QGN18067.1 | W303.asm02.HP0_G0008260 | OG0003550 | chrIV | 353209 | 355710 | CP015061.1 | 143148 | 145424 | 43.30143541 | 833 | 758 | 1.098944591 | 1 | 1 | 1 | FIM1_4385 | YDL056W | YDL056W | MBP1 | n |
| Kmar_QGN16260.1 | W303.asm02.HP0_G0025210 | OG0003551 | chrVIII | 492928 | 493764 | CP015057.1 | 1148905 | 1149726 | 31.42857143 | 278 | 273 | 1.018315018 | 1 | 1 | 1 | FIM1_2965 | YHR192W | YHR192W | LNP1 | n |
| Kmar_QGN16259.1 | W303.asm02.HP0_G0025220 | OG0003552 | chrVIII | 493814 | 494338 | CP015057.1 | 1148238 | 1148759 | 62.64367816 | 174 | 173 | 1.005780347 | 1 | 1 | 1 | FIM1_2964 | YHR193C | YHR193C | EGD2 | n |
| Kmar_QGN17089.1 | W303.asm02.HP0_G0046830 | OG0003553 | chrXIV | 659585 | 661090 | CP015059.1 | 212950 | 214326 | 60.55776892 | 501 | 458 | 1.093886463 | 1 | 1 | 1 | FIM1_3820 | YNR012W | YNR012W | URK1 | n |
| Kmar_QGN17808.1 | W303.asm02.HP0_G0045760 | OG0003554 | chrXIV | 431742 | 436148 | CP015060.1 | 525283 | 529584 | 61.47596479 | 1468 | 1433 | 1.024424285 | 1 | 1 | 1 | FIM1_5017 | YNL102W | YNL102W | POL1 | n |
| Kmar_QGN17853.1 | W303.asm02.HP0_G0021840 | OG0003555 | chrVII | 929455 | 930861 | CP015060.1 | 619971 | 621293 | 25.42735043 | 468 | 440 | 1.063636364 | 1 | 1 | 1 | FIM1_5062 | YGR212W | YGR212W | SLI1 | n |
| Kmar_QGN13482.1 | W303.asm02.HP0_G0029950 | OG0003556 | chrX | 516277 | 518616 | CP015054.1 | 231058 | 233340 | 50.69708492 | 779 | 760 | 1.025 | 1 | 1 | 1 | FIM1_120 | YJR040W | YJR040W | GEF1 | n |
| Kmar_QGN13483.1 | W303.asm02.HP0_G0029960 | OG0003557 | chrX | 518770 | 522294 | CP015054.1 | 233380 | 236814 | 28.98913952 | 1174 | 1144 | 1.026223776 | 1 | 1 | 1 | FIM1_121 | YJR041C | YJR041C | URB2 | n |
| Kmar_QGN13481.1 | W303.asm02.HP0_G0014220 | OG0003558 | chrV | 103126 | 103506 | CP015054.1 | 230091 | 230471 | 96.03174603 | 126 | 126 | 1 | 1 | 1 | 1 | FIM1_119 | YEL026W | YEL026W | SNU13 | n |
| Kmar_QGN13484.1 | W303.asm02.HP0_G0029970 | OG0003559 | chrX | 522586 | 524820 | CP015054.1 | 237436 | 239601 | 42.32309746 | 744 | 721 | 1.031900139 | 1 | 1 | 1 | FIM1_122 | YJR042W | YJR042W | NUP85 | n |
| Kmar_QGN14510.1 | W303.asm02.HP0_G0042850 | OG0003560 | chrXIII | 763755 | 764822 | CP015055.1 | 658280 | 659344 | 59.5567867 | 355 | 354 | 1.002824859 | 1 | 1 | 1 | FIM1_1171 | YMR244W | YMR244W | | n |
| Kmar_QGN18116.1 | W303.asm02.HP0_G0000940 | OG0003561 | chrII | 18481 | 21597 | CP015061.1 | 247049 | 249940 | 58.88568684 | 1038 | 963 | 1.07788162 | 1 | 1 | 1 | FIM1_4436 | YBL104C | YBL104C | SEA4 | n |
| Kmar_QGN18115.1 | W303.asm02.HP0_G0000930 | OG0003562 | chrII | 15782 | 18000 | CP015061.1 | 242994 | 246446 | 38.79310345 | 709 | 1150 | 0.616521739 | 1 | 1 | 1 | FIM1_4435 | YBL105C | YBL105C | PKC1 | n |
| Kmar_QGN17276.1 | W303.asm02.HP0_G0051820 | OG0003563 | chrXV | 882684 | 883991 | CP015059.1 | 599419 | 600705 | 41.94260486 | 435 | 428 | 1.01635514 | 1 | 1 | 1 | FIM1_4007 | YOR301W | YOR301W | RAX1 | n |
| Kmar_QGN18336.1 | W303.asm02.HP0_G0001630 | OG0003564 | chrII | 169830 | 170150 | CP015061.1 | 717718 | 718056 | 58.03571429 | 106 | 112 | 0.946428571 | 1 | 1 | 1 | FIM1_4662 | YBL028C | YBL028C | | n |
| Kmar_QGN14783.1 | W303.asm02.HP0_G0050250 | OG0003565 | chrXV | 565351 | 567066 | CP015055.1 | 1218915 | 1220624 | 70.92819615 | 537 | 569 | 0.943760984 | 1 | 1 | 1 | FIM1_1453 | YOR128C | YOR128C | ADE2 | n |
| Kmar_QGN18117.1 | W303.asm02.HP0_G0055700 | OG0003566 | chrXVI | 638012 | 640258 | CP015061.1 | 250137 | 252305 | 40.89709763 | 748 | 722 | 1.03601108 | 1 | 1 | 1 | FIM1_4437 | YPR031W | YPR031W | NTO1 | n |
| Kmar_QGN18340.1 | W303.asm02.HP0_G0009270 | OG0003567 | chrIV | 574798 | 577875 | CP015061.1 | 728288 | 731368 | 54.75285171 | 1025 | 1026 | 0.999025341 | 1 | 1 | 1 | FIM1_4667 | YDR060W | YDR060W | MAK21 | n |
| Kmar_QGN14780.1 | W303.asm02.HP0_G0050260 | OG0003568 | chrXV | 567773 | 570433 | CP015055.1 | 1211930 | 1214695 | 30.11542497 | 886 | 921 | 0.961997828 | 1 | 1 | 1 | FIM1_1450 | YOR129C | YOR129C | AFI1 | n |
| Kmar_QGN14778.1 | W303.asm02.HP0_G0012330 | OG0003569 | chrIV | 1243905 | 1244443 | CP015055.1 | 1208584 | 1208907 | 50.87719298 | 114 | 107 | 1.065420561 | 1 | 1 | 1 | FIM1_1448 | YDR381C-A | YDR381C-A | COI1 | n |
| Kmar_QGN16972.1 | W303.asm02.HP0_G0006600 | OG0003570 | chrIV | 30732 | 31904 | CP015058.1 | 1284253 | 1285479 | 35.33653846 | 390 | 408 | 0.955882353 | 1 | 1 | 1 | FIM1_3699 | YDL237W | YDL237W | AIM6 | n |
| Kmar_QGN16932.1 | W303.asm02.HP0_G0010530 | OG0003571 | chrIV | 847284 | 848576 | CP015058.1 | 1212056 | 1213777 | 24.78336222 | 430 | 573 | 0.7504363 | 1 | 1 | 1 | FIM1_3659 | YDR192C | YDR192C | NUP42 | n |
| Kmar_QGN16933.1 | W303.asm02.HP0_G0037220 | OG0003572 | chrXII | 618148 | 619113 | CP015058.1 | 1214109 | 1215020 | 39.19753086 | 321 | 303 | 1.059405941 | 1 | 1 | 1 | FIM1_3660 | YLR242C | YLR242C | ARV1 | n |
| Kmar_QGN15965.1 | W303.asm02.HP0_G0037150 | OG0003573 | chrXII | 605013 | 606983 | CP015057.1 | 496177 | 498183 | 53.63372093 | 656 | 668 | 0.982035928 | 1 | 1 | 1 | FIM1_2663 | YLR234W | YLR234W | TOP3 | n |
| Kmar_QGN16563.1 | W303.asm02.HP0_G0038920 | OG0003574 | chrXII | 990103 | 996798 | CP015058.1 | 380512 | 386514 | 42.44572441 | 2231 | 2000 | 1.1155 | 1 | 1 | 1 | FIM1_3277 | YLR430W | YLR430W | SEN1 | n |
| Kmar_QGN16542.1 | W303.asm02.HP0_G0039860 | OG0003576 | chrXIII | 132090 | 133484 | CP015058.1 | 338781 | 340148 | 36.8869936 | 464 | 455 | 1.01978022 | 1 | 1 | 1 | FIM1_3255 | YML068W | YML068W | ITT1 | n |
| Kmar_QGN17510.1 | W303.asm02.HP0_G0024790 | OG0003577 | chrVIII | 403357 | 405096 | CP015059.1 | 1060708 | 1062360 | 32.13058419 | 579 | 550 | 1.052727273 | 1 | 1 | 1 | FIM1_4246 | YHR150W | YHR150W | PEX28 | n |
| Kmar_QGN16902.1 | W303.asm02.HP0_G0010300 | OG0003578 | chrIV | 799907 | 805936 | CP015058.1 | 1137377 | 1142968 | 56.74074074 | 2009 | 1863 | 1.078368223 | 1 | 1 | 1 | FIM1_3629 | YDR170C | YDR170C | SEC7 | n |
| Kmar_QGN18218.1 | W303.asm02.HP0_G0007370 | OG0003579 | chrIV | 188361 | 190793 | CP015061.1 | 474972 | 477458 | 52.68817204 | 810 | 828 | 0.97826087 | 1 | 1 | 1 | FIM1_4543 | YDL148C | YDL148C | NOP14 | n |
| Kmar_QGN17516.1 | W303.asm02.HP0_G0022010 | OG0003580 | chrVII | 966375 | 967061 | CP015059.1 | 1070979 | 1071674 | 51.30434783 | 228 | 231 | 0.987012987 | 1 | 1 | 1 | FIM1_4252 | YGR232W | YGR232W | NAS6 | n |
| Kmar_QGN17050.1 | W303.asm02.HP0_G0007000 | OG0003581 | chrIV | 114746 | 115873 | CP015059.1 | 125381 | 126430 | 49.47089947 | 375 | 349 | 1.074498567 | 1 | 1 | 1 | FIM1_3779 | YDL193W | YDL193W | NUS1 | n |
| Kmar_QGN17052.1 | W303.asm02.HP0_G0007030 | OG0003582 | chrIV | 118781 | 121666 | CP015059.1 | 131085 | 133958 | 60.97308489 | 961 | 957 | 1.004179728 | 1 | 1 | 1 | FIM1_3782 | YDL190C | YDL190C | UFD2 | n |
| Kmar_QGN17053.1 | W303.asm02.HP0_G0007040 | OG0003583 | chrIV | 122290 | 123663 | CP015059.1 | 134880 | 136238 | 20.65217391 | 457 | 452 | 1.011061947 | 1 | 1 | 1 | FIM1_3783 | YDL189W | YDL189W | RBS1 | n |
| Kmar_QGN17054.1 | W303.asm02.HP0_G0007500 | OG0003584 | chrIV | 218886 | 219494 | CP015059.1 | 136440 | 137039 | 80.1980198 | 202 | 199 | 1.015075377 | 1 | 1 | 1 | FIM1_3784 | YDL135C | YDL135C | RDI1 | n |
| Kmar_QGN17511.1 | W303.asm02.HP0_G0024800 | OG0003585 | chrVIII | 405374 | 406954 | CP015059.1 | 1062476 | 1064251 | 31.56146179 | 526 | 591 | 0.89001692 | 1 | 1 | 1 | FIM1_4247 | YHR151C | YHR151C | MTC6 | n |
| Kmar_QGN17515.1 | W303.asm02.HP0_G0022000 | OG0003586 | chrVII | 964963 | 965895 | CP015059.1 | 1069475 | 1070398 | 77.85016287 | 310 | 307 | 1.009771987 | 1 | 1 | 1 | FIM1_4251 | YGR231C | YGR231C | PHB2 | n |
| Kmar_QGN18219.1 | W303.asm02.HP0_G0007360 | OG0003587 | chrIV | 185132 | 188125 | CP015061.1 | 477503 | 480223 | 44.83450351 | 997 | 906 | 1.100441501 | 1 | 1 | 1 | FIM1_4544 | YDL149W | YDL149W | ATG9 | n |
| Kmar_QGN18168.1 | W303.asm02.HP0_G0015130 | OG0003588 | chrV | 279968 | 281296 | CP015061.1 | 362736 | 364031 | 44.14414414 | 442 | 431 | 1.025522042 | 1 | 1 | 1 | FIM1_4491 | YER061C | YER061C | CEM1 | n |
| Kmar_QGN14004.1 | W303.asm02.HP0_G0048370 | OG0003589 | chrXV | 194287 | 195255 | CP015054.1 | 1310084 | 1310920 | 37.5 | 322 | 278 | 1.158273381 | 1 | 1 | 1 | FIM1_655 | YOL073C | YOL073C | DSC2 | n |
| Kmar_QGN16381.1 | W303.asm02.HP0_G0021170 | OG0003590 | chrVII | 792126 | 793106 | CP015057.1 | 1402216 | 1403193 | 75.2293578 | 326 | 325 | 1.003076923 | 1 | 1 | 1 | FIM1_3089 | YGR144W | YGR144W | THI4 | n |
| Kmar_QGN17823.1 | W303.asm02.HP0_G0047880 | OG0003591 | chrXV | 85796 | 87097 | CP015060.1 | 566548 | 567861 | 72.04545455 | 433 | 437 | 0.990846682 | 1 | 1 | 1 | FIM1_5032 | YOL124C | YOL124C | TRM11 | n |
| Kmar_QGN17287.1 | W303.asm02.HP0_G0008460 | OG0003592 | chrIV | 394260 | 397247 | CP015059.1 | 622816 | 625761 | 69.50852558 | 995 | 981 | 1.014271152 | 1 | 1 | 1 | FIM1_4018 | YDL031W | YDL031W | DBP10 | n |
| Kmar_QGN13642.1 | W303.asm02.HP0_G0005790 | OG0003593 | chrIII | 179284 | 179913 | CP015054.1 | 569310 | 569882 | 39.53488372 | 209 | 190 | 1.1 | 1 | 1 | 1 | FIM1_284 | YCR027C | YCR027C | RHB1 | n |
| Kmar_QGN13877.1 | W303.asm02.HP0_G0033770 | OG0003594 | chrXI | 545313 | 557591 | CP015054.1 | 1039613 | 1051789 | 34.60510328 | 4092 | 4058 | 1.008378512 | 1 | 1 | 1 | FIM1_524 | YKR054C | YKR054C | DYN1 | n |
| Kmar_QGN14627.1 | W303.asm02.HP0_G0034620 | OG0003595 | chrXII | 74279 | 76756 | CP015055.1 | 893709 | 895883 | 25.84541063 | 825 | 724 | 1.139502762 | 1 | 1 | 1 | FIM1_1291 | YLL032C | YLL032C | | n |
| Kmar_QGN15195.1 | W303.asm02.HP0_G0048430 | OG0003596 | chrXV | 202440 | 202973 | CP015056.1 | 407067 | 407648 | 39.53488372 | 177 | 193 | 0.917098446 | 1 | 1 | 1 | FIM1_1883 | YOL067C | YOL067C | RTG1 | n |
| Kmar_QGN16036.1 | W303.asm02.HP0_G0008390 | OG0003597 | chrIV | 376566 | 377366 | CP015057.1 | 675763 | 676524 | 49.04214559 | 266 | 253 | 1.051383399 | 1 | 1 | 1 | FIM1_2736 | YDL043C | YDL043C | PRP11 | n |
| Kmar_QGN14124.1 | W303.asm02.HP0_G0036690 | OG0003598 | chrXII | 518069 | 520480 | CP015054.1 | 1587191 | 1589572 | 37.96407186 | 803 | 793 | 1.01261034 | 1 | 1 | 1 | FIM1_777 | YLR182W | YLR182W | SWI6 | n |
| Kmar_QGN14795.1 | W303.asm02.HP0_G0012220 | OG0003599 | chrIV | 1228352 | 1228924 | CP015055.1 | 1244178 | 1244750 | 78.42105263 | 190 | 190 | 1 | 1 | 1 | 1 | FIM1_1466 | YDR373W | YDR373W | FRQ1 | n |
| Kmar_QGN18378.1 | W303.asm02.HP0_G0048390 | OG0003600 | chrXV | 196962 | 197450 | CP015061.1 | 808006 | 808578 | 54.21052632 | 162 | 190 | 0.852631579 | 1 | 1 | 1 | FIM1_4705 | YOL071W | YOL071W | SDH5 | n |
| Kmar_QGN17733.1 | W303.asm02.HP0_G0026170 | OG0003601 | chrIX | 141952 | 145203 | CP015060.1 | 347281 | 350328 | 34.71596032 | 1083 | 1015 | 1.066995074 | 1 | 1 | 1 | FIM1_4942 | YIL112W | YIL112W | HOS4 | n |
| Kmar_QGN17730.1 | W303.asm02.HP0_G0046240 | OG0003602 | chrXIV | 543842 | 545053 | CP015060.1 | 342627 | 343811 | 36.47342995 | 403 | 394 | 1.02284264 | 1 | 1 | 1 | FIM1_4939 | YNL051W | YNL051W | COG5 | n |
| Kmar_QGN16053.1 | W303.asm02.HP0_G0018470 | OG0003603 | chrVII | 199189 | 200301 | CP015057.1 | 709271 | 710293 | 38.87399464 | 370 | 340 | 1.088235294 | 1 | 1 | 1 | FIM1_2753 | YGL159W | YGL159W | | n |
| Kmar_QGN14108.1 | W303.asm02.HP0_G0050480 | OG0003604 | chrXV | 625604 | 627367 | CP015054.1 | 1547588 | 1549288 | 38.49840256 | 587 | 566 | 1.037102473 | 1 | 1 | 1 | FIM1_759 | YOR154W | YOR154W | SLP1 | n |
| Kmar_QGN16936.1 | W303.asm02.HP0_G0010540 | OG0003605 | chrIV | 849667 | 851661 | CP015058.1 | 1220815 | 1222827 | 40.91580502 | 664 | 670 | 0.991044776 | 1 | 1 | 1 | FIM1_3663 | YDR194C | YDR194C | MSS116 | n |
| Kmar_QGN13935.1 | W303.asm02.HP0_G0025180 | OG0003606 | chrVIII | 490130 | 490702 | CP015054.1 | 1174260 | 1174811 | 36.31578947 | 190 | 183 | 1.038251366 | 1 | 1 | 1 | FIM1_583 | YHR189W | YHR189W | PTH1 | n |
| Kmar_QGN18087.1 | W303.asm02.HP0_G0030620 | OG0003607 | chrX | 643388 | 645454 | CP015061.1 | 179769 | 181682 | 47.11815562 | 688 | 637 | 1.080062794 | 1 | 1 | 1 | FIM1_4406 | YJR110W | YJR110W | YMR1 | n |
| Kmar_QGN18255.1 | W303.asm02.HP0_G0056920 | OG0003608 | chrXVI | 884456 | 886000 | CP015061.1 | 561817 | 563382 | 45.12635379 | 514 | 521 | 0.986564299 | 1 | 1 | 1 | FIM1_4580 | YPR169W | YPR169W | JIP5 | n |
| Kmar_QGN18296.1 | W303.asm02.HP0_G0014610 | OG0003609 | chrV | 183747 | 185366 | CP015061.1 | 641366 | 642979 | 48.42883549 | 539 | 537 | 1.003724395 | 1 | 1 | 1 | FIM1_4622 | YER014W | YER014W | HEM14 | n |
| Kmar_QGN18295.1 | W303.asm02.HP0_G0027310 | OG0003610 | chrIX | 365693 | 366028 | CP015061.1 | 640772 | 641203 | 38.19444444 | 111 | 143 | 0.776223776 | 1 | 1 | 1 | FIM1_4621 | YIR009W | YIR009W | MSL1 | n |
| Kmar_QGN17702.1 | W303.asm02.HP0_G0026350 | OG0003611 | chrIX | 178485 | 179279 | CP015060.1 | 280053 | 280835 | 57.57575758 | 264 | 260 | 1.015384615 | 1 | 1 | 1 | FIM1_4909 | YIL093C | YIL093C | RSM25 | n |
| Kmar_QGN17726.1 | W303.asm02.HP0_G0026220 | OG0003612 | chrIX | 153636 | 156119 | CP015060.1 | 333010 | 335355 | 50.90252708 | 827 | 781 | 1.058898848 | 1 | 1 | 1 | FIM1_4935 | YIL107C | YIL107C | PFK26 | n |
| Kmar_QGN16987.1 | W303.asm02.HP0_G0044560 | OG0003613 | chrXIV | 205884 | 207263 | CP015058.1 | 1316984 | 1318273 | 57.29847495 | 459 | 429 | 1.06993007 | 1 | 1 | 1 | FIM1_3714 | YNL237W | YNL237W | YTP1 | n |
| Kmar_QGN16989.1 | W303.asm02.HP0_G0044530 | OG0003614 | chrXIV | 199055 | 200673 | CP015058.1 | 1321880 | 1323271 | 48.57723577 | 491 | 463 | 1.060475162 | 1 | 1 | 1 | FIM1_3716 | YNL240C | YNL240C | NAR1 | n |
| Kmar_QGN14541.1 | W303.asm02.HP0_G0000630 | OG0003615 | chrI | 137680 | 138276 | CP015055.1 | 722226 | 722744 | 28 | 198 | 172 | 1.151162791 | 1 | 1 | 1 | FIM1_1203 | YAL008W | YAL008W | FUN14 | n |
| Kmar_QGN17216.1 | W303.asm02.HP0_G0019050 | OG0003616 | chrVII | 328070 | 331417 | CP015059.1 | 468131 | 471571 | 51.08601216 | 1115 | 1146 | 0.972949389 | 1 | 1 | 1 | FIM1_3947 | YGL094C | YGL094C | PAN2 | n |
| Kmar_QGN14220.1 | W303.asm02.HP0_G0009390 | OG0003617 | chrIV | 594037 | 595620 | CP015055.1 | 58489 | 60126 | 44.48462929 | 527 | 545 | 0.966972477 | 1 | 1 | 1 | FIM1_876 | YDR072C | YDR072C | IPT1 | n |
| Kmar_QGN18135.1 | W303.asm02.HP0_G0015730 | OG0003618 | chrV | 399974 | 401077 | CP015061.1 | 297127 | 298209 | 47.15447154 | 367 | 360 | 1.019444444 | 1 | 1 | 1 | FIM1_4457 | YER118C | YER118C | SHO1 | n |
| Kmar_QGN13551.1 | W303.asm02.HP0_G0019240 | OG0003619 | chrVII | 365706 | 368207 | CP015054.1 | 400374 | 402401 | 27.6546091 | 833 | 675 | 1.234074074 | 1 | 1 | 1 | FIM1_192 | YGL073W | YGL073W | HSF1 | n |
| Kmar_QGN15197.1 | W303.asm02.HP0_G0008660 | OG0003620 | chrIV | 439568 | 440413 | CP015056.1 | 409925 | 410830 | 73.97260274 | 281 | 301 | 0.933554817 | 1 | 1 | 1 | FIM1_1885 | YDL006W | YDL006W | PTC1 | n |
| Kmar_QGN18156.1 | W303.asm02.HP0_G0015210 | OG0003621 | chrV | 297080 | 299671 | CP015061.1 | 341452 | 344031 | 75.54535017 | 863 | 859 | 1.004656577 | 1 | 1 | 1 | FIM1_4479 | YER069W | YER069W | ARG56 | n |
| Kmar_QGN15463.1 | W303.asm02.HP0_G0051350 | OG0003622 | chrXV | 805390 | 805926 | CP015056.1 | 1003919 | 1004494 | 62.92134831 | 178 | 191 | 0.931937173 | 1 | 1 | 1 | FIM1_2154 | YOR252W | YOR252W | TMA16 | n |
| Kmar_QGN13897.1 | W303.asm02.HP0_G0051590 | OG0003623 | chrXV | 843052 | 843537 | CP015054.1 | 1104743 | 1105288 | 46.90721649 | 161 | 181 | 0.889502762 | 1 | 1 | 1 | FIM1_544 | YOR276W | YOR276W | CAF20 | n |
| Kmar_QGN16181.1 | W303.asm02.HP0_G0042520 | OG0003624 | chrXIII | 691274 | 693622 | CP015057.1 | 974320 | 976533 | 44.98094028 | 782 | 737 | 1.061058345 | 1 | 1 | 1 | FIM1_2883 | YMR212C | YMR212C | EFR3 | n |
| Kmar_QGN16182.1 | W303.asm02.HP0_G0042530 | OG0003625 | chrXIII | 693960 | 695732 | CP015057.1 | 977197 | 978813 | 40.76539101 | 590 | 538 | 1.096654275 | 1 | 1 | 1 | FIM1_2884 | YMR213W | YMR213W | CEF1 | n |
| Kmar_QGN16180.1 | W303.asm02.HP0_G0051580 | OG0003626 | chrXV | 840803 | 842788 | CP015057.1 | 972165 | 974117 | 34.08748115 | 661 | 650 | 1.016923077 | 1 | 1 | 1 | FIM1_2882 | YOR275C | YOR275C | RIM20 | n |
| Kmar_QGN16179.1 | W303.asm02.HP0_G0051570 | OG0003627 | chrXV | 839393 | 840679 | CP015057.1 | 970886 | 972163 | 60.13986014 | 428 | 425 | 1.007058824 | 1 | 1 | 1 | FIM1_2881 | YOR274W | YOR274W | MOD5 | n |
| Kmar_QGN14258.1 | W303.asm02.HP0_G0039530 | OG0003628 | chrXIII | 57140 | 62098 | CP015055.1 | 127074 | 131903 | 35.93099346 | 1652 | 1609 | 1.026724674 | 1 | 1 | 1 | FIM1_915 | YML103C | YML103C | NUP188 | n |
| Kmar_QGN16173.1 | W303.asm02.HP0_G0042480 | OG0003629 | chrXIII | 685046 | 686377 | CP015057.1 | 958399 | 959658 | 49.44320713 | 443 | 419 | 1.057279236 | 1 | 1 | 1 | FIM1_2875 | YMR208W | YMR208W | ERG12 | n |
| Kmar_QGN16174.1 | W303.asm02.HP0_G0051540 | OG0003630 | chrXV | 832778 | 833761 | CP015057.1 | 959963 | 960946 | 81.65137615 | 327 | 327 | 1 | 1 | 1 | 1 | FIM1_2876 | YOR271C | YOR271C | FSF1 | n |
| Kmar_QGN14393.1 | W303.asm02.HP0_G0053970 | OG0003631 | chrXVI | 278228 | 278761 | CP015055.1 | 428866 | 429489 | 28.50678733 | 177 | 207 | 0.855072464 | 1 | 1 | 1 | FIM1_1053 | YPL148C | YPL148C | PPT2 | n |
| Kmar_QGN13902.1 | W303.asm02.HP0_G0042570 | OG0003632 | chrXIII | 702369 | 703946 | CP015054.1 | 1116316 | 1117890 | 87.23809524 | 525 | 524 | 1.001908397 | 1 | 1 | 1 | FIM1_550 | YMR217W | YMR217W | GUA1 | n |
| Kmar_QGN16175.1 | W303.asm02.HP0_G0051550 | OG0003633 | chrXV | 834532 | 835914 | CP015057.1 | 962520 | 963854 | 79.34782609 | 460 | 444 | 1.036036036 | 1 | 1 | 1 | FIM1_2877 | YOR272W | YOR272W | YTM1 | n |
| Kmar_QGN16176.1 | W303.asm02.HP0_G0042490 | OG0003634 | chrXIII | 686490 | 687863 | CP015057.1 | 963992 | 965275 | 39.95633188 | 457 | 427 | 1.070257611 | 1 | 1 | 1 | FIM1_2878 | YMR209C | YMR209C | | n |
| Kmar_QGN17415.1 | W303.asm02.HP0_G0033070 | OG0003635 | chrXI | 411660 | 412610 | CP015059.1 | 867228 | 868175 | 50.7788162 | 316 | 315 | 1.003174603 | 1 | 1 | 1 | FIM1_4147 | YKL019W | YKL019W | RAM2 | n |
| Kmar_QGN17715.1 | W303.asm02.HP0_G0026300 | OG0003636 | chrIX | 170130 | 170597 | CP015060.1 | 303504 | 303989 | 35.40372671 | 155 | 161 | 0.962732919 | 1 | 1 | 1 | FIM1_4922 | YIL098C | YIL098C | FMC1 | n |
| Kmar_QGN16970.1 | W303.asm02.HP0_G0044700 | OG0003637 | chrXIV | 235109 | 236410 | CP015058.1 | 1278219 | 1279523 | 84.06466513 | 433 | 434 | 0.997695853 | 1 | 1 | 1 | FIM1_3697 | YNL220W | YNL220W | ADE12 | n |
| Kmar_QGN15042.1 | W303.asm02.HP0_G0031230 | OG0003638 | chrXI | 32982 | 35129 | CP015056.1 | 76738 | 78915 | 53.06957708 | 715 | 725 | 0.986206897 | 1 | 1 | 1 | FIM1_1727 | YKL213C | YKL213C | DOA1 | n |
| Kmar_QGN15489.1 | W303.asm02.HP0_G0015360 | OG0003639 | chrV | 325942 | 327606 | CP015056.1 | 1050285 | 1051961 | 72.27191413 | 554 | 558 | 0.992831541 | 1 | 1 | 1 | FIM1_2180 | YER082C | YER082C | UTP7 | n |

| | | | | | | | | | | | | | | | | | | | | |
|---|---|---|---|---|---|---|---|---|---|---|---|---|---|---|---|---|---|---|---|---|
| Kmar_QGN15492.1 | W303.asm02.HP0_G0015370 | OG0003640 | chrV | 327844 | 328701 | CP015056.1 | 1055820 | 1056833 | 37.38317757 | 285 | 337 | 0.845697329 | 1 | 1 | 1 | FIM1_2183 | YER083C | YER083C | GET2 | n |
| Kmar_QGN18289.1 | W303.asm02.HP0_G0020880 | OG0003641 | chrVII | 717141 | 718310 | CP015061.1 | 624069 | 625175 | 53.58974359 | 389 | 368 | 1.057065217 | 1 | 1 | 1 | FIM1_4615 | YGR112W | YGR112W | SHY1 | n |
| Kmar_QGN17496.1 | W303.asm02.HP0_G0035380 | OG0003642 | chrXII | 247160 | 249220 | CP015059.1 | 1030422 | 1032596 | 47.05075446 | 686 | 724 | 0.947513812 | 1 | 1 | 1 | FIM1_4232 | YLR047C | YLR047C | FRE8 | n |
| Kmar_QGN17494.1 | W303.asm02.HP0_G0021820 | OG0003644 | chrVII | 925918 | 927153 | CP015059.1 | 1027204 | 1028433 | 85.15815085 | 411 | 409 | 1.004889976 | 1 | 1 | 1 | FIM1_4230 | YGR210C | YGR210C | | n |
| Kmar_QGN16229.1 | W303.asm02.HP0_G0057110 | OG0003645 | chrXVI | 918496 | 922794 | CP015057.1 | 1078973 | 1083175 | 44.0027894 | 1432 | 1400 | 1.022857143 | 1 | 1 | 1 | FIM1_2933 | YPR189W | YPR189W | SKI3 | n |
| Kmar_QGN16832.1 | W303.asm02.HP0_G0027330 | OG0003646 | chrIX | 368455 | 369414 | CP015058.1 | 987606 | 988673 | 34.34065934 | 319 | 355 | 0.898591549 | 1 | 1 | 1 | FIM1_3556 | YIR011C | YIR011C | STS1 | n |
| Kmar_QGN16779.1 | W303.asm02.HP0_G0052890 | OG0003647 | chrXVI | 49852 | 51507 | CP015058.1 | 867660 | 869456 | 63.51575456 | 551 | 598 | 0.921404682 | 1 | 1 | 1 | FIM1_3501 | YPL260W | YPL260W | CUB1 | n |
| Kmar_QGN17226.1 | W303.asm02.HP0_G0032970 | OG0003648 | chrXI | 394850 | 396298 | CP015059.1 | 489810 | 491333 | 47.86821705 | 482 | 507 | 0.950690335 | 1 | 1 | 1 | FIM1_3957 | YKL028W | YKL028W | TFA1 | n |
| Kmar_QGN15509.1 | W303.asm02.HP0_G0006080 | OG0003649 | chrIII | 252291 | 253985 | CP015056.1 | 1089807 | 1091504 | 30.70607553 | 564 | 565 | 0.998230088 | 1 | 1 | 1 | FIM1_2200 | YCR065W | YCR065W | HCM1 | n |
| Kmar_QGN15508.1 | W303.asm02.HP0_G0006070 | OG0003650 | chrIII | 251299 | 251772 | CP015056.1 | 1088332 | 1088805 | 66.87898089 | 157 | 157 | 1 | 1 | 1 | 1 | FIM1_2199 | YCR063W | YCR063W | BUD31 | n |
| Kmar_QGN15379.1 | W303.asm02.HP0_G0009480 | OG0003651 | chrIV | 608284 | 611262 | CP015056.1 | 840679 | 843309 | 40.32258065 | 992 | 876 | 1.132420091 | 1 | 1 | 1 | FIM1_2069 | YDR080W | YDR080W | VPS41 | n |
| Kmar_QGN17360.1 | W303.asm02.HP0_G0009520 | OG0003652 | chrIV | 617682 | 618281 | CP015059.1 | 770128 | 770715 | 55.2238806 | 199 | 195 | 1.020512821 | 1 | 1 | 1 | FIM1_4092 | YDR084C | YDR084C | TVP23 | n |
| Kmar_QGN15378.1 | W303.asm02.HP0_G0009490 | OG0003653 | chrIV | 611581 | 614358 | CP015056.1 | 838225 | 840570 | 40.9287257 | 925 | 781 | 1.184379001 | 1 | 1 | 1 | FIM1_2068 | YDR081C | YDR081C | PDC2 | n |
| Kmar_QGN16591.1 | W303.asm02.HP0_G0032320 | OG0003654 | chrXI | 271825 | 272766 | CP015058.1 | 452309 | 453259 | 49.53271028 | 313 | 316 | 0.990506329 | 1 | 1 | 1 | FIM1_3307 | YKL094W | YKL094W | YJU3 | n |
| Kmar_QGN16634.1 | W303.asm02.HP0_G0027980 | OG0003655 | chrX | 66334 | 68793 | CP015058.1 | 550057 | 552318 | 37.10462287 | 819 | 753 | 1.087649402 | 1 | 1 | 1 | FIM1_3351 | YJL187C | YJL187C | SWE1 | n |
| Kmar_QGN15519.1 | W303.asm02.HP0_G0019200 | OG0003656 | chrVII | 357240 | 358811 | CP015056.1 | 1109524 | 1111074 | 77.29007634 | 523 | 516 | 1.013565891 | 1 | 1 | 1 | FIM1_2210 | YGL078C | YGL078C | DBP3 | n |
| Kmar_QGN15349.1 | W303.asm02.HP0_G0013210 | OG0003657 | chrIV | 1426125 | 1426532 | CP015056.1 | 767730 | 768038 | 38.97058824 | 135 | 102 | 1.323529412 | 1 | 1 | 1 | FIM1_2039 | YDR482C | YDR482C | CWC21 | n |
| Kmar_QGN16745.1 | W303.asm02.HP0_G0027250 | OG0003658 | chrIX | 351939 | 353978 | CP015058.1 | 803762 | 805729 | 19.62209302 | 679 | 655 | 1.036641221 | 1 | 1 | 1 | FIM1_3467 | YIR003W | YIR003W | AIM21 | n |
| Kmar_QGN13861.1 | W303.asm02.HP0_G0033850 | OG0003659 | chrXI | 569327 | 570313 | CP015054.1 | 1014343 | 1015194 | 52.27963526 | 328 | 283 | 1.159010601 | 1 | 1 | 1 | FIM1_507 | YKR062W | YKR062W | TFA2 | n |
| Kmar_QGN15041.1 | W303.asm02.HP0_G0030310 | OG0003660 | chrX | 586542 | 587477 | CP015056.1 | 73715 | 74638 | 81.99356913 | 311 | 307 | 1.013029316 | 1 | 1 | 1 | FIM1_1725 | YJR077C | YJR077C | MIR1 | n |
| Kmar_QGN16743.1 | W303.asm02.HP0_G0014570 | OG0003661 | chrV | 173817 | 174521 | CP015058.1 | 801393 | 802115 | 57.67634855 | 234 | 240 | 0.975 | 1 | 1 | 1 | FIM1_3465 | YER010C | YER010C | | n |
| Kmar_QGN13860.1 | W303.asm02.HP0_G0028320 | OG0003662 | chrX | 129965 | 130666 | CP015054.1 | 1013663 | 1014238 | 33.05785124 | 233 | 191 | 1.219895288 | 1 | 1 | 1 | FIM1_506 | YJL148W | YJL148W | RPA34 | n |
| Kmar_QGN13864.1 | W303.asm02.HP0_G0028350 | OG0003663 | chrX | 134688 | 135572 | CP015054.1 | 1019202 | 1020044 | 49.49494949 | 294 | 280 | 1.05 | 1 | 1 | 1 | FIM1_510 | YJL145W | YJL145W | SFH5 | n |
| Kmar_QGN17621.1 | W303.asm02.HP0_G0043200 | OG0003664 | chrXIII | 839261 | 840175 | CP015060.1 | 122663 | 123460 | 35.29411765 | 304 | 265 | 1.147169811 | 1 | 1 | 1 | FIM1_4828 | YMR281W | YMR281W | GPI12 | n |
| Kmar_QGN15893.1 | W303.asm02.HP0_G0035740 | OG0003665 | chrXII | 308626 | 309942 | CP015057.1 | 316620 | 317906 | 57.8713969 | 438 | 428 | 1.023364486 | 1 | 1 | 1 | FIM1_2591 | YLR085C | YLR085C | ARP6 | n |
| Kmar_QGN15507.1 | W303.asm02.HP0_G0006060 | OG0003666 | chrIII | 248544 | 250439 | CP015056.1 | 1085340 | 1087328 | 50.37369208 | 631 | 662 | 0.953172205 | 1 | 1 | 1 | FIM1_2198 | YCR061W | YCR061W | TVS1 | n |
| Kmar_QGN14945.1 | W303.asm02.HP0_G0045230 | OG0003667 | chrXIV | 333289 | 335559 | CP015055.1 | 1545950 | 1548211 | 68.14044213 | 756 | 753 | 1.003984064 | 1 | 1 | 1 | FIM1_1621 | YNL161W | YNL161W | CBK1 | n |
| Kmar_QGN13678.1 | W303.asm02.HP0_G0002360 | OG0003668 | chrII | 339046 | 339573 | CP015054.1 | 648680 | 649216 | 43.71584699 | 175 | 178 | 0.983146067 | 1 | 0 | 0 | FIM1_321 | YBR047W | YBR047W | FMP23 | n |
| Kmar_QGN13701.1 | W303.asm02.HP0_G0043380 | OG0003669 | chrXIII | 870739 | 871191 | CP015054.1 | 688683 | 689141 | 49.375 | 150 | 152 | 0.986842105 | 1 | 0 | 1 | FIM1_344 | YMR298W | YMR298W | LIP1 | n |
| Kmar_QGN14000.1 | W303.asm02.HP0_G0037540 | OG0003670 | chrXII | 694650 | 698675 | CP015054.1 | 1299112 | 1302801 | 43.53932584 | 1341 | 1229 | 1.091131001 | 1 | 0 | 1 | FIM1_651 | YLR278C | YLR278C | | n |
| Kmar_QGN14026.1 | W303.asm02.HP0_G0048540 | OG0003671 | chrXV | 223723 | 224634 | CP015054.1 | 1359213 | 1360076 | 56.47840532 | 303 | 287 | 1.055749129 | 1 | 0 | 0 | FIM1_677 | YOL056W | YOL056W | GPM3 | n |
| Kmar_QGN14038.1 | W303.asm02.HP0_G0048190 | OG0003672 | chrXV | 145789 | 147618 | CP015054.1 | 1387658 | 1389541 | 14.28571429 | 609 | 627 | 0.971291866 | 1 | 0 | 1 | FIM1_689 | YOL091W | YOL091W | SPO21 | n |
| Kmar_QGN14262.1 | W303.asm02.HP0_G0043040 | OG0003673 | chrXIII | 803041 | 804426 | CP015055.1 | 138597 | 140111 | 23.47140039 | 461 | 504 | 0.91468254 | 1 | 0 | 0 | FIM1_919 | YMR265C | YMR265C | | n |
| Kmar_QGN14735.1 | W303.asm02.HP0_G0021150 | OG0003674 | chrVII | 784181 | 785413 | CP015055.1 | 1123689 | 1124810 | 27.66990291 | 410 | 373 | 1.09919571 | 1 | 0 | 1 | FIM1_1404 | YGR142W | YGR142W | BTN2 | n |
| Kmar_QGN15205.1 | W303.asm02.HP0_G0021430 | OG0003675 | chrVII | 848168 | 848446 | CP015056.1 | 426920 | 427192 | 73.33333333 | 92 | 90 | 1.022222222 | 1 | 0 | 0 | FIM1_1893 | YGR169C-A | YGR169C-A | LSO2 | n |
| Kmar_QGN15215.1 | W303.asm02.HP0_G0052530 | OG0003676 | chrXV | 1048507 | 1050084 | CP015056.1 | 447015 | 448562 | 34.84848485 | 525 | 515 | 1.019417476 | 1 | 0 | 0 | FIM1_1903 | YOR377W | YOR377W | ATF1 | n |
| Kmar_QGN15302.1 | W303.asm02.HP0_G0016140 | OG0003677 | chrV | 525918 | 528473 | CP015056.1 | 642186 | 644567 | 15.57377049 | 851 | 793 | 1.073139975 | 1 | 0 | 1 | FIM1_1991 | YER167W | YER167W | BCK2 | n |
| Kmar_QGN15335.1 | W303.asm02.HP0_G0004400 | OG0003678 | chrII | 742388 | 743959 | CP015056.1 | 735695 | 737365 | 21.76258993 | 523 | 556 | 0.940647482 | 1 | 0 | 0 | FIM1_2024 | YBR250W | YBR250W | SPO23 | n |
| Kmar_QGN15387.1 | W303.asm02.HP0_G0001810 | OG0003679 | chrII | 209479 | 211509 | CP015056.1 | 853476 | 855704 | 26.87338501 | 676 | 742 | 0.911051213 | 1 | 0 | 0 | FIM1_2077 | YBL009W | YBL009W | ALK2 | n |
| Kmar_QGN15481.1 | W303.asm02.HP0_G0051460 | OG0003680 | chrXV | 820589 | 821392 | CP015056.1 | 1031751 | 1033073 | 20.22727273 | 267 | 440 | 0.606818182 | 1 | 0 | 0 | FIM1_2172 | YOR264W | YOR264W | DSE3 | n |
| Kmar_QGN15550.1 | W303.asm02.HP0_G0011610 | OG0003681 | chrIV | 1083548 | 1084699 | CP015056.1 | 1166427 | 1167548 | 35.23809524 | 383 | 373 | 1.026809651 | 1 | 0 | 0 | FIM1_2241 | YDR309C | YDR309C | GIC2 | n |
| Kmar_QGN15553.1 | W303.asm02.HP0_G0011620 | OG0003682 | chrIV | 1085628 | 1088946 | CP015056.1 | 1170594 | 1173437 | 28.55910267 | 1045 | 947 | 1.103484688 | 1 | 0 | 1 | FIM1_2244 | YDR310C | YDR310C | SUM1 | n |
| Kmar_QGN15576.1 | W303.asm02.HP0_G0032160 | OG0003683 | chrXI | 234429 | 236624 | CP015056.1 | 1220457 | 1221947 | 31.0626703 | 731 | 496 | 1.473790323 | 1 | 0 | 1 | FIM1_2267 | YKL112W | YKL112W | ABF1 | n |
| Kmar_QGN15968.1 | W303.asm02.HP0_G0044890 | OG0003684 | chrXIV | 265626 | 267516 | CP015057.1 | 502181 | 503311 | 24.12060302 | 595 | 376 | 1.582446809 | 1 | 0 | 1 | FIM1_2666 | YNL199C | YNL199C | GCR2 | n |
| Kmar_QGN16038.1 | W303.asm02.HP0_G0029480 | OG0003685 | chrX | 398546 | 400594 | CP015057.1 | 679108 | 681144 | 34.25797503 | 682 | 678 | 1.005899705 | 1 | 0 | 1 | FIM1_2738 | YJL019W | YJL019W | MPS3 | n |
| Kmar_QGN16133.1 | W303.asm02.HP0_G0025930 | OG0003686 | chrIX | 84096 | 85406 | CP015057.1 | 882962 | 884524 | 27.90262172 | 436 | 520 | 0.838461538 | 1 | 0 | 0 | FIM1_2834 | YIL135C | YIL135C | VHS2 | n |
| Kmar_QGN16378.1 | W303.asm02.HP0_G0018850 | OG0003687 | chrVII | 285465 | 286262 | CP015057.1 | 1395786 | 1396634 | 17.89473684 | 265 | 282 | 0.939716312 | 1 | 0 | 1 | FIM1_3086 | YGL117W | YGL117W | ARO5 | n |
| Kmar_QGN16474.1 | W303.asm02.HP0_G0029250 | OG0003688 | chrX | 339599 | 342127 | CP015058.1 | 183792 | 185903 | 17.84869976 | 842 | 703 | 1.19772404 | 1 | 0 | 1 | FIM1_3187 | YJL047C | YJL047C | RTT101 | n |
| Kmar_QGN16592.1 | W303.asm02.HP0_G0032330 | OG0003689 | chrXI | 273265 | 274284 | CP015058.1 | 455017 | 456189 | 21.22762148 | 339 | 390 | 0.869230769 | 1 | 0 | 0 | FIM1_3308 | YKL093W | YKL093W | MBR1 | n |
| Kmar_QGN16849.1 | W303.asm02.HP0_G0021050 | OG0003690 | chrVII | 763054 | 765570 | CP015058.1 | 1022738 | 1025398 | 26.99662542 | 662 | 886 | 0.74717833 | 1 | 0 | 1 | FIM1_3574 | YGR130C | YGR130C | | n |
| Kmar_QGN16890.1 | W303.asm02.HP0_G0036310 | OG0003691 | chrXII | 441915 | 442103 | CP015058.1 | 1112140 | 1112373 | 29.62962963 | 62 | 77 | 0.805194805 | 1 | 0 | 0 | FIM1_3617 | YLR146W-A | YLR146W-A | | n |
| Kmar_QGN16977.1 | W303.asm02.HP0_G0044650 | OG0003692 | chrXIV | 223420 | 225165 | CP015058.1 | 1294196 | 1296790 | 20.11560694 | 581 | 864 | 0.672453704 | 1 | 0 | 1 | FIM1_3704 | YNL225C | YNL225C | CNM67 | n |
| Kmar_QGN17035.1 | W303.asm02.HP0_G0032840 | OG0003693 | chrXI | 367078 | 368169 | CP015059.1 | 89064 | 90035 | 20.27027027 | 363 | 323 | 1.123839009 | 1 | 0 | 1 | FIM1_3764 | YKL042W | YKL042W | SPC42 | n |
| Kmar_QGN17154.1 | W303.asm02.HP0_G0017610 | OG0003694 | chrVII | 21088 | 22854 | CP015059.1 | 352653 | 354434 | 69.02356902 | 588 | 593 | 0.991568297 | 1 | 0 | 1 | FIM1_3885 | YGL252C | YGL252C | RTG2 | n |
| Kmar_QGN17158.1 | W303.asm02.HP0_G0054920 | OG0003695 | chrXVI | 468841 | 470199 | CP015059.1 | 358748 | 359884 | 16.81415929 | 452 | 378 | 1.195767196 | 1 | 0 | 0 | FIM1_3889 | YPL049C | YPL049C | DIG1 | n |
| Kmar_QGN17230.1 | W303.asm02.HP0_G0041140 | OG0003696 | chrXIII | 409658 | 411130 | CP015059.1 | 502438 | 503832 | 25.05050505 | 490 | 464 | 1.056034483 | 1 | 0 | 1 | FIM1_3961 | YMR070W | YMR070W | MOT3 | n |
| Kmar_QGN17361.1 | W303.asm02.HP0_G0009530 | OG0003697 | chrIV | 618565 | 620427 | CP015059.1 | 771257 | 773263 | 22.33576642 | 620 | 668 | 0.928143713 | 1 | 0 | 0 | FIM1_4093 | YDR085C | YDR085C | AFR1 | n |
| Kmar_QGN17375.1 | W303.asm02.HP0_G0032490 | OG0003698 | chrXI | 301432 | 302784 | CP015059.1 | 798100 | 799233 | 49.22394678 | 450 | 377 | 1.193633952 | 1 | 0 | 1 | FIM1_4107 | YKL075C | YKL075C | AAN1 | n |
| Kmar_QGN17514.1 | W303.asm02.HP0_G0021990 | OG0003699 | chrVII | 964309 | 964722 | CP015059.1 | 1068821 | 1069099 | 30.65693431 | 137 | 92 | 1.489130435 | 1 | 0 | 1 | FIM1_4250 | YGR230W | YGR230W | BNS1 | n |
| Kmar_QGN17520.1 | W303.asm02.HP0_G0022050 | OG0003700 | chrVII | 974945 | 975232 | CP015059.1 | 1081238 | 1081510 | 16.84210526 | 95 | 90 | 1.055555556 | 1 | 0 | 1 | FIM1_4256 | YGR236C | YGR236C | SPG1 | n |
| Kmar_QGN17525.1 | W303.asm02.HP0_G0022080 | OG0003701 | chrVII | 981602 | 982468 | CP015059.1 | 1091962 | 1092939 | 25.65789474 | 288 | 325 | 0.886153846 | 1 | 0 | 1 | FIM1_4261 | YGR239C | YGR239C | PEX21 | n |
| Kmar_QGN18204.1 | W303.asm02.HP0_G0053180 | OG0003702 | chrXVI | 120877 | 122052 | CP015061.1 | 443907 | 445016 | 25.44080605 | 391 | 369 | 1.059620596 | 1 | 0 | 0 | FIM1_4528 | YPL230W | YPL230W | USV1 | n |
| Kmar_QGN18205.1 | W303.asm02.HP0_G0053190 | OG0003703 | chrXVI | 122632 | 123252 | CP015061.1 | 445893 | 446459 | 28.7804878 | 206 | 188 | 1.095744681 | 1 | 0 | 0 | FIM1_4529 | YPL229W | YPL229W | | n |
| Kmar_QGN18226.1 | W303.asm02.HP0_G0012430 | OG0003704 | chrIV | 1263083 | 1263781 | CP015061.1 | 496508 | 497236 | 47.10743802 | 232 | 242 | 0.958677686 | 1 | 0 | 0 | FIM1_4551 | YDR391C | YDR391C | | n |
| Kmar_QGN18367.1 | W303.asm02.HP0_G0049840 | OG0003705 | chrXV | 480268 | 481155 | CP015061.1 | 788237 | 789496 | 18.37708831 | 295 | 419 | 0.704057279 | 1 | 0 | 0 | FIM1_4694 | YOR083W | YOR083W | WHI5 | n |
| Kmar_QGN18334.1 | W303.asm02.HP0_G0001610 | OG0003714 | chrII | 166790 | 167074 | CP015061.1 | 714718 | 714990 | 56.52173913 | 94 | 90 | 1.044444444 | 1 | 0 | 1 | FIM1_4660 | YBL029C-A | YBL029C-A | | n |
| Kmar_QGN15486.1 | W303.asm02.HP0_G0026510 | OG0003718 | chrIX | 213432 | 216449 | CP015056.1 | 1041742 | 1044692 | 33.09572301 | 975 | 956 | 1.019874477 | 1 | 0 | 1 | FIM1_2177 | YIL073C | YIL073C | SPO22 | n |
| Kmar_QGN15380.1 | W303.asm02.HP0_G0009470 | OG0003719 | chrIV | 607869 | 608087 | CP015056.1 | 843524 | 843742 | 77.77777778 | 72 | 72 | 1 | 1 | 0 | 1 | FIM1_2070 | YDR079C-A | YDR079C-A | TFB5 | n |
| Kmar_QGN15235.1 | W303.asm02.HP0_G0034850 | OG0003724 | chrXII | 132390 | 132599 | CP015056.1 | 491851 | 492048 | 61.53846154 | 69 | 65 | 1.061538462 | 1 | 0 | 1 | FIM1_1924 | YLL009C | YLL009C | COX17 | n |
| Kmar_QGN17153.1 | W303.asm02.HP0_G0017620 | OG0003727 | chrVII | 23291 | 27006 | CP015059.1 | 348816 | 352313 | 43.2998325 | 1187 | 1165 | 1.01888412 | 1 | 0 | 1 | FIM1_3884 | YGL251C | YGL251C | HFM1 | n |
| Kmar_QGN16221.1 | W303.asm02.HP0_G0038580 | OG0003729 | chrXII | 904620 | 906068 | CP015057.1 | 1062959 | 1063933 | 23.7006237 | 482 | 324 | 1.487654321 | 1 | 0 | 1 | FIM1_2925 | YLR394W | YLR394W | CST9 | n |
| Kmar_QGN17533.1 | W303.asm02.HP0_G0028530 | OG0003730 | chrX | 171568 | 171726 | CP015059.1 | 1118358 | 1118510 | 58.49056604 | 52 | 50 | 1.04 | 1 | 0 | 1 | FIM1_4269 | YJL127C-B | YJL127C-B | MCO6 | n |
| Kmar_QGN17198.1 | W303.asm02.HP0_G0053570 | OG0003733 | chrXVI | 193859 | 194065 | CP015059.1 | 435144 | 435341 | 49.27536232 | 68 | 65 | 1.046153846 | 1 | 0 | 1 | FIM1_3929 | YPL189C-A | YPL189C-A | COA2 | n |
| Kmar_QGN13503.1 | W303.asm02.HP0_G0008000 | OG0003736 | chrIV | 305567 | 306907 | CP015054.1 | 280054 | 281367 | 87.89237668 | 446 | 437 | 1.020594966 | 1 | 0 | 1 | FIM1_142 | YDL084W | YDL084W | SUB2 | n |
| Kmar_QGN18222.1 | W303.asm02.HP0_G0007330 | OG0003737 | chrIV | 178546 | 181251 | CP015061.1 | 484197 | 486767 | 35.76881134 | 901 | 856 | 1.052570093 | 1 | 0 | 1 | FIM1_4547 | YDL154W | YDL154W | MSH5 | n |
| Kmar_QGN17973.1 | W303.asm02.HP0_G0012130 | OG0003740 | chrIV | 1213644 | 1213913 | CP015060.1 | 858489 | 858686 | 40.4494382 | 89 | 65 | 1.369230769 | 1 | 0 | 1 | FIM1_5182 | YDR363W-A | YDR363W-A | SEM1 | n |
| Kmar_QGN13646.1 | W303.asm02.HP0_G0005710 | OG0003743 | chrIII | 166744 | 167010 | CP015054.1 | 577018 | 577293 | 39.56043956 | 88 | 91 | 0.967032967 | 1 | 0 | 1 | FIM1_288 | YCR020C-A | YCR020C-A | MAK31 | n |
| Kmar_QGN14289.1 | W303.asm02.HP0_G0026260 | OG0003747 | chrIX | 163723 | 163950 | CP015055.1 | 195227 | 195442 | 36 | 75 | 71 | 1.056338028 | 1 | 0 | 1 | FIM1_946 | YIL102C-A | YIL102C-A | | n |
| Kmar_QGN17597.1 | W303.asm02.HP0_G0052790 | OG0003750 | chrXVI | 30631 | 30819 | CP015060.1 | 70458 | 70643 | 43.5483871 | 62 | 61 | 1.016393443 | 1 | 0 | 1 | FIM1_4804 | YPL271W | YPL271W | ATP15 | n |
| Kmar_QGN17029.1 | W303.asm02.HP0_G0046060 | OG0003754 | chrXIV | 495552 | 495734 | CP015059.1 | 76185 | 76367 | 83.33333333 | 60 | 60 | 1 | 1 | 0 | 1 | FIM1_3758 | YNL070W | YNL070W | TOM7 | n |
| Kmar_QGN15384.1 | W303.asm02.HP0_G0016100 | OG0003765 | chrV | 510786 | 511484 | CP015056.1 | 848385 | 849137 | 52.99145299 | 232 | 250 | 0.928 | 1 | 0 | 1 | FIM1_2074 | YER163C | YER163C | GCG1 | n |
| Kmar_QGN17115.1 | W303.asm02.HP0_G0014720 | OG0003772 | chrV | 203331 | 206102 | CP015059.1 | 266945 | 269527 | 51.56418554 | 923 | 860 | 1.073255814 | 1 | 0 | 1 | FIM1_3846 | YER024W | YER024W | YAT2 | n |
| Kmar_QGN18104.1 | W303.asm02.HP0_G0055760 | OG0003774 | chrXVI | 652447 | 652650 | CP015061.1 | 223897 | 224022 | 49.25373134 | 67 | 41 | 1.634146341 | 1 | 0 | 1 | FIM1_4424 | YPR036W-A | YPR036W-A | SPO24 | n |
| Kmar_QGN13492.1 | W303.asm02.HP0_G0014170 | OG0003780 | chrV | 90616 | 94263 | CP015054.1 | 252828 | 256448 | 75.10339123 | 1215 | 1206 | 1.007462687 | 1 | 0 | 1 | FIM1_130 | YEL031W | YEL031W | SPF1 | n |
| Kmar_QGN17166.1 | W303.asm02.HP0_G0011110 | OG0003781 | chrIV | 969614 | 970990 | CP015059.1 | 370556 | 371755 | 25.86956522 | 458 | 399 | 1.147869674 | 1 | 0 | 1 | FIM1_3897 | YDR254W | YDR254W | CHL4 | n |
| Kmar_QGN18166.1 | W303.asm02.HP0_G0015150 | OG0003786 | chrV | 283380 | 284036 | CP015061.1 | 359307 | 359906 | 34.40366972 | 218 | 199 | 1.095477387 | 1 | 0 | 1 | FIM1_4489 | YER063W | YER063W | THO1 | n |
| Kmar_QGN14089.1 | W303.asm02.HP0_G0024850 | OG0003788 | chrVIII | 417490 | 418512 | CP015054.1 | 1494311 | 1495258 | 27.05882353 | 340 | 315 | 1.079365079 | 1 | 0 | 1 | FIM1_740 | YHR156C | YHR156C | LIN1 | n |
| Kmar_QGN14397.1 | W303.asm02.HP0_G0053950 | OG0003790 | chrXVI | 274122 | 276827 | CP015055.1 | 434140 | 436683 | 42.3580786 | 901 | 847 | 1.063754427 | 1 | 0 | 1 | FIM1_1057 | YPL150W | YPL150W | | n |
| Kmar_QGN18043.1 | W303.asm02.HP0_G0035540 | OG0003791 | chrXII | 273407 | 274228 | CP015061.1 | 86438 | 87280 | 35.19163763 | 273 | 280 | 0.975 | 1 | 0 | 1 | FIM1_4359 | YLR064W | YLR064W | PER33 | n |
| Kmar_QGN18041.1 | W303.asm02.HP0_G0035550 | OG0003792 | chrXII | 274324 | 274869 | CP015061.1 | 84476 | 85018 | 39.89361702 | 181 | 180 | 1.005555556 | 1 | 0 | 1 | FIM1_4357 | YLR065C | YLR065C | SND2 | n |
| Kmar_QGN14022.1 | W303.asm02.HP0_G0000840 | OG0003793 | chrI | 191393 | 193456 | CP015054.1 | 1347865 | 1349961 | 59.49367089 | 687 | 698 | 0.984240688 | 1 | 0 | 1 | FIM1_673 | YAR035W | YAR035W | YAT1 | n |
| Kmar_QGN17385.1 | W303.asm02.HP0_G0034420 | OG0003794 | chrXII | 25627 | 26865 | CP015059.1 | 816585 | 817757 | 64.37054632 | 412 | 390 | 1.056410256 | 1 | 0 | 1 | FIM1_4117 | YLL057C | YLL057C | JLP1 | n |
| Kmar_QGN15944.1 | W303.asm02.HP0_G0037470 | OG0003796 | chrXII | 677387 | 678211 | CP015057.1 | 444162 | 444986 | 21.01449275 | 274 | 274 | 1 | 1 | 0 | 1 | FIM1_2642 | YLR271W | YLR271W | CMG1 | n |
| Kmar_QGN14505.1 | W303.asm02.HP0_G0035470 | OG0003802 | chrXII | 261674 | 262771 | CP015055.1 | 650578 | 651636 | 58.03814714 | 365 | 352 | 1.036931818 | 1 | 0 | 1 | FIM1_1166 | YLR056W | YLR056W | ERG3 | n |
| Kmar_QGN18066.1 | W303.asm02.HP0_G0008250 | OG0003808 | chrIV | 351766 | 352752 | CP015061.1 | 140714 | 142576 | 24.66793169 | 328 | 620 | 0.529032258 | 1 | 0 | 1 | FIM1_4384 | YDL057W | YDL057W | | n |
| Kmar_QGN14272.1 | W303.asm02.HP0_G0002980 | OG0003809 | chrII | 463995 | 465602 | CP015055.1 | 155302 | 156870 | 40.1459854 | 535 | 522 | 1.024904215 | 1 | 0 | 1 | FIM1_929 | YBR103W | YBR103W | SIF2 | n |
| Kmar_QGN16606.1 | W303.asm02.HP0_G0002290 | OG0003810 | chrII | 316977 | 319046 | CP015058.1 | 485777 | 487792 | 54.84330484 | 689 | 671 | 1.026825633 | 1 | 0 | 1 | FIM1_3322 | YBR043C | YBR043C | QDR3 | n |
| Kmar_QGN16214.1 | W303.asm02.HP0_G0014430 | OG0003813 | chrV | 148134 | 149162 | CP015057.1 | 1049923 | 1050918 | 37.5 | 342 | 331 | 1.033232628 | 1 | 0 | 1 | FIM1_2918 | YEL004W | YEL004W | YEA4 | n |
| Kmar_QGN17615.1 | W303.asm02.HP0_G0040490 | OG0003815 | chrXIII | 267680 | 268306 | CP015060.1 | 110338 | 110961 | 86.05769231 | 208 | 207 | 1.004830918 | 1 | 0 | 1 | FIM1_4822 | YML001W | YML001W | YPT7 | n |

| | | | | | | | | | | | | | | | | | | | | |
|---|---|---|---|---|---|---|---|---|---|---|---|---|---|---|---|---|---|---|---|---|
| Kmar_QGN13454.1 | W303.asm02.HP0_G0045410 | OG0003816 | chrXIV | 360429 | 361472 | CP015054.1 | 176056 | 177123 | 65.53672316 | 347 | 355 | 0.977464789 | 1 | 0 | 1 | FIM1_91 | YNL141W | YNL141W | AAH1 | n |
| Kmar_QGN16077.1 | W303.asm02.HP0_G0055340 | OG0003819 | chrXVI | 550988 | 554500 | CP015057.1 | 760189 | 763734 | 57.80590717 | 1170 | 1181 | 0.990685859 | 1 | 0 | 1 | FIM1_2777 | YPL006W | YPL006W | NCR1 | n |
| Kmar_QGN16786.1 | W303.asm02.HP0_G0022900 | OG0003822 | chrVIII | 62666 | 64063 | CP015058.1 | 887854 | 889098 | 33.68869936 | 465 | 414 | 1.123188406 | 1 | 0 | 1 | FIM1_3509 | YHL021C | YHL021C | AIM17 | n |
| Kmar_QGN17901.1 | W303.asm02.HP0_G0042970 | OG0003823 | chrXIII | 789122 | 790783 | CP015060.1 | 703410 | 705062 | 45.38878843 | 553 | 550 | 1.005454545 | 1 | 0 | 1 | FIM1_5110 | YMR258C | YMR258C | ROY1 | n |
| Kmar_QGN14824.1 | W303.asm02.HP0_G0006940 | OG0003826 | chrIV | 100575 | 101141 | CP015055.1 | 1298539 | 1299108 | 35.8974359 | 188 | 189 | 0.994708995 | 1 | 0 | 1 | FIM1_1495 | YDL200C | YDL200C | MGT1 | n |
| Kmar_QGN17992.1 | W303.asm02.HP0_G0057220 | OG0003827 | chrXVI | 945754 | 946968 | CP015060.1 | 916868 | 918139 | 61.22931442 | 404 | 423 | 0.955082742 | 1 | 0 | 1 | FIM1_5202 | YPR201W | YPR201W | ARR3 | n |
| Kmar_QGN13726.1 | W303.asm02.HP0_G0022260 | OG0003828 | chrVII | 1018645 | 1019745 | CP015054.1 | 748093 | 749205 | 58.88594164 | 366 | 370 | 0.989189189 | 1 | 0 | 1 | FIM1_370 | YGR257C | YGR257C | MTM1 | n |
| Kmar_QGN16648.1 | W303.asm02.HP0_G0020480 | OG0003835 | chrVII | 620128 | 622542 | CP015058.1 | 587410 | 590226 | 22.4229543 | 804 | 938 | 0.857142857 | 1 | 0 | 1 | FIM1_3367 | YGR067C | YGR067C | | n |
| Kmar_QGN13819.1 | W303.asm02.HP0_G0024090 | OG0003843 | chrVIII | 268166 | 268527 | CP015054.1 | 936885 | 937453 | 43.95604396 | 91 | 62 | 1.467741935 | 1 | 0 | 1 | FIM1_465 | YHR079C-A | YHR079C-A | SAE3 | n |
| Kmar_QGN16171.1 | W303.asm02.HP0_G0051520 | OG0003845 | chrXV | 828104 | 829588 | CP015057.1 | 953468 | 954802 | 37.77335984 | 494 | 444 | 1.112612613 | 1 | 0 | 1 | FIM1_2873 | YOR269W | YOR269W | PAC1 | n |
| Kmar_QGN15505.1 | W303.asm02.HP0_G0006050 | OG0003846 | chrIII | 247380 | 247715 | CP015056.1 | 1076570 | 1076917 | 45.2173913 | 111 | 115 | 0.965217391 | 1 | 0 | 1 | FIM1_2196 | YCR060W | YCR060W | TAH1 | n |
| Kmar_QGN16101.1 | W303.asm02.HP0_G0015020 | OG0003847 | chrV | 255927 | 257405 | CP015057.1 | 815115 | 816548 | 44.37869822 | 492 | 477 | 1.031446541 | 1 | 0 | 1 | FIM1_2802 | YER051W | YER051W | JHD1 | n |
| Kmar_QGN18210.1 | W303.asm02.HP0_G0042170 | OG0003848 | chrXIII | 619063 | 619887 | CP015061.1 | 457085 | 457942 | 55.0877193 | 274 | 285 | 0.961403509 | 1 | 0 | 1 | FIM1_4534 | YMR178W | YMR178W | FPY1 | n |
| Kmar_QGN14410.1 | W303.asm02.HP0_G0016870 | OG0003855 | chrVI | 103004 | 105640 | CP015055.1 | 459298 | 461955 | 41.54535274 | 878 | 885 | 0.992090395 | 1 | 0 | 1 | FIM1_1070 | YFL003C | YFL003C | MSH4 | n |
| Kmar_QGN18213.1 | W303.asm02.HP0_G0042150 | OG0003858 | chrXIII | 612324 | 616559 | CP015061.1 | 461352 | 465770 | 29.47154472 | 1411 | 1472 | 0.958559783 | 1 | 0 | 1 | FIM1_4537 | YMR176W | YMR176W | ECM5 | n |
| Kmar_QGN15667.1 | W303.asm02.HP0_G0012640 | OG0003859 | chrIV | 1302762 | 1303886 | CP015056.1 | 1420569 | 1421696 | 60.73298429 | 374 | 375 | 0.997333333 | 1 | 0 | 1 | FIM1_2360 | YDR415C | YDR415C | | n |
| Kmar_QGN14211.1 | W303.asm02.HP0_G0055820 | OG0003860 | chrXVI | 661638 | 663050 | CP015055.1 | 42234 | 44141 | 25.54517134 | 470 | 635 | 0.74015748 | 1 | 0 | 1 | FIM1_867 | YPR045C | YPR045C | THP3 | n |
| Kmar_QGN16855.1 | W303.asm02.HP0_G0028290 | OG0003867 | chrX | 123860 | 125461 | CP015058.1 | 1039456 | 1041048 | 73.22097378 | 533 | 530 | 1.005660377 | 1 | 0 | 1 | FIM1_3581 | YJL153C | YJL153C | INO1 | n |
| Kmar_QGN18235.1 | W303.asm02.HP0_G0007270 | OG0003868 | chrIV | 169601 | 169843 | CP015061.1 | 512486 | 512725 | 32.14285714 | 80 | 79 | 1.012658228 | 1 | 0 | 1 | FIM1_4560 | YDL160C-A | YDL160C-A | MHF2 | n |
| Kmar_QGN15329.1 | W303.asm02.HP0_G0018250 | OG0003871 | chrVII | 150128 | 151525 | CP015056.1 | 718021 | 719406 | 59.02335456 | 465 | 461 | 1.00867679 | 1 | 0 | 1 | FIM1_2018 | YGL184C | YGL184C | STR3 | n |
| Kmar_QGN16286.1 | W303.asm02.HP0_G0038780 | OG0003873 | chrXII | 950022 | 950813 | CP015057.1 | 1207324 | 1208136 | 39.70037453 | 263 | 270 | 0.974074074 | 1 | 0 | 1 | FIM1_2991 | YLR414C | YLR414C | PUN1 | n |
| Kmar_QGN17594.1 | W303.asm02.HP0_G0013690 | OG0003875 | chrIV | 1507140 | 1507853 | CP015060.1 | 64424 | 65137 | 70.29288703 | 237 | 237 | 1 | 1 | 0 | 0 | FIM1_4801 | YDR533C | YDR533C | HSP31 | n |
| Kmar_QGN17192.1 | W303.asm02.HP0_G0019180 | OG0003876 | chrVII | 355588 | 355980 | CP015059.1 | 423592 | 423960 | 67.1875 | 130 | 122 | 1.06557377 | 1 | 0 | 1 | FIM1_3923 | YGL080W | YGL080W | MPC1 | n |
| Kmar_QGN17987.1 | W303.asm02.HP0_G0022750 | OG0003881 | chrVIII | 23865 | 25505 | CP015060.1 | 898430 | 900037 | 44.40433213 | 546 | 535 | 1.020560748 | 1 | 0 | 1 | FIM1_5196 | YHL036W | YHL036W | MUP3 | n |
| Kmar_QGN14948.1 | W303.asm02.HP0_G0035410 | OG0003885 | chrXII | 253400 | 253885 | CP015055.1 | 1553812 | 1554339 | 30.72625698 | 161 | 175 | 0.92 | 1 | 0 | 1 | FIM1_1624 | YLR050C | YLR050C | EMA19 | n |
| Kmar_QGN16586.1 | W303.asm02.HP0_G0020090 | OG0003885 | chrVII | 529504 | 530217 | CP015058.1 | 440360 | 441073 | 75.10548523 | 237 | 237 | 1 | 1 | 0 | 1 | FIM1_3302 | YGR024C | YGR024C | THG1 | n |
| Kmar_QGN13680.1 | W303.asm02.HP0_G0032540 | OG0003886 | chrXI | 314687 | 315196 | CP015054.1 | 651353 | 652075 | 53.33333333 | 169 | 240 | 0.704166667 | 1 | 0 | 1 | FIM1_323 | YKL070W | YKL070W | | n |
| Kmar_QGN13725.1 | W303.asm02.HP0_G0025140 | OG0003887 | chrVIII | 480731 | 481444 | CP015054.1 | 747051 | 747824 | 22.67657993 | 237 | 257 | 0.922178988 | 1 | 0 | 1 | FIM1_369 | YHR185C | YHR185C | PFS1 | n |
| Kmar_QGN15501.1 | W303.asm02.HP0_G0006020 | OG0003890 | chrIII | 241357 | 243048 | CP015056.1 | 1069453 | 1071063 | 29.29824561 | 563 | 536 | 1.050373134 | 1 | 0 | 1 | FIM1_2192 | YCR054C | YCR054C | CTR86 | n |
| Kmar_QGN14898.1 | W303.asm02.HP0_G0003230 | OG0003891 | chrII | 510639 | 511625 | CP015055.1 | 1454990 | 1455907 | 29.61876833 | 328 | 305 | 1.075409836 | 1 | 0 | 1 | FIM1_1572 | YBR129C | YBR129C | OPY1 | n |
| Kmar_QGN14650.1 | W303.asm02.HP0_G0055510 | OG0003893 | chrXVI | 589558 | 590538 | CP015055.1 | 948614 | 949549 | 70.55214724 | 326 | 311 | 1.048231511 | 1 | 0 | 1 | FIM1_1314 | YPR011C | YPR011C | MRX21 | n |
| Kmar_QGN14555.1 | W303.asm02.HP0_G0048680 | OG0003894 | chrXV | 251722 | 253101 | CP015055.1 | 747511 | 748857 | 54.33403805 | 459 | 448 | 1.024553571 | 1 | 0 | 1 | FIM1_1217 | YOL041C | YOL041C | NOP12 | n |
| Kmar_QGN16883.1 | W303.asm02.HP0_G0036270 | OG0003896 | chrXII | 435375 | 437432 | CP015058.1 | 1099223 | 1101319 | 48.57549858 | 685 | 698 | 0.981375358 | 1 | 0 | 1 | FIM1_3610 | YLR143W | YLR143W | DPH6 | n |
| Kmar_QGN16723.1 | W303.asm02.HP0_G0011750 | OG0003897 | chrIV | 1121189 | 1122292 | CP015058.1 | 745660 | 746775 | 58.13333333 | 367 | 371 | 0.989218329 | 1 | 0 | 1 | FIM1_3444 | YDR322W | YDR322W | MRPL35 | n |
| Kmar_QGN15582.1 | W303.asm02.HP0_G0028680 | OG0003900 | chrX | 208869 | 210032 | CP015056.1 | 1238136 | 1241045 | 15.94650206 | 387 | 969 | 0.399380805 | 1 | 0 | 1 | FIM1_2274 | YJL107C | YJL107C | | n |
| Kmar_QGN17262.1 | W303.asm02.HP0_G0016690 | OG0003902 | chrVI | 61979 | 63490 | CP015059.1 | 571117 | 572616 | 74.5526839 | 503 | 499 | 1.008016032 | 1 | 0 | 1 | FIM1_3993 | YFL022C | YFL022C | FRS2 | n |
| Kmar_QGN14573.1 | W303.asm02.HP0_G0007970 | OG0003903 | chrIV | 301743 | 302564 | CP015055.1 | 787032 | 787844 | 61.9047619 | 273 | 270 | 1.011111111 | 1 | 0 | 1 | FIM1_1235 | YDL086W | YDL086W | | n |
| Kmar_QGN15407.1 | W303.asm02.HP0_G0011340 | OG0003906 | chrIV | 1028010 | 1029254 | CP015056.1 | 898571 | 899638 | 46.24697337 | 414 | 355 | 1.166197183 | 1 | 0 | 1 | FIM1_2097 | YDR282C | YDR282C | MRX10 | n |
| Kmar_QGN16720.1 | W303.asm02.HP0_G0024040 | OG0003917 | chrVIII | 254413 | 255615 | CP015058.1 | 740231 | 741415 | 48.75 | 400 | 394 | 1.015228426 | 1 | 0 | 1 | FIM1_3441 | YHR075C | YHR075C | PPE1 | n |
| Kmar_QGN15358.1 | W303.asm02.HP0_G0003510 | OG0003918 | chrII | 573326 | 574975 | CP015056.1 | 790093 | 791382 | 24.41229656 | 549 | 429 | 1.27972028 | 1 | 0 | 1 | FIM1_2048 | YBR158W | YBR158W | AMN1 | n |
| Kmar_QGN14798.1 | W303.asm02.HP0_G0012200 | OG0003919 | chrIV | 1225006 | 1226541 | CP015055.1 | 1251079 | 1252521 | 37.80718336 | 511 | 480 | 1.064583333 | 1 | 0 | 1 | FIM1_1469 | YDR371W | YDR371W | CTS2 | n |
| Kmar_QGN17228.1 | W303.asm02.HP0_G0023140 | OG0003921 | chrVIII | 104116 | 106078 | CP015059.1 | 495464 | 496588 | 30.26086957 | 575 | 374 | 1.537433155 | 1 | 0 | 1 | FIM1_3959 | YHR004C | YHR004C | NEM1 | n |
| Kmar_QGN13480.1 | W303.asm02.HP0_G0029930 | OG0003924 | chrX | 509244 | 511922 | CP015054.1 | 227101 | 229554 | 37.62486127 | 892 | 817 | 1.091799266 | 1 | 0 | 1 | FIM1_118 | YJR036C | YJR036C | HUL4 | n |
| Kmar_QGN15255.1 | W303.asm02.HP0_G0031710 | OG0003926 | chrXI | 148909 | 150117 | CP015056.1 | 538400 | 539560 | 30.09708738 | 402 | 386 | 1.041450777 | 1 | 0 | 1 | FIM1_1944 | YKL162C | YKL162C | | n |
| Kmar_QGN17647.1 | W303.asm02.HP0_G0047270 | OG0003928 | chrXIV | 740993 | 742753 | CP015060.1 | 183198 | 184964 | 69.4772344 | 586 | 588 | 0.996598639 | 1 | 0 | 1 | FIM1_4854 | YNR055C | YNR055C | HOL1 | n |
| Kmar_QGN13373.1 | W303.asm02.HP0_G0031200 | OG0003929 | chrXI | 26237 | 27181 | CP015054.1 | 26523 | 27470 | 77.38853503 | 314 | 315 | 0.996825397 | 1 | 0 | 1 | FIM1_10 | YKL216W | YKL216W | URA1 | n |
| Kmar_QGN13384.1 | W303.asm02.HP0_G0003590 | OG0003930 | chrII | 585628 | 586461 | CP015054.1 | 47058 | 47801 | 24.64285714 | 277 | 247 | 1.12145749 | 1 | 0 | 1 | FIM1_21 | YBR165W | YBR165W | UBS1 | n |
| Kmar_QGN13387.1 | W303.asm02.HP0_G0054360 | OG0003931 | chrXVI | 354804 | 355310 | CP015054.1 | 50868 | 51464 | 37.8238342 | 168 | 198 | 0.848484848 | 1 | 0 | 1 | FIM1_24 | YPL108W | YPL108W | | n |
| Kmar_QGN13389.1 | W303.asm02.HP0_G0003620 | OG0003932 | chrII | 589146 | 590387 | CP015054.1 | 52119 | 53270 | 29.73621103 | 413 | 383 | 1.078328982 | 1 | 0 | 1 | FIM1_26 | YBR168W | YBR168W | PEX32 | n |
| Kmar_QGN13403.1 | W303.asm02.HP0_G0054430 | OG0003933 | chrXVI | 368230 | 369720 | CP015054.1 | 74790 | 75974 | 37.4749499 | 496 | 394 | 1.258883249 | 1 | 0 | 1 | FIM1_40 | YPL100W | YPL100W | ATG21 | n |
| Kmar_QGN13404.1 | W303.asm02.HP0_G0054440 | OG0003934 | chrXVI | 369878 | 370426 | CP015054.1 | 76061 | 76549 | 43.40659341 | 182 | 162 | 1.12345679 | 1 | 0 | 1 | FIM1_41 | YPL099C | YPL099C | INA17 | n |
| Kmar_QGN13426.1 | W303.asm02.HP0_G0003790 | OG0003935 | chrII | 617328 | 619135 | CP015054.1 | 124240 | 124714 | 9.219858156 | 564 | 132 | 4.272727273 | 1 | 0 | 1 | FIM1_63 | YBR186W | YBR186W | PCH2 | n |
| Kmar_QGN13435.1 | W303.asm02.HP0_G0003860 | OG0003936 | chrII | 626813 | 627184 | CP015054.1 | 141311 | 141731 | 34.14634146 | 123 | 115 | 1.069565217 | 1 | 0 | 1 | FIM1_72 | YBR194W | YBR194W | AIM4 | n |
| Kmar_QGN13442.1 | W303.asm02.HP0_G0054680 | OG0003937 | chrXVI | 419186 | 421720 | CP015054.1 | 153851 | 156304 | 31.10624315 | 844 | 817 | 1.033047736 | 1 | 0 | 1 | FIM1_79 | YPL075W | YPL075W | GCR1 | n |
| Kmar_QGN13446.1 | W303.asm02.HP0_G0054710 | OG0003938 | chrXVI | 426406 | 426876 | CP015054.1 | 164016 | 164573 | 29.78723404 | 156 | 185 | 0.843243243 | 1 | 0 | 1 | FIM1_83 | YPL071C | YPL071C | | n |
| Kmar_QGN13449.1 | W303.asm02.HP0_G0003950 | OG0003939 | chrII | 640351 | 640986 | CP015054.1 | 168026 | 168595 | 26.39593909 | 211 | 189 | 1.116402116 | 1 | 0 | 1 | FIM1_86 | YBR201W | YBR201W | DER1 | n |
| Kmar_QGN13450.1 | W303.asm02.HP0_G0054740 | OG0003940 | chrXVI | 430573 | 431454 | CP015054.1 | 168710 | 169585 | 19.66666667 | 293 | 291 | 1.006872852 | 1 | 0 | 1 | FIM1_87 | YPL068C | YPL068C | | n |
| Kmar_QGN13486.1 | W303.asm02.HP0_G0029980 | OG0003941 | chrX | 524992 | 526044 | CP015054.1 | 241602 | 242810 | 28.42892768 | 350 | 402 | 0.870646766 | 1 | 0 | 1 | FIM1_124 | YJR043C | YJR043C | POL32 | n |
| Kmar_QGN13491.1 | W303.asm02.HP0_G0030010 | OG0003942 | chrX | 530579 | 532393 | CP015054.1 | 250938 | 252665 | 32.73615635 | 604 | 575 | 1.050434783 | 1 | 0 | 1 | FIM1_129 | YJR046W | YJR046W | TAH11 | n |
| Kmar_QGN13502.1 | W303.asm02.HP0_G0041830 | OG0003943 | chrXIII | 553985 | 555013 | CP015054.1 | 278260 | 279393 | 27.70780856 | 342 | 377 | 0.907161804 | 1 | 0 | 1 | FIM1_141 | YMR144W | YMR144W | FDO1 | n |
| Kmar_QGN13513.1 | W303.asm02.HP0_G0012010 | OG0003944 | chrIV | 1187636 | 1189690 | CP015054.1 | 304072 | 306180 | 23.23943662 | 684 | 702 | 0.974358974 | 1 | 0 | 1 | FIM1_153 | YDR350C | YDR350C | ATP22 | n |
| Kmar_QGN13549.1 | W303.asm02.HP0_G0019230 | OG0003945 | chrVII | 363878 | 365041 | CP015054.1 | 396321 | 397199 | 20.41343669 | 387 | 292 | 1.325342466 | 1 | 0 | 1 | FIM1_190 | YGL075C | YGL075C | MPS2 | n |
| Kmar_QGN13573.1 | W303.asm02.HP0_G0038860 | OG0003946 | chrXII | 968586 | 969839 | CP015054.1 | 442669 | 443934 | 32.70588235 | 417 | 421 | 0.990498812 | 1 | 0 | 1 | FIM1_214 | YLR423C | YLR423C | ATG17 | n |
| Kmar_QGN13605.1 | W303.asm02.HP0_G0021190 | OG0003947 | chrVII | 795950 | 796585 | CP015054.1 | 505364 | 505822 | 25 | 211 | 152 | 1.388157895 | 1 | 0 | 1 | FIM1_247 | YGR146C | YGR146C | ECL1 | n |
| Kmar_QGN13609.1 | W303.asm02.HP0_G0043090 | OG0003948 | chrXIII | 811826 | 812923 | CP015054.1 | 514335 | 515609 | 26.26931567 | 365 | 424 | 0.860849057 | 1 | 0 | 1 | FIM1_251 | YMR270C | YMR270C | RRN9 | n |
| Kmar_QGN13612.1 | W303.asm02.HP0_G0039490 | OG0003949 | chrXIII | 49824 | 50984 | CP015054.1 | 518442 | 519416 | 20.15306122 | 386 | 324 | 1.191358025 | 1 | 0 | 1 | FIM1_254 | YML107C | YML107C | PML39 | n |
| Kmar_QGN13615.1 | W303.asm02.HP0_G0039440 | OG0003950 | chrXIII | 39615 | 40505 | CP015054.1 | 522979 | 523950 | 36.7781155 | 296 | 323 | 0.916408669 | 1 | 0 | 1 | FIM1_257 | YML112W | YML112W | CTK3 | n |
| Kmar_QGN13627.1 | W303.asm02.HP0_G0024560 | OG0003951 | chrVIII | 366886 | 367617 | CP015054.1 | 546827 | 547405 | 23.45679012 | 243 | 192 | 1.265625 | 1 | 0 | 1 | FIM1_269 | YHR127W | YHR127W | | n |
| Kmar_QGN13636.1 | W303.asm02.HP0_G0014310 | OG0003952 | chrV | 122653 | 123492 | CP015054.1 | 560165 | 561058 | 22.74247492 | 279 | 297 | 0.939393939 | 1 | 0 | 1 | FIM1_278 | YEL018W | YEL018W | EAF5 | n |
| Kmar_QGN13647.1 | W303.asm02.HP0_G0005720 | OG0003953 | chrIII | 167234 | 167470 | CP015054.1 | 577665 | 577907 | 33.33333333 | 78 | 80 | 0.975 | 1 | 0 | 1 | FIM1_290 | YCR020W-B | YCR020W-B | HTL1 | n |
| Kmar_QGN13659.1 | W303.asm02.HP0_G0005640 | OG0003954 | chrIII | 144425 | 146173 | CP015054.1 | 610959 | 612491 | 26.16984402 | 582 | 510 | 1.141176471 | 1 | 0 | 1 | FIM1_302 | YCR014C | YCR014C | POL4 | n |
| Kmar_QGN13661.1 | W303.asm02.HP0_G0048820 | OG0003955 | chrXV | 275412 | 277394 | CP015054.1 | 614856 | 616595 | 18.14092954 | 660 | 579 | 1.139896373 | 1 | 0 | 1 | FIM1_304 | YOL025W | YOL025W | LAG2 | n |
| Kmar_QGN13674.1 | W303.asm02.HP0_G0008910 | OG0003956 | chrIV | 487731 | 488321 | CP015054.1 | 641271 | 641690 | 20.91836735 | 196 | 139 | 1.410071942 | 1 | 0 | 1 | FIM1_317 | YDR022C | YDR022C | ATG31 | n |
| Kmar_QGN13683.1 | W303.asm02.HP0_G0025670 | OG0003957 | chrIX | 26415 | 28013 | CP015054.1 | 656383 | 658053 | 51.1627907 | 532 | 556 | 0.956834532 | 1 | 0 | 1 | FIM1_326 | YIL162W | YIL162W | SUC2 | n |
| Kmar_QGN13697.1 | W303.asm02.HP0_G0043390 | OG0003958 | chrXIII | 871332 | 872270 | CP015054.1 | 680463 | 681611 | 20.33898305 | 312 | 382 | 0.816753927 | 1 | 0 | 1 | FIM1_340 | YMR299C | YMR299C | DYN3 | n |
| Kmar_QGN13699.1 | W303.asm02.HP0_G0022420 | OG0003959 | chrVII | 1056620 | 1057156 | CP015054.1 | 684314 | 684859 | 12.43243243 | 178 | 181 | 0.983425414 | 1 | 0 | 1 | FIM1_342 | YGR275W | YGR275W | RTT102 | n |
| Kmar_QGN13712.1 | W303.asm02.HP0_G0043310 | OG0003960 | chrXIII | 859551 | 861311 | CP015054.1 | 713210 | 714919 | 41.03852596 | 586 | 569 | 1.029876977 | 1 | 0 | 1 | FIM1_356 | YMR291W | YMR291W | TDA1 | n |
| Kmar_QGN13727.1 | W303.asm02.HP0_G0025130 | OG0003961 | chrVIII | 478848 | 480563 | CP015054.1 | 749444 | 751423 | 20.3030303 | 571 | 659 | 0.86646434 | 1 | 0 | 1 | FIM1_371 | YHR184W | YHR184W | SSP1 | n |
| Kmar_QGN13744.1 | W303.asm02.HP0_G0019850 | OG0003962 | chrVII | 488236 | 489489 | CP015054.1 | 795711 | 796997 | 29.63800905 | 417 | 428 | 0.974299065 | 1 | 0 | 1 | FIM1_388 | YGL004C | YGL004C | RPN14 | n |
| Kmar_QGN13757.1 | W303.asm02.HP0_G0005470 | OG0003963 | chrIII | 113261 | 114031 | CP015054.1 | 819840 | 820811 | 20.76677316 | 256 | 323 | 0.792569659 | 1 | 0 | 1 | FIM1_402 | YCL005W | YCL005W | LDB16 | n |
| Kmar_QGN13763.1 | W303.asm02.HP0_G0023300 | OG0003964 | chrVIII | 132525 | 133682 | CP015054.1 | 830827 | 831987 | 29.29782082 | 385 | 386 | 0.997409326 | 1 | 0 | 1 | FIM1_408 | YHR017W | YHR017W | YSC83 | n |
| Kmar_QGN13769.1 | W303.asm02.HP0_G0040050 | OG0003965 | chrXIII | 166369 | 167676 | CP015054.1 | 844714 | 846102 | 58.06451613 | 435 | 462 | 0.941558442 | 1 | 0 | 1 | FIM1_414 | YML051W | YML051W | GAL80 | n |
| Kmar_QGN13782.1 | W303.asm02.HP0_G0049750 | OG0003966 | chrXV | 465506 | 467278 | CP015054.1 | 871327 | 873255 | 27.74390244 | 590 | 642 | 0.919003115 | 1 | 0 | 1 | FIM1_428 | YOR073W | YOR073W | SGO1 | n |
| Kmar_QGN13798.1 | W303.asm02.HP0_G0048000 | OG0003967 | chrXV | 108713 | 109351 | CP015054.1 | 896874 | 897521 | 36.95652174 | 212 | 215 | 0.986046512 | 1 | 0 | 1 | FIM1_444 | YOL111C | YOL111C | MDY2 | n |
| Kmar_QGN13801.1 | W303.asm02.HP0_G0044020 | OG0003968 | chrXIV | 77393 | 78967 | CP015054.1 | 901991 | 903478 | 32.82442748 | 524 | 495 | 1.058585859 | 1 | 0 | 1 | FIM1_447 | YNL295W | YNL295W | MRX6 | n |
| Kmar_QGN13808.1 | W303.asm02.HP0_G0047940 | OG0003969 | chrXV | 98006 | 99943 | CP015054.1 | 918845 | 920674 | 22.08955224 | 645 | 609 | 1.0591133 | 1 | 0 | 1 | FIM1_454 | YOL117W | YOL117W | RRI2 | n |
| Kmar_QGN13814.1 | W303.asm02.HP0_G0013170 | OG0003970 | chrIV | 1420269 | 1420865 | CP015054.1 | 929206 | 929778 | 31.84079602 | 198 | 190 | 1.042105263 | 1 | 0 | 1 | FIM1_460 | YDR478W | YDR478W | SNM1 | n |
| Kmar_QGN13838.1 | W303.asm02.HP0_G0028130 | OG0003971 | chrX | 90678 | 91322 | CP015054.1 | 974821 | 975384 | 19.33962264 | 214 | 187 | 1.144385027 | 1 | 0 | 1 | FIM1_484 | YJL170C | YJL170C | ASG7 | n |
| Kmar_QGN13841.1 | W303.asm02.HP0_G0028100 | OG0003972 | chrX | 85691 | 86059 | CP015054.1 | 980534 | 980884 | 47.54098361 | 122 | 116 | 1.051724138 | 1 | 0 | 1 | FIM1_487 | YJL173C | YJL173C | RFA3 | n |
| Kmar_QGN13849.1 | W303.asm02.HP0_G0029870 | OG0003973 | chrX | 492490 | 494727 | CP015054.1 | 994137 | 996590 | 22.7863046 | 745 | 817 | 0.911872705 | 1 | 0 | 0 | FIM1_495 | YJR030C | YJR030C | RBH2 | n |
| Kmar_QGN13853.1 | W303.asm02.HP0_G0005810 | OG0003974 | chrIII | 196036 | 196526 | CP015054.1 | 1001461 | 1001859 | 60.74074074 | 135 | 132 | 1.022727273 | 1 | 0 | 1 | FIM1_499 | YCR028C-A | YCR028C-A | RIM1 | n |
| Kmar_QGN13858.1 | W303.asm02.HP0_G0033870 | OG0003975 | chrXI | 572211 | 574802 | CP015054.1 | 1008632 | 1010596 | 23.8150289 | 863 | 654 | 1.319571865 | 1 | 0 | 1 | FIM1_504 | YKR064W | YKR064W | OAF3 | n |
| Kmar_QGN13862.1 | W303.asm02.HP0_G0028330 | OG0003976 | chrX | 130947 | 132095 | CP015054.1 | 1015889 | 1017001 | 22.64631043 | 382 | 370 | 1.032432432 | 1 | 0 | 1 | FIM1_508 | YJL147C | YJL147C | SMT1 | n |
| Kmar_QGN13863.1 | W303.asm02.HP0_G0028340 | OG0003977 | chrX | 132820 | 134229 | CP015054.1 | 1017641 | 1018747 | 23.76873662 | 469 | 368 | 1.274456522 | 1 | 0 | 1 | FIM1_509 | YJL146W | YJL146W | IDS2 | n |
| Kmar_QGN13879.1 | W303.asm02.HP0_G0028460 | OG0003978 | chrX | 149151 | 149375 | CP015054.1 | 1053836 | 1054084 | 47.56097561 | 74 | 82 | 0.902439024 | 1 | 0 | 1 | FIM1_526 | YJL133C-A | YJL133C-A | DPI8 | n |
| Kmar_QGN13890.1 | W303.asm02.HP0_G0022190 | OG0003979 | chrVII | 1003614 | 1005959 | CP015054.1 | 1078888 | 1081479 | 25.81005587 | 781 | 863 | 0.904982619 | 1 | 0 | 1 | FIM1_537 | YGR250C | YGR250C | RIE1 | n |
| Kmar_QGN13895.1 | W303.asm02.HP0_G0024950 | OG0003980 | chrVIII | 445448 | 446233 | CP015054.1 | 1101922 | 1102713 | 29.88929889 | 261 | 263 | 0.992395437 | 1 | 0 | 1 | FIM1_542 | YHR167W | YHR167W | THP2 | n |
| Kmar_QGN13933.1 | W303.asm02.HP0_G0050410 | OG0003981 | chrXV | 607389 | 609350 | CP015054.1 | 1170621 | 1172375 | 31.68168168 | 653 | 584 | 1.118150685 | 1 | 0 | 1 | FIM1_581 | YOR147W | YOR147W | MDM32 | n |
| Kmar_QGN13939.1 | W303.asm02.HP0_G0039370 | OG0003982 | chrXIII | 25162 | 26235 | CP015054.1 | 1183148 | 1183924 | 20.16806723 | 357 | 258 | 1.38372093 | 1 | 0 | 1 | FIM1_587 | YML119W | YML119W | | n |

| | | | | | | | | | | | | | | | | | | | | |
|---|---|---|---|---|---|---|---|---|---|---|---|---|---|---|---|---|---|---|---|---|
| Kmar_QGN13962.1 | W303.asm02.HP0_G0019340 | OG0003983 | chrVII | 385921 | 386664 | CP015054.1 | 1226910 | 1227521 | 30.73770492 | 247 | 203 | 1.216748768 | 1 | 0 | 1 | FIM1_610 | YGL061C | YGL061C | DUO1 | n |
| Kmar_QGN13966.1 | W303.asm02.HP0_G0019360 | OG0003984 | chrVII | 389176 | 390651 | CP015054.1 | 1232975 | 1234384 | 61.91446029 | 491 | 469 | 1.046908316 | 1 | 0 | 1 | FIM1_615 | YGL059W | YGL059W | PKP2 | n |
| Kmar_QGN13973.1 | W303.asm02.HP0_G0004030 | OG0003985 | chrII | 668444 | 669418 | CP015054.1 | 1248179 | 1249039 | 22.42990654 | 324 | 286 | 1.132867133 | 1 | 0 | 1 | FIM1_622 | YBR211C | YBR211C | AME1 | n |
| Kmar_QGN13977.1 | W303.asm02.HP0_G0037730 | OG0003986 | chrXII | 725594 | 726328 | CP015054.1 | 1253022 | 1253747 | 46.93877551 | 244 | 241 | 1.012448133 | 1 | 0 | 1 | FIM1_626 | YLR301W | YLR301W | HRI1 | n |
| Kmar_QGN14007.1 | W303.asm02.HP0_G0003310 | OG0003987 | chrII | 530048 | 531622 | CP015054.1 | 1315823 | 1317229 | 25.23364486 | 524 | 468 | 1.11965812 | 1 | 0 | 1 | FIM1_658 | YBR138C | YBR138C | | n |
| Kmar_QGN14072.1 | W303.asm02.HP0_G0030490 | OG0003988 | chrX | 622959 | 624929 | CP015054.1 | 1462167 | 1464095 | 48.35329341 | 656 | 642 | 1.021806854 | 1 | 0 | 1 | FIM1_723 | YJR098C | YJR098C | | n |
| Kmar_QGN14093.1 | W303.asm02.HP0_G0018750 | OG0003989 | chrVII | 265761 | 266612 | CP015054.1 | 1505302 | 1506117 | 25.34722222 | 283 | 271 | 1.044280443 | 1 | 0 | 1 | FIM1_744 | YGL128C | YGL128C | CWC23 | n |
| Kmar_QGN14095.1 | W303.asm02.HP0_G0018640 | OG0003990 | chrVII | 237821 | 241480 | CP015054.1 | 1507753 | 1511343 | 51.38436482 | 1219 | 1196 | 1.019230769 | 1 | 0 | 1 | FIM1_746 | YGL140C | YGL140C | | n |
| Kmar_QGN14111.1 | W303.asm02.HP0_G0005930 | OG0003991 | chrIII | 229241 | 229624 | CP015054.1 | 1555861 | 1556247 | 44.53125 | 127 | 128 | 0.9921875 | 1 | 0 | 1 | FIM1_762 | YCR043C | YCR043C | | n |
| Kmar_QGN14165.1 | W303.asm02.HP0_G0013680 | OG0003992 | chrIV | 1505089 | 1506246 | CP015054.1 | 1668931 | 1669986 | 20.20725389 | 385 | 351 | 1.096866097 | 1 | 0 | 1 | FIM1_818 | YDR532C | YDR532C | KRE28 | n |
| Kmar_QGN14182.1 | W303.asm02.HP0_G0004990 | OG0003993 | chrIII | 16943 | 18214 | CP015054.1 | 1700076 | 1701560 | 27.32673267 | 423 | 494 | 0.856275304 | 1 | 0 | 1 | FIM1_835 | YCL063W | YCL063W | VAC17 | n |
| Kmar_QGN14185.1 | W303.asm02.HP0_G0004960 | OG0003994 | chrIII | 12039 | 12671 | CP015054.1 | 1709151 | 1709828 | 21.12068966 | 210 | 225 | 0.933333333 | 1 | 0 | 1 | FIM1_838 | YCL067C/YCR039C | YCR039C | MATALPHA2 | y |
| Kmar_QGN14217.1 | W303.asm02.HP0_G0015950 | OG0003995 | chrV | 470533 | 472407 | CP015055.1 | 51346 | 53307 | 30.60921248 | 624 | 653 | 0.955589587 | 1 | 0 | 1 | FIM1_873 | YER147C | YER147C | SCC4 | n |
| Kmar_QGN14219.1 | W303.asm02.HP0_G0009400 | OG0003996 | chrIV | 596715 | 597224 | CP015055.1 | 56705 | 57202 | 31.21387283 | 169 | 165 | 1.024242424 | 1 | 0 | 1 | FIM1_875 | YDR073W | YDR073W | SNF11 | n |
| Kmar_QGN14223.1 | W303.asm02.HP0_G0009370 | OG0003997 | chrIV | 592375 | 592656 | CP015055.1 | 62767 | 63036 | 40.625 | 93 | 89 | 1.04494382 | 1 | 0 | 1 | FIM1_879 | YDR070C | YDR070C | FMP16 | n |
| Kmar_QGN14252.1 | W303.asm02.HP0_G0019590 | OG0003998 | chrVII | 433656 | 433919 | CP015055.1 | 116191 | 116463 | 31.91489362 | 87 | 90 | 0.966666667 | 1 | 0 | 1 | FIM1_908 | YGL032C | YGL032C | AGA2 | n |
| Kmar_QGN14275.1 | W303.asm02.HP0_G0002950 | OG0003999 | chrII | 457801 | 459876 | CP015055.1 | 160826 | 162955 | 23.29700272 | 691 | 709 | 0.97461213 | 1 | 0 | 1 | FIM1_932 | YBR098W | YBR098W | MMS4 | n |
| Kmar_QGN14277.1 | W303.asm02.HP0_G0022860 | OG0004000 | chrVIII | 53055 | 54053 | CP015055.1 | 166656 | 167525 | 22.12389381 | 332 | 289 | 1.148788927 | 1 | 0 | 1 | FIM1_934 | YHL025W | YHL025W | SNF6 | n |
| Kmar_QGN14317.1 | W303.asm02.HP0_G0010780 | OG0004001 | chrIV | 909956 | 911353 | CP015055.1 | 263059 | 264477 | 18.67219917 | 465 | 472 | 0.985169492 | 1 | 0 | 1 | FIM1_974 | YDR219C | YDR219C | MFB1 | n |
| Kmar_QGN14331.1 | W303.asm02.HP0_G0037270 | OG0004002 | chrXII | 623921 | 628591 | CP015055.1 | 289165 | 293709 | 30.66332916 | 1556 | 1514 | 1.027741083 | 1 | 0 | 1 | FIM1_991 | YLR247C | YLR247C | IRC20 | n |
| Kmar_QGN14341.1 | W303.asm02.HP0_G0041350 | OG0004003 | chrXIII | 456330 | 457766 | CP015055.1 | 312586 | 313758 | 26.25 | 478 | 390 | 1.225641026 | 1 | 0 | 1 | FIM1_1001 | YMR094W | YMR094W | CTF13 | n |
| Kmar_QGN14356.1 | W303.asm02.HP0_G0032270 | OG0004004 | chrXI | 265246 | 266319 | CP015055.1 | 345913 | 346800 | 25.41899441 | 357 | 295 | 1.210169492 | 1 | 0 | 1 | FIM1_1016 | YKL098W | YKL098W | MTC2 | n |
| Kmar_QGN14364.1 | W303.asm02.HP0_G0037020 | OG0004005 | chrXII | 574275 | 576461 | CP015055.1 | 363403 | 365550 | 26.79045093 | 728 | 715 | 1.018181818 | 1 | 0 | 1 | FIM1_1024 | YLR219W | YLR219W | MSC3 | n |
| Kmar_QGN14373.1 | W303.asm02.HP0_G0009710 | OG0004006 | chrIV | 665834 | 666688 | CP015055.1 | 381709 | 382485 | 20.07042254 | 284 | 258 | 1.100775194 | 1 | 0 | 1 | FIM1_1033 | YDR106W | YDR106W | ARP10 | n |
| Kmar_QGN14374.1 | W303.asm02.HP0_G0016270 | OG0004007 | chrV | 557426 | 558229 | CP015055.1 | 382591 | 383502 | 19.07894737 | 267 | 303 | 0.881188119 | 1 | 0 | 1 | FIM1_1034 | YER180C | YER180C | ISC10 | n |
| Kmar_QGN14375.1 | W303.asm02.HP0_G0016280 | OG0004008 | chrV | 558567 | 558824 | CP015055.1 | 383726 | 383929 | 42.35294118 | 85 | 67 | 1.268656716 | 1 | 0 | 1 | FIM1_1035 | YER180C-A | YER180C-A | SLO1 | n |
| Kmar_QGN14385.1 | W303.asm02.HP0_G0002730 | OG0004009 | chrII | 408092 | 408580 | CP015055.1 | 414391 | 414810 | 41.97530864 | 162 | 139 | 1.165467626 | 1 | 0 | 1 | FIM1_1045 | YBR077C | YBR077C | SLM4 | n |
| Kmar_QGN14386.1 | W303.asm02.HP0_G0009990 | OG0004010 | chrIV | 733168 | 735159 | CP015055.1 | 415613 | 417409 | 31.58682635 | 663 | 598 | 1.108695652 | 1 | 0 | 1 | FIM1_1046 | YDR137W | YDR137W | RGP1 | n |
| Kmar_QGN14388.1 | W303.asm02.HP0_G0010000 | OG0004011 | chrIV | 735487 | 737745 | CP015055.1 | 419714 | 421942 | 23.46002621 | 752 | 742 | 1.013477089 | 1 | 0 | 1 | FIM1_1048 | YDR138W | YDR138W | HPR1 | n |
| Kmar_QGN14407.1 | W303.asm02.HP0_G0013270 | OG0004012 | chrIV | 1434882 | 1436483 | CP015055.1 | 453355 | 455061 | 21.24789207 | 533 | 568 | 0.938380282 | 1 | 0 | 1 | FIM1_1067 | YDR488C | YDR488C | PAC11 | n |
| Kmar_QGN14417.1 | W303.asm02.HP0_G0025680 | OG0004013 | chrIX | 28464 | 29171 | CP015055.1 | 476069 | 476665 | 23.40425532 | 235 | 198 | 1.186868687 | 1 | 0 | 1 | FIM1_1078 | YIL161W | YIL161W | SMU2 | n |
| Kmar_QGN14420.1 | W303.asm02.HP0_G0018130 | OG0004014 | chrVII | 125396 | 126682 | CP015055.1 | 484168 | 485526 | 41.03671706 | 428 | 452 | 0.946902655 | 1 | 0 | 1 | FIM1_1081 | YGL196W | YGL196W | DSD1 | n |
| Kmar_QGN14435.1 | W303.asm02.HP0_G0018930 | OG0004015 | chrVII | 300596 | 301018 | CP015055.1 | 512475 | 512789 | 35.43307087 | 140 | 104 | 1.346153846 | 1 | 0 | 1 | FIM1_1096 | YGL108C | YGL108C | | n |
| Kmar_QGN14440.1 | W303.asm02.HP0_G0004300 | OG0004016 | chrII | 723346 | 724698 | CP015055.1 | 520558 | 521658 | 24.83370288 | 450 | 366 | 1.229508197 | 1 | 0 | 1 | FIM1_1101 | YBR240C | YBR240C | THI2 | n |
| Kmar_QGN14470.1 | W303.asm02.HP0_G0004160 | OG0004017 | chrII | 696415 | 697977 | CP015055.1 | 582851 | 584356 | 62.71510516 | 520 | 501 | 1.037924152 | 1 | 0 | 1 | FIM1_1131 | YBR227C | YBR227C | MCX1 | n |
| Kmar_QGN14471.1 | W303.asm02.HP0_G0004150 | OG0004018 | chrII | 693467 | 696172 | CP015055.1 | 584509 | 586371 | 23.94678492 | 901 | 620 | 1.453225806 | 1 | 0 | 1 | FIM1_1132 | YBR225W | YBR225W | | n |
| Kmar_QGN14473.1 | W303.asm02.HP0_G0004140 | OG0004019 | chrII | 690947 | 693137 | CP015055.1 | 590119 | 591597 | 24.18772563 | 544 | 492 | 1.105691057 | 1 | 0 | 1 | FIM1_1134 | YBR223C | YBR223C | TDP1 | n |
| Kmar_QGN14500.1 | W303.asm02.HP0_G0016600 | OG0004020 | chrVI | 45279 | 46436 | CP015055.1 | 642746 | 643858 | 53.92670157 | 385 | 370 | 1.040540541 | 1 | 0 | 1 | FIM1_1161 | YFL030W | YFL030W | AGX1 | n |
| Kmar_QGN14502.1 | W303.asm02.HP0_G0035480 | OG0004021 | chrXII | 263118 | 265667 | CP015055.1 | 646434 | 648572 | 21.96261682 | 849 | 712 | 1.19241573 | 1 | 0 | 1 | FIM1_1163 | YLR057W | YLR057W | MNL2 | n |
| Kmar_QGN14532.1 | W303.asm02.HP0_G0020610 | OG0004022 | chrVII | 645159 | 646178 | CP015055.1 | 709016 | 709972 | 30.94555874 | 339 | 318 | 1.066037736 | 1 | 0 | 1 | FIM1_1194 | YGR084C | YGR084C | MRP13 | n |
| Kmar_QGN14533.1 | W303.asm02.HP0_G0056290 | OG0004023 | chrXVI | 736991 | 737518 | CP015055.1 | 710336 | 710833 | 13.63636364 | 175 | 165 | 1.060606061 | 1 | 0 | 1 | FIM1_1195 | YPR101W | YPR101W | SNT309 | n |
| Kmar_QGN14558.1 | W303.asm02.HP0_G0048670 | OG0004024 | chrXV | 250280 | 251371 | CP015055.1 | 751265 | 752380 | 59.29919137 | 363 | 371 | 0.978436658 | 1 | 0 | 1 | FIM1_1220 | YOL042W | YOL042W | NGL1 | n |
| Kmar_QGN14561.1 | W303.asm02.HP0_G0048650 | OG0004025 | chrXV | 247605 | 248756 | CP015055.1 | 756191 | 757255 | 18.2767624 | 383 | 354 | 1.081920904 | 1 | 0 | 1 | FIM1_1223 | YOL044W | YOL044W | PEX15 | n |
| Kmar_QGN14586.1 | W303.asm02.HP0_G0007910 | OG0004026 | chrIV | 293111 | 293551 | CP015055.1 | 803900 | 804325 | 37.90849673 | 146 | 141 | 1.035460993 | 1 | 0 | 1 | FIM1_1248 | YDL092W | YDL092W | SRP14 | n |
| Kmar_QGN14596.1 | W303.asm02.HP0_G0006230 | OG0004027 | chrIII | 281864 | 282250 | CP015055.1 | 826052 | 826450 | 29.85074627 | 128 | 132 | 0.96969697 | 1 | 0 | 1 | FIM1_1258 | YCR082W | YCR082W | AHC2 | n |
| Kmar_QGN14605.1 | W303.asm02.HP0_G0019010 | OG0004028 | chrVII | 314285 | 315022 | CP015055.1 | 844174 | 844947 | 34.21052632 | 245 | 257 | 0.953307393 | 1 | 0 | 1 | FIM1_1268 | YGL098W | YGL098W | USE1 | n |
| Kmar_QGN14610.1 | W303.asm02.HP0_G0052940 | OG0004029 | chrXVI | 73293 | 74450 | CP015055.1 | 852476 | 853720 | 25.16853933 | 385 | 414 | 0.929951691 | 1 | 0 | 1 | FIM1_1273 | YPL255W | YPL255W | BBP1 | n |
| Kmar_QGN14618.1 | W303.asm02.HP0_G0009430 | OG0004030 | chrIV | 602744 | 603964 | CP015055.1 | 868089 | 869357 | 31.84855234 | 406 | 422 | 0.962085308 | 1 | 0 | 1 | FIM1_1282 | YDR076W | YDR076W | RAD55 | n |
| Kmar_QGN14622.1 | W303.asm02.HP0_G0009450 | OG0004031 | chrIV | 606474 | 607145 | CP015055.1 | 879964 | 880617 | 34.21052632 | 223 | 217 | 1.02764977 | 1 | 0 | 1 | FIM1_1286 | YDR078C | YDR078C | SHU2 | n |
| Kmar_QGN14624.1 | W303.asm02.HP0_G0016010 | OG0004032 | chrV | 481606 | 482370 | CP015055.1 | 882948 | 883772 | 20.41522491 | 254 | 274 | 0.927007299 | 1 | 0 | 1 | FIM1_1288 | YER153C | YER153C | PET122 | n |
| Kmar_QGN14628.1 | W303.asm02.HP0_G0034610 | OG0004033 | chrXII | 73418 | 74110 | CP015055.1 | 895972 | 896538 | 29.56521739 | 230 | 188 | 1.223404255 | 1 | 0 | 1 | FIM1_1292 | YLL033W | YLL033W | IRC19 | n |
| Kmar_QGN14643.1 | W303.asm02.HP0_G0012060 | OG0004034 | chrIV | 1197623 | 1200457 | CP015055.1 | 929074 | 931716 | 22.49208025 | 944 | 880 | 1.072727273 | 1 | 0 | 1 | FIM1_1307 | YDR356W | YDR356W | SPC110 | n |
| Kmar_QGN14662.1 | W303.asm02.HP0_G0017070 | OG0004035 | chrVI | 152038 | 153129 | CP015055.1 | 970853 | 971863 | 36.87002653 | 363 | 336 | 1.080357143 | 1 | 0 | 1 | FIM1_1326 | YFR018C | YFR018C | | n |
| Kmar_QGN14694.1 | W303.asm02.HP0_G0021010 | OG0004036 | chrVII | 757505 | 758197 | CP015055.1 | 1050490 | 1050951 | 25.2173913 | 230 | 153 | 1.503267974 | 1 | 0 | 1 | FIM1_1361 | YGR126W | YGR126W | | n |
| Kmar_QGN14698.1 | W303.asm02.HP0_G0056720 | OG0004037 | chrXVI | 838466 | 839773 | CP015055.1 | 1055040 | 1056302 | 48.65470852 | 435 | 420 | 1.035714286 | 1 | 0 | 1 | FIM1_1365 | YPR148C | YPR148C | | n |
| Kmar_QGN14705.1 | W303.asm02.HP0_G0026810 | OG0004038 | chrIX | 268352 | 268768 | CP015055.1 | 1064320 | 1064769 | 18.83116883 | 138 | 149 | 0.926174497 | 1 | 0 | 1 | FIM1_1372 | YIL040W | YIL040W | APQ12 | n |
| Kmar_QGN14711.1 | W303.asm02.HP0_G0026840 | OG0004039 | chrIX | 273658 | 275628 | CP015055.1 | 1075563 | 1077335 | 19.93957704 | 656 | 590 | 1.111864407 | 1 | 0 | 1 | FIM1_1378 | YIL037C | YIL037C | PRM2 | n |
| Kmar_QGN14726.1 | W303.asm02.HP0_G0056740 | OG0004040 | chrXVI | 842688 | 843308 | CP015055.1 | 1103169 | 1103753 | 32.91139241 | 206 | 194 | 1.06185567 | 1 | 0 | 1 | FIM1_1394 | YPR151C | YPR151C | SUE1 | n |
| Kmar_QGN14728.1 | W303.asm02.HP0_G0056750 | OG0004041 | chrXVI | 843694 | 845091 | CP015055.1 | 1108299 | 1109633 | 30.10526316 | 465 | 444 | 1.047297297 | 1 | 0 | 1 | FIM1_1396 | YPR152C | YPR152C | URN1 | n |
| Kmar_QGN14733.1 | W303.asm02.HP0_G0021130 | OG0004042 | chrVII | 779156 | 782026 | CP015055.1 | 1119468 | 1121750 | 25 | 956 | 760 | 1.257894737 | 1 | 0 | 1 | FIM1_1402 | YGR140W | YGR140W | CBF2 | n |
| Kmar_QGN14738.1 | W303.asm02.HP0_G0013050 | OG0004043 | chrIV | 1394565 | 1398872 | CP015055.1 | 1131199 | 1135488 | 23.99737015 | 1435 | 1429 | 1.00419874 | 1 | 0 | 1 | FIM1_1407 | YDR464W | YDR464W | SPP41 | n |
| Kmar_QGN14766.1 | W303.asm02.HP0_G0012390 | OG0004044 | chrIV | 1253748 | 1255415 | CP015055.1 | 1187911 | 1189668 | 50.99337748 | 555 | 585 | 0.948717949 | 1 | 0 | 1 | FIM1_1436 | YDR387C | YDR387C | CIN10 | n |
| Kmar_QGN14770.1 | W303.asm02.HP0_G0050280 | OG0004045 | chrXV | 573058 | 573714 | CP015055.1 | 1197404 | 1198036 | 56.42201835 | 218 | 210 | 1.038095238 | 1 | 0 | 1 | FIM1_1440 | YOR131C | YOR131C | | n |
| Kmar_QGN14804.1 | W303.asm02.HP0_G0012180 | OG0004046 | chrIV | 1220609 | 1223173 | CP015055.1 | 1260312 | 1262801 | 20.04608295 | 854 | 829 | 1.030156815 | 1 | 0 | 1 | FIM1_1475 | YDR369C | YDR369C | XRS2 | n |
| Kmar_QGN14811.1 | W303.asm02.HP0_G0043730 | OG0004047 | chrXIV | 24480 | 27605 | CP015055.1 | 1274608 | 1276335 | 17.96349664 | 1041 | 575 | 1.810434783 | 1 | 0 | 1 | FIM1_1482 | YNL327W | YNL327W | EGT2 | n |
| Kmar_QGN14825.1 | W303.asm02.HP0_G0047170 | OG0004048 | chrXIV | 718721 | 720190 | CP015055.1 | 1299558 | 1300802 | 25.20491803 | 489 | 414 | 1.18115942 | 1 | 0 | 1 | FIM1_1496 | YNR045W | YNR045W | PET494 | n |
| Kmar_QGN14827.1 | W303.asm02.HP0_G0042010 | OG0004049 | chrXIII | 584879 | 586996 | CP015055.1 | 1303212 | 1305236 | 27.29805014 | 705 | 674 | 1.045994065 | 1 | 0 | 1 | FIM1_1498 | YMR163C | YMR163C | INP2 | n |
| Kmar_QGN14834.1 | W303.asm02.HP0_G0041980 | OG0004050 | chrXIII | 575674 | 578124 | CP015055.1 | 1320174 | 1322330 | 34.27172583 | 816 | 718 | 1.136490251 | 1 | 0 | 1 | FIM1_1505 | YMR160W | YMR160W | CVM1 | n |
| Kmar_QGN14835.1 | W303.asm02.HP0_G0041970 | OG0004051 | chrXIII | 575084 | 575536 | CP015055.1 | 1322702 | 1323082 | 30.66666667 | 150 | 126 | 1.19047619 | 1 | 0 | 1 | FIM1_1506 | YMR159C | YMR159C | ATG16 | n |
| Kmar_QGN14837.1 | W303.asm02.HP0_G0007860 | OG0004052 | chrIV | 283748 | 284773 | CP015055.1 | 1325173 | 1326045 | 35.19061584 | 341 | 290 | 1.175862069 | 1 | 0 | 1 | FIM1_1508 | YDL099W | YDL099W | BUG1 | n |
| Kmar_QGN14860.1 | W303.asm02.HP0_G0042060 | OG0004053 | chrXIII | 597916 | 599742 | CP015055.1 | 1365585 | 1367435 | 29.24528302 | 608 | 616 | 0.987012987 | 1 | 0 | 1 | FIM1_1533 | YMR168C | YMR168C | CEP3 | n |
| Kmar_QGN14876.1 | W303.asm02.HP0_G0021680 | OG0004054 | chrVII | 902148 | 904601 | CP015055.1 | 1401926 | 1404328 | 27.4909964 | 817 | 800 | 1.02125 | 1 | 0 | 1 | FIM1_1549 | YGR196C | YGR196C | FYV8 | n |
| Kmar_QGN14895.1 | W303.asm02.HP0_G0003220 | OG0004055 | chrII | 509367 | 510401 | CP015055.1 | 1450849 | 1451766 | 25.79710145 | 344 | 305 | 1.127868852 | 1 | 0 | 1 | FIM1_1569 | YBR128C | YBR128C | ATG14 | n |
| Kmar_QGN14901.1 | W303.asm02.HP0_G0003250 | OG0004056 | chrII | 513449 | 515563 | CP015055.1 | 1459848 | 1461725 | 28.27004219 | 704 | 625 | 1.1264 | 1 | 0 | 1 | FIM1_1575 | YBR131W | YBR131W | CCZ1 | n |
| Kmar_QGN14903.1 | W303.asm02.HP0_G0042230 | OG0004057 | chrXIII | 628772 | 629368 | CP015055.1 | 1464009 | 1464593 | 33.66336634 | 198 | 194 | 1.020618557 | 1 | 0 | 1 | FIM1_1577 | YMR184W | YMR184W | ADD37 | n |
| Kmar_QGN14915.1 | W303.asm02.HP0_G0044990 | OG0004058 | chrXIV | 287004 | 288305 | CP015055.1 | 1485529 | 1486338 | 19.16859122 | 433 | 269 | 1.609665428 | 1 | 0 | 1 | FIM1_1589 | YNL188W | YNL188W | KAR1 | n |
| Kmar_QGN14917.1 | W303.asm02.HP0_G0045000 | OG0004059 | chrXIV | 288516 | 289763 | CP015055.1 | 1487259 | 1488524 | 29.13752914 | 415 | 421 | 0.985748219 | 1 | 0 | 1 | FIM1_1591 | YNL187W | YNL187W | SWT21 | n |
| Kmar_QGN14924.1 | W303.asm02.HP0_G0045040 | OG0004060 | chrXIV | 296656 | 298323 | CP015055.1 | 1503997 | 1505529 | 41.63763066 | 555 | 510 | 1.088235294 | 1 | 0 | 1 | FIM1_1598 | YNL182C | YNL182C | IPI3 | n |
| Kmar_QGN14940.1 | W303.asm02.HP0_G0024720 | OG0004061 | chrVIII | 391615 | 392592 | CP015055.1 | 1535886 | 1536536 | 31.38461538 | 325 | 216 | 1.50462963 | 1 | 0 | 1 | FIM1_1615 | YHR143W | YHR143W | DSE2 | n |
| Kmar_QGN14941.1 | W303.asm02.HP0_G0045190 | OG0004062 | chrXIV | 325957 | 327012 | CP015055.1 | 1537403 | 1538446 | 19.04761905 | 351 | 347 | 1.011527378 | 1 | 0 | 1 | FIM1_1616 | YNL164C | YNL164C | IBD2 | n |
| Kmar_QGN14960.1 | W303.asm02.HP0_G0054020 | OG0004063 | chrXVI | 286413 | 286859 | CP015055.1 | 1575790 | 1576251 | 19.35483871 | 148 | 153 | 0.967320261 | 1 | 0 | 1 | FIM1_1637 | YPL144W | YPL144W | POC4 | n |
| Kmar_QGN14983.1 | W303.asm02.HP0_G0051020 | OG0004064 | chrXV | 748020 | 748577 | CP015055.1 | 1620798 | 1621358 | 52.40641711 | 185 | 186 | 0.994623656 | 1 | 0 | 1 | FIM1_1661 | YOR215C | YOR215C | AIM41 | n |
| Kmar_QGN15006.1 | W303.asm02.HP0_G0028810 | OG0004065 | chrX | 241837 | 244458 | CP015055.1 | 1671614 | 1673977 | 43.65714286 | 873 | 787 | 1.109275731 | 1 | 0 | 1 | FIM1_1687 | YJL094C | YJL094C | KHA1 | n |
| Kmar_QGN15051.1 | W303.asm02.HP0_G0031280 | OG0004066 | chrXI | 48182 | 48997 | CP015056.1 | 97523 | 98245 | 21.77121771 | 271 | 240 | 1.129166667 | 1 | 0 | 1 | FIM1_1736 | YKL208W | YKL208W | CBT1 | n |
| Kmar_QGN15059.1 | W303.asm02.HP0_G0031320 | OG0004067 | chrXI | 54729 | 56627 | CP015056.1 | 110864 | 112807 | 17.37160121 | 632 | 647 | 0.976816074 | 1 | 0 | 1 | FIM1_1744 | YKL204W | YKL204W | EAP1 | n |
| Kmar_QGN15078.1 | W303.asm02.HP0_G0023050 | OG0004068 | chrVIII | 90704 | 91156 | CP015056.1 | 158500 | 159015 | 25 | 150 | 171 | 0.877192982 | 1 | 0 | 1 | FIM1_1763 | YHL006C | YHL006C | SHU1 | n |
| Kmar_QGN15099.1 | W303.asm02.HP0_G0042730 | OG0004069 | chrXIII | 737504 | 739537 | CP015056.1 | 202064 | 204136 | 28.95863053 | 677 | 690 | 0.98115942 | 1 | 0 | 1 | FIM1_1786 | YMR232W | YMR232W | FUS2 | n |
| Kmar_QGN15118.1 | W303.asm02.HP0_G0041480 | OG0004070 | chrXIII | 483520 | 483867 | CP015056.1 | 241098 | 241403 | 32.47863248 | 115 | 101 | 1.138613861 | 1 | 0 | 1 | FIM1_1805 | YMR107W | YMR107W | SPG4 | n |
| Kmar_QGN15124.1 | W303.asm02.HP0_G0032000 | OG0004071 | chrXI | 196593 | 197333 | CP015056.1 | 254805 | 255554 | 41.015625 | 246 | 249 | 0.987951807 | 1 | 0 | 1 | FIM1_1811 | YKL130C | YKL130C | SHE2 | n |
| Kmar_QGN15144.1 | W303.asm02.HP0_G0009320 | OG0004072 | chrIV | 584836 | 585933 | CP015056.1 | 291004 | 292212 | 28.75 | 365 | 402 | 0.907960199 | 1 | 0 | 1 | FIM1_1831 | YDR065W | YDR065W | RRG1 | n |
| Kmar_QGN15191.1 | W303.asm02.HP0_G0021410 | OG0004073 | chrVII | 845133 | 846263 | CP015056.1 | 400565 | 401611 | 19.94680851 | 376 | 348 | 1.08045977 | 1 | 0 | 1 | FIM1_1879 | YGR168C | YGR168C | PEX35 | n |
| Kmar_QGN15198.1 | W303.asm02.HP0_G0008670 | OG0004074 | chrIV | 440673 | 441968 | CP015056.1 | 411068 | 411910 | 24.12993039 | 431 | 280 | 1.539285714 | 1 | 0 | 1 | FIM1_1886 | YDL005C | YDL005C | MED2 | n |
| Kmar_QGN15204.1 | W303.asm02.HP0_G0029460 | OG0004075 | chrX | 393004 | 394047 | CP015056.1 | 425701 | 426516 | 24.63768116 | 347 | 271 | 1.280442804 | 1 | 0 | 1 | FIM1_1892 | YJL023C | YJL023C | PET130 | n |
| Kmar_QGN15212.1 | W303.asm02.HP0_G0021520 | OG0004076 | chrVII | 865453 | 866673 | CP015056.1 | 440611 | 441744 | 22.13541667 | 406 | 377 | 1.076923077 | 1 | 0 | 1 | FIM1_1900 | YGR179C | YGR179C | OKP1 | n |
| Kmar_QGN15233.1 | W303.asm02.HP0_G0035110 | OG0004077 | chrXII | 186097 | 186996 | CP015056.1 | 488738 | 490051 | 20.77625571 | 299 | 437 | 0.684210526 | 1 | 0 | 1 | FIM1_1922 | YLR018C | YLR018C | POM34 | n |
| Kmar_QGN15274.1 | W303.asm02.HP0_G0049680 | OG0004078 | chrXV | 446675 | 447334 | CP015056.1 | 574793 | 575479 | 31.53526971 | 219 | 228 | 0.960526316 | 1 | 0 | 1 | FIM1_1963 | YOR064C | YOR064C | YNG1 | n |
| Kmar_QGN15350.1 | W303.asm02.HP0_G0054890 | OG0004079 | chrXVI | 465432 | 466037 | CP015056.1 | 768381 | 769013 | 17.43119266 | 201 | 210 | 0.957142857 | 1 | 0 | 1 | FIM1_2040 | YPL052W | YPL052W | OAZ1 | n |
| Kmar_QGN15354.1 | W303.asm02.HP0_G0003490 | OG0004080 | chrII | 567881 | 569977 | CP015056.1 | 776857 | 779112 | 22.40932642 | 698 | 751 | 0.92942743 | 1 | 0 | 1 | FIM1_2044 | YBR156C | YBR156C | SLI15 | n |
| Kmar_QGN15359.1 | W303.asm02.HP0_G0054860 | OG0004081 | chrXVI | 460351 | 461349 | CP015056.1 | 792095 | 792409 | 15.06024096 | 332 | 104 | 3.192307692 | 1 | 0 | 1 | FIM1_2049 | YPL055C | YPL055C | LGE1 | n |

| | | | | | | | | | | | | | | | | | | | | |
|---|---|---|---|---|---|---|---|---|---|---|---|---|---|---|---|---|---|---|---|---|
| Kmar_QGN15372.1 | W303.asm02.HP0_G0037350 | OG0004082 | chrXII | 653510 | 654475 | CP015056.1 | 822999 | 823874 | 31.25 | 321 | 291 | 1.103092784 | 1 | 0 | 1 | FIM1_2062 | YLR257W | YLR257W | | n |
| Kmar_QGN15386.1 | W303.asm02.HP0_G0001800 | OG0004083 | chrII | 208392 | 209234 | CP015056.1 | 851891 | 852724 | 26.04166667 | 280 | 277 | 1.010830325 | 1 | 0 | 1 | FIM1_2076 | YBL010C | YBL010C | LAA2 | n |
| Kmar_QGN15411.1 | W303.asm02.HP0_G0011310 | OG0004084 | chrIV | 1023867 | 1024919 | CP015056.1 | 903268 | 904287 | 35.32763533 | 350 | 339 | 1.032448378 | 1 | 0 | 1 | FIM1_2101 | YDR279W | YDR279W | RNH202 | n |
| Kmar_QGN15453.1 | W303.asm02.HP0_G0053840 | OG0004085 | chrXVI | 250252 | 252153 | CP015056.1 | 985080 | 986882 | 32.76397516 | 633 | 600 | 1.055 | 1 | 0 | 1 | FIM1_2144 | YPL161C | YPL161C | BEM4 | n |
| Kmar_QGN15461.1 | W303.asm02.HP0_G0053800 | OG0004086 | chrXVI | 243271 | 243912 | CP015056.1 | 998355 | 998843 | 41.31455399 | 213 | 162 | 1.314814815 | 1 | 0 | 1 | FIM1_2152 | YPL166W | YPL166W | ATG29 | n |
| Kmar_QGN15464.1 | W303.asm02.HP0_G0053780 | OG0004087 | chrXVI | 237150 | 238442 | CP015056.1 | 1004667 | 1005821 | 22.40566038 | 430 | 384 | 1.119791667 | 1 | 0 | 1 | FIM1_2155 | YPL168W | YPL168W | MRX4 | n |
| Kmar_QGN15468.1 | W303.asm02.HP0_G0051380 | OG0004088 | chrXV | 808994 | 809830 | CP015056.1 | 1011443 | 1012093 | 25.47528517 | 278 | 216 | 1.287037037 | 1 | 0 | 1 | FIM1_2159 | YOR255W | YOR255W | OSW1 | n |
| Kmar_QGN15485.1 | W303.asm02.HP0_G0033380 | OG0004089 | chrXI | 471197 | 472258 | CP015056.1 | 1039983 | 1040888 | 25.20775623 | 353 | 301 | 1.172757475 | 1 | 0 | 1 | FIM1_2176 | YKR011C | YKR011C | FPT1 | n |
| Kmar_QGN15493.1 | W303.asm02.HP0_G0026480 | OG0004090 | chrIX | 207153 | 208043 | CP015056.1 | 1057159 | 1058040 | 46.15384615 | 296 | 293 | 1.010238908 | 1 | 0 | 1 | FIM1_2184 | YIL076W | YIL076W | SEC28 | n |
| Kmar_QGN15525.1 | W303.asm02.HP0_G0010970 | OG0004091 | chrIV | 948175 | 949653 | CP015056.1 | 1119854 | 1121281 | 21.73038229 | 492 | 475 | 1.035789474 | 1 | 0 | 1 | FIM1_2216 | YDR240C | YDR240C | SNU56 | n |
| Kmar_QGN15526.1 | W303.asm02.HP0_G0010960 | OG0004092 | chrIV | 945558 | 947921 | CP015056.1 | 1121631 | 1123850 | 30.19559902 | 787 | 739 | 1.064952639 | 1 | 0 | 1 | FIM1_2217 | YDR239C | YDR239C | | n |
| Kmar_QGN15540.1 | W303.asm02.HP0_G0010870 | OG0004093 | chrIV | 929286 | 930647 | CP015056.1 | 1150255 | 1151670 | 36.82092555 | 453 | 471 | 0.961783439 | 1 | 0 | 1 | FIM1_2231 | YDR229W | YDR229W | IVY1 | n |
| Kmar_QGN15543.1 | W303.asm02.HP0_G0040150 | OG0004094 | chrXIII | 204893 | 205915 | CP015056.1 | 1157280 | 1158401 | 13.13672922 | 340 | 373 | 0.91152815 | 1 | 0 | 1 | FIM1_2234 | YML037C | YML037C | LFT1 | n |
| Kmar_QGN15583.1 | W303.asm02.HP0_G0048220 | OG0004095 | chrXV | 154367 | 155200 | CP015056.1 | 1241465 | 1242334 | 26.44067797 | 277 | 289 | 0.958477509 | 1 | 0 | 1 | FIM1_2275 | YOL088C | YOL088C | MPD2 | n |
| Kmar_QGN15626.1 | W303.asm02.HP0_G0016970 | OG0004096 | chrVI | 129018 | 129683 | CP015056.1 | 1340384 | 1341034 | 26.10619469 | 221 | 216 | 1.023148148 | 1 | 0 | 1 | FIM1_2319 | YFR008W | YFR008W | FAR7 | n |
| Kmar_QGN15629.1 | W303.asm02.HP0_G0033340 | OG0004097 | chrXI | 461002 | 461550 | CP015056.1 | 1345645 | 1346100 | 34.61538462 | 182 | 151 | 1.205298013 | 1 | 0 | 1 | FIM1_2322 | YKR007W | YKR007W | MEH1 | n |
| Kmar_QGN15637.1 | W303.asm02.HP0_G0020340 | OG0004098 | chrVII | 591355 | 592464 | CP015056.1 | 1360254 | 1361507 | 33.10185185 | 369 | 417 | 0.884892086 | 1 | 0 | 1 | FIM1_2330 | YGR052W | YGR052W | FMP48 | n |
| Kmar_QGN15677.1 | W303.asm02.HP0_G0040450 | OG0004099 | chrXIII | 256598 | 258922 | CP015056.1 | 1444793 | 1446394 | 29.16129032 | 774 | 533 | 1.452157598 | 1 | 0 | 1 | FIM1_2370 | YML006C | YML006C | GIS4 | n |
| Kmar_QGN15686.1 | W303.asm02.HP0_G0017940 | OG0004100 | chrVII | 78390 | 79769 | CP015056.1 | 1470211 | 1471899 | 42.0212766 | 459 | 562 | 0.816725979 | 1 | 0 | 1 | FIM1_2379 | YGL219C | YGL219C | MDM34 | n |
| Kmar_QGN15689.1 | W303.asm02.HP0_G0014820 | OG0004101 | chrV | 222985 | 223542 | CP015056.1 | 1473896 | 1474453 | 36.70212766 | 185 | 185 | 1 | 1 | 0 | 1 | FIM1_2382 | YER034W | YER034W | | n |
| Kmar_QGN15692.1 | W303.asm02.HP0_G0017900 | OG0004102 | chrVII | 74484 | 75737 | CP015056.1 | 1479815 | 1480984 | 18.05225653 | 417 | 389 | 1.071979434 | 1 | 0 | 1 | FIM1_2385 | YGL223C | YGL223C | COG1 | n |
| Kmar_QGN15694.1 | W303.asm02.HP0_G0014860 | OG0004103 | chrV | 227997 | 229391 | CP015056.1 | 1482340 | 1483176 | 19.39655172 | 464 | 278 | 1.669064748 | 1 | 0 | 1 | FIM1_2387 | YER038C | YER038C | KRE29 | n |
| Kmar_QGN15699.1 | W303.asm02.HP0_G0017860 | OG0004104 | chrVII | 68712 | 69083 | CP015056.1 | 1491610 | 1492038 | 30.71428571 | 123 | 142 | 0.866197183 | 1 | 0 | 1 | FIM1_2392 | YGL226W | YGL226W | MTC3 | n |
| Kmar_QGN15700.1 | W303.asm02.HP0_G0017850 | OG0004105 | chrVII | 68121 | 68530 | CP015056.1 | 1492341 | 1492730 | 36.08247423 | 86 | 97 | 0.886597938 | 1 | 0 | 1 | FIM1_2393 | YGL226C-A | YGL226C-A | OST5 | n |
| Kmar_QGN15701.1 | W303.asm02.HP0_G0017840 | OG0004106 | chrVII | 65043 | 67919 | CP015056.1 | 1492744 | 1495374 | 38.5492228 | 958 | 876 | 1.093607306 | 1 | 0 | 1 | FIM1_2394 | YGL227W | YGL227W | VID30 | n |
| Kmar_QGN15726.1 | W303.asm02.HP0_G0025830 | OG0004107 | chrIX | 63215 | 64804 | CP015056.1 | 1542318 | 1543859 | 25.58558559 | 529 | 513 | 1.031189084 | 1 | 0 | 1 | FIM1_2419 | YIL146C | YIL146C | ATG32 | n |
| Kmar_QGN15785.1 | W303.asm02.HP0_G0037920 | OG0004109 | chrXII | 767625 | 771989 | CP015057.1 | 93740 | 98137 | 25.03337784 | 1454 | 1465 | 0.992491468 | 1 | 0 | 1 | FIM1_2481 | YLR320W | YLR320W | MMS22 | n |
| Kmar_QGN15801.1 | W303.asm02.HP0_G0020030 | OG0004110 | chrVII | 519795 | 520367 | CP015057.1 | 122559 | 123149 | 34.5177665 | 190 | 196 | 0.969387755 | 1 | 0 | 1 | FIM1_2497 | YGR016W | YGR016W | | n |
| Kmar_QGN15830.1 | W303.asm02.HP0_G0041630 | OG0004111 | chrXIII | 511941 | 512195 | CP015057.1 | 178556 | 179044 | 30.24691358 | 84 | 162 | 0.518518519 | 1 | 0 | 1 | FIM1_2526 | YMR122W-A | YMR122W-A | NCW1 | n |
| Kmar_QGN15842.1 | W303.asm02.HP0_G0026970 | OG0004112 | chrIX | 300015 | 301055 | CP015057.1 | 200016 | 201080 | 34.42622951 | 346 | 354 | 0.97740113 | 1 | 0 | 1 | FIM1_2538 | YIL023C | YIL023C | YKE4 | n |
| Kmar_QGN15885.1 | W303.asm02.HP0_G0002670 | OG0004113 | chrII | 396698 | 397333 | CP015057.1 | 293114 | 293476 | 18.0952381 | 211 | 120 | 1.758333333 | 1 | 0 | 1 | FIM1_2583 | YBR071W | YBR071W | | n |
| Kmar_QGN15897.1 | W303.asm02.HP0_G0009750 | OG0004114 | chrIV | 680853 | 682553 | CP015057.1 | 333521 | 334924 | 30.24475524 | 566 | 467 | 1.211991435 | 1 | 0 | 1 | FIM1_2595 | YDR110W | YDR110W | FOB1 | n |
| Kmar_QGN15900.1 | W303.asm02.HP0_G0009770 | OG0004115 | chrIV | 685403 | 686524 | CP015057.1 | 340400 | 341242 | 27.82152231 | 373 | 280 | 1.332142857 | 1 | 0 | 1 | FIM1_2598 | YDR113C | YDR113C | PDS1 | n |
| Kmar_QGN15901.1 | W303.asm02.HP0_G0035800 | OG0004116 | chrXII | 330244 | 331125 | CP015057.1 | 341818 | 342714 | 25.08038585 | 293 | 298 | 0.983221477 | 1 | 0 | 1 | FIM1_2599 | YLR091W | YLR091W | GEP5 | n |
| Kmar_QGN15911.1 | W303.asm02.HP0_G0035820 | OG0004117 | chrXII | 334459 | 335348 | CP015057.1 | 359541 | 360239 | 42.96577947 | 262 | 232 | 1.129310345 | 1 | 0 | 1 | FIM1_2609 | YLR093C | YLR093C | NYV1 | n |
| Kmar_QGN15913.1 | W303.asm02.HP0_G0035830 | OG0004118 | chrXII | 335687 | 337195 | CP015057.1 | 362745 | 364121 | 30.0589391 | 502 | 458 | 1.096069869 | 1 | 0 | 1 | FIM1_2611 | YLR094C | YLR094C | GIS3 | n |
| Kmar_QGN15916.1 | W303.asm02.HP0_G0009870 | OG0004119 | chrIV | 703463 | 704377 | CP015057.1 | 372866 | 373966 | 23.22404372 | 304 | 366 | 0.830601093 | 1 | 0 | 1 | FIM1_2614 | YDR123C | YDR123C | INO2 | n |
| Kmar_QGN15933.1 | W303.asm02.HP0_G0009940 | OG0004120 | chrIV | 720656 | 721531 | CP015057.1 | 416342 | 417022 | 33.1010453 | 291 | 226 | 1.287610619 | 1 | 0 | 1 | FIM1_2631 | YDR130C | YDR130C | FIN1 | n |
| Kmar_QGN15950.1 | W303.asm02.HP0_G0037420 | OG0004121 | chrXII | 669077 | 670105 | CP015057.1 | 453339 | 454298 | 17.8041543 | 342 | 319 | 1.072100313 | 1 | 0 | 1 | FIM1_2648 | YLR265C | YLR265C | NEJ1 | n |
| Kmar_QGN15953.1 | W303.asm02.HP0_G0037400 | OG0004122 | chrXII | 664993 | 667476 | CP015057.1 | 457109 | 459818 | 24.10313901 | 827 | 872 | 0.948394495 | 1 | 0 | 1 | FIM1_2651 | YLR263W | YLR263W | RED1 | n |
| Kmar_QGN15973.1 | W303.asm02.HP0_G0006740 | OG0004123 | chrIV | 62317 | 65091 | CP015057.1 | 517290 | 519980 | 18.95287958 | 924 | 896 | 1.03125 | 1 | 0 | 1 | FIM1_2672 | YDL220C | YDL220C | CDC13 | n |
| Kmar_QGN15991.1 | W303.asm02.HP0_G0028650 | OG0004124 | chrX | 195321 | 197465 | CP015057.1 | 562283 | 564412 | 43.88586957 | 714 | 709 | 1.007052186 | 1 | 0 | 1 | FIM1_2691 | YJL112W | YJL112W | MDV1 | n |
| Kmar_QGN15992.1 | W303.asm02.HP0_G0033620 | OG0004125 | chrXI | 520605 | 521492 | CP015057.1 | 564746 | 565588 | 36.51315789 | 295 | 280 | 1.053571429 | 1 | 0 | 1 | FIM1_2692 | YKR037C | YKR037C | SPC34 | n |
| Kmar_QGN16027.1 | W303.asm02.HP0_G0019560 | OG0004126 | chrVII | 425687 | 428416 | CP015057.1 | 654696 | 657203 | 25.71428571 | 909 | 835 | 1.088622754 | 1 | 0 | 1 | FIM1_2727 | YGL036W | YGL036W | VIR1 | n |
| Kmar_QGN16034.1 | W303.asm02.HP0_G0018330 | OG0004127 | chrVII | 169753 | 170790 | CP015057.1 | 672975 | 674012 | 29.26829268 | 345 | 345 | 1 | 1 | 0 | 1 | FIM1_2734 | YGL175C | YGL175C | SAE2 | n |
| Kmar_QGN16039.1 | W303.asm02.HP0_G0029490 | OG0004128 | chrX | 401237 | 402922 | CP015057.1 | 681846 | 683483 | 29.28452579 | 561 | 545 | 1.029357798 | 1 | 0 | 1 | FIM1_2739 | YJL016W | YJL016W | TPH3 | n |
| Kmar_QGN16048.1 | W303.asm02.HP0_G0036600 | OG0004129 | chrXII | 502682 | 504508 | CP015057.1 | 698130 | 699911 | 24.59807074 | 608 | 593 | 1.02529511 | 1 | 0 | 1 | FIM1_2748 | YLR173W | YLR173W | TAG1 | n |
| Kmar_QGN16069.1 | W303.asm02.HP0_G0032630 | OG0004130 | chrXI | 329619 | 330122 | CP015057.1 | 742496 | 743134 | 25.70093458 | 167 | 212 | 0.787735849 | 1 | 0 | 1 | FIM1_2769 | YKL063C | YKL063C | | n |
| Kmar_QGN16071.1 | W303.asm02.HP0_G0032780 | OG0004131 | chrXI | 355818 | 357740 | CP015057.1 | 746301 | 747710 | 28.34645669 | 640 | 469 | 1.364605544 | 1 | 0 | 1 | FIM1_2771 | YKL048C | YKL048C | ELM1 | n |
| Kmar_QGN16078.1 | W303.asm02.HP0_G0055330 | OG0004132 | chrXVI | 548439 | 550205 | CP015057.1 | 764043 | 765788 | 35.85858586 | 588 | 581 | 1.012048193 | 1 | 0 | 1 | FIM1_2778 | YPL007C | YPL007C | TFC8 | n |
| Kmar_QGN16092.1 | W303.asm02.HP0_G0006170 | OG0004133 | chrIII | 271014 | 271796 | CP015057.1 | 794099 | 794842 | 44.62151394 | 260 | 247 | 1.052631579 | 1 | 0 | 1 | FIM1_2792 | YCR075C | YCR075C | ERS1 | n |
| Kmar_QGN16105.1 | W303.asm02.HP0_G0017280 | OG0004134 | chrVI | 195868 | 196242 | CP015057.1 | 824930 | 825271 | 27.55905512 | 124 | 113 | 1.097345133 | 1 | 0 | 1 | FIM1_2806 | YFR036W | YFR036W | CDC26 | n |
| Kmar_QGN16114.1 | W303.asm02.HP0_G0045260 | OG0004135 | chrXIV | 340304 | 340900 | CP015057.1 | 838307 | 838882 | 30.62200957 | 198 | 191 | 1.036649215 | 1 | 0 | 1 | FIM1_2815 | YNL158W | YNL158W | PGA1 | n |
| Kmar_QGN16115.1 | W303.asm02.HP0_G0045250 | OG0004136 | chrXIV | 339170 | 340039 | CP015057.1 | 839209 | 839910 | 26.29757785 | 289 | 233 | 1.240343348 | 1 | 0 | 1 | FIM1_2816 | YNL159C | YNL159C | ASI2 | n |
| Kmar_QGN16127.1 | W303.asm02.HP0_G0045980 | OG0004137 | chrXIV | 481395 | 483179 | CP015057.1 | 865193 | 866725 | 19.17808219 | 448 | 510 | 0.878431373 | 1 | 0 | 1 | FIM1_2828 | YNL078W | YNL078W | NIS1 | n |
| Kmar_QGN16130.1 | W303.asm02.HP0_G0025920 | OG0004138 | chrIX | 82650 | 83831 | CP015057.1 | 875162 | 876847 | 27.06766917 | 393 | 561 | 0.700534759 | 1 | 0 | 1 | FIM1_2831 | YIL136W | YIL136W | OM45 | n |
| Kmar_QGN16131.1 | W303.asm02.HP0_G0046000 | OG0004139 | chrXIV | 485743 | 487497 | CP015057.1 | 878875 | 880845 | 29.11208151 | 584 | 656 | 0.890243902 | 1 | 0 | 1 | FIM1_2832 | YNL076W | YNL076W | MKS1 | n |
| Kmar_QGN16139.1 | W303.asm02.HP0_G0032650 | OG0004140 | chrXI | 334375 | 334716 | CP015057.1 | 893476 | 893781 | 29.20353982 | 113 | 101 | 1.118811881 | 1 | 0 | 1 | FIM1_2840 | YKL061W | YKL061W | BLI1 | n |
| Kmar_QGN16142.1 | W303.asm02.HP0_G0040730 | OG0004141 | chrXIII | 323803 | 324690 | CP015057.1 | 896853 | 897464 | 18.36734694 | 295 | 203 | 1.45320197 | 1 | 0 | 1 | FIM1_2843 | YMR025W | YMR025W | CSI1 | n |
| Kmar_QGN16155.1 | W303.asm02.HP0_G0043880 | OG0004142 | chrXIV | 49829 | 52120 | CP015057.1 | 917903 | 919915 | 41.03225806 | 763 | 670 | 1.13880597 | 1 | 0 | 1 | FIM1_2856 | YNL311C | YNL311C | SKP2 | n |
| Kmar_QGN16157.1 | W303.asm02.HP0_G0043900 | OG0004143 | chrXIV | 53094 | 54356 | CP015057.1 | 921330 | 922352 | 24.76415094 | 420 | 340 | 1.235294118 | 1 | 0 | 1 | FIM1_2858 | YNL309W | YNL309W | STB1 | n |
| Kmar_QGN16164.1 | W303.asm02.HP0_G0026530 | OG0004144 | chrIX | 219155 | 220489 | CP015057.1 | 938242 | 939408 | 23.64864865 | 444 | 388 | 1.144329897 | 1 | 0 | 1 | FIM1_2866 | YIL071C | YIL071C | PCI8 | n |
| Kmar_QGN16206.1 | W303.asm02.HP0_G0008120 | OG0004145 | chrIV | 326943 | 329897 | CP015057.1 | 1032122 | 1035238 | 25.7518797 | 984 | 1038 | 0.947976879 | 1 | 0 | 1 | FIM1_2909 | YDL073W | YDL073W | AHK1 | n |
| Kmar_QGN16207.1 | W303.asm02.HP0_G0038660 | OG0004146 | chrXII | 925413 | 926270 | CP015057.1 | 1035515 | 1036219 | 20.35087719 | 285 | 234 | 1.217948718 | 1 | 0 | 1 | FIM1_2911 | YLR404W | YLR404W | SEI1 | n |
| Kmar_QGN16249.1 | W303.asm02.HP0_G0030880 | OG0004147 | chrX | 687211 | 688476 | CP015057.1 | 1129543 | 1130817 | 33.56321839 | 421 | 424 | 0.992924528 | 1 | 0 | 1 | FIM1_2954 | YJR136C | YJR136C | TTI2 | n |
| Kmar_QGN16251.1 | W303.asm02.HP0_G0030860 | OG0004148 | chrX | 685831 | 686550 | CP015057.1 | 1131556 | 1132230 | 25.20661157 | 239 | 224 | 1.066964286 | 1 | 0 | 1 | FIM1_2956 | YJR135C | YJR135C | MCM22 | n |
| Kmar_QGN16271.1 | W303.asm02.HP0_G0039750 | OG0004149 | chrXIII | 104787 | 105392 | CP015057.1 | 1168639 | 1169277 | 49.05660377 | 201 | 212 | 0.948113208 | 1 | 0 | 1 | FIM1_2976 | YML079W | YML079W | CFF1 | n |
| Kmar_QGN16274.1 | W303.asm02.HP0_G0038710 | OG0004150 | chrXII | 930554 | 930922 | CP015057.1 | 1172910 | 1173233 | 37.70491803 | 122 | 107 | 1.140186916 | 1 | 0 | 1 | FIM1_2979 | YLR408C | YLR408C | BLS1 | n |
| Kmar_QGN16329.1 | W303.asm02.HP0_G0040220 | OG0004151 | chrXIII | 217870 | 220386 | CP015057.1 | 1300889 | 1303441 | 26.86733556 | 838 | 850 | 0.985882353 | 1 | 0 | 1 | FIM1_3034 | YML029W | YML029W | USA1 | n |
| Kmar_QGN16339.1 | W303.asm02.HP0_G0012870 | OG0004152 | chrIV | 1355845 | 1357908 | CP015057.1 | 1322302 | 1324212 | 45.32374101 | 687 | 636 | 1.080188679 | 1 | 0 | 1 | FIM1_3045 | YDR444W | YDR444W | | n |
| Kmar_QGN16341.1 | W303.asm02.HP0_G0040280 | OG0004153 | chrXIII | 227502 | 229172 | CP015057.1 | 1328719 | 1330248 | 24.59893048 | 556 | 509 | 1.092337917 | 1 | 0 | 1 | FIM1_3047 | YML023C | YML023C | NSE5 | n |
| Kmar_QGN16346.1 | W303.asm02.HP0_G0012830 | OG0004154 | chrIV | 1345232 | 1346275 | CP015057.1 | 1336890 | 1337588 | 21.32564841 | 347 | 232 | 1.495689655 | 1 | 0 | 1 | FIM1_3052 | YDR439W | YDR439W | LRS4 | n |
| Kmar_QGN16361.1 | W303.asm02.HP0_G0040400 | OG0004155 | chrXIII | 247401 | 247934 | CP015057.1 | 1362573 | 1363100 | 40.11299435 | 177 | 175 | 1.011428571 | 1 | 0 | 1 | FIM1_3067 | YML011C | YML011C | RAD33 | n |
| Kmar_QGN16370.1 | W303.asm02.HP0_G0014800 | OG0004156 | chrV | 216055 | 218832 | CP015057.1 | 1379085 | 1381952 | 25.21994135 | 925 | 955 | 0.968586387 | 1 | 0 | 1 | FIM1_3077 | YER032W | YER032W | FIR1 | n |
| Kmar_QGN16391.1 | W303.asm02.HP0_G0052470 | OG0004157 | chrXV | 1037088 | 1038752 | CP015058.1 | 12291 | 13964 | 18.71750433 | 554 | 557 | 0.994614004 | 1 | 0 | 1 | FIM1_3104 | YOR372C | YOR372C | NDD1 | n |
| Kmar_QGN16408.1 | W303.asm02.HP0_G0050850 | OG0004158 | chrXV | 714161 | 716626 | CP015058.1 | 51449 | 53362 | 24.23873325 | 821 | 637 | 1.288854003 | 1 | 0 | 1 | FIM1_3121 | YOR195W | YOR195W | SLK19 | n |
| Kmar_QGN16433.1 | W303.asm02.HP0_G0004630 | OG0004159 | chrII | 772943 | 774526 | CP015058.1 | 109837 | 111402 | 51.03969754 | 527 | 521 | 1.011516315 | 1 | 0 | 1 | FIM1_3146 | YBR274W | YBR274W | CHK1 | n |
| Kmar_QGN16438.1 | W303.asm02.HP0_G0029210 | OG0004160 | chrX | 329806 | 332274 | CP015058.1 | 120260 | 122455 | 23.69047619 | 822 | 731 | 1.124487004 | 1 | 0 | 1 | FIM1_3151 | YJL051W | YJL051W | IRC8 | n |
| Kmar_QGN16498.1 | W303.asm02.HP0_G0000440 | OG0004161 | chrI | 95450 | 96235 | CP015058.1 | 238814 | 239775 | 32.67973856 | 261 | 293 | 0.890784983 | 1 | 0 | 1 | FIM1_3211 | YAL027W | YAL027W | SAW1 | n |
| Kmar_QGN16537.1 | W303.asm02.HP0_G0001850 | OG0004162 | chrII | 218872 | 219414 | CP015058.1 | 329535 | 329999 | 25.27472527 | 180 | 154 | 1.168831169 | 1 | 0 | 1 | FIM1_3250 | YBL006C | YBL006C | LDB7 | n |
| Kmar_QGN16549.1 | W303.asm02.HP0_G0039920 | OG0004163 | chrXIII | 142045 | 143223 | CP015058.1 | 350974 | 352017 | 23.21428571 | 392 | 347 | 1.129682997 | 1 | 0 | 1 | FIM1_3263 | YML062C | YML062C | MFT1 | n |
| Kmar_QGN16562.1 | W303.asm02.HP0_G0038930 | OG0004164 | chrXII | 997003 | 998364 | CP015058.1 | 378871 | 380199 | 25.38293217 | 453 | 442 | 1.024886878 | 1 | 0 | 1 | FIM1_3276 | YLR431C | YLR431C | ATG23 | n |
| Kmar_QGN16578.1 | W303.asm02.HP0_G0020140 | OG0004165 | chrVII | 543303 | 543779 | CP015058.1 | 421880 | 422371 | 31.3253012 | 158 | 163 | 0.969325153 | 1 | 0 | 1 | FIM1_3293 | YGR030C | YGR030C | POP6 | n |
| Kmar_QGN16603.1 | W303.asm02.HP0_G0002300 | OG0004166 | chrII | 319442 | 321160 | CP015058.1 | 478638 | 480317 | 45.54973822 | 572 | 559 | 1.023255814 | 1 | 0 | 1 | FIM1_3319 | YBR044C | YBR044C | TCM62 | n |
| Kmar_QGN16614.1 | W303.asm02.HP0_G0008830 | OG0004167 | chrIV | 473705 | 475648 | CP015058.1 | 504308 | 506044 | 22.10200927 | 647 | 578 | 1.119377163 | 1 | 0 | 1 | FIM1_3331 | YDR014W | YDR014W | RAD61 | n |
| Kmar_QGN16624.1 | W303.asm02.HP0_G0028200 | OG0004168 | chrX | 103405 | 105156 | CP015058.1 | 522648 | 524597 | 26.60550459 | 583 | 649 | 0.898305085 | 1 | 0 | 1 | FIM1_3341 | YJL162C | YJL162C | JJJ2 | n |
| Kmar_QGN16636.1 | W303.asm02.HP0_G0027940 | OG0004169 | chrX | 62241 | 62945 | CP015058.1 | 557448 | 558152 | 42.19409283 | 234 | 234 | 1 | 1 | 0 | 1 | FIM1_3355 | YJL192C | YJL192C | SOP4 | n |
| Kmar_QGN16646.1 | W303.asm02.HP0_G0038340 | OG0004170 | chrXII | 853720 | 855516 | CP015058.1 | 580793 | 582529 | 19.01639344 | 598 | 578 | 1.034602076 | 1 | 0 | 1 | FIM1_3365 | YLR368W | YLR368W | MDM30 | n |
| Kmar_QGN16650.1 | W303.asm02.HP0_G0020460 | OG0004171 | chrVII | 615833 | 617614 | CP015058.1 | 593539 | 595374 | 70.73170732 | 593 | 611 | 0.970540098 | 1 | 0 | 1 | FIM1_3369 | YGR065C | YGR065C | VHT1 | n |
| Kmar_QGN16651.1 | W303.asm02.HP0_G0038320 | OG0004172 | chrXII | 850242 | 850571 | CP015058.1 | 595411 | 595746 | 54.95495495 | 109 | 111 | 0.981981982 | 1 | 0 | 1 | FIM1_3370 | YLR364W | YLR364W | GRX8 | n |
| Kmar_QGN16711.1 | W303.asm02.HP0_G0053040 | OG0004173 | chrXVI | 91151 | 92515 | CP015058.1 | 719679 | 721007 | 61.62280702 | 454 | 442 | 1.027149321 | 1 | 0 | 1 | FIM1_3431 | YPL245W | YPL245W | | n |
| Kmar_QGN16714.1 | W303.asm02.HP0_G0042260 | OG0004174 | chrXIII | 635272 | 636567 | CP015058.1 | 724692 | 726074 | 27.54880694 | 431 | 460 | 0.936956522 | 1 | 0 | 1 | FIM1_3434 | YMR187C | YMR187C | | n |
| Kmar_QGN16766.1 | W303.asm02.HP0_G0014420 | OG0004175 | chrV | 147090 | 147938 | CP015058.1 | 845376 | 846254 | 20.33898305 | 282 | 292 | 0.965753425 | 1 | 0 | 1 | FIM1_3488 | YEL005C | YEL005C | VAB2 | n |
| Kmar_QGN16776.1 | W303.asm02.HP0_G0027120 | OG0004176 | chrIX | 326560 | 326841 | CP015058.1 | 864206 | 864751 | 23.97260274 | 93 | 181 | 0.513812155 | 1 | 0 | 1 | FIM1_3498 | YIL009C-A | YIL009C-A | EST3 | n |
| Kmar_QGN16790.1 | W303.asm02.HP0_G0020650 | OG0004177 | chrVII | 653104 | 654792 | CP015058.1 | 894064 | 895713 | 66.31393298 | 562 | 549 | 1.023679417 | 1 | 0 | 1 | FIM1_3513 | YGR088W | YGR088W | CTT1 | n |
| Kmar_QGN16794.1 | W303.asm02.HP0_G0056370 | OG0004178 | chrXVI | 751188 | 752072 | CP015058.1 | 904218 | 905054 | 24.03846154 | 294 | 278 | 1.057553957 | 1 | 0 | 1 | FIM1_3517 | YPR109W | YPR109W | GLD1 | n |
| Kmar_QGN16803.1 | W303.asm02.HP0_G0056420 | OG0004179 | chrXVI | 760230 | 761177 | CP015058.1 | 924080 | 925063 | 42.40687679 | 315 | 327 | 0.963302752 | 1 | 0 | 1 | FIM1_3527 | YPR114W | YPR114W | | n |
| Kmar_QGN16804.1 | W303.asm02.HP0_G0020740 | OG0004180 | chrVII | 677166 | 680606 | CP015058.1 | 925507 | 928701 | 38.16793893 | 1146 | 1064 | 1.077067669 | 1 | 0 | 0 | FIM1_3528 | YGR097W | YGR097W | ASK10 | n |
| Kmar_QGN16806.1 | W303.asm02.HP0_G0056440 | OG0004181 | chrXVI | 765577 | 766410 | CP015058.1 | 929782 | 930627 | 26.55172414 | 277 | 281 | 0.985765125 | 1 | 0 | 1 | FIM1_3530 | YPR116W | YPR116W | RRG8 | n |

| | | | | | | | | | | | | | | | | | | | | |
|---|---|---|---|---|---|---|---|---|---|---|---|---|---|---|---|---|---|---|---|---|
| Kmar_QGN16816.1 | W303.asm02.HP0_G0020830 | OG0004182 | chrVII | 697434 | 698255 | CP015058.1 | 954616 | 955410 | 30.37037037 | 273 | 264 | 1.034090909 | 1 | 0 | 1 | FIM1_3540 | YGR106C | YGR106C | VOA1 | n |
| Kmar_QGN16825.1 | W303.asm02.HP0_G0056500 | OG0004183 | chrXVI | 788974 | 792600 | CP015058.1 | 974215 | 977748 | 36.49754501 | 1208 | 1177 | 1.026338148 | 1 | 0 | 1 | FIM1_3549 | YPR122W | YPR122W | AXL1 | n |
| Kmar_QGN16831.1 | W303.asm02.HP0_G0004450 | OG0004184 | chrII | 749974 | 750430 | CP015058.1 | 986377 | 987037 | 68.85245902 | 120 | 121 | 0.991735537 | 1 | 0 | 1 | FIM1_3555 | YBR255C-A | YBR255C-A | RCF3 | n |
| Kmar_QGN16837.1 | W303.asm02.HP0_G0027030 | OG0004185 | chrIX | 308829 | 311594 | CP015058.1 | 995184 | 998093 | 26.24618515 | 921 | 969 | 0.950464396 | 1 | 0 | 1 | FIM1_3561 | YIL017C | YIL017C | VID28 | n |
| Kmar_QGN16858.1 | W303.asm02.HP0_G0036170 | OG0004186 | chrXII | 415236 | 416108 | CP015058.1 | 1047683 | 1048510 | 22.8956229 | 290 | 275 | 1.054545455 | 1 | 0 | 1 | FIM1_3584 | YLR132C | YLR132C | USB1 | n |
| Kmar_QGN16869.1 | W303.asm02.HP0_G0010140 | OG0004187 | chrIV | 770450 | 771685 | CP015058.1 | 1071957 | 1073210 | 50.81585082 | 411 | 417 | 0.985611511 | 1 | 0 | 1 | FIM1_3595 | YDR153C | YDR153C | ENT5 | n |
| Kmar_QGN16880.1 | W303.asm02.HP0_G0036240 | OG0004188 | chrXII | 429588 | 431519 | CP015058.1 | 1092444 | 1094519 | 24.72067039 | 643 | 691 | 0.930535456 | 1 | 0 | 1 | FIM1_3607 | YLR139C | YLR139C | SLS1 | n |
| Kmar_QGN16906.1 | W303.asm02.HP0_G0010310 | OG0004189 | chrIV | 810335 | 811462 | CP015058.1 | 1149742 | 1150926 | 39.8989899 | 375 | 394 | 0.95177665 | 1 | 0 | 1 | FIM1_3633 | YDR171W | YDR171W | HSP42 | n |
| Kmar_QGN16912.1 | W303.asm02.HP0_G0010390 | OG0004190 | chrIV | 822423 | 822911 | CP015058.1 | 1177831 | 1178304 | 29.26829268 | 162 | 157 | 1.031847134 | 1 | 0 | 1 | FIM1_3639 | YDR179C | YDR179C | CSN9 | n |
| Kmar_QGN16913.1 | W303.asm02.HP0_G0010400 | OG0004191 | chrIV | 823148 | 824539 | CP015058.1 | 1178604 | 1179911 | 26.34989201 | 463 | 435 | 1.064367816 | 1 | 0 | 1 | FIM1_3640 | YDR179W-A | YDR179W-A | NVJ3 | n |
| Kmar_QGN16919.1 | W303.asm02.HP0_G0010450 | OG0004192 | chrIV | 833300 | 833992 | CP015058.1 | 1191481 | 1192143 | 36.36363636 | 230 | 220 | 1.045454545 | 1 | 0 | 1 | FIM1_3646 | YDR183W | YDR183W | PLP1 | n |
| Kmar_QGN16921.1 | W303.asm02.HP0_G0010460 | OG0004193 | chrIV | 834344 | 835228 | CP015058.1 | 1193497 | 1194348 | 28.62068966 | 294 | 283 | 1.038869258 | 1 | 0 | 1 | FIM1_3648 | YDR184C | YDR184C | ATC1 | n |
| Kmar_QGN16924.1 | W303.asm02.HP0_G0010480 | OG0004194 | chrIV | 836574 | 839207 | CP015058.1 | 1198098 | 1200164 | 33.97075366 | 877 | 688 | 1.274709302 | 1 | 0 | 1 | FIM1_3651 | YDR186C | YDR186C | SND1 | n |
| Kmar_QGN16930.1 | W303.asm02.HP0_G0010520 | OG0004195 | chrIV | 846052 | 847164 | CP015058.1 | 1209618 | 1210784 | 44.50127877 | 370 | 388 | 0.953608247 | 1 | 0 | 1 | FIM1_3657 | YDR191W | YDR191W | HST4 | n |
| Kmar_QGN16937.1 | W303.asm02.HP0_G0010560 | OG0004196 | chrIV | 852314 | 853915 | CP015058.1 | 1223730 | 1225148 | 35.18850987 | 533 | 472 | 1.129237288 | 1 | 0 | 1 | FIM1_3664 | YDR195W | YDR195W | REF2 | n |
| Kmar_QGN16939.1 | W303.asm02.HP0_G0010580 | OG0004197 | chrIV | 854944 | 856113 | CP015058.1 | 1226863 | 1227975 | 30.76923077 | 389 | 370 | 1.051351351 | 1 | 0 | 1 | FIM1_3666 | YDR197W | YDR197W | CBS2 | n |
| Kmar_QGN16953.1 | W303.asm02.HP0_G0044800 | OG0004198 | chrXIV | 251631 | 252443 | CP015058.1 | 1248987 | 1249805 | 20.37037037 | 270 | 272 | 0.992647059 | 1 | 0 | 1 | FIM1_3680 | YNL210W | YNL210W | MER1 | n |
| Kmar_QGN16955.1 | W303.asm02.HP0_G0006650 | OG0004199 | chrIV | 38940 | 42317 | CP015058.1 | 1252261 | 1255392 | 47.60213144 | 1125 | 1043 | 1.078619367 | 1 | 0 | 1 | FIM1_3682 | YDL231C | YDL231C | BRE4 | n |
| Kmar_QGN16960.1 | W303.asm02.HP0_G0044760 | OG0004200 | chrXIV | 246316 | 246915 | CP015058.1 | 1259623 | 1260210 | 27.45098039 | 199 | 195 | 1.020512821 | 1 | 0 | 1 | FIM1_3687 | YNL214W | YNL214W | PEX17 | n |
| Kmar_QGN16962.1 | W303.asm02.HP0_G0044750 | OG0004201 | chrXIV | 245167 | 246129 | CP015058.1 | 1262887 | 1263816 | 27.02702703 | 320 | 309 | 1.035598706 | 1 | 0 | 1 | FIM1_3689 | YNL215W | YNL215W | IES2 | n |
| Kmar_QGN16985.1 | W303.asm02.HP0_G0043590 | OG0004202 | chrXIII | 920908 | 922566 | CP015058.1 | 1311717 | 1313360 | 50.99099099 | 552 | 547 | 1.009140768 | 1 | 0 | 1 | FIM1_3712 | YMR319C | YMR319C | FET4 | n |
| Kmar_QGN17021.1 | W303.asm02.HP0_G0026020 | OG0004203 | chrIX | 106058 | 106678 | CP015059.1 | 53768 | 54367 | 50.97087379 | 206 | 199 | 1.035175879 | 1 | 0 | 1 | FIM1_3750 | YIL127C | YIL127C | RRT14 | n |
| Kmar_QGN17026.1 | W303.asm02.HP0_G0025970 | OG0004204 | chrIX | 88891 | 89532 | CP015059.1 | 72308 | 72934 | 26.33928571 | 213 | 208 | 1.024038462 | 1 | 0 | 1 | FIM1_3755 | YIL132C | YIL132C | CSM2 | n |
| Kmar_QGN17065.1 | W303.asm02.HP0_G0007570 | OG0004205 | chrIV | 231230 | 232105 | CP015059.1 | 157420 | 158274 | 29.37293729 | 291 | 284 | 1.024647887 | 1 | 0 | 1 | FIM1_3795 | YDL129W | YDL129W | | n |
| Kmar_QGN17066.1 | W303.asm02.HP0_G0007110 | OG0004206 | chrIV | 136116 | 137759 | CP015059.1 | 159218 | 160756 | 32.0855615 | 547 | 512 | 1.068359375 | 1 | 0 | 1 | FIM1_3796 | YDL180W | YDL180W | AIT1 | n |
| Kmar_QGN17072.1 | W303.asm02.HP0_G0007140 | OG0004207 | chrIV | 141429 | 141941 | CP015059.1 | 172694 | 173218 | 51.63043478 | 170 | 174 | 0.977011494 | 1 | 0 | 1 | FIM1_3803 | YDL177C | YDL177C | | n |
| Kmar_QGN17074.1 | W303.asm02.HP0_G0007630 | OG0004208 | chrIV | 241624 | 242046 | CP015059.1 | 175695 | 176105 | 52.77777778 | 140 | 136 | 1.029411765 | 1 | 0 | 1 | FIM1_3805 | YDL123W | YDL123W | SNA4 | n |
| Kmar_QGN17096.1 | W303.asm02.HP0_G0046810 | OG0004209 | chrXIV | 655897 | 656346 | CP015059.1 | 226414 | 226968 | 28.96174863 | 149 | 184 | 0.809782609 | 1 | 0 | 1 | FIM1_3827 | YNR010W | YNR010W | CSE2 | n |
| Kmar_QGN17101.1 | W303.asm02.HP0_G0008290 | OG0004210 | chrIV | 362034 | 362591 | CP015059.1 | 234088 | 234561 | 19.67213115 | 185 | 157 | 1.178343949 | 1 | 0 | 1 | FIM1_3832 | YDL053C | YDL053C | PBP4 | n |
| Kmar_QGN17104.1 | W303.asm02.HP0_G0038410 | OG0004211 | chrXII | 869492 | 870220 | CP015059.1 | 238500 | 239099 | 24.66960352 | 242 | 199 | 1.216080402 | 1 | 0 | 1 | FIM1_3835 | YLR376C | YLR376C | PSY3 | n |
| Kmar_QGN17127.1 | W303.asm02.HP0_G0030570 | OG0004212 | chrX | 634611 | 636788 | CP015059.1 | 292165 | 294144 | 31.94070081 | 725 | 659 | 1.100151745 | 1 | 0 | 1 | FIM1_3858 | YJR106W | YJR106W | ECM27 | n |
| Kmar_QGN17152.1 | W303.asm02.HP0_G0017630 | OG0004213 | chrVII | 27280 | 28005 | CP015059.1 | 347441 | 348370 | 30.64516129 | 241 | 309 | 0.779935275 | 1 | 0 | 1 | FIM1_3883 | YGL250W | YGL250W | RMR1 | n |
| Kmar_QGN17157.1 | W303.asm02.HP0_G0017410 | OG0004214 | chrVI | 217428 | 217799 | CP015059.1 | 357568 | 357951 | 47.24409449 | 123 | 127 | 0.968503937 | 1 | 0 | 1 | FIM1_3888 | YFR049W | YFR049W | KGD4 | n |
| Kmar_QGN17175.1 | W303.asm02.HP0_G0011060 | OG0004215 | chrIV | 963182 | 964303 | CP015059.1 | 389667 | 390788 | 24.05063291 | 373 | 373 | 1 | 1 | 0 | 1 | FIM1_3906 | YDR249C | YDR249C | | n |
| Kmar_QGN17193.1 | W303.asm02.HP0_G0053530 | OG0004216 | chrXVI | 186955 | 188100 | CP015059.1 | 424577 | 425650 | 29.84693878 | 381 | 357 | 1.067226891 | 1 | 0 | 1 | FIM1_3924 | YPL193W | YPL193W | RSA1 | n |
| Kmar_QGN17208.1 | W303.asm02.HP0_G0053610 | OG0004217 | chrXVI | 200063 | 200977 | CP015059.1 | 449573 | 450322 | 26.97368421 | 304 | 249 | 1.220883534 | 1 | 0 | 1 | FIM1_3939 | YPL186C | YPL186C | UIP4 | n |
| Kmar_QGN17211.1 | W303.asm02.HP0_G0019080 | OG0004218 | chrVII | 339008 | 339994 | CP015059.1 | 455795 | 456787 | 81.81818182 | 328 | 330 | 0.993939394 | 1 | 0 | 1 | FIM1_3942 | YGL091C | YGL091C | NBP35 | n |
| Kmar_QGN17243.1 | W303.asm02.HP0_G0032920 | OG0004219 | chrXI | 383104 | 383814 | CP015059.1 | 532513 | 533184 | 57.62711864 | 236 | 223 | 1.058295964 | 1 | 0 | 1 | FIM1_3974 | YKL033W-A | YKL033W-A | | n |
| Kmar_QGN17261.1 | W303.asm02.HP0_G0035430 | OG0004220 | chrXII | 255014 | 255766 | CP015059.1 | 570095 | 570673 | 25.89641434 | 250 | 192 | 1.302083333 | 1 | 0 | 1 | FIM1_3992 | YLR052W | YLR052W | IES3 | n |
| Kmar_QGN17263.1 | W303.asm02.HP0_G0035450 | OG0004221 | chrXII | 256530 | 258791 | CP015059.1 | 572707 | 574695 | 29.73720609 | 724 | 662 | 1.093655589 | 1 | 0 | 1 | FIM1_3994 | YLR054C | YLR054C | OSW2 | n |
| Kmar_QGN17297.1 | W303.asm02.HP0_G0045490 | OG0004222 | chrXIV | 375004 | 375525 | CP015059.1 | 642407 | 642952 | 26.53061224 | 173 | 181 | 0.955801105 | 1 | 0 | 1 | FIM1_4028 | YNL133C | YNL133C | FYV6 | n |
| Kmar_QGN17340.1 | W303.asm02.HP0_G0052270 | OG0004223 | chrXV | 995145 | 997136 | CP015059.1 | 726682 | 728553 | 19.73293769 | 663 | 623 | 1.064205457 | 1 | 0 | 1 | FIM1_4072 | YOR350C | YOR350C | MNE1 | n |
| Kmar_QGN17358.1 | W303.asm02.HP0_G0009500 | OG0004224 | chrIV | 614718 | 616202 | CP015059.1 | 767191 | 768441 | 21.9123506 | 494 | 416 | 1.1875 | 1 | 0 | 1 | FIM1_4090 | YDR082W | YDR082W | STN1 | n |
| Kmar_QGN17383.1 | W303.asm02.HP0_G0041080 | OG0004225 | chrXIII | 398406 | 399962 | CP015059.1 | 813079 | 814554 | 18.72586873 | 518 | 491 | 1.054989817 | 1 | 0 | 1 | FIM1_4115 | YMR064W | YMR064W | AEP1 | n |
| Kmar_QGN17389.1 | W303.asm02.HP0_G0041100 | OG0004226 | chrXIII | 402044 | 404740 | CP015059.1 | 824730 | 827270 | 25.52255226 | 898 | 846 | 1.061465721 | 1 | 0 | 1 | FIM1_4121 | YMR066W | YMR066W | SOV1 | n |
| Kmar_QGN17406.1 | W303.asm02.HP0_G0027480 | OG0004227 | chrIX | 394142 | 394792 | CP015059.1 | 850021 | 850710 | 28.39506173 | 216 | 229 | 0.943231441 | 1 | 0 | 1 | FIM1_4138 | YIR024C | YIR024C | INA22 | n |
| Kmar_QGN17413.1 | W303.asm02.HP0_G0027490 | OG0004228 | chrIX | 394960 | 396066 | CP015059.1 | 864912 | 865742 | 22.07084469 | 368 | 276 | 1.333333333 | 1 | 0 | 1 | FIM1_4145 | YIR025W | YIR025W | MND2 | n |
| Kmar_QGN17465.1 | W303.asm02.HP0_G0011170 | OG0004229 | chrIV | 981218 | 981730 | CP015059.1 | 977926 | 978258 | 21.17647059 | 170 | 110 | 1.545454545 | 1 | 0 | 1 | FIM1_4199 | YDR260C | YDR260C | SWM1 | n |
| Kmar_QGN17485.1 | W303.asm02.HP0_G0021780 | OG0004230 | chrVII | 922847 | 923152 | CP015059.1 | 1011317 | 1011598 | 24.50980392 | 101 | 93 | 1.086021505 | 1 | 0 | 1 | FIM1_4221 | YGR206W | YGR206W | MVB12 | n |
| Kmar_QGN17507.1 | W303.asm02.HP0_G0021930 | OG0004231 | chrVII | 952338 | 953219 | CP015059.1 | 1055606 | 1056433 | 26.51006711 | 293 | 275 | 1.065454545 | 1 | 0 | 1 | FIM1_4243 | YGR222W | YGR222W | PET54 | n |
| Kmar_QGN17524.1 | W303.asm02.HP0_G0024870 | OG0004232 | chrVIII | 423655 | 425169 | CP015059.1 | 1090506 | 1091837 | 17.32283465 | 504 | 443 | 1.137697517 | 1 | 0 | 1 | FIM1_4260 | YHR159W | YHR159W | TDA11 | n |
| Kmar_QGN17541.1 | W303.asm02.HP0_G0033720 | OG0004233 | chrXI | 536518 | 536919 | CP015059.1 | 1132307 | 1132702 | 36.2962963 | 133 | 131 | 1.015267176 | 1 | 0 | 1 | FIM1_4277 | YKR049C | YKR049C | FMP46 | n |
| Kmar_QGN17546.1 | W303.asm02.HP0_G0033700 | OG0004234 | chrXI | 533911 | 534762 | CP015059.1 | 1139327 | 1140217 | 32.2147651 | 283 | 296 | 0.956081081 | 1 | 0 | 1 | FIM1_4282 | YKR046C | YKR046C | PLN1 | n |
| Kmar_QGN17554.1 | W303.asm02.HP0_G0018590 | OG0004235 | chrVII | 226711 | 228816 | CP015059.1 | 1153531 | 1155570 | 30.79056865 | 701 | 679 | 1.032400589 | 1 | 0 | 1 | FIM1_4290 | YGL145W | YGL145W | TIP20 | n |
| Kmar_QGN17602.1 | W303.asm02.HP0_G0013900 | OG0004236 | chrV | 42763 | 42987 | CP015060.1 | 80253 | 80471 | 48.64864865 | 74 | 72 | 1.027777778 | 1 | 0 | 1 | FIM1_4809 | YEL059C-A | YEL059C-A | SOM1 | n |
| Kmar_QGN17623.1 | W303.asm02.HP0_G0039390 | OG0004237 | chrXIII | 28794 | 32198 | CP015060.1 | 129802 | 133278 | 28.5012285 | 1134 | 1158 | 0.979274611 | 1 | 0 | 1 | FIM1_4830 | YML117W | YML117W | NAB6 | n |
| Kmar_QGN17654.1 | W303.asm02.HP0_G0035000 | OG0004238 | chrXII | 165817 | 166827 | CP015060.1 | 199514 | 200551 | 28.91891892 | 336 | 345 | 0.973913043 | 1 | 0 | 1 | FIM1_4861 | YLR007W | YLR007W | NSE1 | n |
| Kmar_QGN17664.1 | W303.asm02.HP0_G0007750 | OG0004239 | chrIV | 264718 | 265170 | CP015060.1 | 211885 | 212337 | 39.61038961 | 150 | 150 | 1 | 1 | 0 | 1 | FIM1_4871 | YDL110C | YDL110C | TMA17 | n |
| Kmar_QGN17665.1 | W303.asm02.HP0_G0018580 | OG0004240 | chrVII | 225219 | 226154 | CP015060.1 | 212912 | 213703 | 21.13564669 | 311 | 263 | 1.182509506 | 1 | 0 | 1 | FIM1_4872 | YGL146C | YGL146C | RRT6 | n |
| Kmar_QGN17677.1 | W303.asm02.HP0_G0018540 | OG0004241 | chrVII | 213992 | 217390 | CP015060.1 | 241788 | 245066 | 35.33447684 | 1132 | 1092 | 1.036630037 | 1 | 0 | 1 | FIM1_4884 | YGL151W | YGL151W | NUT1 | n |
| Kmar_QGN17703.1 | W303.asm02.HP0_G0046510 | OG0004242 | chrXIV | 598290 | 599030 | CP015060.1 | 281005 | 281754 | 61.41732283 | 246 | 249 | 0.987951807 | 1 | 0 | 1 | FIM1_4910 | YNL024C | YNL024C | EFM6 | n |
| Kmar_QGN17731.1 | W303.asm02.HP0_G0026190 | OG0004243 | chrIX | 146402 | 147535 | CP015060.1 | 344057 | 345175 | 51.2 | 377 | 372 | 1.01344086 | 1 | 0 | 1 | FIM1_4940 | YIL110W | YIL110W | HPM1 | n |
| Kmar_QGN17798.1 | W303.asm02.HP0_G0049980 | OG0004244 | chrXV | 508683 | 511913 | CP015060.1 | 507212 | 510247 | 22.88058341 | 1076 | 1011 | 1.064292779 | 1 | 0 | 1 | FIM1_5007 | YOR098C | YOR098C | NUP1 | n |
| Kmar_QGN17806.1 | W303.asm02.HP0_G0045780 | OG0004245 | chrXIV | 439268 | 439972 | CP015060.1 | 521255 | 521977 | 30 | 234 | 240 | 0.975 | 1 | 0 | 1 | FIM1_5015 | YNL100W | YNL100W | MIC27 | n |
| Kmar_QGN17811.1 | W303.asm02.HP0_G0050050 | OG0004246 | chrXV | 522088 | 523017 | CP015060.1 | 534338 | 535795 | 13.81443299 | 309 | 485 | 0.637113402 | 1 | 0 | 1 | FIM1_5020 | YOR107W | YOR107W | RGS2 | n |
| Kmar_QGN17815.1 | W303.asm02.HP0_G0052310 | OG0004247 | chrXV | 1003428 | 1005506 | CP015060.1 | 542331 | 544445 | 22.08333333 | 692 | 704 | 0.982954545 | 1 | 0 | 1 | FIM1_5024 | YOR354C | YOR354C | MSC6 | n |
| Kmar_QGN17824.1 | W303.asm02.HP0_G0043950 | OG0004248 | chrXIV | 60731 | 61984 | CP015060.1 | 568137 | 569234 | 39.85680191 | 417 | 365 | 1.142465753 | 1 | 0 | 1 | FIM1_5033 | YNL304W | YNL304W | YPT11 | n |
| Kmar_QGN17838.1 | W303.asm02.HP0_G0040810 | OG0004249 | chrXIII | 335801 | 337810 | CP015060.1 | 593298 | 595208 | 31.68908819 | 669 | 636 | 1.051886792 | 1 | 0 | 1 | FIM1_5047 | YMR032W | YMR032W | HOF1 | n |
| Kmar_QGN17846.1 | W303.asm02.HP0_G0013330 | OG0004250 | chrIV | 1442624 | 1443709 | CP015060.1 | 605961 | 606908 | 31.0626703 | 361 | 315 | 1.146031746 | 1 | 0 | 1 | FIM1_5055 | YDR494W | YDR494W | RSM28 | n |
| Kmar_QGN17851.1 | W303.asm02.HP0_G0048060 | OG0004251 | chrXV | 116851 | 117909 | CP015060.1 | 617116 | 618111 | 19.61325967 | 352 | 331 | 1.063444109 | 1 | 0 | 1 | FIM1_5060 | YOL104C | YOL104C | NDJ1 | n |
| Kmar_QGN17866.1 | W303.asm02.HP0_G0003030 | OG0004252 | chrII | 471108 | 473951 | CP015060.1 | 636912 | 639575 | 24.76586889 | 947 | 887 | 1.067643743 | 1 | 0 | 1 | FIM1_5075 | YBR108W | YBR108W | AIM3 | n |
| Kmar_QGN17871.1 | W303.asm02.HP0_G0056030 | OG0004253 | chrXVI | 692396 | 694096 | CP015060.1 | 645576 | 646877 | 31.21693122 | 566 | 433 | 1.307159353 | 1 | 0 | 1 | FIM1_5080 | YPR070W | YPR070W | MED1 | n |
| Kmar_QGN17886.1 | W303.asm02.HP0_G0003130 | OG0004254 | chrII | 495623 | 496608 | CP015060.1 | 680523 | 681398 | 30.46357616 | 298 | 291 | 1.024054983 | 1 | 0 | 1 | FIM1_5095 | YBR119W | YBR119W | MUD1 | n |
| Kmar_QGN17897.1 | W303.asm02.HP0_G0048570 | OG0004255 | chrXV | 230540 | 231727 | CP015060.1 | 696491 | 697543 | 34.22330097 | 395 | 350 | 1.128571429 | 1 | 0 | 1 | FIM1_5106 | YOL053W | YOL053W | AIM39 | n |
| Kmar_QGN17900.1 | W303.asm02.HP0_G0042960 | OG0004256 | chrXIII | 786131 | 788533 | CP015060.1 | 700539 | 702875 | 22.82878412 | 800 | 778 | 1.028277635 | 1 | 0 | 1 | FIM1_5109 | YMR257C | YMR257C | PET111 | n |
| Kmar_QGN17923.1 | W303.asm02.HP0_G0015480 | OG0004257 | chrV | 344525 | 344902 | CP015060.1 | 746668 | 747021 | 34.12698413 | 125 | 117 | 1.068376068 | 1 | 0 | 1 | FIM1_5132 | YER092W | YER092W | IES5 | n |
| Kmar_QGN17939.1 | W303.asm02.HP0_G0001200 | OG0004258 | chrII | 87875 | 88942 | CP015060.1 | 777891 | 779018 | 31.41361257 | 355 | 375 | 0.946666667 | 1 | 0 | 1 | FIM1_5148 | YBL074C | YBL074C | AAR2 | n |
| Kmar_QGN17947.1 | W303.asm02.HP0_G0001140 | OG0004259 | chrII | 73019 | 74125 | CP015060.1 | 797640 | 798773 | 24.01055409 | 368 | 377 | 0.976127321 | 1 | 0 | 1 | FIM1_5156 | YBL081W | YBL081W | | n |
| Kmar_QGN17969.1 | W303.asm02.HP0_G0050160 | OG0004260 | chrXV | 547732 | 549435 | CP015060.1 | 850722 | 852437 | 47.04861111 | 567 | 571 | 0.992994746 | 1 | 0 | 1 | FIM1_5178 | YOR118W | YOR118W | RTC5 | n |
| Kmar_QGN17980.1 | W303.asm02.HP0_G0034540 | OG0004261 | chrXII | 52096 | 52599 | CP015060.1 | 881343 | 881786 | 26.62721893 | 167 | 147 | 1.136054422 | 1 | 0 | 1 | FIM1_5189 | YLL042C | YLL042C | ATG10 | n |
| Kmar_QGN18009.1 | W303.asm02.HP0_G0053860 | OG0004262 | chrXVI | 256841 | 257602 | CP015061.1 | 26086 | 26817 | 39.68871595 | 253 | 243 | 1.041152263 | 1 | 0 | 1 | FIM1_4325 | YPL159C | YPL159C | PET20 | n |
| Kmar_QGN18011.1 | W303.asm02.HP0_G0053870 | OG0004263 | chrXVI | 257968 | 260244 | CP015061.1 | 28845 | 30656 | 31.58584535 | 758 | 603 | 1.257048093 | 1 | 0 | 1 | FIM1_4327 | YPL158C | YPL158C | AIM44 | n |
| Kmar_QGN18013.1 | W303.asm02.HP0_G0053890 | OG0004264 | chrXVI | 261847 | 262701 | CP015061.1 | 33056 | 33931 | 30.16949153 | 284 | 291 | 0.975945017 | 1 | 0 | 1 | FIM1_4329 | YPL156C | YPL156C | PRM4 | n |
| Kmar_QGN18016.1 | W303.asm02.HP0_G0016410 | OG0004265 | chrVI | 5250 | 7121 | CP015061.1 | 39702 | 41300 | 34.94176373 | 623 | 532 | 1.171052632 | 1 | 0 | 1 | FIM1_4332 | YFL049W | YFL049W | SWP82 | n |
| Kmar_QGN18020.1 | W303.asm02.HP0_G0035680 | OG0004266 | chrXII | 294771 | 295625 | CP015061.1 | 46274 | 47080 | 29.22535211 | 284 | 268 | 1.059701493 | 1 | 0 | 1 | FIM1_4336 | YLR079W | YLR079W | SIC1 | n |
| Kmar_QGN18030.1 | W303.asm02.HP0_G0035630 | OG0004267 | chrXII | 288970 | 289572 | CP015061.1 | 60023 | 60610 | 34.80392157 | 200 | 195 | 1.025641026 | 1 | 0 | 1 | FIM1_4346 | YLR073C | YLR073C | RFU1 | n |
| Kmar_QGN18058.1 | W303.asm02.HP0_G0038490 | OG0004268 | chrXII | 889662 | 890060 | CP015061.1 | 117900 | 118277 | 36.84210526 | 132 | 125 | 1.056 | 1 | 0 | 1 | FIM1_4376 | YLR385C | YLR385C | SWC7 | n |
| Kmar_QGN18072.1 | W303.asm02.HP0_G0055830 | OG0004269 | chrXVI | 663297 | 663842 | CP015061.1 | 153728 | 154222 | 23.20441989 | 181 | 164 | 1.103658537 | 1 | 0 | 1 | FIM1_4390 | YPR046W | YPR046W | MCM16 | n |
| Kmar_QGN18092.1 | W303.asm02.HP0_G0030440 | OG0004270 | chrX | 614342 | 615424 | CP015061.1 | 197300 | 198331 | 17.90633609 | 360 | 343 | 1.049562682 | 1 | 0 | 1 | FIM1_4412 | YJR094C | YJR094C | IME1 | n |
| Kmar_QGN18106.1 | W303.asm02.HP0_G0030350 | OG0004271 | chrX | 591895 | 592824 | CP015061.1 | 226899 | 227555 | 20.86092715 | 309 | 218 | 1.417431193 | 1 | 0 | 1 | FIM1_4426 | YJR083C | YJR083C | ACF4 | n |
| Kmar_QGN18110.1 | W303.asm02.HP0_G0030340 | OG0004272 | chrX | 591201 | 591542 | CP015061.1 | 234758 | 235105 | 36.13445378 | 113 | 115 | 0.982608696 | 1 | 0 | 1 | FIM1_4430 | YJR082C | YJR082C | EAF6 | n |
| Kmar_QGN18138.1 | W303.asm02.HP0_G0015700 | OG0004273 | chrV | 396314 | 396889 | CP015061.1 | 309912 | 310463 | 21.56862745 | 191 | 183 | 1.043715847 | 1 | 0 | 1 | FIM1_4461 | YER115C | YER115C | SPR6 | n |
| Kmar_QGN18164.1 | W303.asm02.HP0_G0026670 | OG0004274 | chrIX | 242671 | 244554 | CP015061.1 | 356916 | 358475 | 21.66934189 | 627 | 519 | 1.208092486 | 1 | 0 | 1 | FIM1_4487 | YIL055C | YIL055C | | n |
| Kmar_QGN18188.1 | W303.asm02.HP0_G0056240 | OG0004275 | chrXVI | 727536 | 731216 | CP015061.1 | 400637 | 403963 | 22.08201893 | 1226 | 1108 | 1.106498195 | 1 | 0 | 1 | FIM1_4512 | YPR095C | YPR095C | SYT1 | n |
| Kmar_QGN18190.1 | W303.asm02.HP0_G0056220 | OG0004276 | chrXVI | 726057 | 726989 | CP015061.1 | 404566 | 405465 | 32.8125 | 310 | 299 | 1.036789298 | 1 | 0 | 1 | FIM1_4514 | YPR093C | YPR093C | ASR1 | n |
| Kmar_QGN18196.1 | W303.asm02.HP0_G0003270 | OG0004277 | chrII | 518090 | 520573 | CP015061.1 | 421036 | 423576 | 41.70403587 | 827 | 846 | 0.977541371 | 1 | 0 | 1 | FIM1_4520 | YBR133C | YBR133C | HSL7 | n |
| Kmar_QGN18256.1 | W303.asm02.HP0_G0056940 | OG0004278 | chrXVI | 889660 | 891390 | CP015061.1 | 564608 | 566233 | 25.70951586 | 576 | 541 | 1.064695009 | 1 | 0 | 1 | FIM1_4582 | YPR171W | YPR171W | BSP1 | n |
| Kmar_QGN18267.1 | W303.asm02.HP0_G0020950 | OG0004279 | chrVII | 731658 | 732866 | CP015061.1 | 583053 | 584225 | 27.64423077 | 402 | 390 | 1.030769231 | 1 | 0 | 1 | FIM1_4593 | YGR122W | YGR122W | | n |
| Kmar_QGN18275.1 | W303.asm02.HP0_G0056600 | OG0004280 | chrXVI | 804980 | 805786 | CP015061.1 | 601237 | 602016 | 19.02985075 | 268 | 259 | 1.034749035 | 1 | 0 | 1 | FIM1_4601 | YPR134W | YPR134W | MSS18 | n |

| | | | | | | | | | | | | | | | | | | | | |
|---|---|---|---|---|---|---|---|---|---|---|---|---|---|---|---|---|---|---|---|---|
| Kmar_QGN18288.1 | W303.asm02.HP0_G0020890 | OG0004281 | chrVII | 718676 | 719707 | CP015061.1 | 622543 | 623589 | 32.41758242 | 343 | 348 | 0.985632184 | 1 | 0 | 1 | FIM1_4614 | YGR113W | YGR113W | DAM1 | n |
| Kmar_QGN18301.1 | W303.asm02.HP0_G0048800 | OG0004282 | chrXV | 272458 | 274179 | CP015061.1 | 652423 | 654219 | 53.65448505 | 573 | 598 | 0.95819398 | 1 | 0 | 1 | FIM1_4627 | YOL027C | YOL027C | MDM38 | n |
| Kmar_QGN18302.1 | W303.asm02.HP0_G0027360 | OG0004283 | chrIX | 372254 | 372982 | CP015061.1 | 654793 | 655422 | 18.18181818 | 242 | 209 | 1.157894737 | 1 | 0 | 1 | FIM1_4628 | YIR014W | YIR014W | VLD1 | n |
| Kmar_QGN18328.1 | W303.asm02.HP0_G0002830 | OG0004284 | chrII | 435193 | 435450 | CP015061.1 | 705997 | 706251 | 59.30232558 | 85 | 84 | 1.011904762 | 1 | 0 | 1 | FIM1_4654 | YBR085C-A | YBR085C-A | | n |
| Kmar_QGN18331.1 | W303.asm02.HP0_G0001590 | OG0004285 | chrII | 164001 | 165017 | CP015061.1 | 710068 | 711276 | 21.39303483 | 338 | 402 | 0.84079602 | 1 | 0 | 1 | FIM1_4657 | YBL031W | YBL031W | SHE1 | n |
| Kmar_QGN18381.1 | W303.asm02.HP0_G0008380 | OG0004286 | chrIV | 374852 | 376174 | CP015061.1 | 810490 | 811710 | 25.39325843 | 440 | 406 | 1.083743842 | 1 | 0 | 1 | FIM1_4708 | YDL044C | YDL044C | MTF2 | n |
| Kmar_QGN14846.1 | W303.asm02.HP0_G0039270 | OG0004296 | chrXIII | 9093 | 9305 | CP015055.1 | 1339714 | 1339911 | 64.28571429 | 70 | 65 | 1.076923077 | 1 | 0 | 1 | FIM1_1518 | YML129C | YML129C | COX14 | n |
| Kmar_QGN17723.1 | W303.asm02.HP0_G0046300 | OG0004311 | chrXIV | 554146 | 556161 | CP015060.1 | 322429 | 324402 | 62.5 | 671 | 657 | 1.02130898 | 1 | 0 | 1 | FIM1_4931 | YNL045W | YNL045W | LAP2 | n |
| Kmar_QGN14394.1 | W303.asm02.HP0_G0051240 | OG0004324 | chrXV | 787379 | 788467 | CP015055.1 | 430395 | 431480 | 68.2320442 | 362 | 361 | 1.002770083 | 1 | 0 | 1 | FIM1_1054 | YOR239W | YOR239W | ABP140 | n |
| Kmar_QGN15613.1 | W303.asm02.HP0_G0030990 | OG0004334 | chrX | 716929 | 718143 | CP015056.1 | 1308610 | 1309695 | 27.00729927 | 404 | 361 | 1.119113573 | 1 | 0 | 1 | FIM1_2306 | YJR149W | YJR149W | | n |
| Kmar_QGN14177.1 | W303.asm02.HP0_G0005040 | OG0004339 | chrIII | 24421 | 26559 | CP015054.1 | 1691341 | 1693479 | 59.527121 | 712 | 712 | 1 | 1 | 0 | 1 | FIM1_830 | YCL057W | YCL057W | PRD1 | n |
| Kmar_QGN13571.1 | W303.asm02.HP0_G0046350 | OG0004340 | chrXIV | 564563 | 565933 | CP015054.1 | 439549 | 440916 | 48.61407249 | 456 | 455 | 1.002197802 | 1 | 0 | 1 | FIM1_212 | YNL040W | YNL040W | | n |
| Kmar_QGN16395.1 | W303.asm02.HP0_G0000150 | OG0004347 | chrI | 38099 | 39607 | CP015058.1 | 21947 | 23827 | 28.39116719 | 502 | 626 | 0.801916933 | 1 | 0 | 1 | FIM1_3108 | YAL058W | YAL058W | CNE1 | n |
| Kmar_QGN14037.1 | W303.asm02.HP0_G0048470 | OG0004356 | chrXV | 207846 | 210719 | CP015054.1 | 1384924 | 1387320 | 19.58333333 | 957 | 798 | 1.19924812 | 1 | 0 | 1 | FIM1_688 | YOL063C | YOL063C | CRT10 | n |
| Kmar_QGN13407.1 | W303.asm02.HP0_G0054470 | OG0004378 | chrXVI | 372887 | 373093 | CP015054.1 | 78916 | 79131 | 32.3943662 | 68 | 71 | 0.957746479 | 1 | 0 | 1 | FIM1_44 | YPL096C-A | YPL096C-A | ERI1 | n |
| Kmar_QGN13668.1 | W303.asm02.HP0_G0048870 | OG0004380 | chrXV | 286627 | 288405 | CP015054.1 | 630201 | 631976 | 70.10135135 | 592 | 591 | 1.001692047 | 1 | 0 | 1 | FIM1_311 | YOL020W | YOL020W | TAT2 | n |
| Kmar_QGN13899.1 | W303.asm02.HP0_G0042550 | OG0004381 | chrXIII | 697376 | 698950 | CP015054.1 | 1107877 | 1109388 | 63.93129771 | 524 | 503 | 1.041749503 | 1 | 0 | 1 | FIM1_546 | YMR215W | YMR215W | GAS3 | n |
| Kmar_QGN13949.1 | W303.asm02.HP0_G0040530 | OG0004383 | chrXIII | 273624 | 274220 | CP015054.1 | 1204210 | 1204983 | 22.96296296 | 198 | 257 | 0.770428016 | 1 | 0 | 1 | FIM1_597 | YMR003W | YMR003W | AIM34 | n |
| Kmar_QGN14127.1 | W303.asm02.HP0_G0013420 | OG0004386 | chrIV | 1460736 | 1461560 | CP015054.1 | 1593215 | 1594048 | 36.42857143 | 274 | 277 | 0.989169675 | 1 | 0 | 1 | FIM1_780 | YDR503C | YDR503C | LPP1 | n |
| Kmar_QGN14130.1 | W303.asm02.HP0_G0013450 | OG0004387 | chrIV | 1465422 | 1467248 | CP015054.1 | 1600083 | 1602017 | 34.32392273 | 608 | 644 | 0.944099379 | 1 | 0 | 1 | FIM1_783 | YDR506C | YDR506C | GMC1 | n |
| Kmar_QGN14160.1 | W303.asm02.HP0_G0013570 | OG0004388 | chrIV | 1486778 | 1489096 | CP015054.1 | 1655770 | 1657710 | 45.73643411 | 772 | 646 | 1.19504644 | 1 | 0 | 1 | FIM1_813 | YDR520C | YDR520C | URC2 | n |
| Kmar_QGN14241.1 | W303.asm02.HP0_G0019660 | OG0004390 | chrVII | 445850 | 447145 | CP015055.1 | 94230 | 95216 | 23.61111111 | 431 | 328 | 1.31402439 | 1 | 0 | 1 | FIM1_897 | YGL025C | YGL025C | PGD1 | n |
| Kmar_QGN14314.1 | W303.asm02.HP0_G0037100 | OG0004391 | chrXII | 591250 | 592731 | CP015055.1 | 258677 | 259774 | 18.38383838 | 493 | 365 | 1.350684932 | 1 | 0 | 1 | FIM1_971 | YLR227C | YLR227C | ADY4 | n |
| Kmar_QGN14773.1 | W303.asm02.HP0_G0012350 | OG0004397 | chrIV | 1245553 | 1246269 | CP015055.1 | 1203906 | 1204538 | 21.25 | 238 | 210 | 1.133333333 | 1 | 0 | 1 | FIM1_1443 | YDR383C | YDR383C | NKP1 | n |
| Kmar_QGN14792.1 | W303.asm02.HP0_G0012240 | OG0004398 | chrIV | 1230350 | 1230619 | CP015055.1 | 1241335 | 1241595 | 22.34042553 | 89 | 86 | 1.034883721 | 1 | 0 | 1 | FIM1_1463 | YDR374W-A | YDR374W-A | WIP1 | n |
| Kmar_QGN14879.1 | W303.asm02.HP0_G0035090 | OG0004399 | chrXII | 184192 | 184806 | CP015055.1 | 1410914 | 1411525 | 51.2195122 | 204 | 203 | 1.004926108 | 1 | 0 | 1 | FIM1_1552 | YLR016C | YLR016C | PML1 | n |
| Kmar_QGN15014.1 | W303.asm02.HP0_G0047280 | OG0004402 | chrXIV | 744185 | 745870 | CP015055.1 | 1691081 | 1692745 | 45.75389948 | 561 | 554 | 1.012635379 | 1 | 0 | 0 | FIM1_1695 | YNR056C | YNR056C | BIO5 | n |
| Kmar_QGN15125.1 | W303.asm02.HP0_G0041520 | OG0004404 | chrXIII | 493029 | 494417 | CP015056.1 | 255587 | 256837 | 31.62393162 | 462 | 416 | 1.110576923 | 1 | 0 | 1 | FIM1_1812 | YMR111C | YMR111C | EUC1 | n |
| Kmar_QGN15136.1 | W303.asm02.HP0_G0031930 | OG0004405 | chrXI | 186107 | 186316 | CP015056.1 | 270990 | 271184 | 37.31343284 | 69 | 64 | 1.078125 | 1 | 0 | 1 | FIM1_1823 | YKL138C-A | YKL138C-A | HSK3 | n |
| Kmar_QGN15157.1 | W303.asm02.HP0_G0018150 | OG0004406 | chrVII | 135232 | 135474 | CP015056.1 | 324081 | 324311 | 24.09638554 | 80 | 76 | 1.052631579 | 1 | 0 | 1 | FIM1_1844 | YGL194C-A | YGL194C-A | | n |
| Kmar_QGN15179.1 | W303.asm02.HP0_G0033480 | OG0004407 | chrXI | 491864 | 492832 | CP015056.1 | 376668 | 377588 | 22.15568862 | 322 | 306 | 1.052287582 | 1 | 0 | 1 | FIM1_1867 | YKR022C | YKR022C | NTR2 | n |
| Kmar_QGN15347.1 | W303.asm02.HP0_G0011690 | OG0004408 | chrIV | 1112250 | 1113494 | CP015056.1 | 760176 | 761306 | 20.75055188 | 414 | 376 | 1.10106383 | 1 | 0 | 1 | FIM1_2037 | YDR317W | YDR317W | HIM1 | n |
| Kmar_QGN15356.1 | W303.asm02.HP0_G0003500 | OG0004409 | chrII | 570320 | 571087 | CP015056.1 | 782346 | 783809 | 17.19745223 | 255 | 487 | 0.523613963 | 1 | 0 | 1 | FIM1_2046 | YBR157C | YBR157C | ICS2 | n |
| Kmar_QGN15420.1 | W303.asm02.HP0_G0011370 | OG0004410 | chrIV | 1036935 | 1039562 | CP015056.1 | 919303 | 921591 | 27.22095672 | 875 | 762 | 1.148293963 | 1 | 0 | 1 | FIM1_2110 | YDR285W | YDR285W | ZIP1 | n |
| Kmar_QGN15706.1 | W303.asm02.HP0_G0017340 | OG0004411 | chrVI | 207362 | 207964 | CP015056.1 | 1505734 | 1506354 | 49.51456311 | 200 | 206 | 0.970873786 | 1 | 0 | 1 | FIM1_2399 | YFR042W | YFR042W | KEG1 | n |
| Kmar_QGN15715.1 | W303.asm02.HP0_G0017380 | OG0004412 | chrVI | 211979 | 213064 | CP015056.1 | 1522266 | 1523060 | 16.02209945 | 361 | 264 | 1.367424242 | 1 | 0 | 1 | FIM1_2408 | YFR046C | YFR046C | CNN1 | n |
| Kmar_QGN15812.1 | W303.asm02.HP0_G0035320 | OG0004413 | chrXII | 236721 | 237395 | CP015057.1 | 139472 | 140083 | 33.48214286 | 224 | 203 | 1.103448276 | 1 | 0 | 1 | FIM1_2508 | YLR040C | YLR040C | AFB1 | n |
| Kmar_QGN15868.1 | W303.asm02.HP0_G0002530 | OG0004414 | chrII | 372611 | 372853 | CP015057.1 | 264468 | 264722 | 39.28571429 | 80 | 84 | 0.952380952 | 1 | 0 | 1 | FIM1_2565 | YBR058C-A | YBR058C-A | TSC3 | n |
| Kmar_QGN15886.1 | W303.asm02.HP0_G0002680 | OG0004415 | chrII | 398317 | 398961 | CP015057.1 | 294565 | 295146 | 52.13270142 | 214 | 193 | 1.10880829 | 1 | 0 | 1 | FIM1_2584 | YBR072W | YBR072W | HSP26 | n |
| Kmar_QGN15908.1 | W303.asm02.HP0_G0009830 | OG0004416 | chrIV | 695923 | 696123 | CP015057.1 | 356010 | 356240 | 27.14285714 | 66 | 76 | 0.868421053 | 1 | 0 | 1 | FIM1_2606 | YDR119W-A | YDR119W-A | COX26 | n |
| Kmar_QGN16059.1 | W303.asm02.HP0_G0032560 | OG0004418 | chrXI | 317682 | 317918 | CP015057.1 | 723692 | 723931 | 27.84810127 | 78 | 79 | 0.987341772 | 1 | 0 | 1 | FIM1_2759 | YKL068W-A | YKL068W-A | | n |
| Kmar_QGN16254.1 | W303.asm02.HP0_G0030840 | OG0004421 | chrX | 682763 | 683392 | CP015057.1 | 1137300 | 1137944 | 75.23364486 | 209 | 214 | 0.976635514 | 1 | 0 | 1 | FIM1_2959 | YJR133W | YJR133W | XPT1 | n |
| Kmar_QGN16387.1 | W303.asm02.HP0_G0000170 | OG0004422 | chrI | 42812 | 43354 | CP015058.1 | 6091 | 6513 | 28.17679558 | 180 | 140 | 1.285714286 | 1 | 0 | 1 | FIM1_3100 | YAL055W | YAL055W | PEX22 | n |
| Kmar_QGN16625.1 | W303.asm02.HP0_G0028210 | OG0004424 | chrX | 106772 | 107314 | CP015058.1 | 525998 | 526525 | 26.04166667 | 180 | 175 | 1.028571429 | 1 | 0 | 1 | FIM1_3342 | YJL161W | YJL161W | FMP33 | n |
| Kmar_QGN16638.1 | W303.asm02.HP0_G0027930 | OG0004425 | chrX | 60897 | 62105 | CP015058.1 | 564399 | 565541 | 46.58536585 | 402 | 380 | 1.057894737 | 1 | 0 | 1 | FIM1_3357 | YJL193W | YJL193W | | n |
| Kmar_QGN16682.1 | W303.asm02.HP0_G0036800 | OG0004426 | chrXII | 540937 | 541701 | CP015058.1 | 662454 | 663044 | 27.95275591 | 254 | 196 | 1.295918367 | 1 | 0 | 1 | FIM1_3401 | YLR194C | YLR194C | NCW2 | n |
| Kmar_QGN16879.1 | W303.asm02.HP0_G0010220 | OG0004428 | chrIV | 785137 | 785664 | CP015058.1 | 1091739 | 1092329 | 32 | 175 | 196 | 0.892857143 | 1 | 0 | 1 | FIM1_3606 | YDR163W | YDR163W | CWC15 | n |
| Kmar_QGN17039.1 | W303.asm02.HP0_G0036980 | OG0004429 | chrXII | 568691 | 570751 | CP015059.1 | 98416 | 100500 | 40.68278805 | 686 | 694 | 0.988472622 | 1 | 0 | 1 | FIM1_3768 | YLR214W | YLR214W | FRE1 | n |
| Kmar_QGN17057.1 | W303.asm02.HP0_G0007060 | OG0004430 | chrIV | 125690 | 126523 | CP015059.1 | 140682 | 141410 | 25.70422535 | 277 | 242 | 1.144628099 | 1 | 0 | 1 | FIM1_3787 | YDL186W | YDL186W | | n |
| Kmar_QGN17073.1 | W303.asm02.HP0_G0007150 | OG0004431 | chrIV | 142317 | 144443 | CP015059.1 | 173492 | 175543 | 30.45897079 | 708 | 683 | 1.036603221 | 1 | 0 | 1 | FIM1_3804 | YDL176W | YDL176W | GID12 | n |
| Kmar_QGN17133.1 | W303.asm02.HP0_G0024650 | OG0004433 | chrVIII | 380627 | 381073 | CP015059.1 | 306294 | 307067 | 21.6 | 148 | 257 | 0.575875486 | 1 | 0 | 1 | FIM1_3864 | YHR136C | YHR136C | SPL2 | n |
| Kmar_QGN17151.1 | W303.asm02.HP0_G0017640 | OG0004434 | chrVII | 28468 | 30582 | CP015059.1 | 345198 | 347291 | 21.17486339 | 704 | 697 | 1.010043042 | 1 | 0 | 1 | FIM1_3882 | YGL249W | YGL249W | ZIP2 | n |
| Kmar_QGN17194.1 | W303.asm02.HP0_G0019170 | OG0004435 | chrVII | 354329 | 355291 | CP015059.1 | 426541 | 427524 | 19.88130564 | 320 | 327 | 0.978593272 | 1 | 0 | 1 | FIM1_3925 | YGL081W | YGL081W | | n |
| Kmar_QGN17196.1 | W303.asm02.HP0_G0052650 | OG0004436 | chrXV | 1076493 | 1078367 | CP015059.1 | 429540 | 431432 | 39.31357254 | 624 | 630 | 0.99047619 | 1 | 0 | 0 | FIM1_3927 | YOR389W | YOR389W | | n |
| Kmar_QGN17326.1 | W303.asm02.HP0_G0050120 | OG0004438 | chrXV | 538445 | 539329 | CP015059.1 | 700291 | 701124 | 30.26315789 | 294 | 277 | 1.061371841 | 1 | 0 | 1 | FIM1_4058 | YOR114W | YOR114W | CIM1 | n |
| Kmar_QGN17482.1 | W303.asm02.HP0_G0029830 | OG0004439 | chrX | 465947 | 466342 | CP015059.1 | 1009535 | 1009831 | 24.24242424 | 131 | 98 | 1.336734694 | 1 | 0 | 1 | FIM1_4218 | W303.asm02.HP0_G0029830 | YJR021C | REC107 | y |
| Kmar_QGN17585.1 | W303.asm02.HP0_G0052620 | OG0004440 | chrXV | 1069120 | 1070817 | CP015060.1 | 43539 | 45257 | 45.7337884 | 565 | 572 | 0.987762238 | 1 | 0 | 1 | FIM1_4792 | YOR386W | YOR386W | PHR1 | n |
| Kmar_QGN17591.1 | W303.asm02.HP0_G0052830 | OG0004441 | chrXVI | 38721 | 39350 | CP015060.1 | 59537 | 60229 | 32.05128205 | 209 | 230 | 0.908695652 | 1 | 0 | 1 | FIM1_4798 | YPL267W | YPL267W | ACM1 | n |
| Kmar_QGN17797.1 | W303.asm02.HP0_G0049970 | OG0004442 | chrXV | 507713 | 508240 | CP015060.1 | 506453 | 506746 | 14.60674157 | 175 | 97 | 1.804123711 | 1 | 0 | 1 | FIM1_5006 | YOR097C | YOR097C | | n |
| Kmar_QGN18075.1 | W303.asm02.HP0_G0001500 | OG0004443 | chrII | 145032 | 145211 | CP015061.1 | 158162 | 158389 | 38.66666667 | 59 | 75 | 0.786666667 | 1 | 0 | 1 | FIM1_4393 | YBL039W-B | YBL039W-B | MIN6 | n |
| Kmar_QGN18145.1 | W303.asm02.HP0_G0043850 | OG0004444 | chrXIV | 44879 | 45646 | CP015061.1 | 320636 | 321454 | 21.24542125 | 255 | 272 | 0.9375 | 1 | 0 | 1 | FIM1_4468 | YNL314W | YNL314W | DAL82 | n |
| Kmar_QGN18365.1 | W303.asm02.HP0_G0042110 | OG0004446 | chrXIII | 609273 | 610565 | CP015061.1 | 786222 | 786443 | 15.27093596 | 430 | 73 | 5.890410959 | 1 | 0 | 1 | FIM1_4692 | YMR173W | YMR173W | DDR48 | n |
| Kmar_QGN18410.1 | W303.asm02.HP0_G0055450 | OG0004447 | chrXVI | 573626 | 575353 | CP015061.1 | 875080 | 876798 | 63.25823224 | 575 | 572 | 1.005244755 | 1 | 0 | 1 | FIM1_4738 | YPR006C | YPR006C | ICL2 | n |
| Kmar_QGN18423.1 | W303.asm02.HP0_G0055410 | OG0004448 | chrXVI | 564742 | 566292 | CP015061.1 | 900568 | 902154 | 71.40151515 | 516 | 528 | 0.977272727 | 1 | 0 | 1 | FIM1_4751 | YPR002W | YPR002W | PDH1 | n |
| Kmar_QGN17787.1 | W303.asm02.HP0_G0045860 | OG0004537 | chrXIV | 452498 | 453700 | CP015060.1 | 483300 | 484514 | 40.93137255 | 400 | 404 | 0.99009901 | 1 | 0 | 1 | FIM1_4996 | YNL092W | YNL092W | | n |
| Kmar_QGN14199.1 | W303.asm02.HP0_G0017570 | OG0004576 | chrVII | 10526 | 11674 | CP015055.1 | 17392 | 18648 | 65.07177033 | 382 | 418 | 0.913875598 | 1 | 0 | 1 | FIM1_855 | YGL256W | YGL256W | ADH4 | n |
| Kmar_QGN15320.1 | W303.asm02.HP0_G0037790 | OG0004581 | chrXII | 742105 | 742368 | CP015056.1 | 697436 | 697684 | 35.63218391 | 87 | 82 | 1.06097561 | 1 | 0 | 1 | FIM1_2009 | YLR307C-A | YLR307C-A | DPA10 | n |
| Kmar_QGN15632.1 | W303.asm02.HP0_G0033320 | OG0004585 | chrXI | 458089 | 459735 | CP015056.1 | 1350710 | 1351810 | 18.40607211 | 525 | 366 | 1.43442623 | 1 | 0 | 1 | FIM1_2325 | YKR005C | YKR005C | | n |
| Kmar_QGN17171.1 | W303.asm02.HP0_G0055010 | OG0004598 | chrXVI | 485585 | 486535 | CP015059.1 | 380998 | 381918 | 23.88888889 | 316 | 306 | 1.032679739 | 1 | 0 | 1 | FIM1_3902 | YPL039W | YPL039W | | n |
| Kmar_QGN17995.1 | W303.asm02.HP0_G0047340 | OG0004603 | chrXIV | 759515 | 761338 | CP015060.1 | 926009 | 927784 | 47.24025974 | 607 | 591 | 1.027072758 | 1 | 0 | 1 | FIM1_5205 | YNR063W | YNR063W | PUL4 | n |
| Kmar_QGN17996.1 | W303.asm02.HP0_G0047330 | OG0004604 | chrXIV | 756925 | 758357 | CP015060.1 | 928698 | 930083 | 60.38961039 | 450 | 461 | 0.976138829 | 1 | 0 | 1 | FIM1_5206 | YNR062C | YNR062C | PUL3 | n |
| Kmar_QGN18337.1 | W303.asm02.HP0_G0001640 | OG0004606 | chrII | 171948 | 172184 | CP015061.1 | 720349 | 720582 | 35.8974359 | 78 | 77 | 1.012987013 | 1 | 0 | 1 | FIM1_4664 | NA | NA | NA | NA |
| Kmar_QGN13685.1 | W303.asm02.HP0_G0031030 | OG0004735 | chrX | 732638 | 733723 | CP015054.1 | 660344 | 661432 | 58.51648352 | 361 | 362 | 0.997237569 | 1 | 0 | 1 | FIM1_328 | YJR153W | YJR153W | PGU1 | n |
| Kmar_QGN13785.1 | W303.asm02.HP0_G0034080 | OG0004736 | chrXI | 616876 | 617213 | CP015054.1 | 878375 | 878653 | 50 | 98 | 92 | 1.065217391 | 1 | 0 | 1 | FIM1_431 | W303.asm02.HP0_G0034080 | YKR084C | HBS1 | y |
| Kmar_QGN17857.1 | W303.asm02.HP0_G0015290 | OG0004763 | chrV | 314260 | 315168 | CP015060.1 | 626435 | 627343 | 20.19230769 | 302 | 302 | 1 | 1 | 0 | 1 | FIM1_5066 | YER076C | YER076C | | n |
